# Deflating the RNA Mg^2+^ bubble. Stereochemistry to the rescue!

**DOI:** 10.1101/2020.04.14.023697

**Authors:** Pascal Auffinger, Eric Ennifar, Luigi D’Ascenzo

## Abstract

Proper evaluation of the ionic structure of biomolecular systems remains challenging in X-ray and cryo-EM techniques but is essential for advancing our understanding of complex structure/activity/solvent relationships. However, numerous studies overestimate the number of Mg^2+^ in the deposited structures and underrate the importance of stereochemical rules to correctly assign these ions. Herein, we re-evaluate the PDBid 6QNR and 6SJ6 models of the ribosome ionic structure and establish that stereochemical principles should always be considered when evaluating ion binding features, even when K^+^ anomalous signals are available as it is the case for 6QNR. Assignment errors can result in misleading conceptions of the solvent structure of ribosomes and other RNA systems and should therefore be avoided. Our analysis resulted in a significant decrease of bound Mg^2+^ ions in the 6QNR structure, suggesting that K^+^ and not Mg^2+^ is the prevalent ion in the ribosome 1^st^ solvation shell. We stress that the use of proper stereochemical guidelines is critical for deflating the current Mg^2+^ bubble witnessed in many ribosome and other RNA structures. Herewith, we would like to draw the attention of the researchers interested in the ionic structure of biomolecular systems on the importance and complementarity of stereochemistry and other ion identification techniques such as those pertaining to the detection of anomalous signals of transition metals and K^+^. We also stress that for the identification of lighter ions such as Mg^2+^, Na^+^, …, stereochemistry coupled with high resolution structures remain the best currently available option.

---

A recent publication (Rozov et al. 2019) reported that long-wavelength X-ray diffraction techniques can now be used to detect the presence of potassium ions (K^+^) in large ribosome structures counting >150.000 heavy atoms. The crystallographic experiments leading to that result involved the detection of the anomalous signal of this biologically essential cation (Epstein 2003; Nierhaus 2014; Auffinger et al. 2016; Danchin and Nikel 2019) through the use of the dedicated I23 *in-vacuum* long-wavelength beamline at the Diamond Light Source facility (Wagner et al. 2016). As a result and in better agreement with experimental coordination distances and geometries, some electron density patterns that were previously attributed to Mg^2+^ (Jenner et al. 2010) could now be associated with K^+^; Interestingly, similar results were described earlier solely based on the use of stereochemical data (Leonarski et al. 2017; Leonarski et al. 2019). Precisely, the authors focused on the role of this monovalent ion at the ribosomal decoding center showing that K^+^ acts as a structural link between the protein uS12, the mRNA codon and the rRNA.

Precise ion identification is critical in biomolecular systems since attribution errors can affect over many years our understanding of the investigated systems (Grime et al. 2020). For instance, the modeling of a weakly coordinating K^+^ (chaotropic) versus a more organizing and stabilizing Mg^2+^ (kosmotropic) ion (Auffinger et al. 2003; Auffinger et al. 2004b; Ball and Hallsworth 2015; Danchin and Nikel 2019) at the decoding center or elsewhere in ribosomes and other RNA systems has profound implications on how we picture the internal dynamics, folding and assembly of these particles. In the ribosomal decoding center where mobility of the different partners is key to the translational process, a monovalent ion should facilitate rather than hinder the shift of tRNAs and mRNAs from one binding site to the other. This can be understood if we consider the respective residency times of water molecules coordinating to K^+^ (<10 ps) and Mg^2+^ (≈µs) in aqueous solution (Bleuzen et al. 1997; Joung and Cheatham 2009). The residency times of Mg^2+^ bound to two or three phosphate groups as observed in the rRNA core (Klein et al. 2004) may even be longer and possibly equal the lifetime of the folded ribosomal subunits. Such, often conserved, Mg^2+^ binding locations are essential for maintaining a stable and functional 3D structure as described for the ribosomal peptidyl transferase center (Hsiao and Williams 2009; Bowman et al. 2012; Bowman et al. 2018; Liebschner et al. 2019). By contrast, the presence of Mg^2+^ in the decoding center is unlikely, given the Mg^2+^ potential for “gripping” the translational machinery by establishing disproportionately strong contacts between uS12, the mRNA codon and the rRNA partners. At such non-rigid locations, the presence of a less structuring ion, such as K^+^, is more probable (Leonarski et al. 2019; Rozov et al. 2019).

The dynamical differences between bound K^+^ and Mg^2+^ mentioned above are deeply rooted in the ion stereochemical features that have been repeatedly described in the literature (Harding et al. 2010; Bowman et al. 2012; Zheng et al. 2014; Leonarski et al. 2017; Bowman et al. 2018; Leonarski et al. 2019). In short, the main distinctive character of these ions, besides their charge, resides in their idiosyncratic coordination distances (Mg^2+^: ≈2.1 Å; K^+^: ≈2.8 Å; **Figure 1**). Such differences make them difficult to misidentify even in medium resolution structures (≈3.0 Å). The second major discriminating factor is related to the strict octahedral coordination of Mg^2+^ that is rarely adopted by K^+^. The latter ion, given its larger ionic radius, prefers binding modes with coordination numbers of 7 to 8, such as for example square anti-prismatic —see UCSF Chimera definition (Pettersen et al. 2004). These binding modes are recurrently observed in potassium channels, nucleic acid quadruplexes and other RNA systems (Zhou et al. 2001; Auffinger et al. 2016; Leonarski et al. 2017).

**Figure 1.**
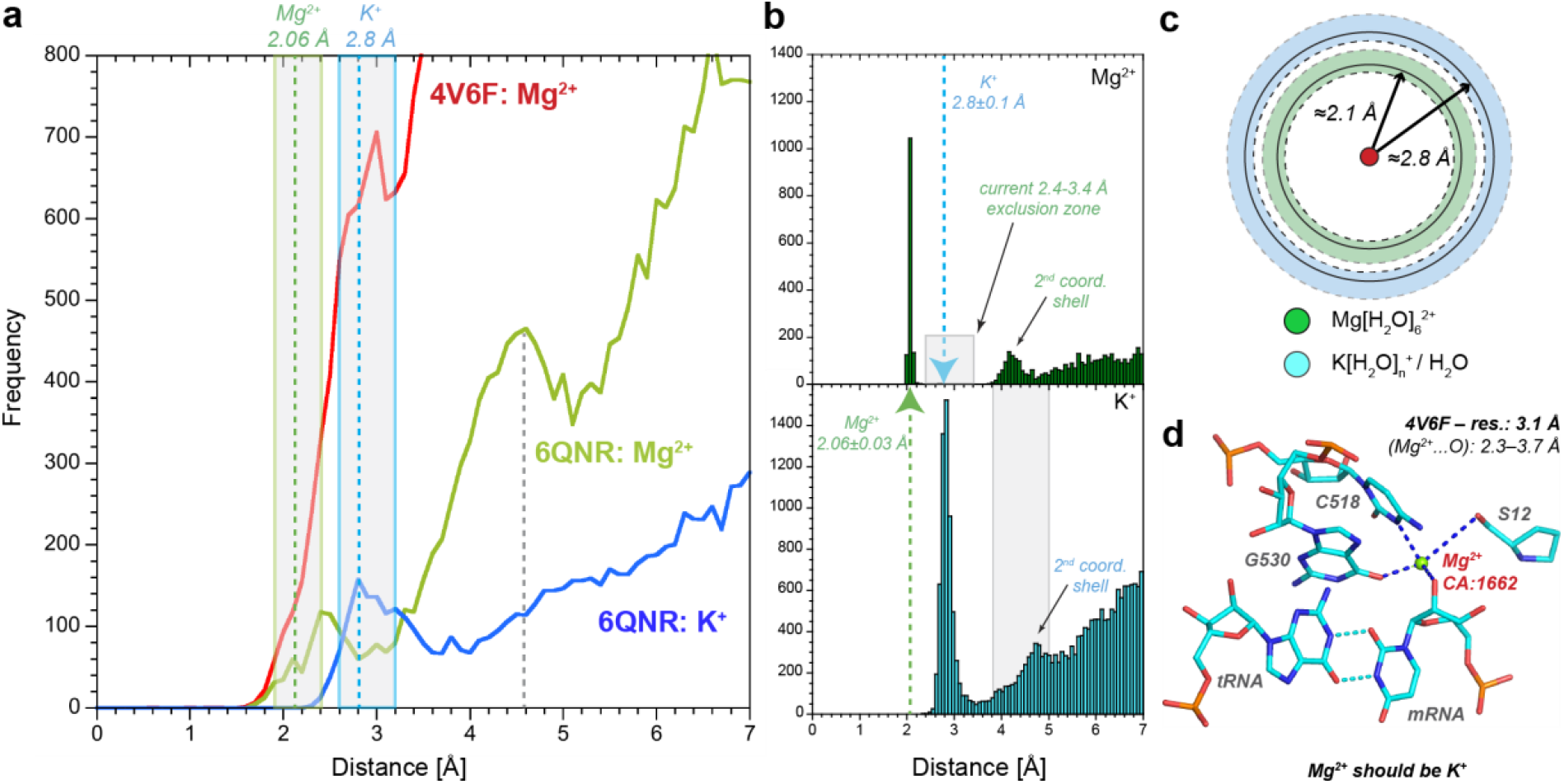
Mg^2+^ and K^+^ coordination distances to oxygen atoms in 4V6F, 6QNR and the CSD. **a** The red, green and blue curves show the number of Mg^2+^ and K^+^ bound to rRNA and r-protein oxygen atoms in the 4V6F (Jenner et al. 2010) and 6QNR (Rozov et al. 2019) structures. Water oxygens are excluded. The colored areas and the green and blue dashed lines highlight two regions where Mg^2+^ (1.9–2.4 Å; favored: 2.06 Å) and K^+^ (2.6–3.2 Å; favored: 2.8 Å) should be present. The large 2^nd^ peak in the green curve (≈4.6 Å) appears in a region where a precise ion assignment is problematic since it could correspond to the 2^nd^ coordination shell of Mg^2+^ or to other solvent molecules, including K^+^. **b** Cambridge Structural Database (CSD; Taylor and Wood 2019) histograms showing the Mg^2+^…H_2_O (green) and K^+^…O (cyan) coordination distances. The 2^nd^ coordination shell for K^+^ and a H_2_O “exclusion zone” for Mg^2+^, where no oxygen atom is expected (grey areas). **c** Schematic representation of the respective 2.1 and 2.8 Å average coordination distances and 1.9–2.4 and 2.6– Å coordination range for Mg^2+^ and K^+^ (see panel **b**). The white area between the green and blue segments (2.4–2.6 Å range) points to improper coordination distances for both Mg^2+^ and K^+^. **d** View of the Mg^2+^ that has been assigned in the 4V6F decoding center (Rozov et al. 2016). An examination of the ion binding features suggests that this ion cannot be Mg^2+^ but is K^+^ as assigned in 6QNR (Rozovet al. 2019) and by us in 5E81 (Leonarski et al. 2019). Density patterns for 6QNR and 5E81 are shown in **Figure S1**.

K^+^ and Mg^2+^ display also distinctive coordination preferences that need to be taken into account for a reliable ion assignment (Leonarski et al. 2017; Leonarski et al. 2019). The primary ligands for monovalent ions in nucleic acids are nucleobase carbonyl groups (Leonarski et al. 2019). As such, the major groove of G•U pairs has persistently been found to constitute an excellent binding location for K^+^ but not for Mg^2+^ ions (Klein et al. 2004; Fan et al. 2005; Wang et al. 2009; Leonarski et al. 2019; Westhof et al. 2019). On the other hand, Mg^2+^ prefers binding to phosphate and carboxyl groups and binds much more rarely to nitrogen atoms or to carbonyl, ester and hydroxyl oxygen atoms (Hsiao et al. 2009; Leonarski et al. 2017; Bowman et al. 2018; Leonarski et al. 2019).

To advance our comprehension of ion binding features and functions in biomolecular systems it is essential to enforce the use of stereochemical rules (Wlodawer 2017) in order to avoid false identifications that would jeopardize our ability to discriminate weakly from strongly coordinating ions and, therefore, affect our understanding of the local structure and dynamics of ribosomal systems. Unfortunately, the assignment of ions in a significant number of biomolecular structures has been based on approximate stereochemical knowledge leading to the deposition in structural databases of locally misinterpreted structures (Williams 2005; Minor et al. 2016; Leonarski et al. 2017; Zheng et al. 2017; Handing et al. 2018; Leonarski et al. 2019; Grime et al. 2020). In order to illustrate important drawbacks associated with non-stereochemically based ion assignments in ribosomal structures, we chose to reexamine the recent 70S elongation complex structure mentioned above (PDBid: 6QNR) that is grounded on the detection of K^+^ anomalous signals. Although the monovalent ion assignments were based on experimental evidence, this was not the case for the Mg^2+^ assignments for which the very weak anomalous signals are currently beyond detection limit (Liu et al. 2013). However, we found a significant number of Mg^2+^ misassignments in 6QNR that question some aspects of the ribosome ionic structure. Such misassignments could have been avoided, in part, by a careful examination of the ion stereochemistry and that of their hydration shell (Leonarski et al. 2019). We believe that it is appropriate and timely to discuss these issues by taking as an example the 6QNR study that is devoted to the analysis of the ionic structure of a *Thermus thermophilus* ribosome where Mg^2+^ misassignments could result in more general interpretation approximations. For instance, based on this structure, one could conclude that Mg^2+^ (1005 assigned ions) and not K^+^ (367 assigned ions) dominates the ribosomal ionic landscape. We note that this study along with that devoted to the first *Haloarcula marismortui* 50S subunit structure (Klein et al. 2004) are the only studies we are aware of that discuss in detail the ribosome ionic structure.

Herein, we provide evidence that the 6QNR ion assignment needs revision and propose that K^+^ and not Mg^2+^ is the prevalent ion in the 1^st^ solvation shell of ribosomes. We also suggest that, in the absence of K^+^ anomalous signals, numerous early incorrect Mg^2+^ assignments in the 4V6F structure (Jenner et al. 2010) could have been avoided by applying appropriate stereochemical rules.

Regarding the K^+^ assignments in 6QNR, a rapid screen of the 3.1 Å resolution structure (**Figure 1a**) shows, as expected, a 1^st^ broad peak centered around 2.8 Å that is indicative of K^+^ coordination. Thus, none of the solvent density peaks with coordination distances around 2.8 Å should have been associated with Mg^2+^ in the initially released 4V6F structure that embeds 5345 assigned Mg^2+^ ions (Jenner et al. 2010). As an illustration, Figure 2 of the Rozov et al. paper, Figure 8 of a related publication that analyzed the parent 5E81 structure (Leonarski et al. 2019) and **Figures 1d/S1** show that stereochemical considerations should have been sufficient to discard an Mg^2+^ assignment for density peaks close to the decoding center and, instead, should have led to the more appropriate placement of K^+^. The assignment of other solvent particles with coordination distances around 2.8 Å is easily discarded since water molecules are usually not observed in structures with resolutions >3.0 Å and would not display high density peaks characteristic of K^+^. Furthermore, in the aforementioned figures, it is clear that the ion coordination geometry is not compatible with a water molecule while the observed non-octahedral coordination pattern disqualifies Mg^2+^ immediately. A tetrahedral NH_4_^+^ establishing bifurcated hydrogen bonds could eventually fit into such a binding pocket, although this seems unlikely since the small density peak associated with this ion would probably be buried into the noise when compared to a significantly denser K^+^ peak. Moreover, it has to be kept in mind that NH_4_^+^ is toxic to mammals (Dasarathy et al. 2017), suggesting that this ion has probably not been selected to be present in the universally conserved decoding center. However, its presence in *in crystallo* conditions cannot be excluded.

**Figure 2.**
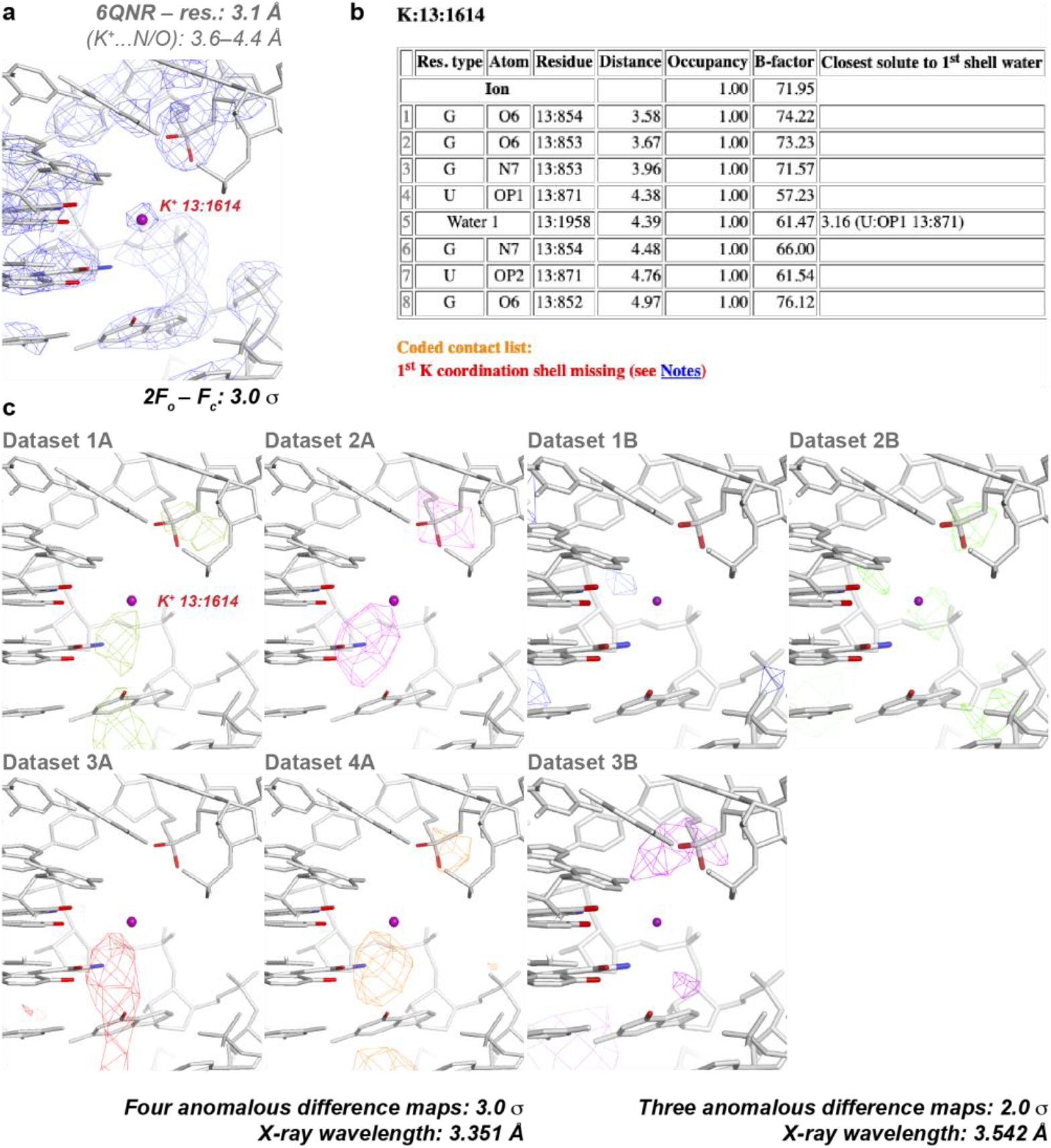
Comparison between 2F_o_-F_c_ and anomalous difference maps around a K^+^ with no 1^st^ coordination shell. **a** 2F_o_-F_c_ density pattern at 3.0 σ. The closest rRNA oxygen and nitrogen atoms are outside the 1^st^ coordination shell of K^+^ (2.8–3.4 Å; **Figure 1b**). **b** Extract of the 6QNR ‘in-house’ diagnosis program output that lists the closest contact distances to the ion shown in **a** —the water molecule mentioned in the table and belonging to a Mg(H_2_O)_6_^2+^ is not shown. **c** Four and three density patterns (four different crystals have been analyzed that differ from the one used to calculate the density maps shown in panel **a**) derived from the 6QNR anomalous diffraction data at the 3.351 and 3.542 Å wavelengths, respectively (Rozov et al. 2019). Note that for K^+^, f≈4.0 e^-^ and f≈0.5 e^-^ at the 3.351 and 3.542 Å wavelengths, respectively. Therefore, a K^+^ anomalous signal should be visible in the “A” dataset but absent in the “B” dataset. The fact that none of the anomalous difference densities overlap with the original assignment underlines the ambiguity of such data and the possibility that the four new and non-isomorphous crystals locally differ from the original crystal obtained 10 years earlier in slightly different conditions even if similar crystallization protocols were used. Note the probable presence of an anomalous signal for a phosphorus atom in crystals 1, 2, and 4 as noted by Rozov et al. (f”=1.75 e^-^ for P at a wavelength of 3.351 Å) and the large amount of densities not directly related to the presence of K^+^.

These observations challenge an assumption made by Rozov et al. suggesting that, at resolutions around 3.0 Å, it is almost impossible to distinguish K^+^ from Mg^2+^ based on average M^n+^…N/O coordination distances. Rather, we suggest that stereochemically based K^+^ and Mg^2+^ assignments, given the large 0.7 Å difference in coordination distance, are possible, provided B-factors are in a reasonable range. To assess our point, we visualized all 6QNR ions by using the Coot and UCSF Chimera programs (Pettersen et al. 2004; Emsley et al. 2010) and checked difference and anomalous maps. We also developed an ‘in-house’ analysis program to screen the stereochemistry of the assigned ions (Leonarski & Auffinger, unpublished). These combined checks confirmed that close to none of the assigned K^+^ in 6QNR could have been mistaken for Mg^2+^ given coordination distances around 2.8 Å and the absence of associated octahedral coordination patterns.

Yet, we isolated a few borderline K^+^ assignments in 6QNR. For instance, 12 K^+^ have no 1^st^ coordination shell, i.e., no ligand coordination distances below 3.4 Å. More precisely, for some density patterns associated with these ions, the presence of an anomalous signal not overlapping with the ion position in the 2F_o_-F_c_ map does not warrant a clear-cut K^+^ assignment (**Figure 2**). Instead, it suggests the occurrence of local structural differences in the solvent and in the ion binding site organization between the 10 years old original crystals and the 2019 batch of four crystals used for the anomalous signal detection, all of these crystals being non-isomorphous (Rozov et al. 2019). As such, the cell dimensions of the four crystals used for gathering the anomalous signals and those of the original 4V6F differ significantly, forbidding multi-crystal averaging. In such instances, we suggest that when the anomalous signal obtained from different crystals does not comply to the expected stereochemistry it should not be used to make an assignment that may be subsequently used as a reference.

While the main focus of the authors that deposited the 6QNR structure was the identification of 367 K^+^, the authors revised also the placement of Mg^2+^ that resulted in the assignment of 1005 Mg^2+^ in 6QNR versus 5345 Mg^2+^ in the original 4V6F structure (Jenner et al. 2010). As for now, it has been almost impossible, besides a single documented exception, to collect anomalous signals for Mg^2+^ in macromolecular systems (Liu et al. 2013). Therefore, precise Mg^2+^ assignments necessitate a resolution around 2.4 Å or better, reasonably low B-factors and appropriate modeling that allows the observation of a well-defined octahedral coordination shell with coordination distances ≈2.1 Å (**Figure S2**). In 6QNR, no Mg^2+^ octahedral density patterns were observed (**Figure 3c**). Nonetheless, a hydration sphere has been modeled for 393 out of 1005 Mg^2+^. Regrettably, these hydration spheres generate a large number of clashes involving Mg^2+^ coordinating water molecules as listed in both the publicly available PDB validation report and the output of our analysis program (**Supplementary Information**). A rapid screen of the water oxygen atom to rRNA and r-protein oxygen atom distances reveals the presence of 564 water contact distances below 2.4 Å, affecting 321 Mg^2+^ (82% of the 393 hydrated Mg^2+^; **Figure 3**). This observation is significant since it has been clearly established in a comment (Kruse et al. 2019) of an article by Qi and Kulik (Qi and Kulik 2019) that hydrogen bonds shorter than 2.4–2.5 Å in crystal structures are artefactual due to inappropriate considerations of alternate conformations in high resolution structures. In 6QNR, such short contacts (<2.4 Å) reveal that Mg^2+^ hydration spheres and sometimes entire ion binding sites, have been poorly modeled and that other mono- or divalent ions or solvent molecules could be present at these locations in the crystal used for obtaining the 4V6F structure. For instance, the ion shown in **Figure 3** is at ≈2.8 Å of an oxygen atom of a phosphate group and is, therefore, more likely K^+^ despite the lack of reported anomalous signal at the 4-sigma level. Rozov et al. mentioned the fact that with the used procedure, they could have missed some K^+^ assignments. Yet, stereochemical considerations should have raised some warning flags and no Mg^2+^ should have been modeled at this location.

**Figure 3.**
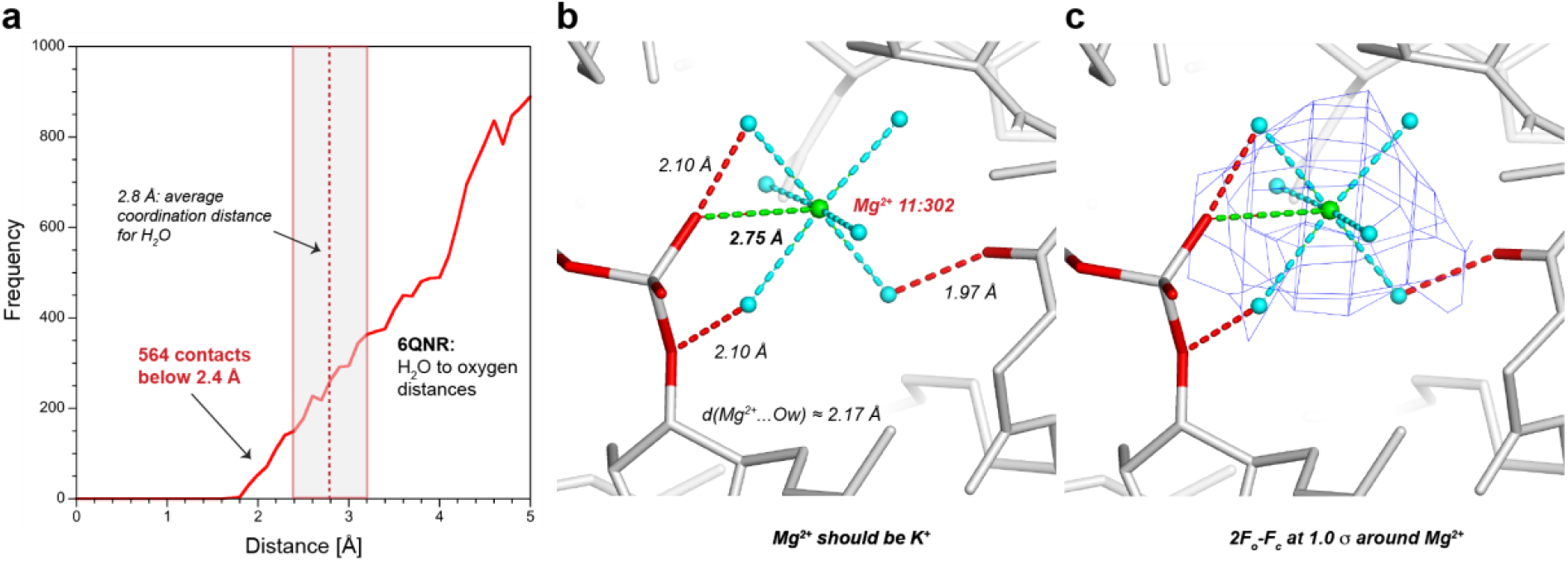
Abnormally short water to rRNA and r-protein contact distances reveal ill-placed Mg^2+^ in 6QNR. **a** The red curve shows the number of contacts between water molecules of the Mg^2+^ first hydration shell and rRNA and r-protein oxygen atoms with respect to distance. The 564 contacts < 2.4 Å reveal the presence of ill-placed Mg^2+^. Note that the forgiving 2.4 Å hydrogen bond limit; a hydrogen bond is only exceptionally shorter than 2.5–2.6 Å (Kruse et al. 2019) and is more generally centered around 2.8 Å (grey box). **b** Example of a Mg^2+^ in 6QNR whose hydration sphere clashes with three oxygen atoms (red dashed lines; see also **Figure S3**). The 2.75 Å distance to a phosphate oxygen atom reveals the probable presence of a K^+^ and unambiguously discards that of Mg^2+^ (green dashed line). Note the average *d*(Mg^2+^…Ow) distance close to 2.17 Å. **c** The corresponding 2F_o_-F_c_ map around the ion reveals no octahedral coordination pattern (**Figure S3**).

In total, we identified 853 out of a total of 1005 Mg^2+^ (≈85%) that do not satisfy all requested stereochemical criteria necessary for unambiguous ion assignments (**Table S1 and Figure 1c**). Among those, 138 and 361 Mg^2+^ display at least one coordination distance that suggests the presence of Na^+^ or K^+^, respectively. In addition, 144 Mg^2+^ have no atoms in their 1^st^ coordination shell rendering any conclusive identification attempt impossible. Indeed, in the absence of stereochemical evidence, some of the latter density spots could correspond to Mg(H_2_O)_6_^2+^ but could as well correspond to buffer solvent molecules of the buffer or noise in the density maps. Therefore, any ion assignment at those locations remains a perilous three-dimensional gambling exercise that is further hampered by locally inaccurate structural modeling. For instance, within medium to low resolution structures, assigning K^+^ or Mg^2+^ with occupancies of 0.5 should clearly be avoided; 24 ions with partial occupancies (0.5) are present in the 6QNR structure.

Less dramatic but not to be neglected, we evidenced through a simple distance check the use of an “idealized” average Mg^2+^…OH_2_ distance of 2.18 ± 0.02 Å (far from the 2.0 Å mentioned in Figure 2 of Rozov et al.) that contrasts with the known coordination distances of water to Mg^2+^ that genuinely turns around 2.06±0.03 Å as shown in **Figures 1 and 4** (see also: Zheng et al. 2015; Leonarski et al. 2017; Zheng et al. 2017; Leonarski et al. 2019)). Such errors can easily become viral (Minor et al. 2016) and may confuse, among others, untrained scientists that lack time to address stereochemical issues.

**Figure 4.**
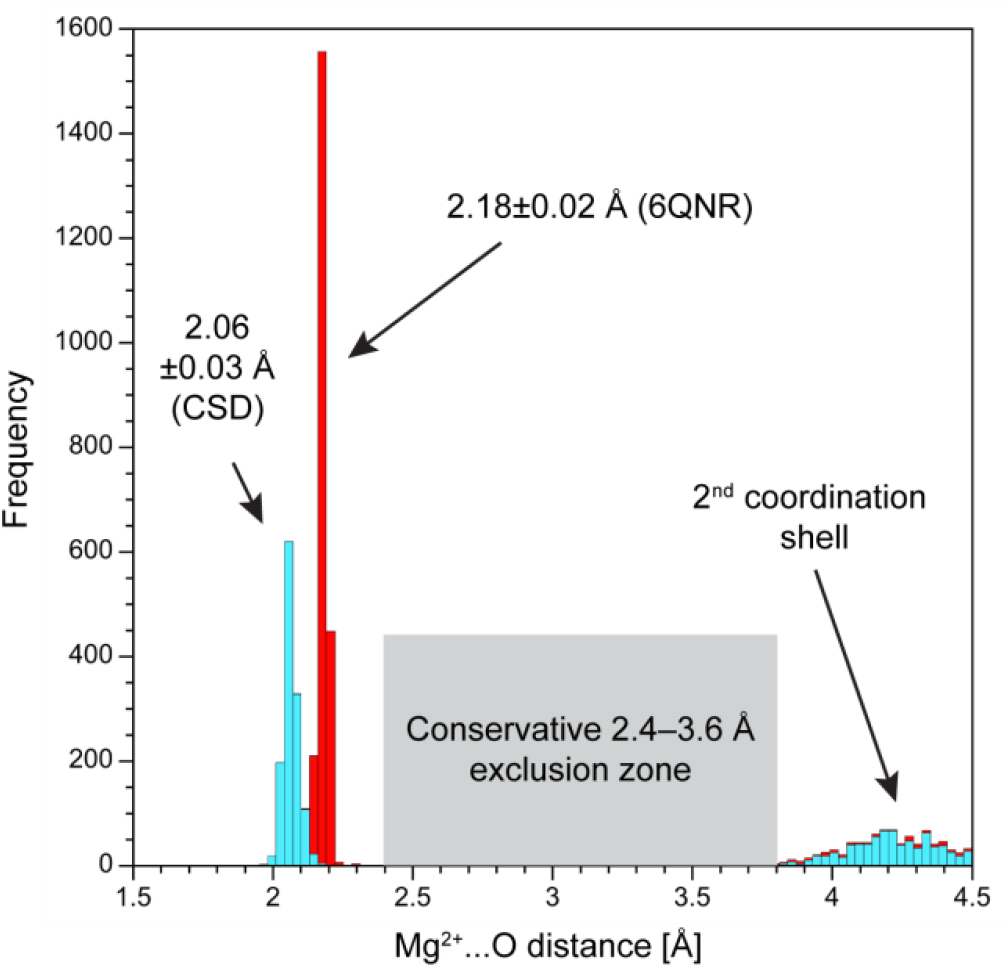
Mg^2+^…O_w_ coordination distance bias towards 2.18 Å. The cyan histogram is derived from the CSD (Groom et al. 2016; Taylor and Wood 2019) and is similar to that in **Figure 1a**. The red histogram highlights the average Mg^2+^…O_w_ distance around 2.18 Å in 6QNR (Rozov et al. 2019). These artefactual distances, common in PDB structures, have been discussed in preceding studies (Zheng et al. 2015; Leonarski et al. 2017; Zheng et al. 2017; Leonarski et al. 2019). For a definition of the exclusion zone see **Table S1** and **Figure 1**.

To summarize, our survey suggests that roughly 80% of the 1005 Mg^2+^ are misassigned or suffer from severe stereochemical issues and that ≈400 of these Mg^2+^ are possibly K^+^ (or eventually Na^+^; see Method section). This number adds to the 361 K^+^ that were assigned based on anomalous signals suggesting the presence of ≈700-800 monovalent ions and ≈200 Mg^2+^ in the 6QNR structure (or ≈400 monovalent ions and ≈100 Mg^2+^ in the 70S ribosomal 1^st^ solvation shell since the 6QNR crystallographic unit comprises two ribosomal particles). NH_4_^+^ ions are certainly also present in the 1^st^ solvation shell since they are mentioned in the crystallization conditions but remain undetectable. At a Å resolution, the numbers we provide can only be qualitative. However, we believe that the proposed orders of magnitude are representative of the balance of mono- and divalent ions in 1^st^ solvation shell of ribosomes. We would like to precise that we did not address issues regarding the ions present in the 2^nd^ solvation shell of ribosomes that are particularly difficult to localize when the resolution does not allow to visualize the coordinating water molecules; i.e. Mg(H_2_O)_6_^2+^. However, given the estimated intracellular Mg^2+^ (2–5 mM) and K^+^ (60–160 mM) concentrations (Nierhaus 2014), it seems unlikely that the Mg(H_2_O)_6_^2+^ concentration may approach that of bound and unbound K^+^ in a ribosome.

At this point, we note that the number of assigned ions is only able to neutralize about 1/5 to 1/4^th^ of the negative charge carried by ribosomes suggesting that the ionic atmosphere is essentially composed of non-coordinating ions and charged particles that remain undetected by X-ray or cryo-EM techniques. Given the larger K^+^ over Mg^2+^ concentration *in cristallo* (Rozov et al. 2019), but also *in vivo* (Nierhaus 2014; Miller-Fleming et al. 2015; Noeske et al. 2015), it is not unreasonable to suggest that K^+^, along with polyamines is responsible for the largest part of the charge neutralization of ribosomes while Mg^2+^ intervenes at a limited number of strategic locations to stabilize intricate structural folds (Bowman et al. 2012). This conclusion is not a surprise if we consider that Mg^2+^ is a regulated resource in the cell that is used with parsimony and for specific structural purposes while K^+^ is more abundant (Nierhaus 2014; Auffinger et al. 2016; Danchin and Nikel 2019). If the Rozov et al. along with the earlier Klein and Steitz structures (Klein et al. 2004) and other studies (Leonarski et al. 2017; Leonarski et al. 2019) add to our knowledge of the monovalent ion binding to nucleic acids and corrects previous stereochemically incorrect assignments (Jenner et al. 2010), the discussed structures continue to mask essential Mg^2+^ binding features.

Although numerous ribosomal structures with deficient ion placements have been deposited to the PDB, none of those studies, with the exception of a seminal study by Klein and Steitz (Klein et al. 2004), placed a strong emphasis on the ribosome ionic structure. The latter were the first authors to identify monovalent ions around a ribosomal subunit. However, in a large number of instances and especially for large structures, ions are placed in X-ray structures by using automatic routines to improve crystallographic statistics (Shabalin et al. 2018). In that respect, a common approach consists in systematically filling the electron density blobs in the immediate vicinity of RNA with Mg^2+^ or water molecules. Such a process helps understand the large and sometimes excessive number of Mg^2+^ present in some deposited ribosome structures (see for example 4V6F with 5345 Mg^2+^ for 9080 nucleotides leading to an apparent excess of 1640 positive charges; note that charged protein and nucleotide residues were not included in this estimate; interestingly, in this structure 4427 Mg^2+^ have no atoms in their 1^st^ coordination shell that is with a coordination distance below 2.4 Å; see **Supplementary Information II**).

Sometimes, similar strategies are used during the refinement of cryo-EM structures leading also to locally deficient ion placements. In a recently published 50S *Staphylococcus aureus* cryo-EM structure at 3.23 Å resolution (PDBid: 6SJ6), the authors reported the use of the *Thermus thermophilus* 6QNR structure discussed herein to model K^+^. They mentioned that densities that can be attributed to solvent molecules have been interpreted as Mg^2+^ (Khusainov et al. 2020). As a result, the 6SJ6 model deposited to the PDB contains 3 K^+^, 56 Mg^2+^ and no water molecules. Surprisingly, the number of K^+^ and Mg^2+^ is well below the number of ions expected for a bacterial 50S subunit although the resolution is close to that of 6QNR. To verify these assignments, we used our in-house diagnosis program on this structure and found that only 14 out of 56 Mg^2+^ satisfy requested stereochemical criteria. In 6SJ6, no octahedral density patterns could be observed around Mg^2+^. A visual inspection of ion positions and density maps led to the conclusion that almost none of these ions were appropriately assigned (**Figure 5**). Therefore, we suggest that in the absence of clear experimental and stereochemical evidence, it may be more suitable to not assign any ions in biomolecular structures with low resolutions since those assignments may subsequently be taken for granted by other groups with the risk of creating unsuitable prior knowledge cases that may affect the outcome of database surveys (Minor et al. 2016; Kowiel et al. 2018).

**Figure 5.**
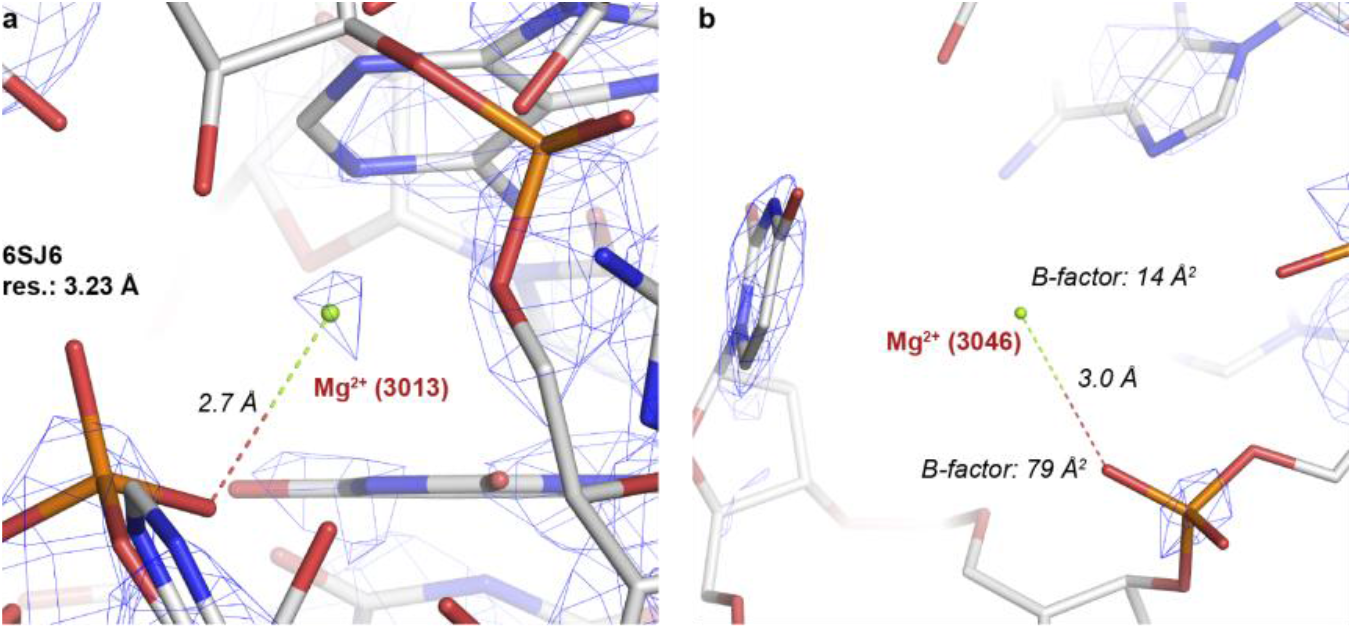
Two examples of improper Mg^2+^ assignments in the 6SJ6 cryo-EM structure at 3.23 Å resolution (Khusainov et al. 2020). **a** The coordination distance to this ion excludes the presence of Mg^2+^ and rather suggests the presence of K^+^. **b** As in (a). Note the large B-factor discrepancy.

It is now widely appreciated that ribosomes like any other biomolecular system cannot fulfill their *in vivo* functions without being immersed in a proper ion and solvent ecosystem. Thus, besides a correct K^+^ and Mg^2+^ identification, the real challenge is to unscramble the structural and functional roles these ions assume. For that, collecting high-resolution X-ray and cryo-EM data coupled with stereochemical knowledge remains the best option for advancing our understanding of mono- and divalent ion binding features since only this combination of techniques allows for a better and often clear-cut identification of the lighter ions (Mg^2+^, Na^+^, NH_4_^+^, …), water and other solutes but also of the heavier ions such as K^+^ and Cl^-^. For instance, it is only at resolutions better than 2.3 Å that Mg^2+^ and Na^+^ can be clearly identified based on their very close but characteristic coordination distances of ≈2.1 and ≈2.4 Å. It has to be noted that for suitable resolutions, density map peak heights provide further clues on the ion identity. Thus, high-resolution data are essential to reduce identification biases and to start the constitution of solid prior-knowledge databases that may significantly improve manual and automatized ion identification strategies in structures at all resolutions (Dauter et al. 2014; Rupp et al. 2016; Kowiel et al. 2018; Brzezinski et al. 2020).

Collecting anomalous signals at short and long-wavelength (Tereshko et al. 2001; Ennifar et al. 2003; Liebschner et al. 2016; Basu et al. 2019; Rozov et al. 2019) is an essential complement to a full structural characterization and should always be attempted given that it displays a few unique benefits. It is a powerful technique to distinguish between light and heavy ions or, more specifically between ions with detectable anomalous signals and the others. In ribosomes, the use of this technique was able to assess the long-overlooked presence of bound K^+^ and allowed to discard a large amount of erroneous Mg^2+^ attributions (Jenner et al. 2010; Rozov et al. 2019). However, it is currently only feasible for heavier ions such as K^+^, transition metals and anions and involves sometimes the use of *in-vacuo* beam lines that are not easily available to all. For instance, it cannot help to distinguish Mg^2+^ from Na^+^ since none of these ions has detectable anomalous signals. However, it can help to distinguish K^+^ from Mg^2+^/Na^+^ or other solvent particles. However, we strongly advocate that ion identification based on anomalous signals has to be associated with ion stereochemistry in order to validate the assignments and we state that Mg^2+^ should not be placed at binding sites where coordination distances exceed 2.4 Å and where coordination is not octahedral. Moreover, at non-atomic resolutions (resolutions > 2.6 Å), the likelihood that the biomolecular part of the system is locally incorrectly modeled is significantly greater. Placing ions in locations where the stereochemistry is not respected may severely jeopardize subsequent efforts to construct a reliable prior-knowledge database (Dauter et al. 2014; Rupp et al. 2016; Kowiel et al. 2018; Brzezinski et al. 2020).

The combined use of all these techniques should contribute to ‘deflate’ the Mg^2+^ bubble currently witnessed in numerous ribosomal structures, a bubble that eventually affects also a significant part of structures shaping the current RNA world (Leonarski et al. 2017; Leonarski et al. 2019). Besides the detection of anomalous signals, it is important to keep in mind that for the light ions present in crystallization buffers (Mg^2+^, Na^+^, NH_4_^+^, …) and for water molecules, anomalous signals cannot be collected. Therefore, to precisely identify them, stereochemistry remains the only viable option.

## METHODS

### Description of our ‘in-house’ ion diagnosis program

To precisely analyse the ion content and ion stereochemistry of 6QNR (Rozov et al. 2019) and related structures, we inspected the output of an ‘in-house’ program (Leonarski and Auffinger, unpublished) that screens PDB structures based on the stereochemistry of the assigned ions as derived from the PDB. This program is based on data from previous investigations (Leonarski et al. 2017; Leonarski et al. 2019) and provides a classification of the coordination features of each ion. It flags those for which ion identity conflicts with prior-knowledge coordination features coming from high-resolution structures (Harding et al. 2010; Bowman et al. 2012; Zheng et al. 2014; Leonarski et al. 2017; Leonarski et al. 2019). The complete ***4V6F_ion_diagnosis***.***html, 6QNR_ion_diagnosis***.***html*** and ***6SJ6_ion_diagnosis***.***html*** outputs (best opened with Firefox or Google Chrome) are given in **Supplementary Information II, III** and **IV**, respectively. This program is, in spirit, similar to the CheckMyMetal program that, unfortunately, does not accept large ‘cif’ files making it unsuitable for the analysis of most ribosomal structures (Zheng et al. 2014; Zheng et al. 2017). This program is also avoiding some of the serious pitfalls of the MgRNA website (Zheng et al. 2015) that had for purpose to classify Mg^2+^ binding sites for RNA structures deposited to the PDB. Those pitfalls are described in (Leonarski et al. 2017; Leonarski et al. 2019) and are essentially associated with the fact that the authors did consider a large amount of idealized (restrained based) Mg^2+^ binding sites derived from structures as low resolutions (sometimes > 4.0 Å). Note that our program utilizes an updated version of the Mg^2+^ binding site classification (Leonarski et al. 2017) originally developed by the MgRNA authors (Zheng et al. 2015).

Given that the 3.1 Å resolution of 6QNR is not ideal for allowing a reliable identification of the various solvent molecules and light ions surrounding the ribosome in the crystal phase, we used the relaxed coordination distance criteria noted in **Table S1** that allowed to take in some way into account crystallographic coordinate errors (Harding et al. 2010). More stringent criteria would have led to the characterization of a larger number of attribution issues. However, we felt that a precise and detailed analysis of the 6QNR ionic structure is not appropriate since clear octahedral coordination shells cannot be observed. For similar reasons, we did not re-analyze the parent 6QNQ initiation complex structure given its lower 3.5 Å resolution (Rozov et al. 2019). We expect that the data provided by this analysis program will be used to correct the ion placement and identification in 6QNR, in related ribosomal structures and in other biomolecular systems. It is important to consider that this program is only able to highlight grey areas where stereochemical criteria are not respected. These areas should be inspected with great care in order to confirm or not the original assignments and eventually correct binding site features. Improved versions of this program are currently under development.

### Notes regarding the possible presence of Na^+^ in ribosome crystals

Na^+^ is not mentioned in the 6QNR crystallization conditions but is probably present in the buffers used during several purification stages. The presence of this ion in 6QNR, although unlikely, cannot be discarded without a precise analysis of the crystal content through X-ray fluorescence or PIXE analysis, for example (Olieric et al. 2016; Grime et al. 2020). This is very challenging for large structures and light ions since Na^+^ and Mg^2+^ are close to the detection limit of these techniques. It is interesting to note that one of the 1^st^ ribosomal structure (PDBid: 4V9F; 2.4 Å resolution) discusses the concomitant presence of both Na^+^ and K^+^ monovalent ions in their structure; Na^+^ and K^+^ are mentioned in the 4V9F crystallization conditions (Klein et al. 2004; Gabdulkhakov et al. 2013).

However, ion coordination distances in the ≈2.4–2.6 Å range that imply the presence of Na^+^ could also find their origin in the improper modeling of some ion binding sites, therefore, leading to short and untrustworthy interatomic distances and rendering any ion assignment at those locations difficult. Inaccurate local structural modeling is not uncommon at resolutions > 3.0 Å and is probably reflected in part in the high 6QNR clashscore of 19 mentioned by the PDB validation report (4Y4O at 2.3 Å resolution, 4V90 at 2.95 Å resolution and 4V6F at 3.1 Å resolution display clashscores of 8, 45 and 52, respectively).

### Notes regarding the detection of anions (Cl^-^, SCN^-^) in 6QNR

The detection of anions in RNA structures is also a matter of concern (Auffinger et al. 2004a) (D’Ascenzo and Auffinger 2016). In 6QNR, a few suspicious ions have been associated with Mg^2+^ while they are contacting amino groups belonging to nucleobases or to amino acids and, therefore, could correspond to Cl^-^, CH_3_COO^-^ or SCN^-^ ions that are mentioned in the crystallographic conditions. For instance, it is highly improbable that the ion shown **Fig. S4** is Mg^2+^. It can be regretted that the authors did not use their technical setup to identify anions featuring atoms with significant anomalous signals (Liu et al. 2013; Panjikar et al. 2015; Liu and Hendrickson 2017) such as Cl^-^ or SCN^-^. This would certainly have been possible since the anomalous signals of phosphorus and sulfur atoms have been detected (**Fig. 2**).

## Supporting information

Spupplemental I

Spupplemental II

Spupplemental III

Spupplemental IV

## Acknowledgment

We thank Alexey Rozov for having provided the anomalous diffraction data and Filip Leonarski for having provided an early version of his “Ion Report” program for the analysis of the ribosomal ionic environment. We thank Philippe Dumas, Quentin Vicens, and Filip Leonarski for critical views on the manuscript.

## Author contributions

PA conceived the project. PA, EE, and LD analyzed the data. LD helped in writing analysis tools. PA wrote the manuscript. All authors discussed the results and conceived the final manuscript.

## Additional information

**Supplementary Information** accompanies this publication.

It consists in two parts: the first contains one table and four figures; the second, third and fourth, a full output of our “Ion_diagnosis” program for the 4V6F, 6QNR and 6SJ6 structures.

## Competing interests

None.

## Notes

### Competing Interest Statement

The authors have declared no competing interest.

### Summary of Updates

We clarified the text; Added Figure 2; added a method section, previously in the SI; added new references; rewrote part of the abstract; the message and the conclusions remain unchanged;

