## Supplementary material for "Deflating the RNA Mg^2+^ bubble. Stereochemistry to the rescue!": Spupplemental I

**Table of content:**

1) **Table S1** Coordination criteria used in this study
2) **Figure S1** Two views of the assigned $Mg^{2+}$ and $K^{+}$ close to G530 in the 5E81 and 6QNR ribosomal decoding center
3) **Figure S2** Illustration of densities corresponding to hexacoordinated $Mg^{2+}$ at high and medium resolutions
4) **Figure S3** Extract of the 6QNR 'in-house' analysis program output related to the $Mg^{2+}$ ion shown in Fig. 3.
5) **Figure S4** $Mg^{2+}$ occupying an anion binding site in 6QNR.
6) **References**

**Table S1. Coordination criteria used in this study.** Some of the values given in the table are derived from earlier studies (D'Ascenzo and Auffinger 2015; Leonarski et al. 2016; Leonarski et al. 2017; Kruse et al. 2019; Leonarski et al. 2019) and from **Figure 1**. Converging views on the coordination of metals to biomolecular systems are provided in the following references (Harding et al. 2010; Zheng et al. 2014). The current reference list is non-exhaustive. The criteria proposed here represent by no means absolute values but point to coordination distance ranges where stereochemistry, ion binding site features and ion identity have to be reassessed. These criteria take in some ways into account crystallographic coordinate errors (Harding et al. 2010).

| | Relaxed coordination criteria (Å)* | Stringent coordination criteria (Å)** | CSD average coordination distance (Å) |
|---|---|---|---|
| **$1^{st}$ coordination shell** | | | |
| $Mg^{2+}$…O | 1.8–2.4 | 1.9–2.3 | 2.06±0.03 |
| $Mg^{2+}$…N | 1.8–2.4 | 1.9–2.4 | 2.10±0.07 |
| $Na^{+}$…O | 2.4–2.6 | 2.3–2.6 | 2.41±0.10 |
| $Na^{+}$…N | 2.4–2.6 | 2.3–2.6 | 2.45±0.05 |
| $K^{+}$…O | 2.6–3.4 | 2.6–3.2 | 2.8±0.1 |
| $K^{+}$…N | 2.6–3.4 | 2.6–3.2 | 2.85±0.06 |
| **$Mg^{2+}$ exclusion zone** | | | |
| | 2.4–3.4 | 2.3–3.8 | 2.2–3.8 |
| **$Na^{+}$ exclusion zone** | | | |
| | 2.8–3.6 | 2.3–3.6 | 2.3–3.6 |
| **$2^{nd}$ coordination shell** | | | |
| $Mg^{2+}$…O | 3.4–4.6 | 3.8–4.6 | 3.8–4.6 |
| $Na^{+}$…O | 3.6–4.9 | 3.6-4.9 | 3.6–4.9 |
| $K^{+}$…O | n.a. | n.a. | n.a. |
| **Hydrogen bond** | | | |
| O…$O_w$ | 2.4–3.0 | 2.5–3.0 | 2.5–3.0 |
| N…$O_w$ | 2.4–3.2 | 2.5–3.2 | 2.5–3.2 |

* criteria suggested for medium to poor resolutions (> 2.6 Å);
** criteria suggested for resolutions ≤ 2.6 Å;

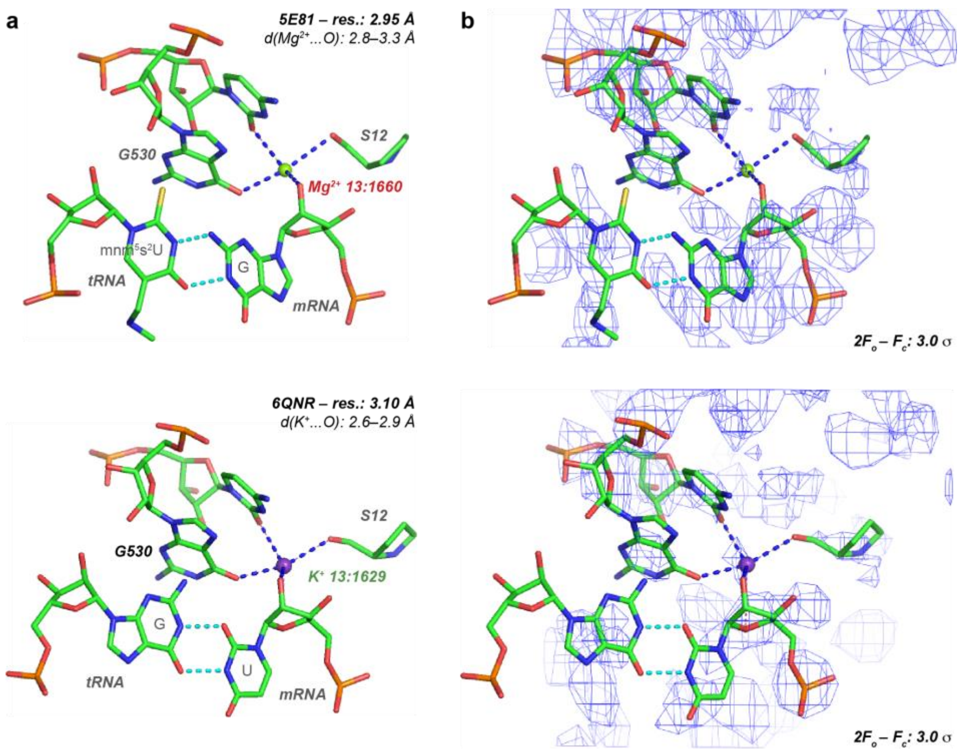


**c**

MG:13:1660 — 5E81

| | Res. type | Atom | Residue | Distance | Occupancy | B-factor | Closest solute to 1st shell water |
|---|---|---|---|---|---|---|---|
| | Ion | | | | 1.00 | 55.65 | |
| 1 | G | O2' | 4K:21 | 2.58 | 1.00 | 73.14 | |
| 2 | G | O6 | 13:530 | 2.66 | 1.00 | 74.53 | |
| 3 | C | O2 | 13:518 | 2.76 | 1.00 | 68.81 | |
| 4 | PRO | O | 31:48 | 3.01 | 1.00 | 76.15 | |
| 5 | C | N3 | 13:518 | 3.14 | 1.00 | 74.22 | |
| 6 | G | O4' | 4K:21 | 3.30 | 1.00 | 78.83 | |
| 7 | ASN | N | 31:49 | 4.25 | 1.00 | 71.35 | |
| 8 | G | N7 | 13:530 | 4.36 | 1.00 | 72.01 | |
| 9 | A | N1 | 13:1492 | 4.38 | 1.00 | 71.14 | |
| 10 | G | N1 | 13:530 | 4.43 | 1.00 | 71.90 | |
| 11 | G | O3' | 4K:21 | 4.65 | 1.00 | 77.83 | |
| 12 | A | N3 | 13:1492 | 4.67 | 1.00 | 69.93 | |
| 13 | C | N1 | 13:518 | 4.70 | 1.00 | 69.89 | |
| 14 | C | N4 | 13:518 | 4.84 | 1.00 | 76.08 | |
| 15 | G | N9 | 4K:21 | 4.97 | 1.00 | 76.83 | |
| 16 | G | O6 | 13:529 | 4.98 | 1.00 | 69.23 | |

Coded contact list:
Exclusion zone contact(s): /N3$_C$/O2/O2'/O4'/O6/O$_{back}$/
Consider K in place of MG (5 valid contact(s) for K vs. 0 for MG)

K:13:1629 — 6QNR

| | Res. type | Atom | Residue | Distance | Occupancy | B-factor | Closest solute to 1st shell water |
|---|---|---|---|---|---|---|---|
| | Ion | | | | 1.00 | 50.77 | |
| 1 | G | O6 | 13:530 | 2.55 | 1.00 | 53.69 | |
| 2 | PRO | O | 31:45 | 2.77 | 1.00 | 52.89 | |
| 3 | C | O2 | 13:518 | 2.79 | 1.00 | 48.94 | |
| 4 | U | O2' | 4K:51 | 2.92 | 1.00 | 64.73 | |
| 5 | C | N3 | 13:518 | 3.16 | 1.00 | 54.68 | |
| 6 | U | O4' | 4K:51 | 3.35 | 1.00 | 51.48 | |
| 7 | U | O3' | 4K:51 | 4.25 | 1.00 | 71.00 | |
| 8 | ASN | N | 31:46 | 4.38 | 1.00 | 48.46 | |
| 9 | G | N7 | 13:530 | 4.48 | 1.00 | 53.80 | |
| 10 | G | N1 | 13:530 | 4.51 | 1.00 | 50.05 | |
| 11 | C | N1 | 13:518 | 4.73 | 1.00 | 49.26 | |
| 12 | C | N4 | 13:518 | 4.78 | 1.00 | 54.67 | |
| 13 | A | N3 | 13:1492 | 4.81 | 1.00 | 46.86 | |
| 14 | ASN | O | 31:46 | 4.93 | 1.00 | 49.73 | |

1st shell ligand angles:
1-2: 164; 1-3: 81; 1-4: 73; 1-5: 81; 1-6: 72; 2-3: 88; 2-4: 106; 2-5: 83; 2-6: 121; 3-4: 124;

Contact(s): /N3$_C$/O2/O2'/O4'/O$_{back}$/
Coded contact list: 2O$_r$O$_b$N$_b$O$_{back}$
Exclusion zone contact(s): /O6/

**Figure S1. Two views of the assigned $Mg^{2+}$ and $K^+$ close to G530 in the 5E81 and 6QNR ribosomal decoding center. a** View of the 5E81 (2016) (Rozov et al. 2016; Leonarski et al. 2019) (top; see **Figure 1d** for the '2010' 4V6F structure (Jenner et al. 2010)) and recent (bottom) 6QNR (Rozov et al. 2019) decoding center structures with modeled $Mg^{2+}$ (5E81/4V6F) and $K^+$ ions (6QNR) displaying very similar coordination distances around 2.8 Å. The text in red and green highlights unfavorable and favorable attributions, respectively. The 5E81 structure (resolution: 2.95 Å) displays a slightly better resolution than the 6QNR and 4V6F (Jenner et al. 2010) structures (resolution: 3.1 Å) and corresponds currently to the best resolution among the *Thermus thermophilus* structures embedding tRNA and mRNA fragments. Note the different tRNA-mRNA pair in 5E81 and 6QNR. **b** Associated $2F_o$-$F_c$ density patterns at 3.0 σ, emphasizing the structural similarity between 5E81 and 6QNR. **a,b** The blue dotted lines show ion coordination distances in the 2.6–3.3 Å range. **c** Extract of the 5E81 and 6QNR 'in-house' diagnosis program output listing the closest contact distance of the ion bound to G530. Red and green colors mark conflicts and agreements between ion identity and stereochemistry, respectively.

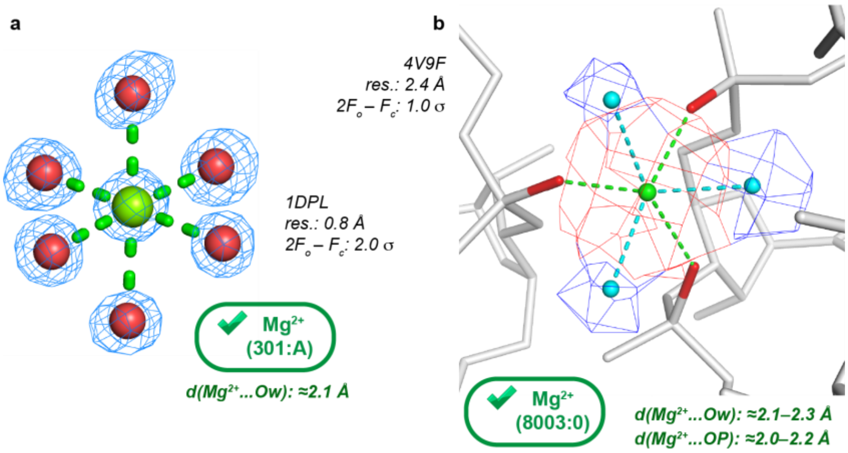


**Figure S2. Illustration of densities corresponding to hexacoordinated $Mg^{2+}$ at high and medium resolutions. a** At resolutions around 1.0 Å, well-separated density blobs allow easy coordination shell placement (Leonarski et al. 2019). **b** At medium resolutions (≈2.4 Å), the density blobs of the coordinating atoms are often fused with that of the ion. Here, the $Mg^{2+}$ ion (red density) is coordinated to three phosphate anionic oxygen atoms (red) and three water molecules (blue densities). For density pattern comparison with resolutions around 3.0 Å, see **Figure 3c**.

**MG:11:302**

| | Res. type | Atom | Residue | Distance | Occupancy | B-factor | Closest solute to 1st shell water |
|---|---|---|---|---|---|---|---|
| Ion | | | | | 1.00 | 14.91 | |
| 1 | Water 1 | | 11:402 | 2.16 | 1.00 | 18.70 | 2.46 (PRO:O 11:232) |
| 2 | Water 2 | | 1H:3661 | 2.16 | 1.00 | 17.30 | 2.10 (A:OP1 1H:2611) |
| 3 | Water 3 | | 1H:3662 | 2.16 | 1.00 | 19.67 | 2.10 (G:O3' 1H:2610) |
| 4 | Water 4 | | 11:407 | 2.17 | 1.00 | 16.52 | 2.91 (ARG:O 11:242) |
| 5 | Water 5 | | 11:401 | 2.17 | 1.00 | 17.06 | 1.97 (ARG:O 11:242) |
| 6 | Water 6 | | 11:404 | 2.19 | 1.00 | 17.01 | 2.68 (PRO:O 11:241) |
| 7 | A | OP1 | 1H:2611 | 2.75 | 1.00 | 13.91 | |
| 8 | ARG | O | 11:242 | 3.18 | 1.00 | 18.42 | |
| 9 | G | O3' | 1H:2610 | 3.91 | 1.00 | 14.64 | |
| 10 | PRO | O | 11:241 | 4.01 | 1.00 | 20.91 | |
| 11 | GLY | O | 11:234 | 4.05 | 1.00 | 17.52 | |
| 12 | HIS | O | 11:233 | 4.23 | 1.00 | 20.82 | |
| 13 | PRO | O | 11:232 | 4.37 | 1.00 | 20.82 | |
| 14 | GLY | N | 11:243 | 4.70 | 1.00 | 14.14 | |
| 15 | A | OP2 | 1H:2611 | 4.81 | 1.00 | 16.73 | |
| 16 | A | O5' | 1H:2611 | 4.91 | 1.00 | 17.13 | |

**1st shell ligand angles:**
1-2: **85**; 1-3: **81**; 1-4: **95**; 1-5: **93**; 1-6: **172**; 2-3: **104**; 2-4: **84**; 2-5: **178**; 2-6: **89**; 3-4: **171**; 3-5: **75**; 3-6: **95**; 4-5: **97**; 4-6: **89**; 5-6: **93**;

**Contact(s): /$O_w$/$O_w$/$O_w$/$O_w$/$O_w$/$O_w$/**
**Coded contact list: 6$O_w$**
**Exclusion zone contact(s): /$O_{back}$/OP1/**
**Consider K in place of MG (2 valid contact(s) for K vs. 0 for MG)**
**MG hydration shell issue: water too close (< 2.4 Angs.)**

**Figure S3. Extract of the 6QNR 'in-house' analysis program output related to the $Mg^{2+}$ ion shown in Fig. 3.** The short 2nd shell contact distances that suggest a poor hydration shell modeling are highlighted in red as well as the two coordination distances to OP1 and O atoms that suggest the presence of $K^+$ instead of the presence of $Mg^{2+}$. Note the $Mg^{2+}$…$O_W$ coordination distances around 2.18 instead of 2.07 Å.

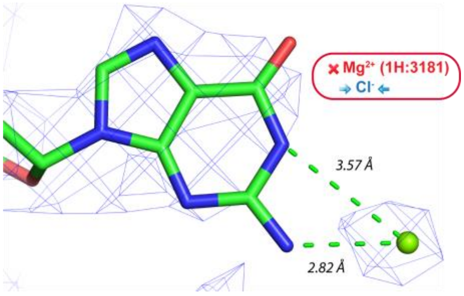


**Figure S4**. **$Mg^{2+}$ occupying an anion binding site in 6QNR.** Anion binding to the guanine Watson-Crick edge has been well documented (Auffinger et al. 2004; D'Ascenzo and Auffinger 2016). Given its position and a ≈2.8 Å coordination distance, the ion shown in this figure is not $Mg^{2+}$ but more likely $Cl^{-}$.
