## Supplementary material for "Deflating the RNA Mg^2+^ bubble. Stereochemistry to the rescue!": Spupplemental II: 4V6F.ion_diagnosis.html

 Ion report: 4V6F  

### Ion report: 4V6F

#### date: 2020-10-01 13:21

Authors: Filip Leonarski and Pascal Auffinger  
\*\*\* beta version: unpublished \*\*\*  
-> reference 1: Leonarski F., D'Ascenzo L., Auffinger P. (2017) NAR, 45, 987-1004  
-> reference 2: Leonarski F., D'Ascenzo L., Auffinger P. (2019) RNA, 25, 173-192  
  
Density maps not assigned - No density checks made in the current version!  
  
\_\_\_\_\_\_\_\_\_\_\_\_\_\_\_\_\_\_\_\_\_\_\_\_\_\_\_\_\_\_\_\_\_\_\_\_\_\_\_\_\_\_\_\_\_\_\_\_\_\_\_\_\_\_\_\_\_\_\_\_\_\_\_\_\_\_\_\_\_\_\_\_\_\_\_\_\_\_\_\_\_  

#### Table of content (ToC):

1) Number of ions of each type and coordination distance ranges used for validation  
2) Ion validation table  
3) Potential issue list  
4) Ion/ligand tables  
5) Sorted coded contact list  
\_\_\_\_\_\_\_\_\_\_\_\_\_\_\_\_\_\_\_\_\_\_\_\_\_\_\_\_\_\_\_\_\_\_\_\_\_\_\_\_\_\_\_\_\_\_\_\_\_\_\_\_\_\_\_\_\_\_\_\_\_\_\_\_\_\_\_\_\_\_\_\_\_\_\_\_\_\_\_\_\_  
  
REMARKS:  
- 1^st^ shell: correct distance range for coordinating atoms;  
- Exclusion zone (excl. zone): a distance range where \*no\* coordinating atoms should be present;  
- Currently coordination distances with sulfur (S) atoms are not considered --> no CD/ZN (cadmium/zinc) validation;  
\_\_\_\_\_\_\_\_\_\_\_\_\_\_\_\_\_\_\_\_\_\_\_\_\_\_\_\_\_\_\_\_\_\_\_\_\_\_\_\_\_\_\_\_\_\_\_\_\_\_\_\_\_\_\_\_\_\_\_\_\_\_\_\_  

#### 1) Number of ions of each categorie and coordination distance ranges used for validation

##### Go to ToC

| Ion type | Number | Ion list | Short excl. zone [Angs] | 1^st^ shell [Angs.] | Long excl. zone [Angs] |
| --- | --- | --- | --- | --- | --- |
| Sodium (NA) | --- |  | <2.4 | 2.4 - 2.6 | 2.6 - 3.6 |
| Potassium (K) | --- |  | <2.6 | 2.6 - 3.4 | --- |
| Caesium (CS) | --- |  | <2.9 | 2.9 - 3.6 | --- |
| Magnesium (MG) | 5345 | A0:201 A1:201 A1:202 A1:203 A1:204 A1:205 A2:201 A3:101 A3:102 A3:103 A3:104 A5:101 A5:102 A5:103 A6:101 A6:102 A7:101 A7:102 A7:103 A7:104 A8:101 A8:102 A8:103 A8:104 AA:3001 AA:3002 AA:3003 AA:3004 AA:3005 AA:3006 AA:3007 AA:3008 AA:3009 AA:3010 AA:3011 AA:3012 AA:3013 AA:3014 AA:3015 AA:3016 AA:3017 AA:3018 AA:3019 AA:3020 AA:3021 AA:3022 AA:3023 AA:3024 AA:3025 AA:3026 AA:3027 AA:3028 AA:3029 AA:3030 AA:3031 AA:3032 AA:3033 AA:3034 AA:3035 AA:3036 AA:3037 AA:3038 AA:3039 AA:3040 AA:3041 AA:3042 AA:3043 AA:3044 AA:3045 AA:3046 AA:3047 AA:3048 AA:3049 AA:3050 AA:3051 AA:3052 AA:3053 AA:3054 AA:3055 AA:3056 AA:3057 AA:3058 AA:3059 AA:3060 AA:3061 AA:3062 AA:3063 AA:3064 AA:3065 AA:3066 AA:3067 AA:3068 AA:3069 AA:3070 AA:3071 AA:3072 AA:3073 AA:3074 AA:3075 AA:3076 AA:3077 AA:3078 AA:3079 AA:3080 AA:3081 AA:3082 AA:3083 AA:3084 AA:3085 AA:3086 AA:3087 AA:3088 AA:3089 AA:3090 AA:3091 AA:3092 AA:3093 AA:3094 AA:3095 AA:3096 AA:3097 AA:3098 AA:3099 AA:3100 AA:3101 AA:3102 AA:3103 AA:3104 AA:3105 AA:3106 AA:3107 AA:3108 AA:3109 AA:3110 AA:3111 AA:3112 AA:3113 AA:3114 AA:3115 AA:3116 AA:3117 AA:3118 AA:3119 AA:3120 AA:3121 AA:3122 AA:3123 AA:3124 AA:3125 AA:3126 AA:3127 AA:3128 AA:3129 AA:3130 AA:3131 AA:3132 AA:3133 AA:3134 AA:3135 AA:3136 AA:3137 AA:3138 AA:3139 AA:3140 AA:3141 AA:3142 AA:3143 AA:3144 AA:3145 AA:3146 AA:3147 AA:3148 AA:3149 AA:3150 AA:3151 AA:3152 AA:3153 AA:3154 AA:3155 AA:3156 AA:3157 AA:3158 AA:3159 AA:3160 AA:3161 AA:3162 AA:3163 AA:3164 AA:3165 AA:3166 AA:3167 AA:3168 AA:3169 AA:3170 AA:3171 AA:3172 AA:3173 AA:3174 AA:3175 AA:3176 AA:3177 AA:3178 AA:3179 AA:3180 AA:3181 AA:3182 AA:3183 AA:3184 AA:3185 AA:3186 AA:3187 AA:3188 AA:3189 AA:3190 AA:3191 AA:3192 AA:3193 AA:3194 AA:3195 AA:3196 AA:3197 AA:3198 AA:3199 AA:3200 AA:3201 AA:3202 AA:3203 AA:3204 AA:3205 AA:3206 AA:3207 AA:3208 AA:3209 AA:3210 AA:3211 AA:3212 AA:3213 AA:3214 AA:3215 AA:3216 AA:3217 AA:3218 AA:3219 AA:3220 AA:3221 AA:3222 AA:3223 AA:3224 AA:3225 AA:3226 AA:3227 AA:3228 AA:3229 AA:3230 AA:3231 AA:3232 AA:3233 AA:3234 AA:3235 AA:3236 AA:3237 AA:3238 AA:3239 AA:3240 AA:3241 AA:3242 AA:3243 AA:3244 AA:3245 AA:3246 AA:3247 AA:3248 AA:3249 AA:3250 AA:3251 AA:3252 AA:3253 AA:3254 AA:3255 AA:3256 AA:3257 AA:3258 AA:3259 AA:3260 AA:3261 AA:3262 AA:3263 AA:3264 AA:3265 AA:3266 AA:3267 AA:3268 AA:3269 AA:3270 AA:3271 AA:3272 AA:3273 AA:3274 AA:3275 AA:3276 AA:3277 AA:3278 AA:3279 AA:3280 AA:3281 AA:3282 AA:3283 AA:3284 AA:3285 AA:3286 AA:3287 AA:3288 AA:3289 AA:3290 AA:3291 AA:3292 AA:3293 AA:3294 AA:3295 AA:3296 AA:3297 AA:3298 AA:3299 AA:3300 AA:3301 AA:3302 AA:3303 AA:3304 AA:3305 AA:3306 AA:3307 AA:3308 AA:3309 AA:3310 AA:3311 AA:3312 AA:3313 AA:3314 AA:3315 AA:3316 AA:3317 AA:3318 AA:3319 AA:3320 AA:3321 AA:3322 AA:3323 AA:3324 AA:3325 AA:3326 AA:3327 AA:3328 AA:3329 AA:3330 AA:3331 AA:3332 AA:3333 AA:3334 AA:3335 AA:3336 AA:3337 AA:3338 AA:3339 AA:3340 AA:3341 AA:3342 AA:3343 AA:3344 AA:3345 AA:3346 AA:3347 AA:3348 AA:3349 AA:3350 AA:3351 AA:3352 AA:3353 AA:3354 AA:3355 AA:3356 AA:3357 AA:3358 AA:3359 AA:3360 AA:3361 AA:3362 AA:3363 AA:3364 AA:3365 AA:3366 AA:3367 AA:3368 AA:3369 AA:3370 AA:3371 AA:3372 AA:3373 AA:3374 AA:3375 AA:3376 AA:3377 AA:3378 AA:3379 AA:3380 AA:3381 AA:3382 AA:3383 AA:3384 AA:3385 AA:3386 AA:3387 AA:3388 AA:3389 AA:3390 AA:3391 AA:3392 AA:3393 AA:3394 AA:3395 AA:3396 AA:3397 AA:3398 AA:3399 AA:3400 AA:3401 AA:3402 AA:3403 AA:3404 AA:3405 AA:3406 AA:3407 AA:3408 AA:3409 AA:3410 AA:3411 AA:3412 AA:3413 AA:3414 AA:3415 AA:3416 AA:3417 AA:3418 AA:3419 AA:3420 AA:3421 AA:3422 AA:3423 AA:3424 AA:3425 AA:3426 AA:3427 AA:3428 AA:3429 AA:3430 AA:3431 AA:3432 AA:3433 AA:3434 AA:3435 AA:3436 AA:3437 AA:3438 AA:3439 AA:3440 AA:3441 AA:3442 AA:3443 AA:3444 AA:3445 AA:3446 AA:3447 AA:3448 AA:3449 AA:3450 AA:3451 AA:3452 AA:3453 AA:3454 AA:3455 AA:3456 AA:3457 AA:3458 AA:3459 AA:3460 AA:3461 AA:3462 AA:3463 AA:3464 AA:3465 AA:3466 AA:3467 AA:3468 AA:3469 AA:3470 AA:3471 AA:3472 AA:3473 AA:3474 AA:3475 AA:3476 AA:3477 AA:3478 AA:3479 AA:3480 AA:3481 AA:3482 AA:3483 AA:3484 AA:3485 AA:3486 AA:3487 AA:3488 AA:3489 AA:3490 AA:3491 AA:3492 AA:3493 AA:3494 AA:3495 AA:3496 AA:3497 AA:3498 AA:3499 AA:3500 AA:3501 AA:3502 AA:3503 AA:3504 AA:3505 AA:3506 AA:3507 AA:3508 AA:3509 AA:3510 AA:3511 AA:3512 AA:3513 AA:3514 AA:3515 AA:3516 AA:3517 AA:3518 AA:3519 AA:3520 AA:3521 AA:3522 AA:3523 AA:3524 AA:3525 AA:3526 AA:3527 AA:3528 AA:3529 AA:3530 AA:3531 AA:3532 AA:3533 AA:3534 AA:3535 AA:3536 AA:3537 AA:3538 AA:3539 AA:3540 AA:3541 AA:3542 AA:3543 AA:3544 AA:3545 AA:3546 AA:3547 AA:3548 AA:3549 AA:3550 AA:3551 AA:3552 AA:3553 AA:3554 AA:3555 AA:3556 AA:3557 AA:3558 AA:3559 AA:3560 AA:3561 AA:3562 AA:3563 AA:3564 AA:3565 AA:3566 AA:3567 AA:3568 AA:3569 AA:3570 AA:3571 AA:3572 AA:3573 AA:3574 AA:3575 AA:3576 AA:3577 AA:3578 AA:3579 AA:3580 AA:3581 AA:3582 AA:3583 AA:3584 AA:3585 AA:3586 AA:3587 AA:3588 AA:3589 AA:3590 AA:3591 AA:3592 AA:3593 AA:3594 AA:3595 AA:3596 AA:3597 AA:3598 AA:3599 AA:3600 AA:3601 AA:3602 AA:3603 AA:3604 AA:3605 AA:3606 AA:3607 AA:3608 AA:3609 AA:3610 AA:3611 AA:3612 AA:3613 AA:3614 AA:3615 AA:3616 AA:3617 AA:3618 AA:3619 AA:3620 AA:3621 AA:3622 AA:3623 AA:3624 AA:3625 AA:3626 AA:3627 AA:3628 AA:3629 AA:3630 AA:3631 AA:3632 AA:3633 AA:3634 AA:3635 AA:3636 AA:3637 AA:3638 AA:3639 AA:3640 AA:3641 AA:3642 AA:3643 AA:3644 AA:3645 AA:3646 AA:3647 AA:3648 AA:3649 AA:3650 AA:3651 AA:3652 AA:3653 AA:3654 AA:3655 AA:3656 AA:3657 AA:3658 AA:3659 AA:3660 AA:3661 AA:3662 AA:3663 AA:3664 AA:3665 AA:3666 AA:3667 AA:3668 AA:3669 AA:3670 AA:3671 AA:3672 AA:3673 AA:3674 AA:3675 AA:3676 AA:3677 AA:3678 AA:3679 AA:3680 AA:3681 AA:3682 AA:3683 AA:3684 AA:3685 AA:3686 AA:3687 AA:3688 AA:3689 AA:3690 AA:3691 AA:3692 AA:3693 AA:3694 AA:3695 AA:3696 AA:3697 AA:3698 AA:3699 AA:3700 AA:3701 AA:3702 AA:3703 AA:3704 AA:3705 AA:3706 AA:3707 AA:3708 AA:3709 AA:3710 AA:3711 AA:3712 AA:3713 AA:3714 AA:3715 AA:3716 AA:3717 AA:3718 AA:3719 AA:3720 AA:3721 AA:3722 AA:3723 AA:3724 AA:3725 AA:3726 AA:3727 AA:3728 AA:3729 AA:3730 AA:3731 AA:3732 AA:3733 AA:3734 AA:3735 AA:3736 AA:3737 AA:3738 AA:3739 AA:3740 AA:3741 AA:3742 AA:3743 AA:3744 AA:3745 AA:3746 AA:3747 AA:3748 AA:3749 AA:3750 AA:3751 AA:3752 AA:3753 AA:3754 AA:3755 AA:3756 AA:3757 AA:3758 AA:3759 AA:3760 AA:3761 AA:3762 AA:3763 AA:3764 AA:3765 AA:3766 AA:3767 AA:3768 AA:3769 AA:3770 AA:3771 AA:3772 AA:3773 AA:3774 AA:3775 AA:3776 AA:3777 AA:3778 AA:3779 AA:3780 AA:3781 AA:3782 AA:3783 AA:3784 AA:3785 AA:3786 AA:3787 AA:3788 AA:3789 AA:3790 AA:3791 AA:3792 AA:3793 AA:3794 AA:3795 AA:3796 AA:3797 AA:3798 AA:3799 AA:3800 AA:3801 AA:3802 AA:3803 AA:3804 AA:3805 AA:3806 AA:3807 AA:3808 AA:3809 AA:3810 AA:3811 AA:3812 AA:3813 AA:3814 AA:3815 AA:3816 AA:3817 AA:3818 AA:3819 AA:3820 AA:3821 AA:3822 AA:3823 AA:3824 AA:3825 AA:3826 AA:3827 AA:3828 AA:3829 AA:3830 AA:3831 AA:3832 AA:3833 AA:3834 AA:3835 AA:3836 AA:3837 AA:3838 AA:3839 AA:3840 AA:3841 AA:3842 AA:3843 AA:3844 AA:3845 AA:3846 AA:3847 AA:3848 AA:3849 AA:3850 AA:3851 AA:3852 AA:3853 AA:3854 AA:3855 AA:3856 AA:3857 AA:3858 AA:3859 AA:3860 AA:3861 AA:3862 AA:3863 AA:3864 AA:3865 AA:3866 AA:3867 AA:3868 AA:3869 AA:3870 AA:3871 AA:3872 AA:3873 AA:3874 AA:3875 AA:3876 AA:3877 AA:3878 AA:3879 AA:3880 AA:3881 AA:3882 AA:3883 AA:3884 AA:3885 AA:3886 AA:3887 AA:3888 AA:3889 AA:3890 AA:3891 AA:3892 AA:3893 AA:3894 AA:3895 AA:3896 AA:3897 AA:3898 AA:3899 AA:3900 AA:3901 AA:3902 AA:3903 AA:3904 AA:3905 AA:3906 AA:3907 AA:3908 AA:3909 AA:3910 AA:3911 AA:3912 AA:3913 AA:3914 AA:3915 AA:3916 AA:3917 AA:3918 AA:3919 AA:3920 AA:3921 AA:3922 AA:3923 AA:3924 AA:3925 AA:3926 AA:3927 AA:3928 AA:3929 AA:3930 AA:3931 AA:3932 AA:3933 AA:3934 AA:3935 AA:3936 AA:3937 AA:3938 AA:3939 AA:3940 AA:3941 AA:3942 AA:3943 AA:3944 AA:3945 AA:3946 AA:3947 AA:3948 AA:3949 AA:3950 AA:3951 AA:3952 AA:3953 AA:3954 AA:3955 AA:3956 AA:3957 AA:3958 AA:3959 AA:3960 AA:3961 AA:3962 AA:3963 AA:3964 AA:3965 AA:3966 AA:3967 AA:3968 AA:3969 AA:3970 AA:3971 AA:3972 AA:3973 AA:3974 AA:3975 AA:3976 AA:3977 AA:3978 AA:3979 AA:3980 AA:3981 AA:3982 AA:3983 AA:3984 AA:3985 AA:3986 AA:3987 AA:3988 AA:3989 AA:3990 AA:3991 AA:3992 AA:3993 AA:3994 AA:3995 AA:3996 AA:3997 AA:3998 AA:3999 AA:4000 AA:4001 AA:4002 AA:4003 AA:4004 AA:4005 AA:4006 AA:4007 AA:4008 AA:4009 AA:4010 AA:4011 AA:4012 AA:4013 AA:4014 AA:4015 AA:4016 AA:4017 AA:4018 AA:4019 AA:4020 AA:4021 AA:4022 AA:4023 AA:4024 AA:4025 AA:4026 AA:4027 AA:4028 AA:4029 AA:4030 AA:4031 AA:4032 AA:4033 AA:4034 AA:4035 AA:4036 AA:4037 AA:4038 AA:4039 AA:4040 AA:4041 AA:4042 AA:4043 AA:4044 AA:4045 AA:4046 AA:4047 AA:4048 AA:4049 AA:4050 AA:4051 AA:4052 AA:4053 AA:4054 AA:4055 AA:4056 AA:4057 AA:4058 AA:4059 AA:4060 AA:4061 AA:4062 AA:4063 AA:4064 AA:4065 AA:4066 AA:4067 AA:4068 AA:4069 AA:4070 AA:4071 AA:4072 AA:4073 AA:4074 AA:4075 AA:4076 AA:4077 AA:4078 AA:4079 AA:4080 AA:4081 AA:4082 AA:4083 AA:4084 AA:4085 AA:4086 AA:4087 AA:4088 AA:4089 AA:4090 AA:4091 AA:4092 AA:4093 AA:4094 AA:4095 AA:4096 AA:4097 AA:4098 AA:4099 AA:4100 AA:4101 AA:4102 AA:4103 AA:4104 AA:4105 AA:4106 AA:4107 AA:4108 AA:4109 AA:4110 AA:4111 AA:4112 AA:4113 AA:4114 AA:4115 AA:4116 AA:4117 AA:4118 AA:4119 AA:4120 AA:4121 AA:4122 AA:4123 AA:4124 AA:4125 AA:4126 AA:4127 AA:4128 AA:4129 AA:4130 AA:4131 AA:4132 AA:4133 AA:4134 AA:4135 AA:4136 AA:4137 AA:4138 AA:4139 AA:4140 AA:4141 AA:4142 AA:4143 AA:4144 AA:4145 AA:4146 AA:4147 AA:4148 AA:4149 AA:4150 AA:4151 AA:4152 AA:4153 AA:4154 AA:4155 AA:4156 AA:4157 AA:4158 AA:4159 AA:4160 AA:4161 AA:4162 AA:4163 AA:4164 AA:4165 AA:4166 AB:201 AB:202 AB:203 AB:204 AB:205 AB:206 AB:207 AB:208 AB:209 AB:210 AB:211 AB:212 AB:213 AB:214 AB:215 AB:216 AB:217 AB:218 AB:219 AB:220 AB:221 AB:222 AB:223 AB:224 AB:225 AB:226 AB:227 AB:228 AB:229 AB:230 AB:231 AB:232 AB:233 AB:234 AB:235 AB:236 AD:301 AD:302 AD:303 AD:304 AD:305 AD:306 AD:307 AD:308 AD:309 AD:310 AD:311 AD:312 AD:313 AE:301 AE:302 AE:303 AE:304 AF:301 AF:302 AF:303 AF:304 AF:305 AF:306 AF:307 AG:201 AG:202 AG:203 AH:201 AK:201 AK:202 AK:203 AN:201 AO:201 AO:202 AO:203 AO:204 AO:205 AO:206 AO:207 AP:201 AQ:201 AQ:202 AQ:203 AQ:204 AQ:205 AR:201 AR:202 AR:203 AR:204 AR:205 AS:201 AS:202 AS:203 AT:101 AT:102 AT:103 AT:104 AU:201 AU:202 AU:203 AU:204 AU:205 AU:206 AV:301 AW:101 AW:102 AZ:101 AZ:102 AZ:103 B1:101 B1:102 B1:103 B1:104 BA:1601 BA:1602 BA:1603 BA:1604 BA:1605 BA:1606 BA:1607 BA:1608 BA:1609 BA:1610 BA:1611 BA:1612 BA:1613 BA:1614 BA:1615 BA:1616 BA:1617 BA:1618 BA:1619 BA:1620 BA:1621 BA:1622 BA:1623 BA:1624 BA:1625 BA:1626 BA:1627 BA:1628 BA:1629 BA:1630 BA:1631 BA:1632 BA:1633 BA:1634 BA:1635 BA:1636 BA:1637 BA:1638 BA:1639 BA:1640 BA:1641 BA:1642 BA:1643 BA:1644 BA:1645 BA:1646 BA:1647 BA:1648 BA:1649 BA:1650 BA:1651 BA:1652 BA:1653 BA:1654 BA:1655 BA:1656 BA:1657 BA:1658 BA:1659 BA:1660 BA:1661 BA:1662 BA:1663 BA:1664 BA:1665 BA:1666 BA:1667 BA:1668 BA:1669 BA:1670 BA:1671 BA:1672 BA:1673 BA:1674 BA:1675 BA:1676 BA:1677 BA:1678 BA:1679 BA:1680 BA:1681 BA:1682 BA:1683 BA:1684 BA:1685 BA:1686 BA:1687 BA:1688 BA:1689 BA:1690 BA:1691 BA:1692 BA:1693 BA:1694 BA:1695 BA:1696 BA:1697 BA:1698 BA:1699 BA:1700 BA:1701 BA:1702 BA:1703 BA:1704 BA:1705 BA:1706 BA:1707 BA:1708 BA:1709 BA:1710 BA:1711 BA:1712 BA:1713 BA:1714 BA:1715 BA:1716 BA:1717 BA:1718 BA:1719 BA:1720 BA:1721 BA:1722 BA:1723 BA:1724 BA:1725 BA:1726 BA:1727 BA:1728 BA:1729 BA:1730 BA:1731 BA:1732 BA:1733 BA:1734 BA:1735 BA:1736 BA:1737 BA:1738 BA:1739 BA:1740 BA:1741 BA:1742 BA:1743 BA:1744 BA:1745 BA:1746 BA:1747 BA:1748 BA:1749 BA:1750 BA:1751 BA:1752 BA:1753 BA:1754 BA:1755 BA:1756 BA:1757 BA:1758 BA:1759 BA:1760 BA:1761 BA:1762 BA:1763 BA:1764 BA:1765 BA:1766 BA:1767 BA:1768 BA:1769 BA:1770 BA:1771 BA:1772 BA:1773 BA:1774 BA:1775 BA:1776 BA:1777 BA:1778 BA:1779 BA:1780 BA:1781 BA:1782 BA:1783 BA:1784 BA:1785 BA:1786 BA:1787 BA:1788 BA:1789 BA:1790 BA:1791 BA:1792 BA:1793 BA:1794 BA:1795 BA:1796 BA:1797 BA:1798 BA:1799 BA:1800 BA:1801 BA:1802 BA:1803 BA:1804 BA:1805 BA:1806 BA:1807 BA:1808 BA:1809 BA:1810 BA:1811 BA:1812 BA:1813 BA:1814 BA:1815 BA:1816 BA:1817 BA:1818 BA:1819 BA:1820 BA:1821 BA:1822 BA:1823 BA:1824 BA:1825 BA:1826 BA:1827 BA:1828 BA:1829 BA:1830 BA:1831 BA:1832 BA:1833 BA:1834 BA:1835 BA:1836 BA:1837 BA:1838 BA:1839 BA:1840 BA:1841 BA:1842 BA:1843 BA:1844 BA:1845 BA:1846 BA:1847 BA:1848 BA:1849 BA:1850 BA:1851 BA:1852 BA:1853 BA:1854 BA:1855 BA:1856 BA:1857 BA:1858 BA:1859 BA:1860 BA:1861 BA:1862 BA:1863 BA:1864 BA:1865 BA:1866 BA:1867 BA:1868 BA:1869 BA:1870 BA:1871 BA:1872 BA:1873 BA:1874 BA:1875 BA:1876 BA:1877 BA:1878 BA:1879 BA:1880 BA:1881 BA:1882 BA:1883 BA:1884 BA:1885 BA:1886 BA:1887 BA:1888 BA:1889 BA:1890 BA:1891 BA:1892 BA:1893 BA:1894 BA:1895 BA:1896 BA:1897 BA:1898 BA:1899 BA:1900 BA:1901 BA:1902 BA:1903 BA:1904 BA:1905 BA:1906 BA:1907 BA:1908 BA:1909 BA:1910 BA:1911 BA:1912 BA:1913 BA:1914 BA:1915 BA:1916 BA:1917 BA:1918 BA:1919 BA:1920 BA:1921 BA:1922 BA:1923 BA:1924 BA:1925 BA:1926 BA:1927 BA:1928 BA:1929 BA:1930 BA:1931 BA:1932 BA:1933 BA:1934 BA:1935 BA:1936 BA:1937 BA:1938 BA:1939 BA:1940 BA:1941 BA:1942 BA:1943 BA:1944 BA:1945 BA:1946 BA:1947 BA:1948 BA:1949 BA:1950 BA:1951 BA:1952 BA:1953 BA:1954 BA:1955 BA:1956 BA:1957 BA:1958 BA:1959 BA:1960 BA:1961 BA:1962 BA:1963 BA:1964 BA:1965 BA:1966 BA:1967 BA:1968 BA:1969 BA:1970 BA:1971 BA:1972 BA:1973 BA:1974 BA:1975 BA:1976 BA:1977 BA:1978 BA:1979 BA:1980 BA:1981 BA:1982 BA:1983 BA:1984 BA:1985 BA:1986 BA:1987 BA:1988 BA:1989 BA:1990 BA:1991 BA:1992 BA:1993 BA:1994 BA:1995 BA:1996 BA:1997 BA:1998 BA:1999 BA:2000 BA:2001 BA:2002 BA:2003 BA:2004 BA:2005 BA:2006 BA:2007 BA:2008 BA:2009 BA:2010 BA:2011 BA:2012 BA:2013 BA:2014 BA:2015 BA:2016 BA:2017 BA:2018 BA:2019 BA:2020 BA:2021 BA:2022 BA:2023 BA:2024 BA:2025 BA:2026 BA:2027 BA:2028 BA:2029 BA:2030 BA:2031 BA:2032 BA:2033 BA:2034 BA:2035 BA:2036 BA:2037 BA:2038 BA:2039 BA:2040 BA:2041 BA:2042 BA:2043 BA:2044 BA:2045 BA:2046 BA:2047 BA:2048 BA:2049 BA:2050 BA:2051 BA:2052 BA:2053 BA:2054 BA:2055 BA:2056 BA:2057 BA:2058 BA:2059 BA:2060 BA:2061 BA:2062 BA:2063 BA:2064 BA:2065 BA:2066 BA:2067 BA:2068 BA:2069 BA:2070 BA:2071 BA:2072 BA:2073 BA:2074 BA:2075 BA:2076 BA:2077 BA:2078 BA:2079 BA:2080 BA:2081 BA:2082 BA:2083 BA:2084 BA:2085 BA:2086 BA:2087 BA:2088 BA:2089 BA:2090 BA:2091 BA:2092 BA:2093 BA:2094 BA:2095 BA:2096 BA:2097 BA:2098 BA:2099 BA:2100 BA:2101 BA:2102 BA:2103 BA:2104 BA:2105 BA:2106 BA:2107 BA:2108 BA:2109 BA:2110 BA:2111 BA:2112 BA:2113 BA:2114 BA:2115 BA:2116 BA:2117 BA:2118 BA:2119 BA:2120 BA:2121 BA:2122 BA:2123 BA:2124 BA:2125 BA:2126 BA:2127 BA:2128 BA:2129 BA:2130 BA:2131 BA:2132 BA:2133 BA:2134 BA:2135 BA:2136 BA:2137 BA:2138 BA:2139 BA:2140 BA:2141 BA:2142 BA:2143 BA:2144 BA:2145 BA:2146 BA:2147 BA:2148 BA:2149 BA:2150 BA:2151 BA:2152 BA:2153 BA:2154 BA:2155 BA:2156 BA:2157 BA:2158 BA:2159 BA:2160 BA:2161 BA:2162 BA:2163 BA:2164 BA:2165 BA:2166 BA:2167 BA:2168 BA:2169 BA:2170 BA:2171 BA:2172 BA:2173 BA:2174 BA:2175 BA:2176 BA:2177 BA:2178 BA:2179 BA:2180 BA:2181 BA:2182 BA:2183 BA:2184 BA:2185 BA:2186 BA:2187 BA:2188 BA:2189 BA:2190 BA:2191 BA:2192 BA:2193 BA:2194 BA:2195 BA:2196 BA:2197 BA:2198 BA:2199 BA:2200 BA:2201 BA:2202 BA:2203 BA:2204 BA:2205 BA:2206 BA:2207 BA:2208 BA:2209 BA:2210 BA:2211 BA:2212 BA:2213 BA:2214 BA:2215 BA:2216 BA:2217 BA:2218 BA:2219 BA:2220 BA:2221 BA:2222 BA:2223 BA:2224 BA:2225 BA:2226 BA:2227 BA:2228 BA:2229 BA:2230 BA:2231 BA:2232 BA:2233 BA:2234 BA:2235 BA:2236 BA:2237 BA:2238 BA:2239 BA:2240 BA:2241 BA:2242 BA:2243 BA:2244 BA:2245 BA:2246 BA:2247 BA:2248 BA:2249 BA:2250 BA:2251 BA:2252 BA:2253 BA:2254 BA:2255 BA:2256 BA:2257 BA:2258 BA:2259 BA:2260 BA:2261 BA:2262 BA:2263 BA:2264 BA:2265 BA:2266 BA:2267 BA:2268 BA:2269 BA:2270 BA:2271 BA:2272 BA:2273 BA:2274 BA:2275 BA:2276 BB:101 BB:102 BB:103 BB:104 BB:105 BB:106 BB:107 BB:108 BB:109 BB:110 BB:111 BB:112 BB:113 BC:101 BC:102 BC:103 BC:104 BC:105 BC:106 BC:107 BC:108 BC:109 BC:110 BC:111 BC:112 BC:113 BC:114 BC:115 BC:116 BD:101 BD:102 BD:103 BD:104 BD:105 BD:106 BD:107 BD:108 BD:109 BD:110 BD:111 BD:112 BD:113 BD:114 BD:115 BD:116 BD:117 BD:118 BD:119 BD:120 BD:121 BD:122 BD:123 BD:124 BD:125 BD:126 BE:301 BE:302 BE:303 BE:304 BE:305 BF:301 BF:302 BG:301 BG:302 BG:303 BG:304 BG:305 BG:306 BG:307 BH:201 BH:202 BH:203 BH:204 BH:205 BI:201 BK:201 BK:202 BK:203 BK:204 BK:205 BK:206 BL:201 BL:202 BM:201 BM:202 BM:203 BO:201 BP:201 BP:202 BQ:101 BQ:102 BQ:103 BS:101 BS:102 BS:103 BS:104 BS:105 BS:106 BS:107 BS:108 BS:109 BT:201 BT:202 BU:1601 BV:101 BW:201 BW:202 BW:203 BW:204 BW:205 BW:206 BW:207 BW:208 BX:101 C1:101 C1:102 C1:103 C1:104 C1:105 C1:106 CA:1601 CA:1602 CA:1603 CA:1604 CA:1605 CA:1606 CA:1607 CA:1608 CA:1609 CA:1610 CA:1611 CA:1612 CA:1613 CA:1614 CA:1615 CA:1616 CA:1617 CA:1618 CA:1619 CA:1620 CA:1621 CA:1622 CA:1623 CA:1624 CA:1625 CA:1626 CA:1627 CA:1628 CA:1629 CA:1630 CA:1631 CA:1632 CA:1633 CA:1634 CA:1635 CA:1636 CA:1637 CA:1638 CA:1639 CA:1640 CA:1641 CA:1642 CA:1643 CA:1644 CA:1645 CA:1646 CA:1647 CA:1648 CA:1649 CA:1650 CA:1651 CA:1652 CA:1653 CA:1654 CA:1655 CA:1656 CA:1657 CA:1658 CA:1659 CA:1660 CA:1661 CA:1662 CA:1663 CA:1664 CA:1665 CA:1666 CA:1667 CA:1668 CA:1669 CA:1670 CA:1671 CA:1672 CA:1673 CA:1674 CA:1675 CA:1676 CA:1677 CA:1678 CA:1679 CA:1680 CA:1681 CA:1682 CA:1683 CA:1684 CA:1685 CA:1686 CA:1687 CA:1688 CA:1689 CA:1690 CA:1691 CA:1692 CA:1693 CA:1694 CA:1695 CA:1696 CA:1697 CA:1698 CA:1699 CA:1700 CA:1701 CA:1702 CA:1703 CA:1704 CA:1705 CA:1706 CA:1707 CA:1708 CA:1709 CA:1710 CA:1711 CA:1712 CA:1713 CA:1714 CA:1715 CA:1716 CA:1717 CA:1718 CA:1719 CA:1720 CA:1721 CA:1722 CA:1723 CA:1724 CA:1725 CA:1726 CA:1727 CA:1728 CA:1729 CA:1730 CA:1731 CA:1732 CA:1733 CA:1734 CA:1735 CA:1736 CA:1737 CA:1738 CA:1739 CA:1740 CA:1741 CA:1742 CA:1743 CA:1744 CA:1745 CA:1746 CA:1747 CA:1748 CA:1749 CA:1750 CA:1751 CA:1752 CA:1753 CA:1754 CA:1755 CA:1756 CA:1757 CA:1758 CA:1759 CA:1760 CA:1761 CA:1762 CA:1763 CA:1764 CA:1765 CA:1766 CA:1767 CA:1768 CA:1769 CA:1770 CA:1771 CA:1772 CA:1773 CA:1774 CA:1775 CA:1776 CA:1777 CA:1778 CA:1779 CA:1780 CA:1781 CA:1782 CA:1783 CA:1784 CA:1785 CA:1786 CA:1787 CA:1788 CA:1789 CA:1790 CA:1791 CA:1792 CA:1793 CA:1794 CA:1795 CA:1796 CA:1797 CA:1798 CA:1799 CA:1800 CA:1801 CA:1802 CA:1803 CA:1804 CA:1805 CA:1806 CA:1807 CA:1808 CA:1809 CA:1810 CA:1811 CA:1812 CA:1813 CA:1814 CA:1815 CA:1816 CA:1817 CA:1818 CA:1819 CA:1820 CA:1821 CA:1822 CA:1823 CA:1824 CA:1825 CA:1826 CA:1827 CA:1828 CA:1829 CA:1830 CA:1831 CA:1832 CA:1833 CA:1834 CA:1835 CA:1836 CA:1837 CA:1838 CA:1839 CA:1840 CA:1841 CA:1842 CA:1843 CA:1844 CA:1845 CA:1846 CA:1847 CA:1848 CA:1849 CA:1850 CA:1851 CA:1852 CA:1853 CA:1854 CA:1855 CA:1856 CA:1857 CA:1858 CA:1859 CA:1860 CA:1861 CA:1862 CA:1863 CA:1864 CA:1865 CA:1866 CA:1867 CA:1868 CA:1869 CA:1870 CA:1871 CA:1872 CA:1873 CA:1874 CA:1875 CA:1876 CA:1877 CA:1878 CA:1879 CA:1880 CA:1881 CA:1882 CA:1883 CA:1884 CA:1885 CA:1886 CA:1887 CA:1888 CA:1889 CA:1890 CA:1891 CA:1892 CA:1893 CA:1894 CA:1895 CA:1896 CA:1897 CA:1898 CA:1899 CA:1900 CA:1901 CA:1902 CA:1903 CA:1904 CA:1905 CA:1906 CA:1907 CA:1908 CA:1909 CA:1910 CA:1911 CA:1912 CA:1913 CA:1914 CA:1915 CA:1916 CA:1917 CA:1918 CA:1919 CA:1920 CA:1921 CA:1922 CA:1923 CA:1924 CA:1925 CA:1926 CA:1927 CA:1928 CA:1929 CA:1930 CA:1931 CA:1932 CA:1933 CA:1934 CA:1935 CA:1936 CA:1937 CA:1938 CA:1939 CA:1940 CA:1941 CA:1942 CA:1943 CA:1944 CA:1945 CA:1946 CA:1947 CA:1948 CA:1949 CA:1950 CA:1951 CA:1952 CA:1953 CA:1954 CA:1955 CA:1956 CA:1957 CA:1958 CA:1959 CA:1960 CA:1961 CA:1962 CA:1963 CA:1964 CA:1965 CA:1966 CA:1967 CA:1968 CA:1969 CA:1970 CA:1971 CA:1972 CA:1973 CA:1974 CA:1975 CA:1976 CA:1977 CA:1978 CA:1979 CA:1980 CA:1981 CA:1982 CA:1983 CA:1984 CA:1985 CA:1986 CA:1987 CA:1988 CA:1989 CA:1990 CA:1991 CA:1992 CA:1993 CA:1994 CA:1995 CA:1996 CA:1997 CA:1998 CA:1999 CA:2000 CA:2001 CA:2002 CA:2003 CA:2004 CA:2005 CA:2006 CA:2007 CA:2008 CA:2009 CA:2010 CA:2011 CA:2012 CA:2013 CA:2014 CA:2015 CA:2016 CA:2017 CA:2018 CA:2019 CA:2020 CA:2021 CA:2022 CA:2023 CA:2024 CA:2025 CA:2026 CA:2027 CA:2028 CA:2029 CA:2030 CA:2031 CA:2032 CA:2033 CA:2034 CA:2035 CA:2036 CA:2037 CA:2038 CA:2039 CA:2040 CA:2041 CA:2042 CA:2043 CA:2044 CA:2045 CA:2046 CA:2047 CA:2048 CA:2049 CA:2050 CA:2051 CA:2052 CA:2053 CA:2054 CA:2055 CA:2056 CA:2057 CA:2058 CA:2059 CA:2060 CA:2061 CA:2062 CA:2063 CA:2064 CA:2065 CA:2066 CA:2067 CA:2068 CA:2069 CA:2070 CA:2071 CA:2072 CA:2073 CA:2074 CA:2075 CA:2076 CA:2077 CA:2078 CA:2079 CA:2080 CA:2081 CA:2082 CA:2083 CA:2084 CA:2085 CA:2086 CA:2087 CA:2088 CA:2089 CA:2090 CA:2091 CA:2092 CA:2093 CA:2094 CA:2095 CA:2096 CA:2097 CA:2098 CA:2099 CA:2100 CA:2101 CA:2102 CA:2103 CA:2104 CA:2105 CA:2106 CA:2107 CA:2108 CA:2109 CA:2110 CA:2111 CA:2112 CA:2113 CA:2114 CA:2115 CA:2116 CA:2117 CA:2118 CA:2119 CA:2120 CA:2121 CA:2122 CA:2123 CA:2124 CA:2125 CA:2126 CA:2127 CA:2128 CA:2129 CA:2130 CA:2131 CA:2132 CA:2133 CA:2134 CA:2135 CA:2136 CA:2137 CA:2138 CA:2139 CA:2140 CA:2141 CA:2142 CA:2143 CA:2144 CA:2145 CA:2146 CA:2147 CA:2148 CA:2149 CA:2150 CA:2151 CA:2152 CA:2153 CA:2154 CA:2155 CA:2156 CA:2157 CA:2158 CA:2159 CA:2160 CA:2161 CA:2162 CA:2163 CA:2164 CA:2165 CA:2166 CA:2167 CA:2168 CA:2169 CA:2170 CA:2171 CA:2172 CA:2173 CA:2174 CA:2175 CA:2176 CA:2177 CA:2178 CA:2179 CA:2180 CA:2181 CA:2182 CA:2183 CA:2184 CA:2185 CA:2186 CA:2187 CA:2188 CA:2189 CA:2190 CA:2191 CA:2192 CA:2193 CA:2194 CA:2195 CA:2196 CA:2197 CA:2198 CA:2199 CA:2200 CA:2201 CA:2202 CA:2203 CA:2204 CA:2205 CA:2206 CA:2207 CA:2208 CA:2209 CA:2210 CA:2211 CA:2212 CA:2213 CA:2214 CA:2215 CA:2216 CA:2217 CA:2218 CA:2219 CA:2220 CA:2221 CA:2222 CA:2223 CA:2224 CA:2225 CA:2226 CA:2227 CA:2228 CA:2229 CA:2230 CA:2231 CA:2232 CA:2233 CA:2234 CA:2235 CA:2236 CA:2237 CA:2238 CA:2239 CA:2240 CA:2241 CA:2242 CA:2243 CA:2244 CA:2245 CA:2246 CA:2247 CA:2248 CA:2249 CA:2250 CA:2251 CA:2252 CA:2253 CA:2254 CA:2255 CA:2256 CA:2257 CA:2258 CA:2259 CA:2260 CA:2261 CA:2262 CA:2263 CA:2264 CA:2265 CA:2266 CA:2267 CA:2268 CA:2269 CA:2270 CA:2271 CA:2272 CA:2273 CA:2274 CA:2275 CA:2276 CA:2277 CA:2278 CA:2279 CA:2280 CA:2281 CA:2282 CA:2283 CA:2284 CA:2285 CA:2286 CA:2287 CA:2288 CA:2289 CA:2290 CA:2291 CA:2292 CA:2293 CA:2294 CA:2295 CA:2296 CA:2297 CA:2298 CA:2299 CA:2300 CA:2301 CA:2302 CA:2303 CA:2304 CA:2305 CA:2306 CA:2307 CA:2308 CA:2309 CA:2310 CA:2311 CA:2312 CA:2313 CA:2314 CA:2315 CA:2316 CA:2317 CA:2318 CA:2319 CA:2320 CA:2321 CA:2322 CA:2323 CA:2324 CB:101 CB:102 CB:103 CB:104 CB:105 CB:106 CB:107 CB:108 CB:109 CB:110 CB:111 CB:112 CB:113 CB:114 CB:115 CB:116 CB:117 CB:118 CB:119 CB:120 CB:121 CC:101 CC:102 CC:103 CC:104 CC:105 CC:106 CC:107 CC:108 CC:109 CC:110 CC:111 CC:112 CC:113 CC:114 CC:115 CC:116 CC:117 CC:118 CC:119 CC:120 CC:121 CC:122 CC:123 CC:124 CC:125 CC:126 CC:127 CD:101 CD:102 CD:103 CD:104 CD:105 CD:106 CD:107 CD:108 CD:109 CD:110 CD:111 CD:112 CD:113 CD:114 CD:115 CD:116 CD:117 CD:118 CD:119 CD:120 CD:121 CD:122 CD:123 CD:124 CD:125 CD:126 CD:127 CD:128 CD:129 CD:130 CE:301 CE:302 CE:303 CE:304 CE:305 CE:306 CF:301 CF:302 CF:303 CG:302 CG:303 CG:304 CG:305 CG:306 CG:307 CG:308 CG:309 CG:310 CG:311 CG:312 CH:201 CH:202 CH:203 CH:204 CH:205 CH:206 CI:201 CJ:201 CK:201 CK:202 CK:203 CK:204 CK:205 CK:206 CK:207 CK:208 CK:209 CK:210 CK:211 CL:201 CM:201 CM:202 CM:203 CN:201 CN:202 CP:201 CP:202 CP:203 CQ:102 CQ:103 CQ:104 CR:101 CR:102 CR:103 CS:101 CS:102 CS:103 CS:104 CS:105 CS:106 CT:201 CT:202 CT:203 CV:101 CV:102 CV:103 CV:104 CW:201 D0:201 D0:202 D0:203 D0:204 D0:205 D0:206 D0:207 D0:208 D0:209 D0:210 D0:211 D1:201 D1:202 D1:203 D1:204 D1:205 D1:206 D1:207 D1:208 D1:209 D1:210 D1:211 D1:212 D2:201 D2:202 D2:203 D2:204 D2:205 D2:206 D2:207 D2:208 D2:209 D2:210 D3:101 D3:102 D3:103 D3:104 D3:105 D3:106 D3:107 D4:101 D4:102 D5:101 D5:102 D5:103 D5:104 D5:105 D5:106 D5:107 D5:108 D6:101 D6:102 D6:103 D7:101 D7:102 D7:103 D7:104 D7:105 D8:101 D8:102 D8:103 D8:104 D8:105 D8:106 D8:107 D8:108 D8:109 DA:3001 DA:3002 DA:3003 DA:3004 DA:3005 DA:3006 DA:3007 DA:3008 DA:3009 DA:3010 DA:3011 DA:3012 DA:3013 DA:3014 DA:3015 DA:3016 DA:3017 DA:3018 DA:3019 DA:3020 DA:3021 DA:3022 DA:3023 DA:3024 DA:3025 DA:3026 DA:3027 DA:3028 DA:3029 DA:3030 DA:3031 DA:3032 DA:3033 DA:3034 DA:3035 DA:3036 DA:3037 DA:3038 DA:3039 DA:3040 DA:3041 DA:3042 DA:3043 DA:3044 DA:3045 DA:3046 DA:3047 DA:3048 DA:3049 DA:3050 DA:3051 DA:3052 DA:3053 DA:3054 DA:3055 DA:3056 DA:3057 DA:3058 DA:3059 DA:3060 DA:3061 DA:3062 DA:3063 DA:3064 DA:3065 DA:3066 DA:3067 DA:3068 DA:3069 DA:3070 DA:3071 DA:3072 DA:3073 DA:3074 DA:3075 DA:3076 DA:3077 DA:3078 DA:3079 DA:3080 DA:3081 DA:3082 DA:3083 DA:3084 DA:3085 DA:3086 DA:3087 DA:3088 DA:3089 DA:3090 DA:3091 DA:3092 DA:3093 DA:3094 DA:3095 DA:3096 DA:3097 DA:3098 DA:3099 DA:3100 DA:3101 DA:3102 DA:3103 DA:3104 DA:3105 DA:3106 DA:3107 DA:3108 DA:3109 DA:3110 DA:3111 DA:3112 DA:3113 DA:3114 DA:3115 DA:3116 DA:3117 DA:3118 DA:3119 DA:3120 DA:3121 DA:3122 DA:3123 DA:3124 DA:3125 DA:3126 DA:3127 DA:3128 DA:3129 DA:3130 DA:3131 DA:3132 DA:3133 DA:3134 DA:3135 DA:3136 DA:3137 DA:3138 DA:3139 DA:3140 DA:3141 DA:3142 DA:3143 DA:3144 DA:3145 DA:3146 DA:3147 DA:3148 DA:3149 DA:3150 DA:3151 DA:3152 DA:3153 DA:3154 DA:3155 DA:3156 DA:3157 DA:3158 DA:3159 DA:3160 DA:3161 DA:3162 DA:3163 DA:3164 DA:3165 DA:3166 DA:3167 DA:3168 DA:3169 DA:3170 DA:3171 DA:3172 DA:3173 DA:3174 DA:3175 DA:3176 DA:3177 DA:3178 DA:3179 DA:3180 DA:3181 DA:3182 DA:3183 DA:3184 DA:3185 DA:3186 DA:3187 DA:3188 DA:3189 DA:3190 DA:3191 DA:3192 DA:3193 DA:3194 DA:3195 DA:3196 DA:3197 DA:3198 DA:3199 DA:3200 DA:3201 DA:3202 DA:3203 DA:3204 DA:3205 DA:3206 DA:3207 DA:3208 DA:3209 DA:3210 DA:3211 DA:3212 DA:3213 DA:3214 DA:3215 DA:3216 DA:3217 DA:3218 DA:3219 DA:3220 DA:3221 DA:3222 DA:3223 DA:3224 DA:3225 DA:3226 DA:3227 DA:3228 DA:3229 DA:3230 DA:3231 DA:3232 DA:3233 DA:3234 DA:3235 DA:3236 DA:3237 DA:3238 DA:3239 DA:3240 DA:3241 DA:3242 DA:3243 DA:3244 DA:3245 DA:3246 DA:3247 DA:3248 DA:3249 DA:3250 DA:3251 DA:3252 DA:3253 DA:3254 DA:3255 DA:3256 DA:3257 DA:3258 DA:3259 DA:3260 DA:3261 DA:3262 DA:3263 DA:3264 DA:3265 DA:3266 DA:3267 DA:3268 DA:3269 DA:3270 DA:3271 DA:3272 DA:3273 DA:3274 DA:3275 DA:3276 DA:3277 DA:3278 DA:3279 DA:3280 DA:3281 DA:3282 DA:3283 DA:3284 DA:3285 DA:3286 DA:3287 DA:3288 DA:3289 DA:3290 DA:3291 DA:3292 DA:3293 DA:3294 DA:3295 DA:3296 DA:3297 DA:3298 DA:3299 DA:3300 DA:3301 DA:3302 DA:3303 DA:3304 DA:3305 DA:3306 DA:3307 DA:3308 DA:3309 DA:3310 DA:3311 DA:3312 DA:3313 DA:3314 DA:3315 DA:3316 DA:3317 DA:3318 DA:3319 DA:3320 DA:3321 DA:3322 DA:3323 DA:3324 DA:3325 DA:3326 DA:3327 DA:3328 DA:3329 DA:3330 DA:3331 DA:3332 DA:3333 DA:3334 DA:3335 DA:3336 DA:3337 DA:3338 DA:3339 DA:3340 DA:3341 DA:3342 DA:3343 DA:3344 DA:3345 DA:3346 DA:3347 DA:3348 DA:3349 DA:3350 DA:3351 DA:3352 DA:3353 DA:3354 DA:3355 DA:3356 DA:3357 DA:3358 DA:3359 DA:3360 DA:3361 DA:3362 DA:3363 DA:3364 DA:3365 DA:3366 DA:3367 DA:3368 DA:3369 DA:3370 DA:3371 DA:3372 DA:3373 DA:3374 DA:3375 DA:3376 DA:3377 DA:3378 DA:3379 DA:3380 DA:3381 DA:3382 DA:3383 DA:3384 DA:3385 DA:3386 DA:3387 DA:3388 DA:3389 DA:3390 DA:3391 DA:3392 DA:3393 DA:3394 DA:3395 DA:3396 DA:3397 DA:3398 DA:3399 DA:3400 DA:3401 DA:3402 DA:3403 DA:3404 DA:3405 DA:3406 DA:3407 DA:3408 DA:3409 DA:3410 DA:3411 DA:3412 DA:3413 DA:3414 DA:3415 DA:3416 DA:3417 DA:3418 DA:3419 DA:3420 DA:3421 DA:3422 DA:3423 DA:3424 DA:3425 DA:3426 DA:3427 DA:3428 DA:3429 DA:3430 DA:3431 DA:3432 DA:3433 DA:3434 DA:3435 DA:3436 DA:3437 DA:3438 DA:3439 DA:3440 DA:3441 DA:3442 DA:3443 DA:3444 DA:3445 DA:3446 DA:3447 DA:3448 DA:3449 DA:3450 DA:3451 DA:3452 DA:3453 DA:3454 DA:3455 DA:3456 DA:3457 DA:3458 DA:3459 DA:3460 DA:3461 DA:3462 DA:3463 DA:3464 DA:3465 DA:3466 DA:3467 DA:3468 DA:3469 DA:3470 DA:3471 DA:3472 DA:3473 DA:3474 DA:3475 DA:3476 DA:3477 DA:3478 DA:3479 DA:3480 DA:3481 DA:3482 DA:3483 DA:3484 DA:3485 DA:3486 DA:3487 DA:3488 DA:3489 DA:3490 DA:3491 DA:3492 DA:3493 DA:3494 DA:3495 DA:3496 DA:3497 DA:3498 DA:3499 DA:3500 DA:3501 DA:3502 DA:3503 DA:3504 DA:3505 DA:3506 DA:3507 DA:3508 DA:3509 DA:3510 DA:3511 DA:3512 DA:3513 DA:3514 DA:3515 DA:3516 DA:3517 DA:3518 DA:3519 DA:3520 DA:3521 DA:3522 DA:3523 DA:3524 DA:3525 DA:3526 DA:3527 DA:3528 DA:3529 DA:3530 DA:3531 DA:3532 DA:3533 DA:3534 DA:3535 DA:3536 DA:3537 DA:3538 DA:3539 DA:3540 DA:3541 DA:3542 DA:3543 DA:3544 DA:3545 DA:3546 DA:3547 DA:3548 DA:3549 DA:3550 DA:3551 DA:3552 DA:3553 DA:3554 DA:3555 DA:3556 DA:3557 DA:3558 DA:3559 DA:3560 DA:3561 DA:3562 DA:3563 DA:3564 DA:3565 DA:3566 DA:3567 DA:3568 DA:3569 DA:3570 DA:3571 DA:3572 DA:3573 DA:3574 DA:3575 DA:3576 DA:3577 DA:3578 DA:3579 DA:3580 DA:3581 DA:3582 DA:3583 DA:3584 DA:3585 DA:3586 DA:3587 DA:3588 DA:3589 DA:3590 DA:3591 DA:3592 DA:3593 DA:3594 DA:3595 DA:3596 DA:3597 DA:3598 DA:3599 DA:3600 DA:3601 DA:3602 DA:3603 DA:3604 DA:3605 DA:3606 DA:3607 DA:3608 DA:3609 DA:3610 DA:3611 DA:3612 DA:3613 DA:3614 DA:3615 DA:3616 DA:3617 DA:3618 DA:3619 DA:3620 DA:3621 DA:3622 DA:3623 DA:3624 DA:3625 DA:3626 DA:3627 DA:3628 DA:3629 DA:3630 DA:3631 DA:3632 DA:3633 DA:3634 DA:3635 DA:3636 DA:3637 DA:3638 DA:3639 DA:3640 DA:3641 DA:3642 DA:3643 DA:3644 DA:3645 DA:3646 DA:3647 DA:3648 DA:3649 DA:3650 DA:3651 DA:3652 DA:3653 DA:3654 DA:3655 DA:3656 DA:3657 DA:3658 DA:3659 DA:3660 DA:3661 DA:3662 DA:3663 DA:3664 DA:3665 DA:3666 DA:3667 DA:3668 DA:3669 DA:3670 DA:3671 DA:3672 DA:3673 DA:3674 DA:3675 DA:3676 DA:3677 DA:3678 DA:3679 DA:3680 DA:3681 DA:3682 DA:3683 DA:3684 DA:3685 DA:3686 DA:3687 DA:3688 DA:3689 DA:3690 DA:3691 DA:3692 DA:3693 DA:3694 DA:3695 DA:3696 DA:3697 DA:3698 DA:3699 DA:3700 DA:3701 DA:3702 DA:3703 DA:3704 DA:3705 DA:3706 DA:3707 DA:3708 DA:3709 DA:3710 DA:3711 DA:3712 DA:3713 DA:3714 DA:3715 DA:3716 DA:3717 DA:3718 DA:3719 DA:3720 DA:3721 DA:3722 DA:3723 DA:3724 DA:3725 DA:3726 DA:3727 DA:3728 DA:3729 DA:3730 DA:3731 DA:3732 DA:3733 DA:3734 DA:3735 DA:3736 DA:3737 DA:3738 DA:3739 DA:3740 DA:3741 DA:3742 DA:3743 DA:3744 DA:3745 DA:3746 DA:3747 DA:3748 DA:3749 DA:3750 DA:3751 DA:3752 DA:3753 DA:3754 DA:3755 DA:3756 DA:3757 DA:3758 DA:3759 DA:3760 DA:3761 DA:3762 DA:3763 DA:3764 DA:3765 DA:3766 DA:3767 DA:3768 DA:3769 DA:3770 DA:3771 DA:3772 DA:3773 DA:3774 DA:3775 DA:3776 DA:3777 DA:3778 DA:3779 DA:3780 DA:3781 DA:3782 DA:3783 DA:3784 DA:3785 DA:3786 DA:3787 DA:3788 DA:3789 DA:3790 DA:3791 DA:3792 DA:3793 DA:3794 DA:3795 DA:3796 DA:3797 DA:3798 DA:3799 DA:3800 DA:3801 DA:3802 DA:3803 DA:3804 DA:3805 DA:3806 DA:3807 DA:3808 DA:3809 DA:3810 DA:3811 DA:3812 DA:3813 DA:3814 DA:3815 DA:3816 DA:3817 DA:3818 DA:3819 DA:3820 DA:3821 DA:3822 DA:3823 DA:3824 DA:3825 DA:3826 DA:3827 DA:3828 DA:3829 DA:3830 DA:3831 DA:3832 DA:3833 DA:3834 DA:3835 DA:3836 DA:3837 DA:3838 DA:3839 DA:3840 DA:3841 DA:3842 DA:3843 DA:3844 DA:3845 DA:3846 DA:3847 DA:3848 DA:3849 DA:3850 DA:3851 DA:3852 DA:3853 DA:3854 DA:3855 DA:3856 DA:3857 DA:3858 DA:3859 DA:3860 DA:3861 DA:3862 DA:3863 DA:3864 DA:3865 DA:3866 DA:3867 DA:3868 DA:3869 DA:3870 DA:3871 DA:3872 DA:3873 DA:3874 DA:3875 DA:3876 DA:3877 DA:3878 DA:3879 DA:3880 DA:3881 DA:3882 DA:3883 DA:3884 DA:3885 DA:3886 DA:3887 DA:3888 DA:3889 DA:3890 DA:3891 DA:3892 DA:3893 DA:3894 DA:3895 DA:3896 DA:3897 DA:3898 DA:3899 DA:3900 DA:3901 DA:3902 DA:3903 DA:3904 DA:3905 DA:3906 DA:3907 DA:3908 DA:3909 DA:3910 DA:3911 DA:3912 DA:3913 DA:3914 DA:3915 DA:3916 DA:3917 DA:3918 DA:3919 DA:3920 DA:3921 DA:3922 DA:3923 DA:3924 DA:3925 DA:3926 DA:3927 DA:3928 DA:3929 DA:3930 DA:3931 DA:3932 DA:3933 DA:3934 DA:3935 DA:3936 DA:3937 DA:3938 DA:3939 DA:3940 DA:3941 DA:3942 DA:3943 DA:3944 DA:3945 DA:3946 DA:3947 DA:3948 DA:3949 DA:3950 DA:3951 DA:3952 DA:3953 DA:3954 DA:3955 DA:3956 DA:3957 DA:3958 DA:3959 DA:3960 DA:3961 DA:3962 DA:3963 DA:3964 DA:3965 DA:3966 DA:3967 DA:3968 DA:3969 DA:3970 DA:3971 DA:3972 DA:3973 DA:3974 DA:3975 DA:3976 DA:3977 DA:3978 DA:3979 DA:3980 DA:3981 DA:3982 DA:3983 DA:3984 DA:3985 DA:3986 DA:3987 DA:3988 DA:3989 DA:3990 DA:3991 DA:3992 DA:3993 DA:3994 DA:3995 DA:3996 DA:3997 DA:3998 DA:3999 DA:4000 DA:4001 DA:4002 DA:4003 DA:4004 DA:4005 DA:4006 DA:4007 DA:4008 DA:4009 DA:4010 DA:4011 DA:4012 DA:4013 DA:4014 DA:4015 DA:4016 DA:4017 DA:4018 DA:4019 DA:4020 DA:4021 DA:4022 DA:4023 DA:4024 DA:4025 DA:4026 DA:4027 DA:4028 DA:4029 DA:4030 DA:4031 DA:4032 DA:4033 DA:4034 DA:4035 DA:4036 DA:4037 DA:4038 DA:4039 DA:4040 DA:4041 DA:4042 DA:4043 DA:4044 DA:4045 DA:4046 DA:4047 DA:4048 DA:4049 DA:4050 DA:4051 DA:4052 DA:4053 DA:4054 DA:4055 DA:4056 DA:4057 DA:4058 DA:4059 DA:4060 DA:4061 DA:4062 DA:4063 DA:4064 DA:4065 DA:4066 DA:4067 DA:4068 DA:4069 DA:4070 DA:4071 DA:4072 DA:4073 DA:4074 DA:4075 DA:4076 DA:4077 DA:4078 DA:4079 DA:4080 DA:4081 DA:4082 DA:4083 DA:4084 DA:4085 DA:4086 DA:4087 DA:4088 DA:4089 DA:4090 DA:4091 DA:4092 DA:4093 DA:4094 DA:4095 DA:4096 DA:4097 DA:4098 DA:4099 DA:4100 DA:4101 DA:4102 DA:4103 DA:4104 DA:4105 DA:4106 DA:4107 DA:4108 DA:4109 DA:4110 DA:4111 DA:4112 DA:4113 DA:4114 DA:4115 DA:4116 DA:4117 DA:4118 DA:4119 DA:4120 DA:4121 DA:4122 DA:4123 DA:4124 DA:4125 DA:4126 DA:4127 DA:4128 DA:4129 DA:4130 DA:4131 DA:4132 DA:4133 DA:4134 DA:4135 DA:4136 DA:4137 DA:4138 DA:4139 DA:4140 DA:4141 DA:4142 DA:4143 DA:4144 DA:4145 DA:4146 DA:4147 DA:4148 DA:4149 DA:4150 DA:4151 DA:4152 DA:4153 DA:4154 DA:4155 DA:4156 DA:4157 DA:4158 DA:4159 DA:4160 DA:4161 DA:4162 DA:4163 DA:4164 DA:4165 DA:4166 DA:4167 DA:4168 DA:4169 DA:4170 DA:4171 DA:4172 DA:4173 DA:4174 DA:4175 DA:4176 DA:4177 DA:4178 DA:4179 DA:4180 DA:4181 DA:4182 DA:4183 DA:4184 DA:4185 DA:4186 DA:4187 DA:4188 DA:4189 DA:4190 DA:4191 DA:4192 DA:4193 DA:4194 DA:4195 DA:4196 DA:4197 DA:4198 DA:4199 DA:4200 DA:4201 DA:4202 DA:4203 DA:4204 DA:4205 DA:4206 DA:4207 DA:4208 DA:4209 DA:4210 DA:4211 DA:4212 DA:4213 DA:4214 DA:4215 DA:4216 DA:4217 DA:4218 DA:4219 DA:4220 DA:4221 DA:4222 DA:4223 DA:4224 DA:4225 DA:4226 DA:4227 DA:4228 DA:4229 DA:4230 DA:4231 DA:4232 DA:4233 DA:4234 DA:4235 DA:4236 DA:4237 DA:4238 DA:4239 DA:4240 DA:4241 DA:4242 DA:4243 DA:4244 DA:4245 DA:4246 DA:4247 DA:4248 DA:4249 DA:4250 DA:4251 DA:4252 DA:4253 DA:4254 DA:4255 DA:4256 DA:4257 DA:4258 DA:4259 DA:4260 DA:4261 DA:4262 DA:4263 DA:4264 DA:4265 DA:4266 DA:4267 DA:4268 DA:4269 DA:4270 DA:4271 DA:4272 DA:4273 DA:4274 DA:4275 DA:4276 DA:4277 DA:4278 DA:4279 DA:4280 DA:4281 DA:4282 DA:4283 DA:4284 DA:4285 DA:4286 DA:4287 DA:4288 DA:4289 DA:4290 DA:4291 DA:4292 DA:4293 DA:4294 DA:4295 DA:4296 DA:4297 DA:4298 DA:4299 DA:4300 DA:4301 DA:4302 DA:4303 DA:4304 DA:4305 DA:4306 DA:4307 DA:4308 DA:4309 DA:4310 DA:4311 DA:4312 DA:4313 DA:4314 DA:4315 DA:4316 DA:4317 DA:4318 DA:4319 DA:4320 DA:4321 DA:4322 DA:4323 DA:4324 DA:4325 DA:4326 DA:4327 DA:4328 DA:4329 DA:4330 DA:4331 DA:4332 DA:4333 DA:4334 DA:4335 DA:4336 DA:4337 DA:4338 DA:4339 DA:4340 DA:4341 DA:4342 DA:4343 DA:4344 DA:4345 DA:4346 DA:4347 DA:4348 DA:4349 DA:4350 DA:4351 DA:4352 DA:4353 DA:4354 DA:4355 DA:4356 DA:4357 DA:4358 DA:4359 DA:4360 DA:4361 DA:4362 DA:4363 DA:4364 DA:4365 DA:4366 DA:4367 DA:4368 DA:4369 DA:4370 DA:4371 DA:4372 DA:4373 DA:4374 DA:4375 DA:4376 DA:4377 DA:4378 DA:4379 DA:4380 DA:4381 DA:4382 DA:4383 DA:4384 DA:4385 DA:4386 DA:4387 DA:4388 DA:4389 DA:4390 DA:4391 DA:4392 DA:4393 DA:4394 DA:4395 DA:4396 DA:4397 DA:4398 DA:4399 DA:4400 DA:4401 DA:4402 DA:4403 DA:4404 DA:4405 DA:4406 DA:4407 DA:4408 DA:4409 DA:4410 DA:4411 DA:4412 DA:4413 DA:4414 DA:4415 DA:4416 DA:4417 DA:4418 DA:4419 DA:4420 DA:4421 DA:4422 DA:4423 DA:4424 DA:4425 DA:4426 DA:4427 DA:4428 DA:4429 DA:4430 DA:4431 DA:4432 DA:4433 DA:4434 DA:4435 DA:4436 DA:4437 DA:4438 DA:4439 DA:4440 DA:4441 DA:4442 DA:4443 DA:4444 DA:4445 DA:4446 DA:4447 DA:4448 DA:4449 DA:4450 DA:4451 DA:4452 DA:4453 DA:4454 DA:4455 DA:4456 DA:4457 DA:4458 DA:4459 DA:4460 DA:4461 DA:4462 DA:4463 DA:4464 DA:4465 DA:4466 DA:4467 DA:4468 DA:4469 DA:4470 DA:4471 DA:4472 DA:4473 DA:4474 DA:4475 DA:4476 DA:4477 DA:4478 DA:4479 DA:4480 DA:4481 DA:4482 DA:4483 DA:4484 DA:4485 DA:4486 DA:4487 DA:4488 DA:4489 DA:4490 DA:4491 DA:4492 DA:4493 DA:4494 DA:4495 DA:4496 DA:4497 DA:4498 DA:4499 DA:4500 DA:4501 DA:4502 DA:4503 DA:4504 DA:4505 DA:4506 DA:4507 DA:4508 DA:4509 DA:4510 DA:4511 DA:4512 DA:4513 DA:4514 DA:4515 DA:4516 DA:4517 DA:4518 DA:4519 DA:4520 DA:4521 DA:4522 DA:4523 DA:4524 DA:4525 DA:4526 DA:4527 DA:4528 DA:4529 DA:4530 DA:4531 DA:4532 DA:4533 DA:4534 DA:4535 DA:4536 DA:4537 DA:4538 DA:4539 DA:4540 DA:4541 DA:4542 DA:4543 DA:4544 DA:4545 DA:4546 DA:4547 DA:4548 DA:4549 DA:4550 DA:4551 DA:4552 DA:4553 DA:4554 DA:4555 DA:4556 DA:4557 DA:4558 DA:4559 DA:4560 DA:4561 DA:4562 DA:4563 DA:4564 DA:4565 DA:4566 DA:4567 DA:4568 DA:4569 DA:4570 DA:4571 DA:4572 DA:4573 DA:4574 DA:4575 DA:4576 DA:4577 DA:4578 DA:4579 DA:4580 DA:4581 DA:4582 DA:4583 DA:4584 DA:4585 DA:4586 DA:4587 DA:4588 DA:4589 DA:4590 DA:4591 DA:4592 DA:4593 DA:4594 DA:4595 DA:4596 DA:4597 DA:4598 DA:4599 DA:4600 DA:4601 DA:4602 DA:4603 DA:4604 DA:4605 DA:4606 DA:4607 DA:4608 DA:4609 DA:4610 DA:4611 DA:4612 DA:4613 DA:4614 DA:4615 DA:4616 DA:4617 DA:4618 DA:4619 DA:4620 DA:4621 DA:4622 DA:4623 DA:4624 DA:4625 DA:4626 DA:4627 DA:4628 DA:4629 DA:4630 DA:4631 DA:4632 DA:4633 DA:4634 DA:4635 DA:4636 DA:4637 DA:4638 DA:4639 DA:4640 DA:4641 DA:4642 DA:4643 DA:4644 DA:4645 DA:4646 DA:4647 DA:4648 DA:4649 DA:4650 DA:4651 DA:4652 DA:4653 DA:4654 DA:4655 DA:4656 DA:4657 DA:4658 DA:4659 DA:4660 DA:4661 DA:4662 DA:4663 DA:4664 DA:4665 DA:4666 DA:4667 DA:4668 DA:4669 DA:4670 DA:4671 DA:4672 DA:4673 DA:4674 DA:4675 DA:4676 DA:4677 DA:4678 DA:4679 DA:4680 DA:4681 DA:4682 DA:4683 DA:4684 DA:4685 DA:4686 DA:4687 DA:4688 DA:4689 DA:4690 DA:4691 DA:4692 DA:4693 DA:4694 DA:4695 DA:4696 DA:4697 DA:4698 DA:4699 DA:4700 DA:4701 DA:4702 DA:4703 DA:4704 DA:4705 DA:4706 DA:4707 DA:4708 DA:4709 DA:4710 DA:4711 DA:4712 DA:4713 DA:4714 DA:4715 DA:4716 DA:4717 DA:4718 DA:4719 DA:4720 DA:4721 DA:4722 DA:4723 DA:4724 DA:4725 DA:4726 DA:4727 DA:4728 DA:4729 DA:4730 DA:4731 DA:4732 DA:4733 DA:4734 DA:4735 DA:4736 DA:4737 DA:4738 DA:4739 DA:4740 DA:4741 DA:4742 DA:4743 DA:4744 DA:4745 DA:4746 DA:4747 DA:4748 DA:4749 DA:4750 DA:4751 DA:4752 DA:4753 DA:4754 DA:4755 DA:4756 DA:4757 DA:4758 DA:4759 DA:4760 DA:4761 DA:4762 DA:4763 DA:4764 DA:4765 DA:4766 DA:4767 DA:4768 DA:4769 DA:4770 DA:4771 DA:4772 DA:4773 DA:4774 DA:4775 DA:4776 DA:4777 DA:4778 DA:4779 DA:4780 DA:4781 DA:4782 DA:4783 DA:4784 DA:4785 DA:4786 DA:4787 DA:4788 DA:4789 DA:4790 DA:4791 DA:4792 DA:4793 DA:4794 DA:4795 DA:4796 DA:4797 DA:4798 DA:4799 DA:4800 DA:4801 DA:4802 DA:4803 DA:4804 DA:4805 DA:4806 DA:4807 DA:4808 DA:4809 DA:4810 DA:4811 DA:4812 DA:4813 DA:4814 DA:4815 DA:4816 DA:4817 DA:4818 DA:4819 DA:4820 DA:4821 DA:4822 DA:4823 DA:4824 DA:4825 DA:4826 DA:4827 DA:4828 DA:4829 DA:4830 DA:4831 DA:4832 DA:4833 DA:4834 DA:4835 DA:4836 DA:4837 DA:4838 DA:4839 DA:4840 DA:4841 DA:4842 DA:4843 DA:4844 DA:4845 DA:4846 DA:4847 DA:4848 DA:4849 DA:4850 DA:4851 DA:4852 DA:4853 DA:4854 DA:4855 DA:4856 DA:4857 DA:4858 DA:4859 DA:4860 DA:4861 DA:4862 DA:4863 DA:4864 DA:4865 DA:4866 DA:4867 DA:4868 DA:4869 DA:4870 DA:4871 DA:4872 DA:4873 DA:4874 DA:4875 DA:4876 DA:4877 DA:4878 DA:4879 DA:4880 DA:4881 DA:4882 DA:4883 DA:4884 DA:4885 DA:4886 DA:4887 DA:4888 DA:4889 DA:4890 DA:4891 DA:4892 DA:4893 DA:4894 DA:4895 DA:4896 DA:4897 DA:4898 DA:4899 DA:4900 DA:4901 DA:4902 DA:4903 DA:4904 DA:4905 DA:4906 DA:4907 DA:4908 DA:4909 DA:4910 DA:4911 DA:4912 DA:4913 DA:4914 DA:4915 DA:4916 DA:4917 DA:4918 DA:4919 DA:4920 DA:4921 DA:4922 DA:4923 DA:4924 DA:4925 DA:4926 DA:4927 DA:4928 DA:4929 DA:4930 DA:4931 DA:4932 DA:4933 DA:4934 DA:4935 DA:4936 DA:4937 DA:4938 DA:4939 DA:4940 DA:4941 DA:4942 DA:4943 DA:4944 DA:4945 DA:4946 DA:4947 DA:4948 DA:4949 DA:4950 DA:4951 DA:4952 DA:4953 DA:4954 DA:4955 DA:4956 DA:4957 DA:4958 DA:4959 DA:4960 DA:4961 DA:4962 DA:4963 DA:4964 DA:4965 DA:4966 DA:4967 DA:4968 DA:4969 DA:4970 DA:4971 DA:4972 DA:4973 DA:4974 DA:4975 DA:4976 DA:4977 DA:4978 DA:4979 DA:4980 DA:4981 DA:4982 DA:4983 DA:4984 DA:4985 DA:4986 DA:4987 DA:4988 DA:4989 DA:4990 DA:4991 DA:4992 DA:4993 DA:4994 DA:4995 DA:4996 DA:4997 DA:4998 DA:4999 DA:5000 DA:5001 DA:5002 DA:5003 DA:5004 DA:5005 DA:5006 DA:5007 DA:5008 DA:5009 DA:5010 DA:5011 DA:5012 DA:5013 DA:5014 DA:5015 DA:5016 DA:5017 DA:5018 DA:5019 DA:5020 DA:5021 DA:5022 DA:5023 DA:5024 DA:5025 DA:5026 DA:5027 DA:5028 DA:5029 DA:5030 DA:5031 DA:5032 DA:5033 DA:5034 DA:5035 DA:5036 DA:5037 DA:5038 DA:5039 DA:5040 DA:5041 DA:5042 DA:5043 DA:5044 DA:5045 DA:5046 DA:5047 DA:5048 DA:5049 DA:5050 DA:5051 DA:5052 DA:5053 DA:5054 DA:5055 DA:5056 DA:5057 DA:5058 DA:5059 DA:5060 DA:5061 DA:5062 DA:5063 DA:5064 DA:5065 DA:5066 DA:5067 DA:5068 DA:5069 DA:5070 DA:5071 DA:5072 DA:5073 DA:5074 DA:5075 DA:5076 DA:5077 DB:201 DB:202 DB:203 DB:204 DB:205 DB:206 DB:207 DB:208 DB:209 DB:210 DB:211 DB:212 DB:213 DB:214 DB:215 DB:216 DB:217 DB:218 DB:219 DB:220 DB:221 DB:222 DB:223 DB:224 DB:225 DB:226 DB:227 DB:228 DB:229 DB:230 DB:231 DB:232 DB:233 DB:234 DB:235 DB:236 DB:237 DB:238 DB:239 DB:240 DB:241 DB:242 DB:243 DB:244 DB:245 DB:246 DB:247 DB:248 DB:249 DB:250 DB:251 DB:252 DB:253 DB:254 DB:255 DB:256 DB:257 DB:258 DB:259 DB:260 DB:261 DB:262 DB:263 DB:264 DB:265 DB:266 DB:267 DB:268 DB:269 DB:270 DB:271 DB:272 DB:273 DB:274 DB:275 DB:276 DD:301 DD:302 DD:303 DD:304 DD:305 DD:306 DD:307 DD:308 DD:309 DD:310 DD:311 DD:312 DD:313 DD:314 DE:301 DE:302 DE:303 DE:304 DE:305 DE:306 DE:307 DE:308 DE:309 DE:310 DE:311 DE:312 DE:313 DE:314 DE:315 DF:301 DF:302 DF:303 DF:304 DF:305 DF:306 DF:307 DF:308 DF:309 DF:310 DF:311 DF:312 DF:313 DF:314 DF:315 DF:316 DF:317 DF:318 DF:319 DF:320 DF:321 DF:322 DF:323 DF:324 DF:325 DG:201 DG:202 DG:203 DG:204 DG:205 DH:201 DH:202 DH:203 DH:204 DH:205 DK:201 DK:202 DL:201 DM:201 DM:202 DM:203 DM:204 DM:205 DM:206 DN:201 DN:202 DO:201 DO:202 DO:203 DO:204 DO:205 DO:206 DO:207 DO:208 DO:209 DO:210 DO:211 DO:212 DO:213 DO:214 DO:215 DO:216 DO:217 DO:218 DP:201 DP:202 DP:203 DP:204 DQ:201 DQ:202 DR:201 DR:202 DR:203 DR:204 DS:201 DS:202 DS:203 DS:204 DS:205 DS:206 DS:207 DS:208 DT:101 DT:102 DT:103 DT:104 DT:105 DT:106 DT:107 DU:201 DU:202 DU:203 DU:204 DU:205 DU:206 DU:207 DU:208 DU:209 DU:210 DU:211 DU:212 DU:213 DU:214 DU:215 DU:216 DU:217 DU:218 DU:219 DV:301 DV:302 DV:303 DV:304 DV:305 DW:101 DW:102 DW:103 DW:104 DW:105 DW:106 DW:107 DX:101 DX:102 DY:201 DY:202 DY:203 DY:204 DZ:101 DZ:102 DZ:103 DZ:104 DZ:105 | <1.8 | 1.8 - 2.4 | 2.4 - 3.4 |
| Calcium (CA) | --- |  | <2.2 | 2.2 - 2.8 | 2.8 - 3.6 |
| Strontium (SR) | --- |  | <2.4 | 2.4 - 2.8 | 2.8 - 4.0 |
| Barium (BA) | --- |  | <2.6 | 2.6 - 3.2 | --- |
| Manganese (MN) | --- |  | <2.0 | 2.0 - 2.3 | 2.3 - 3.6 |
| Zinc (ZN) | 6 | A4:101 BG:308 BQ:104 CG:301 CQ:101 D4:103 | <1.9 | 1.9 - 2.3 | 2.3 - 3.4 |
| Cadmium (CD) | --- |  | <2.2 | 2.2 - 2.5 | 2.5 - 3.6 |
| Cobalt (CO) | --- |  | <1.9 | 1.9 - 2.3 | 2.3 - 3.5 |
| Iridium (IRI) | --- |  | <2.0 | 2.0 - 2.3 | 2.3 - 3.6 |
| Anion (Anion) | --- |  | <2.8 | 2.8 - 3.4 | --- |
| Chloride (CL) | --- |  | <2.8 | 2.8 - 3.4 | --- |
| Sulphate (SO4) | --- |  | <2.8 | 2.8 - 3.4 | --- |
| Phosphate (PO4) | --- |  | <2.8 | 2.8 - 3.4 | --- |
| **Total** | **5351** |  |  |  |  |

---

\_\_\_\_\_\_\_\_\_\_\_\_\_\_\_\_\_\_\_\_\_\_\_\_\_\_\_\_\_\_\_\_\_\_\_\_\_\_\_\_\_\_\_\_\_\_\_\_\_\_\_\_\_\_\_\_\_\_\_\_\_\_\_\_  

#### 2) Ion validation table

##### Go to ToC

NOTES:

  
- O_ph_: OP1 or OP2 anionic phosphate oxygens;  
- O_r_: O2', O3', O4' or O5' hydroxyl and ester oxygens;  
- O_b_: O2, O4 or O6 carbonyl oxygens ;  
- N_b_: N1_A_, N3_C_, N3_G_, N3_A_ or N7 non-protonated nitrogens;  
- O_back_: protein backbone carbonyl group;  
- O_coo_: ASP/GLU carboxyl group;  
- O_cno_: ASN/GLN carbonyl group;  
- OH_Prot_: SER/THR/TYR hydroxyl group;  
- O_w_: water molecule oxygen;  
- Anion: NH group (imino, amino and amonium groups) - suggests potential anion contact or poor binding site modeling;  
  
"Large octahedral discrepancy" means a cumulated ligand-ion-ligand angle deviation > 10 deg.;  
-> the ions that are currently checked for octahedral coordination are Mg^2+^, Mn^2+^ and Na^+^;  
-> could be a hint to the presence of a polyatomic ligand instead of a monoatomic ion;  
1^st^ coordination shell missing:   
1) if Mg^2+^ or Na^+^, consider adding a hydration shell and optimize the water to solute distances;   
2) if preceding step is not conclusive, consider the presence of another solvent molecule (check crystallisation conditions);   
Exclusion atoms:   
-> if O3'-O5' atoms are mentioned, this points to a defficient modelling of the RNA backbone;  
  

| Ion | Occup. | B-factor | Ligands (too close) | Ligands (1^st^ shell) | Ligands (excl. zone) | Contacts & potential issues |
| --- | --- | --- | --- | --- | --- | --- |
| MG:A0:201 | 1.00 | 61.86 |  | 0 | 3 | - Exclusion zone contact(s): /N_back_/O_back_/OD2_Asp_/ Consider K in place of MG (2 valid contact(s) for K vs. 0 for MG) |
| MG:A1:201 | 1.00 | 60.48 |  | 1 | 1 | O_ph_ - Exclusion zone contact(s): /OG1_Thr_/ |
| MG:A1:202 | 1.00 | 61.95 |  | 1 | 4 | Anion - Exclusion zone contact(s): /N_back_/O_back_/O_back_/OP1/ Consider K in place of MG (4 valid contact(s) for K vs. 1 for MG) |
| MG:A1:203 | 1.00 | 77.24 |  | 0 | 2 | - Exclusion zone contact(s): /NE_Arg_/OP2/ Consider K in place of MG (2 valid contact(s) for K vs. 0 for MG) |
| MG:A1:204 | 1.00 | 65.58 |  | 0 | 3 | - Exclusion zone contact(s): /NH2_Arg_/OG1_Thr_/OP2/ Consider K in place of MG (3 valid contact(s) for K vs. 0 for MG) |
| MG:A1:205 | 1.00 | 79.96 |  | 0 |  | 1^st^ MG coordination shell missing (see Notes) |
| MG:A2:201 | 1.00 | 66.09 |  | 0 | 6 | - Exclusion zone contact(s): /N_back_/N_back_/N_back_/NE2_Gln_/O_back_/OP1/ Consider K in place of MG (4 valid contact(s) for K vs. 0 for MG) |
| MG:A3:101 | 1.00 | 40.68 |  | 0 | 1 | - Exclusion zone contact(s): /NH1_Arg_/ Consider K in place of MG (1 valid contact(s) for K vs. 0 for MG) |
| MG:A3:102 | 1.00 | 83.35 |  | 1 | 1 | O_b_ - Exclusion zone contact(s): /O2/ |
| MG:A3:103 | 1.00 | 55.67 |  | 0 |  | 1^st^ MG coordination shell missing (see Notes) |
| MG:A3:104 | 1.00 | 83.91 |  | 0 | 2 | - Exclusion zone contact(s): /OG_Ser_/OP2/ Consider K in place of MG (2 valid contact(s) for K vs. 0 for MG) |
| ZN:A4:101 | 1.00 | 200.41 |  | 0 | 3 | - Exclusion zone contact(s): /N_back_/N_back_/O_back_/ |
| MG:A5:101 | 1.00 | 23.78 |  | 1 | 1 | His - Exclusion zone contact(s): /OP2/ |
| MG:A5:102 | 1.00 | 28.56 |  | 0 | 2 | - Exclusion zone contact(s): /NE2_His_/O4/ Consider NA in place of MG (1 valid contact(s) for NA vs. 0 for MG) |
| MG:A5:103 | 1.00 | 54.74 |  | 0 | 3 | - Exclusion zone contact(s): /N_back_/N_back_/O_back_/ Consider K in place of MG (3 valid contact(s) for K vs. 0 for MG) |
| MG:A6:101 | 1.00 | 55.92 |  | 0 | 2 | - Exclusion zone contact(s): /O_back_/OG1_Thr_/ Consider NA in place of MG (1 valid contact(s) for NA vs. 0 for MG) |
| MG:A6:102 | 1.00 | 60.62 |  | 0 |  | 1^st^ MG coordination shell missing (see Notes) |
| MG:A7:101 | 1.00 | 77.44 |  | 0 | 1 | - Exclusion zone contact(s): /N7/ Consider K in place of MG (1 valid contact(s) for K vs. 0 for MG) |
| MG:A7:102 | 1.00 | 59.24 |  | 0 | 1 | - Exclusion zone contact(s): /OP1/ Consider K in place of MG (1 valid contact(s) for K vs. 0 for MG) |
| MG:A7:103 | 1.00 | 45.57 |  | 0 | 2 | - Exclusion zone contact(s): /N7/NH1_Arg_/ Consider K in place of MG (2 valid contact(s) for K vs. 0 for MG) |
| MG:A7:104 | 1.00 | 88.06 |  | 0 |  | 1^st^ MG coordination shell missing (see Notes) |
| MG:A8:101 | 1.00 | 45.46 |  | 0 | 1 | - Exclusion zone contact(s): /OP1/ Consider K in place of MG (1 valid contact(s) for K vs. 0 for MG) |
| MG:A8:102 | 1.00 | 68.55 |  | 0 | 1 | - Exclusion zone contact(s): /NZ_Lys_/ Consider K in place of MG (1 valid contact(s) for K vs. 0 for MG) |
| MG:A8:103 | 1.00 | 62.00 |  | 0 | 1 | - Exclusion zone contact(s): /NZ_Lys_/ Consider K in place of MG (1 valid contact(s) for K vs. 0 for MG) |
| MG:A8:104 | 1.00 | 68.27 |  | 0 | 4 | - Exclusion zone contact(s): /N_back_/NH2_Arg_/O2'/OP2/ Consider K in place of MG (3 valid contact(s) for K vs. 0 for MG) |
| MG:AA:3001 | 1.00 | 21.73 |  | 1 | 2 | O_ph_ - Exclusion zone contact(s): /OP2/OP2/ Consider NA in place of MG (2 valid contact(s) for NA vs. 1 for MG) |
| MG:AA:3002 | 1.00 | 14.47 |  | 1 |  | O_ph_ |
| MG:AA:3003 | 1.00 | 12.87 |  | 0 | 1 | - Exclusion zone contact(s): /OP2/ Consider NA in place of MG (1 valid contact(s) for NA vs. 0 for MG) |
| MG:AA:3004 | 1.00 | 7.98 |  | 1 | 2 | N_b_ - Exclusion zone contact(s): /N7/O6/ |
| MG:AA:3005 | 1.00 | 17.63 |  | 1 | 1 | O_ph_ - Exclusion zone contact(s): /O5'/ |
| MG:AA:3006 | 1.00 | 14.41 |  | 2 |  | 2O_ph_ |
| MG:AA:3007 | 1.00 | 24.26 |  | 1 | 1 | O_ph_ - Exclusion zone contact(s): /OP1/ |
| MG:AA:3008 | 1.00 | 31.20 |  | 1 | 1 | O_ph_ - Exclusion zone contact(s): /OP2/ |
| MG:AA:3009 | 1.00 | 18.07 | 1 | 2 | 2 | 2O_ph_ - Exclusion zone contact(s): /O5'/O5'/OP2/ |
| MG:AA:3010 | 1.00 | 19.48 |  | 2 |  | O_ph_O_b_ |
| MG:AA:3011 | 1.00 | 17.42 |  | 0 | 3 | - Exclusion zone contact(s): /OP1/OP2/OP2/ Consider K in place of MG (3 valid contact(s) for K vs. 0 for MG) |
| MG:AA:3012 | 1.00 | 21.99 |  | 1 |  | O_ph_ |
| MG:AA:3013 | 1.00 | 49.26 |  | 1 |  | O_ph_ |
| MG:AA:3014 | 1.00 | 20.98 | 1 | 1 |  | O_ph_ - Exclusion zone contact(s): /OP1/ |
| MG:AA:3015 | 1.00 | 19.07 |  | 2 | 1 | 2O_ph_ - Exclusion zone contact(s): /OP2/ |
| MG:AA:3016 | 1.00 | 9.25 |  | 0 | 4 | - Exclusion zone contact(s): /O6/O_back_/OP1/OP2/ Consider NA in place of MG (2 valid contact(s) for NA vs. 0 for MG) |
| MG:AA:3017 | 1.00 | 29.20 |  | 0 |  | 1^st^ MG coordination shell missing (see Notes) |
| MG:AA:3018 | 1.00 | 22.01 |  | 0 | 2 | - Exclusion zone contact(s): /OP1/OP2/ Consider NA in place of MG (1 valid contact(s) for NA vs. 0 for MG) |
| MG:AA:3019 | 1.00 | 31.43 |  | 0 | 1 | - Exclusion zone contact(s): /O4/ Consider K in place of MG (1 valid contact(s) for K vs. 0 for MG) |
| MG:AA:3020 | 1.00 | 27.39 |  | 0 |  | 1^st^ MG coordination shell missing (see Notes) |
| MG:AA:3021 | 1.00 | 26.52 |  | 0 | 1 | - Exclusion zone contact(s): /OP1/ Consider NA in place of MG (1 valid contact(s) for NA vs. 0 for MG) |
| MG:AA:3022 | 1.00 | 63.57 |  | 1 | 1 | O_b_ - Exclusion zone contact(s): /OP2/ |
| MG:AA:3023 | 1.00 | 59.18 |  | 1 |  | O_ph_ |
| MG:AA:3024 | 1.00 | 61.85 |  | 0 | 3 | - Exclusion zone contact(s): /O3'/O5'/OP1/ Consider K in place of MG (3 valid contact(s) for K vs. 0 for MG) |
| MG:AA:3025 | 1.00 | 38.77 |  | 0 |  | 1^st^ MG coordination shell missing (see Notes) |
| MG:AA:3026 | 1.00 | 38.61 |  | 0 |  | 1^st^ MG coordination shell missing (see Notes) |
| MG:AA:3027 | 1.00 | 60.92 |  | 0 | 2 | - Exclusion zone contact(s): /O6/OP2/ Consider NA in place of MG (1 valid contact(s) for NA vs. 0 for MG) |
| MG:AA:3028 | 1.00 | 15.00 |  | 1 | 2 | O_ph_ - Exclusion zone contact(s): /OP1/OP1/ |
| MG:AA:3029 | 1.00 | 46.59 |  | 0 | 4 | - Exclusion zone contact(s): /O3'/OP1/OP1/OP2/ Consider K in place of MG (4 valid contact(s) for K vs. 0 for MG) |
| MG:AA:3030 | 1.00 | 45.62 |  | 0 |  | 1^st^ MG coordination shell missing (see Notes) |
| MG:AA:3031 | 1.00 | 34.94 |  | 0 |  | 1^st^ MG coordination shell missing (see Notes) |
| MG:AA:3032 | 1.00 | 28.73 |  | 0 |  | 1^st^ MG coordination shell missing (see Notes) |
| MG:AA:3033 | 1.00 | 41.61 |  | 2 |  | O_ph_O_b_ |
| MG:AA:3034 | 1.00 | 41.23 |  | 0 | 3 | - Exclusion zone contact(s): /N7/N7/O6/ Consider K in place of MG (3 valid contact(s) for K vs. 0 for MG) |
| MG:AA:3035 | 1.00 | 17.28 |  | 0 | 5 | - Exclusion zone contact(s): /O2/O2'/O4/O4/O6/ Consider K in place of MG (4 valid contact(s) for K vs. 0 for MG) |
| MG:AA:3036 | 1.00 | 36.10 |  | 0 | 3 | - Exclusion zone contact(s): /O3'/O4/OP2/ Consider K in place of MG (3 valid contact(s) for K vs. 0 for MG) |
| MG:AA:3037 | 1.00 | 107.45 |  | 0 | 1 | - Exclusion zone contact(s): /OP1/ Consider NA in place of MG (1 valid contact(s) for NA vs. 0 for MG) |
| MG:AA:3038 | 1.00 | 68.62 |  | 0 | 1 | - Exclusion zone contact(s): /OP2/ Consider K in place of MG (1 valid contact(s) for K vs. 0 for MG) |
| MG:AA:3039 | 1.00 | 30.45 |  | 0 |  | 1^st^ MG coordination shell missing (see Notes) |
| MG:AA:3040 | 1.00 | 37.88 |  | 2 | 1 | 2O_ph_ - Exclusion zone contact(s): /OP1/ |
| MG:AA:3041 | 1.00 | 81.30 |  | 0 | 1 | - Exclusion zone contact(s): /OP1/ Consider NA in place of MG (1 valid contact(s) for NA vs. 0 for MG) |
| MG:AA:3042 | 1.00 | 35.53 |  | 0 | 1 | - Exclusion zone contact(s): /OP2/ Consider K in place of MG (1 valid contact(s) for K vs. 0 for MG) |
| MG:AA:3043 | 1.00 | 42.74 |  | 1 | 3 | O_b_ - Exclusion zone contact(s): /O6/OP1/OP2/ Consider K in place of MG (2 valid contact(s) for K vs. 1 for MG) |
| MG:AA:3044 | 1.00 | 36.57 |  | 1 |  | O_ph_ |
| MG:AA:3045 | 1.00 | 51.58 |  | 1 | 1 | O_ph_ - Exclusion zone contact(s): /OP1/ |
| MG:AA:3046 | 1.00 | 38.54 |  | 0 |  | 1^st^ MG coordination shell missing (see Notes) |
| MG:AA:3047 | 1.00 | 28.73 |  | 1 | 4 | O_b_ - Exclusion zone contact(s): /N3_A_/N3_C_/O2/O5'/ Consider K in place of MG (3 valid contact(s) for K vs. 1 for MG) |
| MG:AA:3048 | 1.00 | 41.14 |  | 1 |  | O_ph_ |
| MG:AA:3049 | 1.00 | 48.97 |  | 0 |  | 1^st^ MG coordination shell missing (see Notes) |
| MG:AA:3050 | 1.00 | 59.18 |  | 0 | 2 | - Exclusion zone contact(s): /O2'/OP1/ Consider K in place of MG (2 valid contact(s) for K vs. 0 for MG) |
| MG:AA:3051 | 1.00 | 62.78 |  | 0 | 1 | - Exclusion zone contact(s): /O6/ Consider K in place of MG (1 valid contact(s) for K vs. 0 for MG) |
| MG:AA:3052 | 1.00 | 60.28 |  | 1 | 1 | O_b_ - Exclusion zone contact(s): /N1_G_/ |
| MG:AA:3053 | 1.00 | 30.47 |  | 2 |  | 2O_ph_ |
| MG:AA:3054 | 1.00 | 44.92 |  | 0 | 5 | - Exclusion zone contact(s): /O2'/O3'/OP1/OP1/OP2/ Consider K in place of MG (4 valid contact(s) for K vs. 0 for MG) |
| MG:AA:3055 | 1.00 | 33.23 |  | 0 |  | 1^st^ MG coordination shell missing (see Notes) |
| MG:AA:3056 | 1.00 | 63.39 |  | 0 | 2 | - Exclusion zone contact(s): /OP2/OP2/ Consider K in place of MG (2 valid contact(s) for K vs. 0 for MG) |
| MG:AA:3057 | 1.00 | 49.18 |  | 1 | 1 | O_ph_ - Exclusion zone contact(s): /OP1/ |
| MG:AA:3058 | 1.00 | 25.75 |  | 0 | 2 | - Exclusion zone contact(s): /OP1/OP1/ Consider NA in place of MG (1 valid contact(s) for NA vs. 0 for MG) |
| MG:AA:3059 | 1.00 | 68.56 |  | 0 |  | 1^st^ MG coordination shell missing (see Notes) |
| MG:AA:3060 | 1.00 | 76.02 |  | 0 |  | 1^st^ MG coordination shell missing (see Notes) |
| MG:AA:3061 | 1.00 | 35.87 |  | 3 |  | 3O_ph_ |
| MG:AA:3062 | 1.00 | 29.65 |  | 0 | 1 | - Exclusion zone contact(s): /OP2/ Consider NA in place of MG (1 valid contact(s) for NA vs. 0 for MG) |
| MG:AA:3063 | 1.00 | 27.18 |  | 1 | 2 | O_ph_ - Exclusion zone contact(s): /OP2/OP2/ |
| MG:AA:3064 | 1.00 | 90.30 |  | 0 | 1 | - Exclusion zone contact(s): /OP1/ Consider K in place of MG (1 valid contact(s) for K vs. 0 for MG) |
| MG:AA:3065 | 1.00 | 60.64 |  | 0 |  | 1^st^ MG coordination shell missing (see Notes) |
| MG:AA:3066 | 1.00 | 17.99 |  | 1 | 2 | O_ph_ - Exclusion zone contact(s): /O_back_/OP2/ Consider K in place of MG (2 valid contact(s) for K vs. 1 for MG) |
| MG:AA:3067 | 1.00 | 37.76 |  | 0 | 3 | - Exclusion zone contact(s): /OP1/OP2/OP2/ Consider K in place of MG (3 valid contact(s) for K vs. 0 for MG) |
| MG:AA:3068 | 1.00 | 40.72 |  | 0 | 2 | - Exclusion zone contact(s): /O2'/OP1/ Consider K in place of MG (2 valid contact(s) for K vs. 0 for MG) |
| MG:AA:3069 | 1.00 | 29.85 |  | 1 | 1 | O_ph_ - Exclusion zone contact(s): /O3'/ |
| MG:AA:3070 | 1.00 | 35.50 |  | 3 | 1 | 3O_ph_ - Exclusion zone contact(s): /OP2/ |
| MG:AA:3071 | 1.00 | 16.11 |  | 0 |  | 1^st^ MG coordination shell missing (see Notes) |
| MG:AA:3072 | 1.00 | 21.02 |  | 0 | 1 | - Exclusion zone contact(s): /OP1/ Consider K in place of MG (1 valid contact(s) for K vs. 0 for MG) |
| MG:AA:3073 | 1.00 | 71.67 |  | 0 |  | 1^st^ MG coordination shell missing (see Notes) |
| MG:AA:3074 | 1.00 | 51.27 |  | 0 | 1 | - Exclusion zone contact(s): /OP2/ Consider NA in place of MG (1 valid contact(s) for NA vs. 0 for MG) |
| MG:AA:3075 | 1.00 | 33.44 |  | 0 | 4 | - Exclusion zone contact(s): /O4/O6/O6/O6/ Consider K in place of MG (3 valid contact(s) for K vs. 0 for MG) |
| MG:AA:3076 | 1.00 | 35.41 |  | 0 | 5 | - Exclusion zone contact(s): /N1_G_/O2/O6/O6/O6/ Consider K in place of MG (5 valid contact(s) for K vs. 0 for MG) |
| MG:AA:3077 | 1.00 | 63.33 |  | 0 | 1 | - Exclusion zone contact(s): /O6/ Consider K in place of MG (1 valid contact(s) for K vs. 0 for MG) |
| MG:AA:3078 | 1.00 | 44.30 |  | 2 |  | O_ph_O_b_ |
| MG:AA:3079 | 1.00 | 40.87 |  | 2 | 3 | 2O_ph_ - Exclusion zone contact(s): /OP1/OP1/OP2/ Consider K in place of MG (3 valid contact(s) for K vs. 2 for MG) |
| MG:AA:3080 | 1.00 | 41.06 |  | 0 | 2 | - Exclusion zone contact(s): /OP1/OP2/ Consider K in place of MG (2 valid contact(s) for K vs. 0 for MG) |
| MG:AA:3081 | 1.00 | 61.80 | 1 | 1 |  | O_ph_ - Exclusion zone contact(s): /OP1/ |
| MG:AA:3082 | 1.00 | 48.08 |  | 0 | 1 | - Exclusion zone contact(s): /OP2/ Consider NA in place of MG (1 valid contact(s) for NA vs. 0 for MG) |
| MG:AA:3083 | 1.00 | 33.56 |  | 0 | 1 | - Exclusion zone contact(s): /OP2/ Consider NA in place of MG (1 valid contact(s) for NA vs. 0 for MG) |
| MG:AA:3084 | 1.00 | 55.41 |  | 2 | 1 | 2O_ph_ - Exclusion zone contact(s): /OP1/ |
| MG:AA:3085 | 1.00 | 29.02 |  | 0 | 2 | - Exclusion zone contact(s): /N7/O6/ Consider NA in place of MG (1 valid contact(s) for NA vs. 0 for MG) |
| MG:AA:3086 | 1.00 | 35.16 |  | 1 |  | O_ph_ |
| MG:AA:3087 | 1.00 | 20.77 |  | 0 | 1 | - Exclusion zone contact(s): /N3_C_/ Consider K in place of MG (1 valid contact(s) for K vs. 0 for MG) |
| MG:AA:3088 | 1.00 | 51.18 |  | 0 | 2 | - Exclusion zone contact(s): /O4/O6/ Consider NA in place of MG (1 valid contact(s) for NA vs. 0 for MG) |
| MG:AA:3089 | 1.00 | 70.32 |  | 1 |  | O_ph_ |
| MG:AA:3090 | 1.00 | 42.81 |  | 1 | 1 | O_ph_ - Exclusion zone contact(s): /N7/ |
| MG:AA:3091 | 1.00 | 59.86 |  | 0 |  | 1^st^ MG coordination shell missing (see Notes) |
| MG:AA:3092 | 1.00 | 16.75 |  | 0 | 1 | - Exclusion zone contact(s): /OP2/ Consider K in place of MG (1 valid contact(s) for K vs. 0 for MG) |
| MG:AA:3093 | 1.00 | 39.84 |  | 0 | 3 | - Exclusion zone contact(s): /O4/O4/OP2/ Consider K in place of MG (2 valid contact(s) for K vs. 0 for MG) |
| MG:AA:3094 | 1.00 | 21.10 |  | 3 |  | 3O_ph_ |
| MG:AA:3095 | 1.00 | 18.72 |  | 0 | 2 | - Exclusion zone contact(s): /O_back_/OP1/ Consider NA in place of MG (1 valid contact(s) for NA vs. 0 for MG) |
| MG:AA:3096 | 1.00 | 67.17 |  | 0 | 1 | - Exclusion zone contact(s): /O6/ Consider K in place of MG (1 valid contact(s) for K vs. 0 for MG) |
| MG:AA:3097 | 1.00 | 73.92 |  | 2 | 1 | 2O_b_ - Exclusion zone contact(s): /N3_U_/ |
| MG:AA:3098 | 1.00 | 45.87 |  | 0 |  | 1^st^ MG coordination shell missing (see Notes) |
| MG:AA:3099 | 1.00 | 33.22 |  | 0 | 4 | - Exclusion zone contact(s): /O2'/O3'/OP1/OP2/ Consider K in place of MG (3 valid contact(s) for K vs. 0 for MG) |
| MG:AA:3100 | 1.00 | 34.80 |  | 0 | 1 | - Exclusion zone contact(s): /O2'/ Consider K in place of MG (1 valid contact(s) for K vs. 0 for MG) |
| MG:AA:3101 | 1.00 | 38.07 |  | 0 | 2 | - Exclusion zone contact(s): /OP1/OP2/ Consider NA in place of MG (1 valid contact(s) for NA vs. 0 for MG) |
| MG:AA:3102 | 1.00 | 65.55 |  | 1 | 1 | O_ph_ - Exclusion zone contact(s): /OP2/ |
| MG:AA:3103 | 1.00 | 45.85 |  | 0 | 2 | - Exclusion zone contact(s): /O2'/OP1/ Consider K in place of MG (2 valid contact(s) for K vs. 0 for MG) |
| MG:AA:3104 | 1.00 | 54.99 |  | 0 |  | 1^st^ MG coordination shell missing (see Notes) |
| MG:AA:3105 | 1.00 | 50.42 |  | 0 | 1 | - Exclusion zone contact(s): /OP2/ Consider K in place of MG (1 valid contact(s) for K vs. 0 for MG) |
| MG:AA:3106 | 1.00 | 17.86 |  | 0 | 1 | - Exclusion zone contact(s): /OP2/ Consider NA in place of MG (1 valid contact(s) for NA vs. 0 for MG) |
| MG:AA:3107 | 1.00 | 32.30 |  | 1 |  | O_ph_ |
| MG:AA:3108 | 1.00 | 41.31 |  | 0 | 1 | - Exclusion zone contact(s): /OP1/ Consider NA in place of MG (1 valid contact(s) for NA vs. 0 for MG) |
| MG:AA:3109 | 1.00 | 40.20 |  | 0 |  | 1^st^ MG coordination shell missing (see Notes) |
| MG:AA:3110 | 1.00 | 59.01 |  | 0 | 2 | - Exclusion zone contact(s): /N7/N7/ Consider K in place of MG (2 valid contact(s) for K vs. 0 for MG) |
| MG:AA:3111 | 1.00 | 46.03 |  | 1 | 3 | O_ph_ - Exclusion zone contact(s): /O2'/O4'/O5'/ Consider K in place of MG (3 valid contact(s) for K vs. 1 for MG) |
| MG:AA:3112 | 1.00 | 102.42 | 1 | 1 | 4 | - Exclusion zone contact(s): /N_back_/N_back_/O3'/O5'/OP1/ Consider K in place of MG (3 valid contact(s) for K vs. 0 for MG) |
| MG:AA:3113 | 1.00 | 85.32 |  | 0 | 3 | - Exclusion zone contact(s): /N7/O6/OP2/ Consider K in place of MG (3 valid contact(s) for K vs. 0 for MG) |
| MG:AA:3114 | 1.00 | 97.62 |  | 0 | 1 | - Exclusion zone contact(s): /OP1/ Consider K in place of MG (1 valid contact(s) for K vs. 0 for MG) |
| MG:AA:3115 | 1.00 | 76.64 |  | 0 |  | 1^st^ MG coordination shell missing (see Notes) |
| MG:AA:3116 | 1.00 | 85.76 |  | 0 | 1 | - Exclusion zone contact(s): /OP1/ Consider K in place of MG (1 valid contact(s) for K vs. 0 for MG) |
| MG:AA:3117 | 1.00 | 72.13 |  | 0 |  | 1^st^ MG coordination shell missing (see Notes) |
| MG:AA:3118 | 1.00 | 52.24 |  | 0 | 3 | - Exclusion zone contact(s): /N4/O6/O6/ Consider K in place of MG (2 valid contact(s) for K vs. 0 for MG) |
| MG:AA:3119 | 1.00 | 46.78 |  | 0 | 3 | - Exclusion zone contact(s): /N7/O6/OP2/ Consider K in place of MG (3 valid contact(s) for K vs. 0 for MG) |
| MG:AA:3120 | 1.00 | 59.45 |  | 0 |  | 1^st^ MG coordination shell missing (see Notes) |
| MG:AA:3121 | 1.00 | 85.65 |  | 0 |  | 1^st^ MG coordination shell missing (see Notes) |
| MG:AA:3122 | 1.00 | 73.25 |  | 0 | 1 | - Exclusion zone contact(s): /OP1/ Consider NA in place of MG (1 valid contact(s) for NA vs. 0 for MG) |
| MG:AA:3123 | 1.00 | 47.97 |  | 0 |  | 1^st^ MG coordination shell missing (see Notes) |
| MG:AA:3124 | 1.00 | 71.81 |  | 0 | 1 | - Exclusion zone contact(s): /N7/ Consider K in place of MG (1 valid contact(s) for K vs. 0 for MG) |
| MG:AA:3125 | 1.00 | 51.81 |  | 0 | 2 | - Exclusion zone contact(s): /N1_A_/N2/ Consider K in place of MG (2 valid contact(s) for K vs. 0 for MG) |
| MG:AA:3126 | 1.00 | 58.26 |  | 2 |  | O_ph_O_b_ |
| MG:AA:3127 | 1.00 | 46.67 |  | 2 | 3 | O_r_O_b_ - Exclusion zone contact(s): /N7/O3'/OP2/ Consider K in place of MG (3 valid contact(s) for K vs. 2 for MG) |
| MG:AA:3128 | 1.00 | 32.62 |  | 0 | 1 | - Exclusion zone contact(s): /OP2/ Consider NA in place of MG (1 valid contact(s) for NA vs. 0 for MG) |
| MG:AA:3129 | 1.00 | 31.27 |  | 2 | 1 | 2O_ph_ - Exclusion zone contact(s): /O5'/ |
| MG:AA:3130 | 1.00 | 24.59 |  | 1 | 1 | O_ph_ - Exclusion zone contact(s): /OP1/ |
| MG:AA:3131 | 1.00 | 34.15 |  | 0 | 1 | - Exclusion zone contact(s): /N7/ Consider K in place of MG (1 valid contact(s) for K vs. 0 for MG) |
| MG:AA:3132 | 1.00 | 44.59 |  | 0 | 2 | - Exclusion zone contact(s): /O5'/OP1/ Consider K in place of MG (2 valid contact(s) for K vs. 0 for MG) |
| MG:AA:3133 | 1.00 | 25.21 |  | 2 | 2 | O_ph_O_r_ - Exclusion zone contact(s): /O5'/OP2/ |
| MG:AA:3134 | 1.00 | 22.39 |  | 1 | 3 | O_ph_ - Exclusion zone contact(s): /OP1/OP1/OP2/ Consider K in place of MG (2 valid contact(s) for K vs. 1 for MG) |
| MG:AA:3135 | 1.00 | 19.92 |  | 1 | 2 | O_b_ - Exclusion zone contact(s): /O3'/OP1/ Consider K in place of MG (2 valid contact(s) for K vs. 1 for MG) |
| MG:AA:3136 | 1.00 | 38.15 |  | 0 |  | 1^st^ MG coordination shell missing (see Notes) |
| MG:AA:3137 | 1.00 | 25.73 |  | 0 | 1 | - Exclusion zone contact(s): /OP2/ Consider NA in place of MG (1 valid contact(s) for NA vs. 0 for MG) |
| MG:AA:3138 | 1.00 | 19.98 |  | 1 | 2 | O_ph_ - Exclusion zone contact(s): /OP1/OP2/ Consider K in place of MG (2 valid contact(s) for K vs. 1 for MG) |
| MG:AA:3139 | 1.00 | 45.49 |  | 0 | 2 | - Exclusion zone contact(s): /O6/OP1/ Consider K in place of MG (2 valid contact(s) for K vs. 0 for MG) |
| MG:AA:3140 | 1.00 | 57.43 |  | 0 |  | 1^st^ MG coordination shell missing (see Notes) |
| MG:AA:3141 | 1.00 | 66.50 |  | 0 |  | 1^st^ MG coordination shell missing (see Notes) |
| MG:AA:3142 | 1.00 | 12.03 |  | 0 | 7 | - Exclusion zone contact(s): /O2'/O3'/O4'/O5'/OP1/OP2/OP2/ Consider K in place of MG (7 valid contact(s) for K vs. 0 for MG) |
| MG:AA:3143 | 1.00 | 37.95 |  | 1 | 3 | O_b_ - Exclusion zone contact(s): /O4/O6/O6/ Consider K in place of MG (2 valid contact(s) for K vs. 1 for MG) |
| MG:AA:3144 | 1.00 | 69.03 |  | 0 |  | 1^st^ MG coordination shell missing (see Notes) |
| MG:AA:3145 | 1.00 | 61.85 |  | 0 |  | 1^st^ MG coordination shell missing (see Notes) |
| MG:AA:3146 | 1.00 | 78.86 |  | 0 |  | 1^st^ MG coordination shell missing (see Notes) |
| MG:AA:3147 | 1.00 | 60.03 |  | 0 | 2 | - Exclusion zone contact(s): /O6/OP2/ Consider K in place of MG (2 valid contact(s) for K vs. 0 for MG) |
| MG:AA:3148 | 1.00 | 53.91 |  | 0 | 2 | - Exclusion zone contact(s): /O3'/OP1/ Consider K in place of MG (2 valid contact(s) for K vs. 0 for MG) |
| MG:AA:3149 | 1.00 | 61.05 |  | 0 |  | 1^st^ MG coordination shell missing (see Notes) |
| MG:AA:3150 | 1.00 | 29.80 |  | 1 |  | O_ph_ |
| MG:AA:3151 | 1.00 | 60.16 |  | 0 | 3 | - Exclusion zone contact(s): /O5'/OP2/OP2/ Consider K in place of MG (2 valid contact(s) for K vs. 0 for MG) |
| MG:AA:3152 | 1.00 | 28.01 |  | 2 | 1 | O_ph_N_b_ - Exclusion zone contact(s): /OP1/ |
| MG:AA:3153 | 1.00 | 40.38 |  | 0 | 4 | - Exclusion zone contact(s): /O4/O4'/OP2/ Consider K in place of MG (3 valid contact(s) for K vs. 0 for MG) |
| MG:AA:3154 | 1.00 | 59.02 |  | 0 |  | 1^st^ MG coordination shell missing (see Notes) |
| MG:AA:3155 | 1.00 | 46.22 |  | 1 | 1 | O_ph_ - Exclusion zone contact(s): /OP2/ |
| MG:AA:3156 | 1.00 | 55.99 |  | 0 | 1 | - Exclusion zone contact(s): /OP1/ Consider K in place of MG (1 valid contact(s) for K vs. 0 for MG) |
| MG:AA:3157 | 1.00 | 73.63 |  | 0 |  | 1^st^ MG coordination shell missing (see Notes) |
| MG:AA:3158 | 1.00 | 55.35 |  | 1 | 1 | O_ph_ - Exclusion zone contact(s): /O3'/ |
| MG:AA:3159 | 1.00 | 61.32 |  | 0 | 1 | - Exclusion zone contact(s): /OP1/ Consider K in place of MG (1 valid contact(s) for K vs. 0 for MG) |
| MG:AA:3160 | 1.00 | 64.66 |  | 0 | 3 | - Exclusion zone contact(s): /O6/OP1/OP2/ Consider K in place of MG (3 valid contact(s) for K vs. 0 for MG) |
| MG:AA:3161 | 1.00 | 63.32 |  | 0 | 2 | - Exclusion zone contact(s): /O2'/OP2/ Consider NA in place of MG (1 valid contact(s) for NA vs. 0 for MG) |
| MG:AA:3162 | 1.00 | 48.58 |  | 0 | 2 | - Exclusion zone contact(s): /O6/O6/ Consider K in place of MG (2 valid contact(s) for K vs. 0 for MG) |
| MG:AA:3163 | 1.00 | 65.91 |  | 0 |  | 1^st^ MG coordination shell missing (see Notes) |
| MG:AA:3164 | 1.00 | 79.43 |  | 0 |  | 1^st^ MG coordination shell missing (see Notes) |
| MG:AA:3165 | 1.00 | 53.54 |  | 0 | 2 | - Exclusion zone contact(s): /O6/O6/ Consider NA in place of MG (1 valid contact(s) for NA vs. 0 for MG) |
| MG:AA:3166 | 1.00 | 65.28 |  | 0 | 4 | - Exclusion zone contact(s): /O5'/OP1/OP1/OP2/ Consider NA in place of MG (2 valid contact(s) for NA vs. 0 for MG) |
| MG:AA:3167 | 1.00 | 61.71 |  | 0 | 3 | - Exclusion zone contact(s): /O4/O6/O6/ Consider K in place of MG (3 valid contact(s) for K vs. 0 for MG) |
| MG:AA:3168 | 1.00 | 48.90 |  | 0 |  | 1^st^ MG coordination shell missing (see Notes) |
| MG:AA:3169 | 1.00 | 11.78 |  | 1 | 5 | O_ph_ - Exclusion zone contact(s): /N7/ND1_His_/NE2_His_/OP1/OP2/ Consider K in place of MG (4 valid contact(s) for K vs. 1 for MG) |
| MG:AA:3170 | 1.00 | 32.29 |  | 1 | 2 | O_ph_ - Exclusion zone contact(s): /O2/OP2/ |
| MG:AA:3171 | 1.00 | 50.62 |  | 0 | 4 | - Exclusion zone contact(s): /N1_A_/N3_C_/O2/OP2/ Consider K in place of MG (4 valid contact(s) for K vs. 0 for MG) |
| MG:AA:3172 | 1.00 | 67.55 |  | 0 |  | 1^st^ MG coordination shell missing (see Notes) |
| MG:AA:3173 | 1.00 | 14.02 |  | 0 | 6 | - Exclusion zone contact(s): /N1_A_/N3_A_/N7/O2'/O6/O6/ Consider K in place of MG (6 valid contact(s) for K vs. 0 for MG) |
| MG:AA:3174 | 1.00 | 63.65 |  | 0 | 2 | - Exclusion zone contact(s): /O6/O6/ Consider K in place of MG (2 valid contact(s) for K vs. 0 for MG) |
| MG:AA:3175 | 1.00 | 29.69 |  | 0 | 6 | - Exclusion zone contact(s): /O3'/O3'/O5'/OP1/OP1/OP2/ Consider K in place of MG (6 valid contact(s) for K vs. 0 for MG) |
| MG:AA:3176 | 1.00 | 33.86 |  | 1 | 1 | O_ph_ - Exclusion zone contact(s): /O3'/ |
| MG:AA:3177 | 1.00 | 26.54 |  | 1 |  | O_ph_ |
| MG:AA:3178 | 1.00 | 41.32 |  | 0 | 4 | - Exclusion zone contact(s): /O4/O4/O6/O6/ Consider K in place of MG (4 valid contact(s) for K vs. 0 for MG) |
| MG:AA:3179 | 1.00 | 38.96 |  | 0 |  | 1^st^ MG coordination shell missing (see Notes) |
| MG:AA:3180 | 1.00 | 62.14 |  | 0 | 1 | - Exclusion zone contact(s): /OP2/ Consider K in place of MG (1 valid contact(s) for K vs. 0 for MG) |
| MG:AA:3181 | 1.00 | 51.85 |  | 0 |  | 1^st^ MG coordination shell missing (see Notes) |
| MG:AA:3182 | 1.00 | 60.60 |  | 0 | 1 | - Exclusion zone contact(s): /O6/ Consider K in place of MG (1 valid contact(s) for K vs. 0 for MG) |
| MG:AA:3183 | 1.00 | 29.01 |  | 1 |  | O_ph_ |
| MG:AA:3184 | 1.00 | 49.15 |  | 0 | 1 | - Exclusion zone contact(s): /OP1/ Consider K in place of MG (1 valid contact(s) for K vs. 0 for MG) |
| MG:AA:3185 | 1.00 | 29.53 |  | 0 | 2 | - Exclusion zone contact(s): /OP1/OP2/ Consider NA in place of MG (1 valid contact(s) for NA vs. 0 for MG) |
| MG:AA:3186 | 1.00 | 19.62 |  | 1 | 2 | N_b_ - Exclusion zone contact(s): /N6/O6/ |
| MG:AA:3187 | 1.00 | 68.66 |  | 0 | 1 | - Exclusion zone contact(s): /N7/ Consider K in place of MG (1 valid contact(s) for K vs. 0 for MG) |
| MG:AA:3188 | 1.00 | 52.15 |  | 0 |  | 1^st^ MG coordination shell missing (see Notes) |
| MG:AA:3189 | 1.00 | 52.29 |  | 0 |  | 1^st^ MG coordination shell missing (see Notes) |
| MG:AA:3190 | 1.00 | 26.09 |  | 1 | 1 | O_ph_ - Exclusion zone contact(s): /OP2/ |
| MG:AA:3191 | 1.00 | 31.51 |  | 1 |  | O_ph_ |
| MG:AA:3192 | 1.00 | 39.98 |  | 2 | 1 | 2O_ph_ - Exclusion zone contact(s): /OP2/ |
| MG:AA:3193 | 1.00 | 99.99 |  | 0 | 2 | - Exclusion zone contact(s): /OP1/OP1/ Consider NA in place of MG (1 valid contact(s) for NA vs. 0 for MG) |
| MG:AA:3194 | 1.00 | 56.68 |  | 0 | 3 | - Exclusion zone contact(s): /N7/O4/O6/ Consider K in place of MG (3 valid contact(s) for K vs. 0 for MG) |
| MG:AA:3195 | 1.00 | 82.71 |  | 0 |  | 1^st^ MG coordination shell missing (see Notes) |
| MG:AA:3196 | 1.00 | 29.80 |  | 2 |  | 2O_ph_ |
| MG:AA:3197 | 1.00 | 50.52 |  | 0 |  | 1^st^ MG coordination shell missing (see Notes) |
| MG:AA:3198 | 1.00 | 34.91 |  | 0 | 2 | - Exclusion zone contact(s): /O2'/O3'/ Consider NA in place of MG (1 valid contact(s) for NA vs. 0 for MG) |
| MG:AA:3199 | 1.00 | 75.67 |  | 0 |  | 1^st^ MG coordination shell missing (see Notes) |
| MG:AA:3200 | 1.00 | 83.39 |  | 0 |  | 1^st^ MG coordination shell missing (see Notes) |
| MG:AA:3201 | 1.00 | 71.57 |  | 0 |  | 1^st^ MG coordination shell missing (see Notes) |
| MG:AA:3202 | 1.00 | 31.32 |  | 0 | 1 | - Exclusion zone contact(s): /OP1/ Consider NA in place of MG (1 valid contact(s) for NA vs. 0 for MG) |
| MG:AA:3203 | 1.00 | 40.89 |  | 0 | 5 | - Exclusion zone contact(s): /N7/O6/O6/OP1/OP2/ Consider K in place of MG (5 valid contact(s) for K vs. 0 for MG) |
| MG:AA:3204 | 1.00 | 68.05 |  | 1 | 2 | O_ph_ - Exclusion zone contact(s): /O3'/O6/ Consider K in place of MG (2 valid contact(s) for K vs. 1 for MG) |
| MG:AA:3205 | 1.00 | 71.22 |  | 0 |  | 1^st^ MG coordination shell missing (see Notes) |
| MG:AA:3206 | 1.00 | 40.85 | 1 | 1 | 2 | - Exclusion zone contact(s): /O4/O5'/OP2/ Consider K in place of MG (2 valid contact(s) for K vs. 0 for MG) |
| MG:AA:3207 | 1.00 | 30.21 |  | 2 |  | 2O_ph_ |
| MG:AA:3208 | 1.00 | 30.06 |  | 1 | 3 | O_ph_ - Exclusion zone contact(s): /N1_G_/N2/O5'/ Consider K in place of MG (3 valid contact(s) for K vs. 1 for MG) |
| MG:AA:3209 | 1.00 | 77.98 |  | 0 |  | 1^st^ MG coordination shell missing (see Notes) |
| MG:AA:3210 | 1.00 | 39.68 |  | 0 | 4 | - Exclusion zone contact(s): /O4/O6/O6/O6/ Consider K in place of MG (4 valid contact(s) for K vs. 0 for MG) |
| MG:AA:3211 | 1.00 | 56.19 |  | 0 |  | 1^st^ MG coordination shell missing (see Notes) |
| MG:AA:3212 | 1.00 | 78.83 |  | 0 |  | 1^st^ MG coordination shell missing (see Notes) |
| MG:AA:3213 | 1.00 | 51.57 |  | 0 | 3 | - Exclusion zone contact(s): /OP1/OP1/OP2/ Consider K in place of MG (3 valid contact(s) for K vs. 0 for MG) |
| MG:AA:3214 | 1.00 | 56.23 |  | 1 |  | O_ph_ |
| MG:AA:3215 | 1.00 | 72.43 |  | 0 |  | 1^st^ MG coordination shell missing (see Notes) |
| MG:AA:3216 | 1.00 | 40.87 |  | 2 |  | 2O_ph_ |
| MG:AA:3217 | 1.00 | 57.34 |  | 0 | 1 | - Exclusion zone contact(s): /OP2/ Consider K in place of MG (1 valid contact(s) for K vs. 0 for MG) |
| MG:AA:3218 | 1.00 | 65.81 |  | 0 | 3 | - Exclusion zone contact(s): /N7/OP2/OP2/ Consider K in place of MG (2 valid contact(s) for K vs. 0 for MG) |
| MG:AA:3219 | 1.00 | 81.83 |  | 0 | 1 | - Exclusion zone contact(s): /O6/ Consider K in place of MG (1 valid contact(s) for K vs. 0 for MG) |
| MG:AA:3220 | 1.00 | 70.71 |  | 0 | 1 | - Exclusion zone contact(s): /OP2/ Consider K in place of MG (1 valid contact(s) for K vs. 0 for MG) |
| MG:AA:3221 | 1.00 | 63.71 |  | 0 |  | 1^st^ MG coordination shell missing (see Notes) |
| MG:AA:3222 | 1.00 | 28.46 |  | 2 | 1 | 2O_ph_ - Exclusion zone contact(s): /OP1/ |
| MG:AA:3223 | 1.00 | 14.73 |  | 2 | 2 | 2O_ph_ - Exclusion zone contact(s): /OP1/OP2/ |
| MG:AA:3224 | 1.00 | 63.49 |  | 0 | 2 | - Exclusion zone contact(s): /N7/O6/ Consider K in place of MG (2 valid contact(s) for K vs. 0 for MG) |
| MG:AA:3225 | 1.00 | 18.74 |  | 0 | 4 | - Exclusion zone contact(s): /N7/O2'/O3'/OP1/ Consider K in place of MG (3 valid contact(s) for K vs. 0 for MG) |
| MG:AA:3226 | 1.00 | 68.99 |  | 3 |  | O_ph_O_r_Anion |
| MG:AA:3227 | 1.00 | 30.99 |  | 0 | 4 | - Exclusion zone contact(s): /O5'/OP2/OP2/OP2/ Consider K in place of MG (3 valid contact(s) for K vs. 0 for MG) |
| MG:AA:3228 | 1.00 | 53.51 |  | 0 | 3 | - Exclusion zone contact(s): /OP1/OP1/OP2/ Consider K in place of MG (2 valid contact(s) for K vs. 0 for MG) |
| MG:AA:3229 | 1.00 | 56.38 |  | 0 | 2 | - Exclusion zone contact(s): /O2'/OP2/ Consider K in place of MG (2 valid contact(s) for K vs. 0 for MG) |
| MG:AA:3230 | 1.00 | 51.96 |  | 1 | 2 | O_ph_ - Exclusion zone contact(s): /O3'/OP2/ |
| MG:AA:3231 | 1.00 | 43.70 |  | 0 | 3 | - Exclusion zone contact(s): /O4'/O5'/O6/ Consider K in place of MG (3 valid contact(s) for K vs. 0 for MG) |
| MG:AA:3232 | 1.00 | 25.64 |  | 0 | 3 | - Exclusion zone contact(s): /O4'/OP2/ Consider K in place of MG (3 valid contact(s) for K vs. 0 for MG) |
| MG:AA:3233 | 1.00 | 55.28 |  | 0 | 1 | - Exclusion zone contact(s): /OP2/ Consider K in place of MG (1 valid contact(s) for K vs. 0 for MG) |
| MG:AA:3234 | 1.00 | 54.02 |  | 0 | 1 | - Exclusion zone contact(s): /OP1/ Consider NA in place of MG (1 valid contact(s) for NA vs. 0 for MG) |
| MG:AA:3235 | 1.00 | 48.56 |  | 0 |  | 1^st^ MG coordination shell missing (see Notes) |
| MG:AA:3236 | 1.00 | 20.38 |  | 0 | 7 | - Exclusion zone contact(s): /O2'/O2'/O3'/O3'/O4/O5'/OP1/ Consider K in place of MG (6 valid contact(s) for K vs. 0 for MG) |
| MG:AA:3237 | 1.00 | 36.17 |  | 0 | 4 | - Exclusion zone contact(s): /N7/O2/O2'/OP2/ Consider K in place of MG (4 valid contact(s) for K vs. 0 for MG) |
| MG:AA:3238 | 1.00 | 63.30 |  | 0 |  | 1^st^ MG coordination shell missing (see Notes) |
| MG:AA:3239 | 1.00 | 51.41 |  | 0 | 4 | - Exclusion zone contact(s): /N7/O2'/O4'/O6/ Consider K in place of MG (4 valid contact(s) for K vs. 0 for MG) |
| MG:AA:3240 | 1.00 | 34.58 |  | 1 |  | O_ph_ |
| MG:AA:3241 | 1.00 | 59.43 |  | 0 | 1 | - Exclusion zone contact(s): /OP1/ Consider K in place of MG (1 valid contact(s) for K vs. 0 for MG) |
| MG:AA:3242 | 1.00 | 56.12 |  | 0 |  | 1^st^ MG coordination shell missing (see Notes) |
| MG:AA:3243 | 1.00 | 99.25 |  | 0 | 2 | - Exclusion zone contact(s): /N4/N7/ Consider K in place of MG (2 valid contact(s) for K vs. 0 for MG) |
| MG:AA:3244 | 1.00 | 55.42 |  | 0 |  | 1^st^ MG coordination shell missing (see Notes) |
| MG:AA:3245 | 1.00 | 69.33 |  | 0 |  | 1^st^ MG coordination shell missing (see Notes) |
| MG:AA:3246 | 1.00 | 45.91 |  | 0 | 2 | - Exclusion zone contact(s): /N7/O6/ Consider K in place of MG (2 valid contact(s) for K vs. 0 for MG) |
| MG:AA:3247 | 1.00 | 47.48 |  | 0 | 1 | - Exclusion zone contact(s): /N4/ Consider K in place of MG (1 valid contact(s) for K vs. 0 for MG) |
| MG:AA:3248 | 1.00 | 78.91 |  | 1 | 2 | O_r_ - Exclusion zone contact(s): /N3_G_/O4'/ |
| MG:AA:3249 | 1.00 | 41.03 |  | 1 |  | O_b_ |
| MG:AA:3250 | 1.00 | 24.67 |  | 0 | 4 | - Exclusion zone contact(s): /N7/O6/OP2/OP2/ Consider K in place of MG (4 valid contact(s) for K vs. 0 for MG) |
| MG:AA:3251 | 1.00 | 24.39 |  | 2 | 1 | 2O_ph_ - Exclusion zone contact(s): /OP1/ |
| MG:AA:3252 | 1.00 | 38.29 |  | 0 | 2 | - Exclusion zone contact(s): /O6/OP2/ Consider NA in place of MG (1 valid contact(s) for NA vs. 0 for MG) |
| MG:AA:3253 | 1.00 | 50.59 |  | 0 | 4 | - Exclusion zone contact(s): /O4/O6/O6/O6/ Consider K in place of MG (4 valid contact(s) for K vs. 0 for MG) |
| MG:AA:3254 | 1.00 | 76.35 |  | 0 | 1 | - Exclusion zone contact(s): /O4/ Consider K in place of MG (1 valid contact(s) for K vs. 0 for MG) |
| MG:AA:3255 | 1.00 | 46.06 |  | 0 | 3 | - Exclusion zone contact(s): /O3'/OP1/OP2/ Consider K in place of MG (2 valid contact(s) for K vs. 0 for MG) |
| MG:AA:3256 | 1.00 | 62.93 |  | 0 |  | 1^st^ MG coordination shell missing (see Notes) |
| MG:AA:3257 | 1.00 | 36.22 |  | 1 |  | O_ph_ |
| MG:AA:3258 | 1.00 | 81.01 |  | 0 |  | 1^st^ MG coordination shell missing (see Notes) |
| MG:AA:3259 | 1.00 | 57.79 |  | 0 |  | 1^st^ MG coordination shell missing (see Notes) |
| MG:AA:3260 | 1.00 | 44.00 |  | 1 | 2 | O_ph_ - Exclusion zone contact(s): /O3'/OP2/ Consider K in place of MG (2 valid contact(s) for K vs. 1 for MG) |
| MG:AA:3261 | 1.00 | 40.12 |  | 0 | 1 | - Exclusion zone contact(s): /OP2/ Consider NA in place of MG (1 valid contact(s) for NA vs. 0 for MG) |
| MG:AA:3262 | 1.00 | 76.77 |  | 1 | 1 | O_ph_ - Exclusion zone contact(s): /N1_A_/ |
| MG:AA:3263 | 1.00 | 57.44 |  | 0 | 4 | - Exclusion zone contact(s): /N1_G_/N3_C_/N6/O2/ Consider K in place of MG (3 valid contact(s) for K vs. 0 for MG) |
| MG:AA:3264 | 1.00 | 43.64 |  | 0 | 2 | - Exclusion zone contact(s): /N7/OP2/ Consider NA in place of MG (1 valid contact(s) for NA vs. 0 for MG) |
| MG:AA:3265 | 1.00 | 71.03 |  | 0 | 1 | - Exclusion zone contact(s): /OP2/ Consider K in place of MG (1 valid contact(s) for K vs. 0 for MG) |
| MG:AA:3266 | 1.00 | 73.01 |  | 0 |  | 1^st^ MG coordination shell missing (see Notes) |
| MG:AA:3267 | 1.00 | 49.19 |  | 2 | 1 | 2O_r_ - Exclusion zone contact(s): /OP1/ |
| MG:AA:3268 | 1.00 | 60.84 |  | 0 |  | 1^st^ MG coordination shell missing (see Notes) |
| MG:AA:3269 | 1.00 | 61.35 |  | 0 |  | 1^st^ MG coordination shell missing (see Notes) |
| MG:AA:3270 | 1.00 | 72.79 |  | 0 |  | 1^st^ MG coordination shell missing (see Notes) |
| MG:AA:3271 | 1.00 | 83.01 |  | 0 | 1 | - Exclusion zone contact(s): /OP2/ Consider NA in place of MG (1 valid contact(s) for NA vs. 0 for MG) |
| MG:AA:3272 | 1.00 | 54.84 |  | 0 | 3 | - Exclusion zone contact(s): /O2'/O2'/O3'/ Consider K in place of MG (2 valid contact(s) for K vs. 0 for MG) |
| MG:AA:3273 | 1.00 | 30.93 |  | 0 | 4 | - Exclusion zone contact(s): /O4/O6/O6/O6/ Consider K in place of MG (4 valid contact(s) for K vs. 0 for MG) |
| MG:AA:3274 | 1.00 | 70.06 |  | 0 |  | 1^st^ MG coordination shell missing (see Notes) |
| MG:AA:3275 | 1.00 | 58.19 |  | 0 |  | 1^st^ MG coordination shell missing (see Notes) |
| MG:AA:3276 | 1.00 | 46.88 |  | 0 | 3 | - Exclusion zone contact(s): /OP1/OP2/OP2/ Consider K in place of MG (3 valid contact(s) for K vs. 0 for MG) |
| MG:AA:3277 | 1.00 | 42.37 |  | 3 | 1 | 3O_ph_ - Exclusion zone contact(s): /O5'/ |
| MG:AA:3278 | 1.00 | 59.29 |  | 0 |  | 1^st^ MG coordination shell missing (see Notes) |
| MG:AA:3279 | 1.00 | 45.92 |  | 1 |  | O_b_ |
| MG:AA:3280 | 1.00 | 56.98 |  | 0 | 1 | - Exclusion zone contact(s): /O6/ Consider K in place of MG (1 valid contact(s) for K vs. 0 for MG) |
| MG:AA:3281 | 1.00 | 45.04 |  | 0 | 1 | - Exclusion zone contact(s): /O6/ Consider K in place of MG (1 valid contact(s) for K vs. 0 for MG) |
| MG:AA:3282 | 1.00 | 64.96 |  | 0 | 2 | - Exclusion zone contact(s): /N4/O6/ Consider K in place of MG (2 valid contact(s) for K vs. 0 for MG) |
| MG:AA:3283 | 1.00 | 74.16 |  | 0 |  | 1^st^ MG coordination shell missing (see Notes) |
| MG:AA:3284 | 1.00 | 78.15 |  | 0 | 2 | - Exclusion zone contact(s): /N4/O6/ Consider K in place of MG (2 valid contact(s) for K vs. 0 for MG) |
| MG:AA:3285 | 1.00 | 62.48 |  | 0 | 1 | - Exclusion zone contact(s): /OP2/ Consider K in place of MG (1 valid contact(s) for K vs. 0 for MG) |
| MG:AA:3286 | 1.00 | 67.72 |  | 0 |  | 1^st^ MG coordination shell missing (see Notes) |
| MG:AA:3287 | 1.00 | 66.74 |  | 0 | 1 | - Exclusion zone contact(s): /OP2/ Consider K in place of MG (1 valid contact(s) for K vs. 0 for MG) |
| MG:AA:3288 | 1.00 | 51.41 |  | 0 | 1 | - Exclusion zone contact(s): /O6/ Consider K in place of MG (1 valid contact(s) for K vs. 0 for MG) |
| MG:AA:3289 | 1.00 | 90.31 |  | 0 |  | 1^st^ MG coordination shell missing (see Notes) |
| MG:AA:3290 | 1.00 | 78.76 |  | 0 | 2 | - Exclusion zone contact(s): /N7/O6/ Consider K in place of MG (2 valid contact(s) for K vs. 0 for MG) |
| MG:AA:3291 | 1.00 | 34.43 |  | 1 | 2 | O_ph_ - Exclusion zone contact(s): /N7/O6/ Consider K in place of MG (2 valid contact(s) for K vs. 1 for MG) |
| MG:AA:3292 | 1.00 | 79.30 |  | 0 |  | 1^st^ MG coordination shell missing (see Notes) |
| MG:AA:3293 | 1.00 | 45.27 |  | 0 | 2 | - Exclusion zone contact(s): /O2'/OP1/ Consider K in place of MG (2 valid contact(s) for K vs. 0 for MG) |
| MG:AA:3294 | 1.00 | 64.90 |  | 0 | 2 | - Exclusion zone contact(s): /N7/OP2/ Consider K in place of MG (2 valid contact(s) for K vs. 0 for MG) |
| MG:AA:3295 | 1.00 | 62.58 |  | 0 | 3 | - Exclusion zone contact(s): /NH1_Arg_/OP1/OP2/ Consider K in place of MG (3 valid contact(s) for K vs. 0 for MG) |
| MG:AA:3296 | 1.00 | 55.33 |  | 1 |  | O_ph_ |
| MG:AA:3297 | 1.00 | 58.42 |  | 1 |  | O_ph_ |
| MG:AA:3298 | 1.00 | 38.21 |  | 0 | 2 | - Exclusion zone contact(s): /O3'/OP1/ Consider K in place of MG (2 valid contact(s) for K vs. 0 for MG) |
| MG:AA:3299 | 1.00 | 82.58 |  | 0 | 3 | - Exclusion zone contact(s): /O2'/O4'/OP2/ Consider K in place of MG (3 valid contact(s) for K vs. 0 for MG) |
| MG:AA:3300 | 1.00 | 38.92 |  | 0 |  | 1^st^ MG coordination shell missing (see Notes) |
| MG:AA:3301 | 1.00 | 50.46 |  | 0 | 2 | - Exclusion zone contact(s): /OP1/OP1/ Consider K in place of MG (2 valid contact(s) for K vs. 0 for MG) |
| MG:AA:3302 | 1.00 | 32.34 |  | 0 | 4 | - Exclusion zone contact(s): /N3_A_/N6/O2/OP1/ Consider K in place of MG (4 valid contact(s) for K vs. 0 for MG) |
| MG:AA:3303 | 1.00 | 94.20 |  | 0 | 2 | - Exclusion zone contact(s): /OP1/OP2/ Consider K in place of MG (2 valid contact(s) for K vs. 0 for MG) |
| MG:AA:3304 | 1.00 | 36.79 |  | 0 | 2 | - Exclusion zone contact(s): /O3'/OP1/ Consider NA in place of MG (1 valid contact(s) for NA vs. 0 for MG) |
| MG:AA:3305 | 1.00 | 47.04 |  | 0 | 2 | - Exclusion zone contact(s): /O5'/OP2/ Consider K in place of MG (2 valid contact(s) for K vs. 0 for MG) |
| MG:AA:3306 | 1.00 | 49.39 |  | 0 |  | 1^st^ MG coordination shell missing (see Notes) |
| MG:AA:3307 | 1.00 | 50.21 |  | 0 |  | 1^st^ MG coordination shell missing (see Notes) |
| MG:AA:3308 | 1.00 | 38.87 |  | 1 | 4 | O_ph_ - Exclusion zone contact(s): /NZ_Lys_/OP1/OP2/OP2/ Consider K in place of MG (3 valid contact(s) for K vs. 1 for MG) |
| MG:AA:3309 | 1.00 | 54.88 |  | 0 | 1 | - Exclusion zone contact(s): /O6/ Consider K in place of MG (1 valid contact(s) for K vs. 0 for MG) |
| MG:AA:3310 | 1.00 | 47.65 |  | 0 | 1 | - Exclusion zone contact(s): /OP2/ Consider K in place of MG (1 valid contact(s) for K vs. 0 for MG) |
| MG:AA:3311 | 1.00 | 49.75 |  | 0 | 1 | - Exclusion zone contact(s): /OP2/ Consider K in place of MG (1 valid contact(s) for K vs. 0 for MG) |
| MG:AA:3312 | 1.00 | 40.35 |  | 0 | 2 | - Exclusion zone contact(s): /OP1/OP2/ Consider K in place of MG (2 valid contact(s) for K vs. 0 for MG) |
| MG:AA:3313 | 1.00 | 83.02 |  | 0 | 1 | - Exclusion zone contact(s): /OP1/ Consider K in place of MG (1 valid contact(s) for K vs. 0 for MG) |
| MG:AA:3314 | 1.00 | 74.92 |  | 0 | 1 | - Exclusion zone contact(s): /OP1/ Consider K in place of MG (1 valid contact(s) for K vs. 0 for MG) |
| MG:AA:3315 | 1.00 | 49.59 |  | 0 | 5 | - Exclusion zone contact(s): /N4/N4/O4/O6/O6/ Consider K in place of MG (5 valid contact(s) for K vs. 0 for MG) |
| MG:AA:3316 | 1.00 | 47.57 |  | 0 | 4 | - Exclusion zone contact(s): /N1/O3'/O4'/OP2/ Consider K in place of MG (4 valid contact(s) for K vs. 0 for MG) |
| MG:AA:3317 | 1.00 | 57.87 |  | 0 | 2 | - Exclusion zone contact(s): /N7/O6/ Consider K in place of MG (2 valid contact(s) for K vs. 0 for MG) |
| MG:AA:3318 | 1.00 | 65.92 |  | 0 |  | 1^st^ MG coordination shell missing (see Notes) |
| MG:AA:3319 | 1.00 | 85.00 |  | 0 |  | 1^st^ MG coordination shell missing (see Notes) |
| MG:AA:3320 | 1.00 | 69.54 |  | 0 |  | 1^st^ MG coordination shell missing (see Notes) |
| MG:AA:3321 | 1.00 | 69.64 |  | 0 |  | 1^st^ MG coordination shell missing (see Notes) |
| MG:AA:3322 | 1.00 | 58.59 |  | 0 | 1 | - Exclusion zone contact(s): /OP1/ Consider K in place of MG (1 valid contact(s) for K vs. 0 for MG) |
| MG:AA:3323 | 1.00 | 48.97 |  | 0 |  | 1^st^ MG coordination shell missing (see Notes) |
| MG:AA:3324 | 1.00 | 43.15 |  | 1 |  | O_ph_ |
| MG:AA:3325 | 1.00 | 50.00 |  | 0 | 3 | - Exclusion zone contact(s): /N7/N7/O6/ Consider K in place of MG (3 valid contact(s) for K vs. 0 for MG) |
| MG:AA:3326 | 1.00 | 22.91 |  | 1 | 3 | O_r_ - Exclusion zone contact(s): /N7/O2'/OP2/ Consider K in place of MG (2 valid contact(s) for K vs. 1 for MG) |
| MG:AA:3327 | 1.00 | 45.33 |  | 0 | 1 | - Exclusion zone contact(s): /OP2/ Consider K in place of MG (1 valid contact(s) for K vs. 0 for MG) |
| MG:AA:3328 | 1.00 | 32.16 |  | 0 | 1 | - Exclusion zone contact(s): /O6/ Consider K in place of MG (1 valid contact(s) for K vs. 0 for MG) |
| MG:AA:3329 | 1.00 | 66.22 |  | 0 |  | 1^st^ MG coordination shell missing (see Notes) |
| MG:AA:3330 | 1.00 | 111.12 |  | 0 | 1 | - Exclusion zone contact(s): /OP1/ Consider NA in place of MG (1 valid contact(s) for NA vs. 0 for MG) |
| MG:AA:3331 | 1.00 | 41.86 |  | 1 |  | O_ph_ |
| MG:AA:3332 | 1.00 | 83.12 |  | 0 | 1 | - Exclusion zone contact(s): /OP2/ Consider K in place of MG (1 valid contact(s) for K vs. 0 for MG) |
| MG:AA:3333 | 1.00 | 59.71 |  | 0 | 3 | - Exclusion zone contact(s): /N2/O4'/ Consider K in place of MG (3 valid contact(s) for K vs. 0 for MG) |
| MG:AA:3334 | 1.00 | 46.61 |  | 0 | 4 | - Exclusion zone contact(s): /O4/O4/O6/O6/ Consider K in place of MG (3 valid contact(s) for K vs. 0 for MG) |
| MG:AA:3335 | 1.00 | 83.83 |  | 0 | 1 | - Exclusion zone contact(s): /OP2/ Consider K in place of MG (1 valid contact(s) for K vs. 0 for MG) |
| MG:AA:3336 | 1.00 | 122.46 |  | 0 | 2 | - Exclusion zone contact(s): /O2'/O3'/ Consider K in place of MG (2 valid contact(s) for K vs. 0 for MG) |
| MG:AA:3337 | 1.00 | 51.68 |  | 0 | 2 | - Exclusion zone contact(s): /O2'/O4'/ Consider K in place of MG (2 valid contact(s) for K vs. 0 for MG) |
| MG:AA:3338 | 1.00 | 72.27 |  | 0 | 1 | - Exclusion zone contact(s): /OP2/ Consider NA in place of MG (1 valid contact(s) for NA vs. 0 for MG) |
| MG:AA:3339 | 1.00 | 50.98 |  | 1 | 2 | O_b_ - Exclusion zone contact(s): /N7/O6/ Consider K in place of MG (2 valid contact(s) for K vs. 1 for MG) |
| MG:AA:3340 | 1.00 | 90.63 |  | 0 | 2 | - Exclusion zone contact(s): /N7/OP2/ Consider NA in place of MG (1 valid contact(s) for NA vs. 0 for MG) |
| MG:AA:3341 | 1.00 | 57.15 |  | 0 | 2 | - Exclusion zone contact(s): /O4/O6/ Consider K in place of MG (2 valid contact(s) for K vs. 0 for MG) |
| MG:AA:3342 | 1.00 | 63.95 |  | 0 | 3 | - Exclusion zone contact(s): /O6/O6/OP2/ Consider K in place of MG (3 valid contact(s) for K vs. 0 for MG) |
| MG:AA:3343 | 1.00 | 48.48 |  | 0 | 1 | - Exclusion zone contact(s): /O6/ Consider K in place of MG (1 valid contact(s) for K vs. 0 for MG) |
| MG:AA:3344 | 1.00 | 76.02 |  | 0 | 2 | - Exclusion zone contact(s): /O6/O6/ Consider K in place of MG (2 valid contact(s) for K vs. 0 for MG) |
| MG:AA:3345 | 1.00 | 55.65 |  | 1 |  | O_ph_ |
| MG:AA:3346 | 1.00 | 58.56 |  | 0 | 1 | - Exclusion zone contact(s): /OP1/ Consider K in place of MG (1 valid contact(s) for K vs. 0 for MG) |
| MG:AA:3347 | 1.00 | 66.65 |  | 0 | 2 | - Exclusion zone contact(s): /N7/OP1/ Consider K in place of MG (2 valid contact(s) for K vs. 0 for MG) |
| MG:AA:3348 | 1.00 | 26.88 |  | 2 |  | O_ph_N_b_ |
| MG:AA:3349 | 1.00 | 42.88 |  | 1 | 2 | O_ph_ - Exclusion zone contact(s): /O2/O2/ Consider K in place of MG (2 valid contact(s) for K vs. 1 for MG) |
| MG:AA:3350 | 1.00 | 36.45 |  | 1 | 3 | O_r_ - Exclusion zone contact(s): /O2'/O4/OP2/ Consider K in place of MG (3 valid contact(s) for K vs. 1 for MG) |
| MG:AA:3351 | 1.00 | 62.63 |  | 1 | 1 | O_r_ - Exclusion zone contact(s): /OP2/ |
| MG:AA:3352 | 1.00 | 28.23 |  | 0 | 4 | - Exclusion zone contact(s): /O3'/O6/OP1/OP2/ Consider K in place of MG (3 valid contact(s) for K vs. 0 for MG) |
| MG:AA:3353 | 1.00 | 83.67 |  | 0 | 1 | - Exclusion zone contact(s): /N7/ Consider K in place of MG (1 valid contact(s) for K vs. 0 for MG) |
| MG:AA:3354 | 1.00 | 53.42 |  | 1 | 4 | O_b_ - Exclusion zone contact(s): /N4/O4/O6/O6/ Consider K in place of MG (4 valid contact(s) for K vs. 1 for MG) |
| MG:AA:3355 | 1.00 | 48.58 |  | 0 | 1 | - Exclusion zone contact(s): /OP2/ Consider NA in place of MG (1 valid contact(s) for NA vs. 0 for MG) |
| MG:AA:3356 | 1.00 | 62.66 |  | 1 |  | O_b_ |
| MG:AA:3357 | 1.00 | 91.55 |  | 0 | 2 | - Exclusion zone contact(s): /O4/OP1/ Consider NA in place of MG (1 valid contact(s) for NA vs. 0 for MG) |
| MG:AA:3358 | 1.00 | 95.89 |  | 0 | 4 | - Exclusion zone contact(s): /N4/O4/O6/O6/ Consider NA in place of MG (2 valid contact(s) for NA vs. 0 for MG) |
| MG:AA:3359 | 1.00 | 48.03 |  | 0 | 3 | - Exclusion zone contact(s): /O4/O6/O6/ Consider K in place of MG (2 valid contact(s) for K vs. 0 for MG) |
| MG:AA:3360 | 1.00 | 95.65 |  | 0 |  | 1^st^ MG coordination shell missing (see Notes) |
| MG:AA:3361 | 1.00 | 93.68 |  | 0 | 1 | - Exclusion zone contact(s): /OP2/ Consider K in place of MG (1 valid contact(s) for K vs. 0 for MG) |
| MG:AA:3362 | 1.00 | 62.45 |  | 1 |  | O_ph_ |
| MG:AA:3363 | 1.00 | 74.62 |  | 0 | 5 | - Exclusion zone contact(s): /O3'/O4'/O5'/OP1/OP1/ Consider K in place of MG (4 valid contact(s) for K vs. 0 for MG) |
| MG:AA:3364 | 1.00 | 57.12 |  | 0 | 2 | - Exclusion zone contact(s): /O6/O6/ Consider K in place of MG (2 valid contact(s) for K vs. 0 for MG) |
| MG:AA:3365 | 1.00 | 39.04 |  | 0 |  | 1^st^ MG coordination shell missing (see Notes) |
| MG:AA:3366 | 1.00 | 53.51 |  | 0 | 2 | - Exclusion zone contact(s): /N7/O6/ Consider K in place of MG (2 valid contact(s) for K vs. 0 for MG) |
| MG:AA:3367 | 1.00 | 49.16 |  | 0 |  | 1^st^ MG coordination shell missing (see Notes) |
| MG:AA:3368 | 1.00 | 87.31 |  | 0 |  | 1^st^ MG coordination shell missing (see Notes) |
| MG:AA:3369 | 1.00 | 81.22 |  | 1 | 2 | N_b_ - Exclusion zone contact(s): /N4/O6/ Consider K in place of MG (2 valid contact(s) for K vs. 1 for MG) |
| MG:AA:3370 | 1.00 | 64.80 |  | 0 |  | 1^st^ MG coordination shell missing (see Notes) |
| MG:AA:3371 | 1.00 | 63.31 |  | 1 | 2 | O_ph_ - Exclusion zone contact(s): /O5'/OP2/ Consider K in place of MG (2 valid contact(s) for K vs. 1 for MG) |
| MG:AA:3372 | 1.00 | 52.09 |  | 0 | 2 | - Exclusion zone contact(s): /O6/O6/ Consider NA in place of MG (1 valid contact(s) for NA vs. 0 for MG) |
| MG:AA:3373 | 1.00 | 48.32 |  | 0 | 3 | - Exclusion zone contact(s): /N7/OP2/ Consider K in place of MG (3 valid contact(s) for K vs. 0 for MG) |
| MG:AA:3374 | 1.00 | 46.95 |  | 0 | 3 | - Exclusion zone contact(s): /O2/O4'/O6/ Consider K in place of MG (2 valid contact(s) for K vs. 0 for MG) |
| MG:AA:3375 | 1.00 | 61.38 |  | 0 |  | 1^st^ MG coordination shell missing (see Notes) |
| MG:AA:3376 | 1.00 | 116.05 |  | 0 |  | 1^st^ MG coordination shell missing (see Notes) |
| MG:AA:3377 | 1.00 | 19.48 |  | 3 | 1 | 3O_ph_ - Exclusion zone contact(s): /OP1/ |
| MG:AA:3378 | 1.00 | 50.06 |  | 4 |  | 4O_ph_ Large octahedral topology discrepancy!!! |
| MG:AA:3379 | 1.00 | 70.12 |  | 0 | 3 | - Exclusion zone contact(s): /N4/O4/O6/ Consider K in place of MG (2 valid contact(s) for K vs. 0 for MG) |
| MG:AA:3380 | 1.00 | 25.33 |  | 0 | 4 | - Exclusion zone contact(s): /O2'/O3'/OP1/OP2/ Consider K in place of MG (4 valid contact(s) for K vs. 0 for MG) |
| MG:AA:3381 | 1.00 | 32.15 |  | 1 |  | O_ph_ |
| MG:AA:3382 | 1.00 | 83.43 |  | 0 |  | 1^st^ MG coordination shell missing (see Notes) |
| MG:AA:3383 | 1.00 | 24.00 |  | 2 |  | 2O_ph_ |
| MG:AA:3384 | 1.00 | 47.88 |  | 0 |  | 1^st^ MG coordination shell missing (see Notes) |
| MG:AA:3385 | 1.00 | 34.82 |  | 0 | 2 | - Exclusion zone contact(s): /O6/OP2/ Consider NA in place of MG (1 valid contact(s) for NA vs. 0 for MG) |
| MG:AA:3386 | 1.00 | 51.73 |  | 1 | 1 | O_ph_ - Exclusion zone contact(s): /OP2/ |
| MG:AA:3387 | 1.00 | 40.29 |  | 0 | 1 | - Exclusion zone contact(s): /OP1/ Consider NA in place of MG (1 valid contact(s) for NA vs. 0 for MG) |
| MG:AA:3388 | 1.00 | 46.85 |  | 0 | 2 | - Exclusion zone contact(s): /O4/OP2/ Consider K in place of MG (2 valid contact(s) for K vs. 0 for MG) |
| MG:AA:3389 | 1.00 | 67.53 |  | 0 | 3 | - Exclusion zone contact(s): /O3'/OP1/OP1/ Consider K in place of MG (2 valid contact(s) for K vs. 0 for MG) |
| MG:AA:3390 | 1.00 | 46.61 |  | 0 | 1 | - Exclusion zone contact(s): /O4/ Consider K in place of MG (1 valid contact(s) for K vs. 0 for MG) |
| MG:AA:3391 | 1.00 | 39.04 |  | 1 |  | O_ph_ |
| MG:AA:3392 | 1.00 | 63.14 |  | 0 | 2 | - Exclusion zone contact(s): /OP1/OP1/ Consider NA in place of MG (1 valid contact(s) for NA vs. 0 for MG) |
| MG:AA:3393 | 1.00 | 63.46 |  | 0 | 1 | - Exclusion zone contact(s): /O2'/ Consider K in place of MG (1 valid contact(s) for K vs. 0 for MG) |
| MG:AA:3394 | 1.00 | 45.39 |  | 0 | 3 | - Exclusion zone contact(s): /O2/O2'/O4'/ Consider K in place of MG (2 valid contact(s) for K vs. 0 for MG) |
| MG:AA:3395 | 1.00 | 88.32 |  | 0 |  | 1^st^ MG coordination shell missing (see Notes) |
| MG:AA:3396 | 1.00 | 58.83 |  | 0 | 4 | - Exclusion zone contact(s): /N1_A_/N6/O3'/OP1/ Consider K in place of MG (4 valid contact(s) for K vs. 0 for MG) |
| MG:AA:3397 | 1.00 | 56.32 |  | 0 | 2 | - Exclusion zone contact(s): /O6/O6/ Consider NA in place of MG (1 valid contact(s) for NA vs. 0 for MG) |
| MG:AA:3398 | 1.00 | 30.59 |  | 0 |  | 1^st^ MG coordination shell missing (see Notes) |
| MG:AA:3399 | 1.00 | 75.77 |  | 0 | 2 | - Exclusion zone contact(s): /N7/O6/ Consider NA in place of MG (1 valid contact(s) for NA vs. 0 for MG) |
| MG:AA:3400 | 1.00 | 58.33 |  | 1 |  | O_ph_ |
| MG:AA:3401 | 1.00 | 46.18 |  | 0 |  | 1^st^ MG coordination shell missing (see Notes) |
| MG:AA:3402 | 1.00 | 66.57 |  | 0 | 3 | - Exclusion zone contact(s): /N7/O6/O6/ Consider NA in place of MG (2 valid contact(s) for NA vs. 0 for MG) |
| MG:AA:3403 | 1.00 | 60.99 |  | 0 | 1 | - Exclusion zone contact(s): /O6/ Consider K in place of MG (1 valid contact(s) for K vs. 0 for MG) |
| MG:AA:3404 | 1.00 | 77.61 |  | 0 |  | 1^st^ MG coordination shell missing (see Notes) |
| MG:AA:3405 | 1.00 | 73.68 |  | 0 |  | No contacts in 5.0 A |
| MG:AA:3406 | 1.00 | 32.52 |  | 0 | 2 | - Exclusion zone contact(s): /N4/OP2/ Consider K in place of MG (2 valid contact(s) for K vs. 0 for MG) |
| MG:AA:3407 | 1.00 | 78.31 |  | 0 |  | No contacts in 5.0 A |
| MG:AA:3408 | 1.00 | 61.54 |  | 0 | 1 | - Exclusion zone contact(s): /OP2/ Consider K in place of MG (1 valid contact(s) for K vs. 0 for MG) |
| MG:AA:3409 | 1.00 | 56.91 |  | 0 |  | 1^st^ MG coordination shell missing (see Notes) |
| MG:AA:3410 | 1.00 | 84.65 |  | 0 | 2 | - Exclusion zone contact(s): /O2'/O3'/ Consider K in place of MG (2 valid contact(s) for K vs. 0 for MG) |
| MG:AA:3411 | 1.00 | 73.75 |  | 1 |  | O_ph_ |
| MG:AA:3412 | 1.00 | 30.72 |  | 0 |  | 1^st^ MG coordination shell missing (see Notes) |
| MG:AA:3413 | 1.00 | 95.99 |  | 0 |  | No contacts in 5.0 A |
| MG:AA:3414 | 1.00 | 56.43 |  | 0 | 1 | - Exclusion zone contact(s): /N7/ Consider K in place of MG (1 valid contact(s) for K vs. 0 for MG) |
| MG:AA:3415 | 1.00 | 63.16 |  | 0 | 2 | - Exclusion zone contact(s): /N7/OP2/ Consider K in place of MG (2 valid contact(s) for K vs. 0 for MG) |
| MG:AA:3416 | 1.00 | 78.19 |  | 0 |  | 1^st^ MG coordination shell missing (see Notes) |
| MG:AA:3417 | 1.00 | 58.79 |  | 0 | 5 | - Exclusion zone contact(s): /N3_U_/O4/O4/O4/O6/ Consider K in place of MG (5 valid contact(s) for K vs. 0 for MG) |
| MG:AA:3418 | 1.00 | 91.89 |  | 0 |  | No contacts in 5.0 A |
| MG:AA:3419 | 1.00 | 82.10 |  | 0 |  | No contacts in 5.0 A |
| MG:AA:3420 | 1.00 | 68.80 |  | 0 | 1 | - Exclusion zone contact(s): /O6/ Consider K in place of MG (1 valid contact(s) for K vs. 0 for MG) |
| MG:AA:3421 | 1.00 | 43.26 |  | 0 | 1 | - Exclusion zone contact(s): /OP1/ Consider K in place of MG (1 valid contact(s) for K vs. 0 for MG) |
| MG:AA:3422 | 1.00 | 73.31 |  | 0 | 4 | - Exclusion zone contact(s): /N1_G_/O6/O6/OP2/ Consider K in place of MG (4 valid contact(s) for K vs. 0 for MG) |
| MG:AA:3423 | 1.00 | 99.88 |  | 0 |  | No contacts in 5.0 A |
| MG:AA:3424 | 1.00 | 54.22 |  | 1 | 5 | O_ph_ - Exclusion zone contact(s): /O3'/OP1/OP1/OP2/OP2/ Consider K in place of MG (4 valid contact(s) for K vs. 1 for MG) |
| MG:AA:3425 | 1.00 | 44.25 |  | 0 | 3 | - Exclusion zone contact(s): /OP2/OP2/OP2/ Consider K in place of MG (3 valid contact(s) for K vs. 0 for MG) |
| MG:AA:3426 | 1.00 | 64.62 |  | 0 | 2 | - Exclusion zone contact(s): /O2'/OP2/ Consider K in place of MG (2 valid contact(s) for K vs. 0 for MG) |
| MG:AA:3427 | 1.00 | 47.30 |  | 2 | 1 | O_ph_N_b_ - Exclusion zone contact(s): /O6/ |
| MG:AA:3428 | 1.00 | 101.84 |  | 0 |  | 1^st^ MG coordination shell missing (see Notes) |
| MG:AA:3429 | 1.00 | 60.62 |  | 1 | 2 | O_r_ - Exclusion zone contact(s): /OP1/OP2/ Consider K in place of MG (2 valid contact(s) for K vs. 1 for MG) |
| MG:AA:3430 | 1.00 | 81.64 |  | 3 | 1 | O_ph_2Anion - Exclusion zone contact(s): /N2/ |
| MG:AA:3431 | 1.00 | 53.00 |  | 0 | 3 | - Exclusion zone contact(s): /O2'/OP1/OP2/ Consider K in place of MG (2 valid contact(s) for K vs. 0 for MG) |
| MG:AA:3432 | 1.00 | 83.79 |  | 0 |  | 1^st^ MG coordination shell missing (see Notes) |
| MG:AA:3433 | 1.00 | 64.18 |  | 0 | 3 | - Exclusion zone contact(s): /N7/O2'/OP1/ Consider K in place of MG (3 valid contact(s) for K vs. 0 for MG) |
| MG:AA:3434 | 1.00 | 65.07 |  | 0 | 4 | - Exclusion zone contact(s): /N2/O4/O6/O6/ Consider K in place of MG (4 valid contact(s) for K vs. 0 for MG) |
| MG:AA:3435 | 1.00 | 69.66 |  | 0 | 2 | - Exclusion zone contact(s): /OP1/OP2/ Consider K in place of MG (2 valid contact(s) for K vs. 0 for MG) |
| MG:AA:3436 | 1.00 | 127.98 |  | 0 | 2 | - Exclusion zone contact(s): /N3_G_/N6/ Consider K in place of MG (2 valid contact(s) for K vs. 0 for MG) |
| MG:AA:3437 | 1.00 | 51.63 |  | 1 | 1 | O_ph_ - Exclusion zone contact(s): /OP2/ |
| MG:AA:3438 | 1.00 | 99.85 |  | 0 | 1 | - Exclusion zone contact(s): /O6/ Consider NA in place of MG (1 valid contact(s) for NA vs. 0 for MG) |
| MG:AA:3439 | 1.00 | 71.22 |  | 0 | 1 | - Exclusion zone contact(s): /OP1/ Consider K in place of MG (1 valid contact(s) for K vs. 0 for MG) |
| MG:AA:3440 | 1.00 | 75.10 |  | 0 | 1 | - Exclusion zone contact(s): /OP1/ Consider K in place of MG (1 valid contact(s) for K vs. 0 for MG) |
| MG:AA:3441 | 1.00 | 102.51 |  | 0 | 3 | - Exclusion zone contact(s): /O5'/OP2/OP2/ Consider K in place of MG (3 valid contact(s) for K vs. 0 for MG) |
| MG:AA:3442 | 1.00 | 128.35 |  | 2 | 5 | N_b_Anion - Exclusion zone contact(s): /N1_A_/N1_G_/N3_A_/N6/N6/ Consider K in place of MG (4 valid contact(s) for K vs. 2 for MG) |
| MG:AA:3443 | 1.00 | 77.65 |  | 0 | 3 | - Exclusion zone contact(s): /OP1/OP1/OP1/ Consider K in place of MG (3 valid contact(s) for K vs. 0 for MG) |
| MG:AA:3444 | 1.00 | 81.54 |  | 0 |  | 1^st^ MG coordination shell missing (see Notes) |
| MG:AA:3445 | 1.00 | 63.38 |  | 0 | 1 | - Exclusion zone contact(s): /O6/ Consider K in place of MG (1 valid contact(s) for K vs. 0 for MG) |
| MG:AA:3446 | 1.00 | 90.83 |  | 0 |  | No contacts in 5.0 A |
| MG:AA:3447 | 1.00 | 71.17 |  | 0 | 3 | - Exclusion zone contact(s): /O5'/OP1/OP2/ Consider K in place of MG (3 valid contact(s) for K vs. 0 for MG) |
| MG:AA:3448 | 1.00 | 93.83 |  | 0 |  | 1^st^ MG coordination shell missing (see Notes) |
| MG:AA:3449 | 1.00 | 82.59 |  | 0 | 1 | - Exclusion zone contact(s): /O3'/ Consider K in place of MG (1 valid contact(s) for K vs. 0 for MG) |
| MG:AA:3450 | 1.00 | 59.72 |  | 1 |  | O_ph_ |
| MG:AA:3451 | 1.00 | 89.90 |  | 0 | 4 | - Exclusion zone contact(s): /O3'/O5'/OP2/OP2/ Consider K in place of MG (3 valid contact(s) for K vs. 0 for MG) |
| MG:AA:3452 | 1.00 | 97.35 |  | 0 |  | 1^st^ MG coordination shell missing (see Notes) |
| MG:AA:3453 | 1.00 | 123.40 |  | 0 | 2 | - Exclusion zone contact(s): /N1/O2'/ Consider NA in place of MG (1 valid contact(s) for NA vs. 0 for MG) |
| MG:AA:3454 | 1.00 | 67.59 |  | 0 | 1 | - Exclusion zone contact(s): /OP1/ Consider K in place of MG (1 valid contact(s) for K vs. 0 for MG) |
| MG:AA:3455 | 1.00 | 22.33 |  | 0 | 2 | - Exclusion zone contact(s): /N7/N7/ Consider K in place of MG (2 valid contact(s) for K vs. 0 for MG) |
| MG:AA:3456 | 1.00 | 20.84 |  | 1 | 1 | O_ph_ - Exclusion zone contact(s): /OP1/ |
| MG:AA:3457 | 1.00 | 57.61 |  | 1 | 2 | O_ph_ - Exclusion zone contact(s): /O6/OP1/ Consider K in place of MG (2 valid contact(s) for K vs. 1 for MG) |
| MG:AA:3458 | 1.00 | 49.59 |  | 1 |  | O_ph_ |
| MG:AA:3459 | 1.00 | 30.62 |  | 0 |  | 1^st^ MG coordination shell missing (see Notes) |
| MG:AA:3460 | 1.00 | 28.26 |  | 0 | 3 | - Exclusion zone contact(s): /N7/O4/O6/ Consider K in place of MG (3 valid contact(s) for K vs. 0 for MG) |
| MG:AA:3461 | 1.00 | 58.79 |  | 0 |  | 1^st^ MG coordination shell missing (see Notes) |
| MG:AA:3462 | 1.00 | 32.83 |  | 2 | 1 | 2O_ph_ - Exclusion zone contact(s): /OP2/ |
| MG:AA:3463 | 1.00 | 3.22 |  | 0 | 5 | - Exclusion zone contact(s): /O2/O2'/O2'/O6/O6/ Consider K in place of MG (4 valid contact(s) for K vs. 0 for MG) |
| MG:AA:3464 | 1.00 | 61.58 |  | 1 | 1 | O_ph_ - Exclusion zone contact(s): /OP2/ |
| MG:AA:3465 | 1.00 | 28.96 |  | 0 | 2 | - Exclusion zone contact(s): /N7/O6/ Consider K in place of MG (2 valid contact(s) for K vs. 0 for MG) |
| MG:AA:3466 | 1.00 | 37.27 |  | 1 | 1 | O_ph_ - Exclusion zone contact(s): /OP2/ |
| MG:AA:3467 | 1.00 | 55.52 |  | 0 | 2 | - Exclusion zone contact(s): /N6/O2/ Consider K in place of MG (2 valid contact(s) for K vs. 0 for MG) |
| MG:AA:3468 | 1.00 | 22.85 |  | 2 | 2 | 2O_ph_ - Exclusion zone contact(s): /O4/OP2/ |
| MG:AA:3469 | 1.00 | 42.27 |  | 2 |  | 2O_ph_ |
| MG:AA:3470 | 1.00 | 76.29 |  | 0 | 1 | - Exclusion zone contact(s): /OP2/ Consider K in place of MG (1 valid contact(s) for K vs. 0 for MG) |
| MG:AA:3471 | 1.00 | 77.28 |  | 0 | 1 | - Exclusion zone contact(s): /OP1/ Consider K in place of MG (1 valid contact(s) for K vs. 0 for MG) |
| MG:AA:3472 | 1.00 | 59.70 |  | 1 | 1 | O_ph_ - Exclusion zone contact(s): /OP2/ |
| MG:AA:3473 | 1.00 | 63.00 |  | 1 |  | O_ph_ |
| MG:AA:3474 | 1.00 | 65.83 |  | 0 | 1 | - Exclusion zone contact(s): /OP1/ Consider K in place of MG (1 valid contact(s) for K vs. 0 for MG) |
| MG:AA:3475 | 1.00 | 85.32 |  | 0 |  | 1^st^ MG coordination shell missing (see Notes) |
| MG:AA:3476 | 1.00 | 47.13 |  | 0 | 2 | - Exclusion zone contact(s): /OP2/OP2/ Consider NA in place of MG (2 valid contact(s) for NA vs. 0 for MG) |
| MG:AA:3477 | 1.00 | 65.14 |  | 0 | 1 | - Exclusion zone contact(s): /O6/ Consider K in place of MG (1 valid contact(s) for K vs. 0 for MG) |
| MG:AA:3478 | 1.00 | 59.96 |  | 0 | 3 | - Exclusion zone contact(s): /O2/O4/O4/ Consider K in place of MG (3 valid contact(s) for K vs. 0 for MG) |
| MG:AA:3479 | 1.00 | 59.42 |  | 0 | 1 | - Exclusion zone contact(s): /OP1/ Consider NA in place of MG (1 valid contact(s) for NA vs. 0 for MG) |
| MG:AA:3480 | 1.00 | 70.99 |  | 0 |  | 1^st^ MG coordination shell missing (see Notes) |
| MG:AA:3481 | 1.00 | 61.26 |  | 0 | 1 | - Exclusion zone contact(s): /OP1/ Consider K in place of MG (1 valid contact(s) for K vs. 0 for MG) |
| MG:AA:3482 | 1.00 | 58.21 |  | 0 | 3 | - Exclusion zone contact(s): /N7/O5'/OP2/ Consider NA in place of MG (2 valid contact(s) for NA vs. 0 for MG) |
| MG:AA:3483 | 1.00 | 65.58 |  | 0 | 5 | - Exclusion zone contact(s): /O3'/OP1/OP1/OP2/OP2/ Consider K in place of MG (4 valid contact(s) for K vs. 0 for MG) |
| MG:AA:3484 | 1.00 | 66.76 |  | 1 | 3 | O_b_ - Exclusion zone contact(s): /N1_G_/N7/O6/ Consider K in place of MG (3 valid contact(s) for K vs. 1 for MG) |
| MG:AA:3485 | 1.00 | 91.90 |  | 0 | 1 | - Exclusion zone contact(s): /OP1/ Consider K in place of MG (1 valid contact(s) for K vs. 0 for MG) |
| MG:AA:3486 | 1.00 | 60.56 |  | 0 | 3 | - Exclusion zone contact(s): /O2/O2'/OP1/ Consider K in place of MG (3 valid contact(s) for K vs. 0 for MG) |
| MG:AA:3487 | 1.00 | 72.21 |  | 0 |  | 1^st^ MG coordination shell missing (see Notes) |
| MG:AA:3488 | 1.00 | 68.19 |  | 0 |  | 1^st^ MG coordination shell missing (see Notes) |
| MG:AA:3489 | 1.00 | 63.20 |  | 0 | 2 | - Exclusion zone contact(s): /N3_C_/N7/ Consider K in place of MG (2 valid contact(s) for K vs. 0 for MG) |
| MG:AA:3490 | 1.00 | 102.04 |  | 0 | 1 | - Exclusion zone contact(s): /N4/ Consider K in place of MG (1 valid contact(s) for K vs. 0 for MG) |
| MG:AA:3491 | 1.00 | 68.35 |  | 0 | 2 | - Exclusion zone contact(s): /N7/O6/ Consider K in place of MG (2 valid contact(s) for K vs. 0 for MG) |
| MG:AA:3492 | 1.00 | 48.57 |  | 1 | 2 | O_r_ - Exclusion zone contact(s): /O3'/OP2/ Consider K in place of MG (2 valid contact(s) for K vs. 1 for MG) |
| MG:AA:3493 | 1.00 | 76.97 |  | 0 |  | 1^st^ MG coordination shell missing (see Notes) |
| MG:AA:3494 | 1.00 | 69.96 |  | 1 | 2 | O_r_ - Exclusion zone contact(s): /OP2/OP2/ |
| MG:AA:3495 | 1.00 | 56.49 |  | 0 | 4 | - Exclusion zone contact(s): /N7/O6/OP1/OP2/ Consider K in place of MG (4 valid contact(s) for K vs. 0 for MG) |
| MG:AA:3496 | 1.00 | 63.04 |  | 0 | 3 | - Exclusion zone contact(s): /O5'/O5'/OP1/ Consider K in place of MG (3 valid contact(s) for K vs. 0 for MG) |
| MG:AA:3497 | 1.00 | 80.92 |  | 0 |  | 1^st^ MG coordination shell missing (see Notes) |
| MG:AA:3498 | 1.00 | 52.42 |  | 0 | 3 | - Exclusion zone contact(s): /O2/O3'/OP1/ Consider K in place of MG (3 valid contact(s) for K vs. 0 for MG) |
| MG:AA:3499 | 1.00 | 101.49 |  | 1 | 1 | O_b_ - Exclusion zone contact(s): /O6/ |
| MG:AA:3500 | 1.00 | 22.71 |  | 1 | 2 | O_ph_ - Exclusion zone contact(s): /O2'/OP2/ Consider K in place of MG (2 valid contact(s) for K vs. 1 for MG) |
| MG:AA:3501 | 1.00 | 24.65 |  | 1 | 1 | O_ph_ - Exclusion zone contact(s): /O2'/ |
| MG:AA:3502 | 1.00 | 55.35 |  | 0 | 2 | - Exclusion zone contact(s): /O3'/OP1/ Consider NA in place of MG (1 valid contact(s) for NA vs. 0 for MG) |
| MG:AA:3503 | 1.00 | 23.33 |  | 2 | 5 | 2O_ph_ - Exclusion zone contact(s): /O2'/O2'/O3'/OP1/OP2/ Consider K in place of MG (5 valid contact(s) for K vs. 2 for MG) |
| MG:AA:3504 | 1.00 | 31.60 |  | 3 | 1 | 2O_ph_O_b_ - Exclusion zone contact(s): /O3'/ |
| MG:AA:3505 | 1.00 | 55.64 |  | 0 | 1 | - Exclusion zone contact(s): /OP2/ Consider NA in place of MG (1 valid contact(s) for NA vs. 0 for MG) |
| MG:AA:3506 | 1.00 | 27.05 |  | 1 |  | O_ph_ |
| MG:AA:3507 | 1.00 | 30.00 |  | 1 |  | O_ph_ |
| MG:AA:3508 | 1.00 | 61.75 |  | 0 | 2 | - Exclusion zone contact(s): /N1_G_/O4/ Consider K in place of MG (2 valid contact(s) for K vs. 0 for MG) |
| MG:AA:3509 | 1.00 | 19.26 |  | 1 | 1 | O_ph_ - Exclusion zone contact(s): /OP1/ |
| MG:AA:3510 | 1.00 | 26.83 |  | 1 |  | O_ph_ |
| MG:AA:3511 | 1.00 | 26.97 |  | 0 | 1 | - Exclusion zone contact(s): /O6/ Consider K in place of MG (1 valid contact(s) for K vs. 0 for MG) |
| MG:AA:3512 | 1.00 | 31.24 |  | 1 | 2 | O_r_ - Exclusion zone contact(s): /OP2/OP2/ |
| MG:AA:3513 | 1.00 | 28.73 |  | 0 |  | 1^st^ MG coordination shell missing (see Notes) |
| MG:AA:3514 | 1.00 | 49.29 |  | 0 |  | 1^st^ MG coordination shell missing (see Notes) |
| MG:AA:3515 | 1.00 | 20.64 |  | 0 | 2 | - Exclusion zone contact(s): /N7/O4/ Consider NA in place of MG (1 valid contact(s) for NA vs. 0 for MG) |
| MG:AA:3516 | 1.00 | 35.81 |  | 1 | 1 | O_ph_ - Exclusion zone contact(s): /OP1/ |
| MG:AA:3517 | 1.00 | 69.11 |  | 0 | 1 | - Exclusion zone contact(s): /OP1/ Consider NA in place of MG (1 valid contact(s) for NA vs. 0 for MG) |
| MG:AA:3518 | 1.00 | 17.31 |  | 0 | 5 | - Exclusion zone contact(s): /N7/O2'/OP2/OP2/OP2/ Consider K in place of MG (3 valid contact(s) for K vs. 0 for MG) |
| MG:AA:3519 | 1.00 | 63.87 |  | 2 |  | 2O_ph_ |
| MG:AA:3520 | 1.00 | 75.04 |  | 2 | 2 | O_ph_O_r_ - Exclusion zone contact(s): /OP1/OP2/ |
| MG:AA:3521 | 1.00 | 32.54 |  | 0 | 1 | - Exclusion zone contact(s): /O2'/ Consider K in place of MG (1 valid contact(s) for K vs. 0 for MG) |
| MG:AA:3522 | 1.00 | 14.68 |  | 1 | 5 | O_b_ - Exclusion zone contact(s): /N7/O2/O4/O4'/O6/ Consider K in place of MG (3 valid contact(s) for K vs. 1 for MG) |
| MG:AA:3523 | 1.00 | 42.97 |  | 2 |  | O_ph_O_b_ |
| MG:AA:3524 | 1.00 | 50.85 |  | 0 | 2 | - Exclusion zone contact(s): /N7/OP2/ Consider K in place of MG (2 valid contact(s) for K vs. 0 for MG) |
| MG:AA:3525 | 1.00 | 36.01 |  | 0 | 3 | - Exclusion zone contact(s): /N7/N7/OP2/ Consider K in place of MG (2 valid contact(s) for K vs. 0 for MG) |
| MG:AA:3526 | 1.00 | 75.64 |  | 0 | 2 | - Exclusion zone contact(s): /O6/O6/ Consider K in place of MG (2 valid contact(s) for K vs. 0 for MG) |
| MG:AA:3527 | 1.00 | 34.65 |  | 0 | 3 | - Exclusion zone contact(s): /O4/O4/O6/ Consider K in place of MG (3 valid contact(s) for K vs. 0 for MG) |
| MG:AA:3528 | 1.00 | 46.97 |  | 0 | 4 | - Exclusion zone contact(s): /NZ_Lys_/OG1_Thr_/OP1/OP2/ Consider K in place of MG (4 valid contact(s) for K vs. 0 for MG) |
| MG:AA:3529 | 1.00 | 29.10 |  | 0 | 2 | - Exclusion zone contact(s): /O2/OP1/ Consider K in place of MG (2 valid contact(s) for K vs. 0 for MG) |
| MG:AA:3530 | 1.00 | 50.55 |  | 0 | 6 | - Exclusion zone contact(s): /O2'/O3'/O5'/OP1/OP2/OP2/ Consider K in place of MG (6 valid contact(s) for K vs. 0 for MG) |
| MG:AA:3531 | 1.00 | 26.28 |  | 0 |  | 1^st^ MG coordination shell missing (see Notes) |
| MG:AA:3532 | 1.00 | 87.83 |  | 0 |  | 1^st^ MG coordination shell missing (see Notes) |
| MG:AA:3533 | 1.00 | 44.66 |  | 0 | 3 | - Exclusion zone contact(s): /N7/O2/O4/ Consider K in place of MG (2 valid contact(s) for K vs. 0 for MG) |
| MG:AA:3534 | 1.00 | 32.62 |  | 2 |  | 2O_ph_ |
| MG:AA:3535 | 1.00 | 58.79 |  | 0 | 5 | - Exclusion zone contact(s): /N7/N7/O5'/OP2/OP2/ Consider K in place of MG (5 valid contact(s) for K vs. 0 for MG) |
| MG:AA:3536 | 1.00 | 46.21 |  | 0 | 1 | - Exclusion zone contact(s): /OP1/ Consider K in place of MG (1 valid contact(s) for K vs. 0 for MG) |
| MG:AA:3537 | 1.00 | 56.81 |  | 0 | 5 | - Exclusion zone contact(s): /N3_A_/N3_A_/O2'/O4'/ Consider K in place of MG (5 valid contact(s) for K vs. 0 for MG) |
| MG:AA:3538 | 1.00 | 45.04 |  | 0 | 1 | - Exclusion zone contact(s): /O4/ Consider K in place of MG (1 valid contact(s) for K vs. 0 for MG) |
| MG:AA:3539 | 1.00 | 50.12 |  | 0 | 1 | - Exclusion zone contact(s): /OP2/ Consider NA in place of MG (1 valid contact(s) for NA vs. 0 for MG) |
| MG:AA:3540 | 1.00 | 61.54 |  | 0 |  | 1^st^ MG coordination shell missing (see Notes) |
| MG:AA:3541 | 1.00 | 49.41 |  | 0 | 4 | - Exclusion zone contact(s): /O2'/O2'/O3'/O3'/ Consider K in place of MG (3 valid contact(s) for K vs. 0 for MG) |
| MG:AA:3542 | 1.00 | 54.33 |  | 0 | 1 | - Exclusion zone contact(s): /N7/ Consider K in place of MG (1 valid contact(s) for K vs. 0 for MG) |
| MG:AA:3543 | 1.00 | 77.91 |  | 1 | 1 | O_b_ - Exclusion zone contact(s): /O4/ |
| MG:AA:3544 | 1.00 | 62.74 |  | 0 | 2 | - Exclusion zone contact(s): /O3'/O5'/ Consider K in place of MG (2 valid contact(s) for K vs. 0 for MG) |
| MG:AA:3545 | 1.00 | 56.53 |  | 0 | 2 | - Exclusion zone contact(s): /O6/O6/ Consider K in place of MG (2 valid contact(s) for K vs. 0 for MG) |
| MG:AA:3546 | 1.00 | 52.86 |  | 0 | 3 | - Exclusion zone contact(s): /O2'/O3'/OP2/ Consider NA in place of MG (2 valid contact(s) for NA vs. 0 for MG) |
| MG:AA:3547 | 1.00 | 47.83 |  | 0 |  | 1^st^ MG coordination shell missing (see Notes) |
| MG:AA:3548 | 1.00 | 59.41 |  | 0 | 2 | - Exclusion zone contact(s): /O6/O6/ Consider K in place of MG (2 valid contact(s) for K vs. 0 for MG) |
| MG:AA:3549 | 1.00 | 44.99 |  | 0 | 4 | - Exclusion zone contact(s): /N4/N6/N7/O6/ Consider K in place of MG (4 valid contact(s) for K vs. 0 for MG) |
| MG:AA:3550 | 1.00 | 82.65 |  | 0 |  | 1^st^ MG coordination shell missing (see Notes) |
| MG:AA:3551 | 1.00 | 44.31 |  | 0 | 4 | - Exclusion zone contact(s): /O2'/O3'/OP1/OP1/ Consider K in place of MG (3 valid contact(s) for K vs. 0 for MG) |
| MG:AA:3552 | 1.00 | 90.53 |  | 0 |  | 1^st^ MG coordination shell missing (see Notes) |
| MG:AA:3553 | 1.00 | 73.88 |  | 0 |  | 1^st^ MG coordination shell missing (see Notes) |
| MG:AA:3554 | 1.00 | 86.10 |  | 0 | 2 | - Exclusion zone contact(s): /O4/O6/ Consider K in place of MG (2 valid contact(s) for K vs. 0 for MG) |
| MG:AA:3555 | 1.00 | 60.64 |  | 1 | 1 | O_ph_ - Exclusion zone contact(s): /OP1/ |
| MG:AA:3556 | 1.00 | 74.57 |  | 0 |  | 1^st^ MG coordination shell missing (see Notes) |
| MG:AA:3557 | 1.00 | 89.04 |  | 1 |  | O_b_ |
| MG:AA:3558 | 1.00 | 50.22 |  | 0 | 2 | - Exclusion zone contact(s): /OP1/OP1/ Consider K in place of MG (2 valid contact(s) for K vs. 0 for MG) |
| MG:AA:3559 | 1.00 | 50.33 |  | 0 | 3 | - Exclusion zone contact(s): /N3_G_/O2'/O_back_/ Consider K in place of MG (3 valid contact(s) for K vs. 0 for MG) |
| MG:AA:3560 | 1.00 | 37.01 |  | 0 |  | 1^st^ MG coordination shell missing (see Notes) |
| MG:AA:3561 | 1.00 | 45.53 |  | 0 | 2 | - Exclusion zone contact(s): /O2/O2'/ Consider NA in place of MG (2 valid contact(s) for NA vs. 0 for MG) |
| MG:AA:3562 | 1.00 | 55.06 |  | 0 | 1 | - Exclusion zone contact(s): /O2'/ Consider K in place of MG (1 valid contact(s) for K vs. 0 for MG) |
| MG:AA:3563 | 1.00 | 67.56 |  | 0 | 2 | - Exclusion zone contact(s): /OP1/OP1/ Consider K in place of MG (2 valid contact(s) for K vs. 0 for MG) |
| MG:AA:3564 | 1.00 | 67.59 |  | 0 |  | 1^st^ MG coordination shell missing (see Notes) |
| MG:AA:3565 | 1.00 | 89.66 |  | 0 |  | 1^st^ MG coordination shell missing (see Notes) |
| MG:AA:3566 | 1.00 | 65.85 |  | 0 |  | 1^st^ MG coordination shell missing (see Notes) |
| MG:AA:3567 | 1.00 | 82.80 |  | 0 | 2 | - Exclusion zone contact(s): /O6/OP2/ Consider K in place of MG (2 valid contact(s) for K vs. 0 for MG) |
| MG:AA:3568 | 1.00 | 41.52 |  | 0 |  | 1^st^ MG coordination shell missing (see Notes) |
| MG:AA:3569 | 1.00 | 75.09 |  | 0 | 1 | - Exclusion zone contact(s): /OP1/ Consider K in place of MG (1 valid contact(s) for K vs. 0 for MG) |
| MG:AA:3570 | 1.00 | 78.80 |  | 0 | 1 | - Exclusion zone contact(s): /OP1/ Consider K in place of MG (1 valid contact(s) for K vs. 0 for MG) |
| MG:AA:3571 | 1.00 | 57.47 |  | 0 | 2 | - Exclusion zone contact(s): /O6/OP2/ Consider K in place of MG (2 valid contact(s) for K vs. 0 for MG) |
| MG:AA:3572 | 1.00 | 64.96 |  | 0 | 1 | - Exclusion zone contact(s): /O4/ Consider K in place of MG (1 valid contact(s) for K vs. 0 for MG) |
| MG:AA:3573 | 1.00 | 44.53 |  | 0 |  | 1^st^ MG coordination shell missing (see Notes) |
| MG:AA:3574 | 1.00 | 57.81 |  | 0 | 1 | - Exclusion zone contact(s): /OP1/ Consider K in place of MG (1 valid contact(s) for K vs. 0 for MG) |
| MG:AA:3575 | 1.00 | 51.70 |  | 0 |  | 1^st^ MG coordination shell missing (see Notes) |
| MG:AA:3576 | 1.00 | 37.76 |  | 0 | 2 | - Exclusion zone contact(s): /O6/O6/ Consider K in place of MG (2 valid contact(s) for K vs. 0 for MG) |
| MG:AA:3577 | 1.00 | 80.55 |  | 2 |  | 2O_ph_ |
| MG:AA:3578 | 1.00 | 33.69 |  | 2 | 2 | 2O_ph_ - Exclusion zone contact(s): /N7/O3'/ |
| MG:AA:3579 | 1.00 | 83.12 |  | 0 | 2 | - Exclusion zone contact(s): /N7/OP2/ Consider NA in place of MG (1 valid contact(s) for NA vs. 0 for MG) |
| MG:AA:3580 | 1.00 | 50.44 |  | 0 | 1 | - Exclusion zone contact(s): /O6/ Consider K in place of MG (1 valid contact(s) for K vs. 0 for MG) |
| MG:AA:3581 | 1.00 | 55.66 |  | 0 |  | 1^st^ MG coordination shell missing (see Notes) |
| MG:AA:3582 | 1.00 | 54.44 |  | 0 | 2 | - Exclusion zone contact(s): /O6/OP2/ Consider K in place of MG (2 valid contact(s) for K vs. 0 for MG) |
| MG:AA:3583 | 1.00 | 72.55 |  | 0 | 2 | - Exclusion zone contact(s): /OP1/OP2/ Consider K in place of MG (2 valid contact(s) for K vs. 0 for MG) |
| MG:AA:3584 | 1.00 | 85.74 |  | 0 | 1 | - Exclusion zone contact(s): /OP1/ Consider K in place of MG (1 valid contact(s) for K vs. 0 for MG) |
| MG:AA:3585 | 1.00 | 45.14 |  | 0 | 2 | - Exclusion zone contact(s): /N7/O6/ Consider K in place of MG (2 valid contact(s) for K vs. 0 for MG) |
| MG:AA:3586 | 1.00 | 41.72 |  | 1 | 1 | O_ph_ - Exclusion zone contact(s): /O5'/ |
| MG:AA:3587 | 1.00 | 55.53 |  | 0 | 2 | - Exclusion zone contact(s): /OP1/OP2/ Consider K in place of MG (2 valid contact(s) for K vs. 0 for MG) |
| MG:AA:3588 | 1.00 | 82.52 |  | 0 | 1 | - Exclusion zone contact(s): /N7/ Consider K in place of MG (1 valid contact(s) for K vs. 0 for MG) |
| MG:AA:3589 | 1.00 | 139.58 | 1 | 1 | 1 | - Exclusion zone contact(s): /O5'/O6/ Consider K in place of MG (1 valid contact(s) for K vs. 0 for MG) |
| MG:AA:3590 | 1.00 | 78.59 |  | 0 |  | 1^st^ MG coordination shell missing (see Notes) |
| MG:AA:3591 | 1.00 | 106.42 |  | 0 |  | 1^st^ MG coordination shell missing (see Notes) |
| MG:AA:3592 | 1.00 | 63.14 |  | 0 |  | 1^st^ MG coordination shell missing (see Notes) |
| MG:AA:3593 | 1.00 | 80.55 |  | 0 | 1 | - Exclusion zone contact(s): /OP1/ Consider K in place of MG (1 valid contact(s) for K vs. 0 for MG) |
| MG:AA:3594 | 1.00 | 78.36 |  | 0 |  | 1^st^ MG coordination shell missing (see Notes) |
| MG:AA:3595 | 1.00 | 103.78 |  | 0 | 1 | - Exclusion zone contact(s): /OP2/ Consider K in place of MG (1 valid contact(s) for K vs. 0 for MG) |
| MG:AA:3596 | 1.00 | 65.97 |  | 0 | 2 | - Exclusion zone contact(s): /OP1/OP1/ Consider NA in place of MG (1 valid contact(s) for NA vs. 0 for MG) |
| MG:AA:3597 | 1.00 | 63.62 |  | 0 |  | 1^st^ MG coordination shell missing (see Notes) |
| MG:AA:3598 | 1.00 | 79.19 |  | 0 |  | 1^st^ MG coordination shell missing (see Notes) |
| MG:AA:3599 | 1.00 | 110.57 |  | 0 |  | 1^st^ MG coordination shell missing (see Notes) |
| MG:AA:3600 | 1.00 | 46.42 |  | 0 | 1 | - Exclusion zone contact(s): /O6/ Consider K in place of MG (1 valid contact(s) for K vs. 0 for MG) |
| MG:AA:3601 | 1.00 | 73.94 |  | 0 |  | 1^st^ MG coordination shell missing (see Notes) |
| MG:AA:3602 | 1.00 | 102.40 |  | 0 | 2 | - Exclusion zone contact(s): /N7/O6/ Consider K in place of MG (2 valid contact(s) for K vs. 0 for MG) |
| MG:AA:3603 | 1.00 | 80.30 |  | 0 |  | 1^st^ MG coordination shell missing (see Notes) |
| MG:AA:3604 | 1.00 | 74.23 |  | 0 | 2 | - Exclusion zone contact(s): /OP1/OP2/ Consider K in place of MG (2 valid contact(s) for K vs. 0 for MG) |
| MG:AA:3605 | 1.00 | 66.02 |  | 0 | 1 | - Exclusion zone contact(s): /OP1/ Consider K in place of MG (1 valid contact(s) for K vs. 0 for MG) |
| MG:AA:3606 | 1.00 | 97.70 |  | 0 |  | 1^st^ MG coordination shell missing (see Notes) |
| MG:AA:3607 | 1.00 | 94.52 |  | 0 | 2 | - Exclusion zone contact(s): /OP1/OP2/ Consider NA in place of MG (1 valid contact(s) for NA vs. 0 for MG) |
| MG:AA:3608 | 1.00 | 86.86 |  | 0 | 1 | - Exclusion zone contact(s): /OP1/ Consider K in place of MG (1 valid contact(s) for K vs. 0 for MG) |
| MG:AA:3609 | 1.00 | 72.56 |  | 0 | 3 | - Exclusion zone contact(s): /OP1/OP1/OP2/ Consider K in place of MG (2 valid contact(s) for K vs. 0 for MG) |
| MG:AA:3610 | 1.00 | 69.79 |  | 0 | 1 | - Exclusion zone contact(s): /OP1/ Consider K in place of MG (1 valid contact(s) for K vs. 0 for MG) |
| MG:AA:3611 | 1.00 | 70.08 |  | 0 | 1 | - Exclusion zone contact(s): /OP1/ Consider K in place of MG (1 valid contact(s) for K vs. 0 for MG) |
| MG:AA:3612 | 1.00 | 106.33 |  | 0 |  | 1^st^ MG coordination shell missing (see Notes) |
| MG:AA:3613 | 1.00 | 84.73 |  | 0 |  | 1^st^ MG coordination shell missing (see Notes) |
| MG:AA:3614 | 1.00 | 62.01 |  | 0 | 2 | - Exclusion zone contact(s): /O6/O6/ Consider K in place of MG (2 valid contact(s) for K vs. 0 for MG) |
| MG:AA:3615 | 1.00 | 47.56 |  | 0 | 1 | - Exclusion zone contact(s): /N4/ Consider K in place of MG (1 valid contact(s) for K vs. 0 for MG) |
| MG:AA:3616 | 1.00 | 42.39 |  | 0 |  | 1^st^ MG coordination shell missing (see Notes) |
| MG:AA:3617 | 1.00 | 59.63 |  | 0 | 2 | - Exclusion zone contact(s): /O3'/OP1/ Consider K in place of MG (2 valid contact(s) for K vs. 0 for MG) |
| MG:AA:3618 | 1.00 | 89.07 |  | 0 | 2 | - Exclusion zone contact(s): /N4/O6/ Consider K in place of MG (2 valid contact(s) for K vs. 0 for MG) |
| MG:AA:3619 | 1.00 | 64.82 |  | 0 | 2 | - Exclusion zone contact(s): /N7/O6/ Consider K in place of MG (2 valid contact(s) for K vs. 0 for MG) |
| MG:AA:3620 | 1.00 | 55.20 |  | 1 | 1 | O_ph_ - Exclusion zone contact(s): /O5'/ |
| MG:AA:3621 | 1.00 | 46.06 |  | 0 | 1 | - Exclusion zone contact(s): /N7/ Consider NA in place of MG (1 valid contact(s) for NA vs. 0 for MG) |
| MG:AA:3622 | 1.00 | 43.82 |  | 0 | 3 | - Exclusion zone contact(s): /O6/O6/OP2/ Consider K in place of MG (2 valid contact(s) for K vs. 0 for MG) |
| MG:AA:3623 | 1.00 | 50.22 |  | 0 | 4 | - Exclusion zone contact(s): /N4/O4/O6/O6/ Consider K in place of MG (4 valid contact(s) for K vs. 0 for MG) |
| MG:AA:3624 | 1.00 | 38.52 |  | 0 |  | 1^st^ MG coordination shell missing (see Notes) |
| MG:AA:3625 | 1.00 | 64.55 |  | 1 | 1 | O_ph_ - Exclusion zone contact(s): /OP1/ |
| MG:AA:3626 | 1.00 | 48.20 |  | 0 |  | 1^st^ MG coordination shell missing (see Notes) |
| MG:AA:3627 | 1.00 | 37.31 |  | 0 |  | 1^st^ MG coordination shell missing (see Notes) |
| MG:AA:3628 | 1.00 | 62.41 |  | 0 | 2 | - Exclusion zone contact(s): /OP1/OP2/ Consider K in place of MG (2 valid contact(s) for K vs. 0 for MG) |
| MG:AA:3629 | 1.00 | 45.69 |  | 0 |  | 1^st^ MG coordination shell missing (see Notes) |
| MG:AA:3630 | 1.00 | 55.57 |  | 0 |  | 1^st^ MG coordination shell missing (see Notes) |
| MG:AA:3631 | 1.00 | 58.95 |  | 0 | 3 | - Exclusion zone contact(s): /N7/O2'/OP2/ Consider K in place of MG (2 valid contact(s) for K vs. 0 for MG) |
| MG:AA:3632 | 1.00 | 43.60 |  | 0 |  | 1^st^ MG coordination shell missing (see Notes) |
| MG:AA:3633 | 1.00 | 41.74 |  | 0 |  | 1^st^ MG coordination shell missing (see Notes) |
| MG:AA:3634 | 1.00 | 52.50 |  | 0 |  | 1^st^ MG coordination shell missing (see Notes) |
| MG:AA:3635 | 1.00 | 54.50 |  | 2 |  | 2O_ph_ |
| MG:AA:3636 | 1.00 | 48.22 |  | 0 | 2 | - Exclusion zone contact(s): /N7/NE2_His_/ Consider NA in place of MG (2 valid contact(s) for NA vs. 0 for MG) |
| MG:AA:3637 | 1.00 | 54.14 |  | 0 | 3 | - Exclusion zone contact(s): /O6/O6/OP1/ Consider K in place of MG (3 valid contact(s) for K vs. 0 for MG) |
| MG:AA:3638 | 1.00 | 51.96 |  | 0 | 2 | - Exclusion zone contact(s): /O6/O6/ Consider K in place of MG (2 valid contact(s) for K vs. 0 for MG) |
| MG:AA:3639 | 1.00 | 55.74 |  | 0 |  | 1^st^ MG coordination shell missing (see Notes) |
| MG:AA:3640 | 1.00 | 45.16 |  | 0 | 1 | - Exclusion zone contact(s): /OP1/ Consider K in place of MG (1 valid contact(s) for K vs. 0 for MG) |
| MG:AA:3641 | 1.00 | 46.03 |  | 0 | 3 | - Exclusion zone contact(s): /O3'/OP2/OP2/ Consider K in place of MG (2 valid contact(s) for K vs. 0 for MG) |
| MG:AA:3642 | 1.00 | 56.68 |  | 0 | 1 | - Exclusion zone contact(s): /N7/ Consider K in place of MG (1 valid contact(s) for K vs. 0 for MG) |
| MG:AA:3643 | 1.00 | 61.97 |  | 1 |  | O_ph_ |
| MG:AA:3644 | 1.00 | 56.71 |  | 0 | 4 | - Exclusion zone contact(s): /N3_C_/N7/N7/O6/ Consider K in place of MG (3 valid contact(s) for K vs. 0 for MG) |
| MG:AA:3645 | 1.00 | 46.66 |  | 0 |  | 1^st^ MG coordination shell missing (see Notes) |
| MG:AA:3646 | 1.00 | 62.65 |  | 0 | 3 | - Exclusion zone contact(s): /O4/O6/O6/ Consider K in place of MG (2 valid contact(s) for K vs. 0 for MG) |
| MG:AA:3647 | 1.00 | 25.82 |  | 0 | 1 | - Exclusion zone contact(s): /OP2/ Consider K in place of MG (1 valid contact(s) for K vs. 0 for MG) |
| MG:AA:3648 | 1.00 | 43.20 |  | 0 | 2 | - Exclusion zone contact(s): /O2'/OP2/ Consider K in place of MG (2 valid contact(s) for K vs. 0 for MG) |
| MG:AA:3649 | 1.00 | 96.11 |  | 0 | 3 | - Exclusion zone contact(s): /N7/OP2/OP2/ Consider K in place of MG (2 valid contact(s) for K vs. 0 for MG) |
| MG:AA:3650 | 1.00 | 70.97 |  | 1 | 2 | O_ph_ - Exclusion zone contact(s): /OP1/OP2/ |
| MG:AA:3651 | 1.00 | 41.20 |  | 0 | 2 | - Exclusion zone contact(s): /O2'/O4'/ Consider K in place of MG (2 valid contact(s) for K vs. 0 for MG) |
| MG:AA:3652 | 1.00 | 44.81 |  | 0 | 4 | - Exclusion zone contact(s): /O2'/O2'/O6/OP2/ Consider K in place of MG (3 valid contact(s) for K vs. 0 for MG) |
| MG:AA:3653 | 1.00 | 58.07 |  | 0 | 4 | - Exclusion zone contact(s): /O3'/O5'/OP1/OP2/ Consider K in place of MG (3 valid contact(s) for K vs. 0 for MG) |
| MG:AA:3654 | 1.00 | 59.98 |  | 0 |  | 1^st^ MG coordination shell missing (see Notes) |
| MG:AA:3655 | 1.00 | 50.50 |  | 0 | 2 | - Exclusion zone contact(s): /O6/O6/ Consider K in place of MG (2 valid contact(s) for K vs. 0 for MG) |
| MG:AA:3656 | 1.00 | 37.36 |  | 0 | 3 | - Exclusion zone contact(s): /O2/O2/O2'/ Consider K in place of MG (3 valid contact(s) for K vs. 0 for MG) |
| MG:AA:3657 | 1.00 | 89.42 |  | 0 |  | 1^st^ MG coordination shell missing (see Notes) |
| MG:AA:3658 | 1.00 | 64.91 |  | 0 | 2 | - Exclusion zone contact(s): /OP2/OP2/ Consider K in place of MG (2 valid contact(s) for K vs. 0 for MG) |
| MG:AA:3659 | 1.00 | 67.03 |  | 1 | 1 | O_ph_ - Exclusion zone contact(s): /OP2/ |
| MG:AA:3660 | 1.00 | 68.18 |  | 0 |  | 1^st^ MG coordination shell missing (see Notes) |
| MG:AA:3661 | 1.00 | 70.69 |  | 0 | 2 | - Exclusion zone contact(s): /O6/O6/ Consider K in place of MG (2 valid contact(s) for K vs. 0 for MG) |
| MG:AA:3662 | 1.00 | 51.23 |  | 0 | 3 | - Exclusion zone contact(s): /O4'/O6/OP1/ Consider K in place of MG (2 valid contact(s) for K vs. 0 for MG) |
| MG:AA:3663 | 1.00 | 61.61 |  | 0 |  | 1^st^ MG coordination shell missing (see Notes) |
| MG:AA:3664 | 1.00 | 68.34 |  | 0 | 2 | - Exclusion zone contact(s): /OP1/OP1/ Consider NA in place of MG (2 valid contact(s) for NA vs. 0 for MG) |
| MG:AA:3665 | 1.00 | 80.76 |  | 0 |  | 1^st^ MG coordination shell missing (see Notes) |
| MG:AA:3666 | 1.00 | 52.46 |  | 0 |  | 1^st^ MG coordination shell missing (see Notes) |
| MG:AA:3667 | 1.00 | 79.04 |  | 0 |  | 1^st^ MG coordination shell missing (see Notes) |
| MG:AA:3668 | 1.00 | 71.68 |  | 0 | 2 | - Exclusion zone contact(s): /OP1/OP2/ Consider K in place of MG (2 valid contact(s) for K vs. 0 for MG) |
| MG:AA:3669 | 1.00 | 44.20 |  | 0 | 2 | - Exclusion zone contact(s): /OP2/OP2/ Consider NA in place of MG (1 valid contact(s) for NA vs. 0 for MG) |
| MG:AA:3670 | 1.00 | 54.93 |  | 0 | 3 | - Exclusion zone contact(s): /O4/O6/O6/ Consider K in place of MG (2 valid contact(s) for K vs. 0 for MG) |
| MG:AA:3671 | 1.00 | 102.68 |  | 0 | 3 | - Exclusion zone contact(s): /N4/N4/N6/ Consider K in place of MG (3 valid contact(s) for K vs. 0 for MG) |
| MG:AA:3672 | 1.00 | 55.15 |  | 0 | 1 | - Exclusion zone contact(s): /O6/ Consider K in place of MG (1 valid contact(s) for K vs. 0 for MG) |
| MG:AA:3673 | 1.00 | 56.57 |  | 0 |  | 1^st^ MG coordination shell missing (see Notes) |
| MG:AA:3674 | 1.00 | 61.07 |  | 0 |  | 1^st^ MG coordination shell missing (see Notes) |
| MG:AA:3675 | 1.00 | 71.00 |  | 0 | 5 | - Exclusion zone contact(s): /N4/O4/O6/O6/O6/ Consider K in place of MG (5 valid contact(s) for K vs. 0 for MG) |
| MG:AA:3676 | 1.00 | 81.27 |  | 0 |  | 1^st^ MG coordination shell missing (see Notes) |
| MG:AA:3677 | 1.00 | 60.18 |  | 0 | 2 | - Exclusion zone contact(s): /O6/O6/ Consider K in place of MG (2 valid contact(s) for K vs. 0 for MG) |
| MG:AA:3678 | 1.00 | 49.08 |  | 0 | 3 | - Exclusion zone contact(s): /O4/O6/O6/ Consider K in place of MG (3 valid contact(s) for K vs. 0 for MG) |
| MG:AA:3679 | 1.00 | 70.67 |  | 0 |  | 1^st^ MG coordination shell missing (see Notes) |
| MG:AA:3680 | 1.00 | 71.62 |  | 0 | 1 | - Exclusion zone contact(s): /N7/ Consider K in place of MG (1 valid contact(s) for K vs. 0 for MG) |
| MG:AA:3681 | 1.00 | 65.28 |  | 0 | 1 | - Exclusion zone contact(s): /O6/ Consider K in place of MG (1 valid contact(s) for K vs. 0 for MG) |
| MG:AA:3682 | 1.00 | 75.62 |  | 0 | 1 | - Exclusion zone contact(s): /N1/ Consider K in place of MG (1 valid contact(s) for K vs. 0 for MG) |
| MG:AA:3683 | 1.00 | 46.03 |  | 1 | 4 | O_ph_ - Exclusion zone contact(s): /N7/ND1_His_/O_back_/OP2/ Consider K in place of MG (4 valid contact(s) for K vs. 1 for MG) |
| MG:AA:3684 | 1.00 | 58.83 |  | 0 |  | 1^st^ MG coordination shell missing (see Notes) |
| MG:AA:3685 | 1.00 | 65.43 |  | 0 |  | 1^st^ MG coordination shell missing (see Notes) |
| MG:AA:3686 | 1.00 | 49.71 |  | 0 |  | 1^st^ MG coordination shell missing (see Notes) |
| MG:AA:3687 | 1.00 | 43.53 |  | 0 | 4 | - Exclusion zone contact(s): /O3'/O6/OP2/ Consider K in place of MG (4 valid contact(s) for K vs. 0 for MG) |
| MG:AA:3688 | 1.00 | 59.51 |  | 0 |  | 1^st^ MG coordination shell missing (see Notes) |
| MG:AA:3689 | 1.00 | 61.01 |  | 0 | 4 | - Exclusion zone contact(s): /O4/O6/O6/O6/ Consider K in place of MG (4 valid contact(s) for K vs. 0 for MG) |
| MG:AA:3690 | 1.00 | 64.83 |  | 1 | 1 | O_r_ - Exclusion zone contact(s): /O4'/ |
| MG:AA:3691 | 1.00 | 79.96 |  | 0 | 3 | - Exclusion zone contact(s): /N7/O4/O6/ Consider K in place of MG (2 valid contact(s) for K vs. 0 for MG) |
| MG:AA:3692 | 1.00 | 85.75 |  | 0 | 3 | - Exclusion zone contact(s): /O4'/O5'/O_back_/ Consider K in place of MG (2 valid contact(s) for K vs. 0 for MG) |
| MG:AA:3693 | 1.00 | 52.64 |  | 0 | 2 | - Exclusion zone contact(s): /N7/O6/ Consider K in place of MG (2 valid contact(s) for K vs. 0 for MG) |
| MG:AA:3694 | 1.00 | 56.35 |  | 1 | 1 | O_b_ - Exclusion zone contact(s): /OP1/ |
| MG:AA:3695 | 1.00 | 49.89 |  | 0 |  | 1^st^ MG coordination shell missing (see Notes) |
| MG:AA:3696 | 1.00 | 47.13 |  | 0 | 3 | - Exclusion zone contact(s): /OP1/OP1/OP2/ Consider K in place of MG (2 valid contact(s) for K vs. 0 for MG) |
| MG:AA:3697 | 1.00 | 78.81 |  | 0 |  | 1^st^ MG coordination shell missing (see Notes) |
| MG:AA:3698 | 1.00 | 64.99 |  | 0 | 3 | - Exclusion zone contact(s): /O4/O6/O6/ Consider K in place of MG (3 valid contact(s) for K vs. 0 for MG) |
| MG:AA:3699 | 1.00 | 59.06 |  | 0 | 3 | - Exclusion zone contact(s): /O3'/OP1/OP1/ Consider K in place of MG (3 valid contact(s) for K vs. 0 for MG) |
| MG:AA:3700 | 1.00 | 58.97 |  | 0 |  | 1^st^ MG coordination shell missing (see Notes) |
| MG:AA:3701 | 1.00 | 55.59 |  | 0 | 1 | - Exclusion zone contact(s): /O4/ Consider K in place of MG (1 valid contact(s) for K vs. 0 for MG) |
| MG:AA:3702 | 1.00 | 62.01 |  | 0 | 3 | - Exclusion zone contact(s): /OP1/OP1/OP2/ Consider K in place of MG (2 valid contact(s) for K vs. 0 for MG) |
| MG:AA:3703 | 1.00 | 51.31 |  | 0 | 4 | - Exclusion zone contact(s): /O4/O6/O6/O6/ Consider K in place of MG (3 valid contact(s) for K vs. 0 for MG) |
| MG:AA:3704 | 1.00 | 59.72 |  | 0 | 2 | - Exclusion zone contact(s): /OP2/OP2/ Consider K in place of MG (2 valid contact(s) for K vs. 0 for MG) |
| MG:AA:3705 | 1.00 | 87.10 |  | 0 |  | 1^st^ MG coordination shell missing (see Notes) |
| MG:AA:3706 | 1.00 | 44.71 |  | 0 | 4 | - Exclusion zone contact(s): /N1_A_/N6/N7/O4'/ Consider K in place of MG (4 valid contact(s) for K vs. 0 for MG) |
| MG:AA:3707 | 1.00 | 63.66 |  | 0 | 2 | - Exclusion zone contact(s): /OP1/OP1/ Consider K in place of MG (2 valid contact(s) for K vs. 0 for MG) |
| MG:AA:3708 | 1.00 | 49.16 |  | 0 | 5 | - Exclusion zone contact(s): /O2/O2/O2'/O2'/O4'/ Consider K in place of MG (5 valid contact(s) for K vs. 0 for MG) |
| MG:AA:3709 | 1.00 | 66.65 |  | 0 |  | 1^st^ MG coordination shell missing (see Notes) |
| MG:AA:3710 | 1.00 | 76.03 |  | 0 | 1 | - Exclusion zone contact(s): /O6/ Consider K in place of MG (1 valid contact(s) for K vs. 0 for MG) |
| MG:AA:3711 | 1.00 | 69.93 |  | 1 |  | O_ph_ |
| MG:AA:3712 | 1.00 | 57.03 |  | 1 | 1 | O_b_ - Exclusion zone contact(s): /N1_A_/ |
| MG:AA:3713 | 1.00 | 83.25 |  | 1 | 1 | O_ph_ - Exclusion zone contact(s): /OP1/ |
| MG:AA:3714 | 1.00 | 70.93 |  | 0 | 2 | - Exclusion zone contact(s): /O6/O6/ Consider K in place of MG (2 valid contact(s) for K vs. 0 for MG) |
| MG:AA:3715 | 1.00 | 47.51 |  | 0 | 3 | - Exclusion zone contact(s): /O3'/O5'/OP2/ Consider K in place of MG (3 valid contact(s) for K vs. 0 for MG) |
| MG:AA:3716 | 1.00 | 74.12 |  | 0 |  | 1^st^ MG coordination shell missing (see Notes) |
| MG:AA:3717 | 1.00 | 106.73 |  | 0 | 1 | - Exclusion zone contact(s): /O6/ Consider K in place of MG (1 valid contact(s) for K vs. 0 for MG) |
| MG:AA:3718 | 1.00 | 84.39 |  | 0 | 2 | - Exclusion zone contact(s): /N7/OP1/ Consider K in place of MG (2 valid contact(s) for K vs. 0 for MG) |
| MG:AA:3719 | 1.00 | 97.41 |  | 0 |  | No contacts in 5.0 A |
| MG:AA:3720 | 1.00 | 52.18 |  | 0 | 5 | - Exclusion zone contact(s): /N3_G_/N6/N7/O2'/ Consider K in place of MG (5 valid contact(s) for K vs. 0 for MG) |
| MG:AA:3721 | 1.00 | 82.93 |  | 0 | 3 | - Exclusion zone contact(s): /N1_A_/N3_U_/O2/ Consider K in place of MG (3 valid contact(s) for K vs. 0 for MG) |
| MG:AA:3722 | 1.00 | 50.03 |  | 0 |  | 1^st^ MG coordination shell missing (see Notes) |
| MG:AA:3723 | 1.00 | 79.26 |  | 0 |  | No contacts in 5.0 A |
| MG:AA:3724 | 1.00 | 65.32 |  | 0 |  | 1^st^ MG coordination shell missing (see Notes) |
| MG:AA:3725 | 1.00 | 89.64 |  | 0 | 3 | - Exclusion zone contact(s): /N4/O6/OP2/ Consider K in place of MG (3 valid contact(s) for K vs. 0 for MG) |
| MG:AA:3726 | 1.00 | 60.84 |  | 0 | 3 | - Exclusion zone contact(s): /O5'/OP1/OP1/ Consider K in place of MG (3 valid contact(s) for K vs. 0 for MG) |
| MG:AA:3727 | 1.00 | 54.45 |  | 0 |  | 1^st^ MG coordination shell missing (see Notes) |
| MG:AA:3728 | 1.00 | 70.26 |  | 0 | 3 | - Exclusion zone contact(s): /N4/O2'/O3'/ Consider K in place of MG (3 valid contact(s) for K vs. 0 for MG) |
| MG:AA:3729 | 1.00 | 69.11 |  | 0 |  | 1^st^ MG coordination shell missing (see Notes) |
| MG:AA:3730 | 1.00 | 58.12 |  | 0 |  | 1^st^ MG coordination shell missing (see Notes) |
| MG:AA:3731 | 1.00 | 79.73 |  | 2 | 2 | 2O_b_ - Exclusion zone contact(s): /N3_U_/O6/ |
| MG:AA:3732 | 1.00 | 76.20 |  | 0 |  | 1^st^ MG coordination shell missing (see Notes) |
| MG:AA:3733 | 1.00 | 72.01 |  | 0 | 1 | - Exclusion zone contact(s): /O6/ Consider K in place of MG (1 valid contact(s) for K vs. 0 for MG) |
| MG:AA:3734 | 1.00 | 49.07 |  | 0 | 5 | - Exclusion zone contact(s): /N_back_/O6/OG1_Thr_/OP2/ Consider K in place of MG (5 valid contact(s) for K vs. 0 for MG) |
| MG:AA:3735 | 1.00 | 50.13 |  | 0 | 3 | - Exclusion zone contact(s): /OP1/OP1/OP2/ Consider K in place of MG (3 valid contact(s) for K vs. 0 for MG) |
| MG:AA:3736 | 1.00 | 72.03 |  | 0 | 1 | - Exclusion zone contact(s): /OP2/ Consider NA in place of MG (1 valid contact(s) for NA vs. 0 for MG) |
| MG:AA:3737 | 1.00 | 55.25 |  | 0 | 2 | - Exclusion zone contact(s): /O2'/OP2/ Consider K in place of MG (2 valid contact(s) for K vs. 0 for MG) |
| MG:AA:3738 | 1.00 | 66.18 |  | 0 | 2 | - Exclusion zone contact(s): /NZ_Lys_/O2/ Consider K in place of MG (2 valid contact(s) for K vs. 0 for MG) |
| MG:AA:3739 | 1.00 | 86.99 |  | 0 |  | 1^st^ MG coordination shell missing (see Notes) |
| MG:AA:3740 | 1.00 | 60.00 |  | 0 | 1 | - Exclusion zone contact(s): /OP2/ Consider K in place of MG (1 valid contact(s) for K vs. 0 for MG) |
| MG:AA:3741 | 1.00 | 63.24 |  | 0 |  | 1^st^ MG coordination shell missing (see Notes) |
| MG:AA:3742 | 1.00 | 70.52 |  | 0 | 2 | - Exclusion zone contact(s): /O6/O6/ Consider K in place of MG (2 valid contact(s) for K vs. 0 for MG) |
| MG:AA:3743 | 1.00 | 71.16 |  | 0 |  | 1^st^ MG coordination shell missing (see Notes) |
| MG:AA:3744 | 1.00 | 64.63 |  | 0 |  | 1^st^ MG coordination shell missing (see Notes) |
| MG:AA:3745 | 1.00 | 57.89 |  | 0 | 2 | - Exclusion zone contact(s): /N7/O4/ Consider K in place of MG (2 valid contact(s) for K vs. 0 for MG) |
| MG:AA:3746 | 1.00 | 66.04 |  | 0 | 1 | - Exclusion zone contact(s): /OP2/ Consider K in place of MG (1 valid contact(s) for K vs. 0 for MG) |
| MG:AA:3747 | 1.00 | 53.11 |  | 0 | 4 | - Exclusion zone contact(s): /O2'/O3'/O_back_/OP1/ Consider K in place of MG (4 valid contact(s) for K vs. 0 for MG) |
| MG:AA:3748 | 1.00 | 84.86 |  | 0 | 4 | - Exclusion zone contact(s): /N7/N7/O6/O6/ Consider K in place of MG (4 valid contact(s) for K vs. 0 for MG) |
| MG:AA:3749 | 1.00 | 33.11 |  | 1 | 1 | Anion - Exclusion zone contact(s): /OP1/ |
| MG:AA:3750 | 1.00 | 66.86 |  | 0 | 1 | - Exclusion zone contact(s): /OP2/ Consider K in place of MG (1 valid contact(s) for K vs. 0 for MG) |
| MG:AA:3751 | 1.00 | 48.61 |  | 0 | 4 | - Exclusion zone contact(s): /O2'/O3'/OG1_Thr_/OP1/ Consider K in place of MG (4 valid contact(s) for K vs. 0 for MG) |
| MG:AA:3752 | 1.00 | 71.78 |  | 0 | 3 | - Exclusion zone contact(s): /O4/O6/O6/ Consider K in place of MG (2 valid contact(s) for K vs. 0 for MG) |
| MG:AA:3753 | 1.00 | 91.42 |  | 0 |  | 1^st^ MG coordination shell missing (see Notes) |
| MG:AA:3754 | 1.00 | 73.00 |  | 0 | 4 | - Exclusion zone contact(s): /N7/N7/O6/O6/ Consider K in place of MG (4 valid contact(s) for K vs. 0 for MG) |
| MG:AA:3755 | 1.00 | 63.52 |  | 0 |  | 1^st^ MG coordination shell missing (see Notes) |
| MG:AA:3756 | 1.00 | 65.87 |  | 0 | 1 | - Exclusion zone contact(s): /O4/ Consider K in place of MG (1 valid contact(s) for K vs. 0 for MG) |
| MG:AA:3757 | 1.00 | 56.36 |  | 0 | 1 | - Exclusion zone contact(s): /O6/ Consider K in place of MG (1 valid contact(s) for K vs. 0 for MG) |
| MG:AA:3758 | 1.00 | 65.53 |  | 0 | 2 | - Exclusion zone contact(s): /N7/OP2/ Consider K in place of MG (2 valid contact(s) for K vs. 0 for MG) |
| MG:AA:3759 | 1.00 | 59.10 |  | 0 | 4 | - Exclusion zone contact(s): /O3'/O4'/O5'/OP1/ Consider K in place of MG (4 valid contact(s) for K vs. 0 for MG) |
| MG:AA:3760 | 1.00 | 85.32 |  | 0 |  | 1^st^ MG coordination shell missing (see Notes) |
| MG:AA:3761 | 1.00 | 40.50 |  | 0 |  | 1^st^ MG coordination shell missing (see Notes) |
| MG:AA:3762 | 1.00 | 63.08 |  | 0 | 4 | - Exclusion zone contact(s): /O3'/O5'/OP2/OP2/ Consider K in place of MG (4 valid contact(s) for K vs. 0 for MG) |
| MG:AA:3763 | 1.00 | 48.45 |  | 0 | 3 | - Exclusion zone contact(s): /O5'/OP1/OP2/ Consider K in place of MG (2 valid contact(s) for K vs. 0 for MG) |
| MG:AA:3764 | 1.00 | 53.72 |  | 1 |  | O_ph_ |
| MG:AA:3765 | 1.00 | 74.16 |  | 0 |  | 1^st^ MG coordination shell missing (see Notes) |
| MG:AA:3766 | 1.00 | 86.00 |  | 0 | 1 | - Exclusion zone contact(s): /O4/ Consider K in place of MG (1 valid contact(s) for K vs. 0 for MG) |
| MG:AA:3767 | 1.00 | 78.17 |  | 0 | 4 | - Exclusion zone contact(s): /O2'/O3'/OP2/OP2/ Consider K in place of MG (4 valid contact(s) for K vs. 0 for MG) |
| MG:AA:3768 | 1.00 | 79.86 |  | 0 |  | 1^st^ MG coordination shell missing (see Notes) |
| MG:AA:3769 | 1.00 | 80.94 |  | 0 | 3 | - Exclusion zone contact(s): /N7/O5'/OP2/ Consider K in place of MG (3 valid contact(s) for K vs. 0 for MG) |
| MG:AA:3770 | 1.00 | 70.29 |  | 0 |  | 1^st^ MG coordination shell missing (see Notes) |
| MG:AA:3771 | 1.00 | 79.27 |  | 0 |  | 1^st^ MG coordination shell missing (see Notes) |
| MG:AA:3772 | 1.00 | 103.45 |  | 0 | 3 | - Exclusion zone contact(s): /O5'/OP1/OP2/ Consider K in place of MG (3 valid contact(s) for K vs. 0 for MG) |
| MG:AA:3773 | 1.00 | 76.37 |  | 0 |  | 1^st^ MG coordination shell missing (see Notes) |
| MG:AA:3774 | 1.00 | 79.32 |  | 0 |  | 1^st^ MG coordination shell missing (see Notes) |
| MG:AA:3775 | 1.00 | 91.62 |  | 0 | 1 | - Exclusion zone contact(s): /OP2/ Consider K in place of MG (1 valid contact(s) for K vs. 0 for MG) |
| MG:AA:3776 | 1.00 | 84.23 |  | 0 |  | 1^st^ MG coordination shell missing (see Notes) |
| MG:AA:3777 | 1.00 | 58.36 |  | 1 | 3 | O_ph_ - Exclusion zone contact(s): /O3'/O5'/OP1/ Consider K in place of MG (3 valid contact(s) for K vs. 1 for MG) |
| MG:AA:3778 | 1.00 | 53.92 |  | 0 | 3 | - Exclusion zone contact(s): /N7/O5'/OP2/ Consider K in place of MG (2 valid contact(s) for K vs. 0 for MG) |
| MG:AA:3779 | 1.00 | 86.28 |  | 0 | 1 | - Exclusion zone contact(s): /OP2/ Consider NA in place of MG (1 valid contact(s) for NA vs. 0 for MG) |
| MG:AA:3780 | 1.00 | 69.98 |  | 0 | 5 | - Exclusion zone contact(s): /NH1_Arg_/O3'/OP1/OP1/OP2/ Consider K in place of MG (5 valid contact(s) for K vs. 0 for MG) |
| MG:AA:3781 | 1.00 | 104.48 |  | 0 | 2 | - Exclusion zone contact(s): /OP2/OP2/ Consider K in place of MG (2 valid contact(s) for K vs. 0 for MG) |
| MG:AA:3782 | 1.00 | 101.12 |  | 0 |  | 1^st^ MG coordination shell missing (see Notes) |
| MG:AA:3783 | 1.00 | 86.35 |  | 0 |  | 1^st^ MG coordination shell missing (see Notes) |
| MG:AA:3784 | 1.00 | 64.48 |  | 0 | 1 | - Exclusion zone contact(s): /OP1/ Consider K in place of MG (1 valid contact(s) for K vs. 0 for MG) |
| MG:AA:3785 | 1.00 | 89.29 |  | 0 |  | 1^st^ MG coordination shell missing (see Notes) |
| MG:AA:3786 | 1.00 | 107.68 |  | 0 |  | No contacts in 5.0 A |
| MG:AA:3787 | 1.00 | 70.38 |  | 0 |  | 1^st^ MG coordination shell missing (see Notes) |
| MG:AA:3788 | 1.00 | 87.36 |  | 0 | 1 | - Exclusion zone contact(s): /OP2/ Consider NA in place of MG (1 valid contact(s) for NA vs. 0 for MG) |
| MG:AA:3789 | 1.00 | 55.27 |  | 0 |  | 1^st^ MG coordination shell missing (see Notes) |
| MG:AA:3790 | 1.00 | 60.31 |  | 0 | 6 | - Exclusion zone contact(s): /N4/N4/O2'/O5'/OP2/OP2/ Consider K in place of MG (6 valid contact(s) for K vs. 0 for MG) |
| MG:AA:3791 | 1.00 | 65.43 |  | 0 | 3 | - Exclusion zone contact(s): /N3_A_/O2/O2'/ Consider K in place of MG (2 valid contact(s) for K vs. 0 for MG) |
| MG:AA:3792 | 1.00 | 82.48 |  | 0 | 6 | - Exclusion zone contact(s): /N1/N3_G_/O2'/O2'/O4'/O4'/ Consider K in place of MG (6 valid contact(s) for K vs. 0 for MG) |
| MG:AA:3793 | 1.00 | 63.36 |  | 1 | 1 | O_r_ - Exclusion zone contact(s): /OP1/ |
| MG:AA:3794 | 1.00 | 70.71 |  | 0 | 1 | - Exclusion zone contact(s): /OP2/ Consider K in place of MG (1 valid contact(s) for K vs. 0 for MG) |
| MG:AA:3795 | 1.00 | 106.32 |  | 0 | 2 | - Exclusion zone contact(s): /OP1/OP2/ Consider K in place of MG (2 valid contact(s) for K vs. 0 for MG) |
| MG:AA:3796 | 1.00 | 94.27 |  | 0 | 2 | - Exclusion zone contact(s): /N4/O6/ Consider K in place of MG (2 valid contact(s) for K vs. 0 for MG) |
| MG:AA:3797 | 1.00 | 96.58 |  | 0 |  | 1^st^ MG coordination shell missing (see Notes) |
| MG:AA:3798 | 1.00 | 47.59 |  | 0 | 3 | - Exclusion zone contact(s): /O_back_/OP1/OP2/ Consider K in place of MG (3 valid contact(s) for K vs. 0 for MG) |
| MG:AA:3799 | 1.00 | 93.09 |  | 0 |  | 1^st^ MG coordination shell missing (see Notes) |
| MG:AA:3800 | 1.00 | 55.74 |  | 0 | 1 | - Exclusion zone contact(s): /O6/ Consider NA in place of MG (1 valid contact(s) for NA vs. 0 for MG) |
| MG:AA:3801 | 1.00 | 90.88 |  | 0 | 1 | Consider K in place of MG (1 valid contact(s) for K vs. 0 for MG) |
| MG:AA:3802 | 1.00 | 78.71 |  | 0 |  | 1^st^ MG coordination shell missing (see Notes) |
| MG:AA:3803 | 1.00 | 72.56 |  | 0 | 3 | - Exclusion zone contact(s): /N6/O2/OP2/ Consider NA in place of MG (2 valid contact(s) for NA vs. 0 for MG) |
| MG:AA:3804 | 1.00 | 94.08 |  | 0 | 1 | - Exclusion zone contact(s): /OP2/ Consider NA in place of MG (1 valid contact(s) for NA vs. 0 for MG) |
| MG:AA:3805 | 1.00 | 94.75 |  | 0 |  | 1^st^ MG coordination shell missing (see Notes) |
| MG:AA:3806 | 1.00 | 79.78 |  | 1 |  | N_b_ |
| MG:AA:3807 | 1.00 | 87.39 |  | 0 |  | 1^st^ MG coordination shell missing (see Notes) |
| MG:AA:3808 | 1.00 | 75.79 |  | 1 | 1 | O_ph_ - Exclusion zone contact(s): /OP2/ |
| MG:AA:3809 | 1.00 | 84.99 |  | 1 | 1 | O_r_ - Exclusion zone contact(s): /N6/ |
| MG:AA:3810 | 1.00 | 85.98 |  | 0 | 1 | - Exclusion zone contact(s): /O6/ Consider K in place of MG (1 valid contact(s) for K vs. 0 for MG) |
| MG:AA:3811 | 1.00 | 58.11 |  | 0 |  | 1^st^ MG coordination shell missing (see Notes) |
| MG:AA:3812 | 1.00 | 83.76 |  | 0 | 1 | - Exclusion zone contact(s): /OP1/ Consider NA in place of MG (1 valid contact(s) for NA vs. 0 for MG) |
| MG:AA:3813 | 1.00 | 99.41 |  | 0 | 3 | - Exclusion zone contact(s): /OP1/OP1/OP1/ Consider K in place of MG (2 valid contact(s) for K vs. 0 for MG) |
| MG:AA:3814 | 1.00 | 67.14 |  | 0 |  | 1^st^ MG coordination shell missing (see Notes) |
| MG:AA:3815 | 1.00 | 61.20 |  | 0 | 2 | - Exclusion zone contact(s): /OP1/OP1/ Consider K in place of MG (2 valid contact(s) for K vs. 0 for MG) |
| MG:AA:3816 | 1.00 | 73.52 |  | 1 | 2 | O_r_ - Exclusion zone contact(s): /O2'/O2'/ Consider K in place of MG (2 valid contact(s) for K vs. 1 for MG) |
| MG:AA:3817 | 1.00 | 66.72 |  | 0 |  | 1^st^ MG coordination shell missing (see Notes) |
| MG:AA:3818 | 1.00 | 80.24 |  | 0 |  | 1^st^ MG coordination shell missing (see Notes) |
| MG:AA:3819 | 1.00 | 94.83 |  | 0 | 1 | - Exclusion zone contact(s): /N3_C_/ Consider K in place of MG (1 valid contact(s) for K vs. 0 for MG) |
| MG:AA:3820 | 1.00 | 59.34 |  | 0 | 6 | - Exclusion zone contact(s): /N3_A_/N7/O2'/O4/O6/O6/ Consider K in place of MG (5 valid contact(s) for K vs. 0 for MG) |
| MG:AA:3821 | 1.00 | 69.41 |  | 0 | 5 | - Exclusion zone contact(s): /N1_G_/N2/N2/N7/OP2/ Consider K in place of MG (4 valid contact(s) for K vs. 0 for MG) |
| MG:AA:3822 | 1.00 | 77.01 |  | 0 |  | 1^st^ MG coordination shell missing (see Notes) |
| MG:AA:3823 | 1.00 | 89.90 |  | 0 | 1 | - Exclusion zone contact(s): /O4/ Consider K in place of MG (1 valid contact(s) for K vs. 0 for MG) |
| MG:AA:3824 | 1.00 | 94.23 |  | 0 | 1 | - Exclusion zone contact(s): /O6/ Consider K in place of MG (1 valid contact(s) for K vs. 0 for MG) |
| MG:AA:3825 | 1.00 | 81.87 |  | 0 |  | 1^st^ MG coordination shell missing (see Notes) |
| MG:AA:3826 | 1.00 | 92.03 |  | 0 | 3 | - Exclusion zone contact(s): /N1_A_/N6/O2'/ Consider K in place of MG (3 valid contact(s) for K vs. 0 for MG) |
| MG:AA:3827 | 1.00 | 62.83 |  | 0 | 1 | - Exclusion zone contact(s): /O4/ Consider K in place of MG (1 valid contact(s) for K vs. 0 for MG) |
| MG:AA:3828 | 1.00 | 113.11 |  | 0 | 4 | - Exclusion zone contact(s): /N4/O4/O6/O6/ Consider K in place of MG (4 valid contact(s) for K vs. 0 for MG) |
| MG:AA:3829 | 1.00 | 88.38 |  | 0 | 4 | - Exclusion zone contact(s): /N6/N7/N7/O6/ Consider K in place of MG (3 valid contact(s) for K vs. 0 for MG) |
| MG:AA:3830 | 1.00 | 83.58 |  | 0 | 3 | - Exclusion zone contact(s): /N7/O6/O6/ Consider NA in place of MG (2 valid contact(s) for NA vs. 0 for MG) |
| MG:AA:3831 | 1.00 | 77.21 |  | 1 |  | O_ph_ |
| MG:AA:3832 | 1.00 | 63.57 |  | 0 | 4 | - Exclusion zone contact(s): /N7/O5'/OP2/OP2/ Consider K in place of MG (4 valid contact(s) for K vs. 0 for MG) |
| MG:AA:3833 | 1.00 | 80.44 |  | 0 | 2 | - Exclusion zone contact(s): /OP1/OP2/ Consider K in place of MG (2 valid contact(s) for K vs. 0 for MG) |
| MG:AA:3834 | 1.00 | 79.06 |  | 0 |  | 1^st^ MG coordination shell missing (see Notes) |
| MG:AA:3835 | 1.00 | 56.85 |  | 0 | 6 | - Exclusion zone contact(s): /N7/O2'/O3'/O6/OP1/OP2/ Consider K in place of MG (6 valid contact(s) for K vs. 0 for MG) |
| MG:AA:3836 | 1.00 | 62.66 |  | 0 |  | 1^st^ MG coordination shell missing (see Notes) |
| MG:AA:3837 | 1.00 | 116.85 |  | 0 |  | 1^st^ MG coordination shell missing (see Notes) |
| MG:AA:3838 | 1.00 | 102.81 |  | 0 | 3 | - Exclusion zone contact(s): /O2'/O3'/OP2/ Consider K in place of MG (2 valid contact(s) for K vs. 0 for MG) |
| MG:AA:3839 | 1.00 | 63.91 |  | 0 |  | 1^st^ MG coordination shell missing (see Notes) |
| MG:AA:3840 | 1.00 | 77.10 |  | 1 | 3 | O_ph_ - Exclusion zone contact(s): /OP1/OP1/OP2/ Consider K in place of MG (2 valid contact(s) for K vs. 1 for MG) |
| MG:AA:3841 | 1.00 | 85.25 |  | 0 |  | 1^st^ MG coordination shell missing (see Notes) |
| MG:AA:3842 | 1.00 | 67.79 |  | 0 |  | 1^st^ MG coordination shell missing (see Notes) |
| MG:AA:3843 | 1.00 | 81.15 |  | 0 |  | 1^st^ MG coordination shell missing (see Notes) |
| MG:AA:3844 | 1.00 | 94.03 |  | 0 |  | 1^st^ MG coordination shell missing (see Notes) |
| MG:AA:3845 | 1.00 | 54.44 |  | 0 | 1 | - Exclusion zone contact(s): /OP1/ Consider K in place of MG (1 valid contact(s) for K vs. 0 for MG) |
| MG:AA:3846 | 1.00 | 88.18 |  | 0 |  | No contacts in 5.0 A |
| MG:AA:3847 | 1.00 | 84.31 |  | 0 | 2 | - Exclusion zone contact(s): /N7/O6/ Consider K in place of MG (2 valid contact(s) for K vs. 0 for MG) |
| MG:AA:3848 | 1.00 | 90.29 |  | 0 | 2 | - Exclusion zone contact(s): /N7/O6/ Consider K in place of MG (2 valid contact(s) for K vs. 0 for MG) |
| MG:AA:3849 | 1.00 | 59.97 |  | 1 | 2 | O_ph_ - Exclusion zone contact(s): /N7/OP2/ Consider K in place of MG (2 valid contact(s) for K vs. 1 for MG) |
| MG:AA:3850 | 1.00 | 41.85 |  | 1 | 2 | N_b_ - Exclusion zone contact(s): /N2/O2'/ |
| MG:AA:3851 | 1.00 | 60.41 |  | 0 |  | 1^st^ MG coordination shell missing (see Notes) |
| MG:AA:3852 | 1.00 | 67.97 |  | 0 | 2 | - Exclusion zone contact(s): /O5'/OP2/ Consider K in place of MG (2 valid contact(s) for K vs. 0 for MG) |
| MG:AA:3853 | 1.00 | 55.88 |  | 0 | 1 | - Exclusion zone contact(s): /OP1/ Consider NA in place of MG (1 valid contact(s) for NA vs. 0 for MG) |
| MG:AA:3854 | 1.00 | 71.45 |  | 0 |  | 1^st^ MG coordination shell missing (see Notes) |
| MG:AA:3855 | 1.00 | 72.81 |  | 0 | 2 | - Exclusion zone contact(s): /O2'/O3'/ Consider K in place of MG (2 valid contact(s) for K vs. 0 for MG) |
| MG:AA:3856 | 1.00 | 86.43 |  | 0 |  | No contacts in 5.0 A |
| MG:AA:3857 | 1.00 | 65.09 |  | 0 | 1 | - Exclusion zone contact(s): /N4/ Consider K in place of MG (1 valid contact(s) for K vs. 0 for MG) |
| MG:AA:3858 | 1.00 | 76.89 |  | 0 |  | 1^st^ MG coordination shell missing (see Notes) |
| MG:AA:3859 | 1.00 | 80.38 |  | 0 | 7 | - Exclusion zone contact(s): /N1_G_/N3_C_/N4/N7/N7/O6/OP2/ Consider K in place of MG (7 valid contact(s) for K vs. 0 for MG) |
| MG:AA:3860 | 1.00 | 83.00 |  | 0 | 1 | - Exclusion zone contact(s): /O2'/ Consider K in place of MG (1 valid contact(s) for K vs. 0 for MG) |
| MG:AA:3861 | 1.00 | 99.44 |  | 0 | 1 | - Exclusion zone contact(s): /N7/ Consider K in place of MG (1 valid contact(s) for K vs. 0 for MG) |
| MG:AA:3862 | 1.00 | 75.62 |  | 0 |  | 1^st^ MG coordination shell missing (see Notes) |
| MG:AA:3863 | 1.00 | 72.58 |  | 0 | 1 | - Exclusion zone contact(s): /O6/ Consider K in place of MG (1 valid contact(s) for K vs. 0 for MG) |
| MG:AA:3864 | 1.00 | 48.73 |  | 0 | 4 | - Exclusion zone contact(s): /O2/O2'/OP1/OP2/ Consider K in place of MG (4 valid contact(s) for K vs. 0 for MG) |
| MG:AA:3865 | 1.00 | 62.61 |  | 0 |  | 1^st^ MG coordination shell missing (see Notes) |
| MG:AA:3866 | 1.00 | 96.23 |  | 0 | 1 | - Exclusion zone contact(s): /OP2/ Consider K in place of MG (1 valid contact(s) for K vs. 0 for MG) |
| MG:AA:3867 | 1.00 | 41.11 |  | 0 | 4 | - Exclusion zone contact(s): /N7/O2'/OP2/ Consider K in place of MG (4 valid contact(s) for K vs. 0 for MG) |
| MG:AA:3868 | 1.00 | 80.39 |  | 0 | 1 | - Exclusion zone contact(s): /OP2/ Consider K in place of MG (1 valid contact(s) for K vs. 0 for MG) |
| MG:AA:3869 | 1.00 | 68.27 |  | 0 | 3 | - Exclusion zone contact(s): /N1_A_/N7/O6/ Consider K in place of MG (3 valid contact(s) for K vs. 0 for MG) |
| MG:AA:3870 | 1.00 | 69.24 |  | 0 | 2 | - Exclusion zone contact(s): /OP1/OP1/ Consider K in place of MG (2 valid contact(s) for K vs. 0 for MG) |
| MG:AA:3871 | 1.00 | 51.99 |  | 1 | 2 | O_r_ - Exclusion zone contact(s): /OP1/OP1/ |
| MG:AA:3872 | 1.00 | 90.01 |  | 0 |  | 1^st^ MG coordination shell missing (see Notes) |
| MG:AA:3873 | 1.00 | 87.66 |  | 0 |  | 1^st^ MG coordination shell missing (see Notes) |
| MG:AA:3874 | 1.00 | 90.26 |  | 0 | 4 | - Exclusion zone contact(s): /N1_G_/N7/O6/O6/ Consider K in place of MG (4 valid contact(s) for K vs. 0 for MG) |
| MG:AA:3875 | 1.00 | 67.64 |  | 0 | 1 | - Exclusion zone contact(s): /N2/ Consider K in place of MG (1 valid contact(s) for K vs. 0 for MG) |
| MG:AA:3876 | 1.00 | 67.89 |  | 0 | 2 | - Exclusion zone contact(s): /OP1/OP2/ Consider K in place of MG (2 valid contact(s) for K vs. 0 for MG) |
| MG:AA:3877 | 1.00 | 109.97 |  | 0 |  | 1^st^ MG coordination shell missing (see Notes) |
| MG:AA:3878 | 1.00 | 84.76 |  | 0 | 1 | - Exclusion zone contact(s): /N6/ Consider K in place of MG (1 valid contact(s) for K vs. 0 for MG) |
| MG:AA:3879 | 1.00 | 91.56 |  | 0 | 2 | - Exclusion zone contact(s): /O2'/OP1/ Consider NA in place of MG (1 valid contact(s) for NA vs. 0 for MG) |
| MG:AA:3880 | 1.00 | 102.48 |  | 0 | 3 | - Exclusion zone contact(s): /O2/O2'/O6/ Consider K in place of MG (2 valid contact(s) for K vs. 0 for MG) |
| MG:AA:3881 | 1.00 | 72.13 |  | 1 | 1 | O_ph_ - Exclusion zone contact(s): /OP2/ |
| MG:AA:3882 | 1.00 | 77.07 |  | 0 |  | 1^st^ MG coordination shell missing (see Notes) |
| MG:AA:3883 | 1.00 | 100.34 |  | 0 | 1 | - Exclusion zone contact(s): /OP1/ Consider K in place of MG (1 valid contact(s) for K vs. 0 for MG) |
| MG:AA:3884 | 1.00 | 94.75 |  | 0 |  | No contacts in 5.0 A |
| MG:AA:3885 | 1.00 | 51.95 |  | 0 | 2 | - Exclusion zone contact(s): /OP1/OP2/ Consider K in place of MG (2 valid contact(s) for K vs. 0 for MG) |
| MG:AA:3886 | 1.00 | 103.14 |  | 0 |  | 1^st^ MG coordination shell missing (see Notes) |
| MG:AA:3887 | 1.00 | 70.38 |  | 0 | 1 | - Exclusion zone contact(s): /O2'/ Consider K in place of MG (1 valid contact(s) for K vs. 0 for MG) |
| MG:AA:3888 | 1.00 | 79.25 |  | 0 | 3 | - Exclusion zone contact(s): /N4/O6/O6/ Consider K in place of MG (2 valid contact(s) for K vs. 0 for MG) |
| MG:AA:3889 | 1.00 | 73.31 |  | 1 | 1 | O_ph_ - Exclusion zone contact(s): /O5'/ |
| MG:AA:3890 | 1.00 | 103.08 |  | 0 | 1 | - Exclusion zone contact(s): /O6/ Consider K in place of MG (1 valid contact(s) for K vs. 0 for MG) |
| MG:AA:3891 | 1.00 | 105.40 |  | 0 | 1 | - Exclusion zone contact(s): /N7/ Consider K in place of MG (1 valid contact(s) for K vs. 0 for MG) |
| MG:AA:3892 | 1.00 | 84.25 |  | 0 |  | 1^st^ MG coordination shell missing (see Notes) |
| MG:AA:3893 | 1.00 | 67.75 |  | 0 | 2 | - Exclusion zone contact(s): /O2'/O3'/ Consider K in place of MG (2 valid contact(s) for K vs. 0 for MG) |
| MG:AA:3894 | 1.00 | 86.05 |  | 0 | 1 | - Exclusion zone contact(s): /OP2/ Consider K in place of MG (1 valid contact(s) for K vs. 0 for MG) |
| MG:AA:3895 | 1.00 | 51.50 |  | 0 | 4 | - Exclusion zone contact(s): /O2'/O5'/OG_Ser_/OP1/ Consider K in place of MG (4 valid contact(s) for K vs. 0 for MG) |
| MG:AA:3896 | 1.00 | 99.68 |  | 0 | 2 | - Exclusion zone contact(s): /OP2/OP2/ Consider NA in place of MG (1 valid contact(s) for NA vs. 0 for MG) |
| MG:AA:3897 | 1.00 | 103.23 |  | 0 |  | 1^st^ MG coordination shell missing (see Notes) |
| MG:AA:3898 | 1.00 | 112.21 |  | 0 |  | No contacts in 5.0 A |
| MG:AA:3899 | 1.00 | 46.44 |  | 0 | 3 | - Exclusion zone contact(s): /OP1/OP2/OP2/ Consider K in place of MG (3 valid contact(s) for K vs. 0 for MG) |
| MG:AA:3900 | 1.00 | 105.63 |  | 2 | 2 | O_r_N_b_ - Exclusion zone contact(s): /O5'/OP1/ |
| MG:AA:3901 | 1.00 | 52.95 |  | 1 |  | O_ph_ |
| MG:AA:3902 | 1.00 | 111.17 |  | 1 | 2 | N_b_ - Exclusion zone contact(s): /N2/N6/ |
| MG:AA:3903 | 1.00 | 95.58 |  | 0 |  | No contacts in 5.0 A |
| MG:AA:3904 | 1.00 | 77.35 |  | 0 | 1 | - Exclusion zone contact(s): /N7/ Consider K in place of MG (1 valid contact(s) for K vs. 0 for MG) |
| MG:AA:3905 | 1.00 | 86.72 |  | 0 | 1 | - Exclusion zone contact(s): /OP1/ Consider NA in place of MG (1 valid contact(s) for NA vs. 0 for MG) |
| MG:AA:3906 | 1.00 | 67.74 |  | 0 | 4 | - Exclusion zone contact(s): /NZ_Lys_/O5'/OP1/OP2/ Consider K in place of MG (4 valid contact(s) for K vs. 0 for MG) |
| MG:AA:3907 | 1.00 | 57.94 |  | 0 | 3 | - Exclusion zone contact(s): /N3_G_/O2/O2'/ Consider K in place of MG (2 valid contact(s) for K vs. 0 for MG) |
| MG:AA:3908 | 1.00 | 68.16 |  | 0 | 5 | - Exclusion zone contact(s): /O2'/O2'/O4'/OP1/OP2/ Consider K in place of MG (5 valid contact(s) for K vs. 0 for MG) |
| MG:AA:3909 | 1.00 | 90.78 |  | 0 | 1 | - Exclusion zone contact(s): /N4/ Consider NA in place of MG (1 valid contact(s) for NA vs. 0 for MG) |
| MG:AA:3910 | 1.00 | 76.03 |  | 0 |  | 1^st^ MG coordination shell missing (see Notes) |
| MG:AA:3911 | 1.00 | 69.19 |  | 0 | 1 | - Exclusion zone contact(s): /OP1/ Consider K in place of MG (1 valid contact(s) for K vs. 0 for MG) |
| MG:AA:3912 | 1.00 | 91.40 |  | 0 |  | 1^st^ MG coordination shell missing (see Notes) |
| MG:AA:3913 | 1.00 | 38.53 |  | 0 | 3 | - Exclusion zone contact(s): /N1_G_/O6/ Consider K in place of MG (2 valid contact(s) for K vs. 0 for MG) |
| MG:AA:3914 | 1.00 | 78.33 |  | 0 |  | 1^st^ MG coordination shell missing (see Notes) |
| MG:AA:3915 | 1.00 | 105.75 |  | 0 |  | 1^st^ MG coordination shell missing (see Notes) |
| MG:AA:3916 | 1.00 | 93.60 |  | 0 | 3 | - Exclusion zone contact(s): /OP1/OP1/OP1/ Consider K in place of MG (3 valid contact(s) for K vs. 0 for MG) |
| MG:AA:3917 | 1.00 | 98.06 |  | 0 | 2 | - Exclusion zone contact(s): /O6/OP2/ Consider K in place of MG (2 valid contact(s) for K vs. 0 for MG) |
| MG:AA:3918 | 1.00 | 90.64 |  | 0 | 1 | - Exclusion zone contact(s): /OP2/ Consider K in place of MG (1 valid contact(s) for K vs. 0 for MG) |
| MG:AA:3919 | 1.00 | 59.90 |  | 0 | 3 | - Exclusion zone contact(s): /N7/O4/O6/ Consider K in place of MG (2 valid contact(s) for K vs. 0 for MG) |
| MG:AA:3920 | 1.00 | 85.49 |  | 0 | 3 | - Exclusion zone contact(s): /O5'/OP1/OP2/ Consider K in place of MG (2 valid contact(s) for K vs. 0 for MG) |
| MG:AA:3921 | 1.00 | 44.45 |  | 0 | 3 | - Exclusion zone contact(s): /N4/N7/O6/ Consider K in place of MG (3 valid contact(s) for K vs. 0 for MG) |
| MG:AA:3922 | 1.00 | 65.97 |  | 0 | 2 | - Exclusion zone contact(s): /O2'/O3'/ Consider K in place of MG (2 valid contact(s) for K vs. 0 for MG) |
| MG:AA:3923 | 1.00 | 107.43 |  | 0 | 1 | - Exclusion zone contact(s): /O5'/ Consider K in place of MG (1 valid contact(s) for K vs. 0 for MG) |
| MG:AA:3924 | 1.00 | 79.73 |  | 0 | 3 | - Exclusion zone contact(s): /OP1/OP2/OP2/ Consider K in place of MG (2 valid contact(s) for K vs. 0 for MG) |
| MG:AA:3925 | 1.00 | 71.85 |  | 0 | 3 | - Exclusion zone contact(s): /N7/O6/O6/ Consider K in place of MG (3 valid contact(s) for K vs. 0 for MG) |
| MG:AA:3926 | 1.00 | 52.20 |  | 1 | 1 | O_ph_ - Exclusion zone contact(s): /OP2/ |
| MG:AA:3927 | 1.00 | 76.53 |  | 0 | 2 | - Exclusion zone contact(s): /OP1/OP2/ Consider K in place of MG (2 valid contact(s) for K vs. 0 for MG) |
| MG:AA:3928 | 1.00 | 71.30 |  | 0 | 1 | - Exclusion zone contact(s): /OP2/ Consider K in place of MG (1 valid contact(s) for K vs. 0 for MG) |
| MG:AA:3929 | 1.00 | 104.04 |  | 0 | 3 | - Exclusion zone contact(s): /O2'/O4'/O5'/ Consider NA in place of MG (3 valid contact(s) for NA vs. 0 for MG) |
| MG:AA:3930 | 1.00 | 46.11 |  | 0 | 4 | - Exclusion zone contact(s): /N1/O2'/OP2/OP2/ Consider K in place of MG (4 valid contact(s) for K vs. 0 for MG) |
| MG:AA:3931 | 1.00 | 86.33 |  | 0 |  | 1^st^ MG coordination shell missing (see Notes) |
| MG:AA:3932 | 1.00 | 56.46 |  | 0 | 2 | - Exclusion zone contact(s): /N1/O2'/ Consider K in place of MG (2 valid contact(s) for K vs. 0 for MG) |
| MG:AA:3933 | 1.00 | 115.15 |  | 0 |  | 1^st^ MG coordination shell missing (see Notes) |
| MG:AA:3934 | 1.00 | 93.55 |  | 0 |  | 1^st^ MG coordination shell missing (see Notes) |
| MG:AA:3935 | 1.00 | 55.72 |  | 0 | 3 | - Exclusion zone contact(s): /N2/O2/O4'/ Consider K in place of MG (3 valid contact(s) for K vs. 0 for MG) |
| MG:AA:3936 | 1.00 | 83.46 |  | 0 | 1 | - Exclusion zone contact(s): /OP2/ Consider K in place of MG (1 valid contact(s) for K vs. 0 for MG) |
| MG:AA:3937 | 1.00 | 67.42 |  | 0 |  | 1^st^ MG coordination shell missing (see Notes) |
| MG:AA:3938 | 1.00 | 76.17 |  | 0 |  | 1^st^ MG coordination shell missing (see Notes) |
| MG:AA:3939 | 1.00 | 54.32 |  | 0 | 2 | - Exclusion zone contact(s): /O4/O6/ Consider K in place of MG (2 valid contact(s) for K vs. 0 for MG) |
| MG:AA:3940 | 1.00 | 61.52 | 1 | 1 | 3 | - Exclusion zone contact(s): /N1/O2'/O3'/OP2/ Consider K in place of MG (3 valid contact(s) for K vs. 0 for MG) |
| MG:AA:3941 | 1.00 | 57.97 |  | 0 | 1 | - Exclusion zone contact(s): /O6/ Consider K in place of MG (1 valid contact(s) for K vs. 0 for MG) |
| MG:AA:3942 | 1.00 | 85.65 |  | 0 | 1 | - Exclusion zone contact(s): /N7/ Consider K in place of MG (1 valid contact(s) for K vs. 0 for MG) |
| MG:AA:3943 | 1.00 | 65.80 |  | 0 | 2 | - Exclusion zone contact(s): /O2'/O3'/ Consider K in place of MG (2 valid contact(s) for K vs. 0 for MG) |
| MG:AA:3944 | 1.00 | 71.54 |  | 0 | 2 | - Exclusion zone contact(s): /OP1/OP2/ Consider K in place of MG (2 valid contact(s) for K vs. 0 for MG) |
| MG:AA:3945 | 1.00 | 108.04 |  | 0 | 2 | - Exclusion zone contact(s): /OP1/OP2/ Consider NA in place of MG (2 valid contact(s) for NA vs. 0 for MG) |
| MG:AA:3946 | 1.00 | 98.80 |  | 0 | 3 | - Exclusion zone contact(s): /N4/N7/OP2/ Consider K in place of MG (3 valid contact(s) for K vs. 0 for MG) |
| MG:AA:3947 | 1.00 | 84.93 |  | 1 | 3 | - Exclusion zone contact(s): /N3_G_/N7/N7/ Consider K in place of MG (3 valid contact(s) for K vs. 1 for MG) |
| MG:AA:3948 | 1.00 | 72.38 |  | 0 | 1 | - Exclusion zone contact(s): /O6/ Consider NA in place of MG (1 valid contact(s) for NA vs. 0 for MG) |
| MG:AA:3949 | 1.00 | 71.28 |  | 0 | 2 | - Exclusion zone contact(s): /OP1/OP2/ Consider K in place of MG (2 valid contact(s) for K vs. 0 for MG) |
| MG:AA:3950 | 1.00 | 63.16 |  | 1 |  | O_ph_ |
| MG:AA:3951 | 1.00 | 98.61 |  | 0 | 3 | - Exclusion zone contact(s): /N3_U_/O2/OP1/ Consider K in place of MG (3 valid contact(s) for K vs. 0 for MG) |
| MG:AA:3952 | 1.00 | 71.42 |  | 0 | 2 | - Exclusion zone contact(s): /OP1/OP2/ Consider K in place of MG (2 valid contact(s) for K vs. 0 for MG) |
| MG:AA:3953 | 1.00 | 60.13 |  | 0 | 1 | - Exclusion zone contact(s): /O6/ Consider K in place of MG (1 valid contact(s) for K vs. 0 for MG) |
| MG:AA:3954 | 1.00 | 71.89 |  | 0 | 2 | - Exclusion zone contact(s): /O2'/OP2/ Consider K in place of MG (2 valid contact(s) for K vs. 0 for MG) |
| MG:AA:3955 | 1.00 | 66.76 |  | 0 | 4 | - Exclusion zone contact(s): /N1_G_/N6/O4/OP2/ Consider K in place of MG (4 valid contact(s) for K vs. 0 for MG) |
| MG:AA:3956 | 1.00 | 93.66 |  | 0 |  | No contacts in 5.0 A |
| MG:AA:3957 | 1.00 | 74.42 |  | 0 | 2 | - Exclusion zone contact(s): /O4/O6/ Consider K in place of MG (2 valid contact(s) for K vs. 0 for MG) |
| MG:AA:3958 | 1.00 | 94.67 |  | 0 |  | 1^st^ MG coordination shell missing (see Notes) |
| MG:AA:3959 | 1.00 | 96.38 |  | 0 |  | 1^st^ MG coordination shell missing (see Notes) |
| MG:AA:3960 | 1.00 | 101.91 |  | 0 |  | No contacts in 5.0 A |
| MG:AA:3961 | 1.00 | 54.02 |  | 0 | 2 | - Exclusion zone contact(s): /N4/O4/ Consider K in place of MG (2 valid contact(s) for K vs. 0 for MG) |
| MG:AA:3962 | 1.00 | 57.11 |  | 0 | 1 | - Exclusion zone contact(s): /OP1/ Consider K in place of MG (1 valid contact(s) for K vs. 0 for MG) |
| MG:AA:3963 | 1.00 | 62.30 |  | 0 | 1 | - Exclusion zone contact(s): /OP2/ Consider K in place of MG (1 valid contact(s) for K vs. 0 for MG) |
| MG:AA:3964 | 1.00 | 111.74 |  | 0 |  | 1^st^ MG coordination shell missing (see Notes) |
| MG:AA:3965 | 1.00 | 111.41 |  | 0 | 1 | - Exclusion zone contact(s): /OP2/ Consider NA in place of MG (1 valid contact(s) for NA vs. 0 for MG) |
| MG:AA:3966 | 1.00 | 75.23 |  | 0 | 2 | - Exclusion zone contact(s): /O5'/OP1/ Consider K in place of MG (2 valid contact(s) for K vs. 0 for MG) |
| MG:AA:3967 | 1.00 | 63.19 |  | 0 |  | 1^st^ MG coordination shell missing (see Notes) |
| MG:AA:3968 | 1.00 | 77.42 |  | 1 |  | O_ph_ |
| MG:AA:3969 | 1.00 | 60.10 |  | 0 |  | 1^st^ MG coordination shell missing (see Notes) |
| MG:AA:3970 | 1.00 | 47.66 |  | 0 | 1 | - Exclusion zone contact(s): /OP2/ Consider NA in place of MG (1 valid contact(s) for NA vs. 0 for MG) |
| MG:AA:3971 | 1.00 | 85.96 |  | 0 | 1 | - Exclusion zone contact(s): /OP2/ Consider K in place of MG (1 valid contact(s) for K vs. 0 for MG) |
| MG:AA:3972 | 1.00 | 60.39 |  | 0 | 2 | - Exclusion zone contact(s): /OP1/OP2/ Consider K in place of MG (2 valid contact(s) for K vs. 0 for MG) |
| MG:AA:3973 | 1.00 | 86.42 |  | 0 |  | 1^st^ MG coordination shell missing (see Notes) |
| MG:AA:3974 | 1.00 | 65.23 |  | 0 | 2 | - Exclusion zone contact(s): /O4'/ Consider NA in place of MG (1 valid contact(s) for NA vs. 0 for MG) |
| MG:AA:3975 | 1.00 | 76.30 |  | 0 | 2 | - Exclusion zone contact(s): /O4/O6/ Consider K in place of MG (2 valid contact(s) for K vs. 0 for MG) |
| MG:AA:3976 | 1.00 | 86.96 |  | 0 |  | 1^st^ MG coordination shell missing (see Notes) |
| MG:AA:3977 | 1.00 | 56.75 |  | 1 | 2 | O_r_ - Exclusion zone contact(s): /OP1/OP2/ Consider K in place of MG (2 valid contact(s) for K vs. 1 for MG) |
| MG:AA:3978 | 1.00 | 70.56 |  | 0 | 4 | - Exclusion zone contact(s): /OD1_Asn_/OP1/OP2/OP2/ Consider K in place of MG (3 valid contact(s) for K vs. 0 for MG) |
| MG:AA:3979 | 1.00 | 94.97 |  | 0 |  | 1^st^ MG coordination shell missing (see Notes) |
| MG:AA:3980 | 1.00 | 65.05 |  | 1 |  | O_ph_ |
| MG:AA:3981 | 1.00 | 131.22 |  | 0 | 4 | - Exclusion zone contact(s): /N1_G_/N3_G_/N6/N7/ Consider K in place of MG (3 valid contact(s) for K vs. 0 for MG) |
| MG:AA:3982 | 1.00 | 71.81 |  | 0 |  | 1^st^ MG coordination shell missing (see Notes) |
| MG:AA:3983 | 1.00 | 126.38 |  | 0 | 2 | - Exclusion zone contact(s): /N7/O6/ Consider K in place of MG (2 valid contact(s) for K vs. 0 for MG) |
| MG:AA:3984 | 1.00 | 98.94 |  | 0 | 2 | - Exclusion zone contact(s): /OP1/OP1/ Consider K in place of MG (2 valid contact(s) for K vs. 0 for MG) |
| MG:AA:3985 | 1.00 | 97.36 |  | 0 | 2 | - Exclusion zone contact(s): /OP1/OP2/ Consider K in place of MG (2 valid contact(s) for K vs. 0 for MG) |
| MG:AA:3986 | 1.00 | 88.15 |  | 0 | 3 | - Exclusion zone contact(s): /OP1/OP1/OP2/ Consider K in place of MG (3 valid contact(s) for K vs. 0 for MG) |
| MG:AA:3987 | 1.00 | 54.74 |  | 0 | 4 | - Exclusion zone contact(s): /N2/N3_G_/O2'/O4'/ Consider NA in place of MG (2 valid contact(s) for NA vs. 0 for MG) |
| MG:AA:3988 | 1.00 | 64.01 |  | 2 | 1 | O_ph_O_b_ - Exclusion zone contact(s): /O_back_/ |
| MG:AA:3989 | 1.00 | 74.52 |  | 0 | 1 | - Exclusion zone contact(s): /N7/ Consider K in place of MG (1 valid contact(s) for K vs. 0 for MG) |
| MG:AA:3990 | 1.00 | 134.88 |  | 0 |  | 1^st^ MG coordination shell missing (see Notes) |
| MG:AA:3991 | 1.00 | 73.35 |  | 0 | 2 | - Exclusion zone contact(s): /OP1/OP2/ Consider K in place of MG (2 valid contact(s) for K vs. 0 for MG) |
| MG:AA:3992 | 1.00 | 75.20 |  | 0 |  | No contacts in 5.0 A |
| MG:AA:3993 | 1.00 | 91.18 |  | 0 |  | No contacts in 5.0 A |
| MG:AA:3994 | 1.00 | 68.86 |  | 0 | 1 | - Exclusion zone contact(s): /O2/ Consider K in place of MG (1 valid contact(s) for K vs. 0 for MG) |
| MG:AA:3995 | 1.00 | 103.70 |  | 0 | 1 | - Exclusion zone contact(s): /OP1/ Consider K in place of MG (1 valid contact(s) for K vs. 0 for MG) |
| MG:AA:3996 | 1.00 | 49.98 |  | 0 | 2 | - Exclusion zone contact(s): /O3'/O5'/ Consider NA in place of MG (1 valid contact(s) for NA vs. 0 for MG) |
| MG:AA:3997 | 1.00 | 78.52 |  | 0 | 2 | - Exclusion zone contact(s): /O5'/OP1/ Consider K in place of MG (2 valid contact(s) for K vs. 0 for MG) |
| MG:AA:3998 | 1.00 | 86.93 |  | 0 |  | 1^st^ MG coordination shell missing (see Notes) |
| MG:AA:3999 | 1.00 | 57.29 |  | 0 | 1 | - Exclusion zone contact(s): /OP2/ Consider K in place of MG (1 valid contact(s) for K vs. 0 for MG) |
| MG:AA:4000 | 1.00 | 91.29 |  | 0 | 2 | - Exclusion zone contact(s): /N3_G_/O4'/ Consider K in place of MG (2 valid contact(s) for K vs. 0 for MG) |
| MG:AA:4001 | 1.00 | 83.60 |  | 0 | 3 | - Exclusion zone contact(s): /O4'/OP2/ Consider K in place of MG (3 valid contact(s) for K vs. 0 for MG) |
| MG:AA:4002 | 1.00 | 93.11 |  | 0 | 2 | - Exclusion zone contact(s): /OP1/OP2/ Consider K in place of MG (2 valid contact(s) for K vs. 0 for MG) |
| MG:AA:4003 | 1.00 | 109.72 |  | 0 | 8 | - Exclusion zone contact(s): /N1_G_/N1_G_/N3_C_/N3_C_/N4/N4/O6/O6/ Consider K in place of MG (7 valid contact(s) for K vs. 0 for MG) |
| MG:AA:4004 | 1.00 | 78.86 |  | 0 | 1 | - Exclusion zone contact(s): /O4/ Consider K in place of MG (1 valid contact(s) for K vs. 0 for MG) |
| MG:AA:4005 | 1.00 | 63.84 |  | 0 | 4 | - Exclusion zone contact(s): /O5'/O5'/OP1/OP1/ Consider K in place of MG (4 valid contact(s) for K vs. 0 for MG) |
| MG:AA:4006 | 1.00 | 65.19 |  | 0 | 1 | - Exclusion zone contact(s): /OP2/ Consider K in place of MG (1 valid contact(s) for K vs. 0 for MG) |
| MG:AA:4007 | 1.00 | 94.61 |  | 0 | 3 | - Exclusion zone contact(s): /N4/O6/O6/ Consider K in place of MG (2 valid contact(s) for K vs. 0 for MG) |
| MG:AA:4008 | 1.00 | 78.80 |  | 1 | 2 | O_ph_ - Exclusion zone contact(s): /O3'/OP2/ Consider K in place of MG (2 valid contact(s) for K vs. 1 for MG) |
| MG:AA:4009 | 1.00 | 99.54 |  | 0 | 2 | - Exclusion zone contact(s): /N3_U_/O4/ Consider NA in place of MG (1 valid contact(s) for NA vs. 0 for MG) |
| MG:AA:4010 | 1.00 | 74.63 |  | 2 |  | 2O_ph_ |
| MG:AA:4011 | 1.00 | 122.28 |  | 0 | 1 | - Exclusion zone contact(s): /O2'/ Consider NA in place of MG (1 valid contact(s) for NA vs. 0 for MG) |
| MG:AA:4012 | 1.00 | 85.16 |  | 0 | 1 | - Exclusion zone contact(s): /OP1/ Consider K in place of MG (1 valid contact(s) for K vs. 0 for MG) |
| MG:AA:4013 | 1.00 | 119.98 |  | 1 | 1 | O_b_ - Exclusion zone contact(s): /N7/ |
| MG:AA:4014 | 1.00 | 84.51 |  | 0 |  | 1^st^ MG coordination shell missing (see Notes) |
| MG:AA:4015 | 1.00 | 104.85 |  | 0 |  | No contacts in 5.0 A |
| MG:AA:4016 | 1.00 | 111.82 | 1 | 1 | 4 | O_b_ - Exclusion zone contact(s): /N1_A_/N1_G_/N4/N6/N7/ Consider K in place of MG (3 valid contact(s) for K vs. 1 for MG) |
| MG:AA:4017 | 1.00 | 96.09 |  | 0 | 3 | - Exclusion zone contact(s): /N7/O6/OP2/ Consider K in place of MG (3 valid contact(s) for K vs. 0 for MG) |
| MG:AA:4018 | 1.00 | 56.68 |  | 0 | 5 | - Exclusion zone contact(s): /N1_A_/N6/O6/OP1/OP2/ Consider K in place of MG (5 valid contact(s) for K vs. 0 for MG) |
| MG:AA:4019 | 1.00 | 99.01 |  | 0 |  | 1^st^ MG coordination shell missing (see Notes) |
| MG:AA:4020 | 1.00 | 85.69 |  | 0 |  | 1^st^ MG coordination shell missing (see Notes) |
| MG:AA:4021 | 1.00 | 80.24 |  | 0 | 1 | - Exclusion zone contact(s): /OP2/ Consider K in place of MG (1 valid contact(s) for K vs. 0 for MG) |
| MG:AA:4022 | 1.00 | 62.20 |  | 0 | 1 | - Exclusion zone contact(s): /O6/ Consider K in place of MG (1 valid contact(s) for K vs. 0 for MG) |
| MG:AA:4023 | 1.00 | 73.43 |  | 0 | 2 | - Exclusion zone contact(s): /N7/OP2/ Consider K in place of MG (2 valid contact(s) for K vs. 0 for MG) |
| MG:AA:4024 | 1.00 | 95.29 |  | 0 |  | 1^st^ MG coordination shell missing (see Notes) |
| MG:AA:4025 | 1.00 | 86.44 |  | 0 |  | 1^st^ MG coordination shell missing (see Notes) |
| MG:AA:4026 | 1.00 | 53.10 |  | 0 | 4 | - Exclusion zone contact(s): /NZ_Lys_/OP1/OP1/OP2/ Consider K in place of MG (4 valid contact(s) for K vs. 0 for MG) |
| MG:AA:4027 | 1.00 | 72.87 |  | 0 | 2 | - Exclusion zone contact(s): /O5'/OP1/ Consider K in place of MG (2 valid contact(s) for K vs. 0 for MG) |
| MG:AA:4028 | 1.00 | 81.03 |  | 0 |  | 1^st^ MG coordination shell missing (see Notes) |
| MG:AA:4029 | 1.00 | 80.41 |  | 0 | 3 | - Exclusion zone contact(s): /O2'/O3'/OP1/ Consider K in place of MG (3 valid contact(s) for K vs. 0 for MG) |
| MG:AA:4030 | 1.00 | 66.16 |  | 0 | 2 | - Exclusion zone contact(s): /OP2/OP2/ Consider NA in place of MG (1 valid contact(s) for NA vs. 0 for MG) |
| MG:AA:4031 | 1.00 | 84.17 |  | 0 | 1 | - Exclusion zone contact(s): /O6/ Consider K in place of MG (1 valid contact(s) for K vs. 0 for MG) |
| MG:AA:4032 | 1.00 | 102.97 |  | 0 |  | No contacts in 5.0 A |
| MG:AA:4033 | 1.00 | 83.28 |  | 1 | 1 | O_ph_ - Exclusion zone contact(s): /OP2/ |
| MG:AA:4034 | 1.00 | 100.81 |  | 0 |  | 1^st^ MG coordination shell missing (see Notes) |
| MG:AA:4035 | 1.00 | 75.86 |  | 0 |  | 1^st^ MG coordination shell missing (see Notes) |
| MG:AA:4036 | 1.00 | 68.22 |  | 0 |  | 1^st^ MG coordination shell missing (see Notes) |
| MG:AA:4037 | 1.00 | 43.25 |  | 1 | 1 | O_ph_ - Exclusion zone contact(s): /O3'/ |
| MG:AA:4038 | 1.00 | 77.30 |  | 0 | 3 | - Exclusion zone contact(s): /O2'/OP2/OP2/ Consider K in place of MG (3 valid contact(s) for K vs. 0 for MG) |
| MG:AA:4039 | 1.00 | 95.07 |  | 1 |  | O_b_ |
| MG:AA:4040 | 1.00 | 67.78 |  | 0 | 4 | - Exclusion zone contact(s): /N6/OD1_Asn_/OP2/OP2/ Consider K in place of MG (4 valid contact(s) for K vs. 0 for MG) |
| MG:AA:4041 | 1.00 | 101.04 |  | 0 | 1 | - Exclusion zone contact(s): /OP1/ Consider K in place of MG (1 valid contact(s) for K vs. 0 for MG) |
| MG:AA:4042 | 1.00 | 88.39 |  | 0 | 1 | - Exclusion zone contact(s): /O6/ Consider K in place of MG (1 valid contact(s) for K vs. 0 for MG) |
| MG:AA:4043 | 1.00 | 132.06 |  | 1 |  | O_ph_ |
| MG:AA:4044 | 1.00 | 93.13 |  | 0 |  | 1^st^ MG coordination shell missing (see Notes) |
| MG:AA:4045 | 1.00 | 57.96 |  | 0 | 3 | - Exclusion zone contact(s): /O2'/OP2/OP2/ Consider K in place of MG (3 valid contact(s) for K vs. 0 for MG) |
| MG:AA:4046 | 1.00 | 111.36 |  | 0 | 3 | - Exclusion zone contact(s): /N7/N7/O6/ Consider K in place of MG (2 valid contact(s) for K vs. 0 for MG) |
| MG:AA:4047 | 1.00 | 59.78 |  | 0 | 1 | - Exclusion zone contact(s): /OP1/ Consider K in place of MG (1 valid contact(s) for K vs. 0 for MG) |
| MG:AA:4048 | 1.00 | 93.65 |  | 0 | 4 | - Exclusion zone contact(s): /O5'/OP1/OP1/OP2/ Consider K in place of MG (3 valid contact(s) for K vs. 0 for MG) |
| MG:AA:4049 | 1.00 | 115.47 |  | 0 | 2 | - Exclusion zone contact(s): /O2/O2'/ Consider K in place of MG (2 valid contact(s) for K vs. 0 for MG) |
| MG:AA:4050 | 1.00 | 70.65 |  | 0 | 1 | - Exclusion zone contact(s): /O3'/ Consider K in place of MG (1 valid contact(s) for K vs. 0 for MG) |
| MG:AA:4051 | 1.00 | 106.12 |  | 0 | 1 | - Exclusion zone contact(s): /OP1/ Consider K in place of MG (1 valid contact(s) for K vs. 0 for MG) |
| MG:AA:4052 | 1.00 | 73.27 |  | 0 |  | 1^st^ MG coordination shell missing (see Notes) |
| MG:AA:4053 | 1.00 | 63.50 |  | 0 | 2 | - Exclusion zone contact(s): /O6/OP2/ Consider NA in place of MG (1 valid contact(s) for NA vs. 0 for MG) |
| MG:AA:4054 | 1.00 | 74.95 |  | 0 |  | No contacts in 5.0 A |
| MG:AA:4055 | 1.00 | 110.06 |  | 0 |  | 1^st^ MG coordination shell missing (see Notes) |
| MG:AA:4056 | 1.00 | 74.94 |  | 0 | 1 | - Exclusion zone contact(s): /O2'/ Consider K in place of MG (1 valid contact(s) for K vs. 0 for MG) |
| MG:AA:4057 | 1.00 | 74.80 |  | 0 | 2 | - Exclusion zone contact(s): /OP1/OP2/ Consider K in place of MG (2 valid contact(s) for K vs. 0 for MG) |
| MG:AA:4058 | 1.00 | 63.21 |  | 0 |  | 1^st^ MG coordination shell missing (see Notes) |
| MG:AA:4059 | 1.00 | 54.95 |  | 0 | 1 | - Exclusion zone contact(s): /N1_A_/ Consider K in place of MG (1 valid contact(s) for K vs. 0 for MG) |
| MG:AA:4060 | 1.00 | 70.07 |  | 0 | 1 | - Exclusion zone contact(s): /N4/ Consider K in place of MG (1 valid contact(s) for K vs. 0 for MG) |
| MG:AA:4061 | 1.00 | 79.04 |  | 0 |  | No contacts in 5.0 A |
| MG:AA:4062 | 1.00 | 81.18 |  | 0 |  | 1^st^ MG coordination shell missing (see Notes) |
| MG:AA:4063 | 1.00 | 75.20 |  | 0 | 1 | - Exclusion zone contact(s): /OP2/ Consider NA in place of MG (1 valid contact(s) for NA vs. 0 for MG) |
| MG:AA:4064 | 1.00 | 76.99 |  | 0 | 2 | - Exclusion zone contact(s): /N3_G_/O5'/ Consider K in place of MG (2 valid contact(s) for K vs. 0 for MG) |
| MG:AA:4065 | 1.00 | 68.49 |  | 0 |  | 1^st^ MG coordination shell missing (see Notes) |
| MG:AA:4066 | 1.00 | 82.82 |  | 0 |  | 1^st^ MG coordination shell missing (see Notes) |
| MG:AA:4067 | 1.00 | 86.07 |  | 0 | 1 | - Exclusion zone contact(s): /N7/ Consider K in place of MG (1 valid contact(s) for K vs. 0 for MG) |
| MG:AA:4068 | 1.00 | 58.69 |  | 0 | 4 | - Exclusion zone contact(s): /O4/O5'/OP1/OP2/ Consider K in place of MG (4 valid contact(s) for K vs. 0 for MG) |
| MG:AA:4069 | 1.00 | 80.88 |  | 0 |  | 1^st^ MG coordination shell missing (see Notes) |
| MG:AA:4070 | 1.00 | 60.91 |  | 0 | 3 | - Exclusion zone contact(s): /O4/O6/OP2/ Consider K in place of MG (3 valid contact(s) for K vs. 0 for MG) |
| MG:AA:4071 | 1.00 | 70.78 |  | 0 | 2 | - Exclusion zone contact(s): /OP1/OP2/ Consider K in place of MG (2 valid contact(s) for K vs. 0 for MG) |
| MG:AA:4072 | 1.00 | 56.39 |  | 0 | 2 | - Exclusion zone contact(s): /O2'/OP2/ Consider NA in place of MG (1 valid contact(s) for NA vs. 0 for MG) |
| MG:AA:4073 | 1.00 | 84.77 |  | 0 | 2 | - Exclusion zone contact(s): /O2'/OP1/ Consider K in place of MG (2 valid contact(s) for K vs. 0 for MG) |
| MG:AA:4074 | 1.00 | 94.94 |  | 0 | 1 | - Exclusion zone contact(s): /O4/ Consider K in place of MG (1 valid contact(s) for K vs. 0 for MG) |
| MG:AA:4075 | 1.00 | 132.46 |  | 0 |  | 1^st^ MG coordination shell missing (see Notes) |
| MG:AA:4076 | 1.00 | 113.64 |  | 1 | 3 | O_ph_ - Exclusion zone contact(s): /N2/N3_C_/N4/ Consider K in place of MG (3 valid contact(s) for K vs. 1 for MG) |
| MG:AA:4077 | 1.00 | 67.56 |  | 0 |  | 1^st^ MG coordination shell missing (see Notes) |
| MG:AA:4078 | 1.00 | 82.60 |  | 0 | 1 | - Exclusion zone contact(s): /OP2/ Consider K in place of MG (1 valid contact(s) for K vs. 0 for MG) |
| MG:AA:4079 | 1.00 | 92.33 |  | 0 | 3 | - Exclusion zone contact(s): /OP1/OP1/OP2/ Consider K in place of MG (3 valid contact(s) for K vs. 0 for MG) |
| MG:AA:4080 | 1.00 | 71.34 |  | 0 | 1 | - Exclusion zone contact(s): /OP1/ Consider K in place of MG (1 valid contact(s) for K vs. 0 for MG) |
| MG:AA:4081 | 1.00 | 49.24 |  | 0 | 2 | - Exclusion zone contact(s): /O5'/OP2/ Consider K in place of MG (2 valid contact(s) for K vs. 0 for MG) |
| MG:AA:4082 | 1.00 | 75.67 |  | 0 | 3 | - Exclusion zone contact(s): /N1_G_/O4/O6/ Consider K in place of MG (3 valid contact(s) for K vs. 0 for MG) |
| MG:AA:4083 | 1.00 | 74.58 |  | 0 | 2 | - Exclusion zone contact(s): /O3'/OP2/ Consider K in place of MG (2 valid contact(s) for K vs. 0 for MG) |
| MG:AA:4084 | 1.00 | 86.89 |  | 0 | 2 | - Exclusion zone contact(s): /OP2/OP2/ Consider K in place of MG (2 valid contact(s) for K vs. 0 for MG) |
| MG:AA:4085 | 1.00 | 60.90 |  | 0 | 5 | - Exclusion zone contact(s): /O2'/O2'/O3'/O3'/O4'/ Consider K in place of MG (5 valid contact(s) for K vs. 0 for MG) |
| MG:AA:4086 | 1.00 | 76.20 |  | 0 |  | 1^st^ MG coordination shell missing (see Notes) |
| MG:AA:4087 | 1.00 | 89.74 |  | 0 | 1 | - Exclusion zone contact(s): /OP2/ Consider K in place of MG (1 valid contact(s) for K vs. 0 for MG) |
| MG:AA:4088 | 1.00 | 97.02 |  | 0 | 2 | - Exclusion zone contact(s): /O4/O6/ Consider K in place of MG (2 valid contact(s) for K vs. 0 for MG) |
| MG:AA:4089 | 1.00 | 107.82 |  | 0 | 1 | - Exclusion zone contact(s): /O2'/ Consider K in place of MG (1 valid contact(s) for K vs. 0 for MG) |
| MG:AA:4090 | 1.00 | 78.03 |  | 0 |  | 1^st^ MG coordination shell missing (see Notes) |
| MG:AA:4091 | 1.00 | 107.43 |  | 0 |  | 1^st^ MG coordination shell missing (see Notes) |
| MG:AA:4092 | 1.00 | 90.37 |  | 1 | 2 | O_r_ - Exclusion zone contact(s): /N3_G_/O4'/ Consider K in place of MG (2 valid contact(s) for K vs. 1 for MG) |
| MG:AA:4093 | 1.00 | 114.13 |  | 0 |  | No contacts in 5.0 A |
| MG:AA:4094 | 1.00 | 57.39 |  | 0 | 3 | - Exclusion zone contact(s): /N3_A_/O2/O2'/ Consider K in place of MG (2 valid contact(s) for K vs. 0 for MG) |
| MG:AA:4095 | 1.00 | 58.71 |  | 0 | 1 | - Exclusion zone contact(s): /OP1/ Consider K in place of MG (1 valid contact(s) for K vs. 0 for MG) |
| MG:AA:4096 | 1.00 | 114.31 |  | 0 |  | 1^st^ MG coordination shell missing (see Notes) |
| MG:AA:4097 | 1.00 | 109.63 |  | 0 |  | No contacts in 5.0 A |
| MG:AA:4098 | 1.00 | 83.11 |  | 0 | 2 | - Exclusion zone contact(s): /N3_C_/N4/ Consider K in place of MG (2 valid contact(s) for K vs. 0 for MG) |
| MG:AA:4099 | 1.00 | 96.24 |  | 0 | 2 | - Exclusion zone contact(s): /OP1/OP1/ Consider K in place of MG (2 valid contact(s) for K vs. 0 for MG) |
| MG:AA:4100 | 1.00 | 103.15 |  | 0 | 6 | - Exclusion zone contact(s): /N1_A_/N3_A_/O2'/O2'/O4'/ Consider K in place of MG (5 valid contact(s) for K vs. 0 for MG) |
| MG:AA:4101 | 1.00 | 99.83 |  | 0 | 1 | - Exclusion zone contact(s): /OP2/ Consider NA in place of MG (1 valid contact(s) for NA vs. 0 for MG) |
| MG:AA:4102 | 1.00 | 90.52 |  | 0 |  | 1^st^ MG coordination shell missing (see Notes) |
| MG:AA:4103 | 1.00 | 54.32 |  | 0 |  | 1^st^ MG coordination shell missing (see Notes) |
| MG:AA:4104 | 1.00 | 71.98 |  | 0 | 2 | - Exclusion zone contact(s): /N7/OP2/ Consider K in place of MG (2 valid contact(s) for K vs. 0 for MG) |
| MG:AA:4105 | 1.00 | 106.80 |  | 0 | 1 | - Exclusion zone contact(s): /OP2/ Consider K in place of MG (1 valid contact(s) for K vs. 0 for MG) |
| MG:AA:4106 | 1.00 | 88.85 |  | 0 | 2 | - Exclusion zone contact(s): /N7/O6/ Consider NA in place of MG (1 valid contact(s) for NA vs. 0 for MG) |
| MG:AA:4107 | 1.00 | 98.67 |  | 0 | 3 | - Exclusion zone contact(s): /O5'/OP2/OP2/ Consider K in place of MG (3 valid contact(s) for K vs. 0 for MG) |
| MG:AA:4108 | 1.00 | 136.81 |  | 0 | 2 | - Exclusion zone contact(s): /OP1/OP2/ Consider NA in place of MG (1 valid contact(s) for NA vs. 0 for MG) |
| MG:AA:4109 | 1.00 | 81.57 |  | 0 |  | 1^st^ MG coordination shell missing (see Notes) |
| MG:AA:4110 | 1.00 | 102.40 |  | 0 | 1 | - Exclusion zone contact(s): /O6/ Consider K in place of MG (1 valid contact(s) for K vs. 0 for MG) |
| MG:AA:4111 | 1.00 | 108.75 |  | 0 | 3 | - Exclusion zone contact(s): /O5'/OP2/OP2/ Consider K in place of MG (3 valid contact(s) for K vs. 0 for MG) |
| MG:AA:4112 | 1.00 | 89.22 |  | 0 |  | No contacts in 5.0 A |
| MG:AA:4113 | 1.00 | 92.13 |  | 0 | 6 | - Exclusion zone contact(s): /N1_G_/N4/N4/O6/O6/OP2/ Consider K in place of MG (5 valid contact(s) for K vs. 0 for MG) |
| MG:AA:4114 | 1.00 | 92.01 |  | 1 | 1 | O_b_ - Exclusion zone contact(s): /OP2/ |
| MG:AA:4115 | 1.00 | 73.87 |  | 0 |  | 1^st^ MG coordination shell missing (see Notes) |
| MG:AA:4116 | 1.00 | 98.87 |  | 0 | 2 | - Exclusion zone contact(s): /N7/OE1_Glu_/ Consider K in place of MG (2 valid contact(s) for K vs. 0 for MG) |
| MG:AA:4117 | 1.00 | 96.41 |  | 0 | 3 | - Exclusion zone contact(s): /O5'/OP2/OP2/ Consider K in place of MG (3 valid contact(s) for K vs. 0 for MG) |
| MG:AA:4118 | 1.00 | 67.46 |  | 0 |  | 1^st^ MG coordination shell missing (see Notes) |
| MG:AA:4119 | 1.00 | 112.83 |  | 0 | 1 | - Exclusion zone contact(s): /OP1/ Consider K in place of MG (1 valid contact(s) for K vs. 0 for MG) |
| MG:AA:4120 | 1.00 | 87.48 |  | 0 |  | 1^st^ MG coordination shell missing (see Notes) |
| MG:AA:4121 | 1.00 | 66.43 |  | 0 | 1 | - Exclusion zone contact(s): /OP2/ Consider K in place of MG (1 valid contact(s) for K vs. 0 for MG) |
| MG:AA:4122 | 1.00 | 83.11 |  | 0 |  | No contacts in 5.0 A |
| MG:AA:4123 | 1.00 | 55.94 |  | 0 | 3 | - Exclusion zone contact(s): /O3'/O5'/OP2/ Consider K in place of MG (2 valid contact(s) for K vs. 0 for MG) |
| MG:AA:4124 | 1.00 | 107.98 |  | 0 |  | 1^st^ MG coordination shell missing (see Notes) |
| MG:AA:4125 | 1.00 | 69.38 |  | 0 | 3 | - Exclusion zone contact(s): /O3'/OP1/OP1/ Consider K in place of MG (2 valid contact(s) for K vs. 0 for MG) |
| MG:AA:4126 | 1.00 | 87.45 |  | 0 | 1 | - Exclusion zone contact(s): /OP1/ Consider K in place of MG (1 valid contact(s) for K vs. 0 for MG) |
| MG:AA:4127 | 1.00 | 52.24 |  | 0 | 3 | - Exclusion zone contact(s): /O3'/OP1/OP1/ Consider NA in place of MG (2 valid contact(s) for NA vs. 0 for MG) |
| MG:AA:4128 | 1.00 | 116.40 |  | 0 | 1 | - Exclusion zone contact(s): /OP2/ Consider NA in place of MG (1 valid contact(s) for NA vs. 0 for MG) |
| MG:AA:4129 | 1.00 | 86.37 |  | 0 | 2 | - Exclusion zone contact(s): /OP1/OP2/ Consider K in place of MG (2 valid contact(s) for K vs. 0 for MG) |
| MG:AA:4130 | 1.00 | 107.15 |  | 0 | 1 | - Exclusion zone contact(s): /N4/ Consider K in place of MG (1 valid contact(s) for K vs. 0 for MG) |
| MG:AA:4131 | 1.00 | 99.79 |  | 0 |  | 1^st^ MG coordination shell missing (see Notes) |
| MG:AA:4132 | 1.00 | 68.75 |  | 0 | 3 | - Exclusion zone contact(s): /O5'/OP2/OP2/ Consider K in place of MG (2 valid contact(s) for K vs. 0 for MG) |
| MG:AA:4133 | 1.00 | 127.69 |  | 0 |  | No contacts in 5.0 A |
| MG:AA:4134 | 1.00 | 117.37 |  | 0 | 3 | - Exclusion zone contact(s): /N1_A_/N6/O2'/ Consider K in place of MG (2 valid contact(s) for K vs. 0 for MG) |
| MG:AA:4135 | 1.00 | 64.81 |  | 0 |  | 1^st^ MG coordination shell missing (see Notes) |
| MG:AA:4136 | 1.00 | 71.46 |  | 0 |  | No contacts in 5.0 A |
| MG:AA:4137 | 1.00 | 90.25 |  | 0 | 1 | - Exclusion zone contact(s): /O6/ Consider K in place of MG (1 valid contact(s) for K vs. 0 for MG) |
| MG:AA:4138 | 1.00 | 57.32 |  | 1 | 2 | O_ph_ - Exclusion zone contact(s): /N1/OP2/ |
| MG:AA:4139 | 1.00 | 67.30 |  | 0 | 1 | - Exclusion zone contact(s): /OP1/ Consider NA in place of MG (1 valid contact(s) for NA vs. 0 for MG) |
| MG:AA:4140 | 1.00 | 74.41 |  | 0 |  | No contacts in 5.0 A |
| MG:AA:4141 | 1.00 | 88.26 |  | 0 | 1 | - Exclusion zone contact(s): /O4/ Consider K in place of MG (1 valid contact(s) for K vs. 0 for MG) |
| MG:AA:4142 | 1.00 | 104.29 |  | 0 |  | 1^st^ MG coordination shell missing (see Notes) |
| MG:AA:4143 | 1.00 | 112.43 |  | 0 |  | No contacts in 5.0 A |
| MG:AA:4144 | 1.00 | 100.43 |  | 0 | 2 | - Exclusion zone contact(s): /N7/O6/ Consider NA in place of MG (1 valid contact(s) for NA vs. 0 for MG) |
| MG:AA:4145 | 1.00 | 77.33 |  | 0 | 5 | - Exclusion zone contact(s): /OP1/OP1/OP1/OP2/OP2/ Consider K in place of MG (4 valid contact(s) for K vs. 0 for MG) |
| MG:AA:4146 | 1.00 | 90.87 |  | 0 |  | 1^st^ MG coordination shell missing (see Notes) |
| MG:AA:4147 | 1.00 | 66.78 |  | 0 |  | No contacts in 5.0 A |
| MG:AA:4148 | 1.00 | 87.12 |  | 0 |  | 1^st^ MG coordination shell missing (see Notes) |
| MG:AA:4149 | 1.00 | 125.28 |  | 0 | 1 | - Exclusion zone contact(s): /OP1/ Consider K in place of MG (1 valid contact(s) for K vs. 0 for MG) |
| MG:AA:4150 | 1.00 | 82.67 |  | 0 |  | 1^st^ MG coordination shell missing (see Notes) |
| MG:AA:4151 | 1.00 | 108.15 |  | 0 |  | No contacts in 5.0 A |
| MG:AA:4152 | 1.00 | 77.12 |  | 0 | 5 | - Exclusion zone contact(s): /O3'/OP1/OP1/OP1/OP2/ Consider K in place of MG (5 valid contact(s) for K vs. 0 for MG) |
| MG:AA:4153 | 1.00 | 109.56 |  | 0 | 3 | - Exclusion zone contact(s): /OP1/OP1/OP2/ Consider K in place of MG (3 valid contact(s) for K vs. 0 for MG) |
| MG:AA:4154 | 1.00 | 89.55 |  | 0 | 1 | - Exclusion zone contact(s): /N7/ Consider K in place of MG (1 valid contact(s) for K vs. 0 for MG) |
| MG:AA:4155 | 1.00 | 80.32 |  | 0 | 2 | - Exclusion zone contact(s): /O6/O6/ Consider K in place of MG (2 valid contact(s) for K vs. 0 for MG) |
| MG:AA:4156 | 1.00 | 59.33 |  | 1 | 2 | O_ph_ - Exclusion zone contact(s): /O3'/OP1/ Consider NA in place of MG (2 valid contact(s) for NA vs. 1 for MG) |
| MG:AA:4157 | 1.00 | 76.63 |  | 0 | 2 | - Exclusion zone contact(s): /N7/O6/ Consider K in place of MG (2 valid contact(s) for K vs. 0 for MG) |
| MG:AA:4158 | 1.00 | 93.49 |  | 0 | 2 | - Exclusion zone contact(s): /OP1/ Consider K in place of MG (2 valid contact(s) for K vs. 0 for MG) |
| MG:AA:4159 | 1.00 | 67.49 |  | 0 | 2 | - Exclusion zone contact(s): /N1/OP1/ Consider K in place of MG (2 valid contact(s) for K vs. 0 for MG) |
| MG:AA:4160 | 1.00 | 74.52 |  | 0 | 2 | - Exclusion zone contact(s): /O2'/OP1/ Consider K in place of MG (2 valid contact(s) for K vs. 0 for MG) |
| MG:AA:4161 | 1.00 | 68.70 |  | 0 | 1 | - Exclusion zone contact(s): /OP2/ Consider K in place of MG (1 valid contact(s) for K vs. 0 for MG) |
| MG:AA:4162 | 1.00 | 94.39 |  | 0 | 2 | - Exclusion zone contact(s): /O6/OP2/ Consider K in place of MG (2 valid contact(s) for K vs. 0 for MG) |
| MG:AA:4163 | 1.00 | 102.08 |  | 0 | 5 | - Exclusion zone contact(s): /N2/O5'/OP1/OP2/OP2/ Consider K in place of MG (5 valid contact(s) for K vs. 0 for MG) |
| MG:AA:4164 | 1.00 | 80.48 |  | 0 |  | No contacts in 5.0 A |
| MG:AA:4165 | 1.00 | 64.84 |  | 2 | 2 | 2O_ph_ - Exclusion zone contact(s): /O5'/OP1/ |
| MG:AA:4166 | 1.00 | 142.10 |  | 0 | 2 | - Exclusion zone contact(s): /N7/O4/ Consider K in place of MG (2 valid contact(s) for K vs. 0 for MG) |
| MG:AB:201 | 1.00 | 42.46 |  | 1 | 2 | O_ph_ - Exclusion zone contact(s): /O5'/OP2/ Consider K in place of MG (2 valid contact(s) for K vs. 1 for MG) |
| MG:AB:202 | 1.00 | 51.42 |  | 1 | 1 | O_ph_ - Exclusion zone contact(s): /OP2/ |
| MG:AB:203 | 1.00 | 37.57 |  | 0 | 1 | - Exclusion zone contact(s): /N7/ Consider K in place of MG (1 valid contact(s) for K vs. 0 for MG) |
| MG:AB:204 | 1.00 | 54.31 |  | 0 | 8 | - Exclusion zone contact(s): /N1_G_/N3_U_/N3_U_/N4/O2/O4/O6/O6/ Consider K in place of MG (8 valid contact(s) for K vs. 0 for MG) |
| MG:AB:205 | 1.00 | 54.92 |  | 2 | 1 | O_ph_O_r_ - Exclusion zone contact(s): /O5'/ |
| MG:AB:206 | 1.00 | 76.61 |  | 0 | 2 | - Exclusion zone contact(s): /OP2/OP2/ Consider K in place of MG (2 valid contact(s) for K vs. 0 for MG) |
| MG:AB:207 | 1.00 | 74.66 |  | 1 | 2 | O_b_ - Exclusion zone contact(s): /N4/O6/ |
| MG:AB:208 | 1.00 | 57.87 |  | 1 | 2 | O_ph_ - Exclusion zone contact(s): /O3'/OP2/ |
| MG:AB:209 | 1.00 | 104.25 |  | 0 | 1 | - Exclusion zone contact(s): /OP2/ Consider K in place of MG (1 valid contact(s) for K vs. 0 for MG) |
| MG:AB:210 | 1.00 | 81.55 |  | 0 | 2 | - Exclusion zone contact(s): /N7/O6/ Consider K in place of MG (2 valid contact(s) for K vs. 0 for MG) |
| MG:AB:211 | 1.00 | 77.89 |  | 0 |  | 1^st^ MG coordination shell missing (see Notes) |
| MG:AB:212 | 1.00 | 83.08 |  | 0 | 1 | - Exclusion zone contact(s): /OP1/ Consider K in place of MG (1 valid contact(s) for K vs. 0 for MG) |
| MG:AB:213 | 1.00 | 60.74 |  | 0 | 3 | - Exclusion zone contact(s): /N1_G_/N3_G_/N7/ Consider K in place of MG (3 valid contact(s) for K vs. 0 for MG) |
| MG:AB:214 | 1.00 | 105.02 |  | 0 | 2 | - Exclusion zone contact(s): /N7/O6/ Consider NA in place of MG (1 valid contact(s) for NA vs. 0 for MG) |
| MG:AB:215 | 1.00 | 70.55 |  | 0 | 3 | - Exclusion zone contact(s): /O2/O2'/O4'/ Consider K in place of MG (3 valid contact(s) for K vs. 0 for MG) |
| MG:AB:216 | 1.00 | 53.73 |  | 1 | 3 | O_ph_ - Exclusion zone contact(s): /N1/N3_U_/OP1/ Consider K in place of MG (2 valid contact(s) for K vs. 1 for MG) |
| MG:AB:217 | 1.00 | 87.48 |  | 0 |  | 1^st^ MG coordination shell missing (see Notes) |
| MG:AB:218 | 1.00 | 81.77 |  | 0 | 1 | - Exclusion zone contact(s): /O2'/ Consider NA in place of MG (1 valid contact(s) for NA vs. 0 for MG) |
| MG:AB:219 | 1.00 | 75.25 |  | 1 | 2 | O_b_ - Exclusion zone contact(s): /N6/O2'/ Consider K in place of MG (2 valid contact(s) for K vs. 1 for MG) |
| MG:AB:220 | 1.00 | 79.59 |  | 0 | 4 | - Exclusion zone contact(s): /N2/N4/O6/O6/ Consider K in place of MG (4 valid contact(s) for K vs. 0 for MG) |
| MG:AB:221 | 1.00 | 61.81 |  | 0 |  | 1^st^ MG coordination shell missing (see Notes) |
| MG:AB:222 | 1.00 | 85.99 |  | 0 | 1 | - Exclusion zone contact(s): /OP2/ Consider K in place of MG (1 valid contact(s) for K vs. 0 for MG) |
| MG:AB:223 | 1.00 | 112.80 |  | 0 | 1 | - Exclusion zone contact(s): /OP2/ Consider K in place of MG (1 valid contact(s) for K vs. 0 for MG) |
| MG:AB:224 | 1.00 | 85.16 |  | 0 | 2 | - Exclusion zone contact(s): /OP2/OP2/ Consider K in place of MG (2 valid contact(s) for K vs. 0 for MG) |
| MG:AB:225 | 1.00 | 125.89 |  | 1 | 3 | O_r_ - Exclusion zone contact(s): /O2'/O3'/O5'/ Consider K in place of MG (2 valid contact(s) for K vs. 1 for MG) |
| MG:AB:226 | 1.00 | 88.78 |  | 0 |  | 1^st^ MG coordination shell missing (see Notes) |
| MG:AB:227 | 1.00 | 82.99 |  | 0 | 2 | - Exclusion zone contact(s): /O2'/O3'/ Consider NA in place of MG (1 valid contact(s) for NA vs. 0 for MG) |
| MG:AB:228 | 1.00 | 64.94 |  | 1 |  | O_ph_ |
| MG:AB:229 | 1.00 | 67.26 |  | 0 | 3 | - Exclusion zone contact(s): /O2'/O4/O6/ Consider K in place of MG (3 valid contact(s) for K vs. 0 for MG) |
| MG:AB:230 | 1.00 | 76.69 |  | 0 | 5 | - Exclusion zone contact(s): /N1/N7/O2/O6/OP2/ Consider K in place of MG (5 valid contact(s) for K vs. 0 for MG) |
| MG:AB:231 | 1.00 | 94.98 |  | 0 | 1 | - Exclusion zone contact(s): /OP1/ Consider K in place of MG (1 valid contact(s) for K vs. 0 for MG) |
| MG:AB:232 | 1.00 | 70.10 |  | 0 | 2 | - Exclusion zone contact(s): /O4/OP1/ Consider K in place of MG (2 valid contact(s) for K vs. 0 for MG) |
| MG:AB:233 | 1.00 | 86.65 |  | 0 | 2 | - Exclusion zone contact(s): /OP1/OP1/ Consider K in place of MG (2 valid contact(s) for K vs. 0 for MG) |
| MG:AB:234 | 1.00 | 105.93 |  | 0 |  | No contacts in 5.0 A |
| MG:AB:235 | 1.00 | 81.35 |  | 0 |  | No contacts in 5.0 A |
| MG:AB:236 | 1.00 | 64.70 |  | 0 | 1 | - Exclusion zone contact(s): /OP1/ Consider K in place of MG (1 valid contact(s) for K vs. 0 for MG) |
| MG:AD:301 | 1.00 | 36.23 |  | 0 |  | 1^st^ MG coordination shell missing (see Notes) |
| MG:AD:302 | 1.00 | 71.91 |  | 0 | 1 | - Exclusion zone contact(s): /O_back_/ Consider K in place of MG (1 valid contact(s) for K vs. 0 for MG) |
| MG:AD:303 | 1.00 | 62.73 |  | 0 | 4 | - Exclusion zone contact(s): /NE_Arg_/O_back_/OP1/OP2/ Consider K in place of MG (3 valid contact(s) for K vs. 0 for MG) |
| MG:AD:304 | 1.00 | 65.35 |  | 0 |  | 1^st^ MG coordination shell missing (see Notes) |
| MG:AD:305 | 1.00 | 53.88 |  | 0 | 2 | - Exclusion zone contact(s): /N_back_/OD2_Asp_/ Consider K in place of MG (2 valid contact(s) for K vs. 0 for MG) |
| MG:AD:306 | 1.00 | 44.10 |  | 0 | 1 | - Exclusion zone contact(s): /N_back_/ Consider K in place of MG (1 valid contact(s) for K vs. 0 for MG) |
| MG:AD:307 | 1.00 | 51.85 |  | 1 | 1 | O_ph_ - Exclusion zone contact(s): /O_back_/ |
| MG:AD:308 | 1.00 | 58.18 |  | 0 | 6 | - Exclusion zone contact(s): /N_back_/ND2_Asn_/NH1_Arg_/O_back_/OD1_Asp_/OD2_Asp_/ Consider K in place of MG (5 valid contact(s) for K vs. 0 for MG) |
| MG:AD:309 | 1.00 | 48.52 |  | 2 | 1 | O_r_Anion - Exclusion zone contact(s): /OE1_Gln_/ |
| MG:AD:310 | 1.00 | 42.52 |  | 0 | 1 | - Exclusion zone contact(s): /OE1_Glu_/ Consider K in place of MG (1 valid contact(s) for K vs. 0 for MG) |
| MG:AD:311 | 1.00 | 54.52 |  | 0 |  | 1^st^ MG coordination shell missing (see Notes) |
| MG:AD:312 | 1.00 | 79.93 |  | 0 |  | 1^st^ MG coordination shell missing (see Notes) |
| MG:AD:313 | 1.00 | 39.85 |  | 0 | 3 | - Exclusion zone contact(s): /ND2_Asn_/O2'/O_back_/ Consider K in place of MG (3 valid contact(s) for K vs. 0 for MG) |
| MG:AE:301 | 1.00 | 35.28 |  | 1 |  | O_coo_ |
| MG:AE:302 | 1.00 | 74.40 |  | 0 | 1 | - Exclusion zone contact(s): /OP1/ Consider K in place of MG (1 valid contact(s) for K vs. 0 for MG) |
| MG:AE:303 | 1.00 | 39.60 |  | 1 | 1 | His - Exclusion zone contact(s): /OP1/ |
| MG:AE:304 | 1.00 | 83.94 |  | 0 | 1 | - Exclusion zone contact(s): /O_back_/ Consider NA in place of MG (1 valid contact(s) for NA vs. 0 for MG) |
| MG:AF:301 | 1.00 | 73.24 |  | 0 | 4 | - Exclusion zone contact(s): /O4/O_back_/O_back_/O_back_/ Consider K in place of MG (3 valid contact(s) for K vs. 0 for MG) |
| MG:AF:302 | 1.00 | 69.14 |  | 0 |  | No contacts in 5.0 A |
| MG:AF:303 | 1.00 | 68.65 |  | 1 | 3 | Anion - Exclusion zone contact(s): /O2'/O_back_/O_back_/ Consider K in place of MG (3 valid contact(s) for K vs. 1 for MG) |
| MG:AF:304 | 1.00 | 42.11 |  | 1 | 2 | Anion - Exclusion zone contact(s): /O_back_/O_back_/ Consider K in place of MG (2 valid contact(s) for K vs. 1 for MG) |
| MG:AF:305 | 1.00 | 89.91 |  | 0 |  | 1^st^ MG coordination shell missing (see Notes) |
| MG:AF:306 | 1.00 | 77.33 |  | 0 |  | No contacts in 5.0 A |
| MG:AF:307 | 1.00 | 63.43 |  | 0 |  | 1^st^ MG coordination shell missing (see Notes) |
| MG:AG:201 | 1.00 | 86.46 |  | 0 | 1 | - Exclusion zone contact(s): /O_back_/ Consider K in place of MG (1 valid contact(s) for K vs. 0 for MG) |
| MG:AG:202 | 1.00 | 83.37 |  | 0 | 5 | - Exclusion zone contact(s): /ND2_Asn_/O_back_/O_back_/O_back_/O_back_/ Consider K in place of MG (4 valid contact(s) for K vs. 0 for MG) |
| MG:AG:203 | 1.00 | 97.06 |  | 0 | 1 | - Exclusion zone contact(s): /NH1_Arg_/ Consider K in place of MG (1 valid contact(s) for K vs. 0 for MG) |
| MG:AH:201 | 1.00 | 78.75 |  | 0 |  | 1^st^ MG coordination shell missing (see Notes) |
| MG:AK:201 | 1.00 | 64.99 |  | 0 |  | 1^st^ MG coordination shell missing (see Notes) |
| MG:AK:202 | 1.00 | 75.41 |  | 0 |  | 1^st^ MG coordination shell missing (see Notes) |
| MG:AK:203 | 1.00 | 86.03 |  | 0 |  | No contacts in 5.0 A |
| MG:AN:201 | 1.00 | 118.58 |  | 0 | 1 | - Exclusion zone contact(s): /O_back_/ Consider K in place of MG (1 valid contact(s) for K vs. 0 for MG) |
| MG:AO:201 | 1.00 | 50.40 |  | 1 | 2 | Anion - Exclusion zone contact(s): /N_back_/O_back_/ |
| MG:AO:202 | 1.00 | 55.69 |  | 0 | 1 | - Exclusion zone contact(s): /O_back_/ Consider K in place of MG (1 valid contact(s) for K vs. 0 for MG) |
| MG:AO:203 | 1.00 | 47.40 |  | 0 | 1 | - Exclusion zone contact(s): /NH1_Arg_/ Consider K in place of MG (1 valid contact(s) for K vs. 0 for MG) |
| MG:AO:204 | 1.00 | 43.23 |  | 0 | 2 | - Exclusion zone contact(s): /NH2_Arg_/O2'/ Consider K in place of MG (2 valid contact(s) for K vs. 0 for MG) |
| MG:AO:205 | 1.00 | 52.80 |  | 0 | 1 | - Exclusion zone contact(s): /O_back_/ Consider K in place of MG (1 valid contact(s) for K vs. 0 for MG) |
| MG:AO:206 | 1.00 | 53.04 |  | 0 | 3 | - Exclusion zone contact(s): /O_back_/OG_Ser_/OP2/ Consider K in place of MG (3 valid contact(s) for K vs. 0 for MG) |
| MG:AO:207 | 1.00 | 102.25 |  | 0 |  | 1^st^ MG coordination shell missing (see Notes) |
| MG:AP:201 | 1.00 | 95.51 |  | 0 | 4 | - Exclusion zone contact(s): /O_back_/O_back_/O_back_/OP1/ Consider K in place of MG (3 valid contact(s) for K vs. 0 for MG) |
| MG:AQ:201 | 1.00 | 56.70 |  | 0 | 2 | - Exclusion zone contact(s): /O2'/O_back_/ Consider K in place of MG (2 valid contact(s) for K vs. 0 for MG) |
| MG:AQ:202 | 1.00 | 94.24 |  | 0 | 4 | - Exclusion zone contact(s): /N_back_/N_back_/O_back_/O_back_/ Consider NA in place of MG (2 valid contact(s) for NA vs. 0 for MG) |
| MG:AQ:203 | 1.00 | 67.24 |  | 0 | 1 | - Exclusion zone contact(s): /NE2_Gln_/ Consider K in place of MG (1 valid contact(s) for K vs. 0 for MG) |
| MG:AQ:204 | 1.00 | 80.84 |  | 0 | 1 | - Exclusion zone contact(s): /N_back_/ Consider K in place of MG (1 valid contact(s) for K vs. 0 for MG) |
| MG:AQ:205 | 1.00 | 94.27 |  | 0 | 2 | - Exclusion zone contact(s): /NE_Arg_/NH1_Arg_/ Consider NA in place of MG (1 valid contact(s) for NA vs. 0 for MG) |
| MG:AR:201 | 1.00 | 71.40 |  | 0 |  | 1^st^ MG coordination shell missing (see Notes) |
| MG:AR:202 | 1.00 | 64.70 |  | 0 | 1 | - Exclusion zone contact(s): /NH1_Arg_/ Consider K in place of MG (1 valid contact(s) for K vs. 0 for MG) |
| MG:AR:203 | 1.00 | 55.18 |  | 1 | 3 | Anion - Exclusion zone contact(s): /N_back_/O_back_/O_back_/ Consider K in place of MG (2 valid contact(s) for K vs. 1 for MG) |
| MG:AR:204 | 1.00 | 76.23 |  | 0 | 1 | - Exclusion zone contact(s): /O_back_/ Consider NA in place of MG (1 valid contact(s) for NA vs. 0 for MG) |
| MG:AR:205 | 1.00 | 54.17 |  | 0 | 1 | - Exclusion zone contact(s): /OG_Ser_/ Consider K in place of MG (1 valid contact(s) for K vs. 0 for MG) |
| MG:AS:201 | 1.00 | 67.54 |  | 0 | 4 | - Exclusion zone contact(s): /N_back_/ND2_Asn_/O2'/O3'/ Consider K in place of MG (3 valid contact(s) for K vs. 0 for MG) |
| MG:AS:202 | 1.00 | 63.64 |  | 0 |  | 1^st^ MG coordination shell missing (see Notes) |
| MG:AS:203 | 1.00 | 72.80 |  | 0 | 2 | - Exclusion zone contact(s): /NZ_Lys_/OE1_Glu_/ Consider K in place of MG (2 valid contact(s) for K vs. 0 for MG) |
| MG:AT:101 | 1.00 | 70.74 |  | 0 | 1 | - Exclusion zone contact(s): /N_back_/ Consider NA in place of MG (1 valid contact(s) for NA vs. 0 for MG) |
| MG:AT:102 | 1.00 | 57.39 |  | 1 | 3 | O_ph_ - Exclusion zone contact(s): /NE2_His_/NZ_Lys_/O_back_/ Consider K in place of MG (3 valid contact(s) for K vs. 1 for MG) |
| MG:AT:103 | 1.00 | 85.53 |  | 0 |  | 1^st^ MG coordination shell missing (see Notes) |
| MG:AT:104 | 1.00 | 89.02 | 1 | 1 | 3 | - Exclusion zone contact(s): /N_back_/NZ_Lys_/O5'/OP1/ Consider K in place of MG (3 valid contact(s) for K vs. 0 for MG) |
| MG:AU:201 | 1.00 | 51.09 |  | 0 |  | 1^st^ MG coordination shell missing (see Notes) |
| MG:AU:202 | 1.00 | 76.00 |  | 0 | 2 | - Exclusion zone contact(s): /N_back_/O_back_/ Consider K in place of MG (2 valid contact(s) for K vs. 0 for MG) |
| MG:AU:203 | 1.00 | 74.07 |  | 0 | 2 | - Exclusion zone contact(s): /O_back_/O_back_/ Consider K in place of MG (2 valid contact(s) for K vs. 0 for MG) |
| MG:AU:204 | 1.00 | 87.62 |  | 0 |  | 1^st^ MG coordination shell missing (see Notes) |
| MG:AU:205 | 1.00 | 69.78 |  | 0 |  | 1^st^ MG coordination shell missing (see Notes) |
| MG:AU:206 | 1.00 | 75.97 |  | 0 |  | 1^st^ MG coordination shell missing (see Notes) |
| MG:AV:301 | 1.00 | 67.43 |  | 0 |  | No contacts in 5.0 A |
| MG:AW:101 | 1.00 | 97.86 |  | 0 |  | 1^st^ MG coordination shell missing (see Notes) |
| MG:AW:102 | 1.00 | 56.49 |  | 0 | 1 | - Exclusion zone contact(s): /OG_Ser_/ Consider K in place of MG (1 valid contact(s) for K vs. 0 for MG) |
| MG:AZ:101 | 1.00 | 83.78 |  | 0 | 1 | - Exclusion zone contact(s): /OE2_Glu_/ Consider K in place of MG (1 valid contact(s) for K vs. 0 for MG) |
| MG:AZ:102 | 1.00 | 50.38 |  | 0 | 3 | - Exclusion zone contact(s): /NH2_Arg_/O_back_/OP1/ Consider K in place of MG (3 valid contact(s) for K vs. 0 for MG) |
| MG:AZ:103 | 1.00 | 75.57 |  | 0 | 2 | - Exclusion zone contact(s): /N_back_/O_back_/ Consider K in place of MG (2 valid contact(s) for K vs. 0 for MG) |
| MG:B1:101 | 1.00 | 60.13 |  | 0 | 1 | - Exclusion zone contact(s): /OP1/ Consider K in place of MG (1 valid contact(s) for K vs. 0 for MG) |
| MG:B1:102 | 1.00 | 72.04 |  | 0 |  | 1^st^ MG coordination shell missing (see Notes) |
| MG:B1:103 | 1.00 | 67.70 |  | 0 |  | 1^st^ MG coordination shell missing (see Notes) |
| MG:B1:104 | 1.00 | 116.17 |  | 0 | 4 | - Exclusion zone contact(s): /N1_G_/N2/O4/O6/ Consider K in place of MG (3 valid contact(s) for K vs. 0 for MG) |
| MG:BA:1601 | 1.00 | 27.12 |  | 0 |  | 1^st^ MG coordination shell missing (see Notes) |
| MG:BA:1602 | 1.00 | 20.68 |  | 1 | 1 | O_ph_ - Exclusion zone contact(s): /OP2/ |
| MG:BA:1603 | 1.00 | 58.05 |  | 0 |  | 1^st^ MG coordination shell missing (see Notes) |
| MG:BA:1604 | 1.00 | 47.36 |  | 0 | 2 | - Exclusion zone contact(s): /O6/OP2/ Consider K in place of MG (2 valid contact(s) for K vs. 0 for MG) |
| MG:BA:1605 | 1.00 | 56.43 |  | 0 | 3 | - Exclusion zone contact(s): /N7/O6/OP2/ Consider K in place of MG (3 valid contact(s) for K vs. 0 for MG) |
| MG:BA:1606 | 1.00 | 93.22 |  | 1 | 1 | O_b_ - Exclusion zone contact(s): /N7/ |
| MG:BA:1607 | 1.00 | 74.06 |  | 0 | 1 | - Exclusion zone contact(s): /OP2/ Consider K in place of MG (1 valid contact(s) for K vs. 0 for MG) |
| MG:BA:1608 | 1.00 | 47.99 | 1 | 2 | 1 | 2O_ph_ - Exclusion zone contact(s): /O3'/OP1/ |
| MG:BA:1609 | 1.00 | 36.25 |  | 0 | 4 | - Exclusion zone contact(s): /O3'/OP1/OP1/OP1/ Consider K in place of MG (3 valid contact(s) for K vs. 0 for MG) |
| MG:BA:1610 | 1.00 | 25.36 | 2 | 2 | 3 | - Exclusion zone contact(s): /O2'/O2'/OP2/OP2/OP2/ Consider K in place of MG (3 valid contact(s) for K vs. 0 for MG) |
| MG:BA:1611 | 1.00 | 37.48 |  | 2 | 1 | O_ph_O_r_ - Exclusion zone contact(s): /O5'/ |
| MG:BA:1612 | 1.00 | 90.78 |  | 0 | 1 | - Exclusion zone contact(s): /OP1/ Consider K in place of MG (1 valid contact(s) for K vs. 0 for MG) |
| MG:BA:1613 | 1.00 | 84.61 |  | 0 |  | 1^st^ MG coordination shell missing (see Notes) |
| MG:BA:1614 | 1.00 | 73.79 |  | 0 |  | No contacts in 5.0 A |
| MG:BA:1615 | 1.00 | 81.59 |  | 0 |  | 1^st^ MG coordination shell missing (see Notes) |
| MG:BA:1616 | 1.00 | 47.23 |  | 2 | 1 | O_ph_O_b_ - Exclusion zone contact(s): /OP1/ |
| MG:BA:1617 | 1.00 | 44.78 |  | 0 |  | 1^st^ MG coordination shell missing (see Notes) |
| MG:BA:1618 | 1.00 | 34.71 |  | 0 |  | 1^st^ MG coordination shell missing (see Notes) |
| MG:BA:1619 | 1.00 | 31.72 |  | 0 | 1 | - Exclusion zone contact(s): /O6/ Consider K in place of MG (1 valid contact(s) for K vs. 0 for MG) |
| MG:BA:1620 | 1.00 | 50.42 |  | 0 | 1 | - Exclusion zone contact(s): /O6/ Consider K in place of MG (1 valid contact(s) for K vs. 0 for MG) |
| MG:BA:1621 | 1.00 | 120.66 |  | 0 |  | 1^st^ MG coordination shell missing (see Notes) |
| MG:BA:1622 | 1.00 | 64.44 |  | 1 | 3 | O_ph_ - Exclusion zone contact(s): /O3'/OP1/OP2/ Consider K in place of MG (3 valid contact(s) for K vs. 1 for MG) |
| MG:BA:1623 | 1.00 | 53.55 |  | 1 | 4 | O_ph_ - Exclusion zone contact(s): /O5'/O6/OP1/OP2/ Consider K in place of MG (4 valid contact(s) for K vs. 1 for MG) |
| MG:BA:1624 | 1.00 | 84.21 |  | 1 | 1 | O_b_ - Exclusion zone contact(s): /O6/ |
| MG:BA:1625 | 1.00 | 74.07 |  | 0 |  | 1^st^ MG coordination shell missing (see Notes) |
| MG:BA:1626 | 1.00 | 68.61 |  | 2 |  | 2O_ph_ |
| MG:BA:1627 | 1.00 | 60.81 |  | 0 |  | 1^st^ MG coordination shell missing (see Notes) |
| MG:BA:1628 | 1.00 | 49.98 |  | 1 |  | O_ph_ |
| MG:BA:1629 | 1.00 | 76.88 |  | 0 | 1 | - Exclusion zone contact(s): /O6/ Consider K in place of MG (1 valid contact(s) for K vs. 0 for MG) |
| MG:BA:1630 | 1.00 | 78.28 |  | 1 |  | O_ph_ |
| MG:BA:1631 | 1.00 | 48.18 |  | 0 | 2 | - Exclusion zone contact(s): /OP1/OP2/ Consider K in place of MG (2 valid contact(s) for K vs. 0 for MG) |
| MG:BA:1632 | 1.00 | 68.80 |  | 0 |  | 1^st^ MG coordination shell missing (see Notes) |
| MG:BA:1633 | 1.00 | 45.17 |  | 0 | 3 | - Exclusion zone contact(s): /O3'/OP1/OP1/ Consider K in place of MG (3 valid contact(s) for K vs. 0 for MG) |
| MG:BA:1634 | 1.00 | 37.69 |  | 0 |  | 1^st^ MG coordination shell missing (see Notes) |
| MG:BA:1635 | 1.00 | 102.75 |  | 0 | 1 | - Exclusion zone contact(s): /O6/ Consider K in place of MG (1 valid contact(s) for K vs. 0 for MG) |
| MG:BA:1636 | 1.00 | 44.26 |  | 0 | 1 | - Exclusion zone contact(s): /O6/ Consider K in place of MG (1 valid contact(s) for K vs. 0 for MG) |
| MG:BA:1637 | 1.00 | 57.83 |  | 0 |  | 1^st^ MG coordination shell missing (see Notes) |
| MG:BA:1638 | 1.00 | 51.29 |  | 0 | 2 | - Exclusion zone contact(s): /N7/O6/ Consider NA in place of MG (1 valid contact(s) for NA vs. 0 for MG) |
| MG:BA:1639 | 1.00 | 72.00 |  | 0 |  | 1^st^ MG coordination shell missing (see Notes) |
| MG:BA:1640 | 1.00 | 27.94 |  | 0 | 1 | - Exclusion zone contact(s): /OP2/ Consider NA in place of MG (1 valid contact(s) for NA vs. 0 for MG) |
| MG:BA:1641 | 1.00 | 53.27 |  | 1 | 1 | O_ph_ - Exclusion zone contact(s): /OP1/ |
| MG:BA:1642 | 1.00 | 45.30 |  | 0 | 2 | - Exclusion zone contact(s): /O6/O6/ Consider K in place of MG (2 valid contact(s) for K vs. 0 for MG) |
| MG:BA:1643 | 1.00 | 53.55 |  | 0 | 1 | - Exclusion zone contact(s): /O6/ Consider K in place of MG (1 valid contact(s) for K vs. 0 for MG) |
| MG:BA:1644 | 1.00 | 102.10 |  | 0 | 1 | - Exclusion zone contact(s): /OP1/ Consider K in place of MG (1 valid contact(s) for K vs. 0 for MG) |
| MG:BA:1645 | 1.00 | 88.73 |  | 1 |  | O_ph_ |
| MG:BA:1646 | 1.00 | 82.56 |  | 2 | 1 | 2O_r_ - Exclusion zone contact(s): /N1_A_/ |
| MG:BA:1647 | 1.00 | 78.12 |  | 1 | 2 | O_ph_ - Exclusion zone contact(s): /OP1/OP2/ Consider K in place of MG (2 valid contact(s) for K vs. 1 for MG) |
| MG:BA:1648 | 1.00 | 50.78 |  | 1 |  | O_ph_ |
| MG:BA:1649 | 1.00 | 63.77 |  | 0 | 1 | - Exclusion zone contact(s): /OP2/ Consider K in place of MG (1 valid contact(s) for K vs. 0 for MG) |
| MG:BA:1650 | 1.00 | 52.58 |  | 1 | 3 | O_ph_ - Exclusion zone contact(s): /O3'/OP1/OP2/ Consider K in place of MG (2 valid contact(s) for K vs. 1 for MG) |
| MG:BA:1651 | 1.00 | 72.93 |  | 0 | 2 | - Exclusion zone contact(s): /O3'/OP1/ Consider K in place of MG (2 valid contact(s) for K vs. 0 for MG) |
| MG:BA:1652 | 1.00 | 40.15 |  | 0 | 1 | - Exclusion zone contact(s): /O6/ Consider K in place of MG (1 valid contact(s) for K vs. 0 for MG) |
| MG:BA:1653 | 1.00 | 47.98 |  | 0 |  | 1^st^ MG coordination shell missing (see Notes) |
| MG:BA:1654 | 1.00 | 53.79 |  | 0 | 1 | - Exclusion zone contact(s): /OP2/ Consider NA in place of MG (1 valid contact(s) for NA vs. 0 for MG) |
| MG:BA:1655 | 1.00 | 58.30 |  | 0 |  | 1^st^ MG coordination shell missing (see Notes) |
| MG:BA:1656 | 1.00 | 57.99 |  | 0 | 2 | - Exclusion zone contact(s): /O6/O6/ Consider K in place of MG (2 valid contact(s) for K vs. 0 for MG) |
| MG:BA:1657 | 1.00 | 63.66 |  | 0 |  | 1^st^ MG coordination shell missing (see Notes) |
| MG:BA:1658 | 1.00 | 49.58 | 1 | 1 | 1 | - Exclusion zone contact(s): /OP1/OP1/ Consider K in place of MG (1 valid contact(s) for K vs. 0 for MG) |
| MG:BA:1659 | 1.00 | 71.99 |  | 0 | 2 | - Exclusion zone contact(s): /O3'/OP2/ Consider K in place of MG (2 valid contact(s) for K vs. 0 for MG) |
| MG:BA:1660 | 1.00 | 101.83 |  | 2 | 1 | 2O_r_ - Exclusion zone contact(s): /OP2/ |
| MG:BA:1661 | 1.00 | 83.67 |  | 0 | 1 | - Exclusion zone contact(s): /OP1/ Consider NA in place of MG (1 valid contact(s) for NA vs. 0 for MG) |
| MG:BA:1662 | 1.00 | 91.66 |  | 0 |  | 1^st^ MG coordination shell missing (see Notes) |
| MG:BA:1663 | 1.00 | 94.59 |  | 0 | 1 | - Exclusion zone contact(s): /OP2/ Consider K in place of MG (1 valid contact(s) for K vs. 0 for MG) |
| MG:BA:1664 | 1.00 | 58.79 |  | 0 | 1 | - Exclusion zone contact(s): /O6/ Consider K in place of MG (1 valid contact(s) for K vs. 0 for MG) |
| MG:BA:1665 | 1.00 | 61.61 |  | 1 |  | O_ph_ |
| MG:BA:1666 | 1.00 | 43.30 |  | 0 | 5 | - Exclusion zone contact(s): /O2'/O3'/O4'/O5'/OP2/ Consider K in place of MG (5 valid contact(s) for K vs. 0 for MG) |
| MG:BA:1667 | 1.00 | 62.87 |  | 1 |  | O_b_ |
| MG:BA:1668 | 1.00 | 56.38 |  | 1 | 2 | O_ph_ - Exclusion zone contact(s): /O3'/OP1/ Consider K in place of MG (2 valid contact(s) for K vs. 1 for MG) |
| MG:BA:1669 | 1.00 | 32.78 |  | 1 |  | O_ph_ |
| MG:BA:1670 | 1.00 | 50.28 |  | 0 | 1 | - Exclusion zone contact(s): /OP2/ Consider K in place of MG (1 valid contact(s) for K vs. 0 for MG) |
| MG:BA:1671 | 1.00 | 50.29 |  | 0 |  | 1^st^ MG coordination shell missing (see Notes) |
| MG:BA:1672 | 1.00 | 39.43 |  | 0 |  | 1^st^ MG coordination shell missing (see Notes) |
| MG:BA:1673 | 1.00 | 73.97 |  | 0 |  | 1^st^ MG coordination shell missing (see Notes) |
| MG:BA:1674 | 1.00 | 45.27 |  | 0 |  | 1^st^ MG coordination shell missing (see Notes) |
| MG:BA:1675 | 1.00 | 28.83 |  | 0 | 6 | - Exclusion zone contact(s): /N4/O4/O4/O4/O6/O6/ Consider K in place of MG (5 valid contact(s) for K vs. 0 for MG) |
| MG:BA:1676 | 1.00 | 84.49 |  | 0 | 1 | - Exclusion zone contact(s): /OP1/ Consider K in place of MG (1 valid contact(s) for K vs. 0 for MG) |
| MG:BA:1677 | 1.00 | 52.67 |  | 1 |  | O_ph_ |
| MG:BA:1678 | 1.00 | 77.24 |  | 0 | 2 | - Exclusion zone contact(s): /OP1/OP2/ Consider K in place of MG (2 valid contact(s) for K vs. 0 for MG) |
| MG:BA:1679 | 1.00 | 50.22 |  | 0 | 2 | - Exclusion zone contact(s): /OP1/OP1/ Consider K in place of MG (2 valid contact(s) for K vs. 0 for MG) |
| MG:BA:1680 | 1.00 | 55.57 |  | 0 | 1 | - Exclusion zone contact(s): /OP2/ Consider K in place of MG (1 valid contact(s) for K vs. 0 for MG) |
| MG:BA:1681 | 1.00 | 43.61 |  | 0 |  | 1^st^ MG coordination shell missing (see Notes) |
| MG:BA:1682 | 1.00 | 51.82 |  | 0 |  | 1^st^ MG coordination shell missing (see Notes) |
| MG:BA:1683 | 1.00 | 51.44 |  | 0 | 3 | - Exclusion zone contact(s): /O4/O6/O6/ Consider K in place of MG (3 valid contact(s) for K vs. 0 for MG) |
| MG:BA:1684 | 1.00 | 36.68 |  | 1 | 5 | O_ph_ - Exclusion zone contact(s): /N2/N3_G_/N7/O2/O6/ Consider K in place of MG (5 valid contact(s) for K vs. 1 for MG) |
| MG:BA:1685 | 1.00 | 45.60 |  | 0 | 4 | - Exclusion zone contact(s): /O4/O6/O6/O6/ Consider K in place of MG (3 valid contact(s) for K vs. 0 for MG) |
| MG:BA:1686 | 1.00 | 34.88 |  | 1 |  | O_ph_ |
| MG:BA:1687 | 1.00 | 41.03 |  | 0 | 4 | - Exclusion zone contact(s): /O2'/O3'/O5'/OP2/ Consider K in place of MG (3 valid contact(s) for K vs. 0 for MG) |
| MG:BA:1688 | 1.00 | 61.22 |  | 0 | 1 | - Exclusion zone contact(s): /N7/ Consider K in place of MG (1 valid contact(s) for K vs. 0 for MG) |
| MG:BA:1689 | 1.00 | 64.16 |  | 0 | 1 | - Exclusion zone contact(s): /O3'/ Consider K in place of MG (1 valid contact(s) for K vs. 0 for MG) |
| MG:BA:1690 | 1.00 | 66.62 |  | 1 | 2 | O_ph_ - Exclusion zone contact(s): /O5'/OP2/ Consider K in place of MG (2 valid contact(s) for K vs. 1 for MG) |
| MG:BA:1691 | 1.00 | 65.60 |  | 0 | 2 | - Exclusion zone contact(s): /O6/OD2_Asp_/ Consider K in place of MG (2 valid contact(s) for K vs. 0 for MG) |
| MG:BA:1692 | 1.00 | 34.53 | 1 | 1 | 1 | - Exclusion zone contact(s): /O2/OP2/ Consider K in place of MG (1 valid contact(s) for K vs. 0 for MG) |
| MG:BA:1693 | 1.00 | 46.18 |  | 1 |  | O_ph_ |
| MG:BA:1694 | 1.00 | 47.39 |  | 1 |  | O_b_ |
| MG:BA:1695 | 1.00 | 44.03 |  | 0 | 5 | - Exclusion zone contact(s): /O2'/O3'/OP1/OP2/OP2/ Consider K in place of MG (4 valid contact(s) for K vs. 0 for MG) |
| MG:BA:1696 | 1.00 | 73.94 |  | 0 | 3 | - Exclusion zone contact(s): /O4/O6/OP2/ Consider K in place of MG (3 valid contact(s) for K vs. 0 for MG) |
| MG:BA:1697 | 1.00 | 52.48 |  | 0 |  | 1^st^ MG coordination shell missing (see Notes) |
| MG:BA:1698 | 1.00 | 104.35 |  | 0 |  | 1^st^ MG coordination shell missing (see Notes) |
| MG:BA:1699 | 1.00 | 90.06 |  | 1 |  | O_ph_ |
| MG:BA:1700 | 1.00 | 46.91 |  | 0 | 4 | - Exclusion zone contact(s): /N7/N7/OP2/OP2/ Consider K in place of MG (4 valid contact(s) for K vs. 0 for MG) |
| MG:BA:1701 | 1.00 | 59.59 |  | 2 | 1 | 2O_ph_ - Exclusion zone contact(s): /O5'/ |
| MG:BA:1702 | 1.00 | 33.59 |  | 3 | 3 | 3O_ph_ - Exclusion zone contact(s): /O5'/O5'/OP1/ |
| MG:BA:1703 | 1.00 | 40.27 |  | 0 | 1 | - Exclusion zone contact(s): /OP2/ Consider NA in place of MG (1 valid contact(s) for NA vs. 0 for MG) |
| MG:BA:1704 | 1.00 | 67.61 |  | 0 | 1 | - Exclusion zone contact(s): /O6/ Consider K in place of MG (1 valid contact(s) for K vs. 0 for MG) |
| MG:BA:1705 | 1.00 | 52.12 |  | 0 | 1 | - Exclusion zone contact(s): /O2'/ Consider K in place of MG (1 valid contact(s) for K vs. 0 for MG) |
| MG:BA:1706 | 1.00 | 52.80 |  | 0 | 2 | - Exclusion zone contact(s): /OP1/OP2/ Consider K in place of MG (2 valid contact(s) for K vs. 0 for MG) |
| MG:BA:1707 | 1.00 | 59.99 |  | 0 | 1 | - Exclusion zone contact(s): /OP2/ Consider K in place of MG (1 valid contact(s) for K vs. 0 for MG) |
| MG:BA:1708 | 1.00 | 57.28 |  | 1 | 1 | O_ph_ - Exclusion zone contact(s): /O6/ |
| MG:BA:1709 | 1.00 | 53.99 |  | 0 | 1 | - Exclusion zone contact(s): /OP2/ Consider NA in place of MG (1 valid contact(s) for NA vs. 0 for MG) |
| MG:BA:1710 | 1.00 | 90.83 |  | 0 | 2 | - Exclusion zone contact(s): /OP1/OP2/ Consider K in place of MG (2 valid contact(s) for K vs. 0 for MG) |
| MG:BA:1711 | 1.00 | 68.29 |  | 0 | 2 | - Exclusion zone contact(s): /OP1/OP1/ Consider K in place of MG (2 valid contact(s) for K vs. 0 for MG) |
| MG:BA:1712 | 1.00 | 56.78 |  | 0 | 1 | - Exclusion zone contact(s): /OP2/ Consider K in place of MG (1 valid contact(s) for K vs. 0 for MG) |
| MG:BA:1713 | 1.00 | 48.71 |  | 0 |  | 1^st^ MG coordination shell missing (see Notes) |
| MG:BA:1714 | 1.00 | 69.29 |  | 1 |  | O_ph_ |
| MG:BA:1715 | 1.00 | 75.05 |  | 0 | 3 | - Exclusion zone contact(s): /N7/O2/O6/ Consider K in place of MG (3 valid contact(s) for K vs. 0 for MG) |
| MG:BA:1716 | 1.00 | 56.20 |  | 1 |  | O_b_ |
| MG:BA:1717 | 1.00 | 55.06 |  | 0 | 1 | - Exclusion zone contact(s): /O6/ Consider K in place of MG (1 valid contact(s) for K vs. 0 for MG) |
| MG:BA:1718 | 1.00 | 43.47 |  | 0 | 3 | - Exclusion zone contact(s): /O3'/OP1/OP1/ Consider K in place of MG (3 valid contact(s) for K vs. 0 for MG) |
| MG:BA:1719 | 1.00 | 43.41 |  | 0 | 4 | - Exclusion zone contact(s): /O2'/O3'/OP1/OP1/ Consider K in place of MG (3 valid contact(s) for K vs. 0 for MG) |
| MG:BA:1720 | 1.00 | 95.06 |  | 0 |  | 1^st^ MG coordination shell missing (see Notes) |
| MG:BA:1721 | 1.00 | 71.88 |  | 1 | 3 | O_ph_ - Exclusion zone contact(s): /O3'/O5'/OP1/ Consider K in place of MG (3 valid contact(s) for K vs. 1 for MG) |
| MG:BA:1722 | 1.00 | 54.05 |  | 0 | 1 | - Exclusion zone contact(s): /O6/ Consider K in place of MG (1 valid contact(s) for K vs. 0 for MG) |
| MG:BA:1723 | 1.00 | 63.71 |  | 1 | 1 | O_b_ - Exclusion zone contact(s): /O6/ |
| MG:BA:1724 | 1.00 | 100.21 |  | 0 | 1 | - Exclusion zone contact(s): /OP2/ Consider K in place of MG (1 valid contact(s) for K vs. 0 for MG) |
| MG:BA:1725 | 1.00 | 109.37 |  | 0 | 3 | - Exclusion zone contact(s): /N6/O4/O6/ Consider K in place of MG (3 valid contact(s) for K vs. 0 for MG) |
| MG:BA:1726 | 1.00 | 62.36 |  | 0 |  | 1^st^ MG coordination shell missing (see Notes) |
| MG:BA:1727 | 1.00 | 52.68 |  | 2 | 1 | 2O_ph_ - Exclusion zone contact(s): /OP1/ |
| MG:BA:1728 | 1.00 | 47.86 |  | 2 | 1 | O_ph_O_b_ - Exclusion zone contact(s): /OP2/ |
| MG:BA:1729 | 1.00 | 69.75 |  | 0 |  | 1^st^ MG coordination shell missing (see Notes) |
| MG:BA:1730 | 1.00 | 37.47 |  | 0 | 3 | - Exclusion zone contact(s): /N7/N7/O6/ Consider K in place of MG (3 valid contact(s) for K vs. 0 for MG) |
| MG:BA:1731 | 1.00 | 46.36 |  | 0 | 1 | - Exclusion zone contact(s): /O6/ Consider K in place of MG (1 valid contact(s) for K vs. 0 for MG) |
| MG:BA:1732 | 1.00 | 62.42 |  | 0 |  | 1^st^ MG coordination shell missing (see Notes) |
| MG:BA:1733 | 1.00 | 46.37 |  | 2 | 2 | O_b_O_back_ - Exclusion zone contact(s): /O6/O_back_/ |
| MG:BA:1734 | 1.00 | 76.26 |  | 0 | 2 | - Exclusion zone contact(s): /O3'/OP2/ Consider NA in place of MG (1 valid contact(s) for NA vs. 0 for MG) |
| MG:BA:1735 | 1.00 | 40.97 |  | 0 | 7 | - Exclusion zone contact(s): /N3_C_/N4/O2/O4/O6/O6/O6/ Consider K in place of MG (6 valid contact(s) for K vs. 0 for MG) |
| MG:BA:1736 | 1.00 | 53.53 |  | 3 |  | 3O_ph_ |
| MG:BA:1737 | 1.00 | 59.48 |  | 1 |  | O_ph_ |
| MG:BA:1738 | 1.00 | 59.12 |  | 1 |  | O_ph_ |
| MG:BA:1739 | 1.00 | 67.21 |  | 1 | 1 | O_ph_ - Exclusion zone contact(s): /N4/ |
| MG:BA:1740 | 1.00 | 42.89 |  | 0 | 3 | - Exclusion zone contact(s): /O4/O6/O6/ Consider K in place of MG (3 valid contact(s) for K vs. 0 for MG) |
| MG:BA:1741 | 1.00 | 85.49 |  | 0 |  | 1^st^ MG coordination shell missing (see Notes) |
| MG:BA:1742 | 1.00 | 46.25 |  | 0 | 1 | - Exclusion zone contact(s): /OP1/ Consider NA in place of MG (1 valid contact(s) for NA vs. 0 for MG) |
| MG:BA:1743 | 1.00 | 43.40 |  | 1 | 2 | O_b_ - Exclusion zone contact(s): /O4/O4/ Consider K in place of MG (2 valid contact(s) for K vs. 1 for MG) |
| MG:BA:1744 | 1.00 | 54.38 |  | 0 | 2 | - Exclusion zone contact(s): /O2'/OP1/ Consider K in place of MG (2 valid contact(s) for K vs. 0 for MG) |
| MG:BA:1745 | 1.00 | 38.27 |  | 0 | 2 | - Exclusion zone contact(s): /O3'/OP2/ Consider NA in place of MG (1 valid contact(s) for NA vs. 0 for MG) |
| MG:BA:1746 | 1.00 | 40.28 |  | 0 | 5 | - Exclusion zone contact(s): /O2'/O3'/O5'/OP1/OP2/ Consider K in place of MG (3 valid contact(s) for K vs. 0 for MG) |
| MG:BA:1747 | 1.00 | 50.69 | 1 | 1 |  | - Exclusion zone contact(s): /OP1/ 1^st^ MG coordination shell missing (see Notes) |
| MG:BA:1748 | 1.00 | 66.96 |  | 0 | 3 | - Exclusion zone contact(s): /O2'/OP2/OP2/ Consider K in place of MG (2 valid contact(s) for K vs. 0 for MG) |
| MG:BA:1749 | 1.00 | 30.94 |  | 1 | 3 | O_r_ - Exclusion zone contact(s): /O2'/O3'/OP2/ Consider K in place of MG (3 valid contact(s) for K vs. 1 for MG) |
| MG:BA:1750 | 1.00 | 65.64 |  | 0 | 3 | - Exclusion zone contact(s): /O4/O6/O6/ Consider K in place of MG (2 valid contact(s) for K vs. 0 for MG) |
| MG:BA:1751 | 1.00 | 62.84 |  | 1 |  | O_ph_ |
| MG:BA:1752 | 1.00 | 48.61 |  | 1 | 1 | O_b_ - Exclusion zone contact(s): /O6/ |
| MG:BA:1753 | 1.00 | 69.02 |  | 1 | 1 | O_ph_ - Exclusion zone contact(s): /O5'/ |
| MG:BA:1754 | 1.00 | 68.41 |  | 0 | 1 | - Exclusion zone contact(s): /OP2/ Consider NA in place of MG (1 valid contact(s) for NA vs. 0 for MG) |
| MG:BA:1755 | 1.00 | 56.45 |  | 0 |  | 1^st^ MG coordination shell missing (see Notes) |
| MG:BA:1756 | 1.00 | 69.09 |  | 0 | 2 | - Exclusion zone contact(s): /N7/O6/ Consider K in place of MG (2 valid contact(s) for K vs. 0 for MG) |
| MG:BA:1757 | 1.00 | 39.59 |  | 0 | 1 | - Exclusion zone contact(s): /N7/ Consider K in place of MG (1 valid contact(s) for K vs. 0 for MG) |
| MG:BA:1758 | 1.00 | 54.92 |  | 0 | 2 | - Exclusion zone contact(s): /O6/O6/ Consider K in place of MG (2 valid contact(s) for K vs. 0 for MG) |
| MG:BA:1759 | 1.00 | 77.05 |  | 1 | 1 | O_ph_ - Exclusion zone contact(s): /O3'/ |
| MG:BA:1760 | 1.00 | 85.89 |  | 1 | 2 | O_ph_ - Exclusion zone contact(s): /OP1/OP2/ Consider K in place of MG (2 valid contact(s) for K vs. 1 for MG) |
| MG:BA:1761 | 1.00 | 58.50 |  | 0 | 3 | - Exclusion zone contact(s): /N7/O6/OP2/ Consider K in place of MG (2 valid contact(s) for K vs. 0 for MG) |
| MG:BA:1762 | 1.00 | 78.54 |  | 0 |  | 1^st^ MG coordination shell missing (see Notes) |
| MG:BA:1763 | 1.00 | 57.63 |  | 1 |  | O_ph_ |
| MG:BA:1764 | 1.00 | 50.16 |  | 1 | 2 | O_ph_ - Exclusion zone contact(s): /OP2/OP2/ |
| MG:BA:1765 | 1.00 | 82.80 |  | 0 | 2 | - Exclusion zone contact(s): /N7/OP2/ Consider NA in place of MG (1 valid contact(s) for NA vs. 0 for MG) |
| MG:BA:1766 | 1.00 | 74.70 |  | 0 | 2 | - Exclusion zone contact(s): /OP1/OP1/ Consider NA in place of MG (1 valid contact(s) for NA vs. 0 for MG) |
| MG:BA:1767 | 1.00 | 50.51 |  | 0 | 2 | - Exclusion zone contact(s): /N7/O6/ Consider K in place of MG (2 valid contact(s) for K vs. 0 for MG) |
| MG:BA:1768 | 1.00 | 75.29 |  | 0 | 1 | - Exclusion zone contact(s): /OP1/ Consider NA in place of MG (1 valid contact(s) for NA vs. 0 for MG) |
| MG:BA:1769 | 1.00 | 87.04 |  | 0 | 1 | - Exclusion zone contact(s): /OP2/ Consider K in place of MG (1 valid contact(s) for K vs. 0 for MG) |
| MG:BA:1770 | 1.00 | 67.47 |  | 0 | 3 | - Exclusion zone contact(s): /N4/O6/O6/ Consider K in place of MG (3 valid contact(s) for K vs. 0 for MG) |
| MG:BA:1771 | 1.00 | 74.32 |  | 0 |  | 1^st^ MG coordination shell missing (see Notes) |
| MG:BA:1772 | 1.00 | 75.46 |  | 0 | 1 | - Exclusion zone contact(s): /N7/ Consider K in place of MG (1 valid contact(s) for K vs. 0 for MG) |
| MG:BA:1773 | 1.00 | 80.55 |  | 0 |  | 1^st^ MG coordination shell missing (see Notes) |
| MG:BA:1774 | 1.00 | 53.26 |  | 0 | 3 | - Exclusion zone contact(s): /N7/O2'/OP2/ Consider K in place of MG (3 valid contact(s) for K vs. 0 for MG) |
| MG:BA:1775 | 1.00 | 93.15 |  | 0 | 2 | - Exclusion zone contact(s): /OP1/OP2/ Consider K in place of MG (2 valid contact(s) for K vs. 0 for MG) |
| MG:BA:1776 | 1.00 | 79.99 |  | 1 | 2 | N_b_ - Exclusion zone contact(s): /OP1/OP2/ |
| MG:BA:1777 | 1.00 | 57.00 |  | 0 | 2 | - Exclusion zone contact(s): /O6/O6/ Consider K in place of MG (2 valid contact(s) for K vs. 0 for MG) |
| MG:BA:1778 | 1.00 | 49.37 |  | 0 |  | 1^st^ MG coordination shell missing (see Notes) |
| MG:BA:1779 | 1.00 | 45.36 |  | 1 | 2 | O_b_ - Exclusion zone contact(s): /O2'/OP1/ |
| MG:BA:1780 | 1.00 | 85.05 |  | 0 | 1 | - Exclusion zone contact(s): /OP2/ Consider K in place of MG (1 valid contact(s) for K vs. 0 for MG) |
| MG:BA:1781 | 1.00 | 69.81 |  | 0 | 2 | - Exclusion zone contact(s): /O3'/OP1/ Consider NA in place of MG (2 valid contact(s) for NA vs. 0 for MG) |
| MG:BA:1782 | 1.00 | 100.11 |  | 0 |  | 1^st^ MG coordination shell missing (see Notes) |
| MG:BA:1783 | 1.00 | 81.54 |  | 1 |  | O_ph_ |
| MG:BA:1784 | 1.00 | 54.70 |  | 0 | 5 | - Exclusion zone contact(s): /O2'/O3'/O3'/OP1/OP1/ Consider K in place of MG (4 valid contact(s) for K vs. 0 for MG) |
| MG:BA:1785 | 1.00 | 67.16 |  | 0 |  | 1^st^ MG coordination shell missing (see Notes) |
| MG:BA:1786 | 1.00 | 57.70 |  | 0 |  | 1^st^ MG coordination shell missing (see Notes) |
| MG:BA:1787 | 1.00 | 49.87 |  | 0 |  | 1^st^ MG coordination shell missing (see Notes) |
| MG:BA:1788 | 1.00 | 53.90 |  | 0 | 1 | - Exclusion zone contact(s): /O5'/ Consider K in place of MG (1 valid contact(s) for K vs. 0 for MG) |
| MG:BA:1789 | 1.00 | 105.64 |  | 1 | 1 | O_ph_ - Exclusion zone contact(s): /O5'/ |
| MG:BA:1790 | 1.00 | 56.46 |  | 1 | 2 | O_ph_ - Exclusion zone contact(s): /O4/O6/ Consider K in place of MG (2 valid contact(s) for K vs. 1 for MG) |
| MG:BA:1791 | 1.00 | 55.69 |  | 0 | 4 | - Exclusion zone contact(s): /O5'/OD1_Asn_/OP1/OP2/ Consider K in place of MG (4 valid contact(s) for K vs. 0 for MG) |
| MG:BA:1792 | 1.00 | 79.85 |  | 0 |  | 1^st^ MG coordination shell missing (see Notes) |
| MG:BA:1793 | 1.00 | 57.10 |  | 1 | 3 | O_b_ - Exclusion zone contact(s): /O4/O4/O6/ Consider K in place of MG (3 valid contact(s) for K vs. 1 for MG) |
| MG:BA:1794 | 1.00 | 49.72 |  | 1 |  | O_ph_ |
| MG:BA:1795 | 1.00 | 67.08 |  | 0 |  | 1^st^ MG coordination shell missing (see Notes) |
| MG:BA:1796 | 1.00 | 77.78 |  | 0 |  | 1^st^ MG coordination shell missing (see Notes) |
| MG:BA:1797 | 1.00 | 63.63 |  | 0 |  | 1^st^ MG coordination shell missing (see Notes) |
| MG:BA:1798 | 1.00 | 93.68 |  | 0 |  | 1^st^ MG coordination shell missing (see Notes) |
| MG:BA:1799 | 1.00 | 55.22 |  | 0 |  | 1^st^ MG coordination shell missing (see Notes) |
| MG:BA:1800 | 1.00 | 66.51 |  | 1 | 2 | O_ph_ - Exclusion zone contact(s): /OP1/OP1/ Consider K in place of MG (2 valid contact(s) for K vs. 1 for MG) |
| MG:BA:1801 | 1.00 | 40.72 |  | 0 |  | 1^st^ MG coordination shell missing (see Notes) |
| MG:BA:1802 | 1.00 | 75.25 |  | 0 |  | 1^st^ MG coordination shell missing (see Notes) |
| MG:BA:1803 | 1.00 | 90.90 |  | 0 |  | 1^st^ MG coordination shell missing (see Notes) |
| MG:BA:1804 | 1.00 | 105.88 |  | 1 | 3 | O_b_ - Exclusion zone contact(s): /N3_U_/N4/O4/ Consider K in place of MG (3 valid contact(s) for K vs. 1 for MG) |
| MG:BA:1805 | 1.00 | 99.65 |  | 0 |  | 1^st^ MG coordination shell missing (see Notes) |
| MG:BA:1806 | 1.00 | 78.79 |  | 0 |  | 1^st^ MG coordination shell missing (see Notes) |
| MG:BA:1807 | 1.00 | 63.72 | 1 | 1 |  | - Exclusion zone contact(s): /OP2/ 1^st^ MG coordination shell missing (see Notes) |
| MG:BA:1808 | 1.00 | 70.02 |  | 0 | 1 | - Exclusion zone contact(s): /N7/ Consider NA in place of MG (1 valid contact(s) for NA vs. 0 for MG) |
| MG:BA:1809 | 1.00 | 114.37 |  | 1 | 2 | N_b_ - Exclusion zone contact(s): /OP2/ Consider K in place of MG (2 valid contact(s) for K vs. 1 for MG) |
| MG:BA:1810 | 1.00 | 93.74 |  | 1 | 1 | N_b_ - Exclusion zone contact(s): /O6/ |
| MG:BA:1811 | 1.00 | 65.56 |  | 1 |  | O_b_ |
| MG:BA:1812 | 1.00 | 136.01 |  | 0 |  | 1^st^ MG coordination shell missing (see Notes) |
| MG:BA:1813 | 1.00 | 94.76 |  | 1 | 1 | O_ph_ - Exclusion zone contact(s): /OP1/ |
| MG:BA:1814 | 1.00 | 115.79 |  | 0 |  | 1^st^ MG coordination shell missing (see Notes) |
| MG:BA:1815 | 1.00 | 119.44 |  | 0 | 1 | - Exclusion zone contact(s): /O2'/ Consider K in place of MG (1 valid contact(s) for K vs. 0 for MG) |
| MG:BA:1816 | 1.00 | 71.08 |  | 0 | 1 | - Exclusion zone contact(s): /O4'/ Consider K in place of MG (1 valid contact(s) for K vs. 0 for MG) |
| MG:BA:1817 | 1.00 | 63.32 |  | 0 | 1 | - Exclusion zone contact(s): /O6/ Consider K in place of MG (1 valid contact(s) for K vs. 0 for MG) |
| MG:BA:1818 | 1.00 | 57.10 |  | 0 |  | 1^st^ MG coordination shell missing (see Notes) |
| MG:BA:1819 | 1.00 | 61.36 |  | 2 | 3 | O_r_O_b_ - Exclusion zone contact(s): /O4/O4/O6/ Consider K in place of MG (3 valid contact(s) for K vs. 2 for MG) |
| MG:BA:1820 | 1.00 | 84.64 |  | 0 | 1 | - Exclusion zone contact(s): /O6/ Consider K in place of MG (1 valid contact(s) for K vs. 0 for MG) |
| MG:BA:1821 | 1.00 | 93.39 |  | 0 | 1 | - Exclusion zone contact(s): /N7/ Consider K in place of MG (1 valid contact(s) for K vs. 0 for MG) |
| MG:BA:1822 | 1.00 | 97.16 |  | 0 |  | 1^st^ MG coordination shell missing (see Notes) |
| MG:BA:1823 | 1.00 | 84.14 |  | 0 | 1 | - Exclusion zone contact(s): /OP2/ Consider NA in place of MG (1 valid contact(s) for NA vs. 0 for MG) |
| MG:BA:1824 | 1.00 | 107.38 |  | 0 | 1 | - Exclusion zone contact(s): /N3_C_/ Consider K in place of MG (1 valid contact(s) for K vs. 0 for MG) |
| MG:BA:1825 | 1.00 | 121.76 |  | 1 |  | O_ph_ |
| MG:BA:1826 | 1.00 | 75.34 |  | 0 | 1 | - Exclusion zone contact(s): /OP2/ Consider K in place of MG (1 valid contact(s) for K vs. 0 for MG) |
| MG:BA:1827 | 1.00 | 101.25 |  | 0 | 1 | - Exclusion zone contact(s): /O2/ Consider K in place of MG (1 valid contact(s) for K vs. 0 for MG) |
| MG:BA:1828 | 1.00 | 114.78 |  | 0 |  | 1^st^ MG coordination shell missing (see Notes) |
| MG:BA:1829 | 1.00 | 75.24 |  | 0 |  | 1^st^ MG coordination shell missing (see Notes) |
| MG:BA:1830 | 1.00 | 90.52 |  | 0 |  | 1^st^ MG coordination shell missing (see Notes) |
| MG:BA:1831 | 1.00 | 81.02 |  | 0 | 1 | - Exclusion zone contact(s): /O2'/ Consider NA in place of MG (1 valid contact(s) for NA vs. 0 for MG) |
| MG:BA:1832 | 1.00 | 89.12 |  | 0 |  | 1^st^ MG coordination shell missing (see Notes) |
| MG:BA:1833 | 1.00 | 71.41 |  | 0 | 2 | - Exclusion zone contact(s): /OP1/OP2/ Consider K in place of MG (2 valid contact(s) for K vs. 0 for MG) |
| MG:BA:1834 | 1.00 | 70.60 |  | 0 | 1 | - Exclusion zone contact(s): /OP2/ Consider K in place of MG (1 valid contact(s) for K vs. 0 for MG) |
| MG:BA:1835 | 1.00 | 72.95 |  | 0 | 3 | - Exclusion zone contact(s): /OP1/OP2/OP2/ Consider K in place of MG (2 valid contact(s) for K vs. 0 for MG) |
| MG:BA:1836 | 1.00 | 57.71 |  | 0 | 3 | - Exclusion zone contact(s): /O3'/O5'/OP1/ Consider NA in place of MG (2 valid contact(s) for NA vs. 0 for MG) |
| MG:BA:1837 | 1.00 | 88.25 |  | 2 |  | O_ph_O_r_ |
| MG:BA:1838 | 1.00 | 91.53 |  | 0 |  | 1^st^ MG coordination shell missing (see Notes) |
| MG:BA:1839 | 1.00 | 136.72 |  | 0 | 4 | - Exclusion zone contact(s): /N4/N4/O6/OP2/ Consider K in place of MG (4 valid contact(s) for K vs. 0 for MG) |
| MG:BA:1840 | 1.00 | 156.86 |  | 2 | 2 | 2Anion - Exclusion zone contact(s): /O6/O6/ |
| MG:BA:1841 | 1.00 | 108.03 |  | 1 | 1 | O_b_ - Exclusion zone contact(s): /O6/ |
| MG:BA:1842 | 1.00 | 92.20 |  | 0 |  | 1^st^ MG coordination shell missing (see Notes) |
| MG:BA:1843 | 1.00 | 63.47 |  | 0 |  | 1^st^ MG coordination shell missing (see Notes) |
| MG:BA:1844 | 1.00 | 118.88 |  | 1 |  | O_ph_ |
| MG:BA:1845 | 1.00 | 92.53 |  | 1 |  | N_b_ |
| MG:BA:1846 | 1.00 | 63.45 |  | 0 | 1 | - Exclusion zone contact(s): /N7/ Consider K in place of MG (1 valid contact(s) for K vs. 0 for MG) |
| MG:BA:1847 | 1.00 | 74.65 | 1 | 1 |  | - Exclusion zone contact(s): /OP2/ 1^st^ MG coordination shell missing (see Notes) |
| MG:BA:1848 | 1.00 | 56.88 |  | 0 | 3 | - Exclusion zone contact(s): /O4'/OP2/ Consider K in place of MG (3 valid contact(s) for K vs. 0 for MG) |
| MG:BA:1849 | 1.00 | 74.74 |  | 0 | 3 | - Exclusion zone contact(s): /O2'/O3'/OP2/ Consider K in place of MG (3 valid contact(s) for K vs. 0 for MG) |
| MG:BA:1850 | 1.00 | 90.03 |  | 0 | 1 | - Exclusion zone contact(s): /O2'/ Consider NA in place of MG (1 valid contact(s) for NA vs. 0 for MG) |
| MG:BA:1851 | 1.00 | 142.74 |  | 0 |  | 1^st^ MG coordination shell missing (see Notes) |
| MG:BA:1852 | 1.00 | 105.85 |  | 1 |  | O_ph_ |
| MG:BA:1853 | 1.00 | 103.05 |  | 0 |  | No contacts in 5.0 A |
| MG:BA:1854 | 1.00 | 84.55 |  | 0 |  | 1^st^ MG coordination shell missing (see Notes) |
| MG:BA:1855 | 1.00 | 76.90 |  | 0 |  | 1^st^ MG coordination shell missing (see Notes) |
| MG:BA:1856 | 1.00 | 60.63 |  | 0 |  | No contacts in 5.0 A |
| MG:BA:1857 | 1.00 | 122.86 |  | 0 |  | 1^st^ MG coordination shell missing (see Notes) |
| MG:BA:1858 | 1.00 | 141.69 |  | 0 | 2 | - Exclusion zone contact(s): /O2'/O4'/ Consider K in place of MG (2 valid contact(s) for K vs. 0 for MG) |
| MG:BA:1859 | 1.00 | 146.52 |  | 0 |  | No contacts in 5.0 A |
| MG:BA:1860 | 1.00 | 60.80 |  | 0 |  | 1^st^ MG coordination shell missing (see Notes) |
| MG:BA:1861 | 1.00 | 78.55 |  | 0 |  | 1^st^ MG coordination shell missing (see Notes) |
| MG:BA:1862 | 1.00 | 77.67 |  | 0 |  | 1^st^ MG coordination shell missing (see Notes) |
| MG:BA:1863 | 1.00 | 81.19 |  | 0 |  | No contacts in 5.0 A |
| MG:BA:1864 | 1.00 | 95.08 |  | 0 |  | 1^st^ MG coordination shell missing (see Notes) |
| MG:BA:1865 | 1.00 | 75.70 |  | 0 |  | 1^st^ MG coordination shell missing (see Notes) |
| MG:BA:1866 | 1.00 | 111.99 |  | 0 |  | 1^st^ MG coordination shell missing (see Notes) |
| MG:BA:1867 | 1.00 | 121.89 |  | 0 |  | 1^st^ MG coordination shell missing (see Notes) |
| MG:BA:1868 | 1.00 | 65.33 |  | 0 | 1 | - Exclusion zone contact(s): /O4'/ Consider NA in place of MG (1 valid contact(s) for NA vs. 0 for MG) |
| MG:BA:1869 | 1.00 | 73.50 |  | 0 |  | No contacts in 5.0 A |
| MG:BA:1870 | 1.00 | 119.77 |  | 0 |  | No contacts in 5.0 A |
| MG:BA:1871 | 1.00 | 74.42 |  | 0 |  | No contacts in 5.0 A |
| MG:BA:1872 | 1.00 | 103.39 |  | 0 | 1 | - Exclusion zone contact(s): /O5'/ Consider K in place of MG (1 valid contact(s) for K vs. 0 for MG) |
| MG:BA:1873 | 1.00 | 109.83 |  | 0 |  | No contacts in 5.0 A |
| MG:BA:1874 | 1.00 | 93.60 |  | 0 |  | No contacts in 5.0 A |
| MG:BA:1875 | 1.00 | 125.20 |  | 0 |  | No contacts in 5.0 A |
| MG:BA:1876 | 1.00 | 62.77 |  | 0 |  | No contacts in 5.0 A |
| MG:BA:1877 | 1.00 | 75.60 |  | 0 |  | 1^st^ MG coordination shell missing (see Notes) |
| MG:BA:1878 | 1.00 | 75.45 |  | 0 | 1 | - Exclusion zone contact(s): /O2'/ Consider NA in place of MG (1 valid contact(s) for NA vs. 0 for MG) |
| MG:BA:1879 | 1.00 | 46.29 |  | 0 |  | 1^st^ MG coordination shell missing (see Notes) |
| MG:BA:1880 | 1.00 | 60.98 |  | 3 |  | 2O_ph_O_b_ |
| MG:BA:1881 | 1.00 | 46.44 |  | 1 | 1 | O_ph_ - Exclusion zone contact(s): /OP2/ |
| MG:BA:1882 | 1.00 | 42.06 |  | 0 |  | 1^st^ MG coordination shell missing (see Notes) |
| MG:BA:1883 | 1.00 | 64.59 |  | 0 |  | 1^st^ MG coordination shell missing (see Notes) |
| MG:BA:1884 | 1.00 | 67.35 |  | 0 |  | 1^st^ MG coordination shell missing (see Notes) |
| MG:BA:1885 | 1.00 | 44.28 |  | 0 |  | 1^st^ MG coordination shell missing (see Notes) |
| MG:BA:1886 | 1.00 | 64.75 |  | 0 |  | 1^st^ MG coordination shell missing (see Notes) |
| MG:BA:1887 | 1.00 | 56.84 |  | 0 | 3 | - Exclusion zone contact(s): /O4/O6/O6/ Consider K in place of MG (2 valid contact(s) for K vs. 0 for MG) |
| MG:BA:1888 | 1.00 | 67.74 |  | 0 |  | 1^st^ MG coordination shell missing (see Notes) |
| MG:BA:1889 | 1.00 | 37.02 |  | 0 | 5 | - Exclusion zone contact(s): /N3_C_/N7/ND2_Asn_/O6/O_back_/ Consider K in place of MG (5 valid contact(s) for K vs. 0 for MG) |
| MG:BA:1890 | 1.00 | 72.40 |  | 0 |  | 1^st^ MG coordination shell missing (see Notes) |
| MG:BA:1891 | 1.00 | 60.06 |  | 1 |  | O_ph_ |
| MG:BA:1892 | 1.00 | 75.26 |  | 0 | 4 | - Exclusion zone contact(s): /N7/N7/O6/O6/ Consider K in place of MG (3 valid contact(s) for K vs. 0 for MG) |
| MG:BA:1893 | 1.00 | 54.06 |  | 0 | 6 | - Exclusion zone contact(s): /N7/N7/N7/O6/OP2/OP2/ Consider K in place of MG (6 valid contact(s) for K vs. 0 for MG) |
| MG:BA:1894 | 1.00 | 83.05 |  | 0 |  | 1^st^ MG coordination shell missing (see Notes) |
| MG:BA:1895 | 1.00 | 54.54 |  | 0 |  | 1^st^ MG coordination shell missing (see Notes) |
| MG:BA:1896 | 1.00 | 83.10 |  | 0 | 3 | - Exclusion zone contact(s): /N7/O6/OP1/ Consider NA in place of MG (2 valid contact(s) for NA vs. 0 for MG) |
| MG:BA:1897 | 1.00 | 65.25 |  | 0 |  | 1^st^ MG coordination shell missing (see Notes) |
| MG:BA:1898 | 1.00 | 49.78 |  | 0 | 1 | - Exclusion zone contact(s): /OP1/ Consider K in place of MG (1 valid contact(s) for K vs. 0 for MG) |
| MG:BA:1899 | 1.00 | 57.84 |  | 1 | 2 | O_r_ - Exclusion zone contact(s): /N6/O4'/ |
| MG:BA:1900 | 1.00 | 83.25 |  | 0 | 2 | - Exclusion zone contact(s): /OP1/OP1/ Consider NA in place of MG (2 valid contact(s) for NA vs. 0 for MG) |
| MG:BA:1901 | 1.00 | 82.56 |  | 1 | 3 | O_b_ - Exclusion zone contact(s): /N3_G_/N4/O2'/ Consider K in place of MG (2 valid contact(s) for K vs. 1 for MG) |
| MG:BA:1902 | 1.00 | 71.72 |  | 0 |  | 1^st^ MG coordination shell missing (see Notes) |
| MG:BA:1903 | 1.00 | 71.14 |  | 0 | 1 | - Exclusion zone contact(s): /O3'/ Consider K in place of MG (1 valid contact(s) for K vs. 0 for MG) |
| MG:BA:1904 | 1.00 | 38.67 |  | 0 | 1 | - Exclusion zone contact(s): /OP1/ Consider K in place of MG (1 valid contact(s) for K vs. 0 for MG) |
| MG:BA:1905 | 1.00 | 74.46 |  | 0 | 4 | - Exclusion zone contact(s): /O2'/O3'/OP1/OP2/ Consider K in place of MG (3 valid contact(s) for K vs. 0 for MG) |
| MG:BA:1906 | 1.00 | 107.70 |  | 0 | 3 | - Exclusion zone contact(s): /O2/O2'/OP1/ Consider K in place of MG (3 valid contact(s) for K vs. 0 for MG) |
| MG:BA:1907 | 1.00 | 56.42 |  | 0 | 1 | - Exclusion zone contact(s): /OP2/ Consider NA in place of MG (1 valid contact(s) for NA vs. 0 for MG) |
| MG:BA:1908 | 1.00 | 79.54 |  | 1 | 1 | O_ph_ - Exclusion zone contact(s): /OP1/ |
| MG:BA:1909 | 1.00 | 66.41 |  | 0 |  | 1^st^ MG coordination shell missing (see Notes) |
| MG:BA:1910 | 1.00 | 59.17 |  | 0 | 4 | - Exclusion zone contact(s): /N1_G_/N7/O6/ Consider K in place of MG (4 valid contact(s) for K vs. 0 for MG) |
| MG:BA:1911 | 1.00 | 76.75 |  | 0 |  | 1^st^ MG coordination shell missing (see Notes) |
| MG:BA:1912 | 1.00 | 82.35 |  | 0 | 1 | - Exclusion zone contact(s): /OP2/ Consider K in place of MG (1 valid contact(s) for K vs. 0 for MG) |
| MG:BA:1913 | 1.00 | 73.20 |  | 0 | 4 | - Exclusion zone contact(s): /OP1/OP1/OP1/OP2/ Consider K in place of MG (3 valid contact(s) for K vs. 0 for MG) |
| MG:BA:1914 | 1.00 | 55.54 |  | 0 | 2 | - Exclusion zone contact(s): /O3'/OP1/ Consider NA in place of MG (1 valid contact(s) for NA vs. 0 for MG) |
| MG:BA:1915 | 1.00 | 69.56 |  | 0 | 2 | - Exclusion zone contact(s): /N7/O6/ Consider K in place of MG (2 valid contact(s) for K vs. 0 for MG) |
| MG:BA:1916 | 1.00 | 56.47 |  | 0 | 3 | - Exclusion zone contact(s): /O2/O2/O2'/ Consider K in place of MG (3 valid contact(s) for K vs. 0 for MG) |
| MG:BA:1917 | 1.00 | 64.95 |  | 0 | 2 | - Exclusion zone contact(s): /OP1/OP2/ Consider K in place of MG (2 valid contact(s) for K vs. 0 for MG) |
| MG:BA:1918 | 1.00 | 74.66 |  | 1 | 3 | O_r_ - Exclusion zone contact(s): /O5'/OP2/OP2/ Consider K in place of MG (3 valid contact(s) for K vs. 1 for MG) |
| MG:BA:1919 | 1.00 | 59.17 |  | 0 | 1 | - Exclusion zone contact(s): /O2/ Consider K in place of MG (1 valid contact(s) for K vs. 0 for MG) |
| MG:BA:1920 | 1.00 | 86.85 |  | 0 | 1 | - Exclusion zone contact(s): /OP2/ Consider K in place of MG (1 valid contact(s) for K vs. 0 for MG) |
| MG:BA:1921 | 1.00 | 85.90 |  | 0 |  | 1^st^ MG coordination shell missing (see Notes) |
| MG:BA:1922 | 1.00 | 71.88 |  | 0 | 4 | - Exclusion zone contact(s): /N4/N7/O6/O6/ Consider K in place of MG (3 valid contact(s) for K vs. 0 for MG) |
| MG:BA:1923 | 1.00 | 81.36 |  | 0 | 1 | - Exclusion zone contact(s): /O2'/ Consider K in place of MG (1 valid contact(s) for K vs. 0 for MG) |
| MG:BA:1924 | 1.00 | 78.66 |  | 0 |  | 1^st^ MG coordination shell missing (see Notes) |
| MG:BA:1925 | 1.00 | 55.46 |  | 1 | 4 | O_ph_ - Exclusion zone contact(s): /N1_A_/N4/N6/OP1/ Consider K in place of MG (4 valid contact(s) for K vs. 1 for MG) |
| MG:BA:1926 | 1.00 | 53.35 |  | 0 | 4 | - Exclusion zone contact(s): /O4/O4/O6/O6/ Consider NA in place of MG (2 valid contact(s) for NA vs. 0 for MG) |
| MG:BA:1927 | 1.00 | 51.54 |  | 0 | 1 | - Exclusion zone contact(s): /OP2/ Consider K in place of MG (1 valid contact(s) for K vs. 0 for MG) |
| MG:BA:1928 | 1.00 | 67.31 |  | 0 | 3 | - Exclusion zone contact(s): /N7/O2'/O4/ Consider K in place of MG (3 valid contact(s) for K vs. 0 for MG) |
| MG:BA:1929 | 1.00 | 85.11 |  | 0 | 1 | - Exclusion zone contact(s): /O4'/ Consider K in place of MG (1 valid contact(s) for K vs. 0 for MG) |
| MG:BA:1930 | 1.00 | 62.22 |  | 0 | 1 | - Exclusion zone contact(s): /O6/ Consider K in place of MG (1 valid contact(s) for K vs. 0 for MG) |
| MG:BA:1931 | 1.00 | 68.36 |  | 0 | 2 | - Exclusion zone contact(s): /N7/O6/ Consider K in place of MG (2 valid contact(s) for K vs. 0 for MG) |
| MG:BA:1932 | 1.00 | 107.79 |  | 0 | 3 | - Exclusion zone contact(s): /N7/O6/OP2/ Consider K in place of MG (2 valid contact(s) for K vs. 0 for MG) |
| MG:BA:1933 | 1.00 | 63.78 |  | 0 | 1 | - Exclusion zone contact(s): /N1_A_/ Consider K in place of MG (1 valid contact(s) for K vs. 0 for MG) |
| MG:BA:1934 | 1.00 | 84.57 |  | 0 |  | 1^st^ MG coordination shell missing (see Notes) |
| MG:BA:1935 | 1.00 | 95.12 |  | 0 | 3 | - Exclusion zone contact(s): /N7/N7/OP2/ Consider K in place of MG (3 valid contact(s) for K vs. 0 for MG) |
| MG:BA:1936 | 1.00 | 80.71 |  | 0 |  | 1^st^ MG coordination shell missing (see Notes) |
| MG:BA:1937 | 1.00 | 71.93 |  | 0 |  | 1^st^ MG coordination shell missing (see Notes) |
| MG:BA:1938 | 1.00 | 69.22 |  | 0 | 2 | - Exclusion zone contact(s): /OP1/OP2/ Consider K in place of MG (2 valid contact(s) for K vs. 0 for MG) |
| MG:BA:1939 | 1.00 | 81.71 |  | 0 |  | 1^st^ MG coordination shell missing (see Notes) |
| MG:BA:1940 | 1.00 | 79.63 |  | 0 |  | 1^st^ MG coordination shell missing (see Notes) |
| MG:BA:1941 | 1.00 | 58.97 |  | 2 | 2 | O_ph_O_r_ - Exclusion zone contact(s): /OP1/OP2/ |
| MG:BA:1942 | 1.00 | 63.12 |  | 0 | 3 | - Exclusion zone contact(s): /N4/O6/OP2/ Consider K in place of MG (3 valid contact(s) for K vs. 0 for MG) |
| MG:BA:1943 | 1.00 | 72.21 |  | 0 | 2 | - Exclusion zone contact(s): /N1/OP1/ Consider NA in place of MG (1 valid contact(s) for NA vs. 0 for MG) |
| MG:BA:1944 | 1.00 | 71.33 |  | 0 |  | No contacts in 5.0 A |
| MG:BA:1945 | 1.00 | 56.27 |  | 1 |  | Anion |
| MG:BA:1946 | 1.00 | 70.04 |  | 0 | 3 | - Exclusion zone contact(s): /N4/N7/OP1/ Consider K in place of MG (3 valid contact(s) for K vs. 0 for MG) |
| MG:BA:1947 | 1.00 | 53.76 |  | 0 | 1 | - Exclusion zone contact(s): /O4/ Consider K in place of MG (1 valid contact(s) for K vs. 0 for MG) |
| MG:BA:1948 | 1.00 | 75.24 |  | 0 | 3 | - Exclusion zone contact(s): /O2/O3'/OP2/ Consider K in place of MG (3 valid contact(s) for K vs. 0 for MG) |
| MG:BA:1949 | 1.00 | 85.17 |  | 0 | 1 | - Exclusion zone contact(s): /O4/ Consider K in place of MG (1 valid contact(s) for K vs. 0 for MG) |
| MG:BA:1950 | 1.00 | 62.51 |  | 0 | 1 | - Exclusion zone contact(s): /OP2/ Consider K in place of MG (1 valid contact(s) for K vs. 0 for MG) |
| MG:BA:1951 | 1.00 | 78.48 |  | 0 |  | 1^st^ MG coordination shell missing (see Notes) |
| MG:BA:1952 | 1.00 | 70.97 |  | 0 | 1 | - Exclusion zone contact(s): /O6/ Consider K in place of MG (1 valid contact(s) for K vs. 0 for MG) |
| MG:BA:1953 | 1.00 | 68.09 |  | 0 | 6 | - Exclusion zone contact(s): /N7/O2'/O3'/O6/OP1/OP2/ Consider K in place of MG (5 valid contact(s) for K vs. 0 for MG) |
| MG:BA:1954 | 1.00 | 104.12 |  | 0 | 2 | - Exclusion zone contact(s): /O5'/OP2/ Consider NA in place of MG (1 valid contact(s) for NA vs. 0 for MG) |
| MG:BA:1955 | 1.00 | 74.28 |  | 0 |  | 1^st^ MG coordination shell missing (see Notes) |
| MG:BA:1956 | 1.00 | 111.93 |  | 0 |  | 1^st^ MG coordination shell missing (see Notes) |
| MG:BA:1957 | 1.00 | 80.65 |  | 0 | 1 | - Exclusion zone contact(s): /OP1/ Consider K in place of MG (1 valid contact(s) for K vs. 0 for MG) |
| MG:BA:1958 | 1.00 | 80.99 |  | 1 | 2 | O_b_ - Exclusion zone contact(s): /O2/OP1/ |
| MG:BA:1959 | 1.00 | 64.07 |  | 0 |  | 1^st^ MG coordination shell missing (see Notes) |
| MG:BA:1960 | 1.00 | 68.91 |  | 0 | 1 | - Exclusion zone contact(s): /OP2/ Consider K in place of MG (1 valid contact(s) for K vs. 0 for MG) |
| MG:BA:1961 | 1.00 | 49.79 |  | 1 | 5 | O_r_ - Exclusion zone contact(s): /N7/O4'/O5'/O6/OP2/ Consider K in place of MG (3 valid contact(s) for K vs. 1 for MG) |
| MG:BA:1962 | 1.00 | 73.16 |  | 0 |  | 1^st^ MG coordination shell missing (see Notes) |
| MG:BA:1963 | 1.00 | 83.04 |  | 1 |  | O_ph_ |
| MG:BA:1964 | 1.00 | 78.98 |  | 0 |  | 1^st^ MG coordination shell missing (see Notes) |
| MG:BA:1965 | 1.00 | 110.48 |  | 0 | 2 | - Exclusion zone contact(s): /O3'/OP2/ Consider NA in place of MG (1 valid contact(s) for NA vs. 0 for MG) |
| MG:BA:1966 | 1.00 | 82.84 |  | 0 | 3 | - Exclusion zone contact(s): /N7/O5'/OP2/ Consider NA in place of MG (2 valid contact(s) for NA vs. 0 for MG) |
| MG:BA:1967 | 1.00 | 77.15 |  | 0 | 1 | - Exclusion zone contact(s): /OP1/ Consider K in place of MG (1 valid contact(s) for K vs. 0 for MG) |
| MG:BA:1968 | 1.00 | 96.47 |  | 0 |  | 1^st^ MG coordination shell missing (see Notes) |
| MG:BA:1969 | 1.00 | 71.04 |  | 0 | 2 | - Exclusion zone contact(s): /O6/O6/ Consider K in place of MG (2 valid contact(s) for K vs. 0 for MG) |
| MG:BA:1970 | 1.00 | 141.60 |  | 2 | 1 | O_ph_O_b_ - Exclusion zone contact(s): /OP1/ |
| MG:BA:1971 | 1.00 | 98.52 |  | 0 | 2 | - Exclusion zone contact(s): /O6/O6/ Consider NA in place of MG (1 valid contact(s) for NA vs. 0 for MG) |
| MG:BA:1972 | 1.00 | 53.66 |  | 0 | 1 | - Exclusion zone contact(s): /O6/ Consider K in place of MG (1 valid contact(s) for K vs. 0 for MG) |
| MG:BA:1973 | 1.00 | 81.61 |  | 0 | 2 | - Exclusion zone contact(s): /N7/OP2/ Consider NA in place of MG (1 valid contact(s) for NA vs. 0 for MG) |
| MG:BA:1974 | 1.00 | 109.52 |  | 0 | 1 | - Exclusion zone contact(s): /O6/ Consider NA in place of MG (1 valid contact(s) for NA vs. 0 for MG) |
| MG:BA:1975 | 1.00 | 109.30 |  | 1 | 1 | O_b_ - Exclusion zone contact(s): /O2'/ |
| MG:BA:1976 | 1.00 | 147.65 |  | 0 | 3 | - Exclusion zone contact(s): /N7/N7/O6/ Consider NA in place of MG (2 valid contact(s) for NA vs. 0 for MG) |
| MG:BA:1977 | 1.00 | 85.53 |  | 0 | 4 | - Exclusion zone contact(s): /N3_G_/N6/O2'/O4'/ Consider K in place of MG (3 valid contact(s) for K vs. 0 for MG) |
| MG:BA:1978 | 1.00 | 73.00 |  | 0 |  | 1^st^ MG coordination shell missing (see Notes) |
| MG:BA:1979 | 1.00 | 73.52 |  | 1 |  | O_ph_ |
| MG:BA:1980 | 1.00 | 102.40 |  | 1 | 1 | N_b_ - Exclusion zone contact(s): /N7/ |
| MG:BA:1981 | 1.00 | 68.03 |  | 0 | 5 | - Exclusion zone contact(s): /N1/N2/N3_G_/O2'/O3'/ Consider K in place of MG (5 valid contact(s) for K vs. 0 for MG) |
| MG:BA:1982 | 1.00 | 96.72 |  | 0 | 3 | - Exclusion zone contact(s): /N7/O5'/OP2/ Consider K in place of MG (3 valid contact(s) for K vs. 0 for MG) |
| MG:BA:1983 | 1.00 | 80.96 |  | 0 | 1 | - Exclusion zone contact(s): /O5'/ Consider K in place of MG (1 valid contact(s) for K vs. 0 for MG) |
| MG:BA:1984 | 1.00 | 84.82 |  | 0 | 3 | - Exclusion zone contact(s): /N6/N7/N7/ Consider K in place of MG (3 valid contact(s) for K vs. 0 for MG) |
| MG:BA:1985 | 1.00 | 105.98 |  | 0 | 4 | - Exclusion zone contact(s): /N4/O6/O6/OP2/ Consider K in place of MG (3 valid contact(s) for K vs. 0 for MG) |
| MG:BA:1986 | 1.00 | 78.90 |  | 1 | 1 | O_ph_ - Exclusion zone contact(s): /OP2/ |
| MG:BA:1987 | 1.00 | 83.97 |  | 0 | 3 | - Exclusion zone contact(s): /O5'/OP1/OP2/ Consider K in place of MG (2 valid contact(s) for K vs. 0 for MG) |
| MG:BA:1988 | 1.00 | 95.40 |  | 0 | 2 | - Exclusion zone contact(s): /N4/N7/ Consider K in place of MG (2 valid contact(s) for K vs. 0 for MG) |
| MG:BA:1989 | 1.00 | 75.95 |  | 1 | 4 | O_r_ - Exclusion zone contact(s): /O2'/O3'/O5'/OP1/ Consider K in place of MG (3 valid contact(s) for K vs. 1 for MG) |
| MG:BA:1990 | 1.00 | 82.46 |  | 0 | 2 | - Exclusion zone contact(s): /O4/O6/ Consider K in place of MG (2 valid contact(s) for K vs. 0 for MG) |
| MG:BA:1991 | 1.00 | 91.45 |  | 0 | 1 | - Exclusion zone contact(s): /OP2/ Consider K in place of MG (1 valid contact(s) for K vs. 0 for MG) |
| MG:BA:1992 | 1.00 | 95.45 |  | 0 | 2 | - Exclusion zone contact(s): /O5'/OP2/ Consider K in place of MG (2 valid contact(s) for K vs. 0 for MG) |
| MG:BA:1993 | 1.00 | 60.60 |  | 0 | 4 | - Exclusion zone contact(s): /N1/N2/O4'/OP1/ Consider K in place of MG (4 valid contact(s) for K vs. 0 for MG) |
| MG:BA:1994 | 1.00 | 107.21 |  | 0 | 2 | - Exclusion zone contact(s): /O4/O6/ Consider NA in place of MG (1 valid contact(s) for NA vs. 0 for MG) |
| MG:BA:1995 | 1.00 | 84.40 |  | 0 | 1 | - Exclusion zone contact(s): /OP2/ Consider K in place of MG (1 valid contact(s) for K vs. 0 for MG) |
| MG:BA:1996 | 1.00 | 86.13 |  | 0 |  | 1^st^ MG coordination shell missing (see Notes) |
| MG:BA:1997 | 1.00 | 87.44 |  | 0 |  | 1^st^ MG coordination shell missing (see Notes) |
| MG:BA:1998 | 1.00 | 76.83 |  | 1 |  | O_b_ |
| MG:BA:1999 | 1.00 | 77.68 |  | 0 | 4 | - Exclusion zone contact(s): /O2/O4/O6/O6/ Consider K in place of MG (4 valid contact(s) for K vs. 0 for MG) |
| MG:BA:2000 | 1.00 | 93.39 |  | 0 | 1 | - Exclusion zone contact(s): /OP2/ Consider K in place of MG (1 valid contact(s) for K vs. 0 for MG) |
| MG:BA:2001 | 1.00 | 96.42 |  | 0 | 1 | - Exclusion zone contact(s): /OP2/ Consider K in place of MG (1 valid contact(s) for K vs. 0 for MG) |
| MG:BA:2002 | 1.00 | 73.79 |  | 0 | 1 | - Exclusion zone contact(s): /O3'/ Consider K in place of MG (1 valid contact(s) for K vs. 0 for MG) |
| MG:BA:2003 | 1.00 | 98.89 |  | 0 | 3 | - Exclusion zone contact(s): /O6/OP2/OP2/ Consider K in place of MG (3 valid contact(s) for K vs. 0 for MG) |
| MG:BA:2004 | 1.00 | 69.20 |  | 0 | 2 | - Exclusion zone contact(s): /O_back_/OP1/ Consider K in place of MG (2 valid contact(s) for K vs. 0 for MG) |
| MG:BA:2005 | 1.00 | 75.25 |  | 0 |  | 1^st^ MG coordination shell missing (see Notes) |
| MG:BA:2006 | 1.00 | 71.27 |  | 0 |  | 1^st^ MG coordination shell missing (see Notes) |
| MG:BA:2007 | 1.00 | 106.40 |  | 0 | 3 | - Exclusion zone contact(s): /N7/O4/O6/ Consider K in place of MG (2 valid contact(s) for K vs. 0 for MG) |
| MG:BA:2008 | 1.00 | 50.69 |  | 1 | 4 | O_ph_ - Exclusion zone contact(s): /O2'/O3'/OP2/OP2/ Consider K in place of MG (3 valid contact(s) for K vs. 1 for MG) |
| MG:BA:2009 | 1.00 | 63.11 |  | 0 | 3 | - Exclusion zone contact(s): /N7/O6/OP1/ Consider NA in place of MG (2 valid contact(s) for NA vs. 0 for MG) |
| MG:BA:2010 | 1.00 | 61.70 |  | 0 | 2 | - Exclusion zone contact(s): /O2'/OP1/ Consider K in place of MG (2 valid contact(s) for K vs. 0 for MG) |
| MG:BA:2011 | 1.00 | 64.02 |  | 0 |  | 1^st^ MG coordination shell missing (see Notes) |
| MG:BA:2012 | 1.00 | 72.35 |  | 0 |  | 1^st^ MG coordination shell missing (see Notes) |
| MG:BA:2013 | 1.00 | 67.65 |  | 1 | 2 | O_ph_ - Exclusion zone contact(s): /OP1/OP2/ |
| MG:BA:2014 | 1.00 | 64.40 |  | 0 | 4 | - Exclusion zone contact(s): /N6/N7/N7/OP2/ Consider K in place of MG (4 valid contact(s) for K vs. 0 for MG) |
| MG:BA:2015 | 1.00 | 71.08 |  | 0 | 6 | - Exclusion zone contact(s): /N3_C_/NE_Arg_/NH2_Arg_/O2/O2'/O4'/ Consider K in place of MG (5 valid contact(s) for K vs. 0 for MG) |
| MG:BA:2016 | 1.00 | 78.44 |  | 0 | 2 | - Exclusion zone contact(s): /OP1/OP2/ Consider K in place of MG (2 valid contact(s) for K vs. 0 for MG) |
| MG:BA:2017 | 1.00 | 85.84 |  | 0 | 1 | - Exclusion zone contact(s): /OP2/ Consider K in place of MG (1 valid contact(s) for K vs. 0 for MG) |
| MG:BA:2018 | 1.00 | 84.02 |  | 0 | 2 | - Exclusion zone contact(s): /OP1/OP1/ Consider K in place of MG (2 valid contact(s) for K vs. 0 for MG) |
| MG:BA:2019 | 1.00 | 80.65 |  | 0 |  | 1^st^ MG coordination shell missing (see Notes) |
| MG:BA:2020 | 1.00 | 90.58 |  | 0 | 2 | - Exclusion zone contact(s): /OG_Ser_/OP2/ Consider K in place of MG (2 valid contact(s) for K vs. 0 for MG) |
| MG:BA:2021 | 1.00 | 84.39 |  | 1 | 2 | O_b_ - Exclusion zone contact(s): /O6/O6/ Consider K in place of MG (2 valid contact(s) for K vs. 1 for MG) |
| MG:BA:2022 | 1.00 | 104.00 |  | 0 | 3 | - Exclusion zone contact(s): /O6/OP2/OP2/ Consider K in place of MG (2 valid contact(s) for K vs. 0 for MG) |
| MG:BA:2023 | 1.00 | 67.90 |  | 0 |  | 1^st^ MG coordination shell missing (see Notes) |
| MG:BA:2024 | 1.00 | 111.49 |  | 0 | 4 | - Exclusion zone contact(s): /N3_G_/O2'/O4'/ Consider NA in place of MG (2 valid contact(s) for NA vs. 0 for MG) |
| MG:BA:2025 | 1.00 | 90.82 |  | 0 | 1 | - Exclusion zone contact(s): /OP1/ Consider K in place of MG (1 valid contact(s) for K vs. 0 for MG) |
| MG:BA:2026 | 1.00 | 128.76 |  | 0 | 4 | - Exclusion zone contact(s): /N6/N7/N7/O6/ Consider K in place of MG (4 valid contact(s) for K vs. 0 for MG) |
| MG:BA:2027 | 1.00 | 107.37 |  | 0 |  | 1^st^ MG coordination shell missing (see Notes) |
| MG:BA:2028 | 1.00 | 77.13 |  | 1 | 3 | O_b_ - Exclusion zone contact(s): /N1_G_/N3_G_/O2'/ Consider K in place of MG (3 valid contact(s) for K vs. 1 for MG) |
| MG:BA:2029 | 1.00 | 76.24 |  | 0 | 1 | - Exclusion zone contact(s): /O4/ Consider K in place of MG (1 valid contact(s) for K vs. 0 for MG) |
| MG:BA:2030 | 1.00 | 71.76 |  | 0 |  | 1^st^ MG coordination shell missing (see Notes) |
| MG:BA:2031 | 1.00 | 71.49 |  | 2 | 2 | 2O_ph_ - Exclusion zone contact(s): /O2'/ |
| MG:BA:2032 | 1.00 | 70.97 |  | 0 |  | 1^st^ MG coordination shell missing (see Notes) |
| MG:BA:2033 | 1.00 | 94.76 |  | 0 | 1 | - Exclusion zone contact(s): /OP2/ Consider K in place of MG (1 valid contact(s) for K vs. 0 for MG) |
| MG:BA:2034 | 1.00 | 68.55 |  | 0 |  | 1^st^ MG coordination shell missing (see Notes) |
| MG:BA:2035 | 1.00 | 91.48 |  | 1 | 2 | O_ph_ - Exclusion zone contact(s): /O5'/OP2/ Consider K in place of MG (2 valid contact(s) for K vs. 1 for MG) |
| MG:BA:2036 | 1.00 | 86.81 |  | 0 | 2 | - Exclusion zone contact(s): /O6/OP1/ Consider NA in place of MG (1 valid contact(s) for NA vs. 0 for MG) |
| MG:BA:2037 | 1.00 | 132.50 |  | 1 | 3 | O_b_ - Exclusion zone contact(s): /N3_U_/O2/O2'/ Consider K in place of MG (2 valid contact(s) for K vs. 1 for MG) |
| MG:BA:2038 | 1.00 | 80.17 |  | 0 |  | 1^st^ MG coordination shell missing (see Notes) |
| MG:BA:2039 | 1.00 | 138.77 |  | 0 | 2 | - Exclusion zone contact(s): /OP1/OP2/ Consider NA in place of MG (1 valid contact(s) for NA vs. 0 for MG) |
| MG:BA:2040 | 1.00 | 93.57 |  | 1 |  | O_b_ |
| MG:BA:2041 | 1.00 | 91.83 |  | 0 |  | 1^st^ MG coordination shell missing (see Notes) |
| MG:BA:2042 | 1.00 | 95.96 |  | 0 | 6 | - Exclusion zone contact(s): /N4/O2/O2'/O4/O4'/OP1/ Consider K in place of MG (5 valid contact(s) for K vs. 0 for MG) |
| MG:BA:2043 | 1.00 | 109.65 |  | 0 | 2 | - Exclusion zone contact(s): /N6/O2/ Consider NA in place of MG (1 valid contact(s) for NA vs. 0 for MG) |
| MG:BA:2044 | 1.00 | 62.41 |  | 0 | 3 | - Exclusion zone contact(s): /N3_C_/O5'/OP1/ Consider K in place of MG (3 valid contact(s) for K vs. 0 for MG) |
| MG:BA:2045 | 1.00 | 66.61 |  | 0 | 2 | - Exclusion zone contact(s): /N1/N4/ Consider K in place of MG (2 valid contact(s) for K vs. 0 for MG) |
| MG:BA:2046 | 1.00 | 95.23 |  | 0 | 1 | - Exclusion zone contact(s): /N7/ Consider NA in place of MG (1 valid contact(s) for NA vs. 0 for MG) |
| MG:BA:2047 | 1.00 | 107.38 |  | 0 | 2 | - Exclusion zone contact(s): /OP1/OP2/ Consider NA in place of MG (1 valid contact(s) for NA vs. 0 for MG) |
| MG:BA:2048 | 1.00 | 95.18 |  | 0 | 3 | - Exclusion zone contact(s): /O3'/OP1/OP1/ Consider K in place of MG (3 valid contact(s) for K vs. 0 for MG) |
| MG:BA:2049 | 1.00 | 55.18 |  | 0 |  | 1^st^ MG coordination shell missing (see Notes) |
| MG:BA:2050 | 1.00 | 85.65 |  | 0 | 3 | - Exclusion zone contact(s): /N3_G_/O2'/O4'/ Consider K in place of MG (2 valid contact(s) for K vs. 0 for MG) |
| MG:BA:2051 | 1.00 | 108.09 |  | 0 | 1 | - Exclusion zone contact(s): /N7/ Consider K in place of MG (1 valid contact(s) for K vs. 0 for MG) |
| MG:BA:2052 | 1.00 | 128.97 |  | 0 | 3 | - Exclusion zone contact(s): /N4/O6/OP2/ Consider K in place of MG (3 valid contact(s) for K vs. 0 for MG) |
| MG:BA:2053 | 1.00 | 90.41 |  | 0 | 2 | - Exclusion zone contact(s): /O6/OP2/ Consider NA in place of MG (1 valid contact(s) for NA vs. 0 for MG) |
| MG:BA:2054 | 1.00 | 68.71 |  | 0 |  | No contacts in 5.0 A |
| MG:BA:2055 | 1.00 | 89.56 |  | 3 | 1 | 2O_ph_O_b_ - Exclusion zone contact(s): /N3_C_/ |
| MG:BA:2056 | 1.00 | 69.44 |  | 0 |  | 1^st^ MG coordination shell missing (see Notes) |
| MG:BA:2057 | 1.00 | 103.47 |  | 1 | 2 | N_b_ - Exclusion zone contact(s): /O5'/OP2/ Consider K in place of MG (2 valid contact(s) for K vs. 1 for MG) |
| MG:BA:2058 | 1.00 | 104.40 |  | 0 |  | No contacts in 5.0 A |
| MG:BA:2059 | 1.00 | 67.99 |  | 0 |  | 1^st^ MG coordination shell missing (see Notes) |
| MG:BA:2060 | 1.00 | 73.86 |  | 0 |  | 1^st^ MG coordination shell missing (see Notes) |
| MG:BA:2061 | 1.00 | 70.63 |  | 0 |  | 1^st^ MG coordination shell missing (see Notes) |
| MG:BA:2062 | 1.00 | 75.60 |  | 0 | 1 | - Exclusion zone contact(s): /O6/ Consider K in place of MG (1 valid contact(s) for K vs. 0 for MG) |
| MG:BA:2063 | 1.00 | 65.70 |  | 1 |  | O_ph_ |
| MG:BA:2064 | 1.00 | 90.66 |  | 0 | 4 | - Exclusion zone contact(s): /N2/N3_G_/O2'/OP1/ Consider K in place of MG (4 valid contact(s) for K vs. 0 for MG) |
| MG:BA:2065 | 1.00 | 73.30 |  | 0 | 1 | - Exclusion zone contact(s): /O5'/ Consider K in place of MG (1 valid contact(s) for K vs. 0 for MG) |
| MG:BA:2066 | 1.00 | 105.28 |  | 0 |  | 1^st^ MG coordination shell missing (see Notes) |
| MG:BA:2067 | 1.00 | 95.64 |  | 0 | 1 | - Exclusion zone contact(s): /N4/ Consider K in place of MG (1 valid contact(s) for K vs. 0 for MG) |
| MG:BA:2068 | 1.00 | 90.02 |  | 0 |  | 1^st^ MG coordination shell missing (see Notes) |
| MG:BA:2069 | 1.00 | 105.04 |  | 0 |  | No contacts in 5.0 A |
| MG:BA:2070 | 1.00 | 89.01 |  | 0 | 4 | - Exclusion zone contact(s): /N4/O4/O6/O6/ Consider K in place of MG (3 valid contact(s) for K vs. 0 for MG) |
| MG:BA:2071 | 1.00 | 93.74 |  | 0 | 2 | - Exclusion zone contact(s): /O4'/O5'/ Consider K in place of MG (2 valid contact(s) for K vs. 0 for MG) |
| MG:BA:2072 | 1.00 | 72.65 |  | 1 | 1 | O_ph_ - Exclusion zone contact(s): /O2'/ |
| MG:BA:2073 | 1.00 | 81.13 |  | 0 | 1 | - Exclusion zone contact(s): /O2'/ Consider NA in place of MG (1 valid contact(s) for NA vs. 0 for MG) |
| MG:BA:2074 | 1.00 | 77.82 |  | 0 |  | 1^st^ MG coordination shell missing (see Notes) |
| MG:BA:2075 | 1.00 | 101.11 |  | 0 |  | 1^st^ MG coordination shell missing (see Notes) |
| MG:BA:2076 | 1.00 | 85.24 |  | 1 | 4 | O_b_ - Exclusion zone contact(s): /N1_G_/O4/O6/OP2/ Consider K in place of MG (3 valid contact(s) for K vs. 1 for MG) |
| MG:BA:2077 | 1.00 | 74.88 |  | 0 | 1 | - Exclusion zone contact(s): /O4/ Consider K in place of MG (1 valid contact(s) for K vs. 0 for MG) |
| MG:BA:2078 | 1.00 | 79.30 |  | 0 | 2 | - Exclusion zone contact(s): /OP1/OP2/ Consider K in place of MG (2 valid contact(s) for K vs. 0 for MG) |
| MG:BA:2079 | 1.00 | 99.78 |  | 0 | 2 | - Exclusion zone contact(s): /N1/O4'/ Consider NA in place of MG (1 valid contact(s) for NA vs. 0 for MG) |
| MG:BA:2080 | 1.00 | 61.44 |  | 0 | 3 | - Exclusion zone contact(s): /OP1/OP1/OP2/ Consider K in place of MG (3 valid contact(s) for K vs. 0 for MG) |
| MG:BA:2081 | 1.00 | 69.94 |  | 0 | 1 | - Exclusion zone contact(s): /O6/ Consider K in place of MG (1 valid contact(s) for K vs. 0 for MG) |
| MG:BA:2082 | 1.00 | 96.82 |  | 0 | 2 | - Exclusion zone contact(s): /N1_A_/O2/ Consider K in place of MG (2 valid contact(s) for K vs. 0 for MG) |
| MG:BA:2083 | 1.00 | 86.51 |  | 1 | 2 | O_r_ - Exclusion zone contact(s): /N2/ Consider K in place of MG (2 valid contact(s) for K vs. 1 for MG) |
| MG:BA:2084 | 1.00 | 90.01 |  | 1 |  | O_ph_ |
| MG:BA:2085 | 1.00 | 102.57 |  | 0 |  | 1^st^ MG coordination shell missing (see Notes) |
| MG:BA:2086 | 1.00 | 83.42 |  | 1 |  | O_ph_ |
| MG:BA:2087 | 1.00 | 81.46 |  | 0 | 3 | - Exclusion zone contact(s): /O2'/O3'/OP2/ Consider K in place of MG (3 valid contact(s) for K vs. 0 for MG) |
| MG:BA:2088 | 1.00 | 98.45 |  | 0 | 1 | - Exclusion zone contact(s): /O5'/ Consider K in place of MG (1 valid contact(s) for K vs. 0 for MG) |
| MG:BA:2089 | 1.00 | 56.69 |  | 0 | 2 | - Exclusion zone contact(s): /O2'/O3'/ Consider K in place of MG (2 valid contact(s) for K vs. 0 for MG) |
| MG:BA:2090 | 1.00 | 119.46 |  | 0 | 4 | - Exclusion zone contact(s): /N7/N7/O6/O6/ Consider K in place of MG (3 valid contact(s) for K vs. 0 for MG) |
| MG:BA:2091 | 1.00 | 108.11 |  | 0 | 2 | - Exclusion zone contact(s): /N2/O2'/ Consider K in place of MG (2 valid contact(s) for K vs. 0 for MG) |
| MG:BA:2092 | 1.00 | 77.96 |  | 0 | 4 | - Exclusion zone contact(s): /N1_G_/O4/O6/O6/ Consider K in place of MG (4 valid contact(s) for K vs. 0 for MG) |
| MG:BA:2093 | 1.00 | 106.93 |  | 0 | 2 | - Exclusion zone contact(s): /O6/OP1/ Consider NA in place of MG (1 valid contact(s) for NA vs. 0 for MG) |
| MG:BA:2094 | 1.00 | 50.68 |  | 0 | 1 | - Exclusion zone contact(s): /OP2/ Consider NA in place of MG (1 valid contact(s) for NA vs. 0 for MG) |
| MG:BA:2095 | 1.00 | 67.50 |  | 0 |  | 1^st^ MG coordination shell missing (see Notes) |
| MG:BA:2096 | 1.00 | 51.57 |  | 0 | 2 | - Exclusion zone contact(s): /N2/O4'/ Consider K in place of MG (2 valid contact(s) for K vs. 0 for MG) |
| MG:BA:2097 | 1.00 | 94.01 |  | 0 | 2 | - Exclusion zone contact(s): /N6/N7/ Consider K in place of MG (2 valid contact(s) for K vs. 0 for MG) |
| MG:BA:2098 | 1.00 | 73.44 |  | 0 |  | 1^st^ MG coordination shell missing (see Notes) |
| MG:BA:2099 | 1.00 | 67.41 |  | 0 | 3 | - Exclusion zone contact(s): /N2/N7/O5'/ Consider K in place of MG (3 valid contact(s) for K vs. 0 for MG) |
| MG:BA:2100 | 1.00 | 69.32 |  | 0 |  | No contacts in 5.0 A |
| MG:BA:2101 | 1.00 | 124.56 |  | 0 | 3 | - Exclusion zone contact(s): /NH2_Arg_/OP2/OP2/ Consider K in place of MG (3 valid contact(s) for K vs. 0 for MG) |
| MG:BA:2102 | 1.00 | 85.69 |  | 0 | 2 | - Exclusion zone contact(s): /OP1/OP2/ Consider K in place of MG (2 valid contact(s) for K vs. 0 for MG) |
| MG:BA:2103 | 1.00 | 80.19 |  | 0 | 2 | - Exclusion zone contact(s): /OP1/OP2/ Consider NA in place of MG (1 valid contact(s) for NA vs. 0 for MG) |
| MG:BA:2104 | 1.00 | 80.42 |  | 0 |  | 1^st^ MG coordination shell missing (see Notes) |
| MG:BA:2105 | 1.00 | 117.04 |  | 0 | 3 | - Exclusion zone contact(s): /O2'/O3'/OP2/ Consider K in place of MG (2 valid contact(s) for K vs. 0 for MG) |
| MG:BA:2106 | 1.00 | 104.57 |  | 0 | 1 | - Exclusion zone contact(s): /OP1/ Consider K in place of MG (1 valid contact(s) for K vs. 0 for MG) |
| MG:BA:2107 | 1.00 | 83.21 |  | 0 | 1 | - Exclusion zone contact(s): /N4/ Consider K in place of MG (1 valid contact(s) for K vs. 0 for MG) |
| MG:BA:2108 | 1.00 | 154.40 |  | 0 | 7 | - Exclusion zone contact(s): /N1_G_/O4'/O5'/O5'/O6/OP1/ Consider K in place of MG (7 valid contact(s) for K vs. 0 for MG) |
| MG:BA:2109 | 1.00 | 93.88 |  | 0 | 4 | - Exclusion zone contact(s): /N3_A_/N6/O4'/ Consider K in place of MG (4 valid contact(s) for K vs. 0 for MG) |
| MG:BA:2110 | 1.00 | 91.13 |  | 0 |  | No contacts in 5.0 A |
| MG:BA:2111 | 1.00 | 79.36 |  | 0 | 4 | - Exclusion zone contact(s): /O2'/O3'/OP1/OP2/ Consider K in place of MG (4 valid contact(s) for K vs. 0 for MG) |
| MG:BA:2112 | 1.00 | 101.06 |  | 0 | 1 | - Exclusion zone contact(s): /OP1/ Consider K in place of MG (1 valid contact(s) for K vs. 0 for MG) |
| MG:BA:2113 | 1.00 | 92.92 |  | 0 | 1 | - Exclusion zone contact(s): /OP2/ Consider K in place of MG (1 valid contact(s) for K vs. 0 for MG) |
| MG:BA:2114 | 1.00 | 71.95 |  | 1 | 5 | N_b_ - Exclusion zone contact(s): /N1_G_/N2/O6/OP2/ Consider K in place of MG (5 valid contact(s) for K vs. 1 for MG) |
| MG:BA:2115 | 1.00 | 118.70 |  | 0 | 2 | - Exclusion zone contact(s): /OP1/OP2/ Consider K in place of MG (2 valid contact(s) for K vs. 0 for MG) |
| MG:BA:2116 | 1.00 | 54.85 |  | 0 | 3 | - Exclusion zone contact(s): /O4/O4'/OP1/ Consider K in place of MG (3 valid contact(s) for K vs. 0 for MG) |
| MG:BA:2117 | 1.00 | 109.31 |  | 0 |  | 1^st^ MG coordination shell missing (see Notes) |
| MG:BA:2118 | 1.00 | 83.42 |  | 0 |  | 1^st^ MG coordination shell missing (see Notes) |
| MG:BA:2119 | 1.00 | 67.83 |  | 0 |  | 1^st^ MG coordination shell missing (see Notes) |
| MG:BA:2120 | 1.00 | 67.00 |  | 0 | 3 | - Exclusion zone contact(s): /NZ_Lys_/O2'/O4'/ Consider K in place of MG (3 valid contact(s) for K vs. 0 for MG) |
| MG:BA:2121 | 1.00 | 84.64 |  | 0 |  | No contacts in 5.0 A |
| MG:BA:2122 | 1.00 | 107.30 |  | 0 | 1 | Consider K in place of MG (1 valid contact(s) for K vs. 0 for MG) |
| MG:BA:2123 | 1.00 | 90.68 |  | 2 | 2 | O_ph_O_r_ - Exclusion zone contact(s): /N4/O2'/ |
| MG:BA:2124 | 1.00 | 72.85 |  | 0 | 2 | - Exclusion zone contact(s): /O5'/OP2/ Consider K in place of MG (2 valid contact(s) for K vs. 0 for MG) |
| MG:BA:2125 | 1.00 | 65.68 |  | 0 | 3 | - Exclusion zone contact(s): /O2'/O3'/O5'/ Consider NA in place of MG (2 valid contact(s) for NA vs. 0 for MG) |
| MG:BA:2126 | 1.00 | 74.99 |  | 0 | 1 | - Exclusion zone contact(s): /N1/ Consider K in place of MG (1 valid contact(s) for K vs. 0 for MG) |
| MG:BA:2127 | 1.00 | 50.63 |  | 2 | 2 | O_b_O_back_ - Exclusion zone contact(s): /N3_C_/O6/ |
| MG:BA:2128 | 1.00 | 108.25 |  | 0 | 2 | - Exclusion zone contact(s): /N1_G_/O6/ Consider K in place of MG (2 valid contact(s) for K vs. 0 for MG) |
| MG:BA:2129 | 1.00 | 75.18 |  | 0 | 3 | - Exclusion zone contact(s): /O2'/O3'/OP2/ Consider NA in place of MG (2 valid contact(s) for NA vs. 0 for MG) |
| MG:BA:2130 | 1.00 | 86.43 |  | 0 | 2 | - Exclusion zone contact(s): /NH2_Arg_/OP2/ Consider K in place of MG (2 valid contact(s) for K vs. 0 for MG) |
| MG:BA:2131 | 1.00 | 61.05 |  | 0 | 1 | - Exclusion zone contact(s): /O6/ Consider K in place of MG (1 valid contact(s) for K vs. 0 for MG) |
| MG:BA:2132 | 1.00 | 87.18 |  | 0 | 5 | - Exclusion zone contact(s): /O3'/O3'/O5'/OP1/OP2/ Consider K in place of MG (4 valid contact(s) for K vs. 0 for MG) |
| MG:BA:2133 | 1.00 | 122.49 |  | 2 | 3 | O_r_O_b_ - Exclusion zone contact(s): /N7/OP2/OP2/ Consider K in place of MG (3 valid contact(s) for K vs. 2 for MG) |
| MG:BA:2134 | 1.00 | 62.51 |  | 0 | 2 | - Exclusion zone contact(s): /N7/OP2/ Consider K in place of MG (2 valid contact(s) for K vs. 0 for MG) |
| MG:BA:2135 | 1.00 | 77.05 |  | 0 | 3 | - Exclusion zone contact(s): /O5'/OP2/OP2/ Consider K in place of MG (3 valid contact(s) for K vs. 0 for MG) |
| MG:BA:2136 | 1.00 | 117.37 |  | 0 | 2 | - Exclusion zone contact(s): /N4/N7/ Consider NA in place of MG (1 valid contact(s) for NA vs. 0 for MG) |
| MG:BA:2137 | 1.00 | 119.48 |  | 0 | 2 | - Exclusion zone contact(s): /N1_G_/N2/ Consider NA in place of MG (1 valid contact(s) for NA vs. 0 for MG) |
| MG:BA:2138 | 1.00 | 74.87 |  | 0 | 2 | - Exclusion zone contact(s): /N4/O6/ Consider K in place of MG (2 valid contact(s) for K vs. 0 for MG) |
| MG:BA:2139 | 1.00 | 82.15 |  | 0 | 1 | - Exclusion zone contact(s): /OP2/ Consider K in place of MG (1 valid contact(s) for K vs. 0 for MG) |
| MG:BA:2140 | 1.00 | 103.45 |  | 0 |  | 1^st^ MG coordination shell missing (see Notes) |
| MG:BA:2141 | 1.00 | 80.58 |  | 0 |  | 1^st^ MG coordination shell missing (see Notes) |
| MG:BA:2142 | 1.00 | 84.45 |  | 0 | 6 | - Exclusion zone contact(s): /N3_A_/O2'/O2'/O3'/O4'/ Consider K in place of MG (5 valid contact(s) for K vs. 0 for MG) |
| MG:BA:2143 | 1.00 | 138.31 |  | 0 | 5 | - Exclusion zone contact(s): /O2'/O3'/O5'/O5'/OP2/ Consider K in place of MG (4 valid contact(s) for K vs. 0 for MG) |
| MG:BA:2144 | 1.00 | 75.19 |  | 0 |  | 1^st^ MG coordination shell missing (see Notes) |
| MG:BA:2145 | 1.00 | 96.83 |  | 0 | 2 | - Exclusion zone contact(s): /O3'/OP1/ Consider NA in place of MG (1 valid contact(s) for NA vs. 0 for MG) |
| MG:BA:2146 | 1.00 | 81.78 |  | 0 |  | 1^st^ MG coordination shell missing (see Notes) |
| MG:BA:2147 | 1.00 | 107.13 |  | 0 | 1 | - Exclusion zone contact(s): /O6/ Consider K in place of MG (1 valid contact(s) for K vs. 0 for MG) |
| MG:BA:2148 | 1.00 | 110.86 |  | 0 | 2 | - Exclusion zone contact(s): /O2'/O5'/ Consider K in place of MG (2 valid contact(s) for K vs. 0 for MG) |
| MG:BA:2149 | 1.00 | 82.00 |  | 0 | 4 | - Exclusion zone contact(s): /N7/O6/OP2/OP2/ Consider K in place of MG (4 valid contact(s) for K vs. 0 for MG) |
| MG:BA:2150 | 1.00 | 96.84 |  | 0 | 1 | - Exclusion zone contact(s): /N4/ Consider K in place of MG (1 valid contact(s) for K vs. 0 for MG) |
| MG:BA:2151 | 1.00 | 81.70 |  | 0 | 2 | - Exclusion zone contact(s): /O2/OP2/ Consider K in place of MG (2 valid contact(s) for K vs. 0 for MG) |
| MG:BA:2152 | 1.00 | 83.90 |  | 0 | 1 | - Exclusion zone contact(s): /N1_G_/ Consider K in place of MG (1 valid contact(s) for K vs. 0 for MG) |
| MG:BA:2153 | 1.00 | 100.94 |  | 0 | 4 | - Exclusion zone contact(s): /N2/NH2_Arg_/O6/OP2/ Consider K in place of MG (3 valid contact(s) for K vs. 0 for MG) |
| MG:BA:2154 | 1.00 | 73.23 |  | 0 | 2 | - Exclusion zone contact(s): /O6/OP2/ Consider K in place of MG (2 valid contact(s) for K vs. 0 for MG) |
| MG:BA:2155 | 1.00 | 71.78 |  | 0 | 2 | - Exclusion zone contact(s): /OP1/OP2/ Consider NA in place of MG (1 valid contact(s) for NA vs. 0 for MG) |
| MG:BA:2156 | 1.00 | 90.51 |  | 1 |  | O_ph_ |
| MG:BA:2157 | 1.00 | 72.10 |  | 0 | 4 | - Exclusion zone contact(s): /O2'/O3'/O4/OP1/ Consider K in place of MG (4 valid contact(s) for K vs. 0 for MG) |
| MG:BA:2158 | 1.00 | 77.58 |  | 0 |  | 1^st^ MG coordination shell missing (see Notes) |
| MG:BA:2159 | 1.00 | 89.26 |  | 0 | 1 | - Exclusion zone contact(s): /OP1/ Consider K in place of MG (1 valid contact(s) for K vs. 0 for MG) |
| MG:BA:2160 | 1.00 | 95.82 |  | 0 | 3 | - Exclusion zone contact(s): /N2/N3_G_/O2'/ Consider K in place of MG (3 valid contact(s) for K vs. 0 for MG) |
| MG:BA:2161 | 1.00 | 114.71 |  | 0 | 1 | - Exclusion zone contact(s): /N2/ Consider NA in place of MG (1 valid contact(s) for NA vs. 0 for MG) |
| MG:BA:2162 | 1.00 | 115.78 |  | 0 | 3 | - Exclusion zone contact(s): /O3'/O3'/OP1/ Consider K in place of MG (2 valid contact(s) for K vs. 0 for MG) |
| MG:BA:2163 | 1.00 | 105.42 |  | 0 | 2 | - Exclusion zone contact(s): /O2'/O5'/ Consider NA in place of MG (1 valid contact(s) for NA vs. 0 for MG) |
| MG:BA:2164 | 1.00 | 78.52 |  | 1 | 2 | O_ph_ - Exclusion zone contact(s): /OP1/OP2/ |
| MG:BA:2165 | 1.00 | 61.57 |  | 0 |  | 1^st^ MG coordination shell missing (see Notes) |
| MG:BA:2166 | 1.00 | 96.47 |  | 0 | 3 | - Exclusion zone contact(s): /N3_A_/O2'/O4'/ Consider K in place of MG (2 valid contact(s) for K vs. 0 for MG) |
| MG:BA:2167 | 1.00 | 101.31 |  | 1 | 1 | O_ph_ - Exclusion zone contact(s): /OP1/ |
| MG:BA:2168 | 1.00 | 73.90 |  | 0 | 2 | - Exclusion zone contact(s): /N3_G_/ Consider NA in place of MG (1 valid contact(s) for NA vs. 0 for MG) |
| MG:BA:2169 | 1.00 | 98.17 |  | 0 | 2 | - Exclusion zone contact(s): /N4/O6/ Consider K in place of MG (2 valid contact(s) for K vs. 0 for MG) |
| MG:BA:2170 | 1.00 | 151.16 |  | 1 | 2 | O_ph_ - Exclusion zone contact(s): /N2/OP2/ |
| MG:BA:2171 | 1.00 | 132.44 |  | 1 |  | O_ph_ |
| MG:BA:2172 | 1.00 | 75.89 |  | 0 | 1 | - Exclusion zone contact(s): /O4/ Consider K in place of MG (1 valid contact(s) for K vs. 0 for MG) |
| MG:BA:2173 | 1.00 | 98.85 |  | 1 | 1 | O_ph_ - Exclusion zone contact(s): /N7/ |
| MG:BA:2174 | 1.00 | 125.96 |  | 0 |  | 1^st^ MG coordination shell missing (see Notes) |
| MG:BA:2175 | 1.00 | 81.49 |  | 0 | 1 | - Exclusion zone contact(s): /OP2/ Consider K in place of MG (1 valid contact(s) for K vs. 0 for MG) |
| MG:BA:2176 | 1.00 | 99.22 |  | 0 | 1 | - Exclusion zone contact(s): /O4/ Consider K in place of MG (1 valid contact(s) for K vs. 0 for MG) |
| MG:BA:2177 | 1.00 | 67.61 |  | 0 |  | 1^st^ MG coordination shell missing (see Notes) |
| MG:BA:2178 | 1.00 | 141.45 |  | 1 |  | N_b_ |
| MG:BA:2179 | 1.00 | 101.40 |  | 0 |  | No contacts in 5.0 A |
| MG:BA:2180 | 1.00 | 88.73 |  | 0 | 2 | - Exclusion zone contact(s): /O2/O2'/ Consider K in place of MG (2 valid contact(s) for K vs. 0 for MG) |
| MG:BA:2181 | 1.00 | 93.47 |  | 0 | 4 | - Exclusion zone contact(s): /NZ_Lys_/O5'/OP2/OP2/ Consider NA in place of MG (2 valid contact(s) for NA vs. 0 for MG) |
| MG:BA:2182 | 1.00 | 110.66 |  | 0 | 3 | - Exclusion zone contact(s): /N2/N4/OP2/ Consider K in place of MG (3 valid contact(s) for K vs. 0 for MG) |
| MG:BA:2183 | 1.00 | 107.55 |  | 0 |  | 1^st^ MG coordination shell missing (see Notes) |
| MG:BA:2184 | 1.00 | 101.26 |  | 0 | 2 | - Exclusion zone contact(s): /O2/O2'/ Consider K in place of MG (2 valid contact(s) for K vs. 0 for MG) |
| MG:BA:2185 | 1.00 | 81.93 |  | 0 |  | No contacts in 5.0 A |
| MG:BA:2186 | 1.00 | 79.02 |  | 0 | 3 | - Exclusion zone contact(s): /O3'/OP1/OP2/ Consider K in place of MG (3 valid contact(s) for K vs. 0 for MG) |
| MG:BA:2187 | 1.00 | 107.58 |  | 0 | 4 | - Exclusion zone contact(s): /N2/O5'/OP1/OP2/ Consider K in place of MG (3 valid contact(s) for K vs. 0 for MG) |
| MG:BA:2188 | 1.00 | 102.88 |  | 0 |  | 1^st^ MG coordination shell missing (see Notes) |
| MG:BA:2189 | 1.00 | 140.23 |  | 0 | 2 | - Exclusion zone contact(s): /OP2/OP2/ Consider K in place of MG (2 valid contact(s) for K vs. 0 for MG) |
| MG:BA:2190 | 1.00 | 62.62 |  | 1 | 3 | N_b_ - Exclusion zone contact(s): /N1_A_/N1_A_/ Consider K in place of MG (3 valid contact(s) for K vs. 1 for MG) |
| MG:BA:2191 | 1.00 | 73.53 |  | 0 | 2 | - Exclusion zone contact(s): /N7/OP1/ Consider K in place of MG (2 valid contact(s) for K vs. 0 for MG) |
| MG:BA:2192 | 1.00 | 102.73 |  | 0 | 2 | - Exclusion zone contact(s): /OP1/OP2/ Consider K in place of MG (2 valid contact(s) for K vs. 0 for MG) |
| MG:BA:2193 | 1.00 | 76.33 |  | 0 |  | 1^st^ MG coordination shell missing (see Notes) |
| MG:BA:2194 | 1.00 | 40.03 |  | 0 | 3 | - Exclusion zone contact(s): /O3'/O5'/OP1/ Consider K in place of MG (2 valid contact(s) for K vs. 0 for MG) |
| MG:BA:2195 | 1.00 | 67.17 |  | 0 |  | 1^st^ MG coordination shell missing (see Notes) |
| MG:BA:2196 | 1.00 | 65.27 |  | 0 | 3 | - Exclusion zone contact(s): /O2'/O3'/OP1/ Consider K in place of MG (2 valid contact(s) for K vs. 0 for MG) |
| MG:BA:2197 | 1.00 | 136.81 |  | 1 | 1 | O_ph_ - Exclusion zone contact(s): /OP2/ |
| MG:BA:2198 | 1.00 | 139.46 |  | 0 |  | No contacts in 5.0 A |
| MG:BA:2199 | 1.00 | 107.46 |  | 0 | 1 | - Exclusion zone contact(s): /O2'/ Consider NA in place of MG (1 valid contact(s) for NA vs. 0 for MG) |
| MG:BA:2200 | 1.00 | 102.21 |  | 0 | 3 | - Exclusion zone contact(s): /O4'/O5'/OP2/ Consider K in place of MG (2 valid contact(s) for K vs. 0 for MG) |
| MG:BA:2201 | 1.00 | 85.47 |  | 0 |  | 1^st^ MG coordination shell missing (see Notes) |
| MG:BA:2202 | 1.00 | 87.35 |  | 0 |  | No contacts in 5.0 A |
| MG:BA:2203 | 1.00 | 118.36 |  | 0 | 3 | - Exclusion zone contact(s): /O2'/O3'/OP1/ Consider K in place of MG (2 valid contact(s) for K vs. 0 for MG) |
| MG:BA:2204 | 1.00 | 70.85 |  | 1 | 2 | O_ph_ - Exclusion zone contact(s): /O5'/OP1/ Consider K in place of MG (2 valid contact(s) for K vs. 1 for MG) |
| MG:BA:2205 | 1.00 | 109.44 |  | 0 | 3 | - Exclusion zone contact(s): /O5'/OP1/OP1/ Consider K in place of MG (2 valid contact(s) for K vs. 0 for MG) |
| MG:BA:2206 | 1.00 | 106.72 |  | 0 | 2 | - Exclusion zone contact(s): /OP1/OP2/ Consider K in place of MG (2 valid contact(s) for K vs. 0 for MG) |
| MG:BA:2207 | 1.00 | 103.23 |  | 0 |  | 1^st^ MG coordination shell missing (see Notes) |
| MG:BA:2208 | 1.00 | 105.92 |  | 0 | 1 | - Exclusion zone contact(s): /OP1/ Consider K in place of MG (1 valid contact(s) for K vs. 0 for MG) |
| MG:BA:2209 | 1.00 | 56.31 |  | 0 | 3 | - Exclusion zone contact(s): /O5'/OP2/OP2/ Consider K in place of MG (2 valid contact(s) for K vs. 0 for MG) |
| MG:BA:2210 | 1.00 | 78.19 |  | 2 | 1 | O_ph_O_r_ - Exclusion zone contact(s): /OP2/ |
| MG:BA:2211 | 1.00 | 78.66 |  | 0 | 3 | - Exclusion zone contact(s): /O2'/OP1/OP2/ Consider K in place of MG (2 valid contact(s) for K vs. 0 for MG) |
| MG:BA:2212 | 1.00 | 91.92 |  | 0 | 1 | - Exclusion zone contact(s): /O2'/ Consider K in place of MG (1 valid contact(s) for K vs. 0 for MG) |
| MG:BA:2213 | 1.00 | 114.40 |  | 0 | 1 | - Exclusion zone contact(s): /N7/ Consider K in place of MG (1 valid contact(s) for K vs. 0 for MG) |
| MG:BA:2214 | 1.00 | 90.18 |  | 0 |  | 1^st^ MG coordination shell missing (see Notes) |
| MG:BA:2215 | 1.00 | 79.24 |  | 0 | 1 | - Exclusion zone contact(s): /OP2/ Consider K in place of MG (1 valid contact(s) for K vs. 0 for MG) |
| MG:BA:2216 | 1.00 | 91.74 |  | 0 |  | 1^st^ MG coordination shell missing (see Notes) |
| MG:BA:2217 | 1.00 | 108.51 |  | 0 | 5 | - Exclusion zone contact(s): /O5'/OG_Ser_/OP1/OP2/OP2/ Consider K in place of MG (5 valid contact(s) for K vs. 0 for MG) |
| MG:BA:2218 | 1.00 | 80.03 |  | 0 | 3 | - Exclusion zone contact(s): /N6/N7/O6/ Consider K in place of MG (3 valid contact(s) for K vs. 0 for MG) |
| MG:BA:2219 | 1.00 | 60.84 |  | 0 | 3 | - Exclusion zone contact(s): /O4/O6/O6/ Consider K in place of MG (2 valid contact(s) for K vs. 0 for MG) |
| MG:BA:2220 | 1.00 | 86.85 |  | 0 | 1 | - Exclusion zone contact(s): /N7/ Consider K in place of MG (1 valid contact(s) for K vs. 0 for MG) |
| MG:BA:2221 | 1.00 | 75.69 |  | 0 | 2 | - Exclusion zone contact(s): /N1/O2/ Consider K in place of MG (2 valid contact(s) for K vs. 0 for MG) |
| MG:BA:2222 | 1.00 | 102.14 |  | 1 | 2 | Anion - Exclusion zone contact(s): /N3_G_/O5'/ Consider K in place of MG (2 valid contact(s) for K vs. 1 for MG) |
| MG:BA:2223 | 1.00 | 71.84 |  | 0 | 3 | - Exclusion zone contact(s): /O2/O4/O6/ Consider K in place of MG (3 valid contact(s) for K vs. 0 for MG) |
| MG:BA:2224 | 1.00 | 83.20 |  | 0 |  | 1^st^ MG coordination shell missing (see Notes) |
| MG:BA:2225 | 1.00 | 98.46 |  | 0 | 1 | - Exclusion zone contact(s): /OP2/ Consider NA in place of MG (1 valid contact(s) for NA vs. 0 for MG) |
| MG:BA:2226 | 1.00 | 101.88 |  | 0 |  | 1^st^ MG coordination shell missing (see Notes) |
| MG:BA:2227 | 1.00 | 72.42 |  | 0 |  | 1^st^ MG coordination shell missing (see Notes) |
| MG:BA:2228 | 1.00 | 98.81 |  | 0 |  | 1^st^ MG coordination shell missing (see Notes) |
| MG:BA:2229 | 1.00 | 67.00 |  | 0 |  | 1^st^ MG coordination shell missing (see Notes) |
| MG:BA:2230 | 1.00 | 111.96 |  | 0 | 2 | - Exclusion zone contact(s): /NZ_Lys_/OP2/ Consider K in place of MG (2 valid contact(s) for K vs. 0 for MG) |
| MG:BA:2231 | 1.00 | 89.36 |  | 0 | 2 | - Exclusion zone contact(s): /OP1/OP2/ Consider NA in place of MG (1 valid contact(s) for NA vs. 0 for MG) |
| MG:BA:2232 | 1.00 | 103.99 |  | 0 | 3 | - Exclusion zone contact(s): /N2/N3_G_/O2/ Consider K in place of MG (3 valid contact(s) for K vs. 0 for MG) |
| MG:BA:2233 | 1.00 | 100.31 |  | 0 | 4 | - Exclusion zone contact(s): /N7/O5'/OP2/OP2/ Consider K in place of MG (4 valid contact(s) for K vs. 0 for MG) |
| MG:BA:2234 | 1.00 | 108.87 |  | 0 | 1 | - Exclusion zone contact(s): /OP2/ Consider K in place of MG (1 valid contact(s) for K vs. 0 for MG) |
| MG:BA:2235 | 1.00 | 69.80 |  | 1 | 2 | O_ph_ - Exclusion zone contact(s): /O3'/OP1/ Consider K in place of MG (2 valid contact(s) for K vs. 1 for MG) |
| MG:BA:2236 | 1.00 | 82.46 |  | 0 | 4 | - Exclusion zone contact(s): /N7/N7/O6/OP2/ Consider K in place of MG (3 valid contact(s) for K vs. 0 for MG) |
| MG:BA:2237 | 1.00 | 77.59 |  | 0 |  | 1^st^ MG coordination shell missing (see Notes) |
| MG:BA:2238 | 1.00 | 95.45 |  | 0 | 1 | - Exclusion zone contact(s): /O6/ Consider K in place of MG (1 valid contact(s) for K vs. 0 for MG) |
| MG:BA:2239 | 1.00 | 63.98 |  | 0 |  | 1^st^ MG coordination shell missing (see Notes) |
| MG:BA:2240 | 1.00 | 86.87 |  | 0 | 4 | - Exclusion zone contact(s): /N7/O5'/OP2/ Consider K in place of MG (4 valid contact(s) for K vs. 0 for MG) |
| MG:BA:2241 | 1.00 | 78.89 |  | 0 | 1 | - Exclusion zone contact(s): /N6/ Consider K in place of MG (1 valid contact(s) for K vs. 0 for MG) |
| MG:BA:2242 | 1.00 | 57.69 |  | 0 |  | 1^st^ MG coordination shell missing (see Notes) |
| MG:BA:2243 | 1.00 | 114.64 |  | 0 |  | No contacts in 5.0 A |
| MG:BA:2244 | 1.00 | 113.71 |  | 0 |  | No contacts in 5.0 A |
| MG:BA:2245 | 1.00 | 71.61 |  | 0 |  | 1^st^ MG coordination shell missing (see Notes) |
| MG:BA:2246 | 1.00 | 125.23 |  | 0 | 3 | - Exclusion zone contact(s): /O2'/OP1/OP2/ Consider K in place of MG (2 valid contact(s) for K vs. 0 for MG) |
| MG:BA:2247 | 1.00 | 82.09 |  | 0 | 1 | - Exclusion zone contact(s): /O3'/ Consider K in place of MG (1 valid contact(s) for K vs. 0 for MG) |
| MG:BA:2248 | 1.00 | 61.12 |  | 0 | 3 | - Exclusion zone contact(s): /O5'/OP2/OP2/ Consider K in place of MG (2 valid contact(s) for K vs. 0 for MG) |
| MG:BA:2249 | 1.00 | 66.31 |  | 0 |  | 1^st^ MG coordination shell missing (see Notes) |
| MG:BA:2250 | 1.00 | 90.70 |  | 0 | 1 | - Exclusion zone contact(s): /OP1/ Consider K in place of MG (1 valid contact(s) for K vs. 0 for MG) |
| MG:BA:2251 | 1.00 | 89.76 |  | 0 |  | 1^st^ MG coordination shell missing (see Notes) |
| MG:BA:2252 | 1.00 | 70.19 |  | 0 | 1 | - Exclusion zone contact(s): /OP2/ Consider K in place of MG (1 valid contact(s) for K vs. 0 for MG) |
| MG:BA:2253 | 1.00 | 103.70 |  | 1 | 4 | O_ph_ - Exclusion zone contact(s): /NE_Arg_/NH1_Arg_/OP2/OP2/ Consider K in place of MG (3 valid contact(s) for K vs. 1 for MG) |
| MG:BA:2254 | 1.00 | 53.06 |  | 0 | 4 | - Exclusion zone contact(s): /O3'/OP1/OP1/OP2/ Consider NA in place of MG (2 valid contact(s) for NA vs. 0 for MG) |
| MG:BA:2255 | 1.00 | 61.64 |  | 0 | 2 | - Exclusion zone contact(s): /N7/OP2/ Consider K in place of MG (2 valid contact(s) for K vs. 0 for MG) |
| MG:BA:2256 | 1.00 | 76.54 |  | 0 |  | 1^st^ MG coordination shell missing (see Notes) |
| MG:BA:2257 | 1.00 | 85.38 |  | 0 | 2 | - Exclusion zone contact(s): /O2'/O6/ Consider NA in place of MG (1 valid contact(s) for NA vs. 0 for MG) |
| MG:BA:2258 | 1.00 | 77.65 |  | 0 | 2 | - Exclusion zone contact(s): /OP2/OP2/ Consider K in place of MG (2 valid contact(s) for K vs. 0 for MG) |
| MG:BA:2259 | 1.00 | 101.94 |  | 0 | 2 | - Exclusion zone contact(s): /O5'/OP2/ Consider NA in place of MG (1 valid contact(s) for NA vs. 0 for MG) |
| MG:BA:2260 | 1.00 | 77.49 |  | 0 | 4 | - Exclusion zone contact(s): /NE2_Gln_/O5'/OE1_Gln_/OP2/ Consider K in place of MG (3 valid contact(s) for K vs. 0 for MG) |
| MG:BA:2261 | 1.00 | 81.30 |  | 0 | 1 | - Exclusion zone contact(s): /N7/ Consider K in place of MG (1 valid contact(s) for K vs. 0 for MG) |
| MG:BA:2262 | 1.00 | 90.68 |  | 0 | 4 | - Exclusion zone contact(s): /O2'/OP2/OP2/OP2/ Consider K in place of MG (3 valid contact(s) for K vs. 0 for MG) |
| MG:BA:2263 | 1.00 | 71.04 |  | 1 | 3 | O_r_ - Exclusion zone contact(s): /O2'/O3'/OP1/ Consider K in place of MG (3 valid contact(s) for K vs. 1 for MG) |
| MG:BA:2264 | 1.00 | 79.88 |  | 0 | 2 | - Exclusion zone contact(s): /O5'/OP2/ Consider K in place of MG (2 valid contact(s) for K vs. 0 for MG) |
| MG:BA:2265 | 1.00 | 111.90 |  | 0 | 2 | - Exclusion zone contact(s): /O2/O2'/ Consider K in place of MG (2 valid contact(s) for K vs. 0 for MG) |
| MG:BA:2266 | 1.00 | 91.52 |  | 0 | 1 | - Exclusion zone contact(s): /OP2/ Consider K in place of MG (1 valid contact(s) for K vs. 0 for MG) |
| MG:BA:2267 | 1.00 | 91.52 |  | 0 |  | 1^st^ MG coordination shell missing (see Notes) |
| MG:BA:2268 | 1.00 | 87.92 |  | 0 | 2 | - Exclusion zone contact(s): /O5'/OP2/ Consider K in place of MG (2 valid contact(s) for K vs. 0 for MG) |
| MG:BA:2269 | 1.00 | 80.13 |  | 0 | 1 | - Exclusion zone contact(s): /OP2/ Consider K in place of MG (1 valid contact(s) for K vs. 0 for MG) |
| MG:BA:2270 | 1.00 | 100.58 |  | 0 |  | 1^st^ MG coordination shell missing (see Notes) |
| MG:BA:2271 | 1.00 | 106.00 |  | 0 | 3 | - Exclusion zone contact(s): /N1/O2/ Consider K in place of MG (2 valid contact(s) for K vs. 0 for MG) |
| MG:BA:2272 | 1.00 | 75.79 |  | 0 | 1 | - Exclusion zone contact(s): /O6/ Consider K in place of MG (1 valid contact(s) for K vs. 0 for MG) |
| MG:BA:2273 | 1.00 | 94.45 |  | 0 | 4 | - Exclusion zone contact(s): /NZ_Lys_/O3'/OP1/OP2/ Consider K in place of MG (4 valid contact(s) for K vs. 0 for MG) |
| MG:BA:2274 | 1.00 | 82.89 |  | 0 | 4 | - Exclusion zone contact(s): /N1/O2'/OP2/OP2/ Consider K in place of MG (3 valid contact(s) for K vs. 0 for MG) |
| MG:BA:2275 | 1.00 | 145.41 |  | 2 |  | O_r_O_b_ |
| MG:BA:2276 | 1.00 | 128.70 |  | 6 |  | 6O_w_ |
| MG:BB:101 | 1.00 | 76.48 |  | 1 | 2 | O_r_ - Exclusion zone contact(s): /O3'/O5'/ Consider K in place of MG (2 valid contact(s) for K vs. 1 for MG) |
| MG:BB:102 | 1.00 | 55.85 |  | 0 |  | 1^st^ MG coordination shell missing (see Notes) |
| MG:BB:103 | 1.00 | 77.26 |  | 0 |  | 1^st^ MG coordination shell missing (see Notes) |
| MG:BB:104 | 1.00 | 71.34 |  | 0 |  | 1^st^ MG coordination shell missing (see Notes) |
| MG:BB:105 | 1.00 | 85.66 |  | 0 | 3 | - Exclusion zone contact(s): /O4/OP1/OP2/ Consider K in place of MG (2 valid contact(s) for K vs. 0 for MG) |
| MG:BB:106 | 1.00 | 116.37 |  | 0 |  | 1^st^ MG coordination shell missing (see Notes) |
| MG:BB:107 | 1.00 | 83.11 |  | 0 |  | 1^st^ MG coordination shell missing (see Notes) |
| MG:BB:108 | 1.00 | 94.14 |  | 0 |  | 1^st^ MG coordination shell missing (see Notes) |
| MG:BB:109 | 1.00 | 119.55 |  | 1 |  | O_r_ |
| MG:BB:110 | 1.00 | 69.45 |  | 1 | 2 | O_coo_ - Exclusion zone contact(s): /OP2/OP2/ |
| MG:BB:111 | 1.00 | 90.04 |  | 0 | 1 | - Exclusion zone contact(s): /O6/ Consider K in place of MG (1 valid contact(s) for K vs. 0 for MG) |
| MG:BB:112 | 1.00 | 107.10 |  | 0 | 2 | - Exclusion zone contact(s): /O5'/OP2/ Consider NA in place of MG (1 valid contact(s) for NA vs. 0 for MG) |
| MG:BB:113 | 1.00 | 142.38 |  | 0 | 2 | - Exclusion zone contact(s): /OP1/OP2/ Consider NA in place of MG (1 valid contact(s) for NA vs. 0 for MG) |
| MG:BC:101 | 1.00 | 53.08 |  | 0 | 1 | - Exclusion zone contact(s): /OP1/ Consider NA in place of MG (1 valid contact(s) for NA vs. 0 for MG) |
| MG:BC:102 | 1.00 | 78.23 |  | 1 |  | O_ph_ |
| MG:BC:103 | 1.00 | 51.37 |  | 1 | 2 | O_ph_ - Exclusion zone contact(s): /O5'/OP1/ |
| MG:BC:104 | 1.00 | 45.35 |  | 0 | 1 | - Exclusion zone contact(s): /OP2/ Consider K in place of MG (1 valid contact(s) for K vs. 0 for MG) |
| MG:BC:105 | 1.00 | 75.59 |  | 1 |  | O_ph_ |
| MG:BC:106 | 1.00 | 58.59 |  | 0 | 1 | - Exclusion zone contact(s): /N7/ Consider K in place of MG (1 valid contact(s) for K vs. 0 for MG) |
| MG:BC:107 | 1.00 | 65.96 |  | 0 |  | 1^st^ MG coordination shell missing (see Notes) |
| MG:BC:108 | 1.00 | 81.73 |  | 0 |  | 1^st^ MG coordination shell missing (see Notes) |
| MG:BC:109 | 1.00 | 72.52 |  | 1 | 3 | Anion - Exclusion zone contact(s): /O4/O6/O6/ Consider K in place of MG (3 valid contact(s) for K vs. 1 for MG) |
| MG:BC:110 | 1.00 | 56.83 |  | 0 |  | 1^st^ MG coordination shell missing (see Notes) |
| MG:BC:111 | 1.00 | 63.66 |  | 0 | 3 | - Exclusion zone contact(s): /O3'/O4/OP1/ Consider K in place of MG (3 valid contact(s) for K vs. 0 for MG) |
| MG:BC:112 | 1.00 | 57.13 |  | 0 |  | 1^st^ MG coordination shell missing (see Notes) |
| MG:BC:113 | 1.00 | 54.80 |  | 3 |  | 3O_ph_ |
| MG:BC:114 | 1.00 | 58.63 |  | 0 |  | 1^st^ MG coordination shell missing (see Notes) |
| MG:BC:115 | 1.00 | 106.33 |  | 0 | 2 | - Exclusion zone contact(s): /N1_G_/N2/ Consider K in place of MG (2 valid contact(s) for K vs. 0 for MG) |
| MG:BC:116 | 1.00 | 107.99 |  | 0 | 3 | - Exclusion zone contact(s): /N7/O6/O6/ Consider K in place of MG (3 valid contact(s) for K vs. 0 for MG) |
| MG:BD:101 | 1.00 | 86.03 |  | 0 |  | 1^st^ MG coordination shell missing (see Notes) |
| MG:BD:102 | 1.00 | 84.22 |  | 0 | 1 | - Exclusion zone contact(s): /OP1/ Consider K in place of MG (1 valid contact(s) for K vs. 0 for MG) |
| MG:BD:103 | 1.00 | 58.66 |  | 0 |  | 1^st^ MG coordination shell missing (see Notes) |
| MG:BD:104 | 1.00 | 51.34 |  | 0 |  | 1^st^ MG coordination shell missing (see Notes) |
| MG:BD:105 | 1.00 | 87.44 |  | 0 | 1 | - Exclusion zone contact(s): /OP2/ Consider K in place of MG (1 valid contact(s) for K vs. 0 for MG) |
| MG:BD:106 | 1.00 | 91.97 |  | 0 |  | No contacts in 5.0 A |
| MG:BD:107 | 1.00 | 71.75 |  | 0 | 1 | - Exclusion zone contact(s): /OP1/ Consider K in place of MG (1 valid contact(s) for K vs. 0 for MG) |
| MG:BD:108 | 1.00 | 82.60 |  | 0 | 3 | - Exclusion zone contact(s): /O4/O4/OP1/ Consider K in place of MG (2 valid contact(s) for K vs. 0 for MG) |
| MG:BD:109 | 1.00 | 139.83 |  | 0 |  | 1^st^ MG coordination shell missing (see Notes) |
| MG:BD:110 | 1.00 | 81.72 |  | 1 |  | O_ph_ |
| MG:BD:111 | 1.00 | 74.04 |  | 0 | 1 | - Exclusion zone contact(s): /O6/ Consider K in place of MG (1 valid contact(s) for K vs. 0 for MG) |
| MG:BD:112 | 1.00 | 78.21 |  | 0 |  | 1^st^ MG coordination shell missing (see Notes) |
| MG:BD:113 | 1.00 | 78.94 |  | 0 |  | 1^st^ MG coordination shell missing (see Notes) |
| MG:BD:114 | 1.00 | 93.21 |  | 0 |  | 1^st^ MG coordination shell missing (see Notes) |
| MG:BD:115 | 1.00 | 61.76 |  | 0 |  | 1^st^ MG coordination shell missing (see Notes) |
| MG:BD:116 | 1.00 | 129.19 |  | 0 |  | 1^st^ MG coordination shell missing (see Notes) |
| MG:BD:117 | 1.00 | 115.08 |  | 0 | 1 | - Exclusion zone contact(s): /OP2/ Consider K in place of MG (1 valid contact(s) for K vs. 0 for MG) |
| MG:BD:118 | 1.00 | 107.12 |  | 0 |  | 1^st^ MG coordination shell missing (see Notes) |
| MG:BD:119 | 1.00 | 84.48 |  | 0 |  | No contacts in 5.0 A |
| MG:BD:120 | 1.00 | 90.74 |  | 0 |  | 1^st^ MG coordination shell missing (see Notes) |
| MG:BD:121 | 1.00 | 79.68 |  | 0 |  | 1^st^ MG coordination shell missing (see Notes) |
| MG:BD:122 | 1.00 | 100.29 |  | 0 |  | No contacts in 5.0 A |
| MG:BD:123 | 1.00 | 82.83 |  | 0 |  | 1^st^ MG coordination shell missing (see Notes) |
| MG:BD:124 | 1.00 | 104.22 |  | 0 | 1 | - Exclusion zone contact(s): /O4'/ Consider K in place of MG (1 valid contact(s) for K vs. 0 for MG) |
| MG:BD:125 | 1.00 | 105.33 |  | 0 |  | No contacts in 5.0 A |
| MG:BD:126 | 1.00 | 133.39 |  | 0 | 1 | - Exclusion zone contact(s): /OP1/ Consider K in place of MG (1 valid contact(s) for K vs. 0 for MG) |
| MG:BE:301 | 1.00 | 63.40 |  | 0 |  | 1^st^ MG coordination shell missing (see Notes) |
| MG:BE:302 | 1.00 | 59.65 |  | 0 |  | 1^st^ MG coordination shell missing (see Notes) |
| MG:BE:303 | 1.00 | 94.26 |  | 0 | 3 | - Exclusion zone contact(s): /OD1_Asp_/OD2_Asp_/OE2_Glu_/ Consider K in place of MG (2 valid contact(s) for K vs. 0 for MG) |
| MG:BE:304 | 1.00 | 85.55 |  | 0 |  | No contacts in 5.0 A |
| MG:BE:305 | 1.00 | 95.09 |  | 0 |  | 1^st^ MG coordination shell missing (see Notes) |
| MG:BF:301 | 1.00 | 104.46 |  | 0 | 1 | - Exclusion zone contact(s): /NH1_Arg_/ Consider K in place of MG (1 valid contact(s) for K vs. 0 for MG) |
| MG:BF:302 | 1.00 | 105.38 |  | 0 | 2 | - Exclusion zone contact(s): /N_back_/O_back_/ Consider K in place of MG (2 valid contact(s) for K vs. 0 for MG) |
| MG:BG:301 | 1.00 | 67.35 |  | 0 |  | 1^st^ MG coordination shell missing (see Notes) |
| MG:BG:302 | 1.00 | 63.90 |  | 0 |  | 1^st^ MG coordination shell missing (see Notes) |
| MG:BG:303 | 1.00 | 59.75 |  | 1 | 3 | O_back_ - Exclusion zone contact(s): /O_back_/O_back_/OG1_Thr_/ Consider K in place of MG (2 valid contact(s) for K vs. 1 for MG) |
| MG:BG:304 | 1.00 | 79.94 |  | 0 |  | 1^st^ MG coordination shell missing (see Notes) |
| MG:BG:305 | 1.00 | 70.48 |  | 0 | 1 | - Exclusion zone contact(s): /N_back_/ Consider K in place of MG (1 valid contact(s) for K vs. 0 for MG) |
| MG:BG:306 | 1.00 | 108.76 |  | 1 | 2 | Anion - Exclusion zone contact(s): /N3_U_/O4/ Consider K in place of MG (2 valid contact(s) for K vs. 1 for MG) |
| MG:BG:307 | 1.00 | 85.26 |  | 0 |  | 1^st^ MG coordination shell missing (see Notes) |
| ZN:BG:308 | 1.00 | 71.76 |  | 0 | 1 | - Exclusion zone contact(s): /O_back_/ |
| MG:BH:201 | 1.00 | 55.30 |  | 1 | 3 | Anion - Exclusion zone contact(s): /N7/NH1_Arg_/O4/ Consider K in place of MG (3 valid contact(s) for K vs. 1 for MG) |
| MG:BH:202 | 1.00 | 61.92 |  | 0 |  | 1^st^ MG coordination shell missing (see Notes) |
| MG:BH:203 | 1.00 | 89.20 |  | 0 | 2 | - Exclusion zone contact(s): /N_back_/O_back_/ Consider K in place of MG (2 valid contact(s) for K vs. 0 for MG) |
| MG:BH:204 | 1.00 | 72.75 |  | 0 |  | No contacts in 5.0 A |
| MG:BH:205 | 1.00 | 78.38 |  | 1 | 1 | O_coo_ - Exclusion zone contact(s): /OE2_Glu_/ |
| MG:BI:201 | 1.00 | 47.12 |  | 0 | 3 | - Exclusion zone contact(s): /O_back_/O_back_/O_back_/ Consider K in place of MG (2 valid contact(s) for K vs. 0 for MG) |
| MG:BK:201 | 1.00 | 96.67 |  | 0 | 1 | - Exclusion zone contact(s): /N_back_/ Consider K in place of MG (1 valid contact(s) for K vs. 0 for MG) |
| MG:BK:202 | 1.00 | 80.12 |  | 0 |  | 1^st^ MG coordination shell missing (see Notes) |
| MG:BK:203 | 1.00 | 69.81 |  | 0 |  | No contacts in 5.0 A |
| MG:BK:204 | 1.00 | 74.24 |  | 0 |  | 1^st^ MG coordination shell missing (see Notes) |
| MG:BK:205 | 1.00 | 86.26 |  | 0 | 2 | - Exclusion zone contact(s): /O_back_/OE1_Glu_/ Consider NA in place of MG (1 valid contact(s) for NA vs. 0 for MG) |
| MG:BK:206 | 1.00 | 108.66 |  | 0 | 1 | - Exclusion zone contact(s): /NH2_Arg_/ Consider NA in place of MG (1 valid contact(s) for NA vs. 0 for MG) |
| MG:BL:201 | 1.00 | 86.58 |  | 0 | 4 | - Exclusion zone contact(s): /N_back_/N_back_/N_back_/O_back_/ Consider K in place of MG (3 valid contact(s) for K vs. 0 for MG) |
| MG:BL:202 | 1.00 | 64.77 |  | 1 | 1 | O_b_ - Exclusion zone contact(s): /N_back_/ |
| MG:BM:201 | 1.00 | 64.83 |  | 1 | 1 | Anion - Exclusion zone contact(s): /O4/ |
| MG:BM:202 | 1.00 | 81.18 |  | 0 | 2 | - Exclusion zone contact(s): /O6/OG_Ser_/ Consider NA in place of MG (1 valid contact(s) for NA vs. 0 for MG) |
| MG:BM:203 | 1.00 | 91.89 |  | 0 |  | 1^st^ MG coordination shell missing (see Notes) |
| MG:BO:201 | 1.00 | 72.85 |  | 0 |  | 1^st^ MG coordination shell missing (see Notes) |
| MG:BP:201 | 1.00 | 61.99 |  | 0 | 2 | - Exclusion zone contact(s): /O_back_/O_back_/ Consider K in place of MG (2 valid contact(s) for K vs. 0 for MG) |
| MG:BP:202 | 1.00 | 50.53 |  | 0 |  | 1^st^ MG coordination shell missing (see Notes) |
| MG:BQ:101 | 1.00 | 66.73 |  | 0 |  | 1^st^ MG coordination shell missing (see Notes) |
| MG:BQ:102 | 1.00 | 63.48 |  | 0 | 3 | - Exclusion zone contact(s): /N_back_/OG1_Thr_/OP1/ Consider K in place of MG (3 valid contact(s) for K vs. 0 for MG) |
| MG:BQ:103 | 1.00 | 69.22 |  | 0 |  | 1^st^ MG coordination shell missing (see Notes) |
| ZN:BQ:104 | 1.00 | 97.14 |  | 0 |  |  |
| MG:BS:101 | 1.00 | 76.18 |  | 0 |  | 1^st^ MG coordination shell missing (see Notes) |
| MG:BS:102 | 1.00 | 69.46 |  | 0 | 1 | - Exclusion zone contact(s): /NH1_Arg_/ Consider NA in place of MG (1 valid contact(s) for NA vs. 0 for MG) |
| MG:BS:103 | 1.00 | 83.53 |  | 0 | 1 | - Exclusion zone contact(s): /O_back_/ Consider K in place of MG (1 valid contact(s) for K vs. 0 for MG) |
| MG:BS:104 | 1.00 | 76.09 |  | 0 | 2 | - Exclusion zone contact(s): /NE_Arg_/OP1/ Consider K in place of MG (2 valid contact(s) for K vs. 0 for MG) |
| MG:BS:105 | 1.00 | 89.46 |  | 0 | 2 | - Exclusion zone contact(s): /O_back_/OD2_Asp_/ Consider K in place of MG (2 valid contact(s) for K vs. 0 for MG) |
| MG:BS:106 | 1.00 | 100.03 |  | 0 |  | 1^st^ MG coordination shell missing (see Notes) |
| MG:BS:107 | 1.00 | 90.51 |  | 0 | 2 | - Exclusion zone contact(s): /NH2_Arg_/O_back_/ Consider K in place of MG (2 valid contact(s) for K vs. 0 for MG) |
| MG:BS:108 | 1.00 | 91.30 |  | 0 | 2 | - Exclusion zone contact(s): /OD1_Asp_/OD2_Asp_/ Consider K in place of MG (2 valid contact(s) for K vs. 0 for MG) |
| MG:BS:109 | 1.00 | 110.45 |  | 0 | 5 | - Exclusion zone contact(s): /N_back_/NH2_Arg_/O2'/O3'/O_back_/ Consider K in place of MG (4 valid contact(s) for K vs. 0 for MG) |
| MG:BT:201 | 1.00 | 110.38 |  | 1 | 4 | O_coo_ - Exclusion zone contact(s): /O_back_/O_back_/O_back_/OE2_Glu_/ Consider K in place of MG (4 valid contact(s) for K vs. 1 for MG) |
| MG:BT:202 | 1.00 | 76.14 |  | 0 | 1 | - Exclusion zone contact(s): /OP1/ Consider K in place of MG (1 valid contact(s) for K vs. 0 for MG) |
| MG:BU:1601 | 1.00 | 56.37 |  | 0 |  | 1^st^ MG coordination shell missing (see Notes) |
| MG:BV:101 | 1.00 | 73.23 |  | 0 | 4 | - Exclusion zone contact(s): /N_back_/NH1_Arg_/O_back_/OP1/ Consider K in place of MG (3 valid contact(s) for K vs. 0 for MG) |
| MG:BW:201 | 1.00 | 88.42 |  | 0 | 2 | - Exclusion zone contact(s): /N_back_/O_back_/ Consider K in place of MG (2 valid contact(s) for K vs. 0 for MG) |
| MG:BW:202 | 1.00 | 73.04 |  | 0 | 1 | - Exclusion zone contact(s): /NZ_Lys_/ Consider K in place of MG (1 valid contact(s) for K vs. 0 for MG) |
| MG:BW:203 | 1.00 | 125.07 |  | 0 | 2 | - Exclusion zone contact(s): /N7/O_back_/ Consider NA in place of MG (1 valid contact(s) for NA vs. 0 for MG) |
| MG:BW:204 | 1.00 | 90.69 |  | 0 | 2 | - Exclusion zone contact(s): /N_back_/O_back_/ Consider NA in place of MG (1 valid contact(s) for NA vs. 0 for MG) |
| MG:BW:205 | 1.00 | 91.96 |  | 0 | 1 | - Exclusion zone contact(s): /O_back_/ Consider K in place of MG (1 valid contact(s) for K vs. 0 for MG) |
| MG:BW:206 | 1.00 | 56.12 |  | 0 | 3 | - Exclusion zone contact(s): /N_back_/N_back_/OG_Ser_/ Consider K in place of MG (3 valid contact(s) for K vs. 0 for MG) |
| MG:BW:207 | 1.00 | 85.93 |  | 0 | 2 | - Exclusion zone contact(s): /O_back_/OP1/ Consider K in place of MG (2 valid contact(s) for K vs. 0 for MG) |
| MG:BW:208 | 1.00 | 62.29 |  | 0 |  | 1^st^ MG coordination shell missing (see Notes) |
| MG:BX:101 | 1.00 | 74.63 |  | 0 |  | 1^st^ MG coordination shell missing (see Notes) |
| MG:C1:101 | 1.00 | 156.36 |  | 1 | 1 | O_ph_ - Exclusion zone contact(s): /OP2/ |
| MG:C1:102 | 1.00 | 68.03 |  | 0 | 2 | - Exclusion zone contact(s): /OP1/OP2/ Consider K in place of MG (2 valid contact(s) for K vs. 0 for MG) |
| MG:C1:103 | 1.00 | 64.04 |  | 0 |  | 1^st^ MG coordination shell missing (see Notes) |
| MG:C1:104 | 1.00 | 71.04 |  | 0 | 2 | - Exclusion zone contact(s): /O4'/OP1/ Consider K in place of MG (2 valid contact(s) for K vs. 0 for MG) |
| MG:C1:105 | 1.00 | 89.89 |  | 0 | 1 | - Exclusion zone contact(s): /OP1/ Consider K in place of MG (1 valid contact(s) for K vs. 0 for MG) |
| MG:C1:106 | 1.00 | 89.34 |  | 0 | 4 | - Exclusion zone contact(s): /O5'/OP1/OP1/OP2/ Consider K in place of MG (3 valid contact(s) for K vs. 0 for MG) |
| MG:CA:1601 | 1.00 | 41.50 |  | 0 | 2 | - Exclusion zone contact(s): /O6/O6/ Consider K in place of MG (2 valid contact(s) for K vs. 0 for MG) |
| MG:CA:1602 | 1.00 | 89.73 |  | 1 | 1 | O_ph_ - Exclusion zone contact(s): /OP1/ |
| MG:CA:1603 | 1.00 | 80.10 |  | 0 | 1 | - Exclusion zone contact(s): /O3'/ Consider K in place of MG (1 valid contact(s) for K vs. 0 for MG) |
| MG:CA:1604 | 1.00 | 10.18 | 1 | 1 | 1 | - Exclusion zone contact(s): /O2/OP2/ Consider K in place of MG (1 valid contact(s) for K vs. 0 for MG) |
| MG:CA:1605 | 1.00 | 36.44 |  | 1 |  | O_ph_ |
| MG:CA:1606 | 1.00 | 22.86 |  | 2 |  | O_ph_O_b_ |
| MG:CA:1607 | 1.00 | 20.17 |  | 1 |  | O_ph_ |
| MG:CA:1608 | 1.00 | 48.96 |  | 1 | 1 | O_ph_ - Exclusion zone contact(s): /OP1/ |
| MG:CA:1609 | 1.00 | 31.01 | 1 | 1 | 2 | O_ph_ - Exclusion zone contact(s): /OP1/OP2/OP2/ Consider K in place of MG (2 valid contact(s) for K vs. 1 for MG) |
| MG:CA:1610 | 1.00 | 48.83 |  | 0 |  | 1^st^ MG coordination shell missing (see Notes) |
| MG:CA:1611 | 1.00 | 39.35 |  | 0 | 1 | - Exclusion zone contact(s): /N7/ Consider K in place of MG (1 valid contact(s) for K vs. 0 for MG) |
| MG:CA:1612 | 1.00 | 57.74 |  | 0 | 1 | - Exclusion zone contact(s): /O6/ Consider K in place of MG (1 valid contact(s) for K vs. 0 for MG) |
| MG:CA:1613 | 1.00 | 41.51 |  | 1 | 3 | O_b_ - Exclusion zone contact(s): /O2'/O3'/OP2/ Consider NA in place of MG (3 valid contact(s) for NA vs. 1 for MG) |
| MG:CA:1614 | 1.00 | 41.09 |  | 0 | 1 | - Exclusion zone contact(s): /O6/ Consider K in place of MG (1 valid contact(s) for K vs. 0 for MG) |
| MG:CA:1615 | 1.00 | 60.97 |  | 1 |  | O_ph_ |
| MG:CA:1616 | 1.00 | 39.13 |  | 0 |  | 1^st^ MG coordination shell missing (see Notes) |
| MG:CA:1617 | 1.00 | 33.10 |  | 2 | 1 | 2O_ph_ - Exclusion zone contact(s): /OP2/ |
| MG:CA:1618 | 1.00 | 24.99 |  | 1 | 4 | O_ph_ - Exclusion zone contact(s): /O5'/OP1/OP1/OP2/ Consider K in place of MG (3 valid contact(s) for K vs. 1 for MG) |
| MG:CA:1619 | 1.00 | 24.53 |  | 0 | 3 | - Exclusion zone contact(s): /OP1/OP2/OP2/ Consider K in place of MG (2 valid contact(s) for K vs. 0 for MG) |
| MG:CA:1620 | 1.00 | 39.27 |  | 0 | 4 | - Exclusion zone contact(s): /N4/O4/O6/O6/ Consider K in place of MG (4 valid contact(s) for K vs. 0 for MG) |
| MG:CA:1621 | 1.00 | 45.63 |  | 0 | 1 | - Exclusion zone contact(s): /O2'/ Consider K in place of MG (1 valid contact(s) for K vs. 0 for MG) |
| MG:CA:1622 | 1.00 | 51.86 | 1 | 2 | 2 | O_ph_O_b_ - Exclusion zone contact(s): /O6/OP1/OP2/ |
| MG:CA:1623 | 1.00 | 49.42 |  | 0 | 1 | - Exclusion zone contact(s): /OP1/ Consider NA in place of MG (1 valid contact(s) for NA vs. 0 for MG) |
| MG:CA:1624 | 1.00 | 80.79 |  | 0 | 1 | - Exclusion zone contact(s): /OP2/ Consider K in place of MG (1 valid contact(s) for K vs. 0 for MG) |
| MG:CA:1625 | 1.00 | 68.12 |  | 0 | 1 | - Exclusion zone contact(s): /N7/ Consider K in place of MG (1 valid contact(s) for K vs. 0 for MG) |
| MG:CA:1626 | 1.00 | 38.79 |  | 0 | 2 | - Exclusion zone contact(s): /N7/O6/ Consider K in place of MG (2 valid contact(s) for K vs. 0 for MG) |
| MG:CA:1627 | 1.00 | 49.12 |  | 0 | 2 | - Exclusion zone contact(s): /N7/O6/ Consider K in place of MG (2 valid contact(s) for K vs. 0 for MG) |
| MG:CA:1628 | 1.00 | 42.29 |  | 0 |  | 1^st^ MG coordination shell missing (see Notes) |
| MG:CA:1629 | 1.00 | 34.46 |  | 0 |  | 1^st^ MG coordination shell missing (see Notes) |
| MG:CA:1630 | 1.00 | 52.89 |  | 1 |  | O_ph_ |
| MG:CA:1631 | 1.00 | 42.13 |  | 0 |  | 1^st^ MG coordination shell missing (see Notes) |
| MG:CA:1632 | 1.00 | 65.76 |  | 0 |  | 1^st^ MG coordination shell missing (see Notes) |
| MG:CA:1633 | 1.00 | 105.11 |  | 0 |  | 1^st^ MG coordination shell missing (see Notes) |
| MG:CA:1634 | 1.00 | 91.13 |  | 0 |  | 1^st^ MG coordination shell missing (see Notes) |
| MG:CA:1635 | 1.00 | 21.61 |  | 0 | 1 | - Exclusion zone contact(s): /O6/ Consider K in place of MG (1 valid contact(s) for K vs. 0 for MG) |
| MG:CA:1636 | 1.00 | 16.11 |  | 0 | 1 | - Exclusion zone contact(s): /OP2/ Consider K in place of MG (1 valid contact(s) for K vs. 0 for MG) |
| MG:CA:1637 | 1.00 | 45.34 |  | 0 |  | 1^st^ MG coordination shell missing (see Notes) |
| MG:CA:1638 | 1.00 | 51.61 |  | 1 | 3 | O_ph_ - Exclusion zone contact(s): /O2'/O3'/OP2/ Consider K in place of MG (3 valid contact(s) for K vs. 1 for MG) |
| MG:CA:1639 | 1.00 | 43.72 |  | 0 | 2 | - Exclusion zone contact(s): /O6/O6/ Consider NA in place of MG (1 valid contact(s) for NA vs. 0 for MG) |
| MG:CA:1640 | 1.00 | 28.75 |  | 1 | 5 | O_b_ - Exclusion zone contact(s): /N4/O2/O6/O6/O6/ Consider K in place of MG (5 valid contact(s) for K vs. 1 for MG) |
| MG:CA:1641 | 1.00 | 28.79 |  | 0 | 3 | - Exclusion zone contact(s): /O4/O6/O6/ Consider K in place of MG (3 valid contact(s) for K vs. 0 for MG) |
| MG:CA:1642 | 1.00 | 25.06 |  | 0 | 5 | - Exclusion zone contact(s): /N4/O4/O4/O6/O6/ Consider K in place of MG (5 valid contact(s) for K vs. 0 for MG) |
| MG:CA:1643 | 1.00 | 51.70 |  | 1 | 3 | O_ph_ - Exclusion zone contact(s): /O3'/O4/O6/ Consider K in place of MG (3 valid contact(s) for K vs. 1 for MG) |
| MG:CA:1644 | 1.00 | 24.49 |  | 1 | 2 | O_ph_ - Exclusion zone contact(s): /O2'/O4/ |
| MG:CA:1645 | 1.00 | 74.97 |  | 0 |  | 1^st^ MG coordination shell missing (see Notes) |
| MG:CA:1646 | 1.00 | 53.01 |  | 0 | 2 | - Exclusion zone contact(s): /O2/OP2/ Consider NA in place of MG (1 valid contact(s) for NA vs. 0 for MG) |
| MG:CA:1647 | 1.00 | 13.24 |  | 0 | 3 | - Exclusion zone contact(s): /O3'/OP2/OP2/ Consider K in place of MG (2 valid contact(s) for K vs. 0 for MG) |
| MG:CA:1648 | 1.00 | 36.01 |  | 1 | 1 | O_ph_ - Exclusion zone contact(s): /O2/ |
| MG:CA:1649 | 1.00 | 47.63 |  | 1 | 2 | O_r_ - Exclusion zone contact(s): /N3_A_/OP2/ Consider K in place of MG (2 valid contact(s) for K vs. 1 for MG) |
| MG:CA:1650 | 1.00 | 36.05 |  | 0 | 1 | - Exclusion zone contact(s): /OP1/ Consider NA in place of MG (1 valid contact(s) for NA vs. 0 for MG) |
| MG:CA:1651 | 1.00 | 33.30 |  | 0 |  | 1^st^ MG coordination shell missing (see Notes) |
| MG:CA:1652 | 1.00 | 39.32 |  | 0 | 2 | - Exclusion zone contact(s): /N7/OP2/ Consider K in place of MG (2 valid contact(s) for K vs. 0 for MG) |
| MG:CA:1653 | 1.00 | 47.06 |  | 1 |  | O_ph_ |
| MG:CA:1654 | 1.00 | 48.05 |  | 1 |  | N_b_ |
| MG:CA:1655 | 1.00 | 18.89 |  | 2 | 3 | 2O_ph_ - Exclusion zone contact(s): /O2'/OP1/OP2/ Consider K in place of MG (3 valid contact(s) for K vs. 2 for MG) |
| MG:CA:1656 | 1.00 | 43.75 |  | 0 | 1 | - Exclusion zone contact(s): /OP1/ Consider K in place of MG (1 valid contact(s) for K vs. 0 for MG) |
| MG:CA:1657 | 1.00 | 35.77 |  | 0 |  | 1^st^ MG coordination shell missing (see Notes) |
| MG:CA:1658 | 1.00 | 48.92 |  | 0 |  | 1^st^ MG coordination shell missing (see Notes) |
| MG:CA:1659 | 1.00 | 41.78 |  | 0 | 2 | - Exclusion zone contact(s): /O2/OP2/ Consider K in place of MG (2 valid contact(s) for K vs. 0 for MG) |
| MG:CA:1660 | 1.00 | 79.95 |  | 0 |  | 1^st^ MG coordination shell missing (see Notes) |
| MG:CA:1661 | 1.00 | 59.94 |  | 1 | 2 | O_ph_ - Exclusion zone contact(s): /O2'/OP2/ Consider K in place of MG (2 valid contact(s) for K vs. 1 for MG) |
| MG:CA:1662 | 1.00 | 19.30 |  | 1 | 3 | O_b_ - Exclusion zone contact(s): /N3_C_/O2'/O4'/ Consider K in place of MG (2 valid contact(s) for K vs. 1 for MG) |
| MG:CA:1663 | 1.00 | 52.83 |  | 0 | 2 | - Exclusion zone contact(s): /N7/OP2/ Consider K in place of MG (2 valid contact(s) for K vs. 0 for MG) |
| MG:CA:1664 | 1.00 | 22.95 |  | 2 | 2 | 2O_ph_ - Exclusion zone contact(s): /O3'/OP1/ |
| MG:CA:1665 | 1.00 | 31.28 |  | 0 | 4 | - Exclusion zone contact(s): /O3'/OP1/OP1/OP1/ Consider K in place of MG (4 valid contact(s) for K vs. 0 for MG) |
| MG:CA:1666 | 1.00 | 60.42 |  | 0 |  | 1^st^ MG coordination shell missing (see Notes) |
| MG:CA:1667 | 1.00 | 50.91 |  | 0 | 1 | - Exclusion zone contact(s): /N7/ Consider K in place of MG (1 valid contact(s) for K vs. 0 for MG) |
| MG:CA:1668 | 1.00 | 88.11 |  | 0 | 1 | - Exclusion zone contact(s): /OP2/ Consider K in place of MG (1 valid contact(s) for K vs. 0 for MG) |
| MG:CA:1669 | 1.00 | 40.73 |  | 1 |  | O_ph_ |
| MG:CA:1670 | 1.00 | 46.93 |  | 0 |  | 1^st^ MG coordination shell missing (see Notes) |
| MG:CA:1671 | 1.00 | 24.48 |  | 0 |  | 1^st^ MG coordination shell missing (see Notes) |
| MG:CA:1672 | 1.00 | 57.05 |  | 0 | 2 | - Exclusion zone contact(s): /OP2/OP2/ Consider K in place of MG (2 valid contact(s) for K vs. 0 for MG) |
| MG:CA:1673 | 1.00 | 97.25 |  | 1 | 1 | O_ph_ - Exclusion zone contact(s): /OP2/ |
| MG:CA:1674 | 1.00 | 60.26 |  | 0 |  | 1^st^ MG coordination shell missing (see Notes) |
| MG:CA:1675 | 1.00 | 53.98 |  | 0 |  | 1^st^ MG coordination shell missing (see Notes) |
| MG:CA:1676 | 1.00 | 89.52 |  | 0 | 2 | - Exclusion zone contact(s): /N4/N7/ Consider K in place of MG (2 valid contact(s) for K vs. 0 for MG) |
| MG:CA:1677 | 1.00 | 89.43 |  | 0 |  | 1^st^ MG coordination shell missing (see Notes) |
| MG:CA:1678 | 1.00 | 65.66 |  | 0 |  | 1^st^ MG coordination shell missing (see Notes) |
| MG:CA:1679 | 1.00 | 60.80 |  | 0 |  | 1^st^ MG coordination shell missing (see Notes) |
| MG:CA:1680 | 1.00 | 40.43 |  | 0 |  | 1^st^ MG coordination shell missing (see Notes) |
| MG:CA:1681 | 1.00 | 24.64 |  | 1 | 2 | O_ph_ - Exclusion zone contact(s): /OP2/OP2/ |
| MG:CA:1682 | 1.00 | 36.25 |  | 0 | 1 | - Exclusion zone contact(s): /O6/ Consider K in place of MG (1 valid contact(s) for K vs. 0 for MG) |
| MG:CA:1683 | 1.00 | 47.45 |  | 0 | 1 | - Exclusion zone contact(s): /OP2/ Consider K in place of MG (1 valid contact(s) for K vs. 0 for MG) |
| MG:CA:1684 | 1.00 | 103.85 |  | 0 |  | 1^st^ MG coordination shell missing (see Notes) |
| MG:CA:1685 | 1.00 | 80.11 |  | 0 | 1 | - Exclusion zone contact(s): /OP2/ Consider K in place of MG (1 valid contact(s) for K vs. 0 for MG) |
| MG:CA:1686 | 1.00 | 35.08 |  | 0 |  | 1^st^ MG coordination shell missing (see Notes) |
| MG:CA:1687 | 1.00 | 46.64 |  | 2 |  | O_ph_O_b_ |
| MG:CA:1688 | 1.00 | 42.41 |  | 0 | 2 | - Exclusion zone contact(s): /O6/O6/ Consider K in place of MG (2 valid contact(s) for K vs. 0 for MG) |
| MG:CA:1689 | 1.00 | 58.87 |  | 0 | 1 | - Exclusion zone contact(s): /OP1/ Consider NA in place of MG (1 valid contact(s) for NA vs. 0 for MG) |
| MG:CA:1690 | 1.00 | 43.02 |  | 1 | 4 | O_ph_ - Exclusion zone contact(s): /N2/N3_G_/O2/O6/ Consider K in place of MG (4 valid contact(s) for K vs. 1 for MG) |
| MG:CA:1691 | 1.00 | 33.41 |  | 1 |  | O_ph_ |
| MG:CA:1692 | 1.00 | 51.19 |  | 0 | 3 | - Exclusion zone contact(s): /O5'/OP1/OP1/ Consider K in place of MG (3 valid contact(s) for K vs. 0 for MG) |
| MG:CA:1693 | 1.00 | 60.31 |  | 0 |  | 1^st^ MG coordination shell missing (see Notes) |
| MG:CA:1694 | 1.00 | 56.44 |  | 0 | 2 | - Exclusion zone contact(s): /O4/O6/ Consider NA in place of MG (1 valid contact(s) for NA vs. 0 for MG) |
| MG:CA:1695 | 1.00 | 50.62 |  | 0 | 3 | - Exclusion zone contact(s): /O4/O6/O6/ Consider K in place of MG (2 valid contact(s) for K vs. 0 for MG) |
| MG:CA:1696 | 1.00 | 33.42 |  | 1 |  | O_ph_ |
| MG:CA:1697 | 1.00 | 44.85 |  | 1 |  | O_b_ |
| MG:CA:1698 | 1.00 | 101.21 |  | 0 |  | 1^st^ MG coordination shell missing (see Notes) |
| MG:CA:1699 | 1.00 | 40.62 |  | 0 | 2 | - Exclusion zone contact(s): /N7/O6/ Consider K in place of MG (2 valid contact(s) for K vs. 0 for MG) |
| MG:CA:1700 | 1.00 | 39.97 |  | 0 | 2 | - Exclusion zone contact(s): /O2'/OP1/ Consider NA in place of MG (1 valid contact(s) for NA vs. 0 for MG) |
| MG:CA:1701 | 1.00 | 56.25 |  | 0 | 3 | - Exclusion zone contact(s): /O3'/OP1/OP1/ Consider K in place of MG (3 valid contact(s) for K vs. 0 for MG) |
| MG:CA:1702 | 1.00 | 45.86 |  | 0 | 5 | - Exclusion zone contact(s): /O2'/O2'/O3'/OP2/OP2/ Consider K in place of MG (4 valid contact(s) for K vs. 0 for MG) |
| MG:CA:1703 | 1.00 | 49.65 |  | 0 | 3 | - Exclusion zone contact(s): /N6/N7/OP2/ Consider K in place of MG (3 valid contact(s) for K vs. 0 for MG) |
| MG:CA:1704 | 1.00 | 46.27 |  | 1 |  | O_ph_ |
| MG:CA:1705 | 1.00 | 54.72 |  | 0 | 4 | - Exclusion zone contact(s): /N7/N7/O6/OP2/ Consider K in place of MG (4 valid contact(s) for K vs. 0 for MG) |
| MG:CA:1706 | 1.00 | 41.19 |  | 0 | 1 | - Exclusion zone contact(s): /O6/ Consider K in place of MG (1 valid contact(s) for K vs. 0 for MG) |
| MG:CA:1707 | 1.00 | 54.94 |  | 0 |  | 1^st^ MG coordination shell missing (see Notes) |
| MG:CA:1708 | 1.00 | 58.98 |  | 1 | 2 | O_ph_ - Exclusion zone contact(s): /O3'/O4/ Consider K in place of MG (2 valid contact(s) for K vs. 1 for MG) |
| MG:CA:1709 | 1.00 | 55.49 |  | 1 | 1 | O_ph_ - Exclusion zone contact(s): /OP2/ |
| MG:CA:1710 | 1.00 | 30.31 |  | 1 | 5 | O_ph_ - Exclusion zone contact(s): /O2'/O3'/O3'/OP2/OP2/ Consider K in place of MG (4 valid contact(s) for K vs. 1 for MG) |
| MG:CA:1711 | 1.00 | 62.98 |  | 1 | 4 | O_ph_ - Exclusion zone contact(s): /O3'/O5'/OP2/OP2/ Consider NA in place of MG (2 valid contact(s) for NA vs. 1 for MG) |
| MG:CA:1712 | 1.00 | 67.29 |  | 0 |  | 1^st^ MG coordination shell missing (see Notes) |
| MG:CA:1713 | 1.00 | 57.71 |  | 0 |  | 1^st^ MG coordination shell missing (see Notes) |
| MG:CA:1714 | 1.00 | 66.79 |  | 0 | 1 | - Exclusion zone contact(s): /OP1/ Consider K in place of MG (1 valid contact(s) for K vs. 0 for MG) |
| MG:CA:1715 | 1.00 | 62.25 |  | 0 | 2 | - Exclusion zone contact(s): /OP1/OP2/ Consider NA in place of MG (1 valid contact(s) for NA vs. 0 for MG) |
| MG:CA:1716 | 1.00 | 66.44 |  | 0 |  | 1^st^ MG coordination shell missing (see Notes) |
| MG:CA:1717 | 1.00 | 67.35 |  | 0 | 4 | - Exclusion zone contact(s): /O2'/O3'/O6/OP1/ Consider K in place of MG (3 valid contact(s) for K vs. 0 for MG) |
| MG:CA:1718 | 1.00 | 60.75 |  | 0 |  | 1^st^ MG coordination shell missing (see Notes) |
| MG:CA:1719 | 1.00 | 62.66 |  | 1 | 3 | O_ph_ - Exclusion zone contact(s): /OP1/OP1/OP2/ Consider K in place of MG (3 valid contact(s) for K vs. 1 for MG) |
| MG:CA:1720 | 1.00 | 80.08 |  | 0 | 1 | - Exclusion zone contact(s): /OP1/ Consider K in place of MG (1 valid contact(s) for K vs. 0 for MG) |
| MG:CA:1721 | 1.00 | 39.16 |  | 0 | 2 | - Exclusion zone contact(s): /O5'/OP1/ Consider K in place of MG (2 valid contact(s) for K vs. 0 for MG) |
| MG:CA:1722 | 1.00 | 50.14 |  | 0 | 2 | - Exclusion zone contact(s): /O6/O6/ Consider K in place of MG (2 valid contact(s) for K vs. 0 for MG) |
| MG:CA:1723 | 1.00 | 61.13 |  | 1 | 2 | O_ph_ - Exclusion zone contact(s): /O3'/OP2/ Consider K in place of MG (2 valid contact(s) for K vs. 1 for MG) |
| MG:CA:1724 | 1.00 | 45.83 | 1 | 1 |  | - Exclusion zone contact(s): /OP1/ 1^st^ MG coordination shell missing (see Notes) |
| MG:CA:1725 | 1.00 | 55.48 |  | 2 | 1 | 2O_ph_ - Exclusion zone contact(s): /OP1/ |
| MG:CA:1726 | 1.00 | 48.81 |  | 1 | 2 | O_b_ - Exclusion zone contact(s): /O6/O6/ Consider K in place of MG (2 valid contact(s) for K vs. 1 for MG) |
| MG:CA:1727 | 1.00 | 56.63 |  | 0 | 1 | - Exclusion zone contact(s): /OP1/ Consider K in place of MG (1 valid contact(s) for K vs. 0 for MG) |
| MG:CA:1728 | 1.00 | 76.32 |  | 0 | 2 | - Exclusion zone contact(s): /OP1/OP2/ Consider K in place of MG (2 valid contact(s) for K vs. 0 for MG) |
| MG:CA:1729 | 1.00 | 46.83 |  | 0 | 2 | - Exclusion zone contact(s): /O6/O6/ Consider K in place of MG (2 valid contact(s) for K vs. 0 for MG) |
| MG:CA:1730 | 1.00 | 38.09 |  | 0 | 4 | - Exclusion zone contact(s): /O4/O4/O6/O6/ Consider K in place of MG (4 valid contact(s) for K vs. 0 for MG) |
| MG:CA:1731 | 1.00 | 56.72 |  | 0 | 2 | - Exclusion zone contact(s): /N7/O6/ Consider K in place of MG (2 valid contact(s) for K vs. 0 for MG) |
| MG:CA:1732 | 1.00 | 48.61 |  | 0 | 1 | - Exclusion zone contact(s): /O6/ Consider K in place of MG (1 valid contact(s) for K vs. 0 for MG) |
| MG:CA:1733 | 1.00 | 55.12 |  | 0 |  | 1^st^ MG coordination shell missing (see Notes) |
| MG:CA:1734 | 1.00 | 38.46 |  | 1 | 4 | O_r_ - Exclusion zone contact(s): /O3'/O_back_/O_back_/OP2/ Consider K in place of MG (4 valid contact(s) for K vs. 1 for MG) |
| MG:CA:1735 | 1.00 | 42.15 |  | 0 | 3 | - Exclusion zone contact(s): /O5'/OP2/OP2/ Consider K in place of MG (3 valid contact(s) for K vs. 0 for MG) |
| MG:CA:1736 | 1.00 | 30.32 |  | 0 | 1 | - Exclusion zone contact(s): /OP1/ Consider NA in place of MG (1 valid contact(s) for NA vs. 0 for MG) |
| MG:CA:1737 | 1.00 | 77.26 |  | 1 | 1 | O_b_ - Exclusion zone contact(s): /N7/ |
| MG:CA:1738 | 1.00 | 66.63 |  | 0 |  | 1^st^ MG coordination shell missing (see Notes) |
| MG:CA:1739 | 1.00 | 43.68 |  | 2 |  | 2O_ph_ |
| MG:CA:1740 | 1.00 | 48.52 |  | 1 |  | O_ph_ |
| MG:CA:1741 | 1.00 | 66.33 |  | 0 |  | 1^st^ MG coordination shell missing (see Notes) |
| MG:CA:1742 | 1.00 | 40.47 |  | 0 | 2 | - Exclusion zone contact(s): /O3'/OP1/ Consider K in place of MG (2 valid contact(s) for K vs. 0 for MG) |
| MG:CA:1743 | 1.00 | 81.83 |  | 0 | 3 | - Exclusion zone contact(s): /O5'/OP2/OP2/ Consider K in place of MG (3 valid contact(s) for K vs. 0 for MG) |
| MG:CA:1744 | 1.00 | 55.99 |  | 0 | 2 | - Exclusion zone contact(s): /OP1/OP2/ Consider K in place of MG (2 valid contact(s) for K vs. 0 for MG) |
| MG:CA:1745 | 1.00 | 56.00 |  | 0 | 1 | - Exclusion zone contact(s): /N7/ Consider K in place of MG (1 valid contact(s) for K vs. 0 for MG) |
| MG:CA:1746 | 1.00 | 62.27 |  | 0 | 1 | - Exclusion zone contact(s): /OP2/ Consider K in place of MG (1 valid contact(s) for K vs. 0 for MG) |
| MG:CA:1747 | 1.00 | 51.74 |  | 0 | 1 | - Exclusion zone contact(s): /O2'/ Consider K in place of MG (1 valid contact(s) for K vs. 0 for MG) |
| MG:CA:1748 | 1.00 | 27.58 |  | 2 |  | 2O_ph_ |
| MG:CA:1749 | 1.00 | 68.87 |  | 1 | 2 | O_r_ - Exclusion zone contact(s): /O2'/OP2/ Consider K in place of MG (2 valid contact(s) for K vs. 1 for MG) |
| MG:CA:1750 | 1.00 | 82.72 |  | 0 | 2 | - Exclusion zone contact(s): /O2/O6/ Consider K in place of MG (2 valid contact(s) for K vs. 0 for MG) |
| MG:CA:1751 | 1.00 | 34.77 |  | 0 |  | 1^st^ MG coordination shell missing (see Notes) |
| MG:CA:1752 | 1.00 | 53.56 |  | 0 | 3 | - Exclusion zone contact(s): /O4/O6/O6/ Consider K in place of MG (3 valid contact(s) for K vs. 0 for MG) |
| MG:CA:1753 | 1.00 | 56.09 |  | 0 |  | 1^st^ MG coordination shell missing (see Notes) |
| MG:CA:1754 | 1.00 | 80.06 |  | 0 |  | 1^st^ MG coordination shell missing (see Notes) |
| MG:CA:1755 | 1.00 | 75.25 |  | 0 | 4 | - Exclusion zone contact(s): /N7/O5'/O5'/OP2/ Consider K in place of MG (4 valid contact(s) for K vs. 0 for MG) |
| MG:CA:1756 | 1.00 | 53.02 |  | 0 | 2 | - Exclusion zone contact(s): /N7/OP2/ Consider K in place of MG (2 valid contact(s) for K vs. 0 for MG) |
| MG:CA:1757 | 1.00 | 26.86 |  | 0 | 3 | - Exclusion zone contact(s): /O2/O2'/OP1/ Consider K in place of MG (3 valid contact(s) for K vs. 0 for MG) |
| MG:CA:1758 | 1.00 | 52.09 |  | 1 | 2 | O_b_ - Exclusion zone contact(s): /O2'/O2'/ Consider K in place of MG (2 valid contact(s) for K vs. 1 for MG) |
| MG:CA:1759 | 1.00 | 75.93 |  | 0 | 1 | - Exclusion zone contact(s): /O6/ Consider K in place of MG (1 valid contact(s) for K vs. 0 for MG) |
| MG:CA:1760 | 1.00 | 69.93 |  | 0 |  | 1^st^ MG coordination shell missing (see Notes) |
| MG:CA:1761 | 1.00 | 65.36 |  | 0 | 2 | - Exclusion zone contact(s): /N7/O6/ Consider NA in place of MG (1 valid contact(s) for NA vs. 0 for MG) |
| MG:CA:1762 | 1.00 | 49.13 |  | 1 | 3 | O_ph_ - Exclusion zone contact(s): /O3'/OP1/OP2/ Consider K in place of MG (3 valid contact(s) for K vs. 1 for MG) |
| MG:CA:1763 | 1.00 | 39.91 |  | 0 | 4 | - Exclusion zone contact(s): /O2/OP1/OP2/OP2/ Consider K in place of MG (4 valid contact(s) for K vs. 0 for MG) |
| MG:CA:1764 | 1.00 | 48.40 |  | 2 |  | O_ph_O_r_ |
| MG:CA:1765 | 1.00 | 56.77 |  | 0 | 2 | - Exclusion zone contact(s): /O2'/OP2/ Consider K in place of MG (2 valid contact(s) for K vs. 0 for MG) |
| MG:CA:1766 | 1.00 | 59.45 |  | 1 | 1 | O_ph_ - Exclusion zone contact(s): /O2/ |
| MG:CA:1767 | 1.00 | 80.73 |  | 0 | 1 | - Exclusion zone contact(s): /OP1/ Consider K in place of MG (1 valid contact(s) for K vs. 0 for MG) |
| MG:CA:1768 | 1.00 | 48.99 |  | 0 | 3 | - Exclusion zone contact(s): /O3'/OP1/OP2/ Consider K in place of MG (2 valid contact(s) for K vs. 0 for MG) |
| MG:CA:1769 | 1.00 | 42.12 |  | 0 | 4 | - Exclusion zone contact(s): /N7/O3'/O6/O6/ Consider K in place of MG (4 valid contact(s) for K vs. 0 for MG) |
| MG:CA:1770 | 1.00 | 56.39 |  | 1 | 1 | O_ph_ - Exclusion zone contact(s): /OP1/ |
| MG:CA:1771 | 1.00 | 44.95 |  | 0 |  | 1^st^ MG coordination shell missing (see Notes) |
| MG:CA:1772 | 1.00 | 62.50 |  | 0 | 1 | - Exclusion zone contact(s): /OP2/ Consider K in place of MG (1 valid contact(s) for K vs. 0 for MG) |
| MG:CA:1773 | 1.00 | 45.45 |  | 0 | 2 | - Exclusion zone contact(s): /N7/OP1/ Consider K in place of MG (2 valid contact(s) for K vs. 0 for MG) |
| MG:CA:1774 | 1.00 | 78.95 |  | 0 | 2 | - Exclusion zone contact(s): /N7/O4/ Consider K in place of MG (2 valid contact(s) for K vs. 0 for MG) |
| MG:CA:1775 | 1.00 | 51.64 |  | 1 |  | O_ph_ |
| MG:CA:1776 | 1.00 | 62.12 |  | 0 |  | 1^st^ MG coordination shell missing (see Notes) |
| MG:CA:1777 | 1.00 | 49.22 |  | 0 |  | 1^st^ MG coordination shell missing (see Notes) |
| MG:CA:1778 | 1.00 | 69.17 |  | 0 |  | 1^st^ MG coordination shell missing (see Notes) |
| MG:CA:1779 | 1.00 | 38.72 |  | 0 |  | 1^st^ MG coordination shell missing (see Notes) |
| MG:CA:1780 | 1.00 | 56.72 |  | 1 | 1 | O_ph_ - Exclusion zone contact(s): /OP2/ |
| MG:CA:1781 | 1.00 | 45.64 |  | 0 |  | 1^st^ MG coordination shell missing (see Notes) |
| MG:CA:1782 | 1.00 | 50.67 |  | 0 | 1 | - Exclusion zone contact(s): /O2'/ Consider K in place of MG (1 valid contact(s) for K vs. 0 for MG) |
| MG:CA:1783 | 1.00 | 56.80 |  | 1 |  | O_ph_ |
| MG:CA:1784 | 1.00 | 54.22 |  | 0 | 1 | - Exclusion zone contact(s): /N3_C_/ Consider K in place of MG (1 valid contact(s) for K vs. 0 for MG) |
| MG:CA:1785 | 1.00 | 63.90 |  | 0 |  | 1^st^ MG coordination shell missing (see Notes) |
| MG:CA:1786 | 1.00 | 93.01 |  | 0 |  | 1^st^ MG coordination shell missing (see Notes) |
| MG:CA:1787 | 1.00 | 59.29 |  | 0 | 2 | - Exclusion zone contact(s): /N3_C_/O2/ Consider NA in place of MG (1 valid contact(s) for NA vs. 0 for MG) |
| MG:CA:1788 | 1.00 | 79.11 |  | 0 | 1 | - Exclusion zone contact(s): /O2/ Consider NA in place of MG (1 valid contact(s) for NA vs. 0 for MG) |
| MG:CA:1789 | 1.00 | 99.52 |  | 0 | 4 | - Exclusion zone contact(s): /N1_G_/N7/O6/OP2/ Consider K in place of MG (4 valid contact(s) for K vs. 0 for MG) |
| MG:CA:1790 | 1.00 | 81.69 |  | 0 | 1 | - Exclusion zone contact(s): /O6/ Consider K in place of MG (1 valid contact(s) for K vs. 0 for MG) |
| MG:CA:1791 | 1.00 | 87.60 |  | 0 |  | 1^st^ MG coordination shell missing (see Notes) |
| MG:CA:1792 | 1.00 | 58.44 |  | 1 |  | O_b_ |
| MG:CA:1793 | 1.00 | 74.52 |  | 0 | 1 | - Exclusion zone contact(s): /N7/ Consider K in place of MG (1 valid contact(s) for K vs. 0 for MG) |
| MG:CA:1794 | 1.00 | 124.90 |  | 0 | 1 | - Exclusion zone contact(s): /N7/ Consider K in place of MG (1 valid contact(s) for K vs. 0 for MG) |
| MG:CA:1795 | 1.00 | 109.55 |  | 0 | 1 | - Exclusion zone contact(s): /N4/ Consider K in place of MG (1 valid contact(s) for K vs. 0 for MG) |
| MG:CA:1796 | 1.00 | 67.54 |  | 0 |  | 1^st^ MG coordination shell missing (see Notes) |
| MG:CA:1797 | 1.00 | 52.85 |  | 0 |  | 1^st^ MG coordination shell missing (see Notes) |
| MG:CA:1798 | 1.00 | 139.90 |  | 1 |  | O_ph_ |
| MG:CA:1799 | 1.00 | 85.99 |  | 0 | 1 | - Exclusion zone contact(s): /N4/ Consider K in place of MG (1 valid contact(s) for K vs. 0 for MG) |
| MG:CA:1800 | 1.00 | 89.72 |  | 0 | 1 | - Exclusion zone contact(s): /OP2/ Consider K in place of MG (1 valid contact(s) for K vs. 0 for MG) |
| MG:CA:1801 | 1.00 | 44.99 |  | 1 | 2 | O_b_ - Exclusion zone contact(s): /O4/O4/ |
| MG:CA:1802 | 1.00 | 76.97 |  | 0 | 2 | - Exclusion zone contact(s): /N7/OP2/ Consider K in place of MG (2 valid contact(s) for K vs. 0 for MG) |
| MG:CA:1803 | 1.00 | 36.53 |  | 0 | 1 | - Exclusion zone contact(s): /OP1/ Consider K in place of MG (1 valid contact(s) for K vs. 0 for MG) |
| MG:CA:1804 | 1.00 | 67.69 |  | 0 | 2 | - Exclusion zone contact(s): /O3'/OP2/ Consider K in place of MG (2 valid contact(s) for K vs. 0 for MG) |
| MG:CA:1805 | 1.00 | 52.09 |  | 0 |  | 1^st^ MG coordination shell missing (see Notes) |
| MG:CA:1806 | 1.00 | 98.81 |  | 0 |  | 1^st^ MG coordination shell missing (see Notes) |
| MG:CA:1807 | 1.00 | 63.01 |  | 0 | 1 | - Exclusion zone contact(s): /N7/ Consider NA in place of MG (1 valid contact(s) for NA vs. 0 for MG) |
| MG:CA:1808 | 1.00 | 148.65 |  | 0 |  | 1^st^ MG coordination shell missing (see Notes) |
| MG:CA:1809 | 1.00 | 82.99 |  | 0 | 2 | - Exclusion zone contact(s): /O4/O6/ Consider K in place of MG (2 valid contact(s) for K vs. 0 for MG) |
| MG:CA:1810 | 1.00 | 61.30 |  | 0 | 2 | - Exclusion zone contact(s): /N7/O6/ Consider K in place of MG (2 valid contact(s) for K vs. 0 for MG) |
| MG:CA:1811 | 1.00 | 46.19 |  | 0 | 3 | - Exclusion zone contact(s): /O6/O6/O6/ Consider K in place of MG (3 valid contact(s) for K vs. 0 for MG) |
| MG:CA:1812 | 1.00 | 45.95 |  | 0 | 1 | - Exclusion zone contact(s): /O6/ Consider K in place of MG (1 valid contact(s) for K vs. 0 for MG) |
| MG:CA:1813 | 1.00 | 116.94 |  | 0 |  | No contacts in 5.0 A |
| MG:CA:1814 | 1.00 | 42.18 |  | 1 | 2 | O_b_ - Exclusion zone contact(s): /N7/O6/ Consider K in place of MG (2 valid contact(s) for K vs. 1 for MG) |
| MG:CA:1815 | 1.00 | 66.57 |  | 0 | 1 | - Exclusion zone contact(s): /O6/ Consider K in place of MG (1 valid contact(s) for K vs. 0 for MG) |
| MG:CA:1816 | 1.00 | 86.90 |  | 0 |  | No contacts in 5.0 A |
| MG:CA:1817 | 1.00 | 85.77 |  | 0 |  | 1^st^ MG coordination shell missing (see Notes) |
| MG:CA:1818 | 1.00 | 101.57 |  | 0 |  | 1^st^ MG coordination shell missing (see Notes) |
| MG:CA:1819 | 1.00 | 72.71 |  | 0 | 5 | - Exclusion zone contact(s): /N6/O6/O6/OP1/OP1/ Consider K in place of MG (4 valid contact(s) for K vs. 0 for MG) |
| MG:CA:1820 | 1.00 | 67.92 |  | 0 |  | 1^st^ MG coordination shell missing (see Notes) |
| MG:CA:1821 | 1.00 | 51.60 |  | 0 |  | 1^st^ MG coordination shell missing (see Notes) |
| MG:CA:1822 | 1.00 | 56.07 |  | 0 | 1 | - Exclusion zone contact(s): /OP2/ Consider K in place of MG (1 valid contact(s) for K vs. 0 for MG) |
| MG:CA:1823 | 1.00 | 67.02 |  | 0 |  | 1^st^ MG coordination shell missing (see Notes) |
| MG:CA:1824 | 1.00 | 102.82 |  | 0 | 2 | - Exclusion zone contact(s): /OP1/OP2/ Consider K in place of MG (2 valid contact(s) for K vs. 0 for MG) |
| MG:CA:1825 | 1.00 | 90.26 |  | 1 |  | O_ph_ |
| MG:CA:1826 | 1.00 | 83.01 |  | 1 |  | O_ph_ |
| MG:CA:1827 | 1.00 | 101.61 |  | 0 |  | 1^st^ MG coordination shell missing (see Notes) |
| MG:CA:1828 | 1.00 | 100.03 |  | 0 |  | 1^st^ MG coordination shell missing (see Notes) |
| MG:CA:1829 | 1.00 | 61.77 |  | 0 | 2 | - Exclusion zone contact(s): /O4/O6/ Consider K in place of MG (2 valid contact(s) for K vs. 0 for MG) |
| MG:CA:1830 | 1.00 | 59.03 |  | 1 | 3 | O_b_ - Exclusion zone contact(s): /O4/O6/O6/ Consider K in place of MG (2 valid contact(s) for K vs. 1 for MG) |
| MG:CA:1831 | 1.00 | 132.30 |  | 0 | 3 | - Exclusion zone contact(s): /N7/N7/O6/ Consider K in place of MG (2 valid contact(s) for K vs. 0 for MG) |
| MG:CA:1832 | 1.00 | 115.41 |  | 0 | 1 | - Exclusion zone contact(s): /OP2/ Consider K in place of MG (1 valid contact(s) for K vs. 0 for MG) |
| MG:CA:1833 | 1.00 | 57.44 |  | 0 |  | 1^st^ MG coordination shell missing (see Notes) |
| MG:CA:1834 | 1.00 | 67.14 |  | 0 | 4 | - Exclusion zone contact(s): /N1_G_/N7/O6/O6/ Consider K in place of MG (4 valid contact(s) for K vs. 0 for MG) |
| MG:CA:1835 | 1.00 | 114.28 |  | 0 | 3 | - Exclusion zone contact(s): /N1_G_/O2'/O3'/ Consider K in place of MG (3 valid contact(s) for K vs. 0 for MG) |
| MG:CA:1836 | 1.00 | 131.58 |  | 0 | 1 | - Exclusion zone contact(s): /OP2/ Consider NA in place of MG (1 valid contact(s) for NA vs. 0 for MG) |
| MG:CA:1837 | 1.00 | 79.72 |  | 0 | 3 | - Exclusion zone contact(s): /O5'/OP2/OP2/ Consider K in place of MG (2 valid contact(s) for K vs. 0 for MG) |
| MG:CA:1838 | 1.00 | 122.54 |  | 0 |  | 1^st^ MG coordination shell missing (see Notes) |
| MG:CA:1839 | 1.00 | 55.26 | 1 | 1 | 1 | - Exclusion zone contact(s): /O5'/OP2/ Consider NA in place of MG (1 valid contact(s) for NA vs. 0 for MG) |
| MG:CA:1840 | 1.00 | 84.68 |  | 0 |  | 1^st^ MG coordination shell missing (see Notes) |
| MG:CA:1841 | 1.00 | 85.44 |  | 0 |  | 1^st^ MG coordination shell missing (see Notes) |
| MG:CA:1842 | 1.00 | 78.41 |  | 0 | 1 | - Exclusion zone contact(s): /OP1/ Consider NA in place of MG (1 valid contact(s) for NA vs. 0 for MG) |
| MG:CA:1843 | 1.00 | 99.23 |  | 0 | 1 | - Exclusion zone contact(s): /OP1/ Consider K in place of MG (1 valid contact(s) for K vs. 0 for MG) |
| MG:CA:1844 | 1.00 | 57.02 |  | 0 | 1 | - Exclusion zone contact(s): /O6/ Consider K in place of MG (1 valid contact(s) for K vs. 0 for MG) |
| MG:CA:1845 | 1.00 | 108.17 |  | 1 | 2 | O_ph_ - Exclusion zone contact(s): /O5'/OP2/ Consider K in place of MG (2 valid contact(s) for K vs. 1 for MG) |
| MG:CA:1846 | 1.00 | 90.74 |  | 0 | 1 | - Exclusion zone contact(s): /OP2/ Consider K in place of MG (1 valid contact(s) for K vs. 0 for MG) |
| MG:CA:1847 | 1.00 | 95.71 |  | 0 | 1 | - Exclusion zone contact(s): /OP2/ Consider K in place of MG (1 valid contact(s) for K vs. 0 for MG) |
| MG:CA:1848 | 1.00 | 100.86 |  | 0 | 1 | - Exclusion zone contact(s): /OP2/ Consider K in place of MG (1 valid contact(s) for K vs. 0 for MG) |
| MG:CA:1849 | 1.00 | 75.52 |  | 0 |  | 1^st^ MG coordination shell missing (see Notes) |
| MG:CA:1850 | 1.00 | 100.28 |  | 0 |  | 1^st^ MG coordination shell missing (see Notes) |
| MG:CA:1851 | 1.00 | 100.40 |  | 1 | 1 | O_ph_ - Exclusion zone contact(s): /OP2/ |
| MG:CA:1852 | 1.00 | 143.23 |  | 0 |  | 1^st^ MG coordination shell missing (see Notes) |
| MG:CA:1853 | 1.00 | 61.63 |  | 0 |  | 1^st^ MG coordination shell missing (see Notes) |
| MG:CA:1854 | 1.00 | 67.54 |  | 0 | 1 | - Exclusion zone contact(s): /O4/ Consider K in place of MG (1 valid contact(s) for K vs. 0 for MG) |
| MG:CA:1855 | 1.00 | 79.32 |  | 0 | 1 | - Exclusion zone contact(s): /O4/ Consider NA in place of MG (1 valid contact(s) for NA vs. 0 for MG) |
| MG:CA:1856 | 1.00 | 81.02 |  | 0 | 1 | - Exclusion zone contact(s): /N4/ Consider K in place of MG (1 valid contact(s) for K vs. 0 for MG) |
| MG:CA:1857 | 1.00 | 105.63 |  | 0 |  | 1^st^ MG coordination shell missing (see Notes) |
| MG:CA:1858 | 1.00 | 73.76 |  | 0 |  | 1^st^ MG coordination shell missing (see Notes) |
| MG:CA:1859 | 1.00 | 106.51 | 1 | 1 | 1 | - Exclusion zone contact(s): /OP1/OP2/ Consider K in place of MG (1 valid contact(s) for K vs. 0 for MG) |
| MG:CA:1860 | 1.00 | 107.38 |  | 0 |  | 1^st^ MG coordination shell missing (see Notes) |
| MG:CA:1861 | 1.00 | 87.79 |  | 0 | 2 | - Exclusion zone contact(s): /O4/O6/ Consider K in place of MG (2 valid contact(s) for K vs. 0 for MG) |
| MG:CA:1862 | 1.00 | 122.12 |  | 2 | 1 | 2O_b_ - Exclusion zone contact(s): /N1_G_/ |
| MG:CA:1863 | 1.00 | 66.67 |  | 0 | 1 | - Exclusion zone contact(s): /N7/ Consider K in place of MG (1 valid contact(s) for K vs. 0 for MG) |
| MG:CA:1864 | 1.00 | 82.59 |  | 0 | 1 | - Exclusion zone contact(s): /N7/ Consider K in place of MG (1 valid contact(s) for K vs. 0 for MG) |
| MG:CA:1865 | 1.00 | 67.50 | 1 | 1 |  | - Exclusion zone contact(s): /O3'/ 1^st^ MG coordination shell missing (see Notes) |
| MG:CA:1866 | 1.00 | 109.68 | 1 | 1 | 2 | O_r_ - Exclusion zone contact(s): /O5'/OP1/OP2/ |
| MG:CA:1867 | 1.00 | 99.10 |  | 1 | 2 | O_b_ - Exclusion zone contact(s): /O6/O6/ Consider K in place of MG (2 valid contact(s) for K vs. 1 for MG) |
| MG:CA:1868 | 1.00 | 29.76 |  | 0 | 3 | - Exclusion zone contact(s): /O2'/O3'/OP1/ Consider K in place of MG (2 valid contact(s) for K vs. 0 for MG) |
| MG:CA:1869 | 1.00 | 69.45 |  | 0 | 2 | - Exclusion zone contact(s): /OP1/OP1/ Consider NA in place of MG (1 valid contact(s) for NA vs. 0 for MG) |
| MG:CA:1870 | 1.00 | 58.59 |  | 0 | 1 | - Exclusion zone contact(s): /OP1/ Consider K in place of MG (1 valid contact(s) for K vs. 0 for MG) |
| MG:CA:1871 | 1.00 | 98.90 |  | 1 |  | O_ph_ |
| MG:CA:1872 | 1.00 | 74.74 |  | 0 |  | 1^st^ MG coordination shell missing (see Notes) |
| MG:CA:1873 | 1.00 | 51.39 |  | 0 |  | 1^st^ MG coordination shell missing (see Notes) |
| MG:CA:1874 | 1.00 | 55.96 |  | 0 | 5 | - Exclusion zone contact(s): /O3'/O4'/O5'/OP2/ Consider K in place of MG (5 valid contact(s) for K vs. 0 for MG) |
| MG:CA:1875 | 1.00 | 71.16 |  | 1 |  | O_ph_ |
| MG:CA:1876 | 1.00 | 79.38 |  | 2 |  | N_b_Anion |
| MG:CA:1877 | 1.00 | 100.84 |  | 1 | 2 | O_ph_ - Exclusion zone contact(s): /N2/OP1/ Consider K in place of MG (2 valid contact(s) for K vs. 1 for MG) |
| MG:CA:1878 | 1.00 | 26.62 |  | 0 | 4 | - Exclusion zone contact(s): /O4/O4/O6/O6/ Consider K in place of MG (4 valid contact(s) for K vs. 0 for MG) |
| MG:CA:1879 | 1.00 | 58.38 |  | 0 |  | 1^st^ MG coordination shell missing (see Notes) |
| MG:CA:1880 | 1.00 | 35.75 |  | 2 | 4 | O_back_Anion - Exclusion zone contact(s): /N3_C_/N4/N7/O6/ Consider K in place of MG (4 valid contact(s) for K vs. 2 for MG) |
| MG:CA:1881 | 1.00 | 48.23 |  | 0 |  | 1^st^ MG coordination shell missing (see Notes) |
| MG:CA:1882 | 1.00 | 55.24 |  | 0 |  | 1^st^ MG coordination shell missing (see Notes) |
| MG:CA:1883 | 1.00 | 43.83 |  | 0 | 6 | - Exclusion zone contact(s): /N3_A_/N3_G_/N7/O2/O2'/O6/ Consider K in place of MG (5 valid contact(s) for K vs. 0 for MG) |
| MG:CA:1884 | 1.00 | 64.83 |  | 0 | 3 | - Exclusion zone contact(s): /O2'/O5'/OP2/ Consider K in place of MG (2 valid contact(s) for K vs. 0 for MG) |
| MG:CA:1885 | 1.00 | 49.27 |  | 0 | 2 | - Exclusion zone contact(s): /N7/O6/ Consider K in place of MG (2 valid contact(s) for K vs. 0 for MG) |
| MG:CA:1886 | 1.00 | 43.16 |  | 0 | 2 | - Exclusion zone contact(s): /OP2/ Consider K in place of MG (2 valid contact(s) for K vs. 0 for MG) |
| MG:CA:1887 | 1.00 | 92.55 |  | 0 | 1 | - Exclusion zone contact(s): /OP1/ Consider K in place of MG (1 valid contact(s) for K vs. 0 for MG) |
| MG:CA:1888 | 1.00 | 72.47 |  | 0 |  | 1^st^ MG coordination shell missing (see Notes) |
| MG:CA:1889 | 1.00 | 59.65 |  | 0 | 2 | - Exclusion zone contact(s): /O4/O4/ Consider NA in place of MG (1 valid contact(s) for NA vs. 0 for MG) |
| MG:CA:1890 | 1.00 | 69.16 |  | 0 | 2 | - Exclusion zone contact(s): /N3_G_/N6/ Consider K in place of MG (2 valid contact(s) for K vs. 0 for MG) |
| MG:CA:1891 | 1.00 | 56.41 |  | 1 | 1 | N_b_ - Exclusion zone contact(s): /O6/ |
| MG:CA:1892 | 1.00 | 68.58 |  | 0 |  | 1^st^ MG coordination shell missing (see Notes) |
| MG:CA:1893 | 1.00 | 54.17 |  | 1 |  | O_r_ |
| MG:CA:1894 | 1.00 | 60.74 |  | 0 | 2 | - Exclusion zone contact(s): /N6/O6/ Consider K in place of MG (2 valid contact(s) for K vs. 0 for MG) |
| MG:CA:1895 | 1.00 | 58.30 |  | 0 | 4 | - Exclusion zone contact(s): /N2/O4'/O4'/ Consider K in place of MG (4 valid contact(s) for K vs. 0 for MG) |
| MG:CA:1896 | 1.00 | 73.91 |  | 0 | 3 | - Exclusion zone contact(s): /N7/O6/OP2/ Consider K in place of MG (2 valid contact(s) for K vs. 0 for MG) |
| MG:CA:1897 | 1.00 | 71.74 |  | 0 |  | 1^st^ MG coordination shell missing (see Notes) |
| MG:CA:1898 | 1.00 | 59.40 |  | 0 | 6 | - Exclusion zone contact(s): /N7/O4/O5'/OP2/OP2/OP2/ Consider K in place of MG (5 valid contact(s) for K vs. 0 for MG) |
| MG:CA:1899 | 1.00 | 49.07 |  | 0 | 1 | - Exclusion zone contact(s): /O6/ Consider K in place of MG (1 valid contact(s) for K vs. 0 for MG) |
| MG:CA:1900 | 1.00 | 74.09 |  | 1 | 4 | O_ph_ - Exclusion zone contact(s): /N4/NH1_Arg_/NH2_Arg_/O6/ Consider K in place of MG (4 valid contact(s) for K vs. 1 for MG) |
| MG:CA:1901 | 1.00 | 65.66 |  | 0 | 1 | - Exclusion zone contact(s): /O6/ Consider K in place of MG (1 valid contact(s) for K vs. 0 for MG) |
| MG:CA:1902 | 1.00 | 52.50 |  | 0 | 1 | - Exclusion zone contact(s): /OP2/ Consider K in place of MG (1 valid contact(s) for K vs. 0 for MG) |
| MG:CA:1903 | 1.00 | 56.27 |  | 0 | 4 | - Exclusion zone contact(s): /N4/O4/O6/O6/ Consider K in place of MG (4 valid contact(s) for K vs. 0 for MG) |
| MG:CA:1904 | 1.00 | 74.58 |  | 0 |  | 1^st^ MG coordination shell missing (see Notes) |
| MG:CA:1905 | 1.00 | 81.90 |  | 0 | 2 | - Exclusion zone contact(s): /O6/OP2/ Consider K in place of MG (2 valid contact(s) for K vs. 0 for MG) |
| MG:CA:1906 | 1.00 | 50.24 |  | 0 |  | 1^st^ MG coordination shell missing (see Notes) |
| MG:CA:1907 | 1.00 | 66.72 |  | 0 |  | 1^st^ MG coordination shell missing (see Notes) |
| MG:CA:1908 | 1.00 | 61.55 |  | 0 |  | 1^st^ MG coordination shell missing (see Notes) |
| MG:CA:1909 | 1.00 | 69.51 |  | 0 |  | 1^st^ MG coordination shell missing (see Notes) |
| MG:CA:1910 | 1.00 | 54.89 |  | 0 | 3 | - Exclusion zone contact(s): /N3_C_/O2/O_back_/ Consider K in place of MG (3 valid contact(s) for K vs. 0 for MG) |
| MG:CA:1911 | 1.00 | 56.63 |  | 0 | 1 | - Exclusion zone contact(s): /OP1/ Consider K in place of MG (1 valid contact(s) for K vs. 0 for MG) |
| MG:CA:1912 | 1.00 | 81.42 |  | 0 |  | 1^st^ MG coordination shell missing (see Notes) |
| MG:CA:1913 | 1.00 | 52.31 |  | 0 | 3 | - Exclusion zone contact(s): /O2/O2'/OP2/ Consider NA in place of MG (2 valid contact(s) for NA vs. 0 for MG) |
| MG:CA:1914 | 1.00 | 58.51 |  | 0 | 4 | - Exclusion zone contact(s): /NZ_Lys_/O4/O6/O6/ Consider K in place of MG (4 valid contact(s) for K vs. 0 for MG) |
| MG:CA:1915 | 1.00 | 108.81 |  | 0 | 3 | - Exclusion zone contact(s): /N7/O4/O6/ Consider K in place of MG (3 valid contact(s) for K vs. 0 for MG) |
| MG:CA:1916 | 1.00 | 61.34 |  | 0 | 1 | - Exclusion zone contact(s): /OP1/ Consider NA in place of MG (1 valid contact(s) for NA vs. 0 for MG) |
| MG:CA:1917 | 1.00 | 64.29 |  | 0 |  | 1^st^ MG coordination shell missing (see Notes) |
| MG:CA:1918 | 1.00 | 86.79 |  | 0 |  | 1^st^ MG coordination shell missing (see Notes) |
| MG:CA:1919 | 1.00 | 50.35 |  | 0 | 4 | - Exclusion zone contact(s): /O2/O4/O6/O6/ Consider K in place of MG (3 valid contact(s) for K vs. 0 for MG) |
| MG:CA:1920 | 1.00 | 85.33 |  | 0 | 2 | - Exclusion zone contact(s): /O6/OP1/ Consider NA in place of MG (1 valid contact(s) for NA vs. 0 for MG) |
| MG:CA:1921 | 1.00 | 52.61 |  | 0 | 1 | - Exclusion zone contact(s): /N7/ Consider K in place of MG (1 valid contact(s) for K vs. 0 for MG) |
| MG:CA:1922 | 1.00 | 84.83 |  | 0 | 3 | - Exclusion zone contact(s): /O4/OP1/OP2/ Consider K in place of MG (2 valid contact(s) for K vs. 0 for MG) |
| MG:CA:1923 | 1.00 | 48.15 |  | 0 |  | 1^st^ MG coordination shell missing (see Notes) |
| MG:CA:1924 | 1.00 | 88.95 |  | 0 | 2 | - Exclusion zone contact(s): /OP1/OP2/ Consider K in place of MG (2 valid contact(s) for K vs. 0 for MG) |
| MG:CA:1925 | 1.00 | 115.69 |  | 0 | 1 | - Exclusion zone contact(s): /O6/ Consider K in place of MG (1 valid contact(s) for K vs. 0 for MG) |
| MG:CA:1926 | 1.00 | 88.68 |  | 0 | 5 | - Exclusion zone contact(s): /N2/N3_G_/O2'/O6/ Consider K in place of MG (3 valid contact(s) for K vs. 0 for MG) |
| MG:CA:1927 | 1.00 | 62.52 |  | 1 | 1 | O_b_ - Exclusion zone contact(s): /O6/ |
| MG:CA:1928 | 1.00 | 78.47 |  | 0 | 2 | - Exclusion zone contact(s): /O4/O6/ Consider K in place of MG (2 valid contact(s) for K vs. 0 for MG) |
| MG:CA:1929 | 1.00 | 40.97 |  | 1 | 3 | O_ph_ - Exclusion zone contact(s): /O2'/O3'/O3'/ Consider K in place of MG (3 valid contact(s) for K vs. 1 for MG) |
| MG:CA:1930 | 1.00 | 42.30 |  | 0 | 4 | - Exclusion zone contact(s): /N2/N3_G_/OP1/OP2/ Consider NA in place of MG (2 valid contact(s) for NA vs. 0 for MG) |
| MG:CA:1931 | 1.00 | 64.24 |  | 0 |  | 1^st^ MG coordination shell missing (see Notes) |
| MG:CA:1932 | 1.00 | 51.90 |  | 0 |  | 1^st^ MG coordination shell missing (see Notes) |
| MG:CA:1933 | 1.00 | 60.76 |  | 0 | 2 | - Exclusion zone contact(s): /O5'/OP1/ Consider K in place of MG (2 valid contact(s) for K vs. 0 for MG) |
| MG:CA:1934 | 1.00 | 110.36 |  | 0 | 1 | - Exclusion zone contact(s): /N2/ Consider K in place of MG (1 valid contact(s) for K vs. 0 for MG) |
| MG:CA:1935 | 1.00 | 45.14 |  | 0 | 3 | - Exclusion zone contact(s): /O4/O4'/OP1/ Consider K in place of MG (3 valid contact(s) for K vs. 0 for MG) |
| MG:CA:1936 | 1.00 | 55.72 |  | 1 | 2 | O_r_ - Exclusion zone contact(s): /O2'/OP1/ |
| MG:CA:1937 | 1.00 | 73.17 |  | 0 | 3 | - Exclusion zone contact(s): /O_back_/OP1/OP2/ Consider K in place of MG (3 valid contact(s) for K vs. 0 for MG) |
| MG:CA:1938 | 1.00 | 87.67 |  | 0 | 1 | - Exclusion zone contact(s): /OP1/ Consider NA in place of MG (1 valid contact(s) for NA vs. 0 for MG) |
| MG:CA:1939 | 1.00 | 81.41 |  | 0 | 2 | - Exclusion zone contact(s): /N7/N7/ Consider K in place of MG (2 valid contact(s) for K vs. 0 for MG) |
| MG:CA:1940 | 1.00 | 67.63 |  | 0 | 1 | - Exclusion zone contact(s): /N7/ Consider K in place of MG (1 valid contact(s) for K vs. 0 for MG) |
| MG:CA:1941 | 1.00 | 91.87 |  | 0 |  | 1^st^ MG coordination shell missing (see Notes) |
| MG:CA:1942 | 1.00 | 105.81 |  | 0 |  | 1^st^ MG coordination shell missing (see Notes) |
| MG:CA:1943 | 1.00 | 73.55 |  | 0 | 3 | - Exclusion zone contact(s): /N7/O6/O6/ Consider K in place of MG (3 valid contact(s) for K vs. 0 for MG) |
| MG:CA:1944 | 1.00 | 61.29 |  | 1 | 2 | O_ph_ - Exclusion zone contact(s): /O5'/OP2/ Consider K in place of MG (2 valid contact(s) for K vs. 1 for MG) |
| MG:CA:1945 | 1.00 | 78.16 |  | 0 | 1 | - Exclusion zone contact(s): /O4/ Consider K in place of MG (1 valid contact(s) for K vs. 0 for MG) |
| MG:CA:1946 | 1.00 | 112.01 |  | 0 |  | 1^st^ MG coordination shell missing (see Notes) |
| MG:CA:1947 | 1.00 | 82.07 |  | 0 |  | 1^st^ MG coordination shell missing (see Notes) |
| MG:CA:1948 | 1.00 | 97.41 |  | 0 | 2 | - Exclusion zone contact(s): /OP2/OP2/ Consider NA in place of MG (1 valid contact(s) for NA vs. 0 for MG) |
| MG:CA:1949 | 1.00 | 60.65 |  | 0 | 2 | - Exclusion zone contact(s): /O2/O2/ Consider NA in place of MG (2 valid contact(s) for NA vs. 0 for MG) |
| MG:CA:1950 | 1.00 | 86.48 |  | 0 | 6 | - Exclusion zone contact(s): /N1_G_/N2/N3_G_/N3_G_/O2'/O6/ Consider K in place of MG (5 valid contact(s) for K vs. 0 for MG) |
| MG:CA:1951 | 1.00 | 82.60 |  | 0 | 1 | - Exclusion zone contact(s): /OP2/ Consider K in place of MG (1 valid contact(s) for K vs. 0 for MG) |
| MG:CA:1952 | 1.00 | 94.11 |  | 0 | 6 | - Exclusion zone contact(s): /N1/N1_A_/N1_G_/N2/N2/N3_G_/ Consider K in place of MG (6 valid contact(s) for K vs. 0 for MG) |
| MG:CA:1953 | 1.00 | 80.76 |  | 1 |  | O_r_ |
| MG:CA:1954 | 1.00 | 90.06 |  | 0 |  | 1^st^ MG coordination shell missing (see Notes) |
| MG:CA:1955 | 1.00 | 58.97 |  | 0 | 2 | - Exclusion zone contact(s): /O4/O6/ Consider K in place of MG (2 valid contact(s) for K vs. 0 for MG) |
| MG:CA:1956 | 1.00 | 74.66 |  | 0 | 4 | - Exclusion zone contact(s): /N3_U_/O2/OP1/OP1/ Consider K in place of MG (3 valid contact(s) for K vs. 0 for MG) |
| MG:CA:1957 | 1.00 | 56.20 |  | 0 | 1 | - Exclusion zone contact(s): /O2'/ Consider K in place of MG (1 valid contact(s) for K vs. 0 for MG) |
| MG:CA:1958 | 1.00 | 75.36 |  | 0 | 3 | - Exclusion zone contact(s): /O2'/O3'/OP2/ Consider NA in place of MG (2 valid contact(s) for NA vs. 0 for MG) |
| MG:CA:1959 | 1.00 | 56.53 |  | 0 | 1 | - Exclusion zone contact(s): /OP1/ Consider K in place of MG (1 valid contact(s) for K vs. 0 for MG) |
| MG:CA:1960 | 1.00 | 91.28 |  | 0 | 4 | - Exclusion zone contact(s): /N7/O2/O2'/O4'/ Consider K in place of MG (4 valid contact(s) for K vs. 0 for MG) |
| MG:CA:1961 | 1.00 | 70.54 |  | 0 | 1 | - Exclusion zone contact(s): /OP1/ Consider K in place of MG (1 valid contact(s) for K vs. 0 for MG) |
| MG:CA:1962 | 1.00 | 76.90 |  | 0 | 3 | - Exclusion zone contact(s): /O2'/O6/OP2/ Consider K in place of MG (3 valid contact(s) for K vs. 0 for MG) |
| MG:CA:1963 | 1.00 | 71.18 |  | 0 | 3 | - Exclusion zone contact(s): /O3'/O_back_/OP1/ Consider K in place of MG (3 valid contact(s) for K vs. 0 for MG) |
| MG:CA:1964 | 1.00 | 41.26 |  | 0 | 3 | - Exclusion zone contact(s): /N7/O3'/OP2/ Consider K in place of MG (3 valid contact(s) for K vs. 0 for MG) |
| MG:CA:1965 | 1.00 | 81.31 |  | 0 | 3 | - Exclusion zone contact(s): /NE_Arg_/O3'/OP1/ Consider K in place of MG (3 valid contact(s) for K vs. 0 for MG) |
| MG:CA:1966 | 1.00 | 69.38 |  | 0 | 2 | - Exclusion zone contact(s): /N6/OP1/ Consider K in place of MG (2 valid contact(s) for K vs. 0 for MG) |
| MG:CA:1967 | 1.00 | 56.20 |  | 1 |  | O_ph_ |
| MG:CA:1968 | 1.00 | 90.60 |  | 0 |  | 1^st^ MG coordination shell missing (see Notes) |
| MG:CA:1969 | 1.00 | 85.55 |  | 1 | 1 | O_b_ - Exclusion zone contact(s): /N3_U_/ |
| MG:CA:1970 | 1.00 | 100.33 |  | 0 | 2 | - Exclusion zone contact(s): /O4/OP2/ Consider NA in place of MG (1 valid contact(s) for NA vs. 0 for MG) |
| MG:CA:1971 | 1.00 | 54.10 |  | 0 | 1 | - Exclusion zone contact(s): /N7/ Consider K in place of MG (1 valid contact(s) for K vs. 0 for MG) |
| MG:CA:1972 | 1.00 | 68.93 |  | 0 |  | 1^st^ MG coordination shell missing (see Notes) |
| MG:CA:1973 | 1.00 | 65.28 |  | 1 | 2 | Anion - Exclusion zone contact(s): /N4/O6/ |
| MG:CA:1974 | 1.00 | 55.70 |  | 0 | 5 | - Exclusion zone contact(s): /O2'/O3'/O3'/OP1/OP2/ Consider K in place of MG (4 valid contact(s) for K vs. 0 for MG) |
| MG:CA:1975 | 1.00 | 71.28 |  | 0 |  | 1^st^ MG coordination shell missing (see Notes) |
| MG:CA:1976 | 1.00 | 88.44 |  | 0 | 1 | - Exclusion zone contact(s): /O4'/ Consider K in place of MG (1 valid contact(s) for K vs. 0 for MG) |
| MG:CA:1977 | 1.00 | 64.82 |  | 0 | 2 | - Exclusion zone contact(s): /N7/O4/ Consider NA in place of MG (1 valid contact(s) for NA vs. 0 for MG) |
| MG:CA:1978 | 1.00 | 93.29 |  | 0 | 2 | - Exclusion zone contact(s): /N7/O6/ Consider NA in place of MG (2 valid contact(s) for NA vs. 0 for MG) |
| MG:CA:1979 | 1.00 | 58.36 |  | 0 | 2 | - Exclusion zone contact(s): /O6/O6/ Consider NA in place of MG (2 valid contact(s) for NA vs. 0 for MG) |
| MG:CA:1980 | 1.00 | 83.08 |  | 0 | 2 | - Exclusion zone contact(s): /O5'/OP1/ Consider K in place of MG (2 valid contact(s) for K vs. 0 for MG) |
| MG:CA:1981 | 1.00 | 78.94 |  | 0 | 1 | - Exclusion zone contact(s): /O6/ Consider K in place of MG (1 valid contact(s) for K vs. 0 for MG) |
| MG:CA:1982 | 1.00 | 59.17 |  | 0 |  | 1^st^ MG coordination shell missing (see Notes) |
| MG:CA:1983 | 1.00 | 43.69 |  | 0 | 1 | - Exclusion zone contact(s): /OP2/ Consider K in place of MG (1 valid contact(s) for K vs. 0 for MG) |
| MG:CA:1984 | 1.00 | 49.22 |  | 0 |  | 1^st^ MG coordination shell missing (see Notes) |
| MG:CA:1985 | 1.00 | 61.65 |  | 1 | 1 | O_ph_ - Exclusion zone contact(s): /OP2/ |
| MG:CA:1986 | 1.00 | 95.01 |  | 1 | 1 | O_r_ - Exclusion zone contact(s): /OP1/ |
| MG:CA:1987 | 1.00 | 73.76 |  | 1 | 1 | O_ph_ - Exclusion zone contact(s): /N7/ |
| MG:CA:1988 | 1.00 | 57.54 |  | 0 | 1 | - Exclusion zone contact(s): /O6/ Consider K in place of MG (1 valid contact(s) for K vs. 0 for MG) |
| MG:CA:1989 | 1.00 | 60.41 |  | 0 | 1 | - Exclusion zone contact(s): /OP2/ Consider K in place of MG (1 valid contact(s) for K vs. 0 for MG) |
| MG:CA:1990 | 1.00 | 64.13 |  | 0 |  | 1^st^ MG coordination shell missing (see Notes) |
| MG:CA:1991 | 1.00 | 76.23 |  | 1 | 1 | O_ph_ - Exclusion zone contact(s): /OP2/ |
| MG:CA:1992 | 1.00 | 77.22 |  | 0 | 5 | - Exclusion zone contact(s): /O3'/O5'/OP1/OP2/OP2/ Consider K in place of MG (5 valid contact(s) for K vs. 0 for MG) |
| MG:CA:1993 | 1.00 | 91.94 |  | 0 | 5 | - Exclusion zone contact(s): /N1_G_/N2/O6/OP2/ Consider K in place of MG (3 valid contact(s) for K vs. 0 for MG) |
| MG:CA:1994 | 1.00 | 94.31 |  | 0 |  | 1^st^ MG coordination shell missing (see Notes) |
| MG:CA:1995 | 1.00 | 48.34 |  | 0 | 2 | - Exclusion zone contact(s): /O2/O2'/ Consider K in place of MG (2 valid contact(s) for K vs. 0 for MG) |
| MG:CA:1996 | 1.00 | 94.39 |  | 0 | 2 | - Exclusion zone contact(s): /OP1/OP2/ Consider K in place of MG (2 valid contact(s) for K vs. 0 for MG) |
| MG:CA:1997 | 1.00 | 96.53 |  | 0 |  | 1^st^ MG coordination shell missing (see Notes) |
| MG:CA:1998 | 1.00 | 90.02 |  | 1 | 6 | - Exclusion zone contact(s): /N3_G_/N3_G_/N7/N7/O4'/ Consider K in place of MG (6 valid contact(s) for K vs. 1 for MG) |
| MG:CA:1999 | 1.00 | 63.42 |  | 0 | 2 | - Exclusion zone contact(s): /O5'/OP2/ Consider K in place of MG (2 valid contact(s) for K vs. 0 for MG) |
| MG:CA:2000 | 1.00 | 99.05 |  | 0 | 3 | - Exclusion zone contact(s): /O5'/OP1/OP2/ Consider K in place of MG (2 valid contact(s) for K vs. 0 for MG) |
| MG:CA:2001 | 1.00 | 75.04 |  | 0 | 2 | - Exclusion zone contact(s): /OP1/OP2/ Consider K in place of MG (2 valid contact(s) for K vs. 0 for MG) |
| MG:CA:2002 | 1.00 | 112.37 |  | 1 | 4 | O_b_ - Exclusion zone contact(s): /N1_G_/N3_C_/N4/O6/ Consider K in place of MG (4 valid contact(s) for K vs. 1 for MG) |
| MG:CA:2003 | 1.00 | 77.38 |  | 0 |  | 1^st^ MG coordination shell missing (see Notes) |
| MG:CA:2004 | 1.00 | 61.25 |  | 0 | 4 | - Exclusion zone contact(s): /N3_A_/O2'/OH_Tyr_/ Consider K in place of MG (3 valid contact(s) for K vs. 0 for MG) |
| MG:CA:2005 | 1.00 | 55.54 |  | 0 |  | 1^st^ MG coordination shell missing (see Notes) |
| MG:CA:2006 | 1.00 | 67.15 |  | 0 | 2 | - Exclusion zone contact(s): /OP1/OP2/ Consider K in place of MG (2 valid contact(s) for K vs. 0 for MG) |
| MG:CA:2007 | 1.00 | 78.26 |  | 0 | 1 | - Exclusion zone contact(s): /OP1/ Consider K in place of MG (1 valid contact(s) for K vs. 0 for MG) |
| MG:CA:2008 | 1.00 | 56.63 |  | 0 | 2 | - Exclusion zone contact(s): /N7/O4/ Consider K in place of MG (2 valid contact(s) for K vs. 0 for MG) |
| MG:CA:2009 | 1.00 | 61.25 |  | 0 | 2 | - Exclusion zone contact(s): /O5'/OP2/ Consider NA in place of MG (1 valid contact(s) for NA vs. 0 for MG) |
| MG:CA:2010 | 1.00 | 104.61 |  | 0 |  | 1^st^ MG coordination shell missing (see Notes) |
| MG:CA:2011 | 1.00 | 62.87 |  | 0 | 2 | - Exclusion zone contact(s): /N6/O4/ Consider K in place of MG (2 valid contact(s) for K vs. 0 for MG) |
| MG:CA:2012 | 1.00 | 69.45 |  | 0 | 1 | - Exclusion zone contact(s): /OP1/ Consider K in place of MG (1 valid contact(s) for K vs. 0 for MG) |
| MG:CA:2013 | 1.00 | 46.71 |  | 1 | 3 | O_ph_ - Exclusion zone contact(s): /O5'/OP1/OP2/ Consider K in place of MG (2 valid contact(s) for K vs. 1 for MG) |
| MG:CA:2014 | 1.00 | 56.53 |  | 1 | 3 | O_ph_ - Exclusion zone contact(s): /O2'/O3'/OP1/ Consider K in place of MG (2 valid contact(s) for K vs. 1 for MG) |
| MG:CA:2015 | 1.00 | 57.26 |  | 0 | 3 | - Exclusion zone contact(s): /N1_A_/N2/OP1/ Consider K in place of MG (3 valid contact(s) for K vs. 0 for MG) |
| MG:CA:2016 | 1.00 | 80.65 |  | 0 |  | 1^st^ MG coordination shell missing (see Notes) |
| MG:CA:2017 | 1.00 | 57.07 |  | 0 | 3 | - Exclusion zone contact(s): /O4/O6/O6/ Consider K in place of MG (3 valid contact(s) for K vs. 0 for MG) |
| MG:CA:2018 | 1.00 | 106.19 |  | 0 |  | 1^st^ MG coordination shell missing (see Notes) |
| MG:CA:2019 | 1.00 | 68.35 |  | 0 | 1 | - Exclusion zone contact(s): /OP1/ Consider K in place of MG (1 valid contact(s) for K vs. 0 for MG) |
| MG:CA:2020 | 1.00 | 67.87 |  | 0 |  | 1^st^ MG coordination shell missing (see Notes) |
| MG:CA:2021 | 1.00 | 63.10 |  | 0 | 3 | - Exclusion zone contact(s): /O2/O2'/O_back_/ Consider K in place of MG (3 valid contact(s) for K vs. 0 for MG) |
| MG:CA:2022 | 1.00 | 88.19 |  | 0 | 1 | - Exclusion zone contact(s): /N7/ Consider K in place of MG (1 valid contact(s) for K vs. 0 for MG) |
| MG:CA:2023 | 1.00 | 79.19 |  | 0 |  | 1^st^ MG coordination shell missing (see Notes) |
| MG:CA:2024 | 1.00 | 96.92 |  | 0 | 4 | - Exclusion zone contact(s): /N1_G_/O5'/O6/OP2/ Consider K in place of MG (4 valid contact(s) for K vs. 0 for MG) |
| MG:CA:2025 | 1.00 | 95.52 |  | 0 | 2 | - Exclusion zone contact(s): /O5'/OP2/ Consider K in place of MG (2 valid contact(s) for K vs. 0 for MG) |
| MG:CA:2026 | 1.00 | 69.01 |  | 0 | 1 | - Exclusion zone contact(s): /O6/ Consider K in place of MG (1 valid contact(s) for K vs. 0 for MG) |
| MG:CA:2027 | 1.00 | 110.32 |  | 0 |  | 1^st^ MG coordination shell missing (see Notes) |
| MG:CA:2028 | 1.00 | 71.97 |  | 1 | 6 | N_b_ - Exclusion zone contact(s): /N2/N7/N7/O6/O6/O6/ Consider K in place of MG (6 valid contact(s) for K vs. 1 for MG) |
| MG:CA:2029 | 1.00 | 115.83 |  | 1 |  | O_ph_ |
| MG:CA:2030 | 1.00 | 72.95 |  | 0 | 4 | - Exclusion zone contact(s): /N6/N7/O2'/O4/ Consider K in place of MG (4 valid contact(s) for K vs. 0 for MG) |
| MG:CA:2031 | 1.00 | 77.27 |  | 0 | 1 | - Exclusion zone contact(s): /OP2/ Consider K in place of MG (1 valid contact(s) for K vs. 0 for MG) |
| MG:CA:2032 | 1.00 | 69.87 |  | 0 |  | 1^st^ MG coordination shell missing (see Notes) |
| MG:CA:2033 | 1.00 | 73.35 |  | 0 | 2 | - Exclusion zone contact(s): /O2/O4/ Consider K in place of MG (2 valid contact(s) for K vs. 0 for MG) |
| MG:CA:2034 | 1.00 | 61.69 |  | 0 | 1 | - Exclusion zone contact(s): /N7/ Consider K in place of MG (1 valid contact(s) for K vs. 0 for MG) |
| MG:CA:2035 | 1.00 | 68.29 |  | 0 | 2 | - Exclusion zone contact(s): /O5'/OP1/ Consider K in place of MG (2 valid contact(s) for K vs. 0 for MG) |
| MG:CA:2036 | 1.00 | 83.00 |  | 1 | 3 | O_b_ - Exclusion zone contact(s): /N2/N2/O2'/ Consider K in place of MG (3 valid contact(s) for K vs. 1 for MG) |
| MG:CA:2037 | 1.00 | 75.23 |  | 0 |  | 1^st^ MG coordination shell missing (see Notes) |
| MG:CA:2038 | 1.00 | 74.87 |  | 0 | 2 | - Exclusion zone contact(s): /N1/OP2/ Consider K in place of MG (2 valid contact(s) for K vs. 0 for MG) |
| MG:CA:2039 | 1.00 | 66.03 |  | 1 | 2 | O_ph_ - Exclusion zone contact(s): /O3'/OP2/ Consider K in place of MG (2 valid contact(s) for K vs. 1 for MG) |
| MG:CA:2040 | 1.00 | 56.91 |  | 0 | 2 | - Exclusion zone contact(s): /O3'/OP2/ Consider NA in place of MG (1 valid contact(s) for NA vs. 0 for MG) |
| MG:CA:2041 | 1.00 | 64.39 |  | 0 | 2 | - Exclusion zone contact(s): /N1_A_/N6/ Consider K in place of MG (2 valid contact(s) for K vs. 0 for MG) |
| MG:CA:2042 | 1.00 | 57.41 |  | 0 | 1 | - Exclusion zone contact(s): /OP1/ Consider NA in place of MG (1 valid contact(s) for NA vs. 0 for MG) |
| MG:CA:2043 | 1.00 | 100.14 |  | 0 |  | 1^st^ MG coordination shell missing (see Notes) |
| MG:CA:2044 | 1.00 | 65.04 |  | 0 |  | 1^st^ MG coordination shell missing (see Notes) |
| MG:CA:2045 | 1.00 | 66.60 |  | 0 | 2 | - Exclusion zone contact(s): /OP2/OP2/ Consider K in place of MG (2 valid contact(s) for K vs. 0 for MG) |
| MG:CA:2046 | 1.00 | 67.70 |  | 0 |  | 1^st^ MG coordination shell missing (see Notes) |
| MG:CA:2047 | 1.00 | 80.04 |  | 0 | 3 | - Exclusion zone contact(s): /N7/O6/OP2/ Consider K in place of MG (3 valid contact(s) for K vs. 0 for MG) |
| MG:CA:2048 | 1.00 | 77.92 |  | 0 |  | 1^st^ MG coordination shell missing (see Notes) |
| MG:CA:2049 | 1.00 | 113.80 |  | 0 |  | No contacts in 5.0 A |
| MG:CA:2050 | 1.00 | 80.56 |  | 0 |  | 1^st^ MG coordination shell missing (see Notes) |
| MG:CA:2051 | 1.00 | 89.15 |  | 0 | 2 | - Exclusion zone contact(s): /N7/O6/ Consider K in place of MG (2 valid contact(s) for K vs. 0 for MG) |
| MG:CA:2052 | 1.00 | 54.58 |  | 0 | 1 | - Exclusion zone contact(s): /OP2/ Consider NA in place of MG (1 valid contact(s) for NA vs. 0 for MG) |
| MG:CA:2053 | 1.00 | 74.86 |  | 0 | 1 | - Exclusion zone contact(s): /N1/ Consider K in place of MG (1 valid contact(s) for K vs. 0 for MG) |
| MG:CA:2054 | 1.00 | 62.63 |  | 0 | 1 | - Exclusion zone contact(s): /N1/ Consider K in place of MG (1 valid contact(s) for K vs. 0 for MG) |
| MG:CA:2055 | 1.00 | 60.11 |  | 0 | 3 | - Exclusion zone contact(s): /N3_A_/O2'/O3'/ Consider K in place of MG (2 valid contact(s) for K vs. 0 for MG) |
| MG:CA:2056 | 1.00 | 120.86 |  | 0 |  | 1^st^ MG coordination shell missing (see Notes) |
| MG:CA:2057 | 1.00 | 111.72 |  | 0 | 3 | - Exclusion zone contact(s): /N1/O4/O4'/ Consider K in place of MG (3 valid contact(s) for K vs. 0 for MG) |
| MG:CA:2058 | 1.00 | 73.80 |  | 0 | 1 | - Exclusion zone contact(s): /N7/ Consider K in place of MG (1 valid contact(s) for K vs. 0 for MG) |
| MG:CA:2059 | 1.00 | 95.95 |  | 0 | 3 | - Exclusion zone contact(s): /O2'/O3'/OP2/ Consider NA in place of MG (2 valid contact(s) for NA vs. 0 for MG) |
| MG:CA:2060 | 1.00 | 44.57 |  | 0 | 6 | - Exclusion zone contact(s): /N7/N7/N7/O6/OP2/OP2/ Consider K in place of MG (6 valid contact(s) for K vs. 0 for MG) |
| MG:CA:2061 | 1.00 | 50.31 |  | 0 | 2 | - Exclusion zone contact(s): /O5'/OP2/ Consider K in place of MG (2 valid contact(s) for K vs. 0 for MG) |
| MG:CA:2062 | 1.00 | 85.67 |  | 0 | 1 | - Exclusion zone contact(s): /O4/ Consider K in place of MG (1 valid contact(s) for K vs. 0 for MG) |
| MG:CA:2063 | 1.00 | 78.11 |  | 0 |  | 1^st^ MG coordination shell missing (see Notes) |
| MG:CA:2064 | 1.00 | 66.95 |  | 0 |  | 1^st^ MG coordination shell missing (see Notes) |
| MG:CA:2065 | 1.00 | 87.02 |  | 0 |  | 1^st^ MG coordination shell missing (see Notes) |
| MG:CA:2066 | 1.00 | 77.79 |  | 0 |  | 1^st^ MG coordination shell missing (see Notes) |
| MG:CA:2067 | 1.00 | 89.14 |  | 0 | 1 | - Exclusion zone contact(s): /OP2/ Consider K in place of MG (1 valid contact(s) for K vs. 0 for MG) |
| MG:CA:2068 | 1.00 | 89.28 |  | 1 | 1 | O_ph_ - Exclusion zone contact(s): /OP2/ |
| MG:CA:2069 | 1.00 | 74.13 |  | 1 | 1 | O_ph_ - Exclusion zone contact(s): /O3'/ |
| MG:CA:2070 | 1.00 | 68.54 |  | 1 | 2 | O_b_ - Exclusion zone contact(s): /N2/N3_G_/ Consider K in place of MG (2 valid contact(s) for K vs. 1 for MG) |
| MG:CA:2071 | 1.00 | 100.78 |  | 0 |  | No contacts in 5.0 A |
| MG:CA:2072 | 1.00 | 48.20 |  | 0 | 4 | - Exclusion zone contact(s): /O2/O2'/O3'/OP1/ Consider K in place of MG (3 valid contact(s) for K vs. 0 for MG) |
| MG:CA:2073 | 1.00 | 41.74 |  | 0 | 4 | - Exclusion zone contact(s): /N1_A_/O6/OP2/OP2/ Consider K in place of MG (4 valid contact(s) for K vs. 0 for MG) |
| MG:CA:2074 | 1.00 | 61.79 |  | 0 | 4 | - Exclusion zone contact(s): /O2'/OP2/OP2/OP2/ Consider K in place of MG (3 valid contact(s) for K vs. 0 for MG) |
| MG:CA:2075 | 1.00 | 84.77 |  | 0 |  | 1^st^ MG coordination shell missing (see Notes) |
| MG:CA:2076 | 1.00 | 113.39 |  | 0 | 1 | - Exclusion zone contact(s): /O2'/ Consider NA in place of MG (1 valid contact(s) for NA vs. 0 for MG) |
| MG:CA:2077 | 1.00 | 96.38 |  | 1 | 1 | O_ph_ - Exclusion zone contact(s): /OP1/ |
| MG:CA:2078 | 1.00 | 75.98 |  | 0 |  | 1^st^ MG coordination shell missing (see Notes) |
| MG:CA:2079 | 1.00 | 83.00 |  | 0 |  | 1^st^ MG coordination shell missing (see Notes) |
| MG:CA:2080 | 1.00 | 83.09 |  | 0 |  | 1^st^ MG coordination shell missing (see Notes) |
| MG:CA:2081 | 1.00 | 107.58 |  | 0 | 1 | - Exclusion zone contact(s): /N4/ Consider K in place of MG (1 valid contact(s) for K vs. 0 for MG) |
| MG:CA:2082 | 1.00 | 63.34 |  | 1 |  | Anion |
| MG:CA:2083 | 1.00 | 87.74 |  | 1 | 2 | O_ph_ - Exclusion zone contact(s): /O2'/OP1/ Consider K in place of MG (2 valid contact(s) for K vs. 1 for MG) |
| MG:CA:2084 | 1.00 | 77.85 |  | 0 | 1 | - Exclusion zone contact(s): /O6/ Consider K in place of MG (1 valid contact(s) for K vs. 0 for MG) |
| MG:CA:2085 | 1.00 | 58.69 |  | 1 | 4 | N_b_ - Exclusion zone contact(s): /N3_G_/N7/O2'/ Consider K in place of MG (3 valid contact(s) for K vs. 1 for MG) |
| MG:CA:2086 | 1.00 | 83.11 |  | 0 | 2 | - Exclusion zone contact(s): /N1_G_/O6/ Consider K in place of MG (2 valid contact(s) for K vs. 0 for MG) |
| MG:CA:2087 | 1.00 | 95.36 |  | 0 | 4 | - Exclusion zone contact(s): /N6/O4/O4/O6/ Consider K in place of MG (4 valid contact(s) for K vs. 0 for MG) |
| MG:CA:2088 | 1.00 | 65.98 |  | 0 | 2 | - Exclusion zone contact(s): /N3_G_/O2'/ Consider K in place of MG (2 valid contact(s) for K vs. 0 for MG) |
| MG:CA:2089 | 1.00 | 63.91 |  | 0 |  | 1^st^ MG coordination shell missing (see Notes) |
| MG:CA:2090 | 1.00 | 69.67 |  | 0 | 5 | - Exclusion zone contact(s): /N3_G_/O2'/O2'/O3'/OP1/ Consider K in place of MG (3 valid contact(s) for K vs. 0 for MG) |
| MG:CA:2091 | 1.00 | 68.48 |  | 0 | 4 | - Exclusion zone contact(s): /OP1/OP1/OP2/OP2/ Consider K in place of MG (4 valid contact(s) for K vs. 0 for MG) |
| MG:CA:2092 | 1.00 | 94.19 |  | 0 |  | No contacts in 5.0 A |
| MG:CA:2093 | 1.00 | 64.54 |  | 2 | 2 | O_ph_O_r_ - Exclusion zone contact(s): /OP1/OP2/ |
| MG:CA:2094 | 1.00 | 87.90 |  | 0 |  | 1^st^ MG coordination shell missing (see Notes) |
| MG:CA:2095 | 1.00 | 64.76 |  | 0 |  | 1^st^ MG coordination shell missing (see Notes) |
| MG:CA:2096 | 1.00 | 60.19 |  | 0 | 4 | - Exclusion zone contact(s): /N7/O2'/O6/OP2/ Consider K in place of MG (4 valid contact(s) for K vs. 0 for MG) |
| MG:CA:2097 | 1.00 | 82.12 |  | 0 |  | 1^st^ MG coordination shell missing (see Notes) |
| MG:CA:2098 | 1.00 | 64.96 |  | 0 | 4 | - Exclusion zone contact(s): /N4/O4/O6/O6/ Consider K in place of MG (4 valid contact(s) for K vs. 0 for MG) |
| MG:CA:2099 | 1.00 | 94.44 |  | 0 | 1 | - Exclusion zone contact(s): /N7/ Consider K in place of MG (1 valid contact(s) for K vs. 0 for MG) |
| MG:CA:2100 | 1.00 | 40.74 |  | 0 | 6 | - Exclusion zone contact(s): /N1/N1_G_/N3_G_/O2/O3'/O4'/ Consider K in place of MG (5 valid contact(s) for K vs. 0 for MG) |
| MG:CA:2101 | 1.00 | 68.39 |  | 0 | 3 | - Exclusion zone contact(s): /O3'/OP1/OP1/ Consider K in place of MG (3 valid contact(s) for K vs. 0 for MG) |
| MG:CA:2102 | 1.00 | 57.43 |  | 0 |  | 1^st^ MG coordination shell missing (see Notes) |
| MG:CA:2103 | 1.00 | 111.22 |  | 0 | 1 | - Exclusion zone contact(s): /N7/ Consider K in place of MG (1 valid contact(s) for K vs. 0 for MG) |
| MG:CA:2104 | 1.00 | 67.29 |  | 0 | 4 | - Exclusion zone contact(s): /O5'/OP1/OP2/OP2/ Consider K in place of MG (4 valid contact(s) for K vs. 0 for MG) |
| MG:CA:2105 | 1.00 | 94.13 |  | 0 |  | 1^st^ MG coordination shell missing (see Notes) |
| MG:CA:2106 | 1.00 | 83.94 |  | 0 | 3 | - Exclusion zone contact(s): /O3'/O5'/OP2/ Consider K in place of MG (2 valid contact(s) for K vs. 0 for MG) |
| MG:CA:2107 | 1.00 | 73.28 |  | 0 | 1 | - Exclusion zone contact(s): /O2'/ Consider NA in place of MG (1 valid contact(s) for NA vs. 0 for MG) |
| MG:CA:2108 | 1.00 | 78.20 |  | 1 | 1 | O_ph_ - Exclusion zone contact(s): /O6/ |
| MG:CA:2109 | 1.00 | 74.90 |  | 0 | 4 | - Exclusion zone contact(s): /N1_G_/N3_G_/O4/ Consider K in place of MG (4 valid contact(s) for K vs. 0 for MG) |
| MG:CA:2110 | 1.00 | 62.65 |  | 0 |  | 1^st^ MG coordination shell missing (see Notes) |
| MG:CA:2111 | 1.00 | 75.27 |  | 0 | 4 | - Exclusion zone contact(s): /O5'/OP1/OP2/OP2/ Consider NA in place of MG (2 valid contact(s) for NA vs. 0 for MG) |
| MG:CA:2112 | 1.00 | 78.81 |  | 0 | 1 | - Exclusion zone contact(s): /O4/ Consider K in place of MG (1 valid contact(s) for K vs. 0 for MG) |
| MG:CA:2113 | 1.00 | 68.28 |  | 0 |  | 1^st^ MG coordination shell missing (see Notes) |
| MG:CA:2114 | 1.00 | 111.48 |  | 0 |  | 1^st^ MG coordination shell missing (see Notes) |
| MG:CA:2115 | 1.00 | 78.28 |  | 0 |  | 1^st^ MG coordination shell missing (see Notes) |
| MG:CA:2116 | 1.00 | 79.78 |  | 0 | 2 | - Exclusion zone contact(s): /O5'/OP2/ Consider K in place of MG (2 valid contact(s) for K vs. 0 for MG) |
| MG:CA:2117 | 1.00 | 84.72 |  | 0 |  | 1^st^ MG coordination shell missing (see Notes) |
| MG:CA:2118 | 1.00 | 82.08 |  | 0 |  | 1^st^ MG coordination shell missing (see Notes) |
| MG:CA:2119 | 1.00 | 76.41 |  | 0 |  | 1^st^ MG coordination shell missing (see Notes) |
| MG:CA:2120 | 1.00 | 84.06 |  | 0 |  | 1^st^ MG coordination shell missing (see Notes) |
| MG:CA:2121 | 1.00 | 125.81 |  | 0 | 4 | - Exclusion zone contact(s): /O5'/OP1/OP2/OP2/ Consider K in place of MG (4 valid contact(s) for K vs. 0 for MG) |
| MG:CA:2122 | 1.00 | 123.06 |  | 0 | 1 | - Exclusion zone contact(s): /O4/ Consider K in place of MG (1 valid contact(s) for K vs. 0 for MG) |
| MG:CA:2123 | 1.00 | 100.42 |  | 0 | 1 | - Exclusion zone contact(s): /N1_A_/ Consider K in place of MG (1 valid contact(s) for K vs. 0 for MG) |
| MG:CA:2124 | 1.00 | 94.45 |  | 0 |  | 1^st^ MG coordination shell missing (see Notes) |
| MG:CA:2125 | 1.00 | 79.97 |  | 0 | 1 | - Exclusion zone contact(s): /OP2/ Consider K in place of MG (1 valid contact(s) for K vs. 0 for MG) |
| MG:CA:2126 | 1.00 | 59.56 |  | 0 | 1 | - Exclusion zone contact(s): /NH1_Arg_/ Consider K in place of MG (1 valid contact(s) for K vs. 0 for MG) |
| MG:CA:2127 | 1.00 | 88.22 |  | 0 | 3 | - Exclusion zone contact(s): /OP1/OP1/OP2/ Consider K in place of MG (3 valid contact(s) for K vs. 0 for MG) |
| MG:CA:2128 | 1.00 | 101.65 |  | 0 | 4 | - Exclusion zone contact(s): /NH2_Arg_/O3'/O5'/OP2/ Consider K in place of MG (4 valid contact(s) for K vs. 0 for MG) |
| MG:CA:2129 | 1.00 | 75.52 |  | 0 | 2 | - Exclusion zone contact(s): /N7/N7/ Consider K in place of MG (2 valid contact(s) for K vs. 0 for MG) |
| MG:CA:2130 | 1.00 | 117.60 |  | 1 | 2 | O_ph_ - Exclusion zone contact(s): /OP2/OP2/ Consider K in place of MG (2 valid contact(s) for K vs. 1 for MG) |
| MG:CA:2131 | 1.00 | 61.11 |  | 0 |  | 1^st^ MG coordination shell missing (see Notes) |
| MG:CA:2132 | 1.00 | 80.03 |  | 0 | 2 | - Exclusion zone contact(s): /O4/OP2/ Consider K in place of MG (2 valid contact(s) for K vs. 0 for MG) |
| MG:CA:2133 | 1.00 | 67.01 |  | 0 | 2 | - Exclusion zone contact(s): /OP1/OP1/ Consider K in place of MG (2 valid contact(s) for K vs. 0 for MG) |
| MG:CA:2134 | 1.00 | 89.69 |  | 0 | 1 | - Exclusion zone contact(s): /OP1/ Consider NA in place of MG (1 valid contact(s) for NA vs. 0 for MG) |
| MG:CA:2135 | 1.00 | 92.43 |  | 0 | 4 | - Exclusion zone contact(s): /N1_G_/N3_U_/O2/O6/ Consider K in place of MG (3 valid contact(s) for K vs. 0 for MG) |
| MG:CA:2136 | 1.00 | 77.59 |  | 0 | 1 | - Exclusion zone contact(s): /OP2/ Consider K in place of MG (1 valid contact(s) for K vs. 0 for MG) |
| MG:CA:2137 | 1.00 | 88.20 |  | 0 | 2 | - Exclusion zone contact(s): /O2'/O5'/ Consider K in place of MG (2 valid contact(s) for K vs. 0 for MG) |
| MG:CA:2138 | 1.00 | 91.20 |  | 0 |  | 1^st^ MG coordination shell missing (see Notes) |
| MG:CA:2139 | 1.00 | 96.33 |  | 0 | 2 | - Exclusion zone contact(s): /O2'/ Consider K in place of MG (2 valid contact(s) for K vs. 0 for MG) |
| MG:CA:2140 | 1.00 | 110.95 |  | 1 | 4 | O_b_ - Exclusion zone contact(s): /N3_U_/N4/O6/O6/ Consider K in place of MG (4 valid contact(s) for K vs. 1 for MG) |
| MG:CA:2141 | 1.00 | 60.89 |  | 0 | 4 | - Exclusion zone contact(s): /N2/O2/O4/OP2/ Consider K in place of MG (4 valid contact(s) for K vs. 0 for MG) |
| MG:CA:2142 | 1.00 | 110.92 |  | 0 |  | No contacts in 5.0 A |
| MG:CA:2143 | 1.00 | 57.72 |  | 1 | 3 | O_r_ - Exclusion zone contact(s): /N2/O5'/OP1/ Consider K in place of MG (2 valid contact(s) for K vs. 1 for MG) |
| MG:CA:2144 | 1.00 | 61.50 |  | 0 | 3 | - Exclusion zone contact(s): /N7/O6/O6/ Consider K in place of MG (2 valid contact(s) for K vs. 0 for MG) |
| MG:CA:2145 | 1.00 | 80.26 |  | 0 | 2 | - Exclusion zone contact(s): /OP1/OP2/ Consider K in place of MG (2 valid contact(s) for K vs. 0 for MG) |
| MG:CA:2146 | 1.00 | 85.05 |  | 0 |  | 1^st^ MG coordination shell missing (see Notes) |
| MG:CA:2147 | 1.00 | 69.68 |  | 0 |  | 1^st^ MG coordination shell missing (see Notes) |
| MG:CA:2148 | 1.00 | 69.63 |  | 0 |  | 1^st^ MG coordination shell missing (see Notes) |
| MG:CA:2149 | 1.00 | 103.13 |  | 0 |  | 1^st^ MG coordination shell missing (see Notes) |
| MG:CA:2150 | 1.00 | 85.16 |  | 0 | 1 | - Exclusion zone contact(s): /OP1/ Consider NA in place of MG (1 valid contact(s) for NA vs. 0 for MG) |
| MG:CA:2151 | 1.00 | 60.46 |  | 0 |  | 1^st^ MG coordination shell missing (see Notes) |
| MG:CA:2152 | 1.00 | 42.86 |  | 0 | 4 | - Exclusion zone contact(s): /N4/O4/O4/O6/ Consider K in place of MG (3 valid contact(s) for K vs. 0 for MG) |
| MG:CA:2153 | 1.00 | 76.13 |  | 0 | 1 | - Exclusion zone contact(s): /OP1/ Consider NA in place of MG (1 valid contact(s) for NA vs. 0 for MG) |
| MG:CA:2154 | 1.00 | 75.32 |  | 0 |  | 1^st^ MG coordination shell missing (see Notes) |
| MG:CA:2155 | 1.00 | 65.13 |  | 0 | 5 | - Exclusion zone contact(s): /N7/N7/O6/O6/OP2/ Consider K in place of MG (5 valid contact(s) for K vs. 0 for MG) |
| MG:CA:2156 | 1.00 | 92.91 |  | 0 |  | 1^st^ MG coordination shell missing (see Notes) |
| MG:CA:2157 | 1.00 | 79.03 |  | 0 | 4 | - Exclusion zone contact(s): /O2'/OP1/OP2/OP2/ Consider K in place of MG (3 valid contact(s) for K vs. 0 for MG) |
| MG:CA:2158 | 1.00 | 58.92 |  | 0 |  | 1^st^ MG coordination shell missing (see Notes) |
| MG:CA:2159 | 1.00 | 46.41 |  | 0 | 2 | - Exclusion zone contact(s): /N7/OP2/ Consider K in place of MG (2 valid contact(s) for K vs. 0 for MG) |
| MG:CA:2160 | 1.00 | 96.13 |  | 0 | 2 | - Exclusion zone contact(s): /OP1/OP2/ Consider K in place of MG (2 valid contact(s) for K vs. 0 for MG) |
| MG:CA:2161 | 1.00 | 96.23 |  | 0 | 2 | - Exclusion zone contact(s): /N7/O_back_/ Consider K in place of MG (2 valid contact(s) for K vs. 0 for MG) |
| MG:CA:2162 | 1.00 | 79.51 |  | 0 | 4 | - Exclusion zone contact(s): /N4/O6/O6/OP1/ Consider K in place of MG (3 valid contact(s) for K vs. 0 for MG) |
| MG:CA:2163 | 1.00 | 61.03 |  | 0 |  | 1^st^ MG coordination shell missing (see Notes) |
| MG:CA:2164 | 1.00 | 103.36 |  | 0 | 4 | - Exclusion zone contact(s): /N1_G_/O5'/O6/OP2/ Consider K in place of MG (4 valid contact(s) for K vs. 0 for MG) |
| MG:CA:2165 | 1.00 | 101.37 |  | 0 | 1 | - Exclusion zone contact(s): /OP2/ Consider NA in place of MG (1 valid contact(s) for NA vs. 0 for MG) |
| MG:CA:2166 | 1.00 | 91.06 |  | 0 |  | 1^st^ MG coordination shell missing (see Notes) |
| MG:CA:2167 | 1.00 | 97.85 |  | 0 | 2 | - Exclusion zone contact(s): /N4/N7/ Consider K in place of MG (2 valid contact(s) for K vs. 0 for MG) |
| MG:CA:2168 | 1.00 | 108.86 |  | 0 | 2 | - Exclusion zone contact(s): /O5'/OP2/ Consider NA in place of MG (1 valid contact(s) for NA vs. 0 for MG) |
| MG:CA:2169 | 1.00 | 95.45 |  | 0 | 3 | - Exclusion zone contact(s): /O5'/OP2/OP2/ Consider K in place of MG (3 valid contact(s) for K vs. 0 for MG) |
| MG:CA:2170 | 1.00 | 70.77 |  | 0 | 1 | - Exclusion zone contact(s): /OP2/ Consider NA in place of MG (1 valid contact(s) for NA vs. 0 for MG) |
| MG:CA:2171 | 1.00 | 115.33 |  | 0 | 3 | - Exclusion zone contact(s): /O6/OP1/OP2/ Consider K in place of MG (3 valid contact(s) for K vs. 0 for MG) |
| MG:CA:2172 | 1.00 | 95.29 |  | 0 |  | 1^st^ MG coordination shell missing (see Notes) |
| MG:CA:2173 | 1.00 | 71.14 |  | 0 | 1 | - Exclusion zone contact(s): /OP1/ Consider NA in place of MG (1 valid contact(s) for NA vs. 0 for MG) |
| MG:CA:2174 | 1.00 | 100.91 |  | 0 | 4 | - Exclusion zone contact(s): /N1_G_/N7/O6/O6/ Consider K in place of MG (3 valid contact(s) for K vs. 0 for MG) |
| MG:CA:2175 | 1.00 | 89.72 |  | 0 |  | 1^st^ MG coordination shell missing (see Notes) |
| MG:CA:2176 | 1.00 | 62.11 |  | 1 | 3 | O_r_ - Exclusion zone contact(s): /O2'/O3'/O4'/ Consider K in place of MG (2 valid contact(s) for K vs. 1 for MG) |
| MG:CA:2177 | 1.00 | 116.13 |  | 1 | 2 | O_b_ - Exclusion zone contact(s): /N4/O6/ |
| MG:CA:2178 | 1.00 | 92.25 |  | 0 | 3 | - Exclusion zone contact(s): /N3_G_/O2'/O2'/ Consider K in place of MG (3 valid contact(s) for K vs. 0 for MG) |
| MG:CA:2179 | 1.00 | 80.83 |  | 0 |  | 1^st^ MG coordination shell missing (see Notes) |
| MG:CA:2180 | 1.00 | 66.73 |  | 0 | 2 | - Exclusion zone contact(s): /N7/O6/ Consider K in place of MG (2 valid contact(s) for K vs. 0 for MG) |
| MG:CA:2181 | 1.00 | 73.80 |  | 0 |  | No contacts in 5.0 A |
| MG:CA:2182 | 1.00 | 78.85 |  | 0 |  | No contacts in 5.0 A |
| MG:CA:2183 | 1.00 | 62.83 |  | 0 |  | 1^st^ MG coordination shell missing (see Notes) |
| MG:CA:2184 | 1.00 | 83.42 |  | 0 |  | 1^st^ MG coordination shell missing (see Notes) |
| MG:CA:2185 | 1.00 | 87.83 |  | 0 | 1 | - Exclusion zone contact(s): /N7/ Consider K in place of MG (1 valid contact(s) for K vs. 0 for MG) |
| MG:CA:2186 | 1.00 | 121.77 |  | 0 | 3 | - Exclusion zone contact(s): /N4/N7/O6/ Consider K in place of MG (3 valid contact(s) for K vs. 0 for MG) |
| MG:CA:2187 | 1.00 | 80.10 |  | 0 | 1 | - Exclusion zone contact(s): /O3'/ Consider K in place of MG (1 valid contact(s) for K vs. 0 for MG) |
| MG:CA:2188 | 1.00 | 79.13 |  | 0 |  | 1^st^ MG coordination shell missing (see Notes) |
| MG:CA:2189 | 1.00 | 77.50 |  | 0 | 1 | - Exclusion zone contact(s): /OP1/ Consider K in place of MG (1 valid contact(s) for K vs. 0 for MG) |
| MG:CA:2190 | 1.00 | 62.64 |  | 0 |  | 1^st^ MG coordination shell missing (see Notes) |
| MG:CA:2191 | 1.00 | 99.89 |  | 0 | 3 | - Exclusion zone contact(s): /N1/N2/N3_C_/ Consider K in place of MG (3 valid contact(s) for K vs. 0 for MG) |
| MG:CA:2192 | 1.00 | 60.13 |  | 0 |  | 1^st^ MG coordination shell missing (see Notes) |
| MG:CA:2193 | 1.00 | 86.69 |  | 0 | 1 | - Exclusion zone contact(s): /O6/ Consider K in place of MG (1 valid contact(s) for K vs. 0 for MG) |
| MG:CA:2194 | 1.00 | 101.22 |  | 0 | 2 | - Exclusion zone contact(s): /N7/O6/ Consider K in place of MG (2 valid contact(s) for K vs. 0 for MG) |
| MG:CA:2195 | 1.00 | 76.11 |  | 0 | 2 | - Exclusion zone contact(s): /N6/N7/ Consider K in place of MG (2 valid contact(s) for K vs. 0 for MG) |
| MG:CA:2196 | 1.00 | 62.79 |  | 0 | 1 | - Exclusion zone contact(s): /OP1/ Consider K in place of MG (1 valid contact(s) for K vs. 0 for MG) |
| MG:CA:2197 | 1.00 | 107.52 |  | 0 | 2 | - Exclusion zone contact(s): /O5'/OP2/ Consider K in place of MG (2 valid contact(s) for K vs. 0 for MG) |
| MG:CA:2198 | 1.00 | 81.16 |  | 0 |  | 1^st^ MG coordination shell missing (see Notes) |
| MG:CA:2199 | 1.00 | 106.10 |  | 0 | 2 | - Exclusion zone contact(s): /N7/O6/ Consider K in place of MG (2 valid contact(s) for K vs. 0 for MG) |
| MG:CA:2200 | 1.00 | 73.15 |  | 0 | 3 | - Exclusion zone contact(s): /N3_U_/O2/O2'/ Consider K in place of MG (3 valid contact(s) for K vs. 0 for MG) |
| MG:CA:2201 | 1.00 | 80.83 |  | 0 | 2 | - Exclusion zone contact(s): /OP1/OP1/ Consider K in place of MG (2 valid contact(s) for K vs. 0 for MG) |
| MG:CA:2202 | 1.00 | 47.49 |  | 0 | 4 | - Exclusion zone contact(s): /N3_G_/O2'/O4'/O4'/ Consider K in place of MG (4 valid contact(s) for K vs. 0 for MG) |
| MG:CA:2203 | 1.00 | 84.39 |  | 0 | 4 | - Exclusion zone contact(s): /O5'/OP1/OP2/OP2/ Consider K in place of MG (3 valid contact(s) for K vs. 0 for MG) |
| MG:CA:2204 | 1.00 | 87.46 |  | 0 |  | No contacts in 5.0 A |
| MG:CA:2205 | 1.00 | 70.30 |  | 0 |  | No contacts in 5.0 A |
| MG:CA:2206 | 1.00 | 52.15 |  | 0 | 4 | - Exclusion zone contact(s): /N4/N4/O6/OP2/ Consider K in place of MG (4 valid contact(s) for K vs. 0 for MG) |
| MG:CA:2207 | 1.00 | 70.67 |  | 1 | 2 | O_ph_ - Exclusion zone contact(s): /O3'/OP2/ |
| MG:CA:2208 | 1.00 | 81.03 |  | 0 | 3 | - Exclusion zone contact(s): /O5'/OP2/OP2/ Consider K in place of MG (3 valid contact(s) for K vs. 0 for MG) |
| MG:CA:2209 | 1.00 | 68.44 |  | 0 | 1 | - Exclusion zone contact(s): /OP2/ Consider K in place of MG (1 valid contact(s) for K vs. 0 for MG) |
| MG:CA:2210 | 1.00 | 144.13 |  | 0 | 4 | - Exclusion zone contact(s): /N1_G_/N2/O5'/OP1/ Consider K in place of MG (4 valid contact(s) for K vs. 0 for MG) |
| MG:CA:2211 | 1.00 | 90.45 |  | 0 |  | 1^st^ MG coordination shell missing (see Notes) |
| MG:CA:2212 | 1.00 | 69.03 |  | 0 |  | 1^st^ MG coordination shell missing (see Notes) |
| MG:CA:2213 | 1.00 | 84.56 |  | 0 | 2 | - Exclusion zone contact(s): /N7/NZ_Lys_/ Consider K in place of MG (2 valid contact(s) for K vs. 0 for MG) |
| MG:CA:2214 | 1.00 | 104.98 |  | 1 | 3 | N_b_ - Exclusion zone contact(s): /N2/OP2/ Consider K in place of MG (2 valid contact(s) for K vs. 1 for MG) |
| MG:CA:2215 | 1.00 | 63.18 |  | 0 | 3 | - Exclusion zone contact(s): /NZ_Lys_/OP1/OP2/ Consider K in place of MG (3 valid contact(s) for K vs. 0 for MG) |
| MG:CA:2216 | 1.00 | 89.80 |  | 0 | 1 | - Exclusion zone contact(s): /OP1/ Consider K in place of MG (1 valid contact(s) for K vs. 0 for MG) |
| MG:CA:2217 | 1.00 | 93.49 |  | 0 |  | 1^st^ MG coordination shell missing (see Notes) |
| MG:CA:2218 | 1.00 | 82.03 |  | 0 |  | 1^st^ MG coordination shell missing (see Notes) |
| MG:CA:2219 | 1.00 | 57.80 |  | 0 |  | 1^st^ MG coordination shell missing (see Notes) |
| MG:CA:2220 | 1.00 | 112.97 |  | 0 | 1 | - Exclusion zone contact(s): /O5'/ Consider NA in place of MG (1 valid contact(s) for NA vs. 0 for MG) |
| MG:CA:2221 | 1.00 | 72.74 |  | 0 | 1 | - Exclusion zone contact(s): /OP2/ Consider K in place of MG (1 valid contact(s) for K vs. 0 for MG) |
| MG:CA:2222 | 1.00 | 72.12 |  | 0 |  | 1^st^ MG coordination shell missing (see Notes) |
| MG:CA:2223 | 1.00 | 67.64 |  | 0 | 2 | - Exclusion zone contact(s): /O3'/OP1/ Consider K in place of MG (2 valid contact(s) for K vs. 0 for MG) |
| MG:CA:2224 | 1.00 | 83.42 |  | 0 |  | 1^st^ MG coordination shell missing (see Notes) |
| MG:CA:2225 | 1.00 | 70.73 |  | 0 | 1 | - Exclusion zone contact(s): /N7/ Consider K in place of MG (1 valid contact(s) for K vs. 0 for MG) |
| MG:CA:2226 | 1.00 | 53.48 |  | 0 | 3 | - Exclusion zone contact(s): /N6/O3'/OP1/ Consider K in place of MG (3 valid contact(s) for K vs. 0 for MG) |
| MG:CA:2227 | 1.00 | 78.80 |  | 0 |  | 1^st^ MG coordination shell missing (see Notes) |
| MG:CA:2228 | 1.00 | 75.11 |  | 0 | 1 | - Exclusion zone contact(s): /N6/ Consider K in place of MG (1 valid contact(s) for K vs. 0 for MG) |
| MG:CA:2229 | 1.00 | 109.85 |  | 0 | 3 | - Exclusion zone contact(s): /O5'/OP1/OP2/ Consider K in place of MG (3 valid contact(s) for K vs. 0 for MG) |
| MG:CA:2230 | 1.00 | 92.18 |  | 0 | 2 | - Exclusion zone contact(s): /O_back_/O_back_/ Consider K in place of MG (2 valid contact(s) for K vs. 0 for MG) |
| MG:CA:2231 | 1.00 | 95.49 |  | 0 | 3 | - Exclusion zone contact(s): /O5'/OP1/OP2/ Consider K in place of MG (3 valid contact(s) for K vs. 0 for MG) |
| MG:CA:2232 | 1.00 | 86.42 |  | 0 |  | 1^st^ MG coordination shell missing (see Notes) |
| MG:CA:2233 | 1.00 | 60.70 |  | 0 | 3 | - Exclusion zone contact(s): /O2'/O2'/OP1/ Consider K in place of MG (2 valid contact(s) for K vs. 0 for MG) |
| MG:CA:2234 | 1.00 | 81.48 |  | 0 |  | 1^st^ MG coordination shell missing (see Notes) |
| MG:CA:2235 | 1.00 | 70.12 |  | 0 | 1 | - Exclusion zone contact(s): /OP2/ Consider K in place of MG (1 valid contact(s) for K vs. 0 for MG) |
| MG:CA:2236 | 1.00 | 104.58 |  | 0 |  | 1^st^ MG coordination shell missing (see Notes) |
| MG:CA:2237 | 1.00 | 62.84 |  | 0 |  | 1^st^ MG coordination shell missing (see Notes) |
| MG:CA:2238 | 1.00 | 65.84 |  | 0 | 3 | - Exclusion zone contact(s): /O2'/O3'/OP1/ Consider NA in place of MG (2 valid contact(s) for NA vs. 0 for MG) |
| MG:CA:2239 | 1.00 | 65.51 |  | 0 | 1 | - Exclusion zone contact(s): /OP1/ Consider K in place of MG (1 valid contact(s) for K vs. 0 for MG) |
| MG:CA:2240 | 1.00 | 70.80 |  | 0 | 1 | - Exclusion zone contact(s): /OP2/ Consider NA in place of MG (1 valid contact(s) for NA vs. 0 for MG) |
| MG:CA:2241 | 1.00 | 98.86 |  | 0 |  | 1^st^ MG coordination shell missing (see Notes) |
| MG:CA:2242 | 1.00 | 84.77 |  | 0 | 3 | - Exclusion zone contact(s): /O5'/OP2/OP2/ Consider K in place of MG (3 valid contact(s) for K vs. 0 for MG) |
| MG:CA:2243 | 1.00 | 53.02 |  | 1 | 1 | O_ph_ - Exclusion zone contact(s): /OP1/ |
| MG:CA:2244 | 1.00 | 126.37 |  | 0 | 5 | - Exclusion zone contact(s): /N4/N7/O2/O4/O6/ Consider K in place of MG (4 valid contact(s) for K vs. 0 for MG) |
| MG:CA:2245 | 1.00 | 133.17 |  | 0 | 3 | - Exclusion zone contact(s): /O5'/OP2/OP2/ Consider K in place of MG (2 valid contact(s) for K vs. 0 for MG) |
| MG:CA:2246 | 1.00 | 91.79 |  | 0 |  | 1^st^ MG coordination shell missing (see Notes) |
| MG:CA:2247 | 1.00 | 103.88 |  | 0 | 2 | - Exclusion zone contact(s): /N1/OP2/ Consider K in place of MG (2 valid contact(s) for K vs. 0 for MG) |
| MG:CA:2248 | 1.00 | 72.81 |  | 0 | 1 | - Exclusion zone contact(s): /N3_G_/ Consider K in place of MG (1 valid contact(s) for K vs. 0 for MG) |
| MG:CA:2249 | 1.00 | 68.81 |  | 0 | 1 | - Exclusion zone contact(s): /N7/ Consider K in place of MG (1 valid contact(s) for K vs. 0 for MG) |
| MG:CA:2250 | 1.00 | 88.11 |  | 0 | 1 | - Exclusion zone contact(s): /N7/ Consider K in place of MG (1 valid contact(s) for K vs. 0 for MG) |
| MG:CA:2251 | 1.00 | 93.75 |  | 0 | 1 | - Exclusion zone contact(s): /OP1/ Consider K in place of MG (1 valid contact(s) for K vs. 0 for MG) |
| MG:CA:2252 | 1.00 | 85.08 |  | 0 | 1 | - Exclusion zone contact(s): /O2'/ Consider K in place of MG (1 valid contact(s) for K vs. 0 for MG) |
| MG:CA:2253 | 1.00 | 85.45 |  | 0 | 1 | - Exclusion zone contact(s): /OP1/ Consider K in place of MG (1 valid contact(s) for K vs. 0 for MG) |
| MG:CA:2254 | 1.00 | 56.22 |  | 1 | 4 | O_ph_ - Exclusion zone contact(s): /O3'/OP1/OP1/OP1/ Consider K in place of MG (4 valid contact(s) for K vs. 1 for MG) |
| MG:CA:2255 | 1.00 | 95.52 |  | 0 | 4 | - Exclusion zone contact(s): /N3_G_/O2'/O4/O6/ Consider K in place of MG (4 valid contact(s) for K vs. 0 for MG) |
| MG:CA:2256 | 1.00 | 110.84 |  | 0 | 2 | - Exclusion zone contact(s): /O2'/O4'/ Consider K in place of MG (2 valid contact(s) for K vs. 0 for MG) |
| MG:CA:2257 | 1.00 | 60.30 |  | 0 | 5 | - Exclusion zone contact(s): /N2/N3_G_/N7/O2'/ Consider K in place of MG (4 valid contact(s) for K vs. 0 for MG) |
| MG:CA:2258 | 1.00 | 75.76 |  | 0 | 1 | - Exclusion zone contact(s): /OP2/ Consider K in place of MG (1 valid contact(s) for K vs. 0 for MG) |
| MG:CA:2259 | 1.00 | 67.07 |  | 0 |  | 1^st^ MG coordination shell missing (see Notes) |
| MG:CA:2260 | 1.00 | 67.46 |  | 0 | 1 | - Exclusion zone contact(s): /OP2/ Consider K in place of MG (1 valid contact(s) for K vs. 0 for MG) |
| MG:CA:2261 | 1.00 | 66.68 |  | 1 | 3 | O_r_ - Exclusion zone contact(s): /O2/O2'/O4'/ Consider K in place of MG (3 valid contact(s) for K vs. 1 for MG) |
| MG:CA:2262 | 1.00 | 61.54 |  | 0 | 1 | - Exclusion zone contact(s): /N4/ Consider K in place of MG (1 valid contact(s) for K vs. 0 for MG) |
| MG:CA:2263 | 1.00 | 98.12 |  | 0 |  | 1^st^ MG coordination shell missing (see Notes) |
| MG:CA:2264 | 1.00 | 77.38 |  | 0 | 1 | - Exclusion zone contact(s): /OP1/ Consider K in place of MG (1 valid contact(s) for K vs. 0 for MG) |
| MG:CA:2265 | 1.00 | 64.63 |  | 0 | 2 | - Exclusion zone contact(s): /O3'/O5'/ Consider K in place of MG (2 valid contact(s) for K vs. 0 for MG) |
| MG:CA:2266 | 1.00 | 73.50 |  | 0 |  | No contacts in 5.0 A |
| MG:CA:2267 | 1.00 | 80.04 |  | 0 |  | 1^st^ MG coordination shell missing (see Notes) |
| MG:CA:2268 | 1.00 | 86.81 |  | 0 |  | 1^st^ MG coordination shell missing (see Notes) |
| MG:CA:2269 | 1.00 | 71.14 |  | 0 | 2 | - Exclusion zone contact(s): /O2/O2'/ Consider K in place of MG (2 valid contact(s) for K vs. 0 for MG) |
| MG:CA:2270 | 1.00 | 68.99 |  | 0 | 1 | - Exclusion zone contact(s): /O2'/ Consider K in place of MG (1 valid contact(s) for K vs. 0 for MG) |
| MG:CA:2271 | 1.00 | 55.11 |  | 0 | 3 | - Exclusion zone contact(s): /O3'/OP1/OP2/ Consider K in place of MG (3 valid contact(s) for K vs. 0 for MG) |
| MG:CA:2272 | 1.00 | 147.89 |  | 1 | 5 | O_b_ - Exclusion zone contact(s): /N1/N2/N3_C_/O2/O2'/ Consider K in place of MG (3 valid contact(s) for K vs. 1 for MG) |
| MG:CA:2273 | 1.00 | 65.78 |  | 0 | 1 | - Exclusion zone contact(s): /O2'/ Consider K in place of MG (1 valid contact(s) for K vs. 0 for MG) |
| MG:CA:2274 | 1.00 | 54.76 |  | 0 | 2 | - Exclusion zone contact(s): /OP1/OP1/ Consider K in place of MG (2 valid contact(s) for K vs. 0 for MG) |
| MG:CA:2275 | 1.00 | 42.57 |  | 0 | 5 | - Exclusion zone contact(s): /O2/O3'/O3'/OP1/OP2/ Consider K in place of MG (5 valid contact(s) for K vs. 0 for MG) |
| MG:CA:2276 | 1.00 | 69.93 |  | 0 | 2 | - Exclusion zone contact(s): /N6/OP2/ Consider K in place of MG (2 valid contact(s) for K vs. 0 for MG) |
| MG:CA:2277 | 1.00 | 103.51 |  | 1 |  | O_b_ |
| MG:CA:2278 | 1.00 | 84.49 |  | 0 | 1 | - Exclusion zone contact(s): /O6/ Consider K in place of MG (1 valid contact(s) for K vs. 0 for MG) |
| MG:CA:2279 | 1.00 | 67.16 |  | 0 | 1 | - Exclusion zone contact(s): /O2/ Consider K in place of MG (1 valid contact(s) for K vs. 0 for MG) |
| MG:CA:2280 | 1.00 | 84.31 |  | 1 | 2 | N_b_ - Exclusion zone contact(s): /N1_G_/N3_G_/ Consider K in place of MG (2 valid contact(s) for K vs. 1 for MG) |
| MG:CA:2281 | 1.00 | 100.59 |  | 0 | 1 | - Exclusion zone contact(s): /OP1/ Consider NA in place of MG (1 valid contact(s) for NA vs. 0 for MG) |
| MG:CA:2282 | 1.00 | 128.52 |  | 0 | 2 | - Exclusion zone contact(s): /O2'/O3'/ Consider K in place of MG (2 valid contact(s) for K vs. 0 for MG) |
| MG:CA:2283 | 1.00 | 68.01 |  | 0 | 2 | - Exclusion zone contact(s): /O2/O2'/ Consider K in place of MG (2 valid contact(s) for K vs. 0 for MG) |
| MG:CA:2284 | 1.00 | 85.59 |  | 0 | 2 | - Exclusion zone contact(s): /OP1/OP2/ Consider K in place of MG (2 valid contact(s) for K vs. 0 for MG) |
| MG:CA:2285 | 1.00 | 135.68 |  | 0 | 2 | - Exclusion zone contact(s): /O2'/ Consider K in place of MG (2 valid contact(s) for K vs. 0 for MG) |
| MG:CA:2286 | 1.00 | 67.64 |  | 0 | 2 | - Exclusion zone contact(s): /NH2_Arg_/OP1/ Consider NA in place of MG (1 valid contact(s) for NA vs. 0 for MG) |
| MG:CA:2287 | 1.00 | 119.37 |  | 0 | 1 | - Exclusion zone contact(s): /N7/ Consider K in place of MG (1 valid contact(s) for K vs. 0 for MG) |
| MG:CA:2288 | 1.00 | 74.52 |  | 0 | 1 | - Exclusion zone contact(s): /N7/ Consider K in place of MG (1 valid contact(s) for K vs. 0 for MG) |
| MG:CA:2289 | 1.00 | 75.58 |  | 0 |  | 1^st^ MG coordination shell missing (see Notes) |
| MG:CA:2290 | 1.00 | 103.06 |  | 0 | 2 | - Exclusion zone contact(s): /N4/N6/ Consider K in place of MG (2 valid contact(s) for K vs. 0 for MG) |
| MG:CA:2291 | 1.00 | 80.90 |  | 0 | 1 | - Exclusion zone contact(s): /O6/ Consider K in place of MG (1 valid contact(s) for K vs. 0 for MG) |
| MG:CA:2292 | 1.00 | 67.80 |  | 0 |  | 1^st^ MG coordination shell missing (see Notes) |
| MG:CA:2293 | 1.00 | 80.71 |  | 0 |  | 1^st^ MG coordination shell missing (see Notes) |
| MG:CA:2294 | 1.00 | 80.59 |  | 0 |  | 1^st^ MG coordination shell missing (see Notes) |
| MG:CA:2295 | 1.00 | 110.37 |  | 0 |  | 1^st^ MG coordination shell missing (see Notes) |
| MG:CA:2296 | 1.00 | 68.41 |  | 1 | 2 | O_ph_ - Exclusion zone contact(s): /O4/OP2/ Consider K in place of MG (2 valid contact(s) for K vs. 1 for MG) |
| MG:CA:2297 | 1.00 | 71.56 |  | 0 | 2 | - Exclusion zone contact(s): /N7/O6/ Consider K in place of MG (2 valid contact(s) for K vs. 0 for MG) |
| MG:CA:2298 | 1.00 | 68.58 |  | 0 | 4 | - Exclusion zone contact(s): /O2/O2'/O4'/OP1/ Consider K in place of MG (3 valid contact(s) for K vs. 0 for MG) |
| MG:CA:2299 | 1.00 | 111.85 |  | 0 | 2 | - Exclusion zone contact(s): /OP1/OP2/ Consider NA in place of MG (1 valid contact(s) for NA vs. 0 for MG) |
| MG:CA:2300 | 1.00 | 119.35 |  | 1 | 3 | O_ph_ - Exclusion zone contact(s): /O5'/OP2/OP2/ Consider K in place of MG (2 valid contact(s) for K vs. 1 for MG) |
| MG:CA:2301 | 1.00 | 55.93 |  | 0 | 2 | - Exclusion zone contact(s): /O2'/OP1/ Consider K in place of MG (2 valid contact(s) for K vs. 0 for MG) |
| MG:CA:2302 | 1.00 | 77.66 |  | 0 | 1 | - Exclusion zone contact(s): /N7/ Consider K in place of MG (1 valid contact(s) for K vs. 0 for MG) |
| MG:CA:2303 | 1.00 | 102.50 |  | 0 |  | 1^st^ MG coordination shell missing (see Notes) |
| MG:CA:2304 | 1.00 | 124.74 |  | 0 |  | 1^st^ MG coordination shell missing (see Notes) |
| MG:CA:2305 | 1.00 | 81.88 |  | 0 | 1 | - Exclusion zone contact(s): /O6/ Consider K in place of MG (1 valid contact(s) for K vs. 0 for MG) |
| MG:CA:2306 | 1.00 | 90.41 |  | 0 |  | 1^st^ MG coordination shell missing (see Notes) |
| MG:CA:2307 | 1.00 | 76.06 |  | 0 |  | 1^st^ MG coordination shell missing (see Notes) |
| MG:CA:2308 | 1.00 | 107.27 |  | 0 |  | 1^st^ MG coordination shell missing (see Notes) |
| MG:CA:2309 | 1.00 | 92.14 |  | 0 | 3 | - Exclusion zone contact(s): /O2'/O3'/O4'/ Consider NA in place of MG (2 valid contact(s) for NA vs. 0 for MG) |
| MG:CA:2310 | 1.00 | 127.87 |  | 1 | 1 | O_ph_ - Exclusion zone contact(s): /O6/ |
| MG:CA:2311 | 1.00 | 81.83 |  | 0 |  | 1^st^ MG coordination shell missing (see Notes) |
| MG:CA:2312 | 1.00 | 78.80 |  | 0 | 1 | - Exclusion zone contact(s): /O3'/ Consider K in place of MG (1 valid contact(s) for K vs. 0 for MG) |
| MG:CA:2313 | 1.00 | 128.64 |  | 0 | 2 | - Exclusion zone contact(s): /N7/O6/ Consider NA in place of MG (1 valid contact(s) for NA vs. 0 for MG) |
| MG:CA:2314 | 1.00 | 92.48 |  | 0 | 2 | - Exclusion zone contact(s): /O3'/OP2/ Consider K in place of MG (2 valid contact(s) for K vs. 0 for MG) |
| MG:CA:2315 | 1.00 | 94.30 |  | 0 | 2 | - Exclusion zone contact(s): /O5'/OP1/ Consider K in place of MG (2 valid contact(s) for K vs. 0 for MG) |
| MG:CA:2316 | 1.00 | 75.35 |  | 0 |  | 1^st^ MG coordination shell missing (see Notes) |
| MG:CA:2317 | 1.00 | 104.12 |  | 0 |  | No contacts in 5.0 A |
| MG:CA:2318 | 1.00 | 112.87 |  | 0 | 1 | - Exclusion zone contact(s): /OP2/ Consider NA in place of MG (1 valid contact(s) for NA vs. 0 for MG) |
| MG:CA:2319 | 1.00 | 60.65 |  | 0 |  | 1^st^ MG coordination shell missing (see Notes) |
| MG:CA:2320 | 1.00 | 81.42 |  | 0 | 1 | - Exclusion zone contact(s): /O2'/ Consider K in place of MG (1 valid contact(s) for K vs. 0 for MG) |
| MG:CA:2321 | 1.00 | 90.37 |  | 0 |  | 1^st^ MG coordination shell missing (see Notes) |
| MG:CA:2322 | 1.00 | 88.27 |  | 0 |  | 1^st^ MG coordination shell missing (see Notes) |
| MG:CA:2323 | 1.00 | 88.49 |  | 0 |  | 1^st^ MG coordination shell missing (see Notes) |
| MG:CA:2324 | 1.00 | 89.29 |  | 2 | 1 | O_r_N_b_ - Exclusion zone contact(s): /O6/ |
| MG:CB:101 | 1.00 | 64.56 |  | 0 | 1 | - Exclusion zone contact(s): /OP1/ Consider K in place of MG (1 valid contact(s) for K vs. 0 for MG) |
| MG:CB:102 | 1.00 | 65.04 |  | 1 | 2 | O_ph_ - Exclusion zone contact(s): /O3'/OP2/ Consider K in place of MG (2 valid contact(s) for K vs. 1 for MG) |
| MG:CB:103 | 1.00 | 53.31 |  | 0 | 4 | - Exclusion zone contact(s): /O4/O4/OP1/OP2/ Consider K in place of MG (4 valid contact(s) for K vs. 0 for MG) |
| MG:CB:104 | 1.00 | 51.82 |  | 0 |  | 1^st^ MG coordination shell missing (see Notes) |
| MG:CB:105 | 1.00 | 76.29 |  | 0 | 2 | - Exclusion zone contact(s): /OP2/OP2/ Consider NA in place of MG (1 valid contact(s) for NA vs. 0 for MG) |
| MG:CB:106 | 1.00 | 97.15 |  | 0 | 1 | - Exclusion zone contact(s): /OP1/ Consider K in place of MG (1 valid contact(s) for K vs. 0 for MG) |
| MG:CB:107 | 1.00 | 106.66 |  | 0 | 2 | - Exclusion zone contact(s): /OP1/OP2/ Consider NA in place of MG (1 valid contact(s) for NA vs. 0 for MG) |
| MG:CB:108 | 1.00 | 87.89 |  | 0 | 1 | - Exclusion zone contact(s): /N7/ Consider NA in place of MG (1 valid contact(s) for NA vs. 0 for MG) |
| MG:CB:109 | 1.00 | 60.69 |  | 1 | 1 | O_r_ - Exclusion zone contact(s): /O3'/ |
| MG:CB:110 | 1.00 | 39.18 |  | 1 | 1 | O_ph_ - Exclusion zone contact(s): /OP2/ |
| MG:CB:111 | 1.00 | 54.08 |  | 0 |  | 1^st^ MG coordination shell missing (see Notes) |
| MG:CB:112 | 1.00 | 77.66 |  | 0 | 2 | - Exclusion zone contact(s): /N3_A_/O2/ Consider NA in place of MG (1 valid contact(s) for NA vs. 0 for MG) |
| MG:CB:113 | 1.00 | 72.09 |  | 0 | 1 | - Exclusion zone contact(s): /O2'/ Consider NA in place of MG (1 valid contact(s) for NA vs. 0 for MG) |
| MG:CB:114 | 1.00 | 71.25 |  | 0 | 1 | - Exclusion zone contact(s): /O2'/ Consider K in place of MG (1 valid contact(s) for K vs. 0 for MG) |
| MG:CB:115 | 1.00 | 97.63 |  | 0 |  | 1^st^ MG coordination shell missing (see Notes) |
| MG:CB:116 | 1.00 | 89.95 |  | 0 | 2 | - Exclusion zone contact(s): /NH1_Arg_/OP2/ Consider K in place of MG (2 valid contact(s) for K vs. 0 for MG) |
| MG:CB:117 | 1.00 | 68.12 |  | 0 |  | 1^st^ MG coordination shell missing (see Notes) |
| MG:CB:118 | 1.00 | 73.73 |  | 1 | 1 | O_b_ - Exclusion zone contact(s): /O2'/ |
| MG:CB:119 | 1.00 | 69.23 |  | 1 | 2 | O_r_ - Exclusion zone contact(s): /OP1/OP2/ |
| MG:CB:120 | 1.00 | 100.74 |  | 0 | 1 | - Exclusion zone contact(s): /O5'/ Consider NA in place of MG (1 valid contact(s) for NA vs. 0 for MG) |
| MG:CB:121 | 1.00 | 101.57 |  | 0 | 4 | - Exclusion zone contact(s): /N4/N6/N7/O6/ Consider K in place of MG (4 valid contact(s) for K vs. 0 for MG) |
| MG:CC:101 | 1.00 | 41.17 |  | 0 | 1 | - Exclusion zone contact(s): /OP2/ Consider K in place of MG (1 valid contact(s) for K vs. 0 for MG) |
| MG:CC:102 | 1.00 | 28.56 | 1 | 1 | 1 | O_ph_ - Exclusion zone contact(s): /OP1/OP1/ |
| MG:CC:103 | 1.00 | 39.10 |  | 0 | 6 | - Exclusion zone contact(s): /O2'/O2'/O3'/O3'/OP1/OP1/ Consider K in place of MG (5 valid contact(s) for K vs. 0 for MG) |
| MG:CC:104 | 1.00 | 23.37 |  | 0 | 4 | - Exclusion zone contact(s): /O2'/O2'/O3'/O3'/ Consider K in place of MG (4 valid contact(s) for K vs. 0 for MG) |
| MG:CC:105 | 1.00 | 71.64 |  | 0 | 2 | - Exclusion zone contact(s): /O2'/OP1/ Consider NA in place of MG (1 valid contact(s) for NA vs. 0 for MG) |
| MG:CC:106 | 1.00 | 60.34 |  | 0 | 2 | - Exclusion zone contact(s): /OP1/OP2/ Consider K in place of MG (2 valid contact(s) for K vs. 0 for MG) |
| MG:CC:107 | 1.00 | 56.02 |  | 0 |  | 1^st^ MG coordination shell missing (see Notes) |
| MG:CC:108 | 1.00 | 57.54 |  | 0 | 2 | - Exclusion zone contact(s): /N4/OP1/ Consider K in place of MG (2 valid contact(s) for K vs. 0 for MG) |
| MG:CC:109 | 1.00 | 49.00 |  | 0 | 1 | - Exclusion zone contact(s): /OP2/ Consider K in place of MG (1 valid contact(s) for K vs. 0 for MG) |
| MG:CC:110 | 1.00 | 71.98 |  | 0 | 1 | - Exclusion zone contact(s): /OP1/ Consider K in place of MG (1 valid contact(s) for K vs. 0 for MG) |
| MG:CC:111 | 1.00 | 68.15 |  | 0 | 1 | - Exclusion zone contact(s): /OP2/ Consider K in place of MG (1 valid contact(s) for K vs. 0 for MG) |
| MG:CC:112 | 1.00 | 63.55 |  | 0 | 4 | - Exclusion zone contact(s): /O4/O4/OP1/OP2/ Consider K in place of MG (4 valid contact(s) for K vs. 0 for MG) |
| MG:CC:113 | 1.00 | 85.02 |  | 0 |  | 1^st^ MG coordination shell missing (see Notes) |
| MG:CC:114 | 1.00 | 73.02 |  | 0 |  | 1^st^ MG coordination shell missing (see Notes) |
| MG:CC:115 | 1.00 | 69.69 |  | 0 |  | 1^st^ MG coordination shell missing (see Notes) |
| MG:CC:116 | 1.00 | 66.26 |  | 1 |  | O_ph_ |
| MG:CC:117 | 1.00 | 45.94 |  | 0 | 3 | - Exclusion zone contact(s): /N4/N7/O6/ Consider K in place of MG (2 valid contact(s) for K vs. 0 for MG) |
| MG:CC:118 | 1.00 | 66.22 |  | 0 | 2 | - Exclusion zone contact(s): /O5'/OP2/ Consider K in place of MG (2 valid contact(s) for K vs. 0 for MG) |
| MG:CC:119 | 1.00 | 46.62 |  | 0 | 5 | - Exclusion zone contact(s): /O5'/OP1/OP1/OP1/OP2/ Consider K in place of MG (5 valid contact(s) for K vs. 0 for MG) |
| MG:CC:120 | 1.00 | 44.35 |  | 0 | 4 | - Exclusion zone contact(s): /N3_U_/O4/O6/O6/ Consider K in place of MG (4 valid contact(s) for K vs. 0 for MG) |
| MG:CC:121 | 1.00 | 49.61 |  | 0 |  | 1^st^ MG coordination shell missing (see Notes) |
| MG:CC:122 | 1.00 | 69.62 |  | 1 |  | O_r_ |
| MG:CC:123 | 1.00 | 74.59 |  | 1 | 1 | O_ph_ - Exclusion zone contact(s): /OP2/ |
| MG:CC:124 | 1.00 | 77.80 |  | 0 |  | 1^st^ MG coordination shell missing (see Notes) |
| MG:CC:125 | 1.00 | 52.48 |  | 0 | 1 | - Exclusion zone contact(s): /O2'/ Consider K in place of MG (1 valid contact(s) for K vs. 0 for MG) |
| MG:CC:126 | 1.00 | 55.52 |  | 1 | 4 | O_ph_ - Exclusion zone contact(s): /O3'/O5'/OP1/OP2/ Consider K in place of MG (4 valid contact(s) for K vs. 1 for MG) |
| MG:CC:127 | 1.00 | 89.33 |  | 0 |  | 1^st^ MG coordination shell missing (see Notes) |
| MG:CD:101 | 1.00 | 85.24 |  | 0 | 2 | - Exclusion zone contact(s): /O2'/OP2/ Consider K in place of MG (2 valid contact(s) for K vs. 0 for MG) |
| MG:CD:102 | 1.00 | 64.00 |  | 0 | 2 | - Exclusion zone contact(s): /O2/O2/ Consider K in place of MG (2 valid contact(s) for K vs. 0 for MG) |
| MG:CD:103 | 1.00 | 75.38 |  | 0 | 1 | - Exclusion zone contact(s): /OP2/ Consider K in place of MG (1 valid contact(s) for K vs. 0 for MG) |
| MG:CD:104 | 1.00 | 74.16 |  | 0 |  | 1^st^ MG coordination shell missing (see Notes) |
| MG:CD:105 | 1.00 | 62.04 |  | 0 |  | No contacts in 5.0 A |
| MG:CD:106 | 1.00 | 88.22 |  | 0 | 1 | - Exclusion zone contact(s): /O6/ Consider K in place of MG (1 valid contact(s) for K vs. 0 for MG) |
| MG:CD:107 | 1.00 | 57.04 |  | 0 |  | No contacts in 5.0 A |
| MG:CD:108 | 1.00 | 97.69 |  | 0 |  | 1^st^ MG coordination shell missing (see Notes) |
| MG:CD:109 | 1.00 | 70.17 |  | 0 | 2 | - Exclusion zone contact(s): /OP1/OP2/ Consider NA in place of MG (1 valid contact(s) for NA vs. 0 for MG) |
| MG:CD:110 | 1.00 | 77.05 |  | 0 | 3 | - Exclusion zone contact(s): /N4/O6/OP2/ Consider K in place of MG (3 valid contact(s) for K vs. 0 for MG) |
| MG:CD:111 | 1.00 | 69.47 |  | 0 |  | No contacts in 5.0 A |
| MG:CD:112 | 1.00 | 142.13 |  | 0 | 2 | - Exclusion zone contact(s): /N6/O6/ Consider K in place of MG (2 valid contact(s) for K vs. 0 for MG) |
| MG:CD:113 | 1.00 | 75.86 |  | 1 |  | Anion |
| MG:CD:114 | 1.00 | 79.33 |  | 0 | 1 | - Exclusion zone contact(s): /O5'/ Consider NA in place of MG (1 valid contact(s) for NA vs. 0 for MG) |
| MG:CD:115 | 1.00 | 85.34 |  | 0 |  | No contacts in 5.0 A |
| MG:CD:116 | 1.00 | 78.59 |  | 0 |  | 1^st^ MG coordination shell missing (see Notes) |
| MG:CD:117 | 1.00 | 58.05 |  | 0 |  | 1^st^ MG coordination shell missing (see Notes) |
| MG:CD:118 | 1.00 | 97.95 |  | 0 |  | 1^st^ MG coordination shell missing (see Notes) |
| MG:CD:119 | 1.00 | 60.17 |  | 0 | 1 | - Exclusion zone contact(s): /O2'/ Consider K in place of MG (1 valid contact(s) for K vs. 0 for MG) |
| MG:CD:120 | 1.00 | 98.14 |  | 0 | 4 | - Exclusion zone contact(s): /N4/N4/O6/O6/ Consider K in place of MG (4 valid contact(s) for K vs. 0 for MG) |
| MG:CD:121 | 1.00 | 133.56 |  | 1 | 3 | Anion - Exclusion zone contact(s): /N4/O4/O6/ Consider NA in place of MG (2 valid contact(s) for NA vs. 1 for MG) |
| MG:CD:122 | 1.00 | 84.68 |  | 0 | 1 | - Exclusion zone contact(s): /O2'/ Consider K in place of MG (1 valid contact(s) for K vs. 0 for MG) |
| MG:CD:123 | 1.00 | 120.34 |  | 0 | 3 | - Exclusion zone contact(s): /O2/O6/O6/ Consider K in place of MG (2 valid contact(s) for K vs. 0 for MG) |
| MG:CD:124 | 1.00 | 73.45 |  | 0 |  | No contacts in 5.0 A |
| MG:CD:125 | 1.00 | 88.31 |  | 0 | 1 | - Exclusion zone contact(s): /OP1/ Consider K in place of MG (1 valid contact(s) for K vs. 0 for MG) |
| MG:CD:126 | 1.00 | 88.01 |  | 1 |  | Anion |
| MG:CD:127 | 1.00 | 101.47 |  | 0 |  | No contacts in 5.0 A |
| MG:CD:128 | 1.00 | 66.43 |  | 0 | 2 | - Exclusion zone contact(s): /N3_C_/N4/ Consider K in place of MG (2 valid contact(s) for K vs. 0 for MG) |
| MG:CD:129 | 1.00 | 95.20 |  | 0 | 2 | - Exclusion zone contact(s): /N1_A_/N7/ Consider K in place of MG (2 valid contact(s) for K vs. 0 for MG) |
| MG:CD:130 | 1.00 | 84.66 |  | 0 | 1 | - Exclusion zone contact(s): /O2'/ Consider K in place of MG (1 valid contact(s) for K vs. 0 for MG) |
| MG:CE:301 | 1.00 | 61.52 |  | 0 | 2 | - Exclusion zone contact(s): /N_back_/OG_Ser_/ Consider NA in place of MG (1 valid contact(s) for NA vs. 0 for MG) |
| MG:CE:302 | 1.00 | 98.31 |  | 0 | 1 | - Exclusion zone contact(s): /O_back_/ Consider K in place of MG (1 valid contact(s) for K vs. 0 for MG) |
| MG:CE:303 | 1.00 | 83.18 |  | 0 | 3 | - Exclusion zone contact(s): /N_back_/ND1_His_/O_back_/ Consider K in place of MG (3 valid contact(s) for K vs. 0 for MG) |
| MG:CE:304 | 1.00 | 62.94 |  | 0 | 2 | - Exclusion zone contact(s): /OD1_Asp_/OD1_Asp_/ Consider NA in place of MG (1 valid contact(s) for NA vs. 0 for MG) |
| MG:CE:305 | 1.00 | 95.18 |  | 0 | 1 | - Exclusion zone contact(s): /O_back_/ Consider K in place of MG (1 valid contact(s) for K vs. 0 for MG) |
| MG:CE:306 | 1.00 | 96.44 |  | 0 |  | 1^st^ MG coordination shell missing (see Notes) |
| MG:CF:301 | 1.00 | 75.25 |  | 0 |  | 1^st^ MG coordination shell missing (see Notes) |
| MG:CF:302 | 1.00 | 79.11 |  | 0 |  | No contacts in 5.0 A |
| MG:CF:303 | 1.00 | 79.12 |  | 0 | 3 | - Exclusion zone contact(s): /NZ_Lys_/O3'/O_back_/ Consider K in place of MG (3 valid contact(s) for K vs. 0 for MG) |
| ZN:CG:301 | 1.00 | 74.63 |  | 1 |  | O_back_ |
| MG:CG:302 | 1.00 | 67.42 |  | 0 |  | 1^st^ MG coordination shell missing (see Notes) |
| MG:CG:303 | 1.00 | 73.71 |  | 0 | 1 | Consider K in place of MG (1 valid contact(s) for K vs. 0 for MG) |
| MG:CG:304 | 1.00 | 95.01 |  | 0 | 1 | Consider K in place of MG (1 valid contact(s) for K vs. 0 for MG) |
| MG:CG:305 | 1.00 | 80.17 |  | 0 | 1 | Consider K in place of MG (1 valid contact(s) for K vs. 0 for MG) |
| MG:CG:306 | 1.00 | 124.11 |  | 0 |  | 1^st^ MG coordination shell missing (see Notes) |
| MG:CG:307 | 1.00 | 106.51 |  | 2 | 2 | O_back_OH_Prot_ - Exclusion zone contact(s): /O_back_/O_back_/ |
| MG:CG:308 | 1.00 | 100.16 |  | 0 | 1 | - Exclusion zone contact(s): /N_back_/ Consider K in place of MG (1 valid contact(s) for K vs. 0 for MG) |
| MG:CG:309 | 1.00 | 78.10 |  | 1 | 3 | Anion - Exclusion zone contact(s): /N_back_/N_back_/N_back_/ Consider K in place of MG (3 valid contact(s) for K vs. 1 for MG) |
| MG:CG:310 | 1.00 | 96.96 |  | 0 | 2 | - Exclusion zone contact(s): /N_back_/O_back_/ Consider K in place of MG (2 valid contact(s) for K vs. 0 for MG) |
| MG:CG:311 | 1.00 | 150.41 |  | 1 | 2 | O_back_ - Exclusion zone contact(s): /N_back_/O_back_/ Consider K in place of MG (2 valid contact(s) for K vs. 1 for MG) |
| MG:CG:312 | 1.00 | 104.84 |  | 0 | 2 | - Exclusion zone contact(s): /NZ_Lys_/OP1/ Consider K in place of MG (2 valid contact(s) for K vs. 0 for MG) |
| MG:CH:201 | 1.00 | 55.54 |  | 0 |  | 1^st^ MG coordination shell missing (see Notes) |
| MG:CH:202 | 1.00 | 62.95 |  | 0 |  | 1^st^ MG coordination shell missing (see Notes) |
| MG:CH:203 | 1.00 | 70.99 |  | 0 | 2 | - Exclusion zone contact(s): /OD1_Asp_/OE2_Glu_/ Consider NA in place of MG (2 valid contact(s) for NA vs. 0 for MG) |
| MG:CH:204 | 1.00 | 93.07 |  | 0 | 3 | - Exclusion zone contact(s): /N_back_/N_back_/N_back_/ Consider K in place of MG (3 valid contact(s) for K vs. 0 for MG) |
| MG:CH:205 | 1.00 | 87.80 |  | 0 |  | 1^st^ MG coordination shell missing (see Notes) |
| MG:CH:206 | 1.00 | 70.08 |  | 0 | 4 | - Exclusion zone contact(s): /N_back_/N_back_/N_back_/O_back_/ Consider K in place of MG (3 valid contact(s) for K vs. 0 for MG) |
| MG:CI:201 | 1.00 | 56.10 |  | 0 | 3 | - Exclusion zone contact(s): /O_back_/O_back_/O_back_/ Consider K in place of MG (2 valid contact(s) for K vs. 0 for MG) |
| MG:CJ:201 | 1.00 | 89.00 |  | 0 | 2 | - Exclusion zone contact(s): /N_back_/O_back_/ Consider K in place of MG (2 valid contact(s) for K vs. 0 for MG) |
| MG:CK:201 | 1.00 | 102.93 |  | 1 | 1 | Anion - Exclusion zone contact(s): /OP2/ |
| MG:CK:202 | 1.00 | 79.89 |  | 1 |  | Anion |
| MG:CK:203 | 1.00 | 68.13 |  | 0 | 1 | - Exclusion zone contact(s): /NH1_Arg_/ Consider K in place of MG (1 valid contact(s) for K vs. 0 for MG) |
| MG:CK:204 | 1.00 | 60.63 |  | 0 | 1 | - Exclusion zone contact(s): /NE_Arg_/ Consider K in place of MG (1 valid contact(s) for K vs. 0 for MG) |
| MG:CK:205 | 1.00 | 75.51 |  | 0 |  | 1^st^ MG coordination shell missing (see Notes) |
| MG:CK:206 | 1.00 | 62.04 |  | 0 |  | 1^st^ MG coordination shell missing (see Notes) |
| MG:CK:207 | 1.00 | 80.37 |  | 0 | 4 | - Exclusion zone contact(s): /O2'/O4'/O_back_/O_back_/ Consider K in place of MG (3 valid contact(s) for K vs. 0 for MG) |
| MG:CK:208 | 1.00 | 69.98 |  | 0 | 1 | - Exclusion zone contact(s): /O2'/ Consider K in place of MG (1 valid contact(s) for K vs. 0 for MG) |
| MG:CK:209 | 1.00 | 60.85 |  | 0 | 1 | - Exclusion zone contact(s): /N_back_/ Consider K in place of MG (1 valid contact(s) for K vs. 0 for MG) |
| MG:CK:210 | 1.00 | 71.99 |  | 0 | 2 | - Exclusion zone contact(s): /NE_Arg_/NH2_Arg_/ Consider NA in place of MG (1 valid contact(s) for NA vs. 0 for MG) |
| MG:CK:211 | 1.00 | 57.18 |  | 0 | 1 | - Exclusion zone contact(s): /NH2_Arg_/ Consider K in place of MG (1 valid contact(s) for K vs. 0 for MG) |
| MG:CL:201 | 1.00 | 54.90 |  | 1 | 3 | O_ph_ - Exclusion zone contact(s): /N_back_/O_back_/OP2/ Consider K in place of MG (3 valid contact(s) for K vs. 1 for MG) |
| MG:CM:201 | 1.00 | 74.25 |  | 0 |  | No contacts in 5.0 A |
| MG:CM:202 | 1.00 | 86.14 |  | 0 |  | 1^st^ MG coordination shell missing (see Notes) |
| MG:CM:203 | 1.00 | 89.50 |  | 0 | 1 | - Exclusion zone contact(s): /OD1_Asp_/ Consider K in place of MG (1 valid contact(s) for K vs. 0 for MG) |
| MG:CN:201 | 1.00 | 33.20 |  | 1 | 4 | O_cno_ - Exclusion zone contact(s): /O4/O6/O_back_/O_back_/ Consider K in place of MG (3 valid contact(s) for K vs. 1 for MG) |
| MG:CN:202 | 1.00 | 64.37 |  | 0 |  | 1^st^ MG coordination shell missing (see Notes) |
| MG:CP:201 | 1.00 | 54.39 |  | 2 | 3 | O_ph_O_back_ - Exclusion zone contact(s): /O_back_/O_back_/OP2/ |
| MG:CP:202 | 1.00 | 63.80 |  | 0 |  | 1^st^ MG coordination shell missing (see Notes) |
| MG:CP:203 | 1.00 | 77.67 |  | 0 | 1 | - Exclusion zone contact(s): /N_back_/ Consider K in place of MG (1 valid contact(s) for K vs. 0 for MG) |
| ZN:CQ:101 | 1.00 | 114.41 |  | 0 | 1 | - Exclusion zone contact(s): /N_back_/ |
| MG:CQ:102 | 1.00 | 51.17 |  | 0 | 3 | - Exclusion zone contact(s): /N2/N_back_/NH1_Arg_/ Consider K in place of MG (2 valid contact(s) for K vs. 0 for MG) |
| MG:CQ:103 | 1.00 | 51.51 |  | 0 | 4 | - Exclusion zone contact(s): /N_back_/O_back_/OG1_Thr_/OP2/ Consider K in place of MG (3 valid contact(s) for K vs. 0 for MG) |
| MG:CQ:104 | 1.00 | 56.62 |  | 0 | 3 | - Exclusion zone contact(s): /N_back_/OG1_Thr_/OP1/ Consider K in place of MG (2 valid contact(s) for K vs. 0 for MG) |
| MG:CR:101 | 1.00 | 89.81 |  | 0 | 1 | - Exclusion zone contact(s): /OE1_Glu_/ Consider NA in place of MG (1 valid contact(s) for NA vs. 0 for MG) |
| MG:CR:102 | 1.00 | 43.51 |  | 1 | 4 | Anion - Exclusion zone contact(s): /N_back_/ND1_His_/O2'/O_back_/ Consider NA in place of MG (2 valid contact(s) for NA vs. 1 for MG) |
| MG:CR:103 | 1.00 | 70.95 |  | 0 |  | 1^st^ MG coordination shell missing (see Notes) |
| MG:CS:101 | 1.00 | 61.30 |  | 0 |  | 1^st^ MG coordination shell missing (see Notes) |
| MG:CS:102 | 1.00 | 65.66 |  | 0 | 1 | - Exclusion zone contact(s): /NE1_Trp_/ Consider K in place of MG (1 valid contact(s) for K vs. 0 for MG) |
| MG:CS:103 | 1.00 | 74.34 |  | 0 | 2 | - Exclusion zone contact(s): /N_back_/O_back_/ Consider K in place of MG (2 valid contact(s) for K vs. 0 for MG) |
| MG:CS:104 | 1.00 | 125.81 |  | 1 |  | O_ph_ |
| MG:CS:105 | 1.00 | 69.52 |  | 0 | 3 | - Exclusion zone contact(s): /N3_A_/N_back_/O2'/ Consider K in place of MG (3 valid contact(s) for K vs. 0 for MG) |
| MG:CS:106 | 1.00 | 77.71 |  | 0 | 2 | - Exclusion zone contact(s): /O_back_/OP1/ Consider K in place of MG (2 valid contact(s) for K vs. 0 for MG) |
| MG:CT:201 | 1.00 | 99.68 |  | 0 | 2 | - Exclusion zone contact(s): /NZ_Lys_/O_back_/ Consider K in place of MG (2 valid contact(s) for K vs. 0 for MG) |
| MG:CT:202 | 1.00 | 85.60 |  | 0 | 2 | - Exclusion zone contact(s): /N_back_/N_back_/ Consider K in place of MG (2 valid contact(s) for K vs. 0 for MG) |
| MG:CT:203 | 1.00 | 54.59 |  | 0 | 3 | - Exclusion zone contact(s): /NZ_Lys_/OP1/OP2/ Consider K in place of MG (3 valid contact(s) for K vs. 0 for MG) |
| MG:CV:101 | 1.00 | 53.57 |  | 1 | 3 | Anion - Exclusion zone contact(s): /N_back_/OP1/OP2/ Consider K in place of MG (3 valid contact(s) for K vs. 1 for MG) |
| MG:CV:102 | 1.00 | 63.09 |  | 0 |  | 1^st^ MG coordination shell missing (see Notes) |
| MG:CV:103 | 1.00 | 84.13 |  | 0 | 3 | - Exclusion zone contact(s): /N_back_/O_back_/O_back_/ Consider K in place of MG (3 valid contact(s) for K vs. 0 for MG) |
| MG:CV:104 | 1.00 | 86.68 |  | 1 | 2 | Anion - Exclusion zone contact(s): /N_back_/O_back_/ |
| MG:CW:201 | 1.00 | 89.70 |  | 0 | 1 | - Exclusion zone contact(s): /N_back_/ Consider K in place of MG (1 valid contact(s) for K vs. 0 for MG) |
| MG:D0:201 | 1.00 | 37.93 |  | 0 |  | 1^st^ MG coordination shell missing (see Notes) |
| MG:D0:202 | 1.00 | 62.38 |  | 0 |  | 1^st^ MG coordination shell missing (see Notes) |
| MG:D0:203 | 1.00 | 97.33 |  | 0 |  | 1^st^ MG coordination shell missing (see Notes) |
| MG:D0:204 | 1.00 | 27.90 |  | 3 | 1 | 2O_ph_O_back_ - Exclusion zone contact(s): /ND1_His_/ |
| MG:D0:205 | 1.00 | 41.44 |  | 0 | 1 | - Exclusion zone contact(s): /O_back_/ Consider K in place of MG (1 valid contact(s) for K vs. 0 for MG) |
| MG:D0:206 | 1.00 | 45.81 |  | 0 | 4 | - Exclusion zone contact(s): /NH2_Arg_/NZ_Lys_/OP1/OP1/ Consider NA in place of MG (3 valid contact(s) for NA vs. 0 for MG) |
| MG:D0:207 | 1.00 | 58.49 |  | 0 | 1 | - Exclusion zone contact(s): /N_back_/ Consider K in place of MG (1 valid contact(s) for K vs. 0 for MG) |
| MG:D0:208 | 1.00 | 72.65 |  | 0 | 2 | - Exclusion zone contact(s): /N3_A_/N_back_/ Consider K in place of MG (2 valid contact(s) for K vs. 0 for MG) |
| MG:D0:209 | 1.00 | 91.95 |  | 0 |  | 1^st^ MG coordination shell missing (see Notes) |
| MG:D0:210 | 1.00 | 75.81 |  | 0 | 2 | - Exclusion zone contact(s): /NH2_Arg_/OP1/ Consider NA in place of MG (1 valid contact(s) for NA vs. 0 for MG) |
| MG:D0:211 | 1.00 | 89.41 |  | 0 | 2 | - Exclusion zone contact(s): /N_back_/N_back_/ Consider K in place of MG (2 valid contact(s) for K vs. 0 for MG) |
| MG:D1:201 | 1.00 | 49.79 |  | 0 | 2 | - Exclusion zone contact(s): /OG1_Thr_/OP2/ Consider K in place of MG (2 valid contact(s) for K vs. 0 for MG) |
| MG:D1:202 | 1.00 | 57.61 |  | 0 | 2 | - Exclusion zone contact(s): /NH2_Arg_/OP2/ Consider K in place of MG (2 valid contact(s) for K vs. 0 for MG) |
| MG:D1:203 | 1.00 | 42.27 |  | 0 | 5 | - Exclusion zone contact(s): /N_back_/N_back_/O_back_/O_back_/OP1/ Consider K in place of MG (3 valid contact(s) for K vs. 0 for MG) |
| MG:D1:204 | 1.00 | 59.04 |  | 0 | 3 | - Exclusion zone contact(s): /O2'/O_back_/O_back_/ Consider K in place of MG (3 valid contact(s) for K vs. 0 for MG) |
| MG:D1:205 | 1.00 | 55.50 |  | 0 | 3 | - Exclusion zone contact(s): /NH2_Arg_/O_back_/OP1/ Consider K in place of MG (3 valid contact(s) for K vs. 0 for MG) |
| MG:D1:206 | 1.00 | 68.95 |  | 0 | 2 | - Exclusion zone contact(s): /O_back_/OP1/ Consider K in place of MG (2 valid contact(s) for K vs. 0 for MG) |
| MG:D1:207 | 1.00 | 64.78 |  | 0 | 4 | - Exclusion zone contact(s): /NE_Arg_/NE_Arg_/NH2_Arg_/NH2_Arg_/ Consider K in place of MG (4 valid contact(s) for K vs. 0 for MG) |
| MG:D1:208 | 1.00 | 58.21 |  | 0 | 1 | - Exclusion zone contact(s): /NH2_Arg_/ Consider K in place of MG (1 valid contact(s) for K vs. 0 for MG) |
| MG:D1:209 | 1.00 | 65.60 |  | 0 | 1 | - Exclusion zone contact(s): /O_back_/ Consider K in place of MG (1 valid contact(s) for K vs. 0 for MG) |
| MG:D1:210 | 1.00 | 66.63 |  | 1 | 1 | O_ph_ - Exclusion zone contact(s): /NZ_Lys_/ |
| MG:D1:211 | 1.00 | 49.07 |  | 0 | 2 | - Exclusion zone contact(s): /NE_Arg_/OD2_Asp_/ Consider NA in place of MG (1 valid contact(s) for NA vs. 0 for MG) |
| MG:D1:212 | 1.00 | 53.96 |  | 0 | 3 | - Exclusion zone contact(s): /N6/N7/O_back_/ Consider NA in place of MG (2 valid contact(s) for NA vs. 0 for MG) |
| MG:D2:201 | 1.00 | 48.99 |  | 0 |  | 1^st^ MG coordination shell missing (see Notes) |
| MG:D2:202 | 1.00 | 52.61 |  | 0 |  | 1^st^ MG coordination shell missing (see Notes) |
| MG:D2:203 | 1.00 | 57.68 |  | 0 | 5 | - Exclusion zone contact(s): /N_back_/NE_Arg_/NH2_Arg_/O_back_/OP1/ Consider K in place of MG (4 valid contact(s) for K vs. 0 for MG) |
| MG:D2:204 | 1.00 | 51.62 |  | 0 | 1 | - Exclusion zone contact(s): /ND1_His_/ Consider K in place of MG (1 valid contact(s) for K vs. 0 for MG) |
| MG:D2:205 | 1.00 | 84.09 |  | 0 | 2 | - Exclusion zone contact(s): /N_back_/OP1/ Consider NA in place of MG (2 valid contact(s) for NA vs. 0 for MG) |
| MG:D2:206 | 1.00 | 55.81 |  | 0 | 2 | - Exclusion zone contact(s): /O_back_/OP2/ Consider K in place of MG (2 valid contact(s) for K vs. 0 for MG) |
| MG:D2:207 | 1.00 | 68.64 |  | 0 |  | 1^st^ MG coordination shell missing (see Notes) |
| MG:D2:208 | 1.00 | 71.44 |  | 0 | 2 | - Exclusion zone contact(s): /N_back_/O_back_/ Consider K in place of MG (2 valid contact(s) for K vs. 0 for MG) |
| MG:D2:209 | 1.00 | 74.84 |  | 0 | 1 | - Exclusion zone contact(s): /N_back_/ Consider K in place of MG (1 valid contact(s) for K vs. 0 for MG) |
| MG:D2:210 | 1.00 | 98.60 |  | 0 |  | No contacts in 5.0 A |
| MG:D3:101 | 1.00 | 33.62 | 3 | 3 | 2 | - Exclusion zone contact(s): /NE_Arg_/NH1_Arg_/NH2_Arg_/O2/O6/ Consider K in place of MG (2 valid contact(s) for K vs. 0 for MG) |
| MG:D3:102 | 1.00 | 47.42 |  | 0 | 4 | - Exclusion zone contact(s): /N_back_/O2'/O3'/O_back_/ Consider K in place of MG (3 valid contact(s) for K vs. 0 for MG) |
| MG:D3:103 | 1.00 | 23.80 |  | 0 | 1 | - Exclusion zone contact(s): /OP1/ Consider NA in place of MG (1 valid contact(s) for NA vs. 0 for MG) |
| MG:D3:104 | 1.00 | 32.59 |  | 0 | 1 | - Exclusion zone contact(s): /NH1_Arg_/ Consider K in place of MG (1 valid contact(s) for K vs. 0 for MG) |
| MG:D3:105 | 1.00 | 48.02 |  | 0 | 2 | - Exclusion zone contact(s): /N_back_/OP1/ Consider K in place of MG (2 valid contact(s) for K vs. 0 for MG) |
| MG:D3:106 | 1.00 | 58.34 |  | 0 | 2 | - Exclusion zone contact(s): /N_back_/OP1/ Consider NA in place of MG (1 valid contact(s) for NA vs. 0 for MG) |
| MG:D3:107 | 1.00 | 53.44 |  | 0 | 3 | - Exclusion zone contact(s): /OD1_Asp_/OG_Ser_/OP2/ Consider K in place of MG (3 valid contact(s) for K vs. 0 for MG) |
| MG:D4:101 | 1.00 | 96.49 |  | 0 |  | No contacts in 5.0 A |
| MG:D4:102 | 1.00 | 86.33 |  | 0 | 1 | - Exclusion zone contact(s): /N_back_/ Consider K in place of MG (1 valid contact(s) for K vs. 0 for MG) |
| ZN:D4:103 | 1.00 | 200.41 |  | 0 | 1 | - Exclusion zone contact(s): /N_back_/ |
| MG:D5:101 | 1.00 | 31.63 |  | 1 | 1 | His - Exclusion zone contact(s): /OP2/ |
| MG:D5:102 | 1.00 | 24.45 |  | 0 | 2 | - Exclusion zone contact(s): /NZ_Lys_/OP1/ Consider K in place of MG (2 valid contact(s) for K vs. 0 for MG) |
| MG:D5:103 | 1.00 | 32.87 |  | 0 | 2 | - Exclusion zone contact(s): /NE2_His_/O4/ Consider NA in place of MG (1 valid contact(s) for NA vs. 0 for MG) |
| MG:D5:104 | 1.00 | 38.07 |  | 0 | 1 | - Exclusion zone contact(s): /N_back_/ Consider NA in place of MG (1 valid contact(s) for NA vs. 0 for MG) |
| MG:D5:105 | 1.00 | 63.75 |  | 0 | 2 | - Exclusion zone contact(s): /OH_Tyr_/OP1/ Consider NA in place of MG (1 valid contact(s) for NA vs. 0 for MG) |
| MG:D5:106 | 1.00 | 58.27 |  | 0 | 4 | - Exclusion zone contact(s): /N_back_/O2/O2'/O_back_/ Consider K in place of MG (4 valid contact(s) for K vs. 0 for MG) |
| MG:D5:107 | 1.00 | 59.10 |  | 1 | 3 | Anion - Exclusion zone contact(s): /N_back_/O_back_/O_back_/ Consider K in place of MG (2 valid contact(s) for K vs. 1 for MG) |
| MG:D5:108 | 1.00 | 63.37 |  | 0 | 4 | - Exclusion zone contact(s): /N_back_/O2/O2'/O_back_/ Consider K in place of MG (3 valid contact(s) for K vs. 0 for MG) |
| MG:D6:101 | 1.00 | 63.33 |  | 0 |  | 1^st^ MG coordination shell missing (see Notes) |
| MG:D6:102 | 1.00 | 45.89 |  | 0 |  | 1^st^ MG coordination shell missing (see Notes) |
| MG:D6:103 | 1.00 | 63.95 |  | 0 | 2 | - Exclusion zone contact(s): /NE1_Trp_/O2'/ Consider NA in place of MG (1 valid contact(s) for NA vs. 0 for MG) |
| MG:D7:101 | 1.00 | 52.03 |  | 0 |  | 1^st^ MG coordination shell missing (see Notes) |
| MG:D7:102 | 1.00 | 41.16 |  | 0 | 3 | - Exclusion zone contact(s): /N_back_/NE2_Gln_/O2/ Consider K in place of MG (3 valid contact(s) for K vs. 0 for MG) |
| MG:D7:103 | 1.00 | 45.17 |  | 0 | 2 | - Exclusion zone contact(s): /N_back_/OP1/ Consider K in place of MG (2 valid contact(s) for K vs. 0 for MG) |
| MG:D7:104 | 1.00 | 47.09 |  | 0 | 2 | - Exclusion zone contact(s): /NE_Arg_/OP1/ Consider NA in place of MG (1 valid contact(s) for NA vs. 0 for MG) |
| MG:D7:105 | 1.00 | 52.18 |  | 0 |  | 1^st^ MG coordination shell missing (see Notes) |
| MG:D8:101 | 1.00 | 42.63 |  | 0 | 1 | - Exclusion zone contact(s): /O_back_/ Consider K in place of MG (1 valid contact(s) for K vs. 0 for MG) |
| MG:D8:102 | 1.00 | 37.12 |  | 0 | 4 | - Exclusion zone contact(s): /N_back_/O_back_/O_back_/OP1/ Consider K in place of MG (3 valid contact(s) for K vs. 0 for MG) |
| MG:D8:103 | 1.00 | 76.86 |  | 0 | 1 | - Exclusion zone contact(s): /OP1/ Consider K in place of MG (1 valid contact(s) for K vs. 0 for MG) |
| MG:D8:104 | 1.00 | 50.61 |  | 0 | 3 | - Exclusion zone contact(s): /NH1_Arg_/OP2/OP2/ Consider K in place of MG (2 valid contact(s) for K vs. 0 for MG) |
| MG:D8:105 | 1.00 | 44.55 |  | 0 |  | 1^st^ MG coordination shell missing (see Notes) |
| MG:D8:106 | 1.00 | 41.14 |  | 0 | 2 | - Exclusion zone contact(s): /N_back_/O_back_/ Consider NA in place of MG (1 valid contact(s) for NA vs. 0 for MG) |
| MG:D8:107 | 1.00 | 44.66 |  | 1 | 3 | Anion - Exclusion zone contact(s): /O3'/O_back_/OP1/ Consider K in place of MG (3 valid contact(s) for K vs. 1 for MG) |
| MG:D8:108 | 1.00 | 52.15 |  | 0 | 2 | - Exclusion zone contact(s): /O_back_/OP1/ Consider NA in place of MG (1 valid contact(s) for NA vs. 0 for MG) |
| MG:D8:109 | 1.00 | 69.99 |  | 0 | 1 | - Exclusion zone contact(s): /OP1/ Consider K in place of MG (1 valid contact(s) for K vs. 0 for MG) |
| MG:DA:3001 | 1.00 | 8.09 |  | 1 | 1 | O_ph_ - Exclusion zone contact(s): /OP2/ |
| MG:DA:3002 | 1.00 | 11.80 |  | 1 |  | O_ph_ |
| MG:DA:3003 | 1.00 | 11.59 |  | 0 | 2 | - Exclusion zone contact(s): /O5'/OP1/ Consider NA in place of MG (1 valid contact(s) for NA vs. 0 for MG) |
| MG:DA:3004 | 1.00 | 9.38 |  | 1 | 1 | O_ph_ - Exclusion zone contact(s): /OP2/ |
| MG:DA:3005 | 1.00 | 17.14 |  | 2 |  | 2O_ph_ |
| MG:DA:3006 | 1.00 | 14.11 |  | 0 | 6 | - Exclusion zone contact(s): /N7/O2/O4'/O6/O6/O6/ Consider K in place of MG (6 valid contact(s) for K vs. 0 for MG) |
| MG:DA:3007 | 1.00 | 10.67 |  | 3 | 1 | 2O_ph_N_b_ - Exclusion zone contact(s): /O3'/ |
| MG:DA:3008 | 1.00 | 10.06 |  | 1 |  | O_ph_ |
| MG:DA:3009 | 1.00 | 10.24 |  | 1 | 3 | O_ph_ - Exclusion zone contact(s): /OP1/OP2/OP2/ Consider NA in place of MG (2 valid contact(s) for NA vs. 1 for MG) |
| MG:DA:3010 | 1.00 | 10.22 |  | 1 | 1 | O_ph_ - Exclusion zone contact(s): /O5'/ |
| MG:DA:3011 | 1.00 | 1.53 |  | 1 | 1 | O_ph_ - Exclusion zone contact(s): /OP1/ |
| MG:DA:3012 | 1.00 | 19.61 |  | 0 |  | 1^st^ MG coordination shell missing (see Notes) |
| MG:DA:3013 | 1.00 | 50.76 |  | 0 | 2 | - Exclusion zone contact(s): /N2/O4/ Consider NA in place of MG (1 valid contact(s) for NA vs. 0 for MG) |
| MG:DA:3014 | 1.00 | 7.70 |  | 0 |  | 1^st^ MG coordination shell missing (see Notes) |
| MG:DA:3015 | 1.00 | 30.75 |  | 0 |  | 1^st^ MG coordination shell missing (see Notes) |
| MG:DA:3016 | 1.00 | 4.62 |  | 1 |  | O_ph_ |
| MG:DA:3017 | 1.00 | 7.80 |  | 1 | 3 | O_ph_ - Exclusion zone contact(s): /O2'/O3'/OP1/ Consider K in place of MG (3 valid contact(s) for K vs. 1 for MG) |
| MG:DA:3018 | 1.00 | 20.46 |  | 0 |  | 1^st^ MG coordination shell missing (see Notes) |
| MG:DA:3019 | 1.00 | 20.35 |  | 1 | 1 | O_ph_ - Exclusion zone contact(s): /OP1/ |
| MG:DA:3020 | 1.00 | 21.79 |  | 0 | 3 | - Exclusion zone contact(s): /O4/O4/O6/ Consider K in place of MG (2 valid contact(s) for K vs. 0 for MG) |
| MG:DA:3021 | 1.00 | 26.41 |  | 1 | 1 | O_b_ - Exclusion zone contact(s): /OP2/ |
| MG:DA:3022 | 1.00 | 12.11 |  | 0 | 1 | - Exclusion zone contact(s): /OP1/ Consider K in place of MG (1 valid contact(s) for K vs. 0 for MG) |
| MG:DA:3023 | 1.00 | 57.55 |  | 0 |  | 1^st^ MG coordination shell missing (see Notes) |
| MG:DA:3024 | 1.00 | 9.91 |  | 1 | 1 | O_ph_ - Exclusion zone contact(s): /N7/ |
| MG:DA:3025 | 1.00 | 44.07 |  | 0 | 1 | - Exclusion zone contact(s): /OP2/ Consider K in place of MG (1 valid contact(s) for K vs. 0 for MG) |
| MG:DA:3026 | 1.00 | 17.95 |  | 0 |  | 1^st^ MG coordination shell missing (see Notes) |
| MG:DA:3027 | 1.00 | 15.69 |  | 2 |  | 2O_ph_ |
| MG:DA:3028 | 1.00 | 18.17 |  | 0 | 2 | - Exclusion zone contact(s): /O6/OP1/ Consider NA in place of MG (1 valid contact(s) for NA vs. 0 for MG) |
| MG:DA:3029 | 1.00 | 7.95 |  | 1 | 1 | O_ph_ - Exclusion zone contact(s): /OP2/ |
| MG:DA:3030 | 1.00 | 3.73 |  | 2 |  | 2N_b_ |
| MG:DA:3031 | 1.00 | 3.08 |  | 1 |  | O_ph_ |
| MG:DA:3032 | 1.00 | 6.96 |  | 1 |  | O_ph_ |
| MG:DA:3033 | 1.00 | 13.55 |  | 2 | 1 | 2O_ph_ - Exclusion zone contact(s): /OP2/ |
| MG:DA:3034 | 1.00 | 11.74 |  | 0 |  | 1^st^ MG coordination shell missing (see Notes) |
| MG:DA:3035 | 1.00 | 15.02 |  | 2 |  | 2O_ph_ |
| MG:DA:3036 | 1.00 | 14.08 |  | 0 |  | 1^st^ MG coordination shell missing (see Notes) |
| MG:DA:3037 | 1.00 | 1.53 |  | 1 | 3 | O_b_ - Exclusion zone contact(s): /O_back_/OP1/OP2/ Consider K in place of MG (2 valid contact(s) for K vs. 1 for MG) |
| MG:DA:3038 | 1.00 | 13.06 |  | 0 | 4 | - Exclusion zone contact(s): /O5'/OP1/OP1/OP1/ Consider K in place of MG (3 valid contact(s) for K vs. 0 for MG) |
| MG:DA:3039 | 1.00 | 10.68 |  | 1 | 1 | O_b_ - Exclusion zone contact(s): /OP1/ |
| MG:DA:3040 | 1.00 | 14.58 |  | 2 |  | O_ph_O_b_ |
| MG:DA:3041 | 1.00 | 20.87 |  | 0 | 1 | - Exclusion zone contact(s): /OP1/ Consider NA in place of MG (1 valid contact(s) for NA vs. 0 for MG) |
| MG:DA:3042 | 1.00 | 21.61 |  | 1 | 2 | O_ph_ - Exclusion zone contact(s): /O3'/OP1/ |
| MG:DA:3043 | 1.00 | 23.27 |  | 2 |  | 2O_ph_ |
| MG:DA:3044 | 1.00 | 18.52 |  | 1 |  | O_ph_ |
| MG:DA:3045 | 1.00 | 23.42 |  | 2 | 1 | 2O_ph_ - Exclusion zone contact(s): /OP2/ |
| MG:DA:3046 | 1.00 | 12.79 |  | 1 |  | O_ph_ |
| MG:DA:3047 | 1.00 | 1.53 |  | 1 | 2 | O_ph_ - Exclusion zone contact(s): /OP1/OP2/ Consider K in place of MG (2 valid contact(s) for K vs. 1 for MG) |
| MG:DA:3048 | 1.00 | 15.83 |  | 1 |  | O_ph_ |
| MG:DA:3049 | 1.00 | 25.83 |  | 1 |  | O_ph_ |
| MG:DA:3050 | 1.00 | 46.65 |  | 1 | 2 | O_b_ - Exclusion zone contact(s): /O4/O6/ Consider K in place of MG (2 valid contact(s) for K vs. 1 for MG) |
| MG:DA:3051 | 1.00 | 17.21 |  | 1 | 1 | O_b_ - Exclusion zone contact(s): /OP2/ |
| MG:DA:3052 | 1.00 | 19.03 |  | 1 |  | O_ph_ |
| MG:DA:3053 | 1.00 | 24.76 |  | 0 |  | 1^st^ MG coordination shell missing (see Notes) |
| MG:DA:3054 | 1.00 | 11.36 |  | 1 |  | O_ph_ |
| MG:DA:3055 | 1.00 | 21.38 |  | 0 | 2 | - Exclusion zone contact(s): /O6/OP2/ Consider K in place of MG (2 valid contact(s) for K vs. 0 for MG) |
| MG:DA:3056 | 1.00 | 12.29 |  | 1 | 2 | O_ph_ - Exclusion zone contact(s): /O3'/OP1/ |
| MG:DA:3057 | 1.00 | 6.92 |  | 1 | 2 | O_ph_ - Exclusion zone contact(s): /OP1/OP2/ Consider K in place of MG (2 valid contact(s) for K vs. 1 for MG) |
| MG:DA:3058 | 1.00 | 15.92 |  | 2 |  | 2O_ph_ |
| MG:DA:3059 | 1.00 | 8.02 |  | 0 | 4 | - Exclusion zone contact(s): /O4/O6/O6/O6/ Consider K in place of MG (3 valid contact(s) for K vs. 0 for MG) |
| MG:DA:3060 | 1.00 | 19.50 |  | 1 | 3 | O_ph_ - Exclusion zone contact(s): /OP1/OP2/OP2/ Consider K in place of MG (2 valid contact(s) for K vs. 1 for MG) |
| MG:DA:3061 | 1.00 | 23.21 |  | 3 |  | 3O_ph_ |
| MG:DA:3062 | 1.00 | 19.63 |  | 0 | 1 | - Exclusion zone contact(s): /OP2/ Consider NA in place of MG (1 valid contact(s) for NA vs. 0 for MG) |
| MG:DA:3063 | 1.00 | 23.28 |  | 0 |  | 1^st^ MG coordination shell missing (see Notes) |
| MG:DA:3064 | 1.00 | 35.49 |  | 0 |  | 1^st^ MG coordination shell missing (see Notes) |
| MG:DA:3065 | 1.00 | 6.24 |  | 1 | 5 | O_b_ - Exclusion zone contact(s): /N3_A_/N3_C_/O2/O5'/OP2/ Consider K in place of MG (5 valid contact(s) for K vs. 1 for MG) |
| MG:DA:3066 | 1.00 | 17.79 |  | 2 |  | 2O_ph_ |
| MG:DA:3067 | 1.00 | 14.44 |  | 0 |  | 1^st^ MG coordination shell missing (see Notes) |
| MG:DA:3068 | 1.00 | 15.25 |  | 1 |  | O_ph_ |
| MG:DA:3069 | 1.00 | 16.47 |  | 1 |  | O_ph_ |
| MG:DA:3070 | 1.00 | 24.99 |  | 0 |  | 1^st^ MG coordination shell missing (see Notes) |
| MG:DA:3071 | 1.00 | 13.12 |  | 2 | 1 | 2O_ph_ - Exclusion zone contact(s): /N7/ |
| MG:DA:3072 | 1.00 | 9.96 |  | 1 |  | O_ph_ |
| MG:DA:3073 | 1.00 | 16.89 |  | 0 | 1 | - Exclusion zone contact(s): /O6/ Consider K in place of MG (1 valid contact(s) for K vs. 0 for MG) |
| MG:DA:3074 | 1.00 | 22.64 |  | 1 | 2 | O_ph_ - Exclusion zone contact(s): /N7/O6/ |
| MG:DA:3075 | 1.00 | 19.41 |  | 0 |  | 1^st^ MG coordination shell missing (see Notes) |
| MG:DA:3076 | 1.00 | 13.61 |  | 0 | 2 | - Exclusion zone contact(s): /O_back_/OP1/ Consider NA in place of MG (1 valid contact(s) for NA vs. 0 for MG) |
| MG:DA:3077 | 1.00 | 47.81 |  | 0 | 1 | - Exclusion zone contact(s): /O6/ Consider K in place of MG (1 valid contact(s) for K vs. 0 for MG) |
| MG:DA:3078 | 1.00 | 17.63 |  | 1 | 1 | O_ph_ - Exclusion zone contact(s): /NE2_His_/ |
| MG:DA:3079 | 1.00 | 37.42 |  | 0 |  | 1^st^ MG coordination shell missing (see Notes) |
| MG:DA:3080 | 1.00 | 31.66 |  | 0 |  | 1^st^ MG coordination shell missing (see Notes) |
| MG:DA:3081 | 1.00 | 24.79 |  | 0 |  | 1^st^ MG coordination shell missing (see Notes) |
| MG:DA:3082 | 1.00 | 27.68 |  | 2 |  | O_ph_O_b_ |
| MG:DA:3083 | 1.00 | 9.89 |  | 1 | 2 | O_ph_ - Exclusion zone contact(s): /OP1/OP2/ |
| MG:DA:3084 | 1.00 | 10.90 |  | 0 | 1 | - Exclusion zone contact(s): /N7/ Consider NA in place of MG (1 valid contact(s) for NA vs. 0 for MG) |
| MG:DA:3085 | 1.00 | 43.85 |  | 0 | 1 | - Exclusion zone contact(s): /OP1/ Consider NA in place of MG (1 valid contact(s) for NA vs. 0 for MG) |
| MG:DA:3086 | 1.00 | 17.04 |  | 3 |  | 3O_ph_ |
| MG:DA:3087 | 1.00 | 15.84 |  | 2 | 1 | 2O_ph_ - Exclusion zone contact(s): /OP2/ |
| MG:DA:3088 | 1.00 | 12.53 |  | 1 |  | O_ph_ |
| MG:DA:3089 | 1.00 | 23.46 |  | 2 |  | 2O_ph_ |
| MG:DA:3090 | 1.00 | 13.73 |  | 0 | 5 | - Exclusion zone contact(s): /O2/O2'/O2'/O3'/OP2/ Consider K in place of MG (4 valid contact(s) for K vs. 0 for MG) |
| MG:DA:3091 | 1.00 | 31.97 |  | 1 |  | O_ph_ |
| MG:DA:3092 | 1.00 | 3.88 |  | 0 | 7 | - Exclusion zone contact(s): /N1_A_/N3_A_/N7/N7/O2'/O6/O6/ Consider K in place of MG (6 valid contact(s) for K vs. 0 for MG) |
| MG:DA:3093 | 1.00 | 13.33 |  | 1 |  | O_ph_ |
| MG:DA:3094 | 1.00 | 23.12 |  | 0 | 2 | - Exclusion zone contact(s): /O3'/OP1/ Consider NA in place of MG (1 valid contact(s) for NA vs. 0 for MG) |
| MG:DA:3095 | 1.00 | 27.24 |  | 1 | 1 | O_ph_ - Exclusion zone contact(s): /N7/ |
| MG:DA:3096 | 1.00 | 7.81 |  | 0 | 4 | - Exclusion zone contact(s): /O4/O6/O6/O6/ Consider K in place of MG (4 valid contact(s) for K vs. 0 for MG) |
| MG:DA:3097 | 1.00 | 21.64 |  | 1 | 1 | O_ph_ - Exclusion zone contact(s): /OP2/ |
| MG:DA:3098 | 1.00 | 34.00 |  | 1 | 1 | O_ph_ - Exclusion zone contact(s): /OP1/ |
| MG:DA:3099 | 1.00 | 48.39 |  | 2 |  | 2O_ph_ |
| MG:DA:3100 | 1.00 | 36.61 |  | 0 |  | 1^st^ MG coordination shell missing (see Notes) |
| MG:DA:3101 | 1.00 | 45.43 |  | 0 | 2 | - Exclusion zone contact(s): /O4/OP2/ Consider NA in place of MG (1 valid contact(s) for NA vs. 0 for MG) |
| MG:DA:3102 | 1.00 | 61.08 |  | 0 | 1 | - Exclusion zone contact(s): /O4/ Consider K in place of MG (1 valid contact(s) for K vs. 0 for MG) |
| MG:DA:3103 | 1.00 | 49.69 |  | 0 |  | 1^st^ MG coordination shell missing (see Notes) |
| MG:DA:3104 | 1.00 | 42.40 |  | 1 | 2 | O_ph_ - Exclusion zone contact(s): /O3'/OP1/ Consider K in place of MG (2 valid contact(s) for K vs. 1 for MG) |
| MG:DA:3105 | 1.00 | 16.30 |  | 2 | 2 | 2O_ph_ - Exclusion zone contact(s): /OP1/OP2/ |
| MG:DA:3106 | 1.00 | 56.89 |  | 0 | 2 | - Exclusion zone contact(s): /OP1/OP2/ Consider K in place of MG (2 valid contact(s) for K vs. 0 for MG) |
| MG:DA:3107 | 1.00 | 43.79 |  | 0 | 2 | - Exclusion zone contact(s): /N7/O6/ Consider K in place of MG (2 valid contact(s) for K vs. 0 for MG) |
| MG:DA:3108 | 1.00 | 76.43 |  | 0 |  | 1^st^ MG coordination shell missing (see Notes) |
| MG:DA:3109 | 1.00 | 18.30 |  | 0 | 1 | - Exclusion zone contact(s): /OP1/ Consider NA in place of MG (1 valid contact(s) for NA vs. 0 for MG) |
| MG:DA:3110 | 1.00 | 35.72 |  | 0 | 5 | - Exclusion zone contact(s): /N7/O4/O4/O6/OP2/ Consider K in place of MG (5 valid contact(s) for K vs. 0 for MG) |
| MG:DA:3111 | 1.00 | 64.67 |  | 0 | 3 | - Exclusion zone contact(s): /O2/OP1/OP2/ Consider K in place of MG (3 valid contact(s) for K vs. 0 for MG) |
| MG:DA:3112 | 1.00 | 49.22 |  | 0 |  | 1^st^ MG coordination shell missing (see Notes) |
| MG:DA:3113 | 1.00 | 73.98 |  | 0 | 2 | - Exclusion zone contact(s): /OP1/OP2/ Consider K in place of MG (2 valid contact(s) for K vs. 0 for MG) |
| MG:DA:3114 | 1.00 | 47.42 |  | 0 |  | 1^st^ MG coordination shell missing (see Notes) |
| MG:DA:3115 | 1.00 | 34.12 |  | 0 | 2 | - Exclusion zone contact(s): /N7/O6/ Consider K in place of MG (2 valid contact(s) for K vs. 0 for MG) |
| MG:DA:3116 | 1.00 | 63.03 |  | 0 | 2 | - Exclusion zone contact(s): /N7/O6/ Consider K in place of MG (2 valid contact(s) for K vs. 0 for MG) |
| MG:DA:3117 | 1.00 | 58.41 |  | 0 | 1 | - Exclusion zone contact(s): /O6/ Consider K in place of MG (1 valid contact(s) for K vs. 0 for MG) |
| MG:DA:3118 | 1.00 | 69.17 |  | 0 | 1 | - Exclusion zone contact(s): /O6/ Consider K in place of MG (1 valid contact(s) for K vs. 0 for MG) |
| MG:DA:3119 | 1.00 | 60.04 |  | 0 | 3 | - Exclusion zone contact(s): /N7/O6/OP1/ Consider K in place of MG (2 valid contact(s) for K vs. 0 for MG) |
| MG:DA:3120 | 1.00 | 57.80 |  | 0 | 1 | - Exclusion zone contact(s): /O6/ Consider K in place of MG (1 valid contact(s) for K vs. 0 for MG) |
| MG:DA:3121 | 1.00 | 51.23 |  | 0 | 1 | - Exclusion zone contact(s): /OP1/ Consider K in place of MG (1 valid contact(s) for K vs. 0 for MG) |
| MG:DA:3122 | 1.00 | 75.99 |  | 0 | 2 | - Exclusion zone contact(s): /O6/O6/ Consider K in place of MG (2 valid contact(s) for K vs. 0 for MG) |
| MG:DA:3123 | 1.00 | 65.65 |  | 0 | 1 | - Exclusion zone contact(s): /O6/ Consider K in place of MG (1 valid contact(s) for K vs. 0 for MG) |
| MG:DA:3124 | 1.00 | 36.79 |  | 1 | 2 | O_ph_ - Exclusion zone contact(s): /O5'/OP1/ Consider K in place of MG (2 valid contact(s) for K vs. 1 for MG) |
| MG:DA:3125 | 1.00 | 37.38 |  | 0 |  | 1^st^ MG coordination shell missing (see Notes) |
| MG:DA:3126 | 1.00 | 63.03 |  | 0 | 3 | - Exclusion zone contact(s): /O6/OP1/OP2/ Consider NA in place of MG (2 valid contact(s) for NA vs. 0 for MG) |
| MG:DA:3127 | 1.00 | 32.57 |  | 1 | 3 | O_r_ - Exclusion zone contact(s): /N7/O6/OP2/ Consider K in place of MG (2 valid contact(s) for K vs. 1 for MG) |
| MG:DA:3128 | 1.00 | 31.86 |  | 0 | 3 | - Exclusion zone contact(s): /O2'/O4/OP2/ Consider NA in place of MG (2 valid contact(s) for NA vs. 0 for MG) |
| MG:DA:3129 | 1.00 | 14.25 |  | 0 | 4 | - Exclusion zone contact(s): /O3'/OP1/OP1/OP2/ Consider K in place of MG (4 valid contact(s) for K vs. 0 for MG) |
| MG:DA:3130 | 1.00 | 73.53 |  | 1 | 1 | O_ph_ - Exclusion zone contact(s): /OP1/ |
| MG:DA:3131 | 1.00 | 32.40 |  | 0 | 3 | - Exclusion zone contact(s): /N6/N7/OP2/ Consider K in place of MG (3 valid contact(s) for K vs. 0 for MG) |
| MG:DA:3132 | 1.00 | 34.67 |  | 0 | 1 | - Exclusion zone contact(s): /OP2/ Consider NA in place of MG (1 valid contact(s) for NA vs. 0 for MG) |
| MG:DA:3133 | 1.00 | 45.41 |  | 1 |  | O_ph_ |
| MG:DA:3134 | 1.00 | 44.31 |  | 0 |  | 1^st^ MG coordination shell missing (see Notes) |
| MG:DA:3135 | 1.00 | 35.86 |  | 0 | 4 | - Exclusion zone contact(s): /N4/N7/O6/O6/ Consider K in place of MG (4 valid contact(s) for K vs. 0 for MG) |
| MG:DA:3136 | 1.00 | 63.82 |  | 0 |  | 1^st^ MG coordination shell missing (see Notes) |
| MG:DA:3137 | 1.00 | 31.91 |  | 0 | 1 | - Exclusion zone contact(s): /OP2/ Consider K in place of MG (1 valid contact(s) for K vs. 0 for MG) |
| MG:DA:3138 | 1.00 | 44.35 |  | 0 | 1 | - Exclusion zone contact(s): /N7/ Consider K in place of MG (1 valid contact(s) for K vs. 0 for MG) |
| MG:DA:3139 | 1.00 | 45.09 |  | 0 |  | 1^st^ MG coordination shell missing (see Notes) |
| MG:DA:3140 | 1.00 | 26.06 |  | 1 |  | O_b_ |
| MG:DA:3141 | 1.00 | 42.85 |  | 1 |  | O_ph_ |
| MG:DA:3142 | 1.00 | 73.16 |  | 0 | 1 | - Exclusion zone contact(s): /O5'/ Consider K in place of MG (1 valid contact(s) for K vs. 0 for MG) |
| MG:DA:3143 | 1.00 | 61.44 |  | 1 |  | O_ph_ |
| MG:DA:3144 | 1.00 | 36.67 |  | 2 |  | 2O_ph_ |
| MG:DA:3145 | 1.00 | 39.54 |  | 0 |  | 1^st^ MG coordination shell missing (see Notes) |
| MG:DA:3146 | 1.00 | 55.67 |  | 0 |  | 1^st^ MG coordination shell missing (see Notes) |
| MG:DA:3147 | 1.00 | 45.07 |  | 0 | 1 | - Exclusion zone contact(s): /OP2/ Consider K in place of MG (1 valid contact(s) for K vs. 0 for MG) |
| MG:DA:3148 | 1.00 | 67.05 |  | 0 |  | 1^st^ MG coordination shell missing (see Notes) |
| MG:DA:3149 | 1.00 | 18.84 |  | 0 | 4 | - Exclusion zone contact(s): /O4/O4/O6/O6/ Consider K in place of MG (4 valid contact(s) for K vs. 0 for MG) |
| MG:DA:3150 | 1.00 | 39.09 |  | 0 |  | 1^st^ MG coordination shell missing (see Notes) |
| MG:DA:3151 | 1.00 | 49.53 |  | 0 | 1 | - Exclusion zone contact(s): /OP2/ Consider K in place of MG (1 valid contact(s) for K vs. 0 for MG) |
| MG:DA:3152 | 1.00 | 52.18 |  | 0 | 1 | - Exclusion zone contact(s): /O6/ Consider K in place of MG (1 valid contact(s) for K vs. 0 for MG) |
| MG:DA:3153 | 1.00 | 26.74 |  | 0 | 1 | - Exclusion zone contact(s): /N7/ Consider K in place of MG (1 valid contact(s) for K vs. 0 for MG) |
| MG:DA:3154 | 1.00 | 46.47 |  | 0 | 2 | - Exclusion zone contact(s): /O6/O6/ Consider K in place of MG (2 valid contact(s) for K vs. 0 for MG) |
| MG:DA:3155 | 1.00 | 32.34 |  | 1 | 1 | O_ph_ - Exclusion zone contact(s): /N7/ |
| MG:DA:3156 | 1.00 | 27.47 |  | 1 |  | O_ph_ |
| MG:DA:3157 | 1.00 | 58.96 |  | 0 |  | 1^st^ MG coordination shell missing (see Notes) |
| MG:DA:3158 | 1.00 | 28.82 |  | 0 | 4 | - Exclusion zone contact(s): /O4/O6/O6/O6/ Consider K in place of MG (3 valid contact(s) for K vs. 0 for MG) |
| MG:DA:3159 | 1.00 | 47.37 |  | 0 |  | 1^st^ MG coordination shell missing (see Notes) |
| MG:DA:3160 | 1.00 | 48.88 |  | 0 | 2 | - Exclusion zone contact(s): /O6/O6/ Consider K in place of MG (2 valid contact(s) for K vs. 0 for MG) |
| MG:DA:3161 | 1.00 | 55.75 |  | 0 |  | 1^st^ MG coordination shell missing (see Notes) |
| MG:DA:3162 | 1.00 | 45.82 |  | 0 | 3 | - Exclusion zone contact(s): /O4/O6/O6/ Consider K in place of MG (3 valid contact(s) for K vs. 0 for MG) |
| MG:DA:3163 | 1.00 | 39.58 |  | 0 | 1 | - Exclusion zone contact(s): /OP1/ Consider K in place of MG (1 valid contact(s) for K vs. 0 for MG) |
| MG:DA:3164 | 1.00 | 53.48 |  | 0 | 1 | - Exclusion zone contact(s): /OP2/ Consider K in place of MG (1 valid contact(s) for K vs. 0 for MG) |
| MG:DA:3165 | 1.00 | 46.57 |  | 0 | 2 | - Exclusion zone contact(s): /O3'/OP1/ Consider K in place of MG (2 valid contact(s) for K vs. 0 for MG) |
| MG:DA:3166 | 1.00 | 56.66 |  | 0 | 1 | - Exclusion zone contact(s): /OP1/ Consider K in place of MG (1 valid contact(s) for K vs. 0 for MG) |
| MG:DA:3167 | 1.00 | 48.37 |  | 0 | 2 | - Exclusion zone contact(s): /O3'/OP1/ Consider K in place of MG (2 valid contact(s) for K vs. 0 for MG) |
| MG:DA:3168 | 1.00 | 40.17 |  | 0 |  | 1^st^ MG coordination shell missing (see Notes) |
| MG:DA:3169 | 1.00 | 59.10 |  | 0 | 1 | - Exclusion zone contact(s): /O6/ Consider NA in place of MG (1 valid contact(s) for NA vs. 0 for MG) |
| MG:DA:3170 | 1.00 | 16.15 |  | 0 |  | 1^st^ MG coordination shell missing (see Notes) |
| MG:DA:3171 | 1.00 | 36.56 |  | 0 | 1 | - Exclusion zone contact(s): /O6/ Consider K in place of MG (1 valid contact(s) for K vs. 0 for MG) |
| MG:DA:3172 | 1.00 | 37.57 |  | 0 | 4 | - Exclusion zone contact(s): /O2'/O3'/OP1/OP2/ Consider K in place of MG (4 valid contact(s) for K vs. 0 for MG) |
| MG:DA:3173 | 1.00 | 36.41 |  | 0 | 1 | - Exclusion zone contact(s): /OP2/ Consider K in place of MG (1 valid contact(s) for K vs. 0 for MG) |
| MG:DA:3174 | 1.00 | 30.62 |  | 0 | 2 | - Exclusion zone contact(s): /N4/O6/ Consider K in place of MG (2 valid contact(s) for K vs. 0 for MG) |
| MG:DA:3175 | 1.00 | 21.74 |  | 3 |  | 3O_ph_ |
| MG:DA:3176 | 1.00 | 39.33 |  | 0 |  | 1^st^ MG coordination shell missing (see Notes) |
| MG:DA:3177 | 1.00 | 39.74 |  | 0 | 1 | - Exclusion zone contact(s): /O6/ Consider K in place of MG (1 valid contact(s) for K vs. 0 for MG) |
| MG:DA:3178 | 1.00 | 49.03 |  | 0 | 1 | - Exclusion zone contact(s): /OP2/ Consider K in place of MG (1 valid contact(s) for K vs. 0 for MG) |
| MG:DA:3179 | 1.00 | 52.71 |  | 0 |  | 1^st^ MG coordination shell missing (see Notes) |
| MG:DA:3180 | 1.00 | 77.17 |  | 1 | 1 | O_b_ - Exclusion zone contact(s): /N7/ |
| MG:DA:3181 | 1.00 | 40.96 |  | 0 | 1 | - Exclusion zone contact(s): /OP1/ Consider K in place of MG (1 valid contact(s) for K vs. 0 for MG) |
| MG:DA:3182 | 1.00 | 65.97 |  | 0 | 1 | - Exclusion zone contact(s): /OP2/ Consider K in place of MG (1 valid contact(s) for K vs. 0 for MG) |
| MG:DA:3183 | 1.00 | 97.39 |  | 0 |  | 1^st^ MG coordination shell missing (see Notes) |
| MG:DA:3184 | 1.00 | 40.69 |  | 0 | 5 | - Exclusion zone contact(s): /O2'/O3'/OP1/OP1/OP2/ Consider K in place of MG (5 valid contact(s) for K vs. 0 for MG) |
| MG:DA:3185 | 1.00 | 54.05 |  | 0 | 1 | - Exclusion zone contact(s): /OP1/ Consider NA in place of MG (1 valid contact(s) for NA vs. 0 for MG) |
| MG:DA:3186 | 1.00 | 39.66 |  | 0 | 4 | - Exclusion zone contact(s): /OP1/OP1/OP2/OP2/ Consider K in place of MG (4 valid contact(s) for K vs. 0 for MG) |
| MG:DA:3187 | 1.00 | 45.70 |  | 0 | 3 | - Exclusion zone contact(s): /N7/ND2_Asn_/O6/ Consider K in place of MG (2 valid contact(s) for K vs. 0 for MG) |
| MG:DA:3188 | 1.00 | 60.04 |  | 0 | 2 | - Exclusion zone contact(s): /OP1/OP2/ Consider K in place of MG (2 valid contact(s) for K vs. 0 for MG) |
| MG:DA:3189 | 1.00 | 26.93 |  | 0 | 4 | - Exclusion zone contact(s): /O4/O4/O6/O6/ Consider K in place of MG (4 valid contact(s) for K vs. 0 for MG) |
| MG:DA:3190 | 1.00 | 60.58 |  | 0 | 3 | - Exclusion zone contact(s): /N7/O6/OP2/ Consider K in place of MG (3 valid contact(s) for K vs. 0 for MG) |
| MG:DA:3191 | 1.00 | 60.09 |  | 0 | 3 | - Exclusion zone contact(s): /O_back_/OP2/OP2/ Consider K in place of MG (3 valid contact(s) for K vs. 0 for MG) |
| MG:DA:3192 | 1.00 | 95.39 |  | 1 | 2 | O_ph_ - Exclusion zone contact(s): /O5'/OP2/ Consider K in place of MG (2 valid contact(s) for K vs. 1 for MG) |
| MG:DA:3193 | 1.00 | 71.54 |  | 1 | 2 | O_ph_ - Exclusion zone contact(s): /O5'/O5'/ Consider K in place of MG (2 valid contact(s) for K vs. 1 for MG) |
| MG:DA:3194 | 1.00 | 57.84 |  | 0 |  | 1^st^ MG coordination shell missing (see Notes) |
| MG:DA:3195 | 1.00 | 50.63 |  | 0 | 1 | - Exclusion zone contact(s): /OP2/ Consider NA in place of MG (1 valid contact(s) for NA vs. 0 for MG) |
| MG:DA:3196 | 1.00 | 43.97 |  | 0 | 1 | - Exclusion zone contact(s): /OP2/ Consider K in place of MG (1 valid contact(s) for K vs. 0 for MG) |
| MG:DA:3197 | 1.00 | 19.50 |  | 0 | 4 | - Exclusion zone contact(s): /O4/O6/OP2/OP2/ Consider K in place of MG (4 valid contact(s) for K vs. 0 for MG) |
| MG:DA:3198 | 1.00 | 21.72 |  | 0 | 4 | - Exclusion zone contact(s): /O4/O6/O6/O6/ Consider NA in place of MG (2 valid contact(s) for NA vs. 0 for MG) |
| MG:DA:3199 | 1.00 | 29.54 |  | 0 | 1 | - Exclusion zone contact(s): /OP2/ Consider K in place of MG (1 valid contact(s) for K vs. 0 for MG) |
| MG:DA:3200 | 1.00 | 67.68 |  | 0 |  | 1^st^ MG coordination shell missing (see Notes) |
| MG:DA:3201 | 1.00 | 45.53 |  | 0 |  | 1^st^ MG coordination shell missing (see Notes) |
| MG:DA:3202 | 1.00 | 37.24 |  | 0 | 2 | - Exclusion zone contact(s): /OP2/OP2/ Consider K in place of MG (2 valid contact(s) for K vs. 0 for MG) |
| MG:DA:3203 | 1.00 | 59.68 |  | 0 | 1 | - Exclusion zone contact(s): /N7/ Consider K in place of MG (1 valid contact(s) for K vs. 0 for MG) |
| MG:DA:3204 | 1.00 | 50.36 |  | 1 | 1 | O_b_ - Exclusion zone contact(s): /O2'/ |
| MG:DA:3205 | 1.00 | 65.00 |  | 0 | 2 | - Exclusion zone contact(s): /OP1/OP2/ Consider K in place of MG (2 valid contact(s) for K vs. 0 for MG) |
| MG:DA:3206 | 1.00 | 76.01 |  | 0 | 2 | - Exclusion zone contact(s): /O2'/O3'/ Consider K in place of MG (2 valid contact(s) for K vs. 0 for MG) |
| MG:DA:3207 | 1.00 | 45.87 |  | 0 |  | 1^st^ MG coordination shell missing (see Notes) |
| MG:DA:3208 | 1.00 | 72.01 |  | 0 |  | 1^st^ MG coordination shell missing (see Notes) |
| MG:DA:3209 | 1.00 | 33.21 |  | 0 |  | 1^st^ MG coordination shell missing (see Notes) |
| MG:DA:3210 | 1.00 | 32.28 |  | 0 | 4 | - Exclusion zone contact(s): /O4'/O5'/O6/OP2/ Consider K in place of MG (4 valid contact(s) for K vs. 0 for MG) |
| MG:DA:3211 | 1.00 | 43.23 |  | 0 |  | 1^st^ MG coordination shell missing (see Notes) |
| MG:DA:3212 | 1.00 | 30.04 |  | 1 | 2 | O_ph_ - Exclusion zone contact(s): /O5'/OP2/ Consider K in place of MG (2 valid contact(s) for K vs. 1 for MG) |
| MG:DA:3213 | 1.00 | 53.13 |  | 1 | 3 | Anion - Exclusion zone contact(s): /N7/O6/O6/ Consider K in place of MG (3 valid contact(s) for K vs. 1 for MG) |
| MG:DA:3214 | 1.00 | 57.74 |  | 0 | 2 | - Exclusion zone contact(s): /OP2/OP2/ Consider K in place of MG (2 valid contact(s) for K vs. 0 for MG) |
| MG:DA:3215 | 1.00 | 84.43 |  | 0 | 1 | - Exclusion zone contact(s): /O2'/ Consider NA in place of MG (1 valid contact(s) for NA vs. 0 for MG) |
| MG:DA:3216 | 1.00 | 54.82 |  | 0 | 2 | - Exclusion zone contact(s): /OP1/OP2/ Consider K in place of MG (2 valid contact(s) for K vs. 0 for MG) |
| MG:DA:3217 | 1.00 | 28.22 |  | 0 | 2 | - Exclusion zone contact(s): /N1_G_/N2/ Consider K in place of MG (2 valid contact(s) for K vs. 0 for MG) |
| MG:DA:3218 | 1.00 | 47.67 |  | 0 | 1 | - Exclusion zone contact(s): /O2/ Consider K in place of MG (1 valid contact(s) for K vs. 0 for MG) |
| MG:DA:3219 | 1.00 | 48.49 |  | 0 | 3 | - Exclusion zone contact(s): /O2/O2'/O4/ Consider K in place of MG (3 valid contact(s) for K vs. 0 for MG) |
| MG:DA:3220 | 1.00 | 37.49 |  | 0 | 2 | - Exclusion zone contact(s): /OP1/OP2/ Consider K in place of MG (2 valid contact(s) for K vs. 0 for MG) |
| MG:DA:3221 | 1.00 | 43.67 |  | 0 |  | 1^st^ MG coordination shell missing (see Notes) |
| MG:DA:3222 | 1.00 | 42.42 |  | 0 | 3 | - Exclusion zone contact(s): /O4/O6/O6/ Consider K in place of MG (2 valid contact(s) for K vs. 0 for MG) |
| MG:DA:3223 | 1.00 | 45.62 |  | 0 |  | 1^st^ MG coordination shell missing (see Notes) |
| MG:DA:3224 | 1.00 | 31.97 |  | 0 | 5 | - Exclusion zone contact(s): /O3'/O4'/O5'/O6/OP1/ Consider K in place of MG (4 valid contact(s) for K vs. 0 for MG) |
| MG:DA:3225 | 1.00 | 39.70 |  | 0 | 3 | - Exclusion zone contact(s): /N2/O4/OP2/ Consider K in place of MG (2 valid contact(s) for K vs. 0 for MG) |
| MG:DA:3226 | 1.00 | 44.55 |  | 0 | 3 | - Exclusion zone contact(s): /N6/N7/O6/ Consider NA in place of MG (2 valid contact(s) for NA vs. 0 for MG) |
| MG:DA:3227 | 1.00 | 52.88 |  | 0 |  | 1^st^ MG coordination shell missing (see Notes) |
| MG:DA:3228 | 1.00 | 36.61 |  | 0 | 3 | - Exclusion zone contact(s): /N7/O4/O6/ Consider K in place of MG (3 valid contact(s) for K vs. 0 for MG) |
| MG:DA:3229 | 1.00 | 33.22 |  | 1 | 4 | O_b_ - Exclusion zone contact(s): /N7/O3'/O6/OP2/ Consider K in place of MG (4 valid contact(s) for K vs. 1 for MG) |
| MG:DA:3230 | 1.00 | 43.81 |  | 0 | 1 | - Exclusion zone contact(s): /OP2/ Consider K in place of MG (1 valid contact(s) for K vs. 0 for MG) |
| MG:DA:3231 | 1.00 | 9.84 |  | 0 | 3 | - Exclusion zone contact(s): /O2/O2'/OP2/ Consider K in place of MG (3 valid contact(s) for K vs. 0 for MG) |
| MG:DA:3232 | 1.00 | 61.55 |  | 0 | 1 | - Exclusion zone contact(s): /OP2/ Consider K in place of MG (1 valid contact(s) for K vs. 0 for MG) |
| MG:DA:3233 | 1.00 | 26.50 |  | 0 | 4 | - Exclusion zone contact(s): /N7/O2/O2'/OP2/ Consider K in place of MG (4 valid contact(s) for K vs. 0 for MG) |
| MG:DA:3234 | 1.00 | 17.22 |  | 1 |  | O_b_ |
| MG:DA:3235 | 1.00 | 25.04 |  | 0 | 3 | - Exclusion zone contact(s): /O6/O6/OP1/ Consider NA in place of MG (3 valid contact(s) for NA vs. 0 for MG) |
| MG:DA:3236 | 1.00 | 20.62 |  | 0 | 1 | - Exclusion zone contact(s): /N7/ Consider K in place of MG (1 valid contact(s) for K vs. 0 for MG) |
| MG:DA:3237 | 1.00 | 13.56 |  | 0 | 3 | - Exclusion zone contact(s): /O_back_/OP1/OP2/ Consider K in place of MG (3 valid contact(s) for K vs. 0 for MG) |
| MG:DA:3238 | 1.00 | 18.88 |  | 1 | 1 | O_ph_ - Exclusion zone contact(s): /OP1/ |
| MG:DA:3239 | 1.00 | 37.27 |  | 0 | 1 | - Exclusion zone contact(s): /OP1/ Consider NA in place of MG (1 valid contact(s) for NA vs. 0 for MG) |
| MG:DA:3240 | 1.00 | 32.44 |  | 1 |  | O_ph_ |
| MG:DA:3241 | 1.00 | 20.14 |  | 1 | 2 | N_b_ - Exclusion zone contact(s): /N6/OP1/ |
| MG:DA:3242 | 1.00 | 18.42 |  | 1 | 2 | Anion - Exclusion zone contact(s): /O6/OP1/ |
| MG:DA:3243 | 1.00 | 48.79 |  | 0 | 2 | - Exclusion zone contact(s): /O3'/OP1/ Consider K in place of MG (2 valid contact(s) for K vs. 0 for MG) |
| MG:DA:3244 | 1.00 | 97.76 |  | 1 | 1 | O_ph_ - Exclusion zone contact(s): /OP2/ |
| MG:DA:3245 | 1.00 | 41.74 |  | 1 | 2 | O_b_ - Exclusion zone contact(s): /O4/O6/ Consider K in place of MG (2 valid contact(s) for K vs. 1 for MG) |
| MG:DA:3246 | 1.00 | 34.51 |  | 0 | 1 | - Exclusion zone contact(s): /O6/ Consider K in place of MG (1 valid contact(s) for K vs. 0 for MG) |
| MG:DA:3247 | 1.00 | 72.50 |  | 0 | 2 | - Exclusion zone contact(s): /OP1/OP2/ Consider K in place of MG (2 valid contact(s) for K vs. 0 for MG) |
| MG:DA:3248 | 1.00 | 70.09 |  | 0 |  | 1^st^ MG coordination shell missing (see Notes) |
| MG:DA:3249 | 1.00 | 13.25 |  | 1 | 4 | O_r_ - Exclusion zone contact(s): /N7/O2'/O3'/OP2/ Consider NA in place of MG (2 valid contact(s) for NA vs. 1 for MG) |
| MG:DA:3250 | 1.00 | 13.21 |  | 1 | 2 | O_ph_ - Exclusion zone contact(s): /O2'/OP1/ Consider K in place of MG (2 valid contact(s) for K vs. 1 for MG) |
| MG:DA:3251 | 1.00 | 12.46 |  | 0 | 4 | - Exclusion zone contact(s): /N3_A_/N6/O2/OP1/ Consider K in place of MG (3 valid contact(s) for K vs. 0 for MG) |
| MG:DA:3252 | 1.00 | 29.51 |  | 1 | 2 | O_b_ - Exclusion zone contact(s): /N7/O6/ Consider K in place of MG (2 valid contact(s) for K vs. 1 for MG) |
| MG:DA:3253 | 1.00 | 15.11 |  | 0 | 5 | - Exclusion zone contact(s): /O3'/O6/OP1/OP1/OP2/ Consider K in place of MG (4 valid contact(s) for K vs. 0 for MG) |
| MG:DA:3254 | 1.00 | 25.65 |  | 0 | 3 | - Exclusion zone contact(s): /O2/O2/OP1/ Consider K in place of MG (3 valid contact(s) for K vs. 0 for MG) |
| MG:DA:3255 | 1.00 | 72.05 |  | 0 |  | 1^st^ MG coordination shell missing (see Notes) |
| MG:DA:3256 | 1.00 | 50.29 |  | 0 | 1 | - Exclusion zone contact(s): /N7/ Consider K in place of MG (1 valid contact(s) for K vs. 0 for MG) |
| MG:DA:3257 | 1.00 | 66.79 |  | 0 |  | 1^st^ MG coordination shell missing (see Notes) |
| MG:DA:3258 | 1.00 | 83.05 |  | 0 | 3 | - Exclusion zone contact(s): /N4/O6/O6/ Consider K in place of MG (3 valid contact(s) for K vs. 0 for MG) |
| MG:DA:3259 | 1.00 | 60.30 |  | 0 |  | 1^st^ MG coordination shell missing (see Notes) |
| MG:DA:3260 | 1.00 | 60.22 |  | 0 |  | 1^st^ MG coordination shell missing (see Notes) |
| MG:DA:3261 | 1.00 | 34.19 |  | 2 | 1 | 2O_ph_ - Exclusion zone contact(s): /O5'/ |
| MG:DA:3262 | 1.00 | 72.20 |  | 0 | 3 | - Exclusion zone contact(s): /N7/O6/O6/ Consider K in place of MG (3 valid contact(s) for K vs. 0 for MG) |
| MG:DA:3263 | 1.00 | 24.79 |  | 0 | 3 | - Exclusion zone contact(s): /O4/O6/O6/ Consider K in place of MG (3 valid contact(s) for K vs. 0 for MG) |
| MG:DA:3264 | 1.00 | 28.42 |  | 1 | 1 | O_ph_ - Exclusion zone contact(s): /OP2/ |
| MG:DA:3265 | 1.00 | 47.61 |  | 0 |  | 1^st^ MG coordination shell missing (see Notes) |
| MG:DA:3266 | 1.00 | 57.57 |  | 0 | 3 | - Exclusion zone contact(s): /N1/O3'/OP1/ Consider K in place of MG (3 valid contact(s) for K vs. 0 for MG) |
| MG:DA:3267 | 1.00 | 72.30 |  | 0 |  | 1^st^ MG coordination shell missing (see Notes) |
| MG:DA:3268 | 1.00 | 50.59 |  | 0 | 1 | - Exclusion zone contact(s): /O4/ Consider K in place of MG (1 valid contact(s) for K vs. 0 for MG) |
| MG:DA:3269 | 1.00 | 36.97 |  | 0 | 4 | - Exclusion zone contact(s): /O2'/O3'/OP1/OP2/ Consider K in place of MG (4 valid contact(s) for K vs. 0 for MG) |
| MG:DA:3270 | 1.00 | 90.43 |  | 0 |  | 1^st^ MG coordination shell missing (see Notes) |
| MG:DA:3271 | 1.00 | 33.48 |  | 0 | 3 | - Exclusion zone contact(s): /N7/O6/OP2/ Consider NA in place of MG (2 valid contact(s) for NA vs. 0 for MG) |
| MG:DA:3272 | 1.00 | 29.68 |  | 0 |  | 1^st^ MG coordination shell missing (see Notes) |
| MG:DA:3273 | 1.00 | 18.63 |  | 0 | 4 | - Exclusion zone contact(s): /O2'/O2'/O6/OP2/ Consider K in place of MG (4 valid contact(s) for K vs. 0 for MG) |
| MG:DA:3274 | 1.00 | 67.23 |  | 0 | 2 | - Exclusion zone contact(s): /O6/O6/ Consider K in place of MG (2 valid contact(s) for K vs. 0 for MG) |
| MG:DA:3275 | 1.00 | 64.60 |  | 0 |  | 1^st^ MG coordination shell missing (see Notes) |
| MG:DA:3276 | 1.00 | 34.96 |  | 1 |  | O_ph_ |
| MG:DA:3277 | 1.00 | 67.26 |  | 0 | 1 | - Exclusion zone contact(s): /OP2/ Consider K in place of MG (1 valid contact(s) for K vs. 0 for MG) |
| MG:DA:3278 | 1.00 | 62.71 |  | 0 | 2 | - Exclusion zone contact(s): /N7/O6/ Consider K in place of MG (2 valid contact(s) for K vs. 0 for MG) |
| MG:DA:3279 | 1.00 | 45.06 |  | 0 | 1 | - Exclusion zone contact(s): /OP2/ Consider K in place of MG (1 valid contact(s) for K vs. 0 for MG) |
| MG:DA:3280 | 1.00 | 32.77 |  | 0 | 3 | - Exclusion zone contact(s): /O6/OP1/OP2/ Consider K in place of MG (3 valid contact(s) for K vs. 0 for MG) |
| MG:DA:3281 | 1.00 | 38.91 |  | 0 | 5 | - Exclusion zone contact(s): /N3_A_/O2'/OP1/OP1/OP2/ Consider K in place of MG (5 valid contact(s) for K vs. 0 for MG) |
| MG:DA:3282 | 1.00 | 12.97 |  | 1 | 4 | O_r_ - Exclusion zone contact(s): /N3_A_/N7/N7/O6/ Consider NA in place of MG (2 valid contact(s) for NA vs. 1 for MG) |
| MG:DA:3283 | 1.00 | 87.73 |  | 0 |  | 1^st^ MG coordination shell missing (see Notes) |
| MG:DA:3284 | 1.00 | 27.46 |  | 0 | 2 | - Exclusion zone contact(s): /OP1/OP2/ Consider K in place of MG (2 valid contact(s) for K vs. 0 for MG) |
| MG:DA:3285 | 1.00 | 56.43 |  | 0 |  | 1^st^ MG coordination shell missing (see Notes) |
| MG:DA:3286 | 1.00 | 54.74 |  | 0 | 1 | - Exclusion zone contact(s): /N7/ Consider NA in place of MG (1 valid contact(s) for NA vs. 0 for MG) |
| MG:DA:3287 | 1.00 | 78.16 |  | 1 | 2 | O_ph_ - Exclusion zone contact(s): /OP1/OP2/ Consider K in place of MG (2 valid contact(s) for K vs. 1 for MG) |
| MG:DA:3288 | 1.00 | 52.02 |  | 0 |  | 1^st^ MG coordination shell missing (see Notes) |
| MG:DA:3289 | 1.00 | 27.80 |  | 1 | 3 | O_ph_ - Exclusion zone contact(s): /O3'/O5'/OP2/ Consider K in place of MG (3 valid contact(s) for K vs. 1 for MG) |
| MG:DA:3290 | 1.00 | 33.76 |  | 0 |  | 1^st^ MG coordination shell missing (see Notes) |
| MG:DA:3291 | 1.00 | 59.16 |  | 0 | 1 | - Exclusion zone contact(s): /O6/ Consider K in place of MG (1 valid contact(s) for K vs. 0 for MG) |
| MG:DA:3292 | 1.00 | 48.29 |  | 3 | 1 | O_b_N_b_Anion - Exclusion zone contact(s): /N3_U_/ |
| MG:DA:3293 | 1.00 | 35.19 |  | 0 | 5 | - Exclusion zone contact(s): /N4/O4/O6/O6/O6/ Consider K in place of MG (4 valid contact(s) for K vs. 0 for MG) |
| MG:DA:3294 | 1.00 | 28.67 |  | 0 | 2 | - Exclusion zone contact(s): /O6/OP2/ Consider K in place of MG (2 valid contact(s) for K vs. 0 for MG) |
| MG:DA:3295 | 1.00 | 14.71 |  | 1 |  | O_ph_ |
| MG:DA:3296 | 1.00 | 19.92 |  | 0 | 2 | - Exclusion zone contact(s): /N7/OP2/ Consider K in place of MG (2 valid contact(s) for K vs. 0 for MG) |
| MG:DA:3297 | 1.00 | 27.57 |  | 1 | 1 | O_ph_ - Exclusion zone contact(s): /O_back_/ |
| MG:DA:3298 | 1.00 | 97.73 |  | 0 | 3 | - Exclusion zone contact(s): /N1_G_/O4/O6/ Consider K in place of MG (3 valid contact(s) for K vs. 0 for MG) |
| MG:DA:3299 | 1.00 | 33.02 |  | 0 | 4 | - Exclusion zone contact(s): /O4/O6/O6/O6/ Consider K in place of MG (4 valid contact(s) for K vs. 0 for MG) |
| MG:DA:3300 | 1.00 | 27.31 |  | 1 | 1 | O_ph_ - Exclusion zone contact(s): /O3'/ |
| MG:DA:3301 | 1.00 | 29.80 |  | 0 | 4 | - Exclusion zone contact(s): /O2/O2'/O6/OP1/ Consider NA in place of MG (2 valid contact(s) for NA vs. 0 for MG) |
| MG:DA:3302 | 1.00 | 50.25 |  | 0 |  | 1^st^ MG coordination shell missing (see Notes) |
| MG:DA:3303 | 1.00 | 46.25 |  | 0 | 1 | - Exclusion zone contact(s): /O6/ Consider K in place of MG (1 valid contact(s) for K vs. 0 for MG) |
| MG:DA:3304 | 1.00 | 43.15 |  | 0 | 2 | - Exclusion zone contact(s): /O6/O6/ Consider K in place of MG (2 valid contact(s) for K vs. 0 for MG) |
| MG:DA:3305 | 1.00 | 21.20 |  | 0 | 2 | - Exclusion zone contact(s): /O6/O6/ Consider K in place of MG (2 valid contact(s) for K vs. 0 for MG) |
| MG:DA:3306 | 1.00 | 61.41 |  | 0 | 1 | - Exclusion zone contact(s): /OP2/ Consider K in place of MG (1 valid contact(s) for K vs. 0 for MG) |
| MG:DA:3307 | 1.00 | 59.64 |  | 0 | 1 | - Exclusion zone contact(s): /O6/ Consider K in place of MG (1 valid contact(s) for K vs. 0 for MG) |
| MG:DA:3308 | 1.00 | 75.34 |  | 0 | 1 | - Exclusion zone contact(s): /O6/ Consider K in place of MG (1 valid contact(s) for K vs. 0 for MG) |
| MG:DA:3309 | 1.00 | 56.27 |  | 0 | 1 | - Exclusion zone contact(s): /OP1/ Consider K in place of MG (1 valid contact(s) for K vs. 0 for MG) |
| MG:DA:3310 | 1.00 | 30.86 |  | 0 | 1 | - Exclusion zone contact(s): /N7/ Consider K in place of MG (1 valid contact(s) for K vs. 0 for MG) |
| MG:DA:3311 | 1.00 | 65.10 |  | 0 |  | 1^st^ MG coordination shell missing (see Notes) |
| MG:DA:3312 | 1.00 | 49.66 |  | 0 | 2 | - Exclusion zone contact(s): /N7/OP2/ Consider NA in place of MG (1 valid contact(s) for NA vs. 0 for MG) |
| MG:DA:3313 | 1.00 | 48.05 |  | 1 | 1 | O_ph_ - Exclusion zone contact(s): /OP2/ |
| MG:DA:3314 | 1.00 | 83.32 |  | 0 | 4 | - Exclusion zone contact(s): /N7/N7/O6/O6/ Consider K in place of MG (4 valid contact(s) for K vs. 0 for MG) |
| MG:DA:3315 | 1.00 | 28.95 |  | 0 |  | 1^st^ MG coordination shell missing (see Notes) |
| MG:DA:3316 | 1.00 | 43.58 |  | 1 | 3 | O_ph_ - Exclusion zone contact(s): /NH2_Arg_/O2'/O2'/ Consider K in place of MG (3 valid contact(s) for K vs. 1 for MG) |
| MG:DA:3317 | 1.00 | 86.38 |  | 0 | 3 | - Exclusion zone contact(s): /N1/N3_U_/O2'/ Consider K in place of MG (3 valid contact(s) for K vs. 0 for MG) |
| MG:DA:3318 | 1.00 | 72.40 |  | 0 | 1 | - Exclusion zone contact(s): /OP1/ Consider K in place of MG (1 valid contact(s) for K vs. 0 for MG) |
| MG:DA:3319 | 1.00 | 46.33 |  | 0 |  | 1^st^ MG coordination shell missing (see Notes) |
| MG:DA:3320 | 1.00 | 14.49 |  | 0 | 5 | - Exclusion zone contact(s): /O2'/O4'/O5'/OP1/OP2/ Consider K in place of MG (3 valid contact(s) for K vs. 0 for MG) |
| MG:DA:3321 | 1.00 | 34.09 |  | 0 | 2 | - Exclusion zone contact(s): /O6/O6/ Consider K in place of MG (2 valid contact(s) for K vs. 0 for MG) |
| MG:DA:3322 | 1.00 | 36.94 |  | 0 | 4 | - Exclusion zone contact(s): /N7/NZ_Lys_/OP1/OP2/ Consider K in place of MG (4 valid contact(s) for K vs. 0 for MG) |
| MG:DA:3323 | 1.00 | 41.06 |  | 0 | 5 | - Exclusion zone contact(s): /N2/O3'/O5'/OP1/OP2/ Consider K in place of MG (3 valid contact(s) for K vs. 0 for MG) |
| MG:DA:3324 | 1.00 | 42.90 |  | 1 | 1 | O_ph_ - Exclusion zone contact(s): /OP1/ |
| MG:DA:3325 | 1.00 | 62.19 |  | 0 |  | 1^st^ MG coordination shell missing (see Notes) |
| MG:DA:3326 | 1.00 | 63.12 |  | 0 |  | 1^st^ MG coordination shell missing (see Notes) |
| MG:DA:3327 | 1.00 | 48.00 |  | 0 | 2 | - Exclusion zone contact(s): /OP1/OP1/ Consider K in place of MG (2 valid contact(s) for K vs. 0 for MG) |
| MG:DA:3328 | 1.00 | 56.16 |  | 1 | 1 | O_ph_ - Exclusion zone contact(s): /O5'/ |
| MG:DA:3329 | 1.00 | 57.00 |  | 0 |  | 1^st^ MG coordination shell missing (see Notes) |
| MG:DA:3330 | 1.00 | 70.43 |  | 0 | 5 | - Exclusion zone contact(s): /O2'/O2'/O3'/OP1/OP2/ Consider K in place of MG (5 valid contact(s) for K vs. 0 for MG) |
| MG:DA:3331 | 1.00 | 68.43 |  | 0 | 4 | - Exclusion zone contact(s): /N7/O6/O6/OP2/ Consider K in place of MG (4 valid contact(s) for K vs. 0 for MG) |
| MG:DA:3332 | 1.00 | 82.81 |  | 0 |  | 1^st^ MG coordination shell missing (see Notes) |
| MG:DA:3333 | 1.00 | 78.86 |  | 1 |  | O_ph_ |
| MG:DA:3334 | 1.00 | 40.49 |  | 0 |  | 1^st^ MG coordination shell missing (see Notes) |
| MG:DA:3335 | 1.00 | 32.23 | 1 | 1 | 3 | - Exclusion zone contact(s): /O3'/OP1/OP2/OP2/ Consider K in place of MG (3 valid contact(s) for K vs. 0 for MG) |
| MG:DA:3336 | 1.00 | 36.70 |  | 2 | 4 | O_ph_O_r_ - Exclusion zone contact(s): /N2/N3_G_/N7/O6/ Consider K in place of MG (3 valid contact(s) for K vs. 2 for MG) |
| MG:DA:3337 | 1.00 | 44.84 |  | 1 | 2 | O_b_ - Exclusion zone contact(s): /O6/O6/ Consider K in place of MG (2 valid contact(s) for K vs. 1 for MG) |
| MG:DA:3338 | 1.00 | 49.35 |  | 0 | 2 | - Exclusion zone contact(s): /O2'/OP1/ Consider K in place of MG (2 valid contact(s) for K vs. 0 for MG) |
| MG:DA:3339 | 1.00 | 68.61 |  | 0 |  | 1^st^ MG coordination shell missing (see Notes) |
| MG:DA:3340 | 1.00 | 37.51 |  | 0 | 2 | - Exclusion zone contact(s): /N7/O6/ Consider K in place of MG (2 valid contact(s) for K vs. 0 for MG) |
| MG:DA:3341 | 1.00 | 49.39 |  | 0 |  | 1^st^ MG coordination shell missing (see Notes) |
| MG:DA:3342 | 1.00 | 55.55 |  | 0 |  | 1^st^ MG coordination shell missing (see Notes) |
| MG:DA:3343 | 1.00 | 58.33 |  | 0 |  | 1^st^ MG coordination shell missing (see Notes) |
| MG:DA:3344 | 1.00 | 90.26 |  | 0 |  | 1^st^ MG coordination shell missing (see Notes) |
| MG:DA:3345 | 1.00 | 80.69 |  | 1 |  | O_b_ |
| MG:DA:3346 | 1.00 | 49.40 |  | 0 |  | 1^st^ MG coordination shell missing (see Notes) |
| MG:DA:3347 | 1.00 | 29.26 |  | 0 | 1 | - Exclusion zone contact(s): /O6/ Consider NA in place of MG (1 valid contact(s) for NA vs. 0 for MG) |
| MG:DA:3348 | 1.00 | 54.29 |  | 0 | 1 | - Exclusion zone contact(s): /O6/ Consider NA in place of MG (1 valid contact(s) for NA vs. 0 for MG) |
| MG:DA:3349 | 1.00 | 56.91 |  | 0 | 1 | - Exclusion zone contact(s): /O6/ Consider K in place of MG (1 valid contact(s) for K vs. 0 for MG) |
| MG:DA:3350 | 1.00 | 50.65 |  | 1 | 4 | Anion - Exclusion zone contact(s): /N2/O4/O6/OP2/ Consider K in place of MG (4 valid contact(s) for K vs. 1 for MG) |
| MG:DA:3351 | 1.00 | 39.34 |  | 0 | 2 | - Exclusion zone contact(s): /O6/OP1/ Consider K in place of MG (2 valid contact(s) for K vs. 0 for MG) |
| MG:DA:3352 | 1.00 | 80.19 |  | 0 | 2 | - Exclusion zone contact(s): /OP1/OP1/ Consider K in place of MG (2 valid contact(s) for K vs. 0 for MG) |
| MG:DA:3353 | 1.00 | 22.10 |  | 0 | 5 | - Exclusion zone contact(s): /N7/O2/O2'/O4'/O6/ Consider K in place of MG (4 valid contact(s) for K vs. 0 for MG) |
| MG:DA:3354 | 1.00 | 66.41 |  | 1 | 1 | O_ph_ - Exclusion zone contact(s): /O2'/ |
| MG:DA:3355 | 1.00 | 64.47 |  | 0 |  | 1^st^ MG coordination shell missing (see Notes) |
| MG:DA:3356 | 1.00 | 58.55 |  | 0 |  | 1^st^ MG coordination shell missing (see Notes) |
| MG:DA:3357 | 1.00 | 59.10 |  | 0 | 2 | - Exclusion zone contact(s): /N7/OP2/ Consider K in place of MG (2 valid contact(s) for K vs. 0 for MG) |
| MG:DA:3358 | 1.00 | 23.27 |  | 1 |  | O_ph_ |
| MG:DA:3359 | 1.00 | 43.53 |  | 0 |  | 1^st^ MG coordination shell missing (see Notes) |
| MG:DA:3360 | 1.00 | 36.16 |  | 0 |  | 1^st^ MG coordination shell missing (see Notes) |
| MG:DA:3361 | 1.00 | 50.15 |  | 0 | 2 | - Exclusion zone contact(s): /OP1/OP2/ Consider K in place of MG (2 valid contact(s) for K vs. 0 for MG) |
| MG:DA:3362 | 1.00 | 35.30 |  | 2 |  | 2O_ph_ |
| MG:DA:3363 | 1.00 | 36.54 |  | 1 | 3 | O_ph_ - Exclusion zone contact(s): /N7/O6/O6/ Consider K in place of MG (3 valid contact(s) for K vs. 1 for MG) |
| MG:DA:3364 | 1.00 | 54.53 |  | 0 |  | 1^st^ MG coordination shell missing (see Notes) |
| MG:DA:3365 | 1.00 | 45.78 |  | 0 | 2 | - Exclusion zone contact(s): /O6/O6/ Consider K in place of MG (2 valid contact(s) for K vs. 0 for MG) |
| MG:DA:3366 | 1.00 | 35.30 |  | 1 | 3 | O_b_ - Exclusion zone contact(s): /N4/O4/O6/ Consider K in place of MG (3 valid contact(s) for K vs. 1 for MG) |
| MG:DA:3367 | 1.00 | 91.13 |  | 0 |  | 1^st^ MG coordination shell missing (see Notes) |
| MG:DA:3368 | 1.00 | 31.31 |  | 0 | 4 | - Exclusion zone contact(s): /O4/O6/O6/O6/ Consider K in place of MG (4 valid contact(s) for K vs. 0 for MG) |
| MG:DA:3369 | 1.00 | 16.06 |  | 2 |  | 2O_ph_ |
| MG:DA:3370 | 1.00 | 73.40 |  | 0 | 1 | - Exclusion zone contact(s): /O2/ Consider K in place of MG (1 valid contact(s) for K vs. 0 for MG) |
| MG:DA:3371 | 1.00 | 73.74 |  | 1 |  | O_ph_ |
| MG:DA:3372 | 1.00 | 54.02 |  | 0 | 2 | - Exclusion zone contact(s): /O5'/OP1/ Consider K in place of MG (2 valid contact(s) for K vs. 0 for MG) |
| MG:DA:3373 | 1.00 | 40.10 |  | 0 |  | 1^st^ MG coordination shell missing (see Notes) |
| MG:DA:3374 | 1.00 | 84.64 |  | 0 | 1 | - Exclusion zone contact(s): /OP2/ Consider K in place of MG (1 valid contact(s) for K vs. 0 for MG) |
| MG:DA:3375 | 1.00 | 64.37 |  | 0 | 3 | - Exclusion zone contact(s): /O2/O2'/OP1/ Consider K in place of MG (2 valid contact(s) for K vs. 0 for MG) |
| MG:DA:3376 | 1.00 | 18.18 |  | 0 | 5 | - Exclusion zone contact(s): /O2/O2/O2'/O2'/O4'/ Consider K in place of MG (4 valid contact(s) for K vs. 0 for MG) |
| MG:DA:3377 | 1.00 | 83.94 |  | 1 |  | O_ph_ |
| MG:DA:3378 | 1.00 | 55.66 |  | 0 |  | 1^st^ MG coordination shell missing (see Notes) |
| MG:DA:3379 | 1.00 | 36.32 |  | 1 | 1 | O_ph_ - Exclusion zone contact(s): /OP1/ |
| MG:DA:3380 | 1.00 | 55.92 |  | 2 | 1 | 2O_ph_ - Exclusion zone contact(s): /OP2/ |
| MG:DA:3381 | 1.00 | 66.93 |  | 0 |  | 1^st^ MG coordination shell missing (see Notes) |
| MG:DA:3382 | 1.00 | 47.05 |  | 0 | 2 | - Exclusion zone contact(s): /O4/O6/ Consider K in place of MG (2 valid contact(s) for K vs. 0 for MG) |
| MG:DA:3383 | 1.00 | 49.18 |  | 0 | 3 | - Exclusion zone contact(s): /O2'/O2'/O4'/ Consider K in place of MG (2 valid contact(s) for K vs. 0 for MG) |
| MG:DA:3384 | 1.00 | 56.99 |  | 0 |  | 1^st^ MG coordination shell missing (see Notes) |
| MG:DA:3385 | 1.00 | 85.92 |  | 1 | 1 | O_ph_ - Exclusion zone contact(s): /OP2/ |
| MG:DA:3386 | 1.00 | 69.13 |  | 0 |  | 1^st^ MG coordination shell missing (see Notes) |
| MG:DA:3387 | 1.00 | 47.77 |  | 1 |  | O_ph_ |
| MG:DA:3388 | 1.00 | 33.73 |  | 0 | 4 | - Exclusion zone contact(s): /N3_A_/N7/O2'/O6/ Consider K in place of MG (3 valid contact(s) for K vs. 0 for MG) |
| MG:DA:3389 | 1.00 | 48.35 |  | 0 | 1 | - Exclusion zone contact(s): /OP1/ Consider K in place of MG (1 valid contact(s) for K vs. 0 for MG) |
| MG:DA:3390 | 1.00 | 83.92 |  | 0 | 2 | - Exclusion zone contact(s): /N4/N7/ Consider NA in place of MG (2 valid contact(s) for NA vs. 0 for MG) |
| MG:DA:3391 | 1.00 | 84.29 |  | 0 |  | 1^st^ MG coordination shell missing (see Notes) |
| MG:DA:3392 | 1.00 | 76.32 |  | 0 | 1 | - Exclusion zone contact(s): /O2'/ Consider K in place of MG (1 valid contact(s) for K vs. 0 for MG) |
| MG:DA:3393 | 1.00 | 88.25 |  | 0 |  | 1^st^ MG coordination shell missing (see Notes) |
| MG:DA:3394 | 1.00 | 43.11 |  | 0 |  | 1^st^ MG coordination shell missing (see Notes) |
| MG:DA:3395 | 1.00 | 42.79 |  | 0 |  | 1^st^ MG coordination shell missing (see Notes) |
| MG:DA:3396 | 1.00 | 46.07 |  | 0 | 2 | - Exclusion zone contact(s): /O2'/O4'/ Consider K in place of MG (2 valid contact(s) for K vs. 0 for MG) |
| MG:DA:3397 | 1.00 | 65.31 |  | 1 | 2 | O_b_ - Exclusion zone contact(s): /O6/O6/ Consider K in place of MG (2 valid contact(s) for K vs. 1 for MG) |
| MG:DA:3398 | 1.00 | 75.14 |  | 0 | 3 | - Exclusion zone contact(s): /O5'/OP1/OP2/ Consider K in place of MG (2 valid contact(s) for K vs. 0 for MG) |
| MG:DA:3399 | 1.00 | 52.80 | 1 | 1 | 2 | - Exclusion zone contact(s): /O3'/OP2/OP2/ Consider K in place of MG (2 valid contact(s) for K vs. 0 for MG) |
| MG:DA:3400 | 1.00 | 48.26 |  | 0 | 1 | - Exclusion zone contact(s): /O4/ Consider K in place of MG (1 valid contact(s) for K vs. 0 for MG) |
| MG:DA:3401 | 1.00 | 45.70 |  | 0 | 2 | - Exclusion zone contact(s): /O6/OP1/ Consider K in place of MG (2 valid contact(s) for K vs. 0 for MG) |
| MG:DA:3402 | 1.00 | 24.49 |  | 0 | 1 | - Exclusion zone contact(s): /OP1/ Consider K in place of MG (1 valid contact(s) for K vs. 0 for MG) |
| MG:DA:3403 | 1.00 | 89.01 |  | 0 | 2 | - Exclusion zone contact(s): /OP1/OP2/ Consider K in place of MG (2 valid contact(s) for K vs. 0 for MG) |
| MG:DA:3404 | 1.00 | 65.84 |  | 0 | 1 | - Exclusion zone contact(s): /N7/ Consider K in place of MG (1 valid contact(s) for K vs. 0 for MG) |
| MG:DA:3405 | 1.00 | 43.05 |  | 0 | 2 | - Exclusion zone contact(s): /N6/N7/ Consider K in place of MG (2 valid contact(s) for K vs. 0 for MG) |
| MG:DA:3406 | 1.00 | 59.81 |  | 0 | 3 | - Exclusion zone contact(s): /O6/OP2/OP2/ Consider K in place of MG (2 valid contact(s) for K vs. 0 for MG) |
| MG:DA:3407 | 1.00 | 70.97 |  | 0 | 1 | - Exclusion zone contact(s): /OP1/ Consider K in place of MG (1 valid contact(s) for K vs. 0 for MG) |
| MG:DA:3408 | 1.00 | 41.41 |  | 0 | 3 | - Exclusion zone contact(s): /O4/O4/OP2/ Consider NA in place of MG (2 valid contact(s) for NA vs. 0 for MG) |
| MG:DA:3409 | 1.00 | 44.85 |  | 0 |  | 1^st^ MG coordination shell missing (see Notes) |
| MG:DA:3410 | 1.00 | 64.33 |  | 3 | 3 | N_b_O_cno_Anion - Exclusion zone contact(s): /N1_G_/N6/O6/ |
| MG:DA:3411 | 1.00 | 56.87 |  | 0 | 3 | - Exclusion zone contact(s): /N1_G_/N2/ Consider K in place of MG (3 valid contact(s) for K vs. 0 for MG) |
| MG:DA:3412 | 1.00 | 68.39 |  | 0 | 1 | - Exclusion zone contact(s): /OP2/ Consider NA in place of MG (1 valid contact(s) for NA vs. 0 for MG) |
| MG:DA:3413 | 1.00 | 45.05 |  | 1 | 1 | O_ph_ - Exclusion zone contact(s): /OP2/ |
| MG:DA:3414 | 1.00 | 45.01 |  | 0 |  | 1^st^ MG coordination shell missing (see Notes) |
| MG:DA:3415 | 1.00 | 49.25 |  | 0 |  | 1^st^ MG coordination shell missing (see Notes) |
| MG:DA:3416 | 1.00 | 62.81 |  | 0 | 1 | - Exclusion zone contact(s): /OP2/ Consider K in place of MG (1 valid contact(s) for K vs. 0 for MG) |
| MG:DA:3417 | 1.00 | 62.75 |  | 1 | 2 | O_ph_ - Exclusion zone contact(s): /O4'/OP2/ Consider K in place of MG (2 valid contact(s) for K vs. 1 for MG) |
| MG:DA:3418 | 1.00 | 60.84 |  | 0 | 2 | - Exclusion zone contact(s): /O3'/O5'/ Consider K in place of MG (2 valid contact(s) for K vs. 0 for MG) |
| MG:DA:3419 | 1.00 | 32.12 |  | 0 | 4 | - Exclusion zone contact(s): /O4/O6/O6/O6/ Consider K in place of MG (4 valid contact(s) for K vs. 0 for MG) |
| MG:DA:3420 | 1.00 | 51.89 |  | 0 |  | 1^st^ MG coordination shell missing (see Notes) |
| MG:DA:3421 | 1.00 | 56.96 |  | 0 | 4 | - Exclusion zone contact(s): /N7/O2/O4/O6/ Consider K in place of MG (3 valid contact(s) for K vs. 0 for MG) |
| MG:DA:3422 | 1.00 | 41.11 |  | 0 | 2 | - Exclusion zone contact(s): /N7/O6/ Consider K in place of MG (2 valid contact(s) for K vs. 0 for MG) |
| MG:DA:3423 | 1.00 | 58.69 |  | 0 | 3 | - Exclusion zone contact(s): /N7/O6/OP2/ Consider K in place of MG (2 valid contact(s) for K vs. 0 for MG) |
| MG:DA:3424 | 1.00 | 47.52 |  | 0 | 3 | - Exclusion zone contact(s): /O4/O6/O6/ Consider K in place of MG (3 valid contact(s) for K vs. 0 for MG) |
| MG:DA:3425 | 1.00 | 63.56 |  | 0 |  | 1^st^ MG coordination shell missing (see Notes) |
| MG:DA:3426 | 1.00 | 39.56 |  | 0 |  | 1^st^ MG coordination shell missing (see Notes) |
| MG:DA:3427 | 1.00 | 44.61 |  | 2 | 2 | O_ph_O_r_ - Exclusion zone contact(s): /OP1/OP2/ |
| MG:DA:3428 | 1.00 | 38.21 |  | 1 |  | O_ph_ |
| MG:DA:3429 | 1.00 | 30.92 |  | 0 | 1 | - Exclusion zone contact(s): /OP2/ Consider K in place of MG (1 valid contact(s) for K vs. 0 for MG) |
| MG:DA:3430 | 1.00 | 60.01 |  | 0 | 2 | - Exclusion zone contact(s): /OP1/OP2/ Consider K in place of MG (2 valid contact(s) for K vs. 0 for MG) |
| MG:DA:3431 | 1.00 | 61.11 |  | 0 | 2 | - Exclusion zone contact(s): /N7/O6/ Consider NA in place of MG (1 valid contact(s) for NA vs. 0 for MG) |
| MG:DA:3432 | 1.00 | 46.30 |  | 1 |  | O_ph_ |
| MG:DA:3433 | 1.00 | 43.87 |  | 1 |  | O_ph_ |
| MG:DA:3434 | 1.00 | 41.92 |  | 0 | 2 | - Exclusion zone contact(s): /N4/N7/ Consider NA in place of MG (1 valid contact(s) for NA vs. 0 for MG) |
| MG:DA:3435 | 1.00 | 49.20 |  | 1 |  | O_ph_ |
| MG:DA:3436 | 1.00 | 48.45 | 1 | 1 |  | O_ph_ - Exclusion zone contact(s): /OP2/ |
| MG:DA:3437 | 1.00 | 47.08 |  | 1 | 2 | O_ph_ - Exclusion zone contact(s): /O3'/OP1/ |
| MG:DA:3438 | 1.00 | 32.80 |  | 0 | 2 | - Exclusion zone contact(s): /OP1/OP1/ Consider K in place of MG (2 valid contact(s) for K vs. 0 for MG) |
| MG:DA:3439 | 1.00 | 86.28 |  | 0 | 3 | - Exclusion zone contact(s): /O4'/O6/OP2/ Consider K in place of MG (3 valid contact(s) for K vs. 0 for MG) |
| MG:DA:3440 | 1.00 | 57.71 |  | 1 |  | O_ph_ |
| MG:DA:3441 | 1.00 | 43.89 |  | 2 |  | 2O_ph_ |
| MG:DA:3442 | 1.00 | 45.32 |  | 0 | 1 | - Exclusion zone contact(s): /OP1/ Consider K in place of MG (1 valid contact(s) for K vs. 0 for MG) |
| MG:DA:3443 | 1.00 | 54.39 |  | 1 | 1 | O_ph_ - Exclusion zone contact(s): /O5'/ |
| MG:DA:3444 | 1.00 | 56.04 |  | 0 | 3 | - Exclusion zone contact(s): /O5'/OP2/OP2/ Consider K in place of MG (3 valid contact(s) for K vs. 0 for MG) |
| MG:DA:3445 | 1.00 | 59.40 |  | 0 |  | 1^st^ MG coordination shell missing (see Notes) |
| MG:DA:3446 | 1.00 | 66.00 |  | 0 | 1 | - Exclusion zone contact(s): /OP2/ Consider NA in place of MG (1 valid contact(s) for NA vs. 0 for MG) |
| MG:DA:3447 | 1.00 | 48.99 |  | 0 | 2 | - Exclusion zone contact(s): /O6/O6/ Consider K in place of MG (2 valid contact(s) for K vs. 0 for MG) |
| MG:DA:3448 | 1.00 | 40.65 |  | 0 |  | 1^st^ MG coordination shell missing (see Notes) |
| MG:DA:3449 | 1.00 | 50.94 |  | 1 | 3 | O_r_ - Exclusion zone contact(s): /N2/O2/O4'/ Consider K in place of MG (3 valid contact(s) for K vs. 1 for MG) |
| MG:DA:3450 | 1.00 | 75.68 |  | 0 | 1 | - Exclusion zone contact(s): /OP1/ Consider K in place of MG (1 valid contact(s) for K vs. 0 for MG) |
| MG:DA:3451 | 1.00 | 76.20 |  | 0 | 2 | - Exclusion zone contact(s): /N4/N4/ Consider K in place of MG (2 valid contact(s) for K vs. 0 for MG) |
| MG:DA:3452 | 1.00 | 68.92 |  | 1 | 2 | O_r_ - Exclusion zone contact(s): /O2'/OP1/ Consider K in place of MG (2 valid contact(s) for K vs. 1 for MG) |
| MG:DA:3453 | 1.00 | 56.70 |  | 0 | 2 | - Exclusion zone contact(s): /O6/O6/ Consider K in place of MG (2 valid contact(s) for K vs. 0 for MG) |
| MG:DA:3454 | 1.00 | 51.29 |  | 0 | 1 | - Exclusion zone contact(s): /OP2/ Consider K in place of MG (1 valid contact(s) for K vs. 0 for MG) |
| MG:DA:3455 | 1.00 | 58.80 |  | 0 | 1 | - Exclusion zone contact(s): /O6/ Consider K in place of MG (1 valid contact(s) for K vs. 0 for MG) |
| MG:DA:3456 | 1.00 | 60.72 |  | 0 | 2 | - Exclusion zone contact(s): /O6/OP2/ Consider K in place of MG (2 valid contact(s) for K vs. 0 for MG) |
| MG:DA:3457 | 1.00 | 76.27 |  | 0 | 1 | - Exclusion zone contact(s): /OP1/ Consider K in place of MG (1 valid contact(s) for K vs. 0 for MG) |
| MG:DA:3458 | 1.00 | 54.35 |  | 0 | 4 | - Exclusion zone contact(s): /N1_A_/N3_C_/O2/OP2/ Consider K in place of MG (4 valid contact(s) for K vs. 0 for MG) |
| MG:DA:3459 | 1.00 | 63.70 |  | 1 | 1 | O_ph_ - Exclusion zone contact(s): /O3'/ |
| MG:DA:3460 | 1.00 | 46.90 |  | 0 | 2 | - Exclusion zone contact(s): /O5'/O5'/ Consider K in place of MG (2 valid contact(s) for K vs. 0 for MG) |
| MG:DA:3461 | 1.00 | 34.05 |  | 0 | 2 | - Exclusion zone contact(s): /O2'/OP1/ Consider NA in place of MG (1 valid contact(s) for NA vs. 0 for MG) |
| MG:DA:3462 | 1.00 | 56.60 |  | 0 | 2 | - Exclusion zone contact(s): /N7/O6/ Consider K in place of MG (2 valid contact(s) for K vs. 0 for MG) |
| MG:DA:3463 | 1.00 | 61.04 |  | 0 |  | 1^st^ MG coordination shell missing (see Notes) |
| MG:DA:3464 | 1.00 | 53.27 |  | 0 |  | 1^st^ MG coordination shell missing (see Notes) |
| MG:DA:3465 | 1.00 | 56.51 |  | 0 | 1 | - Exclusion zone contact(s): /O2/ Consider K in place of MG (1 valid contact(s) for K vs. 0 for MG) |
| MG:DA:3466 | 1.00 | 79.14 |  | 1 | 1 | O_ph_ - Exclusion zone contact(s): /O5'/ |
| MG:DA:3467 | 1.00 | 75.49 |  | 0 |  | 1^st^ MG coordination shell missing (see Notes) |
| MG:DA:3468 | 1.00 | 49.96 |  | 1 | 2 | O_ph_ - Exclusion zone contact(s): /O2'/O3'/ Consider K in place of MG (2 valid contact(s) for K vs. 1 for MG) |
| MG:DA:3469 | 1.00 | 63.70 |  | 1 | 1 | O_ph_ - Exclusion zone contact(s): /OP1/ |
| MG:DA:3470 | 1.00 | 52.25 |  | 1 | 3 | O_ph_ - Exclusion zone contact(s): /NH2_Arg_/O5'/OP2/ Consider K in place of MG (2 valid contact(s) for K vs. 1 for MG) |
| MG:DA:3471 | 1.00 | 27.62 |  | 0 | 3 | - Exclusion zone contact(s): /O4/O6/O6/ Consider K in place of MG (3 valid contact(s) for K vs. 0 for MG) |
| MG:DA:3472 | 1.00 | 55.77 |  | 0 | 1 | - Exclusion zone contact(s): /O4/ Consider K in place of MG (1 valid contact(s) for K vs. 0 for MG) |
| MG:DA:3473 | 1.00 | 44.35 |  | 0 | 1 | - Exclusion zone contact(s): /O6/ Consider K in place of MG (1 valid contact(s) for K vs. 0 for MG) |
| MG:DA:3474 | 1.00 | 41.07 |  | 0 | 1 | - Exclusion zone contact(s): /OP1/ Consider K in place of MG (1 valid contact(s) for K vs. 0 for MG) |
| MG:DA:3475 | 1.00 | 51.31 |  | 0 | 4 | - Exclusion zone contact(s): /N7/O2'/O4/O6/ Consider K in place of MG (3 valid contact(s) for K vs. 0 for MG) |
| MG:DA:3476 | 1.00 | 27.74 |  | 0 | 3 | - Exclusion zone contact(s): /O6/O_back_/OP2/ Consider K in place of MG (2 valid contact(s) for K vs. 0 for MG) |
| MG:DA:3477 | 1.00 | 171.85 |  | 1 | 6 | O_ph_ - Exclusion zone contact(s): /N3_U_/NE_Arg_/NH1_Arg_/O2/O5'/OP1/ Consider K in place of MG (6 valid contact(s) for K vs. 1 for MG) |
| MG:DA:3478 | 1.00 | 18.88 |  | 0 |  | 1^st^ MG coordination shell missing (see Notes) |
| MG:DA:3479 | 1.00 | 74.38 |  | 0 |  | 1^st^ MG coordination shell missing (see Notes) |
| MG:DA:3480 | 1.00 | 46.88 |  | 1 | 2 | O_b_ - Exclusion zone contact(s): /O6/O6/ Consider K in place of MG (2 valid contact(s) for K vs. 1 for MG) |
| MG:DA:3481 | 1.00 | 93.43 |  | 0 | 1 | - Exclusion zone contact(s): /N7/ Consider K in place of MG (1 valid contact(s) for K vs. 0 for MG) |
| MG:DA:3482 | 1.00 | 39.07 | 1 | 1 | 2 | - Exclusion zone contact(s): /N7/O3'/OP1/ Consider K in place of MG (2 valid contact(s) for K vs. 0 for MG) |
| MG:DA:3483 | 1.00 | 61.59 |  | 0 | 3 | - Exclusion zone contact(s): /N6/O6/OP1/ Consider K in place of MG (2 valid contact(s) for K vs. 0 for MG) |
| MG:DA:3484 | 1.00 | 76.15 |  | 1 | 1 | O_ph_ - Exclusion zone contact(s): /OP2/ |
| MG:DA:3485 | 1.00 | 77.33 |  | 0 |  | 1^st^ MG coordination shell missing (see Notes) |
| MG:DA:3486 | 1.00 | 58.87 |  | 0 | 3 | - Exclusion zone contact(s): /O5'/OP1/OP2/ Consider K in place of MG (3 valid contact(s) for K vs. 0 for MG) |
| MG:DA:3487 | 1.00 | 62.28 |  | 0 | 2 | - Exclusion zone contact(s): /O5'/OP2/ Consider K in place of MG (2 valid contact(s) for K vs. 0 for MG) |
| MG:DA:3488 | 1.00 | 37.65 |  | 0 | 1 | - Exclusion zone contact(s): /N7/ Consider K in place of MG (1 valid contact(s) for K vs. 0 for MG) |
| MG:DA:3489 | 1.00 | 41.26 |  | 0 |  | 1^st^ MG coordination shell missing (see Notes) |
| MG:DA:3490 | 1.00 | 64.19 |  | 0 | 1 | - Exclusion zone contact(s): /O6/ Consider K in place of MG (1 valid contact(s) for K vs. 0 for MG) |
| MG:DA:3491 | 1.00 | 78.88 |  | 0 | 1 | - Exclusion zone contact(s): /O6/ Consider K in place of MG (1 valid contact(s) for K vs. 0 for MG) |
| MG:DA:3492 | 1.00 | 53.99 |  | 0 | 1 | - Exclusion zone contact(s): /N1_A_/ Consider K in place of MG (1 valid contact(s) for K vs. 0 for MG) |
| MG:DA:3493 | 1.00 | 50.44 |  | 0 | 3 | - Exclusion zone contact(s): /NH2_Arg_/OP1/OP2/ Consider K in place of MG (2 valid contact(s) for K vs. 0 for MG) |
| MG:DA:3494 | 1.00 | 59.40 |  | 0 | 1 | - Exclusion zone contact(s): /O4/ Consider K in place of MG (1 valid contact(s) for K vs. 0 for MG) |
| MG:DA:3495 | 1.00 | 51.46 |  | 0 |  | 1^st^ MG coordination shell missing (see Notes) |
| MG:DA:3496 | 1.00 | 47.28 |  | 0 | 4 | - Exclusion zone contact(s): /O2'/O3'/O5'/OP1/ Consider K in place of MG (4 valid contact(s) for K vs. 0 for MG) |
| MG:DA:3497 | 1.00 | 38.42 |  | 0 |  | 1^st^ MG coordination shell missing (see Notes) |
| MG:DA:3498 | 1.00 | 44.39 |  | 0 |  | 1^st^ MG coordination shell missing (see Notes) |
| MG:DA:3499 | 1.00 | 70.78 |  | 0 | 1 | - Exclusion zone contact(s): /OP1/ Consider K in place of MG (1 valid contact(s) for K vs. 0 for MG) |
| MG:DA:3500 | 1.00 | 57.13 |  | 0 | 2 | - Exclusion zone contact(s): /N7/OP2/ Consider K in place of MG (2 valid contact(s) for K vs. 0 for MG) |
| MG:DA:3501 | 1.00 | 51.57 |  | 0 | 2 | - Exclusion zone contact(s): /N7/O6/ Consider K in place of MG (2 valid contact(s) for K vs. 0 for MG) |
| MG:DA:3502 | 1.00 | 74.70 |  | 0 |  | 1^st^ MG coordination shell missing (see Notes) |
| MG:DA:3503 | 1.00 | 71.91 |  | 0 | 3 | - Exclusion zone contact(s): /N1_A_/O4/O6/ Consider K in place of MG (3 valid contact(s) for K vs. 0 for MG) |
| MG:DA:3504 | 1.00 | 30.94 |  | 0 |  | 1^st^ MG coordination shell missing (see Notes) |
| MG:DA:3505 | 1.00 | 51.60 |  | 0 | 5 | - Exclusion zone contact(s): /N3_G_/O2'/O4'/O_back_/OG_Ser_/ Consider K in place of MG (4 valid contact(s) for K vs. 0 for MG) |
| MG:DA:3506 | 1.00 | 46.34 |  | 1 | 3 | O_ph_ - Exclusion zone contact(s): /N6/O3'/O6/ Consider K in place of MG (2 valid contact(s) for K vs. 1 for MG) |
| MG:DA:3507 | 1.00 | 66.38 |  | 0 | 4 | - Exclusion zone contact(s): /O2'/O2'/O3'/O3'/ Consider K in place of MG (4 valid contact(s) for K vs. 0 for MG) |
| MG:DA:3508 | 1.00 | 56.78 |  | 0 | 2 | - Exclusion zone contact(s): /O4/O6/ Consider K in place of MG (2 valid contact(s) for K vs. 0 for MG) |
| MG:DA:3509 | 1.00 | 47.59 |  | 0 |  | 1^st^ MG coordination shell missing (see Notes) |
| MG:DA:3510 | 1.00 | 56.63 |  | 0 |  | 1^st^ MG coordination shell missing (see Notes) |
| MG:DA:3511 | 1.00 | 42.58 |  | 0 | 4 | - Exclusion zone contact(s): /O4/O4/O6/OP2/ Consider K in place of MG (4 valid contact(s) for K vs. 0 for MG) |
| MG:DA:3512 | 1.00 | 66.71 |  | 0 | 1 | - Exclusion zone contact(s): /O6/ Consider K in place of MG (1 valid contact(s) for K vs. 0 for MG) |
| MG:DA:3513 | 1.00 | 50.48 |  | 0 |  | 1^st^ MG coordination shell missing (see Notes) |
| MG:DA:3514 | 1.00 | 94.78 |  | 0 | 2 | - Exclusion zone contact(s): /OP1/OP2/ Consider K in place of MG (2 valid contact(s) for K vs. 0 for MG) |
| MG:DA:3515 | 1.00 | 28.19 |  | 0 | 4 | - Exclusion zone contact(s): /NH1_Arg_/O2'/O3'/OP1/ Consider K in place of MG (4 valid contact(s) for K vs. 0 for MG) |
| MG:DA:3516 | 1.00 | 53.34 |  | 1 | 2 | O_ph_ - Exclusion zone contact(s): /O3'/O5'/ Consider K in place of MG (2 valid contact(s) for K vs. 1 for MG) |
| MG:DA:3517 | 1.00 | 29.50 |  | 0 | 5 | - Exclusion zone contact(s): /O2/O2'/O2'/O4'/O4'/ Consider K in place of MG (4 valid contact(s) for K vs. 0 for MG) |
| MG:DA:3518 | 1.00 | 65.43 |  | 0 | 2 | - Exclusion zone contact(s): /O3'/OP1/ Consider K in place of MG (2 valid contact(s) for K vs. 0 for MG) |
| MG:DA:3519 | 1.00 | 36.04 |  | 0 | 1 | - Exclusion zone contact(s): /OP2/ Consider K in place of MG (1 valid contact(s) for K vs. 0 for MG) |
| MG:DA:3520 | 1.00 | 95.62 |  | 0 | 2 | - Exclusion zone contact(s): /N4/O4/ Consider NA in place of MG (1 valid contact(s) for NA vs. 0 for MG) |
| MG:DA:3521 | 1.00 | 61.30 |  | 0 |  | 1^st^ MG coordination shell missing (see Notes) |
| MG:DA:3522 | 1.00 | 63.54 |  | 1 | 1 | N_b_ - Exclusion zone contact(s): /O4/ |
| MG:DA:3523 | 1.00 | 82.04 |  | 1 |  | O_ph_ |
| MG:DA:3524 | 1.00 | 32.11 |  | 0 | 4 | - Exclusion zone contact(s): /O4/O4/OP1/OP2/ Consider K in place of MG (4 valid contact(s) for K vs. 0 for MG) |
| MG:DA:3525 | 1.00 | 43.90 |  | 0 | 3 | - Exclusion zone contact(s): /N6/N7/OP1/ Consider K in place of MG (3 valid contact(s) for K vs. 0 for MG) |
| MG:DA:3526 | 1.00 | 46.30 |  | 0 | 3 | - Exclusion zone contact(s): /O2'/O3'/OP1/ Consider K in place of MG (3 valid contact(s) for K vs. 0 for MG) |
| MG:DA:3527 | 1.00 | 46.31 |  | 0 | 4 | - Exclusion zone contact(s): /O3'/O5'/OP2/OP2/ Consider K in place of MG (3 valid contact(s) for K vs. 0 for MG) |
| MG:DA:3528 | 1.00 | 45.83 |  | 0 | 3 | - Exclusion zone contact(s): /O2'/O6/OP2/ Consider K in place of MG (3 valid contact(s) for K vs. 0 for MG) |
| MG:DA:3529 | 1.00 | 84.92 |  | 0 |  | 1^st^ MG coordination shell missing (see Notes) |
| MG:DA:3530 | 1.00 | 73.01 |  | 0 | 2 | - Exclusion zone contact(s): /N4/O6/ Consider K in place of MG (2 valid contact(s) for K vs. 0 for MG) |
| MG:DA:3531 | 1.00 | 53.17 |  | 0 | 1 | - Exclusion zone contact(s): /OP1/ Consider NA in place of MG (1 valid contact(s) for NA vs. 0 for MG) |
| MG:DA:3532 | 1.00 | 49.94 |  | 0 |  | 1^st^ MG coordination shell missing (see Notes) |
| MG:DA:3533 | 1.00 | 47.49 |  | 0 | 1 | - Exclusion zone contact(s): /O6/ Consider K in place of MG (1 valid contact(s) for K vs. 0 for MG) |
| MG:DA:3534 | 1.00 | 49.26 |  | 0 | 1 | - Exclusion zone contact(s): /OP2/ Consider K in place of MG (1 valid contact(s) for K vs. 0 for MG) |
| MG:DA:3535 | 1.00 | 69.17 |  | 1 |  | O_ph_ |
| MG:DA:3536 | 1.00 | 48.58 |  | 0 | 4 | - Exclusion zone contact(s): /O2'/O3'/O6/OP2/ Consider K in place of MG (4 valid contact(s) for K vs. 0 for MG) |
| MG:DA:3537 | 1.00 | 43.50 |  | 1 |  | O_r_ |
| MG:DA:3538 | 1.00 | 79.77 |  | 0 | 2 | - Exclusion zone contact(s): /O2'/OP1/ Consider NA in place of MG (1 valid contact(s) for NA vs. 0 for MG) |
| MG:DA:3539 | 1.00 | 93.11 |  | 0 | 3 | - Exclusion zone contact(s): /N7/O6/O6/ Consider K in place of MG (3 valid contact(s) for K vs. 0 for MG) |
| MG:DA:3540 | 1.00 | 74.61 |  | 1 | 2 | O_r_ - Exclusion zone contact(s): /O3'/OP1/ |
| MG:DA:3541 | 1.00 | 19.23 |  | 0 | 2 | - Exclusion zone contact(s): /OP1/OP2/ Consider K in place of MG (2 valid contact(s) for K vs. 0 for MG) |
| MG:DA:3542 | 1.00 | 51.84 |  | 0 | 1 | - Exclusion zone contact(s): /O6/ Consider K in place of MG (1 valid contact(s) for K vs. 0 for MG) |
| MG:DA:3543 | 1.00 | 62.52 |  | 1 |  | O_ph_ |
| MG:DA:3544 | 1.00 | 51.13 |  | 1 |  | O_ph_ |
| MG:DA:3545 | 1.00 | 36.75 |  | 0 | 3 | - Exclusion zone contact(s): /O2'/O4'/OP1/ Consider K in place of MG (3 valid contact(s) for K vs. 0 for MG) |
| MG:DA:3546 | 1.00 | 29.61 |  | 0 |  | 1^st^ MG coordination shell missing (see Notes) |
| MG:DA:3547 | 1.00 | 68.85 |  | 0 | 1 | - Exclusion zone contact(s): /OP1/ Consider K in place of MG (1 valid contact(s) for K vs. 0 for MG) |
| MG:DA:3548 | 1.00 | 58.28 |  | 0 |  | No contacts in 5.0 A |
| MG:DA:3549 | 1.00 | 40.00 |  | 0 | 3 | - Exclusion zone contact(s): /O2'/OP1/OP2/ Consider K in place of MG (3 valid contact(s) for K vs. 0 for MG) |
| MG:DA:3550 | 1.00 | 96.72 |  | 0 |  | No contacts in 5.0 A |
| MG:DA:3551 | 1.00 | 29.78 |  | 0 |  | 1^st^ MG coordination shell missing (see Notes) |
| MG:DA:3552 | 1.00 | 45.14 |  | 0 | 2 | - Exclusion zone contact(s): /O4'/OP2/ Consider K in place of MG (2 valid contact(s) for K vs. 0 for MG) |
| MG:DA:3553 | 1.00 | 89.82 |  | 0 |  | 1^st^ MG coordination shell missing (see Notes) |
| MG:DA:3554 | 1.00 | 51.36 |  | 0 |  | 1^st^ MG coordination shell missing (see Notes) |
| MG:DA:3555 | 1.00 | 49.77 |  | 0 | 3 | - Exclusion zone contact(s): /O3'/OP1/OP2/ Consider K in place of MG (3 valid contact(s) for K vs. 0 for MG) |
| MG:DA:3556 | 1.00 | 107.46 |  | 0 | 1 | - Exclusion zone contact(s): /OP2/ Consider K in place of MG (1 valid contact(s) for K vs. 0 for MG) |
| MG:DA:3557 | 1.00 | 55.39 |  | 0 | 2 | - Exclusion zone contact(s): /O6/O6/ Consider K in place of MG (2 valid contact(s) for K vs. 0 for MG) |
| MG:DA:3558 | 1.00 | 64.63 |  | 0 |  | 1^st^ MG coordination shell missing (see Notes) |
| MG:DA:3559 | 1.00 | 23.74 |  | 0 | 5 | - Exclusion zone contact(s): /N7/O2/O2'/O4/O6/ Consider K in place of MG (4 valid contact(s) for K vs. 0 for MG) |
| MG:DA:3560 | 1.00 | 36.31 |  | 0 | 4 | - Exclusion zone contact(s): /N2/N6/OP1/OP2/ Consider K in place of MG (4 valid contact(s) for K vs. 0 for MG) |
| MG:DA:3561 | 1.00 | 51.75 |  | 0 |  | 1^st^ MG coordination shell missing (see Notes) |
| MG:DA:3562 | 1.00 | 73.22 |  | 0 |  | 1^st^ MG coordination shell missing (see Notes) |
| MG:DA:3563 | 1.00 | 44.94 |  | 0 | 5 | - Exclusion zone contact(s): /O2'/O3'/OP1/OP1/OP2/ Consider K in place of MG (3 valid contact(s) for K vs. 0 for MG) |
| MG:DA:3564 | 1.00 | 83.92 |  | 0 |  | 1^st^ MG coordination shell missing (see Notes) |
| MG:DA:3565 | 1.00 | 55.13 |  | 0 |  | 1^st^ MG coordination shell missing (see Notes) |
| MG:DA:3566 | 1.00 | 21.00 |  | 1 | 3 | O_r_ - Exclusion zone contact(s): /O3'/OP1/OP2/ Consider K in place of MG (2 valid contact(s) for K vs. 1 for MG) |
| MG:DA:3567 | 1.00 | 47.86 |  | 0 | 2 | - Exclusion zone contact(s): /O2'/O3'/ Consider NA in place of MG (1 valid contact(s) for NA vs. 0 for MG) |
| MG:DA:3568 | 1.00 | 54.45 |  | 0 |  | 1^st^ MG coordination shell missing (see Notes) |
| MG:DA:3569 | 1.00 | 21.69 |  | 0 |  | 1^st^ MG coordination shell missing (see Notes) |
| MG:DA:3570 | 1.00 | 71.27 |  | 0 | 1 | - Exclusion zone contact(s): /OP2/ Consider K in place of MG (1 valid contact(s) for K vs. 0 for MG) |
| MG:DA:3571 | 1.00 | 93.17 |  | 1 | 2 | O_ph_ - Exclusion zone contact(s): /O5'/OP2/ Consider K in place of MG (2 valid contact(s) for K vs. 1 for MG) |
| MG:DA:3572 | 1.00 | 51.21 |  | 0 | 2 | - Exclusion zone contact(s): /O2'/O3'/ Consider K in place of MG (2 valid contact(s) for K vs. 0 for MG) |
| MG:DA:3573 | 1.00 | 84.30 |  | 0 | 2 | - Exclusion zone contact(s): /N7/O6/ Consider K in place of MG (2 valid contact(s) for K vs. 0 for MG) |
| MG:DA:3574 | 1.00 | 71.40 |  | 0 | 5 | - Exclusion zone contact(s): /N3_U_/O4/O4/O6/O6/ Consider K in place of MG (4 valid contact(s) for K vs. 0 for MG) |
| MG:DA:3575 | 1.00 | 112.43 |  | 0 |  | 1^st^ MG coordination shell missing (see Notes) |
| MG:DA:3576 | 1.00 | 102.80 |  | 0 |  | No contacts in 5.0 A |
| MG:DA:3577 | 1.00 | 101.08 |  | 0 | 1 | - Exclusion zone contact(s): /O6/ Consider K in place of MG (1 valid contact(s) for K vs. 0 for MG) |
| MG:DA:3578 | 1.00 | 62.78 |  | 0 |  | 1^st^ MG coordination shell missing (see Notes) |
| MG:DA:3579 | 1.00 | 21.01 |  | 1 | 1 | His - Exclusion zone contact(s): /OP1/ |
| MG:DA:3580 | 1.00 | 58.34 |  | 0 |  | 1^st^ MG coordination shell missing (see Notes) |
| MG:DA:3581 | 1.00 | 91.17 |  | 0 | 1 | - Exclusion zone contact(s): /N7/ Consider K in place of MG (1 valid contact(s) for K vs. 0 for MG) |
| MG:DA:3582 | 1.00 | 63.00 |  | 0 | 1 | - Exclusion zone contact(s): /O6/ Consider K in place of MG (1 valid contact(s) for K vs. 0 for MG) |
| MG:DA:3583 | 1.00 | 3.25 |  | 2 |  | 2O_ph_ |
| MG:DA:3584 | 1.00 | 63.39 |  | 0 | 1 | - Exclusion zone contact(s): /O2'/ Consider K in place of MG (1 valid contact(s) for K vs. 0 for MG) |
| MG:DA:3585 | 1.00 | 10.53 |  | 0 |  | 1^st^ MG coordination shell missing (see Notes) |
| MG:DA:3586 | 1.00 | 83.67 |  | 0 | 2 | - Exclusion zone contact(s): /O2'/OP2/ Consider K in place of MG (2 valid contact(s) for K vs. 0 for MG) |
| MG:DA:3587 | 1.00 | 21.22 |  | 2 |  | 2O_ph_ |
| MG:DA:3588 | 1.00 | 54.26 |  | 0 | 1 | - Exclusion zone contact(s): /OP2/ Consider K in place of MG (1 valid contact(s) for K vs. 0 for MG) |
| MG:DA:3589 | 1.00 | 7.15 |  | 2 | 2 | 2O_ph_ - Exclusion zone contact(s): /OP1/OP2/ |
| MG:DA:3590 | 1.00 | 46.43 |  | 0 | 2 | - Exclusion zone contact(s): /N4/N7/ Consider K in place of MG (2 valid contact(s) for K vs. 0 for MG) |
| MG:DA:3591 | 1.00 | 16.84 |  | 2 |  | 2O_ph_ |
| MG:DA:3592 | 1.00 | 81.77 |  | 0 | 2 | - Exclusion zone contact(s): /O6/O6/ Consider K in place of MG (2 valid contact(s) for K vs. 0 for MG) |
| MG:DA:3593 | 1.00 | 29.50 |  | 2 |  | 2O_ph_ |
| MG:DA:3594 | 1.00 | 49.73 |  | 0 | 1 | - Exclusion zone contact(s): /N4/ Consider K in place of MG (1 valid contact(s) for K vs. 0 for MG) |
| MG:DA:3595 | 1.00 | 38.68 |  | 0 | 2 | - Exclusion zone contact(s): /N1_G_/N2/ Consider K in place of MG (2 valid contact(s) for K vs. 0 for MG) |
| MG:DA:3596 | 1.00 | 76.02 |  | 0 |  | 1^st^ MG coordination shell missing (see Notes) |
| MG:DA:3597 | 1.00 | 23.20 |  | 0 |  | 1^st^ MG coordination shell missing (see Notes) |
| MG:DA:3598 | 1.00 | 116.31 |  | 0 |  | 1^st^ MG coordination shell missing (see Notes) |
| MG:DA:3599 | 1.00 | 6.61 |  | 0 | 2 | - Exclusion zone contact(s): /N7/N7/ Consider K in place of MG (2 valid contact(s) for K vs. 0 for MG) |
| MG:DA:3600 | 1.00 | 59.58 |  | 0 |  | 1^st^ MG coordination shell missing (see Notes) |
| MG:DA:3601 | 1.00 | 1.67 |  | 1 | 4 | O_b_ - Exclusion zone contact(s): /O2/O2'/O2'/O6/ Consider K in place of MG (4 valid contact(s) for K vs. 1 for MG) |
| MG:DA:3602 | 1.00 | 61.70 |  | 0 |  | 1^st^ MG coordination shell missing (see Notes) |
| MG:DA:3603 | 1.00 | 20.32 |  | 0 | 3 | - Exclusion zone contact(s): /N7/O2/O6/ Consider K in place of MG (2 valid contact(s) for K vs. 0 for MG) |
| MG:DA:3604 | 1.00 | 57.93 |  | 0 | 2 | - Exclusion zone contact(s): /N7/O6/ Consider K in place of MG (2 valid contact(s) for K vs. 0 for MG) |
| MG:DA:3605 | 1.00 | 29.45 |  | 0 | 4 | - Exclusion zone contact(s): /O4/O6/O6/O6/ Consider K in place of MG (4 valid contact(s) for K vs. 0 for MG) |
| MG:DA:3606 | 1.00 | 88.52 |  | 0 |  | 1^st^ MG coordination shell missing (see Notes) |
| MG:DA:3607 | 1.00 | 79.99 |  | 0 |  | 1^st^ MG coordination shell missing (see Notes) |
| MG:DA:3608 | 1.00 | 60.62 |  | 1 |  | O_ph_ |
| MG:DA:3609 | 1.00 | 76.98 |  | 0 |  | 1^st^ MG coordination shell missing (see Notes) |
| MG:DA:3610 | 1.00 | 45.08 |  | 0 | 2 | - Exclusion zone contact(s): /OP1/OP2/ Consider NA in place of MG (1 valid contact(s) for NA vs. 0 for MG) |
| MG:DA:3611 | 1.00 | 115.27 |  | 0 |  | 1^st^ MG coordination shell missing (see Notes) |
| MG:DA:3612 | 1.00 | 64.73 |  | 0 |  | 1^st^ MG coordination shell missing (see Notes) |
| MG:DA:3613 | 1.00 | 68.30 |  | 0 | 3 | - Exclusion zone contact(s): /O2'/O3'/OP1/ Consider K in place of MG (3 valid contact(s) for K vs. 0 for MG) |
| MG:DA:3614 | 1.00 | 53.31 |  | 1 | 1 | O_ph_ - Exclusion zone contact(s): /OP1/ |
| MG:DA:3615 | 1.00 | 112.34 |  | 0 | 2 | - Exclusion zone contact(s): /O2'/O3'/ Consider NA in place of MG (1 valid contact(s) for NA vs. 0 for MG) |
| MG:DA:3616 | 1.00 | 64.03 |  | 0 | 1 | - Exclusion zone contact(s): /OP1/ Consider K in place of MG (1 valid contact(s) for K vs. 0 for MG) |
| MG:DA:3617 | 1.00 | 82.99 |  | 0 | 1 | - Exclusion zone contact(s): /OP1/ Consider NA in place of MG (1 valid contact(s) for NA vs. 0 for MG) |
| MG:DA:3618 | 1.00 | 14.91 |  | 1 | 3 | O_b_ - Exclusion zone contact(s): /OP1/OP2/OP2/ Consider NA in place of MG (2 valid contact(s) for NA vs. 1 for MG) |
| MG:DA:3619 | 1.00 | 59.15 |  | 0 | 1 | - Exclusion zone contact(s): /N7/ Consider K in place of MG (1 valid contact(s) for K vs. 0 for MG) |
| MG:DA:3620 | 1.00 | 58.21 |  | 0 | 1 | - Exclusion zone contact(s): /OP2/ Consider K in place of MG (1 valid contact(s) for K vs. 0 for MG) |
| MG:DA:3621 | 1.00 | 51.01 |  | 0 | 2 | - Exclusion zone contact(s): /O6/O6/ Consider K in place of MG (2 valid contact(s) for K vs. 0 for MG) |
| MG:DA:3622 | 1.00 | 69.12 |  | 0 |  | 1^st^ MG coordination shell missing (see Notes) |
| MG:DA:3623 | 1.00 | 89.30 |  | 0 | 1 | - Exclusion zone contact(s): /O3'/ Consider K in place of MG (1 valid contact(s) for K vs. 0 for MG) |
| MG:DA:3624 | 1.00 | 74.87 |  | 0 | 1 | - Exclusion zone contact(s): /O4/ Consider K in place of MG (1 valid contact(s) for K vs. 0 for MG) |
| MG:DA:3625 | 1.00 | 19.42 |  | 0 | 4 | - Exclusion zone contact(s): /N3_U_/N4/O6/OP2/ Consider K in place of MG (4 valid contact(s) for K vs. 0 for MG) |
| MG:DA:3626 | 1.00 | 84.77 |  | 0 | 2 | - Exclusion zone contact(s): /O4/O6/ Consider K in place of MG (2 valid contact(s) for K vs. 0 for MG) |
| MG:DA:3627 | 1.00 | 40.58 |  | 1 | 3 | O_r_ - Exclusion zone contact(s): /O4'/OP1/OP2/ Consider K in place of MG (3 valid contact(s) for K vs. 1 for MG) |
| MG:DA:3628 | 1.00 | 92.28 |  | 0 |  | 1^st^ MG coordination shell missing (see Notes) |
| MG:DA:3629 | 1.00 | 29.32 |  | 0 | 2 | - Exclusion zone contact(s): /O4/O6/ Consider NA in place of MG (1 valid contact(s) for NA vs. 0 for MG) |
| MG:DA:3630 | 1.00 | 71.40 |  | 0 |  | 1^st^ MG coordination shell missing (see Notes) |
| MG:DA:3631 | 1.00 | 60.82 |  | 0 | 3 | - Exclusion zone contact(s): /OP1/OP2/OP2/ Consider K in place of MG (3 valid contact(s) for K vs. 0 for MG) |
| MG:DA:3632 | 1.00 | 52.37 |  | 1 | 3 | O_r_ - Exclusion zone contact(s): /O2'/O5'/OP1/ Consider K in place of MG (2 valid contact(s) for K vs. 1 for MG) |
| MG:DA:3633 | 1.00 | 48.67 |  | 0 | 1 | - Exclusion zone contact(s): /O4/ Consider K in place of MG (1 valid contact(s) for K vs. 0 for MG) |
| MG:DA:3634 | 1.00 | 54.89 |  | 0 | 1 | - Exclusion zone contact(s): /OP2/ Consider K in place of MG (1 valid contact(s) for K vs. 0 for MG) |
| MG:DA:3635 | 1.00 | 33.32 |  | 0 | 1 | - Exclusion zone contact(s): /OP1/ Consider K in place of MG (1 valid contact(s) for K vs. 0 for MG) |
| MG:DA:3636 | 1.00 | 46.71 |  | 0 |  | 1^st^ MG coordination shell missing (see Notes) |
| MG:DA:3637 | 1.00 | 63.70 |  | 0 | 1 | - Exclusion zone contact(s): /OP1/ Consider K in place of MG (1 valid contact(s) for K vs. 0 for MG) |
| MG:DA:3638 | 1.00 | 78.72 |  | 1 | 2 | O_ph_ - Exclusion zone contact(s): /O5'/OP1/ Consider K in place of MG (2 valid contact(s) for K vs. 1 for MG) |
| MG:DA:3639 | 1.00 | 66.13 |  | 0 | 1 | - Exclusion zone contact(s): /O6/ Consider K in place of MG (1 valid contact(s) for K vs. 0 for MG) |
| MG:DA:3640 | 1.00 | 54.66 |  | 0 | 1 | - Exclusion zone contact(s): /OP1/ Consider NA in place of MG (1 valid contact(s) for NA vs. 0 for MG) |
| MG:DA:3641 | 1.00 | 60.01 |  | 0 | 3 | - Exclusion zone contact(s): /N7/O2'/OP1/ Consider K in place of MG (2 valid contact(s) for K vs. 0 for MG) |
| MG:DA:3642 | 1.00 | 42.21 |  | 0 | 1 | - Exclusion zone contact(s): /OP2/ Consider K in place of MG (1 valid contact(s) for K vs. 0 for MG) |
| MG:DA:3643 | 1.00 | 62.16 |  | 0 | 1 | - Exclusion zone contact(s): /N7/ Consider K in place of MG (1 valid contact(s) for K vs. 0 for MG) |
| MG:DA:3644 | 1.00 | 84.43 |  | 0 |  | No contacts in 5.0 A |
| MG:DA:3645 | 1.00 | 61.58 |  | 0 | 2 | - Exclusion zone contact(s): /N7/O6/ Consider K in place of MG (2 valid contact(s) for K vs. 0 for MG) |
| MG:DA:3646 | 1.00 | 53.17 |  | 0 |  | 1^st^ MG coordination shell missing (see Notes) |
| MG:DA:3647 | 1.00 | 108.50 |  | 0 |  | 1^st^ MG coordination shell missing (see Notes) |
| MG:DA:3648 | 1.00 | 51.92 |  | 1 | 3 | OH_Prot_ - Exclusion zone contact(s): /O2'/O3'/OP1/ Consider K in place of MG (3 valid contact(s) for K vs. 1 for MG) |
| MG:DA:3649 | 1.00 | 71.43 |  | 0 | 2 | - Exclusion zone contact(s): /O5'/OP2/ Consider K in place of MG (2 valid contact(s) for K vs. 0 for MG) |
| MG:DA:3650 | 1.00 | 37.71 |  | 0 | 5 | - Exclusion zone contact(s): /O3'/O5'/OP1/OP1/OP2/ Consider K in place of MG (5 valid contact(s) for K vs. 0 for MG) |
| MG:DA:3651 | 1.00 | 40.23 |  | 0 | 4 | - Exclusion zone contact(s): /N1_G_/N7/O6/O6/ Consider K in place of MG (4 valid contact(s) for K vs. 0 for MG) |
| MG:DA:3652 | 1.00 | 93.87 |  | 0 | 2 | - Exclusion zone contact(s): /OP1/OP2/ Consider NA in place of MG (1 valid contact(s) for NA vs. 0 for MG) |
| MG:DA:3653 | 1.00 | 61.46 |  | 0 | 2 | - Exclusion zone contact(s): /N7/O6/ Consider K in place of MG (2 valid contact(s) for K vs. 0 for MG) |
| MG:DA:3654 | 1.00 | 40.74 |  | 0 | 5 | - Exclusion zone contact(s): /N3_C_/O2/O4/O4/O4/ Consider K in place of MG (5 valid contact(s) for K vs. 0 for MG) |
| MG:DA:3655 | 1.00 | 86.43 |  | 0 |  | 1^st^ MG coordination shell missing (see Notes) |
| MG:DA:3656 | 1.00 | 81.61 |  | 0 |  | 1^st^ MG coordination shell missing (see Notes) |
| MG:DA:3657 | 1.00 | 74.77 |  | 0 | 1 | - Exclusion zone contact(s): /OP2/ Consider K in place of MG (1 valid contact(s) for K vs. 0 for MG) |
| MG:DA:3658 | 1.00 | 60.75 |  | 0 | 1 | - Exclusion zone contact(s): /OP1/ Consider K in place of MG (1 valid contact(s) for K vs. 0 for MG) |
| MG:DA:3659 | 1.00 | 61.60 |  | 0 | 1 | - Exclusion zone contact(s): /N7/ Consider K in place of MG (1 valid contact(s) for K vs. 0 for MG) |
| MG:DA:3660 | 1.00 | 46.98 |  | 0 | 3 | - Exclusion zone contact(s): /O2'/O2'/O3'/ Consider K in place of MG (2 valid contact(s) for K vs. 0 for MG) |
| MG:DA:3661 | 1.00 | 68.31 |  | 0 | 2 | - Exclusion zone contact(s): /O6/O6/ Consider NA in place of MG (1 valid contact(s) for NA vs. 0 for MG) |
| MG:DA:3662 | 1.00 | 73.55 |  | 0 |  | 1^st^ MG coordination shell missing (see Notes) |
| MG:DA:3663 | 1.00 | 79.04 |  | 1 | 2 | O_b_ - Exclusion zone contact(s): /O4/O6/ Consider NA in place of MG (2 valid contact(s) for NA vs. 1 for MG) |
| MG:DA:3664 | 1.00 | 60.38 |  | 0 | 1 | - Exclusion zone contact(s): /OP1/ Consider K in place of MG (1 valid contact(s) for K vs. 0 for MG) |
| MG:DA:3665 | 1.00 | 110.47 |  | 0 | 1 | - Exclusion zone contact(s): /O3'/ Consider K in place of MG (1 valid contact(s) for K vs. 0 for MG) |
| MG:DA:3666 | 1.00 | 53.57 |  | 0 |  | 1^st^ MG coordination shell missing (see Notes) |
| MG:DA:3667 | 1.00 | 74.55 |  | 0 |  | 1^st^ MG coordination shell missing (see Notes) |
| MG:DA:3668 | 1.00 | 80.79 |  | 0 | 1 | - Exclusion zone contact(s): /N2/ Consider K in place of MG (1 valid contact(s) for K vs. 0 for MG) |
| MG:DA:3669 | 1.00 | 68.14 |  | 0 | 1 | - Exclusion zone contact(s): /OP2/ Consider K in place of MG (1 valid contact(s) for K vs. 0 for MG) |
| MG:DA:3670 | 1.00 | 78.74 |  | 0 | 2 | - Exclusion zone contact(s): /OP1/OP2/ Consider NA in place of MG (1 valid contact(s) for NA vs. 0 for MG) |
| MG:DA:3671 | 1.00 | 40.26 |  | 0 | 2 | - Exclusion zone contact(s): /N4/N7/ Consider K in place of MG (2 valid contact(s) for K vs. 0 for MG) |
| MG:DA:3672 | 1.00 | 76.52 |  | 0 |  | 1^st^ MG coordination shell missing (see Notes) |
| MG:DA:3673 | 1.00 | 61.73 |  | 0 |  | 1^st^ MG coordination shell missing (see Notes) |
| MG:DA:3674 | 1.00 | 62.15 |  | 0 | 2 | - Exclusion zone contact(s): /OP1/OP2/ Consider K in place of MG (2 valid contact(s) for K vs. 0 for MG) |
| MG:DA:3675 | 1.00 | 44.95 |  | 0 | 3 | - Exclusion zone contact(s): /N6/N7/OP2/ Consider K in place of MG (3 valid contact(s) for K vs. 0 for MG) |
| MG:DA:3676 | 1.00 | 70.59 |  | 0 | 3 | - Exclusion zone contact(s): /O5'/OP2/OP2/ Consider K in place of MG (3 valid contact(s) for K vs. 0 for MG) |
| MG:DA:3677 | 1.00 | 78.03 |  | 0 | 1 | - Exclusion zone contact(s): /OP1/ Consider NA in place of MG (1 valid contact(s) for NA vs. 0 for MG) |
| MG:DA:3678 | 1.00 | 123.47 |  | 0 | 2 | - Exclusion zone contact(s): /OP2/OP2/ Consider NA in place of MG (1 valid contact(s) for NA vs. 0 for MG) |
| MG:DA:3679 | 1.00 | 69.73 |  | 0 | 3 | - Exclusion zone contact(s): /O4/O6/O6/ Consider K in place of MG (2 valid contact(s) for K vs. 0 for MG) |
| MG:DA:3680 | 1.00 | 76.38 |  | 0 |  | 1^st^ MG coordination shell missing (see Notes) |
| MG:DA:3681 | 1.00 | 73.45 |  | 1 | 1 | O_ph_ - Exclusion zone contact(s): /OP1/ |
| MG:DA:3682 | 1.00 | 83.46 |  | 0 | 2 | - Exclusion zone contact(s): /O2'/O4'/ Consider K in place of MG (2 valid contact(s) for K vs. 0 for MG) |
| MG:DA:3683 | 1.00 | 56.72 |  | 0 | 5 | - Exclusion zone contact(s): /OP1/OP1/OP1/OP2/OP2/ Consider K in place of MG (4 valid contact(s) for K vs. 0 for MG) |
| MG:DA:3684 | 1.00 | 56.85 |  | 0 | 1 | - Exclusion zone contact(s): /OP1/ Consider K in place of MG (1 valid contact(s) for K vs. 0 for MG) |
| MG:DA:3685 | 1.00 | 59.33 |  | 0 | 1 | - Exclusion zone contact(s): /OP2/ Consider K in place of MG (1 valid contact(s) for K vs. 0 for MG) |
| MG:DA:3686 | 1.00 | 70.16 |  | 0 |  | 1^st^ MG coordination shell missing (see Notes) |
| MG:DA:3687 | 1.00 | 41.75 |  | 1 | 1 | O_ph_ - Exclusion zone contact(s): /O3'/ |
| MG:DA:3688 | 1.00 | 42.48 |  | 0 | 2 | - Exclusion zone contact(s): /O6/OP2/ Consider K in place of MG (2 valid contact(s) for K vs. 0 for MG) |
| MG:DA:3689 | 1.00 | 67.94 |  | 0 | 2 | - Exclusion zone contact(s): /O2/OP1/ Consider K in place of MG (2 valid contact(s) for K vs. 0 for MG) |
| MG:DA:3690 | 1.00 | 111.16 |  | 0 |  | 1^st^ MG coordination shell missing (see Notes) |
| MG:DA:3691 | 1.00 | 69.66 |  | 0 |  | 1^st^ MG coordination shell missing (see Notes) |
| MG:DA:3692 | 1.00 | 60.66 |  | 0 | 1 | - Exclusion zone contact(s): /N7/ Consider K in place of MG (1 valid contact(s) for K vs. 0 for MG) |
| MG:DA:3693 | 1.00 | 108.43 |  | 0 |  | 1^st^ MG coordination shell missing (see Notes) |
| MG:DA:3694 | 1.00 | 41.57 |  | 0 | 4 | - Exclusion zone contact(s): /N7/O5'/OP2/OP2/ Consider K in place of MG (3 valid contact(s) for K vs. 0 for MG) |
| MG:DA:3695 | 1.00 | 131.01 |  | 1 |  | Anion |
| MG:DA:3696 | 1.00 | 19.33 |  | 0 | 1 | - Exclusion zone contact(s): /OP2/ Consider K in place of MG (1 valid contact(s) for K vs. 0 for MG) |
| MG:DA:3697 | 1.00 | 6.76 |  | 1 | 2 | O_ph_ - Exclusion zone contact(s): /O2'/OP1/ Consider NA in place of MG (2 valid contact(s) for NA vs. 1 for MG) |
| MG:DA:3698 | 1.00 | 35.96 |  | 0 |  | 1^st^ MG coordination shell missing (see Notes) |
| MG:DA:3699 | 1.00 | 1.53 |  | 2 |  | 2O_ph_ |
| MG:DA:3700 | 1.00 | 39.59 |  | 0 | 2 | - Exclusion zone contact(s): /N4/O6/ Consider NA in place of MG (1 valid contact(s) for NA vs. 0 for MG) |
| MG:DA:3701 | 1.00 | 10.34 |  | 1 |  | O_ph_ |
| MG:DA:3702 | 1.00 | 36.40 |  | 0 | 4 | - Exclusion zone contact(s): /N4/O6/O6/OP1/ Consider K in place of MG (3 valid contact(s) for K vs. 0 for MG) |
| MG:DA:3703 | 1.00 | 12.74 |  | 1 |  | O_b_ |
| MG:DA:3704 | 1.00 | 33.00 |  | 0 |  | 1^st^ MG coordination shell missing (see Notes) |
| MG:DA:3705 | 1.00 | 1.53 |  | 1 | 5 | O_r_ - Exclusion zone contact(s): /N7/O5'/OP2/OP2/OP2/ Consider K in place of MG (4 valid contact(s) for K vs. 1 for MG) |
| MG:DA:3706 | 1.00 | 20.06 |  | 2 |  | O_ph_O_b_ |
| MG:DA:3707 | 1.00 | 28.50 |  | 0 | 4 | - Exclusion zone contact(s): /N7/O5'/O6/OP1/ Consider K in place of MG (4 valid contact(s) for K vs. 0 for MG) |
| MG:DA:3708 | 1.00 | 9.26 |  | 0 | 6 | - Exclusion zone contact(s): /O3'/O3'/OP1/OP1/OP1/OP2/ Consider K in place of MG (5 valid contact(s) for K vs. 0 for MG) |
| MG:DA:3709 | 1.00 | 46.34 |  | 0 | 2 | - Exclusion zone contact(s): /N7/OP2/ Consider K in place of MG (2 valid contact(s) for K vs. 0 for MG) |
| MG:DA:3710 | 1.00 | 14.87 |  | 1 |  | O_ph_ |
| MG:DA:3711 | 1.00 | 85.20 |  | 0 | 2 | - Exclusion zone contact(s): /O2'/O3'/ Consider K in place of MG (2 valid contact(s) for K vs. 0 for MG) |
| MG:DA:3712 | 1.00 | 25.57 |  | 3 | 3 | 3O_ph_ - Exclusion zone contact(s): /O2'/O2'/O3'/ |
| MG:DA:3713 | 1.00 | 70.28 |  | 0 |  | 1^st^ MG coordination shell missing (see Notes) |
| MG:DA:3714 | 1.00 | 21.64 |  | 0 | 4 | - Exclusion zone contact(s): /O2'/OE1_Glu_/OP2/OP2/ Consider K in place of MG (3 valid contact(s) for K vs. 0 for MG) |
| MG:DA:3715 | 1.00 | 75.27 |  | 0 | 1 | - Exclusion zone contact(s): /O6/ Consider K in place of MG (1 valid contact(s) for K vs. 0 for MG) |
| MG:DA:3716 | 1.00 | 13.40 |  | 0 | 2 | - Exclusion zone contact(s): /OP1/OP2/ Consider K in place of MG (2 valid contact(s) for K vs. 0 for MG) |
| MG:DA:3717 | 1.00 | 1.53 |  | 1 | 6 | O_b_ - Exclusion zone contact(s): /N7/N7/O2/O4/O4'/O6/ Consider K in place of MG (5 valid contact(s) for K vs. 1 for MG) |
| MG:DA:3718 | 1.00 | 25.18 |  | 0 | 4 | - Exclusion zone contact(s): /O2/O2'/O4/O6/ Consider K in place of MG (3 valid contact(s) for K vs. 0 for MG) |
| MG:DA:3719 | 1.00 | 29.59 |  | 0 | 2 | - Exclusion zone contact(s): /OP2/OP2/ Consider K in place of MG (2 valid contact(s) for K vs. 0 for MG) |
| MG:DA:3720 | 1.00 | 46.31 |  | 0 | 1 | - Exclusion zone contact(s): /O6/ Consider K in place of MG (1 valid contact(s) for K vs. 0 for MG) |
| MG:DA:3721 | 1.00 | 51.04 |  | 0 |  | 1^st^ MG coordination shell missing (see Notes) |
| MG:DA:3722 | 1.00 | 83.95 | 1 | 1 | 1 | - Exclusion zone contact(s): /O2'/OP2/ Consider K in place of MG (1 valid contact(s) for K vs. 0 for MG) |
| MG:DA:3723 | 1.00 | 19.73 |  | 0 | 2 | - Exclusion zone contact(s): /OP2/OP2/ Consider NA in place of MG (1 valid contact(s) for NA vs. 0 for MG) |
| MG:DA:3724 | 1.00 | 45.62 |  | 0 |  | 1^st^ MG coordination shell missing (see Notes) |
| MG:DA:3725 | 1.00 | 22.85 |  | 0 | 3 | - Exclusion zone contact(s): /O4/O4/O6/ Consider K in place of MG (3 valid contact(s) for K vs. 0 for MG) |
| MG:DA:3726 | 1.00 | 1.53 |  | 2 | 3 | 2O_r_ - Exclusion zone contact(s): /O2'/O5'/OP2/ Consider K in place of MG (3 valid contact(s) for K vs. 2 for MG) |
| MG:DA:3727 | 1.00 | 49.54 |  | 0 |  | 1^st^ MG coordination shell missing (see Notes) |
| MG:DA:3728 | 1.00 | 42.21 |  | 0 |  | 1^st^ MG coordination shell missing (see Notes) |
| MG:DA:3729 | 1.00 | 52.36 |  | 1 | 3 | O_ph_ - Exclusion zone contact(s): /O5'/OP1/OP2/ Consider K in place of MG (2 valid contact(s) for K vs. 1 for MG) |
| MG:DA:3730 | 1.00 | 47.07 |  | 0 |  | 1^st^ MG coordination shell missing (see Notes) |
| MG:DA:3731 | 1.00 | 35.69 |  | 0 | 2 | - Exclusion zone contact(s): /O2'/OP2/ Consider K in place of MG (2 valid contact(s) for K vs. 0 for MG) |
| MG:DA:3732 | 1.00 | 60.48 |  | 0 | 2 | - Exclusion zone contact(s): /O4/O6/ Consider K in place of MG (2 valid contact(s) for K vs. 0 for MG) |
| MG:DA:3733 | 1.00 | 40.47 |  | 0 | 3 | - Exclusion zone contact(s): /OP1/OP1/OP2/ Consider K in place of MG (3 valid contact(s) for K vs. 0 for MG) |
| MG:DA:3734 | 1.00 | 55.15 |  | 1 |  | O_ph_ |
| MG:DA:3735 | 1.00 | 76.96 |  | 0 | 2 | - Exclusion zone contact(s): /N7/OP2/ Consider K in place of MG (2 valid contact(s) for K vs. 0 for MG) |
| MG:DA:3736 | 1.00 | 100.72 |  | 0 |  | 1^st^ MG coordination shell missing (see Notes) |
| MG:DA:3737 | 1.00 | 36.90 |  | 0 | 3 | - Exclusion zone contact(s): /N6/O6/O6/ Consider K in place of MG (3 valid contact(s) for K vs. 0 for MG) |
| MG:DA:3738 | 1.00 | 45.23 |  | 1 | 1 | O_ph_ - Exclusion zone contact(s): /OP1/ |
| MG:DA:3739 | 1.00 | 51.12 |  | 0 | 4 | - Exclusion zone contact(s): /O2'/O3'/O5'/OP1/ Consider K in place of MG (3 valid contact(s) for K vs. 0 for MG) |
| MG:DA:3740 | 1.00 | 49.05 |  | 0 | 2 | - Exclusion zone contact(s): /OP1/OP1/ Consider NA in place of MG (1 valid contact(s) for NA vs. 0 for MG) |
| MG:DA:3741 | 1.00 | 76.58 |  | 0 | 2 | - Exclusion zone contact(s): /N7/O6/ Consider K in place of MG (2 valid contact(s) for K vs. 0 for MG) |
| MG:DA:3742 | 1.00 | 65.95 |  | 1 |  | O_ph_ |
| MG:DA:3743 | 1.00 | 65.57 |  | 0 |  | 1^st^ MG coordination shell missing (see Notes) |
| MG:DA:3744 | 1.00 | 58.74 |  | 0 | 4 | - Exclusion zone contact(s): /O3'/OP1/OP1/OP2/ Consider K in place of MG (3 valid contact(s) for K vs. 0 for MG) |
| MG:DA:3745 | 1.00 | 49.93 |  | 0 | 3 | - Exclusion zone contact(s): /N7/O4/O6/ Consider K in place of MG (2 valid contact(s) for K vs. 0 for MG) |
| MG:DA:3746 | 1.00 | 36.95 |  | 0 |  | 1^st^ MG coordination shell missing (see Notes) |
| MG:DA:3747 | 1.00 | 75.26 |  | 0 | 1 | - Exclusion zone contact(s): /OP2/ Consider NA in place of MG (1 valid contact(s) for NA vs. 0 for MG) |
| MG:DA:3748 | 1.00 | 66.11 |  | 0 | 2 | - Exclusion zone contact(s): /OP1/OP2/ Consider K in place of MG (2 valid contact(s) for K vs. 0 for MG) |
| MG:DA:3749 | 1.00 | 92.26 |  | 0 |  | 1^st^ MG coordination shell missing (see Notes) |
| MG:DA:3750 | 1.00 | 101.92 |  | 0 | 2 | - Exclusion zone contact(s): /OP1/OP1/ Consider K in place of MG (2 valid contact(s) for K vs. 0 for MG) |
| MG:DA:3751 | 1.00 | 97.20 |  | 0 | 2 | - Exclusion zone contact(s): /N7/N7/ Consider K in place of MG (2 valid contact(s) for K vs. 0 for MG) |
| MG:DA:3752 | 1.00 | 103.38 |  | 0 |  | 1^st^ MG coordination shell missing (see Notes) |
| MG:DA:3753 | 1.00 | 62.48 |  | 1 | 1 | O_b_ - Exclusion zone contact(s): /O6/ |
| MG:DA:3754 | 1.00 | 107.29 |  | 0 |  | 1^st^ MG coordination shell missing (see Notes) |
| MG:DA:3755 | 1.00 | 70.46 |  | 0 | 1 | - Exclusion zone contact(s): /O6/ Consider K in place of MG (1 valid contact(s) for K vs. 0 for MG) |
| MG:DA:3756 | 1.00 | 126.64 |  | 0 | 2 | - Exclusion zone contact(s): /N7/O6/ Consider NA in place of MG (1 valid contact(s) for NA vs. 0 for MG) |
| MG:DA:3757 | 1.00 | 128.29 |  | 0 | 1 | - Exclusion zone contact(s): /OP2/ Consider K in place of MG (1 valid contact(s) for K vs. 0 for MG) |
| MG:DA:3758 | 1.00 | 100.64 |  | 0 |  | No contacts in 5.0 A |
| MG:DA:3759 | 1.00 | 84.29 |  | 0 |  | 1^st^ MG coordination shell missing (see Notes) |
| MG:DA:3760 | 1.00 | 78.40 |  | 0 | 1 | - Exclusion zone contact(s): /OP2/ Consider K in place of MG (1 valid contact(s) for K vs. 0 for MG) |
| MG:DA:3761 | 1.00 | 90.04 |  | 0 |  | 1^st^ MG coordination shell missing (see Notes) |
| MG:DA:3762 | 1.00 | 89.35 |  | 0 |  | 1^st^ MG coordination shell missing (see Notes) |
| MG:DA:3763 | 1.00 | 8.33 |  | 0 |  | 1^st^ MG coordination shell missing (see Notes) |
| MG:DA:3764 | 1.00 | 98.03 |  | 1 | 1 | O_ph_ - Exclusion zone contact(s): /OP2/ |
| MG:DA:3765 | 1.00 | 61.43 |  | 0 | 1 | - Exclusion zone contact(s): /O4/ Consider K in place of MG (1 valid contact(s) for K vs. 0 for MG) |
| MG:DA:3766 | 1.00 | 77.04 |  | 0 |  | 1^st^ MG coordination shell missing (see Notes) |
| MG:DA:3767 | 1.00 | 88.51 |  | 0 |  | 1^st^ MG coordination shell missing (see Notes) |
| MG:DA:3768 | 1.00 | 77.71 |  | 0 |  | 1^st^ MG coordination shell missing (see Notes) |
| MG:DA:3769 | 1.00 | 72.06 |  | 0 |  | 1^st^ MG coordination shell missing (see Notes) |
| MG:DA:3770 | 1.00 | 88.78 |  | 0 |  | 1^st^ MG coordination shell missing (see Notes) |
| MG:DA:3771 | 1.00 | 92.12 |  | 1 | 2 | O_r_ - Exclusion zone contact(s): /O3'/O4'/ |
| MG:DA:3772 | 1.00 | 65.83 |  | 1 | 3 | O_ph_ - Exclusion zone contact(s): /O5'/OP1/OP2/ Consider K in place of MG (2 valid contact(s) for K vs. 1 for MG) |
| MG:DA:3773 | 1.00 | 70.48 |  | 0 |  | 1^st^ MG coordination shell missing (see Notes) |
| MG:DA:3774 | 1.00 | 37.89 |  | 3 | 1 | 2O_ph_O_b_ - Exclusion zone contact(s): /O4/ |
| MG:DA:3775 | 1.00 | 87.53 |  | 0 | 1 | - Exclusion zone contact(s): /OP2/ Consider K in place of MG (1 valid contact(s) for K vs. 0 for MG) |
| MG:DA:3776 | 1.00 | 46.86 |  | 2 | 2 | 2O_ph_ - Exclusion zone contact(s): /O6/OP2/ |
| MG:DA:3777 | 1.00 | 107.66 |  | 0 | 1 | - Exclusion zone contact(s): /OP2/ Consider K in place of MG (1 valid contact(s) for K vs. 0 for MG) |
| MG:DA:3778 | 1.00 | 18.15 |  | 0 | 5 | - Exclusion zone contact(s): /N6/O3'/O5'/OP1/OP1/ Consider K in place of MG (5 valid contact(s) for K vs. 0 for MG) |
| MG:DA:3779 | 1.00 | 155.99 |  | 0 | 2 | - Exclusion zone contact(s): /OP1/OP2/ Consider K in place of MG (2 valid contact(s) for K vs. 0 for MG) |
| MG:DA:3780 | 1.00 | 32.87 |  | 1 | 2 | O_ph_ - Exclusion zone contact(s): /O6/OP1/ Consider K in place of MG (2 valid contact(s) for K vs. 1 for MG) |
| MG:DA:3781 | 1.00 | 73.94 |  | 0 | 3 | - Exclusion zone contact(s): /OP1/OP2/OP2/ Consider K in place of MG (2 valid contact(s) for K vs. 0 for MG) |
| MG:DA:3782 | 1.00 | 38.78 |  | 1 | 2 | O_ph_ - Exclusion zone contact(s): /O5'/OP1/ |
| MG:DA:3783 | 1.00 | 63.19 |  | 2 | 1 | 2O_ph_ - Exclusion zone contact(s): /OP2/ |
| MG:DA:3784 | 1.00 | 52.53 |  | 0 | 2 | - Exclusion zone contact(s): /OP1/OP2/ Consider K in place of MG (2 valid contact(s) for K vs. 0 for MG) |
| MG:DA:3785 | 1.00 | 88.72 |  | 0 | 3 | - Exclusion zone contact(s): /O2'/O3'/OP1/ Consider K in place of MG (2 valid contact(s) for K vs. 0 for MG) |
| MG:DA:3786 | 1.00 | 68.26 |  | 0 | 2 | - Exclusion zone contact(s): /O2'/OP2/ Consider K in place of MG (2 valid contact(s) for K vs. 0 for MG) |
| MG:DA:3787 | 1.00 | 114.70 |  | 0 | 2 | - Exclusion zone contact(s): /OP1/OP2/ Consider NA in place of MG (1 valid contact(s) for NA vs. 0 for MG) |
| MG:DA:3788 | 1.00 | 50.10 |  | 0 | 2 | - Exclusion zone contact(s): /O2'/OP2/ Consider NA in place of MG (1 valid contact(s) for NA vs. 0 for MG) |
| MG:DA:3789 | 1.00 | 45.77 |  | 0 | 1 | - Exclusion zone contact(s): /O6/ Consider K in place of MG (1 valid contact(s) for K vs. 0 for MG) |
| MG:DA:3790 | 1.00 | 66.90 |  | 0 | 1 | - Exclusion zone contact(s): /OP1/ Consider K in place of MG (1 valid contact(s) for K vs. 0 for MG) |
| MG:DA:3791 | 1.00 | 109.99 |  | 0 | 1 | - Exclusion zone contact(s): /N7/ Consider K in place of MG (1 valid contact(s) for K vs. 0 for MG) |
| MG:DA:3792 | 1.00 | 92.42 |  | 0 | 3 | - Exclusion zone contact(s): /OP1/OP2/OP2/ Consider K in place of MG (3 valid contact(s) for K vs. 0 for MG) |
| MG:DA:3793 | 1.00 | 81.65 |  | 0 |  | 1^st^ MG coordination shell missing (see Notes) |
| MG:DA:3794 | 1.00 | 38.63 |  | 2 |  | 2O_ph_ |
| MG:DA:3795 | 1.00 | 88.49 |  | 0 | 1 | - Exclusion zone contact(s): /N4/ Consider K in place of MG (1 valid contact(s) for K vs. 0 for MG) |
| MG:DA:3796 | 1.00 | 26.19 |  | 0 | 4 | - Exclusion zone contact(s): /O3'/OP1/OP1/OP2/ Consider K in place of MG (4 valid contact(s) for K vs. 0 for MG) |
| MG:DA:3797 | 1.00 | 95.70 |  | 0 |  | 1^st^ MG coordination shell missing (see Notes) |
| MG:DA:3798 | 1.00 | 30.13 |  | 0 | 2 | - Exclusion zone contact(s): /N1_A_/OP1/ Consider K in place of MG (2 valid contact(s) for K vs. 0 for MG) |
| MG:DA:3799 | 1.00 | 83.07 |  | 0 |  | No contacts in 5.0 A |
| MG:DA:3800 | 1.00 | 33.95 |  | 1 | 1 | O_r_ - Exclusion zone contact(s): /OP1/ |
| MG:DA:3801 | 1.00 | 96.02 |  | 0 | 2 | - Exclusion zone contact(s): /N1_G_/OP2/ Consider K in place of MG (2 valid contact(s) for K vs. 0 for MG) |
| MG:DA:3802 | 1.00 | 88.60 |  | 1 | 2 | N_b_ - Exclusion zone contact(s): /O6/O6/ |
| MG:DA:3803 | 1.00 | 54.85 |  | 0 |  | 1^st^ MG coordination shell missing (see Notes) |
| MG:DA:3804 | 1.00 | 36.23 |  | 0 | 4 | - Exclusion zone contact(s): /N2/O5'/OP2/OP2/ Consider K in place of MG (3 valid contact(s) for K vs. 0 for MG) |
| MG:DA:3805 | 1.00 | 66.93 |  | 0 |  | 1^st^ MG coordination shell missing (see Notes) |
| MG:DA:3806 | 1.00 | 55.95 |  | 0 | 3 | - Exclusion zone contact(s): /O4/OP1/OP2/ Consider K in place of MG (2 valid contact(s) for K vs. 0 for MG) |
| MG:DA:3807 | 1.00 | 59.11 |  | 0 | 2 | - Exclusion zone contact(s): /N4/O6/ Consider NA in place of MG (1 valid contact(s) for NA vs. 0 for MG) |
| MG:DA:3808 | 1.00 | 77.36 |  | 1 | 2 | N_b_ - Exclusion zone contact(s): /N7/O6/ Consider K in place of MG (2 valid contact(s) for K vs. 1 for MG) |
| MG:DA:3809 | 1.00 | 82.75 |  | 1 | 2 | O_ph_ - Exclusion zone contact(s): /N4/N4/ Consider K in place of MG (2 valid contact(s) for K vs. 1 for MG) |
| MG:DA:3810 | 1.00 | 39.38 |  | 0 | 3 | - Exclusion zone contact(s): /N7/OP1/OP2/ Consider K in place of MG (2 valid contact(s) for K vs. 0 for MG) |
| MG:DA:3811 | 1.00 | 101.14 |  | 0 | 2 | - Exclusion zone contact(s): /O5'/OP2/ Consider K in place of MG (2 valid contact(s) for K vs. 0 for MG) |
| MG:DA:3812 | 1.00 | 73.77 |  | 0 | 5 | - Exclusion zone contact(s): /N1_A_/N3_A_/N6/N7/ Consider K in place of MG (4 valid contact(s) for K vs. 0 for MG) |
| MG:DA:3813 | 1.00 | 96.95 |  | 0 |  | 1^st^ MG coordination shell missing (see Notes) |
| MG:DA:3814 | 1.00 | 67.26 |  | 0 | 1 | - Exclusion zone contact(s): /OP2/ Consider K in place of MG (1 valid contact(s) for K vs. 0 for MG) |
| MG:DA:3815 | 1.00 | 83.58 |  | 1 |  | O_ph_ |
| MG:DA:3816 | 1.00 | 89.23 |  | 1 | 1 | O_ph_ - Exclusion zone contact(s): /O6/ |
| MG:DA:3817 | 1.00 | 46.99 |  | 0 | 5 | - Exclusion zone contact(s): /N7/N7/O5'/OP2/OP2/ Consider K in place of MG (5 valid contact(s) for K vs. 0 for MG) |
| MG:DA:3818 | 1.00 | 71.79 |  | 0 | 1 | - Exclusion zone contact(s): /O6/ Consider K in place of MG (1 valid contact(s) for K vs. 0 for MG) |
| MG:DA:3819 | 1.00 | 68.58 |  | 0 |  | 1^st^ MG coordination shell missing (see Notes) |
| MG:DA:3820 | 1.00 | 97.14 |  | 0 | 1 | - Exclusion zone contact(s): /OP2/ Consider K in place of MG (1 valid contact(s) for K vs. 0 for MG) |
| MG:DA:3821 | 1.00 | 81.57 |  | 0 | 2 | - Exclusion zone contact(s): /OP1/OP2/ Consider K in place of MG (2 valid contact(s) for K vs. 0 for MG) |
| MG:DA:3822 | 1.00 | 55.49 |  | 0 | 1 | - Exclusion zone contact(s): /OP1/ Consider K in place of MG (1 valid contact(s) for K vs. 0 for MG) |
| MG:DA:3823 | 1.00 | 81.32 |  | 0 |  | 1^st^ MG coordination shell missing (see Notes) |
| MG:DA:3824 | 1.00 | 73.24 |  | 0 | 2 | - Exclusion zone contact(s): /O4/O4/ Consider NA in place of MG (1 valid contact(s) for NA vs. 0 for MG) |
| MG:DA:3825 | 1.00 | 77.56 |  | 0 | 1 | - Exclusion zone contact(s): /OP2/ Consider K in place of MG (1 valid contact(s) for K vs. 0 for MG) |
| MG:DA:3826 | 1.00 | 33.42 |  | 0 | 4 | - Exclusion zone contact(s): /NH1_Arg_/O5'/OP1/OP1/ Consider K in place of MG (3 valid contact(s) for K vs. 0 for MG) |
| MG:DA:3827 | 1.00 | 97.30 |  | 0 |  | 1^st^ MG coordination shell missing (see Notes) |
| MG:DA:3828 | 1.00 | 70.25 |  | 0 | 2 | - Exclusion zone contact(s): /N6/N7/ Consider K in place of MG (2 valid contact(s) for K vs. 0 for MG) |
| MG:DA:3829 | 1.00 | 66.18 |  | 0 |  | 1^st^ MG coordination shell missing (see Notes) |
| MG:DA:3830 | 1.00 | 39.90 |  | 0 | 4 | - Exclusion zone contact(s): /N_back_/OG1_Thr_/OP1/OP2/ Consider K in place of MG (3 valid contact(s) for K vs. 0 for MG) |
| MG:DA:3831 | 1.00 | 61.65 |  | 0 |  | 1^st^ MG coordination shell missing (see Notes) |
| MG:DA:3832 | 1.00 | 55.72 |  | 0 | 1 | - Exclusion zone contact(s): /OP1/ Consider K in place of MG (1 valid contact(s) for K vs. 0 for MG) |
| MG:DA:3833 | 1.00 | 65.98 |  | 1 | 2 | N_b_ - Exclusion zone contact(s): /N7/OP2/ Consider K in place of MG (2 valid contact(s) for K vs. 1 for MG) |
| MG:DA:3834 | 1.00 | 57.30 |  | 0 | 4 | - Exclusion zone contact(s): /O2'/OP1/OP2/OP2/ Consider K in place of MG (4 valid contact(s) for K vs. 0 for MG) |
| MG:DA:3835 | 1.00 | 61.34 |  | 0 | 2 | - Exclusion zone contact(s): /N7/O6/ Consider K in place of MG (2 valid contact(s) for K vs. 0 for MG) |
| MG:DA:3836 | 1.00 | 85.18 |  | 0 |  | 1^st^ MG coordination shell missing (see Notes) |
| MG:DA:3837 | 1.00 | 61.45 |  | 0 | 1 | - Exclusion zone contact(s): /O6/ Consider K in place of MG (1 valid contact(s) for K vs. 0 for MG) |
| MG:DA:3838 | 1.00 | 80.38 |  | 0 | 1 | - Exclusion zone contact(s): /N4/ Consider K in place of MG (1 valid contact(s) for K vs. 0 for MG) |
| MG:DA:3839 | 1.00 | 54.15 |  | 0 | 1 | - Exclusion zone contact(s): /OP2/ Consider NA in place of MG (1 valid contact(s) for NA vs. 0 for MG) |
| MG:DA:3840 | 1.00 | 58.19 |  | 0 | 2 | - Exclusion zone contact(s): /O2'/OP2/ Consider NA in place of MG (1 valid contact(s) for NA vs. 0 for MG) |
| MG:DA:3841 | 1.00 | 50.90 |  | 1 | 3 | O_b_ - Exclusion zone contact(s): /O2'/O2'/OP2/ Consider K in place of MG (3 valid contact(s) for K vs. 1 for MG) |
| MG:DA:3842 | 1.00 | 54.20 |  | 0 | 1 | - Exclusion zone contact(s): /O2'/ Consider K in place of MG (1 valid contact(s) for K vs. 0 for MG) |
| MG:DA:3843 | 1.00 | 53.71 |  | 0 |  | 1^st^ MG coordination shell missing (see Notes) |
| MG:DA:3844 | 1.00 | 109.00 |  | 0 |  | 1^st^ MG coordination shell missing (see Notes) |
| MG:DA:3845 | 1.00 | 54.37 |  | 0 |  | 1^st^ MG coordination shell missing (see Notes) |
| MG:DA:3846 | 1.00 | 64.86 |  | 0 |  | 1^st^ MG coordination shell missing (see Notes) |
| MG:DA:3847 | 1.00 | 81.89 |  | 0 | 2 | - Exclusion zone contact(s): /OP1/OP2/ Consider K in place of MG (2 valid contact(s) for K vs. 0 for MG) |
| MG:DA:3848 | 1.00 | 35.80 |  | 0 | 5 | - Exclusion zone contact(s): /N6/O5'/OP1/OP2/OP2/ Consider K in place of MG (4 valid contact(s) for K vs. 0 for MG) |
| MG:DA:3849 | 1.00 | 63.77 |  | 0 | 2 | - Exclusion zone contact(s): /OP1/OP2/ Consider K in place of MG (2 valid contact(s) for K vs. 0 for MG) |
| MG:DA:3850 | 1.00 | 51.65 |  | 0 |  | 1^st^ MG coordination shell missing (see Notes) |
| MG:DA:3851 | 1.00 | 72.38 |  | 1 | 1 | O_ph_ - Exclusion zone contact(s): /OP2/ |
| MG:DA:3852 | 1.00 | 55.66 |  | 0 | 3 | - Exclusion zone contact(s): /O5'/OP1/OP1/ Consider NA in place of MG (2 valid contact(s) for NA vs. 0 for MG) |
| MG:DA:3853 | 1.00 | 71.82 |  | 1 | 2 | N_b_ - Exclusion zone contact(s): /N7/ Consider K in place of MG (2 valid contact(s) for K vs. 1 for MG) |
| MG:DA:3854 | 1.00 | 72.32 |  | 0 | 3 | - Exclusion zone contact(s): /OP1/OP1/OP2/ Consider K in place of MG (3 valid contact(s) for K vs. 0 for MG) |
| MG:DA:3855 | 1.00 | 85.52 |  | 0 | 1 | - Exclusion zone contact(s): /N7/ Consider K in place of MG (1 valid contact(s) for K vs. 0 for MG) |
| MG:DA:3856 | 1.00 | 70.22 |  | 0 | 3 | - Exclusion zone contact(s): /O4/O6/O6/ Consider K in place of MG (3 valid contact(s) for K vs. 0 for MG) |
| MG:DA:3857 | 1.00 | 78.40 |  | 0 |  | 1^st^ MG coordination shell missing (see Notes) |
| MG:DA:3858 | 1.00 | 49.71 |  | 0 |  | 1^st^ MG coordination shell missing (see Notes) |
| MG:DA:3859 | 1.00 | 92.38 |  | 0 | 1 | - Exclusion zone contact(s): /N4/ Consider K in place of MG (1 valid contact(s) for K vs. 0 for MG) |
| MG:DA:3860 | 1.00 | 55.21 |  | 0 | 1 | - Exclusion zone contact(s): /OP1/ Consider K in place of MG (1 valid contact(s) for K vs. 0 for MG) |
| MG:DA:3861 | 1.00 | 77.38 |  | 0 |  | 1^st^ MG coordination shell missing (see Notes) |
| MG:DA:3862 | 1.00 | 71.58 |  | 0 |  | 1^st^ MG coordination shell missing (see Notes) |
| MG:DA:3863 | 1.00 | 69.99 |  | 0 |  | 1^st^ MG coordination shell missing (see Notes) |
| MG:DA:3864 | 1.00 | 69.32 |  | 0 | 2 | - Exclusion zone contact(s): /O5'/OP1/ Consider NA in place of MG (1 valid contact(s) for NA vs. 0 for MG) |
| MG:DA:3865 | 1.00 | 87.49 |  | 0 | 3 | - Exclusion zone contact(s): /O3'/O5'/OP1/ Consider K in place of MG (3 valid contact(s) for K vs. 0 for MG) |
| MG:DA:3866 | 1.00 | 59.48 |  | 0 | 4 | - Exclusion zone contact(s): /N4/N6/N7/O6/ Consider K in place of MG (3 valid contact(s) for K vs. 0 for MG) |
| MG:DA:3867 | 1.00 | 100.72 |  | 0 | 1 | - Exclusion zone contact(s): /OP1/ Consider K in place of MG (1 valid contact(s) for K vs. 0 for MG) |
| MG:DA:3868 | 1.00 | 72.43 |  | 2 | 3 | O_ph_O_r_ - Exclusion zone contact(s): /O2'/O5'/O_back_/ Consider K in place of MG (3 valid contact(s) for K vs. 2 for MG) |
| MG:DA:3869 | 1.00 | 83.14 |  | 0 |  | 1^st^ MG coordination shell missing (see Notes) |
| MG:DA:3870 | 1.00 | 67.68 |  | 0 |  | 1^st^ MG coordination shell missing (see Notes) |
| MG:DA:3871 | 1.00 | 49.18 |  | 1 |  | O_ph_ |
| MG:DA:3872 | 1.00 | 75.25 |  | 0 |  | 1^st^ MG coordination shell missing (see Notes) |
| MG:DA:3873 | 1.00 | 119.46 |  | 0 |  | 1^st^ MG coordination shell missing (see Notes) |
| MG:DA:3874 | 1.00 | 69.73 |  | 0 | 2 | - Exclusion zone contact(s): /N7/OP2/ Consider K in place of MG (2 valid contact(s) for K vs. 0 for MG) |
| MG:DA:3875 | 1.00 | 116.54 |  | 1 | 2 | O_ph_ - Exclusion zone contact(s): /N2/OP2/ Consider NA in place of MG (2 valid contact(s) for NA vs. 1 for MG) |
| MG:DA:3876 | 1.00 | 52.47 |  | 0 | 3 | - Exclusion zone contact(s): /O5'/OP2/OP2/ Consider K in place of MG (2 valid contact(s) for K vs. 0 for MG) |
| MG:DA:3877 | 1.00 | 37.39 |  | 0 | 3 | - Exclusion zone contact(s): /O4/O6/O6/ Consider K in place of MG (2 valid contact(s) for K vs. 0 for MG) |
| MG:DA:3878 | 1.00 | 60.63 |  | 0 | 4 | - Exclusion zone contact(s): /O3'/OP1/OP1/OP2/ Consider K in place of MG (4 valid contact(s) for K vs. 0 for MG) |
| MG:DA:3879 | 1.00 | 83.13 |  | 0 | 1 | - Exclusion zone contact(s): /O4/ Consider NA in place of MG (1 valid contact(s) for NA vs. 0 for MG) |
| MG:DA:3880 | 1.00 | 44.93 |  | 0 | 4 | - Exclusion zone contact(s): /N2/O2'/O2'/O4'/ Consider K in place of MG (4 valid contact(s) for K vs. 0 for MG) |
| MG:DA:3881 | 1.00 | 86.63 |  | 0 | 3 | - Exclusion zone contact(s): /N7/O4/OP2/ Consider K in place of MG (3 valid contact(s) for K vs. 0 for MG) |
| MG:DA:3882 | 1.00 | 43.81 |  | 0 | 2 | - Exclusion zone contact(s): /O2/O2'/ Consider NA in place of MG (1 valid contact(s) for NA vs. 0 for MG) |
| MG:DA:3883 | 1.00 | 74.73 |  | 0 | 2 | - Exclusion zone contact(s): /O6/OP2/ Consider K in place of MG (2 valid contact(s) for K vs. 0 for MG) |
| MG:DA:3884 | 1.00 | 60.46 |  | 0 | 1 | - Exclusion zone contact(s): /O4'/ Consider K in place of MG (1 valid contact(s) for K vs. 0 for MG) |
| MG:DA:3885 | 1.00 | 57.46 |  | 0 | 1 | - Exclusion zone contact(s): /OP2/ Consider K in place of MG (1 valid contact(s) for K vs. 0 for MG) |
| MG:DA:3886 | 1.00 | 48.49 |  | 0 | 2 | - Exclusion zone contact(s): /N7/O6/ Consider NA in place of MG (1 valid contact(s) for NA vs. 0 for MG) |
| MG:DA:3887 | 1.00 | 68.65 |  | 0 | 2 | - Exclusion zone contact(s): /O4'/O5'/ Consider K in place of MG (2 valid contact(s) for K vs. 0 for MG) |
| MG:DA:3888 | 1.00 | 55.89 | 1 | 1 | 2 | - Exclusion zone contact(s): /O6/OP1/OP2/ Consider K in place of MG (2 valid contact(s) for K vs. 0 for MG) |
| MG:DA:3889 | 1.00 | 80.71 |  | 0 | 2 | - Exclusion zone contact(s): /N4/N4/ Consider K in place of MG (2 valid contact(s) for K vs. 0 for MG) |
| MG:DA:3890 | 1.00 | 86.51 |  | 0 | 5 | - Exclusion zone contact(s): /N1/N7/O2'/O4'/O6/ Consider K in place of MG (5 valid contact(s) for K vs. 0 for MG) |
| MG:DA:3891 | 1.00 | 82.13 |  | 0 | 1 | - Exclusion zone contact(s): /O4/ Consider K in place of MG (1 valid contact(s) for K vs. 0 for MG) |
| MG:DA:3892 | 1.00 | 46.32 |  | 0 | 4 | - Exclusion zone contact(s): /N1_A_/N6/O2'/O6/ Consider K in place of MG (3 valid contact(s) for K vs. 0 for MG) |
| MG:DA:3893 | 1.00 | 7.15 |  | 3 | 1 | 3O_ph_ - Exclusion zone contact(s): /O3'/ |
| MG:DA:3894 | 1.00 | 101.66 |  | 1 | 1 | O_ph_ - Exclusion zone contact(s): /O4/ |
| MG:DA:3895 | 1.00 | 14.64 |  | 1 | 1 | O_ph_ - Exclusion zone contact(s): /OP2/ |
| MG:DA:3896 | 1.00 | 69.21 |  | 0 |  | 1^st^ MG coordination shell missing (see Notes) |
| MG:DA:3897 | 1.00 | 27.88 |  | 1 |  | O_ph_ |
| MG:DA:3898 | 1.00 | 130.59 |  | 0 |  | 1^st^ MG coordination shell missing (see Notes) |
| MG:DA:3899 | 1.00 | 8.04 |  | 3 | 2 | 3O_ph_ - Exclusion zone contact(s): /O5'/OP1/ |
| MG:DA:3900 | 1.00 | 143.14 |  | 0 |  | 1^st^ MG coordination shell missing (see Notes) |
| MG:DA:3901 | 1.00 | 30.65 |  | 0 | 3 | - Exclusion zone contact(s): /O2'/OP1/OP1/ Consider K in place of MG (2 valid contact(s) for K vs. 0 for MG) |
| MG:DA:3902 | 1.00 | 64.40 |  | 0 |  | 1^st^ MG coordination shell missing (see Notes) |
| MG:DA:3903 | 1.00 | 42.98 |  | 1 |  | O_ph_ |
| MG:DA:3904 | 1.00 | 105.29 |  | 0 | 3 | - Exclusion zone contact(s): /N1_G_/O6/OP1/ Consider NA in place of MG (2 valid contact(s) for NA vs. 0 for MG) |
| MG:DA:3905 | 1.00 | 61.65 |  | 0 |  | 1^st^ MG coordination shell missing (see Notes) |
| MG:DA:3906 | 1.00 | 90.27 |  | 0 | 1 | - Exclusion zone contact(s): /O6/ Consider K in place of MG (1 valid contact(s) for K vs. 0 for MG) |
| MG:DA:3907 | 1.00 | 83.64 |  | 1 |  | Anion |
| MG:DA:3908 | 1.00 | 124.31 |  | 0 |  | 1^st^ MG coordination shell missing (see Notes) |
| MG:DA:3909 | 1.00 | 69.46 |  | 0 | 2 | - Exclusion zone contact(s): /OP1/OP2/ Consider K in place of MG (2 valid contact(s) for K vs. 0 for MG) |
| MG:DA:3910 | 1.00 | 143.39 |  | 0 |  | 1^st^ MG coordination shell missing (see Notes) |
| MG:DA:3911 | 1.00 | 115.14 |  | 0 | 1 | - Exclusion zone contact(s): /N7/ Consider K in place of MG (1 valid contact(s) for K vs. 0 for MG) |
| MG:DA:3912 | 1.00 | 89.24 |  | 0 | 1 | - Exclusion zone contact(s): /O6/ Consider NA in place of MG (1 valid contact(s) for NA vs. 0 for MG) |
| MG:DA:3913 | 1.00 | 120.11 |  | 0 |  | 1^st^ MG coordination shell missing (see Notes) |
| MG:DA:3914 | 1.00 | 36.96 |  | 1 | 3 | O_b_ - Exclusion zone contact(s): /O4/O4/O6/ Consider K in place of MG (3 valid contact(s) for K vs. 1 for MG) |
| MG:DA:3915 | 1.00 | 28.02 |  | 1 | 2 | O_ph_ - Exclusion zone contact(s): /O2'/OP2/ |
| MG:DA:3916 | 1.00 | 36.74 |  | 0 |  | 1^st^ MG coordination shell missing (see Notes) |
| MG:DA:3917 | 1.00 | 77.57 |  | 2 | 2 | N_b_Anion - Exclusion zone contact(s): /N3_U_/O4/ |
| MG:DA:3918 | 1.00 | 50.85 |  | 0 | 1 | - Exclusion zone contact(s): /O4/ Consider K in place of MG (1 valid contact(s) for K vs. 0 for MG) |
| MG:DA:3919 | 1.00 | 76.34 |  | 0 | 6 | - Exclusion zone contact(s): /N3_U_/O4/O5'/OP1/OP1/OP2/ Consider K in place of MG (4 valid contact(s) for K vs. 0 for MG) |
| MG:DA:3920 | 1.00 | 50.82 |  | 0 |  | 1^st^ MG coordination shell missing (see Notes) |
| MG:DA:3921 | 1.00 | 24.14 |  | 0 | 1 | - Exclusion zone contact(s): /N7/ Consider NA in place of MG (1 valid contact(s) for NA vs. 0 for MG) |
| MG:DA:3922 | 1.00 | 69.27 |  | 0 |  | 1^st^ MG coordination shell missing (see Notes) |
| MG:DA:3923 | 1.00 | 70.36 |  | 0 | 2 | - Exclusion zone contact(s): /N3_A_/O4'/ Consider K in place of MG (2 valid contact(s) for K vs. 0 for MG) |
| MG:DA:3924 | 1.00 | 48.94 |  | 0 | 2 | - Exclusion zone contact(s): /N3_A_/N6/ Consider K in place of MG (2 valid contact(s) for K vs. 0 for MG) |
| MG:DA:3925 | 1.00 | 25.93 |  | 0 | 2 | - Exclusion zone contact(s): /O2'/OP1/ Consider NA in place of MG (1 valid contact(s) for NA vs. 0 for MG) |
| MG:DA:3926 | 1.00 | 49.26 |  | 0 | 2 | - Exclusion zone contact(s): /N3_A_/O2'/ Consider K in place of MG (2 valid contact(s) for K vs. 0 for MG) |
| MG:DA:3927 | 1.00 | 66.63 |  | 2 | 3 | O_ph_O_b_ - Exclusion zone contact(s): /N1_G_/O5'/OP2/ Consider K in place of MG (3 valid contact(s) for K vs. 2 for MG) |
| MG:DA:3928 | 1.00 | 49.84 |  | 0 | 1 | - Exclusion zone contact(s): /O6/ Consider K in place of MG (1 valid contact(s) for K vs. 0 for MG) |
| MG:DA:3929 | 1.00 | 73.31 |  | 0 |  | 1^st^ MG coordination shell missing (see Notes) |
| MG:DA:3930 | 1.00 | 72.98 |  | 1 | 3 | O_b_ - Exclusion zone contact(s): /N7/N7/O6/ Consider K in place of MG (3 valid contact(s) for K vs. 1 for MG) |
| MG:DA:3931 | 1.00 | 83.78 |  | 0 |  | No contacts in 5.0 A |
| MG:DA:3932 | 1.00 | 53.67 |  | 0 | 1 | - Exclusion zone contact(s): /OP2/ Consider K in place of MG (1 valid contact(s) for K vs. 0 for MG) |
| MG:DA:3933 | 1.00 | 113.78 |  | 0 |  | 1^st^ MG coordination shell missing (see Notes) |
| MG:DA:3934 | 1.00 | 45.20 |  | 0 | 1 | - Exclusion zone contact(s): /OP2/ Consider K in place of MG (1 valid contact(s) for K vs. 0 for MG) |
| MG:DA:3935 | 1.00 | 75.49 |  | 0 | 2 | - Exclusion zone contact(s): /OP1/OP2/ Consider NA in place of MG (1 valid contact(s) for NA vs. 0 for MG) |
| MG:DA:3936 | 1.00 | 61.73 | 1 | 1 | 2 | - Exclusion zone contact(s): /N7/O6/OP2/ Consider K in place of MG (2 valid contact(s) for K vs. 0 for MG) |
| MG:DA:3937 | 1.00 | 67.42 |  | 0 |  | 1^st^ MG coordination shell missing (see Notes) |
| MG:DA:3938 | 1.00 | 96.71 |  | 1 | 2 | N_b_ - Exclusion zone contact(s): /O6/O6/ |
| MG:DA:3939 | 1.00 | 47.42 |  | 0 |  | 1^st^ MG coordination shell missing (see Notes) |
| MG:DA:3940 | 1.00 | 64.55 |  | 0 | 2 | - Exclusion zone contact(s): /O5'/OP2/ Consider NA in place of MG (1 valid contact(s) for NA vs. 0 for MG) |
| MG:DA:3941 | 1.00 | 61.61 |  | 0 | 4 | - Exclusion zone contact(s): /N7/N7/O6/O6/ Consider K in place of MG (4 valid contact(s) for K vs. 0 for MG) |
| MG:DA:3942 | 1.00 | 64.42 |  | 0 | 1 | - Exclusion zone contact(s): /OP2/ Consider NA in place of MG (1 valid contact(s) for NA vs. 0 for MG) |
| MG:DA:3943 | 1.00 | 47.90 |  | 0 | 1 | - Exclusion zone contact(s): /O6/ Consider K in place of MG (1 valid contact(s) for K vs. 0 for MG) |
| MG:DA:3944 | 1.00 | 72.08 |  | 0 | 3 | - Exclusion zone contact(s): /O5'/OP1/OP2/ Consider NA in place of MG (2 valid contact(s) for NA vs. 0 for MG) |
| MG:DA:3945 | 1.00 | 58.34 |  | 0 | 2 | - Exclusion zone contact(s): /O6/O6/ Consider K in place of MG (2 valid contact(s) for K vs. 0 for MG) |
| MG:DA:3946 | 1.00 | 82.23 |  | 1 |  | O_ph_ |
| MG:DA:3947 | 1.00 | 72.13 |  | 1 |  | O_b_ |
| MG:DA:3948 | 1.00 | 127.88 |  | 0 | 1 | - Exclusion zone contact(s): /OP1/ Consider K in place of MG (1 valid contact(s) for K vs. 0 for MG) |
| MG:DA:3949 | 1.00 | 132.61 |  | 0 |  | 1^st^ MG coordination shell missing (see Notes) |
| MG:DA:3950 | 1.00 | 117.91 |  | 1 | 1 | O_ph_ - Exclusion zone contact(s): /O3'/ |
| MG:DA:3951 | 1.00 | 136.81 |  | 0 | 2 | - Exclusion zone contact(s): /OP1/OP2/ Consider K in place of MG (2 valid contact(s) for K vs. 0 for MG) |
| MG:DA:3952 | 1.00 | 46.47 |  | 1 | 1 | O_ph_ - Exclusion zone contact(s): /OP2/ |
| MG:DA:3953 | 1.00 | 84.78 |  | 0 |  | 1^st^ MG coordination shell missing (see Notes) |
| MG:DA:3954 | 1.00 | 52.97 |  | 0 |  | 1^st^ MG coordination shell missing (see Notes) |
| MG:DA:3955 | 1.00 | 120.28 |  | 0 |  | 1^st^ MG coordination shell missing (see Notes) |
| MG:DA:3956 | 1.00 | 45.35 |  | 0 |  | 1^st^ MG coordination shell missing (see Notes) |
| MG:DA:3957 | 1.00 | 62.07 |  | 0 |  | 1^st^ MG coordination shell missing (see Notes) |
| MG:DA:3958 | 1.00 | 55.80 |  | 0 |  | 1^st^ MG coordination shell missing (see Notes) |
| MG:DA:3959 | 1.00 | 59.77 |  | 0 |  | 1^st^ MG coordination shell missing (see Notes) |
| MG:DA:3960 | 1.00 | 54.12 |  | 0 | 1 | - Exclusion zone contact(s): /O6/ Consider NA in place of MG (1 valid contact(s) for NA vs. 0 for MG) |
| MG:DA:3961 | 1.00 | 41.86 |  | 0 | 3 | - Exclusion zone contact(s): /N3_A_/O4/O4'/ Consider K in place of MG (3 valid contact(s) for K vs. 0 for MG) |
| MG:DA:3962 | 1.00 | 81.33 |  | 0 |  | 1^st^ MG coordination shell missing (see Notes) |
| MG:DA:3963 | 1.00 | 59.00 |  | 0 | 2 | - Exclusion zone contact(s): /OP2/OP2/ Consider NA in place of MG (1 valid contact(s) for NA vs. 0 for MG) |
| MG:DA:3964 | 1.00 | 68.10 |  | 0 | 1 | - Exclusion zone contact(s): /OP2/ Consider NA in place of MG (1 valid contact(s) for NA vs. 0 for MG) |
| MG:DA:3965 | 1.00 | 78.93 |  | 1 | 2 | O_ph_ - Exclusion zone contact(s): /OP1/OP2/ Consider K in place of MG (2 valid contact(s) for K vs. 1 for MG) |
| MG:DA:3966 | 1.00 | 41.18 |  | 1 | 1 | O_ph_ - Exclusion zone contact(s): /OP1/ |
| MG:DA:3967 | 1.00 | 80.53 |  | 1 | 3 | O_ph_ - Exclusion zone contact(s): /O3'/OP1/OP2/ Consider K in place of MG (3 valid contact(s) for K vs. 1 for MG) |
| MG:DA:3968 | 1.00 | 55.45 |  | 0 | 2 | - Exclusion zone contact(s): /O4/O6/ Consider K in place of MG (2 valid contact(s) for K vs. 0 for MG) |
| MG:DA:3969 | 1.00 | 48.05 |  | 1 | 1 | O_ph_ - Exclusion zone contact(s): /OP1/ |
| MG:DA:3970 | 1.00 | 50.89 |  | 0 |  | 1^st^ MG coordination shell missing (see Notes) |
| MG:DA:3971 | 1.00 | 66.37 |  | 0 | 1 | - Exclusion zone contact(s): /OP1/ Consider K in place of MG (1 valid contact(s) for K vs. 0 for MG) |
| MG:DA:3972 | 1.00 | 41.68 |  | 0 | 3 | - Exclusion zone contact(s): /O2'/OP1/OP1/ Consider K in place of MG (3 valid contact(s) for K vs. 0 for MG) |
| MG:DA:3973 | 1.00 | 60.34 |  | 1 | 2 | O_ph_ - Exclusion zone contact(s): /O3'/O5'/ Consider K in place of MG (2 valid contact(s) for K vs. 1 for MG) |
| MG:DA:3974 | 1.00 | 61.52 |  | 0 |  | 1^st^ MG coordination shell missing (see Notes) |
| MG:DA:3975 | 1.00 | 50.74 |  | 0 | 4 | - Exclusion zone contact(s): /O2'/O3'/OP1/OP1/ Consider K in place of MG (3 valid contact(s) for K vs. 0 for MG) |
| MG:DA:3976 | 1.00 | 37.45 |  | 0 | 6 | - Exclusion zone contact(s): /N1/N1_A_/N6/N6/N7/O2/ Consider K in place of MG (6 valid contact(s) for K vs. 0 for MG) |
| MG:DA:3977 | 1.00 | 71.29 |  | 1 |  | O_ph_ |
| MG:DA:3978 | 1.00 | 82.22 |  | 0 |  | 1^st^ MG coordination shell missing (see Notes) |
| MG:DA:3979 | 1.00 | 82.66 |  | 0 |  | 1^st^ MG coordination shell missing (see Notes) |
| MG:DA:3980 | 1.00 | 65.08 |  | 0 | 2 | - Exclusion zone contact(s): /N4/N4/ Consider K in place of MG (2 valid contact(s) for K vs. 0 for MG) |
| MG:DA:3981 | 1.00 | 87.30 |  | 0 |  | 1^st^ MG coordination shell missing (see Notes) |
| MG:DA:3982 | 1.00 | 106.27 |  | 1 |  | O_b_ |
| MG:DA:3983 | 1.00 | 57.53 |  | 0 | 2 | - Exclusion zone contact(s): /O4/O6/ Consider NA in place of MG (1 valid contact(s) for NA vs. 0 for MG) |
| MG:DA:3984 | 1.00 | 117.60 |  | 0 |  | 1^st^ MG coordination shell missing (see Notes) |
| MG:DA:3985 | 1.00 | 81.97 |  | 0 | 1 | - Exclusion zone contact(s): /OP1/ Consider K in place of MG (1 valid contact(s) for K vs. 0 for MG) |
| MG:DA:3986 | 1.00 | 51.00 |  | 0 | 2 | - Exclusion zone contact(s): /N7/O6/ Consider K in place of MG (2 valid contact(s) for K vs. 0 for MG) |
| MG:DA:3987 | 1.00 | 53.64 |  | 0 | 3 | - Exclusion zone contact(s): /O3'/O5'/OP1/ Consider K in place of MG (3 valid contact(s) for K vs. 0 for MG) |
| MG:DA:3988 | 1.00 | 59.08 |  | 0 | 1 | - Exclusion zone contact(s): /OP1/ Consider K in place of MG (1 valid contact(s) for K vs. 0 for MG) |
| MG:DA:3989 | 1.00 | 110.00 |  | 0 | 4 | - Exclusion zone contact(s): /N1_G_/N2/N7/OP2/ Consider K in place of MG (4 valid contact(s) for K vs. 0 for MG) |
| MG:DA:3990 | 1.00 | 31.43 |  | 0 | 2 | - Exclusion zone contact(s): /O2'/OP2/ Consider NA in place of MG (1 valid contact(s) for NA vs. 0 for MG) |
| MG:DA:3991 | 1.00 | 96.13 |  | 0 | 2 | - Exclusion zone contact(s): /N7/OP2/ Consider NA in place of MG (1 valid contact(s) for NA vs. 0 for MG) |
| MG:DA:3992 | 1.00 | 61.54 |  | 0 | 1 | - Exclusion zone contact(s): /O6/ Consider K in place of MG (1 valid contact(s) for K vs. 0 for MG) |
| MG:DA:3993 | 1.00 | 51.60 |  | 0 | 2 | - Exclusion zone contact(s): /O2'/O3'/ Consider K in place of MG (2 valid contact(s) for K vs. 0 for MG) |
| MG:DA:3994 | 1.00 | 48.06 |  | 0 | 2 | - Exclusion zone contact(s): /O6/O6/ Consider K in place of MG (2 valid contact(s) for K vs. 0 for MG) |
| MG:DA:3995 | 1.00 | 71.64 |  | 0 |  | 1^st^ MG coordination shell missing (see Notes) |
| MG:DA:3996 | 1.00 | 63.50 |  | 0 | 3 | - Exclusion zone contact(s): /O2'/O3'/OP1/ Consider K in place of MG (3 valid contact(s) for K vs. 0 for MG) |
| MG:DA:3997 | 1.00 | 57.58 |  | 0 | 2 | - Exclusion zone contact(s): /OP2/OP2/ Consider K in place of MG (2 valid contact(s) for K vs. 0 for MG) |
| MG:DA:3998 | 1.00 | 46.21 |  | 0 | 6 | - Exclusion zone contact(s): /N2/N3_G_/N6/N6/N7/O2'/ Consider K in place of MG (6 valid contact(s) for K vs. 0 for MG) |
| MG:DA:3999 | 1.00 | 83.87 |  | 1 |  | O_ph_ |
| MG:DA:4000 | 1.00 | 50.87 |  | 0 | 3 | - Exclusion zone contact(s): /N1/N7/O6/ Consider K in place of MG (3 valid contact(s) for K vs. 0 for MG) |
| MG:DA:4001 | 1.00 | 121.15 |  | 0 | 1 | - Exclusion zone contact(s): /O6/ Consider K in place of MG (1 valid contact(s) for K vs. 0 for MG) |
| MG:DA:4002 | 1.00 | 42.51 |  | 0 | 3 | - Exclusion zone contact(s): /O3'/O4'/OP2/ Consider K in place of MG (3 valid contact(s) for K vs. 0 for MG) |
| MG:DA:4003 | 1.00 | 39.86 |  | 0 | 3 | - Exclusion zone contact(s): /N7/OP2/ Consider K in place of MG (3 valid contact(s) for K vs. 0 for MG) |
| MG:DA:4004 | 1.00 | 90.11 |  | 0 | 2 | - Exclusion zone contact(s): /OP2/OP2/ Consider K in place of MG (2 valid contact(s) for K vs. 0 for MG) |
| MG:DA:4005 | 1.00 | 84.59 |  | 0 |  | 1^st^ MG coordination shell missing (see Notes) |
| MG:DA:4006 | 1.00 | 71.29 |  | 0 | 1 | - Exclusion zone contact(s): /OP1/ Consider K in place of MG (1 valid contact(s) for K vs. 0 for MG) |
| MG:DA:4007 | 1.00 | 110.19 |  | 0 | 2 | - Exclusion zone contact(s): /N7/O2'/ Consider NA in place of MG (1 valid contact(s) for NA vs. 0 for MG) |
| MG:DA:4008 | 1.00 | 49.51 |  | 0 | 1 | - Exclusion zone contact(s): /O6/ Consider K in place of MG (1 valid contact(s) for K vs. 0 for MG) |
| MG:DA:4009 | 1.00 | 98.15 |  | 0 | 3 | - Exclusion zone contact(s): /N7/O4/O6/ Consider K in place of MG (3 valid contact(s) for K vs. 0 for MG) |
| MG:DA:4010 | 1.00 | 91.40 |  | 1 | 2 | O_b_ - Exclusion zone contact(s): /N4/O6/ Consider K in place of MG (2 valid contact(s) for K vs. 1 for MG) |
| MG:DA:4011 | 1.00 | 52.20 |  | 1 | 2 | O_r_ - Exclusion zone contact(s): /O2'/OP2/ |
| MG:DA:4012 | 1.00 | 63.07 |  | 1 | 3 | O_ph_ - Exclusion zone contact(s): /O5'/OP1/OP2/ Consider K in place of MG (3 valid contact(s) for K vs. 1 for MG) |
| MG:DA:4013 | 1.00 | 42.17 |  | 0 | 4 | - Exclusion zone contact(s): /O3'/O6/O_back_/OP2/ Consider K in place of MG (4 valid contact(s) for K vs. 0 for MG) |
| MG:DA:4014 | 1.00 | 50.92 |  | 0 | 4 | - Exclusion zone contact(s): /O5'/OP1/OP2/OP2/ Consider K in place of MG (3 valid contact(s) for K vs. 0 for MG) |
| MG:DA:4015 | 1.00 | 57.81 |  | 0 | 3 | - Exclusion zone contact(s): /NZ_Lys_/OP1/OP2/ Consider K in place of MG (3 valid contact(s) for K vs. 0 for MG) |
| MG:DA:4016 | 1.00 | 40.57 |  | 0 | 2 | - Exclusion zone contact(s): /N7/O6/ Consider K in place of MG (2 valid contact(s) for K vs. 0 for MG) |
| MG:DA:4017 | 1.00 | 69.29 |  | 0 |  | 1^st^ MG coordination shell missing (see Notes) |
| MG:DA:4018 | 1.00 | 75.42 |  | 0 | 1 | - Exclusion zone contact(s): /OP2/ Consider K in place of MG (1 valid contact(s) for K vs. 0 for MG) |
| MG:DA:4019 | 1.00 | 91.13 |  | 0 | 4 | - Exclusion zone contact(s): /N1_G_/O2'/O4'/O6/ Consider K in place of MG (4 valid contact(s) for K vs. 0 for MG) |
| MG:DA:4020 | 1.00 | 153.71 |  | 0 | 4 | - Exclusion zone contact(s): /N7/O5'/OP2/ Consider K in place of MG (3 valid contact(s) for K vs. 0 for MG) |
| MG:DA:4021 | 1.00 | 76.30 |  | 0 | 2 | - Exclusion zone contact(s): /O4/OP2/ Consider K in place of MG (2 valid contact(s) for K vs. 0 for MG) |
| MG:DA:4022 | 1.00 | 41.32 |  | 0 | 4 | - Exclusion zone contact(s): /N_back_/OG_Ser_/OP1/OP2/ Consider K in place of MG (3 valid contact(s) for K vs. 0 for MG) |
| MG:DA:4023 | 1.00 | 41.75 |  | 1 |  | O_ph_ |
| MG:DA:4024 | 1.00 | 47.19 |  | 0 | 4 | - Exclusion zone contact(s): /N7/N7/O5'/OP2/ Consider K in place of MG (4 valid contact(s) for K vs. 0 for MG) |
| MG:DA:4025 | 1.00 | 48.49 |  | 0 | 2 | - Exclusion zone contact(s): /O6/OP2/ Consider K in place of MG (2 valid contact(s) for K vs. 0 for MG) |
| MG:DA:4026 | 1.00 | 55.91 |  | 0 | 1 | - Exclusion zone contact(s): /OP1/ Consider K in place of MG (1 valid contact(s) for K vs. 0 for MG) |
| MG:DA:4027 | 1.00 | 59.87 |  | 0 | 3 | - Exclusion zone contact(s): /O3'/OP1/OP1/ Consider K in place of MG (3 valid contact(s) for K vs. 0 for MG) |
| MG:DA:4028 | 1.00 | 48.45 |  | 1 | 1 | Anion - Exclusion zone contact(s): /N7/ |
| MG:DA:4029 | 1.00 | 65.32 |  | 0 | 1 | - Exclusion zone contact(s): /O5'/ Consider K in place of MG (1 valid contact(s) for K vs. 0 for MG) |
| MG:DA:4030 | 1.00 | 84.30 |  | 0 | 1 | - Exclusion zone contact(s): /OP2/ Consider NA in place of MG (1 valid contact(s) for NA vs. 0 for MG) |
| MG:DA:4031 | 1.00 | 79.20 |  | 0 |  | No contacts in 5.0 A |
| MG:DA:4032 | 1.00 | 86.64 |  | 0 | 2 | - Exclusion zone contact(s): /O3'/OP1/ Consider NA in place of MG (1 valid contact(s) for NA vs. 0 for MG) |
| MG:DA:4033 | 1.00 | 103.82 |  | 0 | 2 | - Exclusion zone contact(s): /O3'/OP1/ Consider NA in place of MG (1 valid contact(s) for NA vs. 0 for MG) |
| MG:DA:4034 | 1.00 | 71.18 |  | 0 | 3 | - Exclusion zone contact(s): /N3_A_/N6/NZ_Lys_/ Consider NA in place of MG (2 valid contact(s) for NA vs. 0 for MG) |
| MG:DA:4035 | 1.00 | 52.44 |  | 0 | 7 | - Exclusion zone contact(s): /N1_A_/N6/N6/O2'/O4'/O_back_/O_back_/ Consider K in place of MG (5 valid contact(s) for K vs. 0 for MG) |
| MG:DA:4036 | 1.00 | 55.01 |  | 0 | 1 | - Exclusion zone contact(s): /O4/ Consider K in place of MG (1 valid contact(s) for K vs. 0 for MG) |
| MG:DA:4037 | 1.00 | 81.04 |  | 0 | 2 | - Exclusion zone contact(s): /O2'/O3'/ Consider K in place of MG (2 valid contact(s) for K vs. 0 for MG) |
| MG:DA:4038 | 1.00 | 103.45 |  | 0 |  | No contacts in 5.0 A |
| MG:DA:4039 | 1.00 | 75.47 |  | 0 | 2 | - Exclusion zone contact(s): /O3'/OP1/ Consider K in place of MG (2 valid contact(s) for K vs. 0 for MG) |
| MG:DA:4040 | 1.00 | 59.87 |  | 0 | 4 | - Exclusion zone contact(s): /N2/O2/O2/O2'/ Consider K in place of MG (4 valid contact(s) for K vs. 0 for MG) |
| MG:DA:4041 | 1.00 | 101.43 |  | 0 | 1 | - Exclusion zone contact(s): /O2'/ Consider K in place of MG (1 valid contact(s) for K vs. 0 for MG) |
| MG:DA:4042 | 1.00 | 71.46 |  | 0 | 2 | - Exclusion zone contact(s): /N7/O6/ Consider K in place of MG (2 valid contact(s) for K vs. 0 for MG) |
| MG:DA:4043 | 1.00 | 82.66 |  | 1 | 3 | O_ph_ - Exclusion zone contact(s): /O2'/OP1/OP2/ Consider K in place of MG (2 valid contact(s) for K vs. 1 for MG) |
| MG:DA:4044 | 1.00 | 107.10 |  | 0 | 3 | - Exclusion zone contact(s): /N6/N6/N7/ Consider K in place of MG (2 valid contact(s) for K vs. 0 for MG) |
| MG:DA:4045 | 1.00 | 134.82 |  | 1 | 4 | O_ph_ - Exclusion zone contact(s): /N1_G_/N7/O6/OP1/ Consider K in place of MG (3 valid contact(s) for K vs. 1 for MG) |
| MG:DA:4046 | 1.00 | 52.12 |  | 0 | 1 | - Exclusion zone contact(s): /O6/ Consider K in place of MG (1 valid contact(s) for K vs. 0 for MG) |
| MG:DA:4047 | 1.00 | 44.23 |  | 0 | 1 | - Exclusion zone contact(s): /O6/ Consider K in place of MG (1 valid contact(s) for K vs. 0 for MG) |
| MG:DA:4048 | 1.00 | 75.78 |  | 0 | 1 | - Exclusion zone contact(s): /O6/ Consider NA in place of MG (1 valid contact(s) for NA vs. 0 for MG) |
| MG:DA:4049 | 1.00 | 33.67 |  | 0 |  | 1^st^ MG coordination shell missing (see Notes) |
| MG:DA:4050 | 1.00 | 66.17 |  | 0 | 1 | - Exclusion zone contact(s): /O6/ Consider K in place of MG (1 valid contact(s) for K vs. 0 for MG) |
| MG:DA:4051 | 1.00 | 94.89 |  | 0 | 4 | - Exclusion zone contact(s): /N1_G_/O2'/O6/O6/ Consider K in place of MG (4 valid contact(s) for K vs. 0 for MG) |
| MG:DA:4052 | 1.00 | 178.89 |  | 1 |  | O_b_ |
| MG:DA:4053 | 1.00 | 75.59 |  | 0 | 1 | - Exclusion zone contact(s): /OP2/ Consider K in place of MG (1 valid contact(s) for K vs. 0 for MG) |
| MG:DA:4054 | 1.00 | 108.77 |  | 0 | 2 | - Exclusion zone contact(s): /N7/OP2/ Consider K in place of MG (2 valid contact(s) for K vs. 0 for MG) |
| MG:DA:4055 | 1.00 | 68.13 |  | 0 | 1 | - Exclusion zone contact(s): /OP2/ Consider K in place of MG (1 valid contact(s) for K vs. 0 for MG) |
| MG:DA:4056 | 1.00 | 62.79 |  | 1 | 3 | O_r_ - Exclusion zone contact(s): /O3'/OP2/OP2/ Consider K in place of MG (3 valid contact(s) for K vs. 1 for MG) |
| MG:DA:4057 | 1.00 | 87.72 |  | 0 |  | 1^st^ MG coordination shell missing (see Notes) |
| MG:DA:4058 | 1.00 | 81.30 |  | 1 | 3 | O_r_ - Exclusion zone contact(s): /O3'/OP2/OP2/ Consider K in place of MG (2 valid contact(s) for K vs. 1 for MG) |
| MG:DA:4059 | 1.00 | 103.63 |  | 1 | 1 | O_ph_ - Exclusion zone contact(s): /OP1/ |
| MG:DA:4060 | 1.00 | 52.57 |  | 0 |  | 1^st^ MG coordination shell missing (see Notes) |
| MG:DA:4061 | 1.00 | 54.71 |  | 0 | 1 | - Exclusion zone contact(s): /OP2/ Consider K in place of MG (1 valid contact(s) for K vs. 0 for MG) |
| MG:DA:4062 | 1.00 | 49.47 |  | 0 | 3 | - Exclusion zone contact(s): /O4'/O5'/OP1/ Consider K in place of MG (2 valid contact(s) for K vs. 0 for MG) |
| MG:DA:4063 | 1.00 | 52.62 |  | 0 | 4 | - Exclusion zone contact(s): /O2'/O3'/OP1/OP1/ Consider K in place of MG (4 valid contact(s) for K vs. 0 for MG) |
| MG:DA:4064 | 1.00 | 56.44 |  | 1 |  | O_ph_ |
| MG:DA:4065 | 1.00 | 58.66 |  | 0 | 1 | - Exclusion zone contact(s): /OP2/ Consider K in place of MG (1 valid contact(s) for K vs. 0 for MG) |
| MG:DA:4066 | 1.00 | 59.82 |  | 0 | 1 | - Exclusion zone contact(s): /OP1/ Consider K in place of MG (1 valid contact(s) for K vs. 0 for MG) |
| MG:DA:4067 | 1.00 | 39.94 |  | 0 | 4 | - Exclusion zone contact(s): /OP1/OP1/OP2/OP2/ Consider K in place of MG (4 valid contact(s) for K vs. 0 for MG) |
| MG:DA:4068 | 1.00 | 50.95 |  | 1 | 2 | O_ph_ - Exclusion zone contact(s): /OP1/OP1/ Consider K in place of MG (2 valid contact(s) for K vs. 1 for MG) |
| MG:DA:4069 | 1.00 | 24.51 |  | 3 | 2 | 3O_ph_ - Exclusion zone contact(s): /O5'/OP1/ |
| MG:DA:4070 | 1.00 | 45.78 |  | 1 | 2 | O_r_ - Exclusion zone contact(s): /OP1/OP2/ Consider K in place of MG (2 valid contact(s) for K vs. 1 for MG) |
| MG:DA:4071 | 1.00 | 55.82 |  | 0 | 5 | - Exclusion zone contact(s): /N1_G_/N7/O2'/O5'/O6/ Consider K in place of MG (5 valid contact(s) for K vs. 0 for MG) |
| MG:DA:4072 | 1.00 | 48.35 |  | 0 | 2 | - Exclusion zone contact(s): /N7/O6/ Consider K in place of MG (2 valid contact(s) for K vs. 0 for MG) |
| MG:DA:4073 | 1.00 | 106.94 |  | 0 | 1 | - Exclusion zone contact(s): /N7/ Consider K in place of MG (1 valid contact(s) for K vs. 0 for MG) |
| MG:DA:4074 | 1.00 | 54.07 |  | 0 | 1 | - Exclusion zone contact(s): /O6/ Consider K in place of MG (1 valid contact(s) for K vs. 0 for MG) |
| MG:DA:4075 | 1.00 | 25.95 |  | 0 |  | 1^st^ MG coordination shell missing (see Notes) |
| MG:DA:4076 | 1.00 | 27.93 |  | 1 | 3 | O_ph_ - Exclusion zone contact(s): /O2'/O5'/OP2/ Consider K in place of MG (3 valid contact(s) for K vs. 1 for MG) |
| MG:DA:4077 | 1.00 | 51.42 |  | 0 |  | 1^st^ MG coordination shell missing (see Notes) |
| MG:DA:4078 | 1.00 | 65.22 |  | 0 | 4 | - Exclusion zone contact(s): /N6/N7/N7/O6/ Consider K in place of MG (4 valid contact(s) for K vs. 0 for MG) |
| MG:DA:4079 | 1.00 | 66.25 |  | 0 |  | 1^st^ MG coordination shell missing (see Notes) |
| MG:DA:4080 | 1.00 | 89.02 |  | 0 |  | 1^st^ MG coordination shell missing (see Notes) |
| MG:DA:4081 | 1.00 | 67.73 |  | 0 | 1 | - Exclusion zone contact(s): /O6/ Consider K in place of MG (1 valid contact(s) for K vs. 0 for MG) |
| MG:DA:4082 | 1.00 | 75.10 |  | 0 |  | 1^st^ MG coordination shell missing (see Notes) |
| MG:DA:4083 | 1.00 | 52.72 |  | 0 | 3 | - Exclusion zone contact(s): /O5'/OP1/OP2/ Consider K in place of MG (2 valid contact(s) for K vs. 0 for MG) |
| MG:DA:4084 | 1.00 | 79.67 |  | 0 | 1 | - Exclusion zone contact(s): /OP1/ Consider K in place of MG (1 valid contact(s) for K vs. 0 for MG) |
| MG:DA:4085 | 1.00 | 60.81 |  | 0 | 1 | - Exclusion zone contact(s): /OP2/ Consider K in place of MG (1 valid contact(s) for K vs. 0 for MG) |
| MG:DA:4086 | 1.00 | 98.03 |  | 1 | 1 | O_ph_ - Exclusion zone contact(s): /OP1/ |
| MG:DA:4087 | 1.00 | 45.89 |  | 0 | 1 | - Exclusion zone contact(s): /OP2/ Consider K in place of MG (1 valid contact(s) for K vs. 0 for MG) |
| MG:DA:4088 | 1.00 | 83.88 |  | 0 |  | 1^st^ MG coordination shell missing (see Notes) |
| MG:DA:4089 | 1.00 | 63.13 |  | 0 | 2 | - Exclusion zone contact(s): /OP2/OP2/ Consider NA in place of MG (1 valid contact(s) for NA vs. 0 for MG) |
| MG:DA:4090 | 1.00 | 52.05 |  | 0 | 3 | - Exclusion zone contact(s): /OP1/OP2/OP2/ Consider K in place of MG (2 valid contact(s) for K vs. 0 for MG) |
| MG:DA:4091 | 1.00 | 42.96 |  | 1 | 5 | O_ph_ - Exclusion zone contact(s): /O3'/OP1/OP1/OP2/OP2/ Consider K in place of MG (5 valid contact(s) for K vs. 1 for MG) |
| MG:DA:4092 | 1.00 | 99.15 |  | 0 | 2 | - Exclusion zone contact(s): /OP1/OP1/ Consider K in place of MG (2 valid contact(s) for K vs. 0 for MG) |
| MG:DA:4093 | 1.00 | 85.06 |  | 0 | 1 | - Exclusion zone contact(s): /OP1/ Consider K in place of MG (1 valid contact(s) for K vs. 0 for MG) |
| MG:DA:4094 | 1.00 | 41.02 |  | 1 | 2 | O_b_ - Exclusion zone contact(s): /O2/O2'/ Consider K in place of MG (2 valid contact(s) for K vs. 1 for MG) |
| MG:DA:4095 | 1.00 | 115.57 |  | 1 | 1 | O_ph_ - Exclusion zone contact(s): /OP2/ |
| MG:DA:4096 | 1.00 | 73.91 |  | 0 |  | 1^st^ MG coordination shell missing (see Notes) |
| MG:DA:4097 | 1.00 | 64.73 |  | 0 |  | 1^st^ MG coordination shell missing (see Notes) |
| MG:DA:4098 | 1.00 | 43.90 |  | 0 | 2 | - Exclusion zone contact(s): /O6/O6/ Consider K in place of MG (2 valid contact(s) for K vs. 0 for MG) |
| MG:DA:4099 | 1.00 | 55.31 |  | 0 |  | 1^st^ MG coordination shell missing (see Notes) |
| MG:DA:4100 | 1.00 | 72.68 |  | 0 | 1 | - Exclusion zone contact(s): /OP2/ Consider K in place of MG (1 valid contact(s) for K vs. 0 for MG) |
| MG:DA:4101 | 1.00 | 82.30 |  | 0 |  | 1^st^ MG coordination shell missing (see Notes) |
| MG:DA:4102 | 1.00 | 70.56 |  | 0 | 1 | - Exclusion zone contact(s): /O6/ Consider K in place of MG (1 valid contact(s) for K vs. 0 for MG) |
| MG:DA:4103 | 1.00 | 85.24 |  | 0 |  | 1^st^ MG coordination shell missing (see Notes) |
| MG:DA:4104 | 1.00 | 76.24 |  | 0 | 1 | - Exclusion zone contact(s): /N4/ Consider K in place of MG (1 valid contact(s) for K vs. 0 for MG) |
| MG:DA:4105 | 1.00 | 62.44 |  | 0 | 1 | - Exclusion zone contact(s): /OP2/ Consider K in place of MG (1 valid contact(s) for K vs. 0 for MG) |
| MG:DA:4106 | 1.00 | 65.65 |  | 0 | 1 | - Exclusion zone contact(s): /OP1/ Consider K in place of MG (1 valid contact(s) for K vs. 0 for MG) |
| MG:DA:4107 | 1.00 | 49.39 |  | 0 | 1 | - Exclusion zone contact(s): /OP1/ Consider K in place of MG (1 valid contact(s) for K vs. 0 for MG) |
| MG:DA:4108 | 1.00 | 74.33 |  | 0 | 1 | - Exclusion zone contact(s): /OP1/ Consider K in place of MG (1 valid contact(s) for K vs. 0 for MG) |
| MG:DA:4109 | 1.00 | 61.04 |  | 0 |  | 1^st^ MG coordination shell missing (see Notes) |
| MG:DA:4110 | 1.00 | 51.12 |  | 0 | 2 | - Exclusion zone contact(s): /O2'/OE2_Glu_/ Consider K in place of MG (2 valid contact(s) for K vs. 0 for MG) |
| MG:DA:4111 | 1.00 | 54.40 |  | 0 | 1 | - Exclusion zone contact(s): /N7/ Consider K in place of MG (1 valid contact(s) for K vs. 0 for MG) |
| MG:DA:4112 | 1.00 | 45.48 |  | 0 |  | 1^st^ MG coordination shell missing (see Notes) |
| MG:DA:4113 | 1.00 | 45.31 |  | 0 |  | 1^st^ MG coordination shell missing (see Notes) |
| MG:DA:4114 | 1.00 | 61.85 |  | 0 |  | No contacts in 5.0 A |
| MG:DA:4115 | 1.00 | 81.37 |  | 0 |  | 1^st^ MG coordination shell missing (see Notes) |
| MG:DA:4116 | 1.00 | 78.22 |  | 0 | 1 | - Exclusion zone contact(s): /O6/ Consider K in place of MG (1 valid contact(s) for K vs. 0 for MG) |
| MG:DA:4117 | 1.00 | 58.21 |  | 0 | 2 | - Exclusion zone contact(s): /N7/O4/ Consider K in place of MG (2 valid contact(s) for K vs. 0 for MG) |
| MG:DA:4118 | 1.00 | 96.26 |  | 0 | 1 | - Exclusion zone contact(s): /OP1/ Consider K in place of MG (1 valid contact(s) for K vs. 0 for MG) |
| MG:DA:4119 | 1.00 | 47.18 |  | 0 |  | 1^st^ MG coordination shell missing (see Notes) |
| MG:DA:4120 | 1.00 | 48.16 |  | 0 | 1 | - Exclusion zone contact(s): /OP1/ Consider K in place of MG (1 valid contact(s) for K vs. 0 for MG) |
| MG:DA:4121 | 1.00 | 77.71 |  | 0 | 2 | - Exclusion zone contact(s): /OP1/OP1/ Consider K in place of MG (2 valid contact(s) for K vs. 0 for MG) |
| MG:DA:4122 | 1.00 | 105.94 |  | 0 |  | 1^st^ MG coordination shell missing (see Notes) |
| MG:DA:4123 | 1.00 | 75.85 |  | 0 |  | 1^st^ MG coordination shell missing (see Notes) |
| MG:DA:4124 | 1.00 | 58.67 |  | 1 | 1 | O_r_ - Exclusion zone contact(s): /O3'/ |
| MG:DA:4125 | 1.00 | 76.31 |  | 0 |  | 1^st^ MG coordination shell missing (see Notes) |
| MG:DA:4126 | 1.00 | 57.50 |  | 0 | 1 | - Exclusion zone contact(s): /OP2/ Consider K in place of MG (1 valid contact(s) for K vs. 0 for MG) |
| MG:DA:4127 | 1.00 | 70.81 |  | 0 |  | 1^st^ MG coordination shell missing (see Notes) |
| MG:DA:4128 | 1.00 | 85.00 |  | 0 |  | 1^st^ MG coordination shell missing (see Notes) |
| MG:DA:4129 | 1.00 | 109.98 |  | 1 | 2 | O_ph_ - Exclusion zone contact(s): /O2'/OP1/ |
| MG:DA:4130 | 1.00 | 29.07 |  | 0 | 1 | - Exclusion zone contact(s): /OP1/ Consider NA in place of MG (1 valid contact(s) for NA vs. 0 for MG) |
| MG:DA:4131 | 1.00 | 34.82 |  | 0 | 2 | - Exclusion zone contact(s): /O6/O6/ Consider NA in place of MG (1 valid contact(s) for NA vs. 0 for MG) |
| MG:DA:4132 | 1.00 | 40.03 |  | 0 | 2 | - Exclusion zone contact(s): /O3'/OP1/ Consider K in place of MG (2 valid contact(s) for K vs. 0 for MG) |
| MG:DA:4133 | 1.00 | 47.58 |  | 0 | 3 | - Exclusion zone contact(s): /O4/O6/O6/ Consider K in place of MG (3 valid contact(s) for K vs. 0 for MG) |
| MG:DA:4134 | 1.00 | 34.29 |  | 0 | 1 | - Exclusion zone contact(s): /O6/ Consider K in place of MG (1 valid contact(s) for K vs. 0 for MG) |
| MG:DA:4135 | 1.00 | 59.93 |  | 0 | 1 | - Exclusion zone contact(s): /OP2/ Consider NA in place of MG (1 valid contact(s) for NA vs. 0 for MG) |
| MG:DA:4136 | 1.00 | 47.90 |  | 0 | 1 | - Exclusion zone contact(s): /OP2/ Consider K in place of MG (1 valid contact(s) for K vs. 0 for MG) |
| MG:DA:4137 | 1.00 | 39.99 |  | 0 | 4 | - Exclusion zone contact(s): /N1_A_/N6/O3'/OP1/ Consider K in place of MG (4 valid contact(s) for K vs. 0 for MG) |
| MG:DA:4138 | 1.00 | 54.80 |  | 0 | 3 | - Exclusion zone contact(s): /OP1/OP1/OP2/ Consider NA in place of MG (2 valid contact(s) for NA vs. 0 for MG) |
| MG:DA:4139 | 1.00 | 36.42 |  | 0 | 3 | - Exclusion zone contact(s): /N2/O4/O6/ Consider K in place of MG (2 valid contact(s) for K vs. 0 for MG) |
| MG:DA:4140 | 1.00 | 47.37 |  | 0 |  | 1^st^ MG coordination shell missing (see Notes) |
| MG:DA:4141 | 1.00 | 66.31 |  | 0 |  | 1^st^ MG coordination shell missing (see Notes) |
| MG:DA:4142 | 1.00 | 29.80 |  | 0 | 1 | - Exclusion zone contact(s): /OP1/ Consider K in place of MG (1 valid contact(s) for K vs. 0 for MG) |
| MG:DA:4143 | 1.00 | 63.31 |  | 0 | 2 | - Exclusion zone contact(s): /O6/O6/ Consider K in place of MG (2 valid contact(s) for K vs. 0 for MG) |
| MG:DA:4144 | 1.00 | 45.00 |  | 0 |  | 1^st^ MG coordination shell missing (see Notes) |
| MG:DA:4145 | 1.00 | 44.40 |  | 0 | 2 | - Exclusion zone contact(s): /OP2/OP2/ Consider K in place of MG (2 valid contact(s) for K vs. 0 for MG) |
| MG:DA:4146 | 1.00 | 72.99 |  | 0 | 1 | - Exclusion zone contact(s): /OP2/ Consider K in place of MG (1 valid contact(s) for K vs. 0 for MG) |
| MG:DA:4147 | 1.00 | 42.99 |  | 0 |  | 1^st^ MG coordination shell missing (see Notes) |
| MG:DA:4148 | 1.00 | 63.55 |  | 0 |  | 1^st^ MG coordination shell missing (see Notes) |
| MG:DA:4149 | 1.00 | 70.20 |  | 0 | 5 | - Exclusion zone contact(s): /O2'/O3'/O5'/OP2/OP2/ Consider K in place of MG (5 valid contact(s) for K vs. 0 for MG) |
| MG:DA:4150 | 1.00 | 47.29 |  | 0 | 3 | - Exclusion zone contact(s): /O6/OP1/OP1/ Consider K in place of MG (2 valid contact(s) for K vs. 0 for MG) |
| MG:DA:4151 | 1.00 | 66.81 |  | 0 | 5 | - Exclusion zone contact(s): /O3'/O4'/O5'/OP1/OP1/ Consider K in place of MG (5 valid contact(s) for K vs. 0 for MG) |
| MG:DA:4152 | 1.00 | 54.74 |  | 0 | 3 | - Exclusion zone contact(s): /O2/OP1/OP2/ Consider K in place of MG (3 valid contact(s) for K vs. 0 for MG) |
| MG:DA:4153 | 1.00 | 48.59 |  | 0 |  | 1^st^ MG coordination shell missing (see Notes) |
| MG:DA:4154 | 1.00 | 74.08 |  | 0 |  | 1^st^ MG coordination shell missing (see Notes) |
| MG:DA:4155 | 1.00 | 89.53 |  | 1 | 1 | O_ph_ - Exclusion zone contact(s): /OP2/ |
| MG:DA:4156 | 1.00 | 47.07 |  | 0 | 4 | - Exclusion zone contact(s): /N4/N7/O6/OP2/ Consider K in place of MG (4 valid contact(s) for K vs. 0 for MG) |
| MG:DA:4157 | 1.00 | 37.86 |  | 0 | 5 | - Exclusion zone contact(s): /O3'/O5'/OP1/OP2/OP2/ Consider K in place of MG (5 valid contact(s) for K vs. 0 for MG) |
| MG:DA:4158 | 1.00 | 47.74 |  | 0 | 2 | - Exclusion zone contact(s): /O2'/ Consider K in place of MG (2 valid contact(s) for K vs. 0 for MG) |
| MG:DA:4159 | 1.00 | 62.69 |  | 0 | 2 | - Exclusion zone contact(s): /N7/O6/ Consider K in place of MG (2 valid contact(s) for K vs. 0 for MG) |
| MG:DA:4160 | 1.00 | 53.85 |  | 0 |  | 1^st^ MG coordination shell missing (see Notes) |
| MG:DA:4161 | 1.00 | 61.60 |  | 0 |  | 1^st^ MG coordination shell missing (see Notes) |
| MG:DA:4162 | 1.00 | 59.41 |  | 1 |  | O_ph_ |
| MG:DA:4163 | 1.00 | 55.61 |  | 0 | 2 | - Exclusion zone contact(s): /N7/OP2/ Consider K in place of MG (2 valid contact(s) for K vs. 0 for MG) |
| MG:DA:4164 | 1.00 | 29.54 |  | 1 | 3 | O_b_ - Exclusion zone contact(s): /O4/O6/O6/ Consider K in place of MG (3 valid contact(s) for K vs. 1 for MG) |
| MG:DA:4165 | 1.00 | 35.49 |  | 1 | 2 | O_r_ - Exclusion zone contact(s): /O3'/OP2/ Consider NA in place of MG (2 valid contact(s) for NA vs. 1 for MG) |
| MG:DA:4166 | 1.00 | 41.03 |  | 0 | 5 | - Exclusion zone contact(s): /N2/N3_G_/N6/O2'/O6/ Consider K in place of MG (5 valid contact(s) for K vs. 0 for MG) |
| MG:DA:4167 | 1.00 | 44.85 |  | 0 | 3 | - Exclusion zone contact(s): /O6/OP1/OP2/ Consider K in place of MG (3 valid contact(s) for K vs. 0 for MG) |
| MG:DA:4168 | 1.00 | 33.78 |  | 0 | 5 | - Exclusion zone contact(s): /O3'/O3'/OP1/OP1/OP2/ Consider K in place of MG (5 valid contact(s) for K vs. 0 for MG) |
| MG:DA:4169 | 1.00 | 71.69 |  | 0 | 3 | - Exclusion zone contact(s): /N1/NH1_Arg_/OP2/ Consider K in place of MG (3 valid contact(s) for K vs. 0 for MG) |
| MG:DA:4170 | 1.00 | 46.75 |  | 0 |  | 1^st^ MG coordination shell missing (see Notes) |
| MG:DA:4171 | 1.00 | 45.49 |  | 0 | 1 | - Exclusion zone contact(s): /OP2/ Consider K in place of MG (1 valid contact(s) for K vs. 0 for MG) |
| MG:DA:4172 | 1.00 | 41.20 |  | 1 | 4 | O_ph_ - Exclusion zone contact(s): /N3_U_/O3'/O4/OP1/ Consider K in place of MG (3 valid contact(s) for K vs. 1 for MG) |
| MG:DA:4173 | 1.00 | 60.01 |  | 0 | 2 | - Exclusion zone contact(s): /OP1/OP1/ Consider K in place of MG (2 valid contact(s) for K vs. 0 for MG) |
| MG:DA:4174 | 1.00 | 49.09 |  | 0 |  | 1^st^ MG coordination shell missing (see Notes) |
| MG:DA:4175 | 1.00 | 59.11 |  | 0 | 3 | - Exclusion zone contact(s): /OP1/OP1/OP2/ Consider K in place of MG (3 valid contact(s) for K vs. 0 for MG) |
| MG:DA:4176 | 1.00 | 53.17 |  | 0 | 2 | - Exclusion zone contact(s): /OP1/OP2/ Consider NA in place of MG (2 valid contact(s) for NA vs. 0 for MG) |
| MG:DA:4177 | 1.00 | 52.95 |  | 0 | 1 | - Exclusion zone contact(s): /OP1/ Consider K in place of MG (1 valid contact(s) for K vs. 0 for MG) |
| MG:DA:4178 | 1.00 | 61.49 |  | 0 |  | 1^st^ MG coordination shell missing (see Notes) |
| MG:DA:4179 | 1.00 | 57.75 |  | 1 | 3 | O_ph_ - Exclusion zone contact(s): /O5'/OP2/OP2/ Consider K in place of MG (3 valid contact(s) for K vs. 1 for MG) |
| MG:DA:4180 | 1.00 | 66.49 |  | 0 | 3 | - Exclusion zone contact(s): /O2'/O_back_/OP2/ Consider K in place of MG (3 valid contact(s) for K vs. 0 for MG) |
| MG:DA:4181 | 1.00 | 81.42 |  | 0 | 1 | - Exclusion zone contact(s): /O6/ Consider K in place of MG (1 valid contact(s) for K vs. 0 for MG) |
| MG:DA:4182 | 1.00 | 82.19 |  | 0 |  | 1^st^ MG coordination shell missing (see Notes) |
| MG:DA:4183 | 1.00 | 53.24 |  | 0 | 3 | - Exclusion zone contact(s): /O3'/OP1/OP1/ Consider K in place of MG (3 valid contact(s) for K vs. 0 for MG) |
| MG:DA:4184 | 1.00 | 61.35 |  | 0 |  | 1^st^ MG coordination shell missing (see Notes) |
| MG:DA:4185 | 1.00 | 88.49 |  | 0 | 1 | - Exclusion zone contact(s): /N7/ Consider K in place of MG (1 valid contact(s) for K vs. 0 for MG) |
| MG:DA:4186 | 1.00 | 45.01 |  | 0 | 4 | - Exclusion zone contact(s): /N7/O6/OP2/OP2/ Consider K in place of MG (4 valid contact(s) for K vs. 0 for MG) |
| MG:DA:4187 | 1.00 | 46.37 |  | 0 | 1 | - Exclusion zone contact(s): /OP2/ Consider K in place of MG (1 valid contact(s) for K vs. 0 for MG) |
| MG:DA:4188 | 1.00 | 39.69 |  | 0 | 2 | - Exclusion zone contact(s): /O4/O6/ Consider K in place of MG (2 valid contact(s) for K vs. 0 for MG) |
| MG:DA:4189 | 1.00 | 60.79 |  | 1 | 3 | O_ph_ - Exclusion zone contact(s): /O5'/OP1/OP2/ Consider K in place of MG (3 valid contact(s) for K vs. 1 for MG) |
| MG:DA:4190 | 1.00 | 70.13 |  | 1 | 4 | O_ph_ - Exclusion zone contact(s): /N1_G_/N3_C_/N4/N7/ Consider K in place of MG (4 valid contact(s) for K vs. 1 for MG) |
| MG:DA:4191 | 1.00 | 48.91 |  | 1 |  | N_b_ |
| MG:DA:4192 | 1.00 | 65.32 |  | 0 |  | 1^st^ MG coordination shell missing (see Notes) |
| MG:DA:4193 | 1.00 | 78.81 |  | 0 |  | 1^st^ MG coordination shell missing (see Notes) |
| MG:DA:4194 | 1.00 | 37.93 |  | 0 | 2 | - Exclusion zone contact(s): /O2'/OP2/ Consider K in place of MG (2 valid contact(s) for K vs. 0 for MG) |
| MG:DA:4195 | 1.00 | 66.80 |  | 0 |  | 1^st^ MG coordination shell missing (see Notes) |
| MG:DA:4196 | 1.00 | 43.60 |  | 0 | 4 | - Exclusion zone contact(s): /O3'/O5'/OP2/OP2/ Consider K in place of MG (3 valid contact(s) for K vs. 0 for MG) |
| MG:DA:4197 | 1.00 | 51.47 |  | 1 |  | O_b_ |
| MG:DA:4198 | 1.00 | 45.02 |  | 0 | 4 | - Exclusion zone contact(s): /O2'/OP1/OP1/OP2/ Consider K in place of MG (4 valid contact(s) for K vs. 0 for MG) |
| MG:DA:4199 | 1.00 | 48.53 |  | 0 | 4 | - Exclusion zone contact(s): /N7/N_back_/O2'/OP2/ Consider K in place of MG (4 valid contact(s) for K vs. 0 for MG) |
| MG:DA:4200 | 1.00 | 61.30 |  | 0 | 1 | - Exclusion zone contact(s): /OP2/ Consider K in place of MG (1 valid contact(s) for K vs. 0 for MG) |
| MG:DA:4201 | 1.00 | 57.19 |  | 0 | 2 | - Exclusion zone contact(s): /OP1/OP2/ Consider K in place of MG (2 valid contact(s) for K vs. 0 for MG) |
| MG:DA:4202 | 1.00 | 69.65 |  | 0 |  | 1^st^ MG coordination shell missing (see Notes) |
| MG:DA:4203 | 1.00 | 50.23 |  | 1 | 3 | O_b_ - Exclusion zone contact(s): /O3'/O4/OP1/ Consider K in place of MG (2 valid contact(s) for K vs. 1 for MG) |
| MG:DA:4204 | 1.00 | 45.75 |  | 0 | 3 | - Exclusion zone contact(s): /O4/OP1/OP2/ Consider K in place of MG (3 valid contact(s) for K vs. 0 for MG) |
| MG:DA:4205 | 1.00 | 37.93 |  | 0 | 6 | - Exclusion zone contact(s): /N7/O2'/O3'/O5'/OP2/OP2/ Consider K in place of MG (6 valid contact(s) for K vs. 0 for MG) |
| MG:DA:4206 | 1.00 | 45.37 |  | 0 | 3 | - Exclusion zone contact(s): /OP1/OP2/OP2/ Consider NA in place of MG (2 valid contact(s) for NA vs. 0 for MG) |
| MG:DA:4207 | 1.00 | 84.27 |  | 0 | 1 | - Exclusion zone contact(s): /OP1/ Consider K in place of MG (1 valid contact(s) for K vs. 0 for MG) |
| MG:DA:4208 | 1.00 | 74.14 |  | 1 |  | O_ph_ |
| MG:DA:4209 | 1.00 | 83.85 |  | 0 | 3 | - Exclusion zone contact(s): /NH1_Arg_/O3'/O5'/ Consider K in place of MG (3 valid contact(s) for K vs. 0 for MG) |
| MG:DA:4210 | 1.00 | 53.36 |  | 0 | 4 | - Exclusion zone contact(s): /N3_A_/N4/O2'/O4'/ Consider K in place of MG (3 valid contact(s) for K vs. 0 for MG) |
| MG:DA:4211 | 1.00 | 47.89 |  | 0 | 2 | - Exclusion zone contact(s): /O3'/OP1/ Consider K in place of MG (2 valid contact(s) for K vs. 0 for MG) |
| MG:DA:4212 | 1.00 | 54.55 |  | 0 | 3 | - Exclusion zone contact(s): /OP1/OP2/OP2/ Consider K in place of MG (2 valid contact(s) for K vs. 0 for MG) |
| MG:DA:4213 | 1.00 | 49.05 |  | 0 | 2 | - Exclusion zone contact(s): /OP1/OP2/ Consider NA in place of MG (1 valid contact(s) for NA vs. 0 for MG) |
| MG:DA:4214 | 1.00 | 50.65 |  | 0 | 2 | - Exclusion zone contact(s): /OP2/OP2/ Consider K in place of MG (2 valid contact(s) for K vs. 0 for MG) |
| MG:DA:4215 | 1.00 | 53.12 |  | 0 | 3 | - Exclusion zone contact(s): /O2'/OP1/OP2/ Consider K in place of MG (3 valid contact(s) for K vs. 0 for MG) |
| MG:DA:4216 | 1.00 | 52.79 |  | 0 | 3 | - Exclusion zone contact(s): /O5'/OP1/OP2/ Consider K in place of MG (2 valid contact(s) for K vs. 0 for MG) |
| MG:DA:4217 | 1.00 | 57.65 |  | 0 |  | 1^st^ MG coordination shell missing (see Notes) |
| MG:DA:4218 | 1.00 | 51.40 |  | 0 | 3 | - Exclusion zone contact(s): /O2/O2'/O4'/ Consider K in place of MG (2 valid contact(s) for K vs. 0 for MG) |
| MG:DA:4219 | 1.00 | 60.92 |  | 0 | 2 | - Exclusion zone contact(s): /O4/OP2/ Consider K in place of MG (2 valid contact(s) for K vs. 0 for MG) |
| MG:DA:4220 | 1.00 | 40.84 |  | 1 | 4 | O_r_ - Exclusion zone contact(s): /NZ_Lys_/O3'/OP1/OP2/ Consider K in place of MG (4 valid contact(s) for K vs. 1 for MG) |
| MG:DA:4221 | 1.00 | 49.81 |  | 0 | 1 | - Exclusion zone contact(s): /OP1/ Consider K in place of MG (1 valid contact(s) for K vs. 0 for MG) |
| MG:DA:4222 | 1.00 | 54.54 |  | 0 |  | 1^st^ MG coordination shell missing (see Notes) |
| MG:DA:4223 | 1.00 | 60.01 |  | 0 |  | 1^st^ MG coordination shell missing (see Notes) |
| MG:DA:4224 | 1.00 | 90.68 |  | 0 | 3 | - Exclusion zone contact(s): /N7/O3'/OP1/ Consider K in place of MG (2 valid contact(s) for K vs. 0 for MG) |
| MG:DA:4225 | 1.00 | 83.83 |  | 0 |  | 1^st^ MG coordination shell missing (see Notes) |
| MG:DA:4226 | 1.00 | 45.20 |  | 0 | 3 | - Exclusion zone contact(s): /O2'/O2'/O4'/ Consider K in place of MG (3 valid contact(s) for K vs. 0 for MG) |
| MG:DA:4227 | 1.00 | 51.16 |  | 0 | 1 | - Exclusion zone contact(s): /OP2/ Consider NA in place of MG (1 valid contact(s) for NA vs. 0 for MG) |
| MG:DA:4228 | 1.00 | 50.65 |  | 0 | 2 | - Exclusion zone contact(s): /OP1/OP1/ Consider NA in place of MG (1 valid contact(s) for NA vs. 0 for MG) |
| MG:DA:4229 | 1.00 | 48.27 |  | 0 | 2 | - Exclusion zone contact(s): /N2/O4'/ Consider K in place of MG (2 valid contact(s) for K vs. 0 for MG) |
| MG:DA:4230 | 1.00 | 41.24 |  | 0 | 3 | - Exclusion zone contact(s): /O2'/O5'/OP2/ Consider K in place of MG (3 valid contact(s) for K vs. 0 for MG) |
| MG:DA:4231 | 1.00 | 42.77 |  | 0 | 5 | - Exclusion zone contact(s): /NZ_Lys_/O5'/OG1_Thr_/OP1/OP2/ Consider K in place of MG (4 valid contact(s) for K vs. 0 for MG) |
| MG:DA:4232 | 1.00 | 68.37 |  | 0 | 2 | - Exclusion zone contact(s): /N7/O6/ Consider K in place of MG (2 valid contact(s) for K vs. 0 for MG) |
| MG:DA:4233 | 1.00 | 69.91 |  | 0 | 1 | - Exclusion zone contact(s): /OP1/ Consider NA in place of MG (1 valid contact(s) for NA vs. 0 for MG) |
| MG:DA:4234 | 1.00 | 37.76 |  | 1 | 3 | Anion - Exclusion zone contact(s): /O2'/O4'/OP1/ Consider K in place of MG (3 valid contact(s) for K vs. 1 for MG) |
| MG:DA:4235 | 1.00 | 44.87 |  | 0 | 4 | - Exclusion zone contact(s): /O2'/O3'/O_back_/OP1/ Consider K in place of MG (4 valid contact(s) for K vs. 0 for MG) |
| MG:DA:4236 | 1.00 | 52.63 |  | 0 | 2 | - Exclusion zone contact(s): /N1_A_/N2/ Consider K in place of MG (2 valid contact(s) for K vs. 0 for MG) |
| MG:DA:4237 | 1.00 | 78.00 |  | 0 | 2 | - Exclusion zone contact(s): /N2/OP1/ Consider NA in place of MG (2 valid contact(s) for NA vs. 0 for MG) |
| MG:DA:4238 | 1.00 | 52.72 |  | 0 | 4 | - Exclusion zone contact(s): /N7/O2/OP1/OP2/ Consider K in place of MG (4 valid contact(s) for K vs. 0 for MG) |
| MG:DA:4239 | 1.00 | 62.64 |  | 1 |  | O_ph_ |
| MG:DA:4240 | 1.00 | 70.44 |  | 0 | 1 | - Exclusion zone contact(s): /O6/ Consider K in place of MG (1 valid contact(s) for K vs. 0 for MG) |
| MG:DA:4241 | 1.00 | 55.22 |  | 0 | 2 | - Exclusion zone contact(s): /O5'/OP1/ Consider K in place of MG (2 valid contact(s) for K vs. 0 for MG) |
| MG:DA:4242 | 1.00 | 49.38 |  | 0 | 5 | - Exclusion zone contact(s): /N1_A_/O2'/O3'/OP1/OP1/ Consider K in place of MG (4 valid contact(s) for K vs. 0 for MG) |
| MG:DA:4243 | 1.00 | 59.18 |  | 1 | 1 | O_r_ - Exclusion zone contact(s): /O2'/ |
| MG:DA:4244 | 1.00 | 44.75 |  | 0 | 2 | - Exclusion zone contact(s): /N7/O6/ Consider K in place of MG (2 valid contact(s) for K vs. 0 for MG) |
| MG:DA:4245 | 1.00 | 73.01 |  | 0 | 2 | - Exclusion zone contact(s): /N4/O4/ Consider K in place of MG (2 valid contact(s) for K vs. 0 for MG) |
| MG:DA:4246 | 1.00 | 64.28 |  | 0 | 2 | - Exclusion zone contact(s): /OP2/OP2/ Consider NA in place of MG (1 valid contact(s) for NA vs. 0 for MG) |
| MG:DA:4247 | 1.00 | 48.19 |  | 0 | 2 | - Exclusion zone contact(s): /N1_A_/OP2/ Consider NA in place of MG (1 valid contact(s) for NA vs. 0 for MG) |
| MG:DA:4248 | 1.00 | 89.32 |  | 0 | 1 | - Exclusion zone contact(s): /O2'/ Consider NA in place of MG (1 valid contact(s) for NA vs. 0 for MG) |
| MG:DA:4249 | 1.00 | 55.01 |  | 0 | 2 | - Exclusion zone contact(s): /N3_A_/O2'/ Consider K in place of MG (2 valid contact(s) for K vs. 0 for MG) |
| MG:DA:4250 | 1.00 | 49.20 |  | 0 | 3 | - Exclusion zone contact(s): /O2'/O3'/OP1/ Consider K in place of MG (3 valid contact(s) for K vs. 0 for MG) |
| MG:DA:4251 | 1.00 | 51.42 |  | 2 | 2 | O_ph_N_b_ - Exclusion zone contact(s): /O6/OP2/ |
| MG:DA:4252 | 1.00 | 60.69 |  | 0 | 1 | - Exclusion zone contact(s): /O5'/ Consider K in place of MG (1 valid contact(s) for K vs. 0 for MG) |
| MG:DA:4253 | 1.00 | 62.26 |  | 1 | 3 | O_r_ - Exclusion zone contact(s): /N1_A_/O4'/O4'/ Consider K in place of MG (3 valid contact(s) for K vs. 1 for MG) |
| MG:DA:4254 | 1.00 | 83.64 |  | 0 |  | 1^st^ MG coordination shell missing (see Notes) |
| MG:DA:4255 | 1.00 | 54.93 |  | 0 |  | 1^st^ MG coordination shell missing (see Notes) |
| MG:DA:4256 | 1.00 | 115.49 |  | 0 |  | 1^st^ MG coordination shell missing (see Notes) |
| MG:DA:4257 | 1.00 | 59.71 |  | 0 |  | 1^st^ MG coordination shell missing (see Notes) |
| MG:DA:4258 | 1.00 | 64.35 |  | 0 | 2 | - Exclusion zone contact(s): /N7/N7/ Consider K in place of MG (2 valid contact(s) for K vs. 0 for MG) |
| MG:DA:4259 | 1.00 | 54.92 |  | 0 | 1 | - Exclusion zone contact(s): /N2/ Consider K in place of MG (1 valid contact(s) for K vs. 0 for MG) |
| MG:DA:4260 | 1.00 | 56.08 |  | 0 | 1 | - Exclusion zone contact(s): /N7/ Consider K in place of MG (1 valid contact(s) for K vs. 0 for MG) |
| MG:DA:4261 | 1.00 | 65.02 |  | 0 | 2 | - Exclusion zone contact(s): /OP1/OP1/ Consider NA in place of MG (1 valid contact(s) for NA vs. 0 for MG) |
| MG:DA:4262 | 1.00 | 47.65 |  | 0 | 1 | - Exclusion zone contact(s): /O6/ Consider K in place of MG (1 valid contact(s) for K vs. 0 for MG) |
| MG:DA:4263 | 1.00 | 66.40 |  | 0 | 2 | - Exclusion zone contact(s): /O4/O6/ Consider K in place of MG (2 valid contact(s) for K vs. 0 for MG) |
| MG:DA:4264 | 1.00 | 58.59 |  | 0 | 3 | - Exclusion zone contact(s): /O6/OP2/ Consider K in place of MG (2 valid contact(s) for K vs. 0 for MG) |
| MG:DA:4265 | 1.00 | 95.74 |  | 0 | 1 | - Exclusion zone contact(s): /OP2/ Consider NA in place of MG (1 valid contact(s) for NA vs. 0 for MG) |
| MG:DA:4266 | 1.00 | 64.91 |  | 0 | 2 | - Exclusion zone contact(s): /N1_G_/O6/ Consider K in place of MG (2 valid contact(s) for K vs. 0 for MG) |
| MG:DA:4267 | 1.00 | 72.81 |  | 0 |  | 1^st^ MG coordination shell missing (see Notes) |
| MG:DA:4268 | 1.00 | 58.67 |  | 1 | 1 | O_r_ - Exclusion zone contact(s): /N1/ |
| MG:DA:4269 | 1.00 | 72.00 |  | 0 |  | 1^st^ MG coordination shell missing (see Notes) |
| MG:DA:4270 | 1.00 | 46.51 |  | 0 | 2 | - Exclusion zone contact(s): /OP2/OP2/ Consider K in place of MG (2 valid contact(s) for K vs. 0 for MG) |
| MG:DA:4271 | 1.00 | 53.34 |  | 0 | 3 | - Exclusion zone contact(s): /O2'/O3'/O_back_/ Consider K in place of MG (3 valid contact(s) for K vs. 0 for MG) |
| MG:DA:4272 | 1.00 | 56.70 |  | 0 | 1 | - Exclusion zone contact(s): /OP1/ Consider K in place of MG (1 valid contact(s) for K vs. 0 for MG) |
| MG:DA:4273 | 1.00 | 74.45 |  | 0 | 2 | - Exclusion zone contact(s): /OP2/OP2/ Consider K in place of MG (2 valid contact(s) for K vs. 0 for MG) |
| MG:DA:4274 | 1.00 | 41.10 |  | 0 | 3 | - Exclusion zone contact(s): /N1/O2/O6/ Consider K in place of MG (3 valid contact(s) for K vs. 0 for MG) |
| MG:DA:4275 | 1.00 | 78.73 |  | 1 | 1 | O_ph_ - Exclusion zone contact(s): /OP2/ |
| MG:DA:4276 | 1.00 | 75.10 |  | 0 | 6 | - Exclusion zone contact(s): /N3_G_/NH2_Arg_/O2'/O3'/O5'/OP1/ Consider K in place of MG (6 valid contact(s) for K vs. 0 for MG) |
| MG:DA:4277 | 1.00 | 66.53 |  | 0 | 1 | - Exclusion zone contact(s): /O6/ Consider K in place of MG (1 valid contact(s) for K vs. 0 for MG) |
| MG:DA:4278 | 1.00 | 62.93 |  | 0 | 5 | - Exclusion zone contact(s): /N1_G_/N2/ND2_Asn_/NZ_Lys_/O6/ Consider K in place of MG (5 valid contact(s) for K vs. 0 for MG) |
| MG:DA:4279 | 1.00 | 72.70 |  | 2 | 1 | O_ph_Anion - Exclusion zone contact(s): /OP2/ |
| MG:DA:4280 | 1.00 | 91.61 |  | 1 | 2 | N_b_ - Exclusion zone contact(s): /O6/O6/ Consider K in place of MG (2 valid contact(s) for K vs. 1 for MG) |
| MG:DA:4281 | 1.00 | 51.35 |  | 0 | 2 | - Exclusion zone contact(s): /O4'/OH_Tyr_/ Consider K in place of MG (2 valid contact(s) for K vs. 0 for MG) |
| MG:DA:4282 | 1.00 | 60.27 |  | 0 |  | 1^st^ MG coordination shell missing (see Notes) |
| MG:DA:4283 | 1.00 | 63.07 |  | 1 | 4 | O_ph_ - Exclusion zone contact(s): /N2/N7/O6/OP2/ Consider K in place of MG (4 valid contact(s) for K vs. 1 for MG) |
| MG:DA:4284 | 1.00 | 52.58 |  | 0 | 1 | - Exclusion zone contact(s): /O6/ Consider K in place of MG (1 valid contact(s) for K vs. 0 for MG) |
| MG:DA:4285 | 1.00 | 70.87 |  | 0 |  | 1^st^ MG coordination shell missing (see Notes) |
| MG:DA:4286 | 1.00 | 49.36 |  | 0 | 2 | - Exclusion zone contact(s): /O3'/OP2/ Consider NA in place of MG (1 valid contact(s) for NA vs. 0 for MG) |
| MG:DA:4287 | 1.00 | 73.01 |  | 0 | 1 | - Exclusion zone contact(s): /O6/ Consider NA in place of MG (1 valid contact(s) for NA vs. 0 for MG) |
| MG:DA:4288 | 1.00 | 72.71 |  | 0 | 1 | - Exclusion zone contact(s): /O2'/ Consider NA in place of MG (1 valid contact(s) for NA vs. 0 for MG) |
| MG:DA:4289 | 1.00 | 53.37 |  | 0 |  | 1^st^ MG coordination shell missing (see Notes) |
| MG:DA:4290 | 1.00 | 74.43 |  | 0 |  | 1^st^ MG coordination shell missing (see Notes) |
| MG:DA:4291 | 1.00 | 58.01 |  | 0 | 3 | - Exclusion zone contact(s): /O5'/OP1/OP2/ Consider K in place of MG (3 valid contact(s) for K vs. 0 for MG) |
| MG:DA:4292 | 1.00 | 62.02 |  | 0 | 2 | - Exclusion zone contact(s): /OP1/OP1/ Consider K in place of MG (2 valid contact(s) for K vs. 0 for MG) |
| MG:DA:4293 | 1.00 | 54.01 |  | 0 | 1 | - Exclusion zone contact(s): /N7/ Consider K in place of MG (1 valid contact(s) for K vs. 0 for MG) |
| MG:DA:4294 | 1.00 | 83.98 |  | 2 |  | O_ph_O_r_ |
| MG:DA:4295 | 1.00 | 48.55 |  | 0 | 1 | - Exclusion zone contact(s): /OP2/ Consider K in place of MG (1 valid contact(s) for K vs. 0 for MG) |
| MG:DA:4296 | 1.00 | 105.95 |  | 0 |  | 1^st^ MG coordination shell missing (see Notes) |
| MG:DA:4297 | 1.00 | 100.57 |  | 0 |  | 1^st^ MG coordination shell missing (see Notes) |
| MG:DA:4298 | 1.00 | 62.12 |  | 0 | 6 | - Exclusion zone contact(s): /O3'/O5'/OP1/OP1/OP2/OP2/ Consider K in place of MG (6 valid contact(s) for K vs. 0 for MG) |
| MG:DA:4299 | 1.00 | 72.16 |  | 0 | 4 | - Exclusion zone contact(s): /O4/O6/O6/O6/ Consider NA in place of MG (2 valid contact(s) for NA vs. 0 for MG) |
| MG:DA:4300 | 1.00 | 45.81 |  | 0 | 2 | - Exclusion zone contact(s): /N3_G_/O2'/ Consider NA in place of MG (1 valid contact(s) for NA vs. 0 for MG) |
| MG:DA:4301 | 1.00 | 37.41 |  | 0 | 3 | - Exclusion zone contact(s): /N1_A_/N6/O2'/ Consider K in place of MG (3 valid contact(s) for K vs. 0 for MG) |
| MG:DA:4302 | 1.00 | 102.81 |  | 0 | 2 | - Exclusion zone contact(s): /N7/O5'/ Consider K in place of MG (2 valid contact(s) for K vs. 0 for MG) |
| MG:DA:4303 | 1.00 | 47.98 |  | 0 | 1 | - Exclusion zone contact(s): /OP2/ Consider K in place of MG (1 valid contact(s) for K vs. 0 for MG) |
| MG:DA:4304 | 1.00 | 94.98 |  | 1 | 3 | O_ph_ - Exclusion zone contact(s): /N7/O2'/O3'/ Consider K in place of MG (3 valid contact(s) for K vs. 1 for MG) |
| MG:DA:4305 | 1.00 | 46.08 |  | 0 | 1 | - Exclusion zone contact(s): /N1_G_/ Consider K in place of MG (1 valid contact(s) for K vs. 0 for MG) |
| MG:DA:4306 | 1.00 | 45.56 |  | 1 | 3 | O_ph_ - Exclusion zone contact(s): /O2'/O3'/O3'/ Consider K in place of MG (3 valid contact(s) for K vs. 1 for MG) |
| MG:DA:4307 | 1.00 | 57.32 |  | 0 | 1 | - Exclusion zone contact(s): /N7/ Consider K in place of MG (1 valid contact(s) for K vs. 0 for MG) |
| MG:DA:4308 | 1.00 | 71.57 |  | 0 | 1 | - Exclusion zone contact(s): /OP2/ Consider K in place of MG (1 valid contact(s) for K vs. 0 for MG) |
| MG:DA:4309 | 1.00 | 36.30 |  | 0 | 1 | - Exclusion zone contact(s): /OP1/ Consider K in place of MG (1 valid contact(s) for K vs. 0 for MG) |
| MG:DA:4310 | 1.00 | 56.61 |  | 2 | 3 | O_ph_O_r_ - Exclusion zone contact(s): /O3'/O4/OP1/ |
| MG:DA:4311 | 1.00 | 83.40 |  | 0 | 1 | - Exclusion zone contact(s): /N7/ Consider K in place of MG (1 valid contact(s) for K vs. 0 for MG) |
| MG:DA:4312 | 1.00 | 49.06 |  | 0 | 4 | - Exclusion zone contact(s): /OP1/OP2/OP2/OP2/ Consider K in place of MG (4 valid contact(s) for K vs. 0 for MG) |
| MG:DA:4313 | 1.00 | 56.90 |  | 0 | 4 | - Exclusion zone contact(s): /O3'/OP1/OP1/OP2/ Consider K in place of MG (3 valid contact(s) for K vs. 0 for MG) |
| MG:DA:4314 | 1.00 | 67.23 |  | 0 | 2 | - Exclusion zone contact(s): /OP1/OP2/ Consider K in place of MG (2 valid contact(s) for K vs. 0 for MG) |
| MG:DA:4315 | 1.00 | 67.64 |  | 1 | 1 | Anion - Exclusion zone contact(s): /N4/ |
| MG:DA:4316 | 1.00 | 67.89 |  | 0 | 1 | - Exclusion zone contact(s): /N7/ Consider K in place of MG (1 valid contact(s) for K vs. 0 for MG) |
| MG:DA:4317 | 1.00 | 61.86 |  | 0 | 2 | - Exclusion zone contact(s): /O2/O2'/ Consider K in place of MG (2 valid contact(s) for K vs. 0 for MG) |
| MG:DA:4318 | 1.00 | 50.06 |  | 0 | 5 | - Exclusion zone contact(s): /N1_A_/N6/O2'/O4'/ Consider K in place of MG (5 valid contact(s) for K vs. 0 for MG) |
| MG:DA:4319 | 1.00 | 67.39 |  | 0 |  | 1^st^ MG coordination shell missing (see Notes) |
| MG:DA:4320 | 1.00 | 50.83 |  | 1 |  | O_ph_ |
| MG:DA:4321 | 1.00 | 70.28 |  | 0 | 2 | - Exclusion zone contact(s): /O6/OP2/ Consider K in place of MG (2 valid contact(s) for K vs. 0 for MG) |
| MG:DA:4322 | 1.00 | 65.62 |  | 0 | 3 | - Exclusion zone contact(s): /N1/N1/O2/ Consider K in place of MG (3 valid contact(s) for K vs. 0 for MG) |
| MG:DA:4323 | 1.00 | 56.35 |  | 0 | 3 | - Exclusion zone contact(s): /O2'/O3'/OP1/ Consider K in place of MG (2 valid contact(s) for K vs. 0 for MG) |
| MG:DA:4324 | 1.00 | 49.93 |  | 0 | 3 | - Exclusion zone contact(s): /N3_A_/O2'/O4'/ Consider K in place of MG (2 valid contact(s) for K vs. 0 for MG) |
| MG:DA:4325 | 1.00 | 45.30 |  | 1 | 6 | Anion - Exclusion zone contact(s): /N2/N6/N7/O6/O6/OP2/ Consider K in place of MG (6 valid contact(s) for K vs. 1 for MG) |
| MG:DA:4326 | 1.00 | 56.08 |  | 0 | 1 | - Exclusion zone contact(s): /N7/ Consider K in place of MG (1 valid contact(s) for K vs. 0 for MG) |
| MG:DA:4327 | 1.00 | 51.48 |  | 0 | 1 | - Exclusion zone contact(s): /O2/ Consider K in place of MG (1 valid contact(s) for K vs. 0 for MG) |
| MG:DA:4328 | 1.00 | 59.80 |  | 0 | 1 | - Exclusion zone contact(s): /O2/ Consider K in place of MG (1 valid contact(s) for K vs. 0 for MG) |
| MG:DA:4329 | 1.00 | 63.13 |  | 0 | 1 | - Exclusion zone contact(s): /OP1/ Consider K in place of MG (1 valid contact(s) for K vs. 0 for MG) |
| MG:DA:4330 | 1.00 | 56.77 |  | 0 |  | 1^st^ MG coordination shell missing (see Notes) |
| MG:DA:4331 | 1.00 | 63.02 |  | 1 | 3 | N_b_ - Exclusion zone contact(s): /N7/O4'/ Consider K in place of MG (3 valid contact(s) for K vs. 1 for MG) |
| MG:DA:4332 | 1.00 | 53.05 |  | 0 | 4 | - Exclusion zone contact(s): /N3_A_/O2/O3'/OP1/ Consider K in place of MG (4 valid contact(s) for K vs. 0 for MG) |
| MG:DA:4333 | 1.00 | 52.65 |  | 0 | 1 | - Exclusion zone contact(s): /O2'/ Consider K in place of MG (1 valid contact(s) for K vs. 0 for MG) |
| MG:DA:4334 | 1.00 | 70.68 |  | 0 | 3 | - Exclusion zone contact(s): /O2'/O3'/OP1/ Consider K in place of MG (3 valid contact(s) for K vs. 0 for MG) |
| MG:DA:4335 | 1.00 | 50.64 |  | 0 |  | 1^st^ MG coordination shell missing (see Notes) |
| MG:DA:4336 | 1.00 | 73.32 |  | 0 | 1 | - Exclusion zone contact(s): /O6/ Consider K in place of MG (1 valid contact(s) for K vs. 0 for MG) |
| MG:DA:4337 | 1.00 | 58.47 |  | 0 | 1 | - Exclusion zone contact(s): /OP1/ Consider K in place of MG (1 valid contact(s) for K vs. 0 for MG) |
| MG:DA:4338 | 1.00 | 80.81 |  | 0 | 3 | - Exclusion zone contact(s): /N2/O3'/OP1/ Consider K in place of MG (3 valid contact(s) for K vs. 0 for MG) |
| MG:DA:4339 | 1.00 | 71.52 |  | 0 | 2 | - Exclusion zone contact(s): /O3'/OP1/ Consider K in place of MG (2 valid contact(s) for K vs. 0 for MG) |
| MG:DA:4340 | 1.00 | 60.24 |  | 0 | 1 | - Exclusion zone contact(s): /OP2/ Consider K in place of MG (1 valid contact(s) for K vs. 0 for MG) |
| MG:DA:4341 | 1.00 | 59.28 |  | 0 | 2 | - Exclusion zone contact(s): /O6/OP1/ Consider NA in place of MG (1 valid contact(s) for NA vs. 0 for MG) |
| MG:DA:4342 | 1.00 | 93.72 |  | 0 | 1 | - Exclusion zone contact(s): /N1_A_/ Consider NA in place of MG (1 valid contact(s) for NA vs. 0 for MG) |
| MG:DA:4343 | 1.00 | 58.31 |  | 0 |  | 1^st^ MG coordination shell missing (see Notes) |
| MG:DA:4344 | 1.00 | 63.74 |  | 0 |  | 1^st^ MG coordination shell missing (see Notes) |
| MG:DA:4345 | 1.00 | 58.58 |  | 0 | 3 | - Exclusion zone contact(s): /O2/O3'/OP1/ Consider K in place of MG (3 valid contact(s) for K vs. 0 for MG) |
| MG:DA:4346 | 1.00 | 44.76 |  | 0 | 4 | - Exclusion zone contact(s): /N7/O5'/OP2/OP2/ Consider K in place of MG (4 valid contact(s) for K vs. 0 for MG) |
| MG:DA:4347 | 1.00 | 58.09 |  | 0 | 4 | - Exclusion zone contact(s): /O4/O5'/OP1/OP2/ Consider K in place of MG (4 valid contact(s) for K vs. 0 for MG) |
| MG:DA:4348 | 1.00 | 62.84 |  | 0 |  | 1^st^ MG coordination shell missing (see Notes) |
| MG:DA:4349 | 1.00 | 77.14 |  | 0 | 1 | - Exclusion zone contact(s): /O6/ Consider K in place of MG (1 valid contact(s) for K vs. 0 for MG) |
| MG:DA:4350 | 1.00 | 54.14 |  | 1 | 2 | O_r_ - Exclusion zone contact(s): /O2/O4'/ Consider K in place of MG (2 valid contact(s) for K vs. 1 for MG) |
| MG:DA:4351 | 1.00 | 43.61 |  | 0 | 1 | - Exclusion zone contact(s): /O2/ Consider K in place of MG (1 valid contact(s) for K vs. 0 for MG) |
| MG:DA:4352 | 1.00 | 69.63 |  | 0 | 1 | - Exclusion zone contact(s): /N3_C_/ Consider K in place of MG (1 valid contact(s) for K vs. 0 for MG) |
| MG:DA:4353 | 1.00 | 45.78 |  | 0 | 3 | - Exclusion zone contact(s): /O4'/OD1_Asp_/ Consider K in place of MG (3 valid contact(s) for K vs. 0 for MG) |
| MG:DA:4354 | 1.00 | 65.40 |  | 0 |  | 1^st^ MG coordination shell missing (see Notes) |
| MG:DA:4355 | 1.00 | 79.38 |  | 0 |  | 1^st^ MG coordination shell missing (see Notes) |
| MG:DA:4356 | 1.00 | 76.36 |  | 0 |  | 1^st^ MG coordination shell missing (see Notes) |
| MG:DA:4357 | 1.00 | 53.63 |  | 0 | 1 | - Exclusion zone contact(s): /N4/ Consider K in place of MG (1 valid contact(s) for K vs. 0 for MG) |
| MG:DA:4358 | 1.00 | 56.63 |  | 1 | 3 | O_r_ - Exclusion zone contact(s): /O2'/OP1/OP2/ Consider K in place of MG (3 valid contact(s) for K vs. 1 for MG) |
| MG:DA:4359 | 1.00 | 47.75 |  | 0 | 1 | - Exclusion zone contact(s): /O2/ Consider K in place of MG (1 valid contact(s) for K vs. 0 for MG) |
| MG:DA:4360 | 1.00 | 57.24 |  | 1 | 1 | O_b_ - Exclusion zone contact(s): /N4/ |
| MG:DA:4361 | 1.00 | 47.57 |  | 0 | 3 | - Exclusion zone contact(s): /O5'/OP2/OP2/ Consider K in place of MG (3 valid contact(s) for K vs. 0 for MG) |
| MG:DA:4362 | 1.00 | 74.32 |  | 0 | 2 | - Exclusion zone contact(s): /O3'/OP1/ Consider K in place of MG (2 valid contact(s) for K vs. 0 for MG) |
| MG:DA:4363 | 1.00 | 42.57 |  | 0 | 3 | - Exclusion zone contact(s): /O2/OP1/OP2/ Consider K in place of MG (3 valid contact(s) for K vs. 0 for MG) |
| MG:DA:4364 | 1.00 | 90.75 |  | 0 | 2 | - Exclusion zone contact(s): /O2'/O5'/ Consider K in place of MG (2 valid contact(s) for K vs. 0 for MG) |
| MG:DA:4365 | 1.00 | 61.96 |  | 0 | 1 | - Exclusion zone contact(s): /N7/ Consider K in place of MG (1 valid contact(s) for K vs. 0 for MG) |
| MG:DA:4366 | 1.00 | 62.45 |  | 0 | 2 | - Exclusion zone contact(s): /OP1/OP2/ Consider K in place of MG (2 valid contact(s) for K vs. 0 for MG) |
| MG:DA:4367 | 1.00 | 65.78 |  | 0 | 3 | - Exclusion zone contact(s): /O3'/O5'/OP1/ Consider K in place of MG (3 valid contact(s) for K vs. 0 for MG) |
| MG:DA:4368 | 1.00 | 51.05 |  | 0 |  | 1^st^ MG coordination shell missing (see Notes) |
| MG:DA:4369 | 1.00 | 78.53 |  | 0 | 1 | - Exclusion zone contact(s): /O4'/ Consider K in place of MG (1 valid contact(s) for K vs. 0 for MG) |
| MG:DA:4370 | 1.00 | 74.49 |  | 1 |  | O_ph_ |
| MG:DA:4371 | 1.00 | 57.96 |  | 0 | 2 | - Exclusion zone contact(s): /O2/O_back_/ Consider K in place of MG (2 valid contact(s) for K vs. 0 for MG) |
| MG:DA:4372 | 1.00 | 53.44 |  | 0 |  | 1^st^ MG coordination shell missing (see Notes) |
| MG:DA:4373 | 1.00 | 62.16 |  | 1 | 2 | O_ph_ - Exclusion zone contact(s): /N7/O6/ Consider K in place of MG (2 valid contact(s) for K vs. 1 for MG) |
| MG:DA:4374 | 1.00 | 77.60 |  | 0 |  | 1^st^ MG coordination shell missing (see Notes) |
| MG:DA:4375 | 1.00 | 52.18 |  | 0 | 2 | - Exclusion zone contact(s): /OP1/OP2/ Consider K in place of MG (2 valid contact(s) for K vs. 0 for MG) |
| MG:DA:4376 | 1.00 | 60.09 |  | 0 | 2 | - Exclusion zone contact(s): /O2/OP2/ Consider K in place of MG (2 valid contact(s) for K vs. 0 for MG) |
| MG:DA:4377 | 1.00 | 46.89 |  | 0 | 2 | - Exclusion zone contact(s): /N2/OP1/ Consider K in place of MG (2 valid contact(s) for K vs. 0 for MG) |
| MG:DA:4378 | 1.00 | 48.71 |  | 0 | 1 | - Exclusion zone contact(s): /O2'/ Consider K in place of MG (1 valid contact(s) for K vs. 0 for MG) |
| MG:DA:4379 | 1.00 | 74.40 |  | 1 | 3 | O_ph_ - Exclusion zone contact(s): /N4/O5'/OP2/ Consider K in place of MG (3 valid contact(s) for K vs. 1 for MG) |
| MG:DA:4380 | 1.00 | 61.64 |  | 0 |  | 1^st^ MG coordination shell missing (see Notes) |
| MG:DA:4381 | 1.00 | 88.08 |  | 0 | 2 | - Exclusion zone contact(s): /O4/O6/ Consider K in place of MG (2 valid contact(s) for K vs. 0 for MG) |
| MG:DA:4382 | 1.00 | 67.36 |  | 0 |  | 1^st^ MG coordination shell missing (see Notes) |
| MG:DA:4383 | 1.00 | 85.52 |  | 0 |  | 1^st^ MG coordination shell missing (see Notes) |
| MG:DA:4384 | 1.00 | 64.53 |  | 0 | 2 | - Exclusion zone contact(s): /O2'/O3'/ Consider K in place of MG (2 valid contact(s) for K vs. 0 for MG) |
| MG:DA:4385 | 1.00 | 55.37 |  | 0 | 2 | - Exclusion zone contact(s): /OP1/OP1/ Consider K in place of MG (2 valid contact(s) for K vs. 0 for MG) |
| MG:DA:4386 | 1.00 | 49.24 |  | 0 | 2 | - Exclusion zone contact(s): /O4/O6/ Consider K in place of MG (2 valid contact(s) for K vs. 0 for MG) |
| MG:DA:4387 | 1.00 | 57.61 |  | 0 | 1 | - Exclusion zone contact(s): /O2'/ Consider K in place of MG (1 valid contact(s) for K vs. 0 for MG) |
| MG:DA:4388 | 1.00 | 80.36 |  | 0 | 2 | - Exclusion zone contact(s): /O2'/O4'/ Consider K in place of MG (2 valid contact(s) for K vs. 0 for MG) |
| MG:DA:4389 | 1.00 | 60.37 |  | 0 |  | 1^st^ MG coordination shell missing (see Notes) |
| MG:DA:4390 | 1.00 | 53.22 |  | 0 |  | 1^st^ MG coordination shell missing (see Notes) |
| MG:DA:4391 | 1.00 | 52.21 |  | 0 | 4 | - Exclusion zone contact(s): /O2'/O3'/O5'/OP1/ Consider K in place of MG (4 valid contact(s) for K vs. 0 for MG) |
| MG:DA:4392 | 1.00 | 52.61 |  | 0 | 1 | - Exclusion zone contact(s): /O2'/ Consider NA in place of MG (1 valid contact(s) for NA vs. 0 for MG) |
| MG:DA:4393 | 1.00 | 53.57 |  | 0 | 3 | - Exclusion zone contact(s): /N7/O2'/O3'/ Consider K in place of MG (3 valid contact(s) for K vs. 0 for MG) |
| MG:DA:4394 | 1.00 | 69.38 |  | 0 |  | 1^st^ MG coordination shell missing (see Notes) |
| MG:DA:4395 | 1.00 | 46.97 |  | 0 |  | 1^st^ MG coordination shell missing (see Notes) |
| MG:DA:4396 | 1.00 | 85.63 |  | 0 |  | 1^st^ MG coordination shell missing (see Notes) |
| MG:DA:4397 | 1.00 | 56.47 |  | 0 | 1 | - Exclusion zone contact(s): /OP1/ Consider K in place of MG (1 valid contact(s) for K vs. 0 for MG) |
| MG:DA:4398 | 1.00 | 46.33 |  | 0 | 1 | - Exclusion zone contact(s): /OP2/ Consider K in place of MG (1 valid contact(s) for K vs. 0 for MG) |
| MG:DA:4399 | 1.00 | 47.07 |  | 0 | 3 | - Exclusion zone contact(s): /O3'/OG1_Thr_/OP2/ Consider K in place of MG (3 valid contact(s) for K vs. 0 for MG) |
| MG:DA:4400 | 1.00 | 83.04 |  | 0 | 3 | - Exclusion zone contact(s): /OP1/OP2/OP2/ Consider K in place of MG (2 valid contact(s) for K vs. 0 for MG) |
| MG:DA:4401 | 1.00 | 87.52 |  | 0 | 2 | - Exclusion zone contact(s): /OP1/OP2/ Consider K in place of MG (2 valid contact(s) for K vs. 0 for MG) |
| MG:DA:4402 | 1.00 | 35.48 |  | 0 | 2 | - Exclusion zone contact(s): /NH1_Arg_/OP1/ Consider K in place of MG (2 valid contact(s) for K vs. 0 for MG) |
| MG:DA:4403 | 1.00 | 83.38 |  | 0 | 1 | - Exclusion zone contact(s): /OP1/ Consider K in place of MG (1 valid contact(s) for K vs. 0 for MG) |
| MG:DA:4404 | 1.00 | 68.99 |  | 0 | 2 | - Exclusion zone contact(s): /O2'/O6/ Consider NA in place of MG (1 valid contact(s) for NA vs. 0 for MG) |
| MG:DA:4405 | 1.00 | 40.82 |  | 1 | 3 | O_b_ - Exclusion zone contact(s): /O4/O4/O6/ Consider K in place of MG (3 valid contact(s) for K vs. 1 for MG) |
| MG:DA:4406 | 1.00 | 54.35 |  | 0 | 2 | - Exclusion zone contact(s): /O2'/O3'/ Consider K in place of MG (2 valid contact(s) for K vs. 0 for MG) |
| MG:DA:4407 | 1.00 | 85.26 |  | 0 |  | 1^st^ MG coordination shell missing (see Notes) |
| MG:DA:4408 | 1.00 | 54.35 |  | 0 |  | 1^st^ MG coordination shell missing (see Notes) |
| MG:DA:4409 | 1.00 | 75.75 |  | 0 | 2 | - Exclusion zone contact(s): /OP1/OP1/ Consider NA in place of MG (1 valid contact(s) for NA vs. 0 for MG) |
| MG:DA:4410 | 1.00 | 59.56 |  | 0 | 4 | - Exclusion zone contact(s): /N3_G_/O2'/O4'/OD1_Asn_/ Consider K in place of MG (3 valid contact(s) for K vs. 0 for MG) |
| MG:DA:4411 | 1.00 | 42.73 |  | 0 | 3 | - Exclusion zone contact(s): /OP1/OP1/OP2/ Consider K in place of MG (2 valid contact(s) for K vs. 0 for MG) |
| MG:DA:4412 | 1.00 | 82.86 |  | 0 | 2 | - Exclusion zone contact(s): /N4/N7/ Consider K in place of MG (2 valid contact(s) for K vs. 0 for MG) |
| MG:DA:4413 | 1.00 | 65.24 |  | 0 | 1 | - Exclusion zone contact(s): /N6/ Consider K in place of MG (1 valid contact(s) for K vs. 0 for MG) |
| MG:DA:4414 | 1.00 | 59.69 |  | 0 | 3 | - Exclusion zone contact(s): /O4/O6/O6/ Consider K in place of MG (3 valid contact(s) for K vs. 0 for MG) |
| MG:DA:4415 | 1.00 | 74.98 |  | 0 | 3 | - Exclusion zone contact(s): /O5'/OP2/OP2/ Consider K in place of MG (2 valid contact(s) for K vs. 0 for MG) |
| MG:DA:4416 | 1.00 | 74.81 |  | 0 | 1 | - Exclusion zone contact(s): /OP2/ Consider NA in place of MG (1 valid contact(s) for NA vs. 0 for MG) |
| MG:DA:4417 | 1.00 | 64.17 |  | 0 | 2 | - Exclusion zone contact(s): /OP2/OP2/ Consider K in place of MG (2 valid contact(s) for K vs. 0 for MG) |
| MG:DA:4418 | 1.00 | 96.36 |  | 0 | 1 | - Exclusion zone contact(s): /OP1/ Consider K in place of MG (1 valid contact(s) for K vs. 0 for MG) |
| MG:DA:4419 | 1.00 | 57.16 |  | 0 |  | 1^st^ MG coordination shell missing (see Notes) |
| MG:DA:4420 | 1.00 | 40.68 |  | 0 | 2 | - Exclusion zone contact(s): /O3'/OP1/ Consider K in place of MG (2 valid contact(s) for K vs. 0 for MG) |
| MG:DA:4421 | 1.00 | 64.19 |  | 0 |  | 1^st^ MG coordination shell missing (see Notes) |
| MG:DA:4422 | 1.00 | 71.15 |  | 1 | 1 | O_ph_ - Exclusion zone contact(s): /N7/ |
| MG:DA:4423 | 1.00 | 73.20 |  | 0 | 1 | - Exclusion zone contact(s): /OP2/ Consider K in place of MG (1 valid contact(s) for K vs. 0 for MG) |
| MG:DA:4424 | 1.00 | 46.98 |  | 0 | 1 | - Exclusion zone contact(s): /OP2/ Consider K in place of MG (1 valid contact(s) for K vs. 0 for MG) |
| MG:DA:4425 | 1.00 | 109.22 |  | 0 | 3 | - Exclusion zone contact(s): /NH2_Arg_/O3'/O_back_/ Consider K in place of MG (2 valid contact(s) for K vs. 0 for MG) |
| MG:DA:4426 | 1.00 | 75.25 |  | 0 | 4 | - Exclusion zone contact(s): /O2'/O3'/O4/OP2/ Consider NA in place of MG (2 valid contact(s) for NA vs. 0 for MG) |
| MG:DA:4427 | 1.00 | 53.31 |  | 0 | 1 | - Exclusion zone contact(s): /OP1/ Consider K in place of MG (1 valid contact(s) for K vs. 0 for MG) |
| MG:DA:4428 | 1.00 | 64.11 |  | 0 |  | 1^st^ MG coordination shell missing (see Notes) |
| MG:DA:4429 | 1.00 | 55.93 |  | 0 | 3 | - Exclusion zone contact(s): /O5'/OP2/OP2/ Consider K in place of MG (3 valid contact(s) for K vs. 0 for MG) |
| MG:DA:4430 | 1.00 | 83.90 |  | 0 | 3 | - Exclusion zone contact(s): /N2/N3_G_/O2'/ Consider K in place of MG (2 valid contact(s) for K vs. 0 for MG) |
| MG:DA:4431 | 1.00 | 87.89 |  | 0 | 1 | - Exclusion zone contact(s): /N6/ Consider K in place of MG (1 valid contact(s) for K vs. 0 for MG) |
| MG:DA:4432 | 1.00 | 58.75 |  | 0 | 1 | - Exclusion zone contact(s): /O6/ Consider K in place of MG (1 valid contact(s) for K vs. 0 for MG) |
| MG:DA:4433 | 1.00 | 90.56 |  | 0 | 2 | - Exclusion zone contact(s): /O5'/OP2/ Consider K in place of MG (2 valid contact(s) for K vs. 0 for MG) |
| MG:DA:4434 | 1.00 | 59.61 |  | 0 | 3 | - Exclusion zone contact(s): /N6/OP1/OP2/ Consider K in place of MG (2 valid contact(s) for K vs. 0 for MG) |
| MG:DA:4435 | 1.00 | 53.40 |  | 2 | 2 | 2O_ph_ - Exclusion zone contact(s): /N7/O2'/ |
| MG:DA:4436 | 1.00 | 76.49 |  | 0 | 2 | - Exclusion zone contact(s): /N1_G_/N7/ Consider NA in place of MG (1 valid contact(s) for NA vs. 0 for MG) |
| MG:DA:4437 | 1.00 | 27.67 |  | 0 | 4 | - Exclusion zone contact(s): /O2/O2'/O4'/ Consider K in place of MG (3 valid contact(s) for K vs. 0 for MG) |
| MG:DA:4438 | 1.00 | 89.43 |  | 0 |  | 1^st^ MG coordination shell missing (see Notes) |
| MG:DA:4439 | 1.00 | 58.15 |  | 0 | 1 | - Exclusion zone contact(s): /OP1/ Consider K in place of MG (1 valid contact(s) for K vs. 0 for MG) |
| MG:DA:4440 | 1.00 | 54.62 |  | 0 | 3 | - Exclusion zone contact(s): /N3_G_/O2/O2'/ Consider K in place of MG (3 valid contact(s) for K vs. 0 for MG) |
| MG:DA:4441 | 1.00 | 62.16 |  | 0 | 2 | - Exclusion zone contact(s): /O3'/OP1/ Consider K in place of MG (2 valid contact(s) for K vs. 0 for MG) |
| MG:DA:4442 | 1.00 | 49.35 |  | 0 | 4 | - Exclusion zone contact(s): /N1/N1_A_/OG1_Thr_/OP2/ Consider K in place of MG (4 valid contact(s) for K vs. 0 for MG) |
| MG:DA:4443 | 1.00 | 36.48 |  | 0 | 4 | - Exclusion zone contact(s): /N2/N3_G_/O2/O2'/ Consider K in place of MG (4 valid contact(s) for K vs. 0 for MG) |
| MG:DA:4444 | 1.00 | 51.76 |  | 1 | 1 | O_ph_ - Exclusion zone contact(s): /O3'/ |
| MG:DA:4445 | 1.00 | 21.28 |  | 0 | 2 | - Exclusion zone contact(s): /OP1/OP1/ Consider K in place of MG (2 valid contact(s) for K vs. 0 for MG) |
| MG:DA:4446 | 1.00 | 54.52 |  | 0 | 2 | - Exclusion zone contact(s): /O6/O6/ Consider K in place of MG (2 valid contact(s) for K vs. 0 for MG) |
| MG:DA:4447 | 1.00 | 63.61 |  | 0 | 3 | - Exclusion zone contact(s): /N3_G_/O2'/O4'/ Consider K in place of MG (3 valid contact(s) for K vs. 0 for MG) |
| MG:DA:4448 | 1.00 | 95.98 |  | 0 | 1 | - Exclusion zone contact(s): /O6/ Consider NA in place of MG (1 valid contact(s) for NA vs. 0 for MG) |
| MG:DA:4449 | 1.00 | 64.95 |  | 0 |  | 1^st^ MG coordination shell missing (see Notes) |
| MG:DA:4450 | 1.00 | 68.59 |  | 0 | 2 | - Exclusion zone contact(s): /N6/O2'/ Consider K in place of MG (2 valid contact(s) for K vs. 0 for MG) |
| MG:DA:4451 | 1.00 | 63.24 |  | 0 |  | 1^st^ MG coordination shell missing (see Notes) |
| MG:DA:4452 | 1.00 | 50.55 |  | 0 | 3 | - Exclusion zone contact(s): /N3_A_/O2'/OP2/ Consider K in place of MG (2 valid contact(s) for K vs. 0 for MG) |
| MG:DA:4453 | 1.00 | 49.66 |  | 1 | 4 | O_ph_ - Exclusion zone contact(s): /N1/O4'/O5'/OP1/ Consider K in place of MG (4 valid contact(s) for K vs. 1 for MG) |
| MG:DA:4454 | 1.00 | 51.97 |  | 0 | 3 | - Exclusion zone contact(s): /O2/O3'/OP1/ Consider K in place of MG (3 valid contact(s) for K vs. 0 for MG) |
| MG:DA:4455 | 1.00 | 45.93 |  | 0 | 1 | - Exclusion zone contact(s): /O2'/ Consider K in place of MG (1 valid contact(s) for K vs. 0 for MG) |
| MG:DA:4456 | 1.00 | 59.52 |  | 0 | 2 | - Exclusion zone contact(s): /N1_G_/N2/ Consider K in place of MG (2 valid contact(s) for K vs. 0 for MG) |
| MG:DA:4457 | 1.00 | 75.37 |  | 1 | 1 | O_r_ - Exclusion zone contact(s): /O3'/ |
| MG:DA:4458 | 1.00 | 51.82 |  | 0 | 4 | - Exclusion zone contact(s): /N2/N3_G_/O2'/O4'/ Consider K in place of MG (3 valid contact(s) for K vs. 0 for MG) |
| MG:DA:4459 | 1.00 | 35.63 |  | 0 | 3 | - Exclusion zone contact(s): /N3_A_/O2'/O2'/ Consider K in place of MG (3 valid contact(s) for K vs. 0 for MG) |
| MG:DA:4460 | 1.00 | 61.26 |  | 1 | 2 | O_ph_ - Exclusion zone contact(s): /O3'/OP1/ |
| MG:DA:4461 | 1.00 | 88.52 |  | 0 | 1 | - Exclusion zone contact(s): /N7/ Consider K in place of MG (1 valid contact(s) for K vs. 0 for MG) |
| MG:DA:4462 | 1.00 | 74.54 |  | 0 | 3 | - Exclusion zone contact(s): /O6/OP2/OP2/ Consider K in place of MG (3 valid contact(s) for K vs. 0 for MG) |
| MG:DA:4463 | 1.00 | 62.63 |  | 0 | 4 | - Exclusion zone contact(s): /O2'/O3'/O5'/OP2/ Consider K in place of MG (4 valid contact(s) for K vs. 0 for MG) |
| MG:DA:4464 | 1.00 | 51.43 |  | 0 |  | 1^st^ MG coordination shell missing (see Notes) |
| MG:DA:4465 | 1.00 | 48.90 |  | 0 | 2 | - Exclusion zone contact(s): /OP2/ Consider K in place of MG (2 valid contact(s) for K vs. 0 for MG) |
| MG:DA:4466 | 1.00 | 73.81 |  | 0 |  | 1^st^ MG coordination shell missing (see Notes) |
| MG:DA:4467 | 1.00 | 50.62 |  | 0 | 5 | - Exclusion zone contact(s): /O2'/O3'/O4'/O5'/OP1/ Consider K in place of MG (5 valid contact(s) for K vs. 0 for MG) |
| MG:DA:4468 | 1.00 | 60.87 |  | 0 |  | 1^st^ MG coordination shell missing (see Notes) |
| MG:DA:4469 | 1.00 | 52.12 |  | 0 | 3 | - Exclusion zone contact(s): /N2/N6/N7/ Consider K in place of MG (3 valid contact(s) for K vs. 0 for MG) |
| MG:DA:4470 | 1.00 | 41.23 |  | 0 |  | 1^st^ MG coordination shell missing (see Notes) |
| MG:DA:4471 | 1.00 | 46.76 |  | 0 | 2 | - Exclusion zone contact(s): /N_back_/OP1/ Consider K in place of MG (2 valid contact(s) for K vs. 0 for MG) |
| MG:DA:4472 | 1.00 | 56.25 |  | 0 | 2 | - Exclusion zone contact(s): /O3'/OP1/ Consider K in place of MG (2 valid contact(s) for K vs. 0 for MG) |
| MG:DA:4473 | 1.00 | 60.69 |  | 0 | 2 | - Exclusion zone contact(s): /N6/N7/ Consider K in place of MG (2 valid contact(s) for K vs. 0 for MG) |
| MG:DA:4474 | 1.00 | 54.47 |  | 0 | 4 | - Exclusion zone contact(s): /N2/N3_G_/O2'/O4'/ Consider K in place of MG (4 valid contact(s) for K vs. 0 for MG) |
| MG:DA:4475 | 1.00 | 50.67 |  | 0 |  | 1^st^ MG coordination shell missing (see Notes) |
| MG:DA:4476 | 1.00 | 64.90 |  | 0 | 2 | - Exclusion zone contact(s): /OP1/OP1/ Consider K in place of MG (2 valid contact(s) for K vs. 0 for MG) |
| MG:DA:4477 | 1.00 | 61.05 |  | 0 | 1 | - Exclusion zone contact(s): /O6/ Consider NA in place of MG (1 valid contact(s) for NA vs. 0 for MG) |
| MG:DA:4478 | 1.00 | 65.14 |  | 0 | 2 | - Exclusion zone contact(s): /OP1/OP2/ Consider NA in place of MG (1 valid contact(s) for NA vs. 0 for MG) |
| MG:DA:4479 | 1.00 | 50.15 |  | 0 | 1 | - Exclusion zone contact(s): /OP2/ Consider NA in place of MG (1 valid contact(s) for NA vs. 0 for MG) |
| MG:DA:4480 | 1.00 | 77.68 |  | 1 | 1 | O_ph_ - Exclusion zone contact(s): /O5'/ |
| MG:DA:4481 | 1.00 | 62.58 |  | 0 | 2 | - Exclusion zone contact(s): /N1/N7/ Consider K in place of MG (2 valid contact(s) for K vs. 0 for MG) |
| MG:DA:4482 | 1.00 | 80.23 |  | 0 |  | 1^st^ MG coordination shell missing (see Notes) |
| MG:DA:4483 | 1.00 | 64.09 |  | 0 | 1 | - Exclusion zone contact(s): /OP1/ Consider K in place of MG (1 valid contact(s) for K vs. 0 for MG) |
| MG:DA:4484 | 1.00 | 50.53 |  | 0 | 5 | - Exclusion zone contact(s): /N7/NH2_Arg_/OG1_Thr_/OP2/OP2/ Consider K in place of MG (4 valid contact(s) for K vs. 0 for MG) |
| MG:DA:4485 | 1.00 | 138.55 |  | 2 | 4 | O_b_Anion - Exclusion zone contact(s): /N3_U_/O6/OP1/OP2/ Consider K in place of MG (3 valid contact(s) for K vs. 2 for MG) |
| MG:DA:4486 | 1.00 | 63.93 |  | 0 | 2 | - Exclusion zone contact(s): /O2'/O4'/ Consider K in place of MG (2 valid contact(s) for K vs. 0 for MG) |
| MG:DA:4487 | 1.00 | 94.79 |  | 0 | 1 | - Exclusion zone contact(s): /OP1/ Consider K in place of MG (1 valid contact(s) for K vs. 0 for MG) |
| MG:DA:4488 | 1.00 | 89.38 |  | 0 | 4 | - Exclusion zone contact(s): /N1_A_/N3_A_/N6/ Consider K in place of MG (4 valid contact(s) for K vs. 0 for MG) |
| MG:DA:4489 | 1.00 | 42.83 |  | 1 | 3 | O_ph_ - Exclusion zone contact(s): /O3'/OP1/OP1/ Consider K in place of MG (3 valid contact(s) for K vs. 1 for MG) |
| MG:DA:4490 | 1.00 | 125.49 |  | 1 | 3 | Anion - Exclusion zone contact(s): /O4/O4'/O6/ Consider NA in place of MG (2 valid contact(s) for NA vs. 1 for MG) |
| MG:DA:4491 | 1.00 | 66.66 |  | 0 | 1 | - Exclusion zone contact(s): /N2/ Consider K in place of MG (1 valid contact(s) for K vs. 0 for MG) |
| MG:DA:4492 | 1.00 | 66.37 |  | 0 | 2 | - Exclusion zone contact(s): /O3'/OP2/ Consider K in place of MG (2 valid contact(s) for K vs. 0 for MG) |
| MG:DA:4493 | 1.00 | 107.70 |  | 1 |  | O_ph_ |
| MG:DA:4494 | 1.00 | 84.43 |  | 0 | 1 | - Exclusion zone contact(s): /OP2/ Consider K in place of MG (1 valid contact(s) for K vs. 0 for MG) |
| MG:DA:4495 | 1.00 | 111.62 |  | 0 | 1 | - Exclusion zone contact(s): /O6/ Consider K in place of MG (1 valid contact(s) for K vs. 0 for MG) |
| MG:DA:4496 | 1.00 | 81.44 |  | 0 | 3 | - Exclusion zone contact(s): /O2/O2'/O3'/ Consider K in place of MG (3 valid contact(s) for K vs. 0 for MG) |
| MG:DA:4497 | 1.00 | 43.18 |  | 0 | 6 | - Exclusion zone contact(s): /N2/N6/O3'/O4'/O4'/O5'/ Consider K in place of MG (6 valid contact(s) for K vs. 0 for MG) |
| MG:DA:4498 | 1.00 | 82.70 |  | 0 | 4 | - Exclusion zone contact(s): /O2/O2'/OP2/OP2/ Consider K in place of MG (3 valid contact(s) for K vs. 0 for MG) |
| MG:DA:4499 | 1.00 | 51.82 |  | 0 | 2 | - Exclusion zone contact(s): /OP1/OP2/ Consider K in place of MG (2 valid contact(s) for K vs. 0 for MG) |
| MG:DA:4500 | 1.00 | 53.60 |  | 0 | 2 | - Exclusion zone contact(s): /N7/N7/ Consider K in place of MG (2 valid contact(s) for K vs. 0 for MG) |
| MG:DA:4501 | 1.00 | 54.00 |  | 0 | 2 | - Exclusion zone contact(s): /O5'/OP2/ Consider NA in place of MG (1 valid contact(s) for NA vs. 0 for MG) |
| MG:DA:4502 | 1.00 | 62.17 |  | 0 | 3 | - Exclusion zone contact(s): /N1/N1_A_/N6/ Consider K in place of MG (3 valid contact(s) for K vs. 0 for MG) |
| MG:DA:4503 | 1.00 | 96.06 |  | 0 | 1 | Consider K in place of MG (1 valid contact(s) for K vs. 0 for MG) |
| MG:DA:4504 | 1.00 | 82.55 |  | 0 |  | 1^st^ MG coordination shell missing (see Notes) |
| MG:DA:4505 | 1.00 | 97.97 |  | 0 | 1 | - Exclusion zone contact(s): /OP1/ Consider NA in place of MG (1 valid contact(s) for NA vs. 0 for MG) |
| MG:DA:4506 | 1.00 | 65.04 |  | 0 |  | 1^st^ MG coordination shell missing (see Notes) |
| MG:DA:4507 | 1.00 | 88.96 |  | 0 | 1 | - Exclusion zone contact(s): /OP2/ Consider K in place of MG (1 valid contact(s) for K vs. 0 for MG) |
| MG:DA:4508 | 1.00 | 58.55 |  | 0 | 2 | - Exclusion zone contact(s): /O2'/O4'/ Consider K in place of MG (2 valid contact(s) for K vs. 0 for MG) |
| MG:DA:4509 | 1.00 | 64.17 |  | 0 | 1 | - Exclusion zone contact(s): /N7/ Consider K in place of MG (1 valid contact(s) for K vs. 0 for MG) |
| MG:DA:4510 | 1.00 | 78.82 |  | 0 | 1 | - Exclusion zone contact(s): /OP1/ Consider K in place of MG (1 valid contact(s) for K vs. 0 for MG) |
| MG:DA:4511 | 1.00 | 90.26 |  | 0 | 3 | - Exclusion zone contact(s): /N6/N7/OP2/ Consider K in place of MG (3 valid contact(s) for K vs. 0 for MG) |
| MG:DA:4512 | 1.00 | 100.63 |  | 0 |  | 1^st^ MG coordination shell missing (see Notes) |
| MG:DA:4513 | 1.00 | 75.81 |  | 1 | 1 | O_ph_ - Exclusion zone contact(s): /N7/ |
| MG:DA:4514 | 1.00 | 80.33 |  | 1 | 3 | O_b_ - Exclusion zone contact(s): /N7/N7/OP2/ Consider K in place of MG (2 valid contact(s) for K vs. 1 for MG) |
| MG:DA:4515 | 1.00 | 60.11 |  | 0 | 3 | - Exclusion zone contact(s): /O2'/O2'/O3'/ Consider K in place of MG (3 valid contact(s) for K vs. 0 for MG) |
| MG:DA:4516 | 1.00 | 53.92 |  | 1 | 3 | O_ph_ - Exclusion zone contact(s): /NZ_Lys_/O4'/OP2/ Consider K in place of MG (3 valid contact(s) for K vs. 1 for MG) |
| MG:DA:4517 | 1.00 | 97.25 |  | 1 | 2 | O_ph_ - Exclusion zone contact(s): /OP1/OP1/ |
| MG:DA:4518 | 1.00 | 49.81 |  | 0 | 2 | - Exclusion zone contact(s): /O5'/OP2/ Consider K in place of MG (2 valid contact(s) for K vs. 0 for MG) |
| MG:DA:4519 | 1.00 | 118.70 |  | 0 | 2 | - Exclusion zone contact(s): /N1_A_/O2'/ Consider NA in place of MG (1 valid contact(s) for NA vs. 0 for MG) |
| MG:DA:4520 | 1.00 | 84.38 |  | 0 | 2 | - Exclusion zone contact(s): /O3'/OP1/ Consider K in place of MG (2 valid contact(s) for K vs. 0 for MG) |
| MG:DA:4521 | 1.00 | 47.22 |  | 0 | 2 | - Exclusion zone contact(s): /O4/O4/ Consider K in place of MG (2 valid contact(s) for K vs. 0 for MG) |
| MG:DA:4522 | 1.00 | 64.14 |  | 0 | 1 | - Exclusion zone contact(s): /O_back_/ Consider K in place of MG (1 valid contact(s) for K vs. 0 for MG) |
| MG:DA:4523 | 1.00 | 51.34 |  | 0 | 1 | - Exclusion zone contact(s): /OP1/ Consider K in place of MG (1 valid contact(s) for K vs. 0 for MG) |
| MG:DA:4524 | 1.00 | 50.89 |  | 0 | 3 | - Exclusion zone contact(s): /O2'/O4'/O5'/ Consider K in place of MG (2 valid contact(s) for K vs. 0 for MG) |
| MG:DA:4525 | 1.00 | 108.79 |  | 0 | 1 | - Exclusion zone contact(s): /OP1/ Consider NA in place of MG (1 valid contact(s) for NA vs. 0 for MG) |
| MG:DA:4526 | 1.00 | 53.74 |  | 0 | 4 | - Exclusion zone contact(s): /N4/O4/O6/O6/ Consider K in place of MG (4 valid contact(s) for K vs. 0 for MG) |
| MG:DA:4527 | 1.00 | 53.10 |  | 0 | 1 | - Exclusion zone contact(s): /O6/ Consider K in place of MG (1 valid contact(s) for K vs. 0 for MG) |
| MG:DA:4528 | 1.00 | 65.49 |  | 0 | 2 | - Exclusion zone contact(s): /N7/O6/ Consider NA in place of MG (1 valid contact(s) for NA vs. 0 for MG) |
| MG:DA:4529 | 1.00 | 75.82 |  | 0 |  | 1^st^ MG coordination shell missing (see Notes) |
| MG:DA:4530 | 1.00 | 72.93 |  | 0 | 1 | - Exclusion zone contact(s): /OP1/ Consider NA in place of MG (1 valid contact(s) for NA vs. 0 for MG) |
| MG:DA:4531 | 1.00 | 57.66 |  | 0 | 2 | - Exclusion zone contact(s): /N7/O6/ Consider K in place of MG (2 valid contact(s) for K vs. 0 for MG) |
| MG:DA:4532 | 1.00 | 45.13 |  | 0 | 3 | - Exclusion zone contact(s): /O4'/O5'/OP2/ Consider K in place of MG (2 valid contact(s) for K vs. 0 for MG) |
| MG:DA:4533 | 1.00 | 88.56 |  | 0 | 2 | - Exclusion zone contact(s): /OP2/OP2/ Consider K in place of MG (2 valid contact(s) for K vs. 0 for MG) |
| MG:DA:4534 | 1.00 | 118.23 |  | 1 |  | O_b_ |
| MG:DA:4535 | 1.00 | 89.40 |  | 0 | 1 | - Exclusion zone contact(s): /O6/ Consider K in place of MG (1 valid contact(s) for K vs. 0 for MG) |
| MG:DA:4536 | 1.00 | 64.94 |  | 0 | 1 | - Exclusion zone contact(s): /OP1/ Consider K in place of MG (1 valid contact(s) for K vs. 0 for MG) |
| MG:DA:4537 | 1.00 | 58.23 |  | 0 | 3 | - Exclusion zone contact(s): /NH1_Arg_/OP2/OP2/ Consider K in place of MG (2 valid contact(s) for K vs. 0 for MG) |
| MG:DA:4538 | 1.00 | 92.99 |  | 0 | 1 | - Exclusion zone contact(s): /N7/ Consider K in place of MG (1 valid contact(s) for K vs. 0 for MG) |
| MG:DA:4539 | 1.00 | 85.33 |  | 0 | 3 | - Exclusion zone contact(s): /N7/N7/O6/ Consider K in place of MG (3 valid contact(s) for K vs. 0 for MG) |
| MG:DA:4540 | 1.00 | 88.38 |  | 0 | 1 | - Exclusion zone contact(s): /O6/ Consider K in place of MG (1 valid contact(s) for K vs. 0 for MG) |
| MG:DA:4541 | 1.00 | 58.86 |  | 0 |  | 1^st^ MG coordination shell missing (see Notes) |
| MG:DA:4542 | 1.00 | 37.25 |  | 0 | 3 | - Exclusion zone contact(s): /N3_A_/O2'/O4'/ Consider K in place of MG (3 valid contact(s) for K vs. 0 for MG) |
| MG:DA:4543 | 1.00 | 60.36 |  | 0 | 1 | - Exclusion zone contact(s): /OP2/ Consider NA in place of MG (1 valid contact(s) for NA vs. 0 for MG) |
| MG:DA:4544 | 1.00 | 76.92 |  | 0 |  | 1^st^ MG coordination shell missing (see Notes) |
| MG:DA:4545 | 1.00 | 90.13 |  | 1 | 3 | Anion - Exclusion zone contact(s): /O2/O4'/OP1/ Consider K in place of MG (2 valid contact(s) for K vs. 1 for MG) |
| MG:DA:4546 | 1.00 | 67.99 |  | 2 | 2 | O_r_O_b_ - Exclusion zone contact(s): /N1/O2'/ |
| MG:DA:4547 | 1.00 | 46.28 |  | 0 | 2 | - Exclusion zone contact(s): /O2/O2'/ Consider NA in place of MG (1 valid contact(s) for NA vs. 0 for MG) |
| MG:DA:4548 | 1.00 | 53.40 |  | 0 | 1 | - Exclusion zone contact(s): /N3_A_/ Consider K in place of MG (1 valid contact(s) for K vs. 0 for MG) |
| MG:DA:4549 | 1.00 | 98.23 |  | 1 | 1 | O_b_ - Exclusion zone contact(s): /O4'/ |
| MG:DA:4550 | 1.00 | 61.42 |  | 0 | 2 | - Exclusion zone contact(s): /N1/O4/ Consider K in place of MG (2 valid contact(s) for K vs. 0 for MG) |
| MG:DA:4551 | 1.00 | 118.95 |  | 1 | 5 | O_ph_ - Exclusion zone contact(s): /NH1_Arg_/NH2_Arg_/O5'/OP1/OP2/ Consider K in place of MG (4 valid contact(s) for K vs. 1 for MG) |
| MG:DA:4552 | 1.00 | 83.52 |  | 0 | 1 | - Exclusion zone contact(s): /OP2/ Consider K in place of MG (1 valid contact(s) for K vs. 0 for MG) |
| MG:DA:4553 | 1.00 | 69.49 |  | 0 | 4 | - Exclusion zone contact(s): /NZ_Lys_/O2'/O3'/OP1/ Consider K in place of MG (3 valid contact(s) for K vs. 0 for MG) |
| MG:DA:4554 | 1.00 | 81.58 |  | 0 | 1 | - Exclusion zone contact(s): /OP1/ Consider K in place of MG (1 valid contact(s) for K vs. 0 for MG) |
| MG:DA:4555 | 1.00 | 43.15 |  | 0 | 2 | - Exclusion zone contact(s): /OP2/OP2/ Consider K in place of MG (2 valid contact(s) for K vs. 0 for MG) |
| MG:DA:4556 | 1.00 | 41.43 |  | 1 | 2 | O_ph_ - Exclusion zone contact(s): /O2'/O3'/ Consider K in place of MG (2 valid contact(s) for K vs. 1 for MG) |
| MG:DA:4557 | 1.00 | 56.11 |  | 1 | 5 | O_ph_ - Exclusion zone contact(s): /N1/N3_U_/O2'/O3'/O4'/ Consider K in place of MG (4 valid contact(s) for K vs. 1 for MG) |
| MG:DA:4558 | 1.00 | 86.36 |  | 0 | 2 | - Exclusion zone contact(s): /O6/OP2/ Consider NA in place of MG (1 valid contact(s) for NA vs. 0 for MG) |
| MG:DA:4559 | 1.00 | 52.97 |  | 0 | 1 | - Exclusion zone contact(s): /O2'/ Consider K in place of MG (1 valid contact(s) for K vs. 0 for MG) |
| MG:DA:4560 | 1.00 | 61.36 |  | 0 |  | 1^st^ MG coordination shell missing (see Notes) |
| MG:DA:4561 | 1.00 | 43.91 |  | 0 | 3 | - Exclusion zone contact(s): /N3_A_/O2'/O2'/ Consider K in place of MG (2 valid contact(s) for K vs. 0 for MG) |
| MG:DA:4562 | 1.00 | 46.07 |  | 0 | 1 | - Exclusion zone contact(s): /O3'/ Consider K in place of MG (1 valid contact(s) for K vs. 0 for MG) |
| MG:DA:4563 | 1.00 | 77.35 |  | 0 | 1 | - Exclusion zone contact(s): /N1_G_/ Consider K in place of MG (1 valid contact(s) for K vs. 0 for MG) |
| MG:DA:4564 | 1.00 | 39.19 |  | 0 | 5 | - Exclusion zone contact(s): /O3'/O_back_/OP1/OP1/OP2/ Consider K in place of MG (5 valid contact(s) for K vs. 0 for MG) |
| MG:DA:4565 | 1.00 | 119.15 |  | 0 | 5 | - Exclusion zone contact(s): /N1_G_/O4/O6/OP1/OP2/ Consider K in place of MG (3 valid contact(s) for K vs. 0 for MG) |
| MG:DA:4566 | 1.00 | 64.68 |  | 0 | 2 | - Exclusion zone contact(s): /O6/OP1/ Consider NA in place of MG (1 valid contact(s) for NA vs. 0 for MG) |
| MG:DA:4567 | 1.00 | 51.59 |  | 0 | 3 | - Exclusion zone contact(s): /O5'/OP2/OP2/ Consider K in place of MG (3 valid contact(s) for K vs. 0 for MG) |
| MG:DA:4568 | 1.00 | 55.71 |  | 0 | 2 | - Exclusion zone contact(s): /O2'/O5'/ Consider K in place of MG (2 valid contact(s) for K vs. 0 for MG) |
| MG:DA:4569 | 1.00 | 37.75 |  | 0 | 3 | - Exclusion zone contact(s): /N6/O2'/O3'/ Consider K in place of MG (3 valid contact(s) for K vs. 0 for MG) |
| MG:DA:4570 | 1.00 | 59.12 |  | 0 | 2 | - Exclusion zone contact(s): /N7/O6/ Consider K in place of MG (2 valid contact(s) for K vs. 0 for MG) |
| MG:DA:4571 | 1.00 | 61.04 |  | 0 | 1 | - Exclusion zone contact(s): /N4/ Consider K in place of MG (1 valid contact(s) for K vs. 0 for MG) |
| MG:DA:4572 | 1.00 | 76.86 |  | 0 | 4 | - Exclusion zone contact(s): /N3_A_/O2'/O4'/OH_Tyr_/ Consider K in place of MG (4 valid contact(s) for K vs. 0 for MG) |
| MG:DA:4573 | 1.00 | 102.01 |  | 0 | 1 | - Exclusion zone contact(s): /OP2/ Consider K in place of MG (1 valid contact(s) for K vs. 0 for MG) |
| MG:DA:4574 | 1.00 | 73.06 |  | 0 |  | 1^st^ MG coordination shell missing (see Notes) |
| MG:DA:4575 | 1.00 | 68.79 |  | 0 |  | 1^st^ MG coordination shell missing (see Notes) |
| MG:DA:4576 | 1.00 | 87.53 |  | 0 |  | 1^st^ MG coordination shell missing (see Notes) |
| MG:DA:4577 | 1.00 | 57.75 |  | 0 | 1 | - Exclusion zone contact(s): /OP2/ Consider K in place of MG (1 valid contact(s) for K vs. 0 for MG) |
| MG:DA:4578 | 1.00 | 95.97 |  | 0 | 2 | - Exclusion zone contact(s): /O2'/O4'/ Consider K in place of MG (2 valid contact(s) for K vs. 0 for MG) |
| MG:DA:4579 | 1.00 | 81.58 |  | 1 | 2 | O_r_ - Exclusion zone contact(s): /O2'/OP1/ Consider K in place of MG (2 valid contact(s) for K vs. 1 for MG) |
| MG:DA:4580 | 1.00 | 69.78 |  | 0 | 2 | - Exclusion zone contact(s): /OP1/OP2/ Consider K in place of MG (2 valid contact(s) for K vs. 0 for MG) |
| MG:DA:4581 | 1.00 | 55.21 |  | 0 | 3 | - Exclusion zone contact(s): /O2'/O3'/OP1/ Consider K in place of MG (2 valid contact(s) for K vs. 0 for MG) |
| MG:DA:4582 | 1.00 | 69.02 |  | 0 | 4 | - Exclusion zone contact(s): /N3_A_/N4/O2'/OP1/ Consider K in place of MG (3 valid contact(s) for K vs. 0 for MG) |
| MG:DA:4583 | 1.00 | 51.46 |  | 0 | 1 | - Exclusion zone contact(s): /O2'/ Consider K in place of MG (1 valid contact(s) for K vs. 0 for MG) |
| MG:DA:4584 | 1.00 | 40.62 |  | 0 | 2 | - Exclusion zone contact(s): /O6/O6/ Consider K in place of MG (2 valid contact(s) for K vs. 0 for MG) |
| MG:DA:4585 | 1.00 | 76.05 |  | 0 |  | 1^st^ MG coordination shell missing (see Notes) |
| MG:DA:4586 | 1.00 | 78.42 |  | 0 | 1 | - Exclusion zone contact(s): /N7/ Consider K in place of MG (1 valid contact(s) for K vs. 0 for MG) |
| MG:DA:4587 | 1.00 | 71.04 |  | 0 |  | 1^st^ MG coordination shell missing (see Notes) |
| MG:DA:4588 | 1.00 | 41.29 |  | 1 | 4 | Anion - Exclusion zone contact(s): /N1_A_/N2/N3_G_/N6/ Consider K in place of MG (4 valid contact(s) for K vs. 1 for MG) |
| MG:DA:4589 | 1.00 | 58.32 |  | 0 |  | 1^st^ MG coordination shell missing (see Notes) |
| MG:DA:4590 | 1.00 | 81.70 |  | 0 | 4 | - Exclusion zone contact(s): /O2'/O3'/O5'/OP1/ Consider K in place of MG (4 valid contact(s) for K vs. 0 for MG) |
| MG:DA:4591 | 1.00 | 107.95 |  | 0 | 1 | - Exclusion zone contact(s): /OP2/ Consider K in place of MG (1 valid contact(s) for K vs. 0 for MG) |
| MG:DA:4592 | 1.00 | 77.34 |  | 0 | 1 | - Exclusion zone contact(s): /N7/ Consider K in place of MG (1 valid contact(s) for K vs. 0 for MG) |
| MG:DA:4593 | 1.00 | 65.77 |  | 0 | 2 | - Exclusion zone contact(s): /O4/O6/ Consider K in place of MG (2 valid contact(s) for K vs. 0 for MG) |
| MG:DA:4594 | 1.00 | 62.99 |  | 0 | 2 | - Exclusion zone contact(s): /N2/N3_A_/ Consider NA in place of MG (1 valid contact(s) for NA vs. 0 for MG) |
| MG:DA:4595 | 1.00 | 84.68 |  | 0 |  | 1^st^ MG coordination shell missing (see Notes) |
| MG:DA:4596 | 1.00 | 63.87 |  | 0 | 2 | - Exclusion zone contact(s): /OP2/OP2/ Consider K in place of MG (2 valid contact(s) for K vs. 0 for MG) |
| MG:DA:4597 | 1.00 | 59.07 |  | 0 | 2 | - Exclusion zone contact(s): /O3'/OP1/ Consider K in place of MG (2 valid contact(s) for K vs. 0 for MG) |
| MG:DA:4598 | 1.00 | 71.77 |  | 0 |  | No contacts in 5.0 A |
| MG:DA:4599 | 1.00 | 50.73 |  | 0 | 1 | - Exclusion zone contact(s): /OP1/ Consider K in place of MG (1 valid contact(s) for K vs. 0 for MG) |
| MG:DA:4600 | 1.00 | 57.36 |  | 0 | 3 | - Exclusion zone contact(s): /O2'/O3'/OP1/ Consider K in place of MG (3 valid contact(s) for K vs. 0 for MG) |
| MG:DA:4601 | 1.00 | 88.47 |  | 0 | 1 | - Exclusion zone contact(s): /OP2/ Consider K in place of MG (1 valid contact(s) for K vs. 0 for MG) |
| MG:DA:4602 | 1.00 | 71.47 |  | 0 |  | 1^st^ MG coordination shell missing (see Notes) |
| MG:DA:4603 | 1.00 | 55.89 |  | 0 | 2 | - Exclusion zone contact(s): /O2'/O3'/ Consider K in place of MG (2 valid contact(s) for K vs. 0 for MG) |
| MG:DA:4604 | 1.00 | 111.06 |  | 0 | 3 | - Exclusion zone contact(s): /N3_G_/O2'/O4'/ Consider K in place of MG (2 valid contact(s) for K vs. 0 for MG) |
| MG:DA:4605 | 1.00 | 70.14 |  | 0 | 2 | - Exclusion zone contact(s): /OP1/OP2/ Consider K in place of MG (2 valid contact(s) for K vs. 0 for MG) |
| MG:DA:4606 | 1.00 | 41.62 |  | 0 | 5 | - Exclusion zone contact(s): /N3_A_/N3_G_/N3_U_/O2/O2'/ Consider K in place of MG (4 valid contact(s) for K vs. 0 for MG) |
| MG:DA:4607 | 1.00 | 64.69 |  | 0 |  | 1^st^ MG coordination shell missing (see Notes) |
| MG:DA:4608 | 1.00 | 47.93 |  | 0 | 1 | - Exclusion zone contact(s): /OP2/ Consider K in place of MG (1 valid contact(s) for K vs. 0 for MG) |
| MG:DA:4609 | 1.00 | 48.07 |  | 0 |  | 1^st^ MG coordination shell missing (see Notes) |
| MG:DA:4610 | 1.00 | 114.14 |  | 2 | 3 | 2O_r_ - Exclusion zone contact(s): /O3'/O5'/O5'/ |
| MG:DA:4611 | 1.00 | 82.14 |  | 0 |  | 1^st^ MG coordination shell missing (see Notes) |
| MG:DA:4612 | 1.00 | 79.43 |  | 0 |  | 1^st^ MG coordination shell missing (see Notes) |
| MG:DA:4613 | 1.00 | 64.08 |  | 0 | 4 | - Exclusion zone contact(s): /N1_A_/O5'/O6/OP2/ Consider K in place of MG (3 valid contact(s) for K vs. 0 for MG) |
| MG:DA:4614 | 1.00 | 78.49 |  | 0 | 1 | - Exclusion zone contact(s): /OP2/ Consider K in place of MG (1 valid contact(s) for K vs. 0 for MG) |
| MG:DA:4615 | 1.00 | 123.02 |  | 0 | 2 | - Exclusion zone contact(s): /O6/OP2/ Consider NA in place of MG (2 valid contact(s) for NA vs. 0 for MG) |
| MG:DA:4616 | 1.00 | 72.68 |  | 0 | 5 | - Exclusion zone contact(s): /N7/O3'/O5'/OP2/ Consider K in place of MG (5 valid contact(s) for K vs. 0 for MG) |
| MG:DA:4617 | 1.00 | 54.58 |  | 1 | 1 | O_b_ - Exclusion zone contact(s): /O4'/ |
| MG:DA:4618 | 1.00 | 61.81 |  | 0 | 3 | - Exclusion zone contact(s): /OP1/OP2/OP2/ Consider K in place of MG (3 valid contact(s) for K vs. 0 for MG) |
| MG:DA:4619 | 1.00 | 49.30 |  | 0 | 3 | - Exclusion zone contact(s): /N3_G_/O2'/O4'/ Consider K in place of MG (2 valid contact(s) for K vs. 0 for MG) |
| MG:DA:4620 | 1.00 | 84.18 |  | 0 |  | No contacts in 5.0 A |
| MG:DA:4621 | 1.00 | 92.50 |  | 0 |  | 1^st^ MG coordination shell missing (see Notes) |
| MG:DA:4622 | 1.00 | 54.64 |  | 0 | 5 | - Exclusion zone contact(s): /N3_C_/N3_G_/N3_U_/NH1_Arg_/O2/ Consider K in place of MG (4 valid contact(s) for K vs. 0 for MG) |
| MG:DA:4623 | 1.00 | 47.93 |  | 0 | 3 | - Exclusion zone contact(s): /O2/O3'/OP1/ Consider K in place of MG (3 valid contact(s) for K vs. 0 for MG) |
| MG:DA:4624 | 1.00 | 68.60 |  | 0 | 1 | - Exclusion zone contact(s): /OP1/ Consider NA in place of MG (1 valid contact(s) for NA vs. 0 for MG) |
| MG:DA:4625 | 1.00 | 67.41 |  | 0 | 3 | - Exclusion zone contact(s): /N7/OP2/OP2/ Consider K in place of MG (2 valid contact(s) for K vs. 0 for MG) |
| MG:DA:4626 | 1.00 | 56.11 |  | 0 | 1 | - Exclusion zone contact(s): /O6/ Consider K in place of MG (1 valid contact(s) for K vs. 0 for MG) |
| MG:DA:4627 | 1.00 | 80.48 |  | 0 | 1 | - Exclusion zone contact(s): /O2'/ Consider K in place of MG (1 valid contact(s) for K vs. 0 for MG) |
| MG:DA:4628 | 1.00 | 94.45 |  | 0 | 1 | - Exclusion zone contact(s): /OP1/ Consider K in place of MG (1 valid contact(s) for K vs. 0 for MG) |
| MG:DA:4629 | 1.00 | 107.67 |  | 0 | 1 | - Exclusion zone contact(s): /N7/ Consider K in place of MG (1 valid contact(s) for K vs. 0 for MG) |
| MG:DA:4630 | 1.00 | 89.39 |  | 0 | 1 | - Exclusion zone contact(s): /O6/ Consider K in place of MG (1 valid contact(s) for K vs. 0 for MG) |
| MG:DA:4631 | 1.00 | 67.59 |  | 1 | 1 | O_r_ - Exclusion zone contact(s): /OP1/ |
| MG:DA:4632 | 1.00 | 71.06 |  | 1 |  | O_ph_ |
| MG:DA:4633 | 1.00 | 66.52 |  | 0 |  | No contacts in 5.0 A |
| MG:DA:4634 | 1.00 | 82.53 |  | 0 |  | 1^st^ MG coordination shell missing (see Notes) |
| MG:DA:4635 | 1.00 | 71.62 |  | 0 | 1 | - Exclusion zone contact(s): /OP2/ Consider K in place of MG (1 valid contact(s) for K vs. 0 for MG) |
| MG:DA:4636 | 1.00 | 64.32 |  | 0 |  | 1^st^ MG coordination shell missing (see Notes) |
| MG:DA:4637 | 1.00 | 80.83 |  | 0 | 1 | - Exclusion zone contact(s): /OP2/ Consider K in place of MG (1 valid contact(s) for K vs. 0 for MG) |
| MG:DA:4638 | 1.00 | 56.06 |  | 0 | 2 | - Exclusion zone contact(s): /O4/O6/ Consider K in place of MG (2 valid contact(s) for K vs. 0 for MG) |
| MG:DA:4639 | 1.00 | 75.78 |  | 0 | 3 | - Exclusion zone contact(s): /O2'/OP1/OP2/ Consider K in place of MG (2 valid contact(s) for K vs. 0 for MG) |
| MG:DA:4640 | 1.00 | 66.88 |  | 1 |  | O_b_ |
| MG:DA:4641 | 1.00 | 134.55 |  | 0 | 2 | - Exclusion zone contact(s): /N7/N7/ Consider K in place of MG (2 valid contact(s) for K vs. 0 for MG) |
| MG:DA:4642 | 1.00 | 57.60 |  | 0 | 2 | - Exclusion zone contact(s): /N7/O6/ Consider K in place of MG (2 valid contact(s) for K vs. 0 for MG) |
| MG:DA:4643 | 1.00 | 59.10 |  | 0 | 2 | - Exclusion zone contact(s): /N7/OP1/ Consider K in place of MG (2 valid contact(s) for K vs. 0 for MG) |
| MG:DA:4644 | 1.00 | 64.18 |  | 0 |  | 1^st^ MG coordination shell missing (see Notes) |
| MG:DA:4645 | 1.00 | 59.89 |  | 0 |  | 1^st^ MG coordination shell missing (see Notes) |
| MG:DA:4646 | 1.00 | 69.88 |  | 0 |  | 1^st^ MG coordination shell missing (see Notes) |
| MG:DA:4647 | 1.00 | 84.05 |  | 0 | 2 | - Exclusion zone contact(s): /N1_G_/O6/ Consider K in place of MG (2 valid contact(s) for K vs. 0 for MG) |
| MG:DA:4648 | 1.00 | 61.14 |  | 0 |  | 1^st^ MG coordination shell missing (see Notes) |
| MG:DA:4649 | 1.00 | 41.17 |  | 0 | 1 | - Exclusion zone contact(s): /OP2/ Consider NA in place of MG (1 valid contact(s) for NA vs. 0 for MG) |
| MG:DA:4650 | 1.00 | 83.10 |  | 0 |  | 1^st^ MG coordination shell missing (see Notes) |
| MG:DA:4651 | 1.00 | 66.46 |  | 0 | 3 | - Exclusion zone contact(s): /N7/N7/O6/ Consider K in place of MG (3 valid contact(s) for K vs. 0 for MG) |
| MG:DA:4652 | 1.00 | 71.54 |  | 0 | 3 | - Exclusion zone contact(s): /N7/O6/OP2/ Consider K in place of MG (3 valid contact(s) for K vs. 0 for MG) |
| MG:DA:4653 | 1.00 | 69.10 |  | 0 | 3 | - Exclusion zone contact(s): /N1_A_/O6/OP2/ Consider K in place of MG (3 valid contact(s) for K vs. 0 for MG) |
| MG:DA:4654 | 1.00 | 69.32 |  | 0 | 1 | - Exclusion zone contact(s): /OP2/ Consider K in place of MG (1 valid contact(s) for K vs. 0 for MG) |
| MG:DA:4655 | 1.00 | 79.69 |  | 0 | 1 | - Exclusion zone contact(s): /OP1/ Consider NA in place of MG (1 valid contact(s) for NA vs. 0 for MG) |
| MG:DA:4656 | 1.00 | 71.13 |  | 0 |  | 1^st^ MG coordination shell missing (see Notes) |
| MG:DA:4657 | 1.00 | 58.37 |  | 0 | 2 | - Exclusion zone contact(s): /O2'/O4'/ Consider K in place of MG (2 valid contact(s) for K vs. 0 for MG) |
| MG:DA:4658 | 1.00 | 76.81 |  | 0 | 4 | - Exclusion zone contact(s): /N1/O5'/OH_Tyr_/OP1/ Consider K in place of MG (4 valid contact(s) for K vs. 0 for MG) |
| MG:DA:4659 | 1.00 | 68.86 |  | 0 | 1 | - Exclusion zone contact(s): /O6/ Consider K in place of MG (1 valid contact(s) for K vs. 0 for MG) |
| MG:DA:4660 | 1.00 | 73.87 |  | 0 | 3 | - Exclusion zone contact(s): /O5'/OP1/OP2/ Consider K in place of MG (3 valid contact(s) for K vs. 0 for MG) |
| MG:DA:4661 | 1.00 | 76.93 |  | 0 |  | 1^st^ MG coordination shell missing (see Notes) |
| MG:DA:4662 | 1.00 | 101.68 |  | 0 | 2 | - Exclusion zone contact(s): /O2'/OP1/ Consider NA in place of MG (1 valid contact(s) for NA vs. 0 for MG) |
| MG:DA:4663 | 1.00 | 69.09 |  | 0 | 1 | - Exclusion zone contact(s): /OP2/ Consider K in place of MG (1 valid contact(s) for K vs. 0 for MG) |
| MG:DA:4664 | 1.00 | 66.62 |  | 0 |  | 1^st^ MG coordination shell missing (see Notes) |
| MG:DA:4665 | 1.00 | 76.47 |  | 0 | 3 | - Exclusion zone contact(s): /N7/N7/O6/ Consider K in place of MG (3 valid contact(s) for K vs. 0 for MG) |
| MG:DA:4666 | 1.00 | 60.39 |  | 0 | 3 | - Exclusion zone contact(s): /OP1/OP2/OP2/ Consider K in place of MG (2 valid contact(s) for K vs. 0 for MG) |
| MG:DA:4667 | 1.00 | 46.96 |  | 0 | 2 | - Exclusion zone contact(s): /O_back_/OP1/ Consider NA in place of MG (1 valid contact(s) for NA vs. 0 for MG) |
| MG:DA:4668 | 1.00 | 72.80 |  | 0 | 3 | - Exclusion zone contact(s): /N3_U_/O2/O4'/ Consider K in place of MG (3 valid contact(s) for K vs. 0 for MG) |
| MG:DA:4669 | 1.00 | 71.13 |  | 1 | 3 | O_ph_ - Exclusion zone contact(s): /N_back_/OG1_Thr_/OP1/ Consider K in place of MG (3 valid contact(s) for K vs. 1 for MG) |
| MG:DA:4670 | 1.00 | 61.51 |  | 0 | 1 | - Exclusion zone contact(s): /OP1/ Consider K in place of MG (1 valid contact(s) for K vs. 0 for MG) |
| MG:DA:4671 | 1.00 | 72.25 |  | 0 |  | 1^st^ MG coordination shell missing (see Notes) |
| MG:DA:4672 | 1.00 | 95.71 |  | 0 | 1 | - Exclusion zone contact(s): /OP1/ Consider K in place of MG (1 valid contact(s) for K vs. 0 for MG) |
| MG:DA:4673 | 1.00 | 91.34 |  | 0 |  | 1^st^ MG coordination shell missing (see Notes) |
| MG:DA:4674 | 1.00 | 93.63 |  | 0 |  | No contacts in 5.0 A |
| MG:DA:4675 | 1.00 | 73.33 |  | 0 |  | 1^st^ MG coordination shell missing (see Notes) |
| MG:DA:4676 | 1.00 | 82.74 |  | 0 |  | 1^st^ MG coordination shell missing (see Notes) |
| MG:DA:4677 | 1.00 | 61.04 |  | 0 | 2 | - Exclusion zone contact(s): /OG_Ser_/OP1/ Consider NA in place of MG (1 valid contact(s) for NA vs. 0 for MG) |
| MG:DA:4678 | 1.00 | 68.26 |  | 0 | 1 | - Exclusion zone contact(s): /OP1/ Consider K in place of MG (1 valid contact(s) for K vs. 0 for MG) |
| MG:DA:4679 | 1.00 | 47.94 |  | 0 | 2 | - Exclusion zone contact(s): /O2'/OP1/ Consider K in place of MG (2 valid contact(s) for K vs. 0 for MG) |
| MG:DA:4680 | 1.00 | 71.29 |  | 0 | 2 | - Exclusion zone contact(s): /O5'/OP2/ Consider K in place of MG (2 valid contact(s) for K vs. 0 for MG) |
| MG:DA:4681 | 1.00 | 118.10 |  | 0 |  | 1^st^ MG coordination shell missing (see Notes) |
| MG:DA:4682 | 1.00 | 63.78 |  | 0 | 3 | - Exclusion zone contact(s): /O5'/OP2/OP2/ Consider NA in place of MG (2 valid contact(s) for NA vs. 0 for MG) |
| MG:DA:4683 | 1.00 | 100.69 |  | 0 | 2 | - Exclusion zone contact(s): /O5'/OP1/ Consider K in place of MG (2 valid contact(s) for K vs. 0 for MG) |
| MG:DA:4684 | 1.00 | 36.20 |  | 1 | 3 | O_ph_ - Exclusion zone contact(s): /O2'/O4'/O5'/ Consider K in place of MG (3 valid contact(s) for K vs. 1 for MG) |
| MG:DA:4685 | 1.00 | 64.85 |  | 0 | 2 | - Exclusion zone contact(s): /N4/OP2/ Consider K in place of MG (2 valid contact(s) for K vs. 0 for MG) |
| MG:DA:4686 | 1.00 | 59.29 |  | 0 | 2 | - Exclusion zone contact(s): /N3_A_/O4'/ Consider NA in place of MG (1 valid contact(s) for NA vs. 0 for MG) |
| MG:DA:4687 | 1.00 | 76.82 |  | 0 | 2 | - Exclusion zone contact(s): /O3'/OP1/ Consider K in place of MG (2 valid contact(s) for K vs. 0 for MG) |
| MG:DA:4688 | 1.00 | 83.99 |  | 0 |  | 1^st^ MG coordination shell missing (see Notes) |
| MG:DA:4689 | 1.00 | 67.97 |  | 0 | 1 | - Exclusion zone contact(s): /N7/ Consider NA in place of MG (1 valid contact(s) for NA vs. 0 for MG) |
| MG:DA:4690 | 1.00 | 86.74 |  | 0 | 1 | - Exclusion zone contact(s): /N2/ Consider K in place of MG (1 valid contact(s) for K vs. 0 for MG) |
| MG:DA:4691 | 1.00 | 114.96 |  | 0 | 2 | - Exclusion zone contact(s): /OP1/OP2/ Consider K in place of MG (2 valid contact(s) for K vs. 0 for MG) |
| MG:DA:4692 | 1.00 | 74.92 |  | 0 |  | 1^st^ MG coordination shell missing (see Notes) |
| MG:DA:4693 | 1.00 | 62.39 |  | 0 |  | 1^st^ MG coordination shell missing (see Notes) |
| MG:DA:4694 | 1.00 | 56.92 |  | 1 | 1 | O_ph_ - Exclusion zone contact(s): /O5'/ |
| MG:DA:4695 | 1.00 | 52.69 |  | 0 | 4 | - Exclusion zone contact(s): /N3_G_/N7/O4'/O6/ Consider NA in place of MG (2 valid contact(s) for NA vs. 0 for MG) |
| MG:DA:4696 | 1.00 | 57.93 |  | 0 | 1 | - Exclusion zone contact(s): /OP1/ Consider K in place of MG (1 valid contact(s) for K vs. 0 for MG) |
| MG:DA:4697 | 1.00 | 76.94 |  | 1 | 2 | O_r_ - Exclusion zone contact(s): /OP2/OP2/ Consider K in place of MG (2 valid contact(s) for K vs. 1 for MG) |
| MG:DA:4698 | 1.00 | 104.88 |  | 0 |  | 1^st^ MG coordination shell missing (see Notes) |
| MG:DA:4699 | 1.00 | 92.74 |  | 0 |  | 1^st^ MG coordination shell missing (see Notes) |
| MG:DA:4700 | 1.00 | 75.58 |  | 0 | 1 | - Exclusion zone contact(s): /N2/ Consider K in place of MG (1 valid contact(s) for K vs. 0 for MG) |
| MG:DA:4701 | 1.00 | 64.03 |  | 0 | 2 | - Exclusion zone contact(s): /N3_U_/O6/ Consider K in place of MG (2 valid contact(s) for K vs. 0 for MG) |
| MG:DA:4702 | 1.00 | 48.45 |  | 0 | 1 | - Exclusion zone contact(s): /OP1/ Consider K in place of MG (1 valid contact(s) for K vs. 0 for MG) |
| MG:DA:4703 | 1.00 | 49.33 |  | 0 | 3 | - Exclusion zone contact(s): /O2'/O3'/OG_Ser_/ Consider K in place of MG (3 valid contact(s) for K vs. 0 for MG) |
| MG:DA:4704 | 1.00 | 62.64 |  | 0 |  | 1^st^ MG coordination shell missing (see Notes) |
| MG:DA:4705 | 1.00 | 93.35 |  | 0 |  | 1^st^ MG coordination shell missing (see Notes) |
| MG:DA:4706 | 1.00 | 89.53 |  | 0 | 1 | - Exclusion zone contact(s): /OP2/ Consider K in place of MG (1 valid contact(s) for K vs. 0 for MG) |
| MG:DA:4707 | 1.00 | 86.84 |  | 0 |  | 1^st^ MG coordination shell missing (see Notes) |
| MG:DA:4708 | 1.00 | 46.46 |  | 0 |  | 1^st^ MG coordination shell missing (see Notes) |
| MG:DA:4709 | 1.00 | 81.87 |  | 0 |  | 1^st^ MG coordination shell missing (see Notes) |
| MG:DA:4710 | 1.00 | 102.13 |  | 0 | 2 | - Exclusion zone contact(s): /O2'/OP2/ Consider NA in place of MG (1 valid contact(s) for NA vs. 0 for MG) |
| MG:DA:4711 | 1.00 | 94.69 |  | 1 | 2 | O_ph_ - Exclusion zone contact(s): /O5'/OP1/ Consider K in place of MG (2 valid contact(s) for K vs. 1 for MG) |
| MG:DA:4712 | 1.00 | 43.87 |  | 0 | 5 | - Exclusion zone contact(s): /N1_A_/N2/O2/O2'/O4'/ Consider K in place of MG (5 valid contact(s) for K vs. 0 for MG) |
| MG:DA:4713 | 1.00 | 75.53 |  | 0 | 1 | - Exclusion zone contact(s): /N7/ Consider K in place of MG (1 valid contact(s) for K vs. 0 for MG) |
| MG:DA:4714 | 1.00 | 87.34 |  | 0 | 1 | - Exclusion zone contact(s): /OP1/ Consider NA in place of MG (1 valid contact(s) for NA vs. 0 for MG) |
| MG:DA:4715 | 1.00 | 58.19 |  | 0 |  | 1^st^ MG coordination shell missing (see Notes) |
| MG:DA:4716 | 1.00 | 77.79 |  | 1 | 1 | O_r_ - Exclusion zone contact(s): /OP2/ |
| MG:DA:4717 | 1.00 | 80.11 |  | 0 |  | 1^st^ MG coordination shell missing (see Notes) |
| MG:DA:4718 | 1.00 | 69.08 |  | 0 |  | 1^st^ MG coordination shell missing (see Notes) |
| MG:DA:4719 | 1.00 | 54.85 |  | 0 | 2 | - Exclusion zone contact(s): /O6/OP2/ Consider K in place of MG (2 valid contact(s) for K vs. 0 for MG) |
| MG:DA:4720 | 1.00 | 88.77 |  | 1 | 1 | O_ph_ - Exclusion zone contact(s): /N7/ |
| MG:DA:4721 | 1.00 | 75.95 |  | 0 |  | 1^st^ MG coordination shell missing (see Notes) |
| MG:DA:4722 | 1.00 | 109.31 |  | 1 | 2 | Anion - Exclusion zone contact(s): /N7/O6/ Consider K in place of MG (2 valid contact(s) for K vs. 1 for MG) |
| MG:DA:4723 | 1.00 | 71.46 |  | 0 | 3 | - Exclusion zone contact(s): /N2/N3_G_/N6/ Consider K in place of MG (3 valid contact(s) for K vs. 0 for MG) |
| MG:DA:4724 | 1.00 | 89.42 |  | 1 |  | O_ph_ |
| MG:DA:4725 | 1.00 | 56.87 |  | 1 | 2 | N_b_ - Exclusion zone contact(s): /OP2/OP2/ Consider K in place of MG (2 valid contact(s) for K vs. 1 for MG) |
| MG:DA:4726 | 1.00 | 71.51 |  | 0 | 3 | - Exclusion zone contact(s): /O2/O2'/O_back_/ Consider K in place of MG (3 valid contact(s) for K vs. 0 for MG) |
| MG:DA:4727 | 1.00 | 69.65 |  | 0 | 2 | - Exclusion zone contact(s): /O3'/O5'/ Consider NA in place of MG (1 valid contact(s) for NA vs. 0 for MG) |
| MG:DA:4728 | 1.00 | 86.29 |  | 0 | 2 | - Exclusion zone contact(s): /O3'/OP1/ Consider K in place of MG (2 valid contact(s) for K vs. 0 for MG) |
| MG:DA:4729 | 1.00 | 39.45 |  | 0 | 3 | - Exclusion zone contact(s): /N1_A_/O5'/OP2/ Consider K in place of MG (2 valid contact(s) for K vs. 0 for MG) |
| MG:DA:4730 | 1.00 | 57.56 |  | 0 | 1 | - Exclusion zone contact(s): /OP2/ Consider K in place of MG (1 valid contact(s) for K vs. 0 for MG) |
| MG:DA:4731 | 1.00 | 63.94 |  | 0 | 2 | - Exclusion zone contact(s): /O6/O6/ Consider K in place of MG (2 valid contact(s) for K vs. 0 for MG) |
| MG:DA:4732 | 1.00 | 71.25 |  | 0 | 1 | - Exclusion zone contact(s): /O6/ Consider K in place of MG (1 valid contact(s) for K vs. 0 for MG) |
| MG:DA:4733 | 1.00 | 84.54 |  | 0 |  | 1^st^ MG coordination shell missing (see Notes) |
| MG:DA:4734 | 1.00 | 71.75 |  | 0 |  | No contacts in 5.0 A |
| MG:DA:4735 | 1.00 | 88.90 |  | 0 | 1 | - Exclusion zone contact(s): /N7/ Consider K in place of MG (1 valid contact(s) for K vs. 0 for MG) |
| MG:DA:4736 | 1.00 | 77.24 |  | 0 | 2 | - Exclusion zone contact(s): /O2/O2'/ Consider K in place of MG (2 valid contact(s) for K vs. 0 for MG) |
| MG:DA:4737 | 1.00 | 85.82 |  | 0 | 2 | - Exclusion zone contact(s): /N4/O6/ Consider K in place of MG (2 valid contact(s) for K vs. 0 for MG) |
| MG:DA:4738 | 1.00 | 55.86 |  | 0 | 1 | - Exclusion zone contact(s): /OP2/ Consider K in place of MG (1 valid contact(s) for K vs. 0 for MG) |
| MG:DA:4739 | 1.00 | 77.61 |  | 0 |  | 1^st^ MG coordination shell missing (see Notes) |
| MG:DA:4740 | 1.00 | 78.20 |  | 0 | 1 | - Exclusion zone contact(s): /OP1/ Consider K in place of MG (1 valid contact(s) for K vs. 0 for MG) |
| MG:DA:4741 | 1.00 | 62.43 |  | 0 | 1 | - Exclusion zone contact(s): /OP1/ Consider K in place of MG (1 valid contact(s) for K vs. 0 for MG) |
| MG:DA:4742 | 1.00 | 128.69 |  | 1 |  | O_ph_ |
| MG:DA:4743 | 1.00 | 96.58 |  | 0 | 4 | - Exclusion zone contact(s): /NH2_Arg_/NZ_Lys_/OP1/OP2/ Consider K in place of MG (3 valid contact(s) for K vs. 0 for MG) |
| MG:DA:4744 | 1.00 | 70.18 |  | 0 | 2 | - Exclusion zone contact(s): /O2/OP1/ Consider K in place of MG (2 valid contact(s) for K vs. 0 for MG) |
| MG:DA:4745 | 1.00 | 53.19 |  | 0 | 4 | - Exclusion zone contact(s): /N2/O4'/OP2/OP2/ Consider K in place of MG (4 valid contact(s) for K vs. 0 for MG) |
| MG:DA:4746 | 1.00 | 98.77 |  | 0 | 1 | - Exclusion zone contact(s): /OP2/ Consider K in place of MG (1 valid contact(s) for K vs. 0 for MG) |
| MG:DA:4747 | 1.00 | 71.85 |  | 0 | 5 | - Exclusion zone contact(s): /O2/O2'/O2'/O3'/OP2/ Consider K in place of MG (5 valid contact(s) for K vs. 0 for MG) |
| MG:DA:4748 | 1.00 | 83.88 |  | 0 | 3 | - Exclusion zone contact(s): /O3'/O5'/OP2/ Consider K in place of MG (2 valid contact(s) for K vs. 0 for MG) |
| MG:DA:4749 | 1.00 | 78.15 |  | 0 | 2 | - Exclusion zone contact(s): /O3'/OP1/ Consider K in place of MG (2 valid contact(s) for K vs. 0 for MG) |
| MG:DA:4750 | 1.00 | 68.58 |  | 1 |  | O_r_ |
| MG:DA:4751 | 1.00 | 121.25 |  | 0 | 2 | - Exclusion zone contact(s): /O3'/OP1/ Consider K in place of MG (2 valid contact(s) for K vs. 0 for MG) |
| MG:DA:4752 | 1.00 | 84.97 |  | 0 | 2 | - Exclusion zone contact(s): /N4/O6/ Consider K in place of MG (2 valid contact(s) for K vs. 0 for MG) |
| MG:DA:4753 | 1.00 | 68.74 |  | 0 | 3 | - Exclusion zone contact(s): /O3'/O5'/OP1/ Consider K in place of MG (3 valid contact(s) for K vs. 0 for MG) |
| MG:DA:4754 | 1.00 | 71.03 |  | 0 | 2 | - Exclusion zone contact(s): /O2/O2'/ Consider NA in place of MG (1 valid contact(s) for NA vs. 0 for MG) |
| MG:DA:4755 | 1.00 | 80.48 |  | 0 | 3 | - Exclusion zone contact(s): /OP1/OP1/OP2/ Consider K in place of MG (3 valid contact(s) for K vs. 0 for MG) |
| MG:DA:4756 | 1.00 | 57.66 |  | 0 | 1 | - Exclusion zone contact(s): /N2/ Consider K in place of MG (1 valid contact(s) for K vs. 0 for MG) |
| MG:DA:4757 | 1.00 | 46.34 |  | 1 | 1 | O_r_ - Exclusion zone contact(s): /O2'/ |
| MG:DA:4758 | 1.00 | 71.14 |  | 0 |  | 1^st^ MG coordination shell missing (see Notes) |
| MG:DA:4759 | 1.00 | 91.98 |  | 0 |  | 1^st^ MG coordination shell missing (see Notes) |
| MG:DA:4760 | 1.00 | 55.94 |  | 0 | 2 | - Exclusion zone contact(s): /OP1/OP2/ Consider K in place of MG (2 valid contact(s) for K vs. 0 for MG) |
| MG:DA:4761 | 1.00 | 78.21 |  | 0 | 1 | - Exclusion zone contact(s): /OP1/ Consider K in place of MG (1 valid contact(s) for K vs. 0 for MG) |
| MG:DA:4762 | 1.00 | 77.27 |  | 0 | 1 | - Exclusion zone contact(s): /O2'/ Consider K in place of MG (1 valid contact(s) for K vs. 0 for MG) |
| MG:DA:4763 | 1.00 | 51.37 |  | 0 | 4 | - Exclusion zone contact(s): /N1_A_/O2/O2'/O4/ Consider K in place of MG (4 valid contact(s) for K vs. 0 for MG) |
| MG:DA:4764 | 1.00 | 85.56 |  | 0 |  | 1^st^ MG coordination shell missing (see Notes) |
| MG:DA:4765 | 1.00 | 120.55 |  | 0 | 4 | - Exclusion zone contact(s): /N3_A_/N6/N7/O2'/ Consider K in place of MG (3 valid contact(s) for K vs. 0 for MG) |
| MG:DA:4766 | 1.00 | 67.32 |  | 0 | 1 | - Exclusion zone contact(s): /OP1/ Consider K in place of MG (1 valid contact(s) for K vs. 0 for MG) |
| MG:DA:4767 | 1.00 | 52.07 |  | 0 | 2 | - Exclusion zone contact(s): /O2'/O4'/ Consider K in place of MG (2 valid contact(s) for K vs. 0 for MG) |
| MG:DA:4768 | 1.00 | 64.73 |  | 0 |  | 1^st^ MG coordination shell missing (see Notes) |
| MG:DA:4769 | 1.00 | 68.14 |  | 0 | 3 | - Exclusion zone contact(s): /O6/OP1/OP2/ Consider K in place of MG (3 valid contact(s) for K vs. 0 for MG) |
| MG:DA:4770 | 1.00 | 93.34 |  | 1 | 3 | O_ph_ - Exclusion zone contact(s): /O3'/O5'/OP2/ Consider K in place of MG (2 valid contact(s) for K vs. 1 for MG) |
| MG:DA:4771 | 1.00 | 92.65 |  | 0 | 3 | - Exclusion zone contact(s): /NZ_Lys_/O5'/OP2/ Consider K in place of MG (3 valid contact(s) for K vs. 0 for MG) |
| MG:DA:4772 | 1.00 | 46.99 |  | 0 | 3 | - Exclusion zone contact(s): /OP1/OP2/OP2/ Consider K in place of MG (3 valid contact(s) for K vs. 0 for MG) |
| MG:DA:4773 | 1.00 | 81.57 |  | 0 | 1 | - Exclusion zone contact(s): /N7/ Consider K in place of MG (1 valid contact(s) for K vs. 0 for MG) |
| MG:DA:4774 | 1.00 | 70.52 |  | 0 |  | 1^st^ MG coordination shell missing (see Notes) |
| MG:DA:4775 | 1.00 | 86.58 |  | 0 | 2 | - Exclusion zone contact(s): /OP1/OP1/ Consider K in place of MG (2 valid contact(s) for K vs. 0 for MG) |
| MG:DA:4776 | 1.00 | 72.45 |  | 1 | 1 | O_ph_ - Exclusion zone contact(s): /O4/ |
| MG:DA:4777 | 1.00 | 96.71 |  | 1 | 1 | O_r_ - Exclusion zone contact(s): /OP1/ |
| MG:DA:4778 | 1.00 | 66.18 |  | 0 | 1 | - Exclusion zone contact(s): /OP1/ Consider K in place of MG (1 valid contact(s) for K vs. 0 for MG) |
| MG:DA:4779 | 1.00 | 92.38 |  | 0 | 2 | - Exclusion zone contact(s): /OP2/OP2/ Consider K in place of MG (2 valid contact(s) for K vs. 0 for MG) |
| MG:DA:4780 | 1.00 | 62.98 |  | 0 | 1 | - Exclusion zone contact(s): /N7/ Consider K in place of MG (1 valid contact(s) for K vs. 0 for MG) |
| MG:DA:4781 | 1.00 | 105.56 |  | 0 |  | 1^st^ MG coordination shell missing (see Notes) |
| MG:DA:4782 | 1.00 | 83.64 |  | 0 | 2 | - Exclusion zone contact(s): /N7/O6/ Consider K in place of MG (2 valid contact(s) for K vs. 0 for MG) |
| MG:DA:4783 | 1.00 | 93.76 |  | 0 | 1 | - Exclusion zone contact(s): /O6/ Consider K in place of MG (1 valid contact(s) for K vs. 0 for MG) |
| MG:DA:4784 | 1.00 | 74.19 |  | 0 | 1 | - Exclusion zone contact(s): /O5'/ Consider K in place of MG (1 valid contact(s) for K vs. 0 for MG) |
| MG:DA:4785 | 1.00 | 75.54 |  | 0 | 2 | - Exclusion zone contact(s): /OP1/OP2/ Consider NA in place of MG (1 valid contact(s) for NA vs. 0 for MG) |
| MG:DA:4786 | 1.00 | 81.21 |  | 0 | 2 | - Exclusion zone contact(s): /O2/O2'/ Consider NA in place of MG (1 valid contact(s) for NA vs. 0 for MG) |
| MG:DA:4787 | 1.00 | 85.88 |  | 0 |  | No contacts in 5.0 A |
| MG:DA:4788 | 1.00 | 97.38 |  | 0 | 1 | - Exclusion zone contact(s): /O2/ Consider K in place of MG (1 valid contact(s) for K vs. 0 for MG) |
| MG:DA:4789 | 1.00 | 46.45 |  | 0 | 2 | - Exclusion zone contact(s): /O4'/ Consider K in place of MG (2 valid contact(s) for K vs. 0 for MG) |
| MG:DA:4790 | 1.00 | 53.78 |  | 0 |  | 1^st^ MG coordination shell missing (see Notes) |
| MG:DA:4791 | 1.00 | 69.67 |  | 0 |  | 1^st^ MG coordination shell missing (see Notes) |
| MG:DA:4792 | 1.00 | 113.41 |  | 0 | 2 | - Exclusion zone contact(s): /O2'/O3'/ Consider K in place of MG (2 valid contact(s) for K vs. 0 for MG) |
| MG:DA:4793 | 1.00 | 86.35 |  | 0 | 1 | - Exclusion zone contact(s): /OP1/ Consider K in place of MG (1 valid contact(s) for K vs. 0 for MG) |
| MG:DA:4794 | 1.00 | 86.07 |  | 0 |  | 1^st^ MG coordination shell missing (see Notes) |
| MG:DA:4795 | 1.00 | 66.50 |  | 0 | 1 | - Exclusion zone contact(s): /OP2/ Consider K in place of MG (1 valid contact(s) for K vs. 0 for MG) |
| MG:DA:4796 | 1.00 | 45.64 |  | 0 | 1 | - Exclusion zone contact(s): /N4/ Consider K in place of MG (1 valid contact(s) for K vs. 0 for MG) |
| MG:DA:4797 | 1.00 | 88.81 |  | 0 | 1 | - Exclusion zone contact(s): /O4/ Consider K in place of MG (1 valid contact(s) for K vs. 0 for MG) |
| MG:DA:4798 | 1.00 | 72.78 |  | 0 |  | 1^st^ MG coordination shell missing (see Notes) |
| MG:DA:4799 | 1.00 | 98.80 |  | 0 | 3 | - Exclusion zone contact(s): /O5'/OP2/OP2/ Consider NA in place of MG (2 valid contact(s) for NA vs. 0 for MG) |
| MG:DA:4800 | 1.00 | 49.06 |  | 0 | 2 | - Exclusion zone contact(s): /N4/N7/ Consider K in place of MG (2 valid contact(s) for K vs. 0 for MG) |
| MG:DA:4801 | 1.00 | 54.59 |  | 0 | 4 | - Exclusion zone contact(s): /N4/O6/O6/OP1/ Consider K in place of MG (4 valid contact(s) for K vs. 0 for MG) |
| MG:DA:4802 | 1.00 | 75.63 |  | 0 |  | 1^st^ MG coordination shell missing (see Notes) |
| MG:DA:4803 | 1.00 | 82.08 |  | 0 |  | 1^st^ MG coordination shell missing (see Notes) |
| MG:DA:4804 | 1.00 | 69.97 |  | 0 |  | No contacts in 5.0 A |
| MG:DA:4805 | 1.00 | 74.36 |  | 0 |  | 1^st^ MG coordination shell missing (see Notes) |
| MG:DA:4806 | 1.00 | 59.99 |  | 0 | 2 | - Exclusion zone contact(s): /OP1/OP2/ Consider K in place of MG (2 valid contact(s) for K vs. 0 for MG) |
| MG:DA:4807 | 1.00 | 68.50 |  | 0 | 5 | - Exclusion zone contact(s): /N7/O5'/OP2/OP2/OP2/ Consider K in place of MG (5 valid contact(s) for K vs. 0 for MG) |
| MG:DA:4808 | 1.00 | 61.90 |  | 1 | 2 | O_ph_ - Exclusion zone contact(s): /NH2_Arg_/O3'/ Consider K in place of MG (2 valid contact(s) for K vs. 1 for MG) |
| MG:DA:4809 | 1.00 | 59.74 |  | 0 | 2 | - Exclusion zone contact(s): /N7/O2'/ Consider K in place of MG (2 valid contact(s) for K vs. 0 for MG) |
| MG:DA:4810 | 1.00 | 62.33 |  | 0 | 1 | - Exclusion zone contact(s): /OP2/ Consider K in place of MG (1 valid contact(s) for K vs. 0 for MG) |
| MG:DA:4811 | 1.00 | 97.64 |  | 0 | 2 | - Exclusion zone contact(s): /OP2/OP2/ Consider NA in place of MG (1 valid contact(s) for NA vs. 0 for MG) |
| MG:DA:4812 | 1.00 | 87.06 |  | 0 | 1 | - Exclusion zone contact(s): /O4'/ Consider K in place of MG (1 valid contact(s) for K vs. 0 for MG) |
| MG:DA:4813 | 1.00 | 78.98 |  | 0 |  | 1^st^ MG coordination shell missing (see Notes) |
| MG:DA:4814 | 1.00 | 114.78 |  | 0 | 1 | - Exclusion zone contact(s): /OP2/ Consider K in place of MG (1 valid contact(s) for K vs. 0 for MG) |
| MG:DA:4815 | 1.00 | 53.06 |  | 0 | 1 | - Exclusion zone contact(s): /N3_A_/ Consider K in place of MG (1 valid contact(s) for K vs. 0 for MG) |
| MG:DA:4816 | 1.00 | 66.57 |  | 0 | 3 | - Exclusion zone contact(s): /O5'/OP2/OP2/ Consider K in place of MG (2 valid contact(s) for K vs. 0 for MG) |
| MG:DA:4817 | 1.00 | 78.75 |  | 0 | 1 | - Exclusion zone contact(s): /O6/ Consider K in place of MG (1 valid contact(s) for K vs. 0 for MG) |
| MG:DA:4818 | 1.00 | 80.39 |  | 0 | 1 | - Exclusion zone contact(s): /N7/ Consider K in place of MG (1 valid contact(s) for K vs. 0 for MG) |
| MG:DA:4819 | 1.00 | 64.92 |  | 1 |  | O_ph_ |
| MG:DA:4820 | 1.00 | 56.56 |  | 0 | 1 | - Exclusion zone contact(s): /OP2/ Consider K in place of MG (1 valid contact(s) for K vs. 0 for MG) |
| MG:DA:4821 | 1.00 | 54.35 |  | 0 |  | 1^st^ MG coordination shell missing (see Notes) |
| MG:DA:4822 | 1.00 | 39.53 |  | 0 |  | No contacts in 5.0 A |
| MG:DA:4823 | 1.00 | 71.81 |  | 0 | 2 | - Exclusion zone contact(s): /N1/O_back_/ Consider K in place of MG (2 valid contact(s) for K vs. 0 for MG) |
| MG:DA:4824 | 1.00 | 71.23 |  | 0 | 4 | - Exclusion zone contact(s): /NE_Arg_/O2'/O3'/OP1/ Consider K in place of MG (3 valid contact(s) for K vs. 0 for MG) |
| MG:DA:4825 | 1.00 | 63.98 |  | 0 | 3 | - Exclusion zone contact(s): /O4/OP1/OP2/ Consider K in place of MG (3 valid contact(s) for K vs. 0 for MG) |
| MG:DA:4826 | 1.00 | 71.97 |  | 0 | 1 | - Exclusion zone contact(s): /OP2/ Consider K in place of MG (1 valid contact(s) for K vs. 0 for MG) |
| MG:DA:4827 | 1.00 | 87.43 |  | 0 | 2 | - Exclusion zone contact(s): /OP1/OP2/ Consider K in place of MG (2 valid contact(s) for K vs. 0 for MG) |
| MG:DA:4828 | 1.00 | 103.07 |  | 0 | 1 | - Exclusion zone contact(s): /O2'/ Consider K in place of MG (1 valid contact(s) for K vs. 0 for MG) |
| MG:DA:4829 | 1.00 | 106.23 |  | 0 | 2 | - Exclusion zone contact(s): /O5'/OP2/ Consider K in place of MG (2 valid contact(s) for K vs. 0 for MG) |
| MG:DA:4830 | 1.00 | 64.02 |  | 0 | 5 | - Exclusion zone contact(s): /N2/N3_G_/O2'/O4'/OP1/ Consider K in place of MG (4 valid contact(s) for K vs. 0 for MG) |
| MG:DA:4831 | 1.00 | 82.82 |  | 0 |  | 1^st^ MG coordination shell missing (see Notes) |
| MG:DA:4832 | 1.00 | 86.83 |  | 0 |  | 1^st^ MG coordination shell missing (see Notes) |
| MG:DA:4833 | 1.00 | 76.90 |  | 0 | 1 | - Exclusion zone contact(s): /OP1/ Consider K in place of MG (1 valid contact(s) for K vs. 0 for MG) |
| MG:DA:4834 | 1.00 | 78.53 |  | 0 | 2 | - Exclusion zone contact(s): /N1/OP1/ Consider K in place of MG (2 valid contact(s) for K vs. 0 for MG) |
| MG:DA:4835 | 1.00 | 65.12 |  | 0 | 2 | - Exclusion zone contact(s): /N7/OP2/ Consider K in place of MG (2 valid contact(s) for K vs. 0 for MG) |
| MG:DA:4836 | 1.00 | 41.74 |  | 0 | 1 | - Exclusion zone contact(s): /O4/ Consider K in place of MG (1 valid contact(s) for K vs. 0 for MG) |
| MG:DA:4837 | 1.00 | 55.22 |  | 0 | 1 | - Exclusion zone contact(s): /OP1/ Consider NA in place of MG (1 valid contact(s) for NA vs. 0 for MG) |
| MG:DA:4838 | 1.00 | 67.09 |  | 0 | 3 | - Exclusion zone contact(s): /O5'/OP2/OP2/ Consider K in place of MG (2 valid contact(s) for K vs. 0 for MG) |
| MG:DA:4839 | 1.00 | 75.67 |  | 0 |  | 1^st^ MG coordination shell missing (see Notes) |
| MG:DA:4840 | 1.00 | 111.09 |  | 0 |  | 1^st^ MG coordination shell missing (see Notes) |
| MG:DA:4841 | 1.00 | 80.16 |  | 0 |  | 1^st^ MG coordination shell missing (see Notes) |
| MG:DA:4842 | 1.00 | 65.39 |  | 0 | 1 | - Exclusion zone contact(s): /OP2/ Consider K in place of MG (1 valid contact(s) for K vs. 0 for MG) |
| MG:DA:4843 | 1.00 | 72.48 |  | 0 | 3 | - Exclusion zone contact(s): /O2'/O3'/O_back_/ Consider K in place of MG (3 valid contact(s) for K vs. 0 for MG) |
| MG:DA:4844 | 1.00 | 58.24 |  | 0 |  | 1^st^ MG coordination shell missing (see Notes) |
| MG:DA:4845 | 1.00 | 43.76 |  | 0 | 3 | - Exclusion zone contact(s): /N1/N7/O5'/ Consider K in place of MG (3 valid contact(s) for K vs. 0 for MG) |
| MG:DA:4846 | 1.00 | 73.63 |  | 0 | 1 | - Exclusion zone contact(s): /OP1/ Consider NA in place of MG (1 valid contact(s) for NA vs. 0 for MG) |
| MG:DA:4847 | 1.00 | 117.94 |  | 0 | 1 | - Exclusion zone contact(s): /O2'/ Consider K in place of MG (1 valid contact(s) for K vs. 0 for MG) |
| MG:DA:4848 | 1.00 | 67.09 |  | 0 | 1 | - Exclusion zone contact(s): /OP1/ Consider K in place of MG (1 valid contact(s) for K vs. 0 for MG) |
| MG:DA:4849 | 1.00 | 55.86 |  | 0 | 4 | - Exclusion zone contact(s): /N1/N1/N3_C_/O2/ Consider K in place of MG (4 valid contact(s) for K vs. 0 for MG) |
| MG:DA:4850 | 1.00 | 38.40 |  | 0 | 4 | - Exclusion zone contact(s): /OG_Ser_/OP2/OP2/OP2/ Consider NA in place of MG (3 valid contact(s) for NA vs. 0 for MG) |
| MG:DA:4851 | 1.00 | 139.59 |  | 0 | 2 | - Exclusion zone contact(s): /OP2/OP2/ Consider K in place of MG (2 valid contact(s) for K vs. 0 for MG) |
| MG:DA:4852 | 1.00 | 142.82 |  | 0 | 2 | - Exclusion zone contact(s): /N7/O6/ Consider K in place of MG (2 valid contact(s) for K vs. 0 for MG) |
| MG:DA:4853 | 1.00 | 80.29 |  | 0 | 2 | - Exclusion zone contact(s): /OP2/OP2/ Consider K in place of MG (2 valid contact(s) for K vs. 0 for MG) |
| MG:DA:4854 | 1.00 | 66.95 |  | 1 | 1 | N_b_ - Exclusion zone contact(s): /OP1/ |
| MG:DA:4855 | 1.00 | 74.23 |  | 0 | 4 | - Exclusion zone contact(s): /O5'/OP1/OP2/OP2/ Consider K in place of MG (4 valid contact(s) for K vs. 0 for MG) |
| MG:DA:4856 | 1.00 | 49.45 |  | 0 | 4 | - Exclusion zone contact(s): /N1_G_/N2/OP2/OP2/ Consider K in place of MG (4 valid contact(s) for K vs. 0 for MG) |
| MG:DA:4857 | 1.00 | 56.90 |  | 0 | 3 | - Exclusion zone contact(s): /O5'/OP2/OP2/ Consider K in place of MG (3 valid contact(s) for K vs. 0 for MG) |
| MG:DA:4858 | 1.00 | 69.52 |  | 0 | 2 | - Exclusion zone contact(s): /O2'/O3'/ Consider K in place of MG (2 valid contact(s) for K vs. 0 for MG) |
| MG:DA:4859 | 1.00 | 91.48 |  | 0 |  | No contacts in 5.0 A |
| MG:DA:4860 | 1.00 | 64.44 |  | 0 | 2 | - Exclusion zone contact(s): /N3_G_/O2'/ Consider NA in place of MG (1 valid contact(s) for NA vs. 0 for MG) |
| MG:DA:4861 | 1.00 | 74.59 |  | 0 | 1 | - Exclusion zone contact(s): /O6/ Consider NA in place of MG (1 valid contact(s) for NA vs. 0 for MG) |
| MG:DA:4862 | 1.00 | 66.59 |  | 0 | 2 | - Exclusion zone contact(s): /OP1/OP2/ Consider NA in place of MG (1 valid contact(s) for NA vs. 0 for MG) |
| MG:DA:4863 | 1.00 | 83.11 |  | 0 | 1 | - Exclusion zone contact(s): /O4/ Consider K in place of MG (1 valid contact(s) for K vs. 0 for MG) |
| MG:DA:4864 | 1.00 | 93.46 |  | 0 |  | 1^st^ MG coordination shell missing (see Notes) |
| MG:DA:4865 | 1.00 | 67.55 |  | 0 | 1 | - Exclusion zone contact(s): /O6/ Consider K in place of MG (1 valid contact(s) for K vs. 0 for MG) |
| MG:DA:4866 | 1.00 | 53.22 |  | 0 | 1 | - Exclusion zone contact(s): /OP1/ Consider NA in place of MG (1 valid contact(s) for NA vs. 0 for MG) |
| MG:DA:4867 | 1.00 | 62.27 |  | 0 | 3 | - Exclusion zone contact(s): /O5'/OP2/OP2/ Consider K in place of MG (2 valid contact(s) for K vs. 0 for MG) |
| MG:DA:4868 | 1.00 | 70.15 |  | 0 |  | 1^st^ MG coordination shell missing (see Notes) |
| MG:DA:4869 | 1.00 | 44.15 |  | 0 | 2 | - Exclusion zone contact(s): /O2'/O5'/ Consider K in place of MG (2 valid contact(s) for K vs. 0 for MG) |
| MG:DA:4870 | 1.00 | 82.85 |  | 0 |  | No contacts in 5.0 A |
| MG:DA:4871 | 1.00 | 86.63 |  | 0 | 3 | - Exclusion zone contact(s): /OP1/OP2/OP2/ Consider K in place of MG (3 valid contact(s) for K vs. 0 for MG) |
| MG:DA:4872 | 1.00 | 106.79 |  | 0 | 1 | - Exclusion zone contact(s): /OP2/ Consider K in place of MG (1 valid contact(s) for K vs. 0 for MG) |
| MG:DA:4873 | 1.00 | 49.25 |  | 0 | 4 | - Exclusion zone contact(s): /N7/N7/O6/OP2/ Consider K in place of MG (4 valid contact(s) for K vs. 0 for MG) |
| MG:DA:4874 | 1.00 | 84.54 |  | 0 | 1 | - Exclusion zone contact(s): /OP1/ Consider K in place of MG (1 valid contact(s) for K vs. 0 for MG) |
| MG:DA:4875 | 1.00 | 88.96 |  | 1 | 2 | O_ph_ - Exclusion zone contact(s): /O3'/OP2/ Consider K in place of MG (2 valid contact(s) for K vs. 1 for MG) |
| MG:DA:4876 | 1.00 | 61.49 |  | 0 | 1 | - Exclusion zone contact(s): /O4/ Consider K in place of MG (1 valid contact(s) for K vs. 0 for MG) |
| MG:DA:4877 | 1.00 | 77.95 |  | 0 | 1 | - Exclusion zone contact(s): /O4/ Consider K in place of MG (1 valid contact(s) for K vs. 0 for MG) |
| MG:DA:4878 | 1.00 | 50.09 |  | 0 | 1 | - Exclusion zone contact(s): /OP1/ Consider K in place of MG (1 valid contact(s) for K vs. 0 for MG) |
| MG:DA:4879 | 1.00 | 62.63 |  | 0 | 1 | - Exclusion zone contact(s): /N6/ Consider K in place of MG (1 valid contact(s) for K vs. 0 for MG) |
| MG:DA:4880 | 1.00 | 110.74 |  | 0 | 2 | - Exclusion zone contact(s): /OP1/OP2/ Consider K in place of MG (2 valid contact(s) for K vs. 0 for MG) |
| MG:DA:4881 | 1.00 | 85.88 |  | 0 |  | 1^st^ MG coordination shell missing (see Notes) |
| MG:DA:4882 | 1.00 | 74.01 |  | 0 | 2 | - Exclusion zone contact(s): /OP1/OP2/ Consider K in place of MG (2 valid contact(s) for K vs. 0 for MG) |
| MG:DA:4883 | 1.00 | 96.71 |  | 1 | 2 | O_ph_ - Exclusion zone contact(s): /O4'/O5'/ |
| MG:DA:4884 | 1.00 | 73.64 |  | 0 | 2 | - Exclusion zone contact(s): /O2/O2'/ Consider K in place of MG (2 valid contact(s) for K vs. 0 for MG) |
| MG:DA:4885 | 1.00 | 92.27 |  | 0 |  | No contacts in 5.0 A |
| MG:DA:4886 | 1.00 | 60.64 |  | 0 | 2 | - Exclusion zone contact(s): /O3'/OP1/ Consider K in place of MG (2 valid contact(s) for K vs. 0 for MG) |
| MG:DA:4887 | 1.00 | 87.76 |  | 1 | 1 | N_b_ - Exclusion zone contact(s): /O6/ |
| MG:DA:4888 | 1.00 | 76.07 |  | 0 |  | 1^st^ MG coordination shell missing (see Notes) |
| MG:DA:4889 | 1.00 | 75.52 |  | 0 |  | 1^st^ MG coordination shell missing (see Notes) |
| MG:DA:4890 | 1.00 | 96.99 |  | 0 | 2 | - Exclusion zone contact(s): /N1/O4'/ Consider K in place of MG (2 valid contact(s) for K vs. 0 for MG) |
| MG:DA:4891 | 1.00 | 86.90 |  | 0 | 4 | - Exclusion zone contact(s): /N7/O5'/OP2/OP2/ Consider K in place of MG (4 valid contact(s) for K vs. 0 for MG) |
| MG:DA:4892 | 1.00 | 133.09 |  | 0 | 4 | - Exclusion zone contact(s): /O2'/O2'/O3'/O4'/ Consider NA in place of MG (2 valid contact(s) for NA vs. 0 for MG) |
| MG:DA:4893 | 1.00 | 99.30 |  | 1 | 1 | O_ph_ - Exclusion zone contact(s): /OP2/ |
| MG:DA:4894 | 1.00 | 63.32 |  | 0 |  | 1^st^ MG coordination shell missing (see Notes) |
| MG:DA:4895 | 1.00 | 77.51 |  | 0 | 1 | - Exclusion zone contact(s): /OP1/ Consider K in place of MG (1 valid contact(s) for K vs. 0 for MG) |
| MG:DA:4896 | 1.00 | 74.76 |  | 0 |  | 1^st^ MG coordination shell missing (see Notes) |
| MG:DA:4897 | 1.00 | 76.46 |  | 0 | 1 | - Exclusion zone contact(s): /OP1/ Consider NA in place of MG (1 valid contact(s) for NA vs. 0 for MG) |
| MG:DA:4898 | 1.00 | 58.22 |  | 0 | 4 | - Exclusion zone contact(s): /N3_A_/N6/O2'/O2'/ Consider K in place of MG (4 valid contact(s) for K vs. 0 for MG) |
| MG:DA:4899 | 1.00 | 65.73 |  | 0 |  | 1^st^ MG coordination shell missing (see Notes) |
| MG:DA:4900 | 1.00 | 76.15 |  | 0 |  | 1^st^ MG coordination shell missing (see Notes) |
| MG:DA:4901 | 1.00 | 84.85 |  | 0 | 2 | - Exclusion zone contact(s): /N3_A_/O2'/ Consider K in place of MG (2 valid contact(s) for K vs. 0 for MG) |
| MG:DA:4902 | 1.00 | 82.44 |  | 1 | 3 | O_ph_ - Exclusion zone contact(s): /O5'/OP1/OP2/ Consider K in place of MG (3 valid contact(s) for K vs. 1 for MG) |
| MG:DA:4903 | 1.00 | 69.60 |  | 0 | 1 | - Exclusion zone contact(s): /OP1/ Consider K in place of MG (1 valid contact(s) for K vs. 0 for MG) |
| MG:DA:4904 | 1.00 | 71.24 |  | 0 | 2 | - Exclusion zone contact(s): /O4'/OP2/ Consider K in place of MG (2 valid contact(s) for K vs. 0 for MG) |
| MG:DA:4905 | 1.00 | 90.44 |  | 0 |  | 1^st^ MG coordination shell missing (see Notes) |
| MG:DA:4906 | 1.00 | 96.48 |  | 0 |  | 1^st^ MG coordination shell missing (see Notes) |
| MG:DA:4907 | 1.00 | 57.69 |  | 1 | 2 | O_r_ - Exclusion zone contact(s): /O2/O2'/ Consider K in place of MG (2 valid contact(s) for K vs. 1 for MG) |
| MG:DA:4908 | 1.00 | 64.63 |  | 0 |  | 1^st^ MG coordination shell missing (see Notes) |
| MG:DA:4909 | 1.00 | 70.62 |  | 0 |  | No contacts in 5.0 A |
| MG:DA:4910 | 1.00 | 67.30 |  | 0 | 2 | - Exclusion zone contact(s): /O4/O6/ Consider K in place of MG (2 valid contact(s) for K vs. 0 for MG) |
| MG:DA:4911 | 1.00 | 85.84 |  | 0 | 2 | - Exclusion zone contact(s): /N7/OP1/ Consider NA in place of MG (1 valid contact(s) for NA vs. 0 for MG) |
| MG:DA:4912 | 1.00 | 42.97 |  | 0 | 4 | - Exclusion zone contact(s): /O2'/O3'/OP2/OP2/ Consider K in place of MG (3 valid contact(s) for K vs. 0 for MG) |
| MG:DA:4913 | 1.00 | 74.31 |  | 0 | 4 | - Exclusion zone contact(s): /O2'/O4'/OP1/OP2/ Consider K in place of MG (3 valid contact(s) for K vs. 0 for MG) |
| MG:DA:4914 | 1.00 | 44.64 |  | 0 | 1 | - Exclusion zone contact(s): /OP2/ Consider K in place of MG (1 valid contact(s) for K vs. 0 for MG) |
| MG:DA:4915 | 1.00 | 64.47 |  | 0 | 2 | - Exclusion zone contact(s): /N4/O6/ Consider NA in place of MG (1 valid contact(s) for NA vs. 0 for MG) |
| MG:DA:4916 | 1.00 | 88.16 |  | 0 | 3 | - Exclusion zone contact(s): /N6/O3'/OP1/ Consider NA in place of MG (2 valid contact(s) for NA vs. 0 for MG) |
| MG:DA:4917 | 1.00 | 61.22 |  | 0 |  | 1^st^ MG coordination shell missing (see Notes) |
| MG:DA:4918 | 1.00 | 48.71 |  | 0 | 3 | - Exclusion zone contact(s): /O2'/O4'/O5'/ Consider K in place of MG (3 valid contact(s) for K vs. 0 for MG) |
| MG:DA:4919 | 1.00 | 58.14 |  | 0 | 4 | - Exclusion zone contact(s): /N6/O5'/OP2/OP2/ Consider K in place of MG (3 valid contact(s) for K vs. 0 for MG) |
| MG:DA:4920 | 1.00 | 61.60 |  | 0 | 2 | - Exclusion zone contact(s): /N7/OP2/ Consider NA in place of MG (1 valid contact(s) for NA vs. 0 for MG) |
| MG:DA:4921 | 1.00 | 67.82 |  | 0 |  | 1^st^ MG coordination shell missing (see Notes) |
| MG:DA:4922 | 1.00 | 99.63 |  | 0 |  | 1^st^ MG coordination shell missing (see Notes) |
| MG:DA:4923 | 1.00 | 74.10 |  | 0 | 1 | - Exclusion zone contact(s): /OP1/ Consider NA in place of MG (1 valid contact(s) for NA vs. 0 for MG) |
| MG:DA:4924 | 1.00 | 75.40 |  | 0 | 1 | - Exclusion zone contact(s): /OP1/ Consider K in place of MG (1 valid contact(s) for K vs. 0 for MG) |
| MG:DA:4925 | 1.00 | 73.81 |  | 0 |  | 1^st^ MG coordination shell missing (see Notes) |
| MG:DA:4926 | 1.00 | 46.69 |  | 0 |  | 1^st^ MG coordination shell missing (see Notes) |
| MG:DA:4927 | 1.00 | 160.21 |  | 0 | 3 | - Exclusion zone contact(s): /N1/N2/N3_U_/ Consider K in place of MG (3 valid contact(s) for K vs. 0 for MG) |
| MG:DA:4928 | 1.00 | 78.62 |  | 0 | 2 | - Exclusion zone contact(s): /N3_A_/O4'/ Consider K in place of MG (2 valid contact(s) for K vs. 0 for MG) |
| MG:DA:4929 | 1.00 | 52.90 |  | 0 | 3 | - Exclusion zone contact(s): /N1_A_/NH2_Arg_/O2'/ Consider K in place of MG (3 valid contact(s) for K vs. 0 for MG) |
| MG:DA:4930 | 1.00 | 73.00 |  | 0 | 3 | - Exclusion zone contact(s): /N2/OP1/OP2/ Consider K in place of MG (3 valid contact(s) for K vs. 0 for MG) |
| MG:DA:4931 | 1.00 | 41.43 |  | 0 | 3 | - Exclusion zone contact(s): /O4/O4/O6/ Consider K in place of MG (2 valid contact(s) for K vs. 0 for MG) |
| MG:DA:4932 | 1.00 | 69.93 |  | 0 | 1 | - Exclusion zone contact(s): /OP2/ Consider K in place of MG (1 valid contact(s) for K vs. 0 for MG) |
| MG:DA:4933 | 1.00 | 64.91 |  | 0 | 1 | - Exclusion zone contact(s): /OP1/ Consider K in place of MG (1 valid contact(s) for K vs. 0 for MG) |
| MG:DA:4934 | 1.00 | 61.55 |  | 0 |  | 1^st^ MG coordination shell missing (see Notes) |
| MG:DA:4935 | 1.00 | 105.99 |  | 0 |  | 1^st^ MG coordination shell missing (see Notes) |
| MG:DA:4936 | 1.00 | 52.53 |  | 0 | 3 | - Exclusion zone contact(s): /O2'/O2'/O2'/ Consider K in place of MG (3 valid contact(s) for K vs. 0 for MG) |
| MG:DA:4937 | 1.00 | 78.34 |  | 0 | 3 | - Exclusion zone contact(s): /O5'/OP2/OP2/ Consider NA in place of MG (2 valid contact(s) for NA vs. 0 for MG) |
| MG:DA:4938 | 1.00 | 55.54 |  | 0 | 2 | - Exclusion zone contact(s): /OP1/OP2/ Consider NA in place of MG (1 valid contact(s) for NA vs. 0 for MG) |
| MG:DA:4939 | 1.00 | 57.29 |  | 0 | 3 | - Exclusion zone contact(s): /N3_G_/O2'/OH_Tyr_/ Consider K in place of MG (2 valid contact(s) for K vs. 0 for MG) |
| MG:DA:4940 | 1.00 | 69.05 |  | 0 | 2 | - Exclusion zone contact(s): /N4/N6/ Consider K in place of MG (2 valid contact(s) for K vs. 0 for MG) |
| MG:DA:4941 | 1.00 | 88.89 |  | 0 |  | No contacts in 5.0 A |
| MG:DA:4942 | 1.00 | 82.94 |  | 0 | 3 | - Exclusion zone contact(s): /O5'/OP2/OP2/ Consider K in place of MG (2 valid contact(s) for K vs. 0 for MG) |
| MG:DA:4943 | 1.00 | 81.38 |  | 0 | 1 | - Exclusion zone contact(s): /OP2/ Consider K in place of MG (1 valid contact(s) for K vs. 0 for MG) |
| MG:DA:4944 | 1.00 | 56.69 |  | 0 | 1 | - Exclusion zone contact(s): /OP1/ Consider K in place of MG (1 valid contact(s) for K vs. 0 for MG) |
| MG:DA:4945 | 1.00 | 78.66 |  | 0 |  | 1^st^ MG coordination shell missing (see Notes) |
| MG:DA:4946 | 1.00 | 62.75 |  | 1 | 3 | O_r_ - Exclusion zone contact(s): /N3_A_/NZ_Lys_/OH_Tyr_/ Consider K in place of MG (2 valid contact(s) for K vs. 1 for MG) |
| MG:DA:4947 | 1.00 | 87.02 |  | 1 | 3 | O_ph_ - Exclusion zone contact(s): /O5'/OP1/OP2/ Consider K in place of MG (3 valid contact(s) for K vs. 1 for MG) |
| MG:DA:4948 | 1.00 | 126.48 |  | 0 |  | 1^st^ MG coordination shell missing (see Notes) |
| MG:DA:4949 | 1.00 | 71.84 |  | 0 | 2 | - Exclusion zone contact(s): /N1_A_/N7/ Consider K in place of MG (2 valid contact(s) for K vs. 0 for MG) |
| MG:DA:4950 | 1.00 | 125.66 |  | 0 |  | 1^st^ MG coordination shell missing (see Notes) |
| MG:DA:4951 | 1.00 | 102.42 |  | 0 | 2 | - Exclusion zone contact(s): /N3_G_/O2'/ Consider K in place of MG (2 valid contact(s) for K vs. 0 for MG) |
| MG:DA:4952 | 1.00 | 86.81 |  | 0 | 3 | - Exclusion zone contact(s): /O4'/O6/ Consider K in place of MG (3 valid contact(s) for K vs. 0 for MG) |
| MG:DA:4953 | 1.00 | 63.03 |  | 0 | 1 | - Exclusion zone contact(s): /OP2/ Consider K in place of MG (1 valid contact(s) for K vs. 0 for MG) |
| MG:DA:4954 | 1.00 | 73.76 |  | 0 |  | No contacts in 5.0 A |
| MG:DA:4955 | 1.00 | 82.36 |  | 0 |  | 1^st^ MG coordination shell missing (see Notes) |
| MG:DA:4956 | 1.00 | 118.08 |  | 0 | 3 | - Exclusion zone contact(s): /O3'/OP1/OP2/ Consider K in place of MG (2 valid contact(s) for K vs. 0 for MG) |
| MG:DA:4957 | 1.00 | 86.74 |  | 0 | 3 | - Exclusion zone contact(s): /N7/OP1/OP2/ Consider K in place of MG (3 valid contact(s) for K vs. 0 for MG) |
| MG:DA:4958 | 1.00 | 64.15 |  | 0 | 3 | - Exclusion zone contact(s): /O4'/O5'/OP1/ Consider K in place of MG (3 valid contact(s) for K vs. 0 for MG) |
| MG:DA:4959 | 1.00 | 45.78 |  | 0 | 5 | - Exclusion zone contact(s): /N1_A_/N3_C_/N7/O2/OP1/ Consider K in place of MG (5 valid contact(s) for K vs. 0 for MG) |
| MG:DA:4960 | 1.00 | 77.75 |  | 0 |  | 1^st^ MG coordination shell missing (see Notes) |
| MG:DA:4961 | 1.00 | 68.05 |  | 0 | 1 | - Exclusion zone contact(s): /OP1/ Consider K in place of MG (1 valid contact(s) for K vs. 0 for MG) |
| MG:DA:4962 | 1.00 | 77.46 |  | 0 | 2 | - Exclusion zone contact(s): /N6/N7/ Consider K in place of MG (2 valid contact(s) for K vs. 0 for MG) |
| MG:DA:4963 | 1.00 | 63.69 |  | 0 |  | 1^st^ MG coordination shell missing (see Notes) |
| MG:DA:4964 | 1.00 | 73.04 |  | 0 |  | 1^st^ MG coordination shell missing (see Notes) |
| MG:DA:4965 | 1.00 | 61.68 |  | 0 | 2 | - Exclusion zone contact(s): /N3_U_/OP2/ Consider K in place of MG (2 valid contact(s) for K vs. 0 for MG) |
| MG:DA:4966 | 1.00 | 64.00 |  | 0 | 1 | - Exclusion zone contact(s): /O2'/ Consider K in place of MG (1 valid contact(s) for K vs. 0 for MG) |
| MG:DA:4967 | 1.00 | 74.26 |  | 0 | 1 | - Exclusion zone contact(s): /O4/ Consider K in place of MG (1 valid contact(s) for K vs. 0 for MG) |
| MG:DA:4968 | 1.00 | 65.64 |  | 0 | 3 | - Exclusion zone contact(s): /O3'/O5'/OP1/ Consider K in place of MG (2 valid contact(s) for K vs. 0 for MG) |
| MG:DA:4969 | 1.00 | 75.21 |  | 0 | 1 | - Exclusion zone contact(s): /OP2/ Consider NA in place of MG (1 valid contact(s) for NA vs. 0 for MG) |
| MG:DA:4970 | 1.00 | 87.49 |  | 0 |  | 1^st^ MG coordination shell missing (see Notes) |
| MG:DA:4971 | 1.00 | 79.38 |  | 0 |  | 1^st^ MG coordination shell missing (see Notes) |
| MG:DA:4972 | 1.00 | 80.13 |  | 0 | 1 | - Exclusion zone contact(s): /OP1/ Consider NA in place of MG (1 valid contact(s) for NA vs. 0 for MG) |
| MG:DA:4973 | 1.00 | 115.03 |  | 0 | 3 | - Exclusion zone contact(s): /N2/N3_G_/O2/ Consider K in place of MG (3 valid contact(s) for K vs. 0 for MG) |
| MG:DA:4974 | 1.00 | 57.40 |  | 0 | 1 | - Exclusion zone contact(s): /OP2/ Consider K in place of MG (1 valid contact(s) for K vs. 0 for MG) |
| MG:DA:4975 | 1.00 | 78.85 |  | 0 | 2 | - Exclusion zone contact(s): /O5'/OP2/ Consider K in place of MG (2 valid contact(s) for K vs. 0 for MG) |
| MG:DA:4976 | 1.00 | 98.03 |  | 0 | 2 | - Exclusion zone contact(s): /N1_G_/O6/ Consider NA in place of MG (1 valid contact(s) for NA vs. 0 for MG) |
| MG:DA:4977 | 1.00 | 94.43 |  | 2 | 3 | O_ph_O_r_ - Exclusion zone contact(s): /N4/O3'/O4'/ |
| MG:DA:4978 | 1.00 | 57.59 |  | 0 | 1 | - Exclusion zone contact(s): /OP2/ Consider K in place of MG (1 valid contact(s) for K vs. 0 for MG) |
| MG:DA:4979 | 1.00 | 47.68 |  | 0 | 3 | - Exclusion zone contact(s): /O3'/O5'/OP2/ Consider K in place of MG (3 valid contact(s) for K vs. 0 for MG) |
| MG:DA:4980 | 1.00 | 79.59 |  | 0 |  | 1^st^ MG coordination shell missing (see Notes) |
| MG:DA:4981 | 1.00 | 41.31 |  | 0 | 4 | - Exclusion zone contact(s): /N2/O2'/O3'/OP1/ Consider K in place of MG (3 valid contact(s) for K vs. 0 for MG) |
| MG:DA:4982 | 1.00 | 53.72 |  | 0 | 4 | - Exclusion zone contact(s): /NH1_Arg_/NH2_Arg_/OP1/OP2/ Consider K in place of MG (4 valid contact(s) for K vs. 0 for MG) |
| MG:DA:4983 | 1.00 | 53.84 |  | 1 | 2 | O_ph_ - Exclusion zone contact(s): /N7/O5'/ Consider K in place of MG (2 valid contact(s) for K vs. 1 for MG) |
| MG:DA:4984 | 1.00 | 90.29 |  | 0 |  | 1^st^ MG coordination shell missing (see Notes) |
| MG:DA:4985 | 1.00 | 88.52 |  | 0 |  | 1^st^ MG coordination shell missing (see Notes) |
| MG:DA:4986 | 1.00 | 63.13 |  | 0 | 6 | - Exclusion zone contact(s): /N3_C_/N4/N4/N6/N7/O2/ Consider K in place of MG (6 valid contact(s) for K vs. 0 for MG) |
| MG:DA:4987 | 1.00 | 37.21 |  | 0 | 4 | - Exclusion zone contact(s): /O2'/O2'/O3'/O5'/ Consider K in place of MG (4 valid contact(s) for K vs. 0 for MG) |
| MG:DA:4988 | 1.00 | 64.05 |  | 0 | 3 | - Exclusion zone contact(s): /O2'/O3'/OP1/ Consider K in place of MG (2 valid contact(s) for K vs. 0 for MG) |
| MG:DA:4989 | 1.00 | 66.63 |  | 0 |  | No contacts in 5.0 A |
| MG:DA:4990 | 1.00 | 70.77 |  | 0 |  | 1^st^ MG coordination shell missing (see Notes) |
| MG:DA:4991 | 1.00 | 123.99 |  | 0 | 3 | - Exclusion zone contact(s): /N4/OP1/OP2/ Consider NA in place of MG (2 valid contact(s) for NA vs. 0 for MG) |
| MG:DA:4992 | 1.00 | 88.32 |  | 0 | 5 | - Exclusion zone contact(s): /O5'/OP1/OP1/OP2/OP2/ Consider K in place of MG (5 valid contact(s) for K vs. 0 for MG) |
| MG:DA:4993 | 1.00 | 139.68 |  | 1 | 3 | Anion - Exclusion zone contact(s): /N6/N7/OD1_Asn_/ Consider K in place of MG (3 valid contact(s) for K vs. 1 for MG) |
| MG:DA:4994 | 1.00 | 98.91 |  | 0 | 2 | - Exclusion zone contact(s): /O2'/O3'/ Consider K in place of MG (2 valid contact(s) for K vs. 0 for MG) |
| MG:DA:4995 | 1.00 | 121.74 |  | 1 | 2 | O_ph_ - Exclusion zone contact(s): /N7/O6/ Consider NA in place of MG (2 valid contact(s) for NA vs. 1 for MG) |
| MG:DA:4996 | 1.00 | 85.75 |  | 0 |  | 1^st^ MG coordination shell missing (see Notes) |
| MG:DA:4997 | 1.00 | 50.07 |  | 0 | 1 | - Exclusion zone contact(s): /OP1/ Consider K in place of MG (1 valid contact(s) for K vs. 0 for MG) |
| MG:DA:4998 | 1.00 | 110.04 |  | 0 | 2 | - Exclusion zone contact(s): /O2'/O3'/ Consider NA in place of MG (1 valid contact(s) for NA vs. 0 for MG) |
| MG:DA:4999 | 1.00 | 74.46 |  | 0 | 3 | - Exclusion zone contact(s): /O5'/OP2/OP2/ Consider K in place of MG (2 valid contact(s) for K vs. 0 for MG) |
| MG:DA:5000 | 1.00 | 71.90 |  | 0 | 4 | - Exclusion zone contact(s): /O2'/O3'/O_back_/OP1/ Consider K in place of MG (3 valid contact(s) for K vs. 0 for MG) |
| MG:DA:5001 | 1.00 | 75.86 |  | 0 | 1 | - Exclusion zone contact(s): /OP1/ Consider K in place of MG (1 valid contact(s) for K vs. 0 for MG) |
| MG:DA:5002 | 1.00 | 89.77 |  | 0 |  | 1^st^ MG coordination shell missing (see Notes) |
| MG:DA:5003 | 1.00 | 76.42 |  | 0 | 2 | - Exclusion zone contact(s): /OP1/OP2/ Consider K in place of MG (2 valid contact(s) for K vs. 0 for MG) |
| MG:DA:5004 | 1.00 | 80.28 |  | 0 | 2 | - Exclusion zone contact(s): /O2'/O3'/ Consider K in place of MG (2 valid contact(s) for K vs. 0 for MG) |
| MG:DA:5005 | 1.00 | 75.52 |  | 0 |  | 1^st^ MG coordination shell missing (see Notes) |
| MG:DA:5006 | 1.00 | 61.37 |  | 0 | 2 | - Exclusion zone contact(s): /OP1/OP1/ Consider K in place of MG (2 valid contact(s) for K vs. 0 for MG) |
| MG:DA:5007 | 1.00 | 82.49 |  | 0 | 5 | - Exclusion zone contact(s): /N1_A_/O2'/O3'/O5'/ Consider K in place of MG (5 valid contact(s) for K vs. 0 for MG) |
| MG:DA:5008 | 1.00 | 57.65 |  | 0 | 3 | - Exclusion zone contact(s): /NH1_Arg_/O4'/OP1/ Consider K in place of MG (3 valid contact(s) for K vs. 0 for MG) |
| MG:DA:5009 | 1.00 | 66.98 |  | 2 | 4 | O_r_Anion - Exclusion zone contact(s): /N1_G_/O3'/O4'/OP1/ Consider K in place of MG (3 valid contact(s) for K vs. 2 for MG) |
| MG:DA:5010 | 1.00 | 68.90 |  | 0 |  | 1^st^ MG coordination shell missing (see Notes) |
| MG:DA:5011 | 1.00 | 76.17 |  | 0 |  | 1^st^ MG coordination shell missing (see Notes) |
| MG:DA:5012 | 1.00 | 100.24 |  | 0 | 2 | - Exclusion zone contact(s): /O2'/O3'/ Consider K in place of MG (2 valid contact(s) for K vs. 0 for MG) |
| MG:DA:5013 | 1.00 | 54.27 |  | 0 | 1 | - Exclusion zone contact(s): /N7/ Consider K in place of MG (1 valid contact(s) for K vs. 0 for MG) |
| MG:DA:5014 | 1.00 | 64.40 |  | 0 | 2 | - Exclusion zone contact(s): /OP2/OP2/ Consider K in place of MG (2 valid contact(s) for K vs. 0 for MG) |
| MG:DA:5015 | 1.00 | 102.48 |  | 0 | 3 | - Exclusion zone contact(s): /N7/O6/O6/ Consider K in place of MG (3 valid contact(s) for K vs. 0 for MG) |
| MG:DA:5016 | 1.00 | 106.80 |  | 1 | 4 | O_r_ - Exclusion zone contact(s): /O3'/OP1/OP1/OP2/ Consider K in place of MG (4 valid contact(s) for K vs. 1 for MG) |
| MG:DA:5017 | 1.00 | 55.35 |  | 0 | 2 | - Exclusion zone contact(s): /OP1/OP2/ Consider NA in place of MG (1 valid contact(s) for NA vs. 0 for MG) |
| MG:DA:5018 | 1.00 | 89.26 |  | 0 |  | No contacts in 5.0 A |
| MG:DA:5019 | 1.00 | 62.14 |  | 0 | 1 | - Exclusion zone contact(s): /OP2/ Consider K in place of MG (1 valid contact(s) for K vs. 0 for MG) |
| MG:DA:5020 | 1.00 | 63.41 |  | 0 | 1 | - Exclusion zone contact(s): /O2'/ Consider K in place of MG (1 valid contact(s) for K vs. 0 for MG) |
| MG:DA:5021 | 1.00 | 112.74 |  | 0 | 2 | - Exclusion zone contact(s): /OP1/OP2/ Consider NA in place of MG (1 valid contact(s) for NA vs. 0 for MG) |
| MG:DA:5022 | 1.00 | 82.72 |  | 0 | 2 | - Exclusion zone contact(s): /O3'/OP1/ Consider K in place of MG (2 valid contact(s) for K vs. 0 for MG) |
| MG:DA:5023 | 1.00 | 62.96 |  | 1 | 1 | O_r_ - Exclusion zone contact(s): /OP1/ |
| MG:DA:5024 | 1.00 | 43.03 |  | 0 | 4 | - Exclusion zone contact(s): /O2'/O3'/O3'/OP1/ Consider K in place of MG (4 valid contact(s) for K vs. 0 for MG) |
| MG:DA:5025 | 1.00 | 100.39 |  | 0 | 2 | - Exclusion zone contact(s): /OP1/OP2/ Consider K in place of MG (2 valid contact(s) for K vs. 0 for MG) |
| MG:DA:5026 | 1.00 | 42.08 |  | 0 | 4 | - Exclusion zone contact(s): /N2/N3_G_/O2'/O4'/ Consider K in place of MG (4 valid contact(s) for K vs. 0 for MG) |
| MG:DA:5027 | 1.00 | 85.65 |  | 0 |  | 1^st^ MG coordination shell missing (see Notes) |
| MG:DA:5028 | 1.00 | 66.10 |  | 0 | 2 | - Exclusion zone contact(s): /O2/OP1/ Consider K in place of MG (2 valid contact(s) for K vs. 0 for MG) |
| MG:DA:5029 | 1.00 | 39.30 |  | 1 | 2 | O_ph_ - Exclusion zone contact(s): /N6/O5'/ Consider K in place of MG (2 valid contact(s) for K vs. 1 for MG) |
| MG:DA:5030 | 1.00 | 93.67 |  | 0 |  | 1^st^ MG coordination shell missing (see Notes) |
| MG:DA:5031 | 1.00 | 68.65 |  | 0 | 2 | - Exclusion zone contact(s): /N4/O4/ Consider K in place of MG (2 valid contact(s) for K vs. 0 for MG) |
| MG:DA:5032 | 1.00 | 82.55 |  | 0 |  | 1^st^ MG coordination shell missing (see Notes) |
| MG:DA:5033 | 1.00 | 75.21 |  | 0 | 4 | - Exclusion zone contact(s): /N7/O2'/O3'/O6/ Consider K in place of MG (4 valid contact(s) for K vs. 0 for MG) |
| MG:DA:5034 | 1.00 | 76.88 |  | 0 |  | No contacts in 5.0 A |
| MG:DA:5035 | 1.00 | 51.30 |  | 0 | 1 | - Exclusion zone contact(s): /N7/ Consider K in place of MG (1 valid contact(s) for K vs. 0 for MG) |
| MG:DA:5036 | 1.00 | 52.56 |  | 0 | 2 | - Exclusion zone contact(s): /OP2/OP2/ Consider K in place of MG (2 valid contact(s) for K vs. 0 for MG) |
| MG:DA:5037 | 1.00 | 84.92 |  | 0 |  | 1^st^ MG coordination shell missing (see Notes) |
| MG:DA:5038 | 1.00 | 84.88 |  | 0 | 2 | - Exclusion zone contact(s): /O5'/OP2/ Consider K in place of MG (2 valid contact(s) for K vs. 0 for MG) |
| MG:DA:5039 | 1.00 | 75.34 |  | 0 |  | 1^st^ MG coordination shell missing (see Notes) |
| MG:DA:5040 | 1.00 | 90.08 |  | 0 | 4 | - Exclusion zone contact(s): /OP1/OP2/OP2/OP2/ Consider K in place of MG (4 valid contact(s) for K vs. 0 for MG) |
| MG:DA:5041 | 1.00 | 57.80 |  | 0 | 2 | - Exclusion zone contact(s): /O6/OP1/ Consider K in place of MG (2 valid contact(s) for K vs. 0 for MG) |
| MG:DA:5042 | 1.00 | 106.60 |  | 0 | 1 | - Exclusion zone contact(s): /O2'/ Consider K in place of MG (1 valid contact(s) for K vs. 0 for MG) |
| MG:DA:5043 | 1.00 | 73.32 |  | 0 | 3 | - Exclusion zone contact(s): /O2'/O6/O6/ Consider K in place of MG (3 valid contact(s) for K vs. 0 for MG) |
| MG:DA:5044 | 1.00 | 93.76 |  | 0 |  | 1^st^ MG coordination shell missing (see Notes) |
| MG:DA:5045 | 1.00 | 72.83 |  | 1 | 2 | O_r_ - Exclusion zone contact(s): /OP2/OP2/ |
| MG:DA:5046 | 1.00 | 77.75 |  | 0 | 1 | - Exclusion zone contact(s): /O2'/ Consider K in place of MG (1 valid contact(s) for K vs. 0 for MG) |
| MG:DA:5047 | 1.00 | 79.59 |  | 0 |  | 1^st^ MG coordination shell missing (see Notes) |
| MG:DA:5048 | 1.00 | 64.03 |  | 0 | 3 | - Exclusion zone contact(s): /N7/O5'/OP2/ Consider K in place of MG (3 valid contact(s) for K vs. 0 for MG) |
| MG:DA:5049 | 1.00 | 80.50 |  | 0 |  | 1^st^ MG coordination shell missing (see Notes) |
| MG:DA:5050 | 1.00 | 83.52 |  | 0 |  | No contacts in 5.0 A |
| MG:DA:5051 | 1.00 | 103.59 |  | 0 | 1 | - Exclusion zone contact(s): /OP1/ Consider K in place of MG (1 valid contact(s) for K vs. 0 for MG) |
| MG:DA:5052 | 1.00 | 49.05 |  | 0 | 2 | - Exclusion zone contact(s): /O3'/OP2/ Consider K in place of MG (2 valid contact(s) for K vs. 0 for MG) |
| MG:DA:5053 | 1.00 | 101.14 |  | 0 |  | No contacts in 5.0 A |
| MG:DA:5054 | 1.00 | 67.50 |  | 0 | 3 | - Exclusion zone contact(s): /O2'/O3'/OP1/ Consider K in place of MG (2 valid contact(s) for K vs. 0 for MG) |
| MG:DA:5055 | 1.00 | 69.14 |  | 0 | 1 | - Exclusion zone contact(s): /OP2/ Consider NA in place of MG (1 valid contact(s) for NA vs. 0 for MG) |
| MG:DA:5056 | 1.00 | 98.75 |  | 0 |  | 1^st^ MG coordination shell missing (see Notes) |
| MG:DA:5057 | 1.00 | 97.43 |  | 0 |  | 1^st^ MG coordination shell missing (see Notes) |
| MG:DA:5058 | 1.00 | 103.22 |  | 0 | 1 | - Exclusion zone contact(s): /OP1/ Consider K in place of MG (1 valid contact(s) for K vs. 0 for MG) |
| MG:DA:5059 | 1.00 | 66.98 |  | 0 |  | No contacts in 5.0 A |
| MG:DA:5060 | 1.00 | 64.82 |  | 0 |  | 1^st^ MG coordination shell missing (see Notes) |
| MG:DA:5061 | 1.00 | 71.35 |  | 0 | 1 | - Exclusion zone contact(s): /N7/ Consider K in place of MG (1 valid contact(s) for K vs. 0 for MG) |
| MG:DA:5062 | 1.00 | 94.88 |  | 0 | 4 | - Exclusion zone contact(s): /NH2_Arg_/OP1/OP2/OP2/ Consider K in place of MG (4 valid contact(s) for K vs. 0 for MG) |
| MG:DA:5063 | 1.00 | 52.57 |  | 0 | 3 | - Exclusion zone contact(s): /N3_A_/O2'/O4'/ Consider K in place of MG (2 valid contact(s) for K vs. 0 for MG) |
| MG:DA:5064 | 1.00 | 61.51 |  | 0 |  | 1^st^ MG coordination shell missing (see Notes) |
| MG:DA:5065 | 1.00 | 89.97 |  | 0 |  | 1^st^ MG coordination shell missing (see Notes) |
| MG:DA:5066 | 1.00 | 61.31 |  | 0 | 3 | - Exclusion zone contact(s): /O2/O2'/O4'/ Consider K in place of MG (3 valid contact(s) for K vs. 0 for MG) |
| MG:DA:5067 | 1.00 | 119.99 |  | 0 | 2 | - Exclusion zone contact(s): /O4'/OP1/ Consider NA in place of MG (1 valid contact(s) for NA vs. 0 for MG) |
| MG:DA:5068 | 1.00 | 64.43 |  | 0 | 1 | - Exclusion zone contact(s): /OP2/ Consider K in place of MG (1 valid contact(s) for K vs. 0 for MG) |
| MG:DA:5069 | 1.00 | 66.88 |  | 0 | 4 | - Exclusion zone contact(s): /N1_G_/N7/O6/O6/ Consider K in place of MG (3 valid contact(s) for K vs. 0 for MG) |
| MG:DA:5070 | 1.00 | 51.41 |  | 0 | 1 | - Exclusion zone contact(s): /N4/ Consider K in place of MG (1 valid contact(s) for K vs. 0 for MG) |
| MG:DA:5071 | 1.00 | 101.71 |  | 0 | 3 | - Exclusion zone contact(s): /O2'/OP1/OP2/ Consider K in place of MG (3 valid contact(s) for K vs. 0 for MG) |
| MG:DA:5072 | 1.00 | 58.59 |  | 0 |  | 1^st^ MG coordination shell missing (see Notes) |
| MG:DA:5073 | 1.00 | 127.11 |  | 0 | 3 | - Exclusion zone contact(s): /O5'/OP2/OP2/ Consider K in place of MG (2 valid contact(s) for K vs. 0 for MG) |
| MG:DA:5074 | 1.00 | 67.49 |  | 0 | 2 | - Exclusion zone contact(s): /N1/OP1/ Consider K in place of MG (2 valid contact(s) for K vs. 0 for MG) |
| MG:DA:5075 | 1.00 | 50.76 |  | 0 | 3 | - Exclusion zone contact(s): /O5'/OP1/OP2/ Consider K in place of MG (3 valid contact(s) for K vs. 0 for MG) |
| MG:DA:5076 | 1.00 | 85.88 |  | 0 |  | 1^st^ MG coordination shell missing (see Notes) |
| MG:DA:5077 | 1.00 | 102.58 |  | 0 | 1 | - Exclusion zone contact(s): /O6/ Consider K in place of MG (1 valid contact(s) for K vs. 0 for MG) |
| MG:DB:201 | 1.00 | 25.79 |  | 1 |  | O_ph_ |
| MG:DB:202 | 1.00 | 48.37 |  | 0 |  | 1^st^ MG coordination shell missing (see Notes) |
| MG:DB:203 | 1.00 | 46.77 |  | 0 |  | 1^st^ MG coordination shell missing (see Notes) |
| MG:DB:204 | 1.00 | 23.27 |  | 1 | 2 | O_ph_ - Exclusion zone contact(s): /O3'/OP2/ Consider K in place of MG (2 valid contact(s) for K vs. 1 for MG) |
| MG:DB:205 | 1.00 | 27.88 |  | 0 | 6 | - Exclusion zone contact(s): /N1_G_/N2/N3_U_/N4/O4/O6/ Consider K in place of MG (6 valid contact(s) for K vs. 0 for MG) |
| MG:DB:206 | 1.00 | 34.29 |  | 1 | 3 | O_ph_ - Exclusion zone contact(s): /N7/O4'/O6/ Consider K in place of MG (3 valid contact(s) for K vs. 1 for MG) |
| MG:DB:207 | 1.00 | 64.54 |  | 1 |  | O_b_ |
| MG:DB:208 | 1.00 | 46.62 |  | 0 | 2 | - Exclusion zone contact(s): /O4/O6/ Consider K in place of MG (2 valid contact(s) for K vs. 0 for MG) |
| MG:DB:209 | 1.00 | 77.49 |  | 0 |  | 1^st^ MG coordination shell missing (see Notes) |
| MG:DB:210 | 1.00 | 89.31 |  | 0 | 2 | - Exclusion zone contact(s): /N7/O6/ Consider K in place of MG (2 valid contact(s) for K vs. 0 for MG) |
| MG:DB:211 | 1.00 | 90.24 |  | 0 | 1 | - Exclusion zone contact(s): /OP1/ Consider K in place of MG (1 valid contact(s) for K vs. 0 for MG) |
| MG:DB:212 | 1.00 | 71.15 |  | 0 | 1 | - Exclusion zone contact(s): /OP2/ Consider K in place of MG (1 valid contact(s) for K vs. 0 for MG) |
| MG:DB:213 | 1.00 | 64.95 |  | 0 |  | 1^st^ MG coordination shell missing (see Notes) |
| MG:DB:214 | 1.00 | 76.63 |  | 0 |  | 1^st^ MG coordination shell missing (see Notes) |
| MG:DB:215 | 1.00 | 66.91 |  | 0 | 1 | - Exclusion zone contact(s): /O6/ Consider K in place of MG (1 valid contact(s) for K vs. 0 for MG) |
| MG:DB:216 | 1.00 | 43.49 |  | 0 | 3 | - Exclusion zone contact(s): /O2'/O4/O6/ Consider K in place of MG (3 valid contact(s) for K vs. 0 for MG) |
| MG:DB:217 | 1.00 | 46.23 |  | 1 | 1 | O_ph_ - Exclusion zone contact(s): /O3'/ |
| MG:DB:218 | 1.00 | 44.86 |  | 0 | 1 | - Exclusion zone contact(s): /OP1/ Consider K in place of MG (1 valid contact(s) for K vs. 0 for MG) |
| MG:DB:219 | 1.00 | 70.05 |  | 0 |  | 1^st^ MG coordination shell missing (see Notes) |
| MG:DB:220 | 1.00 | 79.48 |  | 1 | 3 | O_b_ - Exclusion zone contact(s): /N4/N7/O6/ Consider K in place of MG (3 valid contact(s) for K vs. 1 for MG) |
| MG:DB:221 | 1.00 | 35.87 |  | 0 | 3 | - Exclusion zone contact(s): /N2/O6/O6/ Consider K in place of MG (3 valid contact(s) for K vs. 0 for MG) |
| MG:DB:222 | 1.00 | 43.17 |  | 0 | 1 | - Exclusion zone contact(s): /O6/ Consider K in place of MG (1 valid contact(s) for K vs. 0 for MG) |
| MG:DB:223 | 1.00 | 58.31 |  | 2 | 1 | O_ph_O_r_ - Exclusion zone contact(s): /OP2/ |
| MG:DB:224 | 1.00 | 51.89 |  | 1 | 3 | O_ph_ - Exclusion zone contact(s): /O2/O2'/O4'/ Consider K in place of MG (3 valid contact(s) for K vs. 1 for MG) |
| MG:DB:225 | 1.00 | 76.15 |  | 2 |  | 2O_ph_ |
| MG:DB:226 | 1.00 | 40.35 |  | 0 | 3 | - Exclusion zone contact(s): /N3_A_/O2'/O4'/ Consider K in place of MG (3 valid contact(s) for K vs. 0 for MG) |
| MG:DB:227 | 1.00 | 59.60 |  | 0 | 4 | - Exclusion zone contact(s): /O4/O4/O6/O6/ Consider NA in place of MG (2 valid contact(s) for NA vs. 0 for MG) |
| MG:DB:228 | 1.00 | 91.16 |  | 0 |  | 1^st^ MG coordination shell missing (see Notes) |
| MG:DB:229 | 1.00 | 83.08 |  | 0 | 1 | - Exclusion zone contact(s): /OP2/ Consider NA in place of MG (1 valid contact(s) for NA vs. 0 for MG) |
| MG:DB:230 | 1.00 | 87.31 |  | 0 |  | 1^st^ MG coordination shell missing (see Notes) |
| MG:DB:231 | 1.00 | 83.11 |  | 0 |  | 1^st^ MG coordination shell missing (see Notes) |
| MG:DB:232 | 1.00 | 54.62 |  | 0 |  | 1^st^ MG coordination shell missing (see Notes) |
| MG:DB:233 | 1.00 | 45.67 |  | 0 | 2 | - Exclusion zone contact(s): /N7/O4/ Consider K in place of MG (2 valid contact(s) for K vs. 0 for MG) |
| MG:DB:234 | 1.00 | 64.59 |  | 0 | 2 | - Exclusion zone contact(s): /OP2/OP2/ Consider K in place of MG (2 valid contact(s) for K vs. 0 for MG) |
| MG:DB:235 | 1.00 | 65.95 |  | 0 | 1 | - Exclusion zone contact(s): /OP2/ Consider K in place of MG (1 valid contact(s) for K vs. 0 for MG) |
| MG:DB:236 | 1.00 | 49.26 |  | 0 | 2 | - Exclusion zone contact(s): /O5'/OP2/ Consider K in place of MG (2 valid contact(s) for K vs. 0 for MG) |
| MG:DB:237 | 1.00 | 57.19 |  | 0 | 3 | - Exclusion zone contact(s): /O2'/O4/OP2/ Consider K in place of MG (2 valid contact(s) for K vs. 0 for MG) |
| MG:DB:238 | 1.00 | 100.66 |  | 0 |  | 1^st^ MG coordination shell missing (see Notes) |
| MG:DB:239 | 1.00 | 81.68 |  | 0 |  | 1^st^ MG coordination shell missing (see Notes) |
| MG:DB:240 | 1.00 | 84.21 |  | 0 |  | 1^st^ MG coordination shell missing (see Notes) |
| MG:DB:241 | 1.00 | 66.33 |  | 1 | 1 | O_ph_ - Exclusion zone contact(s): /N2/ |
| MG:DB:242 | 1.00 | 61.59 |  | 0 |  | 1^st^ MG coordination shell missing (see Notes) |
| MG:DB:243 | 1.00 | 53.38 |  | 0 | 3 | - Exclusion zone contact(s): /O2'/OP2/ Consider K in place of MG (3 valid contact(s) for K vs. 0 for MG) |
| MG:DB:244 | 1.00 | 68.48 |  | 0 | 2 | - Exclusion zone contact(s): /O2'/O4'/ Consider K in place of MG (2 valid contact(s) for K vs. 0 for MG) |
| MG:DB:245 | 1.00 | 59.97 |  | 0 | 2 | - Exclusion zone contact(s): /O5'/OP2/ Consider K in place of MG (2 valid contact(s) for K vs. 0 for MG) |
| MG:DB:246 | 1.00 | 52.56 |  | 0 | 3 | - Exclusion zone contact(s): /O4/O4/O6/ Consider K in place of MG (3 valid contact(s) for K vs. 0 for MG) |
| MG:DB:247 | 1.00 | 61.13 |  | 1 | 5 | O_r_ - Exclusion zone contact(s): /N1/N1_A_/N6/O5'/OP2/ Consider K in place of MG (3 valid contact(s) for K vs. 1 for MG) |
| MG:DB:248 | 1.00 | 76.44 |  | 0 | 2 | - Exclusion zone contact(s): /N4/O6/ Consider NA in place of MG (2 valid contact(s) for NA vs. 0 for MG) |
| MG:DB:249 | 1.00 | 84.85 |  | 0 | 2 | - Exclusion zone contact(s): /N1/N7/ Consider K in place of MG (2 valid contact(s) for K vs. 0 for MG) |
| MG:DB:250 | 1.00 | 74.60 |  | 0 | 5 | - Exclusion zone contact(s): /N1_A_/N1_G_/N2/N2/O2/ Consider K in place of MG (4 valid contact(s) for K vs. 0 for MG) |
| MG:DB:251 | 1.00 | 49.76 |  | 1 | 1 | O_ph_ - Exclusion zone contact(s): /O3'/ |
| MG:DB:252 | 1.00 | 101.77 |  | 0 | 2 | - Exclusion zone contact(s): /O5'/OP1/ Consider K in place of MG (2 valid contact(s) for K vs. 0 for MG) |
| MG:DB:253 | 1.00 | 74.02 |  | 0 | 2 | - Exclusion zone contact(s): /OP2/ Consider K in place of MG (2 valid contact(s) for K vs. 0 for MG) |
| MG:DB:254 | 1.00 | 66.54 |  | 0 |  | 1^st^ MG coordination shell missing (see Notes) |
| MG:DB:255 | 1.00 | 63.45 |  | 0 | 4 | - Exclusion zone contact(s): /N3_A_/O2'/O3'/OP1/ Consider K in place of MG (4 valid contact(s) for K vs. 0 for MG) |
| MG:DB:256 | 1.00 | 65.03 |  | 0 | 2 | - Exclusion zone contact(s): /O5'/OP2/ Consider K in place of MG (2 valid contact(s) for K vs. 0 for MG) |
| MG:DB:257 | 1.00 | 57.85 |  | 0 |  | 1^st^ MG coordination shell missing (see Notes) |
| MG:DB:258 | 1.00 | 77.68 |  | 0 | 1 | - Exclusion zone contact(s): /N7/ Consider K in place of MG (1 valid contact(s) for K vs. 0 for MG) |
| MG:DB:259 | 1.00 | 69.91 |  | 0 |  | 1^st^ MG coordination shell missing (see Notes) |
| MG:DB:260 | 1.00 | 63.44 |  | 0 | 1 | - Exclusion zone contact(s): /O2'/ Consider K in place of MG (1 valid contact(s) for K vs. 0 for MG) |
| MG:DB:261 | 1.00 | 91.22 |  | 0 | 1 | - Exclusion zone contact(s): /N7/ Consider K in place of MG (1 valid contact(s) for K vs. 0 for MG) |
| MG:DB:262 | 1.00 | 89.46 |  | 0 |  | 1^st^ MG coordination shell missing (see Notes) |
| MG:DB:263 | 1.00 | 80.57 |  | 0 | 2 | - Exclusion zone contact(s): /N2/OE1_Glu_/ Consider K in place of MG (2 valid contact(s) for K vs. 0 for MG) |
| MG:DB:264 | 1.00 | 44.55 |  | 0 | 1 | - Exclusion zone contact(s): /OP2/ Consider NA in place of MG (1 valid contact(s) for NA vs. 0 for MG) |
| MG:DB:265 | 1.00 | 81.74 |  | 0 | 2 | - Exclusion zone contact(s): /O3'/OP1/ Consider K in place of MG (2 valid contact(s) for K vs. 0 for MG) |
| MG:DB:266 | 1.00 | 51.28 |  | 0 | 3 | - Exclusion zone contact(s): /N1/N4/O4'/ Consider K in place of MG (3 valid contact(s) for K vs. 0 for MG) |
| MG:DB:267 | 1.00 | 73.38 |  | 0 | 3 | - Exclusion zone contact(s): /O2/O2'/O4'/ Consider K in place of MG (3 valid contact(s) for K vs. 0 for MG) |
| MG:DB:268 | 1.00 | 81.93 |  | 0 | 2 | - Exclusion zone contact(s): /O2'/O3'/ Consider K in place of MG (2 valid contact(s) for K vs. 0 for MG) |
| MG:DB:269 | 1.00 | 91.24 |  | 0 | 1 | - Exclusion zone contact(s): /OP2/ Consider K in place of MG (1 valid contact(s) for K vs. 0 for MG) |
| MG:DB:270 | 1.00 | 61.03 |  | 1 | 4 | O_r_ - Exclusion zone contact(s): /O3'/O5'/OP2/OP2/ Consider K in place of MG (3 valid contact(s) for K vs. 1 for MG) |
| MG:DB:271 | 1.00 | 72.16 |  | 0 | 5 | - Exclusion zone contact(s): /O2'/O2'/O3'/O4'/OH_Tyr_/ Consider K in place of MG (5 valid contact(s) for K vs. 0 for MG) |
| MG:DB:272 | 1.00 | 81.02 |  | 0 | 2 | - Exclusion zone contact(s): /O5'/OP2/ Consider K in place of MG (2 valid contact(s) for K vs. 0 for MG) |
| MG:DB:273 | 1.00 | 100.88 |  | 0 | 2 | - Exclusion zone contact(s): /OP1/OP1/ Consider NA in place of MG (1 valid contact(s) for NA vs. 0 for MG) |
| MG:DB:274 | 1.00 | 79.51 |  | 0 | 2 | - Exclusion zone contact(s): /O4/O6/ Consider K in place of MG (2 valid contact(s) for K vs. 0 for MG) |
| MG:DB:275 | 1.00 | 50.55 |  | 0 | 2 | - Exclusion zone contact(s): /N7/ Consider NA in place of MG (1 valid contact(s) for NA vs. 0 for MG) |
| MG:DB:276 | 1.00 | 66.27 |  | 0 | 4 | - Exclusion zone contact(s): /N4/N4/OP1/OP2/ Consider K in place of MG (4 valid contact(s) for K vs. 0 for MG) |
| MG:DD:301 | 1.00 | 45.04 |  | 0 |  | 1^st^ MG coordination shell missing (see Notes) |
| MG:DD:302 | 1.00 | 58.25 |  | 0 |  | 1^st^ MG coordination shell missing (see Notes) |
| MG:DD:303 | 1.00 | 16.01 |  | 0 | 2 | - Exclusion zone contact(s): /ND2_Asn_/O_back_/ Consider NA in place of MG (1 valid contact(s) for NA vs. 0 for MG) |
| MG:DD:304 | 1.00 | 66.05 |  | 0 | 2 | - Exclusion zone contact(s): /O_back_/OP1/ Consider NA in place of MG (1 valid contact(s) for NA vs. 0 for MG) |
| MG:DD:305 | 1.00 | 78.84 |  | 0 | 5 | - Exclusion zone contact(s): /N_back_/N_back_/N_back_/NZ_Lys_/O_back_/ Consider K in place of MG (4 valid contact(s) for K vs. 0 for MG) |
| MG:DD:306 | 1.00 | 51.55 |  | 0 | 1 | - Exclusion zone contact(s): /O_back_/ Consider K in place of MG (1 valid contact(s) for K vs. 0 for MG) |
| MG:DD:307 | 1.00 | 50.38 |  | 0 | 4 | - Exclusion zone contact(s): /N_back_/O_back_/O_back_/O_back_/ Consider NA in place of MG (2 valid contact(s) for NA vs. 0 for MG) |
| MG:DD:308 | 1.00 | 53.77 |  | 0 | 2 | - Exclusion zone contact(s): /NE2_His_/O_back_/ Consider NA in place of MG (2 valid contact(s) for NA vs. 0 for MG) |
| MG:DD:309 | 1.00 | 57.62 |  | 0 | 3 | - Exclusion zone contact(s): /N_back_/O_back_/OP1/ Consider K in place of MG (3 valid contact(s) for K vs. 0 for MG) |
| MG:DD:310 | 1.00 | 64.61 | 2 | 2 | 2 | - Exclusion zone contact(s): /N_back_/O_back_/OE1_Glu_/OE2_Glu_/ Consider K in place of MG (2 valid contact(s) for K vs. 0 for MG) |
| MG:DD:311 | 1.00 | 65.61 |  | 0 | 1 | - Exclusion zone contact(s): /O2'/ Consider K in place of MG (1 valid contact(s) for K vs. 0 for MG) |
| MG:DD:312 | 1.00 | 37.70 |  | 0 |  | 1^st^ MG coordination shell missing (see Notes) |
| MG:DD:313 | 1.00 | 64.23 |  | 0 |  | 1^st^ MG coordination shell missing (see Notes) |
| MG:DD:314 | 1.00 | 32.11 |  | 1 | 4 | O_back_ - Exclusion zone contact(s): /N_back_/N_back_/N_back_/O2'/ Consider K in place of MG (4 valid contact(s) for K vs. 1 for MG) |
| MG:DE:301 | 1.00 | 19.36 |  | 0 | 5 | - Exclusion zone contact(s): /N_back_/N_back_/O5'/O_back_/OP1/ Consider K in place of MG (5 valid contact(s) for K vs. 0 for MG) |
| MG:DE:302 | 1.00 | 8.59 |  | 1 | 1 | O_coo_ - Exclusion zone contact(s): /O2'/ |
| MG:DE:303 | 1.00 | 52.68 |  | 0 |  | 1^st^ MG coordination shell missing (see Notes) |
| MG:DE:304 | 1.00 | 47.61 |  | 0 | 2 | - Exclusion zone contact(s): /O2'/OP1/ Consider K in place of MG (2 valid contact(s) for K vs. 0 for MG) |
| MG:DE:305 | 1.00 | 44.93 |  | 0 | 3 | - Exclusion zone contact(s): /O_back_/O_back_/OH_Tyr_/ Consider K in place of MG (2 valid contact(s) for K vs. 0 for MG) |
| MG:DE:306 | 1.00 | 45.81 |  | 0 | 3 | - Exclusion zone contact(s): /N_back_/O2'/O_back_/ Consider K in place of MG (2 valid contact(s) for K vs. 0 for MG) |
| MG:DE:307 | 1.00 | 62.46 |  | 0 |  | No contacts in 5.0 A |
| MG:DE:308 | 1.00 | 83.97 |  | 0 | 5 | - Exclusion zone contact(s): /NZ_Lys_/O2'/O_back_/O_back_/OG1_Thr_/ Consider K in place of MG (5 valid contact(s) for K vs. 0 for MG) |
| MG:DE:309 | 1.00 | 49.85 |  | 0 | 1 | - Exclusion zone contact(s): /O_back_/ Consider K in place of MG (1 valid contact(s) for K vs. 0 for MG) |
| MG:DE:310 | 1.00 | 52.11 |  | 0 | 3 | - Exclusion zone contact(s): /N_back_/O_back_/O_back_/ Consider K in place of MG (3 valid contact(s) for K vs. 0 for MG) |
| MG:DE:311 | 1.00 | 42.10 |  | 0 |  | 1^st^ MG coordination shell missing (see Notes) |
| MG:DE:312 | 1.00 | 74.47 |  | 1 |  | O_back_ |
| MG:DE:313 | 1.00 | 73.80 |  | 0 |  | 1^st^ MG coordination shell missing (see Notes) |
| MG:DE:314 | 1.00 | 67.23 |  | 0 | 4 | - Exclusion zone contact(s): /N_back_/NH1_Arg_/O_back_/OD1_Asp_/ Consider K in place of MG (3 valid contact(s) for K vs. 0 for MG) |
| MG:DE:315 | 1.00 | 77.90 |  | 1 | 4 | O_back_ - Exclusion zone contact(s): /N_back_/O_back_/O_back_/O_back_/ Consider K in place of MG (4 valid contact(s) for K vs. 1 for MG) |
| MG:DF:301 | 1.00 | 28.13 |  | 0 | 4 | - Exclusion zone contact(s): /O4/O_back_/O_back_/O_back_/ Consider K in place of MG (3 valid contact(s) for K vs. 0 for MG) |
| MG:DF:302 | 1.00 | 46.99 |  | 0 | 3 | - Exclusion zone contact(s): /N_back_/O2'/OP1/ Consider K in place of MG (2 valid contact(s) for K vs. 0 for MG) |
| MG:DF:303 | 1.00 | 28.99 |  | 0 | 3 | - Exclusion zone contact(s): /N_back_/O2'/O_back_/ Consider K in place of MG (2 valid contact(s) for K vs. 0 for MG) |
| MG:DF:304 | 1.00 | 29.58 |  | 0 | 2 | - Exclusion zone contact(s): /N_back_/OP1/ Consider K in place of MG (2 valid contact(s) for K vs. 0 for MG) |
| MG:DF:305 | 1.00 | 54.53 |  | 0 | 1 | - Exclusion zone contact(s): /NH1_Arg_/ Consider K in place of MG (1 valid contact(s) for K vs. 0 for MG) |
| MG:DF:306 | 1.00 | 46.45 |  | 0 |  | 1^st^ MG coordination shell missing (see Notes) |
| MG:DF:307 | 1.00 | 27.16 |  | 0 | 5 | - Exclusion zone contact(s): /N_back_/N_back_/NZ_Lys_/O_back_/O_back_/ Consider K in place of MG (4 valid contact(s) for K vs. 0 for MG) |
| MG:DF:308 | 1.00 | 30.59 |  | 0 | 5 | - Exclusion zone contact(s): /N_back_/N_back_/O3'/O_back_/OP1/ Consider K in place of MG (4 valid contact(s) for K vs. 0 for MG) |
| MG:DF:309 | 1.00 | 46.34 |  | 0 | 1 | - Exclusion zone contact(s): /O_back_/ Consider K in place of MG (1 valid contact(s) for K vs. 0 for MG) |
| MG:DF:310 | 1.00 | 49.13 |  | 0 | 1 | - Exclusion zone contact(s): /NH1_Arg_/ Consider K in place of MG (1 valid contact(s) for K vs. 0 for MG) |
| MG:DF:311 | 1.00 | 57.60 |  | 0 |  | 1^st^ MG coordination shell missing (see Notes) |
| MG:DF:312 | 1.00 | 75.42 |  | 0 | 2 | - Exclusion zone contact(s): /N_back_/O_back_/ Consider NA in place of MG (1 valid contact(s) for NA vs. 0 for MG) |
| MG:DF:313 | 1.00 | 68.86 |  | 0 | 3 | - Exclusion zone contact(s): /NH1_Arg_/OD1_Asp_/OD2_Asp_/ Consider NA in place of MG (2 valid contact(s) for NA vs. 0 for MG) |
| MG:DF:314 | 1.00 | 58.46 |  | 0 | 1 | - Exclusion zone contact(s): /O_back_/ Consider K in place of MG (1 valid contact(s) for K vs. 0 for MG) |
| MG:DF:315 | 1.00 | 76.11 |  | 0 | 1 | - Exclusion zone contact(s): /OG_Ser_/ Consider K in place of MG (1 valid contact(s) for K vs. 0 for MG) |
| MG:DF:316 | 1.00 | 84.66 |  | 0 | 4 | - Exclusion zone contact(s): /N_back_/ND2_Asn_/O5'/OP2/ Consider K in place of MG (4 valid contact(s) for K vs. 0 for MG) |
| MG:DF:317 | 1.00 | 64.36 |  | 0 |  | 1^st^ MG coordination shell missing (see Notes) |
| MG:DF:318 | 1.00 | 69.09 |  | 0 | 1 | - Exclusion zone contact(s): /O_back_/ Consider K in place of MG (1 valid contact(s) for K vs. 0 for MG) |
| MG:DF:319 | 1.00 | 67.97 |  | 0 |  | No contacts in 5.0 A |
| MG:DF:320 | 1.00 | 58.78 |  | 0 | 2 | - Exclusion zone contact(s): /NZ_Lys_/O3'/ Consider K in place of MG (2 valid contact(s) for K vs. 0 for MG) |
| MG:DF:321 | 1.00 | 55.96 |  | 1 | 2 | Anion - Exclusion zone contact(s): /OP1/OP2/ Consider K in place of MG (2 valid contact(s) for K vs. 1 for MG) |
| MG:DF:322 | 1.00 | 40.51 |  | 0 | 3 | - Exclusion zone contact(s): /NZ_Lys_/O5'/OP1/ Consider K in place of MG (3 valid contact(s) for K vs. 0 for MG) |
| MG:DF:323 | 1.00 | 37.74 |  | 0 | 3 | - Exclusion zone contact(s): /N_back_/O_back_/O_back_/ Consider K in place of MG (3 valid contact(s) for K vs. 0 for MG) |
| MG:DF:324 | 1.00 | 76.37 |  | 0 |  | 1^st^ MG coordination shell missing (see Notes) |
| MG:DF:325 | 1.00 | 47.77 |  | 0 | 1 | - Exclusion zone contact(s): /OE2_Glu_/ Consider K in place of MG (1 valid contact(s) for K vs. 0 for MG) |
| MG:DG:201 | 1.00 | 68.61 |  | 1 | 2 | O_back_ - Exclusion zone contact(s): /N_back_/O_back_/ |
| MG:DG:202 | 1.00 | 47.93 |  | 0 | 4 | - Exclusion zone contact(s): /ND2_Asn_/O_back_/O_back_/O_back_/ Consider K in place of MG (3 valid contact(s) for K vs. 0 for MG) |
| MG:DG:203 | 1.00 | 64.09 |  | 0 |  | 1^st^ MG coordination shell missing (see Notes) |
| MG:DG:204 | 1.00 | 104.59 |  | 0 |  | 1^st^ MG coordination shell missing (see Notes) |
| MG:DG:205 | 1.00 | 64.41 |  | 0 | 1 | - Exclusion zone contact(s): /O_back_/ Consider K in place of MG (1 valid contact(s) for K vs. 0 for MG) |
| MG:DH:201 | 1.00 | 48.37 |  | 0 | 2 | - Exclusion zone contact(s): /NZ_Lys_/O_back_/ Consider NA in place of MG (1 valid contact(s) for NA vs. 0 for MG) |
| MG:DH:202 | 1.00 | 59.71 |  | 0 |  | 1^st^ MG coordination shell missing (see Notes) |
| MG:DH:203 | 1.00 | 78.40 |  | 0 | 1 | - Exclusion zone contact(s): /NH2_Arg_/ Consider NA in place of MG (1 valid contact(s) for NA vs. 0 for MG) |
| MG:DH:204 | 1.00 | 87.67 |  | 0 |  | No contacts in 5.0 A |
| MG:DH:205 | 1.00 | 77.65 |  | 0 | 1 | - Exclusion zone contact(s): /O_back_/ Consider K in place of MG (1 valid contact(s) for K vs. 0 for MG) |
| MG:DK:201 | 1.00 | 71.45 |  | 2 | 1 | 2Anion - Exclusion zone contact(s): /N_back_/ |
| MG:DK:202 | 1.00 | 66.29 |  | 0 | 1 | - Exclusion zone contact(s): /N_back_/ Consider K in place of MG (1 valid contact(s) for K vs. 0 for MG) |
| MG:DL:201 | 1.00 | 94.93 |  | 0 | 2 | - Exclusion zone contact(s): /O_back_/OP1/ Consider NA in place of MG (1 valid contact(s) for NA vs. 0 for MG) |
| MG:DM:201 | 1.00 | 29.67 |  | 0 | 1 | - Exclusion zone contact(s): /OH_Tyr_/ Consider K in place of MG (1 valid contact(s) for K vs. 0 for MG) |
| MG:DM:202 | 1.00 | 48.68 |  | 0 | 3 | - Exclusion zone contact(s): /O_back_/O_back_/O_back_/ Consider K in place of MG (3 valid contact(s) for K vs. 0 for MG) |
| MG:DM:203 | 1.00 | 59.33 |  | 0 |  | 1^st^ MG coordination shell missing (see Notes) |
| MG:DM:204 | 1.00 | 67.24 |  | 0 | 5 | - Exclusion zone contact(s): /N_back_/NH1_Arg_/O_back_/O_back_/O_back_/ Consider K in place of MG (5 valid contact(s) for K vs. 0 for MG) |
| MG:DM:205 | 1.00 | 51.54 |  | 0 | 2 | - Exclusion zone contact(s): /N_back_/N_back_/ Consider K in place of MG (2 valid contact(s) for K vs. 0 for MG) |
| MG:DM:206 | 1.00 | 85.47 |  | 0 | 1 | - Exclusion zone contact(s): /NE_Arg_/ Consider K in place of MG (1 valid contact(s) for K vs. 0 for MG) |
| MG:DN:201 | 1.00 | 50.03 |  | 0 |  | 1^st^ MG coordination shell missing (see Notes) |
| MG:DN:202 | 1.00 | 65.87 |  | 0 |  | 1^st^ MG coordination shell missing (see Notes) |
| MG:DO:201 | 1.00 | 32.30 |  | 0 | 2 | - Exclusion zone contact(s): /NH2_Arg_/O_back_/ Consider K in place of MG (2 valid contact(s) for K vs. 0 for MG) |
| MG:DO:202 | 1.00 | 35.21 |  | 0 |  | 1^st^ MG coordination shell missing (see Notes) |
| MG:DO:203 | 1.00 | 32.67 |  | 0 | 5 | - Exclusion zone contact(s): /N_back_/N_back_/N_back_/N_back_/O_back_/ Consider K in place of MG (4 valid contact(s) for K vs. 0 for MG) |
| MG:DO:204 | 1.00 | 44.69 |  | 0 | 3 | - Exclusion zone contact(s): /N7/ND1_His_/NE2_His_/ Consider K in place of MG (2 valid contact(s) for K vs. 0 for MG) |
| MG:DO:205 | 1.00 | 52.84 |  | 0 |  | 1^st^ MG coordination shell missing (see Notes) |
| MG:DO:206 | 1.00 | 51.89 |  | 0 |  | 1^st^ MG coordination shell missing (see Notes) |
| MG:DO:207 | 1.00 | 37.50 |  | 0 | 2 | - Exclusion zone contact(s): /OG1_Thr_/OP1/ Consider K in place of MG (2 valid contact(s) for K vs. 0 for MG) |
| MG:DO:208 | 1.00 | 59.39 |  | 0 | 3 | - Exclusion zone contact(s): /NZ_Lys_/O_back_/OP1/ Consider K in place of MG (3 valid contact(s) for K vs. 0 for MG) |
| MG:DO:209 | 1.00 | 60.26 |  | 0 | 3 | - Exclusion zone contact(s): /O_back_/OP1/OP2/ Consider K in place of MG (3 valid contact(s) for K vs. 0 for MG) |
| MG:DO:210 | 1.00 | 53.93 |  | 0 | 4 | - Exclusion zone contact(s): /NH2_Arg_/O_back_/O_back_/OP1/ Consider K in place of MG (4 valid contact(s) for K vs. 0 for MG) |
| MG:DO:211 | 1.00 | 53.95 |  | 0 | 1 | - Exclusion zone contact(s): /O_back_/ Consider K in place of MG (1 valid contact(s) for K vs. 0 for MG) |
| MG:DO:212 | 1.00 | 44.26 |  | 0 | 2 | - Exclusion zone contact(s): /ND2_Asn_/OP1/ Consider K in place of MG (2 valid contact(s) for K vs. 0 for MG) |
| MG:DO:213 | 1.00 | 96.62 |  | 0 | 4 | - Exclusion zone contact(s): /ND2_Asn_/O5'/OD1_Asn_/OP1/ Consider K in place of MG (4 valid contact(s) for K vs. 0 for MG) |
| MG:DO:214 | 1.00 | 61.18 |  | 0 | 5 | - Exclusion zone contact(s): /N2/NH1_Arg_/NZ_Lys_/O2'/OE1_Glu_/ Consider K in place of MG (5 valid contact(s) for K vs. 0 for MG) |
| MG:DO:215 | 1.00 | 41.63 |  | 0 | 1 | - Exclusion zone contact(s): /O_back_/ Consider K in place of MG (1 valid contact(s) for K vs. 0 for MG) |
| MG:DO:216 | 1.00 | 58.55 |  | 0 | 2 | - Exclusion zone contact(s): /NH1_Arg_/OH_Tyr_/ Consider NA in place of MG (1 valid contact(s) for NA vs. 0 for MG) |
| MG:DO:217 | 1.00 | 52.44 |  | 0 | 3 | - Exclusion zone contact(s): /O_back_/O_back_/OG_Ser_/ Consider K in place of MG (2 valid contact(s) for K vs. 0 for MG) |
| MG:DO:218 | 1.00 | 74.08 |  | 0 |  | 1^st^ MG coordination shell missing (see Notes) |
| MG:DP:201 | 1.00 | 29.54 |  | 0 | 3 | - Exclusion zone contact(s): /O_back_/O_back_/O_back_/ Consider K in place of MG (3 valid contact(s) for K vs. 0 for MG) |
| MG:DP:202 | 1.00 | 53.06 |  | 0 |  | 1^st^ MG coordination shell missing (see Notes) |
| MG:DP:203 | 1.00 | 92.11 |  | 0 | 3 | - Exclusion zone contact(s): /NE_Arg_/O2'/O2'/ Consider K in place of MG (2 valid contact(s) for K vs. 0 for MG) |
| MG:DP:204 | 1.00 | 51.28 |  | 0 | 1 | - Exclusion zone contact(s): /OP1/ Consider K in place of MG (1 valid contact(s) for K vs. 0 for MG) |
| MG:DQ:201 | 1.00 | 51.24 |  | 0 |  | No contacts in 5.0 A |
| MG:DQ:202 | 1.00 | 42.67 |  | 1 | 2 | O_back_ - Exclusion zone contact(s): /N7/N_back_/ |
| MG:DR:201 | 1.00 | 72.82 |  | 0 | 1 | - Exclusion zone contact(s): /O_back_/ Consider K in place of MG (1 valid contact(s) for K vs. 0 for MG) |
| MG:DR:202 | 1.00 | 60.55 |  | 0 | 2 | - Exclusion zone contact(s): /N_back_/O_back_/ Consider NA in place of MG (1 valid contact(s) for NA vs. 0 for MG) |
| MG:DR:203 | 1.00 | 84.68 |  | 0 |  | 1^st^ MG coordination shell missing (see Notes) |
| MG:DR:204 | 1.00 | 40.30 |  | 0 | 4 | - Exclusion zone contact(s): /N_back_/O2'/O5'/O_back_/ Consider K in place of MG (4 valid contact(s) for K vs. 0 for MG) |
| MG:DS:201 | 1.00 | 88.50 |  | 0 |  | 1^st^ MG coordination shell missing (see Notes) |
| MG:DS:202 | 1.00 | 71.13 |  | 0 | 1 | - Exclusion zone contact(s): /OP1/ Consider K in place of MG (1 valid contact(s) for K vs. 0 for MG) |
| MG:DS:203 | 1.00 | 54.35 |  | 1 | 3 | Anion - Exclusion zone contact(s): /NH2_Arg_/O_back_/OP1/ Consider K in place of MG (3 valid contact(s) for K vs. 1 for MG) |
| MG:DS:204 | 1.00 | 75.24 |  | 0 |  | 1^st^ MG coordination shell missing (see Notes) |
| MG:DS:205 | 1.00 | 58.33 |  | 0 |  | 1^st^ MG coordination shell missing (see Notes) |
| MG:DS:206 | 1.00 | 60.46 |  | 0 | 2 | - Exclusion zone contact(s): /NE_Arg_/OH_Tyr_/ Consider K in place of MG (2 valid contact(s) for K vs. 0 for MG) |
| MG:DS:207 | 1.00 | 109.84 |  | 0 | 2 | - Exclusion zone contact(s): /NZ_Lys_/OP1/ Consider K in place of MG (2 valid contact(s) for K vs. 0 for MG) |
| MG:DS:208 | 1.00 | 54.88 |  | 0 |  | 1^st^ MG coordination shell missing (see Notes) |
| MG:DT:101 | 1.00 | 54.13 |  | 0 | 3 | - Exclusion zone contact(s): /ND2_Asn_/O_back_/OP2/ Consider K in place of MG (3 valid contact(s) for K vs. 0 for MG) |
| MG:DT:102 | 1.00 | 60.63 |  | 0 | 1 | - Exclusion zone contact(s): /N_back_/ Consider K in place of MG (1 valid contact(s) for K vs. 0 for MG) |
| MG:DT:103 | 1.00 | 40.69 |  | 1 | 2 | Anion - Exclusion zone contact(s): /N_back_/N_back_/ Consider K in place of MG (2 valid contact(s) for K vs. 1 for MG) |
| MG:DT:104 | 1.00 | 74.76 |  | 1 | 2 | O_back_ - Exclusion zone contact(s): /N_back_/N_back_/ Consider K in place of MG (2 valid contact(s) for K vs. 1 for MG) |
| MG:DT:105 | 1.00 | 84.04 |  | 0 | 2 | - Exclusion zone contact(s): /N_back_/O_back_/ Consider K in place of MG (2 valid contact(s) for K vs. 0 for MG) |
| MG:DT:106 | 1.00 | 69.55 |  | 0 | 1 | - Exclusion zone contact(s): /O_back_/ Consider K in place of MG (1 valid contact(s) for K vs. 0 for MG) |
| MG:DT:107 | 1.00 | 73.10 |  | 0 | 1 | - Exclusion zone contact(s): /OG1_Thr_/ Consider NA in place of MG (1 valid contact(s) for NA vs. 0 for MG) |
| MG:DU:201 | 1.00 | 36.85 |  | 1 | 1 | Anion - Exclusion zone contact(s): /OP1/ |
| MG:DU:202 | 1.00 | 17.37 |  | 0 |  | 1^st^ MG coordination shell missing (see Notes) |
| MG:DU:203 | 1.00 | 45.15 |  | 0 | 1 | - Exclusion zone contact(s): /OE1_Glu_/ Consider K in place of MG (1 valid contact(s) for K vs. 0 for MG) |
| MG:DU:204 | 1.00 | 62.95 |  | 1 | 2 | O_back_ - Exclusion zone contact(s): /O_back_/O_back_/ Consider NA in place of MG (2 valid contact(s) for NA vs. 1 for MG) |
| MG:DU:205 | 1.00 | 75.28 |  | 0 | 1 | - Exclusion zone contact(s): /OP2/ Consider K in place of MG (1 valid contact(s) for K vs. 0 for MG) |
| MG:DU:206 | 1.00 | 47.11 |  | 2 | 2 | 2O_back_ - Exclusion zone contact(s): /N_back_/NH1_Arg_/ |
| MG:DU:207 | 1.00 | 97.73 |  | 0 |  | 1^st^ MG coordination shell missing (see Notes) |
| MG:DU:208 | 1.00 | 52.32 |  | 0 |  | 1^st^ MG coordination shell missing (see Notes) |
| MG:DU:209 | 1.00 | 96.57 |  | 0 |  | 1^st^ MG coordination shell missing (see Notes) |
| MG:DU:210 | 1.00 | 107.86 |  | 0 | 4 | - Exclusion zone contact(s): /N_back_/N_back_/OH_Tyr_/OP1/ Consider K in place of MG (3 valid contact(s) for K vs. 0 for MG) |
| MG:DU:211 | 1.00 | 116.70 |  | 0 | 1 | - Exclusion zone contact(s): /NZ_Lys_/ Consider K in place of MG (1 valid contact(s) for K vs. 0 for MG) |
| MG:DU:212 | 1.00 | 75.79 |  | 0 | 1 | - Exclusion zone contact(s): /O_back_/ Consider K in place of MG (1 valid contact(s) for K vs. 0 for MG) |
| MG:DU:213 | 1.00 | 70.77 |  | 0 |  | 1^st^ MG coordination shell missing (see Notes) |
| MG:DU:214 | 1.00 | 62.04 |  | 0 |  | 1^st^ MG coordination shell missing (see Notes) |
| MG:DU:215 | 1.00 | 76.66 |  | 0 | 1 | - Exclusion zone contact(s): /NH1_Arg_/ Consider K in place of MG (1 valid contact(s) for K vs. 0 for MG) |
| MG:DU:216 | 1.00 | 80.44 |  | 0 | 3 | - Exclusion zone contact(s): /N_back_/N_back_/O_back_/ Consider K in place of MG (3 valid contact(s) for K vs. 0 for MG) |
| MG:DU:217 | 1.00 | 110.22 |  | 0 |  | 1^st^ MG coordination shell missing (see Notes) |
| MG:DU:218 | 1.00 | 73.68 |  | 0 | 3 | - Exclusion zone contact(s): /NZ_Lys_/O_back_/OH_Tyr_/ Consider K in place of MG (3 valid contact(s) for K vs. 0 for MG) |
| MG:DU:219 | 1.00 | 72.08 |  | 0 |  | 1^st^ MG coordination shell missing (see Notes) |
| MG:DV:301 | 1.00 | 108.25 |  | 0 |  | 1^st^ MG coordination shell missing (see Notes) |
| MG:DV:302 | 1.00 | 87.42 |  | 0 | 1 | - Exclusion zone contact(s): /N_back_/ Consider K in place of MG (1 valid contact(s) for K vs. 0 for MG) |
| MG:DV:303 | 1.00 | 67.25 |  | 0 |  | No contacts in 5.0 A |
| MG:DV:304 | 1.00 | 77.52 |  | 0 |  | 1^st^ MG coordination shell missing (see Notes) |
| MG:DV:305 | 1.00 | 86.38 |  | 0 | 2 | - Exclusion zone contact(s): /N_back_/O_back_/ Consider K in place of MG (2 valid contact(s) for K vs. 0 for MG) |
| MG:DW:101 | 1.00 | 49.50 |  | 0 |  | 1^st^ MG coordination shell missing (see Notes) |
| MG:DW:102 | 1.00 | 37.44 |  | 0 | 2 | - Exclusion zone contact(s): /N_back_/OG_Ser_/ Consider K in place of MG (2 valid contact(s) for K vs. 0 for MG) |
| MG:DW:103 | 1.00 | 78.22 |  | 1 | 2 | O_coo_ - Exclusion zone contact(s): /OG_Ser_/OP1/ |
| MG:DW:104 | 1.00 | 49.84 |  | 0 | 1 | - Exclusion zone contact(s): /NE2_Gln_/ Consider K in place of MG (1 valid contact(s) for K vs. 0 for MG) |
| MG:DW:105 | 1.00 | 81.34 |  | 0 | 1 | - Exclusion zone contact(s): /NZ_Lys_/ Consider K in place of MG (1 valid contact(s) for K vs. 0 for MG) |
| MG:DW:106 | 1.00 | 61.24 |  | 0 | 1 | - Exclusion zone contact(s): /O_back_/ Consider K in place of MG (1 valid contact(s) for K vs. 0 for MG) |
| MG:DW:107 | 1.00 | 70.81 |  | 0 | 2 | - Exclusion zone contact(s): /O_back_/OP2/ Consider K in place of MG (2 valid contact(s) for K vs. 0 for MG) |
| MG:DX:101 | 1.00 | 59.75 |  | 1 | 3 | O_ph_ - Exclusion zone contact(s): /NE2_Gln_/NH1_Arg_/O3'/ Consider K in place of MG (3 valid contact(s) for K vs. 1 for MG) |
| MG:DX:102 | 1.00 | 62.89 |  | 0 | 1 | - Exclusion zone contact(s): /NE2_Gln_/ Consider K in place of MG (1 valid contact(s) for K vs. 0 for MG) |
| MG:DY:201 | 1.00 | 100.92 | 2 | 1 |  | Anion - Exclusion zone contact(s): /O_back_/OE1_Gln_/ |
| MG:DY:202 | 1.00 | 91.74 |  | 1 | 1 | Anion - Exclusion zone contact(s): /OP1/ |
| MG:DY:203 | 1.00 | 75.50 |  | 0 | 1 | - Exclusion zone contact(s): /N_back_/ Consider K in place of MG (1 valid contact(s) for K vs. 0 for MG) |
| MG:DY:204 | 1.00 | 85.10 |  | 0 | 1 | - Exclusion zone contact(s): /O_back_/ Consider K in place of MG (1 valid contact(s) for K vs. 0 for MG) |
| MG:DZ:101 | 1.00 | 87.84 |  | 0 |  | 1^st^ MG coordination shell missing (see Notes) |
| MG:DZ:102 | 1.00 | 49.21 |  | 0 |  | 1^st^ MG coordination shell missing (see Notes) |
| MG:DZ:103 | 1.00 | 87.75 |  | 0 | 1 | - Exclusion zone contact(s): /NH1_Arg_/ Consider K in place of MG (1 valid contact(s) for K vs. 0 for MG) |
| MG:DZ:104 | 1.00 | 59.55 |  | 0 | 1 | - Exclusion zone contact(s): /O6/ Consider K in place of MG (1 valid contact(s) for K vs. 0 for MG) |
| MG:DZ:105 | 1.00 | 54.15 |  | 0 | 3 | - Exclusion zone contact(s): /N_back_/N_back_/O_back_/ Consider K in place of MG (2 valid contact(s) for K vs. 0 for MG) |

---

\_\_\_\_\_\_\_\_\_\_\_\_\_\_\_\_\_\_\_\_\_\_\_\_\_\_\_\_\_\_\_\_\_\_\_\_\_\_\_\_\_\_\_\_\_\_\_\_\_\_\_\_\_\_\_\_\_\_\_\_\_\_\_\_  

#### 3) Potential issue list

##### Go to ToC

|  |  |  |
| --- | --- | --- |
| **Problem** | **Number** | **Affected ions** |
| **1) MG:  could be K** | 2845 | A0:201  A1:202  A1:203  A1:204  A2:201  A3:101  A3:104  A5:103  A7:101  A7:102  A7:103  A8:101  A8:102  A8:103  A8:104  AA:3011  AA:3019  AA:3024  AA:3029  AA:3034  AA:3035  AA:3036  AA:3038  AA:3042  AA:3043  AA:3047  AA:3050  AA:3051  AA:3054  AA:3056  AA:3064  AA:3066  AA:3067  AA:3068  AA:3072  AA:3075  AA:3076  AA:3077  AA:3079  AA:3080  AA:3087  AA:3092  AA:3093  AA:3096  AA:3099  AA:3100  AA:3103  AA:3105  AA:3110  AA:3111  AA:3112  AA:3113  AA:3114  AA:3116  AA:3118  AA:3119  AA:3124  AA:3125  AA:3127  AA:3131  AA:3132  AA:3134  AA:3135  AA:3138  AA:3139  AA:3142  AA:3143  AA:3147  AA:3148  AA:3151  AA:3153  AA:3156  AA:3159  AA:3160  AA:3162  AA:3167  AA:3169  AA:3171  AA:3173  AA:3174  AA:3175  AA:3178  AA:3180  AA:3182  AA:3184  AA:3187  AA:3194  AA:3203  AA:3204  AA:3206  AA:3208  AA:3210  AA:3213  AA:3217  AA:3218  AA:3219  AA:3220  AA:3224  AA:3225  AA:3227  AA:3228  AA:3229  AA:3231  AA:3232  AA:3233  AA:3236  AA:3237  AA:3239  AA:3241  AA:3243  AA:3246  AA:3247  AA:3250  AA:3253  AA:3254  AA:3255  AA:3260  AA:3263  AA:3265  AA:3272  AA:3273  AA:3276  AA:3280  AA:3281  AA:3282  AA:3284  AA:3285  AA:3287  AA:3288  AA:3290  AA:3291  AA:3293  AA:3294  AA:3295  AA:3298  AA:3299  AA:3301  AA:3302  AA:3303  AA:3305  AA:3308  AA:3309  AA:3310  AA:3311  AA:3312  AA:3313  AA:3314  AA:3315  AA:3316  AA:3317  AA:3322  AA:3325  AA:3326  AA:3327  AA:3328  AA:3332  AA:3333  AA:3334  AA:3335  AA:3336  AA:3337  AA:3339  AA:3341  AA:3342  AA:3343  AA:3344  AA:3346  AA:3347  AA:3349  AA:3350  AA:3352  AA:3353  AA:3354  AA:3359  AA:3361  AA:3363  AA:3364  AA:3366  AA:3369  AA:3371  AA:3373  AA:3374  AA:3379  AA:3380  AA:3388  AA:3389  AA:3390  AA:3393  AA:3394  AA:3396  AA:3403  AA:3406  AA:3408  AA:3410  AA:3414  AA:3415  AA:3417  AA:3420  AA:3421  AA:3422  AA:3424  AA:3425  AA:3426  AA:3429  AA:3431  AA:3433  AA:3434  AA:3435  AA:3436  AA:3439  AA:3440  AA:3441  AA:3442  AA:3443  AA:3445  AA:3447  AA:3449  AA:3451  AA:3454  AA:3455  AA:3457  AA:3460  AA:3463  AA:3465  AA:3467  AA:3470  AA:3471  AA:3474  AA:3477  AA:3478  AA:3481  AA:3483  AA:3484  AA:3485  AA:3486  AA:3489  AA:3490  AA:3491  AA:3492  AA:3495  AA:3496  AA:3498  AA:3500  AA:3503  AA:3508  AA:3511  AA:3518  AA:3521  AA:3522  AA:3524  AA:3525  AA:3526  AA:3527  AA:3528  AA:3529  AA:3530  AA:3533  AA:3535  AA:3536  AA:3537  AA:3538  AA:3541  AA:3542  AA:3544  AA:3545  AA:3548  AA:3549  AA:3551  AA:3554  AA:3558  AA:3559  AA:3562  AA:3563  AA:3567  AA:3569  AA:3570  AA:3571  AA:3572  AA:3574  AA:3576  AA:3580  AA:3582  AA:3583  AA:3584  AA:3585  AA:3587  AA:3588  AA:3589  AA:3593  AA:3595  AA:3600  AA:3602  AA:3604  AA:3605  AA:3608  AA:3609  AA:3610  AA:3611  AA:3614  AA:3615  AA:3617  AA:3618  AA:3619  AA:3622  AA:3623  AA:3628  AA:3631  AA:3637  AA:3638  AA:3640  AA:3641  AA:3642  AA:3644  AA:3646  AA:3647  AA:3648  AA:3649  AA:3651  AA:3652  AA:3653  AA:3655  AA:3656  AA:3658  AA:3661  AA:3662  AA:3668  AA:3670  AA:3671  AA:3672  AA:3675  AA:3677  AA:3678  AA:3680  AA:3681  AA:3682  AA:3683  AA:3687  AA:3689  AA:3691  AA:3692  AA:3693  AA:3696  AA:3698  AA:3699  AA:3701  AA:3702  AA:3703  AA:3704  AA:3706  AA:3707  AA:3708  AA:3710  AA:3714  AA:3715  AA:3717  AA:3718  AA:3720  AA:3721  AA:3725  AA:3726  AA:3728  AA:3733  AA:3734  AA:3735  AA:3737  AA:3738  AA:3740  AA:3742  AA:3745  AA:3746  AA:3747  AA:3748  AA:3750  AA:3751  AA:3752  AA:3754  AA:3756  AA:3757  AA:3758  AA:3759  AA:3762  AA:3763  AA:3766  AA:3767  AA:3769  AA:3772  AA:3775  AA:3777  AA:3778  AA:3780  AA:3781  AA:3784  AA:3790  AA:3791  AA:3792  AA:3794  AA:3795  AA:3796  AA:3798  AA:3801  AA:3810  AA:3813  AA:3815  AA:3816  AA:3819  AA:3820  AA:3821  AA:3823  AA:3824  AA:3826  AA:3827  AA:3828  AA:3829  AA:3832  AA:3833  AA:3835  AA:3838  AA:3840  AA:3845  AA:3847  AA:3848  AA:3849  AA:3852  AA:3855  AA:3857  AA:3859  AA:3860  AA:3861  AA:3863  AA:3864  AA:3866  AA:3867  AA:3868  AA:3869  AA:3870  AA:3874  AA:3875  AA:3876  AA:3878  AA:3880  AA:3883  AA:3885  AA:3887  AA:3888  AA:3890  AA:3891  AA:3893  AA:3894  AA:3895  AA:3899  AA:3904  AA:3906  AA:3907  AA:3908  AA:3911  AA:3913  AA:3916  AA:3917  AA:3918  AA:3919  AA:3920  AA:3921  AA:3922  AA:3923  AA:3924  AA:3925  AA:3927  AA:3928  AA:3930  AA:3932  AA:3935  AA:3936  AA:3939  AA:3940  AA:3941  AA:3942  AA:3943  AA:3944  AA:3946  AA:3947  AA:3949  AA:3951  AA:3952  AA:3953  AA:3954  AA:3955  AA:3957  AA:3961  AA:3962  AA:3963  AA:3966  AA:3971  AA:3972  AA:3975  AA:3977  AA:3978  AA:3981  AA:3983  AA:3984  AA:3985  AA:3986  AA:3989  AA:3991  AA:3994  AA:3995  AA:3997  AA:3999  AA:4000  AA:4001  AA:4002  AA:4003  AA:4004  AA:4005  AA:4006  AA:4007  AA:4008  AA:4012  AA:4016  AA:4017  AA:4018  AA:4021  AA:4022  AA:4023  AA:4026  AA:4027  AA:4029  AA:4031  AA:4038  AA:4040  AA:4041  AA:4042  AA:4045  AA:4046  AA:4047  AA:4048  AA:4049  AA:4050  AA:4051  AA:4056  AA:4057  AA:4059  AA:4060  AA:4064  AA:4067  AA:4068  AA:4070  AA:4071  AA:4073  AA:4074  AA:4076  AA:4078  AA:4079  AA:4080  AA:4081  AA:4082  AA:4083  AA:4084  AA:4085  AA:4087  AA:4088  AA:4089  AA:4092  AA:4094  AA:4095  AA:4098  AA:4099  AA:4100  AA:4104  AA:4105  AA:4107  AA:4110  AA:4111  AA:4113  AA:4116  AA:4117  AA:4119  AA:4121  AA:4123  AA:4125  AA:4126  AA:4129  AA:4130  AA:4132  AA:4134  AA:4137  AA:4141  AA:4145  AA:4149  AA:4152  AA:4153  AA:4154  AA:4155  AA:4157  AA:4158  AA:4159  AA:4160  AA:4161  AA:4162  AA:4163  AA:4166  AB:201  AB:203  AB:204  AB:206  AB:209  AB:210  AB:212  AB:213  AB:215  AB:216  AB:219  AB:220  AB:222  AB:223  AB:224  AB:225  AB:229  AB:230  AB:231  AB:232  AB:233  AB:236  AD:302  AD:303  AD:305  AD:306  AD:308  AD:310  AD:313  AE:302  AF:301  AF:303  AF:304  AG:201  AG:202  AG:203  AN:201  AO:202  AO:203  AO:204  AO:205  AO:206  AP:201  AQ:201  AQ:203  AQ:204  AR:202  AR:203  AR:205  AS:201  AS:203  AT:102  AT:104  AU:202  AU:203  AW:102  AZ:101  AZ:102  AZ:103  B1:101  B1:104  BA:1604  BA:1605  BA:1607  BA:1609  BA:1610  BA:1612  BA:1619  BA:1620  BA:1622  BA:1623  BA:1629  BA:1631  BA:1633  BA:1635  BA:1636  BA:1642  BA:1643  BA:1644  BA:1647  BA:1649  BA:1650  BA:1651  BA:1652  BA:1656  BA:1658  BA:1659  BA:1663  BA:1664  BA:1666  BA:1668  BA:1670  BA:1675  BA:1676  BA:1678  BA:1679  BA:1680  BA:1683  BA:1684  BA:1685  BA:1687  BA:1688  BA:1689  BA:1690  BA:1691  BA:1692  BA:1695  BA:1696  BA:1700  BA:1704  BA:1705  BA:1706  BA:1707  BA:1710  BA:1711  BA:1712  BA:1715  BA:1717  BA:1718  BA:1719  BA:1721  BA:1722  BA:1724  BA:1725  BA:1730  BA:1731  BA:1735  BA:1740  BA:1743  BA:1744  BA:1746  BA:1748  BA:1749  BA:1750  BA:1756  BA:1757  BA:1758  BA:1760  BA:1761  BA:1767  BA:1769  BA:1770  BA:1772  BA:1774  BA:1775  BA:1777  BA:1780  BA:1784  BA:1788  BA:1790  BA:1791  BA:1793  BA:1800  BA:1804  BA:1809  BA:1815  BA:1816  BA:1817  BA:1819  BA:1820  BA:1821  BA:1824  BA:1826  BA:1827  BA:1833  BA:1834  BA:1835  BA:1839  BA:1846  BA:1848  BA:1849  BA:1858  BA:1872  BA:1887  BA:1889  BA:1892  BA:1893  BA:1898  BA:1901  BA:1903  BA:1904  BA:1905  BA:1906  BA:1910  BA:1912  BA:1913  BA:1915  BA:1916  BA:1917  BA:1918  BA:1919  BA:1920  BA:1922  BA:1923  BA:1925  BA:1927  BA:1928  BA:1929  BA:1930  BA:1931  BA:1932  BA:1933  BA:1935  BA:1938  BA:1942  BA:1946  BA:1947  BA:1948  BA:1949  BA:1950  BA:1952  BA:1953  BA:1957  BA:1960  BA:1961  BA:1967  BA:1969  BA:1972  BA:1977  BA:1981  BA:1982  BA:1983  BA:1984  BA:1985  BA:1987  BA:1988  BA:1989  BA:1990  BA:1991  BA:1992  BA:1993  BA:1995  BA:1999  BA:2000  BA:2001  BA:2002  BA:2003  BA:2004  BA:2007  BA:2008  BA:2010  BA:2014  BA:2015  BA:2016  BA:2017  BA:2018  BA:2020  BA:2021  BA:2022  BA:2025  BA:2026  BA:2028  BA:2029  BA:2033  BA:2035  BA:2037  BA:2042  BA:2044  BA:2045  BA:2048  BA:2050  BA:2051  BA:2052  BA:2057  BA:2062  BA:2064  BA:2065  BA:2067  BA:2070  BA:2071  BA:2076  BA:2077  BA:2078  BA:2080  BA:2081  BA:2082  BA:2083  BA:2087  BA:2088  BA:2089  BA:2090  BA:2091  BA:2092  BA:2096  BA:2097  BA:2099  BA:2101  BA:2102  BA:2105  BA:2106  BA:2107  BA:2108  BA:2109  BA:2111  BA:2112  BA:2113  BA:2114  BA:2115  BA:2116  BA:2120  BA:2122  BA:2124  BA:2126  BA:2128  BA:2130  BA:2131  BA:2132  BA:2133  BA:2134  BA:2135  BA:2138  BA:2139  BA:2142  BA:2143  BA:2147  BA:2148  BA:2149  BA:2150  BA:2151  BA:2152  BA:2153  BA:2154  BA:2157  BA:2159  BA:2160  BA:2162  BA:2166  BA:2169  BA:2172  BA:2175  BA:2176  BA:2180  BA:2182  BA:2184  BA:2186  BA:2187  BA:2189  BA:2190  BA:2191  BA:2192  BA:2194  BA:2196  BA:2200  BA:2203  BA:2204  BA:2205  BA:2206  BA:2208  BA:2209  BA:2211  BA:2212  BA:2213  BA:2215  BA:2217  BA:2218  BA:2219  BA:2220  BA:2221  BA:2222  BA:2223  BA:2230  BA:2232  BA:2233  BA:2234  BA:2235  BA:2236  BA:2238  BA:2240  BA:2241  BA:2246  BA:2247  BA:2248  BA:2250  BA:2252  BA:2253  BA:2255  BA:2258  BA:2260  BA:2261  BA:2262  BA:2263  BA:2264  BA:2265  BA:2266  BA:2268  BA:2269  BA:2271  BA:2272  BA:2273  BA:2274  BB:101  BB:105  BB:111  BC:104  BC:106  BC:109  BC:111  BC:115  BC:116  BD:102  BD:105  BD:107  BD:108  BD:111  BD:117  BD:124  BD:126  BE:303  BF:301  BF:302  BG:303  BG:305  BG:306  BH:201  BH:203  BI:201  BK:201  BL:201  BP:201  BQ:102  BS:103  BS:104  BS:105  BS:107  BS:108  BS:109  BT:201  BT:202  BV:101  BW:201  BW:202  BW:205  BW:206  BW:207  C1:102  C1:104  C1:105  C1:106  CA:1601  CA:1603  CA:1604  CA:1609  CA:1611  CA:1612  CA:1614  CA:1618  CA:1619  CA:1620  CA:1621  CA:1624  CA:1625  CA:1626  CA:1627  CA:1635  CA:1636  CA:1638  CA:1640  CA:1641  CA:1642  CA:1643  CA:1647  CA:1649  CA:1652  CA:1655  CA:1656  CA:1659  CA:1661  CA:1662  CA:1663  CA:1665  CA:1667  CA:1668  CA:1672  CA:1676  CA:1682  CA:1683  CA:1685  CA:1688  CA:1690  CA:1692  CA:1695  CA:1699  CA:1701  CA:1702  CA:1703  CA:1705  CA:1706  CA:1708  CA:1710  CA:1714  CA:1717  CA:1719  CA:1720  CA:1721  CA:1722  CA:1723  CA:1726  CA:1727  CA:1728  CA:1729  CA:1730  CA:1731  CA:1732  CA:1734  CA:1735  CA:1742  CA:1743  CA:1744  CA:1745  CA:1746  CA:1747  CA:1749  CA:1750  CA:1752  CA:1755  CA:1756  CA:1757  CA:1758  CA:1759  CA:1762  CA:1763  CA:1765  CA:1767  CA:1768  CA:1769  CA:1772  CA:1773  CA:1774  CA:1782  CA:1784  CA:1789  CA:1790  CA:1793  CA:1794  CA:1795  CA:1799  CA:1800  CA:1802  CA:1803  CA:1804  CA:1809  CA:1810  CA:1811  CA:1812  CA:1814  CA:1815  CA:1819  CA:1822  CA:1824  CA:1829  CA:1830  CA:1831  CA:1832  CA:1834  CA:1835  CA:1837  CA:1843  CA:1844  CA:1845  CA:1846  CA:1847  CA:1848  CA:1854  CA:1856  CA:1859  CA:1861  CA:1863  CA:1864  CA:1867  CA:1868  CA:1870  CA:1874  CA:1877  CA:1878  CA:1880  CA:1883  CA:1884  CA:1885  CA:1886  CA:1887  CA:1890  CA:1894  CA:1895  CA:1896  CA:1898  CA:1899  CA:1900  CA:1901  CA:1902  CA:1903  CA:1905  CA:1910  CA:1911  CA:1914  CA:1915  CA:1919  CA:1921  CA:1922  CA:1924  CA:1925  CA:1926  CA:1928  CA:1929  CA:1933  CA:1934  CA:1935  CA:1937  CA:1939  CA:1940  CA:1943  CA:1944  CA:1945  CA:1950  CA:1951  CA:1952  CA:1955  CA:1956  CA:1957  CA:1959  CA:1960  CA:1961  CA:1962  CA:1963  CA:1964  CA:1965  CA:1966  CA:1971  CA:1974  CA:1976  CA:1980  CA:1981  CA:1983  CA:1988  CA:1989  CA:1992  CA:1993  CA:1995  CA:1996  CA:1998  CA:1999  CA:2000  CA:2001  CA:2002  CA:2004  CA:2006  CA:2007  CA:2008  CA:2011  CA:2012  CA:2013  CA:2014  CA:2015  CA:2017  CA:2019  CA:2021  CA:2022  CA:2024  CA:2025  CA:2026  CA:2028  CA:2030  CA:2031  CA:2033  CA:2034  CA:2035  CA:2036  CA:2038  CA:2039  CA:2041  CA:2045  CA:2047  CA:2051  CA:2053  CA:2054  CA:2055  CA:2057  CA:2058  CA:2060  CA:2061  CA:2062  CA:2067  CA:2070  CA:2072  CA:2073  CA:2074  CA:2081  CA:2083  CA:2084  CA:2085  CA:2086  CA:2087  CA:2088  CA:2090  CA:2091  CA:2096  CA:2098  CA:2099  CA:2100  CA:2101  CA:2103  CA:2104  CA:2106  CA:2109  CA:2112  CA:2116  CA:2121  CA:2122  CA:2123  CA:2125  CA:2126  CA:2127  CA:2128  CA:2129  CA:2130  CA:2132  CA:2133  CA:2135  CA:2136  CA:2137  CA:2139  CA:2140  CA:2141  CA:2143  CA:2144  CA:2145  CA:2152  CA:2155  CA:2157  CA:2159  CA:2160  CA:2161  CA:2162  CA:2164  CA:2167  CA:2169  CA:2171  CA:2174  CA:2176  CA:2178  CA:2180  CA:2185  CA:2186  CA:2187  CA:2189  CA:2191  CA:2193  CA:2194  CA:2195  CA:2196  CA:2197  CA:2199  CA:2200  CA:2201  CA:2202  CA:2203  CA:2206  CA:2208  CA:2209  CA:2210  CA:2213  CA:2214  CA:2215  CA:2216  CA:2221  CA:2223  CA:2225  CA:2226  CA:2228  CA:2229  CA:2230  CA:2231  CA:2233  CA:2235  CA:2239  CA:2242  CA:2244  CA:2245  CA:2247  CA:2248  CA:2249  CA:2250  CA:2251  CA:2252  CA:2253  CA:2254  CA:2255  CA:2256  CA:2257  CA:2258  CA:2260  CA:2261  CA:2262  CA:2264  CA:2265  CA:2269  CA:2270  CA:2271  CA:2272  CA:2273  CA:2274  CA:2275  CA:2276  CA:2278  CA:2279  CA:2280  CA:2282  CA:2283  CA:2284  CA:2285  CA:2287  CA:2288  CA:2290  CA:2291  CA:2296  CA:2297  CA:2298  CA:2300  CA:2301  CA:2302  CA:2305  CA:2312  CA:2314  CA:2315  CA:2320  CB:101  CB:102  CB:103  CB:106  CB:114  CB:116  CB:121  CC:101  CC:103  CC:104  CC:106  CC:108  CC:109  CC:110  CC:111  CC:112  CC:117  CC:118  CC:119  CC:120  CC:125  CC:126  CD:101  CD:102  CD:103  CD:106  CD:110  CD:112  CD:119  CD:120  CD:122  CD:123  CD:125  CD:128  CD:129  CD:130  CE:302  CE:303  CE:305  CF:303  CG:303  CG:304  CG:305  CG:308  CG:309  CG:310  CG:311  CG:312  CH:204  CH:206  CI:201  CJ:201  CK:203  CK:204  CK:207  CK:208  CK:209  CK:211  CL:201  CM:203  CN:201  CP:203  CQ:102  CQ:103  CQ:104  CS:102  CS:103  CS:105  CS:106  CT:201  CT:202  CT:203  CV:101  CV:103  CW:201  D0:205  D0:207  D0:208  D0:211  D1:201  D1:202  D1:203  D1:204  D1:205  D1:206  D1:207  D1:208  D1:209  D2:203  D2:204  D2:206  D2:208  D2:209  D3:101  D3:102  D3:104  D3:105  D3:107  D4:102  D5:102  D5:106  D5:107  D5:108  D7:102  D7:103  D8:101  D8:102  D8:103  D8:104  D8:107  D8:109  DA:3006  DA:3017  DA:3020  DA:3022  DA:3025  DA:3037  DA:3038  DA:3047  DA:3050  DA:3055  DA:3057  DA:3059  DA:3060  DA:3065  DA:3073  DA:3077  DA:3090  DA:3092  DA:3096  DA:3102  DA:3104  DA:3106  DA:3107  DA:3110  DA:3111  DA:3113  DA:3115  DA:3116  DA:3117  DA:3118  DA:3119  DA:3120  DA:3121  DA:3122  DA:3123  DA:3124  DA:3127  DA:3129  DA:3131  DA:3135  DA:3137  DA:3138  DA:3142  DA:3147  DA:3149  DA:3151  DA:3152  DA:3153  DA:3154  DA:3158  DA:3160  DA:3162  DA:3163  DA:3164  DA:3165  DA:3166  DA:3167  DA:3171  DA:3172  DA:3173  DA:3174  DA:3177  DA:3178  DA:3181  DA:3182  DA:3184  DA:3186  DA:3187  DA:3188  DA:3189  DA:3190  DA:3191  DA:3192  DA:3193  DA:3196  DA:3197  DA:3199  DA:3202  DA:3203  DA:3205  DA:3206  DA:3210  DA:3212  DA:3213  DA:3214  DA:3216  DA:3217  DA:3218  DA:3219  DA:3220  DA:3222  DA:3224  DA:3225  DA:3228  DA:3229  DA:3230  DA:3231  DA:3232  DA:3233  DA:3236  DA:3237  DA:3243  DA:3245  DA:3246  DA:3247  DA:3250  DA:3251  DA:3252  DA:3253  DA:3254  DA:3256  DA:3258  DA:3262  DA:3263  DA:3266  DA:3268  DA:3269  DA:3273  DA:3274  DA:3277  DA:3278  DA:3279  DA:3280  DA:3281  DA:3284  DA:3287  DA:3289  DA:3291  DA:3293  DA:3294  DA:3296  DA:3298  DA:3299  DA:3303  DA:3304  DA:3305  DA:3306  DA:3307  DA:3308  DA:3309  DA:3310  DA:3314  DA:3316  DA:3317  DA:3318  DA:3320  DA:3321  DA:3322  DA:3323  DA:3327  DA:3330  DA:3331  DA:3335  DA:3336  DA:3337  DA:3338  DA:3340  DA:3349  DA:3350  DA:3351  DA:3352  DA:3353  DA:3357  DA:3361  DA:3363  DA:3365  DA:3366  DA:3368  DA:3370  DA:3372  DA:3374  DA:3375  DA:3376  DA:3382  DA:3383  DA:3388  DA:3389  DA:3392  DA:3396  DA:3397  DA:3398  DA:3399  DA:3400  DA:3401  DA:3402  DA:3403  DA:3404  DA:3405  DA:3406  DA:3407  DA:3411  DA:3416  DA:3417  DA:3418  DA:3419  DA:3421  DA:3422  DA:3423  DA:3424  DA:3429  DA:3430  DA:3438  DA:3439  DA:3442  DA:3444  DA:3447  DA:3449  DA:3450  DA:3451  DA:3452  DA:3453  DA:3454  DA:3455  DA:3456  DA:3457  DA:3458  DA:3460  DA:3462  DA:3465  DA:3468  DA:3470  DA:3471  DA:3472  DA:3473  DA:3474  DA:3475  DA:3476  DA:3477  DA:3480  DA:3481  DA:3482  DA:3483  DA:3486  DA:3487  DA:3488  DA:3490  DA:3491  DA:3492  DA:3493  DA:3494  DA:3496  DA:3499  DA:3500  DA:3501  DA:3503  DA:3505  DA:3506  DA:3507  DA:3508  DA:3511  DA:3512  DA:3514  DA:3515  DA:3516  DA:3517  DA:3518  DA:3519  DA:3524  DA:3525  DA:3526  DA:3527  DA:3528  DA:3530  DA:3533  DA:3534  DA:3536  DA:3539  DA:3541  DA:3542  DA:3545  DA:3547  DA:3549  DA:3552  DA:3555  DA:3556  DA:3557  DA:3559  DA:3560  DA:3563  DA:3566  DA:3570  DA:3571  DA:3572  DA:3573  DA:3574  DA:3577  DA:3581  DA:3582  DA:3584  DA:3586  DA:3588  DA:3590  DA:3592  DA:3594  DA:3595  DA:3599  DA:3601  DA:3603  DA:3604  DA:3605  DA:3613  DA:3616  DA:3619  DA:3620  DA:3621  DA:3623  DA:3624  DA:3625  DA:3626  DA:3627  DA:3631  DA:3632  DA:3633  DA:3634  DA:3635  DA:3637  DA:3638  DA:3639  DA:3641  DA:3642  DA:3643  DA:3645  DA:3648  DA:3649  DA:3650  DA:3651  DA:3653  DA:3654  DA:3657  DA:3658  DA:3659  DA:3660  DA:3664  DA:3665  DA:3668  DA:3669  DA:3671  DA:3674  DA:3675  DA:3676  DA:3679  DA:3682  DA:3683  DA:3684  DA:3685  DA:3688  DA:3689  DA:3692  DA:3694  DA:3696  DA:3702  DA:3705  DA:3707  DA:3708  DA:3709  DA:3711  DA:3714  DA:3715  DA:3716  DA:3717  DA:3718  DA:3719  DA:3720  DA:3722  DA:3725  DA:3726  DA:3729  DA:3731  DA:3732  DA:3733  DA:3735  DA:3737  DA:3739  DA:3741  DA:3744  DA:3745  DA:3748  DA:3750  DA:3751  DA:3755  DA:3757  DA:3760  DA:3765  DA:3772  DA:3775  DA:3777  DA:3778  DA:3779  DA:3780  DA:3781  DA:3784  DA:3785  DA:3786  DA:3789  DA:3790  DA:3791  DA:3792  DA:3795  DA:3796  DA:3798  DA:3801  DA:3804  DA:3806  DA:3808  DA:3809  DA:3810  DA:3811  DA:3812  DA:3814  DA:3817  DA:3818  DA:3820  DA:3821  DA:3822  DA:3825  DA:3826  DA:3828  DA:3830  DA:3832  DA:3833  DA:3834  DA:3835  DA:3837  DA:3838  DA:3841  DA:3842  DA:3847  DA:3848  DA:3849  DA:3853  DA:3854  DA:3855  DA:3856  DA:3859  DA:3860  DA:3865  DA:3866  DA:3867  DA:3868  DA:3874  DA:3876  DA:3877  DA:3878  DA:3880  DA:3881  DA:3883  DA:3884  DA:3885  DA:3887  DA:3888  DA:3889  DA:3890  DA:3891  DA:3892  DA:3901  DA:3906  DA:3909  DA:3911  DA:3914  DA:3918  DA:3919  DA:3923  DA:3924  DA:3926  DA:3927  DA:3928  DA:3930  DA:3932  DA:3934  DA:3936  DA:3941  DA:3943  DA:3945  DA:3948  DA:3951  DA:3961  DA:3965  DA:3967  DA:3968  DA:3971  DA:3972  DA:3973  DA:3975  DA:3976  DA:3980  DA:3985  DA:3986  DA:3987  DA:3988  DA:3989  DA:3992  DA:3993  DA:3994  DA:3996  DA:3997  DA:3998  DA:4000  DA:4001  DA:4002  DA:4003  DA:4004  DA:4006  DA:4008  DA:4009  DA:4010  DA:4012  DA:4013  DA:4014  DA:4015  DA:4016  DA:4018  DA:4019  DA:4020  DA:4021  DA:4022  DA:4024  DA:4025  DA:4026  DA:4027  DA:4029  DA:4035  DA:4036  DA:4037  DA:4039  DA:4040  DA:4041  DA:4042  DA:4043  DA:4044  DA:4045  DA:4046  DA:4047  DA:4050  DA:4051  DA:4053  DA:4054  DA:4055  DA:4056  DA:4058  DA:4061  DA:4062  DA:4063  DA:4065  DA:4066  DA:4067  DA:4068  DA:4070  DA:4071  DA:4072  DA:4073  DA:4074  DA:4076  DA:4078  DA:4081  DA:4083  DA:4084  DA:4085  DA:4087  DA:4090  DA:4091  DA:4092  DA:4093  DA:4094  DA:4098  DA:4100  DA:4102  DA:4104  DA:4105  DA:4106  DA:4107  DA:4108  DA:4110  DA:4111  DA:4116  DA:4117  DA:4118  DA:4120  DA:4121  DA:4126  DA:4132  DA:4133  DA:4134  DA:4136  DA:4137  DA:4139  DA:4142  DA:4143  DA:4145  DA:4146  DA:4149  DA:4150  DA:4151  DA:4152  DA:4156  DA:4157  DA:4158  DA:4159  DA:4163  DA:4164  DA:4166  DA:4167  DA:4168  DA:4169  DA:4171  DA:4172  DA:4173  DA:4175  DA:4177  DA:4179  DA:4180  DA:4181  DA:4183  DA:4185  DA:4186  DA:4187  DA:4188  DA:4189  DA:4190  DA:4194  DA:4196  DA:4198  DA:4199  DA:4200  DA:4201  DA:4203  DA:4204  DA:4205  DA:4207  DA:4209  DA:4210  DA:4211  DA:4212  DA:4214  DA:4215  DA:4216  DA:4218  DA:4219  DA:4220  DA:4221  DA:4224  DA:4226  DA:4229  DA:4230  DA:4231  DA:4232  DA:4234  DA:4235  DA:4236  DA:4238  DA:4240  DA:4241  DA:4242  DA:4244  DA:4245  DA:4249  DA:4250  DA:4252  DA:4253  DA:4258  DA:4259  DA:4260  DA:4262  DA:4263  DA:4264  DA:4266  DA:4270  DA:4271  DA:4272  DA:4273  DA:4274  DA:4276  DA:4277  DA:4278  DA:4280  DA:4281  DA:4283  DA:4284  DA:4291  DA:4292  DA:4293  DA:4295  DA:4298  DA:4301  DA:4302  DA:4303  DA:4304  DA:4305  DA:4306  DA:4307  DA:4308  DA:4309  DA:4311  DA:4312  DA:4313  DA:4314  DA:4316  DA:4317  DA:4318  DA:4321  DA:4322  DA:4323  DA:4324  DA:4325  DA:4326  DA:4327  DA:4328  DA:4329  DA:4331  DA:4332  DA:4333  DA:4334  DA:4336  DA:4337  DA:4338  DA:4339  DA:4340  DA:4345  DA:4346  DA:4347  DA:4349  DA:4350  DA:4351  DA:4352  DA:4353  DA:4357  DA:4358  DA:4359  DA:4361  DA:4362  DA:4363  DA:4364  DA:4365  DA:4366  DA:4367  DA:4369  DA:4371  DA:4373  DA:4375  DA:4376  DA:4377  DA:4378  DA:4379  DA:4381  DA:4384  DA:4385  DA:4386  DA:4387  DA:4388  DA:4391  DA:4393  DA:4397  DA:4398  DA:4399  DA:4400  DA:4401  DA:4402  DA:4403  DA:4405  DA:4406  DA:4410  DA:4411  DA:4412  DA:4413  DA:4414  DA:4415  DA:4417  DA:4418  DA:4420  DA:4423  DA:4424  DA:4425  DA:4427  DA:4429  DA:4430  DA:4431  DA:4432  DA:4433  DA:4434  DA:4437  DA:4439  DA:4440  DA:4441  DA:4442  DA:4443  DA:4445  DA:4446  DA:4447  DA:4450  DA:4452  DA:4453  DA:4454  DA:4455  DA:4456  DA:4458  DA:4459  DA:4461  DA:4462  DA:4463  DA:4465  DA:4467  DA:4469  DA:4471  DA:4472  DA:4473  DA:4474  DA:4476  DA:4481  DA:4483  DA:4484  DA:4485  DA:4486  DA:4487  DA:4488  DA:4489  DA:4491  DA:4492  DA:4494  DA:4495  DA:4496  DA:4497  DA:4498  DA:4499  DA:4500  DA:4502  DA:4503  DA:4507  DA:4508  DA:4509  DA:4510  DA:4511  DA:4514  DA:4515  DA:4516  DA:4518  DA:4520  DA:4521  DA:4522  DA:4523  DA:4524  DA:4526  DA:4527  DA:4531  DA:4532  DA:4533  DA:4535  DA:4536  DA:4537  DA:4538  DA:4539  DA:4540  DA:4542  DA:4545  DA:4548  DA:4550  DA:4551  DA:4552  DA:4553  DA:4554  DA:4555  DA:4556  DA:4557  DA:4559  DA:4561  DA:4562  DA:4563  DA:4564  DA:4565  DA:4567  DA:4568  DA:4569  DA:4570  DA:4571  DA:4572  DA:4573  DA:4577  DA:4578  DA:4579  DA:4580  DA:4581  DA:4582  DA:4583  DA:4584  DA:4586  DA:4588  DA:4590  DA:4591  DA:4592  DA:4593  DA:4596  DA:4597  DA:4599  DA:4600  DA:4601  DA:4603  DA:4604  DA:4605  DA:4606  DA:4608  DA:4613  DA:4614  DA:4616  DA:4618  DA:4619  DA:4622  DA:4623  DA:4625  DA:4626  DA:4627  DA:4628  DA:4629  DA:4630  DA:4635  DA:4637  DA:4638  DA:4639  DA:4641  DA:4642  DA:4643  DA:4647  DA:4651  DA:4652  DA:4653  DA:4654  DA:4657  DA:4658  DA:4659  DA:4660  DA:4663  DA:4665  DA:4666  DA:4668  DA:4669  DA:4670  DA:4672  DA:4678  DA:4679  DA:4680  DA:4683  DA:4684  DA:4685  DA:4687  DA:4690  DA:4691  DA:4696  DA:4697  DA:4700  DA:4701  DA:4702  DA:4703  DA:4706  DA:4711  DA:4712  DA:4713  DA:4719  DA:4722  DA:4723  DA:4725  DA:4726  DA:4728  DA:4729  DA:4730  DA:4731  DA:4732  DA:4735  DA:4736  DA:4737  DA:4738  DA:4740  DA:4741  DA:4743  DA:4744  DA:4745  DA:4746  DA:4747  DA:4748  DA:4749  DA:4751  DA:4752  DA:4753  DA:4755  DA:4756  DA:4760  DA:4761  DA:4762  DA:4763  DA:4765  DA:4766  DA:4767  DA:4769  DA:4770  DA:4771  DA:4772  DA:4773  DA:4775  DA:4778  DA:4779  DA:4780  DA:4782  DA:4783  DA:4784  DA:4788  DA:4789  DA:4792  DA:4793  DA:4795  DA:4796  DA:4797  DA:4800  DA:4801  DA:4806  DA:4807  DA:4808  DA:4809  DA:4810  DA:4812  DA:4814  DA:4815  DA:4816  DA:4817  DA:4818  DA:4820  DA:4823  DA:4824  DA:4825  DA:4826  DA:4827  DA:4828  DA:4829  DA:4830  DA:4833  DA:4834  DA:4835  DA:4836  DA:4838  DA:4842  DA:4843  DA:4845  DA:4847  DA:4848  DA:4849  DA:4851  DA:4852  DA:4853  DA:4855  DA:4856  DA:4857  DA:4858  DA:4863  DA:4865  DA:4867  DA:4869  DA:4871  DA:4872  DA:4873  DA:4874  DA:4875  DA:4876  DA:4877  DA:4878  DA:4879  DA:4880  DA:4882  DA:4884  DA:4886  DA:4890  DA:4891  DA:4895  DA:4898  DA:4901  DA:4902  DA:4903  DA:4904  DA:4907  DA:4910  DA:4912  DA:4913  DA:4914  DA:4918  DA:4919  DA:4924  DA:4927  DA:4928  DA:4929  DA:4930  DA:4931  DA:4932  DA:4933  DA:4936  DA:4939  DA:4940  DA:4942  DA:4943  DA:4944  DA:4946  DA:4947  DA:4949  DA:4951  DA:4952  DA:4953  DA:4956  DA:4957  DA:4958  DA:4959  DA:4961  DA:4962  DA:4965  DA:4966  DA:4967  DA:4968  DA:4973  DA:4974  DA:4975  DA:4978  DA:4979  DA:4981  DA:4982  DA:4983  DA:4986  DA:4987  DA:4988  DA:4992  DA:4993  DA:4994  DA:4997  DA:4999  DA:5000  DA:5001  DA:5003  DA:5004  DA:5006  DA:5007  DA:5008  DA:5009  DA:5012  DA:5013  DA:5014  DA:5015  DA:5016  DA:5019  DA:5020  DA:5022  DA:5024  DA:5025  DA:5026  DA:5028  DA:5029  DA:5031  DA:5033  DA:5035  DA:5036  DA:5038  DA:5040  DA:5041  DA:5042  DA:5043  DA:5046  DA:5048  DA:5051  DA:5052  DA:5054  DA:5058  DA:5061  DA:5062  DA:5063  DA:5066  DA:5068  DA:5069  DA:5070  DA:5071  DA:5073  DA:5074  DA:5075  DA:5077  DB:204  DB:205  DB:206  DB:208  DB:210  DB:211  DB:212  DB:215  DB:216  DB:218  DB:220  DB:221  DB:222  DB:224  DB:226  DB:233  DB:234  DB:235  DB:236  DB:237  DB:243  DB:244  DB:245  DB:246  DB:247  DB:249  DB:250  DB:252  DB:253  DB:255  DB:256  DB:258  DB:260  DB:261  DB:263  DB:265  DB:266  DB:267  DB:268  DB:269  DB:270  DB:271  DB:272  DB:274  DB:276  DD:305  DD:306  DD:309  DD:310  DD:311  DD:314  DE:301  DE:304  DE:305  DE:306  DE:308  DE:309  DE:310  DE:314  DE:315  DF:301  DF:302  DF:303  DF:304  DF:305  DF:307  DF:308  DF:309  DF:310  DF:314  DF:315  DF:316  DF:318  DF:320  DF:321  DF:322  DF:323  DF:325  DG:202  DG:205  DH:205  DK:202  DM:201  DM:202  DM:204  DM:205  DM:206  DO:201  DO:203  DO:204  DO:207  DO:208  DO:209  DO:210  DO:211  DO:212  DO:213  DO:214  DO:215  DO:217  DP:201  DP:203  DP:204  DR:201  DR:204  DS:202  DS:203  DS:206  DS:207  DT:101  DT:102  DT:103  DT:104  DT:105  DT:106  DU:203  DU:205  DU:210  DU:211  DU:212  DU:215  DU:216  DU:218  DV:302  DV:305  DW:102  DW:104  DW:105  DW:106  DW:107  DX:101  DX:102  DY:203  DY:204  DZ:103  DZ:104  DZ:105 |
| **1) MG:  could be NA** | 475 | A5:102  A6:101  AA:3001  AA:3003  AA:3016  AA:3018  AA:3021  AA:3027  AA:3037  AA:3041  AA:3058  AA:3062  AA:3074  AA:3082  AA:3083  AA:3085  AA:3088  AA:3095  AA:3101  AA:3106  AA:3108  AA:3122  AA:3128  AA:3137  AA:3161  AA:3165  AA:3166  AA:3185  AA:3193  AA:3198  AA:3202  AA:3234  AA:3252  AA:3261  AA:3264  AA:3271  AA:3304  AA:3330  AA:3338  AA:3340  AA:3355  AA:3357  AA:3358  AA:3372  AA:3385  AA:3387  AA:3392  AA:3397  AA:3399  AA:3402  AA:3438  AA:3453  AA:3476  AA:3479  AA:3482  AA:3502  AA:3505  AA:3515  AA:3517  AA:3539  AA:3546  AA:3561  AA:3579  AA:3596  AA:3607  AA:3621  AA:3636  AA:3664  AA:3669  AA:3736  AA:3779  AA:3788  AA:3800  AA:3803  AA:3804  AA:3812  AA:3830  AA:3853  AA:3879  AA:3896  AA:3905  AA:3909  AA:3929  AA:3945  AA:3948  AA:3965  AA:3970  AA:3974  AA:3987  AA:3996  AA:4009  AA:4011  AA:4030  AA:4053  AA:4063  AA:4072  AA:4101  AA:4106  AA:4108  AA:4127  AA:4128  AA:4139  AA:4144  AA:4156  AB:214  AB:218  AB:227  AE:304  AQ:202  AQ:205  AR:204  AT:101  BA:1638  BA:1640  BA:1654  BA:1661  BA:1703  BA:1709  BA:1734  BA:1742  BA:1745  BA:1754  BA:1765  BA:1766  BA:1768  BA:1781  BA:1808  BA:1823  BA:1831  BA:1836  BA:1850  BA:1868  BA:1878  BA:1896  BA:1900  BA:1907  BA:1914  BA:1926  BA:1943  BA:1954  BA:1965  BA:1966  BA:1971  BA:1973  BA:1974  BA:1976  BA:1994  BA:2009  BA:2024  BA:2036  BA:2039  BA:2043  BA:2046  BA:2047  BA:2053  BA:2073  BA:2079  BA:2093  BA:2094  BA:2103  BA:2125  BA:2129  BA:2136  BA:2137  BA:2145  BA:2155  BA:2161  BA:2163  BA:2168  BA:2181  BA:2199  BA:2225  BA:2231  BA:2254  BA:2257  BA:2259  BB:112  BB:113  BC:101  BK:205  BK:206  BM:202  BS:102  BW:203  BW:204  CA:1613  CA:1623  CA:1639  CA:1646  CA:1650  CA:1689  CA:1694  CA:1700  CA:1711  CA:1715  CA:1736  CA:1761  CA:1787  CA:1788  CA:1807  CA:1836  CA:1839  CA:1842  CA:1855  CA:1869  CA:1889  CA:1913  CA:1916  CA:1920  CA:1930  CA:1938  CA:1948  CA:1949  CA:1958  CA:1970  CA:1977  CA:1978  CA:1979  CA:2009  CA:2040  CA:2042  CA:2052  CA:2059  CA:2076  CA:2107  CA:2111  CA:2134  CA:2150  CA:2153  CA:2165  CA:2168  CA:2170  CA:2173  CA:2220  CA:2238  CA:2240  CA:2281  CA:2286  CA:2299  CA:2309  CA:2313  CA:2318  CB:105  CB:107  CB:108  CB:112  CB:113  CB:120  CC:105  CD:109  CD:114  CD:121  CE:301  CE:304  CH:203  CK:210  CR:101  CR:102  D0:206  D0:210  D1:211  D1:212  D2:205  D3:103  D3:106  D5:103  D5:104  D5:105  D6:103  D7:104  D8:106  D8:108  DA:3003  DA:3009  DA:3013  DA:3028  DA:3041  DA:3062  DA:3076  DA:3084  DA:3085  DA:3094  DA:3101  DA:3109  DA:3126  DA:3128  DA:3132  DA:3169  DA:3185  DA:3195  DA:3198  DA:3215  DA:3226  DA:3235  DA:3239  DA:3249  DA:3271  DA:3282  DA:3286  DA:3301  DA:3312  DA:3347  DA:3348  DA:3390  DA:3408  DA:3412  DA:3431  DA:3434  DA:3446  DA:3461  DA:3520  DA:3531  DA:3538  DA:3567  DA:3610  DA:3615  DA:3617  DA:3618  DA:3629  DA:3640  DA:3652  DA:3661  DA:3663  DA:3670  DA:3677  DA:3678  DA:3697  DA:3700  DA:3723  DA:3740  DA:3747  DA:3756  DA:3787  DA:3788  DA:3807  DA:3824  DA:3839  DA:3840  DA:3852  DA:3864  DA:3875  DA:3879  DA:3882  DA:3886  DA:3904  DA:3912  DA:3921  DA:3925  DA:3935  DA:3940  DA:3942  DA:3944  DA:3960  DA:3963  DA:3964  DA:3983  DA:3990  DA:3991  DA:4007  DA:4030  DA:4032  DA:4033  DA:4034  DA:4048  DA:4089  DA:4130  DA:4131  DA:4135  DA:4138  DA:4165  DA:4176  DA:4206  DA:4213  DA:4227  DA:4228  DA:4233  DA:4237  DA:4246  DA:4247  DA:4248  DA:4261  DA:4265  DA:4286  DA:4287  DA:4288  DA:4299  DA:4300  DA:4341  DA:4342  DA:4392  DA:4404  DA:4409  DA:4416  DA:4426  DA:4436  DA:4448  DA:4477  DA:4478  DA:4479  DA:4490  DA:4501  DA:4505  DA:4519  DA:4525  DA:4528  DA:4530  DA:4543  DA:4547  DA:4558  DA:4566  DA:4594  DA:4615  DA:4624  DA:4649  DA:4655  DA:4662  DA:4667  DA:4677  DA:4682  DA:4686  DA:4689  DA:4695  DA:4710  DA:4714  DA:4727  DA:4754  DA:4785  DA:4786  DA:4799  DA:4811  DA:4837  DA:4846  DA:4850  DA:4860  DA:4861  DA:4862  DA:4866  DA:4892  DA:4897  DA:4911  DA:4915  DA:4916  DA:4920  DA:4923  DA:4937  DA:4938  DA:4969  DA:4972  DA:4976  DA:4991  DA:4995  DA:4998  DA:5017  DA:5021  DA:5055  DA:5067  DB:227  DB:229  DB:248  DB:264  DB:273  DB:275  DD:303  DD:304  DD:307  DD:308  DF:312  DF:313  DH:201  DH:203  DL:201  DO:216  DR:202  DT:107  DU:204 |
| **2) MG: 1^st^ shell missing** | 1283 | A1:205  A3:103  A6:102  A7:104  AA:3017  AA:3020  AA:3025  AA:3026  AA:3030  AA:3031  AA:3032  AA:3039  AA:3046  AA:3049  AA:3055  AA:3059  AA:3060  AA:3065  AA:3071  AA:3073  AA:3091  AA:3098  AA:3104  AA:3109  AA:3115  AA:3117  AA:3120  AA:3121  AA:3123  AA:3136  AA:3140  AA:3141  AA:3144  AA:3145  AA:3146  AA:3149  AA:3154  AA:3157  AA:3163  AA:3164  AA:3168  AA:3172  AA:3179  AA:3181  AA:3188  AA:3189  AA:3195  AA:3197  AA:3199  AA:3200  AA:3201  AA:3205  AA:3209  AA:3211  AA:3212  AA:3215  AA:3221  AA:3235  AA:3238  AA:3242  AA:3244  AA:3245  AA:3256  AA:3258  AA:3259  AA:3266  AA:3268  AA:3269  AA:3270  AA:3274  AA:3275  AA:3278  AA:3283  AA:3286  AA:3289  AA:3292  AA:3300  AA:3306  AA:3307  AA:3318  AA:3319  AA:3320  AA:3321  AA:3323  AA:3329  AA:3360  AA:3365  AA:3367  AA:3368  AA:3370  AA:3375  AA:3376  AA:3382  AA:3384  AA:3395  AA:3398  AA:3401  AA:3404  AA:3409  AA:3412  AA:3416  AA:3428  AA:3432  AA:3444  AA:3448  AA:3452  AA:3459  AA:3461  AA:3475  AA:3480  AA:3487  AA:3488  AA:3493  AA:3497  AA:3513  AA:3514  AA:3531  AA:3532  AA:3540  AA:3547  AA:3550  AA:3552  AA:3553  AA:3556  AA:3560  AA:3564  AA:3565  AA:3566  AA:3568  AA:3573  AA:3575  AA:3581  AA:3590  AA:3591  AA:3592  AA:3594  AA:3597  AA:3598  AA:3599  AA:3601  AA:3603  AA:3606  AA:3612  AA:3613  AA:3616  AA:3624  AA:3626  AA:3627  AA:3629  AA:3630  AA:3632  AA:3633  AA:3634  AA:3639  AA:3645  AA:3654  AA:3657  AA:3660  AA:3663  AA:3665  AA:3666  AA:3667  AA:3673  AA:3674  AA:3676  AA:3679  AA:3684  AA:3685  AA:3686  AA:3688  AA:3695  AA:3697  AA:3700  AA:3705  AA:3709  AA:3716  AA:3722  AA:3724  AA:3727  AA:3729  AA:3730  AA:3732  AA:3739  AA:3741  AA:3743  AA:3744  AA:3753  AA:3755  AA:3760  AA:3761  AA:3765  AA:3768  AA:3770  AA:3771  AA:3773  AA:3774  AA:3776  AA:3782  AA:3783  AA:3785  AA:3787  AA:3789  AA:3797  AA:3799  AA:3802  AA:3805  AA:3807  AA:3811  AA:3814  AA:3817  AA:3818  AA:3822  AA:3825  AA:3834  AA:3836  AA:3837  AA:3839  AA:3841  AA:3842  AA:3843  AA:3844  AA:3851  AA:3854  AA:3858  AA:3862  AA:3865  AA:3872  AA:3873  AA:3877  AA:3882  AA:3886  AA:3892  AA:3897  AA:3910  AA:3912  AA:3914  AA:3915  AA:3931  AA:3933  AA:3934  AA:3937  AA:3938  AA:3958  AA:3959  AA:3964  AA:3967  AA:3969  AA:3973  AA:3976  AA:3979  AA:3982  AA:3990  AA:3998  AA:4014  AA:4019  AA:4020  AA:4024  AA:4025  AA:4028  AA:4034  AA:4035  AA:4036  AA:4044  AA:4052  AA:4055  AA:4058  AA:4062  AA:4065  AA:4066  AA:4069  AA:4075  AA:4077  AA:4086  AA:4090  AA:4091  AA:4096  AA:4102  AA:4103  AA:4109  AA:4115  AA:4118  AA:4120  AA:4124  AA:4131  AA:4135  AA:4142  AA:4146  AA:4148  AA:4150  AB:211  AB:217  AB:221  AB:226  AD:301  AD:304  AD:311  AD:312  AF:305  AF:307  AH:201  AK:201  AK:202  AO:207  AR:201  AS:202  AT:103  AU:201  AU:204  AU:205  AU:206  AW:101  B1:102  B1:103  BA:1601  BA:1603  BA:1613  BA:1615  BA:1617  BA:1618  BA:1621  BA:1625  BA:1627  BA:1632  BA:1634  BA:1637  BA:1639  BA:1653  BA:1655  BA:1657  BA:1662  BA:1671  BA:1672  BA:1673  BA:1674  BA:1681  BA:1682  BA:1697  BA:1698  BA:1713  BA:1720  BA:1726  BA:1729  BA:1732  BA:1741  BA:1747  BA:1755  BA:1762  BA:1771  BA:1773  BA:1778  BA:1782  BA:1785  BA:1786  BA:1787  BA:1792  BA:1795  BA:1796  BA:1797  BA:1798  BA:1799  BA:1801  BA:1802  BA:1803  BA:1805  BA:1806  BA:1807  BA:1812  BA:1814  BA:1818  BA:1822  BA:1828  BA:1829  BA:1830  BA:1832  BA:1838  BA:1842  BA:1843  BA:1847  BA:1851  BA:1854  BA:1855  BA:1857  BA:1860  BA:1861  BA:1862  BA:1864  BA:1865  BA:1866  BA:1867  BA:1877  BA:1879  BA:1882  BA:1883  BA:1884  BA:1885  BA:1886  BA:1888  BA:1890  BA:1894  BA:1895  BA:1897  BA:1902  BA:1909  BA:1911  BA:1921  BA:1924  BA:1934  BA:1936  BA:1937  BA:1939  BA:1940  BA:1951  BA:1955  BA:1956  BA:1959  BA:1962  BA:1964  BA:1968  BA:1978  BA:1996  BA:1997  BA:2005  BA:2006  BA:2011  BA:2012  BA:2019  BA:2023  BA:2027  BA:2030  BA:2032  BA:2034  BA:2038  BA:2041  BA:2049  BA:2056  BA:2059  BA:2060  BA:2061  BA:2066  BA:2068  BA:2074  BA:2075  BA:2085  BA:2095  BA:2098  BA:2104  BA:2117  BA:2118  BA:2119  BA:2140  BA:2141  BA:2144  BA:2146  BA:2158  BA:2165  BA:2174  BA:2177  BA:2183  BA:2188  BA:2193  BA:2195  BA:2201  BA:2207  BA:2214  BA:2216  BA:2224  BA:2226  BA:2227  BA:2228  BA:2229  BA:2237  BA:2239  BA:2242  BA:2245  BA:2249  BA:2251  BA:2256  BA:2267  BA:2270  BB:102  BB:103  BB:104  BB:106  BB:107  BB:108  BC:107  BC:108  BC:110  BC:112  BC:114  BD:101  BD:103  BD:104  BD:109  BD:112  BD:113  BD:114  BD:115  BD:116  BD:118  BD:120  BD:121  BD:123  BE:301  BE:302  BE:305  BG:301  BG:302  BG:304  BG:307  BH:202  BK:202  BK:204  BM:203  BO:201  BP:202  BQ:101  BQ:103  BS:101  BS:106  BU:1601  BW:208  BX:101  C1:103  CA:1610  CA:1616  CA:1628  CA:1629  CA:1631  CA:1632  CA:1633  CA:1634  CA:1637  CA:1645  CA:1651  CA:1657  CA:1658  CA:1660  CA:1666  CA:1670  CA:1671  CA:1674  CA:1675  CA:1677  CA:1678  CA:1679  CA:1680  CA:1684  CA:1686  CA:1693  CA:1698  CA:1707  CA:1712  CA:1713  CA:1716  CA:1718  CA:1724  CA:1733  CA:1738  CA:1741  CA:1751  CA:1753  CA:1754  CA:1760  CA:1771  CA:1776  CA:1777  CA:1778  CA:1779  CA:1781  CA:1785  CA:1786  CA:1791  CA:1796  CA:1797  CA:1805  CA:1806  CA:1808  CA:1817  CA:1818  CA:1820  CA:1821  CA:1823  CA:1827  CA:1828  CA:1833  CA:1838  CA:1840  CA:1841  CA:1849  CA:1850  CA:1852  CA:1853  CA:1857  CA:1858  CA:1860  CA:1865  CA:1872  CA:1873  CA:1879  CA:1881  CA:1882  CA:1888  CA:1892  CA:1897  CA:1904  CA:1906  CA:1907  CA:1908  CA:1909  CA:1912  CA:1917  CA:1918  CA:1923  CA:1931  CA:1932  CA:1941  CA:1942  CA:1946  CA:1947  CA:1954  CA:1968  CA:1972  CA:1975  CA:1982  CA:1984  CA:1990  CA:1994  CA:1997  CA:2003  CA:2005  CA:2010  CA:2016  CA:2018  CA:2020  CA:2023  CA:2027  CA:2032  CA:2037  CA:2043  CA:2044  CA:2046  CA:2048  CA:2050  CA:2056  CA:2063  CA:2064  CA:2065  CA:2066  CA:2075  CA:2078  CA:2079  CA:2080  CA:2089  CA:2094  CA:2095  CA:2097  CA:2102  CA:2105  CA:2110  CA:2113  CA:2114  CA:2115  CA:2117  CA:2118  CA:2119  CA:2120  CA:2124  CA:2131  CA:2138  CA:2146  CA:2147  CA:2148  CA:2149  CA:2151  CA:2154  CA:2156  CA:2158  CA:2163  CA:2166  CA:2172  CA:2175  CA:2179  CA:2183  CA:2184  CA:2188  CA:2190  CA:2192  CA:2198  CA:2211  CA:2212  CA:2217  CA:2218  CA:2219  CA:2222  CA:2224  CA:2227  CA:2232  CA:2234  CA:2236  CA:2237  CA:2241  CA:2246  CA:2259  CA:2263  CA:2267  CA:2268  CA:2289  CA:2292  CA:2293  CA:2294  CA:2295  CA:2303  CA:2304  CA:2306  CA:2307  CA:2308  CA:2311  CA:2316  CA:2319  CA:2321  CA:2322  CA:2323  CB:104  CB:111  CB:115  CB:117  CC:107  CC:113  CC:114  CC:115  CC:121  CC:124  CC:127  CD:104  CD:108  CD:116  CD:117  CD:118  CE:306  CF:301  CG:302  CG:306  CH:201  CH:202  CH:205  CK:205  CK:206  CM:202  CN:202  CP:202  CR:103  CS:101  CV:102  D0:201  D0:202  D0:203  D0:209  D2:201  D2:202  D2:207  D6:101  D6:102  D7:101  D7:105  D8:105  DA:3012  DA:3014  DA:3015  DA:3018  DA:3023  DA:3026  DA:3034  DA:3036  DA:3053  DA:3063  DA:3064  DA:3067  DA:3070  DA:3075  DA:3079  DA:3080  DA:3081  DA:3100  DA:3103  DA:3108  DA:3112  DA:3114  DA:3125  DA:3134  DA:3136  DA:3139  DA:3145  DA:3146  DA:3148  DA:3150  DA:3157  DA:3159  DA:3161  DA:3168  DA:3170  DA:3176  DA:3179  DA:3183  DA:3194  DA:3200  DA:3201  DA:3207  DA:3208  DA:3209  DA:3211  DA:3221  DA:3223  DA:3227  DA:3248  DA:3255  DA:3257  DA:3259  DA:3260  DA:3265  DA:3267  DA:3270  DA:3272  DA:3275  DA:3283  DA:3285  DA:3288  DA:3290  DA:3302  DA:3311  DA:3315  DA:3319  DA:3325  DA:3326  DA:3329  DA:3332  DA:3334  DA:3339  DA:3341  DA:3342  DA:3343  DA:3344  DA:3346  DA:3355  DA:3356  DA:3359  DA:3360  DA:3364  DA:3367  DA:3373  DA:3378  DA:3381  DA:3384  DA:3386  DA:3391  DA:3393  DA:3394  DA:3395  DA:3409  DA:3414  DA:3415  DA:3420  DA:3425  DA:3426  DA:3445  DA:3448  DA:3463  DA:3464  DA:3467  DA:3478  DA:3479  DA:3485  DA:3489  DA:3495  DA:3497  DA:3498  DA:3502  DA:3504  DA:3509  DA:3510  DA:3513  DA:3521  DA:3529  DA:3532  DA:3546  DA:3551  DA:3553  DA:3554  DA:3558  DA:3561  DA:3562  DA:3564  DA:3565  DA:3568  DA:3569  DA:3575  DA:3578  DA:3580  DA:3585  DA:3596  DA:3597  DA:3598  DA:3600  DA:3602  DA:3606  DA:3607  DA:3609  DA:3611  DA:3612  DA:3622  DA:3628  DA:3630  DA:3636  DA:3646  DA:3647  DA:3655  DA:3656  DA:3662  DA:3666  DA:3667  DA:3672  DA:3673  DA:3680  DA:3686  DA:3690  DA:3691  DA:3693  DA:3698  DA:3704  DA:3713  DA:3721  DA:3724  DA:3727  DA:3728  DA:3730  DA:3736  DA:3743  DA:3746  DA:3749  DA:3752  DA:3754  DA:3759  DA:3761  DA:3762  DA:3763  DA:3766  DA:3767  DA:3768  DA:3769  DA:3770  DA:3773  DA:3793  DA:3797  DA:3803  DA:3805  DA:3813  DA:3819  DA:3823  DA:3827  DA:3829  DA:3831  DA:3836  DA:3843  DA:3844  DA:3845  DA:3846  DA:3850  DA:3857  DA:3858  DA:3861  DA:3862  DA:3863  DA:3869  DA:3870  DA:3872  DA:3873  DA:3896  DA:3898  DA:3900  DA:3902  DA:3905  DA:3908  DA:3910  DA:3913  DA:3916  DA:3920  DA:3922  DA:3929  DA:3933  DA:3937  DA:3939  DA:3949  DA:3953  DA:3954  DA:3955  DA:3956  DA:3957  DA:3958  DA:3959  DA:3962  DA:3970  DA:3974  DA:3978  DA:3979  DA:3981  DA:3984  DA:3995  DA:4005  DA:4017  DA:4049  DA:4057  DA:4060  DA:4075  DA:4077  DA:4079  DA:4080  DA:4082  DA:4088  DA:4096  DA:4097  DA:4099  DA:4101  DA:4103  DA:4109  DA:4112  DA:4113  DA:4115  DA:4119  DA:4122  DA:4123  DA:4125  DA:4127  DA:4128  DA:4140  DA:4141  DA:4144  DA:4147  DA:4148  DA:4153  DA:4154  DA:4160  DA:4161  DA:4170  DA:4174  DA:4178  DA:4182  DA:4184  DA:4192  DA:4193  DA:4195  DA:4202  DA:4217  DA:4222  DA:4223  DA:4225  DA:4254  DA:4255  DA:4256  DA:4257  DA:4267  DA:4269  DA:4282  DA:4285  DA:4289  DA:4290  DA:4296  DA:4297  DA:4319  DA:4330  DA:4335  DA:4343  DA:4344  DA:4348  DA:4354  DA:4355  DA:4356  DA:4368  DA:4372  DA:4374  DA:4380  DA:4382  DA:4383  DA:4389  DA:4390  DA:4394  DA:4395  DA:4396  DA:4407  DA:4408  DA:4419  DA:4421  DA:4428  DA:4438  DA:4449  DA:4451  DA:4464  DA:4466  DA:4468  DA:4470  DA:4475  DA:4482  DA:4504  DA:4506  DA:4512  DA:4529  DA:4541  DA:4544  DA:4560  DA:4574  DA:4575  DA:4576  DA:4585  DA:4587  DA:4589  DA:4595  DA:4602  DA:4607  DA:4609  DA:4611  DA:4612  DA:4621  DA:4634  DA:4636  DA:4644  DA:4645  DA:4646  DA:4648  DA:4650  DA:4656  DA:4661  DA:4664  DA:4671  DA:4673  DA:4675  DA:4676  DA:4681  DA:4688  DA:4692  DA:4693  DA:4698  DA:4699  DA:4704  DA:4705  DA:4707  DA:4708  DA:4709  DA:4715  DA:4717  DA:4718  DA:4721  DA:4733  DA:4739  DA:4758  DA:4759  DA:4764  DA:4768  DA:4774  DA:4781  DA:4790  DA:4791  DA:4794  DA:4798  DA:4802  DA:4803  DA:4805  DA:4813  DA:4821  DA:4831  DA:4832  DA:4839  DA:4840  DA:4841  DA:4844  DA:4864  DA:4868  DA:4881  DA:4888  DA:4889  DA:4894  DA:4896  DA:4899  DA:4900  DA:4905  DA:4906  DA:4908  DA:4917  DA:4921  DA:4922  DA:4925  DA:4926  DA:4934  DA:4935  DA:4945  DA:4948  DA:4950  DA:4955  DA:4960  DA:4963  DA:4964  DA:4970  DA:4971  DA:4980  DA:4984  DA:4985  DA:4990  DA:4996  DA:5002  DA:5005  DA:5010  DA:5011  DA:5027  DA:5030  DA:5032  DA:5037  DA:5039  DA:5044  DA:5047  DA:5049  DA:5056  DA:5057  DA:5060  DA:5064  DA:5065  DA:5072  DA:5076  DB:202  DB:203  DB:209  DB:213  DB:214  DB:219  DB:228  DB:230  DB:231  DB:232  DB:238  DB:239  DB:240  DB:242  DB:254  DB:257  DB:259  DB:262  DD:301  DD:302  DD:312  DD:313  DE:303  DE:311  DE:313  DF:306  DF:311  DF:317  DF:324  DG:203  DG:204  DH:202  DM:203  DN:201  DN:202  DO:202  DO:205  DO:206  DO:218  DP:202  DR:203  DS:201  DS:204  DS:205  DS:208  DU:202  DU:207  DU:208  DU:209  DU:213  DU:214  DU:217  DU:219  DV:301  DV:304  DW:101  DZ:101  DZ:102 |
| **3) MG/N1_A_ contacts** | 4 | AA:3902  BA:1776  CA:2085  DA:4331 |
| **3) MG/N1_G_ contacts** | 4 | AA:3430  DA:3242  DA:3350  DA:4325 |
| **3) MG/N2 contacts** | 10 | AA:3226  AA:3430  AA:3442  BA:2222  CA:1876  DA:3292  DA:3907  DA:4279  DA:4545  DA:5009 |
| **3) MG/N3_A_ contacts** | 4 | AA:3900  BA:2057  BA:2178  BA:2190 |
| **3) MG/N3_G_ contacts** | 4 | AA:3850  CA:1876  CA:2214  DA:3292 |
| **3) MG/N4 contacts** | 11 | BA:1840  BA:1840  BC:109  CA:2082  CD:113  DA:3695  DA:4028  DA:4315  DA:4485  DA:4490  DA:4722 |
| **3) MG/N6 contacts** | 4 | BA:1945  CD:121  CD:126  DA:3917 |
| **3) MG/N7 contacts** | 36 | AA:3004  AA:3152  AA:3186  AA:3348  AA:3369  AA:3427  AA:3442  AA:3806  BA:1809  BA:1810  BA:1845  BA:1980  BA:2114  CA:1654  CA:1891  CA:2028  CA:2280  CA:2324  DA:3007  DA:3030  DA:3030  DA:3241  DA:3410  DA:3522  DA:3802  DA:3808  DA:3833  DA:3853  DA:3917  DA:3938  DA:4191  DA:4251  DA:4280  DA:4725  DA:4854  DA:4887 |
| **3) MG/N_back_ contacts** | 17 | A1:202  AD:309  AF:303  AF:304  AR:203  BG:306  CG:309  CR:102  CV:101  CV:104  D5:107  D8:107  DF:321  DK:201  DK:201  DS:203  DT:103 |
| **3) MG/ND1_His_ contacts** | 2 | A5:101  D5:101 |
| **3) MG/ND2_Asn_ contacts** | 4 | CA:1880  DA:3410  DA:4993  DU:201 |
| **3) MG/NE2_Gln_ contacts** | 2 | CA:1973  DY:201 |
| **3) MG/NE2_His_ contacts** | 2 | AE:303  DA:3579 |
| **3) MG/NE_Arg_ contacts** | 1 | AO:201 |
| **3) MG/NH1_Arg_ contacts** | 2 | AA:3749  DA:4588 |
| **3) MG/NH2_Arg_ contacts** | 3 | BH:201  CK:201  CK:202 |
| **3) MG/NZ_Lys_ contacts** | 4 | BM:201  DA:3213  DA:4234  DY:202 |
| **3) MG/O2 contacts** | 22 | AA:3557  BA:1819  BA:1970  BA:1975  BA:2037  BA:2055  BA:2127  BA:2275  BL:202  CA:1758  CA:1801  CA:1927  CA:1969  CA:2002  CA:2036  CA:2070  CA:2272  CB:118  DA:3234  DA:4094  DA:4546  DA:4617 |
| **3) MG/O2' contacts** | 53 | AA:3127  AA:3248  AA:3326  AA:3512  AA:3690  AA:3793  AA:3809  AA:3816  AA:4092  AB:225  AD:309  BA:1646  BA:1660  BA:1749  BA:1819  BA:1961  BA:2133  BA:2275  BB:109  CA:1649  CA:1734  CA:1953  CA:1986  CA:2261  CA:2324  CB:119  DA:3127  DA:3249  DA:3282  DA:3336  DA:3449  DA:3540  DA:3566  DA:3627  DA:3705  DA:3726  DA:3726  DA:3771  DA:4070  DA:4124  DA:4165  DA:4253  DA:4268  DA:4310  DA:4350  DA:4457  DA:4697  DA:4716  DA:4750  DA:4757  DA:4777  DA:4946  DB:270 |
| **3) MG/O3' contacts** | 28 | AA:3133  AA:3267  AA:3350  AA:3977  AB:205  BA:1611  BA:1660  BA:1899  BA:2123  CA:1749  CA:1866  CA:1936  CA:2143  DA:3452  DA:3537  DA:3632  DA:3868  DA:4011  DA:4243  DA:4294  DA:4358  DA:4579  DA:4610  DA:4631  DA:4907  DA:5016  DA:5023  DB:223 |
| **3) MG/O4 contacts** | 43 | AA:3033  AA:3043  AA:3047  AA:3052  AA:3097  AA:3135  AA:3523  AA:3712  AA:3731  AA:3988  AB:219  BA:1723  BA:1733  BA:1752  BA:1779  BA:1804  BA:1880  BA:1958  BA:1998  BA:2021  BA:2040  BA:2133  CA:1613  CA:1622  CA:1640  CA:1726  CA:1862  CA:2140  DA:3039  DA:3040  DA:3065  DA:3082  DA:3292  DA:3337  DA:3397  DA:3618  DA:3703  DA:3706  DA:3914  DA:3982  DA:4485  DA:4534  DA:4549 |
| **3) MG/O4' contacts** | 10 | AA:3900  BA:1646  BA:1918  BA:1989  BA:2083  BB:101  CB:109  DA:4546  DA:4610  DB:247 |
| **3) MG/O5' contacts** | 25 | AA:3226  AA:3267  AA:3351  AA:3429  AA:3492  AA:3494  AA:3520  AA:3871  BA:1837  BA:1941  BA:2210  BA:2263  CA:1764  CA:1893  CA:2093  CA:2176  CC:122  DA:3427  DA:3800  DA:4056  DA:4058  DA:4220  DA:4977  DA:5009  DA:5045 |
| **3) MG/O6 contacts** | 84 | A3:102  AA:3010  AA:3022  AA:3078  AA:3097  AA:3126  AA:3127  AA:3143  AA:3249  AA:3279  AA:3339  AA:3354  AA:3356  AA:3484  AA:3499  AA:3504  AA:3522  AA:3543  AA:3694  AA:3731  AA:4013  AA:4016  AA:4039  AA:4114  AB:207  BA:1606  BA:1616  BA:1624  BA:1667  BA:1694  BA:1716  BA:1728  BA:1743  BA:1793  BA:1811  BA:1841  BA:1901  BA:2028  BA:2076  CA:1606  CA:1662  CA:1687  CA:1697  CA:1737  CA:1792  CA:1814  CA:1830  CA:1862  CA:1867  CA:2177  CA:2277  DA:3021  DA:3037  DA:3050  DA:3051  DA:3140  DA:3180  DA:3204  DA:3229  DA:3245  DA:3252  DA:3345  DA:3366  DA:3480  DA:3601  DA:3663  DA:3717  DA:3753  DA:3774  DA:3841  DA:3927  DA:3930  DA:3947  DA:4010  DA:4052  DA:4164  DA:4197  DA:4203  DA:4360  DA:4405  DA:4514  DA:4640  DB:207  DB:220 |
| **3) MG/O_back_ contacts** | 17 | BA:1733  BA:2127  BG:303  CA:1880  CG:307  CG:311  CP:201  D0:204  DD:314  DE:312  DE:315  DG:201  DQ:202  DT:104  DU:204  DU:206  DU:206 |
| **3) MG/O_w_ contacts** | 6 | BA:2276  BA:2276  BA:2276  BA:2276  BA:2276  BA:2276 |
| **3) MG/OD1_Asn_ contacts** | 2 | CN:201  DA:3410 |
| **3) MG/OD1_Asp_ contacts** | 1 | BH:205 |
| **3) MG/OD2_Asp_ contacts** | 2 | AE:301  DE:302 |
| **3) MG/OE1_Glu_ contacts** | 2 | BB:110  BT:201 |
| **3) MG/OE2_Glu_ contacts** | 1 | DW:103 |
| **3) MG/OG1_Thr_ contacts** | 2 | CG:307  DA:3648 |
| **3) MG/OP1 contacts** | 329 | AA:3005  AA:3006  AA:3007  AA:3008  AA:3009  AA:3009  AA:3010  AA:3014  AA:3015  AA:3015  AA:3028  AA:3040  AA:3040  AA:3045  AA:3053  AA:3057  AA:3061  AA:3061  AA:3069  AA:3070  AA:3070  AA:3089  AA:3094  AA:3102  AA:3107  AA:3111  AA:3150  AA:3158  AA:3169  AA:3177  AA:3183  AA:3190  AA:3192  AA:3192  AA:3196  AA:3196  AA:3204  AA:3207  AA:3207  AA:3208  AA:3214  AA:3216  AA:3216  AA:3223  AA:3230  AA:3251  AA:3257  AA:3277  AA:3277  AA:3291  AA:3296  AA:3324  AA:3331  AA:3348  AA:3349  AA:3362  AA:3377  AA:3378  AA:3378  AA:3378  AA:3378  AA:3381  AA:3391  AA:3411  AA:3424  AA:3450  AA:3456  AA:3458  AA:3462  AA:3462  AA:3464  AA:3466  AA:3468  AA:3469  AA:3504  AA:3504  AA:3506  AA:3507  AA:3509  AA:3516  AA:3519  AA:3519  AA:3577  AA:3577  AA:3578  AA:3586  AA:3620  AA:3625  AA:3635  AA:3635  AA:3764  AA:3840  AA:3849  AA:3889  AA:3950  AA:3980  AA:4008  AA:4010  AA:4037  AA:4156  AD:307  AT:102  BA:1602  BA:1608  BA:1608  BA:1616  BA:1647  BA:1665  BA:1669  BA:1677  BA:1684  BA:1686  BA:1701  BA:1702  BA:1702  BA:1714  BA:1721  BA:1727  BA:1737  BA:1753  BA:1759  BA:1763  BA:1764  BA:1789  BA:1790  BA:1837  BA:1852  BA:1881  BA:1925  BA:2031  BA:2055  BA:2072  BA:2086  BA:2167  BA:2170  BA:2197  BA:2235  BA:2253  BC:105  BC:113  BC:113  C1:101  CA:1602  CA:1605  CA:1606  CA:1607  CA:1617  CA:1617  CA:1618  CA:1622  CA:1630  CA:1661  CA:1664  CA:1664  CA:1669  CA:1673  CA:1690  CA:1696  CA:1704  CA:1709  CA:1711  CA:1723  CA:1739  CA:1748  CA:1770  CA:1775  CA:1783  CA:1825  CA:1871  CA:1900  CA:1929  CA:1967  CA:1985  CA:1991  CA:2014  CA:2039  CA:2077  CA:2130  CA:2207  CA:2296  CA:2300  CB:102  CB:110  CC:123  CL:201  CP:201  DA:3001  DA:3004  DA:3005  DA:3007  DA:3010  DA:3011  DA:3016  DA:3024  DA:3027  DA:3029  DA:3033  DA:3042  DA:3043  DA:3043  DA:3045  DA:3048  DA:3049  DA:3056  DA:3057  DA:3058  DA:3058  DA:3060  DA:3061  DA:3061  DA:3066  DA:3066  DA:3069  DA:3071  DA:3078  DA:3086  DA:3086  DA:3088  DA:3089  DA:3089  DA:3093  DA:3098  DA:3099  DA:3099  DA:3105  DA:3130  DA:3156  DA:3175  DA:3175  DA:3192  DA:3212  DA:3238  DA:3240  DA:3244  DA:3287  DA:3297  DA:3316  DA:3324  DA:3333  DA:3369  DA:3369  DA:3371  DA:3380  DA:3417  DA:3432  DA:3433  DA:3437  DA:3440  DA:3443  DA:3466  DA:3469  DA:3523  DA:3535  DA:3544  DA:3583  DA:3583  DA:3587  DA:3589  DA:3589  DA:3687  DA:3699  DA:3699  DA:3710  DA:3729  DA:3734  DA:3738  DA:3742  DA:3764  DA:3772  DA:3774  DA:3774  DA:3776  DA:3776  DA:3780  DA:3783  DA:3783  DA:3794  DA:3794  DA:3815  DA:3851  DA:3868  DA:3893  DA:3897  DA:3899  DA:3899  DA:3899  DA:3915  DA:3946  DA:3950  DA:3965  DA:3966  DA:3967  DA:3973  DA:4043  DA:4069  DA:4069  DA:4091  DA:4129  DA:4155  DA:4162  DA:4179  DA:4189  DA:4251  DA:4275  DA:4279  DA:4294  DA:4306  DA:4310  DA:4370  DA:4379  DA:4435  DA:4444  DA:4460  DA:4480  DA:4489  DA:4516  DA:4557  DA:4742  DA:4776  DA:4808  DA:4819  DA:4875  DA:4883  DA:4893  DA:4947  DB:223  DB:224  DB:251  DX:101 |
| **3) MG/OP2 contacts** | 369 | A1:201  AA:3001  AA:3002  AA:3006  AA:3012  AA:3013  AA:3023  AA:3033  AA:3044  AA:3048  AA:3053  AA:3061  AA:3063  AA:3066  AA:3070  AA:3078  AA:3079  AA:3079  AA:3081  AA:3084  AA:3084  AA:3086  AA:3090  AA:3094  AA:3094  AA:3126  AA:3129  AA:3129  AA:3130  AA:3133  AA:3134  AA:3138  AA:3152  AA:3155  AA:3170  AA:3176  AA:3191  AA:3222  AA:3222  AA:3223  AA:3226  AA:3240  AA:3251  AA:3260  AA:3262  AA:3277  AA:3297  AA:3308  AA:3345  AA:3371  AA:3377  AA:3377  AA:3383  AA:3383  AA:3386  AA:3400  AA:3427  AA:3430  AA:3437  AA:3457  AA:3468  AA:3469  AA:3472  AA:3473  AA:3500  AA:3501  AA:3503  AA:3503  AA:3510  AA:3520  AA:3523  AA:3534  AA:3534  AA:3555  AA:3578  AA:3643  AA:3650  AA:3659  AA:3683  AA:3711  AA:3713  AA:3777  AA:3808  AA:3831  AA:3881  AA:3901  AA:3926  AA:3968  AA:3988  AA:4010  AA:4033  AA:4043  AA:4076  AA:4138  AA:4165  AA:4165  AB:201  AB:202  AB:205  AB:208  AB:216  AB:228  BA:1611  BA:1622  BA:1623  BA:1626  BA:1626  BA:1628  BA:1630  BA:1641  BA:1645  BA:1648  BA:1650  BA:1668  BA:1690  BA:1693  BA:1699  BA:1701  BA:1702  BA:1708  BA:1727  BA:1728  BA:1736  BA:1736  BA:1736  BA:1738  BA:1739  BA:1751  BA:1760  BA:1783  BA:1794  BA:1800  BA:1813  BA:1825  BA:1844  BA:1880  BA:1880  BA:1891  BA:1908  BA:1941  BA:1963  BA:1970  BA:1979  BA:1986  BA:2008  BA:2013  BA:2031  BA:2035  BA:2055  BA:2063  BA:2084  BA:2123  BA:2156  BA:2164  BA:2171  BA:2173  BA:2204  BA:2210  BC:102  BC:103  BC:113  BD:110  CA:1608  CA:1609  CA:1615  CA:1638  CA:1643  CA:1644  CA:1648  CA:1653  CA:1655  CA:1655  CA:1681  CA:1687  CA:1691  CA:1708  CA:1710  CA:1719  CA:1725  CA:1725  CA:1739  CA:1740  CA:1748  CA:1762  CA:1764  CA:1766  CA:1780  CA:1798  CA:1826  CA:1845  CA:1851  CA:1875  CA:1877  CA:1944  CA:1987  CA:2013  CA:2029  CA:2068  CA:2069  CA:2083  CA:2093  CA:2108  CA:2243  CA:2254  CA:2310  CC:102  CC:116  CC:126  CS:104  D0:204  D0:204  D1:210  DA:3002  DA:3005  DA:3007  DA:3008  DA:3009  DA:3017  DA:3019  DA:3027  DA:3031  DA:3032  DA:3033  DA:3035  DA:3035  DA:3040  DA:3044  DA:3045  DA:3046  DA:3047  DA:3052  DA:3054  DA:3061  DA:3068  DA:3071  DA:3072  DA:3074  DA:3082  DA:3083  DA:3086  DA:3087  DA:3087  DA:3091  DA:3095  DA:3097  DA:3104  DA:3105  DA:3124  DA:3133  DA:3141  DA:3143  DA:3144  DA:3144  DA:3155  DA:3175  DA:3193  DA:3250  DA:3261  DA:3261  DA:3264  DA:3276  DA:3289  DA:3295  DA:3300  DA:3313  DA:3328  DA:3336  DA:3354  DA:3358  DA:3362  DA:3362  DA:3363  DA:3377  DA:3379  DA:3380  DA:3385  DA:3387  DA:3413  DA:3427  DA:3428  DA:3435  DA:3436  DA:3441  DA:3441  DA:3459  DA:3468  DA:3470  DA:3477  DA:3484  DA:3506  DA:3516  DA:3543  DA:3571  DA:3587  DA:3591  DA:3591  DA:3593  DA:3593  DA:3608  DA:3614  DA:3638  DA:3681  DA:3697  DA:3701  DA:3706  DA:3712  DA:3712  DA:3712  DA:3782  DA:3809  DA:3816  DA:3871  DA:3875  DA:3893  DA:3893  DA:3894  DA:3895  DA:3903  DA:3927  DA:3952  DA:3969  DA:3977  DA:3999  DA:4012  DA:4023  DA:4045  DA:4059  DA:4064  DA:4068  DA:4069  DA:4076  DA:4086  DA:4095  DA:4172  DA:4190  DA:4208  DA:4239  DA:4283  DA:4304  DA:4320  DA:4373  DA:4422  DA:4435  DA:4453  DA:4493  DA:4513  DA:4517  DA:4551  DA:4556  DA:4632  DA:4669  DA:4684  DA:4694  DA:4711  DA:4720  DA:4724  DA:4770  DA:4902  DA:4977  DA:4983  DA:4995  DA:5029  DB:201  DB:204  DB:206  DB:217  DB:225  DB:225  DB:241 |
| **3) ZN/O_back_ contacts** | 1 | CG:301 |
| **3) MG/N1 excl. contacts** | 58 | AA:3316  AA:3453  AA:3682  AA:3792  AA:3930  AA:3932  AA:3940  AA:4138  AA:4159  AB:216  AB:230  BA:1943  BA:1981  BA:1993  BA:2045  BA:2079  BA:2126  BA:2221  BA:2271  BA:2274  CA:1952  CA:2038  CA:2053  CA:2054  CA:2057  CA:2100  CA:2191  CA:2247  CA:2272  DA:3266  DA:3317  DA:3890  DA:3976  DA:4000  DA:4169  DA:4268  DA:4274  DA:4322  DA:4322  DA:4442  DA:4453  DA:4481  DA:4502  DA:4546  DA:4550  DA:4557  DA:4658  DA:4823  DA:4834  DA:4845  DA:4849  DA:4849  DA:4890  DA:4927  DA:5074  DB:247  DB:249  DB:266 |
| **3) MG/N1_A_ excl. contacts** | 61 | AA:3125  AA:3171  AA:3173  AA:3262  AA:3396  AA:3442  AA:3706  AA:3712  AA:3721  AA:3826  AA:3869  AA:4016  AA:4018  AA:4059  AA:4100  AA:4134  BA:1646  BA:1925  BA:1933  BA:2082  BA:2190  BA:2190  CA:1952  CA:2015  CA:2041  CA:2073  CA:2123  CD:129  DA:3092  DA:3458  DA:3492  DA:3503  DA:3798  DA:3812  DA:3892  DA:3976  DA:4035  DA:4137  DA:4236  DA:4242  DA:4247  DA:4253  DA:4301  DA:4318  DA:4342  DA:4442  DA:4488  DA:4502  DA:4519  DA:4588  DA:4613  DA:4653  DA:4712  DA:4729  DA:4763  DA:4929  DA:4949  DA:4959  DA:5007  DB:247  DB:250 |
| **3) MG/N1_G_ excl. contacts** | 78 | AA:3052  AA:3076  AA:3208  AA:3263  AA:3422  AA:3442  AA:3484  AA:3508  AA:3821  AA:3859  AA:3874  AA:3913  AA:3955  AA:3981  AA:4003  AA:4003  AA:4016  AA:4082  AA:4113  AB:204  AB:213  B1:104  BA:1910  BA:2028  BA:2076  BA:2092  BA:2108  BA:2114  BA:2128  BA:2137  BA:2152  BC:115  CA:1789  CA:1834  CA:1835  CA:1862  CA:1950  CA:1952  CA:1993  CA:2002  CA:2024  CA:2086  CA:2100  CA:2109  CA:2135  CA:2164  CA:2174  CA:2210  CA:2280  DA:3217  DA:3298  DA:3410  DA:3411  DA:3595  DA:3651  DA:3801  DA:3904  DA:3927  DA:3989  DA:4019  DA:4045  DA:4051  DA:4071  DA:4190  DA:4266  DA:4278  DA:4305  DA:4436  DA:4456  DA:4563  DA:4565  DA:4647  DA:4856  DA:4976  DA:5009  DA:5069  DB:205  DB:250 |
| **3) MG/N2 excl. contacts** | 115 | AA:3125  AA:3208  AA:3333  AA:3430  AA:3434  AA:3821  AA:3821  AA:3850  AA:3875  AA:3902  AA:3935  AA:3987  AA:4076  AA:4163  AB:220  B1:104  BA:1684  BA:1981  BA:1993  BA:2064  BA:2083  BA:2091  BA:2096  BA:2099  BA:2114  BA:2137  BA:2153  BA:2160  BA:2161  BA:2170  BA:2182  BA:2187  BA:2232  BC:115  CA:1690  CA:1877  CA:1895  CA:1926  CA:1930  CA:1934  CA:1950  CA:1952  CA:1952  CA:1993  CA:2015  CA:2028  CA:2036  CA:2036  CA:2070  CA:2141  CA:2143  CA:2191  CA:2210  CA:2214  CA:2257  CA:2272  CQ:102  DA:3013  DA:3217  DA:3225  DA:3323  DA:3336  DA:3350  DA:3411  DA:3449  DA:3560  DA:3595  DA:3668  DA:3804  DA:3875  DA:3880  DA:3989  DA:3998  DA:4040  DA:4139  DA:4166  DA:4229  DA:4236  DA:4237  DA:4259  DA:4278  DA:4283  DA:4325  DA:4338  DA:4377  DA:4430  DA:4443  DA:4456  DA:4458  DA:4469  DA:4474  DA:4491  DA:4497  DA:4588  DA:4594  DA:4690  DA:4700  DA:4712  DA:4723  DA:4745  DA:4756  DA:4830  DA:4856  DA:4927  DA:4930  DA:4973  DA:4981  DA:5026  DB:205  DB:221  DB:241  DB:250  DB:250  DB:263  DO:214 |
| **3) MG/N3_A_ excl. contacts** | 56 | AA:3047  AA:3173  AA:3302  AA:3442  AA:3537  AA:3537  AA:3791  AA:3820  AA:4094  AA:4100  BA:2109  BA:2142  BA:2166  CA:1649  CA:1883  CA:2004  CA:2055  CB:112  CS:105  D0:208  DA:3065  DA:3092  DA:3251  DA:3281  DA:3282  DA:3388  DA:3812  DA:3923  DA:3924  DA:3926  DA:3961  DA:4034  DA:4210  DA:4249  DA:4324  DA:4332  DA:4452  DA:4459  DA:4488  DA:4542  DA:4548  DA:4561  DA:4572  DA:4582  DA:4594  DA:4606  DA:4686  DA:4765  DA:4815  DA:4898  DA:4901  DA:4928  DA:4946  DA:5063  DB:226  DB:255 |
| **3) MG/N3_C_ excl. contacts** | 37 | AA:3047  AA:3087  AA:3171  AA:3263  AA:3489  AA:3644  AA:3819  AA:3859  AA:4003  AA:4003  AA:4076  AA:4098  BA:1735  BA:1824  BA:1889  BA:2015  BA:2044  BA:2055  BA:2127  CA:1662  CA:1784  CA:1787  CA:1880  CA:1910  CA:2002  CA:2191  CA:2272  CD:128  DA:3065  DA:3458  DA:3654  DA:4190  DA:4352  DA:4622  DA:4849  DA:4959  DA:4986 |
| **3) MG/N3_G_ excl. contacts** | 73 | AA:3248  AA:3436  AA:3559  AA:3720  AA:3792  AA:3907  AA:3947  AA:3981  AA:3987  AA:4000  AA:4064  AA:4092  AB:213  BA:1684  BA:1901  BA:1977  BA:1981  BA:2024  BA:2028  BA:2050  BA:2064  BA:2160  BA:2168  BA:2222  BA:2232  CA:1690  CA:1883  CA:1890  CA:1926  CA:1930  CA:1950  CA:1950  CA:1952  CA:1998  CA:1998  CA:2070  CA:2085  CA:2088  CA:2090  CA:2100  CA:2109  CA:2178  CA:2202  CA:2248  CA:2255  CA:2257  CA:2280  DA:3336  DA:3505  DA:3998  DA:4166  DA:4276  DA:4300  DA:4410  DA:4430  DA:4440  DA:4443  DA:4447  DA:4458  DA:4474  DA:4588  DA:4604  DA:4606  DA:4619  DA:4622  DA:4695  DA:4723  DA:4830  DA:4860  DA:4939  DA:4951  DA:4973  DA:5026 |
| **3) MG/N3_U_ excl. contacts** | 35 | AA:3097  AA:3417  AA:3721  AA:3731  AA:3951  AA:4009  AB:204  AB:204  AB:216  BA:1804  BA:2037  BG:306  CA:1956  CA:1969  CA:2135  CA:2140  CA:2200  CC:120  DA:3292  DA:3317  DA:3477  DA:3574  DA:3625  DA:3917  DA:3919  DA:4172  DA:4485  DA:4557  DA:4606  DA:4622  DA:4668  DA:4701  DA:4927  DA:4965  DB:205 |
| **3) MG/N4 excl. contacts** | 167 | AA:3118  AA:3243  AA:3247  AA:3282  AA:3284  AA:3315  AA:3315  AA:3354  AA:3358  AA:3369  AA:3379  AA:3406  AA:3490  AA:3549  AA:3615  AA:3618  AA:3623  AA:3671  AA:3671  AA:3675  AA:3725  AA:3728  AA:3790  AA:3790  AA:3796  AA:3828  AA:3857  AA:3859  AA:3888  AA:3909  AA:3921  AA:3946  AA:3961  AA:4003  AA:4003  AA:4007  AA:4016  AA:4060  AA:4076  AA:4098  AA:4113  AA:4113  AA:4130  AB:204  AB:207  AB:220  BA:1675  BA:1735  BA:1739  BA:1770  BA:1804  BA:1839  BA:1839  BA:1901  BA:1922  BA:1925  BA:1942  BA:1946  BA:1985  BA:1988  BA:2042  BA:2045  BA:2052  BA:2067  BA:2070  BA:2107  BA:2123  BA:2136  BA:2138  BA:2150  BA:2169  BA:2182  CA:1620  CA:1640  CA:1642  CA:1676  CA:1795  CA:1799  CA:1856  CA:1880  CA:1900  CA:1903  CA:1973  CA:2002  CA:2081  CA:2098  CA:2140  CA:2152  CA:2162  CA:2167  CA:2177  CA:2186  CA:2206  CA:2206  CA:2244  CA:2262  CA:2290  CB:121  CC:108  CC:117  CD:110  CD:120  CD:120  CD:121  CD:128  DA:3135  DA:3174  DA:3258  DA:3293  DA:3366  DA:3390  DA:3434  DA:3451  DA:3451  DA:3520  DA:3530  DA:3590  DA:3594  DA:3625  DA:3671  DA:3700  DA:3702  DA:3795  DA:3807  DA:3809  DA:3809  DA:3838  DA:3859  DA:3866  DA:3889  DA:3889  DA:3980  DA:3980  DA:4010  DA:4104  DA:4156  DA:4190  DA:4210  DA:4245  DA:4315  DA:4357  DA:4360  DA:4379  DA:4412  DA:4526  DA:4571  DA:4582  DA:4685  DA:4737  DA:4752  DA:4796  DA:4800  DA:4801  DA:4915  DA:4940  DA:4977  DA:4986  DA:4986  DA:4991  DA:5031  DA:5070  DB:205  DB:220  DB:248  DB:266  DB:276  DB:276 |
| **3) MG/N6 excl. contacts** | 112 | AA:3186  AA:3263  AA:3302  AA:3396  AA:3436  AA:3442  AA:3442  AA:3467  AA:3549  AA:3671  AA:3706  AA:3720  AA:3803  AA:3809  AA:3826  AA:3829  AA:3878  AA:3902  AA:3955  AA:3981  AA:4016  AA:4018  AA:4040  AA:4134  AB:219  BA:1725  BA:1899  BA:1925  BA:1977  BA:1984  BA:2014  BA:2026  BA:2043  BA:2097  BA:2109  BA:2218  BA:2241  CA:1703  CA:1819  CA:1890  CA:1894  CA:1966  CA:2011  CA:2030  CA:2041  CA:2087  CA:2195  CA:2226  CA:2228  CA:2276  CA:2290  CB:121  CD:112  D1:212  DA:3131  DA:3226  DA:3241  DA:3251  DA:3405  DA:3410  DA:3483  DA:3506  DA:3525  DA:3560  DA:3675  DA:3737  DA:3778  DA:3812  DA:3828  DA:3848  DA:3866  DA:3892  DA:3924  DA:3976  DA:3976  DA:3998  DA:3998  DA:4034  DA:4035  DA:4035  DA:4044  DA:4044  DA:4078  DA:4137  DA:4166  DA:4301  DA:4318  DA:4325  DA:4413  DA:4431  DA:4434  DA:4450  DA:4469  DA:4473  DA:4488  DA:4497  DA:4502  DA:4511  DA:4569  DA:4588  DA:4723  DA:4765  DA:4879  DA:4898  DA:4916  DA:4919  DA:4940  DA:4962  DA:4986  DA:4993  DA:5029  DB:247 |
| **3) MG/N7 excl. contacts** | 576 | A7:101  A7:103  AA:3004  AA:3034  AA:3034  AA:3085  AA:3090  AA:3110  AA:3110  AA:3113  AA:3119  AA:3124  AA:3127  AA:3131  AA:3169  AA:3173  AA:3187  AA:3194  AA:3203  AA:3218  AA:3224  AA:3225  AA:3237  AA:3239  AA:3243  AA:3246  AA:3250  AA:3264  AA:3290  AA:3291  AA:3294  AA:3317  AA:3325  AA:3325  AA:3326  AA:3339  AA:3340  AA:3347  AA:3353  AA:3366  AA:3373  AA:3399  AA:3402  AA:3414  AA:3415  AA:3433  AA:3455  AA:3455  AA:3460  AA:3465  AA:3482  AA:3484  AA:3489  AA:3491  AA:3495  AA:3515  AA:3518  AA:3522  AA:3524  AA:3525  AA:3525  AA:3533  AA:3535  AA:3535  AA:3542  AA:3549  AA:3578  AA:3579  AA:3585  AA:3588  AA:3602  AA:3619  AA:3621  AA:3631  AA:3636  AA:3642  AA:3644  AA:3644  AA:3649  AA:3680  AA:3683  AA:3691  AA:3693  AA:3706  AA:3718  AA:3720  AA:3745  AA:3748  AA:3748  AA:3754  AA:3754  AA:3758  AA:3769  AA:3778  AA:3820  AA:3821  AA:3829  AA:3829  AA:3830  AA:3832  AA:3835  AA:3847  AA:3848  AA:3849  AA:3859  AA:3859  AA:3861  AA:3867  AA:3869  AA:3874  AA:3891  AA:3904  AA:3919  AA:3921  AA:3925  AA:3942  AA:3946  AA:3947  AA:3947  AA:3981  AA:3983  AA:3989  AA:4013  AA:4016  AA:4017  AA:4023  AA:4046  AA:4046  AA:4067  AA:4104  AA:4106  AA:4116  AA:4144  AA:4154  AA:4157  AA:4166  AB:203  AB:210  AB:213  AB:214  AB:230  BA:1605  BA:1606  BA:1638  BA:1684  BA:1688  BA:1700  BA:1700  BA:1715  BA:1730  BA:1730  BA:1756  BA:1757  BA:1761  BA:1765  BA:1767  BA:1772  BA:1774  BA:1808  BA:1821  BA:1846  BA:1889  BA:1892  BA:1892  BA:1893  BA:1893  BA:1893  BA:1896  BA:1910  BA:1915  BA:1922  BA:1928  BA:1931  BA:1932  BA:1935  BA:1935  BA:1946  BA:1953  BA:1961  BA:1966  BA:1973  BA:1976  BA:1976  BA:1980  BA:1982  BA:1984  BA:1984  BA:1988  BA:2007  BA:2009  BA:2014  BA:2014  BA:2026  BA:2026  BA:2046  BA:2051  BA:2090  BA:2090  BA:2097  BA:2099  BA:2133  BA:2134  BA:2136  BA:2149  BA:2173  BA:2191  BA:2213  BA:2218  BA:2220  BA:2233  BA:2236  BA:2236  BA:2240  BA:2255  BA:2261  BC:106  BC:116  BH:201  BW:203  CA:1611  CA:1625  CA:1626  CA:1627  CA:1652  CA:1663  CA:1667  CA:1676  CA:1699  CA:1703  CA:1705  CA:1705  CA:1731  CA:1737  CA:1745  CA:1755  CA:1756  CA:1761  CA:1769  CA:1773  CA:1774  CA:1789  CA:1793  CA:1794  CA:1802  CA:1807  CA:1810  CA:1814  CA:1831  CA:1831  CA:1834  CA:1863  CA:1864  CA:1880  CA:1883  CA:1885  CA:1896  CA:1898  CA:1915  CA:1921  CA:1939  CA:1939  CA:1940  CA:1943  CA:1960  CA:1964  CA:1971  CA:1977  CA:1978  CA:1987  CA:1998  CA:1998  CA:2008  CA:2022  CA:2028  CA:2028  CA:2030  CA:2034  CA:2047  CA:2051  CA:2058  CA:2060  CA:2060  CA:2060  CA:2085  CA:2096  CA:2099  CA:2103  CA:2129  CA:2129  CA:2144  CA:2155  CA:2155  CA:2159  CA:2161  CA:2167  CA:2174  CA:2180  CA:2185  CA:2186  CA:2194  CA:2195  CA:2199  CA:2213  CA:2225  CA:2244  CA:2249  CA:2250  CA:2257  CA:2287  CA:2288  CA:2297  CA:2302  CA:2313  CB:108  CB:121  CC:117  CD:129  D1:212  DA:3006  DA:3024  DA:3071  DA:3074  DA:3084  DA:3092  DA:3092  DA:3095  DA:3107  DA:3110  DA:3115  DA:3116  DA:3119  DA:3127  DA:3131  DA:3135  DA:3138  DA:3153  DA:3155  DA:3180  DA:3187  DA:3190  DA:3203  DA:3213  DA:3226  DA:3228  DA:3229  DA:3233  DA:3236  DA:3249  DA:3252  DA:3256  DA:3262  DA:3271  DA:3278  DA:3282  DA:3282  DA:3286  DA:3296  DA:3310  DA:3312  DA:3314  DA:3314  DA:3322  DA:3331  DA:3336  DA:3340  DA:3353  DA:3357  DA:3363  DA:3388  DA:3390  DA:3404  DA:3405  DA:3421  DA:3422  DA:3423  DA:3431  DA:3434  DA:3462  DA:3475  DA:3481  DA:3482  DA:3488  DA:3500  DA:3501  DA:3525  DA:3539  DA:3559  DA:3573  DA:3581  DA:3590  DA:3599  DA:3599  DA:3603  DA:3604  DA:3619  DA:3641  DA:3643  DA:3645  DA:3651  DA:3653  DA:3659  DA:3671  DA:3675  DA:3692  DA:3694  DA:3705  DA:3707  DA:3709  DA:3717  DA:3717  DA:3735  DA:3741  DA:3745  DA:3751  DA:3751  DA:3756  DA:3791  DA:3808  DA:3810  DA:3812  DA:3817  DA:3817  DA:3828  DA:3833  DA:3835  DA:3853  DA:3855  DA:3866  DA:3874  DA:3881  DA:3886  DA:3890  DA:3911  DA:3921  DA:3930  DA:3930  DA:3936  DA:3941  DA:3941  DA:3976  DA:3986  DA:3989  DA:3991  DA:3998  DA:4000  DA:4003  DA:4007  DA:4009  DA:4016  DA:4020  DA:4024  DA:4024  DA:4028  DA:4042  DA:4044  DA:4045  DA:4054  DA:4071  DA:4072  DA:4073  DA:4078  DA:4078  DA:4111  DA:4117  DA:4156  DA:4159  DA:4163  DA:4185  DA:4186  DA:4190  DA:4199  DA:4205  DA:4224  DA:4232  DA:4238  DA:4244  DA:4258  DA:4258  DA:4260  DA:4283  DA:4293  DA:4302  DA:4304  DA:4307  DA:4311  DA:4316  DA:4325  DA:4326  DA:4331  DA:4346  DA:4365  DA:4373  DA:4393  DA:4412  DA:4422  DA:4435  DA:4436  DA:4461  DA:4469  DA:4473  DA:4481  DA:4484  DA:4500  DA:4500  DA:4509  DA:4511  DA:4513  DA:4514  DA:4514  DA:4528  DA:4531  DA:4538  DA:4539  DA:4539  DA:4570  DA:4586  DA:4592  DA:4616  DA:4625  DA:4629  DA:4641  DA:4641  DA:4642  DA:4643  DA:4651  DA:4651  DA:4652  DA:4665  DA:4665  DA:4689  DA:4695  DA:4713  DA:4720  DA:4722  DA:4735  DA:4765  DA:4773  DA:4780  DA:4782  DA:4800  DA:4807  DA:4809  DA:4818  DA:4835  DA:4845  DA:4852  DA:4873  DA:4873  DA:4891  DA:4911  DA:4920  DA:4949  DA:4957  DA:4959  DA:4962  DA:4983  DA:4986  DA:4993  DA:4995  DA:5013  DA:5015  DA:5033  DA:5035  DA:5048  DA:5061  DA:5069  DB:206  DB:210  DB:220  DB:233  DB:249  DB:258  DB:261  DB:275  DO:204  DQ:202 |
| **3) MG/N_back_ excl. contacts** | 151 | A0:201  A1:202  A2:201  A2:201  A2:201  A5:103  A5:103  A8:104  AA:3112  AA:3112  AA:3734  AD:305  AD:306  AD:308  AO:201  AQ:202  AQ:202  AQ:204  AR:203  AS:201  AT:101  AT:104  AU:202  AZ:103  BF:302  BG:305  BH:203  BK:201  BL:201  BL:201  BL:201  BL:202  BQ:102  BS:109  BV:101  BW:201  BW:204  BW:206  BW:206  CE:301  CE:303  CG:308  CG:309  CG:309  CG:309  CG:310  CG:311  CH:204  CH:204  CH:204  CH:206  CH:206  CH:206  CJ:201  CK:209  CL:201  CP:203  CQ:102  CQ:103  CQ:104  CR:102  CS:103  CS:105  CT:202  CT:202  CV:101  CV:103  CV:104  CW:201  D0:207  D0:208  D0:211  D0:211  D1:203  D1:203  D2:203  D2:205  D2:208  D2:209  D3:102  D3:105  D3:106  D4:102  D5:104  D5:106  D5:107  D5:108  D7:102  D7:103  D8:102  D8:106  DA:3830  DA:4022  DA:4199  DA:4471  DA:4669  DD:305  DD:305  DD:305  DD:307  DD:309  DD:310  DD:314  DD:314  DD:314  DE:301  DE:301  DE:306  DE:310  DE:314  DE:315  DF:302  DF:303  DF:304  DF:307  DF:307  DF:308  DF:308  DF:312  DF:316  DF:323  DG:201  DK:201  DK:202  DM:204  DM:205  DM:205  DO:203  DO:203  DO:203  DO:203  DQ:202  DR:202  DR:204  DT:102  DT:103  DT:103  DT:104  DT:104  DT:105  DU:206  DU:210  DU:210  DU:216  DU:216  DV:302  DV:305  DW:102  DY:203  DZ:105  DZ:105 |
| **3) MG/ND1_His_ excl. contacts** | 7 | AA:3169  AA:3683  CE:303  CR:102  D0:204  D2:204  DO:204 |
| **3) MG/ND2_Asn_ excl. contacts** | 13 | AD:308  AD:313  AG:202  AS:201  BA:1889  DA:3187  DA:4278  DD:303  DF:316  DG:202  DO:212  DO:213  DT:101 |
| **3) MG/NE1_Trp_ excl. contacts** | 2 | CS:102  D6:103 |
| **3) MG/NE2_Gln_ excl. contacts** | 7 | A2:201  AQ:203  BA:2260  D7:102  DW:104  DX:101  DX:102 |
| **3) MG/NE2_His_ excl. contacts** | 8 | A5:102  AA:3169  AA:3636  AT:102  D5:103  DA:3078  DD:308  DO:204 |
| **3) MG/NE_Arg_ excl. contacts** | 20 | A1:203  AD:303  AQ:205  BA:2015  BA:2253  BS:104  CA:1965  CK:204  CK:210  D1:207  D1:207  D1:211  D2:203  D3:101  D7:104  DA:3477  DA:4824  DM:206  DP:203  DS:206 |
| **3) MG/NH1_Arg_ excl. contacts** | 44 | A3:101  A7:103  AA:3295  AA:3780  AD:308  AG:203  AO:203  AQ:205  AR:202  BA:2253  BF:301  BH:201  BS:102  BV:101  CA:1900  CA:2126  CB:116  CK:203  CQ:102  D3:101  D3:104  D8:104  DA:3477  DA:3515  DA:3826  DA:4169  DA:4209  DA:4402  DA:4537  DA:4551  DA:4622  DA:4982  DA:5008  DE:314  DF:305  DF:310  DF:313  DM:204  DO:214  DO:216  DU:206  DU:215  DX:101  DZ:103 |
| **3) MG/NH2_Arg_ excl. contacts** | 41 | A1:204  A8:104  AO:204  AZ:102  BA:2015  BA:2101  BA:2130  BA:2153  BK:206  BS:107  BS:109  CA:1900  CA:2128  CA:2286  CK:210  CK:211  D0:206  D0:210  D1:202  D1:205  D1:207  D1:207  D1:208  D2:203  D3:101  DA:3316  DA:3470  DA:3493  DA:4276  DA:4425  DA:4484  DA:4551  DA:4743  DA:4808  DA:4929  DA:4982  DA:5062  DH:203  DO:201  DO:210  DS:203 |
| **3) MG/NZ_Lys_ excl. contacts** | 48 | A8:102  A8:103  AA:3308  AA:3528  AA:3738  AA:3906  AA:4026  AS:203  AT:102  AT:104  BA:2120  BA:2181  BA:2230  BA:2273  BW:202  CA:1914  CA:2213  CA:2215  CF:303  CG:312  CT:201  CT:203  D0:206  D1:210  D5:102  DA:3322  DA:4015  DA:4034  DA:4220  DA:4231  DA:4278  DA:4516  DA:4553  DA:4743  DA:4771  DA:4946  DD:305  DE:308  DF:307  DF:320  DF:322  DH:201  DO:208  DO:214  DS:207  DU:211  DU:218  DW:105 |
| **3) MG/O2 excl. contacts** | 193 | A3:102  AA:3035  AA:3047  AA:3076  AA:3170  AA:3171  AA:3237  AA:3263  AA:3302  AA:3349  AA:3349  AA:3374  AA:3394  AA:3463  AA:3467  AA:3478  AA:3486  AA:3498  AA:3522  AA:3529  AA:3533  AA:3561  AA:3656  AA:3656  AA:3708  AA:3708  AA:3721  AA:3738  AA:3791  AA:3803  AA:3864  AA:3880  AA:3907  AA:3935  AA:3951  AA:3994  AA:4049  AA:4094  AB:204  AB:215  AB:230  BA:1684  BA:1692  BA:1715  BA:1735  BA:1827  BA:1906  BA:1916  BA:1916  BA:1919  BA:1948  BA:1958  BA:1999  BA:2015  BA:2037  BA:2042  BA:2043  BA:2082  BA:2151  BA:2180  BA:2184  BA:2221  BA:2223  BA:2232  BA:2265  BA:2271  CA:1604  CA:1640  CA:1646  CA:1648  CA:1659  CA:1690  CA:1750  CA:1757  CA:1763  CA:1766  CA:1787  CA:1788  CA:1883  CA:1910  CA:1913  CA:1919  CA:1949  CA:1949  CA:1956  CA:1960  CA:1995  CA:2021  CA:2033  CA:2072  CA:2100  CA:2135  CA:2141  CA:2200  CA:2244  CA:2261  CA:2269  CA:2272  CA:2275  CA:2279  CA:2283  CA:2298  CB:112  CD:102  CD:102  CD:123  D3:101  D5:106  D5:108  D7:102  DA:3006  DA:3065  DA:3090  DA:3111  DA:3218  DA:3219  DA:3231  DA:3233  DA:3251  DA:3254  DA:3254  DA:3301  DA:3353  DA:3370  DA:3375  DA:3376  DA:3376  DA:3421  DA:3449  DA:3458  DA:3465  DA:3477  DA:3517  DA:3559  DA:3601  DA:3603  DA:3654  DA:3689  DA:3717  DA:3718  DA:3882  DA:3976  DA:4040  DA:4040  DA:4094  DA:4152  DA:4218  DA:4238  DA:4274  DA:4317  DA:4322  DA:4327  DA:4328  DA:4332  DA:4345  DA:4350  DA:4351  DA:4359  DA:4363  DA:4371  DA:4376  DA:4437  DA:4440  DA:4443  DA:4454  DA:4496  DA:4498  DA:4545  DA:4547  DA:4606  DA:4622  DA:4623  DA:4668  DA:4712  DA:4726  DA:4736  DA:4744  DA:4747  DA:4754  DA:4763  DA:4786  DA:4788  DA:4849  DA:4884  DA:4907  DA:4959  DA:4973  DA:4986  DA:5028  DA:5066  DB:224  DB:250  DB:267 |
| **3) MG/O2' excl. contacts** | 603 | A8:104  AA:3035  AA:3050  AA:3054  AA:3068  AA:3099  AA:3100  AA:3103  AA:3111  AA:3142  AA:3161  AA:3173  AA:3198  AA:3225  AA:3229  AA:3236  AA:3236  AA:3237  AA:3239  AA:3272  AA:3272  AA:3293  AA:3299  AA:3326  AA:3336  AA:3337  AA:3350  AA:3380  AA:3393  AA:3394  AA:3410  AA:3426  AA:3431  AA:3433  AA:3453  AA:3463  AA:3463  AA:3486  AA:3500  AA:3501  AA:3503  AA:3503  AA:3518  AA:3521  AA:3530  AA:3537  AA:3541  AA:3541  AA:3546  AA:3551  AA:3559  AA:3561  AA:3562  AA:3631  AA:3648  AA:3651  AA:3652  AA:3652  AA:3656  AA:3708  AA:3708  AA:3720  AA:3728  AA:3737  AA:3747  AA:3751  AA:3767  AA:3790  AA:3791  AA:3792  AA:3792  AA:3816  AA:3816  AA:3820  AA:3826  AA:3835  AA:3838  AA:3850  AA:3855  AA:3860  AA:3864  AA:3867  AA:3879  AA:3880  AA:3887  AA:3893  AA:3895  AA:3907  AA:3908  AA:3908  AA:3922  AA:3929  AA:3930  AA:3932  AA:3940  AA:3943  AA:3954  AA:3987  AA:4011  AA:4029  AA:4038  AA:4045  AA:4049  AA:4056  AA:4072  AA:4073  AA:4085  AA:4085  AA:4089  AA:4094  AA:4100  AA:4100  AA:4134  AA:4160  AB:215  AB:218  AB:219  AB:225  AB:227  AB:229  AD:313  AF:303  AO:204  AQ:201  AS:201  BA:1610  BA:1610  BA:1666  BA:1687  BA:1695  BA:1705  BA:1719  BA:1744  BA:1746  BA:1748  BA:1749  BA:1774  BA:1779  BA:1784  BA:1815  BA:1831  BA:1849  BA:1850  BA:1858  BA:1878  BA:1901  BA:1905  BA:1906  BA:1916  BA:1923  BA:1928  BA:1953  BA:1975  BA:1977  BA:1981  BA:1989  BA:2008  BA:2010  BA:2015  BA:2024  BA:2028  BA:2031  BA:2037  BA:2042  BA:2050  BA:2064  BA:2072  BA:2073  BA:2087  BA:2089  BA:2091  BA:2105  BA:2111  BA:2120  BA:2123  BA:2125  BA:2129  BA:2142  BA:2142  BA:2143  BA:2148  BA:2157  BA:2160  BA:2163  BA:2166  BA:2180  BA:2184  BA:2196  BA:2199  BA:2203  BA:2211  BA:2212  BA:2246  BA:2257  BA:2262  BA:2263  BA:2265  BA:2274  BS:109  CA:1613  CA:1621  CA:1638  CA:1644  CA:1655  CA:1661  CA:1662  CA:1700  CA:1702  CA:1702  CA:1710  CA:1717  CA:1747  CA:1749  CA:1757  CA:1758  CA:1758  CA:1765  CA:1782  CA:1835  CA:1868  CA:1883  CA:1884  CA:1913  CA:1926  CA:1929  CA:1936  CA:1950  CA:1957  CA:1958  CA:1960  CA:1962  CA:1974  CA:1995  CA:2004  CA:2014  CA:2021  CA:2030  CA:2036  CA:2055  CA:2059  CA:2072  CA:2074  CA:2076  CA:2083  CA:2085  CA:2088  CA:2090  CA:2090  CA:2096  CA:2107  CA:2137  CA:2139  CA:2157  CA:2176  CA:2178  CA:2178  CA:2200  CA:2202  CA:2233  CA:2233  CA:2238  CA:2252  CA:2255  CA:2256  CA:2257  CA:2261  CA:2269  CA:2270  CA:2272  CA:2273  CA:2282  CA:2283  CA:2285  CA:2298  CA:2301  CA:2309  CA:2320  CB:113  CB:114  CB:118  CC:103  CC:103  CC:104  CC:104  CC:105  CC:125  CD:101  CD:119  CD:122  CD:130  CK:207  CK:208  CR:102  CS:105  D1:204  D3:102  D5:106  D5:108  D6:103  DA:3017  DA:3090  DA:3090  DA:3092  DA:3128  DA:3172  DA:3184  DA:3204  DA:3206  DA:3215  DA:3219  DA:3231  DA:3233  DA:3249  DA:3250  DA:3269  DA:3273  DA:3273  DA:3281  DA:3301  DA:3316  DA:3316  DA:3317  DA:3320  DA:3330  DA:3330  DA:3338  DA:3353  DA:3354  DA:3375  DA:3376  DA:3376  DA:3383  DA:3383  DA:3388  DA:3392  DA:3396  DA:3452  DA:3461  DA:3468  DA:3475  DA:3496  DA:3505  DA:3507  DA:3507  DA:3515  DA:3517  DA:3517  DA:3526  DA:3528  DA:3536  DA:3538  DA:3545  DA:3549  DA:3559  DA:3563  DA:3567  DA:3572  DA:3584  DA:3586  DA:3601  DA:3601  DA:3613  DA:3615  DA:3632  DA:3641  DA:3648  DA:3660  DA:3660  DA:3682  DA:3697  DA:3711  DA:3712  DA:3712  DA:3714  DA:3718  DA:3722  DA:3726  DA:3731  DA:3739  DA:3785  DA:3786  DA:3788  DA:3834  DA:3840  DA:3841  DA:3841  DA:3842  DA:3868  DA:3880  DA:3880  DA:3882  DA:3890  DA:3892  DA:3901  DA:3915  DA:3925  DA:3926  DA:3972  DA:3975  DA:3990  DA:3993  DA:3996  DA:3998  DA:4007  DA:4011  DA:4019  DA:4035  DA:4037  DA:4040  DA:4041  DA:4043  DA:4051  DA:4063  DA:4071  DA:4076  DA:4094  DA:4110  DA:4129  DA:4149  DA:4158  DA:4166  DA:4180  DA:4194  DA:4198  DA:4199  DA:4205  DA:4210  DA:4215  DA:4218  DA:4226  DA:4226  DA:4230  DA:4234  DA:4235  DA:4242  DA:4243  DA:4248  DA:4249  DA:4250  DA:4271  DA:4276  DA:4288  DA:4300  DA:4301  DA:4304  DA:4306  DA:4317  DA:4318  DA:4323  DA:4324  DA:4333  DA:4334  DA:4358  DA:4364  DA:4378  DA:4384  DA:4387  DA:4388  DA:4391  DA:4392  DA:4393  DA:4404  DA:4406  DA:4410  DA:4426  DA:4430  DA:4435  DA:4437  DA:4440  DA:4443  DA:4447  DA:4450  DA:4452  DA:4455  DA:4458  DA:4459  DA:4459  DA:4463  DA:4467  DA:4474  DA:4486  DA:4496  DA:4498  DA:4508  DA:4515  DA:4515  DA:4519  DA:4524  DA:4542  DA:4546  DA:4547  DA:4553  DA:4556  DA:4557  DA:4559  DA:4561  DA:4561  DA:4568  DA:4569  DA:4572  DA:4578  DA:4579  DA:4581  DA:4582  DA:4583  DA:4590  DA:4600  DA:4603  DA:4604  DA:4606  DA:4619  DA:4627  DA:4639  DA:4657  DA:4662  DA:4679  DA:4684  DA:4703  DA:4710  DA:4712  DA:4726  DA:4736  DA:4747  DA:4747  DA:4754  DA:4757  DA:4762  DA:4763  DA:4765  DA:4767  DA:4786  DA:4792  DA:4809  DA:4824  DA:4828  DA:4830  DA:4843  DA:4847  DA:4858  DA:4860  DA:4869  DA:4884  DA:4892  DA:4892  DA:4898  DA:4898  DA:4901  DA:4907  DA:4912  DA:4913  DA:4918  DA:4929  DA:4936  DA:4936  DA:4936  DA:4939  DA:4951  DA:4966  DA:4981  DA:4987  DA:4987  DA:4988  DA:4994  DA:4998  DA:5000  DA:5004  DA:5007  DA:5012  DA:5020  DA:5024  DA:5026  DA:5033  DA:5042  DA:5043  DA:5046  DA:5054  DA:5063  DA:5066  DA:5071  DB:216  DB:224  DB:226  DB:237  DB:243  DB:244  DB:255  DB:260  DB:267  DB:268  DB:271  DB:271  DD:311  DD:314  DE:302  DE:304  DE:306  DE:308  DF:302  DF:303  DO:214  DP:203  DP:203  DR:204 |
| **3) MG/O3' excl. contacts** | 433 | AA:3024  AA:3029  AA:3036  AA:3054  AA:3069  AA:3099  AA:3112  AA:3127  AA:3135  AA:3142  AA:3148  AA:3158  AA:3175  AA:3175  AA:3176  AA:3198  AA:3204  AA:3225  AA:3230  AA:3236  AA:3236  AA:3255  AA:3260  AA:3272  AA:3298  AA:3304  AA:3316  AA:3336  AA:3352  AA:3363  AA:3380  AA:3389  AA:3396  AA:3410  AA:3424  AA:3449  AA:3451  AA:3483  AA:3492  AA:3498  AA:3502  AA:3503  AA:3504  AA:3530  AA:3541  AA:3541  AA:3544  AA:3546  AA:3551  AA:3578  AA:3617  AA:3641  AA:3653  AA:3687  AA:3699  AA:3715  AA:3728  AA:3747  AA:3751  AA:3759  AA:3762  AA:3767  AA:3777  AA:3780  AA:3835  AA:3838  AA:3855  AA:3893  AA:3922  AA:3940  AA:3943  AA:3996  AA:4008  AA:4029  AA:4037  AA:4050  AA:4083  AA:4085  AA:4085  AA:4123  AA:4125  AA:4127  AA:4152  AA:4156  AB:208  AB:225  AB:227  AS:201  BA:1608  BA:1609  BA:1622  BA:1633  BA:1650  BA:1651  BA:1659  BA:1666  BA:1668  BA:1687  BA:1689  BA:1695  BA:1718  BA:1719  BA:1721  BA:1734  BA:1745  BA:1746  BA:1749  BA:1759  BA:1781  BA:1784  BA:1784  BA:1836  BA:1849  BA:1903  BA:1905  BA:1914  BA:1948  BA:1953  BA:1965  BA:1981  BA:1989  BA:2002  BA:2008  BA:2048  BA:2087  BA:2089  BA:2105  BA:2111  BA:2125  BA:2129  BA:2132  BA:2132  BA:2142  BA:2143  BA:2145  BA:2157  BA:2162  BA:2162  BA:2186  BA:2194  BA:2196  BA:2203  BA:2235  BA:2247  BA:2254  BA:2263  BA:2273  BB:101  BC:111  BS:109  CA:1603  CA:1613  CA:1638  CA:1643  CA:1647  CA:1664  CA:1665  CA:1701  CA:1702  CA:1708  CA:1710  CA:1710  CA:1711  CA:1717  CA:1723  CA:1734  CA:1742  CA:1762  CA:1768  CA:1769  CA:1804  CA:1835  CA:1865  CA:1868  CA:1874  CA:1929  CA:1929  CA:1958  CA:1963  CA:1964  CA:1965  CA:1974  CA:1974  CA:1992  CA:2014  CA:2039  CA:2040  CA:2055  CA:2059  CA:2069  CA:2072  CA:2090  CA:2100  CA:2101  CA:2106  CA:2128  CA:2176  CA:2187  CA:2207  CA:2223  CA:2226  CA:2238  CA:2254  CA:2265  CA:2271  CA:2275  CA:2275  CA:2282  CA:2309  CA:2312  CA:2314  CB:102  CB:109  CC:103  CC:103  CC:104  CC:104  CC:126  CF:303  D3:102  D8:107  DA:3007  DA:3017  DA:3042  DA:3056  DA:3090  DA:3094  DA:3104  DA:3129  DA:3165  DA:3167  DA:3172  DA:3184  DA:3206  DA:3224  DA:3229  DA:3243  DA:3249  DA:3253  DA:3266  DA:3269  DA:3289  DA:3300  DA:3323  DA:3330  DA:3335  DA:3399  DA:3418  DA:3437  DA:3459  DA:3468  DA:3482  DA:3496  DA:3506  DA:3507  DA:3507  DA:3515  DA:3516  DA:3518  DA:3526  DA:3527  DA:3536  DA:3540  DA:3555  DA:3563  DA:3566  DA:3567  DA:3572  DA:3613  DA:3615  DA:3623  DA:3648  DA:3650  DA:3660  DA:3665  DA:3687  DA:3708  DA:3708  DA:3711  DA:3712  DA:3739  DA:3744  DA:3771  DA:3778  DA:3785  DA:3796  DA:3865  DA:3878  DA:3893  DA:3950  DA:3967  DA:3973  DA:3975  DA:3987  DA:3993  DA:3996  DA:4002  DA:4013  DA:4027  DA:4032  DA:4033  DA:4037  DA:4039  DA:4056  DA:4058  DA:4063  DA:4091  DA:4124  DA:4132  DA:4137  DA:4149  DA:4151  DA:4157  DA:4165  DA:4168  DA:4168  DA:4172  DA:4183  DA:4196  DA:4203  DA:4205  DA:4209  DA:4211  DA:4220  DA:4224  DA:4235  DA:4242  DA:4250  DA:4271  DA:4276  DA:4286  DA:4298  DA:4304  DA:4306  DA:4306  DA:4310  DA:4313  DA:4323  DA:4332  DA:4334  DA:4338  DA:4339  DA:4345  DA:4362  DA:4367  DA:4384  DA:4391  DA:4393  DA:4399  DA:4406  DA:4420  DA:4425  DA:4426  DA:4441  DA:4444  DA:4454  DA:4457  DA:4460  DA:4463  DA:4467  DA:4472  DA:4489  DA:4492  DA:4496  DA:4497  DA:4515  DA:4520  DA:4553  DA:4556  DA:4557  DA:4562  DA:4564  DA:4569  DA:4581  DA:4590  DA:4597  DA:4600  DA:4603  DA:4610  DA:4616  DA:4623  DA:4687  DA:4703  DA:4727  DA:4728  DA:4747  DA:4748  DA:4749  DA:4751  DA:4753  DA:4770  DA:4792  DA:4808  DA:4824  DA:4843  DA:4858  DA:4875  DA:4886  DA:4892  DA:4912  DA:4916  DA:4956  DA:4968  DA:4977  DA:4979  DA:4981  DA:4987  DA:4988  DA:4994  DA:4998  DA:5000  DA:5004  DA:5007  DA:5009  DA:5012  DA:5016  DA:5022  DA:5024  DA:5024  DA:5033  DA:5052  DA:5054  DB:204  DB:217  DB:251  DB:255  DB:265  DB:268  DB:270  DB:271  DF:308  DF:320  DX:101 |
| **3) MG/O4 excl. contacts** | 346 | A5:102  AA:3019  AA:3035  AA:3035  AA:3036  AA:3075  AA:3088  AA:3093  AA:3093  AA:3143  AA:3153  AA:3167  AA:3178  AA:3178  AA:3194  AA:3206  AA:3210  AA:3236  AA:3253  AA:3254  AA:3273  AA:3315  AA:3334  AA:3334  AA:3341  AA:3350  AA:3354  AA:3357  AA:3358  AA:3359  AA:3379  AA:3388  AA:3390  AA:3417  AA:3417  AA:3417  AA:3434  AA:3460  AA:3468  AA:3478  AA:3478  AA:3508  AA:3515  AA:3522  AA:3527  AA:3527  AA:3533  AA:3538  AA:3543  AA:3554  AA:3572  AA:3623  AA:3646  AA:3670  AA:3675  AA:3678  AA:3689  AA:3691  AA:3698  AA:3701  AA:3703  AA:3745  AA:3752  AA:3756  AA:3766  AA:3820  AA:3823  AA:3827  AA:3828  AA:3919  AA:3939  AA:3955  AA:3957  AA:3961  AA:3975  AA:4004  AA:4009  AA:4068  AA:4070  AA:4074  AA:4082  AA:4088  AA:4141  AA:4166  AB:204  AB:229  AB:232  AF:301  B1:104  BA:1675  BA:1675  BA:1675  BA:1683  BA:1685  BA:1696  BA:1725  BA:1735  BA:1740  BA:1743  BA:1743  BA:1750  BA:1790  BA:1793  BA:1793  BA:1804  BA:1819  BA:1819  BA:1887  BA:1926  BA:1926  BA:1928  BA:1947  BA:1949  BA:1990  BA:1994  BA:1999  BA:2007  BA:2029  BA:2042  BA:2070  BA:2076  BA:2077  BA:2092  BA:2116  BA:2157  BA:2172  BA:2176  BA:2219  BA:2223  BB:105  BC:109  BC:111  BD:108  BD:108  BG:306  BH:201  BM:201  CA:1620  CA:1641  CA:1642  CA:1642  CA:1643  CA:1644  CA:1694  CA:1695  CA:1708  CA:1730  CA:1730  CA:1752  CA:1774  CA:1801  CA:1801  CA:1809  CA:1829  CA:1830  CA:1854  CA:1855  CA:1861  CA:1878  CA:1878  CA:1889  CA:1889  CA:1898  CA:1903  CA:1914  CA:1915  CA:1919  CA:1922  CA:1928  CA:1935  CA:1945  CA:1955  CA:1970  CA:1977  CA:2008  CA:2011  CA:2017  CA:2030  CA:2033  CA:2057  CA:2062  CA:2087  CA:2087  CA:2098  CA:2109  CA:2112  CA:2122  CA:2132  CA:2141  CA:2152  CA:2152  CA:2244  CA:2255  CA:2296  CB:103  CB:103  CC:112  CC:112  CC:120  CD:121  CN:201  D5:103  DA:3013  DA:3020  DA:3020  DA:3050  DA:3059  DA:3096  DA:3101  DA:3102  DA:3110  DA:3110  DA:3128  DA:3149  DA:3149  DA:3158  DA:3162  DA:3189  DA:3189  DA:3197  DA:3198  DA:3219  DA:3222  DA:3225  DA:3228  DA:3245  DA:3263  DA:3268  DA:3293  DA:3298  DA:3299  DA:3350  DA:3366  DA:3368  DA:3382  DA:3400  DA:3408  DA:3408  DA:3419  DA:3421  DA:3424  DA:3471  DA:3472  DA:3475  DA:3494  DA:3503  DA:3508  DA:3511  DA:3511  DA:3520  DA:3522  DA:3524  DA:3524  DA:3559  DA:3574  DA:3574  DA:3605  DA:3624  DA:3626  DA:3629  DA:3633  DA:3654  DA:3654  DA:3654  DA:3663  DA:3679  DA:3717  DA:3718  DA:3725  DA:3725  DA:3732  DA:3745  DA:3765  DA:3774  DA:3806  DA:3824  DA:3824  DA:3856  DA:3877  DA:3879  DA:3881  DA:3891  DA:3894  DA:3914  DA:3914  DA:3917  DA:3918  DA:3919  DA:3961  DA:3968  DA:3983  DA:4009  DA:4021  DA:4036  DA:4117  DA:4133  DA:4139  DA:4164  DA:4172  DA:4188  DA:4203  DA:4204  DA:4219  DA:4245  DA:4263  DA:4299  DA:4310  DA:4347  DA:4381  DA:4386  DA:4405  DA:4405  DA:4414  DA:4426  DA:4490  DA:4521  DA:4521  DA:4526  DA:4550  DA:4565  DA:4593  DA:4638  DA:4763  DA:4776  DA:4797  DA:4825  DA:4836  DA:4863  DA:4876  DA:4877  DA:4910  DA:4931  DA:4931  DA:4967  DA:5031  DB:205  DB:208  DB:216  DB:227  DB:227  DB:233  DB:237  DB:246  DB:246  DB:274  DF:301 |
| **3) MG/O4' excl. contacts** | 184 | AA:3111  AA:3142  AA:3153  AA:3231  AA:3232  AA:3239  AA:3248  AA:3299  AA:3316  AA:3333  AA:3337  AA:3363  AA:3374  AA:3394  AA:3522  AA:3537  AA:3651  AA:3662  AA:3690  AA:3692  AA:3706  AA:3708  AA:3759  AA:3792  AA:3792  AA:3908  AA:3929  AA:3935  AA:3974  AA:3987  AA:4000  AA:4001  AA:4085  AA:4092  AA:4100  AB:215  BA:1666  BA:1816  BA:1848  BA:1858  BA:1868  BA:1899  BA:1929  BA:1961  BA:1977  BA:1993  BA:2015  BA:2024  BA:2042  BA:2050  BA:2071  BA:2079  BA:2096  BA:2108  BA:2109  BA:2116  BA:2120  BA:2142  BA:2166  BA:2200  BD:124  C1:104  CA:1662  CA:1874  CA:1895  CA:1895  CA:1935  CA:1960  CA:1976  CA:1998  CA:2057  CA:2100  CA:2176  CA:2202  CA:2202  CA:2256  CA:2261  CA:2298  CA:2309  CK:207  DA:3006  DA:3210  DA:3224  DA:3320  DA:3353  DA:3376  DA:3383  DA:3396  DA:3417  DA:3439  DA:3449  DA:3505  DA:3517  DA:3517  DA:3545  DA:3552  DA:3627  DA:3682  DA:3717  DA:3771  DA:3880  DA:3884  DA:3887  DA:3890  DA:3923  DA:3961  DA:4002  DA:4019  DA:4035  DA:4062  DA:4151  DA:4210  DA:4218  DA:4226  DA:4229  DA:4234  DA:4253  DA:4253  DA:4281  DA:4318  DA:4324  DA:4331  DA:4350  DA:4353  DA:4369  DA:4388  DA:4410  DA:4437  DA:4447  DA:4453  DA:4458  DA:4467  DA:4474  DA:4486  DA:4490  DA:4497  DA:4497  DA:4508  DA:4516  DA:4524  DA:4532  DA:4542  DA:4545  DA:4549  DA:4557  DA:4572  DA:4578  DA:4604  DA:4617  DA:4619  DA:4657  DA:4668  DA:4684  DA:4686  DA:4695  DA:4712  DA:4745  DA:4767  DA:4789  DA:4812  DA:4830  DA:4883  DA:4890  DA:4892  DA:4904  DA:4913  DA:4918  DA:4928  DA:4952  DA:4958  DA:4977  DA:5008  DA:5009  DA:5026  DA:5063  DA:5066  DA:5067  DB:206  DB:224  DB:226  DB:244  DB:266  DB:267  DB:271 |
| **3) MG/O5' excl. contacts** | 372 | AA:3005  AA:3009  AA:3009  AA:3024  AA:3047  AA:3111  AA:3112  AA:3129  AA:3132  AA:3133  AA:3142  AA:3151  AA:3166  AA:3175  AA:3206  AA:3208  AA:3227  AA:3231  AA:3236  AA:3277  AA:3305  AA:3363  AA:3371  AA:3441  AA:3447  AA:3451  AA:3482  AA:3496  AA:3496  AA:3530  AA:3535  AA:3544  AA:3586  AA:3589  AA:3620  AA:3653  AA:3692  AA:3715  AA:3726  AA:3759  AA:3762  AA:3763  AA:3769  AA:3772  AA:3777  AA:3778  AA:3790  AA:3832  AA:3852  AA:3889  AA:3895  AA:3900  AA:3906  AA:3920  AA:3923  AA:3929  AA:3966  AA:3996  AA:3997  AA:4005  AA:4005  AA:4027  AA:4048  AA:4064  AA:4068  AA:4081  AA:4107  AA:4111  AA:4117  AA:4123  AA:4132  AA:4163  AA:4165  AB:201  AB:205  AB:225  AT:104  BA:1611  BA:1623  BA:1666  BA:1687  BA:1690  BA:1701  BA:1702  BA:1702  BA:1721  BA:1746  BA:1753  BA:1788  BA:1789  BA:1791  BA:1836  BA:1872  BA:1918  BA:1954  BA:1961  BA:1966  BA:1982  BA:1983  BA:1987  BA:1989  BA:1992  BA:2035  BA:2044  BA:2057  BA:2065  BA:2071  BA:2088  BA:2099  BA:2108  BA:2108  BA:2124  BA:2125  BA:2132  BA:2135  BA:2143  BA:2143  BA:2148  BA:2163  BA:2181  BA:2187  BA:2194  BA:2200  BA:2204  BA:2205  BA:2209  BA:2217  BA:2222  BA:2233  BA:2240  BA:2248  BA:2259  BA:2260  BA:2264  BA:2268  BB:101  BB:112  BC:103  C1:106  CA:1618  CA:1692  CA:1711  CA:1721  CA:1735  CA:1743  CA:1755  CA:1755  CA:1837  CA:1839  CA:1845  CA:1866  CA:1874  CA:1884  CA:1898  CA:1933  CA:1944  CA:1980  CA:1992  CA:1999  CA:2000  CA:2009  CA:2013  CA:2024  CA:2025  CA:2035  CA:2061  CA:2104  CA:2106  CA:2111  CA:2116  CA:2121  CA:2128  CA:2137  CA:2143  CA:2164  CA:2168  CA:2169  CA:2197  CA:2203  CA:2208  CA:2210  CA:2220  CA:2229  CA:2231  CA:2242  CA:2245  CA:2265  CA:2300  CA:2315  CB:120  CC:118  CC:119  CC:126  CD:114  DA:3003  DA:3010  DA:3038  DA:3065  DA:3124  DA:3142  DA:3192  DA:3193  DA:3193  DA:3210  DA:3212  DA:3224  DA:3261  DA:3289  DA:3320  DA:3323  DA:3328  DA:3372  DA:3398  DA:3418  DA:3443  DA:3444  DA:3460  DA:3460  DA:3466  DA:3470  DA:3477  DA:3486  DA:3487  DA:3496  DA:3516  DA:3527  DA:3571  DA:3632  DA:3638  DA:3649  DA:3650  DA:3676  DA:3694  DA:3705  DA:3707  DA:3726  DA:3729  DA:3739  DA:3772  DA:3778  DA:3782  DA:3804  DA:3811  DA:3817  DA:3826  DA:3848  DA:3852  DA:3864  DA:3865  DA:3868  DA:3876  DA:3887  DA:3899  DA:3919  DA:3927  DA:3940  DA:3944  DA:3973  DA:3987  DA:4012  DA:4014  DA:4020  DA:4024  DA:4029  DA:4062  DA:4069  DA:4071  DA:4076  DA:4083  DA:4149  DA:4151  DA:4157  DA:4179  DA:4189  DA:4196  DA:4205  DA:4209  DA:4216  DA:4230  DA:4231  DA:4241  DA:4252  DA:4276  DA:4291  DA:4298  DA:4302  DA:4346  DA:4347  DA:4361  DA:4364  DA:4367  DA:4379  DA:4391  DA:4415  DA:4429  DA:4433  DA:4453  DA:4463  DA:4467  DA:4480  DA:4497  DA:4501  DA:4518  DA:4524  DA:4532  DA:4551  DA:4567  DA:4568  DA:4590  DA:4610  DA:4610  DA:4613  DA:4616  DA:4658  DA:4660  DA:4680  DA:4682  DA:4683  DA:4684  DA:4694  DA:4711  DA:4727  DA:4729  DA:4748  DA:4753  DA:4770  DA:4771  DA:4784  DA:4799  DA:4807  DA:4816  DA:4829  DA:4838  DA:4845  DA:4855  DA:4857  DA:4867  DA:4869  DA:4883  DA:4891  DA:4902  DA:4918  DA:4919  DA:4937  DA:4942  DA:4947  DA:4958  DA:4968  DA:4975  DA:4979  DA:4983  DA:4987  DA:4992  DA:4999  DA:5007  DA:5029  DA:5038  DA:5048  DA:5073  DA:5075  DB:236  DB:245  DB:247  DB:252  DB:256  DB:270  DB:272  DE:301  DF:316  DF:322  DO:213  DR:204 |
| **3) MG/O6 excl. contacts** | 1014 | AA:3004  AA:3016  AA:3027  AA:3034  AA:3035  AA:3043  AA:3051  AA:3075  AA:3075  AA:3075  AA:3076  AA:3076  AA:3076  AA:3077  AA:3085  AA:3088  AA:3096  AA:3113  AA:3118  AA:3118  AA:3119  AA:3139  AA:3143  AA:3143  AA:3147  AA:3160  AA:3162  AA:3162  AA:3165  AA:3165  AA:3167  AA:3167  AA:3173  AA:3173  AA:3174  AA:3174  AA:3178  AA:3178  AA:3182  AA:3186  AA:3194  AA:3203  AA:3203  AA:3204  AA:3210  AA:3210  AA:3210  AA:3219  AA:3224  AA:3231  AA:3239  AA:3246  AA:3250  AA:3252  AA:3253  AA:3253  AA:3253  AA:3273  AA:3273  AA:3273  AA:3280  AA:3281  AA:3282  AA:3284  AA:3288  AA:3290  AA:3291  AA:3309  AA:3315  AA:3315  AA:3317  AA:3325  AA:3328  AA:3334  AA:3334  AA:3339  AA:3341  AA:3342  AA:3342  AA:3343  AA:3344  AA:3344  AA:3352  AA:3354  AA:3354  AA:3358  AA:3358  AA:3359  AA:3359  AA:3364  AA:3364  AA:3366  AA:3369  AA:3372  AA:3372  AA:3374  AA:3379  AA:3385  AA:3397  AA:3397  AA:3399  AA:3402  AA:3402  AA:3403  AA:3417  AA:3420  AA:3422  AA:3422  AA:3427  AA:3434  AA:3434  AA:3438  AA:3445  AA:3457  AA:3460  AA:3463  AA:3463  AA:3465  AA:3477  AA:3484  AA:3491  AA:3495  AA:3499  AA:3511  AA:3522  AA:3526  AA:3526  AA:3527  AA:3545  AA:3545  AA:3548  AA:3548  AA:3549  AA:3554  AA:3567  AA:3571  AA:3576  AA:3576  AA:3580  AA:3582  AA:3585  AA:3589  AA:3600  AA:3602  AA:3614  AA:3614  AA:3618  AA:3619  AA:3622  AA:3622  AA:3623  AA:3623  AA:3637  AA:3637  AA:3638  AA:3638  AA:3644  AA:3646  AA:3646  AA:3652  AA:3655  AA:3655  AA:3661  AA:3661  AA:3662  AA:3670  AA:3670  AA:3672  AA:3675  AA:3675  AA:3675  AA:3677  AA:3677  AA:3678  AA:3678  AA:3681  AA:3687  AA:3689  AA:3689  AA:3689  AA:3691  AA:3693  AA:3698  AA:3698  AA:3703  AA:3703  AA:3703  AA:3710  AA:3714  AA:3714  AA:3717  AA:3725  AA:3731  AA:3733  AA:3734  AA:3742  AA:3742  AA:3748  AA:3748  AA:3752  AA:3752  AA:3754  AA:3754  AA:3757  AA:3796  AA:3800  AA:3810  AA:3820  AA:3820  AA:3824  AA:3828  AA:3828  AA:3829  AA:3830  AA:3830  AA:3835  AA:3847  AA:3848  AA:3859  AA:3863  AA:3869  AA:3874  AA:3874  AA:3880  AA:3888  AA:3888  AA:3890  AA:3913  AA:3917  AA:3919  AA:3921  AA:3925  AA:3925  AA:3939  AA:3941  AA:3948  AA:3953  AA:3957  AA:3975  AA:3983  AA:4003  AA:4003  AA:4007  AA:4007  AA:4017  AA:4018  AA:4022  AA:4031  AA:4042  AA:4046  AA:4053  AA:4070  AA:4082  AA:4088  AA:4106  AA:4110  AA:4113  AA:4113  AA:4137  AA:4144  AA:4155  AA:4155  AA:4157  AA:4162  AB:204  AB:204  AB:207  AB:210  AB:214  AB:220  AB:220  AB:229  AB:230  B1:104  BA:1604  BA:1605  BA:1619  BA:1620  BA:1623  BA:1624  BA:1629  BA:1635  BA:1636  BA:1638  BA:1642  BA:1642  BA:1643  BA:1652  BA:1656  BA:1656  BA:1664  BA:1675  BA:1675  BA:1683  BA:1683  BA:1684  BA:1685  BA:1685  BA:1685  BA:1691  BA:1696  BA:1704  BA:1708  BA:1715  BA:1717  BA:1722  BA:1723  BA:1725  BA:1730  BA:1731  BA:1733  BA:1735  BA:1735  BA:1735  BA:1740  BA:1740  BA:1750  BA:1750  BA:1752  BA:1756  BA:1758  BA:1758  BA:1761  BA:1767  BA:1770  BA:1770  BA:1777  BA:1777  BA:1790  BA:1793  BA:1810  BA:1817  BA:1819  BA:1820  BA:1839  BA:1840  BA:1840  BA:1841  BA:1887  BA:1887  BA:1889  BA:1892  BA:1892  BA:1893  BA:1896  BA:1910  BA:1915  BA:1922  BA:1922  BA:1926  BA:1926  BA:1930  BA:1931  BA:1932  BA:1942  BA:1952  BA:1953  BA:1961  BA:1969  BA:1969  BA:1971  BA:1971  BA:1972  BA:1974  BA:1976  BA:1985  BA:1985  BA:1990  BA:1994  BA:1999  BA:1999  BA:2003  BA:2007  BA:2009  BA:2021  BA:2021  BA:2022  BA:2026  BA:2036  BA:2052  BA:2053  BA:2062  BA:2070  BA:2070  BA:2076  BA:2081  BA:2090  BA:2090  BA:2092  BA:2092  BA:2093  BA:2108  BA:2114  BA:2127  BA:2128  BA:2131  BA:2138  BA:2147  BA:2149  BA:2153  BA:2154  BA:2169  BA:2218  BA:2219  BA:2219  BA:2223  BA:2236  BA:2238  BA:2257  BA:2272  BB:111  BC:109  BC:109  BC:116  BC:116  BD:111  BM:202  CA:1601  CA:1601  CA:1612  CA:1614  CA:1620  CA:1620  CA:1622  CA:1626  CA:1627  CA:1635  CA:1639  CA:1639  CA:1640  CA:1640  CA:1640  CA:1641  CA:1641  CA:1642  CA:1642  CA:1643  CA:1682  CA:1688  CA:1688  CA:1690  CA:1694  CA:1695  CA:1695  CA:1699  CA:1705  CA:1706  CA:1717  CA:1722  CA:1722  CA:1726  CA:1726  CA:1729  CA:1729  CA:1730  CA:1730  CA:1731  CA:1732  CA:1750  CA:1752  CA:1752  CA:1759  CA:1761  CA:1769  CA:1769  CA:1789  CA:1790  CA:1809  CA:1810  CA:1811  CA:1811  CA:1811  CA:1812  CA:1814  CA:1815  CA:1819  CA:1819  CA:1829  CA:1830  CA:1830  CA:1831  CA:1834  CA:1834  CA:1844  CA:1861  CA:1867  CA:1867  CA:1878  CA:1878  CA:1880  CA:1883  CA:1885  CA:1891  CA:1894  CA:1896  CA:1899  CA:1900  CA:1901  CA:1903  CA:1903  CA:1905  CA:1914  CA:1914  CA:1915  CA:1919  CA:1919  CA:1920  CA:1925  CA:1926  CA:1927  CA:1928  CA:1943  CA:1943  CA:1950  CA:1955  CA:1962  CA:1973  CA:1978  CA:1979  CA:1979  CA:1981  CA:1988  CA:1993  CA:2002  CA:2017  CA:2017  CA:2024  CA:2026  CA:2028  CA:2028  CA:2028  CA:2047  CA:2051  CA:2060  CA:2073  CA:2084  CA:2086  CA:2087  CA:2096  CA:2098  CA:2098  CA:2108  CA:2135  CA:2140  CA:2140  CA:2144  CA:2144  CA:2152  CA:2155  CA:2155  CA:2162  CA:2162  CA:2164  CA:2171  CA:2174  CA:2174  CA:2177  CA:2180  CA:2186  CA:2193  CA:2194  CA:2199  CA:2206  CA:2244  CA:2255  CA:2278  CA:2291  CA:2297  CA:2305  CA:2310  CA:2313  CA:2324  CB:121  CC:117  CC:120  CC:120  CD:106  CD:110  CD:112  CD:120  CD:120  CD:121  CD:123  CD:123  CN:201  D3:101  DA:3006  DA:3006  DA:3006  DA:3020  DA:3028  DA:3050  DA:3055  DA:3059  DA:3059  DA:3059  DA:3073  DA:3074  DA:3077  DA:3092  DA:3092  DA:3096  DA:3096  DA:3096  DA:3107  DA:3110  DA:3115  DA:3116  DA:3117  DA:3118  DA:3119  DA:3120  DA:3122  DA:3122  DA:3123  DA:3126  DA:3127  DA:3135  DA:3135  DA:3149  DA:3149  DA:3152  DA:3154  DA:3154  DA:3158  DA:3158  DA:3158  DA:3160  DA:3160  DA:3162  DA:3162  DA:3169  DA:3171  DA:3174  DA:3177  DA:3187  DA:3189  DA:3189  DA:3190  DA:3197  DA:3198  DA:3198  DA:3198  DA:3210  DA:3213  DA:3213  DA:3222  DA:3222  DA:3224  DA:3226  DA:3228  DA:3229  DA:3235  DA:3235  DA:3242  DA:3245  DA:3246  DA:3252  DA:3253  DA:3258  DA:3258  DA:3262  DA:3262  DA:3263  DA:3263  DA:3271  DA:3273  DA:3274  DA:3274  DA:3278  DA:3280  DA:3282  DA:3291  DA:3293  DA:3293  DA:3293  DA:3294  DA:3298  DA:3299  DA:3299  DA:3299  DA:3301  DA:3303  DA:3304  DA:3304  DA:3305  DA:3305  DA:3307  DA:3308  DA:3314  DA:3314  DA:3321  DA:3321  DA:3331  DA:3331  DA:3336  DA:3337  DA:3337  DA:3340  DA:3347  DA:3348  DA:3349  DA:3350  DA:3351  DA:3353  DA:3363  DA:3363  DA:3365  DA:3365  DA:3366  DA:3368  DA:3368  DA:3368  DA:3382  DA:3388  DA:3397  DA:3397  DA:3401  DA:3406  DA:3410  DA:3419  DA:3419  DA:3419  DA:3421  DA:3422  DA:3423  DA:3424  DA:3424  DA:3431  DA:3439  DA:3447  DA:3447  DA:3453  DA:3453  DA:3455  DA:3456  DA:3462  DA:3471  DA:3471  DA:3473  DA:3475  DA:3476  DA:3480  DA:3480  DA:3483  DA:3490  DA:3491  DA:3501  DA:3503  DA:3506  DA:3508  DA:3511  DA:3512  DA:3528  DA:3530  DA:3533  DA:3536  DA:3539  DA:3539  DA:3542  DA:3557  DA:3557  DA:3559  DA:3573  DA:3574  DA:3574  DA:3577  DA:3582  DA:3592  DA:3592  DA:3601  DA:3603  DA:3604  DA:3605  DA:3605  DA:3605  DA:3621  DA:3621  DA:3625  DA:3626  DA:3629  DA:3639  DA:3645  DA:3651  DA:3651  DA:3653  DA:3661  DA:3661  DA:3663  DA:3679  DA:3679  DA:3688  DA:3700  DA:3702  DA:3702  DA:3707  DA:3715  DA:3717  DA:3718  DA:3720  DA:3725  DA:3732  DA:3737  DA:3737  DA:3741  DA:3745  DA:3753  DA:3755  DA:3756  DA:3776  DA:3780  DA:3789  DA:3802  DA:3802  DA:3807  DA:3808  DA:3816  DA:3818  DA:3835  DA:3837  DA:3856  DA:3856  DA:3866  DA:3877  DA:3877  DA:3883  DA:3886  DA:3888  DA:3890  DA:3892  DA:3904  DA:3906  DA:3912  DA:3914  DA:3928  DA:3930  DA:3936  DA:3938  DA:3938  DA:3941  DA:3941  DA:3943  DA:3945  DA:3945  DA:3960  DA:3968  DA:3983  DA:3986  DA:3992  DA:3994  DA:3994  DA:4000  DA:4001  DA:4008  DA:4009  DA:4010  DA:4013  DA:4016  DA:4019  DA:4025  DA:4042  DA:4045  DA:4046  DA:4047  DA:4048  DA:4050  DA:4051  DA:4051  DA:4071  DA:4072  DA:4074  DA:4078  DA:4081  DA:4098  DA:4098  DA:4102  DA:4116  DA:4131  DA:4131  DA:4133  DA:4133  DA:4134  DA:4139  DA:4143  DA:4143  DA:4150  DA:4156  DA:4159  DA:4164  DA:4164  DA:4166  DA:4167  DA:4181  DA:4186  DA:4188  DA:4232  DA:4240  DA:4244  DA:4251  DA:4262  DA:4263  DA:4264  DA:4266  DA:4274  DA:4277  DA:4278  DA:4280  DA:4280  DA:4283  DA:4284  DA:4287  DA:4299  DA:4299  DA:4299  DA:4321  DA:4325  DA:4325  DA:4336  DA:4341  DA:4349  DA:4373  DA:4381  DA:4386  DA:4404  DA:4405  DA:4414  DA:4414  DA:4432  DA:4446  DA:4446  DA:4448  DA:4462  DA:4477  DA:4485  DA:4490  DA:4495  DA:4526  DA:4526  DA:4527  DA:4528  DA:4531  DA:4535  DA:4539  DA:4540  DA:4558  DA:4565  DA:4566  DA:4570  DA:4584  DA:4584  DA:4593  DA:4613  DA:4615  DA:4626  DA:4630  DA:4638  DA:4642  DA:4647  DA:4651  DA:4652  DA:4653  DA:4659  DA:4665  DA:4695  DA:4701  DA:4719  DA:4722  DA:4731  DA:4731  DA:4732  DA:4737  DA:4752  DA:4769  DA:4782  DA:4783  DA:4801  DA:4801  DA:4817  DA:4852  DA:4861  DA:4865  DA:4873  DA:4887  DA:4910  DA:4915  DA:4931  DA:4952  DA:4976  DA:4995  DA:5015  DA:5015  DA:5033  DA:5041  DA:5043  DA:5043  DA:5069  DA:5069  DA:5077  DB:205  DB:206  DB:208  DB:210  DB:215  DB:216  DB:220  DB:221  DB:221  DB:222  DB:227  DB:227  DB:246  DB:248  DB:274  DZ:104 |
| **3) MG/O_back_ excl. contacts** | 244 | A0:201  A1:202  A1:202  A2:201  A5:103  A6:101  AA:3016  AA:3066  AA:3095  AA:3559  AA:3683  AA:3692  AA:3747  AA:3798  AA:3988  AD:302  AD:303  AD:307  AD:308  AD:313  AE:304  AF:301  AF:301  AF:301  AF:303  AF:303  AF:304  AF:304  AG:201  AG:202  AG:202  AG:202  AG:202  AN:201  AO:201  AO:202  AO:205  AO:206  AP:201  AP:201  AP:201  AQ:201  AQ:202  AQ:202  AR:203  AR:203  AR:204  AT:102  AU:202  AU:203  AU:203  AZ:102  AZ:103  BA:1733  BA:1889  BA:2004  BF:302  BG:303  BG:303  BH:203  BI:201  BI:201  BI:201  BK:205  BL:201  BP:201  BP:201  BS:103  BS:105  BS:107  BS:109  BT:201  BT:201  BT:201  BV:101  BW:201  BW:203  BW:204  BW:205  BW:207  CA:1734  CA:1734  CA:1910  CA:1937  CA:1963  CA:2021  CA:2161  CA:2230  CA:2230  CE:302  CE:303  CE:305  CF:303  CG:307  CG:307  CG:310  CG:311  CH:206  CI:201  CI:201  CI:201  CJ:201  CK:207  CK:207  CL:201  CN:201  CN:201  CP:201  CP:201  CQ:103  CR:102  CS:103  CS:106  CT:201  CV:103  CV:103  CV:104  D0:205  D1:203  D1:203  D1:204  D1:204  D1:205  D1:206  D1:209  D1:212  D2:203  D2:206  D2:208  D3:102  D5:106  D5:107  D5:107  D5:108  D8:101  D8:102  D8:102  D8:106  D8:107  D8:108  DA:3037  DA:3076  DA:3191  DA:3237  DA:3297  DA:3476  DA:3505  DA:3868  DA:4013  DA:4035  DA:4035  DA:4180  DA:4235  DA:4271  DA:4371  DA:4425  DA:4522  DA:4564  DA:4667  DA:4726  DA:4823  DA:4843  DA:5000  DD:303  DD:304  DD:305  DD:306  DD:307  DD:307  DD:307  DD:308  DD:309  DD:310  DE:301  DE:305  DE:305  DE:306  DE:308  DE:308  DE:309  DE:310  DE:310  DE:314  DE:315  DE:315  DE:315  DF:301  DF:301  DF:301  DF:303  DF:307  DF:307  DF:308  DF:309  DF:312  DF:314  DF:318  DF:323  DF:323  DG:201  DG:202  DG:202  DG:202  DG:205  DH:201  DH:205  DL:201  DM:202  DM:202  DM:202  DM:204  DM:204  DM:204  DO:201  DO:203  DO:208  DO:209  DO:210  DO:210  DO:211  DO:215  DO:217  DO:217  DP:201  DP:201  DP:201  DR:201  DR:202  DR:204  DS:203  DT:101  DT:105  DT:106  DU:204  DU:204  DU:212  DU:216  DU:218  DV:305  DW:106  DW:107  DY:201  DY:204  DZ:105 |
| **3) MG/OD1_Asn_ excl. contacts** | 6 | AA:3978  AA:4040  BA:1791  DA:4410  DA:4993  DO:213 |
| **3) MG/OD1_Asp_ excl. contacts** | 11 | AD:308  BE:303  BS:108  CE:304  CE:304  CH:203  CM:203  D3:107  DA:4353  DE:314  DF:313 |
| **3) MG/OD2_Asp_ excl. contacts** | 9 | A0:201  AD:305  AD:308  BA:1691  BE:303  BS:105  BS:108  D1:211  DF:313 |
| **3) MG/OE1_Gln_ excl. contacts** | 3 | AD:309  BA:2260  DY:201 |
| **3) MG/OE1_Glu_ excl. contacts** | 10 | AA:4116  AD:310  AS:203  BK:205  CR:101  DA:3714  DB:263  DD:310  DO:214  DU:203 |
| **3) MG/OE2_Glu_ excl. contacts** | 8 | AZ:101  BE:303  BH:205  BT:201  CH:203  DA:4110  DD:310  DF:325 |
| **3) MG/OG1_Thr_ excl. contacts** | 20 | A1:201  A1:204  A6:101  AA:3528  AA:3734  AA:3751  BG:303  BQ:102  CQ:103  CQ:104  D1:201  DA:3830  DA:4231  DA:4399  DA:4442  DA:4484  DA:4669  DE:308  DO:207  DT:107 |
| **3) MG/OG_Ser_ excl. contacts** | 20 | A3:104  AA:3895  AO:206  AR:205  AW:102  BA:2020  BA:2217  BM:202  BW:206  CE:301  D3:107  DA:3505  DA:4022  DA:4677  DA:4703  DA:4850  DF:315  DO:217  DW:102  DW:103 |
| **3) MG/OH_Tyr_ excl. contacts** | 14 | CA:2004  D5:105  DA:4281  DA:4572  DA:4658  DA:4939  DA:4946  DB:271  DE:305  DM:201  DO:216  DS:206  DU:210  DU:218 |
| **3) MG/OP1 excl. contacts** | 1266 | A1:202  A2:201  A7:102  A8:101  AA:3007  AA:3011  AA:3014  AA:3016  AA:3018  AA:3021  AA:3024  AA:3028  AA:3028  AA:3029  AA:3029  AA:3037  AA:3040  AA:3041  AA:3043  AA:3045  AA:3050  AA:3054  AA:3054  AA:3057  AA:3058  AA:3058  AA:3064  AA:3067  AA:3068  AA:3072  AA:3079  AA:3079  AA:3080  AA:3081  AA:3084  AA:3095  AA:3099  AA:3101  AA:3103  AA:3108  AA:3112  AA:3114  AA:3116  AA:3122  AA:3130  AA:3132  AA:3134  AA:3134  AA:3135  AA:3138  AA:3139  AA:3142  AA:3148  AA:3152  AA:3156  AA:3159  AA:3160  AA:3166  AA:3166  AA:3169  AA:3175  AA:3175  AA:3184  AA:3185  AA:3193  AA:3193  AA:3202  AA:3203  AA:3213  AA:3213  AA:3222  AA:3223  AA:3225  AA:3228  AA:3228  AA:3234  AA:3236  AA:3241  AA:3251  AA:3255  AA:3267  AA:3276  AA:3293  AA:3295  AA:3298  AA:3301  AA:3301  AA:3302  AA:3303  AA:3304  AA:3308  AA:3312  AA:3313  AA:3314  AA:3322  AA:3330  AA:3346  AA:3347  AA:3352  AA:3357  AA:3363  AA:3363  AA:3377  AA:3380  AA:3387  AA:3389  AA:3389  AA:3392  AA:3392  AA:3396  AA:3421  AA:3424  AA:3424  AA:3429  AA:3431  AA:3433  AA:3435  AA:3439  AA:3440  AA:3443  AA:3443  AA:3443  AA:3447  AA:3454  AA:3456  AA:3457  AA:3471  AA:3474  AA:3479  AA:3481  AA:3483  AA:3483  AA:3485  AA:3486  AA:3495  AA:3496  AA:3498  AA:3502  AA:3503  AA:3509  AA:3516  AA:3517  AA:3520  AA:3528  AA:3529  AA:3530  AA:3536  AA:3551  AA:3551  AA:3555  AA:3558  AA:3558  AA:3563  AA:3563  AA:3569  AA:3570  AA:3574  AA:3583  AA:3584  AA:3587  AA:3593  AA:3596  AA:3596  AA:3604  AA:3605  AA:3607  AA:3608  AA:3609  AA:3609  AA:3610  AA:3611  AA:3617  AA:3625  AA:3628  AA:3637  AA:3640  AA:3650  AA:3653  AA:3662  AA:3664  AA:3664  AA:3668  AA:3694  AA:3696  AA:3696  AA:3699  AA:3699  AA:3702  AA:3702  AA:3707  AA:3707  AA:3713  AA:3718  AA:3726  AA:3726  AA:3735  AA:3735  AA:3747  AA:3749  AA:3751  AA:3759  AA:3763  AA:3772  AA:3777  AA:3780  AA:3780  AA:3784  AA:3793  AA:3795  AA:3798  AA:3812  AA:3813  AA:3813  AA:3813  AA:3815  AA:3815  AA:3833  AA:3835  AA:3840  AA:3840  AA:3845  AA:3853  AA:3864  AA:3870  AA:3870  AA:3871  AA:3871  AA:3876  AA:3879  AA:3883  AA:3885  AA:3895  AA:3899  AA:3900  AA:3905  AA:3906  AA:3908  AA:3911  AA:3916  AA:3916  AA:3916  AA:3920  AA:3924  AA:3927  AA:3944  AA:3945  AA:3949  AA:3951  AA:3952  AA:3962  AA:3966  AA:3972  AA:3977  AA:3978  AA:3984  AA:3984  AA:3985  AA:3986  AA:3986  AA:3991  AA:3995  AA:3997  AA:4002  AA:4005  AA:4005  AA:4012  AA:4018  AA:4026  AA:4026  AA:4027  AA:4029  AA:4041  AA:4047  AA:4048  AA:4048  AA:4051  AA:4057  AA:4068  AA:4071  AA:4073  AA:4079  AA:4079  AA:4080  AA:4095  AA:4099  AA:4099  AA:4108  AA:4119  AA:4125  AA:4125  AA:4126  AA:4127  AA:4127  AA:4129  AA:4139  AA:4145  AA:4145  AA:4145  AA:4149  AA:4152  AA:4152  AA:4152  AA:4153  AA:4153  AA:4156  AA:4158  AA:4159  AA:4160  AA:4163  AA:4165  AB:212  AB:216  AB:231  AB:232  AB:233  AB:233  AB:236  AD:303  AE:302  AE:303  AP:201  AT:104  AZ:102  B1:101  BA:1608  BA:1609  BA:1609  BA:1609  BA:1612  BA:1616  BA:1622  BA:1623  BA:1631  BA:1633  BA:1633  BA:1641  BA:1644  BA:1647  BA:1650  BA:1651  BA:1658  BA:1658  BA:1661  BA:1668  BA:1676  BA:1678  BA:1679  BA:1679  BA:1695  BA:1702  BA:1706  BA:1710  BA:1711  BA:1711  BA:1718  BA:1718  BA:1719  BA:1719  BA:1721  BA:1727  BA:1742  BA:1744  BA:1746  BA:1747  BA:1760  BA:1766  BA:1766  BA:1768  BA:1775  BA:1776  BA:1779  BA:1781  BA:1784  BA:1784  BA:1791  BA:1800  BA:1800  BA:1813  BA:1833  BA:1835  BA:1836  BA:1896  BA:1898  BA:1900  BA:1900  BA:1904  BA:1905  BA:1906  BA:1908  BA:1913  BA:1913  BA:1913  BA:1914  BA:1917  BA:1925  BA:1938  BA:1941  BA:1943  BA:1946  BA:1953  BA:1957  BA:1958  BA:1967  BA:1970  BA:1987  BA:1989  BA:1993  BA:2004  BA:2009  BA:2010  BA:2013  BA:2016  BA:2018  BA:2018  BA:2025  BA:2036  BA:2039  BA:2042  BA:2044  BA:2047  BA:2048  BA:2048  BA:2064  BA:2078  BA:2080  BA:2080  BA:2093  BA:2102  BA:2103  BA:2106  BA:2108  BA:2111  BA:2112  BA:2115  BA:2116  BA:2132  BA:2145  BA:2155  BA:2157  BA:2159  BA:2162  BA:2164  BA:2167  BA:2186  BA:2187  BA:2191  BA:2192  BA:2194  BA:2196  BA:2203  BA:2204  BA:2205  BA:2205  BA:2206  BA:2208  BA:2211  BA:2217  BA:2231  BA:2235  BA:2246  BA:2250  BA:2254  BA:2254  BA:2263  BA:2273  BB:105  BB:113  BC:101  BC:103  BC:111  BD:102  BD:107  BD:108  BD:126  BQ:102  BS:104  BT:202  BV:101  BW:207  C1:102  C1:104  C1:105  C1:106  C1:106  CA:1602  CA:1608  CA:1609  CA:1618  CA:1618  CA:1619  CA:1622  CA:1623  CA:1650  CA:1655  CA:1656  CA:1664  CA:1665  CA:1665  CA:1665  CA:1689  CA:1692  CA:1692  CA:1700  CA:1701  CA:1701  CA:1714  CA:1715  CA:1717  CA:1719  CA:1719  CA:1720  CA:1721  CA:1724  CA:1725  CA:1727  CA:1728  CA:1736  CA:1742  CA:1744  CA:1757  CA:1762  CA:1763  CA:1767  CA:1768  CA:1770  CA:1773  CA:1803  CA:1819  CA:1819  CA:1824  CA:1842  CA:1843  CA:1859  CA:1866  CA:1868  CA:1869  CA:1869  CA:1870  CA:1877  CA:1887  CA:1911  CA:1916  CA:1920  CA:1922  CA:1924  CA:1930  CA:1933  CA:1935  CA:1936  CA:1937  CA:1938  CA:1956  CA:1956  CA:1959  CA:1961  CA:1963  CA:1965  CA:1966  CA:1974  CA:1980  CA:1986  CA:1992  CA:1996  CA:2000  CA:2001  CA:2006  CA:2007  CA:2012  CA:2013  CA:2014  CA:2015  CA:2019  CA:2035  CA:2042  CA:2072  CA:2077  CA:2083  CA:2090  CA:2091  CA:2091  CA:2093  CA:2101  CA:2101  CA:2104  CA:2111  CA:2121  CA:2127  CA:2127  CA:2133  CA:2133  CA:2134  CA:2143  CA:2145  CA:2150  CA:2153  CA:2157  CA:2160  CA:2162  CA:2171  CA:2173  CA:2189  CA:2196  CA:2201  CA:2201  CA:2203  CA:2210  CA:2215  CA:2216  CA:2223  CA:2226  CA:2229  CA:2231  CA:2233  CA:2238  CA:2239  CA:2243  CA:2251  CA:2253  CA:2254  CA:2254  CA:2254  CA:2264  CA:2271  CA:2274  CA:2274  CA:2275  CA:2281  CA:2284  CA:2286  CA:2298  CA:2299  CA:2301  CA:2315  CB:101  CB:103  CB:106  CB:107  CB:119  CC:102  CC:102  CC:103  CC:103  CC:105  CC:106  CC:108  CC:110  CC:112  CC:119  CC:119  CC:119  CC:126  CD:109  CD:125  CG:312  CQ:104  CS:106  CT:203  CV:101  D0:206  D0:206  D0:210  D1:203  D1:205  D1:206  D2:203  D2:205  D3:103  D3:105  D3:106  D5:102  D5:105  D7:103  D7:104  D8:102  D8:103  D8:107  D8:108  D8:109  DA:3003  DA:3009  DA:3011  DA:3017  DA:3019  DA:3022  DA:3028  DA:3037  DA:3038  DA:3038  DA:3038  DA:3039  DA:3041  DA:3042  DA:3047  DA:3056  DA:3057  DA:3060  DA:3076  DA:3083  DA:3085  DA:3094  DA:3098  DA:3104  DA:3105  DA:3106  DA:3109  DA:3111  DA:3113  DA:3119  DA:3121  DA:3124  DA:3126  DA:3129  DA:3129  DA:3130  DA:3163  DA:3165  DA:3166  DA:3167  DA:3172  DA:3181  DA:3184  DA:3184  DA:3185  DA:3186  DA:3186  DA:3188  DA:3205  DA:3216  DA:3220  DA:3224  DA:3235  DA:3237  DA:3238  DA:3239  DA:3241  DA:3242  DA:3243  DA:3247  DA:3250  DA:3251  DA:3253  DA:3253  DA:3254  DA:3266  DA:3269  DA:3280  DA:3281  DA:3281  DA:3284  DA:3287  DA:3301  DA:3309  DA:3318  DA:3320  DA:3322  DA:3323  DA:3324  DA:3327  DA:3327  DA:3330  DA:3335  DA:3338  DA:3351  DA:3352  DA:3352  DA:3361  DA:3372  DA:3375  DA:3379  DA:3389  DA:3398  DA:3401  DA:3402  DA:3403  DA:3407  DA:3427  DA:3430  DA:3437  DA:3438  DA:3438  DA:3442  DA:3450  DA:3452  DA:3457  DA:3461  DA:3469  DA:3474  DA:3477  DA:3482  DA:3483  DA:3486  DA:3493  DA:3496  DA:3499  DA:3514  DA:3515  DA:3518  DA:3524  DA:3525  DA:3526  DA:3531  DA:3538  DA:3540  DA:3541  DA:3545  DA:3547  DA:3549  DA:3555  DA:3560  DA:3563  DA:3563  DA:3566  DA:3579  DA:3589  DA:3610  DA:3613  DA:3614  DA:3616  DA:3617  DA:3618  DA:3627  DA:3631  DA:3632  DA:3635  DA:3637  DA:3638  DA:3640  DA:3641  DA:3648  DA:3650  DA:3650  DA:3652  DA:3658  DA:3664  DA:3670  DA:3674  DA:3677  DA:3681  DA:3683  DA:3683  DA:3683  DA:3684  DA:3689  DA:3697  DA:3702  DA:3707  DA:3708  DA:3708  DA:3708  DA:3716  DA:3729  DA:3733  DA:3733  DA:3738  DA:3739  DA:3740  DA:3740  DA:3744  DA:3744  DA:3748  DA:3750  DA:3750  DA:3772  DA:3778  DA:3778  DA:3779  DA:3780  DA:3781  DA:3782  DA:3784  DA:3785  DA:3787  DA:3790  DA:3792  DA:3796  DA:3796  DA:3798  DA:3800  DA:3806  DA:3810  DA:3821  DA:3822  DA:3826  DA:3826  DA:3830  DA:3832  DA:3834  DA:3847  DA:3848  DA:3849  DA:3852  DA:3852  DA:3854  DA:3854  DA:3860  DA:3864  DA:3865  DA:3867  DA:3878  DA:3878  DA:3888  DA:3899  DA:3901  DA:3901  DA:3904  DA:3909  DA:3919  DA:3919  DA:3925  DA:3935  DA:3944  DA:3948  DA:3951  DA:3965  DA:3966  DA:3967  DA:3969  DA:3971  DA:3972  DA:3972  DA:3975  DA:3975  DA:3985  DA:3987  DA:3988  DA:3996  DA:4006  DA:4012  DA:4014  DA:4015  DA:4022  DA:4026  DA:4027  DA:4027  DA:4032  DA:4033  DA:4039  DA:4043  DA:4045  DA:4059  DA:4062  DA:4063  DA:4063  DA:4066  DA:4067  DA:4067  DA:4068  DA:4068  DA:4069  DA:4070  DA:4083  DA:4084  DA:4086  DA:4090  DA:4091  DA:4091  DA:4092  DA:4092  DA:4093  DA:4106  DA:4107  DA:4108  DA:4118  DA:4120  DA:4121  DA:4121  DA:4129  DA:4130  DA:4132  DA:4137  DA:4138  DA:4138  DA:4142  DA:4150  DA:4150  DA:4151  DA:4151  DA:4152  DA:4157  DA:4167  DA:4168  DA:4168  DA:4172  DA:4173  DA:4173  DA:4175  DA:4175  DA:4176  DA:4177  DA:4183  DA:4183  DA:4189  DA:4198  DA:4198  DA:4201  DA:4203  DA:4204  DA:4206  DA:4207  DA:4211  DA:4212  DA:4213  DA:4215  DA:4216  DA:4220  DA:4221  DA:4224  DA:4228  DA:4228  DA:4231  DA:4233  DA:4234  DA:4235  DA:4237  DA:4238  DA:4241  DA:4242  DA:4242  DA:4250  DA:4261  DA:4261  DA:4272  DA:4276  DA:4291  DA:4292  DA:4292  DA:4298  DA:4298  DA:4309  DA:4310  DA:4312  DA:4313  DA:4313  DA:4314  DA:4323  DA:4329  DA:4332  DA:4334  DA:4337  DA:4338  DA:4339  DA:4341  DA:4345  DA:4347  DA:4358  DA:4362  DA:4363  DA:4366  DA:4367  DA:4375  DA:4377  DA:4385  DA:4385  DA:4391  DA:4397  DA:4400  DA:4401  DA:4402  DA:4403  DA:4409  DA:4409  DA:4411  DA:4411  DA:4418  DA:4420  DA:4427  DA:4434  DA:4439  DA:4441  DA:4445  DA:4445  DA:4453  DA:4454  DA:4460  DA:4467  DA:4471  DA:4472  DA:4476  DA:4476  DA:4478  DA:4483  DA:4485  DA:4487  DA:4489  DA:4489  DA:4499  DA:4505  DA:4510  DA:4517  DA:4517  DA:4520  DA:4523  DA:4525  DA:4530  DA:4536  DA:4545  DA:4551  DA:4553  DA:4554  DA:4564  DA:4564  DA:4565  DA:4566  DA:4579  DA:4580  DA:4581  DA:4582  DA:4590  DA:4597  DA:4599  DA:4600  DA:4605  DA:4618  DA:4623  DA:4624  DA:4628  DA:4631  DA:4639  DA:4643  DA:4655  DA:4658  DA:4660  DA:4662  DA:4666  DA:4667  DA:4669  DA:4670  DA:4672  DA:4677  DA:4678  DA:4679  DA:4683  DA:4687  DA:4691  DA:4696  DA:4702  DA:4711  DA:4714  DA:4728  DA:4740  DA:4741  DA:4743  DA:4744  DA:4749  DA:4751  DA:4753  DA:4755  DA:4755  DA:4760  DA:4761  DA:4766  DA:4769  DA:4772  DA:4775  DA:4775  DA:4777  DA:4778  DA:4785  DA:4793  DA:4801  DA:4806  DA:4824  DA:4825  DA:4827  DA:4830  DA:4833  DA:4834  DA:4837  DA:4846  DA:4848  DA:4854  DA:4855  DA:4862  DA:4866  DA:4871  DA:4874  DA:4878  DA:4880  DA:4882  DA:4886  DA:4895  DA:4897  DA:4902  DA:4903  DA:4911  DA:4913  DA:4916  DA:4923  DA:4924  DA:4930  DA:4933  DA:4938  DA:4944  DA:4947  DA:4956  DA:4957  DA:4958  DA:4959  DA:4961  DA:4968  DA:4972  DA:4981  DA:4982  DA:4988  DA:4991  DA:4992  DA:4992  DA:4997  DA:5000  DA:5001  DA:5003  DA:5006  DA:5006  DA:5008  DA:5009  DA:5016  DA:5016  DA:5017  DA:5021  DA:5022  DA:5023  DA:5024  DA:5025  DA:5028  DA:5040  DA:5041  DA:5051  DA:5054  DA:5058  DA:5062  DA:5067  DA:5071  DA:5074  DA:5075  DB:211  DB:218  DB:252  DB:255  DB:265  DB:273  DB:273  DB:276  DD:304  DD:309  DE:301  DE:304  DF:302  DF:304  DF:308  DF:321  DF:322  DL:201  DO:207  DO:208  DO:209  DO:210  DO:212  DO:213  DP:204  DS:202  DS:203  DS:207  DU:201  DU:210  DW:103  DY:202 |
| **3) MG/OP2 excl. contacts** | 1489 | A1:203  A1:204  A3:104  A5:101  A8:104  AA:3001  AA:3001  AA:3003  AA:3008  AA:3009  AA:3011  AA:3011  AA:3015  AA:3016  AA:3018  AA:3022  AA:3027  AA:3029  AA:3036  AA:3038  AA:3042  AA:3043  AA:3054  AA:3056  AA:3056  AA:3062  AA:3063  AA:3063  AA:3066  AA:3067  AA:3067  AA:3070  AA:3074  AA:3079  AA:3080  AA:3082  AA:3083  AA:3092  AA:3093  AA:3099  AA:3101  AA:3102  AA:3105  AA:3106  AA:3113  AA:3119  AA:3127  AA:3128  AA:3133  AA:3134  AA:3137  AA:3138  AA:3142  AA:3142  AA:3147  AA:3151  AA:3151  AA:3153  AA:3155  AA:3160  AA:3161  AA:3166  AA:3169  AA:3170  AA:3171  AA:3175  AA:3180  AA:3185  AA:3190  AA:3192  AA:3203  AA:3206  AA:3213  AA:3217  AA:3218  AA:3218  AA:3220  AA:3223  AA:3227  AA:3227  AA:3227  AA:3228  AA:3229  AA:3230  AA:3232  AA:3233  AA:3237  AA:3250  AA:3250  AA:3252  AA:3255  AA:3260  AA:3261  AA:3264  AA:3265  AA:3271  AA:3276  AA:3276  AA:3285  AA:3287  AA:3294  AA:3295  AA:3299  AA:3303  AA:3305  AA:3308  AA:3308  AA:3310  AA:3311  AA:3312  AA:3316  AA:3326  AA:3327  AA:3332  AA:3335  AA:3338  AA:3340  AA:3342  AA:3350  AA:3351  AA:3352  AA:3355  AA:3361  AA:3371  AA:3373  AA:3380  AA:3385  AA:3386  AA:3388  AA:3406  AA:3408  AA:3415  AA:3422  AA:3424  AA:3424  AA:3425  AA:3425  AA:3425  AA:3426  AA:3429  AA:3431  AA:3435  AA:3437  AA:3441  AA:3441  AA:3447  AA:3451  AA:3451  AA:3462  AA:3464  AA:3466  AA:3468  AA:3470  AA:3472  AA:3476  AA:3476  AA:3482  AA:3483  AA:3483  AA:3492  AA:3494  AA:3494  AA:3495  AA:3500  AA:3503  AA:3505  AA:3512  AA:3512  AA:3518  AA:3518  AA:3518  AA:3520  AA:3524  AA:3525  AA:3528  AA:3530  AA:3530  AA:3535  AA:3535  AA:3539  AA:3546  AA:3567  AA:3571  AA:3579  AA:3582  AA:3583  AA:3587  AA:3595  AA:3604  AA:3607  AA:3609  AA:3622  AA:3628  AA:3631  AA:3641  AA:3641  AA:3647  AA:3648  AA:3649  AA:3649  AA:3650  AA:3652  AA:3653  AA:3658  AA:3658  AA:3659  AA:3668  AA:3669  AA:3669  AA:3683  AA:3687  AA:3696  AA:3702  AA:3704  AA:3704  AA:3715  AA:3725  AA:3734  AA:3735  AA:3736  AA:3737  AA:3740  AA:3746  AA:3750  AA:3758  AA:3762  AA:3762  AA:3763  AA:3767  AA:3767  AA:3769  AA:3772  AA:3775  AA:3778  AA:3779  AA:3780  AA:3781  AA:3781  AA:3788  AA:3790  AA:3790  AA:3794  AA:3795  AA:3798  AA:3803  AA:3804  AA:3808  AA:3821  AA:3832  AA:3832  AA:3833  AA:3835  AA:3838  AA:3840  AA:3849  AA:3852  AA:3859  AA:3864  AA:3866  AA:3867  AA:3868  AA:3876  AA:3881  AA:3885  AA:3894  AA:3896  AA:3896  AA:3899  AA:3899  AA:3906  AA:3908  AA:3917  AA:3918  AA:3920  AA:3924  AA:3924  AA:3926  AA:3927  AA:3928  AA:3930  AA:3930  AA:3936  AA:3940  AA:3944  AA:3945  AA:3946  AA:3949  AA:3952  AA:3954  AA:3955  AA:3963  AA:3965  AA:3970  AA:3971  AA:3972  AA:3977  AA:3978  AA:3978  AA:3985  AA:3986  AA:3991  AA:3999  AA:4001  AA:4002  AA:4006  AA:4008  AA:4017  AA:4018  AA:4021  AA:4023  AA:4026  AA:4030  AA:4030  AA:4033  AA:4038  AA:4038  AA:4040  AA:4040  AA:4045  AA:4045  AA:4048  AA:4053  AA:4057  AA:4063  AA:4068  AA:4070  AA:4071  AA:4072  AA:4078  AA:4079  AA:4081  AA:4083  AA:4084  AA:4084  AA:4087  AA:4101  AA:4104  AA:4105  AA:4107  AA:4107  AA:4108  AA:4111  AA:4111  AA:4113  AA:4114  AA:4117  AA:4117  AA:4121  AA:4123  AA:4128  AA:4129  AA:4132  AA:4132  AA:4138  AA:4145  AA:4145  AA:4152  AA:4153  AA:4161  AA:4162  AA:4163  AA:4163  AB:201  AB:202  AB:206  AB:206  AB:208  AB:209  AB:222  AB:223  AB:224  AB:224  AB:230  AD:303  AO:206  BA:1602  BA:1604  BA:1605  BA:1607  BA:1610  BA:1610  BA:1610  BA:1622  BA:1623  BA:1631  BA:1640  BA:1647  BA:1649  BA:1650  BA:1654  BA:1659  BA:1660  BA:1663  BA:1666  BA:1670  BA:1678  BA:1680  BA:1687  BA:1690  BA:1692  BA:1695  BA:1695  BA:1696  BA:1700  BA:1700  BA:1703  BA:1706  BA:1707  BA:1709  BA:1710  BA:1712  BA:1724  BA:1728  BA:1734  BA:1745  BA:1746  BA:1748  BA:1748  BA:1749  BA:1754  BA:1760  BA:1761  BA:1764  BA:1764  BA:1765  BA:1769  BA:1774  BA:1775  BA:1776  BA:1780  BA:1791  BA:1807  BA:1809  BA:1823  BA:1826  BA:1833  BA:1834  BA:1835  BA:1835  BA:1839  BA:1847  BA:1848  BA:1849  BA:1881  BA:1893  BA:1893  BA:1905  BA:1907  BA:1912  BA:1913  BA:1917  BA:1918  BA:1918  BA:1920  BA:1927  BA:1932  BA:1935  BA:1938  BA:1941  BA:1942  BA:1948  BA:1950  BA:1953  BA:1954  BA:1960  BA:1961  BA:1965  BA:1966  BA:1973  BA:1982  BA:1985  BA:1986  BA:1987  BA:1991  BA:1992  BA:1995  BA:2000  BA:2001  BA:2003  BA:2003  BA:2008  BA:2008  BA:2013  BA:2014  BA:2016  BA:2017  BA:2020  BA:2022  BA:2022  BA:2033  BA:2035  BA:2039  BA:2047  BA:2052  BA:2053  BA:2057  BA:2076  BA:2078  BA:2080  BA:2087  BA:2094  BA:2101  BA:2101  BA:2102  BA:2103  BA:2105  BA:2111  BA:2113  BA:2114  BA:2115  BA:2124  BA:2129  BA:2130  BA:2132  BA:2133  BA:2133  BA:2134  BA:2135  BA:2135  BA:2139  BA:2143  BA:2149  BA:2149  BA:2151  BA:2153  BA:2154  BA:2155  BA:2164  BA:2170  BA:2175  BA:2181  BA:2181  BA:2182  BA:2186  BA:2187  BA:2189  BA:2189  BA:2192  BA:2197  BA:2200  BA:2206  BA:2209  BA:2209  BA:2210  BA:2211  BA:2215  BA:2217  BA:2217  BA:2225  BA:2230  BA:2231  BA:2233  BA:2233  BA:2234  BA:2236  BA:2240  BA:2246  BA:2248  BA:2248  BA:2252  BA:2253  BA:2253  BA:2254  BA:2255  BA:2258  BA:2258  BA:2259  BA:2260  BA:2262  BA:2262  BA:2262  BA:2264  BA:2266  BA:2268  BA:2269  BA:2273  BA:2274  BA:2274  BB:105  BB:110  BB:110  BB:112  BB:113  BC:104  BD:105  BD:117  C1:101  C1:102  C1:106  CA:1604  CA:1609  CA:1609  CA:1613  CA:1617  CA:1618  CA:1619  CA:1619  CA:1622  CA:1624  CA:1636  CA:1638  CA:1646  CA:1647  CA:1647  CA:1649  CA:1652  CA:1655  CA:1659  CA:1661  CA:1663  CA:1668  CA:1672  CA:1672  CA:1673  CA:1681  CA:1681  CA:1683  CA:1685  CA:1702  CA:1702  CA:1703  CA:1705  CA:1709  CA:1710  CA:1710  CA:1711  CA:1711  CA:1715  CA:1719  CA:1723  CA:1728  CA:1734  CA:1735  CA:1735  CA:1743  CA:1743  CA:1744  CA:1746  CA:1749  CA:1755  CA:1756  CA:1762  CA:1763  CA:1763  CA:1765  CA:1768  CA:1772  CA:1780  CA:1789  CA:1800  CA:1802  CA:1804  CA:1822  CA:1824  CA:1832  CA:1836  CA:1837  CA:1837  CA:1839  CA:1845  CA:1846  CA:1847  CA:1848  CA:1851  CA:1859  CA:1866  CA:1874  CA:1884  CA:1886  CA:1896  CA:1898  CA:1898  CA:1898  CA:1902  CA:1905  CA:1913  CA:1922  CA:1924  CA:1930  CA:1937  CA:1944  CA:1948  CA:1948  CA:1951  CA:1958  CA:1962  CA:1964  CA:1970  CA:1974  CA:1983  CA:1985  CA:1989  CA:1991  CA:1992  CA:1992  CA:1993  CA:1996  CA:1999  CA:2000  CA:2001  CA:2006  CA:2009  CA:2013  CA:2024  CA:2025  CA:2031  CA:2038  CA:2039  CA:2040  CA:2045  CA:2045  CA:2047  CA:2052  CA:2059  CA:2060  CA:2060  CA:2061  CA:2067  CA:2068  CA:2073  CA:2073  CA:2074  CA:2074  CA:2074  CA:2091  CA:2091  CA:2093  CA:2096  CA:2104  CA:2104  CA:2106  CA:2111  CA:2111  CA:2116  CA:2121  CA:2121  CA:2125  CA:2127  CA:2128  CA:2130  CA:2130  CA:2132  CA:2136  CA:2141  CA:2145  CA:2155  CA:2157  CA:2157  CA:2159  CA:2160  CA:2164  CA:2165  CA:2168  CA:2169  CA:2169  CA:2170  CA:2171  CA:2197  CA:2203  CA:2203  CA:2206  CA:2207  CA:2208  CA:2208  CA:2209  CA:2214  CA:2215  CA:2221  CA:2229  CA:2231  CA:2235  CA:2240  CA:2242  CA:2242  CA:2245  CA:2245  CA:2247  CA:2258  CA:2260  CA:2271  CA:2275  CA:2276  CA:2284  CA:2296  CA:2299  CA:2300  CA:2300  CA:2314  CA:2318  CB:102  CB:103  CB:105  CB:105  CB:107  CB:110  CB:116  CB:119  CC:101  CC:106  CC:109  CC:111  CC:112  CC:118  CC:119  CC:123  CC:126  CD:101  CD:103  CD:109  CD:110  CK:201  CL:201  CP:201  CQ:103  CT:203  CV:101  D1:201  D1:202  D2:206  D3:107  D5:101  D8:104  D8:104  DA:3001  DA:3004  DA:3009  DA:3009  DA:3021  DA:3025  DA:3029  DA:3033  DA:3037  DA:3045  DA:3047  DA:3051  DA:3055  DA:3057  DA:3060  DA:3060  DA:3062  DA:3065  DA:3083  DA:3087  DA:3090  DA:3097  DA:3101  DA:3105  DA:3106  DA:3110  DA:3111  DA:3113  DA:3126  DA:3127  DA:3128  DA:3129  DA:3131  DA:3132  DA:3137  DA:3147  DA:3151  DA:3164  DA:3172  DA:3173  DA:3178  DA:3182  DA:3184  DA:3186  DA:3186  DA:3188  DA:3190  DA:3191  DA:3191  DA:3192  DA:3195  DA:3196  DA:3197  DA:3197  DA:3199  DA:3202  DA:3202  DA:3205  DA:3210  DA:3212  DA:3214  DA:3214  DA:3216  DA:3220  DA:3225  DA:3229  DA:3230  DA:3231  DA:3232  DA:3233  DA:3237  DA:3244  DA:3247  DA:3249  DA:3253  DA:3264  DA:3269  DA:3271  DA:3273  DA:3277  DA:3279  DA:3280  DA:3281  DA:3284  DA:3287  DA:3289  DA:3294  DA:3296  DA:3306  DA:3312  DA:3313  DA:3320  DA:3322  DA:3323  DA:3330  DA:3331  DA:3335  DA:3335  DA:3350  DA:3357  DA:3361  DA:3374  DA:3380  DA:3385  DA:3398  DA:3399  DA:3399  DA:3403  DA:3406  DA:3406  DA:3408  DA:3412  DA:3413  DA:3416  DA:3417  DA:3423  DA:3427  DA:3429  DA:3430  DA:3436  DA:3439  DA:3444  DA:3444  DA:3446  DA:3454  DA:3456  DA:3458  DA:3470  DA:3476  DA:3484  DA:3486  DA:3487  DA:3493  DA:3500  DA:3511  DA:3514  DA:3519  DA:3524  DA:3527  DA:3527  DA:3528  DA:3534  DA:3536  DA:3541  DA:3549  DA:3552  DA:3555  DA:3556  DA:3560  DA:3563  DA:3566  DA:3570  DA:3571  DA:3586  DA:3588  DA:3589  DA:3610  DA:3618  DA:3618  DA:3620  DA:3625  DA:3627  DA:3631  DA:3631  DA:3634  DA:3642  DA:3649  DA:3650  DA:3652  DA:3657  DA:3669  DA:3670  DA:3674  DA:3675  DA:3676  DA:3676  DA:3678  DA:3678  DA:3683  DA:3683  DA:3685  DA:3688  DA:3694  DA:3694  DA:3696  DA:3705  DA:3705  DA:3705  DA:3708  DA:3709  DA:3714  DA:3714  DA:3716  DA:3719  DA:3719  DA:3722  DA:3723  DA:3723  DA:3726  DA:3729  DA:3731  DA:3733  DA:3735  DA:3744  DA:3747  DA:3748  DA:3757  DA:3760  DA:3764  DA:3772  DA:3775  DA:3776  DA:3777  DA:3779  DA:3781  DA:3781  DA:3783  DA:3784  DA:3786  DA:3787  DA:3788  DA:3792  DA:3792  DA:3796  DA:3801  DA:3804  DA:3804  DA:3806  DA:3810  DA:3811  DA:3814  DA:3817  DA:3817  DA:3820  DA:3821  DA:3825  DA:3830  DA:3833  DA:3834  DA:3834  DA:3839  DA:3840  DA:3841  DA:3847  DA:3848  DA:3848  DA:3849  DA:3851  DA:3854  DA:3874  DA:3875  DA:3876  DA:3876  DA:3878  DA:3881  DA:3883  DA:3885  DA:3888  DA:3895  DA:3909  DA:3915  DA:3919  DA:3927  DA:3932  DA:3934  DA:3935  DA:3936  DA:3940  DA:3942  DA:3944  DA:3951  DA:3952  DA:3963  DA:3963  DA:3964  DA:3965  DA:3967  DA:3989  DA:3990  DA:3991  DA:3997  DA:3997  DA:4002  DA:4003  DA:4004  DA:4004  DA:4011  DA:4012  DA:4013  DA:4014  DA:4014  DA:4015  DA:4018  DA:4020  DA:4021  DA:4022  DA:4024  DA:4025  DA:4030  DA:4043  DA:4053  DA:4054  DA:4055  DA:4056  DA:4056  DA:4058  DA:4058  DA:4061  DA:4065  DA:4067  DA:4067  DA:4070  DA:4076  DA:4083  DA:4085  DA:4087  DA:4089  DA:4089  DA:4090  DA:4090  DA:4091  DA:4091  DA:4095  DA:4100  DA:4105  DA:4126  DA:4135  DA:4136  DA:4138  DA:4145  DA:4145  DA:4146  DA:4149  DA:4149  DA:4152  DA:4155  DA:4156  DA:4157  DA:4157  DA:4163  DA:4165  DA:4167  DA:4168  DA:4169  DA:4171  DA:4175  DA:4176  DA:4179  DA:4179  DA:4180  DA:4186  DA:4186  DA:4187  DA:4189  DA:4194  DA:4196  DA:4196  DA:4198  DA:4199  DA:4200  DA:4201  DA:4204  DA:4205  DA:4205  DA:4206  DA:4206  DA:4212  DA:4212  DA:4213  DA:4214  DA:4214  DA:4215  DA:4216  DA:4219  DA:4220  DA:4227  DA:4230  DA:4231  DA:4238  DA:4246  DA:4246  DA:4247  DA:4251  DA:4264  DA:4265  DA:4270  DA:4270  DA:4273  DA:4273  DA:4275  DA:4279  DA:4283  DA:4286  DA:4291  DA:4295  DA:4298  DA:4298  DA:4303  DA:4308  DA:4312  DA:4312  DA:4312  DA:4313  DA:4314  DA:4321  DA:4325  DA:4340  DA:4346  DA:4346  DA:4347  DA:4358  DA:4361  DA:4361  DA:4363  DA:4366  DA:4375  DA:4376  DA:4379  DA:4398  DA:4399  DA:4400  DA:4400  DA:4401  DA:4411  DA:4415  DA:4415  DA:4416  DA:4417  DA:4417  DA:4423  DA:4424  DA:4426  DA:4429  DA:4429  DA:4433  DA:4434  DA:4442  DA:4452  DA:4462  DA:4462  DA:4463  DA:4465  DA:4478  DA:4479  DA:4484  DA:4484  DA:4485  DA:4492  DA:4494  DA:4498  DA:4498  DA:4499  DA:4501  DA:4507  DA:4511  DA:4514  DA:4516  DA:4518  DA:4532  DA:4533  DA:4533  DA:4537  DA:4537  DA:4543  DA:4551  DA:4552  DA:4555  DA:4555  DA:4558  DA:4564  DA:4565  DA:4567  DA:4567  DA:4573  DA:4577  DA:4580  DA:4591  DA:4596  DA:4596  DA:4601  DA:4605  DA:4608  DA:4613  DA:4614  DA:4615  DA:4616  DA:4618  DA:4618  DA:4625  DA:4625  DA:4635  DA:4637  DA:4639  DA:4649  DA:4652  DA:4653  DA:4654  DA:4660  DA:4663  DA:4666  DA:4666  DA:4680  DA:4682  DA:4682  DA:4685  DA:4691  DA:4697  DA:4697  DA:4706  DA:4710  DA:4716  DA:4719  DA:4725  DA:4725  DA:4729  DA:4730  DA:4738  DA:4743  DA:4745  DA:4745  DA:4746  DA:4747  DA:4748  DA:4755  DA:4760  DA:4769  DA:4770  DA:4771  DA:4772  DA:4772  DA:4779  DA:4779  DA:4785  DA:4795  DA:4799  DA:4799  DA:4806  DA:4807  DA:4807  DA:4807  DA:4810  DA:4811  DA:4811  DA:4814  DA:4816  DA:4816  DA:4820  DA:4825  DA:4826  DA:4827  DA:4829  DA:4835  DA:4838  DA:4838  DA:4842  DA:4850  DA:4850  DA:4850  DA:4851  DA:4851  DA:4853  DA:4853  DA:4855  DA:4855  DA:4856  DA:4856  DA:4857  DA:4857  DA:4862  DA:4867  DA:4867  DA:4871  DA:4871  DA:4872  DA:4873  DA:4875  DA:4880  DA:4882  DA:4891  DA:4891  DA:4893  DA:4902  DA:4904  DA:4912  DA:4912  DA:4913  DA:4914  DA:4919  DA:4919  DA:4920  DA:4930  DA:4932  DA:4937  DA:4937  DA:4938  DA:4942  DA:4942  DA:4943  DA:4947  DA:4953  DA:4956  DA:4957  DA:4965  DA:4969  DA:4974  DA:4975  DA:4978  DA:4979  DA:4982  DA:4991  DA:4992  DA:4992  DA:4999  DA:4999  DA:5003  DA:5014  DA:5014  DA:5016  DA:5017  DA:5019  DA:5021  DA:5025  DA:5036  DA:5036  DA:5038  DA:5040  DA:5040  DA:5040  DA:5045  DA:5045  DA:5048  DA:5052  DA:5055  DA:5062  DA:5062  DA:5068  DA:5071  DA:5073  DA:5073  DA:5075  DB:204  DB:212  DB:223  DB:229  DB:234  DB:234  DB:235  DB:236  DB:237  DB:243  DB:245  DB:247  DB:253  DB:256  DB:264  DB:269  DB:270  DB:270  DB:272  DB:276  DF:316  DF:321  DO:209  DT:101  DU:205  DW:107 |
| **3) ZN/N_back_ excl. contacts** | 4 | A4:101  A4:101  CQ:101  D4:103 |
| **3) ZN/O_back_ excl. contacts** | 2 | A4:101  BG:308 |
| **4) Large octahedral topology discrepancy** | 1 | AA:3378 |
| **No contacts in 5.0 A** | 129 | AA:3405 AA:3407 AA:3413 AA:3418 AA:3419 AA:3423 AA:3446 AA:3719 AA:3723 AA:3786 AA:3846 AA:3856 AA:3884 AA:3898 AA:3903 AA:3956 AA:3960 AA:3992 AA:3993 AA:4015 AA:4032 AA:4054 AA:4061 AA:4093 AA:4097 AA:4112 AA:4122 AA:4133 AA:4136 AA:4140 AA:4143 AA:4147 AA:4151 AA:4164 AB:234 AB:235 AF:302 AF:306 AK:203 AV:301 BA:1614 BA:1853 BA:1856 BA:1859 BA:1863 BA:1869 BA:1870 BA:1871 BA:1873 BA:1874 BA:1875 BA:1876 BA:1944 BA:2054 BA:2058 BA:2069 BA:2100 BA:2110 BA:2121 BA:2179 BA:2185 BA:2198 BA:2202 BA:2243 BA:2244 BD:106 BD:119 BD:122 BD:125 BE:304 BH:204 BK:203 CA:1813 CA:1816 CA:2049 CA:2071 CA:2092 CA:2142 CA:2181 CA:2182 CA:2204 CA:2205 CA:2266 CA:2317 CD:105 CD:107 CD:111 CD:115 CD:124 CD:127 CF:302 CM:201 D2:210 D4:101 DA:3548 DA:3550 DA:3576 DA:3644 DA:3758 DA:3799 DA:3931 DA:4031 DA:4038 DA:4114 DA:4598 DA:4620 DA:4633 DA:4674 DA:4734 DA:4787 DA:4804 DA:4822 DA:4859 DA:4870 DA:4885 DA:4909 DA:4941 DA:4954 DA:4989 DA:5018 DA:5034 DA:5050 DA:5053 DA:5059 DE:307 DF:319 DH:204 DQ:201 DV:303 |

---

\_\_\_\_\_\_\_\_\_\_\_\_\_\_\_\_\_\_\_\_\_\_\_\_\_\_\_\_\_\_\_\_\_\_\_\_\_\_\_\_\_\_\_\_\_\_\_\_\_\_\_\_\_\_\_\_\_\_\_\_\_\_\_\_  

#### 4) Ion/ligand tables: all contacts up to 5.0 Angs. are listed

##### Go to ToC

### MG:A0:201

|  | Res. type | Atom | Residue | Distance | Occupancy | B-factor | Closest solute to 1^st^ shell water |
| --- | --- | --- | --- | --- | --- | --- | --- |
| **Ion** | | | |  | 1.00 | 61.86 || 1 | VAL | N | A0:73 | 2.59 | 1.00 | 64.47 |
| 2 | GLN | O | A0:71 | 2.66 | 1.00 | 67.15 |  |
| 3 | ASP | OD2 | A0:72 | 3.38 | 1.00 | 71.07 |  |
| 4 | ASP | N | A0:72 | 3.70 | 1.00 | 67.69 |  |
| 5 | C | O2' | AA:2701 | 4.12 | 1.00 | 59.49 |  |
| 6 | ASP | O | A0:72 | 4.43 | 1.00 | 65.87 |  |
| 7 | LYS | N | A0:74 | 4.56 | 1.00 | 62.75 |  |
  |

  
 Coded contact list:   
 Exclusion zone contact(s): /N_back_/O_back_/OD2_Asp_/  
Consider K in place of MG (2 valid contact(s) for K vs. 0 for MG)  

### MG:A1:201

|  | Res. type | Atom | Residue | Distance | Occupancy | B-factor | Closest solute to 1^st^ shell water |
| --- | --- | --- | --- | --- | --- | --- | --- |
| **Ion** | | | |  | 1.00 | 60.48 || 1 | G | OP2 | AA:585 | 2.33 | 1.00 | 49.31 |
| 2 | THR | OG1 | A1:6 | 3.17 | 1.00 | 49.14 |  |
| 3 | G | O5' | AA:585 | 4.25 | 1.00 | 47.63 |  |
| 4 | C | OP2 | AA:584 | 4.30 | 1.00 | 47.30 |  |
| 5 | ARG | NH2 | A1:10 | 4.33 | 1.00 | 47.50 |  |
| 6 | THR | N | A1:6 | 4.41 | 1.00 | 50.29 |  |
| 7 | G | OP1 | AA:585 | 4.47 | 1.00 | 49.04 |  |
| 8 | G | OP1 | AA:1250 | 4.50 | 1.00 | 50.33 |  |
| 9 | C | O5' | AA:584 | 4.60 | 1.00 | 48.97 |  |
| 10 | C | OP1 | AA:584 | 4.61 | 1.00 | 48.15 |  |
| 11 | G | N7 | AA:585 | 4.62 | 1.00 | 48.10 |  |
| 12 | C | O3' | AA:584 | 4.79 | 1.00 | 48.99 |  |
| 13 | ALA | O | A1:4 | 4.95 | 1.00 | 51.47 |  |
| 14 | U | O3' | AA:1249 | 4.95 | 1.00 | 52.68 |  |
  |

  
 Contact(s): /OP2/  
 Coded contact list: O_ph_  
 Exclusion zone contact(s): /OG1_Thr_/  

### MG:A1:202

|  | Res. type | Atom | Residue | Distance | Occupancy | B-factor | Closest solute to 1^st^ shell water |
| --- | --- | --- | --- | --- | --- | --- | --- |
| **Ion** | | | |  | 1.00 | 61.95 || 1 | ARG | N | A1:11 | 2.37 | 1.00 | 47.66 |
| 2 | G | OP1 | AA:583 | 2.72 | 1.00 | 43.37 |  |
| 3 | GLY | O | A1:7 | 2.84 | 1.00 | 49.76 |  |
| 4 | ARG | N | A1:10 | 2.89 | 1.00 | 45.25 |  |
| 5 | VAL | O | A1:8 | 3.38 | 1.00 | 48.38 |  |
| 6 | VAL | N | A1:9 | 3.80 | 1.00 | 46.77 |  |
| 7 | ARG | O | A1:10 | 4.34 | 1.00 | 45.13 |  |
| 8 | VAL | O | A1:9 | 4.34 | 1.00 | 42.97 |  |
| 9 | VAL | N | A1:8 | 4.35 | 1.00 | 49.63 |  |
| 10 | G | O5' | AA:583 | 4.63 | 1.00 | 43.05 |  |
| 11 | ARG | N | A1:12 | 4.72 | 1.00 | 50.52 |  |
| 12 | G | OP2 | AA:583 | 4.86 | 1.00 | 43.29 |  |
  |

  
 Contact(s): /N_back_/  
 Coded contact list: Anion  
 Exclusion zone contact(s): /N_back_/O_back_/O_back_/OP1/  
Consider K in place of MG (4 valid contact(s) for K vs. 1 for MG)  

### MG:A1:203

|  | Res. type | Atom | Residue | Distance | Occupancy | B-factor | Closest solute to 1^st^ shell water |
| --- | --- | --- | --- | --- | --- | --- | --- |
| **Ion** | | | |  | 1.00 | 77.24 || 1 | C | OP2 | AA:998 | 2.61 | 1.00 | 83.90 |
| 2 | ARG | NE | A1:58 | 2.71 | 1.00 | 90.84 |  |
| 3 | LYS | NZ | A1:93 | 3.46 | 1.00 | 91.03 |  |
| 4 | G | OP2 | AA:1154 | 3.79 | 1.00 | 84.53 |  |
| 5 | G | OP2 | AA:997 | 3.82 | 1.00 | 84.17 |  |
| 6 | C | OP1 | AA:1153 | 4.09 | 1.00 | 88.96 |  |
| 7 | ARG | NH2 | A1:58 | 4.18 | 1.00 | 92.28 |  |
| 8 | G | OP1 | AA:997 | 4.37 | 1.00 | 84.08 |  |
| 9 | G | O3' | AA:997 | 4.64 | 1.00 | 82.86 |  |
| 10 | G | O5' | AA:997 | 4.74 | 1.00 | 83.17 |  |
| 11 | C | OP1 | AA:998 | 4.77 | 1.00 | 84.00 |  |
| 12 | ARG | NH1 | A1:58 | 4.86 | 1.00 | 91.58 |  |
  |

  
 Coded contact list:   
 Exclusion zone contact(s): /NE_Arg_/OP2/  
Consider K in place of MG (2 valid contact(s) for K vs. 0 for MG)  

### MG:A1:204

|  | Res. type | Atom | Residue | Distance | Occupancy | B-factor | Closest solute to 1^st^ shell water |
| --- | --- | --- | --- | --- | --- | --- | --- |
| **Ion** | | | |  | 1.00 | 65.58 || 1 | C | OP2 | AA:1251 | 2.61 | 1.00 | 53.29 |
| 2 | THR | OG1 | A1:6 | 2.91 | 1.00 | 49.14 |  |
| 3 | ARG | NH2 | A1:10 | 2.97 | 1.00 | 47.50 |  |
| 4 | G | O3' | AA:1250 | 3.57 | 1.00 | 54.22 |  |
| 5 | G | O5' | AA:1250 | 4.14 | 1.00 | 52.15 |  |
| 6 | ARG | NE | A1:10 | 4.22 | 1.00 | 46.85 |  |
| 7 | C | O5' | AA:1251 | 4.47 | 1.00 | 52.69 |  |
| 8 | U | O2 | AA:811 | 4.53 | 1.00 | 53.70 |  |
| 9 | G | OP1 | AA:1250 | 4.70 | 1.00 | 50.33 |  |
| 10 | C | OP2 | AA:584 | 4.71 | 1.00 | 47.30 |  |
| 11 | G | O4' | AA:1250 | 4.87 | 1.00 | 55.45 |  |
| 12 | G | N7 | AA:585 | 4.92 | 1.00 | 48.10 |  |
| 13 | C | OP1 | AA:1251 | 4.96 | 1.00 | 53.24 |  |
| 14 | ARG | NH1 | A1:10 | 4.96 | 1.00 | 50.06 |  |
  |

  
 Coded contact list:   
 Exclusion zone contact(s): /NH2_Arg_/OG1_Thr_/OP2/  
Consider K in place of MG (3 valid contact(s) for K vs. 0 for MG)  

### MG:A1:205

|  | Res. type | Atom | Residue | Distance | Occupancy | B-factor | Closest solute to 1^st^ shell water |
| --- | --- | --- | --- | --- | --- | --- | --- |
| **Ion** | | | |  | 1.00 | 79.96 || 1 | A | OP1 | AA:515 | 4.64 | 1.00 | 51.03 |
  |

  
 Coded contact list:   
1^st^ MG coordination shell missing (see Notes)  

### MG:A2:201

|  | Res. type | Atom | Residue | Distance | Occupancy | B-factor | Closest solute to 1^st^ shell water |
| --- | --- | --- | --- | --- | --- | --- | --- |
| **Ion** | | | |  | 1.00 | 66.09 || 1 | LYS | O | A2:76 | 2.43 | 1.00 | 45.68 |
| 2 | GLN | N | A2:80 | 2.59 | 1.00 | 52.79 |  |
| 3 | C | OP1 | AA:565 | 2.86 | 1.00 | 41.10 |  |
| 4 | VAL | N | A2:79 | 2.94 | 1.00 | 43.88 |  |
| 5 | GLN | NE2 | A2:80 | 3.31 | 1.00 | 53.48 |  |
| 6 | LYS | N | A2:78 | 3.37 | 1.00 | 45.46 |  |
| 7 | GLN | O | A2:80 | 3.80 | 1.00 | 51.78 |  |
| 8 | ALA | N | A2:77 | 4.29 | 1.00 | 46.03 |  |
| 9 | VAL | O | A2:79 | 4.32 | 1.00 | 49.95 |  |
| 10 | ALA | O | A2:77 | 4.37 | 1.00 | 47.45 |  |
| 11 | TYR | N | A2:81 | 4.52 | 1.00 | 57.52 |  |
| 12 | C | OP2 | AA:565 | 4.73 | 1.00 | 40.02 |  |
| 13 | LYS | N | A2:76 | 4.80 | 1.00 | 53.24 |  |
| 14 | TYR | OH | A2:81 | 4.94 | 1.00 | 59.44 |  |
| 15 | LYS | O | A2:78 | 4.99 | 1.00 | 41.39 |  |
  |

  
 Coded contact list:   
 Exclusion zone contact(s): /N_back_/N_back_/N_back_/NE2_Gln_/O_back_/OP1/  
Consider K in place of MG (4 valid contact(s) for K vs. 0 for MG)  

### MG:A3:101

|  | Res. type | Atom | Residue | Distance | Occupancy | B-factor | Closest solute to 1^st^ shell water |
| --- | --- | --- | --- | --- | --- | --- | --- |
| **Ion** | | | |  | 1.00 | 40.68 || 1 | ARG | NH1 | A3:14 | 3.18 | 1.00 | 50.18 |
| 2 | G | N7 | AA:2280 | 4.36 | 1.00 | 61.45 |  |
| 3 | C | N4 | AA:2281 | 4.50 | 1.00 | 65.37 |  |
| 4 | G | O6 | AA:2280 | 4.82 | 1.00 | 59.68 |  |
| 5 | ARG | NH2 | A3:14 | 4.97 | 1.00 | 48.19 |  |
  |

  
 Coded contact list:   
 Exclusion zone contact(s): /NH1_Arg_/  
Consider K in place of MG (1 valid contact(s) for K vs. 0 for MG)  

### MG:A3:102

|  | Res. type | Atom | Residue | Distance | Occupancy | B-factor | Closest solute to 1^st^ shell water |
| --- | --- | --- | --- | --- | --- | --- | --- |
| **Ion** | | | |  | 1.00 | 83.35 || 1 | G | O6 | AA:2334 | 2.34 | 1.00 | 67.23 |
| 2 | C | O2 | AB:12 | 2.84 | 1.00 | 96.16 |  |
| 3 | ARG | NE | A3:74 | 4.03 | 1.00 | 89.36 |  |
| 4 | C | N3 | AB:12 | 4.10 | 1.00 | 98.27 |  |
| 5 | G | N1 | AA:2334 | 4.31 | 1.00 | 65.52 |  |
| 6 | C | N1 | AB:12 | 4.78 | 1.00 | 98.29 |  |
| 7 | C | O2' | AB:12 | 4.80 | 1.00 | 98.62 |  |
| 8 | G | N7 | AA:2334 | 4.89 | 1.00 | 66.85 |  |
  |

  
 Contact(s): /O6/  
 Coded contact list: O_b_  
 Exclusion zone contact(s): /O2/  

### MG:A3:103

|  | Res. type | Atom | Residue | Distance | Occupancy | B-factor | Closest solute to 1^st^ shell water |
| --- | --- | --- | --- | --- | --- | --- | --- |
| **Ion** | | | |  | 1.00 | 55.67 || 1 | GLU | OE2 | A3:27 | 3.71 | 1.00 | 82.89 |
| 2 | GLU | OE1 | A3:27 | 4.21 | 1.00 | 83.28 |  |
| 3 | C | OP1 | AA:856 | 4.22 | 1.00 | 85.76 |  |
| 4 | PHE | O | A3:69 | 4.57 | 1.00 | 85.76 |  |
| 5 | G | O3' | AA:855 | 4.69 | 1.00 | 85.83 |  |
| 6 | C | O5' | AA:856 | 4.83 | 1.00 | 85.01 |  |
  |

  
 Coded contact list:   
1^st^ MG coordination shell missing (see Notes)  

### MG:A3:104

|  | Res. type | Atom | Residue | Distance | Occupancy | B-factor | Closest solute to 1^st^ shell water |
| --- | --- | --- | --- | --- | --- | --- | --- |
| **Ion** | | | |  | 1.00 | 83.91 || 1 | C | OP2 | AA:2263 | 2.67 | 1.00 | 60.20 |
| 2 | SER | OG | A3:16 | 3.24 | 1.00 | 61.93 |  |
| 3 | SER | O | A3:16 | 3.54 | 1.00 | 62.01 |  |
| 4 | U | OP2 | AA:2262 | 4.01 | 1.00 | 55.28 |  |
| 5 | ASP | OD1 | A3:15 | 4.12 | 1.00 | 60.98 |  |
| 6 | GLN | O | A3:17 | 4.13 | 1.00 | 66.54 |  |
| 7 | ASP | OD2 | A3:15 | 4.14 | 1.00 | 61.18 |  |
| 8 | SER | N | A3:16 | 4.49 | 1.00 | 57.52 |  |
| 9 | U | O5' | AA:2262 | 4.68 | 1.00 | 53.96 |  |
| 10 | C | O5' | AA:2263 | 4.74 | 1.00 | 57.70 |  |
  |

  
 Coded contact list:   
 Exclusion zone contact(s): /OG_Ser_/OP2/  
Consider K in place of MG (2 valid contact(s) for K vs. 0 for MG)  

### ZN:A4:101

|  | Res. type | Atom | Residue | Distance | Occupancy | B-factor | Closest solute to 1^st^ shell water |
| --- | --- | --- | --- | --- | --- | --- | --- |
| **Ion** | | | |  | 1.00 | 200.41 || 1 | CYS | O | A4:36 | 2.37 | 1.00 | 156.68 |
| 2 | CYS | N | A4:36 | 2.99 | 1.00 | 156.86 |  |
| 3 | SER | N | A4:37 | 3.32 | 1.00 | 157.36 |  |
| 4 | GLY | N | A4:17 | 3.45 | 1.00 | 153.55 |  |
| 5 | VAL | O | A4:35 | 3.81 | 1.00 | 156.11 |  |
| 6 | GLU | O | A4:34 | 4.32 | 1.00 | 155.06 |  |
| 7 | CYS | N | A4:18 | 4.36 | 1.00 | 154.21 |  |
| 8 | CYS | N | A4:16 | 4.63 | 1.00 | 151.95 |  |
| 9 | CYS | N | A4:39 | 4.66 | 1.00 | 156.75 |  |
| 10 | CYS | O | A4:16 | 4.82 | 1.00 | 153.29 |  |
| 11 | LYS | N | A4:38 | 4.94 | 1.00 | 156.60 |  |
  |

  
 Coded contact list:   
 Exclusion zone contact(s): /N_back_/N_back_/O_back_/  

### MG:A5:101

|  | Res. type | Atom | Residue | Distance | Occupancy | B-factor | Closest solute to 1^st^ shell water |
| --- | --- | --- | --- | --- | --- | --- | --- |
| **Ion** | | | |  | 1.00 | 23.78 || 1 | HIS | ND1 | A5:4 | 2.32 | 1.00 | 42.77 |
| 2 | C | OP2 | AA:2612 | 2.97 | 1.00 | 51.00 |  |
| 3 | C | O5' | AA:2612 | 3.97 | 1.00 | 51.63 |  |
| 4 | HIS | NE2 | A5:4 | 3.99 | 1.00 | 41.55 |  |
| 5 | A | O2' | AA:2577 | 4.27 | 1.00 | 39.51 |  |
| 6 | A | O3' | AA:2577 | 4.73 | 1.00 | 41.34 |  |
| 7 | U | O3' | AA:2611 | 4.79 | 1.00 | 48.43 |  |
| 8 | C | OP1 | AA:2579 | 4.81 | 1.00 | 36.86 |  |
| 9 | U | OP1 | AA:2611 | 4.86 | 1.00 | 43.66 |  |
  |

  
 Contact(s): /ND1_His_/  
 Coded contact list: His  
 Exclusion zone contact(s): /OP2/  

### MG:A5:102

|  | Res. type | Atom | Residue | Distance | Occupancy | B-factor | Closest solute to 1^st^ shell water |
| --- | --- | --- | --- | --- | --- | --- | --- |
| **Ion** | | | |  | 1.00 | 28.56 || 1 | HIS | NE2 | A5:4 | 2.54 | 1.00 | 41.55 |
| 2 | U | O4 | AA:2611 | 3.24 | 1.00 | 42.54 |  |
| 3 | A | O4' | AA:2577 | 3.46 | 1.00 | 43.18 |  |
| 4 | A | OP2 | AA:2577 | 3.59 | 1.00 | 43.39 |  |
| 5 | G | O6 | AA:2056 | 3.95 | 1.00 | 40.78 |  |
| 6 | A | N6 | AA:2057 | 4.19 | 1.00 | 43.68 |  |
| 7 | HIS | ND1 | A5:4 | 4.19 | 1.00 | 42.77 |  |
| 8 | A | O5' | AA:2577 | 4.41 | 1.00 | 42.97 |  |
| 9 | U | N3 | AA:2611 | 4.51 | 1.00 | 43.24 |  |
  |

  
 Coded contact list:   
 Exclusion zone contact(s): /NE2_His_/O4/  
Consider NA in place of MG (1 valid contact(s) for NA vs. 0 for MG)  

### MG:A5:103

|  | Res. type | Atom | Residue | Distance | Occupancy | B-factor | Closest solute to 1^st^ shell water |
| --- | --- | --- | --- | --- | --- | --- | --- |
| **Ion** | | | |  | 1.00 | 54.74 || 1 | GLY | O | A5:54 | 2.61 | 1.00 | 129.14 |
| 2 | TYR | N | A5:52 | 3.25 | 1.00 | 119.52 |  |
| 3 | TYR | N | A5:51 | 3.36 | 1.00 | 113.77 |  |
| 4 | GLY | N | A5:54 | 3.50 | 1.00 | 127.57 |  |
| 5 | TYR | O | A5:51 | 3.63 | 1.00 | 117.43 |  |
| 6 | ALA | N | A5:53 | 3.63 | 1.00 | 124.24 |  |
| 7 | ARG | N | A5:55 | 4.20 | 1.00 | 131.05 |  |
  |

  
 Coded contact list:   
 Exclusion zone contact(s): /N_back_/N_back_/O_back_/  
Consider K in place of MG (3 valid contact(s) for K vs. 0 for MG)  

### MG:A6:101

|  | Res. type | Atom | Residue | Distance | Occupancy | B-factor | Closest solute to 1^st^ shell water |
| --- | --- | --- | --- | --- | --- | --- | --- |
| **Ion** | | | |  | 1.00 | 55.92 || 1 | THR | OG1 | A6:23 | 2.45 | 1.00 | 152.67 |
| 2 | LYS | O | A8:36 | 2.73 | 1.00 | 85.13 |  |
| 3 | TRP | O | A8:34 | 3.45 | 1.00 | 92.52 |  |
| 4 | THR | O | A6:23 | 4.27 | 1.00 | 152.64 |  |
| 5 | SER | N | A8:37 | 4.45 | 1.00 | 82.11 |  |
| 6 | GLY | N | A8:38 | 4.46 | 1.00 | 77.32 |  |
| 7 | GLU | N | A6:24 | 4.79 | 1.00 | 152.16 |  |
| 8 | LYS | N | A8:36 | 4.86 | 1.00 | 88.43 |  |
| 9 | C | OP2 | AA:2347 | 4.95 | 1.00 | 73.92 |  |
  |

  
 Coded contact list:   
 Exclusion zone contact(s): /O_back_/OG1_Thr_/  
Consider NA in place of MG (1 valid contact(s) for NA vs. 0 for MG)  

### MG:A6:102

|  | Res. type | Atom | Residue | Distance | Occupancy | B-factor | Closest solute to 1^st^ shell water |
| --- | --- | --- | --- | --- | --- | --- | --- |
| **Ion** | | | |  | 1.00 | 60.62 || 1 | THR | OG1 | A6:47 | 3.42 | 1.00 | 133.20 |
| 2 | LYS | NZ | A6:45 | 4.70 | 1.00 | 121.68 |  |
| 3 | HIS | O | A6:46 | 4.88 | 1.00 | 133.93 |  |
  |

  
 Coded contact list:   
1^st^ MG coordination shell missing (see Notes)  

### MG:A7:101

|  | Res. type | Atom | Residue | Distance | Occupancy | B-factor | Closest solute to 1^st^ shell water |
| --- | --- | --- | --- | --- | --- | --- | --- |
| **Ion** | | | |  | 1.00 | 77.44 || 1 | A | N7 | AA:126 | 2.93 | 1.00 | 41.65 |
| 2 | THR | N | A7:43 | 3.71 | 1.00 | 56.29 |  |
| 3 | A | N6 | AA:127 | 3.78 | 1.00 | 47.68 |  |
| 4 | THR | OG1 | A7:43 | 3.85 | 1.00 | 56.47 |  |
| 5 | A | N6 | AA:126 | 3.98 | 1.00 | 38.06 |  |
| 6 | THR | O | A7:43 | 4.11 | 1.00 | 62.55 |  |
  |

  
 Coded contact list:   
 Exclusion zone contact(s): /N7/  
Consider K in place of MG (1 valid contact(s) for K vs. 0 for MG)  

### MG:A7:102

|  | Res. type | Atom | Residue | Distance | Occupancy | B-factor | Closest solute to 1^st^ shell water |
| --- | --- | --- | --- | --- | --- | --- | --- |
| **Ion** | | | |  | 1.00 | 59.24 || 1 | G | OP1 | AA:770 | 3.31 | 1.00 | 39.03 |
| 2 | ASN | ND2 | A7:8 | 3.80 | 1.00 | 39.15 |  |
| 3 | ASN | OD1 | A7:8 | 4.43 | 1.00 | 39.34 |  |
| 4 | G | O3' | AA:769 | 4.43 | 1.00 | 35.15 |  |
| 5 | ARG | NE | A7:10 | 4.49 | 1.00 | 41.85 |  |
| 6 | ARG | N | A7:10 | 4.62 | 1.00 | 34.63 |  |
| 7 | G | OP1 | AA:1309 | 4.71 | 1.00 | 28.06 |  |
  |

  
 Coded contact list:   
 Exclusion zone contact(s): /OP1/  
Consider K in place of MG (1 valid contact(s) for K vs. 0 for MG)  

### MG:A7:103

|  | Res. type | Atom | Residue | Distance | Occupancy | B-factor | Closest solute to 1^st^ shell water |
| --- | --- | --- | --- | --- | --- | --- | --- |
| **Ion** | | | |  | 1.00 | 45.57 || 1 | ARG | NH1 | A7:47 | 3.11 | 1.00 | 107.57 |
| 2 | G | N7 | AA:1311 | 3.39 | 1.00 | 58.63 |  |
| 3 | ARG | O | A7:47 | 3.81 | 1.00 | 108.11 |  |
| 4 | ARG | NH2 | A7:9 | 4.16 | 1.00 | 50.66 |  |
| 5 | ARG | NH2 | A7:47 | 4.26 | 1.00 | 105.93 |  |
| 6 | ARG | N | A7:47 | 4.38 | 1.00 | 103.54 |  |
| 7 | ARG | NE | A7:47 | 4.69 | 1.00 | 107.21 |  |
| 8 | G | O6 | AA:1311 | 4.73 | 1.00 | 64.71 |  |
| 9 | G | N9 | AA:1311 | 4.92 | 1.00 | 55.93 |  |
| 10 | G | OP2 | AA:1311 | 4.99 | 1.00 | 37.84 |  |
  |

  
 Coded contact list:   
 Exclusion zone contact(s): /N7/NH1_Arg_/  
Consider K in place of MG (2 valid contact(s) for K vs. 0 for MG)  

### MG:A7:104

|  | Res. type | Atom | Residue | Distance | Occupancy | B-factor | Closest solute to 1^st^ shell water |
| --- | --- | --- | --- | --- | --- | --- | --- |
| **Ion** | | | |  | 1.00 | 88.06 || 1 | TRP | NE1 | A7:40 | 4.43 | 1.00 | 46.01 |
| 2 | U | OP1 | AA:67 | 4.53 | 1.00 | 54.61 |  |
  |

  
 Coded contact list:   
1^st^ MG coordination shell missing (see Notes)  

### MG:A8:101

|  | Res. type | Atom | Residue | Distance | Occupancy | B-factor | Closest solute to 1^st^ shell water |
| --- | --- | --- | --- | --- | --- | --- | --- |
| **Ion** | | | |  | 1.00 | 45.46 || 1 | A | OP1 | AA:631 | 3.34 | 1.00 | 57.67 |
| 2 | LYS | O | AO:64 | 3.85 | 1.00 | 78.24 |  |
| 3 | C | OP1 | AA:2417 | 4.19 | 1.00 | 69.82 |  |
| 4 | C | O3' | AA:2416 | 4.75 | 1.00 | 70.27 |  |
| 5 | C | OP1 | AA:2416 | 4.84 | 1.00 | 68.95 |  |
| 6 | C | OP2 | AA:2417 | 4.86 | 1.00 | 71.42 |  |
| 7 | A | O5' | AA:631 | 4.93 | 1.00 | 57.11 |  |
| 8 | A | O4' | AA:631 | 4.98 | 1.00 | 57.51 |  |
  |

  
 Coded contact list:   
 Exclusion zone contact(s): /OP1/  
Consider K in place of MG (1 valid contact(s) for K vs. 0 for MG)  

### MG:A8:102

|  | Res. type | Atom | Residue | Distance | Occupancy | B-factor | Closest solute to 1^st^ shell water |
| --- | --- | --- | --- | --- | --- | --- | --- |
| **Ion** | | | |  | 1.00 | 68.55 || 1 | LYS | NZ | A8:15 | 2.69 | 1.00 | 66.37 |
| 2 | G | N7 | AA:630 | 3.91 | 1.00 | 60.26 |  |
| 3 | G | O6 | AA:630 | 4.52 | 1.00 | 61.78 |  |
| 4 | G | N7 | AA:629 | 4.57 | 1.00 | 64.88 |  |
  |

  
 Coded contact list:   
 Exclusion zone contact(s): /NZ_Lys_/  
Consider K in place of MG (1 valid contact(s) for K vs. 0 for MG)  

### MG:A8:103

|  | Res. type | Atom | Residue | Distance | Occupancy | B-factor | Closest solute to 1^st^ shell water |
| --- | --- | --- | --- | --- | --- | --- | --- |
| **Ion** | | | |  | 1.00 | 62.00 || 1 | LYS | NZ | A8:15 | 3.38 | 1.00 | 66.37 |
| 2 | A | OP1 | AA:632 | 3.81 | 1.00 | 58.94 |  |
| 3 | A | OP2 | AA:632 | 4.39 | 1.00 | 58.79 |  |
| 4 | G | N7 | AA:630 | 4.59 | 1.00 | 60.26 |  |
| 5 | A | O3' | AA:631 | 4.82 | 1.00 | 58.89 |  |
  |

  
 Coded contact list:   
 Exclusion zone contact(s): /NZ_Lys_/  
Consider K in place of MG (1 valid contact(s) for K vs. 0 for MG)  

### MG:A8:104

|  | Res. type | Atom | Residue | Distance | Occupancy | B-factor | Closest solute to 1^st^ shell water |
| --- | --- | --- | --- | --- | --- | --- | --- |
| **Ion** | | | |  | 1.00 | 68.27 || 1 | U | OP2 | AA:2348 | 2.54 | 1.00 | 69.26 |
| 2 | ARG | NH2 | A8:42 | 2.70 | 1.00 | 62.50 |  |
| 3 | G | O2' | AA:2382 | 2.70 | 1.00 | 79.52 |  |
| 4 | LYS | N | A8:39 | 3.09 | 1.00 | 76.78 |  |
| 5 | GLY | N | A8:38 | 3.88 | 1.00 | 77.32 |  |
| 6 | GLY | O | A8:38 | 4.36 | 1.00 | 75.82 |  |
| 7 | U | OP1 | AA:2348 | 4.42 | 1.00 | 69.92 |  |
| 8 | G | N3 | AA:2382 | 4.48 | 1.00 | 76.07 |  |
| 9 | U | O5' | AA:2348 | 4.65 | 1.00 | 68.90 |  |
| 10 | ARG | NE | A8:42 | 4.74 | 1.00 | 62.75 |  |
| 11 | ARG | NH1 | A8:42 | 4.79 | 1.00 | 62.01 |  |
| 12 | G | O3' | AA:2382 | 4.84 | 1.00 | 77.78 |  |
| 13 | C | O3' | AA:2347 | 4.99 | 1.00 | 70.77 |  |
  |

  
 Coded contact list:   
 Exclusion zone contact(s): /N_back_/NH2_Arg_/O2'/OP2/  
Consider K in place of MG (3 valid contact(s) for K vs. 0 for MG)  

### MG:AA:3001

|  | Res. type | Atom | Residue | Distance | Occupancy | B-factor | Closest solute to 1^st^ shell water |
| --- | --- | --- | --- | --- | --- | --- | --- |
| **Ion** | | | |  | 1.00 | 21.73 || 1 | A | OP2 | AA:2711 | 2.29 | 1.00 | 45.22 |
| 2 | G | OP2 | AA:2714 | 2.41 | 1.00 | 58.17 |  |
| 3 | A | OP2 | AA:2712A | 2.47 | 1.00 | 45.06 |  |
| 4 | C | OP1 | AA:2710 | 3.71 | 1.00 | 49.23 |  |
| 5 | G | OP1 | AA:2714 | 3.73 | 1.00 | 56.15 |  |
| 6 | SER | OG | A0:15 | 3.78 | 1.00 | 50.88 |  |
| 7 | C | OP2 | AA:2710 | 3.99 | 1.00 | 49.67 |  |
| 8 | A | OP1 | AA:2711 | 4.06 | 1.00 | 46.41 |  |
| 9 | G | O5' | AA:2714 | 4.08 | 1.00 | 57.09 |  |
| 10 | A | OP2 | AA:2713 | 4.09 | 1.00 | 46.80 |  |
| 11 | C | O5' | AA:2710 | 4.12 | 1.00 | 49.74 |  |
| 12 | A | OP1 | AA:2712A | 4.15 | 1.00 | 44.70 |  |
| 13 | C | O3' | AA:2710 | 4.26 | 1.00 | 47.28 |  |
| 14 | U | O3' | AA:2712 | 4.71 | 1.00 | 47.68 |  |
| 15 | A | O5' | AA:2711 | 4.81 | 1.00 | 46.17 |  |
| 16 | A | O5' | AA:2712A | 4.84 | 1.00 | 42.14 |  |
| 17 | A | O3' | AA:2713 | 4.85 | 1.00 | 54.15 |  |
  |

  
 Contact(s): /OP2/  
 Coded contact list: O_ph_  
 Exclusion zone contact(s): /OP2/OP2/  
Consider NA in place of MG (2 valid contact(s) for NA vs. 1 for MG)  

### MG:AA:3002

|  | Res. type | Atom | Residue | Distance | Occupancy | B-factor | Closest solute to 1^st^ shell water |
| --- | --- | --- | --- | --- | --- | --- | --- |
| **Ion** | | | |  | 1.00 | 14.47 || 1 | G | OP2 | AA:1776 | 1.98 | 1.00 | 43.90 |
| 2 | U | OP2 | AA:1775 | 3.72 | 1.00 | 41.22 |  |
| 3 | G | N7 | AA:1776 | 3.75 | 1.00 | 37.24 |  |
| 4 | G | OP1 | AA:1776 | 3.99 | 1.00 | 44.37 |  |
| 5 | U | OP1 | AA:1775 | 4.14 | 1.00 | 41.03 |  |
| 6 | U | O3' | AA:1775 | 4.17 | 1.00 | 43.07 |  |
| 7 | A | OP1 | AA:1981 | 4.37 | 1.00 | 40.05 |  |
| 8 | G | O5' | AA:1776 | 4.44 | 1.00 | 41.21 |  |
| 9 | U | O5' | AA:1775 | 4.64 | 1.00 | 40.29 |  |
  |

  
 Contact(s): /OP2/  
 Coded contact list: O_ph_  

### MG:AA:3003

|  | Res. type | Atom | Residue | Distance | Occupancy | B-factor | Closest solute to 1^st^ shell water |
| --- | --- | --- | --- | --- | --- | --- | --- |
| **Ion** | | | |  | 1.00 | 12.87 || 1 | A | OP2 | AA:1970 | 2.44 | 1.00 | 39.18 |
| 2 | A | OP2 | AA:1900 | 3.87 | 1.00 | 47.86 |  |
| 3 | A | OP1 | AA:1970 | 4.07 | 1.00 | 38.39 |  |
| 4 | C | N4 | AA:1836 | 4.36 | 1.00 | 50.31 |  |
| 5 | G | OP2 | AA:1835 | 4.43 | 1.00 | 46.23 |  |
| 6 | G | N7 | AA:1835 | 4.55 | 1.00 | 47.00 |  |
| 7 | A | O5' | AA:1970 | 4.60 | 1.00 | 36.12 |  |
| 8 | G | O6 | AA:1835 | 4.63 | 1.00 | 45.11 |  |
| 9 | G | O2' | AA:1899 | 4.66 | 1.00 | 51.35 |  |
| 10 | A | O3' | AA:1969 | 4.81 | 1.00 | 37.41 |  |
| 11 | U | O2' | AA:1834 | 4.86 | 1.00 | 45.35 |  |
  |

  
 Coded contact list:   
 Exclusion zone contact(s): /OP2/  
Consider NA in place of MG (1 valid contact(s) for NA vs. 0 for MG)  

### MG:AA:3004

|  | Res. type | Atom | Residue | Distance | Occupancy | B-factor | Closest solute to 1^st^ shell water |
| --- | --- | --- | --- | --- | --- | --- | --- |
| **Ion** | | | |  | 1.00 | 7.98 || 1 | G | N7 | AA:733 | 2.19 | 1.00 | 41.88 |
| 2 | A | N7 | AA:761 | 2.47 | 1.00 | 42.79 |  |
| 3 | G | O6 | AA:733 | 3.24 | 1.00 | 45.77 |  |
| 4 | G | OP2 | AA:733 | 3.44 | 1.00 | 41.17 |  |
| 5 | A | OP2 | AA:761 | 3.85 | 1.00 | 38.67 |  |
| 6 | A | N6 | AA:761 | 4.03 | 1.00 | 46.73 |  |
| 7 | C | OP2 | AA:732 | 4.12 | 1.00 | 41.80 |  |
| 8 | G | N9 | AA:733 | 4.40 | 1.00 | 40.11 |  |
| 9 | A | N9 | AA:761 | 4.48 | 1.00 | 41.08 |  |
| 10 | C | O5' | AA:732 | 4.71 | 1.00 | 42.40 |  |
| 11 | G | OP2 | AA:760 | 4.82 | 1.00 | 40.28 |  |
| 12 | G | O5' | AA:733 | 4.88 | 1.00 | 42.03 |  |
| 13 | G | N1 | AA:733 | 4.93 | 1.00 | 41.21 |  |
  |

  
 Contact(s): /N7/  
 Coded contact list: N_b_  
 Exclusion zone contact(s): /N7/O6/  

### MG:AA:3005

|  | Res. type | Atom | Residue | Distance | Occupancy | B-factor | Closest solute to 1^st^ shell water |
| --- | --- | --- | --- | --- | --- | --- | --- |
| **Ion** | | | |  | 1.00 | 17.63 || 1 | G | OP1 | AA:1828 | 1.83 | 1.00 | 28.05 |
| 2 | G | O5' | AA:1828 | 3.20 | 1.00 | 26.56 |  |
| 3 | A | OP2 | AA:1773 | 3.47 | 1.00 | 39.04 |  |
| 4 | G | OP2 | AA:1828 | 3.64 | 1.00 | 25.52 |  |
| 5 | G | OP1 | AA:1772 | 4.15 | 1.00 | 38.79 |  |
| 6 | C | O3' | AA:1827 | 4.21 | 1.00 | 25.12 |  |
| 7 | A | OP2 | AA:1972 | 4.46 | 1.00 | 43.61 |  |
| 8 | G | OP2 | AA:1973 | 4.59 | 1.00 | 34.83 |  |
  |

  
 Contact(s): /OP1/  
 Coded contact list: O_ph_  
 Exclusion zone contact(s): /O5'/  

### MG:AA:3006

|  | Res. type | Atom | Residue | Distance | Occupancy | B-factor | Closest solute to 1^st^ shell water |
| --- | --- | --- | --- | --- | --- | --- | --- |
| **Ion** | | | |  | 1.00 | 14.41 || 1 | U | OP1 | AA:787 | 1.86 | 1.00 | 32.04 |
| 2 | C | OP2 | AA:790 | 1.93 | 1.00 | 44.16 |  |
| 3 | C | O3' | AA:786 | 3.78 | 1.00 | 31.85 |  |
| 4 | U | OP2 | AA:787 | 3.80 | 1.00 | 31.77 |  |
| 5 | A | N6 | AA:1780 | 3.95 | 1.00 | 55.54 |  |
| 6 | C | O5' | AA:790 | 4.05 | 1.00 | 43.39 |  |
| 7 | C | OP1 | AA:790 | 4.10 | 1.00 | 42.03 |  |
| 8 | A | N7 | AA:1780 | 4.16 | 1.00 | 49.88 |  |
| 9 | U | O5' | AA:787 | 4.39 | 1.00 | 29.49 |  |
| 10 | A | O3' | AA:789 | 4.44 | 1.00 | 47.01 |  |
| 11 | A | O3' | AA:788 | 4.79 | 1.00 | 40.33 |  |
  |

  
**1^st^ shell ligand angles:**   
1-2: 154;   
  
 Contact(s): /OP1/OP2/  
 Coded contact list: 2O_ph_  

### MG:AA:3007

|  | Res. type | Atom | Residue | Distance | Occupancy | B-factor | Closest solute to 1^st^ shell water |
| --- | --- | --- | --- | --- | --- | --- | --- |
| **Ion** | | | |  | 1.00 | 24.26 || 1 | A | OP1 | AA:1970 | 2.11 | 1.00 | 38.39 |
| 2 | A | OP1 | AA:1971 | 2.40 | 1.00 | 37.15 |  |
| 3 | A | O5' | AA:1970 | 3.69 | 1.00 | 36.12 |  |
| 4 | A | OP1 | AA:1969 | 4.02 | 1.00 | 36.60 |  |
| 5 | A | O3' | AA:1969 | 4.26 | 1.00 | 37.41 |  |
| 6 | A | O5' | AA:1971 | 4.51 | 1.00 | 39.24 |  |
| 7 | A | OP2 | AA:1970 | 4.58 | 1.00 | 39.18 |  |
| 8 | A | OP2 | AA:1971 | 4.68 | 1.00 | 37.24 |  |
| 9 | A | O3' | AA:1970 | 4.84 | 1.00 | 33.45 |  |
  |

  
 Contact(s): /OP1/  
 Coded contact list: O_ph_  
 Exclusion zone contact(s): /OP1/  

### MG:AA:3008

|  | Res. type | Atom | Residue | Distance | Occupancy | B-factor | Closest solute to 1^st^ shell water |
| --- | --- | --- | --- | --- | --- | --- | --- |
| **Ion** | | | |  | 1.00 | 31.20 || 1 | A | OP1 | AA:2005 | 1.92 | 1.00 | 48.37 |
| 2 | A | OP2 | AA:1269 | 2.51 | 1.00 | 44.25 |  |
| 3 | C | OP2 | AA:2006 | 3.64 | 1.00 | 48.20 |  |
| 4 | A | O5' | AA:2005 | 4.01 | 1.00 | 49.73 |  |
| 5 | A | OP2 | AA:2005 | 4.17 | 1.00 | 47.71 |  |
| 6 | A | OP2 | AA:1268 | 4.33 | 1.00 | 49.16 |  |
| 7 | A | OP1 | AA:1269 | 4.35 | 1.00 | 43.86 |  |
| 8 | G | O3' | AA:2004 | 4.39 | 1.00 | 48.08 |  |
| 9 | A | N7 | AA:1269 | 4.50 | 1.00 | 44.84 |  |
| 10 | A | O3' | AA:1268 | 4.53 | 1.00 | 43.39 |  |
| 11 | A | O5' | AA:1268 | 4.87 | 1.00 | 48.77 |  |
| 12 | A | OP1 | AA:1268 | 4.96 | 1.00 | 48.98 |  |
| 13 | A | O5' | AA:1269 | 4.99 | 1.00 | 40.34 |  |
  |

  
 Contact(s): /OP1/  
 Coded contact list: O_ph_  
 Exclusion zone contact(s): /OP2/  

### MG:AA:3009

|  | Res. type | Atom | Residue | Distance | Occupancy | B-factor | Closest solute to 1^st^ shell water |
| --- | --- | --- | --- | --- | --- | --- | --- |
| **Ion** | | | |  | 1.00 | 18.07 || 1 | C | OP2 | AA:1315 | 1.71 | 1.00 | 46.02 |
| 2 | C | OP1 | AA:1314 | 1.98 | 1.00 | 40.52 |  |
| 3 | G | OP1 | AA:1332 | 2.15 | 1.00 | 53.63 |  |
| 4 | C | O5' | AA:1314 | 3.24 | 1.00 | 41.13 |  |
| 5 | G | O5' | AA:1332 | 3.37 | 1.00 | 54.28 |  |
| 6 | C | OP1 | AA:1315 | 3.70 | 1.00 | 46.93 |  |
| 7 | C | O3' | AA:1314 | 3.95 | 1.00 | 44.97 |  |
| 8 | C | OP2 | AA:1314 | 4.03 | 1.00 | 41.16 |  |
| 9 | C | O5' | AA:1315 | 4.17 | 1.00 | 47.67 |  |
| 10 | U | O3' | AA:1313 | 4.31 | 1.00 | 41.21 |  |
| 11 | A | O3' | AA:1331 | 4.34 | 1.00 | 50.29 |  |
| 12 | G | OP2 | AA:1332 | 4.41 | 1.00 | 52.77 |  |
  |

  
**1^st^ shell ligand angles:**   
1-2: 111; 1-3: 147; 2-3: 90;   
  
 Contact(s): /OP1/OP1/  
 Coded contact list: 2O_ph_  
 Exclusion zone contact(s): /O5'/O5'/OP2/  

### MG:AA:3010

|  | Res. type | Atom | Residue | Distance | Occupancy | B-factor | Closest solute to 1^st^ shell water |
| --- | --- | --- | --- | --- | --- | --- | --- |
| **Ion** | | | |  | 1.00 | 19.48 || 1 | G | O6 | AA:450 | 2.12 | 1.00 | 45.14 |
| 2 | C | OP1 | AA:453 | 2.12 | 1.00 | 43.11 |  |
| 3 | C | O5' | AA:453 | 3.84 | 1.00 | 43.86 |  |
| 4 | G | N7 | AA:450 | 3.93 | 1.00 | 44.83 |  |
| 5 | G | N1 | AA:450 | 4.38 | 1.00 | 44.24 |  |
| 6 | C | OP2 | AA:453 | 4.40 | 1.00 | 42.27 |  |
| 7 | A | N7 | AA:449 | 4.44 | 1.00 | 40.36 |  |
| 8 | A | OP1 | AA:454 | 4.56 | 1.00 | 44.39 |  |
| 9 | G | O3' | AA:452 | 4.56 | 1.00 | 41.50 |  |
| 10 | A | N6 | AA:449 | 4.59 | 1.00 | 38.01 |  |
| 11 | A | OP1 | AA:471 | 4.92 | 1.00 | 38.10 |  |
  |

  
**1^st^ shell ligand angles:**   
1-2: 75;   
  
 Contact(s): /O6/OP1/  
 Coded contact list: O_ph_O_b_  

### MG:AA:3011

|  | Res. type | Atom | Residue | Distance | Occupancy | B-factor | Closest solute to 1^st^ shell water |
| --- | --- | --- | --- | --- | --- | --- | --- |
| **Ion** | | | |  | 1.00 | 17.42 || 1 | U | OP1 | AA:2074 | 2.63 | 1.00 | 26.02 |
| 2 | G | OP2 | AA:2239 | 3.19 | 1.00 | 34.50 |  |
| 3 | C | OP2 | AA:2240 | 3.39 | 1.00 | 39.91 |  |
| 4 | G | O3' | AA:2238 | 3.55 | 1.00 | 35.58 |  |
| 5 | G | O2' | AA:2238 | 3.88 | 1.00 | 38.20 |  |
| 6 | G | O5' | AA:2239 | 4.03 | 1.00 | 35.93 |  |
| 7 | ARG | NH2 | AD:244 | 4.30 | 1.00 | 44.20 |  |
| 8 | U | OP2 | AA:2075 | 4.32 | 1.00 | 32.48 |  |
| 9 | PRO | O | AD:228 | 4.44 | 1.00 | 32.52 |  |
| 10 | U | O5' | AA:2074 | 4.62 | 1.00 | 26.91 |  |
| 11 | U | OP2 | AA:2074 | 4.65 | 1.00 | 26.90 |  |
| 12 | C | OP1 | AA:2240 | 4.89 | 1.00 | 40.35 |  |
| 13 | G | O3' | AA:2239 | 4.97 | 1.00 | 39.88 |  |
  |

  
 Coded contact list:   
 Exclusion zone contact(s): /OP1/OP2/OP2/  
Consider K in place of MG (3 valid contact(s) for K vs. 0 for MG)  

### MG:AA:3012

|  | Res. type | Atom | Residue | Distance | Occupancy | B-factor | Closest solute to 1^st^ shell water |
| --- | --- | --- | --- | --- | --- | --- | --- |
| **Ion** | | | |  | 1.00 | 21.99 || 1 | C | OP2 | AA:1327 | 2.10 | 1.00 | 39.55 |
| 2 | U | OP1 | AA:1326 | 3.94 | 1.00 | 39.88 |  |
| 3 | C | O5' | AA:1327 | 4.07 | 1.00 | 43.17 |  |
| 4 | G | N7 | AA:1324 | 4.12 | 1.00 | 54.16 |  |
| 5 | G | N7 | AA:1328 | 4.20 | 1.00 | 58.33 |  |
| 6 | C | OP1 | AA:1327 | 4.27 | 1.00 | 40.27 |  |
| 7 | U | OP2 | AA:1326 | 4.58 | 1.00 | 41.19 |  |
| 8 | G | OP2 | AA:1328 | 4.60 | 1.00 | 48.23 |  |
| 9 | U | O3' | AA:1326 | 4.65 | 1.00 | 39.67 |  |
| 10 | G | O6 | AA:1328 | 4.84 | 1.00 | 58.10 |  |
  |

  
 Contact(s): /OP2/  
 Coded contact list: O_ph_  

### MG:AA:3013

|  | Res. type | Atom | Residue | Distance | Occupancy | B-factor | Closest solute to 1^st^ shell water |
| --- | --- | --- | --- | --- | --- | --- | --- |
| **Ion** | | | |  | 1.00 | 49.26 || 1 | C | OP2 | AA:1830 | 2.17 | 1.00 | 35.58 |
| 2 | G | O2' | AA:1792 | 4.32 | 1.00 | 37.28 |  |
| 3 | A | O3' | AA:1829 | 4.37 | 1.00 | 38.69 |  |
| 4 | C | O5' | AA:1830 | 4.37 | 1.00 | 36.99 |  |
| 5 | A | O5' | AA:1829 | 4.46 | 1.00 | 37.34 |  |
| 6 | G | N2 | AA:1792 | 4.52 | 1.00 | 24.40 |  |
| 7 | G | N3 | AA:1792 | 4.56 | 1.00 | 29.39 |  |
| 8 | A | OP1 | AA:1829 | 4.62 | 1.00 | 37.92 |  |
| 9 | C | O4' | AA:1793 | 4.66 | 1.00 | 31.73 |  |
| 10 | C | OP1 | AA:1830 | 4.66 | 1.00 | 37.15 |  |
| 11 | A | N1 | AA:1970 | 4.68 | 1.00 | 31.71 |  |
| 12 | G | O3' | AA:1828 | 4.83 | 1.00 | 35.80 |  |
  |

  
 Contact(s): /OP2/  
 Coded contact list: O_ph_  

### MG:AA:3014

|  | Res. type | Atom | Residue | Distance | Occupancy | B-factor | Closest solute to 1^st^ shell water |
| --- | --- | --- | --- | --- | --- | --- | --- |
| **Ion** | | | |  | 1.00 | 20.98 || 1 | C | OP1 | AA:192 | 1.79 | 1.00 | 36.21 |
| 2 | U | OP1 | AA:2243 | 2.39 | 1.00 | 30.71 |  |
| 3 | C | O5' | AA:192 | 3.60 | 1.00 | 35.55 |  |
| 4 | C | OP2 | AA:192 | 3.89 | 1.00 | 35.85 |  |
| 5 | U | O5' | AA:2243 | 4.00 | 1.00 | 32.87 |  |
| 6 | G | O3' | AA:2242 | 4.22 | 1.00 | 30.47 |  |
| 7 | A | O3' | AA:191 | 4.30 | 1.00 | 32.85 |  |
| 8 | A | OP1 | AA:199 | 4.35 | 1.00 | 40.89 |  |
| 9 | U | OP2 | AA:193 | 4.38 | 1.00 | 39.21 |  |
| 10 | A | OP2 | AA:199 | 4.80 | 1.00 | 41.52 |  |
| 11 | U | OP2 | AA:2243 | 4.94 | 1.00 | 31.15 |  |
  |

  
**1^st^ shell ligand angles:**   
1-2: 93;   
  
 Contact(s): /OP1/  
 Coded contact list: O_ph_  
 Exclusion zone contact(s): /OP1/  

### MG:AA:3015

|  | Res. type | Atom | Residue | Distance | Occupancy | B-factor | Closest solute to 1^st^ shell water |
| --- | --- | --- | --- | --- | --- | --- | --- |
| **Ion** | | | |  | 1.00 | 19.07 || 1 | A | OP1 | AA:2503 | 2.12 | 1.00 | 39.95 |
| 2 | U | OP1 | AA:576 | 2.21 | 1.00 | 47.22 |  |
| 3 | A | OP2 | AA:2503 | 3.16 | 1.00 | 37.97 |  |
| 4 | A | O5' | AA:2503 | 3.55 | 1.00 | 38.68 |  |
| 5 | U | OP2 | AA:2504 | 3.75 | 1.00 | 42.06 |  |
| 6 | A | O3' | AA:575 | 4.03 | 1.00 | 48.35 |  |
| 7 | U | OP2 | AA:576 | 4.24 | 1.00 | 46.55 |  |
| 8 | G | O3' | AA:2502 | 4.38 | 1.00 | 36.89 |  |
| 9 | U | O5' | AA:576 | 4.71 | 1.00 | 44.78 |  |
| 10 | G | OP1 | AA:2056 | 4.75 | 1.00 | 43.67 |  |
| 11 | G | O2' | AA:2502 | 4.85 | 1.00 | 39.42 |  |
| 12 | A | O3' | AA:2503 | 4.97 | 1.00 | 41.19 |  |
  |

  
**1^st^ shell ligand angles:**   
1-2: 74;   
  
 Contact(s): /OP1/OP1/  
 Coded contact list: 2O_ph_  
 Exclusion zone contact(s): /OP2/  

### MG:AA:3016

|  | Res. type | Atom | Residue | Distance | Occupancy | B-factor | Closest solute to 1^st^ shell water |
| --- | --- | --- | --- | --- | --- | --- | --- |
| **Ion** | | | |  | 1.00 | 9.25 || 1 | HIS | O | AD:220 | 2.43 | 1.00 | 38.43 |
| 2 | A | OP1 | AA:1789 | 2.56 | 1.00 | 36.53 |  |
| 3 | C | OP2 | AA:1790 | 2.68 | 1.00 | 34.82 |  |
| 4 | G | O6 | AA:1828 | 2.96 | 1.00 | 39.46 |  |
| 5 | ARG | N | AD:222 | 3.55 | 1.00 | 31.36 |  |
| 6 | A | O5' | AA:1789 | 3.70 | 1.00 | 35.95 |  |
| 7 | G | N1 | AA:1828 | 3.93 | 1.00 | 34.80 |  |
| 8 | A | OP2 | AA:1789 | 4.07 | 1.00 | 36.56 |  |
| 9 | VAL | N | AD:221 | 4.25 | 1.00 | 34.95 |  |
| 10 | ARG | NE | AD:222 | 4.38 | 1.00 | 29.07 |  |
| 11 | A | O3' | AA:1789 | 4.46 | 1.00 | 36.17 |  |
| 12 | C | OP1 | AA:1790 | 4.54 | 1.00 | 34.99 |  |
| 13 | C | N4 | AA:1827 | 4.63 | 1.00 | 28.18 |  |
| 14 | VAL | O | AD:221 | 4.92 | 1.00 | 35.33 |  |
| 15 | C | O3' | AA:1788 | 4.95 | 1.00 | 36.32 |  |
  |

  
 Coded contact list:   
 Exclusion zone contact(s): /O6/O_back_/OP1/OP2/  
Consider NA in place of MG (2 valid contact(s) for NA vs. 0 for MG)  

### MG:AA:3017

|  | Res. type | Atom | Residue | Distance | Occupancy | B-factor | Closest solute to 1^st^ shell water |
| --- | --- | --- | --- | --- | --- | --- | --- |
| **Ion** | | | |  | 1.00 | 29.20 || 1 | C | OP2 | AA:1333 | 3.55 | 1.00 | 55.88 |
| 2 | U | O4 | AA:1335 | 4.16 | 1.00 | 60.03 |  |
| 3 | G | OP2 | AA:1332 | 4.24 | 1.00 | 52.77 |  |
| 4 | G | OP2 | AA:1334 | 4.31 | 1.00 | 53.21 |  |
| 5 | U | OP2 | AA:1313 | 4.34 | 1.00 | 43.84 |  |
| 6 | G | N7 | AA:1334 | 4.68 | 1.00 | 58.92 |  |
| 7 | G | O6 | AA:1334 | 4.87 | 1.00 | 61.56 |  |
| 8 | A | N6 | AA:1336 | 4.88 | 1.00 | 54.73 |  |
| 9 | U | O4 | AA:1316 | 4.99 | 1.00 | 60.66 |  |
  |

  
 Coded contact list:   
1^st^ MG coordination shell missing (see Notes)  

### MG:AA:3018

|  | Res. type | Atom | Residue | Distance | Occupancy | B-factor | Closest solute to 1^st^ shell water |
| --- | --- | --- | --- | --- | --- | --- | --- |
| **Ion** | | | |  | 1.00 | 22.01 || 1 | A | OP2 | AA:586 | 2.50 | 1.00 | 46.70 |
| 2 | U | OP1 | AA:1249 | 2.63 | 1.00 | 53.65 |  |
| 3 | A | OP1 | AA:586 | 3.58 | 1.00 | 44.38 |  |
| 4 | U | O5' | AA:1249 | 3.65 | 1.00 | 52.84 |  |
| 5 | G | O3' | AA:1248 | 3.79 | 1.00 | 51.44 |  |
| 6 | A | O5' | AA:586 | 3.87 | 1.00 | 44.85 |  |
| 7 | G | OP1 | AA:585 | 4.26 | 1.00 | 49.04 |  |
| 8 | G | OP1 | AA:1250 | 4.41 | 1.00 | 50.33 |  |
| 9 | U | O4 | AA:811 | 4.60 | 1.00 | 52.02 |  |
| 10 | G | O2' | AA:1248 | 4.74 | 1.00 | 52.27 |  |
| 11 | C | OP1 | AA:587 | 4.81 | 1.00 | 40.32 |  |
| 12 | G | O3' | AA:585 | 4.82 | 1.00 | 44.96 |  |
| 13 | U | OP2 | AA:1249 | 4.99 | 1.00 | 53.61 |  |
  |

  
 Coded contact list:   
 Exclusion zone contact(s): /OP1/OP2/  
Consider NA in place of MG (1 valid contact(s) for NA vs. 0 for MG)  

### MG:AA:3019

|  | Res. type | Atom | Residue | Distance | Occupancy | B-factor | Closest solute to 1^st^ shell water |
| --- | --- | --- | --- | --- | --- | --- | --- |
| **Ion** | | | |  | 1.00 | 31.43 || 1 | U | O4 | AA:2243 | 3.37 | 1.00 | 33.32 |
| 2 | G | O6 | AA:2242 | 3.83 | 1.00 | 31.64 |  |
| 3 | G | N7 | AA:2242 | 4.15 | 1.00 | 28.35 |  |
| 4 | U | O4 | AA:2074 | 4.34 | 1.00 | 30.11 |  |
| 5 | A | N6 | AA:2077 | 4.39 | 1.00 | 35.50 |  |
| 6 | U | O4 | AA:2075 | 4.42 | 1.00 | 35.29 |  |
| 7 | A | N7 | AA:2241 | 4.44 | 1.00 | 31.39 |  |
| 8 | G | N7 | AA:2238 | 4.51 | 1.00 | 38.77 |  |
| 9 | A | N6 | AA:2241 | 4.84 | 1.00 | 33.60 |  |
  |

  
 Coded contact list:   
 Exclusion zone contact(s): /O4/  
Consider K in place of MG (1 valid contact(s) for K vs. 0 for MG)  

### MG:AA:3020

|  | Res. type | Atom | Residue | Distance | Occupancy | B-factor | Closest solute to 1^st^ shell water |
| --- | --- | --- | --- | --- | --- | --- | --- |
| **Ion** | | | |  | 1.00 | 27.39 || 1 | G | O6 | AA:2009 | 3.57 | 1.00 | 47.48 |
| 2 | G | O6 | AA:2010 | 3.64 | 1.00 | 49.94 |  |
| 3 | G | N7 | AA:2009 | 3.77 | 1.00 | 49.28 |  |
| 4 | G | N7 | AA:2010 | 3.78 | 1.00 | 49.08 |  |
| 5 | U | O4 | AA:2011 | 4.14 | 1.00 | 43.80 |  |
| 6 | U | OP2 | AA:1267 | 4.36 | 1.00 | 45.51 |  |
| 7 | C | N4 | AA:2008 | 4.37 | 1.00 | 47.05 |  |
  |

  
 Coded contact list:   
1^st^ MG coordination shell missing (see Notes)  

### MG:AA:3021

|  | Res. type | Atom | Residue | Distance | Occupancy | B-factor | Closest solute to 1^st^ shell water |
| --- | --- | --- | --- | --- | --- | --- | --- |
| **Ion** | | | |  | 1.00 | 26.52 || 1 | G | OP1 | AA:741 | 2.55 | 1.00 | 43.39 |
| 2 | C | OP1 | AA:1983 | 3.59 | 1.00 | 38.96 |  |
| 3 | A | OP1 | AA:1784 | 3.65 | 1.00 | 45.43 |  |
| 4 | A | OP1 | AA:1785 | 3.68 | 1.00 | 46.06 |  |
| 5 | U | O3' | AA:740 | 3.84 | 1.00 | 41.16 |  |
| 6 | G | O3' | AA:2607 | 4.24 | 1.00 | 38.83 |  |
| 7 | C | O3' | AA:1982 | 4.52 | 1.00 | 39.84 |  |
| 8 | G | O5' | AA:741 | 4.71 | 1.00 | 41.00 |  |
| 9 | G | OP1 | AA:2608 | 4.75 | 1.00 | 38.19 |  |
| 10 | G | OP2 | AA:741 | 4.85 | 1.00 | 43.04 |  |
| 11 | G | O2' | AA:2607 | 4.91 | 1.00 | 36.54 |  |
  |

  
 Coded contact list:   
 Exclusion zone contact(s): /OP1/  
Consider NA in place of MG (1 valid contact(s) for NA vs. 0 for MG)  

### MG:AA:3022

|  | Res. type | Atom | Residue | Distance | Occupancy | B-factor | Closest solute to 1^st^ shell water |
| --- | --- | --- | --- | --- | --- | --- | --- |
| **Ion** | | | |  | 1.00 | 63.57 || 1 | G | O6 | AA:1542 | 2.24 | 1.00 | 103.33 |
| 2 | A | OP2 | AA:1528 | 2.93 | 1.00 | 84.53 |  |
| 3 | A | N6 | AA:1543 | 3.59 | 1.00 | 103.66 |  |
| 4 | A | OP1 | AA:1528 | 3.64 | 1.00 | 84.62 |  |
| 5 | G | N1 | AA:1542 | 3.65 | 1.00 | 104.13 |  |
| 6 | G | O3' | AA:1527 | 4.43 | 1.00 | 84.94 |  |
| 7 | G | O5' | AA:1527 | 4.51 | 1.00 | 83.34 |  |
| 8 | U | O4 | AA:1541 | 4.61 | 1.00 | 108.81 |  |
| 9 | G | OP2 | AA:1527 | 4.88 | 1.00 | 81.81 |  |
  |

  
 Contact(s): /O6/  
 Coded contact list: O_b_  
 Exclusion zone contact(s): /OP2/  

### MG:AA:3023

|  | Res. type | Atom | Residue | Distance | Occupancy | B-factor | Closest solute to 1^st^ shell water |
| --- | --- | --- | --- | --- | --- | --- | --- |
| **Ion** | | | |  | 1.00 | 59.18 || 1 | A | OP2 | AA:311 | 1.87 | 1.00 | 65.21 |
| 2 | A | O5' | AA:311 | 4.13 | 1.00 | 66.06 |  |
| 3 | A | O3' | AA:310 | 4.14 | 1.00 | 62.01 |  |
| 4 | U | O4 | AA:303 | 4.19 | 1.00 | 69.95 |  |
| 5 | G | O6 | AA:304 | 4.20 | 1.00 | 65.68 |  |
| 6 | A | OP1 | AA:311 | 4.32 | 1.00 | 62.78 |  |
| 7 | U | O4 | AA:305 | 4.42 | 1.00 | 56.60 |  |
| 8 | G | N7 | AA:304 | 4.83 | 1.00 | 66.23 |  |
| 9 | A | O2' | AA:310 | 4.89 | 1.00 | 61.13 |  |
  |

  
 Contact(s): /OP2/  
 Coded contact list: O_ph_  

### MG:AA:3024

|  | Res. type | Atom | Residue | Distance | Occupancy | B-factor | Closest solute to 1^st^ shell water |
| --- | --- | --- | --- | --- | --- | --- | --- |
| **Ion** | | | |  | 1.00 | 61.85 || 1 | A | O3' | AA:311 | 2.62 | 1.00 | 68.24 |
| 2 | G | OP1 | AA:312 | 2.87 | 1.00 | 69.28 |  |
| 3 | G | O5' | AA:312 | 3.33 | 1.00 | 65.29 |  |
| 4 | G | OP2 | AA:312 | 4.54 | 1.00 | 69.20 |  |
| 5 | A | O2' | AA:311 | 4.72 | 1.00 | 72.47 |  |
| 6 | A | O4' | AA:332 | 4.72 | 1.00 | 55.45 |  |
  |

  
 Coded contact list:   
 Exclusion zone contact(s): /O3'/O5'/OP1/  
Consider K in place of MG (3 valid contact(s) for K vs. 0 for MG)  

### MG:AA:3025

|  | Res. type | Atom | Residue | Distance | Occupancy | B-factor | Closest solute to 1^st^ shell water |
| --- | --- | --- | --- | --- | --- | --- | --- |
| **Ion** | | | |  | 1.00 | 38.77 || 1 | G | O6 | AA:301 | 3.87 | 1.00 | 66.93 |
| 2 | G | N7 | AA:301 | 4.04 | 1.00 | 65.31 |  |
| 3 | C | N4 | AA:302 | 4.41 | 1.00 | 68.85 |  |
| 4 | C | N4 | AA:334 | 4.68 | 1.00 | 62.85 |  |
  |

  
 Coded contact list:   
1^st^ MG coordination shell missing (see Notes)  

### MG:AA:3026

|  | Res. type | Atom | Residue | Distance | Occupancy | B-factor | Closest solute to 1^st^ shell water |
| --- | --- | --- | --- | --- | --- | --- | --- |
| **Ion** | | | |  | 1.00 | 38.61 || 1 | G | O6 | AA:82 | 3.76 | 1.00 | 65.02 |
| 2 | G | N7 | AA:81 | 4.21 | 1.00 | 67.21 |  |
| 3 | G | O6 | AA:81 | 4.22 | 1.00 | 67.03 |  |
| 4 | G | N7 | AA:80 | 4.24 | 1.00 | 69.46 |  |
| 5 | G | O6 | AA:80 | 4.36 | 1.00 | 69.43 |  |
| 6 | A | OP1 | AA:103 | 4.53 | 1.00 | 63.75 |  |
  |

  
 Coded contact list:   
1^st^ MG coordination shell missing (see Notes)  

### MG:AA:3027

|  | Res. type | Atom | Residue | Distance | Occupancy | B-factor | Closest solute to 1^st^ shell water |
| --- | --- | --- | --- | --- | --- | --- | --- |
| **Ion** | | | |  | 1.00 | 60.92 || 1 | G | O6 | AA:1470 | 2.40 | 1.00 | 83.82 |
| 2 | U | OP2 | AA:1520 | 2.63 | 1.00 | 71.58 |  |
| 3 | G | N7 | AA:1470 | 3.62 | 1.00 | 82.21 |  |
| 4 | U | OP1 | AA:1520 | 4.19 | 1.00 | 72.04 |  |
| 5 | U | O5' | AA:1520 | 4.24 | 1.00 | 74.60 |  |
| 6 | G | N7 | AA:1521 | 4.29 | 1.00 | 91.70 |  |
| 7 | G | N1 | AA:1470 | 4.58 | 1.00 | 84.01 |  |
| 8 | A | N7 | AA:1469 | 4.72 | 1.00 | 82.92 |  |
  |

  
 Coded contact list:   
 Exclusion zone contact(s): /O6/OP2/  
Consider NA in place of MG (1 valid contact(s) for NA vs. 0 for MG)  

### MG:AA:3028

|  | Res. type | Atom | Residue | Distance | Occupancy | B-factor | Closest solute to 1^st^ shell water |
| --- | --- | --- | --- | --- | --- | --- | --- |
| **Ion** | | | |  | 1.00 | 15.00 || 1 | A | OP1 | AA:2711 | 2.38 | 1.00 | 46.41 |
| 2 | A | OP1 | AA:2712A | 2.50 | 1.00 | 44.70 |  |
| 3 | C | OP1 | AA:1297 | 3.07 | 1.00 | 49.97 |  |
| 4 | C | O3' | AA:2710 | 3.42 | 1.00 | 47.28 |  |
| 5 | U | O3' | AA:2712 | 3.95 | 1.00 | 47.68 |  |
| 6 | A | OP2 | AA:2711 | 4.49 | 1.00 | 45.22 |  |
| 7 | A | O5' | AA:2711 | 4.57 | 1.00 | 46.17 |  |
| 8 | A | OP2 | AA:2712A | 4.58 | 1.00 | 45.06 |  |
| 9 | C | O5' | AA:1297 | 4.73 | 1.00 | 48.68 |  |
| 10 | G | O3' | AA:1296 | 4.84 | 1.00 | 51.43 |  |
| 11 | A | O5' | AA:2712A | 4.93 | 1.00 | 42.14 |  |
| 12 | C | O3' | AA:1297 | 4.93 | 1.00 | 49.30 |  |
  |

  
 Contact(s): /OP1/  
 Coded contact list: O_ph_  
 Exclusion zone contact(s): /OP1/OP1/  

### MG:AA:3029

|  | Res. type | Atom | Residue | Distance | Occupancy | B-factor | Closest solute to 1^st^ shell water |
| --- | --- | --- | --- | --- | --- | --- | --- |
| **Ion** | | | |  | 1.00 | 46.59 || 1 | G | O3' | AA:452 | 2.61 | 1.00 | 41.50 |
| 2 | C | OP1 | AA:453 | 2.94 | 1.00 | 43.11 |  |
| 3 | A | OP1 | AA:471 | 2.95 | 1.00 | 38.10 |  |
| 4 | A | OP2 | AA:471 | 3.38 | 1.00 | 39.67 |  |
| 5 | G | O2' | AA:452 | 3.57 | 1.00 | 33.50 |  |
| 6 | C | O5' | AA:453 | 3.60 | 1.00 | 43.86 |  |
| 7 | TYR | OH | AF:59 | 3.94 | 1.00 | 50.32 |  |
| 8 | A | O3' | AA:470 | 3.94 | 1.00 | 37.70 |  |
| 9 | C | OP2 | AA:453 | 4.64 | 1.00 | 42.27 |  |
| 10 | A | O5' | AA:470 | 4.72 | 1.00 | 30.95 |  |
| 11 | A | OP1 | AA:470 | 4.76 | 1.00 | 31.55 |  |
| 12 | A | O5' | AA:471 | 5.00 | 1.00 | 38.73 |  |
  |

  
 Coded contact list:   
 Exclusion zone contact(s): /O3'/OP1/OP1/OP2/  
Consider K in place of MG (4 valid contact(s) for K vs. 0 for MG)  

### MG:AA:3030

|  | Res. type | Atom | Residue | Distance | Occupancy | B-factor | Closest solute to 1^st^ shell water |
| --- | --- | --- | --- | --- | --- | --- | --- |
| **Ion** | | | |  | 1.00 | 45.62 || 1 | G | O6 | AA:83 | 3.88 | 1.00 | 55.30 |
| 2 | G | N7 | AA:83 | 4.16 | 1.00 | 52.92 |  |
| 3 | G | N7 | AA:82 | 4.17 | 1.00 | 64.87 |  |
| 4 | G | OP2 | AA:82 | 4.44 | 1.00 | 66.75 |  |
| 5 | GLU | OE1 | AU:91 | 4.98 | 1.00 | 103.57 |  |
  |

  
 Coded contact list:   
1^st^ MG coordination shell missing (see Notes)  

### MG:AA:3031

|  | Res. type | Atom | Residue | Distance | Occupancy | B-factor | Closest solute to 1^st^ shell water |
| --- | --- | --- | --- | --- | --- | --- | --- |
| **Ion** | | | |  | 1.00 | 34.94 || 1 | C | OP1 | AA:76 | 4.20 | 1.00 | 62.57 |
| 2 | C | OP2 | AA:77 | 4.27 | 1.00 | 57.38 |  |
| 3 | U | OP1 | AA:104 | 4.28 | 1.00 | 66.60 |  |
| 4 | ARG | NH1 | AW:59 | 4.39 | 1.00 | 43.08 |  |
| 5 | C | OP1 | AA:77 | 4.49 | 1.00 | 57.12 |  |
  |

  
 Coded contact list:   
1^st^ MG coordination shell missing (see Notes)  

### MG:AA:3032

|  | Res. type | Atom | Residue | Distance | Occupancy | B-factor | Closest solute to 1^st^ shell water |
| --- | --- | --- | --- | --- | --- | --- | --- |
| **Ion** | | | |  | 1.00 | 28.73 || 1 | G | OP2 | AA:570 | 3.43 | 1.00 | 47.09 |
| 2 | A | N1 | AA:821 | 3.47 | 1.00 | 60.67 |  |
| 3 | G | O2' | AA:972 | 3.74 | 1.00 | 62.10 |  |
| 4 | U | O5' | AA:569 | 3.75 | 1.00 | 48.30 |  |
| 5 | U | OP2 | AA:569 | 3.76 | 1.00 | 47.59 |  |
| 6 | G | OP1 | AA:570 | 3.91 | 1.00 | 47.33 |  |
| 7 | U | O3' | AA:569 | 4.41 | 1.00 | 46.55 |  |
| 8 | A | N1 | AA:820 | 4.74 | 1.00 | 54.66 |  |
| 9 | A | N6 | AA:820 | 4.83 | 1.00 | 55.52 |  |
| 10 | A | N6 | AA:821 | 4.91 | 1.00 | 62.28 |  |
  |

  
 Coded contact list:   
1^st^ MG coordination shell missing (see Notes)  

### MG:AA:3033

|  | Res. type | Atom | Residue | Distance | Occupancy | B-factor | Closest solute to 1^st^ shell water |
| --- | --- | --- | --- | --- | --- | --- | --- |
| **Ion** | | | |  | 1.00 | 41.61 || 1 | A | OP2 | AA:973 | 1.87 | 1.00 | 64.50 |
| 2 | U | O4 | AA:568 | 1.94 | 1.00 | 53.74 |  |
| 3 | G | O3' | AA:972 | 3.65 | 1.00 | 63.57 |  |
| 4 | U | N3 | AA:568 | 3.70 | 1.00 | 51.76 |  |
| 5 | A | O5' | AA:973 | 4.07 | 1.00 | 64.69 |  |
| 6 | G | OP1 | AA:570 | 4.09 | 1.00 | 47.33 |  |
| 7 | A | OP1 | AA:571 | 4.17 | 1.00 | 48.97 |  |
| 8 | A | OP2 | AA:571 | 4.21 | 1.00 | 49.11 |  |
| 9 | A | OP1 | AA:973 | 4.38 | 1.00 | 63.50 |  |
| 10 | A | N6 | AA:567 | 4.80 | 1.00 | 42.68 |  |
| 11 | LYS | O | A2:78 | 4.98 | 1.00 | 41.39 |  |
  |

  
**1^st^ shell ligand angles:**   
1-2: 103;   
  
 Contact(s): /O4/OP2/  
 Coded contact list: O_ph_O_b_  

### MG:AA:3034

|  | Res. type | Atom | Residue | Distance | Occupancy | B-factor | Closest solute to 1^st^ shell water |
| --- | --- | --- | --- | --- | --- | --- | --- |
| **Ion** | | | |  | 1.00 | 41.23 || 1 | G | N7 | AA:2837 | 3.12 | 1.00 | 69.52 |
| 2 | G | N7 | AA:2838 | 3.20 | 1.00 | 68.35 |  |
| 3 | G | O6 | AA:2838 | 3.25 | 1.00 | 67.81 |  |
| 4 | G | O6 | AA:2837 | 3.53 | 1.00 | 69.24 |  |
| 5 | G | O6 | AA:2839 | 3.67 | 1.00 | 68.68 |  |
| 6 | A | N3 | AA:2835 | 3.79 | 1.00 | 71.44 |  |
| 7 | U | O4 | AA:2878 | 3.83 | 1.00 | 71.07 |  |
| 8 | A | O2' | AA:2835 | 4.46 | 1.00 | 78.12 |  |
| 9 | G | N9 | AA:2837 | 4.78 | 1.00 | 67.43 |  |
| 10 | G | N1 | AA:2837 | 4.81 | 1.00 | 68.81 |  |
| 11 | G | N7 | AA:2839 | 4.82 | 1.00 | 66.59 |  |
| 12 | A | N1 | AA:2835 | 4.85 | 1.00 | 69.72 |  |
| 13 | U | N3 | AA:2878 | 4.90 | 1.00 | 70.53 |  |
| 14 | G | OP2 | AA:2837 | 4.96 | 1.00 | 70.49 |  |
  |

  
 Coded contact list:   
 Exclusion zone contact(s): /N7/N7/O6/  
Consider K in place of MG (3 valid contact(s) for K vs. 0 for MG)  

### MG:AA:3035

|  | Res. type | Atom | Residue | Distance | Occupancy | B-factor | Closest solute to 1^st^ shell water |
| --- | --- | --- | --- | --- | --- | --- | --- |
| **Ion** | | | |  | 1.00 | 17.28 || 1 | G | O6 | AA:1842 | 2.54 | 1.00 | 44.70 |
| 2 | U | O4 | AA:1841 | 2.84 | 1.00 | 48.82 |  |
| 3 | C | O2' | AA:1838 | 2.88 | 1.00 | 64.48 |  |
| 4 | U | O4 | AA:1898 | 2.92 | 1.00 | 42.21 |  |
| 5 | C | O2 | AA:1838 | 3.06 | 1.00 | 59.90 |  |
| 6 | U | N3 | AA:1898 | 3.73 | 1.00 | 42.53 |  |
| 7 | A | N6 | AA:1901 | 3.96 | 1.00 | 46.25 |  |
| 8 | G | O6 | AA:1897 | 4.11 | 1.00 | 43.09 |  |
| 9 | G | N7 | AA:1842 | 4.20 | 1.00 | 39.30 |  |
| 10 | G | N1 | AA:1842 | 4.65 | 1.00 | 40.49 |  |
| 11 | C | N1 | AA:1838 | 4.66 | 1.00 | 61.43 |  |
| 12 | U | N3 | AA:1841 | 4.67 | 1.00 | 47.06 |  |
| 13 | G | OP2 | AA:1840 | 4.85 | 1.00 | 55.24 |  |
| 14 | C | N3 | AA:1838 | 4.91 | 1.00 | 60.73 |  |
  |

  
 Coded contact list:   
 Exclusion zone contact(s): /O2/O2'/O4/O4/O6/  
Consider K in place of MG (4 valid contact(s) for K vs. 0 for MG)  

### MG:AA:3036

|  | Res. type | Atom | Residue | Distance | Occupancy | B-factor | Closest solute to 1^st^ shell water |
| --- | --- | --- | --- | --- | --- | --- | --- |
| **Ion** | | | |  | 1.00 | 36.10 || 1 | U | O4 | AA:1898 | 2.89 | 1.00 | 42.21 |
| 2 | G | OP2 | AA:1839 | 2.95 | 1.00 | 61.68 |  |
| 3 | C | O3' | AA:1838 | 3.15 | 1.00 | 60.91 |  |
| 4 | C | O2' | AA:1838 | 3.54 | 1.00 | 64.48 |  |
| 5 | G | N7 | AA:1897 | 3.61 | 1.00 | 49.52 |  |
| 6 | G | OP2 | AA:1897 | 4.10 | 1.00 | 57.24 |  |
| 7 | G | OP1 | AA:1839 | 4.65 | 1.00 | 61.41 |  |
| 8 | G | O5' | AA:1839 | 4.80 | 1.00 | 58.66 |  |
| 9 | G | O6 | AA:1897 | 4.86 | 1.00 | 43.09 |  |
  |

  
 Coded contact list:   
 Exclusion zone contact(s): /O3'/O4/OP2/  
Consider K in place of MG (3 valid contact(s) for K vs. 0 for MG)  

### MG:AA:3037

|  | Res. type | Atom | Residue | Distance | Occupancy | B-factor | Closest solute to 1^st^ shell water |
| --- | --- | --- | --- | --- | --- | --- | --- |
| **Ion** | | | |  | 1.00 | 107.45 || 1 | A | OP1 | AA:2366 | 2.41 | 1.00 | 71.75 |
| 2 | C | OP1 | AA:2381 | 3.81 | 1.00 | 70.04 |  |
| 3 | G | OP1 | AA:2365 | 3.94 | 1.00 | 64.63 |  |
| 4 | A | OP2 | AA:2366 | 4.34 | 1.00 | 72.84 |  |
| 5 | G | O3' | AA:2365 | 4.69 | 1.00 | 70.76 |  |
| 6 | G | OP2 | AA:2382 | 4.69 | 1.00 | 74.15 |  |
| 7 | A | O5' | AA:2366 | 4.78 | 1.00 | 72.94 |  |
| 8 | G | O5' | AA:2365 | 4.87 | 1.00 | 63.67 |  |
  |

  
 Coded contact list:   
 Exclusion zone contact(s): /OP1/  
Consider NA in place of MG (1 valid contact(s) for NA vs. 0 for MG)  

### MG:AA:3038

|  | Res. type | Atom | Residue | Distance | Occupancy | B-factor | Closest solute to 1^st^ shell water |
| --- | --- | --- | --- | --- | --- | --- | --- |
| **Ion** | | | |  | 1.00 | 68.62 || 1 | A | OP2 | AA:2366 | 2.76 | 1.00 | 72.84 |
| 2 | A | O5' | AA:2366 | 3.48 | 1.00 | 72.94 |  |
| 3 | A | OP1 | AA:2366 | 3.80 | 1.00 | 71.75 |  |
| 4 | G | N7 | AA:2382 | 3.95 | 1.00 | 79.59 |  |
| 5 | G | N7 | AA:2367 | 4.05 | 1.00 | 71.23 |  |
| 6 | G | OP2 | AA:2367 | 4.08 | 1.00 | 74.10 |  |
| 7 | G | O6 | AA:2367 | 4.50 | 1.00 | 70.49 |  |
| 8 | G | O6 | AA:2382 | 4.97 | 1.00 | 77.90 |  |
| 9 | G | O3' | AA:2365 | 4.98 | 1.00 | 70.76 |  |
  |

  
 Coded contact list:   
 Exclusion zone contact(s): /OP2/  
Consider K in place of MG (1 valid contact(s) for K vs. 0 for MG)  

### MG:AA:3039

|  | Res. type | Atom | Residue | Distance | Occupancy | B-factor | Closest solute to 1^st^ shell water |
| --- | --- | --- | --- | --- | --- | --- | --- |
| **Ion** | | | |  | 1.00 | 30.45 || 1 | A | N6 | AA:300 | 3.48 | 1.00 | 62.16 |
| 2 | G | O6 | AA:338 | 4.08 | 1.00 | 63.29 |  |
| 3 | A | OP2 | AA:299 | 4.32 | 1.00 | 67.11 |  |
| 4 | C | OP2 | AA:336 | 4.35 | 1.00 | 58.61 |  |
| 5 | A | N7 | AA:300 | 4.36 | 1.00 | 58.49 |  |
| 6 | A | N7 | AA:299 | 4.39 | 1.00 | 60.91 |  |
| 7 | G | N7 | AA:338 | 4.64 | 1.00 | 62.73 |  |
  |

  
 Coded contact list:   
1^st^ MG coordination shell missing (see Notes)  

### MG:AA:3040

|  | Res. type | Atom | Residue | Distance | Occupancy | B-factor | Closest solute to 1^st^ shell water |
| --- | --- | --- | --- | --- | --- | --- | --- |
| **Ion** | | | |  | 1.00 | 37.88 || 1 | A | OP1 | AA:526 | 2.05 | 1.00 | 64.24 |
| 2 | C | OP1 | AA:527 | 2.15 | 1.00 | 58.74 |  |
| 3 | A | OP1 | AA:528 | 3.27 | 1.00 | 68.07 |  |
| 4 | A | O5' | AA:526 | 3.66 | 1.00 | 63.27 |  |
| 5 | C | OP2 | AA:527 | 4.04 | 1.00 | 59.49 |  |
| 6 | A | OP2 | AA:526 | 4.06 | 1.00 | 64.47 |  |
| 7 | C | O5' | AA:527 | 4.12 | 1.00 | 59.36 |  |
| 8 | U | O3' | AA:525 | 4.53 | 1.00 | 64.05 |  |
| 9 | A | O3' | AA:526 | 4.63 | 1.00 | 58.80 |  |
  |

  
**1^st^ shell ligand angles:**   
1-2: 87;   
  
 Contact(s): /OP1/OP1/  
 Coded contact list: 2O_ph_  
 Exclusion zone contact(s): /OP1/  

### MG:AA:3041

|  | Res. type | Atom | Residue | Distance | Occupancy | B-factor | Closest solute to 1^st^ shell water |
| --- | --- | --- | --- | --- | --- | --- | --- |
| **Ion** | | | |  | 1.00 | 81.30 || 1 | G | OP1 | AA:2694 | 2.48 | 1.00 | 60.19 |
| 2 | A | O3' | AA:2693 | 3.97 | 1.00 | 59.40 |  |
| 3 | A | O3' | AA:2850 | 4.15 | 1.00 | 64.66 |  |
| 4 | A | OP1 | AA:2851 | 4.68 | 1.00 | 66.20 |  |
| 5 | G | OP2 | AA:2694 | 4.71 | 1.00 | 59.32 |  |
| 6 | G | O5' | AA:2694 | 4.78 | 1.00 | 59.57 |  |
  |

  
 Coded contact list:   
 Exclusion zone contact(s): /OP1/  
Consider NA in place of MG (1 valid contact(s) for NA vs. 0 for MG)  

### MG:AA:3042

|  | Res. type | Atom | Residue | Distance | Occupancy | B-factor | Closest solute to 1^st^ shell water |
| --- | --- | --- | --- | --- | --- | --- | --- |
| **Ion** | | | |  | 1.00 | 35.53 || 1 | A | OP2 | AA:2850 | 3.15 | 1.00 | 66.80 |
| 2 | U | O4 | AA:2865 | 3.45 | 1.00 | 75.75 |  |
| 3 | U | O3' | AA:2849 | 3.61 | 1.00 | 67.63 |  |
| 4 | A | OP1 | AA:2850 | 3.88 | 1.00 | 65.60 |  |
| 5 | U | N3 | AA:2866 | 4.73 | 1.00 | 79.90 |  |
| 6 | U | O2' | AA:2849 | 4.86 | 1.00 | 72.64 |  |
| 7 | G | OP2 | AA:2864 | 5.00 | 1.00 | 78.93 |  |
  |

  
 Coded contact list:   
 Exclusion zone contact(s): /OP2/  
Consider K in place of MG (1 valid contact(s) for K vs. 0 for MG)  

### MG:AA:3043

|  | Res. type | Atom | Residue | Distance | Occupancy | B-factor | Closest solute to 1^st^ shell water |
| --- | --- | --- | --- | --- | --- | --- | --- |
| **Ion** | | | |  | 1.00 | 42.74 || 1 | U | O4 | AA:2866 | 1.97 | 1.00 | 78.20 |
| 2 | G | O6 | AA:2852 | 2.60 | 1.00 | 70.71 |  |
| 3 | A | OP2 | AA:2850 | 2.87 | 1.00 | 66.80 |  |
| 4 | A | OP1 | AA:2850 | 3.24 | 1.00 | 65.60 |  |
| 5 | A | N7 | AA:2851 | 3.44 | 1.00 | 69.07 |  |
| 6 | U | O4 | AA:2865 | 3.45 | 1.00 | 75.75 |  |
| 7 | A | OP2 | AA:2851 | 3.55 | 1.00 | 66.90 |  |
| 8 | U | N3 | AA:2866 | 3.71 | 1.00 | 79.90 |  |
| 9 | A | O5' | AA:2850 | 3.82 | 1.00 | 65.31 |  |
| 10 | U | N3 | AA:2865 | 3.98 | 1.00 | 78.24 |  |
| 11 | G | N7 | AA:2852 | 4.31 | 1.00 | 70.68 |  |
| 12 | A | N6 | AA:2851 | 4.35 | 1.00 | 68.40 |  |
| 13 | G | N1 | AA:2852 | 4.59 | 1.00 | 70.67 |  |
| 14 | U | O2' | AA:2849 | 4.82 | 1.00 | 72.64 |  |
| 15 | U | O3' | AA:2849 | 4.91 | 1.00 | 67.63 |  |
| 16 | A | N7 | AA:2850 | 4.92 | 1.00 | 65.93 |  |
| 17 | C | N4 | AA:2853 | 4.93 | 1.00 | 71.00 |  |
  |

  
 Contact(s): /O4/  
 Coded contact list: O_b_  
 Exclusion zone contact(s): /O6/OP1/OP2/  
Consider K in place of MG (2 valid contact(s) for K vs. 1 for MG)  

### MG:AA:3044

|  | Res. type | Atom | Residue | Distance | Occupancy | B-factor | Closest solute to 1^st^ shell water |
| --- | --- | --- | --- | --- | --- | --- | --- |
| **Ion** | | | |  | 1.00 | 36.57 || 1 | G | OP2 | AA:446 | 2.39 | 1.00 | 41.54 |
| 2 | G | OP1 | AA:446 | 3.94 | 1.00 | 39.75 |  |
| 3 | C | OP1 | AA:445 | 4.08 | 1.00 | 43.40 |  |
| 4 | U | O2' | AA:1199 | 4.24 | 1.00 | 51.63 |  |
| 5 | C | OP2 | AA:445 | 4.31 | 1.00 | 45.19 |  |
| 6 | U | O3' | AA:1199 | 4.33 | 1.00 | 49.85 |  |
| 7 | G | O5' | AA:446 | 4.52 | 1.00 | 43.95 |  |
| 8 | G | N7 | AA:446 | 4.66 | 1.00 | 60.16 |  |
| 9 | C | O3' | AA:445 | 4.74 | 1.00 | 40.57 |  |
| 10 | C | OP1 | AA:1200 | 4.77 | 1.00 | 48.33 |  |
| 11 | C | O5' | AA:445 | 5.00 | 1.00 | 43.31 |  |
  |

  
 Contact(s): /OP2/  
 Coded contact list: O_ph_  

### MG:AA:3045

|  | Res. type | Atom | Residue | Distance | Occupancy | B-factor | Closest solute to 1^st^ shell water |
| --- | --- | --- | --- | --- | --- | --- | --- |
| **Ion** | | | |  | 1.00 | 51.58 || 1 | C | OP1 | AA:31 | 2.02 | 1.00 | 54.04 |
| 2 | G | OP1 | AA:1239 | 3.26 | 1.00 | 56.88 |  |
| 3 | C | OP1 | AA:1200 | 3.58 | 1.00 | 48.33 |  |
| 4 | C | OP2 | AA:31 | 4.32 | 1.00 | 54.39 |  |
| 5 | U | OP2 | AA:1240 | 4.32 | 1.00 | 57.41 |  |
| 6 | G | O3' | AA:30 | 4.32 | 1.00 | 51.92 |  |
| 7 | C | OP2 | AA:1200 | 4.35 | 1.00 | 47.99 |  |
| 8 | C | O5' | AA:31 | 4.38 | 1.00 | 51.72 |  |
| 9 | U | OP1 | AA:1199 | 4.81 | 1.00 | 58.50 |  |
| 10 | G | O5' | AA:1239 | 4.89 | 1.00 | 56.47 |  |
| 11 | U | OP1 | AA:1240 | 4.90 | 1.00 | 58.00 |  |
| 12 | G | O3' | AA:1239 | 4.96 | 1.00 | 58.80 |  |
  |

  
 Contact(s): /OP1/  
 Coded contact list: O_ph_  
 Exclusion zone contact(s): /OP1/  

### MG:AA:3046

|  | Res. type | Atom | Residue | Distance | Occupancy | B-factor | Closest solute to 1^st^ shell water |
| --- | --- | --- | --- | --- | --- | --- | --- |
| **Ion** | | | |  | 1.00 | 38.54 || 1 | A | OP2 | AA:1609 | 3.69 | 1.00 | 50.36 |
| 2 | C | OP1 | AA:1617 | 3.96 | 1.00 | 43.84 |  |
| 3 | G | OP2 | AA:1619 | 4.36 | 1.00 | 43.37 |  |
| 4 | G | O6 | AA:1619 | 4.41 | 1.00 | 44.10 |  |
| 5 | G | N7 | AA:1619 | 4.50 | 1.00 | 43.23 |  |
| 6 | G | O6 | AA:1613 | 4.68 | 1.00 | 46.81 |  |
| 7 | C | OP2 | AA:1617 | 4.81 | 1.00 | 44.27 |  |
| 8 | A | O3' | AA:1608 | 4.83 | 1.00 | 47.78 |  |
  |

  
 Coded contact list:   
1^st^ MG coordination shell missing (see Notes)  

### MG:AA:3047

|  | Res. type | Atom | Residue | Distance | Occupancy | B-factor | Closest solute to 1^st^ shell water |
| --- | --- | --- | --- | --- | --- | --- | --- |
| **Ion** | | | |  | 1.00 | 28.73 || 1 | U | O4 | AA:1940 | 2.20 | 1.00 | 56.91 |
| 2 | C | O2 | AA:1962 | 2.56 | 1.00 | 47.41 |  |
| 3 | A | N3 | AA:1936 | 2.70 | 1.00 | 52.70 |  |
| 4 | C | N3 | AA:1962 | 2.75 | 1.00 | 52.55 |  |
| 5 | A | O5' | AA:1936 | 2.92 | 1.00 | 44.29 |  |
| 6 | A | OP2 | AA:1936 | 3.43 | 1.00 | 44.99 |  |
| 7 | A | O4' | AA:1936 | 3.82 | 1.00 | 49.78 |  |
| 8 | U | N3 | AA:1940 | 3.83 | 1.00 | 55.64 |  |
| 9 | G | O2' | AA:1935 | 3.88 | 1.00 | 41.98 |  |
| 10 | C | N4 | AA:1961 | 3.89 | 1.00 | 53.37 |  |
| 11 | G | O3' | AA:1935 | 4.06 | 1.00 | 44.47 |  |
| 12 | A | N1 | AA:1936 | 4.08 | 1.00 | 53.58 |  |
| 13 | G | N3 | AA:1935 | 4.30 | 1.00 | 44.87 |  |
| 14 | C | N4 | AA:1962 | 4.39 | 1.00 | 53.68 |  |
| 15 | C | N1 | AA:1962 | 4.40 | 1.00 | 53.55 |  |
| 16 | A | N9 | AA:1936 | 4.52 | 1.00 | 51.83 |  |
| 17 | G | N2 | AA:1935 | 4.72 | 1.00 | 46.80 |  |
| 18 | G | O6 | AA:1945 | 4.79 | 1.00 | 54.07 |  |
  |

  
 Contact(s): /O4/  
 Coded contact list: O_b_  
 Exclusion zone contact(s): /N3_A_/N3_C_/O2/O5'/  
Consider K in place of MG (3 valid contact(s) for K vs. 1 for MG)  

### MG:AA:3048

|  | Res. type | Atom | Residue | Distance | Occupancy | B-factor | Closest solute to 1^st^ shell water |
| --- | --- | --- | --- | --- | --- | --- | --- |
| **Ion** | | | |  | 1.00 | 41.14 || 1 | G | OP2 | AA:1187 | 2.16 | 1.00 | 63.08 |
| 2 | C | OP2 | AA:817 | 3.42 | 1.00 | 62.35 |  |
| 3 | C | OP2 | AA:816 | 3.50 | 1.00 | 59.22 |  |
| 4 | G | OP1 | AA:1187 | 4.31 | 1.00 | 63.89 |  |
| 5 | G | O5' | AA:1187 | 4.39 | 1.00 | 63.24 |  |
| 6 | C | O2' | AA:1185 | 4.42 | 1.00 | 72.96 |  |
| 7 | G | O6 | AA:818 | 4.47 | 1.00 | 66.71 |  |
| 8 | G | N7 | AA:1187 | 4.57 | 1.00 | 64.60 |  |
| 9 | G | O3' | AA:1186 | 4.60 | 1.00 | 64.99 |  |
| 10 | C | OP1 | AA:816 | 4.71 | 1.00 | 59.70 |  |
| 11 | C | O5' | AA:816 | 4.74 | 1.00 | 60.51 |  |
| 12 | C | O3' | AA:1185 | 4.78 | 1.00 | 68.50 |  |
  |

  
 Contact(s): /OP2/  
 Coded contact list: O_ph_  

### MG:AA:3049

|  | Res. type | Atom | Residue | Distance | Occupancy | B-factor | Closest solute to 1^st^ shell water |
| --- | --- | --- | --- | --- | --- | --- | --- |
| **Ion** | | | |  | 1.00 | 48.97 || 1 | A | OP1 | AA:1936 | 3.82 | 1.00 | 43.75 |
| 2 | A | OP1 | AA:1937 | 4.27 | 1.00 | 51.43 |  |
| 3 | A | O3' | AA:1936 | 4.38 | 1.00 | 52.23 |  |
| 4 | C | OP2 | AA:1958 | 4.65 | 1.00 | 40.33 |  |
| 5 | A | O5' | AA:1937 | 4.78 | 1.00 | 49.95 |  |
| 6 | G | OP2 | AA:1959 | 4.82 | 1.00 | 41.01 |  |
| 7 | G | N7 | AA:1959 | 4.97 | 1.00 | 39.29 |  |
  |

  
 Coded contact list:   
1^st^ MG coordination shell missing (see Notes)  

### MG:AA:3050

|  | Res. type | Atom | Residue | Distance | Occupancy | B-factor | Closest solute to 1^st^ shell water |
| --- | --- | --- | --- | --- | --- | --- | --- |
| **Ion** | | | |  | 1.00 | 59.18 || 1 | A | O2' | AA:1936 | 3.21 | 1.00 | 52.20 |
| 2 | G | OP1 | AA:1945 | 3.38 | 1.00 | 56.48 |  |
| 3 | A | O3' | AA:1936 | 3.76 | 1.00 | 52.23 |  |
| 4 | U | O4 | AA:1946 | 4.05 | 1.00 | 51.48 |  |
| 5 | G | N7 | AA:1945 | 4.19 | 1.00 | 53.99 |  |
| 6 | A | OP1 | AA:1937 | 4.70 | 1.00 | 51.43 |  |
| 7 | U | OP2 | AA:1956 | 4.76 | 1.00 | 49.18 |  |
| 8 | G | O6 | AA:1945 | 4.76 | 1.00 | 54.07 |  |
  |

  
 Coded contact list:   
 Exclusion zone contact(s): /O2'/OP1/  
Consider K in place of MG (2 valid contact(s) for K vs. 0 for MG)  

### MG:AA:3051

|  | Res. type | Atom | Residue | Distance | Occupancy | B-factor | Closest solute to 1^st^ shell water |
| --- | --- | --- | --- | --- | --- | --- | --- |
| **Ion** | | | |  | 1.00 | 62.78 || 1 | G | O6 | AA:1949 | 3.32 | 1.00 | 37.26 |
| 2 | G | O6 | AA:1948 | 3.74 | 1.00 | 36.19 |  |
| 3 | G | N7 | AA:1949 | 4.13 | 1.00 | 39.68 |  |
| 4 | G | N7 | AA:1948 | 4.32 | 1.00 | 37.51 |  |
| 5 | U | OP2 | AA:1955 | 4.62 | 1.00 | 53.75 |  |
| 6 | G | O6 | AA:1950 | 4.72 | 1.00 | 45.73 |  |
| 7 | C | N4 | AA:1957 | 4.74 | 1.00 | 38.99 |  |
  |

  
 Coded contact list:   
 Exclusion zone contact(s): /O6/  
Consider K in place of MG (1 valid contact(s) for K vs. 0 for MG)  

### MG:AA:3052

|  | Res. type | Atom | Residue | Distance | Occupancy | B-factor | Closest solute to 1^st^ shell water |
| --- | --- | --- | --- | --- | --- | --- | --- |
| **Ion** | | | |  | 1.00 | 60.28 || 1 | U | O4 | AA:1951 | 2.28 | 1.00 | 56.03 |
| 2 | G | N1 | AA:1950 | 2.91 | 1.00 | 46.65 |  |
| 3 | G | N2 | AA:1950 | 3.43 | 1.00 | 50.39 |  |
| 4 | G | O6 | AA:1950 | 3.82 | 1.00 | 45.73 |  |
| 5 | U | N3 | AA:1951 | 3.88 | 1.00 | 55.14 |  |
| 6 | G | OP2 | AA:1954 | 3.92 | 1.00 | 59.86 |  |
| 7 | G | N7 | AA:1949 | 4.56 | 1.00 | 39.68 |  |
| 8 | G | N3 | AA:1950 | 4.94 | 1.00 | 45.43 |  |
  |

  
 Contact(s): /O4/  
 Coded contact list: O_b_  
 Exclusion zone contact(s): /N1_G_/  

### MG:AA:3053

|  | Res. type | Atom | Residue | Distance | Occupancy | B-factor | Closest solute to 1^st^ shell water |
| --- | --- | --- | --- | --- | --- | --- | --- |
| **Ion** | | | |  | 1.00 | 30.47 || 1 | A | OP2 | AA:1665 | 2.03 | 1.00 | 52.02 |
| 2 | A | OP1 | AA:1664 | 2.34 | 1.00 | 45.27 |  |
| 3 | A | O5' | AA:1664 | 3.95 | 1.00 | 46.68 |  |
| 4 | A | OP1 | AA:1665 | 4.08 | 1.00 | 53.05 |  |
| 5 | A | OP2 | AA:1664 | 4.09 | 1.00 | 46.87 |  |
| 6 | A | O5' | AA:1665 | 4.16 | 1.00 | 51.66 |  |
| 7 | A | O3' | AA:1664 | 4.49 | 1.00 | 50.77 |  |
| 8 | G | OP2 | AA:1666 | 4.63 | 1.00 | 53.83 |  |
| 9 | C | O3' | AA:1663 | 4.78 | 1.00 | 47.47 |  |
  |

  
**1^st^ shell ligand angles:**   
1-2: 86;   
  
 Contact(s): /OP1/OP2/  
 Coded contact list: 2O_ph_  

### MG:AA:3054

|  | Res. type | Atom | Residue | Distance | Occupancy | B-factor | Closest solute to 1^st^ shell water |
| --- | --- | --- | --- | --- | --- | --- | --- |
| **Ion** | | | |  | 1.00 | 44.92 || 1 | C | OP1 | AA:393 | 2.53 | 1.00 | 54.89 |
| 2 | C | OP1 | AA:375 | 2.84 | 1.00 | 60.19 |  |
| 3 | G | O2' | AA:408 | 2.99 | 1.00 | 64.22 |  |
| 4 | G | O3' | AA:408 | 3.08 | 1.00 | 64.17 |  |
| 5 | C | OP2 | AA:393 | 3.16 | 1.00 | 53.47 |  |
| 6 | C | OP1 | AA:409 | 4.13 | 1.00 | 63.58 |  |
| 7 | C | O5' | AA:393 | 4.40 | 1.00 | 54.45 |  |
| 8 | C | O3' | AA:392 | 4.45 | 1.00 | 53.47 |  |
| 9 | A | N7 | AA:223 | 4.64 | 1.00 | 64.59 |  |
| 10 | A | O3' | AA:374 | 4.66 | 1.00 | 59.86 |  |
| 11 | C | O5' | AA:409 | 4.67 | 1.00 | 61.93 |  |
| 12 | C | OP2 | AA:375 | 4.68 | 1.00 | 59.73 |  |
| 13 | G | O4' | AA:408 | 4.73 | 1.00 | 63.92 |  |
  |

  
 Coded contact list:   
 Exclusion zone contact(s): /O2'/O3'/OP1/OP1/OP2/  
Consider K in place of MG (4 valid contact(s) for K vs. 0 for MG)  

### MG:AA:3055

|  | Res. type | Atom | Residue | Distance | Occupancy | B-factor | Closest solute to 1^st^ shell water |
| --- | --- | --- | --- | --- | --- | --- | --- |
| **Ion** | | | |  | 1.00 | 33.23 || 1 | C | OP2 | AA:1990 | 3.90 | 1.00 | 48.79 |
| 2 | G | OP2 | AA:1989 | 4.06 | 1.00 | 50.28 |  |
| 3 | G | OP1 | AA:1989 | 4.57 | 1.00 | 51.04 |  |
| 4 | G | OP2 | AA:1667 | 4.69 | 1.00 | 54.35 |  |
  |

  
 Coded contact list:   
1^st^ MG coordination shell missing (see Notes)  

### MG:AA:3056

|  | Res. type | Atom | Residue | Distance | Occupancy | B-factor | Closest solute to 1^st^ shell water |
| --- | --- | --- | --- | --- | --- | --- | --- |
| **Ion** | | | |  | 1.00 | 63.39 || 1 | G | OP2 | AA:410 | 2.62 | 1.00 | 55.33 |
| 2 | C | OP2 | AA:413 | 2.93 | 1.00 | 62.20 |  |
| 3 | G | O2' | AA:391 | 3.89 | 1.00 | 51.42 |  |
| 4 | G | O3' | AA:391 | 4.08 | 1.00 | 53.64 |  |
| 5 | C | OP1 | AA:413 | 4.09 | 1.00 | 61.55 |  |
| 6 | G | OP1 | AA:410 | 4.37 | 1.00 | 55.04 |  |
| 7 | C | OP1 | AA:392 | 4.39 | 1.00 | 53.79 |  |
| 8 | A | O3' | AA:412 | 4.49 | 1.00 | 61.57 |  |
| 9 | C | O3' | AA:409 | 4.86 | 1.00 | 56.34 |  |
| 10 | C | OP1 | AA:409 | 4.93 | 1.00 | 63.58 |  |
| 11 | G | O5' | AA:410 | 4.98 | 1.00 | 56.07 |  |
  |

  
 Coded contact list:   
 Exclusion zone contact(s): /OP2/OP2/  
Consider K in place of MG (2 valid contact(s) for K vs. 0 for MG)  

### MG:AA:3057

|  | Res. type | Atom | Residue | Distance | Occupancy | B-factor | Closest solute to 1^st^ shell water |
| --- | --- | --- | --- | --- | --- | --- | --- |
| **Ion** | | | |  | 1.00 | 49.18 || 1 | C | OP1 | AA:409 | 2.14 | 1.00 | 63.58 |
| 2 | C | OP1 | AA:392 | 2.53 | 1.00 | 53.79 |  |
| 3 | C | OP1 | AA:375 | 3.66 | 1.00 | 60.19 |  |
| 4 | G | O3' | AA:408 | 3.80 | 1.00 | 64.17 |  |
| 5 | C | OP2 | AA:376 | 3.92 | 1.00 | 55.39 |  |
| 6 | C | OP2 | AA:392 | 3.99 | 1.00 | 53.70 |  |
| 7 | C | OP2 | AA:409 | 4.17 | 1.00 | 64.29 |  |
| 8 | C | OP2 | AA:393 | 4.39 | 1.00 | 53.47 |  |
| 9 | C | OP1 | AA:376 | 4.45 | 1.00 | 54.43 |  |
| 10 | C | O5' | AA:392 | 4.59 | 1.00 | 53.88 |  |
| 11 | C | O5' | AA:375 | 4.62 | 1.00 | 59.07 |  |
| 12 | C | O5' | AA:409 | 4.62 | 1.00 | 61.93 |  |
| 13 | G | O3' | AA:391 | 4.88 | 1.00 | 53.64 |  |
| 14 | C | O3' | AA:375 | 4.97 | 1.00 | 55.03 |  |
  |

  
 Contact(s): /OP1/  
 Coded contact list: O_ph_  
 Exclusion zone contact(s): /OP1/  

### MG:AA:3058

|  | Res. type | Atom | Residue | Distance | Occupancy | B-factor | Closest solute to 1^st^ shell water |
| --- | --- | --- | --- | --- | --- | --- | --- |
| **Ion** | | | |  | 1.00 | 25.75 || 1 | U | OP1 | AA:1394 | 2.56 | 1.00 | 52.93 |
| 2 | C | OP1 | AA:1604 | 2.66 | 1.00 | 34.85 |  |
| 3 | A | O3' | AA:1393 | 3.89 | 1.00 | 53.62 |  |
| 4 | C | O5' | AA:1604 | 4.34 | 1.00 | 36.94 |  |
| 5 | U | OP2 | AA:1394 | 4.60 | 1.00 | 55.47 |  |
| 6 | A | O3' | AA:1603 | 4.67 | 1.00 | 34.35 |  |
| 7 | A | OP2 | AA:1395 | 4.85 | 1.00 | 52.14 |  |
| 8 | U | O5' | AA:1394 | 4.92 | 1.00 | 52.27 |  |
  |

  
 Coded contact list:   
 Exclusion zone contact(s): /OP1/OP1/  
Consider NA in place of MG (1 valid contact(s) for NA vs. 0 for MG)  

### MG:AA:3059

|  | Res. type | Atom | Residue | Distance | Occupancy | B-factor | Closest solute to 1^st^ shell water |
| --- | --- | --- | --- | --- | --- | --- | --- |
| **Ion** | | | |  | 1.00 | 68.56 || 1 | G | O6 | AA:1849 | 3.73 | 1.00 | 53.87 |
| 2 | A | N7 | AA:1848 | 3.84 | 1.00 | 68.79 |  |
| 3 | G | N1 | AA:1846 | 4.16 | 1.00 | 58.69 |  |
| 4 | A | N6 | AA:1848 | 4.36 | 1.00 | 69.05 |  |
| 5 | G | N3 | AA:1846 | 4.50 | 1.00 | 60.17 |  |
| 6 | A | OP2 | AA:1848 | 4.50 | 1.00 | 85.87 |  |
| 7 | G | N7 | AA:1849 | 4.67 | 1.00 | 56.16 |  |
| 8 | G | O6 | AA:1846 | 4.74 | 1.00 | 59.25 |  |
| 9 | G | N2 | AA:1846 | 4.83 | 1.00 | 57.55 |  |
| 10 | G | N7 | AA:1846 | 5.00 | 1.00 | 63.69 |  |
  |

  
 Coded contact list:   
1^st^ MG coordination shell missing (see Notes)  

### MG:AA:3060

|  | Res. type | Atom | Residue | Distance | Occupancy | B-factor | Closest solute to 1^st^ shell water |
| --- | --- | --- | --- | --- | --- | --- | --- |
| **Ion** | | | |  | 1.00 | 76.02 || 1 | G | OP1 | AA:1891 | 4.07 | 1.00 | 56.03 |
| 2 | G | OP2 | AA:1891 | 4.20 | 1.00 | 55.00 |  |
  |

  
 Coded contact list:   
1^st^ MG coordination shell missing (see Notes)  

### MG:AA:3061

|  | Res. type | Atom | Residue | Distance | Occupancy | B-factor | Closest solute to 1^st^ shell water |
| --- | --- | --- | --- | --- | --- | --- | --- |
| **Ion** | | | |  | 1.00 | 35.87 || 1 | A | OP1 | AA:1603 | 2.22 | 1.00 | 45.20 |
| 2 | A | OP1 | AA:1395 | 2.32 | 1.00 | 52.83 |  |
| 3 | C | OP2 | AA:1604 | 2.35 | 1.00 | 34.59 |  |
| 4 | C | OP1 | AA:1604 | 3.92 | 1.00 | 34.85 |  |
| 5 | U | O3' | AA:1394 | 4.37 | 1.00 | 53.46 |  |
| 6 | U | O3' | AA:1602 | 4.44 | 1.00 | 48.36 |  |
| 7 | C | O5' | AA:1604 | 4.50 | 1.00 | 36.94 |  |
| 8 | A | O5' | AA:1603 | 4.50 | 1.00 | 45.11 |  |
| 9 | A | OP2 | AA:1603 | 4.58 | 1.00 | 45.36 |  |
| 10 | A | OP2 | AA:1395 | 4.66 | 1.00 | 52.14 |  |
| 11 | A | O3' | AA:1603 | 4.68 | 1.00 | 34.35 |  |
| 12 | A | O5' | AA:1395 | 4.69 | 1.00 | 53.98 |  |
  |

  
**1^st^ shell ligand angles:**   
1-2: 96; 1-3: 62; 2-3: 89;   
  
 Contact(s): /OP1/OP1/OP2/  
 Coded contact list: 3O_ph_  

### MG:AA:3062

|  | Res. type | Atom | Residue | Distance | Occupancy | B-factor | Closest solute to 1^st^ shell water |
| --- | --- | --- | --- | --- | --- | --- | --- |
| **Ion** | | | |  | 1.00 | 29.65 || 1 | G | OP2 | AA:1975 | 2.45 | 1.00 | 34.18 |
| 2 | C | OP1 | AA:1771 | 3.71 | 1.00 | 38.27 |  |
| 3 | C | O5' | AA:1974 | 3.87 | 1.00 | 39.30 |  |
| 4 | C | OP2 | AA:1974 | 4.25 | 1.00 | 41.79 |  |
| 5 | C | O3' | AA:1974 | 4.31 | 1.00 | 35.88 |  |
| 6 | G | OP1 | AA:1975 | 4.38 | 1.00 | 34.52 |  |
| 7 | C | OP2 | AA:1771 | 4.63 | 1.00 | 36.98 |  |
| 8 | C | OP1 | AA:1974 | 4.83 | 1.00 | 40.98 |  |
| 9 | G | O5' | AA:1975 | 4.96 | 1.00 | 33.28 |  |
  |

  
 Coded contact list:   
 Exclusion zone contact(s): /OP2/  
Consider NA in place of MG (1 valid contact(s) for NA vs. 0 for MG)  

### MG:AA:3063

|  | Res. type | Atom | Residue | Distance | Occupancy | B-factor | Closest solute to 1^st^ shell water |
| --- | --- | --- | --- | --- | --- | --- | --- |
| **Ion** | | | |  | 1.00 | 27.18 || 1 | U | OP2 | AA:1639 | 2.09 | 1.00 | 44.86 |
| 2 | C | OP2 | AA:1298 | 2.55 | 1.00 | 49.54 |  |
| 3 | C | OP2 | AA:1297 | 3.24 | 1.00 | 49.86 |  |
| 4 | U | OP1 | AA:1639 | 3.77 | 1.00 | 45.48 |  |
| 5 | C | O3' | AA:1638 | 4.02 | 1.00 | 43.70 |  |
| 6 | C | OP1 | AA:1298 | 4.18 | 1.00 | 48.98 |  |
| 7 | C | O5' | AA:1297 | 4.18 | 1.00 | 48.68 |  |
| 8 | C | O3' | AA:1297 | 4.22 | 1.00 | 49.30 |  |
| 9 | C | OP1 | AA:1297 | 4.25 | 1.00 | 49.97 |  |
| 10 | C | O5' | AA:1638 | 4.44 | 1.00 | 44.77 |  |
| 11 | C | OP1 | AA:1638 | 4.54 | 1.00 | 45.61 |  |
| 12 | U | O5' | AA:1639 | 4.55 | 1.00 | 42.99 |  |
| 13 | C | O2' | AA:2710 | 4.57 | 1.00 | 48.86 |  |
| 14 | G | O6 | AA:1299 | 4.92 | 1.00 | 39.49 |  |
  |

  
 Contact(s): /OP2/  
 Coded contact list: O_ph_  
 Exclusion zone contact(s): /OP2/OP2/  

### MG:AA:3064

|  | Res. type | Atom | Residue | Distance | Occupancy | B-factor | Closest solute to 1^st^ shell water |
| --- | --- | --- | --- | --- | --- | --- | --- |
| **Ion** | | | |  | 1.00 | 90.30 || 1 | G | OP1 | AA:1770 | 3.15 | 1.00 | 44.39 |
| 2 | G | OP1 | AA:1975 | 3.67 | 1.00 | 34.52 |  |
| 3 | G | OP1 | AA:1935 | 4.13 | 1.00 | 47.12 |  |
| 4 | G | OP1 | AA:1959 | 4.61 | 1.00 | 41.17 |  |
| 5 | G | O3' | AA:1769 | 4.70 | 1.00 | 43.42 |  |
  |

  
 Coded contact list:   
 Exclusion zone contact(s): /OP1/  
Consider K in place of MG (1 valid contact(s) for K vs. 0 for MG)  

### MG:AA:3065

|  | Res. type | Atom | Residue | Distance | Occupancy | B-factor | Closest solute to 1^st^ shell water |
| --- | --- | --- | --- | --- | --- | --- | --- |
| **Ion** | | | |  | 1.00 | 60.64 || 1 | C | N4 | AA:1771 | 4.13 | 1.00 | 36.49 |
| 2 | G | O6 | AA:1772 | 4.57 | 1.00 | 41.59 |  |
| 3 | A | OP2 | AA:1977 | 4.71 | 1.00 | 38.15 |  |
| 4 | U | O4 | AA:1976 | 4.94 | 1.00 | 33.87 |  |
| 5 | G | OP2 | AA:1770 | 4.98 | 1.00 | 42.41 |  |
| 6 | U | OP2 | AA:1976 | 4.98 | 1.00 | 36.80 |  |
  |

  
 Coded contact list:   
1^st^ MG coordination shell missing (see Notes)  

### MG:AA:3066

|  | Res. type | Atom | Residue | Distance | Occupancy | B-factor | Closest solute to 1^st^ shell water |
| --- | --- | --- | --- | --- | --- | --- | --- |
| **Ion** | | | |  | 1.00 | 17.99 || 1 | G | OP2 | AA:450 | 2.15 | 1.00 | 40.85 |
| 2 | A | OP2 | AA:449 | 2.95 | 1.00 | 34.02 |  |
| 3 | GLY | O | AF:86 | 3.06 | 1.00 | 37.98 |  |
| 4 | GLY | O | AF:85 | 3.72 | 1.00 | 29.43 |  |
| 5 | G | OP1 | AA:450 | 4.25 | 1.00 | 41.17 |  |
| 6 | VAL | O | AF:84 | 4.27 | 1.00 | 40.78 |  |
| 7 | G | O5' | AA:450 | 4.45 | 1.00 | 40.46 |  |
| 8 | A | OP1 | AA:449 | 4.47 | 1.00 | 34.38 |  |
| 9 | A | O3' | AA:449 | 4.52 | 1.00 | 39.43 |  |
| 10 | G | OP2 | AA:452 | 4.60 | 1.00 | 38.78 |  |
| 11 | A | O5' | AA:449 | 4.65 | 1.00 | 35.04 |  |
  |

  
 Contact(s): /OP2/  
 Coded contact list: O_ph_  
 Exclusion zone contact(s): /O_back_/OP2/  
Consider K in place of MG (2 valid contact(s) for K vs. 1 for MG)  

### MG:AA:3067

|  | Res. type | Atom | Residue | Distance | Occupancy | B-factor | Closest solute to 1^st^ shell water |
| --- | --- | --- | --- | --- | --- | --- | --- |
| **Ion** | | | |  | 1.00 | 37.76 || 1 | G | OP2 | AA:2447 | 2.88 | 1.00 | 38.02 |
| 2 | G | OP1 | AA:2446 | 3.21 | 1.00 | 40.01 |  |
| 3 | G | OP2 | AA:2446 | 3.28 | 1.00 | 37.51 |  |
| 4 | G | OP1 | AA:2502 | 3.70 | 1.00 | 40.75 |  |
| 5 | G | N7 | AA:2502 | 3.91 | 1.00 | 41.92 |  |
| 6 | G | O5' | AA:2446 | 4.39 | 1.00 | 38.37 |  |
| 7 | G | OP1 | AA:2447 | 4.50 | 1.00 | 38.71 |  |
  |

  
 Coded contact list:   
 Exclusion zone contact(s): /OP1/OP2/OP2/  
Consider K in place of MG (3 valid contact(s) for K vs. 0 for MG)  

### MG:AA:3068

|  | Res. type | Atom | Residue | Distance | Occupancy | B-factor | Closest solute to 1^st^ shell water |
| --- | --- | --- | --- | --- | --- | --- | --- |
| **Ion** | | | |  | 1.00 | 40.72 || 1 | G | OP1 | AA:2447 | 2.67 | 1.00 | 38.71 |
| 2 | A | O2' | AA:567 | 3.11 | 1.00 | 37.73 |  |
| 3 | G | OP2 | AA:2447 | 3.65 | 1.00 | 38.02 |  |
| 4 | G | OP1 | AA:2446 | 3.94 | 1.00 | 40.01 |  |
| 5 | G | O5' | AA:2447 | 4.62 | 1.00 | 37.11 |  |
| 6 | G | OP1 | AA:808 | 4.76 | 1.00 | 45.91 |  |
| 7 | G | O3' | AA:2446 | 4.85 | 1.00 | 37.37 |  |
  |

  
 Coded contact list:   
 Exclusion zone contact(s): /O2'/OP1/  
Consider K in place of MG (2 valid contact(s) for K vs. 0 for MG)  

### MG:AA:3069

|  | Res. type | Atom | Residue | Distance | Occupancy | B-factor | Closest solute to 1^st^ shell water |
| --- | --- | --- | --- | --- | --- | --- | --- |
| **Ion** | | | |  | 1.00 | 29.85 || 1 | A | OP1 | AA:567 | 1.92 | 1.00 | 42.45 |
| 2 | U | O3' | AA:566 | 3.37 | 1.00 | 43.20 |  |
| 3 | A | O5' | AA:567 | 3.91 | 1.00 | 42.30 |  |
| 4 | G | OP2 | AA:808 | 3.95 | 1.00 | 45.44 |  |
| 5 | U | O4 | AA:810 | 4.19 | 1.00 | 52.54 |  |
| 6 | G | OP2 | AA:809 | 4.26 | 1.00 | 43.89 |  |
| 7 | A | OP2 | AA:567 | 4.39 | 1.00 | 41.45 |  |
| 8 | G | N7 | AA:809 | 4.41 | 1.00 | 47.65 |  |
| 9 | G | O5' | AA:808 | 4.95 | 1.00 | 47.13 |  |
| 10 | U | O2' | AA:566 | 4.99 | 1.00 | 44.08 |  |
  |

  
 Contact(s): /OP1/  
 Coded contact list: O_ph_  
 Exclusion zone contact(s): /O3'/  

### MG:AA:3070

|  | Res. type | Atom | Residue | Distance | Occupancy | B-factor | Closest solute to 1^st^ shell water |
| --- | --- | --- | --- | --- | --- | --- | --- |
| **Ion** | | | |  | 1.00 | 35.50 || 1 | G | OP1 | AA:948 | 1.86 | 1.00 | 55.90 |
| 2 | U | OP2 | AA:963 | 1.96 | 1.00 | 57.28 |  |
| 3 | G | OP1 | AA:962 | 2.02 | 1.00 | 58.60 |  |
| 4 | G | OP2 | AA:948 | 3.11 | 1.00 | 54.88 |  |
| 5 | G | O3' | AA:947 | 3.86 | 1.00 | 56.25 |  |
| 6 | G | OP2 | AA:962 | 3.90 | 1.00 | 60.12 |  |
| 7 | U | OP1 | AA:963 | 4.02 | 1.00 | 58.34 |  |
| 8 | G | O5' | AA:948 | 4.10 | 1.00 | 54.89 |  |
| 9 | C | O3' | AA:961 | 4.12 | 1.00 | 60.95 |  |
| 10 | G | O3' | AA:962 | 4.17 | 1.00 | 56.80 |  |
| 11 | U | O5' | AA:963 | 4.40 | 1.00 | 57.72 |  |
| 12 | G | O5' | AA:962 | 4.46 | 1.00 | 59.64 |  |
  |

  
**1^st^ shell ligand angles:**   
1-2: 157; 1-3: 99; 2-3: 73;   
  
 Contact(s): /OP1/OP1/OP2/  
 Coded contact list: 3O_ph_  
 Exclusion zone contact(s): /OP2/  

### MG:AA:3071

|  | Res. type | Atom | Residue | Distance | Occupancy | B-factor | Closest solute to 1^st^ shell water |
| --- | --- | --- | --- | --- | --- | --- | --- |
| **Ion** | | | |  | 1.00 | 16.11 || 1 | G | OP2 | AA:946 | 3.77 | 1.00 | 55.35 |
| 2 | G | OP1 | AA:946 | 3.82 | 1.00 | 56.37 |  |
| 3 | A | OP2 | AA:945 | 4.04 | 1.00 | 53.09 |  |
| 4 | G | O3' | AA:944 | 4.05 | 1.00 | 52.41 |  |
| 5 | G | O2' | AA:944 | 4.18 | 1.00 | 56.38 |  |
| 6 | A | O2' | AA:821 | 4.26 | 1.00 | 56.34 |  |
| 7 | A | O3' | AA:821 | 4.70 | 1.00 | 53.18 |  |
| 8 | U | OP2 | AA:822 | 4.73 | 1.00 | 51.39 |  |
  |

  
 Coded contact list:   
1^st^ MG coordination shell missing (see Notes)  

### MG:AA:3072

|  | Res. type | Atom | Residue | Distance | Occupancy | B-factor | Closest solute to 1^st^ shell water |
| --- | --- | --- | --- | --- | --- | --- | --- |
| **Ion** | | | |  | 1.00 | 21.02 || 1 | A | OP1 | AA:945 | 2.95 | 1.00 | 53.13 |
| 2 | G | N7 | AA:831 | 3.57 | 1.00 | 46.22 |  |
| 3 | G | O2' | AA:944 | 3.72 | 1.00 | 56.38 |  |
| 4 | G | OP2 | AA:831 | 3.98 | 1.00 | 43.32 |  |
| 5 | G | O3' | AA:944 | 4.06 | 1.00 | 52.41 |  |
| 6 | G | N2 | AA:944 | 4.11 | 1.00 | 56.75 |  |
| 7 | A | OP2 | AA:945 | 4.13 | 1.00 | 53.09 |  |
| 8 | G | N3 | AA:944 | 4.13 | 1.00 | 55.44 |  |
| 9 | G | O6 | AA:831 | 4.23 | 1.00 | 46.46 |  |
| 10 | G | N7 | AA:832 | 4.68 | 1.00 | 42.63 |  |
| 11 | G | O6 | AA:832 | 4.84 | 1.00 | 46.22 |  |
  |

  
 Coded contact list:   
 Exclusion zone contact(s): /OP1/  
Consider K in place of MG (1 valid contact(s) for K vs. 0 for MG)  

### MG:AA:3073

|  | Res. type | Atom | Residue | Distance | Occupancy | B-factor | Closest solute to 1^st^ shell water |
| --- | --- | --- | --- | --- | --- | --- | --- |
| **Ion** | | | |  | 1.00 | 71.67 || 1 | G | O3' | AA:946 | 4.51 | 1.00 | 53.01 |
| 2 | C | OP1 | AA:964 | 4.62 | 1.00 | 52.92 |  |
| 3 | U | OP2 | AA:822 | 4.86 | 1.00 | 51.39 |  |
  |

  
 Coded contact list:   
1^st^ MG coordination shell missing (see Notes)  

### MG:AA:3074

|  | Res. type | Atom | Residue | Distance | Occupancy | B-factor | Closest solute to 1^st^ shell water |
| --- | --- | --- | --- | --- | --- | --- | --- |
| **Ion** | | | |  | 1.00 | 51.27 || 1 | C | OP2 | AA:2196 | 2.52 | 1.00 | 71.48 |
| 2 | G | OP2 | AA:2224 | 3.95 | 1.00 | 46.73 |  |
| 3 | C | OP1 | AA:2196 | 4.44 | 1.00 | 71.16 |  |
| 4 | C | O5' | AA:2196 | 4.75 | 1.00 | 69.77 |  |
| 5 | C | O5' | AA:2195 | 4.85 | 1.00 | 84.01 |  |
| 6 | C | O3' | AA:2195 | 4.90 | 1.00 | 71.70 |  |
  |

  
 Coded contact list:   
 Exclusion zone contact(s): /OP2/  
Consider NA in place of MG (1 valid contact(s) for NA vs. 0 for MG)  

### MG:AA:3075

|  | Res. type | Atom | Residue | Distance | Occupancy | B-factor | Closest solute to 1^st^ shell water |
| --- | --- | --- | --- | --- | --- | --- | --- |
| **Ion** | | | |  | 1.00 | 33.44 || 1 | G | O6 | AA:948 | 2.54 | 1.00 | 59.21 |
| 2 | G | O6 | AA:968 | 2.93 | 1.00 | 64.37 |  |
| 3 | U | O4 | AA:969 | 2.95 | 1.00 | 59.29 |  |
| 4 | G | O6 | AA:947 | 3.31 | 1.00 | 62.99 |  |
| 5 | U | N3 | AA:969 | 3.86 | 1.00 | 59.57 |  |
| 6 | C | N4 | AA:949 | 3.91 | 1.00 | 61.83 |  |
| 7 | G | N7 | AA:948 | 4.25 | 1.00 | 56.16 |  |
| 8 | C | N4 | AA:970 | 4.40 | 1.00 | 60.35 |  |
| 9 | G | N1 | AA:947 | 4.62 | 1.00 | 63.05 |  |
| 10 | G | N7 | AA:947 | 4.66 | 1.00 | 62.38 |  |
| 11 | G | N1 | AA:968 | 4.76 | 1.00 | 64.38 |  |
| 12 | G | N1 | AA:948 | 4.80 | 1.00 | 57.23 |  |
| 13 | G | N7 | AA:968 | 4.96 | 1.00 | 65.49 |  |
  |

  
 Coded contact list:   
 Exclusion zone contact(s): /O4/O6/O6/O6/  
Consider K in place of MG (3 valid contact(s) for K vs. 0 for MG)  

### MG:AA:3076

|  | Res. type | Atom | Residue | Distance | Occupancy | B-factor | Closest solute to 1^st^ shell water |
| --- | --- | --- | --- | --- | --- | --- | --- |
| **Ion** | | | |  | 1.00 | 35.41 || 1 | G | O6 | AA:1840 | 2.67 | 1.00 | 44.94 |
| 2 | G | O6 | AA:1839 | 2.94 | 1.00 | 65.06 |  |
| 3 | C | O2 | AA:1838 | 3.22 | 1.00 | 59.90 |  |
| 4 | G | O6 | AA:1899 | 3.29 | 1.00 | 54.07 |  |
| 5 | G | N1 | AA:1899 | 3.33 | 1.00 | 53.94 |  |
| 6 | G | N7 | AA:1839 | 3.42 | 1.00 | 64.82 |  |
| 7 | C | O4' | AA:1838 | 3.63 | 1.00 | 59.29 |  |
| 8 | C | N1 | AA:1838 | 3.71 | 1.00 | 61.43 |  |
| 9 | C | O2' | AA:1838 | 3.90 | 1.00 | 64.48 |  |
| 10 | C | N4 | AA:1902 | 4.06 | 1.00 | 42.85 |  |
| 11 | G | N7 | AA:1840 | 4.18 | 1.00 | 49.55 |  |
| 12 | U | O4 | AA:1841 | 4.33 | 1.00 | 48.82 |  |
| 13 | G | N2 | AA:1903 | 4.35 | 1.00 | 47.43 |  |
| 14 | A | N6 | AA:1901 | 4.47 | 1.00 | 46.25 |  |
| 15 | G | N2 | AA:1899 | 4.59 | 1.00 | 58.16 |  |
| 16 | C | N3 | AA:1838 | 4.70 | 1.00 | 60.73 |  |
| 17 | G | N1 | AA:1839 | 4.71 | 1.00 | 63.05 |  |
| 18 | G | N1 | AA:1840 | 4.92 | 1.00 | 46.70 |  |
  |

  
 Coded contact list:   
 Exclusion zone contact(s): /N1_G_/O2/O6/O6/O6/  
Consider K in place of MG (5 valid contact(s) for K vs. 0 for MG)  

### MG:AA:3077

|  | Res. type | Atom | Residue | Distance | Occupancy | B-factor | Closest solute to 1^st^ shell water |
| --- | --- | --- | --- | --- | --- | --- | --- |
| **Ion** | | | |  | 1.00 | 63.33 || 1 | G | O6 | AA:946 | 3.37 | 1.00 | 58.78 |
| 2 | G | O6 | AA:947 | 3.72 | 1.00 | 62.99 |  |
| 3 | A | OP2 | AA:821 | 4.20 | 1.00 | 56.94 |  |
| 4 | G | N7 | AA:946 | 4.39 | 1.00 | 58.82 |  |
| 5 | C | N4 | AA:970 | 4.61 | 1.00 | 60.35 |  |
| 6 | G | N2 | AA:972 | 4.73 | 1.00 | 66.17 |  |
| 7 | U | O4 | AA:969 | 4.87 | 1.00 | 59.29 |  |
  |

  
 Coded contact list:   
 Exclusion zone contact(s): /O6/  
Consider K in place of MG (1 valid contact(s) for K vs. 0 for MG)  

### MG:AA:3078

|  | Res. type | Atom | Residue | Distance | Occupancy | B-factor | Closest solute to 1^st^ shell water |
| --- | --- | --- | --- | --- | --- | --- | --- |
| **Ion** | | | |  | 1.00 | 44.30 || 1 | G | O6 | AA:1187 | 2.12 | 1.00 | 64.03 |
| 2 | G | OP2 | AA:818 | 2.25 | 1.00 | 59.18 |  |
| 3 | C | OP1 | AA:817 | 4.03 | 1.00 | 61.98 |  |
| 4 | G | N7 | AA:1187 | 4.16 | 1.00 | 64.60 |  |
| 5 | G | N7 | AA:818 | 4.16 | 1.00 | 68.04 |  |
| 6 | C | OP2 | AA:817 | 4.18 | 1.00 | 62.35 |  |
| 7 | C | O3' | AA:817 | 4.19 | 1.00 | 60.08 |  |
| 8 | C | O5' | AA:817 | 4.21 | 1.00 | 62.33 |  |
| 9 | G | OP1 | AA:818 | 4.25 | 1.00 | 59.27 |  |
| 10 | G | N1 | AA:1187 | 4.38 | 1.00 | 65.03 |  |
| 11 | G | OP1 | AA:1186 | 4.45 | 1.00 | 67.82 |  |
| 12 | G | N7 | AA:1186 | 4.53 | 1.00 | 71.60 |  |
| 13 | G | O5' | AA:818 | 4.75 | 1.00 | 59.85 |  |
  |

  
**1^st^ shell ligand angles:**   
1-2: 73;   
  
 Contact(s): /O6/OP2/  
 Coded contact list: O_ph_O_b_  

### MG:AA:3079

|  | Res. type | Atom | Residue | Distance | Occupancy | B-factor | Closest solute to 1^st^ shell water |
| --- | --- | --- | --- | --- | --- | --- | --- |
| **Ion** | | | |  | 1.00 | 40.87 || 1 | A | OP2 | AA:990 | 1.83 | 1.00 | 73.82 |
| 2 | G | OP2 | AA:1186 | 2.24 | 1.00 | 68.26 |  |
| 3 | C | OP2 | AA:991 | 2.77 | 1.00 | 73.77 |  |
| 4 | A | OP1 | AA:990 | 3.13 | 1.00 | 75.19 |  |
| 5 | C | OP1 | AA:1185 | 3.37 | 1.00 | 74.00 |  |
| 6 | G | OP1 | AA:1186 | 3.76 | 1.00 | 67.82 |  |
| 7 | C | OP2 | AA:1185 | 3.84 | 1.00 | 75.01 |  |
| 8 | C | O5' | AA:1185 | 3.85 | 1.00 | 73.47 |  |
| 9 | A | O5' | AA:990 | 3.88 | 1.00 | 74.86 |  |
| 10 | G | O3' | AA:989 | 4.09 | 1.00 | 76.04 |  |
| 11 | G | O5' | AA:1186 | 4.39 | 1.00 | 67.68 |  |
| 12 | C | O3' | AA:1185 | 4.55 | 1.00 | 68.50 |  |
| 13 | C | OP1 | AA:991 | 4.76 | 1.00 | 73.67 |  |
| 14 | C | O5' | AA:991 | 4.99 | 1.00 | 72.21 |  |
  |

  
**1^st^ shell ligand angles:**   
1-2: 102;   
  
 Contact(s): /OP2/OP2/  
 Coded contact list: 2O_ph_  
 Exclusion zone contact(s): /OP1/OP1/OP2/  
Consider K in place of MG (3 valid contact(s) for K vs. 2 for MG)  

### MG:AA:3080

|  | Res. type | Atom | Residue | Distance | Occupancy | B-factor | Closest solute to 1^st^ shell water |
| --- | --- | --- | --- | --- | --- | --- | --- |
| **Ion** | | | |  | 1.00 | 41.06 || 1 | A | OP2 | AA:1938 | 2.96 | 1.00 | 50.95 |
| 2 | C | OP1 | AA:1957 | 3.10 | 1.00 | 45.58 |  |
| 3 | A | O3' | AA:1937 | 3.60 | 1.00 | 51.26 |  |
| 4 | C | OP2 | AA:1958 | 3.61 | 1.00 | 40.33 |  |
| 5 | C | OP1 | AA:1958 | 4.08 | 1.00 | 40.94 |  |
| 6 | C | O5' | AA:1957 | 4.39 | 1.00 | 42.97 |  |
| 7 | A | OP1 | AA:1937 | 4.86 | 1.00 | 51.43 |  |
| 8 | A | O5' | AA:1938 | 4.86 | 1.00 | 51.40 |  |
| 9 | C | O3' | AA:1957 | 4.91 | 1.00 | 39.13 |  |
  |

  
 Coded contact list:   
 Exclusion zone contact(s): /OP1/OP2/  
Consider K in place of MG (2 valid contact(s) for K vs. 0 for MG)  

### MG:AA:3081

|  | Res. type | Atom | Residue | Distance | Occupancy | B-factor | Closest solute to 1^st^ shell water |
| --- | --- | --- | --- | --- | --- | --- | --- |
| **Ion** | | | |  | 1.00 | 61.80 || 1 | G | OP1 | AA:1157 | 1.76 | 1.00 | 78.20 |
| 2 | C | OP2 | AA:1158 | 2.03 | 1.00 | 77.64 |  |
| 3 | G | O3' | AA:1157 | 3.63 | 1.00 | 78.14 |  |
| 4 | G | O5' | AA:1157 | 3.95 | 1.00 | 79.26 |  |
| 5 | A | O5' | AA:990 | 3.96 | 1.00 | 74.86 |  |
| 6 | C | OP1 | AA:1158 | 4.00 | 1.00 | 76.75 |  |
| 7 | A | O3' | AA:1156 | 4.02 | 1.00 | 79.57 |  |
| 8 | G | OP2 | AA:1157 | 4.21 | 1.00 | 78.80 |  |
| 9 | A | OP1 | AA:990 | 4.33 | 1.00 | 75.19 |  |
| 10 | C | O5' | AA:1158 | 4.49 | 1.00 | 78.12 |  |
  |

  
**1^st^ shell ligand angles:**   
1-2: 105;   
  
 Contact(s): /OP2/  
 Coded contact list: O_ph_  
 Exclusion zone contact(s): /OP1/  

### MG:AA:3082

|  | Res. type | Atom | Residue | Distance | Occupancy | B-factor | Closest solute to 1^st^ shell water |
| --- | --- | --- | --- | --- | --- | --- | --- |
| **Ion** | | | |  | 1.00 | 48.08 || 1 | A | OP2 | AA:1156 | 2.54 | 1.00 | 75.83 |
| 2 | G | O6 | AA:1157 | 3.44 | 1.00 | 85.42 |  |
| 3 | C | N4 | AA:998 | 3.95 | 1.00 | 86.74 |  |
| 4 | ARG | NH2 | A1:58 | 4.08 | 1.00 | 92.28 |  |
| 5 | A | O3' | AA:1155 | 4.42 | 1.00 | 74.51 |  |
| 6 | A | OP2 | AA:1155 | 4.50 | 1.00 | 81.80 |  |
| 7 | A | O5' | AA:1156 | 4.63 | 1.00 | 76.57 |  |
| 8 | G | O6 | AA:997 | 4.69 | 1.00 | 78.08 |  |
| 9 | A | O5' | AA:1155 | 4.91 | 1.00 | 80.00 |  |
| 10 | C | N4 | AA:1158 | 4.97 | 1.00 | 78.71 |  |
  |

  
 Coded contact list:   
 Exclusion zone contact(s): /OP2/  
Consider NA in place of MG (1 valid contact(s) for NA vs. 0 for MG)  

### MG:AA:3083

|  | Res. type | Atom | Residue | Distance | Occupancy | B-factor | Closest solute to 1^st^ shell water |
| --- | --- | --- | --- | --- | --- | --- | --- |
| **Ion** | | | |  | 1.00 | 33.56 || 1 | A | OP2 | AA:1932 | 2.56 | 1.00 | 57.86 |
| 2 | G | N7 | AA:1933 | 3.46 | 1.00 | 43.21 |  |
| 3 | G | O6 | AA:1933 | 3.50 | 1.00 | 44.52 |  |
| 4 | A | O5' | AA:1932 | 3.90 | 1.00 | 55.40 |  |
| 5 | A | OP1 | AA:1932 | 4.12 | 1.00 | 58.20 |  |
| 6 | C | N4 | AA:1934 | 4.20 | 1.00 | 47.11 |  |
| 7 | U | OP1 | AA:1931 | 4.70 | 1.00 | 55.66 |  |
| 8 | G | N2 | AA:1968 | 4.82 | 1.00 | 36.28 |  |
| 9 | G | OP2 | AA:1933 | 4.85 | 1.00 | 50.18 |  |
| 10 | U | O3' | AA:1931 | 4.96 | 1.00 | 56.14 |  |
| 11 | U | OP2 | AA:1931 | 4.97 | 1.00 | 54.36 |  |
  |

  
 Coded contact list:   
 Exclusion zone contact(s): /OP2/  
Consider NA in place of MG (1 valid contact(s) for NA vs. 0 for MG)  

### MG:AA:3084

|  | Res. type | Atom | Residue | Distance | Occupancy | B-factor | Closest solute to 1^st^ shell water |
| --- | --- | --- | --- | --- | --- | --- | --- |
| **Ion** | | | |  | 1.00 | 55.41 || 1 | U | OP2 | AA:999 | 2.10 | 1.00 | 81.88 |
| 2 | C | OP2 | AA:1153 | 2.15 | 1.00 | 89.49 |  |
| 3 | C | OP1 | AA:998 | 3.34 | 1.00 | 84.00 |  |
| 4 | C | O5' | AA:998 | 3.59 | 1.00 | 83.77 |  |
| 5 | C | O3' | AA:998 | 3.72 | 1.00 | 81.13 |  |
| 6 | U | OP1 | AA:999 | 3.77 | 1.00 | 81.34 |  |
| 7 | C | OP1 | AA:1153 | 3.83 | 1.00 | 88.96 |  |
| 8 | C | OP2 | AA:998 | 4.06 | 1.00 | 83.90 |  |
| 9 | C | O3' | AA:1152 | 4.22 | 1.00 | 90.33 |  |
| 10 | C | O5' | AA:1153 | 4.56 | 1.00 | 89.04 |  |
| 11 | U | O5' | AA:999 | 4.58 | 1.00 | 81.82 |  |
| 12 | G | O6 | AA:1154 | 4.84 | 1.00 | 87.80 |  |
  |

  
**1^st^ shell ligand angles:**   
1-2: 125;   
  
 Contact(s): /OP2/OP2/  
 Coded contact list: 2O_ph_  
 Exclusion zone contact(s): /OP1/  

### MG:AA:3085

|  | Res. type | Atom | Residue | Distance | Occupancy | B-factor | Closest solute to 1^st^ shell water |
| --- | --- | --- | --- | --- | --- | --- | --- |
| **Ion** | | | |  | 1.00 | 29.02 || 1 | G | O6 | AA:397 | 2.54 | 1.00 | 46.77 |
| 2 | G | N7 | AA:397 | 2.97 | 1.00 | 45.28 |  |
| 3 | G | O6 | AA:396 | 3.83 | 1.00 | 48.96 |  |
| 4 | G | O6 | AA:398 | 4.03 | 1.00 | 56.01 |  |
| 5 | G | N7 | AA:396 | 4.28 | 1.00 | 47.27 |  |
| 6 | G | N1 | AA:397 | 4.51 | 1.00 | 45.65 |  |
| 7 | G | N7 | AA:398 | 4.75 | 1.00 | 54.03 |  |
| 8 | C | N4 | AA:377 | 4.76 | 1.00 | 46.00 |  |
| 9 | G | N1 | AA:396 | 4.84 | 1.00 | 49.40 |  |
  |

  
 Coded contact list:   
 Exclusion zone contact(s): /N7/O6/  
Consider NA in place of MG (1 valid contact(s) for NA vs. 0 for MG)  

### MG:AA:3086

|  | Res. type | Atom | Residue | Distance | Occupancy | B-factor | Closest solute to 1^st^ shell water |
| --- | --- | --- | --- | --- | --- | --- | --- |
| **Ion** | | | |  | 1.00 | 35.16 || 1 | G | OP2 | AA:399 | 1.95 | 1.00 | 57.34 |
| 2 | G | O5' | AA:399 | 3.61 | 1.00 | 57.25 |  |
| 3 | G | O6 | AA:400 | 3.68 | 1.00 | 66.30 |  |
| 4 | G | N7 | AA:400 | 3.90 | 1.00 | 63.80 |  |
| 5 | G | OP1 | AA:399 | 4.24 | 1.00 | 57.89 |  |
| 6 | G | N7 | AA:399 | 4.38 | 1.00 | 56.69 |  |
| 7 | G | O3' | AA:398 | 4.39 | 1.00 | 54.89 |  |
| 8 | G | N7 | AA:372 | 4.87 | 1.00 | 66.42 |  |
  |

  
 Contact(s): /OP2/  
 Coded contact list: O_ph_  

### MG:AA:3087

|  | Res. type | Atom | Residue | Distance | Occupancy | B-factor | Closest solute to 1^st^ shell water |
| --- | --- | --- | --- | --- | --- | --- | --- |
| **Ion** | | | |  | 1.00 | 20.77 || 1 | C | N3 | AA:2871 | 3.23 | 1.00 | 73.09 |
| 2 | C | N4 | AA:2871 | 3.54 | 1.00 | 75.29 |  |
| 3 | G | O6 | AA:2845 | 4.33 | 1.00 | 70.14 |  |
| 4 | C | O2 | AA:2871 | 4.36 | 1.00 | 72.64 |  |
| 5 | G | O6 | AA:2844 | 4.68 | 1.00 | 72.71 |  |
| 6 | G | N7 | AA:2843 | 4.90 | 1.00 | 67.55 |  |
| 7 | G | N1 | AA:2845 | 4.97 | 1.00 | 70.25 |  |
  |

  
 Coded contact list:   
 Exclusion zone contact(s): /N3_C_/  
Consider K in place of MG (1 valid contact(s) for K vs. 0 for MG)  

### MG:AA:3088

|  | Res. type | Atom | Residue | Distance | Occupancy | B-factor | Closest solute to 1^st^ shell water |
| --- | --- | --- | --- | --- | --- | --- | --- |
| **Ion** | | | |  | 1.00 | 51.18 || 1 | G | O6 | AA:1975 | 2.47 | 1.00 | 28.88 |
| 2 | U | O4 | AA:1976 | 3.17 | 1.00 | 33.87 |  |
| 3 | G | N7 | AA:1975 | 3.79 | 1.00 | 32.54 |  |
| 4 | C | N4 | AA:1830 | 4.25 | 1.00 | 25.87 |  |
| 5 | C | N4 | AA:1974 | 4.28 | 1.00 | 31.83 |  |
| 6 | U | N3 | AA:1976 | 4.44 | 1.00 | 34.19 |  |
| 7 | G | N1 | AA:1975 | 4.61 | 1.00 | 29.06 |  |
| 8 | A | OP2 | AA:1773 | 4.63 | 1.00 | 39.04 |  |
| 9 | A | OP1 | AA:1829 | 4.89 | 1.00 | 37.92 |  |
| 10 | G | OP2 | AA:1772 | 4.90 | 1.00 | 39.61 |  |
  |

  
 Coded contact list:   
 Exclusion zone contact(s): /O4/O6/  
Consider NA in place of MG (1 valid contact(s) for NA vs. 0 for MG)  

### MG:AA:3089

|  | Res. type | Atom | Residue | Distance | Occupancy | B-factor | Closest solute to 1^st^ shell water |
| --- | --- | --- | --- | --- | --- | --- | --- |
| **Ion** | | | |  | 1.00 | 70.32 || 1 | G | OP1 | AA:1627 | 2.39 | 1.00 | 48.28 |
| 2 | G | OP2 | AA:1627 | 3.74 | 1.00 | 47.27 |  |
| 3 | G | O5' | AA:1627 | 4.19 | 1.00 | 47.41 |  |
| 4 | U | O4 | AA:1300 | 4.22 | 1.00 | 46.58 |  |
| 5 | A | OP1 | AA:1634 | 4.65 | 1.00 | 45.89 |  |
| 6 | G | O5' | AA:1626 | 4.76 | 1.00 | 49.69 |  |
| 7 | G | O3' | AA:1626 | 4.77 | 1.00 | 46.83 |  |
| 8 | G | OP1 | AA:1626 | 4.83 | 1.00 | 52.97 |  |
| 9 | G | OP2 | AA:1626 | 4.89 | 1.00 | 52.04 |  |
  |

  
 Contact(s): /OP1/  
 Coded contact list: O_ph_  

### MG:AA:3090

|  | Res. type | Atom | Residue | Distance | Occupancy | B-factor | Closest solute to 1^st^ shell water |
| --- | --- | --- | --- | --- | --- | --- | --- |
| **Ion** | | | |  | 1.00 | 42.81 || 1 | A | OP2 | AA:1189 | 1.97 | 1.00 | 52.21 |
| 2 | G | N7 | AA:1190 | 3.12 | 1.00 | 63.30 |  |
| 3 | A | OP1 | AA:1189 | 3.44 | 1.00 | 53.22 |  |
| 4 | A | O5' | AA:1189 | 4.02 | 1.00 | 52.52 |  |
| 5 | G | O6 | AA:1190 | 4.24 | 1.00 | 63.46 |  |
| 6 | U | O3' | AA:1188 | 4.31 | 1.00 | 51.71 |  |
| 7 | G | OP2 | AA:1190 | 4.35 | 1.00 | 52.59 |  |
| 8 | GLY | O | AO:26 | 4.35 | 1.00 | 62.02 |  |
| 9 | U | OP1 | AA:1188 | 4.52 | 1.00 | 51.61 |  |
| 10 | G | N2 | AA:818 | 4.73 | 1.00 | 66.10 |  |
| 11 | U | OP2 | AA:1188 | 4.82 | 1.00 | 53.53 |  |
  |

  
 Contact(s): /OP2/  
 Coded contact list: O_ph_  
 Exclusion zone contact(s): /N7/  

### MG:AA:3091

|  | Res. type | Atom | Residue | Distance | Occupancy | B-factor | Closest solute to 1^st^ shell water |
| --- | --- | --- | --- | --- | --- | --- | --- |
| **Ion** | | | |  | 1.00 | 59.86 || 1 | G | O6 | AA:110 | 3.73 | 1.00 | 57.19 |
| 2 | G | N7 | AA:110 | 4.09 | 1.00 | 58.82 |  |
| 3 | G | O6 | AA:109 | 4.48 | 1.00 | 61.72 |  |
| 4 | G | N7 | AA:109 | 4.63 | 1.00 | 61.37 |  |
  |

  
 Coded contact list:   
1^st^ MG coordination shell missing (see Notes)  

### MG:AA:3092

|  | Res. type | Atom | Residue | Distance | Occupancy | B-factor | Closest solute to 1^st^ shell water |
| --- | --- | --- | --- | --- | --- | --- | --- |
| **Ion** | | | |  | 1.00 | 16.75 || 1 | G | OP2 | AA:1828 | 2.73 | 1.00 | 25.52 |
| 2 | A | OP2 | AA:1972 | 3.87 | 1.00 | 43.61 |  |
| 3 | C | O5' | AA:1827 | 4.09 | 1.00 | 30.62 |  |
| 4 | G | N7 | AA:1828 | 4.18 | 1.00 | 35.41 |  |
| 5 | ARG | NH1 | AD:222 | 4.47 | 1.00 | 27.33 |  |
| 6 | C | OP1 | AA:1827 | 4.51 | 1.00 | 32.40 |  |
| 7 | G | OP1 | AA:1828 | 4.61 | 1.00 | 28.05 |  |
| 8 | C | O3' | AA:1827 | 4.63 | 1.00 | 25.12 |  |
| 9 | C | OP2 | AA:1827 | 4.67 | 1.00 | 34.51 |  |
| 10 | ARG | NE | AD:239 | 4.73 | 1.00 | 40.31 |  |
| 11 | ARG | NH2 | AD:222 | 4.95 | 1.00 | 30.28 |  |
  |

  
 Coded contact list:   
 Exclusion zone contact(s): /OP2/  
Consider K in place of MG (1 valid contact(s) for K vs. 0 for MG)  

### MG:AA:3093

|  | Res. type | Atom | Residue | Distance | Occupancy | B-factor | Closest solute to 1^st^ shell water |
| --- | --- | --- | --- | --- | --- | --- | --- |
| **Ion** | | | |  | 1.00 | 39.84 || 1 | U | O4 | AA:202 | 2.41 | 1.00 | 44.18 |
| 2 | U | O4 | AA:193 | 3.09 | 1.00 | 37.44 |  |
| 3 | C | OP2 | AA:192 | 3.23 | 1.00 | 35.85 |  |
| 4 | U | N3 | AA:202 | 4.07 | 1.00 | 43.77 |  |
| 5 | C | OP1 | AA:192 | 4.51 | 1.00 | 36.21 |  |
| 6 | C | N4 | AA:201 | 4.56 | 1.00 | 38.03 |  |
| 7 | A | OP2 | AA:199 | 4.82 | 1.00 | 41.52 |  |
| 8 | C | N3 | AA:203 | 4.87 | 1.00 | 42.58 |  |
| 9 | G | O6 | AA:194 | 4.89 | 1.00 | 37.38 |  |
  |

  
 Coded contact list:   
 Exclusion zone contact(s): /O4/O4/OP2/  
Consider K in place of MG (2 valid contact(s) for K vs. 0 for MG)  

### MG:AA:3094

|  | Res. type | Atom | Residue | Distance | Occupancy | B-factor | Closest solute to 1^st^ shell water |
| --- | --- | --- | --- | --- | --- | --- | --- |
| **Ion** | | | |  | 1.00 | 21.10 || 1 | A | OP2 | AA:783 | 2.21 | 1.00 | 37.57 |
| 2 | A | OP1 | AA:2589 | 2.23 | 1.00 | 33.98 |  |
| 3 | A | OP2 | AA:784 | 2.32 | 1.00 | 38.65 |  |
| 4 | A | O3' | AA:782 | 3.87 | 1.00 | 39.04 |  |
| 5 | A | O5' | AA:783 | 3.94 | 1.00 | 40.16 |  |
| 6 | ASN | ND2 | AD:227 | 4.17 | 1.00 | 32.73 |  |
| 7 | A | OP2 | AA:2589 | 4.39 | 1.00 | 32.60 |  |
| 8 | A | OP1 | AA:784 | 4.41 | 1.00 | 35.96 |  |
| 9 | G | O3' | AA:2588 | 4.43 | 1.00 | 31.49 |  |
| 10 | A | O5' | AA:784 | 4.59 | 1.00 | 37.42 |  |
| 11 | A | O5' | AA:2589 | 4.63 | 1.00 | 31.62 |  |
| 12 | ASN | OD1 | AD:227 | 4.66 | 1.00 | 35.79 |  |
| 13 | A | O3' | AA:783 | 4.71 | 1.00 | 38.25 |  |
| 14 | ALA | O | AD:225 | 4.71 | 1.00 | 36.08 |  |
| 15 | A | O2' | AA:782 | 4.71 | 1.00 | 44.35 |  |
| 16 | A | OP1 | AA:783 | 4.77 | 1.00 | 41.00 |  |
| 17 | MET | O | AD:226 | 4.97 | 1.00 | 34.42 |  |
  |

  
**1^st^ shell ligand angles:**   
1-2: 74; 1-3: 89; 2-3: 73;   
  
 Contact(s): /OP1/OP2/OP2/  
 Coded contact list: 3O_ph_  

### MG:AA:3095

|  | Res. type | Atom | Residue | Distance | Occupancy | B-factor | Closest solute to 1^st^ shell water |
| --- | --- | --- | --- | --- | --- | --- | --- |
| **Ion** | | | |  | 1.00 | 18.72 || 1 | A | OP1 | AA:2598 | 2.55 | 1.00 | 32.36 |
| 2 | ARG | O | AD:242 | 2.79 | 1.00 | 32.21 |  |
| 3 | G | O3' | AA:2597 | 3.57 | 1.00 | 31.37 |  |
| 4 | PRO | O | AD:241 | 3.63 | 1.00 | 32.26 |  |
| 5 | GLY | N | AD:234 | 4.14 | 1.00 | 33.73 |  |
| 6 | PRO | O | AD:232 | 4.36 | 1.00 | 28.25 |  |
| 7 | A | OP2 | AA:2598 | 4.65 | 1.00 | 32.55 |  |
| 8 | A | O5' | AA:2598 | 4.80 | 1.00 | 31.67 |  |
| 9 | GLY | N | AD:243 | 4.87 | 1.00 | 30.68 |  |
| 10 | GLY | O | AD:235 | 4.95 | 1.00 | 41.73 |  |
  |

  
 Coded contact list:   
 Exclusion zone contact(s): /O_back_/OP1/  
Consider NA in place of MG (1 valid contact(s) for NA vs. 0 for MG)  

### MG:AA:3096

|  | Res. type | Atom | Residue | Distance | Occupancy | B-factor | Closest solute to 1^st^ shell water |
| --- | --- | --- | --- | --- | --- | --- | --- |
| **Ion** | | | |  | 1.00 | 67.17 || 1 | G | O6 | AA:1717 | 3.12 | 1.00 | 111.63 |
| 2 | G | O6 | AA:1718 | 3.73 | 1.00 | 106.21 |  |
| 3 | C | N4 | AA:1742 | 4.17 | 1.00 | 110.60 |  |
| 4 | G | N7 | AA:1717 | 4.66 | 1.00 | 113.04 |  |
| 5 | G | N7 | AA:1718 | 4.82 | 1.00 | 108.68 |  |
| 6 | G | O6 | AA:1743 | 4.84 | 1.00 | 105.42 |  |
| 7 | U | O4 | AA:1716 | 4.95 | 1.00 | 106.71 |  |
  |

  
 Coded contact list:   
 Exclusion zone contact(s): /O6/  
Consider K in place of MG (1 valid contact(s) for K vs. 0 for MG)  

### MG:AA:3097

|  | Res. type | Atom | Residue | Distance | Occupancy | B-factor | Closest solute to 1^st^ shell water |
| --- | --- | --- | --- | --- | --- | --- | --- |
| **Ion** | | | |  | 1.00 | 73.92 || 1 | G | O6 | AA:1682 | 2.30 | 1.00 | 65.03 |
| 2 | U | O4 | AA:1706 | 2.38 | 1.00 | 63.42 |  |
| 3 | U | N3 | AA:1706 | 3.14 | 1.00 | 65.07 |  |
| 4 | U | OP1 | AA:1757 | 3.84 | 1.00 | 63.67 |  |
| 5 | G | O6 | AA:1705 | 4.01 | 1.00 | 59.67 |  |
| 6 | G | N7 | AA:1682 | 4.42 | 1.00 | 61.68 |  |
| 7 | G | N1 | AA:1682 | 4.47 | 1.00 | 62.16 |  |
| 8 | C | N4 | AA:1683 | 4.64 | 1.00 | 61.77 |  |
| 9 | U | O5' | AA:1757 | 4.71 | 1.00 | 64.65 |  |
| 10 | U | O2 | AA:1706 | 4.88 | 1.00 | 64.42 |  |
  |

  
**1^st^ shell ligand angles:**   
1-2: 102;   
  
 Contact(s): /O4/O6/  
 Coded contact list: 2O_b_  
 Exclusion zone contact(s): /N3_U_/  

### MG:AA:3098

|  | Res. type | Atom | Residue | Distance | Occupancy | B-factor | Closest solute to 1^st^ shell water |
| --- | --- | --- | --- | --- | --- | --- | --- |
| **Ion** | | | |  | 1.00 | 45.87 || 1 | G | OP2 | AA:1756 | 3.71 | 1.00 | 57.28 |
| 2 | G | O6 | AA:1756 | 3.81 | 1.00 | 68.95 |  |
| 3 | G | N7 | AA:1756 | 4.09 | 1.00 | 67.20 |  |
| 4 | G | O6 | AA:1753 | 4.16 | 1.00 | 66.61 |  |
| 5 | U | OP2 | AA:1706 | 4.42 | 1.00 | 61.95 |  |
| 6 | ARG | NH1 | AR:112 | 4.98 | 1.00 | 100.10 |  |
| 7 | U | OP1 | AA:1706 | 4.99 | 1.00 | 63.96 |  |
  |

  
 Coded contact list:   
1^st^ MG coordination shell missing (see Notes)  

### MG:AA:3099

|  | Res. type | Atom | Residue | Distance | Occupancy | B-factor | Closest solute to 1^st^ shell water |
| --- | --- | --- | --- | --- | --- | --- | --- |
| **Ion** | | | |  | 1.00 | 33.22 || 1 | G | OP1 | AA:1975 | 2.47 | 1.00 | 34.52 |
| 2 | G | OP2 | AA:1975 | 2.83 | 1.00 | 34.18 |  |
| 3 | C | O3' | AA:1974 | 2.90 | 1.00 | 35.88 |  |
| 4 | C | O2' | AA:1934 | 3.19 | 1.00 | 46.06 |  |
| 5 | C | O3' | AA:1934 | 3.74 | 1.00 | 46.58 |  |
| 6 | C | O5' | AA:1974 | 3.92 | 1.00 | 39.30 |  |
| 7 | G | O5' | AA:1975 | 4.34 | 1.00 | 33.28 |  |
| 8 | C | OP1 | AA:1974 | 4.59 | 1.00 | 40.98 |  |
| 9 | C | O4' | AA:1934 | 4.81 | 1.00 | 44.88 |  |
| 10 | G | OP1 | AA:1935 | 4.96 | 1.00 | 47.12 |  |
  |

  
 Coded contact list:   
 Exclusion zone contact(s): /O2'/O3'/OP1/OP2/  
Consider K in place of MG (3 valid contact(s) for K vs. 0 for MG)  

### MG:AA:3100

|  | Res. type | Atom | Residue | Distance | Occupancy | B-factor | Closest solute to 1^st^ shell water |
| --- | --- | --- | --- | --- | --- | --- | --- |
| **Ion** | | | |  | 1.00 | 34.80 || 1 | G | O2' | AA:1899 | 3.24 | 1.00 | 51.35 |
| 2 | A | OP1 | AA:1970 | 4.06 | 1.00 | 38.39 |  |
| 3 | A | OP2 | AA:1970 | 4.19 | 1.00 | 39.18 |  |
| 4 | G | O6 | AA:1903 | 4.26 | 1.00 | 46.12 |  |
| 5 | A | OP2 | AA:1900 | 4.34 | 1.00 | 47.86 |  |
| 6 | G | N7 | AA:1903 | 4.98 | 1.00 | 47.86 |  |
| 7 | G | O6 | AA:1904 | 4.99 | 1.00 | 46.22 |  |
  |

  
 Coded contact list:   
 Exclusion zone contact(s): /O2'/  
Consider K in place of MG (1 valid contact(s) for K vs. 0 for MG)  

### MG:AA:3101

|  | Res. type | Atom | Residue | Distance | Occupancy | B-factor | Closest solute to 1^st^ shell water |
| --- | --- | --- | --- | --- | --- | --- | --- |
| **Ion** | | | |  | 1.00 | 38.07 || 1 | U | OP1 | AA:2017 | 2.44 | 1.00 | 40.66 |
| 2 | A | OP2 | AA:578 | 3.32 | 1.00 | 46.41 |  |
| 3 | G | OP2 | AA:2018 | 4.13 | 1.00 | 41.14 |  |
| 4 | U | O5' | AA:2017 | 4.39 | 1.00 | 42.63 |  |
| 5 | U | OP2 | AA:2017 | 4.43 | 1.00 | 41.38 |  |
| 6 | G | OP2 | AA:577 | 4.59 | 1.00 | 44.81 |  |
| 7 | A | OP1 | AA:2058 | 4.95 | 1.00 | 43.88 |  |
| 8 | U | O3' | AA:2016 | 4.99 | 1.00 | 41.60 |  |
  |

  
 Coded contact list:   
 Exclusion zone contact(s): /OP1/OP2/  
Consider NA in place of MG (1 valid contact(s) for NA vs. 0 for MG)  

### MG:AA:3102

|  | Res. type | Atom | Residue | Distance | Occupancy | B-factor | Closest solute to 1^st^ shell water |
| --- | --- | --- | --- | --- | --- | --- | --- |
| **Ion** | | | |  | 1.00 | 65.55 || 1 | U | OP1 | AA:2016 | 2.34 | 1.00 | 43.64 |
| 2 | U | OP2 | AA:2017 | 3.33 | 1.00 | 41.38 |  |
| 3 | A | OP1 | AA:2058 | 3.41 | 1.00 | 43.88 |  |
| 4 | U | O5' | AA:2016 | 3.57 | 1.00 | 42.58 |  |
| 5 | U | OP2 | AA:2016 | 4.24 | 1.00 | 42.93 |  |
| 6 | U | OP1 | AA:2017 | 4.70 | 1.00 | 40.66 |  |
| 7 | A | O3' | AA:2015 | 4.71 | 1.00 | 42.34 |  |
| 8 | U | O3' | AA:2016 | 4.82 | 1.00 | 41.60 |  |
| 9 | A | OP2 | AA:2059 | 4.88 | 1.00 | 45.39 |  |
| 10 | A | OP1 | AA:2059 | 4.93 | 1.00 | 46.42 |  |
  |

  
 Contact(s): /OP1/  
 Coded contact list: O_ph_  
 Exclusion zone contact(s): /OP2/  

### MG:AA:3103

|  | Res. type | Atom | Residue | Distance | Occupancy | B-factor | Closest solute to 1^st^ shell water |
| --- | --- | --- | --- | --- | --- | --- | --- |
| **Ion** | | | |  | 1.00 | 45.85 || 1 | U | O2' | AA:1255 | 3.21 | 1.00 | 44.93 |
| 2 | G | OP1 | AA:1256 | 3.39 | 1.00 | 47.50 |  |
| 3 | U | O3' | AA:1255 | 3.66 | 1.00 | 45.38 |  |
| 4 | A | OP1 | AA:578 | 4.18 | 1.00 | 46.98 |  |
| 5 | G | OP2 | AA:1256 | 4.60 | 1.00 | 45.51 |  |
| 6 | GLY | O | AF:71 | 4.82 | 1.00 | 48.95 |  |
  |

  
 Coded contact list:   
 Exclusion zone contact(s): /O2'/OP1/  
Consider K in place of MG (2 valid contact(s) for K vs. 0 for MG)  

### MG:AA:3104

|  | Res. type | Atom | Residue | Distance | Occupancy | B-factor | Closest solute to 1^st^ shell water |
| --- | --- | --- | --- | --- | --- | --- | --- |
| **Ion** | | | |  | 1.00 | 54.99 || 1 | G | O6 | AA:1260 | 3.66 | 1.00 | 52.47 |
| 2 | G | O6 | AA:1259 | 4.10 | 1.00 | 49.58 |  |
| 3 | G | N7 | AA:1259 | 4.60 | 1.00 | 49.95 |  |
| 4 | G | O6 | AA:579 | 4.64 | 1.00 | 47.29 |  |
| 5 | C | N4 | AA:580 | 4.70 | 1.00 | 50.60 |  |
| 6 | G | N7 | AA:1260 | 4.96 | 1.00 | 50.52 |  |
  |

  
 Coded contact list:   
1^st^ MG coordination shell missing (see Notes)  

### MG:AA:3105

|  | Res. type | Atom | Residue | Distance | Occupancy | B-factor | Closest solute to 1^st^ shell water |
| --- | --- | --- | --- | --- | --- | --- | --- |
| **Ion** | | | |  | 1.00 | 50.42 || 1 | G | OP2 | AA:469 | 3.20 | 1.00 | 32.16 |
| 2 | G | O6 | AA:458 | 3.61 | 1.00 | 33.02 |  |
| 3 | ALA | O | AF:58 | 3.62 | 1.00 | 41.86 |  |
| 4 | G | N1 | AA:458 | 3.82 | 1.00 | 33.95 |  |
| 5 | TYR | N | AF:59 | 4.41 | 1.00 | 43.62 |  |
| 6 | G | O5' | AA:469 | 4.77 | 1.00 | 32.09 |  |
| 7 | G | N7 | AA:469 | 4.78 | 1.00 | 25.24 |  |
| 8 | G | OP1 | AA:468 | 4.84 | 1.00 | 32.26 |  |
| 9 | LYS | NZ | A7:37 | 4.93 | 1.00 | 41.57 |  |
  |

  
 Coded contact list:   
 Exclusion zone contact(s): /OP2/  
Consider K in place of MG (1 valid contact(s) for K vs. 0 for MG)  

### MG:AA:3106

|  | Res. type | Atom | Residue | Distance | Occupancy | B-factor | Closest solute to 1^st^ shell water |
| --- | --- | --- | --- | --- | --- | --- | --- |
| **Ion** | | | |  | 1.00 | 17.86 || 1 | G | OP2 | AA:801 | 2.55 | 1.00 | 36.34 |
| 2 | A | O3' | AA:800 | 3.66 | 1.00 | 38.84 |  |
| 3 | GLY | O | AF:55 | 4.05 | 1.00 | 36.66 |  |
| 4 | G | OP1 | AA:798 | 4.17 | 1.00 | 40.21 |  |
| 5 | G | OP2 | AA:799 | 4.18 | 1.00 | 40.08 |  |
| 6 | A | O2' | AA:800 | 4.21 | 1.00 | 45.16 |  |
| 7 | GLY | N | AF:61 | 4.21 | 1.00 | 50.13 |  |
| 8 | TYR | O | AF:59 | 4.57 | 1.00 | 46.15 |  |
| 9 | G | OP1 | AA:801 | 4.71 | 1.00 | 35.90 |  |
| 10 | G | O5' | AA:801 | 4.71 | 1.00 | 37.30 |  |
  |

  
 Coded contact list:   
 Exclusion zone contact(s): /OP2/  
Consider NA in place of MG (1 valid contact(s) for NA vs. 0 for MG)  

### MG:AA:3107

|  | Res. type | Atom | Residue | Distance | Occupancy | B-factor | Closest solute to 1^st^ shell water |
| --- | --- | --- | --- | --- | --- | --- | --- |
| **Ion** | | | |  | 1.00 | 32.30 || 1 | A | OP1 | AA:800 | 2.35 | 1.00 | 37.83 |
| 2 | C | N3 | AA:192 | 3.84 | 1.00 | 39.60 |  |
| 3 | A | OP2 | AA:800 | 4.15 | 1.00 | 37.20 |  |
| 4 | A | N1 | AA:191 | 4.17 | 1.00 | 34.56 |  |
| 5 | G | O2' | AA:799 | 4.23 | 1.00 | 42.46 |  |
| 6 | A | N7 | AA:800 | 4.33 | 1.00 | 47.00 |  |
| 7 | C | O2 | AA:192 | 4.33 | 1.00 | 39.93 |  |
| 8 | G | O3' | AA:799 | 4.35 | 1.00 | 38.43 |  |
| 9 | C | N4 | AA:192 | 4.65 | 1.00 | 36.01 |  |
| 10 | A | O5' | AA:800 | 4.82 | 1.00 | 37.80 |  |
| 11 | C | O2' | AA:203 | 4.92 | 1.00 | 42.88 |  |
  |

  
 Contact(s): /OP1/  
 Coded contact list: O_ph_  

### MG:AA:3108

|  | Res. type | Atom | Residue | Distance | Occupancy | B-factor | Closest solute to 1^st^ shell water |
| --- | --- | --- | --- | --- | --- | --- | --- |
| **Ion** | | | |  | 1.00 | 41.31 || 1 | A | OP1 | AA:802 | 2.41 | 1.00 | 37.46 |
| 2 | G | O3' | AA:801 | 3.51 | 1.00 | 38.46 |  |
| 3 | G | OP2 | AA:674 | 3.57 | 1.00 | 36.14 |  |
| 4 | A | OP2 | AA:675 | 3.92 | 1.00 | 37.76 |  |
| 5 | ARG | NH2 | AF:54 | 3.95 | 1.00 | 38.79 |  |
| 6 | A | OP2 | AA:802 | 4.43 | 1.00 | 39.35 |  |
| 7 | G | OP1 | AA:674 | 4.51 | 1.00 | 37.93 |  |
| 8 | A | O5' | AA:802 | 4.68 | 1.00 | 39.38 |  |
| 9 | ARG | NH1 | AF:54 | 4.76 | 1.00 | 37.75 |  |
| 10 | G | O5' | AA:674 | 4.81 | 1.00 | 36.70 |  |
| 11 | U | O4 | AA:803 | 4.99 | 1.00 | 37.98 |  |
  |

  
 Coded contact list:   
 Exclusion zone contact(s): /OP1/  
Consider NA in place of MG (1 valid contact(s) for NA vs. 0 for MG)  

### MG:AA:3109

|  | Res. type | Atom | Residue | Distance | Occupancy | B-factor | Closest solute to 1^st^ shell water |
| --- | --- | --- | --- | --- | --- | --- | --- |
| **Ion** | | | |  | 1.00 | 40.20 || 1 | G | O6 | AA:798 | 4.44 | 1.00 | 36.93 |
| 2 | G | O6 | AA:799 | 4.52 | 1.00 | 39.87 |  |
| 3 | G | N7 | AA:798 | 4.60 | 1.00 | 40.86 |  |
| 4 | ARG | N | AF:62 | 4.96 | 1.00 | 48.72 |  |
  |

  
 Coded contact list:   
1^st^ MG coordination shell missing (see Notes)  

### MG:AA:3110

|  | Res. type | Atom | Residue | Distance | Occupancy | B-factor | Closest solute to 1^st^ shell water |
| --- | --- | --- | --- | --- | --- | --- | --- |
| **Ion** | | | |  | 1.00 | 59.01 || 1 | G | N7 | AA:35 | 3.32 | 1.00 | 48.30 |
| 2 | G | N7 | AA:36 | 3.38 | 1.00 | 45.32 |  |
| 3 | C | O2 | AA:34 | 3.45 | 1.00 | 87.40 |  |
| 4 | G | O6 | AA:35 | 3.51 | 1.00 | 45.68 |  |
| 5 | G | O6 | AA:36 | 3.64 | 1.00 | 42.47 |  |
| 6 | C | O2' | AA:34 | 4.58 | 1.00 | 79.11 |  |
| 7 | C | N1 | AA:34 | 4.84 | 1.00 | 84.82 |  |
| 8 | C | N3 | AA:34 | 4.90 | 1.00 | 87.68 |  |
  |

  
 Coded contact list:   
 Exclusion zone contact(s): /N7/N7/  
Consider K in place of MG (2 valid contact(s) for K vs. 0 for MG)  

### MG:AA:3111

|  | Res. type | Atom | Residue | Distance | Occupancy | B-factor | Closest solute to 1^st^ shell water |
| --- | --- | --- | --- | --- | --- | --- | --- |
| **Ion** | | | |  | 1.00 | 46.03 || 1 | C | OP1 | AA:444 | 2.05 | 1.00 | 50.68 |
| 2 | G | O2' | AA:442 | 2.91 | 1.00 | 61.15 |  |
| 3 | C | O5' | AA:444 | 3.10 | 1.00 | 50.21 |  |
| 4 | C | O4' | AA:444 | 3.19 | 1.00 | 47.33 |  |
| 5 | THR | N | AF:48 | 3.58 | 1.00 | 53.42 |  |
| 6 | ALA | N | AF:49 | 3.61 | 1.00 | 52.00 |  |
| 7 | C | OP2 | AA:444 | 3.82 | 1.00 | 50.98 |  |
| 8 | G | O3' | AA:442 | 4.06 | 1.00 | 57.98 |  |
| 9 | THR | OG1 | AF:48 | 4.15 | 1.00 | 52.70 |  |
| 10 | GLY | N | AF:47 | 4.19 | 1.00 | 52.88 |  |
| 11 | A | O5' | AA:443 | 4.23 | 1.00 | 55.27 |  |
| 12 | A | O3' | AA:443 | 4.31 | 1.00 | 52.68 |  |
| 13 | THR | O | AF:48 | 4.73 | 1.00 | 53.77 |  |
| 14 | A | OP1 | AA:443 | 4.81 | 1.00 | 57.00 |  |
  |

  
 Contact(s): /OP1/  
 Coded contact list: O_ph_  
 Exclusion zone contact(s): /O2'/O4'/O5'/  
Consider K in place of MG (3 valid contact(s) for K vs. 1 for MG)  

### MG:AA:3112

|  | Res. type | Atom | Residue | Distance | Occupancy | B-factor | Closest solute to 1^st^ shell water |
| --- | --- | --- | --- | --- | --- | --- | --- |
| **Ion** | | | |  | 1.00 | 102.42 || 1 | A | OP1 | AA:443 | 1.62 | 1.00 | 57.00 |
| 2 | ARG | N | AF:46 | 2.59 | 1.00 | 52.87 |  |
| 3 | A | O5' | AA:443 | 2.78 | 1.00 | 55.27 |  |
| 4 | GLY | N | AF:47 | 3.02 | 1.00 | 52.88 |  |
| 5 | G | O3' | AA:442 | 3.28 | 1.00 | 57.98 |  |
| 6 | G | O2' | AA:442 | 3.89 | 1.00 | 61.15 |  |
| 7 | A | OP2 | AA:443 | 3.97 | 1.00 | 58.78 |  |
| 8 | C | OP1 | AA:444 | 4.06 | 1.00 | 50.68 |  |
| 9 | G | N2 | AA:615 | 4.26 | 1.00 | 98.29 |  |
| 10 | ARG | NE | AF:45 | 4.71 | 1.00 | 58.25 |  |
| 11 | ARG | O | AF:46 | 4.71 | 1.00 | 52.90 |  |
| 12 | ARG | O | AF:45 | 4.72 | 1.00 | 52.60 |  |
| 13 | THR | N | AF:48 | 4.72 | 1.00 | 53.42 |  |
| 14 | THR | OG1 | AF:48 | 4.80 | 1.00 | 52.70 |  |
| 15 | G | O4' | AA:442 | 4.96 | 1.00 | 59.65 |  |
  |

  
 Coded contact list:   
 Exclusion zone contact(s): /N_back_/N_back_/O3'/O5'/OP1/  
Consider K in place of MG (3 valid contact(s) for K vs. 0 for MG)  

### MG:AA:3113

|  | Res. type | Atom | Residue | Distance | Occupancy | B-factor | Closest solute to 1^st^ shell water |
| --- | --- | --- | --- | --- | --- | --- | --- |
| **Ion** | | | |  | 1.00 | 85.32 || 1 | G | O6 | AA:1243 | 2.95 | 1.00 | 60.55 |
| 2 | G | N7 | AA:1243 | 3.04 | 1.00 | 64.32 |  |
| 3 | A | OP2 | AA:1242 | 3.07 | 1.00 | 65.84 |  |
| 4 | G | O6 | AA:1244 | 3.60 | 1.00 | 47.77 |  |
| 5 | G | N7 | AA:1244 | 3.74 | 1.00 | 51.45 |  |
| 6 | G | N2 | AA:1203 | 4.46 | 1.00 | 60.18 |  |
| 7 | A | OP1 | AA:1241 | 4.51 | 1.00 | 56.89 |  |
| 8 | G | N1 | AA:1243 | 4.83 | 1.00 | 58.79 |  |
| 9 | A | OP1 | AA:1242 | 4.95 | 1.00 | 65.37 |  |
  |

  
 Coded contact list:   
 Exclusion zone contact(s): /N7/O6/OP2/  
Consider K in place of MG (3 valid contact(s) for K vs. 0 for MG)  

### MG:AA:3114

|  | Res. type | Atom | Residue | Distance | Occupancy | B-factor | Closest solute to 1^st^ shell water |
| --- | --- | --- | --- | --- | --- | --- | --- |
| **Ion** | | | |  | 1.00 | 97.62 || 1 | U | OP1 | AA:1240 | 2.94 | 1.00 | 58.00 |
| 2 | G | O3' | AA:1239 | 4.09 | 1.00 | 58.80 |  |
| 3 | U | OP2 | AA:1199 | 4.59 | 1.00 | 58.09 |  |
| 4 | G | OP1 | AA:1215 | 4.71 | 1.00 | 64.79 |  |
| 5 | U | OP1 | AA:1199 | 4.74 | 1.00 | 58.50 |  |
| 6 | U | O5' | AA:1240 | 4.84 | 1.00 | 57.56 |  |
| 7 | A | O3' | AA:1214 | 4.88 | 1.00 | 63.96 |  |
  |

  
 Coded contact list:   
 Exclusion zone contact(s): /OP1/  
Consider K in place of MG (1 valid contact(s) for K vs. 0 for MG)  

### MG:AA:3115

|  | Res. type | Atom | Residue | Distance | Occupancy | B-factor | Closest solute to 1^st^ shell water |
| --- | --- | --- | --- | --- | --- | --- | --- |
| **Ion** | | | |  | 1.00 | 76.64 || 1 | C | OP2 | AA:32 | 3.99 | 1.00 | 45.63 |
| 2 | C | OP1 | AA:32 | 4.31 | 1.00 | 45.59 |  |
| 3 | G | N7 | AA:446 | 4.69 | 1.00 | 60.16 |  |
| 4 | U | OP2 | AA:33 | 4.72 | 1.00 | 58.57 |  |
| 5 | G | O6 | AA:446 | 4.82 | 1.00 | 62.86 |  |
  |

  
 Coded contact list:   
1^st^ MG coordination shell missing (see Notes)  

### MG:AA:3116

|  | Res. type | Atom | Residue | Distance | Occupancy | B-factor | Closest solute to 1^st^ shell water |
| --- | --- | --- | --- | --- | --- | --- | --- |
| **Ion** | | | |  | 1.00 | 85.76 || 1 | C | OP1 | AA:291 | 3.01 | 1.00 | 86.40 |
| 2 | C | OP2 | AA:291 | 3.95 | 1.00 | 85.83 |  |
| 3 | C | O5' | AA:291 | 4.12 | 1.00 | 85.04 |  |
| 4 | C | OP2 | AA:292 | 4.56 | 1.00 | 80.04 |  |
| 5 | G | O6 | AA:344 | 4.67 | 1.00 | 72.66 |  |
| 6 | G | N7 | AA:344 | 4.88 | 1.00 | 73.17 |  |
  |

  
 Coded contact list:   
 Exclusion zone contact(s): /OP1/  
Consider K in place of MG (1 valid contact(s) for K vs. 0 for MG)  

### MG:AA:3117

|  | Res. type | Atom | Residue | Distance | Occupancy | B-factor | Closest solute to 1^st^ shell water |
| --- | --- | --- | --- | --- | --- | --- | --- |
| **Ion** | | | |  | 1.00 | 72.13 || 1 | G | OP2 | AA:341 | 4.21 | 1.00 | 69.89 |
| 2 | G | N7 | AA:342 | 4.57 | 1.00 | 76.66 |  |
| 3 | G | N7 | AA:341 | 4.97 | 1.00 | 77.74 |  |
  |

  
 Coded contact list:   
1^st^ MG coordination shell missing (see Notes)  

### MG:AA:3118

|  | Res. type | Atom | Residue | Distance | Occupancy | B-factor | Closest solute to 1^st^ shell water |
| --- | --- | --- | --- | --- | --- | --- | --- |
| **Ion** | | | |  | 1.00 | 52.24 || 1 | G | O6 | AA:342 | 2.44 | 1.00 | 76.47 |
| 2 | G | O6 | AA:341 | 2.76 | 1.00 | 74.60 |  |
| 3 | C | N4 | AA:297 | 3.31 | 1.00 | 73.75 |  |
| 4 | C | N4 | AA:296 | 3.69 | 1.00 | 73.79 |  |
| 5 | G | N2 | AA:298 | 4.03 | 1.00 | 71.32 |  |
| 6 | G | N7 | AA:342 | 4.22 | 1.00 | 76.66 |  |
| 7 | G | N1 | AA:342 | 4.55 | 1.00 | 76.60 |  |
| 8 | U | OP2 | AA:339 | 4.57 | 1.00 | 61.32 |  |
| 9 | G | N7 | AA:341 | 4.79 | 1.00 | 77.74 |  |
| 10 | G | N1 | AA:341 | 4.80 | 1.00 | 76.16 |  |
| 11 | U | OP1 | AA:339 | 4.80 | 1.00 | 61.59 |  |
  |

  
 Coded contact list:   
 Exclusion zone contact(s): /N4/O6/O6/  
Consider K in place of MG (2 valid contact(s) for K vs. 0 for MG)  

### MG:AA:3119

|  | Res. type | Atom | Residue | Distance | Occupancy | B-factor | Closest solute to 1^st^ shell water |
| --- | --- | --- | --- | --- | --- | --- | --- |
| **Ion** | | | |  | 1.00 | 46.78 || 1 | G | N7 | AA:298 | 2.76 | 1.00 | 70.12 |
| 2 | G | OP2 | AA:338 | 3.03 | 1.00 | 59.67 |  |
| 3 | G | O6 | AA:298 | 3.22 | 1.00 | 72.95 |  |
| 4 | G | OP1 | AA:338 | 4.70 | 1.00 | 58.16 |  |
| 5 | G | OP2 | AA:298 | 4.73 | 1.00 | 68.01 |  |
| 6 | G | N9 | AA:298 | 4.91 | 1.00 | 69.58 |  |
| 7 | C | OP2 | AA:297 | 4.97 | 1.00 | 70.55 |  |
  |

  
 Coded contact list:   
 Exclusion zone contact(s): /N7/O6/OP2/  
Consider K in place of MG (3 valid contact(s) for K vs. 0 for MG)  

### MG:AA:3120

|  | Res. type | Atom | Residue | Distance | Occupancy | B-factor | Closest solute to 1^st^ shell water |
| --- | --- | --- | --- | --- | --- | --- | --- |
| **Ion** | | | |  | 1.00 | 59.45 || 1 | U | OP2 | AA:434 | 3.78 | 1.00 | 58.30 |
| 2 | C | OP1 | AA:433 | 3.96 | 1.00 | 46.70 |  |
| 3 | C | OP1 | AA:39 | 4.47 | 1.00 | 46.18 |  |
| 4 | U | OP1 | AA:434 | 4.51 | 1.00 | 57.98 |  |
| 5 | C | O5' | AA:433 | 4.78 | 1.00 | 47.95 |  |
| 6 | C | OP2 | AA:40 | 4.94 | 1.00 | 60.91 |  |
  |

  
 Coded contact list:   
1^st^ MG coordination shell missing (see Notes)  

### MG:AA:3121

|  | Res. type | Atom | Residue | Distance | Occupancy | B-factor | Closest solute to 1^st^ shell water |
| --- | --- | --- | --- | --- | --- | --- | --- |
| **Ion** | | | |  | 1.00 | 85.65 || 1 | U | OP1 | AA:434 | 3.92 | 1.00 | 57.98 |
| 2 | C | OP2 | AA:39 | 4.38 | 1.00 | 47.35 |  |
| 3 | A | OP1 | AA:38 | 4.43 | 1.00 | 44.34 |  |
| 4 | C | OP1 | AA:39 | 4.48 | 1.00 | 46.18 |  |
  |

  
 Coded contact list:   
1^st^ MG coordination shell missing (see Notes)  

### MG:AA:3122

|  | Res. type | Atom | Residue | Distance | Occupancy | B-factor | Closest solute to 1^st^ shell water |
| --- | --- | --- | --- | --- | --- | --- | --- |
| **Ion** | | | |  | 1.00 | 73.25 || 1 | A | OP1 | AA:432 | 2.56 | 1.00 | 45.91 |
| 2 | C | OP2 | AA:433 | 3.43 | 1.00 | 46.40 |  |
| 3 | A | OP2 | AA:432 | 3.88 | 1.00 | 46.12 |  |
| 4 | A | O5' | AA:432 | 4.16 | 1.00 | 45.71 |  |
| 5 | C | OP1 | AA:433 | 4.82 | 1.00 | 46.70 |  |
| 6 | U | O3' | AA:431 | 4.95 | 1.00 | 48.21 |  |
  |

  
 Coded contact list:   
 Exclusion zone contact(s): /OP1/  
Consider NA in place of MG (1 valid contact(s) for NA vs. 0 for MG)  

### MG:AA:3123

|  | Res. type | Atom | Residue | Distance | Occupancy | B-factor | Closest solute to 1^st^ shell water |
| --- | --- | --- | --- | --- | --- | --- | --- |
| **Ion** | | | |  | 1.00 | 47.97 || 1 | U | O4 | AA:963 | 4.15 | 1.00 | 56.06 |
| 2 | C | OP2 | AA:961 | 4.55 | 1.00 | 60.10 |  |
| 3 | G | N7 | AA:962 | 4.62 | 1.00 | 60.41 |  |
| 4 | G | OP2 | AA:962 | 4.72 | 1.00 | 60.12 |  |
| 5 | C | OP2 | AA:949 | 4.79 | 1.00 | 56.75 |  |
  |

  
 Coded contact list:   
1^st^ MG coordination shell missing (see Notes)  

### MG:AA:3124

|  | Res. type | Atom | Residue | Distance | Occupancy | B-factor | Closest solute to 1^st^ shell water |
| --- | --- | --- | --- | --- | --- | --- | --- |
| **Ion** | | | |  | 1.00 | 71.81 || 1 | G | N7 | AA:842 | 3.22 | 1.00 | 69.72 |
| 2 | G | O6 | AA:842 | 4.13 | 1.00 | 70.55 |  |
| 3 | G | OP2 | AA:842 | 4.37 | 1.00 | 70.32 |  |
| 4 | G | N7 | AA:843 | 4.44 | 1.00 | 77.58 |  |
| 5 | G | O6 | AA:843 | 4.49 | 1.00 | 74.19 |  |
| 6 | A | N7 | AA:841 | 4.87 | 1.00 | 58.02 |  |
  |

  
 Coded contact list:   
 Exclusion zone contact(s): /N7/  
Consider K in place of MG (1 valid contact(s) for K vs. 0 for MG)  

### MG:AA:3125

|  | Res. type | Atom | Residue | Distance | Occupancy | B-factor | Closest solute to 1^st^ shell water |
| --- | --- | --- | --- | --- | --- | --- | --- |
| **Ion** | | | |  | 1.00 | 51.81 || 1 | G | N2 | AA:845 | 2.73 | 1.00 | 92.53 |
| 2 | A | N1 | AA:933 | 3.09 | 1.00 | 83.54 |  |
| 3 | G | N1 | AA:845 | 3.90 | 1.00 | 93.95 |  |
| 4 | G | O6 | AA:934 | 4.10 | 1.00 | 79.47 |  |
| 5 | A | N6 | AA:933 | 4.37 | 1.00 | 82.22 |  |
| 6 | G | O6 | AA:931 | 4.57 | 1.00 | 77.16 |  |
| 7 | C | N4 | AA:844 | 4.85 | 1.00 | 80.76 |  |
| 8 | A | N3 | AA:933 | 4.96 | 1.00 | 81.12 |  |
| 9 | G | N3 | AA:845 | 4.98 | 1.00 | 91.43 |  |
  |

  
 Coded contact list:   
 Exclusion zone contact(s): /N1_A_/N2/  
Consider K in place of MG (2 valid contact(s) for K vs. 0 for MG)  

### MG:AA:3126

|  | Res. type | Atom | Residue | Distance | Occupancy | B-factor | Closest solute to 1^st^ shell water |
| --- | --- | --- | --- | --- | --- | --- | --- |
| **Ion** | | | |  | 1.00 | 58.26 || 1 | U | OP2 | AA:847 | 2.01 | 1.00 | 73.77 |
| 2 | G | O6 | AA:929 | 2.03 | 1.00 | 87.88 |  |
| 3 | U | O5' | AA:847 | 3.63 | 1.00 | 74.46 |  |
| 4 | G | N1 | AA:929 | 3.73 | 1.00 | 87.69 |  |
| 5 | U | OP1 | AA:847 | 4.19 | 1.00 | 74.70 |  |
| 6 | C | O2' | AA:846 | 4.38 | 1.00 | 79.03 |  |
| 7 | C | O3' | AA:846 | 4.46 | 1.00 | 75.69 |  |
| 8 | G | N7 | AA:928 | 4.50 | 1.00 | 80.05 |  |
| 9 | G | N7 | AA:929 | 4.65 | 1.00 | 88.84 |  |
| 10 | G | OP2 | AA:928 | 4.80 | 1.00 | 85.46 |  |
  |

  
**1^st^ shell ligand angles:**   
1-2: 81;   
  
 Contact(s): /O6/OP2/  
 Coded contact list: O_ph_O_b_  

### MG:AA:3127

|  | Res. type | Atom | Residue | Distance | Occupancy | B-factor | Closest solute to 1^st^ shell water |
| --- | --- | --- | --- | --- | --- | --- | --- |
| **Ion** | | | |  | 1.00 | 46.67 || 1 | G | O6 | AA:929 | 2.33 | 1.00 | 87.88 |
| 2 | C | O2' | AA:846 | 2.33 | 1.00 | 79.03 |  |
| 3 | G | N7 | AA:929 | 2.67 | 1.00 | 88.84 |  |
| 4 | U | OP2 | AA:847 | 3.03 | 1.00 | 73.77 |  |
| 5 | C | O3' | AA:846 | 3.22 | 1.00 | 75.69 |  |
| 6 | U | O5' | AA:847 | 3.93 | 1.00 | 74.46 |  |
| 7 | G | N1 | AA:929 | 3.99 | 1.00 | 87.69 |  |
| 8 | U | O4 | AA:930 | 4.37 | 1.00 | 88.06 |  |
| 9 | G | N9 | AA:929 | 4.59 | 1.00 | 87.69 |  |
| 10 | U | N3 | AA:930 | 4.78 | 1.00 | 87.53 |  |
| 11 | C | O2 | AA:846 | 4.80 | 1.00 | 76.99 |  |
| 12 | G | OP2 | AA:928 | 4.81 | 1.00 | 85.46 |  |
  |

  
**1^st^ shell ligand angles:**   
1-2: 86;   
  
 Contact(s): /O2'/O6/  
 Coded contact list: O_r_O_b_  
 Exclusion zone contact(s): /N7/O3'/OP2/  
Consider K in place of MG (3 valid contact(s) for K vs. 2 for MG)  

### MG:AA:3128

|  | Res. type | Atom | Residue | Distance | Occupancy | B-factor | Closest solute to 1^st^ shell water |
| --- | --- | --- | --- | --- | --- | --- | --- |
| **Ion** | | | |  | 1.00 | 32.62 || 1 | A | OP2 | AA:1664 | 2.59 | 1.00 | 46.87 |
| 2 | G | N7 | AA:1992 | 3.41 | 1.00 | 43.75 |  |
| 3 | G | O6 | AA:1992 | 3.77 | 1.00 | 46.35 |  |
| 4 | G | OP2 | AA:1992 | 3.96 | 1.00 | 47.67 |  |
| 5 | G | OP1 | AA:1992 | 4.02 | 1.00 | 46.48 |  |
| 6 | A | N6 | AA:1665 | 4.04 | 1.00 | 46.60 |  |
| 7 | G | O6 | AA:1666 | 4.35 | 1.00 | 39.75 |  |
| 8 | A | N7 | AA:1664 | 4.43 | 1.00 | 47.90 |  |
| 9 | A | N7 | AA:1665 | 4.45 | 1.00 | 48.47 |  |
| 10 | A | O5' | AA:1664 | 4.75 | 1.00 | 46.68 |  |
| 11 | C | O3' | AA:1663 | 4.84 | 1.00 | 47.47 |  |
| 12 | C | O2' | AA:1663 | 4.85 | 1.00 | 47.42 |  |
  |

  
 Coded contact list:   
 Exclusion zone contact(s): /OP2/  
Consider NA in place of MG (1 valid contact(s) for NA vs. 0 for MG)  

### MG:AA:3129

|  | Res. type | Atom | Residue | Distance | Occupancy | B-factor | Closest solute to 1^st^ shell water |
| --- | --- | --- | --- | --- | --- | --- | --- |
| **Ion** | | | |  | 1.00 | 31.27 || 1 | U | OP2 | AA:1352 | 2.22 | 1.00 | 44.43 |
| 2 | C | OP2 | AA:1376 | 2.33 | 1.00 | 49.43 |  |
| 3 | U | O5' | AA:1352 | 3.39 | 1.00 | 44.42 |  |
| 4 | U | OP1 | AA:1352 | 3.63 | 1.00 | 46.66 |  |
| 5 | C | OP2 | AA:1375 | 3.78 | 1.00 | 56.77 |  |
| 6 | C | O5' | AA:1375 | 3.86 | 1.00 | 55.38 |  |
| 7 | C | OP1 | AA:1375 | 3.94 | 1.00 | 56.60 |  |
| 8 | C | O3' | AA:1375 | 4.38 | 1.00 | 49.33 |  |
| 9 | C | OP1 | AA:1376 | 4.44 | 1.00 | 50.34 |  |
| 10 | C | O3' | AA:1351 | 4.56 | 1.00 | 43.28 |  |
| 11 | G | O6 | AA:1377 | 4.61 | 1.00 | 52.27 |  |
| 12 | C | O5' | AA:1376 | 4.79 | 1.00 | 49.21 |  |
| 13 | C | O5' | AA:1351 | 4.90 | 1.00 | 44.29 |  |
| 14 | C | OP1 | AA:1351 | 4.92 | 1.00 | 43.97 |  |
| 15 | C | OP2 | AA:1351 | 4.94 | 1.00 | 45.27 |  |
  |

  
**1^st^ shell ligand angles:**   
1-2: 135;   
  
 Contact(s): /OP2/OP2/  
 Coded contact list: 2O_ph_  
 Exclusion zone contact(s): /O5'/  

### MG:AA:3130

|  | Res. type | Atom | Residue | Distance | Occupancy | B-factor | Closest solute to 1^st^ shell water |
| --- | --- | --- | --- | --- | --- | --- | --- |
| **Ion** | | | |  | 1.00 | 24.59 || 1 | G | OP2 | AA:1647 | 2.24 | 1.00 | 48.36 |
| 2 | G | OP1 | AA:1647 | 2.76 | 1.00 | 49.60 |  |
| 3 | C | O2' | AA:1646 | 3.80 | 1.00 | 48.53 |  |
| 4 | A | O2' | AA:1616 | 3.98 | 1.00 | 50.86 |  |
| 5 | C | O3' | AA:1646 | 4.03 | 1.00 | 48.42 |  |
| 6 | G | O5' | AA:1647 | 4.10 | 1.00 | 46.53 |  |
| 7 | G | OP2 | AA:1325 | 4.80 | 1.00 | 50.31 |  |
| 8 | A | OP2 | AA:1272 | 4.87 | 1.00 | 48.76 |  |
| 9 | G | OP1 | AA:1325 | 4.93 | 1.00 | 50.00 |  |
  |

  
 Contact(s): /OP2/  
 Coded contact list: O_ph_  
 Exclusion zone contact(s): /OP1/  

### MG:AA:3131

|  | Res. type | Atom | Residue | Distance | Occupancy | B-factor | Closest solute to 1^st^ shell water |
| --- | --- | --- | --- | --- | --- | --- | --- |
| **Ion** | | | |  | 1.00 | 34.15 || 1 | G | N7 | AA:2002 | 2.63 | 1.00 | 40.25 |
| 2 | G | O6 | AA:2002 | 3.69 | 1.00 | 38.73 |  |
| 3 | G | OP2 | AA:2002 | 3.85 | 1.00 | 43.08 |  |
| 4 | G | O6 | AA:2003 | 4.58 | 1.00 | 45.98 |  |
| 5 | A | N7 | AA:2001 | 4.60 | 1.00 | 36.20 |  |
| 6 | G | N9 | AA:2002 | 4.67 | 1.00 | 40.67 |  |
| 7 | G | N7 | AA:2003 | 4.82 | 1.00 | 48.29 |  |
  |

  
 Coded contact list:   
 Exclusion zone contact(s): /N7/  
Consider K in place of MG (1 valid contact(s) for K vs. 0 for MG)  

### MG:AA:3132

|  | Res. type | Atom | Residue | Distance | Occupancy | B-factor | Closest solute to 1^st^ shell water |
| --- | --- | --- | --- | --- | --- | --- | --- |
| **Ion** | | | |  | 1.00 | 44.59 || 1 | U | OP1 | AA:2406 | 2.65 | 1.00 | 70.14 |
| 2 | U | O5' | AA:2406 | 2.74 | 1.00 | 70.91 |  |
| 3 | U | OP2 | AA:2408 | 3.88 | 1.00 | 69.34 |  |
| 4 | G | N7 | AA:2409 | 4.02 | 1.00 | 70.00 |  |
| 5 | U | O4 | AA:2408 | 4.08 | 1.00 | 66.28 |  |
| 6 | G | O6 | AA:2409 | 4.09 | 1.00 | 67.44 |  |
| 7 | G | O6 | AA:2410 | 4.26 | 1.00 | 67.96 |  |
| 8 | U | OP2 | AA:2406 | 4.30 | 1.00 | 69.59 |  |
| 9 | G | N7 | AA:2410 | 4.42 | 1.00 | 70.28 |  |
| 10 | G | O3' | AA:2405 | 4.47 | 1.00 | 68.91 |  |
| 11 | G | OP1 | AA:2407 | 4.57 | 1.00 | 69.54 |  |
| 12 | U | N3 | AA:2408 | 4.88 | 1.00 | 66.39 |  |
| 13 | G | OP2 | AA:2409 | 4.97 | 1.00 | 67.22 |  |
  |

  
 Coded contact list:   
 Exclusion zone contact(s): /O5'/OP1/  
Consider K in place of MG (2 valid contact(s) for K vs. 0 for MG)  

### MG:AA:3133

|  | Res. type | Atom | Residue | Distance | Occupancy | B-factor | Closest solute to 1^st^ shell water |
| --- | --- | --- | --- | --- | --- | --- | --- |
| **Ion** | | | |  | 1.00 | 25.21 || 1 | G | OP2 | AA:1456 | 2.29 | 1.00 | 79.27 |
| 2 | U | O3' | AA:1454 | 2.32 | 1.00 | 70.30 |  |
| 3 | G | OP2 | AA:1455 | 2.75 | 1.00 | 68.93 |  |
| 4 | G | O5' | AA:1455 | 2.92 | 1.00 | 69.79 |  |
| 5 | U | O2' | AA:1454 | 4.09 | 1.00 | 73.89 |  |
| 6 | G | O3' | AA:1455 | 4.20 | 1.00 | 77.56 |  |
| 7 | G | OP1 | AA:1455 | 4.30 | 1.00 | 68.48 |  |
| 8 | G | OP1 | AA:1456 | 4.38 | 1.00 | 79.78 |  |
| 9 | A | OP2 | AA:1453 | 4.70 | 1.00 | 80.36 |  |
| 10 | G | O5' | AA:1456 | 4.77 | 1.00 | 80.46 |  |
  |

  
**1^st^ shell ligand angles:**   
1-2: 135;   
  
 Contact(s): /O3'/OP2/  
 Coded contact list: O_ph_O_r_  
 Exclusion zone contact(s): /O5'/OP2/  

### MG:AA:3134

|  | Res. type | Atom | Residue | Distance | Occupancy | B-factor | Closest solute to 1^st^ shell water |
| --- | --- | --- | --- | --- | --- | --- | --- |
| **Ion** | | | |  | 1.00 | 22.39 || 1 | C | OP2 | AA:730 | 2.27 | 1.00 | 38.43 |
| 2 | C | OP2 | AA:731 | 2.59 | 1.00 | 42.66 |  |
| 3 | G | OP1 | AA:763 | 2.65 | 1.00 | 37.73 |  |
| 4 | A | OP1 | AA:761 | 2.75 | 1.00 | 37.95 |  |
| 5 | U | OP1 | AA:762 | 3.98 | 1.00 | 40.39 |  |
| 6 | G | O3' | AA:729 | 3.98 | 1.00 | 39.11 |  |
| 7 | A | O5' | AA:761 | 4.06 | 1.00 | 39.69 |  |
| 8 | C | O3' | AA:730 | 4.23 | 1.00 | 41.42 |  |
| 9 | C | O5' | AA:730 | 4.50 | 1.00 | 38.86 |  |
| 10 | G | O5' | AA:763 | 4.51 | 1.00 | 39.00 |  |
| 11 | C | OP1 | AA:730 | 4.70 | 1.00 | 39.61 |  |
| 12 | A | OP2 | AA:761 | 4.92 | 1.00 | 38.67 |  |
| 13 | C | OP1 | AA:731 | 4.93 | 1.00 | 40.60 |  |
| 14 | C | O5' | AA:731 | 4.96 | 1.00 | 40.64 |  |
| 15 | G | OP2 | AA:763 | 4.99 | 1.00 | 36.62 |  |
| 16 | U | O5' | AA:762 | 4.99 | 1.00 | 39.46 |  |
  |

  
 Contact(s): /OP2/  
 Coded contact list: O_ph_  
 Exclusion zone contact(s): /OP1/OP1/OP2/  
Consider K in place of MG (2 valid contact(s) for K vs. 1 for MG)  

### MG:AA:3135

|  | Res. type | Atom | Residue | Distance | Occupancy | B-factor | Closest solute to 1^st^ shell water |
| --- | --- | --- | --- | --- | --- | --- | --- |
| **Ion** | | | |  | 1.00 | 19.92 || 1 | U | O4 | AA:2028 | 2.34 | 1.00 | 38.88 |
| 2 | A | OP1 | AA:2033 | 2.92 | 1.00 | 57.74 |  |
| 3 | G | O3' | AA:2032 | 3.11 | 1.00 | 58.70 |  |
| 4 | G | O6 | AA:2029 | 3.63 | 1.00 | 30.76 |  |
| 5 | A | O5' | AA:2033 | 3.75 | 1.00 | 57.07 |  |
| 6 | G | N7 | AA:2027 | 4.51 | 1.00 | 40.80 |  |
| 7 | U | N3 | AA:2028 | 4.59 | 1.00 | 39.81 |  |
| 8 | G | O6 | AA:2027 | 4.68 | 1.00 | 41.46 |  |
| 9 | G | N1 | AA:2029 | 4.76 | 1.00 | 32.90 |  |
| 10 | G | O2' | AA:2032 | 4.77 | 1.00 | 60.51 |  |
| 11 | A | N7 | AA:2033 | 4.85 | 1.00 | 53.25 |  |
| 12 | A | OP2 | AA:2033 | 4.97 | 1.00 | 57.73 |  |
  |

  
 Contact(s): /O4/  
 Coded contact list: O_b_  
 Exclusion zone contact(s): /O3'/OP1/  
Consider K in place of MG (2 valid contact(s) for K vs. 1 for MG)  

### MG:AA:3136

|  | Res. type | Atom | Residue | Distance | Occupancy | B-factor | Closest solute to 1^st^ shell water |
| --- | --- | --- | --- | --- | --- | --- | --- |
| **Ion** | | | |  | 1.00 | 38.15 || 1 | G | OP1 | AA:1987 | 3.74 | 1.00 | 42.27 |
| 2 | A | OP1 | AA:1986 | 3.75 | 1.00 | 44.66 |  |
| 3 | C | OP1 | AA:2551 | 4.29 | 1.00 | 49.75 |  |
| 4 | G | OP2 | AA:1987 | 4.56 | 1.00 | 44.45 |  |
| 5 | A | OP1 | AA:1669 | 4.62 | 1.00 | 48.24 |  |
| 6 | G | N1 | AA:1954 | 4.76 | 1.00 | 56.99 |  |
| 7 | A | OP2 | AA:1669 | 4.83 | 1.00 | 47.14 |  |
| 8 | G | O6 | AA:1954 | 4.96 | 1.00 | 60.81 |  |
  |

  
 Coded contact list:   
1^st^ MG coordination shell missing (see Notes)  

### MG:AA:3137

|  | Res. type | Atom | Residue | Distance | Occupancy | B-factor | Closest solute to 1^st^ shell water |
| --- | --- | --- | --- | --- | --- | --- | --- |
| **Ion** | | | |  | 1.00 | 25.73 || 1 | C | OP2 | AA:1636 | 2.47 | 1.00 | 49.07 |
| 2 | A | OP2 | AA:1637 | 3.80 | 1.00 | 44.64 |  |
| 3 | C | O5' | AA:1636 | 3.86 | 1.00 | 46.84 |  |
| 4 | U | OP1 | AA:1300 | 4.08 | 1.00 | 45.13 |  |
| 5 | G | OP1 | AA:1299 | 4.22 | 1.00 | 47.34 |  |
| 6 | C | OP1 | AA:1636 | 4.36 | 1.00 | 48.58 |  |
| 7 | G | O5' | AA:1635 | 4.66 | 1.00 | 44.24 |  |
| 8 | U | O3' | AA:1300 | 4.69 | 1.00 | 46.42 |  |
| 9 | G | O3' | AA:1635 | 4.90 | 1.00 | 46.68 |  |
  |

  
 Coded contact list:   
 Exclusion zone contact(s): /OP2/  
Consider NA in place of MG (1 valid contact(s) for NA vs. 0 for MG)  

### MG:AA:3138

|  | Res. type | Atom | Residue | Distance | Occupancy | B-factor | Closest solute to 1^st^ shell water |
| --- | --- | --- | --- | --- | --- | --- | --- |
| **Ion** | | | |  | 1.00 | 19.98 || 1 | A | OP2 | AA:1265 | 2.02 | 1.00 | 46.57 |
| 2 | U | OP2 | AA:2615 | 3.05 | 1.00 | 49.76 |  |
| 3 | U | OP1 | AA:2615 | 3.09 | 1.00 | 50.94 |  |
| 4 | A | OP1 | AA:1265 | 3.66 | 1.00 | 46.04 |  |
| 5 | A | O5' | AA:1265 | 3.97 | 1.00 | 47.14 |  |
| 6 | U | O2' | AA:1267 | 4.12 | 1.00 | 51.71 |  |
| 7 | G | O3' | AA:1264 | 4.45 | 1.00 | 45.90 |  |
| 8 | A | OP1 | AA:2614 | 4.57 | 1.00 | 46.25 |  |
| 9 | A | O3' | AA:2614 | 4.66 | 1.00 | 50.83 |  |
| 10 | U | O5' | AA:2615 | 4.73 | 1.00 | 49.46 |  |
  |

  
 Contact(s): /OP2/  
 Coded contact list: O_ph_  
 Exclusion zone contact(s): /OP1/OP2/  
Consider K in place of MG (2 valid contact(s) for K vs. 1 for MG)  

### MG:AA:3139

|  | Res. type | Atom | Residue | Distance | Occupancy | B-factor | Closest solute to 1^st^ shell water |
| --- | --- | --- | --- | --- | --- | --- | --- |
| **Ion** | | | |  | 1.00 | 45.49 || 1 | G | OP1 | AA:2035 | 2.72 | 1.00 | 53.42 |
| 2 | G | O6 | AA:2037 | 3.16 | 1.00 | 38.58 |  |
| 3 | G | O6 | AA:2038 | 3.44 | 1.00 | 56.10 |  |
| 4 | G | N7 | AA:2037 | 3.60 | 1.00 | 41.39 |  |
| 5 | C | N4 | AA:2036 | 3.70 | 1.00 | 40.86 |  |
| 6 | C | N4 | AA:2025 | 3.89 | 1.00 | 55.76 |  |
| 7 | C | N3 | AA:2036 | 4.46 | 1.00 | 41.57 |  |
| 8 | G | O6 | AA:2024 | 4.73 | 1.00 | 55.29 |  |
| 9 | G | OP2 | AA:2035 | 4.73 | 1.00 | 53.51 |  |
| 10 | U | O3' | AA:2034 | 4.91 | 1.00 | 53.87 |  |
  |

  
 Coded contact list:   
 Exclusion zone contact(s): /O6/OP1/  
Consider K in place of MG (2 valid contact(s) for K vs. 0 for MG)  

### MG:AA:3140

|  | Res. type | Atom | Residue | Distance | Occupancy | B-factor | Closest solute to 1^st^ shell water |
| --- | --- | --- | --- | --- | --- | --- | --- |
| **Ion** | | | |  | 1.00 | 57.43 || 1 | G | O6 | AA:1824 | 4.26 | 1.00 | 28.33 |
| 2 | G | N7 | AA:1824 | 4.31 | 1.00 | 25.25 |  |
| 3 | G | N7 | AA:1823 | 4.63 | 1.00 | 29.87 |  |
| 4 | G | OP2 | AA:1823 | 4.67 | 1.00 | 30.58 |  |
  |

  
 Coded contact list:   
1^st^ MG coordination shell missing (see Notes)  

### MG:AA:3141

|  | Res. type | Atom | Residue | Distance | Occupancy | B-factor | Closest solute to 1^st^ shell water |
| --- | --- | --- | --- | --- | --- | --- | --- |
| **Ion** | | | |  | 1.00 | 66.50 || 1 | G | O6 | AA:2857 | 3.68 | 1.00 | 89.21 |
| 2 | G | N7 | AA:2857 | 3.70 | 1.00 | 89.18 |  |
| 3 | G | OP1 | AA:2859 | 4.09 | 1.00 | 84.33 |  |
| 4 | G | OP2 | AA:2859 | 4.10 | 1.00 | 83.78 |  |
  |

  
 Coded contact list:   
1^st^ MG coordination shell missing (see Notes)  

### MG:AA:3142

|  | Res. type | Atom | Residue | Distance | Occupancy | B-factor | Closest solute to 1^st^ shell water |
| --- | --- | --- | --- | --- | --- | --- | --- |
| **Ion** | | | |  | 1.00 | 12.03 || 1 | G | O3' | AA:1980 | 2.65 | 1.00 | 42.90 |
| 2 | G | O2' | AA:1980 | 2.67 | 1.00 | 47.89 |  |
| 3 | C | O5' | AA:1982 | 2.72 | 1.00 | 42.17 |  |
| 4 | A | OP2 | AA:1785 | 2.84 | 1.00 | 44.30 |  |
| 5 | A | OP1 | AA:1981 | 3.08 | 1.00 | 40.05 |  |
| 6 | C | O4' | AA:1982 | 3.19 | 1.00 | 40.09 |  |
| 7 | C | OP2 | AA:1982 | 3.32 | 1.00 | 41.60 |  |
| 8 | A | O5' | AA:1981 | 3.47 | 1.00 | 42.58 |  |
| 9 | A | O3' | AA:1981 | 4.45 | 1.00 | 42.98 |  |
| 10 | A | O5' | AA:1785 | 4.54 | 1.00 | 45.50 |  |
| 11 | C | N1 | AA:1982 | 4.60 | 1.00 | 39.00 |  |
| 12 | A | OP2 | AA:1981 | 4.78 | 1.00 | 41.02 |  |
| 13 | C | OP1 | AA:1982 | 4.80 | 1.00 | 42.40 |  |
| 14 | A | O3' | AA:1784 | 4.94 | 1.00 | 45.81 |  |
| 15 | A | O3' | AA:1785 | 4.98 | 1.00 | 43.01 |  |
  |

  
 Coded contact list:   
 Exclusion zone contact(s): /O2'/O3'/O4'/O5'/OP1/OP2/OP2/  
Consider K in place of MG (7 valid contact(s) for K vs. 0 for MG)  

### MG:AA:3143

|  | Res. type | Atom | Residue | Distance | Occupancy | B-factor | Closest solute to 1^st^ shell water |
| --- | --- | --- | --- | --- | --- | --- | --- |
| **Ion** | | | |  | 1.00 | 37.95 || 1 | G | O6 | AA:2709 | 2.34 | 1.00 | 49.51 |
| 2 | U | O4 | AA:2698 | 2.51 | 1.00 | 54.06 |  |
| 3 | G | O6 | AA:2708 | 3.01 | 1.00 | 47.55 |  |
| 4 | G | O6 | AA:2697 | 3.03 | 1.00 | 52.28 |  |
| 5 | U | N3 | AA:2698 | 3.44 | 1.00 | 49.62 |  |
| 6 | C | N4 | AA:2710 | 3.62 | 1.00 | 51.62 |  |
| 7 | C | N4 | AA:2699 | 3.90 | 1.00 | 48.00 |  |
| 8 | G | N7 | AA:2709 | 4.26 | 1.00 | 48.11 |  |
| 9 | G | N1 | AA:2708 | 4.46 | 1.00 | 46.52 |  |
| 10 | G | N1 | AA:2709 | 4.60 | 1.00 | 49.88 |  |
| 11 | G | N1 | AA:2697 | 4.69 | 1.00 | 53.57 |  |
| 12 | G | N7 | AA:2708 | 4.80 | 1.00 | 49.14 |  |
  |

  
 Contact(s): /O6/  
 Coded contact list: O_b_  
 Exclusion zone contact(s): /O4/O6/O6/  
Consider K in place of MG (2 valid contact(s) for K vs. 1 for MG)  

### MG:AA:3144

|  | Res. type | Atom | Residue | Distance | Occupancy | B-factor | Closest solute to 1^st^ shell water |
| --- | --- | --- | --- | --- | --- | --- | --- |
| **Ion** | | | |  | 1.00 | 69.03 || 1 | A | OP2 | AA:2860 | 3.79 | 1.00 | 77.28 |
| 2 | G | N7 | AA:2861 | 4.16 | 1.00 | 78.41 |  |
| 3 | G | O6 | AA:2861 | 4.24 | 1.00 | 79.94 |  |
| 4 | G | O6 | AA:2857 | 4.52 | 1.00 | 89.21 |  |
| 5 | G | N1 | AA:2857 | 4.77 | 1.00 | 89.55 |  |
  |

  
 Coded contact list:   
1^st^ MG coordination shell missing (see Notes)  

### MG:AA:3145

|  | Res. type | Atom | Residue | Distance | Occupancy | B-factor | Closest solute to 1^st^ shell water |
| --- | --- | --- | --- | --- | --- | --- | --- |
| **Ion** | | | |  | 1.00 | 61.85 || 1 | G | N7 | AA:2714 | 3.67 | 1.00 | 62.83 |
| 2 | G | O6 | AA:2714 | 4.07 | 1.00 | 62.42 |  |
| 3 | C | N4 | AA:2690 | 4.19 | 1.00 | 59.07 |  |
| 4 | C | N3 | AA:2690 | 4.20 | 1.00 | 58.29 |  |
| 5 | G | OP1 | AA:2714 | 4.84 | 1.00 | 56.15 |  |
| 6 | C | N4 | AA:2715 | 4.87 | 1.00 | 58.09 |  |
  |

  
 Coded contact list:   
1^st^ MG coordination shell missing (see Notes)  

### MG:AA:3146

|  | Res. type | Atom | Residue | Distance | Occupancy | B-factor | Closest solute to 1^st^ shell water |
| --- | --- | --- | --- | --- | --- | --- | --- |
| **Ion** | | | |  | 1.00 | 78.86 || 1 | A | OP2 | AA:2693 | 3.87 | 1.00 | 60.01 |
| 2 | A | N7 | AA:2693 | 4.56 | 1.00 | 60.10 |  |
  |

  
 Coded contact list:   
1^st^ MG coordination shell missing (see Notes)  

### MG:AA:3147

|  | Res. type | Atom | Residue | Distance | Occupancy | B-factor | Closest solute to 1^st^ shell water |
| --- | --- | --- | --- | --- | --- | --- | --- |
| **Ion** | | | |  | 1.00 | 60.03 || 1 | A | OP2 | AA:1970 | 2.93 | 1.00 | 39.18 |
| 2 | G | O6 | AA:1835 | 3.02 | 1.00 | 45.11 |  |
| 3 | A | OP1 | AA:1970 | 3.59 | 1.00 | 38.39 |  |
| 4 | C | N4 | AA:1836 | 3.87 | 1.00 | 50.31 |  |
| 5 | G | N7 | AA:1835 | 4.10 | 1.00 | 47.00 |  |
| 6 | G | O6 | AA:1904 | 4.37 | 1.00 | 46.22 |  |
| 7 | A | O5' | AA:1969 | 4.43 | 1.00 | 34.98 |  |
| 8 | C | N4 | AA:1905 | 4.45 | 1.00 | 46.10 |  |
| 9 | A | O3' | AA:1969 | 4.46 | 1.00 | 37.41 |  |
  |

  
 Coded contact list:   
 Exclusion zone contact(s): /O6/OP2/  
Consider K in place of MG (2 valid contact(s) for K vs. 0 for MG)  

### MG:AA:3148

|  | Res. type | Atom | Residue | Distance | Occupancy | B-factor | Closest solute to 1^st^ shell water |
| --- | --- | --- | --- | --- | --- | --- | --- |
| **Ion** | | | |  | 1.00 | 53.91 || 1 | G | OP1 | AA:2707 | 3.35 | 1.00 | 60.66 |
| 2 | A | O3' | AA:2850 | 3.35 | 1.00 | 64.66 |  |
| 3 | A | OP1 | AA:2851 | 3.59 | 1.00 | 66.20 |  |
| 4 | A | O2' | AA:2850 | 3.65 | 1.00 | 63.86 |  |
| 5 | G | OP2 | AA:2707 | 4.09 | 1.00 | 62.15 |  |
| 6 | G | OP1 | AA:2694 | 4.16 | 1.00 | 60.19 |  |
| 7 | A | O5' | AA:2851 | 4.67 | 1.00 | 66.70 |  |
| 8 | G | OP2 | AA:2694 | 4.69 | 1.00 | 59.32 |  |
| 9 | G | O3' | AA:2706 | 4.79 | 1.00 | 61.04 |  |
  |

  
 Coded contact list:   
 Exclusion zone contact(s): /O3'/OP1/  
Consider K in place of MG (2 valid contact(s) for K vs. 0 for MG)  

### MG:AA:3149

|  | Res. type | Atom | Residue | Distance | Occupancy | B-factor | Closest solute to 1^st^ shell water |
| --- | --- | --- | --- | --- | --- | --- | --- |
| **Ion** | | | |  | 1.00 | 61.05 || 1 | G | OP1 | AA:2706 | 4.27 | 1.00 | 62.35 |
| 2 | C | OP2 | AA:2695 | 4.45 | 1.00 | 55.81 |  |
| 3 | G | OP2 | AA:2707 | 4.76 | 1.00 | 62.15 |  |
  |

  
 Coded contact list:   
1^st^ MG coordination shell missing (see Notes)  

### MG:AA:3150

|  | Res. type | Atom | Residue | Distance | Occupancy | B-factor | Closest solute to 1^st^ shell water |
| --- | --- | --- | --- | --- | --- | --- | --- |
| **Ion** | | | |  | 1.00 | 29.80 || 1 | C | OP1 | AA:1774 | 2.02 | 1.00 | 39.07 |
| 2 | G | N3 | AA:1772 | 3.52 | 1.00 | 38.97 |  |
| 3 | A | OP1 | AA:1786 | 3.71 | 1.00 | 42.06 |  |
| 4 | A | OP2 | AA:1786 | 3.75 | 1.00 | 43.16 |  |
| 5 | G | O2' | AA:1772 | 3.81 | 1.00 | 38.87 |  |
| 6 | G | N2 | AA:1772 | 3.86 | 1.00 | 40.37 |  |
| 7 | C | OP2 | AA:1774 | 4.23 | 1.00 | 38.12 |  |
| 8 | C | O5' | AA:1774 | 4.33 | 1.00 | 38.23 |  |
| 9 | A | O3' | AA:1773 | 4.41 | 1.00 | 39.67 |  |
| 10 | G | O6 | AA:1776 | 4.66 | 1.00 | 36.38 |  |
| 11 | U | OP2 | AA:1775 | 4.76 | 1.00 | 41.22 |  |
| 12 | G | N9 | AA:1772 | 4.90 | 1.00 | 37.09 |  |
| 13 | A | N6 | AA:1789 | 4.96 | 1.00 | 42.23 |  |
| 14 | A | O5' | AA:1786 | 4.97 | 1.00 | 43.86 |  |
  |

  
 Contact(s): /OP1/  
 Coded contact list: O_ph_  

### MG:AA:3151

|  | Res. type | Atom | Residue | Distance | Occupancy | B-factor | Closest solute to 1^st^ shell water |
| --- | --- | --- | --- | --- | --- | --- | --- |
| **Ion** | | | |  | 1.00 | 60.16 || 1 | A | OP2 | AA:1970 | 2.54 | 1.00 | 39.18 |
| 2 | A | OP2 | AA:1900 | 2.98 | 1.00 | 47.86 |  |
| 3 | A | O5' | AA:1900 | 3.30 | 1.00 | 47.99 |  |
| 4 | A | O5' | AA:1970 | 3.57 | 1.00 | 36.12 |  |
| 5 | A | O4' | AA:1900 | 4.00 | 1.00 | 41.32 |  |
| 6 | G | O2' | AA:1899 | 4.01 | 1.00 | 51.35 |  |
| 7 | A | OP1 | AA:1970 | 4.20 | 1.00 | 38.39 |  |
| 8 | A | OP1 | AA:1900 | 4.71 | 1.00 | 47.27 |  |
| 9 | A | O3' | AA:1969 | 4.83 | 1.00 | 37.41 |  |
| 10 | G | O3' | AA:1899 | 4.96 | 1.00 | 48.88 |  |
  |

  
 Coded contact list:   
 Exclusion zone contact(s): /O5'/OP2/OP2/  
Consider K in place of MG (2 valid contact(s) for K vs. 0 for MG)  

### MG:AA:3152

|  | Res. type | Atom | Residue | Distance | Occupancy | B-factor | Closest solute to 1^st^ shell water |
| --- | --- | --- | --- | --- | --- | --- | --- |
| **Ion** | | | |  | 1.00 | 28.01 || 1 | C | OP2 | AA:198 | 2.33 | 1.00 | 41.69 |
| 2 | A | N7 | AA:195 | 2.38 | 1.00 | 48.61 |  |
| 3 | A | OP1 | AA:197 | 2.46 | 1.00 | 43.92 |  |
| 4 | A | N6 | AA:195 | 3.84 | 1.00 | 43.72 |  |
| 5 | C | OP1 | AA:198 | 3.94 | 1.00 | 42.76 |  |
| 6 | G | OP2 | AA:194 | 4.00 | 1.00 | 42.62 |  |
| 7 | G | N7 | AA:194 | 4.03 | 1.00 | 40.59 |  |
| 8 | A | O5' | AA:197 | 4.19 | 1.00 | 42.30 |  |
| 9 | A | OP2 | AA:195 | 4.21 | 1.00 | 48.18 |  |
| 10 | A | N9 | AA:195 | 4.47 | 1.00 | 46.23 |  |
| 11 | C | O5' | AA:198 | 4.56 | 1.00 | 41.90 |  |
| 12 | A | O3' | AA:196 | 4.63 | 1.00 | 43.42 |  |
| 13 | A | O3' | AA:197 | 4.67 | 1.00 | 43.21 |  |
| 14 | G | O5' | AA:194 | 4.77 | 1.00 | 42.67 |  |
| 15 | A | OP2 | AA:197 | 4.90 | 1.00 | 43.73 |  |
  |

  
**1^st^ shell ligand angles:**   
1-2: 85;   
  
 Contact(s): /N7/OP2/  
 Coded contact list: O_ph_N_b_  
 Exclusion zone contact(s): /OP1/  

### MG:AA:3153

|  | Res. type | Atom | Residue | Distance | Occupancy | B-factor | Closest solute to 1^st^ shell water |
| --- | --- | --- | --- | --- | --- | --- | --- |
| **Ion** | | | |  | 1.00 | 40.38 || 1 | U | O4 | AA:1991 | 2.60 | 1.00 | 48.10 |
| 2 | G | N9 | AA:1667 | 3.17 | 1.00 | 56.50 |  |
| 3 | G | O4' | AA:1667 | 3.37 | 1.00 | 55.61 |  |
| 4 | A | OP2 | AA:1668 | 3.40 | 1.00 | 52.04 |  |
| 5 | G | O3' | AA:1667 | 3.74 | 1.00 | 54.81 |  |
| 6 | C | N4 | AA:1990 | 3.96 | 1.00 | 48.76 |  |
| 7 | G | O2' | AA:1667 | 4.14 | 1.00 | 60.56 |  |
| 8 | G | OP2 | AA:1667 | 4.22 | 1.00 | 54.35 |  |
| 9 | G | N7 | AA:1667 | 4.24 | 1.00 | 56.95 |  |
| 10 | G | N3 | AA:1667 | 4.40 | 1.00 | 58.06 |  |
| 11 | U | N3 | AA:1991 | 4.63 | 1.00 | 49.71 |  |
| 12 | G | O5' | AA:1667 | 4.98 | 1.00 | 56.31 |  |
  |

  
 Coded contact list:   
 Exclusion zone contact(s): /O4/O4'/OP2/  
Consider K in place of MG (3 valid contact(s) for K vs. 0 for MG)  

### MG:AA:3154

|  | Res. type | Atom | Residue | Distance | Occupancy | B-factor | Closest solute to 1^st^ shell water |
| --- | --- | --- | --- | --- | --- | --- | --- |
| **Ion** | | | |  | 1.00 | 59.02 || 1 | A | N1 | AA:1690 | 3.46 | 1.00 | 42.46 |
| 2 | G | O2' | AA:1697 | 3.48 | 1.00 | 42.65 |  |
| 3 | A | OP1 | AA:1978 | 3.70 | 1.00 | 38.83 |  |
| 4 | A | OP2 | AA:1978 | 3.99 | 1.00 | 37.69 |  |
| 5 | A | N6 | AA:1690 | 4.42 | 1.00 | 42.81 |  |
  |

  
 Coded contact list:   
1^st^ MG coordination shell missing (see Notes)  

### MG:AA:3155

|  | Res. type | Atom | Residue | Distance | Occupancy | B-factor | Closest solute to 1^st^ shell water |
| --- | --- | --- | --- | --- | --- | --- | --- |
| **Ion** | | | |  | 1.00 | 46.22 || 1 | G | OP2 | AA:370 | 1.95 | 1.00 | 65.16 |
| 2 | A | OP2 | AA:422 | 2.51 | 1.00 | 69.42 |  |
| 3 | A | N7 | AA:422 | 3.65 | 1.00 | 69.23 |  |
| 4 | G | OP1 | AA:370 | 3.66 | 1.00 | 65.49 |  |
| 5 | A | OP1 | AA:422 | 3.84 | 1.00 | 68.44 |  |
| 6 | A | O5' | AA:422 | 3.86 | 1.00 | 68.14 |  |
| 7 | G | O5' | AA:370 | 3.86 | 1.00 | 65.30 |  |
| 8 | A | N7 | AA:423 | 4.09 | 1.00 | 65.81 |  |
| 9 | U | O2' | AA:421 | 4.32 | 1.00 | 73.14 |  |
| 10 | C | O3' | AA:366 | 4.40 | 1.00 | 66.64 |  |
| 11 | A | N6 | AA:423 | 4.41 | 1.00 | 66.01 |  |
| 12 | C | O2' | AA:366 | 4.45 | 1.00 | 68.67 |  |
| 13 | G | N7 | AA:370 | 4.70 | 1.00 | 63.34 |  |
| 14 | U | O3' | AA:421 | 4.92 | 1.00 | 69.74 |  |
| 15 | A | N9 | AA:422 | 4.94 | 1.00 | 67.50 |  |
  |

  
 Contact(s): /OP2/  
 Coded contact list: O_ph_  
 Exclusion zone contact(s): /OP2/  

### MG:AA:3156

|  | Res. type | Atom | Residue | Distance | Occupancy | B-factor | Closest solute to 1^st^ shell water |
| --- | --- | --- | --- | --- | --- | --- | --- |
| **Ion** | | | |  | 1.00 | 55.99 || 1 | A | OP1 | AA:422 | 3.32 | 1.00 | 68.44 |
| 2 | G | O6 | AA:424 | 3.73 | 1.00 | 54.51 |  |
| 3 | G | N7 | AA:424 | 3.79 | 1.00 | 54.27 |  |
| 4 | G | O6 | AA:370 | 3.84 | 1.00 | 63.53 |  |
| 5 | A | OP2 | AA:423 | 3.98 | 1.00 | 65.41 |  |
| 6 | G | N7 | AA:370 | 4.65 | 1.00 | 63.34 |  |
  |

  
 Coded contact list:   
 Exclusion zone contact(s): /OP1/  
Consider K in place of MG (1 valid contact(s) for K vs. 0 for MG)  

### MG:AA:3157

|  | Res. type | Atom | Residue | Distance | Occupancy | B-factor | Closest solute to 1^st^ shell water |
| --- | --- | --- | --- | --- | --- | --- | --- |
| **Ion** | | | |  | 1.00 | 73.63 || 1 | G | O6 | AA:425 | 4.21 | 1.00 | 63.43 |
| 2 | G | N7 | AA:425 | 4.41 | 1.00 | 60.06 |  |
| 3 | A | OP1 | AA:423 | 4.75 | 1.00 | 65.62 |  |
| 4 | G | O6 | AA:424 | 4.78 | 1.00 | 54.51 |  |
| 5 | A | OP2 | AA:223 | 4.91 | 1.00 | 59.52 |  |
  |

  
 Coded contact list:   
1^st^ MG coordination shell missing (see Notes)  

### MG:AA:3158

|  | Res. type | Atom | Residue | Distance | Occupancy | B-factor | Closest solute to 1^st^ shell water |
| --- | --- | --- | --- | --- | --- | --- | --- |
| **Ion** | | | |  | 1.00 | 55.35 || 1 | A | OP1 | AA:222 | 2.10 | 1.00 | 65.57 |
| 2 | A | O3' | AA:221 | 2.98 | 1.00 | 67.39 |  |
| 3 | A | O5' | AA:222 | 4.04 | 1.00 | 62.92 |  |
| 4 | A | OP2 | AA:222 | 4.32 | 1.00 | 65.76 |  |
| 5 | C | OP2 | AA:271A | 4.58 | 1.00 | 84.34 |  |
| 6 | C | N4 | AA:267 | 4.91 | 1.00 | 63.08 |  |
  |

  
 Contact(s): /OP1/  
 Coded contact list: O_ph_  
 Exclusion zone contact(s): /O3'/  

### MG:AA:3159

|  | Res. type | Atom | Residue | Distance | Occupancy | B-factor | Closest solute to 1^st^ shell water |
| --- | --- | --- | --- | --- | --- | --- | --- |
| **Ion** | | | |  | 1.00 | 61.32 || 1 | G | OP1 | AA:271B | 3.15 | 1.00 | 85.33 |
| 2 | U | OP1 | AA:421 | 3.59 | 1.00 | 65.97 |  |
| 3 | G | OP2 | AA:271B | 3.68 | 1.00 | 84.77 |  |
| 4 | A | OP2 | AA:222 | 3.74 | 1.00 | 65.76 |  |
| 5 | G | O5' | AA:271B | 4.56 | 1.00 | 85.68 |  |
| 6 | A | OP1 | AA:222 | 4.77 | 1.00 | 65.57 |  |
  |

  
 Coded contact list:   
 Exclusion zone contact(s): /OP1/  
Consider K in place of MG (1 valid contact(s) for K vs. 0 for MG)  

### MG:AA:3160

|  | Res. type | Atom | Residue | Distance | Occupancy | B-factor | Closest solute to 1^st^ shell water |
| --- | --- | --- | --- | --- | --- | --- | --- |
| **Ion** | | | |  | 1.00 | 64.66 || 1 | U | OP2 | AA:271C | 3.02 | 1.00 | 93.61 |
| 2 | C | OP1 | AA:420 | 3.29 | 1.00 | 65.02 |  |
| 3 | G | O6 | AA:232 | 3.29 | 1.00 | 78.80 |  |
| 4 | C | OP2 | AA:420 | 3.87 | 1.00 | 65.21 |  |
| 5 | G | N1 | AA:232 | 3.97 | 1.00 | 79.20 |  |
| 6 | U | OP1 | AA:271C | 4.31 | 1.00 | 94.08 |  |
| 7 | C | O3' | AA:419 | 4.74 | 1.00 | 64.00 |  |
| 8 | G | O3' | AA:271B | 4.80 | 1.00 | 92.78 |  |
  |

  
 Coded contact list:   
 Exclusion zone contact(s): /O6/OP1/OP2/  
Consider K in place of MG (3 valid contact(s) for K vs. 0 for MG)  

### MG:AA:3161

|  | Res. type | Atom | Residue | Distance | Occupancy | B-factor | Closest solute to 1^st^ shell water |
| --- | --- | --- | --- | --- | --- | --- | --- |
| **Ion** | | | |  | 1.00 | 63.32 || 1 | C | OP2 | AA:420 | 2.45 | 1.00 | 65.21 |
| 2 | G | O2' | AA:271B | 3.27 | 1.00 | 90.21 |  |
| 3 | U | OP2 | AA:271C | 3.80 | 1.00 | 93.61 |  |
| 4 | C | OP2 | AA:419 | 4.18 | 1.00 | 68.02 |  |
| 5 | C | O5' | AA:420 | 4.33 | 1.00 | 65.05 |  |
| 6 | G | O3' | AA:271B | 4.52 | 1.00 | 92.78 |  |
| 7 | C | O5' | AA:419 | 4.64 | 1.00 | 66.95 |  |
| 8 | C | OP1 | AA:419 | 4.75 | 1.00 | 67.54 |  |
| 9 | C | O3' | AA:419 | 4.80 | 1.00 | 64.00 |  |
| 10 | C | OP1 | AA:420 | 4.87 | 1.00 | 65.02 |  |
| 11 | U | O5' | AA:271C | 4.99 | 1.00 | 96.31 |  |
  |

  
 Coded contact list:   
 Exclusion zone contact(s): /O2'/OP2/  
Consider NA in place of MG (1 valid contact(s) for NA vs. 0 for MG)  

### MG:AA:3162

|  | Res. type | Atom | Residue | Distance | Occupancy | B-factor | Closest solute to 1^st^ shell water |
| --- | --- | --- | --- | --- | --- | --- | --- |
| **Ion** | | | |  | 1.00 | 48.58 || 1 | G | O6 | AA:406 | 2.84 | 1.00 | 73.54 |
| 2 | G | O6 | AA:407 | 3.08 | 1.00 | 63.93 |  |
| 3 | U | O4 | AA:421 | 3.67 | 1.00 | 72.67 |  |
| 4 | G | N7 | AA:406 | 3.80 | 1.00 | 76.63 |  |
| 5 | G | N2 | AA:271B | 3.99 | 1.00 | 84.69 |  |
| 6 | C | O2 | AA:404 | 4.07 | 1.00 | 83.97 |  |
| 7 | C | N4 | AA:420 | 4.37 | 1.00 | 62.44 |  |
| 8 | G | N7 | AA:407 | 4.42 | 1.00 | 66.04 |  |
| 9 | U | N3 | AA:421 | 4.61 | 1.00 | 73.16 |  |
| 10 | C | O2' | AA:404 | 4.77 | 1.00 | 91.10 |  |
| 11 | G | N1 | AA:407 | 4.87 | 1.00 | 63.28 |  |
  |

  
 Coded contact list:   
 Exclusion zone contact(s): /O6/O6/  
Consider K in place of MG (2 valid contact(s) for K vs. 0 for MG)  

### MG:AA:3163

|  | Res. type | Atom | Residue | Distance | Occupancy | B-factor | Closest solute to 1^st^ shell water |
| --- | --- | --- | --- | --- | --- | --- | --- |
| **Ion** | | | |  | 1.00 | 65.91 || 1 | A | N1 | AA:363F | 3.52 | 1.00 | 99.50 |
| 2 | G | O6 | AA:273 | 4.08 | 1.00 | 83.55 |  |
| 3 | G | O6 | AA:272 | 4.12 | 1.00 | 77.13 |  |
| 4 | G | N7 | AA:273 | 4.19 | 1.00 | 84.62 |  |
| 5 | G | N7 | AA:272 | 4.25 | 1.00 | 76.84 |  |
| 6 | U | O4 | AA:363E | 4.54 | 1.00 | 87.04 |  |
| 7 | A | N3 | AA:363F | 4.67 | 1.00 | 99.23 |  |
  |

  
 Coded contact list:   
1^st^ MG coordination shell missing (see Notes)  

### MG:AA:3164

|  | Res. type | Atom | Residue | Distance | Occupancy | B-factor | Closest solute to 1^st^ shell water |
| --- | --- | --- | --- | --- | --- | --- | --- |
| **Ion** | | | |  | 1.00 | 79.43 || 1 | G | O6 | AA:363D | 3.63 | 1.00 | 93.30 |
| 2 | G | O6 | AA:363C | 4.00 | 1.00 | 102.78 |  |
| 3 | C | N4 | AA:273B | 4.33 | 1.00 | 93.25 |  |
| 4 | G | N7 | AA:363C | 4.71 | 1.00 | 104.18 |  |
| 5 | G | N7 | AA:363D | 4.95 | 1.00 | 96.10 |  |
  |

  
 Coded contact list:   
1^st^ MG coordination shell missing (see Notes)  

### MG:AA:3165

|  | Res. type | Atom | Residue | Distance | Occupancy | B-factor | Closest solute to 1^st^ shell water |
| --- | --- | --- | --- | --- | --- | --- | --- |
| **Ion** | | | |  | 1.00 | 53.54 || 1 | G | O6 | AA:1984 | 2.58 | 1.00 | 46.91 |
| 2 | G | O6 | AA:1769 | 2.72 | 1.00 | 36.74 |  |
| 3 | U | O4 | AA:1768 | 3.48 | 1.00 | 49.42 |  |
| 4 | C | N4 | AA:1983 | 3.60 | 1.00 | 36.75 |  |
| 5 | G | N7 | AA:1984 | 3.89 | 1.00 | 46.06 |  |
| 6 | U | N3 | AA:1768 | 4.28 | 1.00 | 47.19 |  |
| 7 | G | N1 | AA:1769 | 4.34 | 1.00 | 32.20 |  |
| 8 | G | N7 | AA:1769 | 4.45 | 1.00 | 37.19 |  |
| 9 | G | O6 | AA:1985 | 4.50 | 1.00 | 39.55 |  |
| 10 | G | N1 | AA:1984 | 4.79 | 1.00 | 46.32 |  |
| 11 | C | N3 | AA:1983 | 4.90 | 1.00 | 35.22 |  |
  |

  
 Coded contact list:   
 Exclusion zone contact(s): /O6/O6/  
Consider NA in place of MG (1 valid contact(s) for NA vs. 0 for MG)  

### MG:AA:3166

|  | Res. type | Atom | Residue | Distance | Occupancy | B-factor | Closest solute to 1^st^ shell water |
| --- | --- | --- | --- | --- | --- | --- | --- |
| **Ion** | | | |  | 1.00 | 65.28 || 1 | A | OP1 | AA:1543 | 2.52 | 1.00 | 103.24 |
| 2 | A | OP1 | AA:1449 | 2.59 | 1.00 | 87.12 |  |
| 3 | A | O5' | AA:1449 | 3.22 | 1.00 | 88.56 |  |
| 4 | A | OP2 | AA:1543 | 3.24 | 1.00 | 103.33 |  |
| 5 | G | O3' | AA:1448 | 4.34 | 1.00 | 86.41 |  |
| 6 | G | O3' | AA:1542 | 4.37 | 1.00 | 103.72 |  |
| 7 | A | O5' | AA:1543 | 4.54 | 1.00 | 103.35 |  |
| 8 | A | OP2 | AA:1449 | 4.74 | 1.00 | 87.43 |  |
  |

  
 Coded contact list:   
 Exclusion zone contact(s): /O5'/OP1/OP1/OP2/  
Consider NA in place of MG (2 valid contact(s) for NA vs. 0 for MG)  

### MG:AA:3167

|  | Res. type | Atom | Residue | Distance | Occupancy | B-factor | Closest solute to 1^st^ shell water |
| --- | --- | --- | --- | --- | --- | --- | --- |
| **Ion** | | | |  | 1.00 | 61.71 || 1 | G | O6 | AA:1823 | 2.61 | 1.00 | 34.40 |
| 2 | U | O4 | AA:1796 | 3.04 | 1.00 | 33.88 |  |
| 3 | G | O6 | AA:1824 | 3.25 | 1.00 | 28.33 |  |
| 4 | C | N4 | AA:1795 | 3.67 | 1.00 | 27.92 |  |
| 5 | U | N3 | AA:1796 | 3.94 | 1.00 | 33.19 |  |
| 6 | G | O6 | AA:1822 | 3.95 | 1.00 | 32.44 |  |
| 7 | G | N7 | AA:1823 | 4.26 | 1.00 | 29.87 |  |
| 8 | C | N4 | AA:1797 | 4.43 | 1.00 | 32.52 |  |
| 9 | G | N1 | AA:1823 | 4.81 | 1.00 | 32.07 |  |
| 10 | G | N1 | AA:1824 | 4.99 | 1.00 | 23.82 |  |
| 11 | G | N7 | AA:1824 | 4.99 | 1.00 | 25.25 |  |
  |

  
 Coded contact list:   
 Exclusion zone contact(s): /O4/O6/O6/  
Consider K in place of MG (3 valid contact(s) for K vs. 0 for MG)  

### MG:AA:3168

|  | Res. type | Atom | Residue | Distance | Occupancy | B-factor | Closest solute to 1^st^ shell water |
| --- | --- | --- | --- | --- | --- | --- | --- |
| **Ion** | | | |  | 1.00 | 48.90 || 1 | G | O6 | AA:1448 | 3.88 | 1.00 | 79.26 |
| 2 | G | O6 | AA:1447 | 4.04 | 1.00 | 80.43 |  |
| 3 | G | N7 | AA:1448 | 4.20 | 1.00 | 82.12 |  |
| 4 | G | N7 | AA:1447 | 4.65 | 1.00 | 82.15 |  |
  |

  
 Coded contact list:   
1^st^ MG coordination shell missing (see Notes)  

### MG:AA:3169

|  | Res. type | Atom | Residue | Distance | Occupancy | B-factor | Closest solute to 1^st^ shell water |
| --- | --- | --- | --- | --- | --- | --- | --- |
| **Ion** | | | |  | 1.00 | 11.78 || 1 | C | OP1 | AA:2612 | 2.25 | 1.00 | 50.44 |
| 2 | HIS | NE2 | AE:132 | 2.41 | 1.00 | 59.78 |  |
| 3 | A | N7 | AA:746 | 2.96 | 1.00 | 52.76 |  |
| 4 | C | OP2 | AA:2612 | 3.02 | 1.00 | 51.00 |  |
| 5 | U | OP1 | AA:2611 | 3.25 | 1.00 | 43.66 |  |
| 6 | HIS | ND1 | AE:132 | 3.39 | 1.00 | 59.47 |  |
| 7 | U | O3' | AA:2611 | 3.48 | 1.00 | 48.43 |  |
| 8 | U | O5' | AA:2611 | 4.27 | 1.00 | 42.96 |  |
| 9 | C | O5' | AA:2612 | 4.40 | 1.00 | 51.63 |  |
| 10 | A | N6 | AA:746 | 4.71 | 1.00 | 56.53 |  |
| 11 | G | OP2 | AA:745 | 4.75 | 1.00 | 41.95 |  |
| 12 | A | N9 | AA:746 | 4.83 | 1.00 | 51.56 |  |
| 13 | C | O3' | AA:2610 | 4.93 | 1.00 | 44.31 |  |
| 14 | LYS | NZ | AE:133 | 4.93 | 1.00 | 45.37 |  |
  |

  
 Contact(s): /OP1/  
 Coded contact list: O_ph_  
 Exclusion zone contact(s): /N7/ND1_His_/NE2_His_/OP1/OP2/  
Consider K in place of MG (4 valid contact(s) for K vs. 1 for MG)  

### MG:AA:3170

|  | Res. type | Atom | Residue | Distance | Occupancy | B-factor | Closest solute to 1^st^ shell water |
| --- | --- | --- | --- | --- | --- | --- | --- |
| **Ion** | | | |  | 1.00 | 32.29 || 1 | C | OP2 | AA:1345 | 2.37 | 1.00 | 60.02 |
| 2 | G | OP2 | AA:1346 | 2.57 | 1.00 | 49.71 |  |
| 3 | C | O2 | AA:1383 | 3.33 | 1.00 | 72.03 |  |
| 4 | A | N7 | AA:1384 | 3.72 | 1.00 | 68.64 |  |
| 5 | C | OP1 | AA:1345 | 3.80 | 1.00 | 60.08 |  |
| 6 | G | O3' | AA:1344 | 3.84 | 1.00 | 59.23 |  |
| 7 | C | N3 | AA:1383 | 4.29 | 1.00 | 72.57 |  |
| 8 | G | OP1 | AA:1346 | 4.55 | 1.00 | 52.52 |  |
| 9 | A | N6 | AA:1384 | 4.61 | 1.00 | 66.11 |  |
| 10 | C | O3' | AA:1345 | 4.79 | 1.00 | 50.51 |  |
| 11 | C | O5' | AA:1345 | 4.81 | 1.00 | 57.78 |  |
| 12 | G | O5' | AA:1346 | 5.00 | 1.00 | 47.60 |  |
  |

  
 Contact(s): /OP2/  
 Coded contact list: O_ph_  
 Exclusion zone contact(s): /O2/OP2/  

### MG:AA:3171

|  | Res. type | Atom | Residue | Distance | Occupancy | B-factor | Closest solute to 1^st^ shell water |
| --- | --- | --- | --- | --- | --- | --- | --- |
| **Ion** | | | |  | 1.00 | 50.62 || 1 | C | OP2 | AA:1774 | 2.94 | 1.00 | 38.12 |
| 2 | C | N3 | AA:1790 | 3.03 | 1.00 | 26.37 |  |
| 3 | A | N1 | AA:1791 | 3.31 | 1.00 | 33.15 |  |
| 4 | C | O2 | AA:1790 | 3.40 | 1.00 | 30.83 |  |
| 5 | A | N6 | AA:1791 | 3.82 | 1.00 | 33.73 |  |
| 6 | A | OP1 | AA:1773 | 3.92 | 1.00 | 39.40 |  |
| 7 | G | O2' | AA:1828 | 3.97 | 1.00 | 35.18 |  |
| 8 | C | N4 | AA:1790 | 4.19 | 1.00 | 26.00 |  |
| 9 | A | N7 | AA:1773 | 4.24 | 1.00 | 35.92 |  |
| 10 | A | O3' | AA:1773 | 4.35 | 1.00 | 39.67 |  |
| 11 | A | O5' | AA:1773 | 4.39 | 1.00 | 38.02 |  |
| 12 | A | N6 | AA:1789 | 4.50 | 1.00 | 42.23 |  |
| 13 | A | O4' | AA:1773 | 4.56 | 1.00 | 39.44 |  |
| 14 | A | N1 | AA:1789 | 4.69 | 1.00 | 38.21 |  |
| 15 | A | N9 | AA:1773 | 4.78 | 1.00 | 35.55 |  |
| 16 | G | O2' | AA:1772 | 4.78 | 1.00 | 38.87 |  |
| 17 | C | O5' | AA:1774 | 4.85 | 1.00 | 38.23 |  |
| 18 | A | OP2 | AA:1829 | 4.86 | 1.00 | 40.34 |  |
| 19 | G | O3' | AA:1772 | 4.94 | 1.00 | 38.80 |  |
| 20 | C | N1 | AA:1790 | 5.00 | 1.00 | 29.15 |  |
  |

  
 Coded contact list:   
 Exclusion zone contact(s): /N1_A_/N3_C_/O2/OP2/  
Consider K in place of MG (4 valid contact(s) for K vs. 0 for MG)  

### MG:AA:3172

|  | Res. type | Atom | Residue | Distance | Occupancy | B-factor | Closest solute to 1^st^ shell water |
| --- | --- | --- | --- | --- | --- | --- | --- |
| **Ion** | | | |  | 1.00 | 67.55 || 1 | G | N7 | AA:1348 | 3.96 | 1.00 | 51.62 |
| 2 | G | N7 | AA:1347 | 4.22 | 1.00 | 45.44 |  |
| 3 | G | OP2 | AA:1347 | 4.80 | 1.00 | 45.82 |  |
| 4 | C | O2 | AA:1383 | 4.83 | 1.00 | 72.03 |  |
| 5 | G | OP2 | AA:1348 | 4.96 | 1.00 | 47.95 |  |
| 6 | G | O6 | AA:1348 | 4.99 | 1.00 | 54.19 |  |
  |

  
 Coded contact list:   
1^st^ MG coordination shell missing (see Notes)  

### MG:AA:3173

|  | Res. type | Atom | Residue | Distance | Occupancy | B-factor | Closest solute to 1^st^ shell water |
| --- | --- | --- | --- | --- | --- | --- | --- |
| **Ion** | | | |  | 1.00 | 14.02 || 1 | G | N7 | AA:2053 | 2.66 | 1.00 | 41.20 |
| 2 | A | N1 | AA:2614 | 2.70 | 1.00 | 48.96 |  |
| 3 | A | O2' | AA:2051 | 3.01 | 1.00 | 44.77 |  |
| 4 | A | N3 | AA:2051 | 3.23 | 1.00 | 40.83 |  |
| 5 | G | O6 | AA:2053 | 3.31 | 1.00 | 41.08 |  |
| 6 | G | O6 | AA:2052 | 3.39 | 1.00 | 46.90 |  |
| 7 | G | N7 | AA:2052 | 3.56 | 1.00 | 47.97 |  |
| 8 | A | N6 | AA:2577 | 3.77 | 1.00 | 44.69 |  |
| 9 | A | N6 | AA:2614 | 3.94 | 1.00 | 51.06 |  |
| 10 | A | N7 | AA:2577 | 4.42 | 1.00 | 43.50 |  |
| 11 | A | N1 | AA:2577 | 4.45 | 1.00 | 42.47 |  |
| 12 | G | N1 | AA:2052 | 4.46 | 1.00 | 45.54 |  |
| 13 | A | N3 | AA:2614 | 4.56 | 1.00 | 47.87 |  |
| 14 | A | N1 | AA:2051 | 4.78 | 1.00 | 41.05 |  |
| 15 | A | N9 | AA:2051 | 4.84 | 1.00 | 39.57 |  |
| 16 | G | N9 | AA:2053 | 4.88 | 1.00 | 42.90 |  |
| 17 | G | OP2 | AA:2053 | 4.90 | 1.00 | 47.50 |  |
| 18 | G | N9 | AA:2052 | 4.92 | 1.00 | 47.56 |  |
  |

  
 Coded contact list:   
 Exclusion zone contact(s): /N1_A_/N3_A_/N7/O2'/O6/O6/  
Consider K in place of MG (6 valid contact(s) for K vs. 0 for MG)  

### MG:AA:3174

|  | Res. type | Atom | Residue | Distance | Occupancy | B-factor | Closest solute to 1^st^ shell water |
| --- | --- | --- | --- | --- | --- | --- | --- |
| **Ion** | | | |  | 1.00 | 63.65 || 1 | G | O6 | AA:2049 | 3.25 | 1.00 | 32.40 |
| 2 | G | O6 | AA:2048 | 3.30 | 1.00 | 33.06 |  |
| 3 | G | N7 | AA:2048 | 3.42 | 1.00 | 30.68 |  |
| 4 | G | N7 | AA:2049 | 3.54 | 1.00 | 27.09 |  |
| 5 | U | O4 | AA:2047 | 4.43 | 1.00 | 40.98 |  |
  |

  
 Coded contact list:   
 Exclusion zone contact(s): /O6/O6/  
Consider K in place of MG (2 valid contact(s) for K vs. 0 for MG)  

### MG:AA:3175

|  | Res. type | Atom | Residue | Distance | Occupancy | B-factor | Closest solute to 1^st^ shell water |
| --- | --- | --- | --- | --- | --- | --- | --- |
| **Ion** | | | |  | 1.00 | 29.69 || 1 | U | OP2 | AA:2504 | 2.72 | 1.00 | 42.06 |
| 2 | A | O3' | AA:575 | 2.99 | 1.00 | 48.35 |  |
| 3 | U | O3' | AA:2500 | 3.09 | 1.00 | 41.87 |  |
| 4 | C | OP1 | AA:2501 | 3.14 | 1.00 | 42.67 |  |
| 5 | A | O5' | AA:2503 | 3.33 | 1.00 | 38.68 |  |
| 6 | U | OP1 | AA:576 | 3.37 | 1.00 | 47.22 |  |
| 7 | A | OP1 | AA:2503 | 3.51 | 1.00 | 39.95 |  |
| 8 | U | O2' | AA:2500 | 3.60 | 1.00 | 39.51 |  |
| 9 | U | OP1 | AA:2504 | 3.76 | 1.00 | 43.84 |  |
| 10 | A | O2' | AA:575 | 4.45 | 1.00 | 48.44 |  |
| 11 | C | O5' | AA:2501 | 4.72 | 1.00 | 41.63 |  |
| 12 | A | O3' | AA:2503 | 4.75 | 1.00 | 41.19 |  |
| 13 | U | O5' | AA:576 | 4.80 | 1.00 | 44.78 |  |
| 14 | U | O5' | AA:2504 | 4.88 | 1.00 | 41.64 |  |
| 15 | C | OP2 | AA:2501 | 4.91 | 1.00 | 42.25 |  |
| 16 | A | OP2 | AA:2503 | 4.94 | 1.00 | 37.97 |  |
| 17 | U | OP2 | AA:576 | 4.95 | 1.00 | 46.55 |  |
  |

  
 Coded contact list:   
 Exclusion zone contact(s): /O3'/O3'/O5'/OP1/OP1/OP2/  
Consider K in place of MG (6 valid contact(s) for K vs. 0 for MG)  

### MG:AA:3176

|  | Res. type | Atom | Residue | Distance | Occupancy | B-factor | Closest solute to 1^st^ shell water |
| --- | --- | --- | --- | --- | --- | --- | --- |
| **Ion** | | | |  | 1.00 | 33.86 || 1 | G | OP2 | AA:1997 | 2.21 | 1.00 | 46.88 |
| 2 | C | O3' | AA:1996 | 3.33 | 1.00 | 49.02 |  |
| 3 | U | OP2 | AA:1995 | 3.97 | 1.00 | 45.95 |  |
| 4 | GLY | O | AE:125 | 4.03 | 1.00 | 54.10 |  |
| 5 | C | OP1 | AA:1994 | 4.07 | 1.00 | 38.79 |  |
| 6 | ALA | O | AE:123 | 4.32 | 1.00 | 47.12 |  |
| 7 | G | OP1 | AA:1997 | 4.32 | 1.00 | 48.01 |  |
| 8 | ASP | OD1 | AE:127 | 4.35 | 1.00 | 61.02 |  |
| 9 | G | O5' | AA:1997 | 4.46 | 1.00 | 48.79 |  |
| 10 | C | O2' | AA:1996 | 4.63 | 1.00 | 51.41 |  |
| 11 | C | OP2 | AA:1994 | 4.71 | 1.00 | 40.58 |  |
  |

  
 Contact(s): /OP2/  
 Coded contact list: O_ph_  
 Exclusion zone contact(s): /O3'/  

### MG:AA:3177

|  | Res. type | Atom | Residue | Distance | Occupancy | B-factor | Closest solute to 1^st^ shell water |
| --- | --- | --- | --- | --- | --- | --- | --- |
| **Ion** | | | |  | 1.00 | 26.54 || 1 | C | OP1 | AA:574 | 2.23 | 1.00 | 46.60 |
| 2 | G | O6 | AA:2035 | 3.62 | 1.00 | 64.76 |  |
| 3 | G | O6 | AA:2018 | 4.23 | 1.00 | 56.60 |  |
| 4 | G | O3' | AA:573 | 4.33 | 1.00 | 45.19 |  |
| 5 | C | OP2 | AA:574 | 4.51 | 1.00 | 45.56 |  |
| 6 | C | O5' | AA:574 | 4.59 | 1.00 | 47.09 |  |
| 7 | C | N4 | AA:531 | 4.64 | 1.00 | 57.50 |  |
| 8 | G | N7 | AA:2035 | 4.80 | 1.00 | 63.76 |  |
| 9 | C | OP2 | AA:2055 | 4.88 | 1.00 | 44.58 |  |
  |

  
 Contact(s): /OP1/  
 Coded contact list: O_ph_  

### MG:AA:3178

|  | Res. type | Atom | Residue | Distance | Occupancy | B-factor | Closest solute to 1^st^ shell water |
| --- | --- | --- | --- | --- | --- | --- | --- |
| **Ion** | | | |  | 1.00 | 41.32 || 1 | G | O6 | AA:16 | 2.65 | 1.00 | 66.18 |
| 2 | U | O4 | AA:525 | 2.91 | 1.00 | 64.94 |  |
| 3 | G | O6 | AA:15 | 2.91 | 1.00 | 65.33 |  |
| 4 | U | O4 | AA:524 | 3.29 | 1.00 | 64.77 |  |
| 5 | U | N3 | AA:525 | 3.81 | 1.00 | 64.27 |  |
| 6 | G | N7 | AA:16 | 4.03 | 1.00 | 66.81 |  |
| 7 | U | N3 | AA:524 | 4.17 | 1.00 | 66.43 |  |
| 8 | G | N7 | AA:15 | 4.45 | 1.00 | 66.47 |  |
| 9 | A | O2' | AA:13 | 4.87 | 1.00 | 86.65 |  |
| 10 | G | N1 | AA:16 | 4.89 | 1.00 | 66.72 |  |
| 11 | G | N1 | AA:15 | 4.95 | 1.00 | 66.24 |  |
| 12 | G | O6 | AA:17 | 4.97 | 1.00 | 70.11 |  |
  |

  
 Coded contact list:   
 Exclusion zone contact(s): /O4/O4/O6/O6/  
Consider K in place of MG (4 valid contact(s) for K vs. 0 for MG)  

### MG:AA:3179

|  | Res. type | Atom | Residue | Distance | Occupancy | B-factor | Closest solute to 1^st^ shell water |
| --- | --- | --- | --- | --- | --- | --- | --- |
| **Ion** | | | |  | 1.00 | 38.96 || 1 | G | N7 | AA:520 | 3.48 | 1.00 | 63.78 |
| 2 | G | O6 | AA:521 | 3.56 | 1.00 | 70.69 |  |
| 3 | G | O6 | AA:520 | 3.67 | 1.00 | 63.02 |  |
| 4 | G | N7 | AA:521 | 4.23 | 1.00 | 72.45 |  |
| 5 | U | O4 | AA:519 | 4.85 | 1.00 | 56.16 |  |
  |

  
 Coded contact list:   
1^st^ MG coordination shell missing (see Notes)  

### MG:AA:3180

|  | Res. type | Atom | Residue | Distance | Occupancy | B-factor | Closest solute to 1^st^ shell water |
| --- | --- | --- | --- | --- | --- | --- | --- |
| **Ion** | | | |  | 1.00 | 62.14 || 1 | G | OP2 | AA:1839 | 3.31 | 1.00 | 61.68 |
| 2 | G | OP2 | AA:1840 | 3.69 | 1.00 | 55.24 |  |
| 3 | G | OP1 | AA:1840 | 4.06 | 1.00 | 55.72 |  |
| 4 | C | O3' | AA:1838 | 4.36 | 1.00 | 60.91 |  |
| 5 | G | O5' | AA:1839 | 4.50 | 1.00 | 58.66 |  |
  |

  
 Coded contact list:   
 Exclusion zone contact(s): /OP2/  
Consider K in place of MG (1 valid contact(s) for K vs. 0 for MG)  

### MG:AA:3181

|  | Res. type | Atom | Residue | Distance | Occupancy | B-factor | Closest solute to 1^st^ shell water |
| --- | --- | --- | --- | --- | --- | --- | --- |
| **Ion** | | | |  | 1.00 | 51.85 || 1 | U | OP1 | AA:1679 | 3.67 | 1.00 | 55.63 |
| 2 | G | O3' | AA:1678 | 4.61 | 1.00 | 55.48 |  |
| 3 | C | OP2 | AA:1663 | 4.66 | 1.00 | 48.16 |  |
| 4 | C | OP1 | AA:1662 | 4.73 | 1.00 | 46.46 |  |
  |

  
 Coded contact list:   
1^st^ MG coordination shell missing (see Notes)  

### MG:AA:3182

|  | Res. type | Atom | Residue | Distance | Occupancy | B-factor | Closest solute to 1^st^ shell water |
| --- | --- | --- | --- | --- | --- | --- | --- |
| **Ion** | | | |  | 1.00 | 60.60 || 1 | G | O6 | AA:1703 | 3.32 | 1.00 | 50.60 |
| 2 | G | N7 | AA:1703 | 4.31 | 1.00 | 51.22 |  |
| 3 | G | O6 | AA:1702 | 4.50 | 1.00 | 60.61 |  |
| 4 | C | N4 | AA:1685 | 4.76 | 1.00 | 49.97 |  |
| 5 | G | N2 | AA:1699 | 4.83 | 1.00 | 64.97 |  |
  |

  
 Coded contact list:   
 Exclusion zone contact(s): /O6/  
Consider K in place of MG (1 valid contact(s) for K vs. 0 for MG)  

### MG:AA:3183

|  | Res. type | Atom | Residue | Distance | Occupancy | B-factor | Closest solute to 1^st^ shell water |
| --- | --- | --- | --- | --- | --- | --- | --- |
| **Ion** | | | |  | 1.00 | 29.01 || 1 | G | OP1 | AA:739 | 1.96 | 1.00 | 41.52 |
| 2 | A | N1 | AA:1784 | 3.48 | 1.00 | 42.24 |  |
| 3 | U | OP2 | AA:1778 | 3.69 | 1.00 | 41.32 |  |
| 4 | U | OP2 | AA:1779 | 3.85 | 1.00 | 34.13 |  |
| 5 | G | O3' | AA:738 | 3.93 | 1.00 | 41.85 |  |
| 6 | U | OP1 | AA:1778 | 4.04 | 1.00 | 41.92 |  |
| 7 | G | O5' | AA:739 | 4.09 | 1.00 | 40.65 |  |
| 8 | A | N6 | AA:1784 | 4.25 | 1.00 | 42.29 |  |
| 9 | U | O5' | AA:1778 | 4.43 | 1.00 | 40.62 |  |
| 10 | G | OP2 | AA:739 | 4.50 | 1.00 | 39.99 |  |
  |

  
 Contact(s): /OP1/  
 Coded contact list: O_ph_  

### MG:AA:3184

|  | Res. type | Atom | Residue | Distance | Occupancy | B-factor | Closest solute to 1^st^ shell water |
| --- | --- | --- | --- | --- | --- | --- | --- |
| **Ion** | | | |  | 1.00 | 49.15 || 1 | G | OP1 | AA:690 | 3.39 | 1.00 | 36.94 |
| 2 | U | O3' | AA:1777 | 4.10 | 1.00 | 42.34 |  |
| 3 | A | OP1 | AA:781 | 4.12 | 1.00 | 34.96 |  |
| 4 | A | OP1 | AA:782 | 4.12 | 1.00 | 34.74 |  |
| 5 | U | O2' | AA:1777 | 4.20 | 1.00 | 41.56 |  |
| 6 | A | O3' | AA:689 | 4.43 | 1.00 | 36.08 |  |
| 7 | U | OP1 | AA:1778 | 4.44 | 1.00 | 41.92 |  |
| 8 | G | O3' | AA:780 | 4.84 | 1.00 | 29.71 |  |
| 9 | A | N7 | AA:781 | 4.90 | 1.00 | 33.22 |  |
  |

  
 Coded contact list:   
 Exclusion zone contact(s): /OP1/  
Consider K in place of MG (1 valid contact(s) for K vs. 0 for MG)  

### MG:AA:3185

|  | Res. type | Atom | Residue | Distance | Occupancy | B-factor | Closest solute to 1^st^ shell water |
| --- | --- | --- | --- | --- | --- | --- | --- |
| **Ion** | | | |  | 1.00 | 29.53 || 1 | G | OP2 | AA:563 | 2.50 | 1.00 | 58.79 |
| 2 | G | OP1 | AA:563 | 3.30 | 1.00 | 58.10 |  |
| 3 | A | OP1 | AA:572 | 3.45 | 1.00 | 57.30 |  |
| 4 | C | OP2 | AA:564 | 3.51 | 1.00 | 49.98 |  |
| 5 | G | O5' | AA:563 | 3.54 | 1.00 | 56.46 |  |
| 6 | C | OP1 | AA:564 | 3.67 | 1.00 | 50.86 |  |
| 7 | C | O2' | AA:974A | 4.66 | 1.00 | 77.03 |  |
| 8 | U | O3' | AA:562 | 4.67 | 1.00 | 57.80 |  |
| 9 | A | OP2 | AA:572 | 4.94 | 1.00 | 57.56 |  |
| 10 | G | O3' | AA:563 | 4.94 | 1.00 | 48.58 |  |
| 11 | G | OP2 | AA:975 | 4.96 | 1.00 | 75.35 |  |
  |

  
 Coded contact list:   
 Exclusion zone contact(s): /OP1/OP2/  
Consider NA in place of MG (1 valid contact(s) for NA vs. 0 for MG)  

### MG:AA:3186

|  | Res. type | Atom | Residue | Distance | Occupancy | B-factor | Closest solute to 1^st^ shell water |
| --- | --- | --- | --- | --- | --- | --- | --- |
| **Ion** | | | |  | 1.00 | 19.62 || 1 | A | N7 | AA:1632 | 1.94 | 1.00 | 43.94 |
| 2 | G | O6 | AA:1633 | 2.54 | 1.00 | 46.36 |  |
| 3 | A | N6 | AA:1632 | 3.04 | 1.00 | 41.10 |  |
| 4 | A | OP2 | AA:1632 | 3.40 | 1.00 | 54.31 |  |
| 5 | A | N6 | AA:1762 | 3.54 | 1.00 | 69.10 |  |
| 6 | A | N7 | AA:1631 | 4.01 | 1.00 | 45.80 |  |
| 7 | G | N7 | AA:1633 | 4.03 | 1.00 | 44.51 |  |
| 8 | A | OP1 | AA:1631 | 4.12 | 1.00 | 63.55 |  |
| 9 | A | N9 | AA:1632 | 4.17 | 1.00 | 43.68 |  |
| 10 | G | N1 | AA:1633 | 4.44 | 1.00 | 45.36 |  |
| 11 | C | O2 | AA:1630A | 4.64 | 1.00 | 55.99 |  |
| 12 | A | N1 | AA:1632 | 4.71 | 1.00 | 41.80 |  |
| 13 | C | N3 | AA:1630A | 4.82 | 1.00 | 55.75 |  |
| 14 | A | O5' | AA:1631 | 4.87 | 1.00 | 61.51 |  |
| 15 | G | N2 | AA:1635 | 4.90 | 1.00 | 53.02 |  |
| 16 | A | N9 | AA:1631 | 4.96 | 1.00 | 49.38 |  |
  |

  
 Contact(s): /N7/  
 Coded contact list: N_b_  
 Exclusion zone contact(s): /N6/O6/  

### MG:AA:3187

|  | Res. type | Atom | Residue | Distance | Occupancy | B-factor | Closest solute to 1^st^ shell water |
| --- | --- | --- | --- | --- | --- | --- | --- |
| **Ion** | | | |  | 1.00 | 68.66 || 1 | A | N7 | AA:1760 | 3.27 | 1.00 | 41.83 |
| 2 | C | N4 | AA:1761 | 3.67 | 1.00 | 52.32 |  |
| 3 | A | OP2 | AA:1759 | 3.78 | 1.00 | 57.04 |  |
| 4 | A | OP2 | AA:1760 | 4.29 | 1.00 | 49.18 |  |
| 5 | A | N6 | AA:1760 | 4.33 | 1.00 | 40.17 |  |
| 6 | U | N3 | AA:1757 | 4.45 | 1.00 | 68.17 |  |
| 7 | G | OP1 | AA:1758 | 4.46 | 1.00 | 67.99 |  |
| 8 | U | N1 | AA:1757 | 4.81 | 1.00 | 67.88 |  |
| 9 | U | O2 | AA:1757 | 4.85 | 1.00 | 69.82 |  |
| 10 | A | O5' | AA:1759 | 4.86 | 1.00 | 55.23 |  |
  |

  
 Coded contact list:   
 Exclusion zone contact(s): /N7/  
Consider K in place of MG (1 valid contact(s) for K vs. 0 for MG)  

### MG:AA:3188

|  | Res. type | Atom | Residue | Distance | Occupancy | B-factor | Closest solute to 1^st^ shell water |
| --- | --- | --- | --- | --- | --- | --- | --- |
| **Ion** | | | |  | 1.00 | 52.15 || 1 | A | OP2 | AA:1701 | 3.43 | 1.00 | 50.27 |
| 2 | G | N7 | AA:1702 | 3.63 | 1.00 | 56.83 |  |
| 3 | G | O6 | AA:1763 | 3.69 | 1.00 | 59.55 |  |
| 4 | G | OP2 | AA:1702 | 4.21 | 1.00 | 53.16 |  |
| 5 | A | OP1 | AA:1701 | 4.64 | 1.00 | 50.69 |  |
| 6 | G | N1 | AA:1763 | 4.80 | 1.00 | 59.22 |  |
| 7 | G | N7 | AA:1703 | 4.84 | 1.00 | 51.22 |  |
  |

  
 Coded contact list:   
1^st^ MG coordination shell missing (see Notes)  

### MG:AA:3189

|  | Res. type | Atom | Residue | Distance | Occupancy | B-factor | Closest solute to 1^st^ shell water |
| --- | --- | --- | --- | --- | --- | --- | --- |
| **Ion** | | | |  | 1.00 | 52.29 || 1 | G | O6 | AA:759 | 3.40 | 1.00 | 49.95 |
| 2 | A | OP2 | AA:734 | 3.83 | 1.00 | 40.36 |  |
| 3 | G | O6 | AA:760 | 3.87 | 1.00 | 40.93 |  |
| 4 | G | N7 | AA:759 | 3.92 | 1.00 | 49.25 |  |
| 5 | A | N6 | AA:761 | 4.35 | 1.00 | 46.73 |  |
| 6 | G | N7 | AA:760 | 4.53 | 1.00 | 39.41 |  |
  |

  
 Coded contact list:   
1^st^ MG coordination shell missing (see Notes)  

### MG:AA:3190

|  | Res. type | Atom | Residue | Distance | Occupancy | B-factor | Closest solute to 1^st^ shell water |
| --- | --- | --- | --- | --- | --- | --- | --- |
| **Ion** | | | |  | 1.00 | 26.09 || 1 | C | OP1 | AA:974A | 1.99 | 1.00 | 71.72 |
| 2 | C | OP2 | AA:974A | 3.14 | 1.00 | 71.48 |  |
| 3 | G | O2' | AA:974 | 3.97 | 1.00 | 69.97 |  |
| 4 | G | O3' | AA:974 | 4.03 | 1.00 | 69.74 |  |
| 5 | C | O5' | AA:974A | 4.14 | 1.00 | 72.43 |  |
| 6 | C | N4 | AA:976 | 4.44 | 1.00 | 80.15 |  |
| 7 | A | O2' | AA:983 | 4.56 | 1.00 | 55.85 |  |
| 8 | A | O3' | AA:983 | 4.80 | 1.00 | 53.99 |  |
  |

  
 Contact(s): /OP1/  
 Coded contact list: O_ph_  
 Exclusion zone contact(s): /OP2/  

### MG:AA:3191

|  | Res. type | Atom | Residue | Distance | Occupancy | B-factor | Closest solute to 1^st^ shell water |
| --- | --- | --- | --- | --- | --- | --- | --- |
| **Ion** | | | |  | 1.00 | 31.51 || 1 | A | OP2 | AA:1676 | 2.13 | 1.00 | 46.49 |
| 2 | A | OP1 | AA:1676 | 4.03 | 1.00 | 46.97 |  |
| 3 | A | O5' | AA:1676 | 4.13 | 1.00 | 48.18 |  |
| 4 | C | OP2 | AA:1675 | 4.37 | 1.00 | 42.30 |  |
| 5 | A | N7 | AA:1676 | 4.55 | 1.00 | 60.15 |  |
| 6 | A | N7 | AA:1677 | 4.61 | 1.00 | 55.59 |  |
| 7 | C | O3' | AA:1675 | 4.67 | 1.00 | 43.68 |  |
| 8 | A | N6 | AA:1677 | 4.72 | 1.00 | 54.54 |  |
| 9 | A | OP2 | AA:1677 | 4.97 | 1.00 | 51.05 |  |
  |

  
 Contact(s): /OP2/  
 Coded contact list: O_ph_  

### MG:AA:3192

|  | Res. type | Atom | Residue | Distance | Occupancy | B-factor | Closest solute to 1^st^ shell water |
| --- | --- | --- | --- | --- | --- | --- | --- |
| **Ion** | | | |  | 1.00 | 39.98 || 1 | C | OP1 | AA:2036 | 1.85 | 1.00 | 55.69 |
| 2 | C | OP1 | AA:531 | 1.88 | 1.00 | 75.45 |  |
| 3 | C | OP2 | AA:531 | 3.15 | 1.00 | 74.99 |  |
| 4 | C | O5' | AA:2036 | 3.72 | 1.00 | 53.77 |  |
| 5 | C | OP2 | AA:2036 | 3.80 | 1.00 | 55.49 |  |
| 6 | G | O3' | AA:530 | 3.92 | 1.00 | 73.49 |  |
| 7 | C | O5' | AA:531 | 4.09 | 1.00 | 72.71 |  |
| 8 | G | O3' | AA:2035 | 4.35 | 1.00 | 56.79 |  |
| 9 | G | O2' | AA:2035 | 4.36 | 1.00 | 58.40 |  |
| 10 | G | O6 | AA:561 | 4.56 | 1.00 | 70.47 |  |
| 11 | G | N1 | AA:561 | 4.71 | 1.00 | 69.28 |  |
  |

  
**1^st^ shell ligand angles:**   
1-2: 150;   
  
 Contact(s): /OP1/OP1/  
 Coded contact list: 2O_ph_  
 Exclusion zone contact(s): /OP2/  

### MG:AA:3193

|  | Res. type | Atom | Residue | Distance | Occupancy | B-factor | Closest solute to 1^st^ shell water |
| --- | --- | --- | --- | --- | --- | --- | --- |
| **Ion** | | | |  | 1.00 | 99.99 || 1 | G | OP1 | AA:2088 | 2.52 | 1.00 | 50.02 |
| 2 | G | OP1 | AA:2191 | 3.18 | 1.00 | 130.40 |  |
| 3 | U | OP1 | AA:2089 | 3.48 | 1.00 | 55.31 |  |
| 4 | U | OP2 | AA:2089 | 3.62 | 1.00 | 54.33 |  |
| 5 | G | OP2 | AA:2088 | 3.90 | 1.00 | 48.77 |  |
| 6 | G | O3' | AA:2088 | 4.02 | 1.00 | 53.90 |  |
| 7 | G | O5' | AA:2088 | 4.20 | 1.00 | 49.38 |  |
| 8 | G | O5' | AA:2191 | 4.54 | 1.00 | 128.45 |  |
| 9 | G | O3' | AA:2087 | 4.92 | 1.00 | 49.52 |  |
  |

  
 Coded contact list:   
 Exclusion zone contact(s): /OP1/OP1/  
Consider NA in place of MG (1 valid contact(s) for NA vs. 0 for MG)  

### MG:AA:3194

|  | Res. type | Atom | Residue | Distance | Occupancy | B-factor | Closest solute to 1^st^ shell water |
| --- | --- | --- | --- | --- | --- | --- | --- |
| **Ion** | | | |  | 1.00 | 56.68 || 1 | G | N7 | AA:2088 | 2.94 | 1.00 | 52.65 |
| 2 | U | O4 | AA:2089 | 3.03 | 1.00 | 49.91 |  |
| 3 | G | O6 | AA:2088 | 3.21 | 1.00 | 54.29 |  |
| 4 | G | N7 | AA:2087 | 3.82 | 1.00 | 45.80 |  |
| 5 | G | O6 | AA:2087 | 4.23 | 1.00 | 42.69 |  |
| 6 | U | N3 | AA:2089 | 4.64 | 1.00 | 49.50 |  |
| 7 | G | OP2 | AA:2087 | 4.80 | 1.00 | 47.44 |  |
  |

  
 Coded contact list:   
 Exclusion zone contact(s): /N7/O4/O6/  
Consider K in place of MG (3 valid contact(s) for K vs. 0 for MG)  

### MG:AA:3195

|  | Res. type | Atom | Residue | Distance | Occupancy | B-factor | Closest solute to 1^st^ shell water |
| --- | --- | --- | --- | --- | --- | --- | --- |
| **Ion** | | | |  | 1.00 | 82.71 || 1 | C | N4 | AA:2853 | 4.85 | 1.00 | 71.00 |
| 2 | G | OP2 | AA:2852 | 4.85 | 1.00 | 66.27 |  |
  |

  
 Coded contact list:   
1^st^ MG coordination shell missing (see Notes)  

### MG:AA:3196

|  | Res. type | Atom | Residue | Distance | Occupancy | B-factor | Closest solute to 1^st^ shell water |
| --- | --- | --- | --- | --- | --- | --- | --- |
| **Ion** | | | |  | 1.00 | 29.80 || 1 | G | OP1 | AA:805 | 2.12 | 1.00 | 43.42 |
| 2 | A | OP1 | AA:804 | 2.16 | 1.00 | 40.72 |  |
| 3 | A | OP2 | AA:804 | 3.69 | 1.00 | 39.30 |  |
| 4 | G | O5' | AA:805 | 3.98 | 1.00 | 44.99 |  |
| 5 | A | O5' | AA:804 | 4.02 | 1.00 | 37.83 |  |
| 6 | G | OP2 | AA:805 | 4.08 | 1.00 | 44.82 |  |
| 7 | GLY | N | AO:44 | 4.30 | 1.00 | 51.61 |  |
| 8 | GLY | N | AO:43 | 4.33 | 1.00 | 43.78 |  |
| 9 | GLY | O | AO:44 | 4.49 | 1.00 | 52.42 |  |
| 10 | U | O3' | AA:803 | 4.58 | 1.00 | 38.91 |  |
| 11 | A | O3' | AA:804 | 4.63 | 1.00 | 42.15 |  |
  |

  
**1^st^ shell ligand angles:**   
1-2: 78;   
  
 Contact(s): /OP1/OP1/  
 Coded contact list: 2O_ph_  

### MG:AA:3197

|  | Res. type | Atom | Residue | Distance | Occupancy | B-factor | Closest solute to 1^st^ shell water |
| --- | --- | --- | --- | --- | --- | --- | --- |
| **Ion** | | | |  | 1.00 | 50.52 || 1 | A | OP2 | AA:804 | 4.10 | 1.00 | 39.30 |
| 2 | A | N7 | AA:804 | 4.89 | 1.00 | 45.28 |  |
| 3 | G | O6 | AA:674 | 4.97 | 1.00 | 43.86 |  |
| 4 | A | OP1 | AA:670 | 5.00 | 1.00 | 54.86 |  |
  |

  
 Coded contact list:   
1^st^ MG coordination shell missing (see Notes)  

### MG:AA:3198

|  | Res. type | Atom | Residue | Distance | Occupancy | B-factor | Closest solute to 1^st^ shell water |
| --- | --- | --- | --- | --- | --- | --- | --- |
| **Ion** | | | |  | 1.00 | 34.91 || 1 | A | O2' | AA:528 | 2.52 | 1.00 | 68.84 |
| 2 | A | O3' | AA:528 | 3.17 | 1.00 | 75.38 |  |
| 3 | A | O5' | AA:529 | 3.60 | 1.00 | 77.50 |  |
| 4 | A | O4' | AA:529 | 4.09 | 1.00 | 75.42 |  |
| 5 | A | OP1 | AA:529 | 4.62 | 1.00 | 77.94 |  |
| 6 | A | O4' | AA:528 | 4.62 | 1.00 | 69.06 |  |
| 7 | G | OP1 | AA:530 | 4.64 | 1.00 | 78.93 |  |
  |

  
 Coded contact list:   
 Exclusion zone contact(s): /O2'/O3'/  
Consider NA in place of MG (1 valid contact(s) for NA vs. 0 for MG)  

### MG:AA:3199

|  | Res. type | Atom | Residue | Distance | Occupancy | B-factor | Closest solute to 1^st^ shell water |
| --- | --- | --- | --- | --- | --- | --- | --- |
| **Ion** | | | |  | 1.00 | 75.67 || 1 | G | OP1 | AA:530 | 4.40 | 1.00 | 78.93 |
  |

  
 Coded contact list:   
1^st^ MG coordination shell missing (see Notes)  

### MG:AA:3200

|  | Res. type | Atom | Residue | Distance | Occupancy | B-factor | Closest solute to 1^st^ shell water |
| --- | --- | --- | --- | --- | --- | --- | --- |
| **Ion** | | | |  | 1.00 | 83.39 || 1 | U | OP2 | AA:534 | 4.03 | 1.00 | 63.68 |
| 2 | GLY | O | A1:23 | 4.56 | 1.00 | 58.26 |  |
| 3 | U | OP2 | AA:554 | 4.83 | 1.00 | 70.14 |  |
  |

  
 Coded contact list:   
1^st^ MG coordination shell missing (see Notes)  

### MG:AA:3201

|  | Res. type | Atom | Residue | Distance | Occupancy | B-factor | Closest solute to 1^st^ shell water |
| --- | --- | --- | --- | --- | --- | --- | --- |
| **Ion** | | | |  | 1.00 | 71.57 || 1 | C | OP2 | AA:1832 | 3.78 | 1.00 | 45.38 |
| 2 | G | OP1 | BA:771 | 4.05 | 1.00 | 41.25 |  |
| 3 | U | OP1 | BA:772 | 4.87 | 1.00 | 46.71 |  |
  |

  
 Coded contact list:   
1^st^ MG coordination shell missing (see Notes)  

### MG:AA:3202

|  | Res. type | Atom | Residue | Distance | Occupancy | B-factor | Closest solute to 1^st^ shell water |
| --- | --- | --- | --- | --- | --- | --- | --- |
| **Ion** | | | |  | 1.00 | 31.32 || 1 | C | OP1 | AA:624 | 2.48 | 1.00 | 59.23 |
| 2 | G | OP1 | AA:242 | 3.50 | 1.00 | 51.50 |  |
| 3 | G | O3' | AA:623 | 3.80 | 1.00 | 59.47 |  |
| 4 | C | O5' | AA:624 | 4.00 | 1.00 | 57.87 |  |
| 5 | PRO | N | A8:63 | 4.88 | 1.00 | 101.18 |  |
| 6 | C | OP2 | AA:624 | 4.94 | 1.00 | 58.58 |  |
  |

  
 Coded contact list:   
 Exclusion zone contact(s): /OP1/  
Consider NA in place of MG (1 valid contact(s) for NA vs. 0 for MG)  

### MG:AA:3203

|  | Res. type | Atom | Residue | Distance | Occupancy | B-factor | Closest solute to 1^st^ shell water |
| --- | --- | --- | --- | --- | --- | --- | --- |
| **Ion** | | | |  | 1.00 | 40.89 || 1 | U | OP1 | AA:120 | 2.84 | 1.00 | 57.79 |
| 2 | U | OP2 | AA:120 | 2.97 | 1.00 | 57.75 |  |
| 3 | G | N7 | AA:123 | 3.00 | 1.00 | 36.53 |  |
| 4 | G | O6 | AA:123 | 3.10 | 1.00 | 41.69 |  |
| 5 | G | O6 | AA:122 | 3.34 | 1.00 | 44.48 |  |
| 6 | G | O6 | AA:124 | 3.54 | 1.00 | 43.26 |  |
| 7 | G | N7 | AA:122 | 3.98 | 1.00 | 47.05 |  |
| 8 | A | O3' | AA:119 | 4.39 | 1.00 | 58.01 |  |
| 9 | U | O5' | AA:120 | 4.61 | 1.00 | 56.96 |  |
| 10 | G | N7 | AA:124 | 4.78 | 1.00 | 42.32 |  |
| 11 | G | N1 | AA:122 | 4.79 | 1.00 | 45.78 |  |
| 12 | G | N1 | AA:124 | 4.92 | 1.00 | 41.48 |  |
  |

  
 Coded contact list:   
 Exclusion zone contact(s): /N7/O6/O6/OP1/OP2/  
Consider K in place of MG (5 valid contact(s) for K vs. 0 for MG)  

### MG:AA:3204

|  | Res. type | Atom | Residue | Distance | Occupancy | B-factor | Closest solute to 1^st^ shell water |
| --- | --- | --- | --- | --- | --- | --- | --- |
| **Ion** | | | |  | 1.00 | 68.05 || 1 | U | OP1 | AA:120 | 1.95 | 1.00 | 57.79 |
| 2 | G | O6 | AA:124 | 3.09 | 1.00 | 43.26 |  |
| 3 | A | O3' | AA:119 | 3.33 | 1.00 | 58.01 |  |
| 4 | U | O5' | AA:120 | 3.96 | 1.00 | 56.96 |  |
| 5 | G | N7 | AA:124 | 4.01 | 1.00 | 42.32 |  |
| 6 | ARG | NH2 | A7:19 | 4.30 | 1.00 | 37.68 |  |
| 7 | U | OP2 | AA:120 | 4.37 | 1.00 | 57.75 |  |
| 8 | G | N7 | AA:123 | 4.78 | 1.00 | 36.53 |  |
| 9 | G | OP2 | AA:123 | 4.97 | 1.00 | 50.71 |  |
  |

  
 Contact(s): /OP1/  
 Coded contact list: O_ph_  
 Exclusion zone contact(s): /O3'/O6/  
Consider K in place of MG (2 valid contact(s) for K vs. 1 for MG)  

### MG:AA:3205

|  | Res. type | Atom | Residue | Distance | Occupancy | B-factor | Closest solute to 1^st^ shell water |
| --- | --- | --- | --- | --- | --- | --- | --- |
| **Ion** | | | |  | 1.00 | 71.22 || 1 | G | OP2 | AA:1705 | 3.95 | 1.00 | 59.84 |
| 2 | U | O4 | AA:1706 | 4.10 | 1.00 | 63.42 |  |
  |

  
 Coded contact list:   
1^st^ MG coordination shell missing (see Notes)  

### MG:AA:3206

|  | Res. type | Atom | Residue | Distance | Occupancy | B-factor | Closest solute to 1^st^ shell water |
| --- | --- | --- | --- | --- | --- | --- | --- |
| **Ion** | | | |  | 1.00 | 40.85 || 1 | U | O4 | AA:1602 | 1.78 | 1.00 | 53.19 |
| 2 | A | OP2 | AA:1342 | 2.83 | 1.00 | 42.30 |  |
| 3 | A | O5' | AA:1342 | 3.19 | 1.00 | 43.29 |  |
| 4 | U | N3 | AA:1602 | 3.58 | 1.00 | 52.55 |  |
| 5 | G | N7 | AA:1601 | 3.75 | 1.00 | 46.35 |  |
| 6 | G | O6 | AA:1601 | 3.78 | 1.00 | 48.31 |  |
| 7 | A | OP1 | AA:1342 | 4.16 | 1.00 | 42.29 |  |
| 8 | C | N4 | AA:1600 | 4.29 | 1.00 | 40.98 |  |
| 9 | THR | O | AT:56 | 4.66 | 1.00 | 46.82 |  |
| 10 | A | N3 | AA:1342 | 4.82 | 1.00 | 52.67 |  |
| 11 | U | O3' | AA:1341 | 4.89 | 1.00 | 42.53 |  |
  |

  
 Coded contact list:   
 Exclusion zone contact(s): /O4/O5'/OP2/  
Consider K in place of MG (2 valid contact(s) for K vs. 0 for MG)  

### MG:AA:3207

|  | Res. type | Atom | Residue | Distance | Occupancy | B-factor | Closest solute to 1^st^ shell water |
| --- | --- | --- | --- | --- | --- | --- | --- |
| **Ion** | | | |  | 1.00 | 30.21 || 1 | A | OP1 | AA:1268 | 1.99 | 1.00 | 48.98 |
| 2 | C | OP1 | AA:2006 | 2.19 | 1.00 | 48.72 |  |
| 3 | U | O3' | AA:1267 | 3.42 | 1.00 | 48.69 |  |
| 4 | G | OP1 | AA:2048 | 4.15 | 1.00 | 36.67 |  |
| 5 | A | O5' | AA:1268 | 4.28 | 1.00 | 48.77 |  |
| 6 | A | OP2 | AA:1268 | 4.30 | 1.00 | 49.16 |  |
| 7 | C | OP2 | AA:2006 | 4.31 | 1.00 | 48.20 |  |
| 8 | A | O3' | AA:2005 | 4.31 | 1.00 | 48.46 |  |
| 9 | G | OP2 | AA:2049 | 4.55 | 1.00 | 35.18 |  |
| 10 | C | O5' | AA:2006 | 4.65 | 1.00 | 47.17 |  |
  |

  
**1^st^ shell ligand angles:**   
1-2: 85;   
  
 Contact(s): /OP1/OP1/  
 Coded contact list: 2O_ph_  

### MG:AA:3208

|  | Res. type | Atom | Residue | Distance | Occupancy | B-factor | Closest solute to 1^st^ shell water |
| --- | --- | --- | --- | --- | --- | --- | --- |
| **Ion** | | | |  | 1.00 | 30.06 || 1 | U | OP1 | AA:2022 | 2.39 | 1.00 | 53.16 |
| 2 | G | N1 | AA:530 | 2.92 | 1.00 | 69.06 |  |
| 3 | U | O5' | AA:2022 | 2.94 | 1.00 | 52.96 |  |
| 4 | G | N2 | AA:530 | 3.10 | 1.00 | 70.58 |  |
| 5 | G | OP1 | AA:2023 | 3.78 | 1.00 | 53.92 |  |
| 6 | G | O6 | AA:530 | 4.14 | 1.00 | 68.23 |  |
| 7 | C | O3' | AA:2021 | 4.26 | 1.00 | 52.77 |  |
| 8 | U | OP2 | AA:2022 | 4.41 | 1.00 | 52.42 |  |
| 9 | G | OP2 | AA:2023 | 4.51 | 1.00 | 53.70 |  |
| 10 | G | N3 | AA:530 | 4.63 | 1.00 | 69.70 |  |
| 11 | A | N7 | AA:529 | 4.71 | 1.00 | 67.44 |  |
| 12 | C | O2' | AA:2021 | 4.89 | 1.00 | 56.72 |  |
  |

  
 Contact(s): /OP1/  
 Coded contact list: O_ph_  
 Exclusion zone contact(s): /N1_G_/N2/O5'/  
Consider K in place of MG (3 valid contact(s) for K vs. 1 for MG)  

### MG:AA:3209

|  | Res. type | Atom | Residue | Distance | Occupancy | B-factor | Closest solute to 1^st^ shell water |
| --- | --- | --- | --- | --- | --- | --- | --- |
| **Ion** | | | |  | 1.00 | 77.98 || 1 | G | OP2 | AA:1756 | 4.04 | 1.00 | 57.28 |
| 2 | A | O3' | AA:1755 | 4.68 | 1.00 | 58.15 |  |
| 3 | G | OP1 | AA:1756 | 4.95 | 1.00 | 59.61 |  |
  |

  
 Coded contact list:   
1^st^ MG coordination shell missing (see Notes)  

### MG:AA:3210

|  | Res. type | Atom | Residue | Distance | Occupancy | B-factor | Closest solute to 1^st^ shell water |
| --- | --- | --- | --- | --- | --- | --- | --- |
| **Ion** | | | |  | 1.00 | 39.68 || 1 | U | O4 | AA:639 | 2.69 | 1.00 | 72.32 |
| 2 | G | O6 | AA:638 | 2.75 | 1.00 | 80.19 |  |
| 3 | G | O6 | AA:648 | 2.99 | 1.00 | 68.22 |  |
| 4 | G | O6 | AA:649 | 3.03 | 1.00 | 74.58 |  |
| 5 | C | N4 | AA:650 | 3.96 | 1.00 | 79.29 |  |
| 6 | U | N3 | AA:639 | 4.00 | 1.00 | 72.19 |  |
| 7 | C | N4 | AA:640 | 4.38 | 1.00 | 68.19 |  |
| 8 | G | N7 | AA:648 | 4.42 | 1.00 | 67.41 |  |
| 9 | G | N7 | AA:638 | 4.59 | 1.00 | 81.90 |  |
| 10 | G | N7 | AA:649 | 4.66 | 1.00 | 72.24 |  |
| 11 | G | N1 | AA:638 | 4.67 | 1.00 | 80.66 |  |
| 12 | G | N1 | AA:648 | 4.75 | 1.00 | 67.47 |  |
  |

  
 Coded contact list:   
 Exclusion zone contact(s): /O4/O6/O6/O6/  
Consider K in place of MG (4 valid contact(s) for K vs. 0 for MG)  

### MG:AA:3211

|  | Res. type | Atom | Residue | Distance | Occupancy | B-factor | Closest solute to 1^st^ shell water |
| --- | --- | --- | --- | --- | --- | --- | --- |
| **Ion** | | | |  | 1.00 | 56.19 || 1 | G | O6 | AA:1822 | 3.49 | 1.00 | 32.44 |
| 2 | A | N6 | AA:1821 | 3.79 | 1.00 | 38.12 |  |
| 3 | A | N7 | AA:1821 | 4.24 | 1.00 | 38.18 |  |
| 4 | G | N7 | AA:1822 | 4.46 | 1.00 | 31.92 |  |
| 5 | U | O4 | AA:1796 | 4.77 | 1.00 | 33.88 |  |
| 6 | C | N4 | AA:1797 | 4.98 | 1.00 | 32.52 |  |
  |

  
 Coded contact list:   
1^st^ MG coordination shell missing (see Notes)  

### MG:AA:3212

|  | Res. type | Atom | Residue | Distance | Occupancy | B-factor | Closest solute to 1^st^ shell water |
| --- | --- | --- | --- | --- | --- | --- | --- |
| **Ion** | | | |  | 1.00 | 78.83 || 1 | G | OP2 | AA:649 | 3.95 | 1.00 | 71.41 |
| 2 | C | N4 | AA:650 | 4.28 | 1.00 | 79.29 |  |
| 3 | G | O6 | AA:651 | 4.32 | 1.00 | 85.86 |  |
| 4 | G | N7 | AA:651 | 4.81 | 1.00 | 84.13 |  |
  |

  
 Coded contact list:   
1^st^ MG coordination shell missing (see Notes)  

### MG:AA:3213

|  | Res. type | Atom | Residue | Distance | Occupancy | B-factor | Closest solute to 1^st^ shell water |
| --- | --- | --- | --- | --- | --- | --- | --- |
| **Ion** | | | |  | 1.00 | 51.57 || 1 | G | OP1 | AA:760 | 2.63 | 1.00 | 42.36 |
| 2 | C | OP1 | AA:732 | 3.12 | 1.00 | 40.57 |  |
| 3 | C | OP2 | AA:732 | 3.20 | 1.00 | 41.80 |  |
| 4 | C | OP1 | AA:731 | 3.64 | 1.00 | 40.60 |  |
| 5 | C | O3' | AA:731 | 4.23 | 1.00 | 41.60 |  |
| 6 | G | O5' | AA:760 | 4.41 | 1.00 | 39.50 |  |
| 7 | G | OP2 | AA:760 | 4.48 | 1.00 | 40.28 |  |
| 8 | A | OP2 | AA:761 | 4.49 | 1.00 | 38.67 |  |
  |

  
 Coded contact list:   
 Exclusion zone contact(s): /OP1/OP1/OP2/  
Consider K in place of MG (3 valid contact(s) for K vs. 0 for MG)  

### MG:AA:3214

|  | Res. type | Atom | Residue | Distance | Occupancy | B-factor | Closest solute to 1^st^ shell water |
| --- | --- | --- | --- | --- | --- | --- | --- |
| **Ion** | | | |  | 1.00 | 56.23 || 1 | G | OP1 | AA:759 | 2.27 | 1.00 | 46.57 |
| 2 | G | OP2 | AA:760 | 4.03 | 1.00 | 40.28 |  |
| 3 | G | O5' | AA:759 | 4.16 | 1.00 | 44.04 |  |
| 4 | G | OP2 | AA:759 | 4.21 | 1.00 | 45.33 |  |
| 5 | C | OP1 | AA:732 | 4.26 | 1.00 | 40.57 |  |
| 6 | G | OP2 | AA:733 | 4.56 | 1.00 | 41.17 |  |
| 7 | G | OP1 | AA:733 | 4.66 | 1.00 | 42.53 |  |
| 8 | C | O3' | AA:758 | 4.77 | 1.00 | 45.65 |  |
| 9 | G | OP1 | AA:760 | 4.83 | 1.00 | 42.36 |  |
  |

  
 Contact(s): /OP1/  
 Coded contact list: O_ph_  

### MG:AA:3215

|  | Res. type | Atom | Residue | Distance | Occupancy | B-factor | Closest solute to 1^st^ shell water |
| --- | --- | --- | --- | --- | --- | --- | --- |
| **Ion** | | | |  | 1.00 | 72.43 || 1 | G | N7 | AA:2735 | 3.55 | 1.00 | 100.12 |
| 2 | G | O6 | AA:2736 | 4.20 | 1.00 | 97.58 |  |
| 3 | G | N7 | AA:2736 | 4.30 | 1.00 | 95.45 |  |
| 4 | G | O6 | AA:2735 | 4.40 | 1.00 | 96.85 |  |
| 5 | G | O6 | AA:2737 | 4.82 | 1.00 | 85.58 |  |
  |

  
 Coded contact list:   
1^st^ MG coordination shell missing (see Notes)  

### MG:AA:3216

|  | Res. type | Atom | Residue | Distance | Occupancy | B-factor | Closest solute to 1^st^ shell water |
| --- | --- | --- | --- | --- | --- | --- | --- |
| **Ion** | | | |  | 1.00 | 40.87 || 1 | G | OP1 | AA:411 | 1.98 | 1.00 | 60.32 |
| 2 | G | OP1 | AA:2407 | 2.27 | 1.00 | 69.54 |  |
| 3 | G | OP2 | AA:2407 | 3.77 | 1.00 | 69.14 |  |
| 4 | A | OP1 | AA:412 | 3.89 | 1.00 | 56.35 |  |
| 5 | G | O5' | AA:411 | 4.20 | 1.00 | 58.13 |  |
| 6 | G | O3' | AA:410 | 4.32 | 1.00 | 58.03 |  |
| 7 | G | OP2 | AA:411 | 4.38 | 1.00 | 58.08 |  |
| 8 | U | O3' | AA:2406 | 4.45 | 1.00 | 70.37 |  |
| 9 | U | O4 | AA:2408 | 4.62 | 1.00 | 66.28 |  |
| 10 | G | O5' | AA:2407 | 4.62 | 1.00 | 69.17 |  |
| 11 | G | N7 | AA:2407 | 4.71 | 1.00 | 66.12 |  |
| 12 | A | OP2 | AA:412 | 4.77 | 1.00 | 56.22 |  |
  |

  
**1^st^ shell ligand angles:**   
1-2: 88;   
  
 Contact(s): /OP1/OP1/  
 Coded contact list: 2O_ph_  

### MG:AA:3217

|  | Res. type | Atom | Residue | Distance | Occupancy | B-factor | Closest solute to 1^st^ shell water |
| --- | --- | --- | --- | --- | --- | --- | --- |
| **Ion** | | | |  | 1.00 | 57.34 || 1 | G | OP2 | AA:2410 | 3.22 | 1.00 | 70.48 |
| 2 | G | N7 | AA:2405 | 4.03 | 1.00 | 70.23 |  |
| 3 | G | OP1 | AA:2410 | 4.61 | 1.00 | 70.44 |  |
| 4 | G | OP2 | AA:2405 | 4.63 | 1.00 | 66.61 |  |
  |

  
 Coded contact list:   
 Exclusion zone contact(s): /OP2/  
Consider K in place of MG (1 valid contact(s) for K vs. 0 for MG)  

### MG:AA:3218

|  | Res. type | Atom | Residue | Distance | Occupancy | B-factor | Closest solute to 1^st^ shell water |
| --- | --- | --- | --- | --- | --- | --- | --- |
| **Ion** | | | |  | 1.00 | 65.81 || 1 | C | OP2 | AA:2404 | 2.57 | 1.00 | 71.57 |
| 2 | G | OP2 | AA:2405 | 2.74 | 1.00 | 66.61 |  |
| 3 | G | N7 | AA:2405 | 3.38 | 1.00 | 70.23 |  |
| 4 | C | OP1 | AA:2404 | 3.51 | 1.00 | 71.07 |  |
| 5 | C | O5' | AA:2404 | 3.52 | 1.00 | 70.02 |  |
| 6 | C | O3' | AA:2404 | 4.66 | 1.00 | 66.51 |  |
| 7 | G | O5' | AA:2405 | 4.76 | 1.00 | 66.69 |  |
| 8 | C | O3' | AA:2403 | 4.78 | 1.00 | 72.64 |  |
  |

  
 Coded contact list:   
 Exclusion zone contact(s): /N7/OP2/OP2/  
Consider K in place of MG (2 valid contact(s) for K vs. 0 for MG)  

### MG:AA:3219

|  | Res. type | Atom | Residue | Distance | Occupancy | B-factor | Closest solute to 1^st^ shell water |
| --- | --- | --- | --- | --- | --- | --- | --- |
| **Ion** | | | |  | 1.00 | 81.83 || 1 | G | O6 | AA:2405 | 3.30 | 1.00 | 71.28 |
| 2 | C | OP2 | AA:2403 | 3.53 | 1.00 | 86.02 |  |
| 3 | C | OP2 | AA:2404 | 3.78 | 1.00 | 71.57 |  |
| 4 | C | OP1 | AA:2403 | 4.43 | 1.00 | 85.55 |  |
| 5 | G | N7 | AA:2405 | 4.56 | 1.00 | 70.23 |  |
| 6 | C | O5' | AA:2403 | 4.70 | 1.00 | 83.57 |  |
  |

  
 Coded contact list:   
 Exclusion zone contact(s): /O6/  
Consider K in place of MG (1 valid contact(s) for K vs. 0 for MG)  

### MG:AA:3220

|  | Res. type | Atom | Residue | Distance | Occupancy | B-factor | Closest solute to 1^st^ shell water |
| --- | --- | --- | --- | --- | --- | --- | --- |
| **Ion** | | | |  | 1.00 | 70.71 || 1 | G | OP2 | AA:2409 | 3.00 | 1.00 | 67.22 |
| 2 | G | OP1 | AA:2409 | 3.60 | 1.00 | 68.44 |  |
| 3 | G | OP1 | AA:2405 | 3.94 | 1.00 | 66.30 |  |
| 4 | G | OP2 | AA:2405 | 3.98 | 1.00 | 66.61 |  |
| 5 | G | O5' | AA:2405 | 4.14 | 1.00 | 66.69 |  |
| 6 | U | O3' | AA:2408 | 4.51 | 1.00 | 67.24 |  |
| 7 | U | OP1 | AA:2408 | 4.53 | 1.00 | 68.62 |  |
  |

  
 Coded contact list:   
 Exclusion zone contact(s): /OP2/  
Consider K in place of MG (1 valid contact(s) for K vs. 0 for MG)  

### MG:AA:3221

|  | Res. type | Atom | Residue | Distance | Occupancy | B-factor | Closest solute to 1^st^ shell water |
| --- | --- | --- | --- | --- | --- | --- | --- |
| **Ion** | | | |  | 1.00 | 63.71 || 1 | G | O6 | AA:2413 | 3.71 | 1.00 | 67.66 |
| 2 | G | N7 | AA:2413 | 4.00 | 1.00 | 69.37 |  |
| 3 | G | O6 | AA:2414 | 4.13 | 1.00 | 69.49 |  |
| 4 | G | N7 | AA:2414 | 4.22 | 1.00 | 66.77 |  |
| 5 | A | OP2 | AA:2412 | 4.97 | 1.00 | 69.84 |  |
  |

  
 Coded contact list:   
1^st^ MG coordination shell missing (see Notes)  

### MG:AA:3222

|  | Res. type | Atom | Residue | Distance | Occupancy | B-factor | Closest solute to 1^st^ shell water |
| --- | --- | --- | --- | --- | --- | --- | --- |
| **Ion** | | | |  | 1.00 | 28.46 || 1 | U | OP2 | AA:740 | 1.83 | 1.00 | 39.25 |
| 2 | A | OP2 | AA:1784 | 2.29 | 1.00 | 44.62 |  |
| 3 | A | OP1 | AA:1783 | 2.42 | 1.00 | 41.01 |  |
| 4 | U | O5' | AA:740 | 3.75 | 1.00 | 37.33 |  |
| 5 | A | OP1 | AA:1784 | 3.78 | 1.00 | 45.43 |  |
| 6 | U | OP1 | AA:740 | 3.78 | 1.00 | 39.11 |  |
| 7 | G | OP2 | AA:741 | 4.24 | 1.00 | 43.04 |  |
| 8 | G | O3' | AA:739 | 4.39 | 1.00 | 40.55 |  |
| 9 | A | O5' | AA:1784 | 4.49 | 1.00 | 43.61 |  |
| 10 | A | O5' | AA:1783 | 4.54 | 1.00 | 41.08 |  |
| 11 | A | O3' | AA:1783 | 4.58 | 1.00 | 42.40 |  |
| 12 | C | O3' | AA:1782 | 4.76 | 1.00 | 39.19 |  |
| 13 | A | OP2 | AA:1783 | 4.82 | 1.00 | 40.42 |  |
  |

  
**1^st^ shell ligand angles:**   
1-2: 123;   
  
 Contact(s): /OP2/OP2/  
 Coded contact list: 2O_ph_  
 Exclusion zone contact(s): /OP1/  

### MG:AA:3223

|  | Res. type | Atom | Residue | Distance | Occupancy | B-factor | Closest solute to 1^st^ shell water |
| --- | --- | --- | --- | --- | --- | --- | --- |
| **Ion** | | | |  | 1.00 | 14.73 || 1 | A | OP2 | AA:1783 | 1.91 | 1.00 | 40.42 |
| 2 | C | OP1 | AA:1782 | 2.09 | 1.00 | 44.97 |  |
| 3 | A | OP1 | AA:1780 | 2.43 | 1.00 | 38.89 |  |
| 4 | C | OP2 | AA:1782 | 2.84 | 1.00 | 44.39 |  |
| 5 | A | OP2 | AA:1780 | 3.63 | 1.00 | 40.25 |  |
| 6 | C | O5' | AA:1782 | 3.85 | 1.00 | 40.65 |  |
| 7 | U | O3' | AA:1779 | 3.97 | 1.00 | 39.54 |  |
| 8 | A | OP1 | AA:1783 | 4.14 | 1.00 | 41.01 |  |
| 9 | C | O3' | AA:1781 | 4.16 | 1.00 | 44.46 |  |
| 10 | A | O5' | AA:1783 | 4.19 | 1.00 | 41.08 |  |
| 11 | C | O3' | AA:1782 | 4.31 | 1.00 | 39.19 |  |
| 12 | C | OP2 | AA:1781 | 4.37 | 1.00 | 44.08 |  |
| 13 | U | O2' | AA:1779 | 4.58 | 1.00 | 39.28 |  |
| 14 | A | O5' | AA:1780 | 4.81 | 1.00 | 41.64 |  |
| 15 | A | N7 | AA:1783 | 4.94 | 1.00 | 35.67 |  |
  |

  
**1^st^ shell ligand angles:**   
1-2: 88;   
  
 Contact(s): /OP1/OP2/  
 Coded contact list: 2O_ph_  
 Exclusion zone contact(s): /OP1/OP2/  

### MG:AA:3224

|  | Res. type | Atom | Residue | Distance | Occupancy | B-factor | Closest solute to 1^st^ shell water |
| --- | --- | --- | --- | --- | --- | --- | --- |
| **Ion** | | | |  | 1.00 | 63.49 || 1 | G | O6 | AA:398 | 2.70 | 1.00 | 56.01 |
| 2 | G | N7 | AA:398 | 3.24 | 1.00 | 54.03 |  |
| 3 | G | O6 | AA:399 | 3.58 | 1.00 | 53.29 |  |
| 4 | G | O6 | AA:397 | 3.93 | 1.00 | 46.77 |  |
| 5 | U | OP2 | AA:373 | 4.31 | 1.00 | 61.83 |  |
| 6 | G | N7 | AA:399 | 4.35 | 1.00 | 56.69 |  |
| 7 | G | N7 | AA:397 | 4.40 | 1.00 | 45.28 |  |
| 8 | C | N4 | AA:376 | 4.78 | 1.00 | 55.16 |  |
| 9 | G | N1 | AA:398 | 4.78 | 1.00 | 55.96 |  |
| 10 | G | N1 | AA:400 | 4.81 | 1.00 | 64.60 |  |
| 11 | G | O6 | AA:400 | 4.88 | 1.00 | 66.30 |  |
  |

  
 Coded contact list:   
 Exclusion zone contact(s): /N7/O6/  
Consider K in place of MG (2 valid contact(s) for K vs. 0 for MG)  

### MG:AA:3225

|  | Res. type | Atom | Residue | Distance | Occupancy | B-factor | Closest solute to 1^st^ shell water |
| --- | --- | --- | --- | --- | --- | --- | --- |
| **Ion** | | | |  | 1.00 | 18.74 || 1 | A | N7 | AA:1981 | 2.57 | 1.00 | 44.48 |
| 2 | A | O2' | AA:1784 | 2.87 | 1.00 | 48.42 |  |
| 3 | A | OP1 | AA:1981 | 2.96 | 1.00 | 40.05 |  |
| 4 | A | O3' | AA:1784 | 3.24 | 1.00 | 45.81 |  |
| 5 | A | OP2 | AA:1785 | 3.47 | 1.00 | 44.30 |  |
| 6 | A | N6 | AA:1981 | 4.09 | 1.00 | 41.07 |  |
| 7 | A | O5' | AA:1981 | 4.40 | 1.00 | 42.58 |  |
| 8 | A | N9 | AA:1981 | 4.52 | 1.00 | 44.07 |  |
| 9 | A | O5' | AA:1785 | 4.94 | 1.00 | 45.50 |  |
  |

  
 Coded contact list:   
 Exclusion zone contact(s): /N7/O2'/O3'/OP1/  
Consider K in place of MG (3 valid contact(s) for K vs. 0 for MG)  

### MG:AA:3226

|  | Res. type | Atom | Residue | Distance | Occupancy | B-factor | Closest solute to 1^st^ shell water |
| --- | --- | --- | --- | --- | --- | --- | --- |
| **Ion** | | | |  | 1.00 | 68.99 || 1 | G | OP2 | AA:2382 | 1.85 | 1.00 | 74.15 |
| 2 | G | N2 | AA:2345 | 2.17 | 1.00 | 59.36 |  |
| 3 | G | O5' | AA:2382 | 2.24 | 1.00 | 75.11 |  |
| 4 | C | O3' | AA:2381 | 3.40 | 1.00 | 72.51 |  |
| 5 | G | N7 | AA:2382 | 3.49 | 1.00 | 79.59 |  |
| 6 | G | N3 | AA:2345 | 3.51 | 1.00 | 58.83 |  |
| 7 | G | O4' | AA:2382 | 3.69 | 1.00 | 76.60 |  |
| 8 | G | OP1 | AA:2382 | 3.80 | 1.00 | 75.10 |  |
| 9 | G | N9 | AA:2382 | 3.95 | 1.00 | 77.20 |  |
| 10 | G | N1 | AA:2345 | 4.17 | 1.00 | 58.72 |  |
| 11 | C | O5' | AA:2381 | 4.37 | 1.00 | 70.24 |  |
| 12 | G | O2' | AA:2345 | 4.74 | 1.00 | 68.53 |  |
| 13 | C | N1 | AA:2381 | 4.98 | 1.00 | 65.63 |  |
  |

  
**1^st^ shell ligand angles:**   
1-2: 119; 1-3: 74; 2-3: 121;   
  
 Contact(s): /N2/O5'/OP2/  
 Coded contact list: O_ph_O_r_Anion  

### MG:AA:3227

|  | Res. type | Atom | Residue | Distance | Occupancy | B-factor | Closest solute to 1^st^ shell water |
| --- | --- | --- | --- | --- | --- | --- | --- |
| **Ion** | | | |  | 1.00 | 30.99 || 1 | C | OP2 | AA:749 | 2.58 | 1.00 | 44.89 |
| 2 | A | OP2 | AA:750 | 3.26 | 1.00 | 42.72 |  |
| 3 | G | OP2 | AA:748 | 3.31 | 1.00 | 43.49 |  |
| 4 | G | O5' | AA:748 | 3.37 | 1.00 | 44.37 |  |
| 5 | A | OP1 | AA:750 | 3.66 | 1.00 | 40.56 |  |
| 6 | C | O5' | AA:749 | 4.22 | 1.00 | 43.49 |  |
| 7 | ALA | N | AS:89 | 4.33 | 1.00 | 46.24 |  |
| 8 | C | O3' | AA:749 | 4.35 | 1.00 | 41.64 |  |
| 9 | G | O3' | AA:748 | 4.44 | 1.00 | 44.32 |  |
| 10 | G | OP1 | AA:748 | 4.60 | 1.00 | 43.35 |  |
  |

  
 Coded contact list:   
 Exclusion zone contact(s): /O5'/OP2/OP2/OP2/  
Consider K in place of MG (3 valid contact(s) for K vs. 0 for MG)  

### MG:AA:3228

|  | Res. type | Atom | Residue | Distance | Occupancy | B-factor | Closest solute to 1^st^ shell water |
| --- | --- | --- | --- | --- | --- | --- | --- |
| **Ion** | | | |  | 1.00 | 53.51 || 1 | A | OP2 | AA:1665 | 2.54 | 1.00 | 52.02 |
| 2 | A | OP1 | AA:1665 | 3.14 | 1.00 | 53.05 |  |
| 3 | A | OP1 | AA:1664 | 3.25 | 1.00 | 45.27 |  |
| 4 | A | O3' | AA:1664 | 4.11 | 1.00 | 50.77 |  |
| 5 | A | O5' | AA:1665 | 4.62 | 1.00 | 51.66 |  |
| 6 | A | O5' | AA:1664 | 4.62 | 1.00 | 46.68 |  |
  |

  
 Coded contact list:   
 Exclusion zone contact(s): /OP1/OP1/OP2/  
Consider K in place of MG (2 valid contact(s) for K vs. 0 for MG)  

### MG:AA:3229

|  | Res. type | Atom | Residue | Distance | Occupancy | B-factor | Closest solute to 1^st^ shell water |
| --- | --- | --- | --- | --- | --- | --- | --- |
| **Ion** | | | |  | 1.00 | 56.38 || 1 | A | O2' | AA:1269 | 3.12 | 1.00 | 42.70 |
| 2 | C | OP2 | AA:749 | 3.17 | 1.00 | 44.89 |  |
| 3 | G | OP1 | AA:748 | 3.61 | 1.00 | 43.35 |  |
| 4 | G | O6 | AA:1325 | 3.69 | 1.00 | 60.79 |  |
| 5 | G | OP2 | AA:748 | 4.01 | 1.00 | 43.49 |  |
| 6 | G | O5' | AA:748 | 4.04 | 1.00 | 44.37 |  |
| 7 | ARG | NE | AS:88 | 4.37 | 1.00 | 44.18 |  |
| 8 | A | O3' | AA:1269 | 4.65 | 1.00 | 42.15 |  |
| 9 | ARG | NH2 | AS:88 | 4.69 | 1.00 | 45.89 |  |
  |

  
 Coded contact list:   
 Exclusion zone contact(s): /O2'/OP2/  
Consider K in place of MG (2 valid contact(s) for K vs. 0 for MG)  

### MG:AA:3230

|  | Res. type | Atom | Residue | Distance | Occupancy | B-factor | Closest solute to 1^st^ shell water |
| --- | --- | --- | --- | --- | --- | --- | --- |
| **Ion** | | | |  | 1.00 | 51.96 || 1 | C | OP1 | AA:749 | 2.39 | 1.00 | 44.38 |
| 2 | C | OP2 | AA:749 | 2.45 | 1.00 | 44.89 |  |
| 3 | G | O3' | AA:748 | 2.61 | 1.00 | 44.32 |  |
| 4 | C | O5' | AA:749 | 4.05 | 1.00 | 43.49 |  |
| 5 | G | O5' | AA:748 | 4.28 | 1.00 | 44.37 |  |
| 6 | C | N4 | AA:2612 | 4.72 | 1.00 | 62.97 |  |
| 7 | A | O3' | AA:1269 | 4.77 | 1.00 | 42.15 |  |
| 8 | C | OP1 | AA:1270 | 4.81 | 1.00 | 44.18 |  |
| 9 | G | OP1 | AA:748 | 4.91 | 1.00 | 43.35 |  |
  |

  
 Contact(s): /OP1/  
 Coded contact list: O_ph_  
 Exclusion zone contact(s): /O3'/OP2/  

### MG:AA:3231

|  | Res. type | Atom | Residue | Distance | Occupancy | B-factor | Closest solute to 1^st^ shell water |
| --- | --- | --- | --- | --- | --- | --- | --- |
| **Ion** | | | |  | 1.00 | 43.70 || 1 | G | O6 | AA:1271 | 2.80 | 1.00 | 61.23 |
| 2 | G | O4' | AA:1325 | 2.97 | 1.00 | 50.09 |  |
| 3 | G | O5' | AA:1325 | 2.97 | 1.00 | 48.13 |  |
| 4 | G | OP2 | AA:1325 | 3.51 | 1.00 | 50.31 |  |
| 5 | G | N9 | AA:1325 | 4.20 | 1.00 | 53.62 |  |
| 6 | C | O2' | AA:1615 | 4.59 | 1.00 | 50.64 |  |
| 7 | G | O3' | AA:1324 | 4.71 | 1.00 | 50.17 |  |
| 8 | G | N7 | AA:1271 | 4.79 | 1.00 | 57.45 |  |
| 9 | G | N7 | AA:1325 | 4.94 | 1.00 | 57.10 |  |
| 10 | C | O2 | AA:1615 | 4.94 | 1.00 | 52.14 |  |
| 11 | G | OP1 | AA:1324 | 4.98 | 1.00 | 49.13 |  |
  |

  
 Coded contact list:   
 Exclusion zone contact(s): /O4'/O5'/O6/  
Consider K in place of MG (3 valid contact(s) for K vs. 0 for MG)  

### MG:AA:3232

|  | Res. type | Atom | Residue | Distance | Occupancy | B-factor | Closest solute to 1^st^ shell water |
| --- | --- | --- | --- | --- | --- | --- | --- |
| **Ion** | | | |  | 1.00 | 25.64 || 1 | U | OP2 | AA:1326 | 2.72 | 1.00 | 41.19 |
| 2 | G | O4' | AA:1324 | 2.94 | 1.00 | 50.92 |  |
| 3 | G | N9 | AA:1324 | 3.00 | 1.00 | 54.01 |  |
| 4 | G | O3' | AA:1324 | 3.47 | 1.00 | 50.17 |  |
| 5 | G | O5' | AA:1325 | 3.68 | 1.00 | 48.13 |  |
| 6 | G | O2' | AA:1324 | 3.71 | 1.00 | 55.78 |  |
| 7 | G | O6 | AA:1328 | 4.02 | 1.00 | 58.10 |  |
| 8 | G | N7 | AA:1324 | 4.12 | 1.00 | 54.16 |  |
| 9 | G | O5' | AA:1324 | 4.17 | 1.00 | 49.38 |  |
| 10 | G | OP1 | AA:1325 | 4.25 | 1.00 | 50.00 |  |
| 11 | G | O3' | AA:1325 | 4.48 | 1.00 | 44.62 |  |
| 12 | G | OP2 | AA:1324 | 4.48 | 1.00 | 49.28 |  |
| 13 | U | OP1 | AA:1326 | 4.54 | 1.00 | 39.88 |  |
| 14 | G | N3 | AA:1324 | 4.76 | 1.00 | 53.50 |  |
  |

  
 Coded contact list:   
 Exclusion zone contact(s): /O4'/OP2/  
Consider K in place of MG (3 valid contact(s) for K vs. 0 for MG)  

### MG:AA:3233

|  | Res. type | Atom | Residue | Distance | Occupancy | B-factor | Closest solute to 1^st^ shell water |
| --- | --- | --- | --- | --- | --- | --- | --- |
| **Ion** | | | |  | 1.00 | 55.28 || 1 | A | OP2 | AA:1616 | 3.17 | 1.00 | 38.97 |
| 2 | C | OP1 | AA:1615 | 4.57 | 1.00 | 44.90 |  |
| 3 | C | OP2 | AA:1611 | 4.95 | 1.00 | 46.07 |  |
  |

  
 Coded contact list:   
 Exclusion zone contact(s): /OP2/  
Consider K in place of MG (1 valid contact(s) for K vs. 0 for MG)  

### MG:AA:3234

|  | Res. type | Atom | Residue | Distance | Occupancy | B-factor | Closest solute to 1^st^ shell water |
| --- | --- | --- | --- | --- | --- | --- | --- |
| **Ion** | | | |  | 1.00 | 54.02 || 1 | C | OP1 | AA:1611 | 2.58 | 1.00 | 44.83 |
| 2 | C | OP2 | AA:1611 | 4.48 | 1.00 | 46.07 |  |
| 3 | C | O5' | AA:1611 | 4.66 | 1.00 | 44.26 |  |
| 4 | G | O2' | AA:1310 | 4.69 | 1.00 | 36.03 |  |
| 5 | C | OP2 | AA:1612 | 4.83 | 1.00 | 37.50 |  |
  |

  
 Coded contact list:   
 Exclusion zone contact(s): /OP1/  
Consider NA in place of MG (1 valid contact(s) for NA vs. 0 for MG)  

### MG:AA:3235

|  | Res. type | Atom | Residue | Distance | Occupancy | B-factor | Closest solute to 1^st^ shell water |
| --- | --- | --- | --- | --- | --- | --- | --- |
| **Ion** | | | |  | 1.00 | 48.56 || 1 | G | O6 | AA:1613 | 3.42 | 1.00 | 46.81 |
| 2 | G | N7 | AA:1613 | 3.59 | 1.00 | 44.70 |  |
| 3 | C | N4 | AA:1612 | 4.12 | 1.00 | 41.96 |  |
| 4 | C | OP2 | AA:1617 | 4.19 | 1.00 | 44.27 |  |
| 5 | A | OP1 | AA:1610 | 4.31 | 1.00 | 48.69 |  |
| 6 | C | OP2 | AA:1611 | 4.66 | 1.00 | 46.07 |  |
| 7 | C | OP2 | AA:1615 | 4.85 | 1.00 | 44.01 |  |
  |

  
 Coded contact list:   
1^st^ MG coordination shell missing (see Notes)  

### MG:AA:3236

|  | Res. type | Atom | Residue | Distance | Occupancy | B-factor | Closest solute to 1^st^ shell water |
| --- | --- | --- | --- | --- | --- | --- | --- |
| **Ion** | | | |  | 1.00 | 20.38 || 1 | A | O5' | AA:1395 | 2.59 | 1.00 | 53.98 |
| 2 | U | O2' | AA:1602 | 2.73 | 1.00 | 53.30 |  |
| 3 | U | O4 | AA:1397 | 2.77 | 1.00 | 61.01 |  |
| 4 | U | O3' | AA:1394 | 2.89 | 1.00 | 53.46 |  |
| 5 | A | OP1 | AA:1395 | 2.95 | 1.00 | 52.83 |  |
| 6 | U | O2' | AA:1394 | 2.98 | 1.00 | 53.89 |  |
| 7 | U | O3' | AA:1602 | 3.02 | 1.00 | 48.36 |  |
| 8 | A | OP1 | AA:1603 | 3.61 | 1.00 | 45.20 |  |
| 9 | U | N3 | AA:1397 | 4.40 | 1.00 | 61.00 |  |
| 10 | A | O5' | AA:1603 | 4.43 | 1.00 | 45.11 |  |
| 11 | A | OP2 | AA:1395 | 4.48 | 1.00 | 52.14 |  |
| 12 | A | O4' | AA:1395 | 4.81 | 1.00 | 57.21 |  |
| 13 | C | OP2 | AA:1604 | 4.82 | 1.00 | 34.59 |  |
  |

  
 Coded contact list:   
 Exclusion zone contact(s): /O2'/O2'/O3'/O3'/O4/O5'/OP1/  
Consider K in place of MG (6 valid contact(s) for K vs. 0 for MG)  

### MG:AA:3237

|  | Res. type | Atom | Residue | Distance | Occupancy | B-factor | Closest solute to 1^st^ shell water |
| --- | --- | --- | --- | --- | --- | --- | --- |
| **Ion** | | | |  | 1.00 | 36.17 || 1 | C | O2 | AA:419 | 2.73 | 1.00 | 62.31 |
| 2 | C | O2' | AA:419 | 3.11 | 1.00 | 64.17 |  |
| 3 | G | OP2 | AA:224 | 3.12 | 1.00 | 63.50 |  |
| 4 | A | N7 | AA:225 | 3.32 | 1.00 | 76.31 |  |
| 5 | G | N7 | AA:224 | 3.65 | 1.00 | 71.72 |  |
| 6 | C | O4' | AA:420 | 3.73 | 1.00 | 65.59 |  |
| 7 | G | O6 | AA:226 | 3.82 | 1.00 | 86.34 |  |
| 8 | G | N2 | AA:408 | 4.10 | 1.00 | 63.17 |  |
| 9 | C | N1 | AA:419 | 4.19 | 1.00 | 63.75 |  |
| 10 | G | O5' | AA:224 | 4.51 | 1.00 | 64.05 |  |
| 11 | C | O2 | AA:409 | 4.60 | 1.00 | 64.43 |  |
| 12 | C | O4' | AA:419 | 4.67 | 1.00 | 64.81 |  |
| 13 | A | N6 | AA:225 | 4.81 | 1.00 | 82.54 |  |
| 14 | G | O6 | AA:224 | 4.92 | 1.00 | 78.06 |  |
| 15 | G | N9 | AA:224 | 4.92 | 1.00 | 69.71 |  |
| 16 | C | N3 | AA:419 | 4.98 | 1.00 | 62.88 |  |
  |

  
 Coded contact list:   
 Exclusion zone contact(s): /N7/O2/O2'/OP2/  
Consider K in place of MG (4 valid contact(s) for K vs. 0 for MG)  

### MG:AA:3238

|  | Res. type | Atom | Residue | Distance | Occupancy | B-factor | Closest solute to 1^st^ shell water |
| --- | --- | --- | --- | --- | --- | --- | --- |
| **Ion** | | | |  | 1.00 | 63.30 || 1 | G | N7 | AA:2864 | 3.47 | 1.00 | 76.00 |
| 2 | C | N4 | AA:2863 | 3.83 | 1.00 | 78.70 |  |
| 3 | C | OP2 | AA:2863 | 4.02 | 1.00 | 74.38 |  |
| 4 | G | O6 | AA:2864 | 4.04 | 1.00 | 74.35 |  |
| 5 | U | O4 | AA:2865 | 4.12 | 1.00 | 75.75 |  |
| 6 | C | N3 | AA:2863 | 4.13 | 1.00 | 79.28 |  |
| 7 | C | N1 | AA:2863 | 4.17 | 1.00 | 78.88 |  |
| 8 | G | OP2 | AA:2864 | 4.35 | 1.00 | 78.93 |  |
  |

  
 Coded contact list:   
1^st^ MG coordination shell missing (see Notes)  

### MG:AA:3239

|  | Res. type | Atom | Residue | Distance | Occupancy | B-factor | Closest solute to 1^st^ shell water |
| --- | --- | --- | --- | --- | --- | --- | --- |
| **Ion** | | | |  | 1.00 | 51.41 || 1 | G | O6 | AA:226 | 2.60 | 1.00 | 86.34 |
| 2 | C | O2' | AA:409 | 2.88 | 1.00 | 56.90 |  |
| 3 | G | N7 | AA:226 | 3.21 | 1.00 | 84.07 |  |
| 4 | G | O4' | AA:410 | 3.35 | 1.00 | 58.52 |  |
| 5 | C | O2 | AA:409 | 3.69 | 1.00 | 64.43 |  |
| 6 | A | OP2 | AA:225 | 4.08 | 1.00 | 66.69 |  |
| 7 | A | N7 | AA:225 | 4.15 | 1.00 | 76.31 |  |
| 8 | G | OP2 | AA:224 | 4.52 | 1.00 | 63.50 |  |
| 9 | G | N1 | AA:226 | 4.63 | 1.00 | 85.31 |  |
| 10 | C | O2 | AA:419 | 4.66 | 1.00 | 62.31 |  |
| 11 | G | N9 | AA:410 | 4.71 | 1.00 | 60.75 |  |
| 12 | G | N2 | AA:418 | 4.77 | 1.00 | 64.21 |  |
  |

  
 Coded contact list:   
 Exclusion zone contact(s): /N7/O2'/O4'/O6/  
Consider K in place of MG (4 valid contact(s) for K vs. 0 for MG)  

### MG:AA:3240

|  | Res. type | Atom | Residue | Distance | Occupancy | B-factor | Closest solute to 1^st^ shell water |
| --- | --- | --- | --- | --- | --- | --- | --- |
| **Ion** | | | |  | 1.00 | 34.58 || 1 | G | OP2 | AA:987 | 1.80 | 1.00 | 68.36 |
| 2 | C | OP2 | AA:986 | 3.56 | 1.00 | 66.61 |  |
| 3 | C | O5' | AA:986 | 3.64 | 1.00 | 68.09 |  |
| 4 | C | OP1 | AA:986 | 3.90 | 1.00 | 66.63 |  |
| 5 | ILE | O | AX:13 | 3.93 | 1.00 | 65.19 |  |
| 6 | G | O5' | AA:987 | 3.97 | 1.00 | 68.56 |  |
| 7 | C | O3' | AA:986 | 4.17 | 1.00 | 68.76 |  |
| 8 | G | OP1 | AA:987 | 4.22 | 1.00 | 69.20 |  |
| 9 | A | N6 | AA:988 | 4.95 | 1.00 | 71.72 |  |
  |

  
 Contact(s): /OP2/  
 Coded contact list: O_ph_  

### MG:AA:3241

|  | Res. type | Atom | Residue | Distance | Occupancy | B-factor | Closest solute to 1^st^ shell water |
| --- | --- | --- | --- | --- | --- | --- | --- |
| **Ion** | | | |  | 1.00 | 59.43 || 1 | G | OP1 | AA:493 | 3.06 | 1.00 | 53.19 |
| 2 | G | OP1 | AA:508 | 3.49 | 1.00 | 57.25 |  |
| 3 | G | N7 | AA:508 | 3.89 | 1.00 | 69.46 |  |
| 4 | A | OP2 | AA:507 | 4.34 | 1.00 | 50.25 |  |
| 5 | G | O5' | AA:493 | 4.61 | 1.00 | 51.92 |  |
  |

  
 Coded contact list:   
 Exclusion zone contact(s): /OP1/  
Consider K in place of MG (1 valid contact(s) for K vs. 0 for MG)  

### MG:AA:3242

|  | Res. type | Atom | Residue | Distance | Occupancy | B-factor | Closest solute to 1^st^ shell water |
| --- | --- | --- | --- | --- | --- | --- | --- |
| **Ion** | | | |  | 1.00 | 56.12 || 1 | G | OP2 | AA:493 | 3.66 | 1.00 | 52.40 |
| 2 | G | N7 | AA:493 | 4.29 | 1.00 | 48.06 |  |
| 3 | G | OP2 | AA:494 | 4.75 | 1.00 | 52.16 |  |
| 4 | G | N7 | AA:494 | 4.79 | 1.00 | 46.62 |  |
  |

  
 Coded contact list:   
1^st^ MG coordination shell missing (see Notes)  

### MG:AA:3243

|  | Res. type | Atom | Residue | Distance | Occupancy | B-factor | Closest solute to 1^st^ shell water |
| --- | --- | --- | --- | --- | --- | --- | --- |
| **Ion** | | | |  | 1.00 | 99.25 || 1 | G | N7 | AA:1753 | 2.82 | 1.00 | 64.61 |
| 2 | C | N4 | AA:1752 | 3.34 | 1.00 | 69.07 |  |
| 3 | LYS | NZ | AR:113 | 3.58 | 1.00 | 86.01 |  |
| 4 | C | N3 | AA:1752 | 3.59 | 1.00 | 68.83 |  |
| 5 | ARG | NH1 | AR:112 | 3.80 | 1.00 | 100.10 |  |
| 6 | G | O6 | AA:1753 | 3.83 | 1.00 | 66.61 |  |
| 7 | C | N1 | AA:1752 | 4.00 | 1.00 | 66.87 |  |
| 8 | G | OP2 | AA:1753 | 4.86 | 1.00 | 62.04 |  |
| 9 | G | N9 | AA:1753 | 4.97 | 1.00 | 61.66 |  |
| 10 | C | O2 | AA:1752 | 4.98 | 1.00 | 68.17 |  |
  |

  
 Coded contact list:   
 Exclusion zone contact(s): /N4/N7/  
Consider K in place of MG (2 valid contact(s) for K vs. 0 for MG)  

### MG:AA:3244

|  | Res. type | Atom | Residue | Distance | Occupancy | B-factor | Closest solute to 1^st^ shell water |
| --- | --- | --- | --- | --- | --- | --- | --- |
| **Ion** | | | |  | 1.00 | 55.42 || 1 | G | N7 | AA:493 | 3.48 | 1.00 | 48.06 |
| 2 | G | O6 | AA:493 | 3.56 | 1.00 | 48.26 |  |
| 3 | G | N7 | AA:494 | 4.23 | 1.00 | 46.62 |  |
| 4 | G | O6 | AA:494 | 4.28 | 1.00 | 48.44 |  |
| 5 | A | OP2 | AA:492 | 4.38 | 1.00 | 56.90 |  |
| 6 | G | N2 | AA:488 | 4.97 | 1.00 | 51.70 |  |
  |

  
 Coded contact list:   
1^st^ MG coordination shell missing (see Notes)  

### MG:AA:3245

|  | Res. type | Atom | Residue | Distance | Occupancy | B-factor | Closest solute to 1^st^ shell water |
| --- | --- | --- | --- | --- | --- | --- | --- |
| **Ion** | | | |  | 1.00 | 69.33 || 1 | G | OP1 | AA:1839 | 3.75 | 1.00 | 61.41 |
| 2 | C | O4' | AA:1838 | 3.77 | 1.00 | 59.29 |  |
| 3 | C | O3' | AA:1838 | 3.83 | 1.00 | 60.91 |  |
| 4 | U | OP2 | AA:1898 | 3.84 | 1.00 | 51.22 |  |
| 5 | C | N1 | AA:1838 | 4.22 | 1.00 | 61.43 |  |
| 6 | G | OP2 | AA:1897 | 4.23 | 1.00 | 57.24 |  |
| 7 | G | OP2 | AA:1839 | 4.37 | 1.00 | 61.68 |  |
| 8 | U | O4 | AA:1898 | 4.50 | 1.00 | 42.21 |  |
| 9 | G | O5' | AA:1897 | 4.57 | 1.00 | 55.73 |  |
| 10 | C | O5' | AA:1838 | 4.61 | 1.00 | 53.62 |  |
| 11 | C | O2' | AA:1838 | 4.85 | 1.00 | 64.48 |  |
  |

  
 Coded contact list:   
1^st^ MG coordination shell missing (see Notes)  

### MG:AA:3246

|  | Res. type | Atom | Residue | Distance | Occupancy | B-factor | Closest solute to 1^st^ shell water |
| --- | --- | --- | --- | --- | --- | --- | --- |
| **Ion** | | | |  | 1.00 | 45.91 || 1 | G | N7 | AA:495 | 3.18 | 1.00 | 54.19 |
| 2 | G | O6 | AA:495 | 3.24 | 1.00 | 54.60 |  |
| 3 | G | O6 | AA:496 | 3.75 | 1.00 | 61.04 |  |
| 4 | G | N7 | AA:496 | 3.88 | 1.00 | 60.70 |  |
| 5 | A | N6 | AA:497 | 4.36 | 1.00 | 65.47 |  |
| 6 | G | N7 | AA:494 | 4.80 | 1.00 | 46.62 |  |
| 7 | G | O6 | AA:494 | 4.82 | 1.00 | 48.44 |  |
  |

  
 Coded contact list:   
 Exclusion zone contact(s): /N7/O6/  
Consider K in place of MG (2 valid contact(s) for K vs. 0 for MG)  

### MG:AA:3247

|  | Res. type | Atom | Residue | Distance | Occupancy | B-factor | Closest solute to 1^st^ shell water |
| --- | --- | --- | --- | --- | --- | --- | --- |
| **Ion** | | | |  | 1.00 | 47.48 || 1 | C | N4 | AA:486 | 2.82 | 1.00 | 46.69 |
| 2 | G | O6 | AA:488 | 3.65 | 1.00 | 54.95 |  |
| 3 | C | N4 | AA:487 | 4.12 | 1.00 | 49.89 |  |
| 4 | G | O6 | AA:494 | 4.49 | 1.00 | 48.44 |  |
| 5 | C | N4 | AA:485 | 4.54 | 1.00 | 54.99 |  |
| 6 | C | N3 | AA:486 | 4.85 | 1.00 | 49.07 |  |
| 7 | G | N1 | AA:488 | 4.98 | 1.00 | 53.72 |  |
  |

  
 Coded contact list:   
 Exclusion zone contact(s): /N4/  
Consider K in place of MG (1 valid contact(s) for K vs. 0 for MG)  

### MG:AA:3248

|  | Res. type | Atom | Residue | Distance | Occupancy | B-factor | Closest solute to 1^st^ shell water |
| --- | --- | --- | --- | --- | --- | --- | --- |
| **Ion** | | | |  | 1.00 | 78.91 || 1 | G | O2' | AA:723 | 2.33 | 1.00 | 59.18 |
| 2 | G | N3 | AA:723 | 2.56 | 1.00 | 59.01 |  |
| 3 | U | O4' | AA:724 | 3.27 | 1.00 | 60.57 |  |
| 4 | G | N2 | AA:723 | 3.53 | 1.00 | 60.22 |  |
| 5 | G | N9 | AA:723 | 3.74 | 1.00 | 59.32 |  |
| 6 | U | O2 | AA:709 | 4.08 | 1.00 | 70.15 |  |
| 7 | U | N1 | AA:724 | 4.32 | 1.00 | 59.38 |  |
| 8 | G | O4' | AA:723 | 4.42 | 1.00 | 60.73 |  |
| 9 | G | N1 | AA:723 | 4.79 | 1.00 | 59.63 |  |
| 10 | G | O3' | AA:723 | 4.85 | 1.00 | 61.75 |  |
  |

  
 Contact(s): /O2'/  
 Coded contact list: O_r_  
 Exclusion zone contact(s): /N3_G_/O4'/  

### MG:AA:3249

|  | Res. type | Atom | Residue | Distance | Occupancy | B-factor | Closest solute to 1^st^ shell water |
| --- | --- | --- | --- | --- | --- | --- | --- |
| **Ion** | | | |  | 1.00 | 41.03 || 1 | G | O6 | AA:2685 | 1.81 | 1.00 | 60.24 |
| 2 | G | N7 | AA:2685 | 3.65 | 1.00 | 61.40 |  |
| 3 | U | O4 | AA:2684 | 3.68 | 1.00 | 61.03 |  |
| 4 | C | N4 | AA:2724 | 3.70 | 1.00 | 59.59 |  |
| 5 | G | O6 | AA:2686 | 3.91 | 1.00 | 60.80 |  |
| 6 | G | N1 | AA:2685 | 3.92 | 1.00 | 58.99 |  |
| 7 | G | N7 | AA:2686 | 4.36 | 1.00 | 59.24 |  |
  |

  
 Contact(s): /O6/  
 Coded contact list: O_b_  

### MG:AA:3250

|  | Res. type | Atom | Residue | Distance | Occupancy | B-factor | Closest solute to 1^st^ shell water |
| --- | --- | --- | --- | --- | --- | --- | --- |
| **Ion** | | | |  | 1.00 | 24.67 || 1 | G | O6 | AA:2722 | 2.81 | 1.00 | 60.44 |
| 2 | U | OP2 | AA:2720 | 2.97 | 1.00 | 59.81 |  |
| 3 | A | OP2 | AA:2721 | 2.98 | 1.00 | 58.86 |  |
| 4 | A | N7 | AA:2721 | 3.39 | 1.00 | 63.59 |  |
| 5 | U | O5' | AA:2720 | 3.41 | 1.00 | 59.89 |  |
| 6 | U | O4 | AA:2687 | 3.58 | 1.00 | 61.93 |  |
| 7 | U | OP1 | AA:2720 | 3.66 | 1.00 | 59.35 |  |
| 8 | G | N7 | AA:2722 | 4.23 | 1.00 | 62.11 |  |
| 9 | U | N3 | AA:2687 | 4.35 | 1.00 | 60.43 |  |
| 10 | A | O5' | AA:2721 | 4.68 | 1.00 | 58.78 |  |
| 11 | U | O3' | AA:2720 | 4.81 | 1.00 | 57.31 |  |
| 12 | U | N1 | AA:2720 | 4.89 | 1.00 | 59.83 |  |
| 13 | G | N1 | AA:2722 | 4.89 | 1.00 | 60.79 |  |
| 14 | U | O4 | AA:2688 | 4.93 | 1.00 | 65.72 |  |
| 15 | G | O6 | AA:2686 | 4.93 | 1.00 | 60.80 |  |
  |

  
 Coded contact list:   
 Exclusion zone contact(s): /N7/O6/OP2/OP2/  
Consider K in place of MG (4 valid contact(s) for K vs. 0 for MG)  

### MG:AA:3251

|  | Res. type | Atom | Residue | Distance | Occupancy | B-factor | Closest solute to 1^st^ shell water |
| --- | --- | --- | --- | --- | --- | --- | --- |
| **Ion** | | | |  | 1.00 | 24.39 || 1 | A | OP1 | AA:2721 | 1.80 | 1.00 | 58.63 |
| 2 | U | OP2 | AA:2682 | 2.29 | 1.00 | 67.79 |  |
| 3 | U | OP1 | AA:2682 | 2.75 | 1.00 | 68.20 |  |
| 4 | A | O5' | AA:2721 | 3.67 | 1.00 | 58.78 |  |
| 5 | U | O5' | AA:2682 | 3.71 | 1.00 | 65.76 |  |
| 6 | A | OP2 | AA:2721 | 3.89 | 1.00 | 58.86 |  |
| 7 | C | OP2 | AA:2683 | 4.25 | 1.00 | 59.46 |  |
| 8 | U | O3' | AA:2720 | 4.29 | 1.00 | 57.31 |  |
| 9 | C | O3' | AA:2681 | 4.34 | 1.00 | 67.60 |  |
| 10 | G | OP2 | AA:2722 | 4.36 | 1.00 | 57.22 |  |
| 11 | C | O2' | AA:2681 | 4.50 | 1.00 | 69.70 |  |
  |

  
**1^st^ shell ligand angles:**   
1-2: 158;   
  
 Contact(s): /OP1/OP2/  
 Coded contact list: 2O_ph_  
 Exclusion zone contact(s): /OP1/  

### MG:AA:3252

|  | Res. type | Atom | Residue | Distance | Occupancy | B-factor | Closest solute to 1^st^ shell water |
| --- | --- | --- | --- | --- | --- | --- | --- |
| **Ion** | | | |  | 1.00 | 38.29 || 1 | A | OP2 | AA:2705 | 2.42 | 1.00 | 71.23 |
| 2 | G | O6 | AA:2706 | 3.33 | 1.00 | 54.69 |  |
| 3 | G | N7 | AA:2706 | 3.42 | 1.00 | 55.27 |  |
| 4 | A | O5' | AA:2705 | 4.02 | 1.00 | 68.96 |  |
| 5 | A | N7 | AA:2705 | 4.20 | 1.00 | 67.97 |  |
| 6 | C | OP2 | AA:2704 | 4.31 | 1.00 | 70.15 |  |
| 7 | G | OP2 | AA:2706 | 4.60 | 1.00 | 63.03 |  |
| 8 | C | OP1 | AA:2704 | 4.63 | 1.00 | 71.00 |  |
| 9 | A | OP1 | AA:2705 | 4.73 | 1.00 | 69.20 |  |
| 10 | C | O3' | AA:2704 | 4.75 | 1.00 | 70.21 |  |
| 11 | C | O5' | AA:2704 | 4.76 | 1.00 | 70.74 |  |
  |

  
 Coded contact list:   
 Exclusion zone contact(s): /O6/OP2/  
Consider NA in place of MG (1 valid contact(s) for NA vs. 0 for MG)  

### MG:AA:3253

|  | Res. type | Atom | Residue | Distance | Occupancy | B-factor | Closest solute to 1^st^ shell water |
| --- | --- | --- | --- | --- | --- | --- | --- |
| **Ion** | | | |  | 1.00 | 50.59 || 1 | G | O6 | AA:2290 | 2.60 | 1.00 | 73.35 |
| 2 | U | O4 | AA:2291 | 2.81 | 1.00 | 75.25 |  |
| 3 | G | O6 | AA:2341 | 2.92 | 1.00 | 78.84 |  |
| 4 | G | O6 | AA:2340 | 3.34 | 1.00 | 79.36 |  |
| 5 | C | N4 | AA:2342 | 3.64 | 1.00 | 74.69 |  |
| 6 | U | N3 | AA:2291 | 3.87 | 1.00 | 76.00 |  |
| 7 | G | N1 | AA:2290 | 4.38 | 1.00 | 72.51 |  |
| 8 | C | N4 | AA:2292 | 4.39 | 1.00 | 79.54 |  |
| 9 | G | N7 | AA:2290 | 4.48 | 1.00 | 69.81 |  |
| 10 | G | N7 | AA:2341 | 4.60 | 1.00 | 79.83 |  |
| 11 | G | O6 | AA:2289 | 4.80 | 1.00 | 72.97 |  |
| 12 | G | N7 | AA:2340 | 4.99 | 1.00 | 82.12 |  |
  |

  
 Coded contact list:   
 Exclusion zone contact(s): /O4/O6/O6/O6/  
Consider K in place of MG (4 valid contact(s) for K vs. 0 for MG)  

### MG:AA:3254

|  | Res. type | Atom | Residue | Distance | Occupancy | B-factor | Closest solute to 1^st^ shell water |
| --- | --- | --- | --- | --- | --- | --- | --- |
| **Ion** | | | |  | 1.00 | 76.35 || 1 | U | O4 | AA:1911 | 3.03 | 1.00 | 63.74 |
| 2 | A | OP2 | AA:1918 | 3.57 | 1.00 | 67.59 |  |
| 3 | G | O6 | AA:1910 | 4.24 | 1.00 | 64.03 |  |
| 4 | A | O5' | AA:1918 | 4.28 | 1.00 | 68.47 |  |
| 5 | G | N7 | AA:1910 | 4.74 | 1.00 | 61.36 |  |
| 6 | A | OP1 | AA:1918 | 4.88 | 1.00 | 69.05 |  |
  |

  
 Coded contact list:   
 Exclusion zone contact(s): /O4/  
Consider K in place of MG (1 valid contact(s) for K vs. 0 for MG)  

### MG:AA:3255

|  | Res. type | Atom | Residue | Distance | Occupancy | B-factor | Closest solute to 1^st^ shell water |
| --- | --- | --- | --- | --- | --- | --- | --- |
| **Ion** | | | |  | 1.00 | 46.06 || 1 | G | OP1 | AA:1235 | 2.58 | 1.00 | 69.01 |
| 2 | A | OP2 | AA:505 | 2.89 | 1.00 | 70.81 |  |
| 3 | U | O3' | AA:1234 | 3.26 | 1.00 | 69.63 |  |
| 4 | U | O2' | AA:1234 | 4.23 | 1.00 | 75.15 |  |
| 5 | U | O2' | AA:504 | 4.25 | 1.00 | 72.53 |  |
| 6 | U | O3' | AA:504 | 4.30 | 1.00 | 71.71 |  |
| 7 | G | O5' | AA:1235 | 4.39 | 1.00 | 66.34 |  |
| 8 | G | OP2 | AA:512 | 4.52 | 1.00 | 56.20 |  |
| 9 | U | OP1 | AA:511 | 4.71 | 1.00 | 50.71 |  |
| 10 | G | OP2 | AA:1235 | 4.74 | 1.00 | 68.63 |  |
| 11 | A | OP1 | AA:505 | 4.87 | 1.00 | 70.72 |  |
  |

  
 Coded contact list:   
 Exclusion zone contact(s): /O3'/OP1/OP2/  
Consider K in place of MG (2 valid contact(s) for K vs. 0 for MG)  

### MG:AA:3256

|  | Res. type | Atom | Residue | Distance | Occupancy | B-factor | Closest solute to 1^st^ shell water |
| --- | --- | --- | --- | --- | --- | --- | --- |
| **Ion** | | | |  | 1.00 | 62.93 || 1 | U | OP1 | AA:200 | 3.89 | 1.00 | 47.56 |
| 2 | C | OP2 | AA:201 | 4.44 | 1.00 | 45.00 |  |
| 3 | U | OP2 | AA:200 | 4.47 | 1.00 | 49.01 |  |
| 4 | U | O4 | AA:202 | 4.57 | 1.00 | 44.18 |  |
| 5 | A | OP1 | AA:191 | 4.69 | 1.00 | 35.15 |  |
| 6 | A | OP2 | AA:191 | 4.71 | 1.00 | 34.60 |  |
| 7 | C | N4 | AA:203 | 4.94 | 1.00 | 44.65 |  |
  |

  
 Coded contact list:   
1^st^ MG coordination shell missing (see Notes)  

### MG:AA:3257

|  | Res. type | Atom | Residue | Distance | Occupancy | B-factor | Closest solute to 1^st^ shell water |
| --- | --- | --- | --- | --- | --- | --- | --- |
| **Ion** | | | |  | 1.00 | 36.22 || 1 | U | OP1 | AA:1313 | 2.06 | 1.00 | 46.12 |
| 2 | U | OP2 | AA:1313 | 3.48 | 1.00 | 43.84 |  |
| 3 | G | OP1 | AA:1334 | 3.55 | 1.00 | 53.78 |  |
| 4 | G | OP2 | AA:1334 | 3.62 | 1.00 | 53.21 |  |
| 5 | C | OP1 | AA:1333 | 3.91 | 1.00 | 55.41 |  |
| 6 | U | O3' | AA:1312 | 4.11 | 1.00 | 45.58 |  |
| 7 | U | O5' | AA:1313 | 4.37 | 1.00 | 42.95 |  |
  |

  
 Contact(s): /OP1/  
 Coded contact list: O_ph_  

### MG:AA:3258

|  | Res. type | Atom | Residue | Distance | Occupancy | B-factor | Closest solute to 1^st^ shell water |
| --- | --- | --- | --- | --- | --- | --- | --- |
| **Ion** | | | |  | 1.00 | 81.01 || 1 | G | O6 | AA:498 | 4.76 | 1.00 | 56.81 |
| 2 | G | OP2 | AA:496 | 4.85 | 1.00 | 57.01 |  |
  |

  
 Coded contact list:   
1^st^ MG coordination shell missing (see Notes)  

### MG:AA:3259

|  | Res. type | Atom | Residue | Distance | Occupancy | B-factor | Closest solute to 1^st^ shell water |
| --- | --- | --- | --- | --- | --- | --- | --- |
| **Ion** | | | |  | 1.00 | 57.79 || 1 | G | N7 | AA:500 | 4.18 | 1.00 | 65.08 |
| 2 | G | O6 | AA:500 | 4.32 | 1.00 | 68.17 |  |
| 3 | A | OP1 | AA:502 | 4.33 | 1.00 | 56.27 |  |
| 4 | A | OP2 | AA:502 | 4.91 | 1.00 | 55.70 |  |
  |

  
 Coded contact list:   
1^st^ MG coordination shell missing (see Notes)  

### MG:AA:3260

|  | Res. type | Atom | Residue | Distance | Occupancy | B-factor | Closest solute to 1^st^ shell water |
| --- | --- | --- | --- | --- | --- | --- | --- |
| **Ion** | | | |  | 1.00 | 44.00 || 1 | C | OP2 | AA:365 | 1.85 | 1.00 | 75.42 |
| 2 | C | OP2 | AA:364 | 2.65 | 1.00 | 90.99 |  |
| 3 | A | O3' | AA:363F | 3.16 | 1.00 | 92.91 |  |
| 4 | C | OP1 | AA:404 | 3.68 | 1.00 | 85.63 |  |
| 5 | U | O3' | AA:403 | 3.70 | 1.00 | 84.36 |  |
| 6 | C | OP2 | AA:404 | 3.77 | 1.00 | 85.49 |  |
| 7 | C | OP1 | AA:365 | 4.00 | 1.00 | 76.63 |  |
| 8 | C | O5' | AA:365 | 4.14 | 1.00 | 75.71 |  |
| 9 | A | O2' | AA:363F | 4.17 | 1.00 | 97.65 |  |
| 10 | C | O3' | AA:364 | 4.27 | 1.00 | 78.99 |  |
| 11 | C | O5' | AA:364 | 4.32 | 1.00 | 89.19 |  |
| 12 | C | OP1 | AA:364 | 4.77 | 1.00 | 90.92 |  |
  |

  
 Contact(s): /OP2/  
 Coded contact list: O_ph_  
 Exclusion zone contact(s): /O3'/OP2/  
Consider K in place of MG (2 valid contact(s) for K vs. 1 for MG)  

### MG:AA:3261

|  | Res. type | Atom | Residue | Distance | Occupancy | B-factor | Closest solute to 1^st^ shell water |
| --- | --- | --- | --- | --- | --- | --- | --- |
| **Ion** | | | |  | 1.00 | 40.12 || 1 | C | OP2 | AA:574 | 2.54 | 1.00 | 45.56 |
| 2 | U | O4 | AA:576 | 3.50 | 1.00 | 39.55 |  |
| 3 | G | O6 | AA:577 | 3.51 | 1.00 | 45.15 |  |
| 4 | G | OP1 | AA:573 | 3.73 | 1.00 | 49.95 |  |
| 5 | C | N4 | AA:564 | 4.15 | 1.00 | 43.50 |  |
| 6 | G | O3' | AA:573 | 4.24 | 1.00 | 45.19 |  |
| 7 | G | O6 | AA:563 | 4.37 | 1.00 | 44.30 |  |
| 8 | C | OP1 | AA:574 | 4.61 | 1.00 | 46.60 |  |
| 9 | G | O5' | AA:573 | 4.93 | 1.00 | 48.45 |  |
| 10 | A | N6 | AA:578 | 4.94 | 1.00 | 34.92 |  |
| 11 | U | OP2 | AA:576 | 4.95 | 1.00 | 46.55 |  |
  |

  
 Coded contact list:   
 Exclusion zone contact(s): /OP2/  
Consider NA in place of MG (1 valid contact(s) for NA vs. 0 for MG)  

### MG:AA:3262

|  | Res. type | Atom | Residue | Distance | Occupancy | B-factor | Closest solute to 1^st^ shell water |
| --- | --- | --- | --- | --- | --- | --- | --- |
| **Ion** | | | |  | 1.00 | 76.77 || 1 | A | OP2 | AA:1495 | 2.25 | 1.00 | 83.96 |
| 2 | A | N1 | AA:1496 | 3.23 | 1.00 | 81.65 |  |
| 3 | A | O5' | AA:1495 | 3.86 | 1.00 | 82.91 |  |
| 4 | A | N7 | AA:1495 | 4.23 | 1.00 | 74.50 |  |
| 5 | A | OP2 | AA:1496 | 4.33 | 1.00 | 77.97 |  |
| 6 | A | OP1 | AA:1495 | 4.33 | 1.00 | 83.73 |  |
| 7 | A | OP2 | AA:1494 | 4.42 | 1.00 | 88.59 |  |
| 8 | A | N3 | AA:1496 | 4.59 | 1.00 | 80.41 |  |
| 9 | A | O3' | AA:1494 | 4.72 | 1.00 | 84.32 |  |
| 10 | A | O5' | AA:1494 | 4.94 | 1.00 | 88.74 |  |
| 11 | G | N7 | AA:1492 | 4.99 | 1.00 | 72.51 |  |
  |

  
 Contact(s): /OP2/  
 Coded contact list: O_ph_  
 Exclusion zone contact(s): /N1_A_/  

### MG:AA:3263

|  | Res. type | Atom | Residue | Distance | Occupancy | B-factor | Closest solute to 1^st^ shell water |
| --- | --- | --- | --- | --- | --- | --- | --- |
| **Ion** | | | |  | 1.00 | 57.44 || 1 | C | N3 | AA:1774 | 2.41 | 1.00 | 38.87 |
| 2 | A | N6 | AA:1773 | 2.80 | 1.00 | 36.49 |  |
| 3 | C | O2 | AA:1774 | 2.93 | 1.00 | 41.21 |  |
| 4 | G | N1 | AA:729 | 3.22 | 1.00 | 42.47 |  |
| 5 | SER | N | AD:12 | 3.70 | 1.00 | 45.87 |  |
| 6 | C | N4 | AA:1774 | 3.70 | 1.00 | 36.55 |  |
| 7 | G | N2 | AA:729 | 3.70 | 1.00 | 43.20 |  |
| 8 | SER | OG | AD:12 | 3.91 | 1.00 | 47.25 |  |
| 9 | C | N1 | AA:1774 | 3.94 | 1.00 | 39.89 |  |
| 10 | GLY | O | AD:207 | 4.06 | 1.00 | 35.67 |  |
| 11 | G | O6 | AA:729 | 4.13 | 1.00 | 42.89 |  |
| 12 | A | N1 | AA:1773 | 4.54 | 1.00 | 34.06 |  |
| 13 | PRO | O | AD:11 | 4.61 | 1.00 | 46.27 |  |
| 14 | A | N3 | AA:1791 | 4.89 | 1.00 | 31.80 |  |
  |

  
 Coded contact list:   
 Exclusion zone contact(s): /N1_G_/N3_C_/N6/O2/  
Consider K in place of MG (3 valid contact(s) for K vs. 0 for MG)  

### MG:AA:3264

|  | Res. type | Atom | Residue | Distance | Occupancy | B-factor | Closest solute to 1^st^ shell water |
| --- | --- | --- | --- | --- | --- | --- | --- |
| **Ion** | | | |  | 1.00 | 43.64 || 1 | C | OP2 | AA:1417 | 2.51 | 1.00 | 88.22 |
| 2 | G | N7 | AA:1418 | 3.20 | 1.00 | 74.67 |  |
| 3 | U | OP2 | AA:1578 | 3.40 | 1.00 | 77.48 |  |
| 4 | G | O6 | AA:1418 | 3.71 | 1.00 | 74.23 |  |
| 5 | C | O5' | AA:1417 | 3.98 | 1.00 | 85.55 |  |
| 6 | G | OP2 | AA:1418 | 4.35 | 1.00 | 80.32 |  |
| 7 | C | OP1 | AA:1417 | 4.45 | 1.00 | 88.52 |  |
| 8 | C | OP2 | AA:1577 | 4.66 | 1.00 | 75.40 |  |
  |

  
 Coded contact list:   
 Exclusion zone contact(s): /N7/OP2/  
Consider NA in place of MG (1 valid contact(s) for NA vs. 0 for MG)  

### MG:AA:3265

|  | Res. type | Atom | Residue | Distance | Occupancy | B-factor | Closest solute to 1^st^ shell water |
| --- | --- | --- | --- | --- | --- | --- | --- |
| **Ion** | | | |  | 1.00 | 71.03 || 1 | G | OP2 | AA:1906 | 3.38 | 1.00 | 57.32 |
| 2 | G | O3' | AA:1930 | 3.97 | 1.00 | 54.50 |  |
| 3 | G | O5' | AA:1930 | 3.99 | 1.00 | 57.86 |  |
| 4 | G | O5' | AA:1906 | 4.17 | 1.00 | 60.16 |  |
| 5 | G | O4' | AA:1930 | 4.56 | 1.00 | 55.06 |  |
| 6 | G | O3' | AA:1929 | 4.58 | 1.00 | 56.98 |  |
| 7 | G | OP2 | AA:1930 | 4.71 | 1.00 | 57.97 |  |
| 8 | G | OP1 | AA:1906 | 4.79 | 1.00 | 57.72 |  |
| 9 | G | O2' | AA:1929 | 4.97 | 1.00 | 59.54 |  |
  |

  
 Coded contact list:   
 Exclusion zone contact(s): /OP2/  
Consider K in place of MG (1 valid contact(s) for K vs. 0 for MG)  

### MG:AA:3266

|  | Res. type | Atom | Residue | Distance | Occupancy | B-factor | Closest solute to 1^st^ shell water |
| --- | --- | --- | --- | --- | --- | --- | --- |
| **Ion** | | | |  | 1.00 | 73.01 || 1 | A | OP1 | AA:1579 | 3.62 | 1.00 | 85.67 |
| 2 | G | N7 | AA:1416 | 3.88 | 1.00 | 91.86 |  |
| 3 | G | O6 | AA:1416 | 3.93 | 1.00 | 95.77 |  |
  |

  
 Coded contact list:   
1^st^ MG coordination shell missing (see Notes)  

### MG:AA:3267

|  | Res. type | Atom | Residue | Distance | Occupancy | B-factor | Closest solute to 1^st^ shell water |
| --- | --- | --- | --- | --- | --- | --- | --- |
| **Ion** | | | |  | 1.00 | 49.19 || 1 | G | O3' | AA:232 | 2.19 | 1.00 | 79.74 |
| 2 | A | O5' | AA:233 | 2.25 | 1.00 | 75.36 |  |
| 3 | A | OP1 | AA:233 | 3.35 | 1.00 | 78.52 |  |
| 4 | C | OP2 | AA:234 | 3.57 | 1.00 | 61.41 |  |
| 5 | G | O2' | AA:232 | 3.80 | 1.00 | 82.61 |  |
| 6 | A | OP2 | AA:233 | 4.01 | 1.00 | 78.79 |  |
| 7 | A | O3' | AA:233 | 4.24 | 1.00 | 62.82 |  |
| 8 | A | O4' | AA:233 | 4.64 | 1.00 | 66.55 |  |
| 9 | G | O4' | AA:232 | 4.79 | 1.00 | 81.47 |  |
  |

  
**1^st^ shell ligand angles:**   
1-2: 69;   
  
 Contact(s): /O3'/O5'/  
 Coded contact list: 2O_r_  
 Exclusion zone contact(s): /OP1/  

### MG:AA:3268

|  | Res. type | Atom | Residue | Distance | Occupancy | B-factor | Closest solute to 1^st^ shell water |
| --- | --- | --- | --- | --- | --- | --- | --- |
| **Ion** | | | |  | 1.00 | 60.84 || 1 | G | N7 | AA:260 | 3.42 | 1.00 | 65.32 |
| 2 | G | O6 | AA:260 | 3.93 | 1.00 | 64.03 |  |
| 3 | G | N7 | AA:261 | 4.01 | 1.00 | 65.74 |  |
| 4 | G | N7 | AA:259 | 4.68 | 1.00 | 62.18 |  |
| 5 | G | O6 | AA:259 | 4.83 | 1.00 | 60.90 |  |
| 6 | G | O6 | AA:261 | 4.83 | 1.00 | 66.19 |  |
  |

  
 Coded contact list:   
1^st^ MG coordination shell missing (see Notes)  

### MG:AA:3269

|  | Res. type | Atom | Residue | Distance | Occupancy | B-factor | Closest solute to 1^st^ shell water |
| --- | --- | --- | --- | --- | --- | --- | --- |
| **Ion** | | | |  | 1.00 | 61.35 || 1 | G | O6 | AA:259 | 4.49 | 1.00 | 60.90 |
| 2 | G | OP2 | AA:258 | 4.90 | 1.00 | 59.65 |  |
| 3 | G | O6 | AA:260 | 4.94 | 1.00 | 64.03 |  |
  |

  
 Coded contact list:   
1^st^ MG coordination shell missing (see Notes)  

### MG:AA:3270

|  | Res. type | Atom | Residue | Distance | Occupancy | B-factor | Closest solute to 1^st^ shell water |
| --- | --- | --- | --- | --- | --- | --- | --- |
| **Ion** | | | |  | 1.00 | 72.79 || 1 | G | O6 | AA:1747 | 3.47 | 1.00 | 102.39 |
| 2 | C | N4 | AA:1711 | 4.34 | 1.00 | 99.27 |  |
| 3 | G | O6 | AA:1748 | 4.41 | 1.00 | 90.65 |  |
| 4 | C | N4 | AA:1710 | 4.67 | 1.00 | 90.06 |  |
| 5 | G | N7 | AA:1747 | 4.82 | 1.00 | 103.46 |  |
| 6 | G | O6 | AA:1746 | 4.85 | 1.00 | 107.96 |  |
  |

  
 Coded contact list:   
1^st^ MG coordination shell missing (see Notes)  

### MG:AA:3271

|  | Res. type | Atom | Residue | Distance | Occupancy | B-factor | Closest solute to 1^st^ shell water |
| --- | --- | --- | --- | --- | --- | --- | --- |
| **Ion** | | | |  | 1.00 | 83.01 || 1 | G | OP2 | AA:258 | 2.58 | 1.00 | 59.65 |
| 2 | A | OP1 | AA:257 | 3.71 | 1.00 | 54.32 |  |
| 3 | G | OP1 | AA:258 | 3.96 | 1.00 | 59.74 |  |
| 4 | A | O3' | AA:257 | 4.12 | 1.00 | 58.74 |  |
| 5 | A | O5' | AA:257 | 4.76 | 1.00 | 55.29 |  |
| 6 | G | O5' | AA:258 | 4.98 | 1.00 | 57.51 |  |
  |

  
 Coded contact list:   
 Exclusion zone contact(s): /OP2/  
Consider NA in place of MG (1 valid contact(s) for NA vs. 0 for MG)  

### MG:AA:3272

|  | Res. type | Atom | Residue | Distance | Occupancy | B-factor | Closest solute to 1^st^ shell water |
| --- | --- | --- | --- | --- | --- | --- | --- |
| **Ion** | | | |  | 1.00 | 54.84 || 1 | G | O2' | AA:1948 | 2.60 | 1.00 | 43.44 |
| 2 | G | O3' | AA:1948 | 2.89 | 1.00 | 46.87 |  |
| 3 | G | O2' | BA:1482 | 3.38 | 1.00 | 53.32 |  |
| 4 | A | O4' | BA:1483 | 3.47 | 1.00 | 47.12 |  |
| 5 | G | O3' | BA:1482 | 3.75 | 1.00 | 51.19 |  |
| 6 | G | O4' | AA:1948 | 3.98 | 1.00 | 44.64 |  |
| 7 | G | OP1 | AA:1949 | 4.00 | 1.00 | 49.30 |  |
| 8 | A | O5' | BA:1483 | 4.65 | 1.00 | 47.98 |  |
| 9 | G | O5' | AA:1949 | 4.74 | 1.00 | 48.43 |  |
| 10 | A | N6 | BA:1418 | 4.87 | 1.00 | 48.47 |  |
| 11 | A | N1 | BA:1418 | 4.89 | 1.00 | 49.39 |  |
| 12 | G | O5' | AA:1948 | 4.96 | 1.00 | 45.90 |  |
  |

  
 Coded contact list:   
 Exclusion zone contact(s): /O2'/O2'/O3'/  
Consider K in place of MG (2 valid contact(s) for K vs. 0 for MG)  

### MG:AA:3273

|  | Res. type | Atom | Residue | Distance | Occupancy | B-factor | Closest solute to 1^st^ shell water |
| --- | --- | --- | --- | --- | --- | --- | --- |
| **Ion** | | | |  | 1.00 | 30.93 || 1 | U | O4 | AA:594 | 2.61 | 1.00 | 58.33 |
| 2 | G | O6 | AA:662 | 2.70 | 1.00 | 55.86 |  |
| 3 | G | O6 | AA:663 | 2.81 | 1.00 | 56.79 |  |
| 4 | G | O6 | AA:593 | 3.18 | 1.00 | 53.47 |  |
| 5 | C | N4 | AA:595 | 3.65 | 1.00 | 57.95 |  |
| 6 | U | N3 | AA:594 | 3.80 | 1.00 | 57.88 |  |
| 7 | C | N4 | AA:664 | 4.09 | 1.00 | 51.47 |  |
| 8 | G | N1 | AA:662 | 4.48 | 1.00 | 54.18 |  |
| 9 | G | N7 | AA:662 | 4.57 | 1.00 | 52.26 |  |
| 10 | G | N7 | AA:663 | 4.65 | 1.00 | 55.81 |  |
| 11 | C | N4 | AA:661 | 4.92 | 1.00 | 55.41 |  |
| 12 | G | N1 | AA:593 | 4.94 | 1.00 | 52.52 |  |
  |

  
 Coded contact list:   
 Exclusion zone contact(s): /O4/O6/O6/O6/  
Consider K in place of MG (4 valid contact(s) for K vs. 0 for MG)  

### MG:AA:3274

|  | Res. type | Atom | Residue | Distance | Occupancy | B-factor | Closest solute to 1^st^ shell water |
| --- | --- | --- | --- | --- | --- | --- | --- |
| **Ion** | | | |  | 1.00 | 70.06 || 1 | G | O6 | AA:592 | 3.62 | 1.00 | 50.96 |
| 2 | G | O6 | AA:593 | 3.70 | 1.00 | 53.47 |  |
| 3 | G | N7 | AA:592 | 4.57 | 1.00 | 54.47 |  |
| 4 | G | N7 | AA:593 | 4.62 | 1.00 | 55.53 |  |
| 5 | C | N4 | AA:664 | 4.85 | 1.00 | 51.47 |  |
  |

  
 Coded contact list:   
1^st^ MG coordination shell missing (see Notes)  

### MG:AA:3275

|  | Res. type | Atom | Residue | Distance | Occupancy | B-factor | Closest solute to 1^st^ shell water |
| --- | --- | --- | --- | --- | --- | --- | --- |
| **Ion** | | | |  | 1.00 | 58.19 || 1 | A | OP2 | AA:590 | 3.91 | 1.00 | 49.96 |
| 2 | A | OP1 | AA:590 | 4.09 | 1.00 | 51.21 |  |
| 3 | C | OP1 | AA:661 | 4.10 | 1.00 | 57.65 |  |
| 4 | C | OP2 | AA:661 | 4.32 | 1.00 | 57.67 |  |
  |

  
 Coded contact list:   
1^st^ MG coordination shell missing (see Notes)  

### MG:AA:3276

|  | Res. type | Atom | Residue | Distance | Occupancy | B-factor | Closest solute to 1^st^ shell water |
| --- | --- | --- | --- | --- | --- | --- | --- |
| **Ion** | | | |  | 1.00 | 46.88 || 1 | U | OP2 | AA:1273 | 2.76 | 1.00 | 54.76 |
| 2 | A | OP1 | AA:1272 | 2.77 | 1.00 | 49.07 |  |
| 3 | G | OP2 | AA:1271 | 3.05 | 1.00 | 42.52 |  |
| 4 | G | OP1 | AA:1271 | 4.10 | 1.00 | 42.89 |  |
| 5 | A | O5' | AA:1272 | 4.12 | 1.00 | 50.28 |  |
| 6 | G | O3' | AA:1271 | 4.17 | 1.00 | 49.58 |  |
| 7 | G | O5' | AA:1271 | 4.49 | 1.00 | 45.41 |  |
| 8 | U | O5' | AA:1273 | 4.52 | 1.00 | 52.94 |  |
| 9 | A | O3' | AA:1272 | 4.90 | 1.00 | 51.65 |  |
  |

  
 Coded contact list:   
 Exclusion zone contact(s): /OP1/OP2/OP2/  
Consider K in place of MG (3 valid contact(s) for K vs. 0 for MG)  

### MG:AA:3277

|  | Res. type | Atom | Residue | Distance | Occupancy | B-factor | Closest solute to 1^st^ shell water |
| --- | --- | --- | --- | --- | --- | --- | --- |
| **Ion** | | | |  | 1.00 | 42.37 || 1 | G | OP2 | AA:1271 | 1.90 | 1.00 | 42.52 |
| 2 | C | OP1 | AA:1648 | 2.24 | 1.00 | 43.56 |  |
| 3 | A | OP1 | AA:1272 | 2.32 | 1.00 | 49.07 |  |
| 4 | G | O5' | AA:1647 | 2.70 | 1.00 | 46.53 |  |
| 5 | C | O5' | AA:1648 | 3.70 | 1.00 | 42.64 |  |
| 6 | G | OP1 | AA:1647 | 3.82 | 1.00 | 49.60 |  |
| 7 | G | O5' | AA:1271 | 4.00 | 1.00 | 45.41 |  |
| 8 | C | O3' | AA:1270 | 4.08 | 1.00 | 43.39 |  |
| 9 | A | OP2 | AA:1272 | 4.35 | 1.00 | 48.76 |  |
| 10 | G | O3' | AA:1271 | 4.40 | 1.00 | 49.58 |  |
| 11 | G | OP1 | AA:1271 | 4.42 | 1.00 | 42.89 |  |
| 12 | C | OP2 | AA:1648 | 4.49 | 1.00 | 41.95 |  |
| 13 | G | O3' | AA:1647 | 4.61 | 1.00 | 43.20 |  |
| 14 | C | O3' | AA:1646 | 4.78 | 1.00 | 48.42 |  |
| 15 | A | O5' | AA:1272 | 4.79 | 1.00 | 50.28 |  |
| 16 | G | OP2 | AA:1647 | 4.99 | 1.00 | 48.36 |  |
  |

  
**1^st^ shell ligand angles:**   
1-2: 88; 1-3: 97; 2-3: 132;   
  
 Contact(s): /OP1/OP1/OP2/  
 Coded contact list: 3O_ph_  
 Exclusion zone contact(s): /O5'/  

### MG:AA:3278

|  | Res. type | Atom | Residue | Distance | Occupancy | B-factor | Closest solute to 1^st^ shell water |
| --- | --- | --- | --- | --- | --- | --- | --- |
| **Ion** | | | |  | 1.00 | 59.29 || 1 | G | O6 | AA:1651 | 3.74 | 1.00 | 43.09 |
| 2 | ASN | OD1 | A0:11 | 3.87 | 1.00 | 59.10 |  |
| 3 | G | O6 | AA:1650 | 3.88 | 1.00 | 46.78 |  |
| 4 | G | N7 | AA:1651 | 4.30 | 1.00 | 44.50 |  |
| 5 | G | N7 | AA:1650 | 4.31 | 1.00 | 46.92 |  |
| 6 | G | OP2 | AA:2004 | 4.40 | 1.00 | 48.72 |  |
  |

  
 Coded contact list:   
1^st^ MG coordination shell missing (see Notes)  

### MG:AA:3279

|  | Res. type | Atom | Residue | Distance | Occupancy | B-factor | Closest solute to 1^st^ shell water |
| --- | --- | --- | --- | --- | --- | --- | --- |
| **Ion** | | | |  | 1.00 | 45.92 || 1 | G | O6 | AA:1425 | 2.29 | 1.00 | 64.98 |
| 2 | G | O6 | AA:1426 | 3.42 | 1.00 | 61.25 |  |
| 3 | G | N1 | AA:1426 | 3.58 | 1.00 | 58.12 |  |
| 4 | G | N7 | AA:1425 | 4.34 | 1.00 | 62.43 |  |
| 5 | G | N1 | AA:1425 | 4.44 | 1.00 | 63.71 |  |
| 6 | G | N2 | AA:1426 | 4.82 | 1.00 | 57.51 |  |
| 7 | A | OP2 | AA:1571 | 4.98 | 1.00 | 44.33 |  |
  |

  
 Contact(s): /O6/  
 Coded contact list: O_b_  

### MG:AA:3280

|  | Res. type | Atom | Residue | Distance | Occupancy | B-factor | Closest solute to 1^st^ shell water |
| --- | --- | --- | --- | --- | --- | --- | --- |
| **Ion** | | | |  | 1.00 | 56.98 || 1 | G | O6 | AA:1425 | 2.85 | 1.00 | 64.98 |
| 2 | G | O6 | AA:1424 | 3.56 | 1.00 | 65.69 |  |
| 3 | G | N1 | AA:1425 | 4.01 | 1.00 | 63.71 |  |
| 4 | G | N7 | AA:1424 | 4.20 | 1.00 | 67.05 |  |
| 5 | A | OP2 | AA:1572 | 4.44 | 1.00 | 43.49 |  |
| 6 | G | N7 | AA:1423 | 4.54 | 1.00 | 66.21 |  |
| 7 | G | O6 | AA:1423 | 4.56 | 1.00 | 68.40 |  |
  |

  
 Coded contact list:   
 Exclusion zone contact(s): /O6/  
Consider K in place of MG (1 valid contact(s) for K vs. 0 for MG)  

### MG:AA:3281

|  | Res. type | Atom | Residue | Distance | Occupancy | B-factor | Closest solute to 1^st^ shell water |
| --- | --- | --- | --- | --- | --- | --- | --- |
| **Ion** | | | |  | 1.00 | 45.04 || 1 | G | O6 | AA:1421 | 3.09 | 1.00 | 81.72 |
| 2 | U | O4 | AA:1576 | 3.44 | 1.00 | 71.18 |  |
| 3 | G | O6 | AA:1422 | 3.44 | 1.00 | 65.39 |  |
| 4 | G | O6 | AA:1423 | 3.53 | 1.00 | 68.40 |  |
| 5 | C | N4 | AA:1577 | 4.01 | 1.00 | 82.11 |  |
| 6 | C | N4 | AA:1575 | 4.06 | 1.00 | 70.65 |  |
| 7 | G | N1 | AA:1421 | 4.57 | 1.00 | 81.86 |  |
| 8 | G | N7 | AA:1422 | 4.67 | 1.00 | 67.51 |  |
| 9 | U | N3 | AA:1576 | 4.68 | 1.00 | 71.28 |  |
| 10 | G | N7 | AA:1421 | 4.68 | 1.00 | 83.07 |  |
  |

  
 Coded contact list:   
 Exclusion zone contact(s): /O6/  
Consider K in place of MG (1 valid contact(s) for K vs. 0 for MG)  

### MG:AA:3282

|  | Res. type | Atom | Residue | Distance | Occupancy | B-factor | Closest solute to 1^st^ shell water |
| --- | --- | --- | --- | --- | --- | --- | --- |
| **Ion** | | | |  | 1.00 | 64.96 || 1 | G | O6 | AA:1421 | 2.62 | 1.00 | 81.72 |
| 2 | C | N4 | AA:1577 | 2.82 | 1.00 | 82.11 |  |
| 3 | U | O4 | AA:1576 | 3.54 | 1.00 | 71.18 |  |
| 4 | A | OP2 | AA:1419 | 4.06 | 1.00 | 83.38 |  |
| 5 | U | OP2 | AA:1420 | 4.37 | 1.00 | 93.58 |  |
| 6 | U | O4 | AA:1578 | 4.64 | 1.00 | 77.57 |  |
| 7 | G | N1 | AA:1421 | 4.75 | 1.00 | 81.86 |  |
| 8 | C | N3 | AA:1577 | 4.86 | 1.00 | 80.41 |  |
| 9 | G | O6 | AA:1422 | 4.92 | 1.00 | 65.39 |  |
| 10 | A | O4' | AA:1419 | 4.94 | 1.00 | 88.83 |  |
  |

  
 Coded contact list:   
 Exclusion zone contact(s): /N4/O6/  
Consider K in place of MG (2 valid contact(s) for K vs. 0 for MG)  

### MG:AA:3283

|  | Res. type | Atom | Residue | Distance | Occupancy | B-factor | Closest solute to 1^st^ shell water |
| --- | --- | --- | --- | --- | --- | --- | --- |
| **Ion** | | | |  | 1.00 | 74.16 || 1 | U | O4 | AA:1576 | 3.53 | 1.00 | 71.18 |
| 2 | C | N4 | AA:1577 | 4.82 | 1.00 | 82.11 |  |
  |

  
 Coded contact list:   
1^st^ MG coordination shell missing (see Notes)  

### MG:AA:3284

|  | Res. type | Atom | Residue | Distance | Occupancy | B-factor | Closest solute to 1^st^ shell water |
| --- | --- | --- | --- | --- | --- | --- | --- |
| **Ion** | | | |  | 1.00 | 78.15 || 1 | C | N4 | AA:2863 | 3.09 | 1.00 | 78.70 |
| 2 | G | O6 | AA:2862 | 3.38 | 1.00 | 83.94 |  |
| 3 | G | N7 | AA:2862 | 3.62 | 1.00 | 81.62 |  |
| 4 | G | O6 | AA:2854 | 4.63 | 1.00 | 80.72 |  |
| 5 | G | N7 | AA:2861 | 4.74 | 1.00 | 78.41 |  |
| 6 | G | O6 | AA:2861 | 4.78 | 1.00 | 79.94 |  |
  |

  
 Coded contact list:   
 Exclusion zone contact(s): /N4/O6/  
Consider K in place of MG (2 valid contact(s) for K vs. 0 for MG)  

### MG:AA:3285

|  | Res. type | Atom | Residue | Distance | Occupancy | B-factor | Closest solute to 1^st^ shell water |
| --- | --- | --- | --- | --- | --- | --- | --- |
| **Ion** | | | |  | 1.00 | 62.48 || 1 | U | OP2 | AA:1576 | 3.35 | 1.00 | 73.26 |
| 2 | U | OP1 | AA:1576 | 3.68 | 1.00 | 73.85 |  |
| 3 | U | O5' | AA:1576 | 4.77 | 1.00 | 74.03 |  |
| 4 | C | O4' | AA:1589 | 4.89 | 1.00 | 84.68 |  |
| 5 | C | O2' | AA:1588 | 4.98 | 1.00 | 86.58 |  |
  |

  
 Coded contact list:   
 Exclusion zone contact(s): /OP2/  
Consider K in place of MG (1 valid contact(s) for K vs. 0 for MG)  

### MG:AA:3286

|  | Res. type | Atom | Residue | Distance | Occupancy | B-factor | Closest solute to 1^st^ shell water |
| --- | --- | --- | --- | --- | --- | --- | --- |
| **Ion** | | | |  | 1.00 | 67.72 || 1 | C | OP1 | AA:1417 | 3.49 | 1.00 | 88.52 |
| 2 | U | O2' | AA:1415 | 4.22 | 1.00 | 91.36 |  |
| 3 | U | O4' | AA:1415 | 4.45 | 1.00 | 92.82 |  |
| 4 | G | N2 | AA:1414 | 4.81 | 1.00 | 89.60 |  |
  |

  
 Coded contact list:   
1^st^ MG coordination shell missing (see Notes)  

### MG:AA:3287

|  | Res. type | Atom | Residue | Distance | Occupancy | B-factor | Closest solute to 1^st^ shell water |
| --- | --- | --- | --- | --- | --- | --- | --- |
| **Ion** | | | |  | 1.00 | 66.74 || 1 | G | OP2 | AA:1418 | 2.84 | 1.00 | 80.32 |
| 2 | C | OP1 | AA:1417 | 3.55 | 1.00 | 88.52 |  |
| 3 | C | O3' | AA:1417 | 4.08 | 1.00 | 82.01 |  |
| 4 | C | O5' | AA:1417 | 4.26 | 1.00 | 85.55 |  |
| 5 | C | OP2 | AA:1577 | 4.36 | 1.00 | 75.40 |  |
| 6 | C | OP2 | AA:1417 | 4.51 | 1.00 | 88.22 |  |
| 7 | G | OP1 | AA:1418 | 4.57 | 1.00 | 80.06 |  |
| 8 | C | O2' | AA:1588 | 4.80 | 1.00 | 86.58 |  |
  |

  
 Coded contact list:   
 Exclusion zone contact(s): /OP2/  
Consider K in place of MG (1 valid contact(s) for K vs. 0 for MG)  

### MG:AA:3288

|  | Res. type | Atom | Residue | Distance | Occupancy | B-factor | Closest solute to 1^st^ shell water |
| --- | --- | --- | --- | --- | --- | --- | --- |
| **Ion** | | | |  | 1.00 | 51.41 || 1 | G | O6 | AA:2224 | 3.08 | 1.00 | 53.21 |
| 2 | U | OP2 | AA:2197 | 3.41 | 1.00 | 69.44 |  |
| 3 | G | OP2 | AA:2219 | 3.86 | 1.00 | 52.06 |  |
| 4 | G | N7 | AA:2224 | 4.06 | 1.00 | 53.55 |  |
| 5 | G | OP2 | AA:2218 | 4.69 | 1.00 | 60.19 |  |
  |

  
 Coded contact list:   
 Exclusion zone contact(s): /O6/  
Consider K in place of MG (1 valid contact(s) for K vs. 0 for MG)  

### MG:AA:3289

|  | Res. type | Atom | Residue | Distance | Occupancy | B-factor | Closest solute to 1^st^ shell water |
| --- | --- | --- | --- | --- | --- | --- | --- |
| **Ion** | | | |  | 1.00 | 90.31 || 1 | C | O2' | AA:2856 | 4.00 | 1.00 | 82.44 |
| 2 | C | O3' | AA:2856 | 4.99 | 1.00 | 82.59 |  |
  |

  
 Coded contact list:   
1^st^ MG coordination shell missing (see Notes)  

### MG:AA:3290

|  | Res. type | Atom | Residue | Distance | Occupancy | B-factor | Closest solute to 1^st^ shell water |
| --- | --- | --- | --- | --- | --- | --- | --- |
| **Ion** | | | |  | 1.00 | 78.76 || 1 | G | O6 | AA:2217 | 2.76 | 1.00 | 57.29 |
| 2 | G | N7 | AA:2217 | 3.33 | 1.00 | 61.13 |  |
| 3 | G | O6 | AA:2218 | 3.60 | 1.00 | 54.07 |  |
| 4 | G | O6 | AA:2216 | 4.60 | 1.00 | 68.64 |  |
| 5 | G | N7 | AA:2218 | 4.62 | 1.00 | 54.41 |  |
| 6 | C | N4 | AA:2207 | 4.74 | 1.00 | 52.53 |  |
| 7 | G | N1 | AA:2217 | 4.84 | 1.00 | 56.13 |  |
| 8 | G | N7 | AA:2216 | 4.89 | 1.00 | 70.44 |  |
  |

  
 Coded contact list:   
 Exclusion zone contact(s): /N7/O6/  
Consider K in place of MG (2 valid contact(s) for K vs. 0 for MG)  

### MG:AA:3291

|  | Res. type | Atom | Residue | Distance | Occupancy | B-factor | Closest solute to 1^st^ shell water |
| --- | --- | --- | --- | --- | --- | --- | --- |
| **Ion** | | | |  | 1.00 | 34.43 || 1 | A | OP1 | AA:981 | 2.36 | 1.00 | 60.75 |
| 2 | G | N7 | AA:979 | 2.82 | 1.00 | 75.85 |  |
| 3 | G | O6 | AA:979 | 3.24 | 1.00 | 76.58 |  |
| 4 | G | OP2 | AA:978 | 4.00 | 1.00 | 69.60 |  |
| 5 | A | OP2 | AA:981 | 4.14 | 1.00 | 61.18 |  |
| 6 | A | O5' | AA:981 | 4.24 | 1.00 | 59.98 |  |
| 7 | C | OP2 | AA:982 | 4.24 | 1.00 | 60.02 |  |
| 8 | G | N7 | AA:978 | 4.47 | 1.00 | 67.72 |  |
| 9 | G | OP2 | AA:979 | 4.72 | 1.00 | 66.14 |  |
| 10 | A | O3' | AA:980 | 4.84 | 1.00 | 60.41 |  |
  |

  
 Contact(s): /OP1/  
 Coded contact list: O_ph_  
 Exclusion zone contact(s): /N7/O6/  
Consider K in place of MG (2 valid contact(s) for K vs. 1 for MG)  

### MG:AA:3292

|  | Res. type | Atom | Residue | Distance | Occupancy | B-factor | Closest solute to 1^st^ shell water |
| --- | --- | --- | --- | --- | --- | --- | --- |
| **Ion** | | | |  | 1.00 | 79.30 || 1 | G | O5' | AA:312 | 4.52 | 1.00 | 65.29 |
| 2 | A | O2' | AA:331 | 4.62 | 1.00 | 57.24 |  |
| 3 | G | OP2 | AA:333 | 4.66 | 1.00 | 55.30 |  |
| 4 | A | OP1 | AA:332 | 4.72 | 1.00 | 55.26 |  |
| 5 | C | OP2 | AA:313 | 4.80 | 1.00 | 60.00 |  |
  |

  
 Coded contact list:   
1^st^ MG coordination shell missing (see Notes)  

### MG:AA:3293

|  | Res. type | Atom | Residue | Distance | Occupancy | B-factor | Closest solute to 1^st^ shell water |
| --- | --- | --- | --- | --- | --- | --- | --- |
| **Ion** | | | |  | 1.00 | 45.27 || 1 | A | OP1 | AA:332 | 3.23 | 1.00 | 55.26 |
| 2 | A | O2' | AA:331 | 3.33 | 1.00 | 57.24 |  |
| 3 | A | O4' | AA:331 | 4.72 | 1.00 | 57.26 |  |
| 4 | G | O6 | AA:1209 | 4.75 | 1.00 | 54.70 |  |
| 5 | A | N3 | AA:331 | 4.88 | 1.00 | 55.03 |  |
  |

  
 Coded contact list:   
 Exclusion zone contact(s): /O2'/OP1/  
Consider K in place of MG (2 valid contact(s) for K vs. 0 for MG)  

### MG:AA:3294

|  | Res. type | Atom | Residue | Distance | Occupancy | B-factor | Closest solute to 1^st^ shell water |
| --- | --- | --- | --- | --- | --- | --- | --- |
| **Ion** | | | |  | 1.00 | 64.90 || 1 | G | OP2 | AA:1209 | 2.66 | 1.00 | 61.90 |
| 2 | G | N7 | AA:1209 | 3.09 | 1.00 | 55.47 |  |
| 3 | G | O5' | AA:1209 | 3.63 | 1.00 | 63.98 |  |
| 4 | C | OP2 | AA:1208 | 3.88 | 1.00 | 61.30 |  |
| 5 | A | N1 | AA:331 | 4.15 | 1.00 | 54.42 |  |
| 6 | G | N9 | AA:1209 | 4.17 | 1.00 | 57.13 |  |
| 7 | C | O3' | AA:1208 | 4.29 | 1.00 | 61.91 |  |
| 8 | C | N1 | AA:1208 | 4.35 | 1.00 | 58.47 |  |
| 9 | C | O5' | AA:1208 | 4.67 | 1.00 | 61.19 |  |
| 10 | G | O4' | AA:1209 | 4.72 | 1.00 | 60.65 |  |
| 11 | C | O4' | AA:1208 | 4.85 | 1.00 | 60.50 |  |
| 12 | C | N3 | AA:1208 | 4.96 | 1.00 | 56.96 |  |
  |

  
 Coded contact list:   
 Exclusion zone contact(s): /N7/OP2/  
Consider K in place of MG (2 valid contact(s) for K vs. 0 for MG)  

### MG:AA:3295

|  | Res. type | Atom | Residue | Distance | Occupancy | B-factor | Closest solute to 1^st^ shell water |
| --- | --- | --- | --- | --- | --- | --- | --- |
| **Ion** | | | |  | 1.00 | 62.58 || 1 | ARG | NH1 | AR:93 | 2.77 | 1.00 | 86.21 |
| 2 | U | OP1 | AA:2849 | 3.01 | 1.00 | 64.42 |  |
| 3 | U | OP2 | AA:2849 | 3.29 | 1.00 | 63.82 |  |
| 4 | U | O5' | AA:2849 | 3.83 | 1.00 | 64.77 |  |
| 5 | ARG | NE | AR:95 | 4.15 | 1.00 | 62.41 |  |
| 6 | ARG | NH2 | AR:93 | 4.30 | 1.00 | 86.93 |  |
| 7 | ARG | NH1 | AR:95 | 4.36 | 1.00 | 64.13 |  |
| 8 | ARG | NE | AR:93 | 4.44 | 1.00 | 85.08 |  |
| 9 | ARG | NH2 | AR:95 | 4.46 | 1.00 | 63.59 |  |
| 10 | U | O4' | AA:2849 | 4.51 | 1.00 | 68.75 |  |
| 11 | U | N1 | AA:2849 | 4.80 | 1.00 | 71.11 |  |
  |

  
 Coded contact list:   
 Exclusion zone contact(s): /NH1_Arg_/OP1/OP2/  
Consider K in place of MG (3 valid contact(s) for K vs. 0 for MG)  

### MG:AA:3296

|  | Res. type | Atom | Residue | Distance | Occupancy | B-factor | Closest solute to 1^st^ shell water |
| --- | --- | --- | --- | --- | --- | --- | --- |
| **Ion** | | | |  | 1.00 | 55.33 || 1 | U | OP1 | AA:1352 | 2.27 | 1.00 | 46.66 |
| 2 | C | O3' | AA:1351 | 4.06 | 1.00 | 43.28 |  |
| 3 | C | OP1 | AA:1375 | 4.08 | 1.00 | 56.60 |  |
| 4 | U | OP2 | AA:1352 | 4.31 | 1.00 | 44.43 |  |
| 5 | C | OP2 | AA:1375 | 4.44 | 1.00 | 56.77 |  |
| 6 | A | O2' | AA:1571 | 4.62 | 1.00 | 46.59 |  |
| 7 | U | O5' | AA:1352 | 4.76 | 1.00 | 44.42 |  |
  |

  
 Contact(s): /OP1/  
 Coded contact list: O_ph_  

### MG:AA:3297

|  | Res. type | Atom | Residue | Distance | Occupancy | B-factor | Closest solute to 1^st^ shell water |
| --- | --- | --- | --- | --- | --- | --- | --- |
| **Ion** | | | |  | 1.00 | 58.42 || 1 | A | OP2 | AA:1977 | 2.26 | 1.00 | 38.15 |
| 2 | U | O3' | AA:1976 | 4.22 | 1.00 | 37.47 |  |
| 3 | U | N1 | AA:1976 | 4.35 | 1.00 | 35.05 |  |
| 4 | U | O5' | AA:1976 | 4.54 | 1.00 | 37.06 |  |
| 5 | A | OP1 | AA:1977 | 4.60 | 1.00 | 37.65 |  |
| 6 | A | N7 | AA:1977 | 4.61 | 1.00 | 40.33 |  |
| 7 | A | O5' | AA:1977 | 4.65 | 1.00 | 38.36 |  |
| 8 | A | N6 | AA:1978 | 4.84 | 1.00 | 42.04 |  |
| 9 | U | N3 | AA:1976 | 4.96 | 1.00 | 34.19 |  |
  |

  
 Contact(s): /OP2/  
 Coded contact list: O_ph_  

### MG:AA:3298

|  | Res. type | Atom | Residue | Distance | Occupancy | B-factor | Closest solute to 1^st^ shell water |
| --- | --- | --- | --- | --- | --- | --- | --- |
| **Ion** | | | |  | 1.00 | 38.21 || 1 | G | O3' | AA:474 | 2.88 | 1.00 | 50.30 |
| 2 | G | OP1 | AA:474 | 3.00 | 1.00 | 49.29 |  |
| 3 | U | OP2 | AA:475 | 3.61 | 1.00 | 49.04 |  |
| 4 | G | O5' | AA:474 | 4.01 | 1.00 | 48.60 |  |
| 5 | C | N4 | AA:509 | 4.36 | 1.00 | 49.26 |  |
| 6 | U | O5' | AA:475 | 4.61 | 1.00 | 48.19 |  |
| 7 | G | OP2 | AA:474 | 4.66 | 1.00 | 48.27 |  |
  |

  
 Coded contact list:   
 Exclusion zone contact(s): /O3'/OP1/  
Consider K in place of MG (2 valid contact(s) for K vs. 0 for MG)  

### MG:AA:3299

|  | Res. type | Atom | Residue | Distance | Occupancy | B-factor | Closest solute to 1^st^ shell water |
| --- | --- | --- | --- | --- | --- | --- | --- |
| **Ion** | | | |  | 1.00 | 82.58 || 1 | C | O4' | AA:1793 | 2.89 | 1.00 | 31.73 |
| 2 | C | OP2 | AA:1830 | 2.93 | 1.00 | 35.58 |  |
| 3 | G | O2' | AA:1792 | 3.38 | 1.00 | 37.28 |  |
| 4 | C | OP1 | AA:1830 | 4.47 | 1.00 | 37.15 |  |
| 5 | A | N1 | AA:1970 | 4.56 | 1.00 | 31.71 |  |
| 6 | A | N6 | AA:1970 | 4.57 | 1.00 | 32.36 |  |
| 7 | C | N1 | AA:1793 | 4.74 | 1.00 | 31.06 |  |
| 8 | C | O2' | AA:1793 | 4.87 | 1.00 | 31.18 |  |
| 9 | C | O5' | AA:1830 | 4.91 | 1.00 | 36.99 |  |
| 10 | C | O5' | AA:1793 | 4.97 | 1.00 | 34.34 |  |
  |

  
 Coded contact list:   
 Exclusion zone contact(s): /O2'/O4'/OP2/  
Consider K in place of MG (3 valid contact(s) for K vs. 0 for MG)  

### MG:AA:3300

|  | Res. type | Atom | Residue | Distance | Occupancy | B-factor | Closest solute to 1^st^ shell water |
| --- | --- | --- | --- | --- | --- | --- | --- |
| **Ion** | | | |  | 1.00 | 38.92 || 1 | U | OP2 | AA:427 | 3.60 | 1.00 | 59.31 |
| 2 | C | OP2 | AA:426 | 3.74 | 1.00 | 60.64 |  |
| 3 | G | OP2 | AA:219 | 3.84 | 1.00 | 54.58 |  |
| 4 | G | N7 | AA:220 | 4.00 | 1.00 | 66.46 |  |
| 5 | G | O6 | AA:220 | 4.17 | 1.00 | 68.93 |  |
| 6 | C | OP1 | AA:426 | 4.25 | 1.00 | 61.63 |  |
| 7 | C | O5' | AA:426 | 4.54 | 1.00 | 60.45 |  |
  |

  
 Coded contact list:   
1^st^ MG coordination shell missing (see Notes)  

### MG:AA:3301

|  | Res. type | Atom | Residue | Distance | Occupancy | B-factor | Closest solute to 1^st^ shell water |
| --- | --- | --- | --- | --- | --- | --- | --- |
| **Ion** | | | |  | 1.00 | 50.46 || 1 | U | OP1 | AA:1766 | 2.65 | 1.00 | 49.58 |
| 2 | A | OP1 | AA:1698 | 2.95 | 1.00 | 49.91 |  |
| 3 | A | OP2 | AA:1698 | 3.46 | 1.00 | 50.31 |  |
| 4 | U | OP2 | AA:1766 | 4.03 | 1.00 | 47.80 |  |
| 5 | U | O5' | AA:1766 | 4.05 | 1.00 | 48.83 |  |
| 6 | A | O2' | AA:1700 | 4.09 | 1.00 | 50.69 |  |
| 7 | A | N3 | AA:1700 | 4.50 | 1.00 | 45.31 |  |
| 8 | C | OP2 | AA:1767 | 4.53 | 1.00 | 49.70 |  |
| 9 | C | OP1 | AA:1767 | 4.55 | 1.00 | 50.80 |  |
| 10 | G | O3' | AA:1697 | 4.75 | 1.00 | 46.86 |  |
| 11 | A | O5' | AA:1698 | 4.86 | 1.00 | 48.81 |  |
  |

  
 Coded contact list:   
 Exclusion zone contact(s): /OP1/OP1/  
Consider K in place of MG (2 valid contact(s) for K vs. 0 for MG)  

### MG:AA:3302

|  | Res. type | Atom | Residue | Distance | Occupancy | B-factor | Closest solute to 1^st^ shell water |
| --- | --- | --- | --- | --- | --- | --- | --- |
| **Ion** | | | |  | 1.00 | 32.34 || 1 | A | OP1 | AA:783 | 2.65 | 1.00 | 41.00 |
| 2 | A | N3 | AA:1787 | 2.95 | 1.00 | 41.43 |  |
| 3 | U | O2 | AA:1778 | 3.25 | 1.00 | 36.48 |  |
| 4 | A | N6 | AA:1785 | 3.36 | 1.00 | 44.19 |  |
| 5 | C | O4' | AA:1788 | 3.65 | 1.00 | 36.17 |  |
| 6 | A | OP2 | AA:782 | 3.98 | 1.00 | 33.26 |  |
| 7 | C | N1 | AA:1788 | 4.29 | 1.00 | 35.72 |  |
| 8 | A | O5' | AA:783 | 4.32 | 1.00 | 40.16 |  |
| 9 | A | N1 | AA:1787 | 4.34 | 1.00 | 41.42 |  |
| 10 | U | O2 | AA:1777 | 4.35 | 1.00 | 46.42 |  |
| 11 | U | N1 | AA:1778 | 4.36 | 1.00 | 36.21 |  |
| 12 | A | O3' | AA:782 | 4.56 | 1.00 | 39.04 |  |
| 13 | U | N3 | AA:1778 | 4.69 | 1.00 | 39.11 |  |
| 14 | A | O2' | AA:1787 | 4.72 | 1.00 | 40.91 |  |
| 15 | U | O4' | AA:1778 | 4.74 | 1.00 | 38.63 |  |
| 16 | A | O5' | AA:782 | 4.86 | 1.00 | 36.45 |  |
| 17 | U | O2' | AA:1778 | 4.95 | 1.00 | 38.65 |  |
  |

  
 Coded contact list:   
 Exclusion zone contact(s): /N3_A_/N6/O2/OP1/  
Consider K in place of MG (4 valid contact(s) for K vs. 0 for MG)  

### MG:AA:3303

|  | Res. type | Atom | Residue | Distance | Occupancy | B-factor | Closest solute to 1^st^ shell water |
| --- | --- | --- | --- | --- | --- | --- | --- |
| **Ion** | | | |  | 1.00 | 94.20 || 1 | C | OP2 | AA:2856 | 2.74 | 1.00 | 84.06 |
| 2 | C | OP1 | AA:2855 | 3.18 | 1.00 | 81.33 |  |
| 3 | C | O5' | AA:2855 | 3.47 | 1.00 | 81.85 |  |
| 4 | C | OP2 | AA:2855 | 3.55 | 1.00 | 80.82 |  |
| 5 | C | O3' | AA:2855 | 3.99 | 1.00 | 83.33 |  |
| 6 | C | OP1 | AA:2856 | 4.26 | 1.00 | 84.14 |  |
  |

  
 Coded contact list:   
 Exclusion zone contact(s): /OP1/OP2/  
Consider K in place of MG (2 valid contact(s) for K vs. 0 for MG)  

### MG:AA:3304

|  | Res. type | Atom | Residue | Distance | Occupancy | B-factor | Closest solute to 1^st^ shell water |
| --- | --- | --- | --- | --- | --- | --- | --- |
| **Ion** | | | |  | 1.00 | 36.79 || 1 | A | O3' | AA:71 | 2.41 | 1.00 | 51.89 |
| 2 | U | OP1 | AA:72 | 2.85 | 1.00 | 47.29 |  |
| 3 | U | O2' | AA:112 | 3.61 | 1.00 | 49.74 |  |
| 4 | A | O5' | AA:71 | 3.97 | 1.00 | 59.08 |  |
| 5 | G | OP1 | AA:113 | 4.05 | 1.00 | 54.26 |  |
| 6 | A | OP2 | AA:71 | 4.19 | 1.00 | 62.13 |  |
| 7 | U | O3' | AA:112 | 4.23 | 1.00 | 52.77 |  |
| 8 | U | O5' | AA:72 | 4.31 | 1.00 | 47.55 |  |
| 9 | U | OP2 | AA:72 | 4.33 | 1.00 | 47.42 |  |
| 10 | G | O5' | AA:113 | 4.47 | 1.00 | 52.78 |  |
| 11 | A | OP1 | AA:71 | 4.77 | 1.00 | 61.62 |  |
  |

  
 Coded contact list:   
 Exclusion zone contact(s): /O3'/OP1/  
Consider NA in place of MG (1 valid contact(s) for NA vs. 0 for MG)  

### MG:AA:3305

|  | Res. type | Atom | Residue | Distance | Occupancy | B-factor | Closest solute to 1^st^ shell water |
| --- | --- | --- | --- | --- | --- | --- | --- |
| **Ion** | | | |  | 1.00 | 47.04 || 1 | G | OP2 | AA:2000 | 2.82 | 1.00 | 43.72 |
| 2 | G | O5' | AA:2000 | 3.40 | 1.00 | 41.28 |  |
| 3 | A | OP2 | AA:2001 | 3.53 | 1.00 | 44.25 |  |
| 4 | G | OP1 | AA:2000 | 3.82 | 1.00 | 44.35 |  |
| 5 | C | O3' | AA:1999 | 4.95 | 1.00 | 43.24 |  |
  |

  
 Coded contact list:   
 Exclusion zone contact(s): /O5'/OP2/  
Consider K in place of MG (2 valid contact(s) for K vs. 0 for MG)  

### MG:AA:3306

|  | Res. type | Atom | Residue | Distance | Occupancy | B-factor | Closest solute to 1^st^ shell water |
| --- | --- | --- | --- | --- | --- | --- | --- |
| **Ion** | | | |  | 1.00 | 49.39 || 1 | G | O6 | AA:2000 | 4.60 | 1.00 | 41.40 |
| 2 | G | N7 | AA:2000 | 4.87 | 1.00 | 43.19 |  |
  |

  
 Coded contact list:   
1^st^ MG coordination shell missing (see Notes)  

### MG:AA:3307

|  | Res. type | Atom | Residue | Distance | Occupancy | B-factor | Closest solute to 1^st^ shell water |
| --- | --- | --- | --- | --- | --- | --- | --- |
| **Ion** | | | |  | 1.00 | 50.21 || 1 | G | N1 | AA:1332 | 3.49 | 1.00 | 65.61 |
| 2 | G | N2 | AA:1332 | 3.64 | 1.00 | 67.26 |  |
| 3 | C | O3' | AA:1605 | 3.75 | 1.00 | 41.17 |  |
| 4 | A | N7 | AA:1610 | 3.84 | 1.00 | 45.80 |  |
| 5 | C | O2' | AA:1605 | 3.87 | 1.00 | 37.09 |  |
| 6 | G | OP1 | AA:1606 | 4.36 | 1.00 | 43.14 |  |
| 7 | G | O6 | AA:1332 | 4.55 | 1.00 | 64.42 |  |
| 8 | G | N3 | AA:1332 | 4.79 | 1.00 | 62.83 |  |
| 9 | A | OP2 | AA:1610 | 4.97 | 1.00 | 49.32 |  |
  |

  
 Coded contact list:   
1^st^ MG coordination shell missing (see Notes)  

### MG:AA:3308

|  | Res. type | Atom | Residue | Distance | Occupancy | B-factor | Closest solute to 1^st^ shell water |
| --- | --- | --- | --- | --- | --- | --- | --- |
| **Ion** | | | |  | 1.00 | 38.87 || 1 | A | OP2 | AA:1010 | 2.17 | 1.00 | 85.66 |
| 2 | A | OP2 | AA:1009 | 2.41 | 1.00 | 84.39 |  |
| 3 | LYS | NZ | AM:37 | 2.62 | 1.00 | 99.14 |  |
| 4 | C | OP2 | AA:1006 | 3.06 | 1.00 | 85.91 |  |
| 5 | C | OP1 | AA:1006 | 3.30 | 1.00 | 85.75 |  |
| 6 | C | O3' | AA:1008 | 3.58 | 1.00 | 85.48 |  |
| 7 | C | OP2 | AA:1007 | 3.86 | 1.00 | 79.94 |  |
| 8 | A | N7 | AA:1010 | 4.21 | 1.00 | 91.33 |  |
| 9 | A | OP1 | AA:1010 | 4.28 | 1.00 | 85.29 |  |
| 10 | A | O5' | AA:1010 | 4.33 | 1.00 | 87.20 |  |
| 11 | C | O5' | AA:1006 | 4.38 | 1.00 | 85.09 |  |
| 12 | A | O5' | AA:1009 | 4.56 | 1.00 | 84.09 |  |
| 13 | A | O3' | AA:1009 | 4.64 | 1.00 | 86.30 |  |
| 14 | A | OP1 | AA:1009 | 4.65 | 1.00 | 85.24 |  |
| 15 | C | O2' | AA:1008 | 4.89 | 1.00 | 90.99 |  |
  |

  
 Contact(s): /OP2/  
 Coded contact list: O_ph_  
 Exclusion zone contact(s): /NZ_Lys_/OP1/OP2/OP2/  
Consider K in place of MG (3 valid contact(s) for K vs. 1 for MG)  

### MG:AA:3309

|  | Res. type | Atom | Residue | Distance | Occupancy | B-factor | Closest solute to 1^st^ shell water |
| --- | --- | --- | --- | --- | --- | --- | --- |
| **Ion** | | | |  | 1.00 | 54.88 || 1 | G | O6 | AA:1697 | 3.34 | 1.00 | 32.10 |
| 2 | G | N7 | AA:1696 | 3.72 | 1.00 | 37.01 |  |
| 3 | G | O6 | AA:1696 | 4.06 | 1.00 | 37.02 |  |
| 4 | G | N1 | AA:1697 | 4.88 | 1.00 | 32.39 |  |
  |

  
 Coded contact list:   
 Exclusion zone contact(s): /O6/  
Consider K in place of MG (1 valid contact(s) for K vs. 0 for MG)  

### MG:AA:3310

|  | Res. type | Atom | Residue | Distance | Occupancy | B-factor | Closest solute to 1^st^ shell water |
| --- | --- | --- | --- | --- | --- | --- | --- |
| **Ion** | | | |  | 1.00 | 47.65 || 1 | A | OP2 | AA:53 | 2.77 | 1.00 | 45.29 |
| 2 | G | O6 | AA:54 | 3.43 | 1.00 | 37.94 |  |
| 3 | G | N7 | AA:54 | 3.82 | 1.00 | 41.71 |  |
| 4 | A | OP2 | AA:52 | 3.85 | 1.00 | 47.03 |  |
| 5 | A | O5' | AA:52 | 4.02 | 1.00 | 47.01 |  |
| 6 | A | OP1 | AA:52 | 4.43 | 1.00 | 48.58 |  |
| 7 | G | N2 | AA:117 | 4.49 | 1.00 | 37.55 |  |
| 8 | A | OP1 | AA:53 | 4.86 | 1.00 | 48.03 |  |
  |

  
 Coded contact list:   
 Exclusion zone contact(s): /OP2/  
Consider K in place of MG (1 valid contact(s) for K vs. 0 for MG)  

### MG:AA:3311

|  | Res. type | Atom | Residue | Distance | Occupancy | B-factor | Closest solute to 1^st^ shell water |
| --- | --- | --- | --- | --- | --- | --- | --- |
| **Ion** | | | |  | 1.00 | 49.75 || 1 | G | OP2 | AA:770 | 2.92 | 1.00 | 38.76 |
| 2 | G | OP2 | AA:686 | 4.24 | 1.00 | 38.33 |  |
| 3 | G | O5' | AA:770 | 4.44 | 1.00 | 36.19 |  |
| 4 | G | OP2 | AA:771 | 4.59 | 1.00 | 37.40 |  |
| 5 | G | OP1 | AA:770 | 4.80 | 1.00 | 39.03 |  |
| 6 | G | N7 | AA:770 | 4.81 | 1.00 | 38.49 |  |
| 7 | ASN | OD1 | A7:8 | 4.84 | 1.00 | 39.34 |  |
  |

  
 Coded contact list:   
 Exclusion zone contact(s): /OP2/  
Consider K in place of MG (1 valid contact(s) for K vs. 0 for MG)  

### MG:AA:3312

|  | Res. type | Atom | Residue | Distance | Occupancy | B-factor | Closest solute to 1^st^ shell water |
| --- | --- | --- | --- | --- | --- | --- | --- |
| **Ion** | | | |  | 1.00 | 40.35 || 1 | G | OP2 | AA:131 | 3.05 | 1.00 | 51.91 |
| 2 | G | OP1 | AA:142 | 3.38 | 1.00 | 64.81 |  |
| 3 | C | OP1 | AA:130 | 3.92 | 1.00 | 52.04 |  |
| 4 | C | O5' | AA:130 | 3.93 | 1.00 | 50.61 |  |
| 5 | C | OP2 | AA:130 | 4.23 | 1.00 | 53.38 |  |
| 6 | C | O2' | AA:1598 | 4.32 | 1.00 | 51.40 |  |
| 7 | C | O2 | AA:1598 | 4.49 | 1.00 | 52.14 |  |
| 8 | G | O5' | AA:142 | 4.67 | 1.00 | 64.53 |  |
| 9 | C | OP1 | AA:143 | 4.89 | 1.00 | 68.31 |  |
  |

  
 Coded contact list:   
 Exclusion zone contact(s): /OP1/OP2/  
Consider K in place of MG (2 valid contact(s) for K vs. 0 for MG)  

### MG:AA:3313

|  | Res. type | Atom | Residue | Distance | Occupancy | B-factor | Closest solute to 1^st^ shell water |
| --- | --- | --- | --- | --- | --- | --- | --- |
| **Ion** | | | |  | 1.00 | 83.02 || 1 | G | OP1 | AA:131 | 2.84 | 1.00 | 51.97 |
| 2 | G | OP2 | AA:131 | 3.47 | 1.00 | 51.91 |  |
| 3 | G | OP1 | AA:142 | 3.71 | 1.00 | 64.81 |  |
| 4 | G | OP2 | AA:142 | 3.82 | 1.00 | 63.54 |  |
| 5 | C | OP1 | AA:141A | 4.45 | 1.00 | 67.14 |  |
| 6 | A | N9 | AA:1349 | 4.62 | 1.00 | 66.93 |  |
| 7 | A | O2' | AA:1349 | 4.69 | 1.00 | 62.12 |  |
| 8 | C | O3' | AA:130 | 4.70 | 1.00 | 50.87 |  |
| 9 | A | N7 | AA:1349 | 4.77 | 1.00 | 67.56 |  |
| 10 | G | O5' | AA:131 | 4.84 | 1.00 | 51.24 |  |
  |

  
 Coded contact list:   
 Exclusion zone contact(s): /OP1/  
Consider K in place of MG (1 valid contact(s) for K vs. 0 for MG)  

### MG:AA:3314

|  | Res. type | Atom | Residue | Distance | Occupancy | B-factor | Closest solute to 1^st^ shell water |
| --- | --- | --- | --- | --- | --- | --- | --- |
| **Ion** | | | |  | 1.00 | 74.92 || 1 | C | OP1 | AA:1771 | 2.97 | 1.00 | 38.27 |
| 2 | U | O4 | AA:1939 | 3.95 | 1.00 | 48.94 |  |
| 3 | C | OP1 | AA:1974 | 4.05 | 1.00 | 40.98 |  |
| 4 | C | OP2 | AA:1974 | 4.44 | 1.00 | 41.79 |  |
| 5 | C | O5' | AA:1974 | 4.73 | 1.00 | 39.30 |  |
| 6 | G | O3' | AA:1770 | 4.82 | 1.00 | 38.22 |  |
| 7 | G | OP2 | AA:1975 | 4.94 | 1.00 | 34.18 |  |
  |

  
 Coded contact list:   
 Exclusion zone contact(s): /OP1/  
Consider K in place of MG (1 valid contact(s) for K vs. 0 for MG)  

### MG:AA:3315

|  | Res. type | Atom | Residue | Distance | Occupancy | B-factor | Closest solute to 1^st^ shell water |
| --- | --- | --- | --- | --- | --- | --- | --- |
| **Ion** | | | |  | 1.00 | 49.59 || 1 | G | O6 | AA:660 | 2.66 | 1.00 | 56.23 |
| 2 | U | O4 | AA:597 | 3.13 | 1.00 | 56.52 |  |
| 3 | G | O6 | AA:596 | 3.29 | 1.00 | 57.06 |  |
| 4 | C | N4 | AA:659 | 3.31 | 1.00 | 63.27 |  |
| 5 | C | N4 | AA:661 | 3.37 | 1.00 | 55.41 |  |
| 6 | U | N3 | AA:597 | 4.03 | 1.00 | 57.55 |  |
| 7 | G | O6 | AA:598 | 4.09 | 1.00 | 62.39 |  |
| 8 | G | N7 | AA:660 | 4.25 | 1.00 | 56.67 |  |
| 9 | G | N1 | AA:660 | 4.88 | 1.00 | 56.56 |  |
  |

  
 Coded contact list:   
 Exclusion zone contact(s): /N4/N4/O4/O6/O6/  
Consider K in place of MG (5 valid contact(s) for K vs. 0 for MG)  

### MG:AA:3316

|  | Res. type | Atom | Residue | Distance | Occupancy | B-factor | Closest solute to 1^st^ shell water |
| --- | --- | --- | --- | --- | --- | --- | --- |
| **Ion** | | | |  | 1.00 | 47.57 || 1 | G | OP2 | AA:396 | 2.70 | 1.00 | 53.02 |
| 2 | U | O4' | AA:395 | 2.96 | 1.00 | 52.75 |  |
| 3 | U | N1 | AA:395 | 2.98 | 1.00 | 50.33 |  |
| 4 | U | O3' | AA:395 | 3.14 | 1.00 | 51.78 |  |
| 5 | U | O2' | AA:395 | 3.41 | 1.00 | 56.60 |  |
| 6 | U | O5' | AA:395 | 3.48 | 1.00 | 54.67 |  |
| 7 | G | N7 | AA:396 | 3.68 | 1.00 | 47.27 |  |
| 8 | U | O2 | AA:395 | 4.08 | 1.00 | 54.29 |  |
| 9 | G | O5' | AA:396 | 4.45 | 1.00 | 49.99 |  |
| 10 | U | OP2 | AA:395 | 4.47 | 1.00 | 54.18 |  |
| 11 | U | N3 | AA:395 | 4.60 | 1.00 | 50.70 |  |
| 12 | G | OP1 | AA:396 | 4.72 | 1.00 | 52.23 |  |
  |

  
 Coded contact list:   
 Exclusion zone contact(s): /N1/O3'/O4'/OP2/  
Consider K in place of MG (4 valid contact(s) for K vs. 0 for MG)  

### MG:AA:3317

|  | Res. type | Atom | Residue | Distance | Occupancy | B-factor | Closest solute to 1^st^ shell water |
| --- | --- | --- | --- | --- | --- | --- | --- |
| **Ion** | | | |  | 1.00 | 57.87 || 1 | G | O6 | AA:55 | 2.87 | 1.00 | 46.61 |
| 2 | G | N7 | AA:55 | 3.25 | 1.00 | 45.62 |  |
| 3 | A | N6 | AA:56 | 3.94 | 1.00 | 44.37 |  |
| 4 | G | O6 | AA:54 | 4.45 | 1.00 | 37.94 |  |
| 5 | U | O4 | AA:114 | 4.71 | 1.00 | 48.65 |  |
| 6 | G | N1 | AA:55 | 4.92 | 1.00 | 46.19 |  |
  |

  
 Coded contact list:   
 Exclusion zone contact(s): /N7/O6/  
Consider K in place of MG (2 valid contact(s) for K vs. 0 for MG)  

### MG:AA:3318

|  | Res. type | Atom | Residue | Distance | Occupancy | B-factor | Closest solute to 1^st^ shell water |
| --- | --- | --- | --- | --- | --- | --- | --- |
| **Ion** | | | |  | 1.00 | 65.92 || 1 | C | N4 | AA:69 | 4.25 | 1.00 | 56.08 |
  |

  
 Coded contact list:   
1^st^ MG coordination shell missing (see Notes)  

### MG:AA:3319

|  | Res. type | Atom | Residue | Distance | Occupancy | B-factor | Closest solute to 1^st^ shell water |
| --- | --- | --- | --- | --- | --- | --- | --- |
| **Ion** | | | |  | 1.00 | 85.00 || 1 | U | OP2 | AA:1915 | 3.58 | 1.00 | 82.95 |
| 2 | U | O4 | AA:1917 | 4.09 | 1.00 | 66.24 |  |
| 3 | A | OP2 | AA:1912 | 4.11 | 1.00 | 64.47 |  |
| 4 | U | OP1 | AA:1915 | 4.13 | 1.00 | 82.75 |  |
| 5 | A | OP2 | AA:1913 | 4.33 | 1.00 | 73.07 |  |
| 6 | A | O5' | AA:1913 | 4.44 | 1.00 | 75.10 |  |
| 7 | A | N7 | AA:1916 | 4.47 | 1.00 | 75.98 |  |
| 8 | A | O5' | AA:1912 | 4.56 | 1.00 | 64.74 |  |
| 9 | A | OP2 | AA:1916 | 4.95 | 1.00 | 79.95 |  |
  |

  
 Coded contact list:   
1^st^ MG coordination shell missing (see Notes)  

### MG:AA:3320

|  | Res. type | Atom | Residue | Distance | Occupancy | B-factor | Closest solute to 1^st^ shell water |
| --- | --- | --- | --- | --- | --- | --- | --- |
| **Ion** | | | |  | 1.00 | 69.54 || 1 | G | OP2 | AA:68 | 3.52 | 1.00 | 50.58 |
| 2 | G | OP1 | AA:68 | 4.01 | 1.00 | 51.91 |  |
| 3 | U | OP1 | AA:67 | 4.39 | 1.00 | 54.61 |  |
| 4 | U | O3' | AA:67 | 4.91 | 1.00 | 51.98 |  |
  |

  
 Coded contact list:   
1^st^ MG coordination shell missing (see Notes)  

### MG:AA:3321

|  | Res. type | Atom | Residue | Distance | Occupancy | B-factor | Closest solute to 1^st^ shell water |
| --- | --- | --- | --- | --- | --- | --- | --- |
| **Ion** | | | |  | 1.00 | 69.64 || 1 | U | OP2 | AA:1951 | 3.43 | 1.00 | 56.42 |
| 2 | G | OP1 | AA:1950 | 3.74 | 1.00 | 50.32 |  |
| 3 | C | O3' | BA:1420 | 4.55 | 1.00 | 78.84 |  |
| 4 | G | OP1 | BA:1421 | 4.58 | 1.00 | 81.31 |  |
| 5 | U | OP1 | AA:1951 | 4.61 | 1.00 | 54.50 |  |
| 6 | GLU | OE2 | AN:54 | 4.90 | 1.00 | 76.26 |  |
  |

  
 Coded contact list:   
1^st^ MG coordination shell missing (see Notes)  

### MG:AA:3322

|  | Res. type | Atom | Residue | Distance | Occupancy | B-factor | Closest solute to 1^st^ shell water |
| --- | --- | --- | --- | --- | --- | --- | --- |
| **Ion** | | | |  | 1.00 | 58.59 || 1 | U | OP1 | AA:2866 | 2.62 | 1.00 | 77.99 |
| 2 | U | O2' | AA:2865 | 3.78 | 1.00 | 76.13 |  |
| 3 | U | O3' | AA:2865 | 4.47 | 1.00 | 76.74 |  |
| 4 | U | OP2 | AA:2866 | 4.74 | 1.00 | 77.61 |  |
| 5 | A | N6 | AA:2850 | 4.83 | 1.00 | 65.36 |  |
  |

  
 Coded contact list:   
 Exclusion zone contact(s): /OP1/  
Consider K in place of MG (1 valid contact(s) for K vs. 0 for MG)  

### MG:AA:3323

|  | Res. type | Atom | Residue | Distance | Occupancy | B-factor | Closest solute to 1^st^ shell water |
| --- | --- | --- | --- | --- | --- | --- | --- |
| **Ion** | | | |  | 1.00 | 48.97 || 1 | C | N4 | AA:57 | 4.37 | 1.00 | 52.29 |
| 2 | G | O6 | AA:58 | 4.40 | 1.00 | 54.77 |  |
| 3 | C | N4 | AA:69 | 4.54 | 1.00 | 56.08 |  |
| 4 | U | OP2 | AA:67 | 4.62 | 1.00 | 54.38 |  |
| 5 | A | OP2 | AA:56 | 4.76 | 1.00 | 42.92 |  |
  |

  
 Coded contact list:   
1^st^ MG coordination shell missing (see Notes)  

### MG:AA:3324

|  | Res. type | Atom | Residue | Distance | Occupancy | B-factor | Closest solute to 1^st^ shell water |
| --- | --- | --- | --- | --- | --- | --- | --- |
| **Ion** | | | |  | 1.00 | 43.15 || 1 | U | OP1 | AA:1267 | 1.97 | 1.00 | 46.88 |
| 2 | U | OP2 | AA:1267 | 3.68 | 1.00 | 45.51 |  |
| 3 | C | OP2 | AA:2008 | 3.93 | 1.00 | 46.19 |  |
| 4 | U | O5' | AA:1267 | 4.18 | 1.00 | 47.01 |  |
| 5 | G | O3' | AA:1266 | 4.34 | 1.00 | 47.46 |  |
  |

  
 Contact(s): /OP1/  
 Coded contact list: O_ph_  

### MG:AA:3325

|  | Res. type | Atom | Residue | Distance | Occupancy | B-factor | Closest solute to 1^st^ shell water |
| --- | --- | --- | --- | --- | --- | --- | --- |
| **Ion** | | | |  | 1.00 | 50.00 || 1 | G | O6 | AA:1309 | 3.04 | 1.00 | 26.03 |
| 2 | G | N7 | AA:1309 | 3.09 | 1.00 | 23.70 |  |
| 3 | G | N7 | AA:1310 | 3.22 | 1.00 | 33.11 |  |
| 4 | G | O6 | AA:1310 | 3.54 | 1.00 | 40.43 |  |
| 5 | G | O6 | AA:1311 | 4.57 | 1.00 | 64.71 |  |
| 6 | G | N1 | AA:1309 | 4.96 | 1.00 | 23.30 |  |
  |

  
 Coded contact list:   
 Exclusion zone contact(s): /N7/N7/O6/  
Consider K in place of MG (3 valid contact(s) for K vs. 0 for MG)  

### MG:AA:3326

|  | Res. type | Atom | Residue | Distance | Occupancy | B-factor | Closest solute to 1^st^ shell water |
| --- | --- | --- | --- | --- | --- | --- | --- |
| **Ion** | | | |  | 1.00 | 22.91 || 1 | G | O2' | AA:775 | 2.27 | 1.00 | 39.24 |
| 2 | G | OP2 | AA:794 | 2.43 | 1.00 | 38.93 |  |
| 3 | A | O2' | AA:793 | 2.66 | 1.00 | 48.59 |  |
| 4 | G | N7 | AA:776 | 2.94 | 1.00 | 43.25 |  |
| 5 | G | O3' | AA:775 | 3.60 | 1.00 | 41.60 |  |
| 6 | G | OP2 | AA:776 | 4.15 | 1.00 | 42.08 |  |
| 7 | A | O3' | AA:793 | 4.27 | 1.00 | 41.17 |  |
| 8 | G | N3 | AA:775 | 4.36 | 1.00 | 38.35 |  |
| 9 | G | N9 | AA:776 | 4.43 | 1.00 | 40.44 |  |
| 10 | G | OP1 | AA:794 | 4.62 | 1.00 | 39.87 |  |
| 11 | A | N3 | AA:793 | 4.79 | 1.00 | 47.06 |  |
| 12 | G | O5' | AA:794 | 4.86 | 1.00 | 38.95 |  |
| 13 | G | O5' | AA:776 | 4.88 | 1.00 | 41.87 |  |
| 14 | G | O4' | AA:776 | 4.90 | 1.00 | 38.49 |  |
| 15 | G | N2 | AA:775 | 4.97 | 1.00 | 39.62 |  |
  |

  
 Contact(s): /O2'/  
 Coded contact list: O_r_  
 Exclusion zone contact(s): /N7/O2'/OP2/  
Consider K in place of MG (2 valid contact(s) for K vs. 1 for MG)  

### MG:AA:3327

|  | Res. type | Atom | Residue | Distance | Occupancy | B-factor | Closest solute to 1^st^ shell water |
| --- | --- | --- | --- | --- | --- | --- | --- |
| **Ion** | | | |  | 1.00 | 45.33 || 1 | G | OP2 | AA:778 | 2.68 | 1.00 | 33.98 |
| 2 | G | N7 | AA:778 | 3.69 | 1.00 | 30.66 |  |
| 3 | U | O4 | AA:779 | 3.80 | 1.00 | 29.21 |  |
| 4 | G | O5' | AA:778 | 4.81 | 1.00 | 31.78 |  |
| 5 | G | O6 | AA:780 | 4.89 | 1.00 | 36.35 |  |
  |

  
 Coded contact list:   
 Exclusion zone contact(s): /OP2/  
Consider K in place of MG (1 valid contact(s) for K vs. 0 for MG)  

### MG:AA:3328

|  | Res. type | Atom | Residue | Distance | Occupancy | B-factor | Closest solute to 1^st^ shell water |
| --- | --- | --- | --- | --- | --- | --- | --- |
| **Ion** | | | |  | 1.00 | 32.16 || 1 | G | O6 | AA:778 | 3.36 | 1.00 | 32.66 |
| 2 | U | O4 | AA:779 | 3.91 | 1.00 | 29.21 |  |
| 3 | G | N7 | AA:778 | 4.03 | 1.00 | 30.66 |  |
| 4 | A | N1 | AA:793 | 4.69 | 1.00 | 50.44 |  |
| 5 | A | N6 | AA:777 | 4.77 | 1.00 | 27.84 |  |
| 6 | A | N3 | AA:793 | 4.99 | 1.00 | 47.06 |  |
  |

  
 Coded contact list:   
 Exclusion zone contact(s): /O6/  
Consider K in place of MG (1 valid contact(s) for K vs. 0 for MG)  

### MG:AA:3329

|  | Res. type | Atom | Residue | Distance | Occupancy | B-factor | Closest solute to 1^st^ shell water |
| --- | --- | --- | --- | --- | --- | --- | --- |
| **Ion** | | | |  | 1.00 | 66.22 || 1 | G | N2 | AA:1975 | 4.12 | 1.00 | 26.92 |
| 2 | G | O2' | AA:1831 | 4.41 | 1.00 | 36.77 |  |
| 3 | G | O4' | AA:1831 | 4.43 | 1.00 | 39.48 |  |
| 4 | U | O4' | AA:1976 | 4.69 | 1.00 | 38.03 |  |
| 5 | G | N9 | AA:1831 | 4.89 | 1.00 | 36.08 |  |
| 6 | U | O2' | AA:1976 | 4.96 | 1.00 | 35.33 |  |
  |

  
 Coded contact list:   
1^st^ MG coordination shell missing (see Notes)  

### MG:AA:3330

|  | Res. type | Atom | Residue | Distance | Occupancy | B-factor | Closest solute to 1^st^ shell water |
| --- | --- | --- | --- | --- | --- | --- | --- |
| **Ion** | | | |  | 1.00 | 111.12 || 1 | G | OP1 | AA:2869 | 2.40 | 1.00 | 65.58 |
| 2 | A | O3' | AA:2868 | 3.57 | 1.00 | 65.60 |  |
| 3 | G | OP2 | AA:2869 | 4.53 | 1.00 | 63.63 |  |
| 4 | G | O5' | AA:2869 | 4.66 | 1.00 | 63.68 |  |
  |

  
 Coded contact list:   
 Exclusion zone contact(s): /OP1/  
Consider NA in place of MG (1 valid contact(s) for NA vs. 0 for MG)  

### MG:AA:3331

|  | Res. type | Atom | Residue | Distance | Occupancy | B-factor | Closest solute to 1^st^ shell water |
| --- | --- | --- | --- | --- | --- | --- | --- |
| **Ion** | | | |  | 1.00 | 41.86 || 1 | U | OP1 | AA:1639 | 2.20 | 1.00 | 45.48 |
| 2 | G | N3 | AA:2709 | 3.67 | 1.00 | 48.90 |  |
| 3 | G | N2 | AA:2709 | 3.69 | 1.00 | 49.35 |  |
| 4 | G | O2' | AA:2709 | 3.89 | 1.00 | 47.47 |  |
| 5 | C | O3' | AA:1638 | 4.21 | 1.00 | 43.70 |  |
| 6 | U | O5' | AA:1639 | 4.34 | 1.00 | 42.99 |  |
| 7 | C | OP2 | AA:1297 | 4.56 | 1.00 | 49.86 |  |
| 8 | U | OP2 | AA:1639 | 4.73 | 1.00 | 44.86 |  |
| 9 | G | OP1 | AA:1296 | 4.77 | 1.00 | 52.43 |  |
| 10 | C | O4' | AA:2710 | 4.82 | 1.00 | 48.75 |  |
| 11 | G | OP2 | AA:1296 | 4.98 | 1.00 | 51.97 |  |
  |

  
 Contact(s): /OP1/  
 Coded contact list: O_ph_  

### MG:AA:3332

|  | Res. type | Atom | Residue | Distance | Occupancy | B-factor | Closest solute to 1^st^ shell water |
| --- | --- | --- | --- | --- | --- | --- | --- |
| **Ion** | | | |  | 1.00 | 83.12 || 1 | G | OP2 | AA:1470 | 3.06 | 1.00 | 75.20 |
| 2 | G | N7 | AA:1470 | 3.47 | 1.00 | 82.21 |  |
| 3 | G | O6 | AA:1521 | 3.84 | 1.00 | 92.51 |  |
| 4 | G | OP2 | AA:1519 | 4.11 | 1.00 | 70.94 |  |
| 5 | U | OP2 | AA:1520 | 4.26 | 1.00 | 71.58 |  |
| 6 | G | O6 | AA:1470 | 4.74 | 1.00 | 83.82 |  |
| 7 | G | O5' | AA:1470 | 4.79 | 1.00 | 76.41 |  |
| 8 | A | OP2 | AA:1469 | 4.87 | 1.00 | 75.98 |  |
  |

  
 Coded contact list:   
 Exclusion zone contact(s): /OP2/  
Consider K in place of MG (1 valid contact(s) for K vs. 0 for MG)  

### MG:AA:3333

|  | Res. type | Atom | Residue | Distance | Occupancy | B-factor | Closest solute to 1^st^ shell water |
| --- | --- | --- | --- | --- | --- | --- | --- |
| **Ion** | | | |  | 1.00 | 59.71 || 1 | G | N2 | AA:1842 | 2.79 | 1.00 | 41.87 |
| 2 | A | N9 | AA:1901 | 3.18 | 1.00 | 43.42 |  |
| 3 | A | O4' | AA:1901 | 3.30 | 1.00 | 43.46 |  |
| 4 | U | O2' | AA:1898 | 3.75 | 1.00 | 45.65 |  |
| 5 | A | N7 | AA:1901 | 3.87 | 1.00 | 45.38 |  |
| 6 | U | O2 | AA:1898 | 3.96 | 1.00 | 44.05 |  |
| 7 | G | N1 | AA:1842 | 4.56 | 1.00 | 40.49 |  |
| 8 | A | N3 | AA:1901 | 4.87 | 1.00 | 43.18 |  |
  |

  
 Coded contact list:   
 Exclusion zone contact(s): /N2/O4'/  
Consider K in place of MG (3 valid contact(s) for K vs. 0 for MG)  

### MG:AA:3334

|  | Res. type | Atom | Residue | Distance | Occupancy | B-factor | Closest solute to 1^st^ shell water |
| --- | --- | --- | --- | --- | --- | --- | --- |
| **Ion** | | | |  | 1.00 | 46.61 || 1 | G | O6 | AA:68 | 2.58 | 1.00 | 62.03 |
| 2 | G | O6 | AA:58 | 2.86 | 1.00 | 54.77 |  |
| 3 | U | O4 | AA:59 | 2.99 | 1.00 | 61.09 |  |
| 4 | U | O4 | AA:67 | 3.36 | 1.00 | 55.74 |  |
| 5 | U | N3 | AA:59 | 3.97 | 1.00 | 61.06 |  |
| 6 | C | N4 | AA:69 | 4.00 | 1.00 | 56.08 |  |
| 7 | G | N7 | AA:68 | 4.21 | 1.00 | 58.57 |  |
| 8 | G | N1 | AA:58 | 4.42 | 1.00 | 54.43 |  |
| 9 | G | N7 | AA:58 | 4.63 | 1.00 | 57.59 |  |
| 10 | G | N1 | AA:68 | 4.86 | 1.00 | 61.14 |  |
  |

  
 Coded contact list:   
 Exclusion zone contact(s): /O4/O4/O6/O6/  
Consider K in place of MG (3 valid contact(s) for K vs. 0 for MG)  

### MG:AA:3335

|  | Res. type | Atom | Residue | Distance | Occupancy | B-factor | Closest solute to 1^st^ shell water |
| --- | --- | --- | --- | --- | --- | --- | --- |
| **Ion** | | | |  | 1.00 | 83.83 || 1 | G | OP2 | AA:60 | 2.69 | 1.00 | 58.17 |
| 2 | G | N7 | AA:60 | 3.42 | 1.00 | 55.81 |  |
| 3 | G | N9 | AA:60 | 4.11 | 1.00 | 55.51 |  |
| 4 | U | OP2 | AA:59 | 4.50 | 1.00 | 56.16 |  |
| 5 | G | O4' | AA:60 | 4.57 | 1.00 | 55.55 |  |
| 6 | G | OP1 | AA:60 | 4.67 | 1.00 | 59.08 |  |
| 7 | A | OP2 | AA:64 | 4.92 | 1.00 | 69.12 |  |
| 8 | U | O3' | AA:59 | 4.96 | 1.00 | 57.86 |  |
  |

  
 Coded contact list:   
 Exclusion zone contact(s): /OP2/  
Consider K in place of MG (1 valid contact(s) for K vs. 0 for MG)  

### MG:AA:3336

|  | Res. type | Atom | Residue | Distance | Occupancy | B-factor | Closest solute to 1^st^ shell water |
| --- | --- | --- | --- | --- | --- | --- | --- |
| **Ion** | | | |  | 1.00 | 122.46 || 1 | G | O3' | AA:2862 | 2.62 | 1.00 | 75.58 |
| 2 | G | O2' | AA:2862 | 2.78 | 1.00 | 76.57 |  |
| 3 | C | O5' | AA:2863 | 3.43 | 1.00 | 75.09 |  |
| 4 | C | OP1 | AA:2863 | 3.54 | 1.00 | 74.36 |  |
| 5 | C | OP1 | AA:1752 | 4.53 | 1.00 | 68.50 |  |
| 6 | G | O4' | AA:2862 | 4.66 | 1.00 | 77.35 |  |
| 7 | C | OP2 | AA:2863 | 4.82 | 1.00 | 74.38 |  |
  |

  
 Coded contact list:   
 Exclusion zone contact(s): /O2'/O3'/  
Consider K in place of MG (2 valid contact(s) for K vs. 0 for MG)  

### MG:AA:3337

|  | Res. type | Atom | Residue | Distance | Occupancy | B-factor | Closest solute to 1^st^ shell water |
| --- | --- | --- | --- | --- | --- | --- | --- |
| **Ion** | | | |  | 1.00 | 51.68 || 1 | G | O2' | AA:1441 | 2.92 | 1.00 | 53.91 |
| 2 | G | O4' | AA:1442 | 3.11 | 1.00 | 65.05 |  |
| 3 | G | N3 | AA:1441 | 3.64 | 1.00 | 60.11 |  |
| 4 | G | N2 | AA:1441 | 4.13 | 1.00 | 60.61 |  |
| 5 | G | OP1 | AA:1628 | 4.23 | 1.00 | 44.11 |  |
| 6 | G | N9 | AA:1442 | 4.54 | 1.00 | 66.67 |  |
| 7 | U | OP1 | AA:1629 | 4.58 | 1.00 | 51.02 |  |
| 8 | C | O2 | AA:1551 | 4.66 | 1.00 | 55.62 |  |
| 9 | G | N9 | AA:1441 | 4.83 | 1.00 | 57.61 |  |
| 10 | U | OP2 | AA:1629 | 4.85 | 1.00 | 50.86 |  |
| 11 | G | N2 | AA:1440 | 4.93 | 1.00 | 56.96 |  |
  |

  
 Coded contact list:   
 Exclusion zone contact(s): /O2'/O4'/  
Consider K in place of MG (2 valid contact(s) for K vs. 0 for MG)  

### MG:AA:3338

|  | Res. type | Atom | Residue | Distance | Occupancy | B-factor | Closest solute to 1^st^ shell water |
| --- | --- | --- | --- | --- | --- | --- | --- |
| **Ion** | | | |  | 1.00 | 72.27 || 1 | A | OP2 | AA:2851 | 2.54 | 1.00 | 66.90 |
| 2 | C | N4 | AA:2853 | 3.54 | 1.00 | 71.00 |  |
| 3 | G | O6 | AA:2852 | 3.56 | 1.00 | 70.71 |  |
| 4 | G | N7 | AA:2852 | 3.67 | 1.00 | 70.68 |  |
| 5 | A | OP1 | AA:2850 | 3.78 | 1.00 | 65.60 |  |
| 6 | A | O5' | AA:2851 | 4.28 | 1.00 | 66.70 |  |
| 7 | A | OP1 | AA:2851 | 4.43 | 1.00 | 66.20 |  |
| 8 | A | N7 | AA:2851 | 4.83 | 1.00 | 69.07 |  |
  |

  
 Coded contact list:   
 Exclusion zone contact(s): /OP2/  
Consider NA in place of MG (1 valid contact(s) for NA vs. 0 for MG)  

### MG:AA:3339

|  | Res. type | Atom | Residue | Distance | Occupancy | B-factor | Closest solute to 1^st^ shell water |
| --- | --- | --- | --- | --- | --- | --- | --- |
| **Ion** | | | |  | 1.00 | 50.98 || 1 | G | O6 | AA:2373 | 2.07 | 1.00 | 53.31 |
| 2 | G | O6 | AA:2372 | 2.93 | 1.00 | 63.31 |  |
| 3 | G | N7 | AA:2373 | 3.21 | 1.00 | 55.93 |  |
| 4 | C | N4 | AA:2374 | 3.74 | 1.00 | 58.73 |  |
| 5 | G | N1 | AA:2372 | 4.00 | 1.00 | 62.71 |  |
| 6 | C | N4 | AA:2380 | 4.14 | 1.00 | 54.59 |  |
| 7 | G | O6 | AA:2379 | 4.16 | 1.00 | 60.24 |  |
| 8 | G | N7 | AA:2372 | 4.21 | 1.00 | 62.75 |  |
| 9 | G | N1 | AA:2373 | 4.26 | 1.00 | 52.75 |  |
| 10 | C | N4 | AA:2381 | 4.44 | 1.00 | 62.24 |  |
  |

  
 Contact(s): /O6/  
 Coded contact list: O_b_  
 Exclusion zone contact(s): /N7/O6/  
Consider K in place of MG (2 valid contact(s) for K vs. 1 for MG)  

### MG:AA:3340

|  | Res. type | Atom | Residue | Distance | Occupancy | B-factor | Closest solute to 1^st^ shell water |
| --- | --- | --- | --- | --- | --- | --- | --- |
| **Ion** | | | |  | 1.00 | 90.63 || 1 | G | N7 | AA:1681 | 2.44 | 1.00 | 67.16 |
| 2 | U | OP2 | AA:1680 | 2.73 | 1.00 | 60.89 |  |
| 3 | U | O5' | AA:1680 | 3.95 | 1.00 | 59.80 |  |
| 4 | G | O6 | AA:1681 | 3.98 | 1.00 | 68.76 |  |
| 5 | G | N9 | AA:1681 | 4.53 | 1.00 | 64.75 |  |
| 6 | U | OP1 | AA:1680 | 4.55 | 1.00 | 59.73 |  |
| 7 | G | OP2 | AA:1681 | 4.62 | 1.00 | 56.91 |  |
| 8 | U | O5' | AA:1679 | 4.83 | 1.00 | 57.58 |  |
  |

  
 Coded contact list:   
 Exclusion zone contact(s): /N7/OP2/  
Consider NA in place of MG (1 valid contact(s) for NA vs. 0 for MG)  

### MG:AA:3341

|  | Res. type | Atom | Residue | Distance | Occupancy | B-factor | Closest solute to 1^st^ shell water |
| --- | --- | --- | --- | --- | --- | --- | --- |
| **Ion** | | | |  | 1.00 | 57.15 || 1 | U | O4 | AA:464 | 3.15 | 1.00 | 26.40 |
| 2 | G | O6 | AA:465 | 3.37 | 1.00 | 33.46 |  |
| 3 | A | OP2 | AA:789 | 3.88 | 1.00 | 41.63 |  |
| 4 | A | O2' | AA:788 | 4.06 | 1.00 | 43.17 |  |
| 5 | A | N9 | AA:788 | 4.07 | 1.00 | 34.18 |  |
| 6 | G | N1 | AA:465 | 4.35 | 1.00 | 30.66 |  |
| 7 | A | O3' | AA:788 | 4.49 | 1.00 | 40.33 |  |
  |

  
 Coded contact list:   
 Exclusion zone contact(s): /O4/O6/  
Consider K in place of MG (2 valid contact(s) for K vs. 0 for MG)  

### MG:AA:3342

|  | Res. type | Atom | Residue | Distance | Occupancy | B-factor | Closest solute to 1^st^ shell water |
| --- | --- | --- | --- | --- | --- | --- | --- |
| **Ion** | | | |  | 1.00 | 63.95 || 1 | G | O6 | AA:1950 | 2.82 | 1.00 | 45.73 |
| 2 | U | OP2 | AA:1955 | 3.00 | 1.00 | 53.75 |  |
| 3 | G | O6 | AA:1949 | 3.31 | 1.00 | 37.26 |  |
| 4 | G | O3' | AA:1954 | 4.17 | 1.00 | 55.81 |  |
| 5 | G | N1 | AA:1950 | 4.22 | 1.00 | 46.65 |  |
| 6 | C | N4 | AA:1957 | 4.35 | 1.00 | 38.99 |  |
| 7 | U | OP1 | AA:1955 | 4.39 | 1.00 | 54.38 |  |
| 8 | G | OP1 | AA:1954 | 4.70 | 1.00 | 60.78 |  |
| 9 | G | O2' | AA:1954 | 4.75 | 1.00 | 59.51 |  |
| 10 | U | O4 | AA:1956 | 4.86 | 1.00 | 47.58 |  |
| 11 | G | OP2 | AA:1954 | 4.91 | 1.00 | 59.86 |  |
  |

  
 Coded contact list:   
 Exclusion zone contact(s): /O6/O6/OP2/  
Consider K in place of MG (3 valid contact(s) for K vs. 0 for MG)  

### MG:AA:3343

|  | Res. type | Atom | Residue | Distance | Occupancy | B-factor | Closest solute to 1^st^ shell water |
| --- | --- | --- | --- | --- | --- | --- | --- |
| **Ion** | | | |  | 1.00 | 48.48 || 1 | G | O6 | AA:212 | 3.12 | 1.00 | 49.38 |
| 2 | G | N7 | AA:212 | 3.86 | 1.00 | 49.70 |  |
| 3 | A | N6 | AA:213 | 4.56 | 1.00 | 60.30 |  |
| 4 | C | N4 | AA:184 | 4.89 | 1.00 | 46.59 |  |
  |

  
 Coded contact list:   
 Exclusion zone contact(s): /O6/  
Consider K in place of MG (1 valid contact(s) for K vs. 0 for MG)  

### MG:AA:3344

|  | Res. type | Atom | Residue | Distance | Occupancy | B-factor | Closest solute to 1^st^ shell water |
| --- | --- | --- | --- | --- | --- | --- | --- |
| **Ion** | | | |  | 1.00 | 76.02 || 1 | G | O6 | AA:180 | 2.89 | 1.00 | 54.73 |
| 2 | G | O6 | AA:179 | 3.37 | 1.00 | 56.22 |  |
| 3 | A | OP2 | AA:213 | 3.62 | 1.00 | 61.26 |  |
| 4 | G | N7 | AA:179 | 4.29 | 1.00 | 52.28 |  |
| 5 | G | N7 | AA:180 | 4.50 | 1.00 | 53.53 |  |
| 6 | A | N7 | AA:213 | 4.59 | 1.00 | 59.31 |  |
| 7 | C | N4 | AA:46 | 4.85 | 1.00 | 54.69 |  |
| 8 | G | O6 | AA:178 | 4.87 | 1.00 | 58.68 |  |
  |

  
 Coded contact list:   
 Exclusion zone contact(s): /O6/O6/  
Consider K in place of MG (2 valid contact(s) for K vs. 0 for MG)  

### MG:AA:3345

|  | Res. type | Atom | Residue | Distance | Occupancy | B-factor | Closest solute to 1^st^ shell water |
| --- | --- | --- | --- | --- | --- | --- | --- |
| **Ion** | | | |  | 1.00 | 55.65 || 1 | A | OP2 | AA:2518 | 2.30 | 1.00 | 69.97 |
| 2 | A | OP1 | AA:2518 | 3.45 | 1.00 | 69.75 |  |
| 3 | U | OP2 | AA:2519 | 3.57 | 1.00 | 61.71 |  |
| 4 | A | N7 | AA:2541 | 3.61 | 1.00 | 70.99 |  |
| 5 | C | OP2 | AA:2540 | 3.93 | 1.00 | 76.54 |  |
| 6 | A | OP2 | AA:2541 | 4.19 | 1.00 | 75.26 |  |
| 7 | A | O5' | AA:2518 | 4.22 | 1.00 | 68.20 |  |
| 8 | C | O3' | AA:2517 | 4.59 | 1.00 | 69.60 |  |
| 9 | A | N6 | AA:2541 | 4.69 | 1.00 | 75.23 |  |
| 10 | C | O2' | AA:2517 | 4.75 | 1.00 | 74.53 |  |
  |

  
 Contact(s): /OP2/  
 Coded contact list: O_ph_  

### MG:AA:3346

|  | Res. type | Atom | Residue | Distance | Occupancy | B-factor | Closest solute to 1^st^ shell water |
| --- | --- | --- | --- | --- | --- | --- | --- |
| **Ion** | | | |  | 1.00 | 58.56 || 1 | C | OP1 | AA:253 | 2.96 | 1.00 | 45.06 |
| 2 | G | OP2 | AA:254 | 3.41 | 1.00 | 43.51 |  |
| 3 | G | OP1 | AA:254 | 4.46 | 1.00 | 43.53 |  |
| 4 | C | O3' | AA:253 | 4.58 | 1.00 | 43.49 |  |
| 5 | C | OP1 | AA:203 | 4.59 | 1.00 | 44.49 |  |
| 6 | C | O5' | AA:253 | 4.59 | 1.00 | 42.72 |  |
  |

  
 Coded contact list:   
 Exclusion zone contact(s): /OP1/  
Consider K in place of MG (1 valid contact(s) for K vs. 0 for MG)  

### MG:AA:3347

|  | Res. type | Atom | Residue | Distance | Occupancy | B-factor | Closest solute to 1^st^ shell water |
| --- | --- | --- | --- | --- | --- | --- | --- |
| **Ion** | | | |  | 1.00 | 66.65 || 1 | G | OP1 | AA:361 | 2.90 | 1.00 | 108.23 |
| 2 | G | N7 | AA:270Y | 3.20 | 1.00 | 78.36 |  |
| 3 | U | OP1 | AA:269 | 3.77 | 1.00 | 67.91 |  |
| 4 | G | OP2 | AA:270X | 3.81 | 1.00 | 85.64 |  |
| 5 | G | O6 | AA:270Y | 4.14 | 1.00 | 77.65 |  |
| 6 | G | OP2 | AA:270Y | 4.21 | 1.00 | 76.12 |  |
| 7 | U | OP2 | AA:269 | 4.76 | 1.00 | 66.66 |  |
| 8 | G | O3' | AA:360 | 4.97 | 1.00 | 107.66 |  |
  |

  
 Coded contact list:   
 Exclusion zone contact(s): /N7/OP1/  
Consider K in place of MG (2 valid contact(s) for K vs. 0 for MG)  

### MG:AA:3348

|  | Res. type | Atom | Residue | Distance | Occupancy | B-factor | Closest solute to 1^st^ shell water |
| --- | --- | --- | --- | --- | --- | --- | --- |
| **Ion** | | | |  | 1.00 | 26.88 || 1 | U | OP1 | AA:810 | 1.96 | 1.00 | 43.95 |
| 2 | A | N7 | AA:1253 | 2.19 | 1.00 | 46.41 |  |
| 3 | A | N6 | AA:1253 | 3.54 | 1.00 | 44.78 |  |
| 4 | U | OP2 | AA:810 | 3.67 | 1.00 | 44.76 |  |
| 5 | A | N9 | AA:1253 | 4.11 | 1.00 | 44.01 |  |
| 6 | C | O2' | AA:565 | 4.12 | 1.00 | 38.36 |  |
| 7 | G | O3' | AA:809 | 4.17 | 1.00 | 42.80 |  |
| 8 | U | O5' | AA:810 | 4.33 | 1.00 | 43.21 |  |
| 9 | C | O3' | AA:565 | 4.46 | 1.00 | 42.22 |  |
| 10 | A | N1 | AA:1253 | 4.95 | 1.00 | 44.38 |  |
  |

  
**1^st^ shell ligand angles:**   
1-2: 108;   
  
 Contact(s): /N7/OP1/  
 Coded contact list: O_ph_N_b_  

### MG:AA:3349

|  | Res. type | Atom | Residue | Distance | Occupancy | B-factor | Closest solute to 1^st^ shell water |
| --- | --- | --- | --- | --- | --- | --- | --- |
| **Ion** | | | |  | 1.00 | 42.88 || 1 | G | OP1 | AA:809 | 2.37 | 1.00 | 43.17 |
| 2 | C | O2 | AA:565 | 2.72 | 1.00 | 42.61 |  |
| 3 | U | O2 | AA:576 | 3.12 | 1.00 | 41.73 |  |
| 4 | U | O4' | AA:566 | 3.56 | 1.00 | 42.81 |  |
| 5 | G | N3 | AA:577 | 3.67 | 1.00 | 43.11 |  |
| 6 | C | O2' | AA:565 | 4.04 | 1.00 | 38.36 |  |
| 7 | G | N9 | AA:577 | 4.30 | 1.00 | 44.44 |  |
| 8 | G | O3' | AA:808 | 4.36 | 1.00 | 43.40 |  |
| 9 | G | O4' | AA:577 | 4.49 | 1.00 | 44.63 |  |
| 10 | A | O2' | AA:1253 | 4.57 | 1.00 | 43.49 |  |
| 11 | U | N1 | AA:566 | 4.57 | 1.00 | 41.90 |  |
| 12 | U | N3 | AA:576 | 4.58 | 1.00 | 40.41 |  |
| 13 | G | OP2 | AA:809 | 4.58 | 1.00 | 43.89 |  |
| 14 | G | N2 | AA:577 | 4.60 | 1.00 | 43.27 |  |
| 15 | U | O2' | AA:566 | 4.73 | 1.00 | 44.08 |  |
| 16 | C | N3 | AA:565 | 4.79 | 1.00 | 39.91 |  |
| 17 | C | N1 | AA:565 | 4.81 | 1.00 | 37.84 |  |
| 18 | G | O5' | AA:809 | 4.81 | 1.00 | 42.84 |  |
| 19 | U | O2 | AA:566 | 4.87 | 1.00 | 44.30 |  |
  |

  
 Contact(s): /OP1/  
 Coded contact list: O_ph_  
 Exclusion zone contact(s): /O2/O2/  
Consider K in place of MG (2 valid contact(s) for K vs. 1 for MG)  

### MG:AA:3350

|  | Res. type | Atom | Residue | Distance | Occupancy | B-factor | Closest solute to 1^st^ shell water |
| --- | --- | --- | --- | --- | --- | --- | --- |
| **Ion** | | | |  | 1.00 | 36.45 || 1 | A | O3' | AA:670 | 2.37 | 1.00 | 52.71 |
| 2 | U | O4 | AA:588 | 3.09 | 1.00 | 47.93 |  |
| 3 | C | OP2 | AA:671 | 3.13 | 1.00 | 51.80 |  |
| 4 | A | O2' | AA:670 | 3.14 | 1.00 | 56.79 |  |
| 5 | C | O5' | AA:671 | 3.64 | 1.00 | 51.55 |  |
| 6 | C | OP1 | AA:671 | 4.62 | 1.00 | 53.26 |  |
| 7 | C | N4 | AA:589 | 4.81 | 1.00 | 57.50 |  |
| 8 | G | O6 | AA:668 | 4.83 | 1.00 | 58.20 |  |
  |

  
 Contact(s): /O3'/  
 Coded contact list: O_r_  
 Exclusion zone contact(s): /O2'/O4/OP2/  
Consider K in place of MG (3 valid contact(s) for K vs. 1 for MG)  

### MG:AA:3351

|  | Res. type | Atom | Residue | Distance | Occupancy | B-factor | Closest solute to 1^st^ shell water |
| --- | --- | --- | --- | --- | --- | --- | --- |
| **Ion** | | | |  | 1.00 | 62.63 || 1 | G | O5' | AA:1449A | 2.37 | 1.00 | 92.05 |
| 2 | G | OP2 | AA:1449A | 2.72 | 1.00 | 92.65 |  |
| 3 | A | OP2 | AA:1449 | 3.73 | 1.00 | 87.43 |  |
| 4 | A | O3' | AA:1449 | 3.93 | 1.00 | 92.15 |  |
| 5 | A | O5' | AA:1449 | 4.15 | 1.00 | 88.56 |  |
| 6 | G | O4' | AA:1449A | 4.34 | 1.00 | 89.88 |  |
| 7 | G | OP1 | AA:1449A | 4.35 | 1.00 | 92.52 |  |
| 8 | G | N7 | AA:1449A | 4.76 | 1.00 | 87.21 |  |
| 9 | A | O4' | AA:1449 | 4.94 | 1.00 | 91.60 |  |
  |

  
 Contact(s): /O5'/  
 Coded contact list: O_r_  
 Exclusion zone contact(s): /OP2/  

### MG:AA:3352

|  | Res. type | Atom | Residue | Distance | Occupancy | B-factor | Closest solute to 1^st^ shell water |
| --- | --- | --- | --- | --- | --- | --- | --- |
| **Ion** | | | |  | 1.00 | 28.23 || 1 | A | OP2 | AA:793 | 2.55 | 1.00 | 35.84 |
| 2 | G | O6 | AA:776 | 2.68 | 1.00 | 45.38 |  |
| 3 | C | OP1 | AA:678 | 2.71 | 1.00 | 43.05 |  |
| 4 | A | O3' | AA:2071 | 2.96 | 1.00 | 32.30 |  |
| 5 | G | OP1 | AA:2072 | 3.48 | 1.00 | 34.30 |  |
| 6 | A | O2' | AA:2071 | 3.78 | 1.00 | 33.89 |  |
| 7 | A | OP1 | AA:793 | 4.17 | 1.00 | 37.05 |  |
| 8 | A | O3' | AA:677 | 4.33 | 1.00 | 41.75 |  |
| 9 | C | OP2 | AA:678 | 4.43 | 1.00 | 44.13 |  |
| 10 | G | O5' | AA:2072 | 4.49 | 1.00 | 29.83 |  |
| 11 | G | N1 | AA:776 | 4.64 | 1.00 | 44.56 |  |
| 12 | A | O5' | AA:793 | 4.75 | 1.00 | 39.71 |  |
| 13 | G | O3' | AA:792 | 4.86 | 1.00 | 35.91 |  |
  |

  
 Coded contact list:   
 Exclusion zone contact(s): /O3'/O6/OP1/OP2/  
Consider K in place of MG (3 valid contact(s) for K vs. 0 for MG)  

### MG:AA:3353

|  | Res. type | Atom | Residue | Distance | Occupancy | B-factor | Closest solute to 1^st^ shell water |
| --- | --- | --- | --- | --- | --- | --- | --- |
| **Ion** | | | |  | 1.00 | 83.67 || 1 | G | N7 | AA:599 | 2.70 | 1.00 | 62.81 |
| 2 | G | N9 | AA:599 | 4.35 | 1.00 | 62.77 |  |
| 3 | G | OP2 | AA:599 | 4.37 | 1.00 | 63.97 |  |
| 4 | G | N7 | AA:598 | 4.72 | 1.00 | 62.28 |  |
| 5 | G | O6 | AA:599 | 4.81 | 1.00 | 64.57 |  |
| 6 | G | OP2 | AA:598 | 4.86 | 1.00 | 65.38 |  |
  |

  
 Coded contact list:   
 Exclusion zone contact(s): /N7/  
Consider K in place of MG (1 valid contact(s) for K vs. 0 for MG)  

### MG:AA:3354

|  | Res. type | Atom | Residue | Distance | Occupancy | B-factor | Closest solute to 1^st^ shell water |
| --- | --- | --- | --- | --- | --- | --- | --- |
| **Ion** | | | |  | 1.00 | 53.42 || 1 | G | O6 | AA:600 | 2.26 | 1.00 | 60.19 |
| 2 | G | O6 | AA:599 | 2.85 | 1.00 | 64.57 |  |
| 3 | G | O6 | AA:656 | 2.97 | 1.00 | 66.54 |  |
| 4 | U | O4 | AA:657 | 3.04 | 1.00 | 63.57 |  |
| 5 | C | N4 | AA:601 | 3.36 | 1.00 | 68.94 |  |
| 6 | U | N3 | AA:657 | 3.75 | 1.00 | 62.97 |  |
| 7 | G | N7 | AA:600 | 3.86 | 1.00 | 61.24 |  |
| 8 | C | N4 | AA:658 | 3.94 | 1.00 | 62.36 |  |
| 9 | G | N1 | AA:599 | 4.34 | 1.00 | 62.76 |  |
| 10 | G | N1 | AA:600 | 4.50 | 1.00 | 61.79 |  |
| 11 | G | N1 | AA:656 | 4.63 | 1.00 | 64.42 |  |
| 12 | G | N7 | AA:599 | 4.78 | 1.00 | 62.81 |  |
| 13 | C | N3 | AA:601 | 4.82 | 1.00 | 67.56 |  |
  |

  
 Contact(s): /O6/  
 Coded contact list: O_b_  
 Exclusion zone contact(s): /N4/O4/O6/O6/  
Consider K in place of MG (4 valid contact(s) for K vs. 1 for MG)  

### MG:AA:3355

|  | Res. type | Atom | Residue | Distance | Occupancy | B-factor | Closest solute to 1^st^ shell water |
| --- | --- | --- | --- | --- | --- | --- | --- |
| **Ion** | | | |  | 1.00 | 48.58 || 1 | G | OP2 | AA:205 | 2.45 | 1.00 | 44.28 |
| 2 | A | O3' | AA:204 | 3.71 | 1.00 | 44.98 |  |
| 3 | C | OP2 | AA:203 | 3.85 | 1.00 | 44.19 |  |
| 4 | U | OP1 | AA:202 | 3.89 | 1.00 | 42.13 |  |
| 5 | G | OP1 | AA:205 | 4.21 | 1.00 | 46.19 |  |
| 6 | C | O2' | AA:253 | 4.42 | 1.00 | 44.71 |  |
| 7 | A | O2' | AA:204 | 4.45 | 1.00 | 46.92 |  |
| 8 | C | OP1 | AA:203 | 4.68 | 1.00 | 44.49 |  |
| 9 | G | O5' | AA:205 | 4.84 | 1.00 | 44.85 |  |
  |

  
 Coded contact list:   
 Exclusion zone contact(s): /OP2/  
Consider NA in place of MG (1 valid contact(s) for NA vs. 0 for MG)  

### MG:AA:3356

|  | Res. type | Atom | Residue | Distance | Occupancy | B-factor | Closest solute to 1^st^ shell water |
| --- | --- | --- | --- | --- | --- | --- | --- |
| **Ion** | | | |  | 1.00 | 62.66 || 1 | G | O6 | AA:240 | 2.35 | 1.00 | 47.01 |
| 2 | A | N1 | AA:241 | 3.73 | 1.00 | 47.11 |  |
| 3 | G | N1 | AA:240 | 4.25 | 1.00 | 45.96 |  |
| 4 | A | OP2 | AA:256 | 4.60 | 1.00 | 45.38 |  |
| 5 | A | N6 | AA:241 | 4.87 | 1.00 | 51.64 |  |
| 6 | A | N3 | AA:241 | 4.93 | 1.00 | 48.55 |  |
  |

  
 Contact(s): /O6/  
 Coded contact list: O_b_  

### MG:AA:3357

|  | Res. type | Atom | Residue | Distance | Occupancy | B-factor | Closest solute to 1^st^ shell water |
| --- | --- | --- | --- | --- | --- | --- | --- |
| **Ion** | | | |  | 1.00 | 91.55 || 1 | U | O4 | AA:810 | 2.41 | 1.00 | 52.54 |
| 2 | A | OP1 | AA:567 | 3.39 | 1.00 | 42.45 |  |
| 3 | LYS | NZ | AO:29 | 3.87 | 1.00 | 44.47 |  |
| 4 | THR | N | AO:30 | 4.03 | 1.00 | 43.82 |  |
| 5 | U | N3 | AA:810 | 4.29 | 1.00 | 51.54 |  |
| 6 | U | O3' | AA:566 | 4.49 | 1.00 | 43.20 |  |
| 7 | A | OP2 | AA:567 | 4.50 | 1.00 | 41.45 |  |
| 8 | LYS | O | AO:29 | 4.88 | 1.00 | 44.49 |  |
| 9 | G | N7 | AA:809 | 4.97 | 1.00 | 47.65 |  |
  |

  
 Coded contact list:   
 Exclusion zone contact(s): /O4/OP1/  
Consider NA in place of MG (1 valid contact(s) for NA vs. 0 for MG)  

### MG:AA:3358

|  | Res. type | Atom | Residue | Distance | Occupancy | B-factor | Closest solute to 1^st^ shell water |
| --- | --- | --- | --- | --- | --- | --- | --- |
| **Ion** | | | |  | 1.00 | 95.89 || 1 | U | O4 | AA:2102 | 2.45 | 1.00 | 141.72 |
| 2 | G | O6 | AA:2187 | 2.54 | 1.00 | 144.46 |  |
| 3 | G | O6 | AA:2101 | 2.89 | 1.00 | 144.06 |  |
| 4 | C | N4 | AA:2188 | 2.97 | 1.00 | 144.55 |  |
| 5 | U | N3 | AA:2102 | 3.48 | 1.00 | 142.72 |  |
| 6 | G | O6 | AA:2186 | 3.59 | 1.00 | 148.44 |  |
| 7 | C | N4 | AA:2103 | 3.90 | 1.00 | 148.35 |  |
| 8 | G | N1 | AA:2101 | 4.43 | 1.00 | 143.46 |  |
| 9 | G | N7 | AA:2187 | 4.45 | 1.00 | 146.13 |  |
| 10 | C | N3 | AA:2188 | 4.46 | 1.00 | 145.28 |  |
| 11 | G | N1 | AA:2186 | 4.58 | 1.00 | 148.55 |  |
| 12 | G | N1 | AA:2187 | 4.77 | 1.00 | 143.78 |  |
| 13 | C | N3 | AA:2103 | 4.81 | 1.00 | 148.27 |  |
  |

  
 Coded contact list:   
 Exclusion zone contact(s): /N4/O4/O6/O6/  
Consider NA in place of MG (2 valid contact(s) for NA vs. 0 for MG)  

### MG:AA:3359

|  | Res. type | Atom | Residue | Distance | Occupancy | B-factor | Closest solute to 1^st^ shell water |
| --- | --- | --- | --- | --- | --- | --- | --- |
| **Ion** | | | |  | 1.00 | 48.03 || 1 | G | O6 | AA:723 | 2.53 | 1.00 | 58.23 |
| 2 | G | O6 | AA:707 | 2.80 | 1.00 | 54.19 |  |
| 3 | U | O4 | AA:724 | 3.15 | 1.00 | 56.94 |  |
| 4 | C | N4 | AA:708 | 3.53 | 1.00 | 60.44 |  |
| 5 | U | N3 | AA:724 | 4.09 | 1.00 | 57.89 |  |
| 6 | G | N2 | AA:725 | 4.11 | 1.00 | 60.00 |  |
| 7 | G | N7 | AA:707 | 4.27 | 1.00 | 57.36 |  |
| 8 | A | OP2 | AA:706 | 4.31 | 1.00 | 49.80 |  |
| 9 | G | N1 | AA:723 | 4.39 | 1.00 | 59.63 |  |
| 10 | G | N1 | AA:725 | 4.50 | 1.00 | 60.83 |  |
| 11 | G | N7 | AA:723 | 4.67 | 1.00 | 58.59 |  |
| 12 | G | N3 | AA:725 | 4.77 | 1.00 | 61.20 |  |
| 13 | C | N3 | AA:708 | 4.98 | 1.00 | 58.83 |  |
  |

  
 Coded contact list:   
 Exclusion zone contact(s): /O4/O6/O6/  
Consider K in place of MG (2 valid contact(s) for K vs. 0 for MG)  

### MG:AA:3360

|  | Res. type | Atom | Residue | Distance | Occupancy | B-factor | Closest solute to 1^st^ shell water |
| --- | --- | --- | --- | --- | --- | --- | --- |
| **Ion** | | | |  | 1.00 | 95.65 || 1 | U | O4 | AA:2102 | 3.91 | 1.00 | 141.72 |
| 2 | G | O6 | AA:2186 | 4.33 | 1.00 | 148.44 |  |
| 3 | G | O6 | AA:2101 | 4.49 | 1.00 | 144.06 |  |
  |

  
 Coded contact list:   
1^st^ MG coordination shell missing (see Notes)  

### MG:AA:3361

|  | Res. type | Atom | Residue | Distance | Occupancy | B-factor | Closest solute to 1^st^ shell water |
| --- | --- | --- | --- | --- | --- | --- | --- |
| **Ion** | | | |  | 1.00 | 93.68 || 1 | G | OP2 | AA:2100 | 3.37 | 1.00 | 129.34 |
| 2 | G | OP2 | AA:2101 | 3.94 | 1.00 | 133.80 |  |
| 3 | G | O5' | AA:2100 | 4.85 | 1.00 | 129.90 |  |
  |

  
 Coded contact list:   
 Exclusion zone contact(s): /OP2/  
Consider K in place of MG (1 valid contact(s) for K vs. 0 for MG)  

### MG:AA:3362

|  | Res. type | Atom | Residue | Distance | Occupancy | B-factor | Closest solute to 1^st^ shell water |
| --- | --- | --- | --- | --- | --- | --- | --- |
| **Ion** | | | |  | 1.00 | 62.45 || 1 | G | OP1 | AA:1479 | 2.11 | 1.00 | 78.21 |
| 2 | G | OP2 | AA:1479 | 3.61 | 1.00 | 78.89 |  |
| 3 | G | O5' | AA:1479 | 4.19 | 1.00 | 78.22 |  |
| 4 | G | O2' | AA:1560 | 4.42 | 1.00 | 66.36 |  |
| 5 | G | O3' | AA:1478 | 4.46 | 1.00 | 78.23 |  |
  |

  
 Contact(s): /OP1/  
 Coded contact list: O_ph_  

### MG:AA:3363

|  | Res. type | Atom | Residue | Distance | Occupancy | B-factor | Closest solute to 1^st^ shell water |
| --- | --- | --- | --- | --- | --- | --- | --- |
| **Ion** | | | |  | 1.00 | 74.62 || 1 | G | OP1 | AA:1236 | 2.43 | 1.00 | 61.05 |
| 2 | U | OP1 | AA:511 | 2.74 | 1.00 | 50.71 |  |
| 3 | C | O3' | AA:510 | 2.98 | 1.00 | 49.17 |  |
| 4 | A | O5' | AA:505 | 3.15 | 1.00 | 66.73 |  |
| 5 | A | O4' | AA:505 | 3.37 | 1.00 | 58.35 |  |
| 6 | C | O2' | AA:510 | 3.76 | 1.00 | 51.51 |  |
| 7 | A | OP2 | AA:505 | 3.80 | 1.00 | 70.81 |  |
| 8 | G | OP2 | AA:1236 | 4.16 | 1.00 | 59.42 |  |
| 9 | U | O5' | AA:511 | 4.42 | 1.00 | 51.34 |  |
| 10 | U | OP2 | AA:511 | 4.65 | 1.00 | 49.69 |  |
| 11 | G | O5' | AA:1236 | 4.69 | 1.00 | 60.45 |  |
| 12 | G | O3' | AA:1235 | 4.74 | 1.00 | 59.88 |  |
  |

  
 Coded contact list:   
 Exclusion zone contact(s): /O3'/O4'/O5'/OP1/OP1/  
Consider K in place of MG (4 valid contact(s) for K vs. 0 for MG)  

### MG:AA:3364

|  | Res. type | Atom | Residue | Distance | Occupancy | B-factor | Closest solute to 1^st^ shell water |
| --- | --- | --- | --- | --- | --- | --- | --- |
| **Ion** | | | |  | 1.00 | 57.12 || 1 | G | O6 | AA:270S | 2.74 | 1.00 | 113.29 |
| 2 | G | O6 | AA:270T | 3.12 | 1.00 | 107.11 |  |
| 3 | U | O4 | AA:270F | 3.42 | 1.00 | 107.99 |  |
| 4 | C | N4 | AA:270U | 3.68 | 1.00 | 113.35 |  |
| 5 | G | O6 | AA:270E | 3.69 | 1.00 | 111.52 |  |
| 6 | G | N7 | AA:270S | 4.19 | 1.00 | 110.53 |  |
| 7 | G | N7 | AA:270T | 4.22 | 1.00 | 107.17 |  |
| 8 | G | N1 | AA:270S | 4.34 | 1.00 | 112.92 |  |
| 9 | C | N4 | AA:270G | 4.39 | 1.00 | 115.38 |  |
| 10 | U | N3 | AA:270F | 4.59 | 1.00 | 108.83 |  |
  |

  
 Coded contact list:   
 Exclusion zone contact(s): /O6/O6/  
Consider K in place of MG (2 valid contact(s) for K vs. 0 for MG)  

### MG:AA:3365

|  | Res. type | Atom | Residue | Distance | Occupancy | B-factor | Closest solute to 1^st^ shell water |
| --- | --- | --- | --- | --- | --- | --- | --- |
| **Ion** | | | |  | 1.00 | 39.04 || 1 | G | O6 | AA:270S | 3.86 | 1.00 | 113.29 |
| 2 | G | O6 | AA:270R | 4.30 | 1.00 | 118.49 |  |
| 3 | G | N7 | AA:270R | 4.30 | 1.00 | 116.56 |  |
| 4 | C | N4 | AA:270G | 4.63 | 1.00 | 115.38 |  |
| 5 | U | O4 | AA:270F | 4.64 | 1.00 | 107.99 |  |
| 6 | C | N4 | AA:270Q | 4.82 | 1.00 | 128.02 |  |
  |

  
 Coded contact list:   
1^st^ MG coordination shell missing (see Notes)  

### MG:AA:3366

|  | Res. type | Atom | Residue | Distance | Occupancy | B-factor | Closest solute to 1^st^ shell water |
| --- | --- | --- | --- | --- | --- | --- | --- |
| **Ion** | | | |  | 1.00 | 53.51 || 1 | G | O6 | AA:713 | 2.75 | 1.00 | 78.26 |
| 2 | G | N7 | AA:712 | 3.10 | 1.00 | 73.48 |  |
| 3 | G | N7 | AA:713 | 3.85 | 1.00 | 77.10 |  |
| 4 | G | O6 | AA:712 | 4.17 | 1.00 | 71.26 |  |
| 5 | G | N7 | AA:711 | 4.57 | 1.00 | 70.39 |  |
| 6 | G | N1 | AA:713 | 4.71 | 1.00 | 77.57 |  |
| 7 | G | OP2 | AA:712 | 4.87 | 1.00 | 74.79 |  |
  |

  
 Coded contact list:   
 Exclusion zone contact(s): /N7/O6/  
Consider K in place of MG (2 valid contact(s) for K vs. 0 for MG)  

### MG:AA:3367

|  | Res. type | Atom | Residue | Distance | Occupancy | B-factor | Closest solute to 1^st^ shell water |
| --- | --- | --- | --- | --- | --- | --- | --- |
| **Ion** | | | |  | 1.00 | 49.16 || 1 | G | O6 | AA:270V | 3.57 | 1.00 | 100.05 |
| 2 | G | O6 | AA:270W | 3.98 | 1.00 | 99.15 |  |
| 3 | C | N4 | AA:270U | 4.20 | 1.00 | 113.35 |  |
| 4 | A | OP2 | AA:270A | 4.31 | 1.00 | 70.59 |  |
| 5 | G | O6 | AA:270X | 4.32 | 1.00 | 80.11 |  |
| 6 | G | N7 | AA:270V | 4.45 | 1.00 | 99.53 |  |
| 7 | G | N7 | AA:270W | 4.76 | 1.00 | 97.28 |  |
| 8 | G | N1 | AA:270X | 4.93 | 1.00 | 79.60 |  |
  |

  
 Coded contact list:   
1^st^ MG coordination shell missing (see Notes)  

### MG:AA:3368

|  | Res. type | Atom | Residue | Distance | Occupancy | B-factor | Closest solute to 1^st^ shell water |
| --- | --- | --- | --- | --- | --- | --- | --- |
| **Ion** | | | |  | 1.00 | 87.31 || 1 | G | N7 | AA:710 | 4.67 | 1.00 | 70.66 |
| 2 | G | O6 | AA:710 | 4.77 | 1.00 | 68.71 |  |
| 3 | G | O6 | AA:711 | 4.97 | 1.00 | 67.66 |  |
| 4 | U | O4 | AA:709 | 4.99 | 1.00 | 67.70 |  |
  |

  
 Coded contact list:   
1^st^ MG coordination shell missing (see Notes)  

### MG:AA:3369

|  | Res. type | Atom | Residue | Distance | Occupancy | B-factor | Closest solute to 1^st^ shell water |
| --- | --- | --- | --- | --- | --- | --- | --- |
| **Ion** | | | |  | 1.00 | 81.22 || 1 | G | N7 | AA:2379 | 2.27 | 1.00 | 60.36 |
| 2 | G | O6 | AA:2379 | 2.83 | 1.00 | 60.24 |  |
| 3 | C | N4 | AA:2380 | 3.36 | 1.00 | 54.59 |  |
| 4 | G | N9 | AA:2379 | 4.42 | 1.00 | 62.33 |  |
| 5 | G | N1 | AA:2379 | 4.51 | 1.00 | 59.81 |  |
| 6 | G | N1 | AA:2375 | 4.74 | 1.00 | 58.52 |  |
| 7 | G | O6 | AA:2373 | 4.75 | 1.00 | 53.31 |  |
| 8 | A | OP2 | AA:2378 | 4.78 | 1.00 | 71.30 |  |
  |

  
 Contact(s): /N7/  
 Coded contact list: N_b_  
 Exclusion zone contact(s): /N4/O6/  
Consider K in place of MG (2 valid contact(s) for K vs. 1 for MG)  

### MG:AA:3370

|  | Res. type | Atom | Residue | Distance | Occupancy | B-factor | Closest solute to 1^st^ shell water |
| --- | --- | --- | --- | --- | --- | --- | --- |
| **Ion** | | | |  | 1.00 | 64.80 || 1 | G | N7 | AA:2375 | 4.39 | 1.00 | 61.53 |
| 2 | G | O6 | AA:2375 | 4.49 | 1.00 | 59.34 |  |
| 3 | A | OP1 | AA:2377 | 4.86 | 1.00 | 69.07 |  |
| 4 | A | OP2 | AA:2377 | 4.91 | 1.00 | 71.14 |  |
  |

  
 Coded contact list:   
1^st^ MG coordination shell missing (see Notes)  

### MG:AA:3371

|  | Res. type | Atom | Residue | Distance | Occupancy | B-factor | Closest solute to 1^st^ shell water |
| --- | --- | --- | --- | --- | --- | --- | --- |
| **Ion** | | | |  | 1.00 | 63.31 || 1 | G | OP2 | AA:355 | 2.32 | 1.00 | 100.93 |
| 2 | G | O5' | AA:354 | 3.09 | 1.00 | 101.11 |  |
| 3 | G | OP2 | AA:354 | 3.22 | 1.00 | 101.33 |  |
| 4 | G | O3' | AA:354 | 3.55 | 1.00 | 100.46 |  |
| 5 | G | OP1 | AA:354 | 4.12 | 1.00 | 100.76 |  |
| 6 | G | O5' | AA:355 | 4.48 | 1.00 | 99.57 |  |
| 7 | G | N9 | AA:354 | 4.52 | 1.00 | 99.02 |  |
| 8 | G | OP1 | AA:355 | 4.58 | 1.00 | 100.55 |  |
| 9 | G | N7 | AA:354 | 4.63 | 1.00 | 99.30 |  |
| 10 | G | O4' | AA:354 | 4.70 | 1.00 | 100.01 |  |
| 11 | G | N7 | AA:355 | 4.90 | 1.00 | 95.43 |  |
  |

  
 Contact(s): /OP2/  
 Coded contact list: O_ph_  
 Exclusion zone contact(s): /O5'/OP2/  
Consider K in place of MG (2 valid contact(s) for K vs. 1 for MG)  

### MG:AA:3372

|  | Res. type | Atom | Residue | Distance | Occupancy | B-factor | Closest solute to 1^st^ shell water |
| --- | --- | --- | --- | --- | --- | --- | --- |
| **Ion** | | | |  | 1.00 | 52.09 || 1 | G | O6 | AA:94 | 2.48 | 1.00 | 74.48 |
| 2 | G | O6 | AA:95 | 2.99 | 1.00 | 71.64 |  |
| 3 | G | N7 | AA:95 | 3.76 | 1.00 | 71.82 |  |
| 4 | G | N7 | AA:94 | 4.13 | 1.00 | 72.27 |  |
| 5 | G | N1 | AA:94 | 4.51 | 1.00 | 72.58 |  |
| 6 | C | N4 | AA:87 | 4.71 | 1.00 | 71.02 |  |
  |

  
 Coded contact list:   
 Exclusion zone contact(s): /O6/O6/  
Consider NA in place of MG (1 valid contact(s) for NA vs. 0 for MG)  

### MG:AA:3373

|  | Res. type | Atom | Residue | Distance | Occupancy | B-factor | Closest solute to 1^st^ shell water |
| --- | --- | --- | --- | --- | --- | --- | --- |
| **Ion** | | | |  | 1.00 | 48.32 || 1 | U | OP2 | AA:1205 | 3.18 | 1.00 | 64.78 |
| 2 | A | N9 | AA:1204 | 3.20 | 1.00 | 69.90 |  |
| 3 | A | N7 | AA:1204 | 3.28 | 1.00 | 69.91 |  |
| 4 | A | O2' | AA:1204 | 3.75 | 1.00 | 71.29 |  |
| 5 | G | OP2 | AA:1203 | 4.31 | 1.00 | 55.44 |  |
| 6 | A | O4' | AA:1204 | 4.60 | 1.00 | 66.92 |  |
| 7 | A | O3' | AA:1204 | 4.62 | 1.00 | 65.81 |  |
| 8 | C | OP2 | AA:1202 | 4.68 | 1.00 | 47.32 |  |
| 9 | C | OP1 | AA:1202 | 4.96 | 1.00 | 47.02 |  |
  |

  
 Coded contact list:   
 Exclusion zone contact(s): /N7/OP2/  
Consider K in place of MG (3 valid contact(s) for K vs. 0 for MG)  

### MG:AA:3374

|  | Res. type | Atom | Residue | Distance | Occupancy | B-factor | Closest solute to 1^st^ shell water |
| --- | --- | --- | --- | --- | --- | --- | --- |
| **Ion** | | | |  | 1.00 | 46.95 || 1 | G | O6 | AA:1285 | 2.56 | 1.00 | 51.93 |
| 2 | U | O4' | AA:1329 | 3.20 | 1.00 | 48.66 |  |
| 3 | U | O2 | AA:1329 | 3.39 | 1.00 | 42.63 |  |
| 4 | G | O2' | AA:1283 | 3.55 | 1.00 | 59.10 |  |
| 5 | U | N1 | AA:1329 | 3.61 | 1.00 | 45.29 |  |
| 6 | G | N7 | AA:1285 | 4.31 | 1.00 | 50.88 |  |
| 7 | G | O4' | AA:1283 | 4.43 | 1.00 | 58.62 |  |
| 8 | U | N3 | AA:1329 | 4.54 | 1.00 | 43.18 |  |
| 9 | U | O2' | AA:1329 | 4.55 | 1.00 | 51.63 |  |
| 10 | G | O3' | AA:1283 | 4.81 | 1.00 | 59.22 |  |
| 11 | G | N1 | AA:1285 | 4.82 | 1.00 | 50.02 |  |
| 12 | U | O5' | AA:1329 | 4.91 | 1.00 | 49.47 |  |
| 13 | U | O3' | AA:1329 | 4.94 | 1.00 | 50.49 |  |
  |

  
 Coded contact list:   
 Exclusion zone contact(s): /O2/O4'/O6/  
Consider K in place of MG (2 valid contact(s) for K vs. 0 for MG)  

### MG:AA:3375

|  | Res. type | Atom | Residue | Distance | Occupancy | B-factor | Closest solute to 1^st^ shell water |
| --- | --- | --- | --- | --- | --- | --- | --- |
| **Ion** | | | |  | 1.00 | 61.38 || 1 | C | OP1 | AA:1330 | 3.90 | 1.00 | 51.70 |
| 2 | A | OP2 | AA:1331 | 3.97 | 1.00 | 48.84 |  |
| 3 | C | OP2 | AA:1330 | 4.02 | 1.00 | 51.72 |  |
| 4 | C | OP2 | AA:1318 | 4.58 | 1.00 | 57.02 |  |
| 5 | G | N7 | AA:1319 | 4.92 | 1.00 | 62.11 |  |
| 6 | A | OP1 | AA:1331 | 4.93 | 1.00 | 47.68 |  |
| 7 | C | N4 | AA:1318 | 5.00 | 1.00 | 61.15 |  |
  |

  
 Coded contact list:   
1^st^ MG coordination shell missing (see Notes)  

### MG:AA:3376

|  | Res. type | Atom | Residue | Distance | Occupancy | B-factor | Closest solute to 1^st^ shell water |
| --- | --- | --- | --- | --- | --- | --- | --- |
| **Ion** | | | |  | 1.00 | 116.05 || 1 | U | OP2 | AA:1316 | 3.85 | 1.00 | 52.00 |
| 2 | U | OP1 | AA:1316 | 4.23 | 1.00 | 52.30 |  |
| 3 | A | OP2 | AA:1317 | 4.57 | 1.00 | 59.05 |  |
| 4 | A | OP1 | AA:1331 | 5.00 | 1.00 | 47.68 |  |
  |

  
 Coded contact list:   
1^st^ MG coordination shell missing (see Notes)  

### MG:AA:3377

|  | Res. type | Atom | Residue | Distance | Occupancy | B-factor | Closest solute to 1^st^ shell water |
| --- | --- | --- | --- | --- | --- | --- | --- |
| **Ion** | | | |  | 1.00 | 19.48 || 1 | A | OP2 | AA:2430 | 1.87 | 1.00 | 42.75 |
| 2 | U | OP1 | AA:827 | 1.91 | 1.00 | 46.41 |  |
| 3 | G | OP2 | AA:2429 | 2.30 | 1.00 | 44.99 |  |
| 4 | A | OP1 | AA:2430 | 3.27 | 1.00 | 40.59 |  |
| 5 | U | O3' | AA:826 | 3.61 | 1.00 | 46.81 |  |
| 6 | G | O3' | AA:2429 | 3.96 | 1.00 | 42.95 |  |
| 7 | U | OP2 | AA:827 | 4.14 | 1.00 | 45.47 |  |
| 8 | G | O5' | AA:2429 | 4.16 | 1.00 | 45.36 |  |
| 9 | A | O5' | AA:2430 | 4.16 | 1.00 | 40.74 |  |
| 10 | U | O5' | AA:827 | 4.37 | 1.00 | 45.76 |  |
| 11 | G | O3' | AA:2428 | 4.63 | 1.00 | 44.63 |  |
| 12 | G | OP1 | AA:2429 | 4.73 | 1.00 | 44.10 |  |
| 13 | U | O4' | AA:827 | 4.76 | 1.00 | 51.44 |  |
| 14 | GLY | O | AO:54 | 4.92 | 1.00 | 50.78 |  |
  |

  
**1^st^ shell ligand angles:**   
1-2: 167; 1-3: 87; 2-3: 89;   
  
 Contact(s): /OP1/OP2/OP2/  
 Coded contact list: 3O_ph_  
 Exclusion zone contact(s): /OP1/  

### MG:AA:3378

|  | Res. type | Atom | Residue | Distance | Occupancy | B-factor | Closest solute to 1^st^ shell water |
| --- | --- | --- | --- | --- | --- | --- | --- |
| **Ion** | | | |  | 1.00 | 50.06 || 1 | C | OP1 | AA:2427 | 1.96 | 1.00 | 47.65 |
| 2 | G | OP1 | AA:2428 | 2.01 | 1.00 | 46.82 |  |
| 3 | G | OP1 | AA:2429 | 2.29 | 1.00 | 44.10 |  |
| 4 | U | OP1 | AA:826 | 2.38 | 1.00 | 51.76 |  |
| 5 | C | O5' | AA:2427 | 3.56 | 1.00 | 48.52 |  |
| 6 | G | OP2 | AA:2428 | 3.85 | 1.00 | 47.06 |  |
| 7 | U | OP2 | AA:826 | 4.05 | 1.00 | 52.22 |  |
| 8 | G | O5' | AA:2428 | 4.08 | 1.00 | 46.19 |  |
| 9 | A | O3' | AA:2426 | 4.12 | 1.00 | 47.95 |  |
| 10 | C | OP2 | AA:2427 | 4.39 | 1.00 | 49.18 |  |
| 11 | C | O3' | AA:2427 | 4.47 | 1.00 | 47.56 |  |
| 12 | G | O3' | AA:2428 | 4.53 | 1.00 | 44.63 |  |
| 13 | U | O5' | AA:826 | 4.54 | 1.00 | 51.52 |  |
| 14 | G | OP2 | AA:2429 | 4.58 | 1.00 | 44.99 |  |
| 15 | G | O5' | AA:2429 | 4.66 | 1.00 | 45.36 |  |
| 16 | C | O3' | AA:825 | 4.79 | 1.00 | 50.21 |  |
| 17 | U | OP2 | AA:827 | 4.96 | 1.00 | 45.47 |  |
| 18 | C | O2' | AA:2258 | 4.97 | 1.00 | 48.92 |  |
  |

  
**1^st^ shell ligand angles:**   
1-2: 101; 1-3: 114; 1-4: 159; 2-3: 74; 2-4: 98; 3-4: 64;   
  
 Contact(s): /OP1/OP1/OP1/OP1/  
 Coded contact list: 4O_ph_  
Large octahedral topology discrepancy!!!  

### MG:AA:3379

|  | Res. type | Atom | Residue | Distance | Occupancy | B-factor | Closest solute to 1^st^ shell water |
| --- | --- | --- | --- | --- | --- | --- | --- |
| **Ion** | | | |  | 1.00 | 70.12 || 1 | U | O4 | AA:1727 | 2.59 | 1.00 | 102.06 |
| 2 | G | O6 | AA:1733 | 2.83 | 1.00 | 103.76 |  |
| 3 | C | N4 | AA:1734 | 3.26 | 1.00 | 96.89 |  |
| 4 | G | O6 | AA:1726 | 3.49 | 1.00 | 98.62 |  |
| 5 | U | N3 | AA:1727 | 3.81 | 1.00 | 103.01 |  |
| 6 | G | N2 | AA:1731 | 4.04 | 1.00 | 125.80 |  |
| 7 | A | OP2 | AA:1732 | 4.19 | 1.00 | 119.96 |  |
| 8 | G | N7 | AA:1733 | 4.45 | 1.00 | 105.50 |  |
| 9 | G | O6 | AA:1728 | 4.65 | 1.00 | 112.09 |  |
| 10 | C | N3 | AA:1734 | 4.66 | 1.00 | 98.95 |  |
| 11 | G | N7 | AA:1728 | 4.80 | 1.00 | 111.80 |  |
| 12 | G | N1 | AA:1726 | 4.82 | 1.00 | 99.59 |  |
  |

  
 Coded contact list:   
 Exclusion zone contact(s): /N4/O4/O6/  
Consider K in place of MG (2 valid contact(s) for K vs. 0 for MG)  

### MG:AA:3380

|  | Res. type | Atom | Residue | Distance | Occupancy | B-factor | Closest solute to 1^st^ shell water |
| --- | --- | --- | --- | --- | --- | --- | --- |
| **Ion** | | | |  | 1.00 | 25.33 || 1 | A | O2' | AA:2426 | 2.72 | 1.00 | 51.90 |
| 2 | G | OP2 | AA:2429 | 2.94 | 1.00 | 44.99 |  |
| 3 | G | OP1 | AA:2429 | 3.12 | 1.00 | 44.10 |  |
| 4 | A | O3' | AA:2426 | 3.15 | 1.00 | 47.95 |  |
| 5 | C | OP1 | AA:2427 | 3.48 | 1.00 | 47.65 |  |
| 6 | U | OP1 | AA:827 | 3.53 | 1.00 | 46.41 |  |
| 7 | U | OP2 | AA:827 | 3.80 | 1.00 | 45.47 |  |
| 8 | G | O5' | AA:2429 | 3.92 | 1.00 | 45.36 |  |
| 9 | A | OP2 | AA:2430 | 4.60 | 1.00 | 42.75 |  |
| 10 | C | O5' | AA:2427 | 4.78 | 1.00 | 48.52 |  |
| 11 | G | OP1 | AA:2428 | 4.79 | 1.00 | 46.82 |  |
| 12 | A | OP2 | AA:2425 | 4.85 | 1.00 | 43.00 |  |
| 13 | G | O3' | AA:2428 | 4.89 | 1.00 | 44.63 |  |
| 14 | U | OP1 | AA:826 | 4.92 | 1.00 | 51.76 |  |
  |

  
 Coded contact list:   
 Exclusion zone contact(s): /O2'/O3'/OP1/OP2/  
Consider K in place of MG (4 valid contact(s) for K vs. 0 for MG)  

### MG:AA:3381

|  | Res. type | Atom | Residue | Distance | Occupancy | B-factor | Closest solute to 1^st^ shell water |
| --- | --- | --- | --- | --- | --- | --- | --- |
| **Ion** | | | |  | 1.00 | 32.15 || 1 | C | OP1 | AA:249 | 2.23 | 1.00 | 36.41 |
| 2 | C | O2' | AA:2394 | 3.75 | 1.00 | 44.26 |  |
| 3 | G | OP1 | AA:250 | 3.83 | 1.00 | 43.22 |  |
| 4 | C | OP2 | AA:249 | 3.90 | 1.00 | 38.20 |  |
| 5 | C | O5' | AA:249 | 4.14 | 1.00 | 37.47 |  |
| 6 | A | OP1 | AA:2432 | 4.50 | 1.00 | 42.16 |  |
| 7 | G | O3' | AA:248 | 4.69 | 1.00 | 36.89 |  |
| 8 | G | O5' | AA:250 | 4.71 | 1.00 | 43.46 |  |
| 9 | G | O4' | AA:250 | 4.75 | 1.00 | 47.94 |  |
| 10 | U | OP1 | AA:2431 | 4.79 | 1.00 | 38.34 |  |
| 11 | C | O2 | AA:2394 | 4.80 | 1.00 | 45.16 |  |
| 12 | A | N3 | AA:2393 | 4.80 | 1.00 | 52.65 |  |
| 13 | C | O4' | AA:2394 | 4.87 | 1.00 | 43.37 |  |
  |

  
 Contact(s): /OP1/  
 Coded contact list: O_ph_  

### MG:AA:3382

|  | Res. type | Atom | Residue | Distance | Occupancy | B-factor | Closest solute to 1^st^ shell water |
| --- | --- | --- | --- | --- | --- | --- | --- |
| **Ion** | | | |  | 1.00 | 83.43 || 1 | G | O6 | AA:1725 | 3.73 | 1.00 | 99.91 |
| 2 | G | N7 | AA:1725 | 3.83 | 1.00 | 103.28 |  |
| 3 | G | N7 | AA:1718 | 4.89 | 1.00 | 108.68 |  |
| 4 | G | O6 | AA:1718 | 4.97 | 1.00 | 106.21 |  |
  |

  
 Coded contact list:   
1^st^ MG coordination shell missing (see Notes)  

### MG:AA:3383

|  | Res. type | Atom | Residue | Distance | Occupancy | B-factor | Closest solute to 1^st^ shell water |
| --- | --- | --- | --- | --- | --- | --- | --- |
| **Ion** | | | |  | 1.00 | 24.00 || 1 | U | OP2 | AA:2431 | 2.09 | 1.00 | 38.20 |
| 2 | A | OP2 | AA:2432 | 2.17 | 1.00 | 40.23 |  |
| 3 | U | OP1 | AA:2431 | 4.02 | 1.00 | 38.34 |  |
| 4 | A | OP1 | AA:2432 | 4.16 | 1.00 | 42.16 |  |
| 5 | U | O5' | AA:2431 | 4.23 | 1.00 | 37.49 |  |
| 6 | A | O5' | AA:2432 | 4.37 | 1.00 | 42.20 |  |
| 7 | A | O3' | AA:2430 | 4.55 | 1.00 | 38.65 |  |
| 8 | A | OP2 | AA:2433 | 4.64 | 1.00 | 44.64 |  |
| 9 | U | O3' | AA:2431 | 4.64 | 1.00 | 40.57 |  |
| 10 | A | N1 | AA:2393 | 4.93 | 1.00 | 54.40 |  |
  |

  
**1^st^ shell ligand angles:**   
1-2: 80;   
  
 Contact(s): /OP2/OP2/  
 Coded contact list: 2O_ph_  

### MG:AA:3384

|  | Res. type | Atom | Residue | Distance | Occupancy | B-factor | Closest solute to 1^st^ shell water |
| --- | --- | --- | --- | --- | --- | --- | --- |
| **Ion** | | | |  | 1.00 | 47.88 || 1 | G | O6 | AA:1846 | 3.94 | 1.00 | 59.25 |
| 2 | G | O6 | AA:1845 | 4.18 | 1.00 | 56.97 |  |
| 3 | G | N7 | AA:1845 | 4.47 | 1.00 | 56.68 |  |
| 4 | G | N7 | AA:1846 | 4.53 | 1.00 | 63.69 |  |
| 5 | C | OP2 | AA:1892 | 4.74 | 1.00 | 53.53 |  |
| 6 | C | N4 | AA:1844 | 4.88 | 1.00 | 45.39 |  |
  |

  
 Coded contact list:   
1^st^ MG coordination shell missing (see Notes)  

### MG:AA:3385

|  | Res. type | Atom | Residue | Distance | Occupancy | B-factor | Closest solute to 1^st^ shell water |
| --- | --- | --- | --- | --- | --- | --- | --- |
| **Ion** | | | |  | 1.00 | 34.82 || 1 | A | OP2 | AA:1890 | 2.51 | 1.00 | 58.84 |
| 2 | G | O6 | AA:1891 | 2.93 | 1.00 | 60.70 |  |
| 3 | G | N7 | AA:1891 | 3.50 | 1.00 | 56.31 |  |
| 4 | A | N7 | AA:1890 | 3.81 | 1.00 | 55.94 |  |
| 5 | U | O4 | AA:1851 | 4.21 | 1.00 | 63.00 |  |
| 6 | C | N4 | AA:1892 | 4.35 | 1.00 | 47.90 |  |
| 7 | A | O5' | AA:1890 | 4.35 | 1.00 | 57.46 |  |
| 8 | C | O2 | AA:1852 | 4.58 | 1.00 | 71.24 |  |
| 9 | C | N3 | AA:1852 | 4.76 | 1.00 | 71.29 |  |
| 10 | A | O3' | AA:1889 | 4.79 | 1.00 | 60.25 |  |
| 11 | G | O6 | AA:1850 | 4.92 | 1.00 | 54.54 |  |
| 12 | A | OP1 | AA:1890 | 4.96 | 1.00 | 58.70 |  |
| 13 | U | N3 | AA:1851 | 4.98 | 1.00 | 61.70 |  |
  |

  
 Coded contact list:   
 Exclusion zone contact(s): /O6/OP2/  
Consider NA in place of MG (1 valid contact(s) for NA vs. 0 for MG)  

### MG:AA:3386

|  | Res. type | Atom | Residue | Distance | Occupancy | B-factor | Closest solute to 1^st^ shell water |
| --- | --- | --- | --- | --- | --- | --- | --- |
| **Ion** | | | |  | 1.00 | 51.73 || 1 | G | OP2 | AA:2070 | 1.90 | 1.00 | 41.38 |
| 2 | G | OP2 | AA:2069 | 2.87 | 1.00 | 42.46 |  |
| 3 | U | OP2 | AA:2244 | 3.86 | 1.00 | 38.54 |  |
| 4 | G | O5' | AA:2070 | 3.96 | 1.00 | 38.34 |  |
| 5 | G | O3' | AA:2069 | 4.27 | 1.00 | 39.18 |  |
| 6 | G | OP1 | AA:2070 | 4.35 | 1.00 | 40.38 |  |
| 7 | U | OP1 | AA:2244 | 4.53 | 1.00 | 39.66 |  |
  |

  
 Contact(s): /OP2/  
 Coded contact list: O_ph_  
 Exclusion zone contact(s): /OP2/  

### MG:AA:3387

|  | Res. type | Atom | Residue | Distance | Occupancy | B-factor | Closest solute to 1^st^ shell water |
| --- | --- | --- | --- | --- | --- | --- | --- |
| **Ion** | | | |  | 1.00 | 40.29 || 1 | U | OP1 | AA:380 | 2.51 | 1.00 | 51.11 |
| 2 | G | O3' | AA:379 | 3.49 | 1.00 | 50.43 |  |
| 3 | ILE | N | AZ:18 | 3.70 | 1.00 | 45.43 |  |
| 4 | ILE | O | AZ:18 | 3.84 | 1.00 | 44.98 |  |
| 5 | SER | OG | AZ:17 | 4.06 | 1.00 | 43.25 |  |
| 6 | U | O5' | AA:380 | 4.17 | 1.00 | 52.14 |  |
| 7 | ASN | O | AZ:16 | 4.22 | 1.00 | 45.85 |  |
| 8 | G | O2' | AA:379 | 4.40 | 1.00 | 48.51 |  |
| 9 | SER | N | AZ:17 | 4.71 | 1.00 | 43.91 |  |
| 10 | GLN | NE2 | AZ:19 | 4.76 | 1.00 | 46.12 |  |
| 11 | U | OP2 | AA:2233 | 4.81 | 1.00 | 50.88 |  |
| 12 | U | OP2 | AA:380 | 4.89 | 1.00 | 49.29 |  |
| 13 | SER | O | AZ:17 | 4.95 | 1.00 | 46.69 |  |
  |

  
 Coded contact list:   
 Exclusion zone contact(s): /OP1/  
Consider NA in place of MG (1 valid contact(s) for NA vs. 0 for MG)  

### MG:AA:3388

|  | Res. type | Atom | Residue | Distance | Occupancy | B-factor | Closest solute to 1^st^ shell water |
| --- | --- | --- | --- | --- | --- | --- | --- |
| **Ion** | | | |  | 1.00 | 46.85 || 1 | U | O4 | AA:2431 | 2.98 | 1.00 | 43.21 |
| 2 | A | OP2 | AA:2434 | 3.36 | 1.00 | 47.26 |  |
| 3 | A | O5' | AA:2434 | 3.75 | 1.00 | 46.26 |  |
| 4 | U | N3 | AA:2431 | 4.29 | 1.00 | 38.50 |  |
| 5 | A | OP1 | AA:2434 | 4.31 | 1.00 | 47.36 |  |
| 6 | A | OP2 | AA:2435 | 4.86 | 1.00 | 45.11 |  |
| 7 | U | O4 | AA:827 | 4.93 | 1.00 | 53.63 |  |
  |

  
 Coded contact list:   
 Exclusion zone contact(s): /O4/OP2/  
Consider K in place of MG (2 valid contact(s) for K vs. 0 for MG)  

### MG:AA:3389

|  | Res. type | Atom | Residue | Distance | Occupancy | B-factor | Closest solute to 1^st^ shell water |
| --- | --- | --- | --- | --- | --- | --- | --- |
| **Ion** | | | |  | 1.00 | 67.53 || 1 | U | OP1 | AA:2244 | 2.57 | 1.00 | 39.66 |
| 2 | A | O3' | AA:197 | 2.80 | 1.00 | 43.21 |  |
| 3 | C | OP1 | AA:198 | 2.80 | 1.00 | 42.76 |  |
| 4 | G | OP2 | AA:2070 | 3.46 | 1.00 | 41.38 |  |
| 5 | G | OP2 | AA:2069 | 3.68 | 1.00 | 42.46 |  |
| 6 | G | OP1 | AA:2070 | 3.73 | 1.00 | 40.38 |  |
| 7 | A | O2' | AA:197 | 3.76 | 1.00 | 42.87 |  |
| 8 | U | OP2 | AA:2244 | 3.89 | 1.00 | 38.54 |  |
| 9 | C | O5' | AA:198 | 4.01 | 1.00 | 41.90 |  |
| 10 | G | OP1 | AA:2069 | 4.03 | 1.00 | 43.48 |  |
| 11 | U | O3' | AA:2243 | 4.15 | 1.00 | 38.65 |  |
| 12 | C | OP2 | AA:198 | 4.75 | 1.00 | 41.69 |  |
| 13 | U | O5' | AA:2244 | 4.97 | 1.00 | 40.27 |  |
| 14 | G | O3' | AA:2069 | 4.98 | 1.00 | 39.18 |  |
  |

  
 Coded contact list:   
 Exclusion zone contact(s): /O3'/OP1/OP1/  
Consider K in place of MG (2 valid contact(s) for K vs. 0 for MG)  

### MG:AA:3390

|  | Res. type | Atom | Residue | Distance | Occupancy | B-factor | Closest solute to 1^st^ shell water |
| --- | --- | --- | --- | --- | --- | --- | --- |
| **Ion** | | | |  | 1.00 | 46.61 || 1 | U | O4 | AA:2079 | 2.61 | 1.00 | 31.45 |
| 2 | G | O6 | AA:2080 | 3.72 | 1.00 | 40.63 |  |
| 3 | G | O6 | AA:2238 | 3.83 | 1.00 | 37.17 |  |
| 4 | G | N7 | AA:2080 | 4.27 | 1.00 | 40.78 |  |
| 5 | C | N4 | AA:2078 | 4.52 | 1.00 | 31.55 |  |
| 6 | G | N1 | AA:2238 | 4.57 | 1.00 | 37.89 |  |
| 7 | G | N7 | AA:2238 | 4.58 | 1.00 | 38.77 |  |
| 8 | U | N3 | AA:2079 | 4.60 | 1.00 | 33.55 |  |
| 9 | A | N6 | AA:2241 | 4.63 | 1.00 | 33.60 |  |
  |

  
 Coded contact list:   
 Exclusion zone contact(s): /O4/  
Consider K in place of MG (1 valid contact(s) for K vs. 0 for MG)  

### MG:AA:3391

|  | Res. type | Atom | Residue | Distance | Occupancy | B-factor | Closest solute to 1^st^ shell water |
| --- | --- | --- | --- | --- | --- | --- | --- |
| **Ion** | | | |  | 1.00 | 39.04 || 1 | G | OP1 | AA:248 | 2.20 | 1.00 | 38.59 |
| 2 | G | OP2 | AA:248 | 4.12 | 1.00 | 39.58 |  |
| 3 | G | O3' | AA:247 | 4.19 | 1.00 | 38.27 |  |
| 4 | G | O2' | AA:386 | 4.21 | 1.00 | 56.29 |  |
| 5 | LYS | O | AZ:32 | 4.47 | 1.00 | 54.71 |  |
| 6 | G | O5' | AA:248 | 4.67 | 1.00 | 37.54 |  |
| 7 | G | N3 | AA:386 | 4.90 | 1.00 | 48.85 |  |
  |

  
 Contact(s): /OP1/  
 Coded contact list: O_ph_  

### MG:AA:3392

|  | Res. type | Atom | Residue | Distance | Occupancy | B-factor | Closest solute to 1^st^ shell water |
| --- | --- | --- | --- | --- | --- | --- | --- |
| **Ion** | | | |  | 1.00 | 63.14 || 1 | U | OP1 | AA:193 | 2.45 | 1.00 | 38.49 |
| 2 | G | OP1 | AA:2070 | 3.39 | 1.00 | 40.38 |  |
| 3 | G | O3' | AA:2069 | 3.91 | 1.00 | 39.18 |  |
| 4 | U | OP2 | AA:193 | 4.07 | 1.00 | 39.21 |  |
| 5 | U | O5' | AA:193 | 4.31 | 1.00 | 38.38 |  |
| 6 | G | OP2 | AA:194 | 4.70 | 1.00 | 42.62 |  |
| 7 | G | O2' | AA:2069 | 4.80 | 1.00 | 39.15 |  |
| 8 | C | O3' | AA:192 | 4.93 | 1.00 | 38.04 |  |
  |

  
 Coded contact list:   
 Exclusion zone contact(s): /OP1/OP1/  
Consider NA in place of MG (1 valid contact(s) for NA vs. 0 for MG)  

### MG:AA:3393

|  | Res. type | Atom | Residue | Distance | Occupancy | B-factor | Closest solute to 1^st^ shell water |
| --- | --- | --- | --- | --- | --- | --- | --- |
| **Ion** | | | |  | 1.00 | 63.46 || 1 | G | O2' | AA:388 | 2.92 | 1.00 | 50.95 |
| 2 | A | N6 | AA:390 | 3.41 | 1.00 | 44.94 |  |
| 3 | G | OP2 | AA:247 | 3.86 | 1.00 | 37.96 |  |
| 4 | G | O6 | AA:389 | 4.17 | 1.00 | 49.89 |  |
| 5 | C | N4 | AA:249 | 4.24 | 1.00 | 43.84 |  |
| 6 | G | O4' | AA:388 | 4.52 | 1.00 | 50.43 |  |
| 7 | C | OP1 | AA:246 | 4.56 | 1.00 | 42.33 |  |
| 8 | G | N7 | AA:389 | 4.87 | 1.00 | 49.45 |  |
  |

  
 Coded contact list:   
 Exclusion zone contact(s): /O2'/  
Consider K in place of MG (1 valid contact(s) for K vs. 0 for MG)  

### MG:AA:3394

|  | Res. type | Atom | Residue | Distance | Occupancy | B-factor | Closest solute to 1^st^ shell water |
| --- | --- | --- | --- | --- | --- | --- | --- |
| **Ion** | | | |  | 1.00 | 45.39 || 1 | C | O2 | AA:2084 | 2.50 | 1.00 | 40.16 |
| 2 | C | O2' | AA:2084 | 2.78 | 1.00 | 37.62 |  |
| 3 | C | O4' | AA:2085 | 3.29 | 1.00 | 37.00 |  |
| 4 | G | N2 | AA:2235 | 3.85 | 1.00 | 41.48 |  |
| 5 | C | N1 | AA:2084 | 3.98 | 1.00 | 37.39 |  |
| 6 | C | O2 | AA:2236 | 4.48 | 1.00 | 37.18 |  |
| 7 | C | N1 | AA:2085 | 4.55 | 1.00 | 38.12 |  |
| 8 | C | O4' | AA:2084 | 4.59 | 1.00 | 39.55 |  |
| 9 | C | N3 | AA:2084 | 4.74 | 1.00 | 37.66 |  |
| 10 | G | N2 | AA:2083 | 4.74 | 1.00 | 38.36 |  |
  |

  
 Coded contact list:   
 Exclusion zone contact(s): /O2/O2'/O4'/  
Consider K in place of MG (2 valid contact(s) for K vs. 0 for MG)  

### MG:AA:3395

|  | Res. type | Atom | Residue | Distance | Occupancy | B-factor | Closest solute to 1^st^ shell water |
| --- | --- | --- | --- | --- | --- | --- | --- |
| **Ion** | | | |  | 1.00 | 88.32 || 1 | G | O6 | AA:2235 | 3.99 | 1.00 | 41.74 |
| 2 | G | O6 | AA:2234 | 4.09 | 1.00 | 45.54 |  |
| 3 | G | N7 | AA:2235 | 4.18 | 1.00 | 42.19 |  |
| 4 | G | N7 | AA:2234 | 4.25 | 1.00 | 48.16 |  |
  |

  
 Coded contact list:   
1^st^ MG coordination shell missing (see Notes)  

### MG:AA:3396

|  | Res. type | Atom | Residue | Distance | Occupancy | B-factor | Closest solute to 1^st^ shell water |
| --- | --- | --- | --- | --- | --- | --- | --- |
| **Ion** | | | |  | 1.00 | 58.83 || 1 | A | OP1 | AA:501 | 2.94 | 1.00 | 60.84 |
| 2 | G | O3' | AA:500 | 3.10 | 1.00 | 62.54 |  |
| 3 | A | N1 | AA:477 | 3.14 | 1.00 | 47.37 |  |
| 4 | A | N6 | AA:477 | 3.20 | 1.00 | 46.52 |  |
| 5 | G | N1 | AA:329 | 3.41 | 1.00 | 55.03 |  |
| 6 | G | N2 | AA:329 | 3.72 | 1.00 | 56.96 |  |
| 7 | A | OP2 | AA:501 | 3.77 | 1.00 | 62.10 |  |
| 8 | G | O2' | AA:308 | 4.17 | 1.00 | 58.18 |  |
| 9 | G | O6 | AA:329 | 4.28 | 1.00 | 54.70 |  |
| 10 | A | N1 | AA:478 | 4.44 | 1.00 | 51.30 |  |
| 11 | A | O5' | AA:501 | 4.88 | 1.00 | 60.46 |  |
  |

  
 Coded contact list:   
 Exclusion zone contact(s): /N1_A_/N6/O3'/OP1/  
Consider K in place of MG (4 valid contact(s) for K vs. 0 for MG)  

### MG:AA:3397

|  | Res. type | Atom | Residue | Distance | Occupancy | B-factor | Closest solute to 1^st^ shell water |
| --- | --- | --- | --- | --- | --- | --- | --- |
| **Ion** | | | |  | 1.00 | 56.32 || 1 | G | O6 | AA:1380 | 2.57 | 1.00 | 49.61 |
| 2 | G | O6 | AA:1381 | 3.17 | 1.00 | 46.15 |  |
| 3 | G | N7 | AA:1380 | 4.02 | 1.00 | 47.60 |  |
| 4 | G | O6 | AA:1382 | 4.11 | 1.00 | 60.26 |  |
| 5 | C | N4 | AA:1351 | 4.11 | 1.00 | 46.24 |  |
| 6 | A | O2' | AA:1378 | 4.14 | 1.00 | 55.09 |  |
| 7 | C | N4 | AA:1350 | 4.20 | 1.00 | 49.54 |  |
| 8 | G | OP1 | AA:1347 | 4.29 | 1.00 | 47.97 |  |
| 9 | G | N1 | AA:1380 | 4.56 | 1.00 | 47.42 |  |
| 10 | A | O5' | AA:1379 | 4.72 | 1.00 | 60.86 |  |
| 11 | G | N7 | AA:1381 | 4.73 | 1.00 | 46.36 |  |
| 12 | G | OP1 | AA:1348 | 4.74 | 1.00 | 48.33 |  |
  |

  
 Coded contact list:   
 Exclusion zone contact(s): /O6/O6/  
Consider NA in place of MG (1 valid contact(s) for NA vs. 0 for MG)  

### MG:AA:3398

|  | Res. type | Atom | Residue | Distance | Occupancy | B-factor | Closest solute to 1^st^ shell water |
| --- | --- | --- | --- | --- | --- | --- | --- |
| **Ion** | | | |  | 1.00 | 30.59 || 1 | G | N7 | AA:1381 | 3.76 | 1.00 | 46.36 |
| 2 | G | O6 | AA:1382 | 3.86 | 1.00 | 60.26 |  |
| 3 | G | OP2 | AA:1381 | 3.90 | 1.00 | 47.97 |  |
| 4 | A | O3' | AA:1379 | 4.15 | 1.00 | 59.83 |  |
| 5 | G | N7 | AA:1382 | 4.25 | 1.00 | 58.42 |  |
| 6 | G | OP2 | AA:1380 | 4.33 | 1.00 | 62.03 |  |
| 7 | G | O5' | AA:1380 | 4.37 | 1.00 | 58.09 |  |
| 8 | G | N7 | AA:1380 | 4.51 | 1.00 | 47.60 |  |
| 9 | G | O6 | AA:1381 | 4.61 | 1.00 | 46.15 |  |
| 10 | G | OP1 | AA:1347 | 4.61 | 1.00 | 47.97 |  |
| 11 | C | N4 | AA:1383 | 4.94 | 1.00 | 71.85 |  |
  |

  
 Coded contact list:   
1^st^ MG coordination shell missing (see Notes)  

### MG:AA:3399

|  | Res. type | Atom | Residue | Distance | Occupancy | B-factor | Closest solute to 1^st^ shell water |
| --- | --- | --- | --- | --- | --- | --- | --- |
| **Ion** | | | |  | 1.00 | 75.77 || 1 | G | N7 | AA:439 | 2.49 | 1.00 | 70.02 |
| 2 | G | O6 | AA:439 | 3.40 | 1.00 | 68.12 |  |
| 3 | G | N7 | AA:438 | 3.97 | 1.00 | 74.05 |  |
| 4 | G | O6 | AA:438 | 4.21 | 1.00 | 74.27 |  |
| 5 | G | N9 | AA:439 | 4.67 | 1.00 | 72.01 |  |
| 6 | G | O6 | AA:440 | 4.68 | 1.00 | 59.81 |  |
| 7 | G | OP2 | AA:439 | 4.72 | 1.00 | 78.05 |  |
| 8 | G | N1 | AA:438 | 4.93 | 1.00 | 74.08 |  |
  |

  
 Coded contact list:   
 Exclusion zone contact(s): /N7/O6/  
Consider NA in place of MG (1 valid contact(s) for NA vs. 0 for MG)  

### MG:AA:3400

|  | Res. type | Atom | Residue | Distance | Occupancy | B-factor | Closest solute to 1^st^ shell water |
| --- | --- | --- | --- | --- | --- | --- | --- |
| **Ion** | | | |  | 1.00 | 58.33 || 1 | G | OP2 | AA:2782 | 2.03 | 1.00 | 77.71 |
| 2 | G | N7 | AA:2782 | 3.85 | 1.00 | 76.46 |  |
| 3 | G | O5' | AA:2782 | 3.88 | 1.00 | 77.37 |  |
| 4 | G | N7 | AA:2783 | 4.03 | 1.00 | 74.51 |  |
| 5 | G | OP2 | AA:2783 | 4.35 | 1.00 | 80.22 |  |
| 6 | A | O3' | AA:2781 | 4.39 | 1.00 | 76.66 |  |
| 7 | G | OP1 | AA:2782 | 4.42 | 1.00 | 77.32 |  |
| 8 | G | O6 | AA:2783 | 4.72 | 1.00 | 69.99 |  |
| 9 | G | O6 | AA:2782 | 4.99 | 1.00 | 74.66 |  |
  |

  
 Contact(s): /OP2/  
 Coded contact list: O_ph_  

### MG:AA:3401

|  | Res. type | Atom | Residue | Distance | Occupancy | B-factor | Closest solute to 1^st^ shell water |
| --- | --- | --- | --- | --- | --- | --- | --- |
| **Ion** | | | |  | 1.00 | 46.18 || 1 | G | N7 | AA:1855 | 3.73 | 1.00 | 67.95 |
| 2 | G | O6 | AA:1857 | 3.90 | 1.00 | 92.68 |  |
| 3 | G | O6 | AA:1855 | 3.98 | 1.00 | 68.56 |  |
| 4 | G | O6 | AA:1856 | 4.13 | 1.00 | 78.82 |  |
| 5 | G | N1 | AA:1857 | 4.55 | 1.00 | 90.39 |  |
| 6 | A | OP2 | AA:1854 | 4.66 | 1.00 | 57.38 |  |
| 7 | G | N7 | AA:1856 | 4.69 | 1.00 | 79.58 |  |
| 8 | G | OP2 | AA:1855 | 4.93 | 1.00 | 62.89 |  |
  |

  
 Coded contact list:   
1^st^ MG coordination shell missing (see Notes)  

### MG:AA:3402

|  | Res. type | Atom | Residue | Distance | Occupancy | B-factor | Closest solute to 1^st^ shell water |
| --- | --- | --- | --- | --- | --- | --- | --- |
| **Ion** | | | |  | 1.00 | 66.57 || 1 | G | O6 | AA:1857 | 2.51 | 1.00 | 92.68 |
| 2 | G | O6 | AA:1858 | 2.55 | 1.00 | 97.24 |  |
| 3 | G | N7 | AA:1857 | 2.92 | 1.00 | 90.67 |  |
| 4 | G | N1 | AA:1858 | 3.84 | 1.00 | 97.71 |  |
| 5 | G | OP2 | AA:1856 | 4.46 | 1.00 | 67.00 |  |
| 6 | G | N1 | AA:1857 | 4.55 | 1.00 | 90.39 |  |
| 7 | G | N7 | AA:1858 | 4.72 | 1.00 | 97.83 |  |
  |

  
 Coded contact list:   
 Exclusion zone contact(s): /N7/O6/O6/  
Consider NA in place of MG (2 valid contact(s) for NA vs. 0 for MG)  

### MG:AA:3403

|  | Res. type | Atom | Residue | Distance | Occupancy | B-factor | Closest solute to 1^st^ shell water |
| --- | --- | --- | --- | --- | --- | --- | --- |
| **Ion** | | | |  | 1.00 | 60.99 || 1 | G | O6 | AA:1858 | 3.16 | 1.00 | 97.24 |
| 2 | G | N7 | AA:1858 | 3.54 | 1.00 | 97.83 |  |
| 3 | G | N7 | AA:1857 | 4.00 | 1.00 | 90.67 |  |
| 4 | G | OP2 | AA:1857 | 4.24 | 1.00 | 78.58 |  |
| 5 | G | O6 | AA:1883 | 4.50 | 1.00 | 82.01 |  |
  |

  
 Coded contact list:   
 Exclusion zone contact(s): /O6/  
Consider K in place of MG (1 valid contact(s) for K vs. 0 for MG)  

### MG:AA:3404

|  | Res. type | Atom | Residue | Distance | Occupancy | B-factor | Closest solute to 1^st^ shell water |
| --- | --- | --- | --- | --- | --- | --- | --- |
| **Ion** | | | |  | 1.00 | 77.61 || 1 | G | OP1 | AA:1861 | 3.82 | 1.00 | 85.95 |
| 2 | G | O3' | AA:1860 | 4.54 | 1.00 | 86.05 |  |
| 3 | G | OP2 | AA:1861 | 4.96 | 1.00 | 85.65 |  |
  |

  
 Coded contact list:   
1^st^ MG coordination shell missing (see Notes)  

### MG:AA:3406

|  | Res. type | Atom | Residue | Distance | Occupancy | B-factor | Closest solute to 1^st^ shell water |
| --- | --- | --- | --- | --- | --- | --- | --- |
| **Ion** | | | |  | 1.00 | 32.52 || 1 | C | N4 | AA:1790 | 2.66 | 1.00 | 26.00 |
| 2 | A | OP2 | AA:1789 | 2.81 | 1.00 | 36.56 |  |
| 3 | A | N7 | AA:1789 | 3.40 | 1.00 | 35.60 |  |
| 4 | A | O5' | AA:1789 | 3.91 | 1.00 | 35.95 |  |
| 5 | G | O6 | AA:1828 | 3.97 | 1.00 | 39.46 |  |
| 6 | G | N1 | AA:1828 | 4.16 | 1.00 | 34.80 |  |
| 7 | G | N7 | AA:1828 | 4.47 | 1.00 | 35.41 |  |
| 8 | C | OP2 | AA:1788 | 4.52 | 1.00 | 35.89 |  |
| 9 | ARG | NE | AD:222 | 4.57 | 1.00 | 29.07 |  |
| 10 | A | N9 | AA:1789 | 4.63 | 1.00 | 36.03 |  |
| 11 | C | N3 | AA:1790 | 4.67 | 1.00 | 26.37 |  |
| 12 | C | O5' | AA:1788 | 4.82 | 1.00 | 37.42 |  |
| 13 | G | N3 | AA:1828 | 4.91 | 1.00 | 32.75 |  |
| 14 | C | O3' | AA:1788 | 4.94 | 1.00 | 36.32 |  |
| 15 | A | OP1 | AA:1789 | 4.97 | 1.00 | 36.53 |  |
| 16 | ARG | NH2 | AD:222 | 4.99 | 1.00 | 30.28 |  |
  |

  
 Coded contact list:   
 Exclusion zone contact(s): /N4/OP2/  
Consider K in place of MG (2 valid contact(s) for K vs. 0 for MG)  

### MG:AA:3408

|  | Res. type | Atom | Residue | Distance | Occupancy | B-factor | Closest solute to 1^st^ shell water |
| --- | --- | --- | --- | --- | --- | --- | --- |
| **Ion** | | | |  | 1.00 | 61.54 || 1 | C | OP2 | AA:2710 | 2.77 | 1.00 | 49.67 |
| 2 | C | O5' | AA:2710 | 3.85 | 1.00 | 49.74 |  |
| 3 | G | O5' | AA:2709 | 4.18 | 1.00 | 48.80 |  |
| 4 | G | OP1 | AA:2714 | 4.19 | 1.00 | 56.15 |  |
| 5 | G | OP2 | AA:2709 | 4.41 | 1.00 | 51.13 |  |
| 6 | G | O3' | AA:2709 | 4.42 | 1.00 | 48.97 |  |
| 7 | C | N4 | AA:2710 | 4.55 | 1.00 | 51.62 |  |
| 8 | C | N1 | AA:2710 | 4.62 | 1.00 | 49.40 |  |
| 9 | G | N7 | AA:2714 | 4.76 | 1.00 | 62.83 |  |
| 10 | A | N7 | AA:2711 | 4.82 | 1.00 | 56.04 |  |
  |

  
 Coded contact list:   
 Exclusion zone contact(s): /OP2/  
Consider K in place of MG (1 valid contact(s) for K vs. 0 for MG)  

### MG:AA:3409

|  | Res. type | Atom | Residue | Distance | Occupancy | B-factor | Closest solute to 1^st^ shell water |
| --- | --- | --- | --- | --- | --- | --- | --- |
| **Ion** | | | |  | 1.00 | 56.91 || 1 | G | OP1 | AA:2415 | 3.87 | 1.00 | 61.20 |
| 2 | G | N7 | AA:2396 | 4.02 | 1.00 | 61.32 |  |
| 3 | G | OP2 | AA:2396 | 4.05 | 1.00 | 55.23 |  |
| 4 | G | OP2 | AA:2415 | 4.63 | 1.00 | 61.24 |  |
| 5 | C | OP2 | AA:2395 | 4.76 | 1.00 | 47.98 |  |
  |

  
 Coded contact list:   
1^st^ MG coordination shell missing (see Notes)  

### MG:AA:3410

|  | Res. type | Atom | Residue | Distance | Occupancy | B-factor | Closest solute to 1^st^ shell water |
| --- | --- | --- | --- | --- | --- | --- | --- |
| **Ion** | | | |  | 1.00 | 84.65 || 1 | U | O3' | AA:2113 | 2.98 | 1.00 | 194.57 |
| 2 | U | O2' | AA:2113 | 3.07 | 1.00 | 192.25 |  |
| 3 | A | OP1 | AA:2114 | 3.57 | 1.00 | 195.58 |  |
| 4 | A | OP2 | AA:2114 | 4.20 | 1.00 | 195.64 |  |
  |

  
 Coded contact list:   
 Exclusion zone contact(s): /O2'/O3'/  
Consider K in place of MG (2 valid contact(s) for K vs. 0 for MG)  

### MG:AA:3411

|  | Res. type | Atom | Residue | Distance | Occupancy | B-factor | Closest solute to 1^st^ shell water |
| --- | --- | --- | --- | --- | --- | --- | --- |
| **Ion** | | | |  | 1.00 | 73.75 || 1 | C | OP1 | AA:62 | 2.03 | 1.00 | 62.81 |
| 2 | U | OP2 | AA:63 | 3.68 | 1.00 | 68.31 |  |
| 3 | C | O5' | AA:62 | 4.13 | 1.00 | 61.88 |  |
| 4 | C | OP2 | AA:62 | 4.19 | 1.00 | 63.17 |  |
| 5 | G | O3' | AA:61 | 4.48 | 1.00 | 61.89 |  |
| 6 | G | OP1 | AA:60 | 4.83 | 1.00 | 59.08 |  |
  |

  
 Contact(s): /OP1/  
 Coded contact list: O_ph_  

### MG:AA:3412

|  | Res. type | Atom | Residue | Distance | Occupancy | B-factor | Closest solute to 1^st^ shell water |
| --- | --- | --- | --- | --- | --- | --- | --- |
| **Ion** | | | |  | 1.00 | 30.72 || 1 | A | OP2 | AA:2392 | 3.49 | 1.00 | 57.17 |
| 2 | HIS | O | A8:31 | 3.65 | 1.00 | 68.93 |  |
| 3 | A | OP1 | AA:2392 | 3.67 | 1.00 | 58.24 |  |
| 4 | LEU | O | A8:32 | 3.67 | 1.00 | 75.34 |  |
| 5 | ASN | OD1 | A8:33 | 4.38 | 1.00 | 81.08 |  |
| 6 | ASN | ND2 | A8:33 | 4.42 | 1.00 | 79.53 |  |
| 7 | A | N7 | AA:2422 | 4.47 | 1.00 | 54.75 |  |
| 8 | A | N6 | AA:2422 | 4.47 | 1.00 | 54.29 |  |
| 9 | G | O3' | AA:2391 | 4.85 | 1.00 | 57.99 |  |
| 10 | LEU | N | A8:32 | 4.91 | 1.00 | 72.62 |  |
  |

  
 Coded contact list:   
1^st^ MG coordination shell missing (see Notes)  

### MG:AA:3414

|  | Res. type | Atom | Residue | Distance | Occupancy | B-factor | Closest solute to 1^st^ shell water |
| --- | --- | --- | --- | --- | --- | --- | --- |
| **Ion** | | | |  | 1.00 | 56.43 || 1 | G | N7 | AA:2353 | 3.08 | 1.00 | 67.04 |
| 2 | A | OP1 | AA:2352 | 3.69 | 1.00 | 71.59 |  |
| 3 | A | OP2 | AA:2352 | 3.70 | 1.00 | 70.05 |  |
| 4 | G | O6 | AA:2353 | 4.13 | 1.00 | 64.25 |  |
| 5 | G | N7 | AA:2354 | 4.25 | 1.00 | 61.71 |  |
| 6 | A | O5' | AA:2352 | 4.65 | 1.00 | 70.72 |  |
| 7 | G | OP2 | AA:2353 | 4.73 | 1.00 | 68.97 |  |
  |

  
 Coded contact list:   
 Exclusion zone contact(s): /N7/  
Consider K in place of MG (1 valid contact(s) for K vs. 0 for MG)  

### MG:AA:3415

|  | Res. type | Atom | Residue | Distance | Occupancy | B-factor | Closest solute to 1^st^ shell water |
| --- | --- | --- | --- | --- | --- | --- | --- |
| **Ion** | | | |  | 1.00 | 63.16 || 1 | A | OP2 | AA:1859 | 2.64 | 1.00 | 95.53 |
| 2 | G | N7 | AA:1860 | 2.77 | 1.00 | 84.75 |  |
| 3 | G | OP2 | AA:1860 | 3.41 | 1.00 | 88.01 |  |
| 4 | A | OP1 | AA:1859 | 3.47 | 1.00 | 95.23 |  |
| 5 | A | O5' | AA:1859 | 3.61 | 1.00 | 94.12 |  |
| 6 | G | O5' | AA:1860 | 4.58 | 1.00 | 87.08 |  |
| 7 | G | N9 | AA:1860 | 4.59 | 1.00 | 85.26 |  |
| 8 | G | O6 | AA:1860 | 4.63 | 1.00 | 85.00 |  |
| 9 | G | O3' | AA:1858 | 4.84 | 1.00 | 95.33 |  |
  |

  
 Coded contact list:   
 Exclusion zone contact(s): /N7/OP2/  
Consider K in place of MG (2 valid contact(s) for K vs. 0 for MG)  

### MG:AA:3416

|  | Res. type | Atom | Residue | Distance | Occupancy | B-factor | Closest solute to 1^st^ shell water |
| --- | --- | --- | --- | --- | --- | --- | --- |
| **Ion** | | | |  | 1.00 | 78.19 || 1 | C | O2' | AA:2177 | 3.46 | 1.00 | 167.17 |
| 2 | C | O4' | AA:2178 | 4.95 | 1.00 | 160.46 |  |
  |

  
 Coded contact list:   
1^st^ MG coordination shell missing (see Notes)  

### MG:AA:3417

|  | Res. type | Atom | Residue | Distance | Occupancy | B-factor | Closest solute to 1^st^ shell water |
| --- | --- | --- | --- | --- | --- | --- | --- |
| **Ion** | | | |  | 1.00 | 58.79 || 1 | U | O4 | AA:2233 | 2.72 | 1.00 | 43.23 |
| 2 | G | O6 | AA:2087 | 2.78 | 1.00 | 42.69 |  |
| 3 | U | O4 | AA:2086 | 2.94 | 1.00 | 46.83 |  |
| 4 | U | O4 | AA:2232 | 3.13 | 1.00 | 45.34 |  |
| 5 | U | N3 | AA:2086 | 3.30 | 1.00 | 44.02 |  |
| 6 | U | N3 | AA:2232 | 4.16 | 1.00 | 45.31 |  |
| 7 | G | O6 | AA:2088 | 4.19 | 1.00 | 54.29 |  |
| 8 | G | N7 | AA:2087 | 4.22 | 1.00 | 45.80 |  |
| 9 | U | N3 | AA:2233 | 4.38 | 1.00 | 42.68 |  |
| 10 | C | N4 | AA:2231 | 4.66 | 1.00 | 52.31 |  |
| 11 | U | O2 | AA:2086 | 4.83 | 1.00 | 44.40 |  |
  |

  
 Coded contact list:   
 Exclusion zone contact(s): /N3_U_/O4/O4/O4/O6/  
Consider K in place of MG (5 valid contact(s) for K vs. 0 for MG)  

### MG:AA:3420

|  | Res. type | Atom | Residue | Distance | Occupancy | B-factor | Closest solute to 1^st^ shell water |
| --- | --- | --- | --- | --- | --- | --- | --- |
| **Ion** | | | |  | 1.00 | 68.80 || 1 | G | O6 | AA:2415 | 3.34 | 1.00 | 77.80 |
| 2 | G | O6 | AA:2400 | 3.66 | 1.00 | 81.85 |  |
| 3 | G | O6 | AA:2414 | 3.72 | 1.00 | 69.49 |  |
| 4 | G | N7 | AA:2415 | 3.78 | 1.00 | 70.87 |  |
| 5 | G | N7 | AA:2414 | 3.84 | 1.00 | 66.77 |  |
| 6 | U | O4 | AA:2401 | 4.08 | 1.00 | 81.96 |  |
| 7 | C | N4 | AA:2416 | 4.10 | 1.00 | 80.56 |  |
  |

  
 Coded contact list:   
 Exclusion zone contact(s): /O6/  
Consider K in place of MG (1 valid contact(s) for K vs. 0 for MG)  

### MG:AA:3421

|  | Res. type | Atom | Residue | Distance | Occupancy | B-factor | Closest solute to 1^st^ shell water |
| --- | --- | --- | --- | --- | --- | --- | --- |
| **Ion** | | | |  | 1.00 | 43.26 || 1 | C | OP1 | AA:2283 | 3.35 | 1.00 | 65.15 |
| 2 | C | OP2 | AA:2284 | 3.68 | 1.00 | 61.50 |  |
| 3 | C | O3' | AA:2283 | 4.42 | 1.00 | 61.46 |  |
| 4 | C | O5' | AA:2283 | 4.50 | 1.00 | 63.95 |  |
| 5 | C | OP1 | AA:2284 | 4.53 | 1.00 | 61.85 |  |
  |

  
 Coded contact list:   
 Exclusion zone contact(s): /OP1/  
Consider K in place of MG (1 valid contact(s) for K vs. 0 for MG)  

### MG:AA:3422

|  | Res. type | Atom | Residue | Distance | Occupancy | B-factor | Closest solute to 1^st^ shell water |
| --- | --- | --- | --- | --- | --- | --- | --- |
| **Ion** | | | |  | 1.00 | 73.31 || 1 | A | OP2 | AA:270 | 2.68 | 1.00 | 66.76 |
| 2 | G | O6 | AA:270X | 2.75 | 1.00 | 80.11 |  |
| 3 | G | O6 | AA:270Y | 2.82 | 1.00 | 77.65 |  |
| 4 | G | N1 | AA:270Y | 2.92 | 1.00 | 77.75 |  |
| 5 | A | OP1 | AA:270 | 3.73 | 1.00 | 66.41 |  |
| 6 | U | O3' | AA:269 | 4.13 | 1.00 | 66.77 |  |
| 7 | G | N2 | AA:270Y | 4.32 | 1.00 | 79.20 |  |
| 8 | G | N7 | AA:270X | 4.48 | 1.00 | 79.16 |  |
| 9 | U | O5' | AA:269 | 4.63 | 1.00 | 68.02 |  |
| 10 | G | OP2 | AA:270W | 4.90 | 1.00 | 97.77 |  |
| 11 | G | N1 | AA:270X | 4.95 | 1.00 | 79.60 |  |
| 12 | A | O5' | AA:270 | 4.99 | 1.00 | 67.28 |  |
  |

  
 Coded contact list:   
 Exclusion zone contact(s): /N1_G_/O6/O6/OP2/  
Consider K in place of MG (4 valid contact(s) for K vs. 0 for MG)  

### MG:AA:3424

|  | Res. type | Atom | Residue | Distance | Occupancy | B-factor | Closest solute to 1^st^ shell water |
| --- | --- | --- | --- | --- | --- | --- | --- |
| **Ion** | | | |  | 1.00 | 54.22 || 1 | G | OP1 | AA:2069 | 2.36 | 1.00 | 43.48 |
| 2 | G | OP2 | AA:2069 | 2.56 | 1.00 | 42.46 |  |
| 3 | A | O3' | AA:197 | 3.14 | 1.00 | 43.21 |  |
| 4 | G | OP2 | AA:2070 | 3.14 | 1.00 | 41.38 |  |
| 5 | G | OP1 | AA:2070 | 3.31 | 1.00 | 40.38 |  |
| 6 | C | OP1 | AA:198 | 3.35 | 1.00 | 42.76 |  |
| 7 | G | O5' | AA:2069 | 3.66 | 1.00 | 43.34 |  |
| 8 | G | O3' | AA:2069 | 3.74 | 1.00 | 39.18 |  |
| 9 | U | OP1 | AA:2244 | 4.12 | 1.00 | 39.66 |  |
| 10 | A | O2' | AA:197 | 4.13 | 1.00 | 42.87 |  |
| 11 | U | O3' | AA:2068 | 4.24 | 1.00 | 45.50 |  |
| 12 | C | OP2 | AA:198 | 4.84 | 1.00 | 41.69 |  |
  |

  
 Contact(s): /OP1/  
 Coded contact list: O_ph_  
 Exclusion zone contact(s): /O3'/OP1/OP1/OP2/OP2/  
Consider K in place of MG (4 valid contact(s) for K vs. 1 for MG)  

### MG:AA:3425

|  | Res. type | Atom | Residue | Distance | Occupancy | B-factor | Closest solute to 1^st^ shell water |
| --- | --- | --- | --- | --- | --- | --- | --- |
| **Ion** | | | |  | 1.00 | 44.25 || 1 | A | OP2 | AA:2433 | 2.60 | 1.00 | 44.64 |
| 2 | A | OP2 | AA:2432 | 3.21 | 1.00 | 40.23 |  |
| 3 | U | OP2 | AA:2431 | 3.37 | 1.00 | 38.20 |  |
| 4 | A | OP1 | AA:2433 | 3.49 | 1.00 | 43.75 |  |
| 5 | A | O3' | AA:2432 | 4.14 | 1.00 | 43.13 |  |
| 6 | U | O4 | AA:2431 | 4.27 | 1.00 | 43.21 |  |
| 7 | U | N1 | AA:2431 | 4.43 | 1.00 | 38.18 |  |
| 8 | U | N3 | AA:2431 | 4.45 | 1.00 | 38.50 |  |
| 9 | U | O5' | AA:2431 | 4.51 | 1.00 | 37.49 |  |
| 10 | A | O5' | AA:2432 | 4.78 | 1.00 | 42.20 |  |
| 11 | A | O5' | AA:2433 | 4.85 | 1.00 | 44.10 |  |
  |

  
 Coded contact list:   
 Exclusion zone contact(s): /OP2/OP2/OP2/  
Consider K in place of MG (3 valid contact(s) for K vs. 0 for MG)  

### MG:AA:3426

|  | Res. type | Atom | Residue | Distance | Occupancy | B-factor | Closest solute to 1^st^ shell water |
| --- | --- | --- | --- | --- | --- | --- | --- |
| **Ion** | | | |  | 1.00 | 64.62 || 1 | C | O2' | BD:3 | 3.00 | 1.00 | 111.93 |
| 2 | C | OP2 | AA:1852 | 3.20 | 1.00 | 63.83 |  |
| 3 | U | OP1 | AA:1851 | 4.06 | 1.00 | 58.91 |  |
| 4 | C | O4' | BD:3 | 4.59 | 1.00 | 112.75 |  |
| 5 | C | O3' | BD:3 | 4.82 | 1.00 | 115.59 |  |
| 6 | G | O6 | AA:1888 | 4.91 | 1.00 | 90.24 |  |
| 7 | U | O5' | AA:1851 | 4.93 | 1.00 | 59.85 |  |
  |

  
 Coded contact list:   
 Exclusion zone contact(s): /O2'/OP2/  
Consider K in place of MG (2 valid contact(s) for K vs. 0 for MG)  

### MG:AA:3427

|  | Res. type | Atom | Residue | Distance | Occupancy | B-factor | Closest solute to 1^st^ shell water |
| --- | --- | --- | --- | --- | --- | --- | --- |
| **Ion** | | | |  | 1.00 | 47.30 || 1 | C | OP2 | AA:971 | 1.93 | 1.00 | 58.31 |
| 2 | G | N7 | AA:974 | 2.31 | 1.00 | 72.35 |  |
| 3 | G | O6 | AA:974 | 3.12 | 1.00 | 70.93 |  |
| 4 | G | O6 | AA:989 | 3.43 | 1.00 | 81.31 |  |
| 5 | C | OP1 | AA:971 | 3.91 | 1.00 | 59.38 |  |
| 6 | C | O5' | AA:971 | 4.05 | 1.00 | 59.55 |  |
| 7 | G | N1 | AA:989 | 4.32 | 1.00 | 78.48 |  |
| 8 | C | O3' | AA:970 | 4.42 | 1.00 | 59.78 |  |
| 9 | G | N9 | AA:974 | 4.54 | 1.00 | 69.62 |  |
| 10 | G | N7 | AA:972 | 4.69 | 1.00 | 65.35 |  |
| 11 | G | O6 | AA:972 | 4.78 | 1.00 | 69.11 |  |
| 12 | G | N1 | AA:974 | 4.88 | 1.00 | 70.45 |  |
  |

  
**1^st^ shell ligand angles:**   
1-2: 86;   
  
 Contact(s): /N7/OP2/  
 Coded contact list: O_ph_N_b_  
 Exclusion zone contact(s): /O6/  

### MG:AA:3428

|  | Res. type | Atom | Residue | Distance | Occupancy | B-factor | Closest solute to 1^st^ shell water |
| --- | --- | --- | --- | --- | --- | --- | --- |
| **Ion** | | | |  | 1.00 | 101.84 || 1 | G | O2' | CA:1032B | 4.89 | 1.00 | 200.41 |
  |

  
 Coded contact list:   
1^st^ MG coordination shell missing (see Notes)  

### MG:AA:3429

|  | Res. type | Atom | Residue | Distance | Occupancy | B-factor | Closest solute to 1^st^ shell water |
| --- | --- | --- | --- | --- | --- | --- | --- |
| **Ion** | | | |  | 1.00 | 60.62 || 1 | A | O5' | AA:819 | 2.34 | 1.00 | 56.44 |
| 2 | A | OP1 | AA:819 | 3.27 | 1.00 | 57.74 |  |
| 3 | A | OP2 | AA:819 | 3.32 | 1.00 | 56.68 |  |
| 4 | A | N1 | AA:973 | 4.19 | 1.00 | 65.62 |  |
| 5 | A | OP2 | AA:820 | 4.54 | 1.00 | 56.12 |  |
| 6 | G | O3' | AA:818 | 4.54 | 1.00 | 57.96 |  |
| 7 | A | O3' | AA:819 | 4.95 | 1.00 | 54.81 |  |
  |

  
 Contact(s): /O5'/  
 Coded contact list: O_r_  
 Exclusion zone contact(s): /OP1/OP2/  
Consider K in place of MG (2 valid contact(s) for K vs. 1 for MG)  

### MG:AA:3430

|  | Res. type | Atom | Residue | Distance | Occupancy | B-factor | Closest solute to 1^st^ shell water |
| --- | --- | --- | --- | --- | --- | --- | --- |
| **Ion** | | | |  | 1.00 | 81.64 || 1 | G | N1 | AA:1857 | 2.25 | 1.00 | 90.39 |
| 2 | G | N2 | AA:1857 | 2.33 | 1.00 | 89.74 |  |
| 3 | A | OP2 | AA:1884 | 2.33 | 1.00 | 85.91 |  |
| 4 | G | N2 | AA:1858 | 2.47 | 1.00 | 98.04 |  |
| 5 | G | O6 | AA:1857 | 3.80 | 1.00 | 92.68 |  |
| 6 | A | N7 | AA:1884 | 3.86 | 1.00 | 78.20 |  |
| 7 | A | O5' | AA:1884 | 3.96 | 1.00 | 84.58 |  |
| 8 | G | N3 | AA:1857 | 3.96 | 1.00 | 89.25 |  |
| 9 | G | N1 | AA:1858 | 4.00 | 1.00 | 97.71 |  |
| 10 | C | N4 | AA:1886 | 4.22 | 1.00 | 78.65 |  |
| 11 | A | OP1 | AA:1884 | 4.31 | 1.00 | 85.77 |  |
| 12 | G | N3 | AA:1858 | 4.37 | 1.00 | 98.03 |  |
| 13 | A | OP2 | AA:1885 | 4.59 | 1.00 | 81.54 |  |
| 14 | G | O3' | AA:1883 | 4.80 | 1.00 | 85.60 |  |
  |

  
**1^st^ shell ligand angles:**   
1-2: 60; 1-3: 140; 2-3: 157;   
  
 Contact(s): /N1_G_/N2/OP2/  
 Coded contact list: O_ph_2Anion  
 Exclusion zone contact(s): /N2/  

### MG:AA:3431

|  | Res. type | Atom | Residue | Distance | Occupancy | B-factor | Closest solute to 1^st^ shell water |
| --- | --- | --- | --- | --- | --- | --- | --- |
| **Ion** | | | |  | 1.00 | 53.00 || 1 | U | OP1 | AA:588 | 2.53 | 1.00 | 51.41 |
| 2 | C | O2' | AA:587 | 2.83 | 1.00 | 51.66 |  |
| 3 | U | OP2 | AA:588 | 2.86 | 1.00 | 50.83 |  |
| 4 | C | OP2 | AA:664 | 4.17 | 1.00 | 54.13 |  |
| 5 | U | O5' | AA:588 | 4.21 | 1.00 | 51.26 |  |
| 6 | C | O3' | AA:587 | 4.32 | 1.00 | 49.54 |  |
  |

  
 Coded contact list:   
 Exclusion zone contact(s): /O2'/OP1/OP2/  
Consider K in place of MG (2 valid contact(s) for K vs. 0 for MG)  

### MG:AA:3432

|  | Res. type | Atom | Residue | Distance | Occupancy | B-factor | Closest solute to 1^st^ shell water |
| --- | --- | --- | --- | --- | --- | --- | --- |
| **Ion** | | | |  | 1.00 | 83.79 || 1 | G | O2' | AA:2121 | 4.43 | 1.00 | 173.88 |
  |

  
 Coded contact list:   
1^st^ MG coordination shell missing (see Notes)  

### MG:AA:3433

|  | Res. type | Atom | Residue | Distance | Occupancy | B-factor | Closest solute to 1^st^ shell water |
| --- | --- | --- | --- | --- | --- | --- | --- |
| **Ion** | | | |  | 1.00 | 64.18 || 1 | G | N7 | AA:2239 | 3.05 | 1.00 | 37.12 |
| 2 | G | OP1 | AA:2239 | 3.10 | 1.00 | 35.89 |  |
| 3 | G | O2' | AA:2237 | 3.40 | 1.00 | 47.16 |  |
| 4 | G | O3' | AA:2237 | 3.43 | 1.00 | 48.51 |  |
| 5 | G | O5' | AA:2238 | 3.87 | 1.00 | 49.20 |  |
| 6 | G | N9 | AA:2239 | 4.12 | 1.00 | 38.65 |  |
| 7 | G | O5' | AA:2239 | 4.34 | 1.00 | 35.93 |  |
| 8 | G | O4' | AA:2239 | 4.36 | 1.00 | 39.28 |  |
| 9 | GLY | O | AD:251 | 4.59 | 1.00 | 36.20 |  |
  |

  
 Coded contact list:   
 Exclusion zone contact(s): /N7/O2'/OP1/  
Consider K in place of MG (3 valid contact(s) for K vs. 0 for MG)  

### MG:AA:3434

|  | Res. type | Atom | Residue | Distance | Occupancy | B-factor | Closest solute to 1^st^ shell water |
| --- | --- | --- | --- | --- | --- | --- | --- |
| **Ion** | | | |  | 1.00 | 65.07 || 1 | G | O6 | AA:1530 | 2.78 | 1.00 | 116.15 |
| 2 | G | N2 | AA:1542 | 2.83 | 1.00 | 104.83 |  |
| 3 | U | O4 | AA:1541 | 3.00 | 1.00 | 108.81 |  |
| 4 | G | O6 | AA:1540 | 3.13 | 1.00 | 125.66 |  |
| 5 | A | OP2 | AA:1529 | 3.73 | 1.00 | 93.51 |  |
| 6 | G | N1 | AA:1542 | 3.78 | 1.00 | 104.13 |  |
| 7 | C | N4 | AA:1531 | 3.80 | 1.00 | 125.15 |  |
| 8 | U | N3 | AA:1541 | 4.08 | 1.00 | 108.80 |  |
| 9 | G | N7 | AA:1530 | 4.37 | 1.00 | 113.86 |  |
| 10 | G | N3 | AA:1542 | 4.39 | 1.00 | 103.53 |  |
| 11 | G | N7 | AA:1540 | 4.96 | 1.00 | 125.08 |  |
  |

  
 Coded contact list:   
 Exclusion zone contact(s): /N2/O4/O6/O6/  
Consider K in place of MG (4 valid contact(s) for K vs. 0 for MG)  

### MG:AA:3435

|  | Res. type | Atom | Residue | Distance | Occupancy | B-factor | Closest solute to 1^st^ shell water |
| --- | --- | --- | --- | --- | --- | --- | --- |
| **Ion** | | | |  | 1.00 | 69.66 || 1 | G | OP1 | AA:2124 | 2.84 | 1.00 | 186.38 |
| 2 | G | OP2 | AA:2124 | 3.06 | 1.00 | 186.20 |  |
| 3 | G | O5' | AA:2124 | 4.21 | 1.00 | 186.76 |  |
| 4 | G | N7 | AA:2125 | 4.62 | 1.00 | 197.69 |  |
| 5 | G | O3' | AA:2123 | 4.76 | 1.00 | 185.77 |  |
  |

  
 Coded contact list:   
 Exclusion zone contact(s): /OP1/OP2/  
Consider K in place of MG (2 valid contact(s) for K vs. 0 for MG)  

### MG:AA:3436

|  | Res. type | Atom | Residue | Distance | Occupancy | B-factor | Closest solute to 1^st^ shell water |
| --- | --- | --- | --- | --- | --- | --- | --- |
| **Ion** | | | |  | 1.00 | 127.98 || 1 | A | N6 | AA:2169 | 2.63 | 1.00 | 196.36 |
| 2 | G | N3 | AA:2112 | 3.11 | 1.00 | 192.47 |  |
| 3 | A | N1 | AA:2169 | 3.69 | 1.00 | 196.46 |  |
| 4 | G | N9 | AA:2112 | 3.90 | 1.00 | 191.77 |  |
| 5 | G | N2 | AA:2112 | 3.93 | 1.00 | 192.33 |  |
| 6 | G | N1 | AA:2112 | 3.95 | 1.00 | 192.56 |  |
| 7 | A | N7 | AA:2169 | 4.49 | 1.00 | 197.25 |  |
| 8 | A | OP1 | AA:2169 | 4.69 | 1.00 | 199.25 |  |
| 9 | G | N7 | AA:2112 | 4.74 | 1.00 | 191.91 |  |
  |

  
 Coded contact list:   
 Exclusion zone contact(s): /N3_G_/N6/  
Consider K in place of MG (2 valid contact(s) for K vs. 0 for MG)  

### MG:AA:3437

|  | Res. type | Atom | Residue | Distance | Occupancy | B-factor | Closest solute to 1^st^ shell water |
| --- | --- | --- | --- | --- | --- | --- | --- |
| **Ion** | | | |  | 1.00 | 51.63 || 1 | G | OP2 | AA:862 | 2.11 | 1.00 | 74.96 |
| 2 | A | OP2 | AA:861 | 3.30 | 1.00 | 72.00 |  |
| 3 | G | N7 | AA:862 | 3.77 | 1.00 | 78.21 |  |
| 4 | A | OP1 | AA:861 | 3.80 | 1.00 | 70.54 |  |
| 5 | A | N7 | AA:863 | 4.17 | 1.00 | 74.78 |  |
| 6 | A | N6 | AA:863 | 4.22 | 1.00 | 75.00 |  |
| 7 | G | OP1 | AA:862 | 4.26 | 1.00 | 74.26 |  |
| 8 | A | O3' | AA:861 | 4.40 | 1.00 | 73.67 |  |
| 9 | A | O5' | AA:861 | 4.44 | 1.00 | 72.06 |  |
| 10 | G | O5' | AA:862 | 4.48 | 1.00 | 74.21 |  |
| 11 | A | OP1 | AA:910 | 4.54 | 1.00 | 71.27 |  |
  |

  
 Contact(s): /OP2/  
 Coded contact list: O_ph_  
 Exclusion zone contact(s): /OP2/  

### MG:AA:3438

|  | Res. type | Atom | Residue | Distance | Occupancy | B-factor | Closest solute to 1^st^ shell water |
| --- | --- | --- | --- | --- | --- | --- | --- |
| **Ion** | | | |  | 1.00 | 99.85 || 1 | G | O6 | AA:2182 | 2.51 | 1.00 | 150.44 |
| 2 | C | N4 | AA:2107 | 3.66 | 1.00 | 151.08 |  |
| 3 | C | N4 | AA:2183 | 3.97 | 1.00 | 151.57 |  |
| 4 | G | O6 | AA:2181 | 4.11 | 1.00 | 148.04 |  |
| 5 | G | O6 | AA:2106 | 4.15 | 1.00 | 152.03 |  |
| 6 | G | N7 | AA:2182 | 4.36 | 1.00 | 150.10 |  |
| 7 | C | N4 | AA:2108 | 4.52 | 1.00 | 148.48 |  |
| 8 | G | N1 | AA:2182 | 4.73 | 1.00 | 150.93 |  |
  |

  
 Coded contact list:   
 Exclusion zone contact(s): /O6/  
Consider NA in place of MG (1 valid contact(s) for NA vs. 0 for MG)  

### MG:AA:3439

|  | Res. type | Atom | Residue | Distance | Occupancy | B-factor | Closest solute to 1^st^ shell water |
| --- | --- | --- | --- | --- | --- | --- | --- |
| **Ion** | | | |  | 1.00 | 71.22 || 1 | G | OP1 | AA:2168 | 3.25 | 1.00 | 199.85 |
| 2 | G | O5' | AA:2168 | 3.55 | 1.00 | 199.80 |  |
| 3 | U | O3' | AA:2167 | 4.80 | 1.00 | 199.73 |  |
| 4 | G | O3' | AA:2168 | 4.95 | 1.00 | 199.39 |  |
  |

  
 Coded contact list:   
 Exclusion zone contact(s): /OP1/  
Consider K in place of MG (1 valid contact(s) for K vs. 0 for MG)  

### MG:AA:3440

|  | Res. type | Atom | Residue | Distance | Occupancy | B-factor | Closest solute to 1^st^ shell water |
| --- | --- | --- | --- | --- | --- | --- | --- |
| **Ion** | | | |  | 1.00 | 75.10 || 1 | G | OP1 | AA:36 | 2.89 | 1.00 | 51.64 |
| 2 | C | N3 | AA:451 | 3.46 | 1.00 | 39.67 |  |
| 3 | C | O2 | AA:451 | 3.61 | 1.00 | 44.86 |  |
| 4 | C | N4 | AA:453 | 3.78 | 1.00 | 42.29 |  |
| 5 | G | O5' | AA:36 | 3.84 | 1.00 | 49.80 |  |
| 6 | G | O3' | AA:35 | 4.21 | 1.00 | 52.69 |  |
| 7 | C | N4 | AA:451 | 4.67 | 1.00 | 41.10 |  |
| 8 | C | N3 | AA:453 | 4.77 | 1.00 | 42.44 |  |
  |

  
 Coded contact list:   
 Exclusion zone contact(s): /OP1/  
Consider K in place of MG (1 valid contact(s) for K vs. 0 for MG)  

### MG:AA:3441

|  | Res. type | Atom | Residue | Distance | Occupancy | B-factor | Closest solute to 1^st^ shell water |
| --- | --- | --- | --- | --- | --- | --- | --- |
| **Ion** | | | |  | 1.00 | 102.51 || 1 | G | OP2 | AA:1862 | 2.66 | 1.00 | 81.54 |
| 2 | G | O5' | AA:1861 | 2.97 | 1.00 | 85.69 |  |
| 3 | G | OP2 | AA:1861 | 3.19 | 1.00 | 85.65 |  |
| 4 | G | O3' | AA:1861 | 3.93 | 1.00 | 82.30 |  |
| 5 | G | OP1 | AA:1861 | 3.96 | 1.00 | 85.95 |  |
| 6 | G | O4' | AA:1861 | 4.69 | 1.00 | 83.88 |  |
| 7 | G | N9 | AA:1861 | 4.83 | 1.00 | 81.79 |  |
| 8 | G | N7 | AA:1861 | 4.88 | 1.00 | 80.28 |  |
| 9 | G | O5' | AA:1862 | 4.88 | 1.00 | 80.15 |  |
| 10 | G | OP1 | AA:1862 | 4.90 | 1.00 | 81.10 |  |
| 11 | G | O3' | AA:1860 | 5.00 | 1.00 | 86.05 |  |
  |

  
 Coded contact list:   
 Exclusion zone contact(s): /O5'/OP2/OP2/  
Consider K in place of MG (3 valid contact(s) for K vs. 0 for MG)  

### MG:AA:3442

|  | Res. type | Atom | Residue | Distance | Occupancy | B-factor | Closest solute to 1^st^ shell water |
| --- | --- | --- | --- | --- | --- | --- | --- |
| **Ion** | | | |  | 1.00 | 128.35 || 1 | G | N2 | AA:2115 | 2.32 | 1.00 | 200.41 |
| 2 | A | N7 | AA:2119 | 2.33 | 1.00 | 185.96 |  |
| 3 | A | N6 | AA:2114 | 2.51 | 1.00 | 199.77 |  |
| 4 | A | N6 | AA:2119 | 2.99 | 1.00 | 186.20 |  |
| 5 | A | N1 | AA:2114 | 3.12 | 1.00 | 199.36 |  |
| 6 | G | N1 | AA:2115 | 3.30 | 1.00 | 200.41 |  |
| 7 | A | N3 | AA:2171 | 3.32 | 1.00 | 198.98 |  |
| 8 | A | OP1 | AA:2119 | 3.60 | 1.00 | 189.80 |  |
| 9 | G | N3 | AA:2115 | 3.78 | 1.00 | 200.39 |  |
| 10 | A | N6 | AA:2170 | 4.20 | 1.00 | 198.00 |  |
| 11 | A | N1 | AA:2171 | 4.21 | 1.00 | 199.34 |  |
| 12 | A | N9 | AA:2119 | 4.28 | 1.00 | 185.74 |  |
| 13 | A | N1 | AA:2119 | 4.53 | 1.00 | 185.95 |  |
| 14 | U | O4 | AA:2172 | 4.92 | 1.00 | 192.96 |  |
| 15 | U | N3 | AA:2172 | 4.93 | 1.00 | 193.82 |  |
  |

  
**1^st^ shell ligand angles:**   
1-2: 131;   
  
 Contact(s): /N2/N7/  
 Coded contact list: N_b_Anion  
 Exclusion zone contact(s): /N1_A_/N1_G_/N3_A_/N6/N6/  
Consider K in place of MG (4 valid contact(s) for K vs. 2 for MG)  

### MG:AA:3443

|  | Res. type | Atom | Residue | Distance | Occupancy | B-factor | Closest solute to 1^st^ shell water |
| --- | --- | --- | --- | --- | --- | --- | --- |
| **Ion** | | | |  | 1.00 | 77.65 || 1 | U | OP1 | AA:2431 | 2.78 | 1.00 | 38.34 |
| 2 | C | OP1 | AA:249 | 2.94 | 1.00 | 36.41 |  |
| 3 | A | OP1 | AA:2432 | 3.27 | 1.00 | 42.16 |  |
| 4 | A | OP2 | AA:2432 | 4.37 | 1.00 | 40.23 |  |
| 5 | U | OP2 | AA:2431 | 4.42 | 1.00 | 38.20 |  |
| 6 | G | N7 | AA:248 | 4.67 | 1.00 | 37.39 |  |
| 7 | U | O3' | AA:2431 | 4.79 | 1.00 | 40.57 |  |
| 8 | C | OP2 | AA:249 | 5.00 | 1.00 | 38.20 |  |
  |

  
 Coded contact list:   
 Exclusion zone contact(s): /OP1/OP1/OP1/  
Consider K in place of MG (3 valid contact(s) for K vs. 0 for MG)  

### MG:AA:3444

|  | Res. type | Atom | Residue | Distance | Occupancy | B-factor | Closest solute to 1^st^ shell water |
| --- | --- | --- | --- | --- | --- | --- | --- |
| **Ion** | | | |  | 1.00 | 81.54 || 1 | A | OP2 | AA:2425 | 3.76 | 1.00 | 43.00 |
| 2 | C | O3' | AA:2424 | 4.12 | 1.00 | 42.86 |  |
| 3 | A | N3 | AA:2426 | 4.19 | 1.00 | 43.65 |  |
| 4 | C | O2' | AA:2424 | 4.20 | 1.00 | 44.99 |  |
| 5 | A | N1 | AA:2426 | 4.87 | 1.00 | 47.16 |  |
  |

  
 Coded contact list:   
1^st^ MG coordination shell missing (see Notes)  

### MG:AA:3445

|  | Res. type | Atom | Residue | Distance | Occupancy | B-factor | Closest solute to 1^st^ shell water |
| --- | --- | --- | --- | --- | --- | --- | --- |
| **Ion** | | | |  | 1.00 | 63.38 || 1 | G | O6 | AA:1814 | 2.71 | 1.00 | 42.85 |
| 2 | G | OP2 | AA:1813 | 4.56 | 1.00 | 37.45 |  |
| 3 | A | N1 | AA:1815 | 4.66 | 1.00 | 50.85 |  |
| 4 | G | N1 | AA:1814 | 4.79 | 1.00 | 38.85 |  |
  |

  
 Coded contact list:   
 Exclusion zone contact(s): /O6/  
Consider K in place of MG (1 valid contact(s) for K vs. 0 for MG)  

### MG:AA:3447

|  | Res. type | Atom | Residue | Distance | Occupancy | B-factor | Closest solute to 1^st^ shell water |
| --- | --- | --- | --- | --- | --- | --- | --- |
| **Ion** | | | |  | 1.00 | 71.17 || 1 | U | OP1 | AA:475 | 2.79 | 1.00 | 48.11 |
| 2 | G | OP2 | AA:476 | 3.16 | 1.00 | 56.40 |  |
| 3 | U | O5' | AA:475 | 3.20 | 1.00 | 48.19 |  |
| 4 | G | N7 | AA:476 | 3.45 | 1.00 | 59.95 |  |
| 5 | G | O2' | AA:474 | 4.03 | 1.00 | 52.09 |  |
| 6 | U | OP2 | AA:475 | 4.09 | 1.00 | 49.04 |  |
| 7 | G | O5' | AA:476 | 4.50 | 1.00 | 57.56 |  |
| 8 | U | O3' | AA:475 | 4.85 | 1.00 | 56.45 |  |
| 9 | G | O3' | AA:474 | 4.93 | 1.00 | 50.30 |  |
  |

  
 Coded contact list:   
 Exclusion zone contact(s): /O5'/OP1/OP2/  
Consider K in place of MG (3 valid contact(s) for K vs. 0 for MG)  

### MG:AA:3448

|  | Res. type | Atom | Residue | Distance | Occupancy | B-factor | Closest solute to 1^st^ shell water |
| --- | --- | --- | --- | --- | --- | --- | --- |
| **Ion** | | | |  | 1.00 | 93.83 || 1 | G | O6 | AA:2120 | 4.21 | 1.00 | 166.47 |
| 2 | G | O6 | AA:2121 | 4.31 | 1.00 | 169.52 |  |
| 3 | A | N6 | AA:2176 | 4.50 | 1.00 | 177.46 |  |
| 4 | G | N7 | AA:2120 | 4.79 | 1.00 | 170.74 |  |
| 5 | U | O4 | AA:2122 | 4.86 | 1.00 | 178.25 |  |
  |

  
 Coded contact list:   
1^st^ MG coordination shell missing (see Notes)  

### MG:AA:3449

|  | Res. type | Atom | Residue | Distance | Occupancy | B-factor | Closest solute to 1^st^ shell water |
| --- | --- | --- | --- | --- | --- | --- | --- |
| **Ion** | | | |  | 1.00 | 82.59 || 1 | G | O3' | AA:2121 | 3.29 | 1.00 | 174.84 |
| 2 | U | OP1 | AA:2122 | 3.87 | 1.00 | 175.34 |  |
| 3 | G | OP1 | AA:2121 | 4.15 | 1.00 | 175.15 |  |
| 4 | G | O5' | AA:2121 | 4.33 | 1.00 | 174.82 |  |
| 5 | U | OP2 | AA:2122 | 4.40 | 1.00 | 175.11 |  |
  |

  
 Coded contact list:   
 Exclusion zone contact(s): /O3'/  
Consider K in place of MG (1 valid contact(s) for K vs. 0 for MG)  

### MG:AA:3450

|  | Res. type | Atom | Residue | Distance | Occupancy | B-factor | Closest solute to 1^st^ shell water |
| --- | --- | --- | --- | --- | --- | --- | --- |
| **Ion** | | | |  | 1.00 | 59.72 || 1 | G | OP1 | AA:848 | 2.23 | 1.00 | 76.28 |
| 2 | A | OP2 | AA:849 | 3.87 | 1.00 | 74.98 |  |
| 3 | A | OP1 | AA:849 | 4.27 | 1.00 | 75.08 |  |
| 4 | G | O5' | AA:848 | 4.47 | 1.00 | 77.12 |  |
| 5 | G | OP2 | AA:848 | 4.50 | 1.00 | 76.06 |  |
| 6 | U | O3' | AA:847 | 4.58 | 1.00 | 75.02 |  |
  |

  
 Contact(s): /OP1/  
 Coded contact list: O_ph_  

### MG:AA:3451

|  | Res. type | Atom | Residue | Distance | Occupancy | B-factor | Closest solute to 1^st^ shell water |
| --- | --- | --- | --- | --- | --- | --- | --- |
| **Ion** | | | |  | 1.00 | 89.90 || 1 | A | OP2 | AA:390 | 2.45 | 1.00 | 52.24 |
| 2 | G | O5' | AA:389 | 2.97 | 1.00 | 50.60 |  |
| 3 | G | O3' | AA:389 | 2.99 | 1.00 | 51.03 |  |
| 4 | G | OP2 | AA:389 | 3.27 | 1.00 | 50.21 |  |
| 5 | A | OP1 | AA:390 | 3.72 | 1.00 | 52.79 |  |
| 6 | A | O5' | AA:390 | 4.56 | 1.00 | 53.19 |  |
| 7 | G | O3' | AA:388 | 4.70 | 1.00 | 50.99 |  |
| 8 | G | O4' | AA:389 | 4.74 | 1.00 | 48.99 |  |
| 9 | G | OP1 | AA:389 | 4.93 | 1.00 | 51.05 |  |
  |

  
 Coded contact list:   
 Exclusion zone contact(s): /O3'/O5'/OP2/OP2/  
Consider K in place of MG (3 valid contact(s) for K vs. 0 for MG)  

### MG:AA:3452

|  | Res. type | Atom | Residue | Distance | Occupancy | B-factor | Closest solute to 1^st^ shell water |
| --- | --- | --- | --- | --- | --- | --- | --- |
| **Ion** | | | |  | 1.00 | 97.35 || 1 | G | N2 | AA:1860 | 4.19 | 1.00 | 85.08 |
| 2 | G | O4' | AA:1861 | 4.24 | 1.00 | 83.88 |  |
| 3 | G | N3 | AA:1860 | 4.71 | 1.00 | 85.10 |  |
| 4 | G | O2' | AA:1861 | 4.87 | 1.00 | 83.79 |  |
| 5 | G | O2' | AA:1860 | 4.89 | 1.00 | 84.69 |  |
  |

  
 Coded contact list:   
1^st^ MG coordination shell missing (see Notes)  

### MG:AA:3453

|  | Res. type | Atom | Residue | Distance | Occupancy | B-factor | Closest solute to 1^st^ shell water |
| --- | --- | --- | --- | --- | --- | --- | --- |
| **Ion** | | | |  | 1.00 | 123.40 || 1 | G | O2' | AA:2147 | 2.44 | 1.00 | 191.23 |
| 2 | U | N1 | AA:2118 | 2.91 | 1.00 | 193.51 |  |
| 3 | U | N3 | AA:2118 | 3.41 | 1.00 | 193.42 |  |
| 4 | U | O2 | AA:2118 | 3.88 | 1.00 | 193.82 |  |
| 5 | U | O4' | AA:2118 | 3.88 | 1.00 | 193.59 |  |
| 6 | G | O4' | AA:2148 | 4.08 | 1.00 | 190.79 |  |
| 7 | G | O4' | AA:2147 | 4.09 | 1.00 | 191.17 |  |
| 8 | U | O4 | AA:2118 | 4.33 | 1.00 | 193.32 |  |
| 9 | G | N9 | AA:2147 | 4.76 | 1.00 | 192.16 |  |
  |

  
 Coded contact list:   
 Exclusion zone contact(s): /N1/O2'/  
Consider NA in place of MG (1 valid contact(s) for NA vs. 0 for MG)  

### MG:AA:3454

|  | Res. type | Atom | Residue | Distance | Occupancy | B-factor | Closest solute to 1^st^ shell water |
| --- | --- | --- | --- | --- | --- | --- | --- |
| **Ion** | | | |  | 1.00 | 67.59 || 1 | G | OP1 | AA:54 | 3.12 | 1.00 | 45.45 |
| 2 | G | O5' | AA:54 | 3.58 | 1.00 | 45.08 |  |
| 3 | G | OP2 | AA:55 | 3.86 | 1.00 | 43.20 |  |
| 4 | G | OP2 | AA:54 | 3.93 | 1.00 | 45.67 |  |
| 5 | G | OP1 | AA:55 | 4.52 | 1.00 | 45.67 |  |
| 6 | G | O3' | AA:54 | 4.53 | 1.00 | 43.82 |  |
  |

  
 Coded contact list:   
 Exclusion zone contact(s): /OP1/  
Consider K in place of MG (1 valid contact(s) for K vs. 0 for MG)  

### MG:AA:3455

|  | Res. type | Atom | Residue | Distance | Occupancy | B-factor | Closest solute to 1^st^ shell water |
| --- | --- | --- | --- | --- | --- | --- | --- |
| **Ion** | | | |  | 1.00 | 22.33 || 1 | A | N7 | AA:2577 | 2.97 | 1.00 | 43.50 |
| 2 | G | N7 | AA:2056 | 2.98 | 1.00 | 39.82 |  |
| 3 | G | OP2 | AA:2053 | 3.62 | 1.00 | 47.50 |  |
| 4 | G | OP1 | AA:2053 | 4.04 | 1.00 | 45.98 |  |
| 5 | A | N6 | AA:2577 | 4.06 | 1.00 | 44.69 |  |
| 6 | A | OP1 | AA:2577 | 4.13 | 1.00 | 44.36 |  |
| 7 | G | O6 | AA:2056 | 4.36 | 1.00 | 40.78 |  |
| 8 | A | OP2 | AA:2054 | 4.56 | 1.00 | 45.92 |  |
| 9 | G | O5' | AA:2053 | 4.71 | 1.00 | 44.42 |  |
| 10 | A | N9 | AA:2577 | 4.73 | 1.00 | 42.09 |  |
| 11 | G | N9 | AA:2056 | 4.95 | 1.00 | 39.29 |  |
  |

  
 Coded contact list:   
 Exclusion zone contact(s): /N7/N7/  
Consider K in place of MG (2 valid contact(s) for K vs. 0 for MG)  

### MG:AA:3456

|  | Res. type | Atom | Residue | Distance | Occupancy | B-factor | Closest solute to 1^st^ shell water |
| --- | --- | --- | --- | --- | --- | --- | --- |
| **Ion** | | | |  | 1.00 | 20.84 || 1 | A | OP1 | AA:2577 | 2.20 | 1.00 | 44.36 |
| 2 | G | OP1 | AA:2576 | 2.51 | 1.00 | 40.83 |  |
| 3 | G | OP1 | AA:2053 | 3.43 | 1.00 | 45.98 |  |
| 4 | G | OP2 | AA:2053 | 3.70 | 1.00 | 47.50 |  |
| 5 | C | O3' | AA:2575 | 4.07 | 1.00 | 40.81 |  |
| 6 | G | O3' | AA:2576 | 4.43 | 1.00 | 43.10 |  |
| 7 | A | OP2 | AA:2577 | 4.48 | 1.00 | 43.39 |  |
| 8 | G | N2 | AA:2576 | 4.55 | 1.00 | 40.39 |  |
| 9 | A | O5' | AA:2577 | 4.60 | 1.00 | 42.97 |  |
| 10 | G | OP2 | AA:2576 | 4.70 | 1.00 | 39.26 |  |
| 11 | A | N7 | AA:2577 | 4.82 | 1.00 | 43.50 |  |
| 12 | G | OP2 | AA:2052 | 4.83 | 1.00 | 45.55 |  |
| 13 | G | O3' | AA:2052 | 4.86 | 1.00 | 44.88 |  |
| 14 | G | O5' | AA:2576 | 4.94 | 1.00 | 39.82 |  |
  |

  
 Contact(s): /OP1/  
 Coded contact list: O_ph_  
 Exclusion zone contact(s): /OP1/  

### MG:AA:3457

|  | Res. type | Atom | Residue | Distance | Occupancy | B-factor | Closest solute to 1^st^ shell water |
| --- | --- | --- | --- | --- | --- | --- | --- |
| **Ion** | | | |  | 1.00 | 57.61 || 1 | G | OP2 | AA:177 | 2.32 | 1.00 | 56.23 |
| 2 | G | O6 | AA:48 | 2.63 | 1.00 | 60.77 |  |
| 3 | G | OP1 | AA:176 | 3.20 | 1.00 | 74.02 |  |
| 4 | G | OP1 | AA:177 | 3.50 | 1.00 | 56.15 |  |
| 5 | G | O3' | AA:176 | 3.51 | 1.00 | 57.62 |  |
| 6 | G | O5' | AA:176 | 3.58 | 1.00 | 70.98 |  |
| 7 | G | N1 | AA:48 | 3.59 | 1.00 | 59.85 |  |
| 8 | G | N2 | AA:177 | 4.39 | 1.00 | 53.20 |  |
| 9 | G | O5' | AA:177 | 4.64 | 1.00 | 55.41 |  |
| 10 | G | O6 | AA:178 | 4.88 | 1.00 | 58.68 |  |
  |

  
 Contact(s): /OP2/  
 Coded contact list: O_ph_  
 Exclusion zone contact(s): /O6/OP1/  
Consider K in place of MG (2 valid contact(s) for K vs. 1 for MG)  

### MG:AA:3458

|  | Res. type | Atom | Residue | Distance | Occupancy | B-factor | Closest solute to 1^st^ shell water |
| --- | --- | --- | --- | --- | --- | --- | --- |
| **Ion** | | | |  | 1.00 | 49.59 || 1 | C | OP1 | AA:2452 | 2.27 | 1.00 | 50.43 |
| 2 | G | OP2 | AA:2494 | 3.94 | 1.00 | 65.94 |  |
| 3 | A | O5' | BB:76 | 4.23 | 1.00 | 135.02 |  |
| 4 | A | O3' | AA:2451 | 4.42 | 1.00 | 47.67 |  |
| 5 | C | OP2 | AA:2452 | 4.61 | 1.00 | 48.04 |  |
| 6 | C | O5' | AA:2452 | 4.64 | 1.00 | 49.23 |  |
| 7 | A | OP1 | BB:76 | 4.72 | 1.00 | 133.56 |  |
| 8 | A | OP2 | AA:2453 | 4.84 | 1.00 | 54.97 |  |
  |

  
 Contact(s): /OP1/  
 Coded contact list: O_ph_  

### MG:AA:3459

|  | Res. type | Atom | Residue | Distance | Occupancy | B-factor | Closest solute to 1^st^ shell water |
| --- | --- | --- | --- | --- | --- | --- | --- |
| **Ion** | | | |  | 1.00 | 30.62 || 1 | A | N1 | AA:2497 | 4.21 | 1.00 | 44.95 |
| 2 | G | N7 | AA:2454 | 4.22 | 1.00 | 45.73 |  |
| 3 | G | O6 | AA:2455 | 4.24 | 1.00 | 52.73 |  |
| 4 | A | N7 | AA:2453 | 4.25 | 1.00 | 43.43 |  |
| 5 | G | O6 | AA:2454 | 4.31 | 1.00 | 43.26 |  |
| 6 | C | OP2 | AA:2452 | 4.31 | 1.00 | 48.04 |  |
| 7 | A | OP2 | AA:2453 | 4.71 | 1.00 | 54.97 |  |
| 8 | C | N4 | AA:2496 | 4.72 | 1.00 | 51.69 |  |
| 9 | A | OP2 | AA:2451 | 4.99 | 1.00 | 46.52 |  |
  |

  
 Coded contact list:   
1^st^ MG coordination shell missing (see Notes)  

### MG:AA:3460

|  | Res. type | Atom | Residue | Distance | Occupancy | B-factor | Closest solute to 1^st^ shell water |
| --- | --- | --- | --- | --- | --- | --- | --- |
| **Ion** | | | |  | 1.00 | 28.26 || 1 | U | O4 | AA:2580 | 2.72 | 1.00 | 40.90 |
| 2 | G | O6 | AA:2508 | 2.74 | 1.00 | 41.84 |  |
| 3 | G | N7 | AA:2576 | 3.14 | 1.00 | 43.84 |  |
| 4 | C | N4 | AA:2507 | 3.65 | 1.00 | 41.31 |  |
| 5 | U | N3 | AA:2580 | 3.84 | 1.00 | 38.08 |  |
| 6 | G | N7 | AA:2581 | 3.97 | 1.00 | 39.36 |  |
| 7 | G | O6 | AA:2509 | 4.25 | 1.00 | 33.97 |  |
| 8 | G | N9 | AA:2576 | 4.28 | 1.00 | 42.74 |  |
| 9 | G | N7 | AA:2508 | 4.41 | 1.00 | 42.07 |  |
| 10 | C | N4 | AA:2579 | 4.79 | 1.00 | 29.60 |  |
| 11 | G | O6 | AA:2576 | 4.86 | 1.00 | 46.83 |  |
| 12 | G | N1 | AA:2508 | 4.96 | 1.00 | 39.39 |  |
  |

  
 Coded contact list:   
 Exclusion zone contact(s): /N7/O4/O6/  
Consider K in place of MG (3 valid contact(s) for K vs. 0 for MG)  

### MG:AA:3461

|  | Res. type | Atom | Residue | Distance | Occupancy | B-factor | Closest solute to 1^st^ shell water |
| --- | --- | --- | --- | --- | --- | --- | --- |
| **Ion** | | | |  | 1.00 | 58.79 || 1 | G | N7 | AA:308 | 3.69 | 1.00 | 54.44 |
| 2 | U | OP2 | AA:1211 | 3.72 | 1.00 | 72.07 |  |
| 3 | G | O6 | AA:309 | 3.82 | 1.00 | 50.87 |  |
| 4 | G | O6 | AA:1212 | 3.85 | 1.00 | 61.04 |  |
| 5 | G | O2' | AA:307 | 3.97 | 1.00 | 59.57 |  |
| 6 | A | O2' | AA:1210 | 4.25 | 1.00 | 73.87 |  |
| 7 | G | O3' | AA:307 | 4.25 | 1.00 | 61.71 |  |
| 8 | G | O6 | AA:308 | 4.62 | 1.00 | 56.53 |  |
| 9 | G | OP1 | AA:308 | 4.85 | 1.00 | 60.95 |  |
  |

  
 Coded contact list:   
1^st^ MG coordination shell missing (see Notes)  

### MG:AA:3462

|  | Res. type | Atom | Residue | Distance | Occupancy | B-factor | Closest solute to 1^st^ shell water |
| --- | --- | --- | --- | --- | --- | --- | --- |
| **Ion** | | | |  | 1.00 | 32.83 || 1 | G | OP1 | AA:2574 | 2.08 | 1.00 | 55.30 |
| 2 | C | OP1 | AA:2573 | 2.24 | 1.00 | 55.22 |  |
| 3 | C | OP2 | AA:2575 | 2.43 | 1.00 | 47.28 |  |
| 4 | C | O5' | AA:2573 | 3.53 | 1.00 | 56.49 |  |
| 5 | G | OP2 | AA:2574 | 4.00 | 1.00 | 55.11 |  |
| 6 | C | OP2 | AA:2573 | 4.05 | 1.00 | 56.13 |  |
| 7 | C | O3' | AA:2573 | 4.31 | 1.00 | 56.50 |  |
| 8 | C | OP2 | AA:2507 | 4.42 | 1.00 | 58.26 |  |
| 9 | G | O5' | AA:2574 | 4.49 | 1.00 | 52.75 |  |
| 10 | G | O3' | AA:2574 | 4.51 | 1.00 | 47.54 |  |
| 11 | C | OP1 | AA:2575 | 4.57 | 1.00 | 45.05 |  |
| 12 | A | O3' | AA:2572 | 4.65 | 1.00 | 54.06 |  |
| 13 | C | OP1 | AA:2507 | 4.71 | 1.00 | 56.08 |  |
| 14 | G | OP2 | AA:2508 | 4.76 | 1.00 | 48.77 |  |
| 15 | C | O5' | AA:2575 | 4.86 | 1.00 | 44.54 |  |
  |

  
**1^st^ shell ligand angles:**   
1-2: 73;   
  
 Contact(s): /OP1/OP1/  
 Coded contact list: 2O_ph_  
 Exclusion zone contact(s): /OP2/  

### MG:AA:3463

|  | Res. type | Atom | Residue | Distance | Occupancy | B-factor | Closest solute to 1^st^ shell water |
| --- | --- | --- | --- | --- | --- | --- | --- |
| **Ion** | | | |  | 1.00 | 3.22 || 1 | G | O6 | AA:2505 | 2.43 | 1.00 | 46.08 |
| 2 | G | O6 | AA:2581 | 2.72 | 1.00 | 39.21 |  |
| 3 | C | O2' | AA:2610 | 2.89 | 1.00 | 45.87 |  |
| 4 | G | O2' | AA:2576 | 2.92 | 1.00 | 43.65 |  |
| 5 | C | O2 | AA:2610 | 3.00 | 1.00 | 48.00 |  |
| 6 | G | N1 | AA:2505 | 3.84 | 1.00 | 46.79 |  |
| 7 | G | N1 | AA:2581 | 4.00 | 1.00 | 38.01 |  |
| 8 | G | N9 | AA:2576 | 4.07 | 1.00 | 42.74 |  |
| 9 | C | OP2 | AA:2579 | 4.12 | 1.00 | 37.55 |  |
| 10 | U | OP2 | AA:2580 | 4.20 | 1.00 | 36.28 |  |
| 11 | C | N1 | AA:2610 | 4.41 | 1.00 | 46.24 |  |
| 12 | C | OP1 | AA:2579 | 4.49 | 1.00 | 36.86 |  |
| 13 | C | N3 | AA:2610 | 4.72 | 1.00 | 47.27 |  |
| 14 | G | O4' | AA:2576 | 4.80 | 1.00 | 43.81 |  |
| 15 | C | O5' | AA:2579 | 4.90 | 1.00 | 34.77 |  |
| 16 | U | OP2 | AA:2611 | 4.96 | 1.00 | 41.13 |  |
  |

  
 Coded contact list:   
 Exclusion zone contact(s): /O2/O2'/O2'/O6/O6/  
Consider K in place of MG (4 valid contact(s) for K vs. 0 for MG)  

### MG:AA:3464

|  | Res. type | Atom | Residue | Distance | Occupancy | B-factor | Closest solute to 1^st^ shell water |
| --- | --- | --- | --- | --- | --- | --- | --- |
| **Ion** | | | |  | 1.00 | 61.58 || 1 | A | OP1 | AA:324 | 2.23 | 1.00 | 67.89 |
| 2 | G | OP2 | AA:323 | 2.71 | 1.00 | 73.64 |  |
| 3 | A | O4' | AA:324 | 3.64 | 1.00 | 64.08 |  |
| 4 | A | O5' | AA:324 | 3.66 | 1.00 | 66.94 |  |
| 5 | G | O5' | AA:323 | 4.38 | 1.00 | 72.81 |  |
| 6 | A | OP2 | AA:324 | 4.46 | 1.00 | 66.30 |  |
| 7 | G | OP1 | AA:323 | 4.47 | 1.00 | 73.95 |  |
| 8 | G | O3' | AA:323 | 4.59 | 1.00 | 69.29 |  |
  |

  
 Contact(s): /OP1/  
 Coded contact list: O_ph_  
 Exclusion zone contact(s): /OP2/  

### MG:AA:3465

|  | Res. type | Atom | Residue | Distance | Occupancy | B-factor | Closest solute to 1^st^ shell water |
| --- | --- | --- | --- | --- | --- | --- | --- |
| **Ion** | | | |  | 1.00 | 28.96 || 1 | G | N7 | AA:2032 | 2.66 | 1.00 | 49.82 |
| 2 | G | O6 | AA:2032 | 3.08 | 1.00 | 46.04 |  |
| 3 | C | O2 | AA:2055 | 3.42 | 1.00 | 38.12 |  |
| 4 | C | N3 | AA:2055 | 3.43 | 1.00 | 38.27 |  |
| 5 | C | N3 | AA:2499 | 3.75 | 1.00 | 49.88 |  |
| 6 | C | O2 | AA:2499 | 3.93 | 1.00 | 50.92 |  |
| 7 | A | N1 | AA:2572 | 4.12 | 1.00 | 53.43 |  |
| 8 | U | O2 | AA:2500 | 4.19 | 1.00 | 38.41 |  |
| 9 | A | N3 | AA:2453 | 4.63 | 1.00 | 40.86 |  |
| 10 | C | N4 | AA:2055 | 4.74 | 1.00 | 38.41 |  |
| 11 | C | N4 | AA:2499 | 4.83 | 1.00 | 51.17 |  |
| 12 | G | OP1 | AA:2032 | 4.84 | 1.00 | 58.74 |  |
| 13 | G | N9 | AA:2032 | 4.84 | 1.00 | 53.14 |  |
| 14 | G | N1 | AA:2032 | 4.90 | 1.00 | 49.07 |  |
  |

  
 Coded contact list:   
 Exclusion zone contact(s): /N7/O6/  
Consider K in place of MG (2 valid contact(s) for K vs. 0 for MG)  

### MG:AA:3466

|  | Res. type | Atom | Residue | Distance | Occupancy | B-factor | Closest solute to 1^st^ shell water |
| --- | --- | --- | --- | --- | --- | --- | --- |
| **Ion** | | | |  | 1.00 | 37.27 || 1 | U | OP1 | AA:1673 | 2.32 | 1.00 | 49.68 |
| 2 | U | OP2 | AA:2552 | 2.58 | 1.00 | 48.04 |  |
| 3 | U | OP1 | AA:2552 | 3.43 | 1.00 | 47.95 |  |
| 4 | C | O3' | AA:1672 | 3.88 | 1.00 | 48.80 |  |
| 5 | U | O5' | AA:2552 | 4.31 | 1.00 | 48.43 |  |
| 6 | C | OP1 | AA:2551 | 4.36 | 1.00 | 49.75 |  |
| 7 | C | O5' | AA:2551 | 4.49 | 1.00 | 48.96 |  |
| 8 | U | O5' | AA:1673 | 4.50 | 1.00 | 47.71 |  |
| 9 | U | OP2 | AA:1673 | 4.73 | 1.00 | 51.17 |  |
| 10 | C | O3' | AA:2551 | 4.75 | 1.00 | 48.98 |  |
| 11 | C | OP2 | AA:2551 | 4.76 | 1.00 | 48.51 |  |
| 12 | G | OP2 | AA:2553 | 4.85 | 1.00 | 45.56 |  |
| 13 | U | OP1 | AA:2554 | 5.00 | 1.00 | 49.20 |  |
  |

  
 Contact(s): /OP1/  
 Coded contact list: O_ph_  
 Exclusion zone contact(s): /OP2/  

### MG:AA:3467

|  | Res. type | Atom | Residue | Distance | Occupancy | B-factor | Closest solute to 1^st^ shell water |
| --- | --- | --- | --- | --- | --- | --- | --- |
| **Ion** | | | |  | 1.00 | 55.52 || 1 | A | N6 | AA:2361 | 2.99 | 1.00 | 55.81 |
| 2 | U | O2 | AA:2357 | 3.38 | 1.00 | 54.42 |  |
| 3 | A | N7 | AA:2361 | 3.56 | 1.00 | 56.87 |  |
| 4 | A | OP2 | AA:2360 | 4.56 | 1.00 | 51.11 |  |
| 5 | C | N3 | AA:2356 | 4.79 | 1.00 | 57.97 |  |
| 6 | G | O6 | AA:2362 | 4.79 | 1.00 | 56.61 |  |
| 7 | A | N7 | AA:2360 | 4.82 | 1.00 | 57.62 |  |
| 8 | C | O2 | AA:2356 | 4.94 | 1.00 | 59.47 |  |
  |

  
 Coded contact list:   
 Exclusion zone contact(s): /N6/O2/  
Consider K in place of MG (2 valid contact(s) for K vs. 0 for MG)  

### MG:AA:3468

|  | Res. type | Atom | Residue | Distance | Occupancy | B-factor | Closest solute to 1^st^ shell water |
| --- | --- | --- | --- | --- | --- | --- | --- |
| **Ion** | | | |  | 1.00 | 22.85 || 1 | G | OP1 | AA:2550 | 2.17 | 1.00 | 50.30 |
| 2 | C | OP2 | AA:1670 | 2.35 | 1.00 | 47.27 |  |
| 3 | U | O4 | AA:1671 | 2.42 | 1.00 | 50.79 |  |
| 4 | A | OP2 | AA:1669 | 2.75 | 1.00 | 47.14 |  |
| 5 | G | OP2 | AA:2550 | 3.55 | 1.00 | 49.31 |  |
| 6 | C | OP2 | AA:2551 | 3.83 | 1.00 | 48.51 |  |
| 7 | C | OP1 | AA:1670 | 4.10 | 1.00 | 45.50 |  |
| 8 | G | O5' | AA:2550 | 4.15 | 1.00 | 51.02 |  |
| 9 | U | N3 | AA:1671 | 4.43 | 1.00 | 50.49 |  |
| 10 | G | O3' | AA:2549 | 4.52 | 1.00 | 50.19 |  |
| 11 | A | O5' | AA:1669 | 4.52 | 1.00 | 47.02 |  |
| 12 | A | O3' | AA:1669 | 4.61 | 1.00 | 46.39 |  |
| 13 | C | O5' | AA:1670 | 4.69 | 1.00 | 47.53 |  |
| 14 | A | O3' | AA:1668 | 4.83 | 1.00 | 49.05 |  |
  |

  
**1^st^ shell ligand angles:**   
1-2: 77;   
  
 Contact(s): /OP1/OP2/  
 Coded contact list: 2O_ph_  
 Exclusion zone contact(s): /O4/OP2/  

### MG:AA:3469

|  | Res. type | Atom | Residue | Distance | Occupancy | B-factor | Closest solute to 1^st^ shell water |
| --- | --- | --- | --- | --- | --- | --- | --- |
| **Ion** | | | |  | 1.00 | 42.27 || 1 | C | OP1 | AA:1670 | 2.07 | 1.00 | 45.50 |
| 2 | U | OP2 | AA:1671 | 2.27 | 1.00 | 49.47 |  |
| 3 | U | OP1 | AA:1671 | 4.18 | 1.00 | 47.02 |  |
| 4 | G | OP2 | AA:2550 | 4.18 | 1.00 | 49.31 |  |
| 5 | G | OP1 | AA:2549 | 4.32 | 1.00 | 48.03 |  |
| 6 | C | O5' | AA:1670 | 4.34 | 1.00 | 47.53 |  |
| 7 | A | O3' | AA:1669 | 4.35 | 1.00 | 46.39 |  |
| 8 | G | OP2 | AA:2549 | 4.48 | 1.00 | 46.60 |  |
| 9 | C | OP2 | AA:1670 | 4.49 | 1.00 | 47.27 |  |
| 10 | U | O5' | AA:1671 | 4.55 | 1.00 | 48.99 |  |
| 11 | C | O3' | AA:1670 | 4.62 | 1.00 | 46.65 |  |
  |

  
**1^st^ shell ligand angles:**   
1-2: 72;   
  
 Contact(s): /OP1/OP2/  
 Coded contact list: 2O_ph_  

### MG:AA:3470

|  | Res. type | Atom | Residue | Distance | Occupancy | B-factor | Closest solute to 1^st^ shell water |
| --- | --- | --- | --- | --- | --- | --- | --- |
| **Ion** | | | |  | 1.00 | 76.29 || 1 | A | OP2 | AA:497 | 2.88 | 1.00 | 66.44 |
| 2 | G | N7 | AA:498 | 3.55 | 1.00 | 53.57 |  |
| 3 | G | OP2 | AA:498 | 3.66 | 1.00 | 62.89 |  |
| 4 | A | O5' | AA:497 | 3.88 | 1.00 | 65.64 |  |
| 5 | A | N7 | AA:497 | 4.56 | 1.00 | 65.70 |  |
| 6 | A | OP1 | AA:497 | 4.82 | 1.00 | 67.41 |  |
  |

  
 Coded contact list:   
 Exclusion zone contact(s): /OP2/  
Consider K in place of MG (1 valid contact(s) for K vs. 0 for MG)  

### MG:AA:3471

|  | Res. type | Atom | Residue | Distance | Occupancy | B-factor | Closest solute to 1^st^ shell water |
| --- | --- | --- | --- | --- | --- | --- | --- |
| **Ion** | | | |  | 1.00 | 77.28 || 1 | U | OP1 | AA:2492 | 2.61 | 1.00 | 72.27 |
| 2 | C | O4' | AA:2573 | 3.57 | 1.00 | 58.75 |  |
| 3 | G | OP1 | AA:2454 | 4.11 | 1.00 | 47.65 |  |
| 4 | G | OP2 | AA:2454 | 4.12 | 1.00 | 47.39 |  |
| 5 | U | O3' | AA:2491 | 4.13 | 1.00 | 72.71 |  |
| 6 | C | N1 | AA:2573 | 4.50 | 1.00 | 59.46 |  |
| 7 | U | OP2 | AA:2492 | 4.61 | 1.00 | 72.38 |  |
| 8 | A | OP1 | AA:2453 | 4.61 | 1.00 | 54.55 |  |
  |

  
 Coded contact list:   
 Exclusion zone contact(s): /OP1/  
Consider K in place of MG (1 valid contact(s) for K vs. 0 for MG)  

### MG:AA:3472

|  | Res. type | Atom | Residue | Distance | Occupancy | B-factor | Closest solute to 1^st^ shell water |
| --- | --- | --- | --- | --- | --- | --- | --- |
| **Ion** | | | |  | 1.00 | 59.70 || 1 | U | OP2 | AA:2491 | 2.40 | 1.00 | 75.69 |
| 2 | G | OP2 | AA:2455 | 3.37 | 1.00 | 48.65 |  |
| 3 | G | O3' | AA:2490 | 3.46 | 1.00 | 76.34 |  |
| 4 | U | O5' | AA:2491 | 4.17 | 1.00 | 75.72 |  |
| 5 | G | OP1 | AA:2454 | 4.19 | 1.00 | 47.65 |  |
| 6 | G | OP1 | AA:2455 | 4.51 | 1.00 | 50.25 |  |
| 7 | G | O2' | AA:2490 | 4.54 | 1.00 | 78.79 |  |
| 8 | U | OP1 | AA:2491 | 4.74 | 1.00 | 77.50 |  |
  |

  
 Contact(s): /OP2/  
 Coded contact list: O_ph_  
 Exclusion zone contact(s): /OP2/  

### MG:AA:3473

|  | Res. type | Atom | Residue | Distance | Occupancy | B-factor | Closest solute to 1^st^ shell water |
| --- | --- | --- | --- | --- | --- | --- | --- |
| **Ion** | | | |  | 1.00 | 63.00 || 1 | G | OP2 | AA:1581 | 2.20 | 1.00 | 99.37 |
| 2 | A | O5' | AA:1580 | 3.49 | 1.00 | 93.89 |  |
| 3 | A | O3' | AA:1580 | 3.71 | 1.00 | 97.76 |  |
| 4 | G | N7 | AA:1581 | 3.80 | 1.00 | 97.87 |  |
| 5 | A | OP2 | AA:1580 | 3.81 | 1.00 | 94.25 |  |
| 6 | G | O5' | AA:1581 | 4.15 | 1.00 | 100.04 |  |
| 7 | A | OP1 | AA:1580 | 4.22 | 1.00 | 94.09 |  |
| 8 | A | O4' | AA:1580 | 4.50 | 1.00 | 93.93 |  |
| 9 | G | OP1 | AA:1581 | 4.70 | 1.00 | 99.56 |  |
  |

  
 Contact(s): /OP2/  
 Coded contact list: O_ph_  

### MG:AA:3474

|  | Res. type | Atom | Residue | Distance | Occupancy | B-factor | Closest solute to 1^st^ shell water |
| --- | --- | --- | --- | --- | --- | --- | --- |
| **Ion** | | | |  | 1.00 | 65.83 || 1 | G | OP1 | AA:2819 | 2.68 | 1.00 | 51.44 |
| 2 | SER | OG | A0:6 | 4.18 | 1.00 | 69.89 |  |
| 3 | G | O5' | AA:2819 | 4.25 | 1.00 | 49.83 |  |
| 4 | G | O3' | AA:2819 | 4.66 | 1.00 | 51.24 |  |
| 5 | G | O3' | AA:2818 | 4.76 | 1.00 | 53.84 |  |
| 6 | G | O3' | AA:2837 | 4.97 | 1.00 | 66.98 |  |
  |

  
 Coded contact list:   
 Exclusion zone contact(s): /OP1/  
Consider K in place of MG (1 valid contact(s) for K vs. 0 for MG)  

### MG:AA:3475

|  | Res. type | Atom | Residue | Distance | Occupancy | B-factor | Closest solute to 1^st^ shell water |
| --- | --- | --- | --- | --- | --- | --- | --- |
| **Ion** | | | |  | 1.00 | 85.32 || 1 | G | N7 | AA:2876 | 4.11 | 1.00 | 61.64 |
| 2 | G | O6 | AA:2876 | 4.80 | 1.00 | 62.09 |  |
| 3 | G | O6 | AA:2877 | 4.82 | 1.00 | 67.63 |  |
  |

  
 Coded contact list:   
1^st^ MG coordination shell missing (see Notes)  

### MG:AA:3476

|  | Res. type | Atom | Residue | Distance | Occupancy | B-factor | Closest solute to 1^st^ shell water |
| --- | --- | --- | --- | --- | --- | --- | --- |
| **Ion** | | | |  | 1.00 | 47.13 || 1 | G | OP2 | AA:2583 | 2.43 | 1.00 | 48.98 |
| 2 | G | OP2 | AA:2582 | 2.59 | 1.00 | 47.17 |  |
| 3 | G | N7 | AA:2608 | 4.06 | 1.00 | 49.11 |  |
| 4 | G | O6 | AA:2608 | 4.19 | 1.00 | 52.30 |  |
| 5 | G | O3' | AA:2581 | 4.25 | 1.00 | 44.83 |  |
| 6 | G | OP1 | AA:2583 | 4.40 | 1.00 | 47.43 |  |
| 7 | U | O4 | AA:2584 | 4.51 | 1.00 | 71.01 |  |
| 8 | G | O5' | AA:2583 | 4.67 | 1.00 | 47.81 |  |
| 9 | G | O3' | AA:2582 | 4.83 | 1.00 | 46.75 |  |
| 10 | G | O5' | AA:2582 | 4.89 | 1.00 | 46.03 |  |
| 11 | G | O2' | AA:2581 | 4.94 | 1.00 | 46.88 |  |
| 12 | G | OP1 | AA:2582 | 4.96 | 1.00 | 47.00 |  |
  |

  
 Coded contact list:   
 Exclusion zone contact(s): /OP2/OP2/  
Consider NA in place of MG (2 valid contact(s) for NA vs. 0 for MG)  

### MG:AA:3477

|  | Res. type | Atom | Residue | Distance | Occupancy | B-factor | Closest solute to 1^st^ shell water |
| --- | --- | --- | --- | --- | --- | --- | --- |
| **Ion** | | | |  | 1.00 | 65.14 || 1 | G | O6 | AA:2495 | 2.94 | 1.00 | 63.93 |
| 2 | G | O6 | AA:2455 | 3.74 | 1.00 | 52.73 |  |
| 3 | C | N4 | AA:2456 | 3.75 | 1.00 | 62.14 |  |
| 4 | G | O6 | AA:2494 | 3.97 | 1.00 | 66.25 |  |
| 5 | G | N7 | AA:2494 | 4.52 | 1.00 | 66.77 |  |
| 6 | U | O4 | AA:2457 | 4.71 | 1.00 | 65.15 |  |
| 7 | C | N4 | AA:2496 | 4.74 | 1.00 | 51.69 |  |
| 8 | G | N7 | AA:2495 | 4.79 | 1.00 | 61.24 |  |
  |

  
 Coded contact list:   
 Exclusion zone contact(s): /O6/  
Consider K in place of MG (1 valid contact(s) for K vs. 0 for MG)  

### MG:AA:3478

|  | Res. type | Atom | Residue | Distance | Occupancy | B-factor | Closest solute to 1^st^ shell water |
| --- | --- | --- | --- | --- | --- | --- | --- |
| **Ion** | | | |  | 1.00 | 59.96 || 1 | U | O4 | AA:2554 | 2.85 | 1.00 | 51.80 |
| 2 | U | O4 | AA:2555 | 3.01 | 1.00 | 63.44 |  |
| 3 | C | O2 | AA:2556 | 3.29 | 1.00 | 61.17 |  |
| 4 | U | O4 | AA:1955 | 3.84 | 1.00 | 51.32 |  |
| 5 | C | N3 | AA:2556 | 4.15 | 1.00 | 59.60 |  |
| 6 | U | N3 | AA:2554 | 4.67 | 1.00 | 51.47 |  |
  |

  
 Coded contact list:   
 Exclusion zone contact(s): /O2/O4/O4/  
Consider K in place of MG (3 valid contact(s) for K vs. 0 for MG)  

### MG:AA:3479

|  | Res. type | Atom | Residue | Distance | Occupancy | B-factor | Closest solute to 1^st^ shell water |
| --- | --- | --- | --- | --- | --- | --- | --- |
| **Ion** | | | |  | 1.00 | 59.42 || 1 | G | OP1 | AA:2582 | 2.50 | 1.00 | 47.00 |
| 2 | G | OP2 | AA:2608 | 4.11 | 1.00 | 39.47 |  |
| 3 | G | OP2 | AA:2582 | 4.14 | 1.00 | 47.17 |  |
| 4 | G | OP1 | AA:2608 | 4.22 | 1.00 | 38.19 |  |
| 5 | U | OP1 | AA:2609 | 4.43 | 1.00 | 45.47 |  |
| 6 | G | O5' | AA:2582 | 4.76 | 1.00 | 46.03 |  |
| 7 | G | O5' | AA:2608 | 4.78 | 1.00 | 38.21 |  |
| 8 | G | O3' | AA:2581 | 4.83 | 1.00 | 44.83 |  |
| 9 | G | OP1 | AA:742 | 4.95 | 1.00 | 42.30 |  |
  |

  
 Coded contact list:   
 Exclusion zone contact(s): /OP1/  
Consider NA in place of MG (1 valid contact(s) for NA vs. 0 for MG)  

### MG:AA:3480

|  | Res. type | Atom | Residue | Distance | Occupancy | B-factor | Closest solute to 1^st^ shell water |
| --- | --- | --- | --- | --- | --- | --- | --- |
| **Ion** | | | |  | 1.00 | 70.99 || 1 | C | OP2 | AA:2474 | 3.56 | 1.00 | 147.94 |
| 2 | C | OP1 | AA:2474 | 3.66 | 1.00 | 147.78 |  |
| 3 | C | OP2 | AA:2475 | 3.97 | 1.00 | 148.50 |  |
| 4 | C | OP2 | AA:2471 | 4.24 | 1.00 | 126.56 |  |
| 5 | C | O5' | AA:2474 | 4.70 | 1.00 | 147.82 |  |
| 6 | G | N7 | AA:2472 | 4.71 | 1.00 | 134.03 |  |
  |

  
 Coded contact list:   
1^st^ MG coordination shell missing (see Notes)  

### MG:AA:3481

|  | Res. type | Atom | Residue | Distance | Occupancy | B-factor | Closest solute to 1^st^ shell water |
| --- | --- | --- | --- | --- | --- | --- | --- |
| **Ion** | | | |  | 1.00 | 61.26 || 1 | G | OP1 | AA:2583 | 3.00 | 1.00 | 47.43 |
| 2 | U | OP2 | AA:2584 | 3.84 | 1.00 | 54.23 |  |
| 3 | G | N7 | AA:2607 | 4.38 | 1.00 | 36.22 |  |
| 4 | G | O6 | AA:2608 | 4.54 | 1.00 | 52.30 |  |
| 5 | G | OP2 | AA:2583 | 4.72 | 1.00 | 48.98 |  |
| 6 | G | O5' | AA:2583 | 4.83 | 1.00 | 47.81 |  |
| 7 | G | O6 | AA:2607 | 4.98 | 1.00 | 37.61 |  |
  |

  
 Coded contact list:   
 Exclusion zone contact(s): /OP1/  
Consider K in place of MG (1 valid contact(s) for K vs. 0 for MG)  

### MG:AA:3482

|  | Res. type | Atom | Residue | Distance | Occupancy | B-factor | Closest solute to 1^st^ shell water |
| --- | --- | --- | --- | --- | --- | --- | --- |
| **Ion** | | | |  | 1.00 | 58.21 || 1 | G | O5' | AA:488 | 2.54 | 1.00 | 55.71 |
| 2 | G | OP2 | AA:488 | 2.54 | 1.00 | 53.17 |  |
| 3 | G | N7 | AA:488 | 3.36 | 1.00 | 57.00 |  |
| 4 | G | OP2 | AA:489 | 3.41 | 1.00 | 55.47 |  |
| 5 | G | N9 | AA:488 | 3.89 | 1.00 | 55.67 |  |
| 6 | G | OP1 | AA:488 | 4.06 | 1.00 | 54.57 |  |
| 7 | C | O3' | AA:487 | 4.28 | 1.00 | 53.86 |  |
| 8 | G | O4' | AA:488 | 4.32 | 1.00 | 56.85 |  |
| 9 | G | O3' | AA:488 | 4.43 | 1.00 | 54.53 |  |
| 10 | G | O5' | AA:489 | 4.69 | 1.00 | 54.88 |  |
| 11 | G | OP2 | AA:491 | 4.92 | 1.00 | 54.47 |  |
| 12 | C | O5' | AA:487 | 4.95 | 1.00 | 54.41 |  |
  |

  
 Coded contact list:   
 Exclusion zone contact(s): /N7/O5'/OP2/  
Consider NA in place of MG (2 valid contact(s) for NA vs. 0 for MG)  

### MG:AA:3483

|  | Res. type | Atom | Residue | Distance | Occupancy | B-factor | Closest solute to 1^st^ shell water |
| --- | --- | --- | --- | --- | --- | --- | --- |
| **Ion** | | | |  | 1.00 | 65.58 || 1 | U | OP1 | AA:2554 | 2.49 | 1.00 | 49.20 |
| 2 | C | O3' | AA:1672 | 2.84 | 1.00 | 48.80 |  |
| 3 | U | OP2 | AA:2554 | 3.11 | 1.00 | 48.78 |  |
| 4 | U | OP1 | AA:1673 | 3.14 | 1.00 | 49.68 |  |
| 5 | U | OP2 | AA:2552 | 3.30 | 1.00 | 48.04 |  |
| 6 | G | OP2 | AA:2553 | 3.77 | 1.00 | 45.56 |  |
| 7 | G | O3' | AA:2553 | 4.14 | 1.00 | 50.36 |  |
| 8 | U | OP2 | AA:1673 | 4.33 | 1.00 | 51.17 |  |
| 9 | C | O2' | AA:1672 | 4.35 | 1.00 | 51.68 |  |
| 10 | U | O5' | AA:2554 | 4.53 | 1.00 | 48.22 |  |
| 11 | U | N1 | AA:2552 | 4.77 | 1.00 | 45.51 |  |
| 12 | U | O5' | AA:1673 | 4.84 | 1.00 | 47.71 |  |
| 13 | U | O5' | AA:2552 | 4.92 | 1.00 | 48.43 |  |
| 14 | G | O5' | AA:2553 | 4.94 | 1.00 | 48.27 |  |
  |

  
 Coded contact list:   
 Exclusion zone contact(s): /O3'/OP1/OP1/OP2/OP2/  
Consider K in place of MG (4 valid contact(s) for K vs. 0 for MG)  

### MG:AA:3484

|  | Res. type | Atom | Residue | Distance | Occupancy | B-factor | Closest solute to 1^st^ shell water |
| --- | --- | --- | --- | --- | --- | --- | --- |
| **Ion** | | | |  | 1.00 | 66.76 || 1 | G | O6 | AA:2490 | 2.27 | 1.00 | 81.60 |
| 2 | G | O6 | AA:2489 | 2.78 | 1.00 | 71.37 |  |
| 3 | G | N1 | AA:2490 | 3.06 | 1.00 | 81.66 |  |
| 4 | G | N7 | AA:2489 | 3.20 | 1.00 | 69.84 |  |
| 5 | A | N7 | AA:2488 | 3.55 | 1.00 | 69.95 |  |
| 6 | A | N6 | AA:2488 | 4.11 | 1.00 | 69.30 |  |
| 7 | A | OP2 | AA:2459 | 4.41 | 1.00 | 68.56 |  |
| 8 | A | OP2 | AA:2488 | 4.79 | 1.00 | 68.54 |  |
| 9 | G | N7 | AA:2490 | 4.85 | 1.00 | 76.30 |  |
| 10 | G | N2 | AA:2490 | 4.85 | 1.00 | 80.60 |  |
| 11 | G | N1 | AA:2489 | 4.88 | 1.00 | 69.97 |  |
| 12 | C | N4 | AA:2461 | 4.89 | 1.00 | 70.87 |  |
  |

  
 Contact(s): /O6/  
 Coded contact list: O_b_  
 Exclusion zone contact(s): /N1_G_/N7/O6/  
Consider K in place of MG (3 valid contact(s) for K vs. 1 for MG)  

### MG:AA:3485

|  | Res. type | Atom | Residue | Distance | Occupancy | B-factor | Closest solute to 1^st^ shell water |
| --- | --- | --- | --- | --- | --- | --- | --- |
| **Ion** | | | |  | 1.00 | 91.90 || 1 | G | OP1 | AA:491 | 3.20 | 1.00 | 54.39 |
| 2 | G | O6 | AA:488 | 3.84 | 1.00 | 54.95 |  |
| 3 | G | OP2 | AA:491 | 4.13 | 1.00 | 54.47 |  |
| 4 | G | N7 | AA:488 | 4.61 | 1.00 | 57.00 |  |
| 5 | G | N1 | AA:488 | 4.88 | 1.00 | 53.72 |  |
| 6 | G | O3' | AA:489 | 4.95 | 1.00 | 54.46 |  |
  |

  
 Coded contact list:   
 Exclusion zone contact(s): /OP1/  
Consider K in place of MG (1 valid contact(s) for K vs. 0 for MG)  

### MG:AA:3486

|  | Res. type | Atom | Residue | Distance | Occupancy | B-factor | Closest solute to 1^st^ shell water |
| --- | --- | --- | --- | --- | --- | --- | --- |
| **Ion** | | | |  | 1.00 | 60.56 || 1 | U | OP1 | AA:1955 | 2.65 | 1.00 | 54.38 |
| 2 | U | O2' | AA:1944 | 2.68 | 1.00 | 64.12 |  |
| 3 | U | O2 | AA:1944 | 3.06 | 1.00 | 58.39 |  |
| 4 | U | O3' | AA:1944 | 4.56 | 1.00 | 59.47 |  |
| 5 | U | N1 | AA:1944 | 4.70 | 1.00 | 61.37 |  |
| 6 | U | O5' | AA:1955 | 4.71 | 1.00 | 52.97 |  |
| 7 | G | O3' | AA:1954 | 4.73 | 1.00 | 55.81 |  |
  |

  
 Coded contact list:   
 Exclusion zone contact(s): /O2/O2'/OP1/  
Consider K in place of MG (3 valid contact(s) for K vs. 0 for MG)  

### MG:AA:3487

|  | Res. type | Atom | Residue | Distance | Occupancy | B-factor | Closest solute to 1^st^ shell water |
| --- | --- | --- | --- | --- | --- | --- | --- |
| **Ion** | | | |  | 1.00 | 72.21 || 1 | A | OP2 | AA:1916 | 3.68 | 1.00 | 79.95 |
| 2 | A | OP2 | AA:1912 | 4.10 | 1.00 | 64.47 |  |
| 3 | U | O4 | AA:1917 | 4.54 | 1.00 | 66.24 |  |
| 4 | A | N6 | AA:1918 | 4.58 | 1.00 | 69.74 |  |
  |

  
 Coded contact list:   
1^st^ MG coordination shell missing (see Notes)  

### MG:AA:3488

|  | Res. type | Atom | Residue | Distance | Occupancy | B-factor | Closest solute to 1^st^ shell water |
| --- | --- | --- | --- | --- | --- | --- | --- |
| **Ion** | | | |  | 1.00 | 68.19 || 1 | G | O6 | AA:2485 | 3.95 | 1.00 | 77.57 |
| 2 | G | O6 | AA:2484 | 4.52 | 1.00 | 76.63 |  |
| 3 | G | OP1 | AA:2482 | 4.64 | 1.00 | 106.78 |  |
| 4 | C | N4 | AA:2465 | 4.96 | 1.00 | 76.39 |  |
  |

  
 Coded contact list:   
1^st^ MG coordination shell missing (see Notes)  

### MG:AA:3489

|  | Res. type | Atom | Residue | Distance | Occupancy | B-factor | Closest solute to 1^st^ shell water |
| --- | --- | --- | --- | --- | --- | --- | --- |
| **Ion** | | | |  | 1.00 | 63.20 || 1 | C | N3 | AA:2499 | 2.83 | 1.00 | 49.88 |
| 2 | G | N7 | AA:2032 | 3.05 | 1.00 | 49.82 |  |
| 3 | G | N3 | AA:2454 | 3.41 | 1.00 | 44.26 |  |
| 4 | G | O6 | AA:2032 | 3.43 | 1.00 | 46.04 |  |
| 5 | C | N4 | AA:2499 | 3.53 | 1.00 | 51.17 |  |
| 6 | G | OP1 | AA:2032 | 3.61 | 1.00 | 58.74 |  |
| 7 | C | O2 | AA:2499 | 3.75 | 1.00 | 50.92 |  |
| 8 | G | N9 | AA:2454 | 4.09 | 1.00 | 46.78 |  |
| 9 | G | O2' | AA:2454 | 4.29 | 1.00 | 47.63 |  |
| 10 | G | N2 | AA:2454 | 4.44 | 1.00 | 45.07 |  |
| 11 | G | OP2 | AA:2032 | 4.55 | 1.00 | 58.90 |  |
| 12 | G | O4' | AA:2454 | 4.64 | 1.00 | 48.51 |  |
| 13 | A | N3 | AA:2453 | 4.76 | 1.00 | 40.86 |  |
| 14 | A | N1 | AA:2453 | 4.79 | 1.00 | 40.78 |  |
| 15 | C | N1 | AA:2499 | 4.84 | 1.00 | 45.72 |  |
| 16 | U | O2 | AA:2500 | 4.90 | 1.00 | 38.41 |  |
  |

  
 Coded contact list:   
 Exclusion zone contact(s): /N3_C_/N7/  
Consider K in place of MG (2 valid contact(s) for K vs. 0 for MG)  

### MG:AA:3490

|  | Res. type | Atom | Residue | Distance | Occupancy | B-factor | Closest solute to 1^st^ shell water |
| --- | --- | --- | --- | --- | --- | --- | --- |
| **Ion** | | | |  | 1.00 | 102.04 || 1 | C | N4 | AA:893 | 2.65 | 1.00 | 200.41 |
| 2 | U | O4 | AA:895 | 4.27 | 1.00 | 199.38 |  |
| 3 | C | N3 | AA:893 | 4.53 | 1.00 | 200.41 |  |
| 4 | C | N4 | AA:894 | 4.55 | 1.00 | 200.23 |  |
| 5 | G | O6 | AA:883 | 4.87 | 1.00 | 200.41 |  |
  |

  
 Coded contact list:   
 Exclusion zone contact(s): /N4/  
Consider K in place of MG (1 valid contact(s) for K vs. 0 for MG)  

### MG:AA:3491

|  | Res. type | Atom | Residue | Distance | Occupancy | B-factor | Closest solute to 1^st^ shell water |
| --- | --- | --- | --- | --- | --- | --- | --- |
| **Ion** | | | |  | 1.00 | 68.35 || 1 | G | O6 | AA:2548 | 2.92 | 1.00 | 49.84 |
| 2 | G | N7 | AA:2548 | 3.20 | 1.00 | 43.68 |  |
| 3 | U | O4 | AA:2547 | 3.48 | 1.00 | 51.14 |  |
| 4 | G | O6 | AA:2549 | 3.91 | 1.00 | 51.15 |  |
| 5 | G | N7 | AA:2549 | 4.11 | 1.00 | 50.17 |  |
| 6 | U | N3 | AA:2547 | 4.40 | 1.00 | 50.46 |  |
| 7 | G | N1 | AA:2548 | 4.92 | 1.00 | 47.63 |  |
| 8 | A | N6 | AA:2561 | 4.94 | 1.00 | 49.97 |  |
  |

  
 Coded contact list:   
 Exclusion zone contact(s): /N7/O6/  
Consider K in place of MG (2 valid contact(s) for K vs. 0 for MG)  

### MG:AA:3492

|  | Res. type | Atom | Residue | Distance | Occupancy | B-factor | Closest solute to 1^st^ shell water |
| --- | --- | --- | --- | --- | --- | --- | --- |
| **Ion** | | | |  | 1.00 | 48.57 || 1 | C | O5' | AA:2368 | 2.01 | 1.00 | 74.61 |
| 2 | A | OP2 | AA:2369 | 2.64 | 1.00 | 76.13 |  |
| 3 | C | O3' | AA:2368 | 2.93 | 1.00 | 75.49 |  |
| 4 | C | OP2 | AA:2368 | 3.43 | 1.00 | 74.73 |  |
| 5 | C | O4' | AA:2368 | 3.62 | 1.00 | 74.17 |  |
| 6 | C | OP1 | AA:2368 | 3.65 | 1.00 | 75.51 |  |
| 7 | C | N1 | AA:2368 | 4.00 | 1.00 | 69.37 |  |
| 8 | A | O5' | AA:2369 | 4.01 | 1.00 | 75.15 |  |
| 9 | G | O3' | AA:2367 | 4.44 | 1.00 | 74.70 |  |
| 10 | A | OP1 | AA:2369 | 4.65 | 1.00 | 76.66 |  |
| 11 | C | O2' | AA:2368 | 4.78 | 1.00 | 73.91 |  |
  |

  
 Contact(s): /O5'/  
 Coded contact list: O_r_  
 Exclusion zone contact(s): /O3'/OP2/  
Consider K in place of MG (2 valid contact(s) for K vs. 1 for MG)  

### MG:AA:3493

|  | Res. type | Atom | Residue | Distance | Occupancy | B-factor | Closest solute to 1^st^ shell water |
| --- | --- | --- | --- | --- | --- | --- | --- |
| **Ion** | | | |  | 1.00 | 76.97 || 1 | C | OP2 | AA:894 | 4.53 | 1.00 | 200.39 |
  |

  
 Coded contact list:   
1^st^ MG coordination shell missing (see Notes)  

### MG:AA:3494

|  | Res. type | Atom | Residue | Distance | Occupancy | B-factor | Closest solute to 1^st^ shell water |
| --- | --- | --- | --- | --- | --- | --- | --- |
| **Ion** | | | |  | 1.00 | 69.96 || 1 | G | O5' | AA:2370 | 2.21 | 1.00 | 75.58 |
| 2 | G | OP2 | AA:2370 | 2.59 | 1.00 | 75.99 |  |
| 3 | G | OP2 | AA:2371 | 2.93 | 1.00 | 72.62 |  |
| 4 | G | OP1 | AA:2370 | 3.62 | 1.00 | 77.02 |  |
| 5 | G | N7 | AA:2371 | 3.87 | 1.00 | 70.35 |  |
| 6 | G | O4' | AA:2370 | 4.18 | 1.00 | 72.29 |  |
| 7 | G | O3' | AA:2370 | 4.21 | 1.00 | 73.47 |  |
| 8 | A | O3' | AA:2369 | 4.21 | 1.00 | 75.85 |  |
| 9 | G | N9 | AA:2370 | 4.45 | 1.00 | 70.92 |  |
| 10 | G | N7 | AA:2370 | 4.65 | 1.00 | 71.03 |  |
| 11 | G | O5' | AA:2371 | 4.72 | 1.00 | 71.67 |  |
  |

  
 Contact(s): /O5'/  
 Coded contact list: O_r_  
 Exclusion zone contact(s): /OP2/OP2/  

### MG:AA:3495

|  | Res. type | Atom | Residue | Distance | Occupancy | B-factor | Closest solute to 1^st^ shell water |
| --- | --- | --- | --- | --- | --- | --- | --- |
| **Ion** | | | |  | 1.00 | 56.49 || 1 | A | OP1 | AA:2577 | 2.92 | 1.00 | 44.36 |
| 2 | G | O6 | AA:2056 | 2.95 | 1.00 | 40.78 |  |
| 3 | A | N7 | AA:2577 | 3.22 | 1.00 | 43.50 |  |
| 4 | A | OP2 | AA:2577 | 3.31 | 1.00 | 43.39 |  |
| 5 | A | O5' | AA:2577 | 3.57 | 1.00 | 42.97 |  |
| 6 | G | N7 | AA:2056 | 3.60 | 1.00 | 39.82 |  |
| 7 | A | N9 | AA:2577 | 3.89 | 1.00 | 42.09 |  |
| 8 | A | O4' | AA:2577 | 3.99 | 1.00 | 43.18 |  |
| 9 | G | O3' | AA:2576 | 4.90 | 1.00 | 43.10 |  |
  |

  
 Coded contact list:   
 Exclusion zone contact(s): /N7/O6/OP1/OP2/  
Consider K in place of MG (4 valid contact(s) for K vs. 0 for MG)  

### MG:AA:3496

|  | Res. type | Atom | Residue | Distance | Occupancy | B-factor | Closest solute to 1^st^ shell water |
| --- | --- | --- | --- | --- | --- | --- | --- |
| **Ion** | | | |  | 1.00 | 63.04 || 1 | A | O5' | AA:2469 | 2.82 | 1.00 | 122.11 |
| 2 | G | O5' | AA:2470 | 3.07 | 1.00 | 125.99 |  |
| 3 | A | OP1 | AA:2469 | 3.18 | 1.00 | 121.57 |  |
| 4 | A | OP2 | AA:2469 | 3.69 | 1.00 | 121.06 |  |
| 5 | G | N7 | AA:2470 | 3.89 | 1.00 | 122.61 |  |
| 6 | A | O4' | AA:2476 | 4.03 | 1.00 | 145.60 |  |
| 7 | A | O3' | AA:2469 | 4.23 | 1.00 | 125.97 |  |
| 8 | G | O4' | AA:2470 | 4.66 | 1.00 | 124.97 |  |
| 9 | G | OP2 | AA:2470 | 4.69 | 1.00 | 126.97 |  |
| 10 | G | N9 | AA:2470 | 4.83 | 1.00 | 123.54 |  |
| 11 | G | O3' | AA:2468 | 4.87 | 1.00 | 119.47 |  |
  |

  
 Coded contact list:   
 Exclusion zone contact(s): /O5'/O5'/OP1/  
Consider K in place of MG (3 valid contact(s) for K vs. 0 for MG)  

### MG:AA:3497

|  | Res. type | Atom | Residue | Distance | Occupancy | B-factor | Closest solute to 1^st^ shell water |
| --- | --- | --- | --- | --- | --- | --- | --- |
| **Ion** | | | |  | 1.00 | 80.92 || 1 | G | O6 | AA:2371 | 3.61 | 1.00 | 73.80 |
| 2 | G | N7 | AA:2370 | 3.82 | 1.00 | 71.03 |  |
| 3 | G | OP2 | AA:2370 | 3.99 | 1.00 | 75.99 |  |
| 4 | G | N7 | AA:2371 | 4.28 | 1.00 | 70.35 |  |
| 5 | A | OP2 | AA:2369 | 4.31 | 1.00 | 76.13 |  |
  |

  
 Coded contact list:   
1^st^ MG coordination shell missing (see Notes)  

### MG:AA:3498

|  | Res. type | Atom | Residue | Distance | Occupancy | B-factor | Closest solute to 1^st^ shell water |
| --- | --- | --- | --- | --- | --- | --- | --- |
| **Ion** | | | |  | 1.00 | 52.42 || 1 | U | OP1 | AA:2506 | 2.64 | 1.00 | 58.11 |
| 2 | U | O2 | AA:2504 | 2.71 | 1.00 | 43.63 |  |
| 3 | G | O3' | AA:2505 | 2.93 | 1.00 | 55.36 |  |
| 4 | U | O2' | AA:2504 | 3.46 | 1.00 | 45.06 |  |
| 5 | U | OP2 | AA:2506 | 3.68 | 1.00 | 57.69 |  |
| 6 | ARG | NH2 | AE:144 | 3.87 | 1.00 | 52.73 |  |
| 7 | C | O2 | AA:2452 | 3.89 | 1.00 | 52.56 |  |
| 8 | ARG | NH1 | AE:144 | 4.37 | 1.00 | 51.41 |  |
| 9 | A | N3 | AA:2572 | 4.43 | 1.00 | 52.24 |  |
| 10 | U | N1 | AA:2504 | 4.47 | 1.00 | 44.37 |  |
| 11 | U | N3 | AA:2504 | 4.63 | 1.00 | 45.96 |  |
| 12 | U | O5' | AA:2506 | 4.64 | 1.00 | 59.73 |  |
| 13 | A | N1 | AA:2572 | 4.69 | 1.00 | 53.43 |  |
| 14 | U | O3' | AA:2504 | 4.82 | 1.00 | 44.23 |  |
| 15 | G | O5' | AA:2505 | 4.93 | 1.00 | 46.77 |  |
  |

  
 Coded contact list:   
 Exclusion zone contact(s): /O2/O3'/OP1/  
Consider K in place of MG (3 valid contact(s) for K vs. 0 for MG)  

### MG:AA:3499

|  | Res. type | Atom | Residue | Distance | Occupancy | B-factor | Closest solute to 1^st^ shell water |
| --- | --- | --- | --- | --- | --- | --- | --- |
| **Ion** | | | |  | 1.00 | 101.49 || 1 | G | O6 | AA:2371 | 2.18 | 1.00 | 73.80 |
| 2 | G | O6 | AA:2345 | 2.86 | 1.00 | 56.95 |  |
| 3 | G | O6 | AA:2372 | 3.74 | 1.00 | 63.31 |  |
| 4 | G | O6 | AA:2370 | 3.74 | 1.00 | 69.78 |  |
| 5 | G | N1 | AA:2371 | 3.91 | 1.00 | 72.45 |  |
| 6 | G | N7 | AA:2370 | 4.66 | 1.00 | 71.03 |  |
| 7 | C | N4 | AA:2381 | 4.66 | 1.00 | 62.24 |  |
| 8 | G | N1 | AA:2345 | 4.67 | 1.00 | 58.72 |  |
| 9 | G | N7 | AA:2372 | 4.83 | 1.00 | 62.75 |  |
  |

  
 Contact(s): /O6/  
 Coded contact list: O_b_  
 Exclusion zone contact(s): /O6/  

### MG:AA:3500

|  | Res. type | Atom | Residue | Distance | Occupancy | B-factor | Closest solute to 1^st^ shell water |
| --- | --- | --- | --- | --- | --- | --- | --- |
| **Ion** | | | |  | 1.00 | 22.71 || 1 | G | OP2 | AA:2502 | 1.81 | 1.00 | 40.13 |
| 2 | G | OP2 | AA:2061 | 2.88 | 1.00 | 40.30 |  |
| 3 | A | O2' | AA:2060 | 3.05 | 1.00 | 45.21 |  |
| 4 | G | OP1 | AA:2502 | 3.89 | 1.00 | 40.75 |  |
| 5 | G | O5' | AA:2502 | 4.05 | 1.00 | 40.60 |  |
| 6 | C | O2' | AA:2501 | 4.07 | 1.00 | 44.45 |  |
| 7 | G | N7 | AA:2446 | 4.07 | 1.00 | 41.61 |  |
| 8 | G | OP2 | AA:2446 | 4.25 | 1.00 | 37.51 |  |
| 9 | C | O3' | AA:2501 | 4.27 | 1.00 | 40.30 |  |
| 10 | G | N7 | AA:2445 | 4.48 | 1.00 | 31.96 |  |
| 11 | G | OP1 | AA:2061 | 4.65 | 1.00 | 42.19 |  |
| 12 | A | N3 | AA:2060 | 4.98 | 1.00 | 38.75 |  |
  |

  
 Contact(s): /OP2/  
 Coded contact list: O_ph_  
 Exclusion zone contact(s): /O2'/OP2/  
Consider K in place of MG (2 valid contact(s) for K vs. 1 for MG)  

### MG:AA:3501

|  | Res. type | Atom | Residue | Distance | Occupancy | B-factor | Closest solute to 1^st^ shell water |
| --- | --- | --- | --- | --- | --- | --- | --- |
| **Ion** | | | |  | 1.00 | 24.65 || 1 | G | OP2 | AA:785 | 2.36 | 1.00 | 35.38 |
| 2 | C | O2' | AA:2440 | 3.24 | 1.00 | 41.16 |  |
| 3 | G | OP2 | AA:792 | 3.67 | 1.00 | 33.79 |  |
| 4 | G | N7 | AA:792 | 4.17 | 1.00 | 42.77 |  |
| 5 | A | O3' | AA:784 | 4.31 | 1.00 | 36.13 |  |
| 6 | G | OP1 | AA:785 | 4.48 | 1.00 | 36.44 |  |
| 7 | C | O2 | AA:2440 | 4.60 | 1.00 | 44.68 |  |
| 8 | A | O2' | AA:2587 | 4.80 | 1.00 | 40.26 |  |
| 9 | G | O5' | AA:785 | 4.84 | 1.00 | 33.13 |  |
| 10 | C | O3' | AA:791 | 4.87 | 1.00 | 35.98 |  |
  |

  
 Contact(s): /OP2/  
 Coded contact list: O_ph_  
 Exclusion zone contact(s): /O2'/  

### MG:AA:3502

|  | Res. type | Atom | Residue | Distance | Occupancy | B-factor | Closest solute to 1^st^ shell water |
| --- | --- | --- | --- | --- | --- | --- | --- |
| **Ion** | | | |  | 1.00 | 55.35 || 1 | U | OP1 | AA:773 | 2.57 | 1.00 | 34.95 |
| 2 | C | O3' | AA:772 | 3.33 | 1.00 | 37.18 |  |
| 3 | G | OP1 | AA:1356 | 3.52 | 1.00 | 48.31 |  |
| 4 | U | O5' | AA:773 | 4.45 | 1.00 | 37.05 |  |
| 5 | C | OP1 | AA:1370 | 4.59 | 1.00 | 46.86 |  |
| 6 | U | OP2 | AA:773 | 4.74 | 1.00 | 35.58 |  |
  |

  
 Coded contact list:   
 Exclusion zone contact(s): /O3'/OP1/  
Consider NA in place of MG (1 valid contact(s) for NA vs. 0 for MG)  

### MG:AA:3503

|  | Res. type | Atom | Residue | Distance | Occupancy | B-factor | Closest solute to 1^st^ shell water |
| --- | --- | --- | --- | --- | --- | --- | --- |
| **Ion** | | | |  | 1.00 | 23.33 || 1 | A | OP2 | AA:2448 | 2.04 | 1.00 | 42.32 |
| 2 | C | OP2 | AA:2499 | 2.06 | 1.00 | 42.95 |  |
| 3 | C | OP2 | AA:2498 | 2.83 | 1.00 | 43.21 |  |
| 4 | A | O3' | AA:2497 | 2.88 | 1.00 | 44.25 |  |
| 5 | G | O2' | AA:2447 | 2.96 | 1.00 | 34.27 |  |
| 6 | A | OP1 | AA:2448 | 3.32 | 1.00 | 40.10 |  |
| 7 | A | O2' | AA:2497 | 3.34 | 1.00 | 49.84 |  |
| 8 | C | O5' | AA:2498 | 3.84 | 1.00 | 41.23 |  |
| 9 | A | O5' | AA:2448 | 4.13 | 1.00 | 40.15 |  |
| 10 | C | O3' | AA:2498 | 4.14 | 1.00 | 40.63 |  |
| 11 | C | OP1 | AA:2499 | 4.22 | 1.00 | 43.06 |  |
| 12 | U | OP2 | AA:2449 | 4.23 | 1.00 | 50.78 |  |
| 13 | G | O3' | AA:2447 | 4.32 | 1.00 | 38.23 |  |
| 14 | C | O5' | AA:2499 | 4.53 | 1.00 | 43.17 |  |
| 15 | C | OP1 | AA:2498 | 4.81 | 1.00 | 44.33 |  |
  |

  
**1^st^ shell ligand angles:**   
1-2: 89;   
  
 Contact(s): /OP2/OP2/  
 Coded contact list: 2O_ph_  
 Exclusion zone contact(s): /O2'/O2'/O3'/OP1/OP2/  
Consider K in place of MG (5 valid contact(s) for K vs. 2 for MG)  

### MG:AA:3504

|  | Res. type | Atom | Residue | Distance | Occupancy | B-factor | Closest solute to 1^st^ shell water |
| --- | --- | --- | --- | --- | --- | --- | --- |
| **Ion** | | | |  | 1.00 | 31.60 || 1 | G | O6 | AA:570 | 1.93 | 1.00 | 39.82 |
| 2 | C | OP1 | AA:2499 | 2.04 | 1.00 | 43.06 |  |
| 3 | A | OP1 | AA:2448 | 2.15 | 1.00 | 40.10 |  |
| 4 | G | O3' | AA:2447 | 3.32 | 1.00 | 38.23 |  |
| 5 | G | N1 | AA:570 | 3.72 | 1.00 | 34.94 |  |
| 6 | U | O4 | AA:569 | 3.97 | 1.00 | 46.05 |  |
| 7 | U | O2' | AA:568 | 4.07 | 1.00 | 47.08 |  |
| 8 | A | O5' | AA:2448 | 4.13 | 1.00 | 40.15 |  |
| 9 | C | OP2 | AA:2499 | 4.15 | 1.00 | 42.95 |  |
| 10 | C | O3' | AA:2498 | 4.20 | 1.00 | 40.63 |  |
| 11 | U | OP2 | AA:2500 | 4.44 | 1.00 | 42.75 |  |
| 12 | C | O5' | AA:2499 | 4.51 | 1.00 | 43.17 |  |
| 13 | A | OP2 | AA:2448 | 4.52 | 1.00 | 42.32 |  |
| 14 | G | N7 | AA:570 | 4.87 | 1.00 | 38.59 |  |
  |

  
**1^st^ shell ligand angles:**   
1-2: 86; 1-3: 134; 2-3: 74;   
  
 Contact(s): /O6/OP1/OP1/  
 Coded contact list: 2O_ph_O_b_  
 Exclusion zone contact(s): /O3'/  

### MG:AA:3505

|  | Res. type | Atom | Residue | Distance | Occupancy | B-factor | Closest solute to 1^st^ shell water |
| --- | --- | --- | --- | --- | --- | --- | --- |
| **Ion** | | | |  | 1.00 | 55.64 || 1 | G | OP2 | AA:1356 | 2.56 | 1.00 | 46.99 |
| 2 | G | OP1 | AA:1356 | 3.57 | 1.00 | 48.31 |  |
| 3 | G | OP2 | AA:1371 | 3.60 | 1.00 | 53.24 |  |
| 4 | G | O3' | AA:1355 | 4.49 | 1.00 | 46.08 |  |
| 5 | G | OP1 | AA:1371 | 4.63 | 1.00 | 53.14 |  |
| 6 | G | O5' | AA:1356 | 4.76 | 1.00 | 47.90 |  |
  |

  
 Coded contact list:   
 Exclusion zone contact(s): /OP2/  
Consider NA in place of MG (1 valid contact(s) for NA vs. 0 for MG)  

### MG:AA:3506

|  | Res. type | Atom | Residue | Distance | Occupancy | B-factor | Closest solute to 1^st^ shell water |
| --- | --- | --- | --- | --- | --- | --- | --- |
| **Ion** | | | |  | 1.00 | 27.05 || 1 | U | OP1 | AA:963 | 2.35 | 1.00 | 58.34 |
| 2 | C | OP1 | AA:2498 | 3.62 | 1.00 | 44.33 |  |
| 3 | G | OP1 | AA:947 | 4.21 | 1.00 | 52.58 |  |
| 4 | U | O5' | AA:963 | 4.35 | 1.00 | 57.72 |  |
| 5 | A | OP1 | AA:2497 | 4.39 | 1.00 | 44.57 |  |
| 6 | U | OP2 | AA:963 | 4.64 | 1.00 | 57.28 |  |
| 7 | G | O3' | AA:962 | 4.77 | 1.00 | 56.80 |  |
  |

  
 Contact(s): /OP1/  
 Coded contact list: O_ph_  

### MG:AA:3507

|  | Res. type | Atom | Residue | Distance | Occupancy | B-factor | Closest solute to 1^st^ shell water |
| --- | --- | --- | --- | --- | --- | --- | --- |
| **Ion** | | | |  | 1.00 | 30.00 || 1 | A | OP1 | AA:2062 | 1.97 | 1.00 | 50.65 |
| 2 | A | OP2 | AA:2062 | 3.53 | 1.00 | 52.33 |  |
| 3 | G | O3' | AA:2061 | 3.81 | 1.00 | 48.77 |  |
| 4 | G | OP2 | AA:2444 | 3.83 | 1.00 | 38.41 |  |
| 5 | G | OP1 | AA:2061 | 3.89 | 1.00 | 42.19 |  |
| 6 | A | O5' | AA:2062 | 4.38 | 1.00 | 51.57 |  |
| 7 | C | OP2 | AA:2443 | 4.49 | 1.00 | 38.53 |  |
| 8 | G | N7 | AA:2444 | 4.55 | 1.00 | 27.81 |  |
  |

  
 Contact(s): /OP1/  
 Coded contact list: O_ph_  

### MG:AA:3508

|  | Res. type | Atom | Residue | Distance | Occupancy | B-factor | Closest solute to 1^st^ shell water |
| --- | --- | --- | --- | --- | --- | --- | --- |
| **Ion** | | | |  | 1.00 | 61.75 || 1 | U | O4 | AA:1357 | 3.10 | 1.00 | 53.78 |
| 2 | G | N1 | AA:1358 | 3.36 | 1.00 | 68.14 |  |
| 3 | U | OP2 | AA:1372 | 3.61 | 1.00 | 74.81 |  |
| 4 | U | OP1 | AA:1372 | 3.79 | 1.00 | 74.99 |  |
| 5 | G | O6 | AA:1358 | 3.94 | 1.00 | 70.99 |  |
| 6 | G | N2 | AA:1358 | 4.11 | 1.00 | 67.89 |  |
| 7 | G | O6 | AA:1356 | 4.75 | 1.00 | 49.09 |  |
| 8 | G | O6 | AA:1374 | 4.76 | 1.00 | 52.27 |  |
| 9 | A | OP2 | AA:1373 | 4.78 | 1.00 | 78.04 |  |
  |

  
 Coded contact list:   
 Exclusion zone contact(s): /N1_G_/O4/  
Consider K in place of MG (2 valid contact(s) for K vs. 0 for MG)  

### MG:AA:3509

|  | Res. type | Atom | Residue | Distance | Occupancy | B-factor | Closest solute to 1^st^ shell water |
| --- | --- | --- | --- | --- | --- | --- | --- |
| **Ion** | | | |  | 1.00 | 19.26 || 1 | G | OP1 | AA:2588 | 2.36 | 1.00 | 36.17 |
| 2 | A | OP1 | AA:784 | 2.41 | 1.00 | 35.96 |  |
| 3 | G | O6 | AA:792 | 3.79 | 1.00 | 40.22 |  |
| 4 | A | OP2 | AA:2589 | 3.93 | 1.00 | 32.60 |  |
| 5 | ASN | ND2 | AD:227 | 3.93 | 1.00 | 32.73 |  |
| 6 | A | O5' | AA:784 | 4.05 | 1.00 | 37.42 |  |
| 7 | A | OP2 | AA:784 | 4.48 | 1.00 | 38.65 |  |
| 8 | A | O3' | AA:2587 | 4.50 | 1.00 | 37.43 |  |
| 9 | G | OP2 | AA:2588 | 4.56 | 1.00 | 35.91 |  |
| 10 | G | O5' | AA:2588 | 4.76 | 1.00 | 33.02 |  |
| 11 | A | O3' | AA:783 | 4.88 | 1.00 | 38.25 |  |
| 12 | C | OP2 | AA:2440 | 4.93 | 1.00 | 38.80 |  |
  |

  
 Contact(s): /OP1/  
 Coded contact list: O_ph_  
 Exclusion zone contact(s): /OP1/  

### MG:AA:3510

|  | Res. type | Atom | Residue | Distance | Occupancy | B-factor | Closest solute to 1^st^ shell water |
| --- | --- | --- | --- | --- | --- | --- | --- |
| **Ion** | | | |  | 1.00 | 26.83 || 1 | G | OP2 | AA:2588 | 2.19 | 1.00 | 35.91 |
| 2 | A | OP1 | AA:2587 | 3.88 | 1.00 | 45.57 |  |
| 3 | G | O5' | AA:2588 | 4.12 | 1.00 | 33.02 |  |
| 4 | A | N7 | AA:2589 | 4.17 | 1.00 | 25.97 |  |
| 5 | G | OP1 | AA:2588 | 4.21 | 1.00 | 36.17 |  |
| 6 | G | N7 | AA:2588 | 4.47 | 1.00 | 29.01 |  |
| 7 | A | OP2 | AA:2589 | 4.51 | 1.00 | 32.60 |  |
| 8 | A | O2' | AA:2439 | 4.63 | 1.00 | 46.71 |  |
| 9 | A | O3' | AA:2587 | 4.74 | 1.00 | 37.43 |  |
| 10 | A | O5' | AA:2587 | 4.89 | 1.00 | 43.35 |  |
  |

  
 Contact(s): /OP2/  
 Coded contact list: O_ph_  

### MG:AA:3511

|  | Res. type | Atom | Residue | Distance | Occupancy | B-factor | Closest solute to 1^st^ shell water |
| --- | --- | --- | --- | --- | --- | --- | --- |
| **Ion** | | | |  | 1.00 | 26.97 || 1 | G | O6 | AA:1355 | 3.25 | 1.00 | 55.84 |
| 2 | A | OP2 | AA:1354 | 3.61 | 1.00 | 45.86 |  |
| 3 | G | O6 | AA:1356 | 3.92 | 1.00 | 49.09 |  |
| 4 | A | OP1 | AA:1353 | 3.97 | 1.00 | 43.87 |  |
| 5 | A | OP2 | AA:1353 | 4.19 | 1.00 | 44.20 |  |
| 6 | G | N7 | AA:1355 | 4.32 | 1.00 | 52.87 |  |
| 7 | A | OP1 | AA:1354 | 4.49 | 1.00 | 48.14 |  |
| 8 | C | N4 | AA:1375 | 4.54 | 1.00 | 47.62 |  |
| 9 | C | N4 | AA:1376 | 4.72 | 1.00 | 54.62 |  |
  |

  
 Coded contact list:   
 Exclusion zone contact(s): /O6/  
Consider K in place of MG (1 valid contact(s) for K vs. 0 for MG)  

### MG:AA:3512

|  | Res. type | Atom | Residue | Distance | Occupancy | B-factor | Closest solute to 1^st^ shell water |
| --- | --- | --- | --- | --- | --- | --- | --- |
| **Ion** | | | |  | 1.00 | 31.24 || 1 | A | O2' | AA:2439 | 2.31 | 1.00 | 46.71 |
| 2 | C | OP2 | AA:2440 | 2.47 | 1.00 | 38.80 |  |
| 3 | A | OP2 | AA:2600 | 2.74 | 1.00 | 35.23 |  |
| 4 | GLU | OE2 | AD:237 | 3.72 | 1.00 | 48.44 |  |
| 5 | GLU | OE1 | AD:237 | 3.95 | 1.00 | 47.26 |  |
| 6 | A | OP1 | AA:2600 | 4.32 | 1.00 | 37.10 |  |
| 7 | A | O3' | AA:2439 | 4.35 | 1.00 | 41.35 |  |
| 8 | G | OP2 | AA:2588 | 4.49 | 1.00 | 35.91 |  |
| 9 | G | O3' | AA:2599 | 4.62 | 1.00 | 35.24 |  |
| 10 | C | OP1 | AA:2440 | 4.70 | 1.00 | 39.85 |  |
| 11 | C | O5' | AA:2440 | 4.89 | 1.00 | 41.04 |  |
  |

  
 Contact(s): /O2'/  
 Coded contact list: O_r_  
 Exclusion zone contact(s): /OP2/OP2/  

### MG:AA:3513

|  | Res. type | Atom | Residue | Distance | Occupancy | B-factor | Closest solute to 1^st^ shell water |
| --- | --- | --- | --- | --- | --- | --- | --- |
| **Ion** | | | |  | 1.00 | 28.73 || 1 | G | OP1 | AA:2597 | 3.48 | 1.00 | 27.51 |
| 2 | G | N7 | AA:2595 | 3.78 | 1.00 | 34.50 |  |
| 3 | G | O6 | AA:2595 | 4.19 | 1.00 | 39.06 |  |
| 4 | G | OP2 | AA:2597 | 4.27 | 1.00 | 29.59 |  |
| 5 | G | OP1 | AA:1904 | 4.43 | 1.00 | 46.78 |  |
| 6 | U | O3' | AA:2596 | 4.72 | 1.00 | 30.21 |  |
  |

  
 Coded contact list:   
1^st^ MG coordination shell missing (see Notes)  

### MG:AA:3514

|  | Res. type | Atom | Residue | Distance | Occupancy | B-factor | Closest solute to 1^st^ shell water |
| --- | --- | --- | --- | --- | --- | --- | --- |
| **Ion** | | | |  | 1.00 | 49.29 || 1 | G | OP2 | AA:1358 | 3.60 | 1.00 | 59.07 |
| 2 | G | N7 | AA:1358 | 3.87 | 1.00 | 67.06 |  |
| 3 | U | OP2 | AA:1357 | 3.95 | 1.00 | 51.56 |  |
| 4 | G | N7 | AA:1371 | 4.00 | 1.00 | 59.65 |  |
| 5 | C | OP2 | AA:1370 | 4.11 | 1.00 | 47.12 |  |
| 6 | G | O6 | AA:1371 | 4.11 | 1.00 | 55.04 |  |
| 7 | U | O5' | AA:1357 | 4.37 | 1.00 | 53.76 |  |
| 8 | U | OP1 | AA:1357 | 4.99 | 1.00 | 53.25 |  |
  |

  
 Coded contact list:   
1^st^ MG coordination shell missing (see Notes)  

### MG:AA:3515

|  | Res. type | Atom | Residue | Distance | Occupancy | B-factor | Closest solute to 1^st^ shell water |
| --- | --- | --- | --- | --- | --- | --- | --- |
| **Ion** | | | |  | 1.00 | 20.64 || 1 | U | O4 | AA:2593 | 2.53 | 1.00 | 36.40 |
| 2 | G | N7 | AA:2592 | 3.12 | 1.00 | 40.31 |  |
| 3 | G | O6 | AA:2592 | 3.70 | 1.00 | 43.35 |  |
| 4 | GLY | O | AD:238 | 3.72 | 1.00 | 47.89 |  |
| 5 | C | OP2 | AA:2591 | 4.03 | 1.00 | 41.12 |  |
| 6 | G | OP2 | AA:2592 | 4.16 | 1.00 | 44.25 |  |
| 7 | A | N6 | AA:2600 | 4.26 | 1.00 | 35.23 |  |
| 8 | A | N1 | AA:1971 | 4.61 | 1.00 | 48.06 |  |
| 9 | U | N3 | AA:2593 | 4.70 | 1.00 | 35.67 |  |
| 10 | C | N4 | AA:2591 | 4.72 | 1.00 | 42.22 |  |
| 11 | GLY | N | AD:238 | 4.91 | 1.00 | 42.84 |  |
| 12 | C | N4 | AA:2594 | 4.95 | 1.00 | 38.29 |  |
  |

  
 Coded contact list:   
 Exclusion zone contact(s): /N7/O4/  
Consider NA in place of MG (1 valid contact(s) for NA vs. 0 for MG)  

### MG:AA:3516

|  | Res. type | Atom | Residue | Distance | Occupancy | B-factor | Closest solute to 1^st^ shell water |
| --- | --- | --- | --- | --- | --- | --- | --- |
| **Ion** | | | |  | 1.00 | 35.81 || 1 | G | OP1 | AA:2592 | 2.22 | 1.00 | 43.47 |
| 2 | G | OP1 | AA:1968 | 2.75 | 1.00 | 47.44 |  |
| 3 | G | OP1 | AA:1973 | 3.74 | 1.00 | 34.70 |  |
| 4 | C | O3' | AA:2591 | 3.80 | 1.00 | 44.33 |  |
| 5 | C | O3' | AA:1967 | 4.20 | 1.00 | 45.79 |  |
| 6 | G | O5' | AA:1968 | 4.43 | 1.00 | 42.21 |  |
| 7 | G | OP2 | AA:2592 | 4.49 | 1.00 | 44.25 |  |
| 8 | C | O2' | AA:1967 | 4.60 | 1.00 | 45.02 |  |
| 9 | G | O5' | AA:2592 | 4.65 | 1.00 | 44.02 |  |
  |

  
 Contact(s): /OP1/  
 Coded contact list: O_ph_  
 Exclusion zone contact(s): /OP1/  

### MG:AA:3517

|  | Res. type | Atom | Residue | Distance | Occupancy | B-factor | Closest solute to 1^st^ shell water |
| --- | --- | --- | --- | --- | --- | --- | --- |
| **Ion** | | | |  | 1.00 | 69.11 || 1 | G | OP1 | AA:2349 | 2.49 | 1.00 | 62.06 |
| 2 | G | O5' | AA:2349 | 3.51 | 1.00 | 62.12 |  |
| 3 | C | OP2 | AA:2350 | 3.94 | 1.00 | 69.53 |  |
| 4 | G | O3' | AA:2349 | 4.50 | 1.00 | 69.23 |  |
| 5 | U | O3' | AA:2348 | 4.55 | 1.00 | 64.09 |  |
| 6 | C | O3' | AA:2417 | 4.65 | 1.00 | 62.50 |  |
| 7 | G | OP2 | AA:2349 | 4.69 | 1.00 | 60.43 |  |
| 8 | A | OP1 | AA:2418 | 4.73 | 1.00 | 60.88 |  |
  |

  
 Coded contact list:   
 Exclusion zone contact(s): /OP1/  
Consider NA in place of MG (1 valid contact(s) for NA vs. 0 for MG)  

### MG:AA:3518

|  | Res. type | Atom | Residue | Distance | Occupancy | B-factor | Closest solute to 1^st^ shell water |
| --- | --- | --- | --- | --- | --- | --- | --- |
| **Ion** | | | |  | 1.00 | 17.31 || 1 | G | O2' | AA:2250 | 2.53 | 1.00 | 48.64 |
| 2 | G | OP2 | AA:2251 | 2.53 | 1.00 | 41.74 |  |
| 3 | A | N7 | AA:2497 | 2.69 | 1.00 | 46.51 |  |
| 4 | C | OP2 | AA:2496 | 3.09 | 1.00 | 52.74 |  |
| 5 | A | OP2 | AA:2497 | 3.39 | 1.00 | 42.72 |  |
| 6 | G | O3' | AA:2250 | 3.46 | 1.00 | 42.92 |  |
| 7 | C | O5' | AA:2496 | 3.70 | 1.00 | 49.72 |  |
| 8 | C | OP1 | AA:2496 | 3.87 | 1.00 | 52.14 |  |
| 9 | A | N6 | AA:2497 | 4.31 | 1.00 | 46.42 |  |
| 10 | G | OP1 | AA:2251 | 4.46 | 1.00 | 44.08 |  |
| 11 | C | O3' | AA:2496 | 4.47 | 1.00 | 45.23 |  |
| 12 | A | N9 | AA:2497 | 4.67 | 1.00 | 44.35 |  |
| 13 | G | O5' | AA:2251 | 4.72 | 1.00 | 41.57 |  |
  |

  
 Coded contact list:   
 Exclusion zone contact(s): /N7/O2'/OP2/OP2/OP2/  
Consider K in place of MG (3 valid contact(s) for K vs. 0 for MG)  

### MG:AA:3519

|  | Res. type | Atom | Residue | Distance | Occupancy | B-factor | Closest solute to 1^st^ shell water |
| --- | --- | --- | --- | --- | --- | --- | --- |
| **Ion** | | | |  | 1.00 | 63.87 || 1 | G | OP1 | AA:2289 | 2.12 | 1.00 | 73.40 |
| 2 | C | OP1 | AA:2385 | 2.23 | 1.00 | 64.78 |  |
| 3 | G | O3' | AA:2384 | 3.55 | 1.00 | 66.32 |  |
| 4 | G | O5' | AA:2289 | 4.14 | 1.00 | 72.39 |  |
| 5 | C | O5' | AA:2385 | 4.21 | 1.00 | 63.86 |  |
| 6 | G | O2' | AA:2384 | 4.30 | 1.00 | 68.58 |  |
| 7 | G | OP1 | AA:2337 | 4.30 | 1.00 | 77.79 |  |
| 8 | G | OP2 | AA:2289 | 4.48 | 1.00 | 75.05 |  |
| 9 | A | O3' | AA:2288 | 4.52 | 1.00 | 75.10 |  |
| 10 | A | O2' | AA:2288 | 4.53 | 1.00 | 79.40 |  |
| 11 | C | OP2 | AA:2385 | 4.65 | 1.00 | 64.51 |  |
| 12 | G | OP2 | AA:2337 | 4.91 | 1.00 | 77.36 |  |
  |

  
**1^st^ shell ligand angles:**   
1-2: 139;   
  
 Contact(s): /OP1/OP1/  
 Coded contact list: 2O_ph_  

### MG:AA:3520

|  | Res. type | Atom | Residue | Distance | Occupancy | B-factor | Closest solute to 1^st^ shell water |
| --- | --- | --- | --- | --- | --- | --- | --- |
| **Ion** | | | |  | 1.00 | 75.04 || 1 | G | O5' | AA:2337 | 1.93 | 1.00 | 75.33 |
| 2 | G | OP2 | AA:2337 | 2.31 | 1.00 | 77.36 |  |
| 3 | G | OP2 | AA:2338 | 2.73 | 1.00 | 70.85 |  |
| 4 | G | OP1 | AA:2337 | 3.02 | 1.00 | 77.79 |  |
| 5 | A | O3' | AA:2336 | 3.87 | 1.00 | 76.94 |  |
| 6 | G | O3' | AA:2337 | 4.37 | 1.00 | 72.65 |  |
| 7 | G | O4' | AA:2337 | 4.78 | 1.00 | 75.16 |  |
| 8 | C | OP1 | AA:2385 | 4.86 | 1.00 | 64.78 |  |
| 9 | G | OP1 | AA:2338 | 5.00 | 1.00 | 72.33 |  |
  |

  
**1^st^ shell ligand angles:**   
1-2: 72;   
  
 Contact(s): /O5'/OP2/  
 Coded contact list: O_ph_O_r_  
 Exclusion zone contact(s): /OP1/OP2/  

### MG:AA:3521

|  | Res. type | Atom | Residue | Distance | Occupancy | B-factor | Closest solute to 1^st^ shell water |
| --- | --- | --- | --- | --- | --- | --- | --- |
| **Ion** | | | |  | 1.00 | 32.54 || 1 | A | O2' | AA:685 | 2.92 | 1.00 | 41.51 |
| 2 | G | OP2 | AA:686 | 3.47 | 1.00 | 38.33 |  |
| 3 | A | O3' | AA:685 | 3.57 | 1.00 | 38.00 |  |
| 4 | U | O4 | AA:688 | 3.65 | 1.00 | 39.20 |  |
| 5 | G | O6 | AA:690 | 3.98 | 1.00 | 47.64 |  |
| 6 | C | N4 | AA:772 | 4.12 | 1.00 | 44.09 |  |
| 7 | A | N6 | AA:689 | 4.35 | 1.00 | 39.69 |  |
| 8 | G | O5' | AA:686 | 4.73 | 1.00 | 36.78 |  |
| 9 | G | O6 | AA:771 | 4.80 | 1.00 | 39.52 |  |
  |

  
 Coded contact list:   
 Exclusion zone contact(s): /O2'/  
Consider K in place of MG (1 valid contact(s) for K vs. 0 for MG)  

### MG:AA:3522

|  | Res. type | Atom | Residue | Distance | Occupancy | B-factor | Closest solute to 1^st^ shell water |
| --- | --- | --- | --- | --- | --- | --- | --- |
| **Ion** | | | |  | 1.00 | 14.68 || 1 | G | O6 | AA:2447 | 2.13 | 1.00 | 34.74 |
| 2 | G | O6 | AA:2061 | 2.42 | 1.00 | 35.65 |  |
| 3 | C | O4' | AA:2501 | 2.56 | 1.00 | 42.23 |  |
| 4 | G | N7 | AA:2447 | 2.78 | 1.00 | 29.81 |  |
| 5 | U | O4 | AA:2504 | 3.16 | 1.00 | 46.16 |  |
| 6 | C | O2 | AA:2501 | 3.24 | 1.00 | 37.01 |  |
| 7 | G | N7 | AA:2061 | 3.45 | 1.00 | 35.35 |  |
| 8 | C | N1 | AA:2501 | 3.66 | 1.00 | 39.10 |  |
| 9 | G | N1 | AA:2447 | 4.18 | 1.00 | 31.75 |  |
| 10 | A | N6 | AA:2451 | 4.34 | 1.00 | 44.31 |  |
| 11 | C | N3 | AA:2501 | 4.42 | 1.00 | 37.08 |  |
| 12 | C | O5' | AA:2501 | 4.55 | 1.00 | 41.63 |  |
| 13 | G | N1 | AA:2061 | 4.60 | 1.00 | 31.63 |  |
| 14 | G | N9 | AA:2447 | 4.89 | 1.00 | 30.86 |  |
  |

  
 Contact(s): /O6/  
 Coded contact list: O_b_  
 Exclusion zone contact(s): /N7/O2/O4/O4'/O6/  
Consider K in place of MG (3 valid contact(s) for K vs. 1 for MG)  

### MG:AA:3523

|  | Res. type | Atom | Residue | Distance | Occupancy | B-factor | Closest solute to 1^st^ shell water |
| --- | --- | --- | --- | --- | --- | --- | --- |
| **Ion** | | | |  | 1.00 | 42.97 || 1 | U | O4 | AA:2249 | 1.82 | 1.00 | 47.00 |
| 2 | C | OP2 | AA:2248 | 2.39 | 1.00 | 39.70 |  |
| 3 | C | OP1 | AA:2066 | 3.76 | 1.00 | 39.69 |  |
| 4 | U | N3 | AA:2249 | 4.07 | 1.00 | 44.37 |  |
| 5 | G | OP1 | AA:2253 | 4.10 | 1.00 | 43.88 |  |
| 6 | A | O5' | AA:2247 | 4.21 | 1.00 | 41.27 |  |
| 7 | A | OP1 | AA:2247 | 4.32 | 1.00 | 43.75 |  |
| 8 | A | O3' | AA:2247 | 4.43 | 1.00 | 38.96 |  |
| 9 | A | OP2 | AA:2247 | 4.52 | 1.00 | 43.27 |  |
| 10 | C | OP1 | AA:2248 | 4.69 | 1.00 | 41.71 |  |
| 11 | C | O5' | AA:2248 | 4.86 | 1.00 | 40.99 |  |
| 12 | G | O3' | AA:2252 | 4.90 | 1.00 | 43.06 |  |
  |

  
**1^st^ shell ligand angles:**   
1-2: 82;   
  
 Contact(s): /O4/OP2/  
 Coded contact list: O_ph_O_b_  

### MG:AA:3524

|  | Res. type | Atom | Residue | Distance | Occupancy | B-factor | Closest solute to 1^st^ shell water |
| --- | --- | --- | --- | --- | --- | --- | --- |
| **Ion** | | | |  | 1.00 | 50.85 || 1 | G | N7 | AA:768 | 3.08 | 1.00 | 32.72 |
| 2 | G | OP2 | AA:768 | 3.20 | 1.00 | 34.30 |  |
| 3 | U | N3 | AA:767 | 4.15 | 1.00 | 37.59 |  |
| 4 | G | O5' | AA:768 | 4.43 | 1.00 | 33.50 |  |
| 5 | G | N7 | AA:769 | 4.49 | 1.00 | 37.81 |  |
| 6 | U | O4 | AA:767 | 4.55 | 1.00 | 38.03 |  |
| 7 | G | O6 | AA:769 | 4.56 | 1.00 | 37.70 |  |
| 8 | U | N1 | AA:767 | 4.67 | 1.00 | 37.63 |  |
| 9 | G | O6 | AA:768 | 4.76 | 1.00 | 39.07 |  |
| 10 | G | N9 | AA:768 | 4.85 | 1.00 | 33.62 |  |
| 11 | U | O2 | AA:767 | 4.93 | 1.00 | 37.00 |  |
  |

  
 Coded contact list:   
 Exclusion zone contact(s): /N7/OP2/  
Consider K in place of MG (2 valid contact(s) for K vs. 0 for MG)  

### MG:AA:3525

|  | Res. type | Atom | Residue | Distance | Occupancy | B-factor | Closest solute to 1^st^ shell water |
| --- | --- | --- | --- | --- | --- | --- | --- |
| **Ion** | | | |  | 1.00 | 36.01 || 1 | G | OP2 | AA:2255 | 2.51 | 1.00 | 53.35 |
| 2 | G | N7 | AA:2255 | 3.13 | 1.00 | 52.91 |  |
| 3 | G | N7 | AA:2256 | 3.29 | 1.00 | 53.08 |  |
| 4 | G | O5' | AA:2255 | 3.43 | 1.00 | 52.66 |  |
| 5 | G | O6 | AA:2256 | 4.10 | 1.00 | 51.68 |  |
| 6 | C | OP2 | AA:2254 | 4.31 | 1.00 | 48.61 |  |
| 7 | G | N9 | AA:2255 | 4.42 | 1.00 | 52.75 |  |
| 8 | C | O3' | AA:2254 | 4.62 | 1.00 | 51.84 |  |
| 9 | G | OP2 | AA:2256 | 4.65 | 1.00 | 55.41 |  |
| 10 | G | OP1 | AA:2255 | 4.68 | 1.00 | 53.61 |  |
| 11 | C | O5' | AA:2254 | 4.74 | 1.00 | 48.89 |  |
| 12 | U | O4 | AA:2257 | 4.91 | 1.00 | 41.46 |  |
  |

  
 Coded contact list:   
 Exclusion zone contact(s): /N7/N7/OP2/  
Consider K in place of MG (2 valid contact(s) for K vs. 0 for MG)  

### MG:AA:3526

|  | Res. type | Atom | Residue | Distance | Occupancy | B-factor | Closest solute to 1^st^ shell water |
| --- | --- | --- | --- | --- | --- | --- | --- |
| **Ion** | | | |  | 1.00 | 75.64 || 1 | G | O6 | AA:2300 | 2.88 | 1.00 | 95.30 |
| 2 | G | O6 | AA:2299 | 2.95 | 1.00 | 99.14 |  |
| 3 | G | N7 | AA:2299 | 3.77 | 1.00 | 99.54 |  |
| 4 | G | N7 | AA:2300 | 3.86 | 1.00 | 96.49 |  |
| 5 | C | N4 | AA:2317 | 4.43 | 1.00 | 98.18 |  |
| 6 | G | N1 | AA:2299 | 4.50 | 1.00 | 98.71 |  |
| 7 | C | N4 | AA:2316 | 4.54 | 1.00 | 94.61 |  |
| 8 | C | N4 | AA:2301 | 4.59 | 1.00 | 96.81 |  |
| 9 | G | O6 | AA:2315 | 4.86 | 1.00 | 95.40 |  |
  |

  
 Coded contact list:   
 Exclusion zone contact(s): /O6/O6/  
Consider K in place of MG (2 valid contact(s) for K vs. 0 for MG)  

### MG:AA:3527

|  | Res. type | Atom | Residue | Distance | Occupancy | B-factor | Closest solute to 1^st^ shell water |
| --- | --- | --- | --- | --- | --- | --- | --- |
| **Ion** | | | |  | 1.00 | 34.65 || 1 | G | O6 | AA:2436 | 2.62 | 1.00 | 26.01 |
| 2 | U | O4 | AA:2245 | 2.65 | 1.00 | 34.39 |  |
| 3 | U | O4 | AA:2437 | 3.04 | 1.00 | 32.23 |  |
| 4 | U | O4 | AA:2244 | 3.82 | 1.00 | 33.36 |  |
| 5 | C | N4 | AA:2073 | 3.99 | 1.00 | 23.87 |  |
| 6 | G | O6 | AA:2072 | 4.16 | 1.00 | 33.32 |  |
| 7 | G | N7 | AA:2436 | 4.28 | 1.00 | 27.99 |  |
| 8 | A | N6 | AA:2071 | 4.33 | 1.00 | 33.07 |  |
| 9 | U | N3 | AA:2245 | 4.51 | 1.00 | 33.63 |  |
| 10 | U | N3 | AA:2437 | 4.64 | 1.00 | 30.54 |  |
| 11 | G | N1 | AA:2436 | 4.75 | 1.00 | 26.34 |  |
| 12 | A | N6 | AA:2435 | 4.86 | 1.00 | 32.08 |  |
  |

  
 Coded contact list:   
 Exclusion zone contact(s): /O4/O4/O6/  
Consider K in place of MG (3 valid contact(s) for K vs. 0 for MG)  

### MG:AA:3528

|  | Res. type | Atom | Residue | Distance | Occupancy | B-factor | Closest solute to 1^st^ shell water |
| --- | --- | --- | --- | --- | --- | --- | --- |
| **Ion** | | | |  | 1.00 | 46.97 || 1 | THR | OG1 | AT:56 | 2.65 | 1.00 | 47.58 |
| 2 | C | OP1 | AA:1599 | 2.65 | 1.00 | 43.94 |  |
| 3 | LYS | NZ | AT:77 | 2.93 | 1.00 | 47.90 |  |
| 4 | C | OP2 | AA:1600 | 3.00 | 1.00 | 44.49 |  |
| 5 | C | O5' | AA:1599 | 3.55 | 1.00 | 46.76 |  |
| 6 | ALA | O | AT:34 | 3.90 | 1.00 | 61.93 |  |
| 7 | C | OP2 | AA:1599 | 3.97 | 1.00 | 42.07 |  |
| 8 | C | OP1 | AA:1600 | 4.16 | 1.00 | 44.73 |  |
| 9 | LYS | N | AT:36 | 4.19 | 1.00 | 59.10 |  |
| 10 | C | O3' | AA:1599 | 4.51 | 1.00 | 44.12 |  |
| 11 | C | O3' | AA:1598 | 4.91 | 1.00 | 45.23 |  |
  |

  
 Coded contact list:   
 Exclusion zone contact(s): /NZ_Lys_/OG1_Thr_/OP1/OP2/  
Consider K in place of MG (4 valid contact(s) for K vs. 0 for MG)  

### MG:AA:3529

|  | Res. type | Atom | Residue | Distance | Occupancy | B-factor | Closest solute to 1^st^ shell water |
| --- | --- | --- | --- | --- | --- | --- | --- |
| **Ion** | | | |  | 1.00 | 29.10 || 1 | A | OP1 | AA:793 | 3.11 | 1.00 | 37.05 |
| 2 | C | O2 | AA:2441 | 3.29 | 1.00 | 37.61 |  |
| 3 | G | O3' | AA:792 | 3.58 | 1.00 | 35.91 |  |
| 4 | A | O2' | AA:2071 | 3.85 | 1.00 | 33.89 |  |
| 5 | G | O2' | AA:792 | 3.90 | 1.00 | 41.72 |  |
| 6 | C | N3 | AA:2440 | 3.96 | 1.00 | 43.12 |  |
| 7 | G | N2 | AA:2070 | 4.05 | 1.00 | 33.79 |  |
| 8 | C | O2' | AA:2441 | 4.31 | 1.00 | 38.87 |  |
| 9 | C | N4 | AA:2440 | 4.55 | 1.00 | 42.31 |  |
| 10 | C | O2 | AA:2440 | 4.62 | 1.00 | 44.68 |  |
| 11 | C | O4' | AA:2442 | 4.67 | 1.00 | 37.80 |  |
| 12 | A | O5' | AA:793 | 4.93 | 1.00 | 39.71 |  |
| 13 | C | N1 | AA:2441 | 4.96 | 1.00 | 37.88 |  |
  |

  
 Coded contact list:   
 Exclusion zone contact(s): /O2/OP1/  
Consider K in place of MG (2 valid contact(s) for K vs. 0 for MG)  

### MG:AA:3530

|  | Res. type | Atom | Residue | Distance | Occupancy | B-factor | Closest solute to 1^st^ shell water |
| --- | --- | --- | --- | --- | --- | --- | --- |
| **Ion** | | | |  | 1.00 | 50.55 || 1 | A | O2' | AA:447 | 2.64 | 1.00 | 49.03 |
| 2 | A | O3' | AA:447 | 2.80 | 1.00 | 45.82 |  |
| 3 | U | OP2 | AA:448 | 3.14 | 1.00 | 43.19 |  |
| 4 | A | OP1 | AA:472 | 3.26 | 1.00 | 45.16 |  |
| 5 | A | OP2 | AA:472 | 3.33 | 1.00 | 44.99 |  |
| 6 | U | O5' | AA:448 | 3.38 | 1.00 | 42.83 |  |
| 7 | G | OP2 | AA:473 | 3.54 | 1.00 | 50.06 |  |
| 8 | A | O5' | AA:472 | 4.09 | 1.00 | 44.52 |  |
| 9 | G | OP1 | AA:473 | 4.31 | 1.00 | 48.85 |  |
| 10 | U | OP1 | AA:448 | 4.80 | 1.00 | 43.40 |  |
  |

  
 Coded contact list:   
 Exclusion zone contact(s): /O2'/O3'/O5'/OP1/OP2/OP2/  
Consider K in place of MG (6 valid contact(s) for K vs. 0 for MG)  

### MG:AA:3531

|  | Res. type | Atom | Residue | Distance | Occupancy | B-factor | Closest solute to 1^st^ shell water |
| --- | --- | --- | --- | --- | --- | --- | --- |
| **Ion** | | | |  | 1.00 | 26.28 || 1 | U | O4 | AA:29 | 3.51 | 1.00 | 42.74 |
| 2 | G | O6 | AA:474 | 3.94 | 1.00 | 48.23 |  |
| 3 | G | O6 | AA:30 | 4.03 | 1.00 | 54.72 |  |
| 4 | C | N4 | AA:31 | 4.16 | 1.00 | 45.57 |  |
| 5 | A | OP2 | AA:28 | 4.32 | 1.00 | 47.12 |  |
| 6 | G | O6 | AA:473 | 4.60 | 1.00 | 46.22 |  |
| 7 | U | OP2 | AA:29 | 4.72 | 1.00 | 48.44 |  |
| 8 | G | N7 | AA:30 | 4.95 | 1.00 | 55.70 |  |
| 9 | A | O5' | AA:447 | 4.97 | 1.00 | 48.99 |  |
  |

  
 Coded contact list:   
1^st^ MG coordination shell missing (see Notes)  

### MG:AA:3532

|  | Res. type | Atom | Residue | Distance | Occupancy | B-factor | Closest solute to 1^st^ shell water |
| --- | --- | --- | --- | --- | --- | --- | --- |
| **Ion** | | | |  | 1.00 | 87.83 || 1 | G | O6 | AA:512 | 4.35 | 1.00 | 55.32 |
| 2 | C | OP2 | AA:510 | 4.52 | 1.00 | 52.25 |  |
| 3 | U | O4 | AA:511 | 4.52 | 1.00 | 59.72 |  |
| 4 | G | N1 | AA:512 | 4.55 | 1.00 | 53.34 |  |
| 5 | G | OP2 | AA:474 | 4.62 | 1.00 | 48.27 |  |
  |

  
 Coded contact list:   
1^st^ MG coordination shell missing (see Notes)  

### MG:AA:3533

|  | Res. type | Atom | Residue | Distance | Occupancy | B-factor | Closest solute to 1^st^ shell water |
| --- | --- | --- | --- | --- | --- | --- | --- |
| **Ion** | | | |  | 1.00 | 44.66 || 1 | U | O4 | AA:1323 | 2.52 | 1.00 | 53.81 |
| 2 | A | N7 | AA:1322 | 3.10 | 1.00 | 35.30 |  |
| 3 | C | O2 | AA:1320 | 3.16 | 1.00 | 66.36 |  |
| 4 | G | O6 | AA:1324 | 3.76 | 1.00 | 53.69 |  |
| 5 | C | O2' | AA:1320 | 3.79 | 1.00 | 66.17 |  |
| 6 | A | N6 | AA:1322 | 4.01 | 1.00 | 36.20 |  |
| 7 | A | N6 | AA:1331 | 4.06 | 1.00 | 54.93 |  |
| 8 | C | N1 | AA:1320 | 4.14 | 1.00 | 65.27 |  |
| 9 | A | OP2 | AA:1322 | 4.32 | 1.00 | 41.34 |  |
| 10 | C | N3 | AA:1320 | 4.40 | 1.00 | 64.94 |  |
| 11 | A | OP2 | AA:1321 | 4.47 | 1.00 | 59.74 |  |
| 12 | C | N4 | AA:1330 | 4.76 | 1.00 | 53.79 |  |
| 13 | U | N3 | AA:1323 | 4.77 | 1.00 | 52.67 |  |
  |

  
 Coded contact list:   
 Exclusion zone contact(s): /N7/O2/O4/  
Consider K in place of MG (2 valid contact(s) for K vs. 0 for MG)  

### MG:AA:3534

|  | Res. type | Atom | Residue | Distance | Occupancy | B-factor | Closest solute to 1^st^ shell water |
| --- | --- | --- | --- | --- | --- | --- | --- |
| **Ion** | | | |  | 1.00 | 32.62 || 1 | G | OP2 | AA:2056 | 2.30 | 1.00 | 44.12 |
| 2 | A | OP2 | AA:2057 | 2.35 | 1.00 | 42.70 |  |
| 3 | A | N7 | AA:2058 | 4.05 | 1.00 | 46.14 |  |
| 4 | G | O5' | AA:2056 | 4.19 | 1.00 | 44.05 |  |
| 5 | A | N6 | AA:2058 | 4.20 | 1.00 | 44.21 |  |
| 6 | G | OP1 | AA:2056 | 4.22 | 1.00 | 43.67 |  |
| 7 | A | O5' | AA:2057 | 4.25 | 1.00 | 44.30 |  |
| 8 | A | OP1 | AA:2057 | 4.36 | 1.00 | 41.84 |  |
| 9 | A | O3' | AA:2503 | 4.36 | 1.00 | 41.19 |  |
| 10 | A | N7 | AA:2057 | 4.37 | 1.00 | 43.65 |  |
| 11 | G | OP1 | AA:2505 | 4.47 | 1.00 | 44.71 |  |
| 12 | G | O3' | AA:2056 | 4.82 | 1.00 | 43.47 |  |
| 13 | A | O2' | AA:2503 | 4.83 | 1.00 | 39.94 |  |
| 14 | C | O3' | AA:2055 | 4.86 | 1.00 | 43.21 |  |
  |

  
**1^st^ shell ligand angles:**   
1-2: 68;   
  
 Contact(s): /OP2/OP2/  
 Coded contact list: 2O_ph_  

### MG:AA:3535

|  | Res. type | Atom | Residue | Distance | Occupancy | B-factor | Closest solute to 1^st^ shell water |
| --- | --- | --- | --- | --- | --- | --- | --- |
| **Ion** | | | |  | 1.00 | 58.79 || 1 | A | N7 | AA:2589 | 2.70 | 1.00 | 25.97 |
| 2 | A | OP2 | AA:2589 | 3.14 | 1.00 | 32.60 |  |
| 3 | G | N7 | AA:2588 | 3.31 | 1.00 | 29.01 |  |
| 4 | G | OP2 | AA:2588 | 3.32 | 1.00 | 35.91 |  |
| 5 | G | O5' | AA:2588 | 3.34 | 1.00 | 33.02 |  |
| 6 | G | N9 | AA:2588 | 3.44 | 1.00 | 27.95 |  |
| 7 | A | O5' | AA:2589 | 3.63 | 1.00 | 31.62 |  |
| 8 | G | O3' | AA:2588 | 3.94 | 1.00 | 31.49 |  |
| 9 | A | N9 | AA:2589 | 3.97 | 1.00 | 27.56 |  |
| 10 | G | O4' | AA:2588 | 4.31 | 1.00 | 30.00 |  |
| 11 | G | OP1 | AA:2588 | 4.32 | 1.00 | 36.17 |  |
| 12 | A | O4' | AA:2589 | 4.61 | 1.00 | 33.92 |  |
| 13 | G | N3 | AA:2588 | 4.87 | 1.00 | 26.87 |  |
  |

  
 Coded contact list:   
 Exclusion zone contact(s): /N7/N7/O5'/OP2/OP2/  
Consider K in place of MG (5 valid contact(s) for K vs. 0 for MG)  

### MG:AA:3536

|  | Res. type | Atom | Residue | Distance | Occupancy | B-factor | Closest solute to 1^st^ shell water |
| --- | --- | --- | --- | --- | --- | --- | --- |
| **Ion** | | | |  | 1.00 | 46.21 || 1 | C | OP1 | AA:2442 | 3.31 | 1.00 | 41.15 |
| 2 | C | O3' | AA:2441 | 3.64 | 1.00 | 40.19 |  |
| 3 | C | OP1 | AA:2441 | 4.06 | 1.00 | 39.86 |  |
| 4 | C | OP2 | AA:2442 | 4.57 | 1.00 | 40.95 |  |
| 5 | A | N3 | AA:2062 | 4.67 | 1.00 | 77.17 |  |
| 6 | A | O2' | AA:2062 | 4.75 | 1.00 | 62.53 |  |
| 7 | C | OP2 | AA:2441 | 4.80 | 1.00 | 36.53 |  |
  |

  
 Coded contact list:   
 Exclusion zone contact(s): /OP1/  
Consider K in place of MG (1 valid contact(s) for K vs. 0 for MG)  

### MG:AA:3537

|  | Res. type | Atom | Residue | Distance | Occupancy | B-factor | Closest solute to 1^st^ shell water |
| --- | --- | --- | --- | --- | --- | --- | --- |
| **Ion** | | | |  | 1.00 | 56.81 || 1 | A | N3 | AA:2590 | 2.78 | 1.00 | 40.13 |
| 2 | C | O4' | AA:2591 | 2.80 | 1.00 | 43.26 |  |
| 3 | A | O2' | AA:2590 | 2.91 | 1.00 | 41.72 |  |
| 4 | A | N9 | AA:1938 | 2.93 | 1.00 | 51.92 |  |
| 5 | A | N3 | AA:1938 | 3.26 | 1.00 | 50.97 |  |
| 6 | A | N7 | AA:1938 | 3.52 | 1.00 | 53.54 |  |
| 7 | A | O2' | AA:1938 | 3.67 | 1.00 | 57.92 |  |
| 8 | C | N1 | AA:2591 | 3.74 | 1.00 | 42.72 |  |
| 9 | A | N9 | AA:2590 | 4.07 | 1.00 | 39.25 |  |
| 10 | A | N1 | AA:1938 | 4.19 | 1.00 | 51.57 |  |
| 11 | C | O5' | AA:2591 | 4.68 | 1.00 | 42.87 |  |
| 12 | A | O4' | AA:2590 | 4.69 | 1.00 | 40.75 |  |
| 13 | A | O4' | AA:1938 | 4.78 | 1.00 | 52.56 |  |
| 14 | A | N6 | AA:1938 | 4.81 | 1.00 | 54.06 |  |
| 15 | A | N1 | AA:2590 | 4.92 | 1.00 | 40.51 |  |
| 16 | C | O2 | AA:2591 | 4.94 | 1.00 | 47.44 |  |
  |

  
 Coded contact list:   
 Exclusion zone contact(s): /N3_A_/N3_A_/O2'/O4'/  
Consider K in place of MG (5 valid contact(s) for K vs. 0 for MG)  

### MG:AA:3538

|  | Res. type | Atom | Residue | Distance | Occupancy | B-factor | Closest solute to 1^st^ shell water |
| --- | --- | --- | --- | --- | --- | --- | --- |
| **Ion** | | | |  | 1.00 | 45.04 || 1 | U | O4 | AA:2604 | 3.10 | 1.00 | 41.52 |
| 2 | U | O4 | AA:2605 | 4.64 | 1.00 | 29.91 |  |
| 3 | G | N7 | AA:2603 | 4.78 | 1.00 | 49.84 |  |
  |

  
 Coded contact list:   
 Exclusion zone contact(s): /O4/  
Consider K in place of MG (1 valid contact(s) for K vs. 0 for MG)  

### MG:AA:3539

|  | Res. type | Atom | Residue | Distance | Occupancy | B-factor | Closest solute to 1^st^ shell water |
| --- | --- | --- | --- | --- | --- | --- | --- |
| **Ion** | | | |  | 1.00 | 50.12 || 1 | G | OP2 | AA:2308 | 2.59 | 1.00 | 99.37 |
| 2 | G | O3' | AA:2307 | 3.54 | 1.00 | 99.05 |  |
| 3 | G | OP1 | AA:2308 | 3.83 | 1.00 | 100.12 |  |
| 4 | C | OP2 | AA:2306 | 3.93 | 1.00 | 94.57 |  |
| 5 | G | N9 | AA:2307 | 3.99 | 1.00 | 94.77 |  |
| 6 | G | O2' | AA:2307 | 4.33 | 1.00 | 98.27 |  |
| 7 | G | N7 | AA:2307 | 4.37 | 1.00 | 92.44 |  |
| 8 | G | O6 | AA:2304 | 4.38 | 1.00 | 99.33 |  |
| 9 | G | O4' | AA:2307 | 4.42 | 1.00 | 97.20 |  |
| 10 | G | N1 | AA:2304 | 4.64 | 1.00 | 100.39 |  |
| 11 | G | O5' | AA:2308 | 4.84 | 1.00 | 100.33 |  |
  |

  
 Coded contact list:   
 Exclusion zone contact(s): /OP2/  
Consider NA in place of MG (1 valid contact(s) for NA vs. 0 for MG)  

### MG:AA:3540

|  | Res. type | Atom | Residue | Distance | Occupancy | B-factor | Closest solute to 1^st^ shell water |
| --- | --- | --- | --- | --- | --- | --- | --- |
| **Ion** | | | |  | 1.00 | 61.54 || 1 | G | OP1 | AA:2330 | 3.55 | 1.00 | 58.50 |
| 2 | G | O3' | AA:2329 | 3.89 | 1.00 | 58.63 |  |
| 3 | U | OP2 | AA:913 | 4.37 | 1.00 | 76.17 |  |
| 4 | U | OP1 | AA:913 | 4.44 | 1.00 | 76.85 |  |
| 5 | C | OP1 | AA:2264 | 4.73 | 1.00 | 65.26 |  |
| 6 | C | O3' | AA:2263 | 4.84 | 1.00 | 62.40 |  |
  |

  
 Coded contact list:   
1^st^ MG coordination shell missing (see Notes)  

### MG:AA:3541

|  | Res. type | Atom | Residue | Distance | Occupancy | B-factor | Closest solute to 1^st^ shell water |
| --- | --- | --- | --- | --- | --- | --- | --- |
| **Ion** | | | |  | 1.00 | 49.41 || 1 | C | O2' | BC:75 | 2.55 | 1.00 | 56.07 |
| 2 | C | O3' | AA:2063 | 2.80 | 1.00 | 44.64 |  |
| 3 | C | O3' | BC:75 | 3.06 | 1.00 | 60.69 |  |
| 4 | C | O2' | AA:2063 | 3.13 | 1.00 | 46.73 |  |
| 5 | C | OP1 | AA:2064 | 3.56 | 1.00 | 44.76 |  |
| 6 | A | OP1 | BC:76 | 4.12 | 1.00 | 64.23 |  |
| 7 | C | O5' | AA:2064 | 4.70 | 1.00 | 44.63 |  |
| 8 | C | O4' | AA:2063 | 4.77 | 1.00 | 45.46 |  |
| 9 | A | OP2 | BC:76 | 4.89 | 1.00 | 62.87 |  |
| 10 | C | O4' | BC:75 | 4.97 | 1.00 | 54.14 |  |
  |

  
 Coded contact list:   
 Exclusion zone contact(s): /O2'/O2'/O3'/O3'/  
Consider K in place of MG (3 valid contact(s) for K vs. 0 for MG)  

### MG:AA:3542

|  | Res. type | Atom | Residue | Distance | Occupancy | B-factor | Closest solute to 1^st^ shell water |
| --- | --- | --- | --- | --- | --- | --- | --- |
| **Ion** | | | |  | 1.00 | 54.33 || 1 | G | N7 | AA:744 | 3.38 | 1.00 | 42.97 |
| 2 | G | O6 | AA:745 | 3.45 | 1.00 | 39.98 |  |
| 3 | G | O6 | AA:744 | 3.77 | 1.00 | 40.54 |  |
| 4 | G | N7 | AA:743 | 3.95 | 1.00 | 44.94 |  |
| 5 | G | O6 | AA:743 | 4.46 | 1.00 | 43.39 |  |
| 6 | G | N1 | AA:745 | 4.68 | 1.00 | 37.35 |  |
| 7 | G | N7 | AA:745 | 4.88 | 1.00 | 42.33 |  |
  |

  
 Coded contact list:   
 Exclusion zone contact(s): /N7/  
Consider K in place of MG (1 valid contact(s) for K vs. 0 for MG)  

### MG:AA:3543

|  | Res. type | Atom | Residue | Distance | Occupancy | B-factor | Closest solute to 1^st^ shell water |
| --- | --- | --- | --- | --- | --- | --- | --- |
| **Ion** | | | |  | 1.00 | 77.91 || 1 | G | O6 | AA:2323 | 2.32 | 1.00 | 58.99 |
| 2 | U | O4 | AA:2332 | 2.92 | 1.00 | 56.91 |  |
| 3 | G | O6 | AA:2331 | 3.70 | 1.00 | 58.43 |  |
| 4 | U | N3 | AA:2332 | 3.79 | 1.00 | 56.42 |  |
| 5 | G | N7 | AA:2323 | 3.90 | 1.00 | 61.76 |  |
| 6 | G | OP2 | AA:2321 | 3.97 | 1.00 | 90.98 |  |
| 7 | A | OP2 | AA:2322 | 4.01 | 1.00 | 82.55 |  |
| 8 | C | N4 | AA:2324 | 4.01 | 1.00 | 58.51 |  |
| 9 | A | N7 | AA:2322 | 4.53 | 1.00 | 66.84 |  |
| 10 | G | O5' | AA:2321 | 4.54 | 1.00 | 88.70 |  |
| 11 | G | N1 | AA:2323 | 4.58 | 1.00 | 58.62 |  |
  |

  
 Contact(s): /O6/  
 Coded contact list: O_b_  
 Exclusion zone contact(s): /O4/  

### MG:AA:3544

|  | Res. type | Atom | Residue | Distance | Occupancy | B-factor | Closest solute to 1^st^ shell water |
| --- | --- | --- | --- | --- | --- | --- | --- |
| **Ion** | | | |  | 1.00 | 62.74 || 1 | G | O3' | AA:2447 | 3.00 | 1.00 | 38.23 |
| 2 | G | O5' | AA:2447 | 3.06 | 1.00 | 37.11 |  |
| 3 | G | OP1 | AA:2447 | 3.44 | 1.00 | 38.71 |  |
| 4 | U | OP1 | AA:2500 | 3.98 | 1.00 | 42.85 |  |
| 5 | A | OP1 | AA:2448 | 4.08 | 1.00 | 40.10 |  |
| 6 | U | OP2 | AA:2500 | 4.24 | 1.00 | 42.75 |  |
| 7 | C | OP1 | AA:2499 | 4.41 | 1.00 | 43.06 |  |
| 8 | A | O5' | AA:2448 | 4.42 | 1.00 | 40.15 |  |
| 9 | G | O6 | AA:570 | 4.55 | 1.00 | 39.82 |  |
| 10 | G | O3' | AA:2446 | 4.74 | 1.00 | 37.37 |  |
| 11 | U | O2' | AA:568 | 4.86 | 1.00 | 47.08 |  |
| 12 | A | N6 | AA:945 | 4.96 | 1.00 | 55.15 |  |
  |

  
 Coded contact list:   
 Exclusion zone contact(s): /O3'/O5'/  
Consider K in place of MG (2 valid contact(s) for K vs. 0 for MG)  

### MG:AA:3545

|  | Res. type | Atom | Residue | Distance | Occupancy | B-factor | Closest solute to 1^st^ shell water |
| --- | --- | --- | --- | --- | --- | --- | --- |
| **Ion** | | | |  | 1.00 | 56.53 || 1 | G | O6 | AA:2607 | 3.08 | 1.00 | 37.61 |
| 2 | G | O6 | AA:2588 | 3.23 | 1.00 | 28.69 |  |
| 3 | A | N6 | AA:2589 | 3.68 | 1.00 | 31.46 |  |
| 4 | A | OP2 | AA:2587 | 3.77 | 1.00 | 41.55 |  |
| 5 | C | OP1 | AA:2586 | 3.98 | 1.00 | 64.40 |  |
| 6 | G | N1 | AA:2607 | 4.15 | 1.00 | 36.94 |  |
| 7 | C | N4 | AA:2606 | 4.37 | 1.00 | 27.16 |  |
| 8 | U | O4 | AA:2605 | 4.43 | 1.00 | 29.91 |  |
| 9 | G | N7 | AA:2588 | 4.49 | 1.00 | 29.01 |  |
| 10 | A | OP1 | AA:2587 | 4.79 | 1.00 | 45.57 |  |
  |

  
 Coded contact list:   
 Exclusion zone contact(s): /O6/O6/  
Consider K in place of MG (2 valid contact(s) for K vs. 0 for MG)  

### MG:AA:3546

|  | Res. type | Atom | Residue | Distance | Occupancy | B-factor | Closest solute to 1^st^ shell water |
| --- | --- | --- | --- | --- | --- | --- | --- |
| **Ion** | | | |  | 1.00 | 52.86 || 1 | U | O3' | AA:2068 | 2.51 | 1.00 | 45.50 |
| 2 | G | OP2 | AA:2069 | 2.58 | 1.00 | 42.46 |  |
| 3 | U | O2' | AA:2068 | 2.87 | 1.00 | 44.14 |  |
| 4 | G | O5' | AA:2069 | 4.08 | 1.00 | 43.34 |  |
| 5 | G | OP1 | AA:2069 | 4.25 | 1.00 | 43.48 |  |
| 6 | G | OP2 | AA:2070 | 4.37 | 1.00 | 41.38 |  |
| 7 | A | O2' | AA:197 | 4.89 | 1.00 | 42.87 |  |
  |

  
 Coded contact list:   
 Exclusion zone contact(s): /O2'/O3'/OP2/  
Consider NA in place of MG (2 valid contact(s) for NA vs. 0 for MG)  

### MG:AA:3547

|  | Res. type | Atom | Residue | Distance | Occupancy | B-factor | Closest solute to 1^st^ shell water |
| --- | --- | --- | --- | --- | --- | --- | --- |
| **Ion** | | | |  | 1.00 | 47.83 || 1 | G | O6 | AA:741 | 3.65 | 1.00 | 50.96 |
| 2 | G | O2' | AA:739 | 3.69 | 1.00 | 43.93 |  |
| 3 | G | O6 | AA:742 | 4.29 | 1.00 | 46.52 |  |
| 4 | U | O4 | AA:740 | 4.40 | 1.00 | 47.33 |  |
| 5 | G | N7 | AA:741 | 4.43 | 1.00 | 47.39 |  |
| 6 | G | N3 | AA:739 | 4.80 | 1.00 | 52.68 |  |
| 7 | G | N2 | AA:739 | 4.95 | 1.00 | 52.89 |  |
  |

  
 Coded contact list:   
1^st^ MG coordination shell missing (see Notes)  

### MG:AA:3548

|  | Res. type | Atom | Residue | Distance | Occupancy | B-factor | Closest solute to 1^st^ shell water |
| --- | --- | --- | --- | --- | --- | --- | --- |
| **Ion** | | | |  | 1.00 | 59.41 || 1 | G | O6 | AA:2069 | 3.23 | 1.00 | 40.13 |
| 2 | G | O6 | AA:2067 | 3.32 | 1.00 | 39.59 |  |
| 3 | G | N7 | AA:2069 | 3.50 | 1.00 | 35.76 |  |
| 4 | G | N7 | AA:2067 | 3.61 | 1.00 | 41.34 |  |
| 5 | C | N4 | AA:2066 | 4.83 | 1.00 | 31.04 |  |
  |

  
 Coded contact list:   
 Exclusion zone contact(s): /O6/O6/  
Consider K in place of MG (2 valid contact(s) for K vs. 0 for MG)  

### MG:AA:3549

|  | Res. type | Atom | Residue | Distance | Occupancy | B-factor | Closest solute to 1^st^ shell water |
| --- | --- | --- | --- | --- | --- | --- | --- |
| **Ion** | | | |  | 1.00 | 44.99 || 1 | G | O6 | AA:2072 | 2.66 | 1.00 | 33.32 |
| 2 | A | N6 | AA:2071 | 2.74 | 1.00 | 33.07 |  |
| 3 | G | N7 | AA:2072 | 3.33 | 1.00 | 28.51 |  |
| 4 | C | N4 | AA:2073 | 3.39 | 1.00 | 23.87 |  |
| 5 | A | N1 | AA:2071 | 3.74 | 1.00 | 31.57 |  |
| 6 | A | N7 | AA:2071 | 3.79 | 1.00 | 33.52 |  |
| 7 | G | O6 | AA:2436 | 3.89 | 1.00 | 26.01 |  |
| 8 | U | O4 | AA:2437 | 3.98 | 1.00 | 32.23 |  |
| 9 | U | O4 | AA:2438 | 4.54 | 1.00 | 35.31 |  |
| 10 | U | N3 | AA:2437 | 4.67 | 1.00 | 30.54 |  |
| 11 | G | N1 | AA:2072 | 4.81 | 1.00 | 29.91 |  |
| 12 | C | N3 | AA:2073 | 4.98 | 1.00 | 22.83 |  |
  |

  
 Coded contact list:   
 Exclusion zone contact(s): /N4/N6/N7/O6/  
Consider K in place of MG (4 valid contact(s) for K vs. 0 for MG)  

### MG:AA:3550

|  | Res. type | Atom | Residue | Distance | Occupancy | B-factor | Closest solute to 1^st^ shell water |
| --- | --- | --- | --- | --- | --- | --- | --- |
| **Ion** | | | |  | 1.00 | 82.65 || 1 | C | OP2 | AA:754 | 3.49 | 1.00 | 41.56 |
| 2 | G | O6 | AA:739 | 3.65 | 1.00 | 55.66 |  |
| 3 | C | OP1 | AA:754 | 3.92 | 1.00 | 40.96 |  |
| 4 | G | OP1 | AA:1620 | 4.50 | 1.00 | 42.53 |  |
| 5 | C | OP1 | AA:753 | 4.66 | 1.00 | 40.61 |  |
  |

  
 Coded contact list:   
1^st^ MG coordination shell missing (see Notes)  

### MG:AA:3551

|  | Res. type | Atom | Residue | Distance | Occupancy | B-factor | Closest solute to 1^st^ shell water |
| --- | --- | --- | --- | --- | --- | --- | --- |
| **Ion** | | | |  | 1.00 | 44.31 || 1 | U | O3' | AA:2596 | 2.56 | 1.00 | 30.21 |
| 2 | U | O2' | AA:2596 | 3.02 | 1.00 | 31.38 |  |
| 3 | G | OP1 | AA:2597 | 3.27 | 1.00 | 27.51 |  |
| 4 | G | OP1 | AA:1903 | 3.31 | 1.00 | 37.84 |  |
| 5 | G | O5' | AA:1903 | 3.53 | 1.00 | 40.34 |  |
| 6 | C | O2' | AA:1902 | 3.58 | 1.00 | 38.72 |  |
| 7 | GLY | O | AD:243 | 3.80 | 1.00 | 31.68 |  |
| 8 | GLY | N | AD:243 | 4.18 | 1.00 | 30.68 |  |
| 9 | C | O3' | AA:1902 | 4.24 | 1.00 | 37.94 |  |
| 10 | G | O5' | AA:2597 | 4.25 | 1.00 | 29.13 |  |
| 11 | ARG | N | AD:244 | 4.57 | 1.00 | 33.24 |  |
| 12 | G | OP2 | AA:2597 | 4.76 | 1.00 | 29.59 |  |
| 13 | U | O4' | AA:2596 | 4.90 | 1.00 | 31.66 |  |
  |

  
 Coded contact list:   
 Exclusion zone contact(s): /O2'/O3'/OP1/OP1/  
Consider K in place of MG (3 valid contact(s) for K vs. 0 for MG)  

### MG:AA:3552

|  | Res. type | Atom | Residue | Distance | Occupancy | B-factor | Closest solute to 1^st^ shell water |
| --- | --- | --- | --- | --- | --- | --- | --- |
| **Ion** | | | |  | 1.00 | 90.53 || 1 | G | O6 | AA:739 | 4.30 | 1.00 | 55.66 |
| 2 | C | N4 | AA:737 | 4.31 | 1.00 | 46.61 |  |
| 3 | G | N7 | AA:738 | 4.43 | 1.00 | 45.18 |  |
| 4 | G | O6 | AA:738 | 4.66 | 1.00 | 48.46 |  |
  |

  
 Coded contact list:   
1^st^ MG coordination shell missing (see Notes)  

### MG:AA:3553

|  | Res. type | Atom | Residue | Distance | Occupancy | B-factor | Closest solute to 1^st^ shell water |
| --- | --- | --- | --- | --- | --- | --- | --- |
| **Ion** | | | |  | 1.00 | 73.88 || 1 | G | N7 | AA:1968 | 3.60 | 1.00 | 35.47 |
| 2 | G | O6 | AA:1968 | 3.73 | 1.00 | 42.36 |  |
| 3 | G | N7 | AA:1930 | 4.29 | 1.00 | 49.67 |  |
| 4 | G | O6 | AA:1930 | 4.52 | 1.00 | 51.19 |  |
| 5 | C | OP1 | AA:1967 | 4.55 | 1.00 | 51.22 |  |
  |

  
 Coded contact list:   
1^st^ MG coordination shell missing (see Notes)  

### MG:AA:3554

|  | Res. type | Atom | Residue | Distance | Occupancy | B-factor | Closest solute to 1^st^ shell water |
| --- | --- | --- | --- | --- | --- | --- | --- |
| **Ion** | | | |  | 1.00 | 86.10 || 1 | U | O4 | AA:2312 | 3.00 | 1.00 | 100.74 |
| 2 | G | O6 | AA:2304 | 3.32 | 1.00 | 99.33 |  |
| 3 | A | OP1 | AA:2310 | 3.45 | 1.00 | 102.80 |  |
| 4 | A | OP1 | AA:2311 | 3.59 | 1.00 | 99.39 |  |
| 5 | U | N3 | AA:2312 | 4.30 | 1.00 | 99.04 |  |
| 6 | C | N4 | AA:2313 | 4.31 | 1.00 | 90.16 |  |
| 7 | A | OP2 | AA:2311 | 4.33 | 1.00 | 99.01 |  |
| 8 | A | OP2 | AA:2310 | 4.60 | 1.00 | 103.73 |  |
| 9 | G | O6 | AA:2303 | 4.80 | 1.00 | 90.89 |  |
| 10 | G | OP2 | AA:2308 | 4.89 | 1.00 | 99.37 |  |
| 11 | A | O5' | AA:2310 | 4.91 | 1.00 | 103.06 |  |
  |

  
 Coded contact list:   
 Exclusion zone contact(s): /O4/O6/  
Consider K in place of MG (2 valid contact(s) for K vs. 0 for MG)  

### MG:AA:3555

|  | Res. type | Atom | Residue | Distance | Occupancy | B-factor | Closest solute to 1^st^ shell water |
| --- | --- | --- | --- | --- | --- | --- | --- |
| **Ion** | | | |  | 1.00 | 60.64 || 1 | G | OP2 | AA:1622 | 2.06 | 1.00 | 47.56 |
| 2 | G | OP1 | AA:1622 | 3.16 | 1.00 | 48.32 |  |
| 3 | C | N3 | AA:1607 | 3.80 | 1.00 | 53.36 |  |
| 4 | G | O5' | AA:1622 | 3.96 | 1.00 | 46.80 |  |
| 5 | U | OP1 | AA:1621 | 3.98 | 1.00 | 43.36 |  |
| 6 | C | N4 | AA:1607 | 4.01 | 1.00 | 49.78 |  |
| 7 | U | O5' | AA:1621 | 4.10 | 1.00 | 42.85 |  |
| 8 | U | O3' | AA:1621 | 4.30 | 1.00 | 45.51 |  |
| 9 | U | OP2 | AA:1621 | 4.37 | 1.00 | 42.43 |  |
| 10 | C | O2 | AA:1607 | 4.86 | 1.00 | 51.75 |  |
| 11 | C | OP1 | AA:737 | 4.92 | 1.00 | 41.33 |  |
| 12 | C | OP1 | AA:736 | 4.99 | 1.00 | 42.79 |  |
  |

  
 Contact(s): /OP2/  
 Coded contact list: O_ph_  
 Exclusion zone contact(s): /OP1/  

### MG:AA:3556

|  | Res. type | Atom | Residue | Distance | Occupancy | B-factor | Closest solute to 1^st^ shell water |
| --- | --- | --- | --- | --- | --- | --- | --- |
| **Ion** | | | |  | 1.00 | 74.57 || 1 | G | O6 | AA:2325 | 4.13 | 1.00 | 60.78 |
| 2 | G | O6 | AA:2389 | 4.20 | 1.00 | 56.80 |  |
| 3 | G | N1 | AA:2384 | 4.47 | 1.00 | 64.81 |  |
| 4 | G | N3 | AA:2384 | 4.58 | 1.00 | 65.30 |  |
| 5 | G | N7 | AA:2384 | 4.59 | 1.00 | 66.64 |  |
| 6 | C | N4 | AA:2283 | 4.65 | 1.00 | 58.22 |  |
| 7 | C | OP2 | AA:2386 | 4.74 | 1.00 | 61.94 |  |
| 8 | G | O6 | AA:2384 | 4.78 | 1.00 | 64.46 |  |
| 9 | G | N9 | AA:2384 | 4.80 | 1.00 | 66.51 |  |
  |

  
 Coded contact list:   
1^st^ MG coordination shell missing (see Notes)  

### MG:AA:3557

|  | Res. type | Atom | Residue | Distance | Occupancy | B-factor | Closest solute to 1^st^ shell water |
| --- | --- | --- | --- | --- | --- | --- | --- |
| **Ion** | | | |  | 1.00 | 89.04 || 1 | U | O2 | AA:2312 | 2.38 | 1.00 | 98.54 |
| 2 | GLY | N | AG:42 | 3.40 | 1.00 | 103.96 |  |
| 3 | U | N1 | AA:2312 | 3.41 | 1.00 | 97.34 |  |
| 4 | G | O6 | AA:2307 | 3.60 | 1.00 | 90.84 |  |
| 5 | U | O4' | AA:2312 | 3.81 | 1.00 | 96.03 |  |
| 6 | G | N2 | AA:2304 | 3.84 | 1.00 | 102.08 |  |
| 7 | G | N7 | AA:2307 | 3.85 | 1.00 | 92.44 |  |
| 8 | ASN | OD1 | AG:40 | 3.87 | 1.00 | 102.15 |  |
| 9 | U | O2' | AA:2312 | 3.89 | 1.00 | 96.18 |  |
| 10 | C | N4 | AA:2306 | 4.05 | 1.00 | 92.70 |  |
| 11 | U | N3 | AA:2312 | 4.36 | 1.00 | 99.04 |  |
| 12 | G | N1 | AA:2304 | 4.52 | 1.00 | 100.39 |  |
| 13 | LEU | N | AG:43 | 4.83 | 1.00 | 106.76 |  |
| 14 | GLY | O | AG:42 | 4.84 | 1.00 | 105.17 |  |
| 15 | GLN | O | AG:41 | 4.87 | 1.00 | 103.13 |  |
| 16 | ASP | OD2 | AG:156 | 4.92 | 1.00 | 99.14 |  |
| 17 | ASN | ND2 | AG:40 | 4.97 | 1.00 | 101.79 |  |
  |

  
 Contact(s): /O2/  
 Coded contact list: O_b_  

### MG:AA:3558

|  | Res. type | Atom | Residue | Distance | Occupancy | B-factor | Closest solute to 1^st^ shell water |
| --- | --- | --- | --- | --- | --- | --- | --- |
| **Ion** | | | |  | 1.00 | 50.22 || 1 | C | OP1 | AA:737 | 2.73 | 1.00 | 41.33 |
| 2 | U | OP1 | AA:1621 | 2.82 | 1.00 | 43.36 |  |
| 3 | G | O3' | AA:1620 | 4.19 | 1.00 | 41.48 |  |
| 4 | C | O5' | AA:737 | 4.35 | 1.00 | 40.96 |  |
| 5 | C | OP2 | AA:737 | 4.54 | 1.00 | 41.81 |  |
| 6 | G | OP2 | AA:738 | 4.64 | 1.00 | 39.89 |  |
| 7 | G | OP1 | AA:1622 | 4.70 | 1.00 | 48.32 |  |
| 8 | U | O5' | AA:1621 | 4.86 | 1.00 | 42.85 |  |
  |

  
 Coded contact list:   
 Exclusion zone contact(s): /OP1/OP1/  
Consider K in place of MG (2 valid contact(s) for K vs. 0 for MG)  

### MG:AA:3559

|  | Res. type | Atom | Residue | Distance | Occupancy | B-factor | Closest solute to 1^st^ shell water |
| --- | --- | --- | --- | --- | --- | --- | --- |
| **Ion** | | | |  | 1.00 | 50.33 || 1 | G | N3 | AA:2255 | 2.86 | 1.00 | 50.78 |
| 2 | G | O2' | AA:2255 | 3.24 | 1.00 | 53.76 |  |
| 3 | LEU | O | A3:7 | 3.29 | 1.00 | 96.73 |  |
| 4 | G | N9 | AA:2255 | 3.60 | 1.00 | 52.75 |  |
| 5 | C | O2' | BC:2 | 3.89 | 1.00 | 87.34 |  |
| 6 | G | O4' | AA:2255 | 3.95 | 1.00 | 53.34 |  |
| 7 | G | N2 | AA:2255 | 4.05 | 1.00 | 49.00 |  |
| 8 | GLY | N | A3:8 | 4.34 | 1.00 | 94.02 |  |
| 9 | GLY | O | A3:8 | 4.44 | 1.00 | 87.51 |  |
| 10 | C | O3' | BC:2 | 4.55 | 1.00 | 88.81 |  |
| 11 | SER | N | A3:9 | 4.82 | 1.00 | 86.19 |  |
| 12 | GLY | O | A3:6 | 4.97 | 1.00 | 100.73 |  |
  |

  
 Coded contact list:   
 Exclusion zone contact(s): /N3_G_/O2'/O_back_/  
Consider K in place of MG (3 valid contact(s) for K vs. 0 for MG)  

### MG:AA:3560

|  | Res. type | Atom | Residue | Distance | Occupancy | B-factor | Closest solute to 1^st^ shell water |
| --- | --- | --- | --- | --- | --- | --- | --- |
| **Ion** | | | |  | 1.00 | 37.01 || 1 | G | O6 | AA:1623 | 3.81 | 1.00 | 43.66 |
| 2 | G | N7 | AA:1623 | 4.21 | 1.00 | 42.93 |  |
| 3 | C | N4 | AA:1607 | 4.24 | 1.00 | 49.78 |  |
| 4 | C | N1 | AA:1607 | 4.66 | 1.00 | 52.06 |  |
| 5 | G | O6 | AA:1624 | 4.68 | 1.00 | 46.82 |  |
| 6 | C | N3 | AA:1607 | 4.70 | 1.00 | 53.36 |  |
| 7 | C | OP2 | AA:1607 | 4.87 | 1.00 | 48.30 |  |
| 8 | G | O6 | AA:1622 | 4.92 | 1.00 | 46.23 |  |
  |

  
 Coded contact list:   
1^st^ MG coordination shell missing (see Notes)  

### MG:AA:3561

|  | Res. type | Atom | Residue | Distance | Occupancy | B-factor | Closest solute to 1^st^ shell water |
| --- | --- | --- | --- | --- | --- | --- | --- |
| **Ion** | | | |  | 1.00 | 45.53 || 1 | C | O2' | AA:1607 | 2.44 | 1.00 | 52.66 |
| 2 | C | O2 | AA:1607 | 2.58 | 1.00 | 51.75 |  |
| 3 | U | O4 | AA:1621 | 4.22 | 1.00 | 43.03 |  |
| 4 | C | N1 | AA:1607 | 4.38 | 1.00 | 52.06 |  |
| 5 | G | OP2 | AA:1620 | 4.72 | 1.00 | 43.76 |  |
| 6 | C | O3' | AA:1607 | 4.79 | 1.00 | 48.25 |  |
| 7 | C | N3 | AA:1607 | 4.81 | 1.00 | 53.36 |  |
  |

  
 Coded contact list:   
 Exclusion zone contact(s): /O2/O2'/  
Consider NA in place of MG (2 valid contact(s) for NA vs. 0 for MG)  

### MG:AA:3562

|  | Res. type | Atom | Residue | Distance | Occupancy | B-factor | Closest solute to 1^st^ shell water |
| --- | --- | --- | --- | --- | --- | --- | --- |
| **Ion** | | | |  | 1.00 | 55.06 || 1 | C | O2' | AA:1607 | 2.90 | 1.00 | 52.66 |
| 2 | A | OP2 | AA:1302 | 3.72 | 1.00 | 51.00 |  |
| 3 | C | O3' | AA:1607 | 3.93 | 1.00 | 48.25 |  |
| 4 | A | OP2 | AA:1608 | 4.55 | 1.00 | 47.75 |  |
| 5 | A | O5' | AA:1302 | 4.71 | 1.00 | 51.55 |  |
  |

  
 Coded contact list:   
 Exclusion zone contact(s): /O2'/  
Consider K in place of MG (1 valid contact(s) for K vs. 0 for MG)  

### MG:AA:3563

|  | Res. type | Atom | Residue | Distance | Occupancy | B-factor | Closest solute to 1^st^ shell water |
| --- | --- | --- | --- | --- | --- | --- | --- |
| **Ion** | | | |  | 1.00 | 67.56 || 1 | G | OP1 | AA:1606 | 2.66 | 1.00 | 43.14 |
| 2 | C | OP1 | AA:1607 | 3.24 | 1.00 | 48.45 |  |
| 3 | G | OP2 | AA:1606 | 3.60 | 1.00 | 43.42 |  |
| 4 | G | O5' | AA:1606 | 3.68 | 1.00 | 43.33 |  |
| 5 | C | OP2 | AA:1304 | 4.06 | 1.00 | 55.75 |  |
| 6 | G | OP1 | AA:1303 | 4.70 | 1.00 | 55.75 |  |
| 7 | C | O3' | AA:1605 | 4.89 | 1.00 | 41.17 |  |
  |

  
 Coded contact list:   
 Exclusion zone contact(s): /OP1/OP1/  
Consider K in place of MG (2 valid contact(s) for K vs. 0 for MG)  

### MG:AA:3564

|  | Res. type | Atom | Residue | Distance | Occupancy | B-factor | Closest solute to 1^st^ shell water |
| --- | --- | --- | --- | --- | --- | --- | --- |
| **Ion** | | | |  | 1.00 | 67.59 || 1 | C | OP2 | AA:1605 | 3.46 | 1.00 | 36.28 |
| 2 | G | OP2 | AA:1606 | 3.46 | 1.00 | 43.42 |  |
| 3 | G | N7 | AA:1606 | 3.68 | 1.00 | 40.85 |  |
| 4 | C | OP2 | AA:1305 | 3.94 | 1.00 | 52.28 |  |
| 5 | C | OP1 | AA:1605 | 4.07 | 1.00 | 36.90 |  |
| 6 | C | O5' | AA:1605 | 4.22 | 1.00 | 38.37 |  |
| 7 | G | O5' | AA:1606 | 4.87 | 1.00 | 43.33 |  |
| 8 | C | O5' | AA:1305 | 4.98 | 1.00 | 52.77 |  |
  |

  
 Coded contact list:   
1^st^ MG coordination shell missing (see Notes)  

### MG:AA:3565

|  | Res. type | Atom | Residue | Distance | Occupancy | B-factor | Closest solute to 1^st^ shell water |
| --- | --- | --- | --- | --- | --- | --- | --- |
| **Ion** | | | |  | 1.00 | 89.66 || 1 | G | OP2 | AA:1389 | 4.73 | 1.00 | 69.47 |
| 2 | G | N7 | AA:1389 | 4.83 | 1.00 | 66.63 |  |
| 3 | U | O4 | AA:1390 | 4.90 | 1.00 | 72.99 |  |
  |

  
 Coded contact list:   
1^st^ MG coordination shell missing (see Notes)  

### MG:AA:3566

|  | Res. type | Atom | Residue | Distance | Occupancy | B-factor | Closest solute to 1^st^ shell water |
| --- | --- | --- | --- | --- | --- | --- | --- |
| **Ion** | | | |  | 1.00 | 65.85 || 1 | C | OP2 | AA:1387 | 3.47 | 1.00 | 67.41 |
| 2 | G | OP2 | AA:1388 | 3.78 | 1.00 | 69.34 |  |
| 3 | G | N7 | AA:1388 | 4.31 | 1.00 | 70.42 |  |
| 4 | C | O5' | AA:1387 | 4.47 | 1.00 | 67.76 |  |
| 5 | U | O4' | AA:1396 | 4.82 | 1.00 | 64.52 |  |
  |

  
 Coded contact list:   
1^st^ MG coordination shell missing (see Notes)  

### MG:AA:3567

|  | Res. type | Atom | Residue | Distance | Occupancy | B-factor | Closest solute to 1^st^ shell water |
| --- | --- | --- | --- | --- | --- | --- | --- |
| **Ion** | | | |  | 1.00 | 82.80 || 1 | U | OP2 | AA:1397 | 2.77 | 1.00 | 60.02 |
| 2 | G | O6 | AA:1389 | 3.40 | 1.00 | 65.68 |  |
| 3 | C | N4 | AA:1398 | 3.53 | 1.00 | 65.92 |  |
| 4 | A | O2' | AA:1395 | 3.92 | 1.00 | 59.60 |  |
| 5 | G | O6 | AA:1388 | 3.92 | 1.00 | 71.03 |  |
| 6 | U | O5' | AA:1396 | 3.92 | 1.00 | 60.26 |  |
| 7 | U | OP2 | AA:1396 | 4.11 | 1.00 | 58.94 |  |
| 8 | A | O3' | AA:1395 | 4.12 | 1.00 | 58.98 |  |
| 9 | U | O3' | AA:1396 | 4.35 | 1.00 | 62.29 |  |
| 10 | U | O4' | AA:1396 | 4.75 | 1.00 | 64.52 |  |
| 11 | U | OP1 | AA:1397 | 4.86 | 1.00 | 60.85 |  |
| 12 | C | N4 | AA:1399 | 4.96 | 1.00 | 70.12 |  |
  |

  
 Coded contact list:   
 Exclusion zone contact(s): /O6/OP2/  
Consider K in place of MG (2 valid contact(s) for K vs. 0 for MG)  

### MG:AA:3568

|  | Res. type | Atom | Residue | Distance | Occupancy | B-factor | Closest solute to 1^st^ shell water |
| --- | --- | --- | --- | --- | --- | --- | --- |
| **Ion** | | | |  | 1.00 | 41.52 || 1 | G | O6 | AA:1807 | 3.80 | 1.00 | 44.09 |
| 2 | G | N7 | AA:1807 | 4.14 | 1.00 | 44.87 |  |
| 3 | A | OP1 | AA:1809 | 4.52 | 1.00 | 43.83 |  |
| 4 | A | OP2 | AA:1809 | 4.83 | 1.00 | 42.61 |  |
  |

  
 Coded contact list:   
1^st^ MG coordination shell missing (see Notes)  

### MG:AA:3569

|  | Res. type | Atom | Residue | Distance | Occupancy | B-factor | Closest solute to 1^st^ shell water |
| --- | --- | --- | --- | --- | --- | --- | --- |
| **Ion** | | | |  | 1.00 | 75.09 || 1 | C | OP1 | AA:1806 | 2.82 | 1.00 | 37.01 |
| 2 | G | OP2 | AA:1807 | 4.10 | 1.00 | 44.48 |  |
| 3 | C | OP2 | AA:1806 | 4.42 | 1.00 | 35.77 |  |
| 4 | C | O5' | AA:1806 | 4.48 | 1.00 | 37.22 |  |
| 5 | G | OP1 | AA:1807 | 4.84 | 1.00 | 45.58 |  |
| 6 | A | OP2 | AA:777 | 4.90 | 1.00 | 35.82 |  |
  |

  
 Coded contact list:   
 Exclusion zone contact(s): /OP1/  
Consider K in place of MG (1 valid contact(s) for K vs. 0 for MG)  

### MG:AA:3570

|  | Res. type | Atom | Residue | Distance | Occupancy | B-factor | Closest solute to 1^st^ shell water |
| --- | --- | --- | --- | --- | --- | --- | --- |
| **Ion** | | | |  | 1.00 | 78.80 || 1 | C | OP1 | AA:2084 | 2.73 | 1.00 | 36.73 |
| 2 | G | O3' | AA:2083 | 3.99 | 1.00 | 38.87 |  |
| 3 | C | O5' | AA:2084 | 4.61 | 1.00 | 38.07 |  |
  |

  
 Coded contact list:   
 Exclusion zone contact(s): /OP1/  
Consider K in place of MG (1 valid contact(s) for K vs. 0 for MG)  

### MG:AA:3571

|  | Res. type | Atom | Residue | Distance | Occupancy | B-factor | Closest solute to 1^st^ shell water |
| --- | --- | --- | --- | --- | --- | --- | --- |
| **Ion** | | | |  | 1.00 | 57.47 || 1 | A | OP2 | AA:2227 | 3.06 | 1.00 | 48.90 |
| 2 | G | O6 | AA:2228 | 3.30 | 1.00 | 53.26 |  |
| 3 | U | O4 | AA:2091 | 3.76 | 1.00 | 55.86 |  |
| 4 | U | O4 | AA:2092 | 3.87 | 1.00 | 62.16 |  |
| 5 | A | N7 | AA:2227 | 3.88 | 1.00 | 53.92 |  |
| 6 | C | OP2 | AA:2226 | 3.93 | 1.00 | 53.43 |  |
| 7 | G | O6 | AA:2090 | 4.20 | 1.00 | 54.32 |  |
| 8 | G | N7 | AA:2228 | 4.21 | 1.00 | 50.72 |  |
| 9 | C | N4 | AA:2229 | 4.23 | 1.00 | 54.92 |  |
| 10 | U | N3 | AA:2092 | 4.66 | 1.00 | 61.26 |  |
| 11 | C | OP1 | AA:2226 | 4.72 | 1.00 | 52.49 |  |
| 12 | A | O5' | AA:2227 | 4.87 | 1.00 | 48.02 |  |
| 13 | U | N3 | AA:2091 | 4.88 | 1.00 | 54.22 |  |
  |

  
 Coded contact list:   
 Exclusion zone contact(s): /O6/OP2/  
Consider K in place of MG (2 valid contact(s) for K vs. 0 for MG)  

### MG:AA:3572

|  | Res. type | Atom | Residue | Distance | Occupancy | B-factor | Closest solute to 1^st^ shell water |
| --- | --- | --- | --- | --- | --- | --- | --- |
| **Ion** | | | |  | 1.00 | 64.96 || 1 | U | O4 | AA:2091 | 3.08 | 1.00 | 55.86 |
| 2 | U | O2 | AA:2092 | 3.52 | 1.00 | 56.30 |  |
| 3 | G | N7 | AA:2090 | 3.87 | 1.00 | 53.11 |  |
| 4 | G | OP2 | AA:2090 | 4.07 | 1.00 | 52.31 |  |
| 5 | U | N3 | AA:2092 | 4.16 | 1.00 | 61.26 |  |
| 6 | U | O2' | AA:2092 | 4.57 | 1.00 | 57.90 |  |
| 7 | G | O6 | AA:2090 | 4.86 | 1.00 | 54.32 |  |
| 8 | C | OP1 | AA:2226 | 4.95 | 1.00 | 52.49 |  |
| 9 | U | N1 | AA:2092 | 4.97 | 1.00 | 60.06 |  |
  |

  
 Coded contact list:   
 Exclusion zone contact(s): /O4/  
Consider K in place of MG (1 valid contact(s) for K vs. 0 for MG)  

### MG:AA:3573

|  | Res. type | Atom | Residue | Distance | Occupancy | B-factor | Closest solute to 1^st^ shell water |
| --- | --- | --- | --- | --- | --- | --- | --- |
| **Ion** | | | |  | 1.00 | 44.53 || 1 | C | N4 | AA:2073 | 3.64 | 1.00 | 23.87 |
| 2 | G | OP2 | AA:2072 | 4.05 | 1.00 | 32.13 |  |
| 3 | C | OP2 | AA:2073 | 4.75 | 1.00 | 25.31 |  |
| 4 | G | N7 | AA:2072 | 4.81 | 1.00 | 28.51 |  |
| 5 | U | O4 | AA:2243 | 4.86 | 1.00 | 33.32 |  |
  |

  
 Coded contact list:   
1^st^ MG coordination shell missing (see Notes)  

### MG:AA:3574

|  | Res. type | Atom | Residue | Distance | Occupancy | B-factor | Closest solute to 1^st^ shell water |
| --- | --- | --- | --- | --- | --- | --- | --- |
| **Ion** | | | |  | 1.00 | 57.81 || 1 | G | OP1 | AA:794 | 3.18 | 1.00 | 39.87 |
| 2 | C | OP2 | AA:795 | 4.26 | 1.00 | 34.17 |  |
| 3 | G | O5' | AA:794 | 4.29 | 1.00 | 38.95 |  |
| 4 | A | O3' | AA:793 | 4.33 | 1.00 | 41.17 |  |
| 5 | C | OP1 | AA:795 | 4.71 | 1.00 | 33.35 |  |
| 6 | A | OP1 | AA:677 | 4.77 | 1.00 | 39.68 |  |
| 7 | G | O3' | AA:794 | 4.84 | 1.00 | 34.83 |  |
  |

  
 Coded contact list:   
 Exclusion zone contact(s): /OP1/  
Consider K in place of MG (1 valid contact(s) for K vs. 0 for MG)  

### MG:AA:3575

|  | Res. type | Atom | Residue | Distance | Occupancy | B-factor | Closest solute to 1^st^ shell water |
| --- | --- | --- | --- | --- | --- | --- | --- |
| **Ion** | | | |  | 1.00 | 51.70 || 1 | G | OP2 | AA:187 | 3.42 | 1.00 | 41.24 |
| 2 | G | N7 | AA:187 | 3.57 | 1.00 | 37.25 |  |
| 3 | G | N7 | AA:186 | 4.54 | 1.00 | 40.77 |  |
| 4 | G | N7 | AA:188 | 4.66 | 1.00 | 39.08 |  |
| 5 | G | O6 | AA:189 | 4.68 | 1.00 | 36.59 |  |
| 6 | G | OP2 | AA:186 | 4.78 | 1.00 | 45.06 |  |
| 7 | G | O5' | AA:187 | 4.86 | 1.00 | 42.02 |  |
| 8 | G | O5' | AA:186 | 4.91 | 1.00 | 45.28 |  |
  |

  
 Coded contact list:   
1^st^ MG coordination shell missing (see Notes)  

### MG:AA:3576

|  | Res. type | Atom | Residue | Distance | Occupancy | B-factor | Closest solute to 1^st^ shell water |
| --- | --- | --- | --- | --- | --- | --- | --- |
| **Ion** | | | |  | 1.00 | 37.76 || 1 | G | O6 | AA:187 | 2.70 | 1.00 | 35.65 |
| 2 | G | O6 | AA:188 | 3.12 | 1.00 | 38.01 |  |
| 3 | G | O6 | AA:186 | 3.81 | 1.00 | 40.81 |  |
| 4 | G | N7 | AA:187 | 3.96 | 1.00 | 37.25 |  |
| 5 | G | N7 | AA:188 | 4.19 | 1.00 | 39.08 |  |
| 6 | C | N4 | AA:209 | 4.29 | 1.00 | 38.57 |  |
| 7 | C | N4 | AA:208 | 4.41 | 1.00 | 33.87 |  |
| 8 | G | N7 | AA:186 | 4.48 | 1.00 | 40.77 |  |
| 9 | G | N1 | AA:187 | 4.93 | 1.00 | 36.63 |  |
| 10 | G | O6 | AA:189 | 4.96 | 1.00 | 36.59 |  |
| 11 | G | N1 | AA:189 | 4.97 | 1.00 | 33.46 |  |
  |

  
 Coded contact list:   
 Exclusion zone contact(s): /O6/O6/  
Consider K in place of MG (2 valid contact(s) for K vs. 0 for MG)  

### MG:AA:3577

|  | Res. type | Atom | Residue | Distance | Occupancy | B-factor | Closest solute to 1^st^ shell water |
| --- | --- | --- | --- | --- | --- | --- | --- |
| **Ion** | | | |  | 1.00 | 80.55 || 1 | A | OP1 | AA:1439 | 1.86 | 1.00 | 59.45 |
| 2 | C | OP1 | AA:1468 | 2.39 | 1.00 | 77.43 |  |
| 3 | A | OP2 | AA:1439 | 3.43 | 1.00 | 59.38 |  |
| 4 | A | O5' | AA:1439 | 3.62 | 1.00 | 59.42 |  |
| 5 | G | OP2 | AA:1440 | 3.71 | 1.00 | 58.51 |  |
| 6 | U | O3' | AA:1438 | 4.28 | 1.00 | 58.21 |  |
| 7 | C | O5' | AA:1468 | 4.46 | 1.00 | 77.13 |  |
| 8 | C | O3' | AA:1467 | 4.72 | 1.00 | 78.27 |  |
| 9 | C | OP2 | AA:1468 | 4.77 | 1.00 | 77.23 |  |
  |

  
**1^st^ shell ligand angles:**   
1-2: 142;   
  
 Contact(s): /OP1/OP1/  
 Coded contact list: 2O_ph_  

### MG:AA:3578

|  | Res. type | Atom | Residue | Distance | Occupancy | B-factor | Closest solute to 1^st^ shell water |
| --- | --- | --- | --- | --- | --- | --- | --- |
| **Ion** | | | |  | 1.00 | 33.69 || 1 | U | OP2 | AA:511 | 2.15 | 1.00 | 49.69 |
| 2 | C | OP1 | AA:510 | 2.22 | 1.00 | 52.21 |  |
| 3 | G | N7 | AA:512 | 2.63 | 1.00 | 54.25 |  |
| 4 | C | O3' | AA:509 | 3.13 | 1.00 | 52.52 |  |
| 5 | G | O6 | AA:512 | 3.69 | 1.00 | 55.32 |  |
| 6 | C | O3' | AA:510 | 4.08 | 1.00 | 49.17 |  |
| 7 | C | OP2 | AA:510 | 4.27 | 1.00 | 52.25 |  |
| 8 | C | O5' | AA:510 | 4.39 | 1.00 | 52.57 |  |
| 9 | U | O5' | AA:511 | 4.41 | 1.00 | 51.34 |  |
| 10 | U | OP1 | AA:511 | 4.64 | 1.00 | 50.71 |  |
| 11 | C | O2' | AA:509 | 4.65 | 1.00 | 53.94 |  |
| 12 | G | N9 | AA:512 | 4.77 | 1.00 | 55.17 |  |
  |

  
**1^st^ shell ligand angles:**   
1-2: 87;   
  
 Contact(s): /OP1/OP2/  
 Coded contact list: 2O_ph_  
 Exclusion zone contact(s): /N7/O3'/  

### MG:AA:3579

|  | Res. type | Atom | Residue | Distance | Occupancy | B-factor | Closest solute to 1^st^ shell water |
| --- | --- | --- | --- | --- | --- | --- | --- |
| **Ion** | | | |  | 1.00 | 83.12 || 1 | G | N7 | AA:512 | 2.44 | 1.00 | 54.25 |
| 2 | U | OP2 | AA:511 | 3.24 | 1.00 | 49.69 |  |
| 3 | G | O6 | AA:512 | 3.41 | 1.00 | 55.32 |  |
| 4 | C | OP1 | AA:510 | 3.43 | 1.00 | 52.21 |  |
| 5 | C | O3' | AA:509 | 3.98 | 1.00 | 52.52 |  |
| 6 | G | O6 | AA:27 | 4.29 | 1.00 | 55.65 |  |
| 7 | G | N9 | AA:512 | 4.60 | 1.00 | 55.17 |  |
  |

  
 Coded contact list:   
 Exclusion zone contact(s): /N7/OP2/  
Consider NA in place of MG (1 valid contact(s) for NA vs. 0 for MG)  

### MG:AA:3580

|  | Res. type | Atom | Residue | Distance | Occupancy | B-factor | Closest solute to 1^st^ shell water |
| --- | --- | --- | --- | --- | --- | --- | --- |
| **Ion** | | | |  | 1.00 | 50.44 || 1 | G | O6 | AA:27 | 3.25 | 1.00 | 55.65 |
| 2 | G | O6 | AA:26 | 4.05 | 1.00 | 54.47 |  |
| 3 | G | N7 | AA:512 | 4.25 | 1.00 | 54.25 |  |
| 4 | G | N1 | AA:27 | 4.56 | 1.00 | 53.19 |  |
| 5 | G | O6 | AA:512 | 4.82 | 1.00 | 55.32 |  |
  |

  
 Coded contact list:   
 Exclusion zone contact(s): /O6/  
Consider K in place of MG (1 valid contact(s) for K vs. 0 for MG)  

### MG:AA:3581

|  | Res. type | Atom | Residue | Distance | Occupancy | B-factor | Closest solute to 1^st^ shell water |
| --- | --- | --- | --- | --- | --- | --- | --- |
| **Ion** | | | |  | 1.00 | 55.66 || 1 | G | OP1 | AA:506 | 4.43 | 1.00 | 55.09 |
| 2 | U | O2' | AA:504 | 4.60 | 1.00 | 72.53 |  |
| 3 | G | OP1 | AA:24 | 4.94 | 1.00 | 53.73 |  |
  |

  
 Coded contact list:   
1^st^ MG coordination shell missing (see Notes)  

### MG:AA:3582

|  | Res. type | Atom | Residue | Distance | Occupancy | B-factor | Closest solute to 1^st^ shell water |
| --- | --- | --- | --- | --- | --- | --- | --- |
| **Ion** | | | |  | 1.00 | 54.44 || 1 | U | OP2 | AA:25 | 2.72 | 1.00 | 51.04 |
| 2 | G | O6 | AA:26 | 2.88 | 1.00 | 54.47 |  |
| 3 | G | N7 | AA:26 | 4.05 | 1.00 | 53.93 |  |
| 4 | G | O6 | AA:27 | 4.10 | 1.00 | 55.65 |  |
| 5 | G | OP2 | AA:24 | 4.62 | 1.00 | 53.14 |  |
| 6 | U | O5' | AA:25 | 4.64 | 1.00 | 51.97 |  |
| 7 | G | O5' | AA:24 | 4.88 | 1.00 | 52.66 |  |
| 8 | G | N1 | AA:27 | 4.97 | 1.00 | 53.19 |  |
  |

  
 Coded contact list:   
 Exclusion zone contact(s): /O6/OP2/  
Consider K in place of MG (2 valid contact(s) for K vs. 0 for MG)  

### MG:AA:3583

|  | Res. type | Atom | Residue | Distance | Occupancy | B-factor | Closest solute to 1^st^ shell water |
| --- | --- | --- | --- | --- | --- | --- | --- |
| **Ion** | | | |  | 1.00 | 72.55 || 1 | U | OP2 | AA:25 | 2.99 | 1.00 | 51.04 |
| 2 | G | OP1 | AA:24 | 3.22 | 1.00 | 53.73 |  |
| 3 | U | OP1 | AA:25 | 3.91 | 1.00 | 51.70 |  |
| 4 | G | O5' | AA:24 | 4.07 | 1.00 | 52.66 |  |
| 5 | G | O3' | AA:24 | 4.20 | 1.00 | 50.29 |  |
| 6 | G | OP2 | AA:24 | 4.77 | 1.00 | 53.14 |  |
  |

  
 Coded contact list:   
 Exclusion zone contact(s): /OP1/OP2/  
Consider K in place of MG (2 valid contact(s) for K vs. 0 for MG)  

### MG:AA:3584

|  | Res. type | Atom | Residue | Distance | Occupancy | B-factor | Closest solute to 1^st^ shell water |
| --- | --- | --- | --- | --- | --- | --- | --- |
| **Ion** | | | |  | 1.00 | 85.74 || 1 | U | OP1 | AA:25 | 3.12 | 1.00 | 51.70 |
| 2 | G | O2' | AA:508 | 3.44 | 1.00 | 64.07 |  |
| 3 | U | OP2 | AA:25 | 3.46 | 1.00 | 51.04 |  |
| 4 | U | O5' | AA:25 | 4.38 | 1.00 | 51.97 |  |
| 5 | TYR | OH | AS:9 | 4.48 | 1.00 | 53.72 |  |
| 6 | G | OP2 | AA:26 | 4.51 | 1.00 | 49.85 |  |
| 7 | C | OP1 | AA:509 | 4.51 | 1.00 | 55.57 |  |
  |

  
 Coded contact list:   
 Exclusion zone contact(s): /OP1/  
Consider K in place of MG (1 valid contact(s) for K vs. 0 for MG)  

### MG:AA:3585

|  | Res. type | Atom | Residue | Distance | Occupancy | B-factor | Closest solute to 1^st^ shell water |
| --- | --- | --- | --- | --- | --- | --- | --- |
| **Ion** | | | |  | 1.00 | 45.14 || 1 | G | O6 | AA:27 | 3.02 | 1.00 | 55.65 |
| 2 | G | N7 | AA:26 | 3.25 | 1.00 | 53.93 |  |
| 3 | G | OP2 | AA:26 | 3.67 | 1.00 | 49.85 |  |
| 4 | U | OP2 | AA:25 | 4.20 | 1.00 | 51.04 |  |
| 5 | C | OP1 | AA:509 | 4.56 | 1.00 | 55.57 |  |
| 6 | G | N7 | AA:27 | 4.58 | 1.00 | 54.82 |  |
| 7 | G | O6 | AA:26 | 4.66 | 1.00 | 54.47 |  |
| 8 | U | O5' | AA:25 | 4.78 | 1.00 | 51.97 |  |
| 9 | G | O2' | AA:508 | 4.78 | 1.00 | 64.07 |  |
| 10 | G | N1 | AA:27 | 4.83 | 1.00 | 53.19 |  |
  |

  
 Coded contact list:   
 Exclusion zone contact(s): /N7/O6/  
Consider K in place of MG (2 valid contact(s) for K vs. 0 for MG)  

### MG:AA:3586

|  | Res. type | Atom | Residue | Distance | Occupancy | B-factor | Closest solute to 1^st^ shell water |
| --- | --- | --- | --- | --- | --- | --- | --- |
| **Ion** | | | |  | 1.00 | 41.72 || 1 | C | OP1 | AA:509 | 2.23 | 1.00 | 55.57 |
| 2 | C | O5' | AA:509 | 3.24 | 1.00 | 56.30 |  |
| 3 | G | O6 | AA:27 | 3.79 | 1.00 | 55.65 |  |
| 4 | G | O2' | AA:508 | 3.80 | 1.00 | 64.07 |  |
| 5 | C | OP2 | AA:509 | 3.96 | 1.00 | 56.17 |  |
| 6 | G | N7 | AA:27 | 4.04 | 1.00 | 54.82 |  |
| 7 | G | O6 | AA:512 | 4.20 | 1.00 | 55.32 |  |
| 8 | G | O3' | AA:508 | 4.61 | 1.00 | 58.63 |  |
  |

  
 Contact(s): /OP1/  
 Coded contact list: O_ph_  
 Exclusion zone contact(s): /O5'/  

### MG:AA:3587

|  | Res. type | Atom | Residue | Distance | Occupancy | B-factor | Closest solute to 1^st^ shell water |
| --- | --- | --- | --- | --- | --- | --- | --- |
| **Ion** | | | |  | 1.00 | 55.53 || 1 | C | OP1 | AA:509 | 3.20 | 1.00 | 55.57 |
| 2 | G | OP2 | AA:26 | 3.23 | 1.00 | 49.85 |  |
| 3 | G | O2' | AA:508 | 3.61 | 1.00 | 64.07 |  |
| 4 | G | OP1 | AA:26 | 4.73 | 1.00 | 50.03 |  |
| 5 | G | OP2 | AA:27 | 4.92 | 1.00 | 44.87 |  |
| 6 | G | O5' | AA:26 | 4.96 | 1.00 | 49.60 |  |
  |

  
 Coded contact list:   
 Exclusion zone contact(s): /OP1/OP2/  
Consider K in place of MG (2 valid contact(s) for K vs. 0 for MG)  

### MG:AA:3588

|  | Res. type | Atom | Residue | Distance | Occupancy | B-factor | Closest solute to 1^st^ shell water |
| --- | --- | --- | --- | --- | --- | --- | --- |
| **Ion** | | | |  | 1.00 | 82.52 || 1 | G | N7 | AA:27 | 2.96 | 1.00 | 54.82 |
| 2 | C | OP1 | AA:509 | 4.20 | 1.00 | 55.57 |  |
| 3 | G | OP2 | AA:27 | 4.37 | 1.00 | 44.87 |  |
| 4 | G | O6 | AA:512 | 4.75 | 1.00 | 55.32 |  |
| 5 | G | O6 | AA:27 | 4.80 | 1.00 | 55.65 |  |
| 6 | C | OP2 | AA:509 | 4.82 | 1.00 | 56.17 |  |
| 7 | G | N9 | AA:27 | 4.85 | 1.00 | 52.02 |  |
  |

  
 Coded contact list:   
 Exclusion zone contact(s): /N7/  
Consider K in place of MG (1 valid contact(s) for K vs. 0 for MG)  

### MG:AA:3589

|  | Res. type | Atom | Residue | Distance | Occupancy | B-factor | Closest solute to 1^st^ shell water |
| --- | --- | --- | --- | --- | --- | --- | --- |
| **Ion** | | | |  | 1.00 | 139.58 || 1 | G | O6 | AA:512 | 1.76 | 1.00 | 55.32 |
| 2 | C | O5' | AA:509 | 3.14 | 1.00 | 56.30 |  |
| 3 | G | O6 | AA:27 | 3.47 | 1.00 | 55.65 |  |
| 4 | C | OP2 | AA:510 | 3.80 | 1.00 | 52.25 |  |
| 5 | C | O3' | AA:509 | 3.88 | 1.00 | 52.52 |  |
| 6 | G | N1 | AA:512 | 3.92 | 1.00 | 53.34 |  |
| 7 | G | N7 | AA:512 | 3.95 | 1.00 | 54.25 |  |
| 8 | C | OP1 | AA:510 | 3.96 | 1.00 | 52.21 |  |
| 9 | C | OP1 | AA:509 | 4.36 | 1.00 | 55.57 |  |
| 10 | G | N7 | AA:27 | 4.79 | 1.00 | 54.82 |  |
  |

  
 Coded contact list:   
 Exclusion zone contact(s): /O5'/O6/  
Consider K in place of MG (1 valid contact(s) for K vs. 0 for MG)  

### MG:AA:3590

|  | Res. type | Atom | Residue | Distance | Occupancy | B-factor | Closest solute to 1^st^ shell water |
| --- | --- | --- | --- | --- | --- | --- | --- |
| **Ion** | | | |  | 1.00 | 78.59 || 1 | C | OP1 | AA:2700 | 4.86 | 1.00 | 53.08 |
  |

  
 Coded contact list:   
1^st^ MG coordination shell missing (see Notes)  

### MG:AA:3591

|  | Res. type | Atom | Residue | Distance | Occupancy | B-factor | Closest solute to 1^st^ shell water |
| --- | --- | --- | --- | --- | --- | --- | --- |
| **Ion** | | | |  | 1.00 | 106.42 || 1 | C | OP2 | AA:1464 | 4.46 | 1.00 | 83.48 |
| 2 | C | OP1 | AA:1463 | 4.92 | 1.00 | 86.85 |  |
  |

  
 Coded contact list:   
1^st^ MG coordination shell missing (see Notes)  

### MG:AA:3592

|  | Res. type | Atom | Residue | Distance | Occupancy | B-factor | Closest solute to 1^st^ shell water |
| --- | --- | --- | --- | --- | --- | --- | --- |
| **Ion** | | | |  | 1.00 | 63.14 || 1 | G | OP1 | AA:1443 | 3.60 | 1.00 | 67.69 |
  |

  
 Coded contact list:   
1^st^ MG coordination shell missing (see Notes)  

### MG:AA:3593

|  | Res. type | Atom | Residue | Distance | Occupancy | B-factor | Closest solute to 1^st^ shell water |
| --- | --- | --- | --- | --- | --- | --- | --- |
| **Ion** | | | |  | 1.00 | 80.55 || 1 | C | OP1 | AA:1463 | 3.09 | 1.00 | 86.85 |
| 2 | C | OP1 | AA:2703 | 3.75 | 1.00 | 69.14 |  |
  |

  
 Coded contact list:   
 Exclusion zone contact(s): /OP1/  
Consider K in place of MG (1 valid contact(s) for K vs. 0 for MG)  

### MG:AA:3594

|  | Res. type | Atom | Residue | Distance | Occupancy | B-factor | Closest solute to 1^st^ shell water |
| --- | --- | --- | --- | --- | --- | --- | --- |
| **Ion** | | | |  | 1.00 | 78.36 || 1 | C | OP2 | AA:1463 | 4.00 | 1.00 | 86.82 |
| 2 | G | OP1 | AA:1444 | 4.48 | 1.00 | 77.38 |  |
| 3 | C | OP1 | AA:1463 | 4.85 | 1.00 | 86.85 |  |
  |

  
 Coded contact list:   
1^st^ MG coordination shell missing (see Notes)  

### MG:AA:3595

|  | Res. type | Atom | Residue | Distance | Occupancy | B-factor | Closest solute to 1^st^ shell water |
| --- | --- | --- | --- | --- | --- | --- | --- |
| **Ion** | | | |  | 1.00 | 103.78 || 1 | G | OP2 | AA:1444 | 3.38 | 1.00 | 77.22 |
| 2 | G | OP1 | AA:1444 | 4.09 | 1.00 | 77.38 |  |
| 3 | G | O5' | AA:1444 | 4.91 | 1.00 | 78.95 |  |
  |

  
 Coded contact list:   
 Exclusion zone contact(s): /OP2/  
Consider K in place of MG (1 valid contact(s) for K vs. 0 for MG)  

### MG:AA:3596

|  | Res. type | Atom | Residue | Distance | Occupancy | B-factor | Closest solute to 1^st^ shell water |
| --- | --- | --- | --- | --- | --- | --- | --- |
| **Ion** | | | |  | 1.00 | 65.97 || 1 | A | OP1 | AA:1444A | 2.52 | 1.00 | 87.40 |
| 2 | G | OP1 | AA:1444 | 3.30 | 1.00 | 77.38 |  |
| 3 | G | O5' | AA:1444 | 4.03 | 1.00 | 78.95 |  |
| 4 | A | OP2 | AA:1444A | 4.20 | 1.00 | 87.42 |  |
| 5 | C | OP2 | AA:1462 | 4.39 | 1.00 | 93.28 |  |
| 6 | C | OP1 | AA:1462 | 4.54 | 1.00 | 93.95 |  |
| 7 | A | O5' | AA:1444A | 4.78 | 1.00 | 89.95 |  |
| 8 | G | O3' | AA:1444 | 4.84 | 1.00 | 86.82 |  |
| 9 | G | OP2 | AA:1444 | 4.88 | 1.00 | 77.22 |  |
  |

  
 Coded contact list:   
 Exclusion zone contact(s): /OP1/OP1/  
Consider NA in place of MG (1 valid contact(s) for NA vs. 0 for MG)  

### MG:AA:3597

|  | Res. type | Atom | Residue | Distance | Occupancy | B-factor | Closest solute to 1^st^ shell water |
| --- | --- | --- | --- | --- | --- | --- | --- |
| **Ion** | | | |  | 1.00 | 63.62 || 1 | G | O6 | AA:1466 | 3.87 | 1.00 | 86.56 |
| 2 | G | OP2 | AA:1444 | 4.14 | 1.00 | 77.22 |  |
| 3 | G | N7 | AA:1444 | 4.41 | 1.00 | 82.29 |  |
| 4 | G | N7 | AA:1466 | 4.47 | 1.00 | 83.37 |  |
| 5 | G | O6 | AA:1465 | 4.91 | 1.00 | 85.33 |  |
  |

  
 Coded contact list:   
1^st^ MG coordination shell missing (see Notes)  

### MG:AA:3598

|  | Res. type | Atom | Residue | Distance | Occupancy | B-factor | Closest solute to 1^st^ shell water |
| --- | --- | --- | --- | --- | --- | --- | --- |
| **Ion** | | | |  | 1.00 | 79.19 || 1 | G | N7 | AA:1466 | 3.86 | 1.00 | 83.37 |
| 2 | G | N7 | AA:1444 | 3.87 | 1.00 | 82.29 |  |
| 3 | G | O6 | AA:1444 | 4.02 | 1.00 | 81.21 |  |
| 4 | G | O6 | AA:1466 | 4.18 | 1.00 | 86.56 |  |
| 5 | G | N7 | AA:1443 | 4.39 | 1.00 | 71.90 |  |
| 6 | G | O6 | AA:1443 | 4.83 | 1.00 | 73.86 |  |
  |

  
 Coded contact list:   
1^st^ MG coordination shell missing (see Notes)  

### MG:AA:3599

|  | Res. type | Atom | Residue | Distance | Occupancy | B-factor | Closest solute to 1^st^ shell water |
| --- | --- | --- | --- | --- | --- | --- | --- |
| **Ion** | | | |  | 1.00 | 110.57 || 1 | G | OP2 | AA:1442 | 3.78 | 1.00 | 55.97 |
| 2 | G | N7 | AA:1442 | 4.61 | 1.00 | 68.67 |  |
| 3 | G | N7 | AA:1443 | 4.89 | 1.00 | 71.90 |  |
  |

  
 Coded contact list:   
1^st^ MG coordination shell missing (see Notes)  

### MG:AA:3600

|  | Res. type | Atom | Residue | Distance | Occupancy | B-factor | Closest solute to 1^st^ shell water |
| --- | --- | --- | --- | --- | --- | --- | --- |
| **Ion** | | | |  | 1.00 | 46.42 || 1 | G | O6 | AA:1443 | 3.29 | 1.00 | 73.86 |
| 2 | G | O6 | AA:1444 | 3.56 | 1.00 | 81.21 |  |
| 3 | G | N7 | AA:1443 | 4.13 | 1.00 | 71.90 |  |
| 4 | C | N4 | AA:1547 | 4.58 | 1.00 | 80.13 |  |
| 5 | C | N4 | AA:1548 | 4.86 | 1.00 | 75.71 |  |
| 6 | G | O6 | AA:1442 | 4.86 | 1.00 | 71.82 |  |
| 7 | G | N7 | AA:1444 | 4.99 | 1.00 | 82.29 |  |
| 8 | G | N7 | AA:1466 | 4.99 | 1.00 | 83.37 |  |
  |

  
 Coded contact list:   
 Exclusion zone contact(s): /O6/  
Consider K in place of MG (1 valid contact(s) for K vs. 0 for MG)  

### MG:AA:3601

|  | Res. type | Atom | Residue | Distance | Occupancy | B-factor | Closest solute to 1^st^ shell water |
| --- | --- | --- | --- | --- | --- | --- | --- |
| **Ion** | | | |  | 1.00 | 73.94 || 1 | G | O6 | AA:1442 | 3.42 | 1.00 | 71.82 |
| 2 | G | N7 | AA:1442 | 3.83 | 1.00 | 68.67 |  |
| 3 | G | N7 | AA:1441 | 4.52 | 1.00 | 59.19 |  |
| 4 | G | O6 | AA:1441 | 4.68 | 1.00 | 60.81 |  |
| 5 | G | O6 | AA:1443 | 4.97 | 1.00 | 73.86 |  |
  |

  
 Coded contact list:   
1^st^ MG coordination shell missing (see Notes)  

### MG:AA:3602

|  | Res. type | Atom | Residue | Distance | Occupancy | B-factor | Closest solute to 1^st^ shell water |
| --- | --- | --- | --- | --- | --- | --- | --- |
| **Ion** | | | |  | 1.00 | 102.40 || 1 | G | O6 | AA:1441 | 2.71 | 1.00 | 60.81 |
| 2 | G | N7 | AA:1441 | 3.31 | 1.00 | 59.19 |  |
| 3 | G | O6 | AA:1442 | 4.37 | 1.00 | 71.82 |  |
| 4 | G | O6 | AA:1440 | 4.57 | 1.00 | 55.58 |  |
| 5 | C | N4 | AA:1550 | 4.64 | 1.00 | 60.56 |  |
| 6 | G | N1 | AA:1441 | 4.75 | 1.00 | 59.56 |  |
  |

  
 Coded contact list:   
 Exclusion zone contact(s): /N7/O6/  
Consider K in place of MG (2 valid contact(s) for K vs. 0 for MG)  

### MG:AA:3603

|  | Res. type | Atom | Residue | Distance | Occupancy | B-factor | Closest solute to 1^st^ shell water |
| --- | --- | --- | --- | --- | --- | --- | --- |
| **Ion** | | | |  | 1.00 | 80.30 || 1 | G | O6 | AA:1441 | 3.84 | 1.00 | 60.81 |
| 2 | G | O6 | AA:1440 | 3.92 | 1.00 | 55.58 |  |
| 3 | A | OP2 | AA:1439 | 3.98 | 1.00 | 59.38 |  |
| 4 | G | N7 | AA:1440 | 4.22 | 1.00 | 57.08 |  |
| 5 | G | N7 | AA:1441 | 4.50 | 1.00 | 59.19 |  |
  |

  
 Coded contact list:   
1^st^ MG coordination shell missing (see Notes)  

### MG:AA:3604

|  | Res. type | Atom | Residue | Distance | Occupancy | B-factor | Closest solute to 1^st^ shell water |
| --- | --- | --- | --- | --- | --- | --- | --- |
| **Ion** | | | |  | 1.00 | 74.23 || 1 | A | OP2 | AA:1439 | 2.92 | 1.00 | 59.38 |
| 2 | A | OP1 | AA:1439 | 3.31 | 1.00 | 59.45 |  |
| 3 | G | OP2 | AA:1440 | 3.58 | 1.00 | 58.51 |  |
| 4 | C | OP1 | AA:1468 | 3.85 | 1.00 | 77.43 |  |
| 5 | A | O5' | AA:1439 | 3.89 | 1.00 | 59.42 |  |
| 6 | G | N7 | AA:1440 | 4.52 | 1.00 | 57.08 |  |
| 7 | U | O3' | AA:1438 | 4.98 | 1.00 | 58.21 |  |
  |

  
 Coded contact list:   
 Exclusion zone contact(s): /OP1/OP2/  
Consider K in place of MG (2 valid contact(s) for K vs. 0 for MG)  

### MG:AA:3605

|  | Res. type | Atom | Residue | Distance | Occupancy | B-factor | Closest solute to 1^st^ shell water |
| --- | --- | --- | --- | --- | --- | --- | --- |
| **Ion** | | | |  | 1.00 | 66.02 || 1 | C | OP1 | AA:1468 | 3.31 | 1.00 | 77.43 |
| 2 | G | OP2 | AA:1440 | 3.55 | 1.00 | 58.51 |  |
| 3 | C | O3' | AA:1467 | 3.62 | 1.00 | 78.27 |  |
  |

  
 Coded contact list:   
 Exclusion zone contact(s): /OP1/  
Consider K in place of MG (1 valid contact(s) for K vs. 0 for MG)  

### MG:AA:3606

|  | Res. type | Atom | Residue | Distance | Occupancy | B-factor | Closest solute to 1^st^ shell water |
| --- | --- | --- | --- | --- | --- | --- | --- |
| **Ion** | | | |  | 1.00 | 97.70 || 1 | A | OP2 | AA:1439 | 3.96 | 1.00 | 59.38 |
| 2 | U | OP1 | AA:1438 | 4.35 | 1.00 | 58.36 |  |
| 3 | U | OP2 | AA:1438 | 4.54 | 1.00 | 57.10 |  |
| 4 | U | O5' | AA:1438 | 4.76 | 1.00 | 58.20 |  |
  |

  
 Coded contact list:   
1^st^ MG coordination shell missing (see Notes)  

### MG:AA:3607

|  | Res. type | Atom | Residue | Distance | Occupancy | B-factor | Closest solute to 1^st^ shell water |
| --- | --- | --- | --- | --- | --- | --- | --- |
| **Ion** | | | |  | 1.00 | 94.52 || 1 | C | OP1 | AA:1468 | 2.54 | 1.00 | 77.43 |
| 2 | C | OP2 | AA:1468 | 3.16 | 1.00 | 77.23 |  |
| 3 | C | O3' | AA:1467 | 3.65 | 1.00 | 78.27 |  |
| 4 | C | O5' | AA:1468 | 4.65 | 1.00 | 77.13 |  |
| 5 | A | OP1 | AA:1439 | 4.95 | 1.00 | 59.45 |  |
  |

  
 Coded contact list:   
 Exclusion zone contact(s): /OP1/OP2/  
Consider NA in place of MG (1 valid contact(s) for NA vs. 0 for MG)  

### MG:AA:3608

|  | Res. type | Atom | Residue | Distance | Occupancy | B-factor | Closest solute to 1^st^ shell water |
| --- | --- | --- | --- | --- | --- | --- | --- |
| **Ion** | | | |  | 1.00 | 86.86 || 1 | A | OP1 | AA:1439 | 2.98 | 1.00 | 59.45 |
| 2 | C | OP1 | AA:1468 | 3.78 | 1.00 | 77.43 |  |
| 3 | G | OP1 | AA:1519 | 3.97 | 1.00 | 71.64 |  |
| 4 | A | OP2 | AA:1439 | 4.08 | 1.00 | 59.38 |  |
| 5 | U | O3' | AA:1438 | 4.27 | 1.00 | 58.21 |  |
| 6 | C | OP2 | AA:1468 | 4.67 | 1.00 | 77.23 |  |
  |

  
 Coded contact list:   
 Exclusion zone contact(s): /OP1/  
Consider K in place of MG (1 valid contact(s) for K vs. 0 for MG)  

### MG:AA:3609

|  | Res. type | Atom | Residue | Distance | Occupancy | B-factor | Closest solute to 1^st^ shell water |
| --- | --- | --- | --- | --- | --- | --- | --- |
| **Ion** | | | |  | 1.00 | 72.56 || 1 | A | OP1 | AA:1439 | 2.47 | 1.00 | 59.45 |
| 2 | A | OP2 | AA:1469 | 3.27 | 1.00 | 75.98 |  |
| 3 | A | OP1 | AA:1469 | 3.31 | 1.00 | 76.38 |  |
| 4 | U | O3' | AA:1438 | 4.07 | 1.00 | 58.21 |  |
| 5 | C | OP1 | AA:1468 | 4.09 | 1.00 | 77.43 |  |
| 6 | A | O5' | AA:1439 | 4.15 | 1.00 | 59.42 |  |
| 7 | G | OP1 | AA:1519 | 4.44 | 1.00 | 71.64 |  |
| 8 | C | O3' | AA:1468 | 4.45 | 1.00 | 75.62 |  |
| 9 | C | O5' | AA:1468 | 4.59 | 1.00 | 77.13 |  |
  |

  
 Coded contact list:   
 Exclusion zone contact(s): /OP1/OP1/OP2/  
Consider K in place of MG (2 valid contact(s) for K vs. 0 for MG)  

### MG:AA:3610

|  | Res. type | Atom | Residue | Distance | Occupancy | B-factor | Closest solute to 1^st^ shell water |
| --- | --- | --- | --- | --- | --- | --- | --- |
| **Ion** | | | |  | 1.00 | 69.79 || 1 | G | OP1 | AA:1519 | 2.98 | 1.00 | 71.64 |
| 2 | U | O3' | AA:1438 | 3.80 | 1.00 | 58.21 |  |
| 3 | A | OP1 | AA:1439 | 3.92 | 1.00 | 59.45 |  |
| 4 | A | OP1 | AA:1469 | 3.93 | 1.00 | 76.38 |  |
| 5 | G | OP2 | AA:1519 | 4.33 | 1.00 | 70.94 |  |
| 6 | A | OP2 | AA:1469 | 4.40 | 1.00 | 75.98 |  |
| 7 | U | O2' | AA:1438 | 4.46 | 1.00 | 58.03 |  |
| 8 | C | O3' | AA:1518 | 4.54 | 1.00 | 71.63 |  |
| 9 | A | O5' | AA:1439 | 5.00 | 1.00 | 59.42 |  |
  |

  
 Coded contact list:   
 Exclusion zone contact(s): /OP1/  
Consider K in place of MG (1 valid contact(s) for K vs. 0 for MG)  

### MG:AA:3611

|  | Res. type | Atom | Residue | Distance | Occupancy | B-factor | Closest solute to 1^st^ shell water |
| --- | --- | --- | --- | --- | --- | --- | --- |
| **Ion** | | | |  | 1.00 | 70.08 || 1 | C | OP1 | AA:1518 | 3.13 | 1.00 | 65.00 |
| 2 | C | OP2 | AA:1518 | 3.81 | 1.00 | 64.90 |  |
| 3 | C | O5' | AA:1518 | 4.40 | 1.00 | 66.48 |  |
| 4 | G | OP2 | AA:1470 | 4.48 | 1.00 | 75.20 |  |
| 5 | G | OP2 | AA:1519 | 4.52 | 1.00 | 70.94 |  |
| 6 | G | OP1 | AA:1470 | 4.80 | 1.00 | 75.12 |  |
  |

  
 Coded contact list:   
 Exclusion zone contact(s): /OP1/  
Consider K in place of MG (1 valid contact(s) for K vs. 0 for MG)  

### MG:AA:3612

|  | Res. type | Atom | Residue | Distance | Occupancy | B-factor | Closest solute to 1^st^ shell water |
| --- | --- | --- | --- | --- | --- | --- | --- |
| **Ion** | | | |  | 1.00 | 106.33 || 1 | A | OP1 | AA:1471 | 4.35 | 1.00 | 80.65 |
| 2 | G | OP2 | AA:1517 | 4.67 | 1.00 | 67.75 |  |
| 3 | G | OP1 | AA:1517 | 4.68 | 1.00 | 67.66 |  |
| 4 | C | OP2 | AA:1518 | 4.88 | 1.00 | 64.90 |  |
| 5 | G | OP1 | AA:1470 | 4.97 | 1.00 | 75.12 |  |
  |

  
 Coded contact list:   
1^st^ MG coordination shell missing (see Notes)  

### MG:AA:3613

|  | Res. type | Atom | Residue | Distance | Occupancy | B-factor | Closest solute to 1^st^ shell water |
| --- | --- | --- | --- | --- | --- | --- | --- |
| **Ion** | | | |  | 1.00 | 84.73 || 1 | A | OP1 | AA:1472 | 4.11 | 1.00 | 81.14 |
| 2 | A | OP2 | AA:1472 | 4.18 | 1.00 | 81.25 |  |
| 3 | G | OP2 | AA:1517 | 4.75 | 1.00 | 67.75 |  |
| 4 | A | O5' | AA:1471 | 4.98 | 1.00 | 79.73 |  |
  |

  
 Coded contact list:   
1^st^ MG coordination shell missing (see Notes)  

### MG:AA:3614

|  | Res. type | Atom | Residue | Distance | Occupancy | B-factor | Closest solute to 1^st^ shell water |
| --- | --- | --- | --- | --- | --- | --- | --- |
| **Ion** | | | |  | 1.00 | 62.01 || 1 | G | O6 | AA:1401 | 2.62 | 1.00 | 70.96 |
| 2 | G | O6 | AA:1400 | 3.09 | 1.00 | 72.59 |  |
| 3 | G | N7 | AA:1400 | 3.58 | 1.00 | 73.79 |  |
| 4 | G | N7 | AA:1401 | 3.68 | 1.00 | 72.93 |  |
| 5 | G | O6 | AA:1385 | 4.12 | 1.00 | 75.30 |  |
| 6 | C | N4 | AA:1402 | 4.13 | 1.00 | 76.98 |  |
| 7 | C | N4 | AA:1386 | 4.33 | 1.00 | 66.79 |  |
| 8 | C | N4 | AA:1399 | 4.62 | 1.00 | 70.12 |  |
| 9 | G | N1 | AA:1400 | 4.63 | 1.00 | 72.69 |  |
| 10 | G | N1 | AA:1401 | 4.81 | 1.00 | 72.40 |  |
  |

  
 Coded contact list:   
 Exclusion zone contact(s): /O6/O6/  
Consider K in place of MG (2 valid contact(s) for K vs. 0 for MG)  

### MG:AA:3615

|  | Res. type | Atom | Residue | Distance | Occupancy | B-factor | Closest solute to 1^st^ shell water |
| --- | --- | --- | --- | --- | --- | --- | --- |
| **Ion** | | | |  | 1.00 | 47.56 || 1 | C | N4 | AA:1399 | 2.89 | 1.00 | 70.12 |
| 2 | G | O6 | AA:1400 | 3.46 | 1.00 | 72.59 |  |
| 3 | U | OP1 | AA:1397 | 3.57 | 1.00 | 60.85 |  |
| 4 | G | N7 | AA:1400 | 4.05 | 1.00 | 73.79 |  |
| 5 | C | N3 | AA:1399 | 4.41 | 1.00 | 70.80 |  |
| 6 | G | O6 | AA:1401 | 4.53 | 1.00 | 70.96 |  |
| 7 | G | O6 | AA:1388 | 4.90 | 1.00 | 71.03 |  |
| 8 | C | N4 | AA:1386 | 4.94 | 1.00 | 66.79 |  |
  |

  
 Coded contact list:   
 Exclusion zone contact(s): /N4/  
Consider K in place of MG (1 valid contact(s) for K vs. 0 for MG)  

### MG:AA:3616

|  | Res. type | Atom | Residue | Distance | Occupancy | B-factor | Closest solute to 1^st^ shell water |
| --- | --- | --- | --- | --- | --- | --- | --- |
| **Ion** | | | |  | 1.00 | 42.39 || 1 | G | O6 | AA:1385 | 3.96 | 1.00 | 75.30 |
| 2 | G | O6 | AA:1401 | 4.67 | 1.00 | 70.96 |  |
| 3 | C | N4 | AA:1402 | 4.96 | 1.00 | 76.98 |  |
  |

  
 Coded contact list:   
1^st^ MG coordination shell missing (see Notes)  

### MG:AA:3617

|  | Res. type | Atom | Residue | Distance | Occupancy | B-factor | Closest solute to 1^st^ shell water |
| --- | --- | --- | --- | --- | --- | --- | --- |
| **Ion** | | | |  | 1.00 | 59.63 || 1 | U | OP1 | AA:1397 | 3.18 | 1.00 | 60.85 |
| 2 | U | O3' | AA:1397 | 3.39 | 1.00 | 62.92 |  |
| 3 | C | OP2 | AA:1398 | 3.52 | 1.00 | 62.84 |  |
| 4 | C | N4 | AA:1399 | 4.49 | 1.00 | 70.12 |  |
| 5 | C | OP2 | AA:1399 | 4.64 | 1.00 | 62.69 |  |
| 6 | U | O5' | AA:1397 | 4.75 | 1.00 | 60.26 |  |
  |

  
 Coded contact list:   
 Exclusion zone contact(s): /O3'/OP1/  
Consider K in place of MG (2 valid contact(s) for K vs. 0 for MG)  

### MG:AA:3618

|  | Res. type | Atom | Residue | Distance | Occupancy | B-factor | Closest solute to 1^st^ shell water |
| --- | --- | --- | --- | --- | --- | --- | --- |
| **Ion** | | | |  | 1.00 | 89.07 || 1 | G | O6 | AA:1344 | 3.24 | 1.00 | 68.08 |
| 2 | C | N4 | AA:1403 | 3.35 | 1.00 | 66.18 |  |
| 3 | G | O6 | AA:1385 | 3.98 | 1.00 | 75.30 |  |
| 4 | C | N4 | AA:1402 | 4.62 | 1.00 | 76.98 |  |
| 5 | G | O6 | AA:1343 | 4.84 | 1.00 | 61.99 |  |
| 6 | C | N4 | AA:1404 | 4.86 | 1.00 | 63.74 |  |
  |

  
 Coded contact list:   
 Exclusion zone contact(s): /N4/O6/  
Consider K in place of MG (2 valid contact(s) for K vs. 0 for MG)  

### MG:AA:3619

|  | Res. type | Atom | Residue | Distance | Occupancy | B-factor | Closest solute to 1^st^ shell water |
| --- | --- | --- | --- | --- | --- | --- | --- |
| **Ion** | | | |  | 1.00 | 64.82 || 1 | G | O6 | AA:1344 | 2.64 | 1.00 | 68.08 |
| 2 | G | N7 | AA:1344 | 3.09 | 1.00 | 63.59 |  |
| 3 | G | O6 | AA:1343 | 3.74 | 1.00 | 61.99 |  |
| 4 | G | N1 | AA:1343 | 4.69 | 1.00 | 59.53 |  |
| 5 | G | N1 | AA:1344 | 4.71 | 1.00 | 65.90 |  |
| 6 | C | N4 | AA:1403 | 4.73 | 1.00 | 66.18 |  |
| 7 | G | N7 | AA:1385 | 4.76 | 1.00 | 74.34 |  |
| 8 | G | N7 | AA:1343 | 4.89 | 1.00 | 58.15 |  |
| 9 | G | O6 | AA:1385 | 4.97 | 1.00 | 75.30 |  |
  |

  
 Coded contact list:   
 Exclusion zone contact(s): /N7/O6/  
Consider K in place of MG (2 valid contact(s) for K vs. 0 for MG)  

### MG:AA:3620

|  | Res. type | Atom | Residue | Distance | Occupancy | B-factor | Closest solute to 1^st^ shell water |
| --- | --- | --- | --- | --- | --- | --- | --- |
| **Ion** | | | |  | 1.00 | 55.20 || 1 | G | OP1 | AA:1343 | 2.37 | 1.00 | 53.00 |
| 2 | G | O5' | AA:1343 | 3.13 | 1.00 | 52.43 |  |
| 3 | A | O3' | AA:1342 | 3.47 | 1.00 | 51.02 |  |
| 4 | A | O2' | AA:1342 | 3.84 | 1.00 | 54.80 |  |
| 5 | G | N7 | AA:1344 | 3.95 | 1.00 | 63.59 |  |
| 6 | G | OP2 | AA:1344 | 3.99 | 1.00 | 54.22 |  |
| 7 | G | N7 | AA:1343 | 4.40 | 1.00 | 58.15 |  |
| 8 | G | OP2 | AA:1343 | 4.50 | 1.00 | 52.95 |  |
| 9 | G | N9 | AA:1343 | 4.74 | 1.00 | 56.26 |  |
  |

  
 Contact(s): /OP1/  
 Coded contact list: O_ph_  
 Exclusion zone contact(s): /O5'/  

### MG:AA:3621

|  | Res. type | Atom | Residue | Distance | Occupancy | B-factor | Closest solute to 1^st^ shell water |
| --- | --- | --- | --- | --- | --- | --- | --- |
| **Ion** | | | |  | 1.00 | 46.06 || 1 | G | N7 | AA:1344 | 2.59 | 1.00 | 63.59 |
| 2 | G | O6 | AA:1344 | 4.03 | 1.00 | 68.08 |  |
| 3 | G | N9 | AA:1344 | 4.42 | 1.00 | 61.85 |  |
| 4 | G | N7 | AA:1385 | 4.61 | 1.00 | 74.34 |  |
| 5 | G | OP1 | AA:1343 | 4.78 | 1.00 | 53.00 |  |
| 6 | G | OP2 | AA:1344 | 4.87 | 1.00 | 54.22 |  |
  |

  
 Coded contact list:   
 Exclusion zone contact(s): /N7/  
Consider NA in place of MG (1 valid contact(s) for NA vs. 0 for MG)  

### MG:AA:3622

|  | Res. type | Atom | Residue | Distance | Occupancy | B-factor | Closest solute to 1^st^ shell water |
| --- | --- | --- | --- | --- | --- | --- | --- |
| **Ion** | | | |  | 1.00 | 43.82 || 1 | U | OP2 | AA:1300 | 2.57 | 1.00 | 46.66 |
| 2 | G | O6 | AA:1628 | 2.83 | 1.00 | 41.31 |  |
| 3 | G | O6 | AA:1627 | 2.89 | 1.00 | 39.40 |  |
| 4 | G | N7 | AA:1628 | 3.49 | 1.00 | 40.79 |  |
| 5 | G | O3' | AA:1299 | 3.51 | 1.00 | 47.27 |  |
| 6 | G | N7 | AA:1627 | 3.52 | 1.00 | 44.22 |  |
| 7 | U | O5' | AA:1300 | 4.55 | 1.00 | 45.59 |  |
| 8 | U | O4 | AA:1639 | 4.55 | 1.00 | 43.60 |  |
| 9 | G | N1 | AA:1627 | 4.59 | 1.00 | 41.43 |  |
| 10 | G | O2' | AA:1299 | 4.71 | 1.00 | 45.95 |  |
| 11 | C | N4 | AA:1638 | 4.72 | 1.00 | 39.08 |  |
| 12 | U | N3 | AA:1639 | 4.77 | 1.00 | 41.01 |  |
| 13 | G | OP2 | AA:1627 | 4.78 | 1.00 | 47.27 |  |
| 14 | U | OP1 | AA:1300 | 4.80 | 1.00 | 45.13 |  |
| 15 | G | N1 | AA:1628 | 4.93 | 1.00 | 39.68 |  |
  |

  
 Coded contact list:   
 Exclusion zone contact(s): /O6/O6/OP2/  
Consider K in place of MG (2 valid contact(s) for K vs. 0 for MG)  

### MG:AA:3623

|  | Res. type | Atom | Residue | Distance | Occupancy | B-factor | Closest solute to 1^st^ shell water |
| --- | --- | --- | --- | --- | --- | --- | --- |
| **Ion** | | | |  | 1.00 | 50.22 || 1 | G | O6 | AA:1519 | 2.68 | 1.00 | 75.54 |
| 2 | U | O4 | AA:1520 | 2.70 | 1.00 | 81.06 |  |
| 3 | C | N4 | AA:1474 | 3.31 | 1.00 | 77.53 |  |
| 4 | G | O6 | AA:1473 | 3.37 | 1.00 | 81.94 |  |
| 5 | G | N2 | AA:1521 | 3.89 | 1.00 | 92.73 |  |
| 6 | U | N3 | AA:1520 | 4.01 | 1.00 | 80.09 |  |
| 7 | A | OP2 | AA:1472 | 4.29 | 1.00 | 81.25 |  |
| 8 | G | N1 | AA:1521 | 4.48 | 1.00 | 91.80 |  |
| 9 | G | N1 | AA:1519 | 4.58 | 1.00 | 77.07 |  |
| 10 | A | OP2 | AA:1471 | 4.71 | 1.00 | 78.94 |  |
| 11 | G | N7 | AA:1519 | 4.83 | 1.00 | 76.04 |  |
| 12 | G | N3 | AA:1521 | 4.90 | 1.00 | 91.84 |  |
| 13 | C | N3 | AA:1474 | 4.90 | 1.00 | 77.81 |  |
| 14 | G | N7 | AA:1473 | 4.95 | 1.00 | 84.43 |  |
| 15 | G | O6 | AA:1475 | 4.98 | 1.00 | 77.69 |  |
  |

  
 Coded contact list:   
 Exclusion zone contact(s): /N4/O4/O6/O6/  
Consider K in place of MG (4 valid contact(s) for K vs. 0 for MG)  

### MG:AA:3624

|  | Res. type | Atom | Residue | Distance | Occupancy | B-factor | Closest solute to 1^st^ shell water |
| --- | --- | --- | --- | --- | --- | --- | --- |
| **Ion** | | | |  | 1.00 | 38.52 || 1 | U | O4 | AA:2847 | 3.84 | 1.00 | 58.14 |
| 2 | G | N7 | AA:2845 | 4.15 | 1.00 | 69.35 |  |
| 3 | G | N7 | AA:2846 | 4.20 | 1.00 | 65.50 |  |
| 4 | G | O6 | AA:2848 | 4.45 | 1.00 | 54.57 |  |
| 5 | G | OP2 | AA:2845 | 4.56 | 1.00 | 70.50 |  |
| 6 | G | OP2 | AA:2846 | 4.67 | 1.00 | 66.60 |  |
| 7 | G | O6 | AA:2845 | 4.92 | 1.00 | 70.14 |  |
| 8 | G | N1 | AA:2848 | 4.95 | 1.00 | 54.75 |  |
  |

  
 Coded contact list:   
1^st^ MG coordination shell missing (see Notes)  

### MG:AA:3625

|  | Res. type | Atom | Residue | Distance | Occupancy | B-factor | Closest solute to 1^st^ shell water |
| --- | --- | --- | --- | --- | --- | --- | --- |
| **Ion** | | | |  | 1.00 | 64.55 || 1 | A | OP1 | AA:2268 | 2.36 | 1.00 | 70.83 |
| 2 | A | OP1 | AA:2269 | 2.58 | 1.00 | 72.06 |  |
| 3 | A | O3' | AA:2268 | 4.00 | 1.00 | 70.84 |  |
| 4 | G | O6 | AA:2271 | 4.06 | 1.00 | 63.96 |  |
| 5 | A | N3 | AA:2267 | 4.07 | 1.00 | 68.29 |  |
| 6 | A | OP2 | AA:2268 | 4.25 | 1.00 | 69.77 |  |
| 7 | A | O3' | AA:2267 | 4.41 | 1.00 | 70.77 |  |
| 8 | A | OP2 | AA:2269 | 4.81 | 1.00 | 71.27 |  |
| 9 | A | O5' | AA:2268 | 4.82 | 1.00 | 70.99 |  |
| 10 | A | O5' | AA:2269 | 4.92 | 1.00 | 69.53 |  |
  |

  
 Contact(s): /OP1/  
 Coded contact list: O_ph_  
 Exclusion zone contact(s): /OP1/  

### MG:AA:3626

|  | Res. type | Atom | Residue | Distance | Occupancy | B-factor | Closest solute to 1^st^ shell water |
| --- | --- | --- | --- | --- | --- | --- | --- |
| **Ion** | | | |  | 1.00 | 48.20 || 1 | G | O6 | AA:382 | 3.52 | 1.00 | 55.65 |
| 2 | U | O4 | AA:383 | 3.88 | 1.00 | 62.67 |  |
| 3 | G | N7 | AA:382 | 4.21 | 1.00 | 57.26 |  |
| 4 | G | OP2 | AA:386 | 4.35 | 1.00 | 45.63 |  |
| 5 | G | O6 | AA:388 | 4.42 | 1.00 | 52.03 |  |
| 6 | G | O6 | AA:381 | 4.45 | 1.00 | 60.92 |  |
| 7 | G | OP1 | AA:386 | 4.50 | 1.00 | 46.28 |  |
| 8 | G | N7 | AA:381 | 4.59 | 1.00 | 57.99 |  |
| 9 | U | O4 | AA:387 | 4.68 | 1.00 | 52.79 |  |
  |

  
 Coded contact list:   
1^st^ MG coordination shell missing (see Notes)  

### MG:AA:3627

|  | Res. type | Atom | Residue | Distance | Occupancy | B-factor | Closest solute to 1^st^ shell water |
| --- | --- | --- | --- | --- | --- | --- | --- |
| **Ion** | | | |  | 1.00 | 37.31 || 1 | A | N3 | AA:945 | 3.61 | 1.00 | 58.61 |
| 2 | G | OP1 | AA:831 | 3.82 | 1.00 | 41.73 |  |
| 3 | G | OP1 | AA:944 | 4.09 | 1.00 | 47.52 |  |
| 4 | A | O5' | AA:945 | 4.34 | 1.00 | 53.02 |  |
| 5 | A | N6 | AA:2448 | 4.34 | 1.00 | 55.15 |  |
| 6 | U | OP1 | AA:568 | 4.34 | 1.00 | 45.47 |  |
| 7 | A | N1 | AA:2448 | 4.34 | 1.00 | 51.32 |  |
| 8 | G | OP2 | AA:944 | 4.62 | 1.00 | 48.48 |  |
| 9 | A | N1 | AA:945 | 4.83 | 1.00 | 58.28 |  |
  |

  
 Coded contact list:   
1^st^ MG coordination shell missing (see Notes)  

### MG:AA:3628

|  | Res. type | Atom | Residue | Distance | Occupancy | B-factor | Closest solute to 1^st^ shell water |
| --- | --- | --- | --- | --- | --- | --- | --- |
| **Ion** | | | |  | 1.00 | 62.41 || 1 | G | OP2 | AA:382 | 2.94 | 1.00 | 52.23 |
| 2 | G | OP1 | AA:386 | 2.95 | 1.00 | 46.28 |  |
| 3 | G | N7 | AA:382 | 4.71 | 1.00 | 57.26 |  |
| 4 | G | O5' | AA:386 | 4.74 | 1.00 | 46.28 |  |
| 5 | G | O5' | AA:382 | 4.75 | 1.00 | 50.99 |  |
  |

  
 Coded contact list:   
 Exclusion zone contact(s): /OP1/OP2/  
Consider K in place of MG (2 valid contact(s) for K vs. 0 for MG)  

### MG:AA:3629

|  | Res. type | Atom | Residue | Distance | Occupancy | B-factor | Closest solute to 1^st^ shell water |
| --- | --- | --- | --- | --- | --- | --- | --- |
| **Ion** | | | |  | 1.00 | 45.69 || 1 | A | N6 | AA:820 | 3.50 | 1.00 | 55.52 |
| 2 | A | N1 | AA:820 | 3.69 | 1.00 | 54.66 |  |
| 3 | A | OP1 | AA:973 | 3.96 | 1.00 | 63.50 |  |
| 4 | G | O2' | AA:972 | 3.99 | 1.00 | 62.10 |  |
| 5 | A | N1 | AA:821 | 4.13 | 1.00 | 60.67 |  |
| 6 | U | O2 | AA:1188 | 4.72 | 1.00 | 62.95 |  |
| 7 | A | N6 | AA:819 | 4.80 | 1.00 | 52.81 |  |
| 8 | G | OP1 | AA:570 | 4.99 | 1.00 | 47.33 |  |
  |

  
 Coded contact list:   
1^st^ MG coordination shell missing (see Notes)  

### MG:AA:3630

|  | Res. type | Atom | Residue | Distance | Occupancy | B-factor | Closest solute to 1^st^ shell water |
| --- | --- | --- | --- | --- | --- | --- | --- |
| **Ion** | | | |  | 1.00 | 55.57 || 1 | U | O4 | AA:1629 | 3.76 | 1.00 | 45.39 |
| 2 | G | N7 | AA:1628 | 4.12 | 1.00 | 40.79 |  |
| 3 | G | O6 | AA:1628 | 4.43 | 1.00 | 41.31 |  |
| 4 | G | OP2 | AA:1627 | 4.63 | 1.00 | 47.27 |  |
| 5 | A | OP2 | AA:1634 | 4.79 | 1.00 | 44.42 |  |
| 6 | U | OP2 | AA:1300 | 4.81 | 1.00 | 46.66 |  |
| 7 | G | O6 | AA:1630 | 4.92 | 1.00 | 50.33 |  |
| 8 | G | OP2 | AA:1628 | 4.98 | 1.00 | 46.53 |  |
  |

  
 Coded contact list:   
1^st^ MG coordination shell missing (see Notes)  

### MG:AA:3631

|  | Res. type | Atom | Residue | Distance | Occupancy | B-factor | Closest solute to 1^st^ shell water |
| --- | --- | --- | --- | --- | --- | --- | --- |
| **Ion** | | | |  | 1.00 | 58.95 || 1 | A | O2' | AA:2835 | 2.60 | 1.00 | 78.12 |
| 2 | G | OP2 | AA:2837 | 2.64 | 1.00 | 70.49 |  |
| 3 | G | N7 | AA:2837 | 3.00 | 1.00 | 69.52 |  |
| 4 | U | OP2 | AA:2836 | 3.51 | 1.00 | 80.17 |  |
| 5 | U | O5' | AA:2836 | 4.15 | 1.00 | 77.60 |  |
| 6 | A | N3 | AA:2835 | 4.43 | 1.00 | 71.44 |  |
| 7 | G | O5' | AA:2837 | 4.46 | 1.00 | 69.23 |  |
| 8 | G | N9 | AA:2837 | 4.70 | 1.00 | 67.43 |  |
| 9 | U | O4 | AA:2878 | 4.71 | 1.00 | 71.07 |  |
| 10 | A | O3' | AA:2835 | 4.74 | 1.00 | 78.81 |  |
| 11 | U | O3' | AA:2836 | 4.81 | 1.00 | 71.51 |  |
| 12 | U | N1 | AA:2836 | 4.94 | 1.00 | 73.93 |  |
| 13 | G | O6 | AA:2837 | 4.99 | 1.00 | 69.24 |  |
  |

  
 Coded contact list:   
 Exclusion zone contact(s): /N7/O2'/OP2/  
Consider K in place of MG (2 valid contact(s) for K vs. 0 for MG)  

### MG:AA:3632

|  | Res. type | Atom | Residue | Distance | Occupancy | B-factor | Closest solute to 1^st^ shell water |
| --- | --- | --- | --- | --- | --- | --- | --- |
| **Ion** | | | |  | 1.00 | 43.60 || 1 | G | OP2 | AA:582 | 3.59 | 1.00 | 48.93 |
| 2 | G | N7 | AA:582 | 3.97 | 1.00 | 43.91 |  |
| 3 | G | N7 | AA:583 | 4.14 | 1.00 | 49.39 |  |
| 4 | G | O5' | AA:582 | 4.61 | 1.00 | 48.11 |  |
| 5 | G | OP2 | AA:583 | 4.80 | 1.00 | 43.29 |  |
| 6 | G | O3' | AA:1252 | 4.92 | 1.00 | 46.74 |  |
  |

  
 Coded contact list:   
1^st^ MG coordination shell missing (see Notes)  

### MG:AA:3633

|  | Res. type | Atom | Residue | Distance | Occupancy | B-factor | Closest solute to 1^st^ shell water |
| --- | --- | --- | --- | --- | --- | --- | --- |
| **Ion** | | | |  | 1.00 | 41.74 || 1 | G | O6 | AA:2822 | 3.46 | 1.00 | 38.44 |
| 2 | A | OP1 | AA:2820 | 3.57 | 1.00 | 50.26 |  |
| 3 | ARG | O | A0:2 | 3.63 | 1.00 | 49.42 |  |
| 4 | ARG | N | A0:2 | 3.85 | 1.00 | 50.12 |  |
| 5 | LYS | O | A0:5 | 3.88 | 1.00 | 60.75 |  |
| 6 | LYS | N | A0:5 | 4.21 | 1.00 | 55.56 |  |
| 7 | LEU | N | A0:4 | 4.40 | 1.00 | 50.54 |  |
| 8 | HIS | N | A0:3 | 4.66 | 1.00 | 49.64 |  |
| 9 | A | N6 | AA:2821 | 4.76 | 1.00 | 37.43 |  |
| 10 | G | N1 | AA:2822 | 4.88 | 1.00 | 36.06 |  |
| 11 | G | OP2 | AA:1653 | 4.90 | 1.00 | 45.19 |  |
  |

  
 Coded contact list:   
1^st^ MG coordination shell missing (see Notes)  

### MG:AA:3634

|  | Res. type | Atom | Residue | Distance | Occupancy | B-factor | Closest solute to 1^st^ shell water |
| --- | --- | --- | --- | --- | --- | --- | --- |
| **Ion** | | | |  | 1.00 | 52.50 || 1 | G | OP1 | AA:2578 | 3.81 | 1.00 | 41.76 |
| 2 | U | OP1 | AA:2613 | 3.98 | 1.00 | 56.61 |  |
| 3 | G | OP2 | AA:2578 | 4.16 | 1.00 | 40.99 |  |
| 4 | A | N7 | AA:2614 | 4.36 | 1.00 | 49.98 |  |
| 5 | A | N7 | AA:2051 | 4.45 | 1.00 | 40.69 |  |
| 6 | A | OP2 | AA:2051 | 4.60 | 1.00 | 36.67 |  |
| 7 | C | OP2 | AA:2050 | 4.78 | 1.00 | 36.87 |  |
| 8 | A | OP2 | AA:2614 | 4.80 | 1.00 | 46.99 |  |
  |

  
 Coded contact list:   
1^st^ MG coordination shell missing (see Notes)  

### MG:AA:3635

|  | Res. type | Atom | Residue | Distance | Occupancy | B-factor | Closest solute to 1^st^ shell water |
| --- | --- | --- | --- | --- | --- | --- | --- |
| **Ion** | | | |  | 1.00 | 54.50 || 1 | A | OP1 | AA:751 | 1.99 | 1.00 | 45.43 |
| 2 | A | OP1 | AA:1614 | 2.15 | 1.00 | 41.83 |  |
| 3 | A | N3 | AA:1614 | 3.57 | 1.00 | 51.97 |  |
| 4 | A | N1 | AA:1614 | 3.67 | 1.00 | 54.87 |  |
| 5 | A | OP2 | AA:1614 | 3.85 | 1.00 | 40.28 |  |
| 6 | C | N3 | AA:1617 | 3.90 | 1.00 | 48.55 |  |
| 7 | A | OP2 | AA:751 | 3.91 | 1.00 | 44.93 |  |
| 8 | A | O3' | AA:750 | 3.98 | 1.00 | 41.40 |  |
| 9 | A | O5' | AA:1614 | 4.14 | 1.00 | 40.93 |  |
| 10 | C | N4 | AA:1617 | 4.31 | 1.00 | 51.61 |  |
| 11 | A | O5' | AA:751 | 4.44 | 1.00 | 46.19 |  |
| 12 | GLY | N | AS:91 | 4.45 | 1.00 | 46.68 |  |
| 13 | C | N4 | AA:1615 | 4.51 | 1.00 | 52.70 |  |
| 14 | G | O3' | AA:1613 | 4.61 | 1.00 | 39.88 |  |
| 15 | A | N7 | AA:751 | 4.70 | 1.00 | 45.47 |  |
| 16 | C | O2 | AA:1617 | 4.73 | 1.00 | 46.50 |  |
| 17 | G | O2' | AA:1613 | 4.91 | 1.00 | 42.84 |  |
  |

  
**1^st^ shell ligand angles:**   
1-2: 120;   
  
 Contact(s): /OP1/OP1/  
 Coded contact list: 2O_ph_  

### MG:AA:3636

|  | Res. type | Atom | Residue | Distance | Occupancy | B-factor | Closest solute to 1^st^ shell water |
| --- | --- | --- | --- | --- | --- | --- | --- |
| **Ion** | | | |  | 1.00 | 48.22 || 1 | HIS | NE2 | AO:27 | 2.44 | 1.00 | 61.16 |
| 2 | G | N7 | AA:1192 | 2.57 | 1.00 | 63.84 |  |
| 3 | HIS | ND1 | AO:27 | 3.54 | 1.00 | 59.31 |  |
| 4 | G | OP2 | AA:1192 | 3.80 | 1.00 | 56.16 |  |
| 5 | G | O6 | AA:1192 | 3.83 | 1.00 | 65.34 |  |
| 6 | G | N9 | AA:1192 | 4.51 | 1.00 | 60.46 |  |
| 7 | G | N7 | AA:1193 | 4.68 | 1.00 | 64.88 |  |
| 8 | SER | O | AO:25 | 4.71 | 1.00 | 61.22 |  |
| 9 | G | O5' | AA:1192 | 4.88 | 1.00 | 56.95 |  |
| 10 | G | N7 | AA:1191 | 4.96 | 1.00 | 63.84 |  |
| 11 | G | OP2 | AA:1191 | 4.97 | 1.00 | 53.48 |  |
  |

  
 Coded contact list:   
 Exclusion zone contact(s): /N7/NE2_His_/  
Consider NA in place of MG (2 valid contact(s) for NA vs. 0 for MG)  

### MG:AA:3637

|  | Res. type | Atom | Residue | Distance | Occupancy | B-factor | Closest solute to 1^st^ shell water |
| --- | --- | --- | --- | --- | --- | --- | --- |
| **Ion** | | | |  | 1.00 | 54.14 || 1 | A | OP1 | AA:428 | 2.63 | 1.00 | 58.15 |
| 2 | G | O6 | AA:219 | 2.79 | 1.00 | 46.85 |  |
| 3 | G | O6 | AA:220 | 2.79 | 1.00 | 68.93 |  |
| 4 | U | OP2 | AA:427 | 3.83 | 1.00 | 59.31 |  |
| 5 | A | OP2 | AA:218 | 3.91 | 1.00 | 51.86 |  |
| 6 | G | N1 | AA:220 | 4.07 | 1.00 | 68.77 |  |
| 7 | G | N7 | AA:219 | 4.12 | 1.00 | 48.06 |  |
| 8 | U | O5' | AA:427 | 4.37 | 1.00 | 58.38 |  |
| 9 | U | OP1 | AA:427 | 4.41 | 1.00 | 58.78 |  |
| 10 | A | N7 | AA:218 | 4.68 | 1.00 | 48.49 |  |
| 11 | A | OP2 | AA:428 | 4.84 | 1.00 | 56.85 |  |
| 12 | U | O3' | AA:427 | 4.88 | 1.00 | 57.03 |  |
| 13 | A | O5' | AA:428 | 4.97 | 1.00 | 58.50 |  |
  |

  
 Coded contact list:   
 Exclusion zone contact(s): /O6/O6/OP1/  
Consider K in place of MG (3 valid contact(s) for K vs. 0 for MG)  

### MG:AA:3638

|  | Res. type | Atom | Residue | Distance | Occupancy | B-factor | Closest solute to 1^st^ shell water |
| --- | --- | --- | --- | --- | --- | --- | --- |
| **Ion** | | | |  | 1.00 | 51.96 || 1 | G | O6 | AA:327 | 2.65 | 1.00 | 63.92 |
| 2 | G | O6 | AA:326 | 3.36 | 1.00 | 60.71 |  |
| 3 | G | N7 | AA:327 | 3.81 | 1.00 | 60.47 |  |
| 4 | C | N4 | AA:335 | 4.25 | 1.00 | 63.84 |  |
| 5 | G | N7 | AA:326 | 4.38 | 1.00 | 61.17 |  |
| 6 | A | N7 | AA:332 | 4.49 | 1.00 | 58.71 |  |
| 7 | A | N9 | AA:332 | 4.52 | 1.00 | 57.23 |  |
| 8 | U | O4 | AA:328 | 4.58 | 1.00 | 56.25 |  |
| 9 | C | N4 | AA:336 | 4.70 | 1.00 | 61.97 |  |
| 10 | G | N1 | AA:327 | 4.89 | 1.00 | 61.92 |  |
| 11 | A | O2' | AA:332 | 4.89 | 1.00 | 57.27 |  |
| 12 | C | OP2 | AA:334 | 4.94 | 1.00 | 59.39 |  |
| 13 | A | OP2 | AA:332 | 4.95 | 1.00 | 56.99 |  |
| 14 | A | O3' | AA:332 | 4.99 | 1.00 | 56.37 |  |
  |

  
 Coded contact list:   
 Exclusion zone contact(s): /O6/O6/  
Consider K in place of MG (2 valid contact(s) for K vs. 0 for MG)  

### MG:AA:3639

|  | Res. type | Atom | Residue | Distance | Occupancy | B-factor | Closest solute to 1^st^ shell water |
| --- | --- | --- | --- | --- | --- | --- | --- |
| **Ion** | | | |  | 1.00 | 55.74 || 1 | G | N7 | AA:89 | 3.83 | 1.00 | 71.72 |
| 2 | U | OP1 | AA:90 | 4.23 | 1.00 | 88.33 |  |
| 3 | G | O6 | AA:89 | 4.47 | 1.00 | 67.07 |  |
| 4 | G | N9 | AA:89 | 4.48 | 1.00 | 75.13 |  |
| 5 | U | O5' | AA:90 | 4.65 | 1.00 | 92.18 |  |
| 6 | G | N1 | AA:89 | 4.83 | 1.00 | 68.12 |  |
| 7 | U | O4' | AA:90 | 4.83 | 1.00 | 104.44 |  |
| 8 | G | N3 | AA:89 | 4.99 | 1.00 | 72.48 |  |
| 9 | A | N6 | AA:64 | 5.00 | 1.00 | 62.75 |  |
  |

  
 Coded contact list:   
1^st^ MG coordination shell missing (see Notes)  

### MG:AA:3640

|  | Res. type | Atom | Residue | Distance | Occupancy | B-factor | Closest solute to 1^st^ shell water |
| --- | --- | --- | --- | --- | --- | --- | --- |
| **Ion** | | | |  | 1.00 | 45.16 || 1 | U | OP1 | AA:2696 | 3.30 | 1.00 | 54.93 |
| 2 | G | O6 | AA:1758 | 3.42 | 1.00 | 54.86 |  |
| 3 | G | N1 | AA:1758 | 4.19 | 1.00 | 53.81 |  |
| 4 | G | O2' | AA:1707 | 4.22 | 1.00 | 70.64 |  |
| 5 | C | O4' | AA:1708 | 4.40 | 1.00 | 77.22 |  |
| 6 | C | O3' | AA:2695 | 4.45 | 1.00 | 53.30 |  |
| 7 | G | N3 | AA:1707 | 4.91 | 1.00 | 74.30 |  |
| 8 | G | N2 | AA:1756 | 4.92 | 1.00 | 69.10 |  |
  |

  
 Coded contact list:   
 Exclusion zone contact(s): /OP1/  
Consider K in place of MG (1 valid contact(s) for K vs. 0 for MG)  

### MG:AA:3641

|  | Res. type | Atom | Residue | Distance | Occupancy | B-factor | Closest solute to 1^st^ shell water |
| --- | --- | --- | --- | --- | --- | --- | --- |
| **Ion** | | | |  | 1.00 | 46.03 || 1 | G | OP2 | AA:785 | 2.45 | 1.00 | 35.38 |
| 2 | C | O3' | AA:791 | 3.19 | 1.00 | 35.98 |  |
| 3 | G | OP2 | AA:792 | 3.20 | 1.00 | 33.79 |  |
| 4 | A | O3' | AA:784 | 3.82 | 1.00 | 36.13 |  |
| 5 | G | O5' | AA:785 | 4.19 | 1.00 | 33.13 |  |
| 6 | G | O5' | AA:792 | 4.24 | 1.00 | 32.54 |  |
| 7 | C | OP2 | AA:786 | 4.36 | 1.00 | 32.18 |  |
| 8 | A | O2' | AA:784 | 4.56 | 1.00 | 41.21 |  |
| 9 | G | OP1 | AA:785 | 4.92 | 1.00 | 36.44 |  |
  |

  
 Coded contact list:   
 Exclusion zone contact(s): /O3'/OP2/OP2/  
Consider K in place of MG (2 valid contact(s) for K vs. 0 for MG)  

### MG:AA:3642

|  | Res. type | Atom | Residue | Distance | Occupancy | B-factor | Closest solute to 1^st^ shell water |
| --- | --- | --- | --- | --- | --- | --- | --- |
| **Ion** | | | |  | 1.00 | 56.68 || 1 | G | N7 | AA:1256 | 3.06 | 1.00 | 44.85 |
| 2 | A | OP2 | AA:1254 | 3.41 | 1.00 | 44.37 |  |
| 3 | A | O5' | AA:1253 | 4.01 | 1.00 | 43.61 |  |
| 4 | G | O6 | AA:1256 | 4.33 | 1.00 | 47.75 |  |
| 5 | C | N4 | AA:1257 | 4.41 | 1.00 | 46.05 |  |
| 6 | A | O3' | AA:1253 | 4.44 | 1.00 | 44.29 |  |
  |

  
 Coded contact list:   
 Exclusion zone contact(s): /N7/  
Consider K in place of MG (1 valid contact(s) for K vs. 0 for MG)  

### MG:AA:3643

|  | Res. type | Atom | Residue | Distance | Occupancy | B-factor | Closest solute to 1^st^ shell water |
| --- | --- | --- | --- | --- | --- | --- | --- |
| **Ion** | | | |  | 1.00 | 61.97 || 1 | G | OP2 | AA:2270 | 2.37 | 1.00 | 62.98 |
| 2 | U | O4 | AA:2272 | 3.66 | 1.00 | 59.04 |  |
| 3 | A | O2' | AA:2266 | 3.81 | 1.00 | 75.39 |  |
| 4 | A | OP2 | AA:2267 | 3.89 | 1.00 | 69.84 |  |
| 5 | G | N7 | AA:2271 | 4.10 | 1.00 | 65.03 |  |
| 6 | G | O6 | AA:2271 | 4.18 | 1.00 | 63.96 |  |
| 7 | A | O3' | AA:2266 | 4.25 | 1.00 | 72.40 |  |
| 8 | G | O5' | AA:2270 | 4.55 | 1.00 | 63.65 |  |
| 9 | A | O5' | AA:2269 | 4.58 | 1.00 | 69.53 |  |
| 10 | A | O3' | AA:2268 | 4.65 | 1.00 | 70.84 |  |
| 11 | G | OP1 | AA:2270 | 4.73 | 1.00 | 64.25 |  |
| 12 | A | O3' | AA:2269 | 4.75 | 1.00 | 64.66 |  |
| 13 | G | OP2 | AA:2271 | 4.88 | 1.00 | 59.24 |  |
  |

  
 Contact(s): /OP2/  
 Coded contact list: O_ph_  

### MG:AA:3644

|  | Res. type | Atom | Residue | Distance | Occupancy | B-factor | Closest solute to 1^st^ shell water |
| --- | --- | --- | --- | --- | --- | --- | --- |
| **Ion** | | | |  | 1.00 | 56.71 || 1 | G | N7 | AA:274 | 2.40 | 1.00 | 158.23 |
| 2 | G | N7 | AA:275 | 2.60 | 1.00 | 165.65 |  |
| 3 | C | N3 | AA:273F | 2.71 | 1.00 | 136.44 |  |
| 4 | G | O6 | AA:275 | 2.83 | 1.00 | 165.12 |  |
| 5 | G | N1 | AA:363 | 3.53 | 1.00 | 133.10 |  |
| 6 | C | O2 | AA:273F | 3.53 | 1.00 | 135.57 |  |
| 7 | C | N4 | AA:273F | 3.55 | 1.00 | 135.71 |  |
| 8 | G | O6 | AA:274 | 3.78 | 1.00 | 157.41 |  |
| 9 | C | N1 | AA:273F | 3.92 | 1.00 | 139.10 |  |
| 10 | G | O6 | AA:363 | 4.10 | 1.00 | 133.01 |  |
| 11 | G | N2 | AA:363 | 4.18 | 1.00 | 133.28 |  |
| 12 | G | N9 | AA:274 | 4.18 | 1.00 | 158.30 |  |
| 13 | G | N1 | AA:275 | 4.29 | 1.00 | 165.46 |  |
| 14 | G | N9 | AA:275 | 4.43 | 1.00 | 165.32 |  |
  |

  
 Coded contact list:   
 Exclusion zone contact(s): /N3_C_/N7/N7/O6/  
Consider K in place of MG (3 valid contact(s) for K vs. 0 for MG)  

### MG:AA:3645

|  | Res. type | Atom | Residue | Distance | Occupancy | B-factor | Closest solute to 1^st^ shell water |
| --- | --- | --- | --- | --- | --- | --- | --- |
| **Ion** | | | |  | 1.00 | 46.66 || 1 | G | N1 | AA:1681 | 3.83 | 1.00 | 67.38 |
| 2 | G | N3 | AA:1681 | 3.93 | 1.00 | 65.36 |  |
| 3 | G | O6 | AA:1681 | 4.49 | 1.00 | 68.76 |  |
| 4 | G | N2 | AA:1681 | 4.50 | 1.00 | 68.58 |  |
| 5 | G | N9 | AA:1681 | 4.57 | 1.00 | 64.75 |  |
| 6 | G | N7 | AA:1681 | 4.61 | 1.00 | 67.16 |  |
| 7 | C | OP2 | AA:1761 | 4.72 | 1.00 | 46.40 |  |
  |

  
 Coded contact list:   
1^st^ MG coordination shell missing (see Notes)  

### MG:AA:3646

|  | Res. type | Atom | Residue | Distance | Occupancy | B-factor | Closest solute to 1^st^ shell water |
| --- | --- | --- | --- | --- | --- | --- | --- |
| **Ion** | | | |  | 1.00 | 62.65 || 1 | G | O6 | AA:2839 | 2.57 | 1.00 | 68.68 |
| 2 | G | O6 | AA:2877 | 3.09 | 1.00 | 67.63 |  |
| 3 | U | O4 | AA:2878 | 3.23 | 1.00 | 71.07 |  |
| 4 | C | N4 | AA:2840 | 3.70 | 1.00 | 67.19 |  |
| 5 | G | N7 | AA:2839 | 3.81 | 1.00 | 66.59 |  |
| 6 | G | N7 | AA:2838 | 4.08 | 1.00 | 68.35 |  |
| 7 | G | O6 | AA:2838 | 4.12 | 1.00 | 67.81 |  |
| 8 | U | N3 | AA:2878 | 4.17 | 1.00 | 70.53 |  |
| 9 | G | N1 | AA:2877 | 4.80 | 1.00 | 68.19 |  |
| 10 | G | N1 | AA:2839 | 4.83 | 1.00 | 68.52 |  |
  |

  
 Coded contact list:   
 Exclusion zone contact(s): /O4/O6/O6/  
Consider K in place of MG (2 valid contact(s) for K vs. 0 for MG)  

### MG:AA:3647

|  | Res. type | Atom | Residue | Distance | Occupancy | B-factor | Closest solute to 1^st^ shell water |
| --- | --- | --- | --- | --- | --- | --- | --- |
| **Ion** | | | |  | 1.00 | 25.82 || 1 | U | OP2 | AA:807 | 3.30 | 1.00 | 35.81 |
| 2 | U | OP1 | AA:807 | 3.42 | 1.00 | 37.37 |  |
| 3 | U | OP1 | AA:568 | 3.63 | 1.00 | 45.47 |  |
| 4 | LYS | O | AO:36 | 3.69 | 1.00 | 47.53 |  |
| 5 | G | OP2 | AA:808 | 3.96 | 1.00 | 45.44 |  |
| 6 | U | O5' | AA:807 | 4.10 | 1.00 | 39.02 |  |
| 7 | A | O3' | AA:567 | 4.39 | 1.00 | 42.40 |  |
| 8 | ARG | NH1 | AO:41 | 4.42 | 1.00 | 48.65 |  |
| 9 | A | N6 | AA:2448 | 4.99 | 1.00 | 55.15 |  |
  |

  
 Coded contact list:   
 Exclusion zone contact(s): /OP2/  
Consider K in place of MG (1 valid contact(s) for K vs. 0 for MG)  

### MG:AA:3648

|  | Res. type | Atom | Residue | Distance | Occupancy | B-factor | Closest solute to 1^st^ shell water |
| --- | --- | --- | --- | --- | --- | --- | --- |
| **Ion** | | | |  | 1.00 | 43.20 || 1 | G | OP2 | AA:205 | 2.69 | 1.00 | 44.28 |
| 2 | A | O2' | AA:204 | 2.79 | 1.00 | 46.92 |  |
| 3 | U | OP1 | AA:202 | 3.87 | 1.00 | 42.13 |  |
| 4 | C | OP2 | AA:203 | 4.04 | 1.00 | 44.19 |  |
| 5 | A | O3' | AA:204 | 4.18 | 1.00 | 44.98 |  |
| 6 | G | O5' | AA:205 | 4.30 | 1.00 | 44.85 |  |
| 7 | U | OP2 | AA:202 | 4.60 | 1.00 | 43.42 |  |
  |

  
 Coded contact list:   
 Exclusion zone contact(s): /O2'/OP2/  
Consider K in place of MG (2 valid contact(s) for K vs. 0 for MG)  

### MG:AA:3649

|  | Res. type | Atom | Residue | Distance | Occupancy | B-factor | Closest solute to 1^st^ shell water |
| --- | --- | --- | --- | --- | --- | --- | --- |
| **Ion** | | | |  | 1.00 | 96.11 || 1 | G | OP2 | AA:348 | 2.56 | 1.00 | 73.66 |
| 2 | G | N7 | AA:348 | 2.89 | 1.00 | 68.35 |  |
| 3 | A | OP2 | AA:347 | 3.32 | 1.00 | 75.24 |  |
| 4 | A | N7 | AA:347 | 3.48 | 1.00 | 75.29 |  |
| 5 | A | N9 | AA:347 | 3.90 | 1.00 | 74.23 |  |
| 6 | G | O5' | AA:348 | 4.10 | 1.00 | 72.80 |  |
| 7 | A | O3' | AA:347 | 4.27 | 1.00 | 73.45 |  |
| 8 | A | O5' | AA:347 | 4.29 | 1.00 | 75.42 |  |
| 9 | G | N9 | AA:348 | 4.64 | 1.00 | 68.91 |  |
| 10 | G | O6 | AA:348 | 4.95 | 1.00 | 68.02 |  |
  |

  
 Coded contact list:   
 Exclusion zone contact(s): /N7/OP2/OP2/  
Consider K in place of MG (2 valid contact(s) for K vs. 0 for MG)  

### MG:AA:3650

|  | Res. type | Atom | Residue | Distance | Occupancy | B-factor | Closest solute to 1^st^ shell water |
| --- | --- | --- | --- | --- | --- | --- | --- |
| **Ion** | | | |  | 1.00 | 70.97 || 1 | G | OP2 | AA:1561 | 2.28 | 1.00 | 67.40 |
| 2 | G | OP2 | AA:1560 | 2.51 | 1.00 | 78.45 |  |
| 3 | G | OP1 | AA:1560 | 2.80 | 1.00 | 77.81 |  |
| 4 | G | O2' | AA:1559 | 4.19 | 1.00 | 83.66 |  |
| 5 | G | O3' | AA:1559 | 4.23 | 1.00 | 78.71 |  |
| 6 | G | O5' | AA:1560 | 4.27 | 1.00 | 76.37 |  |
| 7 | G | O3' | AA:1560 | 4.53 | 1.00 | 67.10 |  |
| 8 | G | OP1 | AA:1561 | 4.56 | 1.00 | 67.01 |  |
| 9 | G | O5' | AA:1561 | 4.64 | 1.00 | 65.34 |  |
| 10 | G | N7 | AA:1561 | 4.82 | 1.00 | 55.57 |  |
  |

  
 Contact(s): /OP2/  
 Coded contact list: O_ph_  
 Exclusion zone contact(s): /OP1/OP2/  

### MG:AA:3651

|  | Res. type | Atom | Residue | Distance | Occupancy | B-factor | Closest solute to 1^st^ shell water |
| --- | --- | --- | --- | --- | --- | --- | --- |
| **Ion** | | | |  | 1.00 | 41.20 || 1 | G | O4' | AA:831 | 3.26 | 1.00 | 41.03 |
| 2 | G | O2' | AA:805 | 3.38 | 1.00 | 51.54 |  |
| 3 | G | O3' | AA:830 | 3.59 | 1.00 | 43.14 |  |
| 4 | G | O3' | AA:805 | 4.06 | 1.00 | 44.66 |  |
| 5 | A | OP1 | AA:829 | 4.10 | 1.00 | 54.00 |  |
| 6 | G | O2' | AA:830 | 4.15 | 1.00 | 45.13 |  |
| 7 | U | OP2 | AA:2068 | 4.31 | 1.00 | 48.03 |  |
| 8 | A | O5' | AA:829 | 4.58 | 1.00 | 52.27 |  |
| 9 | G | N9 | AA:831 | 4.58 | 1.00 | 43.14 |  |
| 10 | G | N3 | AA:805 | 4.93 | 1.00 | 53.67 |  |
  |

  
 Coded contact list:   
 Exclusion zone contact(s): /O2'/O4'/  
Consider K in place of MG (2 valid contact(s) for K vs. 0 for MG)  

### MG:AA:3652

|  | Res. type | Atom | Residue | Distance | Occupancy | B-factor | Closest solute to 1^st^ shell water |
| --- | --- | --- | --- | --- | --- | --- | --- |
| **Ion** | | | |  | 1.00 | 44.81 || 1 | G | O6 | AA:309 | 2.59 | 1.00 | 50.87 |
| 2 | A | O2' | AA:1210 | 2.62 | 1.00 | 73.87 |  |
| 3 | G | O2' | AA:307 | 2.89 | 1.00 | 59.57 |  |
| 4 | U | OP2 | AA:1211 | 3.02 | 1.00 | 72.07 |  |
| 5 | G | O4' | AA:307 | 4.07 | 1.00 | 61.33 |  |
| 6 | A | O3' | AA:1210 | 4.13 | 1.00 | 72.88 |  |
| 7 | A | N1 | AA:330 | 4.16 | 1.00 | 66.36 |  |
| 8 | G | O3' | AA:307 | 4.31 | 1.00 | 61.71 |  |
| 9 | G | N7 | AA:309 | 4.40 | 1.00 | 53.61 |  |
| 10 | U | OP1 | AA:1211 | 4.51 | 1.00 | 72.89 |  |
| 11 | G | N7 | AA:308 | 4.58 | 1.00 | 54.44 |  |
| 12 | G | N1 | AA:309 | 4.83 | 1.00 | 51.33 |  |
  |

  
 Coded contact list:   
 Exclusion zone contact(s): /O2'/O2'/O6/OP2/  
Consider K in place of MG (3 valid contact(s) for K vs. 0 for MG)  

### MG:AA:3653

|  | Res. type | Atom | Residue | Distance | Occupancy | B-factor | Closest solute to 1^st^ shell water |
| --- | --- | --- | --- | --- | --- | --- | --- |
| **Ion** | | | |  | 1.00 | 58.07 || 1 | C | OP2 | AA:2879 | 2.53 | 1.00 | 80.96 |
| 2 | U | O3' | AA:2878 | 2.94 | 1.00 | 79.44 |  |
| 3 | C | OP1 | AA:2879 | 2.98 | 1.00 | 82.52 |  |
| 4 | U | O5' | AA:2878 | 3.33 | 1.00 | 73.79 |  |
| 5 | U | OP1 | AA:2878 | 4.08 | 1.00 | 72.87 |  |
| 6 | C | O5' | AA:2879 | 4.44 | 1.00 | 81.58 |  |
  |

  
 Coded contact list:   
 Exclusion zone contact(s): /O3'/O5'/OP1/OP2/  
Consider K in place of MG (3 valid contact(s) for K vs. 0 for MG)  

### MG:AA:3654

|  | Res. type | Atom | Residue | Distance | Occupancy | B-factor | Closest solute to 1^st^ shell water |
| --- | --- | --- | --- | --- | --- | --- | --- |
| **Ion** | | | |  | 1.00 | 59.98 || 1 | G | OP1 | AA:2780 | 4.00 | 1.00 | 86.40 |
| 2 | A | N6 | AA:528 | 4.04 | 1.00 | 70.25 |  |
| 3 | C | O2' | AA:527 | 4.44 | 1.00 | 63.38 |  |
| 4 | C | N1 | AA:527 | 4.65 | 1.00 | 59.71 |  |
| 5 | U | OP1 | AA:2779 | 4.97 | 1.00 | 72.53 |  |
  |

  
 Coded contact list:   
1^st^ MG coordination shell missing (see Notes)  

### MG:AA:3655

|  | Res. type | Atom | Residue | Distance | Occupancy | B-factor | Closest solute to 1^st^ shell water |
| --- | --- | --- | --- | --- | --- | --- | --- |
| **Ion** | | | |  | 1.00 | 50.50 || 1 | G | O6 | AA:2718 | 3.05 | 1.00 | 72.03 |
| 2 | G | O6 | AA:2717 | 3.20 | 1.00 | 64.46 |  |
| 3 | G | O6 | AA:2719 | 3.54 | 1.00 | 69.29 |  |
| 4 | G | N7 | AA:2718 | 4.19 | 1.00 | 69.59 |  |
| 5 | C | N4 | AA:2692 | 4.33 | 1.00 | 66.15 |  |
| 6 | G | N7 | AA:2717 | 4.49 | 1.00 | 64.30 |  |
| 7 | C | N4 | AA:2691 | 4.50 | 1.00 | 70.75 |  |
| 8 | U | O4 | AA:2716 | 4.60 | 1.00 | 59.00 |  |
| 9 | G | N1 | AA:2719 | 4.83 | 1.00 | 69.46 |  |
  |

  
 Coded contact list:   
 Exclusion zone contact(s): /O6/O6/  
Consider K in place of MG (2 valid contact(s) for K vs. 0 for MG)  

### MG:AA:3656

|  | Res. type | Atom | Residue | Distance | Occupancy | B-factor | Closest solute to 1^st^ shell water |
| --- | --- | --- | --- | --- | --- | --- | --- |
| **Ion** | | | |  | 1.00 | 37.36 || 1 | U | O2 | AA:2068 | 2.76 | 1.00 | 53.19 |
| 2 | U | O2' | AA:827 | 2.81 | 1.00 | 54.62 |  |
| 3 | U | O2 | AA:827 | 3.08 | 1.00 | 53.97 |  |
| 4 | U | O2' | AA:2068 | 3.43 | 1.00 | 44.14 |  |
| 5 | U | OP1 | AA:2245 | 3.58 | 1.00 | 33.24 |  |
| 6 | A | N1 | AA:2430 | 3.90 | 1.00 | 46.94 |  |
| 7 | U | N1 | AA:2068 | 4.14 | 1.00 | 48.42 |  |
| 8 | G | O3' | AA:2246 | 4.28 | 1.00 | 43.77 |  |
| 9 | U | N1 | AA:827 | 4.46 | 1.00 | 52.27 |  |
| 10 | U | N3 | AA:2068 | 4.77 | 1.00 | 50.82 |  |
| 11 | G | O2' | AA:2246 | 4.78 | 1.00 | 47.02 |  |
| 12 | A | N6 | AA:2430 | 4.85 | 1.00 | 47.93 |  |
| 13 | U | N3 | AA:827 | 4.90 | 1.00 | 53.99 |  |
| 14 | U | O4' | AA:2068 | 4.93 | 1.00 | 46.82 |  |
  |

  
 Coded contact list:   
 Exclusion zone contact(s): /O2/O2/O2'/  
Consider K in place of MG (3 valid contact(s) for K vs. 0 for MG)  

### MG:AA:3657

|  | Res. type | Atom | Residue | Distance | Occupancy | B-factor | Closest solute to 1^st^ shell water |
| --- | --- | --- | --- | --- | --- | --- | --- |
| **Ion** | | | |  | 1.00 | 89.42 || 1 | G | O6 | AA:2303 | 3.41 | 1.00 | 90.89 |
| 2 | G | N7 | AA:2303 | 3.55 | 1.00 | 94.13 |  |
| 3 | G | N7 | AA:2304 | 4.06 | 1.00 | 99.39 |  |
| 4 | G | N1 | AA:2303 | 4.83 | 1.00 | 91.97 |  |
  |

  
 Coded contact list:   
1^st^ MG coordination shell missing (see Notes)  

### MG:AA:3658

|  | Res. type | Atom | Residue | Distance | Occupancy | B-factor | Closest solute to 1^st^ shell water |
| --- | --- | --- | --- | --- | --- | --- | --- |
| **Ion** | | | |  | 1.00 | 64.91 || 1 | U | OP2 | AA:2687 | 2.81 | 1.00 | 52.87 |
| 2 | G | OP2 | AA:2719 | 3.39 | 1.00 | 62.40 |  |
| 3 | U | OP1 | AA:2687 | 3.73 | 1.00 | 52.66 |  |
| 4 | U | O5' | AA:2687 | 4.82 | 1.00 | 53.68 |  |
| 5 | G | O3' | AA:2686 | 4.92 | 1.00 | 52.24 |  |
  |

  
 Coded contact list:   
 Exclusion zone contact(s): /OP2/OP2/  
Consider K in place of MG (2 valid contact(s) for K vs. 0 for MG)  

### MG:AA:3659

|  | Res. type | Atom | Residue | Distance | Occupancy | B-factor | Closest solute to 1^st^ shell water |
| --- | --- | --- | --- | --- | --- | --- | --- |
| **Ion** | | | |  | 1.00 | 67.03 || 1 | G | OP2 | AA:15 | 2.30 | 1.00 | 67.09 |
| 2 | A | OP2 | AA:14 | 2.52 | 1.00 | 74.00 |  |
| 3 | G | OP1 | AA:15 | 4.12 | 1.00 | 66.38 |  |
| 4 | A | O3' | AA:14 | 4.49 | 1.00 | 66.13 |  |
| 5 | A | O5' | AA:14 | 4.67 | 1.00 | 72.33 |  |
| 6 | G | O5' | AA:15 | 4.71 | 1.00 | 65.76 |  |
| 7 | A | OP1 | AA:14 | 4.83 | 1.00 | 75.55 |  |
| 8 | A | O3' | AA:13 | 4.84 | 1.00 | 79.18 |  |
  |

  
 Contact(s): /OP2/  
 Coded contact list: O_ph_  
 Exclusion zone contact(s): /OP2/  

### MG:AA:3660

|  | Res. type | Atom | Residue | Distance | Occupancy | B-factor | Closest solute to 1^st^ shell water |
| --- | --- | --- | --- | --- | --- | --- | --- |
| **Ion** | | | |  | 1.00 | 68.18 || 1 | G | O6 | AA:1436 | 3.83 | 1.00 | 57.47 |
| 2 | G | O6 | AA:1435 | 4.01 | 1.00 | 58.24 |  |
| 3 | G | N7 | AA:1435 | 4.17 | 1.00 | 57.64 |  |
| 4 | A | OP2 | AA:1554 | 4.29 | 1.00 | 64.77 |  |
| 5 | A | N7 | AA:1434 | 4.66 | 1.00 | 56.34 |  |
| 6 | A | OP2 | AA:1434 | 4.70 | 1.00 | 63.66 |  |
| 7 | C | N4 | AA:1556 | 4.72 | 1.00 | 55.30 |  |
  |

  
 Coded contact list:   
1^st^ MG coordination shell missing (see Notes)  

### MG:AA:3661

|  | Res. type | Atom | Residue | Distance | Occupancy | B-factor | Closest solute to 1^st^ shell water |
| --- | --- | --- | --- | --- | --- | --- | --- |
| **Ion** | | | |  | 1.00 | 70.69 || 1 | G | O6 | AA:82 | 3.10 | 1.00 | 65.02 |
| 2 | G | O6 | AA:83 | 3.22 | 1.00 | 55.30 |  |
| 3 | G | N7 | AA:82 | 3.79 | 1.00 | 64.87 |  |
| 4 | G | OP2 | AA:81 | 4.59 | 1.00 | 70.55 |  |
| 5 | G | N1 | AA:83 | 4.87 | 1.00 | 55.48 |  |
  |

  
 Coded contact list:   
 Exclusion zone contact(s): /O6/O6/  
Consider K in place of MG (2 valid contact(s) for K vs. 0 for MG)  

### MG:AA:3662

|  | Res. type | Atom | Residue | Distance | Occupancy | B-factor | Closest solute to 1^st^ shell water |
| --- | --- | --- | --- | --- | --- | --- | --- |
| **Ion** | | | |  | 1.00 | 51.23 || 1 | G | OP1 | AA:1763 | 2.45 | 1.00 | 58.91 |
| 2 | G | O6 | AA:1699 | 2.62 | 1.00 | 65.37 |  |
| 3 | G | O4' | AA:1763 | 3.18 | 1.00 | 55.54 |  |
| 4 | G | O2' | AA:1763 | 4.53 | 1.00 | 56.54 |  |
| 5 | G | N1 | AA:1699 | 4.55 | 1.00 | 63.58 |  |
| 6 | A | O3' | AA:1762 | 4.58 | 1.00 | 59.68 |  |
| 7 | C | O3' | AA:732 | 4.63 | 1.00 | 42.83 |  |
| 8 | G | OP1 | AA:733 | 4.71 | 1.00 | 42.53 |  |
| 9 | G | O5' | AA:1763 | 4.75 | 1.00 | 57.95 |  |
| 10 | G | OP2 | AA:1763 | 4.80 | 1.00 | 59.27 |  |
| 11 | G | OP1 | AA:1682 | 4.84 | 1.00 | 58.05 |  |
| 12 | G | N9 | AA:1763 | 4.85 | 1.00 | 56.38 |  |
  |

  
 Coded contact list:   
 Exclusion zone contact(s): /O4'/O6/OP1/  
Consider K in place of MG (2 valid contact(s) for K vs. 0 for MG)  

### MG:AA:3663

|  | Res. type | Atom | Residue | Distance | Occupancy | B-factor | Closest solute to 1^st^ shell water |
| --- | --- | --- | --- | --- | --- | --- | --- |
| **Ion** | | | |  | 1.00 | 61.61 || 1 | G | O6 | AA:2731 | 3.88 | 1.00 | 62.41 |
| 2 | G | N7 | AA:2731 | 4.19 | 1.00 | 63.91 |  |
| 3 | G | O6 | AA:2732 | 4.43 | 1.00 | 73.31 |  |
  |

  
 Coded contact list:   
1^st^ MG coordination shell missing (see Notes)  

### MG:AA:3664

|  | Res. type | Atom | Residue | Distance | Occupancy | B-factor | Closest solute to 1^st^ shell water |
| --- | --- | --- | --- | --- | --- | --- | --- |
| **Ion** | | | |  | 1.00 | 68.34 || 1 | A | OP1 | AA:1439 | 2.51 | 1.00 | 59.45 |
| 2 | C | OP1 | AA:1468 | 2.56 | 1.00 | 77.43 |  |
| 3 | G | OP2 | AA:1440 | 3.62 | 1.00 | 58.51 |  |
| 4 | C | O5' | AA:1468 | 3.69 | 1.00 | 77.13 |  |
| 5 | A | O5' | AA:1439 | 4.02 | 1.00 | 59.42 |  |
| 6 | A | OP2 | AA:1469 | 4.29 | 1.00 | 75.98 |  |
| 7 | C | O3' | AA:1467 | 4.57 | 1.00 | 78.27 |  |
| 8 | A | OP1 | AA:1469 | 4.63 | 1.00 | 76.38 |  |
| 9 | A | OP2 | AA:1439 | 4.69 | 1.00 | 59.38 |  |
| 10 | A | O3' | AA:1439 | 4.79 | 1.00 | 58.04 |  |
| 11 | C | O3' | AA:1468 | 4.79 | 1.00 | 75.62 |  |
| 12 | C | OP2 | AA:1468 | 4.87 | 1.00 | 77.23 |  |
| 13 | U | O3' | AA:1438 | 4.87 | 1.00 | 58.21 |  |
  |

  
 Coded contact list:   
 Exclusion zone contact(s): /OP1/OP1/  
Consider NA in place of MG (2 valid contact(s) for NA vs. 0 for MG)  

### MG:AA:3665

|  | Res. type | Atom | Residue | Distance | Occupancy | B-factor | Closest solute to 1^st^ shell water |
| --- | --- | --- | --- | --- | --- | --- | --- |
| **Ion** | | | |  | 1.00 | 80.76 || 1 | C | N4 | AA:2676 | 4.31 | 1.00 | 62.02 |
| 2 | G | O6 | AA:2677 | 4.69 | 1.00 | 66.37 |  |
  |

  
 Coded contact list:   
1^st^ MG coordination shell missing (see Notes)  

### MG:AA:3666

|  | Res. type | Atom | Residue | Distance | Occupancy | B-factor | Closest solute to 1^st^ shell water |
| --- | --- | --- | --- | --- | --- | --- | --- |
| **Ion** | | | |  | 1.00 | 52.46 || 1 | U | O4 | AA:1165 | 4.24 | 1.00 | 84.36 |
| 2 | A | N6 | AA:1182 | 4.47 | 1.00 | 108.10 |  |
| 3 | C | N4 | AA:1166 | 4.53 | 1.00 | 96.89 |  |
| 4 | G | O6 | AA:1183 | 4.69 | 1.00 | 97.54 |  |
| 5 | C | N4 | AA:1181 | 5.00 | 1.00 | 125.57 |  |
  |

  
 Coded contact list:   
1^st^ MG coordination shell missing (see Notes)  

### MG:AA:3667

|  | Res. type | Atom | Residue | Distance | Occupancy | B-factor | Closest solute to 1^st^ shell water |
| --- | --- | --- | --- | --- | --- | --- | --- |
| **Ion** | | | |  | 1.00 | 79.04 || 1 | G | N7 | AA:2830 | 3.47 | 1.00 | 58.70 |
| 2 | G | O6 | AA:2830 | 3.67 | 1.00 | 58.26 |  |
| 3 | G | N7 | AA:2831 | 4.01 | 1.00 | 58.85 |  |
| 4 | G | O6 | AA:2831 | 4.22 | 1.00 | 61.47 |  |
  |

  
 Coded contact list:   
1^st^ MG coordination shell missing (see Notes)  

### MG:AA:3668

|  | Res. type | Atom | Residue | Distance | Occupancy | B-factor | Closest solute to 1^st^ shell water |
| --- | --- | --- | --- | --- | --- | --- | --- |
| **Ion** | | | |  | 1.00 | 71.68 || 1 | A | OP1 | AA:1349 | 2.78 | 1.00 | 55.50 |
| 2 | G | OP2 | AA:1377 | 2.87 | 1.00 | 48.62 |  |
| 3 | C | OP1 | AA:1376 | 3.54 | 1.00 | 50.34 |  |
| 4 | C | OP2 | AA:1351 | 3.92 | 1.00 | 45.27 |  |
| 5 | G | O3' | AA:1348 | 4.77 | 1.00 | 52.43 |  |
| 6 | G | OP1 | AA:1377 | 4.81 | 1.00 | 49.60 |  |
| 7 | C | O5' | AA:1376 | 4.82 | 1.00 | 49.21 |  |
| 8 | C | O3' | AA:1376 | 4.87 | 1.00 | 50.11 |  |
| 9 | A | OP2 | AA:1349 | 4.98 | 1.00 | 53.17 |  |
  |

  
 Coded contact list:   
 Exclusion zone contact(s): /OP1/OP2/  
Consider K in place of MG (2 valid contact(s) for K vs. 0 for MG)  

### MG:AA:3669

|  | Res. type | Atom | Residue | Distance | Occupancy | B-factor | Closest solute to 1^st^ shell water |
| --- | --- | --- | --- | --- | --- | --- | --- |
| **Ion** | | | |  | 1.00 | 44.20 || 1 | C | OP2 | AA:1298 | 2.57 | 1.00 | 49.54 |
| 2 | U | OP2 | AA:1639 | 2.71 | 1.00 | 44.86 |  |
| 3 | C | O5' | AA:1638 | 3.56 | 1.00 | 44.77 |  |
| 4 | C | OP2 | AA:1638 | 3.60 | 1.00 | 46.36 |  |
| 5 | C | OP1 | AA:1298 | 3.62 | 1.00 | 48.98 |  |
| 6 | C | OP1 | AA:1638 | 4.13 | 1.00 | 45.61 |  |
| 7 | G | N7 | AA:1299 | 4.35 | 1.00 | 43.07 |  |
| 8 | C | O5' | AA:1298 | 4.49 | 1.00 | 48.34 |  |
| 9 | G | OP2 | AA:1299 | 4.56 | 1.00 | 45.05 |  |
| 10 | C | O3' | AA:1638 | 4.58 | 1.00 | 43.70 |  |
| 11 | C | O3' | AA:1297 | 4.81 | 1.00 | 49.30 |  |
| 12 | G | O6 | AA:1299 | 4.84 | 1.00 | 39.49 |  |
| 13 | U | O5' | AA:1639 | 4.86 | 1.00 | 42.99 |  |
  |

  
 Coded contact list:   
 Exclusion zone contact(s): /OP2/OP2/  
Consider NA in place of MG (1 valid contact(s) for NA vs. 0 for MG)  

### MG:AA:3670

|  | Res. type | Atom | Residue | Distance | Occupancy | B-factor | Closest solute to 1^st^ shell water |
| --- | --- | --- | --- | --- | --- | --- | --- |
| **Ion** | | | |  | 1.00 | 54.93 || 1 | G | O6 | AA:1216 | 2.48 | 1.00 | 69.41 |
| 2 | G | O6 | AA:1215 | 3.16 | 1.00 | 73.39 |  |
| 3 | U | O4 | AA:1234 | 3.32 | 1.00 | 73.94 |  |
| 4 | C | N4 | AA:1233 | 3.89 | 1.00 | 69.30 |  |
| 5 | G | N7 | AA:1216 | 3.94 | 1.00 | 66.90 |  |
| 6 | U | N3 | AA:1234 | 4.09 | 1.00 | 73.78 |  |
| 7 | G | N2 | AA:1235 | 4.22 | 1.00 | 62.37 |  |
| 8 | G | N7 | AA:1215 | 4.50 | 1.00 | 72.04 |  |
| 9 | G | N1 | AA:1235 | 4.54 | 1.00 | 64.00 |  |
| 10 | G | N1 | AA:1216 | 4.65 | 1.00 | 66.84 |  |
| 11 | A | OP2 | AA:1214 | 4.91 | 1.00 | 62.63 |  |
  |

  
 Coded contact list:   
 Exclusion zone contact(s): /O4/O6/O6/  
Consider K in place of MG (2 valid contact(s) for K vs. 0 for MG)  

### MG:AA:3671

|  | Res. type | Atom | Residue | Distance | Occupancy | B-factor | Closest solute to 1^st^ shell water |
| --- | --- | --- | --- | --- | --- | --- | --- |
| **Ion** | | | |  | 1.00 | 102.68 || 1 | C | N4 | AA:1076 | 2.66 | 1.00 | 200.13 |
| 2 | A | N6 | AA:1077 | 3.10 | 1.00 | 198.03 |  |
| 3 | C | N4 | AA:1075 | 3.32 | 1.00 | 200.15 |  |
| 4 | G | O6 | AA:1062 | 3.69 | 1.00 | 200.41 |  |
| 5 | A | N1 | AA:1077 | 4.30 | 1.00 | 198.06 |  |
| 6 | G | OP2 | AA:1089 | 4.33 | 1.00 | 200.33 |  |
| 7 | C | N3 | AA:1076 | 4.94 | 1.00 | 200.26 |  |
  |

  
 Coded contact list:   
 Exclusion zone contact(s): /N4/N4/N6/  
Consider K in place of MG (3 valid contact(s) for K vs. 0 for MG)  

### MG:AA:3672

|  | Res. type | Atom | Residue | Distance | Occupancy | B-factor | Closest solute to 1^st^ shell water |
| --- | --- | --- | --- | --- | --- | --- | --- |
| **Ion** | | | |  | 1.00 | 55.15 || 1 | G | O6 | AA:24 | 2.82 | 1.00 | 52.60 |
| 2 | U | O4 | AA:25 | 3.85 | 1.00 | 58.91 |  |
| 3 | C | N4 | AA:516 | 4.05 | 1.00 | 51.30 |  |
| 4 | G | O6 | AA:23 | 4.28 | 1.00 | 52.92 |  |
| 5 | G | N7 | AA:24 | 4.30 | 1.00 | 47.95 |  |
| 6 | A | OP2 | AA:514 | 4.44 | 1.00 | 58.76 |  |
| 7 | U | N3 | AA:25 | 4.86 | 1.00 | 56.28 |  |
| 8 | G | N7 | AA:23 | 4.97 | 1.00 | 55.36 |  |
| 9 | A | OP1 | AA:513 | 4.98 | 1.00 | 58.29 |  |
  |

  
 Coded contact list:   
 Exclusion zone contact(s): /O6/  
Consider K in place of MG (1 valid contact(s) for K vs. 0 for MG)  

### MG:AA:3673

|  | Res. type | Atom | Residue | Distance | Occupancy | B-factor | Closest solute to 1^st^ shell water |
| --- | --- | --- | --- | --- | --- | --- | --- |
| **Ion** | | | |  | 1.00 | 56.57 || 1 | G | N7 | AA:2428 | 3.48 | 1.00 | 41.85 |
| 2 | G | O6 | AA:2428 | 3.70 | 1.00 | 41.11 |  |
| 3 | C | OP1 | AA:825 | 4.14 | 1.00 | 55.90 |  |
| 4 | C | OP1 | AA:2260 | 4.21 | 1.00 | 56.26 |  |
  |

  
 Coded contact list:   
1^st^ MG coordination shell missing (see Notes)  

### MG:AA:3674

|  | Res. type | Atom | Residue | Distance | Occupancy | B-factor | Closest solute to 1^st^ shell water |
| --- | --- | --- | --- | --- | --- | --- | --- |
| **Ion** | | | |  | 1.00 | 61.07 || 1 | G | O6 | AA:2643 | 3.49 | 1.00 | 74.40 |
| 2 | G | O6 | AA:2642 | 4.01 | 1.00 | 69.25 |  |
| 3 | C | N4 | AA:2771 | 4.38 | 1.00 | 76.30 |  |
| 4 | G | N7 | AA:2642 | 4.47 | 1.00 | 68.16 |  |
| 5 | G | O6 | AA:2644 | 4.49 | 1.00 | 69.50 |  |
| 6 | C | OP2 | AA:2769 | 4.75 | 1.00 | 94.03 |  |
| 7 | G | N7 | AA:2643 | 4.91 | 1.00 | 68.39 |  |
  |

  
 Coded contact list:   
1^st^ MG coordination shell missing (see Notes)  

### MG:AA:3675

|  | Res. type | Atom | Residue | Distance | Occupancy | B-factor | Closest solute to 1^st^ shell water |
| --- | --- | --- | --- | --- | --- | --- | --- |
| **Ion** | | | |  | 1.00 | 71.00 || 1 | G | O6 | AA:1184 | 2.94 | 1.00 | 81.01 |
| 2 | G | O6 | AA:1164 | 3.04 | 1.00 | 76.74 |  |
| 3 | U | O4 | AA:1165 | 3.12 | 1.00 | 84.36 |  |
| 4 | C | N4 | AA:1185 | 3.26 | 1.00 | 77.37 |  |
| 5 | G | O6 | AA:1183 | 3.40 | 1.00 | 97.54 |  |
| 6 | U | N3 | AA:1165 | 4.26 | 1.00 | 84.41 |  |
| 7 | G | N7 | AA:1184 | 4.36 | 1.00 | 80.05 |  |
| 8 | G | N7 | AA:1183 | 4.54 | 1.00 | 96.74 |  |
| 9 | G | N1 | AA:1164 | 4.64 | 1.00 | 76.30 |  |
| 10 | C | N3 | AA:1185 | 4.77 | 1.00 | 76.48 |  |
| 11 | C | N4 | AA:1166 | 4.79 | 1.00 | 96.89 |  |
| 12 | G | N1 | AA:1183 | 4.83 | 1.00 | 97.53 |  |
  |

  
 Coded contact list:   
 Exclusion zone contact(s): /N4/O4/O6/O6/O6/  
Consider K in place of MG (5 valid contact(s) for K vs. 0 for MG)  

### MG:AA:3676

|  | Res. type | Atom | Residue | Distance | Occupancy | B-factor | Closest solute to 1^st^ shell water |
| --- | --- | --- | --- | --- | --- | --- | --- |
| **Ion** | | | |  | 1.00 | 81.27 || 1 | C | OP2 | AA:1551 | 3.41 | 1.00 | 64.68 |
| 2 | G | N7 | AA:1552 | 3.82 | 1.00 | 69.49 |  |
| 3 | G | OP2 | AA:1552 | 4.24 | 1.00 | 64.31 |  |
| 4 | C | OP1 | AA:1551 | 4.68 | 1.00 | 65.85 |  |
| 5 | G | O6 | AA:1552 | 4.75 | 1.00 | 68.85 |  |
| 6 | C | O5' | AA:1551 | 4.99 | 1.00 | 64.61 |  |
  |

  
 Coded contact list:   
1^st^ MG coordination shell missing (see Notes)  

### MG:AA:3677

|  | Res. type | Atom | Residue | Distance | Occupancy | B-factor | Closest solute to 1^st^ shell water |
| --- | --- | --- | --- | --- | --- | --- | --- |
| **Ion** | | | |  | 1.00 | 60.18 || 1 | G | O6 | AA:700 | 2.91 | 1.00 | 44.05 |
| 2 | G | O6 | AA:701 | 3.09 | 1.00 | 47.91 |  |
| 3 | G | N7 | AA:700 | 3.72 | 1.00 | 47.81 |  |
| 4 | G | N7 | AA:701 | 3.87 | 1.00 | 48.73 |  |
| 5 | C | N4 | AA:732 | 4.61 | 1.00 | 43.56 |  |
| 6 | G | N2 | AA:733 | 4.85 | 1.00 | 40.25 |  |
| 7 | A | OP2 | AA:699 | 4.86 | 1.00 | 41.88 |  |
| 8 | C | N4 | AA:731 | 4.90 | 1.00 | 46.10 |  |
| 9 | G | N1 | AA:700 | 4.95 | 1.00 | 43.87 |  |
  |

  
 Coded contact list:   
 Exclusion zone contact(s): /O6/O6/  
Consider K in place of MG (2 valid contact(s) for K vs. 0 for MG)  

### MG:AA:3678

|  | Res. type | Atom | Residue | Distance | Occupancy | B-factor | Closest solute to 1^st^ shell water |
| --- | --- | --- | --- | --- | --- | --- | --- |
| **Ion** | | | |  | 1.00 | 49.08 || 1 | G | O6 | AA:132 | 2.72 | 1.00 | 56.47 |
| 2 | G | O6 | AA:131 | 2.89 | 1.00 | 47.92 |  |
| 3 | U | O4 | AA:147 | 2.90 | 1.00 | 55.00 |  |
| 4 | G | O6 | AA:146 | 3.51 | 1.00 | 62.76 |  |
| 5 | C | N4 | AA:148 | 3.83 | 1.00 | 45.11 |  |
| 6 | U | N3 | AA:147 | 3.95 | 1.00 | 55.43 |  |
| 7 | C | N4 | AA:133 | 4.26 | 1.00 | 60.37 |  |
| 8 | G | N1 | AA:131 | 4.42 | 1.00 | 46.07 |  |
| 9 | G | N7 | AA:132 | 4.46 | 1.00 | 56.23 |  |
| 10 | G | N7 | AA:131 | 4.69 | 1.00 | 44.51 |  |
| 11 | G | N1 | AA:132 | 4.98 | 1.00 | 55.82 |  |
  |

  
 Coded contact list:   
 Exclusion zone contact(s): /O4/O6/O6/  
Consider K in place of MG (3 valid contact(s) for K vs. 0 for MG)  

### MG:AA:3679

|  | Res. type | Atom | Residue | Distance | Occupancy | B-factor | Closest solute to 1^st^ shell water |
| --- | --- | --- | --- | --- | --- | --- | --- |
| **Ion** | | | |  | 1.00 | 70.67 || 1 | U | O4 | AA:63 | 4.09 | 1.00 | 76.46 |
| 2 | G | OP2 | AA:88 | 4.46 | 1.00 | 67.49 |  |
| 3 | G | OP2 | AA:89 | 4.51 | 1.00 | 68.76 |  |
| 4 | G | N7 | AA:89 | 4.52 | 1.00 | 71.72 |  |
| 5 | G | OP1 | AA:88 | 4.60 | 1.00 | 66.07 |  |
| 6 | U | N3 | AA:63 | 4.65 | 1.00 | 74.28 |  |
| 7 | U | O4' | AA:90 | 4.69 | 1.00 | 104.44 |  |
| 8 | G | N7 | AA:88 | 4.85 | 1.00 | 61.56 |  |
  |

  
 Coded contact list:   
1^st^ MG coordination shell missing (see Notes)  

### MG:AA:3680

|  | Res. type | Atom | Residue | Distance | Occupancy | B-factor | Closest solute to 1^st^ shell water |
| --- | --- | --- | --- | --- | --- | --- | --- |
| **Ion** | | | |  | 1.00 | 71.62 || 1 | A | N7 | AA:1046 | 2.69 | 1.00 | 190.80 |
| 2 | A | N6 | AA:1046 | 4.02 | 1.00 | 189.93 |  |
| 3 | A | N9 | AA:1046 | 4.76 | 1.00 | 190.56 |  |
  |

  
 Coded contact list:   
 Exclusion zone contact(s): /N7/  
Consider K in place of MG (1 valid contact(s) for K vs. 0 for MG)  

### MG:AA:3681

|  | Res. type | Atom | Residue | Distance | Occupancy | B-factor | Closest solute to 1^st^ shell water |
| --- | --- | --- | --- | --- | --- | --- | --- |
| **Ion** | | | |  | 1.00 | 65.28 || 1 | G | O6 | AA:1443 | 3.37 | 1.00 | 73.86 |
| 2 | G | O6 | AA:1442 | 3.42 | 1.00 | 71.82 |  |
| 3 | C | N4 | AA:1549 | 4.27 | 1.00 | 71.64 |  |
| 4 | C | N4 | AA:1548 | 4.37 | 1.00 | 75.71 |  |
| 5 | C | OP2 | AA:1467 | 4.53 | 1.00 | 77.22 |  |
| 6 | G | N7 | AA:1443 | 4.70 | 1.00 | 71.90 |  |
| 7 | G | O6 | AA:1444 | 4.76 | 1.00 | 81.21 |  |
| 8 | G | N7 | AA:1442 | 4.84 | 1.00 | 68.67 |  |
  |

  
 Coded contact list:   
 Exclusion zone contact(s): /O6/  
Consider K in place of MG (1 valid contact(s) for K vs. 0 for MG)  

### MG:AA:3682

|  | Res. type | Atom | Residue | Distance | Occupancy | B-factor | Closest solute to 1^st^ shell water |
| --- | --- | --- | --- | --- | --- | --- | --- |
| **Ion** | | | |  | 1.00 | 75.62 || 1 | U | N1 | AA:1454 | 3.23 | 1.00 | 72.81 |
| 2 | A | O3' | AA:1453 | 3.48 | 1.00 | 71.62 |  |
| 3 | A | O2' | AA:1453 | 3.58 | 1.00 | 74.92 |  |
| 4 | U | O4 | AA:2702 | 3.82 | 1.00 | 72.32 |  |
| 5 | U | O5' | AA:1454 | 3.95 | 1.00 | 71.45 |  |
| 6 | U | O2' | AA:1454 | 3.96 | 1.00 | 73.89 |  |
| 7 | U | O3' | AA:1454 | 4.05 | 1.00 | 70.30 |  |
| 8 | U | O4' | AA:1454 | 4.11 | 1.00 | 71.67 |  |
| 9 | A | N9 | AA:1453 | 4.29 | 1.00 | 74.19 |  |
| 10 | U | OP2 | AA:1454 | 4.38 | 1.00 | 71.19 |  |
| 11 | A | O4' | AA:1453 | 4.43 | 1.00 | 73.48 |  |
| 12 | A | O5' | AA:1453 | 4.45 | 1.00 | 78.97 |  |
| 13 | U | N3 | AA:1454 | 4.46 | 1.00 | 73.75 |  |
| 14 | U | O2 | AA:1454 | 4.59 | 1.00 | 72.33 |  |
| 15 | G | O6 | AA:1455 | 4.86 | 1.00 | 75.30 |  |
  |

  
 Coded contact list:   
 Exclusion zone contact(s): /N1/  
Consider K in place of MG (1 valid contact(s) for K vs. 0 for MG)  

### MG:AA:3683

|  | Res. type | Atom | Residue | Distance | Occupancy | B-factor | Closest solute to 1^st^ shell water |
| --- | --- | --- | --- | --- | --- | --- | --- |
| **Ion** | | | |  | 1.00 | 46.03 || 1 | G | OP2 | AA:2822 | 2.04 | 1.00 | 46.61 |
| 2 | G | N7 | AA:2822 | 2.75 | 1.00 | 39.58 |  |
| 3 | GLY | O | AE:110 | 2.99 | 1.00 | 58.34 |  |
| 4 | A | OP2 | AA:2821 | 3.16 | 1.00 | 45.16 |  |
| 5 | HIS | ND1 | A0:3 | 3.27 | 1.00 | 45.10 |  |
| 6 | HIS | NE2 | A0:3 | 3.61 | 1.00 | 44.84 |  |
| 7 | G | O5' | AA:2822 | 3.74 | 1.00 | 45.80 |  |
| 8 | A | O3' | AA:2820 | 4.31 | 1.00 | 45.75 |  |
| 9 | A | O3' | AA:2821 | 4.31 | 1.00 | 48.40 |  |
| 10 | A | O2' | AA:2820 | 4.32 | 1.00 | 48.19 |  |
| 11 | G | OP1 | AA:2822 | 4.47 | 1.00 | 45.57 |  |
| 12 | G | N9 | AA:2822 | 4.52 | 1.00 | 41.02 |  |
| 13 | GLY | N | AE:112 | 4.53 | 1.00 | 59.27 |  |
| 14 | A | O5' | AA:2821 | 4.74 | 1.00 | 45.19 |  |
| 15 | G | O6 | AA:2822 | 4.75 | 1.00 | 38.44 |  |
  |

  
 Contact(s): /OP2/  
 Coded contact list: O_ph_  
 Exclusion zone contact(s): /N7/ND1_His_/O_back_/OP2/  
Consider K in place of MG (4 valid contact(s) for K vs. 1 for MG)  

### MG:AA:3684

|  | Res. type | Atom | Residue | Distance | Occupancy | B-factor | Closest solute to 1^st^ shell water |
| --- | --- | --- | --- | --- | --- | --- | --- |
| **Ion** | | | |  | 1.00 | 58.83 || 1 | G | OP2 | AA:684 | 4.88 | 1.00 | 31.17 |
  |

  
 Coded contact list:   
1^st^ MG coordination shell missing (see Notes)  

### MG:AA:3685

|  | Res. type | Atom | Residue | Distance | Occupancy | B-factor | Closest solute to 1^st^ shell water |
| --- | --- | --- | --- | --- | --- | --- | --- |
| **Ion** | | | |  | 1.00 | 65.43 || 1 | G | O6 | AA:175 | 4.03 | 1.00 | 68.22 |
| 2 | G | O6 | AA:176 | 4.41 | 1.00 | 58.35 |  |
| 3 | G | N7 | AA:175 | 4.47 | 1.00 | 71.45 |  |
  |

  
 Coded contact list:   
1^st^ MG coordination shell missing (see Notes)  

### MG:AA:3686

|  | Res. type | Atom | Residue | Distance | Occupancy | B-factor | Closest solute to 1^st^ shell water |
| --- | --- | --- | --- | --- | --- | --- | --- |
| **Ion** | | | |  | 1.00 | 49.71 || 1 | G | O6 | AA:2004 | 3.51 | 1.00 | 51.80 |
| 2 | A | N6 | AA:2005 | 3.82 | 1.00 | 54.35 |  |
| 3 | G | O6 | AA:2003 | 3.85 | 1.00 | 45.98 |  |
| 4 | G | N7 | AA:2004 | 4.42 | 1.00 | 50.81 |  |
| 5 | G | O6 | AA:2002 | 4.48 | 1.00 | 38.73 |  |
| 6 | A | N1 | AA:2005 | 4.66 | 1.00 | 54.49 |  |
| 7 | G | N7 | AA:2002 | 4.76 | 1.00 | 40.25 |  |
| 8 | G | N2 | AA:1653 | 4.85 | 1.00 | 53.14 |  |
| 9 | C | N4 | AA:1656 | 4.88 | 1.00 | 53.01 |  |
| 10 | C | N4 | AA:1657 | 4.91 | 1.00 | 46.43 |  |
| 11 | G | N7 | AA:2003 | 4.98 | 1.00 | 48.29 |  |
  |

  
 Coded contact list:   
1^st^ MG coordination shell missing (see Notes)  

### MG:AA:3687

|  | Res. type | Atom | Residue | Distance | Occupancy | B-factor | Closest solute to 1^st^ shell water |
| --- | --- | --- | --- | --- | --- | --- | --- |
| **Ion** | | | |  | 1.00 | 43.53 || 1 | A | OP2 | AA:1213 | 3.17 | 1.00 | 63.97 |
| 2 | G | N9 | AA:1212 | 3.27 | 1.00 | 62.23 |  |
| 3 | G | O3' | AA:1212 | 3.30 | 1.00 | 64.87 |  |
| 4 | G | O6 | AA:1235 | 3.30 | 1.00 | 63.63 |  |
| 5 | G | N2 | AA:1236 | 3.51 | 1.00 | 65.28 |  |
| 6 | G | O5' | AA:1212 | 3.52 | 1.00 | 72.80 |  |
| 7 | G | O4' | AA:1212 | 3.74 | 1.00 | 64.95 |  |
| 8 | G | N1 | AA:1236 | 3.75 | 1.00 | 66.05 |  |
| 9 | G | O2' | AA:1212 | 3.92 | 1.00 | 68.62 |  |
| 10 | G | N7 | AA:1212 | 3.93 | 1.00 | 61.27 |  |
| 11 | G | N1 | AA:1235 | 4.20 | 1.00 | 64.00 |  |
| 12 | G | OP2 | AA:1212 | 4.40 | 1.00 | 74.25 |  |
| 13 | A | OP1 | AA:1213 | 4.79 | 1.00 | 63.62 |  |
| 14 | A | O5' | AA:1213 | 5.00 | 1.00 | 62.21 |  |
  |

  
 Coded contact list:   
 Exclusion zone contact(s): /O3'/O6/OP2/  
Consider K in place of MG (4 valid contact(s) for K vs. 0 for MG)  

### MG:AA:3688

|  | Res. type | Atom | Residue | Distance | Occupancy | B-factor | Closest solute to 1^st^ shell water |
| --- | --- | --- | --- | --- | --- | --- | --- |
| **Ion** | | | |  | 1.00 | 59.51 || 1 | U | O2' | AA:362 | 3.45 | 1.00 | 125.99 |
| 2 | U | O3' | AA:362 | 4.80 | 1.00 | 126.96 |  |
| 3 | G | N2 | AA:361 | 4.98 | 1.00 | 113.28 |  |
  |

  
 Coded contact list:   
1^st^ MG coordination shell missing (see Notes)  

### MG:AA:3689

|  | Res. type | Atom | Residue | Distance | Occupancy | B-factor | Closest solute to 1^st^ shell water |
| --- | --- | --- | --- | --- | --- | --- | --- |
| **Ion** | | | |  | 1.00 | 61.01 || 1 | U | O4 | AA:534 | 2.64 | 1.00 | 69.75 |
| 2 | G | O6 | AA:559 | 2.87 | 1.00 | 69.32 |  |
| 3 | G | O6 | AA:558 | 3.21 | 1.00 | 70.11 |  |
| 4 | G | O6 | AA:533 | 3.39 | 1.00 | 67.47 |  |
| 5 | C | N4 | AA:535 | 3.71 | 1.00 | 67.48 |  |
| 6 | U | N3 | AA:534 | 3.85 | 1.00 | 68.14 |  |
| 7 | C | N4 | AA:560 | 4.54 | 1.00 | 71.10 |  |
| 8 | G | N7 | AA:559 | 4.69 | 1.00 | 70.31 |  |
| 9 | G | N7 | AA:533 | 4.71 | 1.00 | 63.78 |  |
| 10 | G | N1 | AA:558 | 4.95 | 1.00 | 70.05 |  |
  |

  
 Coded contact list:   
 Exclusion zone contact(s): /O4/O6/O6/O6/  
Consider K in place of MG (4 valid contact(s) for K vs. 0 for MG)  

### MG:AA:3690

|  | Res. type | Atom | Residue | Distance | Occupancy | B-factor | Closest solute to 1^st^ shell water |
| --- | --- | --- | --- | --- | --- | --- | --- |
| **Ion** | | | |  | 1.00 | 64.83 || 1 | A | O2' | AA:1815 | 2.34 | 1.00 | 48.34 |
| 2 | G | O4' | AA:1816 | 3.11 | 1.00 | 51.57 |  |
| 3 | G | O5' | AA:1816 | 3.96 | 1.00 | 46.95 |  |
| 4 | A | O3' | AA:1815 | 4.25 | 1.00 | 46.42 |  |
| 5 | G | OP2 | AA:1816 | 4.38 | 1.00 | 44.67 |  |
| 6 | G | OP1 | AA:1813 | 4.52 | 1.00 | 37.76 |  |
| 7 | G | N3 | AA:1816 | 4.63 | 1.00 | 65.78 |  |
| 8 | G | OP2 | AA:1814 | 4.77 | 1.00 | 33.57 |  |
| 9 | A | N3 | AA:1815 | 4.96 | 1.00 | 50.54 |  |
  |

  
 Contact(s): /O2'/  
 Coded contact list: O_r_  
 Exclusion zone contact(s): /O4'/  

### MG:AA:3691

|  | Res. type | Atom | Residue | Distance | Occupancy | B-factor | Closest solute to 1^st^ shell water |
| --- | --- | --- | --- | --- | --- | --- | --- |
| **Ion** | | | |  | 1.00 | 79.96 || 1 | G | O6 | AA:1480 | 2.59 | 1.00 | 92.30 |
| 2 | U | O4 | AA:1482 | 2.62 | 1.00 | 93.37 |  |
| 3 | G | N7 | AA:1480 | 3.34 | 1.00 | 90.58 |  |
| 4 | G | O6 | AA:1479 | 4.16 | 1.00 | 87.52 |  |
| 5 | G | N7 | AA:1479 | 4.31 | 1.00 | 84.07 |  |
| 6 | U | N3 | AA:1482 | 4.32 | 1.00 | 94.91 |  |
| 7 | C | N4 | AA:1513 | 4.45 | 1.00 | 94.67 |  |
| 8 | G | O6 | AA:1512 | 4.59 | 1.00 | 106.45 |  |
| 9 | G | N1 | AA:1480 | 4.76 | 1.00 | 92.38 |  |
| 10 | G | OP2 | AA:1479 | 4.88 | 1.00 | 78.89 |  |
  |

  
 Coded contact list:   
 Exclusion zone contact(s): /N7/O4/O6/  
Consider K in place of MG (2 valid contact(s) for K vs. 0 for MG)  

### MG:AA:3692

|  | Res. type | Atom | Residue | Distance | Occupancy | B-factor | Closest solute to 1^st^ shell water |
| --- | --- | --- | --- | --- | --- | --- | --- |
| **Ion** | | | |  | 1.00 | 85.75 || 1 | GLY | O | AF:134 | 2.59 | 1.00 | 77.79 |
| 2 | G | O5' | AA:321 | 2.74 | 1.00 | 72.05 |  |
| 3 | G | O4' | AA:321 | 2.91 | 1.00 | 74.63 |  |
| 4 | G | N9 | AA:321 | 3.54 | 1.00 | 73.89 |  |
| 5 | G | OP1 | AA:321 | 3.68 | 1.00 | 69.66 |  |
| 6 | G | N7 | AA:321 | 3.72 | 1.00 | 75.57 |  |
| 7 | G | OP2 | AA:321 | 3.98 | 1.00 | 70.25 |  |
| 8 | ARG | NH1 | AF:165 | 4.24 | 1.00 | 76.03 |  |
| 9 | LYS | NZ | AF:135 | 4.35 | 1.00 | 78.28 |  |
| 10 | LYS | N | AF:135 | 4.50 | 1.00 | 76.91 |  |
| 11 | ASN | O | AF:133 | 4.86 | 1.00 | 83.76 |  |
  |

  
 Coded contact list:   
 Exclusion zone contact(s): /O4'/O5'/O_back_/  
Consider K in place of MG (2 valid contact(s) for K vs. 0 for MG)  

### MG:AA:3693

|  | Res. type | Atom | Residue | Distance | Occupancy | B-factor | Closest solute to 1^st^ shell water |
| --- | --- | --- | --- | --- | --- | --- | --- |
| **Ion** | | | |  | 1.00 | 52.64 || 1 | G | O6 | AA:2719 | 2.67 | 1.00 | 69.29 |
| 2 | G | N7 | AA:2719 | 3.40 | 1.00 | 68.69 |  |
| 3 | G | N7 | AA:2718 | 3.99 | 1.00 | 69.59 |  |
| 4 | U | OP2 | AA:2688 | 4.26 | 1.00 | 55.88 |  |
| 5 | G | N1 | AA:2719 | 4.78 | 1.00 | 69.46 |  |
| 6 | U | O2 | AA:2688 | 4.85 | 1.00 | 67.96 |  |
  |

  
 Coded contact list:   
 Exclusion zone contact(s): /N7/O6/  
Consider K in place of MG (2 valid contact(s) for K vs. 0 for MG)  

### MG:AA:3694

|  | Res. type | Atom | Residue | Distance | Occupancy | B-factor | Closest solute to 1^st^ shell water |
| --- | --- | --- | --- | --- | --- | --- | --- |
| **Ion** | | | |  | 1.00 | 56.35 || 1 | G | O6 | AA:270Y | 2.24 | 1.00 | 77.65 |
| 2 | U | OP1 | AA:269 | 2.51 | 1.00 | 67.91 |  |
| 3 | G | N7 | AA:270Y | 3.94 | 1.00 | 78.36 |  |
| 4 | G | OP2 | AA:270X | 4.34 | 1.00 | 85.64 |  |
| 5 | U | O5' | AA:269 | 4.36 | 1.00 | 68.02 |  |
| 6 | G | N7 | AA:270X | 4.47 | 1.00 | 79.16 |  |
| 7 | U | OP2 | AA:269 | 4.48 | 1.00 | 66.66 |  |
| 8 | G | N1 | AA:270Y | 4.51 | 1.00 | 77.75 |  |
| 9 | G | OP2 | AA:270W | 4.76 | 1.00 | 97.77 |  |
| 10 | C | O3' | AA:268 | 4.99 | 1.00 | 65.79 |  |
  |

  
 Contact(s): /O6/  
 Coded contact list: O_b_  
 Exclusion zone contact(s): /OP1/  

### MG:AA:3695

|  | Res. type | Atom | Residue | Distance | Occupancy | B-factor | Closest solute to 1^st^ shell water |
| --- | --- | --- | --- | --- | --- | --- | --- |
| **Ion** | | | |  | 1.00 | 49.89 || 1 | G | N7 | AA:2717 | 3.72 | 1.00 | 64.30 |
| 2 | G | O6 | AA:2719 | 3.89 | 1.00 | 69.29 |  |
| 3 | G | N7 | AA:2718 | 3.94 | 1.00 | 69.59 |  |
| 4 | G | O6 | AA:2717 | 4.34 | 1.00 | 64.46 |  |
| 5 | G | O6 | AA:2718 | 4.60 | 1.00 | 72.03 |  |
  |

  
 Coded contact list:   
1^st^ MG coordination shell missing (see Notes)  

### MG:AA:3696

|  | Res. type | Atom | Residue | Distance | Occupancy | B-factor | Closest solute to 1^st^ shell water |
| --- | --- | --- | --- | --- | --- | --- | --- |
| **Ion** | | | |  | 1.00 | 47.13 || 1 | A | OP1 | AA:2503 | 2.48 | 1.00 | 39.95 |
| 2 | U | OP1 | AA:576 | 2.90 | 1.00 | 47.22 |  |
| 3 | A | OP2 | AA:2503 | 3.29 | 1.00 | 37.97 |  |
| 4 | G | O2' | AA:2502 | 3.52 | 1.00 | 39.42 |  |
| 5 | G | OP2 | AA:577 | 3.68 | 1.00 | 44.81 |  |
| 6 | G | OP1 | AA:577 | 4.09 | 1.00 | 43.15 |  |
| 7 | G | O3' | AA:2502 | 4.18 | 1.00 | 36.89 |  |
| 8 | A | O5' | AA:2503 | 4.62 | 1.00 | 38.68 |  |
| 9 | U | OP2 | AA:576 | 4.78 | 1.00 | 46.55 |  |
| 10 | A | OP1 | AA:2057 | 4.88 | 1.00 | 41.84 |  |
| 11 | U | O3' | AA:576 | 4.94 | 1.00 | 43.85 |  |
| 12 | U | O5' | AA:576 | 4.99 | 1.00 | 44.78 |  |
  |

  
 Coded contact list:   
 Exclusion zone contact(s): /OP1/OP1/OP2/  
Consider K in place of MG (2 valid contact(s) for K vs. 0 for MG)  

### MG:AA:3697

|  | Res. type | Atom | Residue | Distance | Occupancy | B-factor | Closest solute to 1^st^ shell water |
| --- | --- | --- | --- | --- | --- | --- | --- |
| **Ion** | | | |  | 1.00 | 78.81 || 1 | C | OP2 | AA:414 | 4.14 | 1.00 | 65.78 |
| 2 | C | OP1 | AA:414 | 4.60 | 1.00 | 64.88 |  |
| 3 | C | OP2 | AA:409 | 4.92 | 1.00 | 64.29 |  |
  |

  
 Coded contact list:   
1^st^ MG coordination shell missing (see Notes)  

### MG:AA:3698

|  | Res. type | Atom | Residue | Distance | Occupancy | B-factor | Closest solute to 1^st^ shell water |
| --- | --- | --- | --- | --- | --- | --- | --- |
| **Ion** | | | |  | 1.00 | 64.99 || 1 | U | O4 | AA:922 | 2.86 | 1.00 | 86.31 |
| 2 | G | O6 | AA:921 | 2.94 | 1.00 | 84.70 |  |
| 3 | G | O6 | AA:855 | 3.00 | 1.00 | 88.97 |  |
| 4 | C | N4 | AA:856 | 3.51 | 1.00 | 83.50 |  |
| 5 | G | O6 | AA:854 | 3.96 | 1.00 | 87.01 |  |
| 6 | U | N3 | AA:922 | 3.98 | 1.00 | 85.67 |  |
| 7 | G | N7 | AA:855 | 4.66 | 1.00 | 89.54 |  |
| 8 | C | N4 | AA:923 | 4.77 | 1.00 | 87.58 |  |
| 9 | G | N7 | AA:921 | 4.89 | 1.00 | 81.94 |  |
| 10 | G | N1 | AA:921 | 4.91 | 1.00 | 83.12 |  |
  |

  
 Coded contact list:   
 Exclusion zone contact(s): /O4/O6/O6/  
Consider K in place of MG (3 valid contact(s) for K vs. 0 for MG)  

### MG:AA:3699

|  | Res. type | Atom | Residue | Distance | Occupancy | B-factor | Closest solute to 1^st^ shell water |
| --- | --- | --- | --- | --- | --- | --- | --- |
| **Ion** | | | |  | 1.00 | 59.06 || 1 | A | O3' | AA:1393 | 2.64 | 1.00 | 53.62 |
| 2 | U | OP1 | AA:1394 | 2.79 | 1.00 | 52.93 |  |
| 3 | C | OP1 | AA:1604 | 2.86 | 1.00 | 34.85 |  |
| 4 | A | O3' | AA:1603 | 3.53 | 1.00 | 34.35 |  |
| 5 | A | O2' | AA:1393 | 3.74 | 1.00 | 57.40 |  |
| 6 | C | O5' | AA:1604 | 3.86 | 1.00 | 36.94 |  |
| 7 | C | O2' | AA:1314 | 3.88 | 1.00 | 40.99 |  |
| 8 | U | O5' | AA:1394 | 4.19 | 1.00 | 52.27 |  |
| 9 | A | O2' | AA:1603 | 4.34 | 1.00 | 36.14 |  |
| 10 | U | OP2 | AA:1394 | 4.49 | 1.00 | 55.47 |  |
  |

  
 Coded contact list:   
 Exclusion zone contact(s): /O3'/OP1/OP1/  
Consider K in place of MG (3 valid contact(s) for K vs. 0 for MG)  

### MG:AA:3700

|  | Res. type | Atom | Residue | Distance | Occupancy | B-factor | Closest solute to 1^st^ shell water |
| --- | --- | --- | --- | --- | --- | --- | --- |
| **Ion** | | | |  | 1.00 | 58.97 || 1 | G | N7 | AA:463 | 3.56 | 1.00 | 40.25 |
| 2 | G | OP1 | AA:465 | 3.75 | 1.00 | 34.60 |  |
| 3 | G | O6 | AA:463 | 3.83 | 1.00 | 44.93 |  |
| 4 | G | OP2 | AA:465 | 4.17 | 1.00 | 30.50 |  |
| 5 | ARG | NH2 | A7:12 | 4.90 | 1.00 | 42.84 |  |
  |

  
 Coded contact list:   
1^st^ MG coordination shell missing (see Notes)  

### MG:AA:3701

|  | Res. type | Atom | Residue | Distance | Occupancy | B-factor | Closest solute to 1^st^ shell water |
| --- | --- | --- | --- | --- | --- | --- | --- |
| **Ion** | | | |  | 1.00 | 55.59 || 1 | U | O4 | AA:943 | 2.65 | 1.00 | 51.99 |
| 2 | G | O6 | AA:836 | 3.65 | 1.00 | 53.17 |  |
| 3 | G | O6 | AA:942 | 3.89 | 1.00 | 49.39 |  |
| 4 | U | N3 | AA:943 | 4.27 | 1.00 | 51.09 |  |
| 5 | A | N7 | AA:835 | 4.72 | 1.00 | 58.78 |  |
| 6 | A | OP2 | AA:835 | 4.89 | 1.00 | 56.99 |  |
| 7 | C | OP2 | AA:834 | 4.91 | 1.00 | 52.22 |  |
  |

  
 Coded contact list:   
 Exclusion zone contact(s): /O4/  
Consider K in place of MG (1 valid contact(s) for K vs. 0 for MG)  

### MG:AA:3702

|  | Res. type | Atom | Residue | Distance | Occupancy | B-factor | Closest solute to 1^st^ shell water |
| --- | --- | --- | --- | --- | --- | --- | --- |
| **Ion** | | | |  | 1.00 | 62.01 || 1 | C | OP1 | AA:2646 | 2.47 | 1.00 | 73.62 |
| 2 | A | OP1 | AA:2733 | 2.78 | 1.00 | 87.33 |  |
| 3 | C | OP2 | AA:2646 | 3.32 | 1.00 | 73.08 |  |
| 4 | A | O2' | AA:2765 | 3.82 | 1.00 | 79.58 |  |
| 5 | U | OP2 | AA:2647 | 3.90 | 1.00 | 67.12 |  |
| 6 | G | O3' | AA:2645 | 4.30 | 1.00 | 73.79 |  |
| 7 | A | O5' | AA:2733 | 4.50 | 1.00 | 90.03 |  |
| 8 | C | O5' | AA:2646 | 4.59 | 1.00 | 73.11 |  |
| 9 | A | OP2 | AA:2733 | 4.80 | 1.00 | 87.93 |  |
  |

  
 Coded contact list:   
 Exclusion zone contact(s): /OP1/OP1/OP2/  
Consider K in place of MG (2 valid contact(s) for K vs. 0 for MG)  

### MG:AA:3703

|  | Res. type | Atom | Residue | Distance | Occupancy | B-factor | Closest solute to 1^st^ shell water |
| --- | --- | --- | --- | --- | --- | --- | --- |
| **Ion** | | | |  | 1.00 | 51.31 || 1 | G | O6 | AA:939 | 2.59 | 1.00 | 51.98 |
| 2 | U | O4 | AA:839 | 2.92 | 1.00 | 53.86 |  |
| 3 | G | O6 | AA:938 | 3.12 | 1.00 | 55.85 |  |
| 4 | G | O6 | AA:940 | 3.29 | 1.00 | 55.33 |  |
| 5 | C | N4 | AA:840 | 3.93 | 1.00 | 56.24 |  |
| 6 | U | N3 | AA:839 | 3.93 | 1.00 | 53.44 |  |
| 7 | G | N7 | AA:939 | 4.25 | 1.00 | 53.22 |  |
| 8 | C | N4 | AA:838 | 4.28 | 1.00 | 53.28 |  |
| 9 | G | N1 | AA:938 | 4.51 | 1.00 | 56.74 |  |
| 10 | G | N1 | AA:939 | 4.86 | 1.00 | 52.56 |  |
| 11 | G | N7 | AA:940 | 4.91 | 1.00 | 56.30 |  |
| 12 | G | N7 | AA:938 | 4.97 | 1.00 | 57.20 |  |
  |

  
 Coded contact list:   
 Exclusion zone contact(s): /O4/O6/O6/O6/  
Consider K in place of MG (3 valid contact(s) for K vs. 0 for MG)  

### MG:AA:3704

|  | Res. type | Atom | Residue | Distance | Occupancy | B-factor | Closest solute to 1^st^ shell water |
| --- | --- | --- | --- | --- | --- | --- | --- |
| **Ion** | | | |  | 1.00 | 59.72 || 1 | G | OP2 | AA:1456 | 2.77 | 1.00 | 79.27 |
| 2 | A | OP2 | AA:1453 | 3.02 | 1.00 | 80.36 |  |
| 3 | G | N7 | AA:1456 | 3.99 | 1.00 | 77.02 |  |
| 4 | U | O3' | AA:1454 | 4.23 | 1.00 | 70.30 |  |
| 5 | A | O5' | AA:1453 | 4.32 | 1.00 | 78.97 |  |
| 6 | G | O5' | AA:1456 | 4.48 | 1.00 | 80.46 |  |
| 7 | C | O3' | AA:1451 | 4.79 | 1.00 | 82.47 |  |
| 8 | A | N7 | AA:1457 | 4.81 | 1.00 | 81.79 |  |
| 9 | G | OP1 | AA:1456 | 4.81 | 1.00 | 79.78 |  |
  |

  
 Coded contact list:   
 Exclusion zone contact(s): /OP2/OP2/  
Consider K in place of MG (2 valid contact(s) for K vs. 0 for MG)  

### MG:AA:3705

|  | Res. type | Atom | Residue | Distance | Occupancy | B-factor | Closest solute to 1^st^ shell water |
| --- | --- | --- | --- | --- | --- | --- | --- |
| **Ion** | | | |  | 1.00 | 87.10 || 1 | G | O6 | AA:1485 | 3.95 | 1.00 | 106.57 |
| 2 | G | N7 | AA:1485 | 4.59 | 1.00 | 107.99 |  |
| 3 | A | N6 | AA:1486 | 4.75 | 1.00 | 105.14 |  |
| 4 | G | O6 | AA:1484 | 4.92 | 1.00 | 112.19 |  |
  |

  
 Coded contact list:   
1^st^ MG coordination shell missing (see Notes)  

### MG:AA:3706

|  | Res. type | Atom | Residue | Distance | Occupancy | B-factor | Closest solute to 1^st^ shell water |
| --- | --- | --- | --- | --- | --- | --- | --- |
| **Ion** | | | |  | 1.00 | 44.71 || 1 | A | N6 | AA:621 | 2.73 | 1.00 | 81.94 |
| 2 | A | N7 | AA:621 | 2.73 | 1.00 | 77.63 |  |
| 3 | A | N1 | AA:621 | 3.11 | 1.00 | 79.42 |  |
| 4 | G | O4' | AA:622 | 3.27 | 1.00 | 59.76 |  |
| 5 | U | O2' | AA:239 | 3.49 | 1.00 | 55.83 |  |
| 6 | U | O4' | AA:239 | 3.62 | 1.00 | 54.93 |  |
| 7 | A | N9 | AA:621 | 3.71 | 1.00 | 74.94 |  |
| 8 | A | N3 | AA:621 | 3.76 | 1.00 | 76.48 |  |
| 9 | G | O2' | AA:622 | 4.02 | 1.00 | 59.86 |  |
| 10 | G | N9 | AA:622 | 4.06 | 1.00 | 60.10 |  |
| 11 | A | N3 | AA:608 | 4.20 | 1.00 | 65.15 |  |
| 12 | G | N2 | AA:259 | 4.26 | 1.00 | 62.49 |  |
| 13 | C | O2 | AA:238 | 4.45 | 1.00 | 60.16 |  |
| 14 | G | N3 | AA:622 | 4.60 | 1.00 | 59.82 |  |
| 15 | C | O2' | AA:238 | 4.63 | 1.00 | 52.01 |  |
| 16 | A | N1 | AA:608 | 4.79 | 1.00 | 65.31 |  |
| 17 | A | N9 | AA:608 | 4.92 | 1.00 | 64.50 |  |
  |

  
 Coded contact list:   
 Exclusion zone contact(s): /N1_A_/N6/N7/O4'/  
Consider K in place of MG (4 valid contact(s) for K vs. 0 for MG)  

### MG:AA:3707

|  | Res. type | Atom | Residue | Distance | Occupancy | B-factor | Closest solute to 1^st^ shell water |
| --- | --- | --- | --- | --- | --- | --- | --- |
| **Ion** | | | |  | 1.00 | 63.66 || 1 | A | OP1 | AA:282 | 2.79 | 1.00 | 86.93 |
| 2 | A | OP1 | AA:283 | 3.32 | 1.00 | 81.09 |  |
| 3 | A | OP2 | AA:283 | 3.73 | 1.00 | 80.63 |  |
| 4 | A | O5' | AA:282 | 4.31 | 1.00 | 85.38 |  |
| 5 | A | O3' | AA:282 | 4.44 | 1.00 | 80.67 |  |
| 6 | A | OP2 | AA:282 | 4.70 | 1.00 | 87.27 |  |
  |

  
 Coded contact list:   
 Exclusion zone contact(s): /OP1/OP1/  
Consider K in place of MG (2 valid contact(s) for K vs. 0 for MG)  

### MG:AA:3708

|  | Res. type | Atom | Residue | Distance | Occupancy | B-factor | Closest solute to 1^st^ shell water |
| --- | --- | --- | --- | --- | --- | --- | --- |
| **Ion** | | | |  | 1.00 | 49.16 || 1 | C | O2' | AA:2616 | 2.66 | 1.00 | 45.82 |
| 2 | C | O2 | AA:2616 | 2.68 | 1.00 | 43.20 |  |
| 3 | U | O2 | AA:2022 | 2.69 | 1.00 | 53.65 |  |
| 4 | C | O4' | AA:2617 | 2.95 | 1.00 | 50.17 |  |
| 5 | U | O2' | AA:2022 | 3.32 | 1.00 | 57.92 |  |
| 6 | G | N2 | AA:2053 | 3.95 | 1.00 | 39.93 |  |
| 7 | C | N1 | AA:2616 | 4.21 | 1.00 | 43.71 |  |
| 8 | C | N1 | AA:2617 | 4.30 | 1.00 | 46.95 |  |
| 9 | U | N3 | AA:2022 | 4.49 | 1.00 | 53.63 |  |
| 10 | U | N1 | AA:2022 | 4.62 | 1.00 | 53.27 |  |
| 11 | C | O4' | AA:2616 | 4.83 | 1.00 | 46.31 |  |
| 12 | C | O5' | AA:2617 | 4.88 | 1.00 | 50.70 |  |
| 13 | C | N3 | AA:2616 | 4.94 | 1.00 | 41.90 |  |
  |

  
 Coded contact list:   
 Exclusion zone contact(s): /O2/O2/O2'/O2'/O4'/  
Consider K in place of MG (5 valid contact(s) for K vs. 0 for MG)  

### MG:AA:3709

|  | Res. type | Atom | Residue | Distance | Occupancy | B-factor | Closest solute to 1^st^ shell water |
| --- | --- | --- | --- | --- | --- | --- | --- |
| **Ion** | | | |  | 1.00 | 66.65 || 1 | G | N7 | AA:2218 | 3.40 | 1.00 | 54.41 |
| 2 | G | OP2 | AA:2218 | 3.95 | 1.00 | 60.19 |  |
| 3 | G | O6 | AA:2224 | 4.26 | 1.00 | 53.21 |  |
| 4 | G | OP2 | AA:2217 | 4.42 | 1.00 | 66.48 |  |
| 5 | G | N7 | AA:2217 | 4.60 | 1.00 | 61.13 |  |
| 6 | A | OP2 | AA:2198 | 4.67 | 1.00 | 69.77 |  |
| 7 | G | O6 | AA:2218 | 4.69 | 1.00 | 54.07 |  |
  |

  
 Coded contact list:   
1^st^ MG coordination shell missing (see Notes)  

### MG:AA:3710

|  | Res. type | Atom | Residue | Distance | Occupancy | B-factor | Closest solute to 1^st^ shell water |
| --- | --- | --- | --- | --- | --- | --- | --- |
| **Ion** | | | |  | 1.00 | 76.03 || 1 | G | O6 | AA:1863 | 3.10 | 1.00 | 75.65 |
| 2 | G | O6 | AA:1862 | 3.51 | 1.00 | 72.20 |  |
| 3 | U | O4 | AA:1864 | 3.94 | 1.00 | 75.98 |  |
| 4 | G | N7 | AA:1863 | 4.37 | 1.00 | 73.38 |  |
| 5 | G | N7 | AA:1862 | 4.43 | 1.00 | 72.91 |  |
| 6 | C | N4 | AA:1879 | 4.43 | 1.00 | 76.58 |  |
| 7 | G | O6 | AA:1878 | 4.82 | 1.00 | 76.89 |  |
| 8 | C | N4 | AA:1880 | 4.98 | 1.00 | 72.69 |  |
  |

  
 Coded contact list:   
 Exclusion zone contact(s): /O6/  
Consider K in place of MG (1 valid contact(s) for K vs. 0 for MG)  

### MG:AA:3711

|  | Res. type | Atom | Residue | Distance | Occupancy | B-factor | Closest solute to 1^st^ shell water |
| --- | --- | --- | --- | --- | --- | --- | --- |
| **Ion** | | | |  | 1.00 | 69.93 || 1 | G | OP2 | AA:217 | 2.38 | 1.00 | 54.45 |
| 2 | A | OP1 | AA:428 | 3.78 | 1.00 | 58.15 |  |
| 3 | G | O6 | AA:219 | 3.85 | 1.00 | 46.85 |  |
| 4 | G | O5' | AA:217 | 4.08 | 1.00 | 53.33 |  |
| 5 | A | N6 | AA:218 | 4.27 | 1.00 | 51.10 |  |
| 6 | A | N7 | AA:218 | 4.30 | 1.00 | 48.49 |  |
| 7 | G | N7 | AA:217 | 4.36 | 1.00 | 51.47 |  |
| 8 | G | OP1 | AA:217 | 4.49 | 1.00 | 54.82 |  |
| 9 | A | O5' | AA:428 | 4.51 | 1.00 | 58.50 |  |
| 10 | A | OP2 | AA:429 | 4.68 | 1.00 | 61.05 |  |
| 11 | A | OP2 | AA:218 | 4.82 | 1.00 | 51.86 |  |
| 12 | A | O3' | AA:216 | 4.88 | 1.00 | 53.82 |  |
  |

  
 Contact(s): /OP2/  
 Coded contact list: O_ph_  

### MG:AA:3712

|  | Res. type | Atom | Residue | Distance | Occupancy | B-factor | Closest solute to 1^st^ shell water |
| --- | --- | --- | --- | --- | --- | --- | --- |
| **Ion** | | | |  | 1.00 | 57.03 || 1 | U | O4 | AA:1341 | 2.22 | 1.00 | 43.76 |
| 2 | A | N1 | AA:1395 | 3.19 | 1.00 | 51.66 |  |
| 3 | LYS | NZ | AT:16 | 3.68 | 1.00 | 50.28 |  |
| 4 | U | N3 | AA:1394 | 3.83 | 1.00 | 62.50 |  |
| 5 | U | O2 | AA:1394 | 3.91 | 1.00 | 58.93 |  |
| 6 | A | N6 | AA:1395 | 3.94 | 1.00 | 51.94 |  |
| 7 | U | N3 | AA:1341 | 4.01 | 1.00 | 42.34 |  |
| 8 | U | O4 | AA:1394 | 4.85 | 1.00 | 63.59 |  |
  |

  
 Contact(s): /O4/  
 Coded contact list: O_b_  
 Exclusion zone contact(s): /N1_A_/  

### MG:AA:3713

|  | Res. type | Atom | Residue | Distance | Occupancy | B-factor | Closest solute to 1^st^ shell water |
| --- | --- | --- | --- | --- | --- | --- | --- |
| **Ion** | | | |  | 1.00 | 83.25 || 1 | U | OP2 | AA:120 | 2.34 | 1.00 | 57.75 |
| 2 | G | OP1 | AA:121 | 3.30 | 1.00 | 59.67 |  |
| 3 | A | O2' | AA:119 | 3.87 | 1.00 | 57.17 |  |
| 4 | G | O6 | AA:122 | 3.97 | 1.00 | 44.48 |  |
| 5 | U | OP1 | AA:120 | 4.13 | 1.00 | 57.79 |  |
| 6 | U | O5' | AA:120 | 4.41 | 1.00 | 56.96 |  |
| 7 | G | N7 | AA:122 | 4.51 | 1.00 | 47.05 |  |
| 8 | U | O3' | AA:120 | 4.60 | 1.00 | 60.28 |  |
| 9 | A | O3' | AA:119 | 4.84 | 1.00 | 58.01 |  |
| 10 | G | N7 | AA:121 | 4.88 | 1.00 | 48.07 |  |
| 11 | G | O6 | AA:123 | 4.95 | 1.00 | 41.69 |  |
  |

  
 Contact(s): /OP2/  
 Coded contact list: O_ph_  
 Exclusion zone contact(s): /OP1/  

### MG:AA:3714

|  | Res. type | Atom | Residue | Distance | Occupancy | B-factor | Closest solute to 1^st^ shell water |
| --- | --- | --- | --- | --- | --- | --- | --- |
| **Ion** | | | |  | 1.00 | 70.93 || 1 | G | O6 | AA:1620 | 2.78 | 1.00 | 40.31 |
| 2 | G | O6 | AA:1619 | 2.91 | 1.00 | 44.10 |  |
| 3 | G | N7 | AA:1619 | 3.56 | 1.00 | 43.23 |  |
| 4 | G | N7 | AA:1620 | 3.73 | 1.00 | 43.26 |  |
| 5 | G | OP2 | AA:1619 | 4.36 | 1.00 | 43.37 |  |
| 6 | C | N4 | AA:1611 | 4.47 | 1.00 | 39.78 |  |
| 7 | G | N1 | AA:1619 | 4.55 | 1.00 | 42.20 |  |
| 8 | U | O4 | AA:1621 | 4.56 | 1.00 | 43.03 |  |
| 9 | C | N4 | AA:1612 | 4.61 | 1.00 | 41.96 |  |
| 10 | G | N1 | AA:1620 | 4.97 | 1.00 | 41.71 |  |
  |

  
 Coded contact list:   
 Exclusion zone contact(s): /O6/O6/  
Consider K in place of MG (2 valid contact(s) for K vs. 0 for MG)  

### MG:AA:3715

|  | Res. type | Atom | Residue | Distance | Occupancy | B-factor | Closest solute to 1^st^ shell water |
| --- | --- | --- | --- | --- | --- | --- | --- |
| **Ion** | | | |  | 1.00 | 47.51 || 1 | G | O3' | AA:668 | 2.77 | 1.00 | 61.49 |
| 2 | G | OP2 | AA:669 | 3.15 | 1.00 | 63.35 |  |
| 3 | G | O5' | AA:669 | 3.24 | 1.00 | 62.09 |  |
| 4 | G | O5' | AA:668 | 3.99 | 1.00 | 55.71 |  |
| 5 | U | OP1 | AA:803 | 4.21 | 1.00 | 41.04 |  |
| 6 | G | O4' | AA:669 | 4.37 | 1.00 | 55.44 |  |
| 7 | G | OP1 | AA:669 | 4.73 | 1.00 | 62.20 |  |
| 8 | A | OP2 | AA:670 | 4.80 | 1.00 | 53.96 |  |
  |

  
 Coded contact list:   
 Exclusion zone contact(s): /O3'/O5'/OP2/  
Consider K in place of MG (3 valid contact(s) for K vs. 0 for MG)  

### MG:AA:3716

|  | Res. type | Atom | Residue | Distance | Occupancy | B-factor | Closest solute to 1^st^ shell water |
| --- | --- | --- | --- | --- | --- | --- | --- |
| **Ion** | | | |  | 1.00 | 74.12 || 1 | C | OP2 | AA:2825 | 4.72 | 1.00 | 51.79 |
| 2 | G | O6 | AA:2819 | 4.84 | 1.00 | 40.33 |  |
  |

  
 Coded contact list:   
1^st^ MG coordination shell missing (see Notes)  

### MG:AA:3717

|  | Res. type | Atom | Residue | Distance | Occupancy | B-factor | Closest solute to 1^st^ shell water |
| --- | --- | --- | --- | --- | --- | --- | --- |
| **Ion** | | | |  | 1.00 | 106.73 || 1 | G | O6 | AA:418 | 3.00 | 1.00 | 63.82 |
| 2 | C | N4 | AA:417 | 3.91 | 1.00 | 65.17 |  |
| 3 | C | N4 | AA:409 | 4.28 | 1.00 | 62.19 |  |
| 4 | G | N7 | AA:418 | 4.33 | 1.00 | 66.10 |  |
  |

  
 Coded contact list:   
 Exclusion zone contact(s): /O6/  
Consider K in place of MG (1 valid contact(s) for K vs. 0 for MG)  

### MG:AA:3718

|  | Res. type | Atom | Residue | Distance | Occupancy | B-factor | Closest solute to 1^st^ shell water |
| --- | --- | --- | --- | --- | --- | --- | --- |
| **Ion** | | | |  | 1.00 | 84.39 || 1 | A | OP1 | AA:1871 | 2.73 | 1.00 | 87.15 |
| 2 | G | N7 | AA:1869 | 3.16 | 1.00 | 80.00 |  |
| 3 | G | O6 | AA:1869 | 3.96 | 1.00 | 78.09 |  |
| 4 | A | OP2 | AA:1871 | 4.22 | 1.00 | 87.40 |  |
| 5 | C | O3' | AA:1870 | 4.71 | 1.00 | 86.29 |  |
| 6 | U | O4 | AA:1864 | 4.87 | 1.00 | 75.98 |  |
  |

  
 Coded contact list:   
 Exclusion zone contact(s): /N7/OP1/  
Consider K in place of MG (2 valid contact(s) for K vs. 0 for MG)  

### MG:AA:3720

|  | Res. type | Atom | Residue | Distance | Occupancy | B-factor | Closest solute to 1^st^ shell water |
| --- | --- | --- | --- | --- | --- | --- | --- |
| **Ion** | | | |  | 1.00 | 52.18 || 1 | G | N9 | AA:1377 | 2.71 | 1.00 | 49.55 |
| 2 | A | N6 | AA:1353 | 2.99 | 1.00 | 49.42 |  |
| 3 | A | N7 | AA:1353 | 3.00 | 1.00 | 49.47 |  |
| 4 | G | N3 | AA:1377 | 3.10 | 1.00 | 49.86 |  |
| 5 | G | O2' | AA:1377 | 3.38 | 1.00 | 49.22 |  |
| 6 | U | O4 | AA:1352 | 3.43 | 1.00 | 44.51 |  |
| 7 | G | N7 | AA:1377 | 3.75 | 1.00 | 51.09 |  |
| 8 | A | N7 | AA:1378 | 4.03 | 1.00 | 48.80 |  |
| 9 | A | N6 | AA:1354 | 4.15 | 1.00 | 43.66 |  |
| 10 | G | O4' | AA:1377 | 4.29 | 1.00 | 49.14 |  |
| 11 | U | N3 | AA:1352 | 4.30 | 1.00 | 42.00 |  |
| 12 | G | N1 | AA:1377 | 4.39 | 1.00 | 51.12 |  |
| 13 | G | O3' | AA:1377 | 4.40 | 1.00 | 47.72 |  |
| 14 | A | O5' | AA:1378 | 4.46 | 1.00 | 49.96 |  |
| 15 | A | N9 | AA:1378 | 4.66 | 1.00 | 50.81 |  |
| 16 | A | O4' | AA:1378 | 4.68 | 1.00 | 54.40 |  |
| 17 | A | OP2 | AA:1378 | 4.75 | 1.00 | 49.21 |  |
| 18 | G | N2 | AA:1377 | 4.77 | 1.00 | 51.41 |  |
  |

  
 Coded contact list:   
 Exclusion zone contact(s): /N3_G_/N6/N7/O2'/  
Consider K in place of MG (5 valid contact(s) for K vs. 0 for MG)  

### MG:AA:3721

|  | Res. type | Atom | Residue | Distance | Occupancy | B-factor | Closest solute to 1^st^ shell water |
| --- | --- | --- | --- | --- | --- | --- | --- |
| **Ion** | | | |  | 1.00 | 82.93 || 1 | A | N1 | AA:2705 | 2.64 | 1.00 | 69.09 |
| 2 | U | N3 | AA:1454 | 2.74 | 1.00 | 73.75 |  |
| 3 | U | O2 | AA:1454 | 3.31 | 1.00 | 72.33 |  |
| 4 | ARG | NE | A0:64 | 3.55 | 1.00 | 58.00 |  |
| 5 | U | O4 | AA:1454 | 3.71 | 1.00 | 74.00 |  |
| 6 | G | N3 | AA:2706 | 3.74 | 1.00 | 53.61 |  |
| 7 | G | N2 | AA:2706 | 4.09 | 1.00 | 55.60 |  |
| 8 | A | N6 | AA:2705 | 4.18 | 1.00 | 67.64 |  |
| 9 | A | N3 | AA:2705 | 4.37 | 1.00 | 67.02 |  |
| 10 | U | O4 | AA:2702 | 4.50 | 1.00 | 72.32 |  |
| 11 | ARG | NH1 | A0:64 | 4.70 | 1.00 | 56.95 |  |
| 12 | U | N1 | AA:1454 | 4.78 | 1.00 | 72.81 |  |
| 13 | G | N9 | AA:2706 | 4.90 | 1.00 | 56.07 |  |
| 14 | G | O6 | AA:1455 | 4.92 | 1.00 | 75.30 |  |
  |

  
 Coded contact list:   
 Exclusion zone contact(s): /N1_A_/N3_U_/O2/  
Consider K in place of MG (3 valid contact(s) for K vs. 0 for MG)  

### MG:AA:3722

|  | Res. type | Atom | Residue | Distance | Occupancy | B-factor | Closest solute to 1^st^ shell water |
| --- | --- | --- | --- | --- | --- | --- | --- |
| **Ion** | | | |  | 1.00 | 50.03 || 1 | A | N1 | AA:2821 | 3.44 | 1.00 | 35.97 |
| 2 | G | N2 | AA:2822 | 3.93 | 1.00 | 35.12 |  |
| 3 | G | N7 | AA:2819 | 4.00 | 1.00 | 40.92 |  |
| 4 | G | O6 | AA:2819 | 4.43 | 1.00 | 40.33 |  |
| 5 | A | N6 | AA:2821 | 4.62 | 1.00 | 37.43 |  |
| 6 | G | N1 | AA:2822 | 4.73 | 1.00 | 36.06 |  |
| 7 | G | OP1 | AA:1653 | 4.78 | 1.00 | 45.74 |  |
| 8 | G | N3 | AA:2822 | 4.79 | 1.00 | 35.60 |  |
  |

  
 Coded contact list:   
1^st^ MG coordination shell missing (see Notes)  

### MG:AA:3724

|  | Res. type | Atom | Residue | Distance | Occupancy | B-factor | Closest solute to 1^st^ shell water |
| --- | --- | --- | --- | --- | --- | --- | --- |
| **Ion** | | | |  | 1.00 | 65.32 || 1 | A | N1 | AA:1634 | 3.68 | 1.00 | 33.29 |
| 2 | G | OP2 | AA:1624 | 4.21 | 1.00 | 42.94 |  |
| 3 | G | O6 | AA:1626 | 4.32 | 1.00 | 48.97 |  |
| 4 | A | O2' | AA:735 | 4.44 | 1.00 | 44.49 |  |
| 5 | A | N3 | AA:1634 | 4.77 | 1.00 | 35.00 |  |
  |

  
 Coded contact list:   
1^st^ MG coordination shell missing (see Notes)  

### MG:AA:3725

|  | Res. type | Atom | Residue | Distance | Occupancy | B-factor | Closest solute to 1^st^ shell water |
| --- | --- | --- | --- | --- | --- | --- | --- |
| **Ion** | | | |  | 1.00 | 89.64 || 1 | A | OP2 | AA:1331 | 2.72 | 1.00 | 48.84 |
| 2 | G | O6 | AA:1319 | 2.84 | 1.00 | 58.88 |  |
| 3 | C | N4 | AA:1318 | 2.93 | 1.00 | 61.15 |  |
| 4 | G | N7 | AA:1319 | 3.50 | 1.00 | 62.11 |  |
| 5 | G | O6 | AA:1334 | 4.42 | 1.00 | 61.56 |  |
| 6 | A | OP1 | AA:1331 | 4.42 | 1.00 | 47.68 |  |
| 7 | C | OP2 | AA:1330 | 4.43 | 1.00 | 51.72 |  |
| 8 | C | N4 | AA:1333 | 4.55 | 1.00 | 58.82 |  |
| 9 | A | O5' | AA:1331 | 4.82 | 1.00 | 48.83 |  |
| 10 | G | N1 | AA:1319 | 4.97 | 1.00 | 58.74 |  |
  |

  
 Coded contact list:   
 Exclusion zone contact(s): /N4/O6/OP2/  
Consider K in place of MG (3 valid contact(s) for K vs. 0 for MG)  

### MG:AA:3726

|  | Res. type | Atom | Residue | Distance | Occupancy | B-factor | Closest solute to 1^st^ shell water |
| --- | --- | --- | --- | --- | --- | --- | --- |
| **Ion** | | | |  | 1.00 | 60.84 || 1 | G | OP1 | AA:2872 | 2.69 | 1.00 | 68.45 |
| 2 | C | O5' | AA:2691 | 3.14 | 1.00 | 65.69 |  |
| 3 | C | OP1 | AA:2691 | 3.36 | 1.00 | 67.50 |  |
| 4 | C | OP2 | AA:2691 | 3.47 | 1.00 | 66.31 |  |
| 5 | C | OP1 | AA:2692 | 3.80 | 1.00 | 66.89 |  |
| 6 | C | O3' | AA:2691 | 4.22 | 1.00 | 65.79 |  |
| 7 | G | O5' | AA:2872 | 4.38 | 1.00 | 67.00 |  |
| 8 | C | OP1 | AA:2871 | 4.39 | 1.00 | 67.36 |  |
| 9 | C | O3' | AA:2871 | 4.40 | 1.00 | 68.68 |  |
| 10 | C | OP2 | AA:2692 | 4.41 | 1.00 | 67.02 |  |
  |

  
 Coded contact list:   
 Exclusion zone contact(s): /O5'/OP1/OP1/  
Consider K in place of MG (3 valid contact(s) for K vs. 0 for MG)  

### MG:AA:3727

|  | Res. type | Atom | Residue | Distance | Occupancy | B-factor | Closest solute to 1^st^ shell water |
| --- | --- | --- | --- | --- | --- | --- | --- |
| **Ion** | | | |  | 1.00 | 54.45 || 1 | G | N7 | AA:98 | 3.59 | 1.00 | 58.73 |
| 2 | G | OP2 | AA:98 | 4.34 | 1.00 | 59.89 |  |
| 3 | C | OP2 | AA:97 | 4.60 | 1.00 | 67.79 |  |
| 4 | C | N4 | AA:97 | 4.69 | 1.00 | 58.57 |  |
| 5 | G | O6 | AA:98 | 4.80 | 1.00 | 57.96 |  |
  |

  
 Coded contact list:   
1^st^ MG coordination shell missing (see Notes)  

### MG:AA:3728

|  | Res. type | Atom | Residue | Distance | Occupancy | B-factor | Closest solute to 1^st^ shell water |
| --- | --- | --- | --- | --- | --- | --- | --- |
| **Ion** | | | |  | 1.00 | 70.26 || 1 | A | O2' | AA:390 | 2.82 | 1.00 | 51.92 |
| 2 | C | N4 | AA:392 | 3.39 | 1.00 | 55.68 |  |
| 3 | A | O3' | AA:390 | 3.39 | 1.00 | 52.46 |  |
| 4 | G | OP2 | AA:391 | 3.86 | 1.00 | 51.00 |  |
| 5 | G | O6 | AA:388 | 4.13 | 1.00 | 52.03 |  |
| 6 | G | N7 | AA:391 | 4.26 | 1.00 | 53.78 |  |
| 7 | G | O6 | AA:382 | 4.72 | 1.00 | 55.65 |  |
| 8 | G | O5' | AA:391 | 4.73 | 1.00 | 51.69 |  |
  |

  
 Coded contact list:   
 Exclusion zone contact(s): /N4/O2'/O3'/  
Consider K in place of MG (3 valid contact(s) for K vs. 0 for MG)  

### MG:AA:3729

|  | Res. type | Atom | Residue | Distance | Occupancy | B-factor | Closest solute to 1^st^ shell water |
| --- | --- | --- | --- | --- | --- | --- | --- |
| **Ion** | | | |  | 1.00 | 69.11 || 1 | U | O2' | AA:1142 | 3.47 | 1.00 | 98.10 |
| 2 | G | N7 | AA:1016 | 4.30 | 1.00 | 116.66 |  |
| 3 | G | O6 | AA:1017 | 4.30 | 1.00 | 108.15 |  |
| 4 | G | N7 | AA:1017 | 4.30 | 1.00 | 111.23 |  |
| 5 | G | O6 | AA:1016 | 4.53 | 1.00 | 116.31 |  |
| 6 | C | N4 | AA:1018 | 4.82 | 1.00 | 104.61 |  |
| 7 | A | OP2 | AA:1142A | 4.93 | 1.00 | 100.08 |  |
  |

  
 Coded contact list:   
1^st^ MG coordination shell missing (see Notes)  

### MG:AA:3730

|  | Res. type | Atom | Residue | Distance | Occupancy | B-factor | Closest solute to 1^st^ shell water |
| --- | --- | --- | --- | --- | --- | --- | --- |
| **Ion** | | | |  | 1.00 | 58.12 || 1 | U | OP1 | AA:1397 | 4.12 | 1.00 | 60.85 |
| 2 | U | O3' | AA:1397 | 4.20 | 1.00 | 62.92 |  |
| 3 | C | OP2 | AA:1398 | 4.42 | 1.00 | 62.84 |  |
| 4 | G | N7 | AA:1344 | 4.86 | 1.00 | 63.59 |  |
  |

  
 Coded contact list:   
1^st^ MG coordination shell missing (see Notes)  

### MG:AA:3731

|  | Res. type | Atom | Residue | Distance | Occupancy | B-factor | Closest solute to 1^st^ shell water |
| --- | --- | --- | --- | --- | --- | --- | --- |
| **Ion** | | | |  | 1.00 | 79.73 || 1 | G | O6 | AA:1512 | 2.37 | 1.00 | 106.45 |
| 2 | U | O4 | AA:1482 | 2.39 | 1.00 | 93.37 |  |
| 3 | G | O6 | AA:1480 | 3.18 | 1.00 | 92.30 |  |
| 4 | U | N3 | AA:1482 | 3.23 | 1.00 | 94.91 |  |
| 5 | C | N4 | AA:1513 | 4.01 | 1.00 | 94.67 |  |
| 6 | G | N7 | AA:1512 | 4.12 | 1.00 | 107.95 |  |
| 7 | A | N7 | AA:1511 | 4.26 | 1.00 | 121.89 |  |
| 8 | A | OP2 | AA:1511 | 4.37 | 1.00 | 131.37 |  |
| 9 | G | N1 | AA:1512 | 4.63 | 1.00 | 107.33 |  |
  |

  
**1^st^ shell ligand angles:**   
1-2: 96;   
  
 Contact(s): /O4/O6/  
 Coded contact list: 2O_b_  
 Exclusion zone contact(s): /N3_U_/O6/  

### MG:AA:3732

|  | Res. type | Atom | Residue | Distance | Occupancy | B-factor | Closest solute to 1^st^ shell water |
| --- | --- | --- | --- | --- | --- | --- | --- |
| **Ion** | | | |  | 1.00 | 76.20 || 1 | C | N4 | AA:291 | 4.57 | 1.00 | 76.62 |
| 2 | A | OP2 | AA:289 | 4.92 | 1.00 | 92.62 |  |
| 3 | A | OP2 | AA:346 | 4.94 | 1.00 | 82.15 |  |
  |

  
 Coded contact list:   
1^st^ MG coordination shell missing (see Notes)  

### MG:AA:3733

|  | Res. type | Atom | Residue | Distance | Occupancy | B-factor | Closest solute to 1^st^ shell water |
| --- | --- | --- | --- | --- | --- | --- | --- |
| **Ion** | | | |  | 1.00 | 72.01 || 1 | G | O6 | AA:1488 | 3.12 | 1.00 | 77.78 |
| 2 | U | O4 | AA:1489 | 3.63 | 1.00 | 78.56 |  |
| 3 | G | O6 | AA:1487 | 3.90 | 1.00 | 93.45 |  |
| 4 | C | N4 | AA:1501 | 4.22 | 1.00 | 74.60 |  |
| 5 | G | N7 | AA:1488 | 4.57 | 1.00 | 81.09 |  |
| 6 | G | N7 | AA:1487 | 4.62 | 1.00 | 94.55 |  |
| 7 | C | OP2 | AA:1498 | 4.79 | 1.00 | 79.02 |  |
| 8 | G | O6 | AA:1500 | 4.84 | 1.00 | 75.74 |  |
  |

  
 Coded contact list:   
 Exclusion zone contact(s): /O6/  
Consider K in place of MG (1 valid contact(s) for K vs. 0 for MG)  

### MG:AA:3734

|  | Res. type | Atom | Residue | Distance | Occupancy | B-factor | Closest solute to 1^st^ shell water |
| --- | --- | --- | --- | --- | --- | --- | --- |
| **Ion** | | | |  | 1.00 | 49.07 || 1 | G | O6 | AA:254 | 2.89 | 1.00 | 36.95 |
| 2 | G | N9 | AA:242 | 3.20 | 1.00 | 46.99 |  |
| 3 | U | OP2 | AA:243 | 3.20 | 1.00 | 52.73 |  |
| 4 | THR | N | A8:6 | 3.21 | 1.00 | 51.15 |  |
| 5 | THR | OG1 | A8:6 | 3.37 | 1.00 | 51.58 |  |
| 6 | MET | O | A8:4 | 3.44 | 1.00 | 49.57 |  |
| 7 | G | O3' | AA:242 | 3.47 | 1.00 | 52.73 |  |
| 8 | G | O2' | AA:242 | 3.48 | 1.00 | 54.09 |  |
| 9 | G | O4' | AA:242 | 3.73 | 1.00 | 49.40 |  |
| 10 | LYS | N | A8:5 | 4.04 | 1.00 | 49.98 |  |
| 11 | G | N7 | AA:242 | 4.13 | 1.00 | 44.77 |  |
| 12 | G | N1 | AA:254 | 4.21 | 1.00 | 37.50 |  |
| 13 | U | OP1 | AA:243 | 4.69 | 1.00 | 55.36 |  |
| 14 | LYS | O | A8:5 | 4.86 | 1.00 | 48.59 |  |
  |

  
 Coded contact list:   
 Exclusion zone contact(s): /N_back_/O6/OG1_Thr_/OP2/  
Consider K in place of MG (5 valid contact(s) for K vs. 0 for MG)  

### MG:AA:3735

|  | Res. type | Atom | Residue | Distance | Occupancy | B-factor | Closest solute to 1^st^ shell water |
| --- | --- | --- | --- | --- | --- | --- | --- |
| **Ion** | | | |  | 1.00 | 50.13 || 1 | U | OP1 | AA:2017 | 2.82 | 1.00 | 40.66 |
| 2 | U | OP2 | AA:2017 | 3.09 | 1.00 | 41.38 |  |
| 3 | A | OP1 | AA:2058 | 3.17 | 1.00 | 43.88 |  |
| 4 | A | OP2 | AA:578 | 3.81 | 1.00 | 46.41 |  |
| 5 | U | O3' | AA:2016 | 4.38 | 1.00 | 41.60 |  |
| 6 | A | OP1 | AA:578 | 4.39 | 1.00 | 46.98 |  |
| 7 | U | O5' | AA:2017 | 4.69 | 1.00 | 42.63 |  |
| 8 | U | O2' | AA:1255 | 4.98 | 1.00 | 44.93 |  |
| 9 | U | OP1 | AA:2016 | 5.00 | 1.00 | 43.64 |  |
  |

  
 Coded contact list:   
 Exclusion zone contact(s): /OP1/OP1/OP2/  
Consider K in place of MG (3 valid contact(s) for K vs. 0 for MG)  

### MG:AA:3736

|  | Res. type | Atom | Residue | Distance | Occupancy | B-factor | Closest solute to 1^st^ shell water |
| --- | --- | --- | --- | --- | --- | --- | --- |
| **Ion** | | | |  | 1.00 | 72.03 || 1 | A | OP2 | AA:2059 | 2.43 | 1.00 | 45.39 |
| 2 | U | O2 | AA:1255 | 3.44 | 1.00 | 47.33 |  |
| 3 | A | OP1 | AA:2059 | 3.61 | 1.00 | 46.42 |  |
| 4 | GLY | O | AF:71 | 4.22 | 1.00 | 48.95 |  |
| 5 | A | OP1 | AA:2058 | 4.26 | 1.00 | 43.88 |  |
| 6 | A | O5' | AA:2059 | 4.26 | 1.00 | 44.59 |  |
| 7 | U | OP1 | AA:2016 | 4.62 | 1.00 | 43.64 |  |
| 8 | U | O2' | AA:1255 | 4.67 | 1.00 | 44.93 |  |
| 9 | A | O5' | AA:2058 | 4.72 | 1.00 | 43.47 |  |
| 10 | A | O3' | AA:2058 | 4.77 | 1.00 | 44.52 |  |
  |

  
 Coded contact list:   
 Exclusion zone contact(s): /OP2/  
Consider NA in place of MG (1 valid contact(s) for NA vs. 0 for MG)  

### MG:AA:3737

|  | Res. type | Atom | Residue | Distance | Occupancy | B-factor | Closest solute to 1^st^ shell water |
| --- | --- | --- | --- | --- | --- | --- | --- |
| **Ion** | | | |  | 1.00 | 55.25 || 1 | G | OP2 | AA:508 | 2.90 | 1.00 | 59.12 |
| 2 | A | O2' | AA:507 | 3.12 | 1.00 | 51.12 |  |
| 3 | A | O3' | AA:507 | 3.47 | 1.00 | 55.36 |  |
| 4 | G | O5' | AA:508 | 3.80 | 1.00 | 58.86 |  |
  |

  
 Coded contact list:   
 Exclusion zone contact(s): /O2'/OP2/  
Consider K in place of MG (2 valid contact(s) for K vs. 0 for MG)  

### MG:AA:3738

|  | Res. type | Atom | Residue | Distance | Occupancy | B-factor | Closest solute to 1^st^ shell water |
| --- | --- | --- | --- | --- | --- | --- | --- |
| **Ion** | | | |  | 1.00 | 66.18 || 1 | U | O2 | AA:1391 | 2.68 | 1.00 | 67.19 |
| 2 | LYS | NZ | AT:16 | 3.30 | 1.00 | 50.28 |  |
| 3 | U | O2 | AA:1394 | 3.46 | 1.00 | 58.93 |  |
| 4 | U | N3 | AA:1394 | 3.48 | 1.00 | 62.50 |  |
| 5 | U | N1 | AA:1394 | 3.66 | 1.00 | 59.22 |  |
| 6 | U | O4' | AA:1394 | 3.68 | 1.00 | 54.85 |  |
| 7 | U | O5' | AA:1394 | 3.99 | 1.00 | 52.27 |  |
| 8 | G | O2' | AA:1339 | 4.02 | 1.00 | 48.69 |  |
| 9 | A | N7 | AA:1393 | 4.38 | 1.00 | 56.32 |  |
| 10 | A | N9 | AA:1393 | 4.53 | 1.00 | 56.20 |  |
| 11 | U | N3 | AA:1391 | 4.55 | 1.00 | 67.72 |  |
| 12 | G | O3' | AA:1339 | 4.66 | 1.00 | 46.10 |  |
| 13 | U | N1 | AA:1391 | 4.70 | 1.00 | 66.96 |  |
| 14 | U | O4 | AA:1394 | 4.73 | 1.00 | 63.59 |  |
| 15 | U | OP2 | AA:1394 | 4.86 | 1.00 | 55.47 |  |
  |

  
 Coded contact list:   
 Exclusion zone contact(s): /NZ_Lys_/O2/  
Consider K in place of MG (2 valid contact(s) for K vs. 0 for MG)  

### MG:AA:3739

|  | Res. type | Atom | Residue | Distance | Occupancy | B-factor | Closest solute to 1^st^ shell water |
| --- | --- | --- | --- | --- | --- | --- | --- |
| **Ion** | | | |  | 1.00 | 86.99 || 1 | C | OP2 | AA:77 | 3.76 | 1.00 | 57.38 |
| 2 | A | N7 | AA:78 | 4.82 | 1.00 | 61.70 |  |
| 3 | A | OP2 | AA:78 | 4.86 | 1.00 | 62.34 |  |
  |

  
 Coded contact list:   
1^st^ MG coordination shell missing (see Notes)  

### MG:AA:3740

|  | Res. type | Atom | Residue | Distance | Occupancy | B-factor | Closest solute to 1^st^ shell water |
| --- | --- | --- | --- | --- | --- | --- | --- |
| **Ion** | | | |  | 1.00 | 60.00 || 1 | C | OP2 | AA:375 | 3.08 | 1.00 | 59.73 |
| 2 | C | OP1 | AA:375 | 4.19 | 1.00 | 60.19 |  |
| 3 | C | OP2 | AA:393 | 4.37 | 1.00 | 53.47 |  |
| 4 | A | OP2 | AA:394 | 4.41 | 1.00 | 51.67 |  |
  |

  
 Coded contact list:   
 Exclusion zone contact(s): /OP2/  
Consider K in place of MG (1 valid contact(s) for K vs. 0 for MG)  

### MG:AA:3741

|  | Res. type | Atom | Residue | Distance | Occupancy | B-factor | Closest solute to 1^st^ shell water |
| --- | --- | --- | --- | --- | --- | --- | --- |
| **Ion** | | | |  | 1.00 | 63.24 || 1 | G | O6 | AA:1229 | 4.57 | 1.00 | 73.11 |
| 2 | G | N7 | AA:1229 | 4.72 | 1.00 | 70.72 |  |
  |

  
 Coded contact list:   
1^st^ MG coordination shell missing (see Notes)  

### MG:AA:3742

|  | Res. type | Atom | Residue | Distance | Occupancy | B-factor | Closest solute to 1^st^ shell water |
| --- | --- | --- | --- | --- | --- | --- | --- |
| **Ion** | | | |  | 1.00 | 70.52 || 1 | G | O6 | AA:259 | 2.85 | 1.00 | 60.90 |
| 2 | G | O6 | AA:258 | 3.04 | 1.00 | 55.00 |  |
| 3 | G | N7 | AA:258 | 3.63 | 1.00 | 55.67 |  |
| 4 | C | N4 | AA:238 | 4.20 | 1.00 | 56.78 |  |
| 5 | G | N7 | AA:259 | 4.21 | 1.00 | 62.18 |  |
| 6 | U | O4 | AA:239 | 4.36 | 1.00 | 56.02 |  |
| 7 | A | OP2 | AA:257 | 4.56 | 1.00 | 54.40 |  |
| 8 | G | N2 | AA:240 | 4.63 | 1.00 | 45.28 |  |
| 9 | U | N3 | AA:239 | 4.94 | 1.00 | 54.13 |  |
| 10 | G | N1 | AA:259 | 4.98 | 1.00 | 60.56 |  |
| 11 | G | N1 | AA:258 | 4.99 | 1.00 | 55.12 |  |
  |

  
 Coded contact list:   
 Exclusion zone contact(s): /O6/O6/  
Consider K in place of MG (2 valid contact(s) for K vs. 0 for MG)  

### MG:AA:3743

|  | Res. type | Atom | Residue | Distance | Occupancy | B-factor | Closest solute to 1^st^ shell water |
| --- | --- | --- | --- | --- | --- | --- | --- |
| **Ion** | | | |  | 1.00 | 71.16 || 1 | G | O6 | AA:1642 | 3.48 | 1.00 | 54.95 |
| 2 | G | N7 | AA:1642 | 3.56 | 1.00 | 58.05 |  |
| 3 | A | OP2 | AA:1641 | 3.56 | 1.00 | 50.32 |  |
| 4 | G | O6 | AA:1643 | 3.70 | 1.00 | 56.44 |  |
| 5 | G | N7 | AA:1643 | 4.19 | 1.00 | 58.65 |  |
  |

  
 Coded contact list:   
1^st^ MG coordination shell missing (see Notes)  

### MG:AA:3744

|  | Res. type | Atom | Residue | Distance | Occupancy | B-factor | Closest solute to 1^st^ shell water |
| --- | --- | --- | --- | --- | --- | --- | --- |
| **Ion** | | | |  | 1.00 | 64.63 || 1 | G | N7 | AA:932 | 3.52 | 1.00 | 82.67 |
| 2 | C | O3' | AA:816 | 3.88 | 1.00 | 62.45 |  |
| 3 | C | O2' | AA:816 | 4.02 | 1.00 | 65.08 |  |
| 4 | G | N3 | AA:1184 | 4.14 | 1.00 | 79.53 |  |
| 5 | G | N2 | AA:1184 | 4.21 | 1.00 | 80.64 |  |
| 6 | C | O2 | AA:1166 | 4.41 | 1.00 | 97.45 |  |
| 7 | C | O2' | AA:1166 | 4.49 | 1.00 | 97.74 |  |
| 8 | G | O2' | AA:1184 | 4.59 | 1.00 | 77.63 |  |
| 9 | C | OP1 | AA:817 | 4.69 | 1.00 | 61.98 |  |
| 10 | G | O6 | AA:932 | 4.93 | 1.00 | 83.54 |  |
  |

  
 Coded contact list:   
1^st^ MG coordination shell missing (see Notes)  

### MG:AA:3745

|  | Res. type | Atom | Residue | Distance | Occupancy | B-factor | Closest solute to 1^st^ shell water |
| --- | --- | --- | --- | --- | --- | --- | --- |
| **Ion** | | | |  | 1.00 | 57.89 || 1 | U | O4 | AA:2075 | 2.95 | 1.00 | 35.29 |
| 2 | G | N7 | AA:2238 | 3.18 | 1.00 | 38.77 |  |
| 3 | G | N9 | AA:2238 | 3.73 | 1.00 | 37.09 |  |
| 4 | U | O4 | AA:2074 | 4.11 | 1.00 | 30.11 |  |
| 5 | A | N6 | AA:2077 | 4.52 | 1.00 | 35.50 |  |
| 6 | G | O2' | AA:2238 | 4.72 | 1.00 | 38.20 |  |
| 7 | U | O4 | AA:2243 | 4.98 | 1.00 | 33.32 |  |
| 8 | U | N3 | AA:2074 | 4.99 | 1.00 | 30.30 |  |
  |

  
 Coded contact list:   
 Exclusion zone contact(s): /N7/O4/  
Consider K in place of MG (2 valid contact(s) for K vs. 0 for MG)  

### MG:AA:3746

|  | Res. type | Atom | Residue | Distance | Occupancy | B-factor | Closest solute to 1^st^ shell water |
| --- | --- | --- | --- | --- | --- | --- | --- |
| **Ion** | | | |  | 1.00 | 66.04 || 1 | G | OP2 | AA:1047 | 2.86 | 1.00 | 187.21 |
| 2 | G | OP1 | AA:1107 | 3.60 | 1.00 | 190.07 |  |
| 3 | G | OP1 | AA:1106 | 4.13 | 1.00 | 194.69 |  |
| 4 | G | OP2 | AA:1107 | 4.46 | 1.00 | 190.05 |  |
| 5 | G | OP1 | AA:1047 | 4.49 | 1.00 | 187.20 |  |
| 6 | G | O5' | AA:1047 | 4.94 | 1.00 | 186.86 |  |
| 7 | G | O3' | AA:1106 | 4.99 | 1.00 | 190.74 |  |
  |

  
 Coded contact list:   
 Exclusion zone contact(s): /OP2/  
Consider K in place of MG (1 valid contact(s) for K vs. 0 for MG)  

### MG:AA:3747

|  | Res. type | Atom | Residue | Distance | Occupancy | B-factor | Closest solute to 1^st^ shell water |
| --- | --- | --- | --- | --- | --- | --- | --- |
| **Ion** | | | |  | 1.00 | 53.11 || 1 | LYS | O | AZ:48 | 2.73 | 1.00 | 55.89 |
| 2 | A | O3' | AA:2199 | 2.82 | 1.00 | 48.54 |  |
| 3 | A | O2' | AA:2199 | 2.90 | 1.00 | 52.01 |  |
| 4 | C | OP1 | AA:2205 | 3.20 | 1.00 | 48.99 |  |
| 5 | U | OP1 | AA:2092 | 4.14 | 1.00 | 54.13 |  |
| 6 | C | O5' | AA:2205 | 4.23 | 1.00 | 48.51 |  |
| 7 | ARG | N | AZ:50 | 4.40 | 1.00 | 59.95 |  |
| 8 | VAL | N | AZ:49 | 4.41 | 1.00 | 55.21 |  |
| 9 | C | OP2 | AA:2205 | 4.95 | 1.00 | 49.15 |  |
  |

  
 Coded contact list:   
 Exclusion zone contact(s): /O2'/O3'/O_back_/OP1/  
Consider K in place of MG (4 valid contact(s) for K vs. 0 for MG)  

### MG:AA:3748

|  | Res. type | Atom | Residue | Distance | Occupancy | B-factor | Closest solute to 1^st^ shell water |
| --- | --- | --- | --- | --- | --- | --- | --- |
| **Ion** | | | |  | 1.00 | 84.86 || 1 | G | O6 | AA:1440 | 2.65 | 1.00 | 55.58 |
| 2 | G | O6 | AA:1441 | 2.74 | 1.00 | 60.81 |  |
| 3 | G | N7 | AA:1441 | 2.76 | 1.00 | 59.19 |  |
| 4 | G | N7 | AA:1440 | 3.19 | 1.00 | 57.08 |  |
| 5 | G | N1 | AA:1440 | 4.15 | 1.00 | 57.40 |  |
| 6 | G | N2 | AA:1552 | 4.61 | 1.00 | 70.63 |  |
| 7 | G | N1 | AA:1441 | 4.61 | 1.00 | 59.56 |  |
| 8 | C | N4 | AA:1551 | 4.68 | 1.00 | 55.95 |  |
| 9 | G | N9 | AA:1440 | 4.87 | 1.00 | 56.93 |  |
| 10 | G | N9 | AA:1441 | 4.87 | 1.00 | 57.61 |  |
  |

  
 Coded contact list:   
 Exclusion zone contact(s): /N7/N7/O6/O6/  
Consider K in place of MG (4 valid contact(s) for K vs. 0 for MG)  

### MG:AA:3749

|  | Res. type | Atom | Residue | Distance | Occupancy | B-factor | Closest solute to 1^st^ shell water |
| --- | --- | --- | --- | --- | --- | --- | --- |
| **Ion** | | | |  | 1.00 | 33.11 || 1 | ARG | NH1 | AD:48 | 2.10 | 1.00 | 47.59 |
| 2 | G | OP1 | AA:778 | 2.69 | 1.00 | 33.95 |  |
| 3 | ARG | NE | AD:48 | 4.18 | 1.00 | 41.09 |  |
| 4 | G | O5' | AA:778 | 4.19 | 1.00 | 31.78 |  |
| 5 | C | O4' | AA:1806 | 4.22 | 1.00 | 38.97 |  |
| 6 | A | O3' | AA:777 | 4.24 | 1.00 | 32.50 |  |
| 7 | ARG | NH2 | AD:48 | 4.30 | 1.00 | 48.20 |  |
| 8 | C | O3' | AA:1806 | 4.33 | 1.00 | 41.28 |  |
| 9 | C | O2' | AA:1806 | 4.63 | 1.00 | 39.85 |  |
| 10 | U | OP2 | AA:779 | 4.79 | 1.00 | 33.29 |  |
| 11 | ILE | O | AD:49 | 4.81 | 1.00 | 40.91 |  |
| 12 | C | O5' | AA:1806 | 4.98 | 1.00 | 37.22 |  |
  |

  
 Contact(s): /NH1_Arg_/  
 Coded contact list: Anion  
 Exclusion zone contact(s): /OP1/  

### MG:AA:3750

|  | Res. type | Atom | Residue | Distance | Occupancy | B-factor | Closest solute to 1^st^ shell water |
| --- | --- | --- | --- | --- | --- | --- | --- |
| **Ion** | | | |  | 1.00 | 66.86 || 1 | G | OP2 | AA:680 | 2.80 | 1.00 | 39.43 |
| 2 | G | OP2 | AA:776 | 4.25 | 1.00 | 42.08 |  |
| 3 | G | N7 | AA:680 | 4.39 | 1.00 | 38.50 |  |
| 4 | G | OP1 | AA:776 | 4.46 | 1.00 | 41.89 |  |
| 5 | G | O3' | AA:775 | 4.72 | 1.00 | 41.60 |  |
| 6 | G | O5' | AA:680 | 4.72 | 1.00 | 37.93 |  |
  |

  
 Coded contact list:   
 Exclusion zone contact(s): /OP2/  
Consider K in place of MG (1 valid contact(s) for K vs. 0 for MG)  

### MG:AA:3751

|  | Res. type | Atom | Residue | Distance | Occupancy | B-factor | Closest solute to 1^st^ shell water |
| --- | --- | --- | --- | --- | --- | --- | --- |
| **Ion** | | | |  | 1.00 | 48.61 || 1 | THR | OG1 | AO:32 | 2.64 | 1.00 | 53.71 |
| 2 | A | O3' | AA:941 | 3.04 | 1.00 | 54.04 |  |
| 3 | G | OP1 | AA:1191 | 3.23 | 1.00 | 52.30 |  |
| 4 | A | O2' | AA:941 | 3.34 | 1.00 | 54.91 |  |
| 5 | THR | O | AO:32 | 3.59 | 1.00 | 49.57 |  |
| 6 | A | O5' | AA:941 | 4.00 | 1.00 | 57.07 |  |
| 7 | G | O3' | AA:1190 | 4.03 | 1.00 | 53.17 |  |
| 8 | G | O2' | AA:940 | 4.14 | 1.00 | 57.76 |  |
| 9 | G | OP1 | AA:942 | 4.22 | 1.00 | 52.09 |  |
| 10 | ARG | NH2 | AO:18 | 4.35 | 1.00 | 84.63 |  |
| 11 | A | O4' | AA:941 | 4.47 | 1.00 | 54.23 |  |
| 12 | G | O3' | AA:940 | 4.66 | 1.00 | 58.56 |  |
  |

  
 Coded contact list:   
 Exclusion zone contact(s): /O2'/O3'/OG1_Thr_/OP1/  
Consider K in place of MG (4 valid contact(s) for K vs. 0 for MG)  

### MG:AA:3752

|  | Res. type | Atom | Residue | Distance | Occupancy | B-factor | Closest solute to 1^st^ shell water |
| --- | --- | --- | --- | --- | --- | --- | --- |
| **Ion** | | | |  | 1.00 | 71.78 || 1 | U | O4 | AA:1240 | 2.54 | 1.00 | 55.26 |
| 2 | G | O6 | AA:1206 | 3.15 | 1.00 | 55.19 |  |
| 3 | G | O6 | AA:1239 | 3.20 | 1.00 | 50.05 |  |
| 4 | U | N3 | AA:1240 | 3.91 | 1.00 | 55.56 |  |
| 5 | C | N4 | AA:1207 | 4.25 | 1.00 | 51.57 |  |
| 6 | G | N7 | AA:1206 | 4.84 | 1.00 | 54.12 |  |
| 7 | G | N7 | AA:1239 | 4.88 | 1.00 | 50.32 |  |
  |

  
 Coded contact list:   
 Exclusion zone contact(s): /O4/O6/O6/  
Consider K in place of MG (2 valid contact(s) for K vs. 0 for MG)  

### MG:AA:3753

|  | Res. type | Atom | Residue | Distance | Occupancy | B-factor | Closest solute to 1^st^ shell water |
| --- | --- | --- | --- | --- | --- | --- | --- |
| **Ion** | | | |  | 1.00 | 91.42 || 1 | G | N7 | AA:1047 | 3.60 | 1.00 | 183.51 |
| 2 | A | O3' | AA:1046 | 3.91 | 1.00 | 188.39 |  |
| 3 | A | O2' | AA:1046 | 4.09 | 1.00 | 189.76 |  |
| 4 | A | O2' | AA:1045 | 4.27 | 1.00 | 193.07 |  |
| 5 | A | O4' | AA:1046 | 4.30 | 1.00 | 190.04 |  |
| 6 | G | N9 | AA:1047 | 4.93 | 1.00 | 184.19 |  |
| 7 | G | O6 | AA:1047 | 4.93 | 1.00 | 182.78 |  |
  |

  
 Coded contact list:   
1^st^ MG coordination shell missing (see Notes)  

### MG:AA:3754

|  | Res. type | Atom | Residue | Distance | Occupancy | B-factor | Closest solute to 1^st^ shell water |
| --- | --- | --- | --- | --- | --- | --- | --- |
| **Ion** | | | |  | 1.00 | 73.00 || 1 | G | O6 | AA:2843 | 2.95 | 1.00 | 64.12 |
| 2 | G | N7 | AA:2843 | 3.01 | 1.00 | 67.55 |  |
| 3 | G | N7 | AA:2844 | 3.21 | 1.00 | 71.23 |  |
| 4 | G | O6 | AA:2844 | 3.23 | 1.00 | 72.71 |  |
| 5 | G | N1 | AA:2843 | 4.50 | 1.00 | 64.35 |  |
| 6 | G | O6 | AA:2845 | 4.72 | 1.00 | 70.14 |  |
| 7 | G | N9 | AA:2843 | 4.89 | 1.00 | 67.98 |  |
  |

  
 Coded contact list:   
 Exclusion zone contact(s): /N7/N7/O6/O6/  
Consider K in place of MG (4 valid contact(s) for K vs. 0 for MG)  

### MG:AA:3755

|  | Res. type | Atom | Residue | Distance | Occupancy | B-factor | Closest solute to 1^st^ shell water |
| --- | --- | --- | --- | --- | --- | --- | --- |
| **Ion** | | | |  | 1.00 | 63.52 || 1 | C | OP1 | AA:1499 | 3.70 | 1.00 | 69.45 |
| 2 | C | OP1 | AA:1498 | 3.72 | 1.00 | 77.12 |  |
| 3 | G | OP2 | AA:1485 | 4.59 | 1.00 | 107.18 |  |
  |

  
 Coded contact list:   
1^st^ MG coordination shell missing (see Notes)  

### MG:AA:3756

|  | Res. type | Atom | Residue | Distance | Occupancy | B-factor | Closest solute to 1^st^ shell water |
| --- | --- | --- | --- | --- | --- | --- | --- |
| **Ion** | | | |  | 1.00 | 65.87 || 1 | U | O4 | AA:1489 | 2.93 | 1.00 | 78.56 |
| 2 | G | N7 | AA:1488 | 3.63 | 1.00 | 81.09 |  |
| 3 | G | OP2 | AA:1488 | 4.09 | 1.00 | 88.40 |  |
| 4 | G | O6 | AA:1488 | 4.88 | 1.00 | 77.78 |  |
| 5 | U | N3 | AA:1489 | 4.96 | 1.00 | 80.75 |  |
  |

  
 Coded contact list:   
 Exclusion zone contact(s): /O4/  
Consider K in place of MG (1 valid contact(s) for K vs. 0 for MG)  

### MG:AA:3757

|  | Res. type | Atom | Residue | Distance | Occupancy | B-factor | Closest solute to 1^st^ shell water |
| --- | --- | --- | --- | --- | --- | --- | --- |
| **Ion** | | | |  | 1.00 | 56.36 || 1 | G | O6 | AA:1283 | 2.98 | 1.00 | 69.17 |
| 2 | G | N1 | AA:1283 | 4.29 | 1.00 | 68.01 |  |
| 3 | G | OP1 | AA:1285 | 4.55 | 1.00 | 54.69 |  |
  |

  
 Coded contact list:   
 Exclusion zone contact(s): /O6/  
Consider K in place of MG (1 valid contact(s) for K vs. 0 for MG)  

### MG:AA:3758

|  | Res. type | Atom | Residue | Distance | Occupancy | B-factor | Closest solute to 1^st^ shell water |
| --- | --- | --- | --- | --- | --- | --- | --- |
| **Ion** | | | |  | 1.00 | 65.53 || 1 | A | OP2 | AA:53 | 2.69 | 1.00 | 45.29 |
| 2 | G | N7 | AA:54 | 3.26 | 1.00 | 41.71 |  |
| 3 | A | OP1 | AA:53 | 3.76 | 1.00 | 48.03 |  |
| 4 | A | O5' | AA:53 | 4.08 | 1.00 | 47.21 |  |
| 5 | G | OP2 | AA:54 | 4.13 | 1.00 | 45.67 |  |
| 6 | G | O6 | AA:54 | 4.68 | 1.00 | 37.94 |  |
  |

  
 Coded contact list:   
 Exclusion zone contact(s): /N7/OP2/  
Consider K in place of MG (2 valid contact(s) for K vs. 0 for MG)  

### MG:AA:3759

|  | Res. type | Atom | Residue | Distance | Occupancy | B-factor | Closest solute to 1^st^ shell water |
| --- | --- | --- | --- | --- | --- | --- | --- |
| **Ion** | | | |  | 1.00 | 59.10 || 1 | C | O3' | AA:404 | 2.70 | 1.00 | 92.23 |
| 2 | U | O5' | AA:405 | 2.72 | 1.00 | 93.87 |  |
| 3 | U | OP1 | AA:405 | 2.87 | 1.00 | 94.34 |  |
| 4 | U | O4' | AA:405 | 3.05 | 1.00 | 96.28 |  |
| 5 | U | N1 | AA:405 | 3.62 | 1.00 | 99.58 |  |
| 6 | U | OP2 | AA:405 | 4.46 | 1.00 | 94.06 |  |
| 7 | C | O2' | AA:404 | 4.60 | 1.00 | 91.10 |  |
| 8 | U | N3 | AA:405 | 4.94 | 1.00 | 101.51 |  |
  |

  
 Coded contact list:   
 Exclusion zone contact(s): /O3'/O4'/O5'/OP1/  
Consider K in place of MG (4 valid contact(s) for K vs. 0 for MG)  

### MG:AA:3760

|  | Res. type | Atom | Residue | Distance | Occupancy | B-factor | Closest solute to 1^st^ shell water |
| --- | --- | --- | --- | --- | --- | --- | --- |
| **Ion** | | | |  | 1.00 | 85.32 || 1 | G | OP2 | AA:1633 | 3.62 | 1.00 | 47.92 |
| 2 | C | N4 | AA:1630A | 4.60 | 1.00 | 57.08 |  |
| 3 | G | OP1 | AA:1633 | 4.73 | 1.00 | 48.65 |  |
| 4 | G | N7 | AA:1630 | 4.82 | 1.00 | 52.15 |  |
| 5 | G | OP2 | AA:1630 | 4.89 | 1.00 | 54.56 |  |
| 6 | A | OP1 | AA:1632 | 4.92 | 1.00 | 56.56 |  |
  |

  
 Coded contact list:   
1^st^ MG coordination shell missing (see Notes)  

### MG:AA:3761

|  | Res. type | Atom | Residue | Distance | Occupancy | B-factor | Closest solute to 1^st^ shell water |
| --- | --- | --- | --- | --- | --- | --- | --- |
| **Ion** | | | |  | 1.00 | 40.50 || 1 | A | OP2 | AA:299 | 3.59 | 1.00 | 67.11 |
| 2 | C | OP2 | AA:337 | 4.58 | 1.00 | 57.66 |  |
| 3 | G | N7 | AA:338 | 4.97 | 1.00 | 62.73 |  |
  |

  
 Coded contact list:   
1^st^ MG coordination shell missing (see Notes)  

### MG:AA:3762

|  | Res. type | Atom | Residue | Distance | Occupancy | B-factor | Closest solute to 1^st^ shell water |
| --- | --- | --- | --- | --- | --- | --- | --- |
| **Ion** | | | |  | 1.00 | 63.08 || 1 | C | OP2 | AA:687 | 2.64 | 1.00 | 34.65 |
| 2 | G | O5' | AA:686 | 2.89 | 1.00 | 36.78 |  |
| 3 | G | OP2 | AA:686 | 2.92 | 1.00 | 38.33 |  |
| 4 | A | O3' | AA:685 | 3.25 | 1.00 | 38.00 |  |
| 5 | U | O4 | AA:688 | 3.99 | 1.00 | 39.20 |  |
| 6 | G | O3' | AA:686 | 4.02 | 1.00 | 36.87 |  |
| 7 | A | O2' | AA:685 | 4.25 | 1.00 | 41.51 |  |
| 8 | C | OP1 | AA:687 | 4.60 | 1.00 | 35.89 |  |
| 9 | G | OP1 | AA:686 | 4.67 | 1.00 | 40.30 |  |
  |

  
 Coded contact list:   
 Exclusion zone contact(s): /O3'/O5'/OP2/OP2/  
Consider K in place of MG (4 valid contact(s) for K vs. 0 for MG)  

### MG:AA:3763

|  | Res. type | Atom | Residue | Distance | Occupancy | B-factor | Closest solute to 1^st^ shell water |
| --- | --- | --- | --- | --- | --- | --- | --- |
| **Ion** | | | |  | 1.00 | 48.45 || 1 | A | O5' | AA:1759 | 2.57 | 1.00 | 55.23 |
| 2 | A | OP1 | AA:1759 | 2.95 | 1.00 | 57.19 |  |
| 3 | A | OP2 | AA:1760 | 3.25 | 1.00 | 49.18 |  |
| 4 | A | OP2 | AA:1759 | 3.47 | 1.00 | 57.04 |  |
| 5 | C | O2' | AA:2715 | 3.84 | 1.00 | 52.60 |  |
| 6 | A | O3' | AA:1759 | 4.33 | 1.00 | 49.35 |  |
| 7 | G | O3' | AA:1758 | 4.63 | 1.00 | 57.76 |  |
  |

  
 Coded contact list:   
 Exclusion zone contact(s): /O5'/OP1/OP2/  
Consider K in place of MG (2 valid contact(s) for K vs. 0 for MG)  

### MG:AA:3764

|  | Res. type | Atom | Residue | Distance | Occupancy | B-factor | Closest solute to 1^st^ shell water |
| --- | --- | --- | --- | --- | --- | --- | --- |
| **Ion** | | | |  | 1.00 | 53.72 || 1 | A | OP1 | AA:2042 | 2.01 | 1.00 | 58.09 |
| 2 | U | O3' | AA:2041 | 4.11 | 1.00 | 58.88 |  |
| 3 | A | OP2 | AA:2042 | 4.31 | 1.00 | 56.96 |  |
| 4 | A | O5' | AA:2042 | 4.46 | 1.00 | 57.87 |  |
| 5 | A | OP2 | AA:2778 | 4.59 | 1.00 | 65.41 |  |
  |

  
 Contact(s): /OP1/  
 Coded contact list: O_ph_  

### MG:AA:3765

|  | Res. type | Atom | Residue | Distance | Occupancy | B-factor | Closest solute to 1^st^ shell water |
| --- | --- | --- | --- | --- | --- | --- | --- |
| **Ion** | | | |  | 1.00 | 74.16 || 1 | C | N3 | AA:435 | 3.44 | 1.00 | 66.47 |
| 2 | C | O2 | AA:435 | 3.62 | 1.00 | 66.80 |  |
| 3 | C | N4 | AA:435 | 4.60 | 1.00 | 66.82 |  |
| 4 | C | O4' | AA:436 | 4.98 | 1.00 | 70.57 |  |
  |

  
 Coded contact list:   
1^st^ MG coordination shell missing (see Notes)  

### MG:AA:3766

|  | Res. type | Atom | Residue | Distance | Occupancy | B-factor | Closest solute to 1^st^ shell water |
| --- | --- | --- | --- | --- | --- | --- | --- |
| **Ion** | | | |  | 1.00 | 86.00 || 1 | U | O4 | AA:1205 | 2.80 | 1.00 | 68.61 |
| 2 | A | OP2 | AA:324 | 3.82 | 1.00 | 66.30 |  |
| 3 | U | N3 | AA:1205 | 4.82 | 1.00 | 67.20 |  |
| 4 | VAL | O | AF:173 | 4.87 | 1.00 | 61.61 |  |
  |

  
 Coded contact list:   
 Exclusion zone contact(s): /O4/  
Consider K in place of MG (1 valid contact(s) for K vs. 0 for MG)  

### MG:AA:3767

|  | Res. type | Atom | Residue | Distance | Occupancy | B-factor | Closest solute to 1^st^ shell water |
| --- | --- | --- | --- | --- | --- | --- | --- |
| **Ion** | | | |  | 1.00 | 78.17 || 1 | G | OP2 | AA:411 | 3.08 | 1.00 | 58.08 |
| 2 | U | O2' | AA:2406 | 3.23 | 1.00 | 73.78 |  |
| 3 | G | OP2 | AA:2407 | 3.32 | 1.00 | 69.14 |  |
| 4 | G | O3' | AA:410 | 3.37 | 1.00 | 58.03 |  |
| 5 | G | O2' | AA:410 | 4.11 | 1.00 | 56.51 |  |
| 6 | U | O3' | AA:2406 | 4.73 | 1.00 | 70.37 |  |
| 7 | A | OP2 | AA:227 | 4.79 | 1.00 | 80.90 |  |
| 8 | G | OP1 | AA:411 | 4.80 | 1.00 | 60.32 |  |
  |

  
 Coded contact list:   
 Exclusion zone contact(s): /O2'/O3'/OP2/OP2/  
Consider K in place of MG (4 valid contact(s) for K vs. 0 for MG)  

### MG:AA:3768

|  | Res. type | Atom | Residue | Distance | Occupancy | B-factor | Closest solute to 1^st^ shell water |
| --- | --- | --- | --- | --- | --- | --- | --- |
| **Ion** | | | |  | 1.00 | 79.86 || 1 | A | N7 | AA:1045 | 3.80 | 1.00 | 194.46 |
| 2 | A | N9 | AA:1045 | 3.82 | 1.00 | 194.10 |  |
| 3 | A | O4' | AA:1046 | 4.15 | 1.00 | 190.04 |  |
| 4 | A | O2' | AA:1045 | 4.45 | 1.00 | 193.07 |  |
| 5 | A | O5' | AA:1046 | 4.54 | 1.00 | 191.65 |  |
| 6 | A | O3' | AA:1046 | 4.95 | 1.00 | 188.39 |  |
  |

  
 Coded contact list:   
1^st^ MG coordination shell missing (see Notes)  

### MG:AA:3769

|  | Res. type | Atom | Residue | Distance | Occupancy | B-factor | Closest solute to 1^st^ shell water |
| --- | --- | --- | --- | --- | --- | --- | --- |
| **Ion** | | | |  | 1.00 | 80.94 || 1 | G | O5' | AA:1039 | 2.72 | 1.00 | 149.83 |
| 2 | G | OP2 | AA:1039 | 2.95 | 1.00 | 148.01 |  |
| 3 | G | N7 | AA:1039 | 3.13 | 1.00 | 154.89 |  |
| 4 | G | N9 | AA:1039 | 3.45 | 1.00 | 155.21 |  |
| 5 | G | O4' | AA:1039 | 3.56 | 1.00 | 155.05 |  |
| 6 | C | O3' | AA:1038 | 4.10 | 1.00 | 146.25 |  |
| 7 | C | N1 | AA:1038 | 4.45 | 1.00 | 141.12 |  |
| 8 | G | OP1 | AA:1039 | 4.73 | 1.00 | 147.94 |  |
| 9 | C | OP2 | AA:1040 | 4.80 | 1.00 | 159.23 |  |
  |

  
 Coded contact list:   
 Exclusion zone contact(s): /N7/O5'/OP2/  
Consider K in place of MG (3 valid contact(s) for K vs. 0 for MG)  

### MG:AA:3770

|  | Res. type | Atom | Residue | Distance | Occupancy | B-factor | Closest solute to 1^st^ shell water |
| --- | --- | --- | --- | --- | --- | --- | --- |
| **Ion** | | | |  | 1.00 | 70.29 || 1 | G | OP1 | AA:1731 | 4.27 | 1.00 | 126.56 |
| 2 | G | OP2 | AA:1725 | 4.44 | 1.00 | 107.87 |  |
| 3 | G | OP2 | AA:1726 | 4.51 | 1.00 | 101.78 |  |
| 4 | U | O4 | AA:1727 | 4.84 | 1.00 | 102.06 |  |
| 5 | G | O6 | AA:1728 | 4.89 | 1.00 | 112.09 |  |
  |

  
 Coded contact list:   
1^st^ MG coordination shell missing (see Notes)  

### MG:AA:3771

|  | Res. type | Atom | Residue | Distance | Occupancy | B-factor | Closest solute to 1^st^ shell water |
| --- | --- | --- | --- | --- | --- | --- | --- |
| **Ion** | | | |  | 1.00 | 79.27 || 1 | G | N7 | AA:135 | 3.86 | 1.00 | 71.01 |
| 2 | G | O6 | AA:135 | 4.13 | 1.00 | 64.63 |  |
| 3 | G | O6 | AA:136 | 4.50 | 1.00 | 70.83 |  |
| 4 | C | N4 | AA:134 | 4.72 | 1.00 | 70.68 |  |
| 5 | G | N7 | AA:136 | 4.95 | 1.00 | 69.84 |  |
  |

  
 Coded contact list:   
1^st^ MG coordination shell missing (see Notes)  

### MG:AA:3772

|  | Res. type | Atom | Residue | Distance | Occupancy | B-factor | Closest solute to 1^st^ shell water |
| --- | --- | --- | --- | --- | --- | --- | --- |
| **Ion** | | | |  | 1.00 | 103.45 || 1 | G | OP2 | AA:361 | 2.69 | 1.00 | 107.54 |
| 2 | G | OP1 | AA:360 | 2.79 | 1.00 | 110.29 |  |
| 3 | G | O5' | AA:360 | 3.26 | 1.00 | 109.13 |  |
| 4 | G | OP1 | AA:361 | 3.47 | 1.00 | 108.23 |  |
| 5 | G | O3' | AA:360 | 3.50 | 1.00 | 107.66 |  |
| 6 | G | OP2 | AA:360 | 4.12 | 1.00 | 110.33 |  |
| 7 | G | O5' | AA:361 | 4.82 | 1.00 | 108.00 |  |
| 8 | G | OP2 | AA:270X | 4.85 | 1.00 | 85.64 |  |
| 9 | A | O3' | AA:359 | 4.91 | 1.00 | 110.27 |  |
  |

  
 Coded contact list:   
 Exclusion zone contact(s): /O5'/OP1/OP2/  
Consider K in place of MG (3 valid contact(s) for K vs. 0 for MG)  

### MG:AA:3773

|  | Res. type | Atom | Residue | Distance | Occupancy | B-factor | Closest solute to 1^st^ shell water |
| --- | --- | --- | --- | --- | --- | --- | --- |
| **Ion** | | | |  | 1.00 | 76.37 || 1 | G | O6 | AA:1524 | 3.84 | 1.00 | 89.49 |
| 2 | U | O4 | AA:1523 | 4.26 | 1.00 | 92.22 |  |
| 3 | G | O6 | AA:1525 | 4.43 | 1.00 | 81.05 |  |
| 4 | G | N7 | AA:1524 | 4.73 | 1.00 | 94.70 |  |
  |

  
 Coded contact list:   
1^st^ MG coordination shell missing (see Notes)  

### MG:AA:3774

|  | Res. type | Atom | Residue | Distance | Occupancy | B-factor | Closest solute to 1^st^ shell water |
| --- | --- | --- | --- | --- | --- | --- | --- |
| **Ion** | | | |  | 1.00 | 79.32 || 1 | G | O6 | AA:1429 | 3.93 | 1.00 | 48.28 |
| 2 | G | N7 | AA:1429 | 4.09 | 1.00 | 50.64 |  |
| 3 | C | N4 | AA:1430 | 4.97 | 1.00 | 46.74 |  |
  |

  
 Coded contact list:   
1^st^ MG coordination shell missing (see Notes)  

### MG:AA:3775

|  | Res. type | Atom | Residue | Distance | Occupancy | B-factor | Closest solute to 1^st^ shell water |
| --- | --- | --- | --- | --- | --- | --- | --- |
| **Ion** | | | |  | 1.00 | 91.62 || 1 | C | OP2 | AA:1605 | 2.94 | 1.00 | 36.28 |
| 2 | C | O5' | AA:1604 | 3.41 | 1.00 | 36.94 |  |
| 3 | G | O6 | AA:1606 | 3.60 | 1.00 | 45.29 |  |
| 4 | C | N1 | AA:1604 | 3.69 | 1.00 | 37.01 |  |
| 5 | C | OP2 | AA:1604 | 3.89 | 1.00 | 34.59 |  |
| 6 | C | O3' | AA:1604 | 4.31 | 1.00 | 38.90 |  |
| 7 | G | N7 | AA:1606 | 4.32 | 1.00 | 40.85 |  |
| 8 | C | N3 | AA:1604 | 4.38 | 1.00 | 35.83 |  |
| 9 | C | O4' | AA:1604 | 4.53 | 1.00 | 38.60 |  |
| 10 | C | N4 | AA:1605 | 4.59 | 1.00 | 32.84 |  |
| 11 | C | O5' | AA:1605 | 4.66 | 1.00 | 38.37 |  |
| 12 | C | N4 | AA:1604 | 4.66 | 1.00 | 34.84 |  |
| 13 | A | OP1 | AA:1603 | 4.80 | 1.00 | 45.20 |  |
  |

  
 Coded contact list:   
 Exclusion zone contact(s): /OP2/  
Consider K in place of MG (1 valid contact(s) for K vs. 0 for MG)  

### MG:AA:3776

|  | Res. type | Atom | Residue | Distance | Occupancy | B-factor | Closest solute to 1^st^ shell water |
| --- | --- | --- | --- | --- | --- | --- | --- |
| **Ion** | | | |  | 1.00 | 84.23 || 1 | G | O6 | AA:348 | 3.74 | 1.00 | 68.02 |
| 2 | G | O6 | AA:349 | 4.05 | 1.00 | 73.75 |  |
| 3 | G | N7 | AA:348 | 4.18 | 1.00 | 68.35 |  |
| 4 | A | OP2 | AA:346 | 4.58 | 1.00 | 82.15 |  |
| 5 | G | N7 | AA:349 | 5.00 | 1.00 | 72.68 |  |
| 6 | A | N6 | AA:347 | 5.00 | 1.00 | 72.13 |  |
  |

  
 Coded contact list:   
1^st^ MG coordination shell missing (see Notes)  

### MG:AA:3777

|  | Res. type | Atom | Residue | Distance | Occupancy | B-factor | Closest solute to 1^st^ shell water |
| --- | --- | --- | --- | --- | --- | --- | --- |
| **Ion** | | | |  | 1.00 | 58.36 || 1 | C | OP2 | AA:1636 | 2.31 | 1.00 | 49.07 |
| 2 | G | O5' | AA:1635 | 2.62 | 1.00 | 44.24 |  |
| 3 | G | OP1 | AA:1635 | 3.06 | 1.00 | 44.22 |  |
| 4 | U | O3' | AA:1300 | 3.22 | 1.00 | 46.42 |  |
| 5 | G | OP2 | AA:1635 | 3.69 | 1.00 | 44.70 |  |
| 6 | G | O3' | AA:1635 | 4.29 | 1.00 | 46.68 |  |
| 7 | A | OP1 | AA:1301 | 4.50 | 1.00 | 46.58 |  |
| 8 | C | OP1 | AA:1636 | 4.57 | 1.00 | 48.58 |  |
| 9 | A | O3' | AA:1634 | 4.69 | 1.00 | 43.40 |  |
| 10 | C | O5' | AA:1636 | 4.76 | 1.00 | 46.84 |  |
| 11 | U | O5' | AA:1300 | 4.84 | 1.00 | 45.59 |  |
| 12 | U | O2' | AA:1300 | 4.98 | 1.00 | 46.27 |  |
  |

  
 Contact(s): /OP2/  
 Coded contact list: O_ph_  
 Exclusion zone contact(s): /O3'/O5'/OP1/  
Consider K in place of MG (3 valid contact(s) for K vs. 1 for MG)  

### MG:AA:3778

|  | Res. type | Atom | Residue | Distance | Occupancy | B-factor | Closest solute to 1^st^ shell water |
| --- | --- | --- | --- | --- | --- | --- | --- |
| **Ion** | | | |  | 1.00 | 53.92 || 1 | G | OP2 | AA:558 | 2.47 | 1.00 | 72.44 |
| 2 | G | N7 | AA:558 | 2.66 | 1.00 | 72.04 |  |
| 3 | G | O5' | AA:558 | 2.70 | 1.00 | 71.68 |  |
| 4 | G | N9 | AA:558 | 3.49 | 1.00 | 71.04 |  |
| 5 | G | OP2 | AA:559 | 3.89 | 1.00 | 71.82 |  |
| 6 | U | O3' | AA:557 | 4.04 | 1.00 | 71.68 |  |
| 7 | G | O4' | AA:558 | 4.11 | 1.00 | 70.88 |  |
| 8 | G | OP1 | AA:558 | 4.40 | 1.00 | 72.01 |  |
| 9 | U | O2' | AA:557 | 4.61 | 1.00 | 73.44 |  |
| 10 | U | N1 | AA:557 | 4.81 | 1.00 | 73.44 |  |
| 11 | G | O3' | AA:558 | 4.82 | 1.00 | 71.95 |  |
  |

  
 Coded contact list:   
 Exclusion zone contact(s): /N7/O5'/OP2/  
Consider K in place of MG (2 valid contact(s) for K vs. 0 for MG)  

### MG:AA:3779

|  | Res. type | Atom | Residue | Distance | Occupancy | B-factor | Closest solute to 1^st^ shell water |
| --- | --- | --- | --- | --- | --- | --- | --- |
| **Ion** | | | |  | 1.00 | 86.28 || 1 | G | OP2 | AA:80 | 2.54 | 1.00 | 70.27 |
| 2 | G | OP2 | AA:81 | 3.57 | 1.00 | 70.55 |  |
| 3 | G | O5' | AA:80 | 3.62 | 1.00 | 70.23 |  |
| 4 | G | OP1 | AA:80 | 4.22 | 1.00 | 71.06 |  |
| 5 | G | N7 | AA:80 | 4.50 | 1.00 | 69.46 |  |
| 6 | G | O3' | AA:79 | 4.91 | 1.00 | 70.21 |  |
  |

  
 Coded contact list:   
 Exclusion zone contact(s): /OP2/  
Consider NA in place of MG (1 valid contact(s) for NA vs. 0 for MG)  

### MG:AA:3780

|  | Res. type | Atom | Residue | Distance | Occupancy | B-factor | Closest solute to 1^st^ shell water |
| --- | --- | --- | --- | --- | --- | --- | --- |
| **Ion** | | | |  | 1.00 | 69.98 || 1 | C | OP1 | AA:76 | 2.72 | 1.00 | 62.57 |
| 2 | ARG | NH1 | AW:59 | 2.79 | 1.00 | 43.08 |  |
| 3 | C | OP2 | AA:77 | 2.97 | 1.00 | 57.38 |  |
| 4 | C | OP1 | AA:77 | 3.22 | 1.00 | 57.12 |  |
| 5 | C | O3' | AA:76 | 3.34 | 1.00 | 58.29 |  |
| 6 | C | O5' | AA:76 | 3.92 | 1.00 | 61.82 |  |
| 7 | C | OP2 | AA:76 | 4.06 | 1.00 | 62.89 |  |
| 8 | ARG | NH2 | AW:59 | 4.69 | 1.00 | 42.17 |  |
| 9 | C | O5' | AA:77 | 4.82 | 1.00 | 57.75 |  |
  |

  
 Coded contact list:   
 Exclusion zone contact(s): /NH1_Arg_/O3'/OP1/OP1/OP2/  
Consider K in place of MG (5 valid contact(s) for K vs. 0 for MG)  

### MG:AA:3781

|  | Res. type | Atom | Residue | Distance | Occupancy | B-factor | Closest solute to 1^st^ shell water |
| --- | --- | --- | --- | --- | --- | --- | --- |
| **Ion** | | | |  | 1.00 | 104.48 || 1 | A | OP2 | AA:347 | 2.66 | 1.00 | 75.24 |
| 2 | A | OP2 | AA:346 | 2.80 | 1.00 | 82.15 |  |
| 3 | A | OP1 | AA:346 | 4.04 | 1.00 | 81.11 |  |
| 4 | A | O5' | AA:346 | 4.16 | 1.00 | 80.36 |  |
| 5 | A | N7 | AA:347 | 4.49 | 1.00 | 75.29 |  |
| 6 | A | O2' | AA:345 | 4.54 | 1.00 | 76.72 |  |
| 7 | A | O3' | AA:346 | 4.57 | 1.00 | 75.89 |  |
  |

  
 Coded contact list:   
 Exclusion zone contact(s): /OP2/OP2/  
Consider K in place of MG (2 valid contact(s) for K vs. 0 for MG)  

### MG:AA:3782

|  | Res. type | Atom | Residue | Distance | Occupancy | B-factor | Closest solute to 1^st^ shell water |
| --- | --- | --- | --- | --- | --- | --- | --- |
| **Ion** | | | |  | 1.00 | 101.12 || 1 | U | O4 | AA:1516 | 4.22 | 1.00 | 77.01 |
| 2 | G | O6 | AA:1517 | 4.77 | 1.00 | 68.59 |  |
  |

  
 Coded contact list:   
1^st^ MG coordination shell missing (see Notes)  

### MG:AA:3783

|  | Res. type | Atom | Residue | Distance | Occupancy | B-factor | Closest solute to 1^st^ shell water |
| --- | --- | --- | --- | --- | --- | --- | --- |
| **Ion** | | | |  | 1.00 | 86.35 || 1 | G | OP1 | AA:352 | 3.50 | 1.00 | 93.95 |
| 2 | G | OP2 | AA:352 | 4.24 | 1.00 | 94.28 |  |
| 3 | G | OP2 | AA:351 | 4.80 | 1.00 | 83.44 |  |
  |

  
 Coded contact list:   
1^st^ MG coordination shell missing (see Notes)  

### MG:AA:3784

|  | Res. type | Atom | Residue | Distance | Occupancy | B-factor | Closest solute to 1^st^ shell water |
| --- | --- | --- | --- | --- | --- | --- | --- |
| **Ion** | | | |  | 1.00 | 64.48 || 1 | C | OP1 | AA:1202 | 3.33 | 1.00 | 47.02 |
| 2 | G | OP2 | AA:1203 | 3.49 | 1.00 | 55.44 |  |
| 3 | U | OP2 | AA:1205 | 4.07 | 1.00 | 64.78 |  |
| 4 | U | OP1 | AA:1205 | 4.46 | 1.00 | 65.78 |  |
| 5 | G | OP1 | AA:1203 | 4.81 | 1.00 | 54.73 |  |
| 6 | C | O5' | AA:1202 | 4.82 | 1.00 | 48.44 |  |
  |

  
 Coded contact list:   
 Exclusion zone contact(s): /OP1/  
Consider K in place of MG (1 valid contact(s) for K vs. 0 for MG)  

### MG:AA:3785

|  | Res. type | Atom | Residue | Distance | Occupancy | B-factor | Closest solute to 1^st^ shell water |
| --- | --- | --- | --- | --- | --- | --- | --- |
| **Ion** | | | |  | 1.00 | 89.29 || 1 | G | O6 | AA:407 | 3.98 | 1.00 | 63.93 |
| 2 | G | N7 | AA:406 | 4.87 | 1.00 | 76.63 |  |
| 3 | G | O6 | AA:406 | 4.98 | 1.00 | 73.54 |  |
  |

  
 Coded contact list:   
1^st^ MG coordination shell missing (see Notes)  

### MG:AA:3787

|  | Res. type | Atom | Residue | Distance | Occupancy | B-factor | Closest solute to 1^st^ shell water |
| --- | --- | --- | --- | --- | --- | --- | --- |
| **Ion** | | | |  | 1.00 | 70.38 || 1 | U | OP1 | AA:104 | 4.45 | 1.00 | 66.60 |
| 2 | A | O3' | AA:103 | 4.48 | 1.00 | 64.08 |  |
  |

  
 Coded contact list:   
1^st^ MG coordination shell missing (see Notes)  

### MG:AA:3788

|  | Res. type | Atom | Residue | Distance | Occupancy | B-factor | Closest solute to 1^st^ shell water |
| --- | --- | --- | --- | --- | --- | --- | --- |
| **Ion** | | | |  | 1.00 | 87.36 || 1 | G | OP2 | AA:325 | 2.59 | 1.00 | 64.98 |
| 2 | G | OP1 | AA:325 | 3.97 | 1.00 | 64.14 |  |
| 3 | G | O5' | AA:325 | 4.38 | 1.00 | 63.21 |  |
  |

  
 Coded contact list:   
 Exclusion zone contact(s): /OP2/  
Consider NA in place of MG (1 valid contact(s) for NA vs. 0 for MG)  

### MG:AA:3789

|  | Res. type | Atom | Residue | Distance | Occupancy | B-factor | Closest solute to 1^st^ shell water |
| --- | --- | --- | --- | --- | --- | --- | --- |
| **Ion** | | | |  | 1.00 | 55.27 || 1 | C | O2' | AA:267 | 4.12 | 1.00 | 64.91 |
| 2 | A | N3 | AA:282 | 4.21 | 1.00 | 82.81 |  |
| 3 | A | N7 | AA:283 | 4.48 | 1.00 | 83.09 |  |
| 4 | A | O2' | AA:282 | 4.69 | 1.00 | 80.10 |  |
| 5 | A | N6 | AA:283 | 4.75 | 1.00 | 82.48 |  |
| 6 | A | N1 | AA:283 | 4.76 | 1.00 | 83.12 |  |
| 7 | A | N9 | AA:283 | 4.93 | 1.00 | 83.31 |  |
| 8 | A | N3 | AA:283 | 4.95 | 1.00 | 83.35 |  |
  |

  
 Coded contact list:   
1^st^ MG coordination shell missing (see Notes)  

### MG:AA:3790

|  | Res. type | Atom | Residue | Distance | Occupancy | B-factor | Closest solute to 1^st^ shell water |
| --- | --- | --- | --- | --- | --- | --- | --- |
| **Ion** | | | |  | 1.00 | 60.31 || 1 | U | OP2 | AA:1313 | 2.69 | 1.00 | 43.84 |
| 2 | U | O2' | AA:1312 | 3.02 | 1.00 | 48.42 |  |
| 3 | U | O5' | AA:1313 | 3.04 | 1.00 | 42.95 |  |
| 4 | C | OP2 | AA:1314 | 3.05 | 1.00 | 41.16 |  |
| 5 | C | N4 | AA:1315 | 3.19 | 1.00 | 50.65 |  |
| 6 | C | N4 | AA:1314 | 3.32 | 1.00 | 35.31 |  |
| 7 | U | O3' | AA:1312 | 3.79 | 1.00 | 45.58 |  |
| 8 | C | N3 | AA:1314 | 4.63 | 1.00 | 37.26 |  |
| 9 | U | OP1 | AA:1313 | 4.80 | 1.00 | 46.12 |  |
| 10 | G | O6 | AA:1337 | 4.81 | 1.00 | 53.08 |  |
| 11 | U | O3' | AA:1313 | 4.86 | 1.00 | 41.21 |  |
  |

  
 Coded contact list:   
 Exclusion zone contact(s): /N4/N4/O2'/O5'/OP2/OP2/  
Consider K in place of MG (6 valid contact(s) for K vs. 0 for MG)  

### MG:AA:3791

|  | Res. type | Atom | Residue | Distance | Occupancy | B-factor | Closest solute to 1^st^ shell water |
| --- | --- | --- | --- | --- | --- | --- | --- |
| **Ion** | | | |  | 1.00 | 65.43 || 1 | A | O2' | AA:2823 | 2.57 | 1.00 | 43.10 |
| 2 | C | O2 | AA:2006 | 2.98 | 1.00 | 41.47 |  |
| 3 | A | N3 | AA:1652 | 3.18 | 1.00 | 39.08 |  |
| 4 | A | O3' | AA:2823 | 3.53 | 1.00 | 45.14 |  |
| 5 | A | O2' | AA:1652 | 3.62 | 1.00 | 44.13 |  |
| 6 | G | N2 | AA:1651 | 4.02 | 1.00 | 38.76 |  |
| 7 | C | O2' | AA:2006 | 4.48 | 1.00 | 41.91 |  |
| 8 | A | N9 | AA:1652 | 4.58 | 1.00 | 40.30 |  |
| 9 | C | N1 | AA:2006 | 4.58 | 1.00 | 38.49 |  |
| 10 | A | O4' | AA:2823 | 4.59 | 1.00 | 42.70 |  |
| 11 | C | N3 | AA:2006 | 4.82 | 1.00 | 38.18 |  |
| 12 | C | OP1 | AA:2824 | 4.84 | 1.00 | 42.75 |  |
  |

  
 Coded contact list:   
 Exclusion zone contact(s): /N3_A_/O2/O2'/  
Consider K in place of MG (2 valid contact(s) for K vs. 0 for MG)  

### MG:AA:3792

|  | Res. type | Atom | Residue | Distance | Occupancy | B-factor | Closest solute to 1^st^ shell water |
| --- | --- | --- | --- | --- | --- | --- | --- |
| **Ion** | | | |  | 1.00 | 82.48 || 1 | A | O2' | AA:1603 | 2.87 | 1.00 | 36.14 |
| 2 | G | O2' | AA:1339 | 2.96 | 1.00 | 48.69 |  |
| 3 | A | O4' | AA:1603 | 3.03 | 1.00 | 38.37 |  |
| 4 | U | O4' | AA:1340 | 3.18 | 1.00 | 47.71 |  |
| 5 | U | N1 | AA:1340 | 3.29 | 1.00 | 50.13 |  |
| 6 | G | N3 | AA:1339 | 3.36 | 1.00 | 48.88 |  |
| 7 | G | N2 | AA:1339 | 3.96 | 1.00 | 48.35 |  |
| 8 | A | O2' | AA:1393 | 4.22 | 1.00 | 57.40 |  |
| 9 | U | N3 | AA:1340 | 4.28 | 1.00 | 52.47 |  |
| 10 | U | O5' | AA:1340 | 4.43 | 1.00 | 47.69 |  |
| 11 | G | N9 | AA:1339 | 4.51 | 1.00 | 45.76 |  |
| 12 | A | O3' | AA:1603 | 4.56 | 1.00 | 34.35 |  |
| 13 | A | N9 | AA:1603 | 4.56 | 1.00 | 38.18 |  |
| 14 | U | O4' | AA:1394 | 4.63 | 1.00 | 54.85 |  |
| 15 | U | O2 | AA:1340 | 4.76 | 1.00 | 49.83 |  |
| 16 | U | O4 | AA:1340 | 4.97 | 1.00 | 56.20 |  |
  |

  
 Coded contact list:   
 Exclusion zone contact(s): /N1/N3_G_/O2'/O2'/O4'/O4'/  
Consider K in place of MG (6 valid contact(s) for K vs. 0 for MG)  

### MG:AA:3793

|  | Res. type | Atom | Residue | Distance | Occupancy | B-factor | Closest solute to 1^st^ shell water |
| --- | --- | --- | --- | --- | --- | --- | --- |
| **Ion** | | | |  | 1.00 | 63.36 || 1 | G | O2' | AA:75 | 2.25 | 1.00 | 62.75 |
| 2 | A | OP1 | AA:73 | 2.53 | 1.00 | 51.15 |  |
| 3 | U | O3' | AA:72 | 3.48 | 1.00 | 48.44 |  |
| 4 | U | O2' | AA:72 | 3.80 | 1.00 | 48.89 |  |
| 5 | G | O3' | AA:75 | 4.08 | 1.00 | 63.11 |  |
| 6 | A | O5' | AA:73 | 4.09 | 1.00 | 52.92 |  |
| 7 | C | O4' | AA:76 | 4.34 | 1.00 | 58.89 |  |
| 8 | C | O5' | AA:76 | 4.35 | 1.00 | 61.82 |  |
| 9 | ARG | N | AW:55 | 4.40 | 1.00 | 56.33 |  |
| 10 | ARG | NE | AW:55 | 4.41 | 1.00 | 64.03 |  |
| 11 | LYS | O | AW:54 | 4.69 | 1.00 | 54.51 |  |
| 12 | A | OP2 | AA:73 | 4.88 | 1.00 | 52.21 |  |
  |

  
 Contact(s): /O2'/  
 Coded contact list: O_r_  
 Exclusion zone contact(s): /OP1/  

### MG:AA:3794

|  | Res. type | Atom | Residue | Distance | Occupancy | B-factor | Closest solute to 1^st^ shell water |
| --- | --- | --- | --- | --- | --- | --- | --- |
| **Ion** | | | |  | 1.00 | 70.71 || 1 | G | OP2 | AA:1209 | 3.36 | 1.00 | 61.90 |
| 2 | A | N1 | AA:331 | 4.16 | 1.00 | 54.42 |  |
| 3 | G | O2' | AA:312 | 4.55 | 1.00 | 58.20 |  |
| 4 | A | OP2 | AA:1210 | 4.65 | 1.00 | 64.00 |  |
| 5 | G | O5' | AA:1209 | 4.68 | 1.00 | 63.98 |  |
| 6 | C | OP1 | AA:313 | 4.93 | 1.00 | 62.40 |  |
| 7 | G | O3' | AA:312 | 4.95 | 1.00 | 59.96 |  |
  |

  
 Coded contact list:   
 Exclusion zone contact(s): /OP2/  
Consider K in place of MG (1 valid contact(s) for K vs. 0 for MG)  

### MG:AA:3795

|  | Res. type | Atom | Residue | Distance | Occupancy | B-factor | Closest solute to 1^st^ shell water |
| --- | --- | --- | --- | --- | --- | --- | --- |
| **Ion** | | | |  | 1.00 | 106.32 || 1 | A | OP2 | AA:78 | 2.67 | 1.00 | 62.34 |
| 2 | C | OP1 | AA:77 | 3.15 | 1.00 | 57.12 |  |
| 3 | C | OP2 | AA:77 | 4.04 | 1.00 | 57.38 |  |
| 4 | C | O5' | AA:77 | 4.37 | 1.00 | 57.75 |  |
| 5 | C | O3' | AA:77 | 4.59 | 1.00 | 61.92 |  |
| 6 | A | OP1 | AA:78 | 4.72 | 1.00 | 63.11 |  |
  |

  
 Coded contact list:   
 Exclusion zone contact(s): /OP1/OP2/  
Consider K in place of MG (2 valid contact(s) for K vs. 0 for MG)  

### MG:AA:3796

|  | Res. type | Atom | Residue | Distance | Occupancy | B-factor | Closest solute to 1^st^ shell water |
| --- | --- | --- | --- | --- | --- | --- | --- |
| **Ion** | | | |  | 1.00 | 94.27 || 1 | G | O6 | AA:70 | 3.27 | 1.00 | 54.36 |
| 2 | C | N4 | AA:57 | 3.33 | 1.00 | 52.29 |  |
| 3 | C | N4 | AA:69 | 4.17 | 1.00 | 56.08 |  |
| 4 | A | N6 | AA:56 | 4.21 | 1.00 | 44.37 |  |
| 5 | A | N7 | AA:56 | 4.32 | 1.00 | 41.91 |  |
| 6 | G | O6 | AA:58 | 4.60 | 1.00 | 54.77 |  |
  |

  
 Coded contact list:   
 Exclusion zone contact(s): /N4/O6/  
Consider K in place of MG (2 valid contact(s) for K vs. 0 for MG)  

### MG:AA:3797

|  | Res. type | Atom | Residue | Distance | Occupancy | B-factor | Closest solute to 1^st^ shell water |
| --- | --- | --- | --- | --- | --- | --- | --- |
| **Ion** | | | |  | 1.00 | 96.58 || 1 | C | N4 | AA:1592 | 4.06 | 1.00 | 87.65 |
| 2 | G | N7 | AA:1591 | 4.59 | 1.00 | 90.47 |  |
| 3 | U | O4 | AA:1590 | 4.75 | 1.00 | 88.15 |  |
  |

  
 Coded contact list:   
1^st^ MG coordination shell missing (see Notes)  

### MG:AA:3798

|  | Res. type | Atom | Residue | Distance | Occupancy | B-factor | Closest solute to 1^st^ shell water |
| --- | --- | --- | --- | --- | --- | --- | --- |
| **Ion** | | | |  | 1.00 | 47.59 || 1 | PRO | O | AE:147 | 2.68 | 1.00 | 48.92 |
| 2 | C | OP1 | AA:2025 | 2.84 | 1.00 | 54.27 |  |
| 3 | C | OP2 | AA:2025 | 2.95 | 1.00 | 54.66 |  |
| 4 | U | O4 | AA:1130 | 4.00 | 1.00 | 76.98 |  |
| 5 | U | N3 | AA:1130 | 4.19 | 1.00 | 75.00 |  |
| 6 | G | O3' | AA:2024 | 4.47 | 1.00 | 54.99 |  |
| 7 | C | O5' | AA:2025 | 4.56 | 1.00 | 54.37 |  |
| 8 | LYS | O | AE:145 | 4.72 | 1.00 | 45.80 |  |
| 9 | GLY | N | AE:148 | 4.82 | 1.00 | 49.50 |  |
  |

  
 Coded contact list:   
 Exclusion zone contact(s): /O_back_/OP1/OP2/  
Consider K in place of MG (3 valid contact(s) for K vs. 0 for MG)  

### MG:AA:3799

|  | Res. type | Atom | Residue | Distance | Occupancy | B-factor | Closest solute to 1^st^ shell water |
| --- | --- | --- | --- | --- | --- | --- | --- |
| **Ion** | | | |  | 1.00 | 93.09 || 1 | U | OP2 | AA:2832 | 3.52 | 1.00 | 66.61 |
| 2 | U | OP1 | AA:2884 | 4.41 | 1.00 | 75.63 |  |
| 3 | U | OP2 | AA:2884 | 4.61 | 1.00 | 75.98 |  |
| 4 | C | OP2 | AA:2885 | 4.65 | 1.00 | 83.49 |  |
| 5 | G | N3 | AA:2831 | 4.79 | 1.00 | 60.35 |  |
  |

  
 Coded contact list:   
1^st^ MG coordination shell missing (see Notes)  

### MG:AA:3800

|  | Res. type | Atom | Residue | Distance | Occupancy | B-factor | Closest solute to 1^st^ shell water |
| --- | --- | --- | --- | --- | --- | --- | --- |
| **Ion** | | | |  | 1.00 | 55.74 || 1 | G | O6 | AA:602 | 2.53 | 1.00 | 69.09 |
| 2 | A | OP2 | AA:654V | 3.41 | 1.00 | 134.81 |  |
| 3 | G | N7 | AA:602 | 3.51 | 1.00 | 67.73 |  |
| 4 | A | N7 | AA:654V | 4.07 | 1.00 | 116.50 |  |
| 5 | G | N1 | AA:602 | 4.73 | 1.00 | 66.95 |  |
| 6 | A | OP2 | AA:654U | 4.75 | 1.00 | 156.49 |  |
| 7 | A | O5' | AA:654U | 4.93 | 1.00 | 153.34 |  |
  |

  
 Coded contact list:   
 Exclusion zone contact(s): /O6/  
Consider NA in place of MG (1 valid contact(s) for NA vs. 0 for MG)  

### MG:AA:3801

|  | Res. type | Atom | Residue | Distance | Occupancy | B-factor | Closest solute to 1^st^ shell water |
| --- | --- | --- | --- | --- | --- | --- | --- |
| **Ion** | | | |  | 1.00 | 90.88 || 1 | A | N9 | AA:1460 | 3.35 | 1.00 | 117.95 |
| 2 | A | N7 | AA:1460 | 3.63 | 1.00 | 119.69 |  |
| 3 | A | O3' | AA:1460 | 3.65 | 1.00 | 109.68 |  |
| 4 | G | O5' | AA:1461 | 3.92 | 1.00 | 105.10 |  |
| 5 | A | O2' | AA:1460 | 4.12 | 1.00 | 115.04 |  |
| 6 | A | O5' | AA:1460 | 4.22 | 1.00 | 104.51 |  |
| 7 | A | O4' | AA:1460 | 4.29 | 1.00 | 113.55 |  |
  |

  
 Coded contact list:   
Consider K in place of MG (1 valid contact(s) for K vs. 0 for MG)  

### MG:AA:3802

|  | Res. type | Atom | Residue | Distance | Occupancy | B-factor | Closest solute to 1^st^ shell water |
| --- | --- | --- | --- | --- | --- | --- | --- |
| **Ion** | | | |  | 1.00 | 78.71 || 1 | G | O6 | AA:609A | 3.60 | 1.00 | 57.36 |
| 2 | C | N4 | AA:610 | 4.38 | 1.00 | 53.43 |  |
| 3 | G | O6 | AA:618 | 4.44 | 1.00 | 54.38 |  |
| 4 | A | OP2 | AA:609 | 4.73 | 1.00 | 66.78 |  |
| 5 | G | N7 | AA:609A | 4.73 | 1.00 | 56.01 |  |
| 6 | C | N4 | AA:618A | 4.83 | 1.00 | 61.66 |  |
  |

  
 Coded contact list:   
1^st^ MG coordination shell missing (see Notes)  

### MG:AA:3803

|  | Res. type | Atom | Residue | Distance | Occupancy | B-factor | Closest solute to 1^st^ shell water |
| --- | --- | --- | --- | --- | --- | --- | --- |
| **Ion** | | | |  | 1.00 | 72.56 || 1 | U | O2 | AA:2702 | 2.42 | 1.00 | 74.47 |
| 2 | C | OP2 | AA:2703 | 2.58 | 1.00 | 69.02 |  |
| 3 | A | N6 | AA:2705 | 3.15 | 1.00 | 67.64 |  |
| 4 | C | N4 | AA:2701 | 3.65 | 1.00 | 56.94 |  |
| 5 | C | N3 | AA:2701 | 3.67 | 1.00 | 56.22 |  |
| 6 | U | N1 | AA:2702 | 3.74 | 1.00 | 70.60 |  |
| 7 | U | O3' | AA:2702 | 3.98 | 1.00 | 68.32 |  |
| 8 | U | N3 | AA:2702 | 4.04 | 1.00 | 72.07 |  |
| 9 | U | O2' | AA:2702 | 4.05 | 1.00 | 71.66 |  |
| 10 | U | O4' | AA:2702 | 4.46 | 1.00 | 69.56 |  |
| 11 | C | OP2 | AA:2704 | 4.50 | 1.00 | 70.15 |  |
| 12 | C | N1 | AA:2701 | 4.54 | 1.00 | 57.31 |  |
| 13 | G | O6 | AA:2706 | 4.55 | 1.00 | 54.69 |  |
| 14 | G | N1 | AA:2706 | 4.58 | 1.00 | 54.91 |  |
| 15 | C | O5' | AA:2703 | 4.74 | 1.00 | 68.71 |  |
| 16 | C | O2 | AA:2701 | 4.82 | 1.00 | 55.61 |  |
| 17 | C | OP1 | AA:2703 | 4.97 | 1.00 | 69.14 |  |
  |

  
 Coded contact list:   
 Exclusion zone contact(s): /N6/O2/OP2/  
Consider NA in place of MG (2 valid contact(s) for NA vs. 0 for MG)  

### MG:AA:3804

|  | Res. type | Atom | Residue | Distance | Occupancy | B-factor | Closest solute to 1^st^ shell water |
| --- | --- | --- | --- | --- | --- | --- | --- |
| **Ion** | | | |  | 1.00 | 94.08 || 1 | C | OP2 | AA:286 | 2.58 | 1.00 | 92.71 |
| 2 | C | OP1 | AA:286 | 4.09 | 1.00 | 93.09 |  |
| 3 | C | O5' | AA:286 | 4.42 | 1.00 | 93.86 |  |
| 4 | C | OP2 | AA:287 | 4.49 | 1.00 | 95.58 |  |
  |

  
 Coded contact list:   
 Exclusion zone contact(s): /OP2/  
Consider NA in place of MG (1 valid contact(s) for NA vs. 0 for MG)  

### MG:AA:3805

|  | Res. type | Atom | Residue | Distance | Occupancy | B-factor | Closest solute to 1^st^ shell water |
| --- | --- | --- | --- | --- | --- | --- | --- |
| **Ion** | | | |  | 1.00 | 94.75 || 1 | A | N1 | AA:1871 | 4.26 | 1.00 | 91.61 |
| 2 | A | N6 | AA:1871 | 4.39 | 1.00 | 91.37 |  |
| 3 | C | N4 | AA:1870 | 4.67 | 1.00 | 84.26 |  |
  |

  
 Coded contact list:   
1^st^ MG coordination shell missing (see Notes)  

### MG:AA:3806

|  | Res. type | Atom | Residue | Distance | Occupancy | B-factor | Closest solute to 1^st^ shell water |
| --- | --- | --- | --- | --- | --- | --- | --- |
| **Ion** | | | |  | 1.00 | 79.78 || 1 | A | N7 | AA:653 | 2.26 | 1.00 | 148.64 |
| 2 | A | N6 | AA:653 | 3.51 | 1.00 | 149.77 |  |
| 3 | C | OP2 | AA:652 | 3.93 | 1.00 | 90.55 |  |
| 4 | A | N9 | AA:653 | 4.39 | 1.00 | 148.15 |  |
| 5 | C | N4 | AA:652 | 4.40 | 1.00 | 105.04 |  |
| 6 | C | N1 | AA:652 | 4.44 | 1.00 | 105.13 |  |
| 7 | A | OP2 | AA:653 | 4.65 | 1.00 | 121.23 |  |
| 8 | C | N3 | AA:652 | 4.82 | 1.00 | 104.80 |  |
  |

  
 Contact(s): /N7/  
 Coded contact list: N_b_  

### MG:AA:3807

|  | Res. type | Atom | Residue | Distance | Occupancy | B-factor | Closest solute to 1^st^ shell water |
| --- | --- | --- | --- | --- | --- | --- | --- |
| **Ion** | | | |  | 1.00 | 87.39 || 1 | C | OP1 | AA:654G | 3.49 | 1.00 | 200.41 |
| 2 | C | O2' | AA:654F | 3.50 | 1.00 | 200.41 |  |
| 3 | C | OP2 | AA:654G | 4.74 | 1.00 | 200.41 |  |
  |

  
 Coded contact list:   
1^st^ MG coordination shell missing (see Notes)  

### MG:AA:3808

|  | Res. type | Atom | Residue | Distance | Occupancy | B-factor | Closest solute to 1^st^ shell water |
| --- | --- | --- | --- | --- | --- | --- | --- |
| **Ion** | | | |  | 1.00 | 75.79 || 1 | C | OP2 | AA:2691 | 2.37 | 1.00 | 66.31 |
| 2 | C | OP2 | AA:2692 | 2.98 | 1.00 | 67.02 |  |
| 3 | C | OP1 | AA:2692 | 3.52 | 1.00 | 66.89 |  |
| 4 | C | O2' | AA:2690 | 4.07 | 1.00 | 63.99 |  |
| 5 | C | O5' | AA:2691 | 4.18 | 1.00 | 65.69 |  |
| 6 | C | OP1 | AA:2691 | 4.32 | 1.00 | 67.50 |  |
| 7 | C | O3' | AA:2691 | 4.67 | 1.00 | 65.79 |  |
| 8 | C | O3' | AA:2690 | 4.91 | 1.00 | 63.69 |  |
  |

  
 Contact(s): /OP2/  
 Coded contact list: O_ph_  
 Exclusion zone contact(s): /OP2/  

### MG:AA:3809

|  | Res. type | Atom | Residue | Distance | Occupancy | B-factor | Closest solute to 1^st^ shell water |
| --- | --- | --- | --- | --- | --- | --- | --- |
| **Ion** | | | |  | 1.00 | 84.99 || 1 | C | O2' | AA:2163 | 2.06 | 1.00 | 197.62 |
| 2 | A | N6 | AA:2126 | 2.91 | 1.00 | 189.47 |  |
| 3 | C | O3' | AA:2163 | 3.89 | 1.00 | 198.37 |  |
| 4 | A | N1 | AA:2126 | 4.28 | 1.00 | 189.68 |  |
| 5 | C | O4' | AA:2163 | 4.37 | 1.00 | 197.82 |  |
| 6 | A | N7 | AA:2126 | 4.58 | 1.00 | 190.72 |  |
  |

  
 Contact(s): /O2'/  
 Coded contact list: O_r_  
 Exclusion zone contact(s): /N6/  

### MG:AA:3810

|  | Res. type | Atom | Residue | Distance | Occupancy | B-factor | Closest solute to 1^st^ shell water |
| --- | --- | --- | --- | --- | --- | --- | --- |
| **Ion** | | | |  | 1.00 | 85.98 || 1 | G | O6 | AA:1526 | 3.22 | 1.00 | 81.19 |
| 2 | G | N1 | AA:1527 | 3.67 | 1.00 | 88.68 |  |
| 3 | G | O6 | AA:1527 | 3.87 | 1.00 | 87.74 |  |
| 4 | G | N7 | AA:1526 | 4.27 | 1.00 | 81.52 |  |
| 5 | C | N4 | AA:1546 | 4.63 | 1.00 | 80.98 |  |
| 6 | G | N2 | AA:1527 | 4.63 | 1.00 | 89.60 |  |
| 7 | G | N7 | AA:1525 | 4.79 | 1.00 | 81.62 |  |
  |

  
 Coded contact list:   
 Exclusion zone contact(s): /O6/  
Consider K in place of MG (1 valid contact(s) for K vs. 0 for MG)  

### MG:AA:3811

|  | Res. type | Atom | Residue | Distance | Occupancy | B-factor | Closest solute to 1^st^ shell water |
| --- | --- | --- | --- | --- | --- | --- | --- |
| **Ion** | | | |  | 1.00 | 58.11 || 1 | G | OP2 | AA:396 | 3.89 | 1.00 | 53.02 |
| 2 | G | N7 | AA:396 | 4.64 | 1.00 | 47.27 |  |
| 3 | G | O6 | AA:397 | 4.88 | 1.00 | 46.77 |  |
| 4 | A | OP2 | AA:374 | 4.90 | 1.00 | 58.32 |  |
| 5 | U | OP1 | AA:373 | 4.94 | 1.00 | 61.67 |  |
  |

  
 Coded contact list:   
1^st^ MG coordination shell missing (see Notes)  

### MG:AA:3812

|  | Res. type | Atom | Residue | Distance | Occupancy | B-factor | Closest solute to 1^st^ shell water |
| --- | --- | --- | --- | --- | --- | --- | --- |
| **Ion** | | | |  | 1.00 | 83.76 || 1 | C | OP1 | AA:1196 | 2.41 | 1.00 | 66.87 |
| 2 | C | OP2 | AA:1196 | 3.61 | 1.00 | 66.36 |  |
| 3 | C | O5' | AA:1196 | 4.55 | 1.00 | 66.15 |  |
| 4 | G | O3' | AA:1195 | 4.58 | 1.00 | 66.35 |  |
  |

  
 Coded contact list:   
 Exclusion zone contact(s): /OP1/  
Consider NA in place of MG (1 valid contact(s) for NA vs. 0 for MG)  

### MG:AA:3813

|  | Res. type | Atom | Residue | Distance | Occupancy | B-factor | Closest solute to 1^st^ shell water |
| --- | --- | --- | --- | --- | --- | --- | --- |
| **Ion** | | | |  | 1.00 | 99.41 || 1 | A | OP1 | AA:1449 | 2.56 | 1.00 | 87.12 |
| 2 | G | OP1 | AA:1448 | 2.74 | 1.00 | 85.73 |  |
| 3 | A | OP1 | AA:1543 | 3.23 | 1.00 | 103.24 |  |
| 4 | G | O5' | AA:1448 | 3.63 | 1.00 | 85.41 |  |
| 5 | G | OP2 | AA:1448 | 4.01 | 1.00 | 84.78 |  |
| 6 | G | O3' | AA:1448 | 4.65 | 1.00 | 86.41 |  |
| 7 | A | O5' | AA:1449 | 4.74 | 1.00 | 88.56 |  |
  |

  
 Coded contact list:   
 Exclusion zone contact(s): /OP1/OP1/OP1/  
Consider K in place of MG (2 valid contact(s) for K vs. 0 for MG)  

### MG:AA:3814

|  | Res. type | Atom | Residue | Distance | Occupancy | B-factor | Closest solute to 1^st^ shell water |
| --- | --- | --- | --- | --- | --- | --- | --- |
| **Ion** | | | |  | 1.00 | 67.14 || 1 | U | OP1 | AA:1090 | 3.59 | 1.00 | 200.25 |
| 2 | G | O2' | AA:1087 | 4.54 | 1.00 | 200.09 |  |
  |

  
 Coded contact list:   
1^st^ MG coordination shell missing (see Notes)  

### MG:AA:3815

|  | Res. type | Atom | Residue | Distance | Occupancy | B-factor | Closest solute to 1^st^ shell water |
| --- | --- | --- | --- | --- | --- | --- | --- |
| **Ion** | | | |  | 1.00 | 61.20 || 1 | A | OP1 | AA:1353 | 2.92 | 1.00 | 43.87 |
| 2 | A | OP1 | AA:1354 | 3.08 | 1.00 | 48.14 |  |
| 3 | A | OP2 | AA:1354 | 3.47 | 1.00 | 45.86 |  |
| 4 | A | OP2 | AA:1353 | 4.48 | 1.00 | 44.20 |  |
| 5 | A | O3' | AA:1353 | 4.80 | 1.00 | 44.58 |  |
| 6 | A | O5' | AA:1353 | 4.85 | 1.00 | 43.94 |  |
| 7 | A | O5' | AA:1354 | 4.99 | 1.00 | 46.75 |  |
  |

  
 Coded contact list:   
 Exclusion zone contact(s): /OP1/OP1/  
Consider K in place of MG (2 valid contact(s) for K vs. 0 for MG)  

### MG:AA:3816

|  | Res. type | Atom | Residue | Distance | Occupancy | B-factor | Closest solute to 1^st^ shell water |
| --- | --- | --- | --- | --- | --- | --- | --- |
| **Ion** | | | |  | 1.00 | 73.52 || 1 | A | O2' | AA:1603 | 2.35 | 1.00 | 36.14 |
| 2 | C | O2' | AA:1314 | 2.68 | 1.00 | 40.99 |  |
| 3 | A | O2' | AA:1393 | 2.76 | 1.00 | 57.40 |  |
| 4 | A | O3' | AA:1603 | 3.42 | 1.00 | 34.35 |  |
| 5 | A | O3' | AA:1393 | 3.81 | 1.00 | 53.62 |  |
| 6 | C | O2 | AA:1314 | 3.85 | 1.00 | 38.86 |  |
| 7 | C | O4' | AA:1314 | 3.86 | 1.00 | 41.60 |  |
| 8 | C | O4' | AA:1604 | 4.02 | 1.00 | 38.60 |  |
| 9 | C | O5' | AA:1604 | 4.15 | 1.00 | 36.94 |  |
| 10 | C | N1 | AA:1314 | 4.18 | 1.00 | 39.68 |  |
| 11 | C | OP1 | AA:1604 | 4.45 | 1.00 | 34.85 |  |
| 12 | U | OP1 | AA:1394 | 4.99 | 1.00 | 52.93 |  |
  |

  
 Contact(s): /O2'/  
 Coded contact list: O_r_  
 Exclusion zone contact(s): /O2'/O2'/  
Consider K in place of MG (2 valid contact(s) for K vs. 1 for MG)  

### MG:AA:3817

|  | Res. type | Atom | Residue | Distance | Occupancy | B-factor | Closest solute to 1^st^ shell water |
| --- | --- | --- | --- | --- | --- | --- | --- |
| **Ion** | | | |  | 1.00 | 66.72 || 1 | G | O6 | AA:1491 | 3.66 | 1.00 | 64.39 |
| 2 | G | O6 | AA:1492 | 3.74 | 1.00 | 72.94 |  |
| 3 | A | OP2 | AA:1490 | 3.98 | 1.00 | 87.40 |  |
| 4 | C | N4 | AA:1498 | 4.25 | 1.00 | 71.95 |  |
| 5 | U | OP2 | AA:1497 | 4.33 | 1.00 | 77.05 |  |
| 6 | A | OP2 | AA:1496 | 4.65 | 1.00 | 77.97 |  |
| 7 | G | N7 | AA:1491 | 4.80 | 1.00 | 70.66 |  |
| 8 | C | N4 | AA:1499 | 4.97 | 1.00 | 61.23 |  |
  |

  
 Coded contact list:   
1^st^ MG coordination shell missing (see Notes)  

### MG:AA:3818

|  | Res. type | Atom | Residue | Distance | Occupancy | B-factor | Closest solute to 1^st^ shell water |
| --- | --- | --- | --- | --- | --- | --- | --- |
| **Ion** | | | |  | 1.00 | 80.24 || 1 | G | N7 | AA:612 | 4.54 | 1.00 | 68.47 |
| 2 | U | O4 | AA:613 | 4.62 | 1.00 | 83.98 |  |
| 3 | G | O6 | AA:612 | 4.94 | 1.00 | 70.86 |  |
| 4 | U | N3 | AA:613 | 4.96 | 1.00 | 82.79 |  |
  |

  
 Coded contact list:   
1^st^ MG coordination shell missing (see Notes)  

### MG:AA:3819

|  | Res. type | Atom | Residue | Distance | Occupancy | B-factor | Closest solute to 1^st^ shell water |
| --- | --- | --- | --- | --- | --- | --- | --- |
| **Ion** | | | |  | 1.00 | 94.83 || 1 | C | N3 | AA:271A | 3.14 | 1.00 | 80.96 |
| 2 | C | N1 | AA:271A | 3.60 | 1.00 | 82.19 |  |
| 3 | G | N7 | AA:271B | 3.84 | 1.00 | 86.46 |  |
| 4 | C | O2 | AA:271A | 3.88 | 1.00 | 83.63 |  |
| 5 | C | N4 | AA:271A | 3.89 | 1.00 | 78.99 |  |
| 6 | U | O3' | AA:421 | 4.29 | 1.00 | 69.74 |  |
| 7 | G | OP2 | AA:271B | 4.43 | 1.00 | 84.77 |  |
  |

  
 Coded contact list:   
 Exclusion zone contact(s): /N3_C_/  
Consider K in place of MG (1 valid contact(s) for K vs. 0 for MG)  

### MG:AA:3820

|  | Res. type | Atom | Residue | Distance | Occupancy | B-factor | Closest solute to 1^st^ shell water |
| --- | --- | --- | --- | --- | --- | --- | --- |
| **Ion** | | | |  | 1.00 | 59.34 || 1 | G | O6 | AA:391 | 2.43 | 1.00 | 58.39 |
| 2 | A | N3 | AA:390 | 3.02 | 1.00 | 43.52 |  |
| 3 | A | O2' | AA:390 | 3.13 | 1.00 | 51.92 |  |
| 4 | G | N7 | AA:391 | 3.14 | 1.00 | 53.78 |  |
| 5 | G | O6 | AA:388 | 3.21 | 1.00 | 52.03 |  |
| 6 | U | O4 | AA:383 | 3.36 | 1.00 | 62.67 |  |
| 7 | C | O2 | AA:385 | 3.64 | 1.00 | 54.68 |  |
| 8 | G | OP2 | AA:386 | 4.07 | 1.00 | 45.63 |  |
| 9 | G | N2 | AA:411 | 4.13 | 1.00 | 54.16 |  |
| 10 | U | N3 | AA:383 | 4.18 | 1.00 | 62.28 |  |
| 11 | C | N4 | AA:392 | 4.19 | 1.00 | 55.68 |  |
| 12 | G | N1 | AA:388 | 4.37 | 1.00 | 48.67 |  |
| 13 | G | O6 | AA:382 | 4.44 | 1.00 | 55.65 |  |
| 14 | G | N1 | AA:391 | 4.59 | 1.00 | 58.28 |  |
| 15 | C | N3 | AA:385 | 4.68 | 1.00 | 52.44 |  |
| 16 | A | N9 | AA:390 | 4.75 | 1.00 | 46.19 |  |
| 17 | A | N1 | AA:390 | 4.87 | 1.00 | 43.76 |  |
  |

  
 Coded contact list:   
 Exclusion zone contact(s): /N3_A_/N7/O2'/O4/O6/O6/  
Consider K in place of MG (5 valid contact(s) for K vs. 0 for MG)  

### MG:AA:3821

|  | Res. type | Atom | Residue | Distance | Occupancy | B-factor | Closest solute to 1^st^ shell water |
| --- | --- | --- | --- | --- | --- | --- | --- |
| **Ion** | | | |  | 1.00 | 69.41 || 1 | G | N1 | AA:1425 | 2.53 | 1.00 | 63.71 |
| 2 | G | N2 | AA:1426 | 2.86 | 1.00 | 57.51 |  |
| 3 | A | OP2 | AA:1572 | 2.90 | 1.00 | 43.49 |  |
| 4 | G | N2 | AA:1425 | 2.98 | 1.00 | 62.35 |  |
| 5 | A | N7 | AA:1572 | 3.22 | 1.00 | 43.93 |  |
| 6 | G | O6 | AA:1425 | 3.67 | 1.00 | 64.98 |  |
| 7 | A | O5' | AA:1572 | 3.75 | 1.00 | 43.74 |  |
| 8 | C | N4 | AA:1574 | 3.93 | 1.00 | 62.36 |  |
| 9 | G | N3 | AA:1426 | 4.15 | 1.00 | 59.78 |  |
| 10 | G | N1 | AA:1426 | 4.28 | 1.00 | 58.12 |  |
| 11 | G | N7 | AA:1573 | 4.41 | 1.00 | 53.84 |  |
| 12 | G | OP2 | AA:1573 | 4.41 | 1.00 | 47.96 |  |
| 13 | G | O6 | AA:1424 | 4.42 | 1.00 | 65.69 |  |
| 14 | G | N3 | AA:1425 | 4.52 | 1.00 | 62.83 |  |
| 15 | A | N9 | AA:1572 | 4.72 | 1.00 | 45.19 |  |
| 16 | A | OP1 | AA:1572 | 4.79 | 1.00 | 43.56 |  |
  |

  
 Coded contact list:   
 Exclusion zone contact(s): /N1_G_/N2/N2/N7/OP2/  
Consider K in place of MG (4 valid contact(s) for K vs. 0 for MG)  

### MG:AA:3822

|  | Res. type | Atom | Residue | Distance | Occupancy | B-factor | Closest solute to 1^st^ shell water |
| --- | --- | --- | --- | --- | --- | --- | --- |
| **Ion** | | | |  | 1.00 | 77.01 || 1 | G | O6 | AA:2843 | 3.49 | 1.00 | 64.12 |
| 2 | G | N7 | AA:2842 | 4.11 | 1.00 | 69.64 |  |
| 3 | G | O6 | AA:2842 | 4.13 | 1.00 | 71.33 |  |
| 4 | G | N7 | AA:2843 | 4.30 | 1.00 | 67.55 |  |
| 5 | C | N4 | AA:2841 | 4.83 | 1.00 | 62.20 |  |
  |

  
 Coded contact list:   
1^st^ MG coordination shell missing (see Notes)  

### MG:AA:3823

|  | Res. type | Atom | Residue | Distance | Occupancy | B-factor | Closest solute to 1^st^ shell water |
| --- | --- | --- | --- | --- | --- | --- | --- |
| **Ion** | | | |  | 1.00 | 89.90 || 1 | U | O4 | AA:59 | 3.10 | 1.00 | 61.09 |
| 2 | C | OP2 | AA:65 | 3.79 | 1.00 | 57.30 |  |
| 3 | C | OP1 | AA:65 | 4.98 | 1.00 | 58.05 |  |
  |

  
 Coded contact list:   
 Exclusion zone contact(s): /O4/  
Consider K in place of MG (1 valid contact(s) for K vs. 0 for MG)  

### MG:AA:3824

|  | Res. type | Atom | Residue | Distance | Occupancy | B-factor | Closest solute to 1^st^ shell water |
| --- | --- | --- | --- | --- | --- | --- | --- |
| **Ion** | | | |  | 1.00 | 94.23 || 1 | G | O6 | AA:1015 | 3.17 | 1.00 | 116.18 |
| 2 | U | O4 | AA:1014 | 3.72 | 1.00 | 115.68 |  |
| 3 | C | N4 | AA:1147 | 3.74 | 1.00 | 116.62 |  |
| 4 | A | N6 | AA:1148 | 3.82 | 1.00 | 115.66 |  |
| 5 | G | O6 | AA:1016 | 4.12 | 1.00 | 116.31 |  |
| 6 | C | N4 | AA:1146 | 4.56 | 1.00 | 116.56 |  |
  |

  
 Coded contact list:   
 Exclusion zone contact(s): /O6/  
Consider K in place of MG (1 valid contact(s) for K vs. 0 for MG)  

### MG:AA:3825

|  | Res. type | Atom | Residue | Distance | Occupancy | B-factor | Closest solute to 1^st^ shell water |
| --- | --- | --- | --- | --- | --- | --- | --- |
| **Ion** | | | |  | 1.00 | 81.87 || 1 | U | O2' | AA:1082 | 3.42 | 1.00 | 200.36 |
| 2 | U | O4' | AA:1082 | 4.93 | 1.00 | 200.41 |  |
  |

  
 Coded contact list:   
1^st^ MG coordination shell missing (see Notes)  

### MG:AA:3826

|  | Res. type | Atom | Residue | Distance | Occupancy | B-factor | Closest solute to 1^st^ shell water |
| --- | --- | --- | --- | --- | --- | --- | --- |
| **Ion** | | | |  | 1.00 | 92.03 || 1 | A | N6 | AA:289 | 2.62 | 1.00 | 84.11 |
| 2 | A | N1 | AA:289 | 2.68 | 1.00 | 85.51 |  |
| 3 | G | O2' | AA:351 | 3.13 | 1.00 | 92.54 |  |
| 4 | G | N9 | AA:353 | 3.51 | 1.00 | 99.03 |  |
| 5 | G | N7 | AA:353 | 3.56 | 1.00 | 96.84 |  |
| 6 | G | N3 | AA:290 | 3.97 | 1.00 | 80.74 |  |
| 7 | G | N3 | AA:353 | 4.31 | 1.00 | 96.43 |  |
| 8 | G | O4' | AA:353 | 4.33 | 1.00 | 101.87 |  |
| 9 | G | N2 | AA:290 | 4.46 | 1.00 | 79.76 |  |
| 10 | G | N1 | AA:290 | 4.78 | 1.00 | 80.15 |  |
| 11 | G | N9 | AA:290 | 4.83 | 1.00 | 84.46 |  |
| 12 | G | N1 | AA:353 | 4.87 | 1.00 | 96.61 |  |
| 13 | G | O6 | AA:353 | 4.96 | 1.00 | 97.95 |  |
  |

  
 Coded contact list:   
 Exclusion zone contact(s): /N1_A_/N6/O2'/  
Consider K in place of MG (3 valid contact(s) for K vs. 0 for MG)  

### MG:AA:3827

|  | Res. type | Atom | Residue | Distance | Occupancy | B-factor | Closest solute to 1^st^ shell water |
| --- | --- | --- | --- | --- | --- | --- | --- |
| **Ion** | | | |  | 1.00 | 62.83 || 1 | U | O4 | AA:1818 | 3.37 | 1.00 | 44.59 |
| 2 | G | OP2 | AA:1801 | 3.54 | 1.00 | 46.43 |  |
| 3 | LYS | NZ | AD:154 | 3.89 | 1.00 | 51.77 |  |
| 4 | G | O6 | AA:1817 | 4.32 | 1.00 | 52.19 |  |
| 5 | G | O5' | AA:1801 | 4.46 | 1.00 | 45.71 |  |
| 6 | C | O2' | AA:1800 | 4.47 | 1.00 | 46.33 |  |
  |

  
 Coded contact list:   
 Exclusion zone contact(s): /O4/  
Consider K in place of MG (1 valid contact(s) for K vs. 0 for MG)  

### MG:AA:3828

|  | Res. type | Atom | Residue | Distance | Occupancy | B-factor | Closest solute to 1^st^ shell water |
| --- | --- | --- | --- | --- | --- | --- | --- |
| **Ion** | | | |  | 1.00 | 113.11 || 1 | G | O6 | AA:290 | 2.65 | 1.00 | 81.32 |
| 2 | C | N4 | AA:291 | 2.91 | 1.00 | 76.62 |  |
| 3 | G | O6 | AA:349 | 3.04 | 1.00 | 73.75 |  |
| 4 | U | O4 | AA:350 | 3.22 | 1.00 | 78.38 |  |
| 5 | G | N2 | AA:351 | 3.69 | 1.00 | 92.01 |  |
| 6 | U | N3 | AA:350 | 3.86 | 1.00 | 78.09 |  |
| 7 | G | N7 | AA:290 | 4.10 | 1.00 | 82.91 |  |
| 8 | G | N3 | AA:351 | 4.51 | 1.00 | 91.95 |  |
| 9 | C | N3 | AA:291 | 4.53 | 1.00 | 76.58 |  |
| 10 | G | N1 | AA:351 | 4.54 | 1.00 | 92.64 |  |
| 11 | G | N1 | AA:349 | 4.70 | 1.00 | 71.80 |  |
| 12 | G | N1 | AA:290 | 4.71 | 1.00 | 80.15 |  |
  |

  
 Coded contact list:   
 Exclusion zone contact(s): /N4/O4/O6/O6/  
Consider K in place of MG (4 valid contact(s) for K vs. 0 for MG)  

### MG:AA:3829

|  | Res. type | Atom | Residue | Distance | Occupancy | B-factor | Closest solute to 1^st^ shell water |
| --- | --- | --- | --- | --- | --- | --- | --- |
| **Ion** | | | |  | 1.00 | 88.38 || 1 | G | O6 | AA:1051 | 2.57 | 1.00 | 184.36 |
| 2 | A | N6 | AA:1050 | 2.77 | 1.00 | 173.03 |  |
| 3 | A | N7 | AA:1050 | 3.05 | 1.00 | 173.02 |  |
| 4 | G | N7 | AA:1051 | 3.20 | 1.00 | 184.13 |  |
| 5 | U | O4 | AA:1108 | 4.10 | 1.00 | 184.27 |  |
| 6 | A | N1 | AA:1050 | 4.40 | 1.00 | 172.65 |  |
| 7 | C | OP2 | AA:1049 | 4.40 | 1.00 | 179.72 |  |
| 8 | G | N1 | AA:1051 | 4.57 | 1.00 | 184.25 |  |
| 9 | U | N3 | AA:1108 | 4.63 | 1.00 | 184.95 |  |
| 10 | C | N4 | AA:1052 | 4.71 | 1.00 | 189.07 |  |
| 11 | A | N9 | AA:1050 | 4.91 | 1.00 | 174.04 |  |
| 12 | C | O2 | AA:1109 | 4.97 | 1.00 | 181.17 |  |
  |

  
 Coded contact list:   
 Exclusion zone contact(s): /N6/N7/N7/O6/  
Consider K in place of MG (3 valid contact(s) for K vs. 0 for MG)  

### MG:AA:3830

|  | Res. type | Atom | Residue | Distance | Occupancy | B-factor | Closest solute to 1^st^ shell water |
| --- | --- | --- | --- | --- | --- | --- | --- |
| **Ion** | | | |  | 1.00 | 83.58 || 1 | G | O6 | AA:270X | 2.42 | 1.00 | 80.11 |
| 2 | G | N7 | AA:270W | 2.55 | 1.00 | 97.28 |  |
| 3 | G | O6 | AA:270W | 2.91 | 1.00 | 99.15 |  |
| 4 | G | N1 | AA:270X | 3.60 | 1.00 | 79.60 |  |
| 5 | G | O6 | AA:270V | 3.92 | 1.00 | 100.05 |  |
| 6 | G | N7 | AA:270X | 4.02 | 1.00 | 79.16 |  |
| 7 | G | N7 | AA:270V | 4.40 | 1.00 | 99.53 |  |
| 8 | G | N9 | AA:270W | 4.76 | 1.00 | 95.63 |  |
| 9 | G | N1 | AA:270W | 4.76 | 1.00 | 98.84 |  |
| 10 | G | N1 | AA:270V | 4.80 | 1.00 | 99.86 |  |
  |

  
 Coded contact list:   
 Exclusion zone contact(s): /N7/O6/O6/  
Consider NA in place of MG (2 valid contact(s) for NA vs. 0 for MG)  

### MG:AA:3831

|  | Res. type | Atom | Residue | Distance | Occupancy | B-factor | Closest solute to 1^st^ shell water |
| --- | --- | --- | --- | --- | --- | --- | --- |
| **Ion** | | | |  | 1.00 | 77.21 || 1 | C | OP2 | AA:1049 | 1.89 | 1.00 | 179.72 |
| 2 | C | OP1 | AA:1049 | 4.07 | 1.00 | 179.81 |  |
| 3 | A | O3' | AA:1048 | 4.10 | 1.00 | 180.13 |  |
| 4 | C | O5' | AA:1049 | 4.32 | 1.00 | 179.36 |  |
| 5 | A | OP2 | AA:1050 | 4.46 | 1.00 | 175.72 |  |
| 6 | A | O5' | AA:1048 | 4.56 | 1.00 | 183.38 |  |
| 7 | U | O4 | AA:1108 | 4.60 | 1.00 | 184.27 |  |
| 8 | A | N7 | AA:1050 | 4.87 | 1.00 | 173.02 |  |
  |

  
 Contact(s): /OP2/  
 Coded contact list: O_ph_  

### MG:AA:3832

|  | Res. type | Atom | Residue | Distance | Occupancy | B-factor | Closest solute to 1^st^ shell water |
| --- | --- | --- | --- | --- | --- | --- | --- |
| **Ion** | | | |  | 1.00 | 63.57 || 1 | G | N7 | AA:298 | 2.72 | 1.00 | 70.12 |
| 2 | C | O5' | AA:297 | 2.94 | 1.00 | 70.78 |  |
| 3 | C | OP2 | AA:297 | 2.98 | 1.00 | 70.55 |  |
| 4 | G | OP2 | AA:298 | 3.19 | 1.00 | 68.01 |  |
| 5 | C | OP1 | AA:297 | 3.96 | 1.00 | 72.15 |  |
| 6 | C | O3' | AA:297 | 4.21 | 1.00 | 67.95 |  |
| 7 | G | O5' | AA:298 | 4.25 | 1.00 | 68.54 |  |
| 8 | G | N9 | AA:298 | 4.47 | 1.00 | 69.58 |  |
| 9 | G | O6 | AA:298 | 4.50 | 1.00 | 72.95 |  |
| 10 | C | N1 | AA:297 | 4.65 | 1.00 | 72.05 |  |
| 11 | C | O4' | AA:297 | 4.79 | 1.00 | 69.75 |  |
| 12 | C | O3' | AA:296 | 4.85 | 1.00 | 71.63 |  |
  |

  
 Coded contact list:   
 Exclusion zone contact(s): /N7/O5'/OP2/OP2/  
Consider K in place of MG (4 valid contact(s) for K vs. 0 for MG)  

### MG:AA:3833

|  | Res. type | Atom | Residue | Distance | Occupancy | B-factor | Closest solute to 1^st^ shell water |
| --- | --- | --- | --- | --- | --- | --- | --- |
| **Ion** | | | |  | 1.00 | 80.44 || 1 | C | OP1 | AA:1605 | 2.92 | 1.00 | 36.90 |
| 2 | G | OP2 | AA:1606 | 2.92 | 1.00 | 43.42 |  |
| 3 | C | OP2 | AA:1305 | 4.16 | 1.00 | 52.28 |  |
| 4 | C | O5' | AA:1605 | 4.17 | 1.00 | 38.37 |  |
| 5 | C | OP2 | AA:1605 | 4.29 | 1.00 | 36.28 |  |
| 6 | G | OP1 | AA:1606 | 5.00 | 1.00 | 43.14 |  |
  |

  
 Coded contact list:   
 Exclusion zone contact(s): /OP1/OP2/  
Consider K in place of MG (2 valid contact(s) for K vs. 0 for MG)  

### MG:AA:3834

|  | Res. type | Atom | Residue | Distance | Occupancy | B-factor | Closest solute to 1^st^ shell water |
| --- | --- | --- | --- | --- | --- | --- | --- |
| **Ion** | | | |  | 1.00 | 79.06 || 1 | A | OP2 | AA:359 | 4.16 | 1.00 | 113.03 |
| 2 | G | N7 | AA:360 | 4.71 | 1.00 | 103.13 |  |
  |

  
 Coded contact list:   
1^st^ MG coordination shell missing (see Notes)  

### MG:AA:3835

|  | Res. type | Atom | Residue | Distance | Occupancy | B-factor | Closest solute to 1^st^ shell water |
| --- | --- | --- | --- | --- | --- | --- | --- |
| **Ion** | | | |  | 1.00 | 56.85 || 1 | G | OP2 | AA:794 | 3.03 | 1.00 | 38.93 |
| 2 | G | N7 | AA:776 | 3.11 | 1.00 | 43.25 |  |
| 3 | A | O2' | AA:793 | 3.15 | 1.00 | 48.59 |  |
| 4 | G | OP1 | AA:794 | 3.30 | 1.00 | 39.87 |  |
| 5 | G | O6 | AA:776 | 3.31 | 1.00 | 45.38 |  |
| 6 | A | O3' | AA:793 | 3.36 | 1.00 | 41.17 |  |
| 7 | A | N9 | AA:793 | 4.35 | 1.00 | 46.03 |  |
| 8 | A | O4' | AA:793 | 4.38 | 1.00 | 44.15 |  |
| 9 | A | OP2 | AA:793 | 4.58 | 1.00 | 35.84 |  |
| 10 | C | OP1 | AA:678 | 4.66 | 1.00 | 43.05 |  |
| 11 | G | O5' | AA:794 | 4.87 | 1.00 | 38.95 |  |
  |

  
 Coded contact list:   
 Exclusion zone contact(s): /N7/O2'/O3'/O6/OP1/OP2/  
Consider K in place of MG (6 valid contact(s) for K vs. 0 for MG)  

### MG:AA:3836

|  | Res. type | Atom | Residue | Distance | Occupancy | B-factor | Closest solute to 1^st^ shell water |
| --- | --- | --- | --- | --- | --- | --- | --- |
| **Ion** | | | |  | 1.00 | 62.66 || 1 | G | O6 | AA:2625 | 3.53 | 1.00 | 40.48 |
| 2 | G | O6 | AA:2777 | 4.00 | 1.00 | 46.87 |  |
| 3 | G | O6 | AA:2624 | 4.50 | 1.00 | 43.88 |  |
| 4 | C | N4 | AA:2043 | 4.54 | 1.00 | 45.35 |  |
| 5 | G | N7 | AA:2625 | 4.90 | 1.00 | 41.65 |  |
| 6 | G | N7 | AA:2624 | 4.96 | 1.00 | 43.65 |  |
  |

  
 Coded contact list:   
1^st^ MG coordination shell missing (see Notes)  

### MG:AA:3837

|  | Res. type | Atom | Residue | Distance | Occupancy | B-factor | Closest solute to 1^st^ shell water |
| --- | --- | --- | --- | --- | --- | --- | --- |
| **Ion** | | | |  | 1.00 | 116.85 || 1 | C | N3 | AA:1870 | 3.91 | 1.00 | 85.01 |
| 2 | C | O2 | AA:1870 | 4.18 | 1.00 | 85.68 |  |
| 3 | C | N4 | AA:1870 | 4.88 | 1.00 | 84.26 |  |
| 4 | C | N1 | AA:1870 | 4.94 | 1.00 | 84.66 |  |
  |

  
 Coded contact list:   
1^st^ MG coordination shell missing (see Notes)  

### MG:AA:3838

|  | Res. type | Atom | Residue | Distance | Occupancy | B-factor | Closest solute to 1^st^ shell water |
| --- | --- | --- | --- | --- | --- | --- | --- |
| **Ion** | | | |  | 1.00 | 102.81 || 1 | U | O3' | AA:405 | 2.49 | 1.00 | 91.99 |
| 2 | U | O2' | AA:405 | 2.67 | 1.00 | 97.30 |  |
| 3 | G | OP2 | AA:406 | 2.85 | 1.00 | 90.13 |  |
| 4 | G | OP1 | AA:406 | 4.04 | 1.00 | 90.46 |  |
| 5 | G | O5' | AA:406 | 4.48 | 1.00 | 87.57 |  |
| 6 | U | O4' | AA:405 | 4.71 | 1.00 | 96.28 |  |
  |

  
 Coded contact list:   
 Exclusion zone contact(s): /O2'/O3'/OP2/  
Consider K in place of MG (2 valid contact(s) for K vs. 0 for MG)  

### MG:AA:3839

|  | Res. type | Atom | Residue | Distance | Occupancy | B-factor | Closest solute to 1^st^ shell water |
| --- | --- | --- | --- | --- | --- | --- | --- |
| **Ion** | | | |  | 1.00 | 63.91 || 1 | G | OP2 | AA:1215 | 4.19 | 1.00 | 63.59 |
| 2 | G | OP2 | AA:1216 | 4.50 | 1.00 | 61.45 |  |
| 3 | G | N7 | AA:1216 | 4.59 | 1.00 | 66.90 |  |
| 4 | C | N4 | AA:1217 | 4.87 | 1.00 | 63.74 |  |
  |

  
 Coded contact list:   
1^st^ MG coordination shell missing (see Notes)  

### MG:AA:3840

|  | Res. type | Atom | Residue | Distance | Occupancy | B-factor | Closest solute to 1^st^ shell water |
| --- | --- | --- | --- | --- | --- | --- | --- |
| **Ion** | | | |  | 1.00 | 77.10 || 1 | C | OP1 | AA:1604 | 2.19 | 1.00 | 34.85 |
| 2 | A | OP2 | AA:1395 | 2.51 | 1.00 | 52.14 |  |
| 3 | U | OP1 | AA:1394 | 2.78 | 1.00 | 52.93 |  |
| 4 | A | OP1 | AA:1395 | 3.14 | 1.00 | 52.83 |  |
| 5 | U | O3' | AA:1394 | 3.83 | 1.00 | 53.46 |  |
| 6 | C | OP2 | AA:1604 | 3.93 | 1.00 | 34.59 |  |
| 7 | A | O3' | AA:1603 | 4.42 | 1.00 | 34.35 |  |
| 8 | C | O5' | AA:1604 | 4.55 | 1.00 | 36.94 |  |
| 9 | A | O5' | AA:1395 | 4.64 | 1.00 | 53.98 |  |
| 10 | U | O5' | AA:1394 | 4.73 | 1.00 | 52.27 |  |
  |

  
 Contact(s): /OP1/  
 Coded contact list: O_ph_  
 Exclusion zone contact(s): /OP1/OP1/OP2/  
Consider K in place of MG (2 valid contact(s) for K vs. 1 for MG)  

### MG:AA:3841

|  | Res. type | Atom | Residue | Distance | Occupancy | B-factor | Closest solute to 1^st^ shell water |
| --- | --- | --- | --- | --- | --- | --- | --- |
| **Ion** | | | |  | 1.00 | 85.25 || 1 | C | O2' | AA:1076 | 4.11 | 1.00 | 199.63 |
| 2 | C | O2 | AA:1076 | 4.67 | 1.00 | 200.11 |  |
| 3 | G | N3 | AA:1063 | 4.99 | 1.00 | 200.33 |  |
  |

  
 Coded contact list:   
1^st^ MG coordination shell missing (see Notes)  

### MG:AA:3842

|  | Res. type | Atom | Residue | Distance | Occupancy | B-factor | Closest solute to 1^st^ shell water |
| --- | --- | --- | --- | --- | --- | --- | --- |
| **Ion** | | | |  | 1.00 | 67.79 || 1 | G | N7 | AA:1443 | 3.41 | 1.00 | 71.90 |
| 2 | G | O6 | AA:1443 | 4.12 | 1.00 | 73.86 |  |
| 3 | G | O6 | AA:1444 | 4.47 | 1.00 | 81.21 |  |
| 4 | G | N7 | AA:1444 | 4.62 | 1.00 | 82.29 |  |
| 5 | G | OP2 | AA:1443 | 4.66 | 1.00 | 68.26 |  |
  |

  
 Coded contact list:   
1^st^ MG coordination shell missing (see Notes)  

### MG:AA:3843

|  | Res. type | Atom | Residue | Distance | Occupancy | B-factor | Closest solute to 1^st^ shell water |
| --- | --- | --- | --- | --- | --- | --- | --- |
| **Ion** | | | |  | 1.00 | 81.15 || 1 | G | OP2 | AA:219 | 3.60 | 1.00 | 54.58 |
| 2 | A | OP1 | AA:218 | 3.70 | 1.00 | 51.58 |  |
| 3 | U | OP2 | AA:427 | 4.25 | 1.00 | 59.31 |  |
| 4 | C | OP1 | AA:426 | 4.45 | 1.00 | 61.63 |  |
| 5 | A | OP2 | AA:218 | 4.61 | 1.00 | 51.86 |  |
| 6 | A | O5' | AA:218 | 4.78 | 1.00 | 51.99 |  |
  |

  
 Coded contact list:   
1^st^ MG coordination shell missing (see Notes)  

### MG:AA:3844

|  | Res. type | Atom | Residue | Distance | Occupancy | B-factor | Closest solute to 1^st^ shell water |
| --- | --- | --- | --- | --- | --- | --- | --- |
| **Ion** | | | |  | 1.00 | 94.03 || 1 | U | O4 | AA:597 | 4.19 | 1.00 | 56.52 |
| 2 | U | O4 | AA:657 | 4.85 | 1.00 | 63.57 |  |
  |

  
 Coded contact list:   
1^st^ MG coordination shell missing (see Notes)  

### MG:AA:3845

|  | Res. type | Atom | Residue | Distance | Occupancy | B-factor | Closest solute to 1^st^ shell water |
| --- | --- | --- | --- | --- | --- | --- | --- |
| **Ion** | | | |  | 1.00 | 54.44 || 1 | A | OP1 | AA:213 | 3.38 | 1.00 | 62.41 |
| 2 | G | OP2 | AA:214 | 4.26 | 1.00 | 69.64 |  |
| 3 | G | OP1 | AA:214 | 4.26 | 1.00 | 70.72 |  |
| 4 | G | OP1 | AA:177 | 4.66 | 1.00 | 56.15 |  |
| 5 | G | O3' | AA:176 | 4.80 | 1.00 | 57.62 |  |
| 6 | G | OP1 | AA:176 | 4.86 | 1.00 | 74.02 |  |
| 7 | G | OP2 | AA:177 | 4.97 | 1.00 | 56.23 |  |
  |

  
 Coded contact list:   
 Exclusion zone contact(s): /OP1/  
Consider K in place of MG (1 valid contact(s) for K vs. 0 for MG)  

### MG:AA:3847

|  | Res. type | Atom | Residue | Distance | Occupancy | B-factor | Closest solute to 1^st^ shell water |
| --- | --- | --- | --- | --- | --- | --- | --- |
| **Ion** | | | |  | 1.00 | 84.31 || 1 | G | N7 | AA:2371 | 2.81 | 1.00 | 70.35 |
| 2 | G | O6 | AA:2371 | 2.82 | 1.00 | 73.80 |  |
| 3 | G | N7 | AA:2372 | 3.54 | 1.00 | 62.75 |  |
| 4 | G | N1 | AA:2371 | 4.37 | 1.00 | 72.45 |  |
| 5 | G | N9 | AA:2371 | 4.66 | 1.00 | 70.57 |  |
| 6 | G | O6 | AA:2372 | 4.99 | 1.00 | 63.31 |  |
  |

  
 Coded contact list:   
 Exclusion zone contact(s): /N7/O6/  
Consider K in place of MG (2 valid contact(s) for K vs. 0 for MG)  

### MG:AA:3848

|  | Res. type | Atom | Residue | Distance | Occupancy | B-factor | Closest solute to 1^st^ shell water |
| --- | --- | --- | --- | --- | --- | --- | --- |
| **Ion** | | | |  | 1.00 | 90.29 || 1 | G | N7 | AA:113 | 3.10 | 1.00 | 50.94 |
| 2 | G | O6 | AA:113 | 3.30 | 1.00 | 49.05 |  |
| 3 | U | OP2 | AA:112 | 3.89 | 1.00 | 57.31 |  |
  |

  
 Coded contact list:   
 Exclusion zone contact(s): /N7/O6/  
Consider K in place of MG (2 valid contact(s) for K vs. 0 for MG)  

### MG:AA:3849

|  | Res. type | Atom | Residue | Distance | Occupancy | B-factor | Closest solute to 1^st^ shell water |
| --- | --- | --- | --- | --- | --- | --- | --- |
| **Ion** | | | |  | 1.00 | 59.97 || 1 | U | OP1 | AA:2832 | 2.30 | 1.00 | 66.63 |
| 2 | U | OP2 | AA:2832 | 3.18 | 1.00 | 66.61 |  |
| 3 | G | N7 | AA:2833 | 3.21 | 1.00 | 106.28 |  |
| 4 | U | O5' | AA:2832 | 4.09 | 1.00 | 69.68 |  |
| 5 | G | O6 | AA:2833 | 4.27 | 1.00 | 108.86 |  |
| 6 | G | O3' | AA:2831 | 4.44 | 1.00 | 66.65 |  |
| 7 | G | O5' | AA:2831 | 4.86 | 1.00 | 63.35 |  |
  |

  
 Contact(s): /OP1/  
 Coded contact list: O_ph_  
 Exclusion zone contact(s): /N7/OP2/  
Consider K in place of MG (2 valid contact(s) for K vs. 1 for MG)  

### MG:AA:3850

|  | Res. type | Atom | Residue | Distance | Occupancy | B-factor | Closest solute to 1^st^ shell water |
| --- | --- | --- | --- | --- | --- | --- | --- |
| **Ion** | | | |  | 1.00 | 41.85 || 1 | G | N3 | AA:2049 | 2.22 | 1.00 | 28.70 |
| 2 | G | O2' | AA:2049 | 2.57 | 1.00 | 32.87 |  |
| 3 | G | N2 | AA:2049 | 3.30 | 1.00 | 28.45 |  |
| 4 | G | N9 | AA:2049 | 3.45 | 1.00 | 30.22 |  |
| 5 | C | O4' | AA:2050 | 3.50 | 1.00 | 36.73 |  |
| 6 | C | O2 | AA:2620 | 3.70 | 1.00 | 32.11 |  |
| 7 | G | N2 | AA:2048 | 3.90 | 1.00 | 32.85 |  |
| 8 | ALA | O | AE:157 | 4.05 | 1.00 | 59.92 |  |
| 9 | G | O4' | AA:2049 | 4.32 | 1.00 | 35.80 |  |
| 10 | G | N1 | AA:2049 | 4.46 | 1.00 | 28.72 |  |
| 11 | C | O5' | AA:2050 | 4.52 | 1.00 | 36.69 |  |
| 12 | ALA | N | AE:157 | 4.68 | 1.00 | 55.56 |  |
| 13 | G | O3' | AA:2049 | 4.77 | 1.00 | 38.24 |  |
| 14 | C | N1 | AA:2050 | 4.85 | 1.00 | 32.94 |  |
| 15 | C | N3 | AA:2620 | 4.89 | 1.00 | 30.95 |  |
| 16 | G | N1 | AA:2048 | 4.96 | 1.00 | 31.26 |  |
  |

  
 Contact(s): /N3_G_/  
 Coded contact list: N_b_  
 Exclusion zone contact(s): /N2/O2'/  

### MG:AA:3851

|  | Res. type | Atom | Residue | Distance | Occupancy | B-factor | Closest solute to 1^st^ shell water |
| --- | --- | --- | --- | --- | --- | --- | --- |
| **Ion** | | | |  | 1.00 | 60.41 || 1 | A | N7 | AA:1321 | 4.78 | 1.00 | 39.85 |
  |

  
 Coded contact list:   
1^st^ MG coordination shell missing (see Notes)  

### MG:AA:3852

|  | Res. type | Atom | Residue | Distance | Occupancy | B-factor | Closest solute to 1^st^ shell water |
| --- | --- | --- | --- | --- | --- | --- | --- |
| **Ion** | | | |  | 1.00 | 67.97 || 1 | C | O5' | AA:850 | 2.83 | 1.00 | 76.11 |
| 2 | U | OP2 | AA:851 | 2.88 | 1.00 | 74.46 |  |
| 3 | C | OP2 | AA:850 | 3.43 | 1.00 | 77.47 |  |
| 4 | C | N1 | AA:850 | 3.54 | 1.00 | 73.25 |  |
| 5 | C | O3' | AA:850 | 3.96 | 1.00 | 73.62 |  |
| 6 | C | O4' | AA:850 | 4.12 | 1.00 | 74.34 |  |
| 7 | U | O5' | AA:851 | 4.32 | 1.00 | 74.92 |  |
| 8 | C | N3 | AA:850 | 4.41 | 1.00 | 73.72 |  |
| 9 | U | O4 | AA:851 | 4.53 | 1.00 | 83.21 |  |
| 10 | C | OP1 | AA:850 | 4.56 | 1.00 | 75.82 |  |
| 11 | A | O3' | AA:849 | 4.92 | 1.00 | 76.81 |  |
| 12 | C | N4 | AA:850 | 4.92 | 1.00 | 72.41 |  |
| 13 | C | O2 | AA:850 | 4.95 | 1.00 | 74.68 |  |
| 14 | U | N1 | AA:851 | 4.95 | 1.00 | 80.67 |  |
  |

  
 Coded contact list:   
 Exclusion zone contact(s): /O5'/OP2/  
Consider K in place of MG (2 valid contact(s) for K vs. 0 for MG)  

### MG:AA:3853

|  | Res. type | Atom | Residue | Distance | Occupancy | B-factor | Closest solute to 1^st^ shell water |
| --- | --- | --- | --- | --- | --- | --- | --- |
| **Ion** | | | |  | 1.00 | 55.88 || 1 | G | OP1 | AA:966 | 2.50 | 1.00 | 56.27 |
| 2 | U | O2' | AA:2272 | 3.58 | 1.00 | 57.82 |  |
| 3 | C | O3' | AA:965 | 3.71 | 1.00 | 55.56 |  |
| 4 | U | O4' | AA:2272 | 3.98 | 1.00 | 58.64 |  |
| 5 | G | OP1 | AA:823 | 4.52 | 1.00 | 56.81 |  |
| 6 | G | O5' | AA:966 | 4.57 | 1.00 | 57.78 |  |
| 7 | G | OP2 | AA:966 | 4.80 | 1.00 | 56.85 |  |
| 8 | U | O3' | AA:2272 | 4.84 | 1.00 | 57.39 |  |
| 9 | U | O3' | AA:822 | 4.97 | 1.00 | 58.00 |  |
  |

  
 Coded contact list:   
 Exclusion zone contact(s): /OP1/  
Consider NA in place of MG (1 valid contact(s) for NA vs. 0 for MG)  

### MG:AA:3854

|  | Res. type | Atom | Residue | Distance | Occupancy | B-factor | Closest solute to 1^st^ shell water |
| --- | --- | --- | --- | --- | --- | --- | --- |
| **Ion** | | | |  | 1.00 | 71.45 || 1 | G | N7 | AA:1162 | 3.82 | 1.00 | 75.67 |
| 2 | G | N7 | AA:1163 | 4.39 | 1.00 | 73.19 |  |
| 3 | G | O6 | AA:1163 | 4.47 | 1.00 | 74.07 |  |
| 4 | G | O6 | AA:1162 | 4.61 | 1.00 | 76.16 |  |
  |

  
 Coded contact list:   
1^st^ MG coordination shell missing (see Notes)  

### MG:AA:3855

|  | Res. type | Atom | Residue | Distance | Occupancy | B-factor | Closest solute to 1^st^ shell water |
| --- | --- | --- | --- | --- | --- | --- | --- |
| **Ion** | | | |  | 1.00 | 72.81 || 1 | C | O3' | AA:277 | 2.75 | 1.00 | 146.24 |
| 2 | C | O2' | AA:277 | 3.01 | 1.00 | 149.71 |  |
| 3 | C | O4' | AA:277 | 3.66 | 1.00 | 150.86 |  |
| 4 | C | O5' | AA:277 | 4.09 | 1.00 | 153.84 |  |
| 5 | A | OP2 | AA:278 | 4.66 | 1.00 | 143.77 |  |
| 6 | A | OP1 | AA:278 | 4.68 | 1.00 | 143.82 |  |
  |

  
 Coded contact list:   
 Exclusion zone contact(s): /O2'/O3'/  
Consider K in place of MG (2 valid contact(s) for K vs. 0 for MG)  

### MG:AA:3857

|  | Res. type | Atom | Residue | Distance | Occupancy | B-factor | Closest solute to 1^st^ shell water |
| --- | --- | --- | --- | --- | --- | --- | --- |
| **Ion** | | | |  | 1.00 | 65.09 || 1 | C | N4 | AA:1663 | 3.30 | 1.00 | 48.02 |
| 2 | G | O2' | AA:1992 | 3.43 | 1.00 | 47.37 |  |
| 3 | G | N9 | AA:1992 | 3.60 | 1.00 | 45.14 |  |
| 4 | U | OP2 | AA:1993 | 3.78 | 1.00 | 47.52 |  |
| 5 | G | O3' | AA:1992 | 3.80 | 1.00 | 45.65 |  |
| 6 | G | O5' | AA:1992 | 4.04 | 1.00 | 45.71 |  |
| 7 | G | O6 | AA:1997 | 4.30 | 1.00 | 49.60 |  |
| 8 | G | O4' | AA:1992 | 4.31 | 1.00 | 47.18 |  |
| 9 | G | N7 | AA:1992 | 4.54 | 1.00 | 43.75 |  |
| 10 | C | N4 | AA:1662 | 4.55 | 1.00 | 44.00 |  |
| 11 | G | OP1 | AA:1992 | 4.99 | 1.00 | 46.48 |  |
  |

  
 Coded contact list:   
 Exclusion zone contact(s): /N4/  
Consider K in place of MG (1 valid contact(s) for K vs. 0 for MG)  

### MG:AA:3858

|  | Res. type | Atom | Residue | Distance | Occupancy | B-factor | Closest solute to 1^st^ shell water |
| --- | --- | --- | --- | --- | --- | --- | --- |
| **Ion** | | | |  | 1.00 | 76.89 || 1 | U | O4 | AA:1165 | 4.21 | 1.00 | 84.36 |
| 2 | G | N7 | AA:1164 | 4.70 | 1.00 | 76.42 |  |
  |

  
 Coded contact list:   
1^st^ MG coordination shell missing (see Notes)  

### MG:AA:3859

|  | Res. type | Atom | Residue | Distance | Occupancy | B-factor | Closest solute to 1^st^ shell water |
| --- | --- | --- | --- | --- | --- | --- | --- |
| **Ion** | | | |  | 1.00 | 80.38 || 1 | C | N3 | AA:1630A | 2.65 | 1.00 | 55.75 |
| 2 | A | OP2 | AA:1632 | 2.73 | 1.00 | 54.31 |  |
| 3 | G | N7 | AA:1633 | 2.74 | 1.00 | 44.51 |  |
| 4 | A | N7 | AA:1632 | 3.16 | 1.00 | 43.94 |  |
| 5 | G | O6 | AA:1633 | 3.16 | 1.00 | 46.36 |  |
| 6 | C | N4 | AA:1630A | 3.25 | 1.00 | 57.08 |  |
| 7 | G | N1 | AA:1635 | 3.33 | 1.00 | 51.39 |  |
| 8 | C | O2 | AA:1630A | 3.72 | 1.00 | 55.99 |  |
| 9 | G | O6 | AA:1635 | 3.79 | 1.00 | 53.97 |  |
| 10 | C | N1 | AA:1630A | 4.00 | 1.00 | 58.34 |  |
| 11 | A | O5' | AA:1632 | 4.17 | 1.00 | 53.53 |  |
| 12 | G | N2 | AA:1635 | 4.19 | 1.00 | 53.02 |  |
| 13 | A | OP1 | AA:1632 | 4.30 | 1.00 | 56.56 |  |
| 14 | A | N9 | AA:1632 | 4.64 | 1.00 | 43.68 |  |
| 15 | G | N3 | AA:1635 | 4.87 | 1.00 | 50.44 |  |
| 16 | G | N9 | AA:1633 | 4.96 | 1.00 | 42.30 |  |
  |

  
 Coded contact list:   
 Exclusion zone contact(s): /N1_G_/N3_C_/N4/N7/N7/O6/OP2/  
Consider K in place of MG (7 valid contact(s) for K vs. 0 for MG)  

### MG:AA:3860

|  | Res. type | Atom | Residue | Distance | Occupancy | B-factor | Closest solute to 1^st^ shell water |
| --- | --- | --- | --- | --- | --- | --- | --- |
| **Ion** | | | |  | 1.00 | 83.00 || 1 | U | O2' | AA:358 | 2.98 | 1.00 | 112.79 |
| 2 | A | O5' | AA:359 | 3.69 | 1.00 | 113.09 |  |
| 3 | U | O3' | AA:358 | 3.71 | 1.00 | 113.10 |  |
| 4 | A | O4' | AA:359 | 3.93 | 1.00 | 110.37 |  |
| 5 | A | OP1 | AA:359 | 4.15 | 1.00 | 113.79 |  |
  |

  
 Coded contact list:   
 Exclusion zone contact(s): /O2'/  
Consider K in place of MG (1 valid contact(s) for K vs. 0 for MG)  

### MG:AA:3861

|  | Res. type | Atom | Residue | Distance | Occupancy | B-factor | Closest solute to 1^st^ shell water |
| --- | --- | --- | --- | --- | --- | --- | --- |
| **Ion** | | | |  | 1.00 | 99.44 || 1 | A | N7 | AA:653 | 3.12 | 1.00 | 148.64 |
| 2 | A | N9 | AA:653 | 4.05 | 1.00 | 148.15 |  |
| 3 | A | OP2 | AA:653 | 4.79 | 1.00 | 121.23 |  |
| 4 | A | O5' | AA:653 | 4.80 | 1.00 | 127.65 |  |
  |

  
 Coded contact list:   
 Exclusion zone contact(s): /N7/  
Consider K in place of MG (1 valid contact(s) for K vs. 0 for MG)  

### MG:AA:3862

|  | Res. type | Atom | Residue | Distance | Occupancy | B-factor | Closest solute to 1^st^ shell water |
| --- | --- | --- | --- | --- | --- | --- | --- |
| **Ion** | | | |  | 1.00 | 75.62 || 1 | A | N1 | AA:340 | 3.95 | 1.00 | 68.50 |
| 2 | G | N7 | AA:321 | 4.66 | 1.00 | 75.57 |  |
| 3 | G | N3 | AA:341 | 4.69 | 1.00 | 77.41 |  |
| 4 | A | N6 | AA:322 | 4.88 | 1.00 | 73.80 |  |
| 5 | G | O2' | AA:341 | 4.89 | 1.00 | 76.13 |  |
  |

  
 Coded contact list:   
1^st^ MG coordination shell missing (see Notes)  

### MG:AA:3863

|  | Res. type | Atom | Residue | Distance | Occupancy | B-factor | Closest solute to 1^st^ shell water |
| --- | --- | --- | --- | --- | --- | --- | --- |
| **Ion** | | | |  | 1.00 | 72.58 || 1 | G | O6 | AA:974 | 3.30 | 1.00 | 70.93 |
| 2 | G | O6 | AA:972 | 4.30 | 1.00 | 69.11 |  |
| 3 | G | O6 | AA:989 | 4.41 | 1.00 | 81.31 |  |
| 4 | C | OP2 | AA:971 | 4.47 | 1.00 | 58.31 |  |
| 5 | G | N7 | AA:974 | 4.63 | 1.00 | 72.35 |  |
  |

  
 Coded contact list:   
 Exclusion zone contact(s): /O6/  
Consider K in place of MG (1 valid contact(s) for K vs. 0 for MG)  

### MG:AA:3864

|  | Res. type | Atom | Residue | Distance | Occupancy | B-factor | Closest solute to 1^st^ shell water |
| --- | --- | --- | --- | --- | --- | --- | --- |
| **Ion** | | | |  | 1.00 | 48.73 || 1 | C | O2 | AA:2464 | 2.70 | 1.00 | 73.97 |
| 2 | C | O2' | AA:2464 | 2.85 | 1.00 | 73.31 |  |
| 3 | A | OP2 | AA:1126 | 3.17 | 1.00 | 71.89 |  |
| 4 | A | OP1 | AA:1126 | 3.26 | 1.00 | 71.90 |  |
| 5 | C | O4' | AA:2465 | 3.42 | 1.00 | 77.58 |  |
| 6 | G | N2 | AA:2486 | 3.98 | 1.00 | 72.96 |  |
| 7 | C | N1 | AA:2464 | 4.21 | 1.00 | 74.24 |  |
| 8 | G | O3' | AA:1125 | 4.35 | 1.00 | 71.71 |  |
| 9 | G | N2 | AA:2487 | 4.37 | 1.00 | 69.38 |  |
| 10 | G | N3 | AA:2487 | 4.63 | 1.00 | 68.45 |  |
| 11 | C | N1 | AA:2465 | 4.67 | 1.00 | 77.51 |  |
| 12 | C | O4' | AA:2464 | 4.76 | 1.00 | 74.27 |  |
| 13 | C | N3 | AA:2464 | 4.96 | 1.00 | 74.09 |  |
| 14 | A | N1 | AA:1028 | 4.99 | 1.00 | 81.60 |  |
  |

  
 Coded contact list:   
 Exclusion zone contact(s): /O2/O2'/OP1/OP2/  
Consider K in place of MG (4 valid contact(s) for K vs. 0 for MG)  

### MG:AA:3865

|  | Res. type | Atom | Residue | Distance | Occupancy | B-factor | Closest solute to 1^st^ shell water |
| --- | --- | --- | --- | --- | --- | --- | --- |
| **Ion** | | | |  | 1.00 | 62.61 || 1 | A | OP1 | AA:52 | 3.67 | 1.00 | 48.58 |
| 2 | U | OP2 | AA:114 | 4.49 | 1.00 | 58.29 |  |
| 3 | G | N7 | AA:113 | 5.00 | 1.00 | 50.94 |  |
  |

  
 Coded contact list:   
1^st^ MG coordination shell missing (see Notes)  

### MG:AA:3866

|  | Res. type | Atom | Residue | Distance | Occupancy | B-factor | Closest solute to 1^st^ shell water |
| --- | --- | --- | --- | --- | --- | --- | --- |
| **Ion** | | | |  | 1.00 | 96.23 || 1 | C | OP2 | AA:1504 | 2.88 | 1.00 | 101.09 |
| 2 | C | N4 | AA:1504 | 3.64 | 1.00 | 104.81 |  |
| 3 | U | O5' | AA:1503 | 3.80 | 1.00 | 95.91 |  |
| 4 | U | N1 | AA:1503 | 3.96 | 1.00 | 101.03 |  |
| 5 | U | OP2 | AA:1503 | 4.00 | 1.00 | 93.19 |  |
| 6 | U | N3 | AA:1503 | 4.40 | 1.00 | 102.36 |  |
| 7 | U | O4 | AA:1503 | 4.40 | 1.00 | 102.51 |  |
| 8 | U | O3' | AA:1503 | 4.59 | 1.00 | 100.09 |  |
| 9 | C | O5' | AA:1504 | 4.81 | 1.00 | 101.98 |  |
| 10 | C | N1 | AA:1504 | 4.81 | 1.00 | 105.16 |  |
| 11 | U | O4' | AA:1503 | 4.81 | 1.00 | 99.38 |  |
| 12 | C | N3 | AA:1504 | 4.86 | 1.00 | 104.80 |  |
  |

  
 Coded contact list:   
 Exclusion zone contact(s): /OP2/  
Consider K in place of MG (1 valid contact(s) for K vs. 0 for MG)  

### MG:AA:3867

|  | Res. type | Atom | Residue | Distance | Occupancy | B-factor | Closest solute to 1^st^ shell water |
| --- | --- | --- | --- | --- | --- | --- | --- |
| **Ion** | | | |  | 1.00 | 41.11 || 1 | A | N9 | AA:2287 | 2.62 | 1.00 | 83.47 |
| 2 | G | N7 | AA:2289 | 2.76 | 1.00 | 70.71 |  |
| 3 | A | O2' | AA:2287 | 3.34 | 1.00 | 84.73 |  |
| 4 | A | OP2 | AA:2288 | 3.35 | 1.00 | 81.07 |  |
| 5 | A | O5' | AA:2288 | 3.47 | 1.00 | 79.32 |  |
| 6 | A | N7 | AA:2287 | 3.48 | 1.00 | 85.36 |  |
| 7 | A | O3' | AA:2287 | 3.60 | 1.00 | 80.13 |  |
| 8 | G | OP2 | AA:2289 | 3.64 | 1.00 | 75.05 |  |
| 9 | A | O4' | AA:2287 | 3.79 | 1.00 | 84.03 |  |
| 10 | G | O6 | AA:2289 | 4.14 | 1.00 | 72.97 |  |
| 11 | A | N3 | AA:2287 | 4.40 | 1.00 | 82.32 |  |
| 12 | A | O3' | AA:2288 | 4.58 | 1.00 | 75.10 |  |
| 13 | G | N9 | AA:2289 | 4.81 | 1.00 | 70.75 |  |
  |

  
 Coded contact list:   
 Exclusion zone contact(s): /N7/O2'/OP2/  
Consider K in place of MG (4 valid contact(s) for K vs. 0 for MG)  

### MG:AA:3868

|  | Res. type | Atom | Residue | Distance | Occupancy | B-factor | Closest solute to 1^st^ shell water |
| --- | --- | --- | --- | --- | --- | --- | --- |
| **Ion** | | | |  | 1.00 | 80.39 || 1 | A | OP2 | AA:64 | 3.10 | 1.00 | 69.12 |
| 2 | U | O5' | AA:63 | 3.44 | 1.00 | 69.62 |  |
| 3 | G | N7 | AA:60 | 3.60 | 1.00 | 55.81 |  |
| 4 | U | O3' | AA:63 | 3.66 | 1.00 | 70.16 |  |
| 5 | U | OP2 | AA:63 | 3.67 | 1.00 | 68.31 |  |
| 6 | U | N1 | AA:63 | 3.74 | 1.00 | 72.28 |  |
| 7 | U | O4' | AA:63 | 3.75 | 1.00 | 72.90 |  |
| 8 | U | O2' | AA:63 | 4.04 | 1.00 | 73.85 |  |
| 9 | A | OP1 | AA:64 | 4.71 | 1.00 | 68.60 |  |
  |

  
 Coded contact list:   
 Exclusion zone contact(s): /OP2/  
Consider K in place of MG (1 valid contact(s) for K vs. 0 for MG)  

### MG:AA:3869

|  | Res. type | Atom | Residue | Distance | Occupancy | B-factor | Closest solute to 1^st^ shell water |
| --- | --- | --- | --- | --- | --- | --- | --- |
| **Ion** | | | |  | 1.00 | 68.27 || 1 | G | N7 | AA:506 | 2.82 | 1.00 | 55.45 |
| 2 | G | O6 | AA:498 | 3.37 | 1.00 | 56.81 |  |
| 3 | A | N1 | AA:482 | 3.38 | 1.00 | 60.64 |  |
| 4 | A | O2' | AA:503 | 3.74 | 1.00 | 59.18 |  |
| 5 | A | N3 | AA:482 | 4.22 | 1.00 | 61.25 |  |
| 6 | G | O6 | AA:506 | 4.30 | 1.00 | 60.86 |  |
| 7 | A | O3' | AA:503 | 4.53 | 1.00 | 60.91 |  |
| 8 | A | N9 | AA:503 | 4.54 | 1.00 | 55.89 |  |
| 9 | G | N1 | AA:498 | 4.73 | 1.00 | 52.25 |  |
| 10 | G | N9 | AA:506 | 4.74 | 1.00 | 55.15 |  |
  |

  
 Coded contact list:   
 Exclusion zone contact(s): /N1_A_/N7/O6/  
Consider K in place of MG (3 valid contact(s) for K vs. 0 for MG)  

### MG:AA:3870

|  | Res. type | Atom | Residue | Distance | Occupancy | B-factor | Closest solute to 1^st^ shell water |
| --- | --- | --- | --- | --- | --- | --- | --- |
| **Ion** | | | |  | 1.00 | 69.24 || 1 | A | OP1 | AA:2851 | 2.91 | 1.00 | 66.20 |
| 2 | G | OP1 | AA:2706 | 3.01 | 1.00 | 62.35 |  |
| 3 | G | OP2 | AA:2707 | 3.41 | 1.00 | 62.15 |  |
| 4 | G | O5' | AA:2706 | 3.91 | 1.00 | 61.15 |  |
| 5 | G | O3' | AA:2706 | 4.20 | 1.00 | 61.04 |  |
| 6 | G | OP1 | AA:2707 | 4.75 | 1.00 | 60.66 |  |
| 7 | A | O5' | AA:2851 | 4.90 | 1.00 | 66.70 |  |
| 8 | A | O3' | AA:2850 | 4.99 | 1.00 | 64.66 |  |
  |

  
 Coded contact list:   
 Exclusion zone contact(s): /OP1/OP1/  
Consider K in place of MG (2 valid contact(s) for K vs. 0 for MG)  

### MG:AA:3871

|  | Res. type | Atom | Residue | Distance | Occupancy | B-factor | Closest solute to 1^st^ shell water |
| --- | --- | --- | --- | --- | --- | --- | --- |
| **Ion** | | | |  | 1.00 | 51.99 || 1 | U | O5' | AA:2243 | 2.38 | 1.00 | 32.87 |
| 2 | U | OP1 | AA:2243 | 2.57 | 1.00 | 30.71 |  |
| 3 | A | OP1 | AA:199 | 3.14 | 1.00 | 40.89 |  |
| 4 | C | O3' | AA:198 | 3.68 | 1.00 | 41.25 |  |
| 5 | G | O3' | AA:2242 | 3.87 | 1.00 | 30.47 |  |
| 6 | A | OP2 | AA:199 | 3.88 | 1.00 | 41.52 |  |
| 7 | C | OP1 | AA:192 | 3.93 | 1.00 | 36.21 |  |
| 8 | U | OP2 | AA:2243 | 4.28 | 1.00 | 31.15 |  |
| 9 | C | OP1 | AA:198 | 4.28 | 1.00 | 42.76 |  |
| 10 | C | O5' | AA:198 | 4.49 | 1.00 | 41.90 |  |
| 11 | U | O3' | AA:2243 | 4.69 | 1.00 | 38.65 |  |
| 12 | G | O2' | AA:2242 | 4.76 | 1.00 | 27.44 |  |
| 13 | U | O4' | AA:2243 | 4.87 | 1.00 | 35.81 |  |
  |

  
 Contact(s): /O5'/  
 Coded contact list: O_r_  
 Exclusion zone contact(s): /OP1/OP1/  

### MG:AA:3872

|  | Res. type | Atom | Residue | Distance | Occupancy | B-factor | Closest solute to 1^st^ shell water |
| --- | --- | --- | --- | --- | --- | --- | --- |
| **Ion** | | | |  | 1.00 | 90.01 || 1 | C | O3' | AA:2177 | 4.85 | 1.00 | 164.98 |
| 2 | C | OP1 | AA:2178 | 4.97 | 1.00 | 163.38 |  |
  |

  
 Coded contact list:   
1^st^ MG coordination shell missing (see Notes)  

### MG:AA:3873

|  | Res. type | Atom | Residue | Distance | Occupancy | B-factor | Closest solute to 1^st^ shell water |
| --- | --- | --- | --- | --- | --- | --- | --- |
| **Ion** | | | |  | 1.00 | 87.66 || 1 | G | O4' | AA:654O | 4.01 | 1.00 | 200.30 |
| 2 | G | O2' | AA:654N | 4.70 | 1.00 | 200.00 |  |
| 3 | G | O2' | AA:654O | 4.84 | 1.00 | 200.23 |  |
  |

  
 Coded contact list:   
1^st^ MG coordination shell missing (see Notes)  

### MG:AA:3874

|  | Res. type | Atom | Residue | Distance | Occupancy | B-factor | Closest solute to 1^st^ shell water |
| --- | --- | --- | --- | --- | --- | --- | --- |
| **Ion** | | | |  | 1.00 | 90.26 || 1 | G | O6 | AA:929 | 2.73 | 1.00 | 87.88 |
| 2 | G | N1 | AA:929 | 2.85 | 1.00 | 87.69 |  |
| 3 | G | N7 | AA:928 | 3.15 | 1.00 | 80.05 |  |
| 4 | G | O6 | AA:928 | 3.34 | 1.00 | 75.45 |  |
| 5 | A | N7 | AA:926 | 3.94 | 1.00 | 86.05 |  |
| 6 | A | N6 | AA:926 | 4.05 | 1.00 | 86.48 |  |
| 7 | U | OP2 | AA:847 | 4.26 | 1.00 | 73.77 |  |
| 8 | G | N2 | AA:929 | 4.35 | 1.00 | 89.07 |  |
| 9 | G | OP2 | AA:848 | 4.46 | 1.00 | 76.06 |  |
| 10 | G | N3 | AA:929 | 4.54 | 1.00 | 87.26 |  |
| 11 | G | N7 | AA:929 | 4.66 | 1.00 | 88.84 |  |
  |

  
 Coded contact list:   
 Exclusion zone contact(s): /N1_G_/N7/O6/O6/  
Consider K in place of MG (4 valid contact(s) for K vs. 0 for MG)  

### MG:AA:3875

|  | Res. type | Atom | Residue | Distance | Occupancy | B-factor | Closest solute to 1^st^ shell water |
| --- | --- | --- | --- | --- | --- | --- | --- |
| **Ion** | | | |  | 1.00 | 67.64 || 1 | G | N2 | AA:654O | 3.23 | 1.00 | 200.20 |
| 2 | G | N3 | AA:654O | 4.34 | 1.00 | 200.41 |  |
| 3 | C | O2 | AA:654F | 4.87 | 1.00 | 200.02 |  |
  |

  
 Coded contact list:   
 Exclusion zone contact(s): /N2/  
Consider K in place of MG (1 valid contact(s) for K vs. 0 for MG)  

### MG:AA:3876

|  | Res. type | Atom | Residue | Distance | Occupancy | B-factor | Closest solute to 1^st^ shell water |
| --- | --- | --- | --- | --- | --- | --- | --- |
| **Ion** | | | |  | 1.00 | 67.89 || 1 | G | OP1 | AA:1630 | 3.22 | 1.00 | 54.56 |
| 2 | G | OP2 | AA:1630 | 3.28 | 1.00 | 54.56 |  |
| 3 | U | O3' | AA:1629 | 4.43 | 1.00 | 52.05 |  |
| 4 | U | OP1 | AA:1629 | 4.54 | 1.00 | 51.02 |  |
  |

  
 Coded contact list:   
 Exclusion zone contact(s): /OP1/OP2/  
Consider K in place of MG (2 valid contact(s) for K vs. 0 for MG)  

### MG:AA:3877

|  | Res. type | Atom | Residue | Distance | Occupancy | B-factor | Closest solute to 1^st^ shell water |
| --- | --- | --- | --- | --- | --- | --- | --- |
| **Ion** | | | |  | 1.00 | 109.97 || 1 | G | O6 | AA:425 | 3.93 | 1.00 | 63.43 |
| 2 | G | O6 | AA:424 | 3.95 | 1.00 | 54.51 |  |
| 3 | C | N4 | AA:267 | 4.72 | 1.00 | 63.08 |  |
| 4 | C | N4 | AA:268 | 4.82 | 1.00 | 57.01 |  |
  |

  
 Coded contact list:   
1^st^ MG coordination shell missing (see Notes)  

### MG:AA:3878

|  | Res. type | Atom | Residue | Distance | Occupancy | B-factor | Closest solute to 1^st^ shell water |
| --- | --- | --- | --- | --- | --- | --- | --- |
| **Ion** | | | |  | 1.00 | 84.76 || 1 | A | N6 | AA:1085 | 2.78 | 1.00 | 200.05 |
| 2 | A | N1 | AA:1085 | 3.51 | 1.00 | 199.79 |  |
| 3 | G | O2' | AA:1055 | 4.14 | 1.00 | 200.06 |  |
| 4 | G | OP1 | AA:1056 | 4.91 | 1.00 | 200.30 |  |
| 5 | A | N1 | AA:1086 | 4.93 | 1.00 | 199.59 |  |
  |

  
 Coded contact list:   
 Exclusion zone contact(s): /N6/  
Consider K in place of MG (1 valid contact(s) for K vs. 0 for MG)  

### MG:AA:3879

|  | Res. type | Atom | Residue | Distance | Occupancy | B-factor | Closest solute to 1^st^ shell water |
| --- | --- | --- | --- | --- | --- | --- | --- |
| **Ion** | | | |  | 1.00 | 91.56 || 1 | C | OP1 | AA:2699 | 2.56 | 1.00 | 53.13 |
| 2 | G | O2' | AA:1628 | 3.13 | 1.00 | 42.99 |  |
| 3 | U | O4' | AA:1629 | 3.52 | 1.00 | 49.29 |  |
| 4 | C | OP2 | AA:2699 | 3.92 | 1.00 | 52.35 |  |
| 5 | C | O5' | AA:2699 | 4.70 | 1.00 | 52.20 |  |
| 6 | U | O3' | AA:2698 | 4.78 | 1.00 | 51.04 |  |
  |

  
 Coded contact list:   
 Exclusion zone contact(s): /O2'/OP1/  
Consider NA in place of MG (1 valid contact(s) for NA vs. 0 for MG)  

### MG:AA:3880

|  | Res. type | Atom | Residue | Distance | Occupancy | B-factor | Closest solute to 1^st^ shell water |
| --- | --- | --- | --- | --- | --- | --- | --- |
| **Ion** | | | |  | 1.00 | 102.48 || 1 | G | O6 | AA:932 | 2.57 | 1.00 | 83.54 |
| 2 | C | O2 | AA:816 | 2.94 | 1.00 | 68.83 |  |
| 3 | C | O2' | AA:816 | 3.37 | 1.00 | 65.08 |  |
| 4 | C | OP1 | AA:840 | 3.50 | 1.00 | 59.27 |  |
| 5 | G | N1 | AA:932 | 4.04 | 1.00 | 82.30 |  |
| 6 | C | N1 | AA:816 | 4.19 | 1.00 | 67.98 |  |
| 7 | C | O4' | AA:816 | 4.42 | 1.00 | 64.09 |  |
| 8 | G | N2 | AA:1191 | 4.57 | 1.00 | 69.45 |  |
| 9 | C | O4' | AA:817 | 4.63 | 1.00 | 60.85 |  |
  |

  
 Coded contact list:   
 Exclusion zone contact(s): /O2/O2'/O6/  
Consider K in place of MG (2 valid contact(s) for K vs. 0 for MG)  

### MG:AA:3881

|  | Res. type | Atom | Residue | Distance | Occupancy | B-factor | Closest solute to 1^st^ shell water |
| --- | --- | --- | --- | --- | --- | --- | --- |
| **Ion** | | | |  | 1.00 | 72.13 || 1 | C | OP2 | AA:105 | 2.24 | 1.00 | 66.05 |
| 2 | C | OP2 | AA:106 | 3.00 | 1.00 | 67.59 |  |
| 3 | C | O5' | AA:105 | 3.72 | 1.00 | 66.08 |  |
| 4 | C | OP1 | AA:105 | 3.73 | 1.00 | 68.51 |  |
| 5 | C | O4' | AA:105 | 4.61 | 1.00 | 67.78 |  |
| 6 | U | O3' | AA:104 | 4.68 | 1.00 | 67.44 |  |
| 7 | C | O3' | AA:105 | 4.77 | 1.00 | 68.00 |  |
| 8 | C | N1 | AA:105 | 4.86 | 1.00 | 66.97 |  |
  |

  
 Contact(s): /OP2/  
 Coded contact list: O_ph_  
 Exclusion zone contact(s): /OP2/  

### MG:AA:3882

|  | Res. type | Atom | Residue | Distance | Occupancy | B-factor | Closest solute to 1^st^ shell water |
| --- | --- | --- | --- | --- | --- | --- | --- |
| **Ion** | | | |  | 1.00 | 77.07 || 1 | G | OP2 | AA:212 | 3.89 | 1.00 | 54.24 |
| 2 | A | OP2 | AA:213 | 4.25 | 1.00 | 61.26 |  |
| 3 | G | OP2 | AA:179 | 4.66 | 1.00 | 49.55 |  |
| 4 | G | O5' | AA:212 | 4.70 | 1.00 | 55.82 |  |
| 5 | G | N7 | AA:179 | 4.91 | 1.00 | 52.28 |  |
| 6 | G | OP2 | AA:178 | 4.93 | 1.00 | 55.91 |  |
  |

  
 Coded contact list:   
1^st^ MG coordination shell missing (see Notes)  

### MG:AA:3883

|  | Res. type | Atom | Residue | Distance | Occupancy | B-factor | Closest solute to 1^st^ shell water |
| --- | --- | --- | --- | --- | --- | --- | --- |
| **Ion** | | | |  | 1.00 | 100.34 || 1 | A | OP1 | AA:103 | 2.98 | 1.00 | 63.75 |
| 2 | A | O5' | AA:103 | 3.84 | 1.00 | 62.87 |  |
| 3 | U | OP2 | AA:104 | 5.00 | 1.00 | 65.24 |  |
  |

  
 Coded contact list:   
 Exclusion zone contact(s): /OP1/  
Consider K in place of MG (1 valid contact(s) for K vs. 0 for MG)  

### MG:AA:3885

|  | Res. type | Atom | Residue | Distance | Occupancy | B-factor | Closest solute to 1^st^ shell water |
| --- | --- | --- | --- | --- | --- | --- | --- |
| **Ion** | | | |  | 1.00 | 51.95 || 1 | A | OP2 | AA:861 | 3.24 | 1.00 | 72.00 |
| 2 | U | OP1 | AA:860 | 3.29 | 1.00 | 77.99 |  |
| 3 | U | O3' | AA:860 | 3.84 | 1.00 | 71.73 |  |
| 4 | U | O4 | AA:913 | 4.12 | 1.00 | 75.14 |  |
| 5 | U | O5' | AA:860 | 4.16 | 1.00 | 76.42 |  |
| 6 | A | OP1 | AA:861 | 4.46 | 1.00 | 70.54 |  |
| 7 | U | OP2 | AA:860 | 4.61 | 1.00 | 78.71 |  |
| 8 | A | OP2 | AA:910 | 4.99 | 1.00 | 72.54 |  |
  |

  
 Coded contact list:   
 Exclusion zone contact(s): /OP1/OP2/  
Consider K in place of MG (2 valid contact(s) for K vs. 0 for MG)  

### MG:AA:3886

|  | Res. type | Atom | Residue | Distance | Occupancy | B-factor | Closest solute to 1^st^ shell water |
| --- | --- | --- | --- | --- | --- | --- | --- |
| **Ion** | | | |  | 1.00 | 103.14 || 1 | U | O4 | AA:350 | 3.73 | 1.00 | 78.38 |
| 2 | G | N7 | AA:349 | 3.93 | 1.00 | 72.68 |  |
| 3 | G | O6 | AA:349 | 4.69 | 1.00 | 73.75 |  |
| 4 | G | OP2 | AA:348 | 4.99 | 1.00 | 73.66 |  |
  |

  
 Coded contact list:   
1^st^ MG coordination shell missing (see Notes)  

### MG:AA:3887

|  | Res. type | Atom | Residue | Distance | Occupancy | B-factor | Closest solute to 1^st^ shell water |
| --- | --- | --- | --- | --- | --- | --- | --- |
| **Ion** | | | |  | 1.00 | 70.38 || 1 | U | O2' | AA:270O | 3.32 | 1.00 | 155.67 |
| 2 | C | OP2 | AA:270P | 4.18 | 1.00 | 151.20 |  |
  |

  
 Coded contact list:   
 Exclusion zone contact(s): /O2'/  
Consider K in place of MG (1 valid contact(s) for K vs. 0 for MG)  

### MG:AA:3888

|  | Res. type | Atom | Residue | Distance | Occupancy | B-factor | Closest solute to 1^st^ shell water |
| --- | --- | --- | --- | --- | --- | --- | --- |
| **Ion** | | | |  | 1.00 | 79.25 || 1 | G | O6 | AA:301 | 2.50 | 1.00 | 66.93 |
| 2 | C | N4 | AA:316 | 3.15 | 1.00 | 71.03 |  |
| 3 | G | O6 | AA:317 | 3.16 | 1.00 | 64.42 |  |
| 4 | G | N7 | AA:317 | 4.09 | 1.00 | 66.72 |  |
| 5 | G | N1 | AA:301 | 4.57 | 1.00 | 66.76 |  |
| 6 | C | N4 | AA:334 | 4.59 | 1.00 | 62.85 |  |
| 7 | G | O6 | AA:315 | 4.63 | 1.00 | 70.04 |  |
| 8 | G | N7 | AA:301 | 4.72 | 1.00 | 65.31 |  |
| 9 | C | N3 | AA:316 | 4.98 | 1.00 | 69.80 |  |
  |

  
 Coded contact list:   
 Exclusion zone contact(s): /N4/O6/O6/  
Consider K in place of MG (2 valid contact(s) for K vs. 0 for MG)  

### MG:AA:3889

|  | Res. type | Atom | Residue | Distance | Occupancy | B-factor | Closest solute to 1^st^ shell water |
| --- | --- | --- | --- | --- | --- | --- | --- |
| **Ion** | | | |  | 1.00 | 73.31 || 1 | G | OP1 | AA:1869 | 2.36 | 1.00 | 74.50 |
| 2 | G | O5' | AA:1869 | 3.39 | 1.00 | 76.10 |  |
| 3 | G | O2' | AA:2409 | 4.12 | 1.00 | 68.19 |  |
| 4 | G | OP2 | AA:1869 | 4.16 | 1.00 | 73.36 |  |
| 5 | G | N3 | AA:2409 | 4.28 | 1.00 | 67.74 |  |
| 6 | C | OP2 | AA:1870 | 4.34 | 1.00 | 82.01 |  |
| 7 | U | O3' | AA:1864 | 4.66 | 1.00 | 73.63 |  |
| 8 | G | N2 | AA:2409 | 4.74 | 1.00 | 69.43 |  |
  |

  
 Contact(s): /OP1/  
 Coded contact list: O_ph_  
 Exclusion zone contact(s): /O5'/  

### MG:AA:3890

|  | Res. type | Atom | Residue | Distance | Occupancy | B-factor | Closest solute to 1^st^ shell water |
| --- | --- | --- | --- | --- | --- | --- | --- |
| **Ion** | | | |  | 1.00 | 103.08 || 1 | G | O6 | AA:45 | 2.60 | 1.00 | 61.12 |
| 2 | C | N4 | AA:433 | 3.93 | 1.00 | 56.65 |  |
| 3 | A | N6 | AA:432 | 4.12 | 1.00 | 43.13 |  |
| 4 | G | N7 | AA:45 | 4.13 | 1.00 | 65.92 |  |
| 5 | A | N6 | AA:44 | 4.50 | 1.00 | 68.60 |  |
| 6 | G | O2' | AA:215 | 4.76 | 1.00 | 62.82 |  |
| 7 | G | N1 | AA:45 | 4.85 | 1.00 | 62.21 |  |
  |

  
 Coded contact list:   
 Exclusion zone contact(s): /O6/  
Consider K in place of MG (1 valid contact(s) for K vs. 0 for MG)  

### MG:AA:3891

|  | Res. type | Atom | Residue | Distance | Occupancy | B-factor | Closest solute to 1^st^ shell water |
| --- | --- | --- | --- | --- | --- | --- | --- |
| **Ion** | | | |  | 1.00 | 105.40 || 1 | G | N7 | AA:2382 | 3.28 | 1.00 | 79.59 |
| 2 | G | N2 | AA:2345 | 3.77 | 1.00 | 59.36 |  |
| 3 | G | OP2 | AA:2382 | 4.33 | 1.00 | 74.15 |  |
| 4 | C | OP2 | AA:2381 | 4.46 | 1.00 | 69.86 |  |
| 5 | G | OP2 | AA:2367 | 4.51 | 1.00 | 74.10 |  |
| 6 | G | O6 | AA:2382 | 4.83 | 1.00 | 77.90 |  |
| 7 | A | OP2 | AA:2366 | 4.88 | 1.00 | 72.84 |  |
| 8 | G | N1 | AA:2345 | 4.95 | 1.00 | 58.72 |  |
  |

  
 Coded contact list:   
 Exclusion zone contact(s): /N7/  
Consider K in place of MG (1 valid contact(s) for K vs. 0 for MG)  

### MG:AA:3892

|  | Res. type | Atom | Residue | Distance | Occupancy | B-factor | Closest solute to 1^st^ shell water |
| --- | --- | --- | --- | --- | --- | --- | --- |
| **Ion** | | | |  | 1.00 | 84.25 || 1 | G | O6 | AA:1449A | 4.13 | 1.00 | 85.34 |
| 2 | G | N7 | AA:1448 | 4.19 | 1.00 | 82.12 |  |
| 3 | G | O6 | AA:1448 | 4.49 | 1.00 | 79.26 |  |
| 4 | G | N7 | AA:1449A | 4.92 | 1.00 | 87.21 |  |
  |

  
 Coded contact list:   
1^st^ MG coordination shell missing (see Notes)  

### MG:AA:3893

|  | Res. type | Atom | Residue | Distance | Occupancy | B-factor | Closest solute to 1^st^ shell water |
| --- | --- | --- | --- | --- | --- | --- | --- |
| **Ion** | | | |  | 1.00 | 67.75 || 1 | G | O3' | AA:1630 | 3.26 | 1.00 | 58.80 |
| 2 | G | O2' | AA:1630 | 3.26 | 1.00 | 55.36 |  |
| 3 | C | OP1 | AA:1630A | 3.67 | 1.00 | 59.40 |  |
| 4 | G | OP1 | AA:2697 | 3.77 | 1.00 | 56.26 |  |
| 5 | C | O5' | AA:1630A | 4.07 | 1.00 | 59.69 |  |
| 6 | C | O4' | AA:1630A | 4.51 | 1.00 | 60.98 |  |
| 7 | G | O5' | AA:2697 | 4.62 | 1.00 | 54.65 |  |
  |

  
 Coded contact list:   
 Exclusion zone contact(s): /O2'/O3'/  
Consider K in place of MG (2 valid contact(s) for K vs. 0 for MG)  

### MG:AA:3894

|  | Res. type | Atom | Residue | Distance | Occupancy | B-factor | Closest solute to 1^st^ shell water |
| --- | --- | --- | --- | --- | --- | --- | --- |
| **Ion** | | | |  | 1.00 | 86.05 || 1 | A | OP2 | AA:1237 | 2.83 | 1.00 | 64.57 |
| 2 | A | OP1 | AA:1237 | 3.95 | 1.00 | 65.26 |  |
| 3 | A | O5' | AA:1237 | 4.49 | 1.00 | 62.69 |  |
| 4 | G | N2 | AA:474 | 4.64 | 1.00 | 47.07 |  |
| 5 | G | OP1 | AA:476 | 4.69 | 1.00 | 56.78 |  |
| 6 | C | O2 | AA:510 | 4.96 | 1.00 | 54.66 |  |
  |

  
 Coded contact list:   
 Exclusion zone contact(s): /OP2/  
Consider K in place of MG (1 valid contact(s) for K vs. 0 for MG)  

### MG:AA:3895

|  | Res. type | Atom | Residue | Distance | Occupancy | B-factor | Closest solute to 1^st^ shell water |
| --- | --- | --- | --- | --- | --- | --- | --- |
| **Ion** | | | |  | 1.00 | 51.50 || 1 | G | O2' | AA:669 | 2.74 | 1.00 | 55.31 |
| 2 | A | OP1 | AA:670 | 2.76 | 1.00 | 54.86 |  |
| 3 | SER | OG | AO:42 | 3.10 | 1.00 | 42.97 |  |
| 4 | A | O5' | AA:670 | 3.39 | 1.00 | 53.87 |  |
| 5 | C | OP2 | AA:672 | 3.75 | 1.00 | 43.74 |  |
| 6 | G | O3' | AA:669 | 4.04 | 1.00 | 55.34 |  |
| 7 | GLY | N | AO:43 | 4.34 | 1.00 | 43.78 |  |
| 8 | A | O4' | AA:670 | 4.94 | 1.00 | 53.73 |  |
| 9 | A | OP2 | AA:670 | 5.00 | 1.00 | 53.96 |  |
  |

  
 Coded contact list:   
 Exclusion zone contact(s): /O2'/O5'/OG_Ser_/OP1/  
Consider K in place of MG (4 valid contact(s) for K vs. 0 for MG)  

### MG:AA:3896

|  | Res. type | Atom | Residue | Distance | Occupancy | B-factor | Closest solute to 1^st^ shell water |
| --- | --- | --- | --- | --- | --- | --- | --- |
| **Ion** | | | |  | 1.00 | 99.68 || 1 | G | OP2 | AA:1187 | 2.59 | 1.00 | 63.08 |
| 2 | C | OP2 | AA:816 | 2.65 | 1.00 | 59.22 |  |
| 3 | C | O2' | AA:1185 | 3.67 | 1.00 | 72.96 |  |
| 4 | G | OP1 | AA:1187 | 3.71 | 1.00 | 63.89 |  |
| 5 | C | OP1 | AA:816 | 3.84 | 1.00 | 59.70 |  |
| 6 | G | O3' | AA:1186 | 4.31 | 1.00 | 64.99 |  |
| 7 | C | O5' | AA:816 | 4.69 | 1.00 | 60.51 |  |
| 8 | C | O3' | AA:1185 | 4.70 | 1.00 | 68.50 |  |
| 9 | C | O3' | AA:815 | 4.89 | 1.00 | 58.88 |  |
| 10 | G | O5' | AA:1187 | 4.91 | 1.00 | 63.24 |  |
| 11 | C | OP2 | AA:817 | 4.93 | 1.00 | 62.35 |  |
  |

  
 Coded contact list:   
 Exclusion zone contact(s): /OP2/OP2/  
Consider NA in place of MG (1 valid contact(s) for NA vs. 0 for MG)  

### MG:AA:3897

|  | Res. type | Atom | Residue | Distance | Occupancy | B-factor | Closest solute to 1^st^ shell water |
| --- | --- | --- | --- | --- | --- | --- | --- |
| **Ion** | | | |  | 1.00 | 103.23 || 1 | G | OP2 | AA:1410 | 3.50 | 1.00 | 77.26 |
| 2 | G | OP1 | AA:1410 | 3.92 | 1.00 | 76.61 |  |
| 3 | G | O5' | AA:1410 | 4.96 | 1.00 | 78.32 |  |
  |

  
 Coded contact list:   
1^st^ MG coordination shell missing (see Notes)  

### MG:AA:3899

|  | Res. type | Atom | Residue | Distance | Occupancy | B-factor | Closest solute to 1^st^ shell water |
| --- | --- | --- | --- | --- | --- | --- | --- |
| **Ion** | | | |  | 1.00 | 46.44 || 1 | A | OP1 | AA:1981 | 2.70 | 1.00 | 40.05 |
| 2 | A | OP2 | AA:1981 | 3.05 | 1.00 | 41.02 |  |
| 3 | G | OP2 | AA:1776 | 3.25 | 1.00 | 43.90 |  |
| 4 | G | OP1 | AA:1776 | 3.80 | 1.00 | 44.37 |  |
| 5 | G | O3' | AA:1980 | 4.10 | 1.00 | 42.90 |  |
| 6 | U | OP1 | AA:1775 | 4.52 | 1.00 | 41.03 |  |
| 7 | G | O2' | AA:759 | 4.60 | 1.00 | 41.16 |  |
| 8 | A | O5' | AA:1981 | 4.67 | 1.00 | 42.58 |  |
| 9 | U | O3' | AA:1775 | 4.72 | 1.00 | 43.07 |  |
  |

  
 Coded contact list:   
 Exclusion zone contact(s): /OP1/OP2/OP2/  
Consider K in place of MG (3 valid contact(s) for K vs. 0 for MG)  

### MG:AA:3900

|  | Res. type | Atom | Residue | Distance | Occupancy | B-factor | Closest solute to 1^st^ shell water |
| --- | --- | --- | --- | --- | --- | --- | --- |
| **Ion** | | | |  | 1.00 | 105.63 || 1 | A | N3 | AA:1204 | 2.14 | 1.00 | 68.46 |
| 2 | A | O4' | AA:1204 | 2.36 | 1.00 | 66.92 |  |
| 3 | A | O5' | AA:1204 | 2.75 | 1.00 | 64.46 |  |
| 4 | A | OP1 | AA:1204 | 3.26 | 1.00 | 65.57 |  |
| 5 | A | N1 | AA:1241 | 3.45 | 1.00 | 64.85 |  |
| 6 | A | N9 | AA:1204 | 3.48 | 1.00 | 69.90 |  |
| 7 | G | N9 | AA:1203 | 3.87 | 1.00 | 58.01 |  |
| 8 | G | O3' | AA:1203 | 3.87 | 1.00 | 61.07 |  |
| 9 | A | N6 | AA:1241 | 3.92 | 1.00 | 62.70 |  |
| 10 | G | N3 | AA:1206 | 3.93 | 1.00 | 52.65 |  |
| 11 | G | N2 | AA:1206 | 4.06 | 1.00 | 52.32 |  |
| 12 | A | N1 | AA:1204 | 4.09 | 1.00 | 67.65 |  |
| 13 | G | O2' | AA:1203 | 4.10 | 1.00 | 57.35 |  |
| 14 | G | N7 | AA:1203 | 4.46 | 1.00 | 59.72 |  |
| 15 | G | N1 | AA:1206 | 4.62 | 1.00 | 54.04 |  |
| 16 | G | O4' | AA:1203 | 4.84 | 1.00 | 57.19 |  |
| 17 | A | OP2 | AA:1204 | 4.89 | 1.00 | 64.57 |  |
  |

  
**1^st^ shell ligand angles:**   
1-2: 80;   
  
 Contact(s): /N3_A_/O4'/  
 Coded contact list: O_r_N_b_  
 Exclusion zone contact(s): /O5'/OP1/  

### MG:AA:3901

|  | Res. type | Atom | Residue | Distance | Occupancy | B-factor | Closest solute to 1^st^ shell water |
| --- | --- | --- | --- | --- | --- | --- | --- |
| **Ion** | | | |  | 1.00 | 52.95 || 1 | C | OP2 | AA:487 | 2.32 | 1.00 | 53.76 |
| 2 | G | N7 | AA:488 | 3.67 | 1.00 | 57.00 |  |
| 3 | C | O5' | AA:487 | 3.93 | 1.00 | 54.41 |  |
| 4 | G | O6 | AA:488 | 4.28 | 1.00 | 54.95 |  |
| 5 | C | OP1 | AA:487 | 4.35 | 1.00 | 54.27 |  |
| 6 | G | OP2 | AA:488 | 4.50 | 1.00 | 53.17 |  |
| 7 | C | O3' | AA:486 | 4.74 | 1.00 | 53.01 |  |
| 8 | C | O5' | AA:486 | 4.90 | 1.00 | 54.03 |  |
  |

  
 Contact(s): /OP2/  
 Coded contact list: O_ph_  

### MG:AA:3902

|  | Res. type | Atom | Residue | Distance | Occupancy | B-factor | Closest solute to 1^st^ shell water |
| --- | --- | --- | --- | --- | --- | --- | --- |
| **Ion** | | | |  | 1.00 | 111.17 || 1 | A | N1 | AA:1069 | 2.38 | 1.00 | 199.17 |
| 2 | G | N2 | AA:1068 | 2.49 | 1.00 | 200.21 |  |
| 3 | A | N6 | AA:1069 | 3.23 | 1.00 | 199.30 |  |
| 4 | G | N1 | AA:1068 | 3.90 | 1.00 | 200.41 |  |
| 5 | G | N3 | AA:1068 | 4.06 | 1.00 | 200.41 |  |
| 6 | A | N3 | AA:1069 | 4.19 | 1.00 | 199.11 |  |
  |

  
 Contact(s): /N1_A_/  
 Coded contact list: N_b_  
 Exclusion zone contact(s): /N2/N6/  

### MG:AA:3904

|  | Res. type | Atom | Residue | Distance | Occupancy | B-factor | Closest solute to 1^st^ shell water |
| --- | --- | --- | --- | --- | --- | --- | --- |
| **Ion** | | | |  | 1.00 | 77.35 || 1 | G | N7 | AA:1478 | 3.09 | 1.00 | 82.71 |
| 2 | G | O6 | AA:1478 | 3.67 | 1.00 | 80.32 |  |
| 3 | A | N7 | AA:1477 | 4.23 | 1.00 | 75.24 |  |
| 4 | A | N6 | AA:1477 | 4.47 | 1.00 | 75.25 |  |
  |

  
 Coded contact list:   
 Exclusion zone contact(s): /N7/  
Consider K in place of MG (1 valid contact(s) for K vs. 0 for MG)  

### MG:AA:3905

|  | Res. type | Atom | Residue | Distance | Occupancy | B-factor | Closest solute to 1^st^ shell water |
| --- | --- | --- | --- | --- | --- | --- | --- |
| **Ion** | | | |  | 1.00 | 86.72 || 1 | G | OP1 | AA:2697 | 2.54 | 1.00 | 56.26 |
| 2 | U | O3' | AA:2696 | 3.49 | 1.00 | 56.50 |  |
| 3 | G | O5' | AA:2697 | 4.31 | 1.00 | 54.65 |  |
| 4 | G | O3' | AA:1707 | 4.53 | 1.00 | 71.96 |  |
| 5 | G | N2 | AA:1758 | 4.68 | 1.00 | 52.09 |  |
| 6 | C | OP1 | AA:1708 | 4.69 | 1.00 | 73.34 |  |
| 7 | G | OP2 | AA:1758 | 4.79 | 1.00 | 67.00 |  |
| 8 | G | OP2 | AA:2697 | 4.86 | 1.00 | 57.05 |  |
  |

  
 Coded contact list:   
 Exclusion zone contact(s): /OP1/  
Consider NA in place of MG (1 valid contact(s) for NA vs. 0 for MG)  

### MG:AA:3906

|  | Res. type | Atom | Residue | Distance | Occupancy | B-factor | Closest solute to 1^st^ shell water |
| --- | --- | --- | --- | --- | --- | --- | --- |
| **Ion** | | | |  | 1.00 | 67.74 || 1 | C | OP2 | AA:687 | 2.66 | 1.00 | 34.65 |
| 2 | C | OP1 | AA:687 | 3.02 | 1.00 | 35.89 |  |
| 3 | C | O5' | AA:687 | 3.09 | 1.00 | 34.79 |  |
| 4 | LYS | NZ | A7:2 | 3.20 | 1.00 | 42.30 |  |
| 5 | U | OP2 | AA:688 | 3.44 | 1.00 | 31.70 |  |
| 6 | G | O3' | AA:686 | 4.62 | 1.00 | 36.87 |  |
| 7 | C | O3' | AA:687 | 4.87 | 1.00 | 33.92 |  |
  |

  
 Coded contact list:   
 Exclusion zone contact(s): /NZ_Lys_/O5'/OP1/OP2/  
Consider K in place of MG (4 valid contact(s) for K vs. 0 for MG)  

### MG:AA:3907

|  | Res. type | Atom | Residue | Distance | Occupancy | B-factor | Closest solute to 1^st^ shell water |
| --- | --- | --- | --- | --- | --- | --- | --- |
| **Ion** | | | |  | 1.00 | 57.94 || 1 | U | O2' | AA:511 | 2.55 | 1.00 | 57.77 |
| 2 | U | O2 | AA:29 | 2.98 | 1.00 | 47.68 |  |
| 3 | G | N3 | AA:30 | 3.27 | 1.00 | 56.21 |  |
| 4 | U | O2 | AA:511 | 3.49 | 1.00 | 62.43 |  |
| 5 | G | O2' | AA:1215 | 3.56 | 1.00 | 64.36 |  |
| 6 | G | N3 | AA:1215 | 3.57 | 1.00 | 73.70 |  |
| 7 | G | N9 | AA:30 | 3.60 | 1.00 | 54.56 |  |
| 8 | A | N1 | AA:1214 | 4.16 | 1.00 | 68.84 |  |
| 9 | G | N9 | AA:1215 | 4.17 | 1.00 | 69.98 |  |
| 10 | G | O4' | AA:1215 | 4.34 | 1.00 | 65.88 |  |
| 11 | G | O4' | AA:30 | 4.38 | 1.00 | 54.22 |  |
| 12 | U | N3 | AA:29 | 4.44 | 1.00 | 44.99 |  |
| 13 | G | N2 | AA:30 | 4.44 | 1.00 | 56.29 |  |
| 14 | G | N1 | AA:30 | 4.47 | 1.00 | 55.39 |  |
| 15 | G | N2 | AA:1215 | 4.68 | 1.00 | 73.44 |  |
| 16 | G | N7 | AA:30 | 4.68 | 1.00 | 55.70 |  |
| 17 | U | N1 | AA:511 | 4.79 | 1.00 | 59.26 |  |
  |

  
 Coded contact list:   
 Exclusion zone contact(s): /N3_G_/O2/O2'/  
Consider K in place of MG (2 valid contact(s) for K vs. 0 for MG)  

### MG:AA:3908

|  | Res. type | Atom | Residue | Distance | Occupancy | B-factor | Closest solute to 1^st^ shell water |
| --- | --- | --- | --- | --- | --- | --- | --- |
| **Ion** | | | |  | 1.00 | 68.16 || 1 | C | O4' | AA:57 | 2.84 | 1.00 | 48.67 |
| 2 | A | O2' | AA:56 | 2.96 | 1.00 | 43.28 |  |
| 3 | C | OP1 | AA:128 | 3.09 | 1.00 | 51.51 |  |
| 4 | A | O2' | AA:127 | 3.28 | 1.00 | 51.26 |  |
| 5 | C | OP2 | AA:128 | 3.35 | 1.00 | 51.36 |  |
| 6 | A | O3' | AA:127 | 3.59 | 1.00 | 51.60 |  |
| 7 | A | N3 | AA:56 | 3.73 | 1.00 | 43.52 |  |
| 8 | C | O2' | AA:57 | 4.39 | 1.00 | 48.35 |  |
| 9 | C | N1 | AA:57 | 4.56 | 1.00 | 50.47 |  |
| 10 | A | N9 | AA:56 | 4.96 | 1.00 | 42.84 |  |
| 11 | C | O2 | AA:115 | 4.97 | 1.00 | 44.10 |  |
| 12 | C | O5' | AA:128 | 4.99 | 1.00 | 52.51 |  |
  |

  
 Coded contact list:   
 Exclusion zone contact(s): /O2'/O2'/O4'/OP1/OP2/  
Consider K in place of MG (5 valid contact(s) for K vs. 0 for MG)  

### MG:AA:3909

|  | Res. type | Atom | Residue | Distance | Occupancy | B-factor | Closest solute to 1^st^ shell water |
| --- | --- | --- | --- | --- | --- | --- | --- |
| **Ion** | | | |  | 1.00 | 90.78 || 1 | C | N4 | AA:1407 | 2.53 | 1.00 | 63.88 |
| 2 | C | N4 | AA:1408 | 3.89 | 1.00 | 65.04 |  |
| 3 | C | N3 | AA:1407 | 4.15 | 1.00 | 63.51 |  |
| 4 | G | O6 | AA:1595 | 4.76 | 1.00 | 65.20 |  |
| 5 | C | OP2 | AA:1407 | 4.88 | 1.00 | 59.94 |  |
| 6 | U | O4 | AA:1406 | 4.91 | 1.00 | 62.02 |  |
  |

  
 Coded contact list:   
 Exclusion zone contact(s): /N4/  
Consider NA in place of MG (1 valid contact(s) for NA vs. 0 for MG)  

### MG:AA:3910

|  | Res. type | Atom | Residue | Distance | Occupancy | B-factor | Closest solute to 1^st^ shell water |
| --- | --- | --- | --- | --- | --- | --- | --- |
| **Ion** | | | |  | 1.00 | 76.03 || 1 | U | OP2 | AA:1431 | 4.59 | 1.00 | 49.69 |
| 2 | C | OP1 | AA:1430 | 4.71 | 1.00 | 48.57 |  |
  |

  
 Coded contact list:   
1^st^ MG coordination shell missing (see Notes)  

### MG:AA:3911

|  | Res. type | Atom | Residue | Distance | Occupancy | B-factor | Closest solute to 1^st^ shell water |
| --- | --- | --- | --- | --- | --- | --- | --- |
| **Ion** | | | |  | 1.00 | 69.19 || 1 | G | OP1 | AA:2035 | 3.09 | 1.00 | 53.42 |
| 2 | G | N7 | AA:2037 | 3.44 | 1.00 | 41.39 |  |
| 3 | C | OP2 | AA:2036 | 3.82 | 1.00 | 55.49 |  |
| 4 | C | N4 | AA:2036 | 4.23 | 1.00 | 40.86 |  |
| 5 | C | N3 | AA:2036 | 4.23 | 1.00 | 41.57 |  |
| 6 | C | N1 | AA:2036 | 4.41 | 1.00 | 44.35 |  |
| 7 | G | OP2 | AA:2037 | 4.50 | 1.00 | 46.24 |  |
| 8 | C | O5' | AA:2036 | 4.53 | 1.00 | 53.77 |  |
| 9 | G | O6 | AA:2037 | 4.66 | 1.00 | 38.58 |  |
| 10 | G | O5' | AA:2035 | 4.72 | 1.00 | 53.50 |  |
| 11 | G | O6 | AA:2038 | 4.92 | 1.00 | 56.10 |  |
  |

  
 Coded contact list:   
 Exclusion zone contact(s): /OP1/  
Consider K in place of MG (1 valid contact(s) for K vs. 0 for MG)  

### MG:AA:3912

|  | Res. type | Atom | Residue | Distance | Occupancy | B-factor | Closest solute to 1^st^ shell water |
| --- | --- | --- | --- | --- | --- | --- | --- |
| **Ion** | | | |  | 1.00 | 91.40 || 1 | A | N7 | AA:1434 | 4.41 | 1.00 | 56.34 |
| 2 | A | OP2 | AA:1554 | 4.52 | 1.00 | 64.77 |  |
| 3 | A | N6 | AA:1434 | 4.58 | 1.00 | 58.10 |  |
| 4 | U | N3 | AA:1433 | 4.64 | 1.00 | 58.57 |  |
| 5 | U | O4 | AA:1433 | 4.74 | 1.00 | 60.47 |  |
| 6 | G | O6 | AA:1560 | 4.90 | 1.00 | 59.60 |  |
  |

  
 Coded contact list:   
1^st^ MG coordination shell missing (see Notes)  

### MG:AA:3913

|  | Res. type | Atom | Residue | Distance | Occupancy | B-factor | Closest solute to 1^st^ shell water |
| --- | --- | --- | --- | --- | --- | --- | --- |
| **Ion** | | | |  | 1.00 | 38.53 || 1 | G | N1 | AA:83 | 2.53 | 1.00 | 55.48 |
| 2 | G | O6 | AA:83 | 2.90 | 1.00 | 55.30 |  |
| 3 | G | N9 | AA:102 | 3.18 | 1.00 | 70.67 |  |
| 4 | G | O3' | AA:102 | 3.42 | 1.00 | 67.83 |  |
| 5 | G | O4' | AA:102 | 3.53 | 1.00 | 72.49 |  |
| 6 | G | O6 | AA:82 | 3.56 | 1.00 | 65.02 |  |
| 7 | A | OP2 | AA:103 | 3.59 | 1.00 | 63.89 |  |
| 8 | A | OP1 | AA:103 | 3.70 | 1.00 | 63.75 |  |
| 9 | G | N2 | AA:83 | 3.77 | 1.00 | 58.42 |  |
| 10 | G | O2' | AA:102 | 3.99 | 1.00 | 72.20 |  |
| 11 | G | N7 | AA:102 | 4.19 | 1.00 | 69.94 |  |
| 12 | G | N3 | AA:102 | 4.44 | 1.00 | 70.66 |  |
| 13 | G | N1 | AA:82 | 4.45 | 1.00 | 64.78 |  |
| 14 | G | N3 | AA:83 | 4.75 | 1.00 | 55.04 |  |
| 15 | A | N6 | AA:84 | 4.87 | 1.00 | 45.50 |  |
  |

  
 Coded contact list:   
 Exclusion zone contact(s): /N1_G_/O6/  
Consider K in place of MG (2 valid contact(s) for K vs. 0 for MG)  

### MG:AA:3914

|  | Res. type | Atom | Residue | Distance | Occupancy | B-factor | Closest solute to 1^st^ shell water |
| --- | --- | --- | --- | --- | --- | --- | --- |
| **Ion** | | | |  | 1.00 | 78.33 || 1 | C | OP1 | AA:1408 | 3.51 | 1.00 | 56.41 |
| 2 | C | OP1 | AA:1588 | 3.83 | 1.00 | 91.20 |  |
| 3 | C | OP2 | AA:1409 | 4.14 | 1.00 | 63.03 |  |
| 4 | C | OP2 | AA:1408 | 4.40 | 1.00 | 55.63 |  |
| 5 | C | O5' | AA:1408 | 4.53 | 1.00 | 58.86 |  |
  |

  
 Coded contact list:   
1^st^ MG coordination shell missing (see Notes)  

### MG:AA:3915

|  | Res. type | Atom | Residue | Distance | Occupancy | B-factor | Closest solute to 1^st^ shell water |
| --- | --- | --- | --- | --- | --- | --- | --- |
| **Ion** | | | |  | 1.00 | 105.75 || 1 | A | N6 | AA:1587 | 4.22 | 1.00 | 93.60 |
| 2 | G | O6 | AA:1414 | 4.40 | 1.00 | 88.45 |  |
| 3 | U | O4 | AA:1415 | 4.71 | 1.00 | 90.71 |  |
  |

  
 Coded contact list:   
1^st^ MG coordination shell missing (see Notes)  

### MG:AA:3916

|  | Res. type | Atom | Residue | Distance | Occupancy | B-factor | Closest solute to 1^st^ shell water |
| --- | --- | --- | --- | --- | --- | --- | --- |
| **Ion** | | | |  | 1.00 | 93.60 || 1 | C | OP1 | AA:409 | 2.96 | 1.00 | 63.58 |
| 2 | C | OP1 | AA:376 | 2.97 | 1.00 | 54.43 |  |
| 3 | C | OP1 | AA:392 | 3.01 | 1.00 | 53.79 |  |
| 4 | C | OP2 | AA:376 | 3.70 | 1.00 | 55.39 |  |
| 5 | C | OP2 | AA:409 | 4.10 | 1.00 | 64.29 |  |
| 6 | C | O3' | AA:375 | 4.29 | 1.00 | 55.03 |  |
| 7 | C | OP2 | AA:392 | 4.56 | 1.00 | 53.70 |  |
| 8 | G | O3' | AA:391 | 4.75 | 1.00 | 53.64 |  |
| 9 | G | O3' | AA:408 | 4.88 | 1.00 | 64.17 |  |
  |

  
 Coded contact list:   
 Exclusion zone contact(s): /OP1/OP1/OP1/  
Consider K in place of MG (3 valid contact(s) for K vs. 0 for MG)  

### MG:AA:3917

|  | Res. type | Atom | Residue | Distance | Occupancy | B-factor | Closest solute to 1^st^ shell water |
| --- | --- | --- | --- | --- | --- | --- | --- |
| **Ion** | | | |  | 1.00 | 98.06 || 1 | G | O6 | AA:500 | 2.71 | 1.00 | 68.17 |
| 2 | A | OP2 | AA:503 | 3.32 | 1.00 | 54.77 |  |
| 3 | A | OP1 | AA:502 | 3.93 | 1.00 | 56.27 |  |
| 4 | G | N1 | AA:500 | 3.96 | 1.00 | 65.21 |  |
| 5 | A | OP1 | AA:503 | 4.79 | 1.00 | 54.53 |  |
  |

  
 Coded contact list:   
 Exclusion zone contact(s): /O6/OP2/  
Consider K in place of MG (2 valid contact(s) for K vs. 0 for MG)  

### MG:AA:3918

|  | Res. type | Atom | Residue | Distance | Occupancy | B-factor | Closest solute to 1^st^ shell water |
| --- | --- | --- | --- | --- | --- | --- | --- |
| **Ion** | | | |  | 1.00 | 90.64 || 1 | A | OP2 | AA:1586 | 2.74 | 1.00 | 115.14 |
| 2 | A | N7 | AA:1587 | 4.41 | 1.00 | 95.55 |  |
| 3 | A | OP1 | AA:1586 | 4.67 | 1.00 | 115.32 |  |
| 4 | A | N6 | AA:1587 | 4.97 | 1.00 | 93.60 |  |
  |

  
 Coded contact list:   
 Exclusion zone contact(s): /OP2/  
Consider K in place of MG (1 valid contact(s) for K vs. 0 for MG)  

### MG:AA:3919

|  | Res. type | Atom | Residue | Distance | Occupancy | B-factor | Closest solute to 1^st^ shell water |
| --- | --- | --- | --- | --- | --- | --- | --- |
| **Ion** | | | |  | 1.00 | 59.90 || 1 | G | O6 | AA:379 | 2.55 | 1.00 | 48.08 |
| 2 | U | O4 | AA:380 | 2.71 | 1.00 | 58.63 |  |
| 3 | G | N7 | AA:379 | 3.02 | 1.00 | 50.60 |  |
| 4 | C | N4 | AA:378 | 3.60 | 1.00 | 51.93 |  |
| 5 | U | O4 | AA:395 | 3.79 | 1.00 | 46.98 |  |
| 6 | U | N3 | AA:380 | 4.18 | 1.00 | 57.23 |  |
| 7 | A | N6 | AA:394 | 4.30 | 1.00 | 57.47 |  |
| 8 | U | N3 | AA:395 | 4.46 | 1.00 | 50.70 |  |
| 9 | G | N1 | AA:379 | 4.64 | 1.00 | 49.74 |  |
| 10 | G | O6 | AA:396 | 4.70 | 1.00 | 48.96 |  |
| 11 | C | N3 | AA:378 | 5.00 | 1.00 | 52.98 |  |
  |

  
 Coded contact list:   
 Exclusion zone contact(s): /N7/O4/O6/  
Consider K in place of MG (2 valid contact(s) for K vs. 0 for MG)  

### MG:AA:3920

|  | Res. type | Atom | Residue | Distance | Occupancy | B-factor | Closest solute to 1^st^ shell water |
| --- | --- | --- | --- | --- | --- | --- | --- |
| **Ion** | | | |  | 1.00 | 85.49 || 1 | G | OP2 | AA:2371 | 2.50 | 1.00 | 72.62 |
| 2 | G | OP1 | AA:2370 | 3.14 | 1.00 | 77.02 |  |
| 3 | G | O5' | AA:2370 | 3.34 | 1.00 | 75.58 |  |
| 4 | G | OP1 | AA:2371 | 3.70 | 1.00 | 73.48 |  |
| 5 | G | O3' | AA:2370 | 3.85 | 1.00 | 73.47 |  |
| 6 | G | OP2 | AA:2370 | 4.66 | 1.00 | 75.99 |  |
| 7 | G | O5' | AA:2371 | 4.85 | 1.00 | 71.67 |  |
  |

  
 Coded contact list:   
 Exclusion zone contact(s): /O5'/OP1/OP2/  
Consider K in place of MG (2 valid contact(s) for K vs. 0 for MG)  

### MG:AA:3921

|  | Res. type | Atom | Residue | Distance | Occupancy | B-factor | Closest solute to 1^st^ shell water |
| --- | --- | --- | --- | --- | --- | --- | --- |
| **Ion** | | | |  | 1.00 | 44.45 || 1 | G | N7 | AA:2709 | 2.92 | 1.00 | 48.11 |
| 2 | G | O6 | AA:2709 | 3.20 | 1.00 | 49.51 |  |
| 3 | C | N4 | AA:2710 | 3.22 | 1.00 | 51.62 |  |
| 4 | G | O6 | AA:2708 | 3.59 | 1.00 | 47.55 |  |
| 5 | G | N7 | AA:2708 | 3.61 | 1.00 | 49.14 |  |
| 6 | G | O6 | AA:2697 | 4.34 | 1.00 | 52.28 |  |
| 7 | G | N1 | AA:2708 | 4.44 | 1.00 | 46.52 |  |
| 8 | G | N9 | AA:2708 | 4.85 | 1.00 | 48.89 |  |
| 9 | U | O4 | AA:2698 | 4.90 | 1.00 | 54.06 |  |
  |

  
 Coded contact list:   
 Exclusion zone contact(s): /N4/N7/O6/  
Consider K in place of MG (3 valid contact(s) for K vs. 0 for MG)  

### MG:AA:3922

|  | Res. type | Atom | Residue | Distance | Occupancy | B-factor | Closest solute to 1^st^ shell water |
| --- | --- | --- | --- | --- | --- | --- | --- |
| **Ion** | | | |  | 1.00 | 65.97 || 1 | C | O2' | AA:2701 | 2.79 | 1.00 | 59.49 |
| 2 | C | O3' | AA:2701 | 3.12 | 1.00 | 60.82 |  |
| 3 | U | OP1 | AA:2702 | 3.60 | 1.00 | 64.71 |  |
| 4 | U | OP2 | AA:2702 | 4.62 | 1.00 | 65.79 |  |
| 5 | U | N1 | AA:2702 | 4.73 | 1.00 | 70.60 |  |
| 6 | VAL | N | A0:73 | 4.77 | 1.00 | 64.47 |  |
| 7 | U | O4' | AA:2702 | 4.79 | 1.00 | 69.56 |  |
  |

  
 Coded contact list:   
 Exclusion zone contact(s): /O2'/O3'/  
Consider K in place of MG (2 valid contact(s) for K vs. 0 for MG)  

### MG:AA:3923

|  | Res. type | Atom | Residue | Distance | Occupancy | B-factor | Closest solute to 1^st^ shell water |
| --- | --- | --- | --- | --- | --- | --- | --- |
| **Ion** | | | |  | 1.00 | 107.43 || 1 | C | O5' | AA:1870 | 2.97 | 1.00 | 82.80 |
| 2 | A | OP2 | AA:1871 | 3.63 | 1.00 | 87.40 |  |
| 3 | G | OP2 | AA:1869 | 3.67 | 1.00 | 73.36 |  |
| 4 | C | O3' | AA:1870 | 3.70 | 1.00 | 86.29 |  |
| 5 | G | N7 | AA:1869 | 3.82 | 1.00 | 80.00 |  |
| 6 | C | OP2 | AA:1870 | 3.87 | 1.00 | 82.01 |  |
| 7 | A | OP1 | AA:1871 | 3.96 | 1.00 | 87.15 |  |
| 8 | G | N9 | AA:1869 | 4.10 | 1.00 | 79.50 |  |
| 9 | G | O5' | AA:1869 | 4.23 | 1.00 | 76.10 |  |
| 10 | G | O3' | AA:1869 | 4.26 | 1.00 | 81.09 |  |
| 11 | G | O4' | AA:1869 | 4.27 | 1.00 | 78.68 |  |
  |

  
 Coded contact list:   
 Exclusion zone contact(s): /O5'/  
Consider K in place of MG (1 valid contact(s) for K vs. 0 for MG)  

### MG:AA:3924

|  | Res. type | Atom | Residue | Distance | Occupancy | B-factor | Closest solute to 1^st^ shell water |
| --- | --- | --- | --- | --- | --- | --- | --- |
| **Ion** | | | |  | 1.00 | 79.73 || 1 | C | OP2 | AA:66 | 2.55 | 1.00 | 54.48 |
| 2 | U | OP2 | AA:67 | 3.12 | 1.00 | 54.38 |  |
| 3 | C | OP1 | AA:66 | 3.13 | 1.00 | 53.38 |  |
| 4 | C | O5' | AA:66 | 3.45 | 1.00 | 53.38 |  |
| 5 | C | O3' | AA:65 | 4.62 | 1.00 | 54.81 |  |
| 6 | C | O3' | AA:66 | 4.83 | 1.00 | 53.51 |  |
  |

  
 Coded contact list:   
 Exclusion zone contact(s): /OP1/OP2/OP2/  
Consider K in place of MG (2 valid contact(s) for K vs. 0 for MG)  

### MG:AA:3925

|  | Res. type | Atom | Residue | Distance | Occupancy | B-factor | Closest solute to 1^st^ shell water |
| --- | --- | --- | --- | --- | --- | --- | --- |
| **Ion** | | | |  | 1.00 | 71.85 || 1 | G | O6 | AA:768 | 3.14 | 1.00 | 39.07 |
| 2 | G | O6 | AA:769 | 3.24 | 1.00 | 37.70 |  |
| 3 | G | N7 | AA:768 | 3.36 | 1.00 | 32.72 |  |
| 4 | U | O4 | AA:767 | 3.79 | 1.00 | 38.03 |  |
| 5 | G | O6 | AA:695 | 3.85 | 1.00 | 39.43 |  |
| 6 | U | N3 | AA:767 | 4.02 | 1.00 | 37.59 |  |
| 7 | U | O4 | AA:694 | 4.29 | 1.00 | 43.82 |  |
| 8 | C | N4 | AA:693 | 4.35 | 1.00 | 33.46 |  |
| 9 | G | N7 | AA:769 | 4.64 | 1.00 | 37.81 |  |
  |

  
 Coded contact list:   
 Exclusion zone contact(s): /N7/O6/O6/  
Consider K in place of MG (3 valid contact(s) for K vs. 0 for MG)  

### MG:AA:3926

|  | Res. type | Atom | Residue | Distance | Occupancy | B-factor | Closest solute to 1^st^ shell water |
| --- | --- | --- | --- | --- | --- | --- | --- |
| **Ion** | | | |  | 1.00 | 52.20 || 1 | C | OP2 | AA:1376 | 2.40 | 1.00 | 49.43 |
| 2 | U | OP2 | AA:1352 | 2.86 | 1.00 | 44.43 |  |
| 3 | C | OP1 | AA:1376 | 3.63 | 1.00 | 50.34 |  |
| 4 | C | OP2 | AA:1351 | 3.84 | 1.00 | 45.27 |  |
| 5 | G | N7 | AA:1377 | 4.06 | 1.00 | 51.09 |  |
| 6 | C | O5' | AA:1376 | 4.14 | 1.00 | 49.21 |  |
| 7 | U | O5' | AA:1352 | 4.19 | 1.00 | 44.42 |  |
| 8 | C | O5' | AA:1351 | 4.33 | 1.00 | 44.29 |  |
| 9 | G | O6 | AA:1377 | 4.41 | 1.00 | 52.27 |  |
| 10 | C | O3' | AA:1375 | 4.75 | 1.00 | 49.33 |  |
| 11 | G | OP2 | AA:1377 | 4.96 | 1.00 | 48.62 |  |
  |

  
 Contact(s): /OP2/  
 Coded contact list: O_ph_  
 Exclusion zone contact(s): /OP2/  

### MG:AA:3927

|  | Res. type | Atom | Residue | Distance | Occupancy | B-factor | Closest solute to 1^st^ shell water |
| --- | --- | --- | --- | --- | --- | --- | --- |
| **Ion** | | | |  | 1.00 | 76.53 || 1 | A | OP2 | AA:2693 | 2.74 | 1.00 | 60.01 |
| 2 | A | OP1 | AA:2693 | 3.16 | 1.00 | 61.27 |  |
| 3 | A | O5' | AA:2693 | 3.85 | 1.00 | 59.30 |  |
| 4 | G | OP2 | AA:2694 | 4.07 | 1.00 | 59.32 |  |
| 5 | ARG | NH1 | A0:68 | 4.57 | 1.00 | 73.76 |  |
| 6 | C | O3' | AA:2692 | 4.78 | 1.00 | 60.79 |  |
  |

  
 Coded contact list:   
 Exclusion zone contact(s): /OP1/OP2/  
Consider K in place of MG (2 valid contact(s) for K vs. 0 for MG)  

### MG:AA:3928

|  | Res. type | Atom | Residue | Distance | Occupancy | B-factor | Closest solute to 1^st^ shell water |
| --- | --- | --- | --- | --- | --- | --- | --- |
| **Ion** | | | |  | 1.00 | 71.30 || 1 | G | OP2 | AA:410 | 2.95 | 1.00 | 55.33 |
| 2 | C | O5' | AA:409 | 3.91 | 1.00 | 61.93 |  |
| 3 | G | N7 | AA:410 | 3.98 | 1.00 | 61.85 |  |
| 4 | C | OP2 | AA:409 | 4.09 | 1.00 | 64.29 |  |
| 5 | C | N1 | AA:409 | 4.48 | 1.00 | 59.37 |  |
| 6 | C | O3' | AA:409 | 4.58 | 1.00 | 56.34 |  |
| 7 | G | O5' | AA:410 | 4.74 | 1.00 | 56.07 |  |
  |

  
 Coded contact list:   
 Exclusion zone contact(s): /OP2/  
Consider K in place of MG (1 valid contact(s) for K vs. 0 for MG)  

### MG:AA:3929

|  | Res. type | Atom | Residue | Distance | Occupancy | B-factor | Closest solute to 1^st^ shell water |
| --- | --- | --- | --- | --- | --- | --- | --- |
| **Ion** | | | |  | 1.00 | 104.04 || 1 | C | O4' | AA:277 | 2.50 | 1.00 | 150.86 |
| 2 | A | O2' | AA:276 | 2.54 | 1.00 | 157.34 |  |
| 3 | C | O5' | AA:277 | 2.58 | 1.00 | 153.84 |  |
| 4 | A | O3' | AA:276 | 3.86 | 1.00 | 156.05 |  |
| 5 | C | OP1 | AA:277 | 4.17 | 1.00 | 154.94 |  |
| 6 | C | N1 | AA:277 | 4.57 | 1.00 | 152.42 |  |
| 7 | C | O2' | AA:277 | 4.73 | 1.00 | 149.71 |  |
| 8 | C | OP2 | AA:277 | 4.98 | 1.00 | 154.82 |  |
  |

  
 Coded contact list:   
 Exclusion zone contact(s): /O2'/O4'/O5'/  
Consider NA in place of MG (3 valid contact(s) for NA vs. 0 for MG)  

### MG:AA:3930

|  | Res. type | Atom | Residue | Distance | Occupancy | B-factor | Closest solute to 1^st^ shell water |
| --- | --- | --- | --- | --- | --- | --- | --- |
| **Ion** | | | |  | 1.00 | 46.11 || 1 | C | OP2 | AA:786 | 3.07 | 1.00 | 32.18 |
| 2 | C | N1 | AA:791 | 3.19 | 1.00 | 34.34 |  |
| 3 | U | OP2 | AA:787 | 3.24 | 1.00 | 31.77 |  |
| 4 | C | O2' | AA:791 | 3.36 | 1.00 | 41.72 |  |
| 5 | C | O5' | AA:786 | 3.76 | 1.00 | 31.14 |  |
| 6 | C | O4' | AA:791 | 3.87 | 1.00 | 36.61 |  |
| 7 | C | O5' | AA:791 | 4.04 | 1.00 | 39.53 |  |
| 8 | C | O3' | AA:791 | 4.07 | 1.00 | 35.98 |  |
| 9 | C | N3 | AA:791 | 4.61 | 1.00 | 32.59 |  |
| 10 | C | O2 | AA:791 | 4.61 | 1.00 | 33.83 |  |
| 11 | C | OP1 | AA:786 | 4.70 | 1.00 | 31.67 |  |
| 12 | C | OP2 | AA:791 | 4.75 | 1.00 | 39.34 |  |
  |

  
 Coded contact list:   
 Exclusion zone contact(s): /N1/O2'/OP2/OP2/  
Consider K in place of MG (4 valid contact(s) for K vs. 0 for MG)  

### MG:AA:3931

|  | Res. type | Atom | Residue | Distance | Occupancy | B-factor | Closest solute to 1^st^ shell water |
| --- | --- | --- | --- | --- | --- | --- | --- |
| **Ion** | | | |  | 1.00 | 86.33 || 1 | G | N7 | AA:1878 | 3.62 | 1.00 | 78.29 |
| 2 | C | N4 | AA:1879 | 4.15 | 1.00 | 76.58 |  |
  |

  
 Coded contact list:   
1^st^ MG coordination shell missing (see Notes)  

### MG:AA:3932

|  | Res. type | Atom | Residue | Distance | Occupancy | B-factor | Closest solute to 1^st^ shell water |
| --- | --- | --- | --- | --- | --- | --- | --- |
| **Ion** | | | |  | 1.00 | 56.46 || 1 | U | N1 | AA:90 | 2.86 | 1.00 | 108.05 |
| 2 | U | O2' | AA:90 | 3.17 | 1.00 | 105.35 |  |
| 3 | U | O2 | AA:90 | 3.55 | 1.00 | 108.53 |  |
| 4 | U | O4' | AA:90 | 3.81 | 1.00 | 104.44 |  |
| 5 | U | N3 | AA:90 | 4.19 | 1.00 | 109.66 |  |
| 6 | G | N7 | AA:92 | 4.73 | 1.00 | 80.53 |  |
| 7 | G | OP2 | AA:92 | 4.92 | 1.00 | 98.85 |  |
| 8 | G | OP1 | AA:88 | 4.95 | 1.00 | 66.07 |  |
  |

  
 Coded contact list:   
 Exclusion zone contact(s): /N1/O2'/  
Consider K in place of MG (2 valid contact(s) for K vs. 0 for MG)  

### MG:AA:3933

|  | Res. type | Atom | Residue | Distance | Occupancy | B-factor | Closest solute to 1^st^ shell water |
| --- | --- | --- | --- | --- | --- | --- | --- |
| **Ion** | | | |  | 1.00 | 115.15 || 1 | C | O2' | AA:1076 | 4.13 | 1.00 | 199.63 |
  |

  
 Coded contact list:   
1^st^ MG coordination shell missing (see Notes)  

### MG:AA:3934

|  | Res. type | Atom | Residue | Distance | Occupancy | B-factor | Closest solute to 1^st^ shell water |
| --- | --- | --- | --- | --- | --- | --- | --- |
| **Ion** | | | |  | 1.00 | 93.55 || 1 | A | OP2 | AA:216 | 3.54 | 1.00 | 58.61 |
| 2 | U | O4 | AA:431 | 3.64 | 1.00 | 50.09 |  |
| 3 | A | N6 | AA:432 | 3.82 | 1.00 | 43.13 |  |
| 4 | G | O3' | AA:215 | 4.44 | 1.00 | 60.45 |  |
| 5 | G | O6 | AA:45 | 4.52 | 1.00 | 61.12 |  |
| 6 | G | O2' | AA:215 | 4.55 | 1.00 | 62.82 |  |
  |

  
 Coded contact list:   
1^st^ MG coordination shell missing (see Notes)  

### MG:AA:3935

|  | Res. type | Atom | Residue | Distance | Occupancy | B-factor | Closest solute to 1^st^ shell water |
| --- | --- | --- | --- | --- | --- | --- | --- |
| **Ion** | | | |  | 1.00 | 55.72 || 1 | G | O4' | AA:438 | 2.99 | 1.00 | 74.83 |
| 2 | G | N2 | AA:43 | 3.14 | 1.00 | 64.69 |  |
| 3 | C | O2 | AA:436 | 3.40 | 1.00 | 66.57 |  |
| 4 | G | N9 | AA:438 | 3.66 | 1.00 | 74.67 |  |
| 5 | C | O2' | AA:436 | 4.00 | 1.00 | 69.58 |  |
| 6 | G | N3 | AA:438 | 4.82 | 1.00 | 74.79 |  |
| 7 | G | N7 | AA:438 | 4.92 | 1.00 | 74.05 |  |
  |

  
 Coded contact list:   
 Exclusion zone contact(s): /N2/O2/O4'/  
Consider K in place of MG (3 valid contact(s) for K vs. 0 for MG)  

### MG:AA:3936

|  | Res. type | Atom | Residue | Distance | Occupancy | B-factor | Closest solute to 1^st^ shell water |
| --- | --- | --- | --- | --- | --- | --- | --- |
| **Ion** | | | |  | 1.00 | 83.46 || 1 | A | OP2 | AA:990 | 3.06 | 1.00 | 73.82 |
| 2 | C | OP1 | AA:1185 | 3.74 | 1.00 | 74.00 |  |
| 3 | G | O2' | AA:989 | 3.76 | 1.00 | 80.46 |  |
| 4 | G | O3' | AA:989 | 3.98 | 1.00 | 76.04 |  |
| 5 | G | OP1 | AA:1186 | 4.04 | 1.00 | 67.82 |  |
| 6 | G | OP2 | AA:1186 | 4.18 | 1.00 | 68.26 |  |
| 7 | G | N2 | AA:974 | 4.23 | 1.00 | 69.63 |  |
| 8 | A | OP1 | AA:990 | 4.35 | 1.00 | 75.19 |  |
| 9 | G | N9 | AA:989 | 4.59 | 1.00 | 77.49 |  |
| 10 | G | N7 | AA:1186 | 4.78 | 1.00 | 71.60 |  |
| 11 | G | O4' | AA:989 | 4.84 | 1.00 | 77.66 |  |
  |

  
 Coded contact list:   
 Exclusion zone contact(s): /OP2/  
Consider K in place of MG (1 valid contact(s) for K vs. 0 for MG)  

### MG:AA:3937

|  | Res. type | Atom | Residue | Distance | Occupancy | B-factor | Closest solute to 1^st^ shell water |
| --- | --- | --- | --- | --- | --- | --- | --- |
| **Ion** | | | |  | 1.00 | 67.42 || 1 | U | O4 | AA:405 | 4.81 | 1.00 | 101.75 |
  |

  
 Coded contact list:   
1^st^ MG coordination shell missing (see Notes)  

### MG:AA:3938

|  | Res. type | Atom | Residue | Distance | Occupancy | B-factor | Closest solute to 1^st^ shell water |
| --- | --- | --- | --- | --- | --- | --- | --- |
| **Ion** | | | |  | 1.00 | 76.17 || 1 | C | OP2 | AA:1630A | 4.01 | 1.00 | 60.21 |
| 2 | C | OP1 | AA:1741 | 4.46 | 1.00 | 95.73 |  |
| 3 | G | OP1 | AA:1630 | 4.85 | 1.00 | 54.56 |  |
  |

  
 Coded contact list:   
1^st^ MG coordination shell missing (see Notes)  

### MG:AA:3939

|  | Res. type | Atom | Residue | Distance | Occupancy | B-factor | Closest solute to 1^st^ shell water |
| --- | --- | --- | --- | --- | --- | --- | --- |
| **Ion** | | | |  | 1.00 | 54.32 || 1 | U | O4 | AA:963 | 2.80 | 1.00 | 56.06 |
| 2 | G | O6 | AA:962 | 2.89 | 1.00 | 62.78 |  |
| 3 | C | N4 | AA:955 | 4.18 | 1.00 | 63.42 |  |
| 4 | G | O6 | AA:954 | 4.25 | 1.00 | 58.22 |  |
| 5 | G | N7 | AA:962 | 4.34 | 1.00 | 60.41 |  |
| 6 | U | N3 | AA:963 | 4.51 | 1.00 | 56.67 |  |
| 7 | G | N1 | AA:956 | 4.77 | 1.00 | 75.04 |  |
  |

  
 Coded contact list:   
 Exclusion zone contact(s): /O4/O6/  
Consider K in place of MG (2 valid contact(s) for K vs. 0 for MG)  

### MG:AA:3940

|  | Res. type | Atom | Residue | Distance | Occupancy | B-factor | Closest solute to 1^st^ shell water |
| --- | --- | --- | --- | --- | --- | --- | --- |
| **Ion** | | | |  | 1.00 | 61.52 || 1 | A | OP2 | AA:457 | 1.75 | 1.00 | 63.28 |
| 2 | C | O2' | AA:456 | 3.08 | 1.00 | 72.18 |  |
| 3 | C | O3' | AA:456 | 3.14 | 1.00 | 65.58 |  |
| 4 | C | N1 | AA:456 | 3.22 | 1.00 | 74.69 |  |
| 5 | A | O5' | AA:457 | 3.86 | 1.00 | 58.29 |  |
| 6 | ARG | NE | AT:68 | 4.06 | 1.00 | 93.48 |  |
| 7 | A | OP1 | AA:457 | 4.11 | 1.00 | 62.58 |  |
| 8 | C | O4' | AA:456 | 4.14 | 1.00 | 72.05 |  |
| 9 | ARG | NH2 | AT:68 | 4.37 | 1.00 | 92.76 |  |
| 10 | C | O2 | AA:456 | 4.71 | 1.00 | 74.73 |  |
| 11 | C | N3 | AA:456 | 4.78 | 1.00 | 76.15 |  |
| 12 | TYR | OH | AT:69 | 4.81 | 1.00 | 95.75 |  |
| 13 | C | O5' | AA:456 | 4.93 | 1.00 | 69.44 |  |
  |

  
 Coded contact list:   
 Exclusion zone contact(s): /N1/O2'/O3'/OP2/  
Consider K in place of MG (3 valid contact(s) for K vs. 0 for MG)  

### MG:AA:3941

|  | Res. type | Atom | Residue | Distance | Occupancy | B-factor | Closest solute to 1^st^ shell water |
| --- | --- | --- | --- | --- | --- | --- | --- |
| **Ion** | | | |  | 1.00 | 57.97 || 1 | G | O6 | AA:2523 | 2.95 | 1.00 | 73.13 |
| 2 | G | N7 | AA:2523 | 3.78 | 1.00 | 72.02 |  |
| 3 | U | OP2 | AA:2522 | 4.26 | 1.00 | 63.97 |  |
| 4 | U | O4 | AA:2522 | 4.61 | 1.00 | 59.88 |  |
| 5 | G | O6 | AA:2524 | 4.66 | 1.00 | 74.40 |  |
| 6 | C | N4 | AA:2540 | 4.69 | 1.00 | 74.24 |  |
| 7 | A | N1 | AA:2541 | 4.81 | 1.00 | 73.58 |  |
  |

  
 Coded contact list:   
 Exclusion zone contact(s): /O6/  
Consider K in place of MG (1 valid contact(s) for K vs. 0 for MG)  

### MG:AA:3942

|  | Res. type | Atom | Residue | Distance | Occupancy | B-factor | Closest solute to 1^st^ shell water |
| --- | --- | --- | --- | --- | --- | --- | --- |
| **Ion** | | | |  | 1.00 | 85.65 || 1 | G | N7 | AA:2215 | 3.37 | 1.00 | 84.59 |
| 2 | G | O6 | AA:2215 | 3.63 | 1.00 | 83.74 |  |
| 3 | G | OP2 | AA:2215 | 4.74 | 1.00 | 102.22 |  |
| 4 | ARG | NH1 | AZ:50 | 4.98 | 1.00 | 72.24 |  |
  |

  
 Coded contact list:   
 Exclusion zone contact(s): /N7/  
Consider K in place of MG (1 valid contact(s) for K vs. 0 for MG)  

### MG:AA:3943

|  | Res. type | Atom | Residue | Distance | Occupancy | B-factor | Closest solute to 1^st^ shell water |
| --- | --- | --- | --- | --- | --- | --- | --- |
| **Ion** | | | |  | 1.00 | 65.80 || 1 | C | O3' | AA:34 | 2.73 | 1.00 | 72.11 |
| 2 | A | O2' | AA:454 | 3.02 | 1.00 | 54.52 |  |
| 3 | C | OP1 | AA:455 | 3.65 | 1.00 | 53.86 |  |
| 4 | G | OP1 | AA:35 | 3.82 | 1.00 | 68.74 |  |
| 5 | G | O5' | AA:35 | 3.93 | 1.00 | 65.43 |  |
| 6 | G | O4' | AA:35 | 4.30 | 1.00 | 54.17 |  |
| 7 | C | O5' | AA:34 | 4.39 | 1.00 | 74.98 |  |
| 8 | C | O2' | AA:34 | 4.69 | 1.00 | 79.11 |  |
| 9 | C | O4' | AA:34 | 4.86 | 1.00 | 79.35 |  |
| 10 | U | O2' | AA:33 | 4.99 | 1.00 | 71.23 |  |
  |

  
 Coded contact list:   
 Exclusion zone contact(s): /O2'/O3'/  
Consider K in place of MG (2 valid contact(s) for K vs. 0 for MG)  

### MG:AA:3944

|  | Res. type | Atom | Residue | Distance | Occupancy | B-factor | Closest solute to 1^st^ shell water |
| --- | --- | --- | --- | --- | --- | --- | --- |
| **Ion** | | | |  | 1.00 | 71.54 || 1 | U | OP2 | AA:67 | 2.89 | 1.00 | 54.38 |
| 2 | U | OP1 | AA:67 | 3.23 | 1.00 | 54.61 |  |
| 3 | U | O5' | AA:67 | 3.95 | 1.00 | 54.69 |  |
| 4 | G | OP2 | AA:68 | 3.95 | 1.00 | 50.58 |  |
| 5 | C | O3' | AA:66 | 4.90 | 1.00 | 53.51 |  |
  |

  
 Coded contact list:   
 Exclusion zone contact(s): /OP1/OP2/  
Consider K in place of MG (2 valid contact(s) for K vs. 0 for MG)  

### MG:AA:3945

|  | Res. type | Atom | Residue | Distance | Occupancy | B-factor | Closest solute to 1^st^ shell water |
| --- | --- | --- | --- | --- | --- | --- | --- |
| **Ion** | | | |  | 1.00 | 108.04 || 1 | G | OP2 | AA:131 | 2.43 | 1.00 | 51.91 |
| 2 | G | OP1 | AA:142 | 2.60 | 1.00 | 64.81 |  |
| 3 | G | OP1 | AA:131 | 3.65 | 1.00 | 51.97 |  |
| 4 | G | OP2 | AA:142 | 4.07 | 1.00 | 63.54 |  |
| 5 | C | O3' | AA:130 | 4.21 | 1.00 | 50.87 |  |
| 6 | A | N9 | AA:1349 | 4.35 | 1.00 | 66.93 |  |
| 7 | C | O2 | AA:1598 | 4.36 | 1.00 | 52.14 |  |
| 8 | A | N7 | AA:1349 | 4.52 | 1.00 | 67.56 |  |
| 9 | G | O5' | AA:131 | 4.76 | 1.00 | 51.24 |  |
| 10 | G | O5' | AA:142 | 4.78 | 1.00 | 64.53 |  |
| 11 | C | O3' | AA:141A | 4.86 | 1.00 | 64.70 |  |
| 12 | C | O5' | AA:130 | 4.90 | 1.00 | 50.61 |  |
  |

  
 Coded contact list:   
 Exclusion zone contact(s): /OP1/OP2/  
Consider NA in place of MG (2 valid contact(s) for NA vs. 0 for MG)  

### MG:AA:3946

|  | Res. type | Atom | Residue | Distance | Occupancy | B-factor | Closest solute to 1^st^ shell water |
| --- | --- | --- | --- | --- | --- | --- | --- |
| **Ion** | | | |  | 1.00 | 98.80 || 1 | G | OP2 | AA:1071 | 2.88 | 1.00 | 200.37 |
| 2 | C | N4 | AA:1072 | 3.12 | 1.00 | 200.07 |  |
| 3 | G | N7 | AA:1071 | 3.39 | 1.00 | 199.62 |  |
| 4 | G | O5' | AA:1071 | 3.99 | 1.00 | 200.41 |  |
| 5 | G | N1 | AA:1089 | 4.05 | 1.00 | 200.41 |  |
| 6 | A | OP1 | AA:1098 | 4.49 | 1.00 | 200.34 |  |
| 7 | G | O6 | AA:1089 | 4.50 | 1.00 | 200.36 |  |
| 8 | A | OP2 | AA:1098 | 4.53 | 1.00 | 200.41 |  |
| 9 | G | N2 | AA:1089 | 4.73 | 1.00 | 200.26 |  |
| 10 | C | N3 | AA:1072 | 4.80 | 1.00 | 200.20 |  |
| 11 | G | N9 | AA:1071 | 4.85 | 1.00 | 199.78 |  |
| 12 | G | OP1 | AA:1071 | 4.85 | 1.00 | 200.41 |  |
  |

  
 Coded contact list:   
 Exclusion zone contact(s): /N4/N7/OP2/  
Consider K in place of MG (3 valid contact(s) for K vs. 0 for MG)  

### MG:AA:3947

|  | Res. type | Atom | Residue | Distance | Occupancy | B-factor | Closest solute to 1^st^ shell water |
| --- | --- | --- | --- | --- | --- | --- | --- |
| **Ion** | | | |  | 1.00 | 84.93 || 1 | G | N9 | AA:1388 | 2.34 | 1.00 | 69.58 |
| 2 | G | N7 | AA:1389 | 2.79 | 1.00 | 66.63 |  |
| 3 | G | N7 | AA:1388 | 3.09 | 1.00 | 70.42 |  |
| 4 | G | N3 | AA:1388 | 3.12 | 1.00 | 71.06 |  |
| 5 | G | O4' | AA:1388 | 3.45 | 1.00 | 68.22 |  |
| 6 | G | OP2 | AA:1389 | 3.61 | 1.00 | 69.47 |  |
| 7 | G | O3' | AA:1388 | 3.72 | 1.00 | 68.43 |  |
| 8 | G | O5' | AA:1389 | 3.89 | 1.00 | 68.57 |  |
| 9 | G | N9 | AA:1389 | 4.05 | 1.00 | 66.53 |  |
| 10 | G | O2' | AA:1388 | 4.15 | 1.00 | 69.84 |  |
| 11 | G | N1 | AA:1388 | 4.18 | 1.00 | 71.36 |  |
| 12 | G | O4' | AA:1389 | 4.40 | 1.00 | 65.56 |  |
| 13 | G | O6 | AA:1388 | 4.80 | 1.00 | 71.03 |  |
| 14 | G | N2 | AA:1388 | 4.90 | 1.00 | 71.50 |  |
| 15 | G | O5' | AA:1388 | 4.94 | 1.00 | 70.47 |  |
| 16 | G | OP2 | AA:1388 | 4.95 | 1.00 | 69.34 |  |
  |

  
 Coded contact list:   
 Exclusion zone contact(s): /N3_G_/N7/N7/  
Consider K in place of MG (3 valid contact(s) for K vs. 1 for MG)  

### MG:AA:3948

|  | Res. type | Atom | Residue | Distance | Occupancy | B-factor | Closest solute to 1^st^ shell water |
| --- | --- | --- | --- | --- | --- | --- | --- |
| **Ion** | | | |  | 1.00 | 72.38 || 1 | G | O6 | AA:142 | 2.56 | 1.00 | 63.99 |
| 2 | C | N4 | AA:143 | 3.62 | 1.00 | 71.65 |  |
| 3 | G | O6 | AA:136 | 3.64 | 1.00 | 70.83 |  |
| 4 | C | N4 | AA:137 | 3.67 | 1.00 | 65.36 |  |
| 5 | C | N4 | AA:141A | 4.36 | 1.00 | 65.77 |  |
| 6 | G | N7 | AA:142 | 4.36 | 1.00 | 64.47 |  |
| 7 | G | O6 | AA:137A | 4.63 | 1.00 | 66.44 |  |
| 8 | G | N1 | AA:142 | 4.72 | 1.00 | 64.36 |  |
| 9 | A | OP1 | AA:141 | 4.82 | 1.00 | 66.95 |  |
  |

  
 Coded contact list:   
 Exclusion zone contact(s): /O6/  
Consider NA in place of MG (1 valid contact(s) for NA vs. 0 for MG)  

### MG:AA:3949

|  | Res. type | Atom | Residue | Distance | Occupancy | B-factor | Closest solute to 1^st^ shell water |
| --- | --- | --- | --- | --- | --- | --- | --- |
| **Ion** | | | |  | 1.00 | 71.28 || 1 | A | OP2 | AA:2670 | 2.65 | 1.00 | 105.69 |
| 2 | A | OP1 | AA:2670 | 3.01 | 1.00 | 105.74 |  |
| 3 | G | O3' | AA:2669 | 3.58 | 1.00 | 106.34 |  |
| 4 | G | OP1 | AA:2669 | 3.75 | 1.00 | 110.04 |  |
| 5 | G | O5' | AA:2669 | 4.09 | 1.00 | 109.60 |  |
| 6 | G | OP2 | AA:2669 | 4.61 | 1.00 | 110.95 |  |
| 7 | A | O5' | AA:2670 | 4.64 | 1.00 | 103.76 |  |
  |

  
 Coded contact list:   
 Exclusion zone contact(s): /OP1/OP2/  
Consider K in place of MG (2 valid contact(s) for K vs. 0 for MG)  

### MG:AA:3950

|  | Res. type | Atom | Residue | Distance | Occupancy | B-factor | Closest solute to 1^st^ shell water |
| --- | --- | --- | --- | --- | --- | --- | --- |
| **Ion** | | | |  | 1.00 | 63.16 || 1 | G | OP1 | AA:1444 | 2.24 | 1.00 | 77.38 |
| 2 | G | O3' | AA:1443 | 4.35 | 1.00 | 75.42 |  |
| 3 | G | O5' | AA:1444 | 4.61 | 1.00 | 78.95 |  |
| 4 | G | OP2 | AA:1444 | 4.61 | 1.00 | 77.22 |  |
  |

  
 Contact(s): /OP1/  
 Coded contact list: O_ph_  

### MG:AA:3951

|  | Res. type | Atom | Residue | Distance | Occupancy | B-factor | Closest solute to 1^st^ shell water |
| --- | --- | --- | --- | --- | --- | --- | --- |
| **Ion** | | | |  | 1.00 | 98.61 || 1 | U | O2 | AA:405 | 3.01 | 1.00 | 101.30 |
| 2 | U | N3 | AA:405 | 3.22 | 1.00 | 101.51 |  |
| 3 | G | OP1 | AA:406 | 3.31 | 1.00 | 90.46 |  |
| 4 | U | N1 | AA:405 | 3.43 | 1.00 | 99.58 |  |
| 5 | G | O5' | AA:406 | 4.06 | 1.00 | 87.57 |  |
| 6 | U | O3' | AA:405 | 4.21 | 1.00 | 91.99 |  |
| 7 | U | O4' | AA:405 | 4.54 | 1.00 | 96.28 |  |
| 8 | U | O4 | AA:405 | 4.65 | 1.00 | 101.75 |  |
| 9 | U | O2' | AA:405 | 4.95 | 1.00 | 97.30 |  |
  |

  
 Coded contact list:   
 Exclusion zone contact(s): /N3_U_/O2/OP1/  
Consider K in place of MG (3 valid contact(s) for K vs. 0 for MG)  

### MG:AA:3952

|  | Res. type | Atom | Residue | Distance | Occupancy | B-factor | Closest solute to 1^st^ shell water |
| --- | --- | --- | --- | --- | --- | --- | --- |
| **Ion** | | | |  | 1.00 | 71.42 || 1 | G | OP1 | AA:1250 | 2.98 | 1.00 | 50.33 |
| 2 | A | OP2 | AA:586 | 3.27 | 1.00 | 46.70 |  |
| 3 | U | O5' | AA:1249 | 3.46 | 1.00 | 52.84 |  |
| 4 | U | O3' | AA:1249 | 3.75 | 1.00 | 52.68 |  |
| 5 | U | OP1 | AA:1249 | 3.96 | 1.00 | 53.65 |  |
| 6 | G | OP1 | AA:585 | 3.97 | 1.00 | 49.04 |  |
| 7 | G | OP2 | AA:585 | 4.13 | 1.00 | 49.31 |  |
| 8 | G | O3' | AA:1248 | 4.27 | 1.00 | 51.44 |  |
| 9 | G | O5' | AA:585 | 4.63 | 1.00 | 47.63 |  |
| 10 | G | O2' | AA:1248 | 4.72 | 1.00 | 52.27 |  |
| 11 | U | N3 | AA:811 | 4.87 | 1.00 | 51.57 |  |
| 12 | U | O4 | AA:811 | 4.88 | 1.00 | 52.02 |  |
| 13 | G | OP2 | AA:1250 | 4.94 | 1.00 | 51.15 |  |
  |

  
 Coded contact list:   
 Exclusion zone contact(s): /OP1/OP2/  
Consider K in place of MG (2 valid contact(s) for K vs. 0 for MG)  

### MG:AA:3953

|  | Res. type | Atom | Residue | Distance | Occupancy | B-factor | Closest solute to 1^st^ shell water |
| --- | --- | --- | --- | --- | --- | --- | --- |
| **Ion** | | | |  | 1.00 | 60.13 || 1 | G | O6 | AA:496 | 3.14 | 1.00 | 61.04 |
| 2 | G | O6 | AA:495 | 3.52 | 1.00 | 54.60 |  |
| 3 | C | N4 | AA:484 | 3.56 | 1.00 | 62.57 |  |
| 4 | A | OP2 | AA:483 | 3.81 | 1.00 | 60.84 |  |
| 5 | C | N4 | AA:485 | 4.30 | 1.00 | 54.99 |  |
| 6 | A | N6 | AA:497 | 4.37 | 1.00 | 65.47 |  |
| 7 | A | N1 | AA:497 | 4.75 | 1.00 | 62.21 |  |
  |

  
 Coded contact list:   
 Exclusion zone contact(s): /O6/  
Consider K in place of MG (1 valid contact(s) for K vs. 0 for MG)  

### MG:AA:3954

|  | Res. type | Atom | Residue | Distance | Occupancy | B-factor | Closest solute to 1^st^ shell water |
| --- | --- | --- | --- | --- | --- | --- | --- |
| **Ion** | | | |  | 1.00 | 71.89 || 1 | A | OP2 | AA:222 | 3.09 | 1.00 | 65.76 |
| 2 | A | O2' | AA:221 | 3.40 | 1.00 | 72.25 |  |
| 3 | A | O5' | AA:222 | 3.81 | 1.00 | 62.92 |  |
| 4 | A | O4' | AA:222 | 4.24 | 1.00 | 58.04 |  |
| 5 | A | N7 | AA:222 | 4.41 | 1.00 | 54.99 |  |
| 6 | U | OP1 | AA:421 | 4.51 | 1.00 | 65.97 |  |
| 7 | A | N9 | AA:222 | 4.57 | 1.00 | 55.07 |  |
| 8 | G | O2' | AA:232 | 4.71 | 1.00 | 82.61 |  |
  |

  
 Coded contact list:   
 Exclusion zone contact(s): /O2'/OP2/  
Consider K in place of MG (2 valid contact(s) for K vs. 0 for MG)  

### MG:AA:3955

|  | Res. type | Atom | Residue | Distance | Occupancy | B-factor | Closest solute to 1^st^ shell water |
| --- | --- | --- | --- | --- | --- | --- | --- |
| **Ion** | | | |  | 1.00 | 66.76 || 1 | U | O4 | AA:2011 | 2.63 | 1.00 | 43.80 |
| 2 | U | OP2 | AA:1267 | 3.04 | 1.00 | 45.51 |  |
| 3 | A | N6 | AA:1269 | 3.24 | 1.00 | 44.36 |  |
| 4 | G | N1 | AA:2012 | 3.28 | 1.00 | 45.88 |  |
| 5 | G | O6 | AA:2010 | 3.79 | 1.00 | 49.94 |  |
| 6 | G | O6 | AA:2012 | 3.96 | 1.00 | 46.41 |  |
| 7 | G | N2 | AA:2012 | 4.00 | 1.00 | 43.49 |  |
| 8 | G | O2' | AA:1266 | 4.27 | 1.00 | 50.09 |  |
| 9 | G | N3 | AA:2012 | 4.72 | 1.00 | 45.54 |  |
| 10 | U | N3 | AA:2011 | 4.81 | 1.00 | 42.87 |  |
| 11 | G | N7 | AA:2010 | 4.94 | 1.00 | 49.08 |  |
| 12 | G | O3' | AA:1266 | 4.97 | 1.00 | 47.46 |  |
  |

  
 Coded contact list:   
 Exclusion zone contact(s): /N1_G_/N6/O4/OP2/  
Consider K in place of MG (4 valid contact(s) for K vs. 0 for MG)  

### MG:AA:3957

|  | Res. type | Atom | Residue | Distance | Occupancy | B-factor | Closest solute to 1^st^ shell water |
| --- | --- | --- | --- | --- | --- | --- | --- |
| **Ion** | | | |  | 1.00 | 74.42 || 1 | U | O4 | AA:2089 | 3.20 | 1.00 | 49.91 |
| 2 | G | O6 | AA:2230 | 3.40 | 1.00 | 47.52 |  |
| 3 | G | O6 | AA:2088 | 3.49 | 1.00 | 54.29 |  |
| 4 | C | N4 | AA:2231 | 3.96 | 1.00 | 52.31 |  |
| 5 | C | N4 | AA:2229 | 3.98 | 1.00 | 54.92 |  |
| 6 | G | O6 | AA:2090 | 4.41 | 1.00 | 54.32 |  |
| 7 | U | N3 | AA:2089 | 4.42 | 1.00 | 49.50 |  |
| 8 | G | N7 | AA:2230 | 4.99 | 1.00 | 46.40 |  |
  |

  
 Coded contact list:   
 Exclusion zone contact(s): /O4/O6/  
Consider K in place of MG (2 valid contact(s) for K vs. 0 for MG)  

### MG:AA:3958

|  | Res. type | Atom | Residue | Distance | Occupancy | B-factor | Closest solute to 1^st^ shell water |
| --- | --- | --- | --- | --- | --- | --- | --- |
| **Ion** | | | |  | 1.00 | 94.67 || 1 | C | OP2 | AA:1290 | 3.91 | 1.00 | 47.87 |
| 2 | C | OP1 | AA:1290 | 4.28 | 1.00 | 47.95 |  |
  |

  
 Coded contact list:   
1^st^ MG coordination shell missing (see Notes)  

### MG:AA:3959

|  | Res. type | Atom | Residue | Distance | Occupancy | B-factor | Closest solute to 1^st^ shell water |
| --- | --- | --- | --- | --- | --- | --- | --- |
| **Ion** | | | |  | 1.00 | 96.38 || 1 | A | N6 | AA:1088 | 4.84 | 1.00 | 199.67 |
  |

  
 Coded contact list:   
1^st^ MG coordination shell missing (see Notes)  

### MG:AA:3961

|  | Res. type | Atom | Residue | Distance | Occupancy | B-factor | Closest solute to 1^st^ shell water |
| --- | --- | --- | --- | --- | --- | --- | --- |
| **Ion** | | | |  | 1.00 | 54.02 || 1 | C | N4 | AA:1432 | 2.81 | 1.00 | 53.85 |
| 2 | U | O4 | AA:1433 | 3.31 | 1.00 | 60.47 |  |
| 3 | U | O4 | AA:1431 | 4.57 | 1.00 | 47.32 |  |
| 4 | C | N3 | AA:1432 | 4.70 | 1.00 | 51.25 |  |
  |

  
 Coded contact list:   
 Exclusion zone contact(s): /N4/O4/  
Consider K in place of MG (2 valid contact(s) for K vs. 0 for MG)  

### MG:AA:3962

|  | Res. type | Atom | Residue | Distance | Occupancy | B-factor | Closest solute to 1^st^ shell water |
| --- | --- | --- | --- | --- | --- | --- | --- |
| **Ion** | | | |  | 1.00 | 57.11 || 1 | G | OP1 | AA:695 | 2.94 | 1.00 | 35.49 |
| 2 | G | O2' | AA:1380 | 3.43 | 1.00 | 49.13 |  |
| 3 | G | O3' | AA:1380 | 3.90 | 1.00 | 49.27 |  |
| 4 | C | O2 | AA:1428 | 3.90 | 1.00 | 62.26 |  |
| 5 | G | OP1 | AA:1381 | 4.06 | 1.00 | 48.73 |  |
| 6 | G | O5' | AA:1381 | 4.66 | 1.00 | 47.12 |  |
| 7 | G | O5' | AA:695 | 4.93 | 1.00 | 34.32 |  |
| 8 | G | OP2 | AA:695 | 4.96 | 1.00 | 35.55 |  |
  |

  
 Coded contact list:   
 Exclusion zone contact(s): /OP1/  
Consider K in place of MG (1 valid contact(s) for K vs. 0 for MG)  

### MG:AA:3963

|  | Res. type | Atom | Residue | Distance | Occupancy | B-factor | Closest solute to 1^st^ shell water |
| --- | --- | --- | --- | --- | --- | --- | --- |
| **Ion** | | | |  | 1.00 | 62.30 || 1 | U | OP2 | AA:606 | 3.38 | 1.00 | 62.21 |
| 2 | G | O6 | AA:620 | 4.21 | 1.00 | 66.91 |  |
| 3 | U | OP2 | AA:607 | 4.56 | 1.00 | 67.53 |  |
| 4 | U | O5' | AA:606 | 4.68 | 1.00 | 64.13 |  |
  |

  
 Coded contact list:   
 Exclusion zone contact(s): /OP2/  
Consider K in place of MG (1 valid contact(s) for K vs. 0 for MG)  

### MG:AA:3964

|  | Res. type | Atom | Residue | Distance | Occupancy | B-factor | Closest solute to 1^st^ shell water |
| --- | --- | --- | --- | --- | --- | --- | --- |
| **Ion** | | | |  | 1.00 | 111.74 || 1 | U | O2' | AA:1082 | 3.43 | 1.00 | 200.36 |
| 2 | U | O2 | AA:1082 | 4.28 | 1.00 | 200.09 |  |
| 3 | A | N1 | AA:1086 | 4.97 | 1.00 | 199.59 |  |
  |

  
 Coded contact list:   
1^st^ MG coordination shell missing (see Notes)  

### MG:AA:3965

|  | Res. type | Atom | Residue | Distance | Occupancy | B-factor | Closest solute to 1^st^ shell water |
| --- | --- | --- | --- | --- | --- | --- | --- |
| **Ion** | | | |  | 1.00 | 111.41 || 1 | G | OP2 | AA:1519 | 2.53 | 1.00 | 70.94 |
| 2 | G | OP2 | AA:1470 | 3.48 | 1.00 | 75.20 |  |
| 3 | G | OP1 | AA:1519 | 4.06 | 1.00 | 71.64 |  |
| 4 | C | O3' | AA:1518 | 4.58 | 1.00 | 71.63 |  |
| 5 | G | O5' | AA:1519 | 4.92 | 1.00 | 71.89 |  |
| 6 | C | OP2 | AA:1518 | 4.97 | 1.00 | 64.90 |  |
  |

  
 Coded contact list:   
 Exclusion zone contact(s): /OP2/  
Consider NA in place of MG (1 valid contact(s) for NA vs. 0 for MG)  

### MG:AA:3966

|  | Res. type | Atom | Residue | Distance | Occupancy | B-factor | Closest solute to 1^st^ shell water |
| --- | --- | --- | --- | --- | --- | --- | --- |
| **Ion** | | | |  | 1.00 | 75.23 || 1 | U | OP1 | AA:2832 | 3.15 | 1.00 | 66.63 |
| 2 | G | O5' | AA:2831 | 3.28 | 1.00 | 63.35 |  |
| 3 | U | OP2 | AA:2832 | 3.69 | 1.00 | 66.61 |  |
| 4 | G | O3' | AA:2831 | 3.96 | 1.00 | 66.65 |  |
| 5 | G | N9 | AA:2831 | 4.01 | 1.00 | 59.43 |  |
| 6 | G | OP2 | AA:2831 | 4.12 | 1.00 | 62.19 |  |
| 7 | G | N7 | AA:2831 | 4.17 | 1.00 | 58.85 |  |
| 8 | G | O4' | AA:2831 | 4.27 | 1.00 | 62.46 |  |
  |

  
 Coded contact list:   
 Exclusion zone contact(s): /O5'/OP1/  
Consider K in place of MG (2 valid contact(s) for K vs. 0 for MG)  

### MG:AA:3967

|  | Res. type | Atom | Residue | Distance | Occupancy | B-factor | Closest solute to 1^st^ shell water |
| --- | --- | --- | --- | --- | --- | --- | --- |
| **Ion** | | | |  | 1.00 | 63.19 || 1 | U | O4 | AA:1273 | 3.67 | 1.00 | 48.90 |
| 2 | U | N3 | AA:1273 | 3.80 | 1.00 | 50.00 |  |
| 3 | U | O2' | AA:1659 | 4.56 | 1.00 | 40.35 |  |
| 4 | U | N1 | AA:1273 | 4.76 | 1.00 | 49.16 |  |
| 5 | U | O4' | AA:1659 | 4.86 | 1.00 | 38.21 |  |
| 6 | A | N1 | AA:2712A | 4.98 | 1.00 | 39.42 |  |
  |

  
 Coded contact list:   
1^st^ MG coordination shell missing (see Notes)  

### MG:AA:3968

|  | Res. type | Atom | Residue | Distance | Occupancy | B-factor | Closest solute to 1^st^ shell water |
| --- | --- | --- | --- | --- | --- | --- | --- |
| **Ion** | | | |  | 1.00 | 77.42 || 1 | A | OP2 | AA:331 | 2.29 | 1.00 | 62.81 |
| 2 | A | N1 | AA:311 | 3.60 | 1.00 | 71.80 |  |
| 3 | U | OP2 | AA:328 | 4.27 | 1.00 | 51.53 |  |
| 4 | A | OP1 | AA:331 | 4.46 | 1.00 | 63.04 |  |
| 5 | A | O5' | AA:331 | 4.63 | 1.00 | 60.98 |  |
| 6 | A | O3' | AA:330 | 4.63 | 1.00 | 62.92 |  |
| 7 | U | O4 | AA:328 | 4.73 | 1.00 | 56.25 |  |
| 8 | A | OP2 | AA:330 | 4.74 | 1.00 | 58.44 |  |
  |

  
 Contact(s): /OP2/  
 Coded contact list: O_ph_  

### MG:AA:3969

|  | Res. type | Atom | Residue | Distance | Occupancy | B-factor | Closest solute to 1^st^ shell water |
| --- | --- | --- | --- | --- | --- | --- | --- |
| **Ion** | | | |  | 1.00 | 60.10 || 1 | G | O2' | AA:1031 | 3.61 | 1.00 | 92.06 |
| 2 | G | O3' | AA:1031 | 3.97 | 1.00 | 97.43 |  |
| 3 | A | OP1 | AA:1032 | 4.83 | 1.00 | 99.54 |  |
  |

  
 Coded contact list:   
1^st^ MG coordination shell missing (see Notes)  

### MG:AA:3970

|  | Res. type | Atom | Residue | Distance | Occupancy | B-factor | Closest solute to 1^st^ shell water |
| --- | --- | --- | --- | --- | --- | --- | --- |
| **Ion** | | | |  | 1.00 | 47.66 || 1 | G | OP2 | AA:452 | 2.54 | 1.00 | 38.78 |
| 2 | G | OP2 | AA:450 | 3.52 | 1.00 | 40.85 |  |
| 3 | A | OP2 | AA:449 | 3.83 | 1.00 | 34.02 |  |
| 4 | GLY | O | AF:85 | 4.02 | 1.00 | 29.43 |  |
| 5 | GLY | O | AF:86 | 4.51 | 1.00 | 37.98 |  |
| 6 | G | O5' | AA:452 | 4.60 | 1.00 | 39.15 |  |
| 7 | C | OP2 | AA:451 | 4.62 | 1.00 | 43.73 |  |
| 8 | G | OP1 | AA:452 | 4.84 | 1.00 | 40.21 |  |
| 9 | G | N7 | AA:450 | 4.93 | 1.00 | 44.83 |  |
| 10 | G | O5' | AA:450 | 4.95 | 1.00 | 40.46 |  |
| 11 | TYR | OH | AF:59 | 4.97 | 1.00 | 50.32 |  |
| 12 | C | O3' | AA:451 | 4.99 | 1.00 | 40.86 |  |
  |

  
 Coded contact list:   
 Exclusion zone contact(s): /OP2/  
Consider NA in place of MG (1 valid contact(s) for NA vs. 0 for MG)  

### MG:AA:3971

|  | Res. type | Atom | Residue | Distance | Occupancy | B-factor | Closest solute to 1^st^ shell water |
| --- | --- | --- | --- | --- | --- | --- | --- |
| **Ion** | | | |  | 1.00 | 85.96 || 1 | G | OP2 | AA:1319 | 3.02 | 1.00 | 60.94 |
| 2 | C | OP2 | AA:1330 | 3.61 | 1.00 | 51.72 |  |
| 3 | U | O3' | AA:1329 | 4.00 | 1.00 | 50.49 |  |
| 4 | C | OP1 | AA:1330 | 4.10 | 1.00 | 51.70 |  |
| 5 | G | N7 | AA:1319 | 4.54 | 1.00 | 62.11 |  |
| 6 | G | O5' | AA:1319 | 4.65 | 1.00 | 61.90 |  |
| 7 | C | OP2 | AA:1320 | 4.85 | 1.00 | 61.43 |  |
  |

  
 Coded contact list:   
 Exclusion zone contact(s): /OP2/  
Consider K in place of MG (1 valid contact(s) for K vs. 0 for MG)  

### MG:AA:3972

|  | Res. type | Atom | Residue | Distance | Occupancy | B-factor | Closest solute to 1^st^ shell water |
| --- | --- | --- | --- | --- | --- | --- | --- |
| **Ion** | | | |  | 1.00 | 60.39 || 1 | G | OP1 | AA:488 | 2.77 | 1.00 | 54.57 |
| 2 | G | OP2 | AA:489 | 2.92 | 1.00 | 55.47 |  |
| 3 | G | O5' | AA:488 | 3.41 | 1.00 | 55.71 |  |
| 4 | G | O3' | AA:488 | 3.76 | 1.00 | 54.53 |  |
| 5 | G | OP1 | AA:489 | 4.05 | 1.00 | 55.80 |  |
| 6 | G | OP2 | AA:488 | 4.50 | 1.00 | 53.17 |  |
| 7 | C | O3' | AA:487 | 4.89 | 1.00 | 53.86 |  |
  |

  
 Coded contact list:   
 Exclusion zone contact(s): /OP1/OP2/  
Consider K in place of MG (2 valid contact(s) for K vs. 0 for MG)  

### MG:AA:3973

|  | Res. type | Atom | Residue | Distance | Occupancy | B-factor | Closest solute to 1^st^ shell water |
| --- | --- | --- | --- | --- | --- | --- | --- |
| **Ion** | | | |  | 1.00 | 86.42 || 1 | G | O6 | AA:1228 | 3.47 | 1.00 | 64.57 |
| 2 | G | O6 | AA:1229 | 3.87 | 1.00 | 73.11 |  |
| 3 | C | N4 | AA:1222 | 4.33 | 1.00 | 75.08 |  |
| 4 | G | N7 | AA:1228 | 4.65 | 1.00 | 63.17 |  |
| 5 | C | N4 | AA:1223 | 4.77 | 1.00 | 65.43 |  |
  |

  
 Coded contact list:   
1^st^ MG coordination shell missing (see Notes)  

### MG:AA:3974

|  | Res. type | Atom | Residue | Distance | Occupancy | B-factor | Closest solute to 1^st^ shell water |
| --- | --- | --- | --- | --- | --- | --- | --- |
| **Ion** | | | |  | 1.00 | 65.23 || 1 | G | N9 | AA:1311 | 2.48 | 1.00 | 55.93 |
| 2 | G | O4' | AA:1311 | 3.23 | 1.00 | 52.42 |  |
| 3 | G | N7 | AA:1311 | 3.46 | 1.00 | 58.63 |  |
| 4 | G | N3 | AA:1311 | 3.79 | 1.00 | 62.98 |  |
| 5 | G | O2' | AA:1311 | 3.90 | 1.00 | 59.56 |  |
| 6 | U | OP2 | AA:1312 | 3.97 | 1.00 | 49.30 |  |
| 7 | G | O3' | AA:1311 | 4.20 | 1.00 | 50.31 |  |
| 8 | G | OP2 | AA:1311 | 4.33 | 1.00 | 37.84 |  |
  |

  
 Coded contact list:   
 Exclusion zone contact(s): /O4'/  
Consider NA in place of MG (1 valid contact(s) for NA vs. 0 for MG)  

### MG:AA:3975

|  | Res. type | Atom | Residue | Distance | Occupancy | B-factor | Closest solute to 1^st^ shell water |
| --- | --- | --- | --- | --- | --- | --- | --- |
| **Ion** | | | |  | 1.00 | 76.30 || 1 | U | O4 | AA:553 | 2.77 | 1.00 | 72.17 |
| 2 | G | O6 | AA:552 | 2.91 | 1.00 | 76.89 |  |
| 3 | G | O6 | AA:540 | 3.91 | 1.00 | 74.40 |  |
| 4 | C | N4 | AA:541 | 4.11 | 1.00 | 78.93 |  |
| 5 | U | N3 | AA:553 | 4.33 | 1.00 | 71.21 |  |
| 6 | G | N7 | AA:552 | 4.54 | 1.00 | 74.39 |  |
| 7 | G | O6 | AA:551 | 4.60 | 1.00 | 74.51 |  |
  |

  
 Coded contact list:   
 Exclusion zone contact(s): /O4/O6/  
Consider K in place of MG (2 valid contact(s) for K vs. 0 for MG)  

### MG:AA:3976

|  | Res. type | Atom | Residue | Distance | Occupancy | B-factor | Closest solute to 1^st^ shell water |
| --- | --- | --- | --- | --- | --- | --- | --- |
| **Ion** | | | |  | 1.00 | 86.96 || 1 | C | N1 | AA:974A | 3.89 | 1.00 | 73.15 |
| 2 | A | N7 | AA:572 | 4.27 | 1.00 | 49.24 |  |
| 3 | C | O3' | AA:982 | 4.40 | 1.00 | 61.46 |  |
| 4 | A | OP1 | AA:983 | 4.46 | 1.00 | 62.84 |  |
| 5 | C | O2' | AA:974A | 4.50 | 1.00 | 77.03 |  |
| 6 | A | O3' | AA:571 | 4.82 | 1.00 | 55.35 |  |
  |

  
 Coded contact list:   
1^st^ MG coordination shell missing (see Notes)  

### MG:AA:3977

|  | Res. type | Atom | Residue | Distance | Occupancy | B-factor | Closest solute to 1^st^ shell water |
| --- | --- | --- | --- | --- | --- | --- | --- |
| **Ion** | | | |  | 1.00 | 56.75 || 1 | A | O3' | AA:1247 | 2.39 | 1.00 | 52.11 |
| 2 | G | OP1 | AA:1248 | 2.61 | 1.00 | 53.17 |  |
| 3 | G | OP2 | AA:1248 | 3.07 | 1.00 | 54.61 |  |
| 4 | PRO | N | A1:2 | 3.45 | 1.00 | 50.05 |  |
| 5 | C | OP2 | AA:444 | 3.51 | 1.00 | 50.98 |  |
| 6 | G | O5' | AA:1248 | 4.28 | 1.00 | 53.78 |  |
| 7 | PRO | O | A1:2 | 4.50 | 1.00 | 49.12 |  |
| 8 | A | O2' | AA:1247 | 4.66 | 1.00 | 52.02 |  |
| 9 | TYR | OH | AF:97 | 4.92 | 1.00 | 42.62 |  |
  |

  
 Contact(s): /O3'/  
 Coded contact list: O_r_  
 Exclusion zone contact(s): /OP1/OP2/  
Consider K in place of MG (2 valid contact(s) for K vs. 1 for MG)  

### MG:AA:3978

|  | Res. type | Atom | Residue | Distance | Occupancy | B-factor | Closest solute to 1^st^ shell water |
| --- | --- | --- | --- | --- | --- | --- | --- |
| **Ion** | | | |  | 1.00 | 70.56 || 1 | G | OP1 | AA:770 | 2.60 | 1.00 | 39.03 |
| 2 | ASN | OD1 | A7:8 | 2.77 | 1.00 | 39.34 |  |
| 3 | G | OP2 | AA:770 | 3.07 | 1.00 | 38.76 |  |
| 4 | G | OP2 | AA:771 | 3.26 | 1.00 | 37.40 |  |
| 5 | G | O5' | AA:770 | 3.51 | 1.00 | 36.19 |  |
| 6 | LYS | N | A7:11 | 4.18 | 1.00 | 33.98 |  |
| 7 | ASN | ND2 | A7:8 | 4.33 | 1.00 | 39.15 |  |
| 8 | G | O3' | AA:769 | 4.63 | 1.00 | 35.15 |  |
| 9 | G | O3' | AA:770 | 4.66 | 1.00 | 35.58 |  |
| 10 | G | OP1 | AA:771 | 4.88 | 1.00 | 37.65 |  |
  |

  
 Coded contact list:   
 Exclusion zone contact(s): /OD1_Asn_/OP1/OP2/OP2/  
Consider K in place of MG (3 valid contact(s) for K vs. 0 for MG)  

### MG:AA:3979

|  | Res. type | Atom | Residue | Distance | Occupancy | B-factor | Closest solute to 1^st^ shell water |
| --- | --- | --- | --- | --- | --- | --- | --- |
| **Ion** | | | |  | 1.00 | 94.97 || 1 | G | OP1 | AA:176 | 4.49 | 1.00 | 74.02 |
| 2 | G | N7 | AA:48 | 4.62 | 1.00 | 58.06 |  |
| 3 | G | O6 | AA:48 | 4.68 | 1.00 | 60.77 |  |
  |

  
 Coded contact list:   
1^st^ MG coordination shell missing (see Notes)  

### MG:AA:3980

|  | Res. type | Atom | Residue | Distance | Occupancy | B-factor | Closest solute to 1^st^ shell water |
| --- | --- | --- | --- | --- | --- | --- | --- |
| **Ion** | | | |  | 1.00 | 65.05 || 1 | G | OP1 | AA:321 | 2.25 | 1.00 | 69.66 |
| 2 | A | N7 | AA:322 | 3.51 | 1.00 | 74.02 |  |
| 3 | G | O5' | AA:321 | 3.57 | 1.00 | 72.05 |  |
| 4 | A | O3' | AA:320 | 3.93 | 1.00 | 70.71 |  |
| 5 | LYS | NZ | AF:135 | 4.17 | 1.00 | 78.28 |  |
| 6 | G | N7 | AA:321 | 4.23 | 1.00 | 75.57 |  |
| 7 | A | N3 | AA:340 | 4.50 | 1.00 | 68.86 |  |
| 8 | G | N9 | AA:321 | 4.65 | 1.00 | 73.89 |  |
| 9 | A | N1 | AA:340 | 4.67 | 1.00 | 68.50 |  |
| 10 | G | OP2 | AA:321 | 4.72 | 1.00 | 70.25 |  |
| 11 | G | O4' | AA:321 | 4.78 | 1.00 | 74.63 |  |
| 12 | A | N6 | AA:322 | 4.82 | 1.00 | 73.80 |  |
  |

  
 Contact(s): /OP1/  
 Coded contact list: O_ph_  

### MG:AA:3981

|  | Res. type | Atom | Residue | Distance | Occupancy | B-factor | Closest solute to 1^st^ shell water |
| --- | --- | --- | --- | --- | --- | --- | --- |
| **Ion** | | | |  | 1.00 | 131.22 || 1 | A | N7 | AA:345 | 2.49 | 1.00 | 71.49 |
| 2 | G | N3 | AA:344 | 3.02 | 1.00 | 72.82 |  |
| 3 | G | N1 | AA:344 | 3.13 | 1.00 | 72.44 |  |
| 4 | A | N6 | AA:345 | 3.28 | 1.00 | 71.43 |  |
| 5 | G | N2 | AA:344 | 3.58 | 1.00 | 72.41 |  |
| 6 | G | N9 | AA:344 | 3.95 | 1.00 | 73.91 |  |
| 7 | G | O6 | AA:344 | 4.20 | 1.00 | 72.66 |  |
| 8 | G | N7 | AA:344 | 4.26 | 1.00 | 73.17 |  |
| 9 | A | OP2 | AA:345 | 4.57 | 1.00 | 77.15 |  |
| 10 | A | N9 | AA:345 | 4.71 | 1.00 | 72.90 |  |
  |

  
 Coded contact list:   
 Exclusion zone contact(s): /N1_G_/N3_G_/N6/N7/  
Consider K in place of MG (3 valid contact(s) for K vs. 0 for MG)  

### MG:AA:3982

|  | Res. type | Atom | Residue | Distance | Occupancy | B-factor | Closest solute to 1^st^ shell water |
| --- | --- | --- | --- | --- | --- | --- | --- |
| **Ion** | | | |  | 1.00 | 71.81 || 1 | C | O2' | AA:652 | 3.48 | 1.00 | 107.34 |
| 2 | C | O3' | AA:652 | 3.50 | 1.00 | 116.20 |  |
| 3 | A | OP1 | AA:653 | 3.85 | 1.00 | 121.64 |  |
| 4 | A | N7 | AA:637 | 3.92 | 1.00 | 93.10 |  |
| 5 | LYS | NZ | AO:86 | 4.25 | 1.00 | 82.61 |  |
| 6 | GLU | OE2 | AO:117 | 4.37 | 1.00 | 94.91 |  |
| 7 | A | OP2 | AA:627 | 4.51 | 1.00 | 70.60 |  |
| 8 | ASN | OD1 | AO:84 | 4.60 | 1.00 | 77.39 |  |
  |

  
 Coded contact list:   
1^st^ MG coordination shell missing (see Notes)  

### MG:AA:3983

|  | Res. type | Atom | Residue | Distance | Occupancy | B-factor | Closest solute to 1^st^ shell water |
| --- | --- | --- | --- | --- | --- | --- | --- |
| **Ion** | | | |  | 1.00 | 126.38 || 1 | G | O6 | AA:1244 | 2.77 | 1.00 | 47.77 |
| 2 | G | N7 | AA:1244 | 3.04 | 1.00 | 51.45 |  |
| 3 | G | O6 | AA:1245 | 4.16 | 1.00 | 42.85 |  |
| 4 | G | O6 | AA:1243 | 4.26 | 1.00 | 60.55 |  |
| 5 | G | N7 | AA:1245 | 4.34 | 1.00 | 47.76 |  |
| 6 | G | N7 | AA:1243 | 4.70 | 1.00 | 64.32 |  |
| 7 | G | N1 | AA:1244 | 4.74 | 1.00 | 47.98 |  |
| 8 | C | N4 | AA:1201 | 4.95 | 1.00 | 45.47 |  |
  |

  
 Coded contact list:   
 Exclusion zone contact(s): /N7/O6/  
Consider K in place of MG (2 valid contact(s) for K vs. 0 for MG)  

### MG:AA:3984

|  | Res. type | Atom | Residue | Distance | Occupancy | B-factor | Closest solute to 1^st^ shell water |
| --- | --- | --- | --- | --- | --- | --- | --- |
| **Ion** | | | |  | 1.00 | 98.94 || 1 | G | OP1 | AA:2308 | 2.68 | 1.00 | 100.12 |
| 2 | C | OP1 | AA:2306 | 3.10 | 1.00 | 94.71 |  |
| 3 | G | O5' | AA:2308 | 3.95 | 1.00 | 100.33 |  |
| 4 | C | OP2 | AA:2306 | 4.52 | 1.00 | 94.57 |  |
| 5 | G | OP2 | AA:2308 | 4.65 | 1.00 | 99.37 |  |
| 6 | A | OP2 | AA:2305 | 4.80 | 1.00 | 93.70 |  |
  |

  
 Coded contact list:   
 Exclusion zone contact(s): /OP1/OP1/  
Consider K in place of MG (2 valid contact(s) for K vs. 0 for MG)  

### MG:AA:3985

|  | Res. type | Atom | Residue | Distance | Occupancy | B-factor | Closest solute to 1^st^ shell water |
| --- | --- | --- | --- | --- | --- | --- | --- |
| **Ion** | | | |  | 1.00 | 97.36 || 1 | U | OP1 | AA:293 | 3.00 | 1.00 | 77.15 |
| 2 | A | OP2 | AA:294 | 3.01 | 1.00 | 71.55 |  |
| 3 | U | OP2 | AA:293 | 4.23 | 1.00 | 76.20 |  |
| 4 | U | O5' | AA:293 | 4.51 | 1.00 | 74.70 |  |
| 5 | A | O5' | AA:294 | 4.99 | 1.00 | 72.32 |  |
  |

  
 Coded contact list:   
 Exclusion zone contact(s): /OP1/OP2/  
Consider K in place of MG (2 valid contact(s) for K vs. 0 for MG)  

### MG:AA:3986

|  | Res. type | Atom | Residue | Distance | Occupancy | B-factor | Closest solute to 1^st^ shell water |
| --- | --- | --- | --- | --- | --- | --- | --- |
| **Ion** | | | |  | 1.00 | 88.15 || 1 | G | OP1 | AA:2373 | 2.62 | 1.00 | 66.05 |
| 2 | C | OP2 | AA:2374 | 2.87 | 1.00 | 66.40 |  |
| 3 | C | OP1 | AA:2374 | 3.05 | 1.00 | 65.54 |  |
| 4 | G | O3' | AA:2373 | 3.40 | 1.00 | 65.81 |  |
| 5 | G | O5' | AA:2373 | 3.75 | 1.00 | 64.85 |  |
| 6 | G | OP2 | AA:2373 | 4.37 | 1.00 | 65.34 |  |
| 7 | C | O5' | AA:2343 | 4.65 | 1.00 | 80.16 |  |
| 8 | C | O5' | AA:2374 | 4.73 | 1.00 | 66.44 |  |
| 9 | C | O3' | AA:2343 | 4.94 | 1.00 | 77.52 |  |
| 10 | G | O3' | AA:2372 | 4.98 | 1.00 | 65.86 |  |
  |

  
 Coded contact list:   
 Exclusion zone contact(s): /OP1/OP1/OP2/  
Consider K in place of MG (3 valid contact(s) for K vs. 0 for MG)  

### MG:AA:3987

|  | Res. type | Atom | Residue | Distance | Occupancy | B-factor | Closest solute to 1^st^ shell water |
| --- | --- | --- | --- | --- | --- | --- | --- |
| **Ion** | | | |  | 1.00 | 54.74 || 1 | G | N3 | AA:2818 | 2.51 | 1.00 | 51.49 |
| 2 | G | O2' | AA:2818 | 2.54 | 1.00 | 54.59 |  |
| 3 | G | O4' | AA:2819 | 3.07 | 1.00 | 48.17 |  |
| 4 | G | N2 | AA:2817 | 3.31 | 1.00 | 60.40 |  |
| 5 | U | O2' | AA:2836 | 3.41 | 1.00 | 71.64 |  |
| 6 | G | N2 | AA:2818 | 3.54 | 1.00 | 51.28 |  |
| 7 | G | N9 | AA:2818 | 3.69 | 1.00 | 52.30 |  |
| 8 | C | O2 | AA:2829 | 3.77 | 1.00 | 61.54 |  |
| 9 | U | O3' | AA:2836 | 4.19 | 1.00 | 71.51 |  |
| 10 | G | N9 | AA:2819 | 4.39 | 1.00 | 42.73 |  |
| 11 | G | O4' | AA:2818 | 4.47 | 1.00 | 54.65 |  |
| 12 | G | O5' | AA:2819 | 4.61 | 1.00 | 49.83 |  |
| 13 | G | N1 | AA:2818 | 4.75 | 1.00 | 51.95 |  |
| 14 | G | N1 | AA:2817 | 4.81 | 1.00 | 58.87 |  |
| 15 | U | O4' | AA:2836 | 4.86 | 1.00 | 73.87 |  |
| 16 | G | N3 | AA:2817 | 4.96 | 1.00 | 56.68 |  |
  |

  
 Coded contact list:   
 Exclusion zone contact(s): /N2/N3_G_/O2'/O4'/  
Consider NA in place of MG (2 valid contact(s) for NA vs. 0 for MG)  

### MG:AA:3988

|  | Res. type | Atom | Residue | Distance | Occupancy | B-factor | Closest solute to 1^st^ shell water |
| --- | --- | --- | --- | --- | --- | --- | --- |
| **Ion** | | | |  | 1.00 | 64.01 || 1 | A | OP2 | AA:1342 | 2.00 | 1.00 | 42.30 |
| 2 | U | O4 | AA:1602 | 2.26 | 1.00 | 53.19 |  |
| 3 | THR | O | AT:56 | 2.71 | 1.00 | 46.82 |  |
| 4 | U | O5' | AA:1341 | 3.89 | 1.00 | 41.39 |  |
| 5 | LEU | N | AT:57 | 4.06 | 1.00 | 45.52 |  |
| 6 | A | O5' | AA:1342 | 4.12 | 1.00 | 43.29 |  |
| 7 | A | OP1 | AA:1342 | 4.23 | 1.00 | 42.29 |  |
| 8 | THR | OG1 | AT:56 | 4.34 | 1.00 | 47.58 |  |
| 9 | U | N3 | AA:1602 | 4.35 | 1.00 | 52.55 |  |
| 10 | U | O3' | AA:1341 | 4.44 | 1.00 | 42.53 |  |
| 11 | G | OP2 | AA:1601 | 4.74 | 1.00 | 40.41 |  |
| 12 | U | OP1 | AA:1341 | 4.79 | 1.00 | 42.74 |  |
| 13 | G | N7 | AA:1601 | 4.87 | 1.00 | 46.35 |  |
  |

  
**1^st^ shell ligand angles:**   
1-2: 71;   
  
 Contact(s): /O4/OP2/  
 Coded contact list: O_ph_O_b_  
 Exclusion zone contact(s): /O_back_/  

### MG:AA:3989

|  | Res. type | Atom | Residue | Distance | Occupancy | B-factor | Closest solute to 1^st^ shell water |
| --- | --- | --- | --- | --- | --- | --- | --- |
| **Ion** | | | |  | 1.00 | 74.52 || 1 | A | N7 | AA:1284 | 2.99 | 1.00 | 47.07 |
| 2 | G | O6 | AA:1285 | 3.62 | 1.00 | 51.93 |  |
| 3 | C | OP1 | AA:1320 | 3.74 | 1.00 | 63.29 |  |
| 4 | A | N6 | AA:1284 | 4.02 | 1.00 | 50.57 |  |
| 5 | C | OP2 | AA:1320 | 4.59 | 1.00 | 61.43 |  |
| 6 | G | N1 | AA:1285 | 4.83 | 1.00 | 50.02 |  |
| 7 | G | O2' | AA:489 | 4.85 | 1.00 | 50.67 |  |
| 8 | G | O2' | AA:1283 | 4.92 | 1.00 | 59.10 |  |
  |

  
 Coded contact list:   
 Exclusion zone contact(s): /N7/  
Consider K in place of MG (1 valid contact(s) for K vs. 0 for MG)  

### MG:AA:3990

|  | Res. type | Atom | Residue | Distance | Occupancy | B-factor | Closest solute to 1^st^ shell water |
| --- | --- | --- | --- | --- | --- | --- | --- |
| **Ion** | | | |  | 1.00 | 134.88 || 1 | U | O4 | AA:922 | 3.46 | 1.00 | 86.31 |
| 2 | G | N7 | AA:921 | 4.92 | 1.00 | 81.94 |  |
| 3 | G | OP2 | AA:921 | 5.00 | 1.00 | 79.38 |  |
  |

  
 Coded contact list:   
1^st^ MG coordination shell missing (see Notes)  

### MG:AA:3991

|  | Res. type | Atom | Residue | Distance | Occupancy | B-factor | Closest solute to 1^st^ shell water |
| --- | --- | --- | --- | --- | --- | --- | --- |
| **Ion** | | | |  | 1.00 | 73.35 || 1 | C | OP2 | AA:982 | 2.82 | 1.00 | 60.02 |
| 2 | C | OP1 | AA:982 | 3.24 | 1.00 | 59.11 |  |
| 3 | C | O5' | AA:982 | 4.23 | 1.00 | 59.76 |  |
| 4 | A | OP2 | AA:983 | 4.25 | 1.00 | 61.18 |  |
| 5 | G | O6 | AA:979 | 4.50 | 1.00 | 76.58 |  |
| 6 | A | O3' | AA:981 | 4.85 | 1.00 | 58.23 |  |
  |

  
 Coded contact list:   
 Exclusion zone contact(s): /OP1/OP2/  
Consider K in place of MG (2 valid contact(s) for K vs. 0 for MG)  

### MG:AA:3994

|  | Res. type | Atom | Residue | Distance | Occupancy | B-factor | Closest solute to 1^st^ shell water |
| --- | --- | --- | --- | --- | --- | --- | --- |
| **Ion** | | | |  | 1.00 | 68.86 || 1 | C | O2 | AA:2612 | 3.26 | 1.00 | 65.11 |
| 2 | A | OP1 | AA:746 | 4.80 | 1.00 | 47.24 |  |
| 3 | C | N3 | AA:2612 | 4.98 | 1.00 | 65.35 |  |
  |

  
 Coded contact list:   
 Exclusion zone contact(s): /O2/  
Consider K in place of MG (1 valid contact(s) for K vs. 0 for MG)  

### MG:AA:3995

|  | Res. type | Atom | Residue | Distance | Occupancy | B-factor | Closest solute to 1^st^ shell water |
| --- | --- | --- | --- | --- | --- | --- | --- |
| **Ion** | | | |  | 1.00 | 103.70 || 1 | G | OP1 | AA:301 | 3.00 | 1.00 | 59.65 |
| 2 | A | O3' | AA:300 | 4.52 | 1.00 | 58.24 |  |
| 3 | C | O4' | AA:318 | 4.93 | 1.00 | 66.78 |  |
  |

  
 Coded contact list:   
 Exclusion zone contact(s): /OP1/  
Consider K in place of MG (1 valid contact(s) for K vs. 0 for MG)  

### MG:AA:3996

|  | Res. type | Atom | Residue | Distance | Occupancy | B-factor | Closest solute to 1^st^ shell water |
| --- | --- | --- | --- | --- | --- | --- | --- |
| **Ion** | | | |  | 1.00 | 49.98 || 1 | G | O5' | AA:2833 | 2.52 | 1.00 | 89.21 |
| 2 | U | O3' | AA:2832 | 2.80 | 1.00 | 84.20 |  |
| 3 | U | O2' | AA:2832 | 3.41 | 1.00 | 77.52 |  |
| 4 | G | OP2 | AA:2833 | 4.04 | 1.00 | 87.21 |  |
| 5 | G | N7 | AA:2834 | 4.18 | 1.00 | 78.96 |  |
| 6 | G | OP1 | AA:2833 | 4.43 | 1.00 | 87.46 |  |
| 7 | G | O4' | AA:2833 | 4.87 | 1.00 | 98.73 |  |
  |

  
 Coded contact list:   
 Exclusion zone contact(s): /O3'/O5'/  
Consider NA in place of MG (1 valid contact(s) for NA vs. 0 for MG)  

### MG:AA:3997

|  | Res. type | Atom | Residue | Distance | Occupancy | B-factor | Closest solute to 1^st^ shell water |
| --- | --- | --- | --- | --- | --- | --- | --- |
| **Ion** | | | |  | 1.00 | 78.52 || 1 | G | OP1 | AA:54 | 2.81 | 1.00 | 45.45 |
| 2 | G | O5' | AA:54 | 3.18 | 1.00 | 45.08 |  |
| 3 | A | O3' | AA:53 | 3.53 | 1.00 | 47.02 |  |
| 4 | A | O2' | AA:53 | 3.97 | 1.00 | 47.04 |  |
| 5 | G | OP2 | AA:54 | 4.87 | 1.00 | 45.67 |  |
  |

  
 Coded contact list:   
 Exclusion zone contact(s): /O5'/OP1/  
Consider K in place of MG (2 valid contact(s) for K vs. 0 for MG)  

### MG:AA:3998

|  | Res. type | Atom | Residue | Distance | Occupancy | B-factor | Closest solute to 1^st^ shell water |
| --- | --- | --- | --- | --- | --- | --- | --- |
| **Ion** | | | |  | 1.00 | 86.93 || 1 | A | N3 | AA:1583 | 4.15 | 1.00 | 108.20 |
| 2 | A | O2' | AA:1583 | 4.50 | 1.00 | 110.61 |  |
| 3 | G | O4' | AA:1416 | 4.55 | 1.00 | 91.51 |  |
| 4 | G | N3 | AA:1416 | 4.82 | 1.00 | 93.19 |  |
| 5 | G | N9 | AA:1416 | 5.00 | 1.00 | 91.98 |  |
  |

  
 Coded contact list:   
1^st^ MG coordination shell missing (see Notes)  

### MG:AA:3999

|  | Res. type | Atom | Residue | Distance | Occupancy | B-factor | Closest solute to 1^st^ shell water |
| --- | --- | --- | --- | --- | --- | --- | --- |
| **Ion** | | | |  | 1.00 | 57.29 || 1 | A | OP2 | AA:1276 | 2.82 | 1.00 | 41.68 |
| 2 | C | OP2 | AA:1646 | 3.41 | 1.00 | 48.99 |  |
| 3 | G | OP1 | AA:1649 | 3.49 | 1.00 | 47.61 |  |
| 4 | A | OP1 | AA:1276 | 3.64 | 1.00 | 41.74 |  |
| 5 | C | OP1 | AA:1646 | 3.97 | 1.00 | 47.88 |  |
| 6 | A | O5' | AA:1276 | 4.60 | 1.00 | 41.64 |  |
| 7 | A | O3' | AA:1275 | 4.95 | 1.00 | 43.06 |  |
  |

  
 Coded contact list:   
 Exclusion zone contact(s): /OP2/  
Consider K in place of MG (1 valid contact(s) for K vs. 0 for MG)  

### MG:AA:4000

|  | Res. type | Atom | Residue | Distance | Occupancy | B-factor | Closest solute to 1^st^ shell water |
| --- | --- | --- | --- | --- | --- | --- | --- |
| **Ion** | | | |  | 1.00 | 91.29 || 1 | C | O4' | AA:1592 | 3.21 | 1.00 | 87.61 |
| 2 | G | N3 | AA:1591 | 3.32 | 1.00 | 88.36 |  |
| 3 | G | O2' | AA:1591 | 3.52 | 1.00 | 91.64 |  |
| 4 | G | N2 | AA:1591 | 3.57 | 1.00 | 88.60 |  |
| 5 | C | N1 | AA:1592 | 4.57 | 1.00 | 87.63 |  |
| 6 | G | N9 | AA:1591 | 4.88 | 1.00 | 89.83 |  |
| 7 | C | O2' | AA:1592 | 4.96 | 1.00 | 85.61 |  |
  |

  
 Coded contact list:   
 Exclusion zone contact(s): /N3_G_/O4'/  
Consider K in place of MG (2 valid contact(s) for K vs. 0 for MG)  

### MG:AA:4001

|  | Res. type | Atom | Residue | Distance | Occupancy | B-factor | Closest solute to 1^st^ shell water |
| --- | --- | --- | --- | --- | --- | --- | --- |
| **Ion** | | | |  | 1.00 | 83.60 || 1 | G | OP2 | AA:1542 | 2.86 | 1.00 | 107.81 |
| 2 | A | N9 | AA:1543 | 3.16 | 1.00 | 106.45 |  |
| 3 | A | O4' | AA:1543 | 3.34 | 1.00 | 105.77 |  |
| 4 | C | OP2 | AA:1544 | 3.60 | 1.00 | 106.64 |  |
| 5 | G | N7 | AA:1542 | 3.73 | 1.00 | 103.91 |  |
| 6 | A | N3 | AA:1543 | 3.85 | 1.00 | 104.81 |  |
| 7 | G | O5' | AA:1542 | 3.88 | 1.00 | 107.13 |  |
| 8 | A | O3' | AA:1543 | 4.12 | 1.00 | 106.60 |  |
| 9 | A | N7 | AA:1543 | 4.14 | 1.00 | 106.40 |  |
| 10 | A | O2' | AA:1543 | 4.72 | 1.00 | 107.26 |  |
| 11 | A | O5' | AA:1543 | 4.74 | 1.00 | 103.35 |  |
| 12 | G | OP1 | AA:1542 | 4.74 | 1.00 | 108.15 |  |
  |

  
 Coded contact list:   
 Exclusion zone contact(s): /O4'/OP2/  
Consider K in place of MG (3 valid contact(s) for K vs. 0 for MG)  

### MG:AA:4002

|  | Res. type | Atom | Residue | Distance | Occupancy | B-factor | Closest solute to 1^st^ shell water |
| --- | --- | --- | --- | --- | --- | --- | --- |
| **Ion** | | | |  | 1.00 | 93.11 || 1 | U | OP2 | AA:1082 | 2.84 | 1.00 | 200.41 |
| 2 | U | OP1 | AA:1082 | 2.97 | 1.00 | 200.38 |  |
| 3 | U | OP1 | AA:1081 | 3.68 | 1.00 | 200.41 |  |
| 4 | U | O3' | AA:1081 | 3.80 | 1.00 | 200.41 |  |
| 5 | U | O5' | AA:1081 | 4.41 | 1.00 | 200.41 |  |
| 6 | U | O5' | AA:1082 | 4.75 | 1.00 | 200.41 |  |
| 7 | U | OP2 | AA:1081 | 4.98 | 1.00 | 200.41 |  |
  |

  
 Coded contact list:   
 Exclusion zone contact(s): /OP1/OP2/  
Consider K in place of MG (2 valid contact(s) for K vs. 0 for MG)  

### MG:AA:4003

|  | Res. type | Atom | Residue | Distance | Occupancy | B-factor | Closest solute to 1^st^ shell water |
| --- | --- | --- | --- | --- | --- | --- | --- |
| **Ion** | | | |  | 1.00 | 109.72 || 1 | G | O6 | AA:2159 | 2.56 | 1.00 | 199.77 |
| 2 | G | N1 | AA:2160 | 2.71 | 1.00 | 200.02 |  |
| 3 | G | O6 | AA:2160 | 2.79 | 1.00 | 199.86 |  |
| 4 | C | N3 | AA:2128 | 2.82 | 1.00 | 200.41 |  |
| 5 | C | N4 | AA:2128 | 2.93 | 1.00 | 200.22 |  |
| 6 | G | N1 | AA:2159 | 2.98 | 1.00 | 200.15 |  |
| 7 | C | N4 | AA:2129 | 2.98 | 1.00 | 198.80 |  |
| 8 | C | N3 | AA:2129 | 3.26 | 1.00 | 199.38 |  |
| 9 | G | N3 | AA:2160 | 4.17 | 1.00 | 200.03 |  |
| 10 | G | N2 | AA:2160 | 4.17 | 1.00 | 199.69 |  |
| 11 | C | O2 | AA:2128 | 4.22 | 1.00 | 200.01 |  |
| 12 | G | N7 | AA:2160 | 4.35 | 1.00 | 200.36 |  |
| 13 | U | O4 | AA:2130 | 4.55 | 1.00 | 198.93 |  |
| 14 | G | N2 | AA:2159 | 4.63 | 1.00 | 199.77 |  |
| 15 | C | O2 | AA:2129 | 4.78 | 1.00 | 199.51 |  |
| 16 | G | N7 | AA:2159 | 4.78 | 1.00 | 200.41 |  |
| 17 | C | N1 | AA:2128 | 4.82 | 1.00 | 200.41 |  |
| 18 | G | N3 | AA:2159 | 4.89 | 1.00 | 200.24 |  |
  |

  
 Coded contact list:   
 Exclusion zone contact(s): /N1_G_/N1_G_/N3_C_/N3_C_/N4/N4/O6/O6/  
Consider K in place of MG (7 valid contact(s) for K vs. 0 for MG)  

### MG:AA:4004

|  | Res. type | Atom | Residue | Distance | Occupancy | B-factor | Closest solute to 1^st^ shell water |
| --- | --- | --- | --- | --- | --- | --- | --- |
| **Ion** | | | |  | 1.00 | 78.86 || 1 | U | O4 | AA:270F | 2.98 | 1.00 | 107.99 |
| 2 | G | N7 | AA:270E | 3.93 | 1.00 | 110.04 |  |
| 3 | G | O6 | AA:270E | 4.82 | 1.00 | 111.52 |  |
  |

  
 Coded contact list:   
 Exclusion zone contact(s): /O4/  
Consider K in place of MG (1 valid contact(s) for K vs. 0 for MG)  

### MG:AA:4005

|  | Res. type | Atom | Residue | Distance | Occupancy | B-factor | Closest solute to 1^st^ shell water |
| --- | --- | --- | --- | --- | --- | --- | --- |
| **Ion** | | | |  | 1.00 | 63.84 || 1 | A | O5' | AA:1543 | 2.67 | 1.00 | 103.35 |
| 2 | A | OP1 | AA:1543 | 3.12 | 1.00 | 103.24 |  |
| 3 | G | OP1 | AA:1448 | 3.31 | 1.00 | 85.73 |  |
| 4 | G | O5' | AA:1448 | 3.38 | 1.00 | 85.41 |  |
| 5 | A | O4' | AA:1543 | 3.44 | 1.00 | 105.77 |  |
| 6 | G | O3' | AA:1447 | 4.20 | 1.00 | 85.98 |  |
| 7 | A | OP2 | AA:1543 | 4.33 | 1.00 | 103.33 |  |
| 8 | A | O3' | AA:1543 | 4.39 | 1.00 | 106.60 |  |
| 9 | G | O2' | AA:1447 | 4.48 | 1.00 | 84.94 |  |
| 10 | A | O2' | AA:1545 | 4.54 | 1.00 | 87.61 |  |
| 11 | A | O2' | AA:1543 | 4.67 | 1.00 | 107.26 |  |
| 12 | G | O3' | AA:1542 | 4.71 | 1.00 | 103.72 |  |
  |

  
 Coded contact list:   
 Exclusion zone contact(s): /O5'/O5'/OP1/OP1/  
Consider K in place of MG (4 valid contact(s) for K vs. 0 for MG)  

### MG:AA:4006

|  | Res. type | Atom | Residue | Distance | Occupancy | B-factor | Closest solute to 1^st^ shell water |
| --- | --- | --- | --- | --- | --- | --- | --- |
| **Ion** | | | |  | 1.00 | 65.19 || 1 | A | OP2 | AA:340 | 2.88 | 1.00 | 61.33 |
| 2 | G | OP2 | AA:341 | 3.47 | 1.00 | 69.89 |  |
| 3 | A | O5' | AA:340 | 3.54 | 1.00 | 64.84 |  |
| 4 | G | N7 | AA:341 | 3.63 | 1.00 | 77.74 |  |
| 5 | A | OP1 | AA:340 | 3.80 | 1.00 | 63.82 |  |
| 6 | G | O6 | AA:341 | 4.84 | 1.00 | 74.60 |  |
  |

  
 Coded contact list:   
 Exclusion zone contact(s): /OP2/  
Consider K in place of MG (1 valid contact(s) for K vs. 0 for MG)  

### MG:AA:4007

|  | Res. type | Atom | Residue | Distance | Occupancy | B-factor | Closest solute to 1^st^ shell water |
| --- | --- | --- | --- | --- | --- | --- | --- |
| **Ion** | | | |  | 1.00 | 94.61 || 1 | G | O6 | AA:2038 | 2.51 | 1.00 | 56.10 |
| 2 | C | N4 | AA:2025 | 2.72 | 1.00 | 55.76 |  |
| 3 | G | O6 | AA:2024 | 2.75 | 1.00 | 55.29 |  |
| 4 | G | O6 | AA:2037 | 3.70 | 1.00 | 38.58 |  |
| 5 | C | N4 | AA:2039 | 3.80 | 1.00 | 58.17 |  |
| 6 | G | OP1 | AA:2035 | 3.92 | 1.00 | 53.42 |  |
| 7 | G | N1 | AA:2024 | 4.39 | 1.00 | 56.12 |  |
| 8 | G | N1 | AA:2038 | 4.60 | 1.00 | 53.25 |  |
| 9 | G | N7 | AA:2024 | 4.67 | 1.00 | 52.51 |  |
| 10 | C | N3 | AA:2025 | 4.69 | 1.00 | 52.60 |  |
| 11 | G | N7 | AA:2037 | 4.84 | 1.00 | 41.39 |  |
  |

  
 Coded contact list:   
 Exclusion zone contact(s): /N4/O6/O6/  
Consider K in place of MG (2 valid contact(s) for K vs. 0 for MG)  

### MG:AA:4008

|  | Res. type | Atom | Residue | Distance | Occupancy | B-factor | Closest solute to 1^st^ shell water |
| --- | --- | --- | --- | --- | --- | --- | --- |
| **Ion** | | | |  | 1.00 | 78.80 || 1 | A | OP1 | AA:28 | 2.40 | 1.00 | 47.53 |
| 2 | A | O3' | AA:447 | 2.94 | 1.00 | 45.82 |  |
| 3 | U | OP2 | AA:448 | 3.10 | 1.00 | 43.19 |  |
| 4 | A | OP2 | AA:28 | 3.49 | 1.00 | 47.12 |  |
| 5 | U | OP1 | AA:448 | 4.11 | 1.00 | 43.40 |  |
| 6 | G | O3' | AA:27 | 4.27 | 1.00 | 47.69 |  |
| 7 | A | O5' | AA:28 | 4.62 | 1.00 | 46.71 |  |
| 8 | U | O5' | AA:448 | 4.95 | 1.00 | 42.83 |  |
  |

  
 Contact(s): /OP1/  
 Coded contact list: O_ph_  
 Exclusion zone contact(s): /O3'/OP2/  
Consider K in place of MG (2 valid contact(s) for K vs. 1 for MG)  

### MG:AA:4009

|  | Res. type | Atom | Residue | Distance | Occupancy | B-factor | Closest solute to 1^st^ shell water |
| --- | --- | --- | --- | --- | --- | --- | --- |
| **Ion** | | | |  | 1.00 | 99.54 || 1 | U | N3 | AA:2132 | 2.50 | 1.00 | 200.41 |
| 2 | U | O4 | AA:2132 | 2.72 | 1.00 | 200.16 |  |
| 3 | U | O2 | AA:2132 | 3.87 | 1.00 | 200.41 |  |
| 4 | U | N1 | AA:2132 | 4.79 | 1.00 | 200.41 |  |
| 5 | U | OP1 | AA:2130 | 4.82 | 1.00 | 200.41 |  |
  |

  
 Coded contact list:   
 Exclusion zone contact(s): /N3_U_/O4/  
Consider NA in place of MG (1 valid contact(s) for NA vs. 0 for MG)  

### MG:AA:4010

|  | Res. type | Atom | Residue | Distance | Occupancy | B-factor | Closest solute to 1^st^ shell water |
| --- | --- | --- | --- | --- | --- | --- | --- |
| **Ion** | | | |  | 1.00 | 74.63 || 1 | C | OP2 | AA:455 | 2.34 | 1.00 | 55.78 |
| 2 | C | OP1 | AA:455 | 2.38 | 1.00 | 53.86 |  |
| 3 | A | O2' | AA:454 | 3.65 | 1.00 | 54.52 |  |
| 4 | C | O5' | AA:455 | 3.94 | 1.00 | 56.68 |  |
| 5 | A | O3' | AA:454 | 3.94 | 1.00 | 54.18 |  |
| 6 | A | O4' | AA:454 | 4.06 | 1.00 | 50.39 |  |
| 7 | G | OP1 | AA:35 | 4.12 | 1.00 | 68.74 |  |
| 8 | G | O4' | AA:35 | 4.30 | 1.00 | 54.17 |  |
| 9 | G | O5' | AA:35 | 4.37 | 1.00 | 65.43 |  |
  |

  
**1^st^ shell ligand angles:**   
1-2: 66;   
  
 Contact(s): /OP1/OP2/  
 Coded contact list: 2O_ph_  

### MG:AA:4011

|  | Res. type | Atom | Residue | Distance | Occupancy | B-factor | Closest solute to 1^st^ shell water |
| --- | --- | --- | --- | --- | --- | --- | --- |
| **Ion** | | | |  | 1.00 | 122.28 || 1 | U | O2' | AA:1090 | 2.50 | 1.00 | 200.41 |
| 2 | U | O2 | AA:1090 | 3.78 | 1.00 | 199.56 |  |
| 3 | G | O4' | AA:1091 | 3.82 | 1.00 | 200.41 |  |
| 4 | U | N1 | AA:1090 | 4.82 | 1.00 | 200.31 |  |
| 5 | G | O2' | AA:1091 | 4.87 | 1.00 | 200.41 |  |
| 6 | U | O4' | AA:1090 | 4.88 | 1.00 | 200.41 |  |
| 7 | G | O5' | AA:1091 | 4.89 | 1.00 | 200.41 |  |
  |

  
 Coded contact list:   
 Exclusion zone contact(s): /O2'/  
Consider NA in place of MG (1 valid contact(s) for NA vs. 0 for MG)  

### MG:AA:4012

|  | Res. type | Atom | Residue | Distance | Occupancy | B-factor | Closest solute to 1^st^ shell water |
| --- | --- | --- | --- | --- | --- | --- | --- |
| **Ion** | | | |  | 1.00 | 85.16 || 1 | G | OP1 | AA:2290 | 3.00 | 1.00 | 70.14 |
| 2 | C | OP2 | AA:2385 | 3.95 | 1.00 | 64.51 |  |
| 3 | U | OP2 | AA:2291 | 4.42 | 1.00 | 67.70 |  |
| 4 | G | OP2 | AA:2290 | 4.61 | 1.00 | 70.06 |  |
  |

  
 Coded contact list:   
 Exclusion zone contact(s): /OP1/  
Consider K in place of MG (1 valid contact(s) for K vs. 0 for MG)  

### MG:AA:4013

|  | Res. type | Atom | Residue | Distance | Occupancy | B-factor | Closest solute to 1^st^ shell water |
| --- | --- | --- | --- | --- | --- | --- | --- |
| **Ion** | | | |  | 1.00 | 119.98 || 1 | G | O6 | AA:1071 | 2.37 | 1.00 | 199.57 |
| 2 | G | N7 | AA:1071 | 2.79 | 1.00 | 199.62 |  |
| 3 | G | N2 | AA:1089 | 3.67 | 1.00 | 200.26 |  |
| 4 | U | O4 | AA:1101 | 3.93 | 1.00 | 200.41 |  |
| 5 | U | O4 | AA:1090 | 4.07 | 1.00 | 200.32 |  |
| 6 | C | N4 | AA:1100 | 4.18 | 1.00 | 199.78 |  |
| 7 | G | O6 | AA:1091 | 4.26 | 1.00 | 200.02 |  |
| 8 | G | N1 | AA:1071 | 4.36 | 1.00 | 199.21 |  |
| 9 | G | N3 | AA:1089 | 4.53 | 1.00 | 200.41 |  |
| 10 | G | O6 | AA:1087 | 4.66 | 1.00 | 200.29 |  |
| 11 | G | N9 | AA:1071 | 4.92 | 1.00 | 199.78 |  |
| 12 | G | N1 | AA:1089 | 4.96 | 1.00 | 200.41 |  |
  |

  
 Contact(s): /O6/  
 Coded contact list: O_b_  
 Exclusion zone contact(s): /N7/  

### MG:AA:4014

|  | Res. type | Atom | Residue | Distance | Occupancy | B-factor | Closest solute to 1^st^ shell water |
| --- | --- | --- | --- | --- | --- | --- | --- |
| **Ion** | | | |  | 1.00 | 84.51 || 1 | C | OP2 | AA:1407 | 3.86 | 1.00 | 59.94 |
| 2 | C | OP2 | AA:1408 | 4.83 | 1.00 | 55.63 |  |
  |

  
 Coded contact list:   
1^st^ MG coordination shell missing (see Notes)  

### MG:AA:4016

|  | Res. type | Atom | Residue | Distance | Occupancy | B-factor | Closest solute to 1^st^ shell water |
| --- | --- | --- | --- | --- | --- | --- | --- |
| **Ion** | | | |  | 1.00 | 111.82 || 1 | A | N6 | AA:2900 | 1.65 | 1.00 | 175.00 |
| 2 | G | O6 | AA:2899 | 1.95 | 1.00 | 170.95 |  |
| 3 | G | N1 | AA:2899 | 2.55 | 1.00 | 170.97 |  |
| 4 | A | N7 | AA:2900 | 2.86 | 1.00 | 175.31 |  |
| 5 | A | N1 | AA:2900 | 3.05 | 1.00 | 174.99 |  |
| 6 | C | N4 | AA:4 | 3.29 | 1.00 | 170.94 |  |
| 7 | C | N3 | AA:4 | 3.54 | 1.00 | 171.02 |  |
| 8 | U | O4 | AA:3 | 3.57 | 1.00 | 175.32 |  |
| 9 | G | N7 | AA:2899 | 3.70 | 1.00 | 171.20 |  |
| 10 | G | N3 | AA:2899 | 4.26 | 1.00 | 171.04 |  |
| 11 | A | N6 | AA:5 | 4.29 | 1.00 | 156.27 |  |
| 12 | A | N3 | AA:2900 | 4.37 | 1.00 | 175.18 |  |
| 13 | U | N3 | AA:3 | 4.43 | 1.00 | 175.64 |  |
| 14 | A | N9 | AA:2900 | 4.43 | 1.00 | 175.48 |  |
| 15 | G | N2 | AA:2899 | 4.49 | 1.00 | 170.30 |  |
| 16 | U | O4 | AA:2898 | 4.54 | 1.00 | 156.17 |  |
| 17 | C | N4 | AA:2901 | 4.70 | 1.00 | 174.52 |  |
| 18 | A | N1 | AA:5 | 4.83 | 1.00 | 156.46 |  |
| 19 | G | N9 | AA:2899 | 4.90 | 1.00 | 171.59 |  |
| 20 | C | O2 | AA:4 | 4.91 | 1.00 | 170.53 |  |
  |

  
**1^st^ shell ligand angles:**   
1-2: 154;   
  
 Contact(s): /O6/  
 Coded contact list: O_b_  
 Exclusion zone contact(s): /N1_A_/N1_G_/N4/N6/N7/  
Consider K in place of MG (3 valid contact(s) for K vs. 1 for MG)  

### MG:AA:4017

|  | Res. type | Atom | Residue | Distance | Occupancy | B-factor | Closest solute to 1^st^ shell water |
| --- | --- | --- | --- | --- | --- | --- | --- |
| **Ion** | | | |  | 1.00 | 96.09 || 1 | G | N7 | AA:2735 | 2.76 | 1.00 | 100.12 |
| 2 | G | O6 | AA:2735 | 2.95 | 1.00 | 96.85 |  |
| 3 | A | OP2 | AA:2734 | 3.29 | 1.00 | 97.70 |  |
| 4 | G | O6 | AA:2736 | 4.65 | 1.00 | 97.58 |  |
| 5 | A | OP1 | AA:2734 | 4.69 | 1.00 | 97.89 |  |
| 6 | G | N1 | AA:2735 | 4.87 | 1.00 | 97.41 |  |
| 7 | G | N9 | AA:2735 | 4.98 | 1.00 | 99.85 |  |
  |

  
 Coded contact list:   
 Exclusion zone contact(s): /N7/O6/OP2/  
Consider K in place of MG (3 valid contact(s) for K vs. 0 for MG)  

### MG:AA:4018

|  | Res. type | Atom | Residue | Distance | Occupancy | B-factor | Closest solute to 1^st^ shell water |
| --- | --- | --- | --- | --- | --- | --- | --- |
| **Ion** | | | |  | 1.00 | 56.68 || 1 | G | O6 | AA:1776 | 2.68 | 1.00 | 36.38 |
| 2 | A | N1 | AA:1789 | 2.72 | 1.00 | 38.21 |  |
| 3 | C | OP1 | AA:1774 | 3.16 | 1.00 | 39.07 |  |
| 4 | C | OP2 | AA:1774 | 3.25 | 1.00 | 38.12 |  |
| 5 | A | N6 | AA:1789 | 3.34 | 1.00 | 42.23 |  |
| 6 | G | N7 | AA:1776 | 3.63 | 1.00 | 37.24 |  |
| 7 | C | O5' | AA:1774 | 3.94 | 1.00 | 38.23 |  |
| 8 | U | OP2 | AA:1775 | 4.07 | 1.00 | 41.22 |  |
| 9 | C | N4 | AA:1788 | 4.34 | 1.00 | 34.34 |  |
| 10 | G | N1 | AA:1776 | 4.45 | 1.00 | 37.94 |  |
| 11 | A | N3 | AA:1789 | 4.80 | 1.00 | 37.83 |  |
| 12 | G | O2' | AA:1772 | 4.85 | 1.00 | 38.87 |  |
  |

  
 Coded contact list:   
 Exclusion zone contact(s): /N1_A_/N6/O6/OP1/OP2/  
Consider K in place of MG (5 valid contact(s) for K vs. 0 for MG)  

### MG:AA:4019

|  | Res. type | Atom | Residue | Distance | Occupancy | B-factor | Closest solute to 1^st^ shell water |
| --- | --- | --- | --- | --- | --- | --- | --- |
| **Ion** | | | |  | 1.00 | 99.01 || 1 | G | OP2 | AA:407 | 4.50 | 1.00 | 83.66 |
| 2 | G | OP2 | AA:406 | 4.82 | 1.00 | 90.13 |  |
  |

  
 Coded contact list:   
1^st^ MG coordination shell missing (see Notes)  

### MG:AA:4020

|  | Res. type | Atom | Residue | Distance | Occupancy | B-factor | Closest solute to 1^st^ shell water |
| --- | --- | --- | --- | --- | --- | --- | --- |
| **Ion** | | | |  | 1.00 | 85.69 || 1 | C | O2' | AA:277 | 5.00 | 1.00 | 149.71 |
  |

  
 Coded contact list:   
1^st^ MG coordination shell missing (see Notes)  

### MG:AA:4021

|  | Res. type | Atom | Residue | Distance | Occupancy | B-factor | Closest solute to 1^st^ shell water |
| --- | --- | --- | --- | --- | --- | --- | --- |
| **Ion** | | | |  | 1.00 | 80.24 || 1 | U | OP2 | AA:1420 | 2.85 | 1.00 | 93.58 |
| 2 | U | O4 | AA:1576 | 4.47 | 1.00 | 71.18 |  |
| 3 | G | O6 | AA:1421 | 4.53 | 1.00 | 81.72 |  |
| 4 | U | O5' | AA:1420 | 4.97 | 1.00 | 94.92 |  |
| 5 | A | O3' | AA:1419 | 4.98 | 1.00 | 92.00 |  |
  |

  
 Coded contact list:   
 Exclusion zone contact(s): /OP2/  
Consider K in place of MG (1 valid contact(s) for K vs. 0 for MG)  

### MG:AA:4022

|  | Res. type | Atom | Residue | Distance | Occupancy | B-factor | Closest solute to 1^st^ shell water |
| --- | --- | --- | --- | --- | --- | --- | --- |
| **Ion** | | | |  | 1.00 | 62.20 || 1 | G | O6 | AA:1645 | 3.32 | 1.00 | 48.66 |
| 2 | G | N7 | AA:1645 | 4.35 | 1.00 | 48.41 |  |
| 3 | C | N4 | AA:1644 | 4.90 | 1.00 | 55.22 |  |
  |

  
 Coded contact list:   
 Exclusion zone contact(s): /O6/  
Consider K in place of MG (1 valid contact(s) for K vs. 0 for MG)  

### MG:AA:4023

|  | Res. type | Atom | Residue | Distance | Occupancy | B-factor | Closest solute to 1^st^ shell water |
| --- | --- | --- | --- | --- | --- | --- | --- |
| **Ion** | | | |  | 1.00 | 73.43 || 1 | A | OP2 | AA:2639 | 2.69 | 1.00 | 79.31 |
| 2 | G | N7 | AA:2640 | 3.04 | 1.00 | 68.34 |  |
| 3 | A | OP1 | AA:2639 | 3.61 | 1.00 | 79.02 |  |
| 4 | G | O6 | AA:2640 | 3.68 | 1.00 | 68.39 |  |
| 5 | A | O5' | AA:2639 | 4.27 | 1.00 | 76.45 |  |
| 6 | G | N7 | AA:2641 | 4.35 | 1.00 | 68.37 |  |
| 7 | G | O6 | AA:2641 | 4.64 | 1.00 | 63.84 |  |
| 8 | G | O3' | AA:2638 | 4.95 | 1.00 | 78.66 |  |
| 9 | G | OP2 | AA:2640 | 4.98 | 1.00 | 73.87 |  |
  |

  
 Coded contact list:   
 Exclusion zone contact(s): /N7/OP2/  
Consider K in place of MG (2 valid contact(s) for K vs. 0 for MG)  

### MG:AA:4024

|  | Res. type | Atom | Residue | Distance | Occupancy | B-factor | Closest solute to 1^st^ shell water |
| --- | --- | --- | --- | --- | --- | --- | --- |
| **Ion** | | | |  | 1.00 | 95.29 || 1 | G | OP1 | AA:1056 | 4.15 | 1.00 | 200.30 |
| 2 | A | O2' | AA:1057 | 4.59 | 1.00 | 200.41 |  |
  |

  
 Coded contact list:   
1^st^ MG coordination shell missing (see Notes)  

### MG:AA:4025

|  | Res. type | Atom | Residue | Distance | Occupancy | B-factor | Closest solute to 1^st^ shell water |
| --- | --- | --- | --- | --- | --- | --- | --- |
| **Ion** | | | |  | 1.00 | 86.44 || 1 | A | OP2 | AA:515 | 4.18 | 1.00 | 52.43 |
| 2 | A | OP1 | AA:515 | 4.84 | 1.00 | 51.03 |  |
| 3 | C | N4 | AA:516 | 4.94 | 1.00 | 51.30 |  |
  |

  
 Coded contact list:   
1^st^ MG coordination shell missing (see Notes)  

### MG:AA:4026

|  | Res. type | Atom | Residue | Distance | Occupancy | B-factor | Closest solute to 1^st^ shell water |
| --- | --- | --- | --- | --- | --- | --- | --- |
| **Ion** | | | |  | 1.00 | 53.10 || 1 | U | OP1 | AA:29 | 2.70 | 1.00 | 49.52 |
| 2 | U | OP2 | AA:29 | 2.97 | 1.00 | 48.44 |  |
| 3 | LYS | NZ | A1:5 | 3.00 | 1.00 | 51.20 |  |
| 4 | A | OP1 | AA:447 | 3.33 | 1.00 | 49.10 |  |
| 5 | U | O5' | AA:29 | 4.06 | 1.00 | 49.90 |  |
| 6 | G | OP2 | AA:30 | 4.65 | 1.00 | 50.11 |  |
| 7 | A | O3' | AA:28 | 4.65 | 1.00 | 48.43 |  |
| 8 | U | OP1 | AA:448 | 4.73 | 1.00 | 43.40 |  |
| 9 | A | O5' | AA:447 | 4.96 | 1.00 | 48.99 |  |
| 10 | G | O2' | AA:583 | 5.00 | 1.00 | 42.31 |  |
  |

  
 Coded contact list:   
 Exclusion zone contact(s): /NZ_Lys_/OP1/OP1/OP2/  
Consider K in place of MG (4 valid contact(s) for K vs. 0 for MG)  

### MG:AA:4027

|  | Res. type | Atom | Residue | Distance | Occupancy | B-factor | Closest solute to 1^st^ shell water |
| --- | --- | --- | --- | --- | --- | --- | --- |
| **Ion** | | | |  | 1.00 | 72.87 || 1 | G | OP1 | AA:51 | 2.80 | 1.00 | 65.04 |
| 2 | G | O5' | AA:51 | 2.80 | 1.00 | 63.30 |  |
| 3 | A | OP1 | AA:49 | 3.85 | 1.00 | 61.48 |  |
| 4 | G | OP2 | AA:51 | 4.19 | 1.00 | 62.13 |  |
| 5 | U | O3' | AA:50 | 4.67 | 1.00 | 65.74 |  |
| 6 | G | O4' | AA:51 | 4.97 | 1.00 | 54.07 |  |
  |

  
 Coded contact list:   
 Exclusion zone contact(s): /O5'/OP1/  
Consider K in place of MG (2 valid contact(s) for K vs. 0 for MG)  

### MG:AA:4028

|  | Res. type | Atom | Residue | Distance | Occupancy | B-factor | Closest solute to 1^st^ shell water |
| --- | --- | --- | --- | --- | --- | --- | --- |
| **Ion** | | | |  | 1.00 | 81.03 || 1 | G | OP2 | AA:599 | 3.77 | 1.00 | 63.97 |
| 2 | G | OP2 | AA:598 | 4.57 | 1.00 | 65.38 |  |
| 3 | G | OP1 | AA:598 | 4.79 | 1.00 | 65.63 |  |
  |

  
 Coded contact list:   
1^st^ MG coordination shell missing (see Notes)  

### MG:AA:4029

|  | Res. type | Atom | Residue | Distance | Occupancy | B-factor | Closest solute to 1^st^ shell water |
| --- | --- | --- | --- | --- | --- | --- | --- |
| **Ion** | | | |  | 1.00 | 80.41 || 1 | G | O2' | AA:399 | 2.68 | 1.00 | 57.62 |
| 2 | G | OP1 | AA:400 | 2.81 | 1.00 | 59.52 |  |
| 3 | G | O3' | AA:399 | 2.89 | 1.00 | 58.72 |  |
| 4 | G | O5' | AA:400 | 3.88 | 1.00 | 60.01 |  |
| 5 | G | OP2 | AA:400 | 4.78 | 1.00 | 59.64 |  |
  |

  
 Coded contact list:   
 Exclusion zone contact(s): /O2'/O3'/OP1/  
Consider K in place of MG (3 valid contact(s) for K vs. 0 for MG)  

### MG:AA:4030

|  | Res. type | Atom | Residue | Distance | Occupancy | B-factor | Closest solute to 1^st^ shell water |
| --- | --- | --- | --- | --- | --- | --- | --- |
| **Ion** | | | |  | 1.00 | 66.16 || 1 | G | OP2 | AA:1807 | 2.58 | 1.00 | 44.48 |
| 2 | C | OP2 | AA:1806 | 2.87 | 1.00 | 35.77 |  |
| 3 | C | O5' | AA:1806 | 3.67 | 1.00 | 37.22 |  |
| 4 | C | OP1 | AA:1806 | 3.76 | 1.00 | 37.01 |  |
| 5 | C | O3' | AA:1806 | 4.83 | 1.00 | 41.28 |  |
| 6 | G | O5' | AA:1807 | 4.86 | 1.00 | 44.61 |  |
| 7 | G | OP1 | AA:1807 | 4.88 | 1.00 | 45.58 |  |
  |

  
 Coded contact list:   
 Exclusion zone contact(s): /OP2/OP2/  
Consider NA in place of MG (1 valid contact(s) for NA vs. 0 for MG)  

### MG:AA:4031

|  | Res. type | Atom | Residue | Distance | Occupancy | B-factor | Closest solute to 1^st^ shell water |
| --- | --- | --- | --- | --- | --- | --- | --- |
| **Ion** | | | |  | 1.00 | 84.17 || 1 | G | O6 | AA:695 | 3.40 | 1.00 | 39.43 |
| 2 | U | O4 | AA:694 | 3.73 | 1.00 | 43.82 |  |
| 3 | U | O4 | AA:767 | 3.85 | 1.00 | 38.03 |  |
| 4 | G | N7 | AA:695 | 4.33 | 1.00 | 34.09 |  |
| 5 | G | O6 | AA:696 | 4.34 | 1.00 | 34.86 |  |
| 6 | C | N4 | AA:766 | 4.70 | 1.00 | 29.72 |  |
| 7 | U | N3 | AA:767 | 4.92 | 1.00 | 37.59 |  |
  |

  
 Coded contact list:   
 Exclusion zone contact(s): /O6/  
Consider K in place of MG (1 valid contact(s) for K vs. 0 for MG)  

### MG:AA:4033

|  | Res. type | Atom | Residue | Distance | Occupancy | B-factor | Closest solute to 1^st^ shell water |
| --- | --- | --- | --- | --- | --- | --- | --- |
| **Ion** | | | |  | 1.00 | 83.28 || 1 | G | OP2 | AA:674 | 2.31 | 1.00 | 36.14 |
| 2 | C | OP2 | AA:673 | 2.98 | 1.00 | 40.03 |  |
| 3 | G | N7 | AA:674 | 3.88 | 1.00 | 40.35 |  |
| 4 | C | O5' | AA:673 | 3.90 | 1.00 | 38.20 |  |
| 5 | G | O5' | AA:674 | 4.04 | 1.00 | 36.70 |  |
| 6 | C | O3' | AA:673 | 4.05 | 1.00 | 35.76 |  |
| 7 | ARG | NH1 | AF:54 | 4.07 | 1.00 | 37.75 |  |
| 8 | C | N1 | AA:673 | 4.28 | 1.00 | 42.60 |  |
| 9 | C | O4' | AA:673 | 4.30 | 1.00 | 39.37 |  |
| 10 | C | OP1 | AA:673 | 4.69 | 1.00 | 39.06 |  |
| 11 | G | OP1 | AA:674 | 4.87 | 1.00 | 37.93 |  |
  |

  
 Contact(s): /OP2/  
 Coded contact list: O_ph_  
 Exclusion zone contact(s): /OP2/  

### MG:AA:4034

|  | Res. type | Atom | Residue | Distance | Occupancy | B-factor | Closest solute to 1^st^ shell water |
| --- | --- | --- | --- | --- | --- | --- | --- |
| **Ion** | | | |  | 1.00 | 100.81 || 1 | U | N1 | AA:405 | 3.45 | 1.00 | 99.58 |
| 2 | U | O4' | AA:405 | 4.33 | 1.00 | 96.28 |  |
| 3 | U | N3 | AA:405 | 4.35 | 1.00 | 101.51 |  |
| 4 | U | O4 | AA:405 | 4.85 | 1.00 | 101.75 |  |
  |

  
 Coded contact list:   
1^st^ MG coordination shell missing (see Notes)  

### MG:AA:4035

|  | Res. type | Atom | Residue | Distance | Occupancy | B-factor | Closest solute to 1^st^ shell water |
| --- | --- | --- | --- | --- | --- | --- | --- |
| **Ion** | | | |  | 1.00 | 75.86 || 1 | U | OP1 | AA:305 | 3.61 | 1.00 | 66.60 |
| 2 | G | O3' | AA:304 | 4.96 | 1.00 | 67.06 |  |
  |

  
 Coded contact list:   
1^st^ MG coordination shell missing (see Notes)  

### MG:AA:4036

|  | Res. type | Atom | Residue | Distance | Occupancy | B-factor | Closest solute to 1^st^ shell water |
| --- | --- | --- | --- | --- | --- | --- | --- |
| **Ion** | | | |  | 1.00 | 68.22 || 1 | C | O5' | AA:1537 | 4.12 | 1.00 | 175.37 |
| 2 | G | OP1 | AA:1538 | 4.30 | 1.00 | 169.59 |  |
| 3 | C | OP2 | AA:1537 | 4.31 | 1.00 | 175.83 |  |
| 4 | C | O3' | AA:1537 | 4.35 | 1.00 | 170.98 |  |
| 5 | C | OP1 | AA:1537 | 4.58 | 1.00 | 176.12 |  |
  |

  
 Coded contact list:   
1^st^ MG coordination shell missing (see Notes)  

### MG:AA:4037

|  | Res. type | Atom | Residue | Distance | Occupancy | B-factor | Closest solute to 1^st^ shell water |
| --- | --- | --- | --- | --- | --- | --- | --- |
| **Ion** | | | |  | 1.00 | 43.25 || 1 | G | OP1 | AA:113 | 2.39 | 1.00 | 54.26 |
| 2 | U | O3' | AA:112 | 3.26 | 1.00 | 52.77 |  |
| 3 | G | O5' | AA:113 | 4.22 | 1.00 | 52.78 |  |
| 4 | U | O2' | AA:112 | 4.55 | 1.00 | 49.74 |  |
| 5 | G | OP2 | AA:113 | 4.60 | 1.00 | 53.98 |  |
  |

  
 Contact(s): /OP1/  
 Coded contact list: O_ph_  
 Exclusion zone contact(s): /O3'/  

### MG:AA:4038

|  | Res. type | Atom | Residue | Distance | Occupancy | B-factor | Closest solute to 1^st^ shell water |
| --- | --- | --- | --- | --- | --- | --- | --- |
| **Ion** | | | |  | 1.00 | 77.30 || 1 | A | OP2 | AA:422 | 2.78 | 1.00 | 69.42 |
| 2 | G | OP2 | AA:370 | 2.85 | 1.00 | 65.16 |  |
| 3 | C | O2' | AA:366 | 3.04 | 1.00 | 68.67 |  |
| 4 | C | N4 | AA:271A | 3.63 | 1.00 | 78.99 |  |
| 5 | A | OP1 | AA:422 | 4.25 | 1.00 | 68.44 |  |
| 6 | C | O2 | AA:366 | 4.30 | 1.00 | 75.99 |  |
| 7 | C | N3 | AA:271A | 4.39 | 1.00 | 80.96 |  |
| 8 | G | O6 | AA:271B | 4.44 | 1.00 | 84.61 |  |
| 9 | C | O3' | AA:366 | 4.53 | 1.00 | 66.64 |  |
| 10 | C | N4 | AA:404 | 4.57 | 1.00 | 84.81 |  |
| 11 | G | OP1 | AA:370 | 4.61 | 1.00 | 65.49 |  |
| 12 | U | O3' | AA:421 | 4.88 | 1.00 | 69.74 |  |
| 13 | G | N7 | AA:370 | 4.93 | 1.00 | 63.34 |  |
  |

  
 Coded contact list:   
 Exclusion zone contact(s): /O2'/OP2/OP2/  
Consider K in place of MG (3 valid contact(s) for K vs. 0 for MG)  

### MG:AA:4039

|  | Res. type | Atom | Residue | Distance | Occupancy | B-factor | Closest solute to 1^st^ shell water |
| --- | --- | --- | --- | --- | --- | --- | --- |
| **Ion** | | | |  | 1.00 | 95.07 || 1 | G | O6 | AA:1385 | 2.23 | 1.00 | 75.30 |
| 2 | C | N4 | AA:1402 | 3.71 | 1.00 | 76.98 |  |
| 3 | G | O6 | AA:1344 | 3.76 | 1.00 | 68.08 |  |
| 4 | C | N4 | AA:1403 | 3.80 | 1.00 | 66.18 |  |
| 5 | G | N7 | AA:1385 | 4.01 | 1.00 | 74.34 |  |
| 6 | G | O6 | AA:1401 | 4.28 | 1.00 | 70.96 |  |
| 7 | G | N1 | AA:1385 | 4.50 | 1.00 | 74.78 |  |
| 8 | C | N4 | AA:1386 | 4.69 | 1.00 | 66.79 |  |
  |

  
 Contact(s): /O6/  
 Coded contact list: O_b_  

### MG:AA:4040

|  | Res. type | Atom | Residue | Distance | Occupancy | B-factor | Closest solute to 1^st^ shell water |
| --- | --- | --- | --- | --- | --- | --- | --- |
| **Ion** | | | |  | 1.00 | 67.78 || 1 | ASN | OD1 | A0:11 | 3.02 | 1.00 | 59.10 |
| 2 | G | OP2 | AA:2004 | 3.07 | 1.00 | 48.72 |  |
| 3 | G | OP2 | AA:2003 | 3.27 | 1.00 | 41.06 |  |
| 4 | A | N6 | AA:1652 | 3.38 | 1.00 | 35.04 |  |
| 5 | G | N2 | AA:1653 | 4.06 | 1.00 | 53.14 |  |
| 6 | ASN | ND2 | A0:11 | 4.12 | 1.00 | 57.51 |  |
| 7 | G | O5' | AA:2003 | 4.25 | 1.00 | 42.86 |  |
| 8 | G | OP1 | AA:2003 | 4.33 | 1.00 | 42.60 |  |
| 9 | G | N1 | AA:1653 | 4.92 | 1.00 | 50.90 |  |
| 10 | G | O5' | AA:2004 | 4.93 | 1.00 | 47.54 |  |
| 11 | A | N1 | AA:1652 | 4.99 | 1.00 | 36.49 |  |
  |

  
 Coded contact list:   
 Exclusion zone contact(s): /N6/OD1_Asn_/OP2/OP2/  
Consider K in place of MG (4 valid contact(s) for K vs. 0 for MG)  

### MG:AA:4041

|  | Res. type | Atom | Residue | Distance | Occupancy | B-factor | Closest solute to 1^st^ shell water |
| --- | --- | --- | --- | --- | --- | --- | --- |
| **Ion** | | | |  | 1.00 | 101.04 || 1 | G | OP1 | AA:348 | 3.11 | 1.00 | 73.92 |
| 2 | G | OP2 | AA:348 | 4.39 | 1.00 | 73.66 |  |
  |

  
 Coded contact list:   
 Exclusion zone contact(s): /OP1/  
Consider K in place of MG (1 valid contact(s) for K vs. 0 for MG)  

### MG:AA:4042

|  | Res. type | Atom | Residue | Distance | Occupancy | B-factor | Closest solute to 1^st^ shell water |
| --- | --- | --- | --- | --- | --- | --- | --- |
| **Ion** | | | |  | 1.00 | 88.39 || 1 | G | O6 | AA:2154 | 3.33 | 1.00 | 200.19 |
| 2 | G | O6 | AA:2153 | 3.67 | 1.00 | 200.00 |  |
| 3 | G | N7 | AA:2153 | 3.94 | 1.00 | 200.00 |  |
| 4 | G | N7 | AA:2154 | 4.29 | 1.00 | 200.15 |  |
| 5 | C | N4 | AA:2137 | 4.78 | 1.00 | 200.41 |  |
  |

  
 Coded contact list:   
 Exclusion zone contact(s): /O6/  
Consider K in place of MG (1 valid contact(s) for K vs. 0 for MG)  

### MG:AA:4043

|  | Res. type | Atom | Residue | Distance | Occupancy | B-factor | Closest solute to 1^st^ shell water |
| --- | --- | --- | --- | --- | --- | --- | --- |
| **Ion** | | | |  | 1.00 | 132.06 || 1 | C | OP2 | AA:286 | 2.39 | 1.00 | 92.71 |
| 2 | C | O5' | AA:286 | 3.71 | 1.00 | 93.86 |  |
| 3 | C | N1 | AA:286 | 3.78 | 1.00 | 95.49 |  |
| 4 | C | O4' | AA:286 | 4.10 | 1.00 | 95.39 |  |
| 5 | C | OP2 | AA:287 | 4.12 | 1.00 | 95.58 |  |
| 6 | C | N4 | AA:286 | 4.52 | 1.00 | 95.26 |  |
| 7 | C | O3' | AA:285 | 4.61 | 1.00 | 92.66 |  |
| 8 | C | N3 | AA:286 | 4.68 | 1.00 | 95.46 |  |
| 9 | C | OP1 | AA:286 | 4.70 | 1.00 | 93.09 |  |
  |

  
 Contact(s): /OP2/  
 Coded contact list: O_ph_  

### MG:AA:4044

|  | Res. type | Atom | Residue | Distance | Occupancy | B-factor | Closest solute to 1^st^ shell water |
| --- | --- | --- | --- | --- | --- | --- | --- |
| **Ion** | | | |  | 1.00 | 93.13 || 1 | A | O2' | AA:1046 | 4.52 | 1.00 | 189.76 |
| 2 | G | OP1 | AA:1107 | 4.85 | 1.00 | 190.07 |  |
  |

  
 Coded contact list:   
1^st^ MG coordination shell missing (see Notes)  

### MG:AA:4045

|  | Res. type | Atom | Residue | Distance | Occupancy | B-factor | Closest solute to 1^st^ shell water |
| --- | --- | --- | --- | --- | --- | --- | --- |
| **Ion** | | | |  | 1.00 | 57.96 || 1 | A | O2' | AA:1265 | 3.00 | 1.00 | 50.65 |
| 2 | U | OP2 | AA:1263 | 3.12 | 1.00 | 46.07 |  |
| 3 | G | OP2 | AA:1264 | 3.34 | 1.00 | 46.80 |  |
| 4 | U | OP1 | AA:1263 | 3.70 | 1.00 | 47.49 |  |
| 5 | ARG | NH2 | A5:20 | 3.76 | 1.00 | 52.88 |  |
| 6 | A | N3 | AA:1265 | 3.96 | 1.00 | 49.91 |  |
| 7 | ARG | NH1 | AS:15 | 4.36 | 1.00 | 53.16 |  |
| 8 | U | O5' | AA:1263 | 4.38 | 1.00 | 48.19 |  |
| 9 | G | N7 | AA:1264 | 4.76 | 1.00 | 49.56 |  |
| 10 | A | N9 | AA:1265 | 4.93 | 1.00 | 49.35 |  |
  |

  
 Coded contact list:   
 Exclusion zone contact(s): /O2'/OP2/OP2/  
Consider K in place of MG (3 valid contact(s) for K vs. 0 for MG)  

### MG:AA:4046

|  | Res. type | Atom | Residue | Distance | Occupancy | B-factor | Closest solute to 1^st^ shell water |
| --- | --- | --- | --- | --- | --- | --- | --- |
| **Ion** | | | |  | 1.00 | 111.36 || 1 | G | O6 | AA:2645 | 2.57 | 1.00 | 76.02 |
| 2 | G | N7 | AA:2644 | 2.74 | 1.00 | 68.18 |  |
| 3 | G | N7 | AA:2645 | 2.94 | 1.00 | 74.90 |  |
| 4 | G | OP2 | AA:2644 | 3.52 | 1.00 | 68.53 |  |
| 5 | G | OP2 | AA:2643 | 4.23 | 1.00 | 65.17 |  |
| 6 | G | O6 | AA:2644 | 4.38 | 1.00 | 69.50 |  |
| 7 | G | OP1 | AA:2544 | 4.44 | 1.00 | 62.24 |  |
| 8 | G | N1 | AA:2645 | 4.50 | 1.00 | 76.60 |  |
| 9 | G | N9 | AA:2644 | 4.67 | 1.00 | 68.87 |  |
| 10 | G | O5' | AA:2644 | 4.83 | 1.00 | 68.56 |  |
  |

  
 Coded contact list:   
 Exclusion zone contact(s): /N7/N7/O6/  
Consider K in place of MG (2 valid contact(s) for K vs. 0 for MG)  

### MG:AA:4047

|  | Res. type | Atom | Residue | Distance | Occupancy | B-factor | Closest solute to 1^st^ shell water |
| --- | --- | --- | --- | --- | --- | --- | --- |
| **Ion** | | | |  | 1.00 | 59.78 || 1 | U | OP1 | AA:2653 | 3.03 | 1.00 | 117.99 |
| 2 | G | OP1 | AA:2655 | 3.66 | 1.00 | 132.44 |  |
| 3 | A | O5' | AA:2654 | 3.92 | 1.00 | 129.11 |  |
| 4 | A | O3' | AA:2654 | 4.36 | 1.00 | 132.23 |  |
| 5 | G | OP2 | AA:2655 | 4.37 | 1.00 | 133.04 |  |
  |

  
 Coded contact list:   
 Exclusion zone contact(s): /OP1/  
Consider K in place of MG (1 valid contact(s) for K vs. 0 for MG)  

### MG:AA:4048

|  | Res. type | Atom | Residue | Distance | Occupancy | B-factor | Closest solute to 1^st^ shell water |
| --- | --- | --- | --- | --- | --- | --- | --- |
| **Ion** | | | |  | 1.00 | 93.65 || 1 | G | OP1 | AA:2289 | 2.58 | 1.00 | 73.40 |
| 2 | G | O5' | AA:2289 | 2.69 | 1.00 | 72.39 |  |
| 3 | G | OP2 | AA:2290 | 2.70 | 1.00 | 70.06 |  |
| 4 | C | OP1 | AA:2385 | 3.27 | 1.00 | 64.78 |  |
| 5 | G | O3' | AA:2384 | 3.53 | 1.00 | 66.32 |  |
| 6 | G | OP2 | AA:2289 | 3.95 | 1.00 | 75.05 |  |
| 7 | G | O3' | AA:2289 | 4.01 | 1.00 | 69.94 |  |
| 8 | G | OP1 | AA:2290 | 4.32 | 1.00 | 70.14 |  |
| 9 | A | O3' | AA:2288 | 4.49 | 1.00 | 75.10 |  |
| 10 | C | OP2 | AA:2385 | 4.56 | 1.00 | 64.51 |  |
| 11 | G | O2' | AA:2384 | 4.94 | 1.00 | 68.58 |  |
  |

  
 Coded contact list:   
 Exclusion zone contact(s): /O5'/OP1/OP1/OP2/  
Consider K in place of MG (3 valid contact(s) for K vs. 0 for MG)  

### MG:AA:4049

|  | Res. type | Atom | Residue | Distance | Occupancy | B-factor | Closest solute to 1^st^ shell water |
| --- | --- | --- | --- | --- | --- | --- | --- |
| **Ion** | | | |  | 1.00 | 115.47 || 1 | C | O2' | AA:316 | 2.68 | 1.00 | 70.99 |
| 2 | C | O2 | AA:316 | 2.97 | 1.00 | 69.07 |  |
| 3 | G | N2 | AA:301 | 3.99 | 1.00 | 66.88 |  |
| 4 | G | O4' | AA:317 | 4.20 | 1.00 | 68.42 |  |
| 5 | C | N1 | AA:316 | 4.57 | 1.00 | 71.63 |  |
| 6 | C | O3' | AA:316 | 4.80 | 1.00 | 72.74 |  |
| 7 | G | N2 | AA:315 | 4.95 | 1.00 | 68.95 |  |
  |

  
 Coded contact list:   
 Exclusion zone contact(s): /O2/O2'/  
Consider K in place of MG (2 valid contact(s) for K vs. 0 for MG)  

### MG:AA:4050

|  | Res. type | Atom | Residue | Distance | Occupancy | B-factor | Closest solute to 1^st^ shell water |
| --- | --- | --- | --- | --- | --- | --- | --- |
| **Ion** | | | |  | 1.00 | 70.65 || 1 | G | O3' | AA:2833 | 2.86 | 1.00 | 94.60 |
| 2 | G | N7 | AA:2834 | 3.81 | 1.00 | 78.96 |  |
| 3 | G | O5' | AA:2834 | 3.81 | 1.00 | 91.28 |  |
| 4 | G | OP2 | AA:2834 | 3.95 | 1.00 | 93.55 |  |
| 5 | G | O5' | AA:2833 | 4.05 | 1.00 | 89.21 |  |
| 6 | G | O4' | AA:2833 | 4.78 | 1.00 | 98.73 |  |
| 7 | G | N9 | AA:2834 | 4.94 | 1.00 | 80.98 |  |
  |

  
 Coded contact list:   
 Exclusion zone contact(s): /O3'/  
Consider K in place of MG (1 valid contact(s) for K vs. 0 for MG)  

### MG:AA:4051

|  | Res. type | Atom | Residue | Distance | Occupancy | B-factor | Closest solute to 1^st^ shell water |
| --- | --- | --- | --- | --- | --- | --- | --- |
| **Ion** | | | |  | 1.00 | 106.12 || 1 | A | OP1 | AA:1103 | 3.21 | 1.00 | 200.37 |
| 2 | A | OP2 | AA:1103 | 3.73 | 1.00 | 200.24 |  |
| 3 | C | OP1 | AA:1102 | 4.14 | 1.00 | 200.26 |  |
| 4 | C | O3' | AA:1102 | 4.66 | 1.00 | 200.41 |  |
| 5 | C | O5' | AA:1102 | 4.92 | 1.00 | 200.37 |  |
  |

  
 Coded contact list:   
 Exclusion zone contact(s): /OP1/  
Consider K in place of MG (1 valid contact(s) for K vs. 0 for MG)  

### MG:AA:4052

|  | Res. type | Atom | Residue | Distance | Occupancy | B-factor | Closest solute to 1^st^ shell water |
| --- | --- | --- | --- | --- | --- | --- | --- |
| **Ion** | | | |  | 1.00 | 73.27 || 1 | G | OP1 | AA:1479 | 4.73 | 1.00 | 78.21 |
  |

  
 Coded contact list:   
1^st^ MG coordination shell missing (see Notes)  

### MG:AA:4053

|  | Res. type | Atom | Residue | Distance | Occupancy | B-factor | Closest solute to 1^st^ shell water |
| --- | --- | --- | --- | --- | --- | --- | --- |
| **Ion** | | | |  | 1.00 | 63.50 || 1 | U | OP2 | AA:2197 | 2.40 | 1.00 | 69.44 |
| 2 | G | O6 | AA:2224 | 3.04 | 1.00 | 53.21 |  |
| 3 | U | O5' | AA:2197 | 3.61 | 1.00 | 69.02 |  |
| 4 | G | N7 | AA:2224 | 4.02 | 1.00 | 53.55 |  |
| 5 | C | O5' | AA:2196 | 4.05 | 1.00 | 69.77 |  |
| 6 | C | O3' | AA:2196 | 4.08 | 1.00 | 67.14 |  |
| 7 | U | N1 | AA:2197 | 4.16 | 1.00 | 69.67 |  |
| 8 | U | O4 | AA:2197 | 4.56 | 1.00 | 70.63 |  |
| 9 | G | N1 | AA:2224 | 4.68 | 1.00 | 53.71 |  |
| 10 | C | OP2 | AA:2196 | 4.77 | 1.00 | 71.48 |  |
| 11 | U | O4' | AA:2197 | 4.78 | 1.00 | 69.53 |  |
| 12 | U | OP1 | AA:2197 | 4.83 | 1.00 | 70.16 |  |
| 13 | C | N1 | AA:2196 | 4.84 | 1.00 | 62.22 |  |
| 14 | U | N3 | AA:2197 | 4.95 | 1.00 | 68.99 |  |
  |

  
 Coded contact list:   
 Exclusion zone contact(s): /O6/OP2/  
Consider NA in place of MG (1 valid contact(s) for NA vs. 0 for MG)  

### MG:AA:4055

|  | Res. type | Atom | Residue | Distance | Occupancy | B-factor | Closest solute to 1^st^ shell water |
| --- | --- | --- | --- | --- | --- | --- | --- |
| **Ion** | | | |  | 1.00 | 110.06 || 1 | G | O2' | AA:1107 | 4.68 | 1.00 | 188.53 |
  |

  
 Coded contact list:   
1^st^ MG coordination shell missing (see Notes)  

### MG:AA:4056

|  | Res. type | Atom | Residue | Distance | Occupancy | B-factor | Closest solute to 1^st^ shell water |
| --- | --- | --- | --- | --- | --- | --- | --- |
| **Ion** | | | |  | 1.00 | 74.94 || 1 | G | O2' | AA:1581 | 2.66 | 1.00 | 102.44 |
| 2 | G | O4' | AA:1581 | 4.19 | 1.00 | 102.45 |  |
| 3 | G | N3 | AA:1581 | 4.58 | 1.00 | 95.65 |  |
| 4 | G | N9 | AA:1581 | 4.65 | 1.00 | 99.83 |  |
| 5 | A | N1 | AA:1580 | 4.83 | 1.00 | 86.83 |  |
  |

  
 Coded contact list:   
 Exclusion zone contact(s): /O2'/  
Consider K in place of MG (1 valid contact(s) for K vs. 0 for MG)  

### MG:AA:4057

|  | Res. type | Atom | Residue | Distance | Occupancy | B-factor | Closest solute to 1^st^ shell water |
| --- | --- | --- | --- | --- | --- | --- | --- |
| **Ion** | | | |  | 1.00 | 74.80 || 1 | C | OP2 | AA:2853 | 2.63 | 1.00 | 67.56 |
| 2 | C | OP1 | AA:2853 | 2.90 | 1.00 | 66.80 |  |
| 3 | G | N2 | AA:1455 | 3.54 | 1.00 | 76.49 |  |
| 4 | G | N3 | AA:1455 | 3.95 | 1.00 | 76.80 |  |
| 5 | G | O3' | AA:2852 | 4.30 | 1.00 | 67.42 |  |
| 6 | G | O4' | AA:1456 | 4.33 | 1.00 | 82.69 |  |
| 7 | C | O5' | AA:2853 | 4.43 | 1.00 | 69.06 |  |
  |

  
 Coded contact list:   
 Exclusion zone contact(s): /OP1/OP2/  
Consider K in place of MG (2 valid contact(s) for K vs. 0 for MG)  

### MG:AA:4058

|  | Res. type | Atom | Residue | Distance | Occupancy | B-factor | Closest solute to 1^st^ shell water |
| --- | --- | --- | --- | --- | --- | --- | --- |
| **Ion** | | | |  | 1.00 | 63.21 || 1 | U | OP2 | AA:2712 | 4.20 | 1.00 | 50.20 |
| 2 | C | OP1 | AA:1761 | 4.28 | 1.00 | 44.30 |  |
  |

  
 Coded contact list:   
1^st^ MG coordination shell missing (see Notes)  

### MG:AA:4059

|  | Res. type | Atom | Residue | Distance | Occupancy | B-factor | Closest solute to 1^st^ shell water |
| --- | --- | --- | --- | --- | --- | --- | --- |
| **Ion** | | | |  | 1.00 | 54.95 || 1 | A | N1 | AA:507 | 3.13 | 1.00 | 39.28 |
| 2 | G | N9 | AA:481 | 3.52 | 1.00 | 57.59 |  |
| 3 | A | OP1 | AA:482 | 3.58 | 1.00 | 59.43 |  |
| 4 | G | O3' | AA:481 | 3.70 | 1.00 | 59.38 |  |
| 5 | G | O4' | AA:481 | 3.80 | 1.00 | 60.84 |  |
| 6 | A | OP2 | AA:482 | 3.94 | 1.00 | 58.51 |  |
| 7 | G | O2' | AA:481 | 4.09 | 1.00 | 64.54 |  |
| 8 | A | N6 | AA:507 | 4.27 | 1.00 | 37.95 |  |
| 9 | G | N7 | AA:481 | 4.28 | 1.00 | 54.13 |  |
  |

  
 Coded contact list:   
 Exclusion zone contact(s): /N1_A_/  
Consider K in place of MG (1 valid contact(s) for K vs. 0 for MG)  

### MG:AA:4060

|  | Res. type | Atom | Residue | Distance | Occupancy | B-factor | Closest solute to 1^st^ shell water |
| --- | --- | --- | --- | --- | --- | --- | --- |
| **Ion** | | | |  | 1.00 | 70.07 || 1 | C | N4 | AA:377 | 3.30 | 1.00 | 46.00 |
| 2 | C | N4 | AA:376 | 3.87 | 1.00 | 55.16 |  |
| 3 | G | O6 | AA:397 | 4.20 | 1.00 | 46.77 |  |
| 4 | U | O4 | AA:395 | 4.21 | 1.00 | 46.98 |  |
| 5 | C | N4 | AA:378 | 4.72 | 1.00 | 51.93 |  |
| 6 | G | O6 | AA:396 | 4.95 | 1.00 | 48.96 |  |
  |

  
 Coded contact list:   
 Exclusion zone contact(s): /N4/  
Consider K in place of MG (1 valid contact(s) for K vs. 0 for MG)  

### MG:AA:4062

|  | Res. type | Atom | Residue | Distance | Occupancy | B-factor | Closest solute to 1^st^ shell water |
| --- | --- | --- | --- | --- | --- | --- | --- |
| **Ion** | | | |  | 1.00 | 81.18 || 1 | A | O4' | AA:1077 | 4.73 | 1.00 | 199.89 |
| 2 | A | O2' | AA:1077 | 5.00 | 1.00 | 199.75 |  |
  |

  
 Coded contact list:   
1^st^ MG coordination shell missing (see Notes)  

### MG:AA:4063

|  | Res. type | Atom | Residue | Distance | Occupancy | B-factor | Closest solute to 1^st^ shell water |
| --- | --- | --- | --- | --- | --- | --- | --- |
| **Ion** | | | |  | 1.00 | 75.20 || 1 | G | OP2 | AA:2677 | 2.47 | 1.00 | 65.84 |
| 2 | C | OP2 | AA:2676 | 4.31 | 1.00 | 60.20 |  |
| 3 | C | O5' | AA:2676 | 4.33 | 1.00 | 61.76 |  |
| 4 | G | OP1 | AA:2677 | 4.58 | 1.00 | 66.29 |  |
| 5 | C | O3' | AA:2676 | 4.71 | 1.00 | 65.50 |  |
| 6 | G | O5' | AA:2677 | 4.87 | 1.00 | 66.81 |  |
| 7 | C | OP1 | AA:2676 | 4.89 | 1.00 | 62.10 |  |
  |

  
 Coded contact list:   
 Exclusion zone contact(s): /OP2/  
Consider NA in place of MG (1 valid contact(s) for NA vs. 0 for MG)  

### MG:AA:4064

|  | Res. type | Atom | Residue | Distance | Occupancy | B-factor | Closest solute to 1^st^ shell water |
| --- | --- | --- | --- | --- | --- | --- | --- |
| **Ion** | | | |  | 1.00 | 76.99 || 1 | A | O5' | AA:1427 | 2.72 | 1.00 | 62.68 |
| 2 | G | N3 | AA:1559 | 3.37 | 1.00 | 91.87 |  |
| 3 | G | O3' | AA:1426 | 3.51 | 1.00 | 62.92 |  |
| 4 | A | OP2 | AA:1427 | 3.64 | 1.00 | 63.45 |  |
| 5 | G | N2 | AA:1559 | 3.95 | 1.00 | 94.34 |  |
| 6 | G | O4' | AA:1559 | 4.05 | 1.00 | 81.54 |  |
| 7 | G | N1 | AA:1559 | 4.28 | 1.00 | 94.35 |  |
| 8 | G | N9 | AA:1559 | 4.32 | 1.00 | 86.39 |  |
| 9 | G | OP1 | AA:1426 | 4.53 | 1.00 | 66.29 |  |
| 10 | G | O5' | AA:1426 | 4.56 | 1.00 | 64.85 |  |
| 11 | A | O3' | AA:1427 | 4.78 | 1.00 | 62.13 |  |
| 12 | G | OP1 | AA:1559 | 4.82 | 1.00 | 80.69 |  |
| 13 | A | OP1 | AA:1427 | 4.91 | 1.00 | 63.70 |  |
  |

  
 Coded contact list:   
 Exclusion zone contact(s): /N3_G_/O5'/  
Consider K in place of MG (2 valid contact(s) for K vs. 0 for MG)  

### MG:AA:4065

|  | Res. type | Atom | Residue | Distance | Occupancy | B-factor | Closest solute to 1^st^ shell water |
| --- | --- | --- | --- | --- | --- | --- | --- |
| **Ion** | | | |  | 1.00 | 68.49 || 1 | C | N4 | AA:691 | 3.91 | 1.00 | 39.77 |
| 2 | G | O6 | AA:690 | 3.94 | 1.00 | 47.64 |  |
| 3 | G | O6 | AA:771 | 4.09 | 1.00 | 39.52 |  |
| 4 | G | N7 | AA:690 | 4.81 | 1.00 | 44.04 |  |
  |

  
 Coded contact list:   
1^st^ MG coordination shell missing (see Notes)  

### MG:AA:4066

|  | Res. type | Atom | Residue | Distance | Occupancy | B-factor | Closest solute to 1^st^ shell water |
| --- | --- | --- | --- | --- | --- | --- | --- |
| **Ion** | | | |  | 1.00 | 82.82 || 1 | C | N4 | AA:654B | 3.75 | 1.00 | 194.08 |
| 2 | A | N7 | AA:654A | 4.03 | 1.00 | 184.93 |  |
| 3 | G | N7 | AA:654C | 4.14 | 1.00 | 200.30 |  |
| 4 | C | OP2 | AA:654B | 4.26 | 1.00 | 192.11 |  |
| 5 | G | OP2 | AA:654C | 4.86 | 1.00 | 198.00 |  |
| 6 | C | N3 | AA:654B | 4.88 | 1.00 | 194.15 |  |
| 7 | C | N1 | AA:654B | 4.93 | 1.00 | 195.16 |  |
| 8 | A | OP2 | AA:654A | 4.96 | 1.00 | 179.77 |  |
  |

  
 Coded contact list:   
1^st^ MG coordination shell missing (see Notes)  

### MG:AA:4067

|  | Res. type | Atom | Residue | Distance | Occupancy | B-factor | Closest solute to 1^st^ shell water |
| --- | --- | --- | --- | --- | --- | --- | --- |
| **Ion** | | | |  | 1.00 | 86.07 || 1 | G | N7 | AA:938 | 2.62 | 1.00 | 57.20 |
| 2 | G | OP2 | AA:938 | 4.21 | 1.00 | 64.49 |  |
| 3 | G | O6 | AA:938 | 4.32 | 1.00 | 55.85 |  |
| 4 | G | N9 | AA:938 | 4.44 | 1.00 | 57.48 |  |
| 5 | G | O5' | AA:938 | 4.71 | 1.00 | 63.94 |  |
  |

  
 Coded contact list:   
 Exclusion zone contact(s): /N7/  
Consider K in place of MG (1 valid contact(s) for K vs. 0 for MG)  

### MG:AA:4068

|  | Res. type | Atom | Residue | Distance | Occupancy | B-factor | Closest solute to 1^st^ shell water |
| --- | --- | --- | --- | --- | --- | --- | --- |
| **Ion** | | | |  | 1.00 | 58.69 || 1 | U | O4 | AA:803 | 2.92 | 1.00 | 37.98 |
| 2 | A | O5' | AA:675 | 2.99 | 1.00 | 36.31 |  |
| 3 | A | OP2 | AA:675 | 3.23 | 1.00 | 37.76 |  |
| 4 | A | OP1 | AA:802 | 3.35 | 1.00 | 37.46 |  |
| 5 | A | N7 | AA:802 | 3.48 | 1.00 | 45.10 |  |
| 6 | A | N6 | AA:804 | 3.80 | 1.00 | 48.74 |  |
| 7 | A | OP1 | AA:675 | 4.39 | 1.00 | 37.01 |  |
| 8 | A | N6 | AA:802 | 4.73 | 1.00 | 48.05 |  |
| 9 | A | O4' | AA:675 | 4.95 | 1.00 | 40.20 |  |
  |

  
 Coded contact list:   
 Exclusion zone contact(s): /O4/O5'/OP1/OP2/  
Consider K in place of MG (4 valid contact(s) for K vs. 0 for MG)  

### MG:AA:4069

|  | Res. type | Atom | Residue | Distance | Occupancy | B-factor | Closest solute to 1^st^ shell water |
| --- | --- | --- | --- | --- | --- | --- | --- |
| **Ion** | | | |  | 1.00 | 80.88 || 1 | C | OP1 | AA:2128 | 4.27 | 1.00 | 198.71 |
| 2 | C | OP1 | AA:2129 | 4.74 | 1.00 | 200.14 |  |
| 3 | A | N1 | AA:2173 | 4.96 | 1.00 | 200.41 |  |
  |

  
 Coded contact list:   
1^st^ MG coordination shell missing (see Notes)  

### MG:AA:4070

|  | Res. type | Atom | Residue | Distance | Occupancy | B-factor | Closest solute to 1^st^ shell water |
| --- | --- | --- | --- | --- | --- | --- | --- |
| **Ion** | | | |  | 1.00 | 60.91 || 1 | A | OP2 | AA:2014 | 2.70 | 1.00 | 42.59 |
| 2 | U | O4 | AA:1263 | 2.96 | 1.00 | 47.35 |  |
| 3 | G | O6 | AA:1264 | 3.02 | 1.00 | 51.66 |  |
| 4 | A | OP1 | AA:2014 | 3.49 | 1.00 | 42.01 |  |
| 5 | G | N1 | AA:1264 | 4.03 | 1.00 | 48.28 |  |
| 6 | A | O5' | AA:2014 | 4.68 | 1.00 | 43.81 |  |
| 7 | A | O3' | AA:2013 | 4.70 | 1.00 | 42.32 |  |
| 8 | A | N6 | AA:1265 | 4.94 | 1.00 | 48.40 |  |
  |

  
 Coded contact list:   
 Exclusion zone contact(s): /O4/O6/OP2/  
Consider K in place of MG (3 valid contact(s) for K vs. 0 for MG)  

### MG:AA:4071

|  | Res. type | Atom | Residue | Distance | Occupancy | B-factor | Closest solute to 1^st^ shell water |
| --- | --- | --- | --- | --- | --- | --- | --- |
| **Ion** | | | |  | 1.00 | 70.78 || 1 | A | OP2 | AA:608 | 2.61 | 1.00 | 68.42 |
| 2 | A | OP1 | AA:608 | 3.09 | 1.00 | 68.22 |  |
| 3 | G | N1 | AA:619 | 3.84 | 1.00 | 64.70 |  |
| 4 | G | N2 | AA:619 | 4.29 | 1.00 | 63.97 |  |
| 5 | A | O5' | AA:608 | 4.34 | 1.00 | 69.09 |  |
| 6 | C | N4 | AA:618A | 4.34 | 1.00 | 61.66 |  |
| 7 | G | O6 | AA:619 | 4.50 | 1.00 | 66.96 |  |
| 8 | U | O3' | AA:607 | 4.55 | 1.00 | 70.84 |  |
| 9 | G | O6 | AA:609A | 4.65 | 1.00 | 57.36 |  |
| 10 | A | OP2 | AA:609 | 4.70 | 1.00 | 66.78 |  |
  |

  
 Coded contact list:   
 Exclusion zone contact(s): /OP1/OP2/  
Consider K in place of MG (2 valid contact(s) for K vs. 0 for MG)  

### MG:AA:4072

|  | Res. type | Atom | Residue | Distance | Occupancy | B-factor | Closest solute to 1^st^ shell water |
| --- | --- | --- | --- | --- | --- | --- | --- |
| **Ion** | | | |  | 1.00 | 56.39 || 1 | A | O2' | AA:800 | 2.41 | 1.00 | 45.16 |
| 2 | G | OP2 | AA:801 | 3.34 | 1.00 | 36.34 |  |
| 3 | A | O3' | AA:800 | 3.72 | 1.00 | 38.84 |  |
| 4 | GLY | N | AF:61 | 4.10 | 1.00 | 50.13 |  |
| 5 | G | O5' | AA:801 | 4.15 | 1.00 | 37.30 |  |
| 6 | G | OP2 | AA:799 | 4.42 | 1.00 | 40.08 |  |
| 7 | ASP | OD2 | AF:77 | 4.66 | 1.00 | 51.96 |  |
| 8 | G | OP2 | AA:798 | 4.87 | 1.00 | 39.13 |  |
  |

  
 Coded contact list:   
 Exclusion zone contact(s): /O2'/OP2/  
Consider NA in place of MG (1 valid contact(s) for NA vs. 0 for MG)  

### MG:AA:4073

|  | Res. type | Atom | Residue | Distance | Occupancy | B-factor | Closest solute to 1^st^ shell water |
| --- | --- | --- | --- | --- | --- | --- | --- |
| **Ion** | | | |  | 1.00 | 84.77 || 1 | G | O2' | AA:1385 | 2.89 | 1.00 | 76.27 |
| 2 | G | OP1 | AA:1385 | 2.95 | 1.00 | 67.23 |  |
| 3 | G | N3 | AA:1385 | 3.61 | 1.00 | 74.69 |  |
| 4 | G | O4' | AA:1385 | 3.89 | 1.00 | 72.95 |  |
| 5 | C | O2 | AA:1403 | 4.16 | 1.00 | 69.77 |  |
| 6 | G | N9 | AA:1385 | 4.30 | 1.00 | 75.39 |  |
| 7 | G | O3' | AA:1385 | 4.35 | 1.00 | 71.58 |  |
| 8 | G | N2 | AA:1344 | 4.47 | 1.00 | 65.83 |  |
| 9 | C | O4' | AA:1386 | 4.52 | 1.00 | 69.73 |  |
| 10 | G | N2 | AA:1385 | 4.56 | 1.00 | 75.45 |  |
| 11 | G | OP2 | AA:1385 | 4.95 | 1.00 | 67.46 |  |
  |

  
 Coded contact list:   
 Exclusion zone contact(s): /O2'/OP1/  
Consider K in place of MG (2 valid contact(s) for K vs. 0 for MG)  

### MG:AA:4074

|  | Res. type | Atom | Residue | Distance | Occupancy | B-factor | Closest solute to 1^st^ shell water |
| --- | --- | --- | --- | --- | --- | --- | --- |
| **Ion** | | | |  | 1.00 | 94.94 || 1 | U | O4 | AA:99 | 3.16 | 1.00 | 91.54 |
| 2 | U | N3 | AA:99 | 3.91 | 1.00 | 93.03 |  |
| 3 | G | N2 | AA:98 | 4.99 | 1.00 | 57.92 |  |
  |

  
 Coded contact list:   
 Exclusion zone contact(s): /O4/  
Consider K in place of MG (1 valid contact(s) for K vs. 0 for MG)  

### MG:AA:4075

|  | Res. type | Atom | Residue | Distance | Occupancy | B-factor | Closest solute to 1^st^ shell water |
| --- | --- | --- | --- | --- | --- | --- | --- |
| **Ion** | | | |  | 1.00 | 132.46 || 1 | G | O2' | AA:654N | 4.88 | 1.00 | 200.00 |
  |

  
 Coded contact list:   
1^st^ MG coordination shell missing (see Notes)  

### MG:AA:4076

|  | Res. type | Atom | Residue | Distance | Occupancy | B-factor | Closest solute to 1^st^ shell water |
| --- | --- | --- | --- | --- | --- | --- | --- |
| **Ion** | | | |  | 1.00 | 113.64 || 1 | A | OP2 | AA:1048 | 1.97 | 1.00 | 184.04 |
| 2 | C | N3 | AA:1109 | 2.74 | 1.00 | 181.26 |  |
| 3 | C | N4 | AA:1109 | 3.17 | 1.00 | 181.43 |  |
| 4 | G | N2 | AA:1110 | 3.18 | 1.00 | 176.51 |  |
| 5 | A | N7 | AA:1048 | 3.73 | 1.00 | 179.36 |  |
| 6 | A | O5' | AA:1048 | 3.74 | 1.00 | 183.38 |  |
| 7 | C | O2 | AA:1109 | 3.94 | 1.00 | 181.17 |  |
| 8 | U | O4 | AA:1108 | 4.06 | 1.00 | 184.27 |  |
| 9 | G | O3' | AA:1047 | 4.25 | 1.00 | 184.49 |  |
| 10 | A | O4' | AA:1048 | 4.30 | 1.00 | 180.84 |  |
| 11 | G | O2' | AA:1047 | 4.31 | 1.00 | 185.41 |  |
| 12 | A | N6 | AA:1050 | 4.41 | 1.00 | 173.03 |  |
| 13 | A | OP1 | AA:1048 | 4.42 | 1.00 | 184.20 |  |
| 14 | A | N9 | AA:1048 | 4.49 | 1.00 | 179.98 |  |
| 15 | G | N3 | AA:1110 | 4.76 | 1.00 | 176.65 |  |
| 16 | C | N4 | AA:1049 | 4.77 | 1.00 | 177.88 |  |
| 17 | G | N1 | AA:1110 | 4.89 | 1.00 | 176.57 |  |
  |

  
 Contact(s): /OP2/  
 Coded contact list: O_ph_  
 Exclusion zone contact(s): /N2/N3_C_/N4/  
Consider K in place of MG (3 valid contact(s) for K vs. 1 for MG)  

### MG:AA:4077

|  | Res. type | Atom | Residue | Distance | Occupancy | B-factor | Closest solute to 1^st^ shell water |
| --- | --- | --- | --- | --- | --- | --- | --- |
| **Ion** | | | |  | 1.00 | 67.56 || 1 | C | N4 | AA:976 | 3.49 | 1.00 | 80.15 |
| 2 | G | O6 | AA:977 | 3.51 | 1.00 | 70.05 |  |
| 3 | G | N7 | AA:977 | 4.36 | 1.00 | 72.09 |  |
| 4 | G | O6 | AA:978 | 4.43 | 1.00 | 66.68 |  |
| 5 | G | O6 | AA:987 | 4.47 | 1.00 | 76.65 |  |
| 6 | C | N3 | AA:976 | 4.77 | 1.00 | 78.64 |  |
| 7 | C | OP2 | AA:974A | 4.87 | 1.00 | 71.48 |  |
  |

  
 Coded contact list:   
1^st^ MG coordination shell missing (see Notes)  

### MG:AA:4078

|  | Res. type | Atom | Residue | Distance | Occupancy | B-factor | Closest solute to 1^st^ shell water |
| --- | --- | --- | --- | --- | --- | --- | --- |
| **Ion** | | | |  | 1.00 | 82.60 || 1 | G | OP2 | AA:1421 | 3.04 | 1.00 | 93.71 |
| 2 | G | O6 | AA:1423 | 4.21 | 1.00 | 68.40 |  |
| 3 | G | OP2 | AA:1422 | 4.66 | 1.00 | 77.24 |  |
| 4 | G | N7 | AA:1422 | 4.70 | 1.00 | 67.51 |  |
| 5 | G | N7 | AA:1421 | 4.79 | 1.00 | 83.07 |  |
| 6 | G | OP1 | AA:1421 | 4.86 | 1.00 | 94.17 |  |
| 7 | G | N7 | AA:1423 | 4.90 | 1.00 | 66.21 |  |
  |

  
 Coded contact list:   
 Exclusion zone contact(s): /OP2/  
Consider K in place of MG (1 valid contact(s) for K vs. 0 for MG)  

### MG:AA:4079

|  | Res. type | Atom | Residue | Distance | Occupancy | B-factor | Closest solute to 1^st^ shell water |
| --- | --- | --- | --- | --- | --- | --- | --- |
| **Ion** | | | |  | 1.00 | 92.33 || 1 | G | OP1 | AA:48 | 2.80 | 1.00 | 58.56 |
| 2 | A | OP1 | AA:49 | 3.07 | 1.00 | 61.48 |  |
| 3 | G | OP2 | AA:48 | 3.13 | 1.00 | 58.16 |  |
| 4 | G | O5' | AA:48 | 3.68 | 1.00 | 59.11 |  |
| 5 | A | OP2 | AA:49 | 3.81 | 1.00 | 60.55 |  |
| 6 | G | O3' | AA:48 | 4.73 | 1.00 | 60.73 |  |
| 7 | C | O3' | AA:47 | 4.81 | 1.00 | 59.09 |  |
  |

  
 Coded contact list:   
 Exclusion zone contact(s): /OP1/OP1/OP2/  
Consider K in place of MG (3 valid contact(s) for K vs. 0 for MG)  

### MG:AA:4080

|  | Res. type | Atom | Residue | Distance | Occupancy | B-factor | Closest solute to 1^st^ shell water |
| --- | --- | --- | --- | --- | --- | --- | --- |
| **Ion** | | | |  | 1.00 | 71.34 || 1 | U | OP1 | AA:1282 | 3.02 | 1.00 | 57.14 |
| 2 | G | O3' | AA:1281 | 3.70 | 1.00 | 56.17 |  |
| 3 | G | O2' | AA:1281 | 3.84 | 1.00 | 55.66 |  |
| 4 | U | O5' | AA:1282 | 4.01 | 1.00 | 56.85 |  |
  |

  
 Coded contact list:   
 Exclusion zone contact(s): /OP1/  
Consider K in place of MG (1 valid contact(s) for K vs. 0 for MG)  

### MG:AA:4081

|  | Res. type | Atom | Residue | Distance | Occupancy | B-factor | Closest solute to 1^st^ shell water |
| --- | --- | --- | --- | --- | --- | --- | --- |
| **Ion** | | | |  | 1.00 | 49.24 || 1 | U | OP2 | AA:740 | 3.03 | 1.00 | 39.25 |
| 2 | U | O5' | AA:740 | 3.27 | 1.00 | 37.33 |  |
| 3 | G | OP2 | AA:741 | 3.40 | 1.00 | 43.04 |  |
| 4 | G | O2' | AA:739 | 3.43 | 1.00 | 43.93 |  |
| 5 | G | O3' | AA:739 | 3.64 | 1.00 | 40.55 |  |
| 6 | G | N7 | AA:741 | 4.51 | 1.00 | 47.39 |  |
| 7 | U | O3' | AA:740 | 4.98 | 1.00 | 41.16 |  |
  |

  
 Coded contact list:   
 Exclusion zone contact(s): /O5'/OP2/  
Consider K in place of MG (2 valid contact(s) for K vs. 0 for MG)  

### MG:AA:4082

|  | Res. type | Atom | Residue | Distance | Occupancy | B-factor | Closest solute to 1^st^ shell water |
| --- | --- | --- | --- | --- | --- | --- | --- |
| **Ion** | | | |  | 1.00 | 75.67 || 1 | G | O6 | AA:240 | 2.60 | 1.00 | 47.01 |
| 2 | U | O4 | AA:239 | 2.91 | 1.00 | 56.02 |  |
| 3 | G | N1 | AA:240 | 3.35 | 1.00 | 45.96 |  |
| 4 | A | OP2 | AA:256 | 4.63 | 1.00 | 45.38 |  |
| 5 | U | N3 | AA:239 | 4.86 | 1.00 | 54.13 |  |
  |

  
 Coded contact list:   
 Exclusion zone contact(s): /N1_G_/O4/O6/  
Consider K in place of MG (3 valid contact(s) for K vs. 0 for MG)  

### MG:AA:4083

|  | Res. type | Atom | Residue | Distance | Occupancy | B-factor | Closest solute to 1^st^ shell water |
| --- | --- | --- | --- | --- | --- | --- | --- |
| **Ion** | | | |  | 1.00 | 74.58 || 1 | G | O3' | AA:352 | 2.82 | 1.00 | 103.71 |
| 2 | G | OP2 | AA:353 | 2.83 | 1.00 | 103.36 |  |
| 3 | G | N7 | AA:354 | 4.06 | 1.00 | 99.30 |  |
| 4 | G | O5' | AA:352 | 4.09 | 1.00 | 97.35 |  |
| 5 | G | OP2 | AA:354 | 4.10 | 1.00 | 101.33 |  |
| 6 | G | O5' | AA:353 | 4.14 | 1.00 | 103.38 |  |
| 7 | G | N7 | AA:353 | 4.14 | 1.00 | 96.84 |  |
| 8 | G | OP1 | AA:353 | 4.74 | 1.00 | 103.41 |  |
| 9 | G | N9 | AA:353 | 4.89 | 1.00 | 99.03 |  |
  |

  
 Coded contact list:   
 Exclusion zone contact(s): /O3'/OP2/  
Consider K in place of MG (2 valid contact(s) for K vs. 0 for MG)  

### MG:AA:4084

|  | Res. type | Atom | Residue | Distance | Occupancy | B-factor | Closest solute to 1^st^ shell water |
| --- | --- | --- | --- | --- | --- | --- | --- |
| **Ion** | | | |  | 1.00 | 86.89 || 1 | G | OP2 | AA:2759 | 2.61 | 1.00 | 129.54 |
| 2 | A | OP2 | AA:2758 | 2.89 | 1.00 | 136.30 |  |
| 3 | U | O2 | AA:2756 | 3.50 | 1.00 | 134.22 |  |
| 4 | G | N7 | AA:2759 | 3.65 | 1.00 | 127.05 |  |
| 5 | U | O2' | AA:2756 | 3.71 | 1.00 | 133.76 |  |
| 6 | G | O5' | AA:2759 | 3.96 | 1.00 | 129.22 |  |
| 7 | A | O5' | AA:2758 | 4.15 | 1.00 | 135.54 |  |
| 8 | A | N7 | AA:2758 | 4.34 | 1.00 | 136.21 |  |
| 9 | A | O3' | AA:2758 | 4.49 | 1.00 | 131.46 |  |
| 10 | A | N9 | AA:2758 | 4.73 | 1.00 | 135.21 |  |
| 11 | G | N9 | AA:2759 | 4.85 | 1.00 | 127.24 |  |
| 12 | G | O4' | AA:2759 | 4.95 | 1.00 | 127.51 |  |
| 13 | A | OP1 | AA:2758 | 4.97 | 1.00 | 136.34 |  |
  |

  
 Coded contact list:   
 Exclusion zone contact(s): /OP2/OP2/  
Consider K in place of MG (2 valid contact(s) for K vs. 0 for MG)  

### MG:AA:4085

|  | Res. type | Atom | Residue | Distance | Occupancy | B-factor | Closest solute to 1^st^ shell water |
| --- | --- | --- | --- | --- | --- | --- | --- |
| **Ion** | | | |  | 1.00 | 60.90 || 1 | G | O2' | AA:2351 | 2.66 | 1.00 | 68.92 |
| 2 | G | O2' | AA:647 | 2.74 | 1.00 | 80.90 |  |
| 3 | G | O3' | AA:647 | 2.83 | 1.00 | 82.18 |  |
| 4 | G | O3' | AA:2351 | 3.28 | 1.00 | 70.63 |  |
| 5 | A | O4' | AA:2352 | 3.35 | 1.00 | 66.51 |  |
| 6 | G | O4' | AA:2351 | 3.61 | 1.00 | 68.95 |  |
| 7 | G | OP1 | AA:648 | 3.88 | 1.00 | 82.41 |  |
| 8 | G | O5' | AA:648 | 4.17 | 1.00 | 80.90 |  |
| 9 | A | O5' | AA:2352 | 4.17 | 1.00 | 70.72 |  |
| 10 | G | O4' | AA:648 | 4.51 | 1.00 | 73.16 |  |
  |

  
 Coded contact list:   
 Exclusion zone contact(s): /O2'/O2'/O3'/O3'/O4'/  
Consider K in place of MG (5 valid contact(s) for K vs. 0 for MG)  

### MG:AA:4086

|  | Res. type | Atom | Residue | Distance | Occupancy | B-factor | Closest solute to 1^st^ shell water |
| --- | --- | --- | --- | --- | --- | --- | --- |
| **Ion** | | | |  | 1.00 | 76.20 || 1 | A | OP2 | AA:429 | 3.41 | 1.00 | 61.05 |
| 2 | A | N6 | AA:218 | 3.68 | 1.00 | 51.10 |  |
| 3 | U | O4 | AA:431 | 3.91 | 1.00 | 50.09 |  |
| 4 | G | N7 | AA:217 | 4.18 | 1.00 | 51.47 |  |
| 5 | U | N3 | AA:431 | 4.36 | 1.00 | 50.08 |  |
| 6 | G | O6 | AA:217 | 4.49 | 1.00 | 57.07 |  |
| 7 | G | N7 | AA:430 | 4.63 | 1.00 | 64.28 |  |
| 8 | G | OP2 | AA:430 | 4.82 | 1.00 | 59.96 |  |
| 9 | G | OP2 | AA:217 | 4.82 | 1.00 | 54.45 |  |
| 10 | A | O5' | AA:429 | 4.94 | 1.00 | 61.37 |  |
| 11 | A | N7 | AA:429 | 5.00 | 1.00 | 63.10 |  |
  |

  
 Coded contact list:   
1^st^ MG coordination shell missing (see Notes)  

### MG:AA:4087

|  | Res. type | Atom | Residue | Distance | Occupancy | B-factor | Closest solute to 1^st^ shell water |
| --- | --- | --- | --- | --- | --- | --- | --- |
| **Ion** | | | |  | 1.00 | 89.74 || 1 | G | OP2 | AA:1283 | 2.92 | 1.00 | 57.40 |
| 2 | G | N7 | AA:1283 | 3.54 | 1.00 | 66.81 |  |
| 3 | G | O5' | AA:1283 | 3.95 | 1.00 | 58.31 |  |
| 4 | U | OP2 | AA:1282 | 4.59 | 1.00 | 56.68 |  |
| 5 | U | O5' | AA:1282 | 4.68 | 1.00 | 56.85 |  |
| 6 | A | OP2 | AA:1284 | 4.74 | 1.00 | 58.38 |  |
| 7 | G | N9 | AA:1283 | 4.83 | 1.00 | 63.03 |  |
| 8 | U | O3' | AA:1282 | 4.96 | 1.00 | 56.38 |  |
  |

  
 Coded contact list:   
 Exclusion zone contact(s): /OP2/  
Consider K in place of MG (1 valid contact(s) for K vs. 0 for MG)  

### MG:AA:4088

|  | Res. type | Atom | Residue | Distance | Occupancy | B-factor | Closest solute to 1^st^ shell water |
| --- | --- | --- | --- | --- | --- | --- | --- |
| **Ion** | | | |  | 1.00 | 97.02 || 1 | G | O6 | AA:2708 | 2.98 | 1.00 | 47.55 |
| 2 | U | O4 | AA:2698 | 3.14 | 1.00 | 54.06 |  |
| 3 | G | O6 | AA:2707 | 3.90 | 1.00 | 56.68 |  |
| 4 | C | N4 | AA:2699 | 3.90 | 1.00 | 48.00 |  |
| 5 | G | O6 | AA:2709 | 4.64 | 1.00 | 49.51 |  |
| 6 | G | N7 | AA:2708 | 4.68 | 1.00 | 49.14 |  |
| 7 | U | N3 | AA:2698 | 4.92 | 1.00 | 49.62 |  |
  |

  
 Coded contact list:   
 Exclusion zone contact(s): /O4/O6/  
Consider K in place of MG (2 valid contact(s) for K vs. 0 for MG)  

### MG:AA:4089

|  | Res. type | Atom | Residue | Distance | Occupancy | B-factor | Closest solute to 1^st^ shell water |
| --- | --- | --- | --- | --- | --- | --- | --- |
| **Ion** | | | |  | 1.00 | 107.82 || 1 | C | O2' | AA:654G | 3.31 | 1.00 | 200.41 |
| 2 | C | O3' | AA:654G | 4.30 | 1.00 | 200.41 |  |
| 3 | C | O4' | AA:654G | 4.44 | 1.00 | 200.41 |  |
| 4 | C | O2 | AA:654G | 4.90 | 1.00 | 200.33 |  |
  |

  
 Coded contact list:   
 Exclusion zone contact(s): /O2'/  
Consider K in place of MG (1 valid contact(s) for K vs. 0 for MG)  

### MG:AA:4090

|  | Res. type | Atom | Residue | Distance | Occupancy | B-factor | Closest solute to 1^st^ shell water |
| --- | --- | --- | --- | --- | --- | --- | --- |
| **Ion** | | | |  | 1.00 | 78.03 || 1 | G | O6 | AA:488 | 4.31 | 1.00 | 54.95 |
  |

  
 Coded contact list:   
1^st^ MG coordination shell missing (see Notes)  

### MG:AA:4091

|  | Res. type | Atom | Residue | Distance | Occupancy | B-factor | Closest solute to 1^st^ shell water |
| --- | --- | --- | --- | --- | --- | --- | --- |
| **Ion** | | | |  | 1.00 | 107.43 || 1 | G | OP2 | AA:1512 | 3.60 | 1.00 | 116.15 |
| 2 | G | OP1 | AA:1512 | 3.68 | 1.00 | 116.05 |  |
| 3 | A | O3' | AA:1511 | 4.48 | 1.00 | 117.96 |  |
| 4 | A | OP1 | AA:1511 | 4.77 | 1.00 | 131.78 |  |
  |

  
 Coded contact list:   
1^st^ MG coordination shell missing (see Notes)  

### MG:AA:4092

|  | Res. type | Atom | Residue | Distance | Occupancy | B-factor | Closest solute to 1^st^ shell water |
| --- | --- | --- | --- | --- | --- | --- | --- |
| **Ion** | | | |  | 1.00 | 90.37 || 1 | G | O2' | AA:622 | 2.36 | 1.00 | 59.86 |
| 2 | G | N3 | AA:622 | 2.79 | 1.00 | 59.82 |  |
| 3 | G | O4' | AA:623 | 2.79 | 1.00 | 60.99 |  |
| 4 | G | N9 | AA:622 | 3.78 | 1.00 | 60.10 |  |
| 5 | U | O3' | AA:239 | 3.80 | 1.00 | 53.81 |  |
| 6 | G | N2 | AA:622 | 3.85 | 1.00 | 57.22 |  |
| 7 | G | N9 | AA:623 | 4.22 | 1.00 | 60.86 |  |
| 8 | U | O2' | AA:239 | 4.31 | 1.00 | 55.83 |  |
| 9 | G | O4' | AA:622 | 4.35 | 1.00 | 59.76 |  |
| 10 | G | O5' | AA:623 | 4.41 | 1.00 | 60.19 |  |
| 11 | G | OP1 | AA:240 | 4.59 | 1.00 | 54.94 |  |
| 12 | A | N1 | AA:621 | 4.89 | 1.00 | 79.42 |  |
| 13 | G | O3' | AA:622 | 4.95 | 1.00 | 59.68 |  |
  |

  
 Contact(s): /O2'/  
 Coded contact list: O_r_  
 Exclusion zone contact(s): /N3_G_/O4'/  
Consider K in place of MG (2 valid contact(s) for K vs. 1 for MG)  

### MG:AA:4094

|  | Res. type | Atom | Residue | Distance | Occupancy | B-factor | Closest solute to 1^st^ shell water |
| --- | --- | --- | --- | --- | --- | --- | --- |
| **Ion** | | | |  | 1.00 | 57.39 || 1 | A | N3 | AA:1596 | 2.45 | 1.00 | 63.48 |
| 2 | C | O2 | AA:1407 | 2.94 | 1.00 | 63.52 |  |
| 3 | A | O2' | AA:141 | 3.35 | 1.00 | 71.26 |  |
| 4 | A | O2' | AA:1596 | 3.43 | 1.00 | 63.38 |  |
| 5 | A | N9 | AA:1596 | 3.70 | 1.00 | 63.22 |  |
| 6 | G | N2 | AA:1595 | 3.79 | 1.00 | 65.16 |  |
| 7 | A | N9 | AA:141 | 3.98 | 1.00 | 73.00 |  |
| 8 | C | O2' | AA:1407 | 4.28 | 1.00 | 59.28 |  |
| 9 | A | O4' | AA:1597 | 4.31 | 1.00 | 60.99 |  |
| 10 | A | O3' | AA:141 | 4.38 | 1.00 | 67.95 |  |
| 11 | A | O4' | AA:1596 | 4.55 | 1.00 | 64.67 |  |
| 12 | A | N1 | AA:1596 | 4.61 | 1.00 | 61.42 |  |
| 13 | A | N7 | AA:141 | 4.69 | 1.00 | 72.79 |  |
| 14 | C | N1 | AA:1407 | 4.75 | 1.00 | 63.00 |  |
| 15 | C | O4' | AA:141A | 4.75 | 1.00 | 64.27 |  |
| 16 | C | N3 | AA:1407 | 4.85 | 1.00 | 63.51 |  |
  |

  
 Coded contact list:   
 Exclusion zone contact(s): /N3_A_/O2/O2'/  
Consider K in place of MG (2 valid contact(s) for K vs. 0 for MG)  

### MG:AA:4095

|  | Res. type | Atom | Residue | Distance | Occupancy | B-factor | Closest solute to 1^st^ shell water |
| --- | --- | --- | --- | --- | --- | --- | --- |
| **Ion** | | | |  | 1.00 | 58.71 || 1 | G | OP1 | AA:642 | 2.69 | 1.00 | 71.06 |
| 2 | C | O3' | AA:641 | 4.14 | 1.00 | 69.68 |  |
| 3 | G | O5' | AA:642 | 4.53 | 1.00 | 72.87 |  |
| 4 | G | OP1 | AA:2349 | 4.78 | 1.00 | 62.06 |  |
| 5 | C | O2' | AA:641 | 4.80 | 1.00 | 67.88 |  |
| 6 | C | O2' | AA:2417 | 4.98 | 1.00 | 64.74 |  |
  |

  
 Coded contact list:   
 Exclusion zone contact(s): /OP1/  
Consider K in place of MG (1 valid contact(s) for K vs. 0 for MG)  

### MG:AA:4096

|  | Res. type | Atom | Residue | Distance | Occupancy | B-factor | Closest solute to 1^st^ shell water |
| --- | --- | --- | --- | --- | --- | --- | --- |
| **Ion** | | | |  | 1.00 | 114.31 || 1 | A | OP1 | AA:216 | 3.57 | 1.00 | 59.28 |
| 2 | A | OP1 | AA:429 | 4.84 | 1.00 | 61.86 |  |
  |

  
 Coded contact list:   
1^st^ MG coordination shell missing (see Notes)  

### MG:AA:4098

|  | Res. type | Atom | Residue | Distance | Occupancy | B-factor | Closest solute to 1^st^ shell water |
| --- | --- | --- | --- | --- | --- | --- | --- |
| **Ion** | | | |  | 1.00 | 83.11 || 1 | C | N4 | AA:376 | 2.93 | 1.00 | 55.16 |
| 2 | C | N3 | AA:375 | 3.36 | 1.00 | 52.61 |  |
| 3 | C | N4 | AA:375 | 3.50 | 1.00 | 51.54 |  |
| 4 | C | N1 | AA:375 | 3.56 | 1.00 | 52.90 |  |
| 5 | C | OP2 | AA:375 | 4.12 | 1.00 | 59.73 |  |
| 6 | C | O5' | AA:375 | 4.33 | 1.00 | 59.07 |  |
| 7 | C | N3 | AA:376 | 4.42 | 1.00 | 53.49 |  |
| 8 | C | OP2 | AA:376 | 4.46 | 1.00 | 55.39 |  |
| 9 | C | O2 | AA:375 | 4.55 | 1.00 | 52.96 |  |
| 10 | C | N4 | AA:377 | 4.61 | 1.00 | 46.00 |  |
| 11 | C | N1 | AA:376 | 4.77 | 1.00 | 53.75 |  |
| 12 | A | OP2 | AA:374 | 4.81 | 1.00 | 58.32 |  |
  |

  
 Coded contact list:   
 Exclusion zone contact(s): /N3_C_/N4/  
Consider K in place of MG (2 valid contact(s) for K vs. 0 for MG)  

### MG:AA:4099

|  | Res. type | Atom | Residue | Distance | Occupancy | B-factor | Closest solute to 1^st^ shell water |
| --- | --- | --- | --- | --- | --- | --- | --- |
| **Ion** | | | |  | 1.00 | 96.24 || 1 | G | OP1 | AA:615 | 2.91 | 1.00 | 104.16 |
| 2 | U | OP1 | AA:614 | 3.19 | 1.00 | 88.94 |  |
| 3 | U | O5' | AA:614 | 3.60 | 1.00 | 92.60 |  |
| 4 | U | O3' | AA:613 | 4.49 | 1.00 | 85.60 |  |
  |

  
 Coded contact list:   
 Exclusion zone contact(s): /OP1/OP1/  
Consider K in place of MG (2 valid contact(s) for K vs. 0 for MG)  

### MG:AA:4100

|  | Res. type | Atom | Residue | Distance | Occupancy | B-factor | Closest solute to 1^st^ shell water |
| --- | --- | --- | --- | --- | --- | --- | --- |
| **Ion** | | | |  | 1.00 | 103.15 || 1 | G | O4' | AA:1044 | 2.50 | 1.00 | 191.79 |
| 2 | G | O2' | AA:1044 | 2.62 | 1.00 | 191.89 |  |
| 3 | A | N9 | AA:1048 | 2.83 | 1.00 | 179.98 |  |
| 4 | A | N3 | AA:1048 | 2.86 | 1.00 | 179.03 |  |
| 5 | G | O2' | AA:1047 | 2.97 | 1.00 | 185.41 |  |
| 6 | A | N1 | AA:1111 | 3.30 | 1.00 | 177.26 |  |
| 7 | A | O4' | AA:1048 | 3.83 | 1.00 | 180.84 |  |
| 8 | G | N9 | AA:1044 | 3.95 | 1.00 | 190.93 |  |
| 9 | A | N7 | AA:1048 | 4.02 | 1.00 | 179.36 |  |
| 10 | A | N6 | AA:1111 | 4.09 | 1.00 | 177.42 |  |
| 11 | A | N1 | AA:1048 | 4.30 | 1.00 | 178.29 |  |
| 12 | G | O4' | AA:1047 | 4.32 | 1.00 | 185.21 |  |
| 13 | A | N3 | AA:1111 | 4.50 | 1.00 | 177.17 |  |
| 14 | G | O3' | AA:1044 | 4.69 | 1.00 | 192.98 |  |
| 15 | G | O3' | AA:1047 | 4.70 | 1.00 | 184.49 |  |
| 16 | G | N3 | AA:1044 | 4.75 | 1.00 | 189.68 |  |
| 17 | A | O2' | AA:1048 | 4.82 | 1.00 | 180.32 |  |
  |

  
 Coded contact list:   
 Exclusion zone contact(s): /N1_A_/N3_A_/O2'/O2'/O4'/  
Consider K in place of MG (5 valid contact(s) for K vs. 0 for MG)  

### MG:AA:4101

|  | Res. type | Atom | Residue | Distance | Occupancy | B-factor | Closest solute to 1^st^ shell water |
| --- | --- | --- | --- | --- | --- | --- | --- |
| **Ion** | | | |  | 1.00 | 99.83 || 1 | G | OP2 | AA:642 | 2.46 | 1.00 | 71.59 |
| 2 | C | OP1 | AA:641 | 4.07 | 1.00 | 65.64 |  |
| 3 | C | OP2 | AA:641 | 4.29 | 1.00 | 65.86 |  |
| 4 | G | O5' | AA:642 | 4.36 | 1.00 | 72.87 |  |
| 5 | C | O2 | AA:2402 | 4.40 | 1.00 | 105.39 |  |
| 6 | G | OP1 | AA:642 | 4.54 | 1.00 | 71.06 |  |
| 7 | G | N7 | AA:642 | 4.73 | 1.00 | 75.77 |  |
| 8 | A | OP2 | AA:643 | 4.86 | 1.00 | 82.18 |  |
| 9 | C | O5' | AA:641 | 4.93 | 1.00 | 65.72 |  |
| 10 | C | O3' | AA:641 | 4.96 | 1.00 | 69.68 |  |
  |

  
 Coded contact list:   
 Exclusion zone contact(s): /OP2/  
Consider NA in place of MG (1 valid contact(s) for NA vs. 0 for MG)  

### MG:AA:4102

|  | Res. type | Atom | Residue | Distance | Occupancy | B-factor | Closest solute to 1^st^ shell water |
| --- | --- | --- | --- | --- | --- | --- | --- |
| **Ion** | | | |  | 1.00 | 90.52 || 1 | A | OP1 | AA:513 | 3.94 | 1.00 | 58.29 |
| 2 | G | OP2 | AA:23 | 4.33 | 1.00 | 60.20 |  |
| 3 | G | N7 | AA:23 | 4.72 | 1.00 | 55.36 |  |
  |

  
 Coded contact list:   
1^st^ MG coordination shell missing (see Notes)  

### MG:AA:4103

|  | Res. type | Atom | Residue | Distance | Occupancy | B-factor | Closest solute to 1^st^ shell water |
| --- | --- | --- | --- | --- | --- | --- | --- |
| **Ion** | | | |  | 1.00 | 54.32 || 1 | G | N3 | AA:98 | 4.16 | 1.00 | 56.91 |
| 2 | U | O4' | AA:99 | 4.46 | 1.00 | 84.25 |  |
| 3 | U | O2 | AA:99 | 4.47 | 1.00 | 90.99 |  |
| 4 | U | OP2 | AA:99 | 4.53 | 1.00 | 70.47 |  |
| 5 | U | N1 | AA:99 | 4.63 | 1.00 | 88.20 |  |
| 6 | U | N3 | AA:99 | 4.66 | 1.00 | 93.03 |  |
| 7 | G | N2 | AA:98 | 4.66 | 1.00 | 57.92 |  |
| 8 | G | N1 | AA:98 | 4.67 | 1.00 | 57.66 |  |
| 9 | U | O5' | AA:99 | 4.88 | 1.00 | 74.14 |  |
| 10 | G | N9 | AA:98 | 4.93 | 1.00 | 58.66 |  |
  |

  
 Coded contact list:   
1^st^ MG coordination shell missing (see Notes)  

### MG:AA:4104

|  | Res. type | Atom | Residue | Distance | Occupancy | B-factor | Closest solute to 1^st^ shell water |
| --- | --- | --- | --- | --- | --- | --- | --- |
| **Ion** | | | |  | 1.00 | 71.98 || 1 | A | OP2 | AA:49 | 3.26 | 1.00 | 60.55 |
| 2 | G | N7 | AA:48 | 3.39 | 1.00 | 58.06 |  |
| 3 | G | N9 | AA:48 | 3.74 | 1.00 | 58.47 |  |
| 4 | G | N3 | AA:48 | 4.19 | 1.00 | 60.00 |  |
| 5 | G | O6 | AA:48 | 4.29 | 1.00 | 60.77 |  |
| 6 | G | N1 | AA:48 | 4.31 | 1.00 | 59.85 |  |
| 7 | G | O3' | AA:48 | 4.73 | 1.00 | 60.73 |  |
| 8 | G | N2 | AA:177 | 4.76 | 1.00 | 53.20 |  |
| 9 | G | OP2 | AA:176 | 4.76 | 1.00 | 72.85 |  |
| 10 | G | OP1 | AA:176 | 4.84 | 1.00 | 74.02 |  |
  |

  
 Coded contact list:   
 Exclusion zone contact(s): /N7/OP2/  
Consider K in place of MG (2 valid contact(s) for K vs. 0 for MG)  

### MG:AA:4105

|  | Res. type | Atom | Residue | Distance | Occupancy | B-factor | Closest solute to 1^st^ shell water |
| --- | --- | --- | --- | --- | --- | --- | --- |
| **Ion** | | | |  | 1.00 | 106.80 || 1 | G | OP2 | AA:2154 | 2.61 | 1.00 | 200.41 |
| 2 | G | O5' | AA:2154 | 4.41 | 1.00 | 200.41 |  |
| 3 | G | OP1 | AA:2154 | 4.52 | 1.00 | 200.41 |  |
| 4 | G | OP2 | AA:2155 | 4.96 | 1.00 | 200.32 |  |
  |

  
 Coded contact list:   
 Exclusion zone contact(s): /OP2/  
Consider K in place of MG (1 valid contact(s) for K vs. 0 for MG)  

### MG:AA:4106

|  | Res. type | Atom | Residue | Distance | Occupancy | B-factor | Closest solute to 1^st^ shell water |
| --- | --- | --- | --- | --- | --- | --- | --- |
| **Ion** | | | |  | 1.00 | 88.85 || 1 | G | O6 | AA:771 | 2.55 | 1.00 | 39.52 |
| 2 | G | N7 | AA:771 | 2.91 | 1.00 | 43.00 |  |
| 3 | G | O6 | AA:770 | 3.54 | 1.00 | 42.39 |  |
| 4 | C | N4 | AA:772 | 4.27 | 1.00 | 44.09 |  |
| 5 | G | OP2 | AA:686 | 4.52 | 1.00 | 38.33 |  |
| 6 | G | N1 | AA:771 | 4.62 | 1.00 | 40.21 |  |
| 7 | G | N7 | AA:770 | 4.63 | 1.00 | 38.49 |  |
| 8 | C | N4 | AA:691 | 4.75 | 1.00 | 39.77 |  |
  |

  
 Coded contact list:   
 Exclusion zone contact(s): /N7/O6/  
Consider NA in place of MG (1 valid contact(s) for NA vs. 0 for MG)  

### MG:AA:4107

|  | Res. type | Atom | Residue | Distance | Occupancy | B-factor | Closest solute to 1^st^ shell water |
| --- | --- | --- | --- | --- | --- | --- | --- |
| **Ion** | | | |  | 1.00 | 98.67 || 1 | G | OP2 | AA:1424 | 2.89 | 1.00 | 65.32 |
| 2 | G | OP2 | AA:1423 | 3.16 | 1.00 | 69.16 |  |
| 3 | G | O5' | AA:1423 | 3.20 | 1.00 | 69.20 |  |
| 4 | G | N7 | AA:1423 | 4.18 | 1.00 | 66.21 |  |
| 5 | G | N7 | AA:1424 | 4.19 | 1.00 | 67.05 |  |
| 6 | G | OP1 | AA:1423 | 4.40 | 1.00 | 68.88 |  |
| 7 | G | N9 | AA:1423 | 4.40 | 1.00 | 66.02 |  |
| 8 | G | O3' | AA:1423 | 4.44 | 1.00 | 65.65 |  |
| 9 | G | O6 | AA:1425 | 4.71 | 1.00 | 64.98 |  |
| 10 | G | O4' | AA:1423 | 4.77 | 1.00 | 66.85 |  |
| 11 | G | O5' | AA:1424 | 4.86 | 1.00 | 64.64 |  |
  |

  
 Coded contact list:   
 Exclusion zone contact(s): /O5'/OP2/OP2/  
Consider K in place of MG (3 valid contact(s) for K vs. 0 for MG)  

### MG:AA:4108

|  | Res. type | Atom | Residue | Distance | Occupancy | B-factor | Closest solute to 1^st^ shell water |
| --- | --- | --- | --- | --- | --- | --- | --- |
| **Ion** | | | |  | 1.00 | 136.81 || 1 | C | OP1 | AA:2196 | 2.50 | 1.00 | 71.16 |
| 2 | U | OP2 | AA:2197 | 2.87 | 1.00 | 69.44 |  |
| 3 | C | O5' | AA:2196 | 3.46 | 1.00 | 69.77 |  |
| 4 | C | OP2 | AA:2196 | 4.32 | 1.00 | 71.48 |  |
| 5 | C | O3' | AA:2196 | 4.33 | 1.00 | 67.14 |  |
| 6 | U | OP1 | AA:2197 | 4.56 | 1.00 | 70.16 |  |
| 7 | C | O3' | AA:2195 | 4.79 | 1.00 | 71.70 |  |
  |

  
 Coded contact list:   
 Exclusion zone contact(s): /OP1/OP2/  
Consider NA in place of MG (1 valid contact(s) for NA vs. 0 for MG)  

### MG:AA:4109

|  | Res. type | Atom | Residue | Distance | Occupancy | B-factor | Closest solute to 1^st^ shell water |
| --- | --- | --- | --- | --- | --- | --- | --- |
| **Ion** | | | |  | 1.00 | 81.57 || 1 | C | OP1 | AA:270Q | 4.42 | 1.00 | 133.83 |
| 2 | C | OP2 | AA:270D | 4.60 | 1.00 | 97.31 |  |
| 3 | C | OP2 | AA:270Q | 4.82 | 1.00 | 134.38 |  |
  |

  
 Coded contact list:   
1^st^ MG coordination shell missing (see Notes)  

### MG:AA:4110

|  | Res. type | Atom | Residue | Distance | Occupancy | B-factor | Closest solute to 1^st^ shell water |
| --- | --- | --- | --- | --- | --- | --- | --- |
| **Ion** | | | |  | 1.00 | 102.40 || 1 | G | O6 | AA:1416 | 2.67 | 1.00 | 95.77 |
| 2 | C | N4 | AA:1582 | 3.75 | 1.00 | 97.67 |  |
| 3 | A | N6 | AA:1583 | 4.36 | 1.00 | 107.80 |  |
| 4 | G | N7 | AA:1416 | 4.42 | 1.00 | 91.86 |  |
| 5 | A | OP1 | AA:1579 | 4.67 | 1.00 | 85.67 |  |
| 6 | G | O6 | AA:1581 | 4.73 | 1.00 | 91.42 |  |
| 7 | G | N1 | AA:1416 | 4.91 | 1.00 | 94.84 |  |
  |

  
 Coded contact list:   
 Exclusion zone contact(s): /O6/  
Consider K in place of MG (1 valid contact(s) for K vs. 0 for MG)  

### MG:AA:4111

|  | Res. type | Atom | Residue | Distance | Occupancy | B-factor | Closest solute to 1^st^ shell water |
| --- | --- | --- | --- | --- | --- | --- | --- |
| **Ion** | | | |  | 1.00 | 108.75 || 1 | G | OP2 | AA:1413 | 2.98 | 1.00 | 90.53 |
| 2 | A | OP2 | AA:1412 | 3.06 | 1.00 | 83.48 |  |
| 3 | A | O5' | AA:1412 | 3.09 | 1.00 | 83.95 |  |
| 4 | A | N7 | AA:1412 | 3.84 | 1.00 | 88.79 |  |
| 5 | G | N7 | AA:1413 | 3.97 | 1.00 | 86.59 |  |
| 6 | A | N9 | AA:1412 | 4.16 | 1.00 | 88.54 |  |
| 7 | A | O3' | AA:1412 | 4.33 | 1.00 | 89.30 |  |
| 8 | A | O4' | AA:1412 | 4.60 | 1.00 | 86.42 |  |
| 9 | A | OP1 | AA:1412 | 4.60 | 1.00 | 83.88 |  |
| 10 | C | O3' | AA:1411 | 4.86 | 1.00 | 83.61 |  |
| 11 | G | O5' | AA:1413 | 4.90 | 1.00 | 90.44 |  |
  |

  
 Coded contact list:   
 Exclusion zone contact(s): /O5'/OP2/OP2/  
Consider K in place of MG (3 valid contact(s) for K vs. 0 for MG)  

### MG:AA:4113

|  | Res. type | Atom | Residue | Distance | Occupancy | B-factor | Closest solute to 1^st^ shell water |
| --- | --- | --- | --- | --- | --- | --- | --- |
| **Ion** | | | |  | 1.00 | 92.13 || 1 | C | N4 | AA:1202 | 2.52 | 1.00 | 54.42 |
| 2 | G | N1 | AA:1203 | 2.74 | 1.00 | 61.53 |  |
| 3 | G | O6 | AA:1203 | 2.92 | 1.00 | 64.99 |  |
| 4 | G | O6 | AA:1243 | 3.03 | 1.00 | 60.55 |  |
| 5 | A | OP2 | AA:1241 | 3.28 | 1.00 | 57.23 |  |
| 6 | C | N4 | AA:1201 | 3.32 | 1.00 | 45.47 |  |
| 7 | A | OP1 | AA:1241 | 3.51 | 1.00 | 56.89 |  |
| 8 | G | O6 | AA:1244 | 3.63 | 1.00 | 47.77 |  |
| 9 | G | N2 | AA:1203 | 4.03 | 1.00 | 60.18 |  |
| 10 | C | N3 | AA:1202 | 4.70 | 1.00 | 54.44 |  |
| 11 | A | O5' | AA:1241 | 4.75 | 1.00 | 59.68 |  |
| 12 | G | N3 | AA:1203 | 4.75 | 1.00 | 59.16 |  |
| 13 | C | N3 | AA:1201 | 4.86 | 1.00 | 45.85 |  |
  |

  
 Coded contact list:   
 Exclusion zone contact(s): /N1_G_/N4/N4/O6/O6/OP2/  
Consider K in place of MG (5 valid contact(s) for K vs. 0 for MG)  

### MG:AA:4114

|  | Res. type | Atom | Residue | Distance | Occupancy | B-factor | Closest solute to 1^st^ shell water |
| --- | --- | --- | --- | --- | --- | --- | --- |
| **Ion** | | | |  | 1.00 | 92.01 || 1 | G | O6 | AA:1203 | 2.36 | 1.00 | 64.99 |
| 2 | A | OP2 | AA:1241 | 2.53 | 1.00 | 57.23 |  |
| 3 | A | N6 | AA:1204 | 3.56 | 1.00 | 68.02 |  |
| 4 | G | N1 | AA:1203 | 3.84 | 1.00 | 61.53 |  |
| 5 | A | OP1 | AA:1241 | 3.85 | 1.00 | 56.89 |  |
| 6 | C | N4 | AA:1202 | 4.19 | 1.00 | 54.42 |  |
| 7 | U | O3' | AA:1240 | 4.24 | 1.00 | 58.67 |  |
| 8 | C | N4 | AA:1201 | 4.35 | 1.00 | 45.47 |  |
| 9 | A | O5' | AA:1241 | 4.93 | 1.00 | 59.68 |  |
  |

  
 Contact(s): /O6/  
 Coded contact list: O_b_  
 Exclusion zone contact(s): /OP2/  

### MG:AA:4115

|  | Res. type | Atom | Residue | Distance | Occupancy | B-factor | Closest solute to 1^st^ shell water |
| --- | --- | --- | --- | --- | --- | --- | --- |
| **Ion** | | | |  | 1.00 | 73.87 || 1 | C | O2' | AA:1318 | 4.49 | 1.00 | 59.91 |
| 2 | G | O2' | AA:1319 | 4.53 | 1.00 | 58.52 |  |
| 3 | C | O2 | AA:1318 | 4.55 | 1.00 | 61.97 |  |
| 4 | G | N2 | AA:1334 | 4.70 | 1.00 | 60.73 |  |
| 5 | G | O4' | AA:1319 | 4.81 | 1.00 | 61.27 |  |
  |

  
 Coded contact list:   
1^st^ MG coordination shell missing (see Notes)  

### MG:AA:4116

|  | Res. type | Atom | Residue | Distance | Occupancy | B-factor | Closest solute to 1^st^ shell water |
| --- | --- | --- | --- | --- | --- | --- | --- |
| **Ion** | | | |  | 1.00 | 98.87 || 1 | GLU | OE1 | AU:91 | 2.84 | 1.00 | 103.57 |
| 2 | G | N7 | AA:83 | 3.23 | 1.00 | 52.92 |  |
| 3 | GLU | OE2 | AU:91 | 3.72 | 1.00 | 103.85 |  |
| 4 | G | O6 | AA:83 | 3.87 | 1.00 | 55.30 |  |
| 5 | G | O6 | AA:102 | 4.61 | 1.00 | 71.96 |  |
  |

  
 Coded contact list:   
 Exclusion zone contact(s): /N7/OE1_Glu_/  
Consider K in place of MG (2 valid contact(s) for K vs. 0 for MG)  

### MG:AA:4117

|  | Res. type | Atom | Residue | Distance | Occupancy | B-factor | Closest solute to 1^st^ shell water |
| --- | --- | --- | --- | --- | --- | --- | --- |
| **Ion** | | | |  | 1.00 | 96.41 || 1 | A | OP2 | AA:1412 | 2.61 | 1.00 | 83.48 |
| 2 | C | O5' | AA:1411 | 2.69 | 1.00 | 81.74 |  |
| 3 | C | OP2 | AA:1411 | 2.98 | 1.00 | 81.66 |  |
| 4 | C | OP1 | AA:1411 | 3.66 | 1.00 | 81.32 |  |
| 5 | C | O3' | AA:1411 | 3.95 | 1.00 | 83.61 |  |
| 6 | A | O5' | AA:1412 | 4.52 | 1.00 | 83.95 |  |
| 7 | G | O3' | AA:1410 | 4.70 | 1.00 | 82.11 |  |
| 8 | C | O4' | AA:1411 | 4.86 | 1.00 | 83.69 |  |
| 9 | C | N1 | AA:1411 | 4.87 | 1.00 | 85.48 |  |
  |

  
 Coded contact list:   
 Exclusion zone contact(s): /O5'/OP2/OP2/  
Consider K in place of MG (3 valid contact(s) for K vs. 0 for MG)  

### MG:AA:4118

|  | Res. type | Atom | Residue | Distance | Occupancy | B-factor | Closest solute to 1^st^ shell water |
| --- | --- | --- | --- | --- | --- | --- | --- |
| **Ion** | | | |  | 1.00 | 67.46 || 1 | A | OP2 | AA:1762 | 3.94 | 1.00 | 57.60 |
| 2 | A | OP1 | AA:1762 | 3.95 | 1.00 | 56.57 |  |
| 3 | C | O3' | AA:1761 | 4.23 | 1.00 | 54.72 |  |
| 4 | U | OP2 | AA:757 | 4.34 | 1.00 | 48.74 |  |
| 5 | U | OP1 | AA:757 | 4.74 | 1.00 | 49.49 |  |
  |

  
 Coded contact list:   
1^st^ MG coordination shell missing (see Notes)  

### MG:AA:4119

|  | Res. type | Atom | Residue | Distance | Occupancy | B-factor | Closest solute to 1^st^ shell water |
| --- | --- | --- | --- | --- | --- | --- | --- |
| **Ion** | | | |  | 1.00 | 112.83 || 1 | C | OP1 | AA:2129 | 2.68 | 1.00 | 200.14 |
| 2 | U | O4 | AA:2132 | 4.27 | 1.00 | 200.16 |  |
| 3 | C | O5' | AA:2129 | 4.47 | 1.00 | 200.27 |  |
| 4 | C | OP2 | AA:2129 | 4.75 | 1.00 | 200.41 |  |
| 5 | C | OP1 | AA:2128 | 4.85 | 1.00 | 198.71 |  |
  |

  
 Coded contact list:   
 Exclusion zone contact(s): /OP1/  
Consider K in place of MG (1 valid contact(s) for K vs. 0 for MG)  

### MG:AA:4120

|  | Res. type | Atom | Residue | Distance | Occupancy | B-factor | Closest solute to 1^st^ shell water |
| --- | --- | --- | --- | --- | --- | --- | --- |
| **Ion** | | | |  | 1.00 | 87.48 || 1 | G | OP2 | AA:2379 | 3.67 | 1.00 | 67.20 |
| 2 | G | N7 | AA:2379 | 3.86 | 1.00 | 60.36 |  |
| 3 | A | OP1 | AA:2378 | 4.27 | 1.00 | 70.56 |  |
| 4 | A | OP2 | AA:2378 | 4.72 | 1.00 | 71.30 |  |
| 5 | A | O5' | AA:2378 | 4.95 | 1.00 | 69.42 |  |
  |

  
 Coded contact list:   
1^st^ MG coordination shell missing (see Notes)  

### MG:AA:4121

|  | Res. type | Atom | Residue | Distance | Occupancy | B-factor | Closest solute to 1^st^ shell water |
| --- | --- | --- | --- | --- | --- | --- | --- |
| **Ion** | | | |  | 1.00 | 66.43 || 1 | U | OP2 | AA:1391 | 2.87 | 1.00 | 68.62 |
| 2 | A | OP2 | AA:1392 | 3.54 | 1.00 | 62.65 |  |
| 3 | U | O5' | AA:1391 | 3.84 | 1.00 | 69.31 |  |
| 4 | A | OP2 | AA:1393 | 3.88 | 1.00 | 60.32 |  |
| 5 | A | OP1 | AA:1393 | 4.31 | 1.00 | 60.74 |  |
| 6 | U | OP1 | AA:1391 | 4.79 | 1.00 | 69.69 |  |
  |

  
 Coded contact list:   
 Exclusion zone contact(s): /OP2/  
Consider K in place of MG (1 valid contact(s) for K vs. 0 for MG)  

### MG:AA:4123

|  | Res. type | Atom | Residue | Distance | Occupancy | B-factor | Closest solute to 1^st^ shell water |
| --- | --- | --- | --- | --- | --- | --- | --- |
| **Ion** | | | |  | 1.00 | 55.94 || 1 | U | OP2 | AA:1454 | 2.57 | 1.00 | 71.19 |
| 2 | A | O5' | AA:1453 | 3.15 | 1.00 | 78.97 |  |
| 3 | U | O3' | AA:1454 | 3.31 | 1.00 | 70.30 |  |
| 4 | A | OP2 | AA:1453 | 3.51 | 1.00 | 80.36 |  |
| 5 | G | OP2 | AA:1455 | 4.15 | 1.00 | 68.93 |  |
| 6 | U | O5' | AA:1454 | 4.42 | 1.00 | 71.45 |  |
| 7 | A | O3' | AA:1453 | 4.47 | 1.00 | 71.62 |  |
| 8 | A | OP1 | AA:1453 | 4.55 | 1.00 | 81.74 |  |
  |

  
 Coded contact list:   
 Exclusion zone contact(s): /O3'/O5'/OP2/  
Consider K in place of MG (2 valid contact(s) for K vs. 0 for MG)  

### MG:AA:4124

|  | Res. type | Atom | Residue | Distance | Occupancy | B-factor | Closest solute to 1^st^ shell water |
| --- | --- | --- | --- | --- | --- | --- | --- |
| **Ion** | | | |  | 1.00 | 107.98 || 1 | G | O6 | AA:2707 | 4.91 | 1.00 | 56.68 |
  |

  
 Coded contact list:   
1^st^ MG coordination shell missing (see Notes)  

### MG:AA:4125

|  | Res. type | Atom | Residue | Distance | Occupancy | B-factor | Closest solute to 1^st^ shell water |
| --- | --- | --- | --- | --- | --- | --- | --- |
| **Ion** | | | |  | 1.00 | 69.38 || 1 | A | OP1 | AA:1603 | 2.47 | 1.00 | 45.20 |
| 2 | U | O3' | AA:1602 | 2.92 | 1.00 | 48.36 |  |
| 3 | A | OP1 | AA:1395 | 3.09 | 1.00 | 52.83 |  |
| 4 | U | O2' | AA:1602 | 3.44 | 1.00 | 53.30 |  |
| 5 | A | OP2 | AA:1603 | 4.05 | 1.00 | 45.36 |  |
| 6 | A | O5' | AA:1395 | 4.18 | 1.00 | 53.98 |  |
| 7 | U | O4 | AA:1397 | 4.26 | 1.00 | 61.01 |  |
| 8 | C | OP2 | AA:1604 | 4.31 | 1.00 | 34.59 |  |
| 9 | A | O5' | AA:1603 | 4.49 | 1.00 | 45.11 |  |
| 10 | U | O3' | AA:1394 | 4.65 | 1.00 | 53.46 |  |
  |

  
 Coded contact list:   
 Exclusion zone contact(s): /O3'/OP1/OP1/  
Consider K in place of MG (2 valid contact(s) for K vs. 0 for MG)  

### MG:AA:4126

|  | Res. type | Atom | Residue | Distance | Occupancy | B-factor | Closest solute to 1^st^ shell water |
| --- | --- | --- | --- | --- | --- | --- | --- |
| **Ion** | | | |  | 1.00 | 87.45 || 1 | C | OP1 | AA:1432 | 2.74 | 1.00 | 45.42 |
| 2 | C | OP2 | AA:1432 | 4.53 | 1.00 | 47.35 |  |
| 3 | C | O5' | AA:1432 | 4.77 | 1.00 | 47.45 |  |
  |

  
 Coded contact list:   
 Exclusion zone contact(s): /OP1/  
Consider K in place of MG (1 valid contact(s) for K vs. 0 for MG)  

### MG:AA:4127

|  | Res. type | Atom | Residue | Distance | Occupancy | B-factor | Closest solute to 1^st^ shell water |
| --- | --- | --- | --- | --- | --- | --- | --- |
| **Ion** | | | |  | 1.00 | 52.24 || 1 | U | OP1 | AA:2243 | 2.55 | 1.00 | 30.71 |
| 2 | C | OP1 | AA:192 | 2.57 | 1.00 | 36.21 |  |
| 3 | A | O3' | AA:191 | 3.08 | 1.00 | 32.85 |  |
| 4 | G | O3' | AA:2242 | 3.43 | 1.00 | 30.47 |  |
| 5 | C | O2' | AA:678 | 3.59 | 1.00 | 41.21 |  |
| 6 | C | O5' | AA:192 | 3.99 | 1.00 | 35.55 |  |
| 7 | U | OP2 | AA:2243 | 4.02 | 1.00 | 31.15 |  |
| 8 | A | O2' | AA:191 | 4.65 | 1.00 | 30.30 |  |
| 9 | C | OP2 | AA:192 | 4.70 | 1.00 | 35.85 |  |
| 10 | U | O5' | AA:2243 | 4.81 | 1.00 | 32.87 |  |
| 11 | C | O3' | AA:678 | 4.90 | 1.00 | 37.74 |  |
| 12 | C | O4' | AA:678 | 4.96 | 1.00 | 39.72 |  |
  |

  
 Coded contact list:   
 Exclusion zone contact(s): /O3'/OP1/OP1/  
Consider NA in place of MG (2 valid contact(s) for NA vs. 0 for MG)  

### MG:AA:4128

|  | Res. type | Atom | Residue | Distance | Occupancy | B-factor | Closest solute to 1^st^ shell water |
| --- | --- | --- | --- | --- | --- | --- | --- |
| **Ion** | | | |  | 1.00 | 116.40 || 1 | C | OP2 | AA:1518 | 2.59 | 1.00 | 64.90 |
| 2 | C | OP1 | AA:1518 | 4.18 | 1.00 | 65.00 |  |
| 3 | G | OP1 | AA:1470 | 4.27 | 1.00 | 75.12 |  |
| 4 | C | O5' | AA:1518 | 4.62 | 1.00 | 66.48 |  |
| 5 | G | OP2 | AA:1470 | 4.90 | 1.00 | 75.20 |  |
  |

  
 Coded contact list:   
 Exclusion zone contact(s): /OP2/  
Consider NA in place of MG (1 valid contact(s) for NA vs. 0 for MG)  

### MG:AA:4129

|  | Res. type | Atom | Residue | Distance | Occupancy | B-factor | Closest solute to 1^st^ shell water |
| --- | --- | --- | --- | --- | --- | --- | --- |
| **Ion** | | | |  | 1.00 | 86.37 || 1 | G | OP2 | AA:327 | 2.87 | 1.00 | 57.27 |
| 2 | G | OP1 | AA:327 | 2.89 | 1.00 | 57.18 |  |
| 3 | G | O5' | AA:327 | 3.81 | 1.00 | 56.59 |  |
| 4 | U | OP2 | AA:328 | 4.64 | 1.00 | 51.53 |  |
| 5 | G | O3' | AA:326 | 4.71 | 1.00 | 56.54 |  |
  |

  
 Coded contact list:   
 Exclusion zone contact(s): /OP1/OP2/  
Consider K in place of MG (2 valid contact(s) for K vs. 0 for MG)  

### MG:AA:4130

|  | Res. type | Atom | Residue | Distance | Occupancy | B-factor | Closest solute to 1^st^ shell water |
| --- | --- | --- | --- | --- | --- | --- | --- |
| **Ion** | | | |  | 1.00 | 107.15 || 1 | C | N4 | AA:277 | 2.80 | 1.00 | 153.23 |
| 2 | A | N1 | AA:276 | 4.07 | 1.00 | 158.81 |  |
| 3 | C | N3 | AA:277 | 4.49 | 1.00 | 153.81 |  |
  |

  
 Coded contact list:   
 Exclusion zone contact(s): /N4/  
Consider K in place of MG (1 valid contact(s) for K vs. 0 for MG)  

### MG:AA:4131

|  | Res. type | Atom | Residue | Distance | Occupancy | B-factor | Closest solute to 1^st^ shell water |
| --- | --- | --- | --- | --- | --- | --- | --- |
| **Ion** | | | |  | 1.00 | 99.79 || 1 | C | N4 | AA:1147 | 4.06 | 1.00 | 116.62 |
| 2 | A | N6 | AA:1148 | 4.13 | 1.00 | 115.66 |  |
| 3 | A | N7 | AA:1148 | 4.69 | 1.00 | 116.14 |  |
| 4 | G | O6 | AA:1149 | 4.92 | 1.00 | 105.73 |  |
  |

  
 Coded contact list:   
1^st^ MG coordination shell missing (see Notes)  

### MG:AA:4132

|  | Res. type | Atom | Residue | Distance | Occupancy | B-factor | Closest solute to 1^st^ shell water |
| --- | --- | --- | --- | --- | --- | --- | --- |
| **Ion** | | | |  | 1.00 | 68.75 || 1 | U | OP2 | AA:2074 | 2.53 | 1.00 | 26.90 |
| 2 | C | O5' | AA:2073 | 2.77 | 1.00 | 23.22 |  |
| 3 | C | OP2 | AA:2073 | 2.93 | 1.00 | 25.31 |  |
| 4 | C | OP1 | AA:2073 | 3.54 | 1.00 | 23.65 |  |
| 5 | C | O3' | AA:2073 | 4.11 | 1.00 | 25.26 |  |
| 6 | C | N1 | AA:2073 | 4.40 | 1.00 | 23.47 |  |
| 7 | C | O4' | AA:2073 | 4.60 | 1.00 | 24.60 |  |
| 8 | G | O3' | AA:2072 | 4.65 | 1.00 | 23.70 |  |
| 9 | U | OP1 | AA:2074 | 4.81 | 1.00 | 26.02 |  |
| 10 | U | O5' | AA:2074 | 4.84 | 1.00 | 26.91 |  |
  |

  
 Coded contact list:   
 Exclusion zone contact(s): /O5'/OP2/OP2/  
Consider K in place of MG (2 valid contact(s) for K vs. 0 for MG)  

### MG:AA:4134

|  | Res. type | Atom | Residue | Distance | Occupancy | B-factor | Closest solute to 1^st^ shell water |
| --- | --- | --- | --- | --- | --- | --- | --- |
| **Ion** | | | |  | 1.00 | 117.37 || 1 | A | N6 | AA:1088 | 2.53 | 1.00 | 199.67 |
| 2 | A | O2' | AA:1077 | 2.75 | 1.00 | 199.75 |  |
| 3 | A | N1 | AA:1088 | 2.99 | 1.00 | 199.98 |  |
| 4 | G | N2 | AA:1062 | 3.55 | 1.00 | 200.04 |  |
| 5 | A | N3 | AA:1077 | 4.19 | 1.00 | 198.41 |  |
| 6 | A | O4' | AA:1077 | 4.31 | 1.00 | 199.89 |  |
| 7 | G | N3 | AA:1062 | 4.36 | 1.00 | 200.41 |  |
| 8 | A | N9 | AA:1077 | 4.57 | 1.00 | 199.32 |  |
| 9 | A | N7 | AA:1088 | 4.84 | 1.00 | 200.25 |  |
| 10 | A | N3 | AA:1088 | 4.94 | 1.00 | 200.24 |  |
  |

  
 Coded contact list:   
 Exclusion zone contact(s): /N1_A_/N6/O2'/  
Consider K in place of MG (2 valid contact(s) for K vs. 0 for MG)  

### MG:AA:4135

|  | Res. type | Atom | Residue | Distance | Occupancy | B-factor | Closest solute to 1^st^ shell water |
| --- | --- | --- | --- | --- | --- | --- | --- |
| **Ion** | | | |  | 1.00 | 64.81 || 1 | G | O4' | AA:187 | 4.14 | 1.00 | 39.63 |
| 2 | G | O2' | AA:187 | 4.66 | 1.00 | 40.28 |  |
| 3 | A | O2' | AA:1365 | 4.99 | 1.00 | 44.61 |  |
  |

  
 Coded contact list:   
1^st^ MG coordination shell missing (see Notes)  

### MG:AA:4137

|  | Res. type | Atom | Residue | Distance | Occupancy | B-factor | Closest solute to 1^st^ shell water |
| --- | --- | --- | --- | --- | --- | --- | --- |
| **Ion** | | | |  | 1.00 | 90.25 || 1 | G | O6 | AA:80 | 2.92 | 1.00 | 69.43 |
| 2 | C | N4 | AA:106 | 3.73 | 1.00 | 67.53 |  |
| 3 | G | O6 | AA:81 | 3.76 | 1.00 | 67.03 |  |
| 4 | G | O6 | AA:79 | 3.95 | 1.00 | 66.83 |  |
| 5 | C | N4 | AA:105 | 3.97 | 1.00 | 67.66 |  |
| 6 | G | N7 | AA:80 | 4.78 | 1.00 | 69.46 |  |
| 7 | C | N4 | AA:107 | 4.79 | 1.00 | 64.43 |  |
| 8 | U | OP2 | AA:104 | 4.98 | 1.00 | 65.24 |  |
  |

  
 Coded contact list:   
 Exclusion zone contact(s): /O6/  
Consider K in place of MG (1 valid contact(s) for K vs. 0 for MG)  

### MG:AA:4138

|  | Res. type | Atom | Residue | Distance | Occupancy | B-factor | Closest solute to 1^st^ shell water |
| --- | --- | --- | --- | --- | --- | --- | --- |
| **Ion** | | | |  | 1.00 | 57.32 || 1 | U | OP2 | AA:1329 | 2.33 | 1.00 | 49.05 |
| 2 | C | N1 | AA:1320 | 2.48 | 1.00 | 65.27 |  |
| 3 | A | OP2 | AA:1321 | 3.39 | 1.00 | 59.74 |  |
| 4 | U | OP1 | AA:1329 | 3.42 | 1.00 | 49.74 |  |
| 5 | C | O2' | AA:1320 | 3.53 | 1.00 | 66.17 |  |
| 6 | C | O4' | AA:1320 | 3.62 | 1.00 | 63.02 |  |
| 7 | U | O5' | AA:1329 | 3.91 | 1.00 | 49.47 |  |
| 8 | C | O3' | AA:1320 | 3.93 | 1.00 | 60.49 |  |
| 9 | C | O2 | AA:1320 | 3.97 | 1.00 | 66.36 |  |
| 10 | C | N3 | AA:1320 | 3.98 | 1.00 | 64.94 |  |
| 11 | C | N4 | AA:1330 | 4.18 | 1.00 | 53.79 |  |
| 12 | C | O5' | AA:1320 | 4.38 | 1.00 | 62.92 |  |
| 13 | G | O3' | AA:1328 | 4.63 | 1.00 | 48.91 |  |
  |

  
 Contact(s): /OP2/  
 Coded contact list: O_ph_  
 Exclusion zone contact(s): /N1/OP2/  

### MG:AA:4139

|  | Res. type | Atom | Residue | Distance | Occupancy | B-factor | Closest solute to 1^st^ shell water |
| --- | --- | --- | --- | --- | --- | --- | --- |
| **Ion** | | | |  | 1.00 | 67.30 || 1 | U | OP1 | AA:1995 | 2.46 | 1.00 | 44.43 |
| 2 | G | OP1 | AA:1997 | 3.42 | 1.00 | 48.01 |  |
| 3 | U | OP1 | AA:2511 | 4.22 | 1.00 | 39.26 |  |
| 4 | C | O5' | AA:1996 | 4.48 | 1.00 | 50.86 |  |
| 5 | U | O5' | AA:1995 | 4.55 | 1.00 | 43.84 |  |
| 6 | U | OP2 | AA:1995 | 4.71 | 1.00 | 45.95 |  |
| 7 | C | O3' | AA:1994 | 4.90 | 1.00 | 42.87 |  |
| 8 | C | OP2 | AA:1996 | 4.96 | 1.00 | 50.14 |  |
  |

  
 Coded contact list:   
 Exclusion zone contact(s): /OP1/  
Consider NA in place of MG (1 valid contact(s) for NA vs. 0 for MG)  

### MG:AA:4141

|  | Res. type | Atom | Residue | Distance | Occupancy | B-factor | Closest solute to 1^st^ shell water |
| --- | --- | --- | --- | --- | --- | --- | --- |
| **Ion** | | | |  | 1.00 | 88.26 || 1 | U | O4 | AA:1482 | 3.34 | 1.00 | 93.37 |
| 2 | A | OP2 | AA:1510 | 3.51 | 1.00 | 148.59 |  |
  |

  
 Coded contact list:   
 Exclusion zone contact(s): /O4/  
Consider K in place of MG (1 valid contact(s) for K vs. 0 for MG)  

### MG:AA:4142

|  | Res. type | Atom | Residue | Distance | Occupancy | B-factor | Closest solute to 1^st^ shell water |
| --- | --- | --- | --- | --- | --- | --- | --- |
| **Ion** | | | |  | 1.00 | 104.29 || 1 | G | O6 | AA:333 | 4.50 | 1.00 | 62.35 |
| 2 | C | N4 | AA:318 | 4.89 | 1.00 | 64.35 |  |
  |

  
 Coded contact list:   
1^st^ MG coordination shell missing (see Notes)  

### MG:AA:4144

|  | Res. type | Atom | Residue | Distance | Occupancy | B-factor | Closest solute to 1^st^ shell water |
| --- | --- | --- | --- | --- | --- | --- | --- |
| **Ion** | | | |  | 1.00 | 100.43 || 1 | G | O6 | AA:361 | 2.48 | 1.00 | 112.98 |
| 2 | G | N7 | AA:361 | 2.74 | 1.00 | 113.04 |  |
| 3 | G | O6 | AA:360 | 4.38 | 1.00 | 101.92 |  |
| 4 | G | N1 | AA:361 | 4.39 | 1.00 | 112.58 |  |
| 5 | U | O4 | AA:362 | 4.81 | 1.00 | 130.86 |  |
| 6 | C | N4 | AA:279 | 4.82 | 1.00 | 113.96 |  |
| 7 | G | N9 | AA:361 | 4.85 | 1.00 | 112.20 |  |
| 8 | G | N7 | AA:360 | 4.92 | 1.00 | 103.13 |  |
  |

  
 Coded contact list:   
 Exclusion zone contact(s): /N7/O6/  
Consider NA in place of MG (1 valid contact(s) for NA vs. 0 for MG)  

### MG:AA:4145

|  | Res. type | Atom | Residue | Distance | Occupancy | B-factor | Closest solute to 1^st^ shell water |
| --- | --- | --- | --- | --- | --- | --- | --- |
| **Ion** | | | |  | 1.00 | 77.33 || 1 | C | OP1 | AA:1648 | 2.58 | 1.00 | 43.56 |
| 2 | A | OP1 | AA:1272 | 2.91 | 1.00 | 49.07 |  |
| 3 | A | OP2 | AA:1272 | 3.12 | 1.00 | 48.76 |  |
| 4 | G | OP2 | AA:1271 | 3.21 | 1.00 | 42.52 |  |
| 5 | G | OP1 | AA:1647 | 3.25 | 1.00 | 49.60 |  |
| 6 | G | O5' | AA:1647 | 3.46 | 1.00 | 46.53 |  |
| 7 | G | O5' | AA:1271 | 3.60 | 1.00 | 45.41 |  |
| 8 | G | O3' | AA:1271 | 3.66 | 1.00 | 49.58 |  |
| 9 | G | N7 | AA:1271 | 4.09 | 1.00 | 57.45 |  |
| 10 | C | O3' | AA:1270 | 4.24 | 1.00 | 43.39 |  |
| 11 | G | N9 | AA:1271 | 4.26 | 1.00 | 52.48 |  |
| 12 | C | OP2 | AA:1648 | 4.33 | 1.00 | 41.95 |  |
| 13 | G | OP2 | AA:1647 | 4.52 | 1.00 | 48.36 |  |
| 14 | G | O3' | AA:1647 | 4.76 | 1.00 | 43.20 |  |
| 15 | G | O4' | AA:1271 | 4.79 | 1.00 | 48.93 |  |
| 16 | A | O5' | AA:1272 | 4.82 | 1.00 | 50.28 |  |
| 17 | C | O5' | AA:1648 | 4.97 | 1.00 | 42.64 |  |
  |

  
 Coded contact list:   
 Exclusion zone contact(s): /OP1/OP1/OP1/OP2/OP2/  
Consider K in place of MG (4 valid contact(s) for K vs. 0 for MG)  

### MG:AA:4146

|  | Res. type | Atom | Residue | Distance | Occupancy | B-factor | Closest solute to 1^st^ shell water |
| --- | --- | --- | --- | --- | --- | --- | --- |
| **Ion** | | | |  | 1.00 | 90.87 || 1 | G | OP1 | AA:1555 | 4.22 | 1.00 | 62.67 |
| 2 | G | OP2 | AA:1555 | 4.38 | 1.00 | 64.69 |  |
| 3 | U | O4 | AA:1433 | 4.64 | 1.00 | 60.47 |  |
  |

  
 Coded contact list:   
1^st^ MG coordination shell missing (see Notes)  

### MG:AA:4148

|  | Res. type | Atom | Residue | Distance | Occupancy | B-factor | Closest solute to 1^st^ shell water |
| --- | --- | --- | --- | --- | --- | --- | --- |
| **Ion** | | | |  | 1.00 | 87.12 || 1 | G | N7 | AA:476 | 4.15 | 1.00 | 59.95 |
| 2 | A | OP1 | AA:478 | 4.66 | 1.00 | 55.58 |  |
| 3 | A | OP2 | AA:478 | 4.74 | 1.00 | 54.84 |  |
| 4 | U | OP1 | AA:475 | 4.83 | 1.00 | 48.11 |  |
  |

  
 Coded contact list:   
1^st^ MG coordination shell missing (see Notes)  

### MG:AA:4149

|  | Res. type | Atom | Residue | Distance | Occupancy | B-factor | Closest solute to 1^st^ shell water |
| --- | --- | --- | --- | --- | --- | --- | --- |
| **Ion** | | | |  | 1.00 | 125.28 || 1 | C | OP1 | AA:1589 | 2.94 | 1.00 | 85.62 |
| 2 | C | OP2 | AA:1589 | 4.33 | 1.00 | 85.74 |  |
| 3 | C | O5' | AA:1589 | 4.55 | 1.00 | 85.86 |  |
| 4 | U | OP2 | AA:1590 | 4.57 | 1.00 | 84.96 |  |
| 5 | C | OP1 | AA:1407 | 4.68 | 1.00 | 61.68 |  |
  |

  
 Coded contact list:   
 Exclusion zone contact(s): /OP1/  
Consider K in place of MG (1 valid contact(s) for K vs. 0 for MG)  

### MG:AA:4150

|  | Res. type | Atom | Residue | Distance | Occupancy | B-factor | Closest solute to 1^st^ shell water |
| --- | --- | --- | --- | --- | --- | --- | --- |
| **Ion** | | | |  | 1.00 | 82.67 || 1 | A | N7 | AA:1020 | 3.45 | 1.00 | 91.12 |
| 2 | A | N6 | AA:1020 | 4.22 | 1.00 | 90.23 |  |
| 3 | A | N9 | AA:1020 | 4.65 | 1.00 | 91.10 |  |
| 4 | ARG | NE | AH:60 | 4.80 | 1.00 | 146.16 |  |
| 5 | A | OP2 | AA:1020 | 4.87 | 1.00 | 95.90 |  |
  |

  
 Coded contact list:   
1^st^ MG coordination shell missing (see Notes)  

### MG:AA:4152

|  | Res. type | Atom | Residue | Distance | Occupancy | B-factor | Closest solute to 1^st^ shell water |
| --- | --- | --- | --- | --- | --- | --- | --- |
| **Ion** | | | |  | 1.00 | 77.12 || 1 | G | OP1 | AA:947 | 2.93 | 1.00 | 52.58 |
| 2 | U | OP2 | AA:963 | 3.22 | 1.00 | 57.28 |  |
| 3 | G | OP1 | AA:948 | 3.32 | 1.00 | 55.90 |  |
| 4 | U | OP1 | AA:963 | 3.34 | 1.00 | 58.34 |  |
| 5 | G | O3' | AA:947 | 3.37 | 1.00 | 56.25 |  |
| 6 | G | OP2 | AA:948 | 3.45 | 1.00 | 54.88 |  |
| 7 | G | O5' | AA:947 | 3.87 | 1.00 | 54.86 |  |
| 8 | G | OP1 | AA:962 | 4.42 | 1.00 | 58.60 |  |
| 9 | G | OP2 | AA:947 | 4.60 | 1.00 | 54.77 |  |
| 10 | U | O5' | AA:963 | 4.74 | 1.00 | 57.72 |  |
  |

  
 Coded contact list:   
 Exclusion zone contact(s): /O3'/OP1/OP1/OP1/OP2/  
Consider K in place of MG (5 valid contact(s) for K vs. 0 for MG)  

### MG:AA:4153

|  | Res. type | Atom | Residue | Distance | Occupancy | B-factor | Closest solute to 1^st^ shell water |
| --- | --- | --- | --- | --- | --- | --- | --- |
| **Ion** | | | |  | 1.00 | 109.56 || 1 | A | OP1 | AA:1210 | 2.91 | 1.00 | 65.32 |
| 2 | G | OP1 | AA:1209 | 3.12 | 1.00 | 63.97 |  |
| 3 | A | OP2 | AA:1210 | 3.17 | 1.00 | 64.00 |  |
| 4 | G | O5' | AA:1209 | 3.51 | 1.00 | 63.98 |  |
| 5 | G | OP2 | AA:1209 | 3.91 | 1.00 | 61.90 |  |
| 6 | G | O3' | AA:1209 | 3.94 | 1.00 | 64.78 |  |
| 7 | G | OP1 | AA:1212 | 4.57 | 1.00 | 76.38 |  |
| 8 | A | O5' | AA:1210 | 4.88 | 1.00 | 67.58 |  |
  |

  
 Coded contact list:   
 Exclusion zone contact(s): /OP1/OP1/OP2/  
Consider K in place of MG (3 valid contact(s) for K vs. 0 for MG)  

### MG:AA:4154

|  | Res. type | Atom | Residue | Distance | Occupancy | B-factor | Closest solute to 1^st^ shell water |
| --- | --- | --- | --- | --- | --- | --- | --- |
| **Ion** | | | |  | 1.00 | 89.55 || 1 | G | N7 | AA:70 | 2.96 | 1.00 | 61.56 |
| 2 | G | O6 | AA:70 | 4.21 | 1.00 | 54.36 |  |
| 3 | G | OP2 | AA:70 | 4.64 | 1.00 | 58.60 |  |
| 4 | C | N4 | AA:69 | 4.67 | 1.00 | 56.08 |  |
| 5 | C | N3 | AA:69 | 4.74 | 1.00 | 55.09 |  |
| 6 | G | N9 | AA:70 | 4.98 | 1.00 | 59.68 |  |
| 7 | C | OP2 | AA:69 | 4.98 | 1.00 | 56.78 |  |
  |

  
 Coded contact list:   
 Exclusion zone contact(s): /N7/  
Consider K in place of MG (1 valid contact(s) for K vs. 0 for MG)  

### MG:AA:4155

|  | Res. type | Atom | Residue | Distance | Occupancy | B-factor | Closest solute to 1^st^ shell water |
| --- | --- | --- | --- | --- | --- | --- | --- |
| **Ion** | | | |  | 1.00 | 80.32 || 1 | G | O6 | AA:1348 | 3.16 | 1.00 | 54.19 |
| 2 | G | O6 | AA:1347 | 3.23 | 1.00 | 47.59 |  |
| 3 | A | N6 | AA:1384 | 4.11 | 1.00 | 66.11 |  |
| 4 | G | N7 | AA:1348 | 4.21 | 1.00 | 51.62 |  |
| 5 | C | N4 | AA:1599 | 4.42 | 1.00 | 46.77 |  |
| 6 | G | O6 | AA:1346 | 4.44 | 1.00 | 40.04 |  |
| 7 | G | N7 | AA:1347 | 4.55 | 1.00 | 45.44 |  |
| 8 | C | N4 | AA:1598 | 4.55 | 1.00 | 52.96 |  |
| 9 | G | OP1 | AA:1344 | 4.58 | 1.00 | 54.62 |  |
| 10 | G | O3' | AA:1343 | 4.70 | 1.00 | 54.07 |  |
| 11 | A | N6 | AA:1349 | 4.93 | 1.00 | 68.89 |  |
| 12 | G | O5' | AA:1344 | 4.96 | 1.00 | 55.85 |  |
  |

  
 Coded contact list:   
 Exclusion zone contact(s): /O6/O6/  
Consider K in place of MG (2 valid contact(s) for K vs. 0 for MG)  

### MG:AA:4156

|  | Res. type | Atom | Residue | Distance | Occupancy | B-factor | Closest solute to 1^st^ shell water |
| --- | --- | --- | --- | --- | --- | --- | --- |
| **Ion** | | | |  | 1.00 | 59.33 || 1 | A | OP1 | AA:2712A | 2.25 | 1.00 | 44.70 |
| 2 | U | O3' | AA:2712 | 2.51 | 1.00 | 47.68 |  |
| 3 | A | OP1 | AA:2711 | 2.53 | 1.00 | 46.41 |  |
| 4 | A | O5' | AA:2712A | 3.82 | 1.00 | 42.14 |  |
| 5 | A | OP2 | AA:2712A | 4.21 | 1.00 | 45.06 |  |
| 6 | C | O3' | AA:2710 | 4.68 | 1.00 | 47.28 |  |
| 7 | A | O5' | AA:2711 | 4.70 | 1.00 | 46.17 |  |
| 8 | U | O2' | AA:2712 | 4.79 | 1.00 | 52.57 |  |
| 9 | A | OP2 | AA:2711 | 4.97 | 1.00 | 45.22 |  |
  |

  
 Contact(s): /OP1/  
 Coded contact list: O_ph_  
 Exclusion zone contact(s): /O3'/OP1/  
Consider NA in place of MG (2 valid contact(s) for NA vs. 1 for MG)  

### MG:AA:4157

|  | Res. type | Atom | Residue | Distance | Occupancy | B-factor | Closest solute to 1^st^ shell water |
| --- | --- | --- | --- | --- | --- | --- | --- |
| **Ion** | | | |  | 1.00 | 76.63 || 1 | G | O6 | AA:2362 | 3.08 | 1.00 | 56.61 |
| 2 | G | N7 | AA:2362 | 3.39 | 1.00 | 58.51 |  |
| 3 | A | N7 | AA:2361 | 4.13 | 1.00 | 56.87 |  |
| 4 | C | N4 | AA:2363 | 4.41 | 1.00 | 58.95 |  |
| 5 | LYS | NZ | A8:26 | 4.45 | 1.00 | 52.60 |  |
| 6 | A | N6 | AA:2361 | 4.97 | 1.00 | 55.81 |  |
  |

  
 Coded contact list:   
 Exclusion zone contact(s): /N7/O6/  
Consider K in place of MG (2 valid contact(s) for K vs. 0 for MG)  

### MG:AA:4158

|  | Res. type | Atom | Residue | Distance | Occupancy | B-factor | Closest solute to 1^st^ shell water |
| --- | --- | --- | --- | --- | --- | --- | --- |
| **Ion** | | | |  | 1.00 | 93.49 || 1 | U | OP1 | AA:2406 | 3.00 | 1.00 | 70.14 |
| 2 | G | N9 | AA:2405 | 3.07 | 1.00 | 69.34 |  |
| 3 | G | O2' | AA:2405 | 3.65 | 1.00 | 70.19 |  |
| 4 | G | N3 | AA:2405 | 3.88 | 1.00 | 68.77 |  |
| 5 | G | O3' | AA:2405 | 3.93 | 1.00 | 68.91 |  |
| 6 | A | N7 | AA:2411 | 4.08 | 1.00 | 69.30 |  |
| 7 | G | N7 | AA:2405 | 4.11 | 1.00 | 70.23 |  |
| 8 | G | O4' | AA:2405 | 4.22 | 1.00 | 68.10 |  |
| 9 | A | N6 | AA:2411 | 4.28 | 1.00 | 66.97 |  |
| 10 | G | N7 | AA:2410 | 4.37 | 1.00 | 70.28 |  |
| 11 | G | O5' | AA:2405 | 4.61 | 1.00 | 66.69 |  |
| 12 | U | OP2 | AA:2406 | 4.85 | 1.00 | 69.59 |  |
  |

  
 Coded contact list:   
 Exclusion zone contact(s): /OP1/  
Consider K in place of MG (2 valid contact(s) for K vs. 0 for MG)  

### MG:AA:4159

|  | Res. type | Atom | Residue | Distance | Occupancy | B-factor | Closest solute to 1^st^ shell water |
| --- | --- | --- | --- | --- | --- | --- | --- |
| **Ion** | | | |  | 1.00 | 67.49 || 1 | G | OP1 | AA:51 | 2.98 | 1.00 | 65.04 |
| 2 | U | N1 | AA:50 | 3.25 | 1.00 | 72.72 |  |
| 3 | U | O2 | AA:50 | 3.40 | 1.00 | 73.27 |  |
| 4 | A | OP1 | AA:49 | 3.43 | 1.00 | 61.48 |  |
| 5 | U | O3' | AA:50 | 3.54 | 1.00 | 65.74 |  |
| 6 | U | O2' | AA:50 | 3.78 | 1.00 | 72.41 |  |
| 7 | U | O4' | AA:50 | 3.89 | 1.00 | 71.31 |  |
| 8 | G | O5' | AA:51 | 4.05 | 1.00 | 63.30 |  |
| 9 | U | N3 | AA:50 | 4.21 | 1.00 | 74.81 |  |
| 10 | A | O5' | AA:49 | 4.58 | 1.00 | 62.36 |  |
  |

  
 Coded contact list:   
 Exclusion zone contact(s): /N1/OP1/  
Consider K in place of MG (2 valid contact(s) for K vs. 0 for MG)  

### MG:AA:4160

|  | Res. type | Atom | Residue | Distance | Occupancy | B-factor | Closest solute to 1^st^ shell water |
| --- | --- | --- | --- | --- | --- | --- | --- |
| **Ion** | | | |  | 1.00 | 74.52 || 1 | G | O2' | AA:348 | 2.69 | 1.00 | 69.95 |
| 2 | G | OP1 | AA:109 | 3.08 | 1.00 | 63.15 |  |
| 3 | G | O3' | AA:348 | 3.75 | 1.00 | 71.39 |  |
| 4 | G | O5' | AA:349 | 4.17 | 1.00 | 71.36 |  |
| 5 | G | O5' | AA:109 | 4.41 | 1.00 | 61.50 |  |
| 6 | G | OP1 | AA:110 | 4.55 | 1.00 | 61.74 |  |
| 7 | G | OP1 | AA:349 | 4.66 | 1.00 | 70.91 |  |
| 8 | G | O3' | AA:109 | 4.85 | 1.00 | 62.56 |  |
  |

  
 Coded contact list:   
 Exclusion zone contact(s): /O2'/OP1/  
Consider K in place of MG (2 valid contact(s) for K vs. 0 for MG)  

### MG:AA:4161

|  | Res. type | Atom | Residue | Distance | Occupancy | B-factor | Closest solute to 1^st^ shell water |
| --- | --- | --- | --- | --- | --- | --- | --- |
| **Ion** | | | |  | 1.00 | 68.70 || 1 | G | OP2 | AA:947 | 2.95 | 1.00 | 54.77 |
| 2 | C | OP2 | AA:965 | 4.50 | 1.00 | 50.48 |  |
| 3 | G | O3' | AA:946 | 4.93 | 1.00 | 53.01 |  |
  |

  
 Coded contact list:   
 Exclusion zone contact(s): /OP2/  
Consider K in place of MG (1 valid contact(s) for K vs. 0 for MG)  

### MG:AA:4162

|  | Res. type | Atom | Residue | Distance | Occupancy | B-factor | Closest solute to 1^st^ shell water |
| --- | --- | --- | --- | --- | --- | --- | --- |
| **Ion** | | | |  | 1.00 | 94.39 || 1 | G | OP2 | AA:2643 | 3.28 | 1.00 | 65.17 |
| 2 | G | O6 | AA:2645 | 3.37 | 1.00 | 76.02 |  |
| 3 | G | O6 | AA:2644 | 4.49 | 1.00 | 69.50 |  |
| 4 | G | N7 | AA:2644 | 4.68 | 1.00 | 68.18 |  |
  |

  
 Coded contact list:   
 Exclusion zone contact(s): /O6/OP2/  
Consider K in place of MG (2 valid contact(s) for K vs. 0 for MG)  

### MG:AA:4163

|  | Res. type | Atom | Residue | Distance | Occupancy | B-factor | Closest solute to 1^st^ shell water |
| --- | --- | --- | --- | --- | --- | --- | --- |
| **Ion** | | | |  | 1.00 | 102.08 || 1 | G | N2 | AA:26 | 2.62 | 1.00 | 53.03 |
| 2 | A | OP2 | AA:514 | 2.96 | 1.00 | 58.76 |  |
| 3 | A | OP1 | AA:513 | 2.98 | 1.00 | 58.29 |  |
| 4 | A | OP2 | AA:513 | 3.14 | 1.00 | 58.08 |  |
| 5 | A | O5' | AA:513 | 3.37 | 1.00 | 57.97 |  |
| 6 | G | N1 | AA:26 | 3.59 | 1.00 | 54.97 |  |
| 7 | U | O4 | AA:25 | 3.61 | 1.00 | 58.91 |  |
| 8 | G | N3 | AA:26 | 4.63 | 1.00 | 54.09 |  |
| 9 | A | O5' | AA:514 | 4.77 | 1.00 | 58.09 |  |
| 10 | G | O3' | AA:512 | 4.80 | 1.00 | 57.15 |  |
| 11 | A | N7 | AA:514 | 4.83 | 1.00 | 59.42 |  |
| 12 | G | N2 | AA:27 | 4.86 | 1.00 | 55.09 |  |
  |

  
 Coded contact list:   
 Exclusion zone contact(s): /N2/O5'/OP1/OP2/OP2/  
Consider K in place of MG (5 valid contact(s) for K vs. 0 for MG)  

### MG:AA:4165

|  | Res. type | Atom | Residue | Distance | Occupancy | B-factor | Closest solute to 1^st^ shell water |
| --- | --- | --- | --- | --- | --- | --- | --- |
| **Ion** | | | |  | 1.00 | 64.84 || 1 | C | OP2 | AA:451 | 2.31 | 1.00 | 43.73 |
| 2 | G | OP2 | AA:450 | 2.34 | 1.00 | 40.85 |  |
| 3 | G | O5' | AA:450 | 2.88 | 1.00 | 40.46 |  |
| 4 | G | OP1 | AA:450 | 3.33 | 1.00 | 41.17 |  |
| 5 | G | OP2 | AA:452 | 3.83 | 1.00 | 38.78 |  |
| 6 | C | O5' | AA:451 | 4.19 | 1.00 | 44.20 |  |
| 7 | G | O3' | AA:450 | 4.32 | 1.00 | 45.06 |  |
| 8 | A | O3' | AA:449 | 4.41 | 1.00 | 39.43 |  |
| 9 | C | OP1 | AA:451 | 4.87 | 1.00 | 43.49 |  |
| 10 | A | OP2 | AA:449 | 4.91 | 1.00 | 34.02 |  |
  |

  
**1^st^ shell ligand angles:**   
1-2: 129;   
  
 Contact(s): /OP2/OP2/  
 Coded contact list: 2O_ph_  
 Exclusion zone contact(s): /O5'/OP1/  

### MG:AA:4166

|  | Res. type | Atom | Residue | Distance | Occupancy | B-factor | Closest solute to 1^st^ shell water |
| --- | --- | --- | --- | --- | --- | --- | --- |
| **Ion** | | | |  | 1.00 | 142.10 || 1 | G | N7 | AA:2131 | 2.63 | 1.00 | 200.14 |
| 2 | U | O4 | AA:2130 | 3.17 | 1.00 | 198.93 |  |
| 3 | U | N3 | AA:2130 | 3.76 | 1.00 | 199.01 |  |
| 4 | G | N9 | AA:2131 | 4.31 | 1.00 | 200.20 |  |
| 5 | U | N1 | AA:2130 | 4.34 | 1.00 | 199.54 |  |
| 6 | G | OP2 | AA:2131 | 4.75 | 1.00 | 199.85 |  |
| 7 | G | O6 | AA:2131 | 4.88 | 1.00 | 200.21 |  |
  |

  
 Coded contact list:   
 Exclusion zone contact(s): /N7/O4/  
Consider K in place of MG (2 valid contact(s) for K vs. 0 for MG)  

### MG:AB:201

|  | Res. type | Atom | Residue | Distance | Occupancy | B-factor | Closest solute to 1^st^ shell water |
| --- | --- | --- | --- | --- | --- | --- | --- |
| **Ion** | | | |  | 1.00 | 42.46 || 1 | C | OP2 | AB:36 | 2.01 | 1.00 | 84.68 |
| 2 | U | OP2 | AB:35 | 2.66 | 1.00 | 91.70 |  |
| 3 | U | O5' | AB:35 | 3.27 | 1.00 | 91.28 |  |
| 4 | U | O3' | AB:35 | 3.58 | 1.00 | 85.46 |  |
| 5 | U | O3' | AB:34 | 3.62 | 1.00 | 92.10 |  |
| 6 | C | OP1 | AB:36 | 3.97 | 1.00 | 84.47 |  |
| 7 | C | O5' | AB:36 | 4.48 | 1.00 | 83.14 |  |
| 8 | U | O2' | AB:34 | 4.69 | 1.00 | 93.31 |  |
| 9 | U | OP1 | AB:35 | 4.86 | 1.00 | 92.46 |  |
  |

  
 Contact(s): /OP2/  
 Coded contact list: O_ph_  
 Exclusion zone contact(s): /O5'/OP2/  
Consider K in place of MG (2 valid contact(s) for K vs. 1 for MG)  

### MG:AB:202

|  | Res. type | Atom | Residue | Distance | Occupancy | B-factor | Closest solute to 1^st^ shell water |
| --- | --- | --- | --- | --- | --- | --- | --- |
| **Ion** | | | |  | 1.00 | 51.42 || 1 | G | OP2 | AB:100 | 2.24 | 1.00 | 79.66 |
| 2 | A | OP2 | AB:73 | 3.29 | 1.00 | 89.73 |  |
| 3 | G | O6 | AB:102 | 4.08 | 1.00 | 76.00 |  |
| 4 | G | O5' | AB:100 | 4.37 | 1.00 | 77.92 |  |
| 5 | G | OP1 | AB:100 | 4.60 | 1.00 | 79.32 |  |
| 6 | A | OP1 | AB:73 | 4.64 | 1.00 | 89.68 |  |
| 7 | A | OP2 | AB:101 | 4.66 | 1.00 | 78.49 |  |
| 8 | A | O3' | AB:99 | 4.68 | 1.00 | 79.18 |  |
| 9 | A | O5' | AB:73 | 4.77 | 1.00 | 89.32 |  |
| 10 | A | N7 | AB:101 | 4.93 | 1.00 | 70.35 |  |
  |

  
 Contact(s): /OP2/  
 Coded contact list: O_ph_  
 Exclusion zone contact(s): /OP2/  

### MG:AB:203

|  | Res. type | Atom | Residue | Distance | Occupancy | B-factor | Closest solute to 1^st^ shell water |
| --- | --- | --- | --- | --- | --- | --- | --- |
| **Ion** | | | |  | 1.00 | 37.57 || 1 | G | N7 | AB:102 | 3.06 | 1.00 | 76.53 |
| 2 | A | OP2 | AB:101 | 3.76 | 1.00 | 78.49 |  |
| 3 | G | O6 | AB:102 | 3.86 | 1.00 | 76.00 |  |
| 4 | G | OP2 | AB:102 | 4.02 | 1.00 | 80.19 |  |
| 5 | U | O4 | AB:103 | 4.41 | 1.00 | 90.61 |  |
| 6 | G | O6 | AB:72 | 4.64 | 1.00 | 93.31 |  |
| 7 | G | N1 | AB:72 | 4.78 | 1.00 | 92.02 |  |
| 8 | A | OP1 | AB:101 | 4.78 | 1.00 | 77.82 |  |
| 9 | G | N7 | AB:72 | 4.81 | 1.00 | 91.85 |  |
| 10 | A | N7 | AB:101 | 4.99 | 1.00 | 70.35 |  |
  |

  
 Coded contact list:   
 Exclusion zone contact(s): /N7/  
Consider K in place of MG (1 valid contact(s) for K vs. 0 for MG)  

### MG:AB:204

|  | Res. type | Atom | Residue | Distance | Occupancy | B-factor | Closest solute to 1^st^ shell water |
| --- | --- | --- | --- | --- | --- | --- | --- |
| **Ion** | | | |  | 1.00 | 54.31 || 1 | G | N1 | AB:81 | 2.75 | 1.00 | 102.14 |
| 2 | C | N4 | AB:94 | 2.88 | 1.00 | 91.74 |  |
| 3 | G | O6 | AB:82 | 2.94 | 1.00 | 89.95 |  |
| 4 | U | O4 | AB:80 | 3.10 | 1.00 | 84.59 |  |
| 5 | U | N3 | AB:80 | 3.17 | 1.00 | 83.12 |  |
| 6 | U | N3 | AB:95 | 3.21 | 1.00 | 103.01 |  |
| 7 | G | O6 | AB:81 | 3.24 | 1.00 | 101.16 |  |
| 8 | U | O2 | AB:95 | 3.32 | 1.00 | 102.50 |  |
| 9 | G | N2 | AB:81 | 3.52 | 1.00 | 103.38 |  |
| 10 | U | O4 | AB:95 | 4.50 | 1.00 | 103.90 |  |
| 11 | C | N3 | AB:94 | 4.52 | 1.00 | 92.23 |  |
| 12 | G | N1 | AB:82 | 4.52 | 1.00 | 90.76 |  |
| 13 | G | O6 | AB:96 | 4.57 | 1.00 | 86.58 |  |
| 14 | U | O2 | AB:80 | 4.62 | 1.00 | 84.27 |  |
| 15 | U | N1 | AB:80 | 4.62 | 1.00 | 81.01 |  |
| 16 | G | N3 | AB:81 | 4.82 | 1.00 | 101.07 |  |
  |

  
 Coded contact list:   
 Exclusion zone contact(s): /N1_G_/N3_U_/N3_U_/N4/O2/O4/O6/O6/  
Consider K in place of MG (8 valid contact(s) for K vs. 0 for MG)  

### MG:AB:205

|  | Res. type | Atom | Residue | Distance | Occupancy | B-factor | Closest solute to 1^st^ shell water |
| --- | --- | --- | --- | --- | --- | --- | --- |
| **Ion** | | | |  | 1.00 | 54.92 || 1 | A | OP2 | AB:25 | 2.12 | 1.00 | 105.44 |
| 2 | G | O3' | AB:24 | 2.40 | 1.00 | 106.19 |  |
| 3 | A | O5' | AB:25 | 2.82 | 1.00 | 104.53 |  |
| 4 | A | OP2 | AB:26 | 3.53 | 1.00 | 99.64 |  |
| 5 | A | O3' | AB:25 | 3.62 | 1.00 | 99.57 |  |
| 6 | G | O2' | AB:24 | 3.65 | 1.00 | 106.40 |  |
| 7 | A | OP1 | AB:25 | 4.02 | 1.00 | 106.36 |  |
| 8 | A | OP1 | AB:26 | 4.49 | 1.00 | 99.63 |  |
  |

  
**1^st^ shell ligand angles:**   
1-2: 67;   
  
 Contact(s): /O3'/OP2/  
 Coded contact list: O_ph_O_r_  
 Exclusion zone contact(s): /O5'/  

### MG:AB:206

|  | Res. type | Atom | Residue | Distance | Occupancy | B-factor | Closest solute to 1^st^ shell water |
| --- | --- | --- | --- | --- | --- | --- | --- |
| **Ion** | | | |  | 1.00 | 76.61 || 1 | U | OP2 | AB:55 | 2.99 | 1.00 | 90.85 |
| 2 | A | OP2 | AB:25 | 3.25 | 1.00 | 105.44 |  |
| 3 | A | OP1 | AB:25 | 4.56 | 1.00 | 106.36 |  |
| 4 | G | O3' | AB:24 | 4.83 | 1.00 | 106.19 |  |
| 5 | U | OP1 | AB:55 | 4.90 | 1.00 | 91.24 |  |
| 6 | U | O5' | AB:55 | 4.95 | 1.00 | 91.47 |  |
  |

  
 Coded contact list:   
 Exclusion zone contact(s): /OP2/OP2/  
Consider K in place of MG (2 valid contact(s) for K vs. 0 for MG)  

### MG:AB:207

|  | Res. type | Atom | Residue | Distance | Occupancy | B-factor | Closest solute to 1^st^ shell water |
| --- | --- | --- | --- | --- | --- | --- | --- |
| **Ion** | | | |  | 1.00 | 74.66 || 1 | G | O6 | AB:21 | 2.01 | 1.00 | 103.66 |
| 2 | C | N4 | AB:62 | 2.43 | 1.00 | 101.40 |  |
| 3 | G | O6 | AB:61 | 2.70 | 1.00 | 105.20 |  |
| 4 | U | O4 | AB:22 | 3.42 | 1.00 | 106.41 |  |
| 5 | G | N1 | AB:21 | 3.89 | 1.00 | 104.63 |  |
| 6 | U | N3 | AB:22 | 4.05 | 1.00 | 106.05 |  |
| 7 | C | N3 | AB:62 | 4.10 | 1.00 | 101.39 |  |
| 8 | G | O6 | AB:63 | 4.18 | 1.00 | 104.99 |  |
| 9 | G | N7 | AB:61 | 4.43 | 1.00 | 103.20 |  |
| 10 | C | N4 | AB:60 | 4.71 | 1.00 | 99.23 |  |
| 11 | G | N1 | AB:61 | 4.72 | 1.00 | 105.03 |  |
| 12 | G | N7 | AB:21 | 4.74 | 1.00 | 106.96 |  |
| 13 | C | N4 | AB:20 | 4.80 | 1.00 | 106.67 |  |
  |

  
 Contact(s): /O6/  
 Coded contact list: O_b_  
 Exclusion zone contact(s): /N4/O6/  

### MG:AB:208

|  | Res. type | Atom | Residue | Distance | Occupancy | B-factor | Closest solute to 1^st^ shell water |
| --- | --- | --- | --- | --- | --- | --- | --- |
| **Ion** | | | |  | 1.00 | 57.87 || 1 | A | OP2 | AB:58 | 2.05 | 1.00 | 96.46 |
| 2 | A | OP2 | AB:57 | 2.60 | 1.00 | 98.13 |  |
| 3 | G | O3' | AB:56 | 3.22 | 1.00 | 97.44 |  |
| 4 | G | O2' | AB:56 | 3.75 | 1.00 | 98.22 |  |
| 5 | G | O6 | AB:23 | 3.79 | 1.00 | 99.09 |  |
| 6 | A | O5' | AB:57 | 3.87 | 1.00 | 97.38 |  |
| 7 | A | O3' | AB:57 | 4.18 | 1.00 | 96.77 |  |
| 8 | A | OP1 | AB:58 | 4.28 | 1.00 | 97.40 |  |
| 9 | C | N4 | AB:60 | 4.28 | 1.00 | 99.23 |  |
| 10 | A | O5' | AB:58 | 4.49 | 1.00 | 96.96 |  |
| 11 | U | O4 | AB:22 | 4.65 | 1.00 | 106.41 |  |
| 12 | A | N6 | AB:59 | 4.69 | 1.00 | 104.51 |  |
| 13 | A | N7 | AB:58 | 4.74 | 1.00 | 88.89 |  |
| 14 | A | N7 | AB:59 | 4.76 | 1.00 | 103.44 |  |
| 15 | A | OP1 | AB:57 | 4.82 | 1.00 | 98.14 |  |
  |

  
 Contact(s): /OP2/  
 Coded contact list: O_ph_  
 Exclusion zone contact(s): /O3'/OP2/  

### MG:AB:209

|  | Res. type | Atom | Residue | Distance | Occupancy | B-factor | Closest solute to 1^st^ shell water |
| --- | --- | --- | --- | --- | --- | --- | --- |
| **Ion** | | | |  | 1.00 | 104.25 || 1 | G | OP2 | AB:54 | 3.31 | 1.00 | 97.08 |
| 2 | G | OP1 | AB:54 | 3.52 | 1.00 | 96.67 |  |
| 3 | A | OP2 | AB:53 | 4.34 | 1.00 | 106.54 |  |
  |

  
 Coded contact list:   
 Exclusion zone contact(s): /OP2/  
Consider K in place of MG (1 valid contact(s) for K vs. 0 for MG)  

### MG:AB:210

|  | Res. type | Atom | Residue | Distance | Occupancy | B-factor | Closest solute to 1^st^ shell water |
| --- | --- | --- | --- | --- | --- | --- | --- |
| **Ion** | | | |  | 1.00 | 81.55 || 1 | G | O6 | AB:76 | 2.96 | 1.00 | 79.17 |
| 2 | G | N7 | AB:76 | 3.10 | 1.00 | 79.96 |  |
| 3 | G | N7 | AB:75 | 4.04 | 1.00 | 89.07 |  |
| 4 | U | O4 | AB:77 | 4.28 | 1.00 | 72.39 |  |
| 5 | A | O5' | AB:99 | 4.34 | 1.00 | 76.83 |  |
  |

  
 Coded contact list:   
 Exclusion zone contact(s): /N7/O6/  
Consider K in place of MG (2 valid contact(s) for K vs. 0 for MG)  

### MG:AB:211

|  | Res. type | Atom | Residue | Distance | Occupancy | B-factor | Closest solute to 1^st^ shell water |
| --- | --- | --- | --- | --- | --- | --- | --- |
| **Ion** | | | |  | 1.00 | 77.89 || 1 | G | N7 | AB:116 | 3.58 | 1.00 | 88.43 |
| 2 | G | OP2 | AB:116 | 4.39 | 1.00 | 91.07 |  |
| 3 | G | N7 | AB:115 | 4.49 | 1.00 | 82.25 |  |
| 4 | G | O6 | AB:116 | 4.69 | 1.00 | 88.39 |  |
| 5 | G | O6 | AB:117 | 4.76 | 1.00 | 104.47 |  |
| 6 | G | N7 | AB:117 | 4.83 | 1.00 | 104.79 |  |
  |

  
 Coded contact list:   
1^st^ MG coordination shell missing (see Notes)  

### MG:AB:212

|  | Res. type | Atom | Residue | Distance | Occupancy | B-factor | Closest solute to 1^st^ shell water |
| --- | --- | --- | --- | --- | --- | --- | --- |
| **Ion** | | | |  | 1.00 | 83.08 || 1 | G | OP1 | AB:33 | 3.30 | 1.00 | 91.96 |
| 2 | G | O5' | AB:33 | 3.93 | 1.00 | 91.35 |  |
  |

  
 Coded contact list:   
 Exclusion zone contact(s): /OP1/  
Consider K in place of MG (1 valid contact(s) for K vs. 0 for MG)  

### MG:AB:213

|  | Res. type | Atom | Residue | Distance | Occupancy | B-factor | Closest solute to 1^st^ shell water |
| --- | --- | --- | --- | --- | --- | --- | --- |
| **Ion** | | | |  | 1.00 | 60.74 || 1 | G | N7 | AB:81 | 2.96 | 1.00 | 98.73 |
| 2 | G | N1 | AB:81 | 3.17 | 1.00 | 102.14 |  |
| 3 | G | N3 | AB:81 | 3.39 | 1.00 | 101.07 |  |
| 4 | G | N9 | AB:82 | 3.42 | 1.00 | 89.31 |  |
| 5 | G | N9 | AB:81 | 3.44 | 1.00 | 95.55 |  |
| 6 | G | N3 | AB:82 | 3.61 | 1.00 | 90.61 |  |
| 7 | G | O6 | AB:81 | 3.68 | 1.00 | 101.16 |  |
| 8 | U | O2 | AB:95 | 3.78 | 1.00 | 102.50 |  |
| 9 | G | N7 | AB:82 | 3.84 | 1.00 | 87.95 |  |
| 10 | G | N1 | AB:82 | 4.34 | 1.00 | 90.76 |  |
| 11 | G | N3 | AB:96 | 4.34 | 1.00 | 88.86 |  |
| 12 | G | O4' | AB:82 | 4.53 | 1.00 | 89.13 |  |
| 13 | G | N2 | AB:81 | 4.53 | 1.00 | 103.38 |  |
| 14 | G | N9 | AB:96 | 4.64 | 1.00 | 89.99 |  |
| 15 | U | O2' | AB:95 | 4.77 | 1.00 | 100.06 |  |
| 16 | G | O6 | AB:82 | 4.85 | 1.00 | 89.95 |  |
  |

  
 Coded contact list:   
 Exclusion zone contact(s): /N1_G_/N3_G_/N7/  
Consider K in place of MG (3 valid contact(s) for K vs. 0 for MG)  

### MG:AB:214

|  | Res. type | Atom | Residue | Distance | Occupancy | B-factor | Closest solute to 1^st^ shell water |
| --- | --- | --- | --- | --- | --- | --- | --- |
| **Ion** | | | |  | 1.00 | 105.02 || 1 | G | N7 | AB:117 | 2.59 | 1.00 | 104.79 |
| 2 | G | O6 | AB:117 | 3.08 | 1.00 | 104.47 |  |
| 3 | G | N7 | AB:118 | 3.96 | 1.00 | 126.00 |  |
| 4 | G | O6 | AB:118 | 4.38 | 1.00 | 126.00 |  |
| 5 | G | N9 | AB:117 | 4.56 | 1.00 | 105.43 |  |
| 6 | G | N1 | AB:117 | 4.61 | 1.00 | 103.58 |  |
| 7 | G | N7 | AB:116 | 4.67 | 1.00 | 88.43 |  |
| 8 | G | O6 | AB:116 | 4.88 | 1.00 | 88.39 |  |
  |

  
 Coded contact list:   
 Exclusion zone contact(s): /N7/O6/  
Consider NA in place of MG (1 valid contact(s) for NA vs. 0 for MG)  

### MG:AB:215

|  | Res. type | Atom | Residue | Distance | Occupancy | B-factor | Closest solute to 1^st^ shell water |
| --- | --- | --- | --- | --- | --- | --- | --- |
| **Ion** | | | |  | 1.00 | 70.55 || 1 | C | O2' | AB:38 | 2.64 | 1.00 | 86.60 |
| 2 | C | O2 | AB:47 | 2.73 | 1.00 | 73.67 |  |
| 3 | A | O4' | AB:39 | 3.02 | 1.00 | 95.61 |  |
| 4 | C | O2' | AB:47 | 3.82 | 1.00 | 76.05 |  |
| 5 | A | O2' | AB:39 | 3.83 | 1.00 | 95.80 |  |
| 6 | A | O4' | AB:48 | 3.98 | 1.00 | 82.38 |  |
| 7 | C | O2 | AB:38 | 3.98 | 1.00 | 84.38 |  |
| 8 | A | O5' | AB:39 | 4.50 | 1.00 | 90.49 |  |
| 9 | A | N9 | AB:39 | 4.54 | 1.00 | 98.15 |  |
| 10 | LYS | NZ | AQ:93 | 4.66 | 1.00 | 81.17 |  |
| 11 | C | N1 | AB:47 | 4.81 | 1.00 | 73.32 |  |
| 12 | C | N3 | AB:47 | 4.82 | 1.00 | 72.94 |  |
| 13 | C | N1 | AB:38 | 4.88 | 1.00 | 85.10 |  |
  |

  
 Coded contact list:   
 Exclusion zone contact(s): /O2/O2'/O4'/  
Consider K in place of MG (3 valid contact(s) for K vs. 0 for MG)  

### MG:AB:216

|  | Res. type | Atom | Residue | Distance | Occupancy | B-factor | Closest solute to 1^st^ shell water |
| --- | --- | --- | --- | --- | --- | --- | --- |
| **Ion** | | | |  | 1.00 | 53.73 || 1 | C | OP2 | AB:43 | 2.28 | 1.00 | 91.76 |
| 2 | U | N1 | AB:40 | 2.59 | 1.00 | 100.46 |  |
| 3 | U | OP1 | AB:41 | 2.88 | 1.00 | 95.44 |  |
| 4 | U | N3 | AB:40 | 3.05 | 1.00 | 102.32 |  |
| 5 | U | O2 | AB:40 | 3.55 | 1.00 | 100.84 |  |
| 6 | U | O3' | AB:40 | 3.83 | 1.00 | 95.68 |  |
| 7 | U | O4' | AB:40 | 4.13 | 1.00 | 98.16 |  |
| 8 | C | O3' | AB:42 | 4.14 | 1.00 | 91.63 |  |
| 9 | U | O2' | AB:40 | 4.18 | 1.00 | 98.71 |  |
| 10 | U | O4 | AB:40 | 4.19 | 1.00 | 104.61 |  |
| 11 | U | O5' | AB:41 | 4.26 | 1.00 | 95.14 |  |
| 12 | C | OP1 | AB:43 | 4.29 | 1.00 | 90.84 |  |
| 13 | U | O5' | AB:40 | 4.31 | 1.00 | 97.07 |  |
| 14 | C | O5' | AB:42 | 4.49 | 1.00 | 93.28 |  |
| 15 | U | O3' | AB:41 | 4.53 | 1.00 | 94.94 |  |
| 16 | C | O5' | AB:43 | 4.75 | 1.00 | 90.33 |  |
| 17 | A | N6 | AB:45 | 4.81 | 1.00 | 84.39 |  |
  |

  
 Contact(s): /OP2/  
 Coded contact list: O_ph_  
 Exclusion zone contact(s): /N1/N3_U_/OP1/  
Consider K in place of MG (2 valid contact(s) for K vs. 1 for MG)  

### MG:AB:217

|  | Res. type | Atom | Residue | Distance | Occupancy | B-factor | Closest solute to 1^st^ shell water |
| --- | --- | --- | --- | --- | --- | --- | --- |
| **Ion** | | | |  | 1.00 | 87.48 || 1 | G | O6 | AB:115 | 3.70 | 1.00 | 79.12 |
| 2 | G | O6 | AB:114 | 4.16 | 1.00 | 75.93 |  |
| 3 | G | N7 | AB:115 | 4.54 | 1.00 | 82.25 |  |
| 4 | G | N7 | AB:114 | 4.85 | 1.00 | 80.07 |  |
  |

  
 Coded contact list:   
1^st^ MG coordination shell missing (see Notes)  

### MG:AB:218

|  | Res. type | Atom | Residue | Distance | Occupancy | B-factor | Closest solute to 1^st^ shell water |
| --- | --- | --- | --- | --- | --- | --- | --- |
| **Ion** | | | |  | 1.00 | 81.77 || 1 | C | O2' | AB:60 | 2.56 | 1.00 | 97.95 |
| 2 | C | O3' | AB:60 | 3.82 | 1.00 | 97.94 |  |
| 3 | G | OP1 | AB:7 | 4.20 | 1.00 | 79.84 |  |
| 4 | C | O4' | AB:60 | 4.66 | 1.00 | 98.12 |  |
| 5 | G | OP2 | AB:7 | 4.81 | 1.00 | 80.37 |  |
  |

  
 Coded contact list:   
 Exclusion zone contact(s): /O2'/  
Consider NA in place of MG (1 valid contact(s) for NA vs. 0 for MG)  

### MG:AB:219

|  | Res. type | Atom | Residue | Distance | Occupancy | B-factor | Closest solute to 1^st^ shell water |
| --- | --- | --- | --- | --- | --- | --- | --- |
| **Ion** | | | |  | 1.00 | 75.25 || 1 | U | O4 | AB:34 | 2.37 | 1.00 | 88.56 |
| 2 | G | O2' | AB:44 | 2.86 | 1.00 | 89.16 |  |
| 3 | A | N6 | AB:48 | 3.20 | 1.00 | 88.03 |  |
| 4 | G | O6 | AB:33 | 3.62 | 1.00 | 83.25 |  |
| 5 | G | O3' | AB:44 | 4.12 | 1.00 | 89.44 |  |
| 6 | G | N7 | AB:33 | 4.32 | 1.00 | 84.43 |  |
| 7 | U | N3 | AB:34 | 4.54 | 1.00 | 88.97 |  |
| 8 | A | N1 | AB:48 | 4.93 | 1.00 | 87.60 |  |
  |

  
 Contact(s): /O4/  
 Coded contact list: O_b_  
 Exclusion zone contact(s): /N6/O2'/  
Consider K in place of MG (2 valid contact(s) for K vs. 1 for MG)  

### MG:AB:220

|  | Res. type | Atom | Residue | Distance | Occupancy | B-factor | Closest solute to 1^st^ shell water |
| --- | --- | --- | --- | --- | --- | --- | --- |
| **Ion** | | | |  | 1.00 | 79.59 || 1 | G | O6 | AB:82 | 2.63 | 1.00 | 89.95 |
| 2 | G | O6 | AB:83 | 2.91 | 1.00 | 89.12 |  |
| 3 | G | N2 | AB:81 | 3.14 | 1.00 | 103.38 |  |
| 4 | C | N4 | AB:94 | 3.36 | 1.00 | 91.74 |  |
| 5 | C | N4 | AB:93 | 4.12 | 1.00 | 89.26 |  |
| 6 | G | N7 | AB:83 | 4.24 | 1.00 | 87.31 |  |
| 7 | G | N1 | AB:81 | 4.43 | 1.00 | 102.14 |  |
| 8 | G | N7 | AB:82 | 4.71 | 1.00 | 87.95 |  |
| 9 | C | N4 | AB:84 | 4.73 | 1.00 | 94.62 |  |
| 10 | G | N1 | AB:82 | 4.79 | 1.00 | 90.76 |  |
| 11 | G | N1 | AB:83 | 4.94 | 1.00 | 88.41 |  |
  |

  
 Coded contact list:   
 Exclusion zone contact(s): /N2/N4/O6/O6/  
Consider K in place of MG (4 valid contact(s) for K vs. 0 for MG)  

### MG:AB:221

|  | Res. type | Atom | Residue | Distance | Occupancy | B-factor | Closest solute to 1^st^ shell water |
| --- | --- | --- | --- | --- | --- | --- | --- |
| **Ion** | | | |  | 1.00 | 61.81 || 1 | C | OP2 | AB:64 | 3.97 | 1.00 | 102.78 |
| 2 | G | OP1 | AB:110 | 4.03 | 1.00 | 109.45 |  |
| 3 | G | OP1 | AB:63 | 4.52 | 1.00 | 101.15 |  |
| 4 | G | O3' | AB:109 | 4.57 | 1.00 | 109.33 |  |
| 5 | G | OP1 | AB:109 | 4.78 | 1.00 | 108.30 |  |
| 6 | C | OP1 | AB:64 | 4.88 | 1.00 | 102.29 |  |
  |

  
 Coded contact list:   
1^st^ MG coordination shell missing (see Notes)  

### MG:AB:222

|  | Res. type | Atom | Residue | Distance | Occupancy | B-factor | Closest solute to 1^st^ shell water |
| --- | --- | --- | --- | --- | --- | --- | --- |
| **Ion** | | | |  | 1.00 | 85.99 || 1 | U | OP2 | AB:35 | 3.27 | 1.00 | 91.70 |
| 2 | C | OP2 | AB:36 | 4.07 | 1.00 | 84.68 |  |
| 3 | U | O3' | AB:35 | 4.17 | 1.00 | 85.46 |  |
| 4 | U | O5' | AB:35 | 4.33 | 1.00 | 91.28 |  |
| 5 | C | OP1 | AB:36 | 4.41 | 1.00 | 84.47 |  |
| 6 | GLU | OE1 | A4:3 | 4.94 | 1.00 | 128.89 |  |
  |

  
 Coded contact list:   
 Exclusion zone contact(s): /OP2/  
Consider K in place of MG (1 valid contact(s) for K vs. 0 for MG)  

### MG:AB:223

|  | Res. type | Atom | Residue | Distance | Occupancy | B-factor | Closest solute to 1^st^ shell water |
| --- | --- | --- | --- | --- | --- | --- | --- |
| **Ion** | | | |  | 1.00 | 112.80 || 1 | A | OP2 | AB:0 | 2.80 | 1.00 | 154.77 |
| 2 | A | O3' | AB:1M | 4.92 | 1.00 | 156.16 |  |
  |

  
 Coded contact list:   
 Exclusion zone contact(s): /OP2/  
Consider K in place of MG (1 valid contact(s) for K vs. 0 for MG)  

### MG:AB:224

|  | Res. type | Atom | Residue | Distance | Occupancy | B-factor | Closest solute to 1^st^ shell water |
| --- | --- | --- | --- | --- | --- | --- | --- |
| **Ion** | | | |  | 1.00 | 85.16 || 1 | C | OP2 | AB:47 | 2.79 | 1.00 | 81.30 |
| 2 | A | OP2 | AB:46 | 2.95 | 1.00 | 82.47 |  |
| 3 | A | O5' | AB:46 | 3.68 | 1.00 | 82.01 |  |
| 4 | A | OP1 | AB:46 | 3.81 | 1.00 | 82.75 |  |
| 5 | A | N7 | AB:48 | 4.27 | 1.00 | 85.73 |  |
| 6 | C | OP1 | AB:47 | 4.50 | 1.00 | 81.66 |  |
| 7 | A | O3' | AB:46 | 4.74 | 1.00 | 81.47 |  |
| 8 | A | N6 | AB:48 | 4.84 | 1.00 | 88.03 |  |
  |

  
 Coded contact list:   
 Exclusion zone contact(s): /OP2/OP2/  
Consider K in place of MG (2 valid contact(s) for K vs. 0 for MG)  

### MG:AB:225

|  | Res. type | Atom | Residue | Distance | Occupancy | B-factor | Closest solute to 1^st^ shell water |
| --- | --- | --- | --- | --- | --- | --- | --- |
| **Ion** | | | |  | 1.00 | 125.89 || 1 | G | O2' | AB:116 | 2.35 | 1.00 | 95.94 |
| 2 | A | O2' | AB:26 | 2.57 | 1.00 | 97.23 |  |
| 3 | A | O3' | AB:26 | 3.11 | 1.00 | 97.36 |  |
| 4 | C | O5' | AB:27 | 3.26 | 1.00 | 96.90 |  |
| 5 | G | O4' | AB:117 | 3.75 | 1.00 | 106.66 |  |
| 6 | C | O4' | AB:27 | 4.11 | 1.00 | 93.19 |  |
| 7 | C | OP1 | AB:27 | 4.32 | 1.00 | 97.16 |  |
| 8 | G | O3' | AB:116 | 4.71 | 1.00 | 98.26 |  |
  |

  
 Contact(s): /O2'/  
 Coded contact list: O_r_  
 Exclusion zone contact(s): /O2'/O3'/O5'/  
Consider K in place of MG (2 valid contact(s) for K vs. 1 for MG)  

### MG:AB:226

|  | Res. type | Atom | Residue | Distance | Occupancy | B-factor | Closest solute to 1^st^ shell water |
| --- | --- | --- | --- | --- | --- | --- | --- |
| **Ion** | | | |  | 1.00 | 88.78 || 1 | A | N3 | AB:119 | 3.56 | 1.00 | 130.17 |
| 2 | A | N9 | AB:119 | 3.78 | 1.00 | 129.33 |  |
| 3 | A | N1 | AB:119 | 4.17 | 1.00 | 130.27 |  |
| 4 | A | N7 | AB:119 | 4.24 | 1.00 | 129.88 |  |
| 5 | A | N1 | AB:0 | 4.63 | 1.00 | 149.15 |  |
| 6 | A | O2' | AB:119 | 4.64 | 1.00 | 128.19 |  |
| 7 | A | O3' | AB:119 | 4.80 | 1.00 | 127.71 |  |
| 8 | A | N6 | AB:119 | 4.98 | 1.00 | 129.85 |  |
  |

  
 Coded contact list:   
1^st^ MG coordination shell missing (see Notes)  

### MG:AB:227

|  | Res. type | Atom | Residue | Distance | Occupancy | B-factor | Closest solute to 1^st^ shell water |
| --- | --- | --- | --- | --- | --- | --- | --- |
| **Ion** | | | |  | 1.00 | 82.99 || 1 | C | O2' | AB:4 | 2.55 | 1.00 | 93.22 |
| 2 | C | O3' | AB:4 | 2.65 | 1.00 | 92.52 |  |
| 3 | C | OP1 | AB:5 | 4.17 | 1.00 | 90.89 |  |
| 4 | C | O5' | AB:5 | 4.33 | 1.00 | 89.70 |  |
| 5 | C | O4' | AB:4 | 4.36 | 1.00 | 94.05 |  |
| 6 | G | N2 | AB:61 | 4.79 | 1.00 | 104.54 |  |
| 7 | G | O2' | AB:23 | 4.90 | 1.00 | 105.70 |  |
| 8 | C | O4' | AB:5 | 4.95 | 1.00 | 82.49 |  |
  |

  
 Coded contact list:   
 Exclusion zone contact(s): /O2'/O3'/  
Consider NA in place of MG (1 valid contact(s) for NA vs. 0 for MG)  

### MG:AB:228

|  | Res. type | Atom | Residue | Distance | Occupancy | B-factor | Closest solute to 1^st^ shell water |
| --- | --- | --- | --- | --- | --- | --- | --- |
| **Ion** | | | |  | 1.00 | 64.94 || 1 | C | OP2 | AB:31 | 2.34 | 1.00 | 87.58 |
| 2 | C | OP1 | AB:31 | 3.50 | 1.00 | 88.03 |  |
| 3 | C | O3' | AB:30 | 3.82 | 1.00 | 87.95 |  |
| 4 | A | OP2 | AB:46 | 4.29 | 1.00 | 82.47 |  |
| 5 | C | O5' | AB:31 | 4.66 | 1.00 | 89.53 |  |
| 6 | C | O5' | AB:30 | 4.94 | 1.00 | 84.83 |  |
  |

  
 Contact(s): /OP2/  
 Coded contact list: O_ph_  

### MG:AB:229

|  | Res. type | Atom | Residue | Distance | Occupancy | B-factor | Closest solute to 1^st^ shell water |
| --- | --- | --- | --- | --- | --- | --- | --- |
| **Ion** | | | |  | 1.00 | 67.26 || 1 | U | O4 | AB:55 | 2.97 | 1.00 | 92.08 |
| 2 | G | O2' | AB:24 | 3.12 | 1.00 | 106.40 |  |
| 3 | G | O6 | AB:56 | 3.26 | 1.00 | 100.53 |  |
| 4 | G | O3' | AB:24 | 4.03 | 1.00 | 106.19 |  |
| 5 | A | N6 | AB:29 | 4.04 | 1.00 | 90.98 |  |
| 6 | C | N4 | AB:28 | 4.31 | 1.00 | 90.62 |  |
| 7 | G | N7 | AB:56 | 4.55 | 1.00 | 97.34 |  |
| 8 | A | OP2 | AB:26 | 4.68 | 1.00 | 99.64 |  |
| 9 | U | N3 | AB:55 | 4.76 | 1.00 | 91.81 |  |
| 10 | C | N4 | AB:27 | 4.95 | 1.00 | 88.75 |  |
  |

  
 Coded contact list:   
 Exclusion zone contact(s): /O2'/O4/O6/  
Consider K in place of MG (3 valid contact(s) for K vs. 0 for MG)  

### MG:AB:230

|  | Res. type | Atom | Residue | Distance | Occupancy | B-factor | Closest solute to 1^st^ shell water |
| --- | --- | --- | --- | --- | --- | --- | --- |
| **Ion** | | | |  | 1.00 | 76.69 || 1 | G | N7 | AB:16 | 2.68 | 1.00 | 92.58 |
| 2 | C | O2 | AB:108 | 2.71 | 1.00 | 106.02 |  |
| 3 | G | O6 | AB:16 | 3.29 | 1.00 | 96.80 |  |
| 4 | C | N1 | AB:108 | 3.37 | 1.00 | 107.06 |  |
| 5 | A | OP2 | AB:15 | 3.40 | 1.00 | 103.56 |  |
| 6 | C | O4' | AB:108 | 3.53 | 1.00 | 105.77 |  |
| 7 | C | O2' | AB:108 | 3.91 | 1.00 | 107.48 |  |
| 8 | A | N1 | AB:66 | 3.95 | 1.00 | 99.89 |  |
| 9 | C | N4 | AB:17 | 4.25 | 1.00 | 96.03 |  |
| 10 | C | N3 | AB:108 | 4.26 | 1.00 | 108.18 |  |
| 11 | U | O2 | AB:107 | 4.36 | 1.00 | 99.81 |  |
| 12 | G | N2 | AB:69 | 4.55 | 1.00 | 92.82 |  |
| 13 | U | O2' | AB:107 | 4.65 | 1.00 | 102.42 |  |
| 14 | A | N3 | AB:66 | 4.73 | 1.00 | 99.28 |  |
| 15 | G | N9 | AB:16 | 4.83 | 1.00 | 94.26 |  |
| 16 | G | N1 | AB:69 | 4.84 | 1.00 | 93.86 |  |
| 17 | A | O5' | AB:15 | 4.97 | 1.00 | 103.26 |  |
  |

  
 Coded contact list:   
 Exclusion zone contact(s): /N1/N7/O2/O6/OP2/  
Consider K in place of MG (5 valid contact(s) for K vs. 0 for MG)  

### MG:AB:231

|  | Res. type | Atom | Residue | Distance | Occupancy | B-factor | Closest solute to 1^st^ shell water |
| --- | --- | --- | --- | --- | --- | --- | --- |
| **Ion** | | | |  | 1.00 | 94.98 || 1 | G | OP1 | AB:63 | 3.19 | 1.00 | 101.15 |
| 2 | G | OP1 | AB:110 | 4.08 | 1.00 | 109.45 |  |
| 3 | G | OP2 | AB:110 | 4.51 | 1.00 | 108.73 |  |
  |

  
 Coded contact list:   
 Exclusion zone contact(s): /OP1/  
Consider K in place of MG (1 valid contact(s) for K vs. 0 for MG)  

### MG:AB:232

|  | Res. type | Atom | Residue | Distance | Occupancy | B-factor | Closest solute to 1^st^ shell water |
| --- | --- | --- | --- | --- | --- | --- | --- |
| **Ion** | | | |  | 1.00 | 70.10 || 1 | U | O4 | AB:40 | 2.79 | 1.00 | 104.61 |
| 2 | C | OP1 | AB:43 | 3.00 | 1.00 | 90.84 |  |
| 3 | LYS | NZ | AG:67 | 3.69 | 1.00 | 113.46 |  |
| 4 | LYS | O | A4:2 | 3.90 | 1.00 | 127.75 |  |
| 5 | C | OP2 | AB:42 | 4.03 | 1.00 | 92.99 |  |
| 6 | C | O3' | AB:42 | 4.22 | 1.00 | 91.63 |  |
| 7 | MET | O | A4:1 | 4.24 | 1.00 | 129.27 |  |
| 8 | C | OP2 | AB:43 | 4.67 | 1.00 | 91.76 |  |
| 9 | U | N3 | AB:40 | 4.72 | 1.00 | 102.32 |  |
| 10 | LYS | N | A4:2 | 4.82 | 1.00 | 128.35 |  |
| 11 | HIS | ND1 | A4:6 | 4.92 | 1.00 | 135.13 |  |
  |

  
 Coded contact list:   
 Exclusion zone contact(s): /O4/OP1/  
Consider K in place of MG (2 valid contact(s) for K vs. 0 for MG)  

### MG:AB:233

|  | Res. type | Atom | Residue | Distance | Occupancy | B-factor | Closest solute to 1^st^ shell water |
| --- | --- | --- | --- | --- | --- | --- | --- |
| **Ion** | | | |  | 1.00 | 86.65 || 1 | A | OP1 | AB:101 | 2.72 | 1.00 | 77.82 |
| 2 | G | OP1 | AB:100 | 3.11 | 1.00 | 79.32 |  |
| 3 | A | OP2 | AB:101 | 3.46 | 1.00 | 78.49 |  |
| 4 | G | OP2 | AB:100 | 3.66 | 1.00 | 79.66 |  |
| 5 | G | O3' | AB:100 | 3.68 | 1.00 | 78.20 |  |
| 6 | G | O5' | AB:100 | 3.84 | 1.00 | 77.92 |  |
| 7 | C | O2' | AA:915 | 4.20 | 1.00 | 75.79 |  |
| 8 | A | O5' | AB:101 | 4.88 | 1.00 | 77.56 |  |
  |

  
 Coded contact list:   
 Exclusion zone contact(s): /OP1/OP1/  
Consider K in place of MG (2 valid contact(s) for K vs. 0 for MG)  

### MG:AB:236

|  | Res. type | Atom | Residue | Distance | Occupancy | B-factor | Closest solute to 1^st^ shell water |
| --- | --- | --- | --- | --- | --- | --- | --- |
| **Ion** | | | |  | 1.00 | 64.70 || 1 | G | OP1 | AB:82 | 2.93 | 1.00 | 84.96 |
| 2 | G | O3' | AB:81 | 3.99 | 1.00 | 86.92 |  |
| 3 | G | OP2 | AB:82 | 4.53 | 1.00 | 84.67 |  |
  |

  
 Coded contact list:   
 Exclusion zone contact(s): /OP1/  
Consider K in place of MG (1 valid contact(s) for K vs. 0 for MG)  

### MG:AD:301

|  | Res. type | Atom | Residue | Distance | Occupancy | B-factor | Closest solute to 1^st^ shell water |
| --- | --- | --- | --- | --- | --- | --- | --- |
| **Ion** | | | |  | 1.00 | 36.23 || 1 | ARG | NH1 | AD:273 | 3.63 | 1.00 | 63.40 |
| 2 | U | OP2 | AA:1798 | 3.86 | 1.00 | 35.91 |  |
| 3 | ARG | NE | AD:273 | 4.43 | 1.00 | 65.20 |  |
| 4 | LYS | NZ | AD:258 | 4.44 | 1.00 | 44.81 |  |
| 5 | C | OP1 | AA:1797 | 4.48 | 1.00 | 35.66 |  |
| 6 | ARG | NH2 | AD:273 | 4.91 | 1.00 | 64.47 |  |
  |

  
 Coded contact list:   
1^st^ MG coordination shell missing (see Notes)  

### MG:AD:302

|  | Res. type | Atom | Residue | Distance | Occupancy | B-factor | Closest solute to 1^st^ shell water |
| --- | --- | --- | --- | --- | --- | --- | --- |
| **Ion** | | | |  | 1.00 | 71.91 || 1 | ARG | O | AD:273 | 3.02 | 1.00 | 64.52 |
  |

  
 Coded contact list:   
 Exclusion zone contact(s): /O_back_/  
Consider K in place of MG (1 valid contact(s) for K vs. 0 for MG)  

### MG:AD:303

|  | Res. type | Atom | Residue | Distance | Occupancy | B-factor | Closest solute to 1^st^ shell water |
| --- | --- | --- | --- | --- | --- | --- | --- |
| **Ion** | | | |  | 1.00 | 62.73 || 1 | ARG | O | AD:157 | 2.51 | 1.00 | 49.30 |
| 2 | U | OP1 | AA:1818 | 2.85 | 1.00 | 43.91 |  |
| 3 | ARG | NE | AD:88 | 3.23 | 1.00 | 38.23 |  |
| 4 | U | OP2 | AA:1818 | 3.25 | 1.00 | 42.45 |  |
| 5 | ALA | N | AD:90 | 4.06 | 1.00 | 40.23 |  |
| 6 | SER | N | AD:89 | 4.40 | 1.00 | 40.66 |  |
| 7 | G | O3' | AA:1817 | 4.52 | 1.00 | 43.33 |  |
| 8 | ALA | N | AD:158 | 4.55 | 1.00 | 48.06 |  |
| 9 | SER | OG | AD:89 | 4.59 | 1.00 | 37.62 |  |
| 10 | U | O5' | AA:1818 | 4.75 | 1.00 | 40.97 |  |
| 11 | ARG | NH2 | AD:88 | 4.89 | 1.00 | 39.98 |  |
  |

  
 Coded contact list:   
 Exclusion zone contact(s): /NE_Arg_/O_back_/OP1/OP2/  
Consider K in place of MG (3 valid contact(s) for K vs. 0 for MG)  

### MG:AD:304

|  | Res. type | Atom | Residue | Distance | Occupancy | B-factor | Closest solute to 1^st^ shell water |
| --- | --- | --- | --- | --- | --- | --- | --- |
| **Ion** | | | |  | 1.00 | 65.35 || 1 | SER | N | AD:89 | 4.62 | 1.00 | 40.66 |
| 2 | G | OP1 | AA:1822 | 4.67 | 1.00 | 36.87 |  |
| 3 | ARG | NH2 | AD:60 | 4.85 | 1.00 | 52.59 |  |
| 4 | PRO | O | AD:86 | 4.96 | 1.00 | 47.42 |  |
  |

  
 Coded contact list:   
1^st^ MG coordination shell missing (see Notes)  

### MG:AD:305

|  | Res. type | Atom | Residue | Distance | Occupancy | B-factor | Closest solute to 1^st^ shell water |
| --- | --- | --- | --- | --- | --- | --- | --- |
| **Ion** | | | |  | 1.00 | 53.88 || 1 | ASP | OD2 | BI:83 | 2.83 | 1.00 | 75.68 |
| 2 | VAL | N | AD:138 | 3.23 | 1.00 | 45.28 |  |
| 3 | ARG | NH2 | AD:168 | 3.98 | 1.00 | 61.81 |  |
| 4 | VAL | O | AD:138 | 4.05 | 1.00 | 46.15 |  |
| 5 | ASP | OD1 | BI:83 | 4.63 | 1.00 | 76.09 |  |
| 6 | PRO | O | AD:137 | 4.66 | 1.00 | 45.19 |  |
| 7 | ILE | O | BI:81 | 4.96 | 1.00 | 65.70 |  |
  |

  
 Coded contact list:   
 Exclusion zone contact(s): /N_back_/OD2_Asp_/  
Consider K in place of MG (2 valid contact(s) for K vs. 0 for MG)  

### MG:AD:306

|  | Res. type | Atom | Residue | Distance | Occupancy | B-factor | Closest solute to 1^st^ shell water |
| --- | --- | --- | --- | --- | --- | --- | --- |
| **Ion** | | | |  | 1.00 | 44.10 || 1 | GLY | N | AD:234 | 2.93 | 1.00 | 33.73 |
| 2 | HIS | O | AD:233 | 3.56 | 1.00 | 32.94 |  |
| 3 | A | OP1 | AA:2598 | 4.24 | 1.00 | 32.36 |  |
| 4 | PRO | O | AD:241 | 4.26 | 1.00 | 32.26 |  |
| 5 | GLY | O | AD:235 | 4.39 | 1.00 | 41.73 |  |
| 6 | ARG | O | AD:242 | 4.41 | 1.00 | 32.21 |  |
| 7 | HIS | N | AD:233 | 4.42 | 1.00 | 29.11 |  |
| 8 | GLY | N | AD:235 | 4.49 | 1.00 | 38.63 |  |
| 9 | PRO | O | AD:232 | 4.75 | 1.00 | 28.25 |  |
| 10 | ALA | N | AD:224 | 4.75 | 1.00 | 35.41 |  |
| 11 | ALA | O | AD:240 | 4.80 | 1.00 | 32.89 |  |
| 12 | GLY | O | AD:234 | 4.83 | 1.00 | 40.96 |  |
  |

  
 Coded contact list:   
 Exclusion zone contact(s): /N_back_/  
Consider K in place of MG (1 valid contact(s) for K vs. 0 for MG)  

### MG:AD:307

|  | Res. type | Atom | Residue | Distance | Occupancy | B-factor | Closest solute to 1^st^ shell water |
| --- | --- | --- | --- | --- | --- | --- | --- |
| **Ion** | | | |  | 1.00 | 51.85 || 1 | U | OP1 | AA:2074 | 2.21 | 1.00 | 26.02 |
| 2 | PRO | O | AD:228 | 3.01 | 1.00 | 32.52 |  |
| 3 | HIS | O | AD:231 | 3.41 | 1.00 | 32.87 |  |
| 4 | U | O5' | AA:2074 | 4.05 | 1.00 | 26.91 |  |
| 5 | C | O3' | AA:2073 | 4.10 | 1.00 | 25.26 |  |
| 6 | PRO | O | AD:232 | 4.36 | 1.00 | 28.25 |  |
| 7 | PRO | N | AD:232 | 4.38 | 1.00 | 31.38 |  |
| 8 | G | OP2 | AA:2239 | 4.74 | 1.00 | 34.50 |  |
| 9 | U | OP2 | AA:2074 | 4.77 | 1.00 | 26.90 |  |
| 10 | U | OP2 | AA:2075 | 4.81 | 1.00 | 32.48 |  |
| 11 | ARG | NH2 | AD:244 | 4.96 | 1.00 | 44.20 |  |
  |

  
 Contact(s): /OP1/  
 Coded contact list: O_ph_  
 Exclusion zone contact(s): /O_back_/  

### MG:AD:308

|  | Res. type | Atom | Residue | Distance | Occupancy | B-factor | Closest solute to 1^st^ shell water |
| --- | --- | --- | --- | --- | --- | --- | --- |
| **Ion** | | | |  | 1.00 | 58.18 || 1 | ARG | NH1 | AD:88 | 2.57 | 1.00 | 39.79 |
| 2 | ARG | N | AD:63 | 2.91 | 1.00 | 39.82 |  |
| 3 | ASP | OD2 | AD:85 | 2.94 | 1.00 | 45.85 |  |
| 4 | ARG | O | AD:63 | 2.95 | 1.00 | 42.74 |  |
| 5 | ASP | OD1 | AD:85 | 3.06 | 1.00 | 44.18 |  |
| 6 | ASN | ND2 | AD:87 | 3.27 | 1.00 | 40.42 |  |
| 7 | ARG | NH2 | AD:88 | 4.53 | 1.00 | 39.98 |  |
| 8 | ILE | N | AD:64 | 4.69 | 1.00 | 43.24 |  |
| 9 | TYR | OH | AD:62 | 4.72 | 1.00 | 33.32 |  |
| 10 | ASN | N | AD:87 | 4.79 | 1.00 | 43.57 |  |
| 11 | ARG | NE | AD:88 | 4.82 | 1.00 | 38.23 |  |
| 12 | G | OP1 | AA:1817 | 4.90 | 1.00 | 43.07 |  |
| 13 | ASN | OD1 | AD:87 | 4.98 | 1.00 | 40.89 |  |
  |

  
 Coded contact list:   
 Exclusion zone contact(s): /N_back_/ND2_Asn_/NH1_Arg_/O_back_/OD1_Asp_/OD2_Asp_/  
Consider K in place of MG (5 valid contact(s) for K vs. 0 for MG)  

### MG:AD:309

|  | Res. type | Atom | Residue | Distance | Occupancy | B-factor | Closest solute to 1^st^ shell water |
| --- | --- | --- | --- | --- | --- | --- | --- |
| **Ion** | | | |  | 1.00 | 48.52 || 1 | A | O2' | AA:1812 | 2.30 | 1.00 | 38.72 |
| 2 | GLN | N | AD:46 | 2.33 | 1.00 | 32.81 |  |
| 3 | GLN | OE1 | AD:46 | 3.27 | 1.00 | 36.13 |  |
| 4 | ASN | O | AD:45 | 3.62 | 1.00 | 32.95 |  |
| 5 | GLN | NE2 | AD:46 | 3.98 | 1.00 | 35.55 |  |
| 6 | A | O3' | AA:1812 | 4.05 | 1.00 | 36.77 |  |
| 7 | ASN | N | AD:45 | 4.06 | 1.00 | 33.52 |  |
| 8 | ASN | O | AD:44 | 4.13 | 1.00 | 36.55 |  |
| 9 | G | O4' | AA:1813 | 4.42 | 1.00 | 31.87 |  |
| 10 | A | O4' | AA:1812 | 4.44 | 1.00 | 38.74 |  |
| 11 | ASN | OD1 | AD:45 | 4.72 | 1.00 | 30.34 |  |
  |

  
**1^st^ shell ligand angles:**   
1-2: 114;   
  
 Contact(s): /N_back_/O2'/  
 Coded contact list: O_r_Anion  
 Exclusion zone contact(s): /OE1_Gln_/  

### MG:AD:310

|  | Res. type | Atom | Residue | Distance | Occupancy | B-factor | Closest solute to 1^st^ shell water |
| --- | --- | --- | --- | --- | --- | --- | --- |
| **Ion** | | | |  | 1.00 | 42.52 || 1 | GLU | OE1 | AD:148 | 3.21 | 1.00 | 55.47 |
| 2 | ARG | NH2 | AD:183 | 3.81 | 1.00 | 49.29 |  |
| 3 | ARG | NH1 | AD:183 | 3.88 | 1.00 | 50.12 |  |
| 4 | C | OP2 | AA:1800 | 4.01 | 1.00 | 48.63 |  |
| 5 | SER | OG | AD:266 | 4.45 | 1.00 | 51.26 |  |
  |

  
 Coded contact list:   
 Exclusion zone contact(s): /OE1_Glu_/  
Consider K in place of MG (1 valid contact(s) for K vs. 0 for MG)  

### MG:AD:311

|  | Res. type | Atom | Residue | Distance | Occupancy | B-factor | Closest solute to 1^st^ shell water |
| --- | --- | --- | --- | --- | --- | --- | --- |
| **Ion** | | | |  | 1.00 | 54.52 || 1 | C | OP1 | AA:1804 | 3.91 | 1.00 | 35.43 |
  |

  
 Coded contact list:   
1^st^ MG coordination shell missing (see Notes)  

### MG:AD:312

|  | Res. type | Atom | Residue | Distance | Occupancy | B-factor | Closest solute to 1^st^ shell water |
| --- | --- | --- | --- | --- | --- | --- | --- |
| **Ion** | | | |  | 1.00 | 79.93 || 1 | G | N7 | AA:1568 | 3.54 | 1.00 | 48.63 |
| 2 | G | O6 | AA:1568 | 3.95 | 1.00 | 46.87 |  |
| 3 | PRO | N | AD:29 | 4.40 | 1.00 | 79.92 |  |
| 4 | G | N7 | AA:1429 | 4.58 | 1.00 | 50.64 |  |
| 5 | G | O6 | AA:1429 | 4.97 | 1.00 | 48.28 |  |
  |

  
 Coded contact list:   
1^st^ MG coordination shell missing (see Notes)  

### MG:AD:313

|  | Res. type | Atom | Residue | Distance | Occupancy | B-factor | Closest solute to 1^st^ shell water |
| --- | --- | --- | --- | --- | --- | --- | --- |
| **Ion** | | | |  | 1.00 | 39.85 || 1 | GLN | O | AD:46 | 3.06 | 1.00 | 27.68 |
| 2 | C | O2' | AA:1806 | 3.28 | 1.00 | 39.85 |  |
| 3 | ASN | ND2 | AD:45 | 3.37 | 1.00 | 30.01 |  |
| 4 | C | O2 | AA:1806 | 3.65 | 1.00 | 44.24 |  |
| 5 | ASN | OD1 | AD:45 | 3.90 | 1.00 | 30.34 |  |
| 6 | G | N2 | AA:1811 | 4.21 | 1.00 | 44.34 |  |
| 7 | G | O4' | AA:1807 | 4.42 | 1.00 | 42.31 |  |
| 8 | GLN | OE1 | AD:46 | 4.49 | 1.00 | 36.13 |  |
| 9 | ARG | N | AD:48 | 4.96 | 1.00 | 38.59 |  |
  |

  
 Coded contact list:   
 Exclusion zone contact(s): /ND2_Asn_/O2'/O_back_/  
Consider K in place of MG (3 valid contact(s) for K vs. 0 for MG)  

### MG:AE:301

|  | Res. type | Atom | Residue | Distance | Occupancy | B-factor | Closest solute to 1^st^ shell water |
| --- | --- | --- | --- | --- | --- | --- | --- |
| **Ion** | | | |  | 1.00 | 35.28 || 1 | ASP | OD2 | AE:127 | 2.01 | 1.00 | 59.81 |
| 2 | G | O2' | AA:1992 | 3.41 | 1.00 | 47.37 |  |
| 3 | G | O6 | AA:1997 | 3.53 | 1.00 | 49.60 |  |
| 4 | G | N7 | AA:1997 | 3.75 | 1.00 | 52.11 |  |
| 5 | ASP | OD1 | AE:127 | 4.06 | 1.00 | 61.02 |  |
| 6 | C | OP2 | AA:1994 | 4.32 | 1.00 | 40.58 |  |
| 7 | U | OP2 | AA:1993 | 4.33 | 1.00 | 47.52 |  |
| 8 | G | N3 | AA:1992 | 4.87 | 1.00 | 41.66 |  |
  |

  
 Contact(s): /OD2_Asp_/  
 Coded contact list: O_coo_  

### MG:AE:302

|  | Res. type | Atom | Residue | Distance | Occupancy | B-factor | Closest solute to 1^st^ shell water |
| --- | --- | --- | --- | --- | --- | --- | --- |
| **Ion** | | | |  | 1.00 | 74.40 || 1 | U | OP1 | AA:2580 | 3.02 | 1.00 | 35.86 |
| 2 | GLY | O | AE:130 | 3.45 | 1.00 | 57.83 |  |
| 3 | G | OP2 | AA:2581 | 3.88 | 1.00 | 42.26 |  |
| 4 | HIS | N | AE:132 | 3.93 | 1.00 | 57.79 |  |
| 5 | G | OP1 | AA:743 | 4.06 | 1.00 | 41.41 |  |
| 6 | U | O5' | AA:2580 | 4.18 | 1.00 | 36.61 |  |
| 7 | G | OP2 | AA:744 | 4.65 | 1.00 | 41.36 |  |
| 8 | C | O3' | AA:2579 | 4.73 | 1.00 | 35.97 |  |
| 9 | G | N2 | AA:2581 | 4.77 | 1.00 | 36.08 |  |
| 10 | ALA | N | AE:131 | 4.80 | 1.00 | 58.51 |  |
| 11 | G | OP2 | AA:743 | 4.84 | 1.00 | 40.79 |  |
| 12 | G | OP1 | AA:2581 | 4.98 | 1.00 | 40.85 |  |
  |

  
 Coded contact list:   
 Exclusion zone contact(s): /OP1/  
Consider K in place of MG (1 valid contact(s) for K vs. 0 for MG)  

### MG:AE:303

|  | Res. type | Atom | Residue | Distance | Occupancy | B-factor | Closest solute to 1^st^ shell water |
| --- | --- | --- | --- | --- | --- | --- | --- |
| **Ion** | | | |  | 1.00 | 39.60 || 1 | HIS | NE2 | AE:135 | 2.21 | 1.00 | 55.18 |
| 2 | C | OP1 | AA:1658 | 2.61 | 1.00 | 48.97 |  |
| 3 | U | OP2 | AA:1659 | 3.89 | 1.00 | 46.58 |  |
| 4 | G | O3' | AA:743 | 3.90 | 1.00 | 40.49 |  |
| 5 | G | OP1 | AA:744 | 3.96 | 1.00 | 41.25 |  |
| 6 | HIS | ND1 | AE:135 | 4.15 | 1.00 | 52.98 |  |
| 7 | G | O2' | AA:743 | 4.42 | 1.00 | 42.49 |  |
| 8 | C | O5' | AA:1658 | 4.45 | 1.00 | 45.90 |  |
| 9 | C | OP2 | AA:1658 | 4.49 | 1.00 | 46.46 |  |
  |

  
 Contact(s): /NE2_His_/  
 Coded contact list: His  
 Exclusion zone contact(s): /OP1/  

### MG:AE:304

|  | Res. type | Atom | Residue | Distance | Occupancy | B-factor | Closest solute to 1^st^ shell water |
| --- | --- | --- | --- | --- | --- | --- | --- |
| **Ion** | | | |  | 1.00 | 83.94 || 1 | LYS | O | AE:118 | 2.53 | 1.00 | 50.49 |
| 2 | ASN | ND2 | AE:121 | 3.43 | 1.00 | 51.44 |  |
| 3 | TYR | OH | AE:160 | 3.75 | 1.00 | 60.58 |  |
| 4 | ASN | OD1 | AE:121 | 3.78 | 1.00 | 50.54 |  |
| 5 | C | OP2 | AA:2678 | 4.39 | 1.00 | 69.57 |  |
| 6 | ARG | O | AE:119 | 4.54 | 1.00 | 50.83 |  |
| 7 | C | OP1 | AA:2678 | 4.59 | 1.00 | 70.79 |  |
| 8 | ASN | N | AE:121 | 4.77 | 1.00 | 46.50 |  |
| 9 | ARG | N | AE:119 | 4.78 | 1.00 | 50.13 |  |
| 10 | A | OP2 | AA:2679 | 4.87 | 1.00 | 63.69 |  |
  |

  
 Coded contact list:   
 Exclusion zone contact(s): /O_back_/  
Consider NA in place of MG (1 valid contact(s) for NA vs. 0 for MG)  

### MG:AF:301

|  | Res. type | Atom | Residue | Distance | Occupancy | B-factor | Closest solute to 1^st^ shell water |
| --- | --- | --- | --- | --- | --- | --- | --- |
| **Ion** | | | |  | 1.00 | 73.24 || 1 | U | O4 | AA:1205 | 2.50 | 1.00 | 68.61 |
| 2 | ARG | O | AF:168 | 2.63 | 1.00 | 68.55 |  |
| 3 | LEU | O | AF:170 | 2.74 | 1.00 | 65.13 |  |
| 4 | VAL | O | AF:173 | 2.82 | 1.00 | 61.61 |  |
| 5 | LEU | N | AF:170 | 4.29 | 1.00 | 63.25 |  |
| 6 | U | N3 | AA:1205 | 4.37 | 1.00 | 67.20 |  |
| 7 | PRO | N | AF:171 | 4.56 | 1.00 | 64.57 |  |
| 8 | ASN | O | AF:169 | 4.62 | 1.00 | 63.67 |  |
| 9 | ASN | N | AF:169 | 4.70 | 1.00 | 67.85 |  |
| 10 | VAL | N | AF:174 | 4.86 | 1.00 | 62.98 |  |
| 11 | VAL | N | AF:173 | 4.99 | 1.00 | 62.86 |  |
| 12 | A | OP2 | AA:324 | 4.99 | 1.00 | 66.30 |  |
| 13 | ALA | O | AF:167 | 4.99 | 1.00 | 70.59 |  |
  |

  
 Coded contact list:   
 Exclusion zone contact(s): /O4/O_back_/O_back_/O_back_/  
Consider K in place of MG (3 valid contact(s) for K vs. 0 for MG)  

### MG:AF:303

|  | Res. type | Atom | Residue | Distance | Occupancy | B-factor | Closest solute to 1^st^ shell water |
| --- | --- | --- | --- | --- | --- | --- | --- |
| **Ion** | | | |  | 1.00 | 68.65 || 1 | VAL | N | AF:84 | 2.36 | 1.00 | 40.32 |
| 2 | PRO | O | AF:81 | 2.84 | 1.00 | 37.57 |  |
| 3 | C | O2' | AA:584 | 3.26 | 1.00 | 52.17 |  |
| 4 | VAL | O | AF:84 | 3.28 | 1.00 | 40.78 |  |
| 5 | PHE | N | AF:83 | 3.61 | 1.00 | 39.40 |  |
| 6 | ILE | O | AF:82 | 4.03 | 1.00 | 38.95 |  |
| 7 | ILE | N | AF:82 | 4.24 | 1.00 | 35.19 |  |
| 8 | GLY | N | AF:86 | 4.25 | 1.00 | 33.84 |  |
| 9 | PHE | O | AF:83 | 4.51 | 1.00 | 43.85 |  |
| 10 | ALA | O | AF:80 | 4.62 | 1.00 | 39.43 |  |
| 11 | GLY | N | AF:85 | 4.63 | 1.00 | 35.41 |  |
| 12 | C | O2 | AA:584 | 4.83 | 1.00 | 48.82 |  |
| 13 | GLY | O | AF:87 | 4.89 | 1.00 | 32.36 |  |
| 14 | GLY | O | AF:86 | 4.96 | 1.00 | 37.98 |  |
  |

  
 Contact(s): /N_back_/  
 Coded contact list: Anion  
 Exclusion zone contact(s): /O2'/O_back_/O_back_/  
Consider K in place of MG (3 valid contact(s) for K vs. 1 for MG)  

### MG:AF:304

|  | Res. type | Atom | Residue | Distance | Occupancy | B-factor | Closest solute to 1^st^ shell water |
| --- | --- | --- | --- | --- | --- | --- | --- |
| **Ion** | | | |  | 1.00 | 42.11 || 1 | HIS | N | AF:75 | 2.32 | 1.00 | 35.40 |
| 2 | HIS | O | AF:75 | 2.91 | 1.00 | 33.66 |  |
| 3 | TRP | O | AF:65 | 3.24 | 1.00 | 40.83 |  |
| 4 | GLN | N | AF:67 | 3.51 | 1.00 | 39.89 |  |
| 5 | LYS | NZ | AF:63 | 3.51 | 1.00 | 45.04 |  |
| 6 | ALA | O | AF:73 | 3.81 | 1.00 | 40.59 |  |
| 7 | ARG | N | AF:74 | 4.24 | 1.00 | 40.36 |  |
| 8 | ARG | O | AF:74 | 4.40 | 1.00 | 37.27 |  |
| 9 | GLN | O | AF:67 | 4.61 | 1.00 | 42.08 |  |
| 10 | TRP | N | AF:65 | 4.62 | 1.00 | 41.86 |  |
| 11 | GLY | N | AF:76 | 4.80 | 1.00 | 34.32 |  |
| 12 | GLN | OE1 | AF:67 | 4.91 | 1.00 | 36.60 |  |
  |

  
 Contact(s): /N_back_/  
 Coded contact list: Anion  
 Exclusion zone contact(s): /O_back_/O_back_/  
Consider K in place of MG (2 valid contact(s) for K vs. 1 for MG)  

### MG:AF:305

|  | Res. type | Atom | Residue | Distance | Occupancy | B-factor | Closest solute to 1^st^ shell water |
| --- | --- | --- | --- | --- | --- | --- | --- |
| **Ion** | | | |  | 1.00 | 89.91 || 1 | THR | O | AF:175 | 4.66 | 1.00 | 67.62 |
| 2 | THR | N | AF:175 | 4.88 | 1.00 | 64.94 |  |
  |

  
 Coded contact list:   
1^st^ MG coordination shell missing (see Notes)  

### MG:AF:307

|  | Res. type | Atom | Residue | Distance | Occupancy | B-factor | Closest solute to 1^st^ shell water |
| --- | --- | --- | --- | --- | --- | --- | --- |
| **Ion** | | | |  | 1.00 | 63.43 || 1 | PRO | O | AF:171 | 4.59 | 1.00 | 64.74 |
| 2 | SER | OG | AF:153 | 4.64 | 1.00 | 69.04 |  |
| 3 | ARG | NH1 | AF:188 | 4.97 | 1.00 | 61.61 |  |
  |

  
 Coded contact list:   
1^st^ MG coordination shell missing (see Notes)  

### MG:AG:201

|  | Res. type | Atom | Residue | Distance | Occupancy | B-factor | Closest solute to 1^st^ shell water |
| --- | --- | --- | --- | --- | --- | --- | --- |
| **Ion** | | | |  | 1.00 | 86.46 || 1 | LEU | O | AG:3 | 2.67 | 1.00 | 112.34 |
| 2 | ASP | N | AG:4 | 3.54 | 1.00 | 111.39 |  |
| 3 | LEU | N | AG:3 | 4.29 | 1.00 | 112.09 |  |
| 4 | ASP | OD2 | AG:4 | 4.81 | 1.00 | 113.42 |  |
  |

  
 Coded contact list:   
 Exclusion zone contact(s): /O_back_/  
Consider K in place of MG (1 valid contact(s) for K vs. 0 for MG)  

### MG:AG:202

|  | Res. type | Atom | Residue | Distance | Occupancy | B-factor | Closest solute to 1^st^ shell water |
| --- | --- | --- | --- | --- | --- | --- | --- |
| **Ion** | | | |  | 1.00 | 83.37 || 1 | ARG | O | AG:128 | 2.43 | 1.00 | 96.53 |
| 2 | THR | O | AG:161 | 2.82 | 1.00 | 94.83 |  |
| 3 | ASN | ND2 | AG:130 | 3.09 | 1.00 | 97.57 |  |
| 4 | ALA | O | AG:163 | 3.16 | 1.00 | 104.58 |  |
| 5 | GLY | O | AG:129 | 3.27 | 1.00 | 95.78 |  |
| 6 | ASN | OD1 | AG:130 | 3.78 | 1.00 | 96.88 |  |
| 7 | ALA | N | AG:163 | 4.35 | 1.00 | 99.53 |  |
| 8 | GLY | N | AG:129 | 4.38 | 1.00 | 95.99 |  |
| 9 | THR | N | AG:162 | 4.61 | 1.00 | 95.31 |  |
| 10 | THR | N | AG:161 | 4.65 | 1.00 | 93.35 |  |
  |

  
 Coded contact list:   
 Exclusion zone contact(s): /ND2_Asn_/O_back_/O_back_/O_back_/O_back_/  
Consider K in place of MG (4 valid contact(s) for K vs. 0 for MG)  

### MG:AG:203

|  | Res. type | Atom | Residue | Distance | Occupancy | B-factor | Closest solute to 1^st^ shell water |
| --- | --- | --- | --- | --- | --- | --- | --- |
| **Ion** | | | |  | 1.00 | 97.06 || 1 | ARG | NH1 | AG:9 | 2.81 | 1.00 | 103.19 |
| 2 | ARG | NH2 | AG:9 | 3.76 | 1.00 | 103.01 |  |
| 3 | ARG | NE | AG:9 | 4.69 | 1.00 | 102.36 |  |
| 4 | ASP | OD1 | AG:4 | 4.89 | 1.00 | 113.90 |  |
  |

  
 Coded contact list:   
 Exclusion zone contact(s): /NH1_Arg_/  
Consider K in place of MG (1 valid contact(s) for K vs. 0 for MG)  

### MG:AH:201

|  | Res. type | Atom | Residue | Distance | Occupancy | B-factor | Closest solute to 1^st^ shell water |
| --- | --- | --- | --- | --- | --- | --- | --- |
| **Ion** | | | |  | 1.00 | 78.75 || 1 | PRO | N | AH:126 | 4.93 | 1.00 | 178.41 |
  |

  
 Coded contact list:   
1^st^ MG coordination shell missing (see Notes)  

### MG:AK:201

|  | Res. type | Atom | Residue | Distance | Occupancy | B-factor | Closest solute to 1^st^ shell water |
| --- | --- | --- | --- | --- | --- | --- | --- |
| **Ion** | | | |  | 1.00 | 64.99 || 1 | C | OP1 | CA:366 | 4.49 | 1.00 | 51.32 |
| 2 | U | OP2 | CA:359 | 4.80 | 1.00 | 58.94 |  |
| 3 | LYS | NZ | AK:87 | 4.92 | 1.00 | 85.25 |  |
  |

  
 Coded contact list:   
1^st^ MG coordination shell missing (see Notes)  

### MG:AK:202

|  | Res. type | Atom | Residue | Distance | Occupancy | B-factor | Closest solute to 1^st^ shell water |
| --- | --- | --- | --- | --- | --- | --- | --- |
| **Ion** | | | |  | 1.00 | 75.41 || 1 | HIS | O | AK:105 | 3.69 | 1.00 | 91.46 |
  |

  
 Coded contact list:   
1^st^ MG coordination shell missing (see Notes)  

### MG:AN:201

|  | Res. type | Atom | Residue | Distance | Occupancy | B-factor | Closest solute to 1^st^ shell water |
| --- | --- | --- | --- | --- | --- | --- | --- |
| **Ion** | | | |  | 1.00 | 118.58 || 1 | GLY | O | AN:110 | 3.09 | 1.00 | 81.54 |
  |

  
 Coded contact list:   
 Exclusion zone contact(s): /O_back_/  
Consider K in place of MG (1 valid contact(s) for K vs. 0 for MG)  

### MG:AO:201

|  | Res. type | Atom | Residue | Distance | Occupancy | B-factor | Closest solute to 1^st^ shell water |
| --- | --- | --- | --- | --- | --- | --- | --- |
| **Ion** | | | |  | 1.00 | 50.40 || 1 | ARG | NE | AO:21 | 2.37 | 1.00 | 40.20 |
| 2 | GLY | O | AO:20 | 2.49 | 1.00 | 62.42 |  |
| 3 | ARG | N | AO:21 | 2.69 | 1.00 | 61.24 |  |
| 4 | U | O2' | AA:810 | 3.45 | 1.00 | 43.49 |  |
| 5 | U | N1 | AA:811 | 3.53 | 1.00 | 50.02 |  |
| 6 | U | OP2 | AA:811 | 3.58 | 1.00 | 45.65 |  |
| 7 | ARG | NH2 | AO:21 | 3.65 | 1.00 | 34.65 |  |
| 8 | ARG | O | AO:21 | 3.80 | 1.00 | 61.22 |  |
| 9 | GLY | N | AO:22 | 4.11 | 1.00 | 58.72 |  |
| 10 | U | O3' | AA:810 | 4.32 | 1.00 | 46.66 |  |
| 11 | U | O4' | AA:811 | 4.35 | 1.00 | 46.07 |  |
| 12 | U | O5' | AA:811 | 4.36 | 1.00 | 46.31 |  |
| 13 | U | N3 | AA:811 | 4.37 | 1.00 | 51.57 |  |
| 14 | GLY | N | AO:20 | 4.41 | 1.00 | 65.23 |  |
| 15 | ARG | NH1 | AO:21 | 4.57 | 1.00 | 37.10 |  |
| 16 | U | O4 | AA:811 | 4.85 | 1.00 | 52.02 |  |
  |

  
 Contact(s): /NE_Arg_/  
 Coded contact list: Anion  
 Exclusion zone contact(s): /N_back_/O_back_/  

### MG:AO:202

|  | Res. type | Atom | Residue | Distance | Occupancy | B-factor | Closest solute to 1^st^ shell water |
| --- | --- | --- | --- | --- | --- | --- | --- |
| **Ion** | | | |  | 1.00 | 55.69 || 1 | LEU | O | AO:59 | 3.02 | 1.00 | 64.33 |
| 2 | ARG | N | AO:61 | 3.59 | 1.00 | 66.59 |  |
| 3 | LEU | N | AO:62 | 3.68 | 1.00 | 67.30 |  |
| 4 | LEU | O | AO:62 | 3.98 | 1.00 | 67.45 |  |
| 5 | ARG | O | AO:61 | 4.33 | 1.00 | 68.13 |  |
| 6 | THR | O | AO:58 | 4.81 | 1.00 | 63.47 |  |
  |

  
 Coded contact list:   
 Exclusion zone contact(s): /O_back_/  
Consider K in place of MG (1 valid contact(s) for K vs. 0 for MG)  

### MG:AO:203

|  | Res. type | Atom | Residue | Distance | Occupancy | B-factor | Closest solute to 1^st^ shell water |
| --- | --- | --- | --- | --- | --- | --- | --- |
| **Ion** | | | |  | 1.00 | 47.40 || 1 | ARG | NH1 | AO:21 | 3.20 | 1.00 | 37.10 |
| 2 | U | N3 | AA:1249 | 3.51 | 1.00 | 63.90 |  |
| 3 | U | O2 | AA:1249 | 3.78 | 1.00 | 64.76 |  |
| 4 | G | OP2 | AA:1250 | 3.96 | 1.00 | 51.15 |  |
| 5 | G | N7 | AA:1250 | 4.03 | 1.00 | 59.59 |  |
| 6 | U | N1 | AA:1249 | 4.05 | 1.00 | 59.07 |  |
| 7 | ARG | NE | AO:21 | 4.51 | 1.00 | 40.20 |  |
| 8 | U | O4 | AA:1249 | 4.61 | 1.00 | 65.43 |  |
  |

  
 Coded contact list:   
 Exclusion zone contact(s): /NH1_Arg_/  
Consider K in place of MG (1 valid contact(s) for K vs. 0 for MG)  

### MG:AO:204

|  | Res. type | Atom | Residue | Distance | Occupancy | B-factor | Closest solute to 1^st^ shell water |
| --- | --- | --- | --- | --- | --- | --- | --- |
| **Ion** | | | |  | 1.00 | 43.23 || 1 | ARG | NH2 | AO:55 | 2.89 | 1.00 | 49.81 |
| 2 | U | O2' | AA:833 | 3.40 | 1.00 | 51.51 |  |
| 3 | THR | N | AO:58 | 3.60 | 1.00 | 64.36 |  |
| 4 | THR | OG1 | AO:58 | 3.74 | 1.00 | 68.57 |  |
| 5 | GLU | N | AO:52 | 3.81 | 1.00 | 52.55 |  |
| 6 | ARG | NE | AO:55 | 4.21 | 1.00 | 51.12 |  |
| 7 | PHE | O | AO:51 | 4.28 | 1.00 | 52.21 |  |
| 8 | U | O4' | AA:833 | 4.85 | 1.00 | 51.79 |  |
| 9 | ARG | NH1 | AO:55 | 4.91 | 1.00 | 51.16 |  |
  |

  
 Coded contact list:   
 Exclusion zone contact(s): /NH2_Arg_/O2'/  
Consider K in place of MG (2 valid contact(s) for K vs. 0 for MG)  

### MG:AO:205

|  | Res. type | Atom | Residue | Distance | Occupancy | B-factor | Closest solute to 1^st^ shell water |
| --- | --- | --- | --- | --- | --- | --- | --- |
| **Ion** | | | |  | 1.00 | 52.80 || 1 | ARG | O | AO:49 | 2.61 | 1.00 | 58.08 |
| 2 | U | O3' | AA:833 | 3.73 | 1.00 | 50.71 |  |
| 3 | U | O2' | AA:833 | 4.01 | 1.00 | 51.51 |  |
| 4 | C | OP1 | AA:834 | 4.07 | 1.00 | 51.04 |  |
| 5 | THR | OG1 | AO:58 | 4.19 | 1.00 | 68.57 |  |
| 6 | PHE | O | AO:51 | 4.55 | 1.00 | 52.21 |  |
| 7 | ARG | N | AO:50 | 4.78 | 1.00 | 55.47 |  |
| 8 | ARG | N | AO:49 | 4.82 | 1.00 | 59.64 |  |
| 9 | PHE | N | AO:51 | 4.91 | 1.00 | 51.06 |  |
  |

  
 Coded contact list:   
 Exclusion zone contact(s): /O_back_/  
Consider K in place of MG (1 valid contact(s) for K vs. 0 for MG)  

### MG:AO:206

|  | Res. type | Atom | Residue | Distance | Occupancy | B-factor | Closest solute to 1^st^ shell water |
| --- | --- | --- | --- | --- | --- | --- | --- |
| **Ion** | | | |  | 1.00 | 53.04 || 1 | U | OP2 | AA:813 | 3.13 | 1.00 | 52.06 |
| 2 | PRO | O | AO:23 | 3.16 | 1.00 | 61.14 |  |
| 3 | SER | OG | AO:25 | 3.38 | 1.00 | 55.90 |  |
| 4 | GLY | O | AO:22 | 3.46 | 1.00 | 59.71 |  |
| 5 | GLY | N | AO:22 | 3.50 | 1.00 | 58.72 |  |
| 6 | C | N1 | AA:812 | 3.66 | 1.00 | 55.95 |  |
| 7 | GLY | O | AO:24 | 3.92 | 1.00 | 60.16 |  |
| 8 | C | O3' | AA:812 | 3.97 | 1.00 | 51.81 |  |
| 9 | C | O5' | AA:812 | 4.09 | 1.00 | 47.10 |  |
| 10 | PRO | N | AO:23 | 4.20 | 1.00 | 57.18 |  |
| 11 | C | O4' | AA:812 | 4.24 | 1.00 | 52.98 |  |
| 12 | C | N3 | AA:812 | 4.54 | 1.00 | 59.43 |  |
| 13 | U | O5' | AA:813 | 4.54 | 1.00 | 52.78 |  |
  |

  
 Coded contact list:   
 Exclusion zone contact(s): /O_back_/OG_Ser_/OP2/  
Consider K in place of MG (3 valid contact(s) for K vs. 0 for MG)  

### MG:AO:207

|  | Res. type | Atom | Residue | Distance | Occupancy | B-factor | Closest solute to 1^st^ shell water |
| --- | --- | --- | --- | --- | --- | --- | --- |
| **Ion** | | | |  | 1.00 | 102.25 || 1 | G | N7 | AA:1190 | 3.77 | 1.00 | 63.30 |
| 2 | GLY | O | AO:26 | 3.83 | 1.00 | 62.02 |  |
| 3 | G | N7 | AA:1191 | 4.18 | 1.00 | 63.84 |  |
| 4 | HIS | N | AO:27 | 4.32 | 1.00 | 57.29 |  |
| 5 | GLY | N | AO:26 | 4.44 | 1.00 | 61.54 |  |
| 6 | HIS | ND1 | AO:27 | 4.44 | 1.00 | 59.31 |  |
| 7 | G | OP2 | AA:1191 | 4.47 | 1.00 | 53.48 |  |
| 8 | A | OP2 | AA:1189 | 4.54 | 1.00 | 52.21 |  |
| 9 | G | O6 | AA:1190 | 4.63 | 1.00 | 63.46 |  |
| 10 | G | O6 | AA:1191 | 4.63 | 1.00 | 66.84 |  |
| 11 | G | N9 | AA:1190 | 4.85 | 1.00 | 59.90 |  |
| 12 | G | OP2 | AA:1190 | 4.87 | 1.00 | 52.59 |  |
  |

  
 Coded contact list:   
1^st^ MG coordination shell missing (see Notes)  

### MG:AP:201

|  | Res. type | Atom | Residue | Distance | Occupancy | B-factor | Closest solute to 1^st^ shell water |
| --- | --- | --- | --- | --- | --- | --- | --- |
| **Ion** | | | |  | 1.00 | 95.51 || 1 | ILE | O | AP:127 | 2.46 | 1.00 | 81.46 |
| 2 | LEU | O | AP:125 | 2.76 | 1.00 | 78.44 |  |
| 3 | GLY | O | AP:122 | 2.87 | 1.00 | 81.43 |  |
| 4 | G | OP1 | AA:1030 | 3.18 | 1.00 | 85.36 |  |
| 5 | A | O3' | AA:1029 | 3.63 | 1.00 | 85.40 |  |
| 6 | A | O2' | AA:1029 | 4.11 | 1.00 | 83.39 |  |
| 7 | ILE | N | AP:127 | 4.17 | 1.00 | 80.07 |  |
| 8 | HIS | N | AP:123 | 4.43 | 1.00 | 80.66 |  |
| 9 | LYS | N | AP:128 | 4.50 | 1.00 | 81.85 |  |
| 10 | LEU | N | AP:125 | 4.61 | 1.00 | 76.87 |  |
| 11 | G | O2' | AA:2485 | 4.73 | 1.00 | 74.10 |  |
| 12 | G | O5' | AA:1030 | 4.81 | 1.00 | 84.13 |  |
| 13 | HIS | O | AP:123 | 4.81 | 1.00 | 79.16 |  |
| 14 | PRO | N | AP:126 | 4.85 | 1.00 | 79.49 |  |
  |

  
 Coded contact list:   
 Exclusion zone contact(s): /O_back_/O_back_/O_back_/OP1/  
Consider K in place of MG (3 valid contact(s) for K vs. 0 for MG)  

### MG:AQ:201

|  | Res. type | Atom | Residue | Distance | Occupancy | B-factor | Closest solute to 1^st^ shell water |
| --- | --- | --- | --- | --- | --- | --- | --- |
| **Ion** | | | |  | 1.00 | 56.70 || 1 | TYR | O | AQ:36 | 2.63 | 1.00 | 88.03 |
| 2 | G | O2' | AB:7 | 2.91 | 1.00 | 85.79 |  |
| 3 | PHE | N | AQ:29 | 3.58 | 1.00 | 78.66 |  |
| 4 | ALA | N | AQ:37 | 3.78 | 1.00 | 89.15 |  |
| 5 | GLN | NE2 | AQ:38 | 4.04 | 1.00 | 89.61 |  |
| 6 | ALA | O | AQ:37 | 4.04 | 1.00 | 89.49 |  |
| 7 | GLN | N | AQ:38 | 4.27 | 1.00 | 90.39 |  |
| 8 | SER | O | AQ:27 | 4.28 | 1.00 | 82.64 |  |
| 9 | TYR | N | AQ:36 | 4.74 | 1.00 | 88.38 |  |
| 10 | PHE | O | AQ:29 | 4.83 | 1.00 | 78.22 |  |
| 11 | VAL | N | AQ:28 | 4.90 | 1.00 | 81.60 |  |
  |

  
 Coded contact list:   
 Exclusion zone contact(s): /O2'/O_back_/  
Consider K in place of MG (2 valid contact(s) for K vs. 0 for MG)  

### MG:AQ:202

|  | Res. type | Atom | Residue | Distance | Occupancy | B-factor | Closest solute to 1^st^ shell water |
| --- | --- | --- | --- | --- | --- | --- | --- |
| **Ion** | | | |  | 1.00 | 94.24 || 1 | LEU | O | AQ:110 | 2.47 | 1.00 | 134.17 |
| 2 | LEU | N | AQ:110 | 2.56 | 1.00 | 136.34 |  |
| 3 | VAL | N | AQ:85 | 2.81 | 1.00 | 90.24 |  |
| 4 | VAL | O | AQ:85 | 3.28 | 1.00 | 85.66 |  |
| 5 | GLY | O | AQ:109 | 3.87 | 1.00 | 136.35 |  |
| 6 | LYS | O | AQ:83 | 3.95 | 1.00 | 100.34 |  |
| 7 | ARG | NH1 | AQ:23 | 4.19 | 1.00 | 98.28 |  |
| 8 | GLU | N | AQ:111 | 4.38 | 1.00 | 135.50 |  |
| 9 | GLN | N | AQ:84 | 4.42 | 1.00 | 97.66 |  |
| 10 | GLY | O | AQ:108 | 4.69 | 1.00 | 135.94 |  |
| 11 | GLN | O | AQ:84 | 4.75 | 1.00 | 91.60 |  |
| 12 | GLY | N | AQ:109 | 4.90 | 1.00 | 136.00 |  |
  |

  
 Coded contact list:   
 Exclusion zone contact(s): /N_back_/N_back_/O_back_/O_back_/  
Consider NA in place of MG (2 valid contact(s) for NA vs. 0 for MG)  

### MG:AQ:203

|  | Res. type | Atom | Residue | Distance | Occupancy | B-factor | Closest solute to 1^st^ shell water |
| --- | --- | --- | --- | --- | --- | --- | --- |
| **Ion** | | | |  | 1.00 | 67.24 || 1 | GLN | NE2 | AQ:68 | 2.71 | 1.00 | 89.53 |
| 2 | GLN | O | AQ:68 | 3.79 | 1.00 | 87.93 |  |
| 3 | GLN | OE1 | AQ:68 | 4.34 | 1.00 | 88.08 |  |
| 4 | GLN | N | AQ:68 | 4.80 | 1.00 | 86.31 |  |
| 5 | VAL | N | AQ:69 | 4.94 | 1.00 | 88.83 |  |
| 6 | ALA | N | AQ:72 | 4.98 | 1.00 | 96.37 |  |
  |

  
 Coded contact list:   
 Exclusion zone contact(s): /NE2_Gln_/  
Consider K in place of MG (1 valid contact(s) for K vs. 0 for MG)  

### MG:AQ:204

|  | Res. type | Atom | Residue | Distance | Occupancy | B-factor | Closest solute to 1^st^ shell water |
| --- | --- | --- | --- | --- | --- | --- | --- |
| **Ion** | | | |  | 1.00 | 80.84 || 1 | ALA | N | AQ:2 | 3.33 | 1.00 | 118.31 |
| 2 | C | OP2 | AA:2317 | 3.57 | 1.00 | 91.85 |  |
| 3 | C | OP2 | AA:2316 | 4.42 | 1.00 | 84.62 |  |
| 4 | G | N1 | AA:2318 | 4.67 | 1.00 | 106.24 |  |
| 5 | G | N2 | AA:2318 | 4.80 | 1.00 | 107.71 |  |
  |

  
 Coded contact list:   
 Exclusion zone contact(s): /N_back_/  
Consider K in place of MG (1 valid contact(s) for K vs. 0 for MG)  

### MG:AQ:205

|  | Res. type | Atom | Residue | Distance | Occupancy | B-factor | Closest solute to 1^st^ shell water |
| --- | --- | --- | --- | --- | --- | --- | --- |
| **Ion** | | | |  | 1.00 | 94.27 || 1 | ARG | NH1 | AQ:71 | 2.57 | 1.00 | 101.35 |
| 2 | ARG | NE | AQ:71 | 3.23 | 1.00 | 102.12 |  |
| 3 | ARG | O | AQ:67 | 4.27 | 1.00 | 84.83 |  |
| 4 | GLN | N | AQ:68 | 4.38 | 1.00 | 86.31 |  |
| 5 | ARG | NE | AQ:67 | 4.40 | 1.00 | 89.62 |  |
| 6 | ARG | NH2 | AQ:71 | 4.52 | 1.00 | 102.35 |  |
  |

  
 Coded contact list:   
 Exclusion zone contact(s): /NE_Arg_/NH1_Arg_/  
Consider NA in place of MG (1 valid contact(s) for NA vs. 0 for MG)  

### MG:AR:201

|  | Res. type | Atom | Residue | Distance | Occupancy | B-factor | Closest solute to 1^st^ shell water |
| --- | --- | --- | --- | --- | --- | --- | --- |
| **Ion** | | | |  | 1.00 | 71.40 || 1 | ARG | NE | AR:115 | 4.17 | 1.00 | 94.38 |
  |

  
 Coded contact list:   
1^st^ MG coordination shell missing (see Notes)  

### MG:AR:202

|  | Res. type | Atom | Residue | Distance | Occupancy | B-factor | Closest solute to 1^st^ shell water |
| --- | --- | --- | --- | --- | --- | --- | --- |
| **Ion** | | | |  | 1.00 | 64.70 || 1 | ARG | NH1 | AR:115 | 3.04 | 1.00 | 94.93 |
| 2 | G | OP2 | AA:1753 | 3.56 | 1.00 | 62.04 |  |
| 3 | ARG | NE | AR:115 | 3.62 | 1.00 | 94.38 |  |
| 4 | C | OP2 | AA:1752 | 3.93 | 1.00 | 68.10 |  |
| 5 | ARG | NH2 | AR:115 | 4.04 | 1.00 | 95.37 |  |
| 6 | G | OP1 | AA:1753 | 4.06 | 1.00 | 62.81 |  |
| 7 | C | OP1 | AA:1752 | 4.06 | 1.00 | 68.50 |  |
| 8 | C | O5' | AA:1752 | 4.20 | 1.00 | 66.88 |  |
| 9 | C | O3' | AA:1752 | 4.67 | 1.00 | 61.68 |  |
| 10 | ARG | O | AR:112 | 4.86 | 1.00 | 85.13 |  |
  |

  
 Coded contact list:   
 Exclusion zone contact(s): /NH1_Arg_/  
Consider K in place of MG (1 valid contact(s) for K vs. 0 for MG)  

### MG:AR:203

|  | Res. type | Atom | Residue | Distance | Occupancy | B-factor | Closest solute to 1^st^ shell water |
| --- | --- | --- | --- | --- | --- | --- | --- |
| **Ion** | | | |  | 1.00 | 55.18 || 1 | ARG | N | AR:13 | 2.29 | 1.00 | 81.94 |
| 2 | VAL | O | AR:10 | 2.46 | 1.00 | 74.31 |  |
| 3 | SER | O | AR:12 | 2.72 | 1.00 | 82.28 |  |
| 4 | SER | N | AR:12 | 3.32 | 1.00 | 79.70 |  |
| 5 | LEU | O | AR:9 | 3.96 | 1.00 | 73.19 |  |
| 6 | GLU | O | AR:11 | 4.12 | 1.00 | 78.92 |  |
| 7 | GLU | N | AR:11 | 4.27 | 1.00 | 75.21 |  |
| 8 | TYR | N | AR:14 | 4.49 | 1.00 | 80.50 |  |
| 9 | VAL | N | AR:10 | 4.98 | 1.00 | 72.62 |  |
  |

  
 Contact(s): /N_back_/  
 Coded contact list: Anion  
 Exclusion zone contact(s): /N_back_/O_back_/O_back_/  
Consider K in place of MG (2 valid contact(s) for K vs. 1 for MG)  

### MG:AR:204

|  | Res. type | Atom | Residue | Distance | Occupancy | B-factor | Closest solute to 1^st^ shell water |
| --- | --- | --- | --- | --- | --- | --- | --- |
| **Ion** | | | |  | 1.00 | 76.23 || 1 | SER | O | AR:12 | 2.58 | 1.00 | 82.28 |
| 2 | ARG | N | AR:13 | 3.89 | 1.00 | 81.94 |  |
| 3 | VAL | N | AR:15 | 4.25 | 1.00 | 78.84 |  |
| 4 | ARG | O | AR:13 | 4.26 | 1.00 | 79.97 |  |
| 5 | TYR | N | AR:14 | 4.30 | 1.00 | 80.50 |  |
| 6 | VAL | O | AR:15 | 4.88 | 1.00 | 78.50 |  |
  |

  
 Coded contact list:   
 Exclusion zone contact(s): /O_back_/  
Consider NA in place of MG (1 valid contact(s) for NA vs. 0 for MG)  

### MG:AR:205

|  | Res. type | Atom | Residue | Distance | Occupancy | B-factor | Closest solute to 1^st^ shell water |
| --- | --- | --- | --- | --- | --- | --- | --- |
| **Ion** | | | |  | 1.00 | 54.17 || 1 | SER | OG | AR:12 | 3.01 | 1.00 | 86.36 |
| 2 | PHE | N | AR:57 | 3.86 | 1.00 | 66.89 |  |
| 3 | LYS | O | AR:8 | 4.13 | 1.00 | 69.23 |  |
| 4 | ASN | N | AR:58 | 4.83 | 1.00 | 64.69 |  |
| 5 | LEU | N | AR:9 | 4.84 | 1.00 | 70.43 |  |
| 6 | ASN | ND2 | AR:58 | 4.89 | 1.00 | 60.45 |  |
| 7 | THR | OG1 | AE:12 | 4.99 | 1.00 | 71.26 |  |
  |

  
 Coded contact list:   
 Exclusion zone contact(s): /OG_Ser_/  
Consider K in place of MG (1 valid contact(s) for K vs. 0 for MG)  

### MG:AS:201

|  | Res. type | Atom | Residue | Distance | Occupancy | B-factor | Closest solute to 1^st^ shell water |
| --- | --- | --- | --- | --- | --- | --- | --- |
| **Ion** | | | |  | 1.00 | 67.54 || 1 | ASN | ND2 | AS:40 | 2.59 | 1.00 | 53.95 |
| 2 | G | O2' | AA:2009 | 2.80 | 1.00 | 45.20 |  |
| 3 | LYS | N | AS:41 | 3.13 | 1.00 | 54.32 |  |
| 4 | G | O3' | AA:2009 | 3.17 | 1.00 | 46.08 |  |
| 5 | ASN | O | AS:40 | 3.48 | 1.00 | 55.20 |  |
| 6 | ARG | N | AS:42 | 3.71 | 1.00 | 53.69 |  |
| 7 | ASN | OD1 | AS:40 | 4.28 | 1.00 | 57.20 |  |
| 8 | LYS | O | AS:41 | 4.31 | 1.00 | 55.16 |  |
| 9 | G | O4' | AA:2009 | 4.42 | 1.00 | 45.89 |  |
| 10 | G | OP1 | AA:2010 | 4.54 | 1.00 | 47.27 |  |
| 11 | G | O5' | AA:2010 | 4.88 | 1.00 | 46.70 |  |
  |

  
 Coded contact list:   
 Exclusion zone contact(s): /N_back_/ND2_Asn_/O2'/O3'/  
Consider K in place of MG (3 valid contact(s) for K vs. 0 for MG)  

### MG:AS:202

|  | Res. type | Atom | Residue | Distance | Occupancy | B-factor | Closest solute to 1^st^ shell water |
| --- | --- | --- | --- | --- | --- | --- | --- |
| **Ion** | | | |  | 1.00 | 63.64 || 1 | ALA | N | AS:56 | 4.50 | 1.00 | 61.62 |
| 2 | GLU | OE2 | AS:66 | 4.91 | 1.00 | 87.41 |  |
  |

  
 Coded contact list:   
1^st^ MG coordination shell missing (see Notes)  

### MG:AS:203

|  | Res. type | Atom | Residue | Distance | Occupancy | B-factor | Closest solute to 1^st^ shell water |
| --- | --- | --- | --- | --- | --- | --- | --- |
| **Ion** | | | |  | 1.00 | 72.80 || 1 | LYS | NZ | AS:72 | 2.98 | 1.00 | 65.22 |
| 2 | GLU | OE1 | AS:2 | 3.34 | 1.00 | 85.94 |  |
| 3 | GLU | OE2 | AS:2 | 4.37 | 1.00 | 84.48 |  |
  |

  
 Coded contact list:   
 Exclusion zone contact(s): /NZ_Lys_/OE1_Glu_/  
Consider K in place of MG (2 valid contact(s) for K vs. 0 for MG)  

### MG:AT:101

|  | Res. type | Atom | Residue | Distance | Occupancy | B-factor | Closest solute to 1^st^ shell water |
| --- | --- | --- | --- | --- | --- | --- | --- |
| **Ion** | | | |  | 1.00 | 70.74 || 1 | GLY | N | AT:61 | 2.52 | 1.00 | 62.49 |
| 2 | LYS | N | AT:62 | 3.51 | 1.00 | 64.30 |  |
| 3 | ARG | N | AT:60 | 3.76 | 1.00 | 58.60 |  |
| 4 | VAL | O | AT:59 | 3.96 | 1.00 | 55.69 |  |
| 5 | ARG | NH1 | AT:76 | 4.34 | 1.00 | 58.26 |  |
| 6 | G | O6 | AA:1339 | 4.54 | 1.00 | 46.64 |  |
| 7 | ARG | O | AT:60 | 4.66 | 1.00 | 62.17 |  |
| 8 | GLY | O | AT:61 | 4.70 | 1.00 | 61.05 |  |
  |

  
 Coded contact list:   
 Exclusion zone contact(s): /N_back_/  
Consider NA in place of MG (1 valid contact(s) for NA vs. 0 for MG)  

### MG:AT:102

|  | Res. type | Atom | Residue | Distance | Occupancy | B-factor | Closest solute to 1^st^ shell water |
| --- | --- | --- | --- | --- | --- | --- | --- |
| **Ion** | | | |  | 1.00 | 57.39 || 1 | C | OP1 | AA:1600 | 2.26 | 1.00 | 44.73 |
| 2 | PRO | O | AT:32 | 2.74 | 1.00 | 60.21 |  |
| 3 | LYS | NZ | AT:77 | 3.05 | 1.00 | 47.90 |  |
| 4 | HIS | NE2 | AT:58 | 3.16 | 1.00 | 50.41 |  |
| 5 | C | O3' | AA:1599 | 3.63 | 1.00 | 44.12 |  |
| 6 | ALA | O | AT:34 | 3.93 | 1.00 | 61.93 |  |
| 7 | ALA | N | AT:34 | 4.35 | 1.00 | 63.12 |  |
| 8 | LYS | N | AT:33 | 4.36 | 1.00 | 61.34 |  |
| 9 | C | OP2 | AA:1600 | 4.40 | 1.00 | 44.49 |  |
| 10 | HIS | ND1 | AT:58 | 4.63 | 1.00 | 50.97 |  |
| 11 | C | O5' | AA:1600 | 4.63 | 1.00 | 43.85 |  |
| 12 | LYS | O | AT:33 | 4.75 | 1.00 | 64.09 |  |
  |

  
 Contact(s): /OP1/  
 Coded contact list: O_ph_  
 Exclusion zone contact(s): /NE2_His_/NZ_Lys_/O_back_/  
Consider K in place of MG (3 valid contact(s) for K vs. 1 for MG)  

### MG:AT:103

|  | Res. type | Atom | Residue | Distance | Occupancy | B-factor | Closest solute to 1^st^ shell water |
| --- | --- | --- | --- | --- | --- | --- | --- |
| **Ion** | | | |  | 1.00 | 85.53 || 1 | U | OP1 | AA:72 | 4.11 | 1.00 | 47.29 |
| 2 | THR | OG1 | AT:3 | 4.11 | 1.00 | 61.30 |  |
| 3 | U | OP2 | AA:72 | 4.36 | 1.00 | 47.42 |  |
| 4 | THR | N | AT:3 | 4.57 | 1.00 | 68.32 |  |
| 5 | LYS | NZ | AW:29 | 4.89 | 1.00 | 70.50 |  |
  |

  
 Coded contact list:   
1^st^ MG coordination shell missing (see Notes)  

### MG:AT:104

|  | Res. type | Atom | Residue | Distance | Occupancy | B-factor | Closest solute to 1^st^ shell water |
| --- | --- | --- | --- | --- | --- | --- | --- |
| **Ion** | | | |  | 1.00 | 89.02 || 1 | A | OP1 | AA:1342 | 1.15 | 1.00 | 42.29 |
| 2 | LYS | NZ | AT:36 | 2.91 | 1.00 | 55.27 |  |
| 3 | A | O5' | AA:1342 | 3.34 | 1.00 | 43.29 |  |
| 4 | THR | N | AT:56 | 3.36 | 1.00 | 46.78 |  |
| 5 | U | O3' | AA:1341 | 3.54 | 1.00 | 42.53 |  |
| 6 | ASN | ND2 | AT:55 | 3.58 | 1.00 | 46.02 |  |
| 7 | A | OP2 | AA:1342 | 3.60 | 1.00 | 42.30 |  |
| 8 | VAL | O | AT:54 | 3.94 | 1.00 | 53.66 |  |
| 9 | THR | OG1 | AT:56 | 4.20 | 1.00 | 47.58 |  |
| 10 | THR | O | AT:56 | 4.20 | 1.00 | 46.82 |  |
| 11 | ASN | N | AT:55 | 4.48 | 1.00 | 51.55 |  |
| 12 | G | OP2 | AA:1343 | 4.81 | 1.00 | 52.95 |  |
| 13 | A | O3' | AA:1342 | 4.85 | 1.00 | 51.02 |  |
| 14 | A | O4' | AA:1342 | 4.98 | 1.00 | 48.76 |  |
  |

  
 Coded contact list:   
 Exclusion zone contact(s): /N_back_/NZ_Lys_/O5'/OP1/  
Consider K in place of MG (3 valid contact(s) for K vs. 0 for MG)  

### MG:AU:201

|  | Res. type | Atom | Residue | Distance | Occupancy | B-factor | Closest solute to 1^st^ shell water |
| --- | --- | --- | --- | --- | --- | --- | --- |
| **Ion** | | | |  | 1.00 | 51.09 || 1 | C | O2' | AA:334 | 3.79 | 1.00 | 61.32 |
| 2 | C | OP2 | AA:335 | 3.89 | 1.00 | 63.33 |  |
| 3 | G | OP2 | AA:301 | 4.04 | 1.00 | 60.34 |  |
| 4 | G | O5' | AA:301 | 4.17 | 1.00 | 59.91 |  |
| 5 | A | N7 | AA:300 | 4.31 | 1.00 | 58.49 |  |
| 6 | C | O3' | AA:334 | 4.58 | 1.00 | 62.16 |  |
| 7 | PRO | N | AU:82 | 4.62 | 1.00 | 84.00 |  |
| 8 | A | OP2 | AA:300 | 4.82 | 1.00 | 58.82 |  |
| 9 | A | O5' | AA:300 | 4.84 | 1.00 | 55.61 |  |
  |

  
 Coded contact list:   
1^st^ MG coordination shell missing (see Notes)  

### MG:AU:202

|  | Res. type | Atom | Residue | Distance | Occupancy | B-factor | Closest solute to 1^st^ shell water |
| --- | --- | --- | --- | --- | --- | --- | --- |
| **Ion** | | | |  | 1.00 | 76.00 || 1 | TYR | N | AU:20 | 2.87 | 1.00 | 79.43 |
| 2 | LYS | O | AU:19 | 3.29 | 1.00 | 79.42 |  |
| 3 | G | O6 | AA:329 | 3.79 | 1.00 | 54.70 |  |
| 4 | LYS | N | AU:19 | 4.49 | 1.00 | 76.99 |  |
  |

  
 Coded contact list:   
 Exclusion zone contact(s): /N_back_/O_back_/  
Consider K in place of MG (2 valid contact(s) for K vs. 0 for MG)  

### MG:AU:203

|  | Res. type | Atom | Residue | Distance | Occupancy | B-factor | Closest solute to 1^st^ shell water |
| --- | --- | --- | --- | --- | --- | --- | --- |
| **Ion** | | | |  | 1.00 | 74.07 || 1 | PRO | O | AU:56 | 2.67 | 1.00 | 136.01 |
| 2 | GLN | O | AU:57 | 3.28 | 1.00 | 133.49 |  |
| 3 | GLY | N | AU:58 | 3.67 | 1.00 | 132.47 |  |
| 4 | GLN | N | AU:57 | 4.17 | 1.00 | 135.32 |  |
| 5 | GLN | NE2 | AU:57 | 4.74 | 1.00 | 132.28 |  |
| 6 | TYR | O | AU:55 | 4.98 | 1.00 | 138.57 |  |
  |

  
 Coded contact list:   
 Exclusion zone contact(s): /O_back_/O_back_/  
Consider K in place of MG (2 valid contact(s) for K vs. 0 for MG)  

### MG:AU:204

|  | Res. type | Atom | Residue | Distance | Occupancy | B-factor | Closest solute to 1^st^ shell water |
| --- | --- | --- | --- | --- | --- | --- | --- |
| **Ion** | | | |  | 1.00 | 87.62 || 1 | LYS | NZ | AU:94 | 3.58 | 1.00 | 71.48 |
| 2 | CYS | O | AU:102 | 3.80 | 1.00 | 122.23 |  |
| 3 | LYS | NZ | AU:8 | 3.95 | 1.00 | 69.34 |  |
| 4 | LYS | N | AU:94 | 4.49 | 1.00 | 80.85 |  |
| 5 | LYS | NZ | AU:101 | 4.57 | 1.00 | 112.08 |  |
| 6 | LYS | O | AU:101 | 4.60 | 1.00 | 118.29 |  |
| 7 | LYS | O | AU:94 | 4.82 | 1.00 | 83.12 |  |
  |

  
 Coded contact list:   
1^st^ MG coordination shell missing (see Notes)  

### MG:AU:205

|  | Res. type | Atom | Residue | Distance | Occupancy | B-factor | Closest solute to 1^st^ shell water |
| --- | --- | --- | --- | --- | --- | --- | --- |
| **Ion** | | | |  | 1.00 | 69.78 || 1 | TYR | OH | AU:20 | 4.83 | 1.00 | 85.63 |
  |

  
 Coded contact list:   
1^st^ MG coordination shell missing (see Notes)  

### MG:AU:206

|  | Res. type | Atom | Residue | Distance | Occupancy | B-factor | Closest solute to 1^st^ shell water |
| --- | --- | --- | --- | --- | --- | --- | --- |
| **Ion** | | | |  | 1.00 | 75.97 || 1 | ILE | N | AU:61 | 4.16 | 1.00 | 121.63 |
| 2 | GLY | N | AU:59 | 4.77 | 1.00 | 129.52 |  |
| 3 | PHE | O | AU:60 | 4.80 | 1.00 | 123.12 |  |
| 4 | PHE | N | AU:60 | 4.90 | 1.00 | 126.51 |  |
| 5 | GLY | N | AU:58 | 4.94 | 1.00 | 132.47 |  |
  |

  
 Coded contact list:   
1^st^ MG coordination shell missing (see Notes)  

### MG:AW:101

|  | Res. type | Atom | Residue | Distance | Occupancy | B-factor | Closest solute to 1^st^ shell water |
| --- | --- | --- | --- | --- | --- | --- | --- |
| **Ion** | | | |  | 1.00 | 97.86 || 1 | LEU | O | AW:44 | 4.33 | 1.00 | 72.48 |
| 2 | SER | N | AW:45 | 4.52 | 1.00 | 70.16 |  |
  |

  
 Coded contact list:   
1^st^ MG coordination shell missing (see Notes)  

### MG:AW:102

|  | Res. type | Atom | Residue | Distance | Occupancy | B-factor | Closest solute to 1^st^ shell water |
| --- | --- | --- | --- | --- | --- | --- | --- |
| **Ion** | | | |  | 1.00 | 56.49 || 1 | SER | OG | AW:4 | 3.23 | 1.00 | 60.21 |
| 2 | SER | N | AW:4 | 3.89 | 1.00 | 62.96 |  |
| 3 | G | OP1 | AA:98 | 3.98 | 1.00 | 59.59 |  |
| 4 | GLU | OE2 | AW:5 | 4.53 | 1.00 | 66.53 |  |
| 5 | ARG | NH2 | AW:7 | 4.66 | 1.00 | 79.48 |  |
| 6 | C | O3' | AA:97 | 4.67 | 1.00 | 61.36 |  |
| 7 | G | O5' | AA:98 | 4.77 | 1.00 | 60.22 |  |
| 8 | A | O4' | AA:103 | 4.83 | 1.00 | 58.87 |  |
| 9 | C | O2' | AA:97 | 4.90 | 1.00 | 58.73 |  |
  |

  
 Coded contact list:   
 Exclusion zone contact(s): /OG_Ser_/  
Consider K in place of MG (1 valid contact(s) for K vs. 0 for MG)  

### MG:AZ:101

|  | Res. type | Atom | Residue | Distance | Occupancy | B-factor | Closest solute to 1^st^ shell water |
| --- | --- | --- | --- | --- | --- | --- | --- |
| **Ion** | | | |  | 1.00 | 83.78 || 1 | GLU | OE2 | AZ:89 | 3.23 | 1.00 | 92.13 |
| 2 | GLU | OE1 | AZ:89 | 4.35 | 1.00 | 93.85 |  |
  |

  
 Coded contact list:   
 Exclusion zone contact(s): /OE2_Glu_/  
Consider K in place of MG (1 valid contact(s) for K vs. 0 for MG)  

### MG:AZ:102

|  | Res. type | Atom | Residue | Distance | Occupancy | B-factor | Closest solute to 1^st^ shell water |
| --- | --- | --- | --- | --- | --- | --- | --- |
| **Ion** | | | |  | 1.00 | 50.38 || 1 | C | OP1 | AA:2231 | 2.76 | 1.00 | 47.86 |
| 2 | ARG | O | AZ:41 | 3.02 | 1.00 | 41.90 |  |
| 3 | ARG | NH2 | AZ:40 | 3.27 | 1.00 | 45.75 |  |
| 4 | C | OP2 | AA:2231 | 3.62 | 1.00 | 46.79 |  |
| 5 | ARG | NE | AZ:40 | 4.13 | 1.00 | 43.59 |  |
| 6 | G | O3' | AA:2230 | 4.71 | 1.00 | 45.37 |  |
| 7 | GLN | N | AZ:42 | 4.73 | 1.00 | 41.94 |  |
| 8 | ARG | N | AZ:41 | 4.79 | 1.00 | 41.86 |  |
| 9 | C | O5' | AA:2231 | 4.83 | 1.00 | 46.02 |  |
| 10 | TYR | N | AZ:43 | 4.94 | 1.00 | 41.44 |  |
  |

  
 Coded contact list:   
 Exclusion zone contact(s): /NH2_Arg_/O_back_/OP1/  
Consider K in place of MG (3 valid contact(s) for K vs. 0 for MG)  

### MG:AZ:103

|  | Res. type | Atom | Residue | Distance | Occupancy | B-factor | Closest solute to 1^st^ shell water |
| --- | --- | --- | --- | --- | --- | --- | --- |
| **Ion** | | | |  | 1.00 | 75.57 || 1 | LEU | O | AZ:85 | 2.98 | 1.00 | 76.39 |
| 2 | LYS | N | AZ:88 | 3.20 | 1.00 | 80.93 |  |
| 3 | SER | N | AZ:86 | 3.61 | 1.00 | 75.92 |  |
| 4 | PRO | N | AZ:87 | 3.79 | 1.00 | 74.80 |  |
| 5 | SER | O | AZ:86 | 3.80 | 1.00 | 72.69 |  |
| 6 | GLU | N | AZ:89 | 4.87 | 1.00 | 85.11 |  |
  |

  
 Coded contact list:   
 Exclusion zone contact(s): /N_back_/O_back_/  
Consider K in place of MG (2 valid contact(s) for K vs. 0 for MG)  

### MG:B1:101

|  | Res. type | Atom | Residue | Distance | Occupancy | B-factor | Closest solute to 1^st^ shell water |
| --- | --- | --- | --- | --- | --- | --- | --- |
| **Ion** | | | |  | 1.00 | 60.13 || 1 | U | OP1 | B1:44 | 2.89 | 1.00 | 114.75 |
| 2 | U | OP2 | B1:45 | 3.75 | 1.00 | 105.95 |  |
| 3 | U | OP1 | B1:45 | 3.80 | 1.00 | 106.43 |  |
| 4 | U | OP2 | BA:1506 | 3.96 | 1.00 | 68.29 |  |
| 5 | U | O5' | B1:44 | 4.01 | 1.00 | 113.90 |  |
| 6 | U | O3' | B1:44 | 4.15 | 1.00 | 108.41 |  |
| 7 | G | O3' | BA:1505 | 4.73 | 1.00 | 68.70 |  |
| 8 | U | O3' | B1:43 | 4.97 | 1.00 | 115.20 |  |
  |

  
 Coded contact list:   
 Exclusion zone contact(s): /OP1/  
Consider K in place of MG (1 valid contact(s) for K vs. 0 for MG)  

### MG:B1:102

|  | Res. type | Atom | Residue | Distance | Occupancy | B-factor | Closest solute to 1^st^ shell water |
| --- | --- | --- | --- | --- | --- | --- | --- |
| **Ion** | | | |  | 1.00 | 72.04 || 1 | C | OP2 | B1:30 | 4.51 | 1.00 | 197.38 |
  |

  
 Coded contact list:   
1^st^ MG coordination shell missing (see Notes)  

### MG:B1:103

|  | Res. type | Atom | Residue | Distance | Occupancy | B-factor | Closest solute to 1^st^ shell water |
| --- | --- | --- | --- | --- | --- | --- | --- |
| **Ion** | | | |  | 1.00 | 67.70 || 1 | U | O2 | B1:54 | 4.15 | 1.00 | 175.50 |
| 2 | U | N3 | B1:54 | 4.41 | 1.00 | 176.01 |  |
| 3 | U | OP1 | BA:1194 | 4.80 | 1.00 | 93.02 |  |
  |

  
 Coded contact list:   
1^st^ MG coordination shell missing (see Notes)  

### MG:B1:104

|  | Res. type | Atom | Residue | Distance | Occupancy | B-factor | Closest solute to 1^st^ shell water |
| --- | --- | --- | --- | --- | --- | --- | --- |
| **Ion** | | | |  | 1.00 | 116.17 || 1 | U | O4 | BA:1542 | 2.53 | 1.00 | 196.27 |
| 2 | G | N2 | B1:28 | 3.14 | 1.00 | 194.99 |  |
| 3 | G | N1 | B1:28 | 3.21 | 1.00 | 195.21 |  |
| 4 | G | O6 | B1:29 | 3.24 | 1.00 | 197.84 |  |
| 5 | G | N1 | B1:29 | 3.44 | 1.00 | 198.04 |  |
| 6 | G | O6 | B1:28 | 4.47 | 1.00 | 194.72 |  |
| 7 | U | N3 | BA:1542 | 4.50 | 1.00 | 196.51 |  |
| 8 | G | N2 | B1:29 | 4.77 | 1.00 | 197.76 |  |
| 9 | G | N3 | B1:28 | 4.95 | 1.00 | 195.35 |  |
  |

  
 Coded contact list:   
 Exclusion zone contact(s): /N1_G_/N2/O4/O6/  
Consider K in place of MG (3 valid contact(s) for K vs. 0 for MG)  

### MG:BA:1601

|  | Res. type | Atom | Residue | Distance | Occupancy | B-factor | Closest solute to 1^st^ shell water |
| --- | --- | --- | --- | --- | --- | --- | --- |
| **Ion** | | | |  | 1.00 | 27.12 || 1 | G | O6 | BA:798 | 4.02 | 1.00 | 54.40 |
| 2 | G | O6 | BA:799 | 4.07 | 1.00 | 55.18 |  |
| 3 | G | N7 | BA:799 | 4.30 | 1.00 | 55.33 |  |
| 4 | A | OP2 | BA:781 | 4.48 | 1.00 | 50.92 |  |
| 5 | G | N7 | BA:798 | 4.55 | 1.00 | 56.74 |  |
| 6 | G | O6 | BA:800 | 4.71 | 1.00 | 60.97 |  |
| 7 | G | N1 | BA:800 | 4.75 | 1.00 | 60.10 |  |
  |

  
 Coded contact list:   
1^st^ MG coordination shell missing (see Notes)  

### MG:BA:1602

|  | Res. type | Atom | Residue | Distance | Occupancy | B-factor | Closest solute to 1^st^ shell water |
| --- | --- | --- | --- | --- | --- | --- | --- |
| **Ion** | | | |  | 1.00 | 20.68 || 1 | C | OP1 | BA:578 | 2.40 | 1.00 | 60.24 |
| 2 | G | OP2 | BA:576 | 2.87 | 1.00 | 57.51 |  |
| 3 | G | O2' | BA:575 | 3.54 | 1.00 | 62.42 |  |
| 4 | G | O3' | BA:577 | 3.73 | 1.00 | 56.75 |  |
| 5 | U | OP1 | BA:820 | 3.84 | 1.00 | 59.74 |  |
| 6 | C | OP2 | BA:817 | 3.98 | 1.00 | 52.41 |  |
| 7 | C | OP1 | BA:817 | 4.53 | 1.00 | 53.87 |  |
| 8 | C | OP2 | BA:578 | 4.57 | 1.00 | 57.44 |  |
| 9 | G | O3' | BA:575 | 4.68 | 1.00 | 59.72 |  |
| 10 | C | O5' | BA:578 | 4.73 | 1.00 | 57.70 |  |
| 11 | G | O5' | BA:576 | 4.90 | 1.00 | 57.00 |  |
| 12 | C | O5' | BA:817 | 4.95 | 1.00 | 53.40 |  |
  |

  
 Contact(s): /OP1/  
 Coded contact list: O_ph_  
 Exclusion zone contact(s): /OP2/  

### MG:BA:1603

|  | Res. type | Atom | Residue | Distance | Occupancy | B-factor | Closest solute to 1^st^ shell water |
| --- | --- | --- | --- | --- | --- | --- | --- |
| **Ion** | | | |  | 1.00 | 58.05 || 1 | U | O4 | BA:125 | 3.59 | 1.00 | 73.92 |
| 2 | G | O6 | BA:126 | 3.67 | 1.00 | 75.03 |  |
| 3 | G | N7 | BA:126 | 4.32 | 1.00 | 75.14 |  |
| 4 | G | O6 | BA:127 | 4.59 | 1.00 | 71.51 |  |
  |

  
 Coded contact list:   
1^st^ MG coordination shell missing (see Notes)  

### MG:BA:1604

|  | Res. type | Atom | Residue | Distance | Occupancy | B-factor | Closest solute to 1^st^ shell water |
| --- | --- | --- | --- | --- | --- | --- | --- |
| **Ion** | | | |  | 1.00 | 47.36 || 1 | A | OP2 | BA:780 | 3.21 | 1.00 | 56.48 |
| 2 | G | O6 | BA:800 | 3.31 | 1.00 | 60.97 |  |
| 3 | C | OP2 | BA:779 | 3.73 | 1.00 | 60.06 |  |
| 4 | G | N7 | BA:800 | 3.92 | 1.00 | 61.15 |  |
| 5 | C | O5' | BA:779 | 4.02 | 1.00 | 57.88 |  |
| 6 | C | OP1 | BA:779 | 4.16 | 1.00 | 58.36 |  |
| 7 | A | N7 | BA:780 | 4.25 | 1.00 | 56.68 |  |
| 8 | G | OP2 | BA:800 | 4.73 | 1.00 | 56.51 |  |
| 9 | G | OP2 | BA:799 | 4.81 | 1.00 | 52.74 |  |
  |

  
 Coded contact list:   
 Exclusion zone contact(s): /O6/OP2/  
Consider K in place of MG (2 valid contact(s) for K vs. 0 for MG)  

### MG:BA:1605

|  | Res. type | Atom | Residue | Distance | Occupancy | B-factor | Closest solute to 1^st^ shell water |
| --- | --- | --- | --- | --- | --- | --- | --- |
| **Ion** | | | |  | 1.00 | 56.43 || 1 | G | N7 | BA:251 | 2.68 | 1.00 | 79.00 |
| 2 | A | OP2 | BA:270 | 3.00 | 1.00 | 82.04 |  |
| 3 | G | O6 | BA:251 | 3.06 | 1.00 | 81.02 |  |
| 4 | G | O6 | BA:266 | 4.47 | 1.00 | 68.91 |  |
| 5 | C | N4 | BA:271 | 4.75 | 1.00 | 76.92 |  |
| 6 | A | N7 | BA:270 | 4.84 | 1.00 | 79.76 |  |
| 7 | G | N9 | BA:251 | 4.90 | 1.00 | 76.87 |  |
| 8 | G | N1 | BA:251 | 4.94 | 1.00 | 78.23 |  |
  |

  
 Coded contact list:   
 Exclusion zone contact(s): /N7/O6/OP2/  
Consider K in place of MG (3 valid contact(s) for K vs. 0 for MG)  

### MG:BA:1606

|  | Res. type | Atom | Residue | Distance | Occupancy | B-factor | Closest solute to 1^st^ shell water |
| --- | --- | --- | --- | --- | --- | --- | --- |
| **Ion** | | | |  | 1.00 | 93.22 || 1 | G | O6 | BA:1178 | 2.29 | 1.00 | 144.58 |
| 2 | G | N7 | BA:1178 | 2.98 | 1.00 | 144.29 |  |
| 3 | A | OP2 | BA:1180 | 3.63 | 1.00 | 140.21 |  |
| 4 | A | OP1 | BA:1180 | 3.77 | 1.00 | 140.20 |  |
| 5 | LYS | NZ | BL:97 | 4.23 | 1.00 | 122.00 |  |
| 6 | G | N1 | BA:1178 | 4.38 | 1.00 | 144.58 |  |
| 7 | A | O5' | BA:1180 | 4.93 | 1.00 | 140.27 |  |
  |

  
 Contact(s): /O6/  
 Coded contact list: O_b_  
 Exclusion zone contact(s): /N7/  

### MG:BA:1607

|  | Res. type | Atom | Residue | Distance | Occupancy | B-factor | Closest solute to 1^st^ shell water |
| --- | --- | --- | --- | --- | --- | --- | --- |
| **Ion** | | | |  | 1.00 | 74.06 || 1 | G | OP2 | BA:476 | 2.84 | 1.00 | 117.73 |
| 2 | G | O5' | BA:476 | 4.38 | 1.00 | 116.95 |  |
| 3 | A | OP1 | BA:453 | 4.54 | 1.00 | 78.76 |  |
| 4 | G | OP1 | BA:476 | 4.61 | 1.00 | 117.65 |  |
  |

  
 Coded contact list:   
 Exclusion zone contact(s): /OP2/  
Consider K in place of MG (1 valid contact(s) for K vs. 0 for MG)  

### MG:BA:1608

|  | Res. type | Atom | Residue | Distance | Occupancy | B-factor | Closest solute to 1^st^ shell water |
| --- | --- | --- | --- | --- | --- | --- | --- |
| **Ion** | | | |  | 1.00 | 47.99 || 1 | G | OP1 | BA:1508 | 1.64 | 1.00 | 55.14 |
| 2 | G | OP1 | BA:1505 | 1.86 | 1.00 | 68.46 |  |
| 3 | A | OP1 | BA:1500 | 2.10 | 1.00 | 56.13 |  |
| 4 | A | O3' | BA:1507 | 3.29 | 1.00 | 55.00 |  |
| 5 | G | O5' | BA:1505 | 3.70 | 1.00 | 68.98 |  |
| 6 | G | OP2 | BA:1505 | 3.74 | 1.00 | 68.10 |  |
| 7 | G | O5' | BA:1508 | 3.91 | 1.00 | 55.02 |  |
| 8 | G | OP2 | BA:1508 | 4.01 | 1.00 | 56.20 |  |
| 9 | A | OP2 | BA:1500 | 4.25 | 1.00 | 56.12 |  |
| 10 | G | O3' | BA:1504 | 4.34 | 1.00 | 67.46 |  |
| 11 | A | O5' | BA:1500 | 4.41 | 1.00 | 55.13 |  |
| 12 | A | O3' | BA:1499 | 4.42 | 1.00 | 55.71 |  |
| 13 | G | O5' | BA:1504 | 4.82 | 1.00 | 67.18 |  |
| 14 | C | OP2 | BA:1501 | 4.84 | 1.00 | 56.95 |  |
  |

  
**1^st^ shell ligand angles:**   
1-2: 150; 1-3: 103; 2-3: 76;   
  
 Contact(s): /OP1/OP1/  
 Coded contact list: 2O_ph_  
 Exclusion zone contact(s): /O3'/OP1/  

### MG:BA:1609

|  | Res. type | Atom | Residue | Distance | Occupancy | B-factor | Closest solute to 1^st^ shell water |
| --- | --- | --- | --- | --- | --- | --- | --- |
| **Ion** | | | |  | 1.00 | 36.25 || 1 | G | OP1 | BA:1521 | 2.41 | 1.00 | 49.51 |
| 2 | A | OP1 | BA:1500 | 2.79 | 1.00 | 56.13 |  |
| 3 | A | O3' | BA:1499 | 3.00 | 1.00 | 55.71 |  |
| 4 | G | OP1 | BA:1508 | 3.19 | 1.00 | 55.14 |  |
| 5 | G | OP2 | BA:1521 | 3.83 | 1.00 | 48.63 |  |
| 6 | A | O2' | BA:1499 | 4.08 | 1.00 | 50.20 |  |
| 7 | A | O5' | BA:1500 | 4.33 | 1.00 | 55.13 |  |
| 8 | G | O3' | BA:1520 | 4.44 | 1.00 | 51.00 |  |
| 9 | G | OP2 | BA:1508 | 4.67 | 1.00 | 56.20 |  |
| 10 | A | OP2 | BA:1500 | 4.75 | 1.00 | 56.12 |  |
| 11 | G | O5' | BA:1521 | 4.76 | 1.00 | 49.27 |  |
| 12 | G | OP1 | BA:1505 | 4.78 | 1.00 | 68.46 |  |
| 13 | G | O5' | BA:1508 | 4.91 | 1.00 | 55.02 |  |
  |

  
 Coded contact list:   
 Exclusion zone contact(s): /O3'/OP1/OP1/OP1/  
Consider K in place of MG (3 valid contact(s) for K vs. 0 for MG)  

### MG:BA:1610

|  | Res. type | Atom | Residue | Distance | Occupancy | B-factor | Closest solute to 1^st^ shell water |
| --- | --- | --- | --- | --- | --- | --- | --- |
| **Ion** | | | |  | 1.00 | 25.36 || 1 | A | OP2 | BA:1499 | 1.74 | 1.00 | 56.28 |
| 2 | A | OP2 | BA:1500 | 1.79 | 1.00 | 56.12 |  |
| 3 | G | O2' | BA:1504 | 3.17 | 1.00 | 69.09 |  |
| 4 | U | O2' | BA:1498 | 3.35 | 1.00 | 56.71 |  |
| 5 | G | OP2 | BA:1505 | 3.38 | 1.00 | 68.10 |  |
| 6 | A | OP1 | BA:1499 | 3.53 | 1.00 | 55.01 |  |
| 7 | A | O5' | BA:1500 | 3.66 | 1.00 | 55.13 |  |
| 8 | G | O3' | BA:1504 | 3.88 | 1.00 | 67.46 |  |
| 9 | A | O5' | BA:1499 | 3.91 | 1.00 | 53.66 |  |
| 10 | A | O3' | BA:1499 | 3.92 | 1.00 | 55.71 |  |
| 11 | A | N7 | BA:1500 | 3.98 | 1.00 | 55.44 |  |
| 12 | U | O3' | BA:1498 | 4.20 | 1.00 | 54.86 |  |
| 13 | A | OP1 | BA:1500 | 4.33 | 1.00 | 56.13 |  |
| 14 | G | OP1 | BA:1505 | 4.69 | 1.00 | 68.46 |  |
  |

  
**1^st^ shell ligand angles:**   
1-2: 104;   
  
 Coded contact list:   
 Exclusion zone contact(s): /O2'/O2'/OP2/OP2/OP2/  
Consider K in place of MG (3 valid contact(s) for K vs. 0 for MG)  

### MG:BA:1611

|  | Res. type | Atom | Residue | Distance | Occupancy | B-factor | Closest solute to 1^st^ shell water |
| --- | --- | --- | --- | --- | --- | --- | --- |
| **Ion** | | | |  | 1.00 | 37.48 || 1 | G | OP2 | BA:1505 | 1.81 | 1.00 | 68.10 |
| 2 | G | O3' | BA:1504 | 2.27 | 1.00 | 67.46 |  |
| 3 | G | O5' | BA:1505 | 3.12 | 1.00 | 68.98 |  |
| 4 | A | OP2 | BA:1500 | 3.41 | 1.00 | 56.12 |  |
| 5 | G | O2' | BA:1504 | 3.52 | 1.00 | 69.09 |  |
| 6 | A | OP1 | BA:1499 | 3.61 | 1.00 | 55.01 |  |
| 7 | A | OP2 | BA:1499 | 3.73 | 1.00 | 56.28 |  |
| 8 | G | N7 | BA:1505 | 3.80 | 1.00 | 69.18 |  |
| 9 | G | OP1 | BA:1505 | 3.81 | 1.00 | 68.46 |  |
| 10 | G | O4' | BA:1505 | 4.15 | 1.00 | 67.97 |  |
| 11 | G | N9 | BA:1505 | 4.23 | 1.00 | 68.98 |  |
| 12 | U | O3' | BA:1498 | 4.82 | 1.00 | 54.86 |  |
| 13 | A | OP1 | BA:1500 | 4.92 | 1.00 | 56.13 |  |
| 14 | U | OP1 | BA:1506 | 4.92 | 1.00 | 67.78 |  |
  |

  
**1^st^ shell ligand angles:**   
1-2: 75;   
  
 Contact(s): /O3'/OP2/  
 Coded contact list: O_ph_O_r_  
 Exclusion zone contact(s): /O5'/  

### MG:BA:1612

|  | Res. type | Atom | Residue | Distance | Occupancy | B-factor | Closest solute to 1^st^ shell water |
| --- | --- | --- | --- | --- | --- | --- | --- |
| **Ion** | | | |  | 1.00 | 90.78 || 1 | G | OP1 | BA:293 | 2.89 | 1.00 | 58.74 |
| 2 | G | OP2 | BA:293 | 4.04 | 1.00 | 57.77 |  |
| 3 | G | O3' | BA:292 | 4.92 | 1.00 | 58.62 |  |
  |

  
 Coded contact list:   
 Exclusion zone contact(s): /OP1/  
Consider K in place of MG (1 valid contact(s) for K vs. 0 for MG)  

### MG:BA:1613

|  | Res. type | Atom | Residue | Distance | Occupancy | B-factor | Closest solute to 1^st^ shell water |
| --- | --- | --- | --- | --- | --- | --- | --- |
| **Ion** | | | |  | 1.00 | 84.61 || 1 | G | O6 | BA:297 | 4.48 | 1.00 | 65.96 |
  |

  
 Coded contact list:   
1^st^ MG coordination shell missing (see Notes)  

### MG:BA:1615

|  | Res. type | Atom | Residue | Distance | Occupancy | B-factor | Closest solute to 1^st^ shell water |
| --- | --- | --- | --- | --- | --- | --- | --- |
| **Ion** | | | |  | 1.00 | 81.59 || 1 | G | O2' | BA:630 | 4.85 | 1.00 | 124.63 |
  |

  
 Coded contact list:   
1^st^ MG coordination shell missing (see Notes)  

### MG:BA:1616

|  | Res. type | Atom | Residue | Distance | Occupancy | B-factor | Closest solute to 1^st^ shell water |
| --- | --- | --- | --- | --- | --- | --- | --- |
| **Ion** | | | |  | 1.00 | 47.23 || 1 | G | OP1 | BA:558 | 1.95 | 1.00 | 63.59 |
| 2 | G | O6 | BA:299 | 2.20 | 1.00 | 62.62 |  |
| 3 | U | OP1 | BA:560 | 3.36 | 1.00 | 73.18 |  |
| 4 | G | O6 | BA:566 | 3.44 | 1.00 | 71.17 |  |
| 5 | G | O3' | BA:557 | 3.79 | 1.00 | 62.96 |  |
| 6 | A | N6 | BA:298 | 3.93 | 1.00 | 72.08 |  |
| 7 | G | N1 | BA:299 | 4.05 | 1.00 | 63.21 |  |
| 8 | G | O2' | BA:297 | 4.11 | 1.00 | 67.28 |  |
| 9 | G | OP2 | BA:558 | 4.12 | 1.00 | 63.65 |  |
| 10 | A | N7 | BA:298 | 4.29 | 1.00 | 71.22 |  |
| 11 | A | N1 | BA:300 | 4.30 | 1.00 | 55.50 |  |
| 12 | G | O5' | BA:558 | 4.47 | 1.00 | 65.11 |  |
| 13 | U | OP2 | BA:560 | 4.74 | 1.00 | 72.92 |  |
| 14 | G | N1 | BA:566 | 4.83 | 1.00 | 69.76 |  |
| 15 | A | N6 | BA:300 | 4.83 | 1.00 | 54.32 |  |
  |

  
**1^st^ shell ligand angles:**   
1-2: 171;   
  
 Contact(s): /O6/OP1/  
 Coded contact list: O_ph_O_b_  
 Exclusion zone contact(s): /OP1/  

### MG:BA:1617

|  | Res. type | Atom | Residue | Distance | Occupancy | B-factor | Closest solute to 1^st^ shell water |
| --- | --- | --- | --- | --- | --- | --- | --- |
| **Ion** | | | |  | 1.00 | 44.78 || 1 | G | O6 | BA:1470 | 4.02 | 1.00 | 68.79 |
| 2 | G | N7 | BA:1469 | 4.09 | 1.00 | 74.55 |  |
| 3 | G | N7 | BA:1470 | 4.35 | 1.00 | 71.70 |  |
| 4 | G | O6 | BA:1471 | 4.65 | 1.00 | 64.35 |  |
| 5 | G | O6 | BA:1469 | 4.79 | 1.00 | 74.12 |  |
  |

  
 Coded contact list:   
1^st^ MG coordination shell missing (see Notes)  

### MG:BA:1618

|  | Res. type | Atom | Residue | Distance | Occupancy | B-factor | Closest solute to 1^st^ shell water |
| --- | --- | --- | --- | --- | --- | --- | --- |
| **Ion** | | | |  | 1.00 | 34.71 || 1 | A | OP2 | BA:1468 | 3.44 | 1.00 | 68.75 |
| 2 | G | N2 | BA:1432 | 3.53 | 1.00 | 68.06 |  |
| 3 | G | O6 | BA:1469 | 3.61 | 1.00 | 74.12 |  |
| 4 | G | N1 | BA:1432 | 4.05 | 1.00 | 69.14 |  |
| 5 | G | OP2 | BA:1467 | 4.54 | 1.00 | 66.61 |  |
| 6 | G | N7 | BA:1469 | 4.72 | 1.00 | 74.55 |  |
| 7 | A | OP1 | BA:1468 | 4.77 | 1.00 | 68.71 |  |
| 8 | G | O5' | BA:1467 | 4.84 | 1.00 | 67.16 |  |
| 9 | C | N4 | BA:1431 | 4.88 | 1.00 | 71.99 |  |
  |

  
 Coded contact list:   
1^st^ MG coordination shell missing (see Notes)  

### MG:BA:1619

|  | Res. type | Atom | Residue | Distance | Occupancy | B-factor | Closest solute to 1^st^ shell water |
| --- | --- | --- | --- | --- | --- | --- | --- |
| **Ion** | | | |  | 1.00 | 31.72 || 1 | G | O6 | BA:410 | 3.33 | 1.00 | 90.19 |
| 2 | A | OP2 | BA:431 | 3.45 | 1.00 | 75.57 |  |
| 3 | A | OP1 | BA:431 | 3.76 | 1.00 | 74.93 |  |
| 4 | G | N1 | BA:410 | 4.53 | 1.00 | 89.54 |  |
| 5 | U | O2' | BA:429 | 4.62 | 1.00 | 82.50 |  |
| 6 | A | N7 | BA:408 | 4.65 | 1.00 | 87.73 |  |
| 7 | A | O3' | BA:430 | 4.72 | 1.00 | 75.12 |  |
| 8 | A | N6 | BA:408 | 4.76 | 1.00 | 89.56 |  |
| 9 | G | O6 | BA:409 | 4.88 | 1.00 | 91.57 |  |
| 10 | G | N7 | BA:409 | 4.97 | 1.00 | 87.15 |  |
  |

  
 Coded contact list:   
 Exclusion zone contact(s): /O6/  
Consider K in place of MG (1 valid contact(s) for K vs. 0 for MG)  

### MG:BA:1620

|  | Res. type | Atom | Residue | Distance | Occupancy | B-factor | Closest solute to 1^st^ shell water |
| --- | --- | --- | --- | --- | --- | --- | --- |
| **Ion** | | | |  | 1.00 | 50.42 || 1 | G | O6 | BA:406 | 2.94 | 1.00 | 86.43 |
| 2 | G | O6 | BA:407 | 3.55 | 1.00 | 85.18 |  |
| 3 | G | N7 | BA:406 | 3.86 | 1.00 | 84.30 |  |
| 4 | C | N4 | BA:435 | 4.10 | 1.00 | 86.90 |  |
| 5 | C | N4 | BA:436 | 4.61 | 1.00 | 88.38 |  |
| 6 | A | O3' | BA:495 | 4.82 | 1.00 | 104.30 |  |
| 7 | G | N1 | BA:406 | 4.91 | 1.00 | 86.05 |  |
  |

  
 Coded contact list:   
 Exclusion zone contact(s): /O6/  
Consider K in place of MG (1 valid contact(s) for K vs. 0 for MG)  

### MG:BA:1621

|  | Res. type | Atom | Residue | Distance | Occupancy | B-factor | Closest solute to 1^st^ shell water |
| --- | --- | --- | --- | --- | --- | --- | --- |
| **Ion** | | | |  | 1.00 | 120.66 || 1 | C | OP2 | BA:169 | 4.02 | 1.00 | 98.40 |
| 2 | C | OP1 | BA:169 | 4.21 | 1.00 | 98.09 |  |
| 3 | G | OP1 | BA:168 | 4.78 | 1.00 | 102.17 |  |
| 4 | A | OP1 | BA:149 | 4.82 | 1.00 | 91.10 |  |
  |

  
 Coded contact list:   
1^st^ MG coordination shell missing (see Notes)  

### MG:BA:1622

|  | Res. type | Atom | Residue | Distance | Occupancy | B-factor | Closest solute to 1^st^ shell water |
| --- | --- | --- | --- | --- | --- | --- | --- |
| **Ion** | | | |  | 1.00 | 64.44 || 1 | A | OP2 | BA:452 | 2.29 | 1.00 | 79.17 |
| 2 | G | OP1 | BA:450 | 3.19 | 1.00 | 81.60 |  |
| 3 | G | OP2 | BA:450 | 3.36 | 1.00 | 81.52 |  |
| 4 | A | O3' | BA:451 | 3.39 | 1.00 | 79.31 |  |
| 5 | A | O2' | BA:451 | 4.00 | 1.00 | 81.44 |  |
| 6 | LYS | NZ | BS:43 | 4.35 | 1.00 | 77.09 |  |
| 7 | A | OP1 | BA:452 | 4.43 | 1.00 | 79.12 |  |
| 8 | A | O5' | BA:452 | 4.46 | 1.00 | 77.13 |  |
| 9 | C | O3' | BA:449 | 4.55 | 1.00 | 81.85 |  |
  |

  
 Contact(s): /OP2/  
 Coded contact list: O_ph_  
 Exclusion zone contact(s): /O3'/OP1/OP2/  
Consider K in place of MG (3 valid contact(s) for K vs. 1 for MG)  

### MG:BA:1623

|  | Res. type | Atom | Residue | Distance | Occupancy | B-factor | Closest solute to 1^st^ shell water |
| --- | --- | --- | --- | --- | --- | --- | --- |
| **Ion** | | | |  | 1.00 | 53.55 || 1 | G | OP2 | BA:450 | 2.13 | 1.00 | 81.52 |
| 2 | C | O5' | BA:449 | 2.67 | 1.00 | 84.29 |  |
| 3 | G | O6 | BA:481 | 2.86 | 1.00 | 82.66 |  |
| 4 | C | OP1 | BA:449 | 3.02 | 1.00 | 85.16 |  |
| 5 | C | OP2 | BA:449 | 3.02 | 1.00 | 85.82 |  |
| 6 | G | N7 | BA:450 | 3.63 | 1.00 | 76.07 |  |
| 7 | G | O5' | BA:450 | 3.79 | 1.00 | 81.50 |  |
| 8 | C | O3' | BA:449 | 4.07 | 1.00 | 81.85 |  |
| 9 | A | O3' | BA:448 | 4.53 | 1.00 | 85.11 |  |
| 10 | G | OP1 | BA:450 | 4.66 | 1.00 | 81.60 |  |
| 11 | G | N1 | BA:481 | 4.94 | 1.00 | 81.79 |  |
| 12 | C | O4' | BA:449 | 4.99 | 1.00 | 83.04 |  |
  |

  
 Contact(s): /OP2/  
 Coded contact list: O_ph_  
 Exclusion zone contact(s): /O5'/O6/OP1/OP2/  
Consider K in place of MG (4 valid contact(s) for K vs. 1 for MG)  

### MG:BA:1624

|  | Res. type | Atom | Residue | Distance | Occupancy | B-factor | Closest solute to 1^st^ shell water |
| --- | --- | --- | --- | --- | --- | --- | --- |
| **Ion** | | | |  | 1.00 | 84.21 || 1 | G | O6 | BA:661 | 2.27 | 1.00 | 79.16 |
| 2 | G | O6 | BA:662 | 2.60 | 1.00 | 78.20 |  |
| 3 | G | N7 | BA:662 | 3.41 | 1.00 | 80.19 |  |
| 4 | G | N1 | BA:661 | 3.69 | 1.00 | 80.44 |  |
| 5 | C | N4 | BA:744 | 3.77 | 1.00 | 77.93 |  |
| 6 | G | N7 | BA:661 | 3.94 | 1.00 | 82.65 |  |
| 7 | U | O4 | BA:743 | 3.94 | 1.00 | 78.15 |  |
| 8 | U | N3 | BA:743 | 4.63 | 1.00 | 75.72 |  |
| 9 | C | N3 | BA:744 | 4.74 | 1.00 | 77.62 |  |
| 10 | G | N1 | BA:662 | 4.74 | 1.00 | 79.10 |  |
  |

  
 Contact(s): /O6/  
 Coded contact list: O_b_  
 Exclusion zone contact(s): /O6/  

### MG:BA:1625

|  | Res. type | Atom | Residue | Distance | Occupancy | B-factor | Closest solute to 1^st^ shell water |
| --- | --- | --- | --- | --- | --- | --- | --- |
| **Ion** | | | |  | 1.00 | 74.07 || 1 | G | O6 | BA:660 | 3.58 | 1.00 | 85.78 |
| 2 | G | N7 | BA:660 | 3.73 | 1.00 | 85.91 |  |
| 3 | G | O6 | BA:661 | 4.18 | 1.00 | 79.16 |  |
| 4 | G | N7 | BA:661 | 4.21 | 1.00 | 82.65 |  |
| 5 | U | O4 | BA:659 | 4.66 | 1.00 | 87.74 |  |
  |

  
 Coded contact list:   
1^st^ MG coordination shell missing (see Notes)  

### MG:BA:1626

|  | Res. type | Atom | Residue | Distance | Occupancy | B-factor | Closest solute to 1^st^ shell water |
| --- | --- | --- | --- | --- | --- | --- | --- |
| **Ion** | | | |  | 1.00 | 68.61 || 1 | G | OP2 | BA:750 | 2.22 | 1.00 | 86.91 |
| 2 | C | OP2 | BA:749 | 2.25 | 1.00 | 92.65 |  |
| 3 | C | OP1 | BA:749 | 3.85 | 1.00 | 92.59 |  |
| 4 | C | O2' | BA:748 | 3.96 | 1.00 | 94.96 |  |
| 5 | G | N7 | BA:750 | 4.26 | 1.00 | 82.76 |  |
| 6 | C | O5' | BA:749 | 4.33 | 1.00 | 91.30 |  |
| 7 | C | O3' | BA:749 | 4.44 | 1.00 | 87.08 |  |
| 8 | G | O5' | BA:750 | 4.49 | 1.00 | 86.51 |  |
| 9 | G | OP1 | BA:750 | 4.64 | 1.00 | 86.42 |  |
| 10 | C | O3' | BA:748 | 4.71 | 1.00 | 93.06 |  |
  |

  
**1^st^ shell ligand angles:**   
1-2: 100;   
  
 Contact(s): /OP2/OP2/  
 Coded contact list: 2O_ph_  

### MG:BA:1627

|  | Res. type | Atom | Residue | Distance | Occupancy | B-factor | Closest solute to 1^st^ shell water |
| --- | --- | --- | --- | --- | --- | --- | --- |
| **Ion** | | | |  | 1.00 | 60.81 || 1 | U | OP2 | BA:652 | 3.87 | 1.00 | 87.69 |
| 2 | U | OP2 | BA:751 | 4.20 | 1.00 | 82.47 |  |
| 3 | G | N7 | BA:752 | 4.21 | 1.00 | 81.77 |  |
| 4 | C | OP1 | BA:651 | 4.43 | 1.00 | 90.92 |  |
| 5 | G | O6 | BA:752 | 4.58 | 1.00 | 82.15 |  |
| 6 | U | OP1 | BA:751 | 4.69 | 1.00 | 82.45 |  |
| 7 | C | OP2 | BA:651 | 4.92 | 1.00 | 90.90 |  |
  |

  
 Coded contact list:   
1^st^ MG coordination shell missing (see Notes)  

### MG:BA:1628

|  | Res. type | Atom | Residue | Distance | Occupancy | B-factor | Closest solute to 1^st^ shell water |
| --- | --- | --- | --- | --- | --- | --- | --- |
| **Ion** | | | |  | 1.00 | 49.98 || 1 | G | OP2 | BA:588 | 2.25 | 1.00 | 75.02 |
| 2 | G | OP1 | BA:588 | 3.65 | 1.00 | 74.29 |  |
| 3 | C | OP2 | BA:645 | 3.84 | 1.00 | 78.53 |  |
| 4 | G | OP1 | BA:644 | 4.14 | 1.00 | 73.78 |  |
| 5 | G | O5' | BA:588 | 4.43 | 1.00 | 74.88 |  |
| 6 | G | O3' | BA:587 | 4.49 | 1.00 | 73.18 |  |
| 7 | C | OP1 | BA:645 | 4.74 | 1.00 | 79.25 |  |
  |

  
 Contact(s): /OP2/  
 Coded contact list: O_ph_  

### MG:BA:1629

|  | Res. type | Atom | Residue | Distance | Occupancy | B-factor | Closest solute to 1^st^ shell water |
| --- | --- | --- | --- | --- | --- | --- | --- |
| **Ion** | | | |  | 1.00 | 76.88 || 1 | G | O6 | BA:595 | 2.89 | 1.00 | 82.93 |
| 2 | G | OP2 | BA:594 | 3.74 | 1.00 | 84.69 |  |
| 3 | G | N7 | BA:595 | 4.17 | 1.00 | 85.41 |  |
| 4 | U | OP2 | BA:641 | 4.72 | 1.00 | 89.07 |  |
| 5 | G | OP2 | BA:593 | 4.88 | 1.00 | 89.41 |  |
  |

  
 Coded contact list:   
 Exclusion zone contact(s): /O6/  
Consider K in place of MG (1 valid contact(s) for K vs. 0 for MG)  

### MG:BA:1630

|  | Res. type | Atom | Residue | Distance | Occupancy | B-factor | Closest solute to 1^st^ shell water |
| --- | --- | --- | --- | --- | --- | --- | --- |
| **Ion** | | | |  | 1.00 | 78.28 || 1 | A | OP2 | BA:608 | 2.03 | 1.00 | 74.53 |
| 2 | A | O5' | BA:608 | 3.80 | 1.00 | 73.14 |  |
| 3 | A | OP1 | BA:607 | 3.92 | 1.00 | 79.88 |  |
| 4 | A | N7 | BA:609 | 4.07 | 1.00 | 73.30 |  |
| 5 | A | N6 | BA:609 | 4.24 | 1.00 | 72.66 |  |
| 6 | A | OP1 | BA:608 | 4.33 | 1.00 | 74.40 |  |
| 7 | A | O3' | BA:607 | 4.46 | 1.00 | 75.15 |  |
| 8 | G | O6 | BA:610 | 4.56 | 1.00 | 66.62 |  |
| 9 | A | N7 | BA:608 | 4.62 | 1.00 | 67.82 |  |
| 10 | A | OP2 | BA:609 | 4.67 | 1.00 | 73.07 |  |
| 11 | G | N2 | BA:631 | 4.68 | 1.00 | 131.73 |  |
| 12 | A | OP2 | BA:607 | 4.73 | 1.00 | 80.10 |  |
  |

  
 Contact(s): /OP2/  
 Coded contact list: O_ph_  

### MG:BA:1631

|  | Res. type | Atom | Residue | Distance | Occupancy | B-factor | Closest solute to 1^st^ shell water |
| --- | --- | --- | --- | --- | --- | --- | --- |
| **Ion** | | | |  | 1.00 | 48.18 || 1 | G | OP1 | BA:292 | 2.67 | 1.00 | 55.52 |
| 2 | G | OP2 | BA:292 | 3.39 | 1.00 | 54.35 |  |
| 3 | G | N7 | BA:293 | 3.76 | 1.00 | 62.64 |  |
| 4 | C | O3' | BA:291 | 4.55 | 1.00 | 56.00 |  |
| 5 | G | O5' | BA:292 | 4.69 | 1.00 | 56.96 |  |
| 6 | G | O6 | BA:293 | 4.82 | 1.00 | 61.02 |  |
| 7 | G | OP2 | BA:293 | 4.89 | 1.00 | 57.77 |  |
  |

  
 Coded contact list:   
 Exclusion zone contact(s): /OP1/OP2/  
Consider K in place of MG (2 valid contact(s) for K vs. 0 for MG)  

### MG:BA:1632

|  | Res. type | Atom | Residue | Distance | Occupancy | B-factor | Closest solute to 1^st^ shell water |
| --- | --- | --- | --- | --- | --- | --- | --- |
| **Ion** | | | |  | 1.00 | 68.80 || 1 | U | OP1 | BA:114 | 3.42 | 1.00 | 58.34 |
| 2 | U | OP2 | BA:114 | 3.69 | 1.00 | 58.36 |  |
| 3 | G | OP1 | BA:113 | 3.84 | 1.00 | 59.18 |  |
| 4 | G | O3' | BA:113 | 4.43 | 1.00 | 59.21 |  |
  |

  
 Coded contact list:   
1^st^ MG coordination shell missing (see Notes)  

### MG:BA:1633

|  | Res. type | Atom | Residue | Distance | Occupancy | B-factor | Closest solute to 1^st^ shell water |
| --- | --- | --- | --- | --- | --- | --- | --- |
| **Ion** | | | |  | 1.00 | 45.17 || 1 | C | OP1 | BA:390 | 2.72 | 1.00 | 66.22 |
| 2 | G | OP1 | BA:113 | 3.10 | 1.00 | 59.18 |  |
| 3 | G | O3' | BA:112 | 3.38 | 1.00 | 59.62 |  |
| 4 | C | OP2 | BA:390 | 3.73 | 1.00 | 65.86 |  |
| 5 | A | O3' | BA:389 | 4.04 | 1.00 | 64.74 |  |
| 6 | G | OP2 | BA:113 | 4.21 | 1.00 | 56.91 |  |
| 7 | C | OP1 | BA:308 | 4.73 | 1.00 | 83.83 |  |
  |

  
 Coded contact list:   
 Exclusion zone contact(s): /O3'/OP1/OP1/  
Consider K in place of MG (3 valid contact(s) for K vs. 0 for MG)  

### MG:BA:1634

|  | Res. type | Atom | Residue | Distance | Occupancy | B-factor | Closest solute to 1^st^ shell water |
| --- | --- | --- | --- | --- | --- | --- | --- |
| **Ion** | | | |  | 1.00 | 37.69 || 1 | G | N7 | BA:392 | 3.79 | 1.00 | 69.38 |
| 2 | G | N7 | BA:391 | 4.00 | 1.00 | 70.87 |  |
| 3 | G | OP2 | BA:391 | 4.47 | 1.00 | 68.05 |  |
| 4 | G | O6 | BA:392 | 4.50 | 1.00 | 70.25 |  |
| 5 | G | O6 | BA:391 | 4.69 | 1.00 | 73.45 |  |
| 6 | C | OP2 | BA:390 | 4.96 | 1.00 | 65.86 |  |
  |

  
 Coded contact list:   
1^st^ MG coordination shell missing (see Notes)  

### MG:BA:1635

|  | Res. type | Atom | Residue | Distance | Occupancy | B-factor | Closest solute to 1^st^ shell water |
| --- | --- | --- | --- | --- | --- | --- | --- |
| **Ion** | | | |  | 1.00 | 102.75 || 1 | G | O6 | BA:394 | 3.31 | 1.00 | 69.91 |
| 2 | C | N3 | BA:366 | 3.66 | 1.00 | 56.24 |  |
| 3 | G | O6 | BA:46 | 3.68 | 1.00 | 69.47 |  |
| 4 | C | O2 | BA:366 | 4.25 | 1.00 | 60.69 |  |
| 5 | U | O4 | BA:367 | 4.34 | 1.00 | 54.15 |  |
| 6 | C | N4 | BA:366 | 4.42 | 1.00 | 54.02 |  |
| 7 | G | N1 | BA:394 | 4.46 | 1.00 | 70.20 |  |
| 8 | G | N7 | BA:46 | 4.59 | 1.00 | 68.22 |  |
| 9 | A | N6 | BA:393 | 4.92 | 1.00 | 55.51 |  |
| 10 | G | N1 | BA:46 | 4.98 | 1.00 | 67.89 |  |
  |

  
 Coded contact list:   
 Exclusion zone contact(s): /O6/  
Consider K in place of MG (1 valid contact(s) for K vs. 0 for MG)  

### MG:BA:1636

|  | Res. type | Atom | Residue | Distance | Occupancy | B-factor | Closest solute to 1^st^ shell water |
| --- | --- | --- | --- | --- | --- | --- | --- |
| **Ion** | | | |  | 1.00 | 44.26 || 1 | G | O6 | BA:357 | 3.25 | 1.00 | 67.05 |
| 2 | C | O2 | BA:352 | 3.71 | 1.00 | 69.29 |  |
| 3 | A | N7 | BA:356 | 3.75 | 1.00 | 70.90 |  |
| 4 | G | N7 | BA:357 | 3.80 | 1.00 | 67.95 |  |
| 5 | C | OP2 | BA:355 | 4.07 | 1.00 | 66.77 |  |
| 6 | C | O2' | BA:352 | 4.71 | 1.00 | 67.43 |  |
| 7 | C | N4 | BA:355 | 4.94 | 1.00 | 75.56 |  |
  |

  
 Coded contact list:   
 Exclusion zone contact(s): /O6/  
Consider K in place of MG (1 valid contact(s) for K vs. 0 for MG)  

### MG:BA:1637

|  | Res. type | Atom | Residue | Distance | Occupancy | B-factor | Closest solute to 1^st^ shell water |
| --- | --- | --- | --- | --- | --- | --- | --- |
| **Ion** | | | |  | 1.00 | 57.83 || 1 | U | OP2 | BA:358 | 3.67 | 1.00 | 64.50 |
| 2 | G | OP2 | BA:357 | 3.90 | 1.00 | 63.14 |  |
| 3 | G | O5' | BA:357 | 4.21 | 1.00 | 63.84 |  |
| 4 | C | OP2 | BA:366 | 4.58 | 1.00 | 55.59 |  |
| 5 | G | N7 | BA:357 | 4.58 | 1.00 | 67.95 |  |
  |

  
 Coded contact list:   
1^st^ MG coordination shell missing (see Notes)  

### MG:BA:1638

|  | Res. type | Atom | Residue | Distance | Occupancy | B-factor | Closest solute to 1^st^ shell water |
| --- | --- | --- | --- | --- | --- | --- | --- |
| **Ion** | | | |  | 1.00 | 51.29 || 1 | G | O6 | BA:362 | 2.55 | 1.00 | 67.38 |
| 2 | G | N7 | BA:362 | 2.95 | 1.00 | 63.65 |  |
| 3 | G | N7 | BA:361 | 4.16 | 1.00 | 64.31 |  |
| 4 | G | OP2 | BA:361 | 4.25 | 1.00 | 59.12 |  |
| 5 | G | N1 | BA:362 | 4.64 | 1.00 | 64.21 |  |
  |

  
 Coded contact list:   
 Exclusion zone contact(s): /N7/O6/  
Consider NA in place of MG (1 valid contact(s) for NA vs. 0 for MG)  

### MG:BA:1639

|  | Res. type | Atom | Residue | Distance | Occupancy | B-factor | Closest solute to 1^st^ shell water |
| --- | --- | --- | --- | --- | --- | --- | --- |
| **Ion** | | | |  | 1.00 | 72.00 || 1 | U | OP2 | BA:56 | 3.42 | 1.00 | 75.07 |
| 2 | A | OP2 | BA:55 | 3.55 | 1.00 | 67.80 |  |
| 3 | A | OP1 | BA:55 | 3.93 | 1.00 | 68.26 |  |
  |

  
 Coded contact list:   
1^st^ MG coordination shell missing (see Notes)  

### MG:BA:1640

|  | Res. type | Atom | Residue | Distance | Occupancy | B-factor | Closest solute to 1^st^ shell water |
| --- | --- | --- | --- | --- | --- | --- | --- |
| **Ion** | | | |  | 1.00 | 27.94 || 1 | C | OP2 | BA:352 | 2.42 | 1.00 | 73.94 |
| 2 | C | O2 | BA:330 | 3.46 | 1.00 | 61.96 |  |
| 3 | A | N1 | BA:59 | 3.74 | 1.00 | 74.47 |  |
| 4 | C | O5' | BA:352 | 3.75 | 1.00 | 72.66 |  |
| 5 | A | N6 | BA:59 | 3.75 | 1.00 | 74.63 |  |
| 6 | G | N7 | BA:354 | 3.99 | 1.00 | 64.62 |  |
| 7 | G | O4' | BA:331 | 4.00 | 1.00 | 65.33 |  |
| 8 | G | OP1 | BA:354 | 4.32 | 1.00 | 61.74 |  |
| 9 | G | O3' | BA:351 | 4.35 | 1.00 | 76.10 |  |
| 10 | C | O4' | BA:352 | 4.46 | 1.00 | 69.59 |  |
| 11 | C | OP1 | BA:352 | 4.87 | 1.00 | 74.06 |  |
| 12 | G | N9 | BA:331 | 4.96 | 1.00 | 64.98 |  |
  |

  
 Coded contact list:   
 Exclusion zone contact(s): /OP2/  
Consider NA in place of MG (1 valid contact(s) for NA vs. 0 for MG)  

### MG:BA:1641

|  | Res. type | Atom | Residue | Distance | Occupancy | B-factor | Closest solute to 1^st^ shell water |
| --- | --- | --- | --- | --- | --- | --- | --- |
| **Ion** | | | |  | 1.00 | 53.27 || 1 | G | OP2 | BA:331 | 2.03 | 1.00 | 63.65 |
| 2 | A | OP1 | BA:109 | 2.49 | 1.00 | 63.50 |  |
| 3 | A | O2' | BA:329 | 3.41 | 1.00 | 75.43 |  |
| 4 | A | OP2 | BA:109 | 3.96 | 1.00 | 63.85 |  |
| 5 | G | O5' | BA:331 | 3.98 | 1.00 | 63.22 |  |
| 6 | G | OP1 | BA:331 | 4.18 | 1.00 | 65.02 |  |
| 7 | C | O3' | BA:330 | 4.54 | 1.00 | 64.22 |  |
| 8 | A | N3 | BA:329 | 4.57 | 1.00 | 74.86 |  |
| 9 | G | N7 | BA:331 | 4.63 | 1.00 | 64.70 |  |
| 10 | A | O5' | BA:109 | 4.65 | 1.00 | 64.00 |  |
| 11 | G | O3' | BA:108 | 4.76 | 1.00 | 63.23 |  |
| 12 | G | O2' | BA:108 | 4.95 | 1.00 | 60.20 |  |
  |

  
 Contact(s): /OP2/  
 Coded contact list: O_ph_  
 Exclusion zone contact(s): /OP1/  

### MG:BA:1642

|  | Res. type | Atom | Residue | Distance | Occupancy | B-factor | Closest solute to 1^st^ shell water |
| --- | --- | --- | --- | --- | --- | --- | --- |
| **Ion** | | | |  | 1.00 | 45.30 || 1 | G | O6 | BA:332 | 2.93 | 1.00 | 69.79 |
| 2 | G | O6 | BA:333 | 2.96 | 1.00 | 64.00 |  |
| 3 | A | OP2 | BA:329 | 3.46 | 1.00 | 74.71 |  |
| 4 | G | N7 | BA:332 | 3.88 | 1.00 | 70.78 |  |
| 5 | A | OP1 | BA:329 | 3.97 | 1.00 | 75.35 |  |
| 6 | G | N7 | BA:333 | 4.17 | 1.00 | 61.95 |  |
| 7 | A | O5' | BA:329 | 4.19 | 1.00 | 74.08 |  |
| 8 | A | N6 | BA:321 | 4.45 | 1.00 | 61.02 |  |
| 9 | C | N4 | BA:320 | 4.48 | 1.00 | 63.05 |  |
| 10 | A | O3' | BA:329 | 4.60 | 1.00 | 72.05 |  |
| 11 | G | N1 | BA:332 | 4.79 | 1.00 | 69.76 |  |
| 12 | C | O2' | BA:328 | 4.88 | 1.00 | 77.97 |  |
  |

  
 Coded contact list:   
 Exclusion zone contact(s): /O6/O6/  
Consider K in place of MG (2 valid contact(s) for K vs. 0 for MG)  

### MG:BA:1643

|  | Res. type | Atom | Residue | Distance | Occupancy | B-factor | Closest solute to 1^st^ shell water |
| --- | --- | --- | --- | --- | --- | --- | --- |
| **Ion** | | | |  | 1.00 | 53.55 || 1 | G | O6 | BA:147 | 3.03 | 1.00 | 93.58 |
| 2 | G | N7 | BA:147 | 4.06 | 1.00 | 96.53 |  |
| 3 | G | O6 | BA:146 | 4.29 | 1.00 | 99.13 |  |
| 4 | C | N4 | BA:175 | 4.59 | 1.00 | 91.70 |  |
| 5 | U | OP2 | BA:173 | 4.95 | 1.00 | 88.19 |  |
  |

  
 Coded contact list:   
 Exclusion zone contact(s): /O6/  
Consider K in place of MG (1 valid contact(s) for K vs. 0 for MG)  

### MG:BA:1644

|  | Res. type | Atom | Residue | Distance | Occupancy | B-factor | Closest solute to 1^st^ shell water |
| --- | --- | --- | --- | --- | --- | --- | --- |
| **Ion** | | | |  | 1.00 | 102.10 || 1 | A | OP1 | BA:171 | 3.14 | 1.00 | 87.04 |
| 2 | A | OP2 | BA:172 | 3.86 | 1.00 | 86.83 |  |
| 3 | A | OP2 | BA:171 | 3.92 | 1.00 | 87.31 |  |
| 4 | G | N7 | BA:148 | 4.02 | 1.00 | 95.79 |  |
| 5 | G | O6 | BA:148 | 4.10 | 1.00 | 98.65 |  |
| 6 | A | O5' | BA:171 | 4.79 | 1.00 | 88.31 |  |
  |

  
 Coded contact list:   
 Exclusion zone contact(s): /OP1/  
Consider K in place of MG (1 valid contact(s) for K vs. 0 for MG)  

### MG:BA:1645

|  | Res. type | Atom | Residue | Distance | Occupancy | B-factor | Closest solute to 1^st^ shell water |
| --- | --- | --- | --- | --- | --- | --- | --- |
| **Ion** | | | |  | 1.00 | 88.73 || 1 | A | OP2 | BA:149 | 2.37 | 1.00 | 90.98 |
| 2 | G | OP2 | BA:148 | 4.09 | 1.00 | 97.35 |  |
| 3 | G | O5' | BA:148 | 4.17 | 1.00 | 97.13 |  |
| 4 | G | O3' | BA:148 | 4.27 | 1.00 | 92.16 |  |
| 5 | A | OP1 | BA:149 | 4.62 | 1.00 | 91.10 |  |
| 6 | A | O5' | BA:149 | 4.78 | 1.00 | 91.55 |  |
| 7 | A | OP2 | BA:171 | 4.79 | 1.00 | 87.31 |  |
| 8 | A | N7 | BA:149 | 4.85 | 1.00 | 91.64 |  |
| 9 | G | N7 | BA:148 | 4.89 | 1.00 | 95.79 |  |
  |

  
 Contact(s): /OP2/  
 Coded contact list: O_ph_  

### MG:BA:1646

|  | Res. type | Atom | Residue | Distance | Occupancy | B-factor | Closest solute to 1^st^ shell water |
| --- | --- | --- | --- | --- | --- | --- | --- |
| **Ion** | | | |  | 1.00 | 82.56 || 1 | A | O4' | BA:149 | 2.17 | 1.00 | 90.85 |
| 2 | G | O2' | BA:148 | 2.35 | 1.00 | 94.14 |  |
| 3 | A | N1 | BA:1451 | 3.29 | 1.00 | 157.56 |  |
| 4 | G | N3 | BA:148 | 3.70 | 1.00 | 95.99 |  |
| 5 | A | N9 | BA:149 | 3.70 | 1.00 | 91.50 |  |
| 6 | A | N6 | BA:1451 | 3.98 | 1.00 | 157.46 |  |
| 7 | A | O5' | BA:149 | 4.30 | 1.00 | 91.55 |  |
| 8 | G | N2 | BA:148 | 4.51 | 1.00 | 96.22 |  |
| 9 | G | N9 | BA:148 | 4.63 | 1.00 | 95.59 |  |
| 10 | A | O2' | BA:149 | 4.65 | 1.00 | 91.18 |  |
| 11 | G | O3' | BA:148 | 4.89 | 1.00 | 92.16 |  |
| 12 | G | O4' | BA:148 | 4.92 | 1.00 | 95.43 |  |
| 13 | C | O2 | BA:175 | 4.98 | 1.00 | 92.55 |  |
  |

  
**1^st^ shell ligand angles:**   
1-2: 85;   
  
 Contact(s): /O2'/O4'/  
 Coded contact list: 2O_r_  
 Exclusion zone contact(s): /N1_A_/  

### MG:BA:1647

|  | Res. type | Atom | Residue | Distance | Occupancy | B-factor | Closest solute to 1^st^ shell water |
| --- | --- | --- | --- | --- | --- | --- | --- |
| **Ion** | | | |  | 1.00 | 78.12 || 1 | C | OP1 | BA:221 | 2.33 | 1.00 | 92.20 |
| 2 | C | OP1 | BA:135 | 3.27 | 1.00 | 72.19 |  |
| 3 | U | OP2 | BA:222 | 3.30 | 1.00 | 84.36 |  |
| 4 | C | O5' | BA:221 | 3.64 | 1.00 | 91.43 |  |
| 5 | C | OP2 | BA:221 | 4.14 | 1.00 | 92.86 |  |
| 6 | U | OP1 | BA:222 | 4.42 | 1.00 | 84.72 |  |
| 7 | C | O3' | BA:221 | 4.48 | 1.00 | 85.58 |  |
| 8 | G | O3' | BA:220 | 4.74 | 1.00 | 93.51 |  |
| 9 | A | O3' | BA:134 | 5.00 | 1.00 | 71.22 |  |
  |

  
 Contact(s): /OP1/  
 Coded contact list: O_ph_  
 Exclusion zone contact(s): /OP1/OP2/  
Consider K in place of MG (2 valid contact(s) for K vs. 1 for MG)  

### MG:BA:1648

|  | Res. type | Atom | Residue | Distance | Occupancy | B-factor | Closest solute to 1^st^ shell water |
| --- | --- | --- | --- | --- | --- | --- | --- |
| **Ion** | | | |  | 1.00 | 50.78 || 1 | A | OP2 | BA:53 | 1.96 | 1.00 | 62.84 |
| 2 | A | OP1 | BA:353 | 3.41 | 1.00 | 64.29 |  |
| 3 | G | O3' | BA:52 | 4.20 | 1.00 | 61.15 |  |
| 4 | A | OP1 | BA:53 | 4.25 | 1.00 | 64.01 |  |
| 5 | A | N7 | BA:353 | 4.25 | 1.00 | 59.79 |  |
| 6 | A | O5' | BA:53 | 4.36 | 1.00 | 61.41 |  |
| 7 | C | O3' | BA:352 | 4.80 | 1.00 | 65.94 |  |
  |

  
 Contact(s): /OP2/  
 Coded contact list: O_ph_  

### MG:BA:1649

|  | Res. type | Atom | Residue | Distance | Occupancy | B-factor | Closest solute to 1^st^ shell water |
| --- | --- | --- | --- | --- | --- | --- | --- |
| **Ion** | | | |  | 1.00 | 63.77 || 1 | A | OP2 | BA:51 | 3.32 | 1.00 | 69.97 |
| 2 | G | N7 | BA:52 | 3.64 | 1.00 | 60.29 |  |
| 3 | G | O6 | BA:52 | 3.89 | 1.00 | 64.42 |  |
| 4 | A | N1 | BA:360 | 4.52 | 1.00 | 59.65 |  |
| 5 | A | N6 | BA:53 | 4.62 | 1.00 | 66.80 |  |
  |

  
 Coded contact list:   
 Exclusion zone contact(s): /OP2/  
Consider K in place of MG (1 valid contact(s) for K vs. 0 for MG)  

### MG:BA:1650

|  | Res. type | Atom | Residue | Distance | Occupancy | B-factor | Closest solute to 1^st^ shell water |
| --- | --- | --- | --- | --- | --- | --- | --- |
| **Ion** | | | |  | 1.00 | 52.58 || 1 | C | OP2 | BA:48 | 2.12 | 1.00 | 71.02 |
| 2 | G | OP1 | BA:115 | 2.54 | 1.00 | 58.74 |  |
| 3 | U | OP2 | BA:49 | 2.64 | 1.00 | 70.16 |  |
| 4 | C | O3' | BA:47 | 3.33 | 1.00 | 69.20 |  |
| 5 | C | O5' | BA:48 | 3.81 | 1.00 | 69.36 |  |
| 6 | C | O2' | BA:47 | 3.90 | 1.00 | 69.58 |  |
| 7 | U | O3' | BA:114 | 4.48 | 1.00 | 57.84 |  |
| 8 | C | OP1 | BA:48 | 4.53 | 1.00 | 71.51 |  |
| 9 | A | OP1 | BA:51 | 4.55 | 1.00 | 69.79 |  |
| 10 | U | OP1 | BA:49 | 4.69 | 1.00 | 70.18 |  |
| 11 | U | O5' | BA:49 | 4.83 | 1.00 | 71.08 |  |
| 12 | G | O5' | BA:115 | 4.86 | 1.00 | 58.40 |  |
| 13 | G | OP2 | BA:115 | 4.96 | 1.00 | 58.19 |  |
  |

  
 Contact(s): /OP2/  
 Coded contact list: O_ph_  
 Exclusion zone contact(s): /O3'/OP1/OP2/  
Consider K in place of MG (2 valid contact(s) for K vs. 1 for MG)  

### MG:BA:1651

|  | Res. type | Atom | Residue | Distance | Occupancy | B-factor | Closest solute to 1^st^ shell water |
| --- | --- | --- | --- | --- | --- | --- | --- |
| **Ion** | | | |  | 1.00 | 72.93 || 1 | G | O3' | BA:306 | 2.84 | 1.00 | 86.26 |
| 2 | C | OP1 | BA:307 | 2.87 | 1.00 | 87.09 |  |
| 3 | G | N2 | BA:31 | 3.73 | 1.00 | 76.60 |  |
| 4 | G | OP2 | BA:115 | 3.86 | 1.00 | 58.19 |  |
| 5 | G | O2' | BA:306 | 4.04 | 1.00 | 86.46 |  |
| 6 | C | OP1 | BA:48 | 4.21 | 1.00 | 71.51 |  |
| 7 | C | O5' | BA:307 | 4.41 | 1.00 | 87.49 |  |
| 8 | G | N1 | BA:31 | 4.50 | 1.00 | 77.60 |  |
| 9 | C | OP2 | BA:307 | 4.66 | 1.00 | 87.01 |  |
| 10 | U | OP1 | BA:114 | 4.83 | 1.00 | 58.34 |  |
  |

  
 Coded contact list:   
 Exclusion zone contact(s): /O3'/OP1/  
Consider K in place of MG (2 valid contact(s) for K vs. 0 for MG)  

### MG:BA:1652

|  | Res. type | Atom | Residue | Distance | Occupancy | B-factor | Closest solute to 1^st^ shell water |
| --- | --- | --- | --- | --- | --- | --- | --- |
| **Ion** | | | |  | 1.00 | 40.15 || 1 | G | O6 | BA:44 | 3.26 | 1.00 | 57.00 |
| 2 | U | O4 | BA:45 | 3.42 | 1.00 | 65.88 |  |
| 3 | G | N7 | BA:44 | 3.63 | 1.00 | 56.05 |  |
| 4 | G | O6 | BA:396 | 3.64 | 1.00 | 56.61 |  |
| 5 | C | N4 | BA:43 | 3.99 | 1.00 | 60.67 |  |
| 6 | U | N3 | BA:45 | 4.32 | 1.00 | 63.53 |  |
| 7 | C | N4 | BA:395 | 4.49 | 1.00 | 67.94 |  |
  |

  
 Coded contact list:   
 Exclusion zone contact(s): /O6/  
Consider K in place of MG (1 valid contact(s) for K vs. 0 for MG)  

### MG:BA:1653

|  | Res. type | Atom | Residue | Distance | Occupancy | B-factor | Closest solute to 1^st^ shell water |
| --- | --- | --- | --- | --- | --- | --- | --- |
| **Ion** | | | |  | 1.00 | 47.98 || 1 | G | N7 | BA:41 | 3.93 | 1.00 | 69.49 |
| 2 | G | O6 | BA:41 | 4.02 | 1.00 | 67.12 |  |
| 3 | G | O6 | BA:42 | 4.49 | 1.00 | 63.77 |  |
| 4 | A | O4' | BA:397 | 4.89 | 1.00 | 59.55 |  |
  |

  
 Coded contact list:   
1^st^ MG coordination shell missing (see Notes)  

### MG:BA:1654

|  | Res. type | Atom | Residue | Distance | Occupancy | B-factor | Closest solute to 1^st^ shell water |
| --- | --- | --- | --- | --- | --- | --- | --- |
| **Ion** | | | |  | 1.00 | 53.79 || 1 | C | OP2 | BA:398 | 2.55 | 1.00 | 58.68 |
| 2 | A | N3 | BA:397 | 4.24 | 1.00 | 60.29 |  |
| 3 | G | O6 | BA:548 | 4.38 | 1.00 | 65.41 |  |
| 4 | U | O4 | BA:37 | 4.49 | 1.00 | 69.52 |  |
| 5 | C | O5' | BA:398 | 4.60 | 1.00 | 59.90 |  |
| 6 | A | N1 | BA:397 | 4.61 | 1.00 | 59.74 |  |
| 7 | G | O6 | BA:35 | 4.74 | 1.00 | 58.78 |  |
| 8 | C | N4 | BA:36 | 4.76 | 1.00 | 62.96 |  |
| 9 | C | OP1 | BA:398 | 4.82 | 1.00 | 59.86 |  |
| 10 | A | O3' | BA:397 | 4.97 | 1.00 | 59.90 |  |
  |

  
 Coded contact list:   
 Exclusion zone contact(s): /OP2/  
Consider NA in place of MG (1 valid contact(s) for NA vs. 0 for MG)  

### MG:BA:1655

|  | Res. type | Atom | Residue | Distance | Occupancy | B-factor | Closest solute to 1^st^ shell water |
| --- | --- | --- | --- | --- | --- | --- | --- |
| **Ion** | | | |  | 1.00 | 58.30 || 1 | G | O6 | BA:38 | 3.64 | 1.00 | 64.98 |
| 2 | G | N7 | BA:38 | 4.27 | 1.00 | 63.71 |  |
| 3 | U | O4 | BA:37 | 4.84 | 1.00 | 69.52 |  |
  |

  
 Coded contact list:   
1^st^ MG coordination shell missing (see Notes)  

### MG:BA:1656

|  | Res. type | Atom | Residue | Distance | Occupancy | B-factor | Closest solute to 1^st^ shell water |
| --- | --- | --- | --- | --- | --- | --- | --- |
| **Ion** | | | |  | 1.00 | 57.99 || 1 | G | O6 | BA:362 | 2.81 | 1.00 | 67.38 |
| 2 | G | O6 | BA:361 | 2.86 | 1.00 | 62.38 |  |
| 3 | G | N7 | BA:361 | 3.46 | 1.00 | 64.31 |  |
| 4 | G | N1 | BA:362 | 3.70 | 1.00 | 64.21 |  |
| 5 | A | N7 | BA:360 | 4.13 | 1.00 | 58.41 |  |
| 6 | A | N6 | BA:360 | 4.41 | 1.00 | 58.57 |  |
| 7 | C | N4 | BA:47 | 4.78 | 1.00 | 63.25 |  |
| 8 | A | OP2 | BA:360 | 4.82 | 1.00 | 63.81 |  |
  |

  
 Coded contact list:   
 Exclusion zone contact(s): /O6/O6/  
Consider K in place of MG (2 valid contact(s) for K vs. 0 for MG)  

### MG:BA:1657

|  | Res. type | Atom | Residue | Distance | Occupancy | B-factor | Closest solute to 1^st^ shell water |
| --- | --- | --- | --- | --- | --- | --- | --- |
| **Ion** | | | |  | 1.00 | 63.66 || 1 | G | N7 | BA:371 | 3.53 | 1.00 | 66.06 |
| 2 | G | O6 | BA:371 | 3.90 | 1.00 | 67.20 |  |
| 3 | G | N7 | BA:388 | 4.51 | 1.00 | 61.36 |  |
| 4 | G | OP2 | BA:388 | 4.76 | 1.00 | 63.04 |  |
| 5 | G | N9 | BA:388 | 4.92 | 1.00 | 60.65 |  |
| 6 | G | OP1 | BA:388 | 4.97 | 1.00 | 61.03 |  |
  |

  
 Coded contact list:   
1^st^ MG coordination shell missing (see Notes)  

### MG:BA:1658

|  | Res. type | Atom | Residue | Distance | Occupancy | B-factor | Closest solute to 1^st^ shell water |
| --- | --- | --- | --- | --- | --- | --- | --- |
| **Ion** | | | |  | 1.00 | 49.58 || 1 | U | OP1 | BA:387 | 1.77 | 1.00 | 76.33 |
| 2 | G | OP1 | BA:388 | 3.20 | 1.00 | 61.03 |  |
| 3 | C | O2' | BA:58 | 3.59 | 1.00 | 69.46 |  |
| 4 | C | O3' | BA:58 | 3.60 | 1.00 | 71.12 |  |
| 5 | A | OP1 | BA:59 | 3.88 | 1.00 | 72.76 |  |
| 6 | U | O5' | BA:387 | 3.92 | 1.00 | 73.26 |  |
| 7 | U | OP2 | BA:387 | 3.96 | 1.00 | 76.09 |  |
| 8 | C | O3' | BA:386 | 4.24 | 1.00 | 77.00 |  |
| 9 | A | O5' | BA:59 | 4.79 | 1.00 | 71.17 |  |
  |

  
 Coded contact list:   
 Exclusion zone contact(s): /OP1/OP1/  
Consider K in place of MG (1 valid contact(s) for K vs. 0 for MG)  

### MG:BA:1659

|  | Res. type | Atom | Residue | Distance | Occupancy | B-factor | Closest solute to 1^st^ shell water |
| --- | --- | --- | --- | --- | --- | --- | --- |
| **Ion** | | | |  | 1.00 | 71.99 || 1 | C | OP2 | BA:201 | 2.73 | 1.00 | 127.53 |
| 2 | G | O3' | BA:200 | 3.27 | 1.00 | 124.20 |  |
| 3 | G | OP1 | BA:200 | 3.77 | 1.00 | 107.64 |  |
| 4 | C | OP1 | BA:201 | 4.20 | 1.00 | 127.39 |  |
| 5 | G | O5' | BA:200 | 4.42 | 1.00 | 110.07 |  |
| 6 | U | OP1 | BA:93 | 4.52 | 1.00 | 135.16 |  |
| 7 | C | O5' | BA:201 | 4.85 | 1.00 | 130.18 |  |
  |

  
 Coded contact list:   
 Exclusion zone contact(s): /O3'/OP2/  
Consider K in place of MG (2 valid contact(s) for K vs. 0 for MG)  

### MG:BA:1660

|  | Res. type | Atom | Residue | Distance | Occupancy | B-factor | Closest solute to 1^st^ shell water |
| --- | --- | --- | --- | --- | --- | --- | --- |
| **Ion** | | | |  | 1.00 | 101.83 || 1 | G | O3' | BA:64 | 2.26 | 1.00 | 87.55 |
| 2 | G | O2' | BA:64 | 2.39 | 1.00 | 88.54 |  |
| 3 | U | OP2 | BA:65 | 3.11 | 1.00 | 88.38 |  |
| 4 | U | O5' | BA:65 | 4.06 | 1.00 | 88.02 |  |
| 5 | U | OP1 | BA:65 | 4.49 | 1.00 | 89.58 |  |
  |

  
**1^st^ shell ligand angles:**   
1-2: 64;   
  
 Contact(s): /O2'/O3'/  
 Coded contact list: 2O_r_  
 Exclusion zone contact(s): /OP2/  

### MG:BA:1661

|  | Res. type | Atom | Residue | Distance | Occupancy | B-factor | Closest solute to 1^st^ shell water |
| --- | --- | --- | --- | --- | --- | --- | --- |
| **Ion** | | | |  | 1.00 | 83.67 || 1 | G | OP1 | BA:95 | 2.58 | 1.00 | 131.62 |
| 2 | U | O3' | BA:93 | 3.69 | 1.00 | 132.01 |  |
| 3 | U | OP2 | BA:65 | 4.26 | 1.00 | 88.38 |  |
| 4 | G | OP2 | BA:95 | 4.31 | 1.00 | 131.46 |  |
| 5 | U | OP1 | BA:93 | 4.77 | 1.00 | 135.16 |  |
| 6 | G | O5' | BA:95 | 4.94 | 1.00 | 130.92 |  |
| 7 | C | OP1 | BA:201 | 4.96 | 1.00 | 127.39 |  |
  |

  
 Coded contact list:   
 Exclusion zone contact(s): /OP1/  
Consider NA in place of MG (1 valid contact(s) for NA vs. 0 for MG)  

### MG:BA:1662

|  | Res. type | Atom | Residue | Distance | Occupancy | B-factor | Closest solute to 1^st^ shell water |
| --- | --- | --- | --- | --- | --- | --- | --- |
| **Ion** | | | |  | 1.00 | 91.66 || 1 | G | N9 | BA:64 | 4.68 | 1.00 | 87.56 |
| 2 | G | OP2 | BA:95 | 4.69 | 1.00 | 131.46 |  |
| 3 | U | O4 | BA:97 | 4.77 | 1.00 | 103.41 |  |
| 4 | G | N7 | BA:64 | 4.83 | 1.00 | 87.46 |  |
| 5 | G | OP2 | BA:96 | 4.84 | 1.00 | 127.20 |  |
  |

  
 Coded contact list:   
1^st^ MG coordination shell missing (see Notes)  

### MG:BA:1663

|  | Res. type | Atom | Residue | Distance | Occupancy | B-factor | Closest solute to 1^st^ shell water |
| --- | --- | --- | --- | --- | --- | --- | --- |
| **Ion** | | | |  | 1.00 | 94.59 || 1 | G | OP2 | BA:95 | 3.21 | 1.00 | 131.46 |
| 2 | G | N7 | BA:95 | 3.69 | 1.00 | 130.59 |  |
| 3 | U | N1 | BA:93 | 3.74 | 1.00 | 132.39 |  |
| 4 | U | N3 | BA:93 | 3.84 | 1.00 | 131.88 |  |
| 5 | U | O4 | BA:93 | 4.09 | 1.00 | 132.02 |  |
| 6 | U | O5' | BA:93 | 4.51 | 1.00 | 134.75 |  |
| 7 | U | OP2 | BA:93 | 4.56 | 1.00 | 135.22 |  |
| 8 | G | N9 | BA:95 | 4.80 | 1.00 | 130.32 |  |
| 9 | G | O4' | BA:95 | 4.84 | 1.00 | 129.28 |  |
| 10 | U | O3' | BA:93 | 4.88 | 1.00 | 132.01 |  |
| 11 | U | O2 | BA:93 | 4.91 | 1.00 | 131.86 |  |
  |

  
 Coded contact list:   
 Exclusion zone contact(s): /OP2/  
Consider K in place of MG (1 valid contact(s) for K vs. 0 for MG)  

### MG:BA:1664

|  | Res. type | Atom | Residue | Distance | Occupancy | B-factor | Closest solute to 1^st^ shell water |
| --- | --- | --- | --- | --- | --- | --- | --- |
| **Ion** | | | |  | 1.00 | 58.79 || 1 | G | O6 | BA:64 | 3.31 | 1.00 | 89.41 |
| 2 | A | N7 | BA:101 | 3.82 | 1.00 | 99.90 |  |
| 3 | A | N6 | BA:101 | 3.99 | 1.00 | 100.13 |  |
| 4 | G | O6 | BA:102 | 4.09 | 1.00 | 81.31 |  |
| 5 | G | N7 | BA:102 | 4.37 | 1.00 | 80.75 |  |
| 6 | C | N4 | BA:99 | 4.77 | 1.00 | 101.11 |  |
| 7 | G | N1 | BA:64 | 4.84 | 1.00 | 87.96 |  |
  |

  
 Coded contact list:   
 Exclusion zone contact(s): /O6/  
Consider K in place of MG (1 valid contact(s) for K vs. 0 for MG)  

### MG:BA:1665

|  | Res. type | Atom | Residue | Distance | Occupancy | B-factor | Closest solute to 1^st^ shell water |
| --- | --- | --- | --- | --- | --- | --- | --- |
| **Ion** | | | |  | 1.00 | 61.61 || 1 | A | OP1 | BA:393 | 2.19 | 1.00 | 64.20 |
| 2 | HIS | NE2 | BS:13 | 3.56 | 1.00 | 63.24 |  |
| 3 | C | OP2 | BA:43 | 3.62 | 1.00 | 64.97 |  |
| 4 | C | OP1 | BA:43 | 3.93 | 1.00 | 65.30 |  |
| 5 | G | OP1 | BA:42 | 3.98 | 1.00 | 70.90 |  |
| 6 | A | OP2 | BA:393 | 4.22 | 1.00 | 64.23 |  |
| 7 | A | O5' | BA:393 | 4.37 | 1.00 | 62.34 |  |
| 8 | LYS | NZ | BS:12 | 4.44 | 1.00 | 69.52 |  |
| 9 | G | O3' | BA:392 | 4.63 | 1.00 | 63.20 |  |
| 10 | G | O3' | BA:42 | 4.95 | 1.00 | 66.04 |  |
  |

  
 Contact(s): /OP1/  
 Coded contact list: O_ph_  

### MG:BA:1666

|  | Res. type | Atom | Residue | Distance | Occupancy | B-factor | Closest solute to 1^st^ shell water |
| --- | --- | --- | --- | --- | --- | --- | --- |
| **Ion** | | | |  | 1.00 | 43.30 || 1 | G | O4' | BA:9 | 2.87 | 1.00 | 67.33 |
| 2 | G | O5' | BA:9 | 3.02 | 1.00 | 65.57 |  |
| 3 | G | OP2 | BA:9 | 3.12 | 1.00 | 67.72 |  |
| 4 | A | O2' | BA:8 | 3.27 | 1.00 | 74.55 |  |
| 5 | A | O3' | BA:8 | 3.27 | 1.00 | 69.41 |  |
| 6 | A | N3 | BA:8 | 3.42 | 1.00 | 70.90 |  |
| 7 | A | N9 | BA:8 | 3.67 | 1.00 | 70.76 |  |
| 8 | A | O4' | BA:8 | 3.86 | 1.00 | 71.97 |  |
| 9 | A | OP1 | BA:8 | 4.20 | 1.00 | 74.27 |  |
| 10 | G | N9 | BA:9 | 4.40 | 1.00 | 70.47 |  |
| 11 | A | O5' | BA:8 | 4.42 | 1.00 | 73.42 |  |
| 12 | G | OP1 | BA:9 | 4.81 | 1.00 | 66.45 |  |
  |

  
 Coded contact list:   
 Exclusion zone contact(s): /O2'/O3'/O4'/O5'/OP2/  
Consider K in place of MG (5 valid contact(s) for K vs. 0 for MG)  

### MG:BA:1667

|  | Res. type | Atom | Residue | Distance | Occupancy | B-factor | Closest solute to 1^st^ shell water |
| --- | --- | --- | --- | --- | --- | --- | --- |
| **Ion** | | | |  | 1.00 | 62.87 || 1 | G | O6 | BA:558 | 2.14 | 1.00 | 64.60 |
| 2 | G | N7 | BA:557 | 3.50 | 1.00 | 60.92 |  |
| 3 | G | O6 | BA:557 | 3.86 | 1.00 | 63.15 |  |
| 4 | G | N1 | BA:558 | 4.12 | 1.00 | 63.99 |  |
| 5 | G | N7 | BA:558 | 4.53 | 1.00 | 64.97 |  |
  |

  
 Contact(s): /O6/  
 Coded contact list: O_b_  

### MG:BA:1668

|  | Res. type | Atom | Residue | Distance | Occupancy | B-factor | Closest solute to 1^st^ shell water |
| --- | --- | --- | --- | --- | --- | --- | --- |
| **Ion** | | | |  | 1.00 | 56.38 || 1 | U | OP2 | BA:560 | 2.17 | 1.00 | 72.92 |
| 2 | C | OP1 | BA:562 | 2.75 | 1.00 | 67.53 |  |
| 3 | A | O3' | BA:559 | 3.16 | 1.00 | 71.85 |  |
| 4 | G | N2 | BA:566 | 3.47 | 1.00 | 68.72 |  |
| 5 | A | O2' | BA:559 | 3.81 | 1.00 | 73.70 |  |
| 6 | G | N1 | BA:566 | 4.10 | 1.00 | 69.76 |  |
| 7 | U | O5' | BA:560 | 4.24 | 1.00 | 71.71 |  |
| 8 | U | OP1 | BA:560 | 4.36 | 1.00 | 73.18 |  |
| 9 | A | OP1 | BA:563 | 4.37 | 1.00 | 66.19 |  |
| 10 | G | OP2 | BA:558 | 4.57 | 1.00 | 63.65 |  |
| 11 | G | OP1 | BA:557 | 4.64 | 1.00 | 59.88 |  |
| 12 | C | OP2 | BA:562 | 4.90 | 1.00 | 66.15 |  |
| 13 | C | O5' | BA:562 | 4.90 | 1.00 | 65.69 |  |
  |

  
 Contact(s): /OP2/  
 Coded contact list: O_ph_  
 Exclusion zone contact(s): /O3'/OP1/  
Consider K in place of MG (2 valid contact(s) for K vs. 1 for MG)  

### MG:BA:1669

|  | Res. type | Atom | Residue | Distance | Occupancy | B-factor | Closest solute to 1^st^ shell water |
| --- | --- | --- | --- | --- | --- | --- | --- |
| **Ion** | | | |  | 1.00 | 32.78 || 1 | C | OP1 | BA:770 | 2.05 | 1.00 | 36.96 |
| 2 | G | O3' | BA:769 | 3.61 | 1.00 | 36.17 |  |
| 3 | C | O5' | BA:770 | 4.00 | 1.00 | 37.80 |  |
| 4 | U | OP1 | AA:1833 | 4.37 | 1.00 | 42.47 |  |
| 5 | C | OP2 | BA:770 | 4.57 | 1.00 | 38.81 |  |
| 6 | C | OP1 | BA:1514 | 4.64 | 1.00 | 53.28 |  |
  |

  
 Contact(s): /OP1/  
 Coded contact list: O_ph_  

### MG:BA:1670

|  | Res. type | Atom | Residue | Distance | Occupancy | B-factor | Closest solute to 1^st^ shell water |
| --- | --- | --- | --- | --- | --- | --- | --- |
| **Ion** | | | |  | 1.00 | 50.28 || 1 | G | OP2 | BA:771 | 3.08 | 1.00 | 41.91 |
| 2 | C | OP1 | BA:770 | 3.42 | 1.00 | 36.96 |  |
| 3 | G | OP1 | BA:771 | 4.31 | 1.00 | 41.25 |  |
| 4 | C | OP2 | BA:770 | 4.39 | 1.00 | 38.81 |  |
| 5 | A | O2' | BA:802 | 4.51 | 1.00 | 52.52 |  |
| 6 | C | O5' | BA:770 | 4.53 | 1.00 | 37.80 |  |
| 7 | G | OP1 | BA:803 | 4.84 | 1.00 | 50.00 |  |
| 8 | C | O3' | BA:770 | 4.87 | 1.00 | 40.10 |  |
  |

  
 Coded contact list:   
 Exclusion zone contact(s): /OP2/  
Consider K in place of MG (1 valid contact(s) for K vs. 0 for MG)  

### MG:BA:1671

|  | Res. type | Atom | Residue | Distance | Occupancy | B-factor | Closest solute to 1^st^ shell water |
| --- | --- | --- | --- | --- | --- | --- | --- |
| **Ion** | | | |  | 1.00 | 50.29 || 1 | G | O6 | BA:774 | 3.68 | 1.00 | 59.90 |
| 2 | G | N7 | BA:774 | 3.82 | 1.00 | 60.70 |  |
| 3 | G | OP1 | BA:803 | 3.97 | 1.00 | 50.00 |  |
| 4 | G | O6 | BA:775 | 3.98 | 1.00 | 64.37 |  |
| 5 | G | O6 | BA:773 | 4.19 | 1.00 | 57.53 |  |
| 6 | G | N7 | BA:773 | 4.28 | 1.00 | 54.60 |  |
| 7 | G | OP2 | BA:803 | 4.41 | 1.00 | 51.98 |  |
  |

  
 Coded contact list:   
1^st^ MG coordination shell missing (see Notes)  

### MG:BA:1672

|  | Res. type | Atom | Residue | Distance | Occupancy | B-factor | Closest solute to 1^st^ shell water |
| --- | --- | --- | --- | --- | --- | --- | --- |
| **Ion** | | | |  | 1.00 | 39.43 || 1 | G | OP1 | BA:803 | 3.73 | 1.00 | 50.00 |
| 2 | U | O4 | BA:772 | 4.29 | 1.00 | 50.11 |  |
| 3 | U | OP2 | BA:804 | 4.42 | 1.00 | 53.04 |  |
| 4 | G | N7 | BA:771 | 4.68 | 1.00 | 43.86 |  |
| 5 | G | O5' | BA:803 | 4.95 | 1.00 | 52.82 |  |
  |

  
 Coded contact list:   
1^st^ MG coordination shell missing (see Notes)  

### MG:BA:1673

|  | Res. type | Atom | Residue | Distance | Occupancy | B-factor | Closest solute to 1^st^ shell water |
| --- | --- | --- | --- | --- | --- | --- | --- |
| **Ion** | | | |  | 1.00 | 73.97 || 1 | G | O6 | BA:809 | 4.29 | 1.00 | 37.62 |
| 2 | C | N4 | BA:810 | 4.42 | 1.00 | 32.63 |  |
| 3 | C | N4 | BA:808 | 4.64 | 1.00 | 49.70 |  |
| 4 | G | N7 | BA:809 | 4.64 | 1.00 | 43.64 |  |
  |

  
 Coded contact list:   
1^st^ MG coordination shell missing (see Notes)  

### MG:BA:1674

|  | Res. type | Atom | Residue | Distance | Occupancy | B-factor | Closest solute to 1^st^ shell water |
| --- | --- | --- | --- | --- | --- | --- | --- |
| **Ion** | | | |  | 1.00 | 45.27 || 1 | U | OP2 | BA:1522 | 3.95 | 1.00 | 51.83 |
| 2 | G | OP2 | BA:1521 | 4.02 | 1.00 | 48.63 |  |
| 3 | U | O4 | BA:1522 | 4.26 | 1.00 | 54.51 |  |
| 4 | U | O4 | BA:1510 | 4.51 | 1.00 | 59.60 |  |
  |

  
 Coded contact list:   
1^st^ MG coordination shell missing (see Notes)  

### MG:BA:1675

|  | Res. type | Atom | Residue | Distance | Occupancy | B-factor | Closest solute to 1^st^ shell water |
| --- | --- | --- | --- | --- | --- | --- | --- |
| **Ion** | | | |  | 1.00 | 28.83 || 1 | G | O6 | BA:1511 | 2.40 | 1.00 | 50.35 |
| 2 | G | O6 | BA:1523 | 2.64 | 1.00 | 56.85 |  |
| 3 | C | N4 | BA:1524 | 3.06 | 1.00 | 50.88 |  |
| 4 | U | O4 | BA:1522 | 3.28 | 1.00 | 54.51 |  |
| 5 | U | O4 | BA:1512 | 3.31 | 1.00 | 58.29 |  |
| 6 | U | O4 | BA:1510 | 3.35 | 1.00 | 59.60 |  |
| 7 | A | N6 | BA:1513 | 3.62 | 1.00 | 55.26 |  |
| 8 | U | N3 | BA:1512 | 4.09 | 1.00 | 54.75 |  |
| 9 | G | N7 | BA:1523 | 4.17 | 1.00 | 56.57 |  |
| 10 | U | N3 | BA:1510 | 4.32 | 1.00 | 57.76 |  |
| 11 | G | O6 | BA:1525 | 4.32 | 1.00 | 57.51 |  |
| 12 | G | N1 | BA:1511 | 4.47 | 1.00 | 52.08 |  |
| 13 | U | N3 | BA:1522 | 4.70 | 1.00 | 55.42 |  |
| 14 | G | N7 | BA:1511 | 4.74 | 1.00 | 52.32 |  |
| 15 | G | N1 | BA:1523 | 4.80 | 1.00 | 56.70 |  |
| 16 | A | N1 | BA:1513 | 4.81 | 1.00 | 54.79 |  |
| 17 | C | N3 | BA:1524 | 4.83 | 1.00 | 51.86 |  |
  |

  
 Coded contact list:   
 Exclusion zone contact(s): /N4/O4/O4/O4/O6/O6/  
Consider K in place of MG (5 valid contact(s) for K vs. 0 for MG)  

### MG:BA:1676

|  | Res. type | Atom | Residue | Distance | Occupancy | B-factor | Closest solute to 1^st^ shell water |
| --- | --- | --- | --- | --- | --- | --- | --- |
| **Ion** | | | |  | 1.00 | 84.49 || 1 | C | OP1 | BA:1509 | 3.29 | 1.00 | 58.96 |
| 2 | G | OP2 | BA:1520 | 3.59 | 1.00 | 54.93 |  |
| 3 | G | OP1 | BA:1520 | 4.06 | 1.00 | 54.94 |  |
| 4 | U | OP2 | BA:1510 | 4.45 | 1.00 | 58.79 |  |
  |

  
 Coded contact list:   
 Exclusion zone contact(s): /OP1/  
Consider K in place of MG (1 valid contact(s) for K vs. 0 for MG)  

### MG:BA:1677

|  | Res. type | Atom | Residue | Distance | Occupancy | B-factor | Closest solute to 1^st^ shell water |
| --- | --- | --- | --- | --- | --- | --- | --- |
| **Ion** | | | |  | 1.00 | 52.67 || 1 | G | OP1 | BA:576 | 2.23 | 1.00 | 56.81 |
| 2 | G | OP1 | BA:575 | 3.58 | 1.00 | 61.96 |  |
| 3 | G | O3' | BA:575 | 3.64 | 1.00 | 59.72 |  |
| 4 | G | OP2 | BA:575 | 3.96 | 1.00 | 61.66 |  |
| 5 | G | O5' | BA:575 | 4.22 | 1.00 | 61.49 |  |
| 6 | G | OP2 | BA:576 | 4.48 | 1.00 | 57.51 |  |
| 7 | G | O5' | BA:576 | 4.54 | 1.00 | 57.00 |  |
| 8 | A | OP1 | BA:815 | 4.78 | 1.00 | 52.56 |  |
  |

  
 Contact(s): /OP1/  
 Coded contact list: O_ph_  

### MG:BA:1678

|  | Res. type | Atom | Residue | Distance | Occupancy | B-factor | Closest solute to 1^st^ shell water |
| --- | --- | --- | --- | --- | --- | --- | --- |
| **Ion** | | | |  | 1.00 | 77.24 || 1 | A | OP2 | BA:815 | 2.99 | 1.00 | 49.28 |
| 2 | C | OP1 | BA:817 | 3.11 | 1.00 | 53.87 |  |
| 3 | G | O4' | BA:577 | 3.63 | 1.00 | 54.67 |  |
| 4 | G | OP1 | BA:576 | 3.80 | 1.00 | 56.81 |  |
| 5 | G | OP2 | BA:577 | 3.95 | 1.00 | 53.01 |  |
| 6 | A | N7 | BA:816 | 4.05 | 1.00 | 53.09 |  |
| 7 | G | O5' | BA:577 | 4.10 | 1.00 | 54.49 |  |
| 8 | A | OP1 | BA:815 | 4.35 | 1.00 | 52.56 |  |
| 9 | G | OP2 | BA:576 | 4.42 | 1.00 | 57.51 |  |
| 10 | A | N9 | BA:816 | 4.73 | 1.00 | 51.82 |  |
| 11 | A | O5' | BA:815 | 4.84 | 1.00 | 52.12 |  |
| 12 | A | O3' | BA:816 | 4.90 | 1.00 | 52.66 |  |
  |

  
 Coded contact list:   
 Exclusion zone contact(s): /OP1/OP2/  
Consider K in place of MG (2 valid contact(s) for K vs. 0 for MG)  

### MG:BA:1679

|  | Res. type | Atom | Residue | Distance | Occupancy | B-factor | Closest solute to 1^st^ shell water |
| --- | --- | --- | --- | --- | --- | --- | --- |
| **Ion** | | | |  | 1.00 | 50.22 || 1 | A | OP1 | BA:815 | 3.31 | 1.00 | 52.56 |
| 2 | A | OP1 | BA:814 | 3.40 | 1.00 | 48.98 |  |
| 3 | A | OP2 | BA:814 | 3.57 | 1.00 | 48.31 |  |
| 4 | A | OP2 | BA:815 | 3.69 | 1.00 | 49.28 |  |
| 5 | A | O5' | BA:814 | 3.99 | 1.00 | 48.28 |  |
| 6 | G | OP2 | BA:577 | 3.99 | 1.00 | 53.01 |  |
| 7 | A | O3' | BA:814 | 4.24 | 1.00 | 50.06 |  |
| 8 | A | N7 | BA:816 | 4.70 | 1.00 | 53.09 |  |
  |

  
 Coded contact list:   
 Exclusion zone contact(s): /OP1/OP1/  
Consider K in place of MG (2 valid contact(s) for K vs. 0 for MG)  

### MG:BA:1680

|  | Res. type | Atom | Residue | Distance | Occupancy | B-factor | Closest solute to 1^st^ shell water |
| --- | --- | --- | --- | --- | --- | --- | --- |
| **Ion** | | | |  | 1.00 | 55.57 || 1 | A | OP2 | BA:814 | 3.36 | 1.00 | 48.31 |
| 2 | C | OP1 | BA:904 | 3.67 | 1.00 | 51.81 |  |
| 3 | A | OP1 | BA:814 | 3.96 | 1.00 | 48.98 |  |
| 4 | U | OP1 | BA:813 | 4.35 | 1.00 | 49.95 |  |
| 5 | G | OP2 | BA:577 | 4.36 | 1.00 | 53.01 |  |
| 6 | U | O3' | BA:813 | 4.65 | 1.00 | 49.73 |  |
| 7 | C | O5' | BA:904 | 4.77 | 1.00 | 50.21 |  |
| 8 | A | N6 | BA:816 | 4.88 | 1.00 | 50.08 |  |
| 9 | U | OP2 | BA:905 | 4.97 | 1.00 | 51.77 |  |
  |

  
 Coded contact list:   
 Exclusion zone contact(s): /OP2/  
Consider K in place of MG (1 valid contact(s) for K vs. 0 for MG)  

### MG:BA:1681

|  | Res. type | Atom | Residue | Distance | Occupancy | B-factor | Closest solute to 1^st^ shell water |
| --- | --- | --- | --- | --- | --- | --- | --- |
| **Ion** | | | |  | 1.00 | 43.61 || 1 | U | OP2 | BA:813 | 3.54 | 1.00 | 49.24 |
| 2 | U | OP1 | BA:813 | 3.60 | 1.00 | 49.95 |  |
| 3 | G | N7 | BA:577 | 3.64 | 1.00 | 56.60 |  |
| 4 | G | OP1 | BA:577 | 4.24 | 1.00 | 53.55 |  |
| 5 | G | O6 | BA:577 | 4.32 | 1.00 | 56.47 |  |
  |

  
 Coded contact list:   
1^st^ MG coordination shell missing (see Notes)  

### MG:BA:1682

|  | Res. type | Atom | Residue | Distance | Occupancy | B-factor | Closest solute to 1^st^ shell water |
| --- | --- | --- | --- | --- | --- | --- | --- |
| **Ion** | | | |  | 1.00 | 51.82 || 1 | G | OP2 | BA:579 | 3.65 | 1.00 | 59.25 |
| 2 | G | N7 | BA:576 | 4.09 | 1.00 | 61.39 |  |
| 3 | G | O6 | BA:576 | 4.27 | 1.00 | 62.75 |  |
| 4 | G | OP1 | BA:758 | 4.63 | 1.00 | 54.03 |  |
| 5 | G | N7 | BA:579 | 4.81 | 1.00 | 59.21 |  |
| 6 | G | N1 | BA:576 | 4.95 | 1.00 | 60.22 |  |
  |

  
 Coded contact list:   
1^st^ MG coordination shell missing (see Notes)  

### MG:BA:1683

|  | Res. type | Atom | Residue | Distance | Occupancy | B-factor | Closest solute to 1^st^ shell water |
| --- | --- | --- | --- | --- | --- | --- | --- |
| **Ion** | | | |  | 1.00 | 51.44 || 1 | G | O6 | BA:761 | 2.96 | 1.00 | 61.48 |
| 2 | G | O6 | BA:579 | 2.98 | 1.00 | 58.65 |  |
| 3 | U | O4 | BA:580 | 3.19 | 1.00 | 65.07 |  |
| 4 | C | N4 | BA:762 | 3.43 | 1.00 | 59.93 |  |
| 5 | A | OP1 | BA:759 | 4.03 | 1.00 | 59.51 |  |
| 6 | G | N1 | BA:581 | 4.19 | 1.00 | 56.78 |  |
| 7 | G | N7 | BA:761 | 4.25 | 1.00 | 63.46 |  |
| 8 | U | N3 | BA:580 | 4.32 | 1.00 | 63.26 |  |
| 9 | G | N2 | BA:581 | 4.57 | 1.00 | 55.63 |  |
| 10 | G | N1 | BA:579 | 4.69 | 1.00 | 57.66 |  |
| 11 | A | OP2 | BA:759 | 4.71 | 1.00 | 59.49 |  |
| 12 | C | N3 | BA:762 | 4.91 | 1.00 | 59.65 |  |
| 13 | G | O6 | BA:581 | 4.93 | 1.00 | 55.00 |  |
| 14 | G | OP2 | BA:760 | 4.97 | 1.00 | 63.31 |  |
  |

  
 Coded contact list:   
 Exclusion zone contact(s): /O4/O6/O6/  
Consider K in place of MG (3 valid contact(s) for K vs. 0 for MG)  

### MG:BA:1684

|  | Res. type | Atom | Residue | Distance | Occupancy | B-factor | Closest solute to 1^st^ shell water |
| --- | --- | --- | --- | --- | --- | --- | --- |
| **Ion** | | | |  | 1.00 | 36.68 || 1 | G | OP1 | BA:758 | 2.20 | 1.00 | 54.03 |
| 2 | G | O6 | BA:576 | 2.94 | 1.00 | 62.75 |  |
| 3 | C | O2 | BA:880 | 3.16 | 1.00 | 60.57 |  |
| 4 | G | N2 | BA:575 | 3.22 | 1.00 | 61.83 |  |
| 5 | G | N7 | BA:576 | 3.24 | 1.00 | 61.39 |  |
| 6 | G | N3 | BA:821 | 3.31 | 1.00 | 59.42 |  |
| 7 | G | N2 | BA:821 | 3.72 | 1.00 | 61.32 |  |
| 8 | G | O5' | BA:758 | 4.18 | 1.00 | 56.17 |  |
| 9 | G | O2' | BA:821 | 4.31 | 1.00 | 60.94 |  |
| 10 | U | O3' | BA:757 | 4.48 | 1.00 | 55.67 |  |
| 11 | G | N9 | BA:821 | 4.65 | 1.00 | 60.78 |  |
| 12 | G | OP2 | BA:758 | 4.69 | 1.00 | 55.72 |  |
| 13 | C | N1 | BA:880 | 4.94 | 1.00 | 58.45 |  |
| 14 | C | N3 | BA:880 | 4.98 | 1.00 | 58.40 |  |
| 15 | C | O2' | BA:880 | 4.99 | 1.00 | 58.43 |  |
  |

  
 Contact(s): /OP1/  
 Coded contact list: O_ph_  
 Exclusion zone contact(s): /N2/N3_G_/N7/O2/O6/  
Consider K in place of MG (5 valid contact(s) for K vs. 1 for MG)  

### MG:BA:1685

|  | Res. type | Atom | Residue | Distance | Occupancy | B-factor | Closest solute to 1^st^ shell water |
| --- | --- | --- | --- | --- | --- | --- | --- |
| **Ion** | | | |  | 1.00 | 45.60 || 1 | U | O4 | BA:12 | 2.41 | 1.00 | 63.27 |
| 2 | G | O6 | BA:22 | 2.75 | 1.00 | 64.33 |  |
| 3 | G | O6 | BA:11 | 3.01 | 1.00 | 62.96 |  |
| 4 | G | O6 | BA:21 | 3.10 | 1.00 | 60.83 |  |
| 5 | U | N3 | BA:12 | 3.65 | 1.00 | 63.99 |  |
| 6 | C | N4 | BA:23 | 3.90 | 1.00 | 63.54 |  |
| 7 | G | N1 | BA:21 | 4.26 | 1.00 | 57.82 |  |
| 8 | G | N7 | BA:22 | 4.55 | 1.00 | 63.34 |  |
| 9 | G | N1 | BA:11 | 4.63 | 1.00 | 62.29 |  |
| 10 | G | N7 | BA:11 | 4.79 | 1.00 | 65.67 |  |
  |

  
 Coded contact list:   
 Exclusion zone contact(s): /O4/O6/O6/O6/  
Consider K in place of MG (3 valid contact(s) for K vs. 0 for MG)  

### MG:BA:1686

|  | Res. type | Atom | Residue | Distance | Occupancy | B-factor | Closest solute to 1^st^ shell water |
| --- | --- | --- | --- | --- | --- | --- | --- |
| **Ion** | | | |  | 1.00 | 34.88 || 1 | G | OP1 | BA:903 | 2.25 | 1.00 | 49.16 |
| 2 | G | O5' | BA:903 | 4.04 | 1.00 | 46.63 |  |
| 3 | C | OP2 | BA:904 | 4.23 | 1.00 | 50.62 |  |
| 4 | U | OP1 | BA:1512 | 4.28 | 1.00 | 55.60 |  |
| 5 | U | OP1 | BA:891 | 4.34 | 1.00 | 57.84 |  |
| 6 | G | OP2 | BA:903 | 4.39 | 1.00 | 48.52 |  |
| 7 | G | O3' | BA:902 | 4.69 | 1.00 | 46.70 |  |
  |

  
 Contact(s): /OP1/  
 Coded contact list: O_ph_  

### MG:BA:1687

|  | Res. type | Atom | Residue | Distance | Occupancy | B-factor | Closest solute to 1^st^ shell water |
| --- | --- | --- | --- | --- | --- | --- | --- |
| **Ion** | | | |  | 1.00 | 41.03 || 1 | U | O2' | BA:65 | 2.51 | 1.00 | 89.12 |
| 2 | G | O5' | BA:66 | 2.69 | 1.00 | 85.68 |  |
| 3 | G | OP2 | BA:66 | 2.87 | 1.00 | 86.09 |  |
| 4 | U | O3' | BA:65 | 2.88 | 1.00 | 88.10 |  |
| 5 | U | O4 | BA:173 | 3.41 | 1.00 | 95.51 |  |
| 6 | G | O4' | BA:66 | 3.75 | 1.00 | 84.92 |  |
| 7 | C | OP1 | BA:381 | 4.17 | 1.00 | 83.62 |  |
| 8 | U | N3 | BA:173 | 4.32 | 1.00 | 95.84 |  |
| 9 | G | OP1 | BA:66 | 4.48 | 1.00 | 87.01 |  |
  |

  
 Coded contact list:   
 Exclusion zone contact(s): /O2'/O3'/O5'/OP2/  
Consider K in place of MG (3 valid contact(s) for K vs. 0 for MG)  

### MG:BA:1688

|  | Res. type | Atom | Residue | Distance | Occupancy | B-factor | Closest solute to 1^st^ shell water |
| --- | --- | --- | --- | --- | --- | --- | --- |
| **Ion** | | | |  | 1.00 | 61.22 || 1 | G | N7 | BA:380 | 3.39 | 1.00 | 86.13 |
| 2 | A | OP2 | BA:382 | 3.98 | 1.00 | 88.48 |  |
| 3 | A | OP1 | BA:382 | 4.12 | 1.00 | 89.50 |  |
| 4 | G | O6 | BA:380 | 4.46 | 1.00 | 85.17 |  |
| 5 | C | O3' | BA:381 | 4.65 | 1.00 | 88.15 |  |
  |

  
 Coded contact list:   
 Exclusion zone contact(s): /N7/  
Consider K in place of MG (1 valid contact(s) for K vs. 0 for MG)  

### MG:BA:1689

|  | Res. type | Atom | Residue | Distance | Occupancy | B-factor | Closest solute to 1^st^ shell water |
| --- | --- | --- | --- | --- | --- | --- | --- |
| **Ion** | | | |  | 1.00 | 64.16 || 1 | C | O3' | BA:135 | 3.06 | 1.00 | 72.78 |
| 2 | C | O2' | BA:135 | 3.47 | 1.00 | 70.78 |  |
| 3 | C | OP1 | BA:136 | 4.02 | 1.00 | 74.28 |  |
| 4 | G | OP2 | BA:380 | 4.83 | 1.00 | 74.14 |  |
| 5 | C | O4' | BA:135 | 4.87 | 1.00 | 70.87 |  |
  |

  
 Coded contact list:   
 Exclusion zone contact(s): /O3'/  
Consider K in place of MG (1 valid contact(s) for K vs. 0 for MG)  

### MG:BA:1690

|  | Res. type | Atom | Residue | Distance | Occupancy | B-factor | Closest solute to 1^st^ shell water |
| --- | --- | --- | --- | --- | --- | --- | --- |
| **Ion** | | | |  | 1.00 | 66.62 || 1 | C | OP2 | BA:379 | 2.23 | 1.00 | 68.72 |
| 2 | G | OP2 | BA:378 | 2.98 | 1.00 | 67.81 |  |
| 3 | G | O5' | BA:378 | 3.38 | 1.00 | 67.46 |  |
| 4 | G | OP1 | BA:378 | 4.08 | 1.00 | 68.25 |  |
| 5 | G | O3' | BA:378 | 4.12 | 1.00 | 67.50 |  |
| 6 | C | O5' | BA:379 | 4.16 | 1.00 | 68.78 |  |
| 7 | G | N7 | BA:378 | 4.42 | 1.00 | 73.39 |  |
| 8 | G | N9 | BA:378 | 4.68 | 1.00 | 71.32 |  |
| 9 | C | OP1 | BA:379 | 4.79 | 1.00 | 68.13 |  |
  |

  
 Contact(s): /OP2/  
 Coded contact list: O_ph_  
 Exclusion zone contact(s): /O5'/OP2/  
Consider K in place of MG (2 valid contact(s) for K vs. 1 for MG)  

### MG:BA:1691

|  | Res. type | Atom | Residue | Distance | Occupancy | B-factor | Closest solute to 1^st^ shell water |
| --- | --- | --- | --- | --- | --- | --- | --- |
| **Ion** | | | |  | 1.00 | 65.60 || 1 | ASP | OD2 | BS:68 | 3.12 | 1.00 | 75.42 |
| 2 | G | O6 | BA:380 | 3.25 | 1.00 | 85.17 |  |
| 3 | ASP | OD1 | BS:68 | 4.29 | 1.00 | 75.67 |  |
| 4 | A | OP1 | BA:382 | 4.49 | 1.00 | 89.50 |  |
| 5 | G | N1 | BA:380 | 4.63 | 1.00 | 85.26 |  |
| 6 | C | O2' | BA:454 | 4.63 | 1.00 | 90.41 |  |
  |

  
 Coded contact list:   
 Exclusion zone contact(s): /O6/OD2_Asp_/  
Consider K in place of MG (2 valid contact(s) for K vs. 0 for MG)  

### MG:BA:1692

|  | Res. type | Atom | Residue | Distance | Occupancy | B-factor | Closest solute to 1^st^ shell water |
| --- | --- | --- | --- | --- | --- | --- | --- |
| **Ion** | | | |  | 1.00 | 34.53 || 1 | A | OP2 | BA:766 | 1.76 | 1.00 | 44.82 |
| 2 | C | O2 | BA:812 | 3.34 | 1.00 | 48.75 |  |
| 3 | G | O3' | BA:765 | 3.94 | 1.00 | 44.06 |  |
| 4 | C | N3 | BA:812 | 4.05 | 1.00 | 49.89 |  |
| 5 | C | N4 | BA:811 | 4.09 | 1.00 | 38.52 |  |
| 6 | A | O5' | BA:766 | 4.09 | 1.00 | 44.46 |  |
| 7 | U | O4 | BA:813 | 4.20 | 1.00 | 47.22 |  |
| 8 | A | OP1 | BA:766 | 4.22 | 1.00 | 43.71 |  |
| 9 | C | N3 | BA:811 | 4.30 | 1.00 | 36.54 |  |
| 10 | A | N7 | BA:767 | 4.88 | 1.00 | 33.99 |  |
| 11 | A | N6 | BA:768 | 4.90 | 1.00 | 34.72 |  |
| 12 | A | OP2 | BA:767 | 4.97 | 1.00 | 49.43 |  |
  |

  
 Coded contact list:   
 Exclusion zone contact(s): /O2/OP2/  
Consider K in place of MG (1 valid contact(s) for K vs. 0 for MG)  

### MG:BA:1693

|  | Res. type | Atom | Residue | Distance | Occupancy | B-factor | Closest solute to 1^st^ shell water |
| --- | --- | --- | --- | --- | --- | --- | --- |
| **Ion** | | | |  | 1.00 | 46.18 || 1 | A | OP2 | BA:768 | 2.15 | 1.00 | 42.08 |
| 2 | A | N7 | BA:768 | 3.98 | 1.00 | 34.92 |  |
| 3 | A | O5' | BA:767 | 4.06 | 1.00 | 45.09 |  |
| 4 | A | OP2 | BA:767 | 4.13 | 1.00 | 49.43 |  |
| 5 | A | O5' | BA:768 | 4.25 | 1.00 | 41.34 |  |
| 6 | A | OP1 | BA:767 | 4.35 | 1.00 | 48.71 |  |
| 7 | U | OP1 | BA:804 | 4.50 | 1.00 | 53.48 |  |
| 8 | A | O3' | BA:767 | 4.54 | 1.00 | 42.07 |  |
| 9 | A | OP1 | BA:768 | 4.56 | 1.00 | 43.42 |  |
| 10 | G | N7 | BA:769 | 4.61 | 1.00 | 27.94 |  |
| 11 | C | OP2 | BA:805 | 4.72 | 1.00 | 54.46 |  |
  |

  
 Contact(s): /OP2/  
 Coded contact list: O_ph_  

### MG:BA:1694

|  | Res. type | Atom | Residue | Distance | Occupancy | B-factor | Closest solute to 1^st^ shell water |
| --- | --- | --- | --- | --- | --- | --- | --- |
| **Ion** | | | |  | 1.00 | 47.39 || 1 | G | O6 | BA:1441 | 2.01 | 1.00 | 81.17 |
| 2 | G | N7 | BA:1441 | 4.10 | 1.00 | 85.47 |  |
| 3 | G | N1 | BA:1441 | 4.24 | 1.00 | 81.52 |  |
| 4 | G | OP1 | BA:1455 | 4.26 | 1.00 | 92.53 |  |
| 5 | G | OP2 | BA:1455 | 4.38 | 1.00 | 91.53 |  |
| 6 | C | N4 | BA:1440 | 4.83 | 1.00 | 84.72 |  |
  |

  
 Contact(s): /O6/  
 Coded contact list: O_b_  

### MG:BA:1695

|  | Res. type | Atom | Residue | Distance | Occupancy | B-factor | Closest solute to 1^st^ shell water |
| --- | --- | --- | --- | --- | --- | --- | --- |
| **Ion** | | | |  | 1.00 | 44.03 || 1 | C | OP2 | BA:1054 | 2.51 | 1.00 | 102.79 |
| 2 | G | OP2 | BA:1197 | 2.84 | 1.00 | 91.12 |  |
| 3 | C | OP1 | BA:1054 | 3.21 | 1.00 | 103.40 |  |
| 4 | G | O2' | BA:1053 | 3.24 | 1.00 | 99.80 |  |
| 5 | G | O3' | BA:1053 | 3.29 | 1.00 | 101.32 |  |
| 6 | U | O4 | BA:1199 | 3.72 | 1.00 | 89.63 |  |
| 7 | G | N7 | BA:1198 | 4.12 | 1.00 | 88.69 |  |
| 8 | G | OP2 | BA:1198 | 4.38 | 1.00 | 85.56 |  |
| 9 | C | O5' | BA:1054 | 4.59 | 1.00 | 103.97 |  |
| 10 | U | O2' | BA:1196 | 4.83 | 1.00 | 98.61 |  |
| 11 | G | OP1 | BA:1197 | 4.84 | 1.00 | 91.39 |  |
| 12 | G | O5' | BA:1197 | 4.96 | 1.00 | 90.65 |  |
  |

  
 Coded contact list:   
 Exclusion zone contact(s): /O2'/O3'/OP1/OP2/OP2/  
Consider K in place of MG (4 valid contact(s) for K vs. 0 for MG)  

### MG:BA:1696

|  | Res. type | Atom | Residue | Distance | Occupancy | B-factor | Closest solute to 1^st^ shell water |
| --- | --- | --- | --- | --- | --- | --- | --- |
| **Ion** | | | |  | 1.00 | 73.94 || 1 | U | O4 | BA:1199 | 3.12 | 1.00 | 89.63 |
| 2 | G | O6 | BA:1058 | 3.27 | 1.00 | 89.37 |  |
| 3 | G | OP2 | BA:1057 | 3.39 | 1.00 | 90.20 |  |
| 4 | G | O6 | BA:1198 | 3.69 | 1.00 | 89.55 |  |
| 5 | G | N2 | BA:1053 | 3.85 | 1.00 | 96.33 |  |
| 6 | C | N4 | BA:1059 | 4.23 | 1.00 | 89.67 |  |
| 7 | U | N3 | BA:1199 | 4.46 | 1.00 | 90.88 |  |
| 8 | G | N7 | BA:1058 | 4.52 | 1.00 | 89.94 |  |
| 9 | G | N3 | BA:1053 | 4.63 | 1.00 | 97.68 |  |
| 10 | G | O2' | BA:1053 | 4.74 | 1.00 | 99.80 |  |
  |

  
 Coded contact list:   
 Exclusion zone contact(s): /O4/O6/OP2/  
Consider K in place of MG (3 valid contact(s) for K vs. 0 for MG)  

### MG:BA:1697

|  | Res. type | Atom | Residue | Distance | Occupancy | B-factor | Closest solute to 1^st^ shell water |
| --- | --- | --- | --- | --- | --- | --- | --- |
| **Ion** | | | |  | 1.00 | 52.48 || 1 | G | O6 | BA:628 | 3.51 | 1.00 | 87.39 |
| 2 | A | OP1 | BA:609 | 4.19 | 1.00 | 73.89 |  |
| 3 | G | O6 | BA:627 | 4.34 | 1.00 | 86.75 |  |
| 4 | G | OP2 | BA:610 | 4.44 | 1.00 | 73.86 |  |
| 5 | G | N7 | BA:627 | 4.57 | 1.00 | 87.28 |  |
| 6 | C | N4 | BA:612 | 4.62 | 1.00 | 86.72 |  |
| 7 | A | N6 | BA:611 | 4.86 | 1.00 | 76.02 |  |
| 8 | G | N7 | BA:628 | 4.89 | 1.00 | 92.62 |  |
| 9 | A | OP2 | BA:609 | 4.91 | 1.00 | 73.07 |  |
  |

  
 Coded contact list:   
1^st^ MG coordination shell missing (see Notes)  

### MG:BA:1698

|  | Res. type | Atom | Residue | Distance | Occupancy | B-factor | Closest solute to 1^st^ shell water |
| --- | --- | --- | --- | --- | --- | --- | --- |
| **Ion** | | | |  | 1.00 | 104.35 || 1 | G | N7 | BA:628 | 3.85 | 1.00 | 92.62 |
| 2 | G | N7 | BA:629 | 4.29 | 1.00 | 110.82 |  |
| 3 | G | OP2 | BA:628 | 4.30 | 1.00 | 91.40 |  |
| 4 | G | O6 | BA:629 | 4.32 | 1.00 | 110.16 |  |
| 5 | G | O6 | BA:628 | 4.98 | 1.00 | 87.39 |  |
  |

  
 Coded contact list:   
1^st^ MG coordination shell missing (see Notes)  

### MG:BA:1699

|  | Res. type | Atom | Residue | Distance | Occupancy | B-factor | Closest solute to 1^st^ shell water |
| --- | --- | --- | --- | --- | --- | --- | --- |
| **Ion** | | | |  | 1.00 | 90.06 || 1 | G | OP2 | BA:631 | 2.14 | 1.00 | 127.90 |
| 2 | G | N7 | BA:631 | 3.50 | 1.00 | 131.12 |  |
| 3 | G | O5' | BA:631 | 3.87 | 1.00 | 127.54 |  |
| 4 | G | O3' | BA:630 | 4.04 | 1.00 | 126.77 |  |
| 5 | G | N1 | BA:606 | 4.33 | 1.00 | 93.16 |  |
| 6 | G | O6 | BA:606 | 4.36 | 1.00 | 92.41 |  |
| 7 | G | OP2 | BA:630 | 4.63 | 1.00 | 117.06 |  |
| 8 | G | O5' | BA:630 | 4.65 | 1.00 | 119.22 |  |
| 9 | G | OP1 | BA:631 | 4.70 | 1.00 | 127.86 |  |
| 10 | G | O4' | BA:630 | 4.91 | 1.00 | 124.31 |  |
| 11 | G | N9 | BA:631 | 4.96 | 1.00 | 129.77 |  |
  |

  
 Contact(s): /OP2/  
 Coded contact list: O_ph_  

### MG:BA:1700

|  | Res. type | Atom | Residue | Distance | Occupancy | B-factor | Closest solute to 1^st^ shell water |
| --- | --- | --- | --- | --- | --- | --- | --- |
| **Ion** | | | |  | 1.00 | 46.91 || 1 | G | N7 | BA:869 | 2.69 | 1.00 | 78.88 |
| 2 | G | N7 | BA:858 | 2.83 | 1.00 | 80.09 |  |
| 3 | C | OP2 | BA:868 | 3.19 | 1.00 | 71.35 |  |
| 4 | G | OP2 | BA:869 | 3.24 | 1.00 | 73.77 |  |
| 5 | G | O6 | BA:858 | 3.41 | 1.00 | 79.76 |  |
| 6 | C | O5' | BA:868 | 3.99 | 1.00 | 70.79 |  |
| 7 | G | OP2 | BA:858 | 4.53 | 1.00 | 83.55 |  |
| 8 | G | N9 | BA:869 | 4.57 | 1.00 | 78.63 |  |
| 9 | G | O6 | BA:869 | 4.60 | 1.00 | 77.66 |  |
| 10 | C | OP2 | BA:857 | 4.61 | 1.00 | 86.24 |  |
| 11 | C | OP1 | BA:868 | 4.65 | 1.00 | 70.91 |  |
| 12 | G | O5' | BA:869 | 4.79 | 1.00 | 73.79 |  |
| 13 | G | N9 | BA:858 | 4.96 | 1.00 | 80.58 |  |
  |

  
 Coded contact list:   
 Exclusion zone contact(s): /N7/N7/OP2/OP2/  
Consider K in place of MG (4 valid contact(s) for K vs. 0 for MG)  

### MG:BA:1701

|  | Res. type | Atom | Residue | Distance | Occupancy | B-factor | Closest solute to 1^st^ shell water |
| --- | --- | --- | --- | --- | --- | --- | --- |
| **Ion** | | | |  | 1.00 | 59.59 || 1 | G | OP2 | BA:1304 | 1.84 | 1.00 | 105.52 |
| 2 | C | OP1 | BA:1303 | 2.28 | 1.00 | 112.66 |  |
| 3 | C | O5' | BA:1303 | 2.61 | 1.00 | 111.90 |  |
| 4 | C | O3' | BA:1303 | 3.86 | 1.00 | 106.30 |  |
| 5 | U | O3' | BA:1302 | 3.91 | 1.00 | 114.27 |  |
| 6 | G | O5' | BA:1304 | 3.96 | 1.00 | 105.25 |  |
| 7 | C | O2' | BA:1242 | 4.07 | 1.00 | 91.66 |  |
| 8 | G | N7 | BA:1304 | 4.11 | 1.00 | 101.55 |  |
| 9 | C | OP2 | BA:1303 | 4.22 | 1.00 | 113.22 |  |
| 10 | G | OP1 | BA:1304 | 4.39 | 1.00 | 105.84 |  |
| 11 | G | N7 | BA:1305 | 4.93 | 1.00 | 100.41 |  |
| 12 | G | O6 | BA:1305 | 4.94 | 1.00 | 101.68 |  |
  |

  
**1^st^ shell ligand angles:**   
1-2: 157;   
  
 Contact(s): /OP1/OP2/  
 Coded contact list: 2O_ph_  
 Exclusion zone contact(s): /O5'/  

### MG:BA:1702

|  | Res. type | Atom | Residue | Distance | Occupancy | B-factor | Closest solute to 1^st^ shell water |
| --- | --- | --- | --- | --- | --- | --- | --- |
| **Ion** | | | |  | 1.00 | 33.59 || 1 | C | OP1 | BA:1054 | 2.06 | 1.00 | 103.40 |
| 2 | G | OP2 | BA:1198 | 2.20 | 1.00 | 85.56 |  |
| 3 | G | OP1 | BA:1197 | 2.25 | 1.00 | 91.39 |  |
| 4 | C | O5' | BA:1054 | 3.05 | 1.00 | 103.97 |  |
| 5 | G | O5' | BA:1197 | 3.31 | 1.00 | 90.65 |  |
| 6 | G | OP1 | BA:1198 | 3.39 | 1.00 | 86.00 |  |
| 7 | G | OP2 | BA:1197 | 3.72 | 1.00 | 91.12 |  |
| 8 | C | OP2 | BA:1054 | 4.01 | 1.00 | 102.79 |  |
| 9 | G | O3' | BA:1197 | 4.22 | 1.00 | 86.76 |  |
| 10 | G | O3' | BA:1053 | 4.26 | 1.00 | 101.32 |  |
| 11 | G | O5' | BA:1198 | 4.41 | 1.00 | 86.90 |  |
| 12 | U | O3' | BA:1196 | 4.58 | 1.00 | 93.62 |  |
| 13 | G | OP1 | BB:34 | 5.00 | 1.00 | 88.63 |  |
  |

  
**1^st^ shell ligand angles:**   
1-2: 71; 1-3: 85; 2-3: 96;   
  
 Contact(s): /OP1/OP1/OP2/  
 Coded contact list: 3O_ph_  
 Exclusion zone contact(s): /O5'/O5'/OP1/  

### MG:BA:1703

|  | Res. type | Atom | Residue | Distance | Occupancy | B-factor | Closest solute to 1^st^ shell water |
| --- | --- | --- | --- | --- | --- | --- | --- |
| **Ion** | | | |  | 1.00 | 40.27 || 1 | U | OP2 | BA:891 | 2.52 | 1.00 | 56.14 |
| 2 | G | N1 | BA:906 | 3.53 | 1.00 | 48.61 |  |
| 3 | U | O4 | BA:905 | 3.61 | 1.00 | 53.22 |  |
| 4 | U | O5' | BA:891 | 3.84 | 1.00 | 55.28 |  |
| 5 | G | N2 | BA:906 | 3.87 | 1.00 | 48.94 |  |
| 6 | U | OP1 | BA:891 | 3.93 | 1.00 | 57.84 |  |
| 7 | G | O2' | BA:890 | 4.37 | 1.00 | 64.33 |  |
| 8 | G | O6 | BA:906 | 4.44 | 1.00 | 47.83 |  |
| 9 | A | OP2 | BA:892 | 4.52 | 1.00 | 55.50 |  |
| 10 | C | N4 | BA:904 | 4.61 | 1.00 | 53.69 |  |
| 11 | G | OP2 | BA:903 | 4.78 | 1.00 | 48.52 |  |
| 12 | G | O3' | BA:890 | 4.99 | 1.00 | 57.60 |  |
  |

  
 Coded contact list:   
 Exclusion zone contact(s): /OP2/  
Consider NA in place of MG (1 valid contact(s) for NA vs. 0 for MG)  

### MG:BA:1704

|  | Res. type | Atom | Residue | Distance | Occupancy | B-factor | Closest solute to 1^st^ shell water |
| --- | --- | --- | --- | --- | --- | --- | --- |
| **Ion** | | | |  | 1.00 | 67.61 || 1 | G | O6 | BA:895 | 2.79 | 1.00 | 53.36 |
| 2 | G | O6 | BA:894 | 3.79 | 1.00 | 54.28 |  |
| 3 | G | N7 | BA:895 | 3.90 | 1.00 | 54.31 |  |
| 4 | G | N7 | BA:894 | 4.06 | 1.00 | 52.57 |  |
| 5 | C | N4 | BA:904 | 4.41 | 1.00 | 53.69 |  |
| 6 | G | O6 | BA:903 | 4.44 | 1.00 | 46.99 |  |
| 7 | C | N4 | BA:896 | 4.48 | 1.00 | 44.54 |  |
  |

  
 Coded contact list:   
 Exclusion zone contact(s): /O6/  
Consider K in place of MG (1 valid contact(s) for K vs. 0 for MG)  

### MG:BA:1705

|  | Res. type | Atom | Residue | Distance | Occupancy | B-factor | Closest solute to 1^st^ shell water |
| --- | --- | --- | --- | --- | --- | --- | --- |
| **Ion** | | | |  | 1.00 | 52.12 || 1 | C | O2' | BA:1399 | 2.61 | 1.00 | 70.02 |
| 2 | G | O6 | BA:926 | 3.84 | 1.00 | 73.31 |  |
| 3 | C | O2 | BA:1399 | 4.27 | 1.00 | 68.72 |  |
| 4 | G | N2 | BA:1504 | 4.36 | 1.00 | 63.74 |  |
| 5 | C | O3' | BA:1399 | 4.41 | 1.00 | 66.21 |  |
| 6 | G | O6 | BA:1505 | 4.44 | 1.00 | 70.11 |  |
| 7 | U | O4 | B1:48 | 4.58 | 1.00 | 78.35 |  |
  |

  
 Coded contact list:   
 Exclusion zone contact(s): /O2'/  
Consider K in place of MG (1 valid contact(s) for K vs. 0 for MG)  

### MG:BA:1706

|  | Res. type | Atom | Residue | Distance | Occupancy | B-factor | Closest solute to 1^st^ shell water |
| --- | --- | --- | --- | --- | --- | --- | --- |
| **Ion** | | | |  | 1.00 | 52.80 || 1 | G | OP2 | BA:1224 | 2.65 | 1.00 | 92.05 |
| 2 | G | OP1 | BA:1224 | 3.16 | 1.00 | 92.75 |  |
| 3 | G | OP1 | BA:951 | 3.70 | 1.00 | 78.60 |  |
| 4 | U | OP2 | BA:952 | 3.91 | 1.00 | 77.57 |  |
| 5 | C | O3' | BA:1223 | 4.46 | 1.00 | 92.31 |  |
| 6 | C | O2' | BA:1223 | 4.54 | 1.00 | 91.78 |  |
| 7 | G | O5' | BA:1224 | 4.56 | 1.00 | 93.26 |  |
| 8 | U | OP1 | BA:952 | 4.60 | 1.00 | 78.03 |  |
| 9 | G | OP2 | BA:951 | 4.78 | 1.00 | 77.60 |  |
| 10 | A | N6 | BA:974 | 4.88 | 1.00 | 94.19 |  |
  |

  
 Coded contact list:   
 Exclusion zone contact(s): /OP1/OP2/  
Consider K in place of MG (2 valid contact(s) for K vs. 0 for MG)  

### MG:BA:1707

|  | Res. type | Atom | Residue | Distance | Occupancy | B-factor | Closest solute to 1^st^ shell water |
| --- | --- | --- | --- | --- | --- | --- | --- |
| **Ion** | | | |  | 1.00 | 59.99 || 1 | A | OP2 | BA:975 | 3.20 | 1.00 | 96.17 |
| 2 | G | O2' | BA:976 | 3.42 | 1.00 | 101.92 |  |
| 3 | A | N7 | BA:1363 | 4.14 | 1.00 | 109.84 |  |
| 4 | A | OP1 | BA:1363 | 4.49 | 1.00 | 108.10 |  |
| 5 | A | O2' | BA:974 | 4.85 | 1.00 | 95.63 |  |
  |

  
 Coded contact list:   
 Exclusion zone contact(s): /OP2/  
Consider K in place of MG (1 valid contact(s) for K vs. 0 for MG)  

### MG:BA:1708

|  | Res. type | Atom | Residue | Distance | Occupancy | B-factor | Closest solute to 1^st^ shell water |
| --- | --- | --- | --- | --- | --- | --- | --- |
| **Ion** | | | |  | 1.00 | 57.28 || 1 | U | OP2 | BA:1083 | 1.87 | 1.00 | 75.58 |
| 2 | G | O6 | BA:1084 | 3.36 | 1.00 | 95.16 |  |
| 3 | G | N7 | BA:1084 | 3.47 | 1.00 | 96.93 |  |
| 4 | U | O5' | BA:1083 | 4.02 | 1.00 | 78.23 |  |
| 5 | G | O3' | BA:1082 | 4.18 | 1.00 | 74.71 |  |
| 6 | U | OP1 | BA:1083 | 4.34 | 1.00 | 76.82 |  |
| 7 | G | OP2 | BA:1082 | 4.65 | 1.00 | 71.10 |  |
| 8 | G | O5' | BA:1082 | 4.67 | 1.00 | 69.33 |  |
| 9 | G | OP1 | BA:1082 | 4.73 | 1.00 | 70.85 |  |
| 10 | U | O2 | BA:1085 | 4.75 | 1.00 | 98.64 |  |
| 11 | U | N3 | BA:1085 | 4.90 | 1.00 | 100.66 |  |
  |

  
 Contact(s): /OP2/  
 Coded contact list: O_ph_  
 Exclusion zone contact(s): /O6/  

### MG:BA:1709

|  | Res. type | Atom | Residue | Distance | Occupancy | B-factor | Closest solute to 1^st^ shell water |
| --- | --- | --- | --- | --- | --- | --- | --- |
| **Ion** | | | |  | 1.00 | 53.99 || 1 | C | OP2 | BA:1069 | 2.52 | 1.00 | 86.14 |
| 2 | U | O2' | BA:1085 | 3.62 | 1.00 | 102.50 |  |
| 3 | U | O2 | BA:1085 | 3.73 | 1.00 | 98.64 |  |
| 4 | G | O6 | BA:1094 | 4.29 | 1.00 | 95.23 |  |
| 5 | C | OP1 | BA:1069 | 4.65 | 1.00 | 86.67 |  |
| 6 | G | O3' | BA:1068 | 4.76 | 1.00 | 87.03 |  |
| 7 | C | OP1 | BA:1389 | 4.81 | 1.00 | 74.09 |  |
| 8 | C | O5' | BA:1069 | 4.90 | 1.00 | 88.33 |  |
| 9 | G | N7 | BA:1094 | 4.93 | 1.00 | 94.38 |  |
  |

  
 Coded contact list:   
 Exclusion zone contact(s): /OP2/  
Consider NA in place of MG (1 valid contact(s) for NA vs. 0 for MG)  

### MG:BA:1710

|  | Res. type | Atom | Residue | Distance | Occupancy | B-factor | Closest solute to 1^st^ shell water |
| --- | --- | --- | --- | --- | --- | --- | --- |
| **Ion** | | | |  | 1.00 | 90.83 || 1 | C | OP1 | BA:1069 | 2.97 | 1.00 | 86.67 |
| 2 | C | OP2 | BA:1069 | 3.31 | 1.00 | 86.14 |  |
| 3 | C | OP1 | BA:1388 | 3.50 | 1.00 | 76.93 |  |
| 4 | G | O3' | BA:1068 | 4.25 | 1.00 | 87.03 |  |
| 5 | GLN | OE1 | BH:20 | 4.32 | 1.00 | 83.50 |  |
| 6 | C | O5' | BA:1388 | 4.47 | 1.00 | 76.49 |  |
| 7 | C | OP2 | BA:1389 | 4.78 | 1.00 | 74.34 |  |
| 8 | C | OP2 | BA:1388 | 4.95 | 1.00 | 77.83 |  |
| 9 | C | O5' | BA:1069 | 4.98 | 1.00 | 88.33 |  |
  |

  
 Coded contact list:   
 Exclusion zone contact(s): /OP1/OP2/  
Consider K in place of MG (2 valid contact(s) for K vs. 0 for MG)  

### MG:BA:1711

|  | Res. type | Atom | Residue | Distance | Occupancy | B-factor | Closest solute to 1^st^ shell water |
| --- | --- | --- | --- | --- | --- | --- | --- |
| **Ion** | | | |  | 1.00 | 68.29 || 1 | C | OP1 | BA:1069 | 2.80 | 1.00 | 86.67 |
| 2 | C | OP1 | BA:1388 | 2.88 | 1.00 | 76.93 |  |
| 3 | GLN | OE1 | BH:20 | 3.47 | 1.00 | 83.50 |  |
| 4 | C | OP2 | BA:1388 | 3.71 | 1.00 | 77.83 |  |
| 5 | G | O3' | BA:1068 | 3.75 | 1.00 | 87.03 |  |
| 6 | C | OP2 | BA:1069 | 4.73 | 1.00 | 86.14 |  |
| 7 | C | O5' | BA:1388 | 4.85 | 1.00 | 76.49 |  |
| 8 | G | OP1 | BA:1387 | 4.92 | 1.00 | 82.15 |  |
| 9 | G | O3' | BA:1387 | 4.94 | 1.00 | 78.41 |  |
  |

  
 Coded contact list:   
 Exclusion zone contact(s): /OP1/OP1/  
Consider K in place of MG (2 valid contact(s) for K vs. 0 for MG)  

### MG:BA:1712

|  | Res. type | Atom | Residue | Distance | Occupancy | B-factor | Closest solute to 1^st^ shell water |
| --- | --- | --- | --- | --- | --- | --- | --- |
| **Ion** | | | |  | 1.00 | 56.78 || 1 | C | OP2 | BA:536 | 3.10 | 1.00 | 74.89 |
| 2 | A | OP1 | BA:535 | 3.57 | 1.00 | 80.04 |  |
| 3 | G | N7 | BA:537 | 4.24 | 1.00 | 73.83 |  |
| 4 | G | O6 | BA:538 | 4.47 | 1.00 | 73.19 |  |
| 5 | G | O6 | BA:537 | 4.77 | 1.00 | 74.34 |  |
| 6 | G | N7 | BA:538 | 4.83 | 1.00 | 71.90 |  |
| 7 | C | O5' | BA:536 | 4.91 | 1.00 | 73.87 |  |
  |

  
 Coded contact list:   
 Exclusion zone contact(s): /OP2/  
Consider K in place of MG (1 valid contact(s) for K vs. 0 for MG)  

### MG:BA:1713

|  | Res. type | Atom | Residue | Distance | Occupancy | B-factor | Closest solute to 1^st^ shell water |
| --- | --- | --- | --- | --- | --- | --- | --- |
| **Ion** | | | |  | 1.00 | 48.71 || 1 | G | OP2 | BA:537 | 3.97 | 1.00 | 69.44 |
| 2 | C | O2' | BA:522 | 4.02 | 1.00 | 66.76 |  |
| 3 | C | O3' | BA:522 | 4.34 | 1.00 | 65.80 |  |
| 4 | C | OP1 | BA:503 | 4.45 | 1.00 | 68.52 |  |
| 5 | G | OP1 | BA:537 | 4.49 | 1.00 | 69.80 |  |
| 6 | C | OP2 | BA:504 | 4.60 | 1.00 | 73.90 |  |
| 7 | ARG | NH2 | BO:113 | 4.64 | 1.00 | 52.33 |  |
| 8 | SER | OG | BO:116 | 4.72 | 1.00 | 50.91 |  |
| 9 | C | OP1 | BA:536 | 4.77 | 1.00 | 73.39 |  |
| 10 | C | O5' | BA:536 | 4.98 | 1.00 | 73.87 |  |
  |

  
 Coded contact list:   
1^st^ MG coordination shell missing (see Notes)  

### MG:BA:1714

|  | Res. type | Atom | Residue | Distance | Occupancy | B-factor | Closest solute to 1^st^ shell water |
| --- | --- | --- | --- | --- | --- | --- | --- |
| **Ion** | | | |  | 1.00 | 69.29 || 1 | C | OP1 | BA:504 | 1.88 | 1.00 | 73.99 |
| 2 | G | N7 | BA:505 | 3.97 | 1.00 | 76.30 |  |
| 3 | C | OP2 | BA:504 | 4.02 | 1.00 | 73.90 |  |
| 4 | C | OP1 | BA:536 | 4.20 | 1.00 | 73.39 |  |
| 5 | C | O5' | BA:504 | 4.21 | 1.00 | 74.97 |  |
| 6 | C | O3' | BA:503 | 4.25 | 1.00 | 72.55 |  |
| 7 | G | OP1 | BA:505 | 4.32 | 1.00 | 73.49 |  |
| 8 | A | O4' | BA:535 | 4.99 | 1.00 | 80.19 |  |
  |

  
 Contact(s): /OP1/  
 Coded contact list: O_ph_  

### MG:BA:1715

|  | Res. type | Atom | Residue | Distance | Occupancy | B-factor | Closest solute to 1^st^ shell water |
| --- | --- | --- | --- | --- | --- | --- | --- |
| **Ion** | | | |  | 1.00 | 75.05 || 1 | G | O6 | BA:31 | 2.71 | 1.00 | 79.78 |
| 2 | C | O2 | BA:48 | 3.19 | 1.00 | 76.04 |  |
| 3 | G | N7 | BA:31 | 3.28 | 1.00 | 76.57 |  |
| 4 | G | O4' | BA:306 | 4.56 | 1.00 | 85.43 |  |
| 5 | C | O2' | BA:48 | 4.62 | 1.00 | 74.84 |  |
| 6 | C | N1 | BA:48 | 4.78 | 1.00 | 76.65 |  |
| 7 | G | N1 | BA:31 | 4.82 | 1.00 | 77.60 |  |
  |

  
 Coded contact list:   
 Exclusion zone contact(s): /N7/O2/O6/  
Consider K in place of MG (3 valid contact(s) for K vs. 0 for MG)  

### MG:BA:1716

|  | Res. type | Atom | Residue | Distance | Occupancy | B-factor | Closest solute to 1^st^ shell water |
| --- | --- | --- | --- | --- | --- | --- | --- |
| **Ion** | | | |  | 1.00 | 56.20 || 1 | G | O6 | BA:1417 | 2.12 | 1.00 | 37.26 |
| 2 | G | O6 | BA:1482 | 3.62 | 1.00 | 52.16 |  |
| 3 | G | N7 | BA:1482 | 4.16 | 1.00 | 48.85 |  |
| 4 | G | N1 | BA:1417 | 4.35 | 1.00 | 35.91 |  |
| 5 | G | N7 | BA:1417 | 4.35 | 1.00 | 38.98 |  |
| 6 | U | O4 | BA:1481 | 4.47 | 1.00 | 59.52 |  |
| 7 | G | N7 | BA:1416 | 4.48 | 1.00 | 52.22 |  |
| 8 | G | OP2 | BA:1416 | 4.73 | 1.00 | 56.78 |  |
  |

  
 Contact(s): /O6/  
 Coded contact list: O_b_  

### MG:BA:1717

|  | Res. type | Atom | Residue | Distance | Occupancy | B-factor | Closest solute to 1^st^ shell water |
| --- | --- | --- | --- | --- | --- | --- | --- |
| **Ion** | | | |  | 1.00 | 55.06 || 1 | G | O6 | BA:1417 | 3.32 | 1.00 | 37.26 |
| 2 | G | O6 | BA:1416 | 3.68 | 1.00 | 53.06 |  |
| 3 | G | N7 | BA:1416 | 3.80 | 1.00 | 52.22 |  |
| 4 | G | N7 | BA:1415 | 4.11 | 1.00 | 50.39 |  |
| 5 | G | O6 | BA:1415 | 4.39 | 1.00 | 47.85 |  |
| 6 | G | OP2 | BA:1415 | 4.54 | 1.00 | 56.65 |  |
| 7 | G | N1 | BA:1417 | 4.60 | 1.00 | 35.91 |  |
  |

  
 Coded contact list:   
 Exclusion zone contact(s): /O6/  
Consider K in place of MG (1 valid contact(s) for K vs. 0 for MG)  

### MG:BA:1718

|  | Res. type | Atom | Residue | Distance | Occupancy | B-factor | Closest solute to 1^st^ shell water |
| --- | --- | --- | --- | --- | --- | --- | --- |
| **Ion** | | | |  | 1.00 | 43.47 || 1 | A | OP1 | BA:1502 | 3.00 | 1.00 | 54.69 |
| 2 | G | OP1 | BA:1530 | 3.11 | 1.00 | 80.25 |  |
| 3 | G | O3' | BA:1392 | 3.28 | 1.00 | 65.46 |  |
| 4 | U | OP1 | BA:1393 | 3.56 | 1.00 | 66.11 |  |
| 5 | G | O2' | BA:1392 | 3.59 | 1.00 | 65.44 |  |
| 6 | U | O5' | BA:1393 | 4.42 | 1.00 | 64.04 |  |
| 7 | A | O5' | BA:1502 | 4.43 | 1.00 | 55.97 |  |
| 8 | G | O3' | BA:1529 | 4.74 | 1.00 | 76.75 |  |
  |

  
 Coded contact list:   
 Exclusion zone contact(s): /O3'/OP1/OP1/  
Consider K in place of MG (3 valid contact(s) for K vs. 0 for MG)  

### MG:BA:1719

|  | Res. type | Atom | Residue | Distance | Occupancy | B-factor | Closest solute to 1^st^ shell water |
| --- | --- | --- | --- | --- | --- | --- | --- |
| **Ion** | | | |  | 1.00 | 43.41 || 1 | G | OP1 | BA:1094 | 2.52 | 1.00 | 93.55 |
| 2 | G | OP1 | BA:1068 | 3.03 | 1.00 | 86.57 |  |
| 3 | G | O2' | BA:1387 | 3.05 | 1.00 | 81.14 |  |
| 4 | A | O3' | BA:1067 | 3.06 | 1.00 | 86.76 |  |
| 5 | A | O3' | BA:1093 | 4.38 | 1.00 | 93.97 |  |
| 6 | A | O5' | BA:1067 | 4.68 | 1.00 | 89.54 |  |
| 7 | G | N3 | BA:1387 | 4.69 | 1.00 | 84.50 |  |
| 8 | G | O5' | BA:1094 | 4.70 | 1.00 | 92.98 |  |
| 9 | G | O5' | BA:1068 | 4.74 | 1.00 | 85.54 |  |
| 10 | C | O4' | BA:1388 | 4.74 | 1.00 | 74.30 |  |
| 11 | G | OP2 | BA:1068 | 4.82 | 1.00 | 85.64 |  |
| 12 | A | OP1 | BA:1067 | 4.90 | 1.00 | 89.88 |  |
  |

  
 Coded contact list:   
 Exclusion zone contact(s): /O2'/O3'/OP1/OP1/  
Consider K in place of MG (3 valid contact(s) for K vs. 0 for MG)  

### MG:BA:1720

|  | Res. type | Atom | Residue | Distance | Occupancy | B-factor | Closest solute to 1^st^ shell water |
| --- | --- | --- | --- | --- | --- | --- | --- |
| **Ion** | | | |  | 1.00 | 95.06 || 1 | G | OP2 | BA:1087 | 3.68 | 1.00 | 103.60 |
| 2 | G | N7 | BA:1088 | 4.05 | 1.00 | 108.66 |  |
| 3 | U | O4 | BA:1090 | 4.17 | 1.00 | 103.39 |  |
| 4 | G | O6 | BA:1088 | 4.32 | 1.00 | 107.00 |  |
| 5 | G | N7 | BA:1089 | 4.85 | 1.00 | 100.43 |  |
| 6 | G | N7 | BA:1087 | 4.85 | 1.00 | 105.95 |  |
| 7 | G | OP2 | BA:1088 | 4.85 | 1.00 | 107.07 |  |
| 8 | G | O6 | BA:1089 | 4.90 | 1.00 | 99.08 |  |
| 9 | G | O5' | BA:1087 | 4.92 | 1.00 | 104.52 |  |
  |

  
 Coded contact list:   
1^st^ MG coordination shell missing (see Notes)  

### MG:BA:1721

|  | Res. type | Atom | Residue | Distance | Occupancy | B-factor | Closest solute to 1^st^ shell water |
| --- | --- | --- | --- | --- | --- | --- | --- |
| **Ion** | | | |  | 1.00 | 71.88 || 1 | G | OP1 | BA:1094 | 2.11 | 1.00 | 93.55 |
| 2 | G | OP1 | BA:1068 | 2.72 | 1.00 | 86.57 |  |
| 3 | G | O5' | BA:1094 | 3.03 | 1.00 | 92.98 |  |
| 4 | A | O3' | BA:1067 | 3.26 | 1.00 | 86.76 |  |
| 5 | G | OP2 | BA:1068 | 3.55 | 1.00 | 85.64 |  |
| 6 | G | OP2 | BA:1094 | 4.15 | 1.00 | 93.74 |  |
| 7 | A | O3' | BA:1093 | 4.25 | 1.00 | 93.97 |  |
| 8 | C | O4' | BA:1388 | 4.75 | 1.00 | 74.30 |  |
| 9 | G | O5' | BA:1068 | 4.81 | 1.00 | 85.54 |  |
| 10 | G | O2' | BA:1387 | 4.81 | 1.00 | 81.14 |  |
  |

  
 Contact(s): /OP1/  
 Coded contact list: O_ph_  
 Exclusion zone contact(s): /O3'/O5'/OP1/  
Consider K in place of MG (3 valid contact(s) for K vs. 1 for MG)  

### MG:BA:1722

|  | Res. type | Atom | Residue | Distance | Occupancy | B-factor | Closest solute to 1^st^ shell water |
| --- | --- | --- | --- | --- | --- | --- | --- |
| **Ion** | | | |  | 1.00 | 54.05 || 1 | G | O6 | BA:1370 | 2.86 | 1.00 | 97.43 |
| 2 | G | N7 | BA:1370 | 3.55 | 1.00 | 97.30 |  |
| 3 | G | O6 | BA:1371 | 3.72 | 1.00 | 95.04 |  |
| 4 | G | N7 | BA:1371 | 3.82 | 1.00 | 96.16 |  |
| 5 | G | N1 | BA:1370 | 4.47 | 1.00 | 96.51 |  |
| 6 | U | O4 | BA:1372 | 4.64 | 1.00 | 89.96 |  |
| 7 | C | N4 | BA:1352 | 4.64 | 1.00 | 97.02 |  |
| 8 | A | N6 | BA:1350 | 4.78 | 1.00 | 89.80 |  |
| 9 | U | O4 | BA:1351 | 4.81 | 1.00 | 95.50 |  |
| 10 | U | OP2 | BA:1348 | 4.82 | 1.00 | 91.29 |  |
  |

  
 Coded contact list:   
 Exclusion zone contact(s): /O6/  
Consider K in place of MG (1 valid contact(s) for K vs. 0 for MG)  

### MG:BA:1723

|  | Res. type | Atom | Residue | Distance | Occupancy | B-factor | Closest solute to 1^st^ shell water |
| --- | --- | --- | --- | --- | --- | --- | --- |
| **Ion** | | | |  | 1.00 | 63.71 || 1 | U | O4 | BA:605 | 2.34 | 1.00 | 90.49 |
| 2 | G | O6 | BA:604 | 2.81 | 1.00 | 90.45 |  |
| 3 | U | O4 | BA:603 | 3.71 | 1.00 | 80.05 |  |
| 4 | G | O6 | BA:633 | 3.75 | 1.00 | 90.16 |  |
| 5 | U | N3 | BA:605 | 3.97 | 1.00 | 89.48 |  |
| 6 | G | N7 | BA:604 | 4.15 | 1.00 | 90.09 |  |
| 7 | C | N4 | BA:634 | 4.16 | 1.00 | 91.99 |  |
| 8 | G | N1 | BA:606 | 4.83 | 1.00 | 93.16 |  |
| 9 | G | O6 | BA:606 | 4.90 | 1.00 | 92.41 |  |
| 10 | U | N3 | BA:603 | 4.93 | 1.00 | 81.28 |  |
| 11 | G | N1 | BA:604 | 4.95 | 1.00 | 89.31 |  |
  |

  
 Contact(s): /O4/  
 Coded contact list: O_b_  
 Exclusion zone contact(s): /O6/  

### MG:BA:1724

|  | Res. type | Atom | Residue | Distance | Occupancy | B-factor | Closest solute to 1^st^ shell water |
| --- | --- | --- | --- | --- | --- | --- | --- |
| **Ion** | | | |  | 1.00 | 100.21 || 1 | A | OP2 | BA:602 | 3.28 | 1.00 | 84.82 |
| 2 | C | OP2 | BA:601 | 3.98 | 1.00 | 81.32 |  |
| 3 | A | N1 | BA:632 | 4.23 | 1.00 | 120.91 |  |
| 4 | A | N6 | BA:632 | 4.36 | 1.00 | 121.88 |  |
| 5 | C | O5' | BA:601 | 4.73 | 1.00 | 82.18 |  |
| 6 | C | OP1 | BA:601 | 4.85 | 1.00 | 80.76 |  |
  |

  
 Coded contact list:   
 Exclusion zone contact(s): /OP2/  
Consider K in place of MG (1 valid contact(s) for K vs. 0 for MG)  

### MG:BA:1725

|  | Res. type | Atom | Residue | Distance | Occupancy | B-factor | Closest solute to 1^st^ shell water |
| --- | --- | --- | --- | --- | --- | --- | --- |
| **Ion** | | | |  | 1.00 | 109.37 || 1 | U | O4 | BA:603 | 2.73 | 1.00 | 80.05 |
| 2 | G | O6 | BA:635 | 2.91 | 1.00 | 83.38 |  |
| 3 | A | N6 | BA:602 | 3.24 | 1.00 | 86.29 |  |
| 4 | G | O6 | BA:604 | 3.54 | 1.00 | 90.45 |  |
| 5 | U | O4 | BA:636 | 3.67 | 1.00 | 84.93 |  |
| 6 | C | N4 | BA:634 | 3.83 | 1.00 | 91.99 |  |
| 7 | U | N3 | BA:603 | 3.91 | 1.00 | 81.28 |  |
| 8 | G | N7 | BA:635 | 4.69 | 1.00 | 85.59 |  |
  |

  
 Coded contact list:   
 Exclusion zone contact(s): /N6/O4/O6/  
Consider K in place of MG (3 valid contact(s) for K vs. 0 for MG)  

### MG:BA:1726

|  | Res. type | Atom | Residue | Distance | Occupancy | B-factor | Closest solute to 1^st^ shell water |
| --- | --- | --- | --- | --- | --- | --- | --- |
| **Ion** | | | |  | 1.00 | 62.36 || 1 | C | OP1 | BA:596 | 3.81 | 1.00 | 80.35 |
  |

  
 Coded contact list:   
1^st^ MG coordination shell missing (see Notes)  

### MG:BA:1727

|  | Res. type | Atom | Residue | Distance | Occupancy | B-factor | Closest solute to 1^st^ shell water |
| --- | --- | --- | --- | --- | --- | --- | --- |
| **Ion** | | | |  | 1.00 | 52.68 || 1 | A | OP1 | BA:782 | 2.00 | 1.00 | 54.82 |
| 2 | A | OP2 | BA:794 | 2.01 | 1.00 | 62.09 |  |
| 3 | A | OP1 | BA:794 | 3.28 | 1.00 | 63.15 |  |
| 4 | A | O5' | BA:782 | 3.91 | 1.00 | 55.86 |  |
| 5 | A | OP2 | BA:782 | 3.98 | 1.00 | 54.01 |  |
| 6 | A | O5' | BA:794 | 4.12 | 1.00 | 63.91 |  |
| 7 | C | OP2 | BA:783 | 4.15 | 1.00 | 56.02 |  |
| 8 | U | O3' | BA:793 | 4.20 | 1.00 | 62.36 |  |
| 9 | G | N2 | BA:1521 | 4.25 | 1.00 | 47.22 |  |
| 10 | G | N3 | BA:1521 | 4.31 | 1.00 | 48.45 |  |
| 11 | G | O2' | BA:1521 | 4.38 | 1.00 | 53.46 |  |
| 12 | A | O3' | BA:781 | 4.53 | 1.00 | 52.80 |  |
| 13 | U | O4' | BA:1522 | 4.57 | 1.00 | 53.38 |  |
  |

  
**1^st^ shell ligand angles:**   
1-2: 156;   
  
 Contact(s): /OP1/OP2/  
 Coded contact list: 2O_ph_  
 Exclusion zone contact(s): /OP1/  

### MG:BA:1728

|  | Res. type | Atom | Residue | Distance | Occupancy | B-factor | Closest solute to 1^st^ shell water |
| --- | --- | --- | --- | --- | --- | --- | --- |
| **Ion** | | | |  | 1.00 | 47.86 || 1 | G | O6 | BA:1108 | 2.14 | 1.00 | 86.27 |
| 2 | U | OP2 | BA:1095 | 2.31 | 1.00 | 90.50 |  |
| 3 | C | OP2 | BA:1096 | 3.36 | 1.00 | 95.94 |  |
| 4 | U | O5' | BA:1095 | 3.63 | 1.00 | 91.45 |  |
| 5 | G | N1 | BA:1108 | 3.63 | 1.00 | 89.21 |  |
| 6 | G | O2' | BA:1094 | 4.05 | 1.00 | 92.96 |  |
| 7 | U | OP1 | BA:1095 | 4.33 | 1.00 | 90.34 |  |
| 8 | C | N3 | BA:1109 | 4.57 | 1.00 | 99.16 |  |
| 9 | C | OP1 | BA:1096 | 4.60 | 1.00 | 95.20 |  |
| 10 | U | O3' | BA:1095 | 4.67 | 1.00 | 94.50 |  |
| 11 | G | O3' | BA:1094 | 4.72 | 1.00 | 91.28 |  |
  |

  
**1^st^ shell ligand angles:**   
1-2: 82;   
  
 Contact(s): /O6/OP2/  
 Coded contact list: O_ph_O_b_  
 Exclusion zone contact(s): /OP2/  

### MG:BA:1729

|  | Res. type | Atom | Residue | Distance | Occupancy | B-factor | Closest solute to 1^st^ shell water |
| --- | --- | --- | --- | --- | --- | --- | --- |
| **Ion** | | | |  | 1.00 | 69.75 || 1 | G | OP2 | BA:1120 | 3.84 | 1.00 | 122.15 |
| 2 | G | N7 | BA:1120 | 4.18 | 1.00 | 124.84 |  |
| 3 | U | O4 | BA:1121 | 4.57 | 1.00 | 124.32 |  |
  |

  
 Coded contact list:   
1^st^ MG coordination shell missing (see Notes)  

### MG:BA:1730

|  | Res. type | Atom | Residue | Distance | Occupancy | B-factor | Closest solute to 1^st^ shell water |
| --- | --- | --- | --- | --- | --- | --- | --- |
| **Ion** | | | |  | 1.00 | 37.47 || 1 | G | N7 | BA:758 | 2.91 | 1.00 | 57.95 |
| 2 | G | N7 | BA:581 | 3.03 | 1.00 | 56.11 |  |
| 3 | G | O6 | BA:758 | 3.12 | 1.00 | 55.08 |  |
| 4 | G | OP2 | BA:581 | 3.62 | 1.00 | 57.87 |  |
| 5 | U | OP2 | BA:580 | 4.38 | 1.00 | 60.78 |  |
| 6 | G | OP2 | BA:758 | 4.45 | 1.00 | 55.72 |  |
| 7 | G | O6 | BA:581 | 4.59 | 1.00 | 55.00 |  |
| 8 | U | OP2 | BA:757 | 4.66 | 1.00 | 58.48 |  |
| 9 | G | N9 | BA:581 | 4.82 | 1.00 | 55.32 |  |
| 10 | G | O5' | BA:581 | 4.89 | 1.00 | 56.81 |  |
  |

  
 Coded contact list:   
 Exclusion zone contact(s): /N7/N7/O6/  
Consider K in place of MG (3 valid contact(s) for K vs. 0 for MG)  

### MG:BA:1731

|  | Res. type | Atom | Residue | Distance | Occupancy | B-factor | Closest solute to 1^st^ shell water |
| --- | --- | --- | --- | --- | --- | --- | --- |
| **Ion** | | | |  | 1.00 | 46.36 || 1 | G | O6 | BA:584 | 3.33 | 1.00 | 59.04 |
| 2 | A | OP2 | BA:583 | 3.77 | 1.00 | 60.67 |  |
| 3 | G | O6 | BA:585 | 3.94 | 1.00 | 61.54 |  |
| 4 | U | O4 | BA:757 | 4.00 | 1.00 | 57.54 |  |
| 5 | G | N7 | BA:584 | 4.07 | 1.00 | 58.91 |  |
| 6 | C | N4 | BA:756 | 4.28 | 1.00 | 61.47 |  |
| 7 | G | N2 | BA:758 | 4.30 | 1.00 | 57.69 |  |
| 8 | U | OP2 | BA:582 | 4.84 | 1.00 | 56.75 |  |
  |

  
 Coded contact list:   
 Exclusion zone contact(s): /O6/  
Consider K in place of MG (1 valid contact(s) for K vs. 0 for MG)  

### MG:BA:1732

|  | Res. type | Atom | Residue | Distance | Occupancy | B-factor | Closest solute to 1^st^ shell water |
| --- | --- | --- | --- | --- | --- | --- | --- |
| **Ion** | | | |  | 1.00 | 62.42 || 1 | U | OP1 | BA:646 | 3.80 | 1.00 | 83.85 |
| 2 | G | N7 | BA:587 | 4.14 | 1.00 | 68.72 |  |
| 3 | C | OP1 | BA:645 | 4.30 | 1.00 | 79.25 |  |
| 4 | U | OP2 | BA:646 | 4.75 | 1.00 | 83.65 |  |
| 5 | C | O3' | BA:645 | 4.91 | 1.00 | 83.23 |  |
  |

  
 Coded contact list:   
1^st^ MG coordination shell missing (see Notes)  

### MG:BA:1733

|  | Res. type | Atom | Residue | Distance | Occupancy | B-factor | Closest solute to 1^st^ shell water |
| --- | --- | --- | --- | --- | --- | --- | --- |
| **Ion** | | | |  | 1.00 | 46.37 || 1 | ASN | O | BN:26 | 2.22 | 1.00 | 78.85 |
| 2 | U | O4 | BA:692 | 2.30 | 1.00 | 63.96 |  |
| 3 | GLY | O | BN:52 | 2.76 | 1.00 | 76.15 |  |
| 4 | G | O6 | BA:691 | 2.91 | 1.00 | 54.34 |  |
| 5 | G | N7 | BA:691 | 3.64 | 1.00 | 51.75 |  |
| 6 | SER | N | BN:53 | 3.76 | 1.00 | 77.75 |  |
| 7 | SER | O | BN:53 | 4.18 | 1.00 | 79.98 |  |
| 8 | LYS | NZ | BN:55 | 4.37 | 1.00 | 77.13 |  |
| 9 | ASN | OD1 | BN:26 | 4.42 | 1.00 | 80.13 |  |
| 10 | ASN | N | BN:27 | 4.45 | 1.00 | 77.54 |  |
| 11 | U | N3 | BA:692 | 4.55 | 1.00 | 61.25 |  |
| 12 | G | N1 | BA:691 | 4.57 | 1.00 | 53.21 |  |
| 13 | GLY | N | BN:56 | 4.67 | 1.00 | 76.16 |  |
| 14 | LYS | N | BN:55 | 4.74 | 1.00 | 76.78 |  |
  |

  
**1^st^ shell ligand angles:**   
1-2: 145;   
  
 Contact(s): /O4/O_back_/  
 Coded contact list: O_b_O_back_  
 Exclusion zone contact(s): /O6/O_back_/  

### MG:BA:1734

|  | Res. type | Atom | Residue | Distance | Occupancy | B-factor | Closest solute to 1^st^ shell water |
| --- | --- | --- | --- | --- | --- | --- | --- |
| **Ion** | | | |  | 1.00 | 76.26 || 1 | U | O3' | BA:793 | 2.53 | 1.00 | 62.36 |
| 2 | A | OP2 | BA:794 | 2.69 | 1.00 | 62.09 |  |
| 3 | U | O2' | BA:793 | 3.53 | 1.00 | 65.14 |  |
| 4 | C | OP2 | BA:783 | 3.91 | 1.00 | 56.02 |  |
| 5 | A | OP1 | BA:794 | 4.10 | 1.00 | 63.15 |  |
| 6 | A | O5' | BA:794 | 4.37 | 1.00 | 63.91 |  |
  |

  
 Coded contact list:   
 Exclusion zone contact(s): /O3'/OP2/  
Consider NA in place of MG (1 valid contact(s) for NA vs. 0 for MG)  

### MG:BA:1735

|  | Res. type | Atom | Residue | Distance | Occupancy | B-factor | Closest solute to 1^st^ shell water |
| --- | --- | --- | --- | --- | --- | --- | --- |
| **Ion** | | | |  | 1.00 | 40.97 || 1 | G | O6 | BA:236 | 2.42 | 1.00 | 71.44 |
| 2 | G | O6 | BA:126 | 2.63 | 1.00 | 75.03 |  |
| 3 | U | O4 | BA:125 | 2.67 | 1.00 | 73.92 |  |
| 4 | C | O2 | BA:121 | 2.80 | 1.00 | 69.96 |  |
| 5 | G | O6 | BA:124 | 2.94 | 1.00 | 67.51 |  |
| 6 | C | N3 | BA:121 | 3.09 | 1.00 | 69.88 |  |
| 7 | C | N4 | BA:235 | 3.09 | 1.00 | 73.01 |  |
| 8 | U | N3 | BA:125 | 3.59 | 1.00 | 73.17 |  |
| 9 | G | N1 | BA:126 | 3.73 | 1.00 | 75.95 |  |
| 10 | C | N4 | BA:237 | 3.74 | 1.00 | 67.64 |  |
| 11 | C | N3 | BA:235 | 4.12 | 1.00 | 74.12 |  |
| 12 | G | N7 | BA:236 | 4.31 | 1.00 | 71.79 |  |
| 13 | G | N1 | BA:236 | 4.68 | 1.00 | 72.07 |  |
| 14 | G | N1 | BA:124 | 4.69 | 1.00 | 66.64 |  |
| 15 | C | N4 | BA:121 | 4.69 | 1.00 | 70.96 |  |
| 16 | C | N1 | BA:121 | 4.72 | 1.00 | 69.43 |  |
| 17 | G | N7 | BA:126 | 4.90 | 1.00 | 75.14 |  |
  |

  
 Coded contact list:   
 Exclusion zone contact(s): /N3_C_/N4/O2/O4/O6/O6/O6/  
Consider K in place of MG (6 valid contact(s) for K vs. 0 for MG)  

### MG:BA:1736

|  | Res. type | Atom | Residue | Distance | Occupancy | B-factor | Closest solute to 1^st^ shell water |
| --- | --- | --- | --- | --- | --- | --- | --- |
| **Ion** | | | |  | 1.00 | 53.53 || 1 | A | OP2 | BA:573 | 1.89 | 1.00 | 62.01 |
| 2 | A | OP2 | BA:572 | 2.19 | 1.00 | 64.58 |  |
| 3 | A | OP2 | BA:574 | 2.23 | 1.00 | 61.47 |  |
| 4 | A | OP1 | BA:572 | 3.53 | 1.00 | 65.83 |  |
| 5 | A | OP1 | BA:573 | 3.75 | 1.00 | 62.38 |  |
| 6 | A | O5' | BA:572 | 3.84 | 1.00 | 65.13 |  |
| 7 | A | O5' | BA:573 | 4.01 | 1.00 | 63.35 |  |
| 8 | A | OP1 | BA:574 | 4.26 | 1.00 | 61.27 |  |
| 9 | A | O3' | BA:572 | 4.34 | 1.00 | 63.48 |  |
| 10 | U | OP1 | BA:571 | 4.35 | 1.00 | 64.09 |  |
| 11 | A | O3' | BA:573 | 4.56 | 1.00 | 61.95 |  |
| 12 | A | O5' | BA:574 | 4.58 | 1.00 | 60.67 |  |
| 13 | U | O3' | BA:571 | 4.58 | 1.00 | 62.57 |  |
| 14 | A | N1 | BA:819 | 4.87 | 1.00 | 54.79 |  |
| 15 | U | O5' | BA:571 | 4.87 | 1.00 | 63.24 |  |
  |

  
**1^st^ shell ligand angles:**   
1-2: 86; 1-3: 90; 2-3: 162;   
  
 Contact(s): /OP2/OP2/OP2/  
 Coded contact list: 3O_ph_  

### MG:BA:1737

|  | Res. type | Atom | Residue | Distance | Occupancy | B-factor | Closest solute to 1^st^ shell water |
| --- | --- | --- | --- | --- | --- | --- | --- |
| **Ion** | | | |  | 1.00 | 59.48 || 1 | A | OP1 | BA:572 | 1.97 | 1.00 | 65.83 |
| 2 | U | O3' | BA:571 | 3.96 | 1.00 | 62.57 |  |
| 3 | A | O5' | BA:572 | 4.01 | 1.00 | 65.13 |  |
| 4 | A | N6 | BA:819 | 4.14 | 1.00 | 52.21 |  |
| 5 | A | N1 | BA:819 | 4.19 | 1.00 | 54.79 |  |
| 6 | A | OP1 | BA:573 | 4.27 | 1.00 | 62.38 |  |
| 7 | A | OP2 | BA:572 | 4.53 | 1.00 | 64.58 |  |
| 8 | G | OP1 | BA:917 | 4.90 | 1.00 | 58.28 |  |
  |

  
 Contact(s): /OP1/  
 Coded contact list: O_ph_  

### MG:BA:1738

|  | Res. type | Atom | Residue | Distance | Occupancy | B-factor | Closest solute to 1^st^ shell water |
| --- | --- | --- | --- | --- | --- | --- | --- |
| **Ion** | | | |  | 1.00 | 59.12 || 1 | C | OP2 | BA:970 | 2.13 | 1.00 | 86.17 |
| 2 | C | OP1 | BA:970 | 3.66 | 1.00 | 85.89 |  |
| 3 | A | OP2 | BA:969 | 3.81 | 1.00 | 82.12 |  |
| 4 | A | OP2 | BA:968 | 3.86 | 1.00 | 75.06 |  |
| 5 | A | O5' | BA:969 | 4.09 | 1.00 | 81.51 |  |
| 6 | A | O3' | BA:969 | 4.22 | 1.00 | 84.54 |  |
| 7 | C | O5' | BA:970 | 4.49 | 1.00 | 85.53 |  |
| 8 | C | OP1 | BA:967 | 4.50 | 1.00 | 74.68 |  |
| 9 | A | O5' | BA:968 | 4.97 | 1.00 | 77.08 |  |
  |

  
 Contact(s): /OP2/  
 Coded contact list: O_ph_  

### MG:BA:1739

|  | Res. type | Atom | Residue | Distance | Occupancy | B-factor | Closest solute to 1^st^ shell water |
| --- | --- | --- | --- | --- | --- | --- | --- |
| **Ion** | | | |  | 1.00 | 67.21 || 1 | A | OP2 | BA:1357 | 1.99 | 1.00 | 100.37 |
| 2 | C | N4 | BA:1359 | 3.28 | 1.00 | 89.71 |  |
| 3 | A | OP1 | BA:1357 | 3.53 | 1.00 | 100.97 |  |
| 4 | C | N3 | BA:1359 | 3.79 | 1.00 | 90.18 |  |
| 5 | G | O3' | BA:1356 | 4.19 | 1.00 | 99.61 |  |
| 6 | A | O5' | BA:1357 | 4.30 | 1.00 | 101.65 |  |
| 7 | U | O4 | BA:1358 | 4.95 | 1.00 | 107.22 |  |
  |

  
 Contact(s): /OP2/  
 Coded contact list: O_ph_  
 Exclusion zone contact(s): /N4/  

### MG:BA:1740

|  | Res. type | Atom | Residue | Distance | Occupancy | B-factor | Closest solute to 1^st^ shell water |
| --- | --- | --- | --- | --- | --- | --- | --- |
| **Ion** | | | |  | 1.00 | 42.89 || 1 | G | O6 | BA:1482 | 2.69 | 1.00 | 52.16 |
| 2 | G | O6 | BA:1419 | 2.75 | 1.00 | 56.69 |  |
| 3 | U | O4 | BA:1481 | 2.82 | 1.00 | 59.52 |  |
| 4 | G | O6 | BA:1480 | 3.78 | 1.00 | 69.27 |  |
| 5 | U | N3 | BA:1481 | 3.90 | 1.00 | 60.84 |  |
| 6 | G | N1 | BA:1482 | 4.10 | 1.00 | 48.41 |  |
| 7 | C | N4 | BA:1420 | 4.27 | 1.00 | 67.63 |  |
| 8 | G | N7 | BA:1419 | 4.46 | 1.00 | 55.81 |  |
  |

  
 Coded contact list:   
 Exclusion zone contact(s): /O4/O6/O6/  
Consider K in place of MG (3 valid contact(s) for K vs. 0 for MG)  

### MG:BA:1741

|  | Res. type | Atom | Residue | Distance | Occupancy | B-factor | Closest solute to 1^st^ shell water |
| --- | --- | --- | --- | --- | --- | --- | --- |
| **Ion** | | | |  | 1.00 | 85.49 || 1 | G | O6 | BA:1423 | 4.06 | 1.00 | 72.47 |
| 2 | G | O6 | BA:1422 | 4.14 | 1.00 | 72.41 |  |
| 3 | G | N7 | BA:1422 | 4.83 | 1.00 | 73.87 |  |
| 4 | C | N4 | BA:1477 | 4.90 | 1.00 | 72.19 |  |
  |

  
 Coded contact list:   
1^st^ MG coordination shell missing (see Notes)  

### MG:BA:1742

|  | Res. type | Atom | Residue | Distance | Occupancy | B-factor | Closest solute to 1^st^ shell water |
| --- | --- | --- | --- | --- | --- | --- | --- |
| **Ion** | | | |  | 1.00 | 46.25 || 1 | C | OP1 | BA:934 | 2.42 | 1.00 | 86.01 |
| 2 | U | OP1 | BA:1345 | 3.76 | 1.00 | 86.18 |  |
| 3 | C | O5' | BA:934 | 4.03 | 1.00 | 86.18 |  |
| 4 | U | OP2 | BA:1345 | 4.31 | 1.00 | 86.76 |  |
| 5 | U | O5' | BA:1345 | 4.40 | 1.00 | 86.62 |  |
| 6 | G | O3' | BA:933 | 4.67 | 1.00 | 86.02 |  |
| 7 | C | OP2 | BA:934 | 4.85 | 1.00 | 85.43 |  |
  |

  
 Coded contact list:   
 Exclusion zone contact(s): /OP1/  
Consider NA in place of MG (1 valid contact(s) for NA vs. 0 for MG)  

### MG:BA:1743

|  | Res. type | Atom | Residue | Distance | Occupancy | B-factor | Closest solute to 1^st^ shell water |
| --- | --- | --- | --- | --- | --- | --- | --- |
| **Ion** | | | |  | 1.00 | 43.40 || 1 | G | O6 | BA:922 | 2.33 | 1.00 | 68.87 |
| 2 | U | O4 | BA:1393 | 2.84 | 1.00 | 68.64 |  |
| 3 | U | O4 | BA:921 | 3.00 | 1.00 | 74.46 |  |
| 4 | A | N6 | BA:923 | 3.55 | 1.00 | 69.39 |  |
| 5 | G | N7 | BA:922 | 3.62 | 1.00 | 70.97 |  |
| 6 | C | N4 | BA:1395 | 4.06 | 1.00 | 65.69 |  |
| 7 | A | N6 | BA:1396 | 4.08 | 1.00 | 73.20 |  |
| 8 | U | N3 | BA:1393 | 4.18 | 1.00 | 67.95 |  |
| 9 | G | N1 | BA:922 | 4.55 | 1.00 | 69.06 |  |
| 10 | U | N3 | BA:921 | 4.60 | 1.00 | 74.57 |  |
| 11 | A | N1 | BA:923 | 4.62 | 1.00 | 69.88 |  |
| 12 | U | O4 | BA:920 | 4.97 | 1.00 | 73.84 |  |
  |

  
 Contact(s): /O6/  
 Coded contact list: O_b_  
 Exclusion zone contact(s): /O4/O4/  
Consider K in place of MG (2 valid contact(s) for K vs. 1 for MG)  

### MG:BA:1744

|  | Res. type | Atom | Residue | Distance | Occupancy | B-factor | Closest solute to 1^st^ shell water |
| --- | --- | --- | --- | --- | --- | --- | --- |
| **Ion** | | | |  | 1.00 | 54.38 || 1 | C | OP1 | BA:1200 | 3.19 | 1.00 | 93.09 |
| 2 | C | O2' | BA:962 | 3.28 | 1.00 | 84.52 |  |
| 3 | G | O4' | BA:963 | 4.27 | 1.00 | 78.27 |  |
| 4 | C | OP2 | BA:1200 | 4.59 | 1.00 | 93.54 |  |
| 5 | A | N7 | BA:1201 | 4.80 | 1.00 | 94.61 |  |
| 6 | U | OP2 | BA:1052 | 4.90 | 1.00 | 95.32 |  |
  |

  
 Coded contact list:   
 Exclusion zone contact(s): /O2'/OP1/  
Consider K in place of MG (2 valid contact(s) for K vs. 0 for MG)  

### MG:BA:1745

|  | Res. type | Atom | Residue | Distance | Occupancy | B-factor | Closest solute to 1^st^ shell water |
| --- | --- | --- | --- | --- | --- | --- | --- |
| **Ion** | | | |  | 1.00 | 38.27 || 1 | A | O3' | BA:509 | 2.56 | 1.00 | 80.86 |
| 2 | A | OP2 | BA:509 | 2.70 | 1.00 | 82.18 |  |
| 3 | C | O3' | BA:508 | 3.50 | 1.00 | 83.06 |  |
| 4 | G | OP1 | BA:506 | 3.54 | 1.00 | 73.16 |  |
| 5 | A | OP2 | BA:510 | 3.57 | 1.00 | 79.68 |  |
| 6 | A | N7 | BA:510 | 3.73 | 1.00 | 77.64 |  |
| 7 | A | O5' | BA:510 | 3.95 | 1.00 | 77.53 |  |
| 8 | C | O2' | BA:508 | 4.02 | 1.00 | 85.17 |  |
| 9 | A | O5' | BA:509 | 4.12 | 1.00 | 82.61 |  |
| 10 | C | OP2 | BA:507 | 4.32 | 1.00 | 74.60 |  |
| 11 | G | OP2 | BA:506 | 4.79 | 1.00 | 71.67 |  |
| 12 | A | OP1 | BA:510 | 4.89 | 1.00 | 80.28 |  |
  |

  
 Coded contact list:   
 Exclusion zone contact(s): /O3'/OP2/  
Consider NA in place of MG (1 valid contact(s) for NA vs. 0 for MG)  

### MG:BA:1746

|  | Res. type | Atom | Residue | Distance | Occupancy | B-factor | Closest solute to 1^st^ shell water |
| --- | --- | --- | --- | --- | --- | --- | --- |
| **Ion** | | | |  | 1.00 | 40.28 || 1 | C | O2' | BA:817 | 2.55 | 1.00 | 52.35 |
| 2 | C | O3' | BA:817 | 2.56 | 1.00 | 50.07 |  |
| 3 | U | OP1 | BA:1528 | 2.77 | 1.00 | 63.93 |  |
| 4 | G | OP2 | BA:818 | 2.86 | 1.00 | 47.45 |  |
| 5 | G | O5' | BA:818 | 3.30 | 1.00 | 50.26 |  |
| 6 | C | O3' | BA:1527 | 3.46 | 1.00 | 63.17 |  |
| 7 | U | O5' | BA:1528 | 4.55 | 1.00 | 64.61 |  |
| 8 | G | OP1 | BA:818 | 4.56 | 1.00 | 48.10 |  |
| 9 | U | OP2 | BA:1528 | 4.98 | 1.00 | 63.30 |  |
  |

  
 Coded contact list:   
 Exclusion zone contact(s): /O2'/O3'/O5'/OP1/OP2/  
Consider K in place of MG (3 valid contact(s) for K vs. 0 for MG)  

### MG:BA:1747

|  | Res. type | Atom | Residue | Distance | Occupancy | B-factor | Closest solute to 1^st^ shell water |
| --- | --- | --- | --- | --- | --- | --- | --- |
| **Ion** | | | |  | 1.00 | 50.69 || 1 | G | OP1 | BA:21 | 1.76 | 1.00 | 55.64 |
| 2 | G | OP2 | BA:21 | 3.56 | 1.00 | 56.93 |  |
| 3 | U | O3' | BA:20 | 3.62 | 1.00 | 56.92 |  |
| 4 | A | N6 | BA:573 | 3.91 | 1.00 | 63.40 |  |
| 5 | A | N1 | BA:573 | 4.05 | 1.00 | 64.69 |  |
| 6 | G | O2' | BA:568 | 4.07 | 1.00 | 57.98 |  |
| 7 | G | N2 | BA:567 | 4.18 | 1.00 | 60.58 |  |
| 8 | G | O5' | BA:21 | 4.23 | 1.00 | 57.30 |  |
| 9 | G | N3 | BA:567 | 4.43 | 1.00 | 60.81 |  |
| 10 | G | O2' | BA:567 | 4.64 | 1.00 | 59.62 |  |
  |

  
 Coded contact list:   
 Exclusion zone contact(s): /OP1/  
1^st^ MG coordination shell missing (see Notes)  

### MG:BA:1748

|  | Res. type | Atom | Residue | Distance | Occupancy | B-factor | Closest solute to 1^st^ shell water |
| --- | --- | --- | --- | --- | --- | --- | --- |
| **Ion** | | | |  | 1.00 | 66.96 || 1 | G | O2' | BA:484 | 2.48 | 1.00 | 82.17 |
| 2 | G | OP2 | BA:485 | 3.30 | 1.00 | 91.84 |  |
| 3 | A | OP2 | BA:482 | 3.37 | 1.00 | 82.26 |  |
| 4 | C | OP2 | BA:483 | 3.90 | 1.00 | 74.75 |  |
| 5 | G | O3' | BA:481 | 3.94 | 1.00 | 83.25 |  |
| 6 | G | O3' | BA:484 | 4.32 | 1.00 | 88.60 |  |
| 7 | G | O5' | BA:485 | 4.45 | 1.00 | 95.46 |  |
| 8 | G | N3 | BA:484 | 4.88 | 1.00 | 76.02 |  |
| 9 | G | O2' | BA:481 | 4.88 | 1.00 | 87.34 |  |
  |

  
 Coded contact list:   
 Exclusion zone contact(s): /O2'/OP2/OP2/  
Consider K in place of MG (2 valid contact(s) for K vs. 0 for MG)  

### MG:BA:1749

|  | Res. type | Atom | Residue | Distance | Occupancy | B-factor | Closest solute to 1^st^ shell water |
| --- | --- | --- | --- | --- | --- | --- | --- |
| **Ion** | | | |  | 1.00 | 30.94 || 1 | G | O2' | BA:266 | 1.82 | 1.00 | 74.64 |
| 2 | C | OP2 | BA:267 | 2.67 | 1.00 | 76.00 |  |
| 3 | G | O2' | BA:251 | 2.79 | 1.00 | 76.40 |  |
| 4 | G | O3' | BA:251 | 3.31 | 1.00 | 72.51 |  |
| 5 | C | OP2 | BA:268 | 3.48 | 1.00 | 70.27 |  |
| 6 | U | OP2 | BA:252 | 3.69 | 1.00 | 70.48 |  |
| 7 | C | OP1 | BA:268 | 3.71 | 1.00 | 70.19 |  |
| 8 | G | O3' | BA:266 | 3.75 | 1.00 | 75.01 |  |
| 9 | C | O5' | BA:267 | 4.27 | 1.00 | 74.34 |  |
| 10 | U | OP2 | BA:253 | 4.40 | 1.00 | 66.30 |  |
| 11 | LYS | NZ | BT:67 | 4.58 | 1.00 | 55.32 |  |
| 12 | C | O3' | BA:267 | 4.61 | 1.00 | 70.98 |  |
| 13 | G | N9 | BA:266 | 4.79 | 1.00 | 72.19 |  |
| 14 | U | O5' | BA:252 | 4.88 | 1.00 | 69.65 |  |
  |

  
 Contact(s): /O2'/  
 Coded contact list: O_r_  
 Exclusion zone contact(s): /O2'/O3'/OP2/  
Consider K in place of MG (3 valid contact(s) for K vs. 1 for MG)  

### MG:BA:1750

|  | Res. type | Atom | Residue | Distance | Occupancy | B-factor | Closest solute to 1^st^ shell water |
| --- | --- | --- | --- | --- | --- | --- | --- |
| **Ion** | | | |  | 1.00 | 65.64 || 1 | G | O6 | BA:105 | 2.51 | 1.00 | 66.32 |
| 2 | G | O6 | BA:61 | 2.93 | 1.00 | 70.24 |  |
| 3 | U | O4 | BA:62 | 3.15 | 1.00 | 71.86 |  |
| 4 | C | N4 | BA:106 | 3.58 | 1.00 | 68.82 |  |
| 5 | U | N3 | BA:62 | 3.93 | 1.00 | 70.46 |  |
| 6 | G | O6 | BA:104 | 3.98 | 1.00 | 72.75 |  |
| 7 | G | N7 | BA:105 | 4.15 | 1.00 | 65.65 |  |
| 8 | C | N4 | BA:63 | 4.43 | 1.00 | 74.27 |  |
| 9 | G | N1 | BA:105 | 4.70 | 1.00 | 67.43 |  |
| 10 | G | N1 | BA:61 | 4.80 | 1.00 | 69.76 |  |
| 11 | G | N7 | BA:61 | 4.96 | 1.00 | 68.71 |  |
  |

  
 Coded contact list:   
 Exclusion zone contact(s): /O4/O6/O6/  
Consider K in place of MG (2 valid contact(s) for K vs. 0 for MG)  

### MG:BA:1751

|  | Res. type | Atom | Residue | Distance | Occupancy | B-factor | Closest solute to 1^st^ shell water |
| --- | --- | --- | --- | --- | --- | --- | --- |
| **Ion** | | | |  | 1.00 | 62.84 || 1 | G | OP2 | BA:128 | 1.87 | 1.00 | 72.49 |
| 2 | G | O5' | BA:128 | 3.78 | 1.00 | 72.78 |  |
| 3 | G | N7 | BA:128 | 3.86 | 1.00 | 68.41 |  |
| 4 | G | O3' | BA:127 | 4.06 | 1.00 | 73.52 |  |
| 5 | G | OP1 | BA:128 | 4.40 | 1.00 | 72.57 |  |
  |

  
 Contact(s): /OP2/  
 Coded contact list: O_ph_  

### MG:BA:1752

|  | Res. type | Atom | Residue | Distance | Occupancy | B-factor | Closest solute to 1^st^ shell water |
| --- | --- | --- | --- | --- | --- | --- | --- |
| **Ion** | | | |  | 1.00 | 48.61 || 1 | U | O4 | BA:129 | 1.97 | 1.00 | 73.91 |
| 2 | G | O6 | BA:232 | 3.13 | 1.00 | 66.73 |  |
| 3 | U | N3 | BA:129 | 3.40 | 1.00 | 74.60 |  |
| 4 | G | O6 | BA:231 | 3.49 | 1.00 | 69.59 |  |
| 5 | G | O6 | BA:128 | 3.73 | 1.00 | 67.09 |  |
| 6 | C | N4 | BA:131 | 3.84 | 1.00 | 69.76 |  |
| 7 | C | N4 | BA:233 | 4.37 | 1.00 | 68.08 |  |
| 8 | G | N7 | BA:232 | 4.89 | 1.00 | 68.92 |  |
  |

  
 Contact(s): /O4/  
 Coded contact list: O_b_  
 Exclusion zone contact(s): /O6/  

### MG:BA:1753

|  | Res. type | Atom | Residue | Distance | Occupancy | B-factor | Closest solute to 1^st^ shell water |
| --- | --- | --- | --- | --- | --- | --- | --- |
| **Ion** | | | |  | 1.00 | 69.02 || 1 | A | OP1 | BA:373 | 2.20 | 1.00 | 77.48 |
| 2 | A | O5' | BA:373 | 2.82 | 1.00 | 76.24 |  |
| 3 | C | O5' | BA:372 | 3.78 | 1.00 | 68.50 |  |
| 4 | C | OP1 | BA:372 | 3.88 | 1.00 | 66.93 |  |
| 5 | G | O3' | BA:371 | 3.99 | 1.00 | 66.29 |  |
| 6 | A | OP2 | BA:373 | 4.14 | 1.00 | 77.21 |  |
| 7 | C | O3' | BA:372 | 4.17 | 1.00 | 76.76 |  |
| 8 | G | O2' | BA:371 | 4.27 | 1.00 | 65.10 |  |
| 9 | A | O4' | BA:373 | 4.32 | 1.00 | 76.35 |  |
| 10 | C | O4' | BA:372 | 4.93 | 1.00 | 74.27 |  |
  |

  
 Contact(s): /OP1/  
 Coded contact list: O_ph_  
 Exclusion zone contact(s): /O5'/  

### MG:BA:1754

|  | Res. type | Atom | Residue | Distance | Occupancy | B-factor | Closest solute to 1^st^ shell water |
| --- | --- | --- | --- | --- | --- | --- | --- |
| **Ion** | | | |  | 1.00 | 68.41 || 1 | G | OP2 | BA:371 | 2.45 | 1.00 | 71.64 |
| 2 | G | O5' | BA:371 | 4.24 | 1.00 | 70.05 |  |
| 3 | C | OP2 | BA:370 | 4.61 | 1.00 | 75.04 |  |
| 4 | G | OP1 | BA:371 | 4.72 | 1.00 | 70.72 |  |
| 5 | C | O3' | BA:370 | 4.89 | 1.00 | 71.64 |  |
| 6 | G | N7 | BA:371 | 4.98 | 1.00 | 66.06 |  |
| 7 | C | OP2 | BA:372 | 4.99 | 1.00 | 67.11 |  |
  |

  
 Coded contact list:   
 Exclusion zone contact(s): /OP2/  
Consider NA in place of MG (1 valid contact(s) for NA vs. 0 for MG)  

### MG:BA:1755

|  | Res. type | Atom | Residue | Distance | Occupancy | B-factor | Closest solute to 1^st^ shell water |
| --- | --- | --- | --- | --- | --- | --- | --- |
| **Ion** | | | |  | 1.00 | 56.45 || 1 | A | OP1 | BA:120 | 3.46 | 1.00 | 70.67 |
| 2 | A | OP2 | BA:120 | 3.58 | 1.00 | 69.83 |  |
| 3 | A | O2' | BA:119 | 3.79 | 1.00 | 68.96 |  |
| 4 | A | O3' | BA:119 | 3.96 | 1.00 | 69.36 |  |
| 5 | C | OP2 | BA:283 | 3.99 | 1.00 | 58.02 |  |
| 6 | A | OP2 | BA:282 | 4.63 | 1.00 | 65.77 |  |
  |

  
 Coded contact list:   
1^st^ MG coordination shell missing (see Notes)  

### MG:BA:1756

|  | Res. type | Atom | Residue | Distance | Occupancy | B-factor | Closest solute to 1^st^ shell water |
| --- | --- | --- | --- | --- | --- | --- | --- |
| **Ion** | | | |  | 1.00 | 69.09 || 1 | G | O6 | BA:284 | 3.15 | 1.00 | 68.78 |
| 2 | G | N7 | BA:284 | 3.35 | 1.00 | 64.72 |  |
| 3 | G | O6 | BA:285 | 3.83 | 1.00 | 70.64 |  |
| 4 | A | OP2 | BA:282 | 3.92 | 1.00 | 65.77 |  |
| 5 | C | N4 | BA:283 | 4.53 | 1.00 | 69.56 |  |
| 6 | G | N7 | BA:285 | 4.61 | 1.00 | 65.79 |  |
  |

  
 Coded contact list:   
 Exclusion zone contact(s): /N7/O6/  
Consider K in place of MG (2 valid contact(s) for K vs. 0 for MG)  

### MG:BA:1757

|  | Res. type | Atom | Residue | Distance | Occupancy | B-factor | Closest solute to 1^st^ shell water |
| --- | --- | --- | --- | --- | --- | --- | --- |
| **Ion** | | | |  | 1.00 | 39.59 || 1 | G | N7 | BA:286 | 3.34 | 1.00 | 68.69 |
| 2 | U | O4 | BA:287 | 3.47 | 1.00 | 72.75 |  |
| 3 | G | O6 | BA:286 | 3.69 | 1.00 | 70.02 |  |
| 4 | G | N7 | BA:285 | 3.91 | 1.00 | 65.79 |  |
| 5 | G | O6 | BA:285 | 4.11 | 1.00 | 70.64 |  |
| 6 | A | OP2 | BA:119 | 4.82 | 1.00 | 67.93 |  |
  |

  
 Coded contact list:   
 Exclusion zone contact(s): /N7/  
Consider K in place of MG (1 valid contact(s) for K vs. 0 for MG)  

### MG:BA:1758

|  | Res. type | Atom | Residue | Distance | Occupancy | B-factor | Closest solute to 1^st^ shell water |
| --- | --- | --- | --- | --- | --- | --- | --- |
| **Ion** | | | |  | 1.00 | 54.92 || 1 | G | O6 | BA:258 | 2.65 | 1.00 | 75.73 |
| 2 | G | O6 | BA:257 | 3.29 | 1.00 | 83.13 |  |
| 3 | G | N7 | BA:258 | 3.51 | 1.00 | 77.52 |  |
| 4 | G | OP2 | BA:266 | 3.66 | 1.00 | 69.42 |  |
| 5 | G | N7 | BA:257 | 4.01 | 1.00 | 81.80 |  |
| 6 | C | N4 | BA:268 | 4.42 | 1.00 | 75.32 |  |
| 7 | G | OP1 | BA:266 | 4.81 | 1.00 | 69.90 |  |
| 8 | G | N1 | BA:258 | 4.84 | 1.00 | 75.04 |  |
| 9 | G | O6 | BA:259 | 4.85 | 1.00 | 69.42 |  |
| 10 | G | N1 | BA:257 | 4.87 | 1.00 | 81.88 |  |
  |

  
 Coded contact list:   
 Exclusion zone contact(s): /O6/O6/  
Consider K in place of MG (2 valid contact(s) for K vs. 0 for MG)  

### MG:BA:1759

|  | Res. type | Atom | Residue | Distance | Occupancy | B-factor | Closest solute to 1^st^ shell water |
| --- | --- | --- | --- | --- | --- | --- | --- |
| **Ion** | | | |  | 1.00 | 77.05 || 1 | U | OP1 | BA:1281 | 2.14 | 1.00 | 110.25 |
| 2 | A | O3' | BA:1280 | 2.84 | 1.00 | 109.87 |  |
| 3 | U | O5' | BA:1281 | 3.79 | 1.00 | 112.21 |  |
| 4 | A | O2' | BA:1280 | 3.84 | 1.00 | 108.02 |  |
| 5 | U | OP2 | BA:1281 | 4.30 | 1.00 | 110.49 |  |
| 6 | C | N4 | BA:1282 | 4.61 | 1.00 | 118.60 |  |
| 7 | G | O6 | BA:1283 | 4.81 | 1.00 | 116.94 |  |
| 8 | G | N7 | BA:1283 | 4.95 | 1.00 | 117.47 |  |
  |

  
 Contact(s): /OP1/  
 Coded contact list: O_ph_  
 Exclusion zone contact(s): /O3'/  

### MG:BA:1760

|  | Res. type | Atom | Residue | Distance | Occupancy | B-factor | Closest solute to 1^st^ shell water |
| --- | --- | --- | --- | --- | --- | --- | --- |
| **Ion** | | | |  | 1.00 | 85.89 || 1 | G | OP2 | BA:183 | 2.24 | 1.00 | 103.42 |
| 2 | U | OP1 | BA:182 | 2.75 | 1.00 | 102.16 |  |
| 3 | U | OP2 | BA:182 | 2.81 | 1.00 | 102.31 |  |
| 4 | G | N7 | BA:183 | 3.57 | 1.00 | 97.96 |  |
| 5 | U | O5' | BA:182 | 3.68 | 1.00 | 101.88 |  |
| 6 | G | OP1 | BA:183 | 4.29 | 1.00 | 103.27 |  |
| 7 | U | O3' | BA:182 | 4.29 | 1.00 | 103.09 |  |
| 8 | G | O3' | BA:181 | 4.62 | 1.00 | 101.93 |  |
| 9 | G | O5' | BA:183 | 4.72 | 1.00 | 102.66 |  |
| 10 | G | N9 | BA:183 | 4.90 | 1.00 | 98.57 |  |
| 11 | G | O2' | BA:181 | 4.91 | 1.00 | 102.43 |  |
  |

  
 Contact(s): /OP2/  
 Coded contact list: O_ph_  
 Exclusion zone contact(s): /OP1/OP2/  
Consider K in place of MG (2 valid contact(s) for K vs. 1 for MG)  

### MG:BA:1761

|  | Res. type | Atom | Residue | Distance | Occupancy | B-factor | Closest solute to 1^st^ shell water |
| --- | --- | --- | --- | --- | --- | --- | --- |
| **Ion** | | | |  | 1.00 | 58.50 || 1 | U | OP2 | BA:182 | 2.60 | 1.00 | 102.31 |
| 2 | G | N7 | BA:183 | 2.75 | 1.00 | 97.96 |  |
| 3 | G | O6 | BA:183 | 2.99 | 1.00 | 96.43 |  |
| 4 | G | O6 | BA:184 | 4.28 | 1.00 | 88.24 |  |
| 5 | U | OP1 | BA:182 | 4.48 | 1.00 | 102.16 |  |
| 6 | G | O3' | BA:181 | 4.63 | 1.00 | 101.93 |  |
| 7 | G | O2' | BA:181 | 4.73 | 1.00 | 102.43 |  |
| 8 | G | N7 | BA:184 | 4.81 | 1.00 | 90.12 |  |
| 9 | G | N1 | BA:183 | 4.93 | 1.00 | 95.56 |  |
| 10 | G | N9 | BA:183 | 4.99 | 1.00 | 98.57 |  |
  |

  
 Coded contact list:   
 Exclusion zone contact(s): /N7/O6/OP2/  
Consider K in place of MG (2 valid contact(s) for K vs. 0 for MG)  

### MG:BA:1762

|  | Res. type | Atom | Residue | Distance | Occupancy | B-factor | Closest solute to 1^st^ shell water |
| --- | --- | --- | --- | --- | --- | --- | --- |
| **Ion** | | | |  | 1.00 | 78.54 || 1 | G | N7 | BA:191E | 3.50 | 1.00 | 103.90 |
| 2 | G | O6 | BA:191 | 4.26 | 1.00 | 91.01 |  |
| 3 | G | O6 | BA:191E | 4.34 | 1.00 | 102.12 |  |
| 4 | C | N4 | BA:186 | 4.72 | 1.00 | 90.29 |  |
| 5 | G | OP2 | BA:191E | 4.95 | 1.00 | 109.72 |  |
| 6 | U | O4 | BA:191F | 4.98 | 1.00 | 103.76 |  |
  |

  
 Coded contact list:   
1^st^ MG coordination shell missing (see Notes)  

### MG:BA:1763

|  | Res. type | Atom | Residue | Distance | Occupancy | B-factor | Closest solute to 1^st^ shell water |
| --- | --- | --- | --- | --- | --- | --- | --- |
| **Ion** | | | |  | 1.00 | 57.63 || 1 | G | OP1 | BA:945 | 1.96 | 1.00 | 80.33 |
| 2 | C | OP2 | BA:1230 | 3.62 | 1.00 | 77.06 |  |
| 3 | G | O3' | BA:944 | 3.88 | 1.00 | 80.44 |  |
| 4 | G | OP2 | BA:945 | 3.93 | 1.00 | 81.15 |  |
| 5 | C | OP1 | BA:1230 | 3.93 | 1.00 | 77.36 |  |
| 6 | ARG | NH1 | BP:114 | 4.06 | 1.00 | 92.32 |  |
| 7 | A | OP2 | BA:946 | 4.19 | 1.00 | 80.68 |  |
| 8 | ARG | NH2 | BP:114 | 4.39 | 1.00 | 92.47 |  |
| 9 | G | O5' | BA:945 | 4.46 | 1.00 | 80.34 |  |
| 10 | C | O5' | BA:1230 | 4.69 | 1.00 | 76.99 |  |
  |

  
 Contact(s): /OP1/  
 Coded contact list: O_ph_  

### MG:BA:1764

|  | Res. type | Atom | Residue | Distance | Occupancy | B-factor | Closest solute to 1^st^ shell water |
| --- | --- | --- | --- | --- | --- | --- | --- |
| **Ion** | | | |  | 1.00 | 50.16 || 1 | G | OP1 | BA:944 | 2.04 | 1.00 | 80.91 |
| 2 | G | OP2 | BA:945 | 2.46 | 1.00 | 81.15 |  |
| 3 | G | OP2 | BA:944 | 3.03 | 1.00 | 81.11 |  |
| 4 | A | N7 | BA:946 | 3.54 | 1.00 | 82.99 |  |
| 5 | G | O5' | BA:944 | 3.56 | 1.00 | 81.47 |  |
| 6 | G | O5' | BA:945 | 4.04 | 1.00 | 80.34 |  |
| 7 | G | N7 | BA:945 | 4.15 | 1.00 | 89.17 |  |
| 8 | U | O3' | BA:943 | 4.25 | 1.00 | 82.27 |  |
| 9 | A | N6 | BA:946 | 4.58 | 1.00 | 84.88 |  |
| 10 | A | OP2 | BA:946 | 4.61 | 1.00 | 80.68 |  |
| 11 | G | OP1 | BA:945 | 4.73 | 1.00 | 80.33 |  |
| 12 | G | O6 | BA:947 | 4.75 | 1.00 | 87.88 |  |
| 13 | G | O3' | BA:944 | 4.81 | 1.00 | 80.44 |  |
| 14 | G | N9 | BA:945 | 5.00 | 1.00 | 87.10 |  |
  |

  
 Contact(s): /OP1/  
 Coded contact list: O_ph_  
 Exclusion zone contact(s): /OP2/OP2/  

### MG:BA:1765

|  | Res. type | Atom | Residue | Distance | Occupancy | B-factor | Closest solute to 1^st^ shell water |
| --- | --- | --- | --- | --- | --- | --- | --- |
| **Ion** | | | |  | 1.00 | 82.80 || 1 | G | N7 | BA:1231 | 2.42 | 1.00 | 72.58 |
| 2 | G | OP2 | BA:1231 | 2.95 | 1.00 | 75.14 |  |
| 3 | G | O5' | BA:1231 | 3.98 | 1.00 | 76.15 |  |
| 4 | C | N3 | BA:1230 | 4.03 | 1.00 | 74.35 |  |
| 5 | G | N9 | BA:1231 | 4.18 | 1.00 | 72.77 |  |
| 6 | C | N1 | BA:1230 | 4.22 | 1.00 | 75.78 |  |
| 7 | U | O4 | BA:1232 | 4.37 | 1.00 | 81.59 |  |
| 8 | C | N4 | BA:1230 | 4.50 | 1.00 | 73.62 |  |
| 9 | G | O6 | BA:1231 | 4.57 | 1.00 | 74.81 |  |
| 10 | C | O3' | BA:1230 | 4.70 | 1.00 | 76.33 |  |
| 11 | C | O2 | BA:1230 | 4.74 | 1.00 | 75.62 |  |
  |

  
 Coded contact list:   
 Exclusion zone contact(s): /N7/OP2/  
Consider NA in place of MG (1 valid contact(s) for NA vs. 0 for MG)  

### MG:BA:1766

|  | Res. type | Atom | Residue | Distance | Occupancy | B-factor | Closest solute to 1^st^ shell water |
| --- | --- | --- | --- | --- | --- | --- | --- |
| **Ion** | | | |  | 1.00 | 74.70 || 1 | G | OP1 | BA:1231 | 2.53 | 1.00 | 76.49 |
| 2 | A | OP1 | BC:31 | 2.81 | 1.00 | 79.38 |  |
| 3 | C | O3' | BA:1230 | 3.70 | 1.00 | 76.33 |  |
| 4 | A | O5' | BC:31 | 3.92 | 1.00 | 77.74 |  |
| 5 | G | O5' | BA:1231 | 4.58 | 1.00 | 76.15 |  |
| 6 | U | OP2 | BC:32 | 4.65 | 1.00 | 72.84 |  |
| 7 | A | O3' | BC:31 | 4.67 | 1.00 | 72.85 |  |
| 8 | G | O3' | BC:30 | 4.68 | 1.00 | 77.32 |  |
| 9 | G | OP2 | BA:1231 | 4.87 | 1.00 | 75.14 |  |
  |

  
 Coded contact list:   
 Exclusion zone contact(s): /OP1/OP1/  
Consider NA in place of MG (1 valid contact(s) for NA vs. 0 for MG)  

### MG:BA:1767

|  | Res. type | Atom | Residue | Distance | Occupancy | B-factor | Closest solute to 1^st^ shell water |
| --- | --- | --- | --- | --- | --- | --- | --- |
| **Ion** | | | |  | 1.00 | 50.51 || 1 | G | N7 | BA:309 | 2.81 | 1.00 | 66.43 |
| 2 | G | O6 | BA:309 | 3.11 | 1.00 | 61.67 |  |
| 3 | G | O6 | BA:310 | 4.31 | 1.00 | 57.88 |  |
| 4 | G | N7 | BA:310 | 4.44 | 1.00 | 60.98 |  |
| 5 | C | N4 | BA:308 | 4.46 | 1.00 | 70.19 |  |
| 6 | C | OP2 | BA:307 | 4.73 | 1.00 | 87.01 |  |
| 7 | C | OP2 | BA:308 | 4.73 | 1.00 | 83.36 |  |
| 8 | G | N1 | BA:309 | 4.80 | 1.00 | 61.83 |  |
| 9 | G | N9 | BA:309 | 4.83 | 1.00 | 65.57 |  |
| 10 | C | N3 | BA:308 | 4.89 | 1.00 | 70.04 |  |
| 11 | G | OP2 | BA:309 | 4.94 | 1.00 | 73.91 |  |
  |

  
 Coded contact list:   
 Exclusion zone contact(s): /N7/O6/  
Consider K in place of MG (2 valid contact(s) for K vs. 0 for MG)  

### MG:BA:1768

|  | Res. type | Atom | Residue | Distance | Occupancy | B-factor | Closest solute to 1^st^ shell water |
| --- | --- | --- | --- | --- | --- | --- | --- |
| **Ion** | | | |  | 1.00 | 75.29 || 1 | U | OP1 | BA:1506 | 2.60 | 1.00 | 67.78 |
| 2 | U | OP2 | BA:1506 | 3.62 | 1.00 | 68.29 |  |
| 3 | G | O3' | BA:1505 | 4.25 | 1.00 | 68.70 |  |
| 4 | U | O5' | BA:1506 | 4.88 | 1.00 | 66.18 |  |
  |

  
 Coded contact list:   
 Exclusion zone contact(s): /OP1/  
Consider NA in place of MG (1 valid contact(s) for NA vs. 0 for MG)  

### MG:BA:1769

|  | Res. type | Atom | Residue | Distance | Occupancy | B-factor | Closest solute to 1^st^ shell water |
| --- | --- | --- | --- | --- | --- | --- | --- |
| **Ion** | | | |  | 1.00 | 87.04 || 1 | G | OP2 | BA:1442 | 3.06 | 1.00 | 94.45 |
| 2 | G | OP1 | BA:1442 | 4.38 | 1.00 | 94.32 |  |
| 3 | G | O3' | BA:1441 | 4.71 | 1.00 | 92.17 |  |
| 4 | G | O6 | BA:1454 | 4.80 | 1.00 | 106.62 |  |
  |

  
 Coded contact list:   
 Exclusion zone contact(s): /OP2/  
Consider K in place of MG (1 valid contact(s) for K vs. 0 for MG)  

### MG:BA:1770

|  | Res. type | Atom | Residue | Distance | Occupancy | B-factor | Closest solute to 1^st^ shell water |
| --- | --- | --- | --- | --- | --- | --- | --- |
| **Ion** | | | |  | 1.00 | 67.47 || 1 | G | O6 | BA:925 | 2.82 | 1.00 | 69.06 |
| 2 | G | O6 | BA:927 | 2.95 | 1.00 | 70.17 |  |
| 3 | C | N4 | BA:924 | 3.36 | 1.00 | 62.68 |  |
| 4 | U | O4 | BA:1390 | 3.45 | 1.00 | 67.31 |  |
| 5 | U | O4 | BA:1391 | 3.75 | 1.00 | 70.16 |  |
| 6 | G | N7 | BA:925 | 3.91 | 1.00 | 67.08 |  |
| 7 | G | N7 | BA:927 | 4.37 | 1.00 | 69.77 |  |
| 8 | U | N3 | BA:1390 | 4.42 | 1.00 | 67.58 |  |
| 9 | U | N3 | BA:1391 | 4.48 | 1.00 | 67.93 |  |
| 10 | C | N3 | BA:924 | 4.65 | 1.00 | 63.41 |  |
| 11 | G | O6 | BA:1392 | 4.87 | 1.00 | 61.66 |  |
| 12 | G | O6 | BA:928 | 4.88 | 1.00 | 74.36 |  |
| 13 | G | N1 | BA:925 | 4.91 | 1.00 | 68.15 |  |
  |

  
 Coded contact list:   
 Exclusion zone contact(s): /N4/O6/O6/  
Consider K in place of MG (3 valid contact(s) for K vs. 0 for MG)  

### MG:BA:1771

|  | Res. type | Atom | Residue | Distance | Occupancy | B-factor | Closest solute to 1^st^ shell water |
| --- | --- | --- | --- | --- | --- | --- | --- |
| **Ion** | | | |  | 1.00 | 74.32 || 1 | G | OP2 | BA:925 | 3.96 | 1.00 | 69.43 |
| 2 | G | N7 | BA:927 | 4.52 | 1.00 | 69.77 |  |
| 3 | G | N7 | BA:925 | 4.55 | 1.00 | 67.08 |  |
| 4 | G | OP2 | BA:926 | 4.66 | 1.00 | 66.46 |  |
| 5 | G | O5' | BA:925 | 4.79 | 1.00 | 68.81 |  |
| 6 | G | OP1 | BA:926 | 4.95 | 1.00 | 66.23 |  |
  |

  
 Coded contact list:   
1^st^ MG coordination shell missing (see Notes)  

### MG:BA:1772

|  | Res. type | Atom | Residue | Distance | Occupancy | B-factor | Closest solute to 1^st^ shell water |
| --- | --- | --- | --- | --- | --- | --- | --- |
| **Ion** | | | |  | 1.00 | 75.46 || 1 | G | N7 | BA:928 | 2.99 | 1.00 | 73.38 |
| 2 | G | OP2 | BA:928 | 3.85 | 1.00 | 70.14 |  |
| 3 | G | O6 | BA:928 | 4.05 | 1.00 | 74.36 |  |
| 4 | G | N7 | BA:927 | 4.38 | 1.00 | 69.77 |  |
| 5 | G | N7 | BA:929 | 4.78 | 1.00 | 74.42 |  |
  |

  
 Coded contact list:   
 Exclusion zone contact(s): /N7/  
Consider K in place of MG (1 valid contact(s) for K vs. 0 for MG)  

### MG:BA:1773

|  | Res. type | Atom | Residue | Distance | Occupancy | B-factor | Closest solute to 1^st^ shell water |
| --- | --- | --- | --- | --- | --- | --- | --- |
| **Ion** | | | |  | 1.00 | 80.55 || 1 | C | OP2 | BA:1382 | 3.77 | 1.00 | 90.49 |
| 2 | C | N4 | BA:931 | 4.39 | 1.00 | 86.88 |  |
  |

  
 Coded contact list:   
1^st^ MG coordination shell missing (see Notes)  

### MG:BA:1774

|  | Res. type | Atom | Residue | Distance | Occupancy | B-factor | Closest solute to 1^st^ shell water |
| --- | --- | --- | --- | --- | --- | --- | --- |
| **Ion** | | | |  | 1.00 | 53.26 || 1 | G | OP2 | BA:107 | 2.81 | 1.00 | 70.20 |
| 2 | A | N7 | BA:325 | 3.13 | 1.00 | 59.67 |  |
| 3 | G | O2' | BA:324 | 3.39 | 1.00 | 72.09 |  |
| 4 | G | O6 | BA:326 | 3.52 | 1.00 | 67.72 |  |
| 5 | G | OP1 | BA:107 | 3.82 | 1.00 | 69.53 |  |
| 6 | G | O3' | BA:324 | 3.96 | 1.00 | 70.32 |  |
| 7 | C | O3' | BA:106 | 4.33 | 1.00 | 68.64 |  |
| 8 | A | OP1 | BA:325 | 4.38 | 1.00 | 69.54 |  |
| 9 | G | N7 | BA:326 | 4.59 | 1.00 | 66.68 |  |
| 10 | C | OP1 | BA:106 | 4.61 | 1.00 | 67.24 |  |
| 11 | A | N6 | BA:325 | 4.92 | 1.00 | 58.41 |  |
| 12 | A | O2' | BA:134 | 4.93 | 1.00 | 66.90 |  |
| 13 | C | O5' | BA:106 | 4.93 | 1.00 | 67.73 |  |
| 14 | A | N9 | BA:325 | 5.00 | 1.00 | 62.78 |  |
  |

  
 Coded contact list:   
 Exclusion zone contact(s): /N7/O2'/OP2/  
Consider K in place of MG (3 valid contact(s) for K vs. 0 for MG)  

### MG:BA:1775

|  | Res. type | Atom | Residue | Distance | Occupancy | B-factor | Closest solute to 1^st^ shell water |
| --- | --- | --- | --- | --- | --- | --- | --- |
| **Ion** | | | |  | 1.00 | 93.15 || 1 | G | OP1 | BA:1361 | 2.86 | 1.00 | 96.19 |
| 2 | G | OP2 | BA:1361 | 3.15 | 1.00 | 97.18 |  |
| 3 | C | O2 | BA:1359 | 3.70 | 1.00 | 89.21 |  |
| 4 | A | O3' | BA:1360 | 4.05 | 1.00 | 95.60 |  |
| 5 | C | N3 | BA:1359 | 4.36 | 1.00 | 90.18 |  |
| 6 | G | O5' | BA:1361 | 4.85 | 1.00 | 97.47 |  |
  |

  
 Coded contact list:   
 Exclusion zone contact(s): /OP1/OP2/  
Consider K in place of MG (2 valid contact(s) for K vs. 0 for MG)  

### MG:BA:1776

|  | Res. type | Atom | Residue | Distance | Occupancy | B-factor | Closest solute to 1^st^ shell water |
| --- | --- | --- | --- | --- | --- | --- | --- |
| **Ion** | | | |  | 1.00 | 79.99 || 1 | A | N1 | BA:452 | 2.38 | 1.00 | 78.15 |
| 2 | A | OP1 | BA:374 | 2.44 | 1.00 | 72.25 |  |
| 3 | U | OP2 | BA:375 | 2.93 | 1.00 | 66.55 |  |
| 4 | A | N1 | BA:453 | 3.79 | 1.00 | 72.83 |  |
| 5 | A | OP2 | BA:374 | 3.93 | 1.00 | 72.40 |  |
| 6 | A | O5' | BA:374 | 3.98 | 1.00 | 70.42 |  |
| 7 | A | N6 | BA:452 | 4.01 | 1.00 | 80.04 |  |
| 8 | A | N3 | BA:453 | 4.03 | 1.00 | 75.16 |  |
| 9 | A | N3 | BA:452 | 4.11 | 1.00 | 77.23 |  |
| 10 | U | OP1 | BA:375 | 4.29 | 1.00 | 65.82 |  |
| 11 | A | O3' | BA:374 | 4.41 | 1.00 | 65.97 |  |
| 12 | A | O3' | BA:373 | 4.86 | 1.00 | 73.46 |  |
  |

  
 Contact(s): /N1_A_/  
 Coded contact list: N_b_  
 Exclusion zone contact(s): /OP1/OP2/  

### MG:BA:1777

|  | Res. type | Atom | Residue | Distance | Occupancy | B-factor | Closest solute to 1^st^ shell water |
| --- | --- | --- | --- | --- | --- | --- | --- |
| **Ion** | | | |  | 1.00 | 57.00 || 1 | G | O6 | BA:829 | 2.66 | 1.00 | 83.84 |
| 2 | G | O6 | BA:830 | 3.11 | 1.00 | 86.60 |  |
| 3 | G | N7 | BA:829 | 3.85 | 1.00 | 86.33 |  |
| 4 | C | N4 | BA:857 | 4.21 | 1.00 | 81.15 |  |
| 5 | C | N4 | BA:856 | 4.46 | 1.00 | 85.65 |  |
| 6 | U | OP2 | BA:870 | 4.49 | 1.00 | 77.99 |  |
| 7 | G | N7 | BA:830 | 4.50 | 1.00 | 87.31 |  |
| 8 | U | OP1 | BA:870 | 4.50 | 1.00 | 77.51 |  |
| 9 | G | N2 | BA:858 | 4.74 | 1.00 | 76.54 |  |
| 10 | G | N1 | BA:829 | 4.78 | 1.00 | 83.26 |  |
| 11 | U | OP1 | BA:871 | 4.92 | 1.00 | 75.59 |  |
  |

  
 Coded contact list:   
 Exclusion zone contact(s): /O6/O6/  
Consider K in place of MG (2 valid contact(s) for K vs. 0 for MG)  

### MG:BA:1778

|  | Res. type | Atom | Residue | Distance | Occupancy | B-factor | Closest solute to 1^st^ shell water |
| --- | --- | --- | --- | --- | --- | --- | --- |
| **Ion** | | | |  | 1.00 | 49.37 || 1 | G | N7 | BA:853 | 4.24 | 1.00 | 78.18 |
| 2 | U | OP1 | BA:871 | 4.33 | 1.00 | 75.59 |  |
| 3 | G | O6 | BA:854 | 4.44 | 1.00 | 79.51 |  |
| 4 | G | N7 | BA:854 | 4.58 | 1.00 | 77.98 |  |
| 5 | U | OP2 | BA:871 | 4.60 | 1.00 | 75.04 |  |
| 6 | G | O6 | BA:853 | 4.90 | 1.00 | 78.18 |  |
  |

  
 Coded contact list:   
1^st^ MG coordination shell missing (see Notes)  

### MG:BA:1779

|  | Res. type | Atom | Residue | Distance | Occupancy | B-factor | Closest solute to 1^st^ shell water |
| --- | --- | --- | --- | --- | --- | --- | --- |
| **Ion** | | | |  | 1.00 | 45.36 || 1 | U | O4 | BA:516 | 1.83 | 1.00 | 85.66 |
| 2 | A | OP1 | BA:533 | 2.41 | 1.00 | 82.63 |  |
| 3 | A | O2' | BA:532 | 2.93 | 1.00 | 94.91 |  |
| 4 | G | N7 | BA:515 | 3.56 | 1.00 | 80.10 |  |
| 5 | G | O6 | BA:515 | 3.91 | 1.00 | 76.48 |  |
| 6 | U | N3 | BA:516 | 3.95 | 1.00 | 86.96 |  |
| 7 | A | O5' | BA:533 | 4.04 | 1.00 | 82.77 |  |
| 8 | G | O6 | BA:517 | 4.26 | 1.00 | 81.77 |  |
| 9 | A | OP2 | BA:533 | 4.63 | 1.00 | 83.54 |  |
| 10 | G | OP2 | BA:515 | 4.82 | 1.00 | 75.65 |  |
| 11 | A | O3' | BA:532 | 4.86 | 1.00 | 87.10 |  |
  |

  
 Contact(s): /O4/  
 Coded contact list: O_b_  
 Exclusion zone contact(s): /O2'/OP1/  

### MG:BA:1780

|  | Res. type | Atom | Residue | Distance | Occupancy | B-factor | Closest solute to 1^st^ shell water |
| --- | --- | --- | --- | --- | --- | --- | --- |
| **Ion** | | | |  | 1.00 | 85.05 || 1 | G | OP2 | BA:111 | 2.70 | 1.00 | 65.07 |
| 2 | C | OP1 | BA:110 | 3.79 | 1.00 | 68.21 |  |
| 3 | G | N7 | BA:112 | 3.80 | 1.00 | 60.69 |  |
| 4 | G | O6 | BA:112 | 4.08 | 1.00 | 58.51 |  |
| 5 | G | N7 | BA:111 | 4.30 | 1.00 | 65.08 |  |
| 6 | C | OP2 | BA:110 | 4.50 | 1.00 | 68.09 |  |
| 7 | C | O5' | BA:110 | 4.69 | 1.00 | 67.95 |  |
| 8 | G | O5' | BA:111 | 4.70 | 1.00 | 65.26 |  |
  |

  
 Coded contact list:   
 Exclusion zone contact(s): /OP2/  
Consider K in place of MG (1 valid contact(s) for K vs. 0 for MG)  

### MG:BA:1781

|  | Res. type | Atom | Residue | Distance | Occupancy | B-factor | Closest solute to 1^st^ shell water |
| --- | --- | --- | --- | --- | --- | --- | --- |
| **Ion** | | | |  | 1.00 | 69.81 || 1 | G | O3' | BA:139 | 2.48 | 1.00 | 91.84 |
| 2 | A | OP1 | BA:140 | 2.55 | 1.00 | 93.00 |  |
| 3 | A | O5' | BA:140 | 3.77 | 1.00 | 93.09 |  |
| 4 | G | O2' | BA:139 | 3.89 | 1.00 | 89.39 |  |
| 5 | A | OP2 | BA:140 | 4.39 | 1.00 | 92.27 |  |
  |

  
 Coded contact list:   
 Exclusion zone contact(s): /O3'/OP1/  
Consider NA in place of MG (2 valid contact(s) for NA vs. 0 for MG)  

### MG:BA:1782

|  | Res. type | Atom | Residue | Distance | Occupancy | B-factor | Closest solute to 1^st^ shell water |
| --- | --- | --- | --- | --- | --- | --- | --- |
| **Ion** | | | |  | 1.00 | 100.11 || 1 | G | N2 | BA:226 | 3.97 | 1.00 | 87.56 |
| 2 | G | O2' | BA:138 | 4.05 | 1.00 | 88.67 |  |
| 3 | G | O4' | BA:138 | 4.07 | 1.00 | 88.99 |  |
| 4 | G | N9 | BA:138 | 4.57 | 1.00 | 90.18 |  |
| 5 | G | N3 | BA:138 | 4.75 | 1.00 | 90.05 |  |
| 6 | G | N3 | BA:226 | 4.84 | 1.00 | 87.24 |  |
| 7 | G | O2' | BA:227 | 4.87 | 1.00 | 81.08 |  |
  |

  
 Coded contact list:   
1^st^ MG coordination shell missing (see Notes)  

### MG:BA:1783

|  | Res. type | Atom | Residue | Distance | Occupancy | B-factor | Closest solute to 1^st^ shell water |
| --- | --- | --- | --- | --- | --- | --- | --- |
| **Ion** | | | |  | 1.00 | 81.54 || 1 | G | OP2 | BA:654 | 2.31 | 1.00 | 82.69 |
| 2 | A | OP2 | BA:653 | 3.88 | 1.00 | 82.97 |  |
| 3 | A | O3' | BA:653 | 4.13 | 1.00 | 83.24 |  |
| 4 | G | OP1 | BA:654 | 4.21 | 1.00 | 83.08 |  |
| 5 | A | O5' | BA:653 | 4.49 | 1.00 | 81.71 |  |
| 6 | A | N6 | BA:655 | 4.57 | 1.00 | 81.51 |  |
| 7 | G | O5' | BA:654 | 4.83 | 1.00 | 82.81 |  |
  |

  
 Contact(s): /OP2/  
 Coded contact list: O_ph_  

### MG:BA:1784

|  | Res. type | Atom | Residue | Distance | Occupancy | B-factor | Closest solute to 1^st^ shell water |
| --- | --- | --- | --- | --- | --- | --- | --- |
| **Ion** | | | |  | 1.00 | 54.70 || 1 | C | O3' | BA:526 | 2.58 | 1.00 | 72.70 |
| 2 | U | O3' | BA:12 | 2.71 | 1.00 | 68.82 |  |
| 3 | U | O2' | BA:12 | 2.78 | 1.00 | 66.21 |  |
| 4 | A | OP1 | BA:914 | 2.93 | 1.00 | 67.57 |  |
| 5 | G | OP1 | BA:527 | 3.24 | 1.00 | 71.60 |  |
| 6 | G | OP2 | BA:527 | 3.43 | 1.00 | 71.42 |  |
| 7 | U | OP1 | BA:13 | 3.63 | 1.00 | 69.73 |  |
| 8 | U | O5' | BA:13 | 3.83 | 1.00 | 70.24 |  |
| 9 | A | OP2 | BA:914 | 3.92 | 1.00 | 66.44 |  |
| 10 | C | O2' | BA:526 | 3.98 | 1.00 | 75.13 |  |
| 11 | U | OP2 | BA:14 | 4.64 | 1.00 | 65.30 |  |
| 12 | G | O5' | BA:527 | 4.70 | 1.00 | 70.92 |  |
| 13 | A | O3' | BA:913 | 4.88 | 1.00 | 66.64 |  |
| 14 | U | O4' | BA:12 | 4.98 | 1.00 | 66.05 |  |
| 15 | U | OP2 | BA:13 | 5.00 | 1.00 | 69.43 |  |
  |

  
 Coded contact list:   
 Exclusion zone contact(s): /O2'/O3'/O3'/OP1/OP1/  
Consider K in place of MG (4 valid contact(s) for K vs. 0 for MG)  

### MG:BA:1785

|  | Res. type | Atom | Residue | Distance | Occupancy | B-factor | Closest solute to 1^st^ shell water |
| --- | --- | --- | --- | --- | --- | --- | --- |
| **Ion** | | | |  | 1.00 | 67.16 || 1 | U | OP1 | BA:13 | 3.49 | 1.00 | 69.73 |
| 2 | U | OP2 | BA:14 | 3.57 | 1.00 | 65.30 |  |
| 3 | A | OP2 | BA:16 | 3.95 | 1.00 | 69.95 |  |
| 4 | A | OP1 | BA:16 | 4.02 | 1.00 | 68.20 |  |
| 5 | G | OP2 | BA:15 | 4.25 | 1.00 | 66.41 |  |
| 6 | U | O4 | BA:14 | 4.27 | 1.00 | 69.72 |  |
| 7 | U | O5' | BA:14 | 4.41 | 1.00 | 66.57 |  |
  |

  
 Coded contact list:   
1^st^ MG coordination shell missing (see Notes)  

### MG:BA:1786

|  | Res. type | Atom | Residue | Distance | Occupancy | B-factor | Closest solute to 1^st^ shell water |
| --- | --- | --- | --- | --- | --- | --- | --- |
| **Ion** | | | |  | 1.00 | 57.70 || 1 | A | OP2 | BA:915 | 3.71 | 1.00 | 60.17 |
| 2 | G | OP2 | BA:916 | 3.99 | 1.00 | 54.84 |  |
| 3 | G | N7 | BA:916 | 4.24 | 1.00 | 54.35 |  |
| 4 | U | O2' | BA:13 | 4.55 | 1.00 | 72.94 |  |
  |

  
 Coded contact list:   
1^st^ MG coordination shell missing (see Notes)  

### MG:BA:1787

|  | Res. type | Atom | Residue | Distance | Occupancy | B-factor | Closest solute to 1^st^ shell water |
| --- | --- | --- | --- | --- | --- | --- | --- |
| **Ion** | | | |  | 1.00 | 49.87 || 1 | A | OP2 | BA:1394 | 3.56 | 1.00 | 62.80 |
| 2 | A | N6 | BA:919 | 3.98 | 1.00 | 66.25 |  |
| 3 | A | OP2 | BA:918 | 4.40 | 1.00 | 66.77 |  |
| 4 | A | N7 | BA:919 | 4.43 | 1.00 | 63.83 |  |
| 5 | A | N7 | BA:918 | 4.51 | 1.00 | 60.25 |  |
| 6 | A | OP1 | BA:1394 | 4.57 | 1.00 | 63.13 |  |
  |

  
 Coded contact list:   
1^st^ MG coordination shell missing (see Notes)  

### MG:BA:1788

|  | Res. type | Atom | Residue | Distance | Occupancy | B-factor | Closest solute to 1^st^ shell water |
| --- | --- | --- | --- | --- | --- | --- | --- |
| **Ion** | | | |  | 1.00 | 53.90 || 1 | C | O5' | BA:328 | 2.82 | 1.00 | 75.79 |
| 2 | C | OP2 | BA:330 | 3.50 | 1.00 | 69.99 |  |
| 3 | C | O3' | BA:328 | 4.08 | 1.00 | 75.72 |  |
| 4 | A | O3' | BA:327 | 4.10 | 1.00 | 74.07 |  |
| 5 | C | OP1 | BA:328 | 4.14 | 1.00 | 75.96 |  |
| 6 | A | O5' | BA:329 | 4.34 | 1.00 | 74.08 |  |
| 7 | A | OP1 | BA:329 | 4.59 | 1.00 | 75.35 |  |
| 8 | A | O2' | BA:327 | 4.62 | 1.00 | 73.89 |  |
| 9 | C | OP2 | BA:314 | 4.72 | 1.00 | 61.23 |  |
| 10 | C | OP1 | BA:330 | 4.80 | 1.00 | 69.09 |  |
  |

  
 Coded contact list:   
 Exclusion zone contact(s): /O5'/  
Consider K in place of MG (1 valid contact(s) for K vs. 0 for MG)  

### MG:BA:1789

|  | Res. type | Atom | Residue | Distance | Occupancy | B-factor | Closest solute to 1^st^ shell water |
| --- | --- | --- | --- | --- | --- | --- | --- |
| **Ion** | | | |  | 1.00 | 105.64 || 1 | A | OP1 | BA:172 | 2.27 | 1.00 | 86.47 |
| 2 | A | O5' | BA:172 | 3.21 | 1.00 | 87.00 |  |
| 3 | A | OP2 | BA:172 | 3.50 | 1.00 | 86.83 |  |
| 4 | U | OP2 | BA:173 | 3.84 | 1.00 | 88.19 |  |
| 5 | A | O3' | BA:171 | 4.52 | 1.00 | 87.11 |  |
| 6 | A | O3' | BA:172 | 4.60 | 1.00 | 87.22 |  |
| 7 | U | OP1 | BA:173 | 4.61 | 1.00 | 88.05 |  |
| 8 | C | N4 | BA:174 | 4.88 | 1.00 | 94.53 |  |
  |

  
 Contact(s): /OP1/  
 Coded contact list: O_ph_  
 Exclusion zone contact(s): /O5'/  

### MG:BA:1790

|  | Res. type | Atom | Residue | Distance | Occupancy | B-factor | Closest solute to 1^st^ shell water |
| --- | --- | --- | --- | --- | --- | --- | --- |
| **Ion** | | | |  | 1.00 | 56.46 || 1 | C | OP1 | BA:980 | 2.28 | 1.00 | 106.25 |
| 2 | U | O4 | BA:1219 | 3.03 | 1.00 | 109.76 |  |
| 3 | G | O6 | BA:1220 | 3.37 | 1.00 | 103.97 |  |
| 4 | G | N7 | BA:1220 | 3.79 | 1.00 | 103.47 |  |
| 5 | C | O3' | BA:979 | 4.40 | 1.00 | 106.35 |  |
| 6 | C | O5' | BA:980 | 4.41 | 1.00 | 106.19 |  |
| 7 | G | O6 | BA:1221 | 4.62 | 1.00 | 99.80 |  |
| 8 | U | N3 | BA:1219 | 4.65 | 1.00 | 109.18 |  |
| 9 | C | OP2 | BA:980 | 4.80 | 1.00 | 106.01 |  |
  |

  
 Contact(s): /OP1/  
 Coded contact list: O_ph_  
 Exclusion zone contact(s): /O4/O6/  
Consider K in place of MG (2 valid contact(s) for K vs. 1 for MG)  

### MG:BA:1791

|  | Res. type | Atom | Residue | Distance | Occupancy | B-factor | Closest solute to 1^st^ shell water |
| --- | --- | --- | --- | --- | --- | --- | --- |
| **Ion** | | | |  | 1.00 | 55.69 || 1 | ASN | OD1 | BM:69 | 2.88 | 1.00 | 114.31 |
| 2 | A | OP2 | BA:1279 | 2.89 | 1.00 | 123.08 |  |
| 3 | A | O5' | BA:1279 | 2.96 | 1.00 | 122.12 |  |
| 4 | A | OP1 | BA:1280 | 3.14 | 1.00 | 115.53 |  |
| 5 | ASN | ND2 | BM:69 | 3.49 | 1.00 | 114.63 |  |
| 6 | A | OP1 | BA:1279 | 3.66 | 1.00 | 123.32 |  |
| 7 | A | OP2 | BA:1280 | 3.77 | 1.00 | 116.32 |  |
| 8 | U | O3' | BA:1278 | 4.79 | 1.00 | 124.28 |  |
| 9 | A | O3' | BA:1279 | 4.95 | 1.00 | 116.29 |  |
  |

  
 Coded contact list:   
 Exclusion zone contact(s): /O5'/OD1_Asn_/OP1/OP2/  
Consider K in place of MG (4 valid contact(s) for K vs. 0 for MG)  

### MG:BA:1792

|  | Res. type | Atom | Residue | Distance | Occupancy | B-factor | Closest solute to 1^st^ shell water |
| --- | --- | --- | --- | --- | --- | --- | --- |
| **Ion** | | | |  | 1.00 | 79.85 || 1 | G | N7 | BA:838 | 3.59 | 1.00 | 128.95 |
| 2 | G | O6 | BA:838 | 3.99 | 1.00 | 131.11 |  |
| 3 | G | N7 | BA:837 | 4.17 | 1.00 | 106.51 |  |
| 4 | G | O6 | BA:837 | 4.77 | 1.00 | 109.41 |  |
  |

  
 Coded contact list:   
1^st^ MG coordination shell missing (see Notes)  

### MG:BA:1793

|  | Res. type | Atom | Residue | Distance | Occupancy | B-factor | Closest solute to 1^st^ shell water |
| --- | --- | --- | --- | --- | --- | --- | --- |
| **Ion** | | | |  | 1.00 | 57.10 || 1 | G | O6 | BA:851 | 2.32 | 1.00 | 89.16 |
| 2 | G | O6 | BA:836 | 2.64 | 1.00 | 97.32 |  |
| 3 | U | O4 | BA:850 | 3.03 | 1.00 | 101.96 |  |
| 4 | U | O4 | BA:835 | 3.33 | 1.00 | 88.00 |  |
| 5 | U | N3 | BA:850 | 3.73 | 1.00 | 101.62 |  |
| 6 | G | N7 | BA:851 | 3.79 | 1.00 | 90.18 |  |
| 7 | G | O6 | BA:852 | 4.01 | 1.00 | 83.08 |  |
| 8 | U | N3 | BA:835 | 4.04 | 1.00 | 87.93 |  |
| 9 | G | N7 | BA:836 | 4.47 | 1.00 | 95.36 |  |
| 10 | G | N1 | BA:851 | 4.54 | 1.00 | 88.14 |  |
| 11 | G | N1 | BA:836 | 4.67 | 1.00 | 98.06 |  |
| 12 | C | N4 | BA:834 | 4.76 | 1.00 | 82.68 |  |
| 13 | G | O6 | BA:837 | 4.99 | 1.00 | 109.41 |  |
  |

  
 Contact(s): /O6/  
 Coded contact list: O_b_  
 Exclusion zone contact(s): /O4/O4/O6/  
Consider K in place of MG (3 valid contact(s) for K vs. 1 for MG)  

### MG:BA:1794

|  | Res. type | Atom | Residue | Distance | Occupancy | B-factor | Closest solute to 1^st^ shell water |
| --- | --- | --- | --- | --- | --- | --- | --- |
| **Ion** | | | |  | 1.00 | 49.72 || 1 | C | OP2 | BA:822 | 2.22 | 1.00 | 65.48 |
| 2 | G | O6 | BA:818 | 3.58 | 1.00 | 56.62 |  |
| 3 | U | O3' | BA:820 | 4.16 | 1.00 | 67.60 |  |
| 4 | G | O5' | BA:821 | 4.32 | 1.00 | 65.99 |  |
| 5 | G | OP1 | BA:821 | 4.41 | 1.00 | 66.39 |  |
| 6 | C | O5' | BA:822 | 4.46 | 1.00 | 62.65 |  |
| 7 | C | OP1 | BA:822 | 4.47 | 1.00 | 63.74 |  |
| 8 | G | N1 | BA:818 | 4.53 | 1.00 | 53.77 |  |
| 9 | U | O2' | BA:820 | 4.59 | 1.00 | 71.56 |  |
| 10 | G | O3' | BA:821 | 4.60 | 1.00 | 63.02 |  |
| 11 | A | OP2 | BA:873 | 4.92 | 1.00 | 72.90 |  |
  |

  
 Contact(s): /OP2/  
 Coded contact list: O_ph_  

### MG:BA:1795

|  | Res. type | Atom | Residue | Distance | Occupancy | B-factor | Closest solute to 1^st^ shell water |
| --- | --- | --- | --- | --- | --- | --- | --- |
| **Ion** | | | |  | 1.00 | 67.08 || 1 | A | OP1 | BA:729 | 3.99 | 1.00 | 59.67 |
| 2 | G | N7 | BA:818 | 4.03 | 1.00 | 54.86 |  |
  |

  
 Coded contact list:   
1^st^ MG coordination shell missing (see Notes)  

### MG:BA:1796

|  | Res. type | Atom | Residue | Distance | Occupancy | B-factor | Closest solute to 1^st^ shell water |
| --- | --- | --- | --- | --- | --- | --- | --- |
| **Ion** | | | |  | 1.00 | 77.78 || 1 | G | N7 | BA:727 | 4.08 | 1.00 | 64.73 |
| 2 | C | OP2 | BA:726 | 4.11 | 1.00 | 66.42 |  |
| 3 | G | OP2 | BA:727 | 4.36 | 1.00 | 64.04 |  |
| 4 | G | O2' | BA:853 | 4.82 | 1.00 | 78.96 |  |
  |

  
 Coded contact list:   
1^st^ MG coordination shell missing (see Notes)  

### MG:BA:1797

|  | Res. type | Atom | Residue | Distance | Occupancy | B-factor | Closest solute to 1^st^ shell water |
| --- | --- | --- | --- | --- | --- | --- | --- |
| **Ion** | | | |  | 1.00 | 63.63 || 1 | G | N7 | BA:818 | 3.93 | 1.00 | 54.86 |
| 2 | G | OP2 | BA:855 | 3.97 | 1.00 | 73.54 |  |
| 3 | G | N9 | BA:818 | 4.03 | 1.00 | 52.06 |  |
| 4 | U | O2 | BA:871 | 4.10 | 1.00 | 81.36 |  |
| 5 | G | O4' | BA:818 | 4.52 | 1.00 | 51.40 |  |
| 6 | G | N3 | BA:818 | 4.95 | 1.00 | 52.69 |  |
  |

  
 Coded contact list:   
1^st^ MG coordination shell missing (see Notes)  

### MG:BA:1798

|  | Res. type | Atom | Residue | Distance | Occupancy | B-factor | Closest solute to 1^st^ shell water |
| --- | --- | --- | --- | --- | --- | --- | --- |
| **Ion** | | | |  | 1.00 | 93.68 || 1 | G | OP1 | BA:869 | 3.46 | 1.00 | 73.35 |
| 2 | C | OP2 | BA:856 | 4.35 | 1.00 | 78.48 |  |
| 3 | C | OP1 | BA:856 | 4.61 | 1.00 | 78.65 |  |
| 4 | C | O3' | BA:868 | 4.95 | 1.00 | 73.02 |  |
  |

  
 Coded contact list:   
1^st^ MG coordination shell missing (see Notes)  

### MG:BA:1799

|  | Res. type | Atom | Residue | Distance | Occupancy | B-factor | Closest solute to 1^st^ shell water |
| --- | --- | --- | --- | --- | --- | --- | --- |
| **Ion** | | | |  | 1.00 | 55.22 || 1 | G | N3 | BA:1338 | 4.61 | 1.00 | 80.84 |
| 2 | G | N9 | BA:1338 | 4.66 | 1.00 | 79.95 |  |
| 3 | G | N1 | BA:1338 | 4.86 | 1.00 | 80.63 |  |
| 4 | G | N7 | BA:1338 | 4.87 | 1.00 | 79.88 |  |
  |

  
 Coded contact list:   
1^st^ MG coordination shell missing (see Notes)  

### MG:BA:1800

|  | Res. type | Atom | Residue | Distance | Occupancy | B-factor | Closest solute to 1^st^ shell water |
| --- | --- | --- | --- | --- | --- | --- | --- |
| **Ion** | | | |  | 1.00 | 66.51 || 1 | C | OP2 | BA:1402 | 2.40 | 1.00 | 57.77 |
| 2 | U | OP1 | B1:49 | 3.24 | 1.00 | 75.03 |  |
| 3 | C | OP1 | BA:1402 | 3.36 | 1.00 | 59.50 |  |
| 4 | G | O3' | BA:1401 | 4.09 | 1.00 | 57.96 |  |
| 5 | G | O5' | BA:1401 | 4.10 | 1.00 | 62.16 |  |
| 6 | G | OP1 | BA:1401 | 4.65 | 1.00 | 63.07 |  |
| 7 | C | O5' | BA:1402 | 4.66 | 1.00 | 56.82 |  |
| 8 | U | OP2 | B1:50 | 4.83 | 1.00 | 77.59 |  |
  |

  
 Contact(s): /OP2/  
 Coded contact list: O_ph_  
 Exclusion zone contact(s): /OP1/OP1/  
Consider K in place of MG (2 valid contact(s) for K vs. 1 for MG)  

### MG:BA:1801

|  | Res. type | Atom | Residue | Distance | Occupancy | B-factor | Closest solute to 1^st^ shell water |
| --- | --- | --- | --- | --- | --- | --- | --- |
| **Ion** | | | |  | 1.00 | 40.72 || 1 | U | O4 | BA:1498 | 3.41 | 1.00 | 51.26 |
| 2 | G | N7 | BA:1497 | 3.48 | 1.00 | 51.71 |  |
| 3 | U | O2' | B1:47 | 4.22 | 1.00 | 65.55 |  |
| 4 | U | O3' | B1:47 | 4.67 | 1.00 | 69.67 |  |
| 5 | G | OP2 | BA:1497 | 4.78 | 1.00 | 48.28 |  |
| 6 | G | O6 | BA:1497 | 4.82 | 1.00 | 48.51 |  |
  |

  
 Coded contact list:   
1^st^ MG coordination shell missing (see Notes)  

### MG:BA:1802

|  | Res. type | Atom | Residue | Distance | Occupancy | B-factor | Closest solute to 1^st^ shell water |
| --- | --- | --- | --- | --- | --- | --- | --- |
| **Ion** | | | |  | 1.00 | 75.25 || 1 | U | O4 | B1:39 | 3.59 | 1.00 | 147.33 |
| 2 | G | N7 | BA:1530 | 4.09 | 1.00 | 86.15 |  |
| 3 | U | O4 | BA:1532 | 4.83 | 1.00 | 118.28 |  |
| 4 | G | O6 | BA:1530 | 4.85 | 1.00 | 85.13 |  |
  |

  
 Coded contact list:   
1^st^ MG coordination shell missing (see Notes)  

### MG:BA:1803

|  | Res. type | Atom | Residue | Distance | Occupancy | B-factor | Closest solute to 1^st^ shell water |
| --- | --- | --- | --- | --- | --- | --- | --- |
| **Ion** | | | |  | 1.00 | 90.90 || 1 | U | O4 | BA:1528 | 3.51 | 1.00 | 61.56 |
| 2 | G | N7 | BA:1530 | 4.54 | 1.00 | 86.15 |  |
| 3 | U | N3 | BA:1528 | 4.75 | 1.00 | 63.46 |  |
  |

  
 Coded contact list:   
1^st^ MG coordination shell missing (see Notes)  

### MG:BA:1804

|  | Res. type | Atom | Residue | Distance | Occupancy | B-factor | Closest solute to 1^st^ shell water |
| --- | --- | --- | --- | --- | --- | --- | --- |
| **Ion** | | | |  | 1.00 | 105.88 || 1 | U | O4 | BA:1532 | 2.20 | 1.00 | 118.28 |
| 2 | U | O4 | B1:39 | 2.62 | 1.00 | 147.33 |  |
| 3 | C | N4 | BA:1533 | 2.97 | 1.00 | 131.04 |  |
| 4 | U | N3 | BA:1532 | 3.40 | 1.00 | 117.45 |  |
| 5 | U | O4 | B1:38 | 3.66 | 1.00 | 149.65 |  |
| 6 | A | N6 | BA:1534 | 4.05 | 1.00 | 149.02 |  |
| 7 | U | N3 | B1:39 | 4.32 | 1.00 | 146.80 |  |
| 8 | U | O4 | B1:40 | 4.73 | 1.00 | 141.66 |  |
| 9 | C | N3 | BA:1533 | 4.96 | 1.00 | 131.21 |  |
  |

  
 Contact(s): /O4/  
 Coded contact list: O_b_  
 Exclusion zone contact(s): /N3_U_/N4/O4/  
Consider K in place of MG (3 valid contact(s) for K vs. 1 for MG)  

### MG:BA:1805

|  | Res. type | Atom | Residue | Distance | Occupancy | B-factor | Closest solute to 1^st^ shell water |
| --- | --- | --- | --- | --- | --- | --- | --- |
| **Ion** | | | |  | 1.00 | 99.65 || 1 | U | O4 | BA:1532 | 4.95 | 1.00 | 118.28 |
  |

  
 Coded contact list:   
1^st^ MG coordination shell missing (see Notes)  

### MG:BA:1806

|  | Res. type | Atom | Residue | Distance | Occupancy | B-factor | Closest solute to 1^st^ shell water |
| --- | --- | --- | --- | --- | --- | --- | --- |
| **Ion** | | | |  | 1.00 | 78.79 || 1 | U | O4 | BA:1537 | 4.08 | 1.00 | 173.23 |
| 2 | C | N4 | BA:1536 | 4.53 | 1.00 | 168.56 |  |
  |

  
 Coded contact list:   
1^st^ MG coordination shell missing (see Notes)  

### MG:BA:1807

|  | Res. type | Atom | Residue | Distance | Occupancy | B-factor | Closest solute to 1^st^ shell water |
| --- | --- | --- | --- | --- | --- | --- | --- |
| **Ion** | | | |  | 1.00 | 63.72 || 1 | U | OP2 | BA:323 | 1.79 | 1.00 | 67.28 |
| 2 | U | OP1 | BA:323 | 3.41 | 1.00 | 67.42 |  |
| 3 | U | O5' | BA:323 | 3.79 | 1.00 | 68.26 |  |
| 4 | C | O3' | BA:322 | 4.20 | 1.00 | 65.93 |  |
| 5 | C | OP1 | BA:322 | 4.39 | 1.00 | 61.25 |  |
| 6 | C | O5' | BA:322 | 4.43 | 1.00 | 61.63 |  |
| 7 | G | OP2 | BA:324 | 4.67 | 1.00 | 66.55 |  |
| 8 | G | N7 | BA:324 | 4.90 | 1.00 | 76.90 |  |
| 9 | C | OP2 | BA:322 | 4.96 | 1.00 | 61.67 |  |
| 10 | ASN | OD1 | BW:26 | 4.97 | 1.00 | 77.75 |  |
  |

  
 Coded contact list:   
 Exclusion zone contact(s): /OP2/  
1^st^ MG coordination shell missing (see Notes)  

### MG:BA:1808

|  | Res. type | Atom | Residue | Distance | Occupancy | B-factor | Closest solute to 1^st^ shell water |
| --- | --- | --- | --- | --- | --- | --- | --- |
| **Ion** | | | |  | 1.00 | 70.02 || 1 | G | N7 | BA:324 | 2.55 | 1.00 | 76.90 |
| 2 | G | OP2 | BA:324 | 3.62 | 1.00 | 66.55 |  |
| 3 | U | OP2 | BA:323 | 3.74 | 1.00 | 67.28 |  |
| 4 | G | O6 | BA:324 | 4.23 | 1.00 | 79.32 |  |
| 5 | U | O5' | BA:323 | 4.48 | 1.00 | 68.26 |  |
| 6 | G | N9 | BA:324 | 4.57 | 1.00 | 75.50 |  |
| 7 | G | O5' | BA:324 | 4.76 | 1.00 | 69.90 |  |
| 8 | THR | O | BW:71 | 4.82 | 1.00 | 84.55 |  |
  |

  
 Coded contact list:   
 Exclusion zone contact(s): /N7/  
Consider NA in place of MG (1 valid contact(s) for NA vs. 0 for MG)  

### MG:BA:1809

|  | Res. type | Atom | Residue | Distance | Occupancy | B-factor | Closest solute to 1^st^ shell water |
| --- | --- | --- | --- | --- | --- | --- | --- |
| **Ion** | | | |  | 1.00 | 114.37 || 1 | G | N7 | BA:324 | 2.28 | 1.00 | 76.90 |
| 2 | G | OP2 | BA:326 | 3.08 | 1.00 | 66.00 |  |
| 3 | G | N9 | BA:324 | 3.39 | 1.00 | 75.50 |  |
| 4 | G | O5' | BA:324 | 3.85 | 1.00 | 69.90 |  |
| 5 | G | OP2 | BA:324 | 3.98 | 1.00 | 66.55 |  |
| 6 | A | OP2 | BA:325 | 4.40 | 1.00 | 70.40 |  |
| 7 | G | OP1 | BA:326 | 4.43 | 1.00 | 66.26 |  |
| 8 | G | O4' | BA:324 | 4.63 | 1.00 | 72.19 |  |
| 9 | G | O6 | BA:324 | 4.66 | 1.00 | 79.32 |  |
| 10 | A | O5' | BA:325 | 4.91 | 1.00 | 69.33 |  |
  |

  
 Contact(s): /N7/  
 Coded contact list: N_b_  
 Exclusion zone contact(s): /OP2/  
Consider K in place of MG (2 valid contact(s) for K vs. 1 for MG)  

### MG:BA:1810

|  | Res. type | Atom | Residue | Distance | Occupancy | B-factor | Closest solute to 1^st^ shell water |
| --- | --- | --- | --- | --- | --- | --- | --- |
| **Ion** | | | |  | 1.00 | 93.74 || 1 | G | N7 | BA:324 | 2.26 | 1.00 | 76.90 |
| 2 | G | O6 | BA:324 | 2.73 | 1.00 | 79.32 |  |
| 3 | G | OP2 | BA:326 | 3.70 | 1.00 | 66.00 |  |
| 4 | G | OP1 | BA:326 | 3.82 | 1.00 | 66.26 |  |
| 5 | G | N9 | BA:324 | 4.31 | 1.00 | 75.50 |  |
| 6 | G | N1 | BA:324 | 4.36 | 1.00 | 77.32 |  |
  |

  
 Contact(s): /N7/  
 Coded contact list: N_b_  
 Exclusion zone contact(s): /O6/  

### MG:BA:1811

|  | Res. type | Atom | Residue | Distance | Occupancy | B-factor | Closest solute to 1^st^ shell water |
| --- | --- | --- | --- | --- | --- | --- | --- |
| **Ion** | | | |  | 1.00 | 65.56 || 1 | G | O6 | BA:324 | 2.16 | 1.00 | 79.32 |
| 2 | G | OP1 | BA:326 | 3.88 | 1.00 | 66.26 |  |
| 3 | G | N1 | BA:324 | 4.18 | 1.00 | 77.32 |  |
| 4 | G | N7 | BA:324 | 4.23 | 1.00 | 76.90 |  |
| 5 | A | OP2 | BA:327 | 4.51 | 1.00 | 68.63 |  |
| 6 | A | OP1 | BA:262 | 4.62 | 1.00 | 76.40 |  |
| 7 | G | OP2 | BA:326 | 4.99 | 1.00 | 66.00 |  |
  |

  
 Contact(s): /O6/  
 Coded contact list: O_b_  

### MG:BA:1812

|  | Res. type | Atom | Residue | Distance | Occupancy | B-factor | Closest solute to 1^st^ shell water |
| --- | --- | --- | --- | --- | --- | --- | --- |
| **Ion** | | | |  | 1.00 | 136.01 || 1 | G | O6 | BA:324 | 4.13 | 1.00 | 79.32 |
| 2 | A | OP1 | BA:262 | 4.58 | 1.00 | 76.40 |  |
| 3 | C | OP2 | BA:322 | 4.92 | 1.00 | 61.67 |  |
  |

  
 Coded contact list:   
1^st^ MG coordination shell missing (see Notes)  

### MG:BA:1813

|  | Res. type | Atom | Residue | Distance | Occupancy | B-factor | Closest solute to 1^st^ shell water |
| --- | --- | --- | --- | --- | --- | --- | --- |
| **Ion** | | | |  | 1.00 | 94.76 || 1 | A | OP2 | BA:327 | 2.38 | 1.00 | 68.63 |
| 2 | A | OP1 | BA:327 | 3.27 | 1.00 | 67.27 |  |
| 3 | G | O6 | BA:324 | 3.52 | 1.00 | 79.32 |  |
| 4 | G | N1 | BA:324 | 3.98 | 1.00 | 77.32 |  |
| 5 | A | O5' | BA:327 | 4.32 | 1.00 | 68.37 |  |
| 6 | G | O3' | BA:326 | 4.46 | 1.00 | 67.71 |  |
| 7 | G | OP1 | BA:326 | 4.46 | 1.00 | 66.26 |  |
| 8 | G | O5' | BA:326 | 4.92 | 1.00 | 67.03 |  |
  |

  
 Contact(s): /OP2/  
 Coded contact list: O_ph_  
 Exclusion zone contact(s): /OP1/  

### MG:BA:1814

|  | Res. type | Atom | Residue | Distance | Occupancy | B-factor | Closest solute to 1^st^ shell water |
| --- | --- | --- | --- | --- | --- | --- | --- |
| **Ion** | | | |  | 1.00 | 115.79 || 1 | A | N7 | BA:262 | 4.30 | 1.00 | 79.05 |
| 2 | U | O3' | BA:261 | 4.58 | 1.00 | 77.51 |  |
| 3 | A | OP2 | BA:327 | 4.66 | 1.00 | 68.63 |  |
| 4 | A | OP1 | BA:327 | 4.69 | 1.00 | 67.27 |  |
| 5 | U | O2' | BA:261 | 4.78 | 1.00 | 78.00 |  |
  |

  
 Coded contact list:   
1^st^ MG coordination shell missing (see Notes)  

### MG:BA:1815

|  | Res. type | Atom | Residue | Distance | Occupancy | B-factor | Closest solute to 1^st^ shell water |
| --- | --- | --- | --- | --- | --- | --- | --- |
| **Ion** | | | |  | 1.00 | 119.44 || 1 | U | O2' | BA:261 | 3.18 | 1.00 | 78.00 |
| 2 | U | O3' | BA:261 | 3.59 | 1.00 | 77.51 |  |
| 3 | A | N7 | BA:262 | 3.74 | 1.00 | 79.05 |  |
| 4 | A | N6 | BA:263 | 4.54 | 1.00 | 68.16 |  |
| 5 | U | O4' | BA:261 | 4.57 | 1.00 | 77.24 |  |
| 6 | A | N6 | BA:262 | 4.91 | 1.00 | 81.05 |  |
  |

  
 Coded contact list:   
 Exclusion zone contact(s): /O2'/  
Consider K in place of MG (1 valid contact(s) for K vs. 0 for MG)  

### MG:BA:1816

|  | Res. type | Atom | Residue | Distance | Occupancy | B-factor | Closest solute to 1^st^ shell water |
| --- | --- | --- | --- | --- | --- | --- | --- |
| **Ion** | | | |  | 1.00 | 71.08 || 1 | U | O4' | BA:261 | 3.38 | 1.00 | 77.24 |
| 2 | G | O2' | BA:260 | 3.76 | 1.00 | 74.34 |  |
| 3 | U | O2' | BA:261 | 4.77 | 1.00 | 78.00 |  |
  |

  
 Coded contact list:   
 Exclusion zone contact(s): /O4'/  
Consider K in place of MG (1 valid contact(s) for K vs. 0 for MG)  

### MG:BA:1817

|  | Res. type | Atom | Residue | Distance | Occupancy | B-factor | Closest solute to 1^st^ shell water |
| --- | --- | --- | --- | --- | --- | --- | --- |
| **Ion** | | | |  | 1.00 | 63.32 || 1 | G | O6 | BA:265 | 3.22 | 1.00 | 69.82 |
| 2 | G | N1 | BA:265 | 4.22 | 1.00 | 69.16 |  |
| 3 | U | O4' | BA:261 | 4.44 | 1.00 | 77.24 |  |
| 4 | C | O2 | BA:267 | 4.56 | 1.00 | 69.66 |  |
| 5 | G | N2 | BA:260 | 4.75 | 1.00 | 72.20 |  |
| 6 | U | O4 | BA:264 | 4.87 | 1.00 | 62.24 |  |
  |

  
 Coded contact list:   
 Exclusion zone contact(s): /O6/  
Consider K in place of MG (1 valid contact(s) for K vs. 0 for MG)  

### MG:BA:1818

|  | Res. type | Atom | Residue | Distance | Occupancy | B-factor | Closest solute to 1^st^ shell water |
| --- | --- | --- | --- | --- | --- | --- | --- |
| **Ion** | | | |  | 1.00 | 57.10 || 1 | G | O6 | BA:377 | 4.11 | 1.00 | 76.60 |
| 2 | G | N7 | BA:376 | 4.25 | 1.00 | 70.45 |  |
| 3 | G | N7 | BA:377 | 4.30 | 1.00 | 72.72 |  |
| 4 | G | O6 | BA:376 | 4.68 | 1.00 | 71.36 |  |
  |

  
 Coded contact list:   
1^st^ MG coordination shell missing (see Notes)  

### MG:BA:1819

|  | Res. type | Atom | Residue | Distance | Occupancy | B-factor | Closest solute to 1^st^ shell water |
| --- | --- | --- | --- | --- | --- | --- | --- |
| **Ion** | | | |  | 1.00 | 61.36 || 1 | C | O2' | BA:372 | 2.27 | 1.00 | 75.52 |
| 2 | C | O2 | BA:372 | 2.35 | 1.00 | 76.60 |  |
| 3 | U | O4 | BA:375 | 2.81 | 1.00 | 67.36 |  |
| 4 | U | O4 | BA:387 | 3.00 | 1.00 | 70.68 |  |
| 5 | G | O6 | BA:376 | 3.15 | 1.00 | 71.36 |  |
| 6 | C | N1 | BA:372 | 4.11 | 1.00 | 75.66 |  |
| 7 | A | N6 | BA:389 | 4.21 | 1.00 | 67.51 |  |
| 8 | U | N3 | BA:387 | 4.24 | 1.00 | 69.95 |  |
| 9 | C | N3 | BA:372 | 4.41 | 1.00 | 77.36 |  |
| 10 | A | N7 | BA:374 | 4.50 | 1.00 | 67.22 |  |
| 11 | A | OP2 | BA:374 | 4.64 | 1.00 | 72.40 |  |
| 12 | C | O3' | BA:372 | 4.66 | 1.00 | 76.76 |  |
| 13 | A | N6 | BA:374 | 4.71 | 1.00 | 70.39 |  |
| 14 | G | N7 | BA:376 | 4.87 | 1.00 | 70.45 |  |
| 15 | U | N3 | BA:375 | 4.99 | 1.00 | 66.22 |  |
  |

  
**1^st^ shell ligand angles:**   
1-2: 80;   
  
 Contact(s): /O2/O2'/  
 Coded contact list: O_r_O_b_  
 Exclusion zone contact(s): /O4/O4/O6/  
Consider K in place of MG (3 valid contact(s) for K vs. 2 for MG)  

### MG:BA:1820

|  | Res. type | Atom | Residue | Distance | Occupancy | B-factor | Closest solute to 1^st^ shell water |
| --- | --- | --- | --- | --- | --- | --- | --- |
| **Ion** | | | |  | 1.00 | 84.64 || 1 | G | O6 | BA:384 | 3.22 | 1.00 | 80.76 |
| 2 | G | O6 | BA:380 | 3.80 | 1.00 | 85.17 |  |
| 3 | G | N1 | BA:380 | 4.37 | 1.00 | 85.26 |  |
| 4 | C | N4 | BA:379 | 4.44 | 1.00 | 80.93 |  |
| 5 | G | N7 | BA:384 | 4.54 | 1.00 | 83.76 |  |
| 6 | ASP | OD2 | BS:68 | 4.91 | 1.00 | 75.42 |  |
| 7 | C | N4 | BA:385 | 4.99 | 1.00 | 73.89 |  |
  |

  
 Coded contact list:   
 Exclusion zone contact(s): /O6/  
Consider K in place of MG (1 valid contact(s) for K vs. 0 for MG)  

### MG:BA:1821

|  | Res. type | Atom | Residue | Distance | Occupancy | B-factor | Closest solute to 1^st^ shell water |
| --- | --- | --- | --- | --- | --- | --- | --- |
| **Ion** | | | |  | 1.00 | 93.39 || 1 | G | N7 | BA:384 | 2.94 | 1.00 | 83.76 |
| 2 | G | O6 | BA:384 | 3.92 | 1.00 | 80.76 |  |
| 3 | C | N4 | BA:385 | 4.04 | 1.00 | 73.89 |  |
| 4 | G | N9 | BA:384 | 4.92 | 1.00 | 81.69 |  |
  |

  
 Coded contact list:   
 Exclusion zone contact(s): /N7/  
Consider K in place of MG (1 valid contact(s) for K vs. 0 for MG)  

### MG:BA:1822

|  | Res. type | Atom | Residue | Distance | Occupancy | B-factor | Closest solute to 1^st^ shell water |
| --- | --- | --- | --- | --- | --- | --- | --- |
| **Ion** | | | |  | 1.00 | 97.16 || 1 | A | OP2 | BA:374 | 4.12 | 1.00 | 72.40 |
| 2 | A | OP1 | BA:374 | 4.52 | 1.00 | 72.25 |  |
| 3 | A | N1 | BA:453 | 4.62 | 1.00 | 72.83 |  |
| 4 | C | O2 | BA:454 | 4.75 | 1.00 | 90.24 |  |
  |

  
 Coded contact list:   
1^st^ MG coordination shell missing (see Notes)  

### MG:BA:1823

|  | Res. type | Atom | Residue | Distance | Occupancy | B-factor | Closest solute to 1^st^ shell water |
| --- | --- | --- | --- | --- | --- | --- | --- |
| **Ion** | | | |  | 1.00 | 84.14 || 1 | G | OP2 | BA:384 | 2.56 | 1.00 | 83.13 |
| 2 | G | O5' | BA:384 | 3.62 | 1.00 | 82.43 |  |
| 3 | G | OP1 | BA:384 | 3.90 | 1.00 | 81.98 |  |
| 4 | C | O2 | BA:454 | 3.92 | 1.00 | 90.24 |  |
| 5 | G | N7 | BA:384 | 4.36 | 1.00 | 83.76 |  |
| 6 | A | O3' | BA:383 | 4.88 | 1.00 | 83.02 |  |
| 7 | C | N3 | BA:454 | 4.97 | 1.00 | 89.27 |  |
  |

  
 Coded contact list:   
 Exclusion zone contact(s): /OP2/  
Consider NA in place of MG (1 valid contact(s) for NA vs. 0 for MG)  

### MG:BA:1824

|  | Res. type | Atom | Residue | Distance | Occupancy | B-factor | Closest solute to 1^st^ shell water |
| --- | --- | --- | --- | --- | --- | --- | --- |
| **Ion** | | | |  | 1.00 | 107.38 || 1 | C | N3 | BA:454 | 3.11 | 1.00 | 89.27 |
| 2 | A | N1 | BA:453 | 3.54 | 1.00 | 72.83 |  |
| 3 | A | N6 | BA:453 | 3.59 | 1.00 | 72.04 |  |
| 4 | C | O2 | BA:454 | 3.64 | 1.00 | 90.24 |  |
| 5 | C | N4 | BA:454 | 4.10 | 1.00 | 87.96 |  |
| 6 | G | OP2 | BA:384 | 4.25 | 1.00 | 83.13 |  |
| 7 | C | O2 | BA:479 | 4.39 | 1.00 | 102.98 |  |
| 8 | G | OP1 | BA:384 | 4.48 | 1.00 | 81.98 |  |
  |

  
 Coded contact list:   
 Exclusion zone contact(s): /N3_C_/  
Consider K in place of MG (1 valid contact(s) for K vs. 0 for MG)  

### MG:BA:1825

|  | Res. type | Atom | Residue | Distance | Occupancy | B-factor | Closest solute to 1^st^ shell water |
| --- | --- | --- | --- | --- | --- | --- | --- |
| **Ion** | | | |  | 1.00 | 121.76 || 1 | A | OP2 | BA:373 | 2.38 | 1.00 | 77.21 |
| 2 | A | OP2 | BA:374 | 3.90 | 1.00 | 72.40 |  |
| 3 | C | O3' | BA:372 | 4.05 | 1.00 | 76.76 |  |
| 4 | A | O3' | BA:373 | 4.48 | 1.00 | 73.46 |  |
| 5 | A | O5' | BA:373 | 4.48 | 1.00 | 76.24 |  |
| 6 | A | OP1 | BA:374 | 4.72 | 1.00 | 72.25 |  |
| 7 | A | OP1 | BA:373 | 4.88 | 1.00 | 77.48 |  |
  |

  
 Contact(s): /OP2/  
 Coded contact list: O_ph_  

### MG:BA:1826

|  | Res. type | Atom | Residue | Distance | Occupancy | B-factor | Closest solute to 1^st^ shell water |
| --- | --- | --- | --- | --- | --- | --- | --- |
| **Ion** | | | |  | 1.00 | 75.34 || 1 | A | OP2 | BA:373 | 3.36 | 1.00 | 77.21 |
| 2 | G | OP1 | BA:384 | 4.80 | 1.00 | 81.98 |  |
  |

  
 Coded contact list:   
 Exclusion zone contact(s): /OP2/  
Consider K in place of MG (1 valid contact(s) for K vs. 0 for MG)  

### MG:BA:1827

|  | Res. type | Atom | Residue | Distance | Occupancy | B-factor | Closest solute to 1^st^ shell water |
| --- | --- | --- | --- | --- | --- | --- | --- |
| **Ion** | | | |  | 1.00 | 101.25 || 1 | C | O2 | BA:479 | 3.21 | 1.00 | 102.98 |
| 2 | G | OP1 | BA:384 | 3.42 | 1.00 | 81.98 |  |
| 3 | C | N4 | BA:454 | 3.83 | 1.00 | 87.96 |  |
| 4 | C | N3 | BA:454 | 4.14 | 1.00 | 89.27 |  |
| 5 | G | OP2 | BA:384 | 4.40 | 1.00 | 83.13 |  |
| 6 | A | O3' | BA:383 | 4.95 | 1.00 | 83.02 |  |
  |

  
 Coded contact list:   
 Exclusion zone contact(s): /O2/  
Consider K in place of MG (1 valid contact(s) for K vs. 0 for MG)  

### MG:BA:1828

|  | Res. type | Atom | Residue | Distance | Occupancy | B-factor | Closest solute to 1^st^ shell water |
| --- | --- | --- | --- | --- | --- | --- | --- |
| **Ion** | | | |  | 1.00 | 114.78 || 1 | C | O2' | BA:455 | 3.97 | 1.00 | 105.88 |
| 2 | G | OP1 | BA:384 | 4.17 | 1.00 | 81.98 |  |
| 3 | A | O3' | BA:383 | 4.28 | 1.00 | 83.02 |  |
| 4 | C | O2 | BA:455 | 4.96 | 1.00 | 108.22 |  |
  |

  
 Coded contact list:   
1^st^ MG coordination shell missing (see Notes)  

### MG:BA:1829

|  | Res. type | Atom | Residue | Distance | Occupancy | B-factor | Closest solute to 1^st^ shell water |
| --- | --- | --- | --- | --- | --- | --- | --- |
| **Ion** | | | |  | 1.00 | 75.24 || 1 | C | OP1 | BA:372 | 3.68 | 1.00 | 66.93 |
| 2 | A | OP1 | BA:373 | 4.90 | 1.00 | 77.48 |  |
  |

  
 Coded contact list:   
1^st^ MG coordination shell missing (see Notes)  

### MG:BA:1830

|  | Res. type | Atom | Residue | Distance | Occupancy | B-factor | Closest solute to 1^st^ shell water |
| --- | --- | --- | --- | --- | --- | --- | --- |
| **Ion** | | | |  | 1.00 | 90.52 || 1 | C | OP1 | BA:385 | 4.45 | 1.00 | 80.64 |
| 2 | C | OP2 | BA:385 | 4.47 | 1.00 | 80.61 |  |
  |

  
 Coded contact list:   
1^st^ MG coordination shell missing (see Notes)  

### MG:BA:1831

|  | Res. type | Atom | Residue | Distance | Occupancy | B-factor | Closest solute to 1^st^ shell water |
| --- | --- | --- | --- | --- | --- | --- | --- |
| **Ion** | | | |  | 1.00 | 81.02 || 1 | A | O2' | BA:382 | 2.50 | 1.00 | 89.64 |
| 2 | A | O4' | BA:383 | 3.61 | 1.00 | 83.56 |  |
| 3 | A | N3 | BA:382 | 4.04 | 1.00 | 88.50 |  |
| 4 | A | O4' | BA:382 | 4.42 | 1.00 | 89.10 |  |
| 5 | A | N9 | BA:382 | 4.51 | 1.00 | 88.58 |  |
  |

  
 Coded contact list:   
 Exclusion zone contact(s): /O2'/  
Consider NA in place of MG (1 valid contact(s) for NA vs. 0 for MG)  

### MG:BA:1832

|  | Res. type | Atom | Residue | Distance | Occupancy | B-factor | Closest solute to 1^st^ shell water |
| --- | --- | --- | --- | --- | --- | --- | --- |
| **Ion** | | | |  | 1.00 | 89.12 || 1 | C | O2 | BA:381 | 3.68 | 1.00 | 85.50 |
| 2 | U | O4 | BA:65 | 4.24 | 1.00 | 87.51 |  |
| 3 | C | N3 | BA:381 | 4.70 | 1.00 | 86.05 |  |
| 4 | A | O4' | BA:382 | 4.82 | 1.00 | 89.10 |  |
  |

  
 Coded contact list:   
1^st^ MG coordination shell missing (see Notes)  

### MG:BA:1833

|  | Res. type | Atom | Residue | Distance | Occupancy | B-factor | Closest solute to 1^st^ shell water |
| --- | --- | --- | --- | --- | --- | --- | --- |
| **Ion** | | | |  | 1.00 | 71.41 || 1 | C | OP1 | BA:456 | 2.65 | 1.00 | 109.46 |
| 2 | C | OP2 | BA:456 | 3.03 | 1.00 | 109.48 |  |
| 3 | C | O3' | BA:455 | 4.16 | 1.00 | 108.26 |  |
| 4 | C | O5' | BA:456 | 4.61 | 1.00 | 110.48 |  |
| 5 | C | OP1 | BA:455 | 4.94 | 1.00 | 96.64 |  |
  |

  
 Coded contact list:   
 Exclusion zone contact(s): /OP1/OP2/  
Consider K in place of MG (2 valid contact(s) for K vs. 0 for MG)  

### MG:BA:1834

|  | Res. type | Atom | Residue | Distance | Occupancy | B-factor | Closest solute to 1^st^ shell water |
| --- | --- | --- | --- | --- | --- | --- | --- |
| **Ion** | | | |  | 1.00 | 70.60 || 1 | C | OP2 | BA:457 | 2.97 | 1.00 | 117.03 |
| 2 | C | OP2 | BA:458 | 3.73 | 1.00 | 125.70 |  |
| 3 | G | O6 | BA:464 | 3.94 | 1.00 | 130.72 |  |
| 4 | C | O5' | BA:457 | 4.08 | 1.00 | 118.69 |  |
| 5 | C | O4' | BA:466 | 4.28 | 1.00 | 137.36 |  |
| 6 | C | OP1 | BA:466 | 4.31 | 1.00 | 137.35 |  |
| 7 | C | OP1 | BA:457 | 4.73 | 1.00 | 117.13 |  |
| 8 | U | O4 | BA:208 | 4.74 | 1.00 | 157.27 |  |
  |

  
 Coded contact list:   
 Exclusion zone contact(s): /OP2/  
Consider K in place of MG (1 valid contact(s) for K vs. 0 for MG)  

### MG:BA:1835

|  | Res. type | Atom | Residue | Distance | Occupancy | B-factor | Closest solute to 1^st^ shell water |
| --- | --- | --- | --- | --- | --- | --- | --- |
| **Ion** | | | |  | 1.00 | 72.95 || 1 | C | OP1 | BA:466 | 2.54 | 1.00 | 137.35 |
| 2 | G | OP2 | BA:464 | 2.69 | 1.00 | 128.70 |  |
| 3 | C | OP2 | BA:466 | 2.92 | 1.00 | 137.62 |  |
| 4 | A | O3' | BA:465 | 3.48 | 1.00 | 137.10 |  |
| 5 | C | OP2 | BA:458 | 3.77 | 1.00 | 125.70 |  |
| 6 | G | N7 | BA:464 | 4.21 | 1.00 | 132.71 |  |
| 7 | C | O5' | BA:458 | 4.43 | 1.00 | 126.37 |  |
| 8 | C | OP1 | BA:458 | 4.45 | 1.00 | 126.07 |  |
| 9 | C | O5' | BA:466 | 4.55 | 1.00 | 137.73 |  |
| 10 | G | O5' | BA:464 | 4.78 | 1.00 | 130.03 |  |
  |

  
 Coded contact list:   
 Exclusion zone contact(s): /OP1/OP2/OP2/  
Consider K in place of MG (2 valid contact(s) for K vs. 0 for MG)  

### MG:BA:1836

|  | Res. type | Atom | Residue | Distance | Occupancy | B-factor | Closest solute to 1^st^ shell water |
| --- | --- | --- | --- | --- | --- | --- | --- |
| **Ion** | | | |  | 1.00 | 57.71 || 1 | A | O5' | BA:453 | 2.46 | 1.00 | 78.80 |
| 2 | A | O3' | BA:452 | 2.59 | 1.00 | 78.18 |  |
| 3 | A | OP1 | BA:453 | 2.64 | 1.00 | 78.76 |  |
| 4 | A | O2' | BA:452 | 3.44 | 1.00 | 80.02 |  |
| 5 | ARG | NE | BS:72 | 3.89 | 1.00 | 77.53 |  |
| 6 | A | OP2 | BA:453 | 4.17 | 1.00 | 77.92 |  |
| 7 | ARG | O | BS:72 | 4.61 | 1.00 | 76.56 |  |
| 8 | A | O4' | BA:453 | 4.78 | 1.00 | 78.84 |  |
| 9 | GLN | NE2 | BS:76 | 4.81 | 1.00 | 87.97 |  |
  |

  
 Coded contact list:   
 Exclusion zone contact(s): /O3'/O5'/OP1/  
Consider NA in place of MG (2 valid contact(s) for NA vs. 0 for MG)  

### MG:BA:1837

|  | Res. type | Atom | Residue | Distance | Occupancy | B-factor | Closest solute to 1^st^ shell water |
| --- | --- | --- | --- | --- | --- | --- | --- |
| **Ion** | | | |  | 1.00 | 88.25 || 1 | A | OP1 | BA:453 | 2.09 | 1.00 | 78.76 |
| 2 | A | O5' | BA:453 | 2.28 | 1.00 | 78.80 |  |
| 3 | A | OP2 | BA:453 | 3.53 | 1.00 | 77.92 |  |
| 4 | A | O3' | BA:452 | 3.96 | 1.00 | 78.18 |  |
| 5 | A | O2' | BA:452 | 4.68 | 1.00 | 80.02 |  |
| 6 | A | O4' | BA:453 | 4.69 | 1.00 | 78.84 |  |
| 7 | A | O3' | BA:453 | 4.90 | 1.00 | 81.66 |  |
  |

  
**1^st^ shell ligand angles:**   
1-2: 69;   
  
 Contact(s): /O5'/OP1/  
 Coded contact list: O_ph_O_r_  

### MG:BA:1838

|  | Res. type | Atom | Residue | Distance | Occupancy | B-factor | Closest solute to 1^st^ shell water |
| --- | --- | --- | --- | --- | --- | --- | --- |
| **Ion** | | | |  | 1.00 | 91.53 || 1 | G | O6 | BA:477 | 3.56 | 1.00 | 119.38 |
| 2 | C | OP2 | BA:454 | 3.61 | 1.00 | 83.74 |  |
| 3 | C | N4 | BA:455 | 3.89 | 1.00 | 106.63 |  |
| 4 | A | OP1 | BA:453 | 4.07 | 1.00 | 78.76 |  |
| 5 | A | O5' | BA:453 | 4.22 | 1.00 | 78.80 |  |
| 6 | G | O6 | BA:476 | 4.25 | 1.00 | 113.58 |  |
| 7 | A | OP2 | BA:453 | 4.56 | 1.00 | 77.92 |  |
| 8 | ARG | NH1 | BS:75 | 4.62 | 1.00 | 88.02 |  |
| 9 | A | O3' | BA:453 | 4.63 | 1.00 | 81.66 |  |
| 10 | C | N4 | BA:456 | 4.96 | 1.00 | 112.72 |  |
  |

  
 Coded contact list:   
1^st^ MG coordination shell missing (see Notes)  

### MG:BA:1839

|  | Res. type | Atom | Residue | Distance | Occupancy | B-factor | Closest solute to 1^st^ shell water |
| --- | --- | --- | --- | --- | --- | --- | --- |
| **Ion** | | | |  | 1.00 | 136.72 || 1 | C | OP2 | BA:454 | 2.92 | 1.00 | 83.74 |
| 2 | C | N4 | BA:456 | 3.02 | 1.00 | 112.72 |  |
| 3 | G | O6 | BA:476 | 3.28 | 1.00 | 113.58 |  |
| 4 | C | N4 | BA:455 | 3.34 | 1.00 | 106.63 |  |
| 5 | G | O6 | BA:477 | 4.03 | 1.00 | 119.38 |  |
| 6 | C | OP1 | BA:454 | 4.16 | 1.00 | 82.73 |  |
| 7 | ARG | NH1 | BS:75 | 4.37 | 1.00 | 88.02 |  |
| 8 | G | O6 | BA:475 | 4.78 | 1.00 | 122.50 |  |
| 9 | C | N4 | BA:457 | 4.79 | 1.00 | 123.06 |  |
| 10 | A | O3' | BA:453 | 4.90 | 1.00 | 81.66 |  |
  |

  
 Coded contact list:   
 Exclusion zone contact(s): /N4/N4/O6/OP2/  
Consider K in place of MG (4 valid contact(s) for K vs. 0 for MG)  

### MG:BA:1840

|  | Res. type | Atom | Residue | Distance | Occupancy | B-factor | Closest solute to 1^st^ shell water |
| --- | --- | --- | --- | --- | --- | --- | --- |
| **Ion** | | | |  | 1.00 | 156.86 || 1 | C | N4 | BA:457 | 2.07 | 1.00 | 123.06 |
| 2 | C | N4 | BA:456 | 2.09 | 1.00 | 112.72 |  |
| 3 | G | O6 | BA:476 | 2.61 | 1.00 | 113.58 |  |
| 4 | G | O6 | BA:475 | 3.28 | 1.00 | 122.50 |  |
| 5 | C | N3 | BA:456 | 3.73 | 1.00 | 112.60 |  |
| 6 | C | N3 | BA:457 | 3.97 | 1.00 | 123.12 |  |
| 7 | G | N1 | BA:476 | 4.08 | 1.00 | 113.93 |  |
| 8 | C | N4 | BA:458 | 4.43 | 1.00 | 125.93 |  |
| 9 | C | N4 | BA:455 | 4.72 | 1.00 | 106.63 |  |
| 10 | G | N1 | BA:475 | 4.73 | 1.00 | 122.27 |  |
| 11 | G | O6 | BA:474 | 4.87 | 1.00 | 124.55 |  |
  |

  
**1^st^ shell ligand angles:**   
1-2: 115;   
  
 Contact(s): /N4/N4/  
 Coded contact list: 2Anion  
 Exclusion zone contact(s): /O6/O6/  

### MG:BA:1841

|  | Res. type | Atom | Residue | Distance | Occupancy | B-factor | Closest solute to 1^st^ shell water |
| --- | --- | --- | --- | --- | --- | --- | --- |
| **Ion** | | | |  | 1.00 | 108.03 || 1 | G | O6 | BA:475 | 2.08 | 1.00 | 122.50 |
| 2 | G | O6 | BA:476 | 2.87 | 1.00 | 113.58 |  |
| 3 | G | N7 | BA:475 | 3.52 | 1.00 | 121.35 |  |
| 4 | C | N4 | BA:457 | 3.79 | 1.00 | 123.06 |  |
| 5 | G | N1 | BA:475 | 3.89 | 1.00 | 122.27 |  |
| 6 | G | N7 | BA:476 | 3.89 | 1.00 | 115.39 |  |
| 7 | G | O6 | BA:474 | 3.89 | 1.00 | 124.55 |  |
| 8 | C | N4 | BA:456 | 4.46 | 1.00 | 112.72 |  |
| 9 | G | N7 | BA:474 | 4.93 | 1.00 | 122.40 |  |
| 10 | C | N4 | BA:458 | 4.94 | 1.00 | 125.93 |  |
| 11 | G | N1 | BA:476 | 4.95 | 1.00 | 113.93 |  |
  |

  
 Contact(s): /O6/  
 Coded contact list: O_b_  
 Exclusion zone contact(s): /O6/  

### MG:BA:1842

|  | Res. type | Atom | Residue | Distance | Occupancy | B-factor | Closest solute to 1^st^ shell water |
| --- | --- | --- | --- | --- | --- | --- | --- |
| **Ion** | | | |  | 1.00 | 92.20 || 1 | G | O6 | BA:476 | 3.99 | 1.00 | 113.58 |
| 2 | ARG | NH1 | BS:75 | 4.11 | 1.00 | 88.02 |  |
| 3 | G | O6 | BA:475 | 4.25 | 1.00 | 122.50 |  |
| 4 | G | N7 | BA:475 | 4.51 | 1.00 | 121.35 |  |
| 5 | G | N7 | BA:476 | 4.75 | 1.00 | 115.39 |  |
  |

  
 Coded contact list:   
1^st^ MG coordination shell missing (see Notes)  

### MG:BA:1843

|  | Res. type | Atom | Residue | Distance | Occupancy | B-factor | Closest solute to 1^st^ shell water |
| --- | --- | --- | --- | --- | --- | --- | --- |
| **Ion** | | | |  | 1.00 | 63.47 || 1 | G | OP2 | BA:475 | 4.15 | 1.00 | 119.00 |
| 2 | G | N7 | BA:475 | 4.45 | 1.00 | 121.35 |  |
| 3 | ARG | NH1 | BS:75 | 4.52 | 1.00 | 88.02 |  |
  |

  
 Coded contact list:   
1^st^ MG coordination shell missing (see Notes)  

### MG:BA:1844

|  | Res. type | Atom | Residue | Distance | Occupancy | B-factor | Closest solute to 1^st^ shell water |
| --- | --- | --- | --- | --- | --- | --- | --- |
| **Ion** | | | |  | 1.00 | 118.88 || 1 | G | OP2 | BA:475 | 1.89 | 1.00 | 119.00 |
| 2 | G | N7 | BA:475 | 3.65 | 1.00 | 121.35 |  |
| 3 | G | O5' | BA:475 | 3.89 | 1.00 | 119.34 |  |
| 4 | G | O3' | BA:474 | 4.22 | 1.00 | 119.56 |  |
| 5 | G | OP1 | BA:475 | 4.33 | 1.00 | 119.14 |  |
| 6 | G | N9 | BA:475 | 4.55 | 1.00 | 120.42 |  |
| 7 | G | O4' | BA:475 | 4.65 | 1.00 | 118.80 |  |
| 8 | G | O5' | BA:474 | 4.79 | 1.00 | 121.44 |  |
| 9 | G | OP2 | BA:476 | 4.80 | 1.00 | 117.73 |  |
  |

  
 Contact(s): /OP2/  
 Coded contact list: O_ph_  

### MG:BA:1845

|  | Res. type | Atom | Residue | Distance | Occupancy | B-factor | Closest solute to 1^st^ shell water |
| --- | --- | --- | --- | --- | --- | --- | --- |
| **Ion** | | | |  | 1.00 | 92.53 || 1 | G | N7 | BA:476 | 1.91 | 1.00 | 115.39 |
| 2 | G | O6 | BA:476 | 3.47 | 1.00 | 113.58 |  |
| 3 | G | N9 | BA:476 | 3.82 | 1.00 | 115.94 |  |
| 4 | G | OP2 | BA:476 | 3.91 | 1.00 | 117.73 |  |
| 5 | G | N7 | BA:477 | 4.20 | 1.00 | 118.98 |  |
| 6 | G | N7 | BA:475 | 4.60 | 1.00 | 121.35 |  |
| 7 | G | O5' | BA:476 | 4.68 | 1.00 | 116.95 |  |
| 8 | G | N1 | BA:476 | 4.74 | 1.00 | 113.93 |  |
| 9 | G | O6 | BA:475 | 4.80 | 1.00 | 122.50 |  |
| 10 | G | N1 | BA:475 | 4.88 | 1.00 | 122.27 |  |
| 11 | G | O6 | BA:477 | 4.98 | 1.00 | 119.38 |  |
  |

  
 Contact(s): /N7/  
 Coded contact list: N_b_  

### MG:BA:1846

|  | Res. type | Atom | Residue | Distance | Occupancy | B-factor | Closest solute to 1^st^ shell water |
| --- | --- | --- | --- | --- | --- | --- | --- |
| **Ion** | | | |  | 1.00 | 63.45 || 1 | G | N7 | BA:477 | 3.22 | 1.00 | 118.98 |
| 2 | A | OP1 | BA:453 | 3.97 | 1.00 | 78.76 |  |
| 3 | G | O6 | BA:477 | 4.01 | 1.00 | 119.38 |  |
| 4 | A | OP2 | BA:453 | 4.53 | 1.00 | 77.92 |  |
| 5 | G | OP2 | BA:477 | 4.56 | 1.00 | 118.10 |  |
  |

  
 Coded contact list:   
 Exclusion zone contact(s): /N7/  
Consider K in place of MG (1 valid contact(s) for K vs. 0 for MG)  

### MG:BA:1847

|  | Res. type | Atom | Residue | Distance | Occupancy | B-factor | Closest solute to 1^st^ shell water |
| --- | --- | --- | --- | --- | --- | --- | --- |
| **Ion** | | | |  | 1.00 | 74.65 || 1 | A | OP2 | BA:453 | 1.65 | 1.00 | 77.92 |
| 2 | A | O3' | BA:452 | 3.53 | 1.00 | 78.18 |  |
| 3 | A | N6 | BA:478 | 3.60 | 1.00 | 115.61 |  |
| 4 | C | N4 | BA:479 | 3.88 | 1.00 | 102.96 |  |
| 5 | A | N7 | BA:453 | 3.95 | 1.00 | 74.32 |  |
| 6 | A | O5' | BA:453 | 4.02 | 1.00 | 78.80 |  |
| 7 | A | OP1 | BA:453 | 4.05 | 1.00 | 78.76 |  |
| 8 | A | O2' | BA:452 | 4.40 | 1.00 | 80.02 |  |
| 9 | A | O4' | BA:453 | 4.58 | 1.00 | 78.84 |  |
| 10 | A | N9 | BA:452 | 4.59 | 1.00 | 78.07 |  |
| 11 | A | N9 | BA:453 | 4.80 | 1.00 | 75.99 |  |
| 12 | G | O6 | BA:477 | 4.95 | 1.00 | 119.38 |  |
  |

  
 Coded contact list:   
 Exclusion zone contact(s): /OP2/  
1^st^ MG coordination shell missing (see Notes)  

### MG:BA:1848

|  | Res. type | Atom | Residue | Distance | Occupancy | B-factor | Closest solute to 1^st^ shell water |
| --- | --- | --- | --- | --- | --- | --- | --- |
| **Ion** | | | |  | 1.00 | 56.88 || 1 | A | N9 | BA:452 | 3.08 | 1.00 | 78.07 |
| 2 | A | O4' | BA:452 | 3.19 | 1.00 | 77.89 |  |
| 3 | A | OP2 | BA:453 | 3.33 | 1.00 | 77.92 |  |
| 4 | A | O3' | BA:452 | 3.43 | 1.00 | 78.18 |  |
| 5 | A | N7 | BA:452 | 3.69 | 1.00 | 78.31 |  |
| 6 | A | O2' | BA:451 | 4.11 | 1.00 | 81.44 |  |
| 7 | C | N4 | BA:479 | 4.19 | 1.00 | 102.96 |  |
| 8 | A | O2' | BA:452 | 4.21 | 1.00 | 80.02 |  |
| 9 | A | N7 | BA:453 | 4.39 | 1.00 | 74.32 |  |
| 10 | U | O4 | BA:480 | 4.40 | 1.00 | 82.83 |  |
| 11 | A | O3' | BA:451 | 4.41 | 1.00 | 79.31 |  |
| 12 | A | O5' | BA:452 | 4.77 | 1.00 | 77.13 |  |
  |

  
 Coded contact list:   
 Exclusion zone contact(s): /O4'/OP2/  
Consider K in place of MG (3 valid contact(s) for K vs. 0 for MG)  

### MG:BA:1849

|  | Res. type | Atom | Residue | Distance | Occupancy | B-factor | Closest solute to 1^st^ shell water |
| --- | --- | --- | --- | --- | --- | --- | --- |
| **Ion** | | | |  | 1.00 | 74.74 || 1 | A | O3' | BA:451 | 2.79 | 1.00 | 79.31 |
| 2 | A | O2' | BA:451 | 3.06 | 1.00 | 81.44 |  |
| 3 | A | OP2 | BA:452 | 3.21 | 1.00 | 79.17 |  |
| 4 | A | O5' | BA:452 | 3.49 | 1.00 | 77.13 |  |
| 5 | A | O4' | BA:452 | 4.16 | 1.00 | 77.89 |  |
| 6 | A | OP1 | BA:452 | 4.84 | 1.00 | 79.12 |  |
  |

  
 Coded contact list:   
 Exclusion zone contact(s): /O2'/O3'/OP2/  
Consider K in place of MG (3 valid contact(s) for K vs. 0 for MG)  

### MG:BA:1850

|  | Res. type | Atom | Residue | Distance | Occupancy | B-factor | Closest solute to 1^st^ shell water |
| --- | --- | --- | --- | --- | --- | --- | --- |
| **Ion** | | | |  | 1.00 | 90.03 || 1 | A | O2' | BA:451 | 2.52 | 1.00 | 81.44 |
| 2 | A | O3' | BA:451 | 3.75 | 1.00 | 79.31 |  |
| 3 | G | OP2 | BA:450 | 4.19 | 1.00 | 81.52 |  |
| 4 | A | OP2 | BA:452 | 4.26 | 1.00 | 79.17 |  |
| 5 | A | N3 | BA:451 | 4.70 | 1.00 | 81.50 |  |
  |

  
 Coded contact list:   
 Exclusion zone contact(s): /O2'/  
Consider NA in place of MG (1 valid contact(s) for NA vs. 0 for MG)  

### MG:BA:1851

|  | Res. type | Atom | Residue | Distance | Occupancy | B-factor | Closest solute to 1^st^ shell water |
| --- | --- | --- | --- | --- | --- | --- | --- |
| **Ion** | | | |  | 1.00 | 142.74 || 1 | C | OP1 | BA:449 | 4.71 | 1.00 | 85.16 |
| 2 | A | O2' | BA:451 | 4.75 | 1.00 | 81.44 |  |
  |

  
 Coded contact list:   
1^st^ MG coordination shell missing (see Notes)  

### MG:BA:1852

|  | Res. type | Atom | Residue | Distance | Occupancy | B-factor | Closest solute to 1^st^ shell water |
| --- | --- | --- | --- | --- | --- | --- | --- |
| **Ion** | | | |  | 1.00 | 105.85 || 1 | C | OP1 | BA:449 | 1.90 | 1.00 | 85.16 |
| 2 | C | OP2 | BA:449 | 3.79 | 1.00 | 85.82 |  |
| 3 | C | O5' | BA:449 | 3.99 | 1.00 | 84.29 |  |
| 4 | G | OP2 | BA:450 | 4.27 | 1.00 | 81.52 |  |
| 5 | A | O3' | BA:448 | 4.34 | 1.00 | 85.11 |  |
  |

  
 Contact(s): /OP1/  
 Coded contact list: O_ph_  

### MG:BA:1854

|  | Res. type | Atom | Residue | Distance | Occupancy | B-factor | Closest solute to 1^st^ shell water |
| --- | --- | --- | --- | --- | --- | --- | --- |
| **Ion** | | | |  | 1.00 | 84.55 || 1 | G | N2 | BA:630 | 3.59 | 1.00 | 125.75 |
| 2 | G | O4' | BA:631 | 4.70 | 1.00 | 128.18 |  |
  |

  
 Coded contact list:   
1^st^ MG coordination shell missing (see Notes)  

### MG:BA:1855

|  | Res. type | Atom | Residue | Distance | Occupancy | B-factor | Closest solute to 1^st^ shell water |
| --- | --- | --- | --- | --- | --- | --- | --- |
| **Ion** | | | |  | 1.00 | 76.90 || 1 | G | N2 | BA:630 | 3.86 | 1.00 | 125.75 |
  |

  
 Coded contact list:   
1^st^ MG coordination shell missing (see Notes)  

### MG:BA:1857

|  | Res. type | Atom | Residue | Distance | Occupancy | B-factor | Closest solute to 1^st^ shell water |
| --- | --- | --- | --- | --- | --- | --- | --- |
| **Ion** | | | |  | 1.00 | 122.86 || 1 | C | O4' | BA:612 | 4.26 | 1.00 | 83.43 |
| 2 | C | O2' | BA:612 | 4.44 | 1.00 | 82.87 |  |
  |

  
 Coded contact list:   
1^st^ MG coordination shell missing (see Notes)  

### MG:BA:1858

|  | Res. type | Atom | Residue | Distance | Occupancy | B-factor | Closest solute to 1^st^ shell water |
| --- | --- | --- | --- | --- | --- | --- | --- |
| **Ion** | | | |  | 1.00 | 141.69 || 1 | C | O4' | BA:612 | 2.95 | 1.00 | 83.43 |
| 2 | C | O2' | BA:612 | 3.24 | 1.00 | 82.87 |  |
| 3 | C | N1 | BA:612 | 3.96 | 1.00 | 85.56 |  |
| 4 | G | N2 | BA:629 | 4.05 | 1.00 | 109.09 |  |
| 5 | C | O2 | BA:612 | 4.33 | 1.00 | 87.34 |  |
| 6 | A | N3 | BA:611 | 4.40 | 1.00 | 76.07 |  |
| 7 | A | O2' | BA:611 | 4.97 | 1.00 | 75.96 |  |
  |

  
 Coded contact list:   
 Exclusion zone contact(s): /O2'/O4'/  
Consider K in place of MG (2 valid contact(s) for K vs. 0 for MG)  

### MG:BA:1860

|  | Res. type | Atom | Residue | Distance | Occupancy | B-factor | Closest solute to 1^st^ shell water |
| --- | --- | --- | --- | --- | --- | --- | --- |
| **Ion** | | | |  | 1.00 | 60.80 || 1 | C | O2' | BA:612 | 4.89 | 1.00 | 82.87 |
  |

  
 Coded contact list:   
1^st^ MG coordination shell missing (see Notes)  

### MG:BA:1861

|  | Res. type | Atom | Residue | Distance | Occupancy | B-factor | Closest solute to 1^st^ shell water |
| --- | --- | --- | --- | --- | --- | --- | --- |
| **Ion** | | | |  | 1.00 | 78.55 || 1 | C | O2' | BA:612 | 4.60 | 1.00 | 82.87 |
  |

  
 Coded contact list:   
1^st^ MG coordination shell missing (see Notes)  

### MG:BA:1862

|  | Res. type | Atom | Residue | Distance | Occupancy | B-factor | Closest solute to 1^st^ shell water |
| --- | --- | --- | --- | --- | --- | --- | --- |
| **Ion** | | | |  | 1.00 | 77.67 || 1 | C | O2' | BA:612 | 4.78 | 1.00 | 82.87 |
  |

  
 Coded contact list:   
1^st^ MG coordination shell missing (see Notes)  

### MG:BA:1864

|  | Res. type | Atom | Residue | Distance | Occupancy | B-factor | Closest solute to 1^st^ shell water |
| --- | --- | --- | --- | --- | --- | --- | --- |
| **Ion** | | | |  | 1.00 | 95.08 || 1 | U | OP1 | BA:5 | 4.02 | 1.00 | 105.78 |
| 2 | U | OP2 | BA:5 | 4.54 | 1.00 | 106.11 |  |
  |

  
 Coded contact list:   
1^st^ MG coordination shell missing (see Notes)  

### MG:BA:1865

|  | Res. type | Atom | Residue | Distance | Occupancy | B-factor | Closest solute to 1^st^ shell water |
| --- | --- | --- | --- | --- | --- | --- | --- |
| **Ion** | | | |  | 1.00 | 75.70 || 1 | U | OP1 | BA:5 | 4.73 | 1.00 | 105.78 |
  |

  
 Coded contact list:   
1^st^ MG coordination shell missing (see Notes)  

### MG:BA:1866

|  | Res. type | Atom | Residue | Distance | Occupancy | B-factor | Closest solute to 1^st^ shell water |
| --- | --- | --- | --- | --- | --- | --- | --- |
| **Ion** | | | |  | 1.00 | 111.99 || 1 | U | OP1 | BA:5 | 4.16 | 1.00 | 105.78 |
| 2 | U | O5' | BA:5 | 4.33 | 1.00 | 104.11 |  |
| 3 | U | OP2 | BA:5 | 4.98 | 1.00 | 106.11 |  |
  |

  
 Coded contact list:   
1^st^ MG coordination shell missing (see Notes)  

### MG:BA:1867

|  | Res. type | Atom | Residue | Distance | Occupancy | B-factor | Closest solute to 1^st^ shell water |
| --- | --- | --- | --- | --- | --- | --- | --- |
| **Ion** | | | |  | 1.00 | 121.89 || 1 | G | O2' | BA:630 | 3.67 | 1.00 | 124.63 |
| 2 | G | N3 | BA:630 | 4.23 | 1.00 | 125.90 |  |
| 3 | G | N2 | BA:630 | 4.26 | 1.00 | 125.75 |  |
  |

  
 Coded contact list:   
1^st^ MG coordination shell missing (see Notes)  

### MG:BA:1868

|  | Res. type | Atom | Residue | Distance | Occupancy | B-factor | Closest solute to 1^st^ shell water |
| --- | --- | --- | --- | --- | --- | --- | --- |
| **Ion** | | | |  | 1.00 | 65.33 || 1 | C | O4' | BA:613 | 2.50 | 1.00 | 86.75 |
| 2 | C | O2' | BA:613 | 3.72 | 1.00 | 86.29 |  |
| 3 | C | N1 | BA:613 | 4.54 | 1.00 | 86.34 |  |
| 4 | C | O2' | BA:612 | 4.55 | 1.00 | 82.87 |  |
| 5 | C | O3' | BA:613 | 4.67 | 1.00 | 86.86 |  |
| 6 | C | O5' | BA:613 | 4.83 | 1.00 | 84.17 |  |
  |

  
 Coded contact list:   
 Exclusion zone contact(s): /O4'/  
Consider NA in place of MG (1 valid contact(s) for NA vs. 0 for MG)  

### MG:BA:1872

|  | Res. type | Atom | Residue | Distance | Occupancy | B-factor | Closest solute to 1^st^ shell water |
| --- | --- | --- | --- | --- | --- | --- | --- |
| **Ion** | | | |  | 1.00 | 103.39 || 1 | U | O5' | BA:4 | 2.92 | 1.00 | 102.46 |
| 2 | U | OP1 | BA:5 | 4.73 | 1.00 | 105.78 |  |
| 3 | U | O4 | BA:5 | 4.77 | 1.00 | 99.64 |  |
| 4 | U | O4' | BA:4 | 4.99 | 1.00 | 104.01 |  |
  |

  
 Coded contact list:   
 Exclusion zone contact(s): /O5'/  
Consider K in place of MG (1 valid contact(s) for K vs. 0 for MG)  

### MG:BA:1877

|  | Res. type | Atom | Residue | Distance | Occupancy | B-factor | Closest solute to 1^st^ shell water |
| --- | --- | --- | --- | --- | --- | --- | --- |
| **Ion** | | | |  | 1.00 | 75.60 || 1 | G | O2' | BA:630 | 3.84 | 1.00 | 124.63 |
| 2 | G | O3' | BA:630 | 4.79 | 1.00 | 126.77 |  |
  |

  
 Coded contact list:   
1^st^ MG coordination shell missing (see Notes)  

### MG:BA:1878

|  | Res. type | Atom | Residue | Distance | Occupancy | B-factor | Closest solute to 1^st^ shell water |
| --- | --- | --- | --- | --- | --- | --- | --- |
| **Ion** | | | |  | 1.00 | 75.45 || 1 | G | O2' | BA:629 | 2.51 | 1.00 | 111.08 |
| 2 | G | O3' | BA:629 | 4.78 | 1.00 | 114.83 |  |
| 3 | G | O4' | BA:630 | 4.92 | 1.00 | 124.31 |  |
  |

  
 Coded contact list:   
 Exclusion zone contact(s): /O2'/  
Consider NA in place of MG (1 valid contact(s) for NA vs. 0 for MG)  

### MG:BA:1879

|  | Res. type | Atom | Residue | Distance | Occupancy | B-factor | Closest solute to 1^st^ shell water |
| --- | --- | --- | --- | --- | --- | --- | --- |
| **Ion** | | | |  | 1.00 | 46.29 || 1 | G | OP2 | BA:617 | 3.96 | 1.00 | 82.32 |
| 2 | A | OP1 | BA:621 | 4.05 | 1.00 | 59.99 |  |
| 3 | G | N7 | BA:617 | 4.18 | 1.00 | 80.10 |  |
| 4 | A | OP2 | BA:621 | 4.23 | 1.00 | 61.06 |  |
| 5 | C | OP1 | BA:620 | 4.50 | 1.00 | 67.48 |  |
| 6 | ARG | NH2 | BG:141 | 4.93 | 1.00 | 87.83 |  |
  |

  
 Coded contact list:   
1^st^ MG coordination shell missing (see Notes)  

### MG:BA:1880

|  | Res. type | Atom | Residue | Distance | Occupancy | B-factor | Closest solute to 1^st^ shell water |
| --- | --- | --- | --- | --- | --- | --- | --- |
| **Ion** | | | |  | 1.00 | 60.98 || 1 | C | OP2 | BA:596 | 2.10 | 1.00 | 80.32 |
| 2 | G | OP2 | BA:597 | 2.12 | 1.00 | 78.48 |  |
| 3 | U | O4 | BA:598 | 2.29 | 1.00 | 83.63 |  |
| 4 | C | OP1 | BA:596 | 3.85 | 1.00 | 80.35 |  |
| 5 | A | N6 | BA:640 | 3.87 | 1.00 | 85.00 |  |
| 6 | G | O2' | BA:595 | 3.98 | 1.00 | 85.35 |  |
| 7 | C | O5' | BA:596 | 4.05 | 1.00 | 79.06 |  |
| 8 | G | OP1 | BA:597 | 4.26 | 1.00 | 78.21 |  |
| 9 | C | N4 | BA:599 | 4.28 | 1.00 | 84.41 |  |
| 10 | C | O3' | BA:596 | 4.42 | 1.00 | 78.22 |  |
| 11 | G | O5' | BA:597 | 4.49 | 1.00 | 77.45 |  |
| 12 | U | N3 | BA:598 | 4.54 | 1.00 | 82.17 |  |
| 13 | G | O3' | BA:595 | 4.61 | 1.00 | 81.11 |  |
| 14 | G | N7 | BA:597 | 4.89 | 1.00 | 76.56 |  |
| 15 | G | O6 | BA:639 | 4.99 | 1.00 | 85.89 |  |
  |

  
**1^st^ shell ligand angles:**   
1-2: 79; 1-3: 122; 2-3: 104;   
  
 Contact(s): /O4/OP2/OP2/  
 Coded contact list: 2O_ph_O_b_  

### MG:BA:1881

|  | Res. type | Atom | Residue | Distance | Occupancy | B-factor | Closest solute to 1^st^ shell water |
| --- | --- | --- | --- | --- | --- | --- | --- |
| **Ion** | | | |  | 1.00 | 46.44 || 1 | C | OP1 | BA:972 | 2.13 | 1.00 | 91.10 |
| 2 | C | OP2 | BA:972 | 2.69 | 1.00 | 90.61 |  |
| 3 | C | O5' | BA:972 | 3.57 | 1.00 | 91.29 |  |
| 4 | G | O3' | BA:971 | 4.15 | 1.00 | 91.34 |  |
| 5 | C | OP1 | BA:1366 | 4.52 | 1.00 | 92.55 |  |
| 6 | ASP | OD1 | BM:58 | 4.65 | 1.00 | 87.46 |  |
| 7 | G | OP1 | BA:971 | 4.97 | 1.00 | 89.40 |  |
  |

  
 Contact(s): /OP1/  
 Coded contact list: O_ph_  
 Exclusion zone contact(s): /OP2/  

### MG:BA:1882

|  | Res. type | Atom | Residue | Distance | Occupancy | B-factor | Closest solute to 1^st^ shell water |
| --- | --- | --- | --- | --- | --- | --- | --- |
| **Ion** | | | |  | 1.00 | 42.06 || 1 | G | O6 | BA:683 | 3.98 | 1.00 | 71.63 |
| 2 | A | OP1 | BA:704 | 4.06 | 1.00 | 77.82 |  |
| 3 | A | N6 | BA:684 | 4.10 | 1.00 | 86.94 |  |
| 4 | G | O6 | BA:682 | 4.26 | 1.00 | 76.55 |  |
| 5 | G | N7 | BA:683 | 4.27 | 1.00 | 77.56 |  |
| 6 | G | N7 | BA:682 | 4.57 | 1.00 | 80.16 |  |
| 7 | A | O5' | BA:704 | 4.76 | 1.00 | 76.59 |  |
  |

  
 Coded contact list:   
1^st^ MG coordination shell missing (see Notes)  

### MG:BA:1883

|  | Res. type | Atom | Residue | Distance | Occupancy | B-factor | Closest solute to 1^st^ shell water |
| --- | --- | --- | --- | --- | --- | --- | --- |
| **Ion** | | | |  | 1.00 | 64.59 || 1 | G | O6 | BA:167 | 3.78 | 1.00 | 111.91 |
| 2 | G | O6 | BA:166 | 4.12 | 1.00 | 125.59 |  |
| 3 | G | N7 | BA:166 | 4.15 | 1.00 | 122.92 |  |
| 4 | A | OP1 | BA:151 | 4.38 | 1.00 | 85.32 |  |
| 5 | G | N7 | BA:167 | 4.42 | 1.00 | 108.27 |  |
| 6 | A | OP2 | BA:152 | 4.44 | 1.00 | 88.00 |  |
| 7 | G | O6 | BA:168 | 4.54 | 1.00 | 105.13 |  |
  |

  
 Coded contact list:   
1^st^ MG coordination shell missing (see Notes)  

### MG:BA:1884

|  | Res. type | Atom | Residue | Distance | Occupancy | B-factor | Closest solute to 1^st^ shell water |
| --- | --- | --- | --- | --- | --- | --- | --- |
| **Ion** | | | |  | 1.00 | 67.35 || 1 | G | O6 | BA:1385 | 4.18 | 1.00 | 93.33 |
| 2 | G | N7 | BA:1386 | 4.20 | 1.00 | 85.23 |  |
| 3 | G | N7 | BA:1385 | 4.22 | 1.00 | 91.16 |  |
| 4 | G | O6 | BA:1386 | 4.32 | 1.00 | 86.41 |  |
  |

  
 Coded contact list:   
1^st^ MG coordination shell missing (see Notes)  

### MG:BA:1885

|  | Res. type | Atom | Residue | Distance | Occupancy | B-factor | Closest solute to 1^st^ shell water |
| --- | --- | --- | --- | --- | --- | --- | --- |
| **Ion** | | | |  | 1.00 | 44.28 || 1 | G | OP1 | BA:550 | 3.61 | 1.00 | 59.33 |
| 2 | G | O2' | BA:502 | 3.72 | 1.00 | 70.37 |  |
| 3 | C | O4' | BA:503 | 4.11 | 1.00 | 71.00 |  |
| 4 | G | N3 | BA:502 | 4.30 | 1.00 | 71.82 |  |
| 5 | G | N2 | BA:502 | 4.61 | 1.00 | 71.40 |  |
| 6 | G | O5' | BA:550 | 4.64 | 1.00 | 60.75 |  |
| 7 | A | N6 | BA:509 | 4.93 | 1.00 | 85.86 |  |
  |

  
 Coded contact list:   
1^st^ MG coordination shell missing (see Notes)  

### MG:BA:1886

|  | Res. type | Atom | Residue | Distance | Occupancy | B-factor | Closest solute to 1^st^ shell water |
| --- | --- | --- | --- | --- | --- | --- | --- |
| **Ion** | | | |  | 1.00 | 64.75 || 1 | G | OP2 | BA:167 | 3.74 | 1.00 | 112.09 |
| 2 | A | OP2 | BA:151 | 4.00 | 1.00 | 85.78 |  |
| 3 | C | OP1 | BA:150 | 4.02 | 1.00 | 87.81 |  |
| 4 | G | OP2 | BA:168 | 4.26 | 1.00 | 102.28 |  |
| 5 | A | OP1 | BA:151 | 4.97 | 1.00 | 85.32 |  |
  |

  
 Coded contact list:   
1^st^ MG coordination shell missing (see Notes)  

### MG:BA:1887

|  | Res. type | Atom | Residue | Distance | Occupancy | B-factor | Closest solute to 1^st^ shell water |
| --- | --- | --- | --- | --- | --- | --- | --- |
| **Ion** | | | |  | 1.00 | 56.84 || 1 | G | O6 | BA:305 | 2.47 | 1.00 | 71.35 |
| 2 | G | O6 | BA:293 | 2.76 | 1.00 | 61.02 |  |
| 3 | U | O4 | BA:304 | 2.99 | 1.00 | 62.03 |  |
| 4 | G | N1 | BA:305 | 3.42 | 1.00 | 71.26 |  |
| 5 | U | N3 | BA:304 | 3.97 | 1.00 | 63.07 |  |
| 6 | G | OP2 | BA:292 | 4.18 | 1.00 | 54.35 |  |
| 7 | G | N7 | BA:293 | 4.29 | 1.00 | 62.64 |  |
| 8 | A | N6 | BA:303 | 4.55 | 1.00 | 69.16 |  |
| 9 | U | O4 | BA:294 | 4.77 | 1.00 | 68.66 |  |
| 10 | G | N7 | BA:305 | 4.88 | 1.00 | 71.31 |  |
| 11 | G | N1 | BA:293 | 4.94 | 1.00 | 61.19 |  |
  |

  
 Coded contact list:   
 Exclusion zone contact(s): /O4/O6/O6/  
Consider K in place of MG (2 valid contact(s) for K vs. 0 for MG)  

### MG:BA:1888

|  | Res. type | Atom | Residue | Distance | Occupancy | B-factor | Closest solute to 1^st^ shell water |
| --- | --- | --- | --- | --- | --- | --- | --- |
| **Ion** | | | |  | 1.00 | 67.74 || 1 | C | N4 | BA:1411 | 4.77 | 1.00 | 64.32 |
| 2 | G | O6 | BA:1410 | 4.89 | 1.00 | 55.78 |  |
  |

  
 Coded contact list:   
1^st^ MG coordination shell missing (see Notes)  

### MG:BA:1889

|  | Res. type | Atom | Residue | Distance | Occupancy | B-factor | Closest solute to 1^st^ shell water |
| --- | --- | --- | --- | --- | --- | --- | --- |
| **Ion** | | | |  | 1.00 | 37.02 || 1 | G | O6 | BA:529 | 2.74 | 1.00 | 65.02 |
| 2 | C | N3 | BA:518 | 2.96 | 1.00 | 65.78 |  |
| 3 | ASN | ND2 | BO:49 | 2.97 | 1.00 | 67.02 |  |
| 4 | G | N7 | BA:529 | 3.09 | 1.00 | 67.75 |  |
| 5 | PRO | O | BO:48 | 3.21 | 1.00 | 69.02 |  |
| 6 | C | N4 | BA:518 | 3.50 | 1.00 | 65.06 |  |
| 7 | C | O2 | BA:518 | 3.98 | 1.00 | 66.15 |  |
| 8 | ASN | N | BO:49 | 4.04 | 1.00 | 66.48 |  |
| 9 | C | N4 | BA:528 | 4.36 | 1.00 | 66.96 |  |
| 10 | C | N1 | BA:518 | 4.54 | 1.00 | 67.39 |  |
| 11 | G | N1 | BA:529 | 4.83 | 1.00 | 63.65 |  |
| 12 | U | O2' | B1:51 | 4.91 | 1.00 | 101.66 |  |
  |

  
 Coded contact list:   
 Exclusion zone contact(s): /N3_C_/N7/ND2_Asn_/O6/O_back_/  
Consider K in place of MG (5 valid contact(s) for K vs. 0 for MG)  

### MG:BA:1890

|  | Res. type | Atom | Residue | Distance | Occupancy | B-factor | Closest solute to 1^st^ shell water |
| --- | --- | --- | --- | --- | --- | --- | --- |
| **Ion** | | | |  | 1.00 | 72.40 || 1 | G | O6 | BA:742 | 3.67 | 1.00 | 72.97 |
| 2 | G | O6 | BA:741 | 3.96 | 1.00 | 63.72 |  |
| 3 | G | N7 | BA:741 | 4.17 | 1.00 | 61.64 |  |
| 4 | G | N7 | BA:742 | 4.29 | 1.00 | 70.61 |  |
| 5 | G | O6 | BA:664 | 4.83 | 1.00 | 68.63 |  |
  |

  
 Coded contact list:   
1^st^ MG coordination shell missing (see Notes)  

### MG:BA:1891

|  | Res. type | Atom | Residue | Distance | Occupancy | B-factor | Closest solute to 1^st^ shell water |
| --- | --- | --- | --- | --- | --- | --- | --- |
| **Ion** | | | |  | 1.00 | 60.06 || 1 | A | OP2 | BA:1110 | 2.34 | 1.00 | 96.38 |
| 2 | A | O5' | BA:1110 | 3.46 | 1.00 | 96.76 |  |
| 3 | A | N7 | BA:1111 | 3.65 | 1.00 | 104.91 |  |
| 4 | C | O2 | BA:1189 | 3.76 | 1.00 | 82.38 |  |
| 5 | A | OP1 | BA:1110 | 3.79 | 1.00 | 96.64 |  |
| 6 | A | N7 | BA:1110 | 3.85 | 1.00 | 96.91 |  |
| 7 | A | OP2 | BA:1111 | 4.13 | 1.00 | 97.10 |  |
| 8 | A | N6 | BA:1111 | 4.15 | 1.00 | 104.81 |  |
| 9 | C | OP2 | BA:1109 | 4.26 | 1.00 | 93.32 |  |
| 10 | C | N4 | BA:1112 | 4.63 | 1.00 | 105.38 |  |
| 11 | C | O3' | BA:1109 | 4.69 | 1.00 | 96.04 |  |
| 12 | A | N6 | BA:1188 | 5.00 | 1.00 | 86.91 |  |
  |

  
 Contact(s): /OP2/  
 Coded contact list: O_ph_  

### MG:BA:1892

|  | Res. type | Atom | Residue | Distance | Occupancy | B-factor | Closest solute to 1^st^ shell water |
| --- | --- | --- | --- | --- | --- | --- | --- |
| **Ion** | | | |  | 1.00 | 75.26 || 1 | G | N7 | BA:474 | 2.49 | 1.00 | 122.40 |
| 2 | G | O6 | BA:474 | 2.97 | 1.00 | 124.55 |  |
| 3 | G | O6 | BA:475 | 3.30 | 1.00 | 122.50 |  |
| 4 | G | N7 | BA:475 | 3.38 | 1.00 | 121.35 |  |
| 5 | G | N9 | BA:474 | 4.38 | 1.00 | 121.58 |  |
| 6 | G | N1 | BA:474 | 4.45 | 1.00 | 123.68 |  |
| 7 | A | OP2 | BA:468 | 4.67 | 1.00 | 122.08 |  |
  |

  
 Coded contact list:   
 Exclusion zone contact(s): /N7/N7/O6/O6/  
Consider K in place of MG (3 valid contact(s) for K vs. 0 for MG)  

### MG:BA:1893

|  | Res. type | Atom | Residue | Distance | Occupancy | B-factor | Closest solute to 1^st^ shell water |
| --- | --- | --- | --- | --- | --- | --- | --- |
| **Ion** | | | |  | 1.00 | 54.06 || 1 | G | N7 | BA:945 | 2.82 | 1.00 | 89.17 |
| 2 | U | OP2 | BA:943 | 2.93 | 1.00 | 84.13 |  |
| 3 | G | OP2 | BA:944 | 2.94 | 1.00 | 81.11 |  |
| 4 | G | N7 | BA:944 | 3.15 | 1.00 | 86.55 |  |
| 5 | G | N7 | BA:1337 | 3.19 | 1.00 | 83.17 |  |
| 6 | G | O6 | BA:945 | 3.34 | 1.00 | 92.30 |  |
| 7 | U | O5' | BA:943 | 3.97 | 1.00 | 84.16 |  |
| 8 | G | O5' | BA:944 | 4.22 | 1.00 | 81.47 |  |
| 9 | G | O6 | BA:1337 | 4.57 | 1.00 | 84.30 |  |
| 10 | U | OP1 | BA:943 | 4.60 | 1.00 | 84.33 |  |
| 11 | U | O3' | BA:943 | 4.67 | 1.00 | 82.27 |  |
| 12 | G | N9 | BA:1337 | 4.69 | 1.00 | 83.26 |  |
| 13 | G | N9 | BA:944 | 4.90 | 1.00 | 84.46 |  |
| 14 | U | N1 | BA:943 | 4.94 | 1.00 | 85.29 |  |
| 15 | G | N9 | BA:945 | 4.97 | 1.00 | 87.10 |  |
  |

  
 Coded contact list:   
 Exclusion zone contact(s): /N7/N7/N7/O6/OP2/OP2/  
Consider K in place of MG (6 valid contact(s) for K vs. 0 for MG)  

### MG:BA:1894

|  | Res. type | Atom | Residue | Distance | Occupancy | B-factor | Closest solute to 1^st^ shell water |
| --- | --- | --- | --- | --- | --- | --- | --- |
| **Ion** | | | |  | 1.00 | 83.05 || 1 | G | O6 | BA:594 | 4.38 | 1.00 | 86.45 |
| 2 | G | N1 | BA:595 | 4.61 | 1.00 | 85.26 |  |
| 3 | G | OP2 | BA:593 | 4.76 | 1.00 | 89.41 |  |
| 4 | U | O3' | BA:641 | 4.79 | 1.00 | 92.73 |  |
| 5 | G | O6 | BA:595 | 4.85 | 1.00 | 82.93 |  |
| 6 | G | N7 | BA:593 | 4.85 | 1.00 | 85.95 |  |
| 7 | G | N7 | BA:592 | 4.87 | 1.00 | 91.72 |  |
| 8 | G | N7 | BA:594 | 5.00 | 1.00 | 86.35 |  |
  |

  
 Coded contact list:   
1^st^ MG coordination shell missing (see Notes)  

### MG:BA:1895

|  | Res. type | Atom | Residue | Distance | Occupancy | B-factor | Closest solute to 1^st^ shell water |
| --- | --- | --- | --- | --- | --- | --- | --- |
| **Ion** | | | |  | 1.00 | 54.54 || 1 | G | O6 | BA:649 | 3.80 | 1.00 | 89.93 |
| 2 | G | O6 | BA:650 | 3.86 | 1.00 | 86.94 |  |
| 3 | G | N7 | BA:650 | 3.93 | 1.00 | 87.50 |  |
| 4 | G | N7 | BA:649 | 4.18 | 1.00 | 92.50 |  |
| 5 | A | OP1 | BA:753 | 4.25 | 1.00 | 79.42 |  |
  |

  
 Coded contact list:   
1^st^ MG coordination shell missing (see Notes)  

### MG:BA:1896

|  | Res. type | Atom | Residue | Distance | Occupancy | B-factor | Closest solute to 1^st^ shell water |
| --- | --- | --- | --- | --- | --- | --- | --- |
| **Ion** | | | |  | 1.00 | 83.10 || 1 | C | OP1 | BA:1303 | 2.49 | 1.00 | 112.66 |
| 2 | G | N7 | BA:1304 | 2.52 | 1.00 | 101.55 |  |
| 3 | G | O6 | BA:1305 | 2.73 | 1.00 | 101.68 |  |
| 4 | C | O5' | BA:1303 | 3.46 | 1.00 | 111.90 |  |
| 5 | C | OP2 | BA:1303 | 3.74 | 1.00 | 113.22 |  |
| 6 | G | OP2 | BA:1304 | 3.82 | 1.00 | 105.52 |  |
| 7 | G | N7 | BA:1305 | 3.86 | 1.00 | 100.41 |  |
| 8 | G | O6 | BA:1304 | 4.03 | 1.00 | 101.70 |  |
| 9 | G | N1 | BA:1305 | 4.49 | 1.00 | 100.59 |  |
| 10 | G | N9 | BA:1304 | 4.59 | 1.00 | 99.99 |  |
| 11 | U | O3' | BA:1302 | 4.77 | 1.00 | 114.27 |  |
| 12 | G | O5' | BA:1304 | 4.95 | 1.00 | 105.25 |  |
  |

  
 Coded contact list:   
 Exclusion zone contact(s): /N7/O6/OP1/  
Consider NA in place of MG (2 valid contact(s) for NA vs. 0 for MG)  

### MG:BA:1897

|  | Res. type | Atom | Residue | Distance | Occupancy | B-factor | Closest solute to 1^st^ shell water |
| --- | --- | --- | --- | --- | --- | --- | --- |
| **Ion** | | | |  | 1.00 | 65.25 || 1 | G | O6 | BA:710 | 3.46 | 1.00 | 68.60 |
| 2 | G | O6 | BA:709 | 3.65 | 1.00 | 79.32 |  |
| 3 | G | N7 | BA:709 | 3.93 | 1.00 | 77.70 |  |
| 4 | G | N7 | BA:710 | 4.15 | 1.00 | 69.91 |  |
| 5 | C | N4 | BA:708 | 4.86 | 1.00 | 75.87 |  |
| 6 | G | O6 | BA:711 | 4.99 | 1.00 | 68.83 |  |
  |

  
 Coded contact list:   
1^st^ MG coordination shell missing (see Notes)  

### MG:BA:1898

|  | Res. type | Atom | Residue | Distance | Occupancy | B-factor | Closest solute to 1^st^ shell water |
| --- | --- | --- | --- | --- | --- | --- | --- |
| **Ion** | | | |  | 1.00 | 49.78 || 1 | G | OP1 | BA:1475 | 3.08 | 1.00 | 65.29 |
| 2 | A | N3 | AA:1689 | 4.37 | 1.00 | 39.93 |  |
| 3 | A | N1 | AA:1700 | 4.72 | 1.00 | 44.20 |  |
| 4 | C | O5' | AA:1767 | 4.77 | 1.00 | 50.83 |  |
| 5 | G | OP2 | BA:1475 | 4.97 | 1.00 | 64.53 |  |
  |

  
 Coded contact list:   
 Exclusion zone contact(s): /OP1/  
Consider K in place of MG (1 valid contact(s) for K vs. 0 for MG)  

### MG:BA:1899

|  | Res. type | Atom | Residue | Distance | Occupancy | B-factor | Closest solute to 1^st^ shell water |
| --- | --- | --- | --- | --- | --- | --- | --- |
| **Ion** | | | |  | 1.00 | 57.84 || 1 | U | O3' | BA:1126 | 2.29 | 1.00 | 135.40 |
| 2 | G | O4' | BA:1127 | 2.53 | 1.00 | 141.86 |  |
| 3 | A | N6 | BA:1280 | 3.22 | 1.00 | 108.12 |  |
| 4 | G | O5' | BA:1127 | 3.53 | 1.00 | 137.66 |  |
| 5 | A | N1 | BA:1280 | 3.87 | 1.00 | 107.25 |  |
| 6 | G | N9 | BA:1127 | 3.90 | 1.00 | 142.13 |  |
| 7 | U | O5' | BA:1126 | 4.08 | 1.00 | 132.51 |  |
| 8 | A | N7 | BA:1280 | 4.34 | 1.00 | 107.77 |  |
| 9 | G | OP2 | BA:1127 | 4.42 | 1.00 | 137.07 |  |
| 10 | G | OP1 | BA:1127 | 4.50 | 1.00 | 136.41 |  |
| 11 | U | O4' | BA:1126 | 4.51 | 1.00 | 133.50 |  |
| 12 | G | N7 | BA:1127 | 4.60 | 1.00 | 142.31 |  |
| 13 | U | O2 | BA:1148 | 4.79 | 1.00 | 114.16 |  |
| 14 | C | O2 | BA:1149 | 4.89 | 1.00 | 122.10 |  |
| 15 | U | N3 | BA:1148 | 4.95 | 1.00 | 113.53 |  |
  |

  
 Contact(s): /O3'/  
 Coded contact list: O_r_  
 Exclusion zone contact(s): /N6/O4'/  

### MG:BA:1900

|  | Res. type | Atom | Residue | Distance | Occupancy | B-factor | Closest solute to 1^st^ shell water |
| --- | --- | --- | --- | --- | --- | --- | --- |
| **Ion** | | | |  | 1.00 | 83.25 || 1 | A | OP1 | BA:547 | 2.54 | 1.00 | 64.03 |
| 2 | G | OP1 | BA:548 | 2.58 | 1.00 | 62.05 |  |
| 3 | C | N4 | BA:403 | 3.80 | 1.00 | 72.03 |  |
| 4 | G | N7 | BA:402 | 4.30 | 1.00 | 69.73 |  |
| 5 | A | O5' | BA:547 | 4.48 | 1.00 | 63.98 |  |
| 6 | G | O5' | BA:548 | 4.49 | 1.00 | 59.35 |  |
| 7 | G | O6 | BA:402 | 4.51 | 1.00 | 70.49 |  |
| 8 | G | O3' | BA:546 | 4.77 | 1.00 | 64.52 |  |
| 9 | A | O3' | BA:547 | 4.88 | 1.00 | 63.38 |  |
  |

  
 Coded contact list:   
 Exclusion zone contact(s): /OP1/OP1/  
Consider NA in place of MG (2 valid contact(s) for NA vs. 0 for MG)  

### MG:BA:1901

|  | Res. type | Atom | Residue | Distance | Occupancy | B-factor | Closest solute to 1^st^ shell water |
| --- | --- | --- | --- | --- | --- | --- | --- |
| **Ion** | | | |  | 1.00 | 82.56 || 1 | G | O6 | BA:1143 | 2.12 | 1.00 | 161.18 |
| 2 | C | N4 | BA:1129 | 2.51 | 1.00 | 162.91 |  |
| 3 | G | O2' | BA:1139 | 2.69 | 1.00 | 150.65 |  |
| 4 | G | N3 | BA:1139 | 2.85 | 1.00 | 151.21 |  |
| 5 | G | N2 | BA:1139 | 3.70 | 1.00 | 151.09 |  |
| 6 | C | N4 | BA:1141 | 3.92 | 1.00 | 152.74 |  |
| 7 | G | N9 | BA:1139 | 4.03 | 1.00 | 150.87 |  |
| 8 | G | N1 | BA:1143 | 4.12 | 1.00 | 160.43 |  |
| 9 | C | N3 | BA:1129 | 4.30 | 1.00 | 163.40 |  |
| 10 | C | N4 | BA:1128 | 4.52 | 1.00 | 154.81 |  |
| 11 | G | O4' | BA:1139 | 4.58 | 1.00 | 150.77 |  |
| 12 | G | N7 | BA:1142 | 4.65 | 1.00 | 154.85 |  |
| 13 | G | N7 | BA:1143 | 4.86 | 1.00 | 157.53 |  |
| 14 | G | O6 | BA:1142 | 4.93 | 1.00 | 156.23 |  |
| 15 | C | OP2 | BA:1141 | 4.95 | 1.00 | 151.75 |  |
| 16 | G | N1 | BA:1139 | 4.99 | 1.00 | 151.17 |  |
  |

  
 Contact(s): /O6/  
 Coded contact list: O_b_  
 Exclusion zone contact(s): /N3_G_/N4/O2'/  
Consider K in place of MG (2 valid contact(s) for K vs. 1 for MG)  

### MG:BA:1902

|  | Res. type | Atom | Residue | Distance | Occupancy | B-factor | Closest solute to 1^st^ shell water |
| --- | --- | --- | --- | --- | --- | --- | --- |
| **Ion** | | | |  | 1.00 | 71.72 || 1 | G | O6 | BA:1276 | 3.51 | 1.00 | 132.33 |
| 2 | G | O6 | BA:1258 | 4.17 | 1.00 | 133.07 |  |
| 3 | C | N4 | BA:1277 | 4.31 | 1.00 | 131.39 |  |
| 4 | C | N4 | BA:1259 | 4.67 | 1.00 | 131.55 |  |
| 5 | G | N7 | BA:1276 | 4.69 | 1.00 | 132.95 |  |
  |

  
 Coded contact list:   
1^st^ MG coordination shell missing (see Notes)  

### MG:BA:1903

|  | Res. type | Atom | Residue | Distance | Occupancy | B-factor | Closest solute to 1^st^ shell water |
| --- | --- | --- | --- | --- | --- | --- | --- |
| **Ion** | | | |  | 1.00 | 71.14 || 1 | U | O3' | BA:86 | 3.28 | 1.00 | 187.40 |
| 2 | U | O4' | BA:86 | 3.51 | 1.00 | 189.81 |  |
| 3 | U | O5' | BA:86 | 3.69 | 1.00 | 193.29 |  |
| 4 | U | N1 | BA:86 | 3.86 | 1.00 | 188.70 |  |
| 5 | U | OP1 | BA:86 | 4.17 | 1.00 | 194.42 |  |
| 6 | A | O4' | BA:87 | 4.24 | 1.00 | 180.09 |  |
| 7 | A | O2' | BA:87 | 4.30 | 1.00 | 179.67 |  |
| 8 | A | OP1 | BA:87 | 4.57 | 1.00 | 186.13 |  |
| 9 | A | O5' | BA:87 | 4.71 | 1.00 | 184.52 |  |
| 10 | U | OP2 | BA:86 | 4.78 | 1.00 | 194.37 |  |
| 11 | U | O2' | BA:86 | 4.96 | 1.00 | 188.72 |  |
  |

  
 Coded contact list:   
 Exclusion zone contact(s): /O3'/  
Consider K in place of MG (1 valid contact(s) for K vs. 0 for MG)  

### MG:BA:1904

|  | Res. type | Atom | Residue | Distance | Occupancy | B-factor | Closest solute to 1^st^ shell water |
| --- | --- | --- | --- | --- | --- | --- | --- |
| **Ion** | | | |  | 1.00 | 38.67 || 1 | C | OP1 | BA:578 | 2.69 | 1.00 | 60.24 |
| 2 | C | OP2 | BA:817 | 4.30 | 1.00 | 52.41 |  |
| 3 | C | O5' | BA:578 | 4.39 | 1.00 | 57.70 |  |
| 4 | G | O3' | BA:821 | 4.47 | 1.00 | 63.02 |  |
| 5 | G | O3' | BA:577 | 4.50 | 1.00 | 56.75 |  |
  |

  
 Coded contact list:   
 Exclusion zone contact(s): /OP1/  
Consider K in place of MG (1 valid contact(s) for K vs. 0 for MG)  

### MG:BA:1905

|  | Res. type | Atom | Residue | Distance | Occupancy | B-factor | Closest solute to 1^st^ shell water |
| --- | --- | --- | --- | --- | --- | --- | --- |
| **Ion** | | | |  | 1.00 | 74.46 || 1 | G | O3' | BA:199 | 2.59 | 1.00 | 105.87 |
| 2 | C | OP2 | BA:67 | 2.75 | 1.00 | 85.60 |  |
| 3 | G | OP1 | BA:200 | 2.88 | 1.00 | 107.64 |  |
| 4 | G | O2' | BA:199 | 3.32 | 1.00 | 104.49 |  |
| 5 | C | OP1 | BA:67 | 3.41 | 1.00 | 85.47 |  |
| 6 | G | O5' | BA:200 | 3.70 | 1.00 | 110.07 |  |
| 7 | G | O3' | BA:66 | 4.23 | 1.00 | 85.44 |  |
| 8 | U | O4' | BA:65 | 4.53 | 1.00 | 87.62 |  |
| 9 | G | OP2 | BA:200 | 4.62 | 1.00 | 107.48 |  |
| 10 | U | O5' | BA:65 | 4.78 | 1.00 | 88.02 |  |
| 11 | C | O5' | BA:67 | 4.88 | 1.00 | 84.74 |  |
| 12 | G | O2' | BA:64 | 4.90 | 1.00 | 88.54 |  |
  |

  
 Coded contact list:   
 Exclusion zone contact(s): /O2'/O3'/OP1/OP2/  
Consider K in place of MG (3 valid contact(s) for K vs. 0 for MG)  

### MG:BA:1906

|  | Res. type | Atom | Residue | Distance | Occupancy | B-factor | Closest solute to 1^st^ shell water |
| --- | --- | --- | --- | --- | --- | --- | --- |
| **Ion** | | | |  | 1.00 | 107.70 || 1 | U | O2' | BA:65 | 2.65 | 1.00 | 89.12 |
| 2 | C | OP1 | BA:381 | 2.72 | 1.00 | 83.62 |  |
| 3 | U | O2 | BA:65 | 2.98 | 1.00 | 89.30 |  |
| 4 | C | O2' | BA:219 | 3.49 | 1.00 | 98.24 |  |
| 5 | C | O2 | BA:219 | 3.71 | 1.00 | 95.60 |  |
| 6 | C | O4' | BA:219 | 4.05 | 1.00 | 99.92 |  |
| 7 | C | O4' | BA:381 | 4.28 | 1.00 | 86.04 |  |
| 8 | C | O5' | BA:381 | 4.35 | 1.00 | 84.74 |  |
| 9 | U | N1 | BA:65 | 4.41 | 1.00 | 88.28 |  |
| 10 | C | N1 | BA:219 | 4.44 | 1.00 | 97.38 |  |
| 11 | G | N2 | BA:199 | 4.59 | 1.00 | 107.18 |  |
| 12 | G | O3' | BA:380 | 4.89 | 1.00 | 81.89 |  |
| 13 | U | O3' | BA:65 | 4.96 | 1.00 | 88.10 |  |
  |

  
 Coded contact list:   
 Exclusion zone contact(s): /O2/O2'/OP1/  
Consider K in place of MG (3 valid contact(s) for K vs. 0 for MG)  

### MG:BA:1907

|  | Res. type | Atom | Residue | Distance | Occupancy | B-factor | Closest solute to 1^st^ shell water |
| --- | --- | --- | --- | --- | --- | --- | --- |
| **Ion** | | | |  | 1.00 | 56.42 || 1 | C | OP2 | BA:123 | 2.53 | 1.00 | 61.31 |
| 2 | C | OP1 | BA:123 | 3.50 | 1.00 | 61.80 |  |
| 3 | C | O5' | BA:123 | 3.95 | 1.00 | 60.91 |  |
| 4 | G | OP2 | BA:124 | 4.08 | 1.00 | 63.93 |  |
| 5 | C | O2' | BA:311 | 4.34 | 1.00 | 68.30 |  |
| 6 | G | O3' | BA:122 | 4.81 | 1.00 | 60.80 |  |
  |

  
 Coded contact list:   
 Exclusion zone contact(s): /OP2/  
Consider NA in place of MG (1 valid contact(s) for NA vs. 0 for MG)  

### MG:BA:1908

|  | Res. type | Atom | Residue | Distance | Occupancy | B-factor | Closest solute to 1^st^ shell water |
| --- | --- | --- | --- | --- | --- | --- | --- |
| **Ion** | | | |  | 1.00 | 79.54 || 1 | C | OP2 | BA:175 | 2.34 | 1.00 | 88.23 |
| 2 | C | OP1 | BA:174 | 3.08 | 1.00 | 92.35 |  |
| 3 | C | O5' | BA:174 | 3.64 | 1.00 | 91.61 |  |
| 4 | C | OP2 | BA:174 | 3.66 | 1.00 | 92.13 |  |
| 5 | C | OP1 | BA:175 | 4.11 | 1.00 | 87.91 |  |
| 6 | C | O3' | BA:174 | 4.23 | 1.00 | 89.14 |  |
| 7 | C | O5' | BA:175 | 4.82 | 1.00 | 88.72 |  |
  |

  
 Contact(s): /OP2/  
 Coded contact list: O_ph_  
 Exclusion zone contact(s): /OP1/  

### MG:BA:1909

|  | Res. type | Atom | Residue | Distance | Occupancy | B-factor | Closest solute to 1^st^ shell water |
| --- | --- | --- | --- | --- | --- | --- | --- |
| **Ion** | | | |  | 1.00 | 66.41 || 1 | U | OP1 | BA:863 | 3.58 | 1.00 | 75.21 |
| 2 | A | OP2 | BA:864 | 3.85 | 1.00 | 66.92 |  |
| 3 | GLY | O | BH:85 | 4.19 | 1.00 | 76.72 |  |
| 4 | GLU | OE1 | BH:83 | 4.57 | 1.00 | 87.28 |  |
| 5 | U | O5' | BA:863 | 4.87 | 1.00 | 74.30 |  |
  |

  
 Coded contact list:   
1^st^ MG coordination shell missing (see Notes)  

### MG:BA:1910

|  | Res. type | Atom | Residue | Distance | Occupancy | B-factor | Closest solute to 1^st^ shell water |
| --- | --- | --- | --- | --- | --- | --- | --- |
| **Ion** | | | |  | 1.00 | 59.17 || 1 | G | O6 | BA:1178 | 2.67 | 1.00 | 144.58 |
| 2 | G | N1 | BA:1178 | 2.93 | 1.00 | 144.58 |  |
| 3 | G | N7 | BA:1178 | 3.28 | 1.00 | 144.29 |  |
| 4 | G | N9 | BA:1177 | 3.37 | 1.00 | 144.70 |  |
| 5 | LYS | NZ | BL:97 | 3.43 | 1.00 | 122.00 |  |
| 6 | G | O5' | BA:1177 | 3.50 | 1.00 | 142.10 |  |
| 7 | G | N7 | BA:1177 | 3.51 | 1.00 | 145.34 |  |
| 8 | G | O4' | BA:1177 | 3.75 | 1.00 | 144.14 |  |
| 9 | G | N3 | BA:1178 | 3.87 | 1.00 | 144.69 |  |
| 10 | G | OP2 | BA:1178 | 3.99 | 1.00 | 145.03 |  |
| 11 | G | N9 | BA:1178 | 4.16 | 1.00 | 143.82 |  |
| 12 | G | OP2 | BA:1177 | 4.16 | 1.00 | 141.42 |  |
| 13 | G | O3' | BA:1177 | 4.30 | 1.00 | 144.72 |  |
| 14 | G | N2 | BA:1178 | 4.57 | 1.00 | 143.99 |  |
| 15 | G | O2' | BA:1177 | 4.77 | 1.00 | 144.18 |  |
  |

  
 Coded contact list:   
 Exclusion zone contact(s): /N1_G_/N7/O6/  
Consider K in place of MG (4 valid contact(s) for K vs. 0 for MG)  

### MG:BA:1911

|  | Res. type | Atom | Residue | Distance | Occupancy | B-factor | Closest solute to 1^st^ shell water |
| --- | --- | --- | --- | --- | --- | --- | --- |
| **Ion** | | | |  | 1.00 | 76.75 || 1 | G | O6 | BA:493 | 4.04 | 1.00 | 93.92 |
| 2 | G | O6 | BA:492 | 4.70 | 1.00 | 88.72 |  |
| 3 | G | N7 | BA:491 | 4.80 | 1.00 | 96.13 |  |
| 4 | G | OP2 | BA:491 | 4.89 | 1.00 | 89.84 |  |
| 5 | A | OP2 | BA:439 | 4.89 | 1.00 | 92.79 |  |
| 6 | G | N7 | BA:492 | 4.93 | 1.00 | 87.46 |  |
  |

  
 Coded contact list:   
1^st^ MG coordination shell missing (see Notes)  

### MG:BA:1912

|  | Res. type | Atom | Residue | Distance | Occupancy | B-factor | Closest solute to 1^st^ shell water |
| --- | --- | --- | --- | --- | --- | --- | --- |
| **Ion** | | | |  | 1.00 | 82.35 || 1 | C | OP2 | BA:1234 | 3.08 | 1.00 | 86.88 |
| 2 | U | OP2 | BA:1235 | 4.42 | 1.00 | 89.86 |  |
| 3 | C | O5' | BA:1234 | 4.43 | 1.00 | 87.24 |  |
| 4 | C | OP1 | BA:1234 | 4.68 | 1.00 | 87.03 |  |
| 5 | U | OP1 | BA:1351 | 4.86 | 1.00 | 88.11 |  |
  |

  
 Coded contact list:   
 Exclusion zone contact(s): /OP2/  
Consider K in place of MG (1 valid contact(s) for K vs. 0 for MG)  

### MG:BA:1913

|  | Res. type | Atom | Residue | Distance | Occupancy | B-factor | Closest solute to 1^st^ shell water |
| --- | --- | --- | --- | --- | --- | --- | --- |
| **Ion** | | | |  | 1.00 | 73.20 || 1 | A | OP1 | BA:964 | 2.42 | 1.00 | 76.19 |
| 2 | U | OP1 | BA:1199 | 2.63 | 1.00 | 87.55 |  |
| 3 | A | OP2 | BA:965 | 3.07 | 1.00 | 77.87 |  |
| 4 | G | OP1 | BA:1198 | 3.09 | 1.00 | 86.00 |  |
| 5 | U | OP2 | BA:1199 | 3.93 | 1.00 | 86.92 |  |
| 6 | G | O3' | BA:1198 | 3.95 | 1.00 | 87.16 |  |
| 7 | G | O5' | BA:1198 | 3.98 | 1.00 | 86.90 |  |
| 8 | A | O5' | BA:964 | 4.18 | 1.00 | 75.15 |  |
| 9 | A | OP2 | BA:964 | 4.70 | 1.00 | 76.12 |  |
| 10 | G | O3' | BA:963 | 4.81 | 1.00 | 77.30 |  |
| 11 | A | OP1 | BA:965 | 4.94 | 1.00 | 77.33 |  |
| 12 | U | O5' | BA:1199 | 5.00 | 1.00 | 88.55 |  |
  |

  
 Coded contact list:   
 Exclusion zone contact(s): /OP1/OP1/OP1/OP2/  
Consider K in place of MG (3 valid contact(s) for K vs. 0 for MG)  

### MG:BA:1914

|  | Res. type | Atom | Residue | Distance | Occupancy | B-factor | Closest solute to 1^st^ shell water |
| --- | --- | --- | --- | --- | --- | --- | --- |
| **Ion** | | | |  | 1.00 | 55.54 || 1 | G | OP1 | BA:1530 | 2.58 | 1.00 | 80.25 |
| 2 | G | O3' | BA:1529 | 2.69 | 1.00 | 76.75 |  |
| 3 | G | O2' | BA:1529 | 3.59 | 1.00 | 72.10 |  |
| 4 | U | OP1 | BA:1393 | 3.62 | 1.00 | 66.11 |  |
| 5 | G | OP2 | BA:1530 | 4.20 | 1.00 | 79.99 |  |
| 6 | G | O5' | BA:1530 | 4.40 | 1.00 | 81.23 |  |
| 7 | A | OP1 | BA:1502 | 4.51 | 1.00 | 54.69 |  |
| 8 | G | O3' | BA:1392 | 4.77 | 1.00 | 65.46 |  |
  |

  
 Coded contact list:   
 Exclusion zone contact(s): /O3'/OP1/  
Consider NA in place of MG (1 valid contact(s) for NA vs. 0 for MG)  

### MG:BA:1915

|  | Res. type | Atom | Residue | Distance | Occupancy | B-factor | Closest solute to 1^st^ shell water |
| --- | --- | --- | --- | --- | --- | --- | --- |
| **Ion** | | | |  | 1.00 | 69.56 || 1 | G | O6 | BA:1077 | 2.95 | 1.00 | 78.03 |
| 2 | G | N7 | BA:1077 | 3.05 | 1.00 | 79.76 |  |
| 3 | G | OP1 | BA:1079 | 3.77 | 1.00 | 78.79 |  |
| 4 | C | N4 | BA:1076 | 4.18 | 1.00 | 72.22 |  |
| 5 | G | OP2 | BA:1079 | 4.65 | 1.00 | 78.52 |  |
| 6 | G | N1 | BA:1077 | 4.92 | 1.00 | 77.88 |  |
  |

  
 Coded contact list:   
 Exclusion zone contact(s): /N7/O6/  
Consider K in place of MG (2 valid contact(s) for K vs. 0 for MG)  

### MG:BA:1916

|  | Res. type | Atom | Residue | Distance | Occupancy | B-factor | Closest solute to 1^st^ shell water |
| --- | --- | --- | --- | --- | --- | --- | --- |
| **Ion** | | | |  | 1.00 | 56.47 || 1 | C | O2 | BA:893 | 2.74 | 1.00 | 56.55 |
| 2 | U | O2 | BA:244 | 2.95 | 1.00 | 56.49 |  |
| 3 | U | O2' | BA:244 | 2.97 | 1.00 | 63.92 |  |
| 4 | G | O4' | BA:894 | 4.21 | 1.00 | 52.43 |  |
| 5 | C | N3 | BA:893 | 4.55 | 1.00 | 56.27 |  |
| 6 | C | O2' | BA:893 | 4.61 | 1.00 | 56.25 |  |
| 7 | U | N1 | BA:244 | 4.81 | 1.00 | 58.32 |  |
| 8 | C | N1 | BA:893 | 4.86 | 1.00 | 54.47 |  |
| 9 | G | N9 | BA:894 | 4.88 | 1.00 | 52.31 |  |
  |

  
 Coded contact list:   
 Exclusion zone contact(s): /O2/O2/O2'/  
Consider K in place of MG (3 valid contact(s) for K vs. 0 for MG)  

### MG:BA:1917

|  | Res. type | Atom | Residue | Distance | Occupancy | B-factor | Closest solute to 1^st^ shell water |
| --- | --- | --- | --- | --- | --- | --- | --- |
| **Ion** | | | |  | 1.00 | 64.95 || 1 | A | OP2 | BA:8 | 2.72 | 1.00 | 74.69 |
| 2 | A | OP1 | BA:298 | 2.82 | 1.00 | 70.46 |  |
| 3 | G | OP1 | BA:7 | 3.43 | 1.00 | 79.26 |  |
| 4 | A | OP2 | BA:298 | 3.74 | 1.00 | 70.32 |  |
| 5 | G | O5' | BA:7 | 4.54 | 1.00 | 79.24 |  |
| 6 | A | OP1 | BA:8 | 4.66 | 1.00 | 74.27 |  |
| 7 | G | O3' | BA:7 | 4.76 | 1.00 | 74.14 |  |
| 8 | G | O3' | BA:297 | 4.85 | 1.00 | 69.90 |  |
| 9 | A | O5' | BA:298 | 4.88 | 1.00 | 69.50 |  |
  |

  
 Coded contact list:   
 Exclusion zone contact(s): /OP1/OP2/  
Consider K in place of MG (2 valid contact(s) for K vs. 0 for MG)  

### MG:BA:1918

|  | Res. type | Atom | Residue | Distance | Occupancy | B-factor | Closest solute to 1^st^ shell water |
| --- | --- | --- | --- | --- | --- | --- | --- |
| **Ion** | | | |  | 1.00 | 74.66 || 1 | C | O4' | BA:1054 | 2.25 | 1.00 | 104.91 |
| 2 | C | OP2 | BA:1054 | 2.74 | 1.00 | 102.79 |  |
| 3 | U | OP2 | BA:1056 | 3.27 | 1.00 | 92.11 |  |
| 4 | C | O5' | BA:1054 | 3.32 | 1.00 | 103.97 |  |
| 5 | U | O2' | BA:1196 | 3.41 | 1.00 | 98.61 |  |
| 6 | C | O2' | BA:1054 | 3.42 | 1.00 | 105.16 |  |
| 7 | G | O3' | BA:1053 | 3.60 | 1.00 | 101.32 |  |
| 8 | U | O2 | BA:1196 | 3.95 | 1.00 | 103.01 |  |
| 9 | C | N1 | BA:1054 | 4.31 | 1.00 | 107.12 |  |
| 10 | C | O3' | BA:1054 | 4.54 | 1.00 | 101.91 |  |
| 11 | G | O2' | BA:1053 | 4.78 | 1.00 | 99.80 |  |
| 12 | G | OP2 | BA:1197 | 4.83 | 1.00 | 91.12 |  |
| 13 | U | OP1 | BA:1056 | 4.90 | 1.00 | 91.94 |  |
| 14 | C | OP1 | BA:1054 | 4.90 | 1.00 | 103.40 |  |
| 15 | G | OP1 | BA:1197 | 4.98 | 1.00 | 91.39 |  |
  |

  
 Contact(s): /O4'/  
 Coded contact list: O_r_  
 Exclusion zone contact(s): /O5'/OP2/OP2/  
Consider K in place of MG (3 valid contact(s) for K vs. 1 for MG)  

### MG:BA:1919

|  | Res. type | Atom | Residue | Distance | Occupancy | B-factor | Closest solute to 1^st^ shell water |
| --- | --- | --- | --- | --- | --- | --- | --- |
| **Ion** | | | |  | 1.00 | 59.17 || 1 | C | O2 | BA:280 | 3.38 | 1.00 | 76.70 |
| 2 | SER | O | BT:39 | 3.60 | 1.00 | 70.64 |  |
| 3 | G | OP1 | BA:281 | 3.76 | 1.00 | 71.06 |  |
| 4 | C | O2' | BA:280 | 4.10 | 1.00 | 73.25 |  |
| 5 | A | OP1 | BA:279 | 4.81 | 1.00 | 76.26 |  |
  |

  
 Coded contact list:   
 Exclusion zone contact(s): /O2/  
Consider K in place of MG (1 valid contact(s) for K vs. 0 for MG)  

### MG:BA:1920

|  | Res. type | Atom | Residue | Distance | Occupancy | B-factor | Closest solute to 1^st^ shell water |
| --- | --- | --- | --- | --- | --- | --- | --- |
| **Ion** | | | |  | 1.00 | 86.85 || 1 | A | OP2 | BA:923 | 2.65 | 1.00 | 72.39 |
| 2 | G | O3' | BA:922 | 3.92 | 1.00 | 72.91 |  |
| 3 | G | OP2 | BA:922 | 4.07 | 1.00 | 75.79 |  |
| 4 | G | O5' | BA:922 | 4.33 | 1.00 | 74.69 |  |
| 5 | A | N7 | BA:923 | 4.41 | 1.00 | 69.10 |  |
| 6 | G | O4' | BA:922 | 4.66 | 1.00 | 72.60 |  |
| 7 | A | O5' | BA:923 | 4.76 | 1.00 | 74.15 |  |
  |

  
 Coded contact list:   
 Exclusion zone contact(s): /OP2/  
Consider K in place of MG (1 valid contact(s) for K vs. 0 for MG)  

### MG:BA:1921

|  | Res. type | Atom | Residue | Distance | Occupancy | B-factor | Closest solute to 1^st^ shell water |
| --- | --- | --- | --- | --- | --- | --- | --- |
| **Ion** | | | |  | 1.00 | 85.90 || 1 | U | N1 | BA:1281 | 3.62 | 1.00 | 116.88 |
| 2 | C | N1 | BA:1282 | 3.65 | 1.00 | 119.32 |  |
| 3 | U | O3' | BA:1281 | 3.92 | 1.00 | 116.20 |  |
| 4 | C | O4' | BA:1282 | 3.94 | 1.00 | 118.50 |  |
| 5 | C | O4' | BA:1277 | 3.97 | 1.00 | 131.65 |  |
| 6 | C | O3' | BA:1277 | 4.20 | 1.00 | 130.88 |  |
| 7 | C | O2' | BA:1277 | 4.29 | 1.00 | 131.46 |  |
| 8 | A | O2' | BA:1279 | 4.34 | 1.00 | 118.05 |  |
| 9 | C | N3 | BA:1282 | 4.53 | 1.00 | 119.79 |  |
| 10 | U | O5' | BA:1281 | 4.55 | 1.00 | 112.21 |  |
| 11 | C | N4 | BA:1282 | 4.68 | 1.00 | 118.60 |  |
| 12 | G | O2' | BA:1276 | 4.72 | 1.00 | 133.57 |  |
| 13 | U | N3 | BA:1281 | 4.74 | 1.00 | 117.26 |  |
| 14 | U | O4' | BA:1281 | 4.74 | 1.00 | 116.00 |  |
| 15 | U | O2' | BA:1281 | 4.83 | 1.00 | 116.88 |  |
| 16 | C | OP1 | BA:1282 | 4.96 | 1.00 | 116.94 |  |
  |

  
 Coded contact list:   
1^st^ MG coordination shell missing (see Notes)  

### MG:BA:1922

|  | Res. type | Atom | Residue | Distance | Occupancy | B-factor | Closest solute to 1^st^ shell water |
| --- | --- | --- | --- | --- | --- | --- | --- |
| **Ion** | | | |  | 1.00 | 71.88 || 1 | G | O6 | BA:73 | 2.52 | 1.00 | 104.35 |
| 2 | C | N4 | BA:74 | 2.82 | 1.00 | 116.52 |  |
| 3 | G | N7 | BA:73 | 3.09 | 1.00 | 105.83 |  |
| 4 | G | O6 | BA:96 | 3.31 | 1.00 | 116.88 |  |
| 5 | G | O6 | BA:69 | 4.00 | 1.00 | 101.74 |  |
| 6 | U | O4 | BA:97 | 4.26 | 1.00 | 103.41 |  |
| 7 | G | N1 | BA:69 | 4.28 | 1.00 | 101.41 |  |
| 8 | G | N1 | BA:73 | 4.46 | 1.00 | 103.61 |  |
| 9 | G | N7 | BA:69 | 4.61 | 1.00 | 101.22 |  |
| 10 | U | N3 | BA:97 | 4.64 | 1.00 | 104.42 |  |
| 11 | C | N3 | BA:74 | 4.73 | 1.00 | 117.43 |  |
| 12 | G | O6 | BA:95 | 4.85 | 1.00 | 130.93 |  |
  |

  
 Coded contact list:   
 Exclusion zone contact(s): /N4/N7/O6/O6/  
Consider K in place of MG (3 valid contact(s) for K vs. 0 for MG)  

### MG:BA:1923

|  | Res. type | Atom | Residue | Distance | Occupancy | B-factor | Closest solute to 1^st^ shell water |
| --- | --- | --- | --- | --- | --- | --- | --- |
| **Ion** | | | |  | 1.00 | 81.36 || 1 | G | O2' | BA:81 | 2.91 | 1.00 | 176.99 |
| 2 | G | O3' | BA:81 | 4.46 | 1.00 | 179.94 |  |
  |

  
 Coded contact list:   
 Exclusion zone contact(s): /O2'/  
Consider K in place of MG (1 valid contact(s) for K vs. 0 for MG)  

### MG:BA:1924

|  | Res. type | Atom | Residue | Distance | Occupancy | B-factor | Closest solute to 1^st^ shell water |
| --- | --- | --- | --- | --- | --- | --- | --- |
| **Ion** | | | |  | 1.00 | 78.66 || 1 | G | O2' | BA:557 | 4.46 | 1.00 | 63.81 |
| 2 | A | OP2 | BA:8 | 4.54 | 1.00 | 74.69 |  |
| 3 | A | OP1 | BA:8 | 4.72 | 1.00 | 74.27 |  |
| 4 | ARG | O | BG:209 | 4.78 | 1.00 | 78.64 |  |
| 5 | ARG | OXT | BG:209 | 4.90 | 1.00 | 79.42 |  |
  |

  
 Coded contact list:   
1^st^ MG coordination shell missing (see Notes)  

### MG:BA:1925

|  | Res. type | Atom | Residue | Distance | Occupancy | B-factor | Closest solute to 1^st^ shell water |
| --- | --- | --- | --- | --- | --- | --- | --- |
| **Ion** | | | |  | 1.00 | 55.46 || 1 | G | OP1 | BA:64 | 2.30 | 1.00 | 79.84 |
| 2 | C | N4 | BA:381 | 2.70 | 1.00 | 86.68 |  |
| 3 | U | OP1 | BA:65 | 2.71 | 1.00 | 89.58 |  |
| 4 | A | N6 | BA:382 | 2.75 | 1.00 | 87.18 |  |
| 5 | A | N1 | BA:382 | 2.82 | 1.00 | 86.77 |  |
| 6 | C | O3' | BA:63 | 3.89 | 1.00 | 78.22 |  |
| 7 | G | O5' | BA:64 | 4.16 | 1.00 | 80.83 |  |
| 8 | A | N3 | BA:382 | 4.70 | 1.00 | 88.50 |  |
| 9 | G | OP2 | BA:64 | 4.83 | 1.00 | 78.27 |  |
| 10 | U | OP2 | BA:65 | 4.84 | 1.00 | 88.38 |  |
| 11 | C | N3 | BA:381 | 4.88 | 1.00 | 86.05 |  |
| 12 | A | N7 | BA:382 | 4.92 | 1.00 | 89.29 |  |
| 13 | U | O5' | BA:65 | 4.97 | 1.00 | 88.02 |  |
  |

  
 Contact(s): /OP1/  
 Coded contact list: O_ph_  
 Exclusion zone contact(s): /N1_A_/N4/N6/OP1/  
Consider K in place of MG (4 valid contact(s) for K vs. 1 for MG)  

### MG:BA:1926

|  | Res. type | Atom | Residue | Distance | Occupancy | B-factor | Closest solute to 1^st^ shell water |
| --- | --- | --- | --- | --- | --- | --- | --- |
| **Ion** | | | |  | 1.00 | 53.35 || 1 | U | O4 | BA:1485 | 2.55 | 1.00 | 59.79 |
| 2 | G | O6 | BA:1486 | 2.55 | 1.00 | 59.06 |  |
| 3 | U | O4 | BA:1414 | 2.87 | 1.00 | 60.41 |  |
| 4 | G | O6 | BA:1415 | 3.13 | 1.00 | 47.85 |  |
| 5 | U | N3 | BA:1485 | 3.84 | 1.00 | 57.05 |  |
| 6 | U | N3 | BA:1414 | 3.85 | 1.00 | 58.42 |  |
| 7 | G | N7 | BA:1486 | 4.17 | 1.00 | 56.40 |  |
| 8 | G | O6 | BA:1487 | 4.52 | 1.00 | 58.21 |  |
| 9 | A | N6 | BA:1413 | 4.79 | 1.00 | 57.19 |  |
| 10 | G | N1 | BA:1486 | 4.80 | 1.00 | 56.32 |  |
| 11 | G | N7 | BA:1415 | 4.95 | 1.00 | 50.39 |  |
| 12 | G | N1 | BA:1415 | 4.97 | 1.00 | 48.40 |  |
  |

  
 Coded contact list:   
 Exclusion zone contact(s): /O4/O4/O6/O6/  
Consider NA in place of MG (2 valid contact(s) for NA vs. 0 for MG)  

### MG:BA:1927

|  | Res. type | Atom | Residue | Distance | Occupancy | B-factor | Closest solute to 1^st^ shell water |
| --- | --- | --- | --- | --- | --- | --- | --- |
| **Ion** | | | |  | 1.00 | 51.54 || 1 | C | OP2 | BA:796 | 2.95 | 1.00 | 56.16 |
| 2 | C | O5' | BA:796 | 3.41 | 1.00 | 57.00 |  |
| 3 | C | OP2 | BA:797 | 3.64 | 1.00 | 56.47 |  |
| 4 | C | N1 | BA:796 | 3.75 | 1.00 | 56.05 |  |
| 5 | C | N4 | BA:796 | 3.97 | 1.00 | 57.23 |  |
| 6 | C | N3 | BA:796 | 4.05 | 1.00 | 55.65 |  |
| 7 | C | OP2 | BA:795 | 4.13 | 1.00 | 59.96 |  |
| 8 | C | O5' | BA:795 | 4.13 | 1.00 | 61.69 |  |
| 9 | C | N4 | BA:797 | 4.15 | 1.00 | 56.03 |  |
| 10 | C | OP1 | BA:795 | 4.55 | 1.00 | 60.22 |  |
| 11 | C | O4' | BA:796 | 4.74 | 1.00 | 58.17 |  |
| 12 | C | O3' | BA:795 | 4.77 | 1.00 | 56.41 |  |
| 13 | C | O3' | BA:796 | 4.77 | 1.00 | 56.97 |  |
| 14 | C | O5' | BA:797 | 4.85 | 1.00 | 55.10 |  |
| 15 | C | OP1 | BA:796 | 4.97 | 1.00 | 55.32 |  |
  |

  
 Coded contact list:   
 Exclusion zone contact(s): /OP2/  
Consider K in place of MG (1 valid contact(s) for K vs. 0 for MG)  

### MG:BA:1928

|  | Res. type | Atom | Residue | Distance | Occupancy | B-factor | Closest solute to 1^st^ shell water |
| --- | --- | --- | --- | --- | --- | --- | --- |
| **Ion** | | | |  | 1.00 | 67.31 || 1 | C | O2' | BA:1200 | 2.95 | 1.00 | 100.40 |
| 2 | A | N7 | BA:1204 | 2.96 | 1.00 | 92.80 |  |
| 3 | U | O4 | BA:1205 | 3.23 | 1.00 | 96.55 |  |
| 4 | C | O2 | BA:1200 | 3.47 | 1.00 | 98.44 |  |
| 5 | A | N6 | BA:1204 | 3.54 | 1.00 | 92.15 |  |
| 6 | A | N9 | BA:1204 | 4.38 | 1.00 | 94.51 |  |
| 7 | A | N1 | BA:1204 | 4.51 | 1.00 | 92.47 |  |
| 8 | C | N1 | BA:1200 | 4.59 | 1.00 | 98.72 |  |
| 9 | U | N3 | BA:1205 | 4.72 | 1.00 | 97.47 |  |
| 10 | A | OP2 | BA:1201 | 4.78 | 1.00 | 99.11 |  |
| 11 | C | O4' | BA:1200 | 4.86 | 1.00 | 98.98 |  |
  |

  
 Coded contact list:   
 Exclusion zone contact(s): /N7/O2'/O4/  
Consider K in place of MG (3 valid contact(s) for K vs. 0 for MG)  

### MG:BA:1929

|  | Res. type | Atom | Residue | Distance | Occupancy | B-factor | Closest solute to 1^st^ shell water |
| --- | --- | --- | --- | --- | --- | --- | --- |
| **Ion** | | | |  | 1.00 | 85.11 || 1 | A | O4' | BA:87 | 3.21 | 1.00 | 180.09 |
| 2 | A | OP2 | BA:87 | 3.44 | 1.00 | 186.21 |  |
| 3 | A | N9 | BA:87 | 3.45 | 1.00 | 179.21 |  |
| 4 | A | N7 | BA:87 | 3.45 | 1.00 | 179.39 |  |
| 5 | U | O3' | BA:86 | 3.50 | 1.00 | 187.40 |  |
| 6 | U | O2' | BA:86 | 4.29 | 1.00 | 188.72 |  |
| 7 | A | O5' | BA:87 | 4.83 | 1.00 | 184.52 |  |
  |

  
 Coded contact list:   
 Exclusion zone contact(s): /O4'/  
Consider K in place of MG (1 valid contact(s) for K vs. 0 for MG)  

### MG:BA:1930

|  | Res. type | Atom | Residue | Distance | Occupancy | B-factor | Closest solute to 1^st^ shell water |
| --- | --- | --- | --- | --- | --- | --- | --- |
| **Ion** | | | |  | 1.00 | 62.22 || 1 | G | O6 | BA:447 | 2.99 | 1.00 | 88.45 |
| 2 | U | OP2 | BA:486 | 3.62 | 1.00 | 99.98 |  |
| 3 | G | N7 | BA:446 | 4.20 | 1.00 | 90.35 |  |
| 4 | G | N1 | BA:447 | 4.22 | 1.00 | 88.54 |  |
| 5 | G | O6 | BA:446 | 4.36 | 1.00 | 86.18 |  |
  |

  
 Coded contact list:   
 Exclusion zone contact(s): /O6/  
Consider K in place of MG (1 valid contact(s) for K vs. 0 for MG)  

### MG:BA:1931

|  | Res. type | Atom | Residue | Distance | Occupancy | B-factor | Closest solute to 1^st^ shell water |
| --- | --- | --- | --- | --- | --- | --- | --- |
| **Ion** | | | |  | 1.00 | 68.36 || 1 | G | N7 | BA:1143 | 3.14 | 1.00 | 157.53 |
| 2 | G | O6 | BA:1143 | 3.27 | 1.00 | 161.18 |  |
| 3 | G | O6 | BA:1144 | 3.87 | 1.00 | 153.27 |  |
| 4 | G | N7 | BA:1144 | 4.44 | 1.00 | 150.41 |  |
| 5 | G | O6 | BA:1127 | 4.78 | 1.00 | 142.48 |  |
| 6 | G | N2 | BA:1139 | 4.78 | 1.00 | 151.09 |  |
| 7 | G | N1 | BA:1143 | 4.79 | 1.00 | 160.43 |  |
| 8 | G | OP2 | BA:1142 | 4.93 | 1.00 | 154.62 |  |
  |

  
 Coded contact list:   
 Exclusion zone contact(s): /N7/O6/  
Consider K in place of MG (2 valid contact(s) for K vs. 0 for MG)  

### MG:BA:1932

|  | Res. type | Atom | Residue | Distance | Occupancy | B-factor | Closest solute to 1^st^ shell water |
| --- | --- | --- | --- | --- | --- | --- | --- |
| **Ion** | | | |  | 1.00 | 107.79 || 1 | A | OP2 | BA:1146 | 2.59 | 1.00 | 130.64 |
| 2 | G | N7 | BA:1124 | 2.72 | 1.00 | 121.15 |  |
| 3 | G | O6 | BA:1124 | 3.18 | 1.00 | 122.15 |  |
| 4 | A | N7 | BA:1123 | 3.79 | 1.00 | 121.75 |  |
| 5 | C | O3' | BA:1145 | 4.06 | 1.00 | 132.77 |  |
| 6 | A | N6 | BA:1123 | 4.21 | 1.00 | 121.58 |  |
| 7 | A | OP1 | BA:1146 | 4.45 | 1.00 | 130.70 |  |
| 8 | C | OP2 | BA:1147 | 4.67 | 1.00 | 118.47 |  |
| 9 | G | N9 | BA:1124 | 4.68 | 1.00 | 122.20 |  |
| 10 | G | N1 | BA:1124 | 4.81 | 1.00 | 121.81 |  |
  |

  
 Coded contact list:   
 Exclusion zone contact(s): /N7/O6/OP2/  
Consider K in place of MG (2 valid contact(s) for K vs. 0 for MG)  

### MG:BA:1933

|  | Res. type | Atom | Residue | Distance | Occupancy | B-factor | Closest solute to 1^st^ shell water |
| --- | --- | --- | --- | --- | --- | --- | --- |
| **Ion** | | | |  | 1.00 | 63.78 || 1 | A | N1 | BA:26 | 3.38 | 1.00 | 72.73 |
| 2 | G | O2' | BA:558 | 3.56 | 1.00 | 65.36 |  |
| 3 | A | OP1 | BA:8 | 4.15 | 1.00 | 74.27 |  |
| 4 | A | N6 | BA:26 | 4.24 | 1.00 | 71.81 |  |
| 5 | G | N7 | BA:9 | 4.44 | 1.00 | 71.79 |  |
| 6 | G | O4' | BA:558 | 4.50 | 1.00 | 65.45 |  |
| 7 | G | N9 | BA:9 | 4.62 | 1.00 | 70.47 |  |
| 8 | ARG | O | BG:209 | 4.85 | 1.00 | 78.64 |  |
| 9 | G | OP2 | BA:9 | 4.90 | 1.00 | 67.72 |  |
  |

  
 Coded contact list:   
 Exclusion zone contact(s): /N1_A_/  
Consider K in place of MG (1 valid contact(s) for K vs. 0 for MG)  

### MG:BA:1934

|  | Res. type | Atom | Residue | Distance | Occupancy | B-factor | Closest solute to 1^st^ shell water |
| --- | --- | --- | --- | --- | --- | --- | --- |
| **Ion** | | | |  | 1.00 | 84.57 || 1 | G | OP2 | BA:191A | 3.90 | 1.00 | 91.14 |
| 2 | A | OP1 | BA:185 | 4.25 | 1.00 | 92.36 |  |
| 3 | C | OP1 | BA:186 | 4.27 | 1.00 | 86.43 |  |
| 4 | G | OP2 | BA:191B | 4.36 | 1.00 | 95.73 |  |
  |

  
 Coded contact list:   
1^st^ MG coordination shell missing (see Notes)  

### MG:BA:1935

|  | Res. type | Atom | Residue | Distance | Occupancy | B-factor | Closest solute to 1^st^ shell water |
| --- | --- | --- | --- | --- | --- | --- | --- |
| **Ion** | | | |  | 1.00 | 95.12 || 1 | G | N7 | BA:1274 | 2.75 | 1.00 | 137.75 |
| 2 | G | OP2 | BA:1273 | 2.90 | 1.00 | 136.55 |  |
| 3 | G | N7 | BA:1273 | 3.38 | 1.00 | 138.15 |  |
| 4 | G | O6 | BA:1274 | 3.60 | 1.00 | 136.47 |  |
| 5 | G | OP2 | BA:1274 | 4.04 | 1.00 | 137.11 |  |
| 6 | G | N9 | BA:1273 | 4.13 | 1.00 | 138.24 |  |
| 7 | G | O4' | BA:1273 | 4.25 | 1.00 | 138.15 |  |
| 8 | G | O5' | BA:1273 | 4.27 | 1.00 | 137.37 |  |
| 9 | G | N9 | BA:1274 | 4.93 | 1.00 | 137.62 |  |
  |

  
 Coded contact list:   
 Exclusion zone contact(s): /N7/N7/OP2/  
Consider K in place of MG (3 valid contact(s) for K vs. 0 for MG)  

### MG:BA:1936

|  | Res. type | Atom | Residue | Distance | Occupancy | B-factor | Closest solute to 1^st^ shell water |
| --- | --- | --- | --- | --- | --- | --- | --- |
| **Ion** | | | |  | 1.00 | 80.71 || 1 | U | OP1 | BA:1150 | 3.80 | 1.00 | 121.23 |
| 2 | C | O3' | BA:1149 | 4.06 | 1.00 | 121.28 |  |
| 3 | U | O5' | BA:1150 | 4.08 | 1.00 | 120.82 |  |
| 4 | A | OP2 | BA:1252 | 4.23 | 1.00 | 97.30 |  |
| 5 | C | O2' | BA:1149 | 4.82 | 1.00 | 121.38 |  |
  |

  
 Coded contact list:   
1^st^ MG coordination shell missing (see Notes)  

### MG:BA:1937

|  | Res. type | Atom | Residue | Distance | Occupancy | B-factor | Closest solute to 1^st^ shell water |
| --- | --- | --- | --- | --- | --- | --- | --- |
| **Ion** | | | |  | 1.00 | 71.93 || 1 | G | O6 | BA:139 | 4.72 | 1.00 | 85.00 |
| 2 | G | N7 | BA:138 | 4.79 | 1.00 | 89.96 |  |
| 3 | G | O6 | BA:138 | 4.85 | 1.00 | 88.85 |  |
  |

  
 Coded contact list:   
1^st^ MG coordination shell missing (see Notes)  

### MG:BA:1938

|  | Res. type | Atom | Residue | Distance | Occupancy | B-factor | Closest solute to 1^st^ shell water |
| --- | --- | --- | --- | --- | --- | --- | --- |
| **Ion** | | | |  | 1.00 | 69.22 || 1 | G | OP2 | BA:588 | 2.73 | 1.00 | 75.02 |
| 2 | G | OP1 | BA:588 | 3.03 | 1.00 | 74.29 |  |
| 3 | G | OP1 | BA:644 | 3.45 | 1.00 | 73.78 |  |
| 4 | G | O3' | BA:587 | 3.82 | 1.00 | 73.18 |  |
| 5 | C | OP2 | BA:645 | 4.06 | 1.00 | 78.53 |  |
| 6 | G | O3' | BA:644 | 4.15 | 1.00 | 78.15 |  |
| 7 | G | O5' | BA:644 | 4.38 | 1.00 | 74.15 |  |
| 8 | G | O5' | BA:587 | 4.44 | 1.00 | 67.39 |  |
| 9 | C | OP1 | BA:645 | 4.54 | 1.00 | 79.25 |  |
| 10 | G | O5' | BA:588 | 4.71 | 1.00 | 74.88 |  |
| 11 | G | OP1 | BA:587 | 4.78 | 1.00 | 67.96 |  |
  |

  
 Coded contact list:   
 Exclusion zone contact(s): /OP1/OP2/  
Consider K in place of MG (2 valid contact(s) for K vs. 0 for MG)  

### MG:BA:1939

|  | Res. type | Atom | Residue | Distance | Occupancy | B-factor | Closest solute to 1^st^ shell water |
| --- | --- | --- | --- | --- | --- | --- | --- |
| **Ion** | | | |  | 1.00 | 81.71 || 1 | C | N3 | BA:187 | 4.19 | 1.00 | 97.24 |
| 2 | C | N4 | BA:187 | 4.35 | 1.00 | 96.67 |  |
| 3 | C | N4 | BA:186F | 4.44 | 1.00 | 98.12 |  |
| 4 | G | O6 | BA:191A | 4.52 | 1.00 | 94.13 |  |
| 5 | G | OP2 | BA:190 | 4.85 | 1.00 | 93.69 |  |
  |

  
 Coded contact list:   
1^st^ MG coordination shell missing (see Notes)  

### MG:BA:1940
