## Supplementary material for "Deflating the RNA Mg^2+^ bubble. Stereochemistry to the rescue!": Spupplemental III: 6QNR.ion_diagnosis.html

 Ion report: 6QNR  

### Ion report: 6QNR

#### date: 2020-02-17 15:38

Authors: Filip Leonarski and Pascal Auffinger  
\*\*\* beta version: unpublished \*\*\*  
-> reference 1: Leonarski F., D'Ascenzo L., Auffinger P. (2017) NAR, 45, 987-1004  
-> reference 2: Leonarski F., D'Ascenzo L., Auffinger P. (2019) RNA, 25, 173-192  
  
Density maps not assigned - No density checks made in the current version!  
  
\_\_\_\_\_\_\_\_\_\_\_\_\_\_\_\_\_\_\_\_\_\_\_\_\_\_\_\_\_\_\_\_\_\_\_\_\_\_\_\_\_\_\_\_\_\_\_\_\_\_\_\_\_\_\_\_\_\_\_\_\_\_\_\_\_\_\_\_\_\_\_\_\_\_\_\_\_\_\_\_\_  

#### Table of content (ToC):

1) Number of ions of each type and coordination distance ranges used for validation  
2) Ion validation table  
3) Potential issue list  
4) Ion/ligand tables  
5) Sorted coded contact list  
\_\_\_\_\_\_\_\_\_\_\_\_\_\_\_\_\_\_\_\_\_\_\_\_\_\_\_\_\_\_\_\_\_\_\_\_\_\_\_\_\_\_\_\_\_\_\_\_\_\_\_\_\_\_\_\_\_\_\_\_\_\_\_\_\_\_\_\_\_\_\_\_\_\_\_\_\_\_\_\_\_  
  
REMARKS:  
- 1^st^ shell: correct distance range for coordinating atoms;  
- Exclusion zone (excl. zone): a distance range where \*no\* coordinating atoms should be present;  
- Currently coordination distances with sulfur (S) atoms are not considered --> no CD/ZN (cadmium/zinc) validation;  
\_\_\_\_\_\_\_\_\_\_\_\_\_\_\_\_\_\_\_\_\_\_\_\_\_\_\_\_\_\_\_\_\_\_\_\_\_\_\_\_\_\_\_\_\_\_\_\_\_\_\_\_\_\_\_\_\_\_\_\_\_\_\_\_  

#### 1) Number of ions of each categorie and coordination distance ranges used for validation

##### Go to ToC

| Ion type | Number | Ion list | Short excl. zone [Angs] | 1^st^ shell [Angs.] | Long excl. zone [Angs] |
| --- | --- | --- | --- | --- | --- |
| Sodium (NA) | --- |  | <2.4 | 2.4 - 2.6 | 2.6 - 3.6 |
| Potassium (K) | 367 | 11:301 13:1601 13:1602 13:1603 13:1604 13:1605 13:1606 13:1607 13:1608 13:1609 13:1610 13:1611 13:1612 13:1613 13:1614 13:1615 13:1616 13:1617 13:1618 13:1619 13:1620 13:1621 13:1622 13:1623 13:1624 13:1625 13:1626 13:1627 13:1628 13:1629 13:1630 13:1631 13:1632 13:1633 13:1634 13:1635 13:1636 13:1637 13:1638 13:1639 13:1640 13:1641 13:1642 13:1643 13:1644 13:1645 13:1646 13:1647 13:1648 13:1649 13:1650 13:1651 14:3001 14:3002 14:3003 14:3004 14:3005 14:3006 14:3007 14:3008 14:3009 14:3010 14:3011 14:3012 14:3013 14:3014 14:3015 14:3016 14:3017 14:3018 14:3019 14:3020 14:3021 14:3022 14:3023 14:3024 14:3025 14:3026 14:3027 14:3028 14:3029 14:3030 14:3031 14:3032 14:3033 14:3034 14:3035 14:3036 14:3037 14:3038 14:3039 14:3040 14:3041 14:3042 14:3043 14:3044 14:3045 14:3046 14:3047 14:3048 14:3049 14:3050 14:3051 14:3052 14:3053 14:3054 14:3055 14:3056 14:3057 14:3058 14:3059 14:3060 14:3061 14:3062 14:3063 14:3064 14:3065 14:3066 14:3067 14:3068 14:3069 14:3070 14:3071 14:3072 14:3073 14:3074 14:3075 14:3076 14:3077 14:3078 14:3079 14:3080 14:3081 14:3082 14:3083 14:3084 14:3085 14:3086 14:3087 14:3088 14:3089 14:3090 14:3091 14:3092 14:3093 14:3094 14:3095 14:3096 14:3097 14:3098 14:3099 14:3100 14:3101 14:3102 14:3103 14:3104 14:3105 14:3106 16:201 16:202 16:203 19:301 1G:1601 1G:1602 1G:1603 1G:1604 1G:1605 1G:1606 1G:1607 1G:1608 1G:1609 1G:1610 1G:1611 1G:1612 1G:1613 1G:1614 1G:1615 1G:1616 1G:1617 1G:1618 1G:1619 1G:1620 1G:1621 1G:1622 1G:1623 1G:1624 1G:1625 1G:1626 1G:1627 1G:1628 1G:1629 1G:1630 1G:1631 1G:1632 1G:1633 1G:1634 1G:1635 1G:1636 1H:3001 1H:3002 1H:3003 1H:3004 1H:3005 1H:3006 1H:3007 1H:3008 1H:3009 1H:3010 1H:3011 1H:3012 1H:3013 1H:3014 1H:3015 1H:3016 1H:3017 1H:3018 1H:3019 1H:3020 1H:3021 1H:3022 1H:3023 1H:3024 1H:3025 1H:3026 1H:3027 1H:3028 1H:3029 1H:3030 1H:3031 1H:3032 1H:3033 1H:3034 1H:3035 1H:3036 1H:3037 1H:3038 1H:3039 1H:3040 1H:3040 1H:3041 1H:3042 1H:3043 1H:3044 1H:3045 1H:3046 1H:3047 1H:3048 1H:3049 1H:3050 1H:3051 1H:3052 1H:3053 1H:3054 1H:3055 1H:3056 1H:3057 1H:3058 1H:3059 1H:3060 1H:3061 1H:3062 1H:3063 1H:3064 1H:3065 1H:3066 1H:3067 1H:3068 1H:3069 1H:3070 1H:3071 1H:3072 1H:3073 1H:3074 1H:3075 1H:3076 1H:3077 1H:3078 1H:3079 1H:3080 1H:3081 1H:3082 1H:3083 1H:3084 1H:3085 1H:3086 1H:3087 1H:3088 1H:3089 1H:3090 1H:3091 1H:3092 1H:3093 1H:3094 1H:3095 1H:3096 1H:3097 1H:3098 1H:3099 1H:3100 1H:3101 1H:3102 1H:3103 1H:3104 1H:3105 1H:3106 1H:3107 1H:3108 1H:3109 1H:3110 1H:3111 1H:3112 1H:3113 1H:3114 1H:3115 1H:3116 1H:3117 1H:3118 1H:3119 1H:3120 1H:3121 1H:3122 1H:3123 1H:3124 1H:3125 1H:3126 1H:3127 1H:3128 1H:3129 1H:3130 1H:3131 1H:3132 1H:3133 1H:3134 1H:3135 1H:3136 1H:3137 1H:3138 1H:3139 1H:3140 1H:3141 1H:3142 1H:3143 1H:3144 1J:201 1K:101 21:301 29:301 2A:201 2K:101 2K:102 2K:103 31:301 31:302 32:301 39:301 39:302 41:201 45:201 49:201 4A:201 4I:201 52:201 5A:101 5E:201 88:201 BA:201 BI:201 | <2.6 | 2.6 - 3.4 | --- |
| Caesium (CS) | --- |  | <2.9 | 2.9 - 3.6 | --- |
| Magnesium (MG) | 1005 | 11:302 13:1652 13:1653 13:1654 13:1655 13:1656 13:1657 13:1658 13:1659 13:1660 13:1661 13:1662 13:1663 13:1664 13:1665 13:1666 13:1667 13:1668 13:1669 13:1670 13:1671 13:1672 13:1673 13:1674 13:1675 13:1676 13:1677 13:1678 13:1679 13:1680 13:1681 13:1682 13:1683 13:1684 13:1685 13:1686 13:1687 13:1688 13:1689 13:1690 13:1691 13:1692 13:1693 13:1694 13:1695 13:1696 13:1697 13:1698 13:1699 13:1700 13:1701 13:1702 13:1703 13:1704 13:1705 13:1706 13:1707 13:1708 13:1709 13:1709 13:1710 13:1711 13:1712 13:1713 13:1714 13:1715 13:1716 13:1717 13:1718 13:1719 13:1720 13:1721 13:1722 13:1723 13:1724 13:1725 13:1726 13:1727 13:1728 13:1729 13:1730 13:1731 13:1732 13:1733 13:1734 13:1735 13:1736 13:1737 13:1738 13:1739 13:1740 13:1741 13:1742 13:1743 13:1744 13:1745 13:1746 13:1747 13:1748 13:1749 13:1750 13:1751 13:1752 13:1753 13:1754 13:1755 13:1756 13:1757 13:1758 13:1759 13:1760 13:1761 13:1762 13:1763 13:1764 13:1765 13:1766 13:1767 13:1768 13:1769 13:1770 14:3107 14:3108 14:3109 14:3110 14:3111 14:3112 14:3113 14:3114 14:3115 14:3116 14:3117 14:3118 14:3119 14:3120 14:3121 14:3122 14:3123 14:3124 14:3125 14:3126 14:3127 14:3128 14:3129 14:3130 14:3131 14:3132 14:3133 14:3134 14:3135 14:3136 14:3137 14:3138 14:3139 14:3140 14:3141 14:3142 14:3143 14:3144 14:3145 14:3146 14:3147 14:3148 14:3149 14:3150 14:3151 14:3152 14:3153 14:3154 14:3155 14:3156 14:3157 14:3158 14:3159 14:3160 14:3161 14:3162 14:3163 14:3164 14:3165 14:3166 14:3167 14:3168 14:3169 14:3170 14:3171 14:3172 14:3173 14:3174 14:3175 14:3175 14:3176 14:3177 14:3178 14:3179 14:3180 14:3181 14:3182 14:3183 14:3184 14:3185 14:3186 14:3187 14:3188 14:3189 14:3190 14:3191 14:3192 14:3193 14:3194 14:3195 14:3196 14:3197 14:3198 14:3199 14:3200 14:3201 14:3202 14:3203 14:3204 14:3205 14:3206 14:3207 14:3208 14:3209 14:3210 14:3211 14:3212 14:3213 14:3214 14:3215 14:3216 14:3217 14:3218 14:3219 14:3220 14:3221 14:3222 14:3223 14:3224 14:3225 14:3226 14:3227 14:3228 14:3229 14:3230 14:3231 14:3232 14:3233 14:3234 14:3235 14:3236 14:3237 14:3238 14:3239 14:3240 14:3241 14:3242 14:3243 14:3244 14:3245 14:3246 14:3247 14:3248 14:3249 14:3250 14:3251 14:3252 14:3253 14:3254 14:3255 14:3256 14:3257 14:3258 14:3259 14:3260 14:3261 14:3262 14:3263 14:3264 14:3265 14:3266 14:3267 14:3268 14:3269 14:3270 14:3271 14:3272 14:3273 14:3274 14:3275 14:3276 14:3277 14:3278 14:3279 14:3280 14:3281 14:3282 14:3283 14:3284 14:3285 14:3286 14:3287 14:3288 14:3289 14:3290 14:3291 14:3292 14:3293 14:3294 14:3295 14:3296 14:3297 14:3298 14:3299 14:3300 14:3301 14:3302 14:3303 14:3304 14:3305 14:3306 14:3307 14:3308 14:3309 14:3310 14:3311 14:3312 14:3313 14:3314 14:3315 14:3316 14:3317 14:3318 14:3319 14:3320 14:3321 14:3322 14:3323 14:3324 14:3325 14:3326 14:3327 14:3328 14:3329 14:3330 14:3331 14:3332 14:3333 14:3334 14:3335 14:3336 14:3337 14:3338 14:3339 14:3340 14:3341 14:3342 14:3343 14:3344 14:3345 14:3346 14:3347 14:3348 14:3349 14:3350 14:3351 14:3352 14:3353 14:3354 14:3355 14:3356 14:3357 14:3358 14:3359 14:3360 14:3361 14:3362 14:3363 14:3364 14:3365 14:3366 14:3367 14:3368 14:3369 14:3370 14:3371 14:3372 14:3373 14:3374 14:3375 14:3376 14:3377 14:3378 14:3379 14:3380 14:3381 14:3382 14:3383 14:3384 14:3385 14:3386 14:3387 14:3388 14:3389 14:3390 14:3391 14:3392 14:3393 14:3394 14:3395 14:3396 14:3397 14:3398 14:3399 14:3400 16:204 16:205 16:206 16:207 16:208 16:209 16:210 16:211 16:212 16:213 16:214 16:215 1G:1637 1G:1638 1G:1639 1G:1640 1G:1641 1G:1642 1G:1643 1G:1644 1G:1645 1G:1646 1G:1647 1G:1648 1G:1649 1G:1650 1G:1651 1G:1652 1G:1653 1G:1654 1G:1655 1G:1656 1G:1657 1G:1658 1G:1659 1G:1660 1G:1661 1G:1662 1G:1663 1G:1663 1G:1664 1G:1665 1G:1666 1G:1667 1G:1668 1G:1669 1G:1670 1G:1671 1G:1672 1G:1673 1G:1674 1G:1675 1G:1676 1G:1677 1G:1678 1G:1679 1G:1680 1G:1681 1G:1682 1G:1683 1G:1683 1G:1684 1G:1685 1G:1686 1G:1687 1G:1688 1G:1689 1G:1690 1G:1691 1G:1692 1G:1693 1G:1694 1G:1695 1G:1696 1G:1697 1G:1698 1G:1699 1G:1700 1G:1701 1G:1702 1G:1703 1G:1704 1G:1705 1G:1706 1G:1707 1G:1708 1G:1709 1G:1710 1G:1711 1G:1712 1G:1713 1G:1714 1G:1715 1G:1716 1G:1717 1G:1718 1G:1719 1G:1720 1G:1721 1G:1722 1G:1723 1G:1724 1G:1725 1G:1726 1G:1727 1G:1728 1G:1729 1G:1730 1G:1731 1G:1732 1G:1733 1G:1734 1G:1735 1G:1736 1H:3145 1H:3146 1H:3147 1H:3148 1H:3149 1H:3150 1H:3151 1H:3152 1H:3153 1H:3154 1H:3155 1H:3155 1H:3156 1H:3157 1H:3158 1H:3159 1H:3160 1H:3161 1H:3162 1H:3163 1H:3164 1H:3165 1H:3166 1H:3167 1H:3168 1H:3169 1H:3170 1H:3171 1H:3172 1H:3173 1H:3174 1H:3175 1H:3176 1H:3176 1H:3177 1H:3178 1H:3179 1H:3180 1H:3181 1H:3182 1H:3183 1H:3184 1H:3185 1H:3186 1H:3187 1H:3188 1H:3189 1H:3190 1H:3191 1H:3192 1H:3193 1H:3194 1H:3195 1H:3196 1H:3197 1H:3198 1H:3199 1H:3200 1H:3201 1H:3202 1H:3203 1H:3204 1H:3205 1H:3206 1H:3207 1H:3208 1H:3209 1H:3210 1H:3211 1H:3212 1H:3213 1H:3214 1H:3215 1H:3216 1H:3217 1H:3218 1H:3219 1H:3220 1H:3221 1H:3222 1H:3223 1H:3224 1H:3225 1H:3226 1H:3227 1H:3228 1H:3229 1H:3230 1H:3231 1H:3232 1H:3233 1H:3234 1H:3235 1H:3236 1H:3237 1H:3238 1H:3239 1H:3240 1H:3241 1H:3242 1H:3243 1H:3244 1H:3245 1H:3246 1H:3247 1H:3248 1H:3249 1H:3250 1H:3251 1H:3252 1H:3253 1H:3254 1H:3255 1H:3256 1H:3257 1H:3258 1H:3259 1H:3260 1H:3261 1H:3262 1H:3263 1H:3264 1H:3265 1H:3266 1H:3267 1H:3268 1H:3269 1H:3270 1H:3271 1H:3272 1H:3273 1H:3274 1H:3275 1H:3276 1H:3277 1H:3278 1H:3279 1H:3280 1H:3281 1H:3282 1H:3283 1H:3284 1H:3285 1H:3286 1H:3287 1H:3288 1H:3289 1H:3290 1H:3291 1H:3292 1H:3293 1H:3294 1H:3295 1H:3296 1H:3297 1H:3298 1H:3299 1H:3300 1H:3301 1H:3302 1H:3303 1H:3304 1H:3305 1H:3306 1H:3307 1H:3308 1H:3309 1H:3310 1H:3311 1H:3312 1H:3313 1H:3314 1H:3315 1H:3316 1H:3317 1H:3318 1H:3319 1H:3320 1H:3320 1H:3321 1H:3322 1H:3323 1H:3324 1H:3325 1H:3326 1H:3327 1H:3328 1H:3328 1H:3329 1H:3330 1H:3331 1H:3332 1H:3333 1H:3334 1H:3335 1H:3336 1H:3337 1H:3338 1H:3339 1H:3340 1H:3341 1H:3342 1H:3343 1H:3343 1H:3344 1H:3345 1H:3345 1H:3346 1H:3347 1H:3348 1H:3349 1H:3350 1H:3351 1H:3352 1H:3353 1H:3354 1H:3355 1H:3356 1H:3357 1H:3358 1H:3359 1H:3360 1H:3361 1H:3362 1H:3363 1H:3364 1H:3365 1H:3366 1H:3367 1H:3368 1H:3369 1H:3370 1H:3371 1H:3372 1H:3373 1H:3374 1H:3375 1H:3376 1H:3377 1H:3378 1H:3379 1H:3380 1H:3381 1H:3382 1H:3383 1H:3384 1H:3385 1H:3386 1H:3387 1H:3388 1H:3389 1H:3390 1H:3391 1H:3392 1H:3393 1H:3394 1H:3395 1H:3396 1H:3397 1H:3398 1H:3399 1H:3400 1H:3401 1H:3402 1H:3403 1H:3404 1H:3405 1H:3406 1H:3407 1H:3408 1H:3409 1H:3410 1H:3411 1H:3412 1H:3413 1H:3414 1H:3415 1H:3416 1H:3417 1H:3418 1H:3419 1H:3420 1H:3421 1H:3422 1H:3423 1H:3424 1H:3425 1H:3426 1H:3427 1H:3428 1H:3429 1H:3430 1H:3431 1H:3432 1H:3433 1H:3434 1H:3435 1H:3436 1H:3437 1H:3438 1H:3439 1H:3440 1H:3441 1H:3442 1H:3443 1H:3444 1H:3445 1H:3446 1H:3447 1H:3448 1H:3449 1H:3450 1H:3451 1H:3452 1H:3453 1H:3454 1H:3455 1H:3456 1H:3457 1H:3458 1H:3459 1H:3460 1H:3461 1H:3462 1H:3463 1H:3464 1H:3465 1H:3466 1H:3467 1H:3468 1H:3469 1H:3470 1H:3471 1H:3472 1H:3473 1H:3474 1H:3475 1H:3476 1H:3477 1H:3478 1H:3479 1H:3480 1H:3481 1H:3482 1H:3483 1H:3484 1H:3485 1H:3486 1H:3487 1H:3488 1H:3489 1H:3490 1H:3491 1H:3492 1H:3493 1H:3494 1H:3495 1H:3496 1H:3497 1H:3498 1H:3499 1H:3500 1H:3501 1H:3502 1H:3503 1H:3504 1H:3505 1H:3506 1H:3507 1H:3508 1H:3509 1H:3510 1H:3511 1H:3512 1H:3513 1H:3514 1H:3515 1H:3516 1H:3517 1H:3518 1H:3519 1H:3520 1H:3521 1H:3522 1H:3523 1H:3524 1H:3525 1H:3526 1H:3527 1H:3528 1H:3529 1H:3530 1H:3531 1H:3532 1H:3533 1H:3534 1H:3535 1H:3536 1H:3537 1H:3538 1H:3539 1H:3540 1H:3541 1H:3542 1H:3543 1H:3544 1H:3545 1H:3546 1H:3547 1H:3548 1H:3549 1H:3550 1H:3551 1H:3552 1H:3553 1H:3554 1H:3555 1H:3556 1H:3557 1H:3558 1H:3559 1H:3560 1H:3561 1H:3562 1H:3563 1H:3564 1H:3565 1H:3566 1H:3567 1H:3568 1H:3569 1H:3570 1H:3571 1H:3572 1H:3573 1H:3574 1H:3575 1H:3576 1H:3577 1J:202 1J:203 1J:204 1J:205 21:302 21:303 29:302 29:303 2K:104 2K:105 2L:101 2L:102 2L:103 31:303 32:302 39:303 3E:301 41:202 42:201 4L:101 6E:201 78:201 78:202 7I:101 C8:201 F8:101 I8:101 J8:101 L8:101 M5:101 M5:102 N8:101 O8:101 P8:101 Q8:101 Q8:101 | <1.8 | 1.8 - 2.4 | 2.4 - 3.4 |
| Calcium (CA) | --- |  | <2.2 | 2.2 - 2.8 | 2.8 - 3.6 |
| Strontium (SR) | --- |  | <2.4 | 2.4 - 2.8 | 2.8 - 4.0 |
| Barium (BA) | --- |  | <2.6 | 2.6 - 3.2 | --- |
| Manganese (MN) | --- |  | <2.0 | 2.0 - 2.3 | 2.3 - 3.6 |
| Zinc (ZN) | 2 | 5A:102 5I:101 | <1.9 | 1.9 - 2.3 | 2.3 - 3.4 |
| Cadmium (CD) | --- |  | <2.2 | 2.2 - 2.5 | 2.5 - 3.6 |
| Cobalt (CO) | --- |  | <1.9 | 1.9 - 2.3 | 2.3 - 3.5 |
| Iridium (IRI) | --- |  | <2.0 | 2.0 - 2.3 | 2.3 - 3.6 |
| Chloride (Anion) | --- |  | <2.8 | 2.8 - 3.4 | --- |
| Sulphate (SO4) | --- |  | <2.8 | 2.8 - 3.4 | --- |
| Phosphate (PO4) | --- |  | <2.8 | 2.8 - 3.4 | --- |
| **Total** | **1374** |  |  |  |  |

---

\_\_\_\_\_\_\_\_\_\_\_\_\_\_\_\_\_\_\_\_\_\_\_\_\_\_\_\_\_\_\_\_\_\_\_\_\_\_\_\_\_\_\_\_\_\_\_\_\_\_\_\_\_\_\_\_\_\_\_\_\_\_\_\_  

#### 2) Ion validation table

##### Go to ToC

NOTES:

  
- O_ph_: OP1 or OP2 anionic phosphate oxygens;  
- O_r_: O2', O3', O4' or O5' hydroxyl and ester oxygens;  
- O_b_: O2, O4 or O6 carbonyl oxygens ;  
- N_b_: N1_A_, N3_C_, N3_G_, N3_A_ or N7 non-protonated nitrogens;  
- O_back_: protein backbone carbonyl group;  
- O_coo_: ASP/GLU carboxyl group;  
- O_cno_: ASN/GLN carbonyl group;  
- OH_Prot_: SER/THR/TYR hydroxyl group;  
- O_w_: water molecule oxygen;  
- Anion: NH group (imino, amino and amonium groups) - suggests potential anion contact or poor binding site modeling;  
  
"Large octahedral discrepancy" means a cumulated ligand-ion-ligand angle deviation > 10 deg.;  
-> the ions that are currently checked for octahedral coordination are Mg^2+^, Mn^2+^ and Na^+^;  
-> could be a hint to the presence of a polyatomic ligand instead of a monoatomic ion;  
1^st^ coordination shell missing:   
1) if Mg^2+^ or Na^+^, consider adding a hydration shell and optimize the water to solute distances;   
2) if preceding step is not conclusive, consider the presence of another solvent molecule (check crystallisation conditions);   
Exclusion atoms:   
-> if O3'-O5' atoms are mentioned, this points to a defficient modelling of the RNA backbone;  
  

| Ion | Occup. | B-factor | Ligands (too close) | Ligands (1^st^ shell) | Ligands (excl. zone) | Contacts & potential issues |
| --- | --- | --- | --- | --- | --- | --- |
| K:11:301 | 1.00 | 31.84 |  | 4 |  | 3O_back_O_w_ |
| MG:11:302 | 1.00 | 14.91 |  | 6 | 2 | 6O_w_ - Exclusion zone contact(s): /O_back_/OP1/ Consider K in place of MG (2 valid contact(s) for K vs. 0 for MG) |
| K:13:1601 | 1.00 | 67.83 |  | 1 |  | N_b_ |
| K:13:1602 | 1.00 | 72.95 |  | 4 |  | 4O_b_ |
| K:13:1603 | 1.00 | 75.94 |  | 3 |  | 2O_r_O_b_ |
| K:13:1604 | 1.00 | 82.90 |  | 3 |  | 2O_b_N_b_ |
| K:13:1605 | 1.00 | 81.86 |  | 3 |  | O_r_O_b_N_b_ |
| K:13:1606 | 1.00 | 57.00 |  | 2 |  | O_ph_O_b_ |
| K:13:1607 | 1.00 | 65.00 |  | 4 |  | 4O_b_ |
| K:13:1608 | 1.00 | 89.51 | 1 | 2 |  | O_b_Anion - Exclusion zone contact(s): /O2/ |
| K:13:1609 | 1.00 | 57.48 | 1 | 2 |  | O_ph_O_r_ - Exclusion zone contact(s): /OP1/ |
| K:13:1610 | 1.00 | 103.84 |  | 4 |  | 3O_b_Anion |
| K:13:1611 | 1.00 | 114.54 |  | 1 |  | O_b_ |
| K:13:1612 | 1.00 | 50.94 |  | 4 |  | 2O_ph_O_r_N_b_ |
| K:13:1613 | 1.00 | 76.59 |  | 2 |  | 2O_b_ |
| K:13:1614 | 1.00 | 71.95 |  | 0 |  | 1^st^ K coordination shell missing (see Notes) |
| K:13:1615 | 1.00 | 90.54 |  | 4 |  | O_ph_2O_b_N_b_ |
| K:13:1616 | 1.00 | 75.54 |  | 3 |  | O_ph_2O_b_ |
| K:13:1617 | 1.00 | 50.54 |  | 1 |  | O_ph_ |
| K:13:1618 | 1.00 | 85.59 |  | 3 |  | 3O_b_ |
| K:13:1619 | 1.00 | 58.24 |  | 4 |  | O_ph_2O_r_O_b_ |
| K:13:1620 | 1.00 | 87.09 |  | 1 |  | O_b_ |
| K:13:1621 | 1.00 | 61.31 |  | 6 |  | 2O_ph_O_b_3N_b_ |
| K:13:1622 | 1.00 | 61.23 |  | 7 |  | 2O_r_2O_b_3N_b_ |
| K:13:1623 | 1.00 | 56.20 |  | 5 |  | 2O_b_N_b_2O_back_ |
| K:13:1624 | 1.00 | 59.27 |  | 5 |  | 4O_b_Anion |
| K:13:1625 | 1.00 | 77.07 |  | 2 |  | O_ph_O_back_ |
| K:13:1626 | 1.00 | 73.63 |  | 4 |  | O_ph_O_r_O_b_N_b_ |
| K:13:1627 | 1.00 | 46.56 |  | 6 |  | 5O_b_Anion |
| K:13:1628 | 1.00 | 51.12 |  | 6 |  | O_b_2N_b_O_back_O_cno_Anion |
| K:13:1629 | 1.00 | 50.77 | 1 | 5 |  | 2O_r_O_b_N_b_O_back_ - Exclusion zone contact(s): /O6/ |
| K:13:1630 | 1.00 | 66.28 |  | 3 |  | O_ph_O_r_O_back_ |
| K:13:1631 | 1.00 | 58.22 |  | 4 |  | 3O_ph_O_r_ |
| K:13:1632 | 1.00 | 79.99 |  | 1 |  | N_b_ |
| K:13:1633 | 1.00 | 55.73 |  | 6 |  | O_ph_2O_b_2N_b_Anion |
| K:13:1634 | 1.00 | 90.67 |  | 3 |  | 3O_b_ |
| K:13:1635 | 1.00 | 95.87 | 1 | 2 |  | 2O_b_ - Exclusion zone contact(s): /O2/ |
| K:13:1636 | 1.00 | 70.96 |  | 4 |  | 2O_ph_2O_r_ |
| K:13:1637 | 1.00 | 64.26 |  | 5 |  | 3O_ph_2O_r_ |
| K:13:1638 | 1.00 | 56.02 |  | 3 |  | 3O_b_ |
| K:13:1639 | 1.00 | 66.33 |  | 3 |  | 3O_b_ |
| K:13:1640 | 1.00 | 60.81 |  | 9 |  | O_ph_4O_r_O_b_2N_b_O_w_ |
| K:13:1641 | 1.00 | 84.82 |  | 1 |  | O_b_ |
| K:13:1642 | 1.00 | 93.47 |  | 1 |  | O_ph_ |
| K:13:1643 | 1.00 | 76.13 |  | 5 |  | 2O_ph_3O_r_ |
| K:13:1644 | 1.00 | 74.58 | 1 | 2 |  | O_ph_O_r_ - Exclusion zone contact(s): /OP1/ |
| K:13:1645 | 1.00 | 72.01 |  | 3 |  | 3O_b_ |
| K:13:1646 | 1.00 | 63.13 |  | 2 |  | 2O_b_ |
| K:13:1647 | 1.00 | 50.65 |  | 4 |  | 4O_b_ |
| K:13:1648 | 1.00 | 71.10 |  | 1 |  | O_b_ |
| K:13:1649 | 1.00 | 81.17 |  | 3 |  | 2O_ph_O_r_ |
| K:13:1650 | 1.00 | 97.83 |  | 3 |  | 2O_ph_O_b_ |
| K:13:1651 | 1.00 | 81.28 | 1 | 2 |  | O_b_N_b_ - Exclusion zone contact(s): /OP2/ |
| MG:13:1652 | 1.00 | 43.51 |  | 0 |  | 1^st^ MG coordination shell missing (see Notes) |
| MG:13:1653 | 1.00 | 67.78 |  | 1 | 1 | O_ph_ - Exclusion zone contact(s): /O2/ |
| MG:13:1654 | 1.00 | 42.74 |  | 0 |  | 1^st^ MG coordination shell missing (see Notes) |
| MG:13:1655 | 1.00 | 51.07 |  | 0 | 1 | - Exclusion zone contact(s): /N7/ Consider K in place of MG (1 valid contact(s) for K vs. 0 for MG) |
| MG:13:1656 | 1.00 | 49.80 |  | 1 | 4 | O_ph_ - Exclusion zone contact(s): /O5'/OP1/OP1/OP2/ Consider K in place of MG (3 valid contact(s) for K vs. 1 for MG) |
| MG:13:1657 | 1.00 | 38.67 |  | 1 | 1 | O_ph_ - Exclusion zone contact(s): /OP1/ |
| MG:13:1658 | 1.00 | 36.87 |  | 0 | 1 | - Exclusion zone contact(s): /N7/ Consider K in place of MG (1 valid contact(s) for K vs. 0 for MG) |
| MG:13:1659 | 1.00 | 56.57 | 1 | 1 | 4 | O_b_ - Exclusion zone contact(s): /O6/O6/OP1/OP1/OP2/ Consider K in place of MG (3 valid contact(s) for K vs. 1 for MG) |
| MG:13:1660 | 1.00 | 39.52 |  | 1 |  | O_ph_ |
| MG:13:1661 | 1.00 | 39.42 |  | 1 | 1 | O_ph_ - Exclusion zone contact(s): /OP1/ |
| MG:13:1662 | 1.00 | 54.19 |  | 0 | 1 | - Exclusion zone contact(s): /OP2/ Consider NA in place of MG (1 valid contact(s) for NA vs. 0 for MG) |
| MG:13:1663 | 1.00 | 47.27 |  | 1 |  | O_ph_ |
| MG:13:1664 | 1.00 | 57.26 |  | 1 | 1 | O_b_ - Exclusion zone contact(s): /OP2/ |
| MG:13:1665 | 1.00 | 51.98 |  | 0 | 1 | - Exclusion zone contact(s): /OP2/ Consider NA in place of MG (1 valid contact(s) for NA vs. 0 for MG) |
| MG:13:1666 | 1.00 | 46.55 |  | 0 | 1 | - Exclusion zone contact(s): /O_w_/ 1^st^ MG coordination shell missing (see Notes) |
| MG:13:1667 | 1.00 | 29.73 |  | 0 | 1 | - Exclusion zone contact(s): /OP1/ Consider K in place of MG (1 valid contact(s) for K vs. 0 for MG) |
| MG:13:1668 | 1.00 | 31.81 |  | 3 | 1 | 3O_ph_ - Exclusion zone contact(s): /O2'/ |
| MG:13:1669 | 1.00 | 34.83 |  | 0 |  | 1^st^ MG coordination shell missing (see Notes) |
| MG:13:1670 | 1.00 | 42.83 |  | 1 |  | O_ph_ |
| MG:13:1671 | 1.00 | 49.91 |  | 0 |  | 1^st^ MG coordination shell missing (see Notes) |
| MG:13:1672 | 1.00 | 49.97 |  | 0 | 2 | - Exclusion zone contact(s): /N7/OP2/ Consider K in place of MG (2 valid contact(s) for K vs. 0 for MG) |
| MG:13:1673 | 1.00 | 55.56 |  | 0 | 1 | - Exclusion zone contact(s): /O6/ Consider K in place of MG (1 valid contact(s) for K vs. 0 for MG) |
| MG:13:1674 | 1.00 | 75.42 |  | 1 | 4 | O_ph_ - Exclusion zone contact(s): /N3_C_/O3'/O4/O6/ Consider K in place of MG (4 valid contact(s) for K vs. 1 for MG) |
| MG:13:1675 | 1.00 | 49.35 |  | 0 |  | 1^st^ MG coordination shell missing (see Notes) |
| MG:13:1676 | 1.00 | 53.19 |  | 0 | 2 | - Exclusion zone contact(s): /O5'/OP1/ Consider K in place of MG (2 valid contact(s) for K vs. 0 for MG) |
| MG:13:1677 | 1.00 | 42.26 |  | 2 |  | 2O_ph_ |
| MG:13:1678 | 1.00 | 38.17 |  | 1 |  | O_ph_ |
| MG:13:1679 | 1.00 | 45.54 |  | 0 |  | 1^st^ MG coordination shell missing (see Notes) |
| MG:13:1680 | 1.00 | 49.72 |  | 0 |  | 1^st^ MG coordination shell missing (see Notes) |
| MG:13:1681 | 1.00 | 44.41 |  | 0 | 1 | - Exclusion zone contact(s): /O6/ Consider K in place of MG (1 valid contact(s) for K vs. 0 for MG) |
| MG:13:1682 | 1.00 | 33.19 |  | 0 |  | 1^st^ MG coordination shell missing (see Notes) |
| MG:13:1683 | 1.00 | 39.19 |  | 2 |  | O_ph_O_b_ |
| MG:13:1684 | 1.00 | 73.76 |  | 1 | 2 | O_ph_ - Exclusion zone contact(s): /OP1/OP2/ Consider K in place of MG (2 valid contact(s) for K vs. 1 for MG) |
| MG:13:1685 | 1.00 | 45.99 |  | 1 | 1 | O_b_ - Exclusion zone contact(s): /OP2/ |
| MG:13:1686 | 1.00 | 34.40 |  | 0 | 1 | - Exclusion zone contact(s): /OP1/ Consider NA in place of MG (1 valid contact(s) for NA vs. 0 for MG) |
| MG:13:1687 | 1.00 | 50.85 |  | 0 | 1 | - Exclusion zone contact(s): /O6/ Consider K in place of MG (1 valid contact(s) for K vs. 0 for MG) |
| MG:13:1688 | 1.00 | 56.75 |  | 0 | 2 | - Exclusion zone contact(s): /O2/OP2/ Consider K in place of MG (2 valid contact(s) for K vs. 0 for MG) |
| MG:13:1689 | 1.00 | 53.63 |  | 0 |  | 1^st^ MG coordination shell missing (see Notes) |
| MG:13:1690 | 1.00 | 51.21 |  | 0 | 3 | - Exclusion zone contact(s): /N7/O6/OP2/ Consider K in place of MG (3 valid contact(s) for K vs. 0 for MG) |
| MG:13:1691 | 1.00 | 50.15 |  | 0 | 2 | - Exclusion zone contact(s): /NZ_Lys_/OP1/ Consider NA in place of MG (1 valid contact(s) for NA vs. 0 for MG) |
| MG:13:1692 | 1.00 | 51.67 |  | 0 | 2 | - Exclusion zone contact(s): /N7/O6/ Consider K in place of MG (2 valid contact(s) for K vs. 0 for MG) |
| MG:13:1693 | 1.00 | 61.12 |  | 1 |  | O_ph_ |
| MG:13:1694 | 1.00 | 57.89 |  | 1 |  | O_b_ |
| MG:13:1695 | 1.00 | 66.98 |  | 1 | 4 | O_b_ - Exclusion zone contact(s): /O2'/O3'/OP2/OP2/ Consider K in place of MG (3 valid contact(s) for K vs. 1 for MG) |
| MG:13:1696 | 1.00 | 64.72 |  | 0 |  | 1^st^ MG coordination shell missing (see Notes) |
| MG:13:1697 | 1.00 | 45.64 |  | 0 | 1 | - Exclusion zone contact(s): /O6/ Consider NA in place of MG (1 valid contact(s) for NA vs. 0 for MG) |
| MG:13:1698 | 1.00 | 84.61 |  | 0 | 1 | - Exclusion zone contact(s): /OP2/ Consider K in place of MG (1 valid contact(s) for K vs. 0 for MG) |
| MG:13:1699 | 1.00 | 43.13 |  | 0 |  | 1^st^ MG coordination shell missing (see Notes) |
| MG:13:1700 | 1.00 | 47.60 |  | 0 | 3 | - Exclusion zone contact(s): /OP1/OP1/OP2/ Consider K in place of MG (3 valid contact(s) for K vs. 0 for MG) |
| MG:13:1701 | 1.00 | 69.77 |  | 0 | 2 | - Exclusion zone contact(s): /O6/O6/ Consider K in place of MG (2 valid contact(s) for K vs. 0 for MG) |
| MG:13:1702 | 1.00 | 60.82 |  | 0 | 2 | - Exclusion zone contact(s): /O6/OP2/ Consider K in place of MG (2 valid contact(s) for K vs. 0 for MG) |
| MG:13:1703 | 1.00 | 49.60 |  | 0 | 1 | - Exclusion zone contact(s): /O6/ Consider K in place of MG (1 valid contact(s) for K vs. 0 for MG) |
| MG:13:1704 | 1.00 | 56.96 |  | 0 | 1 | - Exclusion zone contact(s): /OP1/ Consider K in place of MG (1 valid contact(s) for K vs. 0 for MG) |
| MG:13:1705 | 1.00 | 56.00 |  | 0 | 4 | - Exclusion zone contact(s): /N3_A_/O2'/OP1/OP2/ Consider NA in place of MG (2 valid contact(s) for NA vs. 0 for MG) |
| MG:13:1706 | 1.00 | 46.38 |  | 0 | 2 | - Exclusion zone contact(s): /N7/OP1/ Consider K in place of MG (2 valid contact(s) for K vs. 0 for MG) |
| MG:13:1707 | 1.00 | 58.52 |  | 0 | 1 | - Exclusion zone contact(s): /OP2/ Consider K in place of MG (1 valid contact(s) for K vs. 0 for MG) |
| MG:13:1708 | 1.00 | 60.04 |  | 0 | 4 | - Exclusion zone contact(s): /O4/O6/O6/O6/ Consider K in place of MG (4 valid contact(s) for K vs. 0 for MG) |
| MG:13:1709 | 0.50 | 40.21 |  | 0 | 2 | - Exclusion zone contact(s): /O6/O6/ Consider K in place of MG (2 valid contact(s) for K vs. 0 for MG) |
| MG:13:1709 | 0.50 | 45.05 |  | 0 |  | 1^st^ MG coordination shell missing (see Notes) |
| MG:13:1710 | 1.00 | 52.01 |  | 0 |  | 1^st^ MG coordination shell missing (see Notes) |
| MG:13:1711 | 1.00 | 55.78 |  | 0 | 2 | - Exclusion zone contact(s): /N7/OP2/ Consider K in place of MG (2 valid contact(s) for K vs. 0 for MG) |
| MG:13:1712 | 1.00 | 43.50 |  | 0 | 1 | - Exclusion zone contact(s): /O6/ Consider K in place of MG (1 valid contact(s) for K vs. 0 for MG) |
| MG:13:1713 | 1.00 | 46.36 |  | 0 | 2 | - Exclusion zone contact(s): /O6/O6/ Consider K in place of MG (2 valid contact(s) for K vs. 0 for MG) |
| MG:13:1714 | 1.00 | 83.76 |  | 0 | 1 | - Exclusion zone contact(s): /OP1/ Consider K in place of MG (1 valid contact(s) for K vs. 0 for MG) |
| MG:13:1715 | 1.00 | 60.74 |  | 0 | 1 | - Exclusion zone contact(s): /N7/ Consider K in place of MG (1 valid contact(s) for K vs. 0 for MG) |
| MG:13:1716 | 1.00 | 37.21 |  | 0 |  | 1^st^ MG coordination shell missing (see Notes) |
| MG:13:1717 | 1.00 | 62.06 |  | 0 | 1 | - Exclusion zone contact(s): /OP1/ Consider K in place of MG (1 valid contact(s) for K vs. 0 for MG) |
| MG:13:1718 | 1.00 | 55.93 |  | 0 |  | 1^st^ MG coordination shell missing (see Notes) |
| MG:13:1719 | 1.00 | 63.93 |  | 0 |  | 1^st^ MG coordination shell missing (see Notes) |
| MG:13:1720 | 1.00 | 64.00 |  | 0 | 3 | - Exclusion zone contact(s): /O4/O6/O6/ Consider K in place of MG (3 valid contact(s) for K vs. 0 for MG) |
| MG:13:1721 | 1.00 | 99.67 |  | 0 | 3 | - Exclusion zone contact(s): /O5'/OP2/OP2/ Consider K in place of MG (3 valid contact(s) for K vs. 0 for MG) |
| MG:13:1722 | 1.00 | 52.61 |  | 0 | 3 | - Exclusion zone contact(s): /O4/O4/O6/ Consider K in place of MG (3 valid contact(s) for K vs. 0 for MG) |
| MG:13:1723 | 1.00 | 60.24 |  | 0 |  | 1^st^ MG coordination shell missing (see Notes) |
| MG:13:1724 | 1.00 | 43.19 |  | 1 |  | O_ph_ |
| MG:13:1725 | 1.00 | 58.27 |  | 0 |  | 1^st^ MG coordination shell missing (see Notes) |
| MG:13:1726 | 1.00 | 46.92 |  | 3 | 1 | 3O_ph_ - Exclusion zone contact(s): /O3'/ |
| MG:13:1727 | 1.00 | 60.83 |  | 6 |  | 6O_w_ |
| MG:13:1728 | 1.00 | 60.52 |  | 6 |  | 6O_w_ |
| MG:13:1729 | 1.00 | 61.45 |  | 6 |  | 6O_w_ |
| MG:13:1730 | 1.00 | 76.59 |  | 6 |  | 6O_w_ |
| MG:13:1731 | 1.00 | 38.84 |  | 5 | 2 | 5O_w_ - Exclusion zone contact(s): /OP1/OP2/ Consider K in place of MG (2 valid contact(s) for K vs. 0 for MG) |
| MG:13:1732 | 1.00 | 51.59 |  | 6 |  | 6O_w_ |
| MG:13:1733 | 1.00 | 62.46 |  | 6 |  | 6O_w_ |
| MG:13:1734 | 1.00 | 41.94 |  | 6 |  | 6O_w_ |
| MG:13:1735 | 1.00 | 35.59 |  | 6 | 1 | 6O_w_ - Exclusion zone contact(s): /OP1/ Consider K in place of MG (1 valid contact(s) for K vs. 0 for MG) |
| MG:13:1736 | 1.00 | 44.29 |  | 6 |  | 6O_w_ |
| MG:13:1737 | 1.00 | 65.46 |  | 6 | 2 | 6O_w_ - Exclusion zone contact(s): /OP2/OP2/ Consider K in place of MG (2 valid contact(s) for K vs. 0 for MG)  Possible idealized distances ! |
| MG:13:1738 | 1.00 | 86.34 |  | 6 |  | 6O_w_ |
| MG:13:1739 | 1.00 | 30.03 |  | 6 |  | 6O_w_ |
| MG:13:1740 | 1.00 | 33.83 |  | 6 |  | 6O_w_ |
| MG:13:1741 | 1.00 | 109.99 |  | 6 |  | 6O_w_ |
| MG:13:1742 | 1.00 | 60.34 |  | 6 |  | 6O_w_ |
| MG:13:1743 | 1.00 | 54.10 |  | 6 |  | 6O_w_ |
| MG:13:1744 | 1.00 | 19.82 |  | 5 | 1 | 5O_w_ - Exclusion zone contact(s): /OP2/ Consider NA in place of MG (1 valid contact(s) for NA vs. 0 for MG) |
| MG:13:1745 | 1.00 | 39.18 |  | 5 | 2 | 5O_w_ - Exclusion zone contact(s): /O3'/OP2/ Consider K in place of MG (2 valid contact(s) for K vs. 0 for MG) |
| MG:13:1746 | 1.00 | 37.77 |  | 5 | 1 | 5O_w_ - Exclusion zone contact(s): /OP1/ Consider NA in place of MG (1 valid contact(s) for NA vs. 0 for MG) |
| MG:13:1747 | 1.00 | 32.12 |  | 3 | 4 | 3O_w_ - Exclusion zone contact(s): /OP1/OP2/OP2/OP2/ Consider K in place of MG (3 valid contact(s) for K vs. 0 for MG) |
| MG:13:1748 | 1.00 | 38.60 |  | 5 | 1 | O_ph_4O_w_ - Exclusion zone contact(s): /OP1/ |
| MG:13:1749 | 1.00 | 52.96 |  | 3 | 3 | 3O_w_ - Exclusion zone contact(s): /OP2/OP2/OP2/ Consider K in place of MG (2 valid contact(s) for K vs. 0 for MG) |
| MG:13:1750 | 1.00 | 52.09 |  | 5 | 1 | 5O_w_ - Exclusion zone contact(s): /OP1/ Consider NA in place of MG (1 valid contact(s) for NA vs. 0 for MG) |
| MG:13:1751 | 1.00 | 75.12 |  | 4 | 3 | 4O_w_ - Exclusion zone contact(s): /OP1/OP2/OP2/ Consider K in place of MG (3 valid contact(s) for K vs. 0 for MG)  Possible idealized distances ! |
| MG:13:1752 | 1.00 | 47.91 |  | 5 | 1 | 5O_w_ - Exclusion zone contact(s): /OP1/ Consider K in place of MG (1 valid contact(s) for K vs. 0 for MG)  Possible idealized distances ! |
| MG:13:1753 | 1.00 | 69.21 |  | 5 | 2 | 5O_w_ - Exclusion zone contact(s): /OP1/OP2/ Consider K in place of MG (2 valid contact(s) for K vs. 0 for MG) |
| MG:13:1754 | 1.00 | 35.34 |  | 5 | 1 | 5O_w_ - Exclusion zone contact(s): /OP2/ Consider NA in place of MG (1 valid contact(s) for NA vs. 0 for MG) |
| MG:13:1755 | 1.00 | 83.59 |  | 6 |  | 6O_w_ Possible idealized distances ! |
| MG:13:1756 | 1.00 | 63.53 |  | 6 |  | 6O_w_ |
| MG:13:1757 | 1.00 | 50.95 |  | 6 | 1 | 6O_w_ - Exclusion zone contact(s): /N7/ Consider K in place of MG (1 valid contact(s) for K vs. 0 for MG) |
| MG:13:1758 | 1.00 | 76.99 |  | 6 |  | 6O_w_ Possible idealized distances ! |
| MG:13:1759 | 1.00 | 52.95 |  | 6 |  | 6O_w_ |
| MG:13:1760 | 1.00 | 65.02 |  | 6 |  | 6O_w_ |
| MG:13:1761 | 1.00 | 86.98 |  | 6 | 1 | 6O_w_ - Exclusion zone contact(s): /OP1/ Consider K in place of MG (1 valid contact(s) for K vs. 0 for MG)  Possible idealized distances ! |
| MG:13:1762 | 1.00 | 52.02 |  | 6 |  | 6O_w_ |
| MG:13:1763 | 1.00 | 64.95 |  | 6 | 2 | 6O_w_ - Exclusion zone contact(s): /N7/N7/ Consider K in place of MG (2 valid contact(s) for K vs. 0 for MG) |
| MG:13:1764 | 1.00 | 86.65 |  | 6 |  | 6O_w_ |
| MG:13:1765 | 1.00 | 73.01 |  | 6 |  | 6O_w_ |
| MG:13:1766 | 1.00 | 64.12 |  | 6 |  | 6O_w_ |
| MG:13:1767 | 1.00 | 70.75 |  | 6 |  | 6O_w_ |
| MG:13:1768 | 1.00 | 78.43 |  | 6 |  | 6O_w_ Possible idealized distances ! |
| MG:13:1769 | 1.00 | 54.55 |  | 3 | 4 | 3O_w_ - Exclusion zone contact(s): /O2/O2'/OP1/OP2/ Consider K in place of MG (4 valid contact(s) for K vs. 0 for MG) |
| MG:13:1770 | 1.00 | 71.86 |  | 5 | 1 | 5O_w_ - Exclusion zone contact(s): /N7/ Consider K in place of MG (1 valid contact(s) for K vs. 0 for MG) |
| K:14:3001 | 1.00 | 76.91 |  | 2 |  | O_ph_O_b_ |
| K:14:3002 | 1.00 | 69.11 |  | 3 |  | 2O_ph_O_back_ |
| K:14:3003 | 1.00 | 100.43 |  | 3 |  | 3O_b_ |
| K:14:3004 | 1.00 | 81.83 |  | 2 |  | O_ph_O_b_ |
| K:14:3005 | 1.00 | 82.41 | 1 | 1 |  | O_ph_ - Exclusion zone contact(s): /OP2/ |
| K:14:3006 | 1.00 | 106.03 |  | 2 |  | 2O_b_ |
| K:14:3007 | 1.00 | 64.40 |  | 4 |  | 4O_r_ |
| K:14:3008 | 1.00 | 47.69 | 3 | 3 |  | - Exclusion zone contact(s): /OP2/OP2/OP2/ Consider NA in place of K (3 valid contact(s) for NA vs. 0 for K) |
| K:14:3009 | 1.00 | 64.19 |  | 1 |  | O_ph_ |
| K:14:3010 | 1.00 | 58.07 |  | 2 |  | O_ph_O_r_ |
| K:14:3011 | 1.00 | 80.91 |  | 0 |  | 1^st^ K coordination shell missing (see Notes) |
| K:14:3012 | 1.00 | 117.89 |  | 1 |  | O_b_ |
| K:14:3013 | 1.00 | 84.79 |  | 1 |  | O_r_ |
| K:14:3014 | 1.00 | 80.63 |  | 3 |  | O_b_2N_b_ |
| K:14:3015 | 1.00 | 47.64 |  | 8 |  | 5O_ph_2O_r_O_w_ |
| K:14:3016 | 1.00 | 44.18 |  | 4 |  | O_r_3O_b_ |
| K:14:3017 | 1.00 | 52.00 |  | 4 |  | 3O_r_O_b_ |
| K:14:3018 | 1.00 | 65.33 |  | 5 |  | O_r_O_b_3N_b_ |
| K:14:3019 | 1.00 | 40.24 |  | 5 |  | O_r_4N_b_ |
| K:14:3020 | 1.00 | 64.35 |  | 3 |  | 3O_b_ |
| K:14:3021 | 1.00 | 41.45 | 1 | 5 |  | O_r_3O_b_N_b_ - Exclusion zone contact(s): /O6/ |
| K:14:3022 | 1.00 | 50.85 |  | 6 |  | 2O_ph_O_r_O_coo_2O_w_ |
| K:14:3023 | 1.00 | 60.81 |  | 4 |  | O_ph_O_r_O_b_N_b_ |
| K:14:3024 | 1.00 | 52.53 |  | 5 |  | 2O_ph_2O_r_N_b_ |
| K:14:3025 | 1.00 | 50.23 |  | 5 |  | 2O_ph_2O_r_O_w_ |
| K:14:3026 | 1.00 | 35.89 |  | 6 |  | 3O_ph_3O_r_ |
| K:14:3027 | 1.00 | 54.39 |  | 2 |  | O_ph_N_b_ |
| K:14:3028 | 1.00 | 45.73 | 1 | 2 |  | 2N_b_ - Exclusion zone contact(s): /O_w_/ |
| K:14:3029 | 1.00 | 81.55 |  | 4 |  | 4O_b_ |
| K:14:3030 | 1.00 | 63.16 |  | 4 |  | 4O_b_ |
| K:14:3031 | 1.00 | 29.30 |  | 4 |  | 2O_ph_O_b_O_back_ |
| K:14:3032 | 1.00 | 53.89 | 1 | 3 |  | 2O_r_O_b_ - Exclusion zone contact(s): /O2/ |
| K:14:3033 | 1.00 | 52.89 | 1 | 4 |  | O_r_3O_b_ - Exclusion zone contact(s): /O6/ |
| K:14:3034 | 1.00 | 86.90 |  | 2 |  | O_r_O_b_ |
| K:14:3035 | 1.00 | 83.36 |  | 5 |  | 2O_ph_2O_b_N_b_ |
| K:14:3036 | 1.00 | 41.19 | 1 | 3 |  | O_ph_O_r_O_b_ - Exclusion zone contact(s): /OP2/ |
| K:14:3037 | 1.00 | 60.54 |  | 4 |  | 4O_b_ |
| K:14:3038 | 1.00 | 92.10 |  | 4 |  | 2O_r_O_b_N_b_ |
| K:14:3039 | 1.00 | 61.92 |  | 3 |  | O_b_2N_b_ |
| K:14:3040 | 1.00 | 93.42 |  | 7 |  | 2O_ph_2O_r_2N_b_Anion |
| K:14:3041 | 1.00 | 66.57 |  | 2 |  | O_r_O_back_ |
| K:14:3042 | 1.00 | 51.64 |  | 3 |  | 2O_ph_N_b_ |
| K:14:3043 | 1.00 | 80.73 |  | 4 |  | 4O_b_ |
| K:14:3044 | 1.00 | 69.39 |  | 3 |  | O_ph_2O_b_ |
| K:14:3045 | 1.00 | 83.93 |  | 3 |  | 3O_b_ |
| K:14:3046 | 1.00 | 54.58 |  | 3 |  | 2O_r_O_b_ |
| K:14:3047 | 1.00 | 74.56 |  | 4 |  | 4O_b_ |
| K:14:3048 | 1.00 | 67.81 |  | 4 |  | 4O_b_ |
| K:14:3049 | 1.00 | 49.75 |  | 7 |  | 2O_ph_4O_r_O_b_ |
| K:14:3050 | 1.00 | 63.62 |  | 4 |  | O_ph_O_r_2O_b_ |
| K:14:3051 | 1.00 | 65.57 |  | 4 |  | O_r_3O_b_ |
| K:14:3052 | 1.00 | 76.70 |  | 3 |  | 2O_b_Anion |
| K:14:3053 | 1.00 | 102.84 |  | 1 |  | O_b_ |
| K:14:3054 | 1.00 | 58.19 | 1 | 2 |  | O_ph_O_r_ - Exclusion zone contact(s): /O_w_/ |
| K:14:3055 | 1.00 | 65.71 |  | 5 |  | 2O_ph_2O_r_N_b_ |
| K:14:3056 | 1.00 | 56.10 |  | 4 |  | 4O_b_ |
| K:14:3057 | 1.00 | 54.72 |  | 2 |  | 2O_b_ |
| K:14:3058 | 1.00 | 85.16 |  | 2 |  | O_ph_O_b_ |
| K:14:3059 | 1.00 | 48.52 |  | 4 |  | 2O_b_N_b_ |
| K:14:3060 | 1.00 | 45.97 |  | 5 |  | 4O_b_O_w_ |
| K:14:3061 | 1.00 | 63.49 |  | 2 |  | 2O_b_ |
| K:14:3062 | 1.00 | 76.61 |  | 1 |  | O_b_ |
| K:14:3063 | 1.00 | 57.35 |  | 3 |  | 2O_ph_O_b_ |
| K:14:3064 | 1.00 | 87.97 |  | 1 |  | O_b_ |
| K:14:3065 | 1.00 | 55.80 |  | 6 |  | 5O_ph_O_r_ |
| K:14:3066 | 1.00 | 64.36 |  | 2 |  | O_ph_O_r_ |
| K:14:3067 | 1.00 | 70.59 |  | 1 |  | O_r_ |
| K:14:3068 | 1.00 | 61.15 |  | 3 |  | 2O_r_O_back_ |
| K:14:3069 | 1.00 | 56.45 | 1 | 2 |  | 2O_b_ - Exclusion zone contact(s): /OP1/ |
| K:14:3070 | 1.00 | 92.30 |  | 3 |  | 2O_b_Anion |
| K:14:3071 | 1.00 | 77.65 |  | 2 |  | O_ph_O_b_ |
| K:14:3072 | 1.00 | 85.76 |  | 3 |  | O_ph_O_r_O_b_ |
| K:14:3073 | 1.00 | 60.73 |  | 4 |  | O_r_O_b_2O_w_ |
| K:14:3074 | 1.00 | 73.31 |  | 4 |  | 4O_b_ |
| K:14:3075 | 1.00 | 36.88 |  | 5 |  | O_ph_3O_r_N_b_ |
| K:14:3076 | 1.00 | 86.18 |  | 2 |  | O_b_O_w_ |
| K:14:3077 | 1.00 | 73.04 |  | 2 |  | O_r_O_b_ |
| K:14:3078 | 1.00 | 81.67 | 1 | 3 |  | O_ph_2O_r_ - Exclusion zone contact(s): /OP1/ |
| K:14:3079 | 1.00 | 81.16 |  | 0 |  | 1^st^ K coordination shell missing (see Notes) |
| K:14:3080 | 1.00 | 79.15 |  | 3 |  | 2O_b_Anion |
| K:14:3081 | 1.00 | 54.57 |  | 2 |  | O_ph_O_b_ |
| K:14:3082 | 1.00 | 80.02 |  | 3 |  | 2O_r_O_b_ |
| K:14:3083 | 1.00 | 70.33 |  | 3 |  | O_ph_OH_Prot_O_w_ |
| K:14:3084 | 1.00 | 73.92 |  | 4 |  | 3O_ph_O_w_ |
| K:14:3085 | 1.00 | 62.28 |  | 3 |  | O_ph_O_b_O_w_ |
| K:14:3086 | 1.00 | 65.16 |  | 4 |  | O_ph_3O_r_ |
| K:14:3087 | 1.00 | 74.57 |  | 2 |  | O_ph_O_r_ |
| K:14:3088 | 1.00 | 98.43 |  | 0 |  | 1^st^ K coordination shell missing (see Notes) |
| K:14:3089 | 1.00 | 64.79 |  | 4 |  | 2O_r_O_b_N_b_ |
| K:14:3090 | 1.00 | 64.20 |  | 3 |  | O_ph_2O_r_ |
| K:14:3091 | 1.00 | 78.64 |  | 1 |  | O_b_ |
| K:14:3092 | 1.00 | 106.11 |  | 2 |  | O_b_O_w_ |
| K:14:3093 | 1.00 | 53.26 |  | 2 |  | O_ph_O_r_ |
| K:14:3094 | 1.00 | 86.02 | 1 | 3 |  | 3O_w_ - Exclusion zone contact(s): /O6/ Consider NA in place of K (1 valid contact(s) for NA vs. 0 for K) |
| K:14:3095 | 1.00 | 64.76 |  | 3 |  | 3O_b_ |
| K:14:3096 | 1.00 | 70.82 |  | 2 |  | 2O_b_ |
| K:14:3097 | 1.00 | 91.20 |  | 2 |  | 2O_ph_ |
| K:14:3098 | 1.00 | 67.62 |  | 4 |  | 4O_b_ |
| K:14:3099 | 1.00 | 67.62 |  | 3 |  | 2O_b_N_b_ |
| K:14:3100 | 1.00 | 71.58 |  | 3 |  | O_ph_2O_r_ |
| K:14:3101 | 1.00 | 76.69 |  | 2 |  | 2O_b_ |
| K:14:3102 | 1.00 | 76.27 |  | 3 |  | O_ph_O_r_N_b_ |
| K:14:3103 | 1.00 | 70.70 |  | 3 |  | O_ph_2O_w_ |
| K:14:3104 | 1.00 | 75.78 |  | 4 |  | 4O_ph_ |
| K:14:3105 | 1.00 | 91.10 |  | 1 |  | O_b_ |
| K:14:3106 | 1.00 | 68.56 |  | 2 |  | 2O_b_ |
| MG:14:3107 | 1.00 | 33.71 |  | 2 |  | 2O_ph_ |
| MG:14:3108 | 1.00 | 59.02 |  | 2 | 1 | 2O_ph_ - Exclusion zone contact(s): /OP2/ |
| MG:14:3109 | 1.00 | 41.46 |  | 0 | 3 | - Exclusion zone contact(s): /O6/O_w_/OP2/ Consider NA in place of MG (1 valid contact(s) for NA vs. 0 for MG) |
| MG:14:3110 | 1.00 | 35.26 |  | 1 |  | O_ph_ |
| MG:14:3111 | 1.00 | 32.88 |  | 0 | 1 | - Exclusion zone contact(s): /OP2/ Consider NA in place of MG (1 valid contact(s) for NA vs. 0 for MG) |
| MG:14:3112 | 1.00 | 44.97 |  | 0 | 3 | - Exclusion zone contact(s): /N7/N7/OP2/ Consider K in place of MG (2 valid contact(s) for K vs. 0 for MG) |
| MG:14:3113 | 1.00 | 36.44 |  | 0 |  | 1^st^ MG coordination shell missing (see Notes) |
| MG:14:3114 | 1.00 | 33.13 |  | 1 | 1 | O_ph_ - Exclusion zone contact(s): /OP2/ |
| MG:14:3115 | 1.00 | 42.28 |  | 0 |  | 1^st^ MG coordination shell missing (see Notes) |
| MG:14:3116 | 1.00 | 32.75 |  | 0 |  | 1^st^ MG coordination shell missing (see Notes) |
| MG:14:3117 | 1.00 | 49.61 |  | 0 |  | 1^st^ MG coordination shell missing (see Notes) |
| MG:14:3118 | 1.00 | 58.95 |  | 0 | 2 | - Exclusion zone contact(s): /O6/O6/ Consider K in place of MG (2 valid contact(s) for K vs. 0 for MG) |
| MG:14:3119 | 1.00 | 49.87 |  | 1 |  | O_ph_ |
| MG:14:3120 | 1.00 | 44.07 |  | 0 | 1 | - Exclusion zone contact(s): /OP1/ Consider K in place of MG (1 valid contact(s) for K vs. 0 for MG) |
| MG:14:3121 | 1.00 | 47.51 |  | 0 | 2 | - Exclusion zone contact(s): /N7/O6/ Consider K in place of MG (2 valid contact(s) for K vs. 0 for MG) |
| MG:14:3122 | 1.00 | 49.51 |  | 0 |  | 1^st^ MG coordination shell missing (see Notes) |
| MG:14:3123 | 1.00 | 36.11 |  | 2 | 1 | O_ph_O_w_ - Exclusion zone contact(s): /O_w_/ |
| MG:14:3124 | 1.00 | 58.30 |  | 2 | 2 | 2O_ph_ - Exclusion zone contact(s): /N3_A_/O6/ |
| MG:14:3125 | 1.00 | 60.19 |  | 0 |  | 1^st^ MG coordination shell missing (see Notes) |
| MG:14:3126 | 1.00 | 27.80 |  | 2 | 2 | 2O_ph_ - Exclusion zone contact(s): /OP1/OP1/ |
| MG:14:3127 | 1.00 | 30.64 |  | 1 | 2 | O_ph_ - Exclusion zone contact(s): /O_w_/OP2/ |
| MG:14:3128 | 1.00 | 28.12 |  | 2 | 1 | 2O_ph_ - Exclusion zone contact(s): /OP1/ |
| MG:14:3129 | 1.00 | 34.01 |  | 2 | 1 | 2O_ph_ - Exclusion zone contact(s): /N7/ |
| MG:14:3130 | 1.00 | 38.66 | 1 | 3 |  | 3O_ph_ - Exclusion zone contact(s): /OP1/ Large octahedral topology discrepancy!!! |
| MG:14:3131 | 1.00 | 43.06 |  | 3 | 2 | 2O_ph_His - Exclusion zone contact(s): /N7/O_back_/ |
| MG:14:3132 | 1.00 | 53.28 |  | 2 |  | O_ph_O_b_ |
| MG:14:3133 | 1.00 | 39.95 |  | 1 | 1 | O_ph_ - Exclusion zone contact(s): /OP2/ |
| MG:14:3134 | 1.00 | 34.33 |  | 2 |  | O_ph_O_b_ |
| MG:14:3135 | 1.00 | 47.33 |  | 2 | 3 | O_b_N_b_ - Exclusion zone contact(s): /N6/N6/OP2/ Consider K in place of MG (3 valid contact(s) for K vs. 2 for MG) |
| MG:14:3136 | 1.00 | 59.00 |  | 2 |  | 2O_ph_ |
| MG:14:3137 | 1.00 | 49.77 |  | 0 | 2 | - Exclusion zone contact(s): /OP1/OP2/ Consider K in place of MG (2 valid contact(s) for K vs. 0 for MG) |
| MG:14:3138 | 1.00 | 31.24 |  | 2 |  | 2O_ph_ |
| MG:14:3139 | 1.00 | 41.95 |  | 2 | 1 | 2O_ph_ - Exclusion zone contact(s): /O_w_/ |
| MG:14:3140 | 1.00 | 47.81 |  | 1 | 1 | O_ph_ - Exclusion zone contact(s): /OP1/ |
| MG:14:3141 | 1.00 | 26.09 |  | 0 | 1 | - Exclusion zone contact(s): /O4/ Consider NA in place of MG (1 valid contact(s) for NA vs. 0 for MG) |
| MG:14:3142 | 1.00 | 51.35 |  | 0 |  | 1^st^ MG coordination shell missing (see Notes) |
| MG:14:3143 | 1.00 | 39.18 |  | 0 | 2 | - Exclusion zone contact(s): /OP1/OP2/ Consider K in place of MG (2 valid contact(s) for K vs. 0 for MG) |
| MG:14:3144 | 1.00 | 36.37 |  | 1 | 1 | O_ph_ - Exclusion zone contact(s): /OP2/ |
| MG:14:3145 | 1.00 | 49.74 |  | 0 | 2 | - Exclusion zone contact(s): /OP2/OP2/ Consider K in place of MG (2 valid contact(s) for K vs. 0 for MG) |
| MG:14:3146 | 1.00 | 49.27 | 1 | 1 |  | - Exclusion zone contact(s): /O6/ 1^st^ MG coordination shell missing (see Notes) |
| MG:14:3147 | 1.00 | 40.47 |  | 2 |  | 2O_ph_ |
| MG:14:3148 | 1.00 | 43.75 |  | 0 |  | 1^st^ MG coordination shell missing (see Notes) |
| MG:14:3149 | 1.00 | 37.57 |  | 0 |  | 1^st^ MG coordination shell missing (see Notes) |
| MG:14:3150 | 1.00 | 29.98 |  | 0 | 1 | - Exclusion zone contact(s): /OP1/ Consider NA in place of MG (1 valid contact(s) for NA vs. 0 for MG) |
| MG:14:3151 | 1.00 | 26.48 |  | 1 | 1 | O_ph_ - Exclusion zone contact(s): /OP1/ |
| MG:14:3152 | 1.00 | 32.05 |  | 1 | 2 | O_ph_ - Exclusion zone contact(s): /N1_G_/N2/ |
| MG:14:3153 | 1.00 | 51.45 |  | 2 | 1 | 2O_ph_ - Exclusion zone contact(s): /OP1/ |
| MG:14:3154 | 1.00 | 50.70 |  | 2 |  | 2O_ph_ |
| MG:14:3155 | 1.00 | 38.89 |  | 1 |  | O_ph_ |
| MG:14:3156 | 1.00 | 34.72 |  | 1 |  | O_ph_ |
| MG:14:3157 | 1.00 | 34.56 |  | 2 | 1 | O_ph_N_b_ - Exclusion zone contact(s): /N6/ |
| MG:14:3158 | 1.00 | 49.01 |  | 1 | 1 | O_ph_ - Exclusion zone contact(s): /OP2/ |
| MG:14:3159 | 1.00 | 56.73 |  | 0 |  | 1^st^ MG coordination shell missing (see Notes) |
| MG:14:3160 | 1.00 | 38.36 |  | 3 | 1 | 3O_ph_ - Exclusion zone contact(s): /OP2/ |
| MG:14:3161 | 1.00 | 47.07 |  | 1 |  | O_ph_ |
| MG:14:3162 | 1.00 | 46.76 |  | 2 | 1 | 2O_ph_ - Exclusion zone contact(s): /OP2/ |
| MG:14:3163 | 1.00 | 31.37 |  | 1 | 2 | O_b_ - Exclusion zone contact(s): /O3'/OP1/ Consider K in place of MG (2 valid contact(s) for K vs. 1 for MG) |
| MG:14:3164 | 1.00 | 48.12 |  | 0 |  | 1^st^ MG coordination shell missing (see Notes) |
| MG:14:3165 | 1.00 | 45.19 |  | 0 |  | 1^st^ MG coordination shell missing (see Notes) |
| MG:14:3166 | 1.00 | 58.46 |  | 0 | 3 | - Exclusion zone contact(s): /O4/O_w_/O_w_/ Consider K in place of MG (1 valid contact(s) for K vs. 0 for MG) |
| MG:14:3167 | 1.00 | 38.92 |  | 0 | 2 | - Exclusion zone contact(s): /OP1/OP1/ Consider NA in place of MG (1 valid contact(s) for NA vs. 0 for MG) |
| MG:14:3168 | 1.00 | 37.13 |  | 0 | 1 | - Exclusion zone contact(s): /OP1/ Consider K in place of MG (1 valid contact(s) for K vs. 0 for MG) |
| MG:14:3169 | 1.00 | 33.99 |  | 1 |  | O_ph_ |
| MG:14:3170 | 1.00 | 54.76 |  | 1 | 2 | N_b_ - Exclusion zone contact(s): /O6/OP2/ Consider K in place of MG (2 valid contact(s) for K vs. 1 for MG) |
| MG:14:3171 | 1.00 | 55.65 |  | 2 |  | 2O_ph_ |
| MG:14:3172 | 1.00 | 26.71 |  | 1 |  | O_ph_ |
| MG:14:3173 | 1.00 | 66.48 |  | 0 |  | 1^st^ MG coordination shell missing (see Notes) |
| MG:14:3174 | 1.00 | 38.80 |  | 0 | 3 | - Exclusion zone contact(s): /OP1/OP2/OP2/ Consider K in place of MG (3 valid contact(s) for K vs. 0 for MG) |
| MG:14:3175 | 0.50 | 39.96 |  | 0 | 2 | - Exclusion zone contact(s): /O_w_/OP1/ Consider NA in place of MG (1 valid contact(s) for NA vs. 0 for MG) |
| MG:14:3175 | 0.50 | 36.56 |  | 0 | 2 | - Exclusion zone contact(s): /O3'/OP1/ Consider K in place of MG (2 valid contact(s) for K vs. 0 for MG) |
| MG:14:3176 | 1.00 | 69.09 |  | 0 | 2 | - Exclusion zone contact(s): /N1_A_/OP2/ Consider K in place of MG (2 valid contact(s) for K vs. 0 for MG) |
| MG:14:3177 | 1.00 | 45.32 |  | 1 | 1 | O_ph_ - Exclusion zone contact(s): /OP2/ |
| MG:14:3178 | 1.00 | 53.98 |  | 0 |  | 1^st^ MG coordination shell missing (see Notes) |
| MG:14:3179 | 1.00 | 32.99 |  | 2 |  | O_ph_O_b_ |
| MG:14:3180 | 1.00 | 59.36 |  | 0 | 2 | - Exclusion zone contact(s): /OP1/OP2/ Consider NA in place of MG (1 valid contact(s) for NA vs. 0 for MG) |
| MG:14:3181 | 1.00 | 49.93 |  | 0 | 1 | - Exclusion zone contact(s): /N7/ Consider K in place of MG (1 valid contact(s) for K vs. 0 for MG) |
| MG:14:3182 | 1.00 | 62.18 |  | 0 | 3 | - Exclusion zone contact(s): /N7/O6/OP2/ Consider K in place of MG (3 valid contact(s) for K vs. 0 for MG) |
| MG:14:3183 | 1.00 | 30.71 |  | 0 |  | 1^st^ MG coordination shell missing (see Notes) |
| MG:14:3184 | 1.00 | 48.50 |  | 0 |  | 1^st^ MG coordination shell missing (see Notes) |
| MG:14:3185 | 1.00 | 51.93 |  | 0 | 1 | - Exclusion zone contact(s): /O6/ Consider K in place of MG (1 valid contact(s) for K vs. 0 for MG) |
| MG:14:3186 | 1.00 | 49.52 |  | 0 | 1 | - Exclusion zone contact(s): /O6/ Consider K in place of MG (1 valid contact(s) for K vs. 0 for MG) |
| MG:14:3187 | 1.00 | 51.19 |  | 0 | 1 | - Exclusion zone contact(s): /O6/ Consider K in place of MG (1 valid contact(s) for K vs. 0 for MG) |
| MG:14:3188 | 1.00 | 56.08 |  | 0 |  | 1^st^ MG coordination shell missing (see Notes) |
| MG:14:3189 | 1.00 | 50.60 |  | 1 | 1 | O_ph_ - Exclusion zone contact(s): /OP2/ |
| MG:14:3190 | 1.00 | 77.09 |  | 0 | 2 | - Exclusion zone contact(s): /N6/O6/ Consider K in place of MG (2 valid contact(s) for K vs. 0 for MG) |
| MG:14:3191 | 1.00 | 34.58 |  | 1 | 1 | O_ph_ - Exclusion zone contact(s): /OP1/ |
| MG:14:3192 | 1.00 | 41.32 |  | 1 |  | O_ph_ |
| MG:14:3193 | 1.00 | 40.98 |  | 0 | 1 | - Exclusion zone contact(s): /OP2/ Consider NA in place of MG (1 valid contact(s) for NA vs. 0 for MG) |
| MG:14:3194 | 1.00 | 54.37 |  | 0 |  | 1^st^ MG coordination shell missing (see Notes) |
| MG:14:3195 | 1.00 | 53.77 |  | 0 | 1 | - Exclusion zone contact(s): /OP2/ Consider NA in place of MG (1 valid contact(s) for NA vs. 0 for MG) |
| MG:14:3196 | 1.00 | 59.93 |  | 1 |  | O_ph_ |
| MG:14:3197 | 1.00 | 48.86 |  | 0 | 1 | - Exclusion zone contact(s): /O6/ Consider K in place of MG (1 valid contact(s) for K vs. 0 for MG) |
| MG:14:3198 | 1.00 | 33.62 |  | 1 | 5 | O_ph_ - Exclusion zone contact(s): /O5'/O5'/OP1/OP1/OP1/ Consider K in place of MG (5 valid contact(s) for K vs. 1 for MG) |
| MG:14:3199 | 1.00 | 66.07 |  | 2 | 1 | 2O_ph_ - Exclusion zone contact(s): /O3'/ |
| MG:14:3200 | 1.00 | 52.05 |  | 0 |  | 1^st^ MG coordination shell missing (see Notes) |
| MG:14:3201 | 1.00 | 49.74 |  | 0 | 2 | - Exclusion zone contact(s): /N7/O6/ Consider K in place of MG (2 valid contact(s) for K vs. 0 for MG) |
| MG:14:3202 | 1.00 | 42.42 |  | 0 | 2 | - Exclusion zone contact(s): /O2'/O3'/ Consider K in place of MG (2 valid contact(s) for K vs. 0 for MG) |
| MG:14:3203 | 1.00 | 45.60 |  | 0 |  | 1^st^ MG coordination shell missing (see Notes) |
| MG:14:3204 | 1.00 | 38.65 |  | 0 |  | 1^st^ MG coordination shell missing (see Notes) |
| MG:14:3205 | 1.00 | 51.96 |  | 0 |  | 1^st^ MG coordination shell missing (see Notes) |
| MG:14:3206 | 1.00 | 55.81 |  | 1 |  | O_ph_ |
| MG:14:3207 | 1.00 | 62.53 |  | 0 |  | 1^st^ MG coordination shell missing (see Notes) |
| MG:14:3208 | 1.00 | 58.78 |  | 0 |  | 1^st^ MG coordination shell missing (see Notes) |
| MG:14:3209 | 1.00 | 53.72 |  | 0 | 2 | - Exclusion zone contact(s): /N7/OP1/ Consider NA in place of MG (1 valid contact(s) for NA vs. 0 for MG) |
| MG:14:3210 | 1.00 | 47.34 |  | 0 | 2 | - Exclusion zone contact(s): /OP1/OP2/ Consider NA in place of MG (1 valid contact(s) for NA vs. 0 for MG) |
| MG:14:3211 | 1.00 | 58.20 |  | 1 | 1 | O_ph_ - Exclusion zone contact(s): /OP2/ |
| MG:14:3212 | 1.00 | 44.62 |  | 1 | 1 | O_ph_ - Exclusion zone contact(s): /O6/ |
| MG:14:3213 | 1.00 | 40.32 |  | 0 | 1 | - Exclusion zone contact(s): /OP1/ Consider K in place of MG (1 valid contact(s) for K vs. 0 for MG) |
| MG:14:3214 | 1.00 | 66.69 |  | 2 | 1 | O_ph_O_b_ - Exclusion zone contact(s): /N1_G_/ |
| MG:14:3215 | 1.00 | 45.49 |  | 0 | 1 | - Exclusion zone contact(s): /O6/ Consider NA in place of MG (1 valid contact(s) for NA vs. 0 for MG) |
| MG:14:3216 | 1.00 | 51.13 |  | 1 | 2 | O_ph_ - Exclusion zone contact(s): /N7/O6/ Consider K in place of MG (2 valid contact(s) for K vs. 1 for MG) |
| MG:14:3217 | 1.00 | 69.37 |  | 2 |  | 2O_ph_ |
| MG:14:3218 | 1.00 | 60.28 |  | 1 | 3 | O_ph_ - Exclusion zone contact(s): /NZ_Lys_/OP2/OP2/ Consider K in place of MG (2 valid contact(s) for K vs. 1 for MG) |
| MG:14:3219 | 1.00 | 33.32 |  | 0 |  | 1^st^ MG coordination shell missing (see Notes) |
| MG:14:3220 | 1.00 | 46.61 |  | 0 | 2 | - Exclusion zone contact(s): /OP2/OP2/ Consider NA in place of MG (1 valid contact(s) for NA vs. 0 for MG) |
| MG:14:3221 | 1.00 | 52.42 |  | 0 | 1 | - Exclusion zone contact(s): /OP2/ Consider K in place of MG (1 valid contact(s) for K vs. 0 for MG) |
| MG:14:3222 | 1.00 | 57.98 |  | 1 |  | O_ph_ |
| MG:14:3223 | 1.00 | 40.82 |  | 1 |  | O_ph_ |
| MG:14:3224 | 1.00 | 36.01 |  | 0 |  | 1^st^ MG coordination shell missing (see Notes) |
| MG:14:3225 | 1.00 | 47.68 |  | 0 | 1 | - Exclusion zone contact(s): /OP1/ Consider K in place of MG (1 valid contact(s) for K vs. 0 for MG) |
| MG:14:3226 | 1.00 | 55.65 |  | 0 | 1 | - Exclusion zone contact(s): /OP1/ Consider NA in place of MG (1 valid contact(s) for NA vs. 0 for MG) |
| MG:14:3227 | 1.00 | 38.23 |  | 0 | 1 | - Exclusion zone contact(s): /O4/ Consider K in place of MG (1 valid contact(s) for K vs. 0 for MG) |
| MG:14:3228 | 1.00 | 58.27 |  | 0 | 1 | - Exclusion zone contact(s): /OP1/ Consider NA in place of MG (1 valid contact(s) for NA vs. 0 for MG) |
| MG:14:3229 | 1.00 | 48.75 |  | 1 |  | O_ph_ |
| MG:14:3230 | 1.00 | 54.08 |  | 0 | 2 | - Exclusion zone contact(s): /OP1/OP1/ Consider K in place of MG (2 valid contact(s) for K vs. 0 for MG) |
| MG:14:3231 | 1.00 | 39.22 |  | 0 | 2 | - Exclusion zone contact(s): /O4/OP2/ Consider K in place of MG (2 valid contact(s) for K vs. 0 for MG) |
| MG:14:3232 | 1.00 | 63.94 |  | 0 | 1 | - Exclusion zone contact(s): /N3_C_/ Consider K in place of MG (1 valid contact(s) for K vs. 0 for MG) |
| MG:14:3233 | 1.00 | 52.52 |  | 0 |  | 1^st^ MG coordination shell missing (see Notes) |
| MG:14:3234 | 1.00 | 55.89 |  | 0 | 1 | - Exclusion zone contact(s): /OP2/ Consider NA in place of MG (1 valid contact(s) for NA vs. 0 for MG) |
| MG:14:3235 | 1.00 | 50.44 |  | 0 | 2 | - Exclusion zone contact(s): /N6/O2/ Consider K in place of MG (2 valid contact(s) for K vs. 0 for MG) |
| MG:14:3236 | 1.00 | 50.95 |  | 0 | 2 | - Exclusion zone contact(s): /O2'/O4/ Consider K in place of MG (2 valid contact(s) for K vs. 0 for MG) |
| MG:14:3237 | 1.00 | 96.46 |  | 2 | 3 | O_b_O_coo_ - Exclusion zone contact(s): /N3_C_/N6/OE2_Glu_/ |
| MG:14:3238 | 1.00 | 44.43 |  | 0 |  | 1^st^ MG coordination shell missing (see Notes) |
| MG:14:3239 | 1.00 | 66.98 |  | 0 |  | 1^st^ MG coordination shell missing (see Notes) |
| MG:14:3240 | 1.00 | 57.04 |  | 0 |  | 1^st^ MG coordination shell missing (see Notes) |
| MG:14:3241 | 1.00 | 59.92 |  | 0 | 1 | - Exclusion zone contact(s): /O6/ Consider K in place of MG (1 valid contact(s) for K vs. 0 for MG) |
| MG:14:3242 | 1.00 | 65.51 |  | 0 | 1 | - Exclusion zone contact(s): /OP1/ Consider K in place of MG (1 valid contact(s) for K vs. 0 for MG) |
| MG:14:3243 | 1.00 | 54.53 |  | 0 | 1 | - Exclusion zone contact(s): /O_w_/ 1^st^ MG coordination shell missing (see Notes) |
| MG:14:3244 | 1.00 | 64.51 |  | 0 | 2 | - Exclusion zone contact(s): /OP2/OP2/ Consider K in place of MG (2 valid contact(s) for K vs. 0 for MG) |
| MG:14:3245 | 1.00 | 36.70 |  | 1 |  | O_b_ |
| MG:14:3246 | 1.00 | 35.59 |  | 0 | 2 | - Exclusion zone contact(s): /OP1/OP1/ Consider K in place of MG (2 valid contact(s) for K vs. 0 for MG) |
| MG:14:3247 | 1.00 | 47.65 |  | 1 | 1 | O_ph_ - Exclusion zone contact(s): /O2/ |
| MG:14:3248 | 1.00 | 60.17 |  | 0 | 1 | - Exclusion zone contact(s): /O6/ Consider K in place of MG (1 valid contact(s) for K vs. 0 for MG) |
| MG:14:3249 | 1.00 | 62.31 |  | 0 | 5 | - Exclusion zone contact(s): /N4/O6/O6/O_w_/O_w_/ Consider K in place of MG (3 valid contact(s) for K vs. 0 for MG) |
| MG:14:3250 | 1.00 | 38.34 |  | 0 | 6 | - Exclusion zone contact(s): /N7/O6/O6/O_w_/O_w_/O_w_/ Consider K in place of MG (3 valid contact(s) for K vs. 0 for MG) |
| MG:14:3251 | 1.00 | 45.92 |  | 0 | 1 | - Exclusion zone contact(s): /N7/ Consider K in place of MG (1 valid contact(s) for K vs. 0 for MG) |
| MG:14:3252 | 1.00 | 42.83 |  | 0 |  | 1^st^ MG coordination shell missing (see Notes) |
| MG:14:3253 | 1.00 | 47.77 |  | 1 | 5 | O_w_ - Exclusion zone contact(s): /N7/N7/O6/O_w_/OP2/ Consider K in place of MG (4 valid contact(s) for K vs. 0 for MG) |
| MG:14:3254 | 1.00 | 62.59 |  | 0 | 3 | - Exclusion zone contact(s): /O_w_/O_w_/O_w_/ 1^st^ MG coordination shell missing (see Notes) |
| MG:14:3255 | 1.00 | 63.48 |  | 1 | 2 | O_w_ - Exclusion zone contact(s): /OP2/OP2/ Consider K in place of MG (2 valid contact(s) for K vs. 0 for MG) |
| MG:14:3256 | 1.00 | 67.62 |  | 0 | 1 | - Exclusion zone contact(s): /O6/ Consider K in place of MG (1 valid contact(s) for K vs. 0 for MG) |
| MG:14:3257 | 1.00 | 65.99 |  | 1 |  | O_ph_ |
| MG:14:3258 | 1.00 | 34.70 |  | 2 |  | 2O_ph_ |
| MG:14:3259 | 1.00 | 42.14 |  | 0 | 1 | - Exclusion zone contact(s): /O_w_/ 1^st^ MG coordination shell missing (see Notes) |
| MG:14:3260 | 1.00 | 84.74 |  | 0 | 3 | - Exclusion zone contact(s): /N7/OP1/OP2/ Consider K in place of MG (3 valid contact(s) for K vs. 0 for MG) |
| MG:14:3261 | 1.00 | 43.43 |  | 0 | 2 | - Exclusion zone contact(s): /O_back_/OP1/ Consider K in place of MG (2 valid contact(s) for K vs. 0 for MG) |
| MG:14:3262 | 1.00 | 41.71 |  | 0 |  | 1^st^ MG coordination shell missing (see Notes) |
| MG:14:3263 | 1.00 | 60.81 |  | 0 | 1 | - Exclusion zone contact(s): /O6/ Consider K in place of MG (1 valid contact(s) for K vs. 0 for MG) |
| MG:14:3264 | 1.00 | 45.90 |  | 0 | 1 | - Exclusion zone contact(s): /OP2/ Consider K in place of MG (1 valid contact(s) for K vs. 0 for MG) |
| MG:14:3265 | 1.00 | 34.03 |  | 1 | 1 | O_ph_ - Exclusion zone contact(s): /OP2/ |
| MG:14:3266 | 1.00 | 38.96 |  | 2 | 3 | O_ph_O_r_ - Exclusion zone contact(s): /O3'/O_w_/OP2/ |
| MG:14:3267 | 1.00 | 44.95 |  | 1 | 2 | O_b_ - Exclusion zone contact(s): /O4/O6/ |
| MG:14:3268 | 1.00 | 36.05 |  | 0 | 2 | - Exclusion zone contact(s): /O6/O6/ Consider K in place of MG (2 valid contact(s) for K vs. 0 for MG) |
| MG:14:3269 | 1.00 | 68.48 |  | 0 | 1 | - Exclusion zone contact(s): /OP2/ Consider K in place of MG (1 valid contact(s) for K vs. 0 for MG) |
| MG:14:3270 | 1.00 | 66.52 |  | 0 | 1 | - Exclusion zone contact(s): /OP1/ Consider K in place of MG (1 valid contact(s) for K vs. 0 for MG) |
| MG:14:3271 | 1.00 | 51.23 |  | 0 |  | 1^st^ MG coordination shell missing (see Notes) |
| MG:14:3272 | 1.00 | 61.08 |  | 0 |  | 1^st^ MG coordination shell missing (see Notes) |
| MG:14:3273 | 1.00 | 37.20 |  | 1 | 1 | O_ph_ - Exclusion zone contact(s): /OP1/ |
| MG:14:3274 | 1.00 | 54.81 |  | 0 | 1 | - Exclusion zone contact(s): /O6/ Consider K in place of MG (1 valid contact(s) for K vs. 0 for MG) |
| MG:14:3275 | 1.00 | 42.80 |  | 1 |  | O_b_ |
| MG:14:3276 | 1.00 | 52.22 |  | 0 | 2 | - Exclusion zone contact(s): /N1_A_/O6/ Consider K in place of MG (2 valid contact(s) for K vs. 0 for MG) |
| MG:14:3277 | 1.00 | 61.25 |  | 0 |  | 1^st^ MG coordination shell missing (see Notes) |
| MG:14:3278 | 1.00 | 49.90 |  | 0 | 1 | - Exclusion zone contact(s): /O2/ Consider K in place of MG (1 valid contact(s) for K vs. 0 for MG) |
| MG:14:3279 | 1.00 | 22.92 |  | 4 | 3 | 4O_w_ - Exclusion zone contact(s): /N7/N7/O6/ Consider NA in place of MG (2 valid contact(s) for NA vs. 0 for MG) |
| MG:14:3280 | 1.00 | 45.84 |  | 6 | 1 | 6O_w_ - Exclusion zone contact(s): /OP1/ Consider K in place of MG (1 valid contact(s) for K vs. 0 for MG) |
| MG:14:3281 | 1.00 | 25.82 |  | 5 | 2 | 5O_w_ - Exclusion zone contact(s): /OP1/OP2/ Consider NA in place of MG (1 valid contact(s) for NA vs. 0 for MG) |
| MG:14:3282 | 1.00 | 48.52 |  | 5 | 1 | 5O_w_ - Exclusion zone contact(s): /OP1/ Consider NA in place of MG (1 valid contact(s) for NA vs. 0 for MG) |
| MG:14:3283 | 1.00 | 21.46 |  | 5 | 2 | 5O_w_ - Exclusion zone contact(s): /O5'/OP1/ Consider NA in place of MG (1 valid contact(s) for NA vs. 0 for MG) |
| MG:14:3284 | 1.00 | 34.09 |  | 6 |  | 6O_w_ |
| MG:14:3285 | 1.00 | 30.34 |  | 5 | 1 | 5O_w_ - Exclusion zone contact(s): /OP2/ Consider NA in place of MG (1 valid contact(s) for NA vs. 0 for MG) |
| MG:14:3286 | 1.00 | 25.37 |  | 6 |  | O_ph_5O_w_ Large octahedral topology discrepancy!!! |
| MG:14:3287 | 1.00 | 39.67 |  | 6 |  | 6O_w_ |
| MG:14:3288 | 1.00 | 39.91 |  | 6 |  | 6O_w_ |
| MG:14:3289 | 1.00 | 51.32 |  | 6 | 1 | 6O_w_ - Exclusion zone contact(s): /OP1/ Consider K in place of MG (1 valid contact(s) for K vs. 0 for MG) |
| MG:14:3290 | 1.00 | 36.05 |  | 6 |  | 6O_w_ |
| MG:14:3291 | 1.00 | 58.71 |  | 6 | 1 | 6O_w_ - Exclusion zone contact(s): /O6/ Consider K in place of MG (1 valid contact(s) for K vs. 0 for MG) |
| MG:14:3292 | 1.00 | 29.77 |  | 3 | 4 | 3O_w_ - Exclusion zone contact(s): /O5'/OP1/OP1/OP2/ Consider NA in place of MG (2 valid contact(s) for NA vs. 0 for MG) |
| MG:14:3293 | 1.00 | 60.25 |  | 5 | 1 | 5O_w_ - Exclusion zone contact(s): /OP2/ Consider NA in place of MG (1 valid contact(s) for NA vs. 0 for MG) |
| MG:14:3294 | 1.00 | 51.69 |  | 6 |  | 6O_w_ |
| MG:14:3295 | 1.00 | 50.36 |  | 6 | 1 | 6O_w_ - Exclusion zone contact(s): /OP2/ Consider K in place of MG (1 valid contact(s) for K vs. 0 for MG) |
| MG:14:3296 | 1.00 | 35.72 |  | 6 |  | 6O_w_ |
| MG:14:3297 | 1.00 | 28.36 |  | 6 |  | 6O_w_ |
| MG:14:3298 | 1.00 | 32.74 |  | 6 | 2 | 6O_w_ - Exclusion zone contact(s): /O6/OP1/ Consider K in place of MG (2 valid contact(s) for K vs. 0 for MG) |
| MG:14:3299 | 1.00 | 48.20 |  | 6 |  | 6O_w_ Possible idealized distances ! |
| MG:14:3300 | 1.00 | 61.04 |  | 6 |  | 6O_w_ |
| MG:14:3301 | 1.00 | 38.62 |  | 6 | 3 | 6O_w_ - Exclusion zone contact(s): /N7/OP1/OP2/ Consider K in place of MG (3 valid contact(s) for K vs. 0 for MG) |
| MG:14:3302 | 1.00 | 29.50 |  | 5 | 1 | O_ph_4O_w_ - Exclusion zone contact(s): /OP2/ Large octahedral topology discrepancy!!! |
| MG:14:3303 | 1.00 | 58.59 |  | 6 |  | 6O_w_ |
| MG:14:3304 | 1.00 | 63.58 |  | 6 |  | 6O_w_ |
| MG:14:3305 | 1.00 | 40.11 |  | 4 | 2 | 4O_w_ - Exclusion zone contact(s): /OP1/OP1/ Consider NA in place of MG (1 valid contact(s) for NA vs. 0 for MG) |
| MG:14:3306 | 1.00 | 32.64 |  | 5 | 1 | 5O_w_ - Exclusion zone contact(s): /OP1/ Consider NA in place of MG (1 valid contact(s) for NA vs. 0 for MG) |
| MG:14:3307 | 1.00 | 36.01 |  | 6 |  | 6O_w_ |
| MG:14:3308 | 1.00 | 77.22 |  | 6 |  | 6O_w_ |
| MG:14:3309 | 1.00 | 26.49 |  | 5 | 3 | O_ph_4O_w_ - Exclusion zone contact(s): /OP1/OP2/OP2/ Large octahedral topology discrepancy!!!  Consider K in place of MG (3 valid contact(s) for K vs. 1 for MG) |
| MG:14:3310 | 1.00 | 25.88 |  | 5 | 1 | 5O_w_ - Exclusion zone contact(s): /OP1/ Consider NA in place of MG (1 valid contact(s) for NA vs. 0 for MG) |
| MG:14:3311 | 1.00 | 31.98 |  | 5 | 1 | O_ph_4O_w_ - Exclusion zone contact(s): /OP2/ |
| MG:14:3312 | 1.00 | 38.40 |  | 4 | 3 | 4O_w_ - Exclusion zone contact(s): /OP1/OP1/OP2/ Consider K in place of MG (2 valid contact(s) for K vs. 0 for MG)  Possible idealized distances ! |
| MG:14:3313 | 1.00 | 33.15 |  | 6 |  | O_ph_5O_w_ |
| MG:14:3314 | 1.00 | 28.45 |  | 3 | 5 | 3O_w_ - Exclusion zone contact(s): /O3'/OP1/OP2/OP2/OP2/ Consider K in place of MG (4 valid contact(s) for K vs. 0 for MG) |
| MG:14:3315 | 1.00 | 39.04 |  | 5 | 1 | 5O_w_ - Exclusion zone contact(s): /OP2/ Consider NA in place of MG (1 valid contact(s) for NA vs. 0 for MG) |
| MG:14:3316 | 1.00 | 34.87 |  | 5 | 1 | 5O_w_ - Exclusion zone contact(s): /OP1/ Consider NA in place of MG (1 valid contact(s) for NA vs. 0 for MG) |
| MG:14:3317 | 1.00 | 40.48 |  | 4 | 2 | 4O_w_ - Exclusion zone contact(s): /OP1/OP1/ Consider K in place of MG (2 valid contact(s) for K vs. 0 for MG) |
| MG:14:3318 | 1.00 | 35.50 |  | 5 | 1 | 5O_w_ - Exclusion zone contact(s): /OP2/ Consider NA in place of MG (1 valid contact(s) for NA vs. 0 for MG) |
| MG:14:3319 | 1.00 | 27.35 |  | 5 | 2 | 5O_w_ - Exclusion zone contact(s): /O_back_/OP1/ Consider NA in place of MG (1 valid contact(s) for NA vs. 0 for MG) |
| MG:14:3320 | 1.00 | 25.32 |  | 3 | 4 | 3O_w_ - Exclusion zone contact(s): /O_back_/OP1/OP2/OP2/ Consider NA in place of MG (2 valid contact(s) for NA vs. 0 for MG) |
| MG:14:3321 | 1.00 | 26.92 |  | 5 | 1 | O_ph_4O_w_ - Exclusion zone contact(s): /OP1/ Large octahedral topology discrepancy!!! |
| MG:14:3322 | 1.00 | 41.90 |  | 5 | 1 | 5O_w_ - Exclusion zone contact(s): /OP2/ Consider NA in place of MG (1 valid contact(s) for NA vs. 0 for MG) |
| MG:14:3323 | 1.00 | 56.70 |  | 5 | 1 | 5O_w_ - Exclusion zone contact(s): /OP2/ Consider K in place of MG (1 valid contact(s) for K vs. 0 for MG)  Possible idealized distances ! |
| MG:14:3324 | 1.00 | 43.97 |  | 6 |  | 6O_w_ Possible idealized distances ! |
| MG:14:3325 | 1.00 | 30.58 |  | 5 | 2 | O_ph_4O_w_ - Exclusion zone contact(s): /OP1/OP2/ Large octahedral topology discrepancy!!! |
| MG:14:3326 | 1.00 | 40.35 |  | 5 | 1 | 5O_w_ - Exclusion zone contact(s): /OP1/ Consider K in place of MG (1 valid contact(s) for K vs. 0 for MG) |
| MG:14:3327 | 1.00 | 36.56 |  | 5 | 2 | 5O_w_ - Exclusion zone contact(s): /N7/O6/ Consider K in place of MG (2 valid contact(s) for K vs. 0 for MG) |
| MG:14:3328 | 1.00 | 25.02 |  | 6 |  | 6O_w_ |
| MG:14:3329 | 1.00 | 28.96 |  | 4 | 2 | 4O_w_ - Exclusion zone contact(s): /OP1/OP1/ Consider NA in place of MG (1 valid contact(s) for NA vs. 0 for MG) |
| MG:14:3330 | 1.00 | 39.97 |  | 5 | 1 | 5O_w_ - Exclusion zone contact(s): /OP2/ Consider NA in place of MG (1 valid contact(s) for NA vs. 0 for MG)  Possible idealized distances ! |
| MG:14:3331 | 1.00 | 51.92 |  | 5 | 1 | 5O_w_ - Exclusion zone contact(s): /OP2/ Consider NA in place of MG (1 valid contact(s) for NA vs. 0 for MG) |
| MG:14:3332 | 1.00 | 54.87 |  | 5 | 2 | 5O_w_ - Exclusion zone contact(s): /OP1/OP2/ Consider NA in place of MG (1 valid contact(s) for NA vs. 0 for MG) |
| MG:14:3333 | 1.00 | 62.45 |  | 6 |  | 6O_w_ |
| MG:14:3334 | 1.00 | 33.72 |  | 5 | 1 | 5O_w_ - Exclusion zone contact(s): /OP2/ Consider NA in place of MG (1 valid contact(s) for NA vs. 0 for MG)  Possible idealized distances ! |
| MG:14:3335 | 1.00 | 70.79 |  | 6 |  | 6O_w_ Possible idealized distances ! |
| MG:14:3336 | 1.00 | 78.58 |  | 6 |  | 6O_w_ |
| MG:14:3337 | 1.00 | 83.35 |  | 6 |  | 6O_w_ Possible idealized distances ! |
| MG:14:3338 | 1.00 | 34.23 |  | 6 |  | 6O_w_ |
| MG:14:3339 | 1.00 | 29.35 |  | 6 |  | 6O_w_ |
| MG:14:3340 | 1.00 | 26.06 |  | 3 | 4 | 3O_w_ - Exclusion zone contact(s): /OP1/OP1/OP1/OP2/ Consider NA in place of MG (3 valid contact(s) for NA vs. 0 for MG) |
| MG:14:3341 | 1.00 | 35.98 |  | 4 | 2 | 4O_w_ - Exclusion zone contact(s): /OP1/OP2/ Consider NA in place of MG (2 valid contact(s) for NA vs. 0 for MG) |
| MG:14:3342 | 1.00 | 32.62 |  | 4 | 2 | 4O_w_ - Exclusion zone contact(s): /OP2/OP2/ Consider NA in place of MG (1 valid contact(s) for NA vs. 0 for MG) |
| MG:14:3343 | 1.00 | 50.99 |  | 4 | 2 | 4O_w_ - Exclusion zone contact(s): /OP2/OP2/ Consider NA in place of MG (2 valid contact(s) for NA vs. 0 for MG) |
| MG:14:3344 | 1.00 | 31.71 |  | 4 | 3 | 4O_w_ - Exclusion zone contact(s): /OP1/OP1/OP2/ Consider K in place of MG (2 valid contact(s) for K vs. 0 for MG) |
| MG:14:3345 | 1.00 | 48.73 |  | 6 |  | 6O_w_ |
| MG:14:3346 | 1.00 | 52.96 |  | 5 | 2 | 5O_w_ - Exclusion zone contact(s): /OP1/OP2/ Consider NA in place of MG (1 valid contact(s) for NA vs. 0 for MG)  Possible idealized distances ! |
| MG:14:3347 | 1.00 | 27.52 |  | 3 | 5 | 3O_w_ - Exclusion zone contact(s): /O5'/OP1/OP1/OP2/OP2/ Consider NA in place of MG (3 valid contact(s) for NA vs. 0 for MG) |
| MG:14:3348 | 1.00 | 30.79 |  | 4 | 4 | 4O_w_ - Exclusion zone contact(s): /N7/NE2_His_/OP1/OP2/ Consider K in place of MG (3 valid contact(s) for K vs. 0 for MG) |
| MG:14:3349 | 1.00 | 48.81 |  | 6 |  | 6O_w_ |
| MG:14:3350 | 1.00 | 39.66 |  | 5 | 1 | 5O_w_ - Exclusion zone contact(s): /OP2/ Consider NA in place of MG (1 valid contact(s) for NA vs. 0 for MG) |
| MG:14:3351 | 1.00 | 55.21 |  | 5 | 1 | 5O_w_ - Exclusion zone contact(s): /OP2/ Consider NA in place of MG (1 valid contact(s) for NA vs. 0 for MG) |
| MG:14:3352 | 1.00 | 63.63 |  | 6 |  | 6O_w_ |
| MG:14:3353 | 1.00 | 62.63 |  | 6 |  | 6O_w_ Possible idealized distances ! |
| MG:14:3354 | 1.00 | 30.17 |  | 5 | 3 | 5O_w_ - Exclusion zone contact(s): /O2'/OP1/OP2/ Consider K in place of MG (2 valid contact(s) for K vs. 0 for MG) |
| MG:14:3355 | 1.00 | 39.92 |  | 6 | 2 | 6O_w_ - Exclusion zone contact(s): /OP1/OP2/ Consider K in place of MG (2 valid contact(s) for K vs. 0 for MG) |
| MG:14:3356 | 1.00 | 36.57 |  | 6 |  | O_ph_5O_w_ |
| MG:14:3357 | 1.00 | 48.92 |  | 5 | 3 | 5O_w_ - Exclusion zone contact(s): /N7/OP1/OP2/ Consider K in place of MG (2 valid contact(s) for K vs. 0 for MG)  Possible idealized distances ! |
| MG:14:3358 | 1.00 | 51.03 |  | 3 | 5 | 3O_w_ - Exclusion zone contact(s): /O2'/O3'/OP2/OP2/OP2/ Consider K in place of MG (4 valid contact(s) for K vs. 0 for MG) |
| MG:14:3359 | 1.00 | 33.73 |  | 4 | 2 | 4O_w_ - Exclusion zone contact(s): /O4/OP2/ Consider NA in place of MG (2 valid contact(s) for NA vs. 0 for MG) |
| MG:14:3360 | 1.00 | 37.02 |  | 3 | 4 | 3O_w_ - Exclusion zone contact(s): /O5'/O6/OP1/OP1/ Consider NA in place of MG (2 valid contact(s) for NA vs. 0 for MG) |
| MG:14:3361 | 1.00 | 48.95 |  | 5 | 1 | 5O_w_ - Exclusion zone contact(s): /OP1/ Consider K in place of MG (1 valid contact(s) for K vs. 0 for MG) |
| MG:14:3362 | 1.00 | 68.08 |  | 6 | 1 | 6O_w_ - Exclusion zone contact(s): /O6/ Consider K in place of MG (1 valid contact(s) for K vs. 0 for MG) |
| MG:14:3363 | 1.00 | 26.79 |  | 5 | 1 | 5O_w_ - Exclusion zone contact(s): /OP1/ Consider NA in place of MG (1 valid contact(s) for NA vs. 0 for MG) |
| MG:14:3364 | 1.00 | 36.77 |  | 5 | 1 | 5O_w_ - Exclusion zone contact(s): /OP2/ Consider NA in place of MG (1 valid contact(s) for NA vs. 0 for MG) |
| MG:14:3365 | 1.00 | 28.70 |  | 4 | 3 | 4O_w_ - Exclusion zone contact(s): /OP1/OP2/OP2/ Consider NA in place of MG (2 valid contact(s) for NA vs. 0 for MG) |
| MG:14:3366 | 1.00 | 55.59 |  | 6 | 1 | 6O_w_ - Exclusion zone contact(s): /N6/ Consider K in place of MG (1 valid contact(s) for K vs. 0 for MG) |
| MG:14:3367 | 1.00 | 70.06 |  | 6 |  | 6O_w_ Possible idealized distances ! |
| MG:14:3368 | 1.00 | 29.02 |  | 3 | 5 | 3O_w_ - Exclusion zone contact(s): /O5'/OP1/OP1/OP2/OP2/ Consider K in place of MG (3 valid contact(s) for K vs. 0 for MG) |
| MG:14:3369 | 1.00 | 76.32 |  | 5 | 1 | 5O_w_ - Exclusion zone contact(s): /OP2/ Consider K in place of MG (1 valid contact(s) for K vs. 0 for MG)  Possible idealized distances ! |
| MG:14:3370 | 1.00 | 44.38 |  | 4 | 3 | 4O_w_ - Exclusion zone contact(s): /OP1/OP1/OP2/ Consider K in place of MG (2 valid contact(s) for K vs. 0 for MG)  Possible idealized distances ! |
| MG:14:3371 | 1.00 | 32.30 |  | 4 | 2 | 4O_w_ - Exclusion zone contact(s): /OP1/OP1/ Consider NA in place of MG (1 valid contact(s) for NA vs. 0 for MG) |
| MG:14:3372 | 1.00 | 64.47 |  | 4 | 2 | 4O_w_ - Exclusion zone contact(s): /OP1/OP1/ Consider NA in place of MG (1 valid contact(s) for NA vs. 0 for MG)  Possible idealized distances ! |
| MG:14:3373 | 1.00 | 68.79 |  | 6 |  | 6O_w_ |
| MG:14:3374 | 1.00 | 30.73 |  | 2 | 4 | 2O_w_ - Exclusion zone contact(s): /O4/OP1/OP2/OP2/ Consider K in place of MG (4 valid contact(s) for K vs. 0 for MG) |
| MG:14:3375 | 1.00 | 84.05 |  | 6 |  | 6O_w_ |
| MG:14:3376 | 1.00 | 54.68 |  | 5 | 2 | 5O_w_ - Exclusion zone contact(s): /O3'/OP1/ Consider NA in place of MG (1 valid contact(s) for NA vs. 0 for MG) |
| MG:14:3377 | 1.00 | 59.96 |  | 5 | 1 | 5O_w_ - Exclusion zone contact(s): /OP1/ Consider NA in place of MG (1 valid contact(s) for NA vs. 0 for MG)  Possible idealized distances ! |
| MG:14:3378 | 1.00 | 57.15 |  | 5 | 1 | 5O_w_ - Exclusion zone contact(s): /OP1/ Consider NA in place of MG (1 valid contact(s) for NA vs. 0 for MG) |
| MG:14:3379 | 1.00 | 43.72 |  | 6 |  | 6O_w_ |
| MG:14:3380 | 1.00 | 76.47 |  | 6 |  | 6O_w_ |
| MG:14:3381 | 1.00 | 56.01 |  | 5 | 1 | 5O_w_ - Exclusion zone contact(s): /OP2/ Consider NA in place of MG (1 valid contact(s) for NA vs. 0 for MG)  Possible idealized distances ! |
| MG:14:3382 | 1.00 | 52.65 |  | 6 | 2 | 6O_w_ - Exclusion zone contact(s): /OP1/OP2/ Consider K in place of MG (2 valid contact(s) for K vs. 0 for MG) |
| MG:14:3383 | 1.00 | 51.91 |  | 6 |  | 6O_w_ |
| MG:14:3384 | 1.00 | 47.88 |  | 6 |  | 6O_w_ |
| MG:14:3385 | 1.00 | 58.71 |  | 5 | 2 | 5O_w_ - Exclusion zone contact(s): /O2'/OP2/ Consider NA in place of MG (1 valid contact(s) for NA vs. 0 for MG) |
| MG:14:3386 | 1.00 | 61.45 |  | 6 |  | 6O_w_ |
| MG:14:3387 | 1.00 | 45.54 |  | 6 |  | 6O_w_ |
| MG:14:3388 | 1.00 | 91.23 |  | 6 |  | 6O_w_ Possible idealized distances ! |
| MG:14:3389 | 1.00 | 45.36 |  | 5 | 1 | 5O_w_ - Exclusion zone contact(s): /OP2/ Consider NA in place of MG (1 valid contact(s) for NA vs. 0 for MG) |
| MG:14:3390 | 1.00 | 79.43 |  | 6 |  | 6O_w_ |
| MG:14:3391 | 1.00 | 81.43 |  | 6 |  | 6O_w_ Possible idealized distances ! |
| MG:14:3392 | 1.00 | 75.97 |  | 6 |  | 6O_w_ |
| MG:14:3393 | 1.00 | 63.92 |  | 6 |  | 6O_w_ Possible idealized distances ! |
| MG:14:3394 | 1.00 | 61.39 |  | 6 |  | 6O_w_ |
| MG:14:3395 | 1.00 | 25.67 |  | 5 | 1 | 5O_w_ - Exclusion zone contact(s): /OP2/ Consider NA in place of MG (1 valid contact(s) for NA vs. 0 for MG) |
| MG:14:3396 | 1.00 | 66.06 |  | 6 |  | 6O_w_ |
| MG:14:3397 | 1.00 | 75.21 |  | 5 | 2 | 5O_w_ - Exclusion zone contact(s): /N7/OP2/ Consider K in place of MG (2 valid contact(s) for K vs. 0 for MG) |
| MG:14:3398 | 1.00 | 47.69 |  | 6 |  | 6O_w_ |
| MG:14:3399 | 1.00 | 106.45 |  | 6 |  | 6O_w_ |
| MG:14:3400 | 1.00 | 63.52 |  | 5 | 1 | 5O_w_ - Exclusion zone contact(s): /OP2/ Consider K in place of MG (1 valid contact(s) for K vs. 0 for MG) |
| K:16:201 | 1.00 | 55.47 |  | 6 |  | 2O_ph_O_r_O_b_2N_b_ |
| K:16:202 | 1.00 | 74.09 |  | 5 |  | 4O_b_Anion |
| K:16:203 | 1.00 | 44.38 |  | 7 |  | 3O_b_4Anion |
| MG:16:204 | 1.00 | 39.61 |  | 0 | 1 | - Exclusion zone contact(s): /OP1/ Consider K in place of MG (1 valid contact(s) for K vs. 0 for MG) |
| MG:16:205 | 1.00 | 46.71 |  | 0 | 3 | - Exclusion zone contact(s): /O3'/OP2/OP2/ Consider NA in place of MG (2 valid contact(s) for NA vs. 0 for MG) |
| MG:16:206 | 1.00 | 52.99 |  | 0 | 3 | - Exclusion zone contact(s): /O2'/O4/O6/ Consider K in place of MG (2 valid contact(s) for K vs. 0 for MG) |
| MG:16:207 | 1.00 | 43.94 |  | 0 | 1 | - Exclusion zone contact(s): /O6/ Consider K in place of MG (1 valid contact(s) for K vs. 0 for MG) |
| MG:16:208 | 1.00 | 44.39 |  | 0 | 2 | - Exclusion zone contact(s): /O6/O6/ Consider K in place of MG (2 valid contact(s) for K vs. 0 for MG) |
| MG:16:209 | 1.00 | 59.65 |  | 0 | 1 | - Exclusion zone contact(s): /OP2/ Consider K in place of MG (1 valid contact(s) for K vs. 0 for MG) |
| MG:16:210 | 1.00 | 62.98 |  | 1 | 1 | O_ph_ - Exclusion zone contact(s): /O_w_/ |
| MG:16:211 | 1.00 | 57.73 |  | 5 | 1 | 5O_w_ - Exclusion zone contact(s): /OP1/ Consider K in place of MG (1 valid contact(s) for K vs. 0 for MG)  Possible idealized distances ! |
| MG:16:212 | 1.00 | 36.42 |  | 5 | 1 | 5O_w_ - Exclusion zone contact(s): /OP2/ Consider NA in place of MG (1 valid contact(s) for NA vs. 0 for MG)  Possible idealized distances ! |
| MG:16:213 | 1.00 | 41.74 |  | 6 |  | 6O_w_ |
| MG:16:214 | 1.00 | 57.59 |  | 6 |  | 6O_w_ |
| MG:16:215 | 1.00 | 52.42 |  | 6 |  | 6O_w_ |
| K:19:301 | 1.00 | 52.67 |  | 2 |  | 2O_back_ |
| K:1G:1601 | 1.00 | 115.34 |  | 1 |  | O_ph_ |
| K:1G:1602 | 1.00 | 104.24 |  | 2 |  | 2O_b_ |
| K:1G:1603 | 1.00 | 62.13 | 1 | 2 |  | O_ph_O_r_ - Exclusion zone contact(s): /OP2/ |
| K:1G:1604 | 1.00 | 91.70 |  | 0 |  | 1^st^ K coordination shell missing (see Notes) |
| K:1G:1605 | 1.00 | 92.87 |  | 2 |  | 2O_ph_ |
| K:1G:1606 | 1.00 | 61.36 |  | 1 |  | O_ph_ |
| K:1G:1607 | 1.00 | 87.42 |  | 2 |  | 2O_ph_ |
| K:1G:1608 | 1.00 | 80.69 |  | 4 |  | O_ph_O_b_2N_b_ |
| K:1G:1609 | 1.00 | 91.12 |  | 6 |  | 2O_r_2O_b_2N_b_ |
| K:1G:1610 | 1.00 | 68.23 |  | 8 |  | 5O_b_N_b_2Anion |
| K:1G:1611 | 1.00 | 86.36 |  | 2 |  | O_ph_O_b_ |
| K:1G:1612 | 1.00 | 58.38 |  | 5 |  | 4O_b_Anion |
| K:1G:1613 | 1.00 | 63.22 |  | 4 |  | O_b_2N_b_O_cno_ |
| K:1G:1614 | 1.00 | 86.30 |  | 5 |  | O_r_2O_b_N_b_O_back_ |
| K:1G:1615 | 1.00 | 60.41 |  | 4 |  | 3O_ph_O_r_ |
| K:1G:1616 | 1.00 | 86.73 |  | 1 |  | N_b_ |
| K:1G:1617 | 1.00 | 63.76 |  | 4 |  | O_ph_2O_b_N_b_ |
| K:1G:1618 | 1.00 | 79.66 |  | 4 |  | 3O_b_OH_Prot_ |
| K:1G:1619 | 1.00 | 74.45 | 1 | 5 |  | O_r_3O_b_O_w_ - Exclusion zone contact(s): /O2/ |
| K:1G:1620 | 1.00 | 83.15 |  | 5 |  | 3O_ph_2O_r_ |
| K:1G:1621 | 1.00 | 76.49 |  | 5 |  | 3O_ph_2O_r_ |
| K:1G:1622 | 1.00 | 74.93 |  | 3 |  | 2O_b_Anion |
| K:1G:1623 | 1.00 | 95.15 |  | 1 |  | O_ph_ |
| K:1G:1624 | 1.00 | 60.51 | 1 | 5 |  | 3O_r_O_b_N_b_ - Exclusion zone contact(s): /OP2/ |
| K:1G:1625 | 1.00 | 108.07 | 2 | 2 |  | O_r_O_back_ - Exclusion zone contact(s): /O_back_/OP1/ |
| K:1G:1626 | 1.00 | 67.03 |  | 3 |  | 3O_b_ |
| K:1G:1627 | 1.00 | 92.32 |  | 1 |  | O_ph_ |
| K:1G:1628 | 1.00 | 76.61 | 1 | 3 |  | 2O_ph_O_r_ - Exclusion zone contact(s): /O3'/ |
| K:1G:1629 | 1.00 | 88.10 | 1 | 1 |  | O_ph_ - Exclusion zone contact(s): /O3'/ |
| K:1G:1630 | 1.00 | 80.09 |  | 2 |  | 2O_b_ |
| K:1G:1631 | 1.00 | 86.91 |  | 2 |  | 2O_b_ |
| K:1G:1632 | 1.00 | 78.65 |  | 3 |  | 3O_b_ |
| K:1G:1633 | 1.00 | 76.91 |  | 3 |  | 3O_b_ |
| K:1G:1634 | 1.00 | 62.32 |  | 5 |  | 3O_ph_2O_r_ |
| K:1G:1635 | 1.00 | 91.08 |  | 1 |  | O_b_ |
| K:1G:1636 | 1.00 | 78.06 |  | 3 |  | 3O_ph_ |
| MG:1G:1637 | 1.00 | 55.69 |  | 3 | 1 | 3O_ph_ - Exclusion zone contact(s): /O3'/ |
| MG:1G:1638 | 1.00 | 46.37 |  | 1 |  | O_ph_ |
| MG:1G:1639 | 1.00 | 69.75 |  | 2 |  | O_ph_O_b_ |
| MG:1G:1640 | 1.00 | 51.11 |  | 0 | 1 | - Exclusion zone contact(s): /OP2/ Consider K in place of MG (1 valid contact(s) for K vs. 0 for MG) |
| MG:1G:1641 | 1.00 | 64.55 |  | 1 |  | N_b_ |
| MG:1G:1642 | 1.00 | 45.19 |  | 1 | 2 | O_ph_ - Exclusion zone contact(s): /OP1/OP2/ |
| MG:1G:1643 | 1.00 | 62.23 |  | 1 | 1 | O_ph_ - Exclusion zone contact(s): /OP1/ |
| MG:1G:1644 | 1.00 | 59.26 |  | 3 |  | 3O_ph_ |
| MG:1G:1645 | 1.00 | 62.32 |  | 2 | 1 | 2O_ph_ - Exclusion zone contact(s): /OP1/ |
| MG:1G:1646 | 1.00 | 66.33 |  | 1 | 3 | O_ph_ - Exclusion zone contact(s): /O3'/OP2/OP2/ Consider K in place of MG (3 valid contact(s) for K vs. 1 for MG) |
| MG:1G:1647 | 1.00 | 55.67 |  | 1 |  | O_ph_ |
| MG:1G:1648 | 1.00 | 67.44 |  | 2 | 1 | 2O_ph_ - Exclusion zone contact(s): /O4/ |
| MG:1G:1649 | 1.00 | 74.54 |  | 0 |  | 1^st^ MG coordination shell missing (see Notes) |
| MG:1G:1650 | 1.00 | 68.80 |  | 1 |  | O_ph_ |
| MG:1G:1651 | 1.00 | 43.80 |  | 0 | 1 | - Exclusion zone contact(s): /O6/ Consider K in place of MG (1 valid contact(s) for K vs. 0 for MG) |
| MG:1G:1652 | 1.00 | 17.19 |  | 1 |  | O_ph_ |
| MG:1G:1653 | 1.00 | 53.22 |  | 0 | 1 | - Exclusion zone contact(s): /OP1/ Consider K in place of MG (1 valid contact(s) for K vs. 0 for MG) |
| MG:1G:1654 | 1.00 | 50.31 |  | 1 |  | O_ph_ |
| MG:1G:1655 | 1.00 | 58.86 |  | 1 | 2 | O_ph_ - Exclusion zone contact(s): /O_w_/OP1/ |
| MG:1G:1656 | 1.00 | 58.77 |  | 0 |  | 1^st^ MG coordination shell missing (see Notes) |
| MG:1G:1657 | 1.00 | 53.61 |  | 0 | 1 | - Exclusion zone contact(s): /O6/ Consider K in place of MG (1 valid contact(s) for K vs. 0 for MG) |
| MG:1G:1658 | 1.00 | 46.68 |  | 0 |  | 1^st^ MG coordination shell missing (see Notes) |
| MG:1G:1659 | 1.00 | 64.97 |  | 0 | 1 | - Exclusion zone contact(s): /O6/ Consider K in place of MG (1 valid contact(s) for K vs. 0 for MG) |
| MG:1G:1660 | 1.00 | 46.01 |  | 0 |  | 1^st^ MG coordination shell missing (see Notes) |
| MG:1G:1661 | 1.00 | 79.11 |  | 0 |  | 1^st^ MG coordination shell missing (see Notes) |
| MG:1G:1662 | 1.00 | 51.46 |  | 1 |  | O_ph_ |
| MG:1G:1663 | 0.50 | 29.58 |  | 0 | 1 | - Exclusion zone contact(s): /OP2/ Consider K in place of MG (1 valid contact(s) for K vs. 0 for MG) |
| MG:1G:1663 | 0.50 | 30.72 |  | 0 |  | 1^st^ MG coordination shell missing (see Notes) |
| MG:1G:1664 | 1.00 | 40.37 |  | 0 |  | 1^st^ MG coordination shell missing (see Notes) |
| MG:1G:1665 | 1.00 | 50.20 |  | 0 |  | 1^st^ MG coordination shell missing (see Notes) |
| MG:1G:1666 | 1.00 | 46.58 |  | 0 |  | 1^st^ MG coordination shell missing (see Notes) |
| MG:1G:1667 | 1.00 | 42.81 |  | 0 | 1 | - Exclusion zone contact(s): /OP2/ Consider K in place of MG (1 valid contact(s) for K vs. 0 for MG) |
| MG:1G:1668 | 1.00 | 69.14 |  | 1 | 2 | O_ph_ - Exclusion zone contact(s): /OP1/OP2/ Consider K in place of MG (2 valid contact(s) for K vs. 1 for MG) |
| MG:1G:1669 | 1.00 | 60.17 |  | 0 |  | 1^st^ MG coordination shell missing (see Notes) |
| MG:1G:1670 | 1.00 | 57.10 |  | 0 | 2 | - Exclusion zone contact(s): /O6/OP2/ Consider K in place of MG (2 valid contact(s) for K vs. 0 for MG) |
| MG:1G:1671 | 1.00 | 46.87 |  | 0 |  | 1^st^ MG coordination shell missing (see Notes) |
| MG:1G:1672 | 1.00 | 66.82 |  | 1 |  | O_b_ |
| MG:1G:1673 | 1.00 | 50.93 |  | 1 |  | O_ph_ |
| MG:1G:1674 | 1.00 | 122.50 |  | 0 | 3 | - Exclusion zone contact(s): /OP1/OP2/OP2/ Consider K in place of MG (2 valid contact(s) for K vs. 0 for MG) |
| MG:1G:1675 | 1.00 | 65.15 |  | 0 | 1 | - Exclusion zone contact(s): /O6/ Consider K in place of MG (1 valid contact(s) for K vs. 0 for MG) |
| MG:1G:1676 | 1.00 | 71.49 |  | 2 |  | O_ph_O_b_ |
| MG:1G:1677 | 1.00 | 57.14 |  | 0 | 4 | - Exclusion zone contact(s): /N7/N7/OP2/OP2/ Consider K in place of MG (4 valid contact(s) for K vs. 0 for MG) |
| MG:1G:1678 | 1.00 | 54.04 |  | 0 | 2 | - Exclusion zone contact(s): /O4/O6/ Consider K in place of MG (2 valid contact(s) for K vs. 0 for MG) |
| MG:1G:1679 | 1.00 | 62.94 |  | 0 | 3 | - Exclusion zone contact(s): /N1_G_/O6/O6/ Consider K in place of MG (2 valid contact(s) for K vs. 0 for MG) |
| MG:1G:1680 | 1.00 | 50.93 |  | 0 | 1 | - Exclusion zone contact(s): /OP1/ Consider K in place of MG (1 valid contact(s) for K vs. 0 for MG) |
| MG:1G:1681 | 1.00 | 45.14 |  | 0 | 1 | - Exclusion zone contact(s): /OP2/ Consider K in place of MG (1 valid contact(s) for K vs. 0 for MG) |
| MG:1G:1682 | 1.00 | 41.27 |  | 0 | 1 | - Exclusion zone contact(s): /O6/ Consider K in place of MG (1 valid contact(s) for K vs. 0 for MG) |
| MG:1G:1683 | 0.50 | 51.33 |  | 0 |  | 1^st^ MG coordination shell missing (see Notes) |
| MG:1G:1683 | 0.50 | 45.69 |  | 0 | 1 | - Exclusion zone contact(s): /O6/ Consider K in place of MG (1 valid contact(s) for K vs. 0 for MG) |
| MG:1G:1684 | 1.00 | 79.08 |  | 1 | 1 | O_ph_ - Exclusion zone contact(s): /N7/ |
| MG:1G:1685 | 1.00 | 51.49 |  | 0 | 2 | - Exclusion zone contact(s): /N7/O6/ Consider K in place of MG (2 valid contact(s) for K vs. 0 for MG) |
| MG:1G:1686 | 1.00 | 75.73 |  | 0 | 1 | - Exclusion zone contact(s): /O2'/ Consider K in place of MG (1 valid contact(s) for K vs. 0 for MG) |
| MG:1G:1687 | 1.00 | 67.86 |  | 0 | 2 | - Exclusion zone contact(s): /O4/O6/ Consider K in place of MG (2 valid contact(s) for K vs. 0 for MG) |
| MG:1G:1688 | 1.00 | 65.57 |  | 0 | 1 | - Exclusion zone contact(s): /O4/ Consider K in place of MG (1 valid contact(s) for K vs. 0 for MG) |
| MG:1G:1689 | 1.00 | 57.60 |  | 0 |  | 1^st^ MG coordination shell missing (see Notes) |
| MG:1G:1690 | 1.00 | 74.93 |  | 0 | 1 | - Exclusion zone contact(s): /N7/ Consider K in place of MG (1 valid contact(s) for K vs. 0 for MG) |
| MG:1G:1691 | 1.00 | 64.23 |  | 0 | 2 | - Exclusion zone contact(s): /N7/OP2/ Consider K in place of MG (2 valid contact(s) for K vs. 0 for MG) |
| MG:1G:1692 | 1.00 | 72.26 |  | 0 |  | 1^st^ MG coordination shell missing (see Notes) |
| MG:1G:1693 | 1.00 | 40.65 |  | 2 | 3 | 2O_ph_ - Exclusion zone contact(s): /O2'/OP1/OP2/ Consider K in place of MG (3 valid contact(s) for K vs. 2 for MG) |
| MG:1G:1694 | 1.00 | 43.85 |  | 0 | 4 | - Exclusion zone contact(s): /O2/O2'/O3'/OP1/ Consider K in place of MG (3 valid contact(s) for K vs. 0 for MG) |
| MG:1G:1695 | 1.00 | 72.35 |  | 1 | 2 | O_ph_ - Exclusion zone contact(s): /OP1/OP1/ Consider K in place of MG (2 valid contact(s) for K vs. 1 for MG) |
| MG:1G:1696 | 1.00 | 52.65 |  | 0 |  | 1^st^ MG coordination shell missing (see Notes) |
| MG:1G:1697 | 1.00 | 63.29 |  | 0 |  | 1^st^ MG coordination shell missing (see Notes) |
| MG:1G:1698 | 1.00 | 66.26 |  | 0 |  | 1^st^ MG coordination shell missing (see Notes) |
| MG:1G:1699 | 1.00 | 62.42 |  | 1 | 2 | O_ph_ - Exclusion zone contact(s): /OP1/OP2/ Consider K in place of MG (2 valid contact(s) for K vs. 1 for MG) |
| MG:1G:1700 | 1.00 | 52.41 |  | 0 |  | 1^st^ MG coordination shell missing (see Notes) |
| MG:1G:1701 | 1.00 | 45.05 |  | 6 |  | 6O_w_ |
| MG:1G:1702 | 1.00 | 64.66 |  | 6 |  | 6O_w_ |
| MG:1G:1703 | 1.00 | 81.78 |  | 6 | 1 | 6O_w_ - Exclusion zone contact(s): /OP1/ Consider K in place of MG (1 valid contact(s) for K vs. 0 for MG)  Possible idealized distances ! |
| MG:1G:1704 | 1.00 | 66.29 |  | 6 |  | 6O_w_ |
| MG:1G:1705 | 1.00 | 73.26 |  | 6 |  | 6O_w_ |
| MG:1G:1706 | 1.00 | 43.15 |  | 6 |  | 6O_w_ |
| MG:1G:1707 | 1.00 | 62.66 |  | 6 |  | 6O_w_ |
| MG:1G:1708 | 1.00 | 101.76 |  | 6 |  | 6O_w_ |
| MG:1G:1709 | 1.00 | 73.51 |  | 6 |  | 6O_w_ |
| MG:1G:1710 | 1.00 | 59.25 |  | 6 |  | 6O_w_ |
| MG:1G:1711 | 1.00 | 97.35 |  | 6 | 1 | 6O_w_ - Exclusion zone contact(s): /N3_C_/ Consider K in place of MG (1 valid contact(s) for K vs. 0 for MG)  Possible idealized distances ! |
| MG:1G:1712 | 1.00 | 81.10 |  | 6 |  | 6O_w_ Possible idealized distances ! |
| MG:1G:1713 | 1.00 | 120.09 |  | 3 | 1 | 3O_w_ - Exclusion zone contact(s): /N6/ Consider K in place of MG (1 valid contact(s) for K vs. 0 for MG) |
| MG:1G:1714 | 1.00 | 88.23 |  | 6 |  | 6O_w_ |
| MG:1G:1715 | 1.00 | 39.02 |  | 5 | 1 | 5O_w_ - Exclusion zone contact(s): /OP1/ Consider NA in place of MG (1 valid contact(s) for NA vs. 0 for MG) |
| MG:1G:1716 | 1.00 | 70.48 |  | 6 |  | 6O_w_ Possible idealized distances ! |
| MG:1G:1717 | 1.00 | 67.85 |  | 6 |  | 6O_w_ |
| MG:1G:1718 | 1.00 | 39.02 |  | 5 | 1 | 5O_w_ - Exclusion zone contact(s): /OP2/ Consider NA in place of MG (1 valid contact(s) for NA vs. 0 for MG) |
| MG:1G:1719 | 1.00 | 66.92 |  | 2 | 6 | 2O_w_ - Exclusion zone contact(s): /O5'/O5'/OP1/OP1/OP1/OP2/ Consider K in place of MG (6 valid contact(s) for K vs. 0 for MG) |
| MG:1G:1720 | 1.00 | 73.89 |  | 5 | 2 | 5O_w_ - Exclusion zone contact(s): /OP1/OP2/ Consider K in place of MG (2 valid contact(s) for K vs. 0 for MG)  Possible idealized distances ! |
| MG:1G:1721 | 1.00 | 45.83 |  | 5 | 1 | 5O_w_ - Exclusion zone contact(s): /OP2/ Consider NA in place of MG (1 valid contact(s) for NA vs. 0 for MG) |
| MG:1G:1722 | 1.00 | 46.57 |  | 5 | 1 | 5O_w_ - Exclusion zone contact(s): /OP1/ Consider K in place of MG (1 valid contact(s) for K vs. 0 for MG)  Possible idealized distances ! |
| MG:1G:1723 | 1.00 | 59.32 |  | 4 | 3 | 4O_w_ - Exclusion zone contact(s): /O6/OP1/OP1/ Consider NA in place of MG (2 valid contact(s) for NA vs. 0 for MG) |
| MG:1G:1724 | 1.00 | 82.61 |  | 6 | 1 | 6O_w_ - Exclusion zone contact(s): /OP1/ Consider K in place of MG (1 valid contact(s) for K vs. 0 for MG) |
| MG:1G:1725 | 1.00 | 69.41 |  | 6 |  | 6O_w_ Possible idealized distances ! |
| MG:1G:1726 | 1.00 | 49.62 |  | 5 | 2 | 5O_w_ - Exclusion zone contact(s): /OP1/OP2/ Consider NA in place of MG (1 valid contact(s) for NA vs. 0 for MG) |
| MG:1G:1727 | 1.00 | 59.40 |  | 6 |  | 6O_w_ |
| MG:1G:1728 | 1.00 | 50.89 |  | 6 |  | 6O_w_ |
| MG:1G:1729 | 1.00 | 67.15 |  | 6 |  | 6O_w_ |
| MG:1G:1730 | 1.00 | 78.35 |  | 6 | 2 | 6O_w_ - Exclusion zone contact(s): /OP1/OP1/ Consider K in place of MG (2 valid contact(s) for K vs. 0 for MG) |
| MG:1G:1731 | 1.00 | 81.43 |  | 6 |  | 6O_w_ |
| MG:1G:1732 | 1.00 | 83.20 |  | 6 |  | 6O_w_ Possible idealized distances ! |
| MG:1G:1733 | 1.00 | 81.49 |  | 6 | 1 | 6O_w_ - Exclusion zone contact(s): /N7/ Consider K in place of MG (1 valid contact(s) for K vs. 0 for MG) |
| MG:1G:1734 | 1.00 | 94.86 |  | 6 |  | 6O_w_ Possible idealized distances ! |
| MG:1G:1735 | 1.00 | 88.83 |  | 6 |  | 6O_w_ Possible idealized distances ! |
| MG:1G:1736 | 1.00 | 62.90 |  | 5 | 1 | 5O_w_ - Exclusion zone contact(s): /O6/ Consider K in place of MG (1 valid contact(s) for K vs. 0 for MG) |
| K:1H:3001 | 1.00 | 48.54 |  | 4 |  | O_r_3O_b_ |
| K:1H:3002 | 1.00 | 73.77 |  | 2 |  | 2O_b_ |
| K:1H:3003 | 1.00 | 56.99 |  | 4 |  | 4O_b_ |
| K:1H:3004 | 1.00 | 88.42 |  | 2 |  | 2O_b_ |
| K:1H:3005 | 1.00 | 100.29 |  | 0 |  | 1^st^ K coordination shell missing (see Notes) |
| K:1H:3006 | 1.00 | 44.91 | 1 | 3 |  | 2O_b_O_w_ - Exclusion zone contact(s): /O6/ |
| K:1H:3007 | 1.00 | 55.05 |  | 2 |  | O_b_N_b_ |
| K:1H:3008 | 1.00 | 65.91 | 1 | 1 |  | O_b_ - Exclusion zone contact(s): /O6/ |
| K:1H:3009 | 1.00 | 64.90 | 1 | 2 |  | 2O_b_ - Exclusion zone contact(s): /O6/ |
| K:1H:3010 | 1.00 | 66.49 |  | 4 |  | 2O_b_2O_w_ |
| K:1H:3011 | 1.00 | 59.54 |  | 2 |  | 2O_b_ |
| K:1H:3012 | 1.00 | 51.51 |  | 3 |  | 3O_b_ |
| K:1H:3013 | 1.00 | 81.10 |  | 3 |  | 2O_ph_O_b_ |
| K:1H:3014 | 1.00 | 60.89 |  | 3 |  | 2O_r_O_b_ |
| K:1H:3015 | 1.00 | 44.52 |  | 4 |  | O_ph_2O_r_O_b_ |
| K:1H:3016 | 1.00 | 63.21 |  | 3 |  | O_ph_O_r_Anion |
| K:1H:3017 | 1.00 | 56.93 |  | 2 |  | 2O_b_ |
| K:1H:3018 | 1.00 | 51.71 |  | 4 |  | 3O_r_O_b_ |
| K:1H:3019 | 1.00 | 60.75 |  | 2 |  | 2O_b_ |
| K:1H:3020 | 1.00 | 65.85 |  | 3 |  | O_r_2O_w_ |
| K:1H:3021 | 1.00 | 78.20 |  | 1 |  | O_b_ |
| K:1H:3022 | 1.00 | 66.53 |  | 2 |  | O_ph_O_w_ |
| K:1H:3023 | 1.00 | 63.16 |  | 3 |  | O_ph_O_b_N_b_ |
| K:1H:3024 | 1.00 | 59.66 |  | 4 |  | O_ph_O_r_O_b_N_b_ |
| K:1H:3025 | 1.00 | 53.77 |  | 3 |  | O_ph_2O_r_ |
| K:1H:3026 | 1.00 | 88.40 |  | 0 |  | 1^st^ K coordination shell missing (see Notes) |
| K:1H:3027 | 1.00 | 63.75 |  | 4 |  | O_ph_2O_r_N_b_ |
| K:1H:3028 | 1.00 | 87.72 |  | 1 |  | O_ph_ |
| K:1H:3029 | 1.00 | 85.17 |  | 1 |  | O_b_ |
| K:1H:3030 | 1.00 | 73.54 | 1 | 1 |  | O_b_ - Exclusion zone contact(s): /N2/ |
| K:1H:3031 | 1.00 | 39.70 |  | 3 |  | O_ph_2O_r_ |
| K:1H:3032 | 1.00 | 54.80 |  | 2 |  | 2O_ph_ |
| K:1H:3033 | 1.00 | 36.86 | 2 | 2 |  | - Exclusion zone contact(s): /OP1/OP2/ Consider NA in place of K (2 valid contact(s) for NA vs. 0 for K) |
| K:1H:3034 | 1.00 | 66.51 |  | 2 |  | 2O_b_ |
| K:1H:3035 | 1.00 | 52.07 | 1 | 1 |  | - Exclusion zone contact(s): /O6/ Consider NA in place of K (1 valid contact(s) for NA vs. 0 for K) |
| K:1H:3036 | 1.00 | 60.90 |  | 2 |  | O_r_N_b_ |
| K:1H:3037 | 1.00 | 70.88 |  | 0 |  | 1^st^ K coordination shell missing (see Notes) |
| K:1H:3038 | 1.00 | 56.92 | 1 | 1 |  | - Exclusion zone contact(s): /OP2/ Consider NA in place of K (1 valid contact(s) for NA vs. 0 for K) |
| K:1H:3039 | 1.00 | 54.13 |  | 2 |  | 2Anion |
| K:1H:3040 | 0.50 | 46.91 |  | 0 |  | 1^st^ K coordination shell missing (see Notes) |
| K:1H:3040 | 0.50 | 43.22 |  | 0 |  | 1^st^ K coordination shell missing (see Notes) |
| K:1H:3041 | 1.00 | 68.01 |  | 2 |  | 2O_b_ |
| K:1H:3042 | 1.00 | 54.63 |  | 4 |  | 4O_b_ |
| K:1H:3043 | 1.00 | 58.70 | 1 | 3 |  | 3O_b_ - Exclusion zone contact(s): /O6/ |
| K:1H:3044 | 1.00 | 92.12 |  | 2 |  | 2O_b_ |
| K:1H:3045 | 1.00 | 59.62 |  | 3 |  | O_ph_O_r_O_b_ |
| K:1H:3046 | 1.00 | 72.95 |  | 1 |  | O_b_ |
| K:1H:3047 | 1.00 | 74.12 |  | 5 |  | 3O_b_2O_w_ |
| K:1H:3048 | 1.00 | 72.20 |  | 2 |  | O_ph_O_b_ |
| K:1H:3049 | 1.00 | 55.61 |  | 2 |  | 2O_b_ |
| K:1H:3050 | 1.00 | 53.89 |  | 0 |  | 1^st^ K coordination shell missing (see Notes) |
| K:1H:3051 | 1.00 | 49.61 |  | 3 |  | 3O_b_ |
| K:1H:3052 | 1.00 | 52.89 |  | 2 |  | O_ph_O_r_ |
| K:1H:3053 | 1.00 | 25.61 | 1 | 6 |  | 4O_ph_2O_r_ - Exclusion zone contact(s): /O_w_/ |
| K:1H:3054 | 1.00 | 27.77 |  | 4 |  | O_r_3O_b_ |
| K:1H:3055 | 1.00 | 31.99 |  | 4 |  | 4O_r_ |
| K:1H:3056 | 1.00 | 37.58 |  | 5 |  | O_r_O_b_3N_b_ |
| K:1H:3057 | 1.00 | 23.63 |  | 7 |  | O_r_2O_b_4N_b_ |
| K:1H:3058 | 1.00 | 43.20 |  | 3 |  | 3O_b_ |
| K:1H:3059 | 1.00 | 19.99 | 1 | 6 |  | O_r_3O_b_2N_b_ - Exclusion zone contact(s): /O6/ |
| K:1H:3060 | 1.00 | 34.42 | 1 | 6 |  | O_ph_O_r_O_coo_3O_w_ - Exclusion zone contact(s): /OP2/ |
| K:1H:3061 | 1.00 | 43.18 |  | 5 |  | O_ph_O_r_O_b_2N_b_ |
| K:1H:3062 | 1.00 | 27.71 |  | 5 |  | 2O_ph_2O_r_N_b_ |
| K:1H:3063 | 1.00 | 47.89 | 1 | 2 |  | O_ph_O_r_ - Exclusion zone contact(s): /OP1/ |
| K:1H:3064 | 1.00 | 34.06 |  | 6 |  | 3O_ph_3O_r_ |
| K:1H:3065 | 1.00 | 30.08 | 1 | 3 |  | O_b_N_b_Anion - Exclusion zone contact(s): /OP1/ |
| K:1H:3066 | 1.00 | 27.63 |  | 3 |  | 2N_b_O_w_ |
| K:1H:3067 | 1.00 | 53.25 |  | 3 |  | 3O_b_ |
| K:1H:3068 | 1.00 | 29.26 |  | 5 |  | 4O_b_Anion |
| K:1H:3069 | 1.00 | 16.98 | 2 | 2 |  | 2O_ph_ - Exclusion zone contact(s): /O6/O_back_/ |
| K:1H:3070 | 1.00 | 46.66 | 1 | 3 |  | 2O_r_O_b_ - Exclusion zone contact(s): /O2/ |
| K:1H:3071 | 1.00 | 42.96 | 1 | 5 |  | O_r_3O_b_N_b_ - Exclusion zone contact(s): /O6/ |
| K:1H:3072 | 1.00 | 53.73 |  | 3 |  | O_r_2O_b_ |
| K:1H:3073 | 1.00 | 56.35 |  | 5 |  | O_ph_3O_b_N_b_ |
| K:1H:3074 | 1.00 | 34.33 |  | 4 |  | 2O_ph_O_r_O_b_ |
| K:1H:3075 | 1.00 | 28.19 |  | 4 |  | 4O_b_ |
| K:1H:3076 | 1.00 | 50.60 |  | 5 |  | O_ph_O_r_O_b_N_b_O_w_ |
| K:1H:3077 | 1.00 | 47.67 |  | 4 |  | 2O_b_2N_b_ |
| K:1H:3078 | 1.00 | 45.76 | 2 | 5 |  | O_r_O_b_2N_b_Anion - Exclusion zone contact(s): /O2'/OP2/ |
| K:1H:3079 | 1.00 | 37.86 |  | 5 |  | O_ph_3O_r_O_back_ |
| K:1H:3080 | 1.00 | 45.95 |  | 3 |  | 2O_ph_N_b_ |
| K:1H:3081 | 1.00 | 49.65 |  | 3 |  | 3O_b_ |
| K:1H:3082 | 1.00 | 46.98 |  | 5 |  | O_ph_2O_b_2O_w_ |
| K:1H:3083 | 1.00 | 54.39 |  | 3 |  | 3O_b_ |
| K:1H:3084 | 1.00 | 47.78 |  | 3 |  | O_r_2O_b_ |
| K:1H:3085 | 1.00 | 39.53 | 1 | 3 |  | 3O_b_ - Exclusion zone contact(s): /O6/ |
| K:1H:3086 | 1.00 | 66.80 | 1 | 3 |  | 3O_b_ - Exclusion zone contact(s): /O4/ |
| K:1H:3087 | 1.00 | 31.55 |  | 8 |  | 2O_ph_5O_r_O_b_ |
| K:1H:3088 | 1.00 | 50.79 |  | 5 |  | O_ph_O_r_2O_b_N_b_ |
| K:1H:3089 | 1.00 | 36.46 |  | 5 |  | O_r_4O_b_ |
| K:1H:3090 | 1.00 | 55.06 |  | 4 |  | O_ph_2O_b_Anion |
| K:1H:3091 | 1.00 | 64.49 |  | 4 |  | 4O_b_ |
| K:1H:3092 | 1.00 | 37.38 |  | 5 |  | 3O_ph_O_r_O_w_ |
| K:1H:3093 | 1.00 | 42.07 |  | 3 |  | 2O_ph_O_r_ |
| K:1H:3094 | 1.00 | 51.99 |  | 4 |  | 4O_b_ |
| K:1H:3095 | 1.00 | 58.53 |  | 3 |  | 2O_b_N_b_ |
| K:1H:3096 | 1.00 | 52.15 |  | 4 |  | 2O_ph_O_r_O_b_ |
| K:1H:3097 | 1.00 | 25.66 |  | 6 |  | O_ph_O_r_2O_b_N_b_ |
| K:1H:3098 | 1.00 | 41.07 |  | 3 |  | 3O_b_ |
| K:1H:3099 | 1.00 | 59.79 |  | 1 |  | O_b_ |
| K:1H:3100 | 1.00 | 44.51 |  | 3 |  | 3O_b_ |
| K:1H:3101 | 1.00 | 49.84 |  | 3 |  | 2O_ph_O_b_ |
| K:1H:3102 | 1.00 | 51.58 |  | 3 |  | 3O_b_ |
| K:1H:3103 | 1.00 | 33.06 |  | 7 |  | 5O_ph_O_r_O_w_ |
| K:1H:3104 | 1.00 | 42.03 | 1 | 3 |  | O_ph_O_r_O_w_ - Exclusion zone contact(s): /O_w_/ |
| K:1H:3105 | 1.00 | 50.57 |  | 2 |  | 2O_b_ |
| K:1H:3106 | 1.00 | 59.65 |  | 3 |  | 2O_r_O_back_ |
| K:1H:3107 | 1.00 | 44.66 |  | 3 |  | O_ph_2O_b_ |
| K:1H:3108 | 1.00 | 76.83 |  | 2 |  | 2O_b_ |
| K:1H:3109 | 1.00 | 57.54 |  | 4 |  | O_ph_3O_b_ |
| K:1H:3110 | 1.00 | 60.31 |  | 5 |  | 2O_ph_2O_r_O_b_ |
| K:1H:3111 | 1.00 | 51.93 |  | 3 |  | O_r_O_b_O_w_ |
| K:1H:3112 | 1.00 | 48.71 |  | 4 |  | 4O_b_ |
| K:1H:3113 | 1.00 | 30.67 | 1 | 3 |  | O_ph_O_r_N_b_ - Exclusion zone contact(s): /O2'/ |
| K:1H:3114 | 1.00 | 55.75 |  | 1 |  | O_b_ |
| K:1H:3115 | 1.00 | 52.47 |  | 3 |  | O_r_2O_b_ |
| K:1H:3116 | 1.00 | 61.81 |  | 5 |  | 3O_ph_2O_r_ |
| K:1H:3117 | 1.00 | 57.52 |  | 3 |  | 2O_r_O_b_ |
| K:1H:3118 | 1.00 | 56.64 |  | 3 |  | 3O_b_ |
| K:1H:3119 | 1.00 | 46.13 |  | 3 |  | O_ph_O_b_O_back_ |
| K:1H:3120 | 1.00 | 60.83 |  | 3 |  | 2O_r_O_b_ |
| K:1H:3121 | 1.00 | 62.68 |  | 3 |  | O_ph_OH_Prot_O_w_ |
| K:1H:3122 | 1.00 | 65.63 |  | 3 |  | 3O_ph_ |
| K:1H:3123 | 1.00 | 50.90 |  | 4 |  | O_ph_2O_b_O_w_ |
| K:1H:3124 | 1.00 | 64.99 |  | 2 |  | 2O_r_ |
| K:1H:3125 | 1.00 | 64.60 |  | 1 |  | O_ph_ |
| K:1H:3126 | 1.00 | 86.66 |  | 0 |  | 1^st^ K coordination shell missing (see Notes) |
| K:1H:3127 | 1.00 | 47.09 |  | 4 |  | 2O_r_O_b_N_b_ |
| K:1H:3128 | 1.00 | 64.23 |  | 4 |  | O_ph_2O_r_O_b_ |
| K:1H:3129 | 1.00 | 60.33 |  | 1 |  | O_b_ |
| K:1H:3130 | 1.00 | 61.29 |  | 4 |  | O_ph_2O_b_O_w_ |
| K:1H:3131 | 1.00 | 63.18 |  | 1 |  | O_ph_ |
| K:1H:3132 | 1.00 | 87.33 |  | 3 |  | 2O_b_O_w_ |
| K:1H:3133 | 1.00 | 61.34 |  | 4 |  | 3O_b_N_b_ |
| K:1H:3134 | 1.00 | 54.45 |  | 2 |  | 2O_b_ |
| K:1H:3135 | 1.00 | 82.62 |  | 2 |  | 2O_ph_ |
| K:1H:3136 | 1.00 | 54.76 | 1 | 3 |  | 3O_b_ - Exclusion zone contact(s): /O4/ |
| K:1H:3137 | 1.00 | 63.56 |  | 1 |  | O_b_ |
| K:1H:3138 | 1.00 | 62.69 |  | 4 |  | 2O_ph_2O_r_ |
| K:1H:3139 | 1.00 | 64.30 | 1 | 1 |  | O_b_ - Exclusion zone contact(s): /O6/ |
| K:1H:3140 | 1.00 | 62.74 |  | 1 |  | O_r_ |
| K:1H:3141 | 1.00 | 41.63 |  | 3 |  | 2O_ph_O_r_ |
| K:1H:3142 | 1.00 | 78.67 |  | 3 |  | 3O_ph_ |
| K:1H:3143 | 1.00 | 71.81 |  | 2 |  | 2O_b_ |
| K:1H:3144 | 1.00 | 56.21 |  | 3 |  | 3O_b_ |
| MG:1H:3145 | 1.00 | 17.62 |  | 1 |  | O_ph_ |
| MG:1H:3146 | 1.00 | 38.43 |  | 1 | 2 | O_b_ - Exclusion zone contact(s): /N6/N7/ Consider K in place of MG (2 valid contact(s) for K vs. 1 for MG) |
| MG:1H:3147 | 1.00 | 54.70 |  | 2 |  | 2O_ph_ |
| MG:1H:3148 | 1.00 | 35.60 |  | 0 | 1 | - Exclusion zone contact(s): /N7/ Consider K in place of MG (1 valid contact(s) for K vs. 0 for MG) |
| MG:1H:3149 | 1.00 | 40.15 |  | 2 |  | 2O_ph_ |
| MG:1H:3150 | 1.00 | 25.48 |  | 2 |  | O_ph_O_b_ |
| MG:1H:3151 | 1.00 | 41.59 |  | 0 |  | 1^st^ MG coordination shell missing (see Notes) |
| MG:1H:3152 | 1.00 | 36.87 |  | 1 | 3 | O_w_ - Exclusion zone contact(s): /O_w_/O_w_/OP1/ Consider NA in place of MG (1 valid contact(s) for NA vs. 0 for MG) |
| MG:1H:3153 | 1.00 | 45.53 |  | 0 | 2 | - Exclusion zone contact(s): /O_w_/OP2/ Consider K in place of MG (1 valid contact(s) for K vs. 0 for MG) |
| MG:1H:3154 | 1.00 | 26.87 |  | 1 | 1 | O_ph_ - Exclusion zone contact(s): /O_w_/ |
| MG:1H:3155 | 0.50 | 19.89 |  | 1 | 1 | O_ph_ - Exclusion zone contact(s): /OP1/ |
| MG:1H:3155 | 0.50 | 22.49 |  | 2 | 3 | 2O_ph_ - Exclusion zone contact(s): /O2'/O3'/OP2/ Consider K in place of MG (3 valid contact(s) for K vs. 2 for MG) |
| MG:1H:3156 | 1.00 | 33.38 |  | 0 |  | 1^st^ MG coordination shell missing (see Notes) |
| MG:1H:3157 | 1.00 | 49.73 |  | 0 | 1 | - Exclusion zone contact(s): /OP2/ Consider K in place of MG (1 valid contact(s) for K vs. 0 for MG) |
| MG:1H:3158 | 1.00 | 57.64 |  | 0 | 4 | - Exclusion zone contact(s): /O_w_/O_w_/O_w_/OP2/ Consider K in place of MG (1 valid contact(s) for K vs. 0 for MG) |
| MG:1H:3159 | 1.00 | 44.73 |  | 0 | 1 | - Exclusion zone contact(s): /O6/ Consider K in place of MG (1 valid contact(s) for K vs. 0 for MG) |
| MG:1H:3160 | 1.00 | 29.00 |  | 0 |  | 1^st^ MG coordination shell missing (see Notes) |
| MG:1H:3161 | 1.00 | 55.17 |  | 0 |  | 1^st^ MG coordination shell missing (see Notes) |
| MG:1H:3162 | 1.00 | 58.12 |  | 1 |  | O_ph_ |
| MG:1H:3163 | 1.00 | 55.27 |  | 0 | 2 | - Exclusion zone contact(s): /N7/O6/ Consider K in place of MG (2 valid contact(s) for K vs. 0 for MG) |
| MG:1H:3164 | 1.00 | 58.93 |  | 0 |  | 1^st^ MG coordination shell missing (see Notes) |
| MG:1H:3165 | 1.00 | 40.40 |  | 0 | 1 | - Exclusion zone contact(s): /OP1/ Consider NA in place of MG (1 valid contact(s) for NA vs. 0 for MG) |
| MG:1H:3166 | 1.00 | 41.01 |  | 1 | 1 | O_w_ - Exclusion zone contact(s): /O_w_/ |
| MG:1H:3167 | 1.00 | 20.65 |  | 3 |  | 3O_ph_ |
| MG:1H:3168 | 1.00 | 15.12 |  | 2 | 1 | 2O_ph_ - Exclusion zone contact(s): /N7/ |
| MG:1H:3169 | 1.00 | 15.91 |  | 1 |  | O_ph_ |
| MG:1H:3170 | 1.00 | 18.52 |  | 1 | 3 | O_ph_ - Exclusion zone contact(s): /OP1/OP1/OP2/ Consider NA in place of MG (2 valid contact(s) for NA vs. 1 for MG) |
| MG:1H:3171 | 1.00 | 30.81 |  | 2 | 1 | 2O_ph_ - Exclusion zone contact(s): /O3'/ |
| MG:1H:3172 | 1.00 | 31.94 |  | 0 | 2 | - Exclusion zone contact(s): /OP2/OP2/ Consider NA in place of MG (1 valid contact(s) for NA vs. 0 for MG) |
| MG:1H:3173 | 1.00 | 28.18 |  | 0 |  | 1^st^ MG coordination shell missing (see Notes) |
| MG:1H:3174 | 1.00 | 60.76 |  | 0 |  | 1^st^ MG coordination shell missing (see Notes) |
| MG:1H:3175 | 1.00 | 31.11 | 2 | 2 |  | - Exclusion zone contact(s): /OP1/OP1/ 1^st^ MG coordination shell missing (see Notes) |
| MG:1H:3176 | 0.50 | 21.21 |  | 2 | 3 | 2O_ph_ - Exclusion zone contact(s): /O5'/OP1/OP1/ Consider K in place of MG (3 valid contact(s) for K vs. 2 for MG) |
| MG:1H:3176 | 0.50 | 20.46 |  | 3 | 3 | 3O_ph_ - Exclusion zone contact(s): /O5'/OP1/OP2/ |
| MG:1H:3177 | 1.00 | 29.20 |  | 0 |  | 1^st^ MG coordination shell missing (see Notes) |
| MG:1H:3178 | 1.00 | 17.25 |  | 2 |  | 2O_ph_ |
| MG:1H:3179 | 1.00 | 30.89 |  | 0 |  | 1^st^ MG coordination shell missing (see Notes) |
| MG:1H:3180 | 1.00 | 43.95 |  | 2 |  | 2O_ph_ |
| MG:1H:3181 | 1.00 | 36.24 |  | 0 | 1 | - Exclusion zone contact(s): /N2/ Consider K in place of MG (1 valid contact(s) for K vs. 0 for MG) |
| MG:1H:3182 | 1.00 | 49.32 |  | 0 | 3 | - Exclusion zone contact(s): /O5'/OP1/OP1/ Consider K in place of MG (2 valid contact(s) for K vs. 0 for MG) |
| MG:1H:3183 | 1.00 | 47.98 |  | 0 | 2 | - Exclusion zone contact(s): /O5'/OP1/ Consider K in place of MG (2 valid contact(s) for K vs. 0 for MG) |
| MG:1H:3184 | 1.00 | 40.01 |  | 0 |  | 1^st^ MG coordination shell missing (see Notes) |
| MG:1H:3185 | 1.00 | 40.73 |  | 0 |  | 1^st^ MG coordination shell missing (see Notes) |
| MG:1H:3186 | 1.00 | 55.55 |  | 0 | 2 | - Exclusion zone contact(s): /N7/O6/ Consider K in place of MG (2 valid contact(s) for K vs. 0 for MG) |
| MG:1H:3187 | 1.00 | 30.77 |  | 0 | 1 | - Exclusion zone contact(s): /OP1/ Consider NA in place of MG (1 valid contact(s) for NA vs. 0 for MG) |
| MG:1H:3188 | 1.00 | 20.11 |  | 2 | 2 | 2O_ph_ - Exclusion zone contact(s): /O3'/OP2/ |
| MG:1H:3189 | 1.00 | 41.71 |  | 0 | 2 | - Exclusion zone contact(s): /OP2/OP2/ Consider K in place of MG (2 valid contact(s) for K vs. 0 for MG) |
| MG:1H:3190 | 1.00 | 32.41 |  | 0 |  | 1^st^ MG coordination shell missing (see Notes) |
| MG:1H:3191 | 1.00 | 24.72 |  | 3 | 1 | 3O_ph_ - Exclusion zone contact(s): /O5'/ |
| MG:1H:3192 | 1.00 | 37.96 |  | 1 | 2 | O_ph_ - Exclusion zone contact(s): /O_w_/OP2/ |
| MG:1H:3193 | 1.00 | 40.69 |  | 0 |  | 1^st^ MG coordination shell missing (see Notes) |
| MG:1H:3194 | 1.00 | 14.28 |  | 1 | 1 | O_ph_ - Exclusion zone contact(s): /OP1/ |
| MG:1H:3195 | 1.00 | 37.41 |  | 2 |  | 2O_ph_ |
| MG:1H:3196 | 1.00 | 40.03 |  | 0 |  | 1^st^ MG coordination shell missing (see Notes) |
| MG:1H:3197 | 1.00 | 30.64 |  | 0 | 1 | - Exclusion zone contact(s): /N3_A_/ Consider K in place of MG (1 valid contact(s) for K vs. 0 for MG) |
| MG:1H:3198 | 1.00 | 20.55 |  | 2 | 2 | O_ph_O_b_ - Exclusion zone contact(s): /OP2/OP2/ |
| MG:1H:3199 | 1.00 | 34.81 |  | 1 |  | O_ph_ |
| MG:1H:3200 | 1.00 | 36.98 |  | 1 | 1 | O_ph_ - Exclusion zone contact(s): /O5'/ |
| MG:1H:3201 | 1.00 | 45.84 |  | 0 | 1 | - Exclusion zone contact(s): /O6/ Consider K in place of MG (1 valid contact(s) for K vs. 0 for MG) |
| MG:1H:3202 | 1.00 | 45.21 |  | 0 | 2 | - Exclusion zone contact(s): /O6/O6/ Consider K in place of MG (2 valid contact(s) for K vs. 0 for MG) |
| MG:1H:3203 | 1.00 | 26.98 |  | 1 | 2 | O_ph_ - Exclusion zone contact(s): /N1_G_/O6/ |
| MG:1H:3204 | 1.00 | 29.06 |  | 0 | 1 | - Exclusion zone contact(s): /OP1/ Consider NA in place of MG (1 valid contact(s) for NA vs. 0 for MG) |
| MG:1H:3205 | 1.00 | 33.34 |  | 1 |  | O_ph_ |
| MG:1H:3206 | 1.00 | 25.11 |  | 3 | 1 | 2O_ph_His - Exclusion zone contact(s): /O_back_/ |
| MG:1H:3207 | 1.00 | 38.06 |  | 0 | 2 | - Exclusion zone contact(s): /O2'/O4'/ Consider K in place of MG (2 valid contact(s) for K vs. 0 for MG) |
| MG:1H:3208 | 1.00 | 37.30 |  | 0 | 6 | - Exclusion zone contact(s): /O_back_/O_w_/O_w_/OG_Ser_/OP2/OP2/ Consider K in place of MG (3 valid contact(s) for K vs. 0 for MG) |
| MG:1H:3209 | 1.00 | 52.39 |  | 0 |  | 1^st^ MG coordination shell missing (see Notes) |
| MG:1H:3210 | 1.00 | 38.45 |  | 0 | 1 | - Exclusion zone contact(s): /O6/ Consider K in place of MG (1 valid contact(s) for K vs. 0 for MG) |
| MG:1H:3211 | 1.00 | 56.95 |  | 0 | 1 | - Exclusion zone contact(s): /O6/ Consider K in place of MG (1 valid contact(s) for K vs. 0 for MG) |
| MG:1H:3212 | 1.00 | 44.46 |  | 0 | 1 | - Exclusion zone contact(s): /O6/ Consider K in place of MG (1 valid contact(s) for K vs. 0 for MG) |
| MG:1H:3213 | 1.00 | 25.72 |  | 0 | 1 | - Exclusion zone contact(s): /O2'/ Consider K in place of MG (1 valid contact(s) for K vs. 0 for MG) |
| MG:1H:3214 | 1.00 | 41.74 |  | 0 |  | 1^st^ MG coordination shell missing (see Notes) |
| MG:1H:3215 | 1.00 | 40.71 |  | 1 |  | O_ph_ |
| MG:1H:3216 | 1.00 | 34.63 |  | 0 |  | 1^st^ MG coordination shell missing (see Notes) |
| MG:1H:3217 | 1.00 | 31.09 |  | 0 |  | 1^st^ MG coordination shell missing (see Notes) |
| MG:1H:3218 | 1.00 | 28.01 |  | 1 |  | O_b_ |
| MG:1H:3219 | 1.00 | 31.25 |  | 0 |  | 1^st^ MG coordination shell missing (see Notes) |
| MG:1H:3220 | 1.00 | 50.05 |  | 0 |  | 1^st^ MG coordination shell missing (see Notes) |
| MG:1H:3221 | 1.00 | 34.18 |  | 0 | 3 | - Exclusion zone contact(s): /N7/O6/OP2/ Consider K in place of MG (2 valid contact(s) for K vs. 0 for MG) |
| MG:1H:3222 | 1.00 | 32.11 |  | 0 |  | 1^st^ MG coordination shell missing (see Notes) |
| MG:1H:3223 | 1.00 | 30.47 |  | 0 | 1 | - Exclusion zone contact(s): /OP2/ Consider K in place of MG (1 valid contact(s) for K vs. 0 for MG) |
| MG:1H:3224 | 1.00 | 35.70 |  | 0 |  | 1^st^ MG coordination shell missing (see Notes) |
| MG:1H:3225 | 1.00 | 37.96 |  | 0 | 1 | - Exclusion zone contact(s): /O6/ Consider K in place of MG (1 valid contact(s) for K vs. 0 for MG) |
| MG:1H:3226 | 1.00 | 40.08 |  | 0 |  | 1^st^ MG coordination shell missing (see Notes) |
| MG:1H:3227 | 1.00 | 29.99 |  | 0 | 1 | - Exclusion zone contact(s): /O4/ Consider K in place of MG (1 valid contact(s) for K vs. 0 for MG) |
| MG:1H:3228 | 1.00 | 31.69 |  | 0 |  | 1^st^ MG coordination shell missing (see Notes) |
| MG:1H:3229 | 1.00 | 43.96 |  | 0 | 1 | - Exclusion zone contact(s): /O6/ Consider K in place of MG (1 valid contact(s) for K vs. 0 for MG) |
| MG:1H:3230 | 1.00 | 39.68 |  | 0 | 2 | - Exclusion zone contact(s): /O2/OP1/ Consider K in place of MG (2 valid contact(s) for K vs. 0 for MG) |
| MG:1H:3231 | 1.00 | 43.42 |  | 0 | 1 | - Exclusion zone contact(s): /OP2/ Consider K in place of MG (1 valid contact(s) for K vs. 0 for MG) |
| MG:1H:3232 | 1.00 | 43.75 |  | 2 | 1 | 2O_ph_ - Exclusion zone contact(s): /O2'/ |
| MG:1H:3233 | 1.00 | 51.32 |  | 0 | 1 | - Exclusion zone contact(s): /OP2/ Consider K in place of MG (1 valid contact(s) for K vs. 0 for MG) |
| MG:1H:3234 | 1.00 | 39.11 |  | 0 | 1 | - Exclusion zone contact(s): /OP2/ Consider K in place of MG (1 valid contact(s) for K vs. 0 for MG) |
| MG:1H:3235 | 1.00 | 42.11 |  | 0 |  | 1^st^ MG coordination shell missing (see Notes) |
| MG:1H:3236 | 1.00 | 43.55 |  | 0 | 1 | - Exclusion zone contact(s): /OP1/ Consider K in place of MG (1 valid contact(s) for K vs. 0 for MG) |
| MG:1H:3237 | 1.00 | 38.31 |  | 0 | 1 | - Exclusion zone contact(s): /OP1/ Consider K in place of MG (1 valid contact(s) for K vs. 0 for MG) |
| MG:1H:3238 | 1.00 | 39.74 |  | 0 | 4 | - Exclusion zone contact(s): /O2'/O5'/O_w_/OP2/ Consider K in place of MG (3 valid contact(s) for K vs. 0 for MG) |
| MG:1H:3239 | 1.00 | 26.33 |  | 0 |  | 1^st^ MG coordination shell missing (see Notes) |
| MG:1H:3240 | 1.00 | 39.45 |  | 0 | 3 | - Exclusion zone contact(s): /N7/O6/OP2/ Consider K in place of MG (2 valid contact(s) for K vs. 0 for MG) |
| MG:1H:3241 | 1.00 | 30.18 |  | 0 |  | 1^st^ MG coordination shell missing (see Notes) |
| MG:1H:3242 | 1.00 | 37.37 |  | 0 | 2 | - Exclusion zone contact(s): /OP1/OP2/ Consider K in place of MG (2 valid contact(s) for K vs. 0 for MG) |
| MG:1H:3243 | 1.00 | 41.87 |  | 0 |  | 1^st^ MG coordination shell missing (see Notes) |
| MG:1H:3244 | 1.00 | 34.50 |  | 1 |  | O_ph_ |
| MG:1H:3245 | 1.00 | 50.49 |  | 0 |  | 1^st^ MG coordination shell missing (see Notes) |
| MG:1H:3246 | 1.00 | 35.22 |  | 1 | 1 | O_ph_ - Exclusion zone contact(s): /OP1/ |
| MG:1H:3247 | 1.00 | 21.58 |  | 1 |  | O_b_ |
| MG:1H:3248 | 1.00 | 34.62 |  | 0 | 2 | - Exclusion zone contact(s): /OP1/OP2/ Consider K in place of MG (2 valid contact(s) for K vs. 0 for MG) |
| MG:1H:3249 | 1.00 | 31.13 |  | 0 | 4 | - Exclusion zone contact(s): /O2'/OP1/OP1/OP2/ Consider K in place of MG (4 valid contact(s) for K vs. 0 for MG) |
| MG:1H:3250 | 1.00 | 38.38 |  | 2 |  | 2O_ph_ |
| MG:1H:3251 | 1.00 | 39.20 |  | 0 |  | 1^st^ MG coordination shell missing (see Notes) |
| MG:1H:3252 | 1.00 | 42.85 |  | 0 | 1 | - Exclusion zone contact(s): /N7/ Consider K in place of MG (1 valid contact(s) for K vs. 0 for MG) |
| MG:1H:3253 | 1.00 | 41.75 |  | 0 |  | 1^st^ MG coordination shell missing (see Notes) |
| MG:1H:3254 | 1.00 | 29.20 |  | 0 | 3 | - Exclusion zone contact(s): /O5'/OP1/OP2/ Consider K in place of MG (2 valid contact(s) for K vs. 0 for MG) |
| MG:1H:3255 | 1.00 | 47.48 |  | 1 | 1 | O_ph_ - Exclusion zone contact(s): /OP1/ |
| MG:1H:3256 | 1.00 | 40.50 |  | 0 |  | 1^st^ MG coordination shell missing (see Notes) |
| MG:1H:3257 | 1.00 | 40.13 |  | 2 | 2 | O_ph_O_b_ - Exclusion zone contact(s): /N6/O3'/ |
| MG:1H:3258 | 1.00 | 52.48 |  | 0 |  | 1^st^ MG coordination shell missing (see Notes) |
| MG:1H:3259 | 1.00 | 46.30 |  | 1 |  | O_b_ |
| MG:1H:3260 | 1.00 | 24.57 |  | 0 | 5 | - Exclusion zone contact(s): /O_w_/O_w_/O_w_/OP1/OP1/ Consider K in place of MG (2 valid contact(s) for K vs. 0 for MG) |
| MG:1H:3261 | 1.00 | 44.15 |  | 3 |  | O_ph_2O_w_ |
| MG:1H:3262 | 1.00 | 46.03 |  | 0 |  | 1^st^ MG coordination shell missing (see Notes) |
| MG:1H:3263 | 1.00 | 36.17 |  | 0 | 4 | - Exclusion zone contact(s): /O_w_/O_w_/OP2/OP2/ Consider K in place of MG (2 valid contact(s) for K vs. 0 for MG) |
| MG:1H:3264 | 1.00 | 55.11 |  | 0 | 3 | - Exclusion zone contact(s): /N7/O6/OP2/ Consider K in place of MG (2 valid contact(s) for K vs. 0 for MG) |
| MG:1H:3265 | 1.00 | 48.30 |  | 0 | 4 | - Exclusion zone contact(s): /O3'/O_w_/OP1/OP1/ Consider K in place of MG (3 valid contact(s) for K vs. 0 for MG) |
| MG:1H:3266 | 1.00 | 39.52 |  | 0 | 2 | - Exclusion zone contact(s): /O_w_/O_w_/ 1^st^ MG coordination shell missing (see Notes) |
| MG:1H:3267 | 1.00 | 53.96 |  | 0 | 4 | - Exclusion zone contact(s): /N3_U_/O2/OP1/OP2/ Consider K in place of MG (3 valid contact(s) for K vs. 0 for MG) |
| MG:1H:3268 | 1.00 | 34.84 |  | 1 | 1 | O_w_ - Exclusion zone contact(s): /O_w_/ |
| MG:1H:3269 | 1.00 | 33.07 |  | 1 | 4 | O_ph_ - Exclusion zone contact(s): /O5'/O_w_/OP2/OP2/ Consider K in place of MG (3 valid contact(s) for K vs. 1 for MG) |
| MG:1H:3270 | 1.00 | 42.51 |  | 0 |  | 1^st^ MG coordination shell missing (see Notes) |
| MG:1H:3271 | 1.00 | 47.86 |  | 0 |  | 1^st^ MG coordination shell missing (see Notes) |
| MG:1H:3272 | 1.00 | 55.18 |  | 0 |  | 1^st^ MG coordination shell missing (see Notes) |
| MG:1H:3273 | 1.00 | 61.62 |  | 0 | 2 | - Exclusion zone contact(s): /N7/O6/ Consider K in place of MG (2 valid contact(s) for K vs. 0 for MG) |
| MG:1H:3274 | 1.00 | 31.92 |  | 0 | 4 | - Exclusion zone contact(s): /O_w_/O_w_/OP2/OP2/ Consider K in place of MG (2 valid contact(s) for K vs. 0 for MG) |
| MG:1H:3275 | 1.00 | 44.86 |  | 0 | 1 | - Exclusion zone contact(s): /OP1/ Consider K in place of MG (1 valid contact(s) for K vs. 0 for MG) |
| MG:1H:3276 | 1.00 | 44.72 |  | 0 | 1 | - Exclusion zone contact(s): /OP1/ Consider K in place of MG (1 valid contact(s) for K vs. 0 for MG) |
| MG:1H:3277 | 1.00 | 42.71 |  | 0 |  | 1^st^ MG coordination shell missing (see Notes) |
| MG:1H:3278 | 1.00 | 28.55 |  | 0 | 3 | - Exclusion zone contact(s): /O5'/O_w_/OP1/ Consider K in place of MG (2 valid contact(s) for K vs. 0 for MG) |
| MG:1H:3279 | 1.00 | 32.47 |  | 0 | 5 | - Exclusion zone contact(s): /N6/N7/O_w_/O_w_/OP2/ Consider K in place of MG (3 valid contact(s) for K vs. 0 for MG) |
| MG:1H:3280 | 1.00 | 86.30 |  | 0 | 3 | - Exclusion zone contact(s): /O3'/OP1/OP2/ Consider K in place of MG (2 valid contact(s) for K vs. 0 for MG) |
| MG:1H:3281 | 1.00 | 49.33 |  | 0 | 2 | - Exclusion zone contact(s): /OP1/OP2/ Consider K in place of MG (2 valid contact(s) for K vs. 0 for MG) |
| MG:1H:3282 | 1.00 | 49.16 |  | 0 |  | 1^st^ MG coordination shell missing (see Notes) |
| MG:1H:3283 | 1.00 | 44.22 |  | 0 |  | 1^st^ MG coordination shell missing (see Notes) |
| MG:1H:3284 | 1.00 | 36.02 |  | 1 | 3 | O_w_ - Exclusion zone contact(s): /O5'/O_w_/OP1/ Consider K in place of MG (2 valid contact(s) for K vs. 0 for MG) |
| MG:1H:3285 | 1.00 | 43.85 |  | 0 | 1 | - Exclusion zone contact(s): /OP2/ Consider K in place of MG (1 valid contact(s) for K vs. 0 for MG) |
| MG:1H:3286 | 1.00 | 48.88 |  | 0 | 3 | - Exclusion zone contact(s): /O4/O6/OP2/ Consider K in place of MG (3 valid contact(s) for K vs. 0 for MG) |
| MG:1H:3287 | 1.00 | 37.67 |  | 0 |  | 1^st^ MG coordination shell missing (see Notes) |
| MG:1H:3288 | 1.00 | 44.82 |  | 0 |  | 1^st^ MG coordination shell missing (see Notes) |
| MG:1H:3289 | 1.00 | 35.64 |  | 0 |  | 1^st^ MG coordination shell missing (see Notes) |
| MG:1H:3290 | 1.00 | 38.40 |  | 0 | 2 | - Exclusion zone contact(s): /O2'/OP2/ Consider K in place of MG (2 valid contact(s) for K vs. 0 for MG) |
| MG:1H:3291 | 1.00 | 47.59 |  | 0 | 2 | - Exclusion zone contact(s): /OP2/OP2/ Consider K in place of MG (2 valid contact(s) for K vs. 0 for MG) |
| MG:1H:3292 | 1.00 | 32.65 |  | 0 | 2 | - Exclusion zone contact(s): /O6/O_w_/ Consider K in place of MG (1 valid contact(s) for K vs. 0 for MG) |
| MG:1H:3293 | 1.00 | 40.91 |  | 0 |  | 1^st^ MG coordination shell missing (see Notes) |
| MG:1H:3294 | 1.00 | 59.32 |  | 0 | 1 | - Exclusion zone contact(s): /O_w_/ 1^st^ MG coordination shell missing (see Notes) |
| MG:1H:3295 | 1.00 | 29.53 |  | 0 | 1 | - Exclusion zone contact(s): /OP1/ Consider K in place of MG (1 valid contact(s) for K vs. 0 for MG) |
| MG:1H:3296 | 1.00 | 32.27 |  | 0 |  | 1^st^ MG coordination shell missing (see Notes) |
| MG:1H:3297 | 1.00 | 40.33 |  | 0 | 2 | - Exclusion zone contact(s): /N7/OP2/ Consider K in place of MG (2 valid contact(s) for K vs. 0 for MG) |
| MG:1H:3298 | 1.00 | 28.41 |  | 0 | 3 | - Exclusion zone contact(s): /NZ_Lys_/O2'/O3'/ Consider K in place of MG (2 valid contact(s) for K vs. 0 for MG) |
| MG:1H:3299 | 1.00 | 47.60 |  | 0 | 2 | - Exclusion zone contact(s): /OP1/OP2/ Consider K in place of MG (2 valid contact(s) for K vs. 0 for MG) |
| MG:1H:3300 | 1.00 | 59.51 |  | 0 | 1 | - Exclusion zone contact(s): /OP2/ Consider K in place of MG (1 valid contact(s) for K vs. 0 for MG) |
| MG:1H:3301 | 1.00 | 41.40 |  | 0 |  | 1^st^ MG coordination shell missing (see Notes) |
| MG:1H:3302 | 1.00 | 29.06 |  | 0 | 4 | - Exclusion zone contact(s): /O2'/O3'/O_w_/OP1/ Consider K in place of MG (3 valid contact(s) for K vs. 0 for MG) |
| MG:1H:3303 | 1.00 | 46.67 |  | 0 |  | 1^st^ MG coordination shell missing (see Notes) |
| MG:1H:3304 | 1.00 | 38.84 |  | 0 |  | 1^st^ MG coordination shell missing (see Notes) |
| MG:1H:3305 | 1.00 | 41.66 |  | 0 |  | 1^st^ MG coordination shell missing (see Notes) |
| MG:1H:3306 | 1.00 | 34.50 |  | 0 | 5 | - Exclusion zone contact(s): /N4/N7/O_w_/OP2/OP2/ Consider K in place of MG (4 valid contact(s) for K vs. 0 for MG) |
| MG:1H:3307 | 1.00 | 25.95 |  | 0 | 2 | - Exclusion zone contact(s): /O_w_/O_w_/ 1^st^ MG coordination shell missing (see Notes) |
| MG:1H:3308 | 1.00 | 27.17 |  | 0 | 2 | - Exclusion zone contact(s): /O3'/OP1/ Consider K in place of MG (2 valid contact(s) for K vs. 0 for MG) |
| MG:1H:3309 | 1.00 | 52.87 |  | 1 | 2 | O_w_ - Exclusion zone contact(s): /O_w_/O_w_/ |
| MG:1H:3310 | 1.00 | 18.98 |  | 1 | 2 | O_ph_ - Exclusion zone contact(s): /O_w_/OP2/ |
| MG:1H:3311 | 1.00 | 47.50 |  | 0 | 5 | - Exclusion zone contact(s): /O6/O6/O_w_/O_w_/OP1/ Consider K in place of MG (3 valid contact(s) for K vs. 0 for MG) |
| MG:1H:3312 | 1.00 | 39.80 |  | 0 | 2 | - Exclusion zone contact(s): /O_w_/OP2/ Consider K in place of MG (1 valid contact(s) for K vs. 0 for MG) |
| MG:1H:3313 | 1.00 | 44.89 |  | 1 |  | O_w_ |
| MG:1H:3314 | 1.00 | 41.45 |  | 0 | 1 | - Exclusion zone contact(s): /O_w_/ 1^st^ MG coordination shell missing (see Notes) |
| MG:1H:3315 | 1.00 | 43.67 |  | 0 |  | 1^st^ MG coordination shell missing (see Notes) |
| MG:1H:3316 | 1.00 | 60.58 |  | 0 | 1 | - Exclusion zone contact(s): /O_w_/ 1^st^ MG coordination shell missing (see Notes) |
| MG:1H:3317 | 1.00 | 52.18 |  | 0 |  | 1^st^ MG coordination shell missing (see Notes) |
| MG:1H:3318 | 1.00 | 56.14 |  | 1 | 1 | O_b_ - Exclusion zone contact(s): /O_w_/ |
| MG:1H:3319 | 1.00 | 50.87 |  | 0 | 2 | - Exclusion zone contact(s): /O_w_/OP1/ Consider K in place of MG (1 valid contact(s) for K vs. 0 for MG) |
| MG:1H:3320 | 0.50 | 8.94 |  | 0 | 2 | - Exclusion zone contact(s): /O2'/O4'/ Consider K in place of MG (2 valid contact(s) for K vs. 0 for MG) |
| MG:1H:3320 | 0.50 | 9.02 |  | 0 |  | 1^st^ MG coordination shell missing (see Notes) |
| MG:1H:3321 | 1.00 | 36.81 |  | 0 |  | 1^st^ MG coordination shell missing (see Notes) |
| MG:1H:3322 | 1.00 | 23.61 |  | 0 | 2 | - Exclusion zone contact(s): /OP1/OP2/ Consider NA in place of MG (1 valid contact(s) for NA vs. 0 for MG) |
| MG:1H:3323 | 1.00 | 52.52 |  | 0 | 1 | - Exclusion zone contact(s): /O6/ Consider K in place of MG (1 valid contact(s) for K vs. 0 for MG) |
| MG:1H:3324 | 1.00 | 12.36 |  | 3 | 2 | 3O_ph_ - Exclusion zone contact(s): /O5'/OP1/ |
| MG:1H:3325 | 1.00 | 50.07 |  | 0 | 3 | - Exclusion zone contact(s): /OP1/OP2/OP2/ Consider K in place of MG (3 valid contact(s) for K vs. 0 for MG) |
| MG:1H:3326 | 1.00 | 60.44 |  | 0 |  | 1^st^ MG coordination shell missing (see Notes) |
| MG:1H:3327 | 1.00 | 40.26 |  | 0 | 3 | - Exclusion zone contact(s): /O4/O4/OP2/ Consider K in place of MG (3 valid contact(s) for K vs. 0 for MG) |
| MG:1H:3328 | 0.50 | 22.52 |  | 0 | 2 | - Exclusion zone contact(s): /O4/O6/ Consider NA in place of MG (1 valid contact(s) for NA vs. 0 for MG) |
| MG:1H:3328 | 0.50 | 27.38 |  | 0 | 2 | - Exclusion zone contact(s): /N1_A_/O6/ Consider K in place of MG (2 valid contact(s) for K vs. 0 for MG) |
| MG:1H:3329 | 1.00 | 63.10 |  | 0 | 2 | - Exclusion zone contact(s): /O6/O_w_/ Consider NA in place of MG (1 valid contact(s) for NA vs. 0 for MG) |
| MG:1H:3330 | 1.00 | 45.91 |  | 0 | 2 | - Exclusion zone contact(s): /N7/O4/ Consider NA in place of MG (1 valid contact(s) for NA vs. 0 for MG) |
| MG:1H:3331 | 1.00 | 54.95 |  | 0 | 3 | - Exclusion zone contact(s): /O6/O_w_/O_w_/ Consider K in place of MG (1 valid contact(s) for K vs. 0 for MG) |
| MG:1H:3332 | 1.00 | 48.00 |  | 0 | 3 | - Exclusion zone contact(s): /O_w_/O_w_/O_w_/ 1^st^ MG coordination shell missing (see Notes) |
| MG:1H:3333 | 1.00 | 50.85 |  | 0 | 2 | - Exclusion zone contact(s): /O_w_/OP2/ Consider K in place of MG (1 valid contact(s) for K vs. 0 for MG) |
| MG:1H:3334 | 1.00 | 43.98 |  | 0 | 2 | - Exclusion zone contact(s): /ND1_His_/OP1/ Consider K in place of MG (2 valid contact(s) for K vs. 0 for MG) |
| MG:1H:3335 | 1.00 | 40.77 |  | 0 |  | 1^st^ MG coordination shell missing (see Notes) |
| MG:1H:3336 | 1.00 | 45.88 |  | 0 |  | 1^st^ MG coordination shell missing (see Notes) |
| MG:1H:3337 | 1.00 | 23.20 |  | 0 |  | 1^st^ MG coordination shell missing (see Notes) |
| MG:1H:3338 | 1.00 | 43.36 |  | 0 |  | 1^st^ MG coordination shell missing (see Notes) |
| MG:1H:3339 | 1.00 | 55.37 |  | 0 |  | 1^st^ MG coordination shell missing (see Notes) |
| MG:1H:3340 | 1.00 | 48.29 |  | 0 | 1 | - Exclusion zone contact(s): /N7/ Consider K in place of MG (1 valid contact(s) for K vs. 0 for MG) |
| MG:1H:3341 | 1.00 | 24.92 |  | 0 | 1 | - Exclusion zone contact(s): /O6/ Consider K in place of MG (1 valid contact(s) for K vs. 0 for MG) |
| MG:1H:3342 | 1.00 | 27.95 |  | 0 |  | 1^st^ MG coordination shell missing (see Notes) |
| MG:1H:3343 | 0.50 | 55.96 |  | 0 |  | 1^st^ MG coordination shell missing (see Notes) |
| MG:1H:3343 | 0.50 | 68.21 |  | 0 |  | 1^st^ MG coordination shell missing (see Notes) |
| MG:1H:3344 | 1.00 | 39.26 |  | 0 | 1 | - Exclusion zone contact(s): /OP2/ Consider K in place of MG (1 valid contact(s) for K vs. 0 for MG) |
| MG:1H:3345 | 0.50 | 23.09 |  | 0 | 2 | - Exclusion zone contact(s): /N7/OP2/ Consider K in place of MG (2 valid contact(s) for K vs. 0 for MG) |
| MG:1H:3345 | 0.50 | 24.06 |  | 0 | 3 | - Exclusion zone contact(s): /O_w_/OP2/OP2/ Consider K in place of MG (2 valid contact(s) for K vs. 0 for MG) |
| MG:1H:3346 | 1.00 | 55.33 |  | 0 |  | 1^st^ MG coordination shell missing (see Notes) |
| MG:1H:3347 | 1.00 | 38.40 |  | 0 |  | 1^st^ MG coordination shell missing (see Notes) |
| MG:1H:3348 | 1.00 | 30.67 |  | 0 | 2 | - Exclusion zone contact(s): /N7/O_w_/ Consider K in place of MG (1 valid contact(s) for K vs. 0 for MG) |
| MG:1H:3349 | 1.00 | 48.49 |  | 0 | 3 | - Exclusion zone contact(s): /O4/O6/OP2/ Consider K in place of MG (3 valid contact(s) for K vs. 0 for MG) |
| MG:1H:3350 | 1.00 | 25.47 |  | 3 |  | 2O_ph_O_b_ |
| MG:1H:3351 | 1.00 | 79.08 |  | 0 | 3 | - Exclusion zone contact(s): /O4/O6/O6/ Consider K in place of MG (2 valid contact(s) for K vs. 0 for MG) |
| MG:1H:3352 | 1.00 | 43.03 |  | 0 | 1 | - Exclusion zone contact(s): /N7/ Consider K in place of MG (1 valid contact(s) for K vs. 0 for MG) |
| MG:1H:3353 | 1.00 | 33.41 |  | 0 |  | 1^st^ MG coordination shell missing (see Notes) |
| MG:1H:3354 | 1.00 | 29.95 |  | 0 | 2 | - Exclusion zone contact(s): /O2/O2'/ Consider K in place of MG (2 valid contact(s) for K vs. 0 for MG) |
| MG:1H:3355 | 1.00 | 57.33 |  | 0 | 6 | - Exclusion zone contact(s): /O5'/O_w_/O_w_/O_w_/OP2/OP2/ Consider K in place of MG (3 valid contact(s) for K vs. 0 for MG) |
| MG:1H:3356 | 1.00 | 43.71 |  | 0 | 2 | - Exclusion zone contact(s): /N6/O2/ Consider K in place of MG (2 valid contact(s) for K vs. 0 for MG) |
| MG:1H:3357 | 1.00 | 28.78 |  | 0 | 2 | - Exclusion zone contact(s): /OP2/OP2/ Consider NA in place of MG (1 valid contact(s) for NA vs. 0 for MG) |
| MG:1H:3358 | 1.00 | 41.64 |  | 0 |  | 1^st^ MG coordination shell missing (see Notes) |
| MG:1H:3359 | 1.00 | 62.40 |  | 0 | 1 | - Exclusion zone contact(s): /O6/ Consider K in place of MG (1 valid contact(s) for K vs. 0 for MG) |
| MG:1H:3360 | 1.00 | 28.95 |  | 1 | 1 | O_ph_ - Exclusion zone contact(s): /OP1/ |
| MG:1H:3361 | 1.00 | 57.26 |  | 0 | 2 | - Exclusion zone contact(s): /O2'/O3'/ Consider K in place of MG (2 valid contact(s) for K vs. 0 for MG) |
| MG:1H:3362 | 1.00 | 64.67 |  | 0 |  | 1^st^ MG coordination shell missing (see Notes) |
| MG:1H:3363 | 1.00 | 56.45 |  | 0 | 1 | - Exclusion zone contact(s): /O6/ Consider K in place of MG (1 valid contact(s) for K vs. 0 for MG) |
| MG:1H:3364 | 1.00 | 58.51 |  | 0 |  | 1^st^ MG coordination shell missing (see Notes) |
| MG:1H:3365 | 1.00 | 58.96 |  | 0 | 1 | - Exclusion zone contact(s): /O6/ Consider K in place of MG (1 valid contact(s) for K vs. 0 for MG) |
| MG:1H:3366 | 1.00 | 40.91 |  | 0 | 2 | - Exclusion zone contact(s): /O2'/O4'/ Consider K in place of MG (2 valid contact(s) for K vs. 0 for MG) |
| MG:1H:3367 | 1.00 | 35.85 |  | 0 | 2 | - Exclusion zone contact(s): /O4/OP2/ Consider K in place of MG (2 valid contact(s) for K vs. 0 for MG) |
| MG:1H:3368 | 1.00 | 44.78 |  | 0 | 1 | - Exclusion zone contact(s): /OP2/ Consider NA in place of MG (1 valid contact(s) for NA vs. 0 for MG) |
| MG:1H:3369 | 1.00 | 46.41 |  | 0 | 1 | - Exclusion zone contact(s): /O6/ Consider K in place of MG (1 valid contact(s) for K vs. 0 for MG) |
| MG:1H:3370 | 1.00 | 61.73 |  | 0 |  | 1^st^ MG coordination shell missing (see Notes) |
| MG:1H:3371 | 1.00 | 31.09 |  | 0 | 3 | - Exclusion zone contact(s): /N7/O2'/O3'/ Consider K in place of MG (3 valid contact(s) for K vs. 0 for MG) |
| MG:1H:3372 | 1.00 | 31.43 |  | 0 | 1 | - Exclusion zone contact(s): /O2'/ Consider K in place of MG (1 valid contact(s) for K vs. 0 for MG) |
| MG:1H:3373 | 1.00 | 43.93 |  | 0 | 1 | - Exclusion zone contact(s): /O6/ Consider K in place of MG (1 valid contact(s) for K vs. 0 for MG) |
| MG:1H:3374 | 1.00 | 28.06 |  | 2 | 2 | O_ph_O_w_ - Exclusion zone contact(s): /O_w_/OP1/ |
| MG:1H:3375 | 1.00 | 36.73 |  | 0 | 1 | - Exclusion zone contact(s): /OP2/ Consider K in place of MG (1 valid contact(s) for K vs. 0 for MG) |
| MG:1H:3376 | 1.00 | 54.75 |  | 0 | 1 | - Exclusion zone contact(s): /OP1/ Consider K in place of MG (1 valid contact(s) for K vs. 0 for MG) |
| MG:1H:3377 | 1.00 | 26.37 |  | 0 |  | 1^st^ MG coordination shell missing (see Notes) |
| MG:1H:3378 | 1.00 | 28.95 |  | 0 | 4 | - Exclusion zone contact(s): /O2'/O3'/O_w_/OP2/ Consider K in place of MG (3 valid contact(s) for K vs. 0 for MG) |
| MG:1H:3379 | 1.00 | 47.89 |  | 0 | 1 | - Exclusion zone contact(s): /OP1/ Consider K in place of MG (1 valid contact(s) for K vs. 0 for MG) |
| MG:1H:3380 | 1.00 | 44.48 |  | 0 | 1 | - Exclusion zone contact(s): /OP2/ Consider K in place of MG (1 valid contact(s) for K vs. 0 for MG) |
| MG:1H:3381 | 1.00 | 39.99 |  | 0 | 1 | - Exclusion zone contact(s): /O2/ Consider K in place of MG (1 valid contact(s) for K vs. 0 for MG) |
| MG:1H:3382 | 1.00 | 13.33 |  | 5 | 1 | 5O_w_ - Exclusion zone contact(s): /OP1/ Consider NA in place of MG (1 valid contact(s) for NA vs. 0 for MG) |
| MG:1H:3383 | 1.00 | 18.26 |  | 4 | 2 | 4O_w_ - Exclusion zone contact(s): /OP2/OP2/ Consider NA in place of MG (2 valid contact(s) for NA vs. 0 for MG) |
| MG:1H:3384 | 1.00 | 11.68 |  | 6 | 2 | 6O_w_ - Exclusion zone contact(s): /OP1/OP1/ Consider K in place of MG (2 valid contact(s) for K vs. 0 for MG) |
| MG:1H:3385 | 1.00 | 20.15 |  | 6 | 1 | 6O_w_ - Exclusion zone contact(s): /OP2/ Consider K in place of MG (1 valid contact(s) for K vs. 0 for MG) |
| MG:1H:3386 | 1.00 | 31.80 |  | 4 | 3 | 4O_w_ - Exclusion zone contact(s): /O5'/OP2/OP2/ Consider NA in place of MG (2 valid contact(s) for NA vs. 0 for MG) |
| MG:1H:3387 | 1.00 | 16.11 |  | 6 | 1 | 6O_w_ - Exclusion zone contact(s): /OP2/ Consider K in place of MG (1 valid contact(s) for K vs. 0 for MG) |
| MG:1H:3388 | 1.00 | 15.28 |  | 5 | 1 | 5O_w_ - Exclusion zone contact(s): /O4/ Consider NA in place of MG (1 valid contact(s) for NA vs. 0 for MG) |
| MG:1H:3389 | 1.00 | 13.51 |  | 4 | 3 | 4O_w_ - Exclusion zone contact(s): /NE2_His_/O5'/OP1/ Consider K in place of MG (2 valid contact(s) for K vs. 0 for MG) |
| MG:1H:3390 | 1.00 | 26.24 |  | 6 | 1 | 6O_w_ - Exclusion zone contact(s): /OP2/ Consider K in place of MG (1 valid contact(s) for K vs. 0 for MG) |
| MG:1H:3391 | 1.00 | 20.00 |  | 6 | 2 | O_ph_5O_w_ - Exclusion zone contact(s): /O5'/OP1/ Consider K in place of MG (2 valid contact(s) for K vs. 1 for MG) |
| MG:1H:3392 | 1.00 | 51.19 |  | 6 | 1 | 6O_w_ - Exclusion zone contact(s): /OP1/ Consider K in place of MG (1 valid contact(s) for K vs. 0 for MG) |
| MG:1H:3393 | 1.00 | 17.02 |  | 6 |  | O_ph_5O_w_ |
| MG:1H:3394 | 1.00 | 11.72 |  | 6 | 1 | 6O_w_ - Exclusion zone contact(s): /OP1/ Consider K in place of MG (1 valid contact(s) for K vs. 0 for MG) |
| MG:1H:3395 | 1.00 | 23.11 |  | 6 |  | 6O_w_ |
| MG:1H:3396 | 1.00 | 16.42 |  | 6 |  | 6O_w_ |
| MG:1H:3397 | 1.00 | 17.48 |  | 5 | 1 | 5O_w_ - Exclusion zone contact(s): /OP2/ Consider NA in place of MG (1 valid contact(s) for NA vs. 0 for MG) |
| MG:1H:3398 | 1.00 | 52.68 |  | 6 |  | 6O_w_ |
| MG:1H:3399 | 1.00 | 24.11 |  | 6 | 2 | 6O_w_ - Exclusion zone contact(s): /OP1/OP2/ Consider K in place of MG (2 valid contact(s) for K vs. 0 for MG) |
| MG:1H:3400 | 1.00 | 20.32 |  | 4 | 2 | 4O_w_ - Exclusion zone contact(s): /O6/OP1/ Consider NA in place of MG (2 valid contact(s) for NA vs. 0 for MG) |
| MG:1H:3401 | 1.00 | 16.00 |  | 6 |  | 6O_w_ |
| MG:1H:3402 | 1.00 | 23.67 |  | 6 |  | 6O_w_ |
| MG:1H:3403 | 1.00 | 27.38 |  | 6 |  | 6O_w_ |
| MG:1H:3404 | 1.00 | 43.55 |  | 6 |  | 6O_w_ |
| MG:1H:3405 | 1.00 | 16.64 |  | 6 | 1 | 6O_w_ - Exclusion zone contact(s): /OP1/ Consider K in place of MG (1 valid contact(s) for K vs. 0 for MG) |
| MG:1H:3406 | 1.00 | 14.78 |  | 6 |  | 6O_w_ |
| MG:1H:3407 | 1.00 | 16.59 |  | 6 | 1 | 6O_w_ - Exclusion zone contact(s): /OP2/ Consider K in place of MG (1 valid contact(s) for K vs. 0 for MG) |
| MG:1H:3408 | 1.00 | 29.29 |  | 6 |  | 6O_w_ |
| MG:1H:3409 | 1.00 | 14.54 |  | 4 | 2 | 4O_w_ - Exclusion zone contact(s): /N7/N7/ Consider NA in place of MG (2 valid contact(s) for NA vs. 0 for MG) |
| MG:1H:3410 | 1.00 | 16.18 |  | 6 |  | 6O_w_ |
| MG:1H:3411 | 1.00 | 20.14 |  | 6 | 2 | 6O_w_ - Exclusion zone contact(s): /OP2/OP2/ Consider K in place of MG (2 valid contact(s) for K vs. 0 for MG) |
| MG:1H:3412 | 1.00 | 16.25 |  | 5 | 1 | 5O_w_ - Exclusion zone contact(s): /OP1/ Consider NA in place of MG (1 valid contact(s) for NA vs. 0 for MG) |
| MG:1H:3413 | 1.00 | 49.12 |  | 6 | 2 | 6O_w_ - Exclusion zone contact(s): /N7/OP2/ Consider K in place of MG (2 valid contact(s) for K vs. 0 for MG) |
| MG:1H:3414 | 1.00 | 39.03 |  | 6 |  | 6O_w_ |
| MG:1H:3415 | 1.00 | 16.90 |  | 6 |  | O_ph_5O_w_ |
| MG:1H:3416 | 1.00 | 69.11 |  | 6 |  | 6O_w_ Possible idealized distances ! |
| MG:1H:3417 | 1.00 | 61.68 |  | 6 | 1 | 6O_w_ - Exclusion zone contact(s): /N7/ Consider K in place of MG (1 valid contact(s) for K vs. 0 for MG) |
| MG:1H:3418 | 1.00 | 29.09 |  | 6 |  | 6O_w_ |
| MG:1H:3419 | 1.00 | 27.53 |  | 6 | 2 | 6O_w_ - Exclusion zone contact(s): /OP1/OP1/ Consider K in place of MG (2 valid contact(s) for K vs. 0 for MG) |
| MG:1H:3420 | 1.00 | 20.48 |  | 4 | 3 | 4O_w_ - Exclusion zone contact(s): /OP1/OP1/OP2/ Consider K in place of MG (2 valid contact(s) for K vs. 0 for MG) |
| MG:1H:3421 | 1.00 | 36.78 |  | 6 |  | 6O_w_ |
| MG:1H:3422 | 1.00 | 29.01 |  | 4 | 2 | 4O_w_ - Exclusion zone contact(s): /OP1/OP1/ Consider NA in place of MG (2 valid contact(s) for NA vs. 0 for MG) |
| MG:1H:3423 | 1.00 | 35.39 |  | 4 | 2 | 4O_w_ - Exclusion zone contact(s): /OP2/OP2/ Consider NA in place of MG (1 valid contact(s) for NA vs. 0 for MG) |
| MG:1H:3424 | 1.00 | 42.85 |  | 3 | 4 | 3O_w_ - Exclusion zone contact(s): /O5'/OP1/OP1/OP2/ Consider K in place of MG (3 valid contact(s) for K vs. 0 for MG) |
| MG:1H:3425 | 1.00 | 16.24 |  | 6 |  | 6O_w_ |
| MG:1H:3426 | 1.00 | 12.50 |  | 4 | 2 | 4O_w_ - Exclusion zone contact(s): /OP1/OP1/ Consider NA in place of MG (1 valid contact(s) for NA vs. 0 for MG) |
| MG:1H:3427 | 1.00 | 36.51 |  | 6 |  | 6O_w_ |
| MG:1H:3428 | 1.00 | 23.34 |  | 4 | 3 | 4O_w_ - Exclusion zone contact(s): /O5'/OP1/OP1/ Consider K in place of MG (3 valid contact(s) for K vs. 0 for MG) |
| MG:1H:3429 | 1.00 | 29.95 |  | 4 | 2 | 4O_w_ - Exclusion zone contact(s): /O4/OP2/ Consider K in place of MG (2 valid contact(s) for K vs. 0 for MG) |
| MG:1H:3430 | 1.00 | 54.74 |  | 5 | 1 | 5O_w_ - Exclusion zone contact(s): /OP2/ Consider NA in place of MG (1 valid contact(s) for NA vs. 0 for MG)  Possible idealized distances ! |
| MG:1H:3431 | 1.00 | 46.69 |  | 6 |  | 6O_w_ |
| MG:1H:3432 | 1.00 | 55.09 |  | 6 |  | 6O_w_ |
| MG:1H:3433 | 1.00 | 51.44 |  | 6 | 1 | 6O_w_ - Exclusion zone contact(s): /OP2/ Consider K in place of MG (1 valid contact(s) for K vs. 0 for MG)  Possible idealized distances ! |
| MG:1H:3434 | 1.00 | 46.89 |  | 6 |  | 6O_w_ |
| MG:1H:3435 | 1.00 | 46.29 |  | 6 |  | 6O_w_ |
| MG:1H:3436 | 1.00 | 57.23 |  | 6 |  | 6O_w_ |
| MG:1H:3437 | 1.00 | 51.30 |  | 6 |  | 6O_w_ |
| MG:1H:3438 | 1.00 | 74.52 |  | 6 | 1 | 6O_w_ - Exclusion zone contact(s): /OP1/ Consider K in place of MG (1 valid contact(s) for K vs. 0 for MG) |
| MG:1H:3439 | 1.00 | 53.02 |  | 6 |  | 6O_w_ |
| MG:1H:3440 | 1.00 | 62.00 |  | 6 |  | 6O_w_ Possible idealized distances ! |
| MG:1H:3441 | 1.00 | 61.50 |  | 6 |  | 6O_w_ Possible idealized distances ! |
| MG:1H:3442 | 1.00 | 56.17 |  | 6 |  | 6O_w_ |
| MG:1H:3443 | 1.00 | 42.31 |  | 6 |  | 6O_w_ |
| MG:1H:3444 | 1.00 | 23.03 |  | 6 |  | 6O_w_ |
| MG:1H:3445 | 1.00 | 50.89 |  | 6 |  | 6O_w_ |
| MG:1H:3446 | 1.00 | 48.82 |  | 6 |  | 6O_w_ |
| MG:1H:3447 | 1.00 | 19.06 |  | 6 |  | 6O_w_ |
| MG:1H:3448 | 1.00 | 57.33 |  | 6 |  | 6O_w_ |
| MG:1H:3449 | 1.00 | 15.11 |  | 6 | 1 | 6O_w_ - Exclusion zone contact(s): /OP2/ Consider K in place of MG (1 valid contact(s) for K vs. 0 for MG) |
| MG:1H:3450 | 1.00 | 52.62 |  | 6 |  | 6O_w_ |
| MG:1H:3451 | 1.00 | 82.05 |  | 6 |  | 6O_w_ |
| MG:1H:3452 | 1.00 | 68.79 |  | 6 |  | 6O_w_ |
| MG:1H:3453 | 1.00 | 20.70 |  | 6 | 1 | 6O_w_ - Exclusion zone contact(s): /OP2/ Consider K in place of MG (1 valid contact(s) for K vs. 0 for MG) |
| MG:1H:3454 | 1.00 | 23.38 |  | 6 | 2 | 6O_w_ - Exclusion zone contact(s): /OP1/OP2/ Consider K in place of MG (2 valid contact(s) for K vs. 0 for MG) |
| MG:1H:3455 | 1.00 | 11.67 |  | 3 | 4 | 3O_w_ - Exclusion zone contact(s): /OP1/OP1/OP2/OP2/ Consider NA in place of MG (2 valid contact(s) for NA vs. 0 for MG) |
| MG:1H:3456 | 1.00 | 14.32 |  | 5 | 2 | O_ph_4O_w_ - Exclusion zone contact(s): /OP1/OP2/ Consider K in place of MG (2 valid contact(s) for K vs. 1 for MG) |
| MG:1H:3457 | 1.00 | 25.00 |  | 3 | 4 | 3O_w_ - Exclusion zone contact(s): /OP1/OP1/OP2/OP2/ Consider NA in place of MG (2 valid contact(s) for NA vs. 0 for MG) |
| MG:1H:3458 | 1.00 | 21.71 |  | 4 | 2 | 4O_w_ - Exclusion zone contact(s): /OP2/OP2/ Consider NA in place of MG (2 valid contact(s) for NA vs. 0 for MG) |
| MG:1H:3459 | 1.00 | 15.67 |  | 6 |  | O_ph_5O_w_ Large octahedral topology discrepancy!!! |
| MG:1H:3460 | 1.00 | 11.21 |  | 5 | 1 | 5O_w_ - Exclusion zone contact(s): /OP2/ Consider NA in place of MG (1 valid contact(s) for NA vs. 0 for MG) |
| MG:1H:3461 | 1.00 | 20.17 |  | 6 |  | O_ph_5O_w_ |
| MG:1H:3462 | 1.00 | 27.03 |  | 6 | 1 | 6O_w_ - Exclusion zone contact(s): /OP1/ Consider NA in place of MG (1 valid contact(s) for NA vs. 0 for MG) |
| MG:1H:3463 | 1.00 | 17.66 |  | 6 |  | O_ph_5O_w_ |
| MG:1H:3464 | 1.00 | 23.36 |  | 3 | 4 | 3O_w_ - Exclusion zone contact(s): /OP1/OP1/OP2/OP2/ Consider K in place of MG (3 valid contact(s) for K vs. 0 for MG) |
| MG:1H:3465 | 1.00 | 21.32 |  | 5 | 1 | 5O_w_ - Exclusion zone contact(s): /N7/ Consider NA in place of MG (1 valid contact(s) for NA vs. 0 for MG) |
| MG:1H:3466 | 1.00 | 22.91 |  | 6 | 1 | 6O_w_ - Exclusion zone contact(s): /OP2/ Consider K in place of MG (1 valid contact(s) for K vs. 0 for MG) |
| MG:1H:3467 | 1.00 | 44.20 |  | 6 |  | 6O_w_ |
| MG:1H:3468 | 1.00 | 24.42 |  | 6 | 1 | 6O_w_ - Exclusion zone contact(s): /OP1/ Consider K in place of MG (1 valid contact(s) for K vs. 0 for MG) |
| MG:1H:3469 | 1.00 | 13.47 |  | 5 | 3 | 5O_w_ - Exclusion zone contact(s): /O3'/OP1/OP1/ Consider NA in place of MG (2 valid contact(s) for NA vs. 0 for MG) |
| MG:1H:3470 | 1.00 | 24.36 |  | 5 | 1 | 5O_w_ - Exclusion zone contact(s): /OP2/ Consider NA in place of MG (1 valid contact(s) for NA vs. 0 for MG) |
| MG:1H:3471 | 1.00 | 56.16 |  | 6 |  | 6O_w_ |
| MG:1H:3472 | 1.00 | 17.01 |  | 5 | 2 | 5O_w_ - Exclusion zone contact(s): /O_w_/OP1/ Consider NA in place of MG (1 valid contact(s) for NA vs. 0 for MG) |
| MG:1H:3473 | 1.00 | 19.40 |  | 4 | 2 | 4O_w_ - Exclusion zone contact(s): /O4/OP2/ Consider NA in place of MG (2 valid contact(s) for NA vs. 0 for MG) |
| MG:1H:3474 | 1.00 | 24.96 |  | 6 | 1 | O_ph_5O_w_ - Exclusion zone contact(s): /N7/ |
| MG:1H:3475 | 1.00 | 40.16 |  | 5 | 3 | 5O_w_ - Exclusion zone contact(s): /O6/OP1/OP2/ Consider K in place of MG (2 valid contact(s) for K vs. 0 for MG) |
| MG:1H:3476 | 1.00 | 15.73 |  | 6 | 1 | 6O_w_ - Exclusion zone contact(s): /OP2/ Consider K in place of MG (1 valid contact(s) for K vs. 0 for MG) |
| MG:1H:3477 | 1.00 | 28.25 |  | 4 | 2 | 4O_w_ - Exclusion zone contact(s): /OP2/OP2/ Consider NA in place of MG (1 valid contact(s) for NA vs. 0 for MG) |
| MG:1H:3478 | 1.00 | 44.77 |  | 6 | 1 | 6O_w_ - Exclusion zone contact(s): /OP1/ Consider K in place of MG (1 valid contact(s) for K vs. 0 for MG) |
| MG:1H:3479 | 1.00 | 18.60 |  | 5 | 1 | O_ph_4O_w_ - Exclusion zone contact(s): /OP1/ |
| MG:1H:3480 | 1.00 | 18.27 |  | 5 | 1 | 5O_w_ - Exclusion zone contact(s): /OP2/ Consider NA in place of MG (1 valid contact(s) for NA vs. 0 for MG) |
| MG:1H:3481 | 1.00 | 21.56 |  | 6 |  | O_ph_5O_w_ |
| MG:1H:3482 | 1.00 | 45.14 |  | 6 |  | 6O_w_ |
| MG:1H:3483 | 1.00 | 36.88 |  | 6 |  | 6O_w_ |
| MG:1H:3484 | 1.00 | 16.51 |  | 5 | 1 | O_b_4O_w_ - Exclusion zone contact(s): /OP1/ Large octahedral topology discrepancy!!! |
| MG:1H:3485 | 1.00 | 29.83 |  | 6 |  | 6O_w_ Possible idealized distances ! |
| MG:1H:3486 | 1.00 | 36.26 |  | 5 | 1 | 5O_w_ - Exclusion zone contact(s): /OP2/ Consider NA in place of MG (1 valid contact(s) for NA vs. 0 for MG) |
| MG:1H:3487 | 1.00 | 41.77 |  | 6 | 1 | 6O_w_ - Exclusion zone contact(s): /OP2/ Consider K in place of MG (1 valid contact(s) for K vs. 0 for MG) |
| MG:1H:3488 | 1.00 | 35.94 |  | 6 |  | 6O_w_ |
| MG:1H:3489 | 1.00 | 57.14 |  | 6 | 1 | 6O_w_ - Exclusion zone contact(s): /N7/ Consider K in place of MG (1 valid contact(s) for K vs. 0 for MG) |
| MG:1H:3490 | 1.00 | 32.37 |  | 5 | 1 | 5O_w_ - Exclusion zone contact(s): /OP2/ Consider NA in place of MG (1 valid contact(s) for NA vs. 0 for MG) |
| MG:1H:3491 | 1.00 | 42.97 |  | 6 | 1 | 6O_w_ - Exclusion zone contact(s): /OP1/ Consider K in place of MG (1 valid contact(s) for K vs. 0 for MG) |
| MG:1H:3492 | 1.00 | 63.87 |  | 6 |  | 6O_w_ |
| MG:1H:3493 | 1.00 | 27.17 |  | 6 |  | 6O_w_ |
| MG:1H:3494 | 1.00 | 43.72 |  | 6 |  | 6O_w_ |
| MG:1H:3495 | 1.00 | 48.17 |  | 6 |  | 6O_w_ |
| MG:1H:3496 | 1.00 | 48.29 |  | 6 |  | 6O_w_ |
| MG:1H:3497 | 1.00 | 55.46 |  | 6 |  | 6O_w_ |
| MG:1H:3498 | 1.00 | 24.71 |  | 6 | 1 | 6O_w_ - Exclusion zone contact(s): /OP2/ Consider K in place of MG (1 valid contact(s) for K vs. 0 for MG) |
| MG:1H:3499 | 1.00 | 32.74 |  | 6 |  | O_ph_5O_w_ |
| MG:1H:3500 | 1.00 | 63.92 |  | 4 | 2 | 4O_w_ - Exclusion zone contact(s): /OP2/OP2/ Consider K in place of MG (2 valid contact(s) for K vs. 0 for MG) |
| MG:1H:3501 | 1.00 | 57.41 |  | 6 |  | 6O_w_ |
| MG:1H:3502 | 1.00 | 25.62 |  | 6 | 1 | 6O_w_ - Exclusion zone contact(s): /OP2/ Consider K in place of MG (1 valid contact(s) for K vs. 0 for MG) |
| MG:1H:3503 | 1.00 | 29.08 |  | 4 | 2 | 4O_w_ - Exclusion zone contact(s): /OP1/OP1/ Consider NA in place of MG (1 valid contact(s) for NA vs. 0 for MG) |
| MG:1H:3504 | 1.00 | 20.74 |  | 4 | 3 | O_ph_3O_w_ - Exclusion zone contact(s): /O3'/OP2/OP2/ Large octahedral topology discrepancy!!!  Consider NA in place of MG (2 valid contact(s) for NA vs. 1 for MG) |
| MG:1H:3505 | 1.00 | 16.57 |  | 6 | 2 | 6O_w_ - Exclusion zone contact(s): /OP1/OP2/ Consider K in place of MG (2 valid contact(s) for K vs. 0 for MG) |
| MG:1H:3506 | 1.00 | 15.19 |  | 4 | 2 | O_ph_3O_w_ - Exclusion zone contact(s): /OP1/OP2/ |
| MG:1H:3507 | 1.00 | 17.34 |  | 5 | 3 | 5O_w_ - Exclusion zone contact(s): /OP1/OP1/OP2/ Consider K in place of MG (2 valid contact(s) for K vs. 0 for MG) |
| MG:1H:3508 | 1.00 | 22.54 |  | 4 | 3 | 4O_w_ - Exclusion zone contact(s): /NE2_His_/OP1/OP2/ Consider K in place of MG (2 valid contact(s) for K vs. 0 for MG) |
| MG:1H:3509 | 1.00 | 27.97 |  | 5 | 2 | 5O_w_ - Exclusion zone contact(s): /O3'/OP2/ Consider NA in place of MG (1 valid contact(s) for NA vs. 0 for MG) |
| MG:1H:3510 | 1.00 | 19.65 |  | 6 |  | O_ph_5O_w_ Large octahedral topology discrepancy!!! |
| MG:1H:3511 | 1.00 | 26.78 |  | 6 | 1 | 6O_w_ - Exclusion zone contact(s): /OP1/ Consider K in place of MG (1 valid contact(s) for K vs. 0 for MG) |
| MG:1H:3512 | 1.00 | 39.45 |  | 6 | 1 | O_ph_5O_w_ - Exclusion zone contact(s): /N7/ Large octahedral topology discrepancy!!! |
| MG:1H:3513 | 1.00 | 30.51 |  | 6 | 2 | 6O_w_ - Exclusion zone contact(s): /OP1/OP1/ Consider K in place of MG (2 valid contact(s) for K vs. 0 for MG) |
| MG:1H:3514 | 1.00 | 53.97 |  | 4 | 2 | 4O_w_ - Exclusion zone contact(s): /OP1/OP2/ Consider K in place of MG (2 valid contact(s) for K vs. 0 for MG)  Possible idealized distances ! |
| MG:1H:3515 | 1.00 | 40.85 |  | 4 | 2 | 4O_w_ - Exclusion zone contact(s): /OP2/OP2/ Consider NA in place of MG (1 valid contact(s) for NA vs. 0 for MG) |
| MG:1H:3516 | 1.00 | 21.60 |  | 3 | 4 | 3O_w_ - Exclusion zone contact(s): /OP1/OP1/OP1/OP2/ Consider NA in place of MG (2 valid contact(s) for NA vs. 0 for MG) |
| MG:1H:3517 | 1.00 | 26.04 |  | 3 | 3 | 3O_w_ - Exclusion zone contact(s): /OP2/OP2/OP2/ Consider NA in place of MG (2 valid contact(s) for NA vs. 0 for MG) |
| MG:1H:3518 | 1.00 | 37.63 |  | 5 | 2 | 5O_w_ - Exclusion zone contact(s): /N7/NE2_His_/ Consider K in place of MG (2 valid contact(s) for K vs. 0 for MG) |
| MG:1H:3519 | 1.00 | 25.70 |  | 3 | 4 | 3O_w_ - Exclusion zone contact(s): /N7/O3'/OP1/OP2/ Consider NA in place of MG (2 valid contact(s) for NA vs. 0 for MG) |
| MG:1H:3520 | 1.00 | 37.39 |  | 5 | 1 | 5O_w_ - Exclusion zone contact(s): /OP2/ Consider NA in place of MG (1 valid contact(s) for NA vs. 0 for MG) |
| MG:1H:3521 | 1.00 | 14.49 |  | 5 | 1 | O_ph_4O_w_ - Exclusion zone contact(s): /OP2/ |
| MG:1H:3522 | 1.00 | 49.85 |  | 6 |  | 6O_w_ |
| MG:1H:3523 | 1.00 | 32.94 |  | 5 | 1 | 5O_w_ - Exclusion zone contact(s): /OP1/ Consider NA in place of MG (1 valid contact(s) for NA vs. 0 for MG) |
| MG:1H:3524 | 1.00 | 16.11 |  | 3 | 5 | 3O_w_ - Exclusion zone contact(s): /OP1/OP1/OP1/OP2/OP2/ Consider NA in place of MG (3 valid contact(s) for NA vs. 0 for MG) |
| MG:1H:3525 | 1.00 | 15.77 |  | 6 |  | O_ph_5O_w_ |
| MG:1H:3526 | 1.00 | 21.22 |  | 6 | 1 | 6O_w_ - Exclusion zone contact(s): /OP1/ Consider K in place of MG (1 valid contact(s) for K vs. 0 for MG) |
| MG:1H:3527 | 1.00 | 26.02 |  | 5 | 2 | 5O_w_ - Exclusion zone contact(s): /OP1/OP2/ Consider K in place of MG (2 valid contact(s) for K vs. 0 for MG) |
| MG:1H:3528 | 1.00 | 50.63 |  | 6 | 1 | 6O_w_ - Exclusion zone contact(s): /O4/ Consider K in place of MG (1 valid contact(s) for K vs. 0 for MG) |
| MG:1H:3529 | 1.00 | 54.09 |  | 6 | 1 | 6O_w_ - Exclusion zone contact(s): /O6/ Consider K in place of MG (1 valid contact(s) for K vs. 0 for MG) |
| MG:1H:3530 | 1.00 | 21.54 |  | 3 | 4 | 3O_w_ - Exclusion zone contact(s): /O_w_/O_w_/O_w_/OP2/ Consider K in place of MG (1 valid contact(s) for K vs. 0 for MG) |
| MG:1H:3531 | 1.00 | 21.26 |  | 6 |  | 6O_w_ |
| MG:1H:3532 | 1.00 | 56.05 |  | 6 |  | 6O_w_ |
| MG:1H:3533 | 1.00 | 49.62 |  | 6 |  | 6O_w_ |
| MG:1H:3534 | 1.00 | 27.16 |  | 6 |  | 6O_w_ |
| MG:1H:3535 | 1.00 | 64.84 |  | 6 |  | 6O_w_ |
| MG:1H:3536 | 1.00 | 23.61 |  | 5 | 1 | 5O_w_ - Exclusion zone contact(s): /OP1/ Consider NA in place of MG (1 valid contact(s) for NA vs. 0 for MG) |
| MG:1H:3537 | 1.00 | 54.14 |  | 6 |  | 6O_w_ Possible idealized distances ! |
| MG:1H:3538 | 1.00 | 27.15 |  | 6 | 1 | 6O_w_ - Exclusion zone contact(s): /N7/ Consider K in place of MG (1 valid contact(s) for K vs. 0 for MG) |
| MG:1H:3539 | 1.00 | 66.21 |  | 6 |  | 6O_w_ |
| MG:1H:3540 | 1.00 | 50.95 |  | 6 | 1 | 6O_w_ - Exclusion zone contact(s): /N7/ Consider K in place of MG (1 valid contact(s) for K vs. 0 for MG) |
| MG:1H:3541 | 1.00 | 66.38 |  | 6 |  | 6O_w_ |
| MG:1H:3542 | 1.00 | 33.90 |  | 6 |  | 6O_w_ |
| MG:1H:3543 | 1.00 | 48.71 |  | 4 | 3 | 4O_w_ - Exclusion zone contact(s): /OP1/OP1/OP2/ Consider NA in place of MG (2 valid contact(s) for NA vs. 0 for MG) |
| MG:1H:3544 | 1.00 | 63.80 |  | 5 | 1 | 5O_w_ - Exclusion zone contact(s): /OP2/ Consider K in place of MG (1 valid contact(s) for K vs. 0 for MG)  Possible idealized distances ! |
| MG:1H:3545 | 1.00 | 45.83 |  | 6 |  | 6O_w_ |
| MG:1H:3546 | 1.00 | 61.23 |  | 6 |  | 6O_w_ |
| MG:1H:3547 | 1.00 | 52.95 |  | 6 |  | 6O_w_ |
| MG:1H:3548 | 1.00 | 32.98 |  | 6 | 1 | 6O_w_ - Exclusion zone contact(s): /OP2/ Consider K in place of MG (1 valid contact(s) for K vs. 0 for MG) |
| MG:1H:3549 | 1.00 | 60.73 |  | 6 | 1 | 6O_w_ - Exclusion zone contact(s): /OP2/ Consider K in place of MG (1 valid contact(s) for K vs. 0 for MG) |
| MG:1H:3550 | 1.00 | 64.41 |  | 6 |  | 6O_w_ |
| MG:1H:3551 | 1.00 | 45.39 |  | 5 | 1 | 5O_w_ - Exclusion zone contact(s): /OP1/ Consider NA in place of MG (1 valid contact(s) for NA vs. 0 for MG) |
| MG:1H:3552 | 1.00 | 85.39 |  | 6 |  | 6O_w_ Possible idealized distances ! |
| MG:1H:3553 | 1.00 | 60.79 |  | 6 |  | 6O_w_ |
| MG:1H:3554 | 1.00 | 77.78 |  | 6 |  | 6O_w_ Possible idealized distances ! |
| MG:1H:3555 | 1.00 | 56.62 |  | 6 |  | 6O_w_ |
| MG:1H:3556 | 1.00 | 47.98 |  | 6 |  | 6O_w_ |
| MG:1H:3557 | 1.00 | 22.13 |  | 5 | 2 | 5O_w_ - Exclusion zone contact(s): /O_w_/OP2/ Consider NA in place of MG (1 valid contact(s) for NA vs. 0 for MG) |
| MG:1H:3558 | 1.00 | 56.41 |  | 6 |  | 6O_w_ |
| MG:1H:3559 | 1.00 | 15.75 |  | 6 | 1 | O_ph_5O_w_ - Exclusion zone contact(s): /O3'/ |
| MG:1H:3560 | 1.00 | 13.82 |  | 4 | 2 | 4O_w_ - Exclusion zone contact(s): /O4/OP2/ Consider NA in place of MG (2 valid contact(s) for NA vs. 0 for MG) |
| MG:1H:3561 | 1.00 | 38.91 |  | 6 |  | 6O_w_ |
| MG:1H:3562 | 1.00 | 31.31 |  | 5 | 1 | 5O_w_ - Exclusion zone contact(s): /OP2/ Consider NA in place of MG (1 valid contact(s) for NA vs. 0 for MG) |
| MG:1H:3563 | 1.00 | 75.64 |  | 6 |  | 6O_w_ |
| MG:1H:3564 | 1.00 | 25.47 |  | 4 | 2 | 4O_w_ - Exclusion zone contact(s): /N7/OP1/ Consider NA in place of MG (1 valid contact(s) for NA vs. 0 for MG) |
| MG:1H:3565 | 1.00 | 25.61 |  | 5 | 2 | 5O_w_ - Exclusion zone contact(s): /O3'/OP1/ Consider NA in place of MG (1 valid contact(s) for NA vs. 0 for MG) |
| MG:1H:3566 | 1.00 | 37.50 |  | 5 | 1 | O_ph_4O_w_ - Exclusion zone contact(s): /OP1/ Large octahedral topology discrepancy!!! |
| MG:1H:3567 | 1.00 | 24.37 |  | 5 | 1 | 5O_w_ - Exclusion zone contact(s): /OP1/ Consider NA in place of MG (1 valid contact(s) for NA vs. 0 for MG) |
| MG:1H:3568 | 1.00 | 34.36 |  | 6 |  | 6O_w_ |
| MG:1H:3569 | 1.00 | 62.12 |  | 5 | 1 | 5O_w_ - Exclusion zone contact(s): /OP1/ Consider NA in place of MG (1 valid contact(s) for NA vs. 0 for MG)  Possible idealized distances ! |
| MG:1H:3570 | 1.00 | 50.81 |  | 6 |  | 6O_w_ |
| MG:1H:3571 | 1.00 | 26.33 |  | 5 | 2 | 5O_w_ - Exclusion zone contact(s): /O5'/OP2/ Consider NA in place of MG (1 valid contact(s) for NA vs. 0 for MG) |
| MG:1H:3572 | 1.00 | 25.28 |  | 6 |  | 6O_w_ |
| MG:1H:3573 | 1.00 | 40.19 |  | 4 | 2 | 4O_w_ - Exclusion zone contact(s): /O2/OP2/ Consider K in place of MG (2 valid contact(s) for K vs. 0 for MG) |
| MG:1H:3574 | 1.00 | 24.51 |  | 4 | 3 | 4O_w_ - Exclusion zone contact(s): /O5'/OP2/OP2/ Consider K in place of MG (2 valid contact(s) for K vs. 0 for MG) |
| MG:1H:3575 | 1.00 | 17.73 |  | 6 |  | 2O_ph_4O_w_ Large octahedral topology discrepancy!!! |
| MG:1H:3576 | 1.00 | 69.76 |  | 6 |  | 6O_w_ Possible idealized distances ! |
| MG:1H:3577 | 1.00 | 37.74 |  | 4 | 2 | 4O_w_ - Exclusion zone contact(s): /OP2/OP2/ Consider K in place of MG (2 valid contact(s) for K vs. 0 for MG) |
| K:1J:201 | 1.00 | 81.19 | 1 | 6 |  | 2O_b_4Anion - Exclusion zone contact(s): /O6/ |
| MG:1J:202 | 1.00 | 83.12 |  | 1 | 4 | O_ph_ - Exclusion zone contact(s): /O4/O6/OP1/OP2/ Consider K in place of MG (4 valid contact(s) for K vs. 1 for MG) |
| MG:1J:203 | 1.00 | 48.46 |  | 0 | 1 | - Exclusion zone contact(s): /N7/ Consider K in place of MG (1 valid contact(s) for K vs. 0 for MG) |
| MG:1J:204 | 1.00 | 64.73 |  | 0 | 1 | - Exclusion zone contact(s): /O6/ Consider K in place of MG (1 valid contact(s) for K vs. 0 for MG) |
| MG:1J:205 | 1.00 | 68.34 |  | 5 | 1 | 5O_w_ - Exclusion zone contact(s): /OP2/ Consider K in place of MG (1 valid contact(s) for K vs. 0 for MG) |
| K:1K:101 | 1.00 | 83.86 |  | 3 |  | O_ph_2O_b_ |
| K:21:301 | 1.00 | 62.69 |  | 3 |  | 2O_back_OH_Prot_ |
| MG:21:302 | 1.00 | 14.87 |  | 5 | 1 | 5O_w_ - Exclusion zone contact(s): /OD2_Asp_/ Consider K in place of MG (1 valid contact(s) for K vs. 0 for MG) |
| MG:21:303 | 1.00 | 33.81 |  | 6 | 1 | 6O_w_ - Exclusion zone contact(s): /O_back_/ Consider K in place of MG (1 valid contact(s) for K vs. 0 for MG)  Possible idealized distances ! |
| K:29:301 | 1.00 | 86.83 |  | 3 |  | O_back_OH_Prot_Anion |
| MG:29:302 | 1.00 | 38.35 |  | 1 | 1 | O_coo_ - Exclusion zone contact(s): /O6/ |
| MG:29:303 | 1.00 | 28.21 |  | 4 | 2 | 4O_w_ - Exclusion zone contact(s): /NE2_His_/OP1/ Consider NA in place of MG (2 valid contact(s) for NA vs. 0 for MG) |
| K:2A:201 | 1.00 | 70.49 | 1 | 4 |  | 2O_b_O_back_O_cno_ - Exclusion zone contact(s): /O_back_/ |
| K:2K:101 | 1.00 | 62.35 |  | 5 |  | O_ph_4O_r_ |
| K:2K:102 | 1.00 | 58.55 |  | 5 |  | 2O_b_N_b_2Anion |
| K:2K:103 | 1.00 | 79.58 |  | 3 |  | 2O_b_Anion |
| MG:2K:104 | 1.00 | 48.72 |  | 0 |  | 1^st^ MG coordination shell missing (see Notes) |
| MG:2K:105 | 1.00 | 45.40 |  | 6 | 1 | 6O_w_ - Exclusion zone contact(s): /OP2/ Consider K in place of MG (1 valid contact(s) for K vs. 0 for MG) |
| MG:2L:101 | 1.00 | 58.99 |  | 2 | 1 | 2O_ph_ - Exclusion zone contact(s): /OP1/ |
| MG:2L:102 | 1.00 | 56.51 |  | 0 |  | 1^st^ MG coordination shell missing (see Notes) |
| MG:2L:103 | 1.00 | 45.41 |  | 0 |  | 1^st^ MG coordination shell missing (see Notes) |
| K:31:301 | 1.00 | 48.71 |  | 4 |  | O_b_3O_back_ |
| K:31:302 | 1.00 | 45.34 |  | 4 |  | 3O_back_O_w_ |
| MG:31:303 | 1.00 | 37.64 |  | 0 |  | 1^st^ MG coordination shell missing (see Notes) |
| K:32:301 | 1.00 | 104.22 |  | 3 |  | O_b_O_back_OH_Prot_ |
| MG:32:302 | 1.00 | 79.58 |  | 6 | 1 | 6O_w_ - Exclusion zone contact(s): /NZ_Lys_/ Consider K in place of MG (1 valid contact(s) for K vs. 0 for MG)  Possible idealized distances ! |
| K:39:301 | 1.00 | 75.13 |  | 4 |  | O_b_3O_back_ |
| K:39:302 | 1.00 | 74.54 |  | 4 |  | 2O_back_2O_w_ |
| MG:39:303 | 1.00 | 54.86 |  | 0 | 1 | - Exclusion zone contact(s): /OD2_Asp_/ Consider NA in place of MG (1 valid contact(s) for NA vs. 0 for MG) |
| MG:3E:301 | 1.00 | 82.21 |  | 6 | 1 | 6O_w_ - Exclusion zone contact(s): /NH1_Arg_/ Consider K in place of MG (1 valid contact(s) for K vs. 0 for MG) |
| K:41:201 | 1.00 | 67.82 | 1 | 2 |  | 2O_back_ - Exclusion zone contact(s): /O_back_/ |
| MG:41:202 | 1.00 | 55.21 |  | 2 |  | 2O_back_ |
| MG:42:201 | 1.00 | 70.31 |  | 1 | 3 | Anion - Exclusion zone contact(s): /N7/NH1_Arg_/O4/ Consider K in place of MG (3 valid contact(s) for K vs. 1 for MG) |
| K:45:201 | 1.00 | 90.05 |  | 4 |  | O_ph_3O_back_ |
| K:49:201 | 1.00 | 108.83 |  | 3 |  | 3O_back_ |
| K:4A:201 | 1.00 | 96.06 | 1 | 2 |  | O_back_OH_Prot_ - Exclusion zone contact(s): /OP2/ |
| K:4I:201 | 1.00 | 71.46 | 2 | 2 |  | 2O_back_ - Exclusion zone contact(s): /O_back_/OP1/ |
| MG:4L:101 | 1.00 | 79.35 |  | 0 | 2 | - Exclusion zone contact(s): /OP1/OP2/ Consider NA in place of MG (1 valid contact(s) for NA vs. 0 for MG) |
| K:52:201 | 1.00 | 76.72 |  | 3 |  | 3O_back_ |
| K:5A:101 | 1.00 | 82.78 |  | 2 |  | O_ph_Anion |
| ZN:5A:102 | 1.00 | 82.96 |  | 0 |  |  |
| K:5E:201 | 1.00 | 76.48 |  | 3 |  | 3O_back_ |
| ZN:5I:101 | 1.00 | 70.91 |  | 0 |  |  |
| MG:6E:201 | 1.00 | 62.30 |  | 0 |  | 1^st^ MG coordination shell missing (see Notes) |
| MG:78:201 | 1.00 | 38.55 |  | 0 |  | 1^st^ MG coordination shell missing (see Notes) |
| MG:78:202 | 1.00 | 30.73 |  | 6 |  | 6O_w_ |
| MG:7I:101 | 1.00 | 71.55 |  | 0 |  | 1^st^ MG coordination shell missing (see Notes) |
| K:88:201 | 1.00 | 57.95 |  | 4 |  | O_ph_3O_back_ |
| K:BA:201 | 1.00 | 72.97 |  | 4 |  | 3O_back_Anion |
| K:BI:201 | 1.00 | 108.63 |  | 3 |  | 3O_back_ |
| MG:C8:201 | 1.00 | 44.75 |  | 0 | 3 | - Exclusion zone contact(s): /NH1_Arg_/NH2_Arg_/OP2/ Consider K in place of MG (3 valid contact(s) for K vs. 0 for MG) |
| MG:F8:101 | 1.00 | 32.92 |  | 0 | 3 | - Exclusion zone contact(s): /O_back_/OD1_Asn_/OP2/ Consider K in place of MG (3 valid contact(s) for K vs. 0 for MG) |
| MG:I8:101 | 1.00 | 26.71 |  | 0 | 1 | - Exclusion zone contact(s): /NH1_Arg_/ Consider K in place of MG (1 valid contact(s) for K vs. 0 for MG) |
| MG:J8:101 | 1.00 | 32.85 |  | 0 |  | 1^st^ MG coordination shell missing (see Notes) |
| MG:L8:101 | 1.00 | 38.38 |  | 1 | 3 | O_ph_ - Exclusion zone contact(s): /O_back_/O_w_/O_w_/ |
| MG:M5:101 | 1.00 | 84.23 |  | 6 | 1 | 6O_w_ - Exclusion zone contact(s): /NZ_Lys_/ Consider K in place of MG (1 valid contact(s) for K vs. 0 for MG) |
| MG:M5:102 | 1.00 | 58.06 |  | 6 |  | 6O_w_ |
| MG:N8:101 | 1.00 | 41.27 |  | 0 | 1 | - Exclusion zone contact(s): /OD2_Asp_/ Consider K in place of MG (1 valid contact(s) for K vs. 0 for MG) |
| MG:O8:101 | 1.00 | 60.52 |  | 0 |  | 1^st^ MG coordination shell missing (see Notes) |
| MG:P8:101 | 1.00 | 35.75 |  | 0 |  | 1^st^ MG coordination shell missing (see Notes) |
| MG:Q8:101 | 0.50 | 13.93 |  | 0 |  | 1^st^ MG coordination shell missing (see Notes) |
| MG:Q8:101 | 0.50 | 11.81 |  | 0 |  | 1^st^ MG coordination shell missing (see Notes) |

---

\_\_\_\_\_\_\_\_\_\_\_\_\_\_\_\_\_\_\_\_\_\_\_\_\_\_\_\_\_\_\_\_\_\_\_\_\_\_\_\_\_\_\_\_\_\_\_\_\_\_\_\_\_\_\_\_\_\_\_\_\_\_\_\_  

#### 3) Potential issue list

##### Go to ToC

|  |  |  |
| --- | --- | --- |
| **Problem** | **Number** | **Affected ions** |
| **1) K:  could be NA** | 5 | 14:3008  14:3094  1H:3033  1H:3035  1H:3038 |
| **1) MG:  could be K** | 361 | 11:302  13:1655  13:1656  13:1658  13:1659  13:1667  13:1672  13:1673  13:1674  13:1676  13:1681  13:1684  13:1687  13:1688  13:1690  13:1692  13:1695  13:1698  13:1700  13:1701  13:1702  13:1703  13:1704  13:1706  13:1707  13:1708  13:1709  13:1711  13:1712  13:1713  13:1714  13:1715  13:1717  13:1720  13:1721  13:1722  13:1731  13:1735  13:1737  13:1745  13:1747  13:1749  13:1751  13:1752  13:1753  13:1757  13:1761  13:1763  13:1769  13:1770  14:3112  14:3118  14:3120  14:3121  14:3135  14:3137  14:3143  14:3145  14:3163  14:3166  14:3168  14:3170  14:3174  14:3175  14:3176  14:3181  14:3182  14:3185  14:3186  14:3187  14:3190  14:3197  14:3198  14:3201  14:3202  14:3213  14:3216  14:3218  14:3221  14:3225  14:3227  14:3230  14:3231  14:3232  14:3235  14:3236  14:3241  14:3242  14:3244  14:3246  14:3248  14:3249  14:3250  14:3251  14:3253  14:3255  14:3256  14:3260  14:3261  14:3263  14:3264  14:3268  14:3269  14:3270  14:3274  14:3276  14:3278  14:3280  14:3289  14:3291  14:3295  14:3298  14:3301  14:3309  14:3312  14:3314  14:3317  14:3323  14:3326  14:3327  14:3344  14:3348  14:3354  14:3355  14:3357  14:3358  14:3361  14:3362  14:3366  14:3368  14:3369  14:3370  14:3374  14:3382  14:3397  14:3400  16:204  16:206  16:207  16:208  16:209  16:211  1G:1640  1G:1646  1G:1651  1G:1653  1G:1657  1G:1659  1G:1663  1G:1667  1G:1668  1G:1670  1G:1674  1G:1675  1G:1677  1G:1678  1G:1679  1G:1680  1G:1681  1G:1682  1G:1683  1G:1685  1G:1686  1G:1687  1G:1688  1G:1690  1G:1691  1G:1693  1G:1694  1G:1695  1G:1699  1G:1703  1G:1711  1G:1713  1G:1719  1G:1720  1G:1722  1G:1724  1G:1730  1G:1733  1G:1736  1H:3146  1H:3148  1H:3153  1H:3155  1H:3157  1H:3158  1H:3159  1H:3163  1H:3176  1H:3181  1H:3182  1H:3183  1H:3186  1H:3189  1H:3197  1H:3201  1H:3202  1H:3207  1H:3208  1H:3210  1H:3211  1H:3212  1H:3213  1H:3221  1H:3223  1H:3225  1H:3227  1H:3229  1H:3230  1H:3231  1H:3233  1H:3234  1H:3236  1H:3237  1H:3238  1H:3240  1H:3242  1H:3248  1H:3249  1H:3252  1H:3254  1H:3260  1H:3263  1H:3264  1H:3265  1H:3267  1H:3269  1H:3273  1H:3274  1H:3275  1H:3276  1H:3278  1H:3279  1H:3280  1H:3281  1H:3284  1H:3285  1H:3286  1H:3290  1H:3291  1H:3292  1H:3295  1H:3297  1H:3298  1H:3299  1H:3300  1H:3302  1H:3306  1H:3308  1H:3311  1H:3312  1H:3319  1H:3320  1H:3323  1H:3325  1H:3327  1H:3328  1H:3331  1H:3333  1H:3334  1H:3340  1H:3341  1H:3344  1H:3345  1H:3345  1H:3348  1H:3349  1H:3351  1H:3352  1H:3354  1H:3355  1H:3356  1H:3359  1H:3361  1H:3363  1H:3365  1H:3366  1H:3367  1H:3369  1H:3371  1H:3372  1H:3373  1H:3375  1H:3376  1H:3378  1H:3379  1H:3380  1H:3381  1H:3384  1H:3385  1H:3387  1H:3389  1H:3390  1H:3391  1H:3392  1H:3394  1H:3399  1H:3405  1H:3407  1H:3411  1H:3413  1H:3417  1H:3419  1H:3420  1H:3424  1H:3428  1H:3429  1H:3433  1H:3438  1H:3449  1H:3453  1H:3454  1H:3456  1H:3464  1H:3466  1H:3468  1H:3475  1H:3476  1H:3478  1H:3487  1H:3489  1H:3491  1H:3498  1H:3500  1H:3502  1H:3505  1H:3507  1H:3508  1H:3511  1H:3513  1H:3514  1H:3518  1H:3526  1H:3527  1H:3528  1H:3529  1H:3530  1H:3538  1H:3540  1H:3544  1H:3548  1H:3549  1H:3573  1H:3574  1H:3577  1J:202  1J:203  1J:204  1J:205  21:302  21:303  2K:105  32:302  3E:301  42:201  C8:201  F8:101  I8:101  M5:101  N8:101 |
| **1) MG:  could be NA** | 138 | 13:1662  13:1665  13:1686  13:1691  13:1697  13:1705  13:1744  13:1746  13:1750  13:1754  14:3109  14:3111  14:3141  14:3150  14:3167  14:3175  14:3180  14:3193  14:3195  14:3209  14:3210  14:3215  14:3220  14:3226  14:3228  14:3234  14:3279  14:3281  14:3282  14:3283  14:3285  14:3292  14:3293  14:3305  14:3306  14:3310  14:3315  14:3316  14:3318  14:3319  14:3320  14:3322  14:3329  14:3330  14:3331  14:3332  14:3334  14:3340  14:3341  14:3342  14:3343  14:3346  14:3347  14:3350  14:3351  14:3359  14:3360  14:3363  14:3364  14:3365  14:3371  14:3372  14:3376  14:3377  14:3378  14:3381  14:3385  14:3389  14:3395  16:205  16:212  1G:1715  1G:1718  1G:1721  1G:1723  1G:1726  1H:3152  1H:3165  1H:3170  1H:3172  1H:3187  1H:3204  1H:3322  1H:3328  1H:3329  1H:3330  1H:3357  1H:3368  1H:3382  1H:3383  1H:3386  1H:3388  1H:3397  1H:3400  1H:3409  1H:3412  1H:3422  1H:3423  1H:3426  1H:3430  1H:3455  1H:3457  1H:3458  1H:3460  1H:3462  1H:3465  1H:3469  1H:3470  1H:3472  1H:3473  1H:3477  1H:3480  1H:3486  1H:3490  1H:3503  1H:3504  1H:3509  1H:3515  1H:3516  1H:3517  1H:3519  1H:3520  1H:3523  1H:3524  1H:3536  1H:3543  1H:3551  1H:3557  1H:3560  1H:3562  1H:3564  1H:3565  1H:3567  1H:3569  1H:3571  29:303  39:303  4L:101 |
| **2) K: 1^st^ shell missing** | 12 | 13:1614  14:3011  14:3079  14:3088  1G:1604  1H:3005  1H:3026  1H:3037  1H:3040  1H:3040  1H:3050  1H:3126 |
| **2) MG: 1^st^ shell missing** | 165 | 13:1652  13:1654  13:1666  13:1669  13:1671  13:1675  13:1679  13:1680  13:1682  13:1689  13:1696  13:1699  13:1709  13:1710  13:1716  13:1718  13:1719  13:1723  13:1725  14:3113  14:3115  14:3116  14:3117  14:3122  14:3125  14:3142  14:3146  14:3148  14:3149  14:3159  14:3164  14:3165  14:3173  14:3178  14:3183  14:3184  14:3188  14:3194  14:3200  14:3203  14:3204  14:3205  14:3207  14:3208  14:3219  14:3224  14:3233  14:3238  14:3239  14:3240  14:3243  14:3252  14:3254  14:3259  14:3262  14:3271  14:3272  14:3277  1G:1649  1G:1656  1G:1658  1G:1660  1G:1661  1G:1663  1G:1664  1G:1665  1G:1666  1G:1669  1G:1671  1G:1683  1G:1689  1G:1692  1G:1696  1G:1697  1G:1698  1G:1700  1H:3151  1H:3156  1H:3160  1H:3161  1H:3164  1H:3173  1H:3174  1H:3175  1H:3177  1H:3179  1H:3184  1H:3185  1H:3190  1H:3193  1H:3196  1H:3209  1H:3214  1H:3216  1H:3217  1H:3219  1H:3220  1H:3222  1H:3224  1H:3226  1H:3228  1H:3235  1H:3239  1H:3241  1H:3243  1H:3245  1H:3251  1H:3253  1H:3256  1H:3258  1H:3262  1H:3266  1H:3270  1H:3271  1H:3272  1H:3277  1H:3282  1H:3283  1H:3287  1H:3288  1H:3289  1H:3293  1H:3294  1H:3296  1H:3301  1H:3303  1H:3304  1H:3305  1H:3307  1H:3314  1H:3315  1H:3316  1H:3317  1H:3320  1H:3321  1H:3326  1H:3332  1H:3335  1H:3336  1H:3337  1H:3338  1H:3339  1H:3342  1H:3343  1H:3343  1H:3346  1H:3347  1H:3353  1H:3358  1H:3362  1H:3364  1H:3370  1H:3377  2K:104  2L:102  2L:103  31:303  6E:201  78:201  7I:101  J8:101  O8:101  P8:101  Q8:101  Q8:101 |
| **3) K/N1_A_ contacts** | 4 | 13:1612  14:3019  16:201  1H:3057 |
| **3) K/N1_G_ contacts** | 5 | 14:3052  16:203  1H:3039  1H:3090  1J:201 |
| **3) K/N2 contacts** | 8 | 13:1610  13:1633  14:3040  14:3080  16:203  1H:3039  1H:3078  1J:201 |
| **3) K/N3 contacts** | 2 | 14:3059  1H:3097 |
| **3) K/N3_A_ contacts** | 16 | 13:1605  13:1622  14:3018  14:3019  14:3027  14:3039  14:3055  14:3059  14:3102  1G:1609  1H:3024  1H:3056  1H:3057  1H:3065  1H:3077  1H:3097 |
| **3) K/N3_C_ contacts** | 6 | 13:1628  13:1629  1G:1610  1G:1613  1G:1614  1H:3133 |
| **3) K/N3_G_ contacts** | 6 | 13:1622  13:1633  14:3040  1G:1609  1H:3027  1H:3078 |
| **3) K/N3_U_ contacts** | 5 | 13:1608  16:202  16:203  1G:1610  1J:201 |
| **3) K/N4 contacts** | 13 | 13:1624  13:1627  13:1628  14:3070  16:203  1G:1610  1G:1612  1G:1622  1H:3016  1H:3068  1J:201  2K:102  2K:103 |
| **3) K/N6 contacts** | 2 | 1H:3065  2K:102 |
| **3) K/N7 contacts** | 66 | 13:1601  13:1604  13:1615  13:1621  13:1621  13:1621  13:1622  13:1623  13:1626  13:1628  13:1632  13:1633  13:1640  13:1640  13:1651  14:3014  14:3014  14:3018  14:3018  14:3019  14:3019  14:3021  14:3023  14:3024  14:3028  14:3028  14:3035  14:3038  14:3039  14:3040  14:3042  14:3075  14:3089  14:3099  16:201  1G:1608  1G:1608  1G:1613  1G:1616  1G:1617  1G:1624  1H:3007  1H:3023  1H:3036  1H:3056  1H:3056  1H:3057  1H:3057  1H:3059  1H:3059  1H:3061  1H:3061  1H:3062  1H:3066  1H:3066  1H:3071  1H:3073  1H:3076  1H:3077  1H:3078  1H:3080  1H:3088  1H:3095  1H:3113  1H:3127  2K:102 |
| **3) K/N_back_ contacts** | 2 | 5A:101  BA:201 |
| **3) K/ND2_Asn_ contacts** | 1 | 29:301 |
| **3) K/O2 contacts** | 46 | 13:1602  13:1604  13:1605  13:1619  13:1622  13:1624  13:1629  13:1633  14:3016  14:3017  14:3021  14:3023  14:3032  14:3033  14:3046  14:3058  14:3059  14:3060  14:3069  14:3069  14:3095  1G:1609  1G:1610  1G:1614  1G:1617  1H:3001  1H:3015  1H:3018  1H:3024  1H:3029  1H:3045  1H:3054  1H:3059  1H:3061  1H:3065  1H:3070  1H:3071  1H:3084  1H:3084  1H:3096  1H:3097  1H:3107  1H:3107  1H:3133  1K:101  32:301 |
| **3) K/O2' contacts** | 99 | 13:1603  13:1605  13:1609  13:1619  13:1622  13:1622  13:1626  13:1629  13:1637  13:1640  13:1643  14:3007  14:3007  14:3010  14:3013  14:3016  14:3017  14:3017  14:3017  14:3019  14:3022  14:3023  14:3024  14:3025  14:3032  14:3034  14:3038  14:3040  14:3046  14:3046  14:3049  14:3051  14:3055  14:3066  14:3067  14:3068  14:3073  14:3075  14:3075  14:3077  14:3086  14:3087  14:3089  14:3090  14:3093  14:3100  14:3102  1G:1609  1G:1609  1G:1614  1G:1619  1G:1621  1G:1624  1G:1628  1G:1634  1H:3001  1H:3014  1H:3015  1H:3015  1H:3016  1H:3018  1H:3018  1H:3020  1H:3025  1H:3031  1H:3045  1H:3052  1H:3054  1H:3055  1H:3055  1H:3055  1H:3057  1H:3060  1H:3061  1H:3062  1H:3063  1H:3070  1H:3072  1H:3076  1H:3078  1H:3079  1H:3084  1H:3087  1H:3087  1H:3089  1H:3093  1H:3096  1H:3104  1H:3106  1H:3111  1H:3115  1H:3117  1H:3117  1H:3124  1H:3127  1H:3128  1H:3138  1H:3140  2K:101 |
| **3) K/O3' contacts** | 63 | 13:1603  13:1619  13:1630  13:1631  13:1636  13:1636  13:1637  13:1640  13:1643  13:1643  13:1644  13:1649  14:3007  14:3007  14:3015  14:3015  14:3024  14:3025  14:3026  14:3036  14:3038  14:3040  14:3041  14:3049  14:3049  14:3050  14:3054  14:3065  14:3068  14:3075  14:3086  14:3090  14:3100  1G:1603  1G:1615  1G:1620  1G:1620  1G:1621  1G:1634  1H:3018  1H:3024  1H:3031  1H:3053  1H:3053  1H:3055  1H:3062  1H:3064  1H:3074  1H:3079  1H:3079  1H:3087  1H:3087  1H:3088  1H:3092  1H:3103  1H:3106  1H:3110  1H:3113  1H:3124  1H:3128  1H:3138  2K:101  2K:101 |
| **3) K/O4 contacts** | 137 | 13:1602  13:1606  13:1607  13:1610  13:1610  13:1613  13:1616  13:1618  13:1623  13:1624  13:1627  13:1627  13:1627  13:1634  13:1635  13:1635  13:1638  13:1639  13:1645  13:1646  13:1647  13:1647  13:1648  14:3001  14:3004  14:3006  14:3020  14:3020  14:3021  14:3029  14:3030  14:3035  14:3037  14:3043  14:3045  14:3047  14:3048  14:3049  14:3051  14:3052  14:3056  14:3057  14:3059  14:3060  14:3060  14:3061  14:3070  14:3074  14:3074  14:3080  14:3085  14:3095  14:3095  14:3098  14:3098  14:3098  14:3106  16:202  16:202  16:203  16:203  1G:1610  1G:1612  1G:1612  1G:1618  1G:1619  1G:1619  1G:1622  1G:1626  1G:1631  1G:1632  1G:1632  1G:1633  1H:3001  1H:3002  1H:3003  1H:3004  1H:3006  1H:3006  1H:3008  1H:3011  1H:3012  1H:3021  1H:3030  1H:3034  1H:3042  1H:3043  1H:3047  1H:3049  1H:3051  1H:3058  1H:3058  1H:3059  1H:3067  1H:3068  1H:3073  1H:3073  1H:3075  1H:3077  1H:3083  1H:3085  1H:3087  1H:3089  1H:3089  1H:3090  1H:3091  1H:3094  1H:3095  1H:3097  1H:3098  1H:3098  1H:3100  1H:3101  1H:3102  1H:3105  1H:3108  1H:3109  1H:3109  1H:3112  1H:3112  1H:3115  1H:3118  1H:3123  1H:3123  1H:3128  1H:3133  1H:3133  1H:3136  1H:3136  1H:3144  1J:201  1J:201  1K:101  2A:201  2K:103  31:301  39:301 |
| **3) K/O4' contacts** | 28 | 13:1629  13:1640  14:3018  14:3021  14:3026  14:3032  14:3033  14:3055  14:3072  14:3078  14:3082  14:3086  14:3089  16:201  1G:1624  1H:3014  1H:3025  1H:3027  1H:3056  1H:3059  1H:3064  1H:3070  1H:3071  1H:3097  1H:3110  1H:3116  1H:3120  1H:3127 |
| **3) K/O5' contacts** | 16 | 13:1612  13:1640  14:3026  14:3049  14:3078  14:3082  1G:1624  1G:1625  1H:3027  1H:3036  1H:3064  1H:3087  1H:3116  1H:3120  1H:3141  2K:101 |
| **3) K/O6 contacts** | 277 | 13:1602  13:1602  13:1603  13:1604  13:1607  13:1607  13:1607  13:1608  13:1610  13:1611  13:1613  13:1615  13:1615  13:1616  13:1618  13:1618  13:1620  13:1621  13:1622  13:1623  13:1624  13:1624  13:1626  13:1627  13:1627  13:1628  13:1633  13:1634  13:1634  13:1638  13:1638  13:1639  13:1639  13:1640  13:1641  13:1645  13:1645  13:1646  13:1647  13:1647  13:1650  13:1651  14:3003  14:3003  14:3003  14:3006  14:3012  14:3014  14:3016  14:3016  14:3018  14:3020  14:3021  14:3029  14:3029  14:3029  14:3030  14:3030  14:3030  14:3031  14:3033  14:3033  14:3034  14:3035  14:3036  14:3037  14:3037  14:3037  14:3038  14:3039  14:3043  14:3043  14:3043  14:3044  14:3044  14:3045  14:3045  14:3047  14:3047  14:3047  14:3048  14:3048  14:3048  14:3050  14:3050  14:3051  14:3051  14:3052  14:3053  14:3056  14:3056  14:3056  14:3057  14:3060  14:3061  14:3062  14:3063  14:3064  14:3070  14:3071  14:3072  14:3073  14:3074  14:3074  14:3076  14:3077  14:3080  14:3081  14:3082  14:3089  14:3091  14:3092  14:3096  14:3096  14:3098  14:3099  14:3099  14:3101  14:3101  14:3105  14:3106  16:201  16:202  16:202  16:203  1G:1602  1G:1602  1G:1608  1G:1609  1G:1610  1G:1610  1G:1610  1G:1611  1G:1612  1G:1612  1G:1613  1G:1614  1G:1617  1G:1618  1G:1618  1G:1619  1G:1622  1G:1624  1G:1626  1G:1626  1G:1630  1G:1630  1G:1631  1G:1632  1G:1633  1G:1633  1G:1635  1H:3001  1H:3002  1H:3003  1H:3003  1H:3003  1H:3004  1H:3007  1H:3009  1H:3009  1H:3010  1H:3010  1H:3011  1H:3012  1H:3012  1H:3013  1H:3014  1H:3017  1H:3017  1H:3019  1H:3019  1H:3023  1H:3034  1H:3041  1H:3041  1H:3042  1H:3042  1H:3042  1H:3043  1H:3043  1H:3044  1H:3044  1H:3046  1H:3047  1H:3047  1H:3048  1H:3049  1H:3051  1H:3051  1H:3054  1H:3054  1H:3056  1H:3057  1H:3057  1H:3058  1H:3059  1H:3067  1H:3067  1H:3068  1H:3068  1H:3068  1H:3071  1H:3071  1H:3072  1H:3072  1H:3073  1H:3074  1H:3075  1H:3075  1H:3075  1H:3076  1H:3077  1H:3078  1H:3081  1H:3081  1H:3081  1H:3082  1H:3082  1H:3083  1H:3083  1H:3085  1H:3085  1H:3086  1H:3086  1H:3086  1H:3088  1H:3088  1H:3089  1H:3089  1H:3090  1H:3091  1H:3091  1H:3091  1H:3094  1H:3094  1H:3094  1H:3095  1H:3098  1H:3099  1H:3100  1H:3100  1H:3102  1H:3102  1H:3105  1H:3108  1H:3109  1H:3110  1H:3111  1H:3112  1H:3112  1H:3114  1H:3115  1H:3117  1H:3118  1H:3118  1H:3119  1H:3120  1H:3127  1H:3129  1H:3130  1H:3130  1H:3132  1H:3132  1H:3134  1H:3134  1H:3136  1H:3137  1H:3139  1H:3143  1H:3143  1H:3144  1H:3144  2A:201  2K:102  2K:102  2K:103 |
| **3) K/O_back_ contacts** | 62 | 11:301  11:301  11:301  13:1623  13:1623  13:1625  13:1628  13:1629  13:1630  14:3002  14:3031  14:3041  14:3068  19:301  19:301  1G:1614  1G:1625  1H:3079  1H:3106  1H:3119  21:301  21:301  29:301  2A:201  31:301  31:301  31:301  31:302  31:302  31:302  32:301  39:301  39:301  39:301  39:302  39:302  41:201  41:201  45:201  45:201  45:201  49:201  49:201  49:201  4A:201  4I:201  4I:201  52:201  52:201  52:201  5E:201  5E:201  5E:201  88:201  88:201  88:201  BA:201  BA:201  BA:201  BI:201  BI:201  BI:201 |
| **3) K/O_w_ contacts** | 46 | 11:301  13:1640  14:3015  14:3022  14:3022  14:3025  14:3060  14:3073  14:3073  14:3076  14:3083  14:3084  14:3085  14:3092  14:3094  14:3094  14:3094  14:3103  14:3103  1G:1619  1H:3006  1H:3010  1H:3010  1H:3020  1H:3020  1H:3022  1H:3047  1H:3047  1H:3060  1H:3060  1H:3060  1H:3066  1H:3076  1H:3082  1H:3082  1H:3092  1H:3103  1H:3104  1H:3111  1H:3121  1H:3123  1H:3130  1H:3132  31:302  39:302  39:302 |
| **3) K/OD1_Asn_ contacts** | 3 | 13:1628  1G:1613  2A:201 |
| **3) K/OE1_Glu_ contacts** | 2 | 14:3022  1H:3060 |
| **3) K/OG1_Thr_ contacts** | 3 | 14:3083  1H:3121  32:301 |
| **3) K/OG_Ser_ contacts** | 1 | 1G:1618 |
| **3) K/OH_Tyr_ contacts** | 3 | 21:301  29:301  4A:201 |
| **3) K/OP1 contacts** | 100 | 13:1606  13:1616  13:1619  13:1625  13:1631  13:1631  13:1631  13:1633  13:1636  13:1636  13:1643  13:1650  14:3002  14:3015  14:3015  14:3015  14:3015  14:3025  14:3026  14:3027  14:3031  14:3036  14:3040  14:3042  14:3044  14:3049  14:3049  14:3054  14:3055  14:3058  14:3065  14:3065  14:3065  14:3066  14:3072  14:3078  14:3084  14:3084  14:3093  14:3097  14:3103  14:3104  14:3104  16:201  1G:1601  1G:1603  1G:1607  1G:1615  1G:1615  1G:1615  1G:1617  1G:1620  1G:1620  1G:1628  1G:1629  1G:1636  1H:3024  1H:3025  1H:3028  1H:3032  1H:3032  1H:3045  1H:3048  1H:3053  1H:3053  1H:3053  1H:3063  1H:3064  1H:3069  1H:3074  1H:3079  1H:3080  1H:3082  1H:3087  1H:3087  1H:3092  1H:3092  1H:3093  1H:3096  1H:3103  1H:3103  1H:3103  1H:3103  1H:3104  1H:3107  1H:3110  1H:3110  1H:3116  1H:3116  1H:3122  1H:3122  1H:3131  1H:3135  1H:3141  1H:3141  1H:3142  2K:101  45:201  5A:101  88:201 |
| **3) K/OP2 contacts** | 124 | 13:1609  13:1612  13:1612  13:1615  13:1617  13:1621  13:1621  13:1626  13:1630  13:1637  13:1637  13:1637  13:1640  13:1642  13:1643  13:1644  13:1649  13:1649  13:1650  14:3001  14:3002  14:3004  14:3005  14:3009  14:3010  14:3015  14:3022  14:3022  14:3023  14:3024  14:3024  14:3025  14:3026  14:3026  14:3031  14:3035  14:3035  14:3040  14:3042  14:3050  14:3055  14:3063  14:3063  14:3065  14:3065  14:3071  14:3075  14:3081  14:3083  14:3084  14:3085  14:3086  14:3087  14:3090  14:3097  14:3100  14:3102  14:3104  14:3104  16:201  1G:1605  1G:1605  1G:1606  1G:1607  1G:1608  1G:1611  1G:1620  1G:1621  1G:1621  1G:1621  1G:1623  1G:1627  1G:1628  1G:1634  1G:1634  1G:1634  1G:1636  1G:1636  1H:3013  1H:3013  1H:3015  1H:3016  1H:3022  1H:3023  1H:3027  1H:3031  1H:3052  1H:3053  1H:3060  1H:3061  1H:3062  1H:3062  1H:3064  1H:3064  1H:3069  1H:3073  1H:3074  1H:3076  1H:3080  1H:3088  1H:3090  1H:3092  1H:3093  1H:3096  1H:3097  1H:3101  1H:3101  1H:3103  1H:3109  1H:3113  1H:3116  1H:3119  1H:3121  1H:3122  1H:3123  1H:3125  1H:3128  1H:3130  1H:3135  1H:3138  1H:3138  1H:3142  1H:3142  1K:101 |
| **3) MG/N7 contacts** | 4 | 14:3135  14:3157  14:3170  1G:1641 |
| **3) MG/NE2_His_ contacts** | 2 | 14:3131  1H:3206 |
| **3) MG/NH2_Arg_ contacts** | 1 | 42:201 |
| **3) MG/O2 contacts** | 4 | 13:1664  14:3237  1G:1639  1H:3218 |
| **3) MG/O4 contacts** | 10 | 13:1659  13:1695  14:3134  14:3163  14:3179  14:3245  1H:3198  1H:3247  1H:3318  1H:3484 |
| **3) MG/O5' contacts** | 1 | 14:3266 |
| **3) MG/O6 contacts** | 15 | 13:1683  13:1685  13:1694  14:3132  14:3135  14:3214  14:3267  14:3275  1G:1672  1G:1676  1H:3146  1H:3150  1H:3257  1H:3259  1H:3350 |
| **3) MG/O_back_ contacts** | 2 | 41:202  41:202 |
| **3) MG/O_w_ contacts** | 2207 | 11:302  11:302  11:302  11:302  11:302  11:302  13:1727  13:1727  13:1727  13:1727  13:1727  13:1727  13:1728  13:1728  13:1728  13:1728  13:1728  13:1728  13:1729  13:1729  13:1729  13:1729  13:1729  13:1729  13:1730  13:1730  13:1730  13:1730  13:1730  13:1730  13:1731  13:1731  13:1731  13:1731  13:1731  13:1732  13:1732  13:1732  13:1732  13:1732  13:1732  13:1733  13:1733  13:1733  13:1733  13:1733  13:1733  13:1734  13:1734  13:1734  13:1734  13:1734  13:1734  13:1735  13:1735  13:1735  13:1735  13:1735  13:1735  13:1736  13:1736  13:1736  13:1736  13:1736  13:1736  13:1737  13:1737  13:1737  13:1737  13:1737  13:1737  13:1738  13:1738  13:1738  13:1738  13:1738  13:1738  13:1739  13:1739  13:1739  13:1739  13:1739  13:1739  13:1740  13:1740  13:1740  13:1740  13:1740  13:1740  13:1741  13:1741  13:1741  13:1741  13:1741  13:1741  13:1742  13:1742  13:1742  13:1742  13:1742  13:1742  13:1743  13:1743  13:1743  13:1743  13:1743  13:1743  13:1744  13:1744  13:1744  13:1744  13:1744  13:1745  13:1745  13:1745  13:1745  13:1745  13:1746  13:1746  13:1746  13:1746  13:1746  13:1747  13:1747  13:1747  13:1748  13:1748  13:1748  13:1748  13:1749  13:1749  13:1749  13:1750  13:1750  13:1750  13:1750  13:1750  13:1751  13:1751  13:1751  13:1751  13:1752  13:1752  13:1752  13:1752  13:1752  13:1753  13:1753  13:1753  13:1753  13:1753  13:1754  13:1754  13:1754  13:1754  13:1754  13:1755  13:1755  13:1755  13:1755  13:1755  13:1755  13:1756  13:1756  13:1756  13:1756  13:1756  13:1756  13:1757  13:1757  13:1757  13:1757  13:1757  13:1757  13:1758  13:1758  13:1758  13:1758  13:1758  13:1758  13:1759  13:1759  13:1759  13:1759  13:1759  13:1759  13:1760  13:1760  13:1760  13:1760  13:1760  13:1760  13:1761  13:1761  13:1761  13:1761  13:1761  13:1761  13:1762  13:1762  13:1762  13:1762  13:1762  13:1762  13:1763  13:1763  13:1763  13:1763  13:1763  13:1763  13:1764  13:1764  13:1764  13:1764  13:1764  13:1764  13:1765  13:1765  13:1765  13:1765  13:1765  13:1765  13:1766  13:1766  13:1766  13:1766  13:1766  13:1766  13:1767  13:1767  13:1767  13:1767  13:1767  13:1767  13:1768  13:1768  13:1768  13:1768  13:1768  13:1768  13:1769  13:1769  13:1769  13:1770  13:1770  13:1770  13:1770  13:1770  14:3123  14:3253  14:3255  14:3279  14:3279  14:3279  14:3279  14:3280  14:3280  14:3280  14:3280  14:3280  14:3280  14:3281  14:3281  14:3281  14:3281  14:3281  14:3282  14:3282  14:3282  14:3282  14:3282  14:3283  14:3283  14:3283  14:3283  14:3283  14:3284  14:3284  14:3284  14:3284  14:3284  14:3284  14:3285  14:3285  14:3285  14:3285  14:3285  14:3286  14:3286  14:3286  14:3286  14:3286  14:3287  14:3287  14:3287  14:3287  14:3287  14:3287  14:3288  14:3288  14:3288  14:3288  14:3288  14:3288  14:3289  14:3289  14:3289  14:3289  14:3289  14:3289  14:3290  14:3290  14:3290  14:3290  14:3290  14:3290  14:3291  14:3291  14:3291  14:3291  14:3291  14:3291  14:3292  14:3292  14:3292  14:3293  14:3293  14:3293  14:3293  14:3293  14:3294  14:3294  14:3294  14:3294  14:3294  14:3294  14:3295  14:3295  14:3295  14:3295  14:3295  14:3295  14:3296  14:3296  14:3296  14:3296  14:3296  14:3296  14:3297  14:3297  14:3297  14:3297  14:3297  14:3297  14:3298  14:3298  14:3298  14:3298  14:3298  14:3298  14:3299  14:3299  14:3299  14:3299  14:3299  14:3299  14:3300  14:3300  14:3300  14:3300  14:3300  14:3300  14:3301  14:3301  14:3301  14:3301  14:3301  14:3301  14:3302  14:3302  14:3302  14:3302  14:3303  14:3303  14:3303  14:3303  14:3303  14:3303  14:3304  14:3304  14:3304  14:3304  14:3304  14:3304  14:3305  14:3305  14:3305  14:3305  14:3306  14:3306  14:3306  14:3306  14:3306  14:3307  14:3307  14:3307  14:3307  14:3307  14:3307  14:3308  14:3308  14:3308  14:3308  14:3308  14:3308  14:3309  14:3309  14:3309  14:3309  14:3310  14:3310  14:3310  14:3310  14:3310  14:3311  14:3311  14:3311  14:3311  14:3312  14:3312  14:3312  14:3312  14:3313  14:3313  14:3313  14:3313  14:3313  14:3314  14:3314  14:3314  14:3315  14:3315  14:3315  14:3315  14:3315  14:3316  14:3316  14:3316  14:3316  14:3316  14:3317  14:3317  14:3317  14:3317  14:3318  14:3318  14:3318  14:3318  14:3318  14:3319  14:3319  14:3319  14:3319  14:3319  14:3320  14:3320  14:3320  14:3321  14:3321  14:3321  14:3321  14:3322  14:3322  14:3322  14:3322  14:3322  14:3323  14:3323  14:3323  14:3323  14:3323  14:3324  14:3324  14:3324  14:3324  14:3324  14:3324  14:3325  14:3325  14:3325  14:3325  14:3326  14:3326  14:3326  14:3326  14:3326  14:3327  14:3327  14:3327  14:3327  14:3327  14:3328  14:3328  14:3328  14:3328  14:3328  14:3328  14:3329  14:3329  14:3329  14:3329  14:3330  14:3330  14:3330  14:3330  14:3330  14:3331  14:3331  14:3331  14:3331  14:3331  14:3332  14:3332  14:3332  14:3332  14:3332  14:3333  14:3333  14:3333  14:3333  14:3333  14:3333  14:3334  14:3334  14:3334  14:3334  14:3334  14:3335  14:3335  14:3335  14:3335  14:3335  14:3335  14:3336  14:3336  14:3336  14:3336  14:3336  14:3336  14:3337  14:3337  14:3337  14:3337  14:3337  14:3337  14:3338  14:3338  14:3338  14:3338  14:3338  14:3338  14:3339  14:3339  14:3339  14:3339  14:3339  14:3339  14:3340  14:3340  14:3340  14:3341  14:3341  14:3341  14:3341  14:3342  14:3342  14:3342  14:3342  14:3343  14:3343  14:3343  14:3343  14:3344  14:3344  14:3344  14:3344  14:3345  14:3345  14:3345  14:3345  14:3345  14:3345  14:3346  14:3346  14:3346  14:3346  14:3346  14:3347  14:3347  14:3347  14:3348  14:3348  14:3348  14:3348  14:3349  14:3349  14:3349  14:3349  14:3349  14:3349  14:3350  14:3350  14:3350  14:3350  14:3350  14:3351  14:3351  14:3351  14:3351  14:3351  14:3352  14:3352  14:3352  14:3352  14:3352  14:3352  14:3353  14:3353  14:3353  14:3353  14:3353  14:3353  14:3354  14:3354  14:3354  14:3354  14:3354  14:3355  14:3355  14:3355  14:3355  14:3355  14:3355  14:3356  14:3356  14:3356  14:3356  14:3356  14:3357  14:3357  14:3357  14:3357  14:3357  14:3358  14:3358  14:3358  14:3359  14:3359  14:3359  14:3359  14:3360  14:3360  14:3360  14:3361  14:3361  14:3361  14:3361  14:3361  14:3362  14:3362  14:3362  14:3362  14:3362  14:3362  14:3363  14:3363  14:3363  14:3363  14:3363  14:3364  14:3364  14:3364  14:3364  14:3364  14:3365  14:3365  14:3365  14:3365  14:3366  14:3366  14:3366  14:3366  14:3366  14:3366  14:3367  14:3367  14:3367  14:3367  14:3367  14:3367  14:3368  14:3368  14:3368  14:3369  14:3369  14:3369  14:3369  14:3369  14:3370  14:3370  14:3370  14:3370  14:3371  14:3371  14:3371  14:3371  14:3372  14:3372  14:3372  14:3372  14:3373  14:3373  14:3373  14:3373  14:3373  14:3373  14:3374  14:3374  14:3375  14:3375  14:3375  14:3375  14:3375  14:3375  14:3376  14:3376  14:3376  14:3376  14:3376  14:3377  14:3377  14:3377  14:3377  14:3377  14:3378  14:3378  14:3378  14:3378  14:3378  14:3379  14:3379  14:3379  14:3379  14:3379  14:3379  14:3380  14:3380  14:3380  14:3380  14:3380  14:3380  14:3381  14:3381  14:3381  14:3381  14:3381  14:3382  14:3382  14:3382  14:3382  14:3382  14:3382  14:3383  14:3383  14:3383  14:3383  14:3383  14:3383  14:3384  14:3384  14:3384  14:3384  14:3384  14:3384  14:3385  14:3385  14:3385  14:3385  14:3385  14:3386  14:3386  14:3386  14:3386  14:3386  14:3386  14:3387  14:3387  14:3387  14:3387  14:3387  14:3387  14:3388  14:3388  14:3388  14:3388  14:3388  14:3388  14:3389  14:3389  14:3389  14:3389  14:3389  14:3390  14:3390  14:3390  14:3390  14:3390  14:3390  14:3391  14:3391  14:3391  14:3391  14:3391  14:3391  14:3392  14:3392  14:3392  14:3392  14:3392  14:3392  14:3393  14:3393  14:3393  14:3393  14:3393  14:3393  14:3394  14:3394  14:3394  14:3394  14:3394  14:3394  14:3395  14:3395  14:3395  14:3395  14:3395  14:3396  14:3396  14:3396  14:3396  14:3396  14:3396  14:3397  14:3397  14:3397  14:3397  14:3397  14:3398  14:3398  14:3398  14:3398  14:3398  14:3398  14:3399  14:3399  14:3399  14:3399  14:3399  14:3399  14:3400  14:3400  14:3400  14:3400  14:3400  16:211  16:211  16:211  16:211  16:211  16:212  16:212  16:212  16:212  16:212  16:213  16:213  16:213  16:213  16:213  16:213  16:214  16:214  16:214  16:214  16:214  16:214  16:215  16:215  16:215  16:215  16:215  16:215  1G:1701  1G:1701  1G:1701  1G:1701  1G:1701  1G:1701  1G:1702  1G:1702  1G:1702  1G:1702  1G:1702  1G:1702  1G:1703  1G:1703  1G:1703  1G:1703  1G:1703  1G:1703  1G:1704  1G:1704  1G:1704  1G:1704  1G:1704  1G:1704  1G:1705  1G:1705  1G:1705  1G:1705  1G:1705  1G:1705  1G:1706  1G:1706  1G:1706  1G:1706  1G:1706  1G:1706  1G:1707  1G:1707  1G:1707  1G:1707  1G:1707  1G:1707  1G:1708  1G:1708  1G:1708  1G:1708  1G:1708  1G:1708  1G:1709  1G:1709  1G:1709  1G:1709  1G:1709  1G:1709  1G:1710  1G:1710  1G:1710  1G:1710  1G:1710  1G:1710  1G:1711  1G:1711  1G:1711  1G:1711  1G:1711  1G:1711  1G:1712  1G:1712  1G:1712  1G:1712  1G:1712  1G:1712  1G:1713  1G:1713  1G:1713  1G:1714  1G:1714  1G:1714  1G:1714  1G:1714  1G:1714  1G:1715  1G:1715  1G:1715  1G:1715  1G:1715  1G:1716  1G:1716  1G:1716  1G:1716  1G:1716  1G:1716  1G:1717  1G:1717  1G:1717  1G:1717  1G:1717  1G:1717  1G:1718  1G:1718  1G:1718  1G:1718  1G:1718  1G:1719  1G:1719  1G:1720  1G:1720  1G:1720  1G:1720  1G:1720  1G:1721  1G:1721  1G:1721  1G:1721  1G:1721  1G:1722  1G:1722  1G:1722  1G:1722  1G:1722  1G:1723  1G:1723  1G:1723  1G:1723  1G:1724  1G:1724  1G:1724  1G:1724  1G:1724  1G:1724  1G:1725  1G:1725  1G:1725  1G:1725  1G:1725  1G:1725  1G:1726  1G:1726  1G:1726  1G:1726  1G:1726  1G:1727  1G:1727  1G:1727  1G:1727  1G:1727  1G:1727  1G:1728  1G:1728  1G:1728  1G:1728  1G:1728  1G:1728  1G:1729  1G:1729  1G:1729  1G:1729  1G:1729  1G:1729  1G:1730  1G:1730  1G:1730  1G:1730  1G:1730  1G:1730  1G:1731  1G:1731  1G:1731  1G:1731  1G:1731  1G:1731  1G:1732  1G:1732  1G:1732  1G:1732  1G:1732  1G:1732  1G:1733  1G:1733  1G:1733  1G:1733  1G:1733  1G:1733  1G:1734  1G:1734  1G:1734  1G:1734  1G:1734  1G:1734  1G:1735  1G:1735  1G:1735  1G:1735  1G:1735  1G:1735  1G:1736  1G:1736  1G:1736  1G:1736  1G:1736  1H:3152  1H:3166  1H:3261  1H:3261  1H:3268  1H:3284  1H:3309  1H:3313  1H:3374  1H:3382  1H:3382  1H:3382  1H:3382  1H:3382  1H:3383  1H:3383  1H:3383  1H:3383  1H:3384  1H:3384  1H:3384  1H:3384  1H:3384  1H:3384  1H:3385  1H:3385  1H:3385  1H:3385  1H:3385  1H:3385  1H:3386  1H:3386  1H:3386  1H:3386  1H:3387  1H:3387  1H:3387  1H:3387  1H:3387  1H:3387  1H:3388  1H:3388  1H:3388  1H:3388  1H:3388  1H:3389  1H:3389  1H:3389  1H:3389  1H:3390  1H:3390  1H:3390  1H:3390  1H:3390  1H:3390  1H:3391  1H:3391  1H:3391  1H:3391  1H:3391  1H:3392  1H:3392  1H:3392  1H:3392  1H:3392  1H:3392  1H:3393  1H:3393  1H:3393  1H:3393  1H:3393  1H:3394  1H:3394  1H:3394  1H:3394  1H:3394  1H:3394  1H:3395  1H:3395  1H:3395  1H:3395  1H:3395  1H:3395  1H:3396  1H:3396  1H:3396  1H:3396  1H:3396  1H:3396  1H:3397  1H:3397  1H:3397  1H:3397  1H:3397  1H:3398  1H:3398  1H:3398  1H:3398  1H:3398  1H:3398  1H:3399  1H:3399  1H:3399  1H:3399  1H:3399  1H:3399  1H:3400  1H:3400  1H:3400  1H:3400  1H:3401  1H:3401  1H:3401  1H:3401  1H:3401  1H:3401  1H:3402  1H:3402  1H:3402  1H:3402  1H:3402  1H:3402  1H:3403  1H:3403  1H:3403  1H:3403  1H:3403  1H:3403  1H:3404  1H:3404  1H:3404  1H:3404  1H:3404  1H:3404  1H:3405  1H:3405  1H:3405  1H:3405  1H:3405  1H:3405  1H:3406  1H:3406  1H:3406  1H:3406  1H:3406  1H:3406  1H:3407  1H:3407  1H:3407  1H:3407  1H:3407  1H:3407  1H:3408  1H:3408  1H:3408  1H:3408  1H:3408  1H:3408  1H:3409  1H:3409  1H:3409  1H:3409  1H:3410  1H:3410  1H:3410  1H:3410  1H:3410  1H:3410  1H:3411  1H:3411  1H:3411  1H:3411  1H:3411  1H:3411  1H:3412  1H:3412  1H:3412  1H:3412  1H:3412  1H:3413  1H:3413  1H:3413  1H:3413  1H:3413  1H:3413  1H:3414  1H:3414  1H:3414  1H:3414  1H:3414  1H:3414  1H:3415  1H:3415  1H:3415  1H:3415  1H:3415  1H:3416  1H:3416  1H:3416  1H:3416  1H:3416  1H:3416  1H:3417  1H:3417  1H:3417  1H:3417  1H:3417  1H:3417  1H:3418  1H:3418  1H:3418  1H:3418  1H:3418  1H:3418  1H:3419  1H:3419  1H:3419  1H:3419  1H:3419  1H:3419  1H:3420  1H:3420  1H:3420  1H:3420  1H:3421  1H:3421  1H:3421  1H:3421  1H:3421  1H:3421  1H:3422  1H:3422  1H:3422  1H:3422  1H:3423  1H:3423  1H:3423  1H:3423  1H:3424  1H:3424  1H:3424  1H:3425  1H:3425  1H:3425  1H:3425  1H:3425  1H:3425  1H:3426  1H:3426  1H:3426  1H:3426  1H:3427  1H:3427  1H:3427  1H:3427  1H:3427  1H:3427  1H:3428  1H:3428  1H:3428  1H:3428  1H:3429  1H:3429  1H:3429  1H:3429  1H:3430  1H:3430  1H:3430  1H:3430  1H:3430  1H:3431  1H:3431  1H:3431  1H:3431  1H:3431  1H:3431  1H:3432  1H:3432  1H:3432  1H:3432  1H:3432  1H:3432  1H:3433  1H:3433  1H:3433  1H:3433  1H:3433  1H:3433  1H:3434  1H:3434  1H:3434  1H:3434  1H:3434  1H:3434  1H:3435  1H:3435  1H:3435  1H:3435  1H:3435  1H:3435  1H:3436  1H:3436  1H:3436  1H:3436  1H:3436  1H:3436  1H:3437  1H:3437  1H:3437  1H:3437  1H:3437  1H:3437  1H:3438  1H:3438  1H:3438  1H:3438  1H:3438  1H:3438  1H:3439  1H:3439  1H:3439  1H:3439  1H:3439  1H:3439  1H:3440  1H:3440  1H:3440  1H:3440  1H:3440  1H:3440  1H:3441  1H:3441  1H:3441  1H:3441  1H:3441  1H:3441  1H:3442  1H:3442  1H:3442  1H:3442  1H:3442  1H:3442  1H:3443  1H:3443  1H:3443  1H:3443  1H:3443  1H:3443  1H:3444  1H:3444  1H:3444  1H:3444  1H:3444  1H:3444  1H:3445  1H:3445  1H:3445  1H:3445  1H:3445  1H:3445  1H:3446  1H:3446  1H:3446  1H:3446  1H:3446  1H:3446  1H:3447  1H:3447  1H:3447  1H:3447  1H:3447  1H:3447  1H:3448  1H:3448  1H:3448  1H:3448  1H:3448  1H:3448  1H:3449  1H:3449  1H:3449  1H:3449  1H:3449  1H:3449  1H:3450  1H:3450  1H:3450  1H:3450  1H:3450  1H:3450  1H:3451  1H:3451  1H:3451  1H:3451  1H:3451  1H:3451  1H:3452  1H:3452  1H:3452  1H:3452  1H:3452  1H:3452  1H:3453  1H:3453  1H:3453  1H:3453  1H:3453  1H:3453  1H:3454  1H:3454  1H:3454  1H:3454  1H:3454  1H:3454  1H:3455  1H:3455  1H:3455  1H:3456  1H:3456  1H:3456  1H:3456  1H:3457  1H:3457  1H:3457  1H:3458  1H:3458  1H:3458  1H:3458  1H:3459  1H:3459  1H:3459  1H:3459  1H:3459  1H:3460  1H:3460  1H:3460  1H:3460  1H:3460  1H:3461  1H:3461  1H:3461  1H:3461  1H:3461  1H:3462  1H:3462  1H:3462  1H:3462  1H:3462  1H:3462  1H:3463  1H:3463  1H:3463  1H:3463  1H:3463  1H:3464  1H:3464  1H:3464  1H:3465  1H:3465  1H:3465  1H:3465  1H:3465  1H:3466  1H:3466  1H:3466  1H:3466  1H:3466  1H:3466  1H:3467  1H:3467  1H:3467  1H:3467  1H:3467  1H:3467  1H:3468  1H:3468  1H:3468  1H:3468  1H:3468  1H:3468  1H:3469  1H:3469  1H:3469  1H:3469  1H:3469  1H:3470  1H:3470  1H:3470  1H:3470  1H:3470  1H:3471  1H:3471  1H:3471  1H:3471  1H:3471  1H:3471  1H:3472  1H:3472  1H:3472  1H:3472  1H:3472  1H:3473  1H:3473  1H:3473  1H:3473  1H:3474  1H:3474  1H:3474  1H:3474  1H:3474  1H:3475  1H:3475  1H:3475  1H:3475  1H:3475  1H:3476  1H:3476  1H:3476  1H:3476  1H:3476  1H:3476  1H:3477  1H:3477  1H:3477  1H:3477  1H:3478  1H:3478  1H:3478  1H:3478  1H:3478  1H:3478  1H:3479  1H:3479  1H:3479  1H:3479  1H:3480  1H:3480  1H:3480  1H:3480  1H:3480  1H:3481  1H:3481  1H:3481  1H:3481  1H:3481  1H:3482  1H:3482  1H:3482  1H:3482  1H:3482  1H:3482  1H:3483  1H:3483  1H:3483  1H:3483  1H:3483  1H:3483  1H:3484  1H:3484  1H:3484  1H:3484  1H:3485  1H:3485  1H:3485  1H:3485  1H:3485  1H:3485  1H:3486  1H:3486  1H:3486  1H:3486  1H:3486  1H:3487  1H:3487  1H:3487  1H:3487  1H:3487  1H:3487  1H:3488  1H:3488  1H:3488  1H:3488  1H:3488  1H:3488  1H:3489  1H:3489  1H:3489  1H:3489  1H:3489  1H:3489  1H:3490  1H:3490  1H:3490  1H:3490  1H:3490  1H:3491  1H:3491  1H:3491  1H:3491  1H:3491  1H:3491  1H:3492  1H:3492  1H:3492  1H:3492  1H:3492  1H:3492  1H:3493  1H:3493  1H:3493  1H:3493  1H:3493  1H:3493  1H:3494  1H:3494  1H:3494  1H:3494  1H:3494  1H:3494  1H:3495  1H:3495  1H:3495  1H:3495  1H:3495  1H:3495  1H:3496  1H:3496  1H:3496  1H:3496  1H:3496  1H:3496  1H:3497  1H:3497  1H:3497  1H:3497  1H:3497  1H:3497  1H:3498  1H:3498  1H:3498  1H:3498  1H:3498  1H:3498  1H:3499  1H:3499  1H:3499  1H:3499  1H:3499  1H:3500  1H:3500  1H:3500  1H:3500  1H:3501  1H:3501  1H:3501  1H:3501  1H:3501  1H:3501  1H:3502  1H:3502  1H:3502  1H:3502  1H:3502  1H:3502  1H:3503  1H:3503  1H:3503  1H:3503  1H:3504  1H:3504  1H:3504  1H:3505  1H:3505  1H:3505  1H:3505  1H:3505  1H:3505  1H:3506  1H:3506  1H:3506  1H:3507  1H:3507  1H:3507  1H:3507  1H:3507  1H:3508  1H:3508  1H:3508  1H:3508  1H:3509  1H:3509  1H:3509  1H:3509  1H:3509  1H:3510  1H:3510  1H:3510  1H:3510  1H:3510  1H:3511  1H:3511  1H:3511  1H:3511  1H:3511  1H:3511  1H:3512  1H:3512  1H:3512  1H:3512  1H:3512  1H:3513  1H:3513  1H:3513  1H:3513  1H:3513  1H:3513  1H:3514  1H:3514  1H:3514  1H:3514  1H:3515  1H:3515  1H:3515  1H:3515  1H:3516  1H:3516  1H:3516  1H:3517  1H:3517  1H:3517  1H:3518  1H:3518  1H:3518  1H:3518  1H:3518  1H:3519  1H:3519  1H:3519  1H:3520  1H:3520  1H:3520  1H:3520  1H:3520  1H:3521  1H:3521  1H:3521  1H:3521  1H:3522  1H:3522  1H:3522  1H:3522  1H:3522  1H:3522  1H:3523  1H:3523  1H:3523  1H:3523  1H:3523  1H:3524  1H:3524  1H:3524  1H:3525  1H:3525  1H:3525  1H:3525  1H:3525  1H:3526  1H:3526  1H:3526  1H:3526  1H:3526  1H:3526  1H:3527  1H:3527  1H:3527  1H:3527  1H:3527  1H:3528  1H:3528  1H:3528  1H:3528  1H:3528  1H:3528  1H:3529  1H:3529  1H:3529  1H:3529  1H:3529  1H:3529  1H:3530  1H:3530  1H:3530  1H:3531  1H:3531  1H:3531  1H:3531  1H:3531  1H:3531  1H:3532  1H:3532  1H:3532  1H:3532  1H:3532  1H:3532  1H:3533  1H:3533  1H:3533  1H:3533  1H:3533  1H:3533  1H:3534  1H:3534  1H:3534  1H:3534  1H:3534  1H:3534  1H:3535  1H:3535  1H:3535  1H:3535  1H:3535  1H:3535  1H:3536  1H:3536  1H:3536  1H:3536  1H:3536  1H:3537  1H:3537  1H:3537  1H:3537  1H:3537  1H:3537  1H:3538  1H:3538  1H:3538  1H:3538  1H:3538  1H:3538  1H:3539  1H:3539  1H:3539  1H:3539  1H:3539  1H:3539  1H:3540  1H:3540  1H:3540  1H:3540  1H:3540  1H:3540  1H:3541  1H:3541  1H:3541  1H:3541  1H:3541  1H:3541  1H:3542  1H:3542  1H:3542  1H:3542  1H:3542  1H:3542  1H:3543  1H:3543  1H:3543  1H:3543  1H:3544  1H:3544  1H:3544  1H:3544  1H:3544  1H:3545  1H:3545  1H:3545  1H:3545  1H:3545  1H:3545  1H:3546  1H:3546  1H:3546  1H:3546  1H:3546  1H:3546  1H:3547  1H:3547  1H:3547  1H:3547  1H:3547  1H:3547  1H:3548  1H:3548  1H:3548  1H:3548  1H:3548  1H:3548  1H:3549  1H:3549  1H:3549  1H:3549  1H:3549  1H:3549  1H:3550  1H:3550  1H:3550  1H:3550  1H:3550  1H:3550  1H:3551  1H:3551  1H:3551  1H:3551  1H:3551  1H:3552  1H:3552  1H:3552  1H:3552  1H:3552  1H:3552  1H:3553  1H:3553  1H:3553  1H:3553  1H:3553  1H:3553  1H:3554  1H:3554  1H:3554  1H:3554  1H:3554  1H:3554  1H:3555  1H:3555  1H:3555  1H:3555  1H:3555  1H:3555  1H:3556  1H:3556  1H:3556  1H:3556  1H:3556  1H:3556  1H:3557  1H:3557  1H:3557  1H:3557  1H:3557  1H:3558  1H:3558  1H:3558  1H:3558  1H:3558  1H:3558  1H:3559  1H:3559  1H:3559  1H:3559  1H:3559  1H:3560  1H:3560  1H:3560  1H:3560  1H:3561  1H:3561  1H:3561  1H:3561  1H:3561  1H:3561  1H:3562  1H:3562  1H:3562  1H:3562  1H:3562  1H:3563  1H:3563  1H:3563  1H:3563  1H:3563  1H:3563  1H:3564  1H:3564  1H:3564  1H:3564  1H:3565  1H:3565  1H:3565  1H:3565  1H:3565  1H:3566  1H:3566  1H:3566  1H:3566  1H:3567  1H:3567  1H:3567  1H:3567  1H:3567  1H:3568  1H:3568  1H:3568  1H:3568  1H:3568  1H:3568  1H:3569  1H:3569  1H:3569  1H:3569  1H:3569  1H:3570  1H:3570  1H:3570  1H:3570  1H:3570  1H:3570  1H:3571  1H:3571  1H:3571  1H:3571  1H:3571  1H:3572  1H:3572  1H:3572  1H:3572  1H:3572  1H:3572  1H:3573  1H:3573  1H:3573  1H:3573  1H:3574  1H:3574  1H:3574  1H:3574  1H:3575  1H:3575  1H:3575  1H:3575  1H:3576  1H:3576  1H:3576  1H:3576  1H:3576  1H:3576  1H:3577  1H:3577  1H:3577  1H:3577  1J:205  1J:205  1J:205  1J:205  1J:205  21:302  21:302  21:302  21:302  21:302  21:303  21:303  21:303  21:303  21:303  21:303  29:303  29:303  29:303  29:303  2K:105  2K:105  2K:105  2K:105  2K:105  2K:105  32:302  32:302  32:302  32:302  32:302  32:302  3E:301  3E:301  3E:301  3E:301  3E:301  3E:301  78:202  78:202  78:202  78:202  78:202  78:202  M5:101  M5:101  M5:101  M5:101  M5:101  M5:101  M5:102  M5:102  M5:102  M5:102  M5:102  M5:102 |
| **3) MG/OD2_Asp_ contacts** | 1 | 29:302 |
| **3) MG/OE1_Glu_ contacts** | 1 | 14:3237 |
| **3) MG/OP1 contacts** | 109 | 13:1656  13:1660  13:1663  13:1670  13:1677  13:1678  13:1683  13:1724  13:1726  13:1726  13:1726  13:1748  14:3107  14:3107  14:3110  14:3114  14:3124  14:3124  14:3127  14:3128  14:3128  14:3129  14:3130  14:3130  14:3130  14:3139  14:3139  14:3140  14:3147  14:3151  14:3152  14:3153  14:3153  14:3154  14:3155  14:3157  14:3160  14:3160  14:3161  14:3162  14:3162  14:3169  14:3171  14:3172  14:3191  14:3192  14:3216  14:3223  14:3229  14:3257  14:3258  14:3309  14:3321  14:3325  14:3356  1G:1637  1G:1637  1G:1637  1G:1638  1G:1639  1G:1642  1G:1646  1G:1652  1G:1662  1G:1695  1H:3155  1H:3168  1H:3170  1H:3171  1H:3171  1H:3176  1H:3176  1H:3176  1H:3178  1H:3178  1H:3180  1H:3188  1H:3191  1H:3191  1H:3192  1H:3198  1H:3199  1H:3200  1H:3203  1H:3232  1H:3244  1H:3246  1H:3261  1H:3269  1H:3324  1H:3324  1H:3324  1H:3350  1H:3350  1H:3374  1H:3393  1H:3459  1H:3461  1H:3463  1H:3479  1H:3499  1H:3510  1H:3512  1H:3521  1H:3525  1H:3566  1H:3575  1H:3575  2L:101 |
| **3) MG/OP2 contacts** | 122 | 13:1653  13:1657  13:1661  13:1668  13:1668  13:1668  13:1674  13:1677  13:1684  13:1693  14:3108  14:3108  14:3119  14:3123  14:3126  14:3126  14:3129  14:3131  14:3131  14:3132  14:3133  14:3134  14:3136  14:3136  14:3138  14:3138  14:3144  14:3147  14:3154  14:3156  14:3158  14:3160  14:3171  14:3177  14:3179  14:3189  14:3196  14:3198  14:3199  14:3199  14:3206  14:3211  14:3212  14:3214  14:3217  14:3217  14:3218  14:3222  14:3247  14:3258  14:3265  14:3266  14:3273  14:3286  14:3302  14:3311  14:3313  16:210  1G:1643  1G:1644  1G:1644  1G:1644  1G:1645  1G:1645  1G:1647  1G:1648  1G:1648  1G:1650  1G:1654  1G:1655  1G:1668  1G:1673  1G:1676  1G:1684  1G:1693  1G:1693  1G:1699  1H:3145  1H:3147  1H:3147  1H:3149  1H:3149  1H:3150  1H:3154  1H:3155  1H:3155  1H:3162  1H:3167  1H:3167  1H:3167  1H:3168  1H:3169  1H:3176  1H:3176  1H:3180  1H:3188  1H:3191  1H:3194  1H:3195  1H:3195  1H:3205  1H:3206  1H:3206  1H:3215  1H:3232  1H:3250  1H:3250  1H:3255  1H:3257  1H:3310  1H:3360  1H:3391  1H:3415  1H:3456  1H:3474  1H:3481  1H:3504  1H:3506  1H:3559  1J:202  2L:101  L8:101 |
| **3) K/N2 excl. contacts** | 1 | 1H:3030 |
| **3) K/O2 excl. contacts** | 5 | 13:1608  13:1635  14:3032  1G:1619  1H:3070 |
| **3) K/O2' excl. contacts** | 2 | 1H:3078  1H:3113 |
| **3) K/O3' excl. contacts** | 2 | 1G:1628  1G:1629 |
| **3) K/O4 excl. contacts** | 2 | 1H:3086  1H:3136 |
| **3) K/O6 excl. contacts** | 15 | 13:1629  14:3021  14:3033  14:3094  1H:3006  1H:3008  1H:3009  1H:3035  1H:3043  1H:3059  1H:3069  1H:3071  1H:3085  1H:3139  1J:201 |
| **3) K/O_back_ excl. contacts** | 5 | 1G:1625  1H:3069  2A:201  41:201  4I:201 |
| **3) K/O_w_ excl. contacts** | 4 | 14:3028  14:3054  1H:3053  1H:3104 |
| **3) K/OP1 excl. contacts** | 9 | 13:1609  13:1644  14:3069  14:3078  1G:1625  1H:3033  1H:3063  1H:3065  4I:201 |
| **3) K/OP2 excl. contacts** | 13 | 13:1651  14:3005  14:3008  14:3008  14:3008  14:3036  1G:1603  1G:1624  1H:3033  1H:3038  1H:3060  1H:3078  4A:201 |
| **3) MG/N1_A_ excl. contacts** | 3 | 14:3176  14:3276  1H:3328 |
| **3) MG/N1_G_ excl. contacts** | 4 | 14:3152  14:3214  1G:1679  1H:3203 |
| **3) MG/N2 excl. contacts** | 2 | 14:3152  1H:3181 |
| **3) MG/N3_A_ excl. contacts** | 3 | 13:1705  14:3124  1H:3197 |
| **3) MG/N3_C_ excl. contacts** | 4 | 13:1674  14:3232  14:3237  1G:1711 |
| **3) MG/N3_U_ excl. contacts** | 1 | 1H:3267 |
| **3) MG/N4 excl. contacts** | 2 | 14:3249  1H:3306 |
| **3) MG/N6 excl. contacts** | 12 | 14:3135  14:3135  14:3157  14:3190  14:3235  14:3237  14:3366  1G:1713  1H:3146  1H:3257  1H:3279  1H:3356 |
| **3) MG/N7 excl. contacts** | 75 | 13:1655  13:1658  13:1672  13:1690  13:1692  13:1706  13:1711  13:1715  13:1757  13:1763  13:1763  13:1770  14:3112  14:3112  14:3121  14:3129  14:3131  14:3181  14:3182  14:3201  14:3209  14:3216  14:3250  14:3251  14:3253  14:3253  14:3260  14:3279  14:3279  14:3301  14:3327  14:3348  14:3357  14:3397  1G:1677  1G:1677  1G:1684  1G:1685  1G:1690  1G:1691  1G:1733  1H:3146  1H:3148  1H:3163  1H:3168  1H:3186  1H:3221  1H:3240  1H:3252  1H:3264  1H:3273  1H:3279  1H:3297  1H:3306  1H:3330  1H:3340  1H:3345  1H:3348  1H:3352  1H:3371  1H:3409  1H:3409  1H:3413  1H:3417  1H:3465  1H:3474  1H:3489  1H:3512  1H:3518  1H:3519  1H:3538  1H:3540  1H:3564  1J:203  42:201 |
| **3) MG/ND1_His_ excl. contacts** | 1 | 1H:3334 |
| **3) MG/NE2_His_ excl. contacts** | 5 | 14:3348  1H:3389  1H:3508  1H:3518  29:303 |
| **3) MG/NH1_Arg_ excl. contacts** | 4 | 3E:301  42:201  C8:201  I8:101 |
| **3) MG/NH2_Arg_ excl. contacts** | 1 | C8:201 |
| **3) MG/NZ_Lys_ excl. contacts** | 5 | 13:1691  14:3218  1H:3298  32:302  M5:101 |
| **3) MG/O2 excl. contacts** | 13 | 13:1653  13:1688  13:1769  14:3235  14:3247  14:3278  1G:1694  1H:3230  1H:3267  1H:3354  1H:3356  1H:3381  1H:3573 |
| **3) MG/O2' excl. contacts** | 29 | 13:1668  13:1695  13:1705  13:1769  14:3202  14:3236  14:3354  14:3358  14:3385  16:206  1G:1686  1G:1693  1G:1694  1H:3155  1H:3207  1H:3213  1H:3232  1H:3238  1H:3249  1H:3290  1H:3298  1H:3302  1H:3320  1H:3354  1H:3361  1H:3366  1H:3371  1H:3372  1H:3378 |
| **3) MG/O3' excl. contacts** | 34 | 13:1674  13:1695  13:1726  13:1745  14:3163  14:3175  14:3199  14:3202  14:3266  14:3314  14:3358  14:3376  16:205  1G:1637  1G:1646  1G:1694  1H:3155  1H:3171  1H:3188  1H:3257  1H:3265  1H:3280  1H:3298  1H:3302  1H:3308  1H:3361  1H:3371  1H:3378  1H:3469  1H:3504  1H:3509  1H:3519  1H:3559  1H:3565 |
| **3) MG/O4 excl. contacts** | 34 | 13:1674  13:1708  13:1720  13:1722  13:1722  14:3141  14:3166  14:3227  14:3231  14:3236  14:3267  14:3359  14:3374  16:206  1G:1648  1G:1678  1G:1687  1G:1688  1H:3227  1H:3286  1H:3327  1H:3327  1H:3328  1H:3330  1H:3349  1H:3351  1H:3367  1H:3388  1H:3429  1H:3473  1H:3528  1H:3560  1J:202  42:201 |
| **3) MG/O4' excl. contacts** | 3 | 1H:3207  1H:3320  1H:3366 |
| **3) MG/O5' excl. contacts** | 32 | 13:1656  13:1676  13:1721  14:3198  14:3198  14:3283  14:3292  14:3347  14:3360  14:3368  1G:1719  1G:1719  1H:3176  1H:3176  1H:3182  1H:3183  1H:3191  1H:3200  1H:3238  1H:3254  1H:3269  1H:3278  1H:3284  1H:3324  1H:3355  1H:3386  1H:3389  1H:3391  1H:3424  1H:3428  1H:3571  1H:3574 |
| **3) MG/O6 excl. contacts** | 119 | 13:1659  13:1659  13:1673  13:1674  13:1681  13:1687  13:1690  13:1692  13:1697  13:1701  13:1701  13:1702  13:1703  13:1708  13:1708  13:1708  13:1709  13:1709  13:1712  13:1713  13:1713  13:1720  13:1720  13:1722  14:3109  14:3118  14:3118  14:3121  14:3124  14:3146  14:3170  14:3182  14:3185  14:3186  14:3187  14:3190  14:3197  14:3201  14:3212  14:3215  14:3216  14:3241  14:3248  14:3249  14:3249  14:3250  14:3250  14:3253  14:3256  14:3263  14:3267  14:3268  14:3268  14:3274  14:3276  14:3279  14:3291  14:3298  14:3327  14:3360  14:3362  16:206  16:207  16:208  16:208  1G:1651  1G:1657  1G:1659  1G:1670  1G:1675  1G:1678  1G:1679  1G:1679  1G:1682  1G:1683  1G:1685  1G:1687  1G:1723  1G:1736  1H:3159  1H:3163  1H:3186  1H:3201  1H:3202  1H:3202  1H:3203  1H:3210  1H:3211  1H:3212  1H:3221  1H:3225  1H:3229  1H:3240  1H:3264  1H:3273  1H:3286  1H:3292  1H:3311  1H:3311  1H:3323  1H:3328  1H:3328  1H:3329  1H:3331  1H:3341  1H:3349  1H:3351  1H:3351  1H:3359  1H:3363  1H:3365  1H:3369  1H:3373  1H:3400  1H:3475  1H:3529  1J:202  1J:204  29:302 |
| **3) MG/O_back_ excl. contacts** | 10 | 11:302  14:3131  14:3261  14:3319  14:3320  1H:3206  1H:3208  21:303  F8:101  L8:101 |
| **3) MG/O_w_ excl. contacts** | 87 | 13:1666  14:3109  14:3123  14:3127  14:3139  14:3166  14:3166  14:3175  14:3243  14:3249  14:3249  14:3250  14:3250  14:3250  14:3253  14:3254  14:3254  14:3254  14:3259  14:3266  16:210  1G:1655  1H:3152  1H:3152  1H:3153  1H:3154  1H:3158  1H:3158  1H:3158  1H:3166  1H:3192  1H:3208  1H:3208  1H:3238  1H:3260  1H:3260  1H:3260  1H:3263  1H:3263  1H:3265  1H:3266  1H:3266  1H:3268  1H:3269  1H:3274  1H:3274  1H:3278  1H:3279  1H:3279  1H:3284  1H:3292  1H:3294  1H:3302  1H:3306  1H:3307  1H:3307  1H:3309  1H:3309  1H:3310  1H:3311  1H:3311  1H:3312  1H:3314  1H:3316  1H:3318  1H:3319  1H:3329  1H:3331  1H:3331  1H:3332  1H:3332  1H:3332  1H:3333  1H:3345  1H:3348  1H:3355  1H:3355  1H:3355  1H:3374  1H:3378  1H:3472  1H:3530  1H:3530  1H:3530  1H:3557  L8:101  L8:101 |
| **3) MG/OD1_Asn_ excl. contacts** | 1 | F8:101 |
| **3) MG/OD2_Asp_ excl. contacts** | 3 | 21:302  39:303  N8:101 |
| **3) MG/OE2_Glu_ excl. contacts** | 1 | 14:3237 |
| **3) MG/OG_Ser_ excl. contacts** | 1 | 1H:3208 |
| **3) MG/OP1 excl. contacts** | 282 | 11:302  13:1656  13:1656  13:1657  13:1659  13:1659  13:1661  13:1667  13:1676  13:1684  13:1686  13:1691  13:1700  13:1700  13:1704  13:1705  13:1706  13:1714  13:1717  13:1731  13:1735  13:1746  13:1747  13:1748  13:1750  13:1751  13:1752  13:1753  13:1761  13:1769  14:3120  14:3126  14:3126  14:3128  14:3130  14:3137  14:3140  14:3143  14:3150  14:3151  14:3153  14:3163  14:3167  14:3167  14:3168  14:3174  14:3175  14:3175  14:3180  14:3191  14:3198  14:3198  14:3198  14:3209  14:3210  14:3213  14:3225  14:3226  14:3228  14:3230  14:3230  14:3242  14:3246  14:3246  14:3260  14:3261  14:3270  14:3273  14:3280  14:3281  14:3282  14:3283  14:3289  14:3292  14:3292  14:3298  14:3301  14:3305  14:3305  14:3306  14:3309  14:3310  14:3312  14:3312  14:3314  14:3316  14:3317  14:3317  14:3319  14:3320  14:3321  14:3325  14:3326  14:3329  14:3329  14:3332  14:3340  14:3340  14:3340  14:3341  14:3344  14:3344  14:3346  14:3347  14:3347  14:3348  14:3354  14:3355  14:3357  14:3360  14:3360  14:3361  14:3363  14:3365  14:3368  14:3368  14:3370  14:3370  14:3371  14:3371  14:3372  14:3372  14:3374  14:3376  14:3377  14:3378  14:3382  16:204  16:211  1G:1642  1G:1643  1G:1645  1G:1653  1G:1655  1G:1668  1G:1674  1G:1680  1G:1693  1G:1694  1G:1695  1G:1695  1G:1699  1G:1703  1G:1715  1G:1719  1G:1719  1G:1719  1G:1720  1G:1722  1G:1723  1G:1723  1G:1724  1G:1726  1G:1730  1G:1730  1H:3152  1H:3155  1H:3165  1H:3170  1H:3170  1H:3175  1H:3175  1H:3176  1H:3176  1H:3176  1H:3182  1H:3182  1H:3183  1H:3187  1H:3194  1H:3204  1H:3230  1H:3236  1H:3237  1H:3242  1H:3246  1H:3248  1H:3249  1H:3249  1H:3254  1H:3255  1H:3260  1H:3260  1H:3265  1H:3265  1H:3267  1H:3275  1H:3276  1H:3278  1H:3280  1H:3281  1H:3284  1H:3295  1H:3299  1H:3302  1H:3308  1H:3311  1H:3319  1H:3322  1H:3324  1H:3325  1H:3334  1H:3360  1H:3374  1H:3376  1H:3379  1H:3382  1H:3384  1H:3384  1H:3389  1H:3391  1H:3392  1H:3394  1H:3399  1H:3400  1H:3405  1H:3412  1H:3419  1H:3419  1H:3420  1H:3420  1H:3422  1H:3422  1H:3424  1H:3424  1H:3426  1H:3426  1H:3428  1H:3428  1H:3438  1H:3454  1H:3455  1H:3455  1H:3456  1H:3457  1H:3457  1H:3462  1H:3464  1H:3464  1H:3468  1H:3469  1H:3469  1H:3472  1H:3475  1H:3478  1H:3479  1H:3484  1H:3491  1H:3503  1H:3503  1H:3505  1H:3506  1H:3507  1H:3507  1H:3508  1H:3511  1H:3513  1H:3513  1H:3514  1H:3516  1H:3516  1H:3516  1H:3519  1H:3523  1H:3524  1H:3524  1H:3524  1H:3526  1H:3527  1H:3536  1H:3543  1H:3543  1H:3551  1H:3564  1H:3565  1H:3566  1H:3567  1H:3569  1J:202  29:303  2L:101  4L:101 |
| **3) MG/OP2 excl. contacts** | 324 | 13:1656  13:1659  13:1662  13:1664  13:1665  13:1672  13:1684  13:1685  13:1688  13:1690  13:1695  13:1695  13:1698  13:1700  13:1702  13:1705  13:1707  13:1711  13:1721  13:1721  13:1731  13:1737  13:1737  13:1744  13:1745  13:1747  13:1747  13:1747  13:1749  13:1749  13:1749  13:1751  13:1751  13:1753  13:1754  13:1769  14:3108  14:3109  14:3111  14:3112  14:3114  14:3127  14:3133  14:3135  14:3137  14:3143  14:3144  14:3145  14:3145  14:3158  14:3160  14:3162  14:3170  14:3174  14:3174  14:3176  14:3177  14:3180  14:3182  14:3189  14:3193  14:3195  14:3210  14:3211  14:3218  14:3218  14:3220  14:3220  14:3221  14:3231  14:3234  14:3244  14:3244  14:3253  14:3255  14:3255  14:3260  14:3264  14:3265  14:3266  14:3269  14:3281  14:3285  14:3292  14:3293  14:3295  14:3301  14:3302  14:3309  14:3309  14:3311  14:3312  14:3314  14:3314  14:3314  14:3315  14:3318  14:3320  14:3320  14:3322  14:3323  14:3325  14:3330  14:3331  14:3332  14:3334  14:3340  14:3341  14:3342  14:3342  14:3343  14:3343  14:3344  14:3346  14:3347  14:3347  14:3348  14:3350  14:3351  14:3354  14:3355  14:3357  14:3358  14:3358  14:3358  14:3359  14:3364  14:3365  14:3365  14:3368  14:3368  14:3369  14:3370  14:3374  14:3374  14:3381  14:3382  14:3385  14:3389  14:3395  14:3397  14:3400  16:205  16:205  16:209  16:212  1G:1640  1G:1642  1G:1646  1G:1646  1G:1663  1G:1667  1G:1668  1G:1670  1G:1674  1G:1674  1G:1677  1G:1677  1G:1681  1G:1691  1G:1693  1G:1699  1G:1718  1G:1719  1G:1720  1G:1721  1G:1726  1H:3153  1H:3155  1H:3157  1H:3158  1H:3170  1H:3172  1H:3172  1H:3176  1H:3188  1H:3189  1H:3189  1H:3192  1H:3198  1H:3198  1H:3208  1H:3208  1H:3221  1H:3223  1H:3231  1H:3233  1H:3234  1H:3238  1H:3240  1H:3242  1H:3248  1H:3249  1H:3254  1H:3263  1H:3263  1H:3264  1H:3267  1H:3269  1H:3269  1H:3274  1H:3274  1H:3279  1H:3280  1H:3281  1H:3285  1H:3286  1H:3290  1H:3291  1H:3291  1H:3297  1H:3299  1H:3300  1H:3306  1H:3306  1H:3310  1H:3312  1H:3322  1H:3325  1H:3325  1H:3327  1H:3333  1H:3344  1H:3345  1H:3345  1H:3345  1H:3349  1H:3355  1H:3355  1H:3357  1H:3357  1H:3367  1H:3368  1H:3375  1H:3378  1H:3380  1H:3383  1H:3383  1H:3385  1H:3386  1H:3386  1H:3387  1H:3390  1H:3397  1H:3399  1H:3407  1H:3411  1H:3411  1H:3413  1H:3420  1H:3423  1H:3423  1H:3424  1H:3429  1H:3430  1H:3433  1H:3449  1H:3453  1H:3454  1H:3455  1H:3455  1H:3456  1H:3457  1H:3457  1H:3458  1H:3458  1H:3460  1H:3464  1H:3464  1H:3466  1H:3470  1H:3473  1H:3475  1H:3476  1H:3477  1H:3477  1H:3480  1H:3486  1H:3487  1H:3490  1H:3498  1H:3500  1H:3500  1H:3502  1H:3504  1H:3504  1H:3505  1H:3506  1H:3507  1H:3508  1H:3509  1H:3514  1H:3515  1H:3515  1H:3516  1H:3517  1H:3517  1H:3517  1H:3519  1H:3520  1H:3521  1H:3524  1H:3524  1H:3527  1H:3530  1H:3543  1H:3544  1H:3548  1H:3549  1H:3557  1H:3560  1H:3562  1H:3571  1H:3573  1H:3574  1H:3574  1H:3577  1H:3577  1J:202  1J:205  2K:105  4L:101  C8:201  F8:101 |
| **4) Large octahedral topology discrepancy** | 13 | 14:3130  14:3286  14:3302  14:3309  14:3321  14:3325  1H:3459  1H:3484  1H:3504  1H:3510  1H:3512  1H:3566  1H:3575 |

---

\_\_\_\_\_\_\_\_\_\_\_\_\_\_\_\_\_\_\_\_\_\_\_\_\_\_\_\_\_\_\_\_\_\_\_\_\_\_\_\_\_\_\_\_\_\_\_\_\_\_\_\_\_\_\_\_\_\_\_\_\_\_\_\_  

#### 4) Ion/ligand tables: all contacts up to 5.0 Angs. are listed

##### Go to ToC

### K:11:301

|  | Res. type | Atom | Residue | Distance | Occupancy | B-factor | Closest solute to 1^st^ shell water |
| --- | --- | --- | --- | --- | --- | --- | --- |
| **Ion** | | | |  | 1.00 | 31.84 || 1 | GLY | O | 11:235 | 2.71 | 1.00 | 20.78 |
| 2 | GLU | O | 11:237 | 2.98 | 1.00 | 38.69 |  |
| 3 | Water 1 | | 11:409 | 3.14 | 1.00 | 25.04 | 3.76 (ALA:O 11:224) |
| 4 | GLY | O | 11:236 | 3.30 | 1.00 | 23.87 |  |
| 5 | ARG | N | 11:239 | 3.65 | 1.00 | 20.38 |  |
| 6 | A | OP2 | 1H:2603 | 4.20 | 1.00 | 22.34 |  |
| 7 | A | OP1 | 1H:2603 | 4.45 | 1.00 | 20.25 |  |
| 8 | GLY | N | 11:236 | 4.49 | 1.00 | 20.15 |  |
| 9 | ALA | O | 11:224 | 4.71 | 1.00 | 23.50 |  |
| 10 | GLU | N | 11:237 | 4.78 | 1.00 | 20.42 |  |
| 11 | GLY | N | 11:235 | 4.84 | 1.00 | 17.92 |  |
| 12 | GLY | N | 11:238 | 4.89 | 1.00 | 25.18 |  |
  |

  
**1^st^ shell ligand angles:**   
1-2: 90; 1-3: 96; 1-4: 60; 2-3: 157; 2-4: 53; 3-4: 111;   
  
 Contact(s): /O_back_/O_back_/O_back_/O_w_/  
 Coded contact list: 3O_back_O_w_  

### MG:11:302

|  | Res. type | Atom | Residue | Distance | Occupancy | B-factor | Closest solute to 1^st^ shell water |
| --- | --- | --- | --- | --- | --- | --- | --- |
| **Ion** | | | |  | 1.00 | 14.91 || 1 | Water 1 | | 11:402 | 2.16 | 1.00 | 18.70 | 2.46 (PRO:O 11:232) |
| 2 | Water 2 | | 1H:3661 | 2.16 | 1.00 | 17.30 | 2.10 (A:OP1 1H:2611) |
| 3 | Water 3 | | 1H:3662 | 2.16 | 1.00 | 19.67 | 2.10 (G:O3' 1H:2610) |
| 4 | Water 4 | | 11:407 | 2.17 | 1.00 | 16.52 | 2.91 (ARG:O 11:242) |
| 5 | Water 5 | | 11:401 | 2.17 | 1.00 | 17.06 | 1.97 (ARG:O 11:242) |
| 6 | Water 6 | | 11:404 | 2.19 | 1.00 | 17.01 | 2.68 (PRO:O 11:241) |
| 7 | A | OP1 | 1H:2611 | 2.75 | 1.00 | 13.91 |  |
| 8 | ARG | O | 11:242 | 3.18 | 1.00 | 18.42 |  |
| 9 | G | O3' | 1H:2610 | 3.91 | 1.00 | 14.64 |  |
| 10 | PRO | O | 11:241 | 4.01 | 1.00 | 20.91 |  |
| 11 | GLY | O | 11:234 | 4.05 | 1.00 | 17.52 |  |
| 12 | HIS | O | 11:233 | 4.23 | 1.00 | 20.82 |  |
| 13 | PRO | O | 11:232 | 4.37 | 1.00 | 20.82 |  |
| 14 | GLY | N | 11:243 | 4.70 | 1.00 | 14.14 |  |
| 15 | A | OP2 | 1H:2611 | 4.81 | 1.00 | 16.73 |  |
| 16 | A | O5' | 1H:2611 | 4.91 | 1.00 | 17.13 |  |
  |

  
**1^st^ shell ligand angles:**   
1-2: 85; 1-3: 81; 1-4: 95; 1-5: 93; 1-6: 172; 2-3: 104; 2-4: 84; 2-5: 178; 2-6: 89; 3-4: 171; 3-5: 75; 3-6: 95; 4-5: 97; 4-6: 89; 5-6: 93;   
  
 Contact(s): /O_w_/O_w_/O_w_/O_w_/O_w_/O_w_/  
 Coded contact list: 6O_w_  
 Exclusion zone contact(s): /O_back_/OP1/  
Consider K in place of MG (2 valid contact(s) for K vs. 0 for MG)  
MG hydration shell issue: water too close (< 2.4 Angs.)  

### K:13:1601

|  | Res. type | Atom | Residue | Distance | Occupancy | B-factor | Closest solute to 1^st^ shell water |
| --- | --- | --- | --- | --- | --- | --- | --- |
| **Ion** | | | |  | 1.00 | 67.83 || 1 | G | N7 | 13:1222 | 2.94 | 1.00 | 71.68 |
| 2 | C | N4 | 13:1322 | 3.41 | 1.00 | 73.31 |  |
| 3 | C | OP1 | 13:979 | 3.53 | 1.00 | 64.95 |  |
| 4 | G | OP2 | 13:1222 | 3.58 | 1.00 | 66.99 |  |
| 5 | C | N4 | 13:1223 | 3.70 | 1.00 | 66.02 |  |
| 6 | A | O3' | 13:978 | 3.72 | 1.00 | 68.46 |  |
| 7 | G | O6 | 13:1222 | 3.78 | 1.00 | 69.18 |  |
| 8 | G | N7 | 13:1224 | 4.19 | 1.00 | 74.74 |  |
| 9 | G | N9 | 13:1222 | 4.57 | 1.00 | 71.71 |  |
| 10 | C | N3 | 13:1322 | 4.66 | 1.00 | 73.27 |  |
| 11 | C | OP2 | 13:1223 | 4.79 | 1.00 | 70.64 |  |
| 12 | G | O5' | 13:1222 | 4.85 | 1.00 | 62.52 |  |
| 13 | A | O2' | 13:978 | 4.89 | 1.00 | 69.43 |  |
| 14 | C | OP2 | 13:979 | 4.98 | 1.00 | 65.29 |  |
  |

  
 Contact(s): /N7/  
 Coded contact list: N_b_  

### K:13:1602

|  | Res. type | Atom | Residue | Distance | Occupancy | B-factor | Closest solute to 1^st^ shell water |
| --- | --- | --- | --- | --- | --- | --- | --- |
| **Ion** | | | |  | 1.00 | 72.95 || 1 | 2MG | O6 | 13:1207 | 2.90 | 1.00 | 65.97 |
| 2 | C | O2 | 13:1200 | 2.91 | 1.00 | 63.19 |  |
| 3 | G | O6 | 13:1206 | 2.93 | 1.00 | 64.67 |  |
| 4 | U | O4 | 13:1052 | 3.20 | 1.00 | 64.62 |  |
| 5 | U | O4 | 13:1205 | 3.77 | 1.00 | 63.51 |  |
| 6 | U | N3 | 13:1205 | 3.82 | 1.00 | 58.49 |  |
| 7 | C | N4 | 13:1051 | 3.84 | 1.00 | 62.17 |  |
| 8 | G | N7 | 13:1206 | 3.88 | 1.00 | 65.49 |  |
| 9 | C | O2' | 13:1200 | 4.47 | 1.00 | 66.63 |  |
| 10 | 2MG | N1 | 13:1207 | 4.51 | 1.00 | 62.40 |  |
| 11 | C | N3 | 13:1200 | 4.53 | 1.00 | 63.44 |  |
| 12 | U | N3 | 13:1052 | 4.67 | 1.00 | 66.77 |  |
| 13 | 2MG | N7 | 13:1207 | 4.68 | 1.00 | 64.99 |  |
| 14 | A | N6 | 13:1055 | 4.85 | 1.00 | 63.39 |  |
| 15 | C | N1 | 13:1200 | 4.88 | 1.00 | 55.92 |  |
| 16 | U | O2 | 13:1205 | 4.99 | 1.00 | 59.46 |  |
  |

  
**1^st^ shell ligand angles:**   
1-2: 133; 1-3: 64; 1-4: 81; 2-3: 75; 2-4: 58; 3-4: 60;   
  
 Contact(s): /O2/O4/O6/O6/  
 Coded contact list: 4O_b_  

### K:13:1603

|  | Res. type | Atom | Residue | Distance | Occupancy | B-factor | Closest solute to 1^st^ shell water |
| --- | --- | --- | --- | --- | --- | --- | --- |
| **Ion** | | | |  | 1.00 | 75.94 || 1 | G | O6 | 13:1079 | 2.90 | 1.00 | 57.14 |
| 2 | G | O2' | 13:1077 | 3.05 | 1.00 | 74.54 |  |
| 3 | A | O3' | 13:865 | 3.39 | 1.00 | 48.04 |  |
| 4 | C | OP1 | 13:866 | 3.41 | 1.00 | 49.99 |  |
| 5 | G | O3' | 13:1077 | 4.35 | 1.00 | 59.73 |  |
| 6 | C | OP2 | 13:866 | 4.46 | 1.00 | 46.96 |  |
  |

  
**1^st^ shell ligand angles:**   
1-2: 67; 1-3: 92; 2-3: 155;   
  
 Contact(s): /O2'/O3'/O6/  
 Coded contact list: 2O_r_O_b_  

### K:13:1604

|  | Res. type | Atom | Residue | Distance | Occupancy | B-factor | Closest solute to 1^st^ shell water |
| --- | --- | --- | --- | --- | --- | --- | --- |
| **Ion** | | | |  | 1.00 | 82.90 || 1 | G | O6 | 13:976 | 2.65 | 1.00 | 79.37 |
| 2 | C | O2 | 13:1359 | 3.00 | 1.00 | 74.67 |  |
| 3 | G | N7 | 13:976 | 3.06 | 1.00 | 75.64 |  |
| 4 | U | O4 | 13:1358 | 3.60 | 1.00 | 73.39 |  |
| 5 | C | OP2 | 13:1362 | 4.16 | 1.00 | 71.50 |  |
| 6 | U | N3 | 13:1358 | 4.27 | 1.00 | 71.73 |  |
| 7 | G | N1 | 13:976 | 4.65 | 1.00 | 75.48 |  |
| 8 | C | N3 | 13:1359 | 4.96 | 1.00 | 67.67 |  |
| 9 | C | N1 | 13:1359 | 4.96 | 1.00 | 72.15 |  |
  |

  
**1^st^ shell ligand angles:**   
1-2: 144; 1-3: 67; 2-3: 82;   
  
 Contact(s): /N7/O2/O6/  
 Coded contact list: 2O_b_N_b_  

### K:13:1605

|  | Res. type | Atom | Residue | Distance | Occupancy | B-factor | Closest solute to 1^st^ shell water |
| --- | --- | --- | --- | --- | --- | --- | --- |
| **Ion** | | | |  | 1.00 | 81.86 || 1 | C | O2 | 13:1322 | 2.89 | 1.00 | 74.00 |
| 2 | C | O2' | 13:1322 | 2.96 | 1.00 | 75.40 |  |
| 3 | A | N3 | 13:978 | 3.31 | 1.00 | 76.59 |  |
| 4 | A | O4' | 13:1319 | 3.66 | 1.00 | 74.77 |  |
| 5 | A | O3' | 13:1319 | 3.92 | 1.00 | 82.74 |  |
| 6 | A | N3 | 13:1319 | 3.96 | 1.00 | 80.46 |  |
| 7 | C | N4 | 13:1321 | 4.04 | 1.00 | 73.99 |  |
| 8 | A | O2' | 13:1319 | 4.09 | 1.00 | 78.13 |  |
| 9 | C | N3 | 13:1322 | 4.22 | 1.00 | 73.27 |  |
| 10 | A | N9 | 13:1319 | 4.36 | 1.00 | 78.71 |  |
| 11 | C | OP2 | 13:1321 | 4.50 | 1.00 | 72.89 |  |
| 12 | A | N9 | 13:978 | 4.53 | 1.00 | 72.91 |  |
| 13 | C | N1 | 13:1322 | 4.55 | 1.00 | 73.61 |  |
| 14 | A | O4' | 13:978 | 4.92 | 1.00 | 69.25 |  |
| 15 | A | O2' | 13:978 | 4.95 | 1.00 | 69.43 |  |
| 16 | A | N1 | 13:978 | 4.97 | 1.00 | 81.46 |  |
  |

  
**1^st^ shell ligand angles:**   
1-2: 53; 1-3: 91; 2-3: 134;   
  
 Contact(s): /N3_A_/O2/O2'/  
 Coded contact list: O_r_O_b_N_b_  

### K:13:1606

|  | Res. type | Atom | Residue | Distance | Occupancy | B-factor | Closest solute to 1^st^ shell water |
| --- | --- | --- | --- | --- | --- | --- | --- |
| **Ion** | | | |  | 1.00 | 57.00 || 1 | OMC | OP1 | 1H:1943 | 3.10 | 1.00 | 34.41 |
| 2 | U | O4 | 13:793 | 3.14 | 1.00 | 47.17 |  |
| 3 | G | O4' | 13:1517 | 3.50 | 1.00 | 31.18 |  |
| 4 | G | O5' | 13:1517 | 3.82 | 1.00 | 42.35 |  |
| 5 | OMC | OP2 | 1H:1943 | 4.11 | 1.00 | 36.20 |  |
| 6 | U | N3 | 13:793 | 4.38 | 1.00 | 43.38 |  |
| 7 | G | N9 | 13:1517 | 4.74 | 1.00 | 33.42 |  |
| 8 | A | O3' | 1H:1942 | 4.79 | 1.00 | 30.48 |  |
| 9 | G | OP1 | 13:1517 | 4.97 | 1.00 | 44.23 |  |
  |

  
**1^st^ shell ligand angles:**   
1-2: 83;   
  
 Contact(s): /O4/OP1/  
 Coded contact list: O_ph_O_b_  

### K:13:1607

|  | Res. type | Atom | Residue | Distance | Occupancy | B-factor | Closest solute to 1^st^ shell water |
| --- | --- | --- | --- | --- | --- | --- | --- |
| **Ion** | | | |  | 1.00 | 65.00 || 1 | U | O4 | 13:12 | 2.75 | 1.00 | 60.98 |
| 2 | G | O6 | 13:21 | 3.18 | 1.00 | 73.61 |  |
| 3 | G | O6 | 13:11 | 3.22 | 1.00 | 59.75 |  |
| 4 | G | O6 | 13:22 | 3.35 | 1.00 | 59.32 |  |
| 5 | C | N4 | 13:23 | 3.74 | 1.00 | 58.23 |  |
| 6 | U | N3 | 13:12 | 4.29 | 1.00 | 61.30 |  |
| 7 | G | N1 | 13:21 | 4.40 | 1.00 | 67.73 |  |
| 8 | G | N7 | 13:22 | 4.58 | 1.00 | 58.86 |  |
| 9 | A | N6 | 13:10 | 4.92 | 1.00 | 53.14 |  |
| 10 | G | N1 | 13:11 | 4.97 | 1.00 | 56.79 |  |
  |

  
**1^st^ shell ligand angles:**   
1-2: 85; 1-3: 63; 1-4: 65; 2-3: 142; 2-4: 70; 3-4: 77;   
  
 Contact(s): /O4/O6/O6/O6/  
 Coded contact list: 4O_b_  

### K:13:1608

|  | Res. type | Atom | Residue | Distance | Occupancy | B-factor | Closest solute to 1^st^ shell water |
| --- | --- | --- | --- | --- | --- | --- | --- |
| **Ion** | | | |  | 1.00 | 89.51 || 1 | U | O2 | 13:531 | 2.56 | 1.00 | 72.66 |
| 2 | G | O6 | 13:517 | 3.11 | 1.00 | 62.82 |  |
| 3 | U | N3 | 13:531 | 3.36 | 1.00 | 72.04 |  |
| 4 | Water 1 | | 13:1897 | 3.69 | 1.00 | 70.49 | 3.60 (G:O6 13:517) |
| 5 | PSU | OP2 | 13:516 | 3.88 | 1.00 | 56.02 |  |
| 6 | G | OP2 | 13:515 | 4.48 | 1.00 | 60.48 |  |
| 7 | U | N1 | 13:531 | 4.51 | 1.00 | 62.02 |  |
| 8 | G | N7 | 13:517 | 4.60 | 1.00 | 59.73 |  |
| 9 | G | OP1 | 13:515 | 4.70 | 1.00 | 51.86 |  |
  |

  
**1^st^ shell ligand angles:**   
1-2: 72; 1-3: 43; 2-3: 90;   
  
 Contact(s): /N3_U_/O6/  
 Coded contact list: O_b_Anion  
 Exclusion zone contact(s): /O2/  

### K:13:1609

|  | Res. type | Atom | Residue | Distance | Occupancy | B-factor | Closest solute to 1^st^ shell water |
| --- | --- | --- | --- | --- | --- | --- | --- |
| **Ion** | | | |  | 1.00 | 57.48 || 1 | C | OP1 | 13:1501 | 2.57 | 1.00 | 46.52 |
| 2 | A | O2' | 13:1507 | 2.88 | 1.00 | 41.48 |  |
| 3 | A | OP2 | 13:1502 | 3.34 | 1.00 | 55.64 |  |
| 4 | G | O4' | 13:1508 | 3.50 | 1.00 | 39.69 |  |
| 5 | G | OP2 | 13:1504 | 3.86 | 1.00 | 53.60 |  |
| 6 | C | OP2 | 13:1501 | 4.26 | 1.00 | 34.09 |  |
| 7 | C | O5' | 13:1501 | 4.49 | 1.00 | 38.97 |  |
| 8 | A | O3' | 13:1503 | 4.76 | 1.00 | 57.34 |  |
| 9 | A | O3' | 13:1500 | 4.92 | 1.00 | 41.42 |  |
| 10 | G | O2' | 13:1508 | 4.95 | 1.00 | 41.53 |  |
  |

  
**1^st^ shell ligand angles:**   
1-2: 112; 1-3: 70; 2-3: 143;   
  
 Contact(s): /O2'/OP2/  
 Coded contact list: O_ph_O_r_  
 Exclusion zone contact(s): /OP1/  

### K:13:1610

|  | Res. type | Atom | Residue | Distance | Occupancy | B-factor | Closest solute to 1^st^ shell water |
| --- | --- | --- | --- | --- | --- | --- | --- |
| **Ion** | | | |  | 1.00 | 103.84 || 1 | U | O4 | 13:1073 | 2.66 | 1.00 | 82.08 |
| 2 | U | O4 | 13:1083 | 2.90 | 1.00 | 78.12 |  |
| 3 | G | O6 | 13:1074 | 3.23 | 1.00 | 84.95 |  |
| 4 | G | N2 | 13:1084 | 3.27 | 1.00 | 75.27 |  |
| 5 | G | N1 | 13:1084 | 4.09 | 1.00 | 78.53 |  |
| 6 | G | O6 | 13:1082 | 4.16 | 1.00 | 73.74 |  |
| 7 | U | N3 | 13:1083 | 4.76 | 1.00 | 70.94 |  |
| 8 | U | N3 | 13:1073 | 4.79 | 1.00 | 79.15 |  |
| 9 | C | N4 | 13:1075 | 4.94 | 1.00 | 73.49 |  |
  |

  
**1^st^ shell ligand angles:**   
1-2: 93; 1-3: 55; 1-4: 47; 2-3: 63; 2-4: 65; 3-4: 76;   
  
 Contact(s): /N2/O4/O4/O6/  
 Coded contact list: 3O_b_Anion  

### K:13:1611

|  | Res. type | Atom | Residue | Distance | Occupancy | B-factor | Closest solute to 1^st^ shell water |
| --- | --- | --- | --- | --- | --- | --- | --- |
| **Ion** | | | |  | 1.00 | 114.54 || 1 | G | O6 | 13:475 | 3.14 | 1.00 | 127.04 |
| 2 | G | N7 | 13:474 | 3.70 | 1.00 | 127.57 |  |
| 3 | A | OP2 | 13:468 | 3.70 | 1.00 | 127.25 |  |
| 4 | G | N7 | 13:475 | 4.27 | 1.00 | 127.69 |  |
| 5 | G | O6 | 13:474 | 4.27 | 1.00 | 126.65 |  |
| 6 | G | OP1 | 13:467 | 4.58 | 1.00 | 134.13 |  |
| 7 | ARG | NH2 | 7I:75 | 4.64 | 1.00 | 102.78 |  |
| 8 | ARG | NH1 | 7I:75 | 4.73 | 1.00 | 104.18 |  |
| 9 | G | N2 | 13:464 | 4.78 | 1.00 | 134.68 |  |
| 10 | G | N1 | 13:464 | 4.98 | 1.00 | 141.76 |  |
  |

  
 Contact(s): /O6/  
 Coded contact list: O_b_  

### K:13:1612

|  | Res. type | Atom | Residue | Distance | Occupancy | B-factor | Closest solute to 1^st^ shell water |
| --- | --- | --- | --- | --- | --- | --- | --- |
| **Ion** | | | |  | 1.00 | 50.94 || 1 | U | OP2 | 13:801 | 2.96 | 1.00 | 40.66 |
| 2 | U | O5' | 13:801 | 3.00 | 1.00 | 39.61 |  |
| 3 | A | OP2 | 13:802 | 3.00 | 1.00 | 49.21 |  |
| 4 | A | N1 | 13:780 | 3.15 | 1.00 | 36.34 |  |
| 5 | G | O6 | 13:803 | 3.64 | 1.00 | 41.00 |  |
| 6 | G | N2 | 13:776 | 3.76 | 1.00 | 49.11 |  |
| 7 | A | N6 | 13:780 | 3.77 | 1.00 | 35.17 |  |
| 8 | C | N4 | 13:779 | 4.02 | 1.00 | 38.90 |  |
| 9 | U | OP1 | 13:801 | 4.18 | 1.00 | 35.70 |  |
| 10 | U | O3' | 13:801 | 4.59 | 1.00 | 45.72 |  |
| 11 | G | O3' | 13:800 | 4.93 | 1.00 | 34.38 |  |
  |

  
**1^st^ shell ligand angles:**   
1-2: 48; 1-3: 137; 1-4: 77; 2-3: 89; 2-4: 95; 3-4: 108;   
  
 Contact(s): /N1_A_/O5'/OP2/OP2/  
 Coded contact list: 2O_ph_O_r_N_b_  

### K:13:1613

|  | Res. type | Atom | Residue | Distance | Occupancy | B-factor | Closest solute to 1^st^ shell water |
| --- | --- | --- | --- | --- | --- | --- | --- |
| **Ion** | | | |  | 1.00 | 76.59 || 1 | U | O4 | 13:239 | 2.75 | 1.00 | 63.80 |
| 2 | G | O6 | 13:238 | 3.29 | 1.00 | 58.55 |  |
| 3 | G | O6 | 13:122 | 3.85 | 1.00 | 52.37 |  |
| 4 | U | N3 | 13:239 | 4.16 | 1.00 | 56.22 |  |
| 5 | C | N4 | 13:123 | 4.51 | 1.00 | 52.87 |  |
| 6 | C | O4' | 13:121 | 4.54 | 1.00 | 54.41 |  |
| 7 | C | N1 | 13:121 | 4.73 | 1.00 | 58.16 |  |
| 8 | C | OP2 | 13:121 | 4.84 | 1.00 | 65.91 |  |
| 9 | G | N7 | 13:238 | 4.89 | 1.00 | 60.47 |  |
| 10 | C | O5' | 13:121 | 4.94 | 1.00 | 55.15 |  |
  |

  
**1^st^ shell ligand angles:**   
1-2: 78;   
  
 Contact(s): /O4/O6/  
 Coded contact list: 2O_b_  

### K:13:1614

|  | Res. type | Atom | Residue | Distance | Occupancy | B-factor | Closest solute to 1^st^ shell water |
| --- | --- | --- | --- | --- | --- | --- | --- |
| **Ion** | | | |  | 1.00 | 71.95 || 1 | G | O6 | 13:854 | 3.58 | 1.00 | 74.22 |
| 2 | G | O6 | 13:853 | 3.67 | 1.00 | 73.23 |  |
| 3 | G | N7 | 13:853 | 3.96 | 1.00 | 71.57 |  |
| 4 | U | OP1 | 13:871 | 4.38 | 1.00 | 57.23 |  |
| 5 | Water 1 | | 13:1958 | 4.39 | 1.00 | 61.47 | 3.16 (U:OP1 13:871) |
| 6 | G | N7 | 13:854 | 4.48 | 1.00 | 66.00 |  |
| 7 | U | OP2 | 13:871 | 4.76 | 1.00 | 61.54 |  |
| 8 | G | O6 | 13:852 | 4.97 | 1.00 | 76.12 |  |
  |

  
 Coded contact list:   
1^st^ K coordination shell missing (see Notes)  

### K:13:1615

|  | Res. type | Atom | Residue | Distance | Occupancy | B-factor | Closest solute to 1^st^ shell water |
| --- | --- | --- | --- | --- | --- | --- | --- |
| **Ion** | | | |  | 1.00 | 90.54 || 1 | C | OP2 | 13:1303 | 2.71 | 1.00 | 78.93 |
| 2 | G | O6 | 13:1305 | 2.84 | 1.00 | 84.68 |  |
| 3 | G | O6 | 13:1304 | 3.08 | 1.00 | 91.75 |  |
| 4 | G | N7 | 13:1304 | 3.38 | 1.00 | 79.21 |  |
| 5 | U | O3' | 13:1302 | 4.26 | 1.00 | 93.43 |  |
| 6 | U | O2' | 13:1302 | 4.26 | 1.00 | 99.48 |  |
| 7 | G | N1 | 13:1305 | 4.48 | 1.00 | 78.91 |  |
| 8 | C | O5' | 13:1303 | 4.53 | 1.00 | 77.66 |  |
| 9 | G | O2' | 13:1331 | 4.81 | 1.00 | 74.99 |  |
| 10 | Water 1 | | 1F:101 | 4.98 | 1.00 | 73.24 | 3.80 (G:O6 13:1305) |
  |

  
**1^st^ shell ligand angles:**   
1-2: 100; 1-3: 142; 1-4: 86; 2-3: 78; 2-4: 73; 3-4: 58;   
  
 Contact(s): /N7/O6/O6/OP2/  
 Coded contact list: O_ph_2O_b_N_b_  

### K:13:1616

|  | Res. type | Atom | Residue | Distance | Occupancy | B-factor | Closest solute to 1^st^ shell water |
| --- | --- | --- | --- | --- | --- | --- | --- |
| **Ion** | | | |  | 1.00 | 75.54 || 1 | G | OP1 | 13:944 | 2.67 | 1.00 | 61.41 |
| 2 | G | O6 | 13:1233 | 3.09 | 1.00 | 65.31 |  |
| 3 | U | O4 | 13:1232 | 3.31 | 1.00 | 63.25 |  |
| 4 | G | OP2 | 13:945 | 3.78 | 1.00 | 54.67 |  |
| 5 | G | N7 | 13:1233 | 4.32 | 1.00 | 60.75 |  |
| 6 | U | O3' | 13:943 | 4.38 | 1.00 | 60.98 |  |
| 7 | G | O5' | 13:944 | 4.50 | 1.00 | 56.38 |  |
  |

  
**1^st^ shell ligand angles:**   
1-2: 87; 1-3: 140; 2-3: 65;   
  
 Contact(s): /O4/O6/OP1/  
 Coded contact list: O_ph_2O_b_  

### K:13:1617

|  | Res. type | Atom | Residue | Distance | Occupancy | B-factor | Closest solute to 1^st^ shell water |
| --- | --- | --- | --- | --- | --- | --- | --- |
| **Ion** | | | |  | 1.00 | 50.54 || 1 | A | OP2 | 13:780 | 3.26 | 1.00 | 38.58 |
| 2 | G | O6 | 13:800 | 3.45 | 1.00 | 38.01 |  |
| 3 | G | N7 | 13:800 | 3.78 | 1.00 | 46.32 |  |
| 4 | Water 1 | | 13:1864 | 3.79 | 1.00 | 41.84 | 2.50 (G:O6 13:800) |
| 5 | C | OP2 | 13:779 | 4.19 | 1.00 | 42.10 |  |
| 6 | A | N7 | 13:780 | 4.26 | 1.00 | 43.57 |  |
| 7 | C | OP1 | 13:779 | 4.44 | 1.00 | 40.86 |  |
| 8 | C | O5' | 13:779 | 4.47 | 1.00 | 38.84 |  |
| 9 | G | OP2 | 13:799 | 4.55 | 1.00 | 44.92 |  |
| 10 | G | N7 | 13:799 | 4.80 | 1.00 | 41.35 |  |
  |

  
 Contact(s): /OP2/  
 Coded contact list: O_ph_  

### K:13:1618

|  | Res. type | Atom | Residue | Distance | Occupancy | B-factor | Closest solute to 1^st^ shell water |
| --- | --- | --- | --- | --- | --- | --- | --- |
| **Ion** | | | |  | 1.00 | 85.59 || 1 | U | O4 | 13:427 | 2.69 | 1.00 | 80.59 |
| 2 | G | O6 | 13:416 | 3.07 | 1.00 | 73.20 |  |
| 3 | G | O6 | 13:426 | 3.24 | 1.00 | 74.59 |  |
| 4 | A | OP2 | 13:415 | 3.48 | 1.00 | 77.64 |  |
| 5 | G | O6 | 13:428 | 3.79 | 1.00 | 81.68 |  |
| 6 | A | OP2 | 13:414 | 4.11 | 1.00 | 87.17 |  |
| 7 | G | N1 | 13:428 | 4.17 | 1.00 | 90.30 |  |
| 8 | C | N4 | 13:417 | 4.28 | 1.00 | 75.78 |  |
| 9 | A | OP1 | 13:414 | 4.28 | 1.00 | 93.43 |  |
| 10 | A | N7 | 13:415 | 4.29 | 1.00 | 79.36 |  |
| 11 | G | N7 | 13:416 | 4.46 | 1.00 | 79.05 |  |
| 12 | U | N3 | 13:427 | 4.57 | 1.00 | 74.29 |  |
| 13 | A | O5' | 13:414 | 4.67 | 1.00 | 89.34 |  |
| 14 | G | N1 | 13:426 | 4.96 | 1.00 | 76.57 |  |
  |

  
**1^st^ shell ligand angles:**   
1-2: 57; 1-3: 65; 2-3: 69;   
  
 Contact(s): /O4/O6/O6/  
 Coded contact list: 3O_b_  

### K:13:1619

|  | Res. type | Atom | Residue | Distance | Occupancy | B-factor | Closest solute to 1^st^ shell water |
| --- | --- | --- | --- | --- | --- | --- | --- |
| **Ion** | | | |  | 1.00 | 58.24 || 1 | C | OP1 | 1H:1864 | 2.73 | 1.00 | 32.59 |
| 2 | C | O2 | 13:899 | 3.07 | 1.00 | 30.70 |  |
| 3 | C | O3' | 13:770 | 3.13 | 1.00 | 37.20 |  |
| 4 | C | O2' | 13:770 | 3.20 | 1.00 | 31.93 |  |
| 5 | C | O5' | 1H:1864 | 3.63 | 1.00 | 32.54 |  |
| 6 | G | OP1 | 13:771 | 3.71 | 1.00 | 27.60 |  |
| 7 | C | OP2 | 1H:1864 | 3.95 | 1.00 | 29.93 |  |
| 8 | U | OP2 | 1H:1865 | 4.31 | 1.00 | 38.86 |  |
| 9 | G | O5' | 13:771 | 4.34 | 1.00 | 30.10 |  |
| 10 | C | N3 | 13:899 | 4.50 | 1.00 | 28.79 |  |
| 11 | A | N3 | 13:900 | 4.60 | 1.00 | 32.53 |  |
| 12 | C | O3' | 1H:1864 | 4.74 | 1.00 | 22.51 |  |
| 13 | A | O4' | 13:900 | 4.86 | 1.00 | 31.33 |  |
| 14 | G | O3' | 1H:1863 | 4.91 | 1.00 | 32.90 |  |
  |

  
**1^st^ shell ligand angles:**   
1-2: 59; 1-3: 153; 1-4: 117; 2-3: 104; 2-4: 58; 3-4: 51;   
  
 Contact(s): /O2/O2'/O3'/OP1/  
 Coded contact list: O_ph_2O_r_O_b_  

### K:13:1620

|  | Res. type | Atom | Residue | Distance | Occupancy | B-factor | Closest solute to 1^st^ shell water |
| --- | --- | --- | --- | --- | --- | --- | --- |
| **Ion** | | | |  | 1.00 | 87.09 || 1 | G | O6 | 13:666 | 3.04 | 1.00 | 58.40 |
| 2 | U | O4 | 13:740 | 3.47 | 1.00 | 52.94 |  |
| 3 | G | O6 | 13:664 | 3.47 | 1.00 | 59.61 |  |
| 4 | G | O6 | 13:741 | 3.69 | 1.00 | 60.26 |  |
| 5 | G | N7 | 13:666 | 4.03 | 1.00 | 58.80 |  |
| 6 | U | N3 | 13:740 | 4.31 | 1.00 | 53.07 |  |
| 7 | G | N1 | 13:664 | 4.75 | 1.00 | 58.23 |  |
  |

  
 Contact(s): /O6/  
 Coded contact list: O_b_  

### K:13:1621

|  | Res. type | Atom | Residue | Distance | Occupancy | B-factor | Closest solute to 1^st^ shell water |
| --- | --- | --- | --- | --- | --- | --- | --- |
| **Ion** | | | |  | 1.00 | 61.31 || 1 | G | OP2 | 13:944 | 2.78 | 1.00 | 57.47 |
| 2 | G | N7 | 13:945 | 2.80 | 1.00 | 61.67 |  |
| 3 | U | OP2 | 13:943 | 3.09 | 1.00 | 73.14 |  |
| 4 | G | N7 | 13:944 | 3.11 | 1.00 | 60.52 |  |
| 5 | G | O6 | 13:945 | 3.16 | 1.00 | 69.08 |  |
| 6 | G | N7 | 13:1337 | 3.36 | 1.00 | 68.45 |  |
| 7 | U | O5' | 13:943 | 3.48 | 1.00 | 65.18 |  |
| 8 | U | OP1 | 13:943 | 4.01 | 1.00 | 62.57 |  |
| 9 | G | O5' | 13:944 | 4.19 | 1.00 | 56.38 |  |
| 10 | U | O3' | 13:943 | 4.56 | 1.00 | 60.98 |  |
| 11 | U | N1 | 13:943 | 4.79 | 1.00 | 64.40 |  |
| 12 | G | O6 | 13:1337 | 4.80 | 1.00 | 72.77 |  |
| 13 | G | N9 | 13:944 | 4.88 | 1.00 | 56.42 |  |
| 14 | G | N9 | 13:1337 | 4.98 | 1.00 | 66.02 |  |
| 15 | G | O6 | 13:944 | 5.00 | 1.00 | 61.12 |  |
  |

  
**1^st^ shell ligand angles:**   
1-2: 79; 1-3: 110; 1-4: 105; 1-5: 83; 1-6: 140; 2-3: 149; 2-4: 77; 2-5: 62; 2-6: 67; 3-4: 125; 3-5: 89; 3-6: 91; 4-5: 136; 4-6: 87; 5-6: 63;   
  
 Contact(s): /N7/N7/N7/O6/OP2/OP2/  
 Coded contact list: 2O_ph_O_b_3N_b_  

### K:13:1622

|  | Res. type | Atom | Residue | Distance | Occupancy | B-factor | Closest solute to 1^st^ shell water |
| --- | --- | --- | --- | --- | --- | --- | --- |
| **Ion** | | | |  | 1.00 | 61.23 || 1 | A | N3 | 13:949 | 2.83 | 1.00 | 66.54 |
| 2 | G | N7 | 13:971 | 2.91 | 1.00 | 59.11 |  |
| 3 | G | O2' | 13:1233 | 2.92 | 1.00 | 57.15 |  |
| 4 | G | O6 | 13:971 | 2.98 | 1.00 | 59.44 |  |
| 5 | U | O2 | 13:1364 | 3.02 | 1.00 | 71.02 |  |
| 6 | U | O2' | 13:1364 | 3.07 | 1.00 | 66.67 |  |
| 7 | G | N3 | 13:1233 | 3.35 | 1.00 | 63.46 |  |
| 8 | C | O4' | 13:1234 | 4.13 | 1.00 | 67.04 |  |
| 9 | G | N2 | 13:1233 | 4.23 | 1.00 | 64.27 |  |
| 10 | G | N9 | 13:1233 | 4.30 | 1.00 | 62.09 |  |
| 11 | U | N1 | 13:1364 | 4.41 | 1.00 | 68.53 |  |
| 12 | A | N9 | 13:949 | 4.46 | 1.00 | 66.23 |  |
| 13 | A | O2' | 13:949 | 4.58 | 1.00 | 67.03 |  |
| 14 | A | N1 | 13:949 | 4.64 | 1.00 | 60.23 |  |
| 15 | U | O3' | 13:1364 | 4.70 | 1.00 | 56.74 |  |
| 16 | G | O4' | 13:1233 | 4.76 | 1.00 | 57.79 |  |
| 17 | G | N1 | 13:971 | 4.87 | 1.00 | 55.42 |  |
| 18 | U | N3 | 13:1364 | 4.90 | 1.00 | 73.21 |  |
  |

  
**1^st^ shell ligand angles:**   
1-2: 76; 1-3: 135; 1-4: 76; 1-5: 97; 1-6: 142; 1-7: 63; 2-3: 99; 2-4: 64; 2-5: 150; 2-6: 92; 2-7: 117; 3-4: 142; 3-5: 106; 3-6: 81; 3-7: 82; 4-5: 86; 4-6: 67; 4-7: 136; 5-6: 75; 5-7: 84; 6-7: 148;   
  
 Contact(s): /N3_A_/N3_G_/N7/O2/O2'/O2'/O6/  
 Coded contact list: 2O_r_2O_b_3N_b_  

### K:13:1623

|  | Res. type | Atom | Residue | Distance | Occupancy | B-factor | Closest solute to 1^st^ shell water |
| --- | --- | --- | --- | --- | --- | --- | --- |
| **Ion** | | | |  | 1.00 | 56.20 || 1 | ASN | O | 2I:26 | 2.79 | 1.00 | 58.28 |
| 2 | U | O4 | 13:692 | 2.88 | 1.00 | 55.17 |  |
| 3 | GLY | O | 2I:52 | 2.94 | 1.00 | 60.45 |  |
| 4 | G | O6 | 13:691 | 2.94 | 1.00 | 51.50 |  |
| 5 | G | N7 | 13:691 | 3.31 | 1.00 | 49.41 |  |
| 6 | ASN | OD1 | 2I:26 | 3.79 | 1.00 | 57.61 |  |
| 7 | SER | N | 2I:53 | 4.21 | 1.00 | 60.85 |  |
| 8 | ASN | N | 2I:27 | 4.48 | 1.00 | 54.67 |  |
| 9 | ASN | ND2 | 2I:26 | 4.68 | 1.00 | 47.11 |  |
| 10 | SER | O | 2I:53 | 4.71 | 1.00 | 58.22 |  |
| 11 | GLY | N | 2I:56 | 4.72 | 1.00 | 62.01 |  |
| 12 | G | N1 | 13:691 | 4.79 | 1.00 | 46.38 |  |
| 13 | LYS | NZ | 2I:55 | 4.89 | 1.00 | 53.58 |  |
  |

  
**1^st^ shell ligand angles:**   
1-2: 138; 1-3: 103; 1-4: 153; 1-5: 130; 2-3: 77; 2-4: 66; 2-5: 72; 3-4: 68; 3-5: 125; 4-5: 58;   
  
 Contact(s): /N7/O4/O6/O_back_/O_back_/  
 Coded contact list: 2O_b_N_b_2O_back_  

### K:13:1624

|  | Res. type | Atom | Residue | Distance | Occupancy | B-factor | Closest solute to 1^st^ shell water |
| --- | --- | --- | --- | --- | --- | --- | --- |
| **Ion** | | | |  | 1.00 | 59.27 || 1 | U | O4 | 13:125 | 2.68 | 1.00 | 63.23 |
| 2 | C | O2 | 13:121 | 2.99 | 1.00 | 58.80 |  |
| 3 | G | O6 | 13:124 | 3.01 | 1.00 | 59.35 |  |
| 4 | G | O6 | 13:236 | 3.29 | 1.00 | 58.18 |  |
| 5 | C | N4 | 13:235 | 3.39 | 1.00 | 56.76 |  |
| 6 | C | N3 | 13:121 | 3.46 | 1.00 | 55.85 |  |
| 7 | U | N3 | 13:125 | 3.54 | 1.00 | 54.43 |  |
| 8 | G | O6 | 13:126 | 3.77 | 1.00 | 60.43 |  |
| 9 | G | N7 | 13:124 | 4.14 | 1.00 | 53.83 |  |
| 10 | C | N4 | 13:123 | 4.25 | 1.00 | 52.87 |  |
| 11 | C | N4 | 13:237 | 4.31 | 1.00 | 57.82 |  |
| 12 | C | N3 | 13:235 | 4.58 | 1.00 | 57.70 |  |
| 13 | G | N1 | 13:126 | 4.77 | 1.00 | 57.87 |  |
| 14 | C | N1 | 13:121 | 4.99 | 1.00 | 58.16 |  |
  |

  
**1^st^ shell ligand angles:**   
1-2: 137; 1-3: 88; 1-4: 88; 1-5: 116; 2-3: 93; 2-4: 128; 2-5: 84; 3-4: 60; 3-5: 149; 4-5: 98;   
  
 Contact(s): /N4/O2/O4/O6/O6/  
 Coded contact list: 4O_b_Anion  

### K:13:1625

|  | Res. type | Atom | Residue | Distance | Occupancy | B-factor | Closest solute to 1^st^ shell water |
| --- | --- | --- | --- | --- | --- | --- | --- |
| **Ion** | | | |  | 1.00 | 77.07 || 1 | ALA | O | 5I:30 | 3.02 | 1.00 | 69.86 |
| 2 | C | OP1 | 13:1359 | 3.07 | 1.00 | 72.56 |  |
| 3 | THR | O | 5I:22 | 3.50 | 1.00 | 66.40 |  |
| 4 | THR | OG1 | 5I:22 | 3.66 | 1.00 | 75.67 |  |
| 5 | THR | N | 5I:22 | 4.14 | 1.00 | 65.28 |  |
| 6 | G | OP2 | 13:976 | 4.23 | 1.00 | 73.05 |  |
| 7 | U | O3' | 13:1358 | 4.36 | 1.00 | 78.80 |  |
| 8 | VAL | N | 5I:33 | 4.72 | 1.00 | 77.18 |  |
| 9 | SER | N | 5I:32 | 4.83 | 1.00 | 68.35 |  |
| 10 | ARG | N | 5I:31 | 4.83 | 1.00 | 73.52 |  |
  |

  
**1^st^ shell ligand angles:**   
1-2: 151;   
  
 Contact(s): /O_back_/OP1/  
 Coded contact list: O_ph_O_back_  

### K:13:1626

|  | Res. type | Atom | Residue | Distance | Occupancy | B-factor | Closest solute to 1^st^ shell water |
| --- | --- | --- | --- | --- | --- | --- | --- |
| **Ion** | | | |  | 1.00 | 73.63 || 1 | G | O6 | 13:326 | 2.68 | 1.00 | 58.32 |
| 2 | G | OP2 | 13:107 | 2.78 | 1.00 | 77.64 |  |
| 3 | G | O2' | 13:324 | 2.80 | 1.00 | 69.88 |  |
| 4 | A | N7 | 13:325 | 3.17 | 1.00 | 55.95 |  |
| 5 | G | N7 | 13:326 | 3.68 | 1.00 | 56.10 |  |
| 6 | G | O3' | 13:324 | 3.84 | 1.00 | 70.21 |  |
| 7 | G | OP1 | 13:107 | 3.91 | 1.00 | 62.32 |  |
| 8 | G | N2 | 13:108 | 4.12 | 1.00 | 63.52 |  |
| 9 | ARG | NH2 | BI:15 | 4.27 | 1.00 | 70.76 |  |
| 10 | A | N6 | 13:325 | 4.42 | 1.00 | 53.75 |  |
| 11 | G | N1 | 13:326 | 4.64 | 1.00 | 57.40 |  |
| 12 | G | O5' | 13:107 | 4.85 | 1.00 | 66.51 |  |
| 13 | C | O3' | 13:106 | 4.92 | 1.00 | 76.25 |  |
| 14 | A | N6 | 13:109 | 4.97 | 1.00 | 60.75 |  |
  |

  
**1^st^ shell ligand angles:**   
1-2: 85; 1-3: 80; 1-4: 89; 2-3: 156; 2-4: 101; 3-4: 97;   
  
 Contact(s): /N7/O2'/O6/OP2/  
 Coded contact list: O_ph_O_r_O_b_N_b_  

### K:13:1627

|  | Res. type | Atom | Residue | Distance | Occupancy | B-factor | Closest solute to 1^st^ shell water |
| --- | --- | --- | --- | --- | --- | --- | --- |
| **Ion** | | | |  | 1.00 | 46.56 || 1 | G | O6 | 13:1523 | 2.97 | 1.00 | 33.86 |
| 2 | U | O4 | 13:1522 | 3.13 | 1.00 | 38.99 |  |
| 3 | G | O6 | 13:1511 | 3.15 | 1.00 | 30.71 |  |
| 4 | U | O4 | 13:1510 | 3.16 | 1.00 | 34.94 |  |
| 5 | C | N4 | 13:1524 | 3.17 | 1.00 | 32.30 |  |
| 6 | U | O4 | 13:1512 | 3.25 | 1.00 | 35.97 |  |
| 7 | A | N6 | 13:1513 | 3.88 | 1.00 | 30.56 |  |
| 8 | G | N7 | 13:1523 | 3.96 | 1.00 | 30.77 |  |
| 9 | U | N3 | 13:1512 | 4.43 | 1.00 | 29.27 |  |
| 10 | U | N3 | 13:1522 | 4.52 | 1.00 | 27.47 |  |
| 11 | U | N3 | 13:1510 | 4.62 | 1.00 | 32.41 |  |
| 12 | A | N1 | 13:1513 | 4.86 | 1.00 | 29.56 |  |
| 13 | G | O6 | 13:1525 | 5.00 | 1.00 | 32.95 |  |
| 14 | C | N3 | 13:1524 | 5.00 | 1.00 | 27.64 |  |
  |

  
**1^st^ shell ligand angles:**   
1-2: 96; 1-3: 62; 1-4: 129; 1-5: 59; 1-6: 63; 2-3: 155; 2-4: 134; 2-5: 129; 2-6: 96; 3-4: 70; 3-5: 52; 3-6: 64; 4-5: 79; 4-6: 111; 5-6: 107;   
  
 Contact(s): /N4/O4/O4/O4/O6/O6/  
 Coded contact list: 5O_b_Anion  

### K:13:1628

|  | Res. type | Atom | Residue | Distance | Occupancy | B-factor | Closest solute to 1^st^ shell water |
| --- | --- | --- | --- | --- | --- | --- | --- |
| **Ion** | | | |  | 1.00 | 51.12 || 1 | ASN | OD1 | 3I:46 | 2.74 | 1.00 | 44.72 |
| 2 | C | N3 | 13:518 | 2.80 | 1.00 | 54.68 |  |
| 3 | G | O6 | 13:529 | 2.82 | 1.00 | 46.33 |  |
| 4 | G | N7 | 13:529 | 2.87 | 1.00 | 50.81 |  |
| 5 | PRO | O | 3I:45 | 2.97 | 1.00 | 52.89 |  |
| 6 | C | N4 | 13:518 | 3.21 | 1.00 | 54.67 |  |
| 7 | C | O2 | 13:518 | 3.95 | 1.00 | 48.94 |  |
| 8 | ASN | N | 3I:46 | 4.43 | 1.00 | 48.46 |  |
| 9 | C | N4 | 13:528 | 4.46 | 1.00 | 46.34 |  |
| 10 | C | N1 | 13:518 | 4.49 | 1.00 | 49.26 |  |
| 11 | G | N1 | 13:529 | 4.90 | 1.00 | 43.62 |  |
| 12 | ASN | ND2 | 3I:46 | 4.97 | 1.00 | 46.50 |  |
  |

  
**1^st^ shell ligand angles:**   
1-2: 129; 1-3: 57; 1-4: 97; 1-5: 71; 1-6: 168; 2-3: 78; 2-4: 80; 2-5: 86; 2-6: 44; 3-4: 63; 3-5: 87; 3-6: 112; 4-5: 149; 4-6: 73; 5-6: 115;   
  
 Contact(s): /N3_C_/N4/N7/O6/O_back_/OD1_Asn_/  
 Coded contact list: O_b_2N_b_O_back_O_cno_Anion  

### K:13:1629

|  | Res. type | Atom | Residue | Distance | Occupancy | B-factor | Closest solute to 1^st^ shell water |
| --- | --- | --- | --- | --- | --- | --- | --- |
| **Ion** | | | |  | 1.00 | 50.77 || 1 | G | O6 | 13:530 | 2.55 | 1.00 | 53.69 |
| 2 | PRO | O | 3I:45 | 2.77 | 1.00 | 52.89 |  |
| 3 | C | O2 | 13:518 | 2.79 | 1.00 | 48.94 |  |
| 4 | U | O2' | 4K:51 | 2.92 | 1.00 | 64.73 |  |
| 5 | C | N3 | 13:518 | 3.16 | 1.00 | 54.68 |  |
| 6 | U | O4' | 4K:51 | 3.35 | 1.00 | 51.48 |  |
| 7 | U | O3' | 4K:51 | 4.25 | 1.00 | 71.00 |  |
| 8 | ASN | N | 3I:46 | 4.38 | 1.00 | 48.46 |  |
| 9 | G | N7 | 13:530 | 4.48 | 1.00 | 53.80 |  |
| 10 | G | N1 | 13:530 | 4.51 | 1.00 | 50.05 |  |
| 11 | C | N1 | 13:518 | 4.73 | 1.00 | 49.26 |  |
| 12 | C | N4 | 13:518 | 4.78 | 1.00 | 54.67 |  |
| 13 | A | N3 | 13:1492 | 4.81 | 1.00 | 46.86 |  |
| 14 | ASN | O | 3I:46 | 4.93 | 1.00 | 49.73 |  |
  |

  
**1^st^ shell ligand angles:**   
1-2: 164; 1-3: 81; 1-4: 73; 1-5: 81; 1-6: 72; 2-3: 88; 2-4: 106; 2-5: 83; 2-6: 121; 3-4: 124; 3-5: 43; 3-6: 151; 4-5: 84; 4-6: 58; 5-6: 138;   
  
 Contact(s): /N3_C_/O2/O2'/O4'/O_back_/  
 Coded contact list: 2O_r_O_b_N_b_O_back_  
 Exclusion zone contact(s): /O6/  

### K:13:1630

|  | Res. type | Atom | Residue | Distance | Occupancy | B-factor | Closest solute to 1^st^ shell water |
| --- | --- | --- | --- | --- | --- | --- | --- |
| **Ion** | | | |  | 1.00 | 66.28 || 1 | C | OP2 | 13:1322 | 2.97 | 1.00 | 75.80 |
| 2 | G | O3' | 13:1224 | 2.98 | 1.00 | 68.85 |  |
| 3 | GLY | O | 4I:100 | 3.16 | 1.00 | 78.25 |  |
| 4 | GLN | O | 4I:101 | 3.48 | 1.00 | 69.44 |  |
| 5 | G | O2' | 13:1224 | 3.49 | 1.00 | 73.69 |  |
| 6 | A | OP1 | 13:1225 | 3.55 | 1.00 | 57.05 |  |
| 7 | ARG | N | 4I:102 | 3.64 | 1.00 | 72.60 |  |
| 8 | A | O5' | 13:1225 | 3.82 | 1.00 | 64.64 |  |
| 9 | C | O5' | 13:1322 | 4.10 | 1.00 | 64.84 |  |
| 10 | GLN | N | 4I:101 | 4.47 | 1.00 | 76.87 |  |
| 11 | THR | N | 4I:103 | 4.73 | 1.00 | 62.60 |  |
| 12 | C | OP1 | 13:1322 | 4.78 | 1.00 | 73.29 |  |
| 13 | G | O4' | 13:1224 | 4.89 | 1.00 | 73.94 |  |
  |

  
**1^st^ shell ligand angles:**   
1-2: 93; 1-3: 92; 2-3: 175;   
  
 Contact(s): /O3'/O_back_/OP2/  
 Coded contact list: O_ph_O_r_O_back_  

### K:13:1631

|  | Res. type | Atom | Residue | Distance | Occupancy | B-factor | Closest solute to 1^st^ shell water |
| --- | --- | --- | --- | --- | --- | --- | --- |
| **Ion** | | | |  | 1.00 | 58.22 || 1 | A | O3' | 13:1499 | 2.88 | 1.00 | 40.40 |
| 2 | A | OP1 | 13:1500 | 3.01 | 1.00 | 37.10 |  |
| 3 | G | OP1 | 13:1521 | 3.10 | 1.00 | 31.98 |  |
| 4 | G | OP1 | 13:1508 | 3.15 | 1.00 | 52.94 |  |
| 5 | G | OP2 | 13:1521 | 3.76 | 1.00 | 22.78 |  |
| 6 | A | O2' | 13:1499 | 3.96 | 1.00 | 28.48 |  |
| 7 | A | O5' | 13:1500 | 4.27 | 1.00 | 36.38 |  |
| 8 | G | O3' | 13:1520 | 4.27 | 1.00 | 27.82 |  |
| 9 | G | O5' | 13:1508 | 4.48 | 1.00 | 32.16 |  |
| 10 | G | OP2 | 13:1508 | 4.48 | 1.00 | 39.95 |  |
| 11 | C | OP2 | 13:1509 | 4.73 | 1.00 | 45.00 |  |
| 12 | A | OP2 | 13:1500 | 4.87 | 1.00 | 39.71 |  |
  |

  
**1^st^ shell ligand angles:**   
1-2: 50; 1-3: 93; 1-4: 104; 2-3: 127; 2-4: 54; 3-4: 137;   
  
 Contact(s): /O3'/OP1/OP1/OP1/  
 Coded contact list: 3O_ph_O_r_  

### K:13:1632

|  | Res. type | Atom | Residue | Distance | Occupancy | B-factor | Closest solute to 1^st^ shell water |
| --- | --- | --- | --- | --- | --- | --- | --- |
| **Ion** | | | |  | 1.00 | 79.99 || 1 | A | N7 | 13:790 | 3.37 | 1.00 | 45.48 |
| 2 | U | O2' | 13:789 | 3.42 | 1.00 | 52.57 |  |
| 3 | A | N6 | 13:790 | 4.23 | 1.00 | 42.67 |  |
| 4 | G | O6 | 13:791 | 4.31 | 1.00 | 47.30 |  |
| 5 | U | O3' | 13:789 | 4.56 | 1.00 | 44.73 |  |
| 6 | U | OP1 | 4K:45 | 4.90 | 1.00 | 107.48 |  |
  |

  
 Contact(s): /N7/  
 Coded contact list: N_b_  

### K:13:1633

|  | Res. type | Atom | Residue | Distance | Occupancy | B-factor | Closest solute to 1^st^ shell water |
| --- | --- | --- | --- | --- | --- | --- | --- |
| **Ion** | | | |  | 1.00 | 55.73 || 1 | G | OP1 | 13:758 | 2.73 | 1.00 | 42.05 |
| 2 | G | O6 | 13:576 | 3.08 | 1.00 | 49.16 |  |
| 3 | G | N3 | 13:821 | 3.30 | 1.00 | 49.13 |  |
| 4 | C | O2 | 13:880 | 3.31 | 1.00 | 44.18 |  |
| 5 | G | N2 | 13:575 | 3.39 | 1.00 | 41.95 |  |
| 6 | G | N7 | 13:576 | 3.40 | 1.00 | 46.52 |  |
| 7 | G | O2' | 13:821 | 3.58 | 1.00 | 45.79 |  |
| 8 | G | N2 | 13:821 | 3.93 | 1.00 | 47.23 |  |
| 9 | G | N9 | 13:821 | 4.39 | 1.00 | 40.88 |  |
| 10 | G | N1 | 13:576 | 4.80 | 1.00 | 43.72 |  |
| 11 | G | O4' | 13:821 | 4.87 | 1.00 | 43.21 |  |
| 12 | G | O5' | 13:758 | 4.89 | 1.00 | 46.93 |  |
  |

  
**1^st^ shell ligand angles:**   
1-2: 92; 1-3: 82; 1-4: 85; 1-5: 131; 1-6: 146; 2-3: 115; 2-4: 55; 2-5: 76; 2-6: 55; 3-4: 60; 3-5: 62; 3-6: 114; 4-5: 49; 4-6: 80; 5-6: 52;   
  
 Contact(s): /N2/N3_G_/N7/O2/O6/OP1/  
 Coded contact list: O_ph_2O_b_2N_b_Anion  

### K:13:1634

|  | Res. type | Atom | Residue | Distance | Occupancy | B-factor | Closest solute to 1^st^ shell water |
| --- | --- | --- | --- | --- | --- | --- | --- |
| **Ion** | | | |  | 1.00 | 90.67 || 1 | U | O4 | 13:62 | 2.74 | 1.00 | 79.35 |
| 2 | G | O6 | 13:104 | 3.32 | 1.00 | 87.21 |  |
| 3 | G | O6 | 13:61 | 3.33 | 1.00 | 80.46 |  |
| 4 | C | N4 | 13:106 | 3.52 | 1.00 | 79.26 |  |
| 5 | G | O6 | 13:105 | 3.53 | 1.00 | 83.61 |  |
| 6 | LYS | NZ | BI:14 | 3.66 | 1.00 | 101.45 |  |
| 7 | SER | OG | BI:11 | 3.70 | 1.00 | 81.10 |  |
| 8 | G | N7 | 13:105 | 4.06 | 1.00 | 84.30 |  |
| 9 | U | N3 | 13:62 | 4.19 | 1.00 | 81.85 |  |
| 10 | C | N4 | 13:63 | 4.28 | 1.00 | 87.04 |  |
| 11 | G | N1 | 13:104 | 4.74 | 1.00 | 88.24 |  |
  |

  
**1^st^ shell ligand angles:**   
1-2: 91; 1-3: 66; 2-3: 136;   
  
 Contact(s): /O4/O6/O6/  
 Coded contact list: 3O_b_  

### K:13:1635

|  | Res. type | Atom | Residue | Distance | Occupancy | B-factor | Closest solute to 1^st^ shell water |
| --- | --- | --- | --- | --- | --- | --- | --- |
| **Ion** | | | |  | 1.00 | 95.87 || 1 | C | O2 | 13:372 | 2.55 | 1.00 | 75.94 |
| 2 | U | O4 | 13:387 | 2.91 | 1.00 | 81.43 |  |
| 3 | U | O4 | 13:375 | 3.22 | 1.00 | 73.43 |  |
| 4 | G | O6 | 13:376 | 3.59 | 1.00 | 67.38 |  |
| 5 | Water 1 | | 13:1920 | 3.70 | 1.00 | 79.45 | 3.22 (G:O6 13:376) |
| 6 | C | N3 | 13:372 | 4.01 | 1.00 | 77.16 |  |
| 7 | A | OP2 | 13:374 | 4.29 | 1.00 | 74.33 |  |
| 8 | U | N3 | 13:387 | 4.32 | 1.00 | 74.55 |  |
| 9 | A | N7 | 13:374 | 4.38 | 1.00 | 71.91 |  |
| 10 | A | N6 | 13:374 | 4.55 | 1.00 | 74.49 |  |
| 11 | A | OP2 | 13:373 | 4.58 | 1.00 | 89.19 |  |
| 12 | Water 2 | | 13:2026 | 4.67 | 1.00 | 69.91 | 4.49 (G:O6 13:377) |
| 13 | C | N1 | 13:372 | 4.85 | 1.00 | 78.70 |  |
| 14 | A | N6 | 13:389 | 4.87 | 1.00 | 69.49 |  |
| 15 | G | O6 | 13:377 | 5.00 | 1.00 | 70.66 |  |
  |

  
**1^st^ shell ligand angles:**   
1-2: 63; 1-3: 101; 2-3: 115;   
  
 Contact(s): /O4/O4/  
 Coded contact list: 2O_b_  
 Exclusion zone contact(s): /O2/  

### K:13:1636

|  | Res. type | Atom | Residue | Distance | Occupancy | B-factor | Closest solute to 1^st^ shell water |
| --- | --- | --- | --- | --- | --- | --- | --- |
| **Ion** | | | |  | 1.00 | 70.96 || 1 | A | OP1 | 13:914 | 2.72 | 1.00 | 51.89 |
| 2 | C | O3' | 13:526 | 3.02 | 1.00 | 48.14 |  |
| 3 | U | O3' | 13:12 | 3.20 | 1.00 | 57.98 |  |
| 4 | 7MG | OP1 | 13:527 | 3.21 | 1.00 | 57.52 |  |
| 5 | U | OP1 | 13:13 | 3.73 | 1.00 | 57.40 |  |
| 6 | 7MG | OP2 | 13:527 | 3.75 | 1.00 | 63.62 |  |
| 7 | U | O2' | 13:12 | 3.86 | 1.00 | 55.62 |  |
| 8 | U | O5' | 13:13 | 3.91 | 1.00 | 47.71 |  |
| 9 | Water 1 | | 13:1872 | 4.11 | 1.00 | 47.14 | 2.53 (U:OP1 13:13) |
| 10 | U | OP2 | 13:14 | 4.13 | 1.00 | 54.29 |  |
| 11 | Water 2 | | 13:2023 | 4.43 | 1.00 | 70.54 | 4.18 (U:OP1 13:13) |
| 12 | C | O2' | 13:526 | 4.45 | 1.00 | 48.79 |  |
| 13 | A | O5' | 13:914 | 4.63 | 1.00 | 49.90 |  |
| 14 | A | O3' | 13:913 | 4.90 | 1.00 | 56.19 |  |
| 15 | 7MG | O5' | 13:527 | 4.97 | 1.00 | 44.65 |  |
  |

  
**1^st^ shell ligand angles:**   
1-2: 93; 1-3: 99; 1-4: 109; 2-3: 97; 2-4: 47; 3-4: 134;   
  
 Contact(s): /O3'/O3'/OP1/OP1/  
 Coded contact list: 2O_ph_2O_r_  

### K:13:1637

|  | Res. type | Atom | Residue | Distance | Occupancy | B-factor | Closest solute to 1^st^ shell water |
| --- | --- | --- | --- | --- | --- | --- | --- |
| **Ion** | | | |  | 1.00 | 64.26 || 1 | G | OP2 | 13:567 | 2.75 | 1.00 | 65.95 |
| 2 | U | OP2 | 13:565 | 2.79 | 1.00 | 61.04 |  |
| 3 | C | OP2 | 13:564 | 2.82 | 1.00 | 53.42 |  |
| 4 | A | O2' | 13:563 | 3.12 | 1.00 | 48.06 |  |
| 5 | A | O3' | 13:563 | 3.37 | 1.00 | 47.02 |  |
| 6 | G | O3' | 13:566 | 4.02 | 1.00 | 51.44 |  |
| 7 | U | OP1 | 13:565 | 4.06 | 1.00 | 53.29 |  |
| 8 | C | O5' | 13:564 | 4.12 | 1.00 | 58.56 |  |
| 9 | G | O2' | 13:566 | 4.32 | 1.00 | 57.93 |  |
| 10 | ARG | NH2 | 3I:12 | 4.48 | 1.00 | 54.85 |  |
| 11 | G | O5' | 13:567 | 4.52 | 1.00 | 57.39 |  |
| 12 | C | O3' | 13:564 | 4.63 | 1.00 | 45.00 |  |
  |

  
**1^st^ shell ligand angles:**   
1-2: 87; 1-3: 159; 1-4: 73; 1-5: 117; 2-3: 96; 2-4: 80; 2-5: 67; 3-4: 87; 3-5: 47; 4-5: 48;   
  
 Contact(s): /O2'/O3'/OP2/OP2/OP2/  
 Coded contact list: 3O_ph_2O_r_  

### K:13:1638

|  | Res. type | Atom | Residue | Distance | Occupancy | B-factor | Closest solute to 1^st^ shell water |
| --- | --- | --- | --- | --- | --- | --- | --- |
| **Ion** | | | |  | 1.00 | 56.02 || 1 | G | O6 | 13:927 | 2.91 | 1.00 | 65.90 |
| 2 | G | O6 | 13:925 | 3.01 | 1.00 | 59.69 |  |
| 3 | U | O4 | 13:1390 | 3.12 | 1.00 | 64.90 |  |
| 4 | U | O4 | 13:1391 | 3.71 | 1.00 | 51.36 |  |
| 5 | C | N4 | 13:924 | 3.81 | 1.00 | 54.07 |  |
| 6 | G | N7 | 13:925 | 4.06 | 1.00 | 54.57 |  |
| 7 | G | N7 | 13:927 | 4.22 | 1.00 | 54.28 |  |
| 8 | U | N3 | 13:1391 | 4.26 | 1.00 | 58.24 |  |
| 9 | U | N3 | 13:1390 | 4.35 | 1.00 | 62.29 |  |
| 10 | G | O6 | 13:928 | 4.38 | 1.00 | 59.75 |  |
| 11 | G | N1 | 13:925 | 4.98 | 1.00 | 54.90 |  |
  |

  
**1^st^ shell ligand angles:**   
1-2: 67; 1-3: 67; 2-3: 110;   
  
 Contact(s): /O4/O6/O6/  
 Coded contact list: 3O_b_  

### K:13:1639

|  | Res. type | Atom | Residue | Distance | Occupancy | B-factor | Closest solute to 1^st^ shell water |
| --- | --- | --- | --- | --- | --- | --- | --- |
| **Ion** | | | |  | 1.00 | 66.33 || 1 | G | O6 | 13:1058 | 2.87 | 1.00 | 64.25 |
| 2 | U | O4 | 13:1199 | 3.12 | 1.00 | 62.75 |  |
| 3 | G | O6 | 13:1198 | 3.19 | 1.00 | 55.17 |  |
| 4 | G | OP2 | 13:1057 | 3.50 | 1.00 | 63.09 |  |
| 5 | C | N4 | 13:1059 | 3.75 | 1.00 | 61.27 |  |
| 6 | G | N2 | 13:1053 | 3.76 | 1.00 | 59.69 |  |
| 7 | U | N3 | 13:1199 | 4.25 | 1.00 | 59.71 |  |
| 8 | G | N7 | 13:1058 | 4.34 | 1.00 | 68.13 |  |
| 9 | G | N3 | 13:1053 | 4.56 | 1.00 | 62.53 |  |
| 10 | G | N7 | 13:1057 | 4.86 | 1.00 | 68.89 |  |
  |

  
**1^st^ shell ligand angles:**   
1-2: 72; 1-3: 64; 2-3: 61;   
  
 Contact(s): /O4/O6/O6/  
 Coded contact list: 3O_b_  

### K:13:1640

|  | Res. type | Atom | Residue | Distance | Occupancy | B-factor | Closest solute to 1^st^ shell water |
| --- | --- | --- | --- | --- | --- | --- | --- |
| **Ion** | | | |  | 1.00 | 60.81 || 1 | G | O3' | 13:115 | 2.76 | 1.00 | 57.59 |
| 2 | G | O2' | 13:115 | 2.85 | 1.00 | 50.27 |  |
| 3 | Water 1 | | 13:1834 | 2.85 | 1.00 | 47.48 | 2.29 (A:OP2 13:116) |
| 4 | G | O5' | 13:289 | 3.07 | 1.00 | 56.46 |  |
| 5 | G | O6 | 13:117 | 3.11 | 1.00 | 61.26 |  |
| 6 | G | N7 | 13:117 | 3.12 | 1.00 | 58.30 |  |
| 7 | G | O4' | 13:289 | 3.25 | 1.00 | 57.10 |  |
| 8 | A | N7 | 13:116 | 3.31 | 1.00 | 58.09 |  |
| 9 | A | OP2 | 13:116 | 3.35 | 1.00 | 47.23 |  |
| 10 | U | O4 | 13:118 | 4.29 | 1.00 | 59.28 |  |
| 11 | A | O5' | 13:116 | 4.36 | 1.00 | 49.58 |  |
| 12 | G | OP2 | 13:289 | 4.37 | 1.00 | 59.65 |  |
| 13 | A | N6 | 13:116 | 4.50 | 1.00 | 60.86 |  |
| 14 | G | OP2 | 13:117 | 4.54 | 1.00 | 56.63 |  |
| 15 | G | OP1 | 13:289 | 4.62 | 1.00 | 64.94 |  |
| 16 | G | N3 | 13:115 | 4.66 | 1.00 | 57.63 |  |
| 17 | G | O4' | 13:115 | 4.70 | 1.00 | 58.12 |  |
| 18 | G | N9 | 13:289 | 4.78 | 1.00 | 52.52 |  |
| 19 | A | N9 | 13:116 | 4.84 | 1.00 | 57.22 |  |
| 20 | A | OP1 | 13:116 | 4.89 | 1.00 | 47.56 |  |
| 21 | G | N9 | 13:115 | 4.98 | 1.00 | 61.03 |  |
  |

  
**1^st^ shell ligand angles:**   
1-2: 57; 1-3: 89; 1-4: 95; 1-5: 155; 1-6: 123; 1-7: 137; 1-8: 70; 1-9: 48; 2-3: 120; 2-4: 80; 2-5: 114; 2-6: 159; 2-7: 83; 2-8: 91; 2-9: 95; 3-4: 54; 3-5: 113; 3-6: 80; 3-7: 99; 3-8: 124; 3-9: 42; 4-5: 106; 4-6: 120; 4-7: 59; 4-8: 165; 4-9: 75; 5-6: 57; 5-7: 53; 5-8: 89; 5-9: 150; 6-7: 101; 6-8: 71; 6-9: 97; 7-8: 133; 7-9: 133; 8-9: 94;   
  
 Contact(s): /N7/N7/O2'/O3'/O4'/O5'/O6/O_w_/OP2/  
 Coded contact list: O_ph_4O_r_O_b_2N_b_O_w_  
K hydration shell issue: water too close (< 2.4 Angs.)  

### K:13:1641

|  | Res. type | Atom | Residue | Distance | Occupancy | B-factor | Closest solute to 1^st^ shell water |
| --- | --- | --- | --- | --- | --- | --- | --- |
| **Ion** | | | |  | 1.00 | 84.82 || 1 | G | O6 | 13:305 | 2.84 | 1.00 | 57.83 |
| 2 | G | O6 | 13:293 | 3.48 | 1.00 | 64.48 |  |
| 3 | U | O4 | 13:304 | 3.61 | 1.00 | 63.14 |  |
| 4 | G | OP2 | 13:292 | 4.03 | 1.00 | 58.45 |  |
| 5 | G | N1 | 13:305 | 4.18 | 1.00 | 57.47 |  |
| 6 | A | N6 | 13:303 | 4.49 | 1.00 | 60.67 |  |
| 7 | U | N3 | 13:304 | 4.68 | 1.00 | 54.50 |  |
| 8 | G | N7 | 13:293 | 4.71 | 1.00 | 59.22 |  |
| 9 | G | OP1 | 13:292 | 4.76 | 1.00 | 54.81 |  |
| 10 | C | OP1 | 13:291 | 4.97 | 1.00 | 52.45 |  |
  |

  
 Contact(s): /O6/  
 Coded contact list: O_b_  

### K:13:1642

|  | Res. type | Atom | Residue | Distance | Occupancy | B-factor | Closest solute to 1^st^ shell water |
| --- | --- | --- | --- | --- | --- | --- | --- |
| **Ion** | | | |  | 1.00 | 93.47 || 1 | G | OP2 | 13:731 | 3.24 | 1.00 | 54.60 |
| 2 | G | OP1 | 13:730 | 3.83 | 1.00 | 55.79 |  |
| 3 | A | OP2 | 13:816 | 3.97 | 1.00 | 45.66 |  |
| 4 | C | OP1 | 13:1527 | 4.12 | 1.00 | 52.80 |  |
| 5 | G | O3' | 13:1526 | 4.18 | 1.00 | 38.80 |  |
| 6 | G | O5' | 13:730 | 4.63 | 1.00 | 50.03 |  |
| 7 | G | O2' | 13:1526 | 4.79 | 1.00 | 43.14 |  |
| 8 | G | O3' | 13:730 | 4.85 | 1.00 | 39.62 |  |
  |

  
 Contact(s): /OP2/  
 Coded contact list: O_ph_  

### K:13:1643

|  | Res. type | Atom | Residue | Distance | Occupancy | B-factor | Closest solute to 1^st^ shell water |
| --- | --- | --- | --- | --- | --- | --- | --- |
| **Ion** | | | |  | 1.00 | 76.13 || 1 | U | OP1 | 13:1528 | 2.76 | 1.00 | 53.86 |
| 2 | C | O2' | 13:817 | 2.79 | 1.00 | 54.16 |  |
| 3 | G | OP2 | 13:818 | 2.91 | 1.00 | 45.96 |  |
| 4 | C | O3' | 13:817 | 3.10 | 1.00 | 43.62 |  |
| 5 | C | O3' | 13:1527 | 3.19 | 1.00 | 46.02 |  |
| 6 | G | O5' | 13:818 | 4.21 | 1.00 | 44.91 |  |
| 7 | U | OP2 | 13:1528 | 4.49 | 1.00 | 36.19 |  |
| 8 | U | O5' | 13:1528 | 4.83 | 1.00 | 47.58 |  |
| 9 | G | OP1 | 13:818 | 4.91 | 1.00 | 41.31 |  |
| 10 | C | O2 | 13:817 | 4.94 | 1.00 | 44.60 |  |
  |

  
**1^st^ shell ligand angles:**   
1-2: 112; 1-3: 147; 1-4: 147; 1-5: 49; 2-3: 98; 2-4: 53; 2-5: 72; 3-4: 49; 3-5: 161; 4-5: 125;   
  
 Contact(s): /O2'/O3'/O3'/OP1/OP2/  
 Coded contact list: 2O_ph_3O_r_  

### K:13:1644

|  | Res. type | Atom | Residue | Distance | Occupancy | B-factor | Closest solute to 1^st^ shell water |
| --- | --- | --- | --- | --- | --- | --- | --- |
| **Ion** | | | |  | 1.00 | 74.58 || 1 | A | OP1 | 13:315 | 2.55 | 1.00 | 69.97 |
| 2 | C | O3' | 13:314 | 3.15 | 1.00 | 67.76 |  |
| 3 | G | OP2 | 13:318 | 3.15 | 1.00 | 48.86 |  |
| 4 | A | OP2 | 13:315 | 4.11 | 1.00 | 65.82 |  |
| 5 | G | OP1 | 13:318 | 4.38 | 1.00 | 48.50 |  |
| 6 | G | O5' | 13:318 | 4.46 | 1.00 | 52.65 |  |
| 7 | G | OP2 | 13:319 | 4.55 | 1.00 | 61.48 |  |
| 8 | A | O5' | 13:315 | 4.62 | 1.00 | 63.52 |  |
| 9 | C | O5' | 13:314 | 4.86 | 1.00 | 56.56 |  |
| 10 | C | O2' | 13:314 | 4.94 | 1.00 | 58.23 |  |
  |

  
**1^st^ shell ligand angles:**   
1-2: 52; 1-3: 99; 2-3: 111;   
  
 Contact(s): /O3'/OP2/  
 Coded contact list: O_ph_O_r_  
 Exclusion zone contact(s): /OP1/  

### K:13:1645

|  | Res. type | Atom | Residue | Distance | Occupancy | B-factor | Closest solute to 1^st^ shell water |
| --- | --- | --- | --- | --- | --- | --- | --- |
| **Ion** | | | |  | 1.00 | 72.01 || 1 | G | O6 | 13:579 | 3.02 | 1.00 | 42.43 |
| 2 | U | O4 | 13:580 | 3.18 | 1.00 | 52.38 |  |
| 3 | G | O6 | 13:761 | 3.28 | 1.00 | 51.08 |  |
| 4 | C | N4 | 13:762 | 3.55 | 1.00 | 50.21 |  |
| 5 | A | OP1 | 13:759 | 3.66 | 1.00 | 48.43 |  |
| 6 | A | OP2 | 13:759 | 4.09 | 1.00 | 56.15 |  |
| 7 | G | N1 | 13:581 | 4.11 | 1.00 | 45.09 |  |
| 8 | U | N3 | 13:580 | 4.30 | 1.00 | 46.47 |  |
| 9 | G | N7 | 13:761 | 4.37 | 1.00 | 52.37 |  |
| 10 | G | N2 | 13:576 | 4.59 | 1.00 | 43.98 |  |
| 11 | G | N2 | 13:581 | 4.67 | 1.00 | 48.84 |  |
| 12 | G | O6 | 13:581 | 4.68 | 1.00 | 51.41 |  |
| 13 | G | N7 | 13:579 | 4.98 | 1.00 | 46.04 |  |
  |

  
**1^st^ shell ligand angles:**   
1-2: 66; 1-3: 65; 2-3: 67;   
  
 Contact(s): /O4/O6/O6/  
 Coded contact list: 3O_b_  

### K:13:1646

|  | Res. type | Atom | Residue | Distance | Occupancy | B-factor | Closest solute to 1^st^ shell water |
| --- | --- | --- | --- | --- | --- | --- | --- |
| **Ion** | | | |  | 1.00 | 63.13 || 1 | G | O6 | 13:886 | 2.91 | 1.00 | 35.79 |
| 2 | U | O4 | 13:911 | 3.31 | 1.00 | 40.41 |  |
| 3 | G | O6 | 13:885 | 3.41 | 1.00 | 43.91 |  |
| 4 | G | O6 | 13:887 | 3.67 | 1.00 | 38.69 |  |
| 5 | U | N3 | 13:911 | 4.13 | 1.00 | 34.90 |  |
| 6 | G | N7 | 13:886 | 4.16 | 1.00 | 37.67 |  |
| 7 | C | N4 | 13:910 | 4.20 | 1.00 | 36.65 |  |
| 8 | C | N4 | 13:912 | 4.48 | 1.00 | 33.65 |  |
| 9 | G | N7 | 13:885 | 4.72 | 1.00 | 34.98 |  |
  |

  
**1^st^ shell ligand angles:**   
1-2: 74;   
  
 Contact(s): /O4/O6/  
 Coded contact list: 2O_b_  

### K:13:1647

|  | Res. type | Atom | Residue | Distance | Occupancy | B-factor | Closest solute to 1^st^ shell water |
| --- | --- | --- | --- | --- | --- | --- | --- |
| **Ion** | | | |  | 1.00 | 50.65 || 1 | G | O6 | 13:1486 | 2.79 | 1.00 | 37.62 |
| 2 | G | O6 | 13:1415 | 2.86 | 1.00 | 39.55 |  |
| 3 | U | O4 | 13:1485 | 2.96 | 1.00 | 43.99 |  |
| 4 | U | O4 | 13:1414 | 3.18 | 1.00 | 52.73 |  |
| 5 | U | N3 | 13:1485 | 3.88 | 1.00 | 39.02 |  |
| 6 | G | N7 | 13:1486 | 3.92 | 1.00 | 36.43 |  |
| 7 | U | N3 | 13:1414 | 3.99 | 1.00 | 37.21 |  |
| 8 | G | N7 | 13:1415 | 4.71 | 1.00 | 39.48 |  |
| 9 | G | O6 | 13:1487 | 4.72 | 1.00 | 34.80 |  |
| 10 | G | N1 | 13:1415 | 4.86 | 1.00 | 36.73 |  |
| 11 | G | N1 | 13:1486 | 4.95 | 1.00 | 36.15 |  |
  |

  
**1^st^ shell ligand angles:**   
1-2: 69; 1-3: 132; 1-4: 76; 2-3: 76; 2-4: 119; 3-4: 151;   
  
 Contact(s): /O4/O4/O6/O6/  
 Coded contact list: 4O_b_  

### K:13:1648

|  | Res. type | Atom | Residue | Distance | Occupancy | B-factor | Closest solute to 1^st^ shell water |
| --- | --- | --- | --- | --- | --- | --- | --- |
| **Ion** | | | |  | 1.00 | 71.10 || 1 | U | O4 | 13:1481 | 2.69 | 1.00 | 51.04 |
| 2 | G | O6 | 13:1480 | 3.62 | 1.00 | 40.89 |  |
| 3 | G | O6 | 13:1482 | 4.10 | 1.00 | 40.34 |  |
| 4 | G | O6 | 13:1419 | 4.35 | 1.00 | 43.51 |  |
| 5 | U | N3 | 13:1481 | 4.54 | 1.00 | 40.36 |  |
| 6 | G | N7 | 13:1480 | 4.77 | 1.00 | 36.33 |  |
  |

  
 Contact(s): /O4/  
 Coded contact list: O_b_  

### K:13:1649

|  | Res. type | Atom | Residue | Distance | Occupancy | B-factor | Closest solute to 1^st^ shell water |
| --- | --- | --- | --- | --- | --- | --- | --- |
| **Ion** | | | |  | 1.00 | 81.17 || 1 | C | OP2 | 13:267 | 2.90 | 1.00 | 77.46 |
| 2 | C | OP2 | 13:268 | 2.94 | 1.00 | 65.41 |  |
| 3 | G | O3' | 13:251 | 3.32 | 1.00 | 77.40 |  |
| 4 | U | OP2 | 13:252 | 3.41 | 1.00 | 62.83 |  |
| 5 | G | O2' | 13:251 | 3.62 | 1.00 | 79.42 |  |
| 6 | LYS | NZ | 8I:67 | 3.79 | 1.00 | 55.85 |  |
| 7 | G | O2' | 13:266 | 3.98 | 1.00 | 68.55 |  |
| 8 | C | OP1 | 13:268 | 4.23 | 1.00 | 73.24 |  |
| 9 | U | OP2 | 13:253 | 4.45 | 1.00 | 54.40 |  |
| 10 | C | OP1 | 13:267 | 4.76 | 1.00 | 62.15 |  |
| 11 | C | O3' | 13:267 | 4.82 | 1.00 | 73.04 |  |
| 12 | C | O5' | 13:267 | 4.94 | 1.00 | 63.89 |  |
  |

  
**1^st^ shell ligand angles:**   
1-2: 68; 1-3: 148; 2-3: 105;   
  
 Contact(s): /O3'/OP2/OP2/  
 Coded contact list: 2O_ph_O_r_  

### K:13:1650

|  | Res. type | Atom | Residue | Distance | Occupancy | B-factor | Closest solute to 1^st^ shell water |
| --- | --- | --- | --- | --- | --- | --- | --- |
| **Ion** | | | |  | 1.00 | 97.83 || 1 | G | O6 | 13:258 | 3.12 | 1.00 | 78.42 |
| 2 | G | OP2 | 13:266 | 3.30 | 1.00 | 67.07 |  |
| 3 | G | OP1 | 13:266 | 3.39 | 1.00 | 63.98 |  |
| 4 | G | O6 | 13:257 | 3.46 | 1.00 | 84.07 |  |
| 5 | G | N7 | 13:257 | 4.33 | 1.00 | 84.28 |  |
| 6 | G | N7 | 13:258 | 4.48 | 1.00 | 80.67 |  |
| 7 | C | N4 | 13:267 | 4.80 | 1.00 | 71.15 |  |
| 8 | G | O5' | 13:266 | 4.84 | 1.00 | 73.42 |  |
| 9 | C | N4 | 13:268 | 4.90 | 1.00 | 78.64 |  |
| 10 | U | O4 | 13:256 | 4.96 | 1.00 | 80.41 |  |
  |

  
**1^st^ shell ligand angles:**   
1-2: 87; 1-3: 123; 2-3: 46;   
  
 Contact(s): /O6/OP1/OP2/  
 Coded contact list: 2O_ph_O_b_  

### K:13:1651

|  | Res. type | Atom | Residue | Distance | Occupancy | B-factor | Closest solute to 1^st^ shell water |
| --- | --- | --- | --- | --- | --- | --- | --- |
| **Ion** | | | |  | 1.00 | 81.28 || 1 | C | OP2 | 13:110 | 2.55 | 1.00 | 56.91 |
| 2 | G | N7 | 13:111 | 3.25 | 1.00 | 65.70 |  |
| 3 | G | O6 | 13:111 | 3.31 | 1.00 | 68.99 |  |
| 4 | A | OP2 | 13:109 | 3.90 | 1.00 | 62.59 |  |
| 5 | G | O6 | 13:112 | 3.97 | 1.00 | 62.23 |  |
| 6 | C | O5' | 13:110 | 4.04 | 1.00 | 61.99 |  |
| 7 | C | OP1 | 13:330 | 4.47 | 1.00 | 63.18 |  |
| 8 | A | O3' | 13:109 | 4.87 | 1.00 | 52.21 |  |
| 9 | C | OP1 | 13:110 | 4.91 | 1.00 | 49.76 |  |
  |

  
**1^st^ shell ligand angles:**   
1-2: 115; 1-3: 153; 2-3: 57;   
  
 Contact(s): /N7/O6/  
 Coded contact list: O_b_N_b_  
 Exclusion zone contact(s): /OP2/  

### MG:13:1652

|  | Res. type | Atom | Residue | Distance | Occupancy | B-factor | Closest solute to 1^st^ shell water |
| --- | --- | --- | --- | --- | --- | --- | --- |
| **Ion** | | | |  | 1.00 | 43.51 || 1 | G | N7 | 13:1497 | 3.78 | 1.00 | 31.39 |
| 2 | UR3 | O4 | 13:1498 | 4.04 | 1.00 | 38.06 |  |
| 3 | C | OP2 | 13:1496 | 4.14 | 1.00 | 35.94 |  |
| 4 | G | OP2 | 13:1497 | 4.31 | 1.00 | 38.16 |  |
| 5 | U | O2' | 4K:47 | 4.79 | 1.00 | 41.94 |  |
  |

  
 Coded contact list:   
1^st^ MG coordination shell missing (see Notes)  

### MG:13:1653

|  | Res. type | Atom | Residue | Distance | Occupancy | B-factor | Closest solute to 1^st^ shell water |
| --- | --- | --- | --- | --- | --- | --- | --- |
| **Ion** | | | |  | 1.00 | 67.78 || 1 | A | OP2 | 13:1110 | 1.92 | 1.00 | 69.47 |
| 2 | C | O2 | 13:1189 | 2.99 | 1.00 | 66.42 |  |
| 3 | A | OP1 | 13:1110 | 3.58 | 1.00 | 73.15 |  |
| 4 | A | O5' | 13:1110 | 3.84 | 1.00 | 82.10 |  |
| 5 | C | OP1 | 13:1109 | 3.91 | 1.00 | 61.88 |  |
| 6 | C | OP2 | 13:1109 | 3.97 | 1.00 | 76.06 |  |
| 7 | A | N7 | 13:1110 | 4.25 | 1.00 | 75.61 |  |
| 8 | C | O3' | 13:1109 | 4.38 | 1.00 | 71.40 |  |
| 9 | A | N7 | 13:1111 | 4.46 | 1.00 | 88.06 |  |
| 10 | HIS | NE2 | 2E:176 | 4.56 | 1.00 | 71.16 |  |
| 11 | C | N3 | 13:1189 | 4.60 | 1.00 | 70.85 |  |
| 12 | A | N6 | 13:1111 | 4.79 | 1.00 | 81.44 |  |
| 13 | A | N6 | 13:1188 | 4.81 | 1.00 | 70.13 |  |
| 14 | A | OP2 | 13:1111 | 4.82 | 1.00 | 79.36 |  |
  |

  
 Contact(s): /OP2/  
 Coded contact list: O_ph_  
 Exclusion zone contact(s): /O2/  

### MG:13:1654

|  | Res. type | Atom | Residue | Distance | Occupancy | B-factor | Closest solute to 1^st^ shell water |
| --- | --- | --- | --- | --- | --- | --- | --- |
| **Ion** | | | |  | 1.00 | 42.74 || 1 | G | O6 | 13:1469 | 3.43 | 1.00 | 52.16 |
| 2 | A | OP2 | 13:1468 | 3.72 | 1.00 | 51.32 |  |
| 3 | G | N1 | 13:1432 | 3.85 | 1.00 | 45.72 |  |
| 4 | G | OP2 | 13:1467 | 4.11 | 1.00 | 60.76 |  |
| 5 | G | N2 | 13:1432 | 4.15 | 1.00 | 46.32 |  |
| 6 | A | OP1 | 13:1468 | 4.48 | 1.00 | 50.52 |  |
| 7 | G | O5' | 13:1467 | 4.61 | 1.00 | 53.13 |  |
| 8 | C | N4 | 13:1431 | 4.70 | 1.00 | 43.17 |  |
| 9 | G | O6 | 13:1432 | 4.71 | 1.00 | 47.15 |  |
| 10 | G | N7 | 13:1469 | 4.96 | 1.00 | 47.61 |  |
| 11 | G | OP1 | 13:1467 | 4.98 | 1.00 | 56.84 |  |
  |

  
 Coded contact list:   
1^st^ MG coordination shell missing (see Notes)  

### MG:13:1655

|  | Res. type | Atom | Residue | Distance | Occupancy | B-factor | Closest solute to 1^st^ shell water |
| --- | --- | --- | --- | --- | --- | --- | --- |
| **Ion** | | | |  | 1.00 | 51.07 || 1 | G | N7 | 13:682 | 3.26 | 1.00 | 70.06 |
| 2 | G | O6 | 13:682 | 3.44 | 1.00 | 65.72 |  |
| 3 | G | N7 | 13:683 | 3.94 | 1.00 | 71.30 |  |
| 4 | G | O6 | 13:683 | 4.01 | 1.00 | 70.93 |  |
| 5 | A | N6 | 13:684 | 4.60 | 1.00 | 77.35 |  |
| 6 | G | OP2 | 13:682 | 4.99 | 1.00 | 58.90 |  |
  |

  
 Coded contact list:   
 Exclusion zone contact(s): /N7/  
Consider K in place of MG (1 valid contact(s) for K vs. 0 for MG)  

### MG:13:1656

|  | Res. type | Atom | Residue | Distance | Occupancy | B-factor | Closest solute to 1^st^ shell water |
| --- | --- | --- | --- | --- | --- | --- | --- |
| **Ion** | | | |  | 1.00 | 49.80 || 1 | G | OP1 | 13:1197 | 2.15 | 1.00 | 56.83 |
| 2 | G | OP2 | 13:1198 | 2.42 | 1.00 | 49.23 |  |
| 3 | C | O5' | 13:1054 | 3.01 | 1.00 | 63.86 |  |
| 4 | G | OP1 | 13:1198 | 3.22 | 1.00 | 53.68 |  |
| 5 | C | OP1 | 13:1054 | 3.27 | 1.00 | 64.22 |  |
| 6 | G | O5' | 13:1197 | 3.95 | 1.00 | 48.86 |  |
| 7 | G | O3' | 13:1197 | 4.27 | 1.00 | 50.33 |  |
| 8 | G | OP2 | 13:1197 | 4.44 | 1.00 | 60.27 |  |
| 9 | G | OP1 | 1K:34 | 4.54 | 1.00 | 53.55 |  |
| 10 | G | O5' | 13:1198 | 4.56 | 1.00 | 50.73 |  |
| 11 | U | O3' | 13:1196 | 4.70 | 1.00 | 63.58 |  |
| 12 | C | OP2 | 13:1054 | 4.74 | 1.00 | 61.03 |  |
| 13 | G | O3' | 13:1053 | 4.81 | 1.00 | 66.68 |  |
| 14 | Water 1 | | 13:1832 | 4.97 | 1.00 | 46.63 | 4.26 (G:OP1 13:1198) |
  |

  
 Contact(s): /OP1/  
 Coded contact list: O_ph_  
 Exclusion zone contact(s): /O5'/OP1/OP1/OP2/  
Consider K in place of MG (3 valid contact(s) for K vs. 1 for MG)  

### MG:13:1657

|  | Res. type | Atom | Residue | Distance | Occupancy | B-factor | Closest solute to 1^st^ shell water |
| --- | --- | --- | --- | --- | --- | --- | --- |
| **Ion** | | | |  | 1.00 | 38.67 || 1 | G | OP2 | 13:1197 | 2.32 | 1.00 | 60.27 |
| 2 | C | OP1 | 13:1054 | 2.56 | 1.00 | 64.22 |  |
| 3 | C | OP2 | 13:1054 | 3.44 | 1.00 | 61.03 |  |
| 4 | G | O2' | 13:1053 | 3.59 | 1.00 | 61.50 |  |
| 5 | G | N7 | 13:1198 | 3.80 | 1.00 | 57.45 |  |
| 6 | G | O5' | 13:1197 | 3.82 | 1.00 | 48.86 |  |
| 7 | U | O4 | 13:1199 | 3.84 | 1.00 | 62.75 |  |
| 8 | G | O3' | 13:1053 | 3.93 | 1.00 | 66.68 |  |
| 9 | G | OP2 | 13:1198 | 4.21 | 1.00 | 49.23 |  |
| 10 | G | N7 | 13:1197 | 4.22 | 1.00 | 58.84 |  |
| 11 | G | OP1 | 13:1197 | 4.35 | 1.00 | 56.83 |  |
| 12 | U | O3' | 13:1196 | 4.71 | 1.00 | 63.58 |  |
| 13 | U | O2' | 13:1196 | 4.80 | 1.00 | 70.90 |  |
| 14 | C | O5' | 13:1054 | 4.83 | 1.00 | 63.86 |  |
  |

  
 Contact(s): /OP2/  
 Coded contact list: O_ph_  
 Exclusion zone contact(s): /OP1/  

### MG:13:1658

|  | Res. type | Atom | Residue | Distance | Occupancy | B-factor | Closest solute to 1^st^ shell water |
| --- | --- | --- | --- | --- | --- | --- | --- |
| **Ion** | | | |  | 1.00 | 36.87 || 1 | G | N7 | 13:894 | 3.34 | 1.00 | 37.28 |
| 2 | G | O6 | 13:895 | 3.61 | 1.00 | 38.02 |  |
| 3 | G | N7 | 13:895 | 3.89 | 1.00 | 40.15 |  |
| 4 | G | OP2 | 13:894 | 4.35 | 1.00 | 39.99 |  |
| 5 | G | O6 | 13:894 | 4.75 | 1.00 | 43.46 |  |
| 6 | C | OP2 | 13:893 | 4.81 | 1.00 | 31.16 |  |
  |

  
 Coded contact list:   
 Exclusion zone contact(s): /N7/  
Consider K in place of MG (1 valid contact(s) for K vs. 0 for MG)  

### MG:13:1659

|  | Res. type | Atom | Residue | Distance | Occupancy | B-factor | Closest solute to 1^st^ shell water |
| --- | --- | --- | --- | --- | --- | --- | --- |
| **Ion** | | | |  | 1.00 | 56.57 || 1 | G | O6 | 13:1222 | 1.77 | 1.00 | 69.18 |
| 2 | U | O4 | 13:981 | 2.27 | 1.00 | 61.08 |  |
| 3 | C | OP1 | 13:979 | 2.55 | 1.00 | 64.95 |  |
| 4 | C | OP2 | 13:980 | 2.86 | 1.00 | 59.61 |  |
| 5 | G | O6 | 13:1221 | 3.09 | 1.00 | 73.88 |  |
| 6 | C | OP1 | 13:980 | 3.34 | 1.00 | 73.96 |  |
| 7 | G | N1 | 13:1222 | 3.52 | 1.00 | 64.53 |  |
| 8 | U | O2 | 13:982 | 3.65 | 1.00 | 53.15 |  |
| 9 | C | OP2 | 13:979 | 4.00 | 1.00 | 65.29 |  |
| 10 | C | N4 | 13:1223 | 4.10 | 1.00 | 66.02 |  |
| 11 | C | O5' | 13:979 | 4.18 | 1.00 | 68.14 |  |
| 12 | C | O5' | 13:980 | 4.20 | 1.00 | 61.87 |  |
| 13 | U | N3 | 13:982 | 4.23 | 1.00 | 57.65 |  |
| 14 | U | O2' | 13:982 | 4.32 | 1.00 | 65.57 |  |
| 15 | U | N3 | 13:981 | 4.37 | 1.00 | 61.27 |  |
| 16 | G | N7 | 13:1222 | 4.52 | 1.00 | 71.68 |  |
| 17 | C | N3 | 13:1223 | 4.86 | 1.00 | 65.63 |  |
| 18 | C | O3' | 13:979 | 4.87 | 1.00 | 77.85 |  |
| 19 | C | N4 | 13:984 | 4.89 | 1.00 | 73.21 |  |
| 20 | A | O3' | 13:978 | 4.97 | 1.00 | 68.46 |  |
| 21 | G | N1 | 13:1221 | 5.00 | 1.00 | 76.55 |  |
  |

  
**1^st^ shell ligand angles:**   
1-2: 88;   
  
 Contact(s): /O4/  
 Coded contact list: O_b_  
 Exclusion zone contact(s): /O6/O6/OP1/OP1/OP2/  
Consider K in place of MG (3 valid contact(s) for K vs. 1 for MG)  

### MG:13:1660

|  | Res. type | Atom | Residue | Distance | Occupancy | B-factor | Closest solute to 1^st^ shell water |
| --- | --- | --- | --- | --- | --- | --- | --- |
| **Ion** | | | |  | 1.00 | 39.52 || 1 | G | OP1 | 13:945 | 2.18 | 1.00 | 45.93 |
| 2 | C | OP2 | 13:1230 | 3.57 | 1.00 | 53.26 |  |
| 3 | C | OP1 | 13:1230 | 3.64 | 1.00 | 48.77 |  |
| 4 | G | O3' | 13:944 | 3.93 | 1.00 | 49.01 |  |
| 5 | G | OP2 | 13:945 | 4.27 | 1.00 | 54.67 |  |
| 6 | G | OP2 | 13:1231 | 4.49 | 1.00 | 63.74 |  |
| 7 | C | O5' | 13:1230 | 4.63 | 1.00 | 52.78 |  |
| 8 | G | O5' | 13:945 | 4.68 | 1.00 | 45.16 |  |
| 9 | A | OP2 | 13:946 | 4.80 | 1.00 | 41.68 |  |
  |

  
 Contact(s): /OP1/  
 Coded contact list: O_ph_  

### MG:13:1661

|  | Res. type | Atom | Residue | Distance | Occupancy | B-factor | Closest solute to 1^st^ shell water |
| --- | --- | --- | --- | --- | --- | --- | --- |
| **Ion** | | | |  | 1.00 | 39.42 || 1 | G | OP2 | 13:945 | 2.29 | 1.00 | 54.67 |
| 2 | G | OP1 | 13:944 | 2.41 | 1.00 | 61.41 |  |
| 3 | A | N7 | 13:946 | 3.90 | 1.00 | 70.96 |  |
| 4 | G | OP2 | 13:944 | 3.94 | 1.00 | 57.47 |  |
| 5 | G | O5' | 13:944 | 3.99 | 1.00 | 56.38 |  |
| 6 | G | O5' | 13:945 | 4.11 | 1.00 | 45.16 |  |
| 7 | G | OP1 | 13:945 | 4.14 | 1.00 | 45.93 |  |
| 8 | G | O6 | 13:1233 | 4.29 | 1.00 | 65.31 |  |
| 9 | A | OP2 | 13:946 | 4.42 | 1.00 | 41.68 |  |
| 10 | G | O6 | 13:947 | 4.53 | 1.00 | 68.68 |  |
| 11 | C | N4 | 13:948 | 4.83 | 1.00 | 68.13 |  |
| 12 | G | O3' | 13:944 | 4.85 | 1.00 | 49.01 |  |
| 13 | U | O3' | 13:943 | 4.89 | 1.00 | 60.98 |  |
  |

  
 Contact(s): /OP2/  
 Coded contact list: O_ph_  
 Exclusion zone contact(s): /OP1/  

### MG:13:1662

|  | Res. type | Atom | Residue | Distance | Occupancy | B-factor | Closest solute to 1^st^ shell water |
| --- | --- | --- | --- | --- | --- | --- | --- |
| **Ion** | | | |  | 1.00 | 54.19 || 1 | C | OP2 | 13:970 | 2.45 | 1.00 | 61.79 |
| 2 | A | OP2 | 13:968 | 3.56 | 1.00 | 49.94 |  |
| 3 | 5MC | OP1 | 13:967 | 4.02 | 1.00 | 56.92 |  |
| 4 | A | O5' | 13:968 | 4.26 | 1.00 | 58.08 |  |
| 5 | A | O5' | 13:969 | 4.48 | 1.00 | 47.26 |  |
| 6 | C | OP1 | 13:970 | 4.64 | 1.00 | 67.40 |  |
| 7 | A | O3' | 13:969 | 4.74 | 1.00 | 55.20 |  |
| 8 | C | O5' | 13:970 | 4.91 | 1.00 | 50.29 |  |
  |

  
 Coded contact list:   
 Exclusion zone contact(s): /OP2/  
Consider NA in place of MG (1 valid contact(s) for NA vs. 0 for MG)  

### MG:13:1663

|  | Res. type | Atom | Residue | Distance | Occupancy | B-factor | Closest solute to 1^st^ shell water |
| --- | --- | --- | --- | --- | --- | --- | --- |
| **Ion** | | | |  | 1.00 | 47.27 || 1 | C | OP1 | 13:972 | 1.99 | 1.00 | 46.91 |
| 2 | C | O5' | 13:972 | 3.60 | 1.00 | 46.95 |  |
| 3 | C | OP2 | 13:972 | 4.11 | 1.00 | 61.81 |  |
| 4 | ASP | OD2 | 1I:58 | 4.21 | 1.00 | 67.73 |  |
| 5 | G | O3' | 13:971 | 4.41 | 1.00 | 46.39 |  |
| 6 | G | O2' | 13:971 | 4.57 | 1.00 | 46.75 |  |
| 7 | G | OP1 | 13:971 | 4.89 | 1.00 | 52.40 |  |
  |

  
 Contact(s): /OP1/  
 Coded contact list: O_ph_  

### MG:13:1664

|  | Res. type | Atom | Residue | Distance | Occupancy | B-factor | Closest solute to 1^st^ shell water |
| --- | --- | --- | --- | --- | --- | --- | --- |
| **Ion** | | | |  | 1.00 | 57.26 || 1 | C | O2 | 13:1335 | 2.11 | 1.00 | 88.88 |
| 2 | A | OP2 | 13:1238 | 2.49 | 1.00 | 83.33 |  |
| 3 | A | N1 | 13:1299 | 3.69 | 1.00 | 83.53 |  |
| 4 | C | N3 | 13:1335 | 3.90 | 1.00 | 90.85 |  |
| 5 | G | N2 | 13:1300 | 4.15 | 1.00 | 82.80 |  |
| 6 | U | O4 | 13:1301 | 4.29 | 1.00 | 80.31 |  |
| 7 | A | N3 | 13:1299 | 4.39 | 1.00 | 83.18 |  |
| 8 | C | N1 | 13:1335 | 4.44 | 1.00 | 84.53 |  |
| 9 | C | O3' | 13:1237 | 4.60 | 1.00 | 74.56 |  |
| 10 | A | OP1 | 13:1238 | 4.62 | 1.00 | 83.95 |  |
| 11 | C | O2' | 13:1335 | 4.66 | 1.00 | 86.25 |  |
| 12 | A | OP2 | 13:1299 | 4.78 | 1.00 | 81.05 |  |
| 13 | A | O5' | 13:1238 | 4.94 | 1.00 | 79.34 |  |
  |

  
 Contact(s): /O2/  
 Coded contact list: O_b_  
 Exclusion zone contact(s): /OP2/  

### MG:13:1665

|  | Res. type | Atom | Residue | Distance | Occupancy | B-factor | Closest solute to 1^st^ shell water |
| --- | --- | --- | --- | --- | --- | --- | --- |
| **Ion** | | | |  | 1.00 | 51.98 || 1 | A | OP2 | 13:937 | 2.43 | 1.00 | 77.28 |
| 2 | A | N7 | 13:938 | 3.51 | 1.00 | 72.10 |  |
| 3 | A | O5' | 13:937 | 3.70 | 1.00 | 73.07 |  |
| 4 | A | N6 | 13:938 | 3.72 | 1.00 | 74.69 |  |
| 5 | A | N7 | 13:937 | 3.98 | 1.00 | 75.42 |  |
| 6 | G | N7 | 13:939 | 4.08 | 1.00 | 73.95 |  |
| 7 | G | O6 | 13:939 | 4.22 | 1.00 | 67.21 |  |
| 8 | A | OP2 | 13:938 | 4.33 | 1.00 | 69.46 |  |
| 9 | A | OP1 | 13:937 | 4.48 | 1.00 | 72.65 |  |
| 10 | C | O3' | 13:936 | 4.72 | 1.00 | 63.34 |  |
  |

  
 Coded contact list:   
 Exclusion zone contact(s): /OP2/  
Consider NA in place of MG (1 valid contact(s) for NA vs. 0 for MG)  

### MG:13:1666

|  | Res. type | Atom | Residue | Distance | Occupancy | B-factor | Closest solute to 1^st^ shell water |
| --- | --- | --- | --- | --- | --- | --- | --- |
| **Ion** | | | |  | 1.00 | 46.55 || 1 | Water 1 | | 13:1965 | 3.15 | 1.00 | 41.67 | 3.19 (G:O6 13:773) |
| 2 | G | O6 | 13:774 | 3.71 | 1.00 | 52.58 |  |
| 3 | G | N7 | 13:774 | 3.79 | 1.00 | 55.37 |  |
| 4 | G | N7 | 13:773 | 3.85 | 1.00 | 50.14 |  |
| 5 | G | O6 | 13:775 | 4.37 | 1.00 | 58.07 |  |
| 6 | G | O6 | 13:773 | 4.40 | 1.00 | 46.17 |  |
| 7 | G | OP2 | 13:803 | 4.85 | 1.00 | 35.91 |  |
| 8 | Water 2 | | 13:1809 | 4.86 | 1.00 | 34.65 | 3.86 (G:OP2 13:803) |
  |

  
 Coded contact list:   
 Exclusion zone contact(s): /O_w_/  
1^st^ MG coordination shell missing (see Notes)  

### MG:13:1667

|  | Res. type | Atom | Residue | Distance | Occupancy | B-factor | Closest solute to 1^st^ shell water |
| --- | --- | --- | --- | --- | --- | --- | --- |
| **Ion** | | | |  | 1.00 | 29.73 || 1 | C | OP1 | 13:770 | 2.93 | 1.00 | 31.50 |
| 2 | U | OP2 | 1H:1865 | 3.64 | 1.00 | 38.86 |  |
| 3 | U | OP1 | 1H:1865 | 4.16 | 1.00 | 34.62 |  |
| 4 | G | O3' | 13:769 | 4.23 | 1.00 | 27.27 |  |
| 5 | C | O5' | 13:770 | 4.35 | 1.00 | 24.08 |  |
| 6 | C | OP1 | 13:1514 | 4.47 | 1.00 | 31.20 |  |
  |

  
 Coded contact list:   
 Exclusion zone contact(s): /OP1/  
Consider K in place of MG (1 valid contact(s) for K vs. 0 for MG)  

### MG:13:1668

|  | Res. type | Atom | Residue | Distance | Occupancy | B-factor | Closest solute to 1^st^ shell water |
| --- | --- | --- | --- | --- | --- | --- | --- |
| **Ion** | | | |  | 1.00 | 31.81 || 1 | A | OP2 | 13:1500 | 2.21 | 1.00 | 39.71 |
| 2 | A | OP2 | 13:1499 | 2.35 | 1.00 | 44.19 |  |
| 3 | G | OP2 | 13:1505 | 2.40 | 1.00 | 43.78 |  |
| 4 | UR3 | O2' | 13:1498 | 3.38 | 1.00 | 42.86 |  |
| 5 | A | OP1 | 13:1499 | 3.41 | 1.00 | 40.83 |  |
| 6 | G | O3' | 13:1504 | 3.63 | 1.00 | 45.38 |  |
| 7 | A | N7 | 13:1500 | 4.09 | 1.00 | 39.42 |  |
| 8 | A | O5' | 13:1500 | 4.26 | 1.00 | 36.38 |  |
| 9 | UR3 | O3' | 13:1498 | 4.28 | 1.00 | 43.80 |  |
| 10 | G | OP1 | 13:1505 | 4.35 | 1.00 | 44.96 |  |
| 11 | A | O3' | 13:1499 | 4.47 | 1.00 | 40.40 |  |
| 12 | G | O2' | 13:1504 | 4.49 | 1.00 | 45.95 |  |
| 13 | A | O5' | 13:1499 | 4.59 | 1.00 | 40.09 |  |
| 14 | A | OP1 | 13:1500 | 4.69 | 1.00 | 37.10 |  |
| 15 | G | O5' | 13:1505 | 4.85 | 1.00 | 44.87 |  |
  |

  
**1^st^ shell ligand angles:**   
1-2: 90; 1-3: 73; 2-3: 115;   
  
 Contact(s): /OP2/OP2/OP2/  
 Coded contact list: 3O_ph_  
 Exclusion zone contact(s): /O2'/  

### MG:13:1669

|  | Res. type | Atom | Residue | Distance | Occupancy | B-factor | Closest solute to 1^st^ shell water |
| --- | --- | --- | --- | --- | --- | --- | --- |
| **Ion** | | | |  | 1.00 | 34.83 || 1 | A | OP2 | 13:915 | 3.58 | 1.00 | 52.33 |
| 2 | U | O2' | 13:13 | 4.14 | 1.00 | 52.78 |  |
| 3 | G | OP2 | 13:916 | 4.56 | 1.00 | 44.67 |  |
| 4 | G | N7 | 13:916 | 4.61 | 1.00 | 45.97 |  |
  |

  
 Coded contact list:   
1^st^ MG coordination shell missing (see Notes)  

### MG:13:1670

|  | Res. type | Atom | Residue | Distance | Occupancy | B-factor | Closest solute to 1^st^ shell water |
| --- | --- | --- | --- | --- | --- | --- | --- |
| **Ion** | | | |  | 1.00 | 42.83 || 1 | C | OP1 | 13:504 | 1.83 | 1.00 | 43.82 |
| 2 | G | OP1 | 13:505 | 3.87 | 1.00 | 51.51 |  |
| 3 | C | OP1 | 13:536 | 3.90 | 1.00 | 42.98 |  |
| 4 | G | N7 | 13:505 | 4.04 | 1.00 | 52.82 |  |
| 5 | C | O5' | 13:504 | 4.07 | 1.00 | 49.02 |  |
| 6 | C | OP2 | 13:504 | 4.20 | 1.00 | 52.76 |  |
| 7 | Water 1 | | 13:1993 | 4.22 | 1.00 | 51.19 | 3.45 (C:OP2 13:504) |
| 8 | C | O3' | 13:503 | 4.25 | 1.00 | 46.04 |  |
| 9 | A | O4' | 13:535 | 4.65 | 1.00 | 43.54 |  |
  |

  
 Contact(s): /OP1/  
 Coded contact list: O_ph_  

### MG:13:1671

|  | Res. type | Atom | Residue | Distance | Occupancy | B-factor | Closest solute to 1^st^ shell water |
| --- | --- | --- | --- | --- | --- | --- | --- |
| **Ion** | | | |  | 1.00 | 49.91 || 1 | G | OP1 | 13:550 | 3.57 | 1.00 | 46.49 |
| 2 | G | O2' | 13:502 | 4.00 | 1.00 | 57.44 |  |
| 3 | A | N6 | 13:509 | 4.07 | 1.00 | 57.09 |  |
| 4 | A | N1 | 13:509 | 4.07 | 1.00 | 57.87 |  |
| 5 | C | O4' | 13:503 | 4.20 | 1.00 | 53.83 |  |
| 6 | G | N2 | 13:502 | 4.47 | 1.00 | 52.84 |  |
| 7 | G | N3 | 13:502 | 4.63 | 1.00 | 56.39 |  |
| 8 | C | O2' | 13:503 | 4.74 | 1.00 | 56.25 |  |
| 9 | G | O5' | 13:550 | 4.93 | 1.00 | 54.92 |  |
  |

  
 Coded contact list:   
1^st^ MG coordination shell missing (see Notes)  

### MG:13:1672

|  | Res. type | Atom | Residue | Distance | Occupancy | B-factor | Closest solute to 1^st^ shell water |
| --- | --- | --- | --- | --- | --- | --- | --- |
| **Ion** | | | |  | 1.00 | 49.97 || 1 | A | OP2 | 13:53 | 2.83 | 1.00 | 58.75 |
| 2 | A | N7 | 13:353 | 3.03 | 1.00 | 54.98 |  |
| 3 | A | OP1 | 13:353 | 3.81 | 1.00 | 59.38 |  |
| 4 | A | N6 | 13:353 | 3.96 | 1.00 | 64.14 |  |
| 5 | A | OP1 | 13:53 | 4.07 | 1.00 | 54.13 |  |
| 6 | A | O4' | 13:51 | 4.48 | 1.00 | 53.68 |  |
| 7 | G | O4' | 13:316 | 4.94 | 1.00 | 68.76 |  |
| 8 | A | O5' | 13:53 | 4.95 | 1.00 | 38.57 |  |
  |

  
 Coded contact list:   
 Exclusion zone contact(s): /N7/OP2/  
Consider K in place of MG (2 valid contact(s) for K vs. 0 for MG)  

### MG:13:1673

|  | Res. type | Atom | Residue | Distance | Occupancy | B-factor | Closest solute to 1^st^ shell water |
| --- | --- | --- | --- | --- | --- | --- | --- |
| **Ion** | | | |  | 1.00 | 55.56 || 1 | G | O6 | 13:357 | 3.01 | 1.00 | 58.52 |
| 2 | C | O2 | 13:352 | 3.58 | 1.00 | 62.42 |  |
| 3 | G | N7 | 13:357 | 3.76 | 1.00 | 66.41 |  |
| 4 | A | N7 | 13:356 | 3.87 | 1.00 | 62.51 |  |
| 5 | C | OP2 | 13:355 | 4.17 | 1.00 | 62.88 |  |
| 6 | C | O2' | 13:352 | 4.75 | 1.00 | 54.66 |  |
| 7 | C | N4 | 13:54 | 4.99 | 1.00 | 59.78 |  |
  |

  
 Coded contact list:   
 Exclusion zone contact(s): /O6/  
Consider K in place of MG (1 valid contact(s) for K vs. 0 for MG)  

### MG:13:1674

|  | Res. type | Atom | Residue | Distance | Occupancy | B-factor | Closest solute to 1^st^ shell water |
| --- | --- | --- | --- | --- | --- | --- | --- |
| **Ion** | | | |  | 1.00 | 75.42 || 1 | U | OP2 | 13:1085 | 2.02 | 1.00 | 78.47 |
| 2 | U | O4 | 13:1086 | 2.60 | 1.00 | 89.41 |  |
| 3 | G | O6 | 13:1099 | 2.69 | 1.00 | 91.31 |  |
| 4 | G | O3' | 13:1084 | 3.14 | 1.00 | 77.58 |  |
| 5 | C | N3 | 13:1100 | 3.37 | 1.00 | 88.00 |  |
| 6 | G | N1 | 13:1099 | 3.43 | 1.00 | 90.80 |  |
| 7 | C | O2 | 13:1100 | 3.94 | 1.00 | 88.76 |  |
| 8 | U | O5' | 13:1085 | 4.18 | 1.00 | 83.15 |  |
| 9 | U | OP1 | 13:1085 | 4.18 | 1.00 | 75.05 |  |
| 10 | C | N4 | 13:1100 | 4.24 | 1.00 | 88.01 |  |
| 11 | C | OP2 | 13:1103 | 4.35 | 1.00 | 84.72 |  |
| 12 | U | N3 | 13:1086 | 4.70 | 1.00 | 87.59 |  |
  |

  
 Contact(s): /OP2/  
 Coded contact list: O_ph_  
 Exclusion zone contact(s): /N3_C_/O3'/O4/O6/  
Consider K in place of MG (4 valid contact(s) for K vs. 1 for MG)  

### MG:13:1675

|  | Res. type | Atom | Residue | Distance | Occupancy | B-factor | Closest solute to 1^st^ shell water |
| --- | --- | --- | --- | --- | --- | --- | --- |
| **Ion** | | | |  | 1.00 | 49.35 || 1 | U | O4 | 13:757 | 3.58 | 1.00 | 56.34 |
| 2 | A | OP2 | 13:583 | 3.63 | 1.00 | 43.09 |  |
| 3 | G | O6 | 13:584 | 3.82 | 1.00 | 44.81 |  |
| 4 | C | N4 | 13:756 | 3.97 | 1.00 | 47.95 |  |
| 5 | G | N2 | 13:758 | 4.06 | 1.00 | 40.01 |  |
| 6 | U | OP2 | 13:582 | 4.43 | 1.00 | 48.57 |  |
| 7 | G | O6 | 13:585 | 4.45 | 1.00 | 49.28 |  |
| 8 | G | N7 | 13:584 | 4.63 | 1.00 | 38.02 |  |
| 9 | G | N1 | 13:758 | 4.70 | 1.00 | 41.82 |  |
| 10 | ARG | NH1 | 6I:68 | 4.74 | 1.00 | 63.13 |  |
| 11 | ARG | NH2 | 6I:68 | 4.83 | 1.00 | 69.36 |  |
  |

  
 Coded contact list:   
1^st^ MG coordination shell missing (see Notes)  

### MG:13:1676

|  | Res. type | Atom | Residue | Distance | Occupancy | B-factor | Closest solute to 1^st^ shell water |
| --- | --- | --- | --- | --- | --- | --- | --- |
| **Ion** | | | |  | 1.00 | 53.19 || 1 | C | OP1 | 13:980 | 2.86 | 1.00 | 73.96 |
| 2 | C | O5' | 13:980 | 2.90 | 1.00 | 61.87 |  |
| 3 | G | O6 | 13:1220 | 3.97 | 1.00 | 73.90 |  |
| 4 | C | O3' | 13:979 | 4.28 | 1.00 | 77.85 |  |
| 5 | G | N7 | 13:1220 | 4.31 | 1.00 | 76.66 |  |
| 6 | U | O4 | 13:1219 | 4.42 | 1.00 | 79.57 |  |
| 7 | G | O6 | 13:1221 | 4.44 | 1.00 | 73.88 |  |
| 8 | U | OP2 | 13:981 | 4.54 | 1.00 | 62.16 |  |
| 9 | U | O2' | 13:982 | 4.75 | 1.00 | 65.57 |  |
| 10 | C | OP2 | 13:980 | 4.79 | 1.00 | 59.61 |  |
  |

  
 Coded contact list:   
 Exclusion zone contact(s): /O5'/OP1/  
Consider K in place of MG (2 valid contact(s) for K vs. 0 for MG)  

### MG:13:1677

|  | Res. type | Atom | Residue | Distance | Occupancy | B-factor | Closest solute to 1^st^ shell water |
| --- | --- | --- | --- | --- | --- | --- | --- |
| **Ion** | | | |  | 1.00 | 42.26 || 1 | A | OP1 | 13:782 | 1.98 | 1.00 | 44.61 |
| 2 | A | OP2 | 13:794 | 1.99 | 1.00 | 54.61 |  |
| 3 | A | O5' | 13:794 | 3.49 | 1.00 | 47.89 |  |
| 4 | G | O2' | 13:1521 | 3.92 | 1.00 | 44.85 |  |
| 5 | G | N3 | 13:1521 | 3.98 | 1.00 | 32.84 |  |
| 6 | A | OP2 | 13:782 | 4.01 | 1.00 | 36.56 |  |
| 7 | U | O4' | 13:1522 | 4.10 | 1.00 | 38.70 |  |
| 8 | A | OP1 | 13:794 | 4.11 | 1.00 | 52.91 |  |
| 9 | G | N2 | 13:1521 | 4.14 | 1.00 | 31.69 |  |
| 10 | A | O5' | 13:782 | 4.28 | 1.00 | 38.55 |  |
| 11 | U | O3' | 13:793 | 4.38 | 1.00 | 51.37 |  |
| 12 | A | O3' | 13:781 | 4.39 | 1.00 | 30.99 |  |
| 13 | C | OP2 | 13:783 | 4.44 | 1.00 | 44.22 |  |
  |

  
**1^st^ shell ligand angles:**   
1-2: 138;   
  
 Contact(s): /OP1/OP2/  
 Coded contact list: 2O_ph_  

### MG:13:1678

|  | Res. type | Atom | Residue | Distance | Occupancy | B-factor | Closest solute to 1^st^ shell water |
| --- | --- | --- | --- | --- | --- | --- | --- |
| **Ion** | | | |  | 1.00 | 38.17 || 1 | G | OP1 | 13:576 | 2.40 | 1.00 | 49.63 |
| 2 | G | OP2 | 13:575 | 3.61 | 1.00 | 45.04 |  |
| 3 | G | OP1 | 13:575 | 3.70 | 1.00 | 41.76 |  |
| 4 | G | OP2 | 13:576 | 4.13 | 1.00 | 41.48 |  |
| 5 | G | OP2 | 13:577 | 4.49 | 1.00 | 31.40 |  |
| 6 | G | O5' | 13:575 | 4.51 | 1.00 | 45.03 |  |
| 7 | G | O5' | 13:576 | 4.60 | 1.00 | 33.39 |  |
| 8 | A | OP1 | 13:815 | 4.82 | 1.00 | 27.00 |  |
| 9 | G | O3' | 13:575 | 4.94 | 1.00 | 50.71 |  |
  |

  
 Contact(s): /OP1/  
 Coded contact list: O_ph_  

### MG:13:1679

|  | Res. type | Atom | Residue | Distance | Occupancy | B-factor | Closest solute to 1^st^ shell water |
| --- | --- | --- | --- | --- | --- | --- | --- |
| **Ion** | | | |  | 1.00 | 45.54 || 1 | G | OP1 | 13:666 | 3.65 | 1.00 | 50.93 |
| 2 | G | N7 | 13:666 | 3.97 | 1.00 | 58.80 |  |
| 3 | G | N7 | 13:664 | 4.13 | 1.00 | 61.83 |  |
| 4 | A | OP2 | 13:665 | 4.26 | 1.00 | 58.27 |  |
| 5 | G | N9 | 13:664 | 4.47 | 1.00 | 55.46 |  |
| 6 | A | O3' | 13:665 | 4.58 | 1.00 | 52.83 |  |
| 7 | G | N3 | 13:664 | 4.89 | 1.00 | 56.15 |  |
| 8 | G | O6 | 13:664 | 4.92 | 1.00 | 59.61 |  |
| 9 | G | N1 | 13:664 | 4.97 | 1.00 | 58.23 |  |
  |

  
 Coded contact list:   
1^st^ MG coordination shell missing (see Notes)  

### MG:13:1680

|  | Res. type | Atom | Residue | Distance | Occupancy | B-factor | Closest solute to 1^st^ shell water |
| --- | --- | --- | --- | --- | --- | --- | --- |
| **Ion** | | | |  | 1.00 | 49.72 || 1 | G | O6 | 13:1355 | 3.44 | 1.00 | 72.00 |
| 2 | G | N7 | 13:1355 | 4.17 | 1.00 | 76.31 |  |
| 3 | G | O6 | 13:1356 | 4.38 | 1.00 | 71.42 |  |
| 4 | C | N4 | 13:1354 | 4.42 | 1.00 | 85.60 |  |
| 5 | C | N4 | 13:1366 | 4.95 | 1.00 | 72.72 |  |
  |

  
 Coded contact list:   
1^st^ MG coordination shell missing (see Notes)  

### MG:13:1681

|  | Res. type | Atom | Residue | Distance | Occupancy | B-factor | Closest solute to 1^st^ shell water |
| --- | --- | --- | --- | --- | --- | --- | --- |
| **Ion** | | | |  | 1.00 | 44.41 || 1 | G | O6 | 13:35 | 3.33 | 1.00 | 51.46 |
| 2 | C | OP2 | 13:398 | 3.67 | 1.00 | 58.02 |  |
| 3 | C | N4 | 13:36 | 3.78 | 1.00 | 52.49 |  |
| 4 | C | N4 | 13:34 | 4.20 | 1.00 | 53.60 |  |
| 5 | G | O6 | 13:548 | 4.34 | 1.00 | 57.41 |  |
| 6 | G | N7 | 13:35 | 4.49 | 1.00 | 50.82 |  |
| 7 | U | O4 | 13:37 | 4.74 | 1.00 | 65.47 |  |
  |

  
 Coded contact list:   
 Exclusion zone contact(s): /O6/  
Consider K in place of MG (1 valid contact(s) for K vs. 0 for MG)  

### MG:13:1682

|  | Res. type | Atom | Residue | Distance | Occupancy | B-factor | Closest solute to 1^st^ shell water |
| --- | --- | --- | --- | --- | --- | --- | --- |
| **Ion** | | | |  | 1.00 | 33.19 || 1 | G | N7 | 13:577 | 3.47 | 1.00 | 39.03 |
| 2 | G | OP1 | 13:577 | 3.58 | 1.00 | 40.95 |  |
| 3 | U | OP1 | 13:813 | 3.83 | 1.00 | 38.75 |  |
| 4 | U | OP2 | 13:813 | 3.99 | 1.00 | 42.15 |  |
| 5 | Water 1 | | 13:1895 | 4.07 | 1.00 | 35.81 | 2.70 (G:OP1 13:577) |
| 6 | G | O6 | 13:577 | 4.59 | 1.00 | 37.16 |  |
| 7 | A | N6 | 13:816 | 4.62 | 1.00 | 35.60 |  |
| 8 | G | N2 | 13:765 | 4.75 | 1.00 | 37.56 |  |
  |

  
 Coded contact list:   
1^st^ MG coordination shell missing (see Notes)  

### MG:13:1683

|  | Res. type | Atom | Residue | Distance | Occupancy | B-factor | Closest solute to 1^st^ shell water |
| --- | --- | --- | --- | --- | --- | --- | --- |
| **Ion** | | | |  | 1.00 | 39.19 || 1 | G | OP1 | 13:558 | 1.89 | 1.00 | 53.80 |
| 2 | G | O6 | 13:299 | 2.32 | 1.00 | 59.91 |  |
| 3 | G | O3' | 13:557 | 3.58 | 1.00 | 44.52 |  |
| 4 | A | N7 | 13:298 | 3.81 | 1.00 | 63.13 |  |
| 5 | G | O2' | 13:297 | 3.83 | 1.00 | 48.28 |  |
| 6 | G | OP2 | 13:558 | 4.03 | 1.00 | 52.13 |  |
| 7 | U | OP1 | 13:560 | 4.06 | 1.00 | 50.55 |  |
| 8 | G | O6 | 13:566 | 4.33 | 1.00 | 54.58 |  |
| 9 | G | O5' | 13:558 | 4.44 | 1.00 | 49.36 |  |
| 10 | A | N6 | 13:298 | 4.44 | 1.00 | 64.50 |  |
| 11 | G | N1 | 13:299 | 4.46 | 1.00 | 55.56 |  |
| 12 | A | N6 | 13:300 | 4.48 | 1.00 | 53.10 |  |
| 13 | A | N1 | 13:300 | 4.66 | 1.00 | 57.15 |  |
| 14 | G | N7 | 13:299 | 4.69 | 1.00 | 52.23 |  |
| 15 | G | O3' | 13:297 | 4.95 | 1.00 | 48.26 |  |
  |

  
**1^st^ shell ligand angles:**   
1-2: 154;   
  
 Contact(s): /O6/OP1/  
 Coded contact list: O_ph_O_b_  

### MG:13:1684

|  | Res. type | Atom | Residue | Distance | Occupancy | B-factor | Closest solute to 1^st^ shell water |
| --- | --- | --- | --- | --- | --- | --- | --- |
| **Ion** | | | |  | 1.00 | 73.76 || 1 | G | OP2 | 13:750 | 2.33 | 1.00 | 82.60 |
| 2 | C | OP2 | 13:749 | 3.12 | 1.00 | 88.63 |  |
| 3 | C | OP1 | 13:749 | 3.37 | 1.00 | 85.72 |  |
| 4 | G | N7 | 13:750 | 4.24 | 1.00 | 76.16 |  |
| 5 | G | O5' | 13:750 | 4.48 | 1.00 | 82.71 |  |
| 6 | G | O6 | 13:752 | 4.52 | 1.00 | 81.65 |  |
| 7 | G | OP1 | 13:750 | 4.68 | 1.00 | 91.81 |  |
| 8 | C | O3' | 13:749 | 4.70 | 1.00 | 73.23 |  |
| 9 | C | O3' | 13:748 | 4.84 | 1.00 | 90.09 |  |
| 10 | C | O5' | 13:749 | 4.96 | 1.00 | 75.83 |  |
  |

  
 Contact(s): /OP2/  
 Coded contact list: O_ph_  
 Exclusion zone contact(s): /OP1/OP2/  
Consider K in place of MG (2 valid contact(s) for K vs. 1 for MG)  

### MG:13:1685

|  | Res. type | Atom | Residue | Distance | Occupancy | B-factor | Closest solute to 1^st^ shell water |
| --- | --- | --- | --- | --- | --- | --- | --- |
| **Ion** | | | |  | 1.00 | 45.99 || 1 | G | O6 | 13:1108 | 2.21 | 1.00 | 64.62 |
| 2 | U | OP2 | 13:1095 | 2.59 | 1.00 | 71.08 |  |
| 3 | G | O2' | 13:1094 | 3.82 | 1.00 | 88.01 |  |
| 4 | G | N1 | 13:1108 | 3.93 | 1.00 | 71.45 |  |
| 5 | C | OP2 | 13:1096 | 4.01 | 1.00 | 70.63 |  |
| 6 | U | O5' | 13:1095 | 4.26 | 1.00 | 74.32 |  |
| 7 | C | OP1 | 13:1096 | 4.39 | 1.00 | 68.70 |  |
| 8 | U | OP1 | 13:1095 | 4.57 | 1.00 | 73.49 |  |
| 9 | C | N3 | 13:1109 | 4.79 | 1.00 | 75.17 |  |
| 10 | C | N4 | 13:1107 | 4.90 | 1.00 | 68.75 |  |
  |

  
 Contact(s): /O6/  
 Coded contact list: O_b_  
 Exclusion zone contact(s): /OP2/  

### MG:13:1686

|  | Res. type | Atom | Residue | Distance | Occupancy | B-factor | Closest solute to 1^st^ shell water |
| --- | --- | --- | --- | --- | --- | --- | --- |
| **Ion** | | | |  | 1.00 | 34.40 || 1 | G | OP1 | 13:903 | 2.56 | 1.00 | 39.50 |
| 2 | C | OP2 | 13:904 | 4.02 | 1.00 | 42.95 |  |
| 3 | G | OP2 | 13:903 | 4.41 | 1.00 | 32.05 |  |
| 4 | U | OP1 | 13:891 | 4.43 | 1.00 | 32.03 |  |
| 5 | G | O2' | 13:890 | 4.50 | 1.00 | 38.47 |  |
| 6 | G | O5' | 13:903 | 4.57 | 1.00 | 32.74 |  |
| 7 | U | OP1 | 13:1512 | 4.64 | 1.00 | 38.51 |  |
| 8 | G | O3' | 13:902 | 4.99 | 1.00 | 24.99 |  |
  |

  
 Coded contact list:   
 Exclusion zone contact(s): /OP1/  
Consider NA in place of MG (1 valid contact(s) for NA vs. 0 for MG)  

### MG:13:1687

|  | Res. type | Atom | Residue | Distance | Occupancy | B-factor | Closest solute to 1^st^ shell water |
| --- | --- | --- | --- | --- | --- | --- | --- |
| **Ion** | | | |  | 1.00 | 50.85 || 1 | G | O6 | 13:1370 | 2.63 | 1.00 | 76.46 |
| 2 | G | N7 | 13:1370 | 3.54 | 1.00 | 80.28 |  |
| 3 | G | O6 | 13:1371 | 4.13 | 1.00 | 82.21 |  |
| 4 | C | N4 | 13:1352 | 4.61 | 1.00 | 76.72 |  |
| 5 | G | N1 | 13:1370 | 4.72 | 1.00 | 81.26 |  |
| 6 | LYS | NZ | 8E:118 | 4.78 | 1.00 | 60.59 |  |
| 7 | G | N7 | 13:1371 | 4.88 | 1.00 | 82.72 |  |
| 8 | U | OP2 | 13:1348 | 4.88 | 1.00 | 78.21 |  |
| 9 | C | N4 | 13:1369 | 4.95 | 1.00 | 82.13 |  |
| 10 | U | O4 | 13:1351 | 4.98 | 1.00 | 77.34 |  |
  |

  
 Coded contact list:   
 Exclusion zone contact(s): /O6/  
Consider K in place of MG (1 valid contact(s) for K vs. 0 for MG)  

### MG:13:1688

|  | Res. type | Atom | Residue | Distance | Occupancy | B-factor | Closest solute to 1^st^ shell water |
| --- | --- | --- | --- | --- | --- | --- | --- |
| **Ion** | | | |  | 1.00 | 56.75 || 1 | C | OP2 | 13:1069 | 2.72 | 1.00 | 60.52 |
| 2 | U | O2 | 13:1085 | 2.97 | 1.00 | 87.23 |  |
| 3 | G | O6 | 13:1094 | 4.54 | 1.00 | 79.39 |  |
| 4 | C | OP1 | 13:1069 | 4.75 | 1.00 | 72.35 |  |
| 5 | C | OP1 | 13:1389 | 4.80 | 1.00 | 55.64 |  |
| 6 | U | O2' | 13:1085 | 4.86 | 1.00 | 77.14 |  |
| 7 | C | O5' | 13:1069 | 4.88 | 1.00 | 67.01 |  |
  |

  
 Coded contact list:   
 Exclusion zone contact(s): /O2/OP2/  
Consider K in place of MG (2 valid contact(s) for K vs. 0 for MG)  

### MG:13:1689

|  | Res. type | Atom | Residue | Distance | Occupancy | B-factor | Closest solute to 1^st^ shell water |
| --- | --- | --- | --- | --- | --- | --- | --- |
| **Ion** | | | |  | 1.00 | 53.63 || 1 | G | O6 | 13:1470 | 3.79 | 1.00 | 46.38 |
| 2 | G | N7 | 13:1469 | 4.03 | 1.00 | 47.61 |  |
| 3 | G | N7 | 13:1470 | 4.36 | 1.00 | 44.11 |  |
| 4 | G | O6 | 13:1469 | 4.51 | 1.00 | 52.16 |  |
| 5 | G | O6 | 13:1471 | 4.88 | 1.00 | 49.33 |  |
  |

  
 Coded contact list:   
1^st^ MG coordination shell missing (see Notes)  

### MG:13:1690

|  | Res. type | Atom | Residue | Distance | Occupancy | B-factor | Closest solute to 1^st^ shell water |
| --- | --- | --- | --- | --- | --- | --- | --- |
| **Ion** | | | |  | 1.00 | 51.21 || 1 | G | O6 | 13:1432 | 2.99 | 1.00 | 47.15 |
| 2 | C | OP2 | 13:1466 | 3.04 | 1.00 | 58.03 |  |
| 3 | G | N7 | 13:1467 | 3.33 | 1.00 | 53.59 |  |
| 4 | C | O5' | 13:1466 | 3.68 | 1.00 | 50.60 |  |
| 5 | G | OP2 | 13:1467 | 4.01 | 1.00 | 60.76 |  |
| 6 | C | OP1 | 13:1466 | 4.17 | 1.00 | 56.44 |  |
| 7 | G | O6 | 13:1467 | 4.41 | 1.00 | 53.69 |  |
| 8 | G | N1 | 13:1432 | 4.53 | 1.00 | 45.72 |  |
  |

  
 Coded contact list:   
 Exclusion zone contact(s): /N7/O6/OP2/  
Consider K in place of MG (3 valid contact(s) for K vs. 0 for MG)  

### MG:13:1691

|  | Res. type | Atom | Residue | Distance | Occupancy | B-factor | Closest solute to 1^st^ shell water |
| --- | --- | --- | --- | --- | --- | --- | --- |
| **Ion** | | | |  | 1.00 | 50.15 || 1 | C | OP1 | 13:1352 | 2.52 | 1.00 | 61.54 |
| 2 | LYS | NZ | 1F:3 | 3.22 | 1.00 | 68.85 |  |
| 3 | U | O3' | 13:1351 | 3.76 | 1.00 | 74.23 |  |
| 4 | A | OP1 | 13:1236 | 3.98 | 1.00 | 78.15 |  |
| 5 | A | OP2 | 13:1236 | 4.61 | 1.00 | 76.64 |  |
| 6 | C | OP2 | 13:1352 | 4.76 | 1.00 | 71.31 |  |
| 7 | C | O5' | 13:1352 | 4.81 | 1.00 | 85.78 |  |
  |

  
 Coded contact list:   
 Exclusion zone contact(s): /NZ_Lys_/OP1/  
Consider NA in place of MG (1 valid contact(s) for NA vs. 0 for MG)  

### MG:13:1692

|  | Res. type | Atom | Residue | Distance | Occupancy | B-factor | Closest solute to 1^st^ shell water |
| --- | --- | --- | --- | --- | --- | --- | --- |
| **Ion** | | | |  | 1.00 | 51.67 || 1 | G | O6 | 13:1432 | 3.10 | 1.00 | 47.15 |
| 2 | G | N7 | 13:1432 | 3.18 | 1.00 | 55.53 |  |
| 3 | G | O6 | 13:1467 | 3.51 | 1.00 | 53.69 |  |
| 4 | ARG | NH2 | B8:108 | 4.56 | 1.00 | 66.33 |  |
| 5 | G | N7 | 13:1467 | 4.71 | 1.00 | 53.59 |  |
| 6 | C | OP2 | 13:1466 | 4.75 | 1.00 | 58.03 |  |
| 7 | G | OP2 | 13:1432 | 4.90 | 1.00 | 40.25 |  |
| 8 | G | N1 | 13:1432 | 4.94 | 1.00 | 45.72 |  |
| 9 | C | OP2 | 13:1431 | 4.98 | 1.00 | 45.31 |  |
  |

  
 Coded contact list:   
 Exclusion zone contact(s): /N7/O6/  
Consider K in place of MG (2 valid contact(s) for K vs. 0 for MG)  

### MG:13:1693

|  | Res. type | Atom | Residue | Distance | Occupancy | B-factor | Closest solute to 1^st^ shell water |
| --- | --- | --- | --- | --- | --- | --- | --- |
| **Ion** | | | |  | 1.00 | 61.12 || 1 | A | OP2 | 13:608 | 2.16 | 1.00 | 77.55 |
| 2 | A | OP1 | 13:607 | 3.69 | 1.00 | 88.71 |  |
| 3 | A | O5' | 13:608 | 4.04 | 1.00 | 69.74 |  |
| 4 | A | N6 | 13:609 | 4.08 | 1.00 | 63.00 |  |
| 5 | A | N7 | 13:609 | 4.10 | 1.00 | 69.41 |  |
| 6 | A | OP1 | 13:608 | 4.15 | 1.00 | 78.28 |  |
| 7 | A | N7 | 13:608 | 4.34 | 1.00 | 69.43 |  |
| 8 | G | O6 | 13:610 | 4.46 | 1.00 | 75.55 |  |
| 9 | A | OP2 | 13:607 | 4.51 | 1.00 | 82.75 |  |
| 10 | A | OP2 | 13:609 | 4.63 | 1.00 | 72.65 |  |
| 11 | A | O3' | 13:607 | 4.67 | 1.00 | 67.78 |  |
| 12 | G | N7 | 13:610 | 4.96 | 1.00 | 66.86 |  |
  |

  
 Contact(s): /OP2/  
 Coded contact list: O_ph_  

### MG:13:1694

|  | Res. type | Atom | Residue | Distance | Occupancy | B-factor | Closest solute to 1^st^ shell water |
| --- | --- | --- | --- | --- | --- | --- | --- |
| **Ion** | | | |  | 1.00 | 57.89 || 1 | G | O6 | 13:1441 | 2.09 | 1.00 | 63.34 |
| 2 | G | N7 | 13:1441 | 3.97 | 1.00 | 61.24 |  |
| 3 | G | N1 | 13:1441 | 4.38 | 1.00 | 62.61 |  |
| 4 | C | OP2 | 13:1440 | 4.45 | 1.00 | 64.18 |  |
| 5 | G | OP1 | 13:1455 | 4.61 | 1.00 | 79.90 |  |
| 6 | LYS | NZ | BI:38 | 4.62 | 1.00 | 75.19 |  |
  |

  
 Contact(s): /O6/  
 Coded contact list: O_b_  

### MG:13:1695

|  | Res. type | Atom | Residue | Distance | Occupancy | B-factor | Closest solute to 1^st^ shell water |
| --- | --- | --- | --- | --- | --- | --- | --- |
| **Ion** | | | |  | 1.00 | 66.98 || 1 | U | O4 | 13:598 | 2.37 | 1.00 | 78.66 |
| 2 | G | OP2 | 13:597 | 2.48 | 1.00 | 80.50 |  |
| 3 | G | O2' | 13:595 | 2.73 | 1.00 | 81.29 |  |
| 4 | C | OP2 | 13:596 | 2.88 | 1.00 | 86.90 |  |
| 5 | G | O3' | 13:595 | 3.10 | 1.00 | 83.44 |  |
| 6 | A | N6 | 13:640 | 3.85 | 1.00 | 84.69 |  |
| 7 | G | N7 | 13:597 | 3.98 | 1.00 | 72.74 |  |
| 8 | U | N3 | 13:598 | 4.31 | 1.00 | 76.16 |  |
| 9 | C | O5' | 13:596 | 4.55 | 1.00 | 80.90 |  |
| 10 | G | O5' | 13:597 | 4.61 | 1.00 | 75.54 |  |
| 11 | C | O3' | 13:596 | 4.68 | 1.00 | 77.60 |  |
| 12 | C | OP1 | 13:596 | 4.76 | 1.00 | 83.34 |  |
| 13 | C | N4 | 13:599 | 4.89 | 1.00 | 77.34 |  |
| 14 | G | OP1 | 13:597 | 4.94 | 1.00 | 75.30 |  |
  |

  
 Contact(s): /O4/  
 Coded contact list: O_b_  
 Exclusion zone contact(s): /O2'/O3'/OP2/OP2/  
Consider K in place of MG (3 valid contact(s) for K vs. 1 for MG)  

### MG:13:1696

|  | Res. type | Atom | Residue | Distance | Occupancy | B-factor | Closest solute to 1^st^ shell water |
| --- | --- | --- | --- | --- | --- | --- | --- |
| **Ion** | | | |  | 1.00 | 64.72 || 1 | G | N7 | 13:258 | 3.98 | 1.00 | 80.67 |
| 2 | G | O6 | 13:258 | 4.21 | 1.00 | 78.42 |  |
| 3 | G | OP1 | 13:265 | 4.53 | 1.00 | 57.71 |  |
| 4 | G | OP2 | 13:265 | 4.79 | 1.00 | 59.07 |  |
| 5 | G | O6 | 13:259 | 4.87 | 1.00 | 75.64 |  |
  |

  
 Coded contact list:   
1^st^ MG coordination shell missing (see Notes)  

### MG:13:1697

|  | Res. type | Atom | Residue | Distance | Occupancy | B-factor | Closest solute to 1^st^ shell water |
| --- | --- | --- | --- | --- | --- | --- | --- |
| **Ion** | | | |  | 1.00 | 45.64 || 1 | G | O6 | 13:741 | 2.55 | 1.00 | 60.26 |
| 2 | G | N7 | 13:741 | 3.57 | 1.00 | 60.52 |  |
| 3 | G | O6 | 13:742 | 3.65 | 1.00 | 68.77 |  |
| 4 | G | O6 | 13:664 | 3.73 | 1.00 | 59.61 |  |
| 5 | G | N1 | 13:741 | 4.59 | 1.00 | 56.01 |  |
| 6 | G | N7 | 13:742 | 4.62 | 1.00 | 58.37 |  |
| 7 | G | O6 | 13:666 | 4.93 | 1.00 | 58.40 |  |
  |

  
 Coded contact list:   
 Exclusion zone contact(s): /O6/  
Consider NA in place of MG (1 valid contact(s) for NA vs. 0 for MG)  

### MG:13:1698

|  | Res. type | Atom | Residue | Distance | Occupancy | B-factor | Closest solute to 1^st^ shell water |
| --- | --- | --- | --- | --- | --- | --- | --- |
| **Ion** | | | |  | 1.00 | 84.61 || 1 | A | OP2 | 13:1146 | 2.64 | 1.00 | 107.53 |
| 2 | G | O6 | 13:1124 | 3.53 | 1.00 | 112.67 |  |
| 3 | C | O3' | 13:1145 | 3.90 | 1.00 | 110.99 |  |
| 4 | A | N7 | 13:1123 | 4.09 | 1.00 | 116.25 |  |
| 5 | G | N7 | 13:1124 | 4.38 | 1.00 | 117.19 |  |
| 6 | C | OP2 | 13:1147 | 4.46 | 1.00 | 104.04 |  |
| 7 | A | O5' | 13:1146 | 4.69 | 1.00 | 105.30 |  |
| 8 | A | N6 | 13:1123 | 4.75 | 1.00 | 110.28 |  |
| 9 | U | O4 | 13:1122 | 4.83 | 1.00 | 111.20 |  |
  |

  
 Coded contact list:   
 Exclusion zone contact(s): /OP2/  
Consider K in place of MG (1 valid contact(s) for K vs. 0 for MG)  

### MG:13:1699

|  | Res. type | Atom | Residue | Distance | Occupancy | B-factor | Closest solute to 1^st^ shell water |
| --- | --- | --- | --- | --- | --- | --- | --- |
| **Ion** | | | |  | 1.00 | 43.13 || 1 | G | N7 | 13:11 | 3.46 | 1.00 | 58.78 |
| 2 | A | N7 | 13:10 | 3.59 | 1.00 | 58.56 |  |
| 3 | U | O4 | 13:12 | 3.75 | 1.00 | 60.98 |  |
| 4 | G | OP2 | 13:11 | 3.90 | 1.00 | 45.26 |  |
| 5 | A | OP2 | 13:10 | 4.17 | 1.00 | 56.51 |  |
| 6 | G | O6 | 13:11 | 4.40 | 1.00 | 59.75 |  |
| 7 | A | N9 | 13:10 | 4.57 | 1.00 | 55.60 |  |
  |

  
 Coded contact list:   
1^st^ MG coordination shell missing (see Notes)  

### MG:13:1700

|  | Res. type | Atom | Residue | Distance | Occupancy | B-factor | Closest solute to 1^st^ shell water |
| --- | --- | --- | --- | --- | --- | --- | --- |
| **Ion** | | | |  | 1.00 | 47.60 || 1 | 4OC | OP2 | 13:1402 | 2.84 | 1.00 | 40.91 |
| 2 | U | OP1 | 4K:49 | 3.08 | 1.00 | 41.27 |  |
| 3 | G | OP1 | 13:1401 | 3.34 | 1.00 | 41.24 |  |
| 4 | U | OP2 | 4K:50 | 3.93 | 1.00 | 47.95 |  |
| 5 | 4OC | OP1 | 13:1402 | 4.25 | 1.00 | 46.04 |  |
| 6 | U | OP2 | 4K:49 | 4.60 | 1.00 | 46.82 |  |
| 7 | U | O2' | 4K:48 | 4.90 | 1.00 | 45.19 |  |
  |

  
 Coded contact list:   
 Exclusion zone contact(s): /OP1/OP1/OP2/  
Consider K in place of MG (3 valid contact(s) for K vs. 0 for MG)  

### MG:13:1701

|  | Res. type | Atom | Residue | Distance | Occupancy | B-factor | Closest solute to 1^st^ shell water |
| --- | --- | --- | --- | --- | --- | --- | --- |
| **Ion** | | | |  | 1.00 | 69.77 || 1 | G | O6 | 13:491 | 3.25 | 1.00 | 94.56 |
| 2 | G | O6 | 13:492 | 3.36 | 1.00 | 92.02 |  |
| 3 | G | O6 | 13:493 | 4.18 | 1.00 | 84.81 |  |
| 4 | G | N7 | 13:492 | 4.19 | 1.00 | 96.27 |  |
| 5 | G | N7 | 13:491 | 4.33 | 1.00 | 97.49 |  |
| 6 | G | O6 | 13:490 | 4.40 | 1.00 | 107.10 |  |
| 7 | G | N1 | 13:493 | 4.55 | 1.00 | 91.51 |  |
| 8 | C | N4 | 13:443 | 4.57 | 1.00 | 103.61 |  |
| 9 | C | N4 | 13:442 | 4.96 | 1.00 | 98.57 |  |
  |

  
 Coded contact list:   
 Exclusion zone contact(s): /O6/O6/  
Consider K in place of MG (2 valid contact(s) for K vs. 0 for MG)  

### MG:13:1702

|  | Res. type | Atom | Residue | Distance | Occupancy | B-factor | Closest solute to 1^st^ shell water |
| --- | --- | --- | --- | --- | --- | --- | --- |
| **Ion** | | | |  | 1.00 | 60.82 || 1 | G | O6 | 13:818 | 3.16 | 1.00 | 62.81 |
| 2 | C | OP2 | 13:822 | 3.22 | 1.00 | 42.88 |  |
| 3 | U | O2' | 13:820 | 3.85 | 1.00 | 46.44 |  |
| 4 | G | N1 | 13:818 | 3.91 | 1.00 | 50.23 |  |
| 5 | A | OP1 | 13:873 | 4.04 | 1.00 | 54.95 |  |
| 6 | U | O3' | 13:820 | 4.30 | 1.00 | 40.03 |  |
| 7 | G | O5' | 13:821 | 4.55 | 1.00 | 48.20 |  |
| 8 | A | OP1 | 13:872 | 4.59 | 1.00 | 60.82 |  |
| 9 | G | OP1 | 13:821 | 4.75 | 1.00 | 50.30 |  |
| 10 | A | OP2 | 13:873 | 4.79 | 1.00 | 58.66 |  |
| 11 | C | OP1 | 13:822 | 4.80 | 1.00 | 43.65 |  |
  |

  
 Coded contact list:   
 Exclusion zone contact(s): /O6/OP2/  
Consider K in place of MG (2 valid contact(s) for K vs. 0 for MG)  

### MG:13:1703

|  | Res. type | Atom | Residue | Distance | Occupancy | B-factor | Closest solute to 1^st^ shell water |
| --- | --- | --- | --- | --- | --- | --- | --- |
| **Ion** | | | |  | 1.00 | 49.60 || 1 | G | O6 | 13:558 | 2.62 | 1.00 | 61.88 |
| 2 | G | N7 | 13:557 | 3.73 | 1.00 | 59.48 |  |
| 3 | G | O6 | 13:557 | 3.82 | 1.00 | 62.93 |  |
| 4 | G | N1 | 13:558 | 4.50 | 1.00 | 58.85 |  |
  |

  
 Coded contact list:   
 Exclusion zone contact(s): /O6/  
Consider K in place of MG (1 valid contact(s) for K vs. 0 for MG)  

### MG:13:1704

|  | Res. type | Atom | Residue | Distance | Occupancy | B-factor | Closest solute to 1^st^ shell water |
| --- | --- | --- | --- | --- | --- | --- | --- |
| **Ion** | | | |  | 1.00 | 56.96 || 1 | U | OP1 | 13:1083 | 2.71 | 1.00 | 69.75 |
| 2 | G | OP2 | 13:1084 | 3.70 | 1.00 | 76.80 |  |
| 3 | U | O5' | 13:1083 | 3.70 | 1.00 | 78.39 |  |
| 4 | U | O4' | 13:1086 | 3.88 | 1.00 | 74.92 |  |
| 5 | G | O3' | 13:1082 | 4.15 | 1.00 | 48.87 |  |
| 6 | U | OP1 | 13:1390 | 4.68 | 1.00 | 55.28 |  |
| 7 | U | O2' | 13:1086 | 4.77 | 1.00 | 81.58 |  |
  |

  
 Coded contact list:   
 Exclusion zone contact(s): /OP1/  
Consider K in place of MG (1 valid contact(s) for K vs. 0 for MG)  

### MG:13:1705

|  | Res. type | Atom | Residue | Distance | Occupancy | B-factor | Closest solute to 1^st^ shell water |
| --- | --- | --- | --- | --- | --- | --- | --- |
| **Ion** | | | |  | 1.00 | 56.00 || 1 | A | OP1 | 13:109 | 2.49 | 1.00 | 65.13 |
| 2 | G | OP2 | 13:331 | 2.58 | 1.00 | 63.75 |  |
| 3 | A | O2' | 13:329 | 3.28 | 1.00 | 55.58 |  |
| 4 | A | N3 | 13:329 | 3.36 | 1.00 | 56.43 |  |
| 5 | A | OP2 | 13:109 | 4.00 | 1.00 | 62.59 |  |
| 6 | A | O5' | 13:109 | 4.33 | 1.00 | 55.09 |  |
| 7 | A | O3' | 13:329 | 4.47 | 1.00 | 66.68 |  |
| 8 | G | OP1 | 13:331 | 4.53 | 1.00 | 61.84 |  |
| 9 | G | N7 | 13:332 | 4.64 | 1.00 | 57.06 |  |
| 10 | G | O3' | 13:108 | 4.88 | 1.00 | 61.51 |  |
| 11 | C | O3' | 13:330 | 4.90 | 1.00 | 60.74 |  |
| 12 | G | O5' | 13:331 | 4.91 | 1.00 | 61.13 |  |
| 13 | A | N9 | 13:329 | 4.92 | 1.00 | 53.25 |  |
  |

  
 Coded contact list:   
 Exclusion zone contact(s): /N3_A_/O2'/OP1/OP2/  
Consider NA in place of MG (2 valid contact(s) for NA vs. 0 for MG)  

### MG:13:1706

|  | Res. type | Atom | Residue | Distance | Occupancy | B-factor | Closest solute to 1^st^ shell water |
| --- | --- | --- | --- | --- | --- | --- | --- |
| **Ion** | | | |  | 1.00 | 46.38 || 1 | G | N7 | 13:52 | 3.13 | 1.00 | 47.37 |
| 2 | A | OP1 | 13:51 | 3.28 | 1.00 | 46.87 |  |
| 3 | G | O6 | 13:52 | 3.72 | 1.00 | 47.84 |  |
| 4 | A | N1 | 13:360 | 4.03 | 1.00 | 52.02 |  |
| 5 | A | O2' | 13:50 | 4.08 | 1.00 | 53.70 |  |
| 6 | A | N6 | 13:53 | 4.42 | 1.00 | 53.62 |  |
| 7 | C | O2' | 13:47 | 4.49 | 1.00 | 58.23 |  |
| 8 | A | N6 | 13:360 | 4.54 | 1.00 | 55.66 |  |
| 9 | A | OP2 | 13:51 | 4.60 | 1.00 | 55.11 |  |
  |

  
 Coded contact list:   
 Exclusion zone contact(s): /N7/OP1/  
Consider K in place of MG (2 valid contact(s) for K vs. 0 for MG)  

### MG:13:1707

|  | Res. type | Atom | Residue | Distance | Occupancy | B-factor | Closest solute to 1^st^ shell water |
| --- | --- | --- | --- | --- | --- | --- | --- |
| **Ion** | | | |  | 1.00 | 58.52 || 1 | G | OP2 | 13:22 | 3.04 | 1.00 | 57.55 |
| 2 | G | N7 | 13:21 | 4.13 | 1.00 | 65.70 |  |
| 3 | U | OP1 | 13:561 | 4.20 | 1.00 | 60.89 |  |
| 4 | U | N1 | 13:561 | 4.25 | 1.00 | 62.14 |  |
| 5 | G | N7 | 13:22 | 4.43 | 1.00 | 58.86 |  |
| 6 | U | O4' | 13:561 | 4.55 | 1.00 | 59.15 |  |
| 7 | U | O4 | 13:561 | 4.58 | 1.00 | 61.03 |  |
| 8 | G | O5' | 13:22 | 4.70 | 1.00 | 57.59 |  |
| 9 | U | N3 | 13:561 | 4.87 | 1.00 | 61.00 |  |
| 10 | G | N9 | 13:21 | 4.94 | 1.00 | 60.00 |  |
| 11 | G | OP2 | 13:21 | 4.99 | 1.00 | 46.57 |  |
| 12 | G | O5' | 13:21 | 4.99 | 1.00 | 51.33 |  |
  |

  
 Coded contact list:   
 Exclusion zone contact(s): /OP2/  
Consider K in place of MG (1 valid contact(s) for K vs. 0 for MG)  

### MG:13:1708

|  | Res. type | Atom | Residue | Distance | Occupancy | B-factor | Closest solute to 1^st^ shell water |
| --- | --- | --- | --- | --- | --- | --- | --- |
| **Ion** | | | |  | 1.00 | 60.04 || 1 | G | O6 | 13:592 | 2.67 | 1.00 | 93.70 |
| 2 | G | O6 | 13:593 | 2.81 | 1.00 | 84.53 |  |
| 3 | G | O6 | 13:594 | 3.10 | 1.00 | 87.07 |  |
| 4 | U | O4 | 13:646 | 3.20 | 1.00 | 76.85 |  |
| 5 | C | N4 | 13:645 | 3.67 | 1.00 | 78.27 |  |
| 6 | G | N7 | 13:593 | 4.03 | 1.00 | 90.19 |  |
| 7 | G | N1 | 13:592 | 4.21 | 1.00 | 83.84 |  |
| 8 | U | N3 | 13:646 | 4.30 | 1.00 | 80.33 |  |
| 9 | G | N7 | 13:592 | 4.37 | 1.00 | 86.68 |  |
| 10 | C | N4 | 13:647 | 4.57 | 1.00 | 84.31 |  |
| 11 | G | N1 | 13:594 | 4.71 | 1.00 | 88.35 |  |
  |

  
 Coded contact list:   
 Exclusion zone contact(s): /O4/O6/O6/O6/  
Consider K in place of MG (4 valid contact(s) for K vs. 0 for MG)  

### MG:13:1709

|  | Res. type | Atom | Residue | Distance | Occupancy | B-factor | Closest solute to 1^st^ shell water |
| --- | --- | --- | --- | --- | --- | --- | --- |
| **Ion** | | | |  | 0.50 | 40.21 || 1 | G | O6 | 13:309 | 3.01 | 1.00 | 67.01 |
| 2 | G | O6 | 13:310 | 3.11 | 1.00 | 59.30 |  |
| 3 | C | N4 | 13:308 | 3.93 | 1.00 | 70.86 |  |
| 4 | G | N7 | 13:309 | 4.08 | 1.00 | 60.91 |  |
| 5 | C | N4 | 13:291 | 4.13 | 1.00 | 54.33 |  |
| 6 | G | OP2 | 13:306 | 4.47 | 1.00 | 83.46 |  |
| 7 | C | N4 | 13:290 | 4.68 | 1.00 | 54.87 |  |
| 8 | C | OP2 | 13:307 | 4.76 | 1.00 | 73.90 |  |
| 9 | G | N7 | 13:310 | 4.78 | 1.00 | 59.99 |  |
| 10 | G | N1 | 13:309 | 4.82 | 1.00 | 58.51 |  |
| 11 | C | N4 | 13:311 | 4.91 | 1.00 | 60.12 |  |
  |

  
 Coded contact list:   
 Exclusion zone contact(s): /O6/O6/  
Consider K in place of MG (2 valid contact(s) for K vs. 0 for MG)  

### MG:13:1709

|  | Res. type | Atom | Residue | Distance | Occupancy | B-factor | Closest solute to 1^st^ shell water |
| --- | --- | --- | --- | --- | --- | --- | --- |
| **Ion** | | | |  | 0.50 | 45.05 || 1 | G | N7 | 13:309 | 3.54 | 1.00 | 60.91 |
| 2 | C | OP2 | 13:308 | 4.15 | 1.00 | 75.24 |  |
| 3 | G | OP2 | 13:309 | 4.34 | 1.00 | 64.66 |  |
| 4 | G | O6 | 13:309 | 4.41 | 1.00 | 67.01 |  |
| 5 | C | N4 | 13:308 | 4.58 | 1.00 | 70.86 |  |
| 6 | G | O6 | 13:310 | 4.83 | 1.00 | 59.30 |  |
  |

  
 Coded contact list:   
1^st^ MG coordination shell missing (see Notes)  

### MG:13:1710

|  | Res. type | Atom | Residue | Distance | Occupancy | B-factor | Closest solute to 1^st^ shell water |
| --- | --- | --- | --- | --- | --- | --- | --- |
| **Ion** | | | |  | 1.00 | 52.01 || 1 | G | N7 | 13:886 | 3.75 | 1.00 | 37.67 |
| 2 | G | O6 | 13:888 | 3.98 | 1.00 | 42.29 |  |
| 3 | G | N7 | 13:887 | 4.04 | 1.00 | 29.11 |  |
| 4 | G | O6 | 13:887 | 4.25 | 1.00 | 38.69 |  |
| 5 | G | OP2 | 13:886 | 4.34 | 1.00 | 36.46 |  |
| 6 | G | N1 | 13:888 | 4.86 | 1.00 | 35.33 |  |
| 7 | G | O6 | 13:886 | 4.90 | 1.00 | 35.79 |  |
  |

  
 Coded contact list:   
1^st^ MG coordination shell missing (see Notes)  

### MG:13:1711

|  | Res. type | Atom | Residue | Distance | Occupancy | B-factor | Closest solute to 1^st^ shell water |
| --- | --- | --- | --- | --- | --- | --- | --- |
| **Ion** | | | |  | 1.00 | 55.78 || 1 | G | OP2 | 13:584 | 3.23 | 1.00 | 45.54 |
| 2 | G | N7 | 13:584 | 3.24 | 1.00 | 38.02 |  |
| 3 | A | OP2 | 13:583 | 4.25 | 1.00 | 43.09 |  |
| 4 | A | O5' | 13:583 | 4.43 | 1.00 | 43.80 |  |
| 5 | G | O6 | 13:584 | 4.58 | 1.00 | 44.81 |  |
| 6 | G | N7 | 13:585 | 4.63 | 1.00 | 44.47 |  |
| 7 | G | O5' | 13:584 | 4.66 | 1.00 | 40.48 |  |
| 8 | A | OP1 | 13:583 | 4.86 | 1.00 | 52.42 |  |
| 9 | G | O6 | 13:585 | 4.99 | 1.00 | 49.28 |  |
  |

  
 Coded contact list:   
 Exclusion zone contact(s): /N7/OP2/  
Consider K in place of MG (2 valid contact(s) for K vs. 0 for MG)  

### MG:13:1712

|  | Res. type | Atom | Residue | Distance | Occupancy | B-factor | Closest solute to 1^st^ shell water |
| --- | --- | --- | --- | --- | --- | --- | --- |
| **Ion** | | | |  | 1.00 | 43.50 || 1 | G | O6 | 13:324 | 3.38 | 1.00 | 70.07 |
| 2 | A | OP2 | 13:327 | 3.54 | 1.00 | 62.70 |  |
| 3 | Water 1 | | 13:1951 | 4.13 | 1.00 | 63.81 | 3.09 (G:O6 13:324) |
| 4 | G | OP1 | 13:326 | 4.31 | 1.00 | 68.83 |  |
| 5 | G | N1 | 13:324 | 4.38 | 1.00 | 67.10 |  |
| 6 | A | OP1 | 13:327 | 4.45 | 1.00 | 53.04 |  |
| 7 | A | OP1 | 13:262 | 4.59 | 1.00 | 63.85 |  |
| 8 | G | O5' | 13:326 | 4.85 | 1.00 | 62.86 |  |
| 9 | U | O3' | 13:261 | 4.99 | 1.00 | 74.01 |  |
  |

  
 Coded contact list:   
 Exclusion zone contact(s): /O6/  
Consider K in place of MG (1 valid contact(s) for K vs. 0 for MG)  

### MG:13:1713

|  | Res. type | Atom | Residue | Distance | Occupancy | B-factor | Closest solute to 1^st^ shell water |
| --- | --- | --- | --- | --- | --- | --- | --- |
| **Ion** | | | |  | 1.00 | 46.36 || 1 | G | O6 | 13:317 | 2.69 | 1.00 | 55.94 |
| 2 | G | O6 | 13:318 | 3.06 | 1.00 | 63.93 |  |
| 3 | G | O6 | 13:316 | 3.76 | 1.00 | 65.28 |  |
| 4 | G | N7 | 13:316 | 4.02 | 1.00 | 67.95 |  |
| 5 | G | N7 | 13:317 | 4.21 | 1.00 | 63.45 |  |
| 6 | G | OP1 | 13:316 | 4.42 | 1.00 | 60.82 |  |
| 7 | G | N7 | 13:318 | 4.53 | 1.00 | 59.44 |  |
| 8 | C | N4 | 13:336 | 4.80 | 1.00 | 57.11 |  |
| 9 | C | N4 | 13:335 | 4.84 | 1.00 | 58.55 |  |
| 10 | G | N1 | 13:317 | 4.92 | 1.00 | 53.53 |  |
  |

  
 Coded contact list:   
 Exclusion zone contact(s): /O6/O6/  
Consider K in place of MG (2 valid contact(s) for K vs. 0 for MG)  

### MG:13:1714

|  | Res. type | Atom | Residue | Distance | Occupancy | B-factor | Closest solute to 1^st^ shell water |
| --- | --- | --- | --- | --- | --- | --- | --- |
| **Ion** | | | |  | 1.00 | 83.76 || 1 | C | OP1 | 13:1284 | 3.01 | 1.00 | 95.08 |
| 2 | A | O3' | 13:1285 | 3.59 | 1.00 | 89.60 |  |
| 3 | A | O5' | 13:1285 | 4.20 | 1.00 | 85.39 |  |
| 4 | A | O3' | 13:1261 | 4.20 | 1.00 | 110.13 |  |
| 5 | C | OP2 | 13:1262 | 4.39 | 1.00 | 101.63 |  |
| 6 | C | O5' | 13:1284 | 4.86 | 1.00 | 94.98 |  |
| 7 | A | OP2 | 13:1285 | 4.87 | 1.00 | 89.58 |  |
  |

  
 Coded contact list:   
 Exclusion zone contact(s): /OP1/  
Consider K in place of MG (1 valid contact(s) for K vs. 0 for MG)  

### MG:13:1715

|  | Res. type | Atom | Residue | Distance | Occupancy | B-factor | Closest solute to 1^st^ shell water |
| --- | --- | --- | --- | --- | --- | --- | --- |
| **Ion** | | | |  | 1.00 | 60.74 || 1 | G | N7 | 13:942 | 3.28 | 1.00 | 64.42 |
| 2 | G | N7 | 13:941 | 4.02 | 1.00 | 61.76 |  |
| 3 | G | O6 | 13:941 | 4.11 | 1.00 | 63.47 |  |
| 4 | G | O6 | 13:942 | 4.14 | 1.00 | 62.79 |  |
| 5 | C | N4 | 13:1342 | 4.88 | 1.00 | 63.62 |  |
  |

  
 Coded contact list:   
 Exclusion zone contact(s): /N7/  
Consider K in place of MG (1 valid contact(s) for K vs. 0 for MG)  

### MG:13:1716

|  | Res. type | Atom | Residue | Distance | Occupancy | B-factor | Closest solute to 1^st^ shell water |
| --- | --- | --- | --- | --- | --- | --- | --- |
| **Ion** | | | |  | 1.00 | 37.21 || 1 | C | N4 | 13:808 | 3.76 | 1.00 | 44.44 |
| 2 | G | N7 | 13:809 | 3.83 | 1.00 | 38.19 |  |
| 3 | C | N4 | 13:810 | 4.00 | 1.00 | 32.67 |  |
| 4 | G | O6 | 13:809 | 4.13 | 1.00 | 34.96 |  |
  |

  
 Coded contact list:   
1^st^ MG coordination shell missing (see Notes)  

### MG:13:1717

|  | Res. type | Atom | Residue | Distance | Occupancy | B-factor | Closest solute to 1^st^ shell water |
| --- | --- | --- | --- | --- | --- | --- | --- |
| **Ion** | | | |  | 1.00 | 62.06 || 1 | C | OP1 | 13:308 | 3.31 | 1.00 | 73.06 |
| 2 | G | OP1 | 13:113 | 3.49 | 1.00 | 58.77 |  |
| 3 | C | O3' | 13:307 | 3.99 | 1.00 | 70.87 |  |
| 4 | C | OP2 | 13:308 | 4.39 | 1.00 | 75.24 |  |
| 5 | U | OP2 | 13:114 | 4.91 | 1.00 | 58.05 |  |
| 6 | G | O5' | 13:113 | 4.99 | 1.00 | 61.46 |  |
  |

  
 Coded contact list:   
 Exclusion zone contact(s): /OP1/  
Consider K in place of MG (1 valid contact(s) for K vs. 0 for MG)  

### MG:13:1718

|  | Res. type | Atom | Residue | Distance | Occupancy | B-factor | Closest solute to 1^st^ shell water |
| --- | --- | --- | --- | --- | --- | --- | --- |
| **Ion** | | | |  | 1.00 | 55.93 || 1 | G | OP1 | 13:1079 | 3.65 | 1.00 | 69.48 |
| 2 | G | O6 | 13:1077 | 3.87 | 1.00 | 71.24 |  |
| 3 | G | N7 | 13:1077 | 4.03 | 1.00 | 71.43 |  |
| 4 | G | OP2 | 13:1079 | 4.52 | 1.00 | 62.92 |  |
  |

  
 Coded contact list:   
1^st^ MG coordination shell missing (see Notes)  

### MG:13:1719

|  | Res. type | Atom | Residue | Distance | Occupancy | B-factor | Closest solute to 1^st^ shell water |
| --- | --- | --- | --- | --- | --- | --- | --- |
| **Ion** | | | |  | 1.00 | 63.93 || 1 | G | O6 | 13:230 | 4.12 | 1.00 | 68.16 |
| 2 | G | N7 | 13:231 | 4.15 | 1.00 | 69.16 |  |
| 3 | G | O6 | 13:231 | 4.17 | 1.00 | 71.33 |  |
  |

  
 Coded contact list:   
1^st^ MG coordination shell missing (see Notes)  

### MG:13:1720

|  | Res. type | Atom | Residue | Distance | Occupancy | B-factor | Closest solute to 1^st^ shell water |
| --- | --- | --- | --- | --- | --- | --- | --- |
| **Ion** | | | |  | 1.00 | 64.00 || 1 | G | O6 | 13:232 | 2.87 | 1.00 | 65.70 |
| 2 | G | O6 | 13:231 | 2.92 | 1.00 | 71.33 |  |
| 3 | U | O4 | 13:129 | 3.12 | 1.00 | 67.70 |  |
| 4 | G | N7 | 13:232 | 3.78 | 1.00 | 66.36 |  |
| 5 | C | N4 | 13:233 | 3.91 | 1.00 | 69.29 |  |
| 6 | G | N7 | 13:231 | 4.00 | 1.00 | 69.16 |  |
| 7 | G | O6 | 13:128 | 4.21 | 1.00 | 62.47 |  |
| 8 | U | N3 | 13:129 | 4.31 | 1.00 | 69.23 |  |
| 9 | C | N4 | 13:131 | 4.58 | 1.00 | 63.83 |  |
| 10 | G | N1 | 13:231 | 4.68 | 1.00 | 66.37 |  |
  |

  
 Coded contact list:   
 Exclusion zone contact(s): /O4/O6/O6/  
Consider K in place of MG (3 valid contact(s) for K vs. 0 for MG)  

### MG:13:1721

|  | Res. type | Atom | Residue | Distance | Occupancy | B-factor | Closest solute to 1^st^ shell water |
| --- | --- | --- | --- | --- | --- | --- | --- |
| **Ion** | | | |  | 1.00 | 99.67 || 1 | G | OP2 | 13:474 | 2.90 | 1.00 | 114.27 |
| 2 | G | O5' | 13:474 | 3.06 | 1.00 | 122.29 |  |
| 3 | G | OP2 | 13:475 | 3.19 | 1.00 | 124.55 |  |
| 4 | ARG | NE | 7I:75 | 4.01 | 1.00 | 101.23 |  |
| 5 | G | N7 | 13:475 | 4.01 | 1.00 | 127.69 |  |
| 6 | G | OP1 | 13:474 | 4.20 | 1.00 | 109.22 |  |
| 7 | G | N7 | 13:474 | 4.44 | 1.00 | 127.57 |  |
| 8 | G | N9 | 13:474 | 4.62 | 1.00 | 123.54 |  |
| 9 | G | O3' | 13:474 | 4.70 | 1.00 | 117.44 |  |
| 10 | G | O4' | 13:474 | 4.82 | 1.00 | 119.41 |  |
| 11 | ARG | NH2 | 7I:75 | 4.87 | 1.00 | 102.78 |  |
| 12 | A | O3' | 13:468 | 4.89 | 1.00 | 118.14 |  |
  |

  
 Coded contact list:   
 Exclusion zone contact(s): /O5'/OP2/OP2/  
Consider K in place of MG (3 valid contact(s) for K vs. 0 for MG)  

### MG:13:1722

|  | Res. type | Atom | Residue | Distance | Occupancy | B-factor | Closest solute to 1^st^ shell water |
| --- | --- | --- | --- | --- | --- | --- | --- |
| **Ion** | | | |  | 1.00 | 52.61 || 1 | U | O4 | 13:1393 | 2.71 | 1.00 | 55.86 |
| 2 | G | O6 | 13:922 | 2.87 | 1.00 | 53.45 |  |
| 3 | U | O4 | 13:921 | 3.30 | 1.00 | 55.25 |  |
| 4 | G | N7 | 13:922 | 3.66 | 1.00 | 59.51 |  |
| 5 | A | N6 | 13:923 | 4.30 | 1.00 | 53.56 |  |
| 6 | U | N3 | 13:1393 | 4.55 | 1.00 | 56.83 |  |
  |

  
 Coded contact list:   
 Exclusion zone contact(s): /O4/O4/O6/  
Consider K in place of MG (3 valid contact(s) for K vs. 0 for MG)  

### MG:13:1723

|  | Res. type | Atom | Residue | Distance | Occupancy | B-factor | Closest solute to 1^st^ shell water |
| --- | --- | --- | --- | --- | --- | --- | --- |
| **Ion** | | | |  | 1.00 | 60.24 || 1 | G | N7 | 13:297 | 4.19 | 1.00 | 49.79 |
| 2 | G | O6 | 13:297 | 4.24 | 1.00 | 56.96 |  |
  |

  
 Coded contact list:   
1^st^ MG coordination shell missing (see Notes)  

### MG:13:1724

|  | Res. type | Atom | Residue | Distance | Occupancy | B-factor | Closest solute to 1^st^ shell water |
| --- | --- | --- | --- | --- | --- | --- | --- |
| **Ion** | | | |  | 1.00 | 43.19 || 1 | C | OP1 | 13:934 | 2.19 | 1.00 | 75.86 |
| 2 | U | OP2 | 13:1345 | 3.88 | 1.00 | 62.97 |  |
| 3 | U | OP1 | 13:1345 | 4.06 | 1.00 | 69.09 |  |
| 4 | C | O5' | 13:934 | 4.26 | 1.00 | 68.89 |  |
| 5 | G | O3' | 13:933 | 4.48 | 1.00 | 69.82 |  |
| 6 | U | O2' | 13:1345 | 4.66 | 1.00 | 67.93 |  |
| 7 | C | OP2 | 13:934 | 4.67 | 1.00 | 71.07 |  |
  |

  
 Contact(s): /OP1/  
 Coded contact list: O_ph_  

### MG:13:1725

|  | Res. type | Atom | Residue | Distance | Occupancy | B-factor | Closest solute to 1^st^ shell water |
| --- | --- | --- | --- | --- | --- | --- | --- |
| **Ion** | | | |  | 1.00 | 58.27 || 1 | C | O2 | 13:454 | 3.67 | 1.00 | 105.52 |
| 2 | G | OP2 | 13:384 | 3.77 | 1.00 | 90.78 |  |
| 3 | C | N3 | 13:454 | 4.01 | 1.00 | 107.43 |  |
| 4 | A | N1 | 13:453 | 4.28 | 1.00 | 101.51 |  |
  |

  
 Coded contact list:   
1^st^ MG coordination shell missing (see Notes)  

### MG:13:1726

|  | Res. type | Atom | Residue | Distance | Occupancy | B-factor | Closest solute to 1^st^ shell water |
| --- | --- | --- | --- | --- | --- | --- | --- |
| **Ion** | | | |  | 1.00 | 46.92 || 1 | G | OP1 | 13:1505 | 2.00 | 1.00 | 44.96 |
| 2 | A | OP1 | 13:1500 | 2.07 | 1.00 | 37.10 |  |
| 3 | G | OP1 | 13:1508 | 2.09 | 1.00 | 52.94 |  |
| 4 | A | O3' | 13:1507 | 3.29 | 1.00 | 41.16 |  |
| 5 | U | O2' | 13:1506 | 3.78 | 1.00 | 52.39 |  |
| 6 | G | OP2 | 13:1504 | 3.94 | 1.00 | 53.60 |  |
| 7 | G | O3' | 13:1504 | 3.97 | 1.00 | 45.38 |  |
| 8 | A | OP2 | 13:1500 | 4.08 | 1.00 | 39.71 |  |
| 9 | G | O5' | 13:1504 | 4.18 | 1.00 | 47.36 |  |
| 10 | C | OP2 | 13:1501 | 4.24 | 1.00 | 34.09 |  |
| 11 | G | O5' | 13:1508 | 4.26 | 1.00 | 32.16 |  |
| 12 | G | O5' | 13:1505 | 4.30 | 1.00 | 44.87 |  |
| 13 | A | O5' | 13:1500 | 4.35 | 1.00 | 36.38 |  |
| 14 | G | OP2 | 13:1508 | 4.46 | 1.00 | 39.95 |  |
| 15 | G | OP2 | 13:1505 | 4.48 | 1.00 | 43.78 |  |
| 16 | A | O3' | 13:1499 | 4.53 | 1.00 | 40.40 |  |
| 17 | G | O2' | 13:1504 | 4.62 | 1.00 | 45.95 |  |
  |

  
**1^st^ shell ligand angles:**   
1-2: 81; 1-3: 155; 2-3: 84;   
  
 Contact(s): /OP1/OP1/OP1/  
 Coded contact list: 3O_ph_  
 Exclusion zone contact(s): /O3'/  

### MG:13:1727

|  | Res. type | Atom | Residue | Distance | Occupancy | B-factor | Closest solute to 1^st^ shell water |
| --- | --- | --- | --- | --- | --- | --- | --- |
| **Ion** | | | |  | 1.00 | 60.83 || 1 | Water 1 | | 5I:201 | 2.17 | 1.00 | 58.66 | 2.64 (ALA:O 5I:20) |
| 2 | Water 2 | | 13:1998 | 2.17 | 1.00 | 77.24 | 3.53 (C:O3' 13:1359) |
| 3 | Water 3 | | 13:1933 | 2.17 | 1.00 | 73.78 | 2.93 (A:N7 13:1360) |
| 4 | Water 4 | | 13:1827 | 2.18 | 1.00 | 75.65 | 2.24 (G:O6 13:1361) |
| 5 | Water 5 | | 13:1927 | 2.19 | 1.00 | 75.47 | 2.91 (C:N3 13:980) |
| 6 | Water 6 | | 13:1805 | 2.21 | 1.00 | 78.79 | 1.31 (C:N4 13:979) |
| 7 | C | N4 | 13:979 | 3.52 | 1.00 | 74.46 |  |
| 8 | VAL | O | 5I:18 | 3.77 | 1.00 | 69.43 |  |
| 9 | A | N7 | 13:1360 | 3.92 | 1.00 | 75.25 |  |
| 10 | C | N3 | 13:980 | 4.01 | 1.00 | 68.92 |  |
| 11 | ALA | O | 5I:20 | 4.24 | 1.00 | 71.20 |  |
| 12 | G | O6 | 13:1361 | 4.28 | 1.00 | 70.87 |  |
| 13 | C | O2 | 13:980 | 4.31 | 1.00 | 66.77 |  |
| 14 | C | O3' | 13:1359 | 4.74 | 1.00 | 73.59 |  |
| 15 | A | OP1 | 13:1360 | 4.75 | 1.00 | 63.58 |  |
| 16 | A | N6 | 13:1360 | 4.88 | 1.00 | 73.77 |  |
| 17 | C | N4 | 13:980 | 4.90 | 1.00 | 66.27 |  |
  |

  
**1^st^ shell ligand angles:**   
1-2: 87; 1-3: 90; 1-4: 178; 1-5: 91; 1-6: 93; 2-3: 88; 2-4: 91; 2-5: 90; 2-6: 179; 3-4: 89; 3-5: 177; 3-6: 93; 4-5: 90; 4-6: 89; 5-6: 90;   
  
 Contact(s): /O_w_/O_w_/O_w_/O_w_/O_w_/O_w_/  
 Coded contact list: 6O_w_  
MG hydration shell issue: water too close (< 2.4 Angs.)  

### MG:13:1728

|  | Res. type | Atom | Residue | Distance | Occupancy | B-factor | Closest solute to 1^st^ shell water |
| --- | --- | --- | --- | --- | --- | --- | --- |
| **Ion** | | | |  | 1.00 | 60.52 || 1 | Water 1 | | 13:1901 | 2.17 | 1.00 | 55.08 | 2.74 (A:OP2 13:16) |
| 2 | Water 2 | | 13:1991 | 2.18 | 1.00 | 52.18 | 3.42 (G:OP2 13:15) |
| 3 | Water 3 | | 13:1872 | 2.18 | 1.00 | 47.14 | 2.53 (U:OP1 13:13) |
| 4 | Water 4 | | 13:2023 | 2.19 | 1.00 | 70.54 | 4.18 (U:OP1 13:13) |
| 5 | Water 5 | | 13:2002 | 2.19 | 1.00 | 54.79 | 3.62 (A:OP1 13:16) |
| 6 | Water 6 | | 13:2033 | 2.19 | 1.00 | 58.27 | 4.81 (G:OP2 13:15) |
| 7 | U | OP2 | 13:14 | 4.07 | 1.00 | 54.29 |  |
| 8 | U | OP1 | 13:13 | 4.21 | 1.00 | 57.40 |  |
| 9 | G | OP2 | 13:15 | 4.41 | 1.00 | 56.91 |  |
| 10 | A | OP1 | 13:16 | 4.63 | 1.00 | 47.54 |  |
| 11 | A | OP2 | 13:16 | 4.80 | 1.00 | 52.38 |  |
  |

  
**1^st^ shell ligand angles:**   
1-2: 89; 1-3: 90; 1-4: 176; 1-5: 90; 1-6: 91; 2-3: 94; 2-4: 90; 2-5: 177; 2-6: 85; 3-4: 86; 3-5: 89; 3-6: 178; 4-5: 91; 4-6: 92; 5-6: 92;   
  
 Contact(s): /O_w_/O_w_/O_w_/O_w_/O_w_/O_w_/  
 Coded contact list: 6O_w_  

### MG:13:1729

|  | Res. type | Atom | Residue | Distance | Occupancy | B-factor | Closest solute to 1^st^ shell water |
| --- | --- | --- | --- | --- | --- | --- | --- |
| **Ion** | | | |  | 1.00 | 61.45 || 1 | Water 1 | | 13:1863 | 2.18 | 1.00 | 52.92 | 2.50 (G:O6 13:44) |
| 2 | Water 2 | | 13:1850 | 2.18 | 1.00 | 60.24 | 2.37 (C:N4 13:43) |
| 3 | Water 3 | | 13:1963 | 2.18 | 1.00 | 66.58 | 3.17 (G:N7 13:44) |
| 4 | Water 4 | | 13:2008 | 2.19 | 1.00 | 62.06 | 3.72 (C:N4 13:395) |
| 5 | Water 5 | | 13:1914 | 2.19 | 1.00 | 49.48 | 2.81 (C:N4 13:395) |
| 6 | Water 6 | | 13:2017 | 2.19 | 1.00 | 58.25 | 5.30 (C:N4 13:43) |
| 7 | U | O4 | 13:45 | 3.88 | 1.00 | 63.53 |  |
| 8 | C | N4 | 13:43 | 4.12 | 1.00 | 62.47 |  |
| 9 | G | N7 | 13:44 | 4.13 | 1.00 | 59.15 |  |
| 10 | C | N4 | 13:395 | 4.21 | 1.00 | 51.10 |  |
| 11 | G | O6 | 13:396 | 4.39 | 1.00 | 57.09 |  |
| 12 | G | O6 | 13:44 | 4.60 | 1.00 | 58.81 |  |
  |

  
**1^st^ shell ligand angles:**   
1-2: 87; 1-3: 91; 1-4: 90; 1-5: 92; 1-6: 177; 2-3: 93; 2-4: 88; 2-5: 179; 2-6: 90; 3-4: 179; 3-5: 88; 3-6: 87; 4-5: 92; 4-6: 92; 5-6: 91;   
  
 Contact(s): /O_w_/O_w_/O_w_/O_w_/O_w_/O_w_/  
 Coded contact list: 6O_w_  
MG hydration shell issue: water too close (< 2.4 Angs.)  

### MG:13:1730

|  | Res. type | Atom | Residue | Distance | Occupancy | B-factor | Closest solute to 1^st^ shell water |
| --- | --- | --- | --- | --- | --- | --- | --- |
| **Ion** | | | |  | 1.00 | 76.59 || 1 | Water 1 | | 13:1889 | 2.17 | 1.00 | 78.95 | 2.64 (G:O6 13:409) |
| 2 | Water 2 | | 13:1845 | 2.17 | 1.00 | 86.01 | 2.36 (A:OP1 13:431) |
| 3 | Water 3 | | 13:1973 | 2.17 | 1.00 | 75.71 | 3.26 (G:OP2 13:407) |
| 4 | Water 4 | | 3E:402 | 2.18 | 1.00 | 78.25 | 2.88 (LYS:NZ 3E:22) |
| 5 | Water 5 | | 13:1925 | 2.18 | 1.00 | 83.67 | 2.89 (A:N7 13:408) |
| 6 | Water 6 | | 13:1858 | 2.18 | 1.00 | 76.23 | 2.46 (G:O6 13:410) |
| 7 | G | O6 | 13:410 | 3.62 | 1.00 | 83.76 |  |
| 8 | A | OP2 | 13:431 | 4.12 | 1.00 | 82.20 |  |
| 9 | LYS | NZ | 3E:22 | 4.30 | 1.00 | 74.44 |  |
| 10 | A | N7 | 13:408 | 4.36 | 1.00 | 84.80 |  |
| 11 | A | OP1 | 13:431 | 4.36 | 1.00 | 76.79 |  |
| 12 | G | N7 | 13:409 | 4.42 | 1.00 | 85.21 |  |
| 13 | G | N1 | 13:410 | 4.46 | 1.00 | 80.85 |  |
| 14 | G | O6 | 13:409 | 4.72 | 1.00 | 85.38 |  |
| 15 | A | OP2 | 13:408 | 4.79 | 1.00 | 81.08 |  |
| 16 | G | OP2 | 13:407 | 4.97 | 1.00 | 77.15 |  |
  |

  
**1^st^ shell ligand angles:**   
1-2: 87; 1-3: 178; 1-4: 93; 1-5: 87; 1-6: 92; 2-3: 93; 2-4: 178; 2-5: 89; 2-6: 89; 3-4: 87; 3-5: 91; 3-6: 91; 4-5: 93; 4-6: 90; 5-6: 177;   
  
 Contact(s): /O_w_/O_w_/O_w_/O_w_/O_w_/O_w_/  
 Coded contact list: 6O_w_  
MG hydration shell issue: water too close (< 2.4 Angs.)  

### MG:13:1731

|  | Res. type | Atom | Residue | Distance | Occupancy | B-factor | Closest solute to 1^st^ shell water |
| --- | --- | --- | --- | --- | --- | --- | --- |
| **Ion** | | | |  | 1.00 | 38.84 || 1 | Water 1 | | 13:1892 | 2.16 | 1.00 | 56.96 | 2.68 (C:O2 13:330) |
| 2 | Water 2 | | 13:1885 | 2.17 | 1.00 | 60.22 | 2.62 (A:N1 13:59) |
| 3 | Water 3 | | 13:1987 | 2.17 | 1.00 | 51.72 | 3.36 (G:O4' 13:331) |
| 4 | Water 4 | | 13:1887 | 2.17 | 1.00 | 56.12 | 2.63 (C:OP2 13:352) |
| 5 | Water 5 | | 13:1882 | 2.18 | 1.00 | 54.39 | 2.59 (C:OP1 13:352) |
| 6 | C | OP2 | 13:352 | 2.75 | 1.00 | 63.02 |  |
| 7 | C | OP1 | 13:352 | 3.12 | 1.00 | 55.34 |  |
| 8 | C | O2 | 13:330 | 3.72 | 1.00 | 64.09 |  |
| 9 | G | N7 | 13:354 | 4.06 | 1.00 | 61.04 |  |
| 10 | G | O4' | 13:331 | 4.18 | 1.00 | 60.02 |  |
| 11 | C | O4' | 13:352 | 4.31 | 1.00 | 54.26 |  |
| 12 | G | O3' | 13:351 | 4.40 | 1.00 | 60.81 |  |
| 13 | A | N1 | 13:59 | 4.46 | 1.00 | 59.80 |  |
| 14 | C | O5' | 13:352 | 4.52 | 1.00 | 55.57 |  |
| 15 | G | OP1 | 13:354 | 4.71 | 1.00 | 58.28 |  |
| 16 | A | N6 | 13:59 | 4.87 | 1.00 | 61.69 |  |
  |

  
**1^st^ shell ligand angles:**   
1-2: 90; 1-3: 92; 1-4: 88; 1-5: 176; 2-3: 91; 2-4: 87; 2-5: 95; 3-4: 178; 3-5: 89; 4-5: 92;   
  
 Contact(s): /O_w_/O_w_/O_w_/O_w_/O_w_/  
 Coded contact list: 5O_w_  
 Exclusion zone contact(s): /OP1/OP2/  
Consider K in place of MG (2 valid contact(s) for K vs. 0 for MG)  

### MG:13:1732

|  | Res. type | Atom | Residue | Distance | Occupancy | B-factor | Closest solute to 1^st^ shell water |
| --- | --- | --- | --- | --- | --- | --- | --- |
| **Ion** | | | |  | 1.00 | 51.59 || 1 | Water 1 | | 13:1993 | 2.17 | 1.00 | 51.19 | 3.83 (C:OP1 13:503) |
| 2 | Water 2 | | 13:1886 | 2.17 | 1.00 | 49.45 | 2.62 (C:O2' 13:522) |
| 3 | Water 3 | | 13:1947 | 2.18 | 1.00 | 50.36 | 3.05 (C:OP1 13:503) |
| 4 | Water 4 | | 3I:201 | 2.18 | 1.00 | 43.22 | 2.33 (SER:OG 3I:113) |
| 5 | Water 5 | | 13:1873 | 2.18 | 1.00 | 42.92 | 2.53 (G:OP2 13:537) |
| 6 | Water 6 | | 13:1883 | 2.19 | 1.00 | 51.23 | 2.60 (G:OP1 13:537) |
| 7 | G | OP2 | 13:537 | 3.86 | 1.00 | 52.82 |  |
| 8 | C | OP1 | 13:503 | 4.28 | 1.00 | 44.59 |  |
| 9 | SER | OG | 3I:113 | 4.29 | 1.00 | 37.77 |  |
| 10 | TYR | OH | 3I:117 | 4.33 | 1.00 | 44.36 |  |
| 11 | C | OP1 | 13:536 | 4.39 | 1.00 | 42.98 |  |
| 12 | C | O3' | 13:522 | 4.41 | 1.00 | 43.17 |  |
| 13 | G | OP1 | 13:537 | 4.62 | 1.00 | 49.49 |  |
| 14 | A | OP1 | 13:523 | 4.70 | 1.00 | 43.35 |  |
| 15 | C | O2' | 13:522 | 4.75 | 1.00 | 46.97 |  |
  |

  
**1^st^ shell ligand angles:**   
1-2: 91; 1-3: 85; 1-4: 90; 1-5: 94; 1-6: 179; 2-3: 89; 2-4: 179; 2-5: 89; 2-6: 89; 3-4: 90; 3-5: 177; 3-6: 94; 4-5: 91; 4-6: 90; 5-6: 87;   
  
 Contact(s): /O_w_/O_w_/O_w_/O_w_/O_w_/O_w_/  
 Coded contact list: 6O_w_  
MG hydration shell issue: water too close (< 2.4 Angs.)  

### MG:13:1733

|  | Res. type | Atom | Residue | Distance | Occupancy | B-factor | Closest solute to 1^st^ shell water |
| --- | --- | --- | --- | --- | --- | --- | --- |
| **Ion** | | | |  | 1.00 | 62.46 || 1 | Water 1 | | 13:1902 | 2.17 | 1.00 | 49.09 | 2.74 (C:O2' 13:962) |
| 2 | Water 2 | | 13:1950 | 2.17 | 1.00 | 49.52 | 3.59 (C:OP1 13:1051) |
| 3 | Water 3 | | 13:1968 | 2.17 | 1.00 | 58.75 | 3.30 (C:OP1 13:1200) |
| 4 | Water 4 | | 13:1945 | 2.18 | 1.00 | 65.49 | 3.11 (C:O5' 13:1200) |
| 5 | Water 5 | | 13:1904 | 2.19 | 1.00 | 58.60 | 2.74 (C:O2' 13:962) |
| 6 | Water 6 | | 13:1954 | 2.20 | 1.00 | 53.91 | 3.11 (C:OP1 13:1051) |
| 7 | C | O2' | 13:962 | 3.63 | 1.00 | 50.55 |  |
| 8 | C | OP1 | 13:1200 | 4.14 | 1.00 | 50.83 |  |
| 9 | C | OP1 | 13:1051 | 4.53 | 1.00 | 65.25 |  |
| 10 | C | O5' | 13:1200 | 4.63 | 1.00 | 53.39 |  |
| 11 | A | N7 | 13:1201 | 4.95 | 1.00 | 61.29 |  |
| 12 | G | O4' | 13:963 | 4.96 | 1.00 | 52.24 |  |
| 13 | C | OP2 | 13:1051 | 4.97 | 1.00 | 70.40 |  |
  |

  
**1^st^ shell ligand angles:**   
1-2: 96; 1-3: 90; 1-4: 179; 1-5: 86; 1-6: 89; 2-3: 90; 2-4: 85; 2-5: 178; 2-6: 88; 3-4: 89; 3-5: 89; 3-6: 177; 4-5: 93; 4-6: 92; 5-6: 94;   
  
 Contact(s): /O_w_/O_w_/O_w_/O_w_/O_w_/O_w_/  
 Coded contact list: 6O_w_  

### MG:13:1734

|  | Res. type | Atom | Residue | Distance | Occupancy | B-factor | Closest solute to 1^st^ shell water |
| --- | --- | --- | --- | --- | --- | --- | --- |
| **Ion** | | | |  | 1.00 | 41.94 || 1 | Water 1 | | 13:1895 | 2.17 | 1.00 | 35.81 | 2.70 (G:OP1 13:577) |
| 2 | Water 2 | | 13:1840 | 2.18 | 1.00 | 34.30 | 2.34 (C:OP1 13:904) |
| 3 | Water 3 | | 13:2022 | 2.19 | 1.00 | 37.53 | 4.10 (G:OP2 13:577) |
| 4 | Water 4 | | 13:1949 | 2.19 | 1.00 | 32.67 | 3.20 (U:OP2 13:905) |
| 5 | Water 5 | | 13:1922 | 2.20 | 1.00 | 38.76 | 2.87 (A:OP1 13:814) |
| 6 | Water 6 | | 13:1989 | 2.20 | 1.00 | 43.70 | 3.38 (C:OP1 13:904) |
| 7 | C | OP1 | 13:904 | 3.77 | 1.00 | 41.61 |  |
| 8 | A | OP2 | 13:814 | 3.92 | 1.00 | 33.32 |  |
| 9 | A | OP1 | 13:814 | 4.21 | 1.00 | 31.77 |  |
| 10 | U | OP2 | 13:905 | 4.77 | 1.00 | 35.90 |  |
| 11 | G | OP1 | 13:577 | 4.78 | 1.00 | 40.95 |  |
| 12 | G | OP2 | 13:577 | 4.79 | 1.00 | 31.40 |  |
  |

  
**1^st^ shell ligand angles:**   
1-2: 83; 1-3: 92; 1-4: 93; 1-5: 89; 1-6: 178; 2-3: 175; 2-4: 93; 2-5: 89; 2-6: 95; 3-4: 89; 3-5: 88; 3-6: 89; 4-5: 177; 4-6: 86; 5-6: 92;   
  
 Contact(s): /O_w_/O_w_/O_w_/O_w_/O_w_/O_w_/  
 Coded contact list: 6O_w_  
MG hydration shell issue: water too close (< 2.4 Angs.)  

### MG:13:1735

|  | Res. type | Atom | Residue | Distance | Occupancy | B-factor | Closest solute to 1^st^ shell water |
| --- | --- | --- | --- | --- | --- | --- | --- |
| **Ion** | | | |  | 1.00 | 35.59 || 1 | Water 1 | | 13:1810 | 2.16 | 1.00 | 30.54 | 2.05 (C:OP1 13:578) |
| 2 | Water 2 | | 13:1874 | 2.16 | 1.00 | 43.09 | 2.53 (C:OP2 13:817) |
| 3 | Water 3 | | 13:1811 | 2.17 | 1.00 | 35.88 | 2.05 (C:OP1 13:578) |
| 4 | Water 4 | | 13:1852 | 2.17 | 1.00 | 44.96 | 2.38 (G:O2' 13:575) |
| 5 | Water 5 | | 13:1975 | 2.18 | 1.00 | 46.55 | 3.27 (C:OP1 13:578) |
| 6 | Water 6 | | 13:1997 | 2.19 | 1.00 | 42.15 | 3.48 (U:OP1 13:820) |
| 7 | C | OP1 | 13:578 | 2.73 | 1.00 | 34.16 |  |
| 8 | U | OP1 | 13:820 | 3.96 | 1.00 | 46.76 |  |
| 9 | C | OP2 | 13:817 | 3.98 | 1.00 | 37.60 |  |
| 10 | G | OP2 | 13:576 | 4.10 | 1.00 | 41.48 |  |
| 11 | G | O3' | 13:577 | 4.30 | 1.00 | 37.54 |  |
| 12 | G | O3' | 13:575 | 4.47 | 1.00 | 50.71 |  |
| 13 | G | O2' | 13:575 | 4.47 | 1.00 | 47.97 |  |
| 14 | U | O5' | 13:820 | 4.51 | 1.00 | 44.09 |  |
| 15 | C | O5' | 13:817 | 4.93 | 1.00 | 39.03 |  |
  |

  
**1^st^ shell ligand angles:**   
1-2: 91; 1-3: 95; 1-4: 178; 1-5: 85; 1-6: 89; 2-3: 87; 2-4: 90; 2-5: 175; 2-6: 91; 3-4: 86; 3-5: 91; 3-6: 175; 4-5: 94; 4-6: 89; 5-6: 92;   
  
 Contact(s): /O_w_/O_w_/O_w_/O_w_/O_w_/O_w_/  
 Coded contact list: 6O_w_  
 Exclusion zone contact(s): /OP1/  
Consider K in place of MG (1 valid contact(s) for K vs. 0 for MG)  
MG hydration shell issue: water too close (< 2.4 Angs.)  

### MG:13:1736

|  | Res. type | Atom | Residue | Distance | Occupancy | B-factor | Closest solute to 1^st^ shell water |
| --- | --- | --- | --- | --- | --- | --- | --- |
| **Ion** | | | |  | 1.00 | 44.29 || 1 | Water 1 | | 13:1841 | 2.15 | 1.00 | 49.40 | 2.34 (U:O5' 13:863) |
| 2 | Water 2 | | 13:2025 | 2.18 | 1.00 | 59.40 | 4.52 (U:O5' 13:863) |
| 3 | Water 3 | | 13:1979 | 2.18 | 1.00 | 64.81 | 3.32 (U:OP1 13:863) |
| 4 | Water 4 | | 4E:201 | 2.18 | 1.00 | 57.74 | 3.02 (GLY:O 4E:85) |
| 5 | Water 5 | | 13:1934 | 2.18 | 1.00 | 58.10 | 2.95 (A:OP2 13:864) |
| 6 | Water 6 | | 13:2024 | 2.18 | 1.00 | 50.53 | 5.60 (GLY:O 4E:85) |
| 7 | U | OP1 | 13:863 | 3.96 | 1.00 | 58.85 |  |
| 8 | A | OP2 | 13:864 | 4.00 | 1.00 | 50.74 |  |
| 9 | U | O5' | 13:863 | 4.41 | 1.00 | 55.45 |  |
| 10 | GLY | O | 4E:85 | 4.63 | 1.00 | 48.88 |  |
  |

  
**1^st^ shell ligand angles:**   
1-2: 91; 1-3: 87; 1-4: 90; 1-5: 89; 1-6: 177; 2-3: 90; 2-4: 177; 2-5: 88; 2-6: 91; 3-4: 88; 3-5: 175; 3-6: 95; 4-5: 95; 4-6: 88; 5-6: 89;   
  
 Contact(s): /O_w_/O_w_/O_w_/O_w_/O_w_/O_w_/  
 Coded contact list: 6O_w_  
MG hydration shell issue: water too close (< 2.4 Angs.)  

### MG:13:1737

|  | Res. type | Atom | Residue | Distance | Occupancy | B-factor | Closest solute to 1^st^ shell water |
| --- | --- | --- | --- | --- | --- | --- | --- |
| **Ion** | | | |  | 1.00 | 65.46 || 1 | Water 1 | | 13:1812 | 2.17 | 1.00 | 61.62 | 2.06 (G:OP1 13:588) |
| 2 | Water 2 | | 13:1859 | 2.18 | 1.00 | 62.05 | 2.46 (C:OP2 13:645) |
| 3 | Water 3 | | 13:1910 | 2.18 | 1.00 | 70.55 | 2.80 (G:OP1 13:644) |
| 4 | Water 4 | | 13:1819 | 2.18 | 1.00 | 63.60 | 2.15 (G:OP2 13:588) |
| 5 | Water 5 | | 13:1898 | 2.18 | 1.00 | 63.30 | 2.72 (C:OP2 13:645) |
| 6 | Water 6 | | 13:1842 | 2.19 | 1.00 | 53.38 | 2.34 (G:OP2 13:588) |
| 7 | G | OP2 | 13:588 | 2.92 | 1.00 | 69.54 |  |
| 8 | C | OP2 | 13:645 | 3.38 | 1.00 | 61.13 |  |
| 9 | G | OP1 | 13:644 | 3.75 | 1.00 | 64.36 |  |
| 10 | G | OP1 | 13:588 | 3.75 | 1.00 | 66.90 |  |
| 11 | C | OP1 | 13:645 | 3.98 | 1.00 | 65.34 |  |
| 12 | G | O3' | 13:644 | 4.59 | 1.00 | 62.02 |  |
| 13 | G | O3' | 13:587 | 4.72 | 1.00 | 69.29 |  |
  |

  
**1^st^ shell ligand angles:**   
1-2: 177; 1-3: 90; 1-4: 92; 1-5: 87; 1-6: 90; 2-3: 87; 2-4: 90; 2-5: 92; 2-6: 92; 3-4: 178; 3-5: 93; 3-6: 89; 4-5: 86; 4-6: 92; 5-6: 176;   
  
 Contact(s): /O_w_/O_w_/O_w_/O_w_/O_w_/O_w_/  
 Coded contact list: 6O_w_  
 Exclusion zone contact(s): /OP2/OP2/  
Consider K in place of MG (2 valid contact(s) for K vs. 0 for MG)  
MG hydration shell issue: water too close (< 2.4 Angs.)  

### MG:13:1738

|  | Res. type | Atom | Residue | Distance | Occupancy | B-factor | Closest solute to 1^st^ shell water |
| --- | --- | --- | --- | --- | --- | --- | --- |
| **Ion** | | | |  | 1.00 | 86.34 || 1 | Water 1 | | 13:1826 | 2.17 | 1.00 | 87.17 | 2.23 (G:N7 13:650) |
| 2 | Water 2 | | 13:1994 | 2.17 | 1.00 | 65.73 | 3.45 (G:N7 13:649) |
| 3 | Water 3 | | 13:2029 | 2.18 | 1.00 | 95.78 | 4.60 (A:OP1 13:753) |
| 4 | Water 4 | | 13:1953 | 2.18 | 1.00 | 68.32 | 3.14 (A:OP1 13:753) |
| 5 | Water 5 | | 13:1985 | 2.19 | 1.00 | 84.76 | 3.49 (G:O6 13:649) |
| 6 | Water 6 | | 13:1909 | 2.20 | 1.00 | 80.19 | 2.80 (G:O6 13:650) |
| 7 | G | O6 | 13:650 | 3.89 | 1.00 | 84.12 |  |
| 8 | G | N7 | 13:650 | 4.22 | 1.00 | 89.05 |  |
| 9 | G | O6 | 13:649 | 4.23 | 1.00 | 87.91 |  |
| 10 | G | N7 | 13:649 | 4.26 | 1.00 | 88.52 |  |
| 11 | A | OP1 | 13:753 | 4.99 | 1.00 | 72.92 |  |
  |

  
**1^st^ shell ligand angles:**   
1-2: 91; 1-3: 179; 1-4: 90; 1-5: 89; 1-6: 89; 2-3: 89; 2-4: 90; 2-5: 90; 2-6: 180; 3-4: 90; 3-5: 90; 3-6: 91; 4-5: 180; 4-6: 90; 5-6: 90;   
  
 Contact(s): /O_w_/O_w_/O_w_/O_w_/O_w_/O_w_/  
 Coded contact list: 6O_w_  
MG hydration shell issue: water too close (< 2.4 Angs.)  

### MG:13:1739

|  | Res. type | Atom | Residue | Distance | Occupancy | B-factor | Closest solute to 1^st^ shell water |
| --- | --- | --- | --- | --- | --- | --- | --- |
| **Ion** | | | |  | 1.00 | 30.03 || 1 | Water 1 | | 13:1911 | 2.15 | 1.00 | 30.61 | 2.80 (U:O4 13:772) |
| 2 | Water 2 | | 13:2010 | 2.16 | 1.00 | 42.15 | 4.45 (G:N7 13:771) |
| 3 | Water 3 | | 13:1809 | 2.17 | 1.00 | 34.65 | 2.00 (G:OP1 13:803) |
| 4 | Water 4 | | 13:1974 | 2.17 | 1.00 | 49.99 | 3.26 (G:N7 13:771) |
| 5 | Water 5 | | 13:1930 | 2.17 | 1.00 | 39.73 | 2.91 (U:OP2 13:804) |
| 6 | Water 6 | | 13:1965 | 2.20 | 1.00 | 41.67 | 3.19 (G:O6 13:773) |
| 7 | U | O4 | 13:772 | 3.93 | 1.00 | 41.72 |  |
| 8 | G | OP1 | 13:803 | 4.14 | 1.00 | 31.46 |  |
| 9 | G | N7 | 13:771 | 4.38 | 1.00 | 36.91 |  |
| 10 | U | OP2 | 13:804 | 4.75 | 1.00 | 39.38 |  |
| 11 | G | O6 | 13:773 | 4.85 | 1.00 | 46.17 |  |
| 12 | G | O6 | 13:771 | 4.93 | 1.00 | 35.72 |  |
  |

  
**1^st^ shell ligand angles:**   
1-2: 86; 1-3: 175; 1-4: 90; 1-5: 91; 1-6: 94; 2-3: 90; 2-4: 83; 2-5: 97; 2-6: 178; 3-4: 87; 3-5: 92; 3-6: 90; 4-5: 179; 4-6: 95; 5-6: 85;   
  
 Contact(s): /O_w_/O_w_/O_w_/O_w_/O_w_/O_w_/  
 Coded contact list: 6O_w_  
MG hydration shell issue: water too close (< 2.4 Angs.)  

### MG:13:1740

|  | Res. type | Atom | Residue | Distance | Occupancy | B-factor | Closest solute to 1^st^ shell water |
| --- | --- | --- | --- | --- | --- | --- | --- |
| **Ion** | | | |  | 1.00 | 33.83 || 1 | Water 1 | | 13:1851 | 2.17 | 1.00 | 39.48 | 2.38 (A:OP2 13:781) |
| 2 | Water 2 | | 13:1864 | 2.17 | 1.00 | 41.84 | 2.50 (G:O6 13:800) |
| 3 | Water 3 | | 13:2015 | 2.18 | 1.00 | 48.90 | 4.57 (G:O6 13:800) |
| 4 | Water 4 | | 13:1871 | 2.18 | 1.00 | 45.10 | 2.52 (G:O6 13:799) |
| 5 | Water 5 | | 13:2007 | 2.18 | 1.00 | 44.54 | 3.70 (C:N4 13:797) |
| 6 | Water 6 | | 13:1932 | 2.18 | 1.00 | 43.97 | 2.93 (G:N7 13:798) |
| 7 | G | O6 | 13:800 | 3.79 | 1.00 | 38.01 |  |
| 8 | G | O6 | 13:799 | 4.12 | 1.00 | 37.13 |  |
| 9 | G | N7 | 13:799 | 4.13 | 1.00 | 41.35 |  |
| 10 | A | OP2 | 13:781 | 4.43 | 1.00 | 29.24 |  |
| 11 | G | N1 | 13:800 | 4.47 | 1.00 | 39.35 |  |
| 12 | G | N7 | 13:798 | 4.56 | 1.00 | 43.58 |  |
| 13 | G | O6 | 13:798 | 4.59 | 1.00 | 42.82 |  |
| 14 | C | N4 | 13:797 | 4.95 | 1.00 | 47.11 |  |
  |

  
**1^st^ shell ligand angles:**   
1-2: 89; 1-3: 86; 1-4: 94; 1-5: 89; 1-6: 178; 2-3: 93; 2-4: 90; 2-5: 179; 2-6: 89; 3-4: 177; 3-5: 87; 3-6: 94; 4-5: 90; 4-6: 87; 5-6: 93;   
  
 Contact(s): /O_w_/O_w_/O_w_/O_w_/O_w_/O_w_/  
 Coded contact list: 6O_w_  
MG hydration shell issue: water too close (< 2.4 Angs.)  

### MG:13:1741

|  | Res. type | Atom | Residue | Distance | Occupancy | B-factor | Closest solute to 1^st^ shell water |
| --- | --- | --- | --- | --- | --- | --- | --- |
| **Ion** | | | |  | 1.00 | 109.99 || 1 | Water 1 | | 13:2013 | 2.17 | 1.00 | 96.73 | 4.89 (G:O6 13:102) |
| 2 | Water 2 | | 13:1931 | 2.17 | 1.00 | 114.78 | 2.91 (G:O6 13:64) |
| 3 | Water 3 | | 13:1818 | 2.18 | 1.00 | 106.58 | 2.11 (G:O6 13:102) |
| 4 | Water 4 | | 13:2011 | 2.18 | 1.00 | 117.77 | 3.74 (A:N7 13:101) |
| 5 | Water 5 | | 13:2021 | 2.18 | 1.00 | 108.69 | 4.10 (A:N7 13:101) |
| 6 | Water 6 | | 13:1803 | 2.19 | 1.00 | 120.86 | 1.30 (A:N7 13:101) |
| 7 | A | N7 | 13:101 | 3.43 | 1.00 | 117.14 |  |
| 8 | G | O6 | 13:102 | 4.14 | 1.00 | 114.53 |  |
| 9 | C | N4 | 13:99 | 4.29 | 1.00 | 122.36 |  |
| 10 | G | N7 | 13:102 | 4.35 | 1.00 | 114.98 |  |
| 11 | A | N6 | 13:101 | 4.76 | 1.00 | 120.37 |  |
| 12 | G | O6 | 13:64 | 4.77 | 1.00 | 122.39 |  |
  |

  
**1^st^ shell ligand angles:**   
1-2: 89; 1-3: 85; 1-4: 91; 1-5: 93; 1-6: 178; 2-3: 90; 2-4: 180; 2-5: 90; 2-6: 92; 3-4: 90; 3-5: 178; 3-6: 93; 4-5: 90; 4-6: 88; 5-6: 89;   
  
 Contact(s): /O_w_/O_w_/O_w_/O_w_/O_w_/O_w_/  
 Coded contact list: 6O_w_  
MG hydration shell issue: water too close (< 2.4 Angs.)  

### MG:13:1742

|  | Res. type | Atom | Residue | Distance | Occupancy | B-factor | Closest solute to 1^st^ shell water |
| --- | --- | --- | --- | --- | --- | --- | --- |
| **Ion** | | | |  | 1.00 | 60.34 || 1 | Water 1 | | 13:1877 | 2.17 | 1.00 | 52.71 | 2.56 (G:N7 13:758) |
| 2 | Water 2 | | 13:1833 | 2.18 | 1.00 | 52.31 | 2.28 (G:N7 13:581) |
| 3 | Water 3 | | 13:1870 | 2.19 | 1.00 | 47.78 | 2.52 (G:O6 13:758) |
| 4 | Water 4 | | 13:1868 | 2.19 | 1.00 | 50.19 | 2.52 (U:OP2 13:580) |
| 5 | Water 5 | | 13:1962 | 2.19 | 1.00 | 61.63 | 3.17 (G:OP2 13:758) |
| 6 | Water 6 | | 13:1971 | 2.20 | 1.00 | 46.85 | 3.25 (U:OP2 13:757) |
| 7 | G | N7 | 13:581 | 3.67 | 1.00 | 46.16 |  |
| 8 | G | OP2 | 13:581 | 3.87 | 1.00 | 41.66 |  |
| 9 | G | N7 | 13:758 | 3.88 | 1.00 | 47.39 |  |
| 10 | U | OP2 | 13:580 | 4.15 | 1.00 | 55.16 |  |
| 11 | G | O6 | 13:758 | 4.24 | 1.00 | 41.37 |  |
| 12 | G | OP2 | 13:758 | 4.30 | 1.00 | 45.75 |  |
| 13 | U | OP2 | 13:757 | 4.59 | 1.00 | 54.00 |  |
| 14 | U | O5' | 13:580 | 4.75 | 1.00 | 51.63 |  |
  |

  
**1^st^ shell ligand angles:**   
1-2: 87; 1-3: 89; 1-4: 178; 1-5: 89; 1-6: 91; 2-3: 92; 2-4: 91; 2-5: 87; 2-6: 179; 3-4: 91; 3-5: 178; 3-6: 88; 4-5: 90; 4-6: 90; 5-6: 94;   
  
 Contact(s): /O_w_/O_w_/O_w_/O_w_/O_w_/O_w_/  
 Coded contact list: 6O_w_  
MG hydration shell issue: water too close (< 2.4 Angs.)  

### MG:13:1743

|  | Res. type | Atom | Residue | Distance | Occupancy | B-factor | Closest solute to 1^st^ shell water |
| --- | --- | --- | --- | --- | --- | --- | --- |
| **Ion** | | | |  | 1.00 | 54.10 || 1 | Water 1 | | 13:1829 | 2.17 | 1.00 | 50.89 | 2.26 (C:OP2 13:536) |
| 2 | Water 2 | | 13:1893 | 2.18 | 1.00 | 51.12 | 2.69 (C:OP2 13:536) |
| 3 | Water 3 | | 13:1843 | 2.18 | 1.00 | 50.84 | 2.35 (G:O6 13:538) |
| 4 | Water 4 | | 13:1992 | 2.19 | 1.00 | 53.65 | 3.45 (A:N6 13:539) |
| 5 | Water 5 | | 3I:202 | 2.19 | 1.00 | 48.71 | 3.05 (LYS:NZ 3I:112) |
| 6 | Water 6 | | 13:1869 | 2.20 | 1.00 | 46.85 | 2.52 (A:OP1 13:535) |
| 7 | C | OP2 | 13:536 | 3.46 | 1.00 | 52.62 |  |
| 8 | A | OP1 | 13:535 | 3.85 | 1.00 | 52.04 |  |
| 9 | LYS | NZ | 3I:112 | 4.36 | 1.00 | 57.76 |  |
| 10 | G | N7 | 13:537 | 4.44 | 1.00 | 47.40 |  |
| 11 | G | O6 | 13:538 | 4.46 | 1.00 | 51.91 |  |
| 12 | G | O6 | 13:537 | 4.69 | 1.00 | 49.95 |  |
| 13 | A | N6 | 13:539 | 4.85 | 1.00 | 58.43 |  |
| 14 | G | N7 | 13:538 | 4.86 | 1.00 | 55.16 |  |
  |

  
**1^st^ shell ligand angles:**   
1-2: 89; 1-3: 178; 1-4: 89; 1-5: 87; 1-6: 94; 2-3: 92; 2-4: 179; 2-5: 91; 2-6: 87; 3-4: 90; 3-5: 92; 3-6: 88; 4-5: 89; 4-6: 93; 5-6: 178;   
  
 Contact(s): /O_w_/O_w_/O_w_/O_w_/O_w_/O_w_/  
 Coded contact list: 6O_w_  
MG hydration shell issue: water too close (< 2.4 Angs.)  

### MG:13:1744

|  | Res. type | Atom | Residue | Distance | Occupancy | B-factor | Closest solute to 1^st^ shell water |
| --- | --- | --- | --- | --- | --- | --- | --- |
| **Ion** | | | |  | 1.00 | 19.82 || 1 | Water 1 | | 13:1879 | 2.15 | 1.00 | 27.81 | 2.57 (C:O2 13:812) |
| 2 | Water 2 | | 13:1969 | 2.15 | 1.00 | 34.19 | 3.23 (A:OP2 13:766) |
| 3 | Water 3 | | 13:1846 | 2.15 | 1.00 | 31.62 | 2.36 (A:OP2 13:766) |
| 4 | Water 4 | | 13:1915 | 2.17 | 1.00 | 29.55 | 2.82 (C:N4 13:811) |
| 5 | Water 5 | | 13:1926 | 2.18 | 1.00 | 42.47 | 2.91 (C:N3 13:811) |
| 6 | A | OP2 | 13:766 | 2.41 | 1.00 | 36.33 |  |
| 7 | C | N3 | 13:812 | 3.65 | 1.00 | 29.21 |  |
| 8 | G | O3' | 13:765 | 3.86 | 1.00 | 28.23 |  |
| 9 | C | O2 | 13:812 | 3.90 | 1.00 | 32.77 |  |
| 10 | C | N4 | 13:811 | 3.96 | 1.00 | 31.65 |  |
| 11 | A | OP1 | 13:766 | 4.03 | 1.00 | 36.91 |  |
| 12 | C | N3 | 13:811 | 4.28 | 1.00 | 33.40 |  |
| 13 | A | O5' | 13:766 | 4.58 | 1.00 | 32.34 |  |
| 14 | C | N4 | 13:812 | 4.65 | 1.00 | 31.15 |  |
| 15 | A | N7 | 13:767 | 4.85 | 1.00 | 32.25 |  |
| 16 | A | N6 | 13:768 | 4.94 | 1.00 | 33.55 |  |
| 17 | U | O4 | 13:813 | 5.00 | 1.00 | 33.19 |  |
  |

  
**1^st^ shell ligand angles:**   
1-2: 173; 1-3: 90; 1-4: 91; 1-5: 87; 2-3: 84; 2-4: 96; 2-5: 90; 3-4: 177; 3-5: 89; 4-5: 94;   
  
 Contact(s): /O_w_/O_w_/O_w_/O_w_/O_w_/  
 Coded contact list: 5O_w_  
 Exclusion zone contact(s): /OP2/  
Consider NA in place of MG (1 valid contact(s) for NA vs. 0 for MG)  
MG hydration shell issue: water too close (< 2.4 Angs.)  

### MG:13:1745

|  | Res. type | Atom | Residue | Distance | Occupancy | B-factor | Closest solute to 1^st^ shell water |
| --- | --- | --- | --- | --- | --- | --- | --- |
| **Ion** | | | |  | 1.00 | 39.18 || 1 | Water 1 | | 13:1837 | 2.16 | 1.00 | 52.12 | 2.33 (A:OP2 13:510) |
| 2 | Water 2 | | 13:1822 | 2.17 | 1.00 | 59.37 | 2.16 (G:OP1 13:506) |
| 3 | Water 3 | | 13:1849 | 2.17 | 1.00 | 53.84 | 2.37 (C:O3' 13:508) |
| 4 | Water 4 | | 13:1977 | 2.17 | 1.00 | 55.23 | 3.30 (A:OP2 13:509) |
| 5 | Water 5 | | 13:1836 | 2.18 | 1.00 | 58.43 | 2.33 (A:N7 13:510) |
| 6 | A | OP2 | 13:509 | 2.63 | 1.00 | 52.07 |  |
| 7 | A | O3' | 13:509 | 3.00 | 1.00 | 58.51 |  |
| 8 | A | OP2 | 13:510 | 3.44 | 1.00 | 52.05 |  |
| 9 | C | O3' | 13:508 | 3.75 | 1.00 | 61.79 |  |
| 10 | A | O5' | 13:510 | 4.13 | 1.00 | 46.17 |  |
| 11 | G | OP1 | 13:506 | 4.25 | 1.00 | 64.54 |  |
| 12 | A | N7 | 13:510 | 4.26 | 1.00 | 47.62 |  |
| 13 | C | OP2 | 13:507 | 4.53 | 1.00 | 54.50 |  |
| 14 | A | O5' | 13:509 | 4.53 | 1.00 | 49.82 |  |
| 15 | C | O2' | 13:508 | 4.75 | 1.00 | 52.36 |  |
| 16 | A | OP1 | 13:509 | 4.90 | 1.00 | 57.15 |  |
  |

  
**1^st^ shell ligand angles:**   
1-2: 87; 1-3: 179; 1-4: 93; 1-5: 88; 2-3: 92; 2-4: 91; 2-5: 89; 3-4: 86; 3-5: 92; 4-5: 178;   
  
 Contact(s): /O_w_/O_w_/O_w_/O_w_/O_w_/  
 Coded contact list: 5O_w_  
 Exclusion zone contact(s): /O3'/OP2/  
Consider K in place of MG (2 valid contact(s) for K vs. 0 for MG)  
MG hydration shell issue: water too close (< 2.4 Angs.)  

### MG:13:1746

|  | Res. type | Atom | Residue | Distance | Occupancy | B-factor | Closest solute to 1^st^ shell water |
| --- | --- | --- | --- | --- | --- | --- | --- |
| **Ion** | | | |  | 1.00 | 37.77 || 1 | Water 1 | | 13:1824 | 2.15 | 1.00 | 51.87 | 2.21 (G:N3 13:567) |
| 2 | Water 2 | | 13:1937 | 2.17 | 1.00 | 41.35 | 2.96 (G:O2' 13:568) |
| 3 | Water 3 | | 13:1938 | 2.17 | 1.00 | 57.28 | 2.98 (G:O2' 13:567) |
| 4 | Water 4 | | 13:1935 | 2.18 | 1.00 | 50.87 | 2.95 (A:N6 13:563) |
| 5 | Water 5 | | 13:1862 | 2.18 | 1.00 | 44.89 | 2.49 (A:N1 13:573) |
| 6 | G | OP1 | 13:21 | 2.49 | 1.00 | 46.21 |  |
| 7 | G | OP2 | 13:21 | 3.59 | 1.00 | 46.57 |  |
| 8 | G | N2 | 13:567 | 3.81 | 1.00 | 49.93 |  |
| 9 | G | O2' | 13:567 | 3.92 | 1.00 | 55.45 |  |
| 10 | A | N6 | 13:563 | 4.15 | 1.00 | 47.70 |  |
| 11 | A | N1 | 13:573 | 4.19 | 1.00 | 45.30 |  |
| 12 | G | N3 | 13:567 | 4.20 | 1.00 | 47.57 |  |
| 13 | U | O3' | 13:20 | 4.25 | 1.00 | 44.32 |  |
| 14 | A | N6 | 13:573 | 4.40 | 1.00 | 46.51 |  |
| 15 | G | O2' | 13:568 | 4.64 | 1.00 | 45.08 |  |
| 16 | G | O5' | 13:21 | 4.67 | 1.00 | 51.33 |  |
  |

  
**1^st^ shell ligand angles:**   
1-2: 87; 1-3: 90; 1-4: 96; 1-5: 87; 2-3: 92; 2-4: 177; 2-5: 89; 3-4: 89; 3-5: 177; 4-5: 90;   
  
 Contact(s): /O_w_/O_w_/O_w_/O_w_/O_w_/  
 Coded contact list: 5O_w_  
 Exclusion zone contact(s): /OP1/  
Consider NA in place of MG (1 valid contact(s) for NA vs. 0 for MG)  
MG hydration shell issue: water too close (< 2.4 Angs.)  

### MG:13:1747

|  | Res. type | Atom | Residue | Distance | Occupancy | B-factor | Closest solute to 1^st^ shell water |
| --- | --- | --- | --- | --- | --- | --- | --- |
| **Ion** | | | |  | 1.00 | 32.12 || 1 | Water 1 | | 13:1830 | 2.17 | 1.00 | 44.47 | 2.26 (A:OP2 13:574) |
| 2 | Water 2 | | 13:1900 | 2.17 | 1.00 | 48.83 | 2.74 (A:OP1 13:572) |
| 3 | Water 3 | | 13:1928 | 2.18 | 1.00 | 54.00 | 2.91 (U:OP1 13:571) |
| 4 | A | OP2 | 13:573 | 2.57 | 1.00 | 37.69 |  |
| 5 | A | OP2 | 13:572 | 2.62 | 1.00 | 43.30 |  |
| 6 | A | OP2 | 13:574 | 2.79 | 1.00 | 42.58 |  |
| 7 | A | OP1 | 13:574 | 3.09 | 1.00 | 39.88 |  |
| 8 | A | OP1 | 13:572 | 3.42 | 1.00 | 43.56 |  |
| 9 | U | OP1 | 13:571 | 4.00 | 1.00 | 53.18 |  |
| 10 | A | O5' | 13:573 | 4.04 | 1.00 | 36.35 |  |
| 11 | A | O5' | 13:572 | 4.25 | 1.00 | 42.67 |  |
| 12 | A | O5' | 13:574 | 4.48 | 1.00 | 37.60 |  |
| 13 | U | O5' | 13:571 | 4.49 | 1.00 | 48.13 |  |
| 14 | A | OP1 | 13:573 | 4.50 | 1.00 | 36.56 |  |
| 15 | A | N1 | 13:819 | 4.57 | 1.00 | 41.72 |  |
| 16 | A | N3 | 13:819 | 4.63 | 1.00 | 44.03 |  |
| 17 | A | O3' | 13:573 | 4.65 | 1.00 | 35.49 |  |
| 18 | U | O3' | 13:571 | 4.74 | 1.00 | 38.82 |  |
| 19 | U | OP2 | 13:571 | 4.86 | 1.00 | 44.03 |  |
  |

  
**1^st^ shell ligand angles:**   
1-2: 175; 1-3: 88; 2-3: 96;   
  
 Contact(s): /O_w_/O_w_/O_w_/  
 Coded contact list: 3O_w_  
 Exclusion zone contact(s): /OP1/OP2/OP2/OP2/  
Consider K in place of MG (3 valid contact(s) for K vs. 0 for MG)  
MG hydration shell issue: water too close (< 2.4 Angs.)  

### MG:13:1748

|  | Res. type | Atom | Residue | Distance | Occupancy | B-factor | Closest solute to 1^st^ shell water |
| --- | --- | --- | --- | --- | --- | --- | --- |
| **Ion** | | | |  | 1.00 | 38.60 || 1 | Water 1 | | 13:1832 | 2.16 | 1.00 | 46.63 | 2.27 (U:OP2 13:1199) |
| 2 | Water 2 | | 13:1888 | 2.17 | 1.00 | 54.30 | 2.64 (G:O3' 13:963) |
| 3 | Water 3 | | 13:1876 | 2.17 | 1.00 | 46.52 | 2.55 (G:OP1 13:1198) |
| 4 | Water 4 | | 13:1959 | 2.18 | 1.00 | 53.50 | 3.16 (A:OP1 13:964) |
| 5 | A | OP1 | 13:964 | 2.39 | 1.00 | 46.71 |  |
| 6 | U | OP1 | 13:1199 | 2.59 | 1.00 | 51.05 |  |
| 7 | U | OP2 | 13:1199 | 3.44 | 1.00 | 45.18 |  |
| 8 | G | O3' | 13:963 | 3.59 | 1.00 | 48.16 |  |
| 9 | G | O3' | 13:1198 | 4.01 | 1.00 | 49.56 |  |
| 10 | A | O5' | 13:964 | 4.16 | 1.00 | 45.92 |  |
| 11 | A | OP2 | 13:965 | 4.21 | 1.00 | 44.35 |  |
| 12 | G | OP1 | 13:1198 | 4.42 | 1.00 | 53.68 |  |
| 13 | U | O5' | 13:1199 | 4.71 | 1.00 | 61.30 |  |
| 14 | A | OP2 | 13:964 | 4.72 | 1.00 | 53.53 |  |
| 15 | G | O5' | 13:1198 | 4.80 | 1.00 | 50.73 |  |
  |

  
**1^st^ shell ligand angles:**   
1-2: 92; 1-3: 94; 1-4: 96; 1-5: 175; 2-3: 173; 2-4: 87; 2-5: 91; 3-4: 96; 3-5: 83; 4-5: 88;   
  
 Contact(s): /O_w_/O_w_/O_w_/O_w_/OP1/  
 Coded contact list: O_ph_4O_w_  
 Exclusion zone contact(s): /OP1/  
MG hydration shell issue: water too close (< 2.4 Angs.)  

### MG:13:1749

|  | Res. type | Atom | Residue | Distance | Occupancy | B-factor | Closest solute to 1^st^ shell water |
| --- | --- | --- | --- | --- | --- | --- | --- |
| **Ion** | | | |  | 1.00 | 52.96 || 1 | Water 1 | | 13:1834 | 2.17 | 1.00 | 47.48 | 2.29 (A:OP2 13:116) |
| 2 | Water 2 | | 13:1831 | 2.18 | 1.00 | 57.96 | 2.27 (A:OP2 13:288) |
| 3 | Water 3 | | 13:1918 | 2.19 | 1.00 | 62.62 | 2.84 (G:OP2 13:117) |
| 4 | G | OP2 | 13:289 | 2.56 | 1.00 | 59.65 |  |
| 5 | G | OP2 | 13:117 | 2.67 | 1.00 | 56.63 |  |
| 6 | A | OP2 | 13:116 | 2.74 | 1.00 | 47.23 |  |
| 7 | A | O5' | 13:288 | 3.53 | 1.00 | 57.57 |  |
| 8 | G | OP1 | 13:117 | 3.74 | 1.00 | 56.17 |  |
| 9 | A | OP2 | 13:288 | 3.75 | 1.00 | 45.14 |  |
| 10 | G | O5' | 13:289 | 4.32 | 1.00 | 56.46 |  |
| 11 | A | OP1 | 13:288 | 4.42 | 1.00 | 49.78 |  |
| 12 | G | O5' | 13:117 | 4.62 | 1.00 | 52.16 |  |
| 13 | A | O3' | 13:288 | 4.69 | 1.00 | 55.62 |  |
| 14 | A | O3' | 13:116 | 4.76 | 1.00 | 60.20 |  |
| 15 | A | OP1 | 13:116 | 4.84 | 1.00 | 47.56 |  |
| 16 | A | O5' | 13:116 | 4.91 | 1.00 | 49.58 |  |
| 17 | A | N7 | 13:288 | 4.92 | 1.00 | 60.90 |  |
  |

  
**1^st^ shell ligand angles:**   
1-2: 178; 1-3: 88; 2-3: 91;   
  
 Contact(s): /O_w_/O_w_/O_w_/  
 Coded contact list: 3O_w_  
 Exclusion zone contact(s): /OP2/OP2/OP2/  
Consider K in place of MG (2 valid contact(s) for K vs. 0 for MG)  
MG hydration shell issue: water too close (< 2.4 Angs.)  

### MG:13:1750

|  | Res. type | Atom | Residue | Distance | Occupancy | B-factor | Closest solute to 1^st^ shell water |
| --- | --- | --- | --- | --- | --- | --- | --- |
| **Ion** | | | |  | 1.00 | 52.09 || 1 | Water 1 | | 13:1878 | 2.16 | 1.00 | 49.50 | 2.56 (G:OP1 13:1224) |
| 2 | Water 2 | | 13:1806 | 2.17 | 1.00 | 52.72 | 1.91 (G:OP1 13:951) |
| 3 | Water 3 | | 13:2004 | 2.18 | 1.00 | 46.39 | 3.63 (G:OP1 13:951) |
| 4 | Water 4 | | 13:1855 | 2.18 | 1.00 | 39.31 | 2.42 (G:OP1 13:1224) |
| 5 | Water 5 | | 13:1999 | 2.19 | 1.00 | 71.88 | 3.58 (A:N7 13:977) |
| 6 | G | OP1 | 13:1224 | 2.43 | 1.00 | 51.96 |  |
| 7 | G | OP1 | 13:951 | 3.69 | 1.00 | 51.48 |  |
| 8 | G | OP2 | 13:1224 | 4.03 | 1.00 | 61.44 |  |
| 9 | A | N6 | 13:977 | 4.26 | 1.00 | 68.22 |  |
| 10 | A | N7 | 13:977 | 4.50 | 1.00 | 63.01 |  |
| 11 | G | O5' | 13:1224 | 4.52 | 1.00 | 63.20 |  |
| 12 | U | OP2 | 13:952 | 4.67 | 1.00 | 52.03 |  |
| 13 | C | O3' | 13:1223 | 4.70 | 1.00 | 60.08 |  |
| 14 | C | O2' | 13:1223 | 4.77 | 1.00 | 54.24 |  |
  |

  
**1^st^ shell ligand angles:**   
1-2: 90; 1-3: 93; 1-4: 86; 1-5: 179; 2-3: 94; 2-4: 169; 2-5: 89; 3-4: 96; 3-5: 86; 4-5: 95;   
  
 Contact(s): /O_w_/O_w_/O_w_/O_w_/O_w_/  
 Coded contact list: 5O_w_  
 Exclusion zone contact(s): /OP1/  
Consider NA in place of MG (1 valid contact(s) for NA vs. 0 for MG)  
MG hydration shell issue: water too close (< 2.4 Angs.)  

### MG:13:1751

|  | Res. type | Atom | Residue | Distance | Occupancy | B-factor | Closest solute to 1^st^ shell water |
| --- | --- | --- | --- | --- | --- | --- | --- |
| **Ion** | | | |  | 1.00 | 75.12 || 1 | Water 1 | | 1F:102 | 2.18 | 1.00 | 79.47 | 2.62 (GLY:N 1F:2) |
| 2 | Water 2 | | 13:1857 | 2.18 | 1.00 | 89.97 | 2.44 (C:OP1 13:1303) |
| 3 | Water 3 | | 13:1847 | 2.18 | 1.00 | 82.57 | 2.36 (G:N7 13:1305) |
| 4 | Water 4 | | 1F:101 | 2.18 | 1.00 | 73.24 | 2.44 (ASP:OD2 1F:5) |
| 5 | G | OP2 | 13:1304 | 2.72 | 1.00 | 77.79 |  |
| 6 | C | OP2 | 13:1303 | 2.88 | 1.00 | 78.93 |  |
| 7 | C | OP1 | 13:1303 | 3.21 | 1.00 | 91.95 |  |
| 8 | ASP | OD2 | 1F:5 | 3.63 | 1.00 | 78.23 |  |
| 9 | C | O5' | 13:1303 | 3.67 | 1.00 | 77.66 |  |
| 10 | G | N7 | 13:1305 | 3.93 | 1.00 | 78.77 |  |
| 11 | C | O2' | 13:1242 | 4.45 | 1.00 | 81.37 |  |
| 12 | G | O5' | 13:1304 | 4.50 | 1.00 | 80.66 |  |
| 13 | G | O6 | 13:1305 | 4.62 | 1.00 | 84.68 |  |
| 14 | GLY | N | 1F:2 | 4.64 | 1.00 | 72.55 |  |
| 15 | U | O2' | 13:1302 | 4.85 | 1.00 | 99.48 |  |
| 16 | U | O3' | 13:1302 | 4.90 | 1.00 | 93.43 |  |
| 17 | G | N7 | 13:1304 | 4.99 | 1.00 | 79.21 |  |
  |

  
**1^st^ shell ligand angles:**   
1-2: 90; 1-3: 86; 1-4: 99; 2-3: 176; 2-4: 92; 3-4: 88;   
  
 Contact(s): /O_w_/O_w_/O_w_/O_w_/  
 Coded contact list: 4O_w_  
 Exclusion zone contact(s): /OP1/OP2/OP2/  
Consider K in place of MG (3 valid contact(s) for K vs. 0 for MG)  
MG hydration shell issue: water too close (< 2.4 Angs.)  

### MG:13:1752

|  | Res. type | Atom | Residue | Distance | Occupancy | B-factor | Closest solute to 1^st^ shell water |
| --- | --- | --- | --- | --- | --- | --- | --- |
| **Ion** | | | |  | 1.00 | 47.91 || 1 | Water 1 | | 13:2005 | 2.18 | 1.00 | 63.30 | 3.66 (A:OP1 13:573) |
| 2 | Water 2 | | 13:2030 | 2.18 | 1.00 | 46.56 | 4.64 (A:OP1 13:572) |
| 3 | Water 3 | | 13:1942 | 2.18 | 1.00 | 44.47 | 3.03 (A:OP1 13:572) |
| 4 | Water 4 | | 13:1844 | 2.19 | 1.00 | 46.04 | 2.35 (A:OP1 13:572) |
| 5 | Water 5 | | 13:2009 | 2.19 | 1.00 | 61.92 | 3.72 (A:N1 13:819) |
| 6 | A | OP1 | 13:572 | 2.62 | 1.00 | 43.56 |  |
| 7 | Water 6 | | 13:1900 | 3.70 | 1.00 | 48.83 | 2.74 (A:OP1 13:572) |
| 8 | A | N1 | 13:819 | 4.14 | 1.00 | 41.72 |  |
| 9 | A | OP1 | 13:573 | 4.45 | 1.00 | 36.56 |  |
| 10 | U | O3' | 13:571 | 4.52 | 1.00 | 38.82 |  |
| 11 | A | N6 | 13:819 | 4.64 | 1.00 | 42.06 |  |
| 12 | A | O5' | 13:572 | 4.76 | 1.00 | 42.67 |  |
  |

  
**1^st^ shell ligand angles:**   
1-2: 89; 1-3: 177; 1-4: 88; 1-5: 92; 2-3: 93; 2-4: 94; 2-5: 85; 3-4: 89; 3-5: 91; 4-5: 179;   
  
 Contact(s): /O_w_/O_w_/O_w_/O_w_/O_w_/  
 Coded contact list: 5O_w_  
 Exclusion zone contact(s): /OP1/  
Consider K in place of MG (1 valid contact(s) for K vs. 0 for MG)  
MG hydration shell issue: water too close (< 2.4 Angs.)  

### MG:13:1753

|  | Res. type | Atom | Residue | Distance | Occupancy | B-factor | Closest solute to 1^st^ shell water |
| --- | --- | --- | --- | --- | --- | --- | --- |
| **Ion** | | | |  | 1.00 | 69.21 || 1 | Water 1 | | 13:1807 | 2.17 | 1.00 | 55.94 | 1.91 (G:OP2 13:861) |
| 2 | Water 2 | | 13:1990 | 2.19 | 1.00 | 77.60 | 3.40 (A:OP1 13:859) |
| 3 | Water 3 | | 13:1875 | 2.19 | 1.00 | 54.43 | 2.54 (A:OP2 13:860) |
| 4 | Water 4 | | 13:1980 | 2.19 | 1.00 | 69.57 | 3.33 (A:OP1 13:860) |
| 5 | Water 5 | | 13:2020 | 2.19 | 1.00 | 44.70 | 4.00 (G:N7 13:861) |
| 6 | A | OP2 | 13:860 | 2.81 | 1.00 | 67.93 |  |
| 7 | A | OP1 | 13:860 | 3.24 | 1.00 | 57.92 |  |
| 8 | A | O5' | 13:860 | 3.95 | 1.00 | 55.22 |  |
| 9 | G | OP2 | 13:861 | 3.98 | 1.00 | 57.03 |  |
| 10 | A | OP1 | 13:859 | 4.09 | 1.00 | 56.82 |  |
| 11 | G | N7 | 13:861 | 4.67 | 1.00 | 57.54 |  |
| 12 | A | O3' | 13:859 | 4.89 | 1.00 | 54.64 |  |
  |

  
**1^st^ shell ligand angles:**   
1-2: 178; 1-3: 95; 1-4: 88; 1-5: 86; 2-3: 87; 2-4: 91; 2-5: 96; 3-4: 177; 3-5: 87; 4-5: 94;   
  
 Contact(s): /O_w_/O_w_/O_w_/O_w_/O_w_/  
 Coded contact list: 5O_w_  
 Exclusion zone contact(s): /OP1/OP2/  
Consider K in place of MG (2 valid contact(s) for K vs. 0 for MG)  
MG hydration shell issue: water too close (< 2.4 Angs.)  

### MG:13:1754

|  | Res. type | Atom | Residue | Distance | Occupancy | B-factor | Closest solute to 1^st^ shell water |
| --- | --- | --- | --- | --- | --- | --- | --- |
| **Ion** | | | |  | 1.00 | 35.34 || 1 | Water 1 | | 13:1880 | 2.17 | 1.00 | 42.19 | 2.57 (U:OP1 13:804) |
| 2 | Water 2 | | 13:1906 | 2.17 | 1.00 | 36.18 | 2.78 (G:N7 13:769) |
| 3 | Water 3 | | 13:1816 | 2.18 | 1.00 | 38.23 | 2.10 (A:OP2 13:768) |
| 4 | Water 4 | | 13:1960 | 2.18 | 1.00 | 39.64 | 3.17 (A:OP2 13:767) |
| 5 | Water 5 | | 13:1982 | 2.18 | 1.00 | 26.16 | 4.02 (A:N7 13:768) |
| 6 | A | OP2 | 13:768 | 2.44 | 1.00 | 30.47 |  |
| 7 | U | OP1 | 13:804 | 3.90 | 1.00 | 26.91 |  |
| 8 | A | N7 | 13:768 | 4.32 | 1.00 | 29.42 |  |
| 9 | A | O5' | 13:768 | 4.34 | 1.00 | 34.25 |  |
| 10 | A | OP2 | 13:767 | 4.35 | 1.00 | 36.50 |  |
| 11 | C | OP2 | 13:805 | 4.44 | 1.00 | 45.23 |  |
| 12 | G | N7 | 13:769 | 4.49 | 1.00 | 31.16 |  |
| 13 | A | OP1 | 13:768 | 4.68 | 1.00 | 40.99 |  |
| 14 | A | O3' | 13:767 | 4.82 | 1.00 | 36.83 |  |
| 15 | A | O5' | 13:767 | 4.88 | 1.00 | 34.36 |  |
  |

  
**1^st^ shell ligand angles:**   
1-2: 90; 1-3: 175; 1-4: 92; 1-5: 97; 2-3: 88; 2-4: 175; 2-5: 91; 3-4: 90; 3-5: 88; 4-5: 93;   
  
 Contact(s): /O_w_/O_w_/O_w_/O_w_/O_w_/  
 Coded contact list: 5O_w_  
 Exclusion zone contact(s): /OP2/  
Consider NA in place of MG (1 valid contact(s) for NA vs. 0 for MG)  
MG hydration shell issue: water too close (< 2.4 Angs.)  

### MG:13:1755

|  | Res. type | Atom | Residue | Distance | Occupancy | B-factor | Closest solute to 1^st^ shell water |
| --- | --- | --- | --- | --- | --- | --- | --- |
| **Ion** | | | |  | 1.00 | 83.59 || 1 | Water 1 | | 13:2026 | 2.19 | 1.00 | 69.91 | 4.49 (G:O6 13:377) |
| 2 | Water 2 | | 13:1920 | 2.19 | 1.00 | 79.45 | 2.86 (G:N7 13:376) |
| 3 | Water 3 | | 13:1948 | 2.19 | 1.00 | 85.96 | 3.53 (G:N7 13:377) |
| 4 | Water 4 | | 13:1941 | 2.19 | 1.00 | 68.40 | 3.02 (G:N7 13:377) |
| 5 | Water 5 | | 7I:201 | 2.19 | 1.00 | 68.44 | 5.40 (G:N7 13:377) |
| 6 | Water 6 | | 13:2001 | 2.19 | 1.00 | 81.68 | 4.48 (G:N7 13:376) |
| 7 | G | N7 | 13:377 | 4.06 | 1.00 | 73.30 |  |
| 8 | G | N7 | 13:376 | 4.44 | 1.00 | 72.88 |  |
| 9 | G | O6 | 13:377 | 4.48 | 1.00 | 70.66 |  |
  |

  
**1^st^ shell ligand angles:**   
1-2: 87; 1-3: 179; 1-4: 87; 1-5: 92; 1-6: 95; 2-3: 92; 2-4: 94; 2-5: 179; 2-6: 86; 3-4: 93; 3-5: 89; 3-6: 86; 4-5: 87; 4-6: 179; 5-6: 94;   
  
 Contact(s): /O_w_/O_w_/O_w_/O_w_/O_w_/O_w_/  
 Coded contact list: 6O_w_  

### MG:13:1756

|  | Res. type | Atom | Residue | Distance | Occupancy | B-factor | Closest solute to 1^st^ shell water |
| --- | --- | --- | --- | --- | --- | --- | --- |
| **Ion** | | | |  | 1.00 | 63.53 || 1 | Water 1 | | 13:1961 | 2.18 | 1.00 | 51.95 | 3.17 (C:O2 13:1399) |
| 2 | Water 2 | | 4K:101 | 2.18 | 1.00 | 47.54 | 2.67 (U:O4 4K:48) |
| 3 | Water 3 | | 4K:102 | 2.18 | 1.00 | 64.71 | 3.22 (U:OP2 4K:47) |
| 4 | Water 4 | | 13:2014 | 2.19 | 1.00 | 55.32 | 3.78 (C:O2 13:1399) |
| 5 | Water 5 | | 13:1817 | 2.19 | 1.00 | 60.32 | 2.11 (G:O6 13:926) |
| 6 | Water 6 | | 13:1978 | 2.20 | 1.00 | 59.52 | 3.31 (G:O6 13:926) |
| 7 | G | O6 | 13:926 | 3.69 | 1.00 | 61.49 |  |
| 8 | C | O2 | 13:1399 | 4.63 | 1.00 | 42.78 |  |
| 9 | U | O4 | 4K:48 | 4.67 | 1.00 | 51.56 |  |
| 10 | G | N2 | 13:1504 | 4.75 | 1.00 | 50.54 |  |
| 11 | U | OP2 | 4K:47 | 4.80 | 1.00 | 47.16 |  |
| 12 | 5MC | OP2 | 13:1400 | 4.89 | 1.00 | 48.69 |  |
  |

  
**1^st^ shell ligand angles:**   
1-2: 88; 1-3: 179; 1-4: 89; 1-5: 91; 1-6: 93; 2-3: 93; 2-4: 91; 2-5: 178; 2-6: 89; 3-4: 91; 3-5: 89; 3-6: 88; 4-5: 87; 4-6: 178; 5-6: 93;   
  
 Contact(s): /O_w_/O_w_/O_w_/O_w_/O_w_/O_w_/  
 Coded contact list: 6O_w_  
MG hydration shell issue: water too close (< 2.4 Angs.)  

### MG:13:1757

|  | Res. type | Atom | Residue | Distance | Occupancy | B-factor | Closest solute to 1^st^ shell water |
| --- | --- | --- | --- | --- | --- | --- | --- |
| **Ion** | | | |  | 1.00 | 50.95 || 1 | Water 1 | | 13:1820 | 2.17 | 1.00 | 48.13 | 2.15 (G:N7 13:362) |
| 2 | Water 2 | | 13:1860 | 2.17 | 1.00 | 52.69 | 2.46 (G:N7 13:362) |
| 3 | Water 3 | | 3I:203 | 2.18 | 1.00 | 57.29 | 3.88 (G:N7 13:362) |
| 4 | Water 4 | | 13:1981 | 2.18 | 1.00 | 48.98 | 3.33 (G:N7 13:361) |
| 5 | Water 5 | | 13:2031 | 2.18 | 1.00 | 55.73 | 4.65 (G:O6 13:362) |
| 6 | Water 6 | | 13:1835 | 2.20 | 1.00 | 62.19 | 2.29 (G:OP2 13:361) |
| 7 | G | N7 | 13:362 | 3.19 | 1.00 | 47.62 |  |
| 8 | G | OP2 | 13:361 | 3.96 | 1.00 | 48.82 |  |
| 9 | G | O6 | 13:362 | 4.17 | 1.00 | 49.19 |  |
| 10 | G | N7 | 13:361 | 4.55 | 1.00 | 52.32 |  |
| 11 | G | OP2 | 13:362 | 4.73 | 1.00 | 44.00 |  |
| 12 | G | O5' | 13:361 | 4.82 | 1.00 | 47.34 |  |
  |

  
**1^st^ shell ligand angles:**   
1-2: 92; 1-3: 84; 1-4: 94; 1-5: 89; 1-6: 177; 2-3: 88; 2-4: 91; 2-5: 179; 2-6: 87; 3-4: 179; 3-5: 92; 3-6: 94; 4-5: 88; 4-6: 88; 5-6: 92;   
  
 Contact(s): /O_w_/O_w_/O_w_/O_w_/O_w_/O_w_/  
 Coded contact list: 6O_w_  
 Exclusion zone contact(s): /N7/  
Consider K in place of MG (1 valid contact(s) for K vs. 0 for MG)  
MG hydration shell issue: water too close (< 2.4 Angs.)  

### MG:13:1758

|  | Res. type | Atom | Residue | Distance | Occupancy | B-factor | Closest solute to 1^st^ shell water |
| --- | --- | --- | --- | --- | --- | --- | --- |
| **Ion** | | | |  | 1.00 | 76.99 || 1 | Water 1 | | 13:1946 | 2.18 | 1.00 | 75.34 | 3.05 (C:OP2 13:651) |
| 2 | Water 2 | | 13:1940 | 2.18 | 1.00 | 63.19 | 3.00 (G:OP2 13:752) |
| 3 | Water 3 | | 13:1854 | 2.18 | 1.00 | 75.27 | 2.41 (C:OP1 13:651) |
| 4 | Water 4 | | 13:1967 | 2.19 | 1.00 | 76.37 | 3.22 (G:N7 13:752) |
| 5 | Water 5 | | 13:2028 | 2.19 | 1.00 | 82.29 | 4.57 (C:OP1 13:651) |
| 6 | Water 6 | | 13:2019 | 2.19 | 1.00 | 65.19 | 4.17 (U:OP2 13:652) |
| 7 | C | OP1 | 13:651 | 4.21 | 1.00 | 80.96 |  |
| 8 | U | OP2 | 13:652 | 4.35 | 1.00 | 71.22 |  |
| 9 | C | OP2 | 13:651 | 4.47 | 1.00 | 79.07 |  |
| 10 | G | OP2 | 13:752 | 4.70 | 1.00 | 68.38 |  |
| 11 | G | N7 | 13:752 | 4.74 | 1.00 | 71.24 |  |
| 12 | C | O5' | 13:651 | 4.96 | 1.00 | 82.30 |  |
  |

  
**1^st^ shell ligand angles:**   
1-2: 90; 1-3: 89; 1-4: 89; 1-5: 90; 1-6: 177; 2-3: 179; 2-4: 93; 2-5: 91; 2-6: 87; 3-4: 88; 3-5: 88; 3-6: 93; 4-5: 176; 4-6: 90; 5-6: 91;   
  
 Contact(s): /O_w_/O_w_/O_w_/O_w_/O_w_/O_w_/  
 Coded contact list: 6O_w_  

### MG:13:1759

|  | Res. type | Atom | Residue | Distance | Occupancy | B-factor | Closest solute to 1^st^ shell water |
| --- | --- | --- | --- | --- | --- | --- | --- |
| **Ion** | | | |  | 1.00 | 52.95 || 1 | Water 1 | | 13:1814 | 2.17 | 1.00 | 54.95 | 2.10 (A:N6 13:964) |
| 2 | Water 2 | | 13:1917 | 2.17 | 1.00 | 54.90 | 2.84 (G:OP2 13:963) |
| 3 | Water 3 | | 13:1995 | 2.19 | 1.00 | 57.72 | 3.71 (C:N4 13:962) |
| 4 | Water 4 | | 13:1804 | 2.19 | 1.00 | 56.73 | 1.30 (G:N7 13:963) |
| 5 | Water 5 | | 13:1896 | 2.19 | 1.00 | 61.72 | 2.70 (G:OP1 13:953) |
| 6 | Water 6 | | 13:1923 | 2.20 | 1.00 | 58.23 | 2.87 (G:OP2 13:963) |
| 7 | G | N7 | 13:963 | 3.46 | 1.00 | 54.60 |  |
| 8 | G | OP1 | 13:953 | 3.71 | 1.00 | 53.01 |  |
| 9 | G | OP2 | 13:963 | 4.05 | 1.00 | 57.18 |  |
| 10 | A | N6 | 13:964 | 4.15 | 1.00 | 50.10 |  |
| 11 | G | O6 | 13:963 | 4.54 | 1.00 | 48.72 |  |
| 12 | U | O3' | 13:952 | 4.70 | 1.00 | 53.87 |  |
| 13 | A | N7 | 13:964 | 4.74 | 1.00 | 49.40 |  |
| 14 | G | OP2 | 13:953 | 4.82 | 1.00 | 53.70 |  |
| 15 | C | N4 | 13:962 | 4.89 | 1.00 | 49.64 |  |
  |

  
**1^st^ shell ligand angles:**   
1-2: 178; 1-3: 83; 1-4: 88; 1-5: 93; 1-6: 96; 2-3: 98; 2-4: 94; 2-5: 85; 2-6: 83; 3-4: 89; 3-5: 90; 3-6: 177; 4-5: 178; 4-6: 88; 5-6: 94;   
  
 Contact(s): /O_w_/O_w_/O_w_/O_w_/O_w_/O_w_/  
 Coded contact list: 6O_w_  
MG hydration shell issue: water too close (< 2.4 Angs.)  

### MG:13:1760

|  | Res. type | Atom | Residue | Distance | Occupancy | B-factor | Closest solute to 1^st^ shell water |
| --- | --- | --- | --- | --- | --- | --- | --- |
| **Ion** | | | |  | 1.00 | 65.02 || 1 | Water 1 | | 13:2000 | 2.17 | 1.00 | 52.22 | 3.58 (G:O6 13:587) |
| 2 | Water 2 | | 13:1865 | 2.17 | 1.00 | 64.82 | 2.51 (U:OP1 13:646) |
| 3 | Water 3 | | 13:1815 | 2.18 | 1.00 | 60.74 | 2.10 (G:N7 13:587) |
| 4 | Water 4 | | 13:1970 | 2.19 | 1.00 | 67.92 | 3.60 (C:OP1 13:645) |
| 5 | Water 5 | | 13:2018 | 2.19 | 1.00 | 66.60 | 4.59 (C:N4 13:586) |
| 6 | Water 6 | | 13:1988 | 2.19 | 1.00 | 63.31 | 3.37 (U:OP1 13:646) |
| 7 | U | OP1 | 13:646 | 3.86 | 1.00 | 73.05 |  |
| 8 | G | N7 | 13:587 | 3.90 | 1.00 | 54.57 |  |
| 9 | G | O6 | 13:587 | 4.16 | 1.00 | 51.00 |  |
| 10 | C | OP1 | 13:645 | 4.83 | 1.00 | 65.34 |  |
| 11 | C | N4 | 13:586 | 4.90 | 1.00 | 51.07 |  |
  |

  
**1^st^ shell ligand angles:**   
1-2: 93; 1-3: 87; 1-4: 179; 1-5: 89; 1-6: 89; 2-3: 179; 2-4: 88; 2-5: 90; 2-6: 92; 3-4: 92; 3-5: 90; 3-6: 89; 4-5: 91; 4-6: 91; 5-6: 177;   
  
 Contact(s): /O_w_/O_w_/O_w_/O_w_/O_w_/O_w_/  
 Coded contact list: 6O_w_  
MG hydration shell issue: water too close (< 2.4 Angs.)  

### MG:13:1761

|  | Res. type | Atom | Residue | Distance | Occupancy | B-factor | Closest solute to 1^st^ shell water |
| --- | --- | --- | --- | --- | --- | --- | --- |
| **Ion** | | | |  | 1.00 | 86.98 || 1 | Water 1 | | 13:1966 | 2.18 | 1.00 | 91.09 | 3.19 (A:OP1 13:1285) |
| 2 | Water 2 | | 1F:103 | 2.18 | 1.00 | 93.63 | 5.03 (G:OP1 13:1355) |
| 3 | Water 3 | | 13:1908 | 2.18 | 1.00 | 87.19 | 2.79 (G:OP1 13:1355) |
| 4 | Water 4 | | 13:1881 | 2.18 | 1.00 | 87.63 | 2.59 (G:OP1 13:1355) |
| 5 | Water 5 | | 13:1919 | 2.19 | 1.00 | 77.78 | 2.85 (G:OP1 13:1355) |
| 6 | Water 6 | | 13:1957 | 2.19 | 1.00 | 74.72 | 3.16 (C:O2' 13:1284) |
| 7 | G | OP1 | 13:1355 | 3.34 | 1.00 | 86.02 |  |
| 8 | C | O2' | 13:1284 | 4.18 | 1.00 | 94.31 |  |
| 9 | A | OP1 | 13:1285 | 4.25 | 1.00 | 95.70 |  |
| 10 | A | N1 | 13:1286 | 4.65 | 1.00 | 91.03 |  |
| 11 | G | OP2 | 13:1355 | 4.70 | 1.00 | 79.32 |  |
  |

  
**1^st^ shell ligand angles:**   
1-2: 89; 1-3: 89; 1-4: 91; 1-5: 178; 1-6: 90; 2-3: 90; 2-4: 179; 2-5: 90; 2-6: 90; 3-4: 91; 3-5: 89; 3-6: 179; 4-5: 91; 4-6: 89; 5-6: 92;   
  
 Contact(s): /O_w_/O_w_/O_w_/O_w_/O_w_/O_w_/  
 Coded contact list: 6O_w_  
 Exclusion zone contact(s): /OP1/  
Consider K in place of MG (1 valid contact(s) for K vs. 0 for MG)  

### MG:13:1762

|  | Res. type | Atom | Residue | Distance | Occupancy | B-factor | Closest solute to 1^st^ shell water |
| --- | --- | --- | --- | --- | --- | --- | --- |
| **Ion** | | | |  | 1.00 | 52.02 || 1 | Water 1 | | 13:1848 | 2.15 | 1.00 | 59.90 | 2.37 (G:N7 13:286) |
| 2 | Water 2 | | 13:1825 | 2.18 | 1.00 | 59.10 | 2.23 (U:O4 13:287) |
| 3 | Water 3 | | 13:2003 | 2.18 | 1.00 | 67.45 | 3.64 (G:N7 13:285) |
| 4 | Water 4 | | 13:2006 | 2.19 | 1.00 | 62.81 | 3.66 (G:O6 13:285) |
| 5 | Water 5 | | 13:1939 | 2.19 | 1.00 | 55.02 | 3.22 (A:OP2 13:119) |
| 6 | Water 6 | | 13:1996 | 2.19 | 1.00 | 66.11 | 3.47 (U:O4 13:287) |
| 7 | G | O6 | 13:286 | 3.80 | 1.00 | 60.37 |  |
| 8 | U | O4 | 13:287 | 3.87 | 1.00 | 62.26 |  |
| 9 | G | N7 | 13:285 | 4.04 | 1.00 | 57.89 |  |
| 10 | G | N7 | 13:286 | 4.22 | 1.00 | 55.54 |  |
| 11 | G | O6 | 13:285 | 4.59 | 1.00 | 56.19 |  |
| 12 | A | OP2 | 13:119 | 4.61 | 1.00 | 63.97 |  |
  |

  
**1^st^ shell ligand angles:**   
1-2: 86; 1-3: 93; 1-4: 87; 1-5: 177; 1-6: 89; 2-3: 179; 2-4: 91; 2-5: 91; 2-6: 91; 3-4: 88; 3-5: 90; 3-6: 89; 4-5: 94; 4-6: 175; 5-6: 90;   
  
 Contact(s): /O_w_/O_w_/O_w_/O_w_/O_w_/O_w_/  
 Coded contact list: 6O_w_  
MG hydration shell issue: water too close (< 2.4 Angs.)  

### MG:13:1763

|  | Res. type | Atom | Residue | Distance | Occupancy | B-factor | Closest solute to 1^st^ shell water |
| --- | --- | --- | --- | --- | --- | --- | --- |
| **Ion** | | | |  | 1.00 | 64.95 || 1 | Water 1 | | 13:1823 | 2.17 | 1.00 | 68.96 | 2.20 (C:OP2 13:857) |
| 2 | Water 2 | | 13:1801 | 2.18 | 1.00 | 71.31 | 1.30 (G:N7 13:858) |
| 3 | Water 3 | | 13:1861 | 2.18 | 1.00 | 66.01 | 2.48 (C:OP2 13:868) |
| 4 | Water 4 | | 13:1802 | 2.18 | 1.00 | 61.16 | 1.30 (G:N7 13:869) |
| 5 | Water 5 | | 13:1907 | 2.19 | 1.00 | 60.87 | 2.79 (G:OP2 13:858) |
| 6 | Water 6 | | 13:1913 | 2.19 | 1.00 | 61.25 | 2.81 (G:OP2 13:869) |
| 7 | G | N7 | 13:858 | 3.02 | 1.00 | 68.27 |  |
| 8 | G | N7 | 13:869 | 3.05 | 1.00 | 60.30 |  |
| 9 | G | OP2 | 13:869 | 3.60 | 1.00 | 45.85 |  |
| 10 | C | OP2 | 13:857 | 3.78 | 1.00 | 66.01 |  |
| 11 | G | O6 | 13:858 | 3.96 | 1.00 | 62.82 |  |
| 12 | C | OP2 | 13:868 | 4.12 | 1.00 | 47.83 |  |
| 13 | G | OP2 | 13:858 | 4.16 | 1.00 | 67.04 |  |
| 14 | G | O6 | 13:869 | 4.36 | 1.00 | 64.13 |  |
| 15 | C | O5' | 13:857 | 4.47 | 1.00 | 77.78 |  |
| 16 | C | O5' | 13:868 | 4.89 | 1.00 | 44.95 |  |
| 17 | G | O5' | 13:869 | 4.99 | 1.00 | 53.69 |  |
  |

  
**1^st^ shell ligand angles:**   
1-2: 94; 1-3: 86; 1-4: 176; 1-5: 88; 1-6: 89; 2-3: 179; 2-4: 88; 2-5: 88; 2-6: 92; 3-4: 92; 3-5: 93; 3-6: 87; 4-5: 89; 4-6: 94; 5-6: 178;   
  
 Contact(s): /O_w_/O_w_/O_w_/O_w_/O_w_/O_w_/  
 Coded contact list: 6O_w_  
 Exclusion zone contact(s): /N7/N7/  
Consider K in place of MG (2 valid contact(s) for K vs. 0 for MG)  
MG hydration shell issue: water too close (< 2.4 Angs.)  

### MG:13:1764

|  | Res. type | Atom | Residue | Distance | Occupancy | B-factor | Closest solute to 1^st^ shell water |
| --- | --- | --- | --- | --- | --- | --- | --- |
| **Ion** | | | |  | 1.00 | 86.65 || 1 | Water 1 | | 13:1821 | 2.17 | 1.00 | 59.57 | 2.16 (G:N7 13:1461) |
| 2 | Water 2 | | 13:1972 | 2.18 | 1.00 | 59.77 | 4.40 (A:OP2 13:1460) |
| 3 | Water 3 | | 13:1956 | 2.18 | 1.00 | 62.77 | 3.13 (G:O6 13:1462) |
| 4 | Water 4 | | 13:1976 | 2.18 | 1.00 | 76.48 | 3.29 (G:N7 13:1462) |
| 5 | Water 5 | | 13:1903 | 2.19 | 1.00 | 76.36 | 2.74 (A:OP2 13:1460) |
| 6 | Water 6 | | 13:2032 | 2.20 | 1.00 | 64.21 | 5.62 (G:O6 13:1462) |
| 7 | G | O6 | 13:1461 | 4.19 | 1.00 | 65.66 |  |
| 8 | G | N7 | 13:1461 | 4.24 | 1.00 | 65.64 |  |
| 9 | A | OP2 | 13:1460 | 4.45 | 1.00 | 77.25 |  |
| 10 | G | N7 | 13:1462 | 4.56 | 1.00 | 58.76 |  |
| 11 | G | O6 | 13:1462 | 4.94 | 1.00 | 56.77 |  |
  |

  
**1^st^ shell ligand angles:**   
1-2: 92; 1-3: 87; 1-4: 88; 1-5: 90; 1-6: 176; 2-3: 178; 2-4: 87; 2-5: 90; 2-6: 92; 3-4: 92; 3-5: 91; 3-6: 90; 4-5: 176; 4-6: 91; 5-6: 91;   
  
 Contact(s): /O_w_/O_w_/O_w_/O_w_/O_w_/O_w_/  
 Coded contact list: 6O_w_  
MG hydration shell issue: water too close (< 2.4 Angs.)  

### MG:13:1765

|  | Res. type | Atom | Residue | Distance | Occupancy | B-factor | Closest solute to 1^st^ shell water |
| --- | --- | --- | --- | --- | --- | --- | --- |
| **Ion** | | | |  | 1.00 | 73.01 || 1 | Water 1 | | 13:1808 | 2.16 | 1.00 | 72.46 | 2.00 (G:O6 13:829) |
| 2 | Water 2 | | 13:1944 | 2.18 | 1.00 | 75.68 | 3.04 (U:O5' 13:870) |
| 3 | Water 3 | | 13:1936 | 2.18 | 1.00 | 64.49 | 2.96 (U:O2' 13:870) |
| 4 | Water 4 | | 13:1955 | 2.18 | 1.00 | 76.24 | 3.11 (G:O6 13:830) |
| 5 | Water 5 | | 13:1958 | 2.19 | 1.00 | 61.47 | 3.16 (U:OP1 13:871) |
| 6 | Water 6 | | 13:1921 | 2.19 | 1.00 | 76.65 | 2.86 (C:N4 13:856) |
| 7 | G | O6 | 13:830 | 3.70 | 1.00 | 78.31 |  |
| 8 | G | O6 | 13:829 | 3.98 | 1.00 | 75.43 |  |
| 9 | U | O2' | 13:870 | 4.09 | 1.00 | 64.97 |  |
| 10 | U | OP1 | 13:871 | 4.32 | 1.00 | 57.23 |  |
| 11 | U | OP2 | 13:870 | 4.48 | 1.00 | 60.16 |  |
| 12 | C | N4 | 13:856 | 4.71 | 1.00 | 70.61 |  |
| 13 | G | N7 | 13:829 | 4.80 | 1.00 | 77.01 |  |
| 14 | U | O5' | 13:870 | 4.85 | 1.00 | 61.98 |  |
  |

  
**1^st^ shell ligand angles:**   
1-2: 86; 1-3: 87; 1-4: 91; 1-5: 177; 1-6: 92; 2-3: 91; 2-4: 176; 2-5: 92; 2-6: 89; 3-4: 90; 3-5: 92; 3-6: 179; 4-5: 91; 4-6: 89; 5-6: 89;   
  
 Contact(s): /O_w_/O_w_/O_w_/O_w_/O_w_/O_w_/  
 Coded contact list: 6O_w_  
MG hydration shell issue: water too close (< 2.4 Angs.)  

### MG:13:1766

|  | Res. type | Atom | Residue | Distance | Occupancy | B-factor | Closest solute to 1^st^ shell water |
| --- | --- | --- | --- | --- | --- | --- | --- |
| **Ion** | | | |  | 1.00 | 64.12 || 1 | Water 1 | | 13:1899 | 2.17 | 1.00 | 50.68 | 2.72 (G:N7 13:668) |
| 2 | Water 2 | | 13:2012 | 2.18 | 1.00 | 63.99 | 3.75 (U:O4 13:669) |
| 3 | Water 3 | | 13:1884 | 2.18 | 1.00 | 44.04 | 2.62 (G:O6 13:668) |
| 4 | Water 4 | | 13:2027 | 2.19 | 1.00 | 61.53 | 4.53 (U:O4 13:669) |
| 5 | Water 5 | | 13:2016 | 2.19 | 1.00 | 69.10 | 3.85 (G:OP2 13:667) |
| 6 | Water 6 | | 13:1984 | 2.20 | 1.00 | 33.18 | 3.34 (G:OP2 13:667) |
| 7 | G | N7 | 13:668 | 4.10 | 1.00 | 48.43 |  |
| 8 | G | OP2 | 13:667 | 4.50 | 1.00 | 44.26 |  |
| 9 | U | O4 | 13:669 | 4.52 | 1.00 | 56.43 |  |
| 10 | G | O6 | 13:668 | 4.57 | 1.00 | 51.24 |  |
| 11 | G | OP2 | 13:668 | 4.65 | 1.00 | 41.98 |  |
| 12 | G | N7 | 13:667 | 4.98 | 1.00 | 49.06 |  |
  |

  
**1^st^ shell ligand angles:**   
1-2: 90; 1-3: 89; 1-4: 177; 1-5: 90; 1-6: 90; 2-3: 93; 2-4: 87; 2-5: 177; 2-6: 89; 3-4: 91; 3-5: 84; 3-6: 177; 4-5: 94; 4-6: 90; 5-6: 94;   
  
 Contact(s): /O_w_/O_w_/O_w_/O_w_/O_w_/O_w_/  
 Coded contact list: 6O_w_  

### MG:13:1767

|  | Res. type | Atom | Residue | Distance | Occupancy | B-factor | Closest solute to 1^st^ shell water |
| --- | --- | --- | --- | --- | --- | --- | --- |
| **Ion** | | | |  | 1.00 | 70.75 || 1 | Water 1 | | 13:1866 | 2.17 | 1.00 | 63.43 | 2.51 (A:OP1 13:1324) |
| 2 | Water 2 | | 13:1964 | 2.17 | 1.00 | 79.58 | 4.37 (GLN:N 4I:101) |
| 3 | Water 3 | | 13:1813 | 2.18 | 1.00 | 79.49 | 2.06 (C:O2 13:1362A) |
| 4 | Water 4 | | 4I:301 | 2.18 | 1.00 | 68.81 | 2.57 (GLN:N 4I:101) |
| 5 | Water 5 | | 13:1929 | 2.19 | 1.00 | 72.85 | 2.91 (C:O2 13:1362A) |
| 6 | Water 6 | | 13:1986 | 2.19 | 1.00 | 72.73 | 3.36 (A:OP1 13:1324) |
| 7 | C | O2 | 13:1362A | 3.44 | 1.00 | 69.75 |  |
| 8 | A | OP1 | 13:1324 | 3.52 | 1.00 | 72.52 |  |
| 9 | C | O2' | 13:1362A | 4.21 | 1.00 | 76.30 |  |
| 10 | G | OP1 | 13:1323 | 4.43 | 1.00 | 77.10 |  |
| 11 | GLN | N | 4I:101 | 4.48 | 1.00 | 76.87 |  |
| 12 | A | OP2 | 13:1324 | 4.54 | 1.00 | 59.21 |  |
| 13 | G | N2 | 13:1224 | 4.70 | 1.00 | 71.78 |  |
| 14 | G | O3' | 13:1323 | 4.94 | 1.00 | 70.02 |  |
| 15 | U | OP1 | 13:950 | 4.99 | 1.00 | 65.02 |  |
  |

  
**1^st^ shell ligand angles:**   
1-2: 89; 1-3: 91; 1-4: 178; 1-5: 85; 1-6: 95; 2-3: 180; 2-4: 93; 2-5: 92; 2-6: 87; 3-4: 87; 3-5: 88; 3-6: 93; 4-5: 93; 4-6: 86; 5-6: 178;   
  
 Contact(s): /O_w_/O_w_/O_w_/O_w_/O_w_/O_w_/  
 Coded contact list: 6O_w_  
MG hydration shell issue: water too close (< 2.4 Angs.)  

### MG:13:1768

|  | Res. type | Atom | Residue | Distance | Occupancy | B-factor | Closest solute to 1^st^ shell water |
| --- | --- | --- | --- | --- | --- | --- | --- |
| **Ion** | | | |  | 1.00 | 78.43 || 1 | Water 1 | | 13:1890 | 2.18 | 1.00 | 80.18 | 2.65 (A:N3 13:1092) |
| 2 | Water 2 | | 13:1905 | 2.18 | 1.00 | 78.98 | 2.77 (C:O2' 13:1109) |
| 3 | Water 3 | | 13:1924 | 2.18 | 1.00 | 76.20 | 3.27 (C:OP2 13:1066) |
| 4 | Water 4 | | 13:1853 | 2.18 | 1.00 | 63.62 | 2.39 (U:O2' 13:1065) |
| 5 | Water 5 | | 13:1912 | 2.18 | 1.00 | 75.55 | 2.80 (A:OP1 13:1110) |
| 6 | Water 6 | | 13:1856 | 2.18 | 1.00 | 70.53 | 2.42 (C:OP2 13:1066) |
| 7 | U | O2' | 13:1065 | 3.42 | 1.00 | 67.63 |  |
| 8 | C | OP2 | 13:1066 | 3.86 | 1.00 | 64.21 |  |
| 9 | C | O3' | 13:1109 | 4.30 | 1.00 | 71.40 |  |
| 10 | C | O2' | 13:1109 | 4.45 | 1.00 | 79.92 |  |
| 11 | A | O2' | 13:1092 | 4.45 | 1.00 | 78.30 |  |
| 12 | U | O3' | 13:1065 | 4.52 | 1.00 | 67.97 |  |
| 13 | A | N6 | 13:1183 | 4.62 | 1.00 | 88.45 |  |
| 14 | A | N3 | 13:1092 | 4.70 | 1.00 | 83.29 |  |
| 15 | U | O2 | 13:1065 | 4.82 | 1.00 | 61.84 |  |
| 16 | A | OP1 | 13:1110 | 4.92 | 1.00 | 73.15 |  |
| 17 | A | N7 | 13:1067 | 4.97 | 1.00 | 62.40 |  |
  |

  
**1^st^ shell ligand angles:**   
1-2: 88; 1-3: 90; 1-4: 176; 1-5: 89; 1-6: 92; 2-3: 87; 2-4: 88; 2-5: 95; 2-6: 176; 3-4: 91; 3-5: 178; 3-6: 89; 4-5: 90; 4-6: 92; 5-6: 89;   
  
 Contact(s): /O_w_/O_w_/O_w_/O_w_/O_w_/O_w_/  
 Coded contact list: 6O_w_  
MG hydration shell issue: water too close (< 2.4 Angs.)  

### MG:13:1769

|  | Res. type | Atom | Residue | Distance | Occupancy | B-factor | Closest solute to 1^st^ shell water |
| --- | --- | --- | --- | --- | --- | --- | --- |
| **Ion** | | | |  | 1.00 | 54.55 || 1 | Water 1 | | 13:1838 | 2.18 | 1.00 | 49.58 | 2.34 (PSU:O2 13:516) |
| 2 | Water 2 | | 13:1952 | 2.18 | 1.00 | 63.31 | 3.10 (G:OP2 13:515) |
| 3 | Water 3 | | 13:1897 | 2.19 | 1.00 | 70.49 | 2.70 (A:O2' 13:532) |
| 4 | A | O2' | 13:532 | 2.64 | 1.00 | 73.34 |  |
| 5 | PSU | O2 | 13:516 | 2.82 | 1.00 | 60.98 |  |
| 6 | A | OP2 | 13:533 | 2.91 | 1.00 | 60.57 |  |
| 7 | A | OP1 | 13:533 | 2.99 | 1.00 | 64.59 |  |
| 8 | G | O6 | 13:517 | 3.60 | 1.00 | 62.82 |  |
| 9 | G | N7 | 13:515 | 4.19 | 1.00 | 55.48 |  |
| 10 | PSU | N1 | 13:516 | 4.21 | 1.00 | 55.30 |  |
| 11 | G | N1 | 13:517 | 4.25 | 1.00 | 56.77 |  |
| 12 | G | O6 | 13:515 | 4.34 | 1.00 | 55.33 |  |
| 13 | G | OP2 | 13:515 | 4.43 | 1.00 | 60.48 |  |
| 14 | A | O3' | 13:532 | 4.52 | 1.00 | 61.51 |  |
| 15 | A | O5' | 13:533 | 4.66 | 1.00 | 63.23 |  |
  |

  
**1^st^ shell ligand angles:**   
1-2: 97; 1-3: 177; 2-3: 85;   
  
 Contact(s): /O_w_/O_w_/O_w_/  
 Coded contact list: 3O_w_  
 Exclusion zone contact(s): /O2/O2'/OP1/OP2/  
Consider K in place of MG (4 valid contact(s) for K vs. 0 for MG)  
MG hydration shell issue: water too close (< 2.4 Angs.)  

### MG:13:1770

|  | Res. type | Atom | Residue | Distance | Occupancy | B-factor | Closest solute to 1^st^ shell water |
| --- | --- | --- | --- | --- | --- | --- | --- |
| **Ion** | | | |  | 1.00 | 71.86 || 1 | Water 1 | | 13:1894 | 2.17 | 1.00 | 70.38 | 2.70 (G:N7 13:324) |
| 2 | Water 2 | | BI:301 | 2.18 | 1.00 | 82.75 | 2.82 (THR:O BI:71) |
| 3 | Water 3 | | 13:1891 | 2.19 | 1.00 | 67.40 | 2.68 (U:OP2 13:323) |
| 4 | Water 4 | | 13:1951 | 2.19 | 1.00 | 63.81 | 3.09 (G:O6 13:324) |
| 5 | Water 5 | | 13:1867 | 2.19 | 1.00 | 78.37 | 2.52 (G:OP1 13:326) |
| 6 | G | N7 | 13:324 | 2.74 | 1.00 | 71.15 |  |
| 7 | G | O6 | 13:324 | 3.48 | 1.00 | 70.07 |  |
| 8 | G | OP1 | 13:326 | 4.29 | 1.00 | 68.83 |  |
| 9 | G | OP2 | 13:326 | 4.50 | 1.00 | 62.09 |  |
| 10 | G | OP2 | 13:324 | 4.71 | 1.00 | 67.13 |  |
| 11 | U | OP2 | 13:323 | 4.84 | 1.00 | 53.65 |  |
| 12 | G | N9 | 13:324 | 4.85 | 1.00 | 68.63 |  |
| 13 | THR | O | BI:71 | 4.97 | 1.00 | 82.99 |  |
| 14 | SER | O | BI:70 | 5.00 | 1.00 | 86.30 |  |
  |

  
**1^st^ shell ligand angles:**   
1-2: 90; 1-3: 84; 1-4: 178; 1-5: 92; 2-3: 93; 2-4: 88; 2-5: 89; 3-4: 95; 3-5: 176; 4-5: 88;   
  
 Contact(s): /O_w_/O_w_/O_w_/O_w_/O_w_/  
 Coded contact list: 5O_w_  
 Exclusion zone contact(s): /N7/  
Consider K in place of MG (1 valid contact(s) for K vs. 0 for MG)  

### K:14:3001

|  | Res. type | Atom | Residue | Distance | Occupancy | B-factor | Closest solute to 1^st^ shell water |
| --- | --- | --- | --- | --- | --- | --- | --- |
| **Ion** | | | |  | 1.00 | 76.91 || 1 | U | OP2 | 14:1372 | 2.73 | 1.00 | 53.63 |
| 2 | U | O4 | 14:1357 | 3.28 | 1.00 | 46.43 |  |
| 3 | G | O6 | 14:1358 | 3.48 | 1.00 | 60.74 |  |
| 4 | Water 1 | | 14:3805 | 3.50 | 1.00 | 54.73 | 3.82 (U:O4 14:1357) |
| 5 | Water 2 | | 14:4017 | 3.63 | 1.00 | 58.68 | 3.43 (A:OP2 14:1373) |
| 6 | G | N1 | 14:1358 | 3.66 | 1.00 | 55.59 |  |
| 7 | U | OP1 | 14:1372 | 3.76 | 1.00 | 64.12 |  |
| 8 | Water 3 | | 14:3987 | 4.48 | 1.00 | 52.11 | 5.96 (U:O4 14:1357) |
| 9 | U | O5' | 14:1372 | 4.55 | 1.00 | 70.56 |  |
| 10 | G | N2 | 14:1358 | 4.94 | 1.00 | 59.68 |  |
| 11 | A | OP2 | 14:1373 | 4.97 | 1.00 | 69.92 |  |
  |

  
**1^st^ shell ligand angles:**   
1-2: 97;   
  
 Contact(s): /O4/OP2/  
 Coded contact list: O_ph_O_b_  

### K:14:3002

|  | Res. type | Atom | Residue | Distance | Occupancy | B-factor | Closest solute to 1^st^ shell water |
| --- | --- | --- | --- | --- | --- | --- | --- |
| **Ion** | | | |  | 1.00 | 69.11 || 1 | PRO | O | 29:147 | 2.73 | 1.00 | 49.02 |
| 2 | C | OP2 | 14:2025 | 2.94 | 1.00 | 52.73 |  |
| 3 | C | OP1 | 14:2025 | 3.22 | 1.00 | 50.34 |  |
| 4 | U | O4 | 14:1130 | 4.29 | 1.00 | 49.67 |  |
| 5 | G | O3' | 14:2024 | 4.58 | 1.00 | 54.32 |  |
| 6 | LYS | O | 29:145 | 4.69 | 1.00 | 39.01 |  |
| 7 | C | O5' | 14:2025 | 4.77 | 1.00 | 54.80 |  |
| 8 | GLY | N | 29:148 | 4.88 | 1.00 | 42.91 |  |
| 9 | U | N3 | 14:1130 | 4.94 | 1.00 | 51.38 |  |
  |

  
**1^st^ shell ligand angles:**   
1-2: 125; 1-3: 80; 2-3: 49;   
  
 Contact(s): /O_back_/OP1/OP2/  
 Coded contact list: 2O_ph_O_back_  

### K:14:3003

|  | Res. type | Atom | Residue | Distance | Occupancy | B-factor | Closest solute to 1^st^ shell water |
| --- | --- | --- | --- | --- | --- | --- | --- |
| **Ion** | | | |  | 1.00 | 100.43 || 1 | G | O6 | 14:270S | 2.86 | 1.00 | 105.29 |
| 2 | G | O6 | 14:270T | 2.93 | 1.00 | 103.02 |  |
| 3 | G | O6 | 14:270E | 3.05 | 1.00 | 105.24 |  |
| 4 | C | N4 | 14:270U | 3.72 | 1.00 | 107.77 |  |
| 5 | G | N7 | 14:270S | 3.94 | 1.00 | 107.41 |  |
| 6 | G | N7 | 14:270T | 4.01 | 1.00 | 104.72 |  |
| 7 | U | O4 | 14:270F | 4.17 | 1.00 | 110.05 |  |
| 8 | G | N1 | 14:270S | 4.65 | 1.00 | 104.19 |  |
| 9 | C | N4 | 14:270D | 4.72 | 1.00 | 97.01 |  |
| 10 | Water 1 | | 14:3823 | 4.82 | 1.00 | 103.38 | 2.81 (C:N4 14:270U) |
| 11 | G | N1 | 14:270E | 4.86 | 1.00 | 100.68 |  |
  |

  
**1^st^ shell ligand angles:**   
1-2: 59; 1-3: 107; 2-3: 68;   
  
 Contact(s): /O6/O6/O6/  
 Coded contact list: 3O_b_  

### K:14:3004

|  | Res. type | Atom | Residue | Distance | Occupancy | B-factor | Closest solute to 1^st^ shell water |
| --- | --- | --- | --- | --- | --- | --- | --- |
| **Ion** | | | |  | 1.00 | 81.83 || 1 | A | OP2 | 14:2014 | 3.27 | 1.00 | 43.06 |
| 2 | U | O4 | 14:1263 | 3.30 | 1.00 | 51.54 |  |
| 3 | A | OP1 | 14:2014 | 3.42 | 1.00 | 42.42 |  |
| 4 | G | O6 | 14:1264 | 3.71 | 1.00 | 48.37 |  |
| 5 | G | N1 | 14:1264 | 4.75 | 1.00 | 43.63 |  |
| 6 | A | N7 | 14:1262 | 4.93 | 1.00 | 46.93 |  |
| 7 | A | O3' | 14:2013 | 4.98 | 1.00 | 44.31 |  |
  |

  
**1^st^ shell ligand angles:**   
1-2: 84;   
  
 Contact(s): /O4/OP2/  
 Coded contact list: O_ph_O_b_  

### K:14:3005

|  | Res. type | Atom | Residue | Distance | Occupancy | B-factor | Closest solute to 1^st^ shell water |
| --- | --- | --- | --- | --- | --- | --- | --- |
| **Ion** | | | |  | 1.00 | 82.41 || 1 | U | OP2 | 14:2491 | 2.48 | 1.00 | 60.66 |
| 2 | G | OP2 | 14:2455 | 3.24 | 1.00 | 41.27 |  |
| 3 | G | O3' | 14:2490 | 3.88 | 1.00 | 67.26 |  |
| 4 | G | OP1 | 14:2455 | 4.06 | 1.00 | 52.15 |  |
| 5 | G | OP1 | 14:2454 | 4.20 | 1.00 | 50.48 |  |
| 6 | U | OP1 | 14:2491 | 4.56 | 1.00 | 66.64 |  |
| 7 | G | O2' | 14:2490 | 4.62 | 1.00 | 71.17 |  |
| 8 | G | O5' | 14:2454 | 4.64 | 1.00 | 45.32 |  |
| 9 | U | O5' | 14:2491 | 4.75 | 1.00 | 62.44 |  |
  |

  
**1^st^ shell ligand angles:**   
1-2: 160;   
  
 Contact(s): /OP2/  
 Coded contact list: O_ph_  
 Exclusion zone contact(s): /OP2/  

### K:14:3006

|  | Res. type | Atom | Residue | Distance | Occupancy | B-factor | Closest solute to 1^st^ shell water |
| --- | --- | --- | --- | --- | --- | --- | --- |
| **Ion** | | | |  | 1.00 | 106.03 || 1 | G | O6 | 14:1480 | 2.80 | 1.00 | 96.99 |
| 2 | U | O4 | 14:1482 | 2.85 | 1.00 | 99.78 |  |
| 3 | G | O6 | 14:1512 | 3.95 | 1.00 | 103.13 |  |
| 4 | G | N7 | 14:1480 | 4.06 | 1.00 | 86.39 |  |
| 5 | A | OP2 | 14:1511 | 4.47 | 1.00 | 136.40 |  |
| 6 | G | O6 | 14:1479 | 4.90 | 1.00 | 83.32 |  |
| 7 | U | N3 | 14:1482 | 4.91 | 1.00 | 101.61 |  |
| 8 | A | OP2 | 14:1510 | 4.96 | 1.00 | 157.29 |  |
  |

  
**1^st^ shell ligand angles:**   
1-2: 58;   
  
 Contact(s): /O4/O6/  
 Coded contact list: 2O_b_  

### K:14:3007

|  | Res. type | Atom | Residue | Distance | Occupancy | B-factor | Closest solute to 1^st^ shell water |
| --- | --- | --- | --- | --- | --- | --- | --- |
| **Ion** | | | |  | 1.00 | 64.40 || 1 | C | O2' | 2L:75 | 2.75 | 1.00 | 57.80 |
| 2 | C | O3' | 14:2063 | 2.78 | 1.00 | 40.66 |  |
| 3 | C | O2' | 14:2063 | 3.04 | 1.00 | 51.31 |  |
| 4 | C | O3' | 2L:75 | 3.28 | 1.00 | 56.68 |  |
| 5 | C | OP1 | 14:2064 | 3.43 | 1.00 | 43.58 |  |
| 6 | A | OP1 | 2L:76 | 4.23 | 1.00 | 70.97 |  |
| 7 | C | O5' | 14:2064 | 4.73 | 1.00 | 42.15 |  |
| 8 | C | O4' | 14:2063 | 4.79 | 1.00 | 37.53 |  |
  |

  
**1^st^ shell ligand angles:**   
1-2: 91; 1-3: 75; 1-4: 52; 2-3: 54; 2-4: 143; 3-4: 103;   
  
 Contact(s): /O2'/O2'/O3'/O3'/  
 Coded contact list: 4O_r_  

### K:14:3008

|  | Res. type | Atom | Residue | Distance | Occupancy | B-factor | Closest solute to 1^st^ shell water |
| --- | --- | --- | --- | --- | --- | --- | --- |
| **Ion** | | | |  | 1.00 | 47.69 || 1 | G | OP2 | 14:2714 | 2.40 | 1.00 | 50.91 |
| 2 | A | OP2 | 14:2711 | 2.46 | 1.00 | 39.66 |  |
| 3 | A | OP2 | 14:2712A | 2.52 | 1.00 | 38.41 |  |
| 4 | A | OP2 | 14:2713 | 3.73 | 1.00 | 36.64 |  |
| 5 | SER | OG | 55:15 | 3.87 | 1.00 | 46.23 |  |
| 6 | C | OP1 | 14:2710 | 3.98 | 1.00 | 43.16 |  |
| 7 | A | O3' | 14:2713 | 4.00 | 1.00 | 39.95 |  |
| 8 | U | O2' | 14:2712 | 4.00 | 1.00 | 48.13 |  |
| 9 | G | O5' | 14:2714 | 4.04 | 1.00 | 42.46 |  |
| 10 | A | OP1 | 14:2711 | 4.18 | 1.00 | 42.22 |  |
| 11 | C | OP2 | 14:2710 | 4.28 | 1.00 | 48.78 |  |
| 12 | A | O5' | 14:2712A | 4.57 | 1.00 | 27.48 |  |
| 13 | C | O3' | 14:2710 | 4.58 | 1.00 | 46.29 |  |
| 14 | C | O5' | 14:2710 | 4.59 | 1.00 | 45.90 |  |
| 15 | A | OP1 | 14:2712A | 4.70 | 1.00 | 43.03 |  |
| 16 | U | O3' | 14:2712 | 4.92 | 1.00 | 47.73 |  |
| 17 | HIS | NE2 | 55:13 | 4.93 | 1.00 | 36.62 |  |
| 18 | A | O5' | 14:2711 | 4.93 | 1.00 | 37.06 |  |
| 19 | G | OP1 | 14:2714 | 4.97 | 1.00 | 45.03 |  |
  |

  
**1^st^ shell ligand angles:**   
1-2: 72; 1-3: 109; 2-3: 65;   
  
 Coded contact list:   
 Exclusion zone contact(s): /OP2/OP2/OP2/  
Consider NA in place of K (3 valid contact(s) for NA vs. 0 for K)  

### K:14:3009

|  | Res. type | Atom | Residue | Distance | Occupancy | B-factor | Closest solute to 1^st^ shell water |
| --- | --- | --- | --- | --- | --- | --- | --- |
| **Ion** | | | |  | 1.00 | 64.19 || 1 | C | OP2 | 14:749 | 2.71 | 1.00 | 42.85 |
| 2 | G | OP2 | 14:748 | 3.51 | 1.00 | 40.48 |  |
| 3 | A | OP2 | 14:750 | 3.66 | 1.00 | 40.90 |  |
| 4 | G | O5' | 14:748 | 3.71 | 1.00 | 41.22 |  |
| 5 | A | OP1 | 14:750 | 4.20 | 1.00 | 39.18 |  |
| 6 | C | O5' | 14:749 | 4.47 | 1.00 | 36.68 |  |
| 7 | ALA | N | A5:89 | 4.51 | 1.00 | 42.86 |  |
| 8 | G | OP1 | 14:748 | 4.61 | 1.00 | 42.88 |  |
| 9 | PRO | O | A5:87 | 4.65 | 1.00 | 45.17 |  |
| 10 | C | O3' | 14:749 | 4.75 | 1.00 | 40.35 |  |
| 11 | G | O3' | 14:748 | 4.77 | 1.00 | 44.80 |  |
| 12 | ARG | NE | A5:88 | 4.82 | 1.00 | 40.74 |  |
| 13 | G | O6 | 14:1325 | 4.88 | 1.00 | 57.25 |  |
  |

  
 Contact(s): /OP2/  
 Coded contact list: O_ph_  

### K:14:3010

|  | Res. type | Atom | Residue | Distance | Occupancy | B-factor | Closest solute to 1^st^ shell water |
| --- | --- | --- | --- | --- | --- | --- | --- |
| **Ion** | | | |  | 1.00 | 58.07 || 1 | G | OP2 | 14:205 | 3.00 | 1.00 | 43.88 |
| 2 | A | O2' | 14:204 | 3.22 | 1.00 | 36.84 |  |
| 3 | A | O3' | 14:204 | 3.44 | 1.00 | 38.88 |  |
| 4 | U | OP1 | 14:202 | 3.96 | 1.00 | 36.84 |  |
| 5 | G | O5' | 14:205 | 4.11 | 1.00 | 43.39 |  |
| 6 | C | OP2 | 14:203 | 4.29 | 1.00 | 33.57 |  |
| 7 | U | OP2 | 14:202 | 4.84 | 1.00 | 39.66 |  |
| 8 | A | N3 | 14:204 | 4.94 | 1.00 | 31.15 |  |
  |

  
**1^st^ shell ligand angles:**   
1-2: 86;   
  
 Contact(s): /O2'/OP2/  
 Coded contact list: O_ph_O_r_  

### K:14:3011

|  | Res. type | Atom | Residue | Distance | Occupancy | B-factor | Closest solute to 1^st^ shell water |
| --- | --- | --- | --- | --- | --- | --- | --- |
| **Ion** | | | |  | 1.00 | 80.91 || 1 | U | O4 | 14:2091 | 3.41 | 1.00 | 59.03 |
| 2 | U | O2 | 14:2092 | 3.84 | 1.00 | 58.86 |  |
| 3 | C | OP1 | 14:2226 | 4.08 | 1.00 | 55.68 |  |
| 4 | G | N7 | 14:2090 | 4.27 | 1.00 | 48.91 |  |
| 5 | U | N3 | 14:2092 | 4.36 | 1.00 | 51.15 |  |
| 6 | C | OP2 | 14:2226 | 4.47 | 1.00 | 59.71 |  |
| 7 | G | OP2 | 14:2090 | 4.55 | 1.00 | 47.12 |  |
| 8 | G | O6 | 14:2090 | 4.73 | 1.00 | 50.63 |  |
  |

  
 Coded contact list:   
1^st^ K coordination shell missing (see Notes)  

### K:14:3012

|  | Res. type | Atom | Residue | Distance | Occupancy | B-factor | Closest solute to 1^st^ shell water |
| --- | --- | --- | --- | --- | --- | --- | --- |
| **Ion** | | | |  | 1.00 | 117.89 || 1 | G | O6 | 14:1183 | 2.89 | 1.00 | 94.54 |
| 2 | U | O4 | 14:1165 | 4.01 | 1.00 | 86.94 |  |
| 3 | G | N7 | 14:1183 | 4.19 | 1.00 | 108.37 |  |
| 4 | G | O6 | 14:1164 | 4.20 | 1.00 | 90.32 |  |
| 5 | A | N7 | 14:1182 | 4.99 | 1.00 | 120.15 |  |
  |

  
 Contact(s): /O6/  
 Coded contact list: O_b_  

### K:14:3013

|  | Res. type | Atom | Residue | Distance | Occupancy | B-factor | Closest solute to 1^st^ shell water |
| --- | --- | --- | --- | --- | --- | --- | --- |
| **Ion** | | | |  | 1.00 | 84.79 || 1 | C | O2' | 14:1781 | 3.11 | 1.00 | 52.64 |
| 2 | C | OP2 | 14:1782 | 3.82 | 1.00 | 40.94 |  |
| 3 | C | O3' | 14:1781 | 4.17 | 1.00 | 40.43 |  |
| 4 | C | N1 | 14:1781 | 4.31 | 1.00 | 41.45 |  |
| 5 | A | N6 | 14:752 | 4.60 | 1.00 | 43.23 |  |
| 6 | U | O4 | 14:2609 | 4.62 | 1.00 | 34.87 |  |
  |

  
 Contact(s): /O2'/  
 Coded contact list: O_r_  

### K:14:3014

|  | Res. type | Atom | Residue | Distance | Occupancy | B-factor | Closest solute to 1^st^ shell water |
| --- | --- | --- | --- | --- | --- | --- | --- |
| **Ion** | | | |  | 1.00 | 80.63 || 1 | G | O6 | 14:2838 | 3.24 | 1.00 | 68.30 |
| 2 | G | N7 | 14:2837 | 3.29 | 1.00 | 73.69 |  |
| 3 | G | N7 | 14:2838 | 3.37 | 1.00 | 70.89 |  |
| 4 | U | O4 | 14:2878 | 3.58 | 1.00 | 73.16 |  |
| 5 | G | O6 | 14:2839 | 3.95 | 1.00 | 74.69 |  |
| 6 | A | O2' | 14:2835 | 4.09 | 1.00 | 79.15 |  |
| 7 | G | O6 | 14:2837 | 4.22 | 1.00 | 70.02 |  |
| 8 | A | N3 | 14:2835 | 4.23 | 1.00 | 71.39 |  |
| 9 | G | OP2 | 14:2837 | 4.35 | 1.00 | 76.72 |  |
| 10 | G | O6 | 14:2877 | 4.74 | 1.00 | 71.44 |  |
| 11 | G | N9 | 14:2837 | 4.94 | 1.00 | 65.16 |  |
  |

  
**1^st^ shell ligand angles:**   
1-2: 90; 1-3: 56; 2-3: 63;   
  
 Contact(s): /N7/N7/O6/  
 Coded contact list: O_b_2N_b_  

### K:14:3015

|  | Res. type | Atom | Residue | Distance | Occupancy | B-factor | Closest solute to 1^st^ shell water |
| --- | --- | --- | --- | --- | --- | --- | --- |
| **Ion** | | | |  | 1.00 | 47.64 || 1 | Water 1 | | 14:3722 | 3.00 | 1.00 | 33.10 | 2.63 (U:OP2 14:2504) |
| 2 | C | OP1 | 14:2501 | 3.04 | 1.00 | 43.06 |  |
| 3 | U | OP1 | 14:2504 | 3.08 | 1.00 | 41.00 |  |
| 4 | U | OP2 | 14:2504 | 3.12 | 1.00 | 40.57 |  |
| 5 | A | O3' | 14:575 | 3.17 | 1.00 | 43.32 |  |
| 6 | U | OP1 | 14:576 | 3.23 | 1.00 | 37.93 |  |
| 7 | 2MA | OP1 | 14:2503 | 3.28 | 1.00 | 31.76 |  |
| 8 | U | O3' | 14:2500 | 3.32 | 1.00 | 37.90 |  |
| 9 | 2MA | O5' | 14:2503 | 3.42 | 1.00 | 31.00 |  |
| 10 | U | O2' | 14:2500 | 3.89 | 1.00 | 38.90 |  |
| 11 | Water 2 | | 14:3597 | 4.65 | 1.00 | 45.19 | 2.33 (U:OP1 14:576) |
| 12 | 2MA | OP2 | 14:2503 | 4.78 | 1.00 | 30.67 |  |
| 13 | 2MA | O3' | 14:2503 | 4.78 | 1.00 | 32.74 |  |
| 14 | U | O5' | 14:2504 | 4.78 | 1.00 | 34.75 |  |
| 15 | C | O5' | 14:2501 | 4.79 | 1.00 | 34.82 |  |
| 16 | U | O5' | 14:576 | 4.97 | 1.00 | 28.07 |  |
| 17 | C | OP2 | 14:2501 | 4.99 | 1.00 | 38.49 |  |
  |

  
**1^st^ shell ligand angles:**   
1-2: 152; 1-3: 96; 1-4: 51; 1-5: 99; 1-6: 62; 1-7: 61; 1-8: 161; 2-3: 93; 2-4: 140; 2-5: 80; 2-6: 101; 2-7: 91; 2-8: 45; 3-4: 47; 3-5: 160; 3-6: 155; 3-7: 109; 3-8: 72; 4-5: 140; 4-6: 113; 4-7: 95; 4-8: 113; 5-6: 45; 5-7: 90; 5-8: 90; 6-7: 51; 6-8: 132; 7-8: 136;   
  
 Contact(s): /O3'/O3'/O_w_/OP1/OP1/OP1/OP1/OP2/  
 Coded contact list: 5O_ph_2O_r_O_w_  

### K:14:3016

|  | Res. type | Atom | Residue | Distance | Occupancy | B-factor | Closest solute to 1^st^ shell water |
| --- | --- | --- | --- | --- | --- | --- | --- |
| **Ion** | | | |  | 1.00 | 44.18 || 1 | G | O2' | 14:2576 | 2.63 | 1.00 | 44.24 |
| 2 | G | O6 | 14:2505 | 2.67 | 1.00 | 40.21 |  |
| 3 | C | O2 | 14:2610 | 2.91 | 1.00 | 46.27 |  |
| 4 | G | O6 | 14:2581 | 2.99 | 1.00 | 37.33 |  |
| 5 | C | O2' | 14:2610 | 3.42 | 1.00 | 34.95 |  |
| 6 | G | N1 | 14:2581 | 3.86 | 1.00 | 39.18 |  |
| 7 | U | OP2 | 14:2580 | 4.14 | 1.00 | 36.43 |  |
| 8 | G | N1 | 14:2505 | 4.15 | 1.00 | 42.78 |  |
| 9 | LYS | NZ | J5:3 | 4.29 | 1.00 | 42.43 |  |
| 10 | C | N1 | 14:2610 | 4.31 | 1.00 | 38.95 |  |
| 11 | G | N9 | 14:2576 | 4.45 | 1.00 | 42.13 |  |
| 12 | C | OP2 | 14:2579 | 4.50 | 1.00 | 42.47 |  |
| 13 | C | N3 | 14:2610 | 4.60 | 1.00 | 41.84 |  |
| 14 | C | OP1 | 14:2579 | 4.68 | 1.00 | 39.93 |  |
| 15 | C | O3' | 14:2610 | 4.78 | 1.00 | 40.77 |  |
| 16 | U | OP2 | 14:2611 | 4.87 | 1.00 | 35.82 |  |
| 17 | C | O5' | 14:2579 | 4.90 | 1.00 | 33.15 |  |
  |

  
**1^st^ shell ligand angles:**   
1-2: 73; 1-3: 138; 1-4: 101; 2-3: 66; 2-4: 87; 3-4: 66;   
  
 Contact(s): /O2/O2'/O6/O6/  
 Coded contact list: O_r_3O_b_  

### K:14:3017

|  | Res. type | Atom | Residue | Distance | Occupancy | B-factor | Closest solute to 1^st^ shell water |
| --- | --- | --- | --- | --- | --- | --- | --- |
| **Ion** | | | |  | 1.00 | 52.00 || 1 | 5MU | O2' | 14:1939 | 2.89 | 1.00 | 42.50 |
| 2 | G | O2' | 14:2592 | 3.04 | 1.00 | 41.95 |  |
| 3 | A | O2' | 14:1966 | 3.09 | 1.00 | 42.94 |  |
| 4 | 5MU | O2 | 14:1939 | 3.40 | 1.00 | 40.52 |  |
| 5 | 5MU | O3' | 14:1939 | 3.40 | 1.00 | 34.04 |  |
| 6 | A | N6 | 14:1937 | 3.45 | 1.00 | 39.96 |  |
| 7 | C | O5' | 14:1967 | 3.56 | 1.00 | 50.63 |  |
| 8 | C | OP2 | 14:1967 | 3.81 | 1.00 | 45.28 |  |
| 9 | A | O3' | 14:1966 | 3.95 | 1.00 | 34.25 |  |
| 10 | G | O4' | 14:2592 | 4.13 | 1.00 | 42.27 |  |
| 11 | 5MU | N1 | 14:1939 | 4.14 | 1.00 | 38.92 |  |
| 12 | C | OP1 | 14:1965 | 4.32 | 1.00 | 48.56 |  |
| 13 | U | OP1 | 14:1940 | 4.33 | 1.00 | 43.46 |  |
| 14 | 5MU | O4' | 14:1939 | 4.48 | 1.00 | 40.58 |  |
| 15 | A | N7 | 14:1937 | 4.51 | 1.00 | 45.24 |  |
| 16 | G | O3' | 14:2592 | 4.52 | 1.00 | 41.64 |  |
| 17 | U | O5' | 14:1940 | 4.77 | 1.00 | 46.41 |  |
  |

  
**1^st^ shell ligand angles:**   
1-2: 125; 1-3: 125; 1-4: 84; 2-3: 77; 2-4: 100; 3-4: 147;   
  
 Contact(s): /O2/O2'/O2'/O2'/  
 Coded contact list: 3O_r_O_b_  

### K:14:3018

|  | Res. type | Atom | Residue | Distance | Occupancy | B-factor | Closest solute to 1^st^ shell water |
| --- | --- | --- | --- | --- | --- | --- | --- |
| **Ion** | | | |  | 1.00 | 65.33 || 1 | G | N7 | 14:2544 | 2.73 | 1.00 | 57.14 |
| 2 | A | N3 | 14:2542 | 2.80 | 1.00 | 61.75 |  |
| 3 | G | O6 | 14:2544 | 2.89 | 1.00 | 52.53 |  |
| 4 | A | O4' | 14:2542 | 3.01 | 1.00 | 57.97 |  |
| 5 | G | N7 | 14:2543 | 3.17 | 1.00 | 60.03 |  |
| 6 | A | N9 | 14:2542 | 3.98 | 1.00 | 57.44 |  |
| 7 | G | O6 | 14:2543 | 4.12 | 1.00 | 63.27 |  |
| 8 | A | O3' | 14:2542 | 4.42 | 1.00 | 74.62 |  |
| 9 | G | N9 | 14:2543 | 4.62 | 1.00 | 61.59 |  |
| 10 | G | N1 | 14:2544 | 4.83 | 1.00 | 53.84 |  |
| 11 | G | OP2 | 14:2544 | 4.84 | 1.00 | 64.02 |  |
| 12 | G | N9 | 14:2544 | 4.95 | 1.00 | 60.29 |  |
| 13 | A | N1 | 14:2542 | 4.96 | 1.00 | 55.06 |  |
| 14 | G | O6 | 14:2545 | 4.99 | 1.00 | 53.57 |  |
  |

  
**1^st^ shell ligand angles:**   
1-2: 147; 1-3: 66; 1-4: 142; 1-5: 81; 2-3: 94; 2-4: 70; 2-5: 128; 3-4: 135; 3-5: 96; 4-5: 67;   
  
 Contact(s): /N3_A_/N7/N7/O4'/O6/  
 Coded contact list: O_r_O_b_3N_b_  

### K:14:3019

|  | Res. type | Atom | Residue | Distance | Occupancy | B-factor | Closest solute to 1^st^ shell water |
| --- | --- | --- | --- | --- | --- | --- | --- |
| **Ion** | | | |  | 1.00 | 40.24 || 1 | G | N7 | 14:2053 | 2.72 | 1.00 | 40.10 |
| 2 | A | O2' | 14:2051 | 2.99 | 1.00 | 49.04 |  |
| 3 | A | N1 | 14:2614 | 3.08 | 1.00 | 37.03 |  |
| 4 | A | N3 | 14:2051 | 3.14 | 1.00 | 42.39 |  |
| 5 | G | N7 | 14:2052 | 3.31 | 1.00 | 43.22 |  |
| 6 | G | O6 | 14:2052 | 3.47 | 1.00 | 41.39 |  |
| 7 | G | O6 | 14:2053 | 3.58 | 1.00 | 42.02 |  |
| 8 | A | N6 | 14:2577 | 3.79 | 1.00 | 43.00 |  |
| 9 | A | N6 | 14:2614 | 3.96 | 1.00 | 37.12 |  |
| 10 | A | N7 | 14:2577 | 4.41 | 1.00 | 41.75 |  |
| 11 | G | N1 | 14:2052 | 4.46 | 1.00 | 41.36 |  |
| 12 | A | N1 | 14:2577 | 4.52 | 1.00 | 37.86 |  |
| 13 | G | OP2 | 14:2053 | 4.56 | 1.00 | 46.67 |  |
| 14 | A | N9 | 14:2051 | 4.60 | 1.00 | 39.28 |  |
| 15 | G | N9 | 14:2052 | 4.70 | 1.00 | 46.38 |  |
| 16 | G | N9 | 14:2053 | 4.89 | 1.00 | 39.90 |  |
| 17 | A | O3' | 14:2051 | 4.91 | 1.00 | 51.30 |  |
| 18 | A | N1 | 14:2051 | 4.94 | 1.00 | 40.26 |  |
  |

  
**1^st^ shell ligand angles:**   
1-2: 148; 1-3: 126; 1-4: 126; 1-5: 84; 2-3: 85; 2-4: 73; 2-5: 77; 3-4: 65; 3-5: 138; 4-5: 73;   
  
 Contact(s): /N1_A_/N3_A_/N7/N7/O2'/  
 Coded contact list: O_r_4N_b_  

### K:14:3020

|  | Res. type | Atom | Residue | Distance | Occupancy | B-factor | Closest solute to 1^st^ shell water |
| --- | --- | --- | --- | --- | --- | --- | --- |
| **Ion** | | | |  | 1.00 | 64.35 || 1 | G | O6 | 14:2436 | 2.72 | 1.00 | 30.73 |
| 2 | U | O4 | 14:2245 | 2.75 | 1.00 | 41.73 |  |
| 3 | U | O4 | 14:2437 | 3.38 | 1.00 | 34.86 |  |
| 4 | G | O6 | 14:2072 | 3.70 | 1.00 | 34.70 |  |
| 5 | U | O4 | 14:2244 | 3.90 | 1.00 | 37.41 |  |
| 6 | C | N4 | 14:2073 | 3.99 | 1.00 | 31.47 |  |
| 7 | A | N6 | 14:2071 | 4.02 | 1.00 | 32.82 |  |
| 8 | U | N3 | 14:2245 | 4.77 | 1.00 | 38.13 |  |
| 9 | G | N7 | 14:2436 | 4.80 | 1.00 | 36.11 |  |
| 10 | U | N3 | 14:2437 | 4.85 | 1.00 | 31.91 |  |
| 11 | G | N1 | 14:2436 | 4.87 | 1.00 | 31.16 |  |
  |

  
**1^st^ shell ligand angles:**   
1-2: 85; 1-3: 61; 2-3: 77;   
  
 Contact(s): /O4/O4/O6/  
 Coded contact list: 3O_b_  

### K:14:3021

|  | Res. type | Atom | Residue | Distance | Occupancy | B-factor | Closest solute to 1^st^ shell water |
| --- | --- | --- | --- | --- | --- | --- | --- |
| **Ion** | | | |  | 1.00 | 41.45 || 1 | G | O6 | 14:2447 | 2.57 | 1.00 | 42.16 |
| 2 | G | O6 | 14:2061 | 2.72 | 1.00 | 40.18 |  |
| 3 | C | O4' | 14:2501 | 2.75 | 1.00 | 36.08 |  |
| 4 | G | N7 | 14:2447 | 2.93 | 1.00 | 33.25 |  |
| 5 | U | O4 | 14:2504 | 3.04 | 1.00 | 41.15 |  |
| 6 | C | O2 | 14:2501 | 3.16 | 1.00 | 35.02 |  |
| 7 | G | N7 | 14:2061 | 3.50 | 1.00 | 36.36 |  |
| 8 | C | N1 | 14:2501 | 3.77 | 1.00 | 37.75 |  |
| 9 | A | N6 | 14:2451 | 3.97 | 1.00 | 41.96 |  |
| 10 | C | N3 | 14:2501 | 4.35 | 1.00 | 37.04 |  |
| 11 | G | N1 | 14:2447 | 4.61 | 1.00 | 41.21 |  |
| 12 | G | N1 | 14:2061 | 4.83 | 1.00 | 33.28 |  |
  |

  
**1^st^ shell ligand angles:**   
1-2: 99; 1-3: 120; 1-4: 67; 1-5: 62; 1-6: 86; 2-3: 121; 2-4: 161; 2-5: 92; 2-6: 73; 3-4: 61; 3-5: 145; 3-6: 68; 4-5: 94; 4-6: 91; 5-6: 142;   
  
 Contact(s): /N7/O2/O4/O4'/O6/  
 Coded contact list: O_r_3O_b_N_b_  
 Exclusion zone contact(s): /O6/  

### K:14:3022

|  | Res. type | Atom | Residue | Distance | Occupancy | B-factor | Closest solute to 1^st^ shell water |
| --- | --- | --- | --- | --- | --- | --- | --- |
| **Ion** | | | |  | 1.00 | 50.85 || 1 | C | OP2 | 14:2440 | 2.60 | 1.00 | 30.08 |
| 2 | A | O2' | 14:2439 | 2.71 | 1.00 | 35.20 |  |
| 3 | Water 1 | | 14:3614 | 2.86 | 1.00 | 39.43 | 2.38 (G:OP2 14:2588) |
| 4 | A | OP2 | 14:2600 | 2.93 | 1.00 | 35.54 |  |
| 5 | Water 2 | | 14:3683 | 3.27 | 1.00 | 38.71 | 3.18 (G:OP2 14:2588) |
| 6 | GLU | OE1 | 19:237 | 3.36 | 1.00 | 45.47 |  |
| 7 | GLU | OE2 | 19:237 | 3.56 | 1.00 | 55.72 |  |
| 8 | Water 3 | | 14:3943 | 3.91 | 1.00 | 42.41 | 3.14 (C:OP2 14:2440) |
| 9 | Water 4 | | 19:412 | 4.02 | 1.00 | 33.86 | 3.79 (GLU:OE2 19:237) |
| 10 | G | OP2 | 14:2588 | 4.39 | 1.00 | 32.75 |  |
| 11 | C | OP1 | 14:2440 | 4.57 | 1.00 | 36.47 |  |
| 12 | A | O3' | 14:2439 | 4.60 | 1.00 | 34.88 |  |
| 13 | Water 5 | | 14:3649 | 4.62 | 1.00 | 31.99 | 3.98 (G:OP2 14:2588) |
| 14 | A | OP1 | 14:2600 | 4.66 | 1.00 | 33.98 |  |
| 15 | G | O3' | 14:2599 | 4.73 | 1.00 | 36.66 |  |
  |

  
**1^st^ shell ligand angles:**   
1-2: 52; 1-3: 97; 1-4: 102; 1-5: 93; 1-6: 152; 2-3: 115; 2-4: 68; 2-5: 66; 2-6: 137; 3-4: 156; 3-5: 60; 3-6: 99; 4-5: 105; 4-6: 71; 5-6: 116;   
  
 Contact(s): /O2'/O_w_/O_w_/OE1_Glu_/OP2/OP2/  
 Coded contact list: 2O_ph_O_r_O_coo_2O_w_  
K hydration shell issue: water too close (< 2.4 Angs.)  

### K:14:3023

|  | Res. type | Atom | Residue | Distance | Occupancy | B-factor | Closest solute to 1^st^ shell water |
| --- | --- | --- | --- | --- | --- | --- | --- |
| **Ion** | | | |  | 1.00 | 60.81 || 1 | C | O2 | 14:419 | 2.82 | 1.00 | 60.79 |
| 2 | G | OP2 | 14:224 | 3.03 | 1.00 | 61.51 |  |
| 3 | C | O2' | 14:419 | 3.08 | 1.00 | 60.21 |  |
| 4 | G | N7 | 14:224 | 3.34 | 1.00 | 61.58 |  |
| 5 | A | N7 | 14:225 | 3.41 | 1.00 | 64.82 |  |
| 6 | C | O4' | 14:420 | 3.66 | 1.00 | 66.74 |  |
| 7 | G | N2 | 14:408 | 3.92 | 1.00 | 58.31 |  |
| 8 | G | O6 | 14:226 | 4.34 | 1.00 | 65.56 |  |
| 9 | G | O5' | 14:224 | 4.38 | 1.00 | 60.04 |  |
| 10 | C | O2 | 14:409 | 4.40 | 1.00 | 59.26 |  |
| 11 | C | N1 | 14:419 | 4.43 | 1.00 | 61.16 |  |
| 12 | A | N6 | 14:225 | 4.44 | 1.00 | 68.74 |  |
| 13 | G | N9 | 14:224 | 4.72 | 1.00 | 60.04 |  |
| 14 | G | O6 | 14:224 | 4.86 | 1.00 | 64.77 |  |
| 15 | A | O3' | 14:223 | 4.96 | 1.00 | 54.34 |  |
| 16 | C | O4' | 14:419 | 5.00 | 1.00 | 64.02 |  |
  |

  
**1^st^ shell ligand angles:**   
1-2: 99; 1-3: 78; 1-4: 127; 2-3: 145; 2-4: 95; 3-4: 63;   
  
 Contact(s): /N7/O2/O2'/OP2/  
 Coded contact list: O_ph_O_r_O_b_N_b_  

### K:14:3024

|  | Res. type | Atom | Residue | Distance | Occupancy | B-factor | Closest solute to 1^st^ shell water |
| --- | --- | --- | --- | --- | --- | --- | --- |
| **Ion** | | | |  | 1.00 | 52.53 || 1 | G | O2' | 14:2250 | 2.71 | 1.00 | 41.33 |
| 2 | OMG | OP2 | 14:2251 | 2.88 | 1.00 | 40.94 |  |
| 3 | A | N7 | 14:2497 | 3.03 | 1.00 | 41.43 |  |
| 4 | G | O3' | 14:2250 | 3.11 | 1.00 | 39.91 |  |
| 5 | A | OP2 | 14:2497 | 3.29 | 1.00 | 38.89 |  |
| 6 | C | O5' | 14:2496 | 3.50 | 1.00 | 37.48 |  |
| 7 | C | OP2 | 14:2496 | 3.51 | 1.00 | 52.38 |  |
| 8 | C | OP1 | 14:2496 | 3.79 | 1.00 | 45.06 |  |
| 9 | C | O3' | 14:2496 | 3.92 | 1.00 | 36.37 |  |
| 10 | OMG | OP1 | 14:2251 | 4.49 | 1.00 | 45.43 |  |
| 11 | OMG | O5' | 14:2251 | 4.68 | 1.00 | 44.57 |  |
| 12 | A | N6 | 14:2497 | 4.75 | 1.00 | 46.01 |  |
| 13 | A | O5' | 14:2497 | 4.80 | 1.00 | 45.92 |  |
| 14 | A | N9 | 14:2497 | 4.88 | 1.00 | 36.68 |  |
| 15 | A | OP1 | 14:2450 | 4.90 | 1.00 | 42.57 |  |
| 16 | ARG | NH2 | 45:82 | 4.97 | 1.00 | 51.29 |  |
  |

  
**1^st^ shell ligand angles:**   
1-2: 99; 1-3: 149; 1-4: 55; 1-5: 85; 2-3: 112; 2-4: 51; 2-5: 117; 3-4: 152; 3-5: 84; 4-5: 86;   
  
 Contact(s): /N7/O2'/O3'/OP2/OP2/  
 Coded contact list: 2O_ph_2O_r_N_b_  

### K:14:3025

|  | Res. type | Atom | Residue | Distance | Occupancy | B-factor | Closest solute to 1^st^ shell water |
| --- | --- | --- | --- | --- | --- | --- | --- |
| **Ion** | | | |  | 1.00 | 50.23 || 1 | Water 1 | | 14:3650 | 2.90 | 1.00 | 32.74 | 2.63 (U:OP1 14:827) |
| 2 | G | OP2 | 14:2429 | 3.04 | 1.00 | 48.69 |  |
| 3 | A | O3' | 14:2426 | 3.08 | 1.00 | 38.64 |  |
| 4 | C | OP1 | 14:2427 | 3.28 | 1.00 | 41.65 |  |
| 5 | A | O2' | 14:2426 | 3.31 | 1.00 | 43.93 |  |
| 6 | U | OP1 | 14:827 | 3.57 | 1.00 | 33.10 |  |
| 7 | G | OP1 | 14:2429 | 3.66 | 1.00 | 43.33 |  |
| 8 | U | OP2 | 14:827 | 4.11 | 1.00 | 46.15 |  |
| 9 | G | O5' | 14:2429 | 4.27 | 1.00 | 38.78 |  |
| 10 | C | O5' | 14:2427 | 4.74 | 1.00 | 43.22 |  |
| 11 | A | O4' | 14:2426 | 4.82 | 1.00 | 46.10 |  |
| 12 | A | OP2 | 14:2425 | 4.84 | 1.00 | 40.99 |  |
| 13 | C | OP2 | 14:2427 | 4.88 | 1.00 | 44.97 |  |
| 14 | A | N3 | 14:2426 | 4.90 | 1.00 | 43.54 |  |
  |

  
**1^st^ shell ligand angles:**   
1-2: 67; 1-3: 142; 1-4: 146; 1-5: 92; 2-3: 130; 2-4: 86; 2-5: 134; 3-4: 47; 3-5: 51; 4-5: 92;   
  
 Contact(s): /O2'/O3'/O_w_/OP1/OP2/  
 Coded contact list: 2O_ph_2O_r_O_w_  

### K:14:3026

|  | Res. type | Atom | Residue | Distance | Occupancy | B-factor | Closest solute to 1^st^ shell water |
| --- | --- | --- | --- | --- | --- | --- | --- |
| **Ion** | | | |  | 1.00 | 35.89 || 1 | A | OP2 | 14:1785 | 2.76 | 1.00 | 35.28 |
| 2 | A | OP1 | 14:1981 | 2.87 | 1.00 | 36.63 |  |
| 3 | G | O3' | 14:1980 | 3.00 | 1.00 | 36.96 |  |
| 4 | C | O5' | 14:1982 | 3.01 | 1.00 | 37.12 |  |
| 5 | C | O4' | 14:1982 | 3.18 | 1.00 | 35.18 |  |
| 6 | C | OP2 | 14:1982 | 3.24 | 1.00 | 35.12 |  |
| 7 | A | O5' | 14:1981 | 3.49 | 1.00 | 38.95 |  |
| 8 | G | O2' | 14:1980 | 4.06 | 1.00 | 45.13 |  |
| 9 | Water 1 | | 14:3711 | 4.15 | 1.00 | 42.79 | 2.61 (A:OP1 14:1981) |
| 10 | A | O5' | 14:1785 | 4.32 | 1.00 | 36.78 |  |
| 11 | C | N1 | 14:1982 | 4.54 | 1.00 | 39.35 |  |
| 12 | A | O3' | 14:1981 | 4.59 | 1.00 | 41.05 |  |
| 13 | A | OP2 | 14:1981 | 4.85 | 1.00 | 37.05 |  |
| 14 | A | OP1 | 14:1786 | 4.85 | 1.00 | 37.16 |  |
| 15 | A | O3' | 14:1784 | 4.96 | 1.00 | 38.78 |  |
| 16 | A | O3' | 14:1785 | 4.96 | 1.00 | 35.45 |  |
| 17 | C | OP1 | 14:1982 | 4.98 | 1.00 | 37.67 |  |
| 18 | G | O4' | 14:1980 | 4.99 | 1.00 | 40.25 |  |
  |

  
**1^st^ shell ligand angles:**   
1-2: 90; 1-3: 131; 1-4: 74; 1-5: 77; 1-6: 114; 2-3: 50; 2-4: 125; 2-5: 166; 2-6: 104; 3-4: 104; 3-5: 137; 3-6: 62; 4-5: 47; 4-6: 45; 5-6: 77;   
  
 Contact(s): /O3'/O4'/O5'/OP1/OP2/OP2/  
 Coded contact list: 3O_ph_3O_r_  

### K:14:3027

|  | Res. type | Atom | Residue | Distance | Occupancy | B-factor | Closest solute to 1^st^ shell water |
| --- | --- | --- | --- | --- | --- | --- | --- |
| **Ion** | | | |  | 1.00 | 54.39 || 1 | A | OP1 | 14:783 | 2.65 | 1.00 | 33.85 |
| 2 | A | N3 | 14:1787 | 2.99 | 1.00 | 35.83 |  |
| 3 | A | N6 | 14:1785 | 3.47 | 1.00 | 35.32 |  |
| 4 | U | O2 | 14:1778 | 3.55 | 1.00 | 38.20 |  |
| 5 | C | O4' | 14:1788 | 3.90 | 1.00 | 30.79 |  |
| 6 | A | OP2 | 14:782 | 4.19 | 1.00 | 26.45 |  |
| 7 | A | O5' | 14:783 | 4.34 | 1.00 | 34.17 |  |
| 8 | A | O3' | 14:782 | 4.35 | 1.00 | 27.89 |  |
| 9 | A | N1 | 14:1787 | 4.43 | 1.00 | 35.32 |  |
| 10 | U | O2' | 14:1778 | 4.47 | 1.00 | 33.32 |  |
| 11 | U | N1 | 14:1778 | 4.47 | 1.00 | 30.39 |  |
| 12 | A | O5' | 14:782 | 4.56 | 1.00 | 31.82 |  |
| 13 | C | N1 | 14:1788 | 4.58 | 1.00 | 30.06 |  |
| 14 | U | O2 | 14:1777 | 4.68 | 1.00 | 35.72 |  |
| 15 | U | O4' | 14:1778 | 4.74 | 1.00 | 31.81 |  |
| 16 | A | O4' | 14:783 | 4.98 | 1.00 | 33.03 |  |
  |

  
**1^st^ shell ligand angles:**   
1-2: 96;   
  
 Contact(s): /N3_A_/OP1/  
 Coded contact list: O_ph_N_b_  

### K:14:3028

|  | Res. type | Atom | Residue | Distance | Occupancy | B-factor | Closest solute to 1^st^ shell water |
| --- | --- | --- | --- | --- | --- | --- | --- |
| **Ion** | | | |  | 1.00 | 45.73 || 1 | Water 1 | | 14:3757 | 2.57 | 1.00 | 46.93 | 3.67 (A:N7 14:2577) |
| 2 | G | N7 | 14:2056 | 3.08 | 1.00 | 40.47 |  |
| 3 | A | N7 | 14:2577 | 3.29 | 1.00 | 41.75 |  |
| 4 | Water 2 | | 14:3682 | 3.70 | 1.00 | 52.53 | 2.55 (G:OP2 14:2053) |
| 5 | G | OP2 | 14:2053 | 3.74 | 1.00 | 46.67 |  |
| 6 | G | OP1 | 14:2053 | 3.81 | 1.00 | 43.13 |  |
| 7 | Water 3 | | 14:3581 | 3.81 | 1.00 | 46.02 | 2.28 (G:OP1 14:2053) |
| 8 | A | OP2 | 14:2054 | 3.92 | 1.00 | 42.67 |  |
| 9 | A | N6 | 14:2577 | 3.93 | 1.00 | 43.00 |  |
| 10 | G | O5' | 14:2053 | 4.61 | 1.00 | 37.79 |  |
| 11 | G | O6 | 14:2056 | 4.63 | 1.00 | 41.30 |  |
| 12 | G | N9 | 14:2056 | 4.93 | 1.00 | 36.96 |  |
  |

  
**1^st^ shell ligand angles:**   
1-2: 107; 1-3: 77; 2-3: 70;   
  
 Contact(s): /N7/N7/  
 Coded contact list: 2N_b_  
 Exclusion zone contact(s): /O_w_/  

### K:14:3029

|  | Res. type | Atom | Residue | Distance | Occupancy | B-factor | Closest solute to 1^st^ shell water |
| --- | --- | --- | --- | --- | --- | --- | --- |
| **Ion** | | | |  | 1.00 | 81.55 || 1 | U | O4 | 14:639 | 2.72 | 1.00 | 78.81 |
| 2 | G | O6 | 14:638 | 3.09 | 1.00 | 79.58 |  |
| 3 | G | O6 | 14:648 | 3.12 | 1.00 | 72.22 |  |
| 4 | G | O6 | 14:649 | 3.16 | 1.00 | 76.36 |  |
| 5 | C | N4 | 14:650 | 3.86 | 1.00 | 77.86 |  |
| 6 | U | N3 | 14:639 | 4.11 | 1.00 | 76.21 |  |
| 7 | G | N7 | 14:648 | 4.43 | 1.00 | 66.11 |  |
| 8 | C | N4 | 14:640 | 4.54 | 1.00 | 74.49 |  |
| 9 | G | N7 | 14:649 | 4.57 | 1.00 | 65.77 |  |
| 10 | G | N7 | 14:638 | 4.71 | 1.00 | 82.17 |  |
| 11 | G | O6 | 14:651 | 4.83 | 1.00 | 81.24 |  |
| 12 | G | N1 | 14:648 | 4.85 | 1.00 | 66.82 |  |
  |

  
**1^st^ shell ligand angles:**   
1-2: 68; 1-3: 80; 1-4: 68; 2-3: 127; 2-4: 64; 3-4: 66;   
  
 Contact(s): /O4/O6/O6/O6/  
 Coded contact list: 4O_b_  

### K:14:3030

|  | Res. type | Atom | Residue | Distance | Occupancy | B-factor | Closest solute to 1^st^ shell water |
| --- | --- | --- | --- | --- | --- | --- | --- |
| **Ion** | | | |  | 1.00 | 63.16 || 1 | G | O6 | 14:948 | 2.84 | 1.00 | 57.83 |
| 2 | G | O6 | 14:968 | 2.91 | 1.00 | 58.51 |  |
| 3 | G | O6 | 14:947 | 3.03 | 1.00 | 60.51 |  |
| 4 | U | O4 | 14:969 | 3.18 | 1.00 | 61.17 |  |
| 5 | C | N4 | 14:949 | 3.63 | 1.00 | 54.52 |  |
| 6 | U | N3 | 14:969 | 4.06 | 1.00 | 59.44 |  |
| 7 | G | N7 | 14:948 | 4.09 | 1.00 | 51.36 |  |
| 8 | G | N1 | 14:947 | 4.51 | 1.00 | 58.38 |  |
| 9 | C | N4 | 14:970 | 4.59 | 1.00 | 55.67 |  |
| 10 | G | N7 | 14:947 | 4.73 | 1.00 | 56.47 |  |
| 11 | G | N1 | 14:968 | 4.96 | 1.00 | 60.35 |  |
  |

  
**1^st^ shell ligand angles:**   
1-2: 68; 1-3: 75; 1-4: 75; 2-3: 136; 2-4: 70; 3-4: 78;   
  
 Contact(s): /O4/O6/O6/O6/  
 Coded contact list: 4O_b_  

### K:14:3031

|  | Res. type | Atom | Residue | Distance | Occupancy | B-factor | Closest solute to 1^st^ shell water |
| --- | --- | --- | --- | --- | --- | --- | --- |
| **Ion** | | | |  | 1.00 | 29.30 || 1 | HIS | O | 19:220 | 2.61 | 1.00 | 32.69 |
| 2 | G | O6 | 14:1828 | 2.65 | 1.00 | 28.53 |  |
| 3 | A | OP1 | 14:1789 | 2.68 | 1.00 | 24.69 |  |
| 4 | C | OP2 | 14:1790 | 2.77 | 1.00 | 30.10 |  |
| 5 | G | N1 | 14:1828 | 3.63 | 1.00 | 27.48 |  |
| 6 | ARG | N | 19:222 | 3.85 | 1.00 | 26.30 |  |
| 7 | A | O5' | 14:1789 | 3.94 | 1.00 | 31.52 |  |
| 8 | C | OP1 | 14:1790 | 3.97 | 1.00 | 30.22 |  |
| 9 | A | OP2 | 14:1789 | 4.22 | 1.00 | 32.89 |  |
| 10 | VAL | N | 19:221 | 4.46 | 1.00 | 28.64 |  |
| 11 | A | O3' | 14:1789 | 4.47 | 1.00 | 28.40 |  |
| 12 | C | N4 | 14:1827 | 4.54 | 1.00 | 30.42 |  |
| 13 | ARG | NE | 19:222 | 4.60 | 1.00 | 26.81 |  |
| 14 | HIS | N | 19:220 | 4.91 | 1.00 | 29.83 |  |
  |

  
**1^st^ shell ligand angles:**   
1-2: 173; 1-3: 90; 1-4: 103; 2-3: 86; 2-4: 84; 3-4: 103;   
  
 Contact(s): /O6/O_back_/OP1/OP2/  
 Coded contact list: 2O_ph_O_b_O_back_  

### K:14:3032

|  | Res. type | Atom | Residue | Distance | Occupancy | B-factor | Closest solute to 1^st^ shell water |
| --- | --- | --- | --- | --- | --- | --- | --- |
| **Ion** | | | |  | 1.00 | 53.89 || 1 | U | O2 | 14:2022 | 2.58 | 1.00 | 52.05 |
| 2 | C | O2 | 14:2616 | 2.80 | 1.00 | 49.16 |  |
| 3 | C | O2' | 14:2616 | 2.89 | 1.00 | 45.27 |  |
| 4 | C | O4' | 14:2617 | 3.34 | 1.00 | 43.88 |  |
| 5 | U | O2' | 14:2022 | 3.45 | 1.00 | 54.91 |  |
| 6 | G | N2 | 14:2053 | 3.89 | 1.00 | 39.34 |  |
| 7 | C | N1 | 14:2616 | 4.33 | 1.00 | 42.52 |  |
| 8 | U | N3 | 14:2022 | 4.53 | 1.00 | 47.50 |  |
| 9 | U | N1 | 14:2022 | 4.54 | 1.00 | 49.41 |  |
| 10 | U | O4 | 14:2034 | 4.75 | 1.00 | 45.49 |  |
| 11 | C | N1 | 14:2617 | 4.81 | 1.00 | 39.81 |  |
| 12 | A | N3 | 14:2054 | 4.84 | 1.00 | 41.38 |  |
| 13 | C | O4' | 14:2616 | 4.97 | 1.00 | 41.33 |  |
  |

  
**1^st^ shell ligand angles:**   
1-2: 129; 1-3: 64; 1-4: 131; 2-3: 80; 2-4: 65; 3-4: 76;   
  
 Contact(s): /O2/O2'/O4'/  
 Coded contact list: 2O_r_O_b_  
 Exclusion zone contact(s): /O2/  

### K:14:3033

|  | Res. type | Atom | Residue | Distance | Occupancy | B-factor | Closest solute to 1^st^ shell water |
| --- | --- | --- | --- | --- | --- | --- | --- |
| **Ion** | | | |  | 1.00 | 52.89 || 1 | G | O6 | 14:1840 | 2.42 | 1.00 | 45.52 |
| 2 | G | O6 | 14:1899 | 2.75 | 1.00 | 44.16 |  |
| 3 | C | O2 | 14:1838 | 2.88 | 1.00 | 48.72 |  |
| 4 | C | O4' | 14:1838 | 3.16 | 1.00 | 46.38 |  |
| 5 | G | O6 | 14:1839 | 3.19 | 1.00 | 58.31 |  |
| 6 | G | N7 | 14:1839 | 3.78 | 1.00 | 57.08 |  |
| 7 | G | N1 | 14:1899 | 3.80 | 1.00 | 44.25 |  |
| 8 | C | N1 | 14:1838 | 4.06 | 1.00 | 47.62 |  |
| 9 | G | N7 | 14:1840 | 4.17 | 1.00 | 46.28 |  |
| 10 | C | N4 | 14:1902 | 4.25 | 1.00 | 42.46 |  |
| 11 | C | N3 | 14:1838 | 4.29 | 1.00 | 44.69 |  |
| 12 | A | N6 | 14:1901 | 4.30 | 1.00 | 43.25 |  |
| 13 | G | N2 | 14:1903 | 4.35 | 1.00 | 49.25 |  |
| 14 | U | O4 | 14:1841 | 4.45 | 1.00 | 48.62 |  |
| 15 | G | N1 | 14:1840 | 4.65 | 1.00 | 41.31 |  |
| 16 | C | O2 | 14:1837 | 4.67 | 1.00 | 53.00 |  |
| 17 | G | N1 | 14:1839 | 4.92 | 1.00 | 50.45 |  |
  |

  
**1^st^ shell ligand angles:**   
1-2: 127; 1-3: 102; 1-4: 146; 1-5: 76; 2-3: 88; 2-4: 83; 2-5: 106; 3-4: 59; 3-5: 164; 4-5: 114;   
  
 Contact(s): /O2/O4'/O6/O6/  
 Coded contact list: O_r_3O_b_  
 Exclusion zone contact(s): /O6/  

### K:14:3034

|  | Res. type | Atom | Residue | Distance | Occupancy | B-factor | Closest solute to 1^st^ shell water |
| --- | --- | --- | --- | --- | --- | --- | --- |
| **Ion** | | | |  | 1.00 | 86.90 || 1 | G | O6 | 14:327 | 2.65 | 1.00 | 71.66 |
| 2 | A | O2' | 14:332 | 3.31 | 1.00 | 61.41 |  |
| 3 | G | O6 | 14:326 | 3.57 | 1.00 | 64.31 |  |
| 4 | G | N7 | 14:326 | 3.82 | 1.00 | 66.20 |  |
| 5 | G | N7 | 14:327 | 3.94 | 1.00 | 69.60 |  |
| 6 | G | N1 | 14:327 | 4.79 | 1.00 | 62.37 |  |
| 7 | C | N4 | 14:335 | 4.81 | 1.00 | 63.34 |  |
  |

  
**1^st^ shell ligand angles:**   
1-2: 75;   
  
 Contact(s): /O2'/O6/  
 Coded contact list: O_r_O_b_  

### K:14:3035

|  | Res. type | Atom | Residue | Distance | Occupancy | B-factor | Closest solute to 1^st^ shell water |
| --- | --- | --- | --- | --- | --- | --- | --- |
| **Ion** | | | |  | 1.00 | 83.36 || 1 | G | O6 | 14:2852 | 2.62 | 1.00 | 71.06 |
| 2 | U | O4 | 14:2866 | 3.15 | 1.00 | 69.93 |  |
| 3 | A | OP2 | 14:2851 | 3.23 | 1.00 | 64.01 |  |
| 4 | A | OP2 | 14:2850 | 3.27 | 1.00 | 70.57 |  |
| 5 | G | N7 | 14:2852 | 3.39 | 1.00 | 67.19 |  |
| 6 | U | O4 | 14:2865 | 3.42 | 1.00 | 75.11 |  |
| 7 | A | N7 | 14:2851 | 3.58 | 1.00 | 68.21 |  |
| 8 | C | N4 | 14:2853 | 3.95 | 1.00 | 69.68 |  |
| 9 | A | O5' | 14:2850 | 4.14 | 1.00 | 59.68 |  |
| 10 | G | O6 | 14:2864 | 4.51 | 1.00 | 74.65 |  |
| 11 | U | N3 | 14:2865 | 4.54 | 1.00 | 76.79 |  |
| 12 | A | O5' | 14:2851 | 4.63 | 1.00 | 64.87 |  |
| 13 | A | OP1 | 14:2850 | 4.70 | 1.00 | 71.94 |  |
| 14 | G | N1 | 14:2852 | 4.74 | 1.00 | 71.55 |  |
| 15 | U | N3 | 14:2866 | 4.80 | 1.00 | 72.67 |  |
| 16 | A | N6 | 14:2851 | 4.93 | 1.00 | 68.56 |  |
  |

  
**1^st^ shell ligand angles:**   
1-2: 66; 1-3: 160; 1-4: 109; 1-5: 63; 2-3: 114; 2-4: 50; 2-5: 104; 3-4: 83; 3-5: 99; 4-5: 151;   
  
 Contact(s): /N7/O4/O6/OP2/OP2/  
 Coded contact list: 2O_ph_2O_b_N_b_  

### K:14:3036

|  | Res. type | Atom | Residue | Distance | Occupancy | B-factor | Closest solute to 1^st^ shell water |
| --- | --- | --- | --- | --- | --- | --- | --- |
| **Ion** | | | |  | 1.00 | 41.19 || 1 | A | OP2 | 14:793 | 2.53 | 1.00 | 30.37 |
| 2 | C | OP1 | 14:678 | 2.83 | 1.00 | 36.89 |  |
| 3 | G | O6 | 14:776 | 3.15 | 1.00 | 38.73 |  |
| 4 | A | O3' | 14:2071 | 3.40 | 1.00 | 26.18 |  |
| 5 | A | OP1 | 14:793 | 3.74 | 1.00 | 30.31 |  |
| 6 | A | O2' | 14:2071 | 3.92 | 1.00 | 31.12 |  |
| 7 | G | OP1 | 14:2072 | 4.02 | 1.00 | 36.65 |  |
| 8 | C | OP2 | 14:678 | 4.07 | 1.00 | 32.31 |  |
| 9 | A | O3' | 14:677 | 4.26 | 1.00 | 31.07 |  |
| 10 | A | O5' | 14:793 | 4.28 | 1.00 | 41.99 |  |
| 11 | A | O2' | 14:793 | 4.34 | 1.00 | 37.59 |  |
| 12 | G | O3' | 14:792 | 4.88 | 1.00 | 26.56 |  |
| 13 | G | O5' | 14:2072 | 4.91 | 1.00 | 25.45 |  |
  |

  
**1^st^ shell ligand angles:**   
1-2: 162; 1-3: 84; 1-4: 76; 2-3: 90; 2-4: 87; 3-4: 84;   
  
 Contact(s): /O3'/O6/OP1/  
 Coded contact list: O_ph_O_r_O_b_  
 Exclusion zone contact(s): /OP2/  

### K:14:3037

|  | Res. type | Atom | Residue | Distance | Occupancy | B-factor | Closest solute to 1^st^ shell water |
| --- | --- | --- | --- | --- | --- | --- | --- |
| **Ion** | | | |  | 1.00 | 60.54 || 1 | G | O6 | 14:663 | 2.86 | 1.00 | 51.39 |
| 2 | G | O6 | 14:662 | 3.06 | 1.00 | 51.00 |  |
| 3 | U | O4 | 14:594 | 3.06 | 1.00 | 52.03 |  |
| 4 | G | O6 | 14:593 | 3.33 | 1.00 | 47.80 |  |
| 5 | C | N4 | 14:595 | 3.96 | 1.00 | 52.99 |  |
| 6 | C | N4 | 14:664 | 4.10 | 1.00 | 47.16 |  |
| 7 | G | N7 | 14:662 | 4.19 | 1.00 | 49.37 |  |
| 8 | U | N3 | 14:594 | 4.33 | 1.00 | 46.11 |  |
| 9 | C | N4 | 14:661 | 4.55 | 1.00 | 58.75 |  |
| 10 | G | N7 | 14:663 | 4.58 | 1.00 | 50.82 |  |
| 11 | G | N1 | 14:662 | 4.93 | 1.00 | 51.07 |  |
| 12 | G | N7 | 14:593 | 4.99 | 1.00 | 47.42 |  |
  |

  
**1^st^ shell ligand angles:**   
1-2: 64; 1-3: 75; 1-4: 54; 2-3: 91; 2-4: 116; 3-4: 59;   
  
 Contact(s): /O4/O6/O6/O6/  
 Coded contact list: 4O_b_  

### K:14:3038

|  | Res. type | Atom | Residue | Distance | Occupancy | B-factor | Closest solute to 1^st^ shell water |
| --- | --- | --- | --- | --- | --- | --- | --- |
| **Ion** | | | |  | 1.00 | 92.10 || 1 | G | N7 | 14:929 | 2.65 | 1.00 | 88.42 |
| 2 | C | O2' | 14:846 | 2.68 | 1.00 | 69.05 |  |
| 3 | G | O6 | 14:929 | 2.76 | 1.00 | 76.32 |  |
| 4 | C | O3' | 14:846 | 3.30 | 1.00 | 68.82 |  |
| 5 | U | OP2 | 14:847 | 4.14 | 1.00 | 66.11 |  |
| 6 | C | O2 | 14:846 | 4.16 | 1.00 | 74.64 |  |
| 7 | G | N1 | 14:929 | 4.46 | 1.00 | 77.08 |  |
| 8 | G | OP2 | 14:929 | 4.60 | 1.00 | 82.38 |  |
| 9 | G | N9 | 14:929 | 4.67 | 1.00 | 79.77 |  |
| 10 | U | O5' | 14:847 | 4.69 | 1.00 | 74.60 |  |
| 11 | C | N1 | 14:846 | 4.98 | 1.00 | 76.66 |  |
  |

  
**1^st^ shell ligand angles:**   
1-2: 168; 1-3: 68; 1-4: 133; 2-3: 115; 2-4: 53; 3-4: 66;   
  
 Contact(s): /N7/O2'/O3'/O6/  
 Coded contact list: 2O_r_O_b_N_b_  

### K:14:3039

|  | Res. type | Atom | Residue | Distance | Occupancy | B-factor | Closest solute to 1^st^ shell water |
| --- | --- | --- | --- | --- | --- | --- | --- |
| **Ion** | | | |  | 1.00 | 61.92 || 1 | G | N7 | 14:391 | 2.86 | 1.00 | 50.06 |
| 2 | A | N3 | 14:390 | 3.14 | 1.00 | 46.76 |  |
| 3 | G | O6 | 14:391 | 3.17 | 1.00 | 51.08 |  |
| 4 | G | O6 | 14:388 | 3.56 | 1.00 | 53.19 |  |
| 5 | U | O4 | 14:383 | 3.58 | 1.00 | 54.01 |  |
| 6 | A | O2' | 14:390 | 3.64 | 1.00 | 51.05 |  |
| 7 | C | O2 | 14:385 | 3.75 | 1.00 | 47.91 |  |
| 8 | C | N4 | 14:392 | 3.83 | 1.00 | 52.17 |  |
| 9 | G | N2 | 14:411 | 4.07 | 1.00 | 49.16 |  |
| 10 | G | O6 | 14:382 | 4.41 | 1.00 | 49.97 |  |
| 11 | A | N9 | 14:390 | 4.48 | 1.00 | 46.03 |  |
| 12 | U | N3 | 14:383 | 4.75 | 1.00 | 52.90 |  |
| 13 | A | O3' | 14:390 | 4.77 | 1.00 | 53.64 |  |
| 14 | G | N1 | 14:388 | 4.90 | 1.00 | 45.22 |  |
| 15 | C | N3 | 14:385 | 4.95 | 1.00 | 47.83 |  |
| 16 | C | O2' | 14:385 | 4.98 | 1.00 | 49.67 |  |
  |

  
**1^st^ shell ligand angles:**   
1-2: 96; 1-3: 60; 2-3: 109;   
  
 Contact(s): /N3_A_/N7/O6/  
 Coded contact list: O_b_2N_b_  

### K:14:3040

|  | Res. type | Atom | Residue | Distance | Occupancy | B-factor | Closest solute to 1^st^ shell water |
| --- | --- | --- | --- | --- | --- | --- | --- |
| **Ion** | | | |  | 1.00 | 93.42 || 1 | G | N7 | 14:1024 | 2.68 | 1.00 | 88.31 |
| 2 | U | OP2 | 14:1023 | 2.80 | 1.00 | 85.64 |  |
| 3 | G | O3' | 14:1022 | 2.86 | 1.00 | 78.36 |  |
| 4 | G | N3 | 14:1025 | 2.87 | 1.00 | 94.51 |  |
| 5 | G | O2' | 14:1022 | 2.89 | 1.00 | 82.35 |  |
| 6 | U | OP1 | 14:1026 | 3.18 | 1.00 | 92.77 |  |
| 7 | G | N2 | 14:1025 | 3.38 | 1.00 | 90.83 |  |
| 8 | G | OP2 | 14:1024 | 3.72 | 1.00 | 87.26 |  |
| 9 | G | N1 | 14:1025 | 3.96 | 1.00 | 87.14 |  |
| 10 | G | N9 | 14:1025 | 4.12 | 1.00 | 87.72 |  |
| 11 | G | O6 | 14:1024 | 4.16 | 1.00 | 85.26 |  |
| 12 | U | O5' | 14:1023 | 4.25 | 1.00 | 93.95 |  |
| 13 | G | N9 | 14:1024 | 4.58 | 1.00 | 89.16 |  |
| 14 | U | OP1 | 14:1023 | 4.65 | 1.00 | 82.30 |  |
| 15 | G | O2' | 14:1025 | 4.72 | 1.00 | 92.93 |  |
| 16 | G | O5' | 14:1024 | 4.84 | 1.00 | 87.34 |  |
| 17 | G | O3' | 14:1025 | 4.84 | 1.00 | 100.40 |  |
  |

  
**1^st^ shell ligand angles:**   
1-2: 102; 1-3: 69; 1-4: 99; 1-5: 65; 1-6: 130; 1-7: 113; 2-3: 53; 2-4: 155; 2-5: 105; 2-6: 58; 2-7: 137; 3-4: 151; 3-5: 54; 3-6: 111; 3-7: 117; 4-5: 97; 4-6: 97; 4-7: 42; 5-6: 157; 5-7: 70; 6-7: 111;   
  
 Contact(s): /N2/N3_G_/N7/O2'/O3'/OP1/OP2/  
 Coded contact list: 2O_ph_2O_r_2N_b_Anion  

### K:14:3041

|  | Res. type | Atom | Residue | Distance | Occupancy | B-factor | Closest solute to 1^st^ shell water |
| --- | --- | --- | --- | --- | --- | --- | --- |
| **Ion** | | | |  | 1.00 | 66.57 || 1 | THR | O | 35:32 | 3.08 | 1.00 | 43.27 |
| 2 | A | O3' | 14:941 | 3.38 | 1.00 | 46.43 |  |
| 3 | G | OP1 | 14:1191 | 3.50 | 1.00 | 52.34 |  |
| 4 | A | O2' | 14:941 | 3.72 | 1.00 | 51.61 |  |
| 5 | A | O5' | 14:941 | 3.97 | 1.00 | 49.69 |  |
| 6 | G | OP1 | 14:942 | 4.07 | 1.00 | 49.80 |  |
| 7 | THR | OG1 | 35:32 | 4.12 | 1.00 | 44.07 |  |
| 8 | G | O3' | 14:1190 | 4.25 | 1.00 | 45.62 |  |
| 9 | G | O2' | 14:940 | 4.62 | 1.00 | 51.94 |  |
| 10 | ARG | NH2 | 35:18 | 4.73 | 1.00 | 57.14 |  |
| 11 | G | O3' | 14:940 | 4.74 | 1.00 | 50.35 |  |
| 12 | A | O4' | 14:941 | 4.89 | 1.00 | 46.62 |  |
  |

  
**1^st^ shell ligand angles:**   
1-2: 86;   
  
 Contact(s): /O3'/O_back_/  
 Coded contact list: O_r_O_back_  

### K:14:3042

|  | Res. type | Atom | Residue | Distance | Occupancy | B-factor | Closest solute to 1^st^ shell water |
| --- | --- | --- | --- | --- | --- | --- | --- |
| **Ion** | | | |  | 1.00 | 51.64 || 1 | A | OP1 | 14:1981 | 3.00 | 1.00 | 36.63 |
| 2 | A | N7 | 14:1981 | 3.04 | 1.00 | 38.14 |  |
| 3 | A | OP2 | 14:1785 | 3.35 | 1.00 | 35.28 |  |
| 4 | A | O3' | 14:1784 | 3.43 | 1.00 | 38.78 |  |
| 5 | Water 1 | | 14:3768 | 3.50 | 1.00 | 40.80 | 2.69 (A:OP1 14:1981) |
| 6 | A | O2' | 14:1784 | 3.83 | 1.00 | 41.37 |  |
| 7 | Water 2 | | 14:3994 | 3.98 | 1.00 | 39.10 | 4.59 (A:OP1 14:1981) |
| 8 | Water 3 | | 14:3711 | 4.14 | 1.00 | 42.79 | 2.61 (A:OP1 14:1981) |
| 9 | A | N6 | 14:1981 | 4.58 | 1.00 | 31.71 |  |
| 10 | A | O5' | 14:1981 | 4.68 | 1.00 | 38.95 |  |
| 11 | U | O4 | 14:1778 | 4.93 | 1.00 | 40.48 |  |
| 12 | A | N9 | 14:1981 | 4.94 | 1.00 | 36.74 |  |
  |

  
**1^st^ shell ligand angles:**   
1-2: 90; 1-3: 78; 2-3: 65;   
  
 Contact(s): /N7/OP1/OP2/  
 Coded contact list: 2O_ph_N_b_  

### K:14:3043

|  | Res. type | Atom | Residue | Distance | Occupancy | B-factor | Closest solute to 1^st^ shell water |
| --- | --- | --- | --- | --- | --- | --- | --- |
| **Ion** | | | |  | 1.00 | 80.73 || 1 | G | O6 | 14:558 | 2.66 | 1.00 | 71.58 |
| 2 | G | O6 | 14:559 | 2.75 | 1.00 | 65.24 |  |
| 3 | U | O4 | 14:534 | 2.77 | 1.00 | 64.01 |  |
| 4 | G | O6 | 14:533 | 2.85 | 1.00 | 67.15 |  |
| 5 | U | N3 | 14:534 | 3.88 | 1.00 | 60.16 |  |
| 6 | C | N4 | 14:535 | 3.99 | 1.00 | 62.49 |  |
| 7 | G | N7 | 14:559 | 4.40 | 1.00 | 69.04 |  |
| 8 | G | N1 | 14:533 | 4.49 | 1.00 | 59.27 |  |
| 9 | G | N1 | 14:558 | 4.58 | 1.00 | 60.36 |  |
| 10 | C | N4 | 14:560 | 4.62 | 1.00 | 67.55 |  |
| 11 | G | N7 | 14:533 | 4.69 | 1.00 | 60.66 |  |
| 12 | G | N7 | 14:558 | 4.79 | 1.00 | 69.38 |  |
| 13 | U | O4 | 14:557 | 4.92 | 1.00 | 72.41 |  |
| 14 | G | N1 | 14:559 | 4.94 | 1.00 | 61.60 |  |
  |

  
**1^st^ shell ligand angles:**   
1-2: 77; 1-3: 108; 1-4: 152; 2-3: 78; 2-4: 76; 3-4: 75;   
  
 Contact(s): /O4/O6/O6/O6/  
 Coded contact list: 4O_b_  

### K:14:3044

|  | Res. type | Atom | Residue | Distance | Occupancy | B-factor | Closest solute to 1^st^ shell water |
| --- | --- | --- | --- | --- | --- | --- | --- |
| **Ion** | | | |  | 1.00 | 69.39 || 1 | G | O6 | 14:220 | 2.91 | 1.00 | 57.93 |
| 2 | G | O6 | 14:219 | 2.94 | 1.00 | 58.15 |  |
| 3 | A | OP1 | 14:428 | 2.98 | 1.00 | 56.35 |  |
| 4 | A | OP2 | 14:218 | 3.40 | 1.00 | 45.53 |  |
| 5 | U | OP2 | 14:427 | 4.00 | 1.00 | 60.98 |  |
| 6 | G | N7 | 14:219 | 4.16 | 1.00 | 53.22 |  |
| 7 | U | OP1 | 14:427 | 4.21 | 1.00 | 59.78 |  |
| 8 | G | N1 | 14:220 | 4.36 | 1.00 | 57.05 |  |
| 9 | A | N7 | 14:218 | 4.37 | 1.00 | 50.16 |  |
| 10 | U | O5' | 14:427 | 4.59 | 1.00 | 56.25 |  |
| 11 | G | OP2 | 14:217 | 4.75 | 1.00 | 44.23 |  |
  |

  
**1^st^ shell ligand angles:**   
1-2: 62; 1-3: 72; 2-3: 73;   
  
 Contact(s): /O6/O6/OP1/  
 Coded contact list: O_ph_2O_b_  

### K:14:3045

|  | Res. type | Atom | Residue | Distance | Occupancy | B-factor | Closest solute to 1^st^ shell water |
| --- | --- | --- | --- | --- | --- | --- | --- |
| **Ion** | | | |  | 1.00 | 83.93 || 1 | G | O6 | 14:2290 | 2.99 | 1.00 | 72.14 |
| 2 | G | O6 | 14:2340 | 3.36 | 1.00 | 78.53 |  |
| 3 | U | O4 | 14:2291 | 3.37 | 1.00 | 65.31 |  |
| 4 | G | O6 | 14:2341 | 3.97 | 1.00 | 69.60 |  |
| 5 | G | N7 | 14:2340 | 4.02 | 1.00 | 80.82 |  |
| 6 | C | N4 | 14:2342 | 4.52 | 1.00 | 69.53 |  |
| 7 | G | O6 | 14:2289 | 4.80 | 1.00 | 70.54 |  |
| 8 | G | N7 | 14:2341 | 4.87 | 1.00 | 71.10 |  |
| 9 | U | N3 | 14:2291 | 4.88 | 1.00 | 68.70 |  |
  |

  
**1^st^ shell ligand angles:**   
1-2: 100; 1-3: 56; 2-3: 63;   
  
 Contact(s): /O4/O6/O6/  
 Coded contact list: 3O_b_  

### K:14:3046

|  | Res. type | Atom | Residue | Distance | Occupancy | B-factor | Closest solute to 1^st^ shell water |
| --- | --- | --- | --- | --- | --- | --- | --- |
| **Ion** | | | |  | 1.00 | 54.58 || 1 | U | O2' | 14:827 | 2.80 | 1.00 | 48.06 |
| 2 | U | O2 | 14:2068 | 3.12 | 1.00 | 46.76 |  |
| 3 | U | O2' | 14:2068 | 3.35 | 1.00 | 42.31 |  |
| 4 | U | O2 | 14:827 | 3.44 | 1.00 | 42.94 |  |
| 5 | U | OP1 | 14:2245 | 3.57 | 1.00 | 32.84 |  |
| 6 | A | N1 | 14:2430 | 4.15 | 1.00 | 40.10 |  |
| 7 | U | N1 | 14:2068 | 4.25 | 1.00 | 38.47 |  |
| 8 | G | O3' | 14:2246 | 4.26 | 1.00 | 40.27 |  |
| 9 | U | O4' | 14:2068 | 4.83 | 1.00 | 41.28 |  |
| 10 | U | N1 | 14:827 | 4.86 | 1.00 | 36.50 |  |
  |

  
**1^st^ shell ligand angles:**   
1-2: 70; 1-3: 108; 2-3: 71;   
  
 Contact(s): /O2/O2'/O2'/  
 Coded contact list: 2O_r_O_b_  

### K:14:3047

|  | Res. type | Atom | Residue | Distance | Occupancy | B-factor | Closest solute to 1^st^ shell water |
| --- | --- | --- | --- | --- | --- | --- | --- |
| **Ion** | | | |  | 1.00 | 74.56 || 1 | G | O6 | 14:939 | 2.95 | 1.00 | 51.57 |
| 2 | U | O4 | 14:839 | 3.03 | 1.00 | 50.03 |  |
| 3 | G | O6 | 14:938 | 3.09 | 1.00 | 52.85 |  |
| 4 | G | O6 | 14:940 | 3.22 | 1.00 | 53.50 |  |
| 5 | G | N7 | 14:939 | 3.98 | 1.00 | 54.00 |  |
| 6 | C | N4 | 14:838 | 4.31 | 1.00 | 53.62 |  |
| 7 | C | N4 | 14:840 | 4.48 | 1.00 | 58.95 |  |
| 8 | G | N7 | 14:938 | 4.57 | 1.00 | 56.63 |  |
| 9 | U | N3 | 14:839 | 4.59 | 1.00 | 56.26 |  |
| 10 | G | N1 | 14:938 | 4.79 | 1.00 | 56.36 |  |
| 11 | G | N7 | 14:940 | 4.99 | 1.00 | 51.43 |  |
  |

  
**1^st^ shell ligand angles:**   
1-2: 55; 1-3: 70; 1-4: 60; 2-3: 84; 2-4: 68; 3-4: 131;   
  
 Contact(s): /O4/O6/O6/O6/  
 Coded contact list: 4O_b_  

### K:14:3048

|  | Res. type | Atom | Residue | Distance | Occupancy | B-factor | Closest solute to 1^st^ shell water |
| --- | --- | --- | --- | --- | --- | --- | --- |
| **Ion** | | | |  | 1.00 | 67.81 || 1 | U | O4 | 14:147 | 2.86 | 1.00 | 46.41 |
| 2 | G | O6 | 14:131 | 3.13 | 1.00 | 44.16 |  |
| 3 | G | O6 | 14:146 | 3.17 | 1.00 | 56.59 |  |
| 4 | G | O6 | 14:132 | 3.17 | 1.00 | 47.12 |  |
| 5 | U | N3 | 14:147 | 3.92 | 1.00 | 46.31 |  |
| 6 | C | N4 | 14:133 | 3.99 | 1.00 | 48.81 |  |
| 7 | G | N7 | 14:132 | 4.16 | 1.00 | 42.20 |  |
| 8 | C | N4 | 14:148 | 4.19 | 1.00 | 38.93 |  |
| 9 | G | N1 | 14:131 | 4.59 | 1.00 | 37.86 |  |
| 10 | G | N7 | 14:131 | 4.84 | 1.00 | 35.21 |  |
  |

  
**1^st^ shell ligand angles:**   
1-2: 98; 1-3: 67; 1-4: 74; 2-3: 145; 2-4: 80; 3-4: 65;   
  
 Contact(s): /O4/O6/O6/O6/  
 Coded contact list: 4O_b_  

### K:14:3049

|  | Res. type | Atom | Residue | Distance | Occupancy | B-factor | Closest solute to 1^st^ shell water |
| --- | --- | --- | --- | --- | --- | --- | --- |
| **Ion** | | | |  | 1.00 | 49.75 || 1 | U | O4 | 14:1397 | 2.60 | 1.00 | 48.04 |
| 2 | U | O3' | 14:1394 | 2.83 | 1.00 | 42.05 |  |
| 3 | U | O2' | 14:1602 | 2.94 | 1.00 | 44.44 |  |
| 4 | A | OP1 | 14:1395 | 2.97 | 1.00 | 46.79 |  |
| 5 | A | OP1 | 14:1603 | 2.99 | 1.00 | 33.29 |  |
| 6 | U | O3' | 14:1602 | 3.11 | 1.00 | 42.22 |  |
| 7 | A | O5' | 14:1395 | 3.24 | 1.00 | 46.55 |  |
| 8 | U | O2' | 14:1394 | 3.67 | 1.00 | 41.21 |  |
| 9 | Water 1 | | 14:3712 | 3.69 | 1.00 | 32.12 | 2.61 (A:OP1 14:1603) |
| 10 | A | O5' | 14:1603 | 4.15 | 1.00 | 40.72 |  |
| 11 | U | N3 | 14:1397 | 4.39 | 1.00 | 44.36 |  |
| 12 | C | OP2 | 14:1604 | 4.78 | 1.00 | 31.83 |  |
| 13 | A | OP2 | 14:1395 | 4.79 | 1.00 | 35.30 |  |
| 14 | U | O2 | 14:1602 | 4.86 | 1.00 | 44.82 |  |
| 15 | A | OP2 | 14:1603 | 4.98 | 1.00 | 39.52 |  |
  |

  
**1^st^ shell ligand angles:**   
1-2: 120; 1-3: 80; 1-4: 138; 1-5: 144; 1-6: 108; 1-7: 95; 2-3: 111; 2-4: 49; 2-5: 95; 2-6: 125; 2-7: 49; 3-4: 141; 3-5: 95; 3-6: 52; 3-7: 152; 4-5: 61; 4-6: 108; 4-7: 45; 5-6: 47; 5-7: 105; 6-7: 152;   
  
 Contact(s): /O2'/O3'/O3'/O4/O5'/OP1/OP1/  
 Coded contact list: 2O_ph_4O_r_O_b_  

### K:14:3050

|  | Res. type | Atom | Residue | Distance | Occupancy | B-factor | Closest solute to 1^st^ shell water |
| --- | --- | --- | --- | --- | --- | --- | --- |
| **Ion** | | | |  | 1.00 | 63.62 || 1 | U | OP2 | 14:1300 | 2.86 | 1.00 | 48.04 |
| 2 | G | O6 | 14:1628 | 2.95 | 1.00 | 48.45 |  |
| 3 | G | O6 | 14:1627 | 3.22 | 1.00 | 44.15 |  |
| 4 | G | O3' | 14:1299 | 3.34 | 1.00 | 49.26 |  |
| 5 | G | N7 | 14:1627 | 3.63 | 1.00 | 47.12 |  |
| 6 | G | N7 | 14:1628 | 3.65 | 1.00 | 47.84 |  |
| 7 | U | OP1 | 14:1300 | 4.46 | 1.00 | 36.20 |  |
| 8 | C | N4 | 14:1638 | 4.57 | 1.00 | 42.61 |  |
| 9 | U | O4 | 14:1639 | 4.70 | 1.00 | 48.22 |  |
| 10 | U | N3 | 14:1639 | 4.71 | 1.00 | 42.88 |  |
| 11 | G | O2' | 14:1299 | 4.80 | 1.00 | 41.64 |  |
| 12 | G | N1 | 14:1627 | 4.81 | 1.00 | 43.58 |  |
| 13 | G | OP2 | 14:1627 | 4.91 | 1.00 | 50.57 |  |
| 14 | U | O5' | 14:1300 | 4.92 | 1.00 | 36.67 |  |
  |

  
**1^st^ shell ligand angles:**   
1-2: 113; 1-3: 129; 1-4: 46; 2-3: 72; 2-4: 116; 3-4: 85;   
  
 Contact(s): /O3'/O6/O6/OP2/  
 Coded contact list: O_ph_O_r_2O_b_  

### K:14:3051

|  | Res. type | Atom | Residue | Distance | Occupancy | B-factor | Closest solute to 1^st^ shell water |
| --- | --- | --- | --- | --- | --- | --- | --- |
| **Ion** | | | |  | 1.00 | 65.57 || 1 | G | O6 | 14:16 | 2.63 | 1.00 | 64.06 |
| 2 | G | O6 | 14:15 | 3.01 | 1.00 | 65.66 |  |
| 3 | U | O4 | 14:525 | 3.10 | 1.00 | 68.54 |  |
| 4 | A | O2' | 14:13 | 3.23 | 1.00 | 65.70 |  |
| 5 | U | O4 | 14:524 | 3.42 | 1.00 | 66.55 |  |
| 6 | G | N7 | 14:16 | 4.01 | 1.00 | 63.10 |  |
| 7 | G | N7 | 14:15 | 4.30 | 1.00 | 60.96 |  |
| 8 | U | N3 | 14:524 | 4.36 | 1.00 | 68.88 |  |
| 9 | U | N3 | 14:525 | 4.38 | 1.00 | 65.95 |  |
| 10 | G | N1 | 14:15 | 4.74 | 1.00 | 63.50 |  |
| 11 | G | N1 | 14:16 | 4.92 | 1.00 | 65.05 |  |
| 12 | G | O6 | 14:17 | 4.92 | 1.00 | 66.73 |  |
  |

  
**1^st^ shell ligand angles:**   
1-2: 70; 1-3: 62; 1-4: 119; 2-3: 54; 2-4: 49; 3-4: 85;   
  
 Contact(s): /O2'/O4/O6/O6/  
 Coded contact list: O_r_3O_b_  

### K:14:3052

|  | Res. type | Atom | Residue | Distance | Occupancy | B-factor | Closest solute to 1^st^ shell water |
| --- | --- | --- | --- | --- | --- | --- | --- |
| **Ion** | | | |  | 1.00 | 76.70 || 1 | G | N1 | 14:1950 | 2.84 | 1.00 | 59.21 |
| 2 | U | O4 | 14:1951 | 2.93 | 1.00 | 60.26 |  |
| 3 | G | O6 | 14:1950 | 3.32 | 1.00 | 52.89 |  |
| 4 | G | N2 | 14:1950 | 3.80 | 1.00 | 57.49 |  |
| 5 | G | OP2 | 14:1954 | 3.82 | 1.00 | 46.82 |  |
| 6 | U | N3 | 14:1951 | 4.30 | 1.00 | 49.76 |  |
| 7 | G | OP1 | 14:1954 | 4.75 | 1.00 | 58.21 |  |
  |

  
**1^st^ shell ligand angles:**   
1-2: 56; 1-3: 43; 2-3: 80;   
  
 Contact(s): /N1_G_/O4/O6/  
 Coded contact list: 2O_b_Anion  

### K:14:3053

|  | Res. type | Atom | Residue | Distance | Occupancy | B-factor | Closest solute to 1^st^ shell water |
| --- | --- | --- | --- | --- | --- | --- | --- |
| **Ion** | | | |  | 1.00 | 102.84 || 1 | G | O6 | 14:656 | 2.93 | 1.00 | 62.77 |
| 2 | U | O4 | 14:657 | 3.53 | 1.00 | 59.62 |  |
| 3 | A | OP2 | 14:655 | 3.56 | 1.00 | 102.62 |  |
| 4 | G | O6 | 14:599 | 3.74 | 1.00 | 66.24 |  |
| 5 | C | N4 | 14:601 | 4.07 | 1.00 | 65.54 |  |
| 6 | G | N7 | 14:656 | 4.21 | 1.00 | 69.11 |  |
| 7 | A | OP1 | 14:655 | 4.31 | 1.00 | 106.42 |  |
| 8 | G | O6 | 14:600 | 4.43 | 1.00 | 58.69 |  |
| 9 | U | N3 | 14:657 | 4.52 | 1.00 | 57.92 |  |
| 10 | G | N1 | 14:602 | 4.83 | 1.00 | 77.46 |  |
  |

  
 Contact(s): /O6/  
 Coded contact list: O_b_  

### K:14:3054

|  | Res. type | Atom | Residue | Distance | Occupancy | B-factor | Closest solute to 1^st^ shell water |
| --- | --- | --- | --- | --- | --- | --- | --- |
| **Ion** | | | |  | 1.00 | 58.19 || 1 | Water 1 | | 14:3516 | 2.42 | 1.00 | 40.81 | 1.95 (C:OP1 14:453) |
| 2 | G | O3' | 14:452 | 2.87 | 1.00 | 33.71 |  |
| 3 | A | OP1 | 14:471 | 2.94 | 1.00 | 31.79 |  |
| 4 | C | OP1 | 14:453 | 3.43 | 1.00 | 33.90 |  |
| 5 | Water 2 | | 14:3856 | 3.67 | 1.00 | 49.08 | 3.67 (C:OP1 14:453) |
| 6 | TYR | OH | 39:59 | 3.75 | 1.00 | 40.23 |  |
| 7 | A | O3' | 14:470 | 3.85 | 1.00 | 36.63 |  |
| 8 | G | O2' | 14:452 | 3.95 | 1.00 | 33.15 |  |
| 9 | A | OP2 | 14:471 | 4.31 | 1.00 | 38.36 |  |
| 10 | A | OP1 | 14:470 | 4.51 | 1.00 | 33.07 |  |
| 11 | Water 3 | | 14:3521 | 4.51 | 1.00 | 39.35 | 2.00 (C:OP1 14:453) |
| 12 | Water 4 | | 14:3922 | 4.68 | 1.00 | 30.96 | 4.37 (C:OP1 14:453) |
| 13 | C | O5' | 14:453 | 4.70 | 1.00 | 43.63 |  |
| 14 | Water 5 | | 14:4047 | 4.86 | 1.00 | 45.76 | 3.73 (A:OP1 14:471) |
| 15 | A | O5' | 14:470 | 4.89 | 1.00 | 32.04 |  |
| 16 | C | OP2 | 14:453 | 4.92 | 1.00 | 36.03 |  |
  |

  
**1^st^ shell ligand angles:**   
1-2: 74; 1-3: 72; 2-3: 135;   
  
 Contact(s): /O3'/OP1/  
 Coded contact list: O_ph_O_r_  
 Exclusion zone contact(s): /O_w_/  
K hydration shell issue: water too close (< 2.4 Angs.)  

### K:14:3055

|  | Res. type | Atom | Residue | Distance | Occupancy | B-factor | Closest solute to 1^st^ shell water |
| --- | --- | --- | --- | --- | --- | --- | --- |
| **Ion** | | | |  | 1.00 | 65.71 || 1 | C | OP2 | 14:2573 | 3.07 | 1.00 | 52.31 |
| 2 | U | OP1 | 14:2506 | 3.21 | 1.00 | 55.83 |  |
| 3 | A | O2' | 14:2572 | 3.24 | 1.00 | 54.33 |  |
| 4 | A | N3 | 14:2572 | 3.27 | 1.00 | 41.89 |  |
| 5 | A | O4' | 14:2453 | 3.28 | 1.00 | 44.83 |  |
| 6 | A | O3' | 14:2572 | 3.76 | 1.00 | 43.70 |  |
| 7 | A | O2' | 14:2453 | 3.95 | 1.00 | 44.87 |  |
| 8 | U | O5' | 14:2506 | 4.01 | 1.00 | 50.79 |  |
| 9 | ARG | NH2 | 29:144 | 4.18 | 1.00 | 43.88 |  |
| 10 | A | N9 | 14:2572 | 4.23 | 1.00 | 43.19 |  |
| 11 | C | O2' | 14:2452 | 4.67 | 1.00 | 56.19 |  |
| 12 | C | OP1 | 14:2573 | 4.75 | 1.00 | 47.05 |  |
| 13 | A | O4' | 14:2572 | 4.76 | 1.00 | 45.91 |  |
| 14 | C | O2 | 14:2452 | 4.90 | 1.00 | 51.35 |  |
| 15 | U | OP2 | 14:2506 | 4.92 | 1.00 | 55.32 |  |
  |

  
**1^st^ shell ligand angles:**   
1-2: 131; 1-3: 72; 1-4: 135; 1-5: 83; 2-3: 105; 2-4: 90; 2-5: 121; 3-4: 81; 3-5: 133; 4-5: 90;   
  
 Contact(s): /N3_A_/O2'/O4'/OP1/OP2/  
 Coded contact list: 2O_ph_2O_r_N_b_  

### K:14:3056

|  | Res. type | Atom | Residue | Distance | Occupancy | B-factor | Closest solute to 1^st^ shell water |
| --- | --- | --- | --- | --- | --- | --- | --- |
| **Ion** | | | |  | 1.00 | 56.10 || 1 | U | O4 | 14:2698 | 2.83 | 1.00 | 56.39 |
| 2 | G | O6 | 14:2709 | 2.86 | 1.00 | 44.58 |  |
| 3 | G | O6 | 14:2708 | 2.87 | 1.00 | 48.65 |  |
| 4 | G | O6 | 14:2697 | 3.15 | 1.00 | 54.09 |  |
| 5 | C | N4 | 14:2710 | 3.75 | 1.00 | 48.73 |  |
| 6 | G | N7 | 14:2709 | 3.86 | 1.00 | 46.09 |  |
| 7 | U | N3 | 14:2698 | 3.98 | 1.00 | 42.51 |  |
| 8 | G | N7 | 14:2708 | 4.24 | 1.00 | 48.34 |  |
| 9 | C | N4 | 14:2699 | 4.40 | 1.00 | 43.77 |  |
| 10 | G | N1 | 14:2708 | 4.54 | 1.00 | 41.66 |  |
| 11 | U | O4 | 14:2696 | 4.75 | 1.00 | 59.53 |  |
| 12 | G | N1 | 14:2697 | 4.97 | 1.00 | 47.54 |  |
  |

  
**1^st^ shell ligand angles:**   
1-2: 67; 1-3: 81; 1-4: 67; 2-3: 71; 2-4: 69; 3-4: 136;   
  
 Contact(s): /O4/O6/O6/O6/  
 Coded contact list: 4O_b_  

### K:14:3057

|  | Res. type | Atom | Residue | Distance | Occupancy | B-factor | Closest solute to 1^st^ shell water |
| --- | --- | --- | --- | --- | --- | --- | --- |
| **Ion** | | | |  | 1.00 | 54.72 || 1 | U | O4 | 14:2580 | 2.79 | 1.00 | 47.19 |
| 2 | G | O6 | 14:2508 | 2.93 | 1.00 | 47.98 |  |
| 3 | G | N7 | 14:2576 | 3.58 | 1.00 | 49.79 |  |
| 4 | C | N4 | 14:2507 | 3.94 | 1.00 | 46.99 |  |
| 5 | G | O6 | 14:2509 | 3.95 | 1.00 | 45.82 |  |
| 6 | U | N3 | 14:2580 | 4.14 | 1.00 | 37.49 |  |
| 7 | G | N7 | 14:2508 | 4.15 | 1.00 | 41.91 |  |
| 8 | G | N9 | 14:2576 | 4.25 | 1.00 | 42.13 |  |
| 9 | C | N4 | 14:2579 | 4.70 | 1.00 | 43.12 |  |
| 10 | G | N7 | 14:2581 | 4.78 | 1.00 | 43.58 |  |
| 11 | C | OP2 | 14:2507 | 4.91 | 1.00 | 64.62 |  |
| 12 | G | O4' | 14:2576 | 4.96 | 1.00 | 44.71 |  |
  |

  
**1^st^ shell ligand angles:**   
1-2: 71;   
  
 Contact(s): /O4/O6/  
 Coded contact list: 2O_b_  

### K:14:3058

|  | Res. type | Atom | Residue | Distance | Occupancy | B-factor | Closest solute to 1^st^ shell water |
| --- | --- | --- | --- | --- | --- | --- | --- |
| **Ion** | | | |  | 1.00 | 85.16 || 1 | A | OP1 | 14:1126 | 2.72 | 1.00 | 70.86 |
| 2 | C | O2 | 14:2464 | 3.16 | 1.00 | 68.13 |  |
| 3 | C | O2' | 14:2464 | 3.53 | 1.00 | 65.13 |  |
| 4 | A | OP2 | 14:1126 | 3.58 | 1.00 | 68.66 |  |
| 5 | G | N2 | 14:2486 | 3.93 | 1.00 | 68.90 |  |
| 6 | A | N6 | 14:1028 | 4.16 | 1.00 | 71.48 |  |
| 7 | G | N2 | 14:2487 | 4.21 | 1.00 | 71.37 |  |
| 8 | C | O4' | 14:2465 | 4.30 | 1.00 | 75.39 |  |
| 9 | G | N3 | 14:2487 | 4.37 | 1.00 | 64.55 |  |
| 10 | G | O3' | 14:1125 | 4.46 | 1.00 | 75.55 |  |
| 11 | G | O2' | 14:2487 | 4.57 | 1.00 | 55.29 |  |
| 12 | A | N1 | 14:1028 | 4.62 | 1.00 | 66.62 |  |
| 13 | C | N1 | 14:2464 | 4.70 | 1.00 | 66.89 |  |
| 14 | A | O5' | 14:1126 | 4.80 | 1.00 | 74.11 |  |
| 15 | A | O4' | 14:2488 | 4.89 | 1.00 | 60.93 |  |
  |

  
**1^st^ shell ligand angles:**   
1-2: 157;   
  
 Contact(s): /O2/OP1/  
 Coded contact list: O_ph_O_b_  

### K:14:3059

|  | Res. type | Atom | Residue | Distance | Occupancy | B-factor | Closest solute to 1^st^ shell water |
| --- | --- | --- | --- | --- | --- | --- | --- |
| **Ion** | | | |  | 1.00 | 48.52 || 1 | A | N3 | 14:1936 | 2.67 | 1.00 | 48.74 |
| 2 | 5MC | O2 | 14:1962 | 2.73 | 1.00 | 43.09 |  |
| 3 | U | O4 | 14:1940 | 2.73 | 1.00 | 50.35 |  |
| 4 | 5MC | N3 | 14:1962 | 2.77 | 1.00 | 44.06 |  |
| 5 | A | O5' | 14:1936 | 3.46 | 1.00 | 42.29 |  |
| 6 | A | O4' | 14:1936 | 3.62 | 1.00 | 48.81 |  |
| 7 | C | N4 | 14:1961 | 3.62 | 1.00 | 47.60 |  |
| 8 | U | N3 | 14:1940 | 3.69 | 1.00 | 46.07 |  |
| 9 | A | OP2 | 14:1936 | 3.96 | 1.00 | 43.19 |  |
| 10 | G | O3' | 14:1935 | 4.08 | 1.00 | 45.23 |  |
| 11 | A | N1 | 14:1936 | 4.22 | 1.00 | 47.64 |  |
| 12 | G | O2' | 14:1935 | 4.36 | 1.00 | 45.42 |  |
| 13 | G | N3 | 14:1935 | 4.41 | 1.00 | 46.54 |  |
| 14 | A | N9 | 14:1936 | 4.47 | 1.00 | 51.35 |  |
| 15 | G | N2 | 14:1935 | 4.49 | 1.00 | 45.99 |  |
| 16 | 5MC | N4 | 14:1962 | 4.56 | 1.00 | 48.69 |  |
| 17 | 5MC | N1 | 14:1962 | 4.63 | 1.00 | 46.77 |  |
| 18 | C | N3 | 14:1961 | 4.79 | 1.00 | 47.31 |  |
| 19 | G | O6 | 14:1945 | 4.88 | 1.00 | 44.52 |  |
  |

  
**1^st^ shell ligand angles:**   
1-2: 99; 1-3: 101; 1-4: 126; 2-3: 93; 2-4: 48; 3-4: 119;   
  
 Contact(s): /N3/N3_A_/O2/O4/  
 Coded contact list: 2O_b_N_b_  

### K:14:3060

|  | Res. type | Atom | Residue | Distance | Occupancy | B-factor | Closest solute to 1^st^ shell water |
| --- | --- | --- | --- | --- | --- | --- | --- |
| **Ion** | | | |  | 1.00 | 45.97 || 1 | U | O4 | 14:1898 | 2.74 | 1.00 | 44.69 |
| 2 | G | O6 | 14:1842 | 2.82 | 1.00 | 43.81 |  |
| 3 | U | O4 | 14:1841 | 2.83 | 1.00 | 48.62 |  |
| 4 | Water 1 | | 14:3928 | 3.00 | 1.00 | 49.57 | 3.10 (U:O4 14:1898) |
| 5 | C | O2 | 14:1838 | 3.17 | 1.00 | 48.72 |  |
| 6 | U | N3 | 14:1898 | 3.75 | 1.00 | 42.40 |  |
| 7 | G | O6 | 14:1897 | 3.92 | 1.00 | 44.66 |  |
| 8 | G | N7 | 14:1842 | 4.11 | 1.00 | 41.70 |  |
| 9 | A | N6 | 14:1901 | 4.23 | 1.00 | 43.25 |  |
| 10 | C | N3 | 14:1838 | 4.73 | 1.00 | 44.69 |  |
| 11 | U | N3 | 14:1841 | 4.75 | 1.00 | 44.94 |  |
| 12 | G | OP2 | 14:1840 | 4.77 | 1.00 | 48.25 |  |
| 13 | G | N1 | 14:1842 | 4.84 | 1.00 | 40.28 |  |
| 14 | C | N1 | 14:1838 | 4.89 | 1.00 | 47.62 |  |
| 15 | C | N4 | 14:1843 | 4.94 | 1.00 | 45.18 |  |
  |

  
**1^st^ shell ligand angles:**   
1-2: 76; 1-3: 136; 1-4: 65; 1-5: 65; 2-3: 91; 2-4: 129; 2-5: 98; 3-4: 139; 3-5: 76; 4-5: 94;   
  
 Contact(s): /O2/O4/O4/O6/O_w_/  
 Coded contact list: 4O_b_O_w_  

### K:14:3061

|  | Res. type | Atom | Residue | Distance | Occupancy | B-factor | Closest solute to 1^st^ shell water |
| --- | --- | --- | --- | --- | --- | --- | --- |
| **Ion** | | | |  | 1.00 | 63.49 || 1 | G | O6 | 14:1975 | 2.96 | 1.00 | 33.88 |
| 2 | U | O4 | 14:1976 | 3.38 | 1.00 | 33.91 |  |
| 3 | Water 1 | | 14:3670 | 3.80 | 1.00 | 32.87 | 2.53 (A:OP2 14:1773) |
| 4 | A | OP2 | 14:1773 | 4.02 | 1.00 | 41.58 |  |
| 5 | G | N7 | 14:1975 | 4.25 | 1.00 | 33.51 |  |
| 6 | C | N4 | 14:1830 | 4.41 | 1.00 | 30.31 |  |
| 7 | Water 2 | | 14:3684 | 4.65 | 1.00 | 40.17 | 3.76 (A:OP2 14:1773) |
| 8 | A | OP2 | 14:1829 | 4.69 | 1.00 | 38.30 |  |
| 9 | C | N4 | 14:1974 | 4.71 | 1.00 | 28.89 |  |
| 10 | A | O5' | 14:1773 | 4.76 | 1.00 | 35.17 |  |
| 11 | G | OP2 | 14:1772 | 4.82 | 1.00 | 41.62 |  |
| 12 | U | N3 | 14:1976 | 4.94 | 1.00 | 32.59 |  |
  |

  
**1^st^ shell ligand angles:**   
1-2: 61;   
  
 Contact(s): /O4/O6/  
 Coded contact list: 2O_b_  

### K:14:3062

|  | Res. type | Atom | Residue | Distance | Occupancy | B-factor | Closest solute to 1^st^ shell water |
| --- | --- | --- | --- | --- | --- | --- | --- |
| **Ion** | | | |  | 1.00 | 76.61 || 1 | G | O6 | 14:2569 | 3.28 | 1.00 | 58.08 |
| 2 | G | O6 | 14:2513 | 3.56 | 1.00 | 49.16 |  |
| 3 | G | O6 | 14:2570 | 3.98 | 1.00 | 50.59 |  |
| 4 | U | O4 | 14:2514 | 3.99 | 1.00 | 55.04 |  |
| 5 | C | N4 | 14:2571 | 4.10 | 1.00 | 46.11 |  |
| 6 | G | N7 | 14:2569 | 4.54 | 1.00 | 57.10 |  |
| 7 | G | N7 | 14:2570 | 4.66 | 1.00 | 46.37 |  |
| 8 | C | N4 | 14:2515 | 4.93 | 1.00 | 55.25 |  |
  |

  
 Contact(s): /O6/  
 Coded contact list: O_b_  

### K:14:3063

|  | Res. type | Atom | Residue | Distance | Occupancy | B-factor | Closest solute to 1^st^ shell water |
| --- | --- | --- | --- | --- | --- | --- | --- |
| **Ion** | | | |  | 1.00 | 57.35 || 1 | G | O6 | 14:2722 | 2.82 | 1.00 | 55.63 |
| 2 | U | OP2 | 14:2720 | 2.94 | 1.00 | 48.94 |  |
| 3 | A | OP2 | 14:2721 | 3.04 | 1.00 | 52.90 |  |
| 4 | U | O5' | 14:2720 | 3.43 | 1.00 | 54.36 |  |
| 5 | A | N7 | 14:2721 | 3.49 | 1.00 | 56.66 |  |
| 6 | U | OP1 | 14:2720 | 3.59 | 1.00 | 53.18 |  |
| 7 | U | O4 | 14:2687 | 3.83 | 1.00 | 57.59 |  |
| 8 | G | N7 | 14:2722 | 4.19 | 1.00 | 57.73 |  |
| 9 | U | N3 | 14:2687 | 4.20 | 1.00 | 54.55 |  |
| 10 | G | O6 | 14:2686 | 4.43 | 1.00 | 60.25 |  |
| 11 | A | O5' | 14:2721 | 4.69 | 1.00 | 58.39 |  |
| 12 | U | N1 | 14:2720 | 4.82 | 1.00 | 55.78 |  |
| 13 | U | O3' | 14:2720 | 4.85 | 1.00 | 52.85 |  |
| 14 | G | N1 | 14:2722 | 4.87 | 1.00 | 52.92 |  |
  |

  
**1^st^ shell ligand angles:**   
1-2: 128; 1-3: 120; 2-3: 108;   
  
 Contact(s): /O6/OP2/OP2/  
 Coded contact list: 2O_ph_O_b_  

### K:14:3064

|  | Res. type | Atom | Residue | Distance | Occupancy | B-factor | Closest solute to 1^st^ shell water |
| --- | --- | --- | --- | --- | --- | --- | --- |
| **Ion** | | | |  | 1.00 | 87.97 || 1 | G | O6 | 14:723 | 2.94 | 1.00 | 64.41 |
| 2 | U | O4 | 14:724 | 3.70 | 1.00 | 61.40 |  |
| 3 | G | O6 | 14:707 | 3.94 | 1.00 | 58.61 |  |
| 4 | A | OP2 | 14:706 | 4.00 | 1.00 | 47.71 |  |
| 5 | A | OP2 | 14:705 | 4.40 | 1.00 | 49.67 |  |
| 6 | G | N7 | 14:723 | 4.75 | 1.00 | 63.41 |  |
| 7 | G | N2 | 14:725 | 4.81 | 1.00 | 51.54 |  |
| 8 | C | N4 | 14:708 | 4.86 | 1.00 | 59.90 |  |
  |

  
 Contact(s): /O6/  
 Coded contact list: O_b_  

### K:14:3065

|  | Res. type | Atom | Residue | Distance | Occupancy | B-factor | Closest solute to 1^st^ shell water |
| --- | --- | --- | --- | --- | --- | --- | --- |
| **Ion** | | | |  | 1.00 | 55.80 || 1 | A | O3' | 14:197 | 2.98 | 1.00 | 41.45 |
| 2 | G | OP2 | 14:2069 | 3.08 | 1.00 | 31.97 |  |
| 3 | C | OP1 | 14:198 | 3.09 | 1.00 | 35.32 |  |
| 4 | G | OP1 | 14:2069 | 3.14 | 1.00 | 32.51 |  |
| 5 | U | OP1 | 14:2244 | 3.20 | 1.00 | 39.80 |  |
| 6 | G | OP2 | 14:2070 | 3.34 | 1.00 | 43.89 |  |
| 7 | G | OP1 | 14:2070 | 3.42 | 1.00 | 41.69 |  |
| 8 | A | O2' | 14:197 | 3.71 | 1.00 | 36.39 |  |
| 9 | G | O5' | 14:2069 | 4.20 | 1.00 | 43.55 |  |
| 10 | G | O3' | 14:2069 | 4.54 | 1.00 | 34.76 |  |
| 11 | U | OP2 | 14:2244 | 4.61 | 1.00 | 35.38 |  |
| 12 | C | O5' | 14:198 | 4.61 | 1.00 | 36.06 |  |
| 13 | U | O3' | 14:2068 | 4.93 | 1.00 | 43.60 |  |
| 14 | C | OP2 | 14:198 | 4.96 | 1.00 | 33.93 |  |
  |

  
**1^st^ shell ligand angles:**   
1-2: 120; 1-3: 47; 1-4: 83; 1-5: 69; 1-6: 169; 2-3: 159; 2-4: 51; 2-5: 100; 2-6: 49; 3-4: 108; 3-5: 90; 3-6: 144; 4-5: 120; 4-6: 89; 5-6: 108;   
  
 Contact(s): /O3'/OP1/OP1/OP1/OP2/OP2/  
 Coded contact list: 5O_ph_O_r_  

### K:14:3066

|  | Res. type | Atom | Residue | Distance | Occupancy | B-factor | Closest solute to 1^st^ shell water |
| --- | --- | --- | --- | --- | --- | --- | --- |
| **Ion** | | | |  | 1.00 | 64.36 || 1 | G | OP1 | 14:2447 | 2.98 | 1.00 | 45.55 |
| 2 | A | O2' | 14:567 | 3.10 | 1.00 | 32.62 |  |
| 3 | G | OP2 | 14:2447 | 3.51 | 1.00 | 45.42 |  |
| 4 | G | OP1 | 14:2446 | 3.82 | 1.00 | 37.81 |  |
| 5 | G | O5' | 14:2447 | 4.50 | 1.00 | 36.14 |  |
| 6 | G | OP1 | 14:808 | 4.82 | 1.00 | 39.56 |  |
  |

  
**1^st^ shell ligand angles:**   
1-2: 93;   
  
 Contact(s): /O2'/OP1/  
 Coded contact list: O_ph_O_r_  

### K:14:3067

|  | Res. type | Atom | Residue | Distance | Occupancy | B-factor | Closest solute to 1^st^ shell water |
| --- | --- | --- | --- | --- | --- | --- | --- |
| **Ion** | | | |  | 1.00 | 70.59 || 1 | C | O2' | 14:1320 | 3.09 | 1.00 | 55.29 |
| 2 | C | O3' | 14:1320 | 3.66 | 1.00 | 53.53 |  |
| 3 | U | O4 | 14:1323 | 3.80 | 1.00 | 48.05 |  |
| 4 | A | OP2 | 14:1322 | 3.94 | 1.00 | 38.50 |  |
| 5 | A | N7 | 14:1322 | 4.18 | 1.00 | 42.69 |  |
| 6 | C | O2 | 14:1320 | 4.21 | 1.00 | 55.36 |  |
| 7 | Water 1 | | 14:4004 | 4.28 | 1.00 | 40.07 | 4.07 (U:O4 14:1323) |
| 8 | C | N1 | 14:1320 | 4.48 | 1.00 | 60.24 |  |
| 9 | G | O6 | 14:1324 | 4.49 | 1.00 | 54.06 |  |
| 10 | A | O5' | 14:1321 | 4.56 | 1.00 | 48.53 |  |
| 11 | U | OP2 | 14:1329 | 4.71 | 1.00 | 47.27 |  |
| 12 | A | OP2 | 14:1321 | 4.76 | 1.00 | 50.66 |  |
| 13 | C | N4 | 14:1330 | 4.89 | 1.00 | 50.91 |  |
| 14 | C | O4' | 14:1320 | 4.97 | 1.00 | 52.74 |  |
  |

  
 Contact(s): /O2'/  
 Coded contact list: O_r_  

### K:14:3068

|  | Res. type | Atom | Residue | Distance | Occupancy | B-factor | Closest solute to 1^st^ shell water |
| --- | --- | --- | --- | --- | --- | --- | --- |
| **Ion** | | | |  | 1.00 | 61.15 || 1 | A | O2' | 14:2199 | 2.86 | 1.00 | 54.28 |
| 2 | LYS | O | F5:48 | 2.88 | 1.00 | 57.92 |  |
| 3 | A | O3' | 14:2199 | 3.07 | 1.00 | 48.85 |  |
| 4 | C | OP1 | 14:2205 | 3.41 | 1.00 | 48.56 |  |
| 5 | VAL | N | F5:49 | 4.36 | 1.00 | 53.75 |  |
| 6 | C | O5' | 14:2205 | 4.39 | 1.00 | 45.57 |  |
| 7 | U | OP2 | 14:2092 | 4.53 | 1.00 | 56.90 |  |
| 8 | U | OP1 | 14:2092 | 4.60 | 1.00 | 44.09 |  |
| 9 | ARG | N | F5:50 | 4.64 | 1.00 | 54.77 |  |
| 10 | ASN | OD1 | 69:28 | 4.98 | 1.00 | 56.20 |  |
  |

  
**1^st^ shell ligand angles:**   
1-2: 144; 1-3: 56; 2-3: 123;   
  
 Contact(s): /O2'/O3'/O_back_/  
 Coded contact list: 2O_r_O_back_  

### K:14:3069

|  | Res. type | Atom | Residue | Distance | Occupancy | B-factor | Closest solute to 1^st^ shell water |
| --- | --- | --- | --- | --- | --- | --- | --- |
| **Ion** | | | |  | 1.00 | 56.45 || 1 | G | OP1 | 14:809 | 2.49 | 1.00 | 35.49 |
| 2 | U | O2 | 14:576 | 2.81 | 1.00 | 36.81 |  |
| 3 | C | O2 | 14:565 | 3.23 | 1.00 | 35.56 |  |
| 4 | G | N3 | 14:577 | 3.68 | 1.00 | 34.15 |  |
| 5 | G | N9 | 14:577 | 3.89 | 1.00 | 35.43 |  |
| 6 | G | O4' | 14:577 | 3.99 | 1.00 | 35.55 |  |
| 7 | U | O4' | 14:566 | 4.04 | 1.00 | 37.59 |  |
| 8 | A | O2' | 14:1253 | 4.11 | 1.00 | 47.37 |  |
| 9 | G | O3' | 14:808 | 4.12 | 1.00 | 32.45 |  |
| 10 | U | N3 | 14:576 | 4.45 | 1.00 | 33.52 |  |
| 11 | C | O2' | 14:565 | 4.49 | 1.00 | 35.91 |  |
| 12 | U | O2' | 14:566 | 4.66 | 1.00 | 41.01 |  |
| 13 | G | OP2 | 14:809 | 4.69 | 1.00 | 40.74 |  |
| 14 | U | O2 | 14:566 | 4.89 | 1.00 | 33.02 |  |
| 15 | G | O5' | 14:809 | 4.90 | 1.00 | 34.74 |  |
| 16 | G | N2 | 14:577 | 4.91 | 1.00 | 35.44 |  |
| 17 | U | N1 | 14:566 | 4.93 | 1.00 | 31.89 |  |
| 18 | G | O2' | 14:577 | 4.98 | 1.00 | 43.70 |  |
  |

  
**1^st^ shell ligand angles:**   
1-2: 129; 1-3: 114; 2-3: 73;   
  
 Contact(s): /O2/O2/  
 Coded contact list: 2O_b_  
 Exclusion zone contact(s): /OP1/  

### K:14:3070

|  | Res. type | Atom | Residue | Distance | Occupancy | B-factor | Closest solute to 1^st^ shell water |
| --- | --- | --- | --- | --- | --- | --- | --- |
| **Ion** | | | |  | 1.00 | 92.30 || 1 | U | O4 | 14:1727 | 2.65 | 1.00 | 102.88 |
| 2 | C | N4 | 14:1734 | 3.22 | 1.00 | 100.05 |  |
| 3 | G | O6 | 14:1733 | 3.38 | 1.00 | 106.79 |  |
| 4 | G | O6 | 14:1726 | 3.60 | 1.00 | 103.23 |  |
| 5 | U | N3 | 14:1727 | 3.81 | 1.00 | 108.23 |  |
| 6 | A | OP2 | 14:1732 | 4.16 | 1.00 | 119.64 |  |
| 7 | G | N2 | 14:1731 | 4.36 | 1.00 | 121.74 |  |
| 8 | G | N7 | 14:1733 | 4.42 | 1.00 | 111.24 |  |
| 9 | G | O6 | 14:1728 | 4.55 | 1.00 | 116.28 |  |
| 10 | C | N3 | 14:1734 | 4.56 | 1.00 | 101.28 |  |
| 11 | G | O6 | 14:1725 | 4.84 | 1.00 | 116.23 |  |
| 12 | G | N7 | 14:1728 | 5.00 | 1.00 | 117.66 |  |
  |

  
**1^st^ shell ligand angles:**   
1-2: 101; 1-3: 77; 2-3: 69;   
  
 Contact(s): /N4/O4/O6/  
 Coded contact list: 2O_b_Anion  

### K:14:3071

|  | Res. type | Atom | Residue | Distance | Occupancy | B-factor | Closest solute to 1^st^ shell water |
| --- | --- | --- | --- | --- | --- | --- | --- |
| **Ion** | | | |  | 1.00 | 77.65 || 1 | A | OP2 | 14:2227 | 3.24 | 1.00 | 46.54 |
| 2 | G | O6 | 14:2090 | 3.36 | 1.00 | 50.63 |  |
| 3 | G | O6 | 14:2228 | 3.59 | 1.00 | 48.09 |  |
| 4 | U | O4 | 14:2091 | 3.62 | 1.00 | 59.03 |  |
| 5 | C | N4 | 14:2229 | 3.64 | 1.00 | 52.89 |  |
| 6 | C | OP2 | 14:2226 | 3.66 | 1.00 | 59.71 |  |
| 7 | G | N7 | 14:2228 | 4.37 | 1.00 | 46.26 |  |
| 8 | U | O4 | 14:2092 | 4.43 | 1.00 | 51.55 |  |
| 9 | A | N7 | 14:2227 | 4.53 | 1.00 | 52.58 |  |
| 10 | U | N3 | 14:2091 | 4.77 | 1.00 | 45.29 |  |
| 11 | U | O4 | 14:2089 | 4.90 | 1.00 | 54.22 |  |
  |

  
**1^st^ shell ligand angles:**   
1-2: 169;   
  
 Contact(s): /O6/OP2/  
 Coded contact list: O_ph_O_b_  

### K:14:3072

|  | Res. type | Atom | Residue | Distance | Occupancy | B-factor | Closest solute to 1^st^ shell water |
| --- | --- | --- | --- | --- | --- | --- | --- |
| **Ion** | | | |  | 1.00 | 85.76 || 1 | G | OP1 | 14:1763 | 2.61 | 1.00 | 55.01 |
| 2 | G | O6 | 14:1699 | 3.13 | 1.00 | 61.86 |  |
| 3 | G | O4' | 14:1763 | 3.38 | 1.00 | 56.08 |  |
| 4 | G | OP1 | 14:733 | 4.26 | 1.00 | 37.79 |  |
| 5 | G | OP2 | 14:1763 | 4.36 | 1.00 | 53.62 |  |
| 6 | G | O2' | 14:1763 | 4.42 | 1.00 | 60.73 |  |
| 7 | C | O3' | 14:732 | 4.50 | 1.00 | 35.51 |  |
| 8 | G | OP1 | 14:1682 | 4.56 | 1.00 | 51.75 |  |
| 9 | G | O5' | 14:1763 | 4.81 | 1.00 | 52.42 |  |
| 10 | A | O3' | 14:1762 | 4.97 | 1.00 | 58.80 |  |
  |

  
**1^st^ shell ligand angles:**   
1-2: 99; 1-3: 56; 2-3: 60;   
  
 Contact(s): /O4'/O6/OP1/  
 Coded contact list: O_ph_O_r_O_b_  

### K:14:3073

|  | Res. type | Atom | Residue | Distance | Occupancy | B-factor | Closest solute to 1^st^ shell water |
| --- | --- | --- | --- | --- | --- | --- | --- |
| **Ion** | | | |  | 1.00 | 60.73 || 1 | Water 1 | | 14:3604 | 2.87 | 1.00 | 47.13 | 2.35 (G:O6 14:1381) |
| 2 | G | O6 | 14:1380 | 3.13 | 1.00 | 44.90 |  |
| 3 | Water 2 | | 14:3794 | 3.35 | 1.00 | 56.65 | 3.20 (G:OP1 14:1347) |
| 4 | A | O2' | 14:1378 | 3.40 | 1.00 | 45.74 |  |
| 5 | G | O6 | 14:1381 | 3.57 | 1.00 | 43.25 |  |
| 6 | G | N7 | 14:1380 | 4.04 | 1.00 | 37.09 |  |
| 7 | C | N4 | 14:1351 | 4.12 | 1.00 | 40.52 |  |
| 8 | A | OP2 | 14:1379 | 4.20 | 1.00 | 58.92 |  |
| 9 | G | OP1 | 14:1347 | 4.33 | 1.00 | 54.79 |  |
| 10 | C | N4 | 14:1350 | 4.39 | 1.00 | 44.83 |  |
| 11 | Water 3 | | 14:3633 | 4.55 | 1.00 | 54.47 | 2.42 (G:O6 14:1382) |
| 12 | G | OP1 | 14:1348 | 4.59 | 1.00 | 42.72 |  |
| 13 | G | O6 | 14:1382 | 4.68 | 1.00 | 51.50 |  |
| 14 | G | OP2 | 14:1348 | 5.00 | 1.00 | 39.34 |  |
  |

  
**1^st^ shell ligand angles:**   
1-2: 82; 1-3: 59; 1-4: 113; 2-3: 121; 2-4: 81; 3-4: 77;   
  
 Contact(s): /O2'/O6/O_w_/O_w_/  
 Coded contact list: O_r_O_b_2O_w_  
K hydration shell issue: water too close (< 2.4 Angs.)  

### K:14:3074

|  | Res. type | Atom | Residue | Distance | Occupancy | B-factor | Closest solute to 1^st^ shell water |
| --- | --- | --- | --- | --- | --- | --- | --- |
| **Ion** | | | |  | 1.00 | 73.31 || 1 | G | O6 | 14:68 | 2.83 | 1.00 | 59.91 |
| 2 | G | O6 | 14:58 | 2.84 | 1.00 | 54.09 |  |
| 3 | U | O4 | 14:67 | 3.23 | 1.00 | 59.57 |  |
| 4 | U | O4 | 14:59 | 3.25 | 1.00 | 56.94 |  |
| 5 | U | N3 | 14:59 | 4.17 | 1.00 | 57.59 |  |
| 6 | C | OP2 | 14:66 | 4.68 | 1.00 | 68.18 |  |
| 7 | C | N4 | 14:69 | 4.74 | 1.00 | 59.21 |  |
| 8 | G | N7 | 14:68 | 4.79 | 1.00 | 62.38 |  |
| 9 | G | N7 | 14:58 | 4.79 | 1.00 | 57.73 |  |
| 10 | G | N1 | 14:58 | 4.85 | 1.00 | 54.93 |  |
  |

  
**1^st^ shell ligand angles:**   
1-2: 65; 1-3: 64; 1-4: 88; 2-3: 128; 2-4: 70; 3-4: 99;   
  
 Contact(s): /O4/O4/O6/O6/  
 Coded contact list: 4O_b_  

### K:14:3075

|  | Res. type | Atom | Residue | Distance | Occupancy | B-factor | Closest solute to 1^st^ shell water |
| --- | --- | --- | --- | --- | --- | --- | --- |
| **Ion** | | | |  | 1.00 | 36.88 || 1 | A | O2' | 14:793 | 2.66 | 1.00 | 37.59 |
| 2 | G | OP2 | 14:794 | 2.90 | 1.00 | 35.57 |  |
| 3 | G | O2' | 14:775 | 2.99 | 1.00 | 35.43 |  |
| 4 | G | N7 | 14:776 | 3.00 | 1.00 | 35.30 |  |
| 5 | G | O3' | 14:775 | 3.37 | 1.00 | 38.04 |  |
| 6 | A | O3' | 14:793 | 3.42 | 1.00 | 34.04 |  |
| 7 | G | OP2 | 14:776 | 3.73 | 1.00 | 44.12 |  |
| 8 | G | N3 | 14:775 | 4.22 | 1.00 | 33.68 |  |
| 9 | G | O5' | 14:776 | 4.54 | 1.00 | 36.67 |  |
| 10 | G | O5' | 14:794 | 4.65 | 1.00 | 32.79 |  |
| 11 | A | N3 | 14:793 | 4.70 | 1.00 | 36.58 |  |
| 12 | G | N9 | 14:776 | 4.74 | 1.00 | 33.34 |  |
| 13 | G | N9 | 14:775 | 4.93 | 1.00 | 29.33 |  |
| 14 | G | OP1 | 14:794 | 4.96 | 1.00 | 35.16 |  |
  |

  
**1^st^ shell ligand angles:**   
1-2: 86; 1-3: 162; 1-4: 50; 1-5: 144; 2-3: 82; 2-4: 135; 2-5: 130; 3-4: 143; 3-5: 50; 4-5: 95;   
  
 Contact(s): /N7/O2'/O2'/O3'/OP2/  
 Coded contact list: O_ph_3O_r_N_b_  

### K:14:3076

|  | Res. type | Atom | Residue | Distance | Occupancy | B-factor | Closest solute to 1^st^ shell water |
| --- | --- | --- | --- | --- | --- | --- | --- |
| **Ion** | | | |  | 1.00 | 86.18 || 1 | Water 1 | | 14:3678 | 2.78 | 1.00 | 73.39 | 2.54 (G:O6 14:342) |
| 2 | G | O6 | 14:342 | 3.10 | 1.00 | 70.74 |  |
| 3 | U | OP1 | 14:339 | 3.58 | 1.00 | 59.32 |  |
| 4 | U | OP2 | 14:339 | 3.78 | 1.00 | 63.73 |  |
| 5 | G | O6 | 14:341 | 3.80 | 1.00 | 70.99 |  |
| 6 | C | N4 | 14:296 | 3.97 | 1.00 | 68.76 |  |
| 7 | C | N4 | 14:297 | 4.00 | 1.00 | 67.43 |  |
| 8 | G | N2 | 14:298 | 4.42 | 1.00 | 71.43 |  |
| 9 | Water 2 | | 14:4098 | 4.58 | 1.00 | 73.01 | 4.92 (A:OP2 14:340) |
| 10 | A | OP2 | 14:340 | 4.71 | 1.00 | 57.09 |  |
| 11 | G | N1 | 14:342 | 4.95 | 1.00 | 76.19 |  |
  |

  
**1^st^ shell ligand angles:**   
1-2: 51;   
  
 Contact(s): /O6/O_w_/  
 Coded contact list: O_b_O_w_  

### K:14:3077

|  | Res. type | Atom | Residue | Distance | Occupancy | B-factor | Closest solute to 1^st^ shell water |
| --- | --- | --- | --- | --- | --- | --- | --- |
| **Ion** | | | |  | 1.00 | 73.04 || 1 | G | O6 | 14:465 | 2.73 | 1.00 | 37.95 |
| 2 | A | O2' | 14:788 | 2.85 | 1.00 | 37.11 |  |
| 3 | U | O4 | 14:464 | 3.62 | 1.00 | 35.09 |  |
| 4 | G | N1 | 14:465 | 4.11 | 1.00 | 31.85 |  |
| 5 | A | OP2 | 14:789 | 4.38 | 1.00 | 29.49 |  |
| 6 | G | O2' | 14:463 | 4.49 | 1.00 | 40.26 |  |
  |

  
**1^st^ shell ligand angles:**   
1-2: 134;   
  
 Contact(s): /O2'/O6/  
 Coded contact list: O_r_O_b_  

### K:14:3078

|  | Res. type | Atom | Residue | Distance | Occupancy | B-factor | Closest solute to 1^st^ shell water |
| --- | --- | --- | --- | --- | --- | --- | --- |
| **Ion** | | | |  | 1.00 | 81.67 || 1 | G | OP1 | 14:1236 | 2.55 | 1.00 | 56.77 |
| 2 | A | O5' | 14:505 | 2.91 | 1.00 | 59.26 |  |
| 3 | A | O4' | 14:505 | 3.06 | 1.00 | 55.92 |  |
| 4 | U | OP1 | 14:511 | 3.39 | 1.00 | 54.15 |  |
| 5 | C | O3' | 14:510 | 3.70 | 1.00 | 44.28 |  |
| 6 | A | OP2 | 14:505 | 3.85 | 1.00 | 72.61 |  |
| 7 | G | OP2 | 14:1236 | 4.26 | 1.00 | 56.66 |  |
| 8 | G | O5' | 14:1236 | 4.41 | 1.00 | 53.50 |  |
| 9 | A | N3 | 14:502 | 4.68 | 1.00 | 47.71 |  |
| 10 | A | OP1 | 14:505 | 4.75 | 1.00 | 68.30 |  |
| 11 | C | O2' | 14:510 | 4.84 | 1.00 | 53.16 |  |
  |

  
**1^st^ shell ligand angles:**   
1-2: 133; 1-3: 160; 1-4: 74; 2-3: 62; 2-4: 69; 3-4: 108;   
  
 Contact(s): /O4'/O5'/OP1/  
 Coded contact list: O_ph_2O_r_  
 Exclusion zone contact(s): /OP1/  

### K:14:3079

|  | Res. type | Atom | Residue | Distance | Occupancy | B-factor | Closest solute to 1^st^ shell water |
| --- | --- | --- | --- | --- | --- | --- | --- |
| **Ion** | | | |  | 1.00 | 81.16 || 1 | G | O6 | 14:1212 | 3.41 | 1.00 | 69.53 |
| 2 | G | O6 | 14:309 | 3.45 | 1.00 | 51.84 |  |
| 3 | G | O2' | 14:307 | 3.62 | 1.00 | 56.28 |  |
| 4 | G | N7 | 14:308 | 3.83 | 1.00 | 55.18 |  |
| 5 | U | OP2 | 14:1211 | 4.07 | 1.00 | 71.18 |  |
| 6 | G | O3' | 14:307 | 4.12 | 1.00 | 56.95 |  |
| 7 | U | OP1 | 14:1211 | 4.27 | 1.00 | 69.40 |  |
| 8 | G | O6 | 14:308 | 4.38 | 1.00 | 57.39 |  |
| 9 | A | O2' | 14:1210 | 4.58 | 1.00 | 68.28 |  |
| 10 | G | N1 | 14:1212 | 4.97 | 1.00 | 62.79 |  |
  |

  
 Coded contact list:   
1^st^ K coordination shell missing (see Notes)  

### K:14:3080

|  | Res. type | Atom | Residue | Distance | Occupancy | B-factor | Closest solute to 1^st^ shell water |
| --- | --- | --- | --- | --- | --- | --- | --- |
| **Ion** | | | |  | 1.00 | 79.15 || 1 | G | O6 | 14:1519 | 2.78 | 1.00 | 72.07 |
| 2 | U | O4 | 14:1520 | 2.96 | 1.00 | 81.04 |  |
| 3 | G | N2 | 14:1521 | 3.35 | 1.00 | 86.82 |  |
| 4 | G | O6 | 14:1473 | 3.81 | 1.00 | 77.40 |  |
| 5 | C | N4 | 14:1474 | 3.85 | 1.00 | 77.29 |  |
| 6 | A | OP2 | 14:1472 | 4.26 | 1.00 | 67.63 |  |
| 7 | G | N1 | 14:1521 | 4.28 | 1.00 | 79.15 |  |
| 8 | A | OP2 | 14:1471 | 4.29 | 1.00 | 66.19 |  |
| 9 | U | N3 | 14:1520 | 4.37 | 1.00 | 77.95 |  |
| 10 | C | N4 | 14:1518 | 4.59 | 1.00 | 73.80 |  |
| 11 | G | N3 | 14:1521 | 4.69 | 1.00 | 80.80 |  |
| 12 | G | N7 | 14:1473 | 4.70 | 1.00 | 84.97 |  |
| 13 | G | N1 | 14:1519 | 4.81 | 1.00 | 73.89 |  |
| 14 | G | N7 | 14:1519 | 4.88 | 1.00 | 73.58 |  |
  |

  
**1^st^ shell ligand angles:**   
1-2: 66; 1-3: 123; 2-3: 67;   
  
 Contact(s): /N2/O4/O6/  
 Coded contact list: 2O_b_Anion  

### K:14:3081

|  | Res. type | Atom | Residue | Distance | Occupancy | B-factor | Closest solute to 1^st^ shell water |
| --- | --- | --- | --- | --- | --- | --- | --- |
| **Ion** | | | |  | 1.00 | 54.57 || 1 | G | O6 | 14:458 | 3.06 | 1.00 | 32.02 |
| 2 | G | OP2 | 14:469 | 3.23 | 1.00 | 30.98 |  |
| 3 | ALA | O | 39:58 | 3.67 | 1.00 | 41.57 |  |
| 4 | G | N1 | 14:458 | 4.00 | 1.00 | 34.26 |  |
| 5 | G | O5' | 14:469 | 4.52 | 1.00 | 33.56 |  |
| 6 | G | N7 | 14:469 | 4.58 | 1.00 | 32.65 |  |
| 7 | G | OP1 | 14:468 | 4.63 | 1.00 | 33.92 |  |
| 8 | LYS | NZ | L5:37 | 4.79 | 1.00 | 39.82 |  |
| 9 | G | OP1 | 14:469 | 4.79 | 1.00 | 32.10 |  |
| 10 | TYR | N | 39:59 | 4.81 | 1.00 | 37.15 |  |
| 11 | A | OP2 | 14:470 | 4.89 | 1.00 | 29.72 |  |
  |

  
**1^st^ shell ligand angles:**   
1-2: 145;   
  
 Contact(s): /O6/OP2/  
 Coded contact list: O_ph_O_b_  

### K:14:3082

|  | Res. type | Atom | Residue | Distance | Occupancy | B-factor | Closest solute to 1^st^ shell water |
| --- | --- | --- | --- | --- | --- | --- | --- |
| **Ion** | | | |  | 1.00 | 80.02 || 1 | G | O5' | 14:1325 | 2.95 | 1.00 | 43.12 |
| 2 | G | O6 | 14:1271 | 3.17 | 1.00 | 60.60 |  |
| 3 | G | O4' | 14:1325 | 3.37 | 1.00 | 48.33 |  |
| 4 | G | OP2 | 14:1325 | 3.62 | 1.00 | 45.81 |  |
| 5 | G | OP1 | 14:1324 | 4.45 | 1.00 | 46.73 |  |
| 6 | C | O2 | 14:1615 | 4.72 | 1.00 | 45.76 |  |
| 7 | G | N9 | 14:1325 | 4.77 | 1.00 | 47.59 |  |
| 8 | G | O3' | 14:1324 | 4.84 | 1.00 | 41.34 |  |
  |

  
**1^st^ shell ligand angles:**   
1-2: 120; 1-3: 56; 2-3: 100;   
  
 Contact(s): /O4'/O5'/O6/  
 Coded contact list: 2O_r_O_b_  

### K:14:3083

|  | Res. type | Atom | Residue | Distance | Occupancy | B-factor | Closest solute to 1^st^ shell water |
| --- | --- | --- | --- | --- | --- | --- | --- |
| **Ion** | | | |  | 1.00 | 70.33 || 1 | G | OP2 | 14:585 | 2.80 | 1.00 | 42.44 |
| 2 | THR | OG1 | 85:6 | 3.24 | 1.00 | 50.88 |  |
| 3 | Water 1 | | 14:3796 | 3.37 | 1.00 | 47.33 | 2.74 (G:OP1 14:1250) |
| 4 | Water 2 | | 14:3786 | 4.05 | 1.00 | 46.81 | 2.72 (G:OP1 14:585) |
| 5 | G | OP1 | 14:1250 | 4.10 | 1.00 | 41.71 |  |
| 6 | U | O3' | 14:1249 | 4.29 | 1.00 | 49.30 |  |
| 7 | THR | N | 85:6 | 4.32 | 1.00 | 43.29 |  |
| 8 | G | OP1 | 14:585 | 4.56 | 1.00 | 38.92 |  |
| 9 | ALA | O | 85:4 | 4.58 | 1.00 | 47.65 |  |
| 10 | G | O5' | 14:585 | 4.78 | 1.00 | 38.89 |  |
| 11 | C | OP1 | 14:584 | 4.84 | 1.00 | 45.05 |  |
| 12 | G | O5' | 14:1250 | 4.95 | 1.00 | 42.30 |  |
  |

  
**1^st^ shell ligand angles:**   
1-2: 106; 1-3: 86; 2-3: 154;   
  
 Contact(s): /O_w_/OG1_Thr_/OP2/  
 Coded contact list: O_ph_OH_Prot_O_w_  

### K:14:3084

|  | Res. type | Atom | Residue | Distance | Occupancy | B-factor | Closest solute to 1^st^ shell water |
| --- | --- | --- | --- | --- | --- | --- | --- |
| **Ion** | | | |  | 1.00 | 73.92 || 1 | C | OP1 | 14:393 | 2.66 | 1.00 | 55.06 |
| 2 | C | OP1 | 14:375 | 3.04 | 1.00 | 56.65 |  |
| 3 | Water 1 | | 14:3626 | 3.35 | 1.00 | 52.56 | 2.61 (C:OP1 14:375) |
| 4 | C | OP2 | 14:393 | 3.37 | 1.00 | 46.67 |  |
| 5 | G | O2' | 14:408 | 3.58 | 1.00 | 57.92 |  |
| 6 | G | O3' | 14:408 | 3.89 | 1.00 | 60.70 |  |
| 7 | C | O5' | 14:393 | 3.98 | 1.00 | 50.03 |  |
| 8 | C | OP2 | 14:375 | 4.31 | 1.00 | 56.69 |  |
| 9 | A | N7 | 14:223 | 4.47 | 1.00 | 65.93 |  |
| 10 | A | OP2 | 14:394 | 4.50 | 1.00 | 46.52 |  |
| 11 | A | O3' | 14:374 | 4.51 | 1.00 | 58.75 |  |
| 12 | C | O3' | 14:392 | 4.82 | 1.00 | 44.97 |  |
  |

  
**1^st^ shell ligand angles:**   
1-2: 134; 1-3: 88; 1-4: 49; 2-3: 48; 2-4: 102; 3-4: 59;   
  
 Contact(s): /O_w_/OP1/OP1/OP2/  
 Coded contact list: 3O_ph_O_w_  

### K:14:3085

|  | Res. type | Atom | Residue | Distance | Occupancy | B-factor | Closest solute to 1^st^ shell water |
| --- | --- | --- | --- | --- | --- | --- | --- |
| **Ion** | | | |  | 1.00 | 62.28 || 1 | U | O4 | 14:202 | 3.03 | 1.00 | 34.93 |
| 2 | C | OP2 | 14:192 | 3.17 | 1.00 | 33.66 |  |
| 3 | Water 1 | | 14:3932 | 3.30 | 1.00 | 37.26 | 3.10 (A:OP2 14:199) |
| 4 | U | O4 | 14:193 | 3.68 | 1.00 | 36.69 |  |
| 5 | Water 2 | | 14:3520 | 4.35 | 1.00 | 28.11 | 1.99 (C:OP1 14:192) |
| 6 | A | OP2 | 14:199 | 4.37 | 1.00 | 36.42 |  |
| 7 | C | OP1 | 14:192 | 4.64 | 1.00 | 37.79 |  |
| 8 | C | N4 | 14:201 | 4.77 | 1.00 | 35.67 |  |
| 9 | A | OP1 | 14:199 | 4.79 | 1.00 | 35.67 |  |
| 10 | U | N3 | 14:202 | 4.83 | 1.00 | 31.40 |  |
| 11 | U | OP2 | 14:200 | 4.89 | 1.00 | 43.26 |  |
| 12 | A | O3' | 14:191 | 4.99 | 1.00 | 30.06 |  |
  |

  
**1^st^ shell ligand angles:**   
1-2: 103; 1-3: 108; 2-3: 60;   
  
 Contact(s): /O4/O_w_/OP2/  
 Coded contact list: O_ph_O_b_O_w_  
K hydration shell issue: water too close (< 2.4 Angs.)  

### K:14:3086

|  | Res. type | Atom | Residue | Distance | Occupancy | B-factor | Closest solute to 1^st^ shell water |
| --- | --- | --- | --- | --- | --- | --- | --- |
| **Ion** | | | |  | 1.00 | 65.16 || 1 | G | O4' | 14:1816 | 3.01 | 1.00 | 47.59 |
| 2 | A | O3' | 14:1815 | 3.13 | 1.00 | 34.46 |  |
| 3 | A | O2' | 14:1815 | 3.39 | 1.00 | 35.36 |  |
| 4 | G | OP2 | 14:1816 | 3.39 | 1.00 | 38.45 |  |
| 5 | G | O5' | 14:1816 | 4.10 | 1.00 | 38.77 |  |
| 6 | G | OP2 | 14:1814 | 4.44 | 1.00 | 33.25 |  |
| 7 | G | OP1 | 14:1813 | 4.59 | 1.00 | 42.94 |  |
| 8 | G | N3 | 14:1816 | 4.78 | 1.00 | 76.60 |  |
| 9 | G | O2' | 14:1816 | 4.93 | 1.00 | 48.66 |  |
| 10 | G | N9 | 14:1816 | 4.96 | 1.00 | 60.29 |  |
| 11 | A | N3 | 14:1815 | 4.97 | 1.00 | 41.42 |  |
  |

  
**1^st^ shell ligand angles:**   
1-2: 91; 1-3: 128; 1-4: 85; 2-3: 48; 2-4: 44; 3-4: 85;   
  
 Contact(s): /O2'/O3'/O4'/OP2/  
 Coded contact list: O_ph_3O_r_  

### K:14:3087

|  | Res. type | Atom | Residue | Distance | Occupancy | B-factor | Closest solute to 1^st^ shell water |
| --- | --- | --- | --- | --- | --- | --- | --- |
| **Ion** | | | |  | 1.00 | 74.57 || 1 | U | OP2 | 14:1205 | 3.25 | 1.00 | 59.42 |
| 2 | A | O2' | 14:1204 | 3.28 | 1.00 | 63.84 |  |
| 3 | A | N9 | 14:1204 | 3.55 | 1.00 | 62.04 |  |
| 4 | A | N7 | 14:1204 | 3.66 | 1.00 | 58.43 |  |
| 5 | C | OP1 | 14:1202 | 4.37 | 1.00 | 44.00 |  |
| 6 | G | OP2 | 14:1203 | 4.39 | 1.00 | 51.25 |  |
| 7 | C | OP2 | 14:1202 | 4.42 | 1.00 | 38.04 |  |
  |

  
**1^st^ shell ligand angles:**   
1-2: 50;   
  
 Contact(s): /O2'/OP2/  
 Coded contact list: O_ph_O_r_  

### K:14:3088

|  | Res. type | Atom | Residue | Distance | Occupancy | B-factor | Closest solute to 1^st^ shell water |
| --- | --- | --- | --- | --- | --- | --- | --- |
| **Ion** | | | |  | 1.00 | 98.43 || 1 | U | O4 | 14:2089 | 3.83 | 1.00 | 54.22 |
| 2 | G | O6 | 14:2230 | 4.08 | 1.00 | 45.40 |  |
| 3 | G | O6 | 14:2088 | 4.36 | 1.00 | 48.38 |  |
| 4 | C | N4 | 14:2229 | 4.47 | 1.00 | 52.89 |  |
| 5 | C | N4 | 14:2231 | 4.71 | 1.00 | 45.67 |  |
  |

  
 Coded contact list:   
1^st^ K coordination shell missing (see Notes)  

### K:14:3089

|  | Res. type | Atom | Residue | Distance | Occupancy | B-factor | Closest solute to 1^st^ shell water |
| --- | --- | --- | --- | --- | --- | --- | --- |
| **Ion** | | | |  | 1.00 | 64.79 || 1 | C | O2' | 14:409 | 2.88 | 1.00 | 51.72 |
| 2 | G | O6 | 14:226 | 2.94 | 1.00 | 65.56 |  |
| 3 | G | N7 | 14:226 | 2.96 | 1.00 | 64.30 |  |
| 4 | G | O4' | 14:410 | 3.36 | 1.00 | 55.27 |  |
| 5 | C | O2 | 14:409 | 3.54 | 1.00 | 59.26 |  |
| 6 | A | N7 | 14:225 | 3.89 | 1.00 | 64.82 |  |
| 7 | A | OP2 | 14:225 | 4.28 | 1.00 | 58.34 |  |
| 8 | G | N9 | 14:410 | 4.68 | 1.00 | 56.02 |  |
| 9 | C | O2 | 14:419 | 4.73 | 1.00 | 60.79 |  |
| 10 | G | OP2 | 14:224 | 4.79 | 1.00 | 61.51 |  |
| 11 | G | N1 | 14:226 | 4.88 | 1.00 | 68.75 |  |
| 12 | G | N2 | 14:418 | 4.91 | 1.00 | 59.51 |  |
| 13 | C | N1 | 14:409 | 4.97 | 1.00 | 51.75 |  |
  |

  
**1^st^ shell ligand angles:**   
1-2: 136; 1-3: 131; 1-4: 64; 2-3: 63; 2-4: 84; 3-4: 79;   
  
 Contact(s): /N7/O2'/O4'/O6/  
 Coded contact list: 2O_r_O_b_N_b_  

### K:14:3090

|  | Res. type | Atom | Residue | Distance | Occupancy | B-factor | Closest solute to 1^st^ shell water |
| --- | --- | --- | --- | --- | --- | --- | --- |
| **Ion** | | | |  | 1.00 | 64.20 || 1 | C | OP2 | 14:671 | 2.70 | 1.00 | 52.55 |
| 2 | A | O3' | 14:670 | 2.75 | 1.00 | 40.51 |  |
| 3 | A | O2' | 14:670 | 3.16 | 1.00 | 46.96 |  |
| 4 | U | O4 | 14:588 | 3.46 | 1.00 | 48.12 |  |
| 5 | C | O5' | 14:671 | 4.38 | 1.00 | 49.93 |  |
| 6 | C | OP1 | 14:671 | 4.57 | 1.00 | 38.34 |  |
| 7 | C | N4 | 14:589 | 4.84 | 1.00 | 41.04 |  |
  |

  
**1^st^ shell ligand angles:**   
1-2: 53; 1-3: 110; 2-3: 58;   
  
 Contact(s): /O2'/O3'/OP2/  
 Coded contact list: O_ph_2O_r_  

### K:14:3091

|  | Res. type | Atom | Residue | Distance | Occupancy | B-factor | Closest solute to 1^st^ shell water |
| --- | --- | --- | --- | --- | --- | --- | --- |
| **Ion** | | | |  | 1.00 | 78.64 || 1 | G | O6 | 14:2717 | 3.36 | 1.00 | 63.45 |
| 2 | G | O6 | 14:2719 | 3.56 | 1.00 | 68.81 |  |
| 3 | G | O6 | 14:2718 | 3.85 | 1.00 | 63.86 |  |
| 4 | G | N7 | 14:2718 | 4.06 | 1.00 | 62.55 |  |
| 5 | U | O4 | 14:2716 | 4.28 | 1.00 | 66.76 |  |
| 6 | G | N7 | 14:2717 | 4.28 | 1.00 | 65.78 |  |
| 7 | G | N1 | 14:2719 | 4.99 | 1.00 | 61.25 |  |
  |

  
 Contact(s): /O6/  
 Coded contact list: O_b_  

### K:14:3092

|  | Res. type | Atom | Residue | Distance | Occupancy | B-factor | Closest solute to 1^st^ shell water |
| --- | --- | --- | --- | --- | --- | --- | --- |
| **Ion** | | | |  | 1.00 | 106.11 || 1 | G | O6 | 14:270X | 2.81 | 1.00 | 90.22 |
| 2 | Water 1 | | 14:3660 | 2.90 | 1.00 | 86.97 | 2.51 (G:O6 14:270X) |
| 3 | Water 2 | | 14:4018 | 3.56 | 1.00 | 95.00 | 4.19 (G:OP2 14:270V) |
| 4 | A | OP2 | 14:270 | 3.77 | 1.00 | 76.88 |  |
| 5 | G | OP2 | 14:270W | 3.94 | 1.00 | 101.58 |  |
| 6 | Water 3 | | 14:3738 | 4.08 | 1.00 | 91.42 | 2.66 (G:N7 14:270W) |
| 7 | G | O6 | 14:270Y | 4.17 | 1.00 | 81.04 |  |
| 8 | G | N1 | 14:270Y | 4.18 | 1.00 | 83.81 |  |
| 9 | A | OP1 | 14:270 | 4.22 | 1.00 | 73.28 |  |
| 10 | G | N7 | 14:270W | 4.72 | 1.00 | 94.93 |  |
| 11 | G | N7 | 14:270X | 4.77 | 1.00 | 86.46 |  |
| 12 | G | OP2 | 14:270V | 4.79 | 1.00 | 100.76 |  |
| 13 | U | O3' | 14:269 | 4.92 | 1.00 | 69.15 |  |
| 14 | G | N1 | 14:270X | 4.96 | 1.00 | 87.03 |  |
  |

  
**1^st^ shell ligand angles:**   
1-2: 52;   
  
 Contact(s): /O6/O_w_/  
 Coded contact list: O_b_O_w_  

### K:14:3093

|  | Res. type | Atom | Residue | Distance | Occupancy | B-factor | Closest solute to 1^st^ shell water |
| --- | --- | --- | --- | --- | --- | --- | --- |
| **Ion** | | | |  | 1.00 | 53.26 || 1 | G | O2' | 14:792 | 3.30 | 1.00 | 44.06 |
| 2 | A | OP1 | 14:793 | 3.39 | 1.00 | 30.31 |  |
| 3 | C | O2 | 14:2441 | 3.63 | 1.00 | 37.34 |  |
| 4 | C | O2' | 14:2441 | 3.84 | 1.00 | 38.77 |  |
| 5 | G | O3' | 14:792 | 4.02 | 1.00 | 26.56 |  |
| 6 | G | N2 | 14:2070 | 4.04 | 1.00 | 33.14 |  |
| 7 | A | O2' | 14:2071 | 4.17 | 1.00 | 31.12 |  |
| 8 | C | O4' | 14:2442 | 4.29 | 1.00 | 37.06 |  |
| 9 | C | N3 | 14:2440 | 4.62 | 1.00 | 39.44 |  |
  |

  
**1^st^ shell ligand angles:**   
1-2: 81;   
  
 Contact(s): /O2'/OP1/  
 Coded contact list: O_ph_O_r_  

### K:14:3094

|  | Res. type | Atom | Residue | Distance | Occupancy | B-factor | Closest solute to 1^st^ shell water |
| --- | --- | --- | --- | --- | --- | --- | --- |
| **Ion** | | | |  | 1.00 | 86.02 || 1 | G | O6 | 14:1857 | 2.47 | 1.00 | 83.35 |
| 2 | Water 1 | | 14:3828 | 2.69 | 1.00 | 76.98 | 2.82 (G:OP2 14:1856) |
| 3 | Water 2 | | 14:3575 | 3.08 | 1.00 | 85.23 | 2.27 (G:O6 14:1858) |
| 4 | Water 3 | | 14:3951 | 3.26 | 1.00 | 71.97 | 3.17 (G:O6 14:1857) |
| 5 | Water 4 | | 14:4067 | 3.69 | 1.00 | 66.18 | 4.71 (G:O6 14:1857) |
| 6 | G | O6 | 14:1858 | 3.71 | 1.00 | 86.80 |  |
| 7 | Water 5 | | 14:3930 | 4.41 | 1.00 | 64.28 | 3.10 (G:N1 14:1857) |
| 8 | Water 6 | | 14:3585 | 4.44 | 1.00 | 84.79 | 2.29 (G:N7 14:1857) |
| 9 | G | OP2 | 14:1856 | 4.55 | 1.00 | 71.57 |  |
| 10 | G | N1 | 14:1857 | 4.57 | 1.00 | 79.75 |  |
| 11 | Water 7 | | 14:4078 | 4.62 | 1.00 | 81.27 | 4.31 (G:OP2 14:1856) |
| 12 | G | N1 | 14:1858 | 4.75 | 1.00 | 83.53 |  |
| 13 | G | N7 | 14:1857 | 4.87 | 1.00 | 83.02 |  |
  |

  
**1^st^ shell ligand angles:**   
1-2: 68; 1-3: 85; 1-4: 65; 2-3: 65; 2-4: 80; 3-4: 141;   
  
 Contact(s): /O_w_/O_w_/O_w_/  
 Coded contact list: 3O_w_  
 Exclusion zone contact(s): /O6/  
Consider NA in place of K (1 valid contact(s) for NA vs. 0 for K)  
K hydration shell issue: water too close (< 2.4 Angs.)  

### K:14:3095

|  | Res. type | Atom | Residue | Distance | Occupancy | B-factor | Closest solute to 1^st^ shell water |
| --- | --- | --- | --- | --- | --- | --- | --- |
| **Ion** | | | |  | 1.00 | 64.76 || 1 | U | O4 | 14:2554 | 2.96 | 1.00 | 63.42 |
| 2 | C | O2 | 14:2556 | 3.29 | 1.00 | 63.51 |  |
| 3 | U | O4 | 14:1955 | 3.39 | 1.00 | 53.79 |  |
| 4 | U | O4 | 14:2555 | 3.50 | 1.00 | 68.70 |  |
| 5 | C | N3 | 14:2556 | 3.78 | 1.00 | 55.24 |  |
| 6 | U | N3 | 14:2554 | 4.77 | 1.00 | 65.29 |  |
  |

  
**1^st^ shell ligand angles:**   
1-2: 112; 1-3: 71; 2-3: 92;   
  
 Contact(s): /O2/O4/O4/  
 Coded contact list: 3O_b_  

### K:14:3096

|  | Res. type | Atom | Residue | Distance | Occupancy | B-factor | Closest solute to 1^st^ shell water |
| --- | --- | --- | --- | --- | --- | --- | --- |
| **Ion** | | | |  | 1.00 | 70.82 || 1 | G | O6 | 14:1216 | 2.69 | 1.00 | 65.37 |
| 2 | G | O6 | 14:1215 | 2.92 | 1.00 | 60.54 |  |
| 3 | U | O4 | 14:1234 | 3.74 | 1.00 | 65.19 |  |
| 4 | C | N4 | 14:1233 | 3.78 | 1.00 | 55.34 |  |
| 5 | G | N7 | 14:1216 | 4.11 | 1.00 | 59.60 |  |
| 6 | U | N3 | 14:1234 | 4.13 | 1.00 | 63.15 |  |
| 7 | G | N7 | 14:1215 | 4.24 | 1.00 | 61.68 |  |
| 8 | G | N2 | 14:1235 | 4.50 | 1.00 | 63.86 |  |
| 9 | A | OP2 | 14:1214 | 4.64 | 1.00 | 55.37 |  |
| 10 | G | N1 | 14:1216 | 4.67 | 1.00 | 57.03 |  |
| 11 | G | N1 | 14:1235 | 4.67 | 1.00 | 66.84 |  |
| 12 | G | N1 | 14:1215 | 4.85 | 1.00 | 56.61 |  |
| 13 | A | OP2 | 14:1213 | 4.89 | 1.00 | 52.62 |  |
  |

  
**1^st^ shell ligand angles:**   
1-2: 90;   
  
 Contact(s): /O6/O6/  
 Coded contact list: 2O_b_  

### K:14:3097

|  | Res. type | Atom | Residue | Distance | Occupancy | B-factor | Closest solute to 1^st^ shell water |
| --- | --- | --- | --- | --- | --- | --- | --- |
| **Ion** | | | |  | 1.00 | 91.20 || 1 | A | OP1 | 14:324 | 2.91 | 1.00 | 63.28 |
| 2 | G | OP2 | 14:323 | 3.12 | 1.00 | 66.12 |  |
| 3 | A | O4' | 14:324 | 3.60 | 1.00 | 60.81 |  |
| 4 | G | O5' | 14:323 | 3.98 | 1.00 | 65.48 |  |
| 5 | A | O5' | 14:324 | 4.31 | 1.00 | 65.33 |  |
| 6 | U | O2' | 14:339 | 4.80 | 1.00 | 63.66 |  |
  |

  
**1^st^ shell ligand angles:**   
1-2: 104;   
  
 Contact(s): /OP1/OP2/  
 Coded contact list: 2O_ph_  

### K:14:3098

|  | Res. type | Atom | Residue | Distance | Occupancy | B-factor | Closest solute to 1^st^ shell water |
| --- | --- | --- | --- | --- | --- | --- | --- |
| **Ion** | | | |  | 1.00 | 67.62 || 1 | U | O4 | 14:2232 | 2.70 | 1.00 | 45.35 |
| 2 | U | O4 | 14:2233 | 2.75 | 1.00 | 44.68 |  |
| 3 | G | O6 | 14:2087 | 3.02 | 1.00 | 44.14 |  |
| 4 | U | O4 | 14:2086 | 3.10 | 1.00 | 47.23 |  |
| 5 | U | N3 | 14:2086 | 3.51 | 1.00 | 42.75 |  |
| 6 | U | N3 | 14:2232 | 3.79 | 1.00 | 43.51 |  |
| 7 | C | N4 | 14:2231 | 4.23 | 1.00 | 45.67 |  |
| 8 | G | O6 | 14:2088 | 4.49 | 1.00 | 48.38 |  |
| 9 | U | N3 | 14:2233 | 4.55 | 1.00 | 41.92 |  |
| 10 | G | N7 | 14:2087 | 4.86 | 1.00 | 46.34 |  |
  |

  
**1^st^ shell ligand angles:**   
1-2: 95; 1-3: 86; 1-4: 139; 2-3: 72; 2-4: 77; 3-4: 127;   
  
 Contact(s): /O4/O4/O4/O6/  
 Coded contact list: 4O_b_  

### K:14:3099

|  | Res. type | Atom | Residue | Distance | Occupancy | B-factor | Closest solute to 1^st^ shell water |
| --- | --- | --- | --- | --- | --- | --- | --- |
| **Ion** | | | |  | 1.00 | 67.62 || 1 | G | O6 | 14:179 | 3.02 | 1.00 | 51.72 |
| 2 | G | N7 | 14:179 | 3.11 | 1.00 | 44.57 |  |
| 3 | G | O6 | 14:178 | 3.21 | 1.00 | 52.95 |  |
| 4 | G | N7 | 14:178 | 3.86 | 1.00 | 47.82 |  |
| 5 | A | OP2 | 14:213 | 4.12 | 1.00 | 57.40 |  |
| 6 | G | O6 | 14:180 | 4.19 | 1.00 | 46.23 |  |
| 7 | C | N4 | 14:47 | 4.57 | 1.00 | 50.32 |  |
| 8 | G | OP1 | 14:177 | 4.67 | 1.00 | 54.37 |  |
| 9 | C | N4 | 14:46 | 4.72 | 1.00 | 49.18 |  |
| 10 | G | N1 | 14:178 | 4.77 | 1.00 | 46.00 |  |
| 11 | G | N1 | 14:179 | 4.98 | 1.00 | 41.75 |  |
  |

  
**1^st^ shell ligand angles:**   
1-2: 62; 1-3: 64; 2-3: 66;   
  
 Contact(s): /N7/O6/O6/  
 Coded contact list: 2O_b_N_b_  

### K:14:3100

|  | Res. type | Atom | Residue | Distance | Occupancy | B-factor | Closest solute to 1^st^ shell water |
| --- | --- | --- | --- | --- | --- | --- | --- |
| **Ion** | | | |  | 1.00 | 71.58 || 1 | U | OP2 | 14:448 | 2.88 | 1.00 | 48.00 |
| 2 | A | O2' | 14:447 | 3.06 | 1.00 | 45.87 |  |
| 3 | A | O3' | 14:447 | 3.12 | 1.00 | 43.89 |  |
| 4 | G | OP2 | 14:473 | 3.47 | 1.00 | 49.14 |  |
| 5 | U | O5' | 14:448 | 3.88 | 1.00 | 43.78 |  |
| 6 | A | O5' | 14:472 | 3.95 | 1.00 | 48.65 |  |
| 7 | A | OP1 | 14:472 | 3.96 | 1.00 | 39.09 |  |
| 8 | G | OP1 | 14:473 | 4.03 | 1.00 | 42.60 |  |
| 9 | A | OP2 | 14:472 | 4.07 | 1.00 | 41.16 |  |
| 10 | Water 1 | | 14:3976 | 4.62 | 1.00 | 54.22 | 3.24 (A:O2' 14:447) |
| 11 | U | OP1 | 14:448 | 4.85 | 1.00 | 37.12 |  |
  |

  
**1^st^ shell ligand angles:**   
1-2: 82; 1-3: 51; 2-3: 53;   
  
 Contact(s): /O2'/O3'/OP2/  
 Coded contact list: O_ph_2O_r_  

### K:14:3101

|  | Res. type | Atom | Residue | Distance | Occupancy | B-factor | Closest solute to 1^st^ shell water |
| --- | --- | --- | --- | --- | --- | --- | --- |
| **Ion** | | | |  | 1.00 | 76.69 || 1 | G | O6 | 14:700 | 2.91 | 1.00 | 42.23 |
| 2 | G | O6 | 14:701 | 3.14 | 1.00 | 46.83 |  |
| 3 | G | N7 | 14:700 | 3.82 | 1.00 | 48.62 |  |
| 4 | G | N7 | 14:701 | 4.13 | 1.00 | 41.46 |  |
| 5 | C | N4 | 14:732 | 4.54 | 1.00 | 41.97 |  |
| 6 | G | N2 | 14:733 | 4.61 | 1.00 | 40.25 |  |
| 7 | A | OP2 | 14:699 | 4.72 | 1.00 | 40.71 |  |
| 8 | U | OP2 | 14:762 | 4.85 | 1.00 | 43.19 |  |
| 9 | C | N4 | 14:731 | 4.89 | 1.00 | 38.61 |  |
  |

  
**1^st^ shell ligand angles:**   
1-2: 65;   
  
 Contact(s): /O6/O6/  
 Coded contact list: 2O_b_  

### K:14:3102

|  | Res. type | Atom | Residue | Distance | Occupancy | B-factor | Closest solute to 1^st^ shell water |
| --- | --- | --- | --- | --- | --- | --- | --- |
| **Ion** | | | |  | 1.00 | 76.27 || 1 | A | O2' | 14:1265 | 2.89 | 1.00 | 41.99 |
| 2 | U | OP2 | 14:1263 | 3.20 | 1.00 | 45.33 |  |
| 3 | A | N3 | 14:1265 | 3.33 | 1.00 | 43.09 |  |
| 4 | U | OP1 | 14:1263 | 3.82 | 1.00 | 44.64 |  |
| 5 | G | OP2 | 14:1264 | 3.97 | 1.00 | 42.45 |  |
| 6 | ARG | NH2 | J5:20 | 4.07 | 1.00 | 52.78 |  |
| 7 | ARG | NH1 | A5:15 | 4.12 | 1.00 | 52.87 |  |
| 8 | U | O5' | 14:1263 | 4.46 | 1.00 | 48.32 |  |
| 9 | G | N7 | 14:1264 | 4.64 | 1.00 | 44.26 |  |
| 10 | A | N1 | 14:1265 | 4.80 | 1.00 | 45.38 |  |
| 11 | A | N9 | 14:1265 | 4.82 | 1.00 | 40.26 |  |
  |

  
**1^st^ shell ligand angles:**   
1-2: 136; 1-3: 64; 2-3: 129;   
  
 Contact(s): /N3_A_/O2'/OP2/  
 Coded contact list: O_ph_O_r_N_b_  

### K:14:3103

|  | Res. type | Atom | Residue | Distance | Occupancy | B-factor | Closest solute to 1^st^ shell water |
| --- | --- | --- | --- | --- | --- | --- | --- |
| **Ion** | | | |  | 1.00 | 70.70 || 1 | Water 1 | | 14:3610 | 2.87 | 1.00 | 46.40 | 2.37 (A:O5' 14:2448) |
| 2 | Water 2 | | 14:3526 | 3.07 | 1.00 | 36.37 | 2.63 (G:O6 14:570) |
| 3 | G | OP1 | 14:2447 | 3.39 | 1.00 | 45.55 |  |
| 4 | G | O5' | 14:2447 | 3.48 | 1.00 | 36.14 |  |
| 5 | U | OP1 | 14:2500 | 3.80 | 1.00 | 35.33 |  |
| 6 | G | O3' | 14:2447 | 4.01 | 1.00 | 38.97 |  |
| 7 | U | O2' | 14:568 | 4.45 | 1.00 | 42.41 |  |
| 8 | A | N6 | 14:945 | 4.51 | 1.00 | 33.87 |  |
| 9 | A | O2' | 14:567 | 4.77 | 1.00 | 32.62 |  |
| 10 | G | O6 | 14:570 | 4.85 | 1.00 | 38.87 |  |
| 11 | U | O4' | 14:568 | 4.85 | 1.00 | 39.33 |  |
| 12 | A | O5' | 14:2448 | 4.86 | 1.00 | 34.47 |  |
| 13 | U | OP2 | 14:2500 | 4.97 | 1.00 | 35.38 |  |
  |

  
**1^st^ shell ligand angles:**   
1-2: 60; 1-3: 92; 2-3: 144;   
  
 Contact(s): /O_w_/O_w_/OP1/  
 Coded contact list: O_ph_2O_w_  
K hydration shell issue: water too close (< 2.4 Angs.)  

### K:14:3104

|  | Res. type | Atom | Residue | Distance | Occupancy | B-factor | Closest solute to 1^st^ shell water |
| --- | --- | --- | --- | --- | --- | --- | --- |
| **Ion** | | | |  | 1.00 | 75.78 || 1 | A | OP1 | 14:1272 | 2.79 | 1.00 | 48.09 |
| 2 | G | OP2 | 14:1271 | 2.82 | 1.00 | 41.18 |  |
| 3 | U | OP2 | 14:1273 | 3.23 | 1.00 | 52.36 |  |
| 4 | G | OP1 | 14:1271 | 3.25 | 1.00 | 35.86 |  |
| 5 | G | O3' | 14:1271 | 3.61 | 1.00 | 46.23 |  |
| 6 | G | O5' | 14:1271 | 4.18 | 1.00 | 40.73 |  |
| 7 | C | O3' | 14:1270 | 4.85 | 1.00 | 39.74 |  |
| 8 | A | O5' | 14:1272 | 4.91 | 1.00 | 48.71 |  |
| 9 | A | OP2 | 14:1272 | 4.92 | 1.00 | 43.66 |  |
| 10 | U | OP1 | 14:1273 | 5.00 | 1.00 | 48.87 |  |
  |

  
**1^st^ shell ligand angles:**   
1-2: 81; 1-3: 85; 1-4: 125; 2-3: 149; 2-4: 50; 3-4: 124;   
  
 Contact(s): /OP1/OP1/OP2/OP2/  
 Coded contact list: 4O_ph_  

### K:14:3105

|  | Res. type | Atom | Residue | Distance | Occupancy | B-factor | Closest solute to 1^st^ shell water |
| --- | --- | --- | --- | --- | --- | --- | --- |
| **Ion** | | | |  | 1.00 | 91.10 || 1 | G | O6 | 14:1421 | 3.25 | 1.00 | 82.85 |
| 2 | U | O4 | 14:1576 | 3.68 | 1.00 | 67.66 |  |
| 3 | G | O6 | 14:1422 | 3.73 | 1.00 | 61.52 |  |
| 4 | G | O6 | 14:1423 | 3.84 | 1.00 | 65.11 |  |
| 5 | C | N4 | 14:1575 | 3.93 | 1.00 | 66.14 |  |
| 6 | G | N7 | 14:1421 | 4.71 | 1.00 | 83.97 |  |
| 7 | G | N1 | 14:1421 | 4.81 | 1.00 | 75.98 |  |
| 8 | G | N7 | 14:1422 | 4.87 | 1.00 | 71.75 |  |
| 9 | U | N3 | 14:1576 | 4.90 | 1.00 | 68.49 |  |
  |

  
 Contact(s): /O6/  
 Coded contact list: O_b_  

### K:14:3106

|  | Res. type | Atom | Residue | Distance | Occupancy | B-factor | Closest solute to 1^st^ shell water |
| --- | --- | --- | --- | --- | --- | --- | --- |
| **Ion** | | | |  | 1.00 | 68.56 || 1 | G | O6 | 14:728 | 3.12 | 1.00 | 44.66 |
| 2 | U | O4 | 14:703 | 3.12 | 1.00 | 46.13 |  |
| 3 | G | O6 | 14:702 | 3.79 | 1.00 | 54.85 |  |
| 4 | G | N2 | 14:704 | 3.82 | 1.00 | 48.27 |  |
| 5 | A | OP2 | 14:727 | 3.83 | 1.00 | 42.45 |  |
| 6 | C | N4 | 14:730 | 3.84 | 1.00 | 41.81 |  |
| 7 | G | O2' | 14:726 | 3.91 | 1.00 | 38.00 |  |
| 8 | G | N7 | 14:728 | 4.00 | 1.00 | 45.62 |  |
| 9 | U | N3 | 14:703 | 4.39 | 1.00 | 45.54 |  |
| 10 | A | N7 | 14:727 | 4.40 | 1.00 | 39.35 |  |
| 11 | G | N1 | 14:704 | 4.79 | 1.00 | 51.85 |  |
  |

  
**1^st^ shell ligand angles:**   
1-2: 61;   
  
 Contact(s): /O4/O6/  
 Coded contact list: 2O_b_  

### MG:14:3107

|  | Res. type | Atom | Residue | Distance | Occupancy | B-factor | Closest solute to 1^st^ shell water |
| --- | --- | --- | --- | --- | --- | --- | --- |
| **Ion** | | | |  | 1.00 | 33.71 || 1 | A | OP1 | 14:1614 | 1.94 | 1.00 | 36.83 |
| 2 | A | OP1 | 14:751 | 2.13 | 1.00 | 41.89 |  |
| 3 | A | OP2 | 14:1614 | 3.62 | 1.00 | 31.60 |  |
| 4 | A | N1 | 14:789 | 3.62 | 1.00 | 38.06 |  |
| 5 | G | O2' | 14:1613 | 3.71 | 1.00 | 42.46 |  |
| 6 | A | O3' | 14:750 | 3.94 | 1.00 | 32.28 |  |
| 7 | G | O3' | 14:1613 | 3.99 | 1.00 | 29.85 |  |
| 8 | A | N7 | 14:751 | 4.06 | 1.00 | 40.24 |  |
| 9 | A | N6 | 14:789 | 4.07 | 1.00 | 39.81 |  |
| 10 | A | O5' | 14:751 | 4.09 | 1.00 | 46.78 |  |
| 11 | A | N3 | 14:1614 | 4.39 | 1.00 | 38.83 |  |
| 12 | A | O5' | 14:1614 | 4.40 | 1.00 | 33.13 |  |
| 13 | A | OP2 | 14:751 | 4.70 | 1.00 | 38.15 |  |
| 14 | C | N3 | 14:1617 | 4.79 | 1.00 | 35.02 |  |
| 15 | C | N4 | 14:1617 | 4.80 | 1.00 | 40.62 |  |
| 16 | A | N1 | 14:1614 | 5.00 | 1.00 | 48.46 |  |
  |

  
**1^st^ shell ligand angles:**   
1-2: 138;   
  
 Contact(s): /OP1/OP1/  
 Coded contact list: 2O_ph_  

### MG:14:3108

|  | Res. type | Atom | Residue | Distance | Occupancy | B-factor | Closest solute to 1^st^ shell water |
| --- | --- | --- | --- | --- | --- | --- | --- |
| **Ion** | | | |  | 1.00 | 59.02 || 1 | G | OP2 | 14:1456 | 2.23 | 1.00 | 76.30 |
| 2 | G | OP2 | 14:1455 | 2.35 | 1.00 | 59.07 |  |
| 3 | A | OP2 | 14:1453 | 3.24 | 1.00 | 75.45 |  |
| 4 | G | O5' | 14:1455 | 3.61 | 1.00 | 67.63 |  |
| 5 | U | O3' | 14:1454 | 3.93 | 1.00 | 64.68 |  |
| 6 | U | O2' | 14:1454 | 4.21 | 1.00 | 63.37 |  |
| 7 | G | OP1 | 14:1456 | 4.53 | 1.00 | 91.33 |  |
| 8 | G | O5' | 14:1456 | 4.53 | 1.00 | 79.83 |  |
| 9 | G | O3' | 14:1455 | 4.60 | 1.00 | 69.22 |  |
| 10 | G | N7 | 14:1456 | 4.64 | 1.00 | 68.80 |  |
| 11 | A | O5' | 14:1453 | 4.68 | 1.00 | 69.11 |  |
| 12 | G | OP1 | 14:1455 | 4.89 | 1.00 | 54.74 |  |
  |

  
**1^st^ shell ligand angles:**   
1-2: 96;   
  
 Contact(s): /OP2/OP2/  
 Coded contact list: 2O_ph_  
 Exclusion zone contact(s): /OP2/  

### MG:14:3109

|  | Res. type | Atom | Residue | Distance | Occupancy | B-factor | Closest solute to 1^st^ shell water |
| --- | --- | --- | --- | --- | --- | --- | --- |
| **Ion** | | | |  | 1.00 | 41.46 || 1 | U | OP2 | 14:120 | 2.44 | 1.00 | 49.25 |
| 2 | Water 1 | | 14:3775 | 2.52 | 1.00 | 35.25 | 2.71 (G:O6 14:124) |
| 3 | G | O6 | 14:122 | 2.88 | 1.00 | 30.53 |  |
| 4 | G | O6 | 14:123 | 3.51 | 1.00 | 31.70 |  |
| 5 | G | N7 | 14:123 | 3.61 | 1.00 | 29.67 |  |
| 6 | G | N7 | 14:122 | 3.62 | 1.00 | 37.15 |  |
| 7 | Water 2 | | 14:3975 | 4.29 | 1.00 | 46.28 | 3.81 (U:OP2 14:120) |
| 8 | G | O6 | 14:124 | 4.31 | 1.00 | 36.50 |  |
| 9 | A | O3' | 14:119 | 4.56 | 1.00 | 49.21 |  |
| 10 | U | OP1 | 14:120 | 4.60 | 1.00 | 48.69 |  |
| 11 | G | OP1 | 14:121 | 4.62 | 1.00 | 45.88 |  |
| 12 | G | N1 | 14:122 | 4.73 | 1.00 | 30.94 |  |
| 13 | C | N4 | 14:128 | 4.75 | 1.00 | 27.09 |  |
| 14 | C | N4 | 14:129 | 4.96 | 1.00 | 30.90 |  |
  |

  
 Coded contact list:   
 Exclusion zone contact(s): /O6/O_w_/OP2/  
Consider NA in place of MG (1 valid contact(s) for NA vs. 0 for MG)  

### MG:14:3110

|  | Res. type | Atom | Residue | Distance | Occupancy | B-factor | Closest solute to 1^st^ shell water |
| --- | --- | --- | --- | --- | --- | --- | --- |
| **Ion** | | | |  | 1.00 | 35.26 || 1 | C | OP1 | 14:1774 | 2.04 | 1.00 | 35.28 |
| 2 | G | O2' | 14:1772 | 4.03 | 1.00 | 35.41 |  |
| 3 | G | O6 | 14:1776 | 4.13 | 1.00 | 33.65 |  |
| 4 | C | O5' | 14:1774 | 4.13 | 1.00 | 35.37 |  |
| 5 | U | OP2 | 14:1775 | 4.14 | 1.00 | 34.36 |  |
| 6 | C | OP2 | 14:1774 | 4.15 | 1.00 | 30.11 |  |
| 7 | G | N3 | 14:1772 | 4.17 | 1.00 | 38.37 |  |
| 8 | A | OP2 | 14:1786 | 4.29 | 1.00 | 35.77 |  |
| 9 | G | N2 | 14:1772 | 4.30 | 1.00 | 36.79 |  |
| 10 | A | OP1 | 14:1786 | 4.36 | 1.00 | 37.16 |  |
| 11 | A | O3' | 14:1773 | 4.57 | 1.00 | 39.29 |  |
  |

  
 Contact(s): /OP1/  
 Coded contact list: O_ph_  

### MG:14:3111

|  | Res. type | Atom | Residue | Distance | Occupancy | B-factor | Closest solute to 1^st^ shell water |
| --- | --- | --- | --- | --- | --- | --- | --- |
| **Ion** | | | |  | 1.00 | 32.88 || 1 | G | OP2 | 14:1975 | 2.46 | 1.00 | 36.87 |
| 2 | C | OP1 | 14:1771 | 3.49 | 1.00 | 38.81 |  |
| 3 | G | OP1 | 14:1975 | 4.15 | 1.00 | 36.18 |  |
| 4 | C | OP2 | 14:1771 | 4.36 | 1.00 | 29.49 |  |
| 5 | C | O5' | 14:1974 | 4.42 | 1.00 | 36.51 |  |
| 6 | C | O3' | 14:1974 | 4.56 | 1.00 | 40.55 |  |
| 7 | C | OP2 | 14:1974 | 4.75 | 1.00 | 37.49 |  |
| 8 | G | O5' | 14:1975 | 4.93 | 1.00 | 25.79 |  |
  |

  
 Coded contact list:   
 Exclusion zone contact(s): /OP2/  
Consider NA in place of MG (1 valid contact(s) for NA vs. 0 for MG)  

### MG:14:3112

|  | Res. type | Atom | Residue | Distance | Occupancy | B-factor | Closest solute to 1^st^ shell water |
| --- | --- | --- | --- | --- | --- | --- | --- |
| **Ion** | | | |  | 1.00 | 44.97 || 1 | G | OP2 | 14:2255 | 2.44 | 1.00 | 49.89 |
| 2 | G | N7 | 14:2256 | 3.07 | 1.00 | 48.30 |  |
| 3 | G | N7 | 14:2255 | 3.33 | 1.00 | 47.48 |  |
| 4 | G | O5' | 14:2255 | 3.49 | 1.00 | 48.61 |  |
| 5 | G | O6 | 14:2256 | 4.00 | 1.00 | 43.17 |  |
| 6 | G | OP2 | 14:2256 | 4.09 | 1.00 | 54.86 |  |
| 7 | G | OP1 | 14:2255 | 4.56 | 1.00 | 51.52 |  |
| 8 | G | N9 | 14:2255 | 4.68 | 1.00 | 47.94 |  |
| 9 | U | O4 | 14:2257 | 4.75 | 1.00 | 45.06 |  |
| 10 | C | O3' | 14:2254 | 4.81 | 1.00 | 43.04 |  |
| 11 | C | OP2 | 14:2254 | 4.91 | 1.00 | 51.13 |  |
  |

  
 Coded contact list:   
 Exclusion zone contact(s): /N7/N7/OP2/  
Consider K in place of MG (2 valid contact(s) for K vs. 0 for MG)  

### MG:14:3113

|  | Res. type | Atom | Residue | Distance | Occupancy | B-factor | Closest solute to 1^st^ shell water |
| --- | --- | --- | --- | --- | --- | --- | --- |
| **Ion** | | | |  | 1.00 | 36.44 || 1 | G | O6 | 14:2428 | 3.53 | 1.00 | 41.48 |
| 2 | G | N7 | 14:2428 | 3.63 | 1.00 | 43.32 |  |
| 3 | C | OP1 | 14:2260 | 4.07 | 1.00 | 46.22 |  |
| 4 | C | OP1 | 14:825 | 4.48 | 1.00 | 57.81 |  |
| 5 | G | OP2 | 14:2428 | 4.95 | 1.00 | 41.69 |  |
  |

  
 Coded contact list:   
1^st^ MG coordination shell missing (see Notes)  

### MG:14:3114

|  | Res. type | Atom | Residue | Distance | Occupancy | B-factor | Closest solute to 1^st^ shell water |
| --- | --- | --- | --- | --- | --- | --- | --- |
| **Ion** | | | |  | 1.00 | 33.13 || 1 | U | OP1 | 14:1673 | 2.33 | 1.00 | 53.23 |
| 2 | OMU | OP2 | 14:2552 | 2.52 | 1.00 | 43.04 |  |
| 3 | OMU | OP1 | 14:2552 | 3.47 | 1.00 | 52.12 |  |
| 4 | C | O3' | 14:1672 | 3.60 | 1.00 | 35.23 |  |
| 5 | U | OP1 | 14:2554 | 3.90 | 1.00 | 36.12 |  |
| 6 | OMU | O5' | 14:2552 | 4.24 | 1.00 | 47.66 |  |
| 7 | G | OP2 | 14:2553 | 4.31 | 1.00 | 43.83 |  |
| 8 | C | O5' | 14:2551 | 4.51 | 1.00 | 46.89 |  |
| 9 | U | O5' | 14:1673 | 4.51 | 1.00 | 39.93 |  |
| 10 | U | OP2 | 14:1673 | 4.52 | 1.00 | 46.58 |  |
| 11 | C | O3' | 14:2551 | 4.74 | 1.00 | 45.37 |  |
| 12 | C | OP1 | 14:2551 | 4.79 | 1.00 | 55.29 |  |
| 13 | C | OP2 | 14:2551 | 4.97 | 1.00 | 43.87 |  |
  |

  
 Contact(s): /OP1/  
 Coded contact list: O_ph_  
 Exclusion zone contact(s): /OP2/  

### MG:14:3115

|  | Res. type | Atom | Residue | Distance | Occupancy | B-factor | Closest solute to 1^st^ shell water |
| --- | --- | --- | --- | --- | --- | --- | --- |
| **Ion** | | | |  | 1.00 | 42.28 || 1 | G | OP1 | 14:465 | 3.44 | 1.00 | 40.83 |
| 2 | G | N7 | 14:463 | 3.47 | 1.00 | 36.87 |  |
| 3 | G | O6 | 14:463 | 3.86 | 1.00 | 43.81 |  |
| 4 | G | OP2 | 14:465 | 4.66 | 1.00 | 29.76 |  |
| 5 | ARG | NH2 | L5:12 | 4.87 | 1.00 | 44.80 |  |
  |

  
 Coded contact list:   
1^st^ MG coordination shell missing (see Notes)  

### MG:14:3116

|  | Res. type | Atom | Residue | Distance | Occupancy | B-factor | Closest solute to 1^st^ shell water |
| --- | --- | --- | --- | --- | --- | --- | --- |
| **Ion** | | | |  | 1.00 | 32.75 || 1 | G | OP2 | 14:1987 | 3.67 | 1.00 | 43.93 |
| 2 | A | OP1 | 14:1986 | 3.71 | 1.00 | 54.50 |  |
| 3 | G | OP1 | 14:1987 | 3.94 | 1.00 | 53.40 |  |
| 4 | C | OP1 | 14:2551 | 4.28 | 1.00 | 55.29 |  |
| 5 | G | O6 | 14:1954 | 4.37 | 1.00 | 61.12 |  |
| 6 | A | OP1 | 14:1669 | 4.47 | 1.00 | 38.09 |  |
| 7 | G | N1 | 14:1954 | 4.68 | 1.00 | 47.68 |  |
| 8 | Water 1 | | 14:3529 | 4.78 | 1.00 | 39.32 | 3.67 (C:OP1 14:2551) |
| 9 | A | O5' | 14:1986 | 4.98 | 1.00 | 36.26 |  |
  |

  
 Coded contact list:   
1^st^ MG coordination shell missing (see Notes)  

### MG:14:3117

|  | Res. type | Atom | Residue | Distance | Occupancy | B-factor | Closest solute to 1^st^ shell water |
| --- | --- | --- | --- | --- | --- | --- | --- |
| **Ion** | | | |  | 1.00 | 49.61 || 1 | G | O6 | 14:1824 | 4.16 | 1.00 | 37.05 |
| 2 | G | N7 | 14:1824 | 4.23 | 1.00 | 34.57 |  |
| 3 | G | N7 | 14:1823 | 4.72 | 1.00 | 30.17 |  |
| 4 | A | N6 | 14:1825 | 4.94 | 1.00 | 29.48 |  |
| 5 | G | OP2 | 14:1823 | 4.97 | 1.00 | 28.39 |  |
  |

  
 Coded contact list:   
1^st^ MG coordination shell missing (see Notes)  

### MG:14:3118

|  | Res. type | Atom | Residue | Distance | Occupancy | B-factor | Closest solute to 1^st^ shell water |
| --- | --- | --- | --- | --- | --- | --- | --- |
| **Ion** | | | |  | 1.00 | 58.95 || 1 | G | O6 | 14:1425 | 2.83 | 1.00 | 65.67 |
| 2 | G | O6 | 14:1424 | 3.22 | 1.00 | 66.21 |  |
| 3 | G | N1 | 14:1425 | 4.02 | 1.00 | 55.41 |  |
| 4 | G | N7 | 14:1424 | 4.28 | 1.00 | 62.87 |  |
| 5 | A | OP2 | 14:1572 | 4.35 | 1.00 | 47.17 |  |
| 6 | G | N7 | 14:1423 | 4.49 | 1.00 | 63.08 |  |
| 7 | G | O6 | 14:1423 | 4.64 | 1.00 | 65.11 |  |
  |

  
 Coded contact list:   
 Exclusion zone contact(s): /O6/O6/  
Consider K in place of MG (2 valid contact(s) for K vs. 0 for MG)  

### MG:14:3119

|  | Res. type | Atom | Residue | Distance | Occupancy | B-factor | Closest solute to 1^st^ shell water |
| --- | --- | --- | --- | --- | --- | --- | --- |
| **Ion** | | | |  | 1.00 | 49.87 || 1 | G | OP2 | 14:2270 | 2.02 | 1.00 | 58.53 |
| 2 | A | OP2 | 14:2267 | 3.61 | 1.00 | 57.04 |  |
| 3 | G | N7 | 14:2271 | 3.82 | 1.00 | 61.75 |  |
| 4 | U | O4 | 14:2272 | 3.87 | 1.00 | 67.09 |  |
| 5 | G | O6 | 14:2271 | 3.89 | 1.00 | 64.33 |  |
| 6 | G | O5' | 14:2270 | 4.07 | 1.00 | 63.94 |  |
| 7 | A | OP1 | 14:2269 | 4.34 | 1.00 | 74.20 |  |
| 8 | G | OP1 | 14:2270 | 4.41 | 1.00 | 61.51 |  |
| 9 | A | O3' | 14:2269 | 4.45 | 1.00 | 56.17 |  |
| 10 | A | O5' | 14:2269 | 4.57 | 1.00 | 62.75 |  |
| 11 | A | O3' | 14:2266 | 4.63 | 1.00 | 71.80 |  |
| 12 | A | O3' | 14:2268 | 4.74 | 1.00 | 68.34 |  |
| 13 | G | N7 | 14:2270 | 4.76 | 1.00 | 62.72 |  |
| 14 | G | OP2 | 14:2271 | 4.81 | 1.00 | 53.76 |  |
  |

  
 Contact(s): /OP2/  
 Coded contact list: O_ph_  

### MG:14:3120

|  | Res. type | Atom | Residue | Distance | Occupancy | B-factor | Closest solute to 1^st^ shell water |
| --- | --- | --- | --- | --- | --- | --- | --- |
| **Ion** | | | |  | 1.00 | 44.07 || 1 | U | OP1 | 14:773 | 2.82 | 1.00 | 25.39 |
| 2 | G | OP1 | 14:1356 | 3.50 | 1.00 | 50.48 |  |
| 3 | C | O3' | 14:772 | 3.76 | 1.00 | 32.69 |  |
| 4 | C | OP1 | 14:1370 | 4.06 | 1.00 | 37.81 |  |
| 5 | U | O5' | 14:773 | 4.75 | 1.00 | 32.01 |  |
  |

  
 Coded contact list:   
 Exclusion zone contact(s): /OP1/  
Consider K in place of MG (1 valid contact(s) for K vs. 0 for MG)  

### MG:14:3121

|  | Res. type | Atom | Residue | Distance | Occupancy | B-factor | Closest solute to 1^st^ shell water |
| --- | --- | --- | --- | --- | --- | --- | --- |
| **Ion** | | | |  | 1.00 | 47.51 || 1 | G | O6 | 14:397 | 2.71 | 1.00 | 55.44 |
| 2 | G | N7 | 14:397 | 3.35 | 1.00 | 53.08 |  |
| 3 | G | O6 | 14:396 | 4.19 | 1.00 | 46.90 |  |
| 4 | G | O6 | 14:398 | 4.29 | 1.00 | 58.20 |  |
| 5 | G | N7 | 14:396 | 4.35 | 1.00 | 50.80 |  |
| 6 | G | OP2 | 14:396 | 4.36 | 1.00 | 60.83 |  |
| 7 | G | N1 | 14:397 | 4.80 | 1.00 | 46.44 |  |
  |

  
 Coded contact list:   
 Exclusion zone contact(s): /N7/O6/  
Consider K in place of MG (2 valid contact(s) for K vs. 0 for MG)  

### MG:14:3122

|  | Res. type | Atom | Residue | Distance | Occupancy | B-factor | Closest solute to 1^st^ shell water |
| --- | --- | --- | --- | --- | --- | --- | --- |
| **Ion** | | | |  | 1.00 | 49.51 || 1 | G | N7 | 14:1642 | 3.47 | 1.00 | 51.14 |
| 2 | G | O6 | 14:1643 | 3.59 | 1.00 | 43.67 |  |
| 3 | G | O6 | 14:1642 | 3.70 | 1.00 | 48.30 |  |
| 4 | G | N7 | 14:1643 | 3.80 | 1.00 | 47.07 |  |
| 5 | A | OP2 | 14:1641 | 3.96 | 1.00 | 46.32 |  |
  |

  
 Coded contact list:   
1^st^ MG coordination shell missing (see Notes)  

### MG:14:3123

|  | Res. type | Atom | Residue | Distance | Occupancy | B-factor | Closest solute to 1^st^ shell water |
| --- | --- | --- | --- | --- | --- | --- | --- |
| **Ion** | | | |  | 1.00 | 36.11 || 1 | Water 1 | | 14:3814 | 1.85 | 1.00 | 31.71 | 2.79 (G:OP2 14:778) |
| 2 | G | OP2 | 14:778 | 2.14 | 1.00 | 32.34 |  |
| 3 | Water 2 | | 14:3990 | 2.83 | 1.00 | 40.72 | 4.20 (G:OP2 14:778) |
| 4 | Water 3 | | 19:411 | 3.83 | 1.00 | 40.08 | 3.92 (U:O4 14:779) |
| 5 | Water 4 | | 14:3707 | 4.02 | 1.00 | 35.55 | 2.60 (G:N7 14:778) |
| 6 | G | OP1 | 14:778 | 4.08 | 1.00 | 37.15 |  |
| 7 | A | O3' | 14:777 | 4.43 | 1.00 | 28.27 |  |
| 8 | G | O5' | 14:778 | 4.49 | 1.00 | 30.91 |  |
| 9 | G | N7 | 14:778 | 4.67 | 1.00 | 30.26 |  |
| 10 | U | O4 | 14:779 | 4.68 | 1.00 | 31.60 |  |
  |

  
**1^st^ shell ligand angles:**   
1-2: 88;   
  
 Contact(s): /O_w_/OP2/  
 Coded contact list: O_ph_O_w_  
 Exclusion zone contact(s): /O_w_/  

### MG:14:3124

|  | Res. type | Atom | Residue | Distance | Occupancy | B-factor | Closest solute to 1^st^ shell water |
| --- | --- | --- | --- | --- | --- | --- | --- |
| **Ion** | | | |  | 1.00 | 58.30 || 1 | A | OP1 | 14:2268 | 2.20 | 1.00 | 68.13 |
| 2 | A | OP1 | 14:2269 | 2.38 | 1.00 | 74.20 |  |
| 3 | G | O6 | 14:2271 | 3.31 | 1.00 | 64.33 |  |
| 4 | A | N3 | 14:2267 | 3.33 | 1.00 | 62.37 |  |
| 5 | A | OP2 | 14:2267 | 3.68 | 1.00 | 57.04 |  |
| 6 | A | O5' | 14:2267 | 4.16 | 1.00 | 61.78 |  |
| 7 | A | O3' | 14:2267 | 4.33 | 1.00 | 63.10 |  |
| 8 | A | O3' | 14:2268 | 4.36 | 1.00 | 68.34 |  |
| 9 | A | N1 | 14:2267 | 4.53 | 1.00 | 58.12 |  |
| 10 | A | O5' | 14:2268 | 4.54 | 1.00 | 68.96 |  |
| 11 | A | OP2 | 14:2268 | 4.61 | 1.00 | 64.95 |  |
| 12 | A | OP2 | 14:2269 | 4.66 | 1.00 | 71.99 |  |
| 13 | A | O5' | 14:2269 | 4.79 | 1.00 | 62.75 |  |
| 14 | G | N1 | 14:2271 | 4.99 | 1.00 | 59.97 |  |
  |

  
**1^st^ shell ligand angles:**   
1-2: 74;   
  
 Contact(s): /OP1/OP1/  
 Coded contact list: 2O_ph_  
 Exclusion zone contact(s): /N3_A_/O6/  

### MG:14:3125

|  | Res. type | Atom | Residue | Distance | Occupancy | B-factor | Closest solute to 1^st^ shell water |
| --- | --- | --- | --- | --- | --- | --- | --- |
| **Ion** | | | |  | 1.00 | 60.19 || 1 | G | OP2 | 14:212 | 3.73 | 1.00 | 46.52 |
| 2 | A | OP2 | 14:213 | 3.89 | 1.00 | 57.40 |  |
| 3 | G | OP2 | 14:178 | 3.99 | 1.00 | 53.49 |  |
| 4 | G | N7 | 14:178 | 4.51 | 1.00 | 47.82 |  |
| 5 | G | O5' | 14:212 | 4.56 | 1.00 | 51.39 |  |
| 6 | G | OP1 | 14:212 | 4.70 | 1.00 | 46.83 |  |
| 7 | G | OP1 | 14:177 | 4.76 | 1.00 | 54.37 |  |
| 8 | G | OP2 | 14:179 | 4.78 | 1.00 | 43.49 |  |
  |

  
 Coded contact list:   
1^st^ MG coordination shell missing (see Notes)  

### MG:14:3126

|  | Res. type | Atom | Residue | Distance | Occupancy | B-factor | Closest solute to 1^st^ shell water |
| --- | --- | --- | --- | --- | --- | --- | --- |
| **Ion** | | | |  | 1.00 | 27.80 || 1 | C | OP2 | 14:730 | 2.21 | 1.00 | 30.34 |
| 2 | C | OP2 | 14:731 | 2.40 | 1.00 | 40.01 |  |
| 3 | A | OP1 | 14:761 | 2.48 | 1.00 | 34.86 |  |
| 4 | G | OP1 | 14:763 | 3.03 | 1.00 | 32.00 |  |
| 5 | G | O3' | 14:729 | 4.00 | 1.00 | 26.56 |  |
| 6 | A | O5' | 14:761 | 4.02 | 1.00 | 38.68 |  |
| 7 | C | O3' | 14:730 | 4.11 | 1.00 | 30.19 |  |
| 8 | C | O5' | 14:730 | 4.38 | 1.00 | 34.62 |  |
| 9 | U | OP1 | 14:762 | 4.41 | 1.00 | 40.12 |  |
| 10 | G | O5' | 14:763 | 4.60 | 1.00 | 34.39 |  |
| 11 | C | OP1 | 14:730 | 4.69 | 1.00 | 32.72 |  |
| 12 | A | OP2 | 14:761 | 4.72 | 1.00 | 31.53 |  |
| 13 | C | OP1 | 14:731 | 4.78 | 1.00 | 31.40 |  |
| 14 | C | O5' | 14:731 | 4.81 | 1.00 | 37.19 |  |
| 15 | G | O3' | 14:760 | 4.96 | 1.00 | 29.70 |  |
  |

  
**1^st^ shell ligand angles:**   
1-2: 68;   
  
 Contact(s): /OP2/OP2/  
 Coded contact list: 2O_ph_  
 Exclusion zone contact(s): /OP1/OP1/  

### MG:14:3127

|  | Res. type | Atom | Residue | Distance | Occupancy | B-factor | Closest solute to 1^st^ shell water |
| --- | --- | --- | --- | --- | --- | --- | --- |
| **Ion** | | | |  | 1.00 | 30.64 || 1 | C | OP1 | 14:1670 | 2.21 | 1.00 | 35.40 |
| 2 | U | OP2 | 14:1671 | 2.45 | 1.00 | 48.92 |  |
| 3 | Water 1 | | 14:3644 | 3.31 | 1.00 | 43.80 | 2.48 (G:OP2 14:2550) |
| 4 | G | OP1 | 14:2549 | 3.89 | 1.00 | 44.54 |  |
| 5 | G | OP2 | 14:2550 | 4.08 | 1.00 | 38.94 |  |
| 6 | A | O3' | 14:1669 | 4.49 | 1.00 | 42.56 |  |
| 7 | U | OP1 | 14:1671 | 4.54 | 1.00 | 35.47 |  |
| 8 | C | OP2 | 14:1670 | 4.59 | 1.00 | 37.68 |  |
| 9 | C | O5' | 14:1670 | 4.60 | 1.00 | 46.51 |  |
| 10 | U | O5' | 14:1671 | 4.64 | 1.00 | 37.46 |  |
| 11 | G | OP2 | 14:2549 | 4.73 | 1.00 | 45.91 |  |
| 12 | C | O3' | 14:1670 | 4.87 | 1.00 | 36.42 |  |
  |

  
 Contact(s): /OP1/  
 Coded contact list: O_ph_  
 Exclusion zone contact(s): /O_w_/OP2/  

### MG:14:3128

|  | Res. type | Atom | Residue | Distance | Occupancy | B-factor | Closest solute to 1^st^ shell water |
| --- | --- | --- | --- | --- | --- | --- | --- |
| **Ion** | | | |  | 1.00 | 28.12 || 1 | A | OP1 | 14:2711 | 2.14 | 1.00 | 42.22 |
| 2 | A | OP1 | 14:2712A | 2.30 | 1.00 | 43.03 |  |
| 3 | C | OP1 | 14:1297 | 3.33 | 1.00 | 51.05 |  |
| 4 | C | O3' | 14:2710 | 3.49 | 1.00 | 46.29 |  |
| 5 | U | O3' | 14:2712 | 3.71 | 1.00 | 47.73 |  |
| 6 | A | OP2 | 14:2712A | 3.80 | 1.00 | 38.41 |  |
| 7 | A | OP2 | 14:2711 | 4.36 | 1.00 | 39.66 |  |
| 8 | A | O5' | 14:2711 | 4.50 | 1.00 | 37.06 |  |
| 9 | A | O5' | 14:2712A | 4.80 | 1.00 | 27.48 |  |
  |

  
**1^st^ shell ligand angles:**   
1-2: 81;   
  
 Contact(s): /OP1/OP1/  
 Coded contact list: 2O_ph_  
 Exclusion zone contact(s): /OP1/  

### MG:14:3129

|  | Res. type | Atom | Residue | Distance | Occupancy | B-factor | Closest solute to 1^st^ shell water |
| --- | --- | --- | --- | --- | --- | --- | --- |
| **Ion** | | | |  | 1.00 | 34.01 || 1 | A | OP1 | 14:197 | 2.31 | 1.00 | 41.76 |
| 2 | C | OP2 | 14:198 | 2.32 | 1.00 | 33.93 |  |
| 3 | A | N7 | 14:195 | 2.58 | 1.00 | 35.01 |  |
| 4 | C | OP1 | 14:198 | 3.81 | 1.00 | 35.32 |  |
| 5 | A | OP2 | 14:195 | 3.82 | 1.00 | 38.62 |  |
| 6 | G | OP2 | 14:194 | 3.92 | 1.00 | 36.08 |  |
| 7 | A | N6 | 14:195 | 4.11 | 1.00 | 35.19 |  |
| 8 | G | N7 | 14:194 | 4.11 | 1.00 | 32.38 |  |
| 9 | A | O5' | 14:197 | 4.47 | 1.00 | 27.98 |  |
| 10 | G | O5' | 14:194 | 4.53 | 1.00 | 30.96 |  |
| 11 | A | O3' | 14:196 | 4.56 | 1.00 | 27.25 |  |
| 12 | A | O3' | 14:197 | 4.58 | 1.00 | 41.45 |  |
| 13 | A | OP2 | 14:197 | 4.58 | 1.00 | 34.57 |  |
| 14 | C | O5' | 14:198 | 4.64 | 1.00 | 36.06 |  |
| 15 | A | N9 | 14:195 | 4.66 | 1.00 | 35.21 |  |
| 16 | Water 1 | | 14:3754 | 4.72 | 1.00 | 38.25 | 3.69 (C:OP1 14:198) |
  |

  
**1^st^ shell ligand angles:**   
1-2: 70;   
  
 Contact(s): /OP1/OP2/  
 Coded contact list: 2O_ph_  
 Exclusion zone contact(s): /N7/  

### MG:14:3130

|  | Res. type | Atom | Residue | Distance | Occupancy | B-factor | Closest solute to 1^st^ shell water |
| --- | --- | --- | --- | --- | --- | --- | --- |
| **Ion** | | | |  | 1.00 | 38.66 || 1 | C | OP1 | 14:2427 | 1.77 | 1.00 | 41.65 |
| 2 | U | OP1 | 14:826 | 2.06 | 1.00 | 41.33 |  |
| 3 | G | OP1 | 14:2428 | 2.33 | 1.00 | 40.57 |  |
| 4 | G | OP1 | 14:2429 | 2.38 | 1.00 | 43.33 |  |
| 5 | G | OP2 | 14:2429 | 3.86 | 1.00 | 48.69 |  |
| 6 | C | OP2 | 14:2427 | 3.87 | 1.00 | 44.97 |  |
| 7 | C | O5' | 14:2427 | 3.91 | 1.00 | 43.22 |  |
| 8 | U | O5' | 14:826 | 4.02 | 1.00 | 47.38 |  |
| 9 | G | OP2 | 14:2428 | 4.05 | 1.00 | 41.69 |  |
| 10 | U | OP2 | 14:826 | 4.11 | 1.00 | 43.52 |  |
| 11 | A | O3' | 14:2426 | 4.24 | 1.00 | 38.64 |  |
| 12 | G | O5' | 14:2428 | 4.33 | 1.00 | 39.34 |  |
| 13 | G | O3' | 14:2428 | 4.47 | 1.00 | 37.95 |  |
| 14 | C | O3' | 14:825 | 4.65 | 1.00 | 38.07 |  |
| 15 | C | O3' | 14:2427 | 4.78 | 1.00 | 43.68 |  |
| 16 | G | O5' | 14:2429 | 4.90 | 1.00 | 38.78 |  |
| 17 | U | OP2 | 14:827 | 4.99 | 1.00 | 46.15 |  |
  |

  
**1^st^ shell ligand angles:**   
1-2: 166; 1-3: 77; 1-4: 88; 2-3: 95; 2-4: 78; 3-4: 61;   
  
 Contact(s): /OP1/OP1/OP1/  
 Coded contact list: 3O_ph_  
 Exclusion zone contact(s): /OP1/  
Large octahedral topology discrepancy!!!  

### MG:14:3131

|  | Res. type | Atom | Residue | Distance | Occupancy | B-factor | Closest solute to 1^st^ shell water |
| --- | --- | --- | --- | --- | --- | --- | --- |
| **Ion** | | | |  | 1.00 | 43.06 || 1 | G | OP2 | 14:2822 | 1.86 | 1.00 | 46.25 |
| 2 | A | OP2 | 14:2821 | 2.19 | 1.00 | 46.30 |  |
| 3 | HIS | NE2 | 55:3 | 2.19 | 1.00 | 42.76 |  |
| 4 | GLY | O | 29:110 | 2.73 | 1.00 | 51.31 |  |
| 5 | G | N7 | 14:2822 | 3.19 | 1.00 | 46.35 |  |
| 6 | A | O3' | 14:2820 | 3.67 | 1.00 | 45.63 |  |
| 7 | G | O5' | 14:2822 | 3.97 | 1.00 | 43.44 |  |
| 8 | A | O3' | 14:2821 | 4.05 | 1.00 | 52.91 |  |
| 9 | A | O5' | 14:2821 | 4.30 | 1.00 | 41.20 |  |
| 10 | HIS | ND1 | 55:3 | 4.31 | 1.00 | 39.01 |  |
| 11 | G | OP1 | 14:2822 | 4.42 | 1.00 | 44.28 |  |
| 12 | A | OP1 | 14:2821 | 4.69 | 1.00 | 42.92 |  |
| 13 | ARG | O | 29:111 | 4.78 | 1.00 | 50.49 |  |
| 14 | G | O6 | 14:2822 | 4.87 | 1.00 | 50.88 |  |
| 15 | ARG | N | 29:111 | 4.93 | 1.00 | 51.93 |  |
| 16 | GLY | N | 29:112 | 4.98 | 1.00 | 39.48 |  |
| 17 | GLY | N | 29:110 | 5.00 | 1.00 | 54.04 |  |
  |

  
**1^st^ shell ligand angles:**   
1-2: 78; 1-3: 139; 2-3: 78;   
  
 Contact(s): /NE2_His_/OP2/OP2/  
 Coded contact list: 2O_ph_His  
 Exclusion zone contact(s): /N7/O_back_/  

### MG:14:3132

|  | Res. type | Atom | Residue | Distance | Occupancy | B-factor | Closest solute to 1^st^ shell water |
| --- | --- | --- | --- | --- | --- | --- | --- |
| **Ion** | | | |  | 1.00 | 53.28 || 1 | G | O6 | 14:1187 | 2.02 | 1.00 | 61.80 |
| 2 | G | OP2 | 14:818 | 2.38 | 1.00 | 59.39 |  |
| 3 | C | OP1 | 14:817 | 3.78 | 1.00 | 61.85 |  |
| 4 | G | OP1 | 14:1186 | 4.14 | 1.00 | 66.09 |  |
| 5 | C | OP2 | 14:817 | 4.18 | 1.00 | 64.91 |  |
| 6 | G | N1 | 14:1187 | 4.20 | 1.00 | 64.63 |  |
| 7 | C | O5' | 14:817 | 4.21 | 1.00 | 65.13 |  |
| 8 | G | N7 | 14:1187 | 4.21 | 1.00 | 69.17 |  |
| 9 | G | OP1 | 14:818 | 4.37 | 1.00 | 62.58 |  |
| 10 | G | N7 | 14:1186 | 4.38 | 1.00 | 67.91 |  |
| 11 | G | O5' | 14:818 | 4.51 | 1.00 | 57.49 |  |
| 12 | G | N7 | 14:818 | 4.68 | 1.00 | 66.42 |  |
| 13 | C | O3' | 14:817 | 4.85 | 1.00 | 46.43 |  |
  |

  
**1^st^ shell ligand angles:**   
1-2: 76;   
  
 Contact(s): /O6/OP2/  
 Coded contact list: O_ph_O_b_  

### MG:14:3133

|  | Res. type | Atom | Residue | Distance | Occupancy | B-factor | Closest solute to 1^st^ shell water |
| --- | --- | --- | --- | --- | --- | --- | --- |
| **Ion** | | | |  | 1.00 | 39.95 || 1 | C | OP2 | 14:1345 | 2.22 | 1.00 | 53.86 |
| 2 | G | OP2 | 14:1346 | 2.72 | 1.00 | 43.89 |  |
| 3 | C | O2 | 14:1383 | 3.42 | 1.00 | 60.32 |  |
| 4 | A | N7 | 14:1384 | 3.74 | 1.00 | 63.05 |  |
| 5 | C | N3 | 14:1383 | 4.17 | 1.00 | 55.23 |  |
| 6 | A | N6 | 14:1384 | 4.27 | 1.00 | 55.44 |  |
| 7 | G | O3' | 14:1344 | 4.32 | 1.00 | 55.61 |  |
| 8 | C | OP1 | 14:1345 | 4.41 | 1.00 | 53.37 |  |
| 9 | C | O5' | 14:1345 | 4.70 | 1.00 | 54.64 |  |
| 10 | G | O5' | 14:1346 | 4.89 | 1.00 | 40.07 |  |
| 11 | G | OP1 | 14:1346 | 4.97 | 1.00 | 60.84 |  |
  |

  
 Contact(s): /OP2/  
 Coded contact list: O_ph_  
 Exclusion zone contact(s): /OP2/  

### MG:14:3134

|  | Res. type | Atom | Residue | Distance | Occupancy | B-factor | Closest solute to 1^st^ shell water |
| --- | --- | --- | --- | --- | --- | --- | --- |
| **Ion** | | | |  | 1.00 | 34.33 || 1 | U | O4 | 14:2249 | 2.15 | 1.00 | 41.08 |
| 2 | C | OP2 | 14:2248 | 2.22 | 1.00 | 40.50 |  |
| 3 | C | OP1 | 14:2066 | 3.73 | 1.00 | 40.98 |  |
| 4 | G | OP1 | 14:2253 | 3.93 | 1.00 | 43.21 |  |
| 5 | A | O5' | 14:2247 | 4.02 | 1.00 | 39.93 |  |
| 6 | U | N3 | 14:2249 | 4.40 | 1.00 | 39.07 |  |
| 7 | A | O3' | 14:2247 | 4.40 | 1.00 | 38.21 |  |
| 8 | A | OP1 | 14:2247 | 4.49 | 1.00 | 40.26 |  |
| 9 | C | O5' | 14:2248 | 4.63 | 1.00 | 43.46 |  |
| 10 | C | OP1 | 14:2248 | 4.69 | 1.00 | 40.60 |  |
| 11 | A | OP2 | 14:2247 | 4.93 | 1.00 | 39.32 |  |
| 12 | G | O3' | 14:2252 | 4.95 | 1.00 | 42.60 |  |
  |

  
**1^st^ shell ligand angles:**   
1-2: 89;   
  
 Contact(s): /O4/OP2/  
 Coded contact list: O_ph_O_b_  

### MG:14:3135

|  | Res. type | Atom | Residue | Distance | Occupancy | B-factor | Closest solute to 1^st^ shell water |
| --- | --- | --- | --- | --- | --- | --- | --- |
| **Ion** | | | |  | 1.00 | 47.33 || 1 | A | N7 | 14:1632 | 2.14 | 1.00 | 48.05 |
| 2 | G | O6 | 14:1633 | 2.17 | 1.00 | 48.96 |  |
| 3 | A | OP2 | 14:1632 | 3.20 | 1.00 | 51.47 |  |
| 4 | A | N6 | 14:1632 | 3.27 | 1.00 | 47.38 |  |
| 5 | A | N6 | 14:1762 | 3.31 | 1.00 | 60.95 |  |
| 6 | A | OP1 | 14:1631 | 3.73 | 1.00 | 67.66 |  |
| 7 | G | N7 | 14:1633 | 3.80 | 1.00 | 44.75 |  |
| 8 | G | N1 | 14:1633 | 4.26 | 1.00 | 44.93 |  |
| 9 | C | O2 | 14:1630A | 4.29 | 1.00 | 48.64 |  |
| 10 | C | N3 | 14:1630A | 4.37 | 1.00 | 47.31 |  |
| 11 | A | N9 | 14:1632 | 4.37 | 1.00 | 44.80 |  |
| 12 | A | N7 | 14:1631 | 4.40 | 1.00 | 46.21 |  |
| 13 | G | N2 | 14:1635 | 4.57 | 1.00 | 50.00 |  |
| 14 | G | N1 | 14:1635 | 4.65 | 1.00 | 42.97 |  |
| 15 | A | O5' | 14:1631 | 4.73 | 1.00 | 59.54 |  |
| 16 | C | N1 | 14:1630A | 4.96 | 1.00 | 48.35 |  |
  |

  
**1^st^ shell ligand angles:**   
1-2: 87;   
  
 Contact(s): /N7/O6/  
 Coded contact list: O_b_N_b_  
 Exclusion zone contact(s): /N6/N6/OP2/  
Consider K in place of MG (3 valid contact(s) for K vs. 2 for MG)  

### MG:14:3136

|  | Res. type | Atom | Residue | Distance | Occupancy | B-factor | Closest solute to 1^st^ shell water |
| --- | --- | --- | --- | --- | --- | --- | --- |
| **Ion** | | | |  | 1.00 | 59.00 || 1 | G | OP2 | 14:370 | 2.15 | 1.00 | 62.19 |
| 2 | A | OP2 | 14:422 | 2.31 | 1.00 | 65.26 |  |
| 3 | A | N7 | 14:422 | 3.46 | 1.00 | 68.31 |  |
| 4 | G | OP1 | 14:370 | 3.48 | 1.00 | 64.95 |  |
| 5 | A | O5' | 14:422 | 3.89 | 1.00 | 63.44 |  |
| 6 | G | O5' | 14:370 | 3.89 | 1.00 | 58.38 |  |
| 7 | A | N7 | 14:423 | 4.20 | 1.00 | 60.33 |  |
| 8 | U | O3' | 14:421 | 4.49 | 1.00 | 59.09 |  |
| 9 | C | O3' | 14:366 | 4.50 | 1.00 | 63.68 |  |
| 10 | C | O2' | 14:366 | 4.54 | 1.00 | 69.54 |  |
| 11 | U | O2' | 14:421 | 4.59 | 1.00 | 63.04 |  |
| 12 | G | N7 | 14:370 | 4.66 | 1.00 | 58.80 |  |
| 13 | A | OP1 | 14:422 | 4.74 | 1.00 | 62.84 |  |
| 14 | A | N6 | 14:423 | 4.80 | 1.00 | 62.01 |  |
| 15 | A | OP2 | 14:423 | 4.80 | 1.00 | 58.28 |  |
| 16 | A | N9 | 14:422 | 4.99 | 1.00 | 60.83 |  |
  |

  
**1^st^ shell ligand angles:**   
1-2: 133;   
  
 Contact(s): /OP2/OP2/  
 Coded contact list: 2O_ph_  

### MG:14:3137

|  | Res. type | Atom | Residue | Distance | Occupancy | B-factor | Closest solute to 1^st^ shell water |
| --- | --- | --- | --- | --- | --- | --- | --- |
| **Ion** | | | |  | 1.00 | 49.77 || 1 | G | OP2 | 14:131 | 2.99 | 1.00 | 57.18 |
| 2 | G | OP1 | 14:142 | 3.24 | 1.00 | 59.44 |  |
| 3 | C | O2 | 14:1598 | 4.06 | 1.00 | 55.72 |  |
| 4 | C | OP1 | 14:130 | 4.09 | 1.00 | 49.81 |  |
| 5 | C | O2' | 14:1598 | 4.31 | 1.00 | 55.26 |  |
| 6 | C | O5' | 14:130 | 4.34 | 1.00 | 39.15 |  |
| 7 | G | O5' | 14:142 | 4.35 | 1.00 | 55.65 |  |
| 8 | C | OP2 | 14:130 | 4.51 | 1.00 | 41.83 |  |
| 9 | C | OP2 | 14:143 | 4.72 | 1.00 | 57.85 |  |
| 10 | C | OP1 | 14:143 | 4.99 | 1.00 | 66.55 |  |
  |

  
 Coded contact list:   
 Exclusion zone contact(s): /OP1/OP2/  
Consider K in place of MG (2 valid contact(s) for K vs. 0 for MG)  

### MG:14:3138

|  | Res. type | Atom | Residue | Distance | Occupancy | B-factor | Closest solute to 1^st^ shell water |
| --- | --- | --- | --- | --- | --- | --- | --- |
| **Ion** | | | |  | 1.00 | 31.24 || 1 | U | OP2 | 14:2431 | 1.98 | 1.00 | 35.45 |
| 2 | A | OP2 | 14:2432 | 2.31 | 1.00 | 33.94 |  |
| 3 | Water 1 | | 14:3518 | 3.90 | 1.00 | 36.91 | 1.99 (U:OP1 14:2431) |
| 4 | U | O5' | 14:2431 | 4.14 | 1.00 | 32.18 |  |
| 5 | A | OP2 | 14:2433 | 4.23 | 1.00 | 37.49 |  |
| 6 | U | OP1 | 14:2431 | 4.24 | 1.00 | 30.61 |  |
| 7 | A | OP1 | 14:2432 | 4.44 | 1.00 | 39.37 |  |
| 8 | A | O3' | 14:2430 | 4.47 | 1.00 | 31.92 |  |
| 9 | A | O5' | 14:2432 | 4.58 | 1.00 | 34.83 |  |
| 10 | U | O3' | 14:2431 | 4.72 | 1.00 | 39.10 |  |
| 11 | Water 2 | | 14:3986 | 4.83 | 1.00 | 46.07 | 3.69 (A:OP1 14:2432) |
  |

  
**1^st^ shell ligand angles:**   
1-2: 80;   
  
 Contact(s): /OP2/OP2/  
 Coded contact list: 2O_ph_  
MG hydration shell issue: water too close (< 2.4 Angs.)  

### MG:14:3139

|  | Res. type | Atom | Residue | Distance | Occupancy | B-factor | Closest solute to 1^st^ shell water |
| --- | --- | --- | --- | --- | --- | --- | --- |
| **Ion** | | | |  | 1.00 | 41.95 || 1 | G | OP1 | 14:2407 | 1.93 | 1.00 | 53.10 |
| 2 | G | OP1 | 14:411 | 2.30 | 1.00 | 55.29 |  |
| 3 | Water 1 | | 14:3789 | 3.28 | 1.00 | 63.35 | 2.73 (G:OP1 14:2407) |
| 4 | Water 2 | | 14:3688 | 3.60 | 1.00 | 60.81 | 4.34 (G:OP1 14:2407) |
| 5 | U | O2' | 14:2406 | 3.83 | 1.00 | 45.06 |  |
| 6 | G | OP2 | 14:2407 | 4.13 | 1.00 | 52.94 |  |
| 7 | U | O4 | 14:2408 | 4.13 | 1.00 | 58.75 |  |
| 8 | U | O3' | 14:2406 | 4.23 | 1.00 | 53.20 |  |
| 9 | G | N7 | 14:2407 | 4.31 | 1.00 | 59.10 |  |
| 10 | G | O5' | 14:2407 | 4.34 | 1.00 | 54.15 |  |
| 11 | A | OP1 | 14:412 | 4.38 | 1.00 | 47.09 |  |
| 12 | G | O3' | 14:410 | 4.41 | 1.00 | 51.66 |  |
| 13 | G | OP2 | 14:411 | 4.62 | 1.00 | 47.38 |  |
| 14 | G | O5' | 14:411 | 4.70 | 1.00 | 45.39 |  |
  |

  
**1^st^ shell ligand angles:**   
1-2: 78;   
  
 Contact(s): /OP1/OP1/  
 Coded contact list: 2O_ph_  
 Exclusion zone contact(s): /O_w_/  

### MG:14:3140

|  | Res. type | Atom | Residue | Distance | Occupancy | B-factor | Closest solute to 1^st^ shell water |
| --- | --- | --- | --- | --- | --- | --- | --- |
| **Ion** | | | |  | 1.00 | 47.81 || 1 | A | OP1 | 14:1349 | 2.29 | 1.00 | 57.29 |
| 2 | C | OP1 | 14:1376 | 3.40 | 1.00 | 52.09 |  |
| 3 | C | OP2 | 14:1351 | 3.68 | 1.00 | 36.42 |  |
| 4 | G | OP2 | 14:1377 | 3.91 | 1.00 | 44.35 |  |
| 5 | G | O3' | 14:1348 | 4.40 | 1.00 | 43.22 |  |
| 6 | A | O5' | 14:1349 | 4.52 | 1.00 | 46.53 |  |
| 7 | A | OP2 | 14:1349 | 4.69 | 1.00 | 41.69 |  |
| 8 | C | O5' | 14:1376 | 4.88 | 1.00 | 46.08 |  |
| 9 | Water 1 | | 14:3695 | 4.98 | 1.00 | 43.05 | 3.37 (C:OP1 14:1376) |
  |

  
 Contact(s): /OP1/  
 Coded contact list: O_ph_  
 Exclusion zone contact(s): /OP1/  

### MG:14:3141

|  | Res. type | Atom | Residue | Distance | Occupancy | B-factor | Closest solute to 1^st^ shell water |
| --- | --- | --- | --- | --- | --- | --- | --- |
| **Ion** | | | |  | 1.00 | 26.09 || 1 | U | O4 | 14:2593 | 2.41 | 1.00 | 39.10 |
| 2 | G | N7 | 14:2592 | 3.54 | 1.00 | 37.55 |  |
| 3 | C | OP2 | 14:2591 | 3.99 | 1.00 | 30.93 |  |
| 4 | G | O6 | 14:2592 | 4.04 | 1.00 | 39.88 |  |
| 5 | A | N6 | 14:2600 | 4.15 | 1.00 | 36.47 |  |
| 6 | G | OP2 | 14:2592 | 4.16 | 1.00 | 32.39 |  |
| 7 | U | N3 | 14:2593 | 4.62 | 1.00 | 36.17 |  |
| 8 | A | N1 | 14:1971 | 4.65 | 1.00 | 49.24 |  |
| 9 | C | N4 | 14:2591 | 4.70 | 1.00 | 27.23 |  |
| 10 | GLY | N | 19:238 | 4.76 | 1.00 | 30.78 |  |
| 11 | GLY | O | 19:238 | 4.88 | 1.00 | 36.38 |  |
| 12 | C | N4 | 14:2594 | 4.96 | 1.00 | 37.50 |  |
  |

  
 Coded contact list:   
 Exclusion zone contact(s): /O4/  
Consider NA in place of MG (1 valid contact(s) for NA vs. 0 for MG)  

### MG:14:3142

|  | Res. type | Atom | Residue | Distance | Occupancy | B-factor | Closest solute to 1^st^ shell water |
| --- | --- | --- | --- | --- | --- | --- | --- |
| **Ion** | | | |  | 1.00 | 51.35 || 1 | A | OP1 | 14:1936 | 3.67 | 1.00 | 47.73 |
| 2 | A | OP1 | 14:1937 | 4.12 | 1.00 | 45.78 |  |
| 3 | G | N7 | 14:1959 | 4.36 | 1.00 | 40.67 |  |
| 4 | G | OP2 | 14:1959 | 4.61 | 1.00 | 35.07 |  |
| 5 | A | N7 | 14:1960 | 4.66 | 1.00 | 46.64 |  |
| 6 | A | N6 | 14:1960 | 4.76 | 1.00 | 45.94 |  |
  |

  
 Coded contact list:   
1^st^ MG coordination shell missing (see Notes)  

### MG:14:3143

|  | Res. type | Atom | Residue | Distance | Occupancy | B-factor | Closest solute to 1^st^ shell water |
| --- | --- | --- | --- | --- | --- | --- | --- |
| **Ion** | | | |  | 1.00 | 39.18 || 1 | A | OP2 | 14:1938 | 2.68 | 1.00 | 45.32 |
| 2 | C | OP1 | 14:1957 | 3.34 | 1.00 | 46.59 |  |
| 3 | C | OP2 | 14:1958 | 3.50 | 1.00 | 38.23 |  |
| 4 | C | OP1 | 14:1958 | 3.75 | 1.00 | 45.72 |  |
| 5 | A | OP1 | 14:1938 | 4.54 | 1.00 | 36.60 |  |
| 6 | C | O5' | 14:1957 | 4.76 | 1.00 | 40.99 |  |
| 7 | A | O3' | 14:1937 | 4.80 | 1.00 | 49.33 |  |
| 8 | C | O3' | 14:1957 | 4.80 | 1.00 | 42.05 |  |
  |

  
 Coded contact list:   
 Exclusion zone contact(s): /OP1/OP2/  
Consider K in place of MG (2 valid contact(s) for K vs. 0 for MG)  

### MG:14:3144

|  | Res. type | Atom | Residue | Distance | Occupancy | B-factor | Closest solute to 1^st^ shell water |
| --- | --- | --- | --- | --- | --- | --- | --- |
| **Ion** | | | |  | 1.00 | 36.37 || 1 | G | OP2 | 14:2582 | 2.34 | 1.00 | 45.12 |
| 2 | G | OP2 | 14:2583 | 2.46 | 1.00 | 45.00 |  |
| 3 | G | N7 | 14:2608 | 4.01 | 1.00 | 41.64 |  |
| 4 | G | O3' | 14:2581 | 4.16 | 1.00 | 34.50 |  |
| 5 | G | O6 | 14:2608 | 4.35 | 1.00 | 49.07 |  |
| 6 | G | OP1 | 14:2582 | 4.41 | 1.00 | 46.72 |  |
| 7 | G | OP1 | 14:2583 | 4.57 | 1.00 | 45.53 |  |
| 8 | U | O4 | 14:2584 | 4.72 | 1.00 | 83.75 |  |
| 9 | G | O5' | 14:2583 | 4.72 | 1.00 | 47.34 |  |
| 10 | G | O5' | 14:2582 | 4.85 | 1.00 | 38.84 |  |
| 11 | G | O3' | 14:2582 | 4.87 | 1.00 | 43.49 |  |
| 12 | G | O2' | 14:2581 | 4.90 | 1.00 | 41.57 |  |
  |

  
 Contact(s): /OP2/  
 Coded contact list: O_ph_  
 Exclusion zone contact(s): /OP2/  

### MG:14:3145

|  | Res. type | Atom | Residue | Distance | Occupancy | B-factor | Closest solute to 1^st^ shell water |
| --- | --- | --- | --- | --- | --- | --- | --- |
| **Ion** | | | |  | 1.00 | 49.74 || 1 | G | OP2 | 14:2719 | 2.66 | 1.00 | 57.96 |
| 2 | U | OP2 | 14:2687 | 3.39 | 1.00 | 42.80 |  |
| 3 | G | O3' | 14:2718 | 4.65 | 1.00 | 48.75 |  |
| 4 | G | OP2 | 14:2718 | 4.94 | 1.00 | 54.96 |  |
| 5 | G | O5' | 14:2719 | 4.98 | 1.00 | 57.93 |  |
  |

  
 Coded contact list:   
 Exclusion zone contact(s): /OP2/OP2/  
Consider K in place of MG (2 valid contact(s) for K vs. 0 for MG)  

### MG:14:3146

|  | Res. type | Atom | Residue | Distance | Occupancy | B-factor | Closest solute to 1^st^ shell water |
| --- | --- | --- | --- | --- | --- | --- | --- |
| **Ion** | | | |  | 1.00 | 49.27 || 1 | G | O6 | 14:2685 | 1.70 | 1.00 | 59.59 |
| 2 | G | O6 | 14:2686 | 3.46 | 1.00 | 60.25 |  |
| 3 | C | N4 | 14:2724 | 3.86 | 1.00 | 57.63 |  |
| 4 | G | N1 | 14:2685 | 3.91 | 1.00 | 55.12 |  |
| 5 | U | O4 | 14:2684 | 4.01 | 1.00 | 53.94 |  |
| 6 | G | N7 | 14:2685 | 4.15 | 1.00 | 63.11 |  |
| 7 | G | N7 | 14:2686 | 4.66 | 1.00 | 63.30 |  |
| 8 | C | O2' | 14:2681 | 4.80 | 1.00 | 60.54 |  |
  |

  
 Coded contact list:   
 Exclusion zone contact(s): /O6/  
1^st^ MG coordination shell missing (see Notes)  

### MG:14:3147

|  | Res. type | Atom | Residue | Distance | Occupancy | B-factor | Closest solute to 1^st^ shell water |
| --- | --- | --- | --- | --- | --- | --- | --- |
| **Ion** | | | |  | 1.00 | 40.47 || 1 | A | OP1 | 14:2721 | 1.89 | 1.00 | 52.35 |
| 2 | U | OP2 | 14:2682 | 2.07 | 1.00 | 63.22 |  |
| 3 | A | O5' | 14:2721 | 3.79 | 1.00 | 58.39 |  |
| 4 | C | O3' | 14:2681 | 3.85 | 1.00 | 65.59 |  |
| 5 | A | OP2 | 14:2721 | 3.93 | 1.00 | 52.90 |  |
| 6 | G | OP2 | 14:2722 | 4.06 | 1.00 | 56.90 |  |
| 7 | C | OP2 | 14:2683 | 4.13 | 1.00 | 54.34 |  |
| 8 | U | OP1 | 14:2682 | 4.16 | 1.00 | 61.10 |  |
| 9 | U | O3' | 14:2720 | 4.45 | 1.00 | 52.85 |  |
| 10 | C | O2' | 14:2681 | 4.52 | 1.00 | 60.54 |  |
| 11 | U | O5' | 14:2682 | 4.56 | 1.00 | 59.13 |  |
  |

  
**1^st^ shell ligand angles:**   
1-2: 150;   
  
 Contact(s): /OP1/OP2/  
 Coded contact list: 2O_ph_  

### MG:14:3148

|  | Res. type | Atom | Residue | Distance | Occupancy | B-factor | Closest solute to 1^st^ shell water |
| --- | --- | --- | --- | --- | --- | --- | --- |
| **Ion** | | | |  | 1.00 | 43.75 || 1 | G | O6 | 14:1626 | 3.68 | 1.00 | 48.64 |
| 2 | A | N1 | 14:1634 | 3.90 | 1.00 | 32.18 |  |
| 3 | A | O2' | 14:735 | 4.15 | 1.00 | 44.99 |  |
| 4 | G | OP2 | 14:1624 | 4.42 | 1.00 | 36.76 |  |
| 5 | U | O2 | 14:1300 | 4.56 | 1.00 | 49.74 |  |
| 6 | A | N3 | 14:1634 | 4.66 | 1.00 | 36.87 |  |
  |

  
 Coded contact list:   
1^st^ MG coordination shell missing (see Notes)  

### MG:14:3149

|  | Res. type | Atom | Residue | Distance | Occupancy | B-factor | Closest solute to 1^st^ shell water |
| --- | --- | --- | --- | --- | --- | --- | --- |
| **Ion** | | | |  | 1.00 | 37.57 || 1 | G | O6 | 14:1613 | 3.56 | 1.00 | 36.54 |
| 2 | A | OP1 | 14:1610 | 3.69 | 1.00 | 43.54 |  |
| 3 | G | N7 | 14:1613 | 3.85 | 1.00 | 35.98 |  |
| 4 | C | OP2 | 14:1617 | 4.10 | 1.00 | 33.44 |  |
| 5 | C | N4 | 14:1612 | 4.25 | 1.00 | 36.47 |  |
| 6 | Water 1 | | 14:3884 | 4.28 | 1.00 | 47.17 | 2.98 (G:O6 14:1613) |
| 7 | C | OP2 | 14:1611 | 4.71 | 1.00 | 35.20 |  |
| 8 | A | O3' | 14:1609 | 4.94 | 1.00 | 41.10 |  |
  |

  
 Coded contact list:   
1^st^ MG coordination shell missing (see Notes)  

### MG:14:3150

|  | Res. type | Atom | Residue | Distance | Occupancy | B-factor | Closest solute to 1^st^ shell water |
| --- | --- | --- | --- | --- | --- | --- | --- |
| **Ion** | | | |  | 1.00 | 29.98 || 1 | C | OP1 | 14:1611 | 2.40 | 1.00 | 35.89 |
| 2 | C | OP2 | 14:1611 | 3.63 | 1.00 | 35.20 |  |
| 3 | A | O3' | 14:1610 | 4.35 | 1.00 | 43.77 |  |
| 4 | C | O5' | 14:1611 | 4.69 | 1.00 | 33.68 |  |
| 5 | G | O2' | 14:1310 | 4.69 | 1.00 | 37.02 |  |
| 6 | C | OP2 | 14:1612 | 4.85 | 1.00 | 24.33 |  |
  |

  
 Coded contact list:   
 Exclusion zone contact(s): /OP1/  
Consider NA in place of MG (1 valid contact(s) for NA vs. 0 for MG)  

### MG:14:3151

|  | Res. type | Atom | Residue | Distance | Occupancy | B-factor | Closest solute to 1^st^ shell water |
| --- | --- | --- | --- | --- | --- | --- | --- |
| **Ion** | | | |  | 1.00 | 26.48 || 1 | A | OP1 | 14:1268 | 2.16 | 1.00 | 38.94 |
| 2 | C | OP1 | 14:2006 | 2.43 | 1.00 | 36.04 |  |
| 3 | U | O3' | 14:1267 | 3.59 | 1.00 | 40.04 |  |
| 4 | G | OP1 | 14:2048 | 3.79 | 1.00 | 35.72 |  |
| 5 | G | OP2 | 14:2049 | 4.21 | 1.00 | 28.52 |  |
| 6 | A | O3' | 14:2005 | 4.52 | 1.00 | 33.62 |  |
| 7 | A | OP2 | 14:1268 | 4.53 | 1.00 | 42.00 |  |
| 8 | A | O5' | 14:1268 | 4.55 | 1.00 | 39.06 |  |
| 9 | Water 1 | | 14:3584 | 4.61 | 1.00 | 39.67 | 2.28 (C:OP2 14:2006) |
| 10 | C | OP2 | 14:2006 | 4.75 | 1.00 | 41.14 |  |
| 11 | C | O5' | 14:2006 | 4.91 | 1.00 | 33.99 |  |
| 12 | G | O5' | 14:2048 | 5.00 | 1.00 | 36.62 |  |
  |

  
 Contact(s): /OP1/  
 Coded contact list: O_ph_  
 Exclusion zone contact(s): /OP1/  

### MG:14:3152

|  | Res. type | Atom | Residue | Distance | Occupancy | B-factor | Closest solute to 1^st^ shell water |
| --- | --- | --- | --- | --- | --- | --- | --- |
| **Ion** | | | |  | 1.00 | 32.05 || 1 | U | OP1 | 14:2022 | 2.08 | 1.00 | 43.77 |
| 2 | G | N2 | 14:530 | 2.55 | 1.00 | 56.59 |  |
| 3 | G | N1 | 14:530 | 2.95 | 1.00 | 54.02 |  |
| 4 | G | OP1 | 14:2023 | 3.65 | 1.00 | 61.52 |  |
| 5 | C | O3' | 14:2021 | 4.21 | 1.00 | 44.95 |  |
| 6 | A | N7 | 14:529 | 4.40 | 1.00 | 59.73 |  |
| 7 | U | OP2 | 14:2022 | 4.42 | 1.00 | 46.55 |  |
| 8 | G | N3 | 14:530 | 4.46 | 1.00 | 50.48 |  |
| 9 | U | O5' | 14:2022 | 4.50 | 1.00 | 49.89 |  |
| 10 | G | O6 | 14:530 | 4.57 | 1.00 | 52.88 |  |
  |

  
 Contact(s): /OP1/  
 Coded contact list: O_ph_  
 Exclusion zone contact(s): /N1_G_/N2/  

### MG:14:3153

|  | Res. type | Atom | Residue | Distance | Occupancy | B-factor | Closest solute to 1^st^ shell water |
| --- | --- | --- | --- | --- | --- | --- | --- |
| **Ion** | | | |  | 1.00 | 51.45 || 1 | A | OP1 | 14:526 | 2.05 | 1.00 | 63.24 |
| 2 | C | OP1 | 14:527 | 2.27 | 1.00 | 61.74 |  |
| 3 | A | OP1 | 14:528 | 3.16 | 1.00 | 62.50 |  |
| 4 | C | OP2 | 14:527 | 3.79 | 1.00 | 59.74 |  |
| 5 | A | O5' | 14:526 | 3.85 | 1.00 | 62.59 |  |
| 6 | C | O5' | 14:527 | 4.06 | 1.00 | 53.96 |  |
| 7 | A | OP2 | 14:526 | 4.42 | 1.00 | 65.26 |  |
| 8 | U | O3' | 14:525 | 4.47 | 1.00 | 61.96 |  |
| 9 | A | O3' | 14:526 | 4.75 | 1.00 | 51.95 |  |
  |

  
**1^st^ shell ligand angles:**   
1-2: 79;   
  
 Contact(s): /OP1/OP1/  
 Coded contact list: 2O_ph_  
 Exclusion zone contact(s): /OP1/  

### MG:14:3154

|  | Res. type | Atom | Residue | Distance | Occupancy | B-factor | Closest solute to 1^st^ shell water |
| --- | --- | --- | --- | --- | --- | --- | --- |
| **Ion** | | | |  | 1.00 | 50.70 || 1 | C | OP1 | 14:2036 | 1.88 | 1.00 | 54.11 |
| 2 | C | OP2 | 14:531 | 2.09 | 1.00 | 63.10 |  |
| 3 | C | OP2 | 14:2036 | 3.96 | 1.00 | 56.92 |  |
| 4 | G | O6 | 14:561 | 3.97 | 1.00 | 69.62 |  |
| 5 | G | N1 | 14:561 | 4.00 | 1.00 | 60.16 |  |
| 6 | C | OP1 | 14:531 | 4.09 | 1.00 | 54.62 |  |
| 7 | G | O3' | 14:2035 | 4.24 | 1.00 | 53.74 |  |
| 8 | G | O3' | 14:530 | 4.28 | 1.00 | 55.95 |  |
| 9 | C | O5' | 14:2036 | 4.30 | 1.00 | 55.98 |  |
| 10 | G | N7 | 14:561 | 4.54 | 1.00 | 68.61 |  |
| 11 | C | O5' | 14:531 | 4.55 | 1.00 | 53.60 |  |
| 12 | G | N3 | 14:561 | 4.73 | 1.00 | 64.40 |  |
  |

  
**1^st^ shell ligand angles:**   
1-2: 116;   
  
 Contact(s): /OP1/OP2/  
 Coded contact list: 2O_ph_  

### MG:14:3155

|  | Res. type | Atom | Residue | Distance | Occupancy | B-factor | Closest solute to 1^st^ shell water |
| --- | --- | --- | --- | --- | --- | --- | --- |
| **Ion** | | | |  | 1.00 | 38.89 || 1 | C | OP1 | 14:574 | 2.10 | 1.00 | 45.55 |
| 2 | G | O6 | 14:2035 | 4.14 | 1.00 | 50.50 |  |
| 3 | C | OP2 | 14:574 | 4.26 | 1.00 | 44.12 |  |
| 4 | G | O6 | 14:2018 | 4.29 | 1.00 | 51.81 |  |
| 5 | G | O3' | 14:573 | 4.44 | 1.00 | 40.91 |  |
| 6 | C | O5' | 14:574 | 4.46 | 1.00 | 40.38 |  |
| 7 | C | OP2 | 14:2055 | 4.80 | 1.00 | 36.52 |  |
| 8 | C | N4 | 14:531 | 4.81 | 1.00 | 45.62 |  |
  |

  
 Contact(s): /OP1/  
 Coded contact list: O_ph_  

### MG:14:3156

|  | Res. type | Atom | Residue | Distance | Occupancy | B-factor | Closest solute to 1^st^ shell water |
| --- | --- | --- | --- | --- | --- | --- | --- |
| **Ion** | | | |  | 1.00 | 34.72 || 1 | C | OP2 | 14:574 | 2.06 | 1.00 | 44.12 |
| 2 | G | O6 | 14:577 | 3.76 | 1.00 | 36.61 |  |
| 3 | C | O5' | 14:574 | 4.03 | 1.00 | 40.38 |  |
| 4 | G | OP1 | 14:573 | 4.11 | 1.00 | 44.23 |  |
| 5 | U | O4 | 14:576 | 4.20 | 1.00 | 34.35 |  |
| 6 | C | OP1 | 14:574 | 4.28 | 1.00 | 45.55 |  |
| 7 | U | OP2 | 14:576 | 4.33 | 1.00 | 32.60 |  |
| 8 | G | O3' | 14:573 | 4.58 | 1.00 | 40.91 |  |
| 9 | G | O6 | 14:563 | 4.61 | 1.00 | 41.72 |  |
| 10 | C | N4 | 14:564 | 4.82 | 1.00 | 36.23 |  |
  |

  
 Contact(s): /OP2/  
 Coded contact list: O_ph_  

### MG:14:3157

|  | Res. type | Atom | Residue | Distance | Occupancy | B-factor | Closest solute to 1^st^ shell water |
| --- | --- | --- | --- | --- | --- | --- | --- |
| **Ion** | | | |  | 1.00 | 34.56 || 1 | U | OP1 | 14:810 | 2.01 | 1.00 | 41.06 |
| 2 | A | N7 | 14:1253 | 2.33 | 1.00 | 43.80 |  |
| 3 | A | N6 | 14:1253 | 3.28 | 1.00 | 42.84 |  |
| 4 | G | O2' | 14:1252 | 3.89 | 1.00 | 52.10 |  |
| 5 | U | O5' | 14:810 | 4.23 | 1.00 | 33.10 |  |
| 6 | U | OP2 | 14:810 | 4.24 | 1.00 | 43.47 |  |
| 7 | A | N9 | 14:1253 | 4.35 | 1.00 | 39.53 |  |
| 8 | U | OP1 | 14:811 | 4.49 | 1.00 | 37.41 |  |
| 9 | G | O3' | 14:809 | 4.53 | 1.00 | 33.12 |  |
| 10 | C | O2' | 14:565 | 4.89 | 1.00 | 35.91 |  |
| 11 | C | O3' | 14:565 | 4.95 | 1.00 | 43.59 |  |
| 12 | A | N1 | 14:1253 | 4.99 | 1.00 | 35.63 |  |
  |

  
**1^st^ shell ligand angles:**   
1-2: 97;   
  
 Contact(s): /N7/OP1/  
 Coded contact list: O_ph_N_b_  
 Exclusion zone contact(s): /N6/  

### MG:14:3158

|  | Res. type | Atom | Residue | Distance | Occupancy | B-factor | Closest solute to 1^st^ shell water |
| --- | --- | --- | --- | --- | --- | --- | --- |
| **Ion** | | | |  | 1.00 | 49.01 || 1 | G | OP2 | 14:1187 | 2.11 | 1.00 | 53.06 |
| 2 | C | OP2 | 14:817 | 3.25 | 1.00 | 64.91 |  |
| 3 | C | OP2 | 14:816 | 3.61 | 1.00 | 58.97 |  |
| 4 | G | O6 | 14:818 | 3.62 | 1.00 | 60.98 |  |
| 5 | G | N7 | 14:1187 | 4.23 | 1.00 | 69.17 |  |
| 6 | G | O5' | 14:1187 | 4.27 | 1.00 | 61.17 |  |
| 7 | C | O5' | 14:816 | 4.30 | 1.00 | 60.90 |  |
| 8 | G | OP1 | 14:1187 | 4.36 | 1.00 | 56.67 |  |
| 9 | C | O2' | 14:1185 | 4.41 | 1.00 | 82.02 |  |
| 10 | C | OP1 | 14:816 | 4.44 | 1.00 | 61.10 |  |
| 11 | G | O3' | 14:1186 | 4.64 | 1.00 | 55.15 |  |
| 12 | C | O3' | 14:1185 | 4.69 | 1.00 | 62.04 |  |
  |

  
 Contact(s): /OP2/  
 Coded contact list: O_ph_  
 Exclusion zone contact(s): /OP2/  

### MG:14:3159

|  | Res. type | Atom | Residue | Distance | Occupancy | B-factor | Closest solute to 1^st^ shell water |
| --- | --- | --- | --- | --- | --- | --- | --- |
| **Ion** | | | |  | 1.00 | 56.73 || 1 | G | N7 | 14:842 | 3.84 | 1.00 | 60.18 |
| 2 | G | O6 | 14:842 | 4.19 | 1.00 | 66.87 |  |
| 3 | G | O6 | 14:843 | 4.44 | 1.00 | 73.03 |  |
| 4 | G | N7 | 14:843 | 4.83 | 1.00 | 62.39 |  |
| 5 | G | OP2 | 14:842 | 4.93 | 1.00 | 62.82 |  |
  |

  
 Coded contact list:   
1^st^ MG coordination shell missing (see Notes)  

### MG:14:3160

|  | Res. type | Atom | Residue | Distance | Occupancy | B-factor | Closest solute to 1^st^ shell water |
| --- | --- | --- | --- | --- | --- | --- | --- |
| **Ion** | | | |  | 1.00 | 38.36 || 1 | G | OP1 | 14:948 | 1.92 | 1.00 | 49.50 |
| 2 | U | OP2 | 14:963 | 1.92 | 1.00 | 48.03 |  |
| 3 | G | OP1 | 14:962 | 2.03 | 1.00 | 39.31 |  |
| 4 | G | OP2 | 14:948 | 3.10 | 1.00 | 45.39 |  |
| 5 | G | OP2 | 14:962 | 3.96 | 1.00 | 51.92 |  |
| 6 | G | O5' | 14:948 | 3.96 | 1.00 | 41.89 |  |
| 7 | U | OP1 | 14:963 | 4.01 | 1.00 | 45.85 |  |
| 8 | G | O3' | 14:947 | 4.09 | 1.00 | 52.33 |  |
| 9 | G | O3' | 14:962 | 4.19 | 1.00 | 34.41 |  |
| 10 | G | O5' | 14:962 | 4.26 | 1.00 | 45.45 |  |
| 11 | U | O5' | 14:963 | 4.40 | 1.00 | 49.17 |  |
| 12 | C | O3' | 14:961 | 4.53 | 1.00 | 46.82 |  |
  |

  
**1^st^ shell ligand angles:**   
1-2: 150; 1-3: 86; 2-3: 85;   
  
 Contact(s): /OP1/OP1/OP2/  
 Coded contact list: 3O_ph_  
 Exclusion zone contact(s): /OP2/  

### MG:14:3161

|  | Res. type | Atom | Residue | Distance | Occupancy | B-factor | Closest solute to 1^st^ shell water |
| --- | --- | --- | --- | --- | --- | --- | --- |
| **Ion** | | | |  | 1.00 | 47.07 || 1 | C | OP1 | 14:2452 | 2.27 | 1.00 | 57.79 |
| 2 | C | O5' | 14:2452 | 3.92 | 1.00 | 47.66 |  |
| 3 | A | OP2 | 14:2453 | 3.98 | 1.00 | 50.14 |  |
| 4 | A | O2' | 1L:76 | 4.11 | 1.00 | 83.85 |  |
| 5 | A | O3' | 1L:76 | 4.31 | 1.00 | 95.02 |  |
| 6 | G | OP2 | 14:2494 | 4.33 | 1.00 | 62.23 |  |
| 7 | C | OP2 | 14:2452 | 4.61 | 1.00 | 39.21 |  |
| 8 | A | O3' | 14:2451 | 4.63 | 1.00 | 48.94 |  |
| 9 | A | OP1 | 14:2453 | 4.73 | 1.00 | 49.27 |  |
  |

  
 Contact(s): /OP1/  
 Coded contact list: O_ph_  

### MG:14:3162

|  | Res. type | Atom | Residue | Distance | Occupancy | B-factor | Closest solute to 1^st^ shell water |
| --- | --- | --- | --- | --- | --- | --- | --- |
| **Ion** | | | |  | 1.00 | 46.76 || 1 | G | OP1 | 14:2574 | 2.24 | 1.00 | 47.51 |
| 2 | C | OP1 | 14:2573 | 2.29 | 1.00 | 47.05 |  |
| 3 | C | OP2 | 14:2575 | 2.58 | 1.00 | 47.85 |  |
| 4 | C | OP1 | 14:2507 | 3.60 | 1.00 | 55.13 |  |
| 5 | C | OP2 | 14:2573 | 3.83 | 1.00 | 52.31 |  |
| 6 | C | O5' | 14:2573 | 3.90 | 1.00 | 55.01 |  |
| 7 | G | OP2 | 14:2508 | 4.47 | 1.00 | 50.53 |  |
| 8 | C | O3' | 14:2573 | 4.53 | 1.00 | 54.76 |  |
| 9 | G | OP2 | 14:2574 | 4.57 | 1.00 | 51.68 |  |
| 10 | G | O5' | 14:2574 | 4.60 | 1.00 | 42.26 |  |
| 11 | C | OP1 | 14:2575 | 4.61 | 1.00 | 39.87 |  |
| 12 | C | OP2 | 14:2507 | 4.77 | 1.00 | 64.62 |  |
| 13 | G | O3' | 14:2574 | 4.79 | 1.00 | 41.08 |  |
| 14 | A | O3' | 14:2572 | 4.84 | 1.00 | 43.70 |  |
| 15 | C | O5' | 14:2575 | 4.89 | 1.00 | 44.44 |  |
| 16 | C | O5' | 14:2507 | 4.94 | 1.00 | 54.48 |  |
  |

  
**1^st^ shell ligand angles:**   
1-2: 74;   
  
 Contact(s): /OP1/OP1/  
 Coded contact list: 2O_ph_  
 Exclusion zone contact(s): /OP2/  

### MG:14:3163

|  | Res. type | Atom | Residue | Distance | Occupancy | B-factor | Closest solute to 1^st^ shell water |
| --- | --- | --- | --- | --- | --- | --- | --- |
| **Ion** | | | |  | 1.00 | 31.37 || 1 | U | O4 | 14:2028 | 1.98 | 1.00 | 40.12 |
| 2 | A | OP1 | 14:2033 | 2.71 | 1.00 | 46.49 |  |
| 3 | G | O3' | 14:2032 | 3.30 | 1.00 | 57.80 |  |
| 4 | G | O6 | 14:2029 | 3.72 | 1.00 | 44.44 |  |
| 5 | A | O5' | 14:2033 | 4.01 | 1.00 | 46.40 |  |
| 6 | G | N7 | 14:2027 | 4.12 | 1.00 | 44.89 |  |
| 7 | G | O6 | 14:2027 | 4.16 | 1.00 | 48.72 |  |
| 8 | U | N3 | 14:2028 | 4.30 | 1.00 | 40.80 |  |
  |

  
 Contact(s): /O4/  
 Coded contact list: O_b_  
 Exclusion zone contact(s): /O3'/OP1/  
Consider K in place of MG (2 valid contact(s) for K vs. 1 for MG)  

### MG:14:3164

|  | Res. type | Atom | Residue | Distance | Occupancy | B-factor | Closest solute to 1^st^ shell water |
| --- | --- | --- | --- | --- | --- | --- | --- |
| **Ion** | | | |  | 1.00 | 48.12 || 1 | G | O6 | 14:2819 | 3.85 | 1.00 | 47.55 |
| 2 | G | N7 | 14:2819 | 3.93 | 1.00 | 46.49 |  |
| 3 | A | N1 | 14:2821 | 4.00 | 1.00 | 38.89 |  |
| 4 | G | N2 | 14:2822 | 4.18 | 1.00 | 39.95 |  |
| 5 | G | OP1 | 14:1653 | 4.77 | 1.00 | 44.44 |  |
  |

  
 Coded contact list:   
1^st^ MG coordination shell missing (see Notes)  

### MG:14:3165

|  | Res. type | Atom | Residue | Distance | Occupancy | B-factor | Closest solute to 1^st^ shell water |
| --- | --- | --- | --- | --- | --- | --- | --- |
| **Ion** | | | |  | 1.00 | 45.19 || 1 | G | O6 | 14:592 | 3.62 | 1.00 | 45.33 |
| 2 | G | O6 | 14:593 | 4.14 | 1.00 | 47.80 |  |
| 3 | C | N4 | 14:664 | 4.51 | 1.00 | 47.16 |  |
| 4 | C | N4 | 14:665 | 4.92 | 1.00 | 41.75 |  |
  |

  
 Coded contact list:   
1^st^ MG coordination shell missing (see Notes)  

### MG:14:3166

|  | Res. type | Atom | Residue | Distance | Occupancy | B-factor | Closest solute to 1^st^ shell water |
| --- | --- | --- | --- | --- | --- | --- | --- |
| **Ion** | | | |  | 1.00 | 58.46 || 1 | Water 1 | | 14:4081 | 3.06 | 1.00 | 72.66 | 4.20 (C:OP2 14:238) |
| 2 | Water 2 | | 14:3708 | 3.29 | 1.00 | 46.28 | 3.24 (U:O4 14:239) |
| 3 | U | O4 | 14:239 | 3.37 | 1.00 | 54.41 |  |
| 4 | C | N4 | 14:238 | 3.67 | 1.00 | 51.80 |  |
| 5 | C | N4 | 14:237 | 4.42 | 1.00 | 59.35 |  |
| 6 | C | OP2 | 14:238 | 4.79 | 1.00 | 49.17 |  |
| 7 | C | OP2 | 14:237 | 4.83 | 1.00 | 58.45 |  |
| 8 | Water 3 | | 14:4028 | 4.88 | 1.00 | 54.99 | 6.25 (U:O4 14:239) |
| 9 | Water 4 | | 14:3632 | 4.99 | 1.00 | 48.76 | 4.57 (U:O4 14:239) |
  |

  
 Coded contact list:   
 Exclusion zone contact(s): /O4/O_w_/O_w_/  
Consider K in place of MG (1 valid contact(s) for K vs. 0 for MG)  

### MG:14:3167

|  | Res. type | Atom | Residue | Distance | Occupancy | B-factor | Closest solute to 1^st^ shell water |
| --- | --- | --- | --- | --- | --- | --- | --- |
| **Ion** | | | |  | 1.00 | 38.92 || 1 | G | OP1 | 14:760 | 2.56 | 1.00 | 44.68 |
| 2 | C | OP1 | 14:731 | 3.01 | 1.00 | 31.40 |  |
| 3 | C | OP2 | 14:732 | 3.48 | 1.00 | 36.72 |  |
| 4 | C | OP1 | 14:732 | 3.84 | 1.00 | 27.54 |  |
| 5 | A | OP2 | 14:761 | 3.89 | 1.00 | 31.53 |  |
| 6 | G | O5' | 14:760 | 3.95 | 1.00 | 30.56 |  |
| 7 | Water 1 | | 14:3675 | 4.44 | 1.00 | 35.48 | 2.53 (A:OP2 14:761) |
| 8 | C | O3' | 14:731 | 4.73 | 1.00 | 35.79 |  |
| 9 | G | OP2 | 14:760 | 4.75 | 1.00 | 34.50 |  |
| 10 | C | O5' | 14:731 | 4.78 | 1.00 | 37.19 |  |
| 11 | C | OP2 | 14:731 | 4.90 | 1.00 | 40.01 |  |
  |

  
 Coded contact list:   
 Exclusion zone contact(s): /OP1/OP1/  
Consider NA in place of MG (1 valid contact(s) for NA vs. 0 for MG)  

### MG:14:3168

|  | Res. type | Atom | Residue | Distance | Occupancy | B-factor | Closest solute to 1^st^ shell water |
| --- | --- | --- | --- | --- | --- | --- | --- |
| **Ion** | | | |  | 1.00 | 37.13 || 1 | C | OP1 | 14:1330 | 2.88 | 1.00 | 49.01 |
| 2 | A | OP2 | 14:1331 | 3.87 | 1.00 | 50.41 |  |
| 3 | C | OP2 | 14:1330 | 4.10 | 1.00 | 43.15 |  |
| 4 | C | O5' | 14:1330 | 4.21 | 1.00 | 49.17 |  |
| 5 | C | O3' | 14:1330 | 4.85 | 1.00 | 42.69 |  |
| 6 | A | OP1 | 14:1331 | 4.86 | 1.00 | 35.44 |  |
  |

  
 Coded contact list:   
 Exclusion zone contact(s): /OP1/  
Consider K in place of MG (1 valid contact(s) for K vs. 0 for MG)  

### MG:14:3169

|  | Res. type | Atom | Residue | Distance | Occupancy | B-factor | Closest solute to 1^st^ shell water |
| --- | --- | --- | --- | --- | --- | --- | --- |
| **Ion** | | | |  | 1.00 | 33.99 || 1 | U | OP1 | 14:1313 | 2.16 | 1.00 | 45.08 |
| 2 | G | OP2 | 14:1334 | 3.53 | 1.00 | 44.37 |  |
| 3 | G | OP1 | 14:1334 | 3.61 | 1.00 | 46.95 |  |
| 4 | C | OP1 | 14:1333 | 3.70 | 1.00 | 41.13 |  |
| 5 | U | O5' | 14:1313 | 3.84 | 1.00 | 42.26 |  |
| 6 | U | OP2 | 14:1313 | 4.29 | 1.00 | 46.13 |  |
| 7 | Water 1 | | 14:3603 | 4.49 | 1.00 | 39.27 | 3.09 (U:OP2 14:1313) |
| 8 | U | O3' | 14:1312 | 4.66 | 1.00 | 36.95 |  |
  |

  
 Contact(s): /OP1/  
 Coded contact list: O_ph_  

### MG:14:3170

|  | Res. type | Atom | Residue | Distance | Occupancy | B-factor | Closest solute to 1^st^ shell water |
| --- | --- | --- | --- | --- | --- | --- | --- |
| **Ion** | | | |  | 1.00 | 54.76 || 1 | G | N7 | 14:298 | 2.36 | 1.00 | 67.60 |
| 2 | G | O6 | 14:298 | 3.25 | 1.00 | 72.15 |  |
| 3 | G | OP2 | 14:338 | 3.33 | 1.00 | 60.99 |  |
| 4 | G | OP2 | 14:298 | 4.19 | 1.00 | 65.87 |  |
| 5 | C | OP2 | 14:297 | 4.47 | 1.00 | 59.49 |  |
| 6 | G | OP1 | 14:338 | 4.54 | 1.00 | 58.32 |  |
| 7 | G | N9 | 14:298 | 4.55 | 1.00 | 66.03 |  |
| 8 | Water 1 | | 14:4044 | 4.86 | 1.00 | 42.91 | 5.40 (G:N7 14:298) |
  |

  
 Contact(s): /N7/  
 Coded contact list: N_b_  
 Exclusion zone contact(s): /O6/OP2/  
Consider K in place of MG (2 valid contact(s) for K vs. 1 for MG)  

### MG:14:3171

|  | Res. type | Atom | Residue | Distance | Occupancy | B-factor | Closest solute to 1^st^ shell water |
| --- | --- | --- | --- | --- | --- | --- | --- |
| **Ion** | | | |  | 1.00 | 55.65 || 1 | A | OP2 | 14:505 | 2.16 | 1.00 | 72.61 |
| 2 | G | OP1 | 14:1235 | 2.37 | 1.00 | 70.93 |  |
| 3 | U | O3' | 14:504 | 3.67 | 1.00 | 75.89 |  |
| 4 | U | O3' | 14:1234 | 3.84 | 1.00 | 58.29 |  |
| 5 | U | O2' | 14:504 | 3.97 | 1.00 | 78.59 |  |
| 6 | U | OP1 | 14:511 | 4.05 | 1.00 | 54.15 |  |
| 7 | G | OP2 | 14:512 | 4.39 | 1.00 | 50.58 |  |
| 8 | G | O5' | 14:1235 | 4.41 | 1.00 | 56.15 |  |
| 9 | A | OP1 | 14:505 | 4.42 | 1.00 | 68.30 |  |
| 10 | A | O5' | 14:505 | 4.49 | 1.00 | 59.26 |  |
| 11 | G | OP2 | 14:1235 | 4.84 | 1.00 | 59.66 |  |
  |

  
**1^st^ shell ligand angles:**   
1-2: 85;   
  
 Contact(s): /OP1/OP2/  
 Coded contact list: 2O_ph_  

### MG:14:3172

|  | Res. type | Atom | Residue | Distance | Occupancy | B-factor | Closest solute to 1^st^ shell water |
| --- | --- | --- | --- | --- | --- | --- | --- |
| **Ion** | | | |  | 1.00 | 26.71 || 1 | A | OP1 | 14:2062 | 2.18 | 1.00 | 38.47 |
| 2 | A | OP2 | 14:2062 | 3.63 | 1.00 | 43.28 |  |
| 3 | G | OP2 | 14:2444 | 4.19 | 1.00 | 37.71 |  |
| 4 | C | OP2 | 14:2443 | 4.29 | 1.00 | 35.47 |  |
| 5 | G | N7 | 14:2444 | 4.29 | 1.00 | 34.27 |  |
| 6 | A | O5' | 14:2062 | 4.29 | 1.00 | 40.11 |  |
| 7 | Water 1 | | 14:3903 | 4.36 | 1.00 | 40.94 | 3.78 (G:OP1 14:2061) |
| 8 | G | OP1 | 14:2061 | 4.41 | 1.00 | 35.00 |  |
| 9 | G | O3' | 14:2061 | 4.57 | 1.00 | 34.55 |  |
| 10 | C | O5' | 14:2443 | 4.93 | 1.00 | 32.06 |  |
| 11 | LYS | NZ | 39:68 | 4.96 | 1.00 | 37.59 |  |
  |

  
 Contact(s): /OP1/  
 Coded contact list: O_ph_  

### MG:14:3173

|  | Res. type | Atom | Residue | Distance | Occupancy | B-factor | Closest solute to 1^st^ shell water |
| --- | --- | --- | --- | --- | --- | --- | --- |
| **Ion** | | | |  | 1.00 | 66.48 || 1 | C | OP1 | 14:2601 | 3.59 | 1.00 | 44.21 |
| 2 | A | OP1 | 14:2439 | 4.35 | 1.00 | 37.92 |  |
| 3 | C | O4' | 2L:75 | 4.56 | 1.00 | 49.59 |  |
| 4 | A | O3' | 14:2600 | 4.73 | 1.00 | 38.84 |  |
| 5 | U | OP1 | 14:2438 | 4.80 | 1.00 | 25.80 |  |
| 6 | C | OP1 | 14:2064 | 4.95 | 1.00 | 43.58 |  |
  |

  
 Coded contact list:   
1^st^ MG coordination shell missing (see Notes)  

### MG:14:3174

|  | Res. type | Atom | Residue | Distance | Occupancy | B-factor | Closest solute to 1^st^ shell water |
| --- | --- | --- | --- | --- | --- | --- | --- |
| **Ion** | | | |  | 1.00 | 38.80 || 1 | G | OP2 | 14:2447 | 2.73 | 1.00 | 45.42 |
| 2 | G | OP2 | 14:2446 | 3.33 | 1.00 | 37.34 |  |
| 3 | G | OP1 | 14:2446 | 3.36 | 1.00 | 37.81 |  |
| 4 | G | OP1 | 14:2502 | 3.74 | 1.00 | 36.48 |  |
| 5 | G | N7 | 14:2502 | 3.85 | 1.00 | 44.73 |  |
| 6 | G | O5' | 14:2446 | 4.38 | 1.00 | 42.03 |  |
| 7 | G | OP1 | 14:2447 | 4.56 | 1.00 | 45.55 |  |
| 8 | G | O5' | 14:2447 | 4.79 | 1.00 | 36.14 |  |
  |

  
 Coded contact list:   
 Exclusion zone contact(s): /OP1/OP2/OP2/  
Consider K in place of MG (3 valid contact(s) for K vs. 0 for MG)  

### MG:14:3175

|  | Res. type | Atom | Residue | Distance | Occupancy | B-factor | Closest solute to 1^st^ shell water |
| --- | --- | --- | --- | --- | --- | --- | --- |
| **Ion** | | | |  | 0.50 | 39.96 || 1 | U | OP1 | 14:1352 | 2.47 | 1.00 | 49.28 |
| 2 | Water 1 | | 14:3556 | 2.84 | 1.00 | 54.86 | 2.22 (C:OP1 14:1375) |
| 3 | C | OP1 | 14:1375 | 3.65 | 1.00 | 57.28 |  |
| 4 | U | OP2 | 14:1352 | 3.91 | 1.00 | 37.76 |  |
| 5 | C | O3' | 14:1351 | 4.39 | 1.00 | 42.52 |  |
| 6 | C | OP2 | 14:1375 | 4.46 | 1.00 | 50.26 |  |
| 7 | C | OP1 | 14:1351 | 4.65 | 1.00 | 38.25 |  |
| 8 | U | O5' | 14:1352 | 4.90 | 1.00 | 44.63 |  |
  |

  
 Coded contact list:   
 Exclusion zone contact(s): /O_w_/OP1/  
Consider NA in place of MG (1 valid contact(s) for NA vs. 0 for MG)  
MG hydration shell issue: water too close (< 2.4 Angs.)  

### MG:14:3175

|  | Res. type | Atom | Residue | Distance | Occupancy | B-factor | Closest solute to 1^st^ shell water |
| --- | --- | --- | --- | --- | --- | --- | --- |
| **Ion** | | | |  | 0.50 | 36.56 || 1 | U | OP1 | 14:1352 | 2.70 | 1.00 | 49.28 |
| 2 | A | O3' | 14:1571 | 3.28 | 1.00 | 48.08 |  |
| 3 | A | O2' | 14:1571 | 3.48 | 1.00 | 46.31 |  |
| 4 | A | OP1 | 14:1572 | 3.91 | 1.00 | 36.30 |  |
| 5 | C | O3' | 14:1351 | 4.37 | 1.00 | 42.52 |  |
| 6 | A | O5' | 14:1572 | 4.45 | 1.00 | 44.55 |  |
| 7 | Water 1 | | 14:3556 | 4.96 | 1.00 | 54.86 | 3.08 (U:OP1 14:1352) |
| 8 | U | O5' | 14:1352 | 4.98 | 1.00 | 44.63 |  |
  |

  
 Coded contact list:   
 Exclusion zone contact(s): /O3'/OP1/  
Consider K in place of MG (2 valid contact(s) for K vs. 0 for MG)  

### MG:14:3176

|  | Res. type | Atom | Residue | Distance | Occupancy | B-factor | Closest solute to 1^st^ shell water |
| --- | --- | --- | --- | --- | --- | --- | --- |
| **Ion** | | | |  | 1.00 | 69.09 || 1 | A | OP2 | 14:1495 | 2.74 | 1.00 | 83.45 |
| 2 | A | N1 | 14:1496 | 3.25 | 1.00 | 85.79 |  |
| 3 | A | OP2 | 14:1494 | 3.66 | 1.00 | 82.96 |  |
| 4 | A | OP1 | 14:1495 | 3.98 | 1.00 | 88.92 |  |
| 5 | G | N7 | 14:1492 | 4.26 | 1.00 | 70.03 |  |
| 6 | A | O5' | 14:1495 | 4.62 | 1.00 | 80.79 |  |
| 7 | A | OP2 | 14:1496 | 4.68 | 1.00 | 81.28 |  |
| 8 | G | O6 | 14:1492 | 4.71 | 1.00 | 82.21 |  |
| 9 | A | N7 | 14:1495 | 4.74 | 1.00 | 79.08 |  |
| 10 | A | N3 | 14:1496 | 4.76 | 1.00 | 78.34 |  |
| 11 | A | N6 | 14:1496 | 4.89 | 1.00 | 77.10 |  |
  |

  
 Coded contact list:   
 Exclusion zone contact(s): /N1_A_/OP2/  
Consider K in place of MG (2 valid contact(s) for K vs. 0 for MG)  

### MG:14:3177

|  | Res. type | Atom | Residue | Distance | Occupancy | B-factor | Closest solute to 1^st^ shell water |
| --- | --- | --- | --- | --- | --- | --- | --- |
| **Ion** | | | |  | 1.00 | 45.32 || 1 | G | OP2 | 14:2070 | 1.94 | 1.00 | 43.89 |
| 2 | G | OP2 | 14:2069 | 3.12 | 1.00 | 31.97 |  |
| 3 | U | OP2 | 14:2244 | 3.44 | 1.00 | 35.38 |  |
| 4 | G | O5' | 14:2070 | 3.90 | 1.00 | 37.20 |  |
| 5 | G | OP1 | 14:2070 | 3.94 | 1.00 | 41.69 |  |
| 6 | U | OP1 | 14:2244 | 4.18 | 1.00 | 39.80 |  |
| 7 | G | O3' | 14:2069 | 4.50 | 1.00 | 34.76 |  |
  |

  
 Contact(s): /OP2/  
 Coded contact list: O_ph_  
 Exclusion zone contact(s): /OP2/  

### MG:14:3178

|  | Res. type | Atom | Residue | Distance | Occupancy | B-factor | Closest solute to 1^st^ shell water |
| --- | --- | --- | --- | --- | --- | --- | --- |
| **Ion** | | | |  | 1.00 | 53.98 || 1 | G | N7 | 14:2375 | 4.53 | 1.00 | 53.48 |
  |

  
 Coded contact list:   
1^st^ MG coordination shell missing (see Notes)  

### MG:14:3179

|  | Res. type | Atom | Residue | Distance | Occupancy | B-factor | Closest solute to 1^st^ shell water |
| --- | --- | --- | --- | --- | --- | --- | --- |
| **Ion** | | | |  | 1.00 | 32.99 || 1 | U | O4 | 14:1602 | 1.86 | 1.00 | 38.55 |
| 2 | A | OP2 | 14:1342 | 2.36 | 1.00 | 51.72 |  |
| 3 | G | N7 | 14:1601 | 3.75 | 1.00 | 44.76 |  |
| 4 | U | N3 | 14:1602 | 4.13 | 1.00 | 41.39 |  |
| 5 | A | O5' | 14:1342 | 4.20 | 1.00 | 46.77 |  |
| 6 | G | O6 | 14:1601 | 4.27 | 1.00 | 47.76 |  |
| 7 | THR | O | B5:56 | 4.28 | 1.00 | 44.52 |  |
| 8 | A | OP1 | 14:1342 | 4.43 | 1.00 | 40.84 |  |
| 9 | U | O3' | 14:1341 | 4.81 | 1.00 | 40.03 |  |
| 10 | C | N4 | 14:1600 | 4.83 | 1.00 | 51.40 |  |
| 11 | A | N3 | 14:1342 | 4.85 | 1.00 | 43.66 |  |
  |

  
**1^st^ shell ligand angles:**   
1-2: 89;   
  
 Contact(s): /O4/OP2/  
 Coded contact list: O_ph_O_b_  

### MG:14:3180

|  | Res. type | Atom | Residue | Distance | Occupancy | B-factor | Closest solute to 1^st^ shell water |
| --- | --- | --- | --- | --- | --- | --- | --- |
| **Ion** | | | |  | 1.00 | 59.36 || 1 | G | OP1 | 14:1560 | 2.56 | 1.00 | 60.22 |
| 2 | G | OP2 | 14:1561 | 2.83 | 1.00 | 60.26 |  |
| 3 | G | OP2 | 14:1560 | 3.63 | 1.00 | 71.45 |  |
| 4 | G | N7 | 14:1561 | 4.28 | 1.00 | 50.56 |  |
| 5 | G | O5' | 14:1560 | 4.53 | 1.00 | 66.21 |  |
| 6 | G | O3' | 14:1559 | 4.71 | 1.00 | 54.06 |  |
| 7 | G | O5' | 14:1561 | 4.85 | 1.00 | 54.64 |  |
  |

  
 Coded contact list:   
 Exclusion zone contact(s): /OP1/OP2/  
Consider NA in place of MG (1 valid contact(s) for NA vs. 0 for MG)  

### MG:14:3181

|  | Res. type | Atom | Residue | Distance | Occupancy | B-factor | Closest solute to 1^st^ shell water |
| --- | --- | --- | --- | --- | --- | --- | --- |
| **Ion** | | | |  | 1.00 | 49.93 || 1 | G | N7 | 14:2353 | 3.32 | 1.00 | 70.66 |
| 2 | G | O6 | 14:2353 | 3.78 | 1.00 | 66.88 |  |
| 3 | A | OP1 | 14:2352 | 3.81 | 1.00 | 84.35 |  |
| 4 | A | OP2 | 14:2352 | 3.93 | 1.00 | 64.89 |  |
| 5 | G | N7 | 14:2354 | 4.30 | 1.00 | 58.32 |  |
| 6 | G | O6 | 14:2354 | 4.37 | 1.00 | 66.17 |  |
  |

  
 Coded contact list:   
 Exclusion zone contact(s): /N7/  
Consider K in place of MG (1 valid contact(s) for K vs. 0 for MG)  

### MG:14:3182

|  | Res. type | Atom | Residue | Distance | Occupancy | B-factor | Closest solute to 1^st^ shell water |
| --- | --- | --- | --- | --- | --- | --- | --- |
| **Ion** | | | |  | 1.00 | 62.18 || 1 | U | OP2 | 14:1520 | 2.82 | 1.00 | 59.15 |
| 2 | G | O6 | 14:1470 | 2.96 | 1.00 | 87.67 |  |
| 3 | G | N7 | 14:1470 | 3.20 | 1.00 | 81.34 |  |
| 4 | A | N7 | 14:1469 | 4.36 | 1.00 | 78.77 |  |
| 5 | G | N7 | 14:1521 | 4.53 | 1.00 | 82.49 |  |
| 6 | U | O5' | 14:1520 | 4.68 | 1.00 | 78.72 |  |
| 7 | U | OP1 | 14:1520 | 4.69 | 1.00 | 64.70 |  |
| 8 | G | O6 | 14:1521 | 4.85 | 1.00 | 82.41 |  |
| 9 | G | N1 | 14:1470 | 4.98 | 1.00 | 85.62 |  |
| 10 | A | OP2 | 14:1469 | 4.99 | 1.00 | 70.94 |  |
  |

  
 Coded contact list:   
 Exclusion zone contact(s): /N7/O6/OP2/  
Consider K in place of MG (3 valid contact(s) for K vs. 0 for MG)  

### MG:14:3183

|  | Res. type | Atom | Residue | Distance | Occupancy | B-factor | Closest solute to 1^st^ shell water |
| --- | --- | --- | --- | --- | --- | --- | --- |
| **Ion** | | | |  | 1.00 | 30.71 || 1 | G | O6 | 14:799 | 4.04 | 1.00 | 38.48 |
| 2 | GLY | O | 39:61 | 4.15 | 1.00 | 39.83 |  |
| 3 | G | O6 | 14:798 | 4.17 | 1.00 | 36.96 |  |
| 4 | G | N7 | 14:798 | 4.50 | 1.00 | 33.60 |  |
| 5 | A | N3 | 14:800 | 4.89 | 1.00 | 38.14 |  |
  |

  
 Coded contact list:   
1^st^ MG coordination shell missing (see Notes)  

### MG:14:3184

|  | Res. type | Atom | Residue | Distance | Occupancy | B-factor | Closest solute to 1^st^ shell water |
| --- | --- | --- | --- | --- | --- | --- | --- |
| **Ion** | | | |  | 1.00 | 48.50 || 1 | U | OP1 | 14:2696 | 3.54 | 1.00 | 49.04 |
| 2 | G | O2' | 14:1707 | 4.04 | 1.00 | 64.41 |  |
| 3 | G | O6 | 14:1758 | 4.08 | 1.00 | 61.37 |  |
| 4 | G | N1 | 14:1758 | 4.22 | 1.00 | 54.99 |  |
| 5 | C | O4' | 14:1708 | 4.25 | 1.00 | 66.04 |  |
| 6 | G | N3 | 14:1707 | 4.71 | 1.00 | 70.32 |  |
| 7 | C | O3' | 14:2695 | 4.84 | 1.00 | 47.20 |  |
| 8 | G | N2 | 14:1707 | 4.95 | 1.00 | 69.10 |  |
  |

  
 Coded contact list:   
1^st^ MG coordination shell missing (see Notes)  

### MG:14:3185

|  | Res. type | Atom | Residue | Distance | Occupancy | B-factor | Closest solute to 1^st^ shell water |
| --- | --- | --- | --- | --- | --- | --- | --- |
| **Ion** | | | |  | 1.00 | 51.93 || 1 | G | O6 | 14:495 | 3.15 | 1.00 | 65.12 |
| 2 | G | O6 | 14:496 | 3.68 | 1.00 | 58.41 |  |
| 3 | C | N4 | 14:484 | 4.32 | 1.00 | 66.31 |  |
| 4 | A | N6 | 14:497 | 4.39 | 1.00 | 62.52 |  |
| 5 | A | OP2 | 14:483 | 4.41 | 1.00 | 52.31 |  |
| 6 | C | N4 | 14:485 | 4.72 | 1.00 | 56.81 |  |
| 7 | A | N1 | 14:497 | 4.76 | 1.00 | 53.02 |  |
| 8 | G | N7 | 14:496 | 4.80 | 1.00 | 56.71 |  |
| 9 | G | N7 | 14:495 | 4.81 | 1.00 | 52.14 |  |
| 10 | A | N3 | 14:482 | 4.87 | 1.00 | 53.68 |  |
  |

  
 Coded contact list:   
 Exclusion zone contact(s): /O6/  
Consider K in place of MG (1 valid contact(s) for K vs. 0 for MG)  

### MG:14:3186

|  | Res. type | Atom | Residue | Distance | Occupancy | B-factor | Closest solute to 1^st^ shell water |
| --- | --- | --- | --- | --- | --- | --- | --- |
| **Ion** | | | |  | 1.00 | 49.52 || 1 | G | O6 | 14:493 | 3.38 | 1.00 | 48.46 |
| 2 | G | N7 | 14:493 | 3.49 | 1.00 | 50.65 |  |
| 3 | G | O6 | 14:494 | 4.17 | 1.00 | 53.04 |  |
| 4 | A | OP2 | 14:492 | 4.30 | 1.00 | 51.91 |  |
| 5 | G | N7 | 14:494 | 4.43 | 1.00 | 45.96 |  |
| 6 | G | N2 | 14:488 | 4.84 | 1.00 | 45.71 |  |
  |

  
 Coded contact list:   
 Exclusion zone contact(s): /O6/  
Consider K in place of MG (1 valid contact(s) for K vs. 0 for MG)  

### MG:14:3187

|  | Res. type | Atom | Residue | Distance | Occupancy | B-factor | Closest solute to 1^st^ shell water |
| --- | --- | --- | --- | --- | --- | --- | --- |
| **Ion** | | | |  | 1.00 | 51.19 || 1 | G | O6 | 14:521 | 2.97 | 1.00 | 61.05 |
| 2 | G | N7 | 14:520 | 3.66 | 1.00 | 60.84 |  |
| 3 | G | O6 | 14:520 | 3.87 | 1.00 | 65.00 |  |
| 4 | G | N7 | 14:521 | 4.04 | 1.00 | 66.40 |  |
  |

  
 Coded contact list:   
 Exclusion zone contact(s): /O6/  
Consider K in place of MG (1 valid contact(s) for K vs. 0 for MG)  

### MG:14:3188

|  | Res. type | Atom | Residue | Distance | Occupancy | B-factor | Closest solute to 1^st^ shell water |
| --- | --- | --- | --- | --- | --- | --- | --- |
| **Ion** | | | |  | 1.00 | 56.08 || 1 | A | OP2 | 14:861 | 3.47 | 1.00 | 74.98 |
| 2 | U | O5' | 14:860 | 4.12 | 1.00 | 76.63 |  |
| 3 | U | O4 | 14:913 | 4.23 | 1.00 | 74.14 |  |
| 4 | A | OP1 | 14:911 | 4.33 | 1.00 | 71.13 |  |
| 5 | A | OP2 | 14:910 | 4.34 | 1.00 | 71.96 |  |
| 6 | U | OP2 | 14:860 | 4.45 | 1.00 | 83.47 |  |
| 7 | U | O3' | 14:860 | 4.60 | 1.00 | 64.45 |  |
| 8 | U | OP1 | 14:860 | 4.62 | 1.00 | 70.36 |  |
| 9 | A | OP1 | 14:861 | 4.75 | 1.00 | 58.00 |  |
| 10 | A | O5' | 14:910 | 4.85 | 1.00 | 58.27 |  |
| 11 | A | OP1 | 14:910 | 5.00 | 1.00 | 55.50 |  |
  |

  
 Coded contact list:   
1^st^ MG coordination shell missing (see Notes)  

### MG:14:3189

|  | Res. type | Atom | Residue | Distance | Occupancy | B-factor | Closest solute to 1^st^ shell water |
| --- | --- | --- | --- | --- | --- | --- | --- |
| **Ion** | | | |  | 1.00 | 50.60 || 1 | G | OP2 | 14:862 | 2.14 | 1.00 | 74.88 |
| 2 | A | OP2 | 14:861 | 3.38 | 1.00 | 74.98 |  |
| 3 | G | N7 | 14:862 | 3.84 | 1.00 | 84.81 |  |
| 4 | A | OP1 | 14:861 | 3.88 | 1.00 | 58.00 |  |
| 5 | G | O5' | 14:862 | 3.91 | 1.00 | 70.94 |  |
| 6 | G | OP1 | 14:862 | 4.18 | 1.00 | 66.92 |  |
| 7 | A | N7 | 14:863 | 4.21 | 1.00 | 80.89 |  |
| 8 | A | OP1 | 14:910 | 4.25 | 1.00 | 55.50 |  |
| 9 | A | O5' | 14:861 | 4.25 | 1.00 | 66.58 |  |
| 10 | A | N6 | 14:863 | 4.35 | 1.00 | 79.42 |  |
| 11 | A | O3' | 14:861 | 4.63 | 1.00 | 62.07 |  |
  |

  
 Contact(s): /OP2/  
 Coded contact list: O_ph_  
 Exclusion zone contact(s): /OP2/  

### MG:14:3190

|  | Res. type | Atom | Residue | Distance | Occupancy | B-factor | Closest solute to 1^st^ shell water |
| --- | --- | --- | --- | --- | --- | --- | --- |
| **Ion** | | | |  | 1.00 | 77.09 || 1 | G | O6 | 14:1542 | 2.78 | 1.00 | 103.31 |
| 2 | A | N6 | 14:1543 | 3.30 | 1.00 | 97.54 |  |
| 3 | A | OP2 | 14:1528 | 3.58 | 1.00 | 86.05 |  |
| 4 | A | OP1 | 14:1528 | 3.76 | 1.00 | 82.09 |  |
| 5 | G | N1 | 14:1542 | 4.25 | 1.00 | 112.16 |  |
| 6 | U | O4 | 14:1541 | 4.30 | 1.00 | 123.28 |  |
| 7 | G | O5' | 14:1527 | 4.58 | 1.00 | 87.49 |  |
| 8 | G | O3' | 14:1527 | 4.79 | 1.00 | 87.09 |  |
| 9 | G | OP2 | 14:1527 | 4.97 | 1.00 | 75.94 |  |
  |

  
 Coded contact list:   
 Exclusion zone contact(s): /N6/O6/  
Consider K in place of MG (2 valid contact(s) for K vs. 0 for MG)  

### MG:14:3191

|  | Res. type | Atom | Residue | Distance | Occupancy | B-factor | Closest solute to 1^st^ shell water |
| --- | --- | --- | --- | --- | --- | --- | --- |
| **Ion** | | | |  | 1.00 | 34.58 || 1 | G | OP1 | 14:2592 | 2.00 | 1.00 | 32.71 |
| 2 | G | OP1 | 14:1968 | 2.90 | 1.00 | 43.87 |  |
| 3 | C | O3' | 14:2591 | 3.75 | 1.00 | 41.95 |  |
| 4 | G | OP1 | 14:1973 | 3.89 | 1.00 | 30.07 |  |
| 5 | C | O3' | 14:1967 | 4.01 | 1.00 | 36.70 |  |
| 6 | G | OP2 | 14:2592 | 4.42 | 1.00 | 32.39 |  |
| 7 | G | O5' | 14:1968 | 4.52 | 1.00 | 32.19 |  |
| 8 | G | O5' | 14:2592 | 4.52 | 1.00 | 40.76 |  |
| 9 | C | O2' | 14:1967 | 4.66 | 1.00 | 44.44 |  |
  |

  
 Contact(s): /OP1/  
 Coded contact list: O_ph_  
 Exclusion zone contact(s): /OP1/  

### MG:14:3192

|  | Res. type | Atom | Residue | Distance | Occupancy | B-factor | Closest solute to 1^st^ shell water |
| --- | --- | --- | --- | --- | --- | --- | --- |
| **Ion** | | | |  | 1.00 | 41.32 || 1 | C | OP1 | 14:974A | 1.90 | 1.00 | 60.08 |
| 2 | C | OP2 | 14:974A | 3.50 | 1.00 | 63.55 |  |
| 3 | C | O5' | 14:974A | 4.12 | 1.00 | 65.88 |  |
| 4 | G | O2' | 14:974 | 4.12 | 1.00 | 62.55 |  |
| 5 | G | O3' | 14:974 | 4.30 | 1.00 | 60.24 |  |
| 6 | A | O2' | 14:983 | 4.52 | 1.00 | 55.84 |  |
| 7 | A | O3' | 14:983 | 4.68 | 1.00 | 51.32 |  |
| 8 | A | OP1 | 14:984 | 4.82 | 1.00 | 53.28 |  |
  |

  
 Contact(s): /OP1/  
 Coded contact list: O_ph_  

### MG:14:3193

|  | Res. type | Atom | Residue | Distance | Occupancy | B-factor | Closest solute to 1^st^ shell water |
| --- | --- | --- | --- | --- | --- | --- | --- |
| **Ion** | | | |  | 1.00 | 40.98 || 1 | C | OP2 | 14:2196 | 2.57 | 1.00 | 67.72 |
| 2 | G | OP2 | 14:2224 | 4.09 | 1.00 | 45.79 |  |
| 3 | C | OP1 | 14:2196 | 4.25 | 1.00 | 64.82 |  |
| 4 | C | O5' | 14:2195 | 4.40 | 1.00 | 85.58 |  |
| 5 | C | O5' | 14:2196 | 4.78 | 1.00 | 63.79 |  |
| 6 | C | O3' | 14:2195 | 4.89 | 1.00 | 62.40 |  |
  |

  
 Coded contact list:   
 Exclusion zone contact(s): /OP2/  
Consider NA in place of MG (1 valid contact(s) for NA vs. 0 for MG)  

### MG:14:3194

|  | Res. type | Atom | Residue | Distance | Occupancy | B-factor | Closest solute to 1^st^ shell water |
| --- | --- | --- | --- | --- | --- | --- | --- |
| **Ion** | | | |  | 1.00 | 54.37 || 1 | ASN | ND2 | 55:11 | 3.65 | 1.00 | 51.61 |
| 2 | G | O6 | 14:1651 | 3.98 | 1.00 | 47.73 |  |
| 3 | G | OP2 | 14:2004 | 4.13 | 1.00 | 43.53 |  |
| 4 | G | O6 | 14:1650 | 4.29 | 1.00 | 49.12 |  |
| 5 | G | N7 | 14:1650 | 4.52 | 1.00 | 48.72 |  |
| 6 | G | N7 | 14:1651 | 4.62 | 1.00 | 47.61 |  |
| 7 | G | OP1 | 14:2004 | 4.76 | 1.00 | 52.53 |  |
  |

  
 Coded contact list:   
1^st^ MG coordination shell missing (see Notes)  

### MG:14:3195

|  | Res. type | Atom | Residue | Distance | Occupancy | B-factor | Closest solute to 1^st^ shell water |
| --- | --- | --- | --- | --- | --- | --- | --- |
| **Ion** | | | |  | 1.00 | 53.77 || 1 | A | OP2 | 14:53 | 2.42 | 1.00 | 41.77 |
| 2 | G | O6 | 14:54 | 3.66 | 1.00 | 44.01 |  |
| 3 | G | N7 | 14:54 | 3.70 | 1.00 | 47.39 |  |
| 4 | A | OP1 | 14:53 | 4.21 | 1.00 | 35.68 |  |
| 5 | A | OP2 | 14:52 | 4.32 | 1.00 | 44.06 |  |
| 6 | A | O5' | 14:53 | 4.51 | 1.00 | 45.50 |  |
| 7 | A | O5' | 14:52 | 4.60 | 1.00 | 42.47 |  |
| 8 | A | O3' | 14:52 | 4.81 | 1.00 | 47.93 |  |
| 9 | G | N2 | 14:117 | 4.92 | 1.00 | 40.70 |  |
  |

  
 Coded contact list:   
 Exclusion zone contact(s): /OP2/  
Consider NA in place of MG (1 valid contact(s) for NA vs. 0 for MG)  

### MG:14:3196

|  | Res. type | Atom | Residue | Distance | Occupancy | B-factor | Closest solute to 1^st^ shell water |
| --- | --- | --- | --- | --- | --- | --- | --- |
| **Ion** | | | |  | 1.00 | 59.93 || 1 | G | OP2 | 14:2782 | 1.99 | 1.00 | 74.55 |
| 2 | G | O5' | 14:2782 | 3.91 | 1.00 | 71.61 |  |
| 3 | G | N7 | 14:2782 | 3.91 | 1.00 | 70.41 |  |
| 4 | G | OP1 | 14:2782 | 4.15 | 1.00 | 70.01 |  |
| 5 | G | N7 | 14:2783 | 4.25 | 1.00 | 75.25 |  |
| 6 | A | O3' | 14:2781 | 4.49 | 1.00 | 74.15 |  |
| 7 | G | OP2 | 14:2783 | 4.64 | 1.00 | 73.59 |  |
| 8 | G | O6 | 14:2783 | 4.65 | 1.00 | 64.07 |  |
  |

  
 Contact(s): /OP2/  
 Coded contact list: O_ph_  

### MG:14:3197

|  | Res. type | Atom | Residue | Distance | Occupancy | B-factor | Closest solute to 1^st^ shell water |
| --- | --- | --- | --- | --- | --- | --- | --- |
| **Ion** | | | |  | 1.00 | 48.86 || 1 | G | O6 | 14:1949 | 3.15 | 1.00 | 43.85 |
| 2 | G | O6 | 14:1950 | 3.55 | 1.00 | 52.89 |  |
| 3 | G | O6 | 14:1948 | 3.73 | 1.00 | 43.84 |  |
| 4 | G | N7 | 14:1949 | 3.85 | 1.00 | 42.33 |  |
| 5 | U | OP2 | 14:1955 | 4.22 | 1.00 | 48.53 |  |
| 6 | G | N7 | 14:1948 | 4.50 | 1.00 | 43.60 |  |
| 7 | C | N4 | 14:1957 | 4.56 | 1.00 | 46.00 |  |
| 8 | G | O2' | 14:1954 | 4.62 | 1.00 | 53.03 |  |
| 9 | G | N1 | 14:1950 | 4.63 | 1.00 | 59.21 |  |
  |

  
 Coded contact list:   
 Exclusion zone contact(s): /O6/  
Consider K in place of MG (1 valid contact(s) for K vs. 0 for MG)  

### MG:14:3198

|  | Res. type | Atom | Residue | Distance | Occupancy | B-factor | Closest solute to 1^st^ shell water |
| --- | --- | --- | --- | --- | --- | --- | --- |
| **Ion** | | | |  | 1.00 | 33.62 || 1 | G | OP2 | 14:1271 | 2.07 | 1.00 | 41.18 |
| 2 | C | OP1 | 14:1648 | 2.70 | 1.00 | 42.86 |  |
| 3 | G | O5' | 14:1647 | 2.72 | 1.00 | 43.15 |  |
| 4 | A | OP1 | 14:1272 | 2.92 | 1.00 | 48.09 |  |
| 5 | G | OP1 | 14:1647 | 3.03 | 1.00 | 49.34 |  |
| 6 | C | O5' | 14:1648 | 3.24 | 1.00 | 38.94 |  |
| 7 | A | OP2 | 14:1272 | 3.53 | 1.00 | 43.66 |  |
| 8 | G | O3' | 14:1271 | 4.05 | 1.00 | 46.23 |  |
| 9 | C | O3' | 14:1270 | 4.17 | 1.00 | 39.74 |  |
| 10 | G | O3' | 14:1647 | 4.32 | 1.00 | 40.98 |  |
| 11 | G | O5' | 14:1271 | 4.35 | 1.00 | 40.73 |  |
| 12 | Water 1 | | 14:3820 | 4.40 | 1.00 | 46.38 | 2.81 (G:OP1 14:1647) |
| 13 | G | OP2 | 14:1647 | 4.42 | 1.00 | 37.28 |  |
| 14 | G | OP1 | 14:1271 | 4.47 | 1.00 | 35.86 |  |
| 15 | C | O3' | 14:1646 | 4.69 | 1.00 | 40.91 |  |
| 16 | C | OP2 | 14:1648 | 4.82 | 1.00 | 38.13 |  |
  |

  
 Contact(s): /OP2/  
 Coded contact list: O_ph_  
 Exclusion zone contact(s): /O5'/O5'/OP1/OP1/OP1/  
Consider K in place of MG (5 valid contact(s) for K vs. 1 for MG)  

### MG:14:3199

|  | Res. type | Atom | Residue | Distance | Occupancy | B-factor | Closest solute to 1^st^ shell water |
| --- | --- | --- | --- | --- | --- | --- | --- |
| **Ion** | | | |  | 1.00 | 66.07 || 1 | C | OP2 | 14:365 | 2.09 | 1.00 | 72.68 |
| 2 | C | OP2 | 14:364 | 2.39 | 1.00 | 83.29 |  |
| 3 | U | O3' | 14:403 | 3.03 | 1.00 | 84.65 |  |
| 4 | C | OP1 | 14:365 | 3.71 | 1.00 | 81.79 |  |
| 5 | U | O2' | 14:403 | 3.83 | 1.00 | 70.94 |  |
| 6 | C | OP1 | 14:404 | 4.15 | 1.00 | 71.84 |  |
| 7 | C | O5' | 14:365 | 4.21 | 1.00 | 71.19 |  |
| 8 | A | O2' | 14:363F | 4.29 | 1.00 | 89.58 |  |
| 9 | A | O3' | 14:363F | 4.48 | 1.00 | 83.94 |  |
| 10 | C | O3' | 14:364 | 4.52 | 1.00 | 78.53 |  |
| 11 | C | OP1 | 14:364 | 4.76 | 1.00 | 88.00 |  |
| 12 | C | O5' | 14:364 | 4.82 | 1.00 | 87.05 |  |
| 13 | C | OP2 | 14:404 | 4.99 | 1.00 | 80.16 |  |
  |

  
**1^st^ shell ligand angles:**   
1-2: 72;   
  
 Contact(s): /OP2/OP2/  
 Coded contact list: 2O_ph_  
 Exclusion zone contact(s): /O3'/  

### MG:14:3200

|  | Res. type | Atom | Residue | Distance | Occupancy | B-factor | Closest solute to 1^st^ shell water |
| --- | --- | --- | --- | --- | --- | --- | --- |
| **Ion** | | | |  | 1.00 | 52.05 || 1 | U | O4 | 14:1629 | 3.44 | 1.00 | 41.85 |
| 2 | G | N7 | 14:1628 | 4.23 | 1.00 | 47.84 |  |
| 3 | G | O6 | 14:1630 | 4.41 | 1.00 | 43.65 |  |
| 4 | G | O6 | 14:1628 | 4.44 | 1.00 | 48.45 |  |
| 5 | C | N4 | 14:1636 | 4.70 | 1.00 | 42.03 |  |
| 6 | A | OP2 | 14:1634 | 4.76 | 1.00 | 41.84 |  |
| 7 | G | OP2 | 14:1627 | 4.79 | 1.00 | 50.57 |  |
| 8 | G | OP2 | 14:1628 | 5.00 | 1.00 | 55.88 |  |
  |

  
 Coded contact list:   
1^st^ MG coordination shell missing (see Notes)  

### MG:14:3201

|  | Res. type | Atom | Residue | Distance | Occupancy | B-factor | Closest solute to 1^st^ shell water |
| --- | --- | --- | --- | --- | --- | --- | --- |
| **Ion** | | | |  | 1.00 | 49.74 || 1 | G | N7 | 14:2830 | 3.31 | 1.00 | 50.68 |
| 2 | G | O6 | 14:2830 | 3.35 | 1.00 | 49.54 |  |
| 3 | G | O6 | 14:2831 | 4.19 | 1.00 | 60.34 |  |
| 4 | G | N7 | 14:2831 | 4.23 | 1.00 | 55.93 |  |
  |

  
 Coded contact list:   
 Exclusion zone contact(s): /N7/O6/  
Consider K in place of MG (2 valid contact(s) for K vs. 0 for MG)  

### MG:14:3202

|  | Res. type | Atom | Residue | Distance | Occupancy | B-factor | Closest solute to 1^st^ shell water |
| --- | --- | --- | --- | --- | --- | --- | --- |
| **Ion** | | | |  | 1.00 | 42.42 || 1 | A | O2' | 14:528 | 2.78 | 1.00 | 57.06 |
| 2 | A | O3' | 14:528 | 2.98 | 1.00 | 74.25 |  |
| 3 | A | O5' | 14:529 | 4.14 | 1.00 | 69.31 |  |
| 4 | A | O4' | 14:528 | 4.42 | 1.00 | 58.32 |  |
| 5 | A | OP1 | 14:529 | 4.66 | 1.00 | 73.39 |  |
| 6 | A | O4' | 14:529 | 4.81 | 1.00 | 67.27 |  |
  |

  
 Coded contact list:   
 Exclusion zone contact(s): /O2'/O3'/  
Consider K in place of MG (2 valid contact(s) for K vs. 0 for MG)  

### MG:14:3203

|  | Res. type | Atom | Residue | Distance | Occupancy | B-factor | Closest solute to 1^st^ shell water |
| --- | --- | --- | --- | --- | --- | --- | --- |
| **Ion** | | | |  | 1.00 | 45.60 || 1 | G | O6 | 14:110 | 3.46 | 1.00 | 61.61 |
| 2 | G | O6 | 14:109 | 3.80 | 1.00 | 67.13 |  |
| 3 | G | N7 | 14:110 | 4.44 | 1.00 | 63.69 |  |
| 4 | G | N7 | 14:109 | 4.53 | 1.00 | 67.15 |  |
  |

  
 Coded contact list:   
1^st^ MG coordination shell missing (see Notes)  

### MG:14:3204

|  | Res. type | Atom | Residue | Distance | Occupancy | B-factor | Closest solute to 1^st^ shell water |
| --- | --- | --- | --- | --- | --- | --- | --- |
| **Ion** | | | |  | 1.00 | 38.65 || 1 | G | O6 | 14:1845 | 3.63 | 1.00 | 50.41 |
| 2 | G | O6 | 14:1846 | 3.73 | 1.00 | 48.70 |  |
| 3 | G | N7 | 14:1845 | 4.44 | 1.00 | 53.46 |  |
| 4 | G | N7 | 14:1846 | 4.77 | 1.00 | 57.52 |  |
| 5 | C | OP2 | 14:1892 | 4.84 | 1.00 | 45.21 |  |
| 6 | C | N4 | 14:1844 | 4.93 | 1.00 | 44.95 |  |
  |

  
 Coded contact list:   
1^st^ MG coordination shell missing (see Notes)  

### MG:14:3205

|  | Res. type | Atom | Residue | Distance | Occupancy | B-factor | Closest solute to 1^st^ shell water |
| --- | --- | --- | --- | --- | --- | --- | --- |
| **Ion** | | | |  | 1.00 | 51.96 || 1 | U | OP2 | 14:427 | 3.49 | 1.00 | 60.98 |
| 2 | G | N7 | 14:220 | 3.62 | 1.00 | 53.99 |  |
| 3 | G | OP2 | 14:219 | 3.85 | 1.00 | 63.32 |  |
| 4 | G | O6 | 14:220 | 4.03 | 1.00 | 57.93 |  |
| 5 | C | OP2 | 14:426 | 4.15 | 1.00 | 58.13 |  |
| 6 | C | OP1 | 14:426 | 4.51 | 1.00 | 59.04 |  |
| 7 | C | O5' | 14:426 | 4.67 | 1.00 | 52.79 |  |
| 8 | G | N7 | 14:219 | 4.87 | 1.00 | 53.22 |  |
  |

  
 Coded contact list:   
1^st^ MG coordination shell missing (see Notes)  

### MG:14:3206

|  | Res. type | Atom | Residue | Distance | Occupancy | B-factor | Closest solute to 1^st^ shell water |
| --- | --- | --- | --- | --- | --- | --- | --- |
| **Ion** | | | |  | 1.00 | 55.81 || 1 | G | OP2 | 14:1622 | 2.24 | 1.00 | 44.41 |
| 2 | G | OP1 | 14:1622 | 3.43 | 1.00 | 47.76 |  |
| 3 | U | OP1 | 14:1621 | 4.03 | 1.00 | 43.89 |  |
| 4 | G | O5' | 14:1622 | 4.14 | 1.00 | 44.65 |  |
| 5 | C | N4 | 14:1607 | 4.17 | 1.00 | 40.39 |  |
| 6 | C | N3 | 14:1607 | 4.30 | 1.00 | 48.33 |  |
| 7 | U | O5' | 14:1621 | 4.39 | 1.00 | 36.00 |  |
| 8 | U | O3' | 14:1621 | 4.59 | 1.00 | 37.11 |  |
| 9 | C | OP1 | 14:737 | 4.62 | 1.00 | 34.56 |  |
| 10 | U | OP2 | 14:1621 | 4.76 | 1.00 | 36.60 |  |
| 11 | C | OP1 | 14:736 | 4.84 | 1.00 | 47.00 |  |
| 12 | C | O3' | 14:736 | 4.95 | 1.00 | 35.91 |  |
  |

  
 Contact(s): /OP2/  
 Coded contact list: O_ph_  

### MG:14:3207

|  | Res. type | Atom | Residue | Distance | Occupancy | B-factor | Closest solute to 1^st^ shell water |
| --- | --- | --- | --- | --- | --- | --- | --- |
| **Ion** | | | |  | 1.00 | 62.53 || 1 | U | OP2 | 14:504 | 4.80 | 1.00 | 64.65 |
| 2 | A | O2' | 14:503 | 4.90 | 1.00 | 59.45 |  |
| 3 | G | OP2 | 14:496 | 4.95 | 1.00 | 52.30 |  |
  |

  
 Coded contact list:   
1^st^ MG coordination shell missing (see Notes)  

### MG:14:3208

|  | Res. type | Atom | Residue | Distance | Occupancy | B-factor | Closest solute to 1^st^ shell water |
| --- | --- | --- | --- | --- | --- | --- | --- |
| **Ion** | | | |  | 1.00 | 58.78 || 1 | G | O6 | 14:1260 | 3.59 | 1.00 | 47.38 |
| 2 | G | O6 | 14:1259 | 4.22 | 1.00 | 41.04 |  |
| 3 | G | N7 | 14:1259 | 4.32 | 1.00 | 43.80 |  |
| 4 | G | O6 | 14:579 | 4.83 | 1.00 | 44.35 |  |
| 5 | C | N4 | 14:580 | 4.88 | 1.00 | 49.88 |  |
| 6 | C | N4 | 14:1258 | 4.99 | 1.00 | 41.36 |  |
  |

  
 Coded contact list:   
1^st^ MG coordination shell missing (see Notes)  

### MG:14:3209

|  | Res. type | Atom | Residue | Distance | Occupancy | B-factor | Closest solute to 1^st^ shell water |
| --- | --- | --- | --- | --- | --- | --- | --- |
| **Ion** | | | |  | 1.00 | 53.72 || 1 | C | OP1 | 14:509 | 2.46 | 1.00 | 52.26 |
| 2 | G | N7 | 14:27 | 2.87 | 1.00 | 56.56 |  |
| 3 | G | O6 | 14:512 | 4.12 | 1.00 | 50.47 |  |
| 4 | G | O6 | 14:27 | 4.29 | 1.00 | 56.47 |  |
| 5 | G | O3' | 14:508 | 4.47 | 1.00 | 46.91 |  |
| 6 | C | OP2 | 14:509 | 4.82 | 1.00 | 55.86 |  |
| 7 | C | O5' | 14:509 | 4.84 | 1.00 | 55.57 |  |
| 8 | G | N9 | 14:27 | 4.92 | 1.00 | 47.02 |  |
| 9 | G | OP2 | 14:27 | 4.98 | 1.00 | 40.81 |  |
  |

  
 Coded contact list:   
 Exclusion zone contact(s): /N7/OP1/  
Consider NA in place of MG (1 valid contact(s) for NA vs. 0 for MG)  

### MG:14:3210

|  | Res. type | Atom | Residue | Distance | Occupancy | B-factor | Closest solute to 1^st^ shell water |
| --- | --- | --- | --- | --- | --- | --- | --- |
| **Ion** | | | |  | 1.00 | 47.34 || 1 | A | OP1 | 14:802 | 2.51 | 1.00 | 37.31 |
| 2 | G | OP2 | 14:674 | 3.30 | 1.00 | 35.65 |  |
| 3 | A | OP2 | 14:675 | 3.77 | 1.00 | 33.85 |  |
| 4 | G | OP1 | 14:674 | 4.27 | 1.00 | 35.73 |  |
| 5 | G | O3' | 14:801 | 4.33 | 1.00 | 39.45 |  |
| 6 | G | O5' | 14:674 | 4.39 | 1.00 | 29.23 |  |
| 7 | U | O4 | 14:803 | 4.63 | 1.00 | 40.57 |  |
| 8 | ARG | NH2 | 39:54 | 4.68 | 1.00 | 34.05 |  |
| 9 | A | N6 | 14:804 | 4.80 | 1.00 | 39.36 |  |
| 10 | A | OP2 | 14:802 | 4.84 | 1.00 | 36.76 |  |
| 11 | A | O5' | 14:802 | 4.93 | 1.00 | 34.77 |  |
  |

  
 Coded contact list:   
 Exclusion zone contact(s): /OP1/OP2/  
Consider NA in place of MG (1 valid contact(s) for NA vs. 0 for MG)  

### MG:14:3211

|  | Res. type | Atom | Residue | Distance | Occupancy | B-factor | Closest solute to 1^st^ shell water |
| --- | --- | --- | --- | --- | --- | --- | --- |
| **Ion** | | | |  | 1.00 | 58.20 || 1 | A | OP2 | 14:2518 | 2.22 | 1.00 | 63.35 |
| 2 | U | OP2 | 14:2519 | 3.33 | 1.00 | 57.89 |  |
| 3 | A | N7 | 14:2541 | 3.48 | 1.00 | 65.88 |  |
| 4 | C | OP2 | 14:2540 | 3.91 | 1.00 | 71.25 |  |
| 5 | C | O3' | 14:2517 | 4.01 | 1.00 | 64.31 |  |
| 6 | A | OP2 | 14:2541 | 4.21 | 1.00 | 78.86 |  |
| 7 | C | O2' | 14:2517 | 4.24 | 1.00 | 54.34 |  |
| 8 | A | N6 | 14:2541 | 4.35 | 1.00 | 67.06 |  |
| 9 | A | O5' | 14:2518 | 4.62 | 1.00 | 64.29 |  |
| 10 | A | OP1 | 14:2518 | 4.67 | 1.00 | 56.90 |  |
  |

  
 Contact(s): /OP2/  
 Coded contact list: O_ph_  
 Exclusion zone contact(s): /OP2/  

### MG:14:3212

|  | Res. type | Atom | Residue | Distance | Occupancy | B-factor | Closest solute to 1^st^ shell water |
| --- | --- | --- | --- | --- | --- | --- | --- |
| **Ion** | | | |  | 1.00 | 44.62 || 1 | A | OP2 | 14:1890 | 2.39 | 1.00 | 53.30 |
| 2 | G | O6 | 14:1891 | 3.20 | 1.00 | 62.00 |  |
| 3 | G | N7 | 14:1891 | 3.61 | 1.00 | 53.29 |  |
| 4 | A | N7 | 14:1890 | 3.84 | 1.00 | 54.36 |  |
| 5 | A | O5' | 14:1890 | 4.21 | 1.00 | 48.27 |  |
| 6 | C | N4 | 14:1892 | 4.43 | 1.00 | 50.56 |  |
| 7 | A | O3' | 14:1889 | 4.59 | 1.00 | 48.83 |  |
| 8 | U | O4 | 14:1851 | 4.63 | 1.00 | 69.57 |  |
| 9 | G | O6 | 14:1850 | 4.79 | 1.00 | 68.06 |  |
| 10 | C | N3 | 14:1852 | 4.83 | 1.00 | 72.31 |  |
| 11 | C | O2 | 14:1852 | 4.88 | 1.00 | 62.77 |  |
| 12 | A | OP1 | 14:1890 | 4.89 | 1.00 | 45.96 |  |
  |

  
 Contact(s): /OP2/  
 Coded contact list: O_ph_  
 Exclusion zone contact(s): /O6/  

### MG:14:3213

|  | Res. type | Atom | Residue | Distance | Occupancy | B-factor | Closest solute to 1^st^ shell water |
| --- | --- | --- | --- | --- | --- | --- | --- |
| **Ion** | | | |  | 1.00 | 40.32 || 1 | U | OP1 | 14:380 | 2.94 | 1.00 | 45.17 |
| 2 | G | O3' | 14:379 | 3.72 | 1.00 | 45.89 |  |
| 3 | ILE | N | F5:18 | 3.98 | 1.00 | 45.53 |  |
| 4 | SER | OG | F5:17 | 4.09 | 1.00 | 36.79 |  |
| 5 | ILE | O | F5:18 | 4.10 | 1.00 | 41.89 |  |
| 6 | U | O5' | 14:380 | 4.62 | 1.00 | 50.20 |  |
| 7 | U | OP2 | 14:2233 | 4.64 | 1.00 | 47.57 |  |
| 8 | G | O2' | 14:379 | 4.71 | 1.00 | 44.40 |  |
| 9 | GLN | NE2 | F5:19 | 4.83 | 1.00 | 45.96 |  |
| 10 | ASN | O | F5:16 | 4.92 | 1.00 | 44.00 |  |
| 11 | U | OP1 | 14:2233 | 4.92 | 1.00 | 51.90 |  |
  |

  
 Coded contact list:   
 Exclusion zone contact(s): /OP1/  
Consider K in place of MG (1 valid contact(s) for K vs. 0 for MG)  

### MG:14:3214

|  | Res. type | Atom | Residue | Distance | Occupancy | B-factor | Closest solute to 1^st^ shell water |
| --- | --- | --- | --- | --- | --- | --- | --- |
| **Ion** | | | |  | 1.00 | 66.69 || 1 | U | OP2 | 14:847 | 2.29 | 1.00 | 66.11 |
| 2 | G | O6 | 14:929 | 2.39 | 1.00 | 76.32 |  |
| 3 | G | N1 | 14:929 | 3.35 | 1.00 | 77.08 |  |
| 4 | U | O5' | 14:847 | 3.56 | 1.00 | 74.60 |  |
| 5 | G | N7 | 14:928 | 3.81 | 1.00 | 82.72 |  |
| 6 | G | O6 | 14:928 | 3.90 | 1.00 | 76.97 |  |
| 7 | C | O3' | 14:846 | 4.51 | 1.00 | 68.82 |  |
| 8 | G | N7 | 14:929 | 4.62 | 1.00 | 88.42 |  |
| 9 | A | N7 | 14:926 | 4.65 | 1.00 | 82.35 |  |
| 10 | U | OP1 | 14:847 | 4.66 | 1.00 | 64.42 |  |
| 11 | G | OP2 | 14:928 | 4.95 | 1.00 | 85.19 |  |
  |

  
**1^st^ shell ligand angles:**   
1-2: 68;   
  
 Contact(s): /O6/OP2/  
 Coded contact list: O_ph_O_b_  
 Exclusion zone contact(s): /N1_G_/  

### MG:14:3215

|  | Res. type | Atom | Residue | Distance | Occupancy | B-factor | Closest solute to 1^st^ shell water |
| --- | --- | --- | --- | --- | --- | --- | --- |
| **Ion** | | | |  | 1.00 | 45.49 || 1 | G | O6 | 14:2495 | 2.57 | 1.00 | 57.43 |
| 2 | G | O6 | 14:2455 | 3.47 | 1.00 | 48.13 |  |
| 3 | Water 1 | | 14:4043 | 3.59 | 1.00 | 49.82 | 3.67 (G:O6 14:2455) |
| 4 | G | O6 | 14:2494 | 3.69 | 1.00 | 59.39 |  |
| 5 | C | N4 | 14:2456 | 3.91 | 1.00 | 49.55 |  |
| 6 | G | N7 | 14:2494 | 4.11 | 1.00 | 65.04 |  |
| 7 | U | O4 | 14:2457 | 4.74 | 1.00 | 57.36 |  |
| 8 | G | N1 | 14:2495 | 4.77 | 1.00 | 53.06 |  |
| 9 | G | N7 | 14:2495 | 4.83 | 1.00 | 58.68 |  |
| 10 | C | N4 | 14:2496 | 4.86 | 1.00 | 48.38 |  |
  |

  
 Coded contact list:   
 Exclusion zone contact(s): /O6/  
Consider NA in place of MG (1 valid contact(s) for NA vs. 0 for MG)  

### MG:14:3216

|  | Res. type | Atom | Residue | Distance | Occupancy | B-factor | Closest solute to 1^st^ shell water |
| --- | --- | --- | --- | --- | --- | --- | --- |
| **Ion** | | | |  | 1.00 | 51.13 || 1 | A | OP1 | 14:981 | 2.17 | 1.00 | 53.26 |
| 2 | G | N7 | 14:979 | 2.75 | 1.00 | 68.69 |  |
| 3 | G | O6 | 14:979 | 2.85 | 1.00 | 72.10 |  |
| 4 | A | OP2 | 14:981 | 4.00 | 1.00 | 61.82 |  |
| 5 | G | OP2 | 14:978 | 4.00 | 1.00 | 76.77 |  |
| 6 | G | N7 | 14:978 | 4.26 | 1.00 | 74.48 |  |
| 7 | C | OP2 | 14:982 | 4.34 | 1.00 | 57.53 |  |
| 8 | A | O5' | 14:981 | 4.36 | 1.00 | 60.82 |  |
| 9 | G | OP2 | 14:979 | 4.53 | 1.00 | 63.45 |  |
| 10 | A | O3' | 14:980 | 4.63 | 1.00 | 53.59 |  |
| 11 | G | N1 | 14:979 | 4.71 | 1.00 | 70.10 |  |
| 12 | G | N9 | 14:979 | 4.90 | 1.00 | 66.83 |  |
  |

  
 Contact(s): /OP1/  
 Coded contact list: O_ph_  
 Exclusion zone contact(s): /N7/O6/  
Consider K in place of MG (2 valid contact(s) for K vs. 1 for MG)  

### MG:14:3217

|  | Res. type | Atom | Residue | Distance | Occupancy | B-factor | Closest solute to 1^st^ shell water |
| --- | --- | --- | --- | --- | --- | --- | --- |
| **Ion** | | | |  | 1.00 | 69.37 || 1 | C | OP2 | 14:1153 | 2.11 | 1.00 | 88.79 |
| 2 | U | OP2 | 14:999 | 2.26 | 1.00 | 85.82 |  |
| 3 | C | OP1 | 14:998 | 3.44 | 1.00 | 89.28 |  |
| 4 | C | OP1 | 14:1153 | 3.61 | 1.00 | 83.75 |  |
| 5 | C | O5' | 14:998 | 3.62 | 1.00 | 84.57 |  |
| 6 | U | OP1 | 14:999 | 3.66 | 1.00 | 72.64 |  |
| 7 | C | O3' | 14:998 | 3.75 | 1.00 | 71.02 |  |
| 8 | C | O3' | 14:1152 | 4.16 | 1.00 | 87.81 |  |
| 9 | C | OP2 | 14:998 | 4.44 | 1.00 | 83.56 |  |
| 10 | C | O5' | 14:1153 | 4.47 | 1.00 | 84.70 |  |
| 11 | U | O5' | 14:999 | 4.72 | 1.00 | 71.19 |  |
  |

  
**1^st^ shell ligand angles:**   
1-2: 120;   
  
 Contact(s): /OP2/OP2/  
 Coded contact list: 2O_ph_  

### MG:14:3218

|  | Res. type | Atom | Residue | Distance | Occupancy | B-factor | Closest solute to 1^st^ shell water |
| --- | --- | --- | --- | --- | --- | --- | --- |
| **Ion** | | | |  | 1.00 | 60.28 || 1 | A | OP2 | 14:1010 | 2.31 | 1.00 | 79.61 |
| 2 | A | OP2 | 14:1009 | 2.43 | 1.00 | 77.05 |  |
| 3 | LYS | NZ | 15:37 | 2.68 | 1.00 | 86.87 |  |
| 4 | C | OP2 | 14:1006 | 3.11 | 1.00 | 82.31 |  |
| 5 | C | OP2 | 14:1007 | 3.56 | 1.00 | 75.26 |  |
| 6 | C | OP1 | 14:1006 | 3.65 | 1.00 | 77.34 |  |
| 7 | C | O3' | 14:1008 | 4.08 | 1.00 | 74.51 |  |
| 8 | A | OP1 | 14:1010 | 4.11 | 1.00 | 80.76 |  |
| 9 | A | N7 | 14:1010 | 4.25 | 1.00 | 87.06 |  |
| 10 | A | OP1 | 14:1009 | 4.46 | 1.00 | 77.97 |  |
| 11 | C | O5' | 14:1006 | 4.52 | 1.00 | 83.37 |  |
| 12 | A | O5' | 14:1010 | 4.61 | 1.00 | 83.11 |  |
| 13 | A | O3' | 14:1009 | 4.70 | 1.00 | 81.21 |  |
| 14 | C | O2' | 14:1008 | 4.77 | 1.00 | 79.63 |  |
| 15 | A | O5' | 14:1009 | 4.91 | 1.00 | 67.70 |  |
  |

  
 Contact(s): /OP2/  
 Coded contact list: O_ph_  
 Exclusion zone contact(s): /NZ_Lys_/OP2/OP2/  
Consider K in place of MG (2 valid contact(s) for K vs. 1 for MG)  

### MG:14:3219

|  | Res. type | Atom | Residue | Distance | Occupancy | B-factor | Closest solute to 1^st^ shell water |
| --- | --- | --- | --- | --- | --- | --- | --- |
| **Ion** | | | |  | 1.00 | 33.32 || 1 | G | O2' | 14:1667 | 3.92 | 1.00 | 37.62 |
| 2 | C | OP2 | 14:1990 | 4.10 | 1.00 | 49.15 |  |
| 3 | G | OP2 | 14:1989 | 4.34 | 1.00 | 43.74 |  |
| 4 | G | OP2 | 14:1667 | 4.69 | 1.00 | 42.47 |  |
| 5 | G | O5' | 14:1667 | 4.88 | 1.00 | 45.74 |  |
  |

  
 Coded contact list:   
1^st^ MG coordination shell missing (see Notes)  

### MG:14:3220

|  | Res. type | Atom | Residue | Distance | Occupancy | B-factor | Closest solute to 1^st^ shell water |
| --- | --- | --- | --- | --- | --- | --- | --- |
| **Ion** | | | |  | 1.00 | 46.61 || 1 | G | OP2 | 14:15 | 2.40 | 1.00 | 57.44 |
| 2 | A | OP2 | 14:14 | 2.72 | 1.00 | 57.69 |  |
| 3 | A | O5' | 14:14 | 3.65 | 1.00 | 49.34 |  |
| 4 | A | OP1 | 14:14 | 4.32 | 1.00 | 64.44 |  |
| 5 | G | OP1 | 14:15 | 4.38 | 1.00 | 56.80 |  |
| 6 | A | O3' | 14:14 | 4.69 | 1.00 | 51.30 |  |
| 7 | G | O5' | 14:15 | 4.76 | 1.00 | 57.04 |  |
  |

  
 Coded contact list:   
 Exclusion zone contact(s): /OP2/OP2/  
Consider NA in place of MG (1 valid contact(s) for NA vs. 0 for MG)  

### MG:14:3221

|  | Res. type | Atom | Residue | Distance | Occupancy | B-factor | Closest solute to 1^st^ shell water |
| --- | --- | --- | --- | --- | --- | --- | --- |
| **Ion** | | | |  | 1.00 | 52.42 || 1 | G | OP2 | 14:2000 | 3.01 | 1.00 | 47.08 |
| 2 | Water 1 | | 14:4068 | 3.59 | 1.00 | 55.83 | 3.98 (A:OP2 14:2001) |
| 3 | Water 2 | | 14:4033 | 3.68 | 1.00 | 42.90 | 3.57 (G:OP2 14:2000) |
| 4 | A | OP2 | 14:2001 | 3.83 | 1.00 | 46.13 |  |
| 5 | G | O5' | 14:2000 | 4.11 | 1.00 | 46.84 |  |
| 6 | Water 3 | | 14:4085 | 4.22 | 1.00 | 74.05 | 4.24 (A:OP1 14:1655) |
| 7 | G | OP1 | 14:2000 | 4.67 | 1.00 | 53.28 |  |
| 8 | A | OP1 | 14:1655 | 4.94 | 1.00 | 37.02 |  |
  |

  
 Coded contact list:   
 Exclusion zone contact(s): /OP2/  
Consider K in place of MG (1 valid contact(s) for K vs. 0 for MG)  

### MG:14:3222

|  | Res. type | Atom | Residue | Distance | Occupancy | B-factor | Closest solute to 1^st^ shell water |
| --- | --- | --- | --- | --- | --- | --- | --- |
| **Ion** | | | |  | 1.00 | 57.98 || 1 | A | OP2 | 14:1156 | 2.27 | 1.00 | 74.87 |
| 2 | G | O6 | 14:1157 | 3.58 | 1.00 | 81.71 |  |
| 3 | C | N4 | 14:998 | 4.04 | 1.00 | 83.55 |  |
| 4 | C | N4 | 14:1158 | 4.30 | 1.00 | 84.73 |  |
| 5 | A | O5' | 14:1156 | 4.52 | 1.00 | 61.06 |  |
| 6 | A | O3' | 14:1155 | 4.56 | 1.00 | 61.75 |  |
| 7 | G | O6 | 14:997 | 4.59 | 1.00 | 82.23 |  |
| 8 | A | OP2 | 14:1155 | 4.63 | 1.00 | 86.66 |  |
| 9 | A | OP1 | 14:1156 | 4.67 | 1.00 | 65.11 |  |
| 10 | A | O5' | 14:1155 | 4.75 | 1.00 | 75.85 |  |
  |

  
 Contact(s): /OP2/  
 Coded contact list: O_ph_  

### MG:14:3223

|  | Res. type | Atom | Residue | Distance | Occupancy | B-factor | Closest solute to 1^st^ shell water |
| --- | --- | --- | --- | --- | --- | --- | --- |
| **Ion** | | | |  | 1.00 | 40.82 || 1 | G | OP1 | 14:248 | 2.35 | 1.00 | 38.22 |
| 2 | U | OP1 | 14:387 | 3.97 | 1.00 | 45.75 |  |
| 3 | G | O2' | 14:386 | 4.01 | 1.00 | 44.82 |  |
| 4 | G | O3' | 14:386 | 4.14 | 1.00 | 40.01 |  |
| 5 | G | O3' | 14:247 | 4.32 | 1.00 | 37.47 |  |
| 6 | LYS | O | F5:32 | 4.35 | 1.00 | 49.27 |  |
| 7 | G | OP2 | 14:248 | 4.39 | 1.00 | 39.23 |  |
| 8 | G | N3 | 14:386 | 4.69 | 1.00 | 37.74 |  |
| 9 | G | O5' | 14:248 | 4.84 | 1.00 | 30.73 |  |
  |

  
 Contact(s): /OP1/  
 Coded contact list: O_ph_  

### MG:14:3224

|  | Res. type | Atom | Residue | Distance | Occupancy | B-factor | Closest solute to 1^st^ shell water |
| --- | --- | --- | --- | --- | --- | --- | --- |
| **Ion** | | | |  | 1.00 | 36.01 || 1 | G | OP1 | 1G:1475 | 4.06 | 1.00 | 62.69 |
| 2 | C | O5' | 14:1767 | 4.75 | 1.00 | 59.44 |  |
| 3 | C | O3' | 14:1767 | 4.78 | 1.00 | 48.58 |  |
  |

  
 Coded contact list:   
1^st^ MG coordination shell missing (see Notes)  

### MG:14:3225

|  | Res. type | Atom | Residue | Distance | Occupancy | B-factor | Closest solute to 1^st^ shell water |
| --- | --- | --- | --- | --- | --- | --- | --- |
| **Ion** | | | |  | 1.00 | 47.68 || 1 | C | OP1 | 14:2442 | 3.28 | 1.00 | 43.34 |
| 2 | C | OP1 | 14:2441 | 3.93 | 1.00 | 43.06 |  |
| 3 | C | O3' | 14:2441 | 3.96 | 1.00 | 39.24 |  |
| 4 | C | OP2 | 14:2441 | 4.44 | 1.00 | 33.29 |  |
| 5 | C | OP2 | 14:2442 | 4.62 | 1.00 | 46.43 |  |
| 6 | A | O4' | 14:2062 | 4.75 | 1.00 | 55.56 |  |
  |

  
 Coded contact list:   
 Exclusion zone contact(s): /OP1/  
Consider K in place of MG (1 valid contact(s) for K vs. 0 for MG)  

### MG:14:3226

|  | Res. type | Atom | Residue | Distance | Occupancy | B-factor | Closest solute to 1^st^ shell water |
| --- | --- | --- | --- | --- | --- | --- | --- |
| **Ion** | | | |  | 1.00 | 55.65 || 1 | U | OP1 | 14:72 | 2.42 | 1.00 | 48.83 |
| 2 | G | OP1 | 14:113 | 3.42 | 1.00 | 56.87 |  |
| 3 | A | OP2 | 14:71 | 3.56 | 1.00 | 51.19 |  |
| 4 | U | O3' | 14:112 | 3.71 | 1.00 | 55.84 |  |
| 5 | U | O2' | 14:112 | 4.19 | 1.00 | 40.81 |  |
| 6 | A | O2' | 14:71 | 4.49 | 1.00 | 60.33 |  |
| 7 | A | O3' | 14:71 | 4.59 | 1.00 | 48.60 |  |
| 8 | U | O5' | 14:72 | 4.72 | 1.00 | 47.50 |  |
| 9 | G | O5' | 14:113 | 4.80 | 1.00 | 46.28 |  |
| 10 | MET | N | B5:1 | 4.87 | 1.00 | 59.50 |  |
| 11 | U | OP2 | 14:72 | 4.89 | 1.00 | 54.49 |  |
  |

  
 Coded contact list:   
 Exclusion zone contact(s): /OP1/  
Consider NA in place of MG (1 valid contact(s) for NA vs. 0 for MG)  

### MG:14:3227

|  | Res. type | Atom | Residue | Distance | Occupancy | B-factor | Closest solute to 1^st^ shell water |
| --- | --- | --- | --- | --- | --- | --- | --- |
| **Ion** | | | |  | 1.00 | 38.23 || 1 | U | O4 | 14:2079 | 2.96 | 1.00 | 32.81 |
| 2 | G | O6 | 14:2080 | 3.47 | 1.00 | 42.61 |  |
| 3 | G | O6 | 14:2238 | 4.32 | 1.00 | 38.26 |  |
| 4 | G | N7 | 14:2080 | 4.41 | 1.00 | 38.89 |  |
| 5 | G | OP2 | 14:2238 | 4.57 | 1.00 | 45.55 |  |
| 6 | G | N1 | 14:2238 | 4.59 | 1.00 | 37.43 |  |
| 7 | G | N7 | 14:2238 | 4.77 | 1.00 | 34.07 |  |
| 8 | C | N4 | 14:2078 | 4.89 | 1.00 | 30.93 |  |
| 9 | A | N6 | 14:2241 | 4.89 | 1.00 | 33.45 |  |
| 10 | C | N4 | 14:2240 | 4.98 | 1.00 | 35.02 |  |
| 11 | U | N3 | 14:2079 | 4.99 | 1.00 | 34.78 |  |
  |

  
 Coded contact list:   
 Exclusion zone contact(s): /O4/  
Consider K in place of MG (1 valid contact(s) for K vs. 0 for MG)  

### MG:14:3228

|  | Res. type | Atom | Residue | Distance | Occupancy | B-factor | Closest solute to 1^st^ shell water |
| --- | --- | --- | --- | --- | --- | --- | --- |
| **Ion** | | | |  | 1.00 | 58.27 || 1 | G | OP1 | 14:848 | 2.52 | 1.00 | 78.50 |
| 2 | A | OP2 | 14:849 | 4.29 | 1.00 | 73.00 |  |
| 3 | G | O5' | 14:848 | 4.29 | 1.00 | 79.18 |  |
| 4 | A | OP1 | 14:849 | 4.37 | 1.00 | 79.36 |  |
| 5 | U | O3' | 14:847 | 4.45 | 1.00 | 65.90 |  |
  |

  
 Coded contact list:   
 Exclusion zone contact(s): /OP1/  
Consider NA in place of MG (1 valid contact(s) for NA vs. 0 for MG)  

### MG:14:3229

|  | Res. type | Atom | Residue | Distance | Occupancy | B-factor | Closest solute to 1^st^ shell water |
| --- | --- | --- | --- | --- | --- | --- | --- |
| **Ion** | | | |  | 1.00 | 48.75 || 1 | A | OP1 | 14:2042 | 2.10 | 1.00 | 56.81 |
| 2 | A | OP2 | 14:2778 | 3.77 | 1.00 | 65.80 |  |
| 3 | U | O3' | 14:2041 | 4.25 | 1.00 | 53.79 |  |
| 4 | A | O5' | 14:2042 | 4.36 | 1.00 | 52.83 |  |
| 5 | A | OP2 | 14:2042 | 4.57 | 1.00 | 51.53 |  |
  |

  
 Contact(s): /OP1/  
 Coded contact list: O_ph_  

### MG:14:3230

|  | Res. type | Atom | Residue | Distance | Occupancy | B-factor | Closest solute to 1^st^ shell water |
| --- | --- | --- | --- | --- | --- | --- | --- |
| **Ion** | | | |  | 1.00 | 54.08 || 1 | U | OP1 | 14:2016 | 2.81 | 1.00 | 44.09 |
| 2 | A | OP1 | 14:2058 | 3.32 | 1.00 | 33.68 |  |
| 3 | Water 1 | | 14:3811 | 3.52 | 1.00 | 39.74 | 2.83 (U:OP2 14:2017) |
| 4 | A | OP2 | 14:2059 | 3.69 | 1.00 | 33.77 |  |
| 5 | U | O5' | 14:2016 | 3.74 | 1.00 | 37.42 |  |
| 6 | U | OP2 | 14:2017 | 3.92 | 1.00 | 42.12 |  |
| 7 | U | O2' | 14:1255 | 4.69 | 1.00 | 45.40 |  |
| 8 | A | O5' | 14:2058 | 4.72 | 1.00 | 38.87 |  |
| 9 | U | OP2 | 14:2016 | 4.79 | 1.00 | 42.98 |  |
| 10 | A | OP1 | 14:2059 | 4.89 | 1.00 | 40.80 |  |
| 11 | A | O3' | 14:2015 | 4.98 | 1.00 | 35.27 |  |
  |

  
 Coded contact list:   
 Exclusion zone contact(s): /OP1/OP1/  
Consider K in place of MG (2 valid contact(s) for K vs. 0 for MG)  

### MG:14:3231

|  | Res. type | Atom | Residue | Distance | Occupancy | B-factor | Closest solute to 1^st^ shell water |
| --- | --- | --- | --- | --- | --- | --- | --- |
| **Ion** | | | |  | 1.00 | 39.22 || 1 | U | O4 | 14:2431 | 2.70 | 1.00 | 43.15 |
| 2 | A | OP2 | 14:2434 | 3.23 | 1.00 | 39.50 |  |
| 3 | A | O5' | 14:2434 | 3.83 | 1.00 | 38.86 |  |
| 4 | U | O4 | 14:827 | 4.36 | 1.00 | 44.53 |  |
| 5 | A | OP1 | 14:2434 | 4.40 | 1.00 | 45.24 |  |
| 6 | U | N3 | 14:2431 | 4.42 | 1.00 | 35.57 |  |
| 7 | A | OP2 | 14:2435 | 4.60 | 1.00 | 43.82 |  |
| 8 | A | N7 | 14:2434 | 4.92 | 1.00 | 40.92 |  |
  |

  
 Coded contact list:   
 Exclusion zone contact(s): /O4/OP2/  
Consider K in place of MG (2 valid contact(s) for K vs. 0 for MG)  

### MG:14:3232

|  | Res. type | Atom | Residue | Distance | Occupancy | B-factor | Closest solute to 1^st^ shell water |
| --- | --- | --- | --- | --- | --- | --- | --- |
| **Ion** | | | |  | 1.00 | 63.94 || 1 | C | N3 | 14:435 | 3.23 | 1.00 | 62.29 |
| 2 | C | O2 | 14:435 | 3.60 | 1.00 | 62.49 |  |
| 3 | C | N4 | 14:435 | 4.26 | 1.00 | 66.48 |  |
| 4 | G | O2' | 14:45 | 4.86 | 1.00 | 60.91 |  |
  |

  
 Coded contact list:   
 Exclusion zone contact(s): /N3_C_/  
Consider K in place of MG (1 valid contact(s) for K vs. 0 for MG)  

### MG:14:3233

|  | Res. type | Atom | Residue | Distance | Occupancy | B-factor | Closest solute to 1^st^ shell water |
| --- | --- | --- | --- | --- | --- | --- | --- |
| **Ion** | | | |  | 1.00 | 52.52 || 1 | G | N7 | 14:1229A | 3.84 | 1.00 | 66.46 |
| 2 | G | O6 | 14:1229 | 4.27 | 1.00 | 64.96 |  |
| 3 | G | N7 | 14:1229 | 4.35 | 1.00 | 61.76 |  |
| 4 | G | O6 | 14:1229A | 4.37 | 1.00 | 69.44 |  |
| 5 | G | O6 | 14:1219 | 4.47 | 1.00 | 68.33 |  |
| 6 | C | N4 | 14:1230 | 4.58 | 1.00 | 70.35 |  |
  |

  
 Coded contact list:   
1^st^ MG coordination shell missing (see Notes)  

### MG:14:3234

|  | Res. type | Atom | Residue | Distance | Occupancy | B-factor | Closest solute to 1^st^ shell water |
| --- | --- | --- | --- | --- | --- | --- | --- |
| **Ion** | | | |  | 1.00 | 55.89 || 1 | G | OP2 | 14:217 | 2.53 | 1.00 | 44.23 |
| 2 | A | N7 | 14:218 | 3.46 | 1.00 | 50.16 |  |
| 3 | G | N7 | 14:217 | 3.53 | 1.00 | 48.50 |  |
| 4 | G | O6 | 14:219 | 3.85 | 1.00 | 58.15 |  |
| 5 | A | N6 | 14:218 | 3.88 | 1.00 | 57.60 |  |
| 6 | G | O5' | 14:217 | 4.04 | 1.00 | 52.11 |  |
| 7 | A | OP1 | 14:428 | 4.39 | 1.00 | 56.35 |  |
| 8 | A | OP2 | 14:218 | 4.50 | 1.00 | 45.53 |  |
| 9 | G | OP1 | 14:217 | 4.83 | 1.00 | 62.56 |  |
| 10 | A | O3' | 14:216 | 4.90 | 1.00 | 46.74 |  |
  |

  
 Coded contact list:   
 Exclusion zone contact(s): /OP2/  
Consider NA in place of MG (1 valid contact(s) for NA vs. 0 for MG)  

### MG:14:3235

|  | Res. type | Atom | Residue | Distance | Occupancy | B-factor | Closest solute to 1^st^ shell water |
| --- | --- | --- | --- | --- | --- | --- | --- |
| **Ion** | | | |  | 1.00 | 50.44 || 1 | U | O2 | 14:2357 | 2.99 | 1.00 | 56.22 |
| 2 | A | N6 | 14:2361 | 3.15 | 1.00 | 50.83 |  |
| 3 | A | N7 | 14:2361 | 4.06 | 1.00 | 47.30 |  |
| 4 | A | OP2 | 14:2360 | 4.17 | 1.00 | 52.39 |  |
| 5 | U | N3 | 14:2357 | 4.44 | 1.00 | 52.56 |  |
| 6 | C | OP1 | 14:2359 | 4.57 | 1.00 | 40.49 |  |
| 7 | A | N7 | 14:2360 | 4.83 | 1.00 | 44.91 |  |
| 8 | C | OP2 | 14:2359 | 4.88 | 1.00 | 45.37 |  |
  |

  
 Coded contact list:   
 Exclusion zone contact(s): /N6/O2/  
Consider K in place of MG (2 valid contact(s) for K vs. 0 for MG)  

### MG:14:3236

|  | Res. type | Atom | Residue | Distance | Occupancy | B-factor | Closest solute to 1^st^ shell water |
| --- | --- | --- | --- | --- | --- | --- | --- |
| **Ion** | | | |  | 1.00 | 50.95 || 1 | U | O2' | 14:2849 | 3.30 | 1.00 | 59.39 |
| 2 | U | O4 | 14:2865 | 3.38 | 1.00 | 75.11 |  |
| 3 | A | OP2 | 14:2850 | 3.60 | 1.00 | 70.57 |  |
| 4 | A | OP1 | 14:2850 | 3.84 | 1.00 | 71.94 |  |
| 5 | G | N7 | 14:2864 | 4.30 | 1.00 | 81.28 |  |
| 6 | G | OP2 | 14:2864 | 4.65 | 1.00 | 78.54 |  |
| 7 | U | N3 | 14:2866 | 4.73 | 1.00 | 72.67 |  |
| 8 | U | O3' | 14:2849 | 4.82 | 1.00 | 55.90 |  |
| 9 | G | O6 | 14:2864 | 4.82 | 1.00 | 74.65 |  |
| 10 | U | O2 | 14:2866 | 4.84 | 1.00 | 75.20 |  |
  |

  
 Coded contact list:   
 Exclusion zone contact(s): /O2'/O4/  
Consider K in place of MG (2 valid contact(s) for K vs. 0 for MG)  

### MG:14:3237

|  | Res. type | Atom | Residue | Distance | Occupancy | B-factor | Closest solute to 1^st^ shell water |
| --- | --- | --- | --- | --- | --- | --- | --- |
| **Ion** | | | |  | 1.00 | 96.46 || 1 | C | O2 | 14:2306 | 2.32 | 1.00 | 94.57 |
| 2 | GLU | OE1 | 49:45 | 2.39 | 1.00 | 108.47 |  |
| 3 | GLU | OE2 | 49:45 | 2.43 | 1.00 | 97.74 |  |
| 4 | C | N3 | 14:2306 | 2.81 | 1.00 | 89.79 |  |
| 5 | A | N6 | 14:2305 | 3.38 | 1.00 | 91.83 |  |
| 6 | A | N1 | 14:2305 | 3.69 | 1.00 | 99.14 |  |
| 7 | G | N1 | 14:2307 | 3.72 | 1.00 | 93.54 |  |
| 8 | GLY | O | 49:42 | 3.97 | 1.00 | 89.51 |  |
| 9 | G | N2 | 14:2307 | 4.00 | 1.00 | 93.24 |  |
| 10 | C | N1 | 14:2306 | 4.29 | 1.00 | 96.44 |  |
| 11 | C | N4 | 14:2306 | 4.60 | 1.00 | 90.11 |  |
| 12 | G | O6 | 14:2307 | 4.65 | 1.00 | 92.29 |  |
| 13 | GLY | N | 49:44 | 4.88 | 1.00 | 101.73 |  |
  |

  
**1^st^ shell ligand angles:**   
1-2: 116;   
  
 Contact(s): /O2/OE1_Glu_/  
 Coded contact list: O_b_O_coo_  
 Exclusion zone contact(s): /N3_C_/N6/OE2_Glu_/  

### MG:14:3238

|  | Res. type | Atom | Residue | Distance | Occupancy | B-factor | Closest solute to 1^st^ shell water |
| --- | --- | --- | --- | --- | --- | --- | --- |
| **Ion** | | | |  | 1.00 | 44.43 || 1 | A | N3 | 14:2426 | 3.90 | 1.00 | 43.54 |
| 2 | A | OP1 | 14:2426 | 4.11 | 1.00 | 45.78 |  |
| 3 | A | OP2 | 14:2425 | 4.12 | 1.00 | 40.99 |  |
| 4 | A | O2' | 14:2426 | 4.24 | 1.00 | 43.93 |  |
| 5 | A | N1 | 14:2426 | 4.28 | 1.00 | 41.96 |  |
| 6 | A | N9 | 14:2426 | 4.29 | 1.00 | 38.31 |  |
| 7 | A | N7 | 14:2426 | 4.64 | 1.00 | 43.66 |  |
| 8 | C | O3' | 14:2424 | 4.71 | 1.00 | 39.47 |  |
| 9 | C | OP2 | 14:2424 | 4.77 | 1.00 | 43.61 |  |
  |

  
 Coded contact list:   
1^st^ MG coordination shell missing (see Notes)  

### MG:14:3239

|  | Res. type | Atom | Residue | Distance | Occupancy | B-factor | Closest solute to 1^st^ shell water |
| --- | --- | --- | --- | --- | --- | --- | --- |
| **Ion** | | | |  | 1.00 | 66.98 || 1 | G | OP2 | 14:1630 | 4.02 | 1.00 | 57.47 |
| 2 | G | OP2 | 14:1633 | 4.12 | 1.00 | 43.39 |  |
| 3 | G | N7 | 14:1630 | 4.30 | 1.00 | 52.46 |  |
| 4 | C | N4 | 14:1630A | 4.57 | 1.00 | 49.54 |  |
| 5 | A | OP1 | 14:1632 | 4.91 | 1.00 | 56.69 |  |
  |

  
 Coded contact list:   
1^st^ MG coordination shell missing (see Notes)  

### MG:14:3240

|  | Res. type | Atom | Residue | Distance | Occupancy | B-factor | Closest solute to 1^st^ shell water |
| --- | --- | --- | --- | --- | --- | --- | --- |
| **Ion** | | | |  | 1.00 | 57.04 || 1 | A | OP2 | 14:1701 | 3.49 | 1.00 | 63.59 |
| 2 | G | N7 | 14:1702 | 3.81 | 1.00 | 48.60 |  |
| 3 | G | O6 | 14:1763 | 3.95 | 1.00 | 57.76 |  |
| 4 | G | N2 | 14:1699 | 4.23 | 1.00 | 64.22 |  |
| 5 | G | N7 | 14:1703 | 4.42 | 1.00 | 50.94 |  |
| 6 | G | O6 | 14:1702 | 4.72 | 1.00 | 54.29 |  |
| 7 | G | OP2 | 14:1702 | 4.97 | 1.00 | 52.79 |  |
  |

  
 Coded contact list:   
1^st^ MG coordination shell missing (see Notes)  

### MG:14:3241

|  | Res. type | Atom | Residue | Distance | Occupancy | B-factor | Closest solute to 1^st^ shell water |
| --- | --- | --- | --- | --- | --- | --- | --- |
| **Ion** | | | |  | 1.00 | 59.92 || 1 | G | O6 | 14:406 | 2.78 | 1.00 | 64.62 |
| 2 | G | O6 | 14:407 | 3.55 | 1.00 | 56.59 |  |
| 3 | U | O4 | 14:421 | 3.93 | 1.00 | 66.51 |  |
| 4 | G | N7 | 14:406 | 4.01 | 1.00 | 71.59 |  |
| 5 | C | O2 | 14:404 | 4.03 | 1.00 | 67.78 |  |
| 6 | C | N4 | 14:420 | 4.14 | 1.00 | 57.97 |  |
| 7 | G | N2 | 14:271B | 4.26 | 1.00 | 73.92 |  |
| 8 | U | N3 | 14:421 | 4.84 | 1.00 | 63.11 |  |
| 9 | G | N7 | 14:407 | 4.85 | 1.00 | 64.13 |  |
  |

  
 Coded contact list:   
 Exclusion zone contact(s): /O6/  
Consider K in place of MG (1 valid contact(s) for K vs. 0 for MG)  

### MG:14:3242

|  | Res. type | Atom | Residue | Distance | Occupancy | B-factor | Closest solute to 1^st^ shell water |
| --- | --- | --- | --- | --- | --- | --- | --- |
| **Ion** | | | |  | 1.00 | 65.51 || 1 | G | OP1 | 14:131 | 2.99 | 1.00 | 62.96 |
| 2 | G | OP2 | 14:131 | 4.02 | 1.00 | 57.18 |  |
| 3 | C | OP1 | 14:141A | 4.38 | 1.00 | 66.36 |  |
| 4 | A | N7 | 14:1349 | 4.42 | 1.00 | 63.03 |  |
| 5 | A | O2' | 14:1349 | 4.52 | 1.00 | 59.76 |  |
| 6 | A | N9 | 14:1349 | 4.52 | 1.00 | 68.20 |  |
| 7 | G | OP1 | 14:142 | 4.93 | 1.00 | 59.44 |  |
| 8 | C | O3' | 14:130 | 4.98 | 1.00 | 53.14 |  |
  |

  
 Coded contact list:   
 Exclusion zone contact(s): /OP1/  
Consider K in place of MG (1 valid contact(s) for K vs. 0 for MG)  

### MG:14:3243

|  | Res. type | Atom | Residue | Distance | Occupancy | B-factor | Closest solute to 1^st^ shell water |
| --- | --- | --- | --- | --- | --- | --- | --- |
| **Ion** | | | |  | 1.00 | 54.53 || 1 | Water 1 | | 14:3857 | 2.48 | 1.00 | 56.40 | 2.90 (G:O6 14:1443) |
| 2 | G | O6 | 14:1442 | 3.41 | 1.00 | 68.47 |  |
| 3 | G | O6 | 14:1443 | 3.82 | 1.00 | 67.79 |  |
| 4 | Water 2 | | 14:4021 | 4.09 | 1.00 | 66.15 | 3.69 (G:N7 14:1442) |
| 5 | C | N4 | 14:1549 | 4.22 | 1.00 | 71.21 |  |
| 6 | Water 3 | | 14:4020 | 4.41 | 1.00 | 86.07 | 5.31 (C:OP1 14:1467) |
| 7 | G | N7 | 14:1442 | 4.55 | 1.00 | 63.64 |  |
| 8 | C | N4 | 14:1548 | 4.67 | 1.00 | 74.27 |  |
| 9 | C | OP1 | 14:1467 | 4.79 | 1.00 | 77.36 |  |
| 10 | G | O6 | 14:1441 | 4.95 | 1.00 | 76.85 |  |
  |

  
 Coded contact list:   
 Exclusion zone contact(s): /O_w_/  
1^st^ MG coordination shell missing (see Notes)  

### MG:14:3244

|  | Res. type | Atom | Residue | Distance | Occupancy | B-factor | Closest solute to 1^st^ shell water |
| --- | --- | --- | --- | --- | --- | --- | --- |
| **Ion** | | | |  | 1.00 | 64.51 || 1 | C | OP2 | 14:2692 | 2.69 | 1.00 | 71.59 |
| 2 | C | OP2 | 14:2691 | 3.28 | 1.00 | 77.91 |  |
| 3 | C | O2' | 14:2690 | 3.44 | 1.00 | 63.62 |  |
| 4 | C | O3' | 14:2690 | 3.77 | 1.00 | 55.64 |  |
| 5 | C | OP1 | 14:2692 | 3.88 | 1.00 | 68.35 |  |
| 6 | C | O5' | 14:2691 | 4.17 | 1.00 | 67.41 |  |
| 7 | C | O3' | 14:2691 | 4.75 | 1.00 | 55.83 |  |
| 8 | C | O5' | 14:2692 | 4.99 | 1.00 | 53.76 |  |
  |

  
 Coded contact list:   
 Exclusion zone contact(s): /OP2/OP2/  
Consider K in place of MG (2 valid contact(s) for K vs. 0 for MG)  

### MG:14:3245

|  | Res. type | Atom | Residue | Distance | Occupancy | B-factor | Closest solute to 1^st^ shell water |
| --- | --- | --- | --- | --- | --- | --- | --- |
| **Ion** | | | |  | 1.00 | 36.70 || 1 | U | O4 | 14:2419 | 2.24 | 1.00 | 65.07 |
| 2 | ARG | O | M5:30 | 3.65 | 1.00 | 55.30 |  |
| 3 | U | N3 | 14:2419 | 4.41 | 1.00 | 66.47 |  |
| 4 | C | N4 | 14:2420 | 4.63 | 1.00 | 59.72 |  |
| 5 | LYS | O | M5:29 | 4.66 | 1.00 | 62.74 |  |
| 6 | A | N7 | 14:2418 | 4.77 | 1.00 | 65.75 |  |
  |

  
 Contact(s): /O4/  
 Coded contact list: O_b_  

### MG:14:3246

|  | Res. type | Atom | Residue | Distance | Occupancy | B-factor | Closest solute to 1^st^ shell water |
| --- | --- | --- | --- | --- | --- | --- | --- |
| **Ion** | | | |  | 1.00 | 35.59 || 1 | C | OP1 | 14:737 | 3.30 | 1.00 | 34.56 |
| 2 | U | OP1 | 14:1621 | 3.37 | 1.00 | 43.89 |  |
| 3 | G | O3' | 14:1620 | 4.47 | 1.00 | 31.12 |  |
| 4 | G | OP2 | 14:738 | 4.63 | 1.00 | 28.09 |  |
| 5 | C | O5' | 14:737 | 4.84 | 1.00 | 41.12 |  |
| 6 | G | OP1 | 14:1622 | 4.99 | 1.00 | 47.76 |  |
  |

  
 Coded contact list:   
 Exclusion zone contact(s): /OP1/OP1/  
Consider K in place of MG (2 valid contact(s) for K vs. 0 for MG)  

### MG:14:3247

|  | Res. type | Atom | Residue | Distance | Occupancy | B-factor | Closest solute to 1^st^ shell water |
| --- | --- | --- | --- | --- | --- | --- | --- |
| **Ion** | | | |  | 1.00 | 47.65 || 1 | A | OP2 | 14:2059 | 2.33 | 1.00 | 33.77 |
| 2 | U | O2 | 14:1255 | 2.97 | 1.00 | 41.81 |  |
| 3 | A | O5' | 14:2059 | 3.79 | 1.00 | 31.73 |  |
| 4 | GLY | O | 39:71 | 4.38 | 1.00 | 39.05 |  |
| 5 | A | OP2 | 14:2060 | 4.56 | 1.00 | 31.91 |  |
| 6 | A | OP1 | 14:2059 | 4.65 | 1.00 | 40.80 |  |
| 7 | A | O3' | 14:2058 | 4.68 | 1.00 | 29.77 |  |
| 8 | A | O5' | 14:2060 | 4.70 | 1.00 | 42.61 |  |
| 9 | A | OP1 | 14:2058 | 4.74 | 1.00 | 33.68 |  |
| 10 | A | O5' | 14:2058 | 4.83 | 1.00 | 38.87 |  |
| 11 | U | N3 | 14:1255 | 4.87 | 1.00 | 33.03 |  |
  |

  
 Contact(s): /OP2/  
 Coded contact list: O_ph_  
 Exclusion zone contact(s): /O2/  

### MG:14:3248

|  | Res. type | Atom | Residue | Distance | Occupancy | B-factor | Closest solute to 1^st^ shell water |
| --- | --- | --- | --- | --- | --- | --- | --- |
| **Ion** | | | |  | 1.00 | 60.17 || 1 | G | O6 | 14:424 | 3.40 | 1.00 | 59.56 |
| 2 | G | N7 | 14:424 | 3.46 | 1.00 | 63.30 |  |
| 3 | A | OP2 | 14:423 | 3.85 | 1.00 | 58.28 |  |
| 4 | A | OP1 | 14:422 | 4.18 | 1.00 | 62.84 |  |
| 5 | G | O6 | 14:370 | 4.58 | 1.00 | 59.40 |  |
| 6 | A | OP1 | 14:423 | 4.69 | 1.00 | 69.80 |  |
  |

  
 Coded contact list:   
 Exclusion zone contact(s): /O6/  
Consider K in place of MG (1 valid contact(s) for K vs. 0 for MG)  

### MG:14:3249

|  | Res. type | Atom | Residue | Distance | Occupancy | B-factor | Closest solute to 1^st^ shell water |
| --- | --- | --- | --- | --- | --- | --- | --- |
| **Ion** | | | |  | 1.00 | 62.31 || 1 | Water 1 | | 14:3874 | 2.53 | 1.00 | 55.16 | 2.95 (G:O6 14:301) |
| 2 | G | O6 | 14:301 | 2.81 | 1.00 | 64.79 |  |
| 3 | C | N4 | 14:316 | 3.07 | 1.00 | 71.69 |  |
| 4 | Water 2 | | 14:3835 | 3.10 | 1.00 | 71.90 | 2.83 (C:N4 14:334) |
| 5 | G | O6 | 14:317 | 3.35 | 1.00 | 64.77 |  |
| 6 | Water 3 | | 14:4003 | 4.10 | 1.00 | 55.22 | 5.72 (G:O6 14:301) |
| 7 | G | O6 | 14:315 | 4.37 | 1.00 | 63.41 |  |
| 8 | G | N7 | 14:317 | 4.47 | 1.00 | 64.21 |  |
| 9 | Water 4 | | 14:3750 | 4.76 | 1.00 | 59.99 | 3.93 (G:O6 14:301) |
| 10 | C | N4 | 14:334 | 4.91 | 1.00 | 65.36 |  |
| 11 | G | N1 | 14:301 | 4.92 | 1.00 | 65.42 |  |
  |

  
 Coded contact list:   
 Exclusion zone contact(s): /N4/O6/O6/O_w_/O_w_/  
Consider K in place of MG (3 valid contact(s) for K vs. 0 for MG)  

### MG:14:3250

|  | Res. type | Atom | Residue | Distance | Occupancy | B-factor | Closest solute to 1^st^ shell water |
| --- | --- | --- | --- | --- | --- | --- | --- |
| **Ion** | | | |  | 1.00 | 38.34 || 1 | Water 1 | | 14:3895 | 3.05 | 1.00 | 37.04 | 3.00 (G:O6 14:771) |
| 2 | G | N7 | 14:771 | 3.05 | 1.00 | 36.14 |  |
| 3 | G | O6 | 14:770 | 3.07 | 1.00 | 39.46 |  |
| 4 | Water 2 | | 14:4037 | 3.09 | 1.00 | 35.16 | 3.59 (G:O6 14:771) |
| 5 | Water 3 | | 14:3657 | 3.18 | 1.00 | 32.85 | 2.51 (G:OP2 14:686) |
| 6 | G | O6 | 14:771 | 3.38 | 1.00 | 32.36 |  |
| 7 | G | N7 | 14:770 | 3.79 | 1.00 | 33.90 |  |
| 8 | Water 4 | | 14:4057 | 3.92 | 1.00 | 41.11 | 4.26 (G:OP2 14:686) |
| 9 | Water 5 | | 14:3592 | 4.10 | 1.00 | 35.27 | 4.22 (G:N7 14:770) |
| 10 | Water 6 | | 14:3869 | 4.15 | 1.00 | 30.01 | 3.76 (G:N7 14:770) |
| 11 | G | OP2 | 14:686 | 4.69 | 1.00 | 40.59 |  |
| 12 | G | N1 | 14:770 | 4.98 | 1.00 | 34.67 |  |
  |

  
 Coded contact list:   
 Exclusion zone contact(s): /N7/O6/O6/O_w_/O_w_/O_w_/  
Consider K in place of MG (3 valid contact(s) for K vs. 0 for MG)  

### MG:14:3251

|  | Res. type | Atom | Residue | Distance | Occupancy | B-factor | Closest solute to 1^st^ shell water |
| --- | --- | --- | --- | --- | --- | --- | --- |
| **Ion** | | | |  | 1.00 | 45.92 || 1 | G | N7 | 14:1606 | 3.27 | 1.00 | 42.78 |
| 2 | C | OP2 | 14:1605 | 3.69 | 1.00 | 30.85 |  |
| 3 | G | OP2 | 14:1606 | 3.79 | 1.00 | 48.05 |  |
| 4 | C | OP2 | 14:1305 | 4.08 | 1.00 | 47.54 |  |
| 5 | C | OP1 | 14:1605 | 4.23 | 1.00 | 35.09 |  |
| 6 | C | O5' | 14:1605 | 4.34 | 1.00 | 44.26 |  |
| 7 | G | O5' | 14:1606 | 4.91 | 1.00 | 43.68 |  |
| 8 | C | OP2 | 14:1306 | 4.93 | 1.00 | 61.37 |  |
| 9 | G | N9 | 14:1606 | 4.95 | 1.00 | 36.47 |  |
  |

  
 Coded contact list:   
 Exclusion zone contact(s): /N7/  
Consider K in place of MG (1 valid contact(s) for K vs. 0 for MG)  

### MG:14:3252

|  | Res. type | Atom | Residue | Distance | Occupancy | B-factor | Closest solute to 1^st^ shell water |
| --- | --- | --- | --- | --- | --- | --- | --- |
| **Ion** | | | |  | 1.00 | 42.83 || 1 | G | OP2 | 14:187 | 3.63 | 1.00 | 37.06 |
| 2 | G | N7 | 14:187 | 3.74 | 1.00 | 31.48 |  |
| 3 | G | OP2 | 14:186 | 4.43 | 1.00 | 39.08 |  |
| 4 | G | O6 | 14:189 | 4.66 | 1.00 | 36.41 |  |
| 5 | G | N7 | 14:186 | 4.66 | 1.00 | 33.76 |  |
| 6 | G | N7 | 14:188 | 4.73 | 1.00 | 31.74 |  |
| 7 | G | O5' | 14:186 | 4.85 | 1.00 | 38.72 |  |
  |

  
 Coded contact list:   
1^st^ MG coordination shell missing (see Notes)  

### MG:14:3253

|  | Res. type | Atom | Residue | Distance | Occupancy | B-factor | Closest solute to 1^st^ shell water |
| --- | --- | --- | --- | --- | --- | --- | --- |
| **Ion** | | | |  | 1.00 | 47.77 || 1 | Water 1 | | 14:3609 | 2.33 | 1.00 | 44.73 | 2.36 (G:N7 14:1371) |
| 2 | Water 2 | | 14:3589 | 2.81 | 1.00 | 38.63 | 2.31 (U:OP2 14:1357) |
| 3 | G | N7 | 14:1358 | 3.26 | 1.00 | 47.13 |  |
| 4 | G | N7 | 14:1371 | 3.26 | 1.00 | 47.60 |  |
| 5 | G | O6 | 14:1358 | 3.26 | 1.00 | 60.74 |  |
| 6 | G | OP2 | 14:1371 | 3.27 | 1.00 | 45.53 |  |
| 7 | Water 3 | | 14:3870 | 3.44 | 1.00 | 46.63 | 2.94 (G:N7 14:1358) |
| 8 | G | O5' | 14:1371 | 4.06 | 1.00 | 47.16 |  |
| 9 | U | OP2 | 14:1357 | 4.51 | 1.00 | 41.27 |  |
| 10 | Water 4 | | 14:3594 | 4.76 | 1.00 | 47.25 | 4.86 (U:OP2 14:1357) |
| 11 | G | N9 | 14:1371 | 4.80 | 1.00 | 43.59 |  |
| 12 | U | O4 | 14:1357 | 4.86 | 1.00 | 46.43 |  |
| 13 | C | O3' | 14:1370 | 4.94 | 1.00 | 40.59 |  |
| 14 | Water 5 | | 14:3765 | 4.97 | 1.00 | 41.22 | 4.29 (G:N7 14:1371) |
  |

  
 Contact(s): /O_w_/  
 Coded contact list: O_w_  
 Exclusion zone contact(s): /N7/N7/O6/O_w_/OP2/  
Consider K in place of MG (4 valid contact(s) for K vs. 0 for MG)  
MG hydration shell issue: water too close (< 2.4 Angs.)  

### MG:14:3254

|  | Res. type | Atom | Residue | Distance | Occupancy | B-factor | Closest solute to 1^st^ shell water |
| --- | --- | --- | --- | --- | --- | --- | --- |
| **Ion** | | | |  | 1.00 | 62.59 || 1 | Water 1 | | 14:4081 | 2.54 | 1.00 | 72.66 | 5.01 (C:OP2 14:237) |
| 2 | Water 2 | | 14:4097 | 2.58 | 1.00 | 66.02 | 7.01 (C:OP1 14:237) |
| 3 | Water 3 | | 14:4028 | 3.26 | 1.00 | 54.99 | 6.40 (C:OP2 14:237) |
| 4 | C | OP1 | 14:237 | 4.57 | 1.00 | 53.18 |  |
| 5 | Water 4 | | 14:3632 | 4.67 | 1.00 | 48.76 | 8.06 (C:OP2 14:237) |
| 6 | C | OP2 | 14:237 | 4.89 | 1.00 | 58.45 |  |
| 7 | Water 5 | | 14:3708 | 4.97 | 1.00 | 46.28 | 7.09 (C:OP2 14:237) |
| 8 | Water 6 | | 14:3861 | 5.00 | 1.00 | 56.16 | 8.90 (C:OP2 14:237) |
  |

  
 Coded contact list:   
 Exclusion zone contact(s): /O_w_/O_w_/O_w_/  
1^st^ MG coordination shell missing (see Notes)  

### MG:14:3255

|  | Res. type | Atom | Residue | Distance | Occupancy | B-factor | Closest solute to 1^st^ shell water |
| --- | --- | --- | --- | --- | --- | --- | --- |
| **Ion** | | | |  | 1.00 | 63.48 || 1 | Water 1 | | 14:3944 | 2.22 | 1.00 | 62.46 | 3.15 (G:N7 14:500) |
| 2 | A | OP2 | 14:501 | 3.12 | 1.00 | 56.23 |  |
| 3 | G | OP2 | 14:500 | 3.22 | 1.00 | 53.09 |  |
| 4 | Water 2 | | 14:4101 | 3.62 | 1.00 | 71.70 | 5.75 (G:OP2 14:500) |
| 5 | Water 3 | | 14:3718 | 3.70 | 1.00 | 55.11 | 3.84 (G:OP2 14:500) |
| 6 | G | O5' | 14:500 | 3.90 | 1.00 | 60.57 |  |
| 7 | A | O5' | 14:501 | 4.43 | 1.00 | 55.02 |  |
| 8 | G | N7 | 14:500 | 4.56 | 1.00 | 54.08 |  |
| 9 | G | OP1 | 14:500 | 4.57 | 1.00 | 55.23 |  |
| 10 | A | OP2 | 14:502 | 4.77 | 1.00 | 51.34 |  |
  |

  
 Contact(s): /O_w_/  
 Coded contact list: O_w_  
 Exclusion zone contact(s): /OP2/OP2/  
Consider K in place of MG (2 valid contact(s) for K vs. 0 for MG)  

### MG:14:3256

|  | Res. type | Atom | Residue | Distance | Occupancy | B-factor | Closest solute to 1^st^ shell water |
| --- | --- | --- | --- | --- | --- | --- | --- |
| **Ion** | | | |  | 1.00 | 67.62 || 1 | G | O6 | 14:1466 | 2.75 | 1.00 | 80.43 |
| 2 | Water 1 | | 14:3744 | 3.48 | 1.00 | 74.19 | 2.67 (G:N7 14:1466) |
| 3 | Water 2 | | 14:4087 | 3.70 | 1.00 | 77.26 | 5.12 (G:N7 14:1466) |
| 4 | G | N7 | 14:1444 | 3.72 | 1.00 | 74.04 |  |
| 5 | G | O6 | 14:1465 | 3.86 | 1.00 | 84.63 |  |
| 6 | G | N7 | 14:1466 | 3.89 | 1.00 | 84.97 |  |
| 7 | Water 3 | | 14:3802 | 3.90 | 1.00 | 71.97 | 2.89 (G:N7 14:1444) |
| 8 | C | N4 | 14:1445 | 4.31 | 1.00 | 83.83 |  |
| 9 | G | OP2 | 14:1444 | 4.37 | 1.00 | 77.46 |  |
| 10 | G | O6 | 14:1444 | 4.73 | 1.00 | 74.23 |  |
| 11 | G | N7 | 14:1465 | 4.98 | 1.00 | 87.88 |  |
  |

  
 Coded contact list:   
 Exclusion zone contact(s): /O6/  
Consider K in place of MG (1 valid contact(s) for K vs. 0 for MG)  

### MG:14:3257

|  | Res. type | Atom | Residue | Distance | Occupancy | B-factor | Closest solute to 1^st^ shell water |
| --- | --- | --- | --- | --- | --- | --- | --- |
| **Ion** | | | |  | 1.00 | 65.99 || 1 | A | OP1 | 14:222 | 2.08 | 1.00 | 68.92 |
| 2 | A | O5' | 14:222 | 3.71 | 1.00 | 66.35 |  |
| 3 | C | N4 | 14:267 | 3.86 | 1.00 | 73.46 |  |
| 4 | C | OP2 | 14:271A | 4.14 | 1.00 | 84.05 |  |
| 5 | A | O3' | 14:221 | 4.21 | 1.00 | 70.10 |  |
| 6 | A | OP2 | 14:222 | 4.64 | 1.00 | 73.10 |  |
| 7 | A | O2' | 14:221 | 4.86 | 1.00 | 70.77 |  |
  |

  
 Contact(s): /OP1/  
 Coded contact list: O_ph_  

### MG:14:3258

|  | Res. type | Atom | Residue | Distance | Occupancy | B-factor | Closest solute to 1^st^ shell water |
| --- | --- | --- | --- | --- | --- | --- | --- |
| **Ion** | | | |  | 1.00 | 34.70 || 1 | A | OP2 | 14:1265 | 2.01 | 1.00 | 44.27 |
| 2 | U | OP1 | 14:2615 | 2.18 | 1.00 | 48.35 |  |
| 3 | A | O5' | 14:1265 | 3.75 | 1.00 | 40.60 |  |
| 4 | U | OP2 | 14:2615 | 4.24 | 1.00 | 43.94 |  |
| 5 | U | O5' | 14:2615 | 4.32 | 1.00 | 38.98 |  |
| 6 | A | OP1 | 14:1265 | 4.33 | 1.00 | 37.67 |  |
| 7 | A | OP1 | 14:2614 | 4.33 | 1.00 | 41.65 |  |
| 8 | G | O3' | 14:1264 | 4.53 | 1.00 | 33.94 |  |
| 9 | A | O3' | 14:2614 | 4.58 | 1.00 | 45.99 |  |
| 10 | U | O2' | 14:1267 | 4.84 | 1.00 | 48.84 |  |
  |

  
**1^st^ shell ligand angles:**   
1-2: 85;   
  
 Contact(s): /OP1/OP2/  
 Coded contact list: 2O_ph_  

### MG:14:3259

|  | Res. type | Atom | Residue | Distance | Occupancy | B-factor | Closest solute to 1^st^ shell water |
| --- | --- | --- | --- | --- | --- | --- | --- |
| **Ion** | | | |  | 1.00 | 42.14 || 1 | Water 1 | | 14:3832 | 2.86 | 1.00 | 43.86 | 2.83 (G:N7 14:382) |
| 2 | G | O6 | 14:382 | 3.64 | 1.00 | 49.97 |  |
| 3 | U | O4 | 14:387 | 3.81 | 1.00 | 55.18 |  |
| 4 | G | OP2 | 14:386 | 3.83 | 1.00 | 36.60 |  |
| 5 | U | O4 | 14:383 | 4.03 | 1.00 | 54.01 |  |
| 6 | G | O6 | 14:388 | 4.12 | 1.00 | 53.19 |  |
| 7 | G | O6 | 14:381 | 4.17 | 1.00 | 53.30 |  |
| 8 | Water 2 | | 14:3787 | 4.18 | 1.00 | 51.05 | 3.07 (G:OP2 14:386) |
| 9 | G | N7 | 14:381 | 4.63 | 1.00 | 46.95 |  |
| 10 | G | N7 | 14:382 | 4.68 | 1.00 | 53.59 |  |
| 11 | C | N4 | 14:392 | 4.76 | 1.00 | 52.17 |  |
| 12 | Water 3 | | 14:3598 | 4.97 | 1.00 | 49.33 | 2.96 (G:OP2 14:386) |
  |

  
 Coded contact list:   
 Exclusion zone contact(s): /O_w_/  
1^st^ MG coordination shell missing (see Notes)  

### MG:14:3260

|  | Res. type | Atom | Residue | Distance | Occupancy | B-factor | Closest solute to 1^st^ shell water |
| --- | --- | --- | --- | --- | --- | --- | --- |
| **Ion** | | | |  | 1.00 | 84.74 || 1 | G | OP1 | 14:361 | 2.68 | 1.00 | 114.27 |
| 2 | G | N7 | 14:270Y | 2.84 | 1.00 | 86.96 |  |
| 3 | G | OP2 | 14:270X | 3.34 | 1.00 | 84.68 |  |
| 4 | G | OP2 | 14:270Y | 3.94 | 1.00 | 72.25 |  |
| 5 | U | OP1 | 14:269 | 4.00 | 1.00 | 77.35 |  |
| 6 | G | O6 | 14:270Y | 4.07 | 1.00 | 81.04 |  |
| 7 | G | O3' | 14:360 | 4.82 | 1.00 | 109.64 |  |
| 8 | G | OP2 | 14:361 | 4.87 | 1.00 | 109.87 |  |
| 9 | G | N9 | 14:270Y | 4.97 | 1.00 | 82.37 |  |
  |

  
 Coded contact list:   
 Exclusion zone contact(s): /N7/OP1/OP2/  
Consider K in place of MG (3 valid contact(s) for K vs. 0 for MG)  

### MG:14:3261

|  | Res. type | Atom | Residue | Distance | Occupancy | B-factor | Closest solute to 1^st^ shell water |
| --- | --- | --- | --- | --- | --- | --- | --- |
| **Ion** | | | |  | 1.00 | 43.43 || 1 | VAL | O | 19:34 | 3.02 | 1.00 | 44.39 |
| 2 | A | OP1 | 14:1570 | 3.29 | 1.00 | 36.31 |  |
| 3 | LYS | O | 19:31 | 3.92 | 1.00 | 43.42 |  |
| 4 | LYS | N | 19:35 | 4.25 | 1.00 | 34.51 |  |
| 5 | A | N1 | 14:1427 | 4.87 | 1.00 | 39.66 |  |
| 6 | VAL | N | 19:34 | 4.97 | 1.00 | 37.29 |  |
| 7 | A | O5' | 14:1570 | 4.99 | 1.00 | 35.13 |  |
  |

  
 Coded contact list:   
 Exclusion zone contact(s): /O_back_/OP1/  
Consider K in place of MG (2 valid contact(s) for K vs. 0 for MG)  

### MG:14:3262

|  | Res. type | Atom | Residue | Distance | Occupancy | B-factor | Closest solute to 1^st^ shell water |
| --- | --- | --- | --- | --- | --- | --- | --- |
| **Ion** | | | |  | 1.00 | 41.71 || 1 | G | O6 | 14:1807 | 4.20 | 1.00 | 46.36 |
| 2 | G | N7 | 14:1807 | 4.27 | 1.00 | 39.92 |  |
| 3 | A | OP1 | 14:1809 | 4.47 | 1.00 | 40.40 |  |
| 4 | G | OP1 | 14:2083 | 4.58 | 1.00 | 42.07 |  |
| 5 | A | OP2 | 14:1809 | 4.78 | 1.00 | 37.63 |  |
  |

  
 Coded contact list:   
1^st^ MG coordination shell missing (see Notes)  

### MG:14:3263

|  | Res. type | Atom | Residue | Distance | Occupancy | B-factor | Closest solute to 1^st^ shell water |
| --- | --- | --- | --- | --- | --- | --- | --- |
| **Ion** | | | |  | 1.00 | 60.81 || 1 | G | O6 | 14:270R | 3.27 | 1.00 | 107.65 |
| 2 | G | O6 | 14:270S | 3.91 | 1.00 | 105.29 |  |
| 3 | U | O4 | 14:270F | 4.05 | 1.00 | 110.05 |  |
| 4 | G | N7 | 14:270R | 4.40 | 1.00 | 112.99 |  |
| 5 | C | N4 | 14:270G | 4.55 | 1.00 | 112.04 |  |
  |

  
 Coded contact list:   
 Exclusion zone contact(s): /O6/  
Consider K in place of MG (1 valid contact(s) for K vs. 0 for MG)  

### MG:14:3264

|  | Res. type | Atom | Residue | Distance | Occupancy | B-factor | Closest solute to 1^st^ shell water |
| --- | --- | --- | --- | --- | --- | --- | --- |
| **Ion** | | | |  | 1.00 | 45.90 || 1 | G | OP2 | 14:205 | 2.64 | 1.00 | 43.88 |
| 2 | C | O2' | 14:253 | 3.59 | 1.00 | 43.68 |  |
| 3 | C | OP2 | 14:203 | 3.80 | 1.00 | 33.57 |  |
| 4 | C | O3' | 14:253 | 4.00 | 1.00 | 45.22 |  |
| 5 | U | OP1 | 14:202 | 4.23 | 1.00 | 36.84 |  |
| 6 | G | OP1 | 14:205 | 4.45 | 1.00 | 37.16 |  |
| 7 | A | O3' | 14:204 | 4.60 | 1.00 | 38.88 |  |
| 8 | C | OP1 | 14:203 | 4.71 | 1.00 | 39.27 |  |
| 9 | G | OP1 | 14:254 | 4.78 | 1.00 | 49.90 |  |
  |

  
 Coded contact list:   
 Exclusion zone contact(s): /OP2/  
Consider K in place of MG (1 valid contact(s) for K vs. 0 for MG)  

### MG:14:3265

|  | Res. type | Atom | Residue | Distance | Occupancy | B-factor | Closest solute to 1^st^ shell water |
| --- | --- | --- | --- | --- | --- | --- | --- |
| **Ion** | | | |  | 1.00 | 34.03 || 1 | A | OP2 | 14:945 | 1.97 | 1.00 | 47.29 |
| 2 | G | OP2 | 14:946 | 2.40 | 1.00 | 50.14 |  |
| 3 | G | O3' | 14:944 | 3.46 | 1.00 | 47.46 |  |
| 4 | G | OP1 | 14:946 | 3.51 | 1.00 | 54.02 |  |
| 5 | A | O5' | 14:945 | 3.75 | 1.00 | 48.56 |  |
| 6 | Water 1 | | 14:3988 | 3.92 | 1.00 | 56.06 | 3.29 (A:OP2 14:945) |
| 7 | U | OP2 | 14:822 | 4.34 | 1.00 | 50.32 |  |
| 8 | A | O2' | 14:821 | 4.36 | 1.00 | 53.58 |  |
| 9 | A | O3' | 14:821 | 4.39 | 1.00 | 48.56 |  |
| 10 | A | O3' | 14:945 | 4.46 | 1.00 | 54.20 |  |
| 11 | A | OP1 | 14:945 | 4.47 | 1.00 | 42.69 |  |
| 12 | Water 2 | | 14:3700 | 4.49 | 1.00 | 45.09 | 2.58 (A:OP1 14:945) |
| 13 | G | O5' | 14:946 | 4.53 | 1.00 | 51.02 |  |
  |

  
 Contact(s): /OP2/  
 Coded contact list: O_ph_  
 Exclusion zone contact(s): /OP2/  

### MG:14:3266

|  | Res. type | Atom | Residue | Distance | Occupancy | B-factor | Closest solute to 1^st^ shell water |
| --- | --- | --- | --- | --- | --- | --- | --- |
| **Ion** | | | |  | 1.00 | 38.96 || 1 | G | O5' | 14:686 | 1.92 | 1.00 | 29.53 |
| 2 | C | OP2 | 14:687 | 2.23 | 1.00 | 33.27 |  |
| 3 | G | OP2 | 14:686 | 2.76 | 1.00 | 40.59 |  |
| 4 | Water 1 | | 14:4057 | 2.84 | 1.00 | 41.11 | 3.82 (C:OP2 14:687) |
| 5 | A | O3' | 14:685 | 2.91 | 1.00 | 34.60 |  |
| 6 | Water 2 | | 14:3843 | 3.41 | 1.00 | 39.82 | 2.86 (U:O4 14:688) |
| 7 | Water 3 | | 14:3657 | 3.89 | 1.00 | 32.85 | 2.51 (G:OP2 14:686) |
| 8 | Water 4 | | 14:3895 | 3.93 | 1.00 | 37.04 | 3.36 (G:OP2 14:686) |
| 9 | G | OP1 | 14:686 | 4.03 | 1.00 | 27.58 |  |
| 10 | U | O4 | 14:688 | 4.23 | 1.00 | 34.01 |  |
| 11 | G | O3' | 14:686 | 4.38 | 1.00 | 27.53 |  |
| 12 | C | O5' | 14:687 | 4.56 | 1.00 | 27.56 |  |
| 13 | C | OP1 | 14:687 | 4.57 | 1.00 | 28.03 |  |
| 14 | G | O4' | 14:686 | 4.62 | 1.00 | 33.52 |  |
  |

  
**1^st^ shell ligand angles:**   
1-2: 96;   
  
 Contact(s): /O5'/OP2/  
 Coded contact list: O_ph_O_r_  
 Exclusion zone contact(s): /O3'/O_w_/OP2/  

### MG:14:3267

|  | Res. type | Atom | Residue | Distance | Occupancy | B-factor | Closest solute to 1^st^ shell water |
| --- | --- | --- | --- | --- | --- | --- | --- |
| **Ion** | | | |  | 1.00 | 44.95 || 1 | G | O6 | 14:1984 | 2.14 | 1.00 | 45.99 |
| 2 | G | O6 | 14:1769 | 2.55 | 1.00 | 42.53 |  |
| 3 | U | O4 | 14:1768 | 3.26 | 1.00 | 50.83 |  |
| 4 | U | N3 | 14:1768 | 3.91 | 1.00 | 42.47 |  |
| 5 | G | N7 | 14:1769 | 3.96 | 1.00 | 44.85 |  |
| 6 | C | N4 | 14:1983 | 3.98 | 1.00 | 38.35 |  |
| 7 | G | N7 | 14:1984 | 4.21 | 1.00 | 43.33 |  |
| 8 | G | N1 | 14:1984 | 4.24 | 1.00 | 37.70 |  |
| 9 | G | N1 | 14:1769 | 4.36 | 1.00 | 35.55 |  |
| 10 | G | O6 | 14:1985 | 4.54 | 1.00 | 41.88 |  |
  |

  
 Contact(s): /O6/  
 Coded contact list: O_b_  
 Exclusion zone contact(s): /O4/O6/  

### MG:14:3268

|  | Res. type | Atom | Residue | Distance | Occupancy | B-factor | Closest solute to 1^st^ shell water |
| --- | --- | --- | --- | --- | --- | --- | --- |
| **Ion** | | | |  | 1.00 | 36.05 || 1 | G | O6 | 14:187 | 2.73 | 1.00 | 28.67 |
| 2 | G | O6 | 14:188 | 3.03 | 1.00 | 32.14 |  |
| 3 | G | O6 | 14:186 | 3.78 | 1.00 | 35.51 |  |
| 4 | G | N7 | 14:187 | 3.79 | 1.00 | 31.48 |  |
| 5 | G | N7 | 14:188 | 4.27 | 1.00 | 31.74 |  |
| 6 | C | N4 | 14:208 | 4.36 | 1.00 | 28.90 |  |
| 7 | C | N4 | 14:209 | 4.44 | 1.00 | 28.19 |  |
| 8 | G | N7 | 14:186 | 4.57 | 1.00 | 33.76 |  |
| 9 | G | N1 | 14:187 | 4.96 | 1.00 | 29.49 |  |
| 10 | G | N1 | 14:188 | 4.97 | 1.00 | 27.56 |  |
  |

  
 Coded contact list:   
 Exclusion zone contact(s): /O6/O6/  
Consider K in place of MG (2 valid contact(s) for K vs. 0 for MG)  

### MG:14:3269

|  | Res. type | Atom | Residue | Distance | Occupancy | B-factor | Closest solute to 1^st^ shell water |
| --- | --- | --- | --- | --- | --- | --- | --- |
| **Ion** | | | |  | 1.00 | 68.48 || 1 | C | OP2 | 14:2471 | 3.10 | 1.00 | 129.09 |
| 2 | C | OP2 | 14:2474 | 3.86 | 1.00 | 145.25 |  |
| 3 | G | N7 | 14:2472 | 4.10 | 1.00 | 129.06 |  |
| 4 | C | OP1 | 14:2471 | 4.25 | 1.00 | 134.37 |  |
| 5 | G | OP2 | 14:2472 | 4.30 | 1.00 | 132.18 |  |
| 6 | C | OP2 | 14:2475 | 4.40 | 1.00 | 149.30 |  |
| 7 | C | O5' | 14:2471 | 4.80 | 1.00 | 127.69 |  |
  |

  
 Coded contact list:   
 Exclusion zone contact(s): /OP2/  
Consider K in place of MG (1 valid contact(s) for K vs. 0 for MG)  

### MG:14:3270

|  | Res. type | Atom | Residue | Distance | Occupancy | B-factor | Closest solute to 1^st^ shell water |
| --- | --- | --- | --- | --- | --- | --- | --- |
| **Ion** | | | |  | 1.00 | 66.52 || 1 | C | OP1 | 14:62 | 3.00 | 1.00 | 64.49 |
| 2 | G | OP1 | 14:60 | 3.66 | 1.00 | 56.02 |  |
| 3 | C | OP2 | 14:62 | 3.97 | 1.00 | 63.64 |  |
| 4 | G | OP2 | 14:60 | 4.65 | 1.00 | 57.82 |  |
| 5 | U | OP2 | 14:63 | 4.78 | 1.00 | 65.76 |  |
| 6 | G | O3' | 14:61 | 4.95 | 1.00 | 63.76 |  |
  |

  
 Coded contact list:   
 Exclusion zone contact(s): /OP1/  
Consider K in place of MG (1 valid contact(s) for K vs. 0 for MG)  

### MG:14:3271

|  | Res. type | Atom | Residue | Distance | Occupancy | B-factor | Closest solute to 1^st^ shell water |
| --- | --- | --- | --- | --- | --- | --- | --- |
| **Ion** | | | |  | 1.00 | 51.23 || 1 | G | N7 | 14:2714 | 3.82 | 1.00 | 56.79 |
| 2 | C | N3 | 14:2690 | 4.47 | 1.00 | 45.87 |  |
| 3 | G | O6 | 14:2714 | 4.76 | 1.00 | 57.16 |  |
| 4 | C | N4 | 14:2715 | 4.78 | 1.00 | 53.44 |  |
| 5 | C | N4 | 14:2690 | 4.82 | 1.00 | 49.09 |  |
| 6 | G | OP1 | 14:2714 | 4.93 | 1.00 | 45.03 |  |
  |

  
 Coded contact list:   
1^st^ MG coordination shell missing (see Notes)  

### MG:14:3272

|  | Res. type | Atom | Residue | Distance | Occupancy | B-factor | Closest solute to 1^st^ shell water |
| --- | --- | --- | --- | --- | --- | --- | --- |
| **Ion** | | | |  | 1.00 | 61.08 || 1 | C | OP2 | 14:1290 | 3.97 | 1.00 | 40.15 |
| 2 | C | OP1 | 14:1290 | 4.09 | 1.00 | 38.60 |  |
| 3 | C | OP1 | 14:1289 | 4.63 | 1.00 | 45.19 |  |
  |

  
 Coded contact list:   
1^st^ MG coordination shell missing (see Notes)  

### MG:14:3273

|  | Res. type | Atom | Residue | Distance | Occupancy | B-factor | Closest solute to 1^st^ shell water |
| --- | --- | --- | --- | --- | --- | --- | --- |
| **Ion** | | | |  | 1.00 | 37.20 || 1 | C | OP2 | 14:1830 | 2.18 | 1.00 | 34.87 |
| 2 | A | OP1 | 14:1829 | 3.03 | 1.00 | 38.70 |  |
| 3 | G | O2' | 14:1792 | 3.94 | 1.00 | 45.07 |  |
| 4 | C | O4' | 14:1793 | 4.22 | 1.00 | 34.47 |  |
| 5 | C | OP1 | 14:1830 | 4.29 | 1.00 | 34.77 |  |
| 6 | C | O5' | 14:1830 | 4.40 | 1.00 | 32.01 |  |
| 7 | A | N1 | 14:1970 | 4.40 | 1.00 | 37.56 |  |
| 8 | G | N3 | 14:1792 | 4.46 | 1.00 | 36.65 |  |
| 9 | A | O3' | 14:1829 | 4.59 | 1.00 | 38.76 |  |
| 10 | G | N2 | 14:1792 | 4.63 | 1.00 | 28.80 |  |
| 11 | A | O5' | 14:1829 | 4.72 | 1.00 | 39.43 |  |
| 12 | G | O3' | 14:1828 | 4.96 | 1.00 | 29.15 |  |
  |

  
 Contact(s): /OP2/  
 Coded contact list: O_ph_  
 Exclusion zone contact(s): /OP1/  

### MG:14:3274

|  | Res. type | Atom | Residue | Distance | Occupancy | B-factor | Closest solute to 1^st^ shell water |
| --- | --- | --- | --- | --- | --- | --- | --- |
| **Ion** | | | |  | 1.00 | 54.81 || 1 | G | O6 | 14:2235 | 3.38 | 1.00 | 43.82 |
| 2 | G | O6 | 14:2234 | 4.06 | 1.00 | 48.47 |  |
| 3 | G | N7 | 14:2234 | 4.23 | 1.00 | 48.90 |  |
| 4 | G | N7 | 14:2235 | 4.25 | 1.00 | 42.19 |  |
  |

  
 Coded contact list:   
 Exclusion zone contact(s): /O6/  
Consider K in place of MG (1 valid contact(s) for K vs. 0 for MG)  

### MG:14:3275

|  | Res. type | Atom | Residue | Distance | Occupancy | B-factor | Closest solute to 1^st^ shell water |
| --- | --- | --- | --- | --- | --- | --- | --- |
| **Ion** | | | |  | 1.00 | 42.80 || 1 | G | O6 | 14:1425 | 2.17 | 1.00 | 65.67 |
| 2 | G | N7 | 14:1425 | 3.69 | 1.00 | 56.53 |  |
| 3 | G | N1 | 14:1426 | 3.71 | 1.00 | 46.86 |  |
| 4 | G | O6 | 14:1426 | 3.92 | 1.00 | 53.68 |  |
| 5 | G | N1 | 14:1425 | 4.44 | 1.00 | 55.41 |  |
| 6 | G | N2 | 14:1426 | 4.66 | 1.00 | 45.27 |  |
  |

  
 Contact(s): /O6/  
 Coded contact list: O_b_  

### MG:14:3276

|  | Res. type | Atom | Residue | Distance | Occupancy | B-factor | Closest solute to 1^st^ shell water |
| --- | --- | --- | --- | --- | --- | --- | --- |
| **Ion** | | | |  | 1.00 | 52.22 || 1 | G | O6 | 14:1264 | 2.89 | 1.00 | 48.37 |
| 2 | A | N1 | 14:1265 | 3.14 | 1.00 | 45.38 |  |
| 3 | ARG | NH2 | A5:99 | 4.21 | 1.00 | 61.07 |  |
| 4 | A | OP2 | 14:2014 | 4.27 | 1.00 | 43.06 |  |
| 5 | U | OP2 | 14:1263 | 4.34 | 1.00 | 45.33 |  |
| 6 | G | N7 | 14:1264 | 4.38 | 1.00 | 44.26 |  |
| 7 | A | N3 | 14:1265 | 4.45 | 1.00 | 43.09 |  |
| 8 | A | N6 | 14:1265 | 4.57 | 1.00 | 39.96 |  |
| 9 | U | O4 | 14:1263 | 4.85 | 1.00 | 51.54 |  |
  |

  
 Coded contact list:   
 Exclusion zone contact(s): /N1_A_/O6/  
Consider K in place of MG (2 valid contact(s) for K vs. 0 for MG)  

### MG:14:3277

|  | Res. type | Atom | Residue | Distance | Occupancy | B-factor | Closest solute to 1^st^ shell water |
| --- | --- | --- | --- | --- | --- | --- | --- |
| **Ion** | | | |  | 1.00 | 61.25 || 1 | G | OP2 | 14:245 | 3.60 | 1.00 | 45.97 |
| 2 | G | OP1 | 14:245 | 3.89 | 1.00 | 64.94 |  |
| 3 | GLN | O | 35:68 | 4.34 | 1.00 | 51.23 |  |
| 4 | A | OP1 | 14:244 | 4.39 | 1.00 | 41.58 |  |
| 5 | MET | O | 35:67 | 4.72 | 1.00 | 62.46 |  |
| 6 | A | O5' | 14:244 | 4.84 | 1.00 | 42.23 |  |
| 7 | GLN | OE1 | 35:68 | 4.91 | 1.00 | 55.36 |  |
  |

  
 Coded contact list:   
1^st^ MG coordination shell missing (see Notes)  

### MG:14:3278

|  | Res. type | Atom | Residue | Distance | Occupancy | B-factor | Closest solute to 1^st^ shell water |
| --- | --- | --- | --- | --- | --- | --- | --- |
| **Ion** | | | |  | 1.00 | 49.90 || 1 | C | O2 | 14:1607 | 2.68 | 1.00 | 46.37 |
| 2 | C | O2' | 14:1607 | 3.94 | 1.00 | 54.37 |  |
| 3 | U | O4 | 14:1621 | 4.00 | 1.00 | 39.51 |  |
| 4 | C | N1 | 14:1607 | 4.60 | 1.00 | 41.33 |  |
| 5 | C | O3' | 14:1607 | 4.61 | 1.00 | 34.99 |  |
| 6 | G | OP2 | 14:1620 | 4.85 | 1.00 | 37.39 |  |
| 7 | C | N3 | 14:1607 | 4.92 | 1.00 | 48.33 |  |
  |

  
 Coded contact list:   
 Exclusion zone contact(s): /O2/  
Consider K in place of MG (1 valid contact(s) for K vs. 0 for MG)  

### MG:14:3279

|  | Res. type | Atom | Residue | Distance | Occupancy | B-factor | Closest solute to 1^st^ shell water |
| --- | --- | --- | --- | --- | --- | --- | --- |
| **Ion** | | | |  | 1.00 | 22.92 || 1 | Water 1 | | 14:3535 | 2.16 | 1.00 | 31.22 | 2.09 (G:OP2 14:733) |
| 2 | Water 2 | | 14:3848 | 2.16 | 1.00 | 35.27 | 2.88 (A:N6 14:761) |
| 3 | Water 3 | | 14:3675 | 2.17 | 1.00 | 35.48 | 2.53 (A:OP2 14:761) |
| 4 | Water 4 | | 14:3643 | 2.18 | 1.00 | 40.84 | 2.47 (G:O6 14:733) |
| 5 | A | N7 | 14:761 | 2.50 | 1.00 | 34.70 |  |
| 6 | G | N7 | 14:733 | 2.51 | 1.00 | 38.02 |  |
| 7 | G | O6 | 14:733 | 3.16 | 1.00 | 43.80 |  |
| 8 | G | OP2 | 14:733 | 3.85 | 1.00 | 30.06 |  |
| 9 | A | OP2 | 14:761 | 3.88 | 1.00 | 31.53 |  |
| 10 | A | N6 | 14:761 | 3.98 | 1.00 | 37.73 |  |
| 11 | C | OP2 | 14:732 | 4.40 | 1.00 | 36.72 |  |
| 12 | A | N9 | 14:761 | 4.48 | 1.00 | 31.72 |  |
| 13 | G | N9 | 14:733 | 4.65 | 1.00 | 36.57 |  |
| 14 | G | OP2 | 14:760 | 4.87 | 1.00 | 34.50 |  |
| 15 | A | O5' | 14:761 | 4.87 | 1.00 | 38.68 |  |
| 16 | C | O5' | 14:732 | 4.88 | 1.00 | 36.33 |  |
| 17 | G | N1 | 14:733 | 4.97 | 1.00 | 36.95 |  |
  |

  
**1^st^ shell ligand angles:**   
1-2: 82; 1-3: 100; 1-4: 96; 2-3: 85; 2-4: 178; 3-4: 94;   
  
 Contact(s): /O_w_/O_w_/O_w_/O_w_/  
 Coded contact list: 4O_w_  
 Exclusion zone contact(s): /N7/N7/O6/  
Consider NA in place of MG (2 valid contact(s) for NA vs. 0 for MG)  
MG hydration shell issue: water too close (< 2.4 Angs.)  

### MG:14:3280

|  | Res. type | Atom | Residue | Distance | Occupancy | B-factor | Closest solute to 1^st^ shell water |
| --- | --- | --- | --- | --- | --- | --- | --- |
| **Ion** | | | |  | 1.00 | 45.84 || 1 | Water 1 | | 14:3512 | 2.14 | 1.00 | 37.85 | 1.80 (C:OP1 14:249) |
| 2 | Water 2 | | 14:3986 | 2.18 | 1.00 | 46.07 | 3.28 (A:N1 14:2393) |
| 3 | Water 3 | | 14:3730 | 2.18 | 1.00 | 35.77 | 2.65 (C:OP1 14:249) |
| 4 | Water 4 | | 14:3518 | 2.18 | 1.00 | 36.91 | 1.99 (U:OP1 14:2431) |
| 5 | Water 5 | | 14:4076 | 2.18 | 1.00 | 51.71 | 4.06 (A:N1 14:2393) |
| 6 | Water 6 | | 14:3962 | 2.20 | 1.00 | 36.23 | 3.57 (C:OP1 14:249) |
| 7 | C | OP1 | 14:249 | 2.98 | 1.00 | 36.51 |  |
| 8 | A | OP1 | 14:2432 | 3.83 | 1.00 | 39.37 |  |
| 9 | U | OP1 | 14:2431 | 3.87 | 1.00 | 30.61 |  |
| 10 | A | N1 | 14:2393 | 4.79 | 1.00 | 46.71 |  |
| 11 | C | OP2 | 14:249 | 4.89 | 1.00 | 45.62 |  |
  |

  
**1^st^ shell ligand angles:**   
1-2: 84; 1-3: 95; 1-4: 84; 1-5: 174; 1-6: 92; 2-3: 87; 2-4: 93; 2-5: 93; 2-6: 175; 3-4: 179; 3-5: 90; 3-6: 92; 4-5: 91; 4-6: 88; 5-6: 91;   
  
 Contact(s): /O_w_/O_w_/O_w_/O_w_/O_w_/O_w_/  
 Coded contact list: 6O_w_  
 Exclusion zone contact(s): /OP1/  
Consider K in place of MG (1 valid contact(s) for K vs. 0 for MG)  
MG hydration shell issue: water too close (< 2.4 Angs.)  

### MG:14:3281

|  | Res. type | Atom | Residue | Distance | Occupancy | B-factor | Closest solute to 1^st^ shell water |
| --- | --- | --- | --- | --- | --- | --- | --- |
| **Ion** | | | |  | 1.00 | 25.82 || 1 | Water 1 | | 14:3559 | 2.15 | 1.00 | 29.48 | 2.23 (G:OP2 14:1828) |
| 2 | Water 2 | | 14:3574 | 2.16 | 1.00 | 38.08 | 2.27 (C:O5' 14:1827) |
| 3 | Water 3 | | 14:3913 | 2.17 | 1.00 | 35.07 | 3.05 (C:OP1 14:1788) |
| 4 | Water 4 | | 19:408 | 2.17 | 1.00 | 33.66 | 2.89 (ARG:NH1 19:239) |
| 5 | Water 5 | | 19:401 | 2.21 | 1.00 | 53.03 | 1.31 (ARG:NE 19:239) |
| 6 | G | OP2 | 14:1828 | 2.58 | 1.00 | 31.08 |  |
| 7 | G | OP1 | 14:1828 | 3.32 | 1.00 | 31.51 |  |
| 8 | A | OP2 | 14:1972 | 3.42 | 1.00 | 35.67 |  |
| 9 | ARG | NE | 19:239 | 3.43 | 1.00 | 46.48 |  |
| 10 | ARG | NH1 | 19:239 | 3.72 | 1.00 | 44.72 |  |
| 11 | Water 6 | | 14:3871 | 3.98 | 1.00 | 40.20 | 2.94 (G:OP1 14:1828) |
| 12 | C | O5' | 14:1827 | 4.22 | 1.00 | 28.52 |  |
| 13 | C | O3' | 14:1827 | 4.50 | 1.00 | 27.27 |  |
| 14 | G | O5' | 14:1828 | 4.63 | 1.00 | 25.93 |  |
| 15 | G | N7 | 14:1828 | 4.84 | 1.00 | 35.04 |  |
| 16 | C | OP1 | 14:1788 | 4.88 | 1.00 | 40.48 |  |
| 17 | Water 7 | | 14:3636 | 4.89 | 1.00 | 30.03 | 2.46 (G:OP1 14:1828) |
  |

  
**1^st^ shell ligand angles:**   
1-2: 81; 1-3: 83; 1-4: 94; 1-5: 171; 2-3: 88; 2-4: 175; 2-5: 94; 3-4: 94; 3-5: 89; 4-5: 91;   
  
 Contact(s): /O_w_/O_w_/O_w_/O_w_/O_w_/  
 Coded contact list: 5O_w_  
 Exclusion zone contact(s): /OP1/OP2/  
Consider NA in place of MG (1 valid contact(s) for NA vs. 0 for MG)  
MG hydration shell issue: water too close (< 2.4 Angs.)  

### MG:14:3282

|  | Res. type | Atom | Residue | Distance | Occupancy | B-factor | Closest solute to 1^st^ shell water |
| --- | --- | --- | --- | --- | --- | --- | --- |
| **Ion** | | | |  | 1.00 | 48.52 || 1 | Water 1 | | 14:3770 | 2.17 | 1.00 | 39.11 | 2.70 (G:O2' 14:2709) |
| 2 | Water 2 | | 14:3694 | 2.17 | 1.00 | 51.43 | 2.57 (U:OP1 14:1639) |
| 3 | Water 3 | | 14:3501 | 2.18 | 1.00 | 54.76 | 1.30 (G:N3 14:2709) |
| 4 | Water 4 | | 14:3921 | 2.19 | 1.00 | 49.78 | 3.08 (C:O2 14:2699) |
| 5 | Water 5 | | 14:4026 | 2.20 | 1.00 | 54.40 | 3.50 (G:OP1 14:1296) |
| 6 | U | OP1 | 14:1639 | 2.48 | 1.00 | 40.23 |  |
| 7 | G | N3 | 14:2709 | 3.44 | 1.00 | 47.24 |  |
| 8 | G | N2 | 14:2709 | 3.47 | 1.00 | 45.48 |  |
| 9 | G | O2' | 14:2709 | 4.00 | 1.00 | 44.23 |  |
| 10 | C | O3' | 14:1638 | 4.24 | 1.00 | 41.67 |  |
| 11 | U | O5' | 14:1639 | 4.39 | 1.00 | 39.25 |  |
| 12 | C | O2 | 14:2699 | 4.42 | 1.00 | 43.36 |  |
| 13 | G | OP1 | 14:1296 | 4.73 | 1.00 | 46.93 |  |
| 14 | U | OP2 | 14:1639 | 4.88 | 1.00 | 39.24 |  |
| 15 | G | N2 | 14:2708 | 4.99 | 1.00 | 45.24 |  |
  |

  
**1^st^ shell ligand angles:**   
1-2: 83; 1-3: 91; 1-4: 176; 1-5: 95; 2-3: 172; 2-4: 94; 2-5: 94; 3-4: 92; 3-5: 91; 4-5: 87;   
  
 Contact(s): /O_w_/O_w_/O_w_/O_w_/O_w_/  
 Coded contact list: 5O_w_  
 Exclusion zone contact(s): /OP1/  
Consider NA in place of MG (1 valid contact(s) for NA vs. 0 for MG)  
MG hydration shell issue: water too close (< 2.4 Angs.)  

### MG:14:3283

|  | Res. type | Atom | Residue | Distance | Occupancy | B-factor | Closest solute to 1^st^ shell water |
| --- | --- | --- | --- | --- | --- | --- | --- |
| **Ion** | | | |  | 1.00 | 21.46 || 1 | Water 1 | | 14:3670 | 2.15 | 1.00 | 32.87 | 2.53 (A:OP2 14:1773) |
| 2 | Water 2 | | 14:3920 | 2.17 | 1.00 | 41.49 | 3.08 (G:OP2 14:1973) |
| 3 | Water 3 | | 14:3636 | 2.17 | 1.00 | 30.03 | 2.46 (G:OP1 14:1828) |
| 4 | Water 4 | | 14:3871 | 2.17 | 1.00 | 40.20 | 2.94 (G:OP1 14:1828) |
| 5 | Water 5 | | 14:3684 | 2.19 | 1.00 | 40.17 | 2.55 (G:OP1 14:1772) |
| 6 | G | OP1 | 14:1828 | 2.56 | 1.00 | 31.51 |  |
| 7 | G | O5' | 14:1828 | 3.14 | 1.00 | 25.93 |  |
| 8 | A | OP2 | 14:1773 | 3.94 | 1.00 | 41.58 |  |
| 9 | G | OP1 | 14:1772 | 4.26 | 1.00 | 38.56 |  |
| 10 | G | OP2 | 14:1973 | 4.27 | 1.00 | 32.12 |  |
| 11 | G | OP2 | 14:1828 | 4.46 | 1.00 | 31.08 |  |
| 12 | C | O3' | 14:1827 | 4.49 | 1.00 | 27.27 |  |
| 13 | A | O5' | 14:1972 | 4.49 | 1.00 | 31.43 |  |
| 14 | A | OP2 | 14:1972 | 4.64 | 1.00 | 35.67 |  |
  |

  
**1^st^ shell ligand angles:**   
1-2: 90; 1-3: 175; 1-4: 90; 1-5: 87; 2-3: 89; 2-4: 177; 2-5: 93; 3-4: 92; 3-5: 88; 4-5: 91;   
  
 Contact(s): /O_w_/O_w_/O_w_/O_w_/O_w_/  
 Coded contact list: 5O_w_  
 Exclusion zone contact(s): /O5'/OP1/  
Consider NA in place of MG (1 valid contact(s) for NA vs. 0 for MG)  

### MG:14:3284

|  | Res. type | Atom | Residue | Distance | Occupancy | B-factor | Closest solute to 1^st^ shell water |
| --- | --- | --- | --- | --- | --- | --- | --- |
| **Ion** | | | |  | 1.00 | 34.09 || 1 | Water 1 | | 14:3623 | 2.16 | 1.00 | 35.62 | 2.40 (G:N7 14:2595) |
| 2 | Water 2 | | 14:3879 | 2.17 | 1.00 | 42.92 | 2.96 (G:O6 14:2595) |
| 3 | Water 3 | | 14:4010 | 2.17 | 1.00 | 47.61 | 3.39 (G:OP1 14:1904) |
| 4 | Water 4 | | 14:3613 | 2.17 | 1.00 | 37.88 | 2.38 (G:OP1 14:2597) |
| 5 | Water 5 | | 14:3853 | 2.19 | 1.00 | 43.17 | 2.89 (G:OP1 14:1904) |
| 6 | Water 6 | | 14:3981 | 2.19 | 1.00 | 47.75 | 3.27 (G:OP1 14:1904) |
| 7 | G | OP1 | 14:1904 | 3.90 | 1.00 | 45.95 |  |
| 8 | G | N7 | 14:2595 | 4.17 | 1.00 | 37.70 |  |
| 9 | G | OP1 | 14:2597 | 4.29 | 1.00 | 33.20 |  |
| 10 | G | O6 | 14:2595 | 4.54 | 1.00 | 41.60 |  |
| 11 | G | OP2 | 14:2597 | 4.68 | 1.00 | 33.60 |  |
| 12 | G | O3' | 14:1903 | 4.85 | 1.00 | 40.74 |  |
  |

  
**1^st^ shell ligand angles:**   
1-2: 88; 1-3: 176; 1-4: 90; 1-5: 92; 1-6: 90; 2-3: 89; 2-4: 90; 2-5: 179; 2-6: 91; 3-4: 92; 3-5: 91; 3-6: 87; 4-5: 89; 4-6: 179; 5-6: 90;   
  
 Contact(s): /O_w_/O_w_/O_w_/O_w_/O_w_/O_w_/  
 Coded contact list: 6O_w_  
MG hydration shell issue: water too close (< 2.4 Angs.)  

### MG:14:3285

|  | Res. type | Atom | Residue | Distance | Occupancy | B-factor | Closest solute to 1^st^ shell water |
| --- | --- | --- | --- | --- | --- | --- | --- |
| **Ion** | | | |  | 1.00 | 30.34 || 1 | Water 1 | | 14:3614 | 2.16 | 1.00 | 39.43 | 2.38 (G:OP2 14:2588) |
| 2 | Water 2 | | 14:3704 | 2.16 | 1.00 | 29.43 | 2.59 (A:N7 14:2589) |
| 3 | Water 3 | | 14:3683 | 2.17 | 1.00 | 38.71 | 2.55 (A:OP1 14:2587) |
| 4 | Water 4 | | 14:3916 | 2.18 | 1.00 | 28.84 | 3.06 (G:N7 14:2588) |
| 5 | Water 5 | | 19:412 | 2.18 | 1.00 | 33.86 | 3.90 (A:N7 14:2589) |
| 6 | G | OP2 | 14:2588 | 2.50 | 1.00 | 32.75 |  |
| 7 | A | N7 | 14:2589 | 3.92 | 1.00 | 25.12 |  |
| 8 | A | OP1 | 14:2587 | 3.95 | 1.00 | 42.89 |  |
| 9 | G | N7 | 14:2588 | 4.18 | 1.00 | 30.17 |  |
| 10 | G | O5' | 14:2588 | 4.23 | 1.00 | 30.79 |  |
| 11 | G | OP1 | 14:2588 | 4.37 | 1.00 | 35.82 |  |
| 12 | A | OP2 | 14:2589 | 4.57 | 1.00 | 28.72 |  |
| 13 | Water 6 | | 14:3649 | 4.71 | 1.00 | 31.99 | 2.49 (A:OP2 14:2589) |
| 14 | A | O5' | 14:2587 | 4.72 | 1.00 | 36.49 |  |
| 15 | A | O3' | 14:2587 | 4.95 | 1.00 | 37.49 |  |
| 16 | A | N6 | 14:2589 | 5.00 | 1.00 | 28.78 |  |
  |

  
**1^st^ shell ligand angles:**   
1-2: 88; 1-3: 90; 1-4: 177; 1-5: 85; 2-3: 176; 2-4: 92; 2-5: 89; 3-4: 90; 3-5: 94; 4-5: 98;   
  
 Contact(s): /O_w_/O_w_/O_w_/O_w_/O_w_/  
 Coded contact list: 5O_w_  
 Exclusion zone contact(s): /OP2/  
Consider NA in place of MG (1 valid contact(s) for NA vs. 0 for MG)  
MG hydration shell issue: water too close (< 2.4 Angs.)  

### MG:14:3286

|  | Res. type | Atom | Residue | Distance | Occupancy | B-factor | Closest solute to 1^st^ shell water |
| --- | --- | --- | --- | --- | --- | --- | --- |
| **Ion** | | | |  | 1.00 | 25.37 || 1 | Water 1 | | 14:3669 | 2.15 | 1.00 | 46.97 | 2.53 (C:OP2 14:1327) |
| 2 | Water 2 | | 14:3593 | 2.16 | 1.00 | 36.00 | 2.32 (C:OP2 14:1327) |
| 3 | Water 3 | | 14:3681 | 2.17 | 1.00 | 40.63 | 2.54 (G:O6 14:1328) |
| 4 | Water 4 | | 14:4004 | 2.18 | 1.00 | 40.07 | 3.36 (G:N7 14:1324) |
| 5 | Water 5 | | 14:3918 | 2.18 | 1.00 | 47.49 | 3.07 (U:OP1 14:1326) |
| 6 | C | OP2 | 14:1327 | 2.40 | 1.00 | 30.72 |  |
| 7 | U | OP1 | 14:1326 | 3.92 | 1.00 | 28.04 |  |
| 8 | U | OP2 | 14:1326 | 4.11 | 1.00 | 35.03 |  |
| 9 | C | OP1 | 14:1327 | 4.20 | 1.00 | 37.07 |  |
| 10 | C | O5' | 14:1327 | 4.22 | 1.00 | 45.17 |  |
| 11 | G | N7 | 14:1328 | 4.23 | 1.00 | 48.59 |  |
| 12 | G | N7 | 14:1324 | 4.24 | 1.00 | 46.85 |  |
| 13 | G | O6 | 14:1328 | 4.53 | 1.00 | 47.72 |  |
| 14 | U | O3' | 14:1326 | 4.81 | 1.00 | 35.55 |  |
| 15 | U | O5' | 14:1326 | 4.81 | 1.00 | 32.17 |  |
| 16 | G | OP2 | 14:1328 | 4.87 | 1.00 | 48.55 |  |
  |

  
**1^st^ shell ligand angles:**   
1-2: 80; 1-3: 93; 1-4: 93; 1-5: 175; 1-6: 67; 2-3: 171; 2-4: 94; 2-5: 95; 2-6: 61; 3-4: 92; 3-5: 92; 3-6: 112; 4-5: 88; 4-6: 149; 5-6: 110;   
  
 Contact(s): /O_w_/O_w_/O_w_/O_w_/O_w_/OP2/  
 Coded contact list: O_ph_5O_w_  
Large octahedral topology discrepancy!!!  
MG hydration shell issue: water too close (< 2.4 Angs.)  

### MG:14:3287

|  | Res. type | Atom | Residue | Distance | Occupancy | B-factor | Closest solute to 1^st^ shell water |
| --- | --- | --- | --- | --- | --- | --- | --- |
| **Ion** | | | |  | 1.00 | 39.67 || 1 | Water 1 | | 14:3607 | 2.16 | 1.00 | 46.57 | 2.36 (A:N1 14:821) |
| 2 | Water 2 | | 14:4059 | 2.17 | 1.00 | 50.53 | 3.83 (U:O4 14:568) |
| 3 | Water 3 | | 14:3734 | 2.18 | 1.00 | 38.42 | 2.66 (U:O5' 14:569) |
| 4 | Water 4 | | 14:3964 | 2.18 | 1.00 | 39.85 | 3.20 (A:OP1 14:973) |
| 5 | Water 5 | | 14:3624 | 2.18 | 1.00 | 48.03 | 2.40 (G:OP1 14:570) |
| 6 | Water 6 | | 14:3958 | 2.19 | 1.00 | 44.89 | 3.19 (A:N1 14:820) |
| 7 | G | O2' | 14:972 | 3.85 | 1.00 | 49.70 |  |
| 8 | A | N1 | 14:821 | 3.97 | 1.00 | 48.97 |  |
| 9 | G | OP2 | 14:570 | 4.04 | 1.00 | 44.96 |  |
| 10 | A | OP2 | 14:973 | 4.28 | 1.00 | 49.72 |  |
| 11 | G | OP1 | 14:570 | 4.31 | 1.00 | 44.11 |  |
| 12 | A | N6 | 14:820 | 4.35 | 1.00 | 50.00 |  |
| 13 | A | N1 | 14:820 | 4.40 | 1.00 | 46.70 |  |
| 14 | A | OP1 | 14:973 | 4.69 | 1.00 | 51.69 |  |
| 15 | G | O3' | 14:972 | 4.70 | 1.00 | 53.42 |  |
| 16 | U | O5' | 14:569 | 4.74 | 1.00 | 40.91 |  |
| 17 | U | O4 | 14:568 | 4.74 | 1.00 | 46.16 |  |
| 18 | U | OP2 | 14:569 | 4.84 | 1.00 | 39.44 |  |
  |

  
**1^st^ shell ligand angles:**   
1-2: 178; 1-3: 94; 1-4: 88; 1-5: 88; 1-6: 91; 2-3: 89; 2-4: 90; 2-5: 92; 2-6: 89; 3-4: 178; 3-5: 88; 3-6: 91; 4-5: 92; 4-6: 89; 5-6: 178;   
  
 Contact(s): /O_w_/O_w_/O_w_/O_w_/O_w_/O_w_/  
 Coded contact list: 6O_w_  
MG hydration shell issue: water too close (< 2.4 Angs.)  

### MG:14:3288

|  | Res. type | Atom | Residue | Distance | Occupancy | B-factor | Closest solute to 1^st^ shell water |
| --- | --- | --- | --- | --- | --- | --- | --- |
| **Ion** | | | |  | 1.00 | 39.91 || 1 | Water 1 | | 14:3603 | 2.16 | 1.00 | 39.27 | 2.35 (C:OP2 14:1333) |
| 2 | Water 2 | | 14:3894 | 2.17 | 1.00 | 42.07 | 3.00 (U:O4 14:1316) |
| 3 | Water 3 | | 14:3793 | 2.17 | 1.00 | 51.76 | 2.73 (G:N7 14:1334) |
| 4 | Water 4 | | 14:3771 | 2.17 | 1.00 | 49.08 | 2.70 (U:O4 14:1335) |
| 5 | Water 5 | | 14:3570 | 2.18 | 1.00 | 46.83 | 2.26 (G:OP2 14:1332) |
| 6 | Water 6 | | 14:3714 | 2.19 | 1.00 | 47.04 | 2.61 (G:O6 14:1334) |
| 7 | C | OP2 | 14:1333 | 3.66 | 1.00 | 50.73 |  |
| 8 | U | O4 | 14:1335 | 4.20 | 1.00 | 57.05 |  |
| 9 | G | OP2 | 14:1332 | 4.22 | 1.00 | 43.79 |  |
| 10 | G | O6 | 14:1334 | 4.28 | 1.00 | 53.54 |  |
| 11 | G | N7 | 14:1334 | 4.35 | 1.00 | 49.30 |  |
| 12 | C | N4 | 14:1333 | 4.46 | 1.00 | 52.63 |  |
| 13 | U | OP2 | 14:1313 | 4.50 | 1.00 | 46.13 |  |
| 14 | U | O4 | 14:1316 | 4.57 | 1.00 | 59.46 |  |
| 15 | G | OP2 | 14:1334 | 4.62 | 1.00 | 44.37 |  |
| 16 | A | N6 | 14:1317 | 4.98 | 1.00 | 55.56 |  |
  |

  
**1^st^ shell ligand angles:**   
1-2: 89; 1-3: 93; 1-4: 89; 1-5: 89; 1-6: 179; 2-3: 179; 2-4: 89; 2-5: 92; 2-6: 93; 3-4: 91; 3-5: 87; 3-6: 86; 4-5: 177; 4-6: 92; 5-6: 90;   
  
 Contact(s): /O_w_/O_w_/O_w_/O_w_/O_w_/O_w_/  
 Coded contact list: 6O_w_  
MG hydration shell issue: water too close (< 2.4 Angs.)  

### MG:14:3289

|  | Res. type | Atom | Residue | Distance | Occupancy | B-factor | Closest solute to 1^st^ shell water |
| --- | --- | --- | --- | --- | --- | --- | --- |
| **Ion** | | | |  | 1.00 | 51.32 || 1 | Water 1 | | 14:3590 | 2.14 | 1.00 | 40.98 | 2.31 (A:OP1 14:945) |
| 2 | Water 2 | | 14:3691 | 2.15 | 1.00 | 44.85 | 2.56 (G:N7 14:831) |
| 3 | Water 3 | | 14:3863 | 2.18 | 1.00 | 49.79 | 2.91 (G:O6 14:831) |
| 4 | Water 4 | | 14:3700 | 2.18 | 1.00 | 45.09 | 2.58 (A:OP1 14:945) |
| 5 | Water 5 | | 14:3988 | 2.18 | 1.00 | 56.06 | 3.29 (A:OP2 14:945) |
| 6 | Water 6 | | 14:3902 | 2.19 | 1.00 | 56.50 | 3.02 (A:O4' 14:829) |
| 7 | A | OP1 | 14:945 | 3.32 | 1.00 | 42.69 |  |
| 8 | A | OP2 | 14:945 | 4.03 | 1.00 | 47.29 |  |
| 9 | G | O3' | 14:944 | 4.14 | 1.00 | 47.46 |  |
| 10 | G | O6 | 14:831 | 4.42 | 1.00 | 47.52 |  |
| 11 | G | N7 | 14:831 | 4.46 | 1.00 | 47.15 |  |
| 12 | A | O4' | 14:829 | 4.61 | 1.00 | 41.16 |  |
| 13 | G | O2' | 14:944 | 4.72 | 1.00 | 60.10 |  |
| 14 | G | OP2 | 14:831 | 4.96 | 1.00 | 39.52 |  |
  |

  
**1^st^ shell ligand angles:**   
1-2: 83; 1-3: 87; 1-4: 94; 1-5: 93; 1-6: 177; 2-3: 85; 2-4: 93; 2-5: 176; 2-6: 94; 3-4: 177; 3-5: 94; 3-6: 93; 4-5: 89; 4-6: 85; 5-6: 90;   
  
 Contact(s): /O_w_/O_w_/O_w_/O_w_/O_w_/O_w_/  
 Coded contact list: 6O_w_  
 Exclusion zone contact(s): /OP1/  
Consider K in place of MG (1 valid contact(s) for K vs. 0 for MG)  
MG hydration shell issue: water too close (< 2.4 Angs.)  

### MG:14:3290

|  | Res. type | Atom | Residue | Distance | Occupancy | B-factor | Closest solute to 1^st^ shell water |
| --- | --- | --- | --- | --- | --- | --- | --- |
| **Ion** | | | |  | 1.00 | 36.05 || 1 | Water 1 | | 14:4050 | 2.16 | 1.00 | 31.49 | 3.78 (G:OP2 14:944) |
| 2 | Water 2 | | 14:3548 | 2.17 | 1.00 | 38.11 | 2.18 (U:OP1 14:568) |
| 3 | Water 3 | | 14:3671 | 2.17 | 1.00 | 50.15 | 2.53 (G:OP1 14:831) |
| 4 | Water 4 | | 14:3872 | 2.17 | 1.00 | 41.16 | 2.94 (G:OP1 14:944) |
| 5 | Water 5 | | 14:3998 | 2.18 | 1.00 | 47.02 | 3.32 (A:N6 14:2448) |
| 6 | Water 6 | | 14:3510 | 2.24 | 1.00 | 44.61 | 1.31 (A:N1 14:2448) |
| 7 | A | N1 | 14:2448 | 3.46 | 1.00 | 44.28 |  |
| 8 | A | N3 | 14:945 | 3.83 | 1.00 | 36.36 |  |
| 9 | A | N6 | 14:2448 | 3.87 | 1.00 | 41.24 |  |
| 10 | G | OP1 | 14:831 | 4.07 | 1.00 | 35.17 |  |
| 11 | U | OP1 | 14:568 | 4.32 | 1.00 | 42.44 |  |
| 12 | G | OP1 | 14:944 | 4.58 | 1.00 | 32.72 |  |
| 13 | G | O3' | 14:830 | 4.64 | 1.00 | 35.52 |  |
| 14 | G | OP2 | 14:944 | 4.82 | 1.00 | 43.53 |  |
  |

  
**1^st^ shell ligand angles:**   
1-2: 92; 1-3: 87; 1-4: 87; 1-5: 87; 1-6: 176; 2-3: 178; 2-4: 87; 2-5: 90; 2-6: 92; 3-4: 94; 3-5: 89; 3-6: 89; 4-5: 173; 4-6: 94; 5-6: 92;   
  
 Contact(s): /O_w_/O_w_/O_w_/O_w_/O_w_/O_w_/  
 Coded contact list: 6O_w_  
MG hydration shell issue: water too close (< 2.4 Angs.)  

### MG:14:3291

|  | Res. type | Atom | Residue | Distance | Occupancy | B-factor | Closest solute to 1^st^ shell water |
| --- | --- | --- | --- | --- | --- | --- | --- |
| **Ion** | | | |  | 1.00 | 58.71 || 1 | Water 1 | | 14:3629 | 2.16 | 1.00 | 46.80 | 2.41 (G:OP2 14:1380) |
| 2 | Water 2 | | 14:4038 | 2.18 | 1.00 | 52.21 | 3.62 (G:OP2 14:1380) |
| 3 | Water 3 | | 14:3725 | 2.18 | 1.00 | 55.33 | 2.63 (G:N7 14:1382) |
| 4 | Water 4 | | 14:3794 | 2.19 | 1.00 | 56.65 | 2.73 (G:OP2 14:1380) |
| 5 | Water 5 | | 14:3604 | 2.19 | 1.00 | 47.13 | 2.35 (G:O6 14:1381) |
| 6 | Water 6 | | 14:3633 | 2.20 | 1.00 | 54.47 | 2.42 (G:O6 14:1382) |
| 7 | G | O6 | 14:1382 | 3.27 | 1.00 | 51.50 |  |
| 8 | G | OP2 | 14:1380 | 3.44 | 1.00 | 55.37 |  |
| 9 | G | OP1 | 14:1347 | 4.13 | 1.00 | 54.79 |  |
| 10 | C | N4 | 14:1383 | 4.34 | 1.00 | 58.86 |  |
| 11 | G | N7 | 14:1382 | 4.35 | 1.00 | 43.69 |  |
| 12 | G | N7 | 14:1381 | 4.37 | 1.00 | 41.46 |  |
| 13 | G | O6 | 14:1381 | 4.45 | 1.00 | 43.25 |  |
| 14 | G | OP2 | 14:1381 | 4.91 | 1.00 | 40.13 |  |
| 15 | G | OP2 | 14:1347 | 4.94 | 1.00 | 39.91 |  |
  |

  
**1^st^ shell ligand angles:**   
1-2: 89; 1-3: 85; 1-4: 94; 1-5: 89; 1-6: 176; 2-3: 92; 2-4: 89; 2-5: 178; 2-6: 89; 3-4: 179; 3-5: 90; 3-6: 92; 4-5: 90; 4-6: 90; 5-6: 93;   
  
 Contact(s): /O_w_/O_w_/O_w_/O_w_/O_w_/O_w_/  
 Coded contact list: 6O_w_  
 Exclusion zone contact(s): /O6/  
Consider K in place of MG (1 valid contact(s) for K vs. 0 for MG)  
MG hydration shell issue: water too close (< 2.4 Angs.)  

### MG:14:3292

|  | Res. type | Atom | Residue | Distance | Occupancy | B-factor | Closest solute to 1^st^ shell water |
| --- | --- | --- | --- | --- | --- | --- | --- |
| **Ion** | | | |  | 1.00 | 29.77 || 1 | Water 1 | | 14:3525 | 2.15 | 1.00 | 34.40 | 2.03 (C:OP1 14:1314) |
| 2 | Water 2 | | 14:3621 | 2.18 | 1.00 | 45.60 | 2.39 (C:OP2 14:1315) |
| 3 | Water 3 | | 14:4066 | 2.19 | 1.00 | 55.30 | 3.93 (G:OP1 14:1332) |
| 4 | G | OP1 | 14:1332 | 2.44 | 1.00 | 41.03 |  |
| 5 | C | OP2 | 14:1315 | 2.50 | 1.00 | 35.26 |  |
| 6 | C | O5' | 14:1314 | 3.05 | 1.00 | 40.69 |  |
| 7 | C | OP1 | 14:1314 | 3.07 | 1.00 | 40.80 |  |
| 8 | A | O3' | 14:1331 | 3.88 | 1.00 | 38.71 |  |
| 9 | C | OP1 | 14:1315 | 3.90 | 1.00 | 41.90 |  |
| 10 | G | O5' | 14:1332 | 4.13 | 1.00 | 47.13 |  |
| 11 | U | O3' | 14:1313 | 4.50 | 1.00 | 44.17 |  |
| 12 | C | O3' | 14:1314 | 4.60 | 1.00 | 40.30 |  |
| 13 | C | O5' | 14:1315 | 4.71 | 1.00 | 50.49 |  |
| 14 | G | OP2 | 14:1332 | 4.81 | 1.00 | 43.79 |  |
| 15 | C | OP2 | 14:1314 | 4.89 | 1.00 | 38.94 |  |
  |

  
**1^st^ shell ligand angles:**   
1-2: 83; 1-3: 175; 2-3: 97;   
  
 Contact(s): /O_w_/O_w_/O_w_/  
 Coded contact list: 3O_w_  
 Exclusion zone contact(s): /O5'/OP1/OP1/OP2/  
Consider NA in place of MG (2 valid contact(s) for NA vs. 0 for MG)  
MG hydration shell issue: water too close (< 2.4 Angs.)  

### MG:14:3293

|  | Res. type | Atom | Residue | Distance | Occupancy | B-factor | Closest solute to 1^st^ shell water |
| --- | --- | --- | --- | --- | --- | --- | --- |
| **Ion** | | | |  | 1.00 | 60.25 || 1 | Water 1 | | 14:3699 | 2.16 | 1.00 | 37.60 | 2.58 (G:OP2 14:563) |
| 2 | Water 2 | | 14:3827 | 2.17 | 1.00 | 55.59 | 2.82 (A:OP2 14:572) |
| 3 | Water 3 | | 14:3807 | 2.18 | 1.00 | 55.36 | 2.76 (G:OP1 14:975) |
| 4 | Water 4 | | 14:3851 | 2.18 | 1.00 | 44.75 | 2.89 (G:OP1 14:563) |
| 5 | Water 5 | | 14:3908 | 2.19 | 1.00 | 65.57 | 3.04 (C:O3' 14:974A) |
| 6 | G | OP2 | 14:563 | 2.49 | 1.00 | 47.46 |  |
| 7 | G | OP1 | 14:563 | 3.91 | 1.00 | 48.36 |  |
| 8 | C | O2' | 14:974A | 4.07 | 1.00 | 63.85 |  |
| 9 | G | OP2 | 14:975 | 4.22 | 1.00 | 59.61 |  |
| 10 | U | O2' | 14:562 | 4.37 | 1.00 | 48.88 |  |
| 11 | G | O5' | 14:563 | 4.39 | 1.00 | 48.68 |  |
| 12 | C | O3' | 14:974A | 4.41 | 1.00 | 69.96 |  |
| 13 | A | OP2 | 14:572 | 4.65 | 1.00 | 42.73 |  |
| 14 | C | OP1 | 14:564 | 4.65 | 1.00 | 46.90 |  |
| 15 | G | OP1 | 14:975 | 4.68 | 1.00 | 64.40 |  |
| 16 | U | O3' | 14:562 | 4.73 | 1.00 | 45.22 |  |
| 17 | C | OP2 | 14:564 | 4.87 | 1.00 | 43.20 |  |
  |

  
**1^st^ shell ligand angles:**   
1-2: 84; 1-3: 92; 1-4: 91; 1-5: 176; 2-3: 93; 2-4: 171; 2-5: 92; 3-4: 95; 3-5: 87; 4-5: 92;   
  
 Contact(s): /O_w_/O_w_/O_w_/O_w_/O_w_/  
 Coded contact list: 5O_w_  
 Exclusion zone contact(s): /OP2/  
Consider NA in place of MG (1 valid contact(s) for NA vs. 0 for MG)  

### MG:14:3294

|  | Res. type | Atom | Residue | Distance | Occupancy | B-factor | Closest solute to 1^st^ shell water |
| --- | --- | --- | --- | --- | --- | --- | --- |
| **Ion** | | | |  | 1.00 | 51.69 || 1 | Water 1 | | 14:4019 | 2.16 | 1.00 | 54.82 | 3.46 (C:N4 14:2871) |
| 2 | Water 2 | | 14:3652 | 2.17 | 1.00 | 60.95 | 2.49 (G:O6 14:2843) |
| 3 | Water 3 | | 14:3686 | 2.17 | 1.00 | 59.80 | 2.55 (G:O6 14:2845) |
| 4 | Water 4 | | 14:3900 | 2.18 | 1.00 | 59.49 | 3.01 (G:N7 14:2843) |
| 5 | Water 5 | | 14:3940 | 2.18 | 1.00 | 60.96 | 3.13 (C:N4 14:2871) |
| 6 | Water 6 | | 14:3999 | 2.20 | 1.00 | 69.44 | 3.32 (G:N7 14:2844) |
| 7 | C | N4 | 14:2871 | 4.16 | 1.00 | 66.66 |  |
| 8 | G | N7 | 14:2843 | 4.21 | 1.00 | 67.59 |  |
| 9 | G | O6 | 14:2844 | 4.22 | 1.00 | 64.86 |  |
| 10 | G | O6 | 14:2845 | 4.22 | 1.00 | 59.51 |  |
| 11 | C | N3 | 14:2871 | 4.25 | 1.00 | 67.08 |  |
| 12 | G | O6 | 14:2843 | 4.25 | 1.00 | 58.96 |  |
| 13 | G | N7 | 14:2844 | 4.40 | 1.00 | 60.82 |  |
| 14 | G | N1 | 14:2845 | 4.91 | 1.00 | 64.34 |  |
  |

  
**1^st^ shell ligand angles:**   
1-2: 91; 1-3: 85; 1-4: 94; 1-5: 89; 1-6: 177; 2-3: 92; 2-4: 85; 2-5: 178; 2-6: 91; 3-4: 177; 3-5: 89; 3-6: 93; 4-5: 93; 4-6: 88; 5-6: 90;   
  
 Contact(s): /O_w_/O_w_/O_w_/O_w_/O_w_/O_w_/  
 Coded contact list: 6O_w_  

### MG:14:3295

|  | Res. type | Atom | Residue | Distance | Occupancy | B-factor | Closest solute to 1^st^ shell water |
| --- | --- | --- | --- | --- | --- | --- | --- |
| **Ion** | | | |  | 1.00 | 50.36 || 1 | Water 1 | | 14:3959 | 2.17 | 1.00 | 48.93 | 3.19 (A:OP2 14:1637) |
| 2 | Water 2 | | 14:3740 | 2.17 | 1.00 | 26.45 | 2.66 (C:OP2 14:1636) |
| 3 | Water 3 | | 14:3545 | 2.18 | 1.00 | 46.92 | 2.18 (A:OP2 14:1637) |
| 4 | Water 4 | | 14:3941 | 2.18 | 1.00 | 48.09 | 3.14 (G:OP1 14:1299) |
| 5 | Water 5 | | 14:3985 | 2.19 | 1.00 | 51.69 | 3.28 (G:OP1 14:1299) |
| 6 | Water 6 | | 14:3560 | 2.19 | 1.00 | 40.47 | 2.23 (C:OP2 14:1636) |
| 7 | C | OP2 | 14:1636 | 3.04 | 1.00 | 44.71 |  |
| 8 | A | OP2 | 14:1637 | 3.83 | 1.00 | 42.88 |  |
| 9 | G | OP1 | 14:1299 | 3.83 | 1.00 | 41.01 |  |
| 10 | C | OP1 | 14:1636 | 3.96 | 1.00 | 43.63 |  |
| 11 | C | O5' | 14:1636 | 4.10 | 1.00 | 35.48 |  |
| 12 | G | OP1 | 14:1635 | 4.38 | 1.00 | 38.95 |  |
| 13 | U | OP1 | 14:1300 | 4.60 | 1.00 | 36.20 |  |
  |

  
**1^st^ shell ligand angles:**   
1-2: 88; 1-3: 85; 1-4: 94; 1-5: 91; 1-6: 177; 2-3: 96; 2-4: 84; 2-5: 177; 2-6: 95; 3-4: 179; 3-5: 86; 3-6: 93; 4-5: 93; 4-6: 87; 5-6: 87;   
  
 Contact(s): /O_w_/O_w_/O_w_/O_w_/O_w_/O_w_/  
 Coded contact list: 6O_w_  
 Exclusion zone contact(s): /OP2/  
Consider K in place of MG (1 valid contact(s) for K vs. 0 for MG)  
MG hydration shell issue: water too close (< 2.4 Angs.)  

### MG:14:3296

|  | Res. type | Atom | Residue | Distance | Occupancy | B-factor | Closest solute to 1^st^ shell water |
| --- | --- | --- | --- | --- | --- | --- | --- |
| **Ion** | | | |  | 1.00 | 35.72 || 1 | Water 1 | | 14:3907 | 2.15 | 1.00 | 40.15 | 3.03 (U:OP2 14:1267) |
| 2 | Water 2 | | 14:3978 | 2.16 | 1.00 | 39.58 | 3.24 (G:O6 14:2009) |
| 3 | Water 3 | | 14:3804 | 2.17 | 1.00 | 47.12 | 2.75 (G:N7 14:2009) |
| 4 | Water 4 | | 14:3572 | 2.17 | 1.00 | 39.11 | 2.26 (U:O4 14:2011) |
| 5 | Water 5 | | 14:4061 | 2.19 | 1.00 | 60.54 | 3.85 (G:N7 14:2010) |
| 6 | Water 6 | | 14:3509 | 2.22 | 1.00 | 48.74 | 1.31 (G:N7 14:2010) |
| 7 | G | N7 | 14:2010 | 3.47 | 1.00 | 46.62 |  |
| 8 | U | O4 | 14:2011 | 4.01 | 1.00 | 39.57 |  |
| 9 | G | O6 | 14:2009 | 4.02 | 1.00 | 46.80 |  |
| 10 | G | N7 | 14:2009 | 4.03 | 1.00 | 44.50 |  |
| 11 | G | O6 | 14:2010 | 4.07 | 1.00 | 45.27 |  |
| 12 | U | OP2 | 14:1267 | 4.27 | 1.00 | 41.91 |  |
  |

  
**1^st^ shell ligand angles:**   
1-2: 89; 1-3: 83; 1-4: 90; 1-5: 90; 1-6: 178; 2-3: 93; 2-4: 88; 2-5: 178; 2-6: 92; 3-4: 173; 3-5: 86; 3-6: 96; 4-5: 93; 4-6: 90; 5-6: 89;   
  
 Contact(s): /O_w_/O_w_/O_w_/O_w_/O_w_/O_w_/  
 Coded contact list: 6O_w_  
MG hydration shell issue: water too close (< 2.4 Angs.)  

### MG:14:3297

|  | Res. type | Atom | Residue | Distance | Occupancy | B-factor | Closest solute to 1^st^ shell water |
| --- | --- | --- | --- | --- | --- | --- | --- |
| **Ion** | | | |  | 1.00 | 28.36 || 1 | Water 1 | | 14:3749 | 2.15 | 1.00 | 42.21 | 2.67 (G:O6 14:2454) |
| 2 | Water 2 | | 14:3577 | 2.16 | 1.00 | 41.78 | 2.27 (G:N7 14:2454) |
| 3 | Water 3 | | 14:3942 | 2.17 | 1.00 | 43.38 | 3.14 (A:N1 14:2497) |
| 4 | Water 4 | | 14:4043 | 2.17 | 1.00 | 49.82 | 3.67 (G:O6 14:2455) |
| 5 | Water 5 | | 14:3825 | 2.18 | 1.00 | 40.70 | 2.81 (A:OP2 14:2453) |
| 6 | Water 6 | | 14:3713 | 2.18 | 1.00 | 46.76 | 2.61 (A:N6 14:2453) |
| 7 | A | N7 | 14:2453 | 3.97 | 1.00 | 40.43 |  |
| 8 | G | O6 | 14:2454 | 4.10 | 1.00 | 42.84 |  |
| 9 | G | N7 | 14:2454 | 4.11 | 1.00 | 39.82 |  |
| 10 | G | O6 | 14:2455 | 4.27 | 1.00 | 48.13 |  |
| 11 | A | N1 | 14:2497 | 4.35 | 1.00 | 37.39 |  |
| 12 | C | OP2 | 14:2452 | 4.48 | 1.00 | 39.21 |  |
| 13 | A | N6 | 14:2453 | 4.75 | 1.00 | 35.09 |  |
| 14 | C | N4 | 14:2496 | 4.82 | 1.00 | 48.38 |  |
| 15 | A | OP2 | 14:2453 | 4.84 | 1.00 | 50.14 |  |
  |

  
**1^st^ shell ligand angles:**   
1-2: 90; 1-3: 88; 1-4: 90; 1-5: 179; 1-6: 89; 2-3: 178; 2-4: 91; 2-5: 91; 2-6: 88; 3-4: 90; 3-5: 91; 3-6: 91; 4-5: 90; 4-6: 179; 5-6: 91;   
  
 Contact(s): /O_w_/O_w_/O_w_/O_w_/O_w_/O_w_/  
 Coded contact list: 6O_w_  
MG hydration shell issue: water too close (< 2.4 Angs.)  

### MG:14:3298

|  | Res. type | Atom | Residue | Distance | Occupancy | B-factor | Closest solute to 1^st^ shell water |
| --- | --- | --- | --- | --- | --- | --- | --- |
| **Ion** | | | |  | 1.00 | 32.74 || 1 | Water 1 | | 14:3521 | 2.16 | 1.00 | 39.35 | 2.00 (C:OP1 14:453) |
| 2 | Water 2 | | 14:3578 | 2.16 | 1.00 | 43.13 | 2.27 (A:N7 14:449) |
| 3 | Water 3 | | 14:3517 | 2.17 | 1.00 | 38.16 | 1.98 (G:O6 14:450) |
| 4 | Water 4 | | 14:3516 | 2.17 | 1.00 | 40.81 | 1.95 (C:OP1 14:453) |
| 5 | Water 5 | | 14:4047 | 2.18 | 1.00 | 45.76 | 3.73 (A:OP1 14:471) |
| 6 | Water 6 | | 14:3856 | 2.19 | 1.00 | 49.08 | 2.90 (A:OP2 14:449) |
| 7 | C | OP1 | 14:453 | 2.60 | 1.00 | 33.90 |  |
| 8 | G | O6 | 14:450 | 2.84 | 1.00 | 41.51 |  |
| 9 | G | N7 | 14:450 | 3.97 | 1.00 | 39.45 |  |
| 10 | A | N7 | 14:449 | 4.29 | 1.00 | 40.15 |  |
| 11 | A | OP1 | 14:471 | 4.50 | 1.00 | 31.79 |  |
| 12 | A | OP2 | 14:449 | 4.70 | 1.00 | 29.08 |  |
| 13 | C | O5' | 14:453 | 4.73 | 1.00 | 43.63 |  |
| 14 | G | O3' | 14:452 | 4.83 | 1.00 | 33.71 |  |
| 15 | A | N6 | 14:449 | 4.84 | 1.00 | 37.98 |  |
  |

  
**1^st^ shell ligand angles:**   
1-2: 88; 1-3: 94; 1-4: 92; 1-5: 178; 1-6: 84; 2-3: 89; 2-4: 178; 2-5: 92; 2-6: 93; 3-4: 90; 3-5: 88; 3-6: 177; 4-5: 89; 4-6: 88; 5-6: 94;   
  
 Contact(s): /O_w_/O_w_/O_w_/O_w_/O_w_/O_w_/  
 Coded contact list: 6O_w_  
 Exclusion zone contact(s): /O6/OP1/  
Consider K in place of MG (2 valid contact(s) for K vs. 0 for MG)  
MG hydration shell issue: water too close (< 2.4 Angs.)  

### MG:14:3299

|  | Res. type | Atom | Residue | Distance | Occupancy | B-factor | Closest solute to 1^st^ shell water |
| --- | --- | --- | --- | --- | --- | --- | --- |
| **Ion** | | | |  | 1.00 | 48.20 || 1 | Water 1 | | 14:3609 | 2.17 | 1.00 | 44.73 | 2.36 (G:N7 14:1371) |
| 2 | Water 2 | | 14:3765 | 2.18 | 1.00 | 41.22 | 2.69 (G:OP2 14:1358) |
| 3 | Water 3 | | 14:3870 | 2.18 | 1.00 | 46.63 | 2.94 (G:N7 14:1358) |
| 4 | Water 4 | | 14:3594 | 2.18 | 1.00 | 47.25 | 2.32 (C:OP2 14:1370) |
| 5 | Water 5 | | 14:3589 | 2.18 | 1.00 | 38.63 | 2.31 (U:OP2 14:1357) |
| 6 | Water 6 | | 14:3877 | 2.19 | 1.00 | 43.91 | 2.95 (U:OP1 14:1357) |
| 7 | U | OP2 | 14:1357 | 3.59 | 1.00 | 41.27 |  |
| 8 | G | OP2 | 14:1358 | 3.87 | 1.00 | 43.02 |  |
| 9 | C | OP2 | 14:1370 | 3.98 | 1.00 | 43.80 |  |
| 10 | G | N7 | 14:1371 | 4.29 | 1.00 | 47.60 |  |
| 11 | G | O6 | 14:1371 | 4.44 | 1.00 | 39.49 |  |
| 12 | G | N7 | 14:1358 | 4.45 | 1.00 | 47.13 |  |
| 13 | U | O5' | 14:1357 | 4.62 | 1.00 | 44.14 |  |
| 14 | U | OP1 | 14:1357 | 4.78 | 1.00 | 47.15 |  |
  |

  
**1^st^ shell ligand angles:**   
1-2: 89; 1-3: 92; 1-4: 90; 1-5: 88; 1-6: 176; 2-3: 88; 2-4: 87; 2-5: 176; 2-6: 95; 3-4: 175; 3-5: 90; 3-6: 90; 4-5: 95; 4-6: 88; 5-6: 88;   
  
 Contact(s): /O_w_/O_w_/O_w_/O_w_/O_w_/O_w_/  
 Coded contact list: 6O_w_  
MG hydration shell issue: water too close (< 2.4 Angs.)  

### MG:14:3300

|  | Res. type | Atom | Residue | Distance | Occupancy | B-factor | Closest solute to 1^st^ shell water |
| --- | --- | --- | --- | --- | --- | --- | --- |
| **Ion** | | | |  | 1.00 | 61.04 || 1 | Water 1 | | 14:3645 | 2.17 | 1.00 | 61.04 | 2.48 (G:O6 14:81) |
| 2 | Water 2 | | 14:4063 | 2.18 | 1.00 | 57.81 | 4.60 (G:N7 14:80) |
| 3 | Water 3 | | 14:3938 | 2.18 | 1.00 | 63.36 | 3.12 (G:O6 14:80) |
| 4 | Water 4 | | 14:3830 | 2.18 | 1.00 | 69.83 | 2.83 (A:OP1 14:103) |
| 5 | Water 5 | | 14:3776 | 2.18 | 1.00 | 63.27 | 2.71 (G:N7 14:80) |
| 6 | Water 6 | | 14:3580 | 2.19 | 1.00 | 59.43 | 2.28 (G:O6 14:82) |
| 7 | G | O6 | 14:82 | 3.79 | 1.00 | 60.31 |  |
| 8 | G | N7 | 14:80 | 4.03 | 1.00 | 64.58 |  |
| 9 | G | O6 | 14:80 | 4.06 | 1.00 | 62.06 |  |
| 10 | G | N7 | 14:81 | 4.42 | 1.00 | 58.51 |  |
| 11 | A | OP1 | 14:103 | 4.62 | 1.00 | 55.19 |  |
| 12 | G | O6 | 14:81 | 4.63 | 1.00 | 56.65 |  |
| 13 | G | N1 | 14:82 | 4.92 | 1.00 | 63.98 |  |
  |

  
**1^st^ shell ligand angles:**   
1-2: 178; 1-3: 90; 1-4: 89; 1-5: 93; 1-6: 88; 2-3: 88; 2-4: 91; 2-5: 87; 2-6: 94; 3-4: 88; 3-5: 93; 3-6: 178; 4-5: 178; 4-6: 92; 5-6: 87;   
  
 Contact(s): /O_w_/O_w_/O_w_/O_w_/O_w_/O_w_/  
 Coded contact list: 6O_w_  

### MG:14:3301

|  | Res. type | Atom | Residue | Distance | Occupancy | B-factor | Closest solute to 1^st^ shell water |
| --- | --- | --- | --- | --- | --- | --- | --- |
| **Ion** | | | |  | 1.00 | 38.62 || 1 | Water 1 | | 14:3513 | 2.15 | 1.00 | 46.78 | 1.91 (U:OP2 14:511) |
| 2 | Water 2 | | 14:3502 | 2.16 | 1.00 | 53.95 | 1.30 (G:N7 14:512) |
| 3 | Water 3 | | 14:3622 | 2.17 | 1.00 | 52.17 | 2.40 (C:O3' 14:509) |
| 4 | Water 4 | | 14:3514 | 2.17 | 1.00 | 44.73 | 1.92 (U:OP2 14:511) |
| 5 | Water 5 | | 14:4032 | 2.19 | 1.00 | 51.20 | 3.55 (C:O2' 14:509) |
| 6 | Water 6 | | 14:3858 | 2.20 | 1.00 | 39.72 | 2.90 (G:N7 14:512) |
| 7 | U | OP2 | 14:511 | 2.63 | 1.00 | 46.37 |  |
| 8 | G | N7 | 14:512 | 2.83 | 1.00 | 59.27 |  |
| 9 | C | OP1 | 14:510 | 3.06 | 1.00 | 45.33 |  |
| 10 | C | O3' | 14:509 | 3.50 | 1.00 | 50.29 |  |
| 11 | G | O6 | 14:512 | 4.07 | 1.00 | 50.47 |  |
| 12 | C | O2' | 14:509 | 4.74 | 1.00 | 51.55 |  |
| 13 | U | OP1 | 14:511 | 4.77 | 1.00 | 54.15 |  |
| 14 | U | O5' | 14:511 | 4.83 | 1.00 | 49.40 |  |
| 15 | C | O3' | 14:510 | 4.84 | 1.00 | 44.28 |  |
| 16 | G | N9 | 14:512 | 4.93 | 1.00 | 48.61 |  |
| 17 | C | O5' | 14:510 | 4.96 | 1.00 | 51.48 |  |
  |

  
**1^st^ shell ligand angles:**   
1-2: 88; 1-3: 87; 1-4: 89; 1-5: 87; 1-6: 178; 2-3: 89; 2-4: 89; 2-5: 175; 2-6: 92; 3-4: 175; 3-5: 93; 3-6: 91; 4-5: 88; 4-6: 94; 5-6: 93;   
  
 Contact(s): /O_w_/O_w_/O_w_/O_w_/O_w_/O_w_/  
 Coded contact list: 6O_w_  
 Exclusion zone contact(s): /N7/OP1/OP2/  
Consider K in place of MG (3 valid contact(s) for K vs. 0 for MG)  
MG hydration shell issue: water too close (< 2.4 Angs.)  

### MG:14:3302

|  | Res. type | Atom | Residue | Distance | Occupancy | B-factor | Closest solute to 1^st^ shell water |
| --- | --- | --- | --- | --- | --- | --- | --- |
| **Ion** | | | |  | 1.00 | 29.50 || 1 | Water 1 | | 14:3717 | 2.15 | 1.00 | 36.55 | 2.61 (A:N3 14:2060) |
| 2 | Water 2 | | 14:3742 | 2.18 | 1.00 | 44.47 | 2.66 (G:OP2 14:2446) |
| 3 | Water 3 | | 14:3698 | 2.18 | 1.00 | 28.67 | 2.58 (G:OP2 14:2502) |
| 4 | Water 4 | | 14:3903 | 2.19 | 1.00 | 40.94 | 3.02 (G:OP2 14:2061) |
| 5 | G | OP2 | 14:2502 | 2.38 | 1.00 | 31.53 |  |
| 6 | G | OP2 | 14:2061 | 2.61 | 1.00 | 32.70 |  |
| 7 | G | OP1 | 14:2502 | 3.62 | 1.00 | 36.48 |  |
| 8 | A | O2' | 14:2060 | 4.12 | 1.00 | 42.03 |  |
| 9 | G | O5' | 14:2502 | 4.15 | 1.00 | 38.76 |  |
| 10 | G | N7 | 14:2446 | 4.23 | 1.00 | 38.52 |  |
| 11 | C | O2' | 14:2501 | 4.24 | 1.00 | 37.55 |  |
| 12 | G | OP2 | 14:2446 | 4.41 | 1.00 | 37.34 |  |
| 13 | A | O3' | 14:2060 | 4.42 | 1.00 | 36.65 |  |
| 14 | G | N7 | 14:2445 | 4.47 | 1.00 | 34.07 |  |
| 15 | G | OP1 | 14:2061 | 4.62 | 1.00 | 35.00 |  |
| 16 | A | N3 | 14:2060 | 4.68 | 1.00 | 38.03 |  |
| 17 | C | O3' | 14:2501 | 4.68 | 1.00 | 36.21 |  |
| 18 | G | O5' | 14:2061 | 4.98 | 1.00 | 34.99 |  |
  |

  
**1^st^ shell ligand angles:**   
1-2: 93; 1-3: 174; 1-4: 89; 1-5: 107; 2-3: 93; 2-4: 94; 2-5: 113; 3-4: 93; 3-5: 69; 4-5: 147;   
  
 Contact(s): /O_w_/O_w_/O_w_/O_w_/OP2/  
 Coded contact list: O_ph_4O_w_  
 Exclusion zone contact(s): /OP2/  
Large octahedral topology discrepancy!!!  

### MG:14:3303

|  | Res. type | Atom | Residue | Distance | Occupancy | B-factor | Closest solute to 1^st^ shell water |
| --- | --- | --- | --- | --- | --- | --- | --- |
| **Ion** | | | |  | 1.00 | 58.59 || 1 | Water 1 | | 14:3792 | 2.16 | 1.00 | 70.05 | 2.73 (G:OP2 14:82) |
| 2 | Water 2 | | 14:4100 | 2.18 | 1.00 | 61.76 | 5.33 (G:O6 14:83) |
| 3 | Water 3 | | 14:4084 | 2.18 | 1.00 | 69.92 | 4.22 (G:N7 14:83) |
| 4 | Water 4 | | 14:4055 | 2.18 | 1.00 | 64.44 | 3.82 (G:O6 14:83) |
| 5 | Water 5 | | 14:3710 | 2.19 | 1.00 | 60.58 | 2.61 (G:O6 14:83) |
| 6 | Water 6 | | 14:3507 | 2.22 | 1.00 | 59.91 | 1.31 (G:N7 14:83) |
| 7 | G | N7 | 14:83 | 3.51 | 1.00 | 61.14 |  |
| 8 | G | O6 | 14:83 | 3.74 | 1.00 | 64.54 |  |
| 9 | G | N7 | 14:82 | 4.58 | 1.00 | 61.85 |  |
| 10 | G | OP2 | 14:82 | 4.82 | 1.00 | 60.57 |  |
  |

  
**1^st^ shell ligand angles:**   
1-2: 84; 1-3: 90; 1-4: 176; 1-5: 87; 1-6: 96; 2-3: 88; 2-4: 93; 2-5: 89; 2-6: 179; 3-4: 87; 3-5: 176; 3-6: 93; 4-5: 95; 4-6: 87; 5-6: 91;   
  
 Contact(s): /O_w_/O_w_/O_w_/O_w_/O_w_/O_w_/  
 Coded contact list: 6O_w_  

### MG:14:3304

|  | Res. type | Atom | Residue | Distance | Occupancy | B-factor | Closest solute to 1^st^ shell water |
| --- | --- | --- | --- | --- | --- | --- | --- |
| **Ion** | | | |  | 1.00 | 63.58 || 1 | Water 1 | | 14:4009 | 2.18 | 1.00 | 64.03 | 3.39 (A:OP2 14:1854) |
| 2 | Water 2 | | 14:3568 | 2.18 | 1.00 | 69.63 | 2.25 (G:O6 14:1855) |
| 3 | Water 3 | | 14:3797 | 2.18 | 1.00 | 67.68 | 2.74 (G:N7 14:1855) |
| 4 | Water 4 | | 14:3930 | 2.18 | 1.00 | 64.28 | 3.10 (G:N1 14:1857) |
| 5 | Water 5 | | 14:4067 | 2.19 | 1.00 | 66.18 | 3.95 (G:OP2 14:1855) |
| 6 | Water 6 | | 14:3951 | 2.19 | 1.00 | 71.97 | 3.17 (G:O6 14:1857) |
| 7 | G | O6 | 14:1856 | 4.00 | 1.00 | 75.64 |  |
| 8 | G | N7 | 14:1855 | 4.07 | 1.00 | 68.09 |  |
| 9 | G | O6 | 14:1855 | 4.23 | 1.00 | 72.45 |  |
| 10 | A | OP2 | 14:1854 | 4.35 | 1.00 | 56.99 |  |
| 11 | G | O6 | 14:1857 | 4.50 | 1.00 | 83.35 |  |
| 12 | G | OP2 | 14:1855 | 4.60 | 1.00 | 66.64 |  |
| 13 | G | N1 | 14:1857 | 4.75 | 1.00 | 79.75 |  |
| 14 | G | N7 | 14:1856 | 4.81 | 1.00 | 78.13 |  |
  |

  
**1^st^ shell ligand angles:**   
1-2: 92; 1-3: 86; 1-4: 91; 1-5: 90; 1-6: 177; 2-3: 88; 2-4: 89; 2-5: 178; 2-6: 89; 3-4: 175; 3-5: 92; 3-6: 92; 4-5: 91; 4-6: 92; 5-6: 88;   
  
 Contact(s): /O_w_/O_w_/O_w_/O_w_/O_w_/O_w_/  
 Coded contact list: 6O_w_  
MG hydration shell issue: water too close (< 2.4 Angs.)  

### MG:14:3305

|  | Res. type | Atom | Residue | Distance | Occupancy | B-factor | Closest solute to 1^st^ shell water |
| --- | --- | --- | --- | --- | --- | --- | --- |
| **Ion** | | | |  | 1.00 | 40.11 || 1 | Water 1 | | 14:3543 | 2.17 | 1.00 | 31.74 | 2.17 (G:OP1 14:805) |
| 2 | Water 2 | | 35:202 | 2.18 | 1.00 | 40.09 | 2.35 (SER:O 35:42) |
| 3 | Water 3 | | 35:207 | 2.18 | 1.00 | 50.08 | 3.17 (GLY:O 35:44) |
| 4 | Water 4 | | 14:3881 | 2.19 | 1.00 | 37.61 | 2.96 (A:OP1 14:804) |
| 5 | A | OP1 | 14:804 | 2.49 | 1.00 | 43.91 |  |
| 6 | G | OP1 | 14:805 | 2.81 | 1.00 | 28.57 |  |
| 7 | A | OP2 | 14:804 | 3.58 | 1.00 | 34.42 |  |
| 8 | GLY | O | 35:44 | 4.14 | 1.00 | 42.16 |  |
| 9 | A | O5' | 14:804 | 4.15 | 1.00 | 32.58 |  |
| 10 | SER | O | 35:42 | 4.29 | 1.00 | 36.24 |  |
| 11 | U | O3' | 14:803 | 4.54 | 1.00 | 33.50 |  |
| 12 | GLY | N | 35:44 | 4.62 | 1.00 | 41.95 |  |
| 13 | G | O5' | 14:805 | 4.88 | 1.00 | 35.30 |  |
| 14 | G | OP2 | 14:805 | 4.97 | 1.00 | 38.15 |  |
  |

  
**1^st^ shell ligand angles:**   
1-2: 89; 1-3: 104; 1-4: 91; 2-3: 90; 2-4: 177; 3-4: 87;   
  
 Contact(s): /O_w_/O_w_/O_w_/O_w_/  
 Coded contact list: 4O_w_  
 Exclusion zone contact(s): /OP1/OP1/  
Consider NA in place of MG (1 valid contact(s) for NA vs. 0 for MG)  
MG hydration shell issue: water too close (< 2.4 Angs.)  

### MG:14:3306

|  | Res. type | Atom | Residue | Distance | Occupancy | B-factor | Closest solute to 1^st^ shell water |
| --- | --- | --- | --- | --- | --- | --- | --- |
| **Ion** | | | |  | 1.00 | 32.64 || 1 | Water 1 | | 14:3764 | 2.17 | 1.00 | 26.90 | 2.69 (U:OP2 14:1778) |
| 2 | Water 2 | | 14:3927 | 2.18 | 1.00 | 35.91 | 3.10 (G:OP1 14:739) |
| 3 | Water 3 | | 14:3969 | 2.19 | 1.00 | 38.93 | 3.21 (G:OP2 14:739) |
| 4 | Water 4 | | 14:3524 | 2.19 | 1.00 | 38.44 | 2.03 (U:OP2 14:1779) |
| 5 | Water 5 | | 14:3979 | 2.19 | 1.00 | 34.94 | 3.25 (U:OP1 14:1778) |
| 6 | G | OP1 | 14:739 | 2.41 | 1.00 | 34.10 |  |
| 7 | U | OP2 | 14:1779 | 3.83 | 1.00 | 26.45 |  |
| 8 | G | OP2 | 14:739 | 3.86 | 1.00 | 38.66 |  |
| 9 | G | O3' | 14:738 | 4.07 | 1.00 | 38.94 |  |
| 10 | A | N1 | 14:1784 | 4.24 | 1.00 | 33.65 |  |
| 11 | U | OP1 | 14:1778 | 4.31 | 1.00 | 29.40 |  |
| 12 | G | O5' | 14:739 | 4.60 | 1.00 | 35.17 |  |
| 13 | U | OP2 | 14:1778 | 4.69 | 1.00 | 35.26 |  |
| 14 | U | O5' | 14:1778 | 4.86 | 1.00 | 30.73 |  |
| 15 | A | N6 | 14:1784 | 4.96 | 1.00 | 32.11 |  |
  |

  
**1^st^ shell ligand angles:**   
1-2: 86; 1-3: 174; 1-4: 92; 1-5: 95; 2-3: 92; 2-4: 176; 2-5: 94; 3-4: 89; 3-5: 90; 4-5: 89;   
  
 Contact(s): /O_w_/O_w_/O_w_/O_w_/O_w_/  
 Coded contact list: 5O_w_  
 Exclusion zone contact(s): /OP1/  
Consider NA in place of MG (1 valid contact(s) for NA vs. 0 for MG)  
MG hydration shell issue: water too close (< 2.4 Angs.)  

### MG:14:3307

|  | Res. type | Atom | Residue | Distance | Occupancy | B-factor | Closest solute to 1^st^ shell water |
| --- | --- | --- | --- | --- | --- | --- | --- |
| **Ion** | | | |  | 1.00 | 36.01 || 1 | Water 1 | | 14:3917 | 2.17 | 1.00 | 45.08 | 3.07 (G:O6 14:2280) |
| 2 | Water 2 | | 14:4102 | 2.17 | 1.00 | 47.92 | 5.62 (C:N4 14:2281) |
| 3 | Water 3 | | 14:4053 | 2.17 | 1.00 | 48.94 | 3.80 (G:N7 14:2280) |
| 4 | Water 4 | | 14:3876 | 2.17 | 1.00 | 46.30 | 2.95 (C:N4 14:2281) |
| 5 | Water 5 | | E5:101 | 2.18 | 1.00 | 53.40 | 3.39 (ARG:NH1 E5:14) |
| 6 | Water 6 | | 14:4064 | 2.18 | 1.00 | 55.13 | 3.88 (C:N4 14:2281) |
| 7 | G | N7 | 14:2280 | 4.38 | 1.00 | 51.80 |  |
| 8 | ARG | NH1 | E5:14 | 4.40 | 1.00 | 44.35 |  |
| 9 | C | N4 | 14:2281 | 4.44 | 1.00 | 55.61 |  |
| 10 | G | O6 | 14:2280 | 4.90 | 1.00 | 51.57 |  |
  |

  
**1^st^ shell ligand angles:**   
1-2: 178; 1-3: 90; 1-4: 87; 1-5: 91; 1-6: 92; 2-3: 90; 2-4: 91; 2-5: 90; 2-6: 87; 3-4: 94; 3-5: 86; 3-6: 177; 4-5: 178; 4-6: 87; 5-6: 93;   
  
 Contact(s): /O_w_/O_w_/O_w_/O_w_/O_w_/O_w_/  
 Coded contact list: 6O_w_  

### MG:14:3308

|  | Res. type | Atom | Residue | Distance | Occupancy | B-factor | Closest solute to 1^st^ shell water |
| --- | --- | --- | --- | --- | --- | --- | --- |
| **Ion** | | | |  | 1.00 | 77.22 || 1 | Water 1 | | 14:4080 | 2.17 | 1.00 | 60.63 | 4.17 (G:OP2 14:341) |
| 2 | Water 2 | | 14:3838 | 2.18 | 1.00 | 70.73 | 2.84 (G:OP2 14:341) |
| 3 | Water 3 | | 14:4049 | 2.18 | 1.00 | 56.38 | 3.78 (G:O6 14:342) |
| 4 | Water 4 | | 14:3790 | 2.18 | 1.00 | 73.67 | 2.73 (G:N7 14:342) |
| 5 | Water 5 | | 14:3678 | 2.19 | 1.00 | 73.39 | 2.54 (G:O6 14:342) |
| 6 | Water 6 | | 14:4098 | 2.19 | 1.00 | 73.01 | 4.92 (A:OP2 14:340) |
| 7 | G | O6 | 14:342 | 4.13 | 1.00 | 70.74 |  |
| 8 | G | N7 | 14:342 | 4.19 | 1.00 | 74.52 |  |
| 9 | G | OP2 | 14:341 | 4.33 | 1.00 | 66.20 |  |
| 10 | G | N7 | 14:341 | 4.56 | 1.00 | 73.67 |  |
| 11 | A | OP2 | 14:340 | 4.81 | 1.00 | 57.09 |  |
  |

  
**1^st^ shell ligand angles:**   
1-2: 92; 1-3: 89; 1-4: 86; 1-5: 177; 1-6: 92; 2-3: 178; 2-4: 89; 2-5: 86; 2-6: 91; 3-4: 92; 3-5: 93; 3-6: 88; 4-5: 92; 4-6: 178; 5-6: 91;   
  
 Contact(s): /O_w_/O_w_/O_w_/O_w_/O_w_/O_w_/  
 Coded contact list: 6O_w_  

### MG:14:3309

|  | Res. type | Atom | Residue | Distance | Occupancy | B-factor | Closest solute to 1^st^ shell water |
| --- | --- | --- | --- | --- | --- | --- | --- |
| **Ion** | | | |  | 1.00 | 26.49 || 1 | Water 1 | | 14:3587 | 2.16 | 1.00 | 27.81 | 2.30 (C:OP1 14:1782) |
| 2 | Water 2 | | 14:3852 | 2.19 | 1.00 | 37.67 | 2.89 (A:OP1 14:1783) |
| 3 | Water 3 | | 14:3616 | 2.20 | 1.00 | 36.36 | 2.38 (A:OP2 14:1783) |
| 4 | Water 4 | | 14:3511 | 2.20 | 1.00 | 39.89 | 1.75 (A:OP1 14:1780) |
| 5 | C | OP1 | 14:1782 | 2.32 | 1.00 | 27.88 |  |
| 6 | A | OP2 | 14:1783 | 2.68 | 1.00 | 32.11 |  |
| 7 | A | OP1 | 14:1780 | 3.03 | 1.00 | 30.29 |  |
| 8 | A | OP2 | 14:1780 | 3.22 | 1.00 | 36.60 |  |
| 9 | C | OP2 | 14:1781 | 3.62 | 1.00 | 34.87 |  |
| 10 | A | OP1 | 14:1783 | 3.74 | 1.00 | 33.65 |  |
| 11 | C | O3' | 14:1781 | 4.32 | 1.00 | 40.43 |  |
| 12 | C | O5' | 14:1782 | 4.43 | 1.00 | 29.62 |  |
| 13 | Water 5 | | 14:3697 | 4.49 | 1.00 | 25.93 | 2.60 (A:OP1 14:1783) |
| 14 | U | O3' | 14:1779 | 4.50 | 1.00 | 40.70 |  |
| 15 | C | O3' | 14:1782 | 4.71 | 1.00 | 33.39 |  |
| 16 | A | O5' | 14:1783 | 4.75 | 1.00 | 33.91 |  |
| 17 | C | OP2 | 14:1782 | 4.79 | 1.00 | 40.94 |  |
| 18 | A | O5' | 14:1780 | 4.80 | 1.00 | 36.48 |  |
  |

  
**1^st^ shell ligand angles:**   
1-2: 172; 1-3: 83; 1-4: 101; 1-5: 62; 2-3: 99; 2-4: 88; 2-5: 111; 3-4: 86; 3-5: 121; 4-5: 142;   
  
 Contact(s): /O_w_/O_w_/O_w_/O_w_/OP1/  
 Coded contact list: O_ph_4O_w_  
 Exclusion zone contact(s): /OP1/OP2/OP2/  
Large octahedral topology discrepancy!!!  
Consider K in place of MG (3 valid contact(s) for K vs. 1 for MG)  
MG hydration shell issue: water too close (< 2.4 Angs.)  

### MG:14:3310

|  | Res. type | Atom | Residue | Distance | Occupancy | B-factor | Closest solute to 1^st^ shell water |
| --- | --- | --- | --- | --- | --- | --- | --- |
| **Ion** | | | |  | 1.00 | 25.88 || 1 | Water 1 | | 19:402 | 2.15 | 1.00 | 24.26 | 2.14 (PRO:O 19:228) |
| 2 | Water 2 | | 14:3596 | 2.15 | 1.00 | 31.57 | 2.33 (C:OP2 14:2240) |
| 3 | Water 3 | | 14:3709 | 2.17 | 1.00 | 24.71 | 2.60 (G:OP2 14:2239) |
| 4 | Water 4 | | 14:3631 | 2.18 | 1.00 | 38.51 | 2.42 (U:OP2 14:2075) |
| 5 | Water 5 | | 14:4022 | 2.20 | 1.00 | 29.50 | 3.48 (U:O5' 14:2074) |
| 6 | U | OP1 | 14:2074 | 2.41 | 1.00 | 21.78 |  |
| 7 | U | O5' | 14:2074 | 3.60 | 1.00 | 27.77 |  |
| 8 | G | OP2 | 14:2239 | 3.67 | 1.00 | 29.75 |  |
| 9 | U | OP2 | 14:2075 | 3.97 | 1.00 | 28.87 |  |
| 10 | HIS | O | 19:231 | 4.22 | 1.00 | 27.74 |  |
| 11 | G | O3' | 14:2238 | 4.23 | 1.00 | 30.79 |  |
| 12 | PRO | O | 19:228 | 4.23 | 1.00 | 29.71 |  |
| 13 | G | O2' | 14:2238 | 4.36 | 1.00 | 26.58 |  |
| 14 | ARG | NH1 | 19:244 | 4.42 | 1.00 | 30.89 |  |
| 15 | C | OP2 | 14:2240 | 4.47 | 1.00 | 32.68 |  |
| 16 | C | O3' | 14:2073 | 4.60 | 1.00 | 22.22 |  |
| 17 | U | OP2 | 14:2074 | 4.62 | 1.00 | 25.59 |  |
| 18 | PRO | N | 19:232 | 4.87 | 1.00 | 25.35 |  |
| 19 | G | O5' | 14:2239 | 4.97 | 1.00 | 33.17 |  |
  |

  
**1^st^ shell ligand angles:**   
1-2: 94; 1-3: 85; 1-4: 179; 1-5: 86; 2-3: 87; 2-4: 85; 2-5: 176; 3-4: 95; 3-5: 97; 4-5: 94;   
  
 Contact(s): /O_w_/O_w_/O_w_/O_w_/O_w_/  
 Coded contact list: 5O_w_  
 Exclusion zone contact(s): /OP1/  
Consider NA in place of MG (1 valid contact(s) for NA vs. 0 for MG)  
MG hydration shell issue: water too close (< 2.4 Angs.)  

### MG:14:3311

|  | Res. type | Atom | Residue | Distance | Occupancy | B-factor | Closest solute to 1^st^ shell water |
| --- | --- | --- | --- | --- | --- | --- | --- |
| **Ion** | | | |  | 1.00 | 31.98 || 1 | Water 1 | | 14:3659 | 2.16 | 1.00 | 50.31 | 2.51 (C:OP1 14:1638) |
| 2 | Water 2 | | 14:3679 | 2.16 | 1.00 | 29.62 | 2.54 (C:OP2 14:1297) |
| 3 | Water 3 | | 14:4008 | 2.17 | 1.00 | 35.90 | 3.38 (C:OP2 14:1638) |
| 4 | Water 4 | | 14:3788 | 2.17 | 1.00 | 37.61 | 2.73 (G:O6 14:1299) |
| 5 | C | OP2 | 14:1298 | 2.39 | 1.00 | 40.93 |  |
| 6 | U | OP2 | 14:1639 | 2.43 | 1.00 | 39.24 |  |
| 7 | C | OP1 | 14:1298 | 3.58 | 1.00 | 38.31 |  |
| 8 | C | O5' | 14:1298 | 4.04 | 1.00 | 33.52 |  |
| 9 | C | O5' | 14:1638 | 4.07 | 1.00 | 40.44 |  |
| 10 | U | OP1 | 14:1639 | 4.36 | 1.00 | 40.23 |  |
| 11 | C | OP2 | 14:1638 | 4.48 | 1.00 | 45.19 |  |
| 12 | C | O3' | 14:1638 | 4.48 | 1.00 | 41.67 |  |
| 13 | C | O3' | 14:1297 | 4.49 | 1.00 | 38.85 |  |
| 14 | C | OP1 | 14:1638 | 4.53 | 1.00 | 43.60 |  |
| 15 | C | OP2 | 14:1297 | 4.65 | 1.00 | 38.15 |  |
| 16 | U | O5' | 14:1639 | 4.68 | 1.00 | 39.25 |  |
| 17 | G | O6 | 14:1299 | 4.69 | 1.00 | 39.26 |  |
| 18 | G | N7 | 14:1299 | 4.73 | 1.00 | 42.33 |  |
| 19 | C | O5' | 14:1297 | 4.90 | 1.00 | 41.68 |  |
| 20 | C | O2' | 14:2710 | 4.98 | 1.00 | 44.15 |  |
  |

  
**1^st^ shell ligand angles:**   
1-2: 87; 1-3: 94; 1-4: 175; 1-5: 99; 2-3: 179; 2-4: 92; 2-5: 75; 3-4: 87; 3-5: 105; 4-5: 76;   
  
 Contact(s): /O_w_/O_w_/O_w_/O_w_/OP2/  
 Coded contact list: O_ph_4O_w_  
 Exclusion zone contact(s): /OP2/  

### MG:14:3312

|  | Res. type | Atom | Residue | Distance | Occupancy | B-factor | Closest solute to 1^st^ shell water |
| --- | --- | --- | --- | --- | --- | --- | --- |
| **Ion** | | | |  | 1.00 | 38.40 || 1 | Water 1 | | 14:3696 | 2.17 | 1.00 | 42.85 | 2.57 (A:O5' 14:586) |
| 2 | Water 2 | | 14:3728 | 2.17 | 1.00 | 42.42 | 2.64 (A:OP1 14:586) |
| 3 | Water 3 | | 14:3796 | 2.18 | 1.00 | 47.33 | 2.74 (G:OP1 14:1250) |
| 4 | Water 4 | | 14:3786 | 2.18 | 1.00 | 46.81 | 2.72 (G:OP1 14:585) |
| 5 | U | OP1 | 14:1249 | 2.43 | 1.00 | 45.38 |  |
| 6 | A | OP2 | 14:586 | 2.70 | 1.00 | 42.54 |  |
| 7 | A | OP1 | 14:586 | 3.14 | 1.00 | 36.85 |  |
| 8 | A | O5' | 14:586 | 3.75 | 1.00 | 40.19 |  |
| 9 | G | O3' | 14:1248 | 3.98 | 1.00 | 41.85 |  |
| 10 | G | OP1 | 14:585 | 4.01 | 1.00 | 38.92 |  |
| 11 | U | OP2 | 14:1249 | 4.37 | 1.00 | 46.82 |  |
| 12 | G | OP1 | 14:1250 | 4.61 | 1.00 | 41.71 |  |
| 13 | G | O3' | 14:585 | 4.68 | 1.00 | 35.07 |  |
| 14 | U | O4 | 14:811 | 4.70 | 1.00 | 37.69 |  |
| 15 | G | O5' | 14:585 | 4.72 | 1.00 | 38.89 |  |
| 16 | U | O5' | 14:1249 | 4.73 | 1.00 | 46.38 |  |
| 17 | U | O3' | 14:1249 | 4.77 | 1.00 | 49.30 |  |
| 18 | G | OP2 | 14:585 | 5.00 | 1.00 | 42.44 |  |
  |

  
**1^st^ shell ligand angles:**   
1-2: 87; 1-3: 91; 1-4: 173; 2-3: 178; 2-4: 96; 3-4: 86;   
  
 Contact(s): /O_w_/O_w_/O_w_/O_w_/  
 Coded contact list: 4O_w_  
 Exclusion zone contact(s): /OP1/OP1/OP2/  
Consider K in place of MG (2 valid contact(s) for K vs. 0 for MG)  

### MG:14:3313

|  | Res. type | Atom | Residue | Distance | Occupancy | B-factor | Closest solute to 1^st^ shell water |
| --- | --- | --- | --- | --- | --- | --- | --- |
| **Ion** | | | |  | 1.00 | 33.15 || 1 | Water 1 | | 14:3859 | 2.16 | 1.00 | 39.66 | 2.90 (G:N7 14:792) |
| 2 | Water 2 | | 14:3531 | 2.17 | 1.00 | 37.41 | 2.07 (G:OP2 14:792) |
| 3 | Water 3 | | 14:3860 | 2.18 | 1.00 | 43.97 | 2.90 (G:OP2 14:785) |
| 4 | Water 4 | | 14:3963 | 2.18 | 1.00 | 58.67 | 3.20 (G:OP2 14:785) |
| 5 | Water 5 | | 14:3905 | 2.19 | 1.00 | 37.90 | 3.03 (G:OP2 14:792) |
| 6 | G | OP2 | 14:785 | 2.40 | 1.00 | 32.03 |  |
| 7 | G | OP2 | 14:792 | 3.52 | 1.00 | 29.14 |  |
| 8 | A | O3' | 14:784 | 3.76 | 1.00 | 29.70 |  |
| 9 | C | O2' | 14:2440 | 4.45 | 1.00 | 39.22 |  |
| 10 | G | OP1 | 14:785 | 4.47 | 1.00 | 39.51 |  |
| 11 | G | O5' | 14:785 | 4.55 | 1.00 | 32.23 |  |
| 12 | G | N7 | 14:792 | 4.59 | 1.00 | 35.63 |  |
| 13 | G | O5' | 14:792 | 4.67 | 1.00 | 24.11 |  |
| 14 | C | O3' | 14:791 | 4.98 | 1.00 | 33.87 |  |
| 15 | A | O2' | 14:784 | 5.00 | 1.00 | 38.90 |  |
  |

  
**1^st^ shell ligand angles:**   
1-2: 90; 1-3: 86; 1-4: 176; 1-5: 96; 1-6: 88; 2-3: 94; 2-4: 93; 2-5: 88; 2-6: 172; 3-4: 91; 3-5: 178; 3-6: 79; 4-5: 87; 4-6: 89; 5-6: 100;   
  
 Contact(s): /O_w_/O_w_/O_w_/O_w_/O_w_/OP2/  
 Coded contact list: O_ph_5O_w_  
MG hydration shell issue: water too close (< 2.4 Angs.)  

### MG:14:3314

|  | Res. type | Atom | Residue | Distance | Occupancy | B-factor | Closest solute to 1^st^ shell water |
| --- | --- | --- | --- | --- | --- | --- | --- |
| **Ion** | | | |  | 1.00 | 28.45 || 1 | Water 1 | | 14:3702 | 2.17 | 1.00 | 34.55 | 2.59 (G:O2' 14:2447) |
| 2 | Water 2 | | 14:3690 | 2.18 | 1.00 | 38.91 | 2.56 (A:OP2 14:2448) |
| 3 | Water 3 | | 14:3957 | 2.19 | 1.00 | 42.12 | 3.18 (A:O2' 14:2497) |
| 4 | C | OP2 | 14:2499 | 2.54 | 1.00 | 33.64 |  |
| 5 | C | OP2 | 14:2498 | 2.60 | 1.00 | 44.53 |  |
| 6 | A | OP1 | 14:2448 | 2.85 | 1.00 | 30.44 |  |
| 7 | A | O3' | 14:2497 | 2.91 | 1.00 | 34.73 |  |
| 8 | A | OP2 | 14:2448 | 3.02 | 1.00 | 48.63 |  |
| 9 | A | O2' | 14:2497 | 3.68 | 1.00 | 44.23 |  |
| 10 | C | O5' | 14:2498 | 3.97 | 1.00 | 33.58 |  |
| 11 | U | OP2 | 14:2449 | 4.38 | 1.00 | 45.63 |  |
| 12 | G | O2' | 14:2447 | 4.47 | 1.00 | 40.49 |  |
| 13 | C | OP1 | 14:2499 | 4.54 | 1.00 | 35.23 |  |
| 14 | C | O3' | 14:2498 | 4.56 | 1.00 | 33.62 |  |
| 15 | C | OP1 | 14:2498 | 4.56 | 1.00 | 36.83 |  |
| 16 | G | O3' | 14:2447 | 4.57 | 1.00 | 38.97 |  |
| 17 | A | O5' | 14:2448 | 4.63 | 1.00 | 34.47 |  |
| 18 | C | O5' | 14:2499 | 4.91 | 1.00 | 38.13 |  |
| 19 | A | OP2 | 14:2450 | 4.95 | 1.00 | 41.55 |  |
  |

  
**1^st^ shell ligand angles:**   
1-2: 94; 1-3: 83; 2-3: 94;   
  
 Contact(s): /O_w_/O_w_/O_w_/  
 Coded contact list: 3O_w_  
 Exclusion zone contact(s): /O3'/OP1/OP2/OP2/OP2/  
Consider K in place of MG (4 valid contact(s) for K vs. 0 for MG)  

### MG:14:3315

|  | Res. type | Atom | Residue | Distance | Occupancy | B-factor | Closest solute to 1^st^ shell water |
| --- | --- | --- | --- | --- | --- | --- | --- |
| **Ion** | | | |  | 1.00 | 39.04 || 1 | Water 1 | | 14:3818 | 2.15 | 1.00 | 32.76 | 2.80 (A:OP1 14:1676) |
| 2 | Water 2 | | 14:3661 | 2.16 | 1.00 | 41.00 | 2.51 (A:OP2 14:1676) |
| 3 | Water 3 | | 14:4040 | 2.17 | 1.00 | 48.29 | 4.24 (A:N6 14:1677) |
| 4 | Water 4 | | 14:3617 | 2.17 | 1.00 | 49.20 | 2.38 (A:N6 14:1677) |
| 5 | Water 5 | | 14:3541 | 2.18 | 1.00 | 46.82 | 2.17 (C:OP2 14:1675) |
| 6 | A | OP2 | 14:1676 | 2.49 | 1.00 | 38.70 |  |
| 7 | A | OP1 | 14:1676 | 3.72 | 1.00 | 38.44 |  |
| 8 | C | OP2 | 14:1675 | 3.99 | 1.00 | 34.87 |  |
| 9 | A | N7 | 14:1676 | 4.16 | 1.00 | 49.50 |  |
| 10 | A | N6 | 14:1677 | 4.31 | 1.00 | 53.78 |  |
| 11 | A | O5' | 14:1676 | 4.40 | 1.00 | 43.19 |  |
| 12 | A | N7 | 14:1677 | 4.45 | 1.00 | 49.54 |  |
| 13 | C | O3' | 14:1675 | 4.71 | 1.00 | 36.34 |  |
| 14 | C | O5' | 14:1675 | 4.72 | 1.00 | 35.15 |  |
| 15 | G | O3' | 14:1674 | 4.91 | 1.00 | 40.59 |  |
  |

  
**1^st^ shell ligand angles:**   
1-2: 91; 1-3: 90; 1-4: 176; 1-5: 91; 2-3: 90; 2-4: 85; 2-5: 178; 3-4: 91; 3-5: 91; 4-5: 92;   
  
 Contact(s): /O_w_/O_w_/O_w_/O_w_/O_w_/  
 Coded contact list: 5O_w_  
 Exclusion zone contact(s): /OP2/  
Consider NA in place of MG (1 valid contact(s) for NA vs. 0 for MG)  
MG hydration shell issue: water too close (< 2.4 Angs.)  

### MG:14:3316

|  | Res. type | Atom | Residue | Distance | Occupancy | B-factor | Closest solute to 1^st^ shell water |
| --- | --- | --- | --- | --- | --- | --- | --- |
| **Ion** | | | |  | 1.00 | 34.87 || 1 | Water 1 | | 14:3648 | 2.16 | 1.00 | 31.48 | 2.49 (G:OP1 14:2608) |
| 2 | Water 2 | | 14:3839 | 2.17 | 1.00 | 36.22 | 2.85 (C:OP1 14:1983) |
| 3 | Water 3 | | 14:3539 | 2.17 | 1.00 | 34.53 | 2.12 (A:OP1 14:1784) |
| 4 | Water 4 | | 14:3847 | 2.18 | 1.00 | 48.43 | 2.87 (C:OP1 14:1983) |
| 5 | Water 5 | | 14:3864 | 2.18 | 1.00 | 44.69 | 2.91 (A:OP1 14:1785) |
| 6 | G | OP1 | 14:741 | 2.55 | 1.00 | 36.94 |  |
| 7 | U | O3' | 14:740 | 3.71 | 1.00 | 32.44 |  |
| 8 | C | OP1 | 14:1983 | 3.84 | 1.00 | 36.63 |  |
| 9 | A | OP1 | 14:1784 | 4.16 | 1.00 | 34.08 |  |
| 10 | A | OP1 | 14:1785 | 4.17 | 1.00 | 30.83 |  |
| 11 | G | O3' | 14:2607 | 4.24 | 1.00 | 32.77 |  |
| 12 | G | OP1 | 14:2608 | 4.28 | 1.00 | 27.04 |  |
| 13 | G | O5' | 14:741 | 4.41 | 1.00 | 35.00 |  |
| 14 | G | OP2 | 14:741 | 4.73 | 1.00 | 37.34 |  |
| 15 | Water 6 | | 14:3773 | 4.73 | 1.00 | 42.67 | 2.70 (A:OP1 14:1784) |
| 16 | C | O3' | 14:1982 | 4.79 | 1.00 | 40.28 |  |
  |

  
**1^st^ shell ligand angles:**   
1-2: 95; 1-3: 90; 1-4: 89; 1-5: 172; 2-3: 93; 2-4: 85; 2-5: 93; 3-4: 178; 3-5: 90; 4-5: 91;   
  
 Contact(s): /O_w_/O_w_/O_w_/O_w_/O_w_/  
 Coded contact list: 5O_w_  
 Exclusion zone contact(s): /OP1/  
Consider NA in place of MG (1 valid contact(s) for NA vs. 0 for MG)  
MG hydration shell issue: water too close (< 2.4 Angs.)  

### MG:14:3317

|  | Res. type | Atom | Residue | Distance | Occupancy | B-factor | Closest solute to 1^st^ shell water |
| --- | --- | --- | --- | --- | --- | --- | --- |
| **Ion** | | | |  | 1.00 | 40.48 || 1 | Water 1 | | 14:3519 | 2.17 | 1.00 | 36.73 | 1.99 (A:OP1 14:1970) |
| 2 | Water 2 | | 14:3984 | 2.19 | 1.00 | 46.61 | 3.28 (A:OP1 14:1969) |
| 3 | Water 3 | | 14:3689 | 2.19 | 1.00 | 46.26 | 2.55 (A:OP1 14:1971) |
| 4 | Water 4 | | 14:4073 | 2.19 | 1.00 | 29.14 | 4.04 (A:OP1 14:1970) |
| 5 | A | OP1 | 14:1970 | 2.77 | 1.00 | 34.43 |  |
| 6 | A | OP1 | 14:1971 | 2.98 | 1.00 | 38.67 |  |
| 7 | A | OP1 | 14:1969 | 3.66 | 1.00 | 39.94 |  |
| 8 | A | O3' | 14:1969 | 4.77 | 1.00 | 38.09 |  |
| 9 | A | O5' | 14:1971 | 4.81 | 1.00 | 35.51 |  |
| 10 | Water 5 | | 14:3953 | 4.89 | 1.00 | 48.59 | 3.82 (A:OP1 14:1970) |
  |

  
**1^st^ shell ligand angles:**   
1-2: 179; 1-3: 92; 1-4: 88; 2-3: 89; 2-4: 92; 3-4: 83;   
  
 Contact(s): /O_w_/O_w_/O_w_/O_w_/  
 Coded contact list: 4O_w_  
 Exclusion zone contact(s): /OP1/OP1/  
Consider K in place of MG (2 valid contact(s) for K vs. 0 for MG)  
MG hydration shell issue: water too close (< 2.4 Angs.)  

### MG:14:3318

|  | Res. type | Atom | Residue | Distance | Occupancy | B-factor | Closest solute to 1^st^ shell water |
| --- | --- | --- | --- | --- | --- | --- | --- |
| **Ion** | | | |  | 1.00 | 35.50 || 1 | Water 1 | | 14:3582 | 2.16 | 1.00 | 36.16 | 2.28 (A:OP2 14:1970) |
| 2 | Water 2 | | 14:3638 | 2.16 | 1.00 | 40.74 | 2.47 (A:OP2 14:1900) |
| 3 | Water 3 | | 14:3890 | 2.17 | 1.00 | 43.40 | 2.99 (G:O6 14:1835) |
| 4 | Water 4 | | 14:4005 | 2.18 | 1.00 | 51.00 | 3.36 (C:N4 14:1836) |
| 5 | Water 5 | | 14:3953 | 2.20 | 1.00 | 48.59 | 3.17 (G:O2' 14:1899) |
| 6 | A | OP2 | 14:1970 | 2.45 | 1.00 | 37.52 |  |
| 7 | A | OP1 | 14:1970 | 3.65 | 1.00 | 34.43 |  |
| 8 | A | OP2 | 14:1900 | 4.02 | 1.00 | 41.87 |  |
| 9 | G | O2' | 14:1899 | 4.17 | 1.00 | 55.21 |  |
| 10 | A | O5' | 14:1970 | 4.33 | 1.00 | 37.40 |  |
| 11 | Water 6 | | 14:3519 | 4.44 | 1.00 | 36.73 | 1.99 (A:OP1 14:1970) |
| 12 | C | N4 | 14:1836 | 4.62 | 1.00 | 46.82 |  |
| 13 | A | O3' | 14:1969 | 4.68 | 1.00 | 38.09 |  |
| 14 | G | O6 | 14:1835 | 4.80 | 1.00 | 38.39 |  |
| 15 | G | OP2 | 14:1835 | 4.87 | 1.00 | 40.16 |  |
| 16 | A | O5' | 14:1900 | 4.97 | 1.00 | 50.09 |  |
| 17 | G | N7 | 14:1835 | 4.98 | 1.00 | 44.60 |  |
  |

  
**1^st^ shell ligand angles:**   
1-2: 87; 1-3: 89; 1-4: 92; 1-5: 179; 2-3: 175; 2-4: 91; 2-5: 92; 3-4: 92; 3-5: 92; 4-5: 89;   
  
 Contact(s): /O_w_/O_w_/O_w_/O_w_/O_w_/  
 Coded contact list: 5O_w_  
 Exclusion zone contact(s): /OP2/  
Consider NA in place of MG (1 valid contact(s) for NA vs. 0 for MG)  
MG hydration shell issue: water too close (< 2.4 Angs.)  

### MG:14:3319

|  | Res. type | Atom | Residue | Distance | Occupancy | B-factor | Closest solute to 1^st^ shell water |
| --- | --- | --- | --- | --- | --- | --- | --- |
| **Ion** | | | |  | 1.00 | 27.35 || 1 | Water 1 | | 19:403 | 2.17 | 1.00 | 31.68 | 2.17 (GLY:O 19:234) |
| 2 | Water 2 | | 19:409 | 2.17 | 1.00 | 23.07 | 2.90 (ARG:O 19:242) |
| 3 | Water 3 | | 19:406 | 2.17 | 1.00 | 23.52 | 2.70 (ARG:O 19:242) |
| 4 | Water 4 | | 19:404 | 2.18 | 1.00 | 35.75 | 2.36 (PRO:O 19:241) |
| 5 | Water 5 | | 19:405 | 2.18 | 1.00 | 34.20 | 2.58 (GLY:O 19:235) |
| 6 | A | OP1 | 14:2598 | 2.45 | 1.00 | 26.48 |  |
| 7 | ARG | O | 19:242 | 3.33 | 1.00 | 28.13 |  |
| 8 | G | O3' | 14:2597 | 3.64 | 1.00 | 25.63 |  |
| 9 | HIS | O | 19:233 | 3.70 | 1.00 | 29.21 |  |
| 10 | PRO | O | 19:241 | 3.80 | 1.00 | 33.54 |  |
| 11 | GLY | O | 19:234 | 3.85 | 1.00 | 31.73 |  |
| 12 | PRO | O | 19:232 | 4.32 | 1.00 | 26.36 |  |
| 13 | A | OP2 | 14:2598 | 4.42 | 1.00 | 31.78 |  |
| 14 | GLY | O | 19:235 | 4.45 | 1.00 | 40.10 |  |
| 15 | A | O5' | 14:2598 | 4.70 | 1.00 | 28.25 |  |
| 16 | GLY | N | 19:234 | 4.81 | 1.00 | 24.29 |  |
| 17 | ALA | O | 19:240 | 5.00 | 1.00 | 35.08 |  |
  |

  
**1^st^ shell ligand angles:**   
1-2: 94; 1-3: 86; 1-4: 174; 1-5: 90; 2-3: 101; 2-4: 92; 2-5: 81; 3-4: 91; 3-5: 176; 4-5: 92;   
  
 Contact(s): /O_w_/O_w_/O_w_/O_w_/O_w_/  
 Coded contact list: 5O_w_  
 Exclusion zone contact(s): /O_back_/OP1/  
Consider NA in place of MG (1 valid contact(s) for NA vs. 0 for MG)  
MG hydration shell issue: water too close (< 2.4 Angs.)  

### MG:14:3320

|  | Res. type | Atom | Residue | Distance | Occupancy | B-factor | Closest solute to 1^st^ shell water |
| --- | --- | --- | --- | --- | --- | --- | --- |
| **Ion** | | | |  | 1.00 | 25.32 || 1 | Water 1 | | 14:3533 | 2.17 | 1.00 | 29.45 | 2.08 (A:OP2 14:783) |
| 2 | Water 2 | | 14:3540 | 2.17 | 1.00 | 32.56 | 2.15 (A:OP1 14:2589) |
| 3 | Water 3 | | 19:410 | 2.20 | 1.00 | 51.83 | 3.08 (MET:O 19:226) |
| 4 | A | OP2 | 14:783 | 2.48 | 1.00 | 29.29 |  |
| 5 | A | OP2 | 14:784 | 2.59 | 1.00 | 33.77 |  |
| 6 | A | OP1 | 14:2589 | 3.00 | 1.00 | 31.02 |  |
| 7 | ALA | O | 19:225 | 3.14 | 1.00 | 31.47 |  |
| 8 | ASN | OD1 | 19:227 | 3.54 | 1.00 | 26.59 |  |
| 9 | A | O3' | 14:782 | 3.83 | 1.00 | 27.89 |  |
| 10 | A | O5' | 14:783 | 3.96 | 1.00 | 34.17 |  |
| 11 | MET | O | 19:226 | 4.19 | 1.00 | 28.44 |  |
| 12 | A | O5' | 14:784 | 4.76 | 1.00 | 29.90 |  |
| 13 | MET | N | 19:226 | 4.88 | 1.00 | 30.91 |  |
| 14 | A | O3' | 14:783 | 4.89 | 1.00 | 34.25 |  |
| 15 | A | OP1 | 14:784 | 4.89 | 1.00 | 28.27 |  |
| 16 | ASN | ND2 | 19:227 | 4.90 | 1.00 | 32.06 |  |
| 17 | A | OP1 | 14:783 | 4.93 | 1.00 | 33.85 |  |
| 18 | A | OP2 | 14:2589 | 4.93 | 1.00 | 28.72 |  |
| 19 | Water 4 | | 19:407 | 4.97 | 1.00 | 29.41 | 2.76 (ASN:OD1 19:227) |
| 20 | ASN | N | 19:227 | 5.00 | 1.00 | 28.28 |  |
  |

  
**1^st^ shell ligand angles:**   
1-2: 85; 1-3: 81; 2-3: 94;   
  
 Contact(s): /O_w_/O_w_/O_w_/  
 Coded contact list: 3O_w_  
 Exclusion zone contact(s): /O_back_/OP1/OP2/OP2/  
Consider NA in place of MG (2 valid contact(s) for NA vs. 0 for MG)  
MG hydration shell issue: water too close (< 2.4 Angs.)  

### MG:14:3321

|  | Res. type | Atom | Residue | Distance | Occupancy | B-factor | Closest solute to 1^st^ shell water |
| --- | --- | --- | --- | --- | --- | --- | --- |
| **Ion** | | | |  | 1.00 | 26.92 || 1 | Water 1 | | 14:3649 | 2.16 | 1.00 | 31.99 | 2.49 (A:OP2 14:2589) |
| 2 | Water 2 | | 19:407 | 2.17 | 1.00 | 29.41 | 2.76 (ASN:OD1 19:227) |
| 3 | Water 3 | | 14:3576 | 2.18 | 1.00 | 39.24 | 2.27 (G:O6 14:792) |
| 4 | Water 4 | | 14:3943 | 2.20 | 1.00 | 42.41 | 3.14 (C:OP2 14:2440) |
| 5 | G | OP1 | 14:2588 | 2.37 | 1.00 | 35.82 |  |
| 6 | A | OP1 | 14:784 | 2.64 | 1.00 | 28.27 |  |
| 7 | G | O6 | 14:792 | 3.70 | 1.00 | 30.94 |  |
| 8 | Water 5 | | 14:3614 | 3.97 | 1.00 | 39.43 | 2.38 (G:OP2 14:2588) |
| 9 | A | O3' | 14:2587 | 3.99 | 1.00 | 37.49 |  |
| 10 | G | OP2 | 14:2588 | 4.22 | 1.00 | 32.75 |  |
| 11 | A | OP2 | 14:2589 | 4.31 | 1.00 | 28.72 |  |
| 12 | Water 6 | | 14:3540 | 4.40 | 1.00 | 32.56 | 2.21 (A:OP2 14:784) |
| 13 | A | O5' | 14:784 | 4.50 | 1.00 | 29.90 |  |
| 14 | C | OP2 | 14:2440 | 4.60 | 1.00 | 30.08 |  |
| 15 | G | O5' | 14:2588 | 4.67 | 1.00 | 30.79 |  |
| 16 | A | OP2 | 14:784 | 4.70 | 1.00 | 33.77 |  |
| 17 | ASN | OD1 | 19:227 | 4.84 | 1.00 | 26.59 |  |
| 18 | C | O4' | 14:2440 | 4.96 | 1.00 | 35.34 |  |
  |

  
**1^st^ shell ligand angles:**   
1-2: 88; 1-3: 168; 1-4: 96; 1-5: 66; 2-3: 93; 2-4: 93; 2-5: 133; 3-4: 96; 3-5: 106; 4-5: 126;   
  
 Contact(s): /O_w_/O_w_/O_w_/O_w_/OP1/  
 Coded contact list: O_ph_4O_w_  
 Exclusion zone contact(s): /OP1/  
Large octahedral topology discrepancy!!!  
MG hydration shell issue: water too close (< 2.4 Angs.)  

### MG:14:3322

|  | Res. type | Atom | Residue | Distance | Occupancy | B-factor | Closest solute to 1^st^ shell water |
| --- | --- | --- | --- | --- | --- | --- | --- |
| **Ion** | | | |  | 1.00 | 41.90 || 1 | Water 1 | | 14:3583 | 2.16 | 1.00 | 44.65 | 2.28 (G:N7 14:1992) |
| 2 | Water 2 | | 14:3929 | 2.17 | 1.00 | 42.27 | 3.10 (G:O6 14:1667) |
| 3 | Water 3 | | 14:3544 | 2.18 | 1.00 | 52.60 | 2.18 (G:OP2 14:1992) |
| 4 | Water 4 | | 14:3875 | 2.18 | 1.00 | 37.88 | 2.95 (A:N7 14:1664) |
| 5 | Water 5 | | 14:3763 | 2.18 | 1.00 | 43.57 | 2.69 (A:OP2 14:1664) |
| 6 | A | OP2 | 14:1664 | 2.55 | 1.00 | 40.84 |  |
| 7 | G | OP2 | 14:1992 | 3.73 | 1.00 | 45.95 |  |
| 8 | Water 6 | | 14:3618 | 3.87 | 1.00 | 50.05 | 2.39 (A:OP1 14:1664) |
| 9 | G | O6 | 14:1667 | 3.95 | 1.00 | 53.80 |  |
| 10 | G | N7 | 14:1992 | 4.01 | 1.00 | 36.87 |  |
| 11 | G | OP1 | 14:1992 | 4.13 | 1.00 | 42.31 |  |
| 12 | A | N7 | 14:1665 | 4.25 | 1.00 | 46.15 |  |
| 13 | G | O6 | 14:1666 | 4.29 | 1.00 | 43.50 |  |
| 14 | A | OP1 | 14:1664 | 4.36 | 1.00 | 42.18 |  |
| 15 | C | O3' | 14:1663 | 4.44 | 1.00 | 46.00 |  |
| 16 | A | N6 | 14:1665 | 4.52 | 1.00 | 45.10 |  |
| 17 | A | N7 | 14:1664 | 4.55 | 1.00 | 48.34 |  |
| 18 | G | N1 | 14:1667 | 4.61 | 1.00 | 49.63 |  |
| 19 | G | O6 | 14:1992 | 4.63 | 1.00 | 42.88 |  |
| 20 | G | N7 | 14:1666 | 4.81 | 1.00 | 45.77 |  |
| 21 | A | O5' | 14:1664 | 4.89 | 1.00 | 49.37 |  |
  |

  
**1^st^ shell ligand angles:**   
1-2: 91; 1-3: 84; 1-4: 89; 1-5: 177; 2-3: 88; 2-4: 179; 2-5: 90; 3-4: 93; 3-5: 93; 4-5: 91;   
  
 Contact(s): /O_w_/O_w_/O_w_/O_w_/O_w_/  
 Coded contact list: 5O_w_  
 Exclusion zone contact(s): /OP2/  
Consider NA in place of MG (1 valid contact(s) for NA vs. 0 for MG)  
MG hydration shell issue: water too close (< 2.4 Angs.)  

### MG:14:3323

|  | Res. type | Atom | Residue | Distance | Occupancy | B-factor | Closest solute to 1^st^ shell water |
| --- | --- | --- | --- | --- | --- | --- | --- |
| **Ion** | | | |  | 1.00 | 56.70 || 1 | Water 1 | | 14:3655 | 2.17 | 1.00 | 56.17 | 2.50 (G:N7 14:2706) |
| 2 | Water 2 | | 14:3952 | 2.17 | 1.00 | 63.88 | 3.17 (C:OP1 14:2704) |
| 3 | Water 3 | | 14:3620 | 2.17 | 1.00 | 58.15 | 2.39 (A:OP2 14:2705) |
| 4 | Water 4 | | 14:3554 | 2.18 | 1.00 | 61.69 | 2.21 (G:O6 14:2706) |
| 5 | Water 5 | | 14:4082 | 2.18 | 1.00 | 63.95 | 4.20 (A:OP2 14:2705) |
| 6 | A | OP2 | 14:2705 | 2.63 | 1.00 | 65.11 |  |
| 7 | G | N7 | 14:2706 | 3.95 | 1.00 | 53.10 |  |
| 8 | A | O5' | 14:2705 | 3.96 | 1.00 | 60.70 |  |
| 9 | G | O6 | 14:2706 | 4.24 | 1.00 | 50.96 |  |
| 10 | C | OP1 | 14:2704 | 4.38 | 1.00 | 62.49 |  |
| 11 | A | N7 | 14:2705 | 4.51 | 1.00 | 60.19 |  |
| 12 | A | OP1 | 14:2705 | 4.65 | 1.00 | 63.78 |  |
| 13 | C | OP2 | 14:2704 | 4.69 | 1.00 | 60.89 |  |
| 14 | G | OP2 | 14:2706 | 4.72 | 1.00 | 64.79 |  |
| 15 | C | O3' | 14:2704 | 4.89 | 1.00 | 65.01 |  |
| 16 | C | O5' | 14:2704 | 4.89 | 1.00 | 65.20 |  |
  |

  
**1^st^ shell ligand angles:**   
1-2: 178; 1-3: 91; 1-4: 91; 1-5: 89; 2-3: 90; 2-4: 91; 2-5: 90; 3-4: 87; 3-5: 180; 4-5: 93;   
  
 Contact(s): /O_w_/O_w_/O_w_/O_w_/O_w_/  
 Coded contact list: 5O_w_  
 Exclusion zone contact(s): /OP2/  
Consider K in place of MG (1 valid contact(s) for K vs. 0 for MG)  
MG hydration shell issue: water too close (< 2.4 Angs.)  

### MG:14:3324

|  | Res. type | Atom | Residue | Distance | Occupancy | B-factor | Closest solute to 1^st^ shell water |
| --- | --- | --- | --- | --- | --- | --- | --- |
| **Ion** | | | |  | 1.00 | 43.97 || 1 | Water 1 | | 14:3733 | 2.18 | 1.00 | 40.66 | 2.66 (G:N7 14:1619) |
| 2 | Water 2 | | 14:3615 | 2.18 | 1.00 | 53.26 | 2.38 (A:OP2 14:1609) |
| 3 | Water 3 | | 14:3993 | 2.18 | 1.00 | 44.52 | 3.29 (G:OP2 14:1619) |
| 4 | Water 4 | | 14:3884 | 2.18 | 1.00 | 47.17 | 2.98 (G:O6 14:1613) |
| 5 | Water 5 | | 14:3703 | 2.19 | 1.00 | 36.64 | 2.59 (C:OP1 14:1617) |
| 6 | Water 6 | | 14:3837 | 2.19 | 1.00 | 39.73 | 2.84 (G:O6 14:1620) |
| 7 | G | O6 | 14:1619 | 4.07 | 1.00 | 38.65 |  |
| 8 | A | OP2 | 14:1609 | 4.15 | 1.00 | 44.60 |  |
| 9 | A | O2' | 14:1608 | 4.20 | 1.00 | 37.18 |  |
| 10 | G | N7 | 14:1619 | 4.23 | 1.00 | 39.02 |  |
| 11 | G | O6 | 14:1613 | 4.38 | 1.00 | 36.54 |  |
| 12 | C | OP2 | 14:1617 | 4.47 | 1.00 | 33.44 |  |
| 13 | G | OP2 | 14:1619 | 4.50 | 1.00 | 34.90 |  |
| 14 | G | O6 | 14:1620 | 4.69 | 1.00 | 38.74 |  |
| 15 | C | OP1 | 14:1617 | 4.72 | 1.00 | 31.84 |  |
  |

  
**1^st^ shell ligand angles:**   
1-2: 177; 1-3: 91; 1-4: 91; 1-5: 91; 1-6: 90; 2-3: 88; 2-4: 90; 2-5: 86; 2-6: 93; 3-4: 177; 3-5: 89; 3-6: 93; 4-5: 93; 4-6: 85; 5-6: 178;   
  
 Contact(s): /O_w_/O_w_/O_w_/O_w_/O_w_/O_w_/  
 Coded contact list: 6O_w_  
MG hydration shell issue: water too close (< 2.4 Angs.)  

### MG:14:3325

|  | Res. type | Atom | Residue | Distance | Occupancy | B-factor | Closest solute to 1^st^ shell water |
| --- | --- | --- | --- | --- | --- | --- | --- |
| **Ion** | | | |  | 1.00 | 30.58 || 1 | Water 1 | | 14:3722 | 2.16 | 1.00 | 33.10 | 2.63 (U:OP2 14:2504) |
| 2 | Water 2 | | 14:3597 | 2.17 | 1.00 | 45.19 | 2.33 (U:OP1 14:576) |
| 3 | Water 3 | | 14:3850 | 2.19 | 1.00 | 33.21 | 2.88 (G:OP2 14:577) |
| 4 | Water 4 | | 14:3668 | 2.20 | 1.00 | 45.78 | 2.52 (A:OP1 14:2057) |
| 5 | 2MA | OP1 | 14:2503 | 2.36 | 1.00 | 31.76 |  |
| 6 | U | OP1 | 14:576 | 2.59 | 1.00 | 37.93 |  |
| 7 | 2MA | OP2 | 14:2503 | 3.03 | 1.00 | 30.67 |  |
| 8 | Water 5 | | 14:3891 | 3.70 | 1.00 | 38.79 | 2.99 (2MA:OP2 14:2503) |
| 9 | 2MA | O5' | 14:2503 | 4.16 | 1.00 | 31.00 |  |
| 10 | G | O3' | 14:2502 | 4.24 | 1.00 | 25.66 |  |
| 11 | G | O2' | 14:2502 | 4.33 | 1.00 | 36.98 |  |
| 12 | A | OP1 | 14:2057 | 4.53 | 1.00 | 32.52 |  |
| 13 | U | OP2 | 14:2504 | 4.62 | 1.00 | 40.57 |  |
| 14 | G | OP2 | 14:577 | 4.62 | 1.00 | 35.21 |  |
| 15 | U | OP2 | 14:576 | 4.69 | 1.00 | 32.60 |  |
| 16 | A | O3' | 14:575 | 4.89 | 1.00 | 43.32 |  |
| 17 | U | O5' | 14:576 | 4.89 | 1.00 | 28.07 |  |
| 18 | Water 6 | | 14:3896 | 4.94 | 1.00 | 34.69 | 4.37 (G:OP2 14:577) |
  |

  
**1^st^ shell ligand angles:**   
1-2: 84; 1-3: 171; 1-4: 97; 1-5: 90; 2-3: 94; 2-4: 95; 2-5: 126; 3-4: 92; 3-5: 83; 4-5: 139;   
  
 Contact(s): /O_w_/O_w_/O_w_/O_w_/OP1/  
 Coded contact list: O_ph_4O_w_  
 Exclusion zone contact(s): /OP1/OP2/  
Large octahedral topology discrepancy!!!  
MG hydration shell issue: water too close (< 2.4 Angs.)  

### MG:14:3326

|  | Res. type | Atom | Residue | Distance | Occupancy | B-factor | Closest solute to 1^st^ shell water |
| --- | --- | --- | --- | --- | --- | --- | --- |
| **Ion** | | | |  | 1.00 | 40.35 || 1 | Water 1 | | 14:3625 | 2.16 | 1.00 | 30.62 | 2.40 (U:OP1 14:963) |
| 2 | Water 2 | | 14:4002 | 2.17 | 1.00 | 50.90 | 3.35 (C:OP1 14:2498) |
| 3 | Water 3 | | 14:3960 | 2.18 | 1.00 | 59.90 | 3.19 (U:OP1 14:963) |
| 4 | Water 4 | | 14:3639 | 2.19 | 1.00 | 57.60 | 2.47 (G:OP1 14:947) |
| 5 | Water 5 | | 14:4089 | 2.20 | 1.00 | 44.41 | 4.48 (C:OP1 14:2498) |
| 6 | U | OP1 | 14:963 | 2.70 | 1.00 | 45.85 |  |
| 7 | G | OP1 | 14:947 | 3.79 | 1.00 | 37.36 |  |
| 8 | C | OP1 | 14:2498 | 4.36 | 1.00 | 36.83 |  |
| 9 | U | O5' | 14:963 | 4.40 | 1.00 | 49.17 |  |
| 10 | A | OP1 | 14:2497 | 4.91 | 1.00 | 34.84 |  |
| 11 | U | OP2 | 14:963 | 4.92 | 1.00 | 48.03 |  |
  |

  
**1^st^ shell ligand angles:**   
1-2: 88; 1-3: 90; 1-4: 178; 1-5: 95; 2-3: 178; 2-4: 90; 2-5: 96; 3-4: 92; 3-5: 85; 4-5: 84;   
  
 Contact(s): /O_w_/O_w_/O_w_/O_w_/O_w_/  
 Coded contact list: 5O_w_  
 Exclusion zone contact(s): /OP1/  
Consider K in place of MG (1 valid contact(s) for K vs. 0 for MG)  

### MG:14:3327

|  | Res. type | Atom | Residue | Distance | Occupancy | B-factor | Closest solute to 1^st^ shell water |
| --- | --- | --- | --- | --- | --- | --- | --- |
| **Ion** | | | |  | 1.00 | 36.56 || 1 | Water 1 | | 14:3782 | 2.16 | 1.00 | 43.53 | 2.72 (C:O2 14:2499) |
| 2 | Water 2 | | 14:3542 | 2.16 | 1.00 | 34.97 | 2.17 (C:O2 14:2055) |
| 3 | Water 3 | | 14:3646 | 2.17 | 1.00 | 45.72 | 2.48 (G:N3 14:2454) |
| 4 | Water 4 | | 14:3527 | 2.18 | 1.00 | 36.70 | 2.04 (G:O6 14:2032) |
| 5 | Water 5 | | 14:3756 | 2.18 | 1.00 | 43.19 | 2.68 (C:N3 14:2499) |
| 6 | G | N7 | 14:2032 | 2.61 | 1.00 | 37.15 |  |
| 7 | G | O6 | 14:2032 | 3.06 | 1.00 | 38.38 |  |
| 8 | C | N3 | 14:2499 | 3.93 | 1.00 | 36.72 |  |
| 9 | G | OP1 | 14:2032 | 4.16 | 1.00 | 43.28 |  |
| 10 | C | O2 | 14:2055 | 4.19 | 1.00 | 40.02 |  |
| 11 | C | O2 | 14:2499 | 4.21 | 1.00 | 42.68 |  |
| 12 | A | N3 | 14:2453 | 4.24 | 1.00 | 34.75 |  |
| 13 | C | N3 | 14:2055 | 4.37 | 1.00 | 37.68 |  |
| 14 | A | N1 | 14:2572 | 4.50 | 1.00 | 39.97 |  |
| 15 | G | N3 | 14:2454 | 4.62 | 1.00 | 40.01 |  |
| 16 | U | O2 | 14:2500 | 4.67 | 1.00 | 37.86 |  |
| 17 | G | O2' | 14:2454 | 4.71 | 1.00 | 41.48 |  |
| 18 | G | N9 | 14:2032 | 4.79 | 1.00 | 41.81 |  |
| 19 | G | N9 | 14:2454 | 4.81 | 1.00 | 40.11 |  |
| 20 | C | N4 | 14:2499 | 4.89 | 1.00 | 45.21 |  |
| 21 | G | N1 | 14:2032 | 4.93 | 1.00 | 37.75 |  |
| 22 | A | N1 | 14:2453 | 4.97 | 1.00 | 39.07 |  |
| 23 | G | O4' | 14:2454 | 4.99 | 1.00 | 42.48 |  |
  |

  
**1^st^ shell ligand angles:**   
1-2: 87; 1-3: 91; 1-4: 177; 1-5: 89; 2-3: 176; 2-4: 95; 2-5: 91; 3-4: 88; 3-5: 86; 4-5: 94;   
  
 Contact(s): /O_w_/O_w_/O_w_/O_w_/O_w_/  
 Coded contact list: 5O_w_  
 Exclusion zone contact(s): /N7/O6/  
Consider K in place of MG (2 valid contact(s) for K vs. 0 for MG)  
MG hydration shell issue: water too close (< 2.4 Angs.)  

### MG:14:3328

|  | Res. type | Atom | Residue | Distance | Occupancy | B-factor | Closest solute to 1^st^ shell water |
| --- | --- | --- | --- | --- | --- | --- | --- |
| **Ion** | | | |  | 1.00 | 25.02 || 1 | Water 1 | | 14:3833 | 2.16 | 1.00 | 34.33 | 2.83 (A:N7 14:2241) |
| 2 | Water 2 | | 14:3772 | 2.17 | 1.00 | 32.39 | 2.70 (G:O6 14:2242) |
| 3 | Water 3 | | 14:3619 | 2.18 | 1.00 | 24.74 | 2.39 (U:O4 14:2074) |
| 4 | Water 4 | | 14:3801 | 2.19 | 1.00 | 24.05 | 2.75 (G:N7 14:2242) |
| 5 | Water 5 | | 14:4088 | 2.19 | 1.00 | 34.30 | 4.31 (U:O4 14:2074) |
| 6 | Water 6 | | 14:3781 | 2.19 | 1.00 | 23.90 | 2.71 (U:O4 14:2075) |
| 7 | U | O4 | 14:2074 | 4.02 | 1.00 | 29.30 |  |
| 8 | U | O4 | 14:2243 | 4.07 | 1.00 | 29.57 |  |
| 9 | U | O4 | 14:2075 | 4.12 | 1.00 | 35.19 |  |
| 10 | G | N7 | 14:2238 | 4.24 | 1.00 | 34.07 |  |
| 11 | A | N7 | 14:2241 | 4.36 | 1.00 | 32.32 |  |
| 12 | G | N7 | 14:2242 | 4.47 | 1.00 | 30.03 |  |
| 13 | G | O6 | 14:2242 | 4.52 | 1.00 | 29.85 |  |
| 14 | A | N6 | 14:2077 | 4.62 | 1.00 | 34.89 |  |
| 15 | A | N6 | 14:2241 | 4.70 | 1.00 | 33.45 |  |
  |

  
**1^st^ shell ligand angles:**   
1-2: 89; 1-3: 175; 1-4: 89; 1-5: 94; 1-6: 85; 2-3: 87; 2-4: 90; 2-5: 176; 2-6: 91; 3-4: 94; 3-5: 90; 3-6: 92; 4-5: 88; 4-6: 174; 5-6: 91;   
  
 Contact(s): /O_w_/O_w_/O_w_/O_w_/O_w_/O_w_/  
 Coded contact list: 6O_w_  
MG hydration shell issue: water too close (< 2.4 Angs.)  

### MG:14:3329

|  | Res. type | Atom | Residue | Distance | Occupancy | B-factor | Closest solute to 1^st^ shell water |
| --- | --- | --- | --- | --- | --- | --- | --- |
| **Ion** | | | |  | 1.00 | 28.96 || 1 | Water 1 | | 14:3520 | 2.18 | 1.00 | 28.11 | 1.99 (C:OP1 14:192) |
| 2 | Water 2 | | 14:3882 | 2.19 | 1.00 | 30.28 | 2.97 (U:O5' 14:2243) |
| 3 | Water 3 | | 14:3754 | 2.20 | 1.00 | 38.25 | 2.67 (U:OP2 14:193) |
| 4 | Water 4 | | 14:3932 | 2.20 | 1.00 | 37.26 | 3.10 (A:OP2 14:199) |
| 5 | C | OP1 | 14:192 | 2.51 | 1.00 | 37.79 |  |
| 6 | U | OP1 | 14:2243 | 2.97 | 1.00 | 29.50 |  |
| 7 | C | O5' | 14:192 | 3.71 | 1.00 | 31.00 |  |
| 8 | C | OP2 | 14:192 | 3.78 | 1.00 | 33.66 |  |
| 9 | U | OP2 | 14:193 | 3.78 | 1.00 | 30.87 |  |
| 10 | U | O5' | 14:2243 | 4.31 | 1.00 | 32.45 |  |
| 11 | A | OP1 | 14:199 | 4.44 | 1.00 | 35.67 |  |
| 12 | A | OP2 | 14:199 | 4.53 | 1.00 | 36.42 |  |
| 13 | A | O3' | 14:191 | 4.70 | 1.00 | 30.06 |  |
| 14 | C | OP1 | 14:198 | 4.83 | 1.00 | 35.32 |  |
| 15 | G | O3' | 14:2242 | 4.84 | 1.00 | 24.83 |  |
  |

  
**1^st^ shell ligand angles:**   
1-2: 87; 1-3: 172; 1-4: 89; 2-3: 94; 2-4: 95; 3-4: 83;   
  
 Contact(s): /O_w_/O_w_/O_w_/O_w_/  
 Coded contact list: 4O_w_  
 Exclusion zone contact(s): /OP1/OP1/  
Consider NA in place of MG (1 valid contact(s) for NA vs. 0 for MG)  
MG hydration shell issue: water too close (< 2.4 Angs.)  

### MG:14:3330

|  | Res. type | Atom | Residue | Distance | Occupancy | B-factor | Closest solute to 1^st^ shell water |
| --- | --- | --- | --- | --- | --- | --- | --- |
| **Ion** | | | |  | 1.00 | 39.97 || 1 | Water 1 | | 14:3878 | 2.17 | 1.00 | 52.55 | 2.96 (G:O5' 14:446) |
| 2 | Water 2 | | 14:3724 | 2.17 | 1.00 | 51.91 | 2.63 (U:O2' 14:1199) |
| 3 | Water 3 | | 14:3762 | 2.17 | 1.00 | 36.63 | 2.69 (G:OP2 14:446) |
| 4 | Water 4 | | 14:3873 | 2.18 | 1.00 | 49.42 | 2.95 (C:OP1 14:1200) |
| 5 | Water 5 | | 14:4006 | 2.18 | 1.00 | 46.50 | 3.36 (G:OP1 14:446) |
| 6 | G | OP2 | 14:446 | 2.51 | 1.00 | 39.64 |  |
| 7 | G | OP1 | 14:446 | 3.76 | 1.00 | 42.06 |  |
| 8 | G | O5' | 14:446 | 3.95 | 1.00 | 49.66 |  |
| 9 | C | OP1 | 14:445 | 4.27 | 1.00 | 39.62 |  |
| 10 | U | O2' | 14:1199 | 4.27 | 1.00 | 41.34 |  |
| 11 | U | O3' | 14:1199 | 4.27 | 1.00 | 46.88 |  |
| 12 | C | O3' | 14:445 | 4.72 | 1.00 | 42.07 |  |
| 13 | C | OP1 | 14:1200 | 4.75 | 1.00 | 54.11 |  |
| 14 | C | OP2 | 14:445 | 4.77 | 1.00 | 47.33 |  |
| 15 | ARG | O | 85:3 | 4.88 | 1.00 | 49.63 |  |
  |

  
**1^st^ shell ligand angles:**   
1-2: 178; 1-3: 94; 1-4: 90; 1-5: 85; 2-3: 85; 2-4: 92; 2-5: 96; 3-4: 94; 3-5: 175; 4-5: 92;   
  
 Contact(s): /O_w_/O_w_/O_w_/O_w_/O_w_/  
 Coded contact list: 5O_w_  
 Exclusion zone contact(s): /OP2/  
Consider NA in place of MG (1 valid contact(s) for NA vs. 0 for MG)  

### MG:14:3331

|  | Res. type | Atom | Residue | Distance | Occupancy | B-factor | Closest solute to 1^st^ shell water |
| --- | --- | --- | --- | --- | --- | --- | --- |
| **Ion** | | | |  | 1.00 | 51.92 || 1 | Water 1 | | 14:3808 | 2.16 | 1.00 | 42.54 | 2.77 (G:OP2 14:399) |
| 2 | Water 2 | | 14:3563 | 2.17 | 1.00 | 61.22 | 2.23 (G:O6 14:400) |
| 3 | Water 3 | | 14:3785 | 2.17 | 1.00 | 52.13 | 2.72 (G:OP2 14:399) |
| 4 | Water 4 | | 14:3971 | 2.17 | 1.00 | 56.92 | 3.21 (G:N7 14:400) |
| 5 | Water 5 | | 14:4041 | 2.19 | 1.00 | 60.17 | 3.66 (G:OP2 14:399) |
| 6 | G | OP2 | 14:399 | 2.44 | 1.00 | 55.47 |  |
| 7 | G | O5' | 14:399 | 3.70 | 1.00 | 52.17 |  |
| 8 | G | O6 | 14:400 | 3.95 | 1.00 | 58.15 |  |
| 9 | G | OP1 | 14:399 | 4.11 | 1.00 | 59.99 |  |
| 10 | G | N7 | 14:400 | 4.33 | 1.00 | 53.24 |  |
| 11 | G | N7 | 14:372 | 4.50 | 1.00 | 56.44 |  |
| 12 | G | O3' | 14:398 | 4.70 | 1.00 | 53.66 |  |
| 13 | G | O5' | 14:398 | 4.98 | 1.00 | 46.86 |  |
  |

  
**1^st^ shell ligand angles:**   
1-2: 90; 1-3: 87; 1-4: 90; 1-5: 178; 2-3: 88; 2-4: 94; 2-5: 91; 3-4: 177; 3-5: 91; 4-5: 91;   
  
 Contact(s): /O_w_/O_w_/O_w_/O_w_/O_w_/  
 Coded contact list: 5O_w_  
 Exclusion zone contact(s): /OP2/  
Consider NA in place of MG (1 valid contact(s) for NA vs. 0 for MG)  
MG hydration shell issue: water too close (< 2.4 Angs.)  

### MG:14:3332

|  | Res. type | Atom | Residue | Distance | Occupancy | B-factor | Closest solute to 1^st^ shell water |
| --- | --- | --- | --- | --- | --- | --- | --- |
| **Ion** | | | |  | 1.00 | 54.87 || 1 | Water 1 | | 14:3893 | 2.19 | 1.00 | 48.44 | 3.00 (U:OP1 14:120) |
| 2 | Water 2 | | 14:3775 | 2.19 | 1.00 | 35.25 | 2.71 (G:O6 14:124) |
| 3 | Water 3 | | 14:3561 | 2.19 | 1.00 | 39.78 | 2.23 (U:OP1 14:120) |
| 4 | Water 4 | | 14:3975 | 2.20 | 1.00 | 46.28 | 3.23 (G:OP2 14:123) |
| 5 | Water 5 | | 14:3933 | 2.22 | 1.00 | 38.76 | 3.12 (G:N7 14:124) |
| 6 | U | OP1 | 14:120 | 2.49 | 1.00 | 48.69 |  |
| 7 | U | OP2 | 14:120 | 3.36 | 1.00 | 49.25 |  |
| 8 | G | O6 | 14:124 | 3.64 | 1.00 | 36.50 |  |
| 9 | U | O5' | 14:120 | 3.94 | 1.00 | 41.45 |  |
| 10 | G | N7 | 14:124 | 4.26 | 1.00 | 29.06 |  |
| 11 | G | OP2 | 14:123 | 4.34 | 1.00 | 46.85 |  |
| 12 | G | N7 | 14:123 | 4.37 | 1.00 | 29.67 |  |
| 13 | A | O3' | 14:119 | 4.65 | 1.00 | 49.21 |  |
| 14 | G | OP2 | 14:122 | 4.99 | 1.00 | 48.71 |  |
  |

  
**1^st^ shell ligand angles:**   
1-2: 174; 1-3: 88; 1-4: 92; 1-5: 91; 2-3: 92; 2-4: 88; 2-5: 95; 3-4: 176; 3-5: 90; 4-5: 95;   
  
 Contact(s): /O_w_/O_w_/O_w_/O_w_/O_w_/  
 Coded contact list: 5O_w_  
 Exclusion zone contact(s): /OP1/OP2/  
Consider NA in place of MG (1 valid contact(s) for NA vs. 0 for MG)  
MG hydration shell issue: water too close (< 2.4 Angs.)  

### MG:14:3333

|  | Res. type | Atom | Residue | Distance | Occupancy | B-factor | Closest solute to 1^st^ shell water |
| --- | --- | --- | --- | --- | --- | --- | --- |
| **Ion** | | | |  | 1.00 | 62.45 || 1 | Water 1 | | 14:3750 | 2.17 | 1.00 | 59.99 | 2.67 (G:N7 14:301) |
| 2 | Water 2 | | 14:4003 | 2.17 | 1.00 | 55.22 | 3.36 (G:OP1 14:312) |
| 3 | Water 3 | | 14:3835 | 2.18 | 1.00 | 71.90 | 2.83 (C:N4 14:334) |
| 4 | Water 4 | | 14:3803 | 2.18 | 1.00 | 72.64 | 2.75 (G:OP2 14:312) |
| 5 | Water 5 | | 14:3874 | 2.18 | 1.00 | 55.16 | 2.95 (G:O6 14:301) |
| 6 | Water 6 | | 14:4036 | 2.18 | 1.00 | 67.73 | 3.58 (G:OP1 14:312) |
| 7 | G | OP2 | 14:312 | 4.06 | 1.00 | 66.43 |  |
| 8 | G | O6 | 14:301 | 4.38 | 1.00 | 64.79 |  |
| 9 | G | N7 | 14:301 | 4.49 | 1.00 | 63.52 |  |
| 10 | G | OP1 | 14:312 | 4.59 | 1.00 | 70.15 |  |
| 11 | C | N4 | 14:334 | 4.64 | 1.00 | 65.36 |  |
| 12 | C | N4 | 14:302 | 4.88 | 1.00 | 62.67 |  |
  |

  
**1^st^ shell ligand angles:**   
1-2: 178; 1-3: 85; 1-4: 93; 1-5: 92; 1-6: 89; 2-3: 94; 2-4: 88; 2-5: 89; 2-6: 89; 3-4: 177; 3-5: 89; 3-6: 92; 4-5: 92; 4-6: 87; 5-6: 178;   
  
 Contact(s): /O_w_/O_w_/O_w_/O_w_/O_w_/O_w_/  
 Coded contact list: 6O_w_  

### MG:14:3334

|  | Res. type | Atom | Residue | Distance | Occupancy | B-factor | Closest solute to 1^st^ shell water |
| --- | --- | --- | --- | --- | --- | --- | --- |
| **Ion** | | | |  | 1.00 | 33.72 || 1 | Water 1 | | 39:401 | 2.17 | 1.00 | 38.32 | 2.72 (GLY:O 39:55) |
| 2 | Water 2 | | 14:3911 | 2.18 | 1.00 | 39.33 | 3.05 (G:OP2 14:799) |
| 3 | Water 3 | | 39:402 | 2.18 | 1.00 | 49.59 | 2.78 (ASP:OD2 39:77) |
| 4 | Water 4 | | 14:3949 | 2.18 | 1.00 | 34.04 | 3.16 (A:O2' 14:800) |
| 5 | Water 5 | | 14:3778 | 2.18 | 1.00 | 42.38 | 2.71 (G:OP1 14:798) |
| 6 | G | OP2 | 14:801 | 2.47 | 1.00 | 34.88 |  |
| 7 | A | O3' | 14:800 | 3.73 | 1.00 | 33.24 |  |
| 8 | GLY | O | 39:55 | 4.18 | 1.00 | 43.00 |  |
| 9 | G | OP2 | 14:799 | 4.26 | 1.00 | 32.32 |  |
| 10 | TYR | O | 39:59 | 4.37 | 1.00 | 40.10 |  |
| 11 | G | OP1 | 14:798 | 4.46 | 1.00 | 40.53 |  |
| 12 | GLY | N | 39:61 | 4.47 | 1.00 | 35.74 |  |
| 13 | G | OP1 | 14:801 | 4.54 | 1.00 | 28.73 |  |
| 14 | G | O5' | 14:801 | 4.57 | 1.00 | 31.29 |  |
| 15 | ARG | O | 39:54 | 4.78 | 1.00 | 36.01 |  |
| 16 | A | O2' | 14:800 | 4.86 | 1.00 | 41.54 |  |
| 17 | ASP | OD2 | 39:77 | 4.89 | 1.00 | 46.36 |  |
  |

  
**1^st^ shell ligand angles:**   
1-2: 90; 1-3: 87; 1-4: 174; 1-5: 93; 2-3: 177; 2-4: 94; 2-5: 88; 3-4: 90; 3-5: 92; 4-5: 92;   
  
 Contact(s): /O_w_/O_w_/O_w_/O_w_/O_w_/  
 Coded contact list: 5O_w_  
 Exclusion zone contact(s): /OP2/  
Consider NA in place of MG (1 valid contact(s) for NA vs. 0 for MG)  

### MG:14:3335

|  | Res. type | Atom | Residue | Distance | Occupancy | B-factor | Closest solute to 1^st^ shell water |
| --- | --- | --- | --- | --- | --- | --- | --- |
| **Ion** | | | |  | 1.00 | 70.79 || 1 | Water 1 | | 14:3676 | 2.17 | 1.00 | 59.50 | 2.53 (G:O6 14:1400) |
| 2 | Water 2 | | 14:3558 | 2.18 | 1.00 | 70.33 | 2.22 (G:O6 14:1401) |
| 3 | Water 3 | | 14:4029 | 2.18 | 1.00 | 65.11 | 3.52 (U:OP1 14:1397) |
| 4 | Water 4 | | 14:4071 | 2.18 | 1.00 | 60.02 | 4.00 (G:O6 14:1385) |
| 5 | Water 5 | | 14:4001 | 2.18 | 1.00 | 65.63 | 3.33 (G:N7 14:1400) |
| 6 | Water 6 | | 14:3919 | 2.19 | 1.00 | 62.86 | 4.12 (U:OP1 14:1397) |
| 7 | Water 7 | | 14:3946 | 3.53 | 1.00 | 66.46 | 3.15 (G:O6 14:1385) |
| 8 | Water 8 | | 14:3947 | 3.95 | 1.00 | 49.75 | 4.02 (U:OP1 14:1397) |
| 9 | G | O6 | 14:1400 | 4.17 | 1.00 | 70.76 |  |
| 10 | G | O6 | 14:1401 | 4.19 | 1.00 | 76.04 |  |
| 11 | G | N7 | 14:1400 | 4.57 | 1.00 | 70.58 |  |
| 12 | U | OP1 | 14:1397 | 4.57 | 1.00 | 58.87 |  |
| 13 | G | O6 | 14:1385 | 4.58 | 1.00 | 71.83 |  |
| 14 | C | N4 | 14:1399 | 4.60 | 1.00 | 65.51 |  |
| 15 | Water 9 | | 14:4077 | 4.81 | 1.00 | 81.59 | 6.27 (G:O6 14:1385) |
| 16 | C | N4 | 14:1386 | 4.90 | 1.00 | 63.91 |  |
  |

  
**1^st^ shell ligand angles:**   
1-2: 92; 1-3: 91; 1-4: 179; 1-5: 88; 1-6: 89; 2-3: 90; 2-4: 88; 2-5: 90; 2-6: 180; 3-4: 89; 3-5: 178; 3-6: 90; 4-5: 92; 4-6: 92; 5-6: 91;   
  
 Contact(s): /O_w_/O_w_/O_w_/O_w_/O_w_/O_w_/  
 Coded contact list: 6O_w_  
MG hydration shell issue: water too close (< 2.4 Angs.)  

### MG:14:3336

|  | Res. type | Atom | Residue | Distance | Occupancy | B-factor | Closest solute to 1^st^ shell water |
| --- | --- | --- | --- | --- | --- | --- | --- |
| **Ion** | | | |  | 1.00 | 78.58 || 1 | Water 1 | | 14:3666 | 2.17 | 1.00 | 55.86 | 2.52 (G:N7 14:1344) |
| 2 | Water 2 | | 14:3656 | 2.19 | 1.00 | 56.82 | 2.50 (G:OP1 14:1343) |
| 3 | Water 3 | | 14:3755 | 2.19 | 1.00 | 63.20 | 2.68 (G:O6 14:1344) |
| 4 | Water 4 | | 14:3947 | 2.20 | 1.00 | 49.75 | 3.16 (C:OP2 14:1398) |
| 5 | Water 5 | | 14:3946 | 2.20 | 1.00 | 66.46 | 3.55 (G:O6 14:1344) |
| 6 | Water 6 | | 14:4077 | 2.20 | 1.00 | 81.59 | 4.08 (G:OP1 14:1343) |
| 7 | Water 7 | | 14:4071 | 3.68 | 1.00 | 60.02 | 5.44 (G:O6 14:1344) |
| 8 | Water 8 | | 14:4029 | 3.79 | 1.00 | 65.11 | 4.29 (G:N7 14:1385) |
| 9 | G | N7 | 14:1344 | 3.82 | 1.00 | 60.71 |  |
| 10 | G | O6 | 14:1344 | 3.97 | 1.00 | 66.81 |  |
| 11 | G | OP1 | 14:1343 | 4.33 | 1.00 | 48.18 |  |
| 12 | Water 9 | | 14:3919 | 4.55 | 1.00 | 62.86 | 3.07 (C:OP2 14:1398) |
| 13 | G | N7 | 14:1385 | 4.68 | 1.00 | 64.80 |  |
| 14 | C | OP2 | 14:1398 | 4.78 | 1.00 | 54.72 |  |
  |

  
**1^st^ shell ligand angles:**   
1-2: 88; 1-3: 89; 1-4: 92; 1-5: 90; 1-6: 179; 2-3: 93; 2-4: 86; 2-5: 178; 2-6: 92; 3-4: 179; 3-5: 86; 3-6: 91; 4-5: 95; 4-6: 89; 5-6: 90;   
  
 Contact(s): /O_w_/O_w_/O_w_/O_w_/O_w_/O_w_/  
 Coded contact list: 6O_w_  

### MG:14:3337

|  | Res. type | Atom | Residue | Distance | Occupancy | B-factor | Closest solute to 1^st^ shell water |
| --- | --- | --- | --- | --- | --- | --- | --- |
| **Ion** | | | |  | 1.00 | 83.35 || 1 | Water 1 | | 14:3667 | 2.18 | 1.00 | 71.21 | 2.52 (G:N7 14:2405) |
| 2 | Water 2 | | 14:3914 | 2.18 | 1.00 | 71.33 | 3.06 (G:N7 14:2405) |
| 3 | Water 3 | | 14:4086 | 2.19 | 1.00 | 63.64 | 4.26 (C:OP2 14:2404) |
| 4 | Water 4 | | 14:4034 | 2.19 | 1.00 | 67.81 | 3.58 (G:OP2 14:2410) |
| 5 | Water 5 | | 14:3753 | 2.19 | 1.00 | 71.59 | 2.67 (G:OP2 14:2405) |
| 6 | Water 6 | | 14:3634 | 2.19 | 1.00 | 67.74 | 2.43 (G:OP2 14:2410) |
| 7 | G | N7 | 14:2405 | 3.83 | 1.00 | 64.58 |  |
| 8 | G | OP2 | 14:2410 | 3.95 | 1.00 | 73.22 |  |
| 9 | G | OP2 | 14:2405 | 4.25 | 1.00 | 63.57 |  |
| 10 | C | OP2 | 14:2404 | 4.44 | 1.00 | 69.39 |  |
| 11 | G | OP1 | 14:2410 | 4.99 | 1.00 | 72.71 |  |
  |

  
**1^st^ shell ligand angles:**   
1-2: 91; 1-3: 89; 1-4: 88; 1-5: 90; 1-6: 179; 2-3: 180; 2-4: 91; 2-5: 89; 2-6: 88; 3-4: 89; 3-5: 91; 3-6: 92; 4-5: 177; 4-6: 92; 5-6: 90;   
  
 Contact(s): /O_w_/O_w_/O_w_/O_w_/O_w_/O_w_/  
 Coded contact list: 6O_w_  

### MG:14:3338

|  | Res. type | Atom | Residue | Distance | Occupancy | B-factor | Closest solute to 1^st^ shell water |
| --- | --- | --- | --- | --- | --- | --- | --- |
| **Ion** | | | |  | 1.00 | 34.23 || 1 | Water 1 | | 14:3809 | 2.17 | 1.00 | 59.61 | 2.78 (U:OP2 14:29) |
| 2 | Water 2 | | 14:3747 | 2.17 | 1.00 | 48.22 | 2.67 (G:O6 14:473) |
| 3 | Water 3 | | 14:3601 | 2.17 | 1.00 | 46.39 | 2.34 (U:O4 14:29) |
| 4 | Water 4 | | 14:3910 | 2.18 | 1.00 | 55.15 | 3.04 (G:O6 14:474) |
| 5 | Water 5 | | 14:3726 | 2.18 | 1.00 | 54.66 | 2.64 (C:N4 14:31) |
| 6 | Water 6 | | 14:3976 | 2.19 | 1.00 | 54.22 | 3.23 (A:OP2 14:28) |
| 7 | A | OP2 | 14:28 | 4.01 | 1.00 | 51.38 |  |
| 8 | U | O4 | 14:29 | 4.06 | 1.00 | 43.52 |  |
| 9 | A | OP2 | 14:447 | 4.15 | 1.00 | 47.22 |  |
| 10 | G | O6 | 14:30 | 4.21 | 1.00 | 47.79 |  |
| 11 | G | O6 | 14:474 | 4.24 | 1.00 | 48.05 |  |
| 12 | C | N4 | 14:31 | 4.38 | 1.00 | 48.10 |  |
| 13 | G | O6 | 14:473 | 4.41 | 1.00 | 46.01 |  |
| 14 | U | OP2 | 14:29 | 4.52 | 1.00 | 48.75 |  |
| 15 | A | O2' | 14:447 | 4.65 | 1.00 | 45.87 |  |
  |

  
**1^st^ shell ligand angles:**   
1-2: 89; 1-3: 92; 1-4: 179; 1-5: 90; 1-6: 88; 2-3: 177; 2-4: 92; 2-5: 91; 2-6: 88; 3-4: 87; 3-5: 86; 3-6: 94; 4-5: 91; 4-6: 91; 5-6: 177;   
  
 Contact(s): /O_w_/O_w_/O_w_/O_w_/O_w_/O_w_/  
 Coded contact list: 6O_w_  
MG hydration shell issue: water too close (< 2.4 Angs.)  

### MG:14:3339

|  | Res. type | Atom | Residue | Distance | Occupancy | B-factor | Closest solute to 1^st^ shell water |
| --- | --- | --- | --- | --- | --- | --- | --- |
| **Ion** | | | |  | 1.00 | 29.35 || 1 | Water 1 | | 14:3707 | 2.16 | 1.00 | 35.55 | 2.60 (G:N7 14:778) |
| 2 | Water 2 | | 14:3611 | 2.16 | 1.00 | 30.32 | 2.37 (G:O6 14:778) |
| 3 | Water 3 | | 14:4054 | 2.18 | 1.00 | 27.42 | 3.80 (A:N1 14:793) |
| 4 | Water 4 | | 14:3814 | 2.18 | 1.00 | 31.71 | 2.79 (G:OP2 14:778) |
| 5 | Water 5 | | 14:3990 | 2.19 | 1.00 | 40.72 | 3.29 (A:N1 14:793) |
| 6 | Water 6 | | 19:411 | 2.19 | 1.00 | 40.08 | 3.92 (U:O4 14:779) |
| 7 | U | O4 | 14:779 | 3.63 | 1.00 | 31.60 |  |
| 8 | G | N7 | 14:778 | 3.75 | 1.00 | 30.26 |  |
| 9 | G | O6 | 14:778 | 3.98 | 1.00 | 31.84 |  |
| 10 | G | OP2 | 14:778 | 4.48 | 1.00 | 32.34 |  |
| 11 | A | N1 | 14:793 | 4.52 | 1.00 | 38.29 |  |
  |

  
**1^st^ shell ligand angles:**   
1-2: 96; 1-3: 89; 1-4: 89; 1-5: 86; 1-6: 174; 2-3: 75; 2-4: 100; 2-5: 173; 2-6: 85; 3-4: 174; 3-5: 98; 3-6: 97; 4-5: 87; 4-6: 85; 5-6: 94;   
  
 Contact(s): /O_w_/O_w_/O_w_/O_w_/O_w_/O_w_/  
 Coded contact list: 6O_w_  
MG hydration shell issue: water too close (< 2.4 Angs.)  

### MG:14:3340

|  | Res. type | Atom | Residue | Distance | Occupancy | B-factor | Closest solute to 1^st^ shell water |
| --- | --- | --- | --- | --- | --- | --- | --- |
| **Ion** | | | |  | 1.00 | 26.06 || 1 | Water 1 | | 14:3712 | 2.17 | 1.00 | 32.12 | 2.61 (A:OP1 14:1603) |
| 2 | Water 2 | | 14:3980 | 2.18 | 1.00 | 33.37 | 3.27 (C:OP1 14:1604) |
| 3 | Water 3 | | 14:3973 | 2.20 | 1.00 | 46.14 | 3.23 (A:OP1 14:1603) |
| 4 | A | OP1 | 14:1603 | 2.45 | 1.00 | 33.29 |  |
| 5 | C | OP2 | 14:1604 | 2.48 | 1.00 | 31.83 |  |
| 6 | A | OP1 | 14:1395 | 2.53 | 1.00 | 46.79 |  |
| 7 | C | OP1 | 14:1604 | 3.06 | 1.00 | 36.70 |  |
| 8 | C | O5' | 14:1604 | 3.89 | 1.00 | 39.13 |  |
| 9 | Water 4 | | 14:3553 | 4.06 | 1.00 | 42.60 | 2.20 (C:OP1 14:1604) |
| 10 | A | OP2 | 14:1603 | 4.35 | 1.00 | 39.52 |  |
| 11 | A | O5' | 14:1603 | 4.37 | 1.00 | 40.72 |  |
| 12 | A | OP2 | 14:1395 | 4.41 | 1.00 | 35.30 |  |
| 13 | Water 5 | | 14:3586 | 4.43 | 1.00 | 33.16 | 2.29 (A:OP2 14:1395) |
| 14 | A | O3' | 14:1603 | 4.50 | 1.00 | 33.48 |  |
| 15 | U | O3' | 14:1394 | 4.67 | 1.00 | 42.05 |  |
| 16 | A | O5' | 14:1395 | 4.81 | 1.00 | 46.55 |  |
| 17 | U | O3' | 14:1602 | 4.89 | 1.00 | 42.22 |  |
  |

  
**1^st^ shell ligand angles:**   
1-2: 101; 1-3: 95; 2-3: 101;   
  
 Contact(s): /O_w_/O_w_/O_w_/  
 Coded contact list: 3O_w_  
 Exclusion zone contact(s): /OP1/OP1/OP1/OP2/  
Consider NA in place of MG (3 valid contact(s) for NA vs. 0 for MG)  

### MG:14:3341

|  | Res. type | Atom | Residue | Distance | Occupancy | B-factor | Closest solute to 1^st^ shell water |
| --- | --- | --- | --- | --- | --- | --- | --- |
| **Ion** | | | |  | 1.00 | 35.98 || 1 | Water 1 | | 14:3836 | 2.17 | 1.00 | 33.27 | 2.83 (A:OP1 14:2005) |
| 2 | Water 2 | | 14:3955 | 2.17 | 1.00 | 52.28 | 3.18 (A:OP1 14:2005) |
| 3 | Water 3 | | 14:3584 | 2.17 | 1.00 | 39.67 | 2.28 (C:OP2 14:2006) |
| 4 | Water 4 | | 14:3663 | 2.18 | 1.00 | 40.10 | 2.52 (A:OP2 14:1268) |
| 5 | A | OP2 | 14:1269 | 2.54 | 1.00 | 38.02 |  |
| 6 | A | OP1 | 14:2005 | 2.59 | 1.00 | 28.40 |  |
| 7 | C | OP2 | 14:2006 | 3.71 | 1.00 | 41.14 |  |
| 8 | A | OP2 | 14:1268 | 4.02 | 1.00 | 42.00 |  |
| 9 | A | O5' | 14:2005 | 4.05 | 1.00 | 43.39 |  |
| 10 | A | O5' | 14:1268 | 4.21 | 1.00 | 39.06 |  |
| 11 | A | N7 | 14:1269 | 4.26 | 1.00 | 37.39 |  |
| 12 | A | O5' | 14:1269 | 4.28 | 1.00 | 35.52 |  |
| 13 | A | O3' | 14:1268 | 4.49 | 1.00 | 38.55 |  |
| 14 | A | OP1 | 14:1268 | 4.75 | 1.00 | 38.94 |  |
| 15 | G | O3' | 14:2004 | 4.83 | 1.00 | 34.61 |  |
| 16 | A | OP2 | 14:2005 | 4.98 | 1.00 | 36.32 |  |
  |

  
**1^st^ shell ligand angles:**   
1-2: 90; 1-3: 175; 1-4: 91; 2-3: 94; 2-4: 83; 3-4: 92;   
  
 Contact(s): /O_w_/O_w_/O_w_/O_w_/  
 Coded contact list: 4O_w_  
 Exclusion zone contact(s): /OP1/OP2/  
Consider NA in place of MG (2 valid contact(s) for NA vs. 0 for MG)  
MG hydration shell issue: water too close (< 2.4 Angs.)  

### MG:14:3342

|  | Res. type | Atom | Residue | Distance | Occupancy | B-factor | Closest solute to 1^st^ shell water |
| --- | --- | --- | --- | --- | --- | --- | --- |
| **Ion** | | | |  | 1.00 | 32.62 || 1 | Water 1 | | 14:3743 | 2.16 | 1.00 | 32.76 | 2.67 (A:OP2 14:2057) |
| 2 | Water 2 | | 14:3891 | 2.16 | 1.00 | 38.79 | 2.99 (2MA:OP2 14:2503) |
| 3 | Water 3 | | 14:3595 | 2.17 | 1.00 | 47.31 | 2.33 (G:OP2 14:2056) |
| 4 | Water 4 | | 14:3824 | 2.18 | 1.00 | 30.67 | 2.81 (A:N7 14:2058) |
| 5 | A | OP2 | 14:2057 | 2.43 | 1.00 | 38.38 |  |
| 6 | G | OP2 | 14:2056 | 2.63 | 1.00 | 39.90 |  |
| 7 | G | O5' | 14:2056 | 3.78 | 1.00 | 39.11 |  |
| 8 | G | OP1 | 14:2056 | 4.13 | 1.00 | 37.69 |  |
| 9 | A | OP1 | 14:2057 | 4.13 | 1.00 | 32.52 |  |
| 10 | A | N7 | 14:2058 | 4.17 | 1.00 | 35.85 |  |
| 11 | A | O5' | 14:2057 | 4.21 | 1.00 | 42.33 |  |
| 12 | Water 5 | | 14:3668 | 4.38 | 1.00 | 45.78 | 2.52 (A:OP1 14:2057) |
| 13 | A | N6 | 14:2058 | 4.38 | 1.00 | 33.53 |  |
| 14 | 2MA | O2' | 14:2503 | 4.56 | 1.00 | 36.83 |  |
| 15 | U | OP2 | 14:2504 | 4.65 | 1.00 | 40.57 |  |
| 16 | 2MA | OP2 | 14:2503 | 4.67 | 1.00 | 30.67 |  |
| 17 | A | N7 | 14:2057 | 4.69 | 1.00 | 40.44 |  |
| 18 | Water 6 | | 14:3722 | 4.71 | 1.00 | 33.10 | 2.63 (U:OP2 14:2504) |
| 19 | G | O3' | 14:2056 | 4.78 | 1.00 | 37.38 |  |
| 20 | 2MA | O3' | 14:2503 | 4.84 | 1.00 | 32.74 |  |
  |

  
**1^st^ shell ligand angles:**   
1-2: 173; 1-3: 81; 1-4: 98; 2-3: 96; 2-4: 88; 3-4: 89;   
  
 Contact(s): /O_w_/O_w_/O_w_/O_w_/  
 Coded contact list: 4O_w_  
 Exclusion zone contact(s): /OP2/OP2/  
Consider NA in place of MG (1 valid contact(s) for NA vs. 0 for MG)  
MG hydration shell issue: water too close (< 2.4 Angs.)  

### MG:14:3343

|  | Res. type | Atom | Residue | Distance | Occupancy | B-factor | Closest solute to 1^st^ shell water |
| --- | --- | --- | --- | --- | --- | --- | --- |
| **Ion** | | | |  | 1.00 | 50.99 || 1 | Water 1 | | 14:3886 | 2.17 | 1.00 | 47.06 | 2.98 (C:OP2 14:1376) |
| 2 | Water 2 | | 14:3695 | 2.18 | 1.00 | 43.05 | 2.57 (C:OP2 14:1376) |
| 3 | Water 3 | | 14:3556 | 2.19 | 1.00 | 54.86 | 2.22 (C:OP1 14:1375) |
| 4 | Water 4 | | 14:3630 | 2.20 | 1.00 | 57.93 | 2.42 (C:OP1 14:1351) |
| 5 | U | OP2 | 14:1352 | 2.49 | 1.00 | 37.76 |  |
| 6 | C | OP2 | 14:1376 | 2.54 | 1.00 | 39.28 |  |
| 7 | U | O5' | 14:1352 | 3.75 | 1.00 | 44.63 |  |
| 8 | C | OP1 | 14:1376 | 3.85 | 1.00 | 52.09 |  |
| 9 | C | O5' | 14:1375 | 3.97 | 1.00 | 49.16 |  |
| 10 | C | OP1 | 14:1375 | 3.98 | 1.00 | 57.28 |  |
| 11 | U | OP1 | 14:1352 | 4.05 | 1.00 | 49.28 |  |
| 12 | C | OP1 | 14:1351 | 4.08 | 1.00 | 38.25 |  |
| 13 | C | OP2 | 14:1375 | 4.15 | 1.00 | 50.26 |  |
| 14 | C | O3' | 14:1375 | 4.41 | 1.00 | 42.70 |  |
| 15 | C | OP2 | 14:1351 | 4.51 | 1.00 | 36.42 |  |
| 16 | C | O5' | 14:1351 | 4.61 | 1.00 | 40.81 |  |
| 17 | C | O5' | 14:1376 | 4.81 | 1.00 | 46.08 |  |
| 18 | C | O3' | 14:1351 | 4.84 | 1.00 | 42.52 |  |
  |

  
**1^st^ shell ligand angles:**   
1-2: 88; 1-3: 92; 1-4: 173; 2-3: 167; 2-4: 92; 3-4: 90;   
  
 Contact(s): /O_w_/O_w_/O_w_/O_w_/  
 Coded contact list: 4O_w_  
 Exclusion zone contact(s): /OP2/OP2/  
Consider NA in place of MG (2 valid contact(s) for NA vs. 0 for MG)  
MG hydration shell issue: water too close (< 2.4 Angs.)  

### MG:14:3344

|  | Res. type | Atom | Residue | Distance | Occupancy | B-factor | Closest solute to 1^st^ shell water |
| --- | --- | --- | --- | --- | --- | --- | --- |
| **Ion** | | | |  | 1.00 | 31.71 || 1 | Water 1 | | 14:3567 | 2.15 | 1.00 | 35.30 | 2.25 (C:O3' 14:2575) |
| 2 | Water 2 | | 14:3682 | 2.16 | 1.00 | 52.53 | 2.55 (G:OP2 14:2053) |
| 3 | Water 3 | | 14:3581 | 2.16 | 1.00 | 46.02 | 2.28 (G:OP1 14:2053) |
| 4 | Water 4 | | 14:3757 | 2.18 | 1.00 | 46.93 | 2.68 (A:OP2 14:2577) |
| 5 | A | OP1 | 14:2577 | 2.54 | 1.00 | 44.67 |  |
| 6 | G | OP1 | 14:2576 | 2.65 | 1.00 | 40.68 |  |
| 7 | A | OP2 | 14:2577 | 3.31 | 1.00 | 36.41 |  |
| 8 | G | OP1 | 14:2053 | 3.79 | 1.00 | 43.13 |  |
| 9 | C | O3' | 14:2575 | 4.08 | 1.00 | 39.92 |  |
| 10 | A | O5' | 14:2577 | 4.30 | 1.00 | 44.70 |  |
| 11 | G | OP2 | 14:2053 | 4.30 | 1.00 | 46.67 |  |
| 12 | G | N2 | 14:2576 | 4.31 | 1.00 | 45.28 |  |
| 13 | G | O3' | 14:2576 | 4.50 | 1.00 | 37.90 |  |
| 14 | G | OP2 | 14:2052 | 4.57 | 1.00 | 45.47 |  |
| 15 | A | N7 | 14:2577 | 4.70 | 1.00 | 41.75 |  |
| 16 | G | O5' | 14:2576 | 4.80 | 1.00 | 39.73 |  |
| 17 | G | OP2 | 14:2576 | 4.96 | 1.00 | 44.18 |  |
  |

  
**1^st^ shell ligand angles:**   
1-2: 85; 1-3: 93; 1-4: 172; 2-3: 93; 2-4: 92; 3-4: 95;   
  
 Contact(s): /O_w_/O_w_/O_w_/O_w_/  
 Coded contact list: 4O_w_  
 Exclusion zone contact(s): /OP1/OP1/OP2/  
Consider K in place of MG (2 valid contact(s) for K vs. 0 for MG)  
MG hydration shell issue: water too close (< 2.4 Angs.)  

### MG:14:3345

|  | Res. type | Atom | Residue | Distance | Occupancy | B-factor | Closest solute to 1^st^ shell water |
| --- | --- | --- | --- | --- | --- | --- | --- |
| **Ion** | | | |  | 1.00 | 48.73 || 1 | Water 1 | | 14:3642 | 2.14 | 1.00 | 46.87 | 2.47 (G:O6 14:1355) |
| 2 | Water 2 | | 14:3987 | 2.17 | 1.00 | 52.11 | 3.29 (A:OP2 14:1354) |
| 3 | Water 3 | | 14:3732 | 2.18 | 1.00 | 32.01 | 2.65 (A:OP1 14:1353) |
| 4 | Water 4 | | 14:4051 | 2.19 | 1.00 | 57.89 | 3.78 (C:N4 14:1375) |
| 5 | Water 5 | | 14:3805 | 2.20 | 1.00 | 54.73 | 2.76 (G:O6 14:1356) |
| 6 | Water 6 | | 14:4017 | 2.20 | 1.00 | 58.68 | 5.66 (A:OP1 14:1354) |
| 7 | G | O6 | 14:1355 | 4.10 | 1.00 | 49.35 |  |
| 8 | A | OP2 | 14:1354 | 4.25 | 1.00 | 36.46 |  |
| 9 | A | OP1 | 14:1353 | 4.28 | 1.00 | 38.85 |  |
| 10 | C | N4 | 14:1375 | 4.42 | 1.00 | 49.05 |  |
| 11 | G | O6 | 14:1356 | 4.55 | 1.00 | 44.15 |  |
| 12 | A | OP1 | 14:1354 | 4.76 | 1.00 | 45.00 |  |
| 13 | A | OP2 | 14:1353 | 4.80 | 1.00 | 37.76 |  |
  |

  
**1^st^ shell ligand angles:**   
1-2: 83; 1-3: 86; 1-4: 94; 1-5: 93; 1-6: 178; 2-3: 89; 2-4: 177; 2-5: 90; 2-6: 95; 3-4: 89; 3-5: 178; 3-6: 95; 4-5: 91; 4-6: 88; 5-6: 86;   
  
 Contact(s): /O_w_/O_w_/O_w_/O_w_/O_w_/O_w_/  
 Coded contact list: 6O_w_  

### MG:14:3346

|  | Res. type | Atom | Residue | Distance | Occupancy | B-factor | Closest solute to 1^st^ shell water |
| --- | --- | --- | --- | --- | --- | --- | --- |
| **Ion** | | | |  | 1.00 | 52.96 || 1 | Water 1 | | 14:3789 | 2.17 | 1.00 | 63.35 | 2.73 (G:OP1 14:2407) |
| 2 | Water 2 | | 14:3637 | 2.17 | 1.00 | 51.69 | 2.47 (U:OP1 14:2406) |
| 3 | Water 3 | | 14:3662 | 2.17 | 1.00 | 63.29 | 2.51 (U:OP2 14:2408) |
| 4 | Water 4 | | 14:3688 | 2.17 | 1.00 | 60.81 | 2.55 (U:OP2 14:2406) |
| 5 | Water 5 | | 14:3759 | 2.18 | 1.00 | 56.21 | 2.68 (G:N7 14:2409) |
| 6 | U | OP1 | 14:2406 | 2.51 | 1.00 | 59.15 |  |
| 7 | U | OP2 | 14:2406 | 3.23 | 1.00 | 54.77 |  |
| 8 | U | OP2 | 14:2408 | 3.82 | 1.00 | 58.84 |  |
| 9 | G | O6 | 14:2410 | 4.05 | 1.00 | 60.82 |  |
| 10 | U | O4 | 14:2408 | 4.08 | 1.00 | 58.75 |  |
| 11 | U | O5' | 14:2406 | 4.19 | 1.00 | 63.44 |  |
| 12 | G | O3' | 14:2405 | 4.45 | 1.00 | 61.87 |  |
| 13 | G | N7 | 14:2409 | 4.56 | 1.00 | 62.71 |  |
| 14 | G | O2' | 14:2405 | 4.60 | 1.00 | 62.15 |  |
| 15 | G | OP1 | 14:2407 | 4.63 | 1.00 | 53.10 |  |
| 16 | G | N7 | 14:2410 | 4.79 | 1.00 | 62.14 |  |
| 17 | G | O6 | 14:2409 | 4.91 | 1.00 | 58.18 |  |
| 18 | A | N6 | 14:2411 | 4.99 | 1.00 | 60.88 |  |
  |

  
**1^st^ shell ligand angles:**   
1-2: 177; 1-3: 88; 1-4: 88; 1-5: 98; 2-3: 92; 2-4: 92; 2-5: 85; 3-4: 175; 3-5: 89; 4-5: 95;   
  
 Contact(s): /O_w_/O_w_/O_w_/O_w_/O_w_/  
 Coded contact list: 5O_w_  
 Exclusion zone contact(s): /OP1/OP2/  
Consider NA in place of MG (1 valid contact(s) for NA vs. 0 for MG)  

### MG:14:3347

|  | Res. type | Atom | Residue | Distance | Occupancy | B-factor | Closest solute to 1^st^ shell water |
| --- | --- | --- | --- | --- | --- | --- | --- |
| **Ion** | | | |  | 1.00 | 27.52 || 1 | Water 1 | | 14:3697 | 2.17 | 1.00 | 25.93 | 2.58 (U:OP2 14:740) |
| 2 | Water 2 | | 14:3773 | 2.18 | 1.00 | 42.67 | 2.70 (A:OP1 14:1784) |
| 3 | Water 3 | | 14:3687 | 2.18 | 1.00 | 42.13 | 2.55 (U:OP2 14:740) |
| 4 | A | OP2 | 14:1784 | 2.51 | 1.00 | 39.88 |  |
| 5 | U | OP2 | 14:740 | 2.53 | 1.00 | 34.62 |  |
| 6 | A | OP1 | 14:1783 | 2.59 | 1.00 | 33.65 |  |
| 7 | A | O5' | 14:1783 | 3.17 | 1.00 | 33.91 |  |
| 8 | A | OP1 | 14:1784 | 3.20 | 1.00 | 34.08 |  |
| 9 | A | O3' | 14:1783 | 3.88 | 1.00 | 34.60 |  |
| 10 | G | OP2 | 14:741 | 3.90 | 1.00 | 37.34 |  |
| 11 | U | OP1 | 14:740 | 3.93 | 1.00 | 33.03 |  |
| 12 | U | O5' | 14:740 | 4.15 | 1.00 | 32.75 |  |
| 13 | A | O5' | 14:1784 | 4.42 | 1.00 | 31.63 |  |
| 14 | C | O3' | 14:1782 | 4.42 | 1.00 | 33.39 |  |
| 15 | Water 4 | | 14:3616 | 4.69 | 1.00 | 36.36 | 2.38 (A:OP2 14:1783) |
| 16 | A | OP2 | 14:1783 | 4.76 | 1.00 | 32.11 |  |
| 17 | Water 5 | | 14:3852 | 4.83 | 1.00 | 37.67 | 2.89 (A:OP1 14:1783) |
| 18 | G | O3' | 14:739 | 4.91 | 1.00 | 34.27 |  |
  |

  
**1^st^ shell ligand angles:**   
1-2: 175; 1-3: 92; 2-3: 93;   
  
 Contact(s): /O_w_/O_w_/O_w_/  
 Coded contact list: 3O_w_  
 Exclusion zone contact(s): /O5'/OP1/OP1/OP2/OP2/  
Consider NA in place of MG (3 valid contact(s) for NA vs. 0 for MG)  

### MG:14:3348

|  | Res. type | Atom | Residue | Distance | Occupancy | B-factor | Closest solute to 1^st^ shell water |
| --- | --- | --- | --- | --- | --- | --- | --- |
| **Ion** | | | |  | 1.00 | 30.79 || 1 | Water 1 | | 29:401 | 2.16 | 1.00 | 41.92 | 1.30 (HIS:NE2 29:132) |
| 2 | Water 2 | | 14:3505 | 2.18 | 1.00 | 46.06 | 1.30 (A:N7 14:746) |
| 3 | Water 3 | | 14:3537 | 2.18 | 1.00 | 43.13 | 2.11 (U:OP1 14:2611) |
| 4 | Water 4 | | 14:3569 | 2.18 | 1.00 | 52.09 | 2.25 (C:OP1 14:2612) |
| 5 | HIS | NE2 | 29:132 | 2.54 | 1.00 | 49.24 |  |
| 6 | C | OP2 | 14:2612 | 2.70 | 1.00 | 40.82 |  |
| 7 | C | OP1 | 14:2612 | 3.02 | 1.00 | 48.74 |  |
| 8 | A | N7 | 14:746 | 3.32 | 1.00 | 48.00 |  |
| 9 | U | O3' | 14:2611 | 3.73 | 1.00 | 40.83 |  |
| 10 | U | OP1 | 14:2611 | 4.00 | 1.00 | 37.09 |  |
| 11 | HIS | ND1 | 29:132 | 4.20 | 1.00 | 39.27 |  |
| 12 | G | OP2 | 14:745 | 4.40 | 1.00 | 36.56 |  |
| 13 | U | O5' | 14:2611 | 4.55 | 1.00 | 37.74 |  |
| 14 | C | O5' | 14:2612 | 4.63 | 1.00 | 45.99 |  |
| 15 | U | OP2 | 14:2611 | 4.78 | 1.00 | 35.82 |  |
| 16 | LYS | NZ | 29:133 | 4.90 | 1.00 | 36.24 |  |
| 17 | A | N6 | 14:746 | 5.00 | 1.00 | 45.73 |  |
  |

  
**1^st^ shell ligand angles:**   
1-2: 178; 1-3: 93; 1-4: 87; 2-3: 88; 2-4: 92; 3-4: 176;   
  
 Contact(s): /O_w_/O_w_/O_w_/O_w_/  
 Coded contact list: 4O_w_  
 Exclusion zone contact(s): /N7/NE2_His_/OP1/OP2/  
Consider K in place of MG (3 valid contact(s) for K vs. 0 for MG)  
MG hydration shell issue: water too close (< 2.4 Angs.)  

### MG:14:3349

|  | Res. type | Atom | Residue | Distance | Occupancy | B-factor | Closest solute to 1^st^ shell water |
| --- | --- | --- | --- | --- | --- | --- | --- |
| **Ion** | | | |  | 1.00 | 48.81 || 1 | Water 1 | | 14:3901 | 2.17 | 1.00 | 55.81 | 3.01 (U:OP1 14:2613) |
| 2 | Water 2 | | 14:3791 | 2.18 | 1.00 | 43.33 | 2.73 (G:OP2 14:2578) |
| 3 | Water 3 | | 14:3612 | 2.18 | 1.00 | 47.03 | 2.38 (A:OP2 14:2051) |
| 4 | Water 4 | | 14:3766 | 2.18 | 1.00 | 40.95 | 2.69 (G:OP1 14:2578) |
| 5 | Water 5 | | 14:3961 | 2.19 | 1.00 | 39.06 | 3.20 (A:OP2 14:2614) |
| 6 | Water 6 | | 14:3936 | 2.19 | 1.00 | 47.91 | 3.12 (C:OP2 14:2050) |
| 7 | G | OP1 | 14:2578 | 3.92 | 1.00 | 39.47 |  |
| 8 | A | OP2 | 14:2051 | 3.95 | 1.00 | 40.49 |  |
| 9 | G | OP2 | 14:2578 | 4.19 | 1.00 | 42.11 |  |
| 10 | A | N7 | 14:2051 | 4.38 | 1.00 | 41.57 |  |
| 11 | A | N7 | 14:2614 | 4.45 | 1.00 | 38.67 |  |
| 12 | C | OP2 | 14:2050 | 4.45 | 1.00 | 34.96 |  |
| 13 | U | OP1 | 14:2613 | 4.52 | 1.00 | 51.51 |  |
| 14 | A | OP2 | 14:2614 | 5.00 | 1.00 | 40.38 |  |
  |

  
**1^st^ shell ligand angles:**   
1-2: 90; 1-3: 178; 1-4: 91; 1-5: 92; 1-6: 88; 2-3: 88; 2-4: 99; 2-5: 178; 2-6: 82; 3-4: 88; 3-5: 90; 3-6: 92; 4-5: 83; 4-6: 179; 5-6: 96;   
  
 Contact(s): /O_w_/O_w_/O_w_/O_w_/O_w_/O_w_/  
 Coded contact list: 6O_w_  
MG hydration shell issue: water too close (< 2.4 Angs.)  

### MG:14:3350

|  | Res. type | Atom | Residue | Distance | Occupancy | B-factor | Closest solute to 1^st^ shell water |
| --- | --- | --- | --- | --- | --- | --- | --- |
| **Ion** | | | |  | 1.00 | 39.66 || 1 | Water 1 | | 14:3538 | 2.16 | 1.00 | 44.83 | 2.12 (G:OP2 14:1997) |
| 2 | Water 2 | | 14:3898 | 2.17 | 1.00 | 42.15 | 3.01 (C:O2' 14:1996) |
| 3 | Water 3 | | 29:403 | 2.17 | 1.00 | 38.15 | 2.78 (GLY:O 29:125) |
| 4 | Water 4 | | 14:3605 | 2.18 | 1.00 | 48.96 | 2.35 (C:OP1 14:1994) |
| 5 | Water 5 | | 29:402 | 2.19 | 1.00 | 48.95 | 2.78 (ASP:OD1 29:127) |
| 6 | G | OP2 | 14:1997 | 2.51 | 1.00 | 38.32 |  |
| 7 | C | O3' | 14:1996 | 3.79 | 1.00 | 41.40 |  |
| 8 | GLY | O | 29:125 | 4.10 | 1.00 | 46.08 |  |
| 9 | ASP | OD1 | 29:127 | 4.25 | 1.00 | 52.86 |  |
| 10 | U | OP2 | 14:1995 | 4.36 | 1.00 | 45.02 |  |
| 11 | ALA | O | 29:123 | 4.41 | 1.00 | 43.18 |  |
| 12 | C | OP1 | 14:1994 | 4.43 | 1.00 | 40.75 |  |
| 13 | C | OP2 | 14:1994 | 4.54 | 1.00 | 47.40 |  |
| 14 | G | O5' | 14:1997 | 4.76 | 1.00 | 46.04 |  |
| 15 | G | N7 | 14:1997 | 4.76 | 1.00 | 43.12 |  |
| 16 | G | OP1 | 14:1997 | 4.79 | 1.00 | 40.41 |  |
| 17 | C | O2' | 14:1996 | 4.91 | 1.00 | 39.08 |  |
| 18 | ASP | OD2 | 29:127 | 4.98 | 1.00 | 54.09 |  |
  |

  
**1^st^ shell ligand angles:**   
1-2: 86; 1-3: 90; 1-4: 84; 1-5: 176; 2-3: 173; 2-4: 95; 2-5: 92; 3-4: 91; 3-5: 92; 4-5: 92;   
  
 Contact(s): /O_w_/O_w_/O_w_/O_w_/O_w_/  
 Coded contact list: 5O_w_  
 Exclusion zone contact(s): /OP2/  
Consider NA in place of MG (1 valid contact(s) for NA vs. 0 for MG)  
MG hydration shell issue: water too close (< 2.4 Angs.)  

### MG:14:3351

|  | Res. type | Atom | Residue | Distance | Occupancy | B-factor | Closest solute to 1^st^ shell water |
| --- | --- | --- | --- | --- | --- | --- | --- |
| **Ion** | | | |  | 1.00 | 55.21 || 1 | Water 1 | | 14:3618 | 2.16 | 1.00 | 50.05 | 2.39 (A:OP1 14:1664) |
| 2 | Water 2 | | 14:3673 | 2.18 | 1.00 | 38.20 | 2.53 (A:OP2 14:1665) |
| 3 | Water 3 | | 14:3899 | 2.19 | 1.00 | 58.31 | 3.01 (G:OP2 14:1666) |
| 4 | Water 4 | | 14:4090 | 2.19 | 1.00 | 62.70 | 4.46 (G:O6 14:1667) |
| 5 | Water 5 | | 14:3923 | 2.19 | 1.00 | 62.02 | 3.08 (G:OP2 14:1666) |
| 6 | A | OP2 | 14:1665 | 2.53 | 1.00 | 44.59 |  |
| 7 | A | OP1 | 14:1664 | 3.91 | 1.00 | 42.18 |  |
| 8 | G | OP2 | 14:1666 | 3.97 | 1.00 | 52.00 |  |
| 9 | A | O5' | 14:1665 | 4.00 | 1.00 | 52.41 |  |
| 10 | A | OP1 | 14:1665 | 4.10 | 1.00 | 53.51 |  |
| 11 | A | O5' | 14:1664 | 4.12 | 1.00 | 49.37 |  |
| 12 | Water 6 | | 14:3763 | 4.26 | 1.00 | 43.57 | 2.69 (A:OP2 14:1664) |
| 13 | A | OP2 | 14:1664 | 4.66 | 1.00 | 40.84 |  |
| 14 | G | O6 | 14:1667 | 4.68 | 1.00 | 53.80 |  |
| 15 | A | O3' | 14:1664 | 4.89 | 1.00 | 47.51 |  |
  |

  
**1^st^ shell ligand angles:**   
1-2: 88; 1-3: 91; 1-4: 92; 1-5: 177; 2-3: 177; 2-4: 92; 2-5: 91; 3-4: 90; 3-5: 91; 4-5: 91;   
  
 Contact(s): /O_w_/O_w_/O_w_/O_w_/O_w_/  
 Coded contact list: 5O_w_  
 Exclusion zone contact(s): /OP2/  
Consider NA in place of MG (1 valid contact(s) for NA vs. 0 for MG)  
MG hydration shell issue: water too close (< 2.4 Angs.)  

### MG:14:3352

|  | Res. type | Atom | Residue | Distance | Occupancy | B-factor | Closest solute to 1^st^ shell water |
| --- | --- | --- | --- | --- | --- | --- | --- |
| **Ion** | | | |  | 1.00 | 63.63 || 1 | Water 1 | | 14:4048 | 2.17 | 1.00 | 65.14 | 3.77 (G:N7 14:2845) |
| 2 | Water 2 | | 14:3672 | 2.17 | 1.00 | 56.44 | 2.53 (G:OP2 14:2845) |
| 3 | Water 3 | | 14:3812 | 2.18 | 1.00 | 61.24 | 2.78 (G:N7 14:2846) |
| 4 | Water 4 | | 14:3758 | 2.19 | 1.00 | 58.53 | 2.68 (G:O6 14:2848) |
| 5 | Water 5 | | 75:201 | 2.19 | 1.00 | 78.17 | 2.48 (ARG:NE 75:54) |
| 6 | Water 6 | | 14:3972 | 2.19 | 1.00 | 56.90 | 3.22 (G:OP2 14:2845) |
| 7 | G | O6 | 14:2848 | 4.02 | 1.00 | 66.54 |  |
| 8 | G | OP2 | 14:2845 | 4.09 | 1.00 | 64.30 |  |
| 9 | G | OP2 | 14:2846 | 4.10 | 1.00 | 64.18 |  |
| 10 | ARG | NE | 75:54 | 4.30 | 1.00 | 63.33 |  |
| 11 | G | N7 | 14:2845 | 4.40 | 1.00 | 64.85 |  |
| 12 | G | N7 | 14:2846 | 4.63 | 1.00 | 63.39 |  |
| 13 | U | O4 | 14:2847 | 4.77 | 1.00 | 63.09 |  |
  |

  
**1^st^ shell ligand angles:**   
1-2: 90; 1-3: 84; 1-4: 92; 1-5: 176; 1-6: 96; 2-3: 95; 2-4: 178; 2-5: 91; 2-6: 83; 3-4: 87; 3-5: 92; 3-6: 178; 4-5: 88; 4-6: 95; 5-6: 89;   
  
 Contact(s): /O_w_/O_w_/O_w_/O_w_/O_w_/O_w_/  
 Coded contact list: 6O_w_  

### MG:14:3353

|  | Res. type | Atom | Residue | Distance | Occupancy | B-factor | Closest solute to 1^st^ shell water |
| --- | --- | --- | --- | --- | --- | --- | --- |
| **Ion** | | | |  | 1.00 | 62.63 || 1 | Water 1 | | 14:4069 | 2.18 | 1.00 | 32.11 | 5.37 (C:OP2 14:1607) |
| 2 | Water 2 | | 14:3549 | 2.18 | 1.00 | 48.94 | 2.18 (G:O6 14:1624) |
| 3 | Water 3 | | 14:3995 | 2.19 | 1.00 | 39.62 | 3.30 (G:N7 14:1623) |
| 4 | Water 4 | | 14:3939 | 2.19 | 1.00 | 54.70 | 3.12 (G:O6 14:1624) |
| 5 | Water 5 | | 14:4075 | 2.19 | 1.00 | 61.79 | 4.04 (G:O6 14:1624) |
| 6 | Water 6 | | 14:3840 | 2.20 | 1.00 | 44.81 | 2.86 (C:OP2 14:1607) |
| 7 | G | O6 | 14:1624 | 3.74 | 1.00 | 49.44 |  |
| 8 | G | O6 | 14:1623 | 3.96 | 1.00 | 43.32 |  |
| 9 | G | N7 | 14:1623 | 4.22 | 1.00 | 39.87 |  |
| 10 | C | N4 | 14:1305 | 4.60 | 1.00 | 48.35 |  |
| 11 | C | OP2 | 14:1607 | 4.92 | 1.00 | 45.72 |  |
| 12 | C | N4 | 14:1304 | 4.96 | 1.00 | 45.71 |  |
  |

  
**1^st^ shell ligand angles:**   
1-2: 178; 1-3: 89; 1-4: 91; 1-5: 92; 1-6: 92; 2-3: 94; 2-4: 87; 2-5: 89; 2-6: 88; 3-4: 178; 3-5: 91; 3-6: 88; 4-5: 87; 4-6: 94; 5-6: 177;   
  
 Contact(s): /O_w_/O_w_/O_w_/O_w_/O_w_/O_w_/  
 Coded contact list: 6O_w_  
MG hydration shell issue: water too close (< 2.4 Angs.)  

### MG:14:3354

|  | Res. type | Atom | Residue | Distance | Occupancy | B-factor | Closest solute to 1^st^ shell water |
| --- | --- | --- | --- | --- | --- | --- | --- |
| **Ion** | | | |  | 1.00 | 30.17 || 1 | Water 1 | | 14:3658 | 2.17 | 1.00 | 39.38 | 2.51 (C:O2' 14:1646) |
| 2 | Water 2 | | 14:3536 | 2.17 | 1.00 | 45.79 | 2.10 (A:O2' 14:1616) |
| 3 | Water 3 | | 14:3820 | 2.17 | 1.00 | 46.38 | 2.81 (G:OP1 14:1647) |
| 4 | Water 4 | | 14:3705 | 2.18 | 1.00 | 42.74 | 2.60 (A:O2' 14:1616) |
| 5 | Water 5 | | 14:4000 | 2.19 | 1.00 | 40.70 | 3.32 (G:OP2 14:1647) |
| 6 | G | OP2 | 14:1647 | 2.57 | 1.00 | 37.28 |  |
| 7 | G | OP1 | 14:1647 | 3.34 | 1.00 | 49.34 |  |
| 8 | A | O2' | 14:1616 | 3.37 | 1.00 | 38.31 |  |
| 9 | C | O3' | 14:1646 | 4.19 | 1.00 | 40.91 |  |
| 10 | G | OP1 | 14:1325 | 4.19 | 1.00 | 44.81 |  |
| 11 | C | O2' | 14:1646 | 4.64 | 1.00 | 47.51 |  |
| 12 | G | O5' | 14:1647 | 4.72 | 1.00 | 43.15 |  |
| 13 | A | OP2 | 14:1272 | 4.84 | 1.00 | 43.66 |  |
| 14 | A | N9 | 14:1616 | 4.96 | 1.00 | 50.66 |  |
| 15 | A | O3' | 14:1616 | 4.97 | 1.00 | 41.81 |  |
  |

  
**1^st^ shell ligand angles:**   
1-2: 178; 1-3: 92; 1-4: 90; 1-5: 88; 2-3: 88; 2-4: 87; 2-5: 92; 3-4: 90; 3-5: 177; 4-5: 93;   
  
 Contact(s): /O_w_/O_w_/O_w_/O_w_/O_w_/  
 Coded contact list: 5O_w_  
 Exclusion zone contact(s): /O2'/OP1/OP2/  
Consider K in place of MG (2 valid contact(s) for K vs. 0 for MG)  
MG hydration shell issue: water too close (< 2.4 Angs.)  

### MG:14:3355

|  | Res. type | Atom | Residue | Distance | Occupancy | B-factor | Closest solute to 1^st^ shell water |
| --- | --- | --- | --- | --- | --- | --- | --- |
| **Ion** | | | |  | 1.00 | 39.92 || 1 | Water 1 | | 14:4024 | 2.17 | 1.00 | 43.78 | 3.70 (A:OP2 14:578) |
| 2 | Water 2 | | 14:3896 | 2.17 | 1.00 | 34.69 | 3.00 (U:OP1 14:2017) |
| 3 | Water 3 | | 14:3565 | 2.18 | 1.00 | 46.79 | 2.24 (A:OP2 14:578) |
| 4 | Water 4 | | 14:3522 | 2.18 | 1.00 | 41.56 | 2.00 (U:OP1 14:2017) |
| 5 | Water 5 | | 14:3534 | 2.19 | 1.00 | 46.23 | 2.08 (A:OP2 14:578) |
| 6 | Water 6 | | 14:3811 | 2.20 | 1.00 | 39.74 | 2.78 (U:OP1 14:2017) |
| 7 | U | OP1 | 14:2017 | 2.99 | 1.00 | 40.33 |  |
| 8 | A | OP2 | 14:578 | 3.03 | 1.00 | 36.64 |  |
| 9 | G | OP2 | 14:2018 | 4.01 | 1.00 | 34.43 |  |
| 10 | U | OP2 | 14:2017 | 4.12 | 1.00 | 42.12 |  |
| 11 | G | OP2 | 14:577 | 4.47 | 1.00 | 35.21 |  |
| 12 | U | O5' | 14:2017 | 4.65 | 1.00 | 36.78 |  |
| 13 | A | O5' | 14:578 | 4.84 | 1.00 | 42.29 |  |
  |

  
**1^st^ shell ligand angles:**   
1-2: 93; 1-3: 91; 1-4: 83; 1-5: 89; 1-6: 178; 2-3: 91; 2-4: 90; 2-5: 178; 2-6: 89; 3-4: 174; 3-5: 91; 3-6: 88; 4-5: 88; 4-6: 98; 5-6: 90;   
  
 Contact(s): /O_w_/O_w_/O_w_/O_w_/O_w_/O_w_/  
 Coded contact list: 6O_w_  
 Exclusion zone contact(s): /OP1/OP2/  
Consider K in place of MG (2 valid contact(s) for K vs. 0 for MG)  
MG hydration shell issue: water too close (< 2.4 Angs.)  

### MG:14:3356

|  | Res. type | Atom | Residue | Distance | Occupancy | B-factor | Closest solute to 1^st^ shell water |
| --- | --- | --- | --- | --- | --- | --- | --- |
| **Ion** | | | |  | 1.00 | 36.57 || 1 | Water 1 | | 14:3767 | 2.17 | 1.00 | 40.18 | 2.69 (U:O4 14:810) |
| 2 | Water 2 | | 14:3829 | 2.17 | 1.00 | 42.33 | 2.82 (A:OP1 14:567) |
| 3 | Water 3 | | 14:4012 | 2.19 | 1.00 | 43.42 | 3.39 (G:N7 14:809) |
| 4 | Water 4 | | 14:4013 | 2.19 | 1.00 | 50.10 | 3.39 (U:O4 14:810) |
| 5 | Water 5 | | 14:3834 | 2.19 | 1.00 | 37.81 | 2.83 (U:O3' 14:566) |
| 6 | A | OP1 | 14:567 | 2.33 | 1.00 | 38.54 |  |
| 7 | U | O3' | 14:566 | 3.87 | 1.00 | 44.20 |  |
| 8 | U | O4 | 14:810 | 4.02 | 1.00 | 40.96 |  |
| 9 | G | OP2 | 14:808 | 4.12 | 1.00 | 36.39 |  |
| 10 | A | O5' | 14:567 | 4.39 | 1.00 | 41.68 |  |
| 11 | G | N7 | 14:809 | 4.54 | 1.00 | 39.80 |  |
| 12 | A | OP2 | 14:567 | 4.74 | 1.00 | 41.81 |  |
| 13 | G | OP2 | 14:809 | 4.79 | 1.00 | 40.74 |  |
  |

  
**1^st^ shell ligand angles:**   
1-2: 169; 1-3: 94; 1-4: 90; 1-5: 94; 1-6: 92; 2-3: 97; 2-4: 88; 2-5: 87; 2-6: 77; 3-4: 89; 3-5: 92; 3-6: 168; 4-5: 175; 4-6: 101; 5-6: 78;   
  
 Contact(s): /O_w_/O_w_/O_w_/O_w_/O_w_/OP1/  
 Coded contact list: O_ph_5O_w_  

### MG:14:3357

|  | Res. type | Atom | Residue | Distance | Occupancy | B-factor | Closest solute to 1^st^ shell water |
| --- | --- | --- | --- | --- | --- | --- | --- |
| **Ion** | | | |  | 1.00 | 48.92 || 1 | Water 1 | | 14:3821 | 2.17 | 1.00 | 55.99 | 2.81 (G:O6 14:1190) |
| 2 | Water 2 | | 14:3635 | 2.17 | 1.00 | 60.36 | 2.45 (A:OP1 14:1189) |
| 3 | Water 3 | | 14:3503 | 2.18 | 1.00 | 67.33 | 1.30 (G:N7 14:1190) |
| 4 | Water 4 | | 14:3968 | 2.18 | 1.00 | 64.08 | 3.21 (A:OP1 14:1189) |
| 5 | Water 5 | | 35:204 | 2.18 | 1.00 | 55.17 | 2.84 (GLY:O 35:26) |
| 6 | A | OP2 | 14:1189 | 2.58 | 1.00 | 44.85 |  |
| 7 | A | OP1 | 14:1189 | 3.16 | 1.00 | 60.97 |  |
| 8 | G | N7 | 14:1190 | 3.24 | 1.00 | 63.72 |  |
| 9 | GLY | O | 35:26 | 3.79 | 1.00 | 54.68 |  |
| 10 | A | O5' | 14:1189 | 3.92 | 1.00 | 46.61 |  |
| 11 | G | O6 | 14:1190 | 4.32 | 1.00 | 59.69 |  |
| 12 | U | O3' | 14:1188 | 4.62 | 1.00 | 39.66 |  |
| 13 | G | OP2 | 14:1190 | 4.71 | 1.00 | 46.76 |  |
| 14 | U | OP1 | 14:1188 | 4.76 | 1.00 | 45.20 |  |
| 15 | U | OP2 | 14:1188 | 4.93 | 1.00 | 50.40 |  |
  |

  
**1^st^ shell ligand angles:**   
1-2: 177; 1-3: 91; 1-4: 88; 1-5: 91; 2-3: 88; 2-4: 94; 2-5: 91; 3-4: 178; 3-5: 88; 4-5: 93;   
  
 Contact(s): /O_w_/O_w_/O_w_/O_w_/O_w_/  
 Coded contact list: 5O_w_  
 Exclusion zone contact(s): /N7/OP1/OP2/  
Consider K in place of MG (2 valid contact(s) for K vs. 0 for MG)  
MG hydration shell issue: water too close (< 2.4 Angs.)  

### MG:14:3358

|  | Res. type | Atom | Residue | Distance | Occupancy | B-factor | Closest solute to 1^st^ shell water |
| --- | --- | --- | --- | --- | --- | --- | --- |
| **Ion** | | | |  | 1.00 | 51.03 || 1 | Water 1 | | 14:3555 | 2.17 | 1.00 | 67.70 | 2.21 (G:O2' 14:989) |
| 2 | Water 2 | | 14:3674 | 2.17 | 1.00 | 63.71 | 2.53 (A:OP2 14:990) |
| 3 | Water 3 | | 14:3600 | 2.18 | 1.00 | 67.24 | 2.34 (C:OP1 14:1185) |
| 4 | A | OP2 | 14:990 | 2.51 | 1.00 | 59.02 |  |
| 5 | G | OP2 | 14:1186 | 2.66 | 1.00 | 64.78 |  |
| 6 | G | O3' | 14:989 | 2.92 | 1.00 | 64.68 |  |
| 7 | C | OP2 | 14:991 | 3.04 | 1.00 | 69.90 |  |
| 8 | G | O2' | 14:989 | 3.27 | 1.00 | 69.86 |  |
| 9 | G | OP1 | 14:1186 | 3.68 | 1.00 | 66.09 |  |
| 10 | A | OP1 | 14:990 | 4.03 | 1.00 | 64.58 |  |
| 11 | G | O5' | 14:1186 | 4.11 | 1.00 | 64.16 |  |
| 12 | C | OP1 | 14:1185 | 4.22 | 1.00 | 70.65 |  |
| 13 | A | O5' | 14:990 | 4.48 | 1.00 | 66.84 |  |
| 14 | C | O5' | 14:1185 | 4.64 | 1.00 | 72.59 |  |
| 15 | C | OP1 | 14:991 | 4.73 | 1.00 | 72.71 |  |
| 16 | G | O4' | 14:1186 | 4.82 | 1.00 | 61.57 |  |
| 17 | C | O3' | 14:1185 | 4.90 | 1.00 | 62.04 |  |
| 18 | C | OP2 | 14:1185 | 4.95 | 1.00 | 70.86 |  |
  |

  
**1^st^ shell ligand angles:**   
1-2: 178; 1-3: 92; 2-3: 89;   
  
 Contact(s): /O_w_/O_w_/O_w_/  
 Coded contact list: 3O_w_  
 Exclusion zone contact(s): /O2'/O3'/OP2/OP2/OP2/  
Consider K in place of MG (4 valid contact(s) for K vs. 0 for MG)  
MG hydration shell issue: water too close (< 2.4 Angs.)  

### MG:14:3359

|  | Res. type | Atom | Residue | Distance | Occupancy | B-factor | Closest solute to 1^st^ shell water |
| --- | --- | --- | --- | --- | --- | --- | --- |
| **Ion** | | | |  | 1.00 | 33.73 || 1 | Water 1 | | 14:3855 | 2.16 | 1.00 | 48.47 | 2.90 (U:O4 14:568) |
| 2 | Water 2 | | 14:3654 | 2.16 | 1.00 | 51.74 | 2.50 (G:OP1 14:570) |
| 3 | Water 3 | | 14:3751 | 2.17 | 1.00 | 42.18 | 2.67 (A:OP2 14:571) |
| 4 | Water 4 | | 14:3806 | 2.18 | 1.00 | 53.21 | 2.76 (A:O5' 14:973) |
| 5 | A | OP2 | 14:973 | 2.41 | 1.00 | 49.72 |  |
| 6 | U | O4 | 14:568 | 2.47 | 1.00 | 46.16 |  |
| 7 | A | O5' | 14:973 | 3.73 | 1.00 | 52.26 |  |
| 8 | U | N3 | 14:568 | 3.77 | 1.00 | 41.11 |  |
| 9 | G | O3' | 14:972 | 3.84 | 1.00 | 53.42 |  |
| 10 | A | OP2 | 14:571 | 4.01 | 1.00 | 37.66 |  |
| 11 | Water 5 | | 14:3624 | 4.07 | 1.00 | 48.03 | 2.40 (G:OP1 14:570) |
| 12 | G | OP1 | 14:570 | 4.22 | 1.00 | 44.11 |  |
| 13 | A | OP1 | 14:973 | 4.77 | 1.00 | 51.69 |  |
| 14 | A | OP1 | 14:571 | 4.81 | 1.00 | 43.47 |  |
  |

  
**1^st^ shell ligand angles:**   
1-2: 175; 1-3: 94; 1-4: 89; 2-3: 91; 2-4: 94; 3-4: 92;   
  
 Contact(s): /O_w_/O_w_/O_w_/O_w_/  
 Coded contact list: 4O_w_  
 Exclusion zone contact(s): /O4/OP2/  
Consider NA in place of MG (2 valid contact(s) for NA vs. 0 for MG)  

### MG:14:3360

|  | Res. type | Atom | Residue | Distance | Occupancy | B-factor | Closest solute to 1^st^ shell water |
| --- | --- | --- | --- | --- | --- | --- | --- |
| **Ion** | | | |  | 1.00 | 37.02 || 1 | Water 1 | | 14:3526 | 2.15 | 1.00 | 36.37 | 2.04 (C:OP1 14:2499) |
| 2 | Water 2 | | 14:3610 | 2.19 | 1.00 | 46.40 | 2.37 (A:O5' 14:2448) |
| 3 | Water 3 | | 14:3950 | 2.20 | 1.00 | 47.48 | 3.17 (U:O4 14:569) |
| 4 | G | O6 | 14:570 | 2.52 | 1.00 | 38.87 |  |
| 5 | C | OP1 | 14:2499 | 2.54 | 1.00 | 35.23 |  |
| 6 | A | OP1 | 14:2448 | 2.66 | 1.00 | 30.44 |  |
| 7 | A | O5' | 14:2448 | 3.30 | 1.00 | 34.47 |  |
| 8 | G | O3' | 14:2447 | 3.45 | 1.00 | 38.97 |  |
| 9 | U | O4 | 14:569 | 3.56 | 1.00 | 39.00 |  |
| 10 | U | O2' | 14:568 | 4.03 | 1.00 | 42.41 |  |
| 11 | G | N1 | 14:570 | 4.18 | 1.00 | 31.84 |  |
| 12 | A | N6 | 14:945 | 4.36 | 1.00 | 33.87 |  |
| 13 | C | OP2 | 14:2498 | 4.38 | 1.00 | 44.53 |  |
| 14 | C | O3' | 14:2498 | 4.44 | 1.00 | 33.62 |  |
| 15 | C | OP2 | 14:2499 | 4.47 | 1.00 | 33.64 |  |
| 16 | A | OP2 | 14:2448 | 4.75 | 1.00 | 48.63 |  |
  |

  
**1^st^ shell ligand angles:**   
1-2: 86; 1-3: 174; 2-3: 100;   
  
 Contact(s): /O_w_/O_w_/O_w_/  
 Coded contact list: 3O_w_  
 Exclusion zone contact(s): /O5'/O6/OP1/OP1/  
Consider NA in place of MG (2 valid contact(s) for NA vs. 0 for MG)  
MG hydration shell issue: water too close (< 2.4 Angs.)  

### MG:14:3361

|  | Res. type | Atom | Residue | Distance | Occupancy | B-factor | Closest solute to 1^st^ shell water |
| --- | --- | --- | --- | --- | --- | --- | --- |
| **Ion** | | | |  | 1.00 | 48.95 || 1 | Water 1 | | 14:3602 | 2.16 | 1.00 | 51.15 | 2.35 (G:N7 14:2037) |
| 2 | Water 2 | | 14:3719 | 2.17 | 1.00 | 45.00 | 2.62 (C:N4 14:2025) |
| 3 | Water 3 | | 14:3640 | 2.18 | 1.00 | 56.68 | 2.47 (G:O6 14:2038) |
| 4 | Water 4 | | 14:4058 | 2.19 | 1.00 | 50.87 | 3.87 (G:OP1 14:2035) |
| 5 | Water 5 | | 14:3883 | 2.19 | 1.00 | 48.83 | 2.98 (G:OP1 14:2035) |
| 6 | G | OP1 | 14:2035 | 2.63 | 1.00 | 49.30 |  |
| 7 | G | O6 | 14:2038 | 4.03 | 1.00 | 53.46 |  |
| 8 | G | O6 | 14:2024 | 4.22 | 1.00 | 50.36 |  |
| 9 | C | N4 | 14:2025 | 4.34 | 1.00 | 53.85 |  |
| 10 | G | O6 | 14:2037 | 4.34 | 1.00 | 43.49 |  |
| 11 | U | O3' | 14:2034 | 4.36 | 1.00 | 48.35 |  |
| 12 | G | N7 | 14:2037 | 4.50 | 1.00 | 48.53 |  |
| 13 | C | N4 | 14:2036 | 4.61 | 1.00 | 41.46 |  |
| 14 | G | O5' | 14:2035 | 4.65 | 1.00 | 42.88 |  |
  |

  
**1^st^ shell ligand angles:**   
1-2: 91; 1-3: 89; 1-4: 87; 1-5: 178; 2-3: 87; 2-4: 178; 2-5: 89; 3-4: 92; 3-5: 93; 4-5: 93;   
  
 Contact(s): /O_w_/O_w_/O_w_/O_w_/O_w_/  
 Coded contact list: 5O_w_  
 Exclusion zone contact(s): /OP1/  
Consider K in place of MG (1 valid contact(s) for K vs. 0 for MG)  
MG hydration shell issue: water too close (< 2.4 Angs.)  

### MG:14:3362

|  | Res. type | Atom | Residue | Distance | Occupancy | B-factor | Closest solute to 1^st^ shell water |
| --- | --- | --- | --- | --- | --- | --- | --- |
| **Ion** | | | |  | 1.00 | 68.08 || 1 | Water 1 | | 14:3861 | 2.18 | 1.00 | 56.16 | 2.90 (G:O6 14:240) |
| 2 | Water 2 | | 14:4081 | 2.18 | 1.00 | 72.66 | 4.59 (G:O6 14:240) |
| 3 | Water 3 | | 14:3708 | 2.18 | 1.00 | 46.28 | 2.60 (G:O6 14:240) |
| 4 | Water 4 | | 14:4097 | 2.18 | 1.00 | 66.02 | 4.78 (G:O6 14:240) |
| 5 | Water 5 | | 14:3632 | 2.19 | 1.00 | 48.76 | 2.42 (G:O6 14:240) |
| 6 | Water 6 | | 14:4028 | 2.19 | 1.00 | 54.99 | 3.69 (A:OP2 14:256) |
| 7 | G | O6 | 14:240 | 3.14 | 1.00 | 49.41 |  |
| 8 | A | OP2 | 14:256 | 4.15 | 1.00 | 44.06 |  |
| 9 | A | N1 | 14:241 | 4.76 | 1.00 | 36.69 |  |
| 10 | G | N1 | 14:240 | 4.80 | 1.00 | 42.38 |  |
  |

  
**1^st^ shell ligand angles:**   
1-2: 178; 1-3: 95; 1-4: 88; 1-5: 91; 1-6: 86; 2-3: 86; 2-4: 91; 2-5: 90; 2-6: 93; 3-4: 177; 3-5: 90; 3-6: 90; 4-5: 90; 4-6: 91; 5-6: 177;   
  
 Contact(s): /O_w_/O_w_/O_w_/O_w_/O_w_/O_w_/  
 Coded contact list: 6O_w_  
 Exclusion zone contact(s): /O6/  
Consider K in place of MG (1 valid contact(s) for K vs. 0 for MG)  

### MG:14:3363

|  | Res. type | Atom | Residue | Distance | Occupancy | B-factor | Closest solute to 1^st^ shell water |
| --- | --- | --- | --- | --- | --- | --- | --- |
| **Ion** | | | |  | 1.00 | 26.79 || 1 | Water 1 | | 14:3842 | 2.16 | 1.00 | 38.17 | 2.86 (A:OP2 14:800) |
| 2 | Water 2 | | 14:3887 | 2.17 | 1.00 | 34.63 | 2.98 (A:OP1 14:800) |
| 3 | Water 3 | | 14:3937 | 2.17 | 1.00 | 39.46 | 3.12 (C:N3 14:192) |
| 4 | Water 4 | | 14:3680 | 2.17 | 1.00 | 33.52 | 2.54 (C:N3 14:192) |
| 5 | Water 5 | | 14:3826 | 2.18 | 1.00 | 29.02 | 2.81 (A:N1 14:191) |
| 6 | A | OP1 | 14:800 | 2.47 | 1.00 | 32.64 |  |
| 7 | C | N3 | 14:192 | 3.89 | 1.00 | 31.06 |  |
| 8 | A | OP2 | 14:800 | 3.91 | 1.00 | 24.05 |  |
| 9 | G | O3' | 14:799 | 4.31 | 1.00 | 29.33 |  |
| 10 | A | N1 | 14:191 | 4.31 | 1.00 | 28.38 |  |
| 11 | A | N7 | 14:800 | 4.38 | 1.00 | 36.87 |  |
| 12 | G | O2' | 14:799 | 4.39 | 1.00 | 33.91 |  |
| 13 | C | N4 | 14:192 | 4.57 | 1.00 | 28.52 |  |
| 14 | C | O2 | 14:192 | 4.57 | 1.00 | 34.50 |  |
| 15 | A | O5' | 14:800 | 4.70 | 1.00 | 28.27 |  |
  |

  
**1^st^ shell ligand angles:**   
1-2: 91; 1-3: 92; 1-4: 178; 1-5: 90; 2-3: 89; 2-4: 88; 2-5: 178; 3-4: 89; 3-5: 93; 4-5: 92;   
  
 Contact(s): /O_w_/O_w_/O_w_/O_w_/O_w_/  
 Coded contact list: 5O_w_  
 Exclusion zone contact(s): /OP1/  
Consider NA in place of MG (1 valid contact(s) for NA vs. 0 for MG)  

### MG:14:3364

|  | Res. type | Atom | Residue | Distance | Occupancy | B-factor | Closest solute to 1^st^ shell water |
| --- | --- | --- | --- | --- | --- | --- | --- |
| **Ion** | | | |  | 1.00 | 36.77 || 1 | Water 1 | | 14:3769 | 2.16 | 1.00 | 25.57 | 2.70 (G:OP2 14:450) |
| 2 | Water 2 | | 14:3741 | 2.17 | 1.00 | 32.52 | 2.66 (G:OP2 14:450) |
| 3 | Water 3 | | 14:3706 | 2.17 | 1.00 | 39.80 | 2.60 (G:OP2 14:452) |
| 4 | Water 4 | | 14:3841 | 2.18 | 1.00 | 39.39 | 2.86 (G:OP2 14:452) |
| 5 | Water 5 | | 14:3922 | 2.20 | 1.00 | 30.96 | 3.08 (G:OP2 14:452) |
| 6 | G | OP2 | 14:450 | 2.49 | 1.00 | 34.38 |  |
| 7 | G | OP2 | 14:452 | 3.42 | 1.00 | 27.48 |  |
| 8 | GLY | O | 39:86 | 3.86 | 1.00 | 36.89 |  |
| 9 | A | OP2 | 14:449 | 3.94 | 1.00 | 29.08 |  |
| 10 | G | O5' | 14:450 | 4.07 | 1.00 | 38.04 |  |
| 11 | C | OP2 | 14:451 | 4.27 | 1.00 | 33.87 |  |
| 12 | G | OP1 | 14:450 | 4.32 | 1.00 | 42.25 |  |
| 13 | GLY | O | 39:85 | 4.64 | 1.00 | 26.17 |  |
| 14 | A | O3' | 14:449 | 4.86 | 1.00 | 37.05 |  |
  |

  
**1^st^ shell ligand angles:**   
1-2: 86; 1-3: 171; 1-4: 89; 1-5: 97; 2-3: 85; 2-4: 92; 2-5: 176; 3-4: 95; 3-5: 92; 4-5: 86;   
  
 Contact(s): /O_w_/O_w_/O_w_/O_w_/O_w_/  
 Coded contact list: 5O_w_  
 Exclusion zone contact(s): /OP2/  
Consider NA in place of MG (1 valid contact(s) for NA vs. 0 for MG)  

### MG:14:3365

|  | Res. type | Atom | Residue | Distance | Occupancy | B-factor | Closest solute to 1^st^ shell water |
| --- | --- | --- | --- | --- | --- | --- | --- |
| **Ion** | | | |  | 1.00 | 28.70 || 1 | Water 1 | | 14:3566 | 2.16 | 1.00 | 31.29 | 2.24 (C:OP2 14:790) |
| 2 | Water 2 | | 14:3970 | 2.17 | 1.00 | 32.83 | 3.21 (C:OP2 14:790) |
| 3 | Water 3 | | 14:3795 | 2.18 | 1.00 | 44.10 | 2.74 (A:OP1 14:789) |
| 4 | Water 4 | | 14:3677 | 2.20 | 1.00 | 40.84 | 2.54 (C:OP2 14:791) |
| 5 | U | OP1 | 14:787 | 2.46 | 1.00 | 27.18 |  |
| 6 | C | OP2 | 14:790 | 2.47 | 1.00 | 37.08 |  |
| 7 | U | OP2 | 14:787 | 3.21 | 1.00 | 34.88 |  |
| 8 | C | O5' | 14:790 | 3.43 | 1.00 | 30.50 |  |
| 9 | C | OP2 | 14:791 | 3.85 | 1.00 | 41.32 |  |
| 10 | C | O3' | 14:786 | 4.15 | 1.00 | 28.95 |  |
| 11 | C | OP1 | 14:790 | 4.29 | 1.00 | 43.72 |  |
| 12 | A | OP1 | 14:789 | 4.43 | 1.00 | 39.30 |  |
| 13 | U | O5' | 14:787 | 4.53 | 1.00 | 26.23 |  |
| 14 | A | O3' | 14:789 | 4.66 | 1.00 | 46.33 |  |
| 15 | A | N6 | 14:1780 | 4.81 | 1.00 | 39.22 |  |
| 16 | A | N7 | 14:1780 | 4.89 | 1.00 | 36.27 |  |
  |

  
**1^st^ shell ligand angles:**   
1-2: 81; 1-3: 94; 1-4: 177; 2-3: 174; 2-4: 97; 3-4: 88;   
  
 Contact(s): /O_w_/O_w_/O_w_/O_w_/  
 Coded contact list: 4O_w_  
 Exclusion zone contact(s): /OP1/OP2/OP2/  
Consider NA in place of MG (2 valid contact(s) for NA vs. 0 for MG)  
MG hydration shell issue: water too close (< 2.4 Angs.)  

### MG:14:3366

|  | Res. type | Atom | Residue | Distance | Occupancy | B-factor | Closest solute to 1^st^ shell water |
| --- | --- | --- | --- | --- | --- | --- | --- |
| **Ion** | | | |  | 1.00 | 55.59 || 1 | Water 1 | | 14:3504 | 2.17 | 1.00 | 70.65 | 1.30 (A:N6 14:300) |
| 2 | Water 2 | | 14:3915 | 2.17 | 1.00 | 73.61 | 3.06 (G:N7 14:338) |
| 3 | Water 3 | | 14:3819 | 2.17 | 1.00 | 67.05 | 2.80 (A:N7 14:299) |
| 4 | Water 4 | | 14:3716 | 2.18 | 1.00 | 59.11 | 2.61 (C:OP2 14:336) |
| 5 | Water 5 | | C5:201 | 2.19 | 1.00 | 70.16 | 2.79 (ARG:NH1 C5:84) |
| 6 | Water 6 | | 14:4044 | 2.20 | 1.00 | 42.91 | 3.67 (A:OP2 14:299) |
| 7 | A | N6 | 14:300 | 3.40 | 1.00 | 76.13 |  |
| 8 | A | OP2 | 14:299 | 3.95 | 1.00 | 56.63 |  |
| 9 | G | O6 | 14:338 | 4.36 | 1.00 | 66.55 |  |
| 10 | A | N7 | 14:299 | 4.41 | 1.00 | 67.87 |  |
| 11 | A | N7 | 14:300 | 4.46 | 1.00 | 66.67 |  |
| 12 | ARG | NH1 | C5:84 | 4.78 | 1.00 | 59.68 |  |
| 13 | C | OP2 | 14:336 | 4.78 | 1.00 | 61.76 |  |
| 14 | G | N7 | 14:338 | 4.99 | 1.00 | 63.85 |  |
  |

  
**1^st^ shell ligand angles:**   
1-2: 87; 1-3: 89; 1-4: 91; 1-5: 91; 1-6: 179; 2-3: 89; 2-4: 89; 2-5: 177; 2-6: 92; 3-4: 178; 3-5: 89; 3-6: 91; 4-5: 93; 4-6: 89; 5-6: 90;   
  
 Contact(s): /O_w_/O_w_/O_w_/O_w_/O_w_/O_w_/  
 Coded contact list: 6O_w_  
 Exclusion zone contact(s): /N6/  
Consider K in place of MG (1 valid contact(s) for K vs. 0 for MG)  
MG hydration shell issue: water too close (< 2.4 Angs.)  

### MG:14:3367

|  | Res. type | Atom | Residue | Distance | Occupancy | B-factor | Closest solute to 1^st^ shell water |
| --- | --- | --- | --- | --- | --- | --- | --- |
| **Ion** | | | |  | 1.00 | 70.06 || 1 | Water 1 | | 14:4101 | 2.18 | 1.00 | 71.70 | 5.58 (U:OP2 14:499) |
| 2 | Water 2 | | 14:4091 | 2.19 | 1.00 | 71.09 | 4.46 (U:OP2 14:499) |
| 3 | Water 3 | | 14:3817 | 2.19 | 1.00 | 56.20 | 2.79 (G:O6 14:500) |
| 4 | Water 4 | | 14:3944 | 2.19 | 1.00 | 62.46 | 3.15 (G:N7 14:500) |
| 5 | Water 5 | | 14:3718 | 2.19 | 1.00 | 55.11 | 2.62 (U:OP2 14:499) |
| 6 | Water 6 | | 14:3966 | 2.19 | 1.00 | 43.97 | 3.21 (A:OP1 14:502) |
| 7 | G | N7 | 14:500 | 4.06 | 1.00 | 54.08 |  |
| 8 | U | OP2 | 14:499 | 4.45 | 1.00 | 62.14 |  |
| 9 | G | O6 | 14:500 | 4.59 | 1.00 | 63.30 |  |
| 10 | A | OP1 | 14:502 | 4.69 | 1.00 | 54.04 |  |
  |

  
**1^st^ shell ligand angles:**   
1-2: 92; 1-3: 179; 1-4: 87; 1-5: 91; 1-6: 89; 2-3: 88; 2-4: 179; 2-5: 91; 2-6: 90; 3-4: 93; 3-5: 88; 3-6: 92; 4-5: 89; 4-6: 90; 5-6: 179;   
  
 Contact(s): /O_w_/O_w_/O_w_/O_w_/O_w_/O_w_/  
 Coded contact list: 6O_w_  

### MG:14:3368

|  | Res. type | Atom | Residue | Distance | Occupancy | B-factor | Closest solute to 1^st^ shell water |
| --- | --- | --- | --- | --- | --- | --- | --- |
| **Ion** | | | |  | 1.00 | 29.02 || 1 | Water 1 | | 14:3588 | 2.16 | 1.00 | 44.99 | 2.30 (U:OP1 14:827) |
| 2 | Water 2 | | 14:3650 | 2.17 | 1.00 | 32.74 | 2.49 (A:OP1 14:2430) |
| 3 | Water 3 | | 14:3720 | 2.17 | 1.00 | 48.16 | 2.62 (A:OP2 14:2430) |
| 4 | U | OP1 | 14:827 | 2.48 | 1.00 | 33.10 |  |
| 5 | G | OP2 | 14:2429 | 2.59 | 1.00 | 48.69 |  |
| 6 | A | OP2 | 14:2430 | 2.65 | 1.00 | 33.88 |  |
| 7 | A | OP1 | 14:2430 | 2.76 | 1.00 | 29.64 |  |
| 8 | G | O5' | 14:2429 | 3.31 | 1.00 | 38.78 |  |
| 9 | U | O3' | 14:826 | 3.54 | 1.00 | 40.64 |  |
| 10 | G | O3' | 14:2429 | 3.72 | 1.00 | 46.86 |  |
| 11 | G | O3' | 14:2428 | 4.10 | 1.00 | 37.95 |  |
| 12 | A | O5' | 14:2430 | 4.37 | 1.00 | 38.45 |  |
| 13 | U | OP2 | 14:827 | 4.48 | 1.00 | 46.15 |  |
| 14 | G | OP1 | 14:2429 | 4.71 | 1.00 | 43.33 |  |
| 15 | U | O5' | 14:827 | 4.75 | 1.00 | 43.08 |  |
| 16 | GLY | O | 35:54 | 4.86 | 1.00 | 43.07 |  |
  |

  
**1^st^ shell ligand angles:**   
1-2: 91; 1-3: 90; 2-3: 179;   
  
 Contact(s): /O_w_/O_w_/O_w_/  
 Coded contact list: 3O_w_  
 Exclusion zone contact(s): /O5'/OP1/OP1/OP2/OP2/  
Consider K in place of MG (3 valid contact(s) for K vs. 0 for MG)  
MG hydration shell issue: water too close (< 2.4 Angs.)  

### MG:14:3369

|  | Res. type | Atom | Residue | Distance | Occupancy | B-factor | Closest solute to 1^st^ shell water |
| --- | --- | --- | --- | --- | --- | --- | --- |
| **Ion** | | | |  | 1.00 | 76.32 || 1 | Water 1 | | 14:3866 | 2.17 | 1.00 | 70.73 | 2.92 (C:OP2 14:1577) |
| 2 | Water 2 | | 14:3579 | 2.18 | 1.00 | 70.40 | 2.28 (G:OP2 14:1418) |
| 3 | Water 3 | | 14:3647 | 2.18 | 1.00 | 57.33 | 2.48 (G:O6 14:1418) |
| 4 | Water 4 | | 14:3892 | 2.18 | 1.00 | 79.88 | 2.99 (C:OP1 14:1417) |
| 5 | Water 5 | | 14:3774 | 2.19 | 1.00 | 74.74 | 2.71 (C:OP2 14:1417) |
| 6 | C | OP2 | 14:1417 | 2.69 | 1.00 | 83.11 |  |
| 7 | C | O5' | 14:1417 | 3.82 | 1.00 | 70.45 |  |
| 8 | C | OP1 | 14:1417 | 3.82 | 1.00 | 83.80 |  |
| 9 | U | OP2 | 14:1578 | 4.11 | 1.00 | 70.38 |  |
| 10 | G | N7 | 14:1418 | 4.12 | 1.00 | 70.05 |  |
| 11 | G | OP2 | 14:1418 | 4.22 | 1.00 | 74.49 |  |
| 12 | C | OP2 | 14:1577 | 4.24 | 1.00 | 66.95 |  |
| 13 | G | O6 | 14:1418 | 4.60 | 1.00 | 70.13 |  |
| 14 | C | O5' | 14:1577 | 4.80 | 1.00 | 71.52 |  |
| 15 | G | O3' | 14:1416 | 4.95 | 1.00 | 89.16 |  |
  |

  
**1^st^ shell ligand angles:**   
1-2: 93; 1-3: 95; 1-4: 86; 1-5: 89; 2-3: 92; 2-4: 88; 2-5: 179; 3-4: 179; 3-5: 88; 4-5: 92;   
  
 Contact(s): /O_w_/O_w_/O_w_/O_w_/O_w_/  
 Coded contact list: 5O_w_  
 Exclusion zone contact(s): /OP2/  
Consider K in place of MG (1 valid contact(s) for K vs. 0 for MG)  
MG hydration shell issue: water too close (< 2.4 Angs.)  

### MG:14:3370

|  | Res. type | Atom | Residue | Distance | Occupancy | B-factor | Closest solute to 1^st^ shell water |
| --- | --- | --- | --- | --- | --- | --- | --- |
| **Ion** | | | |  | 1.00 | 44.38 || 1 | Water 1 | | 14:3564 | 2.17 | 1.00 | 52.53 | 2.23 (C:OP2 14:376) |
| 2 | Water 2 | | 14:3889 | 2.17 | 1.00 | 57.12 | 2.99 (C:OP2 14:392) |
| 3 | Water 3 | | 14:3626 | 2.18 | 1.00 | 52.56 | 2.41 (C:OP1 14:409) |
| 4 | Water 4 | | 14:3737 | 2.18 | 1.00 | 53.28 | 2.66 (C:OP1 14:409) |
| 5 | C | OP1 | 14:409 | 2.59 | 1.00 | 48.85 |  |
| 6 | C | OP1 | 14:392 | 2.61 | 1.00 | 52.22 |  |
| 7 | C | OP2 | 14:392 | 3.10 | 1.00 | 49.60 |  |
| 8 | C | OP2 | 14:376 | 4.03 | 1.00 | 48.33 |  |
| 9 | C | OP1 | 14:375 | 4.20 | 1.00 | 56.65 |  |
| 10 | G | O3' | 14:391 | 4.34 | 1.00 | 47.54 |  |
| 11 | G | O3' | 14:408 | 4.36 | 1.00 | 60.70 |  |
| 12 | C | O5' | 14:392 | 4.41 | 1.00 | 47.22 |  |
| 13 | C | OP1 | 14:376 | 4.43 | 1.00 | 42.79 |  |
| 14 | C | OP2 | 14:393 | 4.49 | 1.00 | 46.67 |  |
| 15 | C | OP2 | 14:409 | 4.54 | 1.00 | 57.75 |  |
  |

  
**1^st^ shell ligand angles:**   
1-2: 87; 1-3: 97; 1-4: 92; 2-3: 176; 2-4: 92; 3-4: 86;   
  
 Contact(s): /O_w_/O_w_/O_w_/O_w_/  
 Coded contact list: 4O_w_  
 Exclusion zone contact(s): /OP1/OP1/OP2/  
Consider K in place of MG (2 valid contact(s) for K vs. 0 for MG)  
MG hydration shell issue: water too close (< 2.4 Angs.)  

### MG:14:3371

|  | Res. type | Atom | Residue | Distance | Occupancy | B-factor | Closest solute to 1^st^ shell water |
| --- | --- | --- | --- | --- | --- | --- | --- |
| **Ion** | | | |  | 1.00 | 32.30 || 1 | Water 1 | | 14:4007 | 2.16 | 1.00 | 39.71 | 3.37 (A:O3' 14:1393) |
| 2 | Water 2 | | 14:4072 | 2.17 | 1.00 | 35.28 | 4.00 (U:OP1 14:1394) |
| 3 | Water 3 | | 14:3553 | 2.17 | 1.00 | 42.60 | 2.20 (C:OP1 14:1604) |
| 4 | Water 4 | | 14:3586 | 2.18 | 1.00 | 33.16 | 2.29 (A:OP2 14:1395) |
| 5 | U | OP1 | 14:1394 | 2.55 | 1.00 | 42.65 |  |
| 6 | C | OP1 | 14:1604 | 2.61 | 1.00 | 36.70 |  |
| 7 | A | O3' | 14:1393 | 3.62 | 1.00 | 44.59 |  |
| 8 | A | OP2 | 14:1395 | 3.99 | 1.00 | 35.30 |  |
| 9 | C | O5' | 14:1604 | 4.08 | 1.00 | 39.13 |  |
| 10 | A | O3' | 14:1603 | 4.32 | 1.00 | 33.48 |  |
| 11 | U | OP2 | 14:1394 | 4.49 | 1.00 | 57.03 |  |
| 12 | Water 5 | | 14:3980 | 4.53 | 1.00 | 33.37 | 3.27 (C:OP1 14:1604) |
| 13 | U | O5' | 14:1394 | 4.54 | 1.00 | 41.27 |  |
  |

  
**1^st^ shell ligand angles:**   
1-2: 91; 1-3: 94; 1-4: 176; 2-3: 92; 2-4: 93; 3-4: 84;   
  
 Contact(s): /O_w_/O_w_/O_w_/O_w_/  
 Coded contact list: 4O_w_  
 Exclusion zone contact(s): /OP1/OP1/  
Consider NA in place of MG (1 valid contact(s) for NA vs. 0 for MG)  
MG hydration shell issue: water too close (< 2.4 Angs.)  

### MG:14:3372

|  | Res. type | Atom | Residue | Distance | Occupancy | B-factor | Closest solute to 1^st^ shell water |
| --- | --- | --- | --- | --- | --- | --- | --- |
| **Ion** | | | |  | 1.00 | 64.47 || 1 | Water 1 | | 14:3528 | 2.18 | 1.00 | 66.39 | 2.05 (A:OP1 14:1439) |
| 2 | Water 2 | | 14:3693 | 2.18 | 1.00 | 68.48 | 2.57 (A:OP1 14:1469) |
| 3 | Water 3 | | 14:3831 | 2.18 | 1.00 | 75.90 | 2.83 (A:OP1 14:1439) |
| 4 | Water 4 | | 14:3729 | 2.19 | 1.00 | 76.51 | 2.65 (C:OP1 14:1468) |
| 5 | A | OP1 | 14:1439 | 2.48 | 1.00 | 52.56 |  |
| 6 | C | OP1 | 14:1468 | 2.81 | 1.00 | 77.47 |  |
| 7 | C | O5' | 14:1468 | 3.56 | 1.00 | 72.64 |  |
| 8 | A | OP2 | 14:1469 | 3.65 | 1.00 | 70.94 |  |
| 9 | A | OP1 | 14:1469 | 4.00 | 1.00 | 68.65 |  |
| 10 | G | OP2 | 14:1440 | 4.36 | 1.00 | 60.37 |  |
| 11 | A | O5' | 14:1439 | 4.45 | 1.00 | 59.86 |  |
| 12 | C | O3' | 14:1468 | 4.62 | 1.00 | 56.15 |  |
| 13 | C | O3' | 14:1467 | 4.67 | 1.00 | 76.47 |  |
| 14 | U | O3' | 14:1438 | 4.79 | 1.00 | 51.00 |  |
| 15 | A | OP2 | 14:1439 | 4.80 | 1.00 | 57.36 |  |
| 16 | C | OP2 | 14:1468 | 4.89 | 1.00 | 74.07 |  |
  |

  
**1^st^ shell ligand angles:**   
1-2: 178; 1-3: 89; 1-4: 88; 2-3: 91; 2-4: 92; 3-4: 177;   
  
 Contact(s): /O_w_/O_w_/O_w_/O_w_/  
 Coded contact list: 4O_w_  
 Exclusion zone contact(s): /OP1/OP1/  
Consider NA in place of MG (1 valid contact(s) for NA vs. 0 for MG)  
MG hydration shell issue: water too close (< 2.4 Angs.)  

### MG:14:3373

|  | Res. type | Atom | Residue | Distance | Occupancy | B-factor | Closest solute to 1^st^ shell water |
| --- | --- | --- | --- | --- | --- | --- | --- |
| **Ion** | | | |  | 1.00 | 68.79 || 1 | Water 1 | | 14:3744 | 2.17 | 1.00 | 74.19 | 2.67 (G:N7 14:1466) |
| 2 | Water 2 | | 14:3802 | 2.18 | 1.00 | 71.97 | 2.75 (G:N7 14:1443) |
| 3 | Water 3 | | 14:4021 | 2.18 | 1.00 | 66.15 | 3.47 (G:N7 14:1443) |
| 4 | Water 4 | | 14:4020 | 2.18 | 1.00 | 86.07 | 5.13 (G:N7 14:1466) |
| 5 | Water 5 | | 14:3857 | 2.19 | 1.00 | 56.40 | 2.90 (G:O6 14:1443) |
| 6 | Water 6 | | 14:4087 | 2.19 | 1.00 | 77.26 | 5.09 (G:N7 14:1443) |
| 7 | G | N7 | 14:1443 | 4.10 | 1.00 | 69.36 |  |
| 8 | G | O6 | 14:1443 | 4.19 | 1.00 | 67.79 |  |
| 9 | G | O6 | 14:1444 | 4.38 | 1.00 | 74.23 |  |
| 10 | G | N7 | 14:1444 | 4.74 | 1.00 | 74.04 |  |
| 11 | G | N7 | 14:1466 | 4.83 | 1.00 | 84.97 |  |
  |

  
**1^st^ shell ligand angles:**   
1-2: 90; 1-3: 178; 1-4: 89; 1-5: 92; 1-6: 87; 2-3: 92; 2-4: 178; 2-5: 94; 2-6: 85; 3-4: 89; 3-5: 88; 3-6: 93; 4-5: 88; 4-6: 93; 5-6: 179;   
  
 Contact(s): /O_w_/O_w_/O_w_/O_w_/O_w_/O_w_/  
 Coded contact list: 6O_w_  

### MG:14:3374

|  | Res. type | Atom | Residue | Distance | Occupancy | B-factor | Closest solute to 1^st^ shell water |
| --- | --- | --- | --- | --- | --- | --- | --- |
| **Ion** | | | |  | 1.00 | 30.73 || 1 | Water 1 | | 14:3529 | 2.17 | 1.00 | 39.32 | 2.07 (G:OP1 14:2550) |
| 2 | Water 2 | | 14:3644 | 2.18 | 1.00 | 43.80 | 2.48 (G:OP2 14:2550) |
| 3 | U | O4 | 14:1671 | 2.61 | 1.00 | 50.47 |  |
| 4 | C | OP2 | 14:1670 | 2.76 | 1.00 | 37.68 |  |
| 5 | G | OP1 | 14:2550 | 2.81 | 1.00 | 41.19 |  |
| 6 | A | OP2 | 14:1669 | 2.84 | 1.00 | 36.38 |  |
| 7 | G | OP2 | 14:2550 | 4.09 | 1.00 | 38.94 |  |
| 8 | C | OP2 | 14:2551 | 4.16 | 1.00 | 43.87 |  |
| 9 | C | OP1 | 14:1670 | 4.21 | 1.00 | 35.40 |  |
| 10 | U | N3 | 14:1671 | 4.46 | 1.00 | 41.80 |  |
| 11 | G | O5' | 14:2550 | 4.61 | 1.00 | 48.10 |  |
| 12 | C | O5' | 14:1670 | 4.83 | 1.00 | 46.51 |  |
| 13 | U | OP2 | 14:1673 | 4.85 | 1.00 | 46.58 |  |
| 14 | A | O5' | 14:1669 | 4.89 | 1.00 | 36.95 |  |
| 15 | C | OP1 | 14:2551 | 4.92 | 1.00 | 55.29 |  |
  |

  
**1^st^ shell ligand angles:**   
1-2: 103;   
  
 Contact(s): /O_w_/O_w_/  
 Coded contact list: 2O_w_  
 Exclusion zone contact(s): /O4/OP1/OP2/OP2/  
Consider K in place of MG (4 valid contact(s) for K vs. 0 for MG)  
MG hydration shell issue: water too close (< 2.4 Angs.)  

### MG:14:3375

|  | Res. type | Atom | Residue | Distance | Occupancy | B-factor | Closest solute to 1^st^ shell water |
| --- | --- | --- | --- | --- | --- | --- | --- |
| **Ion** | | | |  | 1.00 | 84.05 || 1 | Water 1 | | 14:4065 | 2.17 | 1.00 | 58.13 | 3.91 (G:OP1 14:2859) |
| 2 | Water 2 | | 14:3888 | 2.17 | 1.00 | 78.02 | 2.99 (G:OP2 14:2859) |
| 3 | Water 3 | | 14:3752 | 2.18 | 1.00 | 76.84 | 2.67 (G:N7 14:2857) |
| 4 | Water 4 | | 14:4099 | 2.18 | 1.00 | 81.77 | 5.06 (G:OP2 14:2857) |
| 5 | Water 5 | | 14:4074 | 2.19 | 1.00 | 76.68 | 5.38 (G:N7 14:2857) |
| 6 | Water 6 | | 14:3822 | 2.19 | 1.00 | 84.93 | 2.81 (G:OP2 14:2857) |
| 7 | G | N7 | 14:2857 | 3.92 | 1.00 | 85.63 |  |
| 8 | G | O6 | 14:2857 | 4.47 | 1.00 | 83.73 |  |
| 9 | G | OP2 | 14:2859 | 4.76 | 1.00 | 79.97 |  |
| 10 | G | OP2 | 14:2857 | 4.80 | 1.00 | 77.42 |  |
| 11 | G | OP1 | 14:2859 | 4.95 | 1.00 | 87.34 |  |
  |

  
**1^st^ shell ligand angles:**   
1-2: 90; 1-3: 92; 1-4: 86; 1-5: 91; 1-6: 179; 2-3: 90; 2-4: 90; 2-5: 179; 2-6: 90; 3-4: 178; 3-5: 90; 3-6: 89; 4-5: 90; 4-6: 93; 5-6: 89;   
  
 Contact(s): /O_w_/O_w_/O_w_/O_w_/O_w_/O_w_/  
 Coded contact list: 6O_w_  

### MG:14:3376

|  | Res. type | Atom | Residue | Distance | Occupancy | B-factor | Closest solute to 1^st^ shell water |
| --- | --- | --- | --- | --- | --- | --- | --- |
| **Ion** | | | |  | 1.00 | 54.68 || 1 | Water 1 | | 14:3965 | 2.18 | 1.00 | 65.92 | 3.20 (G:O3' 14:623) |
| 2 | Water 2 | | 14:3760 | 2.18 | 1.00 | 51.46 | 2.68 (G:OP1 14:242) |
| 3 | Water 3 | | 14:3815 | 2.18 | 1.00 | 60.50 | 2.79 (C:OP1 14:624) |
| 4 | Water 4 | | 14:3723 | 2.19 | 1.00 | 49.72 | 2.63 (C:OP1 14:624) |
| 5 | Water 5 | | 14:3956 | 2.19 | 1.00 | 55.23 | 3.18 (G:OP1 14:242) |
| 6 | C | OP1 | 14:624 | 2.50 | 1.00 | 51.73 |  |
| 7 | G | O3' | 14:623 | 3.24 | 1.00 | 54.58 |  |
| 8 | G | OP1 | 14:242 | 3.68 | 1.00 | 51.97 |  |
| 9 | C | O5' | 14:624 | 4.36 | 1.00 | 48.68 |  |
| 10 | C | OP2 | 14:624 | 4.45 | 1.00 | 52.04 |  |
| 11 | TYR | O | M5:64 | 4.73 | 1.00 | 64.67 |  |
  |

  
**1^st^ shell ligand angles:**   
1-2: 91; 1-3: 176; 1-4: 91; 1-5: 89; 2-3: 94; 2-4: 91; 2-5: 93; 3-4: 89; 3-5: 91; 4-5: 176;   
  
 Contact(s): /O_w_/O_w_/O_w_/O_w_/O_w_/  
 Coded contact list: 5O_w_  
 Exclusion zone contact(s): /O3'/OP1/  
Consider NA in place of MG (1 valid contact(s) for NA vs. 0 for MG)  

### MG:14:3377

|  | Res. type | Atom | Residue | Distance | Occupancy | B-factor | Closest solute to 1^st^ shell water |
| --- | --- | --- | --- | --- | --- | --- | --- |
| **Ion** | | | |  | 1.00 | 59.96 || 1 | Water 1 | | 14:3784 | 2.18 | 1.00 | 52.84 | 2.72 (U:OP1 14:1199) |
| 2 | Water 2 | | 14:3653 | 2.18 | 1.00 | 55.87 | 2.49 (C:OP1 14:31) |
| 3 | Water 3 | | 14:3731 | 2.18 | 1.00 | 56.18 | 2.65 (C:OP1 14:1200) |
| 4 | Water 4 | | 14:3854 | 2.19 | 1.00 | 58.40 | 2.90 (C:OP1 14:1200) |
| 5 | Water 5 | | 14:3515 | 2.19 | 1.00 | 50.93 | 1.93 (C:OP1 14:31) |
| 6 | C | OP1 | 14:31 | 2.55 | 1.00 | 50.62 |  |
| 7 | C | OP1 | 14:1200 | 3.69 | 1.00 | 54.11 |  |
| 8 | G | OP1 | 14:1239 | 3.86 | 1.00 | 50.48 |  |
| 9 | C | OP2 | 14:1200 | 4.10 | 1.00 | 45.35 |  |
| 10 | G | O3' | 14:30 | 4.12 | 1.00 | 38.08 |  |
| 11 | C | OP2 | 14:31 | 4.14 | 1.00 | 52.05 |  |
| 12 | U | OP1 | 14:1199 | 4.46 | 1.00 | 59.26 |  |
| 13 | U | OP1 | 14:1240 | 4.70 | 1.00 | 54.59 |  |
| 14 | U | OP2 | 14:1240 | 4.80 | 1.00 | 45.69 |  |
| 15 | C | O5' | 14:31 | 4.83 | 1.00 | 52.45 |  |
| 16 | U | O3' | 14:1199 | 4.94 | 1.00 | 46.88 |  |
  |

  
**1^st^ shell ligand angles:**   
1-2: 88; 1-3: 92; 1-4: 91; 1-5: 179; 2-3: 177; 2-4: 90; 2-5: 91; 3-4: 93; 3-5: 88; 4-5: 87;   
  
 Contact(s): /O_w_/O_w_/O_w_/O_w_/O_w_/  
 Coded contact list: 5O_w_  
 Exclusion zone contact(s): /OP1/  
Consider NA in place of MG (1 valid contact(s) for NA vs. 0 for MG)  
MG hydration shell issue: water too close (< 2.4 Angs.)  

### MG:14:3378

|  | Res. type | Atom | Residue | Distance | Occupancy | B-factor | Closest solute to 1^st^ shell water |
| --- | --- | --- | --- | --- | --- | --- | --- |
| **Ion** | | | |  | 1.00 | 57.15 || 1 | Water 1 | | 14:3736 | 2.17 | 1.00 | 52.18 | 2.66 (U:O4 14:1300) |
| 2 | Water 2 | | 14:4030 | 2.18 | 1.00 | 58.76 | 3.53 (A:OP1 14:1634) |
| 3 | Water 3 | | 14:3739 | 2.18 | 1.00 | 53.75 | 2.66 (G:OP2 14:1626) |
| 4 | Water 4 | | 14:3982 | 2.19 | 1.00 | 52.93 | 3.28 (G:OP1 14:1627) |
| 5 | Water 5 | | 14:3867 | 2.19 | 1.00 | 46.08 | 2.92 (G:OP2 14:1627) |
| 6 | G | OP1 | 14:1627 | 2.58 | 1.00 | 45.47 |  |
| 7 | G | OP2 | 14:1627 | 3.68 | 1.00 | 50.57 |  |
| 8 | U | O4 | 14:1300 | 4.23 | 1.00 | 39.62 |  |
| 9 | G | O5' | 14:1626 | 4.40 | 1.00 | 42.44 |  |
| 10 | G | O5' | 14:1627 | 4.46 | 1.00 | 49.84 |  |
| 11 | A | OP1 | 14:1634 | 4.60 | 1.00 | 45.15 |  |
| 12 | G | OP1 | 14:1626 | 4.62 | 1.00 | 56.05 |  |
| 13 | G | OP2 | 14:1626 | 4.66 | 1.00 | 46.10 |  |
| 14 | G | O3' | 14:1626 | 4.80 | 1.00 | 41.88 |  |
  |

  
**1^st^ shell ligand angles:**   
1-2: 87; 1-3: 88; 1-4: 178; 1-5: 91; 2-3: 92; 2-4: 92; 2-5: 93; 3-4: 90; 3-5: 175; 4-5: 91;   
  
 Contact(s): /O_w_/O_w_/O_w_/O_w_/O_w_/  
 Coded contact list: 5O_w_  
 Exclusion zone contact(s): /OP1/  
Consider NA in place of MG (1 valid contact(s) for NA vs. 0 for MG)  

### MG:14:3379

|  | Res. type | Atom | Residue | Distance | Occupancy | B-factor | Closest solute to 1^st^ shell water |
| --- | --- | --- | --- | --- | --- | --- | --- |
| **Ion** | | | |  | 1.00 | 43.72 || 1 | Water 1 | | 14:3546 | 2.16 | 1.00 | 43.05 | 2.18 (G:O6 14:2002) |
| 2 | Water 2 | | 14:3641 | 2.17 | 1.00 | 38.68 | 2.47 (G:OP2 14:2002) |
| 3 | Water 3 | | 14:3954 | 2.18 | 1.00 | 43.58 | 3.17 (G:N7 14:2002) |
| 4 | Water 4 | | 14:4068 | 2.19 | 1.00 | 55.83 | 4.70 (G:N7 14:2002) |
| 5 | Water 5 | | 14:4039 | 2.19 | 1.00 | 51.20 | 3.93 (G:N2 14:1653) |
| 6 | Water 6 | | 14:3931 | 2.19 | 1.00 | 33.62 | 3.67 (G:O6 14:2003) |
| 7 | G | N7 | 14:2002 | 3.42 | 1.00 | 40.33 |  |
| 8 | G | O6 | 14:2002 | 4.18 | 1.00 | 41.99 |  |
| 9 | G | OP2 | 14:2002 | 4.39 | 1.00 | 38.84 |  |
| 10 | G | O6 | 14:2003 | 4.64 | 1.00 | 40.05 |  |
| 11 | G | N2 | 14:1653 | 4.95 | 1.00 | 50.40 |  |
  |

  
**1^st^ shell ligand angles:**   
1-2: 88; 1-3: 91; 1-4: 85; 1-5: 176; 1-6: 92; 2-3: 91; 2-4: 89; 2-5: 91; 2-6: 177; 3-4: 176; 3-5: 93; 3-6: 87; 4-5: 91; 4-6: 94; 5-6: 89;   
  
 Contact(s): /O_w_/O_w_/O_w_/O_w_/O_w_/O_w_/  
 Coded contact list: 6O_w_  
MG hydration shell issue: water too close (< 2.4 Angs.)  

### MG:14:3380

|  | Res. type | Atom | Residue | Distance | Occupancy | B-factor | Closest solute to 1^st^ shell water |
| --- | --- | --- | --- | --- | --- | --- | --- |
| **Ion** | | | |  | 1.00 | 76.47 || 1 | Water 1 | | 14:3748 | 2.18 | 1.00 | 78.07 | 2.67 (G:O6 14:175) |
| 2 | Water 2 | | 14:4093 | 2.18 | 1.00 | 62.51 | 4.48 (G:O6 14:175) |
| 3 | Water 3 | | 14:4083 | 2.19 | 1.00 | 68.30 | 4.22 (C:N4 14:150) |
| 4 | Water 4 | | 14:3715 | 2.19 | 1.00 | 68.87 | 2.61 (A:O2' 14:49) |
| 5 | Water 5 | | 14:4011 | 2.20 | 1.00 | 66.27 | 3.39 (A:O2' 14:49) |
| 6 | Water 6 | | 14:3991 | 2.20 | 1.00 | 61.08 | 3.29 (G:O6 14:177) |
| 7 | A | O2' | 14:49 | 3.84 | 1.00 | 59.62 |  |
| 8 | G | O6 | 14:176 | 4.23 | 1.00 | 62.80 |  |
| 9 | G | O6 | 14:177 | 4.54 | 1.00 | 54.62 |  |
| 10 | G | O6 | 14:175 | 4.70 | 1.00 | 74.37 |  |
| 11 | C | N4 | 14:150 | 4.94 | 1.00 | 59.63 |  |
  |

  
**1^st^ shell ligand angles:**   
1-2: 86; 1-3: 89; 1-4: 89; 1-5: 178; 1-6: 93; 2-3: 87; 2-4: 93; 2-5: 92; 2-6: 178; 3-4: 178; 3-5: 92; 3-6: 92; 4-5: 90; 4-6: 89; 5-6: 89;   
  
 Contact(s): /O_w_/O_w_/O_w_/O_w_/O_w_/O_w_/  
 Coded contact list: 6O_w_  

### MG:14:3381

|  | Res. type | Atom | Residue | Distance | Occupancy | B-factor | Closest solute to 1^st^ shell water |
| --- | --- | --- | --- | --- | --- | --- | --- |
| **Ion** | | | |  | 1.00 | 56.01 || 1 | Water 1 | | 14:3562 | 2.17 | 1.00 | 62.26 | 2.23 (C:OP2 14:986) |
| 2 | Water 2 | | 14:3608 | 2.17 | 1.00 | 71.91 | 2.36 (A:N7 14:988) |
| 3 | Water 3 | | 14:3746 | 2.17 | 1.00 | 66.09 | 2.67 (G:OP2 14:987) |
| 4 | Water 4 | | H5:101 | 2.17 | 1.00 | 70.62 | 2.23 (ILE:O H5:13) |
| 5 | Water 5 | | H5:102 | 2.18 | 1.00 | 65.26 | 2.98 (ILE:O H5:13) |
| 6 | G | OP2 | 14:987 | 2.42 | 1.00 | 59.44 |  |
| 7 | ILE | O | H5:13 | 3.46 | 1.00 | 63.97 |  |
| 8 | G | O5' | 14:987 | 3.74 | 1.00 | 63.18 |  |
| 9 | G | OP1 | 14:987 | 4.14 | 1.00 | 70.22 |  |
| 10 | A | OP2 | 14:988 | 4.29 | 1.00 | 73.66 |  |
| 11 | A | N7 | 14:988 | 4.37 | 1.00 | 70.50 |  |
| 12 | C | OP2 | 14:986 | 4.38 | 1.00 | 66.50 |  |
| 13 | C | O3' | 14:986 | 4.68 | 1.00 | 68.37 |  |
| 14 | A | N6 | 14:988 | 4.72 | 1.00 | 66.52 |  |
| 15 | C | O5' | 14:986 | 4.80 | 1.00 | 69.37 |  |
| 16 | C | OP1 | 14:986 | 4.82 | 1.00 | 58.68 |  |
| 17 | G | N7 | 14:987 | 4.95 | 1.00 | 71.27 |  |
| 18 | GLY | N | H5:14 | 5.00 | 1.00 | 61.02 |  |
  |

  
**1^st^ shell ligand angles:**   
1-2: 90; 1-3: 87; 1-4: 176; 1-5: 90; 2-3: 177; 2-4: 92; 2-5: 87; 3-4: 91; 3-5: 92; 4-5: 94;   
  
 Contact(s): /O_w_/O_w_/O_w_/O_w_/O_w_/  
 Coded contact list: 5O_w_  
 Exclusion zone contact(s): /OP2/  
Consider NA in place of MG (1 valid contact(s) for NA vs. 0 for MG)  
MG hydration shell issue: water too close (< 2.4 Angs.)  

### MG:14:3382

|  | Res. type | Atom | Residue | Distance | Occupancy | B-factor | Closest solute to 1^st^ shell water |
| --- | --- | --- | --- | --- | --- | --- | --- |
| **Ion** | | | |  | 1.00 | 52.65 || 1 | Water 1 | | 14:3787 | 2.17 | 1.00 | 51.05 | 2.73 (G:OP1 14:386) |
| 2 | Water 2 | | 14:3727 | 2.17 | 1.00 | 56.01 | 2.64 (G:OP2 14:382) |
| 3 | Water 3 | | 14:3598 | 2.18 | 1.00 | 49.33 | 2.34 (G:OP1 14:386) |
| 4 | Water 4 | | 14:3832 | 2.18 | 1.00 | 43.86 | 2.83 (G:N7 14:382) |
| 5 | Water 5 | | 14:3552 | 2.19 | 1.00 | 55.19 | 2.20 (G:OP2 14:382) |
| 6 | Water 6 | | 14:3865 | 2.19 | 1.00 | 42.52 | 2.91 (G:OP1 14:386) |
| 7 | G | OP1 | 14:386 | 3.04 | 1.00 | 47.97 |  |
| 8 | G | OP2 | 14:382 | 3.18 | 1.00 | 49.71 |  |
| 9 | G | OP2 | 14:386 | 3.90 | 1.00 | 36.60 |  |
| 10 | G | N7 | 14:382 | 4.13 | 1.00 | 53.59 |  |
| 11 | G | OP2 | 14:381 | 4.85 | 1.00 | 49.28 |  |
| 12 | G | O5' | 14:386 | 4.85 | 1.00 | 41.35 |  |
| 13 | G | O5' | 14:382 | 4.90 | 1.00 | 37.04 |  |
  |

  
**1^st^ shell ligand angles:**   
1-2: 83; 1-3: 98; 1-4: 85; 1-5: 177; 1-6: 92; 2-3: 178; 2-4: 94; 2-5: 94; 2-6: 88; 3-4: 84; 3-5: 84; 3-6: 94; 4-5: 94; 4-6: 176; 5-6: 89;   
  
 Contact(s): /O_w_/O_w_/O_w_/O_w_/O_w_/O_w_/  
 Coded contact list: 6O_w_  
 Exclusion zone contact(s): /OP1/OP2/  
Consider K in place of MG (2 valid contact(s) for K vs. 0 for MG)  
MG hydration shell issue: water too close (< 2.4 Angs.)  

### MG:14:3383

|  | Res. type | Atom | Residue | Distance | Occupancy | B-factor | Closest solute to 1^st^ shell water |
| --- | --- | --- | --- | --- | --- | --- | --- |
| **Ion** | | | |  | 1.00 | 51.91 || 1 | Water 1 | | 14:3573 | 2.16 | 1.00 | 66.13 | 2.26 (G:N3 14:1681) |
| 2 | Water 2 | | 14:3651 | 2.18 | 1.00 | 56.57 | 2.49 (A:O2' 14:1762) |
| 3 | Water 3 | | 14:3945 | 2.19 | 1.00 | 60.62 | 3.15 (G:N1 14:1681) |
| 4 | Water 4 | | 14:3862 | 2.19 | 1.00 | 46.08 | 2.91 (U:O4 14:1757) |
| 5 | Water 5 | | 14:3974 | 2.19 | 1.00 | 39.39 | 3.95 (G:N2 14:1681) |
| 6 | Water 6 | | 14:3924 | 2.20 | 1.00 | 75.92 | 3.09 (C:OP2 14:1761) |
| 7 | A | O2' | 14:1762 | 3.90 | 1.00 | 61.21 |  |
| 8 | G | N3 | 14:1681 | 4.20 | 1.00 | 52.89 |  |
| 9 | G | N1 | 14:1681 | 4.25 | 1.00 | 54.83 |  |
| 10 | G | O2' | 14:1681 | 4.27 | 1.00 | 63.45 |  |
| 11 | U | O4 | 14:1757 | 4.28 | 1.00 | 61.17 |  |
| 12 | G | N2 | 14:1681 | 4.58 | 1.00 | 55.08 |  |
| 13 | C | OP2 | 14:1761 | 4.60 | 1.00 | 47.99 |  |
| 14 | G | N9 | 14:1681 | 4.95 | 1.00 | 56.55 |  |
  |

  
**1^st^ shell ligand angles:**   
1-2: 81; 1-3: 97; 1-4: 89; 1-5: 88; 1-6: 173; 2-3: 177; 2-4: 93; 2-5: 91; 2-6: 92; 3-4: 85; 3-5: 91; 3-6: 90; 4-5: 175; 4-6: 92; 5-6: 91;   
  
 Contact(s): /O_w_/O_w_/O_w_/O_w_/O_w_/O_w_/  
 Coded contact list: 6O_w_  
MG hydration shell issue: water too close (< 2.4 Angs.)  

### MG:14:3384

|  | Res. type | Atom | Residue | Distance | Occupancy | B-factor | Closest solute to 1^st^ shell water |
| --- | --- | --- | --- | --- | --- | --- | --- |
| **Ion** | | | |  | 1.00 | 47.88 || 1 | Water 1 | | 14:4042 | 2.18 | 1.00 | 39.46 | 3.66 (G:OP2 14:771) |
| 2 | Water 2 | | 14:3592 | 2.18 | 1.00 | 35.27 | 2.32 (G:OP2 14:770) |
| 3 | Water 3 | | 14:3657 | 2.18 | 1.00 | 32.85 | 2.51 (G:OP2 14:686) |
| 4 | Water 4 | | 14:3869 | 2.18 | 1.00 | 30.01 | 2.94 (G:OP2 14:771) |
| 5 | Water 5 | | L5:101 | 2.20 | 1.00 | 34.62 | 3.93 (ASN:ND2 L5:8) |
| 6 | Water 6 | | 14:3628 | 2.21 | 1.00 | 36.42 | 2.41 (G:OP1 14:770) |
| 7 | G | OP2 | 14:770 | 3.41 | 1.00 | 40.01 |  |
| 8 | G | OP2 | 14:771 | 4.00 | 1.00 | 28.83 |  |
| 9 | G | OP2 | 14:686 | 4.17 | 1.00 | 40.59 |  |
| 10 | G | OP1 | 14:770 | 4.33 | 1.00 | 37.76 |  |
| 11 | G | O5' | 14:770 | 4.35 | 1.00 | 28.81 |  |
| 12 | ASN | ND2 | L5:8 | 4.95 | 1.00 | 35.36 |  |
  |

  
**1^st^ shell ligand angles:**   
1-2: 175; 1-3: 92; 1-4: 88; 1-5: 90; 1-6: 87; 2-3: 84; 2-4: 90; 2-5: 91; 2-6: 98; 3-4: 89; 3-5: 95; 3-6: 178; 4-5: 177; 4-6: 93; 5-6: 84;   
  
 Contact(s): /O_w_/O_w_/O_w_/O_w_/O_w_/O_w_/  
 Coded contact list: 6O_w_  
MG hydration shell issue: water too close (< 2.4 Angs.)  

### MG:14:3385

|  | Res. type | Atom | Residue | Distance | Occupancy | B-factor | Closest solute to 1^st^ shell water |
| --- | --- | --- | --- | --- | --- | --- | --- |
| **Ion** | | | |  | 1.00 | 58.71 || 1 | Water 1 | | 14:3845 | 2.18 | 1.00 | 61.92 | 2.87 (U:O4 14:303) |
| 2 | Water 2 | | 14:3599 | 2.18 | 1.00 | 47.53 | 2.34 (A:OP2 14:311) |
| 3 | Water 3 | | 14:4014 | 2.18 | 1.00 | 55.24 | 3.42 (A:OP2 14:311) |
| 4 | Water 4 | | 14:3557 | 2.18 | 1.00 | 53.63 | 2.22 (G:O6 14:304) |
| 5 | Water 5 | | 14:3967 | 2.20 | 1.00 | 65.95 | 3.21 (G:N7 14:304) |
| 6 | A | OP2 | 14:311 | 2.52 | 1.00 | 59.78 |  |
| 7 | A | O2' | 14:310 | 3.36 | 1.00 | 57.10 |  |
| 8 | U | O4 | 14:303 | 3.89 | 1.00 | 65.55 |  |
| 9 | A | O5' | 14:311 | 4.18 | 1.00 | 62.40 |  |
| 10 | G | O6 | 14:304 | 4.26 | 1.00 | 66.30 |  |
| 11 | U | O4 | 14:305 | 4.30 | 1.00 | 56.83 |  |
| 12 | A | O3' | 14:310 | 4.56 | 1.00 | 49.80 |  |
| 13 | G | N7 | 14:304 | 4.58 | 1.00 | 62.60 |  |
| 14 | A | OP1 | 14:311 | 4.95 | 1.00 | 61.48 |  |
  |

  
**1^st^ shell ligand angles:**   
1-2: 84; 1-3: 176; 1-4: 92; 1-5: 95; 2-3: 93; 2-4: 175; 2-5: 91; 3-4: 91; 3-5: 88; 4-5: 93;   
  
 Contact(s): /O_w_/O_w_/O_w_/O_w_/O_w_/  
 Coded contact list: 5O_w_  
 Exclusion zone contact(s): /O2'/OP2/  
Consider NA in place of MG (1 valid contact(s) for NA vs. 0 for MG)  
MG hydration shell issue: water too close (< 2.4 Angs.)  

### MG:14:3386

|  | Res. type | Atom | Residue | Distance | Occupancy | B-factor | Closest solute to 1^st^ shell water |
| --- | --- | --- | --- | --- | --- | --- | --- |
| **Ion** | | | |  | 1.00 | 61.45 || 1 | Water 1 | | 14:3868 | 2.18 | 1.00 | 39.44 | 3.06 (G:N7 14:2000) |
| 2 | Water 2 | | 14:4085 | 2.18 | 1.00 | 74.05 | 5.10 (G:N7 14:2000) |
| 3 | Water 3 | | 29:407 | 2.18 | 1.00 | 30.10 | 4.29 (ARG:NH1 29:136) |
| 4 | Water 4 | | 14:4033 | 2.19 | 1.00 | 42.90 | 3.97 (G:N7 14:2000) |
| 5 | Water 5 | | 14:4027 | 2.19 | 1.00 | 35.02 | 4.68 (ARG:NH1 29:136) |
| 6 | Water 6 | | 29:405 | 2.20 | 1.00 | 57.91 | 3.40 (ARG:NH1 29:136) |
| 7 | G | N7 | 14:2000 | 4.65 | 1.00 | 41.62 |  |
| 8 | ARG | NH1 | 29:136 | 4.84 | 1.00 | 36.29 |  |
  |

  
**1^st^ shell ligand angles:**   
1-2: 88; 1-3: 88; 1-4: 92; 1-5: 177; 1-6: 91; 2-3: 86; 2-4: 92; 2-5: 90; 2-6: 179; 3-4: 179; 3-5: 94; 3-6: 93; 4-5: 86; 4-6: 88; 5-6: 91;   
  
 Contact(s): /O_w_/O_w_/O_w_/O_w_/O_w_/O_w_/  
 Coded contact list: 6O_w_  

### MG:14:3387

|  | Res. type | Atom | Residue | Distance | Occupancy | B-factor | Closest solute to 1^st^ shell water |
| --- | --- | --- | --- | --- | --- | --- | --- |
| **Ion** | | | |  | 1.00 | 45.54 || 1 | Water 1 | | 14:3977 | 2.17 | 1.00 | 47.44 | 3.24 (A:OP1 14:1253) |
| 2 | Water 2 | | 14:3912 | 2.17 | 1.00 | 37.06 | 3.05 (G:OP2 14:582) |
| 3 | Water 3 | | 14:3948 | 2.20 | 1.00 | 41.94 | 3.16 (A:OP1 14:1253) |
| 4 | Water 4 | | 14:3798 | 2.20 | 1.00 | 36.77 | 2.74 (G:OP2 14:582) |
| 5 | Water 5 | | 14:3701 | 2.20 | 1.00 | 39.76 | 2.58 (G:N7 14:583) |
| 6 | Water 6 | | 14:3506 | 2.22 | 1.00 | 52.28 | 1.30 (G:N7 14:582) |
| 7 | G | N7 | 14:582 | 3.51 | 1.00 | 44.10 |  |
| 8 | G | OP2 | 14:582 | 3.86 | 1.00 | 44.16 |  |
| 9 | G | N7 | 14:583 | 3.95 | 1.00 | 38.81 |  |
| 10 | A | OP1 | 14:1253 | 4.26 | 1.00 | 46.80 |  |
| 11 | ARG | NH1 | 85:10 | 4.26 | 1.00 | 47.80 |  |
| 12 | G | O6 | 14:583 | 4.71 | 1.00 | 36.59 |  |
| 13 | G | O6 | 14:582 | 4.73 | 1.00 | 43.37 |  |
| 14 | G | O5' | 14:582 | 4.81 | 1.00 | 41.15 |  |
  |

  
**1^st^ shell ligand angles:**   
1-2: 90; 1-3: 94; 1-4: 86; 1-5: 86; 1-6: 178; 2-3: 85; 2-4: 94; 2-5: 174; 2-6: 90; 3-4: 179; 3-5: 92; 3-6: 88; 4-5: 89; 4-6: 92; 5-6: 94;   
  
 Contact(s): /O_w_/O_w_/O_w_/O_w_/O_w_/O_w_/  
 Coded contact list: 6O_w_  

### MG:14:3388

|  | Res. type | Atom | Residue | Distance | Occupancy | B-factor | Closest solute to 1^st^ shell water |
| --- | --- | --- | --- | --- | --- | --- | --- |
| **Ion** | | | |  | 1.00 | 91.23 || 1 | Water 1 | | 14:3738 | 2.18 | 1.00 | 91.42 | 2.66 (G:N7 14:270W) |
| 2 | Water 2 | | 14:4015 | 2.18 | 1.00 | 82.21 | 3.43 (A:OP2 14:270A) |
| 3 | Water 3 | | 14:3823 | 2.18 | 1.00 | 103.38 | 2.81 (C:N4 14:270U) |
| 4 | Water 4 | | 14:3989 | 2.18 | 1.00 | 92.33 | 3.29 (A:OP1 14:270A) |
| 5 | Water 5 | | 14:3660 | 2.18 | 1.00 | 86.97 | 2.51 (G:O6 14:270X) |
| 6 | Water 6 | | 14:4018 | 2.19 | 1.00 | 95.00 | 3.45 (G:N7 14:270V) |
| 7 | G | O6 | 14:270V | 3.82 | 1.00 | 99.10 |  |
| 8 | G | N7 | 14:270V | 4.25 | 1.00 | 99.85 |  |
| 9 | G | O6 | 14:270X | 4.26 | 1.00 | 90.22 |  |
| 10 | A | OP2 | 14:270A | 4.38 | 1.00 | 75.72 |  |
| 11 | G | O6 | 14:270W | 4.39 | 1.00 | 94.13 |  |
| 12 | G | N1 | 14:270X | 4.58 | 1.00 | 87.03 |  |
| 13 | C | N4 | 14:270U | 4.77 | 1.00 | 107.77 |  |
| 14 | G | N7 | 14:270W | 4.79 | 1.00 | 94.93 |  |
| 15 | A | OP1 | 14:270A | 4.81 | 1.00 | 75.84 |  |
| 16 | A | OP2 | 14:270 | 4.96 | 1.00 | 76.88 |  |
| 17 | A | OP1 | 14:270 | 4.97 | 1.00 | 73.28 |  |
  |

  
**1^st^ shell ligand angles:**   
1-2: 88; 1-3: 92; 1-4: 179; 1-5: 89; 1-6: 91; 2-3: 89; 2-4: 90; 2-5: 92; 2-6: 179; 3-4: 88; 3-5: 179; 3-6: 90; 4-5: 90; 4-6: 90; 5-6: 89;   
  
 Contact(s): /O_w_/O_w_/O_w_/O_w_/O_w_/O_w_/  
 Coded contact list: 6O_w_  

### MG:14:3389

|  | Res. type | Atom | Residue | Distance | Occupancy | B-factor | Closest solute to 1^st^ shell water |
| --- | --- | --- | --- | --- | --- | --- | --- |
| **Ion** | | | |  | 1.00 | 45.36 || 1 | Water 1 | | 35:201 | 2.17 | 1.00 | 49.38 | 2.25 (HIS:O 35:35) |
| 2 | Water 2 | | 35:203 | 2.17 | 1.00 | 45.01 | 2.77 (LYS:O 35:36) |
| 3 | Water 3 | | 35:206 | 2.18 | 1.00 | 45.73 | 3.11 (LYS:O 35:36) |
| 4 | Water 4 | | 14:3810 | 2.18 | 1.00 | 41.56 | 2.78 (U:OP2 14:943) |
| 5 | Water 5 | | 14:3885 | 2.18 | 1.00 | 51.59 | 2.98 (G:N1 14:944) |
| 6 | U | OP2 | 14:943 | 2.52 | 1.00 | 42.40 |  |
| 7 | LYS | O | 35:36 | 3.69 | 1.00 | 42.61 |  |
| 8 | HIS | O | 35:35 | 4.01 | 1.00 | 53.54 |  |
| 9 | U | OP1 | 14:943 | 4.17 | 1.00 | 51.92 |  |
| 10 | G | OP2 | 14:832 | 4.29 | 1.00 | 37.13 |  |
| 11 | U | O5' | 14:943 | 4.37 | 1.00 | 52.02 |  |
| 12 | G | OP1 | 14:832 | 4.41 | 1.00 | 37.45 |  |
| 13 | G | O3' | 14:942 | 4.86 | 1.00 | 42.49 |  |
| 14 | G | N1 | 14:944 | 4.95 | 1.00 | 43.08 |  |
| 15 | ARG | O | 35:33 | 4.96 | 1.00 | 43.11 |  |
  |

  
**1^st^ shell ligand angles:**   
1-2: 87; 1-3: 94; 1-4: 85; 1-5: 177; 2-3: 88; 2-4: 95; 2-5: 91; 3-4: 176; 3-5: 89; 4-5: 93;   
  
 Contact(s): /O_w_/O_w_/O_w_/O_w_/O_w_/  
 Coded contact list: 5O_w_  
 Exclusion zone contact(s): /OP2/  
Consider NA in place of MG (1 valid contact(s) for NA vs. 0 for MG)  
MG hydration shell issue: water too close (< 2.4 Angs.)  

### MG:14:3390

|  | Res. type | Atom | Residue | Distance | Occupancy | B-factor | Closest solute to 1^st^ shell water |
| --- | --- | --- | --- | --- | --- | --- | --- |
| **Ion** | | | |  | 1.00 | 79.43 || 1 | Water 1 | | 14:3585 | 2.16 | 1.00 | 84.79 | 2.29 (G:N7 14:1857) |
| 2 | Water 2 | | 14:3692 | 2.17 | 1.00 | 77.94 | 2.56 (G:OP2 14:1857) |
| 3 | Water 3 | | 14:4078 | 2.18 | 1.00 | 81.27 | 4.12 (G:OP1 14:1856) |
| 4 | Water 4 | | 14:4060 | 2.18 | 1.00 | 84.63 | 3.85 (G:N7 14:1858) |
| 5 | Water 5 | | 14:3828 | 2.18 | 1.00 | 76.98 | 2.82 (G:OP2 14:1856) |
| 6 | Water 6 | | 14:3575 | 2.19 | 1.00 | 85.23 | 2.27 (G:O6 14:1858) |
| 7 | G | O6 | 14:1858 | 3.44 | 1.00 | 86.80 |  |
| 8 | G | N7 | 14:1857 | 3.74 | 1.00 | 83.02 |  |
| 9 | G | O6 | 14:1857 | 4.00 | 1.00 | 83.35 |  |
| 10 | G | N7 | 14:1858 | 4.22 | 1.00 | 88.05 |  |
| 11 | G | OP2 | 14:1856 | 4.48 | 1.00 | 71.57 |  |
| 12 | G | OP2 | 14:1857 | 4.66 | 1.00 | 72.09 |  |
| 13 | G | O6 | 14:1883 | 4.78 | 1.00 | 81.93 |  |
| 14 | G | OP1 | 14:1856 | 4.94 | 1.00 | 63.19 |  |
  |

  
**1^st^ shell ligand angles:**   
1-2: 85; 1-3: 178; 1-4: 92; 1-5: 87; 1-6: 93; 2-3: 93; 2-4: 87; 2-5: 91; 2-6: 177; 3-4: 88; 3-5: 92; 3-6: 89; 4-5: 178; 4-6: 91; 5-6: 91;   
  
 Contact(s): /O_w_/O_w_/O_w_/O_w_/O_w_/O_w_/  
 Coded contact list: 6O_w_  
MG hydration shell issue: water too close (< 2.4 Angs.)  

### MG:14:3391

|  | Res. type | Atom | Residue | Distance | Occupancy | B-factor | Closest solute to 1^st^ shell water |
| --- | --- | --- | --- | --- | --- | --- | --- |
| **Ion** | | | |  | 1.00 | 81.43 || 1 | Water 1 | | 14:3745 | 2.18 | 1.00 | 76.14 | 2.67 (G:N7 14:2382) |
| 2 | Water 2 | | 14:3800 | 2.18 | 1.00 | 76.40 | 2.75 (A:OP2 14:2366) |
| 3 | Water 3 | | 14:3547 | 2.18 | 1.00 | 67.91 | 2.18 (A:OP2 14:2366) |
| 4 | Water 4 | | 14:4056 | 2.18 | 1.00 | 69.52 | 3.82 (G:N7 14:2382) |
| 5 | Water 5 | | 14:3816 | 2.19 | 1.00 | 79.94 | 2.79 (G:OP2 14:2367) |
| 6 | Water 6 | | 14:3779 | 2.19 | 1.00 | 60.56 | 2.71 (G:N2 14:2345) |
| 7 | A | OP2 | 14:2366 | 3.41 | 1.00 | 71.80 |  |
| 8 | A | O5' | 14:2366 | 3.88 | 1.00 | 74.65 |  |
| 9 | G | N7 | 14:2382 | 4.09 | 1.00 | 69.91 |  |
| 10 | G | OP2 | 14:2367 | 4.19 | 1.00 | 67.87 |  |
| 11 | G | OP2 | 14:2382 | 4.36 | 1.00 | 62.82 |  |
| 12 | A | OP1 | 14:2366 | 4.42 | 1.00 | 63.92 |  |
| 13 | G | N7 | 14:2367 | 4.53 | 1.00 | 67.65 |  |
| 14 | G | N2 | 14:2345 | 4.73 | 1.00 | 55.56 |  |
| 15 | G | O6 | 14:2367 | 4.75 | 1.00 | 70.60 |  |
  |

  
**1^st^ shell ligand angles:**   
1-2: 89; 1-3: 87; 1-4: 90; 1-5: 177; 1-6: 94; 2-3: 91; 2-4: 179; 2-5: 88; 2-6: 92; 3-4: 88; 3-5: 93; 3-6: 177; 4-5: 92; 4-6: 88; 5-6: 87;   
  
 Contact(s): /O_w_/O_w_/O_w_/O_w_/O_w_/O_w_/  
 Coded contact list: 6O_w_  
MG hydration shell issue: water too close (< 2.4 Angs.)  

### MG:14:3392

|  | Res. type | Atom | Residue | Distance | Occupancy | B-factor | Closest solute to 1^st^ shell water |
| --- | --- | --- | --- | --- | --- | --- | --- |
| **Ion** | | | |  | 1.00 | 75.97 || 1 | Water 1 | | 14:4094 | 2.17 | 1.00 | 70.49 | 4.56 (C:OP1 14:77) |
| 2 | Water 2 | | 14:3906 | 2.18 | 1.00 | 47.85 | 3.03 (C:OP1 14:76) |
| 3 | Water 3 | | 14:3926 | 2.19 | 1.00 | 70.84 | 3.09 (U:OP1 14:104) |
| 4 | Water 4 | | 14:4052 | 2.19 | 1.00 | 82.23 | 3.79 (U:OP1 14:104) |
| 5 | Water 5 | | 14:3880 | 2.19 | 1.00 | 75.03 | 2.96 (U:OP1 14:104) |
| 6 | Water 6 | | 14:3735 | 2.19 | 1.00 | 84.68 | 2.66 (C:OP2 14:77) |
| 7 | U | OP1 | 14:104 | 3.99 | 1.00 | 64.62 |  |
| 8 | C | OP2 | 14:77 | 4.18 | 1.00 | 56.53 |  |
| 9 | C | OP1 | 14:77 | 4.46 | 1.00 | 57.73 |  |
| 10 | C | OP1 | 14:76 | 4.72 | 1.00 | 58.49 |  |
| 11 | A | O3' | 14:103 | 4.76 | 1.00 | 68.75 |  |
  |

  
**1^st^ shell ligand angles:**   
1-2: 90; 1-3: 91; 1-4: 88; 1-5: 178; 1-6: 89; 2-3: 89; 2-4: 178; 2-5: 92; 2-6: 90; 3-4: 92; 3-5: 89; 3-6: 179; 4-5: 90; 4-6: 90; 5-6: 90;   
  
 Contact(s): /O_w_/O_w_/O_w_/O_w_/O_w_/O_w_/  
 Coded contact list: 6O_w_  

### MG:14:3393

|  | Res. type | Atom | Residue | Distance | Occupancy | B-factor | Closest solute to 1^st^ shell water |
| --- | --- | --- | --- | --- | --- | --- | --- |
| **Ion** | | | |  | 1.00 | 63.92 || 1 | Water 1 | | 14:3799 | 2.18 | 1.00 | 51.66 | 2.75 (G:O6 14:1209) |
| 2 | Water 2 | | 14:4016 | 2.19 | 1.00 | 59.92 | 3.43 (A:O3' 14:331) |
| 3 | Water 3 | | 14:4045 | 2.19 | 1.00 | 76.83 | 5.23 (A:N3 14:331) |
| 4 | Water 4 | | 14:3627 | 2.19 | 1.00 | 60.52 | 2.41 (A:OP1 14:332) |
| 5 | Water 5 | | 14:4035 | 2.19 | 1.00 | 59.13 | 3.58 (G:OP1 14:1206) |
| 6 | Water 6 | | 14:4046 | 2.20 | 1.00 | 61.30 | 3.72 (G:OP1 14:1206) |
| 7 | A | OP1 | 14:332 | 4.09 | 1.00 | 60.84 |  |
| 8 | A | O3' | 14:331 | 4.70 | 1.00 | 51.91 |  |
| 9 | G | O6 | 14:1209 | 4.74 | 1.00 | 48.94 |  |
| 10 | G | OP1 | 14:1206 | 4.76 | 1.00 | 61.16 |  |
| 11 | A | N3 | 14:331 | 4.97 | 1.00 | 55.09 |  |
  |

  
**1^st^ shell ligand angles:**   
1-2: 91; 1-3: 88; 1-4: 93; 1-5: 88; 1-6: 178; 2-3: 91; 2-4: 89; 2-5: 178; 2-6: 88; 3-4: 179; 3-5: 88; 3-6: 91; 4-5: 92; 4-6: 88; 5-6: 93;   
  
 Contact(s): /O_w_/O_w_/O_w_/O_w_/O_w_/O_w_/  
 Coded contact list: 6O_w_  

### MG:14:3394

|  | Res. type | Atom | Residue | Distance | Occupancy | B-factor | Closest solute to 1^st^ shell water |
| --- | --- | --- | --- | --- | --- | --- | --- |
| **Ion** | | | |  | 1.00 | 61.39 || 1 | Water 1 | | 14:3685 | 2.18 | 1.00 | 38.72 | 2.55 (G:O6 14:690) |
| 2 | Water 2 | | 14:3895 | 2.18 | 1.00 | 37.04 | 3.00 (G:O6 14:771) |
| 3 | Water 3 | | 14:4037 | 2.18 | 1.00 | 35.16 | 3.59 (G:O6 14:771) |
| 4 | Water 4 | | 14:3843 | 2.19 | 1.00 | 39.82 | 4.27 (G:O6 14:690) |
| 5 | Water 5 | | 14:4057 | 2.19 | 1.00 | 41.11 | 4.26 (G:OP2 14:686) |
| 6 | Water 6 | | 14:4095 | 2.20 | 1.00 | 47.58 | 5.33 (C:N4 14:691) |
| 7 | G | O6 | 14:690 | 4.16 | 1.00 | 36.42 |  |
| 8 | G | O6 | 14:771 | 4.19 | 1.00 | 32.36 |  |
| 9 | C | N4 | 14:691 | 4.52 | 1.00 | 33.05 |  |
| 10 | G | OP2 | 14:686 | 4.89 | 1.00 | 40.59 |  |
| 11 | C | N4 | 14:772 | 4.96 | 1.00 | 34.96 |  |
  |

  
**1^st^ shell ligand angles:**   
1-2: 96; 1-3: 90; 1-4: 88; 1-5: 177; 1-6: 84; 2-3: 88; 2-4: 88; 2-5: 87; 2-6: 180; 3-4: 175; 3-5: 90; 3-6: 93; 4-5: 92; 4-6: 92; 5-6: 94;   
  
 Contact(s): /O_w_/O_w_/O_w_/O_w_/O_w_/O_w_/  
 Coded contact list: 6O_w_  

### MG:14:3395

|  | Res. type | Atom | Residue | Distance | Occupancy | B-factor | Closest solute to 1^st^ shell water |
| --- | --- | --- | --- | --- | --- | --- | --- |
| **Ion** | | | |  | 1.00 | 25.67 || 1 | Water 1 | | 14:3904 | 2.16 | 1.00 | 40.65 | 3.03 (G:OP2 14:1776) |
| 2 | Water 2 | | 14:3591 | 2.16 | 1.00 | 24.94 | 2.31 (G:OP2 14:1776) |
| 3 | Water 3 | | 14:3711 | 2.18 | 1.00 | 42.79 | 2.61 (A:OP1 14:1981) |
| 4 | Water 4 | | 14:3994 | 2.18 | 1.00 | 39.10 | 3.45 (G:OP2 14:1776) |
| 5 | Water 5 | | 14:3768 | 2.20 | 1.00 | 40.80 | 2.69 (A:OP1 14:1981) |
| 6 | G | OP2 | 14:1776 | 2.57 | 1.00 | 39.87 |  |
| 7 | G | OP1 | 14:1776 | 3.42 | 1.00 | 39.40 |  |
| 8 | A | OP1 | 14:1981 | 3.74 | 1.00 | 36.63 |  |
| 9 | A | OP2 | 14:1981 | 4.25 | 1.00 | 37.05 |  |
| 10 | U | OP1 | 14:1775 | 4.42 | 1.00 | 30.49 |  |
| 11 | G | N7 | 14:1776 | 4.44 | 1.00 | 29.09 |  |
| 12 | U | O3' | 14:1775 | 4.50 | 1.00 | 31.09 |  |
| 13 | G | O5' | 14:1776 | 4.58 | 1.00 | 29.66 |  |
| 14 | U | OP2 | 14:1775 | 4.74 | 1.00 | 34.36 |  |
  |

  
**1^st^ shell ligand angles:**   
1-2: 92; 1-3: 95; 1-4: 171; 1-5: 87; 2-3: 89; 2-4: 85; 2-5: 177; 3-4: 94; 3-5: 88; 4-5: 95;   
  
 Contact(s): /O_w_/O_w_/O_w_/O_w_/O_w_/  
 Coded contact list: 5O_w_  
 Exclusion zone contact(s): /OP2/  
Consider NA in place of MG (1 valid contact(s) for NA vs. 0 for MG)  
MG hydration shell issue: water too close (< 2.4 Angs.)  

### MG:14:3396

|  | Res. type | Atom | Residue | Distance | Occupancy | B-factor | Closest solute to 1^st^ shell water |
| --- | --- | --- | --- | --- | --- | --- | --- |
| **Ion** | | | |  | 1.00 | 66.06 || 1 | Water 1 | | 14:3523 | 2.17 | 1.00 | 61.81 | 2.02 (G:OP2 14:1756) |
| 2 | Water 2 | | 14:3780 | 2.18 | 1.00 | 59.12 | 2.71 (G:O6 14:1756) |
| 3 | Water 3 | | 75:202 | 2.19 | 1.00 | 76.32 | 5.53 (U:OP2 14:1706) |
| 4 | Water 4 | | 14:3721 | 2.19 | 1.00 | 69.93 | 2.62 (U:OP2 14:1706) |
| 5 | Water 5 | | 14:4062 | 2.19 | 1.00 | 53.02 | 3.87 (G:OP2 14:1756) |
| 6 | Water 6 | | 14:3909 | 2.19 | 1.00 | 76.46 | 3.04 (G:O6 14:1753) |
| 7 | G | OP2 | 14:1756 | 3.88 | 1.00 | 56.21 |  |
| 8 | G | O6 | 14:1753 | 4.17 | 1.00 | 69.36 |  |
| 9 | G | O6 | 14:1756 | 4.21 | 1.00 | 70.58 |  |
| 10 | U | OP2 | 14:1706 | 4.70 | 1.00 | 50.83 |  |
| 11 | G | N7 | 14:1756 | 4.87 | 1.00 | 63.36 |  |
  |

  
**1^st^ shell ligand angles:**   
1-2: 87; 1-3: 178; 1-4: 89; 1-5: 91; 1-6: 92; 2-3: 92; 2-4: 88; 2-5: 177; 2-6: 93; 3-4: 90; 3-5: 90; 3-6: 89; 4-5: 94; 4-6: 179; 5-6: 85;   
  
 Contact(s): /O_w_/O_w_/O_w_/O_w_/O_w_/O_w_/  
 Coded contact list: 6O_w_  
MG hydration shell issue: water too close (< 2.4 Angs.)  

### MG:14:3397

|  | Res. type | Atom | Residue | Distance | Occupancy | B-factor | Closest solute to 1^st^ shell water |
| --- | --- | --- | --- | --- | --- | --- | --- |
| **Ion** | | | |  | 1.00 | 75.21 || 1 | Water 1 | | 14:3530 | 2.17 | 1.00 | 64.73 | 2.07 (C:OP2 14:971) |
| 2 | Water 2 | | 14:3532 | 2.18 | 1.00 | 71.16 | 2.08 (G:O6 14:974) |
| 3 | Water 3 | | 14:3897 | 2.18 | 1.00 | 68.25 | 3.01 (G:O6 14:972) |
| 4 | Water 4 | | 14:3997 | 2.19 | 1.00 | 78.94 | 3.84 (C:O5' 14:970) |
| 5 | Water 5 | | 14:3508 | 2.20 | 1.00 | 71.62 | 1.31 (G:N7 14:974) |
| 6 | C | OP2 | 14:971 | 3.00 | 1.00 | 54.56 |  |
| 7 | G | N7 | 14:974 | 3.18 | 1.00 | 74.58 |  |
| 8 | G | O6 | 14:974 | 3.55 | 1.00 | 74.04 |  |
| 9 | G | O6 | 14:989 | 4.10 | 1.00 | 78.32 |  |
| 10 | G | O6 | 14:972 | 4.16 | 1.00 | 67.98 |  |
| 11 | C | O5' | 14:970 | 4.92 | 1.00 | 63.63 |  |
| 12 | G | N7 | 14:972 | 4.99 | 1.00 | 61.67 |  |
  |

  
**1^st^ shell ligand angles:**   
1-2: 90; 1-3: 178; 1-4: 92; 1-5: 85; 2-3: 88; 2-4: 86; 2-5: 94; 3-4: 87; 3-5: 97; 4-5: 177;   
  
 Contact(s): /O_w_/O_w_/O_w_/O_w_/O_w_/  
 Coded contact list: 5O_w_  
 Exclusion zone contact(s): /N7/OP2/  
Consider K in place of MG (2 valid contact(s) for K vs. 0 for MG)  
MG hydration shell issue: water too close (< 2.4 Angs.)  

### MG:14:3398

|  | Res. type | Atom | Residue | Distance | Occupancy | B-factor | Closest solute to 1^st^ shell water |
| --- | --- | --- | --- | --- | --- | --- | --- |
| **Ion** | | | |  | 1.00 | 47.69 || 1 | Water 1 | | 14:3571 | 2.17 | 1.00 | 38.07 | 2.26 (G:O6 14:1309) |
| 2 | Water 2 | | 14:3606 | 2.17 | 1.00 | 45.96 | 2.35 (G:O6 14:1311) |
| 3 | Water 3 | | 14:3846 | 2.18 | 1.00 | 47.19 | 2.87 (G:N7 14:1309) |
| 4 | Water 4 | | 14:3844 | 2.18 | 1.00 | 39.48 | 2.87 (A:OP2 14:1308) |
| 5 | Water 5 | | 14:4092 | 2.20 | 1.00 | 55.54 | 4.87 (G:O6 14:1311) |
| 6 | Water 6 | | 14:3992 | 2.20 | 1.00 | 40.05 | 3.29 (G:O6 14:1311) |
| 7 | G | O6 | 14:1311 | 3.66 | 1.00 | 46.56 |  |
| 8 | G | N7 | 14:1309 | 4.05 | 1.00 | 33.15 |  |
| 9 | ARG | NH2 | L5:9 | 4.19 | 1.00 | 47.34 |  |
| 10 | G | O6 | 14:1309 | 4.28 | 1.00 | 39.59 |  |
| 11 | G | O6 | 14:1310 | 4.49 | 1.00 | 47.70 |  |
| 12 | G | N7 | 14:1310 | 4.51 | 1.00 | 38.04 |  |
| 13 | A | OP2 | 14:1308 | 4.80 | 1.00 | 37.75 |  |
| 14 | G | N1 | 14:1311 | 4.81 | 1.00 | 55.08 |  |
| 15 | G | OP2 | 14:1309 | 4.89 | 1.00 | 39.99 |  |
  |

  
**1^st^ shell ligand angles:**   
1-2: 85; 1-3: 91; 1-4: 90; 1-5: 92; 1-6: 176; 2-3: 91; 2-4: 175; 2-5: 89; 2-6: 93; 3-4: 89; 3-5: 178; 3-6: 86; 4-5: 91; 4-6: 92; 5-6: 91;   
  
 Contact(s): /O_w_/O_w_/O_w_/O_w_/O_w_/O_w_/  
 Coded contact list: 6O_w_  
MG hydration shell issue: water too close (< 2.4 Angs.)  

### MG:14:3399

|  | Res. type | Atom | Residue | Distance | Occupancy | B-factor | Closest solute to 1^st^ shell water |
| --- | --- | --- | --- | --- | --- | --- | --- |
| **Ion** | | | |  | 1.00 | 106.45 || 1 | Water 1 | | 14:3550 | 2.17 | 1.00 | 96.33 | 2.19 (G:N7 14:2304) |
| 2 | Water 2 | | 14:3935 | 2.19 | 1.00 | 81.66 | 3.12 (G:OP2 14:2303) |
| 3 | Water 3 | | 14:3777 | 2.19 | 1.00 | 90.27 | 2.71 (G:O6 14:2303) |
| 4 | Water 4 | | 14:3664 | 2.19 | 1.00 | 100.86 | 2.52 (G:O6 14:2304) |
| 5 | Water 5 | | 14:4096 | 2.19 | 1.00 | 94.55 | 4.76 (G:N7 14:2304) |
| 6 | Water 6 | | 14:4023 | 2.19 | 1.00 | 101.06 | 5.18 (G:O6 14:2304) |
| 7 | G | O6 | 14:2304 | 3.89 | 1.00 | 102.90 |  |
| 8 | G | N7 | 14:2304 | 4.07 | 1.00 | 98.13 |  |
| 9 | G | N7 | 14:2303 | 4.41 | 1.00 | 99.11 |  |
| 10 | G | O6 | 14:2303 | 4.51 | 1.00 | 91.30 |  |
| 11 | G | OP2 | 14:2303 | 4.94 | 1.00 | 101.39 |  |
  |

  
**1^st^ shell ligand angles:**   
1-2: 91; 1-3: 91; 1-4: 89; 1-5: 87; 1-6: 178; 2-3: 89; 2-4: 179; 2-5: 90; 2-6: 90; 3-4: 91; 3-5: 178; 3-6: 91; 4-5: 90; 4-6: 90; 5-6: 91;   
  
 Contact(s): /O_w_/O_w_/O_w_/O_w_/O_w_/O_w_/  
 Coded contact list: 6O_w_  
MG hydration shell issue: water too close (< 2.4 Angs.)  

### MG:14:3400

|  | Res. type | Atom | Residue | Distance | Occupancy | B-factor | Closest solute to 1^st^ shell water |
| --- | --- | --- | --- | --- | --- | --- | --- |
| **Ion** | | | |  | 1.00 | 63.52 || 1 | Water 1 | | 14:3813 | 2.17 | 1.00 | 41.70 | 2.78 (C:O2' 14:1838) |
| 2 | Water 2 | | 14:3928 | 2.18 | 1.00 | 49.57 | 3.10 (U:O4 14:1898) |
| 3 | Water 3 | | 14:3996 | 2.18 | 1.00 | 55.62 | 3.43 (G:N7 14:1897) |
| 4 | Water 4 | | 14:3783 | 2.19 | 1.00 | 48.29 | 2.72 (G:OP2 14:1839) |
| 5 | Water 5 | | 14:3925 | 2.19 | 1.00 | 48.53 | 3.24 (G:OP2 14:1840) |
| 6 | G | OP2 | 14:1839 | 2.74 | 1.00 | 52.07 |  |
| 7 | C | O3' | 14:1838 | 3.64 | 1.00 | 49.79 |  |
| 8 | G | OP2 | 14:1840 | 4.11 | 1.00 | 48.25 |  |
| 9 | G | O5' | 14:1839 | 4.37 | 1.00 | 42.04 |  |
| 10 | U | O4 | 14:1898 | 4.43 | 1.00 | 44.69 |  |
| 11 | C | O2' | 14:1838 | 4.59 | 1.00 | 43.63 |  |
| 12 | G | N7 | 14:1897 | 4.71 | 1.00 | 50.51 |  |
| 13 | G | OP2 | 14:1897 | 4.79 | 1.00 | 48.39 |  |
| 14 | G | OP1 | 14:1839 | 4.97 | 1.00 | 58.24 |  |
  |

  
**1^st^ shell ligand angles:**   
1-2: 84; 1-3: 90; 1-4: 94; 1-5: 177; 2-3: 92; 2-4: 174; 2-5: 95; 3-4: 94; 3-5: 87; 4-5: 88;   
  
 Contact(s): /O_w_/O_w_/O_w_/O_w_/O_w_/  
 Coded contact list: 5O_w_  
 Exclusion zone contact(s): /OP2/  
Consider K in place of MG (1 valid contact(s) for K vs. 0 for MG)  

### K:16:201

|  | Res. type | Atom | Residue | Distance | Occupancy | B-factor | Closest solute to 1^st^ shell water |
| --- | --- | --- | --- | --- | --- | --- | --- |
| **Ion** | | | |  | 1.00 | 55.47 || 1 | G | O6 | 16:16 | 2.70 | 1.00 | 53.31 |
| 2 | C | O4' | 16:108 | 2.85 | 1.00 | 55.89 |  |
| 3 | A | OP1 | 16:15 | 2.93 | 1.00 | 55.52 |  |
| 4 | G | N7 | 16:16 | 3.01 | 1.00 | 54.20 |  |
| 5 | A | OP2 | 16:15 | 3.22 | 1.00 | 60.77 |  |
| 6 | A | N1 | 16:66 | 3.26 | 1.00 | 57.76 |  |
| 7 | C | O2 | 16:108 | 3.60 | 1.00 | 54.49 |  |
| 8 | U | O2 | 16:107 | 3.67 | 1.00 | 51.10 |  |
| 9 | C | N1 | 16:108 | 3.86 | 1.00 | 55.07 |  |
| 10 | G | N1 | 16:69 | 4.10 | 1.00 | 52.52 |  |
| 11 | A | O5' | 16:15 | 4.13 | 1.00 | 57.63 |  |
| 12 | G | N2 | 16:69 | 4.30 | 1.00 | 56.25 |  |
| 13 | A | N6 | 16:66 | 4.57 | 1.00 | 56.22 |  |
| 14 | C | N3 | 16:108 | 4.58 | 1.00 | 54.51 |  |
| 15 | U | O2' | 16:107 | 4.72 | 1.00 | 55.81 |  |
| 16 | G | N1 | 16:16 | 4.76 | 1.00 | 55.82 |  |
| 17 | C | O5' | 16:108 | 4.79 | 1.00 | 48.71 |  |
| 18 | G | O6 | 16:69 | 4.94 | 1.00 | 53.49 |  |
| 19 | U | O3' | 16:14 | 4.97 | 1.00 | 56.23 |  |
  |

  
**1^st^ shell ligand angles:**   
1-2: 132; 1-3: 148; 1-4: 65; 1-5: 106; 1-6: 79; 2-3: 79; 2-4: 131; 2-5: 118; 2-6: 76; 3-4: 91; 3-5: 48; 3-6: 121; 4-5: 85; 4-6: 144; 5-6: 103;   
  
 Contact(s): /N1_A_/N7/O4'/O6/OP1/OP2/  
 Coded contact list: 2O_ph_O_r_O_b_2N_b_  

### K:16:202

|  | Res. type | Atom | Residue | Distance | Occupancy | B-factor | Closest solute to 1^st^ shell water |
| --- | --- | --- | --- | --- | --- | --- | --- |
| **Ion** | | | |  | 1.00 | 74.09 || 1 | G | O6 | 16:112 | 2.70 | 1.00 | 50.40 |
| 2 | U | O4 | 16:111 | 2.96 | 1.00 | 70.93 |  |
| 3 | U | N3 | 16:111 | 2.96 | 1.00 | 58.74 |  |
| 4 | G | O6 | 16:9 | 3.00 | 1.00 | 55.54 |  |
| 5 | U | O4 | 16:8 | 3.30 | 1.00 | 55.00 |  |
| 6 | G | N7 | 16:112 | 3.59 | 1.00 | 53.67 |  |
| 7 | U | O2 | 16:111 | 4.31 | 1.00 | 58.87 |  |
| 8 | U | N3 | 16:8 | 4.33 | 1.00 | 53.30 |  |
| 9 | G | O6 | 16:7 | 4.33 | 1.00 | 44.83 |  |
| 10 | G | N1 | 16:9 | 4.41 | 1.00 | 53.12 |  |
| 11 | C | N4 | 16:113 | 4.45 | 1.00 | 49.60 |  |
| 12 | G | N7 | 16:9 | 4.76 | 1.00 | 57.26 |  |
| 13 | G | N1 | 16:112 | 4.89 | 1.00 | 50.01 |  |
| 14 | U | N1 | 16:111 | 4.97 | 1.00 | 56.97 |  |
  |

  
**1^st^ shell ligand angles:**   
1-2: 153; 1-3: 115; 1-4: 76; 1-5: 64; 2-3: 44; 2-4: 77; 2-5: 113; 3-4: 55; 3-5: 131; 4-5: 81;   
  
 Contact(s): /N3_U_/O4/O4/O6/O6/  
 Coded contact list: 4O_b_Anion  

### K:16:203

|  | Res. type | Atom | Residue | Distance | Occupancy | B-factor | Closest solute to 1^st^ shell water |
| --- | --- | --- | --- | --- | --- | --- | --- |
| **Ion** | | | |  | 1.00 | 44.38 || 1 | G | O6 | 16:82 | 2.88 | 1.00 | 44.29 |
| 2 | G | N2 | 16:81 | 2.89 | 1.00 | 49.19 |  |
| 3 | U | O4 | 16:95 | 2.97 | 1.00 | 45.89 |  |
| 4 | G | N1 | 16:81 | 3.16 | 1.00 | 46.10 |  |
| 5 | C | N4 | 16:94 | 3.25 | 1.00 | 44.72 |  |
| 6 | U | N3 | 16:80 | 3.35 | 1.00 | 41.92 |  |
| 7 | U | O4 | 16:80 | 3.36 | 1.00 | 44.17 |  |
| 8 | G | O6 | 16:96 | 3.50 | 1.00 | 45.83 |  |
| 9 | U | N3 | 16:95 | 4.36 | 1.00 | 46.16 |  |
| 10 | U | O2 | 16:80 | 4.57 | 1.00 | 43.69 |  |
| 11 | G | O6 | 16:81 | 4.58 | 1.00 | 41.62 |  |
| 12 | U | N1 | 16:80 | 4.63 | 1.00 | 44.29 |  |
| 13 | G | N1 | 16:82 | 4.70 | 1.00 | 44.78 |  |
| 14 | G | N7 | 16:82 | 4.80 | 1.00 | 43.54 |  |
| 15 | C | N3 | 16:94 | 4.83 | 1.00 | 48.63 |  |
| 16 | G | N3 | 16:81 | 4.86 | 1.00 | 34.90 |  |
| 17 | G | N1 | 16:96 | 4.88 | 1.00 | 41.60 |  |
  |

  
**1^st^ shell ligand angles:**   
1-2: 96; 1-3: 80; 1-4: 72; 1-5: 60; 1-6: 156; 1-7: 165; 2-3: 125; 2-4: 44; 2-5: 156; 2-6: 62; 2-7: 99; 3-4: 84; 3-5: 53; 3-6: 102; 3-7: 91; 4-5: 119; 4-6: 85; 4-7: 120; 5-6: 140; 5-7: 104; 6-7: 39;   
  
 Contact(s): /N1_G_/N2/N3_U_/N4/O4/O4/O6/  
 Coded contact list: 3O_b_4Anion  

### MG:16:204

|  | Res. type | Atom | Residue | Distance | Occupancy | B-factor | Closest solute to 1^st^ shell water |
| --- | --- | --- | --- | --- | --- | --- | --- |
| **Ion** | | | |  | 1.00 | 39.61 || 1 | G | OP1 | 16:100 | 2.78 | 1.00 | 39.35 |
| 2 | G | OP1 | 1H:963 | 3.54 | 1.00 | 36.55 |  |
| 3 | C | O3' | 1H:962 | 3.77 | 1.00 | 27.42 |  |
| 4 | Water 1 | | 16:313 | 4.13 | 1.00 | 45.00 | 2.89 (G:OP1 16:100) |
| 5 | C | O2' | 1H:962 | 4.48 | 1.00 | 36.40 |  |
| 6 | G | O5' | 1H:963 | 4.86 | 1.00 | 30.95 |  |
| 7 | G | O5' | 16:100 | 4.89 | 1.00 | 40.06 |  |
  |

  
 Coded contact list:   
 Exclusion zone contact(s): /OP1/  
Consider K in place of MG (1 valid contact(s) for K vs. 0 for MG)  

### MG:16:205

|  | Res. type | Atom | Residue | Distance | Occupancy | B-factor | Closest solute to 1^st^ shell water |
| --- | --- | --- | --- | --- | --- | --- | --- |
| **Ion** | | | |  | 1.00 | 46.71 || 1 | A | OP2 | 16:58 | 2.50 | 1.00 | 50.28 |
| 2 | A | OP2 | 16:57 | 2.56 | 1.00 | 65.23 |  |
| 3 | G | O3' | 16:56 | 3.32 | 1.00 | 53.70 |  |
| 4 | G | O6 | 16:23 | 3.50 | 1.00 | 59.31 |  |
| 5 | G | O2' | 16:56 | 3.61 | 1.00 | 55.40 |  |
| 6 | A | N6 | 16:59 | 3.94 | 1.00 | 57.21 |  |
| 7 | A | N7 | 16:58 | 4.02 | 1.00 | 55.25 |  |
| 8 | A | N7 | 16:59 | 4.05 | 1.00 | 53.68 |  |
| 9 | C | N4 | 16:60 | 4.06 | 1.00 | 54.18 |  |
| 10 | A | O5' | 16:58 | 4.19 | 1.00 | 55.05 |  |
| 11 | A | O5' | 16:57 | 4.51 | 1.00 | 49.13 |  |
| 12 | A | OP1 | 16:57 | 4.65 | 1.00 | 57.53 |  |
| 13 | A | O3' | 16:57 | 4.65 | 1.00 | 55.96 |  |
  |

  
 Coded contact list:   
 Exclusion zone contact(s): /O3'/OP2/OP2/  
Consider NA in place of MG (2 valid contact(s) for NA vs. 0 for MG)  

### MG:16:206

|  | Res. type | Atom | Residue | Distance | Occupancy | B-factor | Closest solute to 1^st^ shell water |
| --- | --- | --- | --- | --- | --- | --- | --- |
| **Ion** | | | |  | 1.00 | 52.99 || 1 | G | O6 | 16:56 | 2.44 | 1.00 | 53.76 |
| 2 | G | O2' | 16:24 | 2.77 | 1.00 | 61.38 |  |
| 3 | U | O4 | 16:55 | 2.92 | 1.00 | 49.34 |  |
| 4 | G | O3' | 16:24 | 3.77 | 1.00 | 67.27 |  |
| 5 | A | OP2 | 16:25 | 3.80 | 1.00 | 57.94 |  |
| 6 | G | N7 | 16:56 | 4.14 | 1.00 | 52.32 |  |
| 7 | U | N3 | 16:55 | 4.20 | 1.00 | 51.84 |  |
| 8 | Water 1 | | 16:306 | 4.34 | 1.00 | 68.27 | 2.42 (G:O2' 16:24) |
| 9 | C | N4 | 16:28 | 4.37 | 1.00 | 53.58 |  |
| 10 | A | N6 | 16:29 | 4.53 | 1.00 | 47.84 |  |
| 11 | G | N1 | 16:56 | 4.70 | 1.00 | 51.53 |  |
| 12 | A | OP2 | 16:26 | 4.92 | 1.00 | 65.16 |  |
  |

  
 Coded contact list:   
 Exclusion zone contact(s): /O2'/O4/O6/  
Consider K in place of MG (2 valid contact(s) for K vs. 0 for MG)  

### MG:16:207

|  | Res. type | Atom | Residue | Distance | Occupancy | B-factor | Closest solute to 1^st^ shell water |
| --- | --- | --- | --- | --- | --- | --- | --- |
| **Ion** | | | |  | 1.00 | 43.94 || 1 | G | O6 | 16:85 | 3.26 | 1.00 | 58.57 |
| 2 | G | O6 | 16:86 | 3.91 | 1.00 | 63.41 |  |
| 3 | G | N7 | 16:85 | 4.16 | 1.00 | 58.68 |  |
| 4 | C | N4 | 16:84 | 4.50 | 1.00 | 53.38 |  |
| 5 | C | OP1 | 1H:997 | 4.68 | 1.00 | 25.81 |  |
| 6 | G | N7 | 16:86 | 4.74 | 1.00 | 62.32 |  |
| 7 | C | N4 | 16:91 | 4.84 | 1.00 | 57.74 |  |
| 8 | G | O3' | 1H:996 | 4.91 | 1.00 | 26.33 |  |
  |

  
 Coded contact list:   
 Exclusion zone contact(s): /O6/  
Consider K in place of MG (1 valid contact(s) for K vs. 0 for MG)  

### MG:16:208

|  | Res. type | Atom | Residue | Distance | Occupancy | B-factor | Closest solute to 1^st^ shell water |
| --- | --- | --- | --- | --- | --- | --- | --- |
| **Ion** | | | |  | 1.00 | 44.39 || 1 | G | O6 | 16:82 | 2.76 | 1.00 | 44.29 |
| 2 | G | O6 | 16:83 | 3.40 | 1.00 | 50.91 |  |
| 3 | G | N7 | 16:82 | 3.46 | 1.00 | 43.54 |  |
| 4 | G | N7 | 16:83 | 3.96 | 1.00 | 49.09 |  |
| 5 | G | N2 | 16:81 | 3.99 | 1.00 | 49.19 |  |
| 6 | G | OP2 | 16:81 | 4.12 | 1.00 | 37.55 |  |
| 7 | G | N1 | 16:81 | 4.64 | 1.00 | 46.10 |  |
| 8 | C | N4 | 16:94 | 4.73 | 1.00 | 44.72 |  |
| 9 | G | N1 | 16:82 | 4.78 | 1.00 | 44.78 |  |
| 10 | C | N4 | 16:93 | 4.82 | 1.00 | 47.33 |  |
  |

  
 Coded contact list:   
 Exclusion zone contact(s): /O6/O6/  
Consider K in place of MG (2 valid contact(s) for K vs. 0 for MG)  

### MG:16:209

|  | Res. type | Atom | Residue | Distance | Occupancy | B-factor | Closest solute to 1^st^ shell water |
| --- | --- | --- | --- | --- | --- | --- | --- |
| **Ion** | | | |  | 1.00 | 59.65 || 1 | U | OP2 | 16:22 | 3.09 | 1.00 | 58.19 |
| 2 | A | OP1 | 16:57 | 3.60 | 1.00 | 57.53 |  |
| 3 | G | OP2 | 16:23 | 4.56 | 1.00 | 49.75 |  |
| 4 | U | O5' | 16:22 | 4.63 | 1.00 | 62.45 |  |
| 5 | U | OP1 | 16:22 | 4.88 | 1.00 | 76.91 |  |
  |

  
 Coded contact list:   
 Exclusion zone contact(s): /OP2/  
Consider K in place of MG (1 valid contact(s) for K vs. 0 for MG)  

### MG:16:210

|  | Res. type | Atom | Residue | Distance | Occupancy | B-factor | Closest solute to 1^st^ shell water |
| --- | --- | --- | --- | --- | --- | --- | --- |
| **Ion** | | | |  | 1.00 | 62.98 || 1 | A | OP2 | 16:25 | 1.98 | 1.00 | 57.94 |
| 2 | Water 1 | | 16:305 | 2.70 | 1.00 | 54.78 | 2.38 (A:OP1 16:25) |
| 3 | A | OP1 | 16:25 | 3.80 | 1.00 | 59.80 |  |
| 4 | G | O3' | 16:24 | 4.12 | 1.00 | 67.27 |  |
| 5 | Water 2 | | 16:306 | 4.28 | 1.00 | 68.27 | 2.42 (G:O2' 16:24) |
| 6 | A | OP2 | 16:26 | 4.30 | 1.00 | 65.16 |  |
| 7 | A | O5' | 16:25 | 4.42 | 1.00 | 56.28 |  |
| 8 | G | O2' | 16:24 | 4.47 | 1.00 | 61.38 |  |
| 9 | Water 3 | | 16:310 | 4.82 | 1.00 | 57.03 | 4.81 (A:OP1 16:25) |
  |

  
 Contact(s): /OP2/  
 Coded contact list: O_ph_  
 Exclusion zone contact(s): /O_w_/  
MG hydration shell issue: water too close (< 2.4 Angs.)  

### MG:16:211

|  | Res. type | Atom | Residue | Distance | Occupancy | B-factor | Closest solute to 1^st^ shell water |
| --- | --- | --- | --- | --- | --- | --- | --- |
| **Ion** | | | |  | 1.00 | 57.73 || 1 | Water 1 | | 16:306 | 2.18 | 1.00 | 68.27 | 2.42 (G:O2' 16:24) |
| 2 | Water 2 | | 16:324 | 2.18 | 1.00 | 65.81 | 3.46 (G:OP2 16:56) |
| 3 | Water 3 | | 16:305 | 2.18 | 1.00 | 54.78 | 2.38 (A:OP1 16:25) |
| 4 | Water 4 | | 16:310 | 2.18 | 1.00 | 57.03 | 2.65 (U:OP2 16:55) |
| 5 | Water 5 | | 16:301 | 2.19 | 1.00 | 62.17 | 2.03 (A:OP1 16:25) |
| 6 | A | OP1 | 16:25 | 2.97 | 1.00 | 59.80 |  |
| 7 | G | O2' | 16:24 | 4.41 | 1.00 | 61.38 |  |
| 8 | A | OP2 | 16:25 | 4.65 | 1.00 | 57.94 |  |
| 9 | G | OP2 | 16:56 | 4.73 | 1.00 | 58.97 |  |
| 10 | U | OP2 | 16:55 | 4.81 | 1.00 | 47.94 |  |
| 11 | G | O3' | 16:24 | 4.83 | 1.00 | 67.27 |  |
  |

  
**1^st^ shell ligand angles:**   
1-2: 90; 1-3: 89; 1-4: 87; 1-5: 92; 2-3: 178; 2-4: 89; 2-5: 89; 3-4: 90; 3-5: 91; 4-5: 178;   
  
 Contact(s): /O_w_/O_w_/O_w_/O_w_/O_w_/  
 Coded contact list: 5O_w_  
 Exclusion zone contact(s): /OP1/  
Consider K in place of MG (1 valid contact(s) for K vs. 0 for MG)  
MG hydration shell issue: water too close (< 2.4 Angs.)  

### MG:16:212

|  | Res. type | Atom | Residue | Distance | Occupancy | B-factor | Closest solute to 1^st^ shell water |
| --- | --- | --- | --- | --- | --- | --- | --- |
| **Ion** | | | |  | 1.00 | 36.42 || 1 | Water 1 | | 16:313 | 2.17 | 1.00 | 45.00 | 2.82 (G:OP2 16:100) |
| 2 | Water 2 | | 16:308 | 2.17 | 1.00 | 45.82 | 2.51 (A:OP2 16:101) |
| 3 | Water 3 | | 16:320 | 2.18 | 1.00 | 45.75 | 3.33 (G:O6 16:102) |
| 4 | Water 4 | | 16:303 | 2.18 | 1.00 | 52.47 | 2.21 (A:OP2 16:73) |
| 5 | Water 5 | | 16:317 | 2.18 | 1.00 | 43.87 | 3.04 (A:O5' 16:73) |
| 6 | G | OP2 | 16:100 | 2.53 | 1.00 | 43.48 |  |
| 7 | A | OP2 | 16:73 | 3.61 | 1.00 | 45.65 |  |
| 8 | Water 6 | | 16:318 | 3.80 | 1.00 | 43.68 | 3.05 (A:OP2 16:101) |
| 9 | A | O5' | 16:73 | 4.15 | 1.00 | 49.24 |  |
| 10 | G | OP1 | 16:100 | 4.20 | 1.00 | 39.35 |  |
| 11 | G | O5' | 16:100 | 4.34 | 1.00 | 40.06 |  |
| 12 | G | O6 | 16:102 | 4.43 | 1.00 | 43.38 |  |
| 13 | A | OP2 | 16:101 | 4.50 | 1.00 | 37.76 |  |
| 14 | A | N7 | 16:101 | 4.65 | 1.00 | 36.82 |  |
| 15 | Water 7 | | 16:315 | 4.82 | 1.00 | 41.24 | 3.06 (G:O6 16:102) |
| 16 | A | O3' | 16:99 | 4.88 | 1.00 | 37.04 |  |
| 17 | Water 8 | | 16:314 | 4.89 | 1.00 | 51.14 | 4.87 (G:OP2 16:100) |
| 18 | A | OP1 | 16:73 | 4.94 | 1.00 | 53.83 |  |
  |

  
**1^st^ shell ligand angles:**   
1-2: 92; 1-3: 179; 1-4: 90; 1-5: 88; 2-3: 88; 2-4: 90; 2-5: 174; 3-4: 90; 3-5: 92; 4-5: 96;   
  
 Contact(s): /O_w_/O_w_/O_w_/O_w_/O_w_/  
 Coded contact list: 5O_w_  
 Exclusion zone contact(s): /OP2/  
Consider NA in place of MG (1 valid contact(s) for NA vs. 0 for MG)  
MG hydration shell issue: water too close (< 2.4 Angs.)  

### MG:16:213

|  | Res. type | Atom | Residue | Distance | Occupancy | B-factor | Closest solute to 1^st^ shell water |
| --- | --- | --- | --- | --- | --- | --- | --- |
| **Ion** | | | |  | 1.00 | 41.74 || 1 | Water 1 | | 16:307 | 2.16 | 1.00 | 45.81 | 2.50 (G:N7 16:102) |
| 2 | Water 2 | | 16:312 | 2.16 | 1.00 | 54.50 | 2.81 (G:N7 16:72) |
| 3 | Water 3 | | 16:323 | 2.17 | 1.00 | 46.47 | 3.30 (A:OP1 16:101) |
| 4 | Water 4 | | 16:315 | 2.18 | 1.00 | 41.24 | 2.97 (U:O4 16:103) |
| 5 | Water 5 | | 16:322 | 2.18 | 1.00 | 44.91 | 3.28 (G:N1 16:72) |
| 6 | Water 6 | | 16:318 | 2.18 | 1.00 | 43.68 | 3.05 (A:OP2 16:101) |
| 7 | Water 7 | | 16:303 | 3.90 | 1.00 | 52.47 | 2.21 (A:OP2 16:73) |
| 8 | G | N7 | 16:102 | 4.01 | 1.00 | 40.70 |  |
| 9 | A | OP2 | 16:101 | 4.07 | 1.00 | 37.76 |  |
| 10 | G | OP2 | 16:102 | 4.17 | 1.00 | 32.54 |  |
| 11 | A | OP2 | 16:73 | 4.35 | 1.00 | 45.65 |  |
| 12 | Water 8 | | 16:308 | 4.42 | 1.00 | 45.82 | 2.51 (A:OP2 16:101) |
| 13 | G | N1 | 16:72 | 4.63 | 1.00 | 48.73 |  |
| 14 | G | O6 | 16:72 | 4.67 | 1.00 | 49.02 |  |
| 15 | G | N7 | 16:72 | 4.69 | 1.00 | 42.68 |  |
| 16 | A | O5' | 16:101 | 4.74 | 1.00 | 30.33 |  |
| 17 | G | O6 | 16:102 | 4.77 | 1.00 | 43.38 |  |
| 18 | A | OP1 | 16:101 | 4.83 | 1.00 | 32.91 |  |
| 19 | U | O4 | 16:103 | 4.93 | 1.00 | 49.53 |  |
| 20 | G | N3 | 16:72 | 4.95 | 1.00 | 45.36 |  |
  |

  
**1^st^ shell ligand angles:**   
1-2: 180; 1-3: 91; 1-4: 92; 1-5: 81; 1-6: 98; 2-3: 89; 2-4: 88; 2-5: 99; 2-6: 82; 3-4: 176; 3-5: 88; 3-6: 91; 4-5: 90; 4-6: 91; 5-6: 179;   
  
 Contact(s): /O_w_/O_w_/O_w_/O_w_/O_w_/O_w_/  
 Coded contact list: 6O_w_  

### MG:16:214

|  | Res. type | Atom | Residue | Distance | Occupancy | B-factor | Closest solute to 1^st^ shell water |
| --- | --- | --- | --- | --- | --- | --- | --- |
| **Ion** | | | |  | 1.00 | 57.59 || 1 | Water 1 | | 16:319 | 2.16 | 1.00 | 47.52 | 3.12 (G:N7 16:75) |
| 2 | Water 2 | | 16:302 | 2.17 | 1.00 | 47.64 | 2.13 (G:N7 16:76) |
| 3 | Water 3 | | 16:316 | 2.18 | 1.00 | 45.40 | 4.17 (U:OP2 16:74) |
| 4 | Water 4 | | 16:321 | 2.19 | 1.00 | 41.11 | 3.26 (U:OP2 16:74) |
| 5 | Water 5 | | 16:314 | 2.19 | 1.00 | 51.14 | 2.82 (G:O6 16:76) |
| 6 | Water 6 | | 16:311 | 2.21 | 1.00 | 46.84 | 2.77 (A:OP2 16:99) |
| 7 | A | O5' | 16:99 | 4.01 | 1.00 | 33.23 |  |
| 8 | A | OP2 | 16:99 | 4.07 | 1.00 | 38.11 |  |
| 9 | G | O6 | 16:76 | 4.08 | 1.00 | 40.22 |  |
| 10 | G | N7 | 16:76 | 4.24 | 1.00 | 45.74 |  |
| 11 | U | OP2 | 16:74 | 4.34 | 1.00 | 48.35 |  |
| 12 | G | N7 | 16:75 | 4.37 | 1.00 | 43.83 |  |
| 13 | U | O4 | 16:77 | 4.77 | 1.00 | 42.85 |  |
| 14 | Water 7 | | 16:317 | 4.87 | 1.00 | 43.87 | 3.74 (U:OP2 16:74) |
| 15 | G | O6 | 16:75 | 4.95 | 1.00 | 48.42 |  |
  |

  
**1^st^ shell ligand angles:**   
1-2: 92; 1-3: 90; 1-4: 86; 1-5: 87; 1-6: 177; 2-3: 90; 2-4: 178; 2-5: 88; 2-6: 88; 3-4: 89; 3-5: 177; 3-6: 93; 4-5: 93; 4-6: 94; 5-6: 90;   
  
 Contact(s): /O_w_/O_w_/O_w_/O_w_/O_w_/O_w_/  
 Coded contact list: 6O_w_  
MG hydration shell issue: water too close (< 2.4 Angs.)  

### MG:16:215

|  | Res. type | Atom | Residue | Distance | Occupancy | B-factor | Closest solute to 1^st^ shell water |
| --- | --- | --- | --- | --- | --- | --- | --- |
| **Ion** | | | |  | 1.00 | 52.42 || 1 | Water 1 | | 16:309 | 2.17 | 1.00 | 53.85 | 2.52 (C:O4' 16:10) |
| 2 | Water 2 | | 16:304 | 2.17 | 1.00 | 54.46 | 2.37 (G:O2' 16:9) |
| 3 | Water 3 | | A8:202 | 2.18 | 1.00 | 54.05 | 3.45 (LYS:O A8:44) |
| 4 | Water 4 | | 16:326 | 2.19 | 1.00 | 31.91 | 4.41 (G:N2 16:9) |
| 5 | Water 5 | | A8:201 | 2.19 | 1.00 | 58.26 | 2.86 (LYS:O A8:44) |
| 6 | Water 6 | | 16:325 | 2.19 | 1.00 | 66.59 | 3.73 (C:O4' 16:10) |
| 7 | G | O2' | 16:9 | 4.03 | 1.00 | 54.67 |  |
| 8 | C | O4' | 16:10 | 4.03 | 1.00 | 59.24 |  |
| 9 | LYS | O | A8:44 | 4.25 | 1.00 | 66.61 |  |
| 10 | GLY | N | A8:45 | 4.63 | 1.00 | 65.01 |  |
| 11 | G | N3 | 16:9 | 4.64 | 1.00 | 52.85 |  |
| 12 | G | N2 | 16:9 | 4.71 | 1.00 | 48.84 |  |
  |

  
**1^st^ shell ligand angles:**   
1-2: 92; 1-3: 84; 1-4: 87; 1-5: 175; 1-6: 94; 2-3: 97; 2-4: 176; 2-5: 86; 2-6: 84; 3-4: 87; 3-5: 92; 3-6: 177; 4-5: 95; 4-6: 91; 5-6: 90;   
  
 Contact(s): /O_w_/O_w_/O_w_/O_w_/O_w_/O_w_/  
 Coded contact list: 6O_w_  
MG hydration shell issue: water too close (< 2.4 Angs.)  

### K:19:301

|  | Res. type | Atom | Residue | Distance | Occupancy | B-factor | Closest solute to 1^st^ shell water |
| --- | --- | --- | --- | --- | --- | --- | --- |
| **Ion** | | | |  | 1.00 | 52.67 || 1 | GLY | O | 19:235 | 2.61 | 1.00 | 40.10 |
| 2 | GLU | O | 19:237 | 2.84 | 1.00 | 48.74 |  |
| 3 | Water 1 | | 19:410 | 3.56 | 1.00 | 51.83 | 3.69 (ALA:O 19:224) |
| 4 | GLY | O | 19:236 | 3.76 | 1.00 | 42.28 |  |
| 5 | GLY | N | 19:236 | 3.94 | 1.00 | 35.78 |  |
| 6 | GLY | O | 19:238 | 4.05 | 1.00 | 36.38 |  |
| 7 | ALA | O | 19:224 | 4.41 | 1.00 | 30.14 |  |
| 8 | Water 2 | | 19:405 | 4.41 | 1.00 | 34.20 | 2.58 (GLY:O 19:235) |
| 9 | ARG | O | 19:239 | 4.48 | 1.00 | 50.63 |  |
| 10 | GLU | N | 19:237 | 4.54 | 1.00 | 34.42 |  |
| 11 | A | OP2 | 14:2590 | 4.67 | 1.00 | 34.43 |  |
| 12 | GLY | N | 19:235 | 4.78 | 1.00 | 30.49 |  |
| 13 | GLY | N | 19:238 | 4.93 | 1.00 | 30.78 |  |
| 14 | A | OP1 | 14:2590 | 4.99 | 1.00 | 31.56 |  |
  |

  
**1^st^ shell ligand angles:**   
1-2: 111;   
  
 Contact(s): /O_back_/O_back_/  
 Coded contact list: 2O_back_  

### K:1G:1601

|  | Res. type | Atom | Residue | Distance | Occupancy | B-factor | Closest solute to 1^st^ shell water |
| --- | --- | --- | --- | --- | --- | --- | --- |
| **Ion** | | | |  | 1.00 | 115.34 || 1 | C | OP1 | 1G:449 | 2.84 | 1.00 | 87.20 |
| 2 | G | OP2 | 1G:450 | 3.80 | 1.00 | 80.46 |  |
| 3 | A | O2' | 1G:451 | 4.08 | 1.00 | 81.62 |  |
| 4 | C | O5' | 1G:449 | 4.25 | 1.00 | 84.83 |  |
| 5 | C | OP2 | 1G:449 | 4.43 | 1.00 | 91.15 |  |
| 6 | A | N3 | 1G:451 | 4.45 | 1.00 | 84.76 |  |
| 7 | G | O6 | 1G:481 | 4.67 | 1.00 | 78.91 |  |
  |

  
 Contact(s): /OP1/  
 Coded contact list: O_ph_  

### K:1G:1602

|  | Res. type | Atom | Residue | Distance | Occupancy | B-factor | Closest solute to 1^st^ shell water |
| --- | --- | --- | --- | --- | --- | --- | --- |
| **Ion** | | | |  | 1.00 | 104.24 || 1 | G | O6 | 1G:69 | 2.71 | 1.00 | 102.76 |
| 2 | G | O6 | 1G:73 | 3.40 | 1.00 | 103.42 |  |
| 3 | G | O6 | 1G:96 | 3.54 | 1.00 | 116.04 |  |
| 4 | U | O4 | 1G:97 | 3.83 | 1.00 | 107.88 |  |
| 5 | G | N7 | 1G:69 | 4.04 | 1.00 | 100.37 |  |
| 6 | C | N4 | 1G:74 | 4.08 | 1.00 | 105.30 |  |
| 7 | G | O6 | 1G:95 | 4.16 | 1.00 | 121.87 |  |
| 8 | U | N3 | 1G:97 | 4.54 | 1.00 | 106.62 |  |
| 9 | G | N1 | 1G:69 | 4.61 | 1.00 | 102.58 |  |
| 10 | G | N7 | 1G:73 | 4.69 | 1.00 | 107.92 |  |
  |

  
**1^st^ shell ligand angles:**   
1-2: 63;   
  
 Contact(s): /O6/O6/  
 Coded contact list: 2O_b_  

### K:1G:1603

|  | Res. type | Atom | Residue | Distance | Occupancy | B-factor | Closest solute to 1^st^ shell water |
| --- | --- | --- | --- | --- | --- | --- | --- |
| **Ion** | | | |  | 1.00 | 62.13 || 1 | A | OP2 | 1G:509 | 2.52 | 1.00 | 71.03 |
| 2 | A | O3' | 1G:509 | 3.12 | 1.00 | 74.89 |  |
| 3 | G | OP1 | 1G:506 | 3.26 | 1.00 | 75.61 |  |
| 4 | A | OP2 | 1G:510 | 3.43 | 1.00 | 76.58 |  |
| 5 | C | O3' | 1G:508 | 3.73 | 1.00 | 79.69 |  |
| 6 | C | OP2 | 1G:507 | 4.05 | 1.00 | 61.92 |  |
| 7 | A | N7 | 1G:510 | 4.12 | 1.00 | 70.74 |  |
| 8 | A | O5' | 1G:510 | 4.29 | 1.00 | 62.52 |  |
| 9 | G | OP2 | 1G:506 | 4.38 | 1.00 | 57.42 |  |
| 10 | G | O5' | 1G:506 | 4.51 | 1.00 | 66.89 |  |
| 11 | C | O2' | 1G:508 | 4.53 | 1.00 | 61.58 |  |
| 12 | A | O5' | 1G:509 | 4.54 | 1.00 | 76.24 |  |
| 13 | A | OP1 | 1G:509 | 4.89 | 1.00 | 69.78 |  |
  |

  
**1^st^ shell ligand angles:**   
1-2: 60; 1-3: 144; 2-3: 156;   
  
 Contact(s): /O3'/OP1/  
 Coded contact list: O_ph_O_r_  
 Exclusion zone contact(s): /OP2/  

### K:1G:1604

|  | Res. type | Atom | Residue | Distance | Occupancy | B-factor | Closest solute to 1^st^ shell water |
| --- | --- | --- | --- | --- | --- | --- | --- |
| **Ion** | | | |  | 1.00 | 91.70 || 1 | SER | O | 8A:39 | 3.52 | 1.00 | 73.44 |
| 2 | C | O2' | 1G:280 | 3.52 | 1.00 | 77.43 |  |
| 3 | C | O2 | 1G:280 | 3.86 | 1.00 | 81.98 |  |
| 4 | A | OP1 | 1G:279 | 4.46 | 1.00 | 76.33 |  |
| 5 | C | O3' | 1G:280 | 4.57 | 1.00 | 63.36 |  |
  |

  
 Coded contact list:   
1^st^ K coordination shell missing (see Notes)  

### K:1G:1605

|  | Res. type | Atom | Residue | Distance | Occupancy | B-factor | Closest solute to 1^st^ shell water |
| --- | --- | --- | --- | --- | --- | --- | --- |
| **Ion** | | | |  | 1.00 | 92.87 || 1 | G | OP2 | 1G:1343 | 2.81 | 1.00 | 87.10 |
| 2 | C | OP2 | 1G:936 | 3.12 | 1.00 | 94.27 |  |
| 3 | A | OP2 | 1G:935 | 3.53 | 1.00 | 87.70 |  |
| 4 | C | O3' | 1G:934 | 3.57 | 1.00 | 85.52 |  |
| 5 | C | O2' | 1G:934 | 3.59 | 1.00 | 79.13 |  |
| 6 | G | OP1 | 1G:1343 | 3.88 | 1.00 | 78.98 |  |
| 7 | A | O5' | 1G:935 | 3.97 | 1.00 | 86.18 |  |
| 8 | G | O5' | 1G:1343 | 4.25 | 1.00 | 85.41 |  |
| 9 | A | O3' | 1G:935 | 4.30 | 1.00 | 85.17 |  |
| 10 | C | OP1 | 1G:936 | 4.31 | 1.00 | 88.71 |  |
| 11 | C | OP2 | 1G:1344 | 4.31 | 1.00 | 80.05 |  |
  |

  
**1^st^ shell ligand angles:**   
1-2: 126;   
  
 Contact(s): /OP2/OP2/  
 Coded contact list: 2O_ph_  

### K:1G:1606

|  | Res. type | Atom | Residue | Distance | Occupancy | B-factor | Closest solute to 1^st^ shell water |
| --- | --- | --- | --- | --- | --- | --- | --- |
| **Ion** | | | |  | 1.00 | 61.36 || 1 | C | OP2 | 1G:352 | 2.60 | 1.00 | 76.24 |
| 2 | C | O2 | 1G:330 | 3.59 | 1.00 | 63.33 |  |
| 3 | A | N1 | 1G:59 | 3.75 | 1.00 | 64.37 |  |
| 4 | A | N6 | 1G:59 | 3.84 | 1.00 | 65.59 |  |
| 5 | C | OP1 | 1G:352 | 3.87 | 1.00 | 70.70 |  |
| 6 | G | N7 | 1G:354 | 4.00 | 1.00 | 61.60 |  |
| 7 | G | OP1 | 1G:354 | 4.26 | 1.00 | 63.98 |  |
| 8 | G | O4' | 1G:331 | 4.34 | 1.00 | 62.33 |  |
| 9 | C | O5' | 1G:352 | 4.46 | 1.00 | 61.65 |  |
| 10 | C | O4' | 1G:352 | 4.50 | 1.00 | 68.70 |  |
| 11 | G | O3' | 1G:351 | 4.99 | 1.00 | 78.18 |  |
| 12 | G | N9 | 1G:331 | 4.99 | 1.00 | 62.52 |  |
  |

  
 Contact(s): /OP2/  
 Coded contact list: O_ph_  

### K:1G:1607

|  | Res. type | Atom | Residue | Distance | Occupancy | B-factor | Closest solute to 1^st^ shell water |
| --- | --- | --- | --- | --- | --- | --- | --- |
| **Ion** | | | |  | 1.00 | 87.42 || 1 | A | OP2 | 1G:452 | 2.71 | 1.00 | 83.11 |
| 2 | G | OP1 | 1G:450 | 2.83 | 1.00 | 78.84 |  |
| 3 | A | O2' | 1G:451 | 3.44 | 1.00 | 81.62 |  |
| 4 | A | O3' | 1G:451 | 3.68 | 1.00 | 77.44 |  |
| 5 | G | OP2 | 1G:450 | 3.82 | 1.00 | 80.46 |  |
| 6 | LYS | NZ | 7A:43 | 3.85 | 1.00 | 77.27 |  |
| 7 | C | O3' | 1G:449 | 4.37 | 1.00 | 81.76 |  |
| 8 | A | OP1 | 1G:452 | 4.57 | 1.00 | 77.66 |  |
| 9 | A | O5' | 1G:452 | 4.96 | 1.00 | 65.71 |  |
  |

  
**1^st^ shell ligand angles:**   
1-2: 72;   
  
 Contact(s): /OP1/OP2/  
 Coded contact list: 2O_ph_  

### K:1G:1608

|  | Res. type | Atom | Residue | Distance | Occupancy | B-factor | Closest solute to 1^st^ shell water |
| --- | --- | --- | --- | --- | --- | --- | --- |
| **Ion** | | | |  | 1.00 | 80.69 || 1 | G | N7 | 1G:945 | 3.07 | 1.00 | 86.73 |
| 2 | G | O6 | 1G:945 | 3.09 | 1.00 | 87.04 |  |
| 3 | G | N7 | 1G:944 | 3.09 | 1.00 | 85.80 |  |
| 4 | G | OP2 | 1G:944 | 3.19 | 1.00 | 81.22 |  |
| 5 | U | O5' | 1G:943 | 3.53 | 1.00 | 78.79 |  |
| 6 | G | N7 | 1G:1337 | 3.53 | 1.00 | 88.56 |  |
| 7 | U | OP2 | 1G:943 | 3.63 | 1.00 | 77.48 |  |
| 8 | G | O5' | 1G:944 | 4.45 | 1.00 | 84.45 |  |
| 9 | G | O6 | 1G:944 | 4.55 | 1.00 | 86.64 |  |
| 10 | U | N1 | 1G:943 | 4.58 | 1.00 | 79.75 |  |
| 11 | G | N9 | 1G:1337 | 4.70 | 1.00 | 87.42 |  |
| 12 | U | O3' | 1G:943 | 4.75 | 1.00 | 75.95 |  |
| 13 | U | OP1 | 1G:943 | 4.93 | 1.00 | 82.98 |  |
| 14 | U | N3 | 1G:943 | 4.95 | 1.00 | 79.13 |  |
| 15 | G | N9 | 1G:944 | 4.99 | 1.00 | 78.74 |  |
  |

  
**1^st^ shell ligand angles:**   
1-2: 59; 1-3: 84; 1-4: 67; 2-3: 142; 2-4: 76; 3-4: 99;   
  
 Contact(s): /N7/N7/O6/OP2/  
 Coded contact list: O_ph_O_b_2N_b_  

### K:1G:1609

|  | Res. type | Atom | Residue | Distance | Occupancy | B-factor | Closest solute to 1^st^ shell water |
| --- | --- | --- | --- | --- | --- | --- | --- |
| **Ion** | | | |  | 1.00 | 91.12 || 1 | U | O2 | 1G:1364 | 2.72 | 1.00 | 92.87 |
| 2 | G | N3 | 1G:1233 | 2.89 | 1.00 | 90.10 |  |
| 3 | G | O2' | 1G:1233 | 2.94 | 1.00 | 91.82 |  |
| 4 | A | N3 | 1G:949 | 3.10 | 1.00 | 87.24 |  |
| 5 | U | O2' | 1G:1364 | 3.11 | 1.00 | 90.69 |  |
| 6 | G | O6 | 1G:971 | 3.15 | 1.00 | 87.62 |  |
| 7 | G | N7 | 1G:971 | 3.54 | 1.00 | 86.14 |  |
| 8 | G | N9 | 1G:1233 | 3.74 | 1.00 | 91.75 |  |
| 9 | C | O4' | 1G:1234 | 3.86 | 1.00 | 87.76 |  |
| 10 | G | N2 | 1G:1233 | 4.05 | 1.00 | 88.05 |  |
| 11 | U | O3' | 1G:1364 | 4.13 | 1.00 | 84.58 |  |
| 12 | U | N1 | 1G:1364 | 4.30 | 1.00 | 93.69 |  |
| 13 | G | O4' | 1G:1233 | 4.34 | 1.00 | 91.85 |  |
| 14 | A | N9 | 1G:949 | 4.61 | 1.00 | 83.89 |  |
| 15 | C | N1 | 1G:1234 | 4.66 | 1.00 | 92.87 |  |
| 16 | A | N1 | 1G:949 | 4.81 | 1.00 | 88.97 |  |
| 17 | A | O2' | 1G:949 | 4.82 | 1.00 | 80.93 |  |
| 18 | U | N3 | 1G:1364 | 4.88 | 1.00 | 94.26 |  |
| 19 | G | N1 | 1G:971 | 4.92 | 1.00 | 89.57 |  |
  |

  
**1^st^ shell ligand angles:**   
1-2: 86; 1-3: 111; 1-4: 87; 1-5: 75; 1-6: 83; 2-3: 91; 2-4: 65; 2-5: 160; 2-6: 129; 3-4: 150; 3-5: 89; 3-6: 139; 4-5: 119; 4-6: 65; 5-6: 56;   
  
 Contact(s): /N3_A_/N3_G_/O2/O2'/O2'/O6/  
 Coded contact list: 2O_r_2O_b_2N_b_  

### K:1G:1610

|  | Res. type | Atom | Residue | Distance | Occupancy | B-factor | Closest solute to 1^st^ shell water |
| --- | --- | --- | --- | --- | --- | --- | --- |
| **Ion** | | | |  | 1.00 | 68.23 || 1 | G | O6 | 1G:236 | 2.66 | 1.00 | 65.00 |
| 2 | G | O6 | 1G:126 | 2.75 | 1.00 | 76.07 |  |
| 3 | C | N3 | 1G:121 | 2.90 | 1.00 | 67.33 |  |
| 4 | U | O4 | 1G:125 | 2.94 | 1.00 | 76.28 |  |
| 5 | G | O6 | 1G:124 | 2.99 | 1.00 | 69.24 |  |
| 6 | C | O2 | 1G:121 | 3.29 | 1.00 | 73.33 |  |
| 7 | U | N3 | 1G:125 | 3.34 | 1.00 | 72.15 |  |
| 8 | C | N4 | 1G:235 | 3.38 | 1.00 | 67.96 |  |
| 9 | G | N1 | 1G:126 | 3.55 | 1.00 | 72.52 |  |
| 10 | C | N4 | 1G:237 | 3.81 | 1.00 | 67.62 |  |
| 11 | G | N7 | 1G:236 | 4.05 | 1.00 | 66.08 |  |
| 12 | C | N4 | 1G:121 | 4.10 | 1.00 | 65.88 |  |
| 13 | C | N3 | 1G:235 | 4.21 | 1.00 | 68.35 |  |
| 14 | C | N3 | 1G:237 | 4.61 | 1.00 | 63.50 |  |
| 15 | G | N1 | 1G:124 | 4.70 | 1.00 | 65.20 |  |
| 16 | U | O2 | 1G:125 | 4.82 | 1.00 | 69.96 |  |
| 17 | G | N7 | 1G:126 | 4.86 | 1.00 | 66.58 |  |
| 18 | C | N1 | 1G:121 | 4.89 | 1.00 | 66.74 |  |
| 19 | G | N1 | 1G:236 | 4.92 | 1.00 | 67.56 |  |
| 20 | C | N4 | 1G:123 | 4.98 | 1.00 | 62.10 |  |
  |

  
**1^st^ shell ligand angles:**   
1-2: 83; 1-3: 125; 1-4: 96; 1-5: 86; 1-6: 158; 1-7: 55; 1-8: 115; 2-3: 129; 2-4: 68; 2-5: 136; 2-6: 118; 2-7: 75; 2-8: 50; 3-4: 134; 3-5: 92; 3-6: 43; 3-7: 155; 3-8: 79; 4-5: 71; 4-6: 92; 4-7: 42; 4-8: 102; 5-6: 78; 5-7: 63; 5-8: 159; 6-7: 126; 6-8: 83; 7-8: 125;   
  
 Contact(s): /N3_C_/N3_U_/N4/O2/O4/O6/O6/O6/  
 Coded contact list: 5O_b_N_b_2Anion  

### K:1G:1611

|  | Res. type | Atom | Residue | Distance | Occupancy | B-factor | Closest solute to 1^st^ shell water |
| --- | --- | --- | --- | --- | --- | --- | --- |
| **Ion** | | | |  | 1.00 | 86.36 || 1 | G | OP2 | 1G:107 | 2.78 | 1.00 | 71.45 |
| 2 | G | O6 | 1G:326 | 3.38 | 1.00 | 68.51 |  |
| 3 | A | N7 | 1G:325 | 3.45 | 1.00 | 60.15 |  |
| 4 | G | O2' | 1G:324 | 3.53 | 1.00 | 70.13 |  |
| 5 | G | OP1 | 1G:107 | 3.80 | 1.00 | 69.89 |  |
| 6 | G | O3' | 1G:324 | 4.16 | 1.00 | 69.96 |  |
| 7 | G | N7 | 1G:326 | 4.53 | 1.00 | 68.57 |  |
| 8 | C | O3' | 1G:106 | 4.54 | 1.00 | 69.22 |  |
| 9 | C | OP1 | 1G:106 | 4.69 | 1.00 | 69.33 |  |
| 10 | C | O5' | 1G:106 | 4.73 | 1.00 | 73.55 |  |
| 11 | A | OP1 | 1G:325 | 4.74 | 1.00 | 65.54 |  |
| 12 | C | OP2 | 1G:106 | 4.90 | 1.00 | 63.60 |  |
| 13 | ARG | NH2 | BA:15 | 5.00 | 1.00 | 67.62 |  |
  |

  
**1^st^ shell ligand angles:**   
1-2: 73;   
  
 Contact(s): /O6/OP2/  
 Coded contact list: O_ph_O_b_  

### K:1G:1612

|  | Res. type | Atom | Residue | Distance | Occupancy | B-factor | Closest solute to 1^st^ shell water |
| --- | --- | --- | --- | --- | --- | --- | --- |
| **Ion** | | | |  | 1.00 | 58.38 || 1 | G | O6 | 1G:1511 | 2.64 | 1.00 | 48.28 |
| 2 | G | O6 | 1G:1523 | 2.87 | 1.00 | 50.89 |  |
| 3 | U | O4 | 1G:1510 | 2.97 | 1.00 | 56.18 |  |
| 4 | U | O4 | 1G:1522 | 3.06 | 1.00 | 57.13 |  |
| 5 | C | N4 | 1G:1524 | 3.31 | 1.00 | 50.48 |  |
| 6 | U | O4 | 1G:1512 | 3.43 | 1.00 | 59.94 |  |
| 7 | A | N6 | 1G:1513 | 3.93 | 1.00 | 53.91 |  |
| 8 | G | N7 | 1G:1523 | 4.31 | 1.00 | 51.20 |  |
| 9 | U | N3 | 1G:1512 | 4.39 | 1.00 | 47.24 |  |
| 10 | U | N3 | 1G:1510 | 4.45 | 1.00 | 47.40 |  |
| 11 | U | N3 | 1G:1522 | 4.70 | 1.00 | 52.08 |  |
| 12 | G | O6 | 1G:1525 | 4.78 | 1.00 | 50.82 |  |
| 13 | G | N1 | 1G:1511 | 4.79 | 1.00 | 49.16 |  |
| 14 | A | N1 | 1G:1513 | 4.84 | 1.00 | 54.06 |  |
| 15 | G | N7 | 1G:1511 | 4.87 | 1.00 | 48.38 |  |
  |

  
**1^st^ shell ligand angles:**   
1-2: 64; 1-3: 77; 1-4: 149; 1-5: 51; 2-3: 136; 2-4: 85; 2-5: 53; 3-4: 132; 3-5: 86; 4-5: 113;   
  
 Contact(s): /N4/O4/O4/O6/O6/  
 Coded contact list: 4O_b_Anion  

### K:1G:1613

|  | Res. type | Atom | Residue | Distance | Occupancy | B-factor | Closest solute to 1^st^ shell water |
| --- | --- | --- | --- | --- | --- | --- | --- |
| **Ion** | | | |  | 1.00 | 63.22 || 1 | C | N3 | 1G:518 | 2.91 | 1.00 | 71.86 |
| 2 | G | O6 | 1G:529 | 2.98 | 1.00 | 60.93 |  |
| 3 | G | N7 | 1G:529 | 3.35 | 1.00 | 67.25 |  |
| 4 | ASN | OD1 | 3A:46 | 3.35 | 1.00 | 65.92 |  |
| 5 | C | N4 | 1G:518 | 3.56 | 1.00 | 67.74 |  |
| 6 | PRO | O | 3A:45 | 3.61 | 1.00 | 73.72 |  |
| 7 | ASN | N | 3A:46 | 3.74 | 1.00 | 61.19 |  |
| 8 | C | O2 | 1G:518 | 3.87 | 1.00 | 65.29 |  |
| 9 | C | N4 | 1G:528 | 4.27 | 1.00 | 60.64 |  |
| 10 | C | N1 | 1G:518 | 4.48 | 1.00 | 67.04 |  |
  |

  
**1^st^ shell ligand angles:**   
1-2: 83; 1-3: 75; 1-4: 151; 2-3: 56; 2-4: 69; 3-4: 94;   
  
 Contact(s): /N3_C_/N7/O6/OD1_Asn_/  
 Coded contact list: O_b_2N_b_O_cno_  

### K:1G:1614

|  | Res. type | Atom | Residue | Distance | Occupancy | B-factor | Closest solute to 1^st^ shell water |
| --- | --- | --- | --- | --- | --- | --- | --- |
| **Ion** | | | |  | 1.00 | 86.30 || 1 | U | O2' | 4L:51 | 2.72 | 1.00 | 96.19 |
| 2 | G | O6 | 1G:530 | 2.91 | 1.00 | 74.92 |  |
| 3 | C | O2 | 1G:518 | 3.12 | 1.00 | 65.29 |  |
| 4 | PRO | O | 3A:45 | 3.17 | 1.00 | 73.72 |  |
| 5 | C | N3 | 1G:518 | 3.32 | 1.00 | 71.86 |  |
| 6 | U | O4' | 4L:51 | 3.45 | 1.00 | 82.70 |  |
| 7 | U | O3' | 4L:51 | 4.38 | 1.00 | 100.81 |  |
| 8 | G | N1 | 1G:530 | 4.41 | 1.00 | 77.96 |  |
| 9 | G | N7 | 1G:530 | 4.61 | 1.00 | 81.36 |  |
| 10 | C | N4 | 1G:518 | 4.83 | 1.00 | 67.74 |  |
  |

  
**1^st^ shell ligand angles:**   
1-2: 62; 1-3: 130; 1-4: 151; 1-5: 96; 2-3: 85; 2-4: 147; 2-5: 85; 3-4: 71; 3-5: 41; 4-5: 91;   
  
 Contact(s): /N3_C_/O2/O2'/O6/O_back_/  
 Coded contact list: O_r_2O_b_N_b_O_back_  

### K:1G:1615

|  | Res. type | Atom | Residue | Distance | Occupancy | B-factor | Closest solute to 1^st^ shell water |
| --- | --- | --- | --- | --- | --- | --- | --- |
| **Ion** | | | |  | 1.00 | 60.41 || 1 | G | OP1 | 1G:1521 | 2.64 | 1.00 | 47.76 |
| 2 | G | OP1 | 1G:1508 | 2.74 | 1.00 | 51.11 |  |
| 3 | A | OP1 | 1G:1500 | 2.78 | 1.00 | 49.94 |  |
| 4 | A | O3' | 1G:1499 | 3.18 | 1.00 | 58.26 |  |
| 5 | G | OP2 | 1G:1508 | 4.21 | 1.00 | 59.55 |  |
| 6 | G | O5' | 1G:1508 | 4.31 | 1.00 | 50.27 |  |
| 7 | G | OP2 | 1G:1521 | 4.48 | 1.00 | 45.65 |  |
| 8 | A | O5' | 1G:1500 | 4.51 | 1.00 | 47.42 |  |
| 9 | G | OP1 | 1G:1505 | 4.68 | 1.00 | 54.17 |  |
| 10 | A | OP2 | 1G:1500 | 4.74 | 1.00 | 49.92 |  |
| 11 | A | O2' | 1G:1499 | 4.78 | 1.00 | 48.92 |  |
| 12 | G | O3' | 1G:1520 | 4.90 | 1.00 | 48.90 |  |
| 13 | U | O2' | 1G:1506 | 4.95 | 1.00 | 54.43 |  |
| 14 | G | O5' | 1G:1521 | 4.98 | 1.00 | 43.84 |  |
  |

  
**1^st^ shell ligand angles:**   
1-2: 167; 1-3: 134; 1-4: 86; 2-3: 59; 2-4: 107; 3-4: 49;   
  
 Contact(s): /O3'/OP1/OP1/OP1/  
 Coded contact list: 3O_ph_O_r_  

### K:1G:1616

|  | Res. type | Atom | Residue | Distance | Occupancy | B-factor | Closest solute to 1^st^ shell water |
| --- | --- | --- | --- | --- | --- | --- | --- |
| **Ion** | | | |  | 1.00 | 86.73 || 1 | A | N7 | 1G:790 | 3.02 | 1.00 | 66.27 |
| 2 | U | O2' | 1G:789 | 3.62 | 1.00 | 67.84 |  |
| 3 | U | O3' | 1G:789 | 3.89 | 1.00 | 73.60 |  |
| 4 | A | N6 | 1G:790 | 3.95 | 1.00 | 60.91 |  |
| 5 | G | O6 | 1G:791 | 4.07 | 1.00 | 60.05 |  |
| 6 | U | OP1 | 4L:45 | 4.81 | 1.00 | 91.25 |  |
| 7 | A | OP1 | 1G:790 | 4.92 | 1.00 | 57.23 |  |
  |

  
 Contact(s): /N7/  
 Coded contact list: N_b_  

### K:1G:1617

|  | Res. type | Atom | Residue | Distance | Occupancy | B-factor | Closest solute to 1^st^ shell water |
| --- | --- | --- | --- | --- | --- | --- | --- |
| **Ion** | | | |  | 1.00 | 63.76 || 1 | G | OP1 | 1G:758 | 2.83 | 1.00 | 46.31 |
| 2 | G | N7 | 1G:576 | 3.14 | 1.00 | 59.01 |  |
| 3 | G | O6 | 1G:576 | 3.37 | 1.00 | 60.16 |  |
| 4 | C | O2 | 1G:880 | 3.39 | 1.00 | 59.05 |  |
| 5 | G | N3 | 1G:821 | 3.63 | 1.00 | 58.37 |  |
| 6 | G | N2 | 1G:575 | 3.66 | 1.00 | 59.12 |  |
| 7 | G | N2 | 1G:821 | 4.18 | 1.00 | 60.23 |  |
| 8 | G | O5' | 1G:758 | 4.42 | 1.00 | 60.10 |  |
| 9 | C | O2' | 1G:880 | 4.60 | 1.00 | 57.92 |  |
| 10 | G | O2' | 1G:821 | 4.70 | 1.00 | 54.26 |  |
| 11 | G | N9 | 1G:821 | 4.77 | 1.00 | 59.23 |  |
  |

  
**1^st^ shell ligand angles:**   
1-2: 164; 1-3: 108; 1-4: 106; 2-3: 57; 2-4: 76; 3-4: 72;   
  
 Contact(s): /N7/O2/O6/OP1/  
 Coded contact list: O_ph_2O_b_N_b_  

### K:1G:1618

|  | Res. type | Atom | Residue | Distance | Occupancy | B-factor | Closest solute to 1^st^ shell water |
| --- | --- | --- | --- | --- | --- | --- | --- |
| **Ion** | | | |  | 1.00 | 79.66 || 1 | SER | OG | BA:11 | 2.81 | 1.00 | 72.13 |
| 2 | U | O4 | 1G:62 | 2.85 | 1.00 | 81.57 |  |
| 3 | G | O6 | 1G:61 | 2.88 | 1.00 | 73.16 |  |
| 4 | G | O6 | 1G:105 | 2.93 | 1.00 | 69.97 |  |
| 5 | LYS | NZ | BA:14 | 3.46 | 1.00 | 85.46 |  |
| 6 | C | N4 | 1G:106 | 3.68 | 1.00 | 64.86 |  |
| 7 | G | O6 | 1G:104 | 3.78 | 1.00 | 78.08 |  |
| 8 | U | N3 | 1G:62 | 3.80 | 1.00 | 69.29 |  |
| 9 | G | N7 | 1G:105 | 4.18 | 1.00 | 71.56 |  |
| 10 | C | N4 | 1G:63 | 4.35 | 1.00 | 75.46 |  |
| 11 | G | N7 | 1G:61 | 4.69 | 1.00 | 67.70 |  |
| 12 | SER | O | BA:11 | 4.82 | 1.00 | 73.27 |  |
| 13 | G | N1 | 1G:61 | 4.82 | 1.00 | 70.29 |  |
  |

  
**1^st^ shell ligand angles:**   
1-2: 69; 1-3: 63; 1-4: 123; 2-3: 80; 2-4: 76; 3-4: 68;   
  
 Contact(s): /O4/O6/O6/OG_Ser_/  
 Coded contact list: 3O_b_OH_Prot_  

### K:1G:1619

|  | Res. type | Atom | Residue | Distance | Occupancy | B-factor | Closest solute to 1^st^ shell water |
| --- | --- | --- | --- | --- | --- | --- | --- |
| **Ion** | | | |  | 1.00 | 74.45 || 1 | C | O2 | 1G:372 | 2.51 | 1.00 | 68.25 |
| 2 | C | O2' | 1G:372 | 2.62 | 1.00 | 68.90 |  |
| 3 | G | O6 | 1G:376 | 2.90 | 1.00 | 74.83 |  |
| 4 | U | O4 | 1G:387 | 2.92 | 1.00 | 71.09 |  |
| 5 | U | O4 | 1G:375 | 2.96 | 1.00 | 66.91 |  |
| 6 | Water 1 | | 1G:1891 | 2.99 | 1.00 | 59.90 | 2.75 (G:O6 1G:376) |
| 7 | C | N1 | 1G:372 | 4.30 | 1.00 | 64.51 |  |
| 8 | A | OP2 | 1G:374 | 4.32 | 1.00 | 78.98 |  |
| 9 | A | N7 | 1G:374 | 4.33 | 1.00 | 68.11 |  |
| 10 | C | N3 | 1G:372 | 4.38 | 1.00 | 66.23 |  |
| 11 | A | N6 | 1G:389 | 4.41 | 1.00 | 66.67 |  |
| 12 | U | N3 | 1G:387 | 4.50 | 1.00 | 66.91 |  |
| 13 | A | N6 | 1G:374 | 4.58 | 1.00 | 68.01 |  |
| 14 | G | N1 | 1G:376 | 4.90 | 1.00 | 66.07 |  |
| 15 | C | O3' | 1G:372 | 4.96 | 1.00 | 71.86 |  |
  |

  
**1^st^ shell ligand angles:**   
1-2: 71; 1-3: 106; 1-4: 78; 1-5: 79; 1-6: 161; 2-3: 140; 2-4: 67; 2-5: 149; 2-6: 124; 3-4: 73; 3-5: 57; 3-6: 55; 4-5: 115; 4-6: 97; 5-6: 87;   
  
 Contact(s): /O2'/O4/O4/O6/O_w_/  
 Coded contact list: O_r_3O_b_O_w_  
 Exclusion zone contact(s): /O2/  

### K:1G:1620

|  | Res. type | Atom | Residue | Distance | Occupancy | B-factor | Closest solute to 1^st^ shell water |
| --- | --- | --- | --- | --- | --- | --- | --- |
| **Ion** | | | |  | 1.00 | 83.15 || 1 | 7MG | OP1 | 1G:527 | 2.76 | 1.00 | 72.59 |
| 2 | C | O3' | 1G:526 | 2.88 | 1.00 | 69.76 |  |
| 3 | A | OP1 | 1G:914 | 3.12 | 1.00 | 78.82 |  |
| 4 | 7MG | OP2 | 1G:527 | 3.24 | 1.00 | 70.98 |  |
| 5 | U | O3' | 1G:12 | 3.34 | 1.00 | 71.23 |  |
| 6 | U | O5' | 1G:13 | 3.49 | 1.00 | 69.47 |  |
| 7 | Water 1 | | 1G:1841 | 3.60 | 1.00 | 65.21 | 2.38 (U:OP2 1G:14) |
| 8 | U | OP1 | 1G:13 | 3.82 | 1.00 | 75.85 |  |
| 9 | U | O2' | 1G:12 | 3.86 | 1.00 | 69.46 |  |
| 10 | U | OP2 | 1G:14 | 4.20 | 1.00 | 63.16 |  |
| 11 | C | O2' | 1G:526 | 4.36 | 1.00 | 72.89 |  |
| 12 | 7MG | O5' | 1G:527 | 4.60 | 1.00 | 60.77 |  |
  |

  
**1^st^ shell ligand angles:**   
1-2: 52; 1-3: 115; 1-4: 50; 1-5: 136; 2-3: 95; 2-4: 49; 2-5: 92; 3-4: 66; 3-5: 90; 4-5: 127;   
  
 Contact(s): /O3'/O3'/OP1/OP1/OP2/  
 Coded contact list: 3O_ph_2O_r_  

### K:1G:1621

|  | Res. type | Atom | Residue | Distance | Occupancy | B-factor | Closest solute to 1^st^ shell water |
| --- | --- | --- | --- | --- | --- | --- | --- |
| **Ion** | | | |  | 1.00 | 76.49 || 1 | G | OP2 | 1G:567 | 2.72 | 1.00 | 64.98 |
| 2 | C | OP2 | 1G:564 | 2.85 | 1.00 | 65.64 |  |
| 3 | U | OP2 | 1G:565 | 2.85 | 1.00 | 65.74 |  |
| 4 | A | O2' | 1G:563 | 3.06 | 1.00 | 59.93 |  |
| 5 | A | O3' | 1G:563 | 3.10 | 1.00 | 58.84 |  |
| 6 | G | O3' | 1G:566 | 3.49 | 1.00 | 68.68 |  |
| 7 | G | O2' | 1G:566 | 3.95 | 1.00 | 66.58 |  |
| 8 | C | O5' | 1G:564 | 4.15 | 1.00 | 59.23 |  |
| 9 | G | O5' | 1G:567 | 4.28 | 1.00 | 64.17 |  |
| 10 | U | OP1 | 1G:565 | 4.37 | 1.00 | 65.36 |  |
| 11 | ARG | NH2 | 3A:12 | 4.47 | 1.00 | 63.39 |  |
| 12 | G | N7 | 1G:567 | 4.89 | 1.00 | 64.41 |  |
| 13 | C | OP1 | 1G:564 | 4.90 | 1.00 | 62.57 |  |
| 14 | G | OP1 | 1G:567 | 4.98 | 1.00 | 71.72 |  |
| 15 | C | O3' | 1G:564 | 4.99 | 1.00 | 64.57 |  |
  |

  
**1^st^ shell ligand angles:**   
1-2: 174; 1-3: 97; 1-4: 80; 1-5: 129; 2-3: 88; 2-4: 97; 2-5: 49; 3-4: 85; 3-5: 72; 4-5: 50;   
  
 Contact(s): /O2'/O3'/OP2/OP2/OP2/  
 Coded contact list: 3O_ph_2O_r_  

### K:1G:1622

|  | Res. type | Atom | Residue | Distance | Occupancy | B-factor | Closest solute to 1^st^ shell water |
| --- | --- | --- | --- | --- | --- | --- | --- |
| **Ion** | | | |  | 1.00 | 74.93 || 1 | G | O6 | 1G:927 | 2.84 | 1.00 | 69.17 |
| 2 | C | N4 | 1G:924 | 3.38 | 1.00 | 66.52 |  |
| 3 | U | O4 | 1G:1390 | 3.40 | 1.00 | 59.30 |  |
| 4 | G | O6 | 1G:925 | 3.47 | 1.00 | 60.16 |  |
| 5 | G | N7 | 1G:925 | 3.83 | 1.00 | 64.14 |  |
| 6 | U | O4 | 1G:1391 | 4.08 | 1.00 | 65.14 |  |
| 7 | G | N7 | 1G:927 | 4.11 | 1.00 | 67.20 |  |
| 8 | U | N3 | 1G:1390 | 4.36 | 1.00 | 64.50 |  |
| 9 | G | O6 | 1G:928 | 4.77 | 1.00 | 71.05 |  |
| 10 | C | N4 | 1G:1389 | 4.84 | 1.00 | 66.75 |  |
| 11 | C | N3 | 1G:924 | 4.85 | 1.00 | 61.96 |  |
| 12 | U | N3 | 1G:1391 | 4.92 | 1.00 | 64.49 |  |
  |

  
**1^st^ shell ligand angles:**   
1-2: 110; 1-3: 67; 2-3: 82;   
  
 Contact(s): /N4/O4/O6/  
 Coded contact list: 2O_b_Anion  

### K:1G:1623

|  | Res. type | Atom | Residue | Distance | Occupancy | B-factor | Closest solute to 1^st^ shell water |
| --- | --- | --- | --- | --- | --- | --- | --- |
| **Ion** | | | |  | 1.00 | 95.15 || 1 | G | OP2 | 1G:1057 | 2.98 | 1.00 | 96.33 |
| 2 | U | O4 | 1G:1199 | 3.44 | 1.00 | 86.15 |  |
| 3 | C | N4 | 1G:1059 | 3.94 | 1.00 | 96.30 |  |
| 4 | G | O6 | 1G:1058 | 3.97 | 1.00 | 95.89 |  |
| 5 | G | O6 | 1G:1198 | 4.08 | 1.00 | 91.80 |  |
| 6 | G | N2 | 1G:1053 | 4.19 | 1.00 | 87.36 |  |
| 7 | G | N7 | 1G:1058 | 4.52 | 1.00 | 94.15 |  |
| 8 | G | O5' | 1G:1057 | 4.91 | 1.00 | 101.91 |  |
  |

  
 Contact(s): /OP2/  
 Coded contact list: O_ph_  

### K:1G:1624

|  | Res. type | Atom | Residue | Distance | Occupancy | B-factor | Closest solute to 1^st^ shell water |
| --- | --- | --- | --- | --- | --- | --- | --- |
| **Ion** | | | |  | 1.00 | 60.51 || 1 | A | OP2 | 1G:116 | 2.55 | 1.00 | 58.78 |
| 2 | G | O4' | 1G:289 | 2.65 | 1.00 | 57.89 |  |
| 3 | G | N7 | 1G:117 | 2.78 | 1.00 | 63.18 |  |
| 4 | G | O2' | 1G:115 | 2.85 | 1.00 | 51.60 |  |
| 5 | G | O5' | 1G:289 | 2.87 | 1.00 | 56.08 |  |
| 6 | G | O6 | 1G:117 | 3.12 | 1.00 | 64.13 |  |
| 7 | U | O4 | 1G:118 | 3.61 | 1.00 | 62.56 |  |
| 8 | A | N7 | 1G:116 | 3.76 | 1.00 | 57.91 |  |
| 9 | G | OP2 | 1G:289 | 4.02 | 1.00 | 59.30 |  |
| 10 | G | N3 | 1G:115 | 4.27 | 1.00 | 58.03 |  |
| 11 | G | N9 | 1G:289 | 4.32 | 1.00 | 54.62 |  |
| 12 | G | O3' | 1G:115 | 4.52 | 1.00 | 67.40 |  |
| 13 | A | OP1 | 1G:116 | 4.55 | 1.00 | 55.93 |  |
| 14 | A | N6 | 1G:116 | 4.60 | 1.00 | 56.53 |  |
| 15 | U | N3 | 1G:118 | 4.76 | 1.00 | 61.28 |  |
| 16 | G | OP2 | 1G:117 | 4.93 | 1.00 | 62.57 |  |
| 17 | G | N1 | 1G:117 | 4.96 | 1.00 | 65.08 |  |
| 18 | G | N9 | 1G:117 | 4.97 | 1.00 | 58.33 |  |
  |

  
**1^st^ shell ligand angles:**   
1-2: 144; 1-3: 92; 1-4: 58; 1-5: 85; 1-6: 140; 2-3: 124; 2-4: 91; 2-5: 65; 2-6: 66; 3-4: 134; 3-5: 153; 3-6: 64; 4-5: 65; 4-6: 117; 5-6: 131;   
  
 Contact(s): /N7/O2'/O4'/O5'/O6/  
 Coded contact list: 3O_r_O_b_N_b_  
 Exclusion zone contact(s): /OP2/  

### K:1G:1625

|  | Res. type | Atom | Residue | Distance | Occupancy | B-factor | Closest solute to 1^st^ shell water |
| --- | --- | --- | --- | --- | --- | --- | --- |
| **Ion** | | | |  | 1.00 | 108.07 || 1 | U | OP1 | 1G:1330 | 2.50 | 1.00 | 109.70 |
| 2 | THR | O | 4A:20 | 2.54 | 1.00 | 107.81 |  |
| 3 | ILE | O | 4A:22 | 2.76 | 1.00 | 110.65 |  |
| 4 | U | O5' | 1G:1330 | 3.37 | 1.00 | 110.65 |  |
| 5 | GLY | N | 4A:26 | 3.81 | 1.00 | 113.25 |  |
| 6 | ILE | N | 4A:25 | 4.07 | 1.00 | 113.22 |  |
| 7 | GLY | N | 4A:24 | 4.13 | 1.00 | 118.59 |  |
| 8 | ILE | N | 4A:22 | 4.16 | 1.00 | 111.13 |  |
| 9 | A | O3' | 1G:1329 | 4.30 | 1.00 | 114.53 |  |
| 10 | TYR | N | 4A:23 | 4.37 | 1.00 | 115.98 |  |
| 11 | U | OP2 | 1G:1330 | 4.69 | 1.00 | 107.88 |  |
| 12 | TYR | N | 4A:21 | 4.72 | 1.00 | 111.14 |  |
| 13 | G | O5' | 1G:1331 | 4.74 | 1.00 | 113.88 |  |
| 14 | G | OP2 | 1G:1331 | 4.74 | 1.00 | 121.28 |  |
  |

  
**1^st^ shell ligand angles:**   
1-2: 125; 1-3: 117; 1-4: 49; 2-3: 80; 2-4: 174; 3-4: 101;   
  
 Contact(s): /O5'/O_back_/  
 Coded contact list: O_r_O_back_  
 Exclusion zone contact(s): /O_back_/OP1/  

### K:1G:1626

|  | Res. type | Atom | Residue | Distance | Occupancy | B-factor | Closest solute to 1^st^ shell water |
| --- | --- | --- | --- | --- | --- | --- | --- |
| **Ion** | | | |  | 1.00 | 67.03 || 1 | G | O6 | 1G:305 | 2.66 | 1.00 | 54.19 |
| 2 | G | O6 | 1G:293 | 2.80 | 1.00 | 54.49 |  |
| 3 | U | O4 | 1G:304 | 2.88 | 1.00 | 58.04 |  |
| 4 | G | N1 | 1G:305 | 3.84 | 1.00 | 53.61 |  |
| 5 | U | N3 | 1G:304 | 3.89 | 1.00 | 55.88 |  |
| 6 | G | N7 | 1G:293 | 4.10 | 1.00 | 54.36 |  |
| 7 | G | OP2 | 1G:292 | 4.20 | 1.00 | 47.01 |  |
| 8 | A | N6 | 1G:303 | 4.39 | 1.00 | 56.55 |  |
| 9 | G | N1 | 1G:293 | 4.86 | 1.00 | 54.22 |  |
| 10 | C | OP1 | 1G:291 | 4.95 | 1.00 | 53.76 |  |
  |

  
**1^st^ shell ligand angles:**   
1-2: 78; 1-3: 77; 2-3: 75;   
  
 Contact(s): /O4/O6/O6/  
 Coded contact list: 3O_b_  

### K:1G:1627

|  | Res. type | Atom | Residue | Distance | Occupancy | B-factor | Closest solute to 1^st^ shell water |
| --- | --- | --- | --- | --- | --- | --- | --- |
| **Ion** | | | |  | 1.00 | 92.32 || 1 | A | OP2 | 1G:816 | 3.23 | 1.00 | 50.22 |
| 2 | G | OP2 | 1G:731 | 3.53 | 1.00 | 66.10 |  |
| 3 | G | O3' | 1G:1526 | 3.99 | 1.00 | 59.96 |  |
| 4 | C | OP1 | 1G:1527 | 4.09 | 1.00 | 66.85 |  |
| 5 | G | OP1 | 1G:730 | 4.18 | 1.00 | 73.81 |  |
| 6 | G | O2' | 1G:1526 | 4.25 | 1.00 | 53.26 |  |
| 7 | A | O5' | 1G:816 | 4.43 | 1.00 | 55.33 |  |
| 8 | C | O2' | 1G:817 | 4.66 | 1.00 | 63.56 |  |
| 9 | A | OP1 | 1G:816 | 4.82 | 1.00 | 58.04 |  |
  |

  
 Contact(s): /OP2/  
 Coded contact list: O_ph_  

### K:1G:1628

|  | Res. type | Atom | Residue | Distance | Occupancy | B-factor | Closest solute to 1^st^ shell water |
| --- | --- | --- | --- | --- | --- | --- | --- |
| **Ion** | | | |  | 1.00 | 76.61 || 1 | C | O3' | 1G:817 | 2.52 | 1.00 | 50.49 |
| 2 | G | OP2 | 1G:818 | 2.67 | 1.00 | 66.14 |  |
| 3 | U | OP1 | 1G:1528 | 2.93 | 1.00 | 55.17 |  |
| 4 | C | O2' | 1G:817 | 2.99 | 1.00 | 63.56 |  |
| 5 | G | O5' | 1G:818 | 3.68 | 1.00 | 60.58 |  |
| 6 | C | O3' | 1G:1527 | 3.78 | 1.00 | 61.91 |  |
| 7 | A | OP2 | 1G:819 | 4.41 | 1.00 | 62.05 |  |
| 8 | C | O2 | 1G:817 | 4.53 | 1.00 | 56.93 |  |
| 9 | G | OP1 | 1G:818 | 4.53 | 1.00 | 54.89 |  |
| 10 | U | OP2 | 1G:1528 | 4.94 | 1.00 | 59.28 |  |
  |

  
**1^st^ shell ligand angles:**   
1-2: 57; 1-3: 161; 1-4: 59; 2-3: 134; 2-4: 103; 3-4: 121;   
  
 Contact(s): /O2'/OP1/OP2/  
 Coded contact list: 2O_ph_O_r_  
 Exclusion zone contact(s): /O3'/  

### K:1G:1629

|  | Res. type | Atom | Residue | Distance | Occupancy | B-factor | Closest solute to 1^st^ shell water |
| --- | --- | --- | --- | --- | --- | --- | --- |
| **Ion** | | | |  | 1.00 | 88.10 || 1 | C | O3' | 1G:314 | 2.56 | 1.00 | 63.28 |
| 2 | A | OP1 | 1G:315 | 2.62 | 1.00 | 79.97 |  |
| 3 | G | OP2 | 1G:318 | 3.52 | 1.00 | 69.10 |  |
| 4 | A | O5' | 1G:315 | 4.15 | 1.00 | 65.04 |  |
| 5 | A | OP2 | 1G:315 | 4.25 | 1.00 | 75.36 |  |
| 6 | C | O2' | 1G:314 | 4.43 | 1.00 | 59.81 |  |
| 7 | G | O5' | 1G:318 | 4.89 | 1.00 | 72.70 |  |
  |

  
**1^st^ shell ligand angles:**   
1-2: 59;   
  
 Contact(s): /OP1/  
 Coded contact list: O_ph_  
 Exclusion zone contact(s): /O3'/  

### K:1G:1630

|  | Res. type | Atom | Residue | Distance | Occupancy | B-factor | Closest solute to 1^st^ shell water |
| --- | --- | --- | --- | --- | --- | --- | --- |
| **Ion** | | | |  | 1.00 | 80.09 || 1 | G | O6 | 1G:579 | 3.08 | 1.00 | 61.68 |
| 2 | G | O6 | 1G:761 | 3.26 | 1.00 | 59.58 |  |
| 3 | U | O4 | 1G:580 | 3.45 | 1.00 | 68.57 |  |
| 4 | C | N4 | 1G:762 | 3.72 | 1.00 | 60.88 |  |
| 5 | A | OP1 | 1G:759 | 3.88 | 1.00 | 60.44 |  |
| 6 | A | OP2 | 1G:759 | 4.05 | 1.00 | 61.84 |  |
| 7 | G | N1 | 1G:581 | 4.24 | 1.00 | 60.75 |  |
| 8 | G | N7 | 1G:761 | 4.28 | 1.00 | 63.74 |  |
| 9 | G | N2 | 1G:581 | 4.31 | 1.00 | 61.03 |  |
| 10 | U | N3 | 1G:580 | 4.49 | 1.00 | 63.61 |  |
| 11 | G | OP2 | 1G:760 | 4.74 | 1.00 | 62.13 |  |
| 12 | G | N1 | 1G:579 | 4.88 | 1.00 | 57.89 |  |
| 13 | G | N2 | 1G:576 | 4.93 | 1.00 | 66.48 |  |
  |

  
**1^st^ shell ligand angles:**   
1-2: 74;   
  
 Contact(s): /O6/O6/  
 Coded contact list: 2O_b_  

### K:1G:1631

|  | Res. type | Atom | Residue | Distance | Occupancy | B-factor | Closest solute to 1^st^ shell water |
| --- | --- | --- | --- | --- | --- | --- | --- |
| **Ion** | | | |  | 1.00 | 86.91 || 1 | G | O6 | 1G:886 | 2.87 | 1.00 | 50.54 |
| 2 | U | O4 | 1G:911 | 3.36 | 1.00 | 45.97 |  |
| 3 | G | O6 | 1G:887 | 3.65 | 1.00 | 42.06 |  |
| 4 | G | O6 | 1G:885 | 3.67 | 1.00 | 65.75 |  |
| 5 | G | N7 | 1G:886 | 3.72 | 1.00 | 53.75 |  |
| 6 | G | N7 | 1G:887 | 4.03 | 1.00 | 49.04 |  |
| 7 | U | N3 | 1G:911 | 4.48 | 1.00 | 46.43 |  |
| 8 | G | N7 | 1G:885 | 4.74 | 1.00 | 65.03 |  |
| 9 | C | N4 | 1G:910 | 4.78 | 1.00 | 44.47 |  |
| 10 | G | N1 | 1G:885 | 4.82 | 1.00 | 54.94 |  |
| 11 | G | N1 | 1G:886 | 4.89 | 1.00 | 42.26 |  |
  |

  
**1^st^ shell ligand angles:**   
1-2: 50;   
  
 Contact(s): /O4/O6/  
 Coded contact list: 2O_b_  

### K:1G:1632

|  | Res. type | Atom | Residue | Distance | Occupancy | B-factor | Closest solute to 1^st^ shell water |
| --- | --- | --- | --- | --- | --- | --- | --- |
| **Ion** | | | |  | 1.00 | 78.65 || 1 | U | O4 | 1G:1485 | 2.73 | 1.00 | 53.20 |
| 2 | U | O4 | 1G:1414 | 2.82 | 1.00 | 62.84 |  |
| 3 | G | O6 | 1G:1486 | 3.20 | 1.00 | 50.71 |  |
| 4 | G | O6 | 1G:1415 | 3.46 | 1.00 | 46.57 |  |
| 5 | U | N3 | 1G:1414 | 3.84 | 1.00 | 50.30 |  |
| 6 | U | N3 | 1G:1485 | 4.14 | 1.00 | 44.40 |  |
| 7 | G | N7 | 1G:1486 | 4.41 | 1.00 | 44.64 |  |
| 8 | A | N6 | 1G:1413 | 4.58 | 1.00 | 51.29 |  |
| 9 | G | O6 | 1G:1487 | 4.64 | 1.00 | 50.55 |  |
  |

  
**1^st^ shell ligand angles:**   
1-2: 136; 1-3: 114; 2-3: 85;   
  
 Contact(s): /O4/O4/O6/  
 Coded contact list: 3O_b_  

### K:1G:1633

|  | Res. type | Atom | Residue | Distance | Occupancy | B-factor | Closest solute to 1^st^ shell water |
| --- | --- | --- | --- | --- | --- | --- | --- |
| **Ion** | | | |  | 1.00 | 76.91 || 1 | G | O6 | 1G:1482 | 2.83 | 1.00 | 56.15 |
| 2 | U | O4 | 1G:1481 | 3.07 | 1.00 | 58.26 |  |
| 3 | G | O6 | 1G:1419 | 3.08 | 1.00 | 49.92 |  |
| 4 | G | O6 | 1G:1480 | 3.81 | 1.00 | 68.22 |  |
| 5 | U | N3 | 1G:1481 | 4.17 | 1.00 | 59.28 |  |
| 6 | G | N1 | 1G:1482 | 4.25 | 1.00 | 46.44 |  |
| 7 | Water 1 | | 1G:1871 | 4.50 | 1.00 | 44.33 | 2.84 (G:O6 1G:1482) |
| 8 | G | N7 | 1G:1419 | 4.52 | 1.00 | 52.81 |  |
| 9 | C | N4 | 1G:1420 | 4.78 | 1.00 | 52.05 |  |
  |

  
**1^st^ shell ligand angles:**   
1-2: 69; 1-3: 65; 2-3: 75;   
  
 Contact(s): /O4/O6/O6/  
 Coded contact list: 3O_b_  

### K:1G:1634

|  | Res. type | Atom | Residue | Distance | Occupancy | B-factor | Closest solute to 1^st^ shell water |
| --- | --- | --- | --- | --- | --- | --- | --- |
| **Ion** | | | |  | 1.00 | 62.32 || 1 | G | O2' | 1G:266 | 2.82 | 1.00 | 72.90 |
| 2 | G | O3' | 1G:251 | 3.17 | 1.00 | 78.86 |  |
| 3 | C | OP2 | 1G:268 | 3.34 | 1.00 | 64.87 |  |
| 4 | C | OP2 | 1G:267 | 3.36 | 1.00 | 73.40 |  |
| 5 | U | OP2 | 1G:252 | 3.40 | 1.00 | 65.05 |  |
| 6 | G | O2' | 1G:251 | 3.42 | 1.00 | 79.76 |  |
| 7 | C | OP1 | 1G:268 | 3.90 | 1.00 | 62.46 |  |
| 8 | G | O3' | 1G:266 | 4.01 | 1.00 | 70.52 |  |
| 9 | LYS | NZ | 8A:67 | 4.10 | 1.00 | 63.14 |  |
| 10 | U | OP2 | 1G:253 | 4.24 | 1.00 | 61.42 |  |
| 11 | U | O5' | 1G:252 | 4.66 | 1.00 | 67.37 |  |
  |

  
**1^st^ shell ligand angles:**   
1-2: 121; 1-3: 82; 1-4: 73; 1-5: 142; 2-3: 105; 2-4: 166; 2-5: 44; 3-4: 75; 3-5: 132; 4-5: 126;   
  
 Contact(s): /O2'/O3'/OP2/OP2/OP2/  
 Coded contact list: 3O_ph_2O_r_  

### K:1G:1635

|  | Res. type | Atom | Residue | Distance | Occupancy | B-factor | Closest solute to 1^st^ shell water |
| --- | --- | --- | --- | --- | --- | --- | --- |
| **Ion** | | | |  | 1.00 | 91.08 || 1 | G | O6 | 1G:258 | 2.97 | 1.00 | 80.05 |
| 2 | G | N7 | 1G:258 | 3.73 | 1.00 | 84.14 |  |
| 3 | G | O6 | 1G:257 | 3.83 | 1.00 | 80.83 |  |
| 4 | G | OP2 | 1G:266 | 4.41 | 1.00 | 66.58 |  |
| 5 | G | N7 | 1G:257 | 4.51 | 1.00 | 84.86 |  |
| 6 | G | OP1 | 1G:266 | 4.67 | 1.00 | 72.46 |  |
| 7 | G | OP1 | 1G:265 | 4.80 | 1.00 | 68.76 |  |
| 8 | G | O6 | 1G:259 | 4.85 | 1.00 | 69.27 |  |
  |

  
 Contact(s): /O6/  
 Coded contact list: O_b_  

### K:1G:1636

|  | Res. type | Atom | Residue | Distance | Occupancy | B-factor | Closest solute to 1^st^ shell water |
| --- | --- | --- | --- | --- | --- | --- | --- |
| **Ion** | | | |  | 1.00 | 78.06 || 1 | C | OP2 | 1G:110 | 2.93 | 1.00 | 59.54 |
| 2 | A | OP2 | 1G:109 | 3.11 | 1.00 | 66.30 |  |
| 3 | C | OP1 | 1G:330 | 3.30 | 1.00 | 65.34 |  |
| 4 | G | O6 | 1G:111 | 3.48 | 1.00 | 62.60 |  |
| 5 | Water 1 | | 1G:1940 | 3.85 | 1.00 | 70.02 | 3.23 (C:OP2 1G:110) |
| 6 | G | O6 | 1G:112 | 3.94 | 1.00 | 65.69 |  |
| 7 | G | N7 | 1G:111 | 4.70 | 1.00 | 64.99 |  |
| 8 | A | N6 | 1G:315 | 4.87 | 1.00 | 56.21 |  |
| 9 | C | O5' | 1G:330 | 4.89 | 1.00 | 56.82 |  |
| 10 | A | O5' | 1G:109 | 4.93 | 1.00 | 55.91 |  |
  |

  
**1^st^ shell ligand angles:**   
1-2: 85; 1-3: 99; 2-3: 108;   
  
 Contact(s): /OP1/OP2/OP2/  
 Coded contact list: 3O_ph_  

### MG:1G:1637

|  | Res. type | Atom | Residue | Distance | Occupancy | B-factor | Closest solute to 1^st^ shell water |
| --- | --- | --- | --- | --- | --- | --- | --- |
| **Ion** | | | |  | 1.00 | 55.69 || 1 | G | OP1 | 1G:1508 | 1.94 | 1.00 | 51.11 |
| 2 | G | OP1 | 1G:1505 | 2.08 | 1.00 | 54.17 |  |
| 3 | A | OP1 | 1G:1500 | 2.24 | 1.00 | 49.94 |  |
| 4 | A | O3' | 1G:1507 | 3.35 | 1.00 | 58.44 |  |
| 5 | U | O2' | 1G:1506 | 3.91 | 1.00 | 54.43 |  |
| 6 | G | O3' | 1G:1504 | 3.99 | 1.00 | 60.69 |  |
| 7 | G | OP1 | 1G:1504 | 4.08 | 1.00 | 59.87 |  |
| 8 | G | O5' | 1G:1505 | 4.15 | 1.00 | 60.01 |  |
| 9 | G | OP2 | 1G:1508 | 4.25 | 1.00 | 59.55 |  |
| 10 | A | OP2 | 1G:1500 | 4.27 | 1.00 | 49.92 |  |
| 11 | G | O5' | 1G:1508 | 4.29 | 1.00 | 50.27 |  |
| 12 | G | OP2 | 1G:1504 | 4.34 | 1.00 | 58.21 |  |
| 13 | A | O5' | 1G:1500 | 4.50 | 1.00 | 47.42 |  |
| 14 | C | OP2 | 1G:1501 | 4.51 | 1.00 | 55.07 |  |
| 15 | A | O3' | 1G:1499 | 4.65 | 1.00 | 58.26 |  |
| 16 | G | OP2 | 1G:1505 | 4.69 | 1.00 | 49.73 |  |
| 17 | G | O2' | 1G:1504 | 4.70 | 1.00 | 54.58 |  |
  |

  
**1^st^ shell ligand angles:**   
1-2: 155; 1-3: 81; 2-3: 77;   
  
 Contact(s): /OP1/OP1/OP1/  
 Coded contact list: 3O_ph_  
 Exclusion zone contact(s): /O3'/  

### MG:1G:1638

|  | Res. type | Atom | Residue | Distance | Occupancy | B-factor | Closest solute to 1^st^ shell water |
| --- | --- | --- | --- | --- | --- | --- | --- |
| **Ion** | | | |  | 1.00 | 46.37 || 1 | A | OP1 | 1G:572 | 2.30 | 1.00 | 67.86 |
| 2 | A | N1 | 1G:819 | 4.13 | 1.00 | 56.23 |  |
| 3 | U | O3' | 1G:571 | 4.40 | 1.00 | 56.01 |  |
| 4 | A | N6 | 1G:819 | 4.47 | 1.00 | 52.03 |  |
| 5 | A | OP1 | 1G:573 | 4.56 | 1.00 | 64.37 |  |
| 6 | A | O5' | 1G:572 | 4.59 | 1.00 | 63.44 |  |
| 7 | A | OP2 | 1G:572 | 4.69 | 1.00 | 63.77 |  |
  |

  
 Contact(s): /OP1/  
 Coded contact list: O_ph_  

### MG:1G:1639

|  | Res. type | Atom | Residue | Distance | Occupancy | B-factor | Closest solute to 1^st^ shell water |
| --- | --- | --- | --- | --- | --- | --- | --- |
| **Ion** | | | |  | 1.00 | 69.75 || 1 | PSU | O2 | 1G:516 | 2.07 | 1.00 | 72.50 |
| 2 | A | OP1 | 1G:533 | 2.25 | 1.00 | 65.78 |  |
| 3 | G | N7 | 1G:515 | 3.41 | 1.00 | 81.42 |  |
| 4 | PSU | N1 | 1G:516 | 3.51 | 1.00 | 77.61 |  |
| 5 | G | O6 | 1G:515 | 3.79 | 1.00 | 75.13 |  |
| 6 | G | O6 | 1G:517 | 4.00 | 1.00 | 77.86 |  |
| 7 | PSU | N3 | 1G:516 | 4.13 | 1.00 | 77.52 |  |
| 8 | A | O5' | 1G:533 | 4.17 | 1.00 | 71.65 |  |
| 9 | A | OP2 | 1G:533 | 4.22 | 1.00 | 68.75 |  |
| 10 | G | OP2 | 1G:515 | 4.59 | 1.00 | 64.11 |  |
| 11 | C | N4 | 1G:514 | 4.69 | 1.00 | 77.60 |  |
| 12 | A | O3' | 1G:532 | 4.79 | 1.00 | 76.97 |  |
| 13 | G | N1 | 1G:517 | 4.85 | 1.00 | 73.33 |  |
  |

  
**1^st^ shell ligand angles:**   
1-2: 78;   
  
 Contact(s): /O2/OP1/  
 Coded contact list: O_ph_O_b_  

### MG:1G:1640

|  | Res. type | Atom | Residue | Distance | Occupancy | B-factor | Closest solute to 1^st^ shell water |
| --- | --- | --- | --- | --- | --- | --- | --- |
| **Ion** | | | |  | 1.00 | 51.11 || 1 | C | OP2 | 1G:536 | 2.82 | 1.00 | 69.55 |
| 2 | A | OP1 | 1G:535 | 3.50 | 1.00 | 68.55 |  |
| 3 | G | N7 | 1G:537 | 3.92 | 1.00 | 70.74 |  |
| 4 | G | O6 | 1G:538 | 3.97 | 1.00 | 80.31 |  |
| 5 | C | O5' | 1G:536 | 4.68 | 1.00 | 66.03 |  |
| 6 | G | O6 | 1G:537 | 4.79 | 1.00 | 77.50 |  |
| 7 | G | N7 | 1G:538 | 4.90 | 1.00 | 73.88 |  |
  |

  
 Coded contact list:   
 Exclusion zone contact(s): /OP2/  
Consider K in place of MG (1 valid contact(s) for K vs. 0 for MG)  

### MG:1G:1641

|  | Res. type | Atom | Residue | Distance | Occupancy | B-factor | Closest solute to 1^st^ shell water |
| --- | --- | --- | --- | --- | --- | --- | --- |
| **Ion** | | | |  | 1.00 | 64.55 || 1 | G | N7 | 1G:324 | 2.03 | 1.00 | 74.77 |
| 2 | G | O6 | 1G:324 | 3.41 | 1.00 | 76.64 |  |
| 3 | G | OP2 | 1G:324 | 3.62 | 1.00 | 65.36 |  |
| 4 | SER | O | BA:70 | 3.94 | 1.00 | 81.15 |  |
| 5 | G | N9 | 1G:324 | 4.23 | 1.00 | 74.18 |  |
| 6 | U | OP2 | 1G:323 | 4.26 | 1.00 | 60.73 |  |
| 7 | U | O5' | 1G:323 | 4.90 | 1.00 | 75.05 |  |
| 8 | G | OP2 | 1G:326 | 4.99 | 1.00 | 66.00 |  |
  |

  
 Contact(s): /N7/  
 Coded contact list: N_b_  

### MG:1G:1642

|  | Res. type | Atom | Residue | Distance | Occupancy | B-factor | Closest solute to 1^st^ shell water |
| --- | --- | --- | --- | --- | --- | --- | --- |
| **Ion** | | | |  | 1.00 | 45.19 || 1 | A | OP1 | 1G:782 | 2.17 | 1.00 | 54.46 |
| 2 | A | OP2 | 1G:794 | 2.52 | 1.00 | 60.14 |  |
| 3 | A | OP1 | 1G:794 | 2.67 | 1.00 | 55.68 |  |
| 4 | A | O5' | 1G:782 | 3.66 | 1.00 | 56.64 |  |
| 5 | U | O3' | 1G:793 | 3.95 | 1.00 | 54.89 |  |
| 6 | C | OP2 | 1G:783 | 4.01 | 1.00 | 61.97 |  |
| 7 | A | OP2 | 1G:782 | 4.28 | 1.00 | 53.09 |  |
| 8 | G | N2 | 1G:1521 | 4.29 | 1.00 | 49.45 |  |
| 9 | G | N3 | 1G:1521 | 4.31 | 1.00 | 53.88 |  |
| 10 | A | O5' | 1G:794 | 4.36 | 1.00 | 53.16 |  |
| 11 | G | O2' | 1G:1521 | 4.48 | 1.00 | 57.54 |  |
| 12 | A | O3' | 1G:781 | 4.55 | 1.00 | 47.82 |  |
| 13 | U | O4' | 1G:1522 | 4.65 | 1.00 | 56.61 |  |
| 14 | C | O2 | 1G:1515 | 4.89 | 1.00 | 55.65 |  |
  |

  
 Contact(s): /OP1/  
 Coded contact list: O_ph_  
 Exclusion zone contact(s): /OP1/OP2/  

### MG:1G:1643

|  | Res. type | Atom | Residue | Distance | Occupancy | B-factor | Closest solute to 1^st^ shell water |
| --- | --- | --- | --- | --- | --- | --- | --- |
| **Ion** | | | |  | 1.00 | 62.23 || 1 | G | OP2 | 1G:331 | 1.98 | 1.00 | 60.60 |
| 2 | A | OP1 | 1G:109 | 2.43 | 1.00 | 71.56 |  |
| 3 | A | O2' | 1G:329 | 3.52 | 1.00 | 63.03 |  |
| 4 | A | N3 | 1G:329 | 3.82 | 1.00 | 68.74 |  |
| 5 | G | OP1 | 1G:331 | 3.96 | 1.00 | 69.29 |  |
| 6 | A | OP2 | 1G:109 | 4.07 | 1.00 | 66.30 |  |
| 7 | G | O5' | 1G:331 | 4.21 | 1.00 | 57.15 |  |
| 8 | A | O3' | 1G:329 | 4.34 | 1.00 | 68.76 |  |
| 9 | A | O5' | 1G:109 | 4.48 | 1.00 | 55.91 |  |
| 10 | C | O3' | 1G:330 | 4.49 | 1.00 | 61.82 |  |
| 11 | G | N7 | 1G:332 | 4.64 | 1.00 | 73.54 |  |
| 12 | G | O3' | 1G:108 | 4.79 | 1.00 | 61.04 |  |
  |

  
 Contact(s): /OP2/  
 Coded contact list: O_ph_  
 Exclusion zone contact(s): /OP1/  

### MG:1G:1644

|  | Res. type | Atom | Residue | Distance | Occupancy | B-factor | Closest solute to 1^st^ shell water |
| --- | --- | --- | --- | --- | --- | --- | --- |
| **Ion** | | | |  | 1.00 | 59.26 || 1 | A | OP2 | 1G:572 | 2.12 | 1.00 | 63.77 |
| 2 | A | OP2 | 1G:573 | 2.18 | 1.00 | 56.29 |  |
| 3 | A | OP2 | 1G:574 | 2.28 | 1.00 | 59.13 |  |
| 4 | U | OP1 | 1G:571 | 3.46 | 1.00 | 56.94 |  |
| 5 | U | OP2 | 1G:571 | 3.92 | 1.00 | 59.26 |  |
| 6 | A | O5' | 1G:572 | 3.98 | 1.00 | 63.44 |  |
| 7 | A | O5' | 1G:573 | 4.31 | 1.00 | 62.38 |  |
| 8 | A | OP1 | 1G:572 | 4.32 | 1.00 | 67.86 |  |
| 9 | A | OP1 | 1G:573 | 4.33 | 1.00 | 64.37 |  |
| 10 | U | O5' | 1G:571 | 4.44 | 1.00 | 60.23 |  |
| 11 | A | OP1 | 1G:574 | 4.47 | 1.00 | 61.45 |  |
| 12 | A | O5' | 1G:574 | 4.48 | 1.00 | 54.06 |  |
| 13 | A | O3' | 1G:572 | 4.61 | 1.00 | 59.01 |  |
| 14 | U | O3' | 1G:571 | 4.63 | 1.00 | 56.01 |  |
| 15 | A | O3' | 1G:573 | 4.75 | 1.00 | 54.53 |  |
| 16 | A | N7 | 1G:574 | 4.76 | 1.00 | 62.98 |  |
  |

  
**1^st^ shell ligand angles:**   
1-2: 82; 1-3: 148; 2-3: 82;   
  
 Contact(s): /OP2/OP2/OP2/  
 Coded contact list: 3O_ph_  

### MG:1G:1645

|  | Res. type | Atom | Residue | Distance | Occupancy | B-factor | Closest solute to 1^st^ shell water |
| --- | --- | --- | --- | --- | --- | --- | --- |
| **Ion** | | | |  | 1.00 | 62.32 || 1 | G | OP2 | 1G:117 | 1.90 | 1.00 | 62.57 |
| 2 | G | OP2 | 1G:289 | 2.15 | 1.00 | 59.30 |  |
| 3 | A | OP1 | 1G:116 | 2.62 | 1.00 | 55.93 |  |
| 4 | A | O5' | 1G:288 | 3.76 | 1.00 | 61.07 |  |
| 5 | A | OP2 | 1G:116 | 3.88 | 1.00 | 58.78 |  |
| 6 | G | OP1 | 1G:117 | 3.96 | 1.00 | 56.46 |  |
| 7 | A | OP2 | 1G:288 | 4.14 | 1.00 | 63.76 |  |
| 8 | A | O3' | 1G:116 | 4.18 | 1.00 | 62.33 |  |
| 9 | G | O5' | 1G:117 | 4.34 | 1.00 | 58.76 |  |
| 10 | G | O5' | 1G:289 | 4.36 | 1.00 | 56.08 |  |
| 11 | G | OP1 | 1G:289 | 4.44 | 1.00 | 51.93 |  |
| 12 | A | OP1 | 1G:288 | 4.50 | 1.00 | 71.56 |  |
| 13 | A | O3' | 1G:288 | 4.54 | 1.00 | 53.43 |  |
| 14 | A | O5' | 1G:116 | 4.57 | 1.00 | 53.19 |  |
| 15 | U | O4 | 1G:118 | 4.71 | 1.00 | 62.56 |  |
| 16 | G | O3' | 1G:115 | 4.93 | 1.00 | 67.40 |  |
  |

  
**1^st^ shell ligand angles:**   
1-2: 142;   
  
 Contact(s): /OP2/OP2/  
 Coded contact list: 2O_ph_  
 Exclusion zone contact(s): /OP1/  

### MG:1G:1646

|  | Res. type | Atom | Residue | Distance | Occupancy | B-factor | Closest solute to 1^st^ shell water |
| --- | --- | --- | --- | --- | --- | --- | --- |
| **Ion** | | | |  | 1.00 | 66.33 || 1 | C | OP1 | 1G:1054 | 2.38 | 1.00 | 96.95 |
| 2 | G | OP2 | 1G:1197 | 2.82 | 1.00 | 89.68 |  |
| 3 | C | OP2 | 1G:1054 | 2.83 | 1.00 | 90.59 |  |
| 4 | G | O3' | 1G:1053 | 3.28 | 1.00 | 94.37 |  |
| 5 | G | O2' | 1G:1053 | 3.61 | 1.00 | 91.19 |  |
| 6 | G | N7 | 1G:1198 | 4.28 | 1.00 | 89.88 |  |
| 7 | G | O5' | 1G:1197 | 4.32 | 1.00 | 80.16 |  |
| 8 | C | O5' | 1G:1054 | 4.41 | 1.00 | 96.03 |  |
| 9 | U | O4 | 1G:1199 | 4.45 | 1.00 | 86.15 |  |
| 10 | G | OP1 | 1G:1197 | 4.54 | 1.00 | 86.09 |  |
| 11 | G | OP2 | 1G:1198 | 4.74 | 1.00 | 84.22 |  |
| 12 | U | O2' | 1G:1196 | 4.77 | 1.00 | 105.56 |  |
  |

  
 Contact(s): /OP1/  
 Coded contact list: O_ph_  
 Exclusion zone contact(s): /O3'/OP2/OP2/  
Consider K in place of MG (3 valid contact(s) for K vs. 1 for MG)  

### MG:1G:1647

|  | Res. type | Atom | Residue | Distance | Occupancy | B-factor | Closest solute to 1^st^ shell water |
| --- | --- | --- | --- | --- | --- | --- | --- |
| **Ion** | | | |  | 1.00 | 55.67 || 1 | A | OP2 | 1G:608 | 2.12 | 1.00 | 67.52 |
| 2 | A | N7 | 1G:609 | 3.56 | 1.00 | 64.06 |  |
| 3 | A | O5' | 1G:608 | 3.59 | 1.00 | 65.26 |  |
| 4 | A | OP1 | 1G:607 | 3.84 | 1.00 | 75.09 |  |
| 5 | A | N6 | 1G:609 | 4.16 | 1.00 | 61.88 |  |
| 6 | A | N7 | 1G:608 | 4.22 | 1.00 | 60.21 |  |
| 7 | A | OP1 | 1G:608 | 4.39 | 1.00 | 73.08 |  |
| 8 | A | OP2 | 1G:609 | 4.42 | 1.00 | 67.62 |  |
| 9 | G | O6 | 1G:610 | 4.44 | 1.00 | 67.21 |  |
| 10 | A | O3' | 1G:607 | 4.58 | 1.00 | 76.48 |  |
| 11 | A | OP2 | 1G:607 | 4.97 | 1.00 | 75.70 |  |
  |

  
 Contact(s): /OP2/  
 Coded contact list: O_ph_  

### MG:1G:1648

|  | Res. type | Atom | Residue | Distance | Occupancy | B-factor | Closest solute to 1^st^ shell water |
| --- | --- | --- | --- | --- | --- | --- | --- |
| **Ion** | | | |  | 1.00 | 67.44 || 1 | C | OP2 | 1G:596 | 2.11 | 1.00 | 83.83 |
| 2 | G | OP2 | 1G:597 | 2.15 | 1.00 | 72.58 |  |
| 3 | U | O4 | 1G:598 | 2.47 | 1.00 | 83.85 |  |
| 4 | C | O5' | 1G:596 | 3.54 | 1.00 | 78.43 |  |
| 5 | G | OP1 | 1G:597 | 3.90 | 1.00 | 73.91 |  |
| 6 | G | O2' | 1G:595 | 4.00 | 1.00 | 85.96 |  |
| 7 | G | O3' | 1G:595 | 4.19 | 1.00 | 82.53 |  |
| 8 | A | N6 | 1G:640 | 4.24 | 1.00 | 86.60 |  |
| 9 | G | O5' | 1G:597 | 4.33 | 1.00 | 75.65 |  |
| 10 | C | O3' | 1G:596 | 4.56 | 1.00 | 68.52 |  |
| 11 | C | OP1 | 1G:596 | 4.59 | 1.00 | 70.37 |  |
| 12 | C | N4 | 1G:599 | 4.61 | 1.00 | 83.00 |  |
| 13 | U | N3 | 1G:598 | 4.65 | 1.00 | 74.17 |  |
| 14 | G | N7 | 1G:597 | 4.96 | 1.00 | 79.32 |  |
  |

  
**1^st^ shell ligand angles:**   
1-2: 95;   
  
 Contact(s): /OP2/OP2/  
 Coded contact list: 2O_ph_  
 Exclusion zone contact(s): /O4/  

### MG:1G:1649

|  | Res. type | Atom | Residue | Distance | Occupancy | B-factor | Closest solute to 1^st^ shell water |
| --- | --- | --- | --- | --- | --- | --- | --- |
| **Ion** | | | |  | 1.00 | 74.54 || 1 | G | O6 | 1G:829 | 3.56 | 1.00 | 85.21 |
| 2 | G | O6 | 1G:830 | 3.86 | 1.00 | 87.11 |  |
| 3 | U | OP1 | 1G:871 | 3.98 | 1.00 | 73.46 |  |
| 4 | Water 1 | | 1G:1976 | 4.20 | 1.00 | 73.62 | 4.08 (U:OP1 1G:871) |
| 5 | U | O2' | 1G:870 | 4.20 | 1.00 | 74.48 |  |
| 6 | G | N7 | 1G:829 | 4.26 | 1.00 | 91.72 |  |
| 7 | U | OP1 | 1G:870 | 4.75 | 1.00 | 76.05 |  |
| 8 | U | O5' | 1G:870 | 4.79 | 1.00 | 71.74 |  |
  |

  
 Coded contact list:   
1^st^ MG coordination shell missing (see Notes)  

### MG:1G:1650

|  | Res. type | Atom | Residue | Distance | Occupancy | B-factor | Closest solute to 1^st^ shell water |
| --- | --- | --- | --- | --- | --- | --- | --- |
| **Ion** | | | |  | 1.00 | 68.80 || 1 | A | OP2 | 1G:195 | 2.12 | 1.00 | 83.48 |
| 2 | U | O4 | 1G:180 | 3.60 | 1.00 | 92.98 |  |
| 3 | A | N7 | 1G:179 | 3.93 | 1.00 | 100.92 |  |
| 4 | A | O5' | 1G:195 | 3.95 | 1.00 | 78.47 |  |
| 5 | A | OP1 | 1G:195 | 4.21 | 1.00 | 80.26 |  |
| 6 | A | OP2 | 1G:179 | 4.49 | 1.00 | 93.90 |  |
| 7 | C | O3' | 1G:194 | 4.59 | 1.00 | 82.08 |  |
| 8 | C | OP2 | 1G:194 | 4.61 | 1.00 | 83.51 |  |
| 9 | A | N6 | 1G:179 | 4.70 | 1.00 | 94.68 |  |
| 10 | C | O5' | 1G:194 | 4.72 | 1.00 | 90.88 |  |
| 11 | G | O6 | 1G:181 | 4.74 | 1.00 | 95.57 |  |
| 12 | C | OP1 | 1G:194 | 4.91 | 1.00 | 84.55 |  |
  |

  
 Contact(s): /OP2/  
 Coded contact list: O_ph_  

### MG:1G:1651

|  | Res. type | Atom | Residue | Distance | Occupancy | B-factor | Closest solute to 1^st^ shell water |
| --- | --- | --- | --- | --- | --- | --- | --- |
| **Ion** | | | |  | 1.00 | 43.80 || 1 | G | O6 | 1G:774 | 3.36 | 1.00 | 58.12 |
| 2 | G | OP1 | 1G:803 | 3.58 | 1.00 | 39.29 |  |
| 3 | Water 1 | | 1G:1925 | 3.60 | 1.00 | 55.81 | 3.79 (G:O6 1G:773) |
| 4 | Water 2 | | 1G:1948 | 3.84 | 1.00 | 36.42 | 3.28 (G:OP1 1G:803) |
| 5 | G | O6 | 1G:773 | 3.88 | 1.00 | 59.08 |  |
| 6 | G | N7 | 1G:774 | 4.03 | 1.00 | 54.49 |  |
| 7 | G | O6 | 1G:775 | 4.07 | 1.00 | 60.95 |  |
| 8 | G | N7 | 1G:773 | 4.16 | 1.00 | 50.63 |  |
| 9 | G | OP2 | 1G:803 | 4.59 | 1.00 | 46.25 |  |
  |

  
 Coded contact list:   
 Exclusion zone contact(s): /O6/  
Consider K in place of MG (1 valid contact(s) for K vs. 0 for MG)  

### MG:1G:1652

|  | Res. type | Atom | Residue | Distance | Occupancy | B-factor | Closest solute to 1^st^ shell water |
| --- | --- | --- | --- | --- | --- | --- | --- |
| **Ion** | | | |  | 1.00 | 17.19 || 1 | C | OP1 | 1G:770 | 2.27 | 1.00 | 25.38 |
| 2 | C | O5' | 1G:770 | 3.77 | 1.00 | 38.84 |  |
| 3 | G | O3' | 1G:769 | 3.82 | 1.00 | 35.49 |  |
| 4 | U | OP1 | 14:1833 | 4.03 | 1.00 | 43.08 |  |
| 5 | C | OP1 | 1G:1514 | 4.67 | 1.00 | 48.03 |  |
| 6 | C | OP2 | 1G:770 | 4.75 | 1.00 | 36.18 |  |
  |

  
 Contact(s): /OP1/  
 Coded contact list: O_ph_  

### MG:1G:1653

|  | Res. type | Atom | Residue | Distance | Occupancy | B-factor | Closest solute to 1^st^ shell water |
| --- | --- | --- | --- | --- | --- | --- | --- |
| **Ion** | | | |  | 1.00 | 53.22 || 1 | U | OP1 | 1G:1506 | 2.80 | 1.00 | 60.10 |
| 2 | A | N1 | 1G:794 | 3.67 | 1.00 | 61.98 |  |
| 3 | U | OP2 | 1G:1506 | 4.07 | 1.00 | 63.32 |  |
| 4 | A | N6 | 1G:792 | 4.61 | 1.00 | 64.13 |  |
| 5 | G | O3' | 1G:1505 | 4.75 | 1.00 | 72.20 |  |
| 6 | C | O2 | 1G:795 | 4.82 | 1.00 | 65.23 |  |
| 7 | A | N3 | 1G:794 | 4.86 | 1.00 | 57.82 |  |
| 8 | A | N1 | 1G:792 | 4.99 | 1.00 | 56.01 |  |
  |

  
 Coded contact list:   
 Exclusion zone contact(s): /OP1/  
Consider K in place of MG (1 valid contact(s) for K vs. 0 for MG)  

### MG:1G:1654

|  | Res. type | Atom | Residue | Distance | Occupancy | B-factor | Closest solute to 1^st^ shell water |
| --- | --- | --- | --- | --- | --- | --- | --- |
| **Ion** | | | |  | 1.00 | 50.31 || 1 | A | OP2 | 1G:53 | 2.16 | 1.00 | 63.46 |
| 2 | A | OP1 | 1G:353 | 3.62 | 1.00 | 59.93 |  |
| 3 | A | OP1 | 1G:53 | 3.91 | 1.00 | 68.02 |  |
| 4 | A | N7 | 1G:353 | 4.18 | 1.00 | 61.93 |  |
| 5 | A | O5' | 1G:53 | 4.34 | 1.00 | 51.46 |  |
| 6 | C | O3' | 1G:352 | 4.47 | 1.00 | 59.85 |  |
| 7 | G | O3' | 1G:52 | 4.60 | 1.00 | 51.63 |  |
| 8 | G | O4' | 1G:316 | 4.73 | 1.00 | 73.68 |  |
  |

  
 Contact(s): /OP2/  
 Coded contact list: O_ph_  

### MG:1G:1655

|  | Res. type | Atom | Residue | Distance | Occupancy | B-factor | Closest solute to 1^st^ shell water |
| --- | --- | --- | --- | --- | --- | --- | --- |
| **Ion** | | | |  | 1.00 | 58.86 || 1 | U | OP2 | 1G:560 | 2.23 | 1.00 | 69.85 |
| 2 | C | OP1 | 1G:562 | 3.24 | 1.00 | 59.55 |  |
| 3 | Water 1 | | 1G:1952 | 3.28 | 1.00 | 64.59 | 3.39 (G:OP2 1G:558) |
| 4 | G | N2 | 1G:566 | 3.87 | 1.00 | 62.02 |  |
| 5 | G | OP1 | 1G:557 | 3.93 | 1.00 | 52.08 |  |
| 6 | A | O3' | 1G:559 | 3.94 | 1.00 | 60.07 |  |
| 7 | U | OP1 | 1G:560 | 4.34 | 1.00 | 65.51 |  |
| 8 | A | OP1 | 1G:563 | 4.49 | 1.00 | 58.34 |  |
| 9 | G | N1 | 1G:566 | 4.53 | 1.00 | 64.92 |  |
| 10 | G | OP2 | 1G:558 | 4.53 | 1.00 | 62.36 |  |
| 11 | U | O5' | 1G:560 | 4.67 | 1.00 | 62.61 |  |
  |

  
 Contact(s): /OP2/  
 Coded contact list: O_ph_  
 Exclusion zone contact(s): /O_w_/OP1/  

### MG:1G:1656

|  | Res. type | Atom | Residue | Distance | Occupancy | B-factor | Closest solute to 1^st^ shell water |
| --- | --- | --- | --- | --- | --- | --- | --- |
| **Ion** | | | |  | 1.00 | 58.77 || 1 | GLY | O | 42:85 | 3.53 | 1.00 | 71.74 |
| 2 | A | OP2 | 1G:864 | 3.57 | 1.00 | 67.40 |  |
| 3 | U | OP1 | 1G:863 | 3.67 | 1.00 | 74.93 |  |
| 4 | GLU | OE1 | 42:83 | 3.99 | 1.00 | 75.47 |  |
| 5 | U | O5' | 1G:863 | 4.70 | 1.00 | 70.29 |  |
  |

  
 Coded contact list:   
1^st^ MG coordination shell missing (see Notes)  

### MG:1G:1657

|  | Res. type | Atom | Residue | Distance | Occupancy | B-factor | Closest solute to 1^st^ shell water |
| --- | --- | --- | --- | --- | --- | --- | --- |
| **Ion** | | | |  | 1.00 | 53.61 || 1 | G | O6 | 1G:650 | 3.30 | 1.00 | 94.10 |
| 2 | G | O6 | 1G:649 | 3.76 | 1.00 | 89.90 |  |
| 3 | A | OP1 | 1G:753 | 4.20 | 1.00 | 76.76 |  |
| 4 | G | N7 | 1G:650 | 4.32 | 1.00 | 91.75 |  |
| 5 | G | N7 | 1G:649 | 4.46 | 1.00 | 97.46 |  |
| 6 | A | OP2 | 1G:753 | 4.71 | 1.00 | 82.59 |  |
  |

  
 Coded contact list:   
 Exclusion zone contact(s): /O6/  
Consider K in place of MG (1 valid contact(s) for K vs. 0 for MG)  

### MG:1G:1658

|  | Res. type | Atom | Residue | Distance | Occupancy | B-factor | Closest solute to 1^st^ shell water |
| --- | --- | --- | --- | --- | --- | --- | --- |
| **Ion** | | | |  | 1.00 | 46.68 || 1 | G | O2' | 1G:502 | 3.54 | 1.00 | 66.10 |
| 2 | G | OP1 | 1G:550 | 3.58 | 1.00 | 51.33 |  |
| 3 | C | O4' | 1G:503 | 4.04 | 1.00 | 63.08 |  |
| 4 | G | N3 | 1G:502 | 4.10 | 1.00 | 70.50 |  |
| 5 | G | O5' | 1G:550 | 4.60 | 1.00 | 56.53 |  |
| 6 | G | N2 | 1G:502 | 4.64 | 1.00 | 68.82 |  |
  |

  
 Coded contact list:   
1^st^ MG coordination shell missing (see Notes)  

### MG:1G:1659

|  | Res. type | Atom | Residue | Distance | Occupancy | B-factor | Closest solute to 1^st^ shell water |
| --- | --- | --- | --- | --- | --- | --- | --- |
| **Ion** | | | |  | 1.00 | 64.97 || 1 | G | O6 | 1G:628 | 3.10 | 1.00 | 76.81 |
| 2 | G | O6 | 1G:627 | 3.87 | 1.00 | 85.90 |  |
| 3 | G | OP2 | 1G:610 | 3.93 | 1.00 | 68.69 |  |
| 4 | A | OP1 | 1G:609 | 4.27 | 1.00 | 72.61 |  |
| 5 | G | N7 | 1G:627 | 4.48 | 1.00 | 89.66 |  |
| 6 | C | N4 | 1G:612 | 4.78 | 1.00 | 78.56 |  |
  |

  
 Coded contact list:   
 Exclusion zone contact(s): /O6/  
Consider K in place of MG (1 valid contact(s) for K vs. 0 for MG)  

### MG:1G:1660

|  | Res. type | Atom | Residue | Distance | Occupancy | B-factor | Closest solute to 1^st^ shell water |
| --- | --- | --- | --- | --- | --- | --- | --- |
| **Ion** | | | |  | 1.00 | 46.01 || 1 | G | N7 | 1G:41 | 3.66 | 1.00 | 62.68 |
| 2 | G | O6 | 1G:41 | 3.79 | 1.00 | 64.39 |  |
| 3 | G | O6 | 1G:42 | 4.57 | 1.00 | 59.12 |  |
| 4 | A | O4' | 1G:397 | 4.97 | 1.00 | 61.86 |  |
  |

  
 Coded contact list:   
1^st^ MG coordination shell missing (see Notes)  

### MG:1G:1661

|  | Res. type | Atom | Residue | Distance | Occupancy | B-factor | Closest solute to 1^st^ shell water |
| --- | --- | --- | --- | --- | --- | --- | --- |
| **Ion** | | | |  | 1.00 | 79.11 || 1 | G | O6 | 1G:447 | 3.46 | 1.00 | 100.18 |
| 2 | G | N7 | 1G:445 | 3.81 | 1.00 | 103.22 |  |
| 3 | G | O6 | 1G:445 | 3.90 | 1.00 | 91.07 |  |
| 4 | G | N7 | 1G:446 | 4.12 | 1.00 | 97.42 |  |
| 5 | U | OP2 | 1G:486 | 4.18 | 1.00 | 95.78 |  |
| 6 | G | O6 | 1G:446 | 4.29 | 1.00 | 87.99 |  |
| 7 | G | N1 | 1G:447 | 4.47 | 1.00 | 97.32 |  |
  |

  
 Coded contact list:   
1^st^ MG coordination shell missing (see Notes)  

### MG:1G:1662

|  | Res. type | Atom | Residue | Distance | Occupancy | B-factor | Closest solute to 1^st^ shell water |
| --- | --- | --- | --- | --- | --- | --- | --- |
| **Ion** | | | |  | 1.00 | 51.46 || 1 | G | OP1 | 1G:21 | 2.01 | 1.00 | 52.54 |
| 2 | G | OP2 | 1G:21 | 3.51 | 1.00 | 61.80 |  |
| 3 | G | N2 | 1G:567 | 3.82 | 1.00 | 63.84 |  |
| 4 | G | O2' | 1G:567 | 3.85 | 1.00 | 64.87 |  |
| 5 | G | N3 | 1G:567 | 3.99 | 1.00 | 63.53 |  |
| 6 | G | O2' | 1G:568 | 4.00 | 1.00 | 55.65 |  |
| 7 | U | O3' | 1G:20 | 4.00 | 1.00 | 63.04 |  |
| 8 | G | O5' | 1G:21 | 4.47 | 1.00 | 54.71 |  |
| 9 | A | N6 | 1G:573 | 4.49 | 1.00 | 63.06 |  |
| 10 | A | N1 | 1G:573 | 4.56 | 1.00 | 61.23 |  |
| 11 | A | N6 | 1G:563 | 4.70 | 1.00 | 58.69 |  |
  |

  
 Contact(s): /OP1/  
 Coded contact list: O_ph_  

### MG:1G:1663

|  | Res. type | Atom | Residue | Distance | Occupancy | B-factor | Closest solute to 1^st^ shell water |
| --- | --- | --- | --- | --- | --- | --- | --- |
| **Ion** | | | |  | 0.50 | 29.58 || 1 | A | OP2 | 1G:814 | 2.91 | 1.00 | 42.90 |
| 2 | C | OP1 | 1G:904 | 3.89 | 1.00 | 47.54 |  |
| 3 | G | OP1 | 1G:577 | 4.00 | 1.00 | 52.20 |  |
| 4 | U | O5' | 1G:813 | 4.21 | 1.00 | 45.14 |  |
| 5 | A | OP1 | 1G:814 | 4.23 | 1.00 | 47.67 |  |
| 6 | A | N6 | 1G:816 | 4.32 | 1.00 | 49.02 |  |
| 7 | G | OP2 | 1G:577 | 4.56 | 1.00 | 48.92 |  |
| 8 | U | OP2 | 1G:813 | 4.70 | 1.00 | 41.73 |  |
| 9 | U | O3' | 1G:813 | 4.73 | 1.00 | 55.43 |  |
  |

  
 Coded contact list:   
 Exclusion zone contact(s): /OP2/  
Consider K in place of MG (1 valid contact(s) for K vs. 0 for MG)  

### MG:1G:1663

|  | Res. type | Atom | Residue | Distance | Occupancy | B-factor | Closest solute to 1^st^ shell water |
| --- | --- | --- | --- | --- | --- | --- | --- |
| **Ion** | | | |  | 0.50 | 30.72 || 1 | U | OP2 | 1G:905 | 3.72 | 1.00 | 46.07 |
| 2 | C | OP1 | 1G:904 | 3.83 | 1.00 | 47.54 |  |
| 3 | U | OP1 | 1G:905 | 4.29 | 1.00 | 55.18 |  |
| 4 | C | O3' | 1G:904 | 4.50 | 1.00 | 53.81 |  |
| 5 | A | OP2 | 1G:814 | 4.74 | 1.00 | 42.90 |  |
| 6 | C | O5' | 1G:904 | 4.76 | 1.00 | 45.21 |  |
| 7 | A | OP1 | 1G:814 | 4.99 | 1.00 | 47.67 |  |
  |

  
 Coded contact list:   
1^st^ MG coordination shell missing (see Notes)  

### MG:1G:1664

|  | Res. type | Atom | Residue | Distance | Occupancy | B-factor | Closest solute to 1^st^ shell water |
| --- | --- | --- | --- | --- | --- | --- | --- |
| **Ion** | | | |  | 1.00 | 40.37 || 1 | A | OP2 | 1G:915 | 3.91 | 1.00 | 67.06 |
| 2 | U | O2' | 1G:13 | 4.01 | 1.00 | 68.71 |  |
| 3 | G | OP2 | 1G:916 | 4.03 | 1.00 | 50.75 |  |
| 4 | G | N7 | 1G:916 | 4.12 | 1.00 | 63.73 |  |
  |

  
 Coded contact list:   
1^st^ MG coordination shell missing (see Notes)  

### MG:1G:1665

|  | Res. type | Atom | Residue | Distance | Occupancy | B-factor | Closest solute to 1^st^ shell water |
| --- | --- | --- | --- | --- | --- | --- | --- |
| **Ion** | | | |  | 1.00 | 50.20 || 1 | C | OP2 | 14:1832 | 3.82 | 1.00 | 38.55 |
| 2 | G | OP1 | 1G:771 | 3.99 | 1.00 | 42.18 |  |
| 3 | U | OP1 | 1G:772 | 4.35 | 1.00 | 47.65 |  |
| 4 | U | OP2 | 1G:772 | 4.66 | 1.00 | 51.31 |  |
  |

  
 Coded contact list:   
1^st^ MG coordination shell missing (see Notes)  

### MG:1G:1666

|  | Res. type | Atom | Residue | Distance | Occupancy | B-factor | Closest solute to 1^st^ shell water |
| --- | --- | --- | --- | --- | --- | --- | --- |
| **Ion** | | | |  | 1.00 | 46.58 || 1 | U | O4 | 1G:45 | 3.62 | 1.00 | 59.58 |
| 2 | C | N4 | 1G:43 | 3.69 | 1.00 | 62.38 |  |
| 3 | G | O6 | 1G:44 | 3.89 | 1.00 | 56.72 |  |
| 4 | G | O6 | 1G:396 | 4.12 | 1.00 | 52.98 |  |
| 5 | G | N7 | 1G:44 | 4.14 | 1.00 | 56.81 |  |
| 6 | C | N4 | 1G:395 | 4.18 | 1.00 | 52.92 |  |
| 7 | U | N3 | 1G:45 | 4.76 | 1.00 | 51.34 |  |
  |

  
 Coded contact list:   
1^st^ MG coordination shell missing (see Notes)  

### MG:1G:1667

|  | Res. type | Atom | Residue | Distance | Occupancy | B-factor | Closest solute to 1^st^ shell water |
| --- | --- | --- | --- | --- | --- | --- | --- |
| **Ion** | | | |  | 1.00 | 42.81 || 1 | U | OP2 | 1G:813 | 3.03 | 1.00 | 41.73 |
| 2 | U | OP1 | 1G:813 | 3.80 | 1.00 | 48.30 |  |
| 3 | G | N7 | 1G:577 | 3.81 | 1.00 | 51.57 |  |
| 4 | G | O6 | 1G:577 | 4.05 | 1.00 | 51.56 |  |
| 5 | G | OP1 | 1G:577 | 4.24 | 1.00 | 52.20 |  |
| 6 | U | O5' | 1G:813 | 4.91 | 1.00 | 45.14 |  |
| 7 | A | N6 | 1G:816 | 4.94 | 1.00 | 49.02 |  |
  |

  
 Coded contact list:   
 Exclusion zone contact(s): /OP2/  
Consider K in place of MG (1 valid contact(s) for K vs. 0 for MG)  

### MG:1G:1668

|  | Res. type | Atom | Residue | Distance | Occupancy | B-factor | Closest solute to 1^st^ shell water |
| --- | --- | --- | --- | --- | --- | --- | --- |
| **Ion** | | | |  | 1.00 | 69.14 || 1 | G | OP2 | 1G:750 | 2.28 | 1.00 | 90.73 |
| 2 | C | OP1 | 1G:749 | 3.06 | 1.00 | 93.47 |  |
| 3 | C | OP2 | 1G:749 | 3.22 | 1.00 | 96.81 |  |
| 4 | G | OP1 | 1G:750 | 4.37 | 1.00 | 84.86 |  |
| 5 | C | O2' | 1G:748 | 4.42 | 1.00 | 88.88 |  |
| 6 | G | O5' | 1G:750 | 4.49 | 1.00 | 88.62 |  |
| 7 | C | O3' | 1G:748 | 4.53 | 1.00 | 98.71 |  |
| 8 | G | O6 | 1G:752 | 4.67 | 1.00 | 86.72 |  |
| 9 | C | O3' | 1G:749 | 4.68 | 1.00 | 90.20 |  |
| 10 | G | N7 | 1G:750 | 4.76 | 1.00 | 88.92 |  |
| 11 | C | O5' | 1G:749 | 4.96 | 1.00 | 90.36 |  |
  |

  
 Contact(s): /OP2/  
 Coded contact list: O_ph_  
 Exclusion zone contact(s): /OP1/OP2/  
Consider K in place of MG (2 valid contact(s) for K vs. 1 for MG)  

### MG:1G:1669

|  | Res. type | Atom | Residue | Distance | Occupancy | B-factor | Closest solute to 1^st^ shell water |
| --- | --- | --- | --- | --- | --- | --- | --- |
| **Ion** | | | |  | 1.00 | 60.17 || 1 | G | O6 | 1G:492 | 3.81 | 1.00 | 87.16 |
| 2 | G | N7 | 1G:491 | 4.13 | 1.00 | 94.01 |  |
| 3 | G | O6 | 1G:491 | 4.23 | 1.00 | 96.22 |  |
| 4 | G | N7 | 1G:492 | 4.44 | 1.00 | 87.69 |  |
| 5 | G | O6 | 1G:493 | 4.69 | 1.00 | 84.55 |  |
| 6 | Water 1 | | 32:403 | 4.80 | 1.00 | 83.22 | 3.80 (G:O6 1G:493) |
  |

  
 Coded contact list:   
1^st^ MG coordination shell missing (see Notes)  

### MG:1G:1670

|  | Res. type | Atom | Residue | Distance | Occupancy | B-factor | Closest solute to 1^st^ shell water |
| --- | --- | --- | --- | --- | --- | --- | --- |
| **Ion** | | | |  | 1.00 | 57.10 || 1 | A | OP2 | 1G:1468 | 3.19 | 1.00 | 71.35 |
| 2 | G | O6 | 1G:1469 | 3.30 | 1.00 | 65.24 |  |
| 3 | G | N2 | 1G:1432 | 3.42 | 1.00 | 66.50 |  |
| 4 | Water 1 | | 1G:1933 | 3.46 | 1.00 | 74.79 | 3.13 (G:O6 1G:1469) |
| 5 | G | N1 | 1G:1432 | 4.02 | 1.00 | 75.88 |  |
| 6 | G | OP2 | 1G:1467 | 4.28 | 1.00 | 68.70 |  |
| 7 | G | O5' | 1G:1467 | 4.35 | 1.00 | 62.03 |  |
| 8 | A | OP1 | 1G:1468 | 4.37 | 1.00 | 68.84 |  |
| 9 | Water 2 | | 1G:1829 | 4.83 | 1.00 | 61.76 | 2.31 (G:N7 1G:1469) |
| 10 | G | N7 | 1G:1469 | 4.90 | 1.00 | 65.90 |  |
| 11 | G | O3' | 1G:1467 | 4.91 | 1.00 | 63.93 |  |
| 12 | G | OP1 | 1G:1467 | 4.92 | 1.00 | 60.59 |  |
  |

  
 Coded contact list:   
 Exclusion zone contact(s): /O6/OP2/  
Consider K in place of MG (2 valid contact(s) for K vs. 0 for MG)  

### MG:1G:1671

|  | Res. type | Atom | Residue | Distance | Occupancy | B-factor | Closest solute to 1^st^ shell water |
| --- | --- | --- | --- | --- | --- | --- | --- |
| **Ion** | | | |  | 1.00 | 46.87 || 1 | G | N7 | 1G:392 | 3.69 | 1.00 | 69.62 |
| 2 | G | O6 | 1G:391 | 3.75 | 1.00 | 68.13 |  |
| 3 | G | N7 | 1G:391 | 3.88 | 1.00 | 68.91 |  |
| 4 | G | O6 | 1G:392 | 4.40 | 1.00 | 63.72 |  |
| 5 | G | OP2 | 1G:391 | 4.90 | 1.00 | 65.02 |  |
  |

  
 Coded contact list:   
1^st^ MG coordination shell missing (see Notes)  

### MG:1G:1672

|  | Res. type | Atom | Residue | Distance | Occupancy | B-factor | Closest solute to 1^st^ shell water |
| --- | --- | --- | --- | --- | --- | --- | --- |
| **Ion** | | | |  | 1.00 | 66.82 || 1 | G | O6 | 1G:1441 | 2.09 | 1.00 | 68.82 |
| 2 | G | OP1 | 1G:1455 | 4.11 | 1.00 | 79.56 |  |
| 3 | G | N1 | 1G:1441 | 4.16 | 1.00 | 80.59 |  |
| 4 | G | N7 | 1G:1441 | 4.57 | 1.00 | 81.48 |  |
| 5 | G | OP2 | 1G:1455 | 4.65 | 1.00 | 81.86 |  |
| 6 | C | N4 | 1G:1440 | 4.72 | 1.00 | 79.04 |  |
| 7 | LYS | NZ | BA:38 | 4.96 | 1.00 | 75.65 |  |
  |

  
 Contact(s): /O6/  
 Coded contact list: O_b_  

### MG:1G:1673

|  | Res. type | Atom | Residue | Distance | Occupancy | B-factor | Closest solute to 1^st^ shell water |
| --- | --- | --- | --- | --- | --- | --- | --- |
| **Ion** | | | |  | 1.00 | 50.93 || 1 | U | OP2 | 1G:891 | 2.07 | 1.00 | 50.67 |
| 2 | G | N1 | 1G:906 | 3.69 | 1.00 | 45.41 |  |
| 3 | G | N2 | 1G:906 | 3.74 | 1.00 | 43.14 |  |
| 4 | U | O5' | 1G:891 | 3.74 | 1.00 | 47.55 |  |
| 5 | U | O4 | 1G:905 | 3.85 | 1.00 | 41.26 |  |
| 6 | U | OP1 | 1G:891 | 4.09 | 1.00 | 50.37 |  |
| 7 | A | OP2 | 1G:892 | 4.23 | 1.00 | 47.14 |  |
| 8 | G | O2' | 1G:890 | 4.49 | 1.00 | 54.77 |  |
| 9 | G | O3' | 1G:890 | 4.60 | 1.00 | 47.73 |  |
| 10 | C | N4 | 1G:904 | 4.63 | 1.00 | 49.42 |  |
| 11 | G | OP2 | 1G:903 | 4.80 | 1.00 | 48.52 |  |
| 12 | G | O6 | 1G:906 | 4.83 | 1.00 | 49.19 |  |
  |

  
 Contact(s): /OP2/  
 Coded contact list: O_ph_  

### MG:1G:1674

|  | Res. type | Atom | Residue | Distance | Occupancy | B-factor | Closest solute to 1^st^ shell water |
| --- | --- | --- | --- | --- | --- | --- | --- |
| **Ion** | | | |  | 1.00 | 122.50 || 1 | C | OP1 | 1G:466 | 2.57 | 1.00 | 144.29 |
| 2 | G | OP2 | 1G:464 | 2.86 | 1.00 | 127.91 |  |
| 3 | C | OP2 | 1G:466 | 3.05 | 1.00 | 136.20 |  |
| 4 | A | O3' | 1G:465 | 4.03 | 1.00 | 131.93 |  |
| 5 | C | OP2 | 1G:458 | 4.34 | 1.00 | 130.05 |  |
| 6 | C | O5' | 1G:458 | 4.52 | 1.00 | 130.20 |  |
| 7 | G | N7 | 1G:464 | 4.52 | 1.00 | 131.43 |  |
| 8 | U | N3 | 1G:208 | 4.63 | 1.00 | 161.82 |  |
| 9 | C | O5' | 1G:466 | 4.65 | 1.00 | 140.79 |  |
| 10 | G | O5' | 1G:464 | 4.69 | 1.00 | 127.23 |  |
| 11 | C | OP1 | 1G:458 | 4.71 | 1.00 | 132.02 |  |
| 12 | A | O5' | 1G:465 | 4.72 | 1.00 | 141.80 |  |
| 13 | U | O4 | 1G:208 | 4.93 | 1.00 | 157.60 |  |
  |

  
 Coded contact list:   
 Exclusion zone contact(s): /OP1/OP2/OP2/  
Consider K in place of MG (2 valid contact(s) for K vs. 0 for MG)  

### MG:1G:1675

|  | Res. type | Atom | Residue | Distance | Occupancy | B-factor | Closest solute to 1^st^ shell water |
| --- | --- | --- | --- | --- | --- | --- | --- |
| **Ion** | | | |  | 1.00 | 65.15 || 1 | G | O6 | 1G:660 | 2.97 | 1.00 | 91.18 |
| 2 | G | O6 | 1G:661 | 3.75 | 1.00 | 84.72 |  |
| 3 | G | N7 | 1G:660 | 3.97 | 1.00 | 90.29 |  |
| 4 | G | N7 | 1G:661 | 4.02 | 1.00 | 93.76 |  |
| 5 | G | N1 | 1G:660 | 4.80 | 1.00 | 90.82 |  |
  |

  
 Coded contact list:   
 Exclusion zone contact(s): /O6/  
Consider K in place of MG (1 valid contact(s) for K vs. 0 for MG)  

### MG:1G:1676

|  | Res. type | Atom | Residue | Distance | Occupancy | B-factor | Closest solute to 1^st^ shell water |
| --- | --- | --- | --- | --- | --- | --- | --- |
| **Ion** | | | |  | 1.00 | 71.49 || 1 | U | OP2 | 1G:1095 | 2.23 | 1.00 | 88.15 |
| 2 | G | O6 | 1G:1108 | 2.34 | 1.00 | 81.23 |  |
| 3 | U | O5' | 1G:1095 | 3.63 | 1.00 | 91.45 |  |
| 4 | C | OP2 | 1G:1096 | 3.63 | 1.00 | 102.29 |  |
| 5 | G | N1 | 1G:1108 | 3.67 | 1.00 | 89.65 |  |
| 6 | G | O2' | 1G:1094 | 3.73 | 1.00 | 101.36 |  |
| 7 | U | OP1 | 1G:1095 | 4.47 | 1.00 | 89.95 |  |
| 8 | C | N3 | 1G:1109 | 4.64 | 1.00 | 92.78 |  |
| 9 | G | O3' | 1G:1094 | 4.70 | 1.00 | 89.44 |  |
| 10 | C | N4 | 1G:1109 | 4.86 | 1.00 | 95.52 |  |
| 11 | C | OP1 | 1G:1096 | 4.91 | 1.00 | 95.12 |  |
| 12 | U | O3' | 1G:1095 | 4.91 | 1.00 | 93.16 |  |
| 13 | G | O6 | 1G:1068 | 4.98 | 1.00 | 91.97 |  |
  |

  
**1^st^ shell ligand angles:**   
1-2: 83;   
  
 Contact(s): /O6/OP2/  
 Coded contact list: O_ph_O_b_  

### MG:1G:1677

|  | Res. type | Atom | Residue | Distance | Occupancy | B-factor | Closest solute to 1^st^ shell water |
| --- | --- | --- | --- | --- | --- | --- | --- |
| **Ion** | | | |  | 1.00 | 57.14 || 1 | G | N7 | 1G:869 | 2.61 | 1.00 | 76.29 |
| 2 | G | N7 | 1G:858 | 3.00 | 1.00 | 79.54 |  |
| 3 | C | OP2 | 1G:868 | 3.07 | 1.00 | 70.89 |  |
| 4 | G | OP2 | 1G:869 | 3.32 | 1.00 | 72.06 |  |
| 5 | G | O6 | 1G:858 | 3.79 | 1.00 | 73.53 |  |
| 6 | C | O5' | 1G:868 | 4.00 | 1.00 | 67.75 |  |
| 7 | G | O6 | 1G:869 | 4.22 | 1.00 | 81.84 |  |
| 8 | C | OP2 | 1G:857 | 4.26 | 1.00 | 78.15 |  |
| 9 | G | OP2 | 1G:858 | 4.31 | 1.00 | 85.81 |  |
| 10 | C | OP1 | 1G:868 | 4.38 | 1.00 | 62.25 |  |
| 11 | G | N9 | 1G:869 | 4.65 | 1.00 | 74.53 |  |
| 12 | G | O5' | 1G:869 | 4.95 | 1.00 | 64.92 |  |
  |

  
 Coded contact list:   
 Exclusion zone contact(s): /N7/N7/OP2/OP2/  
Consider K in place of MG (4 valid contact(s) for K vs. 0 for MG)  

### MG:1G:1678

|  | Res. type | Atom | Residue | Distance | Occupancy | B-factor | Closest solute to 1^st^ shell water |
| --- | --- | --- | --- | --- | --- | --- | --- |
| **Ion** | | | |  | 1.00 | 54.04 || 1 | G | O6 | 1G:126 | 3.26 | 1.00 | 76.07 |
| 2 | U | O4 | 1G:125 | 3.31 | 1.00 | 76.28 |  |
| 3 | G | N7 | 1G:126 | 4.14 | 1.00 | 66.58 |  |
| 4 | G | O6 | 1G:127 | 4.34 | 1.00 | 70.67 |  |
| 5 | C | N4 | 1G:235 | 4.58 | 1.00 | 67.96 |  |
| 6 | G | N7 | 1G:127 | 4.90 | 1.00 | 70.76 |  |
| 7 | U | N3 | 1G:125 | 5.00 | 1.00 | 72.15 |  |
  |

  
 Coded contact list:   
 Exclusion zone contact(s): /O4/O6/  
Consider K in place of MG (2 valid contact(s) for K vs. 0 for MG)  

### MG:1G:1679

|  | Res. type | Atom | Residue | Distance | Occupancy | B-factor | Closest solute to 1^st^ shell water |
| --- | --- | --- | --- | --- | --- | --- | --- |
| **Ion** | | | |  | 1.00 | 62.94 || 1 | G | O6 | 1G:741 | 2.58 | 1.00 | 67.13 |
| 2 | G | O6 | 1G:664 | 3.20 | 1.00 | 72.89 |  |
| 3 | G | N1 | 1G:664 | 3.34 | 1.00 | 70.62 |  |
| 4 | G | O6 | 1G:666 | 3.63 | 1.00 | 67.93 |  |
| 5 | U | O4 | 1G:740 | 4.12 | 1.00 | 69.19 |  |
| 6 | G | N7 | 1G:741 | 4.12 | 1.00 | 67.32 |  |
| 7 | G | O6 | 1G:742 | 4.54 | 1.00 | 73.47 |  |
| 8 | A | N6 | 1G:663 | 4.62 | 1.00 | 73.86 |  |
| 9 | G | N2 | 1G:664 | 4.67 | 1.00 | 66.37 |  |
| 10 | G | N1 | 1G:741 | 4.85 | 1.00 | 63.78 |  |
| 11 | U | N3 | 1G:740 | 4.86 | 1.00 | 65.87 |  |
  |

  
 Coded contact list:   
 Exclusion zone contact(s): /N1_G_/O6/O6/  
Consider K in place of MG (2 valid contact(s) for K vs. 0 for MG)  

### MG:1G:1680

|  | Res. type | Atom | Residue | Distance | Occupancy | B-factor | Closest solute to 1^st^ shell water |
| --- | --- | --- | --- | --- | --- | --- | --- |
| **Ion** | | | |  | 1.00 | 50.93 || 1 | G | OP1 | 1G:292 | 2.72 | 1.00 | 55.16 |
| 2 | G | OP2 | 1G:292 | 3.84 | 1.00 | 47.01 |  |
| 3 | G | N7 | 1G:293 | 3.92 | 1.00 | 54.36 |  |
| 4 | C | O3' | 1G:291 | 4.75 | 1.00 | 55.23 |  |
| 5 | G | O5' | 1G:292 | 4.88 | 1.00 | 52.61 |  |
  |

  
 Coded contact list:   
 Exclusion zone contact(s): /OP1/  
Consider K in place of MG (1 valid contact(s) for K vs. 0 for MG)  

### MG:1G:1681

|  | Res. type | Atom | Residue | Distance | Occupancy | B-factor | Closest solute to 1^st^ shell water |
| --- | --- | --- | --- | --- | --- | --- | --- |
| **Ion** | | | |  | 1.00 | 45.14 || 1 | G | OP2 | 1G:579 | 3.18 | 1.00 | 64.43 |
| 2 | G | N7 | 1G:579 | 3.89 | 1.00 | 64.11 |  |
| 3 | U | O4 | 1G:580 | 4.90 | 1.00 | 68.57 |  |
  |

  
 Coded contact list:   
 Exclusion zone contact(s): /OP2/  
Consider K in place of MG (1 valid contact(s) for K vs. 0 for MG)  

### MG:1G:1682

|  | Res. type | Atom | Residue | Distance | Occupancy | B-factor | Closest solute to 1^st^ shell water |
| --- | --- | --- | --- | --- | --- | --- | --- |
| **Ion** | | | |  | 1.00 | 41.27 || 1 | G | O6 | 1G:357 | 3.22 | 1.00 | 60.71 |
| 2 | C | O2 | 1G:352 | 3.50 | 1.00 | 71.20 |  |
| 3 | A | N7 | 1G:356 | 3.63 | 1.00 | 66.68 |  |
| 4 | G | N7 | 1G:357 | 3.67 | 1.00 | 67.21 |  |
| 5 | C | OP2 | 1G:355 | 4.10 | 1.00 | 55.73 |  |
| 6 | C | O2' | 1G:352 | 4.98 | 1.00 | 59.07 |  |
  |

  
 Coded contact list:   
 Exclusion zone contact(s): /O6/  
Consider K in place of MG (1 valid contact(s) for K vs. 0 for MG)  

### MG:1G:1683

|  | Res. type | Atom | Residue | Distance | Occupancy | B-factor | Closest solute to 1^st^ shell water |
| --- | --- | --- | --- | --- | --- | --- | --- |
| **Ion** | | | |  | 0.50 | 51.33 || 1 | M2G | N7 | 1G:966 | 3.63 | 1.00 | 67.31 |
| 2 | M2G | OP2 | 1G:966 | 4.09 | 1.00 | 77.56 |  |
| 3 | M2G | OP1 | 1G:966 | 4.35 | 1.00 | 77.51 |  |
| 4 | M2G | O6 | 1G:966 | 4.73 | 1.00 | 68.03 |  |
  |

  
 Coded contact list:   
1^st^ MG coordination shell missing (see Notes)  

### MG:1G:1683

|  | Res. type | Atom | Residue | Distance | Occupancy | B-factor | Closest solute to 1^st^ shell water |
| --- | --- | --- | --- | --- | --- | --- | --- |
| **Ion** | | | |  | 0.50 | 45.69 || 1 | M2G | O6 | 1G:966 | 3.18 | 1.00 | 68.03 |
| 2 | 5MC | N3 | 1G:1400 | 3.91 | 1.00 | 71.70 |  |
| 3 | M2G | N7 | 1G:966 | 4.08 | 1.00 | 67.31 |  |
| 4 | 5MC | N4 | 1G:1400 | 4.60 | 1.00 | 72.54 |  |
| 5 | 5MC | N4 | 1G:967 | 4.65 | 1.00 | 68.49 |  |
| 6 | 5MC | O2 | 1G:1400 | 4.77 | 1.00 | 67.23 |  |
  |

  
 Coded contact list:   
 Exclusion zone contact(s): /O6/  
Consider K in place of MG (1 valid contact(s) for K vs. 0 for MG)  

### MG:1G:1684

|  | Res. type | Atom | Residue | Distance | Occupancy | B-factor | Closest solute to 1^st^ shell water |
| --- | --- | --- | --- | --- | --- | --- | --- |
| **Ion** | | | |  | 1.00 | 79.08 || 1 | G | OP2 | 1G:1334 | 2.10 | 1.00 | 89.31 |
| 2 | G | N7 | 1G:1300 | 3.20 | 1.00 | 102.78 |  |
| 3 | G | O6 | 1G:1300 | 3.88 | 1.00 | 103.50 |  |
| 4 | G | OP1 | 1G:1334 | 3.98 | 1.00 | 92.54 |  |
| 5 | A | OP2 | 1G:1333 | 4.07 | 1.00 | 96.97 |  |
| 6 | A | OP1 | 1G:1333 | 4.16 | 1.00 | 110.81 |  |
| 7 | G | N7 | 1G:1334 | 4.19 | 1.00 | 94.10 |  |
| 8 | G | O5' | 1G:1334 | 4.22 | 1.00 | 85.03 |  |
| 9 | A | O3' | 1G:1333 | 4.61 | 1.00 | 88.79 |  |
| 10 | A | O5' | 1G:1333 | 4.94 | 1.00 | 96.30 |  |
  |

  
 Contact(s): /OP2/  
 Coded contact list: O_ph_  
 Exclusion zone contact(s): /N7/  

### MG:1G:1685

|  | Res. type | Atom | Residue | Distance | Occupancy | B-factor | Closest solute to 1^st^ shell water |
| --- | --- | --- | --- | --- | --- | --- | --- |
| **Ion** | | | |  | 1.00 | 51.49 || 1 | G | N7 | 1G:362 | 2.84 | 1.00 | 65.08 |
| 2 | G | O6 | 1G:362 | 3.01 | 1.00 | 77.48 |  |
| 3 | G | OP2 | 1G:361 | 4.52 | 1.00 | 68.20 |  |
| 4 | G | N7 | 1G:361 | 4.78 | 1.00 | 66.69 |  |
| 5 | G | N1 | 1G:362 | 4.99 | 1.00 | 68.44 |  |
  |

  
 Coded contact list:   
 Exclusion zone contact(s): /N7/O6/  
Consider K in place of MG (2 valid contact(s) for K vs. 0 for MG)  

### MG:1G:1686

|  | Res. type | Atom | Residue | Distance | Occupancy | B-factor | Closest solute to 1^st^ shell water |
| --- | --- | --- | --- | --- | --- | --- | --- |
| **Ion** | | | |  | 1.00 | 75.73 || 1 | G | O2' | 1G:976 | 3.12 | 1.00 | 111.29 |
| 2 | A | OP1 | 1G:1363 | 3.81 | 1.00 | 95.96 |  |
| 3 | A | N7 | 1G:1363 | 3.82 | 1.00 | 100.87 |  |
| 4 | A | OP2 | 1G:975 | 3.89 | 1.00 | 79.62 |  |
| 5 | A | N6 | 1G:1363 | 4.28 | 1.00 | 100.41 |  |
| 6 | G | N2 | 1G:971 | 4.76 | 1.00 | 96.53 |  |
  |

  
 Coded contact list:   
 Exclusion zone contact(s): /O2'/  
Consider K in place of MG (1 valid contact(s) for K vs. 0 for MG)  

### MG:1G:1687

|  | Res. type | Atom | Residue | Distance | Occupancy | B-factor | Closest solute to 1^st^ shell water |
| --- | --- | --- | --- | --- | --- | --- | --- |
| **Ion** | | | |  | 1.00 | 67.86 || 1 | G | O6 | 1G:238 | 2.70 | 1.00 | 66.29 |
| 2 | U | O4 | 1G:239 | 3.02 | 1.00 | 70.49 |  |
| 3 | G | O6 | 1G:122 | 3.80 | 1.00 | 69.21 |  |
| 4 | C | N4 | 1G:237 | 4.01 | 1.00 | 67.62 |  |
| 5 | C | O4' | 1G:121 | 4.05 | 1.00 | 64.73 |  |
| 6 | C | N4 | 1G:123 | 4.18 | 1.00 | 62.10 |  |
| 7 | C | N1 | 1G:121 | 4.20 | 1.00 | 66.74 |  |
| 8 | G | N1 | 1G:238 | 4.71 | 1.00 | 63.79 |  |
| 9 | C | N3 | 1G:121 | 4.73 | 1.00 | 67.33 |  |
| 10 | U | N3 | 1G:239 | 4.73 | 1.00 | 65.19 |  |
| 11 | C | O5' | 1G:121 | 4.79 | 1.00 | 73.52 |  |
| 12 | C | OP2 | 1G:121 | 5.00 | 1.00 | 66.06 |  |
  |

  
 Coded contact list:   
 Exclusion zone contact(s): /O4/O6/  
Consider K in place of MG (2 valid contact(s) for K vs. 0 for MG)  

### MG:1G:1688

|  | Res. type | Atom | Residue | Distance | Occupancy | B-factor | Closest solute to 1^st^ shell water |
| --- | --- | --- | --- | --- | --- | --- | --- |
| **Ion** | | | |  | 1.00 | 65.57 || 1 | U | O4 | 1G:239 | 2.94 | 1.00 | 70.49 |
| 2 | G | O6 | 1G:238 | 4.37 | 1.00 | 66.29 |  |
| 3 | G | N7 | 1G:238 | 4.60 | 1.00 | 64.31 |  |
  |

  
 Coded contact list:   
 Exclusion zone contact(s): /O4/  
Consider K in place of MG (1 valid contact(s) for K vs. 0 for MG)  

### MG:1G:1689

|  | Res. type | Atom | Residue | Distance | Occupancy | B-factor | Closest solute to 1^st^ shell water |
| --- | --- | --- | --- | --- | --- | --- | --- |
| **Ion** | | | |  | 1.00 | 57.60 || 1 | A | OP1 | 1G:185 | 3.84 | 1.00 | 94.24 |
| 2 | LYS | NZ | BA:74 | 4.92 | 1.00 | 84.07 |  |
  |

  
 Coded contact list:   
1^st^ MG coordination shell missing (see Notes)  

### MG:1G:1690

|  | Res. type | Atom | Residue | Distance | Occupancy | B-factor | Closest solute to 1^st^ shell water |
| --- | --- | --- | --- | --- | --- | --- | --- |
| **Ion** | | | |  | 1.00 | 74.93 || 1 | G | N7 | 1G:231 | 3.00 | 1.00 | 67.53 |
| 2 | G | O6 | 1G:231 | 4.24 | 1.00 | 71.25 |  |
| 3 | G | N7 | 1G:230 | 4.55 | 1.00 | 70.80 |  |
| 4 | G | O6 | 1G:230 | 4.57 | 1.00 | 68.19 |  |
  |

  
 Coded contact list:   
 Exclusion zone contact(s): /N7/  
Consider K in place of MG (1 valid contact(s) for K vs. 0 for MG)  

### MG:1G:1691

|  | Res. type | Atom | Residue | Distance | Occupancy | B-factor | Closest solute to 1^st^ shell water |
| --- | --- | --- | --- | --- | --- | --- | --- |
| **Ion** | | | |  | 1.00 | 64.23 || 1 | G | OP2 | 1G:953 | 3.19 | 1.00 | 76.84 |
| 2 | G | N7 | 1G:954 | 3.33 | 1.00 | 85.99 |  |
| 3 | G | O6 | 1G:954 | 3.62 | 1.00 | 88.93 |  |
| 4 | ARG | NH1 | 4A:104 | 3.63 | 1.00 | 83.76 |  |
| 5 | A | N6 | 1G:1225 | 3.77 | 1.00 | 86.38 |  |
| 6 | U | O4 | 1G:955 | 3.77 | 1.00 | 90.85 |  |
| 7 | ARG | NH2 | 4A:104 | 4.19 | 1.00 | 84.71 |  |
| 8 | G | N7 | 1G:953 | 4.67 | 1.00 | 86.64 |  |
| 9 | G | OP1 | 1G:953 | 4.74 | 1.00 | 87.10 |  |
| 10 | G | O5' | 1G:953 | 4.88 | 1.00 | 88.31 |  |
  |

  
 Coded contact list:   
 Exclusion zone contact(s): /N7/OP2/  
Consider K in place of MG (2 valid contact(s) for K vs. 0 for MG)  

### MG:1G:1692

|  | Res. type | Atom | Residue | Distance | Occupancy | B-factor | Closest solute to 1^st^ shell water |
| --- | --- | --- | --- | --- | --- | --- | --- |
| **Ion** | | | |  | 1.00 | 72.26 || 1 | G | OP2 | 1G:167 | 4.00 | 1.00 | 111.17 |
| 2 | A | OP2 | 1G:151 | 4.27 | 1.00 | 85.65 |  |
| 3 | G | N7 | 1G:168 | 4.28 | 1.00 | 106.17 |  |
| 4 | G | OP2 | 1G:168 | 4.30 | 1.00 | 103.90 |  |
| 5 | A | OP1 | 1G:151 | 4.34 | 1.00 | 84.82 |  |
| 6 | C | OP1 | 1G:150 | 4.37 | 1.00 | 84.21 |  |
| 7 | Water 1 | | 1G:1918 | 4.62 | 1.00 | 106.22 | 4.83 (G:N7 1G:168) |
| 8 | C | O3' | 1G:150 | 4.99 | 1.00 | 82.44 |  |
  |

  
 Coded contact list:   
1^st^ MG coordination shell missing (see Notes)  

### MG:1G:1693

|  | Res. type | Atom | Residue | Distance | Occupancy | B-factor | Closest solute to 1^st^ shell water |
| --- | --- | --- | --- | --- | --- | --- | --- |
| **Ion** | | | |  | 1.00 | 40.65 || 1 | A | OP2 | 1G:1500 | 1.80 | 1.00 | 49.92 |
| 2 | A | OP2 | 1G:1499 | 2.06 | 1.00 | 50.89 |  |
| 3 | G | OP2 | 1G:1505 | 2.79 | 1.00 | 49.73 |  |
| 4 | A | OP1 | 1G:1499 | 3.00 | 1.00 | 49.52 |  |
| 5 | UR3 | O2' | 1G:1498 | 3.26 | 1.00 | 49.30 |  |
| 6 | G | O3' | 1G:1504 | 3.52 | 1.00 | 60.69 |  |
| 7 | A | O5' | 1G:1500 | 3.86 | 1.00 | 47.42 |  |
| 8 | G | OP1 | 1G:1505 | 4.05 | 1.00 | 54.17 |  |
| 9 | UR3 | O3' | 1G:1498 | 4.06 | 1.00 | 53.15 |  |
| 10 | A | O3' | 1G:1499 | 4.08 | 1.00 | 58.26 |  |
| 11 | A | N7 | 1G:1500 | 4.19 | 1.00 | 52.47 |  |
| 12 | A | O5' | 1G:1499 | 4.20 | 1.00 | 53.57 |  |
| 13 | G | O2' | 1G:1504 | 4.28 | 1.00 | 54.58 |  |
| 14 | A | OP1 | 1G:1500 | 4.30 | 1.00 | 49.94 |  |
| 15 | G | O5' | 1G:1505 | 4.99 | 1.00 | 60.01 |  |
  |

  
**1^st^ shell ligand angles:**   
1-2: 95;   
  
 Contact(s): /OP2/OP2/  
 Coded contact list: 2O_ph_  
 Exclusion zone contact(s): /O2'/OP1/OP2/  
Consider K in place of MG (3 valid contact(s) for K vs. 2 for MG)  

### MG:1G:1694

|  | Res. type | Atom | Residue | Distance | Occupancy | B-factor | Closest solute to 1^st^ shell water |
| --- | --- | --- | --- | --- | --- | --- | --- |
| **Ion** | | | |  | 1.00 | 43.85 || 1 | C | O3' | 1G:770 | 2.53 | 1.00 | 42.48 |
| 2 | C | OP1 | 14:1832 | 2.76 | 1.00 | 43.16 |  |
| 3 | C | O2' | 1G:770 | 2.77 | 1.00 | 35.12 |  |
| 4 | C | O2 | 1G:899 | 3.09 | 1.00 | 42.37 |  |
| 5 | G | OP1 | 1G:771 | 3.81 | 1.00 | 42.18 |  |
| 6 | G | O5' | 1G:771 | 4.12 | 1.00 | 43.49 |  |
| 7 | C | O5' | 14:1832 | 4.28 | 1.00 | 44.29 |  |
| 8 | C | OP2 | 14:1832 | 4.37 | 1.00 | 38.55 |  |
| 9 | C | N3 | 1G:899 | 4.60 | 1.00 | 41.21 |  |
| 10 | U | OP1 | 14:1833 | 4.68 | 1.00 | 43.08 |  |
| 11 | U | OP2 | 14:1833 | 4.70 | 1.00 | 44.24 |  |
| 12 | A | N3 | 1G:900 | 4.71 | 1.00 | 38.96 |  |
| 13 | C | O4' | 1G:770 | 4.71 | 1.00 | 38.17 |  |
| 14 | G | OP2 | 1G:771 | 4.91 | 1.00 | 42.52 |  |
  |

  
 Coded contact list:   
 Exclusion zone contact(s): /O2/O2'/O3'/OP1/  
Consider K in place of MG (3 valid contact(s) for K vs. 0 for MG)  

### MG:1G:1695

|  | Res. type | Atom | Residue | Distance | Occupancy | B-factor | Closest solute to 1^st^ shell water |
| --- | --- | --- | --- | --- | --- | --- | --- |
| **Ion** | | | |  | 1.00 | 72.35 || 1 | A | OP1 | 1G:964 | 2.35 | 1.00 | 79.39 |
| 2 | U | OP1 | 1G:1199 | 2.61 | 1.00 | 82.05 |  |
| 3 | G | OP1 | 1G:1198 | 3.10 | 1.00 | 86.20 |  |
| 4 | G | O5' | 1G:1198 | 3.40 | 1.00 | 89.01 |  |
| 5 | A | OP2 | 1G:965 | 3.47 | 1.00 | 86.64 |  |
| 6 | A | O5' | 1G:964 | 3.79 | 1.00 | 81.92 |  |
| 7 | G | O3' | 1G:1198 | 4.13 | 1.00 | 83.62 |  |
| 8 | U | OP2 | 1G:1199 | 4.21 | 1.00 | 79.47 |  |
| 9 | Water 1 | | 1G:1949 | 4.49 | 1.00 | 74.87 | 3.28 (G:OP1 1G:1198) |
| 10 | G | O3' | 1G:963 | 4.56 | 1.00 | 86.92 |  |
| 11 | A | OP2 | 1G:964 | 4.79 | 1.00 | 76.39 |  |
| 12 | G | OP2 | 1G:1198 | 4.86 | 1.00 | 84.22 |  |
  |

  
 Contact(s): /OP1/  
 Coded contact list: O_ph_  
 Exclusion zone contact(s): /OP1/OP1/  
Consider K in place of MG (2 valid contact(s) for K vs. 1 for MG)  

### MG:1G:1696

|  | Res. type | Atom | Residue | Distance | Occupancy | B-factor | Closest solute to 1^st^ shell water |
| --- | --- | --- | --- | --- | --- | --- | --- |
| **Ion** | | | |  | 1.00 | 52.65 || 1 | G | N7 | 1G:1525 | 3.75 | 1.00 | 49.25 |
| 2 | G | N7 | 1G:1526 | 3.85 | 1.00 | 50.59 |  |
| 3 | G | O6 | 1G:1526 | 3.91 | 1.00 | 55.69 |  |
| 4 | G | O6 | 1G:1525 | 4.40 | 1.00 | 50.82 |  |
  |

  
 Coded contact list:   
1^st^ MG coordination shell missing (see Notes)  

### MG:1G:1697

|  | Res. type | Atom | Residue | Distance | Occupancy | B-factor | Closest solute to 1^st^ shell water |
| --- | --- | --- | --- | --- | --- | --- | --- |
| **Ion** | | | |  | 1.00 | 63.29 || 1 | G | OP1 | 1G:730 | 3.53 | 1.00 | 73.81 |
| 2 | C | OP1 | 1G:1527 | 3.81 | 1.00 | 66.85 |  |
| 3 | G | OP2 | 1G:731 | 4.39 | 1.00 | 66.10 |  |
| 4 | G | OP2 | 1G:818 | 4.52 | 1.00 | 66.14 |  |
| 5 | G | N7 | 1G:731 | 4.62 | 1.00 | 60.87 |  |
  |

  
 Coded contact list:   
1^st^ MG coordination shell missing (see Notes)  

### MG:1G:1698

|  | Res. type | Atom | Residue | Distance | Occupancy | B-factor | Closest solute to 1^st^ shell water |
| --- | --- | --- | --- | --- | --- | --- | --- |
| **Ion** | | | |  | 1.00 | 66.26 || 1 | G | OP1 | 1G:666 | 3.79 | 1.00 | 65.16 |
| 2 | A | OP2 | 1G:665 | 4.48 | 1.00 | 63.98 |  |
| 3 | G | N7 | 1G:664 | 4.59 | 1.00 | 74.50 |  |
| 4 | A | O5' | 1G:665 | 4.88 | 1.00 | 70.67 |  |
| 5 | G | N7 | 1G:666 | 4.89 | 1.00 | 69.63 |  |
| 6 | G | N9 | 1G:664 | 5.00 | 1.00 | 66.67 |  |
| 7 | A | O3' | 1G:665 | 5.00 | 1.00 | 62.07 |  |
  |

  
 Coded contact list:   
1^st^ MG coordination shell missing (see Notes)  

### MG:1G:1699

|  | Res. type | Atom | Residue | Distance | Occupancy | B-factor | Closest solute to 1^st^ shell water |
| --- | --- | --- | --- | --- | --- | --- | --- |
| **Ion** | | | |  | 1.00 | 62.42 || 1 | C | OP2 | 1G:48 | 1.91 | 1.00 | 75.18 |
| 2 | U | OP2 | 1G:49 | 3.09 | 1.00 | 61.01 |  |
| 3 | G | OP1 | 1G:115 | 3.15 | 1.00 | 62.76 |  |
| 4 | C | O3' | 1G:47 | 3.71 | 1.00 | 59.90 |  |
| 5 | A | OP1 | 1G:51 | 3.85 | 1.00 | 63.59 |  |
| 6 | C | OP1 | 1G:48 | 4.21 | 1.00 | 70.39 |  |
| 7 | C | O2' | 1G:47 | 4.28 | 1.00 | 73.21 |  |
| 8 | C | O5' | 1G:48 | 4.31 | 1.00 | 58.97 |  |
| 9 | U | O3' | 1G:114 | 4.49 | 1.00 | 64.77 |  |
| 10 | G | O5' | 1G:115 | 4.99 | 1.00 | 62.96 |  |
  |

  
 Contact(s): /OP2/  
 Coded contact list: O_ph_  
 Exclusion zone contact(s): /OP1/OP2/  
Consider K in place of MG (2 valid contact(s) for K vs. 1 for MG)  

### MG:1G:1700

|  | Res. type | Atom | Residue | Distance | Occupancy | B-factor | Closest solute to 1^st^ shell water |
| --- | --- | --- | --- | --- | --- | --- | --- |
| **Ion** | | | |  | 1.00 | 52.41 || 1 | C | OP1 | 1G:291 | 3.93 | 1.00 | 53.76 |
| 2 | G | O6 | 1G:305 | 4.17 | 1.00 | 54.19 |  |
| 3 | G | N7 | 1G:305 | 4.32 | 1.00 | 51.89 |  |
| 4 | C | OP2 | 1G:291 | 4.60 | 1.00 | 54.18 |  |
| 5 | U | O4 | 1G:304 | 4.94 | 1.00 | 58.04 |  |
  |

  
 Coded contact list:   
1^st^ MG coordination shell missing (see Notes)  

### MG:1G:1701

|  | Res. type | Atom | Residue | Distance | Occupancy | B-factor | Closest solute to 1^st^ shell water |
| --- | --- | --- | --- | --- | --- | --- | --- |
| **Ion** | | | |  | 1.00 | 45.05 || 1 | Water 1 | | 1G:1900 | 2.15 | 1.00 | 27.21 | 2.83 (G:OP1 1G:803) |
| 2 | Water 2 | | 1G:1948 | 2.17 | 1.00 | 36.42 | 3.28 (G:OP1 1G:803) |
| 3 | Water 3 | | 1G:1947 | 2.17 | 1.00 | 43.99 | 3.27 (U:OP2 1G:804) |
| 4 | Water 4 | | 1G:1886 | 2.17 | 1.00 | 51.31 | 2.70 (G:N7 1G:771) |
| 5 | Water 5 | | 1G:1855 | 2.18 | 1.00 | 48.00 | 2.45 (U:O4 1G:772) |
| 6 | Water 6 | | 1G:1925 | 2.19 | 1.00 | 55.81 | 3.07 (U:O4 1G:772) |
| 7 | U | O4 | 1G:772 | 3.78 | 1.00 | 49.31 |  |
| 8 | G | OP1 | 1G:803 | 4.00 | 1.00 | 39.29 |  |
| 9 | G | N7 | 1G:771 | 4.29 | 1.00 | 42.54 |  |
| 10 | G | O6 | 1G:773 | 4.42 | 1.00 | 59.08 |  |
| 11 | U | OP2 | 1G:804 | 4.56 | 1.00 | 41.00 |  |
| 12 | G | O6 | 1G:771 | 4.72 | 1.00 | 48.88 |  |
| 13 | C | N4 | 1G:770 | 4.81 | 1.00 | 43.96 |  |
| 14 | G | O5' | 1G:803 | 4.89 | 1.00 | 48.81 |  |
  |

  
**1^st^ shell ligand angles:**   
1-2: 94; 1-3: 92; 1-4: 84; 1-5: 174; 1-6: 89; 2-3: 89; 2-4: 179; 2-5: 92; 2-6: 89; 3-4: 91; 3-5: 89; 3-6: 177; 4-5: 89; 4-6: 91; 5-6: 91;   
  
 Contact(s): /O_w_/O_w_/O_w_/O_w_/O_w_/O_w_/  
 Coded contact list: 6O_w_  

### MG:1G:1702

|  | Res. type | Atom | Residue | Distance | Occupancy | B-factor | Closest solute to 1^st^ shell water |
| --- | --- | --- | --- | --- | --- | --- | --- |
| **Ion** | | | |  | 1.00 | 64.66 || 1 | Water 1 | | 1G:1822 | 2.17 | 1.00 | 57.45 | 2.23 (G:N7 1G:800) |
| 2 | Water 2 | | 1G:1832 | 2.18 | 1.00 | 56.95 | 2.33 (C:O5' 1G:779) |
| 3 | Water 3 | | 1G:1872 | 2.18 | 1.00 | 52.98 | 2.60 (G:OP2 1G:799) |
| 4 | Water 4 | | 1G:1853 | 2.18 | 1.00 | 54.68 | 2.44 (G:O6 1G:800) |
| 5 | Water 5 | | 1G:1866 | 2.18 | 1.00 | 67.46 | 2.54 (C:OP2 1G:779) |
| 6 | Water 6 | | 1G:1896 | 2.19 | 1.00 | 58.47 | 2.79 (C:OP1 1G:779) |
| 7 | A | OP2 | 1G:780 | 3.54 | 1.00 | 56.82 |  |
| 8 | G | O6 | 1G:800 | 3.60 | 1.00 | 55.28 |  |
| 9 | C | O5' | 1G:779 | 3.90 | 1.00 | 51.08 |  |
| 10 | C | OP2 | 1G:779 | 3.95 | 1.00 | 58.67 |  |
| 11 | C | OP1 | 1G:779 | 4.06 | 1.00 | 53.49 |  |
| 12 | G | N7 | 1G:800 | 4.15 | 1.00 | 60.23 |  |
| 13 | G | OP2 | 1G:799 | 4.42 | 1.00 | 50.90 |  |
| 14 | Water 7 | | 1G:1921 | 4.46 | 1.00 | 59.54 | 2.99 (G:O6 1G:800) |
| 15 | A | N7 | 1G:780 | 4.62 | 1.00 | 54.39 |  |
| 16 | G | OP2 | 1G:800 | 4.67 | 1.00 | 54.00 |  |
  |

  
**1^st^ shell ligand angles:**   
1-2: 87; 1-3: 90; 1-4: 88; 1-5: 91; 1-6: 177; 2-3: 177; 2-4: 89; 2-5: 92; 2-6: 91; 3-4: 91; 3-5: 88; 3-6: 92; 4-5: 179; 4-6: 94; 5-6: 87;   
  
 Contact(s): /O_w_/O_w_/O_w_/O_w_/O_w_/O_w_/  
 Coded contact list: 6O_w_  
MG hydration shell issue: water too close (< 2.4 Angs.)  

### MG:1G:1703

|  | Res. type | Atom | Residue | Distance | Occupancy | B-factor | Closest solute to 1^st^ shell water |
| --- | --- | --- | --- | --- | --- | --- | --- |
| **Ion** | | | |  | 1.00 | 81.78 || 1 | Water 1 | | 1G:1807 | 2.17 | 1.00 | 81.93 | 2.05 (G:OP1 1G:1224) |
| 2 | Water 2 | | 1G:1838 | 2.18 | 1.00 | 92.86 | 2.36 (U:OP2 1G:952) |
| 3 | Water 3 | | 1G:1812 | 2.18 | 1.00 | 79.06 | 2.10 (G:OP1 1G:1224) |
| 4 | Water 4 | | 1G:1920 | 2.18 | 1.00 | 88.27 | 2.98 (A:N7 1G:974) |
| 5 | Water 5 | | 1G:1915 | 2.18 | 1.00 | 79.04 | 2.95 (G:OP1 1G:951) |
| 6 | Water 6 | | 1G:1907 | 2.18 | 1.00 | 82.80 | 2.86 (A:N7 1G:974) |
| 7 | G | OP1 | 1G:1224 | 2.89 | 1.00 | 83.06 |  |
| 8 | G | OP1 | 1G:951 | 3.89 | 1.00 | 93.98 |  |
| 9 | A | N7 | 1G:974 | 3.90 | 1.00 | 83.31 |  |
| 10 | G | OP2 | 1G:1224 | 4.24 | 1.00 | 82.55 |  |
| 11 | U | OP2 | 1G:952 | 4.35 | 1.00 | 70.58 |  |
| 12 | U | OP1 | 1G:952 | 4.69 | 1.00 | 76.88 |  |
| 13 | C | O2' | 1G:1223 | 4.72 | 1.00 | 85.14 |  |
| 14 | C | O3' | 1G:1223 | 4.88 | 1.00 | 85.00 |  |
  |

  
**1^st^ shell ligand angles:**   
1-2: 89; 1-3: 91; 1-4: 91; 1-5: 178; 1-6: 87; 2-3: 89; 2-4: 179; 2-5: 91; 2-6: 90; 3-4: 90; 3-5: 90; 3-6: 178; 4-5: 88; 4-6: 91; 5-6: 92;   
  
 Contact(s): /O_w_/O_w_/O_w_/O_w_/O_w_/O_w_/  
 Coded contact list: 6O_w_  
 Exclusion zone contact(s): /OP1/  
Consider K in place of MG (1 valid contact(s) for K vs. 0 for MG)  
MG hydration shell issue: water too close (< 2.4 Angs.)  

### MG:1G:1704

|  | Res. type | Atom | Residue | Distance | Occupancy | B-factor | Closest solute to 1^st^ shell water |
| --- | --- | --- | --- | --- | --- | --- | --- |
| **Ion** | | | |  | 1.00 | 66.29 || 1 | Water 1 | | 1G:1815 | 2.17 | 1.00 | 72.43 | 2.12 (G:O6 1G:682) |
| 2 | Water 2 | | 1G:1860 | 2.17 | 1.00 | 77.05 | 2.48 (A:N6 1G:684) |
| 3 | Water 3 | | 1G:1951 | 2.17 | 1.00 | 79.20 | 3.30 (A:OP1 1G:704) |
| 4 | Water 4 | | 1G:1979 | 2.18 | 1.00 | 80.64 | 3.84 (G:N7 1G:682) |
| 5 | Water 5 | | 1G:1851 | 2.18 | 1.00 | 75.80 | 2.43 (A:OP1 1G:704) |
| 6 | Water 6 | | 1G:1977 | 2.18 | 1.00 | 70.81 | 3.77 (G:N7 1G:683) |
| 7 | A | OP1 | 1G:704 | 3.44 | 1.00 | 80.85 |  |
| 8 | G | O6 | 1G:683 | 4.10 | 1.00 | 72.43 |  |
| 9 | A | N6 | 1G:684 | 4.23 | 1.00 | 88.59 |  |
| 10 | G | O6 | 1G:682 | 4.27 | 1.00 | 74.16 |  |
| 11 | G | N7 | 1G:682 | 4.50 | 1.00 | 80.44 |  |
| 12 | G | N7 | 1G:683 | 4.71 | 1.00 | 80.32 |  |
  |

  
**1^st^ shell ligand angles:**   
1-2: 89; 1-3: 90; 1-4: 88; 1-5: 179; 1-6: 92; 2-3: 87; 2-4: 177; 2-5: 91; 2-6: 93; 3-4: 92; 3-5: 89; 3-6: 179; 4-5: 92; 4-6: 88; 5-6: 89;   
  
 Contact(s): /O_w_/O_w_/O_w_/O_w_/O_w_/O_w_/  
 Coded contact list: 6O_w_  
MG hydration shell issue: water too close (< 2.4 Angs.)  

### MG:1G:1705

|  | Res. type | Atom | Residue | Distance | Occupancy | B-factor | Closest solute to 1^st^ shell water |
| --- | --- | --- | --- | --- | --- | --- | --- |
| **Ion** | | | |  | 1.00 | 73.26 || 1 | Water 1 | | 1G:1879 | 2.17 | 1.00 | 79.09 | 2.68 (G:N7 1G:617) |
| 2 | Water 2 | | 1G:1922 | 2.18 | 1.00 | 67.60 | 3.00 (A:OP1 1G:621) |
| 3 | Water 3 | | 1G:1897 | 2.18 | 1.00 | 65.12 | 2.80 (A:OP2 1G:621) |
| 4 | Water 4 | | 32:404 | 2.18 | 1.00 | 82.24 | 4.17 (ARG:NH2 32:141) |
| 5 | Water 5 | | 1G:1819 | 2.19 | 1.00 | 77.74 | 2.19 (G:OP2 1G:617) |
| 6 | Water 6 | | 32:402 | 2.19 | 1.00 | 74.89 | 2.46 (ARG:NH2 32:141) |
| 7 | ARG | NH2 | 32:141 | 3.82 | 1.00 | 91.64 |  |
| 8 | G | OP2 | 1G:617 | 3.96 | 1.00 | 73.64 |  |
| 9 | G | N7 | 1G:617 | 4.28 | 1.00 | 72.87 |  |
| 10 | A | OP1 | 1G:621 | 4.28 | 1.00 | 52.25 |  |
| 11 | C | OP1 | 1G:620 | 4.61 | 1.00 | 71.77 |  |
| 12 | A | OP2 | 1G:621 | 4.71 | 1.00 | 61.81 |  |
| 13 | G | O6 | 1G:617 | 4.92 | 1.00 | 75.40 |  |
  |

  
**1^st^ shell ligand angles:**   
1-2: 91; 1-3: 92; 1-4: 179; 1-5: 89; 1-6: 87; 2-3: 88; 2-4: 88; 2-5: 178; 2-6: 92; 3-4: 89; 3-5: 90; 3-6: 179; 4-5: 92; 4-6: 92; 5-6: 90;   
  
 Contact(s): /O_w_/O_w_/O_w_/O_w_/O_w_/O_w_/  
 Coded contact list: 6O_w_  
MG hydration shell issue: water too close (< 2.4 Angs.)  

### MG:1G:1706

|  | Res. type | Atom | Residue | Distance | Occupancy | B-factor | Closest solute to 1^st^ shell water |
| --- | --- | --- | --- | --- | --- | --- | --- |
| **Ion** | | | |  | 1.00 | 43.15 || 1 | Water 1 | | 1G:1980 | 2.16 | 1.00 | 43.60 | 3.89 (C:N4 1G:797) |
| 2 | Water 2 | | 1G:1885 | 2.17 | 1.00 | 53.68 | 2.70 (A:OP2 1G:781) |
| 3 | Water 3 | | 1G:1884 | 2.17 | 1.00 | 47.39 | 2.70 (G:O6 1G:798) |
| 4 | Water 4 | | 1G:1974 | 2.18 | 1.00 | 56.49 | 3.81 (G:N7 1G:798) |
| 5 | Water 5 | | 1G:1921 | 2.18 | 1.00 | 59.54 | 2.99 (G:O6 1G:800) |
| 6 | Water 6 | | 1G:1944 | 2.18 | 1.00 | 55.10 | 4.42 (G:O6 1G:800) |
| 7 | Water 7 | | 1G:1853 | 4.30 | 1.00 | 54.68 | 2.44 (G:O6 1G:800) |
| 8 | G | O6 | 1G:800 | 4.56 | 1.00 | 55.28 |  |
| 9 | G | O6 | 1G:799 | 4.59 | 1.00 | 51.22 |  |
| 10 | G | N7 | 1G:798 | 4.61 | 1.00 | 56.91 |  |
| 11 | G | N7 | 1G:799 | 4.63 | 1.00 | 54.61 |  |
| 12 | A | OP2 | 1G:781 | 4.77 | 1.00 | 47.38 |  |
| 13 | G | O6 | 1G:798 | 4.82 | 1.00 | 56.45 |  |
| 14 | G | N1 | 1G:800 | 4.83 | 1.00 | 56.12 |  |
| 15 | C | N4 | 1G:797 | 4.92 | 1.00 | 61.82 |  |
  |

  
**1^st^ shell ligand angles:**   
1-2: 87; 1-3: 90; 1-4: 90; 1-5: 177; 1-6: 95; 2-3: 92; 2-4: 176; 2-5: 90; 2-6: 90; 3-4: 89; 3-5: 90; 3-6: 175; 4-5: 93; 4-6: 89; 5-6: 85;   
  
 Contact(s): /O_w_/O_w_/O_w_/O_w_/O_w_/O_w_/  
 Coded contact list: 6O_w_  

### MG:1G:1707

|  | Res. type | Atom | Residue | Distance | Occupancy | B-factor | Closest solute to 1^st^ shell water |
| --- | --- | --- | --- | --- | --- | --- | --- |
| **Ion** | | | |  | 1.00 | 62.66 || 1 | Water 1 | | 1G:1845 | 2.17 | 1.00 | 60.23 | 2.41 (G:O6 1G:585) |
| 2 | Water 2 | | 1G:1960 | 2.18 | 1.00 | 47.60 | 3.96 (A:OP2 1G:583) |
| 3 | Water 3 | | 1G:1867 | 2.18 | 1.00 | 69.75 | 2.54 (G:N2 1G:758) |
| 4 | Water 4 | | 1G:1984 | 2.18 | 1.00 | 51.60 | 4.08 (G:O6 1G:585) |
| 5 | Water 5 | | 1G:1890 | 2.19 | 1.00 | 66.33 | 2.74 (A:OP2 1G:583) |
| 6 | Water 6 | | 1G:1927 | 2.20 | 1.00 | 63.66 | 3.08 (G:N7 1G:584) |
| 7 | A | OP2 | 1G:583 | 3.80 | 1.00 | 69.15 |  |
| 8 | G | O6 | 1G:585 | 4.01 | 1.00 | 63.25 |  |
| 9 | G | N7 | 1G:584 | 4.14 | 1.00 | 58.17 |  |
| 10 | G | O6 | 1G:584 | 4.24 | 1.00 | 63.69 |  |
| 11 | C | N4 | 1G:756 | 4.56 | 1.00 | 63.02 |  |
| 12 | G | N2 | 1G:758 | 4.62 | 1.00 | 59.02 |  |
| 13 | U | O4 | 1G:757 | 4.91 | 1.00 | 58.67 |  |
  |

  
**1^st^ shell ligand angles:**   
1-2: 92; 1-3: 88; 1-4: 89; 1-5: 179; 1-6: 90; 2-3: 84; 2-4: 95; 2-5: 88; 2-6: 177; 3-4: 177; 3-5: 91; 3-6: 94; 4-5: 92; 4-6: 87; 5-6: 89;   
  
 Contact(s): /O_w_/O_w_/O_w_/O_w_/O_w_/O_w_/  
 Coded contact list: 6O_w_  

### MG:1G:1708

|  | Res. type | Atom | Residue | Distance | Occupancy | B-factor | Closest solute to 1^st^ shell water |
| --- | --- | --- | --- | --- | --- | --- | --- |
| **Ion** | | | |  | 1.00 | 101.76 || 1 | Water 1 | | 1G:1882 | 2.17 | 1.00 | 102.49 | 2.69 (G:O6 1G:166) |
| 2 | Water 2 | | 1G:1918 | 2.18 | 1.00 | 106.22 | 2.99 (G:O6 1G:167) |
| 3 | Water 3 | | 1G:1959 | 2.18 | 1.00 | 90.49 | 3.42 (A:OP2 1G:152) |
| 4 | Water 4 | | 1G:1941 | 2.18 | 1.00 | 100.33 | 3.23 (A:OP2 1G:152) |
| 5 | Water 5 | | 1G:1914 | 2.18 | 1.00 | 109.40 | 2.94 (G:N7 1G:166) |
| 6 | Water 6 | | 1G:1894 | 2.19 | 1.00 | 84.80 | 2.76 (A:OP1 1G:151) |
| 7 | G | O6 | 1G:166 | 3.72 | 1.00 | 118.24 |  |
| 8 | G | O6 | 1G:167 | 4.06 | 1.00 | 109.12 |  |
| 9 | A | OP1 | 1G:151 | 4.27 | 1.00 | 84.82 |  |
| 10 | A | OP2 | 1G:152 | 4.29 | 1.00 | 92.59 |  |
| 11 | G | N7 | 1G:166 | 4.45 | 1.00 | 118.67 |  |
  |

  
**1^st^ shell ligand angles:**   
1-2: 88; 1-3: 89; 1-4: 90; 1-5: 90; 1-6: 180; 2-3: 88; 2-4: 178; 2-5: 92; 2-6: 92; 3-4: 92; 3-5: 179; 3-6: 91; 4-5: 88; 4-6: 90; 5-6: 90;   
  
 Contact(s): /O_w_/O_w_/O_w_/O_w_/O_w_/O_w_/  
 Coded contact list: 6O_w_  

### MG:1G:1709

|  | Res. type | Atom | Residue | Distance | Occupancy | B-factor | Closest solute to 1^st^ shell water |
| --- | --- | --- | --- | --- | --- | --- | --- |
| **Ion** | | | |  | 1.00 | 73.51 || 1 | Water 1 | | 1G:1916 | 2.17 | 1.00 | 73.15 | 2.95 (U:OP2 1G:871) |
| 2 | Water 2 | | 1G:1964 | 2.18 | 1.00 | 70.70 | 3.55 (U:OP2 1G:871) |
| 3 | Water 3 | | 1G:1806 | 2.18 | 1.00 | 81.47 | 2.00 (G:O6 1G:854) |
| 4 | Water 4 | | 1G:1976 | 2.19 | 1.00 | 73.62 | 3.77 (U:OP2 1G:871) |
| 5 | Water 5 | | 1G:1975 | 2.19 | 1.00 | 71.40 | 3.73 (G:O6 1G:853) |
| 6 | Water 6 | | 1G:1804 | 2.20 | 1.00 | 92.61 | 1.30 (G:N7 1G:853) |
| 7 | G | N7 | 1G:853 | 3.48 | 1.00 | 84.89 |  |
| 8 | G | O6 | 1G:854 | 3.93 | 1.00 | 81.14 |  |
| 9 | U | OP2 | 1G:871 | 4.20 | 1.00 | 68.86 |  |
| 10 | G | O6 | 1G:853 | 4.26 | 1.00 | 86.17 |  |
| 11 | G | N7 | 1G:854 | 4.57 | 1.00 | 79.08 |  |
| 12 | U | OP1 | 1G:871 | 5.00 | 1.00 | 73.46 |  |
  |

  
**1^st^ shell ligand angles:**   
1-2: 89; 1-3: 88; 1-4: 86; 1-5: 178; 1-6: 95; 2-3: 177; 2-4: 91; 2-5: 93; 2-6: 88; 3-4: 88; 3-5: 91; 3-6: 92; 4-5: 94; 4-6: 179; 5-6: 86;   
  
 Contact(s): /O_w_/O_w_/O_w_/O_w_/O_w_/O_w_/  
 Coded contact list: 6O_w_  
MG hydration shell issue: water too close (< 2.4 Angs.)  

### MG:1G:1710

|  | Res. type | Atom | Residue | Distance | Occupancy | B-factor | Closest solute to 1^st^ shell water |
| --- | --- | --- | --- | --- | --- | --- | --- |
| **Ion** | | | |  | 1.00 | 59.25 || 1 | Water 1 | | 3A:202 | 2.17 | 1.00 | 60.30 | 3.52 (C:O3' 1G:522) |
| 2 | Water 2 | | 1G:1857 | 2.17 | 1.00 | 60.13 | 2.46 (C:O2' 1G:522) |
| 3 | Water 3 | | 1G:1887 | 2.17 | 1.00 | 70.58 | 2.71 (C:OP1 1G:503) |
| 4 | Water 4 | | 3A:201 | 2.18 | 1.00 | 55.41 | 3.20 (SER:OG 3A:113) |
| 5 | Water 5 | | 1G:1826 | 2.18 | 1.00 | 64.55 | 2.25 (G:OP2 1G:537) |
| 6 | Water 6 | | 1G:1875 | 2.19 | 1.00 | 60.60 | 2.64 (C:OP2 1G:504) |
| 7 | C | OP1 | 1G:536 | 4.12 | 1.00 | 64.14 |  |
| 8 | C | OP1 | 1G:503 | 4.16 | 1.00 | 58.08 |  |
| 9 | C | O3' | 1G:522 | 4.21 | 1.00 | 59.41 |  |
| 10 | G | OP2 | 1G:537 | 4.32 | 1.00 | 59.86 |  |
| 11 | C | O2' | 1G:522 | 4.37 | 1.00 | 59.23 |  |
| 12 | C | OP2 | 1G:504 | 4.51 | 1.00 | 64.95 |  |
| 13 | A | OP1 | 1G:523 | 4.65 | 1.00 | 58.80 |  |
| 14 | G | OP1 | 1G:537 | 4.86 | 1.00 | 66.78 |  |
| 15 | ARG | NH2 | 3A:110 | 4.88 | 1.00 | 46.03 |  |
| 16 | SER | OG | 3A:113 | 4.98 | 1.00 | 48.90 |  |
  |

  
**1^st^ shell ligand angles:**   
1-2: 91; 1-3: 90; 1-4: 88; 1-5: 88; 1-6: 179; 2-3: 88; 2-4: 179; 2-5: 91; 2-6: 88; 3-4: 91; 3-5: 178; 3-6: 90; 4-5: 89; 4-6: 93; 5-6: 92;   
  
 Contact(s): /O_w_/O_w_/O_w_/O_w_/O_w_/O_w_/  
 Coded contact list: 6O_w_  
MG hydration shell issue: water too close (< 2.4 Angs.)  

### MG:1G:1711

|  | Res. type | Atom | Residue | Distance | Occupancy | B-factor | Closest solute to 1^st^ shell water |
| --- | --- | --- | --- | --- | --- | --- | --- |
| **Ion** | | | |  | 1.00 | 97.35 || 1 | Water 1 | | 1G:1912 | 2.18 | 1.00 | 101.17 | 2.93 (G:O6 1G:1361) |
| 2 | Water 2 | | 1G:1850 | 2.18 | 1.00 | 104.92 | 2.43 (C:O2 1G:980) |
| 3 | Water 3 | | 5A:201 | 2.18 | 1.00 | 95.30 | 3.92 (ALA:O 5A:20) |
| 4 | Water 4 | | 1G:1864 | 2.18 | 1.00 | 102.39 | 2.51 (A:OP1 1G:1360) |
| 5 | Water 5 | | 1G:1803 | 2.18 | 1.00 | 108.92 | 1.30 (C:N3 1G:980) |
| 6 | Water 6 | | 1G:1932 | 2.19 | 1.00 | 95.79 | 3.23 (VAL:O 5A:18) |
| 7 | C | N3 | 1G:980 | 3.12 | 1.00 | 110.05 |  |
| 8 | C | O2 | 1G:980 | 3.76 | 1.00 | 106.78 |  |
| 9 | C | N4 | 1G:980 | 4.04 | 1.00 | 100.73 |  |
| 10 | VAL | O | 5A:18 | 4.14 | 1.00 | 108.31 |  |
| 11 | A | N7 | 1G:1360 | 4.33 | 1.00 | 104.46 |  |
| 12 | ALA | O | 5A:20 | 4.41 | 1.00 | 109.36 |  |
| 13 | C | O3' | 1G:1359 | 4.46 | 1.00 | 106.75 |  |
| 14 | A | OP1 | 1G:1360 | 4.60 | 1.00 | 100.51 |  |
| 15 | G | O6 | 1G:1361 | 4.67 | 1.00 | 105.78 |  |
| 16 | TYR | N | 5A:21 | 4.71 | 1.00 | 97.35 |  |
  |

  
**1^st^ shell ligand angles:**   
1-2: 178; 1-3: 87; 1-4: 89; 1-5: 94; 1-6: 91; 2-3: 91; 2-4: 90; 2-5: 87; 2-6: 91; 3-4: 90; 3-5: 91; 3-6: 178; 4-5: 177; 4-6: 90; 5-6: 88;   
  
 Contact(s): /O_w_/O_w_/O_w_/O_w_/O_w_/O_w_/  
 Coded contact list: 6O_w_  
 Exclusion zone contact(s): /N3_C_/  
Consider K in place of MG (1 valid contact(s) for K vs. 0 for MG)  
MG hydration shell issue: water too close (< 2.4 Angs.)  

### MG:1G:1712

|  | Res. type | Atom | Residue | Distance | Occupancy | B-factor | Closest solute to 1^st^ shell water |
| --- | --- | --- | --- | --- | --- | --- | --- |
| **Ion** | | | |  | 1.00 | 81.10 || 1 | Water 1 | | 1G:1833 | 2.18 | 1.00 | 71.57 | 2.33 (G:N7 1G:587) |
| 2 | Water 2 | | 1G:1987 | 2.18 | 1.00 | 73.45 | 5.13 (G:N7 1G:587) |
| 3 | Water 3 | | 1G:1898 | 2.18 | 1.00 | 78.42 | 2.82 (C:OP1 1G:645) |
| 4 | Water 4 | | 1G:1923 | 2.18 | 1.00 | 70.45 | 3.01 (U:OP1 1G:646) |
| 5 | Water 5 | | 1G:1937 | 2.19 | 1.00 | 73.20 | 4.33 (G:N7 1G:587) |
| 6 | Water 6 | | 1G:1935 | 2.19 | 1.00 | 91.94 | 3.15 (U:OP1 1G:646) |
| 7 | U | OP1 | 1G:646 | 3.97 | 1.00 | 82.36 |  |
| 8 | G | N7 | 1G:587 | 4.35 | 1.00 | 65.87 |  |
| 9 | C | OP1 | 1G:645 | 4.73 | 1.00 | 81.99 |  |
| 10 | U | OP2 | 1G:646 | 4.92 | 1.00 | 77.73 |  |
  |

  
**1^st^ shell ligand angles:**   
1-2: 88; 1-3: 92; 1-4: 92; 1-5: 87; 1-6: 177; 2-3: 178; 2-4: 90; 2-5: 92; 2-6: 90; 3-4: 89; 3-5: 90; 3-6: 90; 4-5: 179; 4-6: 89; 5-6: 91;   
  
 Contact(s): /O_w_/O_w_/O_w_/O_w_/O_w_/O_w_/  
 Coded contact list: 6O_w_  
MG hydration shell issue: water too close (< 2.4 Angs.)  

### MG:1G:1713

|  | Res. type | Atom | Residue | Distance | Occupancy | B-factor | Closest solute to 1^st^ shell water |
| --- | --- | --- | --- | --- | --- | --- | --- |
| **Ion** | | | |  | 1.00 | 120.09 || 1 | Water 1 | | 1G:1801 | 2.18 | 1.00 | 125.51 | 1.30 (A:N6 1G:87) |
| 2 | Water 2 | | 1G:1928 | 2.18 | 1.00 | 124.81 | 3.08 (A:N6 1G:87) |
| 3 | Water 3 | | 1G:1930 | 2.18 | 1.00 | 128.04 | 3.11 (A:N6 1G:87) |
| 4 | A | N6 | 1G:87 | 2.94 | 1.00 | 129.08 |  |
| 5 | A | N7 | 1G:87 | 3.98 | 1.00 | 135.76 |  |
| 6 | A | N1 | 1G:87 | 4.46 | 1.00 | 136.87 |  |
  |

  
**1^st^ shell ligand angles:**   
1-2: 89; 1-3: 90; 2-3: 90;   
  
 Contact(s): /O_w_/O_w_/O_w_/  
 Coded contact list: 3O_w_  
 Exclusion zone contact(s): /N6/  
Consider K in place of MG (1 valid contact(s) for K vs. 0 for MG)  
MG hydration shell issue: water too close (< 2.4 Angs.)  

### MG:1G:1714

|  | Res. type | Atom | Residue | Distance | Occupancy | B-factor | Closest solute to 1^st^ shell water |
| --- | --- | --- | --- | --- | --- | --- | --- |
| **Ion** | | | |  | 1.00 | 88.23 || 1 | Water 1 | | 1G:1992 | 2.18 | 1.00 | 92.10 | 4.85 (G:OP2 1G:1385) |
| 2 | Water 2 | | 1G:1938 | 2.18 | 1.00 | 84.24 | 3.19 (G:O6 1G:1386) |
| 3 | Water 3 | | 1G:1981 | 2.18 | 1.00 | 83.11 | 4.23 (G:N7 1G:1385) |
| 4 | Water 4 | | 1G:1968 | 2.19 | 1.00 | 84.48 | 3.66 (G:N7 1G:1386) |
| 5 | Water 5 | | 1G:1843 | 2.20 | 1.00 | 86.46 | 2.38 (G:OP2 1G:1385) |
| 6 | Water 6 | | 1G:1805 | 2.20 | 1.00 | 91.39 | 1.31 (G:N7 1G:1385) |
| 7 | G | N7 | 1G:1385 | 3.47 | 1.00 | 88.69 |  |
| 8 | G | N7 | 1G:1386 | 4.29 | 1.00 | 82.23 |  |
| 9 | G | OP2 | 1G:1385 | 4.30 | 1.00 | 83.97 |  |
| 10 | G | O6 | 1G:1385 | 4.92 | 1.00 | 84.40 |  |
| 11 | G | O6 | 1G:1386 | 4.95 | 1.00 | 82.13 |  |
  |

  
**1^st^ shell ligand angles:**   
1-2: 87; 1-3: 90; 1-4: 89; 1-5: 90; 1-6: 179; 2-3: 86; 2-4: 91; 2-5: 176; 2-6: 94; 3-4: 178; 3-5: 92; 3-6: 90; 4-5: 90; 4-6: 91; 5-6: 89;   
  
 Contact(s): /O_w_/O_w_/O_w_/O_w_/O_w_/O_w_/  
 Coded contact list: 6O_w_  
MG hydration shell issue: water too close (< 2.4 Angs.)  

### MG:1G:1715

|  | Res. type | Atom | Residue | Distance | Occupancy | B-factor | Closest solute to 1^st^ shell water |
| --- | --- | --- | --- | --- | --- | --- | --- |
| **Ion** | | | |  | 1.00 | 39.02 || 1 | Water 1 | | 1G:1903 | 2.16 | 1.00 | 46.59 | 2.84 (G:O2' 1G:575) |
| 2 | Water 2 | | 1G:1837 | 2.16 | 1.00 | 50.59 | 2.36 (G:OP2 1G:576) |
| 3 | Water 3 | | 1G:1913 | 2.17 | 1.00 | 44.34 | 2.93 (C:OP1 1G:578) |
| 4 | Water 4 | | 1G:1835 | 2.17 | 1.00 | 51.31 | 2.34 (C:OP2 1G:817) |
| 5 | Water 5 | | 1G:1880 | 2.18 | 1.00 | 59.80 | 2.68 (U:OP1 1G:820) |
| 6 | C | OP1 | 1G:578 | 2.55 | 1.00 | 59.93 |  |
| 7 | G | O3' | 1G:577 | 3.63 | 1.00 | 48.27 |  |
| 8 | C | OP2 | 1G:817 | 3.87 | 1.00 | 50.26 |  |
| 9 | U | OP1 | 1G:820 | 4.07 | 1.00 | 55.85 |  |
| 10 | G | OP2 | 1G:576 | 4.09 | 1.00 | 51.46 |  |
| 11 | G | O3' | 1G:575 | 4.44 | 1.00 | 53.62 |  |
| 12 | C | O5' | 1G:578 | 4.55 | 1.00 | 49.37 |  |
| 13 | G | O2' | 1G:575 | 4.59 | 1.00 | 59.75 |  |
| 14 | C | OP2 | 1G:578 | 4.68 | 1.00 | 50.05 |  |
| 15 | C | OP1 | 1G:817 | 4.70 | 1.00 | 58.47 |  |
| 16 | C | O5' | 1G:817 | 4.82 | 1.00 | 57.77 |  |
  |

  
**1^st^ shell ligand angles:**   
1-2: 87; 1-3: 94; 1-4: 174; 1-5: 94; 2-3: 178; 2-4: 93; 2-5: 88; 3-4: 86; 3-5: 90; 4-5: 92;   
  
 Contact(s): /O_w_/O_w_/O_w_/O_w_/O_w_/  
 Coded contact list: 5O_w_  
 Exclusion zone contact(s): /OP1/  
Consider NA in place of MG (1 valid contact(s) for NA vs. 0 for MG)  
MG hydration shell issue: water too close (< 2.4 Angs.)  

### MG:1G:1716

|  | Res. type | Atom | Residue | Distance | Occupancy | B-factor | Closest solute to 1^st^ shell water |
| --- | --- | --- | --- | --- | --- | --- | --- |
| **Ion** | | | |  | 1.00 | 70.48 || 1 | Water 1 | | 1G:1967 | 2.18 | 1.00 | 74.58 | 3.62 (U:OP1 1G:13) |
| 2 | Water 2 | | 1G:1899 | 2.18 | 1.00 | 69.03 | 2.82 (U:O4 1G:14) |
| 3 | Water 3 | | 1G:1823 | 2.18 | 1.00 | 66.79 | 2.23 (G:OP2 1G:15) |
| 4 | Water 4 | | 1G:1841 | 2.18 | 1.00 | 65.21 | 2.38 (U:OP2 1G:14) |
| 5 | Water 5 | | 1G:1858 | 2.19 | 1.00 | 72.50 | 2.46 (A:OP2 1G:16) |
| 6 | Water 6 | | 1G:1988 | 2.19 | 1.00 | 67.94 | 4.48 (G:OP2 1G:15) |
| 7 | A | OP2 | 1G:16 | 3.57 | 1.00 | 66.42 |  |
| 8 | U | OP2 | 1G:14 | 4.01 | 1.00 | 63.16 |  |
| 9 | U | OP1 | 1G:13 | 4.28 | 1.00 | 75.85 |  |
| 10 | G | OP2 | 1G:15 | 4.31 | 1.00 | 62.92 |  |
| 11 | A | OP1 | 1G:16 | 4.37 | 1.00 | 66.32 |  |
| 12 | U | O5' | 1G:14 | 4.65 | 1.00 | 61.35 |  |
| 13 | U | O4 | 1G:14 | 4.80 | 1.00 | 74.59 |  |
  |

  
**1^st^ shell ligand angles:**   
1-2: 89; 1-3: 179; 1-4: 89; 1-5: 91; 1-6: 90; 2-3: 91; 2-4: 91; 2-5: 89; 2-6: 178; 3-4: 91; 3-5: 90; 3-6: 90; 4-5: 180; 4-6: 90; 5-6: 90;   
  
 Contact(s): /O_w_/O_w_/O_w_/O_w_/O_w_/O_w_/  
 Coded contact list: 6O_w_  
MG hydration shell issue: water too close (< 2.4 Angs.)  

### MG:1G:1717

|  | Res. type | Atom | Residue | Distance | Occupancy | B-factor | Closest solute to 1^st^ shell water |
| --- | --- | --- | --- | --- | --- | --- | --- |
| **Ion** | | | |  | 1.00 | 67.85 || 1 | Water 1 | | 1G:1852 | 2.16 | 1.00 | 87.09 | 2.43 (G:N7 1G:409) |
| 2 | Water 2 | | 1G:1840 | 2.17 | 1.00 | 83.43 | 2.37 (A:N7 1G:408) |
| 3 | Water 3 | | 1G:1883 | 2.17 | 1.00 | 65.00 | 2.69 (G:OP2 1G:407) |
| 4 | Water 4 | | 1G:1906 | 2.18 | 1.00 | 72.51 | 2.85 (A:OP2 1G:431) |
| 5 | Water 5 | | 1G:1945 | 2.18 | 1.00 | 78.01 | 3.25 (A:OP2 1G:408) |
| 6 | Water 6 | | 1G:1859 | 2.18 | 1.00 | 63.93 | 2.48 (A:OP1 1G:431) |
| 7 | A | N7 | 1G:408 | 4.05 | 1.00 | 80.16 |  |
| 8 | A | OP1 | 1G:431 | 4.06 | 1.00 | 72.21 |  |
| 9 | A | N6 | 1G:408 | 4.10 | 1.00 | 85.50 |  |
| 10 | G | O6 | 1G:410 | 4.16 | 1.00 | 81.94 |  |
| 11 | A | OP2 | 1G:431 | 4.26 | 1.00 | 79.71 |  |
| 12 | G | N7 | 1G:409 | 4.36 | 1.00 | 81.59 |  |
| 13 | G | O6 | 1G:409 | 4.57 | 1.00 | 85.79 |  |
| 14 | G | OP2 | 1G:407 | 4.76 | 1.00 | 66.50 |  |
| 15 | G | N1 | 1G:410 | 4.82 | 1.00 | 79.33 |  |
| 16 | A | OP2 | 1G:408 | 4.84 | 1.00 | 73.12 |  |
  |

  
**1^st^ shell ligand angles:**   
1-2: 89; 1-3: 177; 1-4: 90; 1-5: 88; 1-6: 91; 2-3: 89; 2-4: 179; 2-5: 87; 2-6: 93; 3-4: 93; 3-5: 90; 3-6: 91; 4-5: 93; 4-6: 86; 5-6: 179;   
  
 Contact(s): /O_w_/O_w_/O_w_/O_w_/O_w_/O_w_/  
 Coded contact list: 6O_w_  
MG hydration shell issue: water too close (< 2.4 Angs.)  

### MG:1G:1718

|  | Res. type | Atom | Residue | Distance | Occupancy | B-factor | Closest solute to 1^st^ shell water |
| --- | --- | --- | --- | --- | --- | --- | --- |
| **Ion** | | | |  | 1.00 | 39.02 || 1 | Water 1 | | 1G:1881 | 2.17 | 1.00 | 44.66 | 2.69 (C:O2 1G:812) |
| 2 | Water 2 | | 1G:1811 | 2.18 | 1.00 | 46.69 | 2.10 (A:OP2 1G:766) |
| 3 | Water 3 | | 1G:1848 | 2.18 | 1.00 | 53.39 | 2.42 (C:N3 1G:812) |
| 4 | Water 4 | | 1G:1877 | 2.19 | 1.00 | 45.76 | 2.66 (C:N3 1G:811) |
| 5 | Water 5 | | 1G:1917 | 2.19 | 1.00 | 53.91 | 2.97 (A:OP2 1G:766) |
| 6 | A | OP2 | 1G:766 | 2.42 | 1.00 | 39.53 |  |
| 7 | C | N4 | 1G:811 | 3.83 | 1.00 | 45.83 |  |
| 8 | A | OP1 | 1G:766 | 4.03 | 1.00 | 43.73 |  |
| 9 | C | N3 | 1G:811 | 4.12 | 1.00 | 39.92 |  |
| 10 | C | N3 | 1G:812 | 4.16 | 1.00 | 47.79 |  |
| 11 | G | O3' | 1G:765 | 4.21 | 1.00 | 45.10 |  |
| 12 | C | O2 | 1G:812 | 4.28 | 1.00 | 44.94 |  |
| 13 | A | O5' | 1G:766 | 4.67 | 1.00 | 43.68 |  |
| 14 | A | N6 | 1G:768 | 4.83 | 1.00 | 42.71 |  |
| 15 | U | O4 | 1G:813 | 4.99 | 1.00 | 49.25 |  |
| 16 | A | N7 | 1G:767 | 5.00 | 1.00 | 40.39 |  |
  |

  
**1^st^ shell ligand angles:**   
1-2: 93; 1-3: 87; 1-4: 92; 1-5: 178; 2-3: 177; 2-4: 82; 2-5: 89; 3-4: 101; 3-5: 91; 4-5: 88;   
  
 Contact(s): /O_w_/O_w_/O_w_/O_w_/O_w_/  
 Coded contact list: 5O_w_  
 Exclusion zone contact(s): /OP2/  
Consider NA in place of MG (1 valid contact(s) for NA vs. 0 for MG)  
MG hydration shell issue: water too close (< 2.4 Angs.)  

### MG:1G:1719

|  | Res. type | Atom | Residue | Distance | Occupancy | B-factor | Closest solute to 1^st^ shell water |
| --- | --- | --- | --- | --- | --- | --- | --- |
| **Ion** | | | |  | 1.00 | 66.92 || 1 | Water 1 | | 1G:1949 | 2.17 | 1.00 | 74.87 | 3.28 (G:OP1 1G:1198) |
| 2 | Water 2 | | 1G:1946 | 2.17 | 1.00 | 88.65 | 3.26 (G:OP1 1G:1198) |
| 3 | G | OP1 | 1G:1197 | 2.64 | 1.00 | 86.09 |  |
| 4 | G | OP2 | 1G:1198 | 2.69 | 1.00 | 84.22 |  |
| 5 | C | O5' | 1G:1054 | 2.72 | 1.00 | 96.03 |  |
| 6 | G | O5' | 1G:1197 | 2.98 | 1.00 | 80.16 |  |
| 7 | C | OP1 | 1G:1054 | 3.03 | 1.00 | 96.95 |  |
| 8 | G | OP1 | 1G:1198 | 3.07 | 1.00 | 86.20 |  |
| 9 | G | O3' | 1G:1197 | 4.10 | 1.00 | 78.38 |  |
| 10 | G | OP2 | 1G:1197 | 4.41 | 1.00 | 89.68 |  |
| 11 | G | O3' | 1G:1053 | 4.47 | 1.00 | 94.37 |  |
| 12 | U | O3' | 1G:1196 | 4.58 | 1.00 | 99.29 |  |
| 13 | C | OP2 | 1G:1054 | 4.62 | 1.00 | 90.59 |  |
| 14 | G | O5' | 1G:1198 | 4.70 | 1.00 | 89.01 |  |
| 15 | G | OP1 | 1L:34 | 4.86 | 1.00 | 97.42 |  |
  |

  
**1^st^ shell ligand angles:**   
1-2: 90;   
  
 Contact(s): /O_w_/O_w_/  
 Coded contact list: 2O_w_  
 Exclusion zone contact(s): /O5'/O5'/OP1/OP1/OP1/OP2/  
Consider K in place of MG (6 valid contact(s) for K vs. 0 for MG)  

### MG:1G:1720

|  | Res. type | Atom | Residue | Distance | Occupancy | B-factor | Closest solute to 1^st^ shell water |
| --- | --- | --- | --- | --- | --- | --- | --- |
| **Ion** | | | |  | 1.00 | 73.89 || 1 | Water 1 | | 1G:1934 | 2.17 | 1.00 | 76.85 | 3.14 (G:OP2 1G:588) |
| 2 | Water 2 | | 1G:1954 | 2.18 | 1.00 | 74.81 | 3.37 (G:OP2 1G:588) |
| 3 | Water 3 | | 1G:1908 | 2.18 | 1.00 | 77.09 | 2.86 (C:OP2 1G:645) |
| 4 | Water 4 | | 1G:1957 | 2.18 | 1.00 | 70.62 | 3.40 (G:OP2 1G:588) |
| 5 | Water 5 | | 1G:1821 | 2.18 | 1.00 | 80.93 | 2.23 (G:OP1 1G:588) |
| 6 | G | OP2 | 1G:588 | 2.70 | 1.00 | 82.91 |  |
| 7 | G | OP1 | 1G:588 | 3.00 | 1.00 | 80.75 |  |
| 8 | G | OP1 | 1G:644 | 3.96 | 1.00 | 81.36 |  |
| 9 | C | OP2 | 1G:645 | 4.28 | 1.00 | 73.46 |  |
| 10 | G | O3' | 1G:587 | 4.36 | 1.00 | 81.01 |  |
| 11 | G | O5' | 1G:588 | 4.40 | 1.00 | 76.37 |  |
| 12 | G | O5' | 1G:644 | 4.95 | 1.00 | 71.03 |  |
  |

  
**1^st^ shell ligand angles:**   
1-2: 89; 1-3: 89; 1-4: 91; 1-5: 178; 2-3: 93; 2-4: 175; 2-5: 89; 3-4: 92; 3-5: 90; 4-5: 91;   
  
 Contact(s): /O_w_/O_w_/O_w_/O_w_/O_w_/  
 Coded contact list: 5O_w_  
 Exclusion zone contact(s): /OP1/OP2/  
Consider K in place of MG (2 valid contact(s) for K vs. 0 for MG)  
MG hydration shell issue: water too close (< 2.4 Angs.)  

### MG:1G:1721

|  | Res. type | Atom | Residue | Distance | Occupancy | B-factor | Closest solute to 1^st^ shell water |
| --- | --- | --- | --- | --- | --- | --- | --- |
| **Ion** | | | |  | 1.00 | 45.83 || 1 | Water 1 | | 1G:1816 | 2.17 | 1.00 | 45.06 | 2.14 (A:N7 1G:768) |
| 2 | Water 2 | | 1G:1878 | 2.17 | 1.00 | 42.79 | 2.68 (A:OP2 1G:767) |
| 3 | Water 3 | | 1G:1869 | 2.18 | 1.00 | 45.03 | 2.57 (C:OP2 1G:805) |
| 4 | Water 4 | | 1G:1961 | 2.18 | 1.00 | 57.57 | 3.48 (G:O6 1G:769) |
| 5 | Water 5 | | 1G:1893 | 2.21 | 1.00 | 45.09 | 2.76 (U:OP1 1G:804) |
| 6 | A | OP2 | 1G:768 | 2.48 | 1.00 | 45.97 |  |
| 7 | U | OP1 | 1G:804 | 3.80 | 1.00 | 41.80 |  |
| 8 | A | O5' | 1G:768 | 3.94 | 1.00 | 41.32 |  |
| 9 | A | N7 | 1G:768 | 4.09 | 1.00 | 43.17 |  |
| 10 | G | N7 | 1G:769 | 4.23 | 1.00 | 39.32 |  |
| 11 | A | OP1 | 1G:768 | 4.29 | 1.00 | 45.57 |  |
| 12 | A | OP2 | 1G:767 | 4.60 | 1.00 | 54.10 |  |
| 13 | C | OP2 | 1G:805 | 4.67 | 1.00 | 48.20 |  |
| 14 | A | O3' | 1G:767 | 4.76 | 1.00 | 46.11 |  |
| 15 | A | O5' | 1G:767 | 4.90 | 1.00 | 41.00 |  |
| 16 | G | O6 | 1G:769 | 4.97 | 1.00 | 40.10 |  |
  |

  
**1^st^ shell ligand angles:**   
1-2: 91; 1-3: 178; 1-4: 87; 1-5: 89; 2-3: 89; 2-4: 85; 2-5: 178; 3-4: 94; 3-5: 91; 4-5: 96;   
  
 Contact(s): /O_w_/O_w_/O_w_/O_w_/O_w_/  
 Coded contact list: 5O_w_  
 Exclusion zone contact(s): /OP2/  
Consider NA in place of MG (1 valid contact(s) for NA vs. 0 for MG)  
MG hydration shell issue: water too close (< 2.4 Angs.)  

### MG:1G:1722

|  | Res. type | Atom | Residue | Distance | Occupancy | B-factor | Closest solute to 1^st^ shell water |
| --- | --- | --- | --- | --- | --- | --- | --- |
| **Ion** | | | |  | 1.00 | 46.57 || 1 | Water 1 | | 1G:1955 | 2.18 | 1.00 | 46.21 | 3.80 (C:OP1 1G:904) |
| 2 | Water 2 | | 1G:1842 | 2.18 | 1.00 | 56.22 | 2.38 (C:OP2 1G:904) |
| 3 | Water 3 | | 1G:1861 | 2.18 | 1.00 | 41.59 | 2.49 (G:OP1 1G:903) |
| 4 | Water 4 | | 1G:1874 | 2.18 | 1.00 | 44.85 | 2.61 (U:OP1 1G:1512) |
| 5 | Water 5 | | 1G:1990 | 2.18 | 1.00 | 52.37 | 4.72 (G:OP1 1G:903) |
| 6 | G | OP1 | 1G:903 | 2.73 | 1.00 | 49.61 |  |
| 7 | G | O5' | 1G:903 | 3.81 | 1.00 | 41.12 |  |
| 8 | C | OP2 | 1G:904 | 4.12 | 1.00 | 45.26 |  |
| 9 | U | OP1 | 1G:1512 | 4.39 | 1.00 | 56.19 |  |
| 10 | U | OP1 | 1G:891 | 4.77 | 1.00 | 50.37 |  |
| 11 | G | OP2 | 1G:903 | 4.78 | 1.00 | 48.52 |  |
| 12 | G | O3' | 1G:902 | 4.80 | 1.00 | 48.57 |  |
| 13 | C | OP1 | 1G:904 | 4.82 | 1.00 | 47.54 |  |
  |

  
**1^st^ shell ligand angles:**   
1-2: 91; 1-3: 177; 1-4: 91; 1-5: 87; 2-3: 92; 2-4: 176; 2-5: 85; 3-4: 87; 3-5: 92; 4-5: 99;   
  
 Contact(s): /O_w_/O_w_/O_w_/O_w_/O_w_/  
 Coded contact list: 5O_w_  
 Exclusion zone contact(s): /OP1/  
Consider K in place of MG (1 valid contact(s) for K vs. 0 for MG)  
MG hydration shell issue: water too close (< 2.4 Angs.)  

### MG:1G:1723

|  | Res. type | Atom | Residue | Distance | Occupancy | B-factor | Closest solute to 1^st^ shell water |
| --- | --- | --- | --- | --- | --- | --- | --- |
| **Ion** | | | |  | 1.00 | 59.32 || 1 | Water 1 | | 1G:1836 | 2.16 | 1.00 | 62.16 | 2.35 (G:O6 1G:299) |
| 2 | Water 2 | | 1G:1870 | 2.17 | 1.00 | 61.73 | 2.58 (G:O6 1G:299) |
| 3 | Water 3 | | 1G:1813 | 2.18 | 1.00 | 57.75 | 2.11 (U:OP1 1G:560) |
| 4 | Water 4 | | 1G:1952 | 2.20 | 1.00 | 64.59 | 3.31 (A:N1 1G:300) |
| 5 | G | OP1 | 1G:558 | 2.43 | 1.00 | 52.73 |  |
| 6 | G | O6 | 1G:299 | 2.55 | 1.00 | 59.29 |  |
| 7 | U | OP1 | 1G:560 | 3.17 | 1.00 | 65.51 |  |
| 8 | G | O6 | 1G:566 | 3.71 | 1.00 | 61.36 |  |
| 9 | G | N1 | 1G:299 | 3.75 | 1.00 | 56.18 |  |
| 10 | G | OP2 | 1G:558 | 3.87 | 1.00 | 62.36 |  |
| 11 | A | N1 | 1G:300 | 3.89 | 1.00 | 56.75 |  |
| 12 | G | O3' | 1G:557 | 4.23 | 1.00 | 62.33 |  |
| 13 | A | N6 | 1G:298 | 4.32 | 1.00 | 59.06 |  |
| 14 | U | OP2 | 1G:560 | 4.50 | 1.00 | 69.85 |  |
| 15 | A | N6 | 1G:300 | 4.52 | 1.00 | 57.31 |  |
| 16 | G | O2' | 1G:297 | 4.56 | 1.00 | 61.85 |  |
| 17 | G | N1 | 1G:566 | 4.59 | 1.00 | 64.92 |  |
| 18 | G | O5' | 1G:558 | 4.76 | 1.00 | 62.87 |  |
  |

  
**1^st^ shell ligand angles:**   
1-2: 84; 1-3: 175; 1-4: 90; 2-3: 91; 2-4: 173; 3-4: 95;   
  
 Contact(s): /O_w_/O_w_/O_w_/O_w_/  
 Coded contact list: 4O_w_  
 Exclusion zone contact(s): /O6/OP1/OP1/  
Consider NA in place of MG (2 valid contact(s) for NA vs. 0 for MG)  
MG hydration shell issue: water too close (< 2.4 Angs.)  

### MG:1G:1724

|  | Res. type | Atom | Residue | Distance | Occupancy | B-factor | Closest solute to 1^st^ shell water |
| --- | --- | --- | --- | --- | --- | --- | --- |
| **Ion** | | | |  | 1.00 | 82.61 || 1 | Water 1 | | 1G:1909 | 2.16 | 1.00 | 71.01 | 2.90 (A:OP1 1G:1346) |
| 2 | Water 2 | | 1G:1865 | 2.17 | 1.00 | 83.04 | 2.53 (C:OP1 1G:934) |
| 3 | Water 3 | | 1G:1827 | 2.18 | 1.00 | 72.20 | 2.28 (U:OP1 1G:1345) |
| 4 | Water 4 | | 1G:1810 | 2.18 | 1.00 | 74.82 | 2.09 (C:OP1 1G:934) |
| 5 | Water 5 | | 1G:1983 | 2.18 | 1.00 | 85.94 | 4.05 (A:OP1 1G:1346) |
| 6 | Water 6 | | 1G:1868 | 2.19 | 1.00 | 78.78 | 2.56 (C:OP1 1G:934) |
| 7 | C | OP1 | 1G:934 | 2.77 | 1.00 | 79.98 |  |
| 8 | U | OP1 | 1G:1345 | 3.89 | 1.00 | 85.88 |  |
| 9 | U | OP2 | 1G:1345 | 4.19 | 1.00 | 78.32 |  |
| 10 | C | O5' | 1G:934 | 4.36 | 1.00 | 75.96 |  |
| 11 | A | OP1 | 1G:1346 | 4.56 | 1.00 | 82.73 |  |
| 12 | G | O3' | 1G:933 | 4.94 | 1.00 | 78.46 |  |
| 13 | U | O2' | 1G:1345 | 4.95 | 1.00 | 78.23 |  |
  |

  
**1^st^ shell ligand angles:**   
1-2: 85; 1-3: 93; 1-4: 87; 1-5: 92; 1-6: 178; 2-3: 89; 2-4: 91; 2-5: 177; 2-6: 93; 3-4: 180; 3-5: 91; 3-6: 87; 4-5: 89; 4-6: 93; 5-6: 90;   
  
 Contact(s): /O_w_/O_w_/O_w_/O_w_/O_w_/O_w_/  
 Coded contact list: 6O_w_  
 Exclusion zone contact(s): /OP1/  
Consider K in place of MG (1 valid contact(s) for K vs. 0 for MG)  
MG hydration shell issue: water too close (< 2.4 Angs.)  

### MG:1G:1725

|  | Res. type | Atom | Residue | Distance | Occupancy | B-factor | Closest solute to 1^st^ shell water |
| --- | --- | --- | --- | --- | --- | --- | --- |
| **Ion** | | | |  | 1.00 | 69.41 || 1 | Water 1 | | 1G:1943 | 2.17 | 1.00 | 62.15 | 3.49 (U:O4 4L:48) |
| 2 | Water 2 | | 1G:1814 | 2.17 | 1.00 | 65.08 | 2.12 (C:O2 1G:1399) |
| 3 | Water 3 | | 4L:203 | 2.18 | 1.00 | 70.66 | 3.58 (U:OP2 4L:47) |
| 4 | Water 4 | | 1G:1911 | 2.18 | 1.00 | 67.93 | 2.92 (G:O6 1G:1505) |
| 5 | Water 5 | | 1G:1839 | 2.18 | 1.00 | 79.11 | 2.36 (G:O6 1G:926) |
| 6 | Water 6 | | 4L:202 | 2.19 | 1.00 | 65.10 | 3.56 (U:O4 4L:48) |
| 7 | C | O2 | 1G:1399 | 3.95 | 1.00 | 59.63 |  |
| 8 | G | O6 | 1G:926 | 4.15 | 1.00 | 76.00 |  |
| 9 | G | O6 | 1G:1505 | 4.33 | 1.00 | 70.57 |  |
| 10 | U | O4 | 4L:48 | 4.35 | 1.00 | 70.86 |  |
| 11 | G | N2 | 1G:1504 | 4.44 | 1.00 | 57.43 |  |
| 12 | U | OP2 | 4L:47 | 4.82 | 1.00 | 56.88 |  |
  |

  
**1^st^ shell ligand angles:**   
1-2: 92; 1-3: 89; 1-4: 176; 1-5: 89; 1-6: 90; 2-3: 179; 2-4: 90; 2-5: 89; 2-6: 89; 3-4: 89; 3-5: 92; 3-6: 90; 4-5: 88; 4-6: 93; 5-6: 178;   
  
 Contact(s): /O_w_/O_w_/O_w_/O_w_/O_w_/O_w_/  
 Coded contact list: 6O_w_  
MG hydration shell issue: water too close (< 2.4 Angs.)  

### MG:1G:1726

|  | Res. type | Atom | Residue | Distance | Occupancy | B-factor | Closest solute to 1^st^ shell water |
| --- | --- | --- | --- | --- | --- | --- | --- |
| **Ion** | | | |  | 1.00 | 49.62 || 1 | Water 1 | | 1G:1873 | 2.17 | 1.00 | 52.79 | 2.60 (G:OP2 1G:576) |
| 2 | Water 2 | | 1G:1931 | 2.18 | 1.00 | 55.88 | 4.69 (G:OP1 1G:576) |
| 3 | Water 3 | | 1G:1924 | 2.18 | 1.00 | 50.04 | 3.02 (G:OP1 1G:576) |
| 4 | Water 4 | | 1G:1905 | 2.19 | 1.00 | 54.68 | 2.85 (G:OP2 1G:575) |
| 5 | Water 5 | | 1G:1919 | 2.20 | 1.00 | 47.50 | 2.98 (G:OP2 1G:576) |
| 6 | G | OP1 | 1G:576 | 2.57 | 1.00 | 49.98 |  |
| 7 | G | OP2 | 1G:576 | 3.31 | 1.00 | 51.46 |  |
| 8 | G | OP1 | 1G:575 | 3.90 | 1.00 | 59.31 |  |
| 9 | G | O3' | 1G:575 | 4.26 | 1.00 | 53.62 |  |
| 10 | G | OP2 | 1G:575 | 4.33 | 1.00 | 57.24 |  |
| 11 | G | O5' | 1G:575 | 4.56 | 1.00 | 53.44 |  |
| 12 | G | O5' | 1G:576 | 4.65 | 1.00 | 52.76 |  |
| 13 | G | OP2 | 1G:577 | 4.98 | 1.00 | 48.92 |  |
  |

  
**1^st^ shell ligand angles:**   
1-2: 93; 1-3: 88; 1-4: 176; 1-5: 90; 2-3: 88; 2-4: 91; 2-5: 92; 3-4: 92; 3-5: 177; 4-5: 91;   
  
 Contact(s): /O_w_/O_w_/O_w_/O_w_/O_w_/  
 Coded contact list: 5O_w_  
 Exclusion zone contact(s): /OP1/OP2/  
Consider NA in place of MG (1 valid contact(s) for NA vs. 0 for MG)  

### MG:1G:1727

|  | Res. type | Atom | Residue | Distance | Occupancy | B-factor | Closest solute to 1^st^ shell water |
| --- | --- | --- | --- | --- | --- | --- | --- |
| **Ion** | | | |  | 1.00 | 59.40 || 1 | Water 1 | | 1G:1829 | 2.17 | 1.00 | 61.76 | 2.31 (G:N7 1G:1469) |
| 2 | Water 2 | | 1G:1994 | 2.17 | 1.00 | 64.25 | 5.26 (G:N7 1G:1469) |
| 3 | Water 3 | | 1G:1830 | 2.17 | 1.00 | 65.43 | 2.31 (G:O6 1G:1470) |
| 4 | Water 4 | | 1G:1989 | 2.18 | 1.00 | 74.88 | 4.55 (G:N7 1G:1470) |
| 5 | Water 5 | | 1G:1942 | 2.18 | 1.00 | 70.13 | 3.23 (G:O6 1G:1471) |
| 6 | Water 6 | | 1G:1933 | 2.19 | 1.00 | 74.79 | 3.13 (G:O6 1G:1469) |
| 7 | G | O6 | 1G:1470 | 4.12 | 1.00 | 62.66 |  |
| 8 | G | O6 | 1G:1469 | 4.29 | 1.00 | 65.24 |  |
| 9 | G | N7 | 1G:1469 | 4.29 | 1.00 | 65.90 |  |
| 10 | G | O6 | 1G:1471 | 4.34 | 1.00 | 63.83 |  |
| 11 | G | N7 | 1G:1470 | 4.36 | 1.00 | 65.03 |  |
  |

  
**1^st^ shell ligand angles:**   
1-2: 90; 1-3: 91; 1-4: 90; 1-5: 179; 1-6: 89; 2-3: 178; 2-4: 87; 2-5: 90; 2-6: 94; 3-4: 91; 3-5: 89; 3-6: 88; 4-5: 91; 4-6: 179; 5-6: 90;   
  
 Contact(s): /O_w_/O_w_/O_w_/O_w_/O_w_/O_w_/  
 Coded contact list: 6O_w_  
MG hydration shell issue: water too close (< 2.4 Angs.)  

### MG:1G:1728

|  | Res. type | Atom | Residue | Distance | Occupancy | B-factor | Closest solute to 1^st^ shell water |
| --- | --- | --- | --- | --- | --- | --- | --- |
| **Ion** | | | |  | 1.00 | 50.89 || 1 | Water 1 | | 1G:1824 | 2.17 | 1.00 | 52.33 | 2.23 (UR3:O4 1G:1498) |
| 2 | Water 2 | | 4L:204 | 2.17 | 1.00 | 66.69 | 3.58 (U:O2' 4L:47) |
| 3 | Water 3 | | 1G:1950 | 2.17 | 1.00 | 54.69 | 3.29 (G:N7 1G:1497) |
| 4 | Water 4 | | 1G:1958 | 2.18 | 1.00 | 57.42 | 3.41 (C:OP2 1G:1496) |
| 5 | Water 5 | | 1G:1834 | 2.18 | 1.00 | 37.65 | 2.33 (G:OP2 1G:1497) |
| 6 | Water 6 | | 4L:201 | 2.19 | 1.00 | 64.88 | 2.79 (U:O2' 4L:47) |
| 7 | G | N7 | 1G:1497 | 4.14 | 1.00 | 48.41 |  |
| 8 | U | O2' | 4L:47 | 4.20 | 1.00 | 57.80 |  |
| 9 | UR3 | O4 | 1G:1498 | 4.29 | 1.00 | 47.64 |  |
| 10 | G | OP2 | 1G:1497 | 4.49 | 1.00 | 48.30 |  |
| 11 | C | OP2 | 1G:1496 | 4.68 | 1.00 | 50.15 |  |
  |

  
**1^st^ shell ligand angles:**   
1-2: 88; 1-3: 89; 1-4: 179; 1-5: 93; 1-6: 90; 2-3: 83; 2-4: 93; 2-5: 179; 2-6: 93; 3-4: 92; 3-5: 95; 3-6: 176; 4-5: 86; 4-6: 89; 5-6: 88;   
  
 Contact(s): /O_w_/O_w_/O_w_/O_w_/O_w_/O_w_/  
 Coded contact list: 6O_w_  
MG hydration shell issue: water too close (< 2.4 Angs.)  

### MG:1G:1729

|  | Res. type | Atom | Residue | Distance | Occupancy | B-factor | Closest solute to 1^st^ shell water |
| --- | --- | --- | --- | --- | --- | --- | --- |
| **Ion** | | | |  | 1.00 | 67.15 || 1 | Water 1 | | 1G:1854 | 2.18 | 1.00 | 64.69 | 2.45 (G:N7 1G:377) |
| 2 | Water 2 | | 1G:1936 | 2.18 | 1.00 | 70.22 | 3.17 (G:O6 1G:377) |
| 3 | Water 3 | | 1G:1891 | 2.19 | 1.00 | 59.90 | 2.75 (G:O6 1G:376) |
| 4 | Water 4 | | 1G:1993 | 2.19 | 1.00 | 61.33 | 4.85 (G:N7 1G:377) |
| 5 | Water 5 | | 1G:1963 | 2.19 | 1.00 | 61.26 | 3.58 (G:N7 1G:376) |
| 6 | Water 6 | | 1G:1991 | 2.20 | 1.00 | 66.28 | 5.76 (G:O6 1G:376) |
| 7 | G | O6 | 1G:376 | 4.20 | 1.00 | 74.83 |  |
| 8 | G | N7 | 1G:376 | 4.22 | 1.00 | 66.35 |  |
| 9 | G | N7 | 1G:377 | 4.46 | 1.00 | 68.64 |  |
| 10 | G | O6 | 1G:377 | 4.49 | 1.00 | 70.07 |  |
  |

  
**1^st^ shell ligand angles:**   
1-2: 92; 1-3: 90; 1-4: 87; 1-5: 90; 1-6: 180; 2-3: 89; 2-4: 89; 2-5: 179; 2-6: 89; 3-4: 176; 3-5: 91; 3-6: 90; 4-5: 92; 4-6: 93; 5-6: 90;   
  
 Contact(s): /O_w_/O_w_/O_w_/O_w_/O_w_/O_w_/  
 Coded contact list: 6O_w_  

### MG:1G:1730

|  | Res. type | Atom | Residue | Distance | Occupancy | B-factor | Closest solute to 1^st^ shell water |
| --- | --- | --- | --- | --- | --- | --- | --- |
| **Ion** | | | |  | 1.00 | 78.35 || 1 | Water 1 | | 1G:1847 | 2.16 | 1.00 | 59.48 | 2.42 (G:O2' 1G:1392) |
| 2 | Water 2 | | 1G:1892 | 2.18 | 1.00 | 55.16 | 2.76 (G:OP1 1G:1530) |
| 3 | Water 3 | | 1G:1818 | 2.18 | 1.00 | 79.78 | 2.18 (A:OP1 1G:1502) |
| 4 | Water 4 | | 1G:1929 | 2.19 | 1.00 | 66.63 | 3.09 (G:O2' 1G:1530) |
| 5 | Water 5 | | 1G:1809 | 2.19 | 1.00 | 67.97 | 2.07 (G:OP1 1G:1530) |
| 6 | Water 6 | | 1G:1902 | 2.20 | 1.00 | 69.12 | 2.84 (G:OP1 1G:1530) |
| 7 | G | OP1 | 1G:1530 | 3.09 | 1.00 | 73.78 |  |
| 8 | A | OP1 | 1G:1502 | 3.37 | 1.00 | 57.59 |  |
| 9 | G | O3' | 1G:1392 | 3.83 | 1.00 | 59.66 |  |
| 10 | U | OP1 | 1G:1393 | 4.10 | 1.00 | 58.06 |  |
| 11 | G | O2' | 1G:1392 | 4.32 | 1.00 | 63.37 |  |
| 12 | G | O3' | 1G:1530 | 4.71 | 1.00 | 81.69 |  |
| 13 | G | O5' | 1G:1530 | 4.72 | 1.00 | 66.91 |  |
| 14 | G | O2' | 1G:1530 | 4.78 | 1.00 | 78.58 |  |
  |

  
**1^st^ shell ligand angles:**   
1-2: 93; 1-3: 89; 1-4: 85; 1-5: 178; 1-6: 91; 2-3: 90; 2-4: 177; 2-5: 87; 2-6: 91; 3-4: 89; 3-5: 89; 3-6: 180; 4-5: 95; 4-6: 91; 5-6: 91;   
  
 Contact(s): /O_w_/O_w_/O_w_/O_w_/O_w_/O_w_/  
 Coded contact list: 6O_w_  
 Exclusion zone contact(s): /OP1/OP1/  
Consider K in place of MG (2 valid contact(s) for K vs. 0 for MG)  
MG hydration shell issue: water too close (< 2.4 Angs.)  

### MG:1G:1731

|  | Res. type | Atom | Residue | Distance | Occupancy | B-factor | Closest solute to 1^st^ shell water |
| --- | --- | --- | --- | --- | --- | --- | --- |
| **Ion** | | | |  | 1.00 | 81.43 || 1 | Water 1 | | 1G:1856 | 2.17 | 1.00 | 70.67 | 2.45 (G:O6 1G:112) |
| 2 | Water 2 | | 1G:1940 | 2.18 | 1.00 | 70.02 | 3.84 (G:OP2 1G:111) |
| 3 | Water 3 | | 1G:1876 | 2.18 | 1.00 | 65.07 | 2.64 (G:N7 1G:112) |
| 4 | Water 4 | | 1G:1970 | 2.18 | 1.00 | 69.28 | 3.64 (G:O6 1G:113) |
| 5 | Water 5 | | 1G:1846 | 2.20 | 1.00 | 65.70 | 2.41 (G:OP2 1G:111) |
| 6 | Water 6 | | 1G:1986 | 2.20 | 1.00 | 69.52 | 4.37 (G:N7 1G:113) |
| 7 | G | OP2 | 1G:111 | 3.52 | 1.00 | 64.00 |  |
| 8 | G | N7 | 1G:112 | 3.95 | 1.00 | 63.20 |  |
| 9 | G | O6 | 1G:112 | 4.23 | 1.00 | 65.69 |  |
| 10 | G | O6 | 1G:113 | 4.63 | 1.00 | 59.86 |  |
| 11 | G | N7 | 1G:111 | 4.83 | 1.00 | 64.99 |  |
| 12 | G | N7 | 1G:113 | 4.97 | 1.00 | 54.11 |  |
  |

  
**1^st^ shell ligand angles:**   
1-2: 89; 1-3: 91; 1-4: 87; 1-5: 176; 1-6: 88; 2-3: 87; 2-4: 94; 2-5: 89; 2-6: 177; 3-4: 178; 3-5: 92; 3-6: 92; 4-5: 91; 4-6: 87; 5-6: 94;   
  
 Contact(s): /O_w_/O_w_/O_w_/O_w_/O_w_/O_w_/  
 Coded contact list: 6O_w_  

### MG:1G:1732

|  | Res. type | Atom | Residue | Distance | Occupancy | B-factor | Closest solute to 1^st^ shell water |
| --- | --- | --- | --- | --- | --- | --- | --- |
| **Ion** | | | |  | 1.00 | 83.20 || 1 | Water 1 | | 1G:1939 | 2.17 | 1.00 | 84.31 | 3.22 (C:OP2 1G:1200) |
| 2 | Water 2 | | 1G:1910 | 2.18 | 1.00 | 91.07 | 2.92 (C:OP1 1G:1051) |
| 3 | Water 3 | | 1G:1889 | 2.18 | 1.00 | 90.26 | 2.74 (C:O2' 1G:962) |
| 4 | Water 4 | | 1G:1817 | 2.18 | 1.00 | 79.51 | 2.15 (C:OP1 1G:1200) |
| 5 | Water 5 | | 1G:1825 | 2.18 | 1.00 | 84.39 | 2.23 (C:O2' 1G:962) |
| 6 | Water 6 | | 1G:1962 | 2.19 | 1.00 | 95.80 | 3.50 (C:OP2 1G:1051) |
| 7 | C | O2' | 1G:962 | 3.51 | 1.00 | 86.05 |  |
| 8 | C | OP1 | 1G:1200 | 3.81 | 1.00 | 78.53 |  |
| 9 | C | OP1 | 1G:1051 | 4.44 | 1.00 | 96.55 |  |
| 10 | U | OP2 | 1G:1052 | 4.52 | 1.00 | 93.40 |  |
| 11 | C | OP2 | 1G:1051 | 4.66 | 1.00 | 93.45 |  |
| 12 | C | OP2 | 1G:1200 | 4.72 | 1.00 | 92.72 |  |
  |

  
**1^st^ shell ligand angles:**   
1-2: 91; 1-3: 180; 1-4: 87; 1-5: 92; 1-6: 89; 2-3: 89; 2-4: 178; 2-5: 91; 2-6: 88; 3-4: 93; 3-5: 88; 3-6: 91; 4-5: 89; 4-6: 92; 5-6: 178;   
  
 Contact(s): /O_w_/O_w_/O_w_/O_w_/O_w_/O_w_/  
 Coded contact list: 6O_w_  
MG hydration shell issue: water too close (< 2.4 Angs.)  

### MG:1G:1733

|  | Res. type | Atom | Residue | Distance | Occupancy | B-factor | Closest solute to 1^st^ shell water |
| --- | --- | --- | --- | --- | --- | --- | --- |
| **Ion** | | | |  | 1.00 | 81.49 || 1 | Water 1 | | 1G:1831 | 2.18 | 1.00 | 79.63 | 2.31 (G:OP1 1G:1079) |
| 2 | Water 2 | | 1G:1978 | 2.18 | 1.00 | 77.10 | 3.78 (G:OP1 1G:1079) |
| 3 | Water 3 | | 1G:1969 | 2.18 | 1.00 | 61.17 | 3.63 (G:O6 1G:1077) |
| 4 | Water 4 | | 1G:1802 | 2.19 | 1.00 | 84.58 | 1.30 (G:N7 1G:1077) |
| 5 | Water 5 | | 1G:1901 | 2.19 | 1.00 | 73.79 | 2.83 (G:OP2 1G:1079) |
| 6 | Water 6 | | 1G:1982 | 2.19 | 1.00 | 67.20 | 4.35 (G:N7 1G:1077) |
| 7 | G | N7 | 1G:1077 | 3.30 | 1.00 | 79.02 |  |
| 8 | G | OP1 | 1G:1079 | 3.68 | 1.00 | 76.33 |  |
| 9 | G | O6 | 1G:1077 | 3.87 | 1.00 | 76.78 |  |
| 10 | G | OP2 | 1G:1079 | 4.22 | 1.00 | 75.21 |  |
  |

  
**1^st^ shell ligand angles:**   
1-2: 89; 1-3: 87; 1-4: 89; 1-5: 92; 1-6: 179; 2-3: 93; 2-4: 177; 2-5: 88; 2-6: 92; 3-4: 88; 3-5: 179; 3-6: 92; 4-5: 91; 4-6: 91; 5-6: 89;   
  
 Contact(s): /O_w_/O_w_/O_w_/O_w_/O_w_/O_w_/  
 Coded contact list: 6O_w_  
 Exclusion zone contact(s): /N7/  
Consider K in place of MG (1 valid contact(s) for K vs. 0 for MG)  
MG hydration shell issue: water too close (< 2.4 Angs.)  

### MG:1G:1734

|  | Res. type | Atom | Residue | Distance | Occupancy | B-factor | Closest solute to 1^st^ shell water |
| --- | --- | --- | --- | --- | --- | --- | --- |
| **Ion** | | | |  | 1.00 | 94.86 || 1 | Water 1 | | 1G:1820 | 2.17 | 1.00 | 96.77 | 2.21 (G:N7 1G:1370) |
| 2 | Water 2 | | 1G:1808 | 2.17 | 1.00 | 89.84 | 2.07 (U:OP1 1G:1348) |
| 3 | Water 3 | | 1G:1844 | 2.18 | 1.00 | 92.41 | 2.40 (G:O6 1G:1371) |
| 4 | Water 4 | | 82:201 | 2.18 | 1.00 | 98.34 | 3.51 (U:O4 1G:1351) |
| 5 | Water 5 | | 1G:1828 | 2.18 | 1.00 | 94.54 | 2.30 (G:O6 1G:1370) |
| 6 | Water 6 | | 1G:1985 | 2.18 | 1.00 | 99.21 | 4.11 (G:N7 1G:1370) |
| 7 | G | O6 | 1G:1370 | 3.81 | 1.00 | 100.92 |  |
| 8 | G | N7 | 1G:1370 | 3.98 | 1.00 | 100.80 |  |
| 9 | U | OP1 | 1G:1348 | 4.02 | 1.00 | 86.79 |  |
| 10 | U | O4 | 1G:1351 | 4.30 | 1.00 | 87.72 |  |
| 11 | G | O6 | 1G:1371 | 4.35 | 1.00 | 92.21 |  |
| 12 | A | N6 | 1G:1350 | 4.85 | 1.00 | 85.81 |  |
  |

  
**1^st^ shell ligand angles:**   
1-2: 90; 1-3: 86; 1-4: 176; 1-5: 93; 1-6: 91; 2-3: 91; 2-4: 86; 2-5: 178; 2-6: 91; 3-4: 92; 3-5: 90; 3-6: 176; 4-5: 91; 4-6: 90; 5-6: 88;   
  
 Contact(s): /O_w_/O_w_/O_w_/O_w_/O_w_/O_w_/  
 Coded contact list: 6O_w_  
MG hydration shell issue: water too close (< 2.4 Angs.)  

### MG:1G:1735

|  | Res. type | Atom | Residue | Distance | Occupancy | B-factor | Closest solute to 1^st^ shell water |
| --- | --- | --- | --- | --- | --- | --- | --- |
| **Ion** | | | |  | 1.00 | 88.83 || 1 | Water 1 | | 1G:1971 | 2.17 | 1.00 | 80.39 | 3.67 (G:OP1 1G:384) |
| 2 | Water 2 | | 1G:1973 | 2.18 | 1.00 | 79.10 | 3.69 (A:N6 1G:453) |
| 3 | Water 3 | | 1G:1904 | 2.18 | 1.00 | 79.66 | 2.85 (G:OP2 1G:384) |
| 4 | Water 4 | | 1G:1895 | 2.19 | 1.00 | 80.94 | 2.77 (C:N3 1G:454) |
| 5 | Water 5 | | 1G:1972 | 2.19 | 1.00 | 78.49 | 5.38 (A:N1 1G:453) |
| 6 | Water 6 | | 1G:1888 | 2.19 | 1.00 | 77.46 | 2.71 (A:N1 1G:453) |
| 7 | A | N1 | 1G:453 | 4.17 | 1.00 | 77.28 |  |
| 8 | G | OP2 | 1G:384 | 4.35 | 1.00 | 80.02 |  |
| 9 | C | O2 | 1G:454 | 4.36 | 1.00 | 92.52 |  |
| 10 | C | N3 | 1G:454 | 4.42 | 1.00 | 86.00 |  |
| 11 | A | N6 | 1G:453 | 4.68 | 1.00 | 76.83 |  |
| 12 | G | OP1 | 1G:384 | 4.81 | 1.00 | 77.49 |  |
  |

  
**1^st^ shell ligand angles:**   
1-2: 88; 1-3: 91; 1-4: 91; 1-5: 88; 1-6: 179; 2-3: 179; 2-4: 92; 2-5: 87; 2-6: 93; 3-4: 88; 3-5: 93; 3-6: 88; 4-5: 178; 4-6: 89; 5-6: 92;   
  
 Contact(s): /O_w_/O_w_/O_w_/O_w_/O_w_/O_w_/  
 Coded contact list: 6O_w_  

### MG:1G:1736

|  | Res. type | Atom | Residue | Distance | Occupancy | B-factor | Closest solute to 1^st^ shell water |
| --- | --- | --- | --- | --- | --- | --- | --- |
| **Ion** | | | |  | 1.00 | 62.90 || 1 | Water 1 | | 1G:1926 | 2.18 | 1.00 | 57.11 | 3.33 (G:O6 1G:1417) |
| 2 | Water 2 | | 1G:1871 | 2.19 | 1.00 | 44.33 | 2.60 (G:O6 1G:1417) |
| 3 | Water 3 | | 1G:1956 | 2.19 | 1.00 | 68.50 | 3.39 (G:O6 1G:1417) |
| 4 | Water 4 | | 1G:1953 | 2.19 | 1.00 | 73.42 | 3.36 (G:OP2 1G:1415) |
| 5 | Water 5 | | 1G:1966 | 2.20 | 1.00 | 45.92 | 3.98 (G:O6 1G:1417) |
| 6 | G | O6 | 1G:1417 | 2.60 | 1.00 | 47.84 |  |
| 7 | G | N7 | 1G:1416 | 4.36 | 1.00 | 45.21 |  |
| 8 | G | N1 | 1G:1417 | 4.57 | 1.00 | 44.92 |  |
| 9 | G | OP2 | 1G:1415 | 4.57 | 1.00 | 47.85 |  |
| 10 | G | O6 | 1G:1416 | 4.65 | 1.00 | 50.11 |  |
| 11 | G | O6 | 1G:1482 | 4.68 | 1.00 | 56.15 |  |
| 12 | G | N7 | 1G:1482 | 4.70 | 1.00 | 49.62 |  |
| 13 | G | N7 | 1G:1417 | 4.92 | 1.00 | 44.64 |  |
  |

  
**1^st^ shell ligand angles:**   
1-2: 85; 1-3: 177; 1-4: 90; 1-5: 96; 2-3: 93; 2-4: 91; 2-5: 176; 3-4: 91; 3-5: 86; 4-5: 93;   
  
 Contact(s): /O_w_/O_w_/O_w_/O_w_/O_w_/  
 Coded contact list: 5O_w_  
 Exclusion zone contact(s): /O6/  
Consider K in place of MG (1 valid contact(s) for K vs. 0 for MG)  

### K:1H:3001

|  | Res. type | Atom | Residue | Distance | Occupancy | B-factor | Closest solute to 1^st^ shell water |
| --- | --- | --- | --- | --- | --- | --- | --- |
| **Ion** | | | |  | 1.00 | 48.54 || 1 | G | O6 | 1H:917 | 2.65 | 1.00 | 36.72 |
| 2 | C | O2 | 1H:913 | 2.69 | 1.00 | 36.28 |  |
| 3 | U | O4 | 1H:916 | 2.96 | 1.00 | 38.65 |  |
| 4 | C | O2' | 1H:913 | 3.22 | 1.00 | 40.93 |  |
| 5 | G | N7 | 1H:917 | 3.98 | 1.00 | 39.19 |  |
| 6 | A | N6 | 1H:918 | 4.08 | 1.00 | 42.24 |  |
| 7 | C | N4 | 1H:955 | 4.14 | 1.00 | 33.77 |  |
| 8 | C | N1 | 1H:913 | 4.33 | 1.00 | 40.78 |  |
| 9 | A | N6 | 1H:956 | 4.36 | 1.00 | 35.47 |  |
| 10 | C | N3 | 1H:913 | 4.39 | 1.00 | 37.45 |  |
| 11 | C | OP2 | 1H:915 | 4.80 | 1.00 | 46.46 |  |
| 12 | G | N1 | 1H:917 | 4.83 | 1.00 | 36.20 |  |
| 13 | C | N4 | 1H:915 | 4.95 | 1.00 | 39.07 |  |
  |

  
**1^st^ shell ligand angles:**   
1-2: 88; 1-3: 61; 1-4: 122; 2-3: 70; 2-4: 72; 3-4: 141;   
  
 Contact(s): /O2/O2'/O4/O6/  
 Coded contact list: O_r_3O_b_  

### K:1H:3002

|  | Res. type | Atom | Residue | Distance | Occupancy | B-factor | Closest solute to 1^st^ shell water |
| --- | --- | --- | --- | --- | --- | --- | --- |
| **Ion** | | | |  | 1.00 | 73.77 || 1 | U | O4 | 1H:2325 | 3.02 | 1.00 | 54.45 |
| 2 | G | O6 | 1H:2317 | 3.13 | 1.00 | 65.59 |  |
| 3 | A | OP2 | 1H:2324 | 3.60 | 1.00 | 75.15 |  |
| 4 | A | OP1 | 1H:2324 | 3.77 | 1.00 | 73.41 |  |
| 5 | A | OP1 | 1H:2323 | 4.12 | 1.00 | 86.76 |  |
| 6 | G | N1 | 1H:2317 | 4.15 | 1.00 | 60.72 |  |
| 7 | G | OP2 | 1H:2321 | 4.38 | 1.00 | 69.50 |  |
| 8 | C | N4 | 1H:2326 | 4.86 | 1.00 | 66.40 |  |
| 9 | A | OP2 | 1H:2323 | 4.90 | 1.00 | 76.47 |  |
| 10 | G | O6 | 1H:2316 | 4.95 | 1.00 | 59.65 |  |
  |

  
**1^st^ shell ligand angles:**   
1-2: 79;   
  
 Contact(s): /O4/O6/  
 Coded contact list: 2O_b_  

### K:1H:3003

|  | Res. type | Atom | Residue | Distance | Occupancy | B-factor | Closest solute to 1^st^ shell water |
| --- | --- | --- | --- | --- | --- | --- | --- |
| **Ion** | | | |  | 1.00 | 56.99 || 1 | U | O4 | 1H:969 | 2.86 | 1.00 | 45.04 |
| 2 | G | O6 | 1H:903 | 2.93 | 1.00 | 44.19 |  |
| 3 | G | O6 | 1H:968 | 2.96 | 1.00 | 43.58 |  |
| 4 | G | O6 | 1H:902 | 3.24 | 1.00 | 50.93 |  |
| 5 | C | N4 | 1H:904 | 3.83 | 1.00 | 39.31 |  |
| 6 | Water 1 | | 1H:4587 | 4.21 | 1.00 | 58.13 | 4.55 (G:O6 1H:902) |
| 7 | Water 2 | | 1H:3859 | 4.36 | 1.00 | 56.82 | 2.94 (G:O6 1H:902) |
| 8 | G | N7 | 1H:903 | 4.40 | 1.00 | 50.35 |  |
| 9 | U | N3 | 1H:969 | 4.51 | 1.00 | 35.87 |  |
| 10 | C | N4 | 1H:970 | 4.67 | 1.00 | 48.05 |  |
| 11 | G | N7 | 1H:902 | 4.85 | 1.00 | 57.01 |  |
| 12 | G | N1 | 1H:968 | 4.93 | 1.00 | 34.88 |  |
| 13 | G | N7 | 1H:968 | 4.97 | 1.00 | 37.32 |  |
  |

  
**1^st^ shell ligand angles:**   
1-2: 63; 1-3: 59; 1-4: 84; 2-3: 70; 2-4: 64; 3-4: 130;   
  
 Contact(s): /O4/O6/O6/O6/  
 Coded contact list: 4O_b_  

### K:1H:3004

|  | Res. type | Atom | Residue | Distance | Occupancy | B-factor | Closest solute to 1^st^ shell water |
| --- | --- | --- | --- | --- | --- | --- | --- |
| **Ion** | | | |  | 1.00 | 88.42 || 1 | G | O6 | 1H:1535 | 2.83 | 1.00 | 74.10 |
| 2 | U | O4 | 1H:1536 | 3.04 | 1.00 | 71.54 |  |
| 3 | G | O6 | 1H:1547 | 3.46 | 1.00 | 65.17 |  |
| 4 | C | N4 | 1H:1548 | 3.97 | 1.00 | 57.94 |  |
| 5 | C | OP2 | 1H:1545 | 4.18 | 1.00 | 57.01 |  |
| 6 | U | N3 | 1H:1536 | 4.43 | 1.00 | 72.44 |  |
| 7 | G | O6 | 1H:1534 | 4.50 | 1.00 | 74.48 |  |
| 8 | G | N1 | 1H:1535 | 4.77 | 1.00 | 65.89 |  |
  |

  
**1^st^ shell ligand angles:**   
1-2: 65;   
  
 Contact(s): /O4/O6/  
 Coded contact list: 2O_b_  

### K:1H:3005

|  | Res. type | Atom | Residue | Distance | Occupancy | B-factor | Closest solute to 1^st^ shell water |
| --- | --- | --- | --- | --- | --- | --- | --- |
| **Ion** | | | |  | 1.00 | 100.29 || 1 | A | OP2 | 1H:2531 | 3.67 | 1.00 | 37.77 |
| 2 | Water 1 | | 1H:4268 | 3.77 | 1.00 | 40.61 | 3.00 (A:OP2 1H:2531) |
| 3 | A | OP1 | 1H:2531 | 4.44 | 1.00 | 37.32 |  |
| 4 | A | OP2 | 1H:2554 | 4.52 | 1.00 | 32.96 |  |
| 5 | A | OP2 | 1H:1174 | 4.55 | 1.00 | 59.11 |  |
| 6 | A | OP1 | 1H:1174 | 4.81 | 1.00 | 50.07 |  |
| 7 | A | O5' | 1H:1174 | 4.90 | 1.00 | 49.20 |  |
  |

  
 Coded contact list:   
1^st^ K coordination shell missing (see Notes)  

### K:1H:3006

|  | Res. type | Atom | Residue | Distance | Occupancy | B-factor | Closest solute to 1^st^ shell water |
| --- | --- | --- | --- | --- | --- | --- | --- |
| **Ion** | | | |  | 1.00 | 44.91 || 1 | G | O6 | 1H:743 | 2.51 | 1.00 | 25.89 |
| 2 | U | O4 | 1H:815 | 2.86 | 1.00 | 29.24 |  |
| 3 | Water 1 | | 1H:4188 | 2.97 | 1.00 | 24.29 | 2.88 (G:O6 1H:816) |
| 4 | U | O4 | 1H:742 | 3.07 | 1.00 | 20.92 |  |
| 5 | U | N3 | 1H:815 | 3.67 | 1.00 | 26.36 |  |
| 6 | G | O6 | 1H:816 | 3.71 | 1.00 | 23.20 |  |
| 7 | G | N7 | 1H:743 | 4.18 | 1.00 | 19.74 |  |
| 8 | Water 2 | | 1H:4539 | 4.53 | 1.00 | 31.25 | 3.90 (U:O4 1H:815) |
| 9 | Water 3 | | 1H:4280 | 4.54 | 1.00 | 23.83 | 3.13 (U:N3 1H:815) |
| 10 | U | N3 | 1H:742 | 4.58 | 1.00 | 21.96 |  |
| 11 | G | N1 | 1H:743 | 4.59 | 1.00 | 20.08 |  |
| 12 | C | N4 | 1H:741 | 4.82 | 1.00 | 18.95 |  |
| 13 | G | O6 | 1H:744 | 4.96 | 1.00 | 25.60 |  |
  |

  
**1^st^ shell ligand angles:**   
1-2: 79; 1-3: 99; 1-4: 107; 2-3: 94; 2-4: 173; 3-4: 83;   
  
 Contact(s): /O4/O4/O_w_/  
 Coded contact list: 2O_b_O_w_  
 Exclusion zone contact(s): /O6/  

### K:1H:3007

|  | Res. type | Atom | Residue | Distance | Occupancy | B-factor | Closest solute to 1^st^ shell water |
| --- | --- | --- | --- | --- | --- | --- | --- |
| **Ion** | | | |  | 1.00 | 55.05 || 1 | G | O6 | 1H:2502 | 3.22 | 1.00 | 35.57 |
| 2 | G | N7 | 1H:2502 | 3.32 | 1.00 | 33.95 |  |
| 3 | G | O6 | 1H:2503 | 3.42 | 1.00 | 36.71 |  |
| 4 | G | N1 | 1H:2503 | 3.56 | 1.00 | 35.70 |  |
| 5 | A | N7 | 1H:2501 | 3.78 | 1.00 | 37.33 |  |
| 6 | G | O2' | 1H:2471 | 3.82 | 1.00 | 33.49 |  |
| 7 | A | OP2 | 1H:2501 | 4.47 | 1.00 | 43.52 |  |
| 8 | A | N6 | 1H:2501 | 4.68 | 1.00 | 32.35 |  |
| 9 | G | N2 | 1H:2503 | 4.79 | 1.00 | 49.96 |  |
  |

  
**1^st^ shell ligand angles:**   
1-2: 56;   
  
 Contact(s): /N7/O6/  
 Coded contact list: O_b_N_b_  

### K:1H:3008

|  | Res. type | Atom | Residue | Distance | Occupancy | B-factor | Closest solute to 1^st^ shell water |
| --- | --- | --- | --- | --- | --- | --- | --- |
| **Ion** | | | |  | 1.00 | 65.91 || 1 | G | O6 | 1H:1081 | 2.55 | 1.00 | 43.63 |
| 2 | U | O4 | 1H:1082 | 2.85 | 1.00 | 51.58 |  |
| 3 | G | O6 | 1H:1167 | 3.53 | 1.00 | 54.44 |  |
| 4 | C | N4 | 1H:1168 | 3.66 | 1.00 | 52.63 |  |
| 5 | C | N4 | 1H:1166 | 3.92 | 1.00 | 69.85 |  |
| 6 | U | N3 | 1H:1082 | 4.10 | 1.00 | 50.24 |  |
| 7 | G | N7 | 1H:1081 | 4.39 | 1.00 | 46.69 |  |
| 8 | G | O6 | 1H:1169 | 4.54 | 1.00 | 38.79 |  |
| 9 | G | N1 | 1H:1081 | 4.72 | 1.00 | 40.35 |  |
| 10 | G | O6 | 1H:1083 | 4.76 | 1.00 | 67.96 |  |
  |

  
**1^st^ shell ligand angles:**   
1-2: 77;   
  
 Contact(s): /O4/  
 Coded contact list: O_b_  
 Exclusion zone contact(s): /O6/  

### K:1H:3009

|  | Res. type | Atom | Residue | Distance | Occupancy | B-factor | Closest solute to 1^st^ shell water |
| --- | --- | --- | --- | --- | --- | --- | --- |
| **Ion** | | | |  | 1.00 | 64.90 || 1 | G | O6 | 1H:2384 | 2.54 | 1.00 | 49.35 |
| 2 | G | O6 | 1H:2383 | 2.72 | 1.00 | 52.06 |  |
| 3 | G | O6 | 1H:2358 | 2.91 | 1.00 | 40.10 |  |
| 4 | G | N1 | 1H:2384 | 3.85 | 1.00 | 44.81 |  |
| 5 | G | N7 | 1H:2383 | 4.14 | 1.00 | 52.96 |  |
| 6 | G | O6 | 1H:2385 | 4.17 | 1.00 | 37.16 |  |
| 7 | C | N4 | 1H:2360 | 4.52 | 1.00 | 44.73 |  |
| 8 | G | N1 | 1H:2358 | 4.56 | 1.00 | 32.58 |  |
| 9 | A | N6 | 1H:2382 | 4.63 | 1.00 | 47.22 |  |
| 10 | G | N1 | 1H:2383 | 4.91 | 1.00 | 46.13 |  |
| 11 | C | N4 | 1H:2394 | 4.95 | 1.00 | 34.29 |  |
  |

  
**1^st^ shell ligand angles:**   
1-2: 72; 1-3: 86; 2-3: 68;   
  
 Contact(s): /O6/O6/  
 Coded contact list: 2O_b_  
 Exclusion zone contact(s): /O6/  

### K:1H:3010

|  | Res. type | Atom | Residue | Distance | Occupancy | B-factor | Closest solute to 1^st^ shell water |
| --- | --- | --- | --- | --- | --- | --- | --- |
| **Ion** | | | |  | 1.00 | 66.49 || 1 | G | O6 | 1H:81 | 2.74 | 1.00 | 49.44 |
| 2 | Water 1 | | 1H:3762 | 2.87 | 1.00 | 43.02 | 2.31 (G:O6 1H:82) |
| 3 | Water 2 | | 1H:4197 | 3.13 | 1.00 | 52.19 | 2.89 (G:O6 1H:81) |
| 4 | G | O6 | 1H:82 | 3.40 | 1.00 | 49.91 |  |
| 5 | Water 3 | | 1H:3861 | 3.56 | 1.00 | 56.98 | 2.46 (G:OP2 1H:80) |
| 6 | Water 4 | | 1H:4512 | 3.57 | 1.00 | 54.55 | 3.75 (G:OP2 1H:80) |
| 7 | Water 5 | | 1H:4334 | 3.80 | 1.00 | 48.26 | 3.74 (G:O6 1H:81) |
| 8 | G | O2' | 1H:100 | 4.53 | 1.00 | 46.52 |  |
| 9 | G | N7 | 1H:81 | 4.62 | 1.00 | 42.01 |  |
| 10 | G | OP2 | 1H:80 | 4.84 | 1.00 | 46.82 |  |
| 11 | G | N1 | 1H:81 | 4.98 | 1.00 | 40.52 |  |
  |

  
**1^st^ shell ligand angles:**   
1-2: 78; 1-3: 58; 1-4: 52; 2-3: 133; 2-4: 42; 3-4: 93;   
  
 Contact(s): /O6/O6/O_w_/O_w_/  
 Coded contact list: 2O_b_2O_w_  
K hydration shell issue: water too close (< 2.4 Angs.)  

### K:1H:3011

|  | Res. type | Atom | Residue | Distance | Occupancy | B-factor | Closest solute to 1^st^ shell water |
| --- | --- | --- | --- | --- | --- | --- | --- |
| **Ion** | | | |  | 1.00 | 59.54 || 1 | U | O4 | 1H:578 | 2.97 | 1.00 | 41.59 |
| 2 | G | O6 | 1H:564 | 3.15 | 1.00 | 41.63 |  |
| 3 | U | O4 | 1H:579 | 3.56 | 1.00 | 35.76 |  |
| 4 | C | N4 | 1H:563 | 4.10 | 1.00 | 43.46 |  |
| 5 | G | O6 | 1H:565 | 4.26 | 1.00 | 38.19 |  |
| 6 | G | N7 | 1H:564 | 4.41 | 1.00 | 47.77 |  |
| 7 | U | N3 | 1H:578 | 4.45 | 1.00 | 35.85 |  |
| 8 | U | N3 | 1H:579 | 4.46 | 1.00 | 37.26 |  |
| 9 | G | O6 | 1H:577 | 4.49 | 1.00 | 40.41 |  |
  |

  
**1^st^ shell ligand angles:**   
1-2: 97;   
  
 Contact(s): /O4/O6/  
 Coded contact list: 2O_b_  

### K:1H:3012

|  | Res. type | Atom | Residue | Distance | Occupancy | B-factor | Closest solute to 1^st^ shell water |
| --- | --- | --- | --- | --- | --- | --- | --- |
| **Ion** | | | |  | 1.00 | 51.51 || 1 | G | O6 | 1H:620 | 2.95 | 1.00 | 29.70 |
| 2 | G | O6 | 1H:708 | 3.15 | 1.00 | 32.92 |  |
| 3 | U | O4 | 1H:621 | 3.26 | 1.00 | 24.78 |  |
| 4 | C | N4 | 1H:707 | 3.63 | 1.00 | 33.07 |  |
| 5 | C | N4 | 1H:709 | 3.80 | 1.00 | 29.22 |  |
| 6 | U | N3 | 1H:621 | 4.42 | 1.00 | 26.57 |  |
| 7 | C | N4 | 1H:619 | 4.60 | 1.00 | 26.61 |  |
| 8 | G | O6 | 1H:622 | 4.62 | 1.00 | 34.46 |  |
| 9 | G | N7 | 1H:708 | 4.66 | 1.00 | 30.60 |  |
  |

  
**1^st^ shell ligand angles:**   
1-2: 70; 1-3: 61; 2-3: 67;   
  
 Contact(s): /O4/O6/O6/  
 Coded contact list: 3O_b_  

### K:1H:3013

|  | Res. type | Atom | Residue | Distance | Occupancy | B-factor | Closest solute to 1^st^ shell water |
| --- | --- | --- | --- | --- | --- | --- | --- |
| **Ion** | | | |  | 1.00 | 81.10 || 1 | U | OP2 | 1H:2768 | 3.22 | 1.00 | 54.70 |
| 2 | G | O6 | 1H:2761 | 3.26 | 1.00 | 55.98 |  |
| 3 | C | OP2 | 1H:2769 | 3.39 | 1.00 | 54.75 |  |
| 4 | G | N7 | 1H:2761 | 3.60 | 1.00 | 59.59 |  |
| 5 | Water 1 | | 1H:4166 | 4.08 | 1.00 | 66.80 | 2.85 (U:OP1 1H:2768) |
| 6 | U | O5' | 1H:2768 | 4.21 | 1.00 | 62.74 |  |
| 7 | U | OP2 | 1H:2760 | 4.23 | 1.00 | 53.27 |  |
| 8 | U | OP1 | 1H:2768 | 4.57 | 1.00 | 60.98 |  |
| 9 | C | OP2 | 1H:2759 | 4.68 | 1.00 | 48.80 |  |
  |

  
**1^st^ shell ligand angles:**   
1-2: 98; 1-3: 76; 2-3: 57;   
  
 Contact(s): /O6/OP2/OP2/  
 Coded contact list: 2O_ph_O_b_  

### K:1H:3014

|  | Res. type | Atom | Residue | Distance | Occupancy | B-factor | Closest solute to 1^st^ shell water |
| --- | --- | --- | --- | --- | --- | --- | --- |
| **Ion** | | | |  | 1.00 | 60.89 || 1 | G | O6 | 1H:1332 | 2.64 | 1.00 | 40.39 |
| 2 | G | O2' | 1H:1330 | 2.96 | 1.00 | 48.36 |  |
| 3 | U | O4' | 1H:1376 | 3.32 | 1.00 | 28.16 |  |
| 4 | U | O2 | 1H:1376 | 3.69 | 1.00 | 41.83 |  |
| 5 | U | N1 | 1H:1376 | 4.23 | 1.00 | 33.14 |  |
| 6 | G | N7 | 1H:1332 | 4.42 | 1.00 | 40.32 |  |
| 7 | A | N7 | 1H:1331 | 4.51 | 1.00 | 49.94 |  |
| 8 | G | O4' | 1H:1330 | 4.51 | 1.00 | 47.44 |  |
| 9 | U | O5' | 1H:1376 | 4.67 | 1.00 | 28.88 |  |
| 10 | G | O3' | 1H:1330 | 4.72 | 1.00 | 50.04 |  |
| 11 | U | N3 | 1H:1376 | 4.84 | 1.00 | 39.48 |  |
| 12 | G | N1 | 1H:1332 | 4.86 | 1.00 | 37.69 |  |
| 13 | C | OP2 | 1H:1367 | 4.90 | 1.00 | 31.21 |  |
  |

  
**1^st^ shell ligand angles:**   
1-2: 84; 1-3: 64; 2-3: 124;   
  
 Contact(s): /O2'/O4'/O6/  
 Coded contact list: 2O_r_O_b_  

### K:1H:3015

|  | Res. type | Atom | Residue | Distance | Occupancy | B-factor | Closest solute to 1^st^ shell water |
| --- | --- | --- | --- | --- | --- | --- | --- |
| **Ion** | | | |  | 1.00 | 44.52 || 1 | A | O2' | 1H:2795 | 2.87 | 1.00 | 39.04 |
| 2 | A | O2' | 1H:2790 | 2.92 | 1.00 | 41.49 |  |
| 3 | G | OP2 | 1H:2796 | 3.03 | 1.00 | 41.61 |  |
| 4 | C | O2 | 1H:2641 | 3.33 | 1.00 | 37.32 |  |
| 5 | G | OP2 | 1H:2791 | 3.85 | 1.00 | 35.67 |  |
| 6 | A | O3' | 1H:2790 | 3.87 | 1.00 | 36.57 |  |
| 7 | A | O3' | 1H:2795 | 3.95 | 1.00 | 41.46 |  |
| 8 | C | N3 | 1H:2641 | 4.33 | 1.00 | 25.57 |  |
| 9 | G | N7 | 1H:2796 | 4.56 | 1.00 | 39.97 |  |
| 10 | G | OP1 | 1H:2796 | 4.61 | 1.00 | 36.41 |  |
| 11 | G | OP1 | 1H:2791 | 4.75 | 1.00 | 35.25 |  |
  |

  
**1^st^ shell ligand angles:**   
1-2: 78; 1-3: 82; 1-4: 67; 2-3: 95; 2-4: 143; 3-4: 91;   
  
 Contact(s): /O2/O2'/O2'/OP2/  
 Coded contact list: O_ph_2O_r_O_b_  

### K:1H:3016

|  | Res. type | Atom | Residue | Distance | Occupancy | B-factor | Closest solute to 1^st^ shell water |
| --- | --- | --- | --- | --- | --- | --- | --- |
| **Ion** | | | |  | 1.00 | 63.21 || 1 | G | O2' | 1H:138 | 2.77 | 1.00 | 45.26 |
| 2 | C | N4 | 1H:135 | 2.81 | 1.00 | 46.08 |  |
| 3 | A | OP2 | 1H:139 | 2.92 | 1.00 | 48.05 |  |
| 4 | G | O6 | 1H:136 | 3.42 | 1.00 | 35.49 |  |
| 5 | Water 1 | | 1H:4524 | 3.48 | 1.00 | 62.79 | 3.82 (A:OP2 1H:139) |
| 6 | G | N7 | 1H:136 | 3.70 | 1.00 | 42.44 |  |
| 7 | C | N4 | 1H:141 | 3.96 | 1.00 | 35.11 |  |
| 8 | G | O6 | 1H:142 | 4.21 | 1.00 | 38.91 |  |
| 9 | G | O3' | 1H:138 | 4.27 | 1.00 | 40.58 |  |
| 10 | C | N3 | 1H:135 | 4.37 | 1.00 | 38.73 |  |
| 11 | Water 2 | | 1H:4125 | 4.39 | 1.00 | 53.65 | 2.80 (A:OP2 1H:139) |
| 12 | A | O5' | 1H:139 | 4.47 | 1.00 | 39.52 |  |
| 13 | Water 3 | | 1H:4151 | 4.67 | 1.00 | 46.93 | 3.80 (C:N4 1H:135) |
| 14 | A | OP2 | 1H:140 | 4.73 | 1.00 | 40.36 |  |
  |

  
**1^st^ shell ligand angles:**   
1-2: 140; 1-3: 72; 2-3: 145;   
  
 Contact(s): /N4/O2'/OP2/  
 Coded contact list: O_ph_O_r_Anion  

### K:1H:3017

|  | Res. type | Atom | Residue | Distance | Occupancy | B-factor | Closest solute to 1^st^ shell water |
| --- | --- | --- | --- | --- | --- | --- | --- |
| **Ion** | | | |  | 1.00 | 56.93 || 1 | G | O6 | 1H:26 | 3.00 | 1.00 | 37.67 |
| 2 | G | O6 | 1H:27 | 3.09 | 1.00 | 38.72 |  |
| 3 | U | OP2 | 1H:25 | 3.97 | 1.00 | 29.22 |  |
| 4 | G | N1 | 1H:27 | 4.46 | 1.00 | 27.49 |  |
| 5 | G | N7 | 1H:26 | 4.47 | 1.00 | 26.99 |  |
| 6 | Water 1 | | 1H:4012 | 4.56 | 1.00 | 39.02 | 4.00 (G:O6 1H:27) |
| 7 | G | O2' | 1H:538 | 4.67 | 1.00 | 32.98 |  |
  |

  
**1^st^ shell ligand angles:**   
1-2: 58;   
  
 Contact(s): /O6/O6/  
 Coded contact list: 2O_b_  

### K:1H:3018

|  | Res. type | Atom | Residue | Distance | Occupancy | B-factor | Closest solute to 1^st^ shell water |
| --- | --- | --- | --- | --- | --- | --- | --- |
| **Ion** | | | |  | 1.00 | 51.71 || 1 | A | O2' | 1H:2834 | 2.75 | 1.00 | 22.92 |
| 2 | C | O2 | 1H:2029 | 3.26 | 1.00 | 25.39 |  |
| 3 | A | O3' | 1H:2834 | 3.27 | 1.00 | 29.32 |  |
| 4 | A | O2' | 1H:1700 | 3.30 | 1.00 | 33.95 |  |
| 5 | A | N3 | 1H:1700 | 3.60 | 1.00 | 28.16 |  |
| 6 | G | N2 | 1H:1699 | 4.11 | 1.00 | 27.53 |  |
| 7 | C | O2' | 1H:2029 | 4.39 | 1.00 | 23.35 |  |
| 8 | C | OP1 | 1H:2835 | 4.44 | 1.00 | 26.73 |  |
| 9 | A | O4' | 1H:2834 | 4.66 | 1.00 | 21.81 |  |
| 10 | C | O5' | 1H:2835 | 4.85 | 1.00 | 25.42 |  |
| 11 | G | O4' | 1H:1701 | 4.86 | 1.00 | 28.23 |  |
| 12 | A | N9 | 1H:1700 | 4.88 | 1.00 | 31.76 |  |
| 13 | C | N1 | 1H:2029 | 4.98 | 1.00 | 21.41 |  |
  |

  
**1^st^ shell ligand angles:**   
1-2: 81; 1-3: 53; 1-4: 138; 2-3: 133; 2-4: 122; 3-4: 91;   
  
 Contact(s): /O2/O2'/O2'/O3'/  
 Coded contact list: 3O_r_O_b_  

### K:1H:3019

|  | Res. type | Atom | Residue | Distance | Occupancy | B-factor | Closest solute to 1^st^ shell water |
| --- | --- | --- | --- | --- | --- | --- | --- |
| **Ion** | | | |  | 1.00 | 60.75 || 1 | G | O6 | 1H:1025 | 3.17 | 1.00 | 31.85 |
| 2 | G | O6 | 1H:1026 | 3.22 | 1.00 | 32.09 |  |
| 3 | Water 1 | | 1H:4264 | 3.48 | 1.00 | 37.35 | 2.99 (G:N7 1H:1025) |
| 4 | Water 2 | | 1H:3812 | 3.53 | 1.00 | 50.17 | 2.39 (G:O6 1H:1026) |
| 5 | G | N7 | 1H:1025 | 3.60 | 1.00 | 36.12 |  |
| 6 | Water 3 | | 1H:4537 | 4.02 | 1.00 | 35.08 | 4.15 (G:O6 1H:1024) |
| 7 | G | N7 | 1H:1024 | 4.31 | 1.00 | 35.52 |  |
| 8 | G | O6 | 1H:1024 | 4.46 | 1.00 | 33.81 |  |
| 9 | G | N1 | 1H:1026 | 4.62 | 1.00 | 36.31 |  |
| 10 | Water 4 | | 1H:3602 | 4.80 | 1.00 | 36.80 | 1.30 (G:N7 1H:1026) |
| 11 | A | OP2 | 1H:1030 | 4.88 | 1.00 | 40.06 |  |
| 12 | Water 5 | | 1H:4431 | 4.98 | 1.00 | 30.72 | 3.99 (A:OP2 1H:1030) |
| 13 | G | N7 | 1H:1026 | 4.99 | 1.00 | 37.75 |  |
  |

  
**1^st^ shell ligand angles:**   
1-2: 67;   
  
 Contact(s): /O6/O6/  
 Coded contact list: 2O_b_  
K hydration shell issue: water too close (< 2.4 Angs.)  

### K:1H:3020

|  | Res. type | Atom | Residue | Distance | Occupancy | B-factor | Closest solute to 1^st^ shell water |
| --- | --- | --- | --- | --- | --- | --- | --- |
| **Ion** | | | |  | 1.00 | 65.85 || 1 | Water 1 | | 1H:4360 | 2.99 | 1.00 | 40.57 | 5.04 (A:OP2 1H:2280) |
| 2 | A | O2' | 1H:2279 | 3.03 | 1.00 | 32.30 |  |
| 3 | Water 2 | | 1H:3676 | 3.13 | 1.00 | 36.27 | 2.15 (U:O4 1H:2285) |
| 4 | A | N3 | 1H:2279 | 3.67 | 1.00 | 37.54 |  |
| 5 | Water 3 | | 1H:4261 | 3.91 | 1.00 | 55.47 | 5.00 (A:N3 1H:2279) |
| 6 | A | O3' | 1H:2279 | 4.01 | 1.00 | 31.03 |  |
| 7 | Water 4 | | 1H:3751 | 4.04 | 1.00 | 39.10 | 2.30 (A:OP2 1H:2280) |
| 8 | U | O4 | 1H:2285 | 4.16 | 1.00 | 34.65 |  |
| 9 | Water 5 | | 1H:4568 | 4.53 | 1.00 | 44.20 | 4.74 (A:N3 1H:2279) |
| 10 | U | OP2 | 1H:2278 | 4.67 | 1.00 | 31.27 |  |
| 11 | A | N9 | 1H:2279 | 4.71 | 1.00 | 33.46 |  |
| 12 | A | OP2 | 1H:2280 | 4.89 | 1.00 | 33.91 |  |
| 13 | Water 6 | | 1H:3736 | 4.97 | 1.00 | 36.11 | 4.28 (U:O4 1H:2285) |
  |

  
**1^st^ shell ligand angles:**   
1-2: 147; 1-3: 61; 2-3: 97;   
  
 Contact(s): /O2'/O_w_/O_w_/  
 Coded contact list: O_r_2O_w_  
K hydration shell issue: water too close (< 2.4 Angs.)  

### K:1H:3021

|  | Res. type | Atom | Residue | Distance | Occupancy | B-factor | Closest solute to 1^st^ shell water |
| --- | --- | --- | --- | --- | --- | --- | --- |
| **Ion** | | | |  | 1.00 | 78.20 || 1 | U | O4 | 1H:266 | 3.15 | 1.00 | 98.08 |
| 2 | G | O6 | 1H:278 | 3.59 | 1.00 | 102.98 |  |
| 3 | G | O6 | 1H:279 | 3.61 | 1.00 | 102.49 |  |
| 4 | G | N7 | 1H:278 | 4.34 | 1.00 | 107.42 |  |
| 5 | C | N4 | 1H:267 | 4.55 | 1.00 | 103.21 |  |
| 6 | C | N4 | 1H:277 | 4.66 | 1.00 | 119.40 |  |
  |

  
 Contact(s): /O4/  
 Coded contact list: O_b_  

### K:1H:3022

|  | Res. type | Atom | Residue | Distance | Occupancy | B-factor | Closest solute to 1^st^ shell water |
| --- | --- | --- | --- | --- | --- | --- | --- |
| **Ion** | | | |  | 1.00 | 66.53 || 1 | Water 1 | | 1H:4153 | 2.99 | 1.00 | 25.37 | 2.83 (A:OP2 1H:1685) |
| 2 | G | OP2 | 1H:1346 | 3.17 | 1.00 | 22.74 |  |
| 3 | A | OP2 | 1H:1685 | 3.52 | 1.00 | 28.53 |  |
| 4 | A | OP1 | 1H:1685 | 3.83 | 1.00 | 39.37 |  |
| 5 | C | OP2 | 1H:1686 | 3.85 | 1.00 | 29.10 |  |
| 6 | G | OP1 | 1H:1346 | 4.04 | 1.00 | 22.65 |  |
| 7 | C | OP1 | 1H:1345 | 4.22 | 1.00 | 24.95 |  |
| 8 | A | O5' | 1H:1685 | 4.23 | 1.00 | 29.89 |  |
| 9 | Water 2 | | 1H:4232 | 4.93 | 1.00 | 40.46 | 3.01 (G:OP1 1H:1346) |
  |

  
**1^st^ shell ligand angles:**   
1-2: 91;   
  
 Contact(s): /O_w_/OP2/  
 Coded contact list: O_ph_O_w_  

### K:1H:3023

|  | Res. type | Atom | Residue | Distance | Occupancy | B-factor | Closest solute to 1^st^ shell water |
| --- | --- | --- | --- | --- | --- | --- | --- |
| **Ion** | | | |  | 1.00 | 63.16 || 1 | C | OP2 | 1H:1017 | 2.98 | 1.00 | 23.08 |
| 2 | G | N7 | 1H:1020 | 3.19 | 1.00 | 35.29 |  |
| 3 | G | O6 | 1H:1020 | 3.25 | 1.00 | 34.51 |  |
| 4 | G | O6 | 1H:1036 | 3.80 | 1.00 | 33.17 |  |
| 5 | G | O6 | 1H:1018 | 4.53 | 1.00 | 34.47 |  |
  |

  
**1^st^ shell ligand angles:**   
1-2: 52; 1-3: 97; 2-3: 58;   
  
 Contact(s): /N7/O6/OP2/  
 Coded contact list: O_ph_O_b_N_b_  

### K:1H:3024

|  | Res. type | Atom | Residue | Distance | Occupancy | B-factor | Closest solute to 1^st^ shell water |
| --- | --- | --- | --- | --- | --- | --- | --- |
| **Ion** | | | |  | 1.00 | 59.66 || 1 | G | OP1 | 1H:248 | 2.90 | 1.00 | 36.97 |
| 2 | A | O3' | 1H:247 | 3.03 | 1.00 | 31.62 |  |
| 3 | A | N3 | 1H:215 | 3.04 | 1.00 | 40.93 |  |
| 4 | C | O2 | 1H:221 | 3.27 | 1.00 | 49.57 |  |
| 5 | A | O2' | 1H:215 | 3.88 | 1.00 | 43.99 |  |
| 6 | A | N6 | 1H:218 | 3.92 | 1.00 | 48.44 |  |
| 7 | A | O2' | 1H:247 | 3.95 | 1.00 | 37.40 |  |
| 8 | G | O5' | 1H:248 | 4.32 | 1.00 | 31.00 |  |
| 9 | G | N2 | 1H:214 | 4.54 | 1.00 | 43.12 |  |
| 10 | A | N1 | 1H:215 | 4.61 | 1.00 | 40.52 |  |
| 11 | U | O2 | 1H:220 | 4.66 | 1.00 | 57.78 |  |
| 12 | G | O4' | 1H:216 | 4.80 | 1.00 | 42.43 |  |
| 13 | G | OP2 | 1H:248 | 4.83 | 1.00 | 37.34 |  |
| 14 | C | O2' | 1H:221 | 4.83 | 1.00 | 57.31 |  |
| 15 | C | N3 | 1H:221 | 4.87 | 1.00 | 44.69 |  |
| 16 | C | N1 | 1H:221 | 4.90 | 1.00 | 50.10 |  |
  |

  
**1^st^ shell ligand angles:**   
1-2: 50; 1-3: 136; 1-4: 81; 2-3: 129; 2-4: 129; 3-4: 73;   
  
 Contact(s): /N3_A_/O2/O3'/OP1/  
 Coded contact list: O_ph_O_r_O_b_N_b_  

### K:1H:3025

|  | Res. type | Atom | Residue | Distance | Occupancy | B-factor | Closest solute to 1^st^ shell water |
| --- | --- | --- | --- | --- | --- | --- | --- |
| **Ion** | | | |  | 1.00 | 53.77 || 1 | G | OP1 | 1H:986 | 2.63 | 1.00 | 33.25 |
| 2 | C | O2' | 1H:712 | 2.78 | 1.00 | 23.08 |  |
| 3 | C | O4' | 1H:713 | 3.31 | 1.00 | 24.17 |  |
| 4 | C | O2 | 1H:712 | 3.60 | 1.00 | 24.97 |  |
| 5 | G | OP2 | 1H:986 | 4.29 | 1.00 | 23.51 |  |
| 6 | G | N2 | 1H:617 | 4.60 | 1.00 | 26.55 |  |
| 7 | A | OP2 | 1H:987 | 4.82 | 1.00 | 28.77 |  |
| 8 | G | O5' | 1H:986 | 4.84 | 1.00 | 20.33 |  |
| 9 | C | N1 | 1H:712 | 4.88 | 1.00 | 25.47 |  |
| 10 | G | O3' | 1H:985 | 4.89 | 1.00 | 26.17 |  |
| 11 | C | O5' | 1H:713 | 4.95 | 1.00 | 25.09 |  |
  |

  
**1^st^ shell ligand angles:**   
1-2: 80; 1-3: 138; 2-3: 66;   
  
 Contact(s): /O2'/O4'/OP1/  
 Coded contact list: O_ph_2O_r_  

### K:1H:3026

|  | Res. type | Atom | Residue | Distance | Occupancy | B-factor | Closest solute to 1^st^ shell water |
| --- | --- | --- | --- | --- | --- | --- | --- |
| **Ion** | | | |  | 1.00 | 88.40 || 1 | G | O6 | 1H:2658 | 3.45 | 1.00 | 50.51 |
| 2 | G | OP2 | 1H:2656 | 3.57 | 1.00 | 31.24 |  |
| 3 | Water 1 | | 1H:4193 | 3.67 | 1.00 | 50.93 | 2.89 (G:N7 1H:2656) |
| 4 | Water 2 | | 1H:4574 | 3.67 | 1.00 | 46.97 | 4.63 (G:O6 1H:2657) |
| 5 | G | O6 | 1H:2657 | 4.00 | 1.00 | 46.60 |  |
| 6 | G | N1 | 1H:2658 | 4.69 | 1.00 | 44.58 |  |
| 7 | G | N7 | 1H:2657 | 4.78 | 1.00 | 44.24 |  |
| 8 | Water 3 | | 1H:4240 | 4.95 | 1.00 | 48.21 | 2.95 (G:O6 1H:2657) |
| 9 | G | N7 | 1H:2656 | 4.98 | 1.00 | 37.03 |  |
  |

  
 Coded contact list:   
1^st^ K coordination shell missing (see Notes)  

### K:1H:3027

|  | Res. type | Atom | Residue | Distance | Occupancy | B-factor | Closest solute to 1^st^ shell water |
| --- | --- | --- | --- | --- | --- | --- | --- |
| **Ion** | | | |  | 1.00 | 63.75 || 1 | G | O5' | 1H:2263 | 2.78 | 1.00 | 22.28 |
| 2 | G | O4' | 1H:2263 | 3.02 | 1.00 | 20.52 |  |
| 3 | G | N3 | 1H:2263 | 3.08 | 1.00 | 21.31 |  |
| 4 | G | OP2 | 1H:2263 | 3.20 | 1.00 | 21.79 |  |
| 5 | G | N2 | 1H:2263 | 3.86 | 1.00 | 25.48 |  |
| 6 | G | N9 | 1H:2263 | 4.06 | 1.00 | 24.79 |  |
| 7 | G | N1 | 1H:2263 | 4.69 | 1.00 | 25.19 |  |
| 8 | U | O3' | 1H:2262 | 4.71 | 1.00 | 20.39 |  |
| 9 | G | OP1 | 1H:2263 | 4.76 | 1.00 | 25.46 |  |
  |

  
**1^st^ shell ligand angles:**   
1-2: 59; 1-3: 75; 1-4: 46; 2-3: 61; 2-4: 105; 3-4: 96;   
  
 Contact(s): /N3_G_/O4'/O5'/OP2/  
 Coded contact list: O_ph_2O_r_N_b_  

### K:1H:3028

|  | Res. type | Atom | Residue | Distance | Occupancy | B-factor | Closest solute to 1^st^ shell water |
| --- | --- | --- | --- | --- | --- | --- | --- |
| **Ion** | | | |  | 1.00 | 87.72 || 1 | C | OP1 | 1H:1516 | 3.14 | 1.00 | 55.16 |
| 2 | Water 1 | | 1H:3933 | 3.49 | 1.00 | 56.73 | 2.55 (C:OP1 1H:1516) |
| 3 | C | O3' | 1H:1515 | 3.61 | 1.00 | 62.57 |  |
| 4 | Water 2 | | 1H:4408 | 3.97 | 1.00 | 46.68 | 4.74 (G:OP2 1H:1488) |
| 5 | Water 3 | | 1H:4567 | 4.07 | 1.00 | 62.13 | 4.18 (C:OP1 1H:1516) |
| 6 | C | O2' | 1H:1515 | 4.14 | 1.00 | 59.02 |  |
| 7 | G | OP2 | 1H:1487 | 4.18 | 1.00 | 48.47 |  |
| 8 | G | OP2 | 1H:1488 | 4.92 | 1.00 | 39.75 |  |
  |

  
 Contact(s): /OP1/  
 Coded contact list: O_ph_  

### K:1H:3029

|  | Res. type | Atom | Residue | Distance | Occupancy | B-factor | Closest solute to 1^st^ shell water |
| --- | --- | --- | --- | --- | --- | --- | --- |
| **Ion** | | | |  | 1.00 | 85.17 || 1 | U | O2 | 1H:2419 | 3.04 | 1.00 | 34.99 |
| 2 | GLU | O | 78:74 | 3.41 | 1.00 | 46.64 |  |
| 3 | PRO | O | 78:72 | 4.50 | 1.00 | 30.18 |  |
| 4 | LYS | N | 78:76 | 4.71 | 1.00 | 49.56 |  |
| 5 | U | O2' | 1H:2419 | 4.78 | 1.00 | 60.36 |  |
| 6 | GLY | O | 78:73 | 4.86 | 1.00 | 35.46 |  |
| 7 | A | OP2 | 1H:217 | 4.87 | 1.00 | 44.93 |  |
  |

  
 Contact(s): /O2/  
 Coded contact list: O_b_  

### K:1H:3030

|  | Res. type | Atom | Residue | Distance | Occupancy | B-factor | Closest solute to 1^st^ shell water |
| --- | --- | --- | --- | --- | --- | --- | --- |
| **Ion** | | | |  | 1.00 | 73.54 || 1 | G | N2 | 1H:1973 | 2.53 | 1.00 | 36.33 |
| 2 | U | O4 | 1H:1974 | 3.05 | 1.00 | 39.36 |  |
| 3 | G | OP2 | 1H:1973 | 4.11 | 1.00 | 27.25 |  |
| 4 | G | OP2 | 1H:1972 | 4.32 | 1.00 | 27.49 |  |
| 5 | G | N1 | 1H:1973 | 4.34 | 1.00 | 39.42 |  |
| 6 | A | OP1 | 1H:1976 | 4.42 | 1.00 | 37.83 |  |
| 7 | G | N3 | 1H:1973 | 4.62 | 1.00 | 27.07 |  |
  |

  
**1^st^ shell ligand angles:**   
1-2: 62;   
  
 Contact(s): /O4/  
 Coded contact list: O_b_  
 Exclusion zone contact(s): /N2/  

### K:1H:3031

|  | Res. type | Atom | Residue | Distance | Occupancy | B-factor | Closest solute to 1^st^ shell water |
| --- | --- | --- | --- | --- | --- | --- | --- |
| **Ion** | | | |  | 1.00 | 39.70 || 1 | A | O3' | 1H:186 | 2.73 | 1.00 | 13.95 |
| 2 | A | O2' | 1H:186 | 2.92 | 1.00 | 16.98 |  |
| 3 | A | OP2 | 1H:185 | 3.39 | 1.00 | 25.14 |  |
| 4 | A | OP1 | 1H:187 | 3.54 | 1.00 | 16.05 |  |
| 5 | A | OP1 | 1H:185 | 4.00 | 1.00 | 18.42 |  |
| 6 | G | OP2 | 1H:184 | 4.23 | 1.00 | 19.80 |  |
| 7 | A | O5' | 1H:187 | 4.37 | 1.00 | 16.08 |  |
| 8 | G | OP1 | 1H:184 | 4.58 | 1.00 | 23.75 |  |
| 9 | A | O4' | 1H:186 | 4.73 | 1.00 | 20.20 |  |
| 10 | A | N9 | 1H:186 | 4.89 | 1.00 | 19.06 |  |
| 11 | G | O5' | 1H:184 | 4.93 | 1.00 | 20.73 |  |
  |

  
**1^st^ shell ligand angles:**   
1-2: 55; 1-3: 75; 2-3: 121;   
  
 Contact(s): /O2'/O3'/OP2/  
 Coded contact list: O_ph_2O_r_  

### K:1H:3032

|  | Res. type | Atom | Residue | Distance | Occupancy | B-factor | Closest solute to 1^st^ shell water |
| --- | --- | --- | --- | --- | --- | --- | --- |
| **Ion** | | | |  | 1.00 | 54.80 || 1 | C | OP1 | 1H:2659 | 2.66 | 1.00 | 43.49 |
| 2 | A | OP1 | 1H:2747 | 2.88 | 1.00 | 45.45 |  |
| 3 | G | O2' | 1H:2746 | 3.55 | 1.00 | 49.70 |  |
| 4 | C | OP2 | 1H:2659 | 3.56 | 1.00 | 44.60 |  |
| 5 | U | OP2 | 1H:2660 | 4.46 | 1.00 | 38.09 |  |
| 6 | G | O3' | 1H:2658 | 4.56 | 1.00 | 48.80 |  |
| 7 | C | O5' | 1H:2659 | 4.68 | 1.00 | 34.33 |  |
| 8 | A | O3' | 1H:2747 | 4.74 | 1.00 | 71.28 |  |
| 9 | G | O3' | 1H:2746 | 4.79 | 1.00 | 52.73 |  |
| 10 | A | O5' | 1H:2747 | 4.88 | 1.00 | 60.52 |  |
  |

  
**1^st^ shell ligand angles:**   
1-2: 69;   
  
 Contact(s): /OP1/OP1/  
 Coded contact list: 2O_ph_  

### K:1H:3033

|  | Res. type | Atom | Residue | Distance | Occupancy | B-factor | Closest solute to 1^st^ shell water |
| --- | --- | --- | --- | --- | --- | --- | --- |
| **Ion** | | | |  | 1.00 | 36.86 || 1 | C | OP1 | 1H:1718 | 2.41 | 1.00 | 27.86 |
| 2 | U | OP2 | 1H:1719 | 2.59 | 1.00 | 23.80 |  |
| 3 | G | OP2 | 1H:2563 | 4.03 | 1.00 | 27.73 |  |
| 4 | C | O5' | 1H:1718 | 4.49 | 1.00 | 24.37 |  |
| 5 | G | OP1 | 1H:2562 | 4.67 | 1.00 | 25.22 |  |
| 6 | U | O5' | 1H:1719 | 4.71 | 1.00 | 23.32 |  |
| 7 | C | OP2 | 1H:1718 | 4.74 | 1.00 | 21.37 |  |
| 8 | A | O3' | 1H:1717 | 4.74 | 1.00 | 22.18 |  |
| 9 | U | OP1 | 1H:1719 | 4.80 | 1.00 | 21.08 |  |
| 10 | G | OP2 | 1H:2562 | 4.91 | 1.00 | 28.61 |  |
  |

  
**1^st^ shell ligand angles:**   
1-2: 73;   
  
 Coded contact list:   
 Exclusion zone contact(s): /OP1/OP2/  
Consider NA in place of K (2 valid contact(s) for NA vs. 0 for K)  

### K:1H:3034

|  | Res. type | Atom | Residue | Distance | Occupancy | B-factor | Closest solute to 1^st^ shell water |
| --- | --- | --- | --- | --- | --- | --- | --- |
| **Ion** | | | |  | 1.00 | 66.51 || 1 | G | O6 | 1H:24 | 3.03 | 1.00 | 31.26 |
| 2 | U | O4 | 1H:25 | 3.14 | 1.00 | 29.64 |  |
| 3 | A | OP2 | 1H:540 | 3.72 | 1.00 | 23.98 |  |
| 4 | A | OP2 | 1H:539 | 3.87 | 1.00 | 28.03 |  |
| 5 | G | N7 | 1H:24 | 4.47 | 1.00 | 25.33 |  |
| 6 | U | N3 | 1H:25 | 4.47 | 1.00 | 27.42 |  |
| 7 | C | N4 | 1H:542 | 4.51 | 1.00 | 27.38 |  |
| 8 | G | N2 | 1H:26 | 4.65 | 1.00 | 24.37 |  |
| 9 | G | O6 | 1H:23 | 4.82 | 1.00 | 26.74 |  |
| 10 | G | N1 | 1H:26 | 4.99 | 1.00 | 27.68 |  |
  |

  
**1^st^ shell ligand angles:**   
1-2: 67;   
  
 Contact(s): /O4/O6/  
 Coded contact list: 2O_b_  

### K:1H:3035

|  | Res. type | Atom | Residue | Distance | Occupancy | B-factor | Closest solute to 1^st^ shell water |
| --- | --- | --- | --- | --- | --- | --- | --- |
| **Ion** | | | |  | 1.00 | 52.07 || 1 | G | O6 | 1H:2698 | 2.45 | 1.00 | 45.42 |
| 2 | G | O6 | 1H:2699 | 3.81 | 1.00 | 36.34 |  |
| 3 | G | N7 | 1H:2698 | 3.92 | 1.00 | 41.87 |  |
| 4 | G | N7 | 1H:2699 | 4.11 | 1.00 | 36.35 |  |
| 5 | U | O4 | 1H:2697 | 4.22 | 1.00 | 39.24 |  |
| 6 | G | N1 | 1H:2698 | 4.40 | 1.00 | 36.54 |  |
| 7 | C | N4 | 1H:2738 | 4.45 | 1.00 | 38.34 |  |
| 8 | U | O4 | 1H:2700 | 4.94 | 1.00 | 34.16 |  |
| 9 | A | OP2 | 1H:2735 | 5.00 | 1.00 | 45.49 |  |
  |

  
 Coded contact list:   
 Exclusion zone contact(s): /O6/  
Consider NA in place of K (1 valid contact(s) for NA vs. 0 for K)  

### K:1H:3036

|  | Res. type | Atom | Residue | Distance | Occupancy | B-factor | Closest solute to 1^st^ shell water |
| --- | --- | --- | --- | --- | --- | --- | --- |
| **Ion** | | | |  | 1.00 | 60.90 || 1 | A | O5' | 1H:1300 | 3.16 | 1.00 | 26.01 |
| 2 | G | N7 | 1H:1303 | 3.33 | 1.00 | 25.61 |  |
| 3 | A | OP1 | 1H:1300 | 3.45 | 1.00 | 26.45 |  |
| 4 | G | O6 | 1H:1303 | 3.63 | 1.00 | 26.26 |  |
| 5 | A | OP2 | 1H:1301 | 3.71 | 1.00 | 27.63 |  |
| 6 | C | N4 | 1H:1304 | 3.91 | 1.00 | 23.78 |  |
| 7 | Water 1 | | 1H:4093 | 3.92 | 1.00 | 25.61 | 2.96 (A:OP1 1H:1300) |
| 8 | G | O3' | 1H:1299 | 4.97 | 1.00 | 27.08 |  |
  |

  
**1^st^ shell ligand angles:**   
1-2: 72;   
  
 Contact(s): /N7/O5'/  
 Coded contact list: O_r_N_b_  

### K:1H:3037

|  | Res. type | Atom | Residue | Distance | Occupancy | B-factor | Closest solute to 1^st^ shell water |
| --- | --- | --- | --- | --- | --- | --- | --- |
| **Ion** | | | |  | 1.00 | 70.88 || 1 | U | O4 | 1H:989 | 3.42 | 1.00 | 26.03 |
| 2 | A | N7 | 1H:883 | 4.09 | 1.00 | 28.77 |  |
| 3 | G | O6 | 1H:988 | 4.22 | 1.00 | 21.00 |  |
| 4 | G | O6 | 1H:884 | 4.24 | 1.00 | 26.35 |  |
| 5 | C | OP2 | 1H:882 | 4.49 | 1.00 | 20.37 |  |
| 6 | A | OP2 | 1H:883 | 4.56 | 1.00 | 23.26 |  |
| 7 | C | N4 | 1H:882 | 4.73 | 1.00 | 22.57 |  |
| 8 | U | N3 | 1H:989 | 4.84 | 1.00 | 18.88 |  |
  |

  
 Coded contact list:   
1^st^ K coordination shell missing (see Notes)  

### K:1H:3038

|  | Res. type | Atom | Residue | Distance | Occupancy | B-factor | Closest solute to 1^st^ shell water |
| --- | --- | --- | --- | --- | --- | --- | --- |
| **Ion** | | | |  | 1.00 | 56.92 || 1 | G | OP2 | 1H:427 | 2.54 | 1.00 | 36.62 |
| 2 | G | O5' | 1H:427 | 3.75 | 1.00 | 36.05 |  |
| 3 | G | O6 | 1H:428 | 3.77 | 1.00 | 44.45 |  |
| 4 | G | N7 | 1H:427 | 4.12 | 1.00 | 34.52 |  |
| 5 | G | N7 | 1H:428 | 4.13 | 1.00 | 45.28 |  |
| 6 | G | O3' | 1H:426 | 4.64 | 1.00 | 28.86 |  |
| 7 | G | OP1 | 1H:427 | 4.89 | 1.00 | 36.70 |  |
| 8 | G | OP2 | 1H:426 | 4.90 | 1.00 | 38.78 |  |
  |

  
 Coded contact list:   
 Exclusion zone contact(s): /OP2/  
Consider NA in place of K (1 valid contact(s) for NA vs. 0 for K)  

### K:1H:3039

|  | Res. type | Atom | Residue | Distance | Occupancy | B-factor | Closest solute to 1^st^ shell water |
| --- | --- | --- | --- | --- | --- | --- | --- |
| **Ion** | | | |  | 1.00 | 54.13 || 1 | G | N1 | 1H:2503 | 3.02 | 1.00 | 35.70 |
| 2 | G | N2 | 1H:2503 | 3.14 | 1.00 | 49.96 |  |
| 3 | G | OP2 | 1H:2471 | 3.41 | 1.00 | 29.86 |  |
| 4 | G | O6 | 1H:2503 | 4.18 | 1.00 | 36.71 |  |
| 5 | G | OP2 | 1H:2502 | 4.41 | 1.00 | 28.98 |  |
| 6 | G | O5' | 1H:2471 | 4.51 | 1.00 | 27.06 |  |
| 7 | G | N7 | 1H:2471 | 4.57 | 1.00 | 35.56 |  |
| 8 | G | N7 | 1H:2502 | 4.66 | 1.00 | 33.95 |  |
| 9 | G | N3 | 1H:2503 | 4.93 | 1.00 | 29.03 |  |
  |

  
**1^st^ shell ligand angles:**   
1-2: 43;   
  
 Contact(s): /N1_G_/N2/  
 Coded contact list: 2Anion  

### K:1H:3040

|  | Res. type | Atom | Residue | Distance | Occupancy | B-factor | Closest solute to 1^st^ shell water |
| --- | --- | --- | --- | --- | --- | --- | --- |
| **Ion** | | | |  | 0.50 | 46.91 || 1 | G | N7 | 1H:2536 | 3.42 | 1.00 | 38.53 |
| 2 | G | O6 | 1H:2536 | 3.43 | 1.00 | 36.14 |  |
| 3 | G | O6 | 1H:2537 | 4.19 | 1.00 | 41.82 |  |
| 4 | U | OP2 | 1H:2535 | 4.25 | 1.00 | 34.51 |  |
| 5 | G | N7 | 1H:2537 | 4.53 | 1.00 | 40.41 |  |
  |

  
 Coded contact list:   
1^st^ K coordination shell missing (see Notes)  

### K:1H:3040

|  | Res. type | Atom | Residue | Distance | Occupancy | B-factor | Closest solute to 1^st^ shell water |
| --- | --- | --- | --- | --- | --- | --- | --- |
| **Ion** | | | |  | 0.50 | 43.22 || 1 | G | N7 | 1H:2537 | 3.92 | 1.00 | 40.41 |
| 2 | G | OP2 | 1H:2536 | 4.38 | 1.00 | 34.30 |  |
| 3 | G | N7 | 1H:2536 | 4.48 | 1.00 | 38.53 |  |
| 4 | G | OP2 | 1H:2537 | 4.62 | 1.00 | 34.18 |  |
| 5 | G | O6 | 1H:2537 | 4.71 | 1.00 | 41.82 |  |
  |

  
 Coded contact list:   
1^st^ K coordination shell missing (see Notes)  

### K:1H:3041

|  | Res. type | Atom | Residue | Distance | Occupancy | B-factor | Closest solute to 1^st^ shell water |
| --- | --- | --- | --- | --- | --- | --- | --- |
| **Ion** | | | |  | 1.00 | 68.01 || 1 | G | O6 | 1H:2857 | 2.79 | 1.00 | 50.60 |
| 2 | G | O6 | 1H:2880 | 3.25 | 1.00 | 44.83 |  |
| 3 | Water 1 | | 1H:4049 | 3.59 | 1.00 | 41.99 | 2.69 (G:N7 1H:2857) |
| 4 | Water 2 | | 1H:4048 | 3.74 | 1.00 | 44.24 | 2.69 (G:N1 1H:2859) |
| 5 | U | O4 | 1H:2858 | 3.76 | 1.00 | 45.17 |  |
| 6 | C | N4 | 1H:2881 | 3.88 | 1.00 | 45.34 |  |
| 7 | G | N2 | 1H:2859 | 3.99 | 1.00 | 48.91 |  |
| 8 | Water 3 | | 1H:4597 | 4.16 | 1.00 | 68.97 | 4.95 (U:O4 1H:2858) |
| 9 | U | N3 | 1H:2858 | 4.22 | 1.00 | 45.05 |  |
| 10 | A | OP2 | 1H:2879 | 4.49 | 1.00 | 48.83 |  |
| 11 | G | N7 | 1H:2880 | 4.53 | 1.00 | 47.02 |  |
| 12 | G | O6 | 1H:2856 | 4.67 | 1.00 | 50.65 |  |
| 13 | G | N1 | 1H:2859 | 4.69 | 1.00 | 48.07 |  |
| 14 | G | N1 | 1H:2857 | 4.91 | 1.00 | 47.72 |  |
| 15 | G | N7 | 1H:2857 | 4.98 | 1.00 | 45.65 |  |
| 16 | G | O2' | 1H:2878 | 4.98 | 1.00 | 53.44 |  |
  |

  
**1^st^ shell ligand angles:**   
1-2: 60;   
  
 Contact(s): /O6/O6/  
 Coded contact list: 2O_b_  

### K:1H:3042

|  | Res. type | Atom | Residue | Distance | Occupancy | B-factor | Closest solute to 1^st^ shell water |
| --- | --- | --- | --- | --- | --- | --- | --- |
| **Ion** | | | |  | 1.00 | 54.63 || 1 | G | O6 | 1H:2549 | 2.70 | 1.00 | 50.51 |
| 2 | U | O4 | 1H:2550 | 2.85 | 1.00 | 42.93 |  |
| 3 | G | O6 | 1H:2539 | 3.08 | 1.00 | 44.88 |  |
| 4 | G | O6 | 1H:2538 | 3.29 | 1.00 | 38.92 |  |
| 5 | C | N4 | 1H:2540 | 3.78 | 1.00 | 50.64 |  |
| 6 | U | N3 | 1H:2550 | 4.34 | 1.00 | 42.89 |  |
| 7 | G | N7 | 1H:2539 | 4.40 | 1.00 | 40.17 |  |
| 8 | C | N4 | 1H:2551 | 4.56 | 1.00 | 41.84 |  |
| 9 | G | N1 | 1H:2549 | 4.58 | 1.00 | 47.39 |  |
| 10 | G | N7 | 1H:2538 | 4.68 | 1.00 | 40.30 |  |
| 11 | G | N1 | 1H:2538 | 4.76 | 1.00 | 38.63 |  |
  |

  
**1^st^ shell ligand angles:**   
1-2: 58; 1-3: 74; 1-4: 135; 2-3: 68; 2-4: 81; 3-4: 73;   
  
 Contact(s): /O4/O6/O6/O6/  
 Coded contact list: 4O_b_  

### K:1H:3043

|  | Res. type | Atom | Residue | Distance | Occupancy | B-factor | Closest solute to 1^st^ shell water |
| --- | --- | --- | --- | --- | --- | --- | --- |
| **Ion** | | | |  | 1.00 | 58.70 || 1 | G | O6 | 1H:1229 | 2.57 | 1.00 | 61.67 |
| 2 | U | O4 | 1H:1212 | 2.86 | 1.00 | 48.87 |  |
| 3 | G | O6 | 1H:1211 | 2.90 | 1.00 | 39.09 |  |
| 4 | G | O6 | 1H:1230 | 3.09 | 1.00 | 51.08 |  |
| 5 | G | N7 | 1H:1230 | 3.87 | 1.00 | 49.08 |  |
| 6 | U | N3 | 1H:1212 | 3.93 | 1.00 | 49.86 |  |
| 7 | G | N7 | 1H:1229 | 4.06 | 1.00 | 55.16 |  |
| 8 | C | N4 | 1H:1231 | 4.17 | 1.00 | 44.92 |  |
| 9 | C | N4 | 1H:1213 | 4.22 | 1.00 | 58.95 |  |
| 10 | G | N1 | 1H:1211 | 4.54 | 1.00 | 43.05 |  |
| 11 | G | N1 | 1H:1229 | 4.66 | 1.00 | 52.28 |  |
| 12 | G | N7 | 1H:1211 | 4.90 | 1.00 | 46.54 |  |
| 13 | A | N6 | 1H:1228 | 4.99 | 1.00 | 69.12 |  |
  |

  
**1^st^ shell ligand angles:**   
1-2: 96; 1-3: 141; 1-4: 75; 2-3: 72; 2-4: 76; 3-4: 66;   
  
 Contact(s): /O4/O6/O6/  
 Coded contact list: 3O_b_  
 Exclusion zone contact(s): /O6/  

### K:1H:3044

|  | Res. type | Atom | Residue | Distance | Occupancy | B-factor | Closest solute to 1^st^ shell water |
| --- | --- | --- | --- | --- | --- | --- | --- |
| **Ion** | | | |  | 1.00 | 92.12 || 1 | G | O6 | 1H:2234 | 3.00 | 1.00 | 44.25 |
| 2 | G | O6 | 1H:2233 | 3.16 | 1.00 | 54.63 |  |
| 3 | G | N7 | 1H:2233 | 3.77 | 1.00 | 61.59 |  |
| 4 | G | N7 | 1H:2234 | 3.94 | 1.00 | 48.53 |  |
| 5 | C | N4 | 1H:2225 | 4.53 | 1.00 | 44.64 |  |
| 6 | U | O4 | 1H:2226 | 4.65 | 1.00 | 45.92 |  |
| 7 | G | O6 | 1H:2232 | 4.87 | 1.00 | 66.02 |  |
| 8 | G | N1 | 1H:2233 | 4.96 | 1.00 | 50.72 |  |
| 9 | G | N7 | 1H:2232 | 4.98 | 1.00 | 68.98 |  |
  |

  
**1^st^ shell ligand angles:**   
1-2: 66;   
  
 Contact(s): /O6/O6/  
 Coded contact list: 2O_b_  

### K:1H:3045

|  | Res. type | Atom | Residue | Distance | Occupancy | B-factor | Closest solute to 1^st^ shell water |
| --- | --- | --- | --- | --- | --- | --- | --- |
| **Ion** | | | |  | 1.00 | 59.62 || 1 | C | OP1 | 1H:838 | 2.79 | 1.00 | 35.33 |
| 2 | C | O2 | 1H:1813 | 2.85 | 1.00 | 20.97 |  |
| 3 | C | O2' | 1H:1813 | 3.18 | 1.00 | 25.24 |  |
| 4 | C | N3 | 1H:1813 | 3.61 | 1.00 | 37.19 |  |
| 5 | C | N1 | 1H:1813 | 3.83 | 1.00 | 24.50 |  |
| 6 | A | O2' | 1H:1812 | 4.35 | 1.00 | 24.81 |  |
| 7 | C | OP2 | 1H:838 | 4.73 | 1.00 | 19.75 |  |
  |

  
**1^st^ shell ligand angles:**   
1-2: 97; 1-3: 162; 2-3: 66;   
  
 Contact(s): /O2/O2'/OP1/  
 Coded contact list: O_ph_O_r_O_b_  

### K:1H:3046

|  | Res. type | Atom | Residue | Distance | Occupancy | B-factor | Closest solute to 1^st^ shell water |
| --- | --- | --- | --- | --- | --- | --- | --- |
| **Ion** | | | |  | 1.00 | 72.95 || 1 | G | O6 | 1H:1472 | 2.96 | 1.00 | 38.09 |
| 2 | G | OP2 | 1H:1471 | 4.45 | 1.00 | 40.09 |  |
| 3 | G | O6 | 1H:1473 | 4.47 | 1.00 | 35.37 |  |
| 4 | G | N7 | 1H:1472 | 4.77 | 1.00 | 32.27 |  |
| 5 | G | N1 | 1H:1473 | 4.83 | 1.00 | 29.06 |  |
| 6 | G | N7 | 1H:1471 | 4.93 | 1.00 | 36.09 |  |
  |

  
 Contact(s): /O6/  
 Coded contact list: O_b_  

### K:1H:3047

|  | Res. type | Atom | Residue | Distance | Occupancy | B-factor | Closest solute to 1^st^ shell water |
| --- | --- | --- | --- | --- | --- | --- | --- |
| **Ion** | | | |  | 1.00 | 74.12 || 1 | U | O4 | 1H:2889 | 2.74 | 1.00 | 57.67 |
| 2 | Water 1 | | 1H:3876 | 2.96 | 1.00 | 48.05 | 2.49 (G:O6 1H:2849) |
| 3 | Water 2 | | 1H:4082 | 2.98 | 1.00 | 49.09 | 2.84 (U:O4 1H:2889) |
| 4 | G | O6 | 1H:2888 | 3.00 | 1.00 | 45.14 |  |
| 5 | G | O6 | 1H:2850 | 3.03 | 1.00 | 48.55 |  |
| 6 | Water 3 | | 1H:3886 | 3.73 | 1.00 | 45.49 | 5.45 (U:O4 1H:2889) |
| 7 | C | N4 | 1H:2851 | 4.03 | 1.00 | 48.25 |  |
| 8 | U | N3 | 1H:2889 | 4.19 | 1.00 | 47.17 |  |
| 9 | G | N7 | 1H:2888 | 4.44 | 1.00 | 51.84 |  |
| 10 | G | O6 | 1H:2849 | 4.66 | 1.00 | 45.83 |  |
| 11 | G | N7 | 1H:2850 | 4.74 | 1.00 | 44.81 |  |
| 12 | G | N1 | 1H:2888 | 4.93 | 1.00 | 53.75 |  |
| 13 | Water 4 | | 1H:4115 | 4.97 | 1.00 | 46.30 | 3.03 (G:O6 1H:2849) |
  |

  
**1^st^ shell ligand angles:**   
1-2: 92; 1-3: 59; 1-4: 79; 1-5: 66; 2-3: 64; 2-4: 138; 2-5: 73; 3-4: 136; 3-5: 106; 4-5: 66;   
  
 Contact(s): /O4/O6/O6/O_w_/O_w_/  
 Coded contact list: 3O_b_2O_w_  

### K:1H:3048

|  | Res. type | Atom | Residue | Distance | Occupancy | B-factor | Closest solute to 1^st^ shell water |
| --- | --- | --- | --- | --- | --- | --- | --- |
| **Ion** | | | |  | 1.00 | 72.20 || 1 | G | O6 | 1H:285 | 2.70 | 1.00 | 46.21 |
| 2 | U | OP1 | 1H:259 | 2.85 | 1.00 | 57.94 |  |
| 3 | G | OP2 | 1H:283 | 3.98 | 1.00 | 73.85 |  |
| 4 | G | N7 | 1H:284 | 4.19 | 1.00 | 58.73 |  |
| 5 | G | O6 | 1H:284 | 4.63 | 1.00 | 53.24 |  |
| 6 | G | N7 | 1H:285 | 4.66 | 1.00 | 50.41 |  |
| 7 | U | O5' | 1H:259 | 4.72 | 1.00 | 42.48 |  |
| 8 | G | OP1 | 1H:283 | 4.87 | 1.00 | 75.32 |  |
| 9 | G | OP2 | 1H:284 | 4.87 | 1.00 | 61.90 |  |
| 10 | U | OP2 | 1H:259 | 4.90 | 1.00 | 38.48 |  |
| 11 | G | N1 | 1H:285 | 4.93 | 1.00 | 47.38 |  |
  |

  
**1^st^ shell ligand angles:**   
1-2: 83;   
  
 Contact(s): /O6/OP1/  
 Coded contact list: O_ph_O_b_  

### K:1H:3049

|  | Res. type | Atom | Residue | Distance | Occupancy | B-factor | Closest solute to 1^st^ shell water |
| --- | --- | --- | --- | --- | --- | --- | --- |
| **Ion** | | | |  | 1.00 | 55.61 || 1 | G | O6 | 1H:1801 | 2.85 | 1.00 | 29.55 |
| 2 | U | O4 | 1H:1800 | 3.40 | 1.00 | 26.51 |  |
| 3 | G | O6 | 1H:2007 | 3.41 | 1.00 | 23.14 |  |
| 4 | C | N4 | 1H:2006 | 3.53 | 1.00 | 23.81 |  |
| 5 | G | N7 | 1H:2007 | 4.17 | 1.00 | 24.00 |  |
| 6 | G | N7 | 1H:1801 | 4.41 | 1.00 | 26.93 |  |
| 7 | U | N3 | 1H:1800 | 4.64 | 1.00 | 24.46 |  |
| 8 | G | N1 | 1H:1801 | 4.90 | 1.00 | 19.92 |  |
| 9 | G | O6 | 1H:2008 | 4.93 | 1.00 | 22.20 |  |
  |

  
**1^st^ shell ligand angles:**   
1-2: 112;   
  
 Contact(s): /O4/O6/  
 Coded contact list: 2O_b_  

### K:1H:3050

|  | Res. type | Atom | Residue | Distance | Occupancy | B-factor | Closest solute to 1^st^ shell water |
| --- | --- | --- | --- | --- | --- | --- | --- |
| **Ion** | | | |  | 1.00 | 53.89 || 1 | G | N7 | 1H:177 | 3.77 | 1.00 | 17.14 |
| 2 | G | O6 | 1H:179 | 3.89 | 1.00 | 27.37 |  |
| 3 | G | N7 | 1H:178 | 4.11 | 1.00 | 18.44 |  |
| 4 | G | O6 | 1H:178 | 4.30 | 1.00 | 20.52 |  |
| 5 | G | O6 | 1H:177 | 4.31 | 1.00 | 16.65 |  |
| 6 | G | OP2 | 1H:177 | 4.44 | 1.00 | 22.66 |  |
| 7 | G | O2' | 1H:195 | 4.70 | 1.00 | 21.36 |  |
  |

  
 Coded contact list:   
1^st^ K coordination shell missing (see Notes)  

### K:1H:3051

|  | Res. type | Atom | Residue | Distance | Occupancy | B-factor | Closest solute to 1^st^ shell water |
| --- | --- | --- | --- | --- | --- | --- | --- |
| **Ion** | | | |  | 1.00 | 49.61 || 1 | U | O4 | 1H:1828 | 2.80 | 1.00 | 19.99 |
| 2 | G | O6 | 1H:1855 | 2.90 | 1.00 | 17.33 |  |
| 3 | G | O6 | 1H:1854 | 3.20 | 1.00 | 19.39 |  |
| 4 | G | O6 | 1H:1856 | 3.76 | 1.00 | 21.10 |  |
| 5 | C | N4 | 1H:1827 | 4.03 | 1.00 | 20.29 |  |
| 6 | G | N7 | 1H:1855 | 4.07 | 1.00 | 17.18 |  |
| 7 | C | N4 | 1H:1829 | 4.15 | 1.00 | 16.63 |  |
| 8 | U | N3 | 1H:1828 | 4.23 | 1.00 | 17.35 |  |
| 9 | G | N7 | 1H:1854 | 4.90 | 1.00 | 17.79 |  |
| 10 | G | N1 | 1H:1854 | 4.93 | 1.00 | 16.00 |  |
  |

  
**1^st^ shell ligand angles:**   
1-2: 63; 1-3: 90; 2-3: 71;   
  
 Contact(s): /O4/O6/O6/  
 Coded contact list: 3O_b_  

### K:1H:3052

|  | Res. type | Atom | Residue | Distance | Occupancy | B-factor | Closest solute to 1^st^ shell water |
| --- | --- | --- | --- | --- | --- | --- | --- |
| **Ion** | | | |  | 1.00 | 52.89 || 1 | G | OP2 | 1H:1871 | 2.96 | 1.00 | 40.08 |
| 2 | C | O2' | 1H:1870 | 3.40 | 1.00 | 35.56 |  |
| 3 | C | O3' | 1H:1870 | 3.59 | 1.00 | 36.80 |  |
| 4 | U | O4 | 1H:1921 | 3.60 | 1.00 | 37.38 |  |
| 5 | G | N7 | 1H:1920 | 3.74 | 1.00 | 38.71 |  |
| 6 | G | OP2 | 1H:1920 | 4.02 | 1.00 | 39.57 |  |
| 7 | G | O6 | 1H:1920 | 4.63 | 1.00 | 39.68 |  |
| 8 | G | OP2 | 1H:1872 | 4.73 | 1.00 | 35.36 |  |
| 9 | G | O5' | 1H:1871 | 4.89 | 1.00 | 33.98 |  |
| 10 | G | N7 | 1H:1919 | 4.96 | 1.00 | 47.23 |  |
| 11 | G | OP1 | 1H:1871 | 5.00 | 1.00 | 38.24 |  |
  |

  
**1^st^ shell ligand angles:**   
1-2: 82;   
  
 Contact(s): /O2'/OP2/  
 Coded contact list: O_ph_O_r_  

### K:1H:3053

|  | Res. type | Atom | Residue | Distance | Occupancy | B-factor | Closest solute to 1^st^ shell water |
| --- | --- | --- | --- | --- | --- | --- | --- |
| **Ion** | | | |  | 1.00 | 25.61 || 1 | Water 1 | | 1H:3635 | 2.45 | 1.00 | 18.98 | 2.01 (U:OP1 1H:600) |
| 2 | U | O3' | 1H:2513 | 2.82 | 1.00 | 16.71 |  |
| 3 | C | OP1 | 1H:2514 | 2.96 | 1.00 | 20.98 |  |
| 4 | U | OP2 | 1H:2517 | 2.96 | 1.00 | 19.86 |  |
| 5 | A | O3' | 1H:599 | 3.14 | 1.00 | 19.41 |  |
| 6 | 2MA | OP1 | 1H:2516 | 3.16 | 1.00 | 15.66 |  |
| 7 | U | OP1 | 1H:2517 | 3.37 | 1.00 | 17.46 |  |
| 8 | U | O2' | 1H:2513 | 3.63 | 1.00 | 19.63 |  |
| 9 | U | OP1 | 1H:600 | 3.87 | 1.00 | 19.93 |  |
| 10 | A | O2' | 1H:599 | 4.02 | 1.00 | 22.61 |  |
| 11 | Water 2 | | 1H:3813 | 4.37 | 1.00 | 18.26 | 2.81 (U:OP2 1H:2517) |
| 12 | C | O5' | 1H:2514 | 4.41 | 1.00 | 19.00 |  |
| 13 | 2MA | O5' | 1H:2516 | 4.61 | 1.00 | 16.05 |  |
| 14 | C | OP2 | 1H:2514 | 4.69 | 1.00 | 22.40 |  |
| 15 | Water 3 | | 1H:4295 | 4.71 | 1.00 | 33.55 | 3.74 (U:OP2 1H:2517) |
| 16 | U | O5' | 1H:2517 | 4.73 | 1.00 | 17.15 |  |
| 17 | 2MA | O3' | 1H:2516 | 4.90 | 1.00 | 17.03 |  |
  |

  
**1^st^ shell ligand angles:**   
1-2: 169; 1-3: 124; 1-4: 72; 1-5: 60; 1-6: 47; 1-7: 116; 2-3: 52; 2-4: 117; 2-5: 109; 2-6: 137; 2-7: 74; 3-4: 142; 3-5: 92; 3-6: 90; 3-7: 101; 4-5: 124; 4-6: 78; 4-7: 46; 5-6: 90; 5-7: 165; 6-7: 98;   
  
 Contact(s): /O3'/O3'/OP1/OP1/OP1/OP2/  
 Coded contact list: 4O_ph_2O_r_  
 Exclusion zone contact(s): /O_w_/  
K hydration shell issue: water too close (< 2.4 Angs.)  

### K:1H:3054

|  | Res. type | Atom | Residue | Distance | Occupancy | B-factor | Closest solute to 1^st^ shell water |
| --- | --- | --- | --- | --- | --- | --- | --- |
| **Ion** | | | |  | 1.00 | 27.77 || 1 | G | O2' | 1H:2589 | 2.67 | 1.00 | 26.79 |
| 2 | G | O6 | 1H:2594 | 2.68 | 1.00 | 24.24 |  |
| 3 | G | O6 | 1H:2518 | 3.34 | 1.00 | 21.70 |  |
| 4 | C | O2 | 1H:2623 | 3.34 | 1.00 | 26.94 |  |
| 5 | C | O2' | 1H:2623 | 3.47 | 1.00 | 25.18 |  |
| 6 | U | OP2 | 1H:2593 | 3.63 | 1.00 | 21.02 |  |
| 7 | G | N1 | 1H:2594 | 3.68 | 1.00 | 24.45 |  |
| 8 | C | OP2 | 1H:2592 | 4.37 | 1.00 | 20.42 |  |
| 9 | G | N1 | 1H:2518 | 4.39 | 1.00 | 26.03 |  |
| 10 | C | N1 | 1H:2623 | 4.39 | 1.00 | 22.20 |  |
| 11 | G | N9 | 1H:2589 | 4.50 | 1.00 | 24.28 |  |
| 12 | C | N3 | 1H:2623 | 4.68 | 1.00 | 24.86 |  |
| 13 | C | O5' | 1H:2592 | 4.79 | 1.00 | 16.45 |  |
| 14 | C | OP1 | 1H:2592 | 4.84 | 1.00 | 20.17 |  |
| 15 | G | O4' | 1H:2589 | 4.97 | 1.00 | 20.54 |  |
  |

  
**1^st^ shell ligand angles:**   
1-2: 111; 1-3: 63; 1-4: 121; 2-3: 95; 2-4: 66; 3-4: 59;   
  
 Contact(s): /O2/O2'/O6/O6/  
 Coded contact list: O_r_3O_b_  

### K:1H:3055

|  | Res. type | Atom | Residue | Distance | Occupancy | B-factor | Closest solute to 1^st^ shell water |
| --- | --- | --- | --- | --- | --- | --- | --- |
| **Ion** | | | |  | 1.00 | 31.99 || 1 | G | O2' | 1H:2605 | 2.64 | 1.00 | 17.78 |
| 2 | A | O2' | 1H:1989 | 2.94 | 1.00 | 29.33 |  |
| 3 | 5MU | O2' | 1H:1962 | 3.06 | 1.00 | 21.32 |  |
| 4 | A | O3' | 1H:1989 | 3.16 | 1.00 | 27.38 |  |
| 5 | C | O5' | 1H:1990 | 3.46 | 1.00 | 29.06 |  |
| 6 | 5MU | O2 | 1H:1962 | 3.48 | 1.00 | 23.06 |  |
| 7 | C | OP1 | 1H:1988 | 3.64 | 1.00 | 28.46 |  |
| 8 | 5MU | O3' | 1H:1962 | 3.70 | 1.00 | 20.80 |  |
| 9 | C | OP2 | 1H:1990 | 3.99 | 1.00 | 28.16 |  |
| 10 | U | OP1 | 1H:1963 | 4.01 | 1.00 | 24.28 |  |
| 11 | A | N6 | 1H:1960 | 4.17 | 1.00 | 29.87 |  |
| 12 | G | O3' | 1H:2605 | 4.43 | 1.00 | 22.93 |  |
| 13 | G | O4' | 1H:2605 | 4.47 | 1.00 | 24.14 |  |
| 14 | A | N3 | 1H:1989 | 4.66 | 1.00 | 28.49 |  |
| 15 | 5MU | N1 | 1H:1962 | 4.75 | 1.00 | 23.65 |  |
| 16 | U | O5' | 1H:1963 | 4.77 | 1.00 | 24.95 |  |
| 17 | A | N7 | 1H:1960 | 4.89 | 1.00 | 25.01 |  |
  |

  
**1^st^ shell ligand angles:**   
1-2: 85; 1-3: 111; 1-4: 88; 2-3: 133; 2-4: 53; 3-4: 160;   
  
 Contact(s): /O2'/O2'/O2'/O3'/  
 Coded contact list: 4O_r_  

### K:1H:3056

|  | Res. type | Atom | Residue | Distance | Occupancy | B-factor | Closest solute to 1^st^ shell water |
| --- | --- | --- | --- | --- | --- | --- | --- |
| **Ion** | | | |  | 1.00 | 37.58 || 1 | G | N7 | 1H:2557 | 2.68 | 1.00 | 30.37 |
| 2 | A | N3 | 1H:2555 | 2.91 | 1.00 | 30.99 |  |
| 3 | G | O6 | 1H:2557 | 3.05 | 1.00 | 28.56 |  |
| 4 | G | N7 | 1H:2556 | 3.05 | 1.00 | 29.69 |  |
| 5 | A | O4' | 1H:2555 | 3.06 | 1.00 | 34.80 |  |
| 6 | G | O6 | 1H:2556 | 3.72 | 1.00 | 33.67 |  |
| 7 | A | N9 | 1H:2555 | 4.08 | 1.00 | 29.37 |  |
| 8 | G | N9 | 1H:2556 | 4.76 | 1.00 | 30.45 |  |
| 9 | A | O3' | 1H:2555 | 4.80 | 1.00 | 36.09 |  |
| 10 | G | N9 | 1H:2557 | 4.81 | 1.00 | 27.92 |  |
| 11 | C | N4 | 1H:2534 | 4.84 | 1.00 | 24.97 |  |
| 12 | G | N1 | 1H:2557 | 4.85 | 1.00 | 23.77 |  |
| 13 | U | N3 | 1H:2532 | 4.91 | 1.00 | 29.07 |  |
| 14 | G | O6 | 1H:2558 | 4.95 | 1.00 | 28.40 |  |
| 15 | A | N1 | 1H:2555 | 4.96 | 1.00 | 29.80 |  |
  |

  
**1^st^ shell ligand angles:**   
1-2: 139; 1-3: 64; 1-4: 80; 1-5: 141; 2-3: 100; 2-4: 141; 2-5: 71; 3-4: 103; 3-5: 149; 4-5: 73;   
  
 Contact(s): /N3_A_/N7/N7/O4'/O6/  
 Coded contact list: O_r_O_b_3N_b_  

### K:1H:3057

|  | Res. type | Atom | Residue | Distance | Occupancy | B-factor | Closest solute to 1^st^ shell water |
| --- | --- | --- | --- | --- | --- | --- | --- |
| **Ion** | | | |  | 1.00 | 23.63 || 1 | G | N7 | 1H:2076 | 2.67 | 1.00 | 22.34 |
| 2 | A | N1 | 1H:2627 | 2.94 | 1.00 | 21.28 |  |
| 3 | A | O2' | 1H:2074 | 3.06 | 1.00 | 19.36 |  |
| 4 | A | N3 | 1H:2074 | 3.07 | 1.00 | 21.13 |  |
| 5 | G | O6 | 1H:2075 | 3.07 | 1.00 | 23.07 |  |
| 6 | G | N7 | 1H:2075 | 3.11 | 1.00 | 21.24 |  |
| 7 | G | O6 | 1H:2076 | 3.26 | 1.00 | 20.49 |  |
| 8 | A | N6 | 1H:2590 | 3.52 | 1.00 | 21.24 |  |
| 9 | A | N7 | 1H:2590 | 4.13 | 1.00 | 19.91 |  |
| 10 | A | N6 | 1H:2627 | 4.29 | 1.00 | 22.10 |  |
| 11 | G | N1 | 1H:2075 | 4.37 | 1.00 | 23.02 |  |
| 12 | A | N1 | 1H:2590 | 4.61 | 1.00 | 21.60 |  |
| 13 | A | N9 | 1H:2074 | 4.64 | 1.00 | 17.84 |  |
| 14 | A | N1 | 1H:2074 | 4.69 | 1.00 | 18.79 |  |
| 15 | G | N9 | 1H:2075 | 4.70 | 1.00 | 22.03 |  |
| 16 | A | N3 | 1H:2627 | 4.72 | 1.00 | 23.68 |  |
| 17 | G | N9 | 1H:2076 | 4.82 | 1.00 | 21.21 |  |
| 18 | G | OP2 | 1H:2076 | 4.88 | 1.00 | 23.64 |  |
  |

  
**1^st^ shell ligand angles:**   
1-2: 131; 1-3: 134; 1-4: 137; 1-5: 77; 1-6: 86; 1-7: 63; 2-3: 87; 2-4: 70; 2-5: 110; 2-6: 140; 2-7: 77; 3-4: 71; 3-5: 116; 3-6: 69; 3-7: 163; 4-5: 60; 4-6: 72; 4-7: 96; 5-6: 58; 5-7: 64; 6-7: 118;   
  
 Contact(s): /N1_A_/N3_A_/N7/N7/O2'/O6/O6/  
 Coded contact list: O_r_2O_b_4N_b_  

### K:1H:3058

|  | Res. type | Atom | Residue | Distance | Occupancy | B-factor | Closest solute to 1^st^ shell water |
| --- | --- | --- | --- | --- | --- | --- | --- |
| **Ion** | | | |  | 1.00 | 43.20 || 1 | U | O4 | 1H:2258 | 2.82 | 1.00 | 25.36 |
| 2 | G | O6 | 1H:2449 | 2.87 | 1.00 | 16.89 |  |
| 3 | U | O4 | 1H:2450 | 3.12 | 1.00 | 16.23 |  |
| 4 | C | N4 | 1H:2096 | 3.70 | 1.00 | 11.49 |  |
| 5 | G | O6 | 1H:2095 | 3.73 | 1.00 | 16.13 |  |
| 6 | U | O4 | 1H:2257 | 4.04 | 1.00 | 19.90 |  |
| 7 | A | N6 | 1H:2094 | 4.17 | 1.00 | 17.13 |  |
| 8 | U | N3 | 1H:2450 | 4.59 | 1.00 | 13.52 |  |
| 9 | U | N3 | 1H:2258 | 4.63 | 1.00 | 16.54 |  |
| 10 | G | N7 | 1H:2449 | 4.79 | 1.00 | 13.80 |  |
| 11 | G | N1 | 1H:2449 | 4.90 | 1.00 | 13.05 |  |
  |

  
**1^st^ shell ligand angles:**   
1-2: 96; 1-3: 79; 2-3: 69;   
  
 Contact(s): /O4/O4/O6/  
 Coded contact list: 3O_b_  

### K:1H:3059

|  | Res. type | Atom | Residue | Distance | Occupancy | B-factor | Closest solute to 1^st^ shell water |
| --- | --- | --- | --- | --- | --- | --- | --- |
| **Ion** | | | |  | 1.00 | 19.99 || 1 | G | O6 | 1H:2460 | 2.46 | 1.00 | 20.93 |
| 2 | C | O2 | 1H:2514 | 2.76 | 1.00 | 20.95 |  |
| 3 | C | O4' | 1H:2514 | 2.81 | 1.00 | 19.94 |  |
| 4 | G | O6 | 1H:2084 | 2.87 | 1.00 | 19.63 |  |
| 5 | G | N7 | 1H:2460 | 2.95 | 1.00 | 20.38 |  |
| 6 | U | O4 | 1H:2517 | 2.99 | 1.00 | 19.72 |  |
| 7 | G | N7 | 1H:2084 | 3.34 | 1.00 | 20.43 |  |
| 8 | C | N1 | 1H:2514 | 4.00 | 1.00 | 18.41 |  |
| 9 | A | N6 | 1H:2464 | 4.22 | 1.00 | 22.55 |  |
| 10 | C | N3 | 1H:2514 | 4.39 | 1.00 | 17.74 |  |
| 11 | G | N1 | 1H:2460 | 4.55 | 1.00 | 18.26 |  |
| 12 | C | O5' | 1H:2514 | 4.70 | 1.00 | 19.00 |  |
| 13 | G | N1 | 1H:2084 | 4.83 | 1.00 | 20.60 |  |
| 14 | U | N3 | 1H:2517 | 4.91 | 1.00 | 17.74 |  |
  |

  
**1^st^ shell ligand angles:**   
1-2: 72; 1-3: 115; 1-4: 86; 1-5: 69; 1-6: 58; 1-7: 133; 2-3: 63; 2-4: 73; 2-5: 82; 2-6: 128; 2-7: 68; 3-4: 120; 3-5: 60; 3-6: 151; 3-7: 65; 4-5: 149; 4-6: 88; 4-7: 61; 5-6: 94; 5-7: 125; 6-7: 142;   
  
 Contact(s): /N7/N7/O2/O4/O4'/O6/  
 Coded contact list: O_r_3O_b_2N_b_  
 Exclusion zone contact(s): /O6/  

### K:1H:3060

|  | Res. type | Atom | Residue | Distance | Occupancy | B-factor | Closest solute to 1^st^ shell water |
| --- | --- | --- | --- | --- | --- | --- | --- |
| **Ion** | | | |  | 1.00 | 34.42 || 1 | C | OP2 | 1H:2453 | 2.35 | 1.00 | 17.51 |
| 2 | Water 1 | | 1H:3654 | 2.67 | 1.00 | 18.81 | 2.08 (G:OP2 1H:2601) |
| 3 | GLU | OE1 | 11:237 | 2.71 | 1.00 | 23.05 |  |
| 4 | A | O2' | 1H:2452 | 2.79 | 1.00 | 23.89 |  |
| 5 | A | OP2 | 1H:2613 | 2.84 | 1.00 | 21.64 |  |
| 6 | Water 2 | | 1H:4047 | 2.88 | 1.00 | 22.52 | 2.98 (A:O2' 1H:2452) |
| 7 | Water 3 | | 1H:4310 | 3.31 | 1.00 | 30.87 | 3.09 (C:OP2 1H:2453) |
| 8 | Water 4 | | 11:406 | 3.78 | 1.00 | 28.89 | 2.81 (GLU:OE1 11:237) |
| 9 | GLU | OE2 | 11:237 | 3.91 | 1.00 | 30.98 |  |
| 10 | C | OP1 | 1H:2453 | 4.03 | 1.00 | 22.02 |  |
| 11 | A | O3' | 1H:2452 | 4.15 | 1.00 | 22.58 |  |
| 12 | G | OP2 | 1H:2601 | 4.16 | 1.00 | 14.99 |  |
| 13 | A | OP1 | 1H:2613 | 4.54 | 1.00 | 19.57 |  |
| 14 | G | O3' | 1H:2612 | 4.61 | 1.00 | 23.28 |  |
| 15 | Water 5 | | 1H:3614 | 4.72 | 1.00 | 21.66 | 1.85 (G:OP2 1H:2601) |
| 16 | C | O5' | 1H:2453 | 4.87 | 1.00 | 19.33 |  |
| 17 | Water 6 | | 11:410 | 4.96 | 1.00 | 35.54 | 4.56 (GLU:OE1 11:237) |
  |

  
**1^st^ shell ligand angles:**   
1-2: 98; 1-3: 163; 1-4: 59; 1-5: 106; 1-6: 94; 1-7: 63; 2-3: 76; 2-4: 125; 2-5: 155; 2-6: 70; 2-7: 61; 3-4: 137; 3-5: 82; 3-6: 98; 3-7: 100; 4-5: 67; 4-6: 63; 4-7: 123; 5-6: 104; 5-7: 134; 6-7: 121;   
  
 Contact(s): /O2'/O_w_/O_w_/O_w_/OE1_Glu_/OP2/  
 Coded contact list: O_ph_O_r_O_coo_3O_w_  
 Exclusion zone contact(s): /OP2/  
K hydration shell issue: water too close (< 2.4 Angs.)  

### K:1H:3061

|  | Res. type | Atom | Residue | Distance | Occupancy | B-factor | Closest solute to 1^st^ shell water |
| --- | --- | --- | --- | --- | --- | --- | --- |
| **Ion** | | | |  | 1.00 | 43.18 || 1 | G | N7 | 1H:214 | 2.68 | 1.00 | 34.88 |
| 2 | C | O2' | 1H:447 | 2.93 | 1.00 | 37.29 |  |
| 3 | C | O2 | 1H:447 | 2.97 | 1.00 | 36.68 |  |
| 4 | G | OP2 | 1H:214 | 3.05 | 1.00 | 27.89 |  |
| 5 | A | N7 | 1H:215 | 3.36 | 1.00 | 33.73 |  |
| 6 | C | O4' | 1H:448 | 3.86 | 1.00 | 34.05 |  |
| 7 | G | O6 | 1H:214 | 4.05 | 1.00 | 37.13 |  |
| 8 | G | N2 | 1H:436 | 4.10 | 1.00 | 33.68 |  |
| 9 | G | O6 | 1H:216 | 4.21 | 1.00 | 38.75 |  |
| 10 | G | N9 | 1H:214 | 4.24 | 1.00 | 31.28 |  |
| 11 | G | O5' | 1H:214 | 4.32 | 1.00 | 31.79 |  |
| 12 | C | O2 | 1H:437 | 4.66 | 1.00 | 32.48 |  |
| 13 | C | N1 | 1H:447 | 4.67 | 1.00 | 35.33 |  |
| 14 | C | O2' | 1H:448 | 4.84 | 1.00 | 38.25 |  |
  |

  
**1^st^ shell ligand angles:**   
1-2: 79; 1-3: 128; 1-4: 91; 1-5: 85; 2-3: 72; 2-4: 147; 2-5: 73; 3-4: 90; 3-5: 125; 4-5: 138;   
  
 Contact(s): /N7/N7/O2/O2'/OP2/  
 Coded contact list: O_ph_O_r_O_b_2N_b_  

### K:1H:3062

|  | Res. type | Atom | Residue | Distance | Occupancy | B-factor | Closest solute to 1^st^ shell water |
| --- | --- | --- | --- | --- | --- | --- | --- |
| **Ion** | | | |  | 1.00 | 27.71 || 1 | OMG | OP2 | 1H:2264 | 2.61 | 1.00 | 25.46 |
| 2 | G | O2' | 1H:2263 | 2.69 | 1.00 | 25.98 |  |
| 3 | A | OP2 | 1H:2510 | 2.91 | 1.00 | 19.86 |  |
| 4 | A | N7 | 1H:2510 | 3.08 | 1.00 | 18.70 |  |
| 5 | G | O3' | 1H:2263 | 3.20 | 1.00 | 22.02 |  |
| 6 | C | O5' | 1H:2509 | 3.42 | 1.00 | 17.83 |  |
| 7 | C | OP2 | 1H:2509 | 3.70 | 1.00 | 26.56 |  |
| 8 | C | OP1 | 1H:2509 | 3.86 | 1.00 | 24.84 |  |
| 9 | OMG | OP1 | 1H:2264 | 4.58 | 1.00 | 24.67 |  |
| 10 | OMG | O5' | 1H:2264 | 4.71 | 1.00 | 20.47 |  |
| 11 | A | N9 | 1H:2510 | 4.74 | 1.00 | 17.04 |  |
| 12 | C | O3' | 1H:2509 | 4.94 | 1.00 | 19.19 |  |
  |

  
**1^st^ shell ligand angles:**   
1-2: 93; 1-3: 135; 1-4: 112; 1-5: 49; 2-3: 83; 2-4: 153; 2-5: 55; 3-4: 86; 3-5: 96; 4-5: 151;   
  
 Contact(s): /N7/O2'/O3'/OP2/OP2/  
 Coded contact list: 2O_ph_2O_r_N_b_  

### K:1H:3063

|  | Res. type | Atom | Residue | Distance | Occupancy | B-factor | Closest solute to 1^st^ shell water |
| --- | --- | --- | --- | --- | --- | --- | --- |
| **Ion** | | | |  | 1.00 | 47.89 || 1 | C | OP1 | 1H:2440 | 2.37 | 1.00 | 18.09 |
| 2 | A | O2' | 1H:2439 | 3.06 | 1.00 | 21.26 |  |
| 3 | G | OP1 | 1H:2442 | 3.32 | 1.00 | 19.80 |  |
| 4 | U | OP1 | 1H:875 | 3.53 | 1.00 | 23.02 |  |
| 5 | G | OP2 | 1H:2442 | 3.57 | 1.00 | 20.29 |  |
| 6 | U | OP2 | 1H:875 | 3.63 | 1.00 | 26.49 |  |
| 7 | A | O3' | 1H:2439 | 4.37 | 1.00 | 24.04 |  |
| 8 | C | O5' | 1H:2440 | 4.51 | 1.00 | 19.58 |  |
| 9 | U | O4 | 1H:876 | 4.59 | 1.00 | 28.12 |  |
| 10 | A | N3 | 1H:2439 | 4.64 | 1.00 | 18.41 |  |
| 11 | G | O5' | 1H:2442 | 4.65 | 1.00 | 18.50 |  |
| 12 | A | O4' | 1H:2439 | 4.78 | 1.00 | 20.50 |  |
| 13 | C | OP2 | 1H:2440 | 4.83 | 1.00 | 17.35 |  |
| 14 | G | N2 | 1H:2259 | 4.85 | 1.00 | 21.68 |  |
| 15 | A | N9 | 1H:2439 | 4.99 | 1.00 | 21.88 |  |
  |

  
**1^st^ shell ligand angles:**   
1-2: 59; 1-3: 66; 2-3: 99;   
  
 Contact(s): /O2'/OP1/  
 Coded contact list: O_ph_O_r_  
 Exclusion zone contact(s): /OP1/  

### K:1H:3064

|  | Res. type | Atom | Residue | Distance | Occupancy | B-factor | Closest solute to 1^st^ shell water |
| --- | --- | --- | --- | --- | --- | --- | --- |
| **Ion** | | | |  | 1.00 | 34.06 || 1 | A | OP1 | 1H:2004 | 2.92 | 1.00 | 24.73 |
| 2 | A | OP2 | 1H:1817 | 3.00 | 1.00 | 17.49 |  |
| 3 | G | O3' | 1H:2003 | 3.17 | 1.00 | 21.23 |  |
| 4 | C | O4' | 1H:2005 | 3.21 | 1.00 | 17.73 |  |
| 5 | C | O5' | 1H:2005 | 3.23 | 1.00 | 20.56 |  |
| 6 | C | OP2 | 1H:2005 | 3.36 | 1.00 | 22.74 |  |
| 7 | A | O5' | 1H:2004 | 4.09 | 1.00 | 20.64 |  |
| 8 | Water 1 | | 1H:4333 | 4.26 | 1.00 | 24.07 | 3.14 (A:OP1 1H:2004) |
| 9 | G | O2' | 1H:2003 | 4.32 | 1.00 | 22.71 |  |
| 10 | C | N1 | 1H:2005 | 4.43 | 1.00 | 20.71 |  |
| 11 | A | O5' | 1H:1817 | 4.46 | 1.00 | 17.29 |  |
| 12 | A | OP1 | 1H:1818 | 4.56 | 1.00 | 20.06 |  |
| 13 | A | O3' | 1H:1817 | 4.83 | 1.00 | 18.43 |  |
| 14 | A | OP2 | 1H:2004 | 4.88 | 1.00 | 20.91 |  |
| 15 | G | O4' | 1H:2003 | 4.91 | 1.00 | 21.19 |  |
| 16 | A | O3' | 1H:2004 | 4.98 | 1.00 | 22.28 |  |
  |

  
**1^st^ shell ligand angles:**   
1-2: 80; 1-3: 50; 1-4: 150; 1-5: 110; 1-6: 95; 2-3: 121; 2-4: 77; 2-5: 72; 2-6: 107; 3-4: 132; 3-5: 94; 3-6: 58; 4-5: 45; 4-6: 75; 5-6: 41;   
  
 Contact(s): /O3'/O4'/O5'/OP1/OP2/OP2/  
 Coded contact list: 3O_ph_3O_r_  

### K:1H:3065

|  | Res. type | Atom | Residue | Distance | Occupancy | B-factor | Closest solute to 1^st^ shell water |
| --- | --- | --- | --- | --- | --- | --- | --- |
| **Ion** | | | |  | 1.00 | 30.08 || 1 | A | OP1 | 1H:831 | 2.57 | 1.00 | 17.27 |
| 2 | A | N3 | 1H:1819 | 3.03 | 1.00 | 15.86 |  |
| 3 | U | O2 | 1H:1810 | 3.13 | 1.00 | 17.95 |  |
| 4 | A | N6 | 1H:1817 | 3.31 | 1.00 | 16.26 |  |
| 5 | C | O4' | 1H:1820 | 3.62 | 1.00 | 15.90 |  |
| 6 | A | O5' | 1H:831 | 3.66 | 1.00 | 20.28 |  |
| 7 | A | O2' | 1H:1819 | 4.19 | 1.00 | 17.07 |  |
| 8 | A | O3' | 1H:830 | 4.24 | 1.00 | 15.92 |  |
| 9 | C | N1 | 1H:1820 | 4.35 | 1.00 | 14.28 |  |
| 10 | A | OP2 | 1H:830 | 4.38 | 1.00 | 17.22 |  |
| 11 | U | N1 | 1H:1810 | 4.48 | 1.00 | 16.00 |  |
| 12 | U | N3 | 1H:1810 | 4.66 | 1.00 | 13.96 |  |
| 13 | U | O2 | 1H:1809 | 4.71 | 1.00 | 17.58 |  |
| 14 | A | N9 | 1H:1819 | 4.76 | 1.00 | 14.59 |  |
| 15 | A | N1 | 1H:1819 | 4.83 | 1.00 | 15.56 |  |
| 16 | A | O5' | 1H:830 | 4.86 | 1.00 | 18.36 |  |
| 17 | A | OP2 | 1H:831 | 4.95 | 1.00 | 15.77 |  |
| 18 | A | N1 | 1H:1817 | 4.99 | 1.00 | 14.45 |  |
  |

  
**1^st^ shell ligand angles:**   
1-2: 110; 1-3: 139; 1-4: 99; 2-3: 82; 2-4: 68; 3-4: 47;   
  
 Contact(s): /N3_A_/N6/O2/  
 Coded contact list: O_b_N_b_Anion  
 Exclusion zone contact(s): /OP1/  

### K:1H:3066

|  | Res. type | Atom | Residue | Distance | Occupancy | B-factor | Closest solute to 1^st^ shell water |
| --- | --- | --- | --- | --- | --- | --- | --- |
| **Ion** | | | |  | 1.00 | 27.63 || 1 | G | N7 | 1H:2079 | 2.86 | 1.00 | 21.23 |
| 2 | A | N7 | 1H:2590 | 3.07 | 1.00 | 19.91 |  |
| 3 | Water 1 | | 1H:3724 | 3.11 | 1.00 | 21.84 | 2.25 (G:OP1 1H:2076) |
| 4 | Water 2 | | 1H:3997 | 3.49 | 1.00 | 24.15 | 2.79 (A:OP1 1H:2590) |
| 5 | Water 3 | | 1H:3639 | 3.70 | 1.00 | 29.59 | 2.02 (A:OP1 1H:2590) |
| 6 | G | O6 | 1H:2079 | 3.81 | 1.00 | 23.33 |  |
| 7 | G | OP2 | 1H:2076 | 4.14 | 1.00 | 23.64 |  |
| 8 | A | OP2 | 1H:2590 | 4.40 | 1.00 | 21.44 |  |
| 9 | A | N9 | 1H:2590 | 4.41 | 1.00 | 24.76 |  |
| 10 | G | OP1 | 1H:2076 | 4.59 | 1.00 | 24.21 |  |
| 11 | A | N6 | 1H:2590 | 4.76 | 1.00 | 21.24 |  |
| 12 | A | OP1 | 1H:2590 | 4.84 | 1.00 | 20.08 |  |
| 13 | G | N9 | 1H:2079 | 4.87 | 1.00 | 18.99 |  |
| 14 | A | OP2 | 1H:2077 | 4.94 | 1.00 | 23.75 |  |
  |

  
**1^st^ shell ligand angles:**   
1-2: 77; 1-3: 161; 2-3: 99;   
  
 Contact(s): /N7/N7/O_w_/  
 Coded contact list: 2N_b_O_w_  
K hydration shell issue: water too close (< 2.4 Angs.)  

### K:1H:3067

|  | Res. type | Atom | Residue | Distance | Occupancy | B-factor | Closest solute to 1^st^ shell water |
| --- | --- | --- | --- | --- | --- | --- | --- |
| **Ion** | | | |  | 1.00 | 53.25 || 1 | G | O6 | 1H:664 | 2.75 | 1.00 | 54.34 |
| 2 | G | O6 | 1H:674 | 2.83 | 1.00 | 47.17 |  |
| 3 | U | O4 | 1H:665 | 3.06 | 1.00 | 56.82 |  |
| 4 | G | O6 | 1H:675 | 3.54 | 1.00 | 49.39 |  |
| 5 | C | N4 | 1H:676 | 3.59 | 1.00 | 51.29 |  |
| 6 | G | N7 | 1H:675 | 3.96 | 1.00 | 50.22 |  |
| 7 | U | N3 | 1H:665 | 4.09 | 1.00 | 52.25 |  |
| 8 | G | N7 | 1H:674 | 4.36 | 1.00 | 54.34 |  |
| 9 | C | N4 | 1H:666 | 4.38 | 1.00 | 48.00 |  |
| 10 | G | N1 | 1H:674 | 4.55 | 1.00 | 54.56 |  |
| 11 | G | N1 | 1H:664 | 4.60 | 1.00 | 53.27 |  |
| 12 | G | N7 | 1H:664 | 4.83 | 1.00 | 57.12 |  |
| 13 | G | O6 | 1H:677 | 4.97 | 1.00 | 67.05 |  |
  |

  
**1^st^ shell ligand angles:**   
1-2: 145; 1-3: 72; 2-3: 87;   
  
 Contact(s): /O4/O6/O6/  
 Coded contact list: 3O_b_  

### K:1H:3068

|  | Res. type | Atom | Residue | Distance | Occupancy | B-factor | Closest solute to 1^st^ shell water |
| --- | --- | --- | --- | --- | --- | --- | --- |
| **Ion** | | | |  | 1.00 | 29.26 || 1 | G | O6 | 1H:994 | 2.66 | 1.00 | 26.80 |
| 2 | G | O6 | 1H:993 | 2.96 | 1.00 | 32.34 |  |
| 3 | G | O6 | 1H:1014 | 3.00 | 1.00 | 28.37 |  |
| 4 | U | O4 | 1H:1015 | 3.12 | 1.00 | 29.48 |  |
| 5 | C | N4 | 1H:995 | 3.38 | 1.00 | 23.29 |  |
| 6 | U | N3 | 1H:1015 | 3.85 | 1.00 | 28.68 |  |
| 7 | G | N7 | 1H:994 | 3.99 | 1.00 | 22.44 |  |
| 8 | G | N1 | 1H:993 | 4.43 | 1.00 | 24.84 |  |
| 9 | G | N7 | 1H:993 | 4.70 | 1.00 | 24.24 |  |
| 10 | C | N3 | 1H:995 | 4.86 | 1.00 | 24.01 |  |
| 11 | C | N4 | 1H:1016 | 4.88 | 1.00 | 26.49 |  |
| 12 | G | N1 | 1H:1014 | 4.91 | 1.00 | 25.99 |  |
| 13 | G | N1 | 1H:994 | 4.93 | 1.00 | 22.90 |  |
  |

  
**1^st^ shell ligand angles:**   
1-2: 72; 1-3: 76; 1-4: 76; 1-5: 66; 2-3: 140; 2-4: 77; 2-5: 131; 3-4: 72; 3-5: 47; 4-5: 113;   
  
 Contact(s): /N4/O4/O6/O6/O6/  
 Coded contact list: 4O_b_Anion  

### K:1H:3069

|  | Res. type | Atom | Residue | Distance | Occupancy | B-factor | Closest solute to 1^st^ shell water |
| --- | --- | --- | --- | --- | --- | --- | --- |
| **Ion** | | | |  | 1.00 | 16.98 || 1 | HIS | O | 11:220 | 2.44 | 1.00 | 16.21 |
| 2 | G | O6 | 1H:1860 | 2.46 | 1.00 | 15.43 |  |
| 3 | C | OP2 | 1H:1822 | 2.95 | 1.00 | 19.01 |  |
| 4 | A | OP1 | 1H:1821 | 3.00 | 1.00 | 16.09 |  |
| 5 | G | N1 | 1H:1860 | 3.56 | 1.00 | 17.44 |  |
| 6 | A | O5' | 1H:1821 | 3.90 | 1.00 | 13.34 |  |
| 7 | ARG | N | 11:222 | 3.94 | 1.00 | 13.66 |  |
| 8 | A | OP2 | 1H:1821 | 3.95 | 1.00 | 14.11 |  |
| 9 | ARG | NE | 11:222 | 4.24 | 1.00 | 15.67 |  |
| 10 | VAL | N | 11:221 | 4.30 | 1.00 | 17.30 |  |
| 11 | C | OP1 | 1H:1822 | 4.41 | 1.00 | 15.60 |  |
| 12 | A | O3' | 1H:1821 | 4.60 | 1.00 | 16.76 |  |
| 13 | C | N4 | 1H:1859 | 4.79 | 1.00 | 17.93 |  |
  |

  
**1^st^ shell ligand angles:**   
1-2: 165; 1-3: 80; 1-4: 80; 2-3: 87; 2-4: 97; 3-4: 105;   
  
 Contact(s): /OP1/OP2/  
 Coded contact list: 2O_ph_  
 Exclusion zone contact(s): /O6/O_back_/  

### K:1H:3070

|  | Res. type | Atom | Residue | Distance | Occupancy | B-factor | Closest solute to 1^st^ shell water |
| --- | --- | --- | --- | --- | --- | --- | --- |
| **Ion** | | | |  | 1.00 | 46.66 || 1 | U | O2 | 1H:2045 | 2.48 | 1.00 | 33.25 |
| 2 | C | O2 | 1H:2629 | 2.88 | 1.00 | 22.69 |  |
| 3 | C | O2' | 1H:2629 | 2.91 | 1.00 | 27.17 |  |
| 4 | C | O4' | 1H:2630 | 3.36 | 1.00 | 29.30 |  |
| 5 | G | N2 | 1H:2076 | 3.80 | 1.00 | 20.80 |  |
| 6 | U | O2' | 1H:2045 | 3.82 | 1.00 | 38.54 |  |
| 7 | U | N3 | 1H:2045 | 4.31 | 1.00 | 31.13 |  |
| 8 | C | N1 | 1H:2629 | 4.52 | 1.00 | 22.70 |  |
| 9 | U | N1 | 1H:2045 | 4.58 | 1.00 | 30.57 |  |
| 10 | A | N3 | 1H:2077 | 4.62 | 1.00 | 23.30 |  |
| 11 | C | N1 | 1H:2630 | 4.65 | 1.00 | 24.96 |  |
| 12 | U | O4 | 1H:2057 | 4.76 | 1.00 | 30.85 |  |
| 13 | U | N3 | 1H:2057 | 4.89 | 1.00 | 28.69 |  |
| 14 | A | O2' | 1H:2077 | 5.00 | 1.00 | 22.72 |  |
  |

  
**1^st^ shell ligand angles:**   
1-2: 122; 1-3: 61; 1-4: 119; 2-3: 78; 2-4: 78; 3-4: 71;   
  
 Contact(s): /O2/O2'/O4'/  
 Coded contact list: 2O_r_O_b_  
 Exclusion zone contact(s): /O2/  

### K:1H:3071

|  | Res. type | Atom | Residue | Distance | Occupancy | B-factor | Closest solute to 1^st^ shell water |
| --- | --- | --- | --- | --- | --- | --- | --- |
| **Ion** | | | |  | 1.00 | 42.96 || 1 | G | O6 | 1H:1922 | 2.55 | 1.00 | 31.37 |
| 2 | G | O6 | 1H:1872 | 2.88 | 1.00 | 30.12 |  |
| 3 | C | O2 | 1H:1870 | 3.01 | 1.00 | 34.11 |  |
| 4 | C | O4' | 1H:1870 | 3.09 | 1.00 | 37.61 |  |
| 5 | G | O6 | 1H:1871 | 3.16 | 1.00 | 36.49 |  |
| 6 | G | N7 | 1H:1871 | 3.36 | 1.00 | 34.78 |  |
| 7 | C | N1 | 1H:1870 | 3.89 | 1.00 | 30.96 |  |
| 8 | G | N7 | 1H:1872 | 3.93 | 1.00 | 30.57 |  |
| 9 | G | N2 | 1H:1926 | 4.08 | 1.00 | 27.56 |  |
| 10 | G | N1 | 1H:1922 | 4.12 | 1.00 | 31.52 |  |
| 11 | C | N3 | 1H:1870 | 4.28 | 1.00 | 33.84 |  |
| 12 | C | N4 | 1H:1925 | 4.46 | 1.00 | 26.83 |  |
| 13 | C | O2 | 1H:1869 | 4.51 | 1.00 | 32.92 |  |
| 14 | A | N6 | 1H:1924 | 4.73 | 1.00 | 30.07 |  |
| 15 | U | O4 | 1H:1873 | 4.82 | 1.00 | 33.85 |  |
  |

  
**1^st^ shell ligand angles:**   
1-2: 109; 1-3: 84; 1-4: 88; 1-5: 126; 1-6: 144; 2-3: 81; 2-4: 139; 2-5: 75; 2-6: 105; 3-4: 64; 3-5: 147; 3-6: 114; 4-5: 125; 4-6: 74; 5-6: 53;   
  
 Contact(s): /N7/O2/O4'/O6/O6/  
 Coded contact list: O_r_3O_b_N_b_  
 Exclusion zone contact(s): /O6/  

### K:1H:3072

|  | Res. type | Atom | Residue | Distance | Occupancy | B-factor | Closest solute to 1^st^ shell water |
| --- | --- | --- | --- | --- | --- | --- | --- |
| **Ion** | | | |  | 1.00 | 53.73 || 1 | A | O2' | 1H:357 | 2.82 | 1.00 | 46.41 |
| 2 | G | O6 | 1H:351 | 2.88 | 1.00 | 42.18 |  |
| 3 | G | O6 | 1H:352 | 3.02 | 1.00 | 44.92 |  |
| 4 | G | N7 | 1H:352 | 3.98 | 1.00 | 41.93 |  |
| 5 | C | N4 | 1H:360 | 4.03 | 1.00 | 39.45 |  |
| 6 | C | OP1 | 1H:359 | 4.12 | 1.00 | 37.80 |  |
| 7 | G | N7 | 1H:351 | 4.41 | 1.00 | 39.51 |  |
| 8 | C | N4 | 1H:361 | 4.44 | 1.00 | 37.04 |  |
| 9 | C | OP2 | 1H:359 | 4.82 | 1.00 | 43.46 |  |
| 10 | G | N1 | 1H:351 | 4.93 | 1.00 | 39.56 |  |
  |

  
**1^st^ shell ligand angles:**   
1-2: 141; 1-3: 81; 2-3: 77;   
  
 Contact(s): /O2'/O6/O6/  
 Coded contact list: O_r_2O_b_  

### K:1H:3073

|  | Res. type | Atom | Residue | Distance | Occupancy | B-factor | Closest solute to 1^st^ shell water |
| --- | --- | --- | --- | --- | --- | --- | --- |
| **Ion** | | | |  | 1.00 | 56.35 || 1 | G | O6 | 1H:2863 | 2.88 | 1.00 | 57.63 |
| 2 | U | O4 | 1H:2877 | 2.95 | 1.00 | 55.81 |  |
| 3 | A | OP2 | 1H:2861 | 3.01 | 1.00 | 56.90 |  |
| 4 | A | N7 | 1H:2862 | 3.14 | 1.00 | 52.93 |  |
| 5 | U | O4 | 1H:2876 | 3.32 | 1.00 | 59.23 |  |
| 6 | A | OP2 | 1H:2862 | 3.42 | 1.00 | 49.73 |  |
| 7 | G | N7 | 1H:2863 | 3.67 | 1.00 | 55.41 |  |
| 8 | A | O5' | 1H:2861 | 3.90 | 1.00 | 45.96 |  |
| 9 | U | N3 | 1H:2876 | 4.08 | 1.00 | 56.05 |  |
| 10 | U | N3 | 1H:2877 | 4.26 | 1.00 | 48.98 |  |
| 11 | G | O6 | 1H:2875 | 4.36 | 1.00 | 61.85 |  |
| 12 | C | N4 | 1H:2864 | 4.44 | 1.00 | 56.10 |  |
| 13 | A | N6 | 1H:2862 | 4.46 | 1.00 | 51.43 |  |
| 14 | A | O5' | 1H:2862 | 4.53 | 1.00 | 50.99 |  |
| 15 | A | OP1 | 1H:2861 | 4.61 | 1.00 | 46.93 |  |
| 16 | A | N9 | 1H:2862 | 4.88 | 1.00 | 52.29 |  |
| 17 | G | N1 | 1H:2863 | 4.94 | 1.00 | 57.05 |  |
  |

  
**1^st^ shell ligand angles:**   
1-2: 65; 1-3: 122; 1-4: 86; 1-5: 71; 2-3: 71; 2-4: 68; 2-5: 63; 3-4: 112; 3-5: 55; 4-5: 131;   
  
 Contact(s): /N7/O4/O4/O6/OP2/  
 Coded contact list: O_ph_3O_b_N_b_  

### K:1H:3074

|  | Res. type | Atom | Residue | Distance | Occupancy | B-factor | Closest solute to 1^st^ shell water |
| --- | --- | --- | --- | --- | --- | --- | --- |
| **Ion** | | | |  | 1.00 | 34.33 || 1 | A | OP2 | 1H:841 | 2.62 | 1.00 | 17.97 |
| 2 | C | OP1 | 1H:726 | 2.79 | 1.00 | 19.98 |  |
| 3 | G | O6 | 1H:824 | 2.97 | 1.00 | 22.19 |  |
| 4 | A | O3' | 1H:2094 | 3.04 | 1.00 | 16.43 |  |
| 5 | G | OP1 | 1H:2095 | 3.47 | 1.00 | 23.48 |  |
| 6 | C | OP2 | 1H:726 | 3.63 | 1.00 | 16.06 |  |
| 7 | A | O2' | 1H:2094 | 3.99 | 1.00 | 20.59 |  |
| 8 | A | O2' | 1H:841 | 4.02 | 1.00 | 22.26 |  |
| 9 | A | O5' | 1H:841 | 4.07 | 1.00 | 17.10 |  |
| 10 | A | O3' | 1H:725 | 4.18 | 1.00 | 14.21 |  |
| 11 | A | OP1 | 1H:841 | 4.37 | 1.00 | 21.19 |  |
| 12 | G | O5' | 1H:2095 | 4.70 | 1.00 | 17.55 |  |
| 13 | G | N1 | 1H:824 | 4.81 | 1.00 | 21.32 |  |
  |

  
**1^st^ shell ligand angles:**   
1-2: 152; 1-3: 96; 1-4: 66; 2-3: 103; 2-4: 91; 3-4: 95;   
  
 Contact(s): /O3'/O6/OP1/OP2/  
 Coded contact list: 2O_ph_O_r_O_b_  

### K:1H:3075

|  | Res. type | Atom | Residue | Distance | Occupancy | B-factor | Closest solute to 1^st^ shell water |
| --- | --- | --- | --- | --- | --- | --- | --- |
| **Ion** | | | |  | 1.00 | 28.19 || 1 | G | O6 | 1H:711 | 2.70 | 1.00 | 22.59 |
| 2 | U | O4 | 1H:618 | 2.84 | 1.00 | 22.33 |  |
| 3 | G | O6 | 1H:617 | 2.89 | 1.00 | 27.11 |  |
| 4 | G | O6 | 1H:710 | 2.90 | 1.00 | 25.35 |  |
| 5 | C | N4 | 1H:712 | 3.68 | 1.00 | 27.49 |  |
| 6 | U | N3 | 1H:618 | 3.90 | 1.00 | 22.43 |  |
| 7 | C | N4 | 1H:619 | 3.99 | 1.00 | 26.61 |  |
| 8 | G | N7 | 1H:711 | 4.18 | 1.00 | 25.52 |  |
| 9 | G | N7 | 1H:710 | 4.53 | 1.00 | 23.36 |  |
| 10 | G | N1 | 1H:710 | 4.54 | 1.00 | 22.85 |  |
| 11 | G | N1 | 1H:617 | 4.75 | 1.00 | 23.19 |  |
| 12 | G | N7 | 1H:617 | 4.79 | 1.00 | 25.52 |  |
| 13 | G | N1 | 1H:711 | 4.91 | 1.00 | 22.54 |  |
  |

  
**1^st^ shell ligand angles:**   
1-2: 75; 1-3: 69; 1-4: 72; 2-3: 72; 2-4: 93; 3-4: 140;   
  
 Contact(s): /O4/O6/O6/O6/  
 Coded contact list: 4O_b_  

### K:1H:3076

|  | Res. type | Atom | Residue | Distance | Occupancy | B-factor | Closest solute to 1^st^ shell water |
| --- | --- | --- | --- | --- | --- | --- | --- |
| **Ion** | | | |  | 1.00 | 50.60 || 1 | U | OP2 | 1H:895 | 2.62 | 1.00 | 35.85 |
| 2 | C | O2' | 1H:894 | 2.88 | 1.00 | 40.98 |  |
| 3 | G | N7 | 1H:975 | 3.15 | 1.00 | 48.86 |  |
| 4 | G | O6 | 1H:975 | 3.28 | 1.00 | 42.58 |  |
| 5 | Water 1 | | 1H:3826 | 3.33 | 1.00 | 40.86 | 2.41 (G:O6 1H:975) |
| 6 | Water 2 | | 1H:3728 | 3.98 | 1.00 | 58.12 | 2.26 (U:OP1 1H:895) |
| 7 | C | O2 | 1H:894 | 4.49 | 1.00 | 41.82 |  |
| 8 | C | O3' | 1H:894 | 4.58 | 1.00 | 53.01 |  |
| 9 | G | OP2 | 1H:975 | 4.69 | 1.00 | 47.46 |  |
| 10 | G | N1 | 1H:975 | 4.79 | 1.00 | 37.49 |  |
| 11 | G | N9 | 1H:975 | 4.91 | 1.00 | 45.42 |  |
| 12 | U | OP1 | 1H:895 | 4.99 | 1.00 | 46.17 |  |
  |

  
**1^st^ shell ligand angles:**   
1-2: 61; 1-3: 113; 1-4: 60; 1-5: 72; 2-3: 165; 2-4: 115; 2-5: 129; 3-4: 55; 3-5: 52; 4-5: 43;   
  
 Contact(s): /N7/O2'/O6/O_w_/OP2/  
 Coded contact list: O_ph_O_r_O_b_N_b_O_w_  
K hydration shell issue: water too close (< 2.4 Angs.)  

### K:1H:3077

|  | Res. type | Atom | Residue | Distance | Occupancy | B-factor | Closest solute to 1^st^ shell water |
| --- | --- | --- | --- | --- | --- | --- | --- |
| **Ion** | | | |  | 1.00 | 47.67 || 1 | G | N7 | 1H:419 | 2.79 | 1.00 | 34.90 |
| 2 | G | O6 | 1H:419 | 2.94 | 1.00 | 32.19 |  |
| 3 | A | N3 | 1H:418 | 3.12 | 1.00 | 35.93 |  |
| 4 | U | O4 | 1H:411 | 3.35 | 1.00 | 33.55 |  |
| 5 | C | N4 | 1H:420 | 3.58 | 1.00 | 36.55 |  |
| 6 | A | O2' | 1H:418 | 3.88 | 1.00 | 33.84 |  |
| 7 | G | O6 | 1H:416 | 4.07 | 1.00 | 39.77 |  |
| 8 | G | O6 | 1H:410 | 4.19 | 1.00 | 32.70 |  |
| 9 | G | N2 | 1H:439 | 4.20 | 1.00 | 33.11 |  |
| 10 | C | O2 | 1H:413 | 4.32 | 1.00 | 36.19 |  |
| 11 | A | N9 | 1H:418 | 4.62 | 1.00 | 33.25 |  |
| 12 | U | N3 | 1H:411 | 4.64 | 1.00 | 31.82 |  |
| 13 | A | O3' | 1H:418 | 4.79 | 1.00 | 38.02 |  |
| 14 | G | N1 | 1H:419 | 4.96 | 1.00 | 32.34 |  |
| 15 | G | N9 | 1H:419 | 4.99 | 1.00 | 30.59 |  |
  |

  
**1^st^ shell ligand angles:**   
1-2: 63; 1-3: 91; 1-4: 128; 2-3: 111; 2-4: 65; 3-4: 109;   
  
 Contact(s): /N3_A_/N7/O4/O6/  
 Coded contact list: 2O_b_2N_b_  

### K:1H:3078

|  | Res. type | Atom | Residue | Distance | Occupancy | B-factor | Closest solute to 1^st^ shell water |
| --- | --- | --- | --- | --- | --- | --- | --- |
| **Ion** | | | |  | 1.00 | 45.76 || 1 | U | OP2 | 1H:1070 | 2.36 | 1.00 | 42.78 |
| 2 | G | O2' | 1H:1069 | 2.52 | 1.00 | 43.13 |  |
| 3 | G | N3 | 1H:1072 | 2.75 | 1.00 | 39.70 |  |
| 4 | G | O2' | 1H:1072 | 2.79 | 1.00 | 45.44 |  |
| 5 | G | O6 | 1H:1071 | 2.85 | 1.00 | 42.13 |  |
| 6 | G | N7 | 1H:1071 | 3.12 | 1.00 | 45.30 |  |
| 7 | G | N2 | 1H:1072 | 3.19 | 1.00 | 39.78 |  |
| 8 | U | OP1 | 1H:1070 | 3.48 | 1.00 | 48.28 |  |
| 9 | G | O3' | 1H:1069 | 3.66 | 1.00 | 43.88 |  |
| 10 | G | N1 | 1H:1072 | 3.94 | 1.00 | 37.06 |  |
| 11 | G | N9 | 1H:1072 | 4.08 | 1.00 | 41.92 |  |
| 12 | U | O5' | 1H:1070 | 4.66 | 1.00 | 45.09 |  |
| 13 | G | OP2 | 1H:1071 | 4.83 | 1.00 | 40.29 |  |
| 14 | G | N1 | 1H:1071 | 4.99 | 1.00 | 38.26 |  |
  |

  
**1^st^ shell ligand angles:**   
1-2: 72; 1-3: 147; 1-4: 90; 1-5: 82; 1-6: 60; 1-7: 155; 2-3: 127; 2-4: 158; 2-5: 72; 2-6: 113; 2-7: 87; 3-4: 76; 3-5: 80; 3-6: 87; 3-7: 44; 4-5: 119; 4-6: 64; 4-7: 114; 5-6: 59; 5-7: 80; 6-7: 123;   
  
 Contact(s): /N2/N3_G_/N7/O2'/O6/  
 Coded contact list: O_r_O_b_2N_b_Anion  
 Exclusion zone contact(s): /O2'/OP2/  

### K:1H:3079

|  | Res. type | Atom | Residue | Distance | Occupancy | B-factor | Closest solute to 1^st^ shell water |
| --- | --- | --- | --- | --- | --- | --- | --- |
| **Ion** | | | |  | 1.00 | 37.86 || 1 | THR | O | 78:32 | 2.67 | 1.00 | 24.62 |
| 2 | G | OP1 | 1H:1237 | 2.83 | 1.00 | 30.03 |  |
| 3 | A | O3' | 1H:987 | 3.27 | 1.00 | 20.22 |  |
| 4 | A | O2' | 1H:987 | 3.28 | 1.00 | 21.33 |  |
| 5 | G | O3' | 1H:1236 | 3.35 | 1.00 | 26.77 |  |
| 6 | THR | OG1 | 78:32 | 3.83 | 1.00 | 33.16 |  |
| 7 | G | OP1 | 1H:988 | 3.95 | 1.00 | 29.10 |  |
| 8 | ARG | N | 78:33 | 4.67 | 1.00 | 24.10 |  |
| 9 | ARG | NH2 | 78:18 | 4.73 | 1.00 | 39.02 |  |
| 10 | G | O5' | 1H:1237 | 4.78 | 1.00 | 25.40 |  |
| 11 | G | OP2 | 1H:1237 | 4.94 | 1.00 | 25.01 |  |
| 12 | G | O5' | 1H:988 | 4.95 | 1.00 | 21.42 |  |
  |

  
**1^st^ shell ligand angles:**   
1-2: 103; 1-3: 99; 1-4: 108; 1-5: 88; 2-3: 133; 2-4: 85; 2-5: 44; 3-4: 48; 3-5: 97; 4-5: 51;   
  
 Contact(s): /O2'/O3'/O3'/O_back_/OP1/  
 Coded contact list: O_ph_3O_r_O_back_  

### K:1H:3080

|  | Res. type | Atom | Residue | Distance | Occupancy | B-factor | Closest solute to 1^st^ shell water |
| --- | --- | --- | --- | --- | --- | --- | --- |
| **Ion** | | | |  | 1.00 | 45.95 || 1 | A | N7 | 1H:2004 | 2.99 | 1.00 | 28.00 |
| 2 | A | OP1 | 1H:2004 | 3.02 | 1.00 | 24.73 |  |
| 3 | A | OP2 | 1H:1817 | 3.12 | 1.00 | 17.49 |  |
| 4 | A | O3' | 1H:1816 | 3.60 | 1.00 | 21.02 |  |
| 5 | Water 1 | | 1H:4418 | 3.90 | 1.00 | 29.52 | 3.53 (A:OP1 1H:2004) |
| 6 | A | O2' | 1H:1816 | 4.07 | 1.00 | 20.43 |  |
| 7 | A | N6 | 1H:2004 | 4.55 | 1.00 | 26.33 |  |
| 8 | Water 2 | | 1H:4367 | 4.56 | 1.00 | 17.79 | 5.04 (A:OP1 1H:2004) |
| 9 | Water 3 | | 1H:4333 | 4.62 | 1.00 | 24.07 | 3.14 (A:OP1 1H:2004) |
| 10 | A | O5' | 1H:2004 | 4.78 | 1.00 | 20.64 |  |
| 11 | A | N9 | 1H:2004 | 4.87 | 1.00 | 19.12 |  |
  |

  
**1^st^ shell ligand angles:**   
1-2: 84; 1-3: 74; 2-3: 76;   
  
 Contact(s): /N7/OP1/OP2/  
 Coded contact list: 2O_ph_N_b_  

### K:1H:3081

|  | Res. type | Atom | Residue | Distance | Occupancy | B-factor | Closest solute to 1^st^ shell water |
| --- | --- | --- | --- | --- | --- | --- | --- |
| **Ion** | | | |  | 1.00 | 49.65 || 1 | G | O6 | 1H:559 | 2.89 | 1.00 | 27.70 |
| 2 | G | O6 | 1H:583 | 3.08 | 1.00 | 31.76 |  |
| 3 | G | O6 | 1H:582 | 3.25 | 1.00 | 34.80 |  |
| 4 | C | N4 | 1H:584 | 3.50 | 1.00 | 26.54 |  |
| 5 | U | O4 | 1H:560 | 3.57 | 1.00 | 31.16 |  |
| 6 | G | N7 | 1H:583 | 3.88 | 1.00 | 32.03 |  |
| 7 | U | N3 | 1H:560 | 4.44 | 1.00 | 28.63 |  |
| 8 | G | N7 | 1H:582 | 4.53 | 1.00 | 37.68 |  |
| 9 | C | N4 | 1H:561 | 4.65 | 1.00 | 34.13 |  |
| 10 | G | N1 | 1H:559 | 4.85 | 1.00 | 26.94 |  |
| 11 | G | N1 | 1H:582 | 4.92 | 1.00 | 35.56 |  |
| 12 | G | N7 | 1H:559 | 4.97 | 1.00 | 27.20 |  |
  |

  
**1^st^ shell ligand angles:**   
1-2: 69; 1-3: 132; 2-3: 68;   
  
 Contact(s): /O6/O6/O6/  
 Coded contact list: 3O_b_  

### K:1H:3082

|  | Res. type | Atom | Residue | Distance | Occupancy | B-factor | Closest solute to 1^st^ shell water |
| --- | --- | --- | --- | --- | --- | --- | --- |
| **Ion** | | | |  | 1.00 | 46.98 || 1 | A | OP1 | 1H:456 | 2.73 | 1.00 | 25.72 |
| 2 | G | O6 | 1H:209 | 2.81 | 1.00 | 39.75 |  |
| 3 | G | O6 | 1H:210 | 2.87 | 1.00 | 33.41 |  |
| 4 | Water 1 | | 1H:3674 | 3.03 | 1.00 | 29.72 | 2.15 (A:OP1 1H:456) |
| 5 | Water 2 | | 1H:4143 | 3.25 | 1.00 | 30.28 | 2.82 (A:N7 1H:208) |
| 6 | A | OP2 | 1H:208 | 3.58 | 1.00 | 26.82 |  |
| 7 | G | N7 | 1H:209 | 3.77 | 1.00 | 32.08 |  |
| 8 | U | OP2 | 1H:455 | 3.88 | 1.00 | 35.30 |  |
| 9 | U | O5' | 1H:455 | 4.42 | 1.00 | 26.11 |  |
| 10 | G | N1 | 1H:210 | 4.44 | 1.00 | 32.81 |  |
| 11 | U | OP1 | 1H:455 | 4.50 | 1.00 | 37.54 |  |
| 12 | G | OP2 | 1H:207 | 4.53 | 1.00 | 27.41 |  |
| 13 | A | OP2 | 1H:456 | 4.60 | 1.00 | 30.57 |  |
| 14 | Water 3 | | 1H:4338 | 4.72 | 1.00 | 35.68 | 3.22 (G:O6 1H:209) |
| 15 | A | N7 | 1H:208 | 4.76 | 1.00 | 33.08 |  |
| 16 | A | O5' | 1H:456 | 4.96 | 1.00 | 26.20 |  |
  |

  
**1^st^ shell ligand angles:**   
1-2: 72; 1-3: 79; 1-4: 43; 1-5: 97; 2-3: 69; 2-4: 79; 2-5: 66; 3-4: 120; 3-5: 135; 4-5: 60;   
  
 Contact(s): /O6/O6/O_w_/O_w_/OP1/  
 Coded contact list: O_ph_2O_b_2O_w_  
K hydration shell issue: water too close (< 2.4 Angs.)  

### K:1H:3083

|  | Res. type | Atom | Residue | Distance | Occupancy | B-factor | Closest solute to 1^st^ shell water |
| --- | --- | --- | --- | --- | --- | --- | --- |
| **Ion** | | | |  | 1.00 | 54.39 || 1 | G | O6 | 1H:2303 | 2.80 | 1.00 | 43.42 |
| 2 | G | O6 | 1H:2354 | 2.80 | 1.00 | 52.99 |  |
| 3 | U | O4 | 1H:2304 | 3.08 | 1.00 | 46.17 |  |
| 4 | G | O6 | 1H:2353 | 3.53 | 1.00 | 51.36 |  |
| 5 | C | N4 | 1H:2355 | 3.90 | 1.00 | 56.77 |  |
| 6 | U | N3 | 1H:2304 | 4.10 | 1.00 | 44.80 |  |
| 7 | G | N7 | 1H:2354 | 4.34 | 1.00 | 56.52 |  |
| 8 | G | N7 | 1H:2353 | 4.37 | 1.00 | 47.45 |  |
| 9 | G | O6 | 1H:2302 | 4.72 | 1.00 | 49.35 |  |
| 10 | G | N1 | 1H:2303 | 4.79 | 1.00 | 43.17 |  |
| 11 | G | N1 | 1H:2353 | 4.97 | 1.00 | 48.45 |  |
| 12 | C | N4 | 1H:2305 | 5.00 | 1.00 | 44.54 |  |
  |

  
**1^st^ shell ligand angles:**   
1-2: 68; 1-3: 67; 2-3: 75;   
  
 Contact(s): /O4/O6/O6/  
 Coded contact list: 3O_b_  

### K:1H:3084

|  | Res. type | Atom | Residue | Distance | Occupancy | B-factor | Closest solute to 1^st^ shell water |
| --- | --- | --- | --- | --- | --- | --- | --- |
| **Ion** | | | |  | 1.00 | 47.78 || 1 | U | O2 | 1H:875 | 2.77 | 1.00 | 26.05 |
| 2 | U | O2 | 1H:2091 | 2.77 | 1.00 | 23.70 |  |
| 3 | U | O2' | 1H:875 | 3.08 | 1.00 | 25.24 |  |
| 4 | U | OP1 | 1H:2258 | 3.40 | 1.00 | 21.95 |  |
| 5 | U | O2' | 1H:2091 | 4.02 | 1.00 | 27.64 |  |
| 6 | U | N1 | 1H:2091 | 4.20 | 1.00 | 17.83 |  |
| 7 | G | O3' | 1H:2259 | 4.33 | 1.00 | 19.22 |  |
| 8 | U | OP2 | 1H:2258 | 4.35 | 1.00 | 22.29 |  |
| 9 | A | N1 | 1H:2443 | 4.59 | 1.00 | 24.10 |  |
| 10 | U | N1 | 1H:875 | 4.62 | 1.00 | 19.74 |  |
| 11 | U | N3 | 1H:2091 | 4.73 | 1.00 | 17.79 |  |
| 12 | U | N3 | 1H:875 | 4.77 | 1.00 | 23.84 |  |
| 13 | G | O2' | 1H:2259 | 4.80 | 1.00 | 26.52 |  |
| 14 | U | O4' | 1H:2091 | 4.89 | 1.00 | 17.72 |  |
  |

  
**1^st^ shell ligand angles:**   
1-2: 93; 1-3: 65; 2-3: 80;   
  
 Contact(s): /O2/O2/O2'/  
 Coded contact list: O_r_2O_b_  

### K:1H:3085

|  | Res. type | Atom | Residue | Distance | Occupancy | B-factor | Closest solute to 1^st^ shell water |
| --- | --- | --- | --- | --- | --- | --- | --- |
| **Ion** | | | |  | 1.00 | 39.53 || 1 | G | O6 | 1H:985 | 2.55 | 1.00 | 24.05 |
| 2 | U | O4 | 1H:887 | 3.02 | 1.00 | 22.83 |  |
| 3 | G | O6 | 1H:984 | 3.14 | 1.00 | 30.95 |  |
| 4 | G | O6 | 1H:986 | 3.27 | 1.00 | 31.89 |  |
| 5 | G | N7 | 1H:985 | 3.48 | 1.00 | 26.77 |  |
| 6 | U | N3 | 1H:887 | 3.97 | 1.00 | 22.15 |  |
| 7 | C | N4 | 1H:888 | 4.09 | 1.00 | 28.52 |  |
| 8 | C | N4 | 1H:886 | 4.26 | 1.00 | 21.47 |  |
| 9 | G | N1 | 1H:984 | 4.59 | 1.00 | 28.16 |  |
| 10 | G | N1 | 1H:985 | 4.69 | 1.00 | 23.54 |  |
| 11 | G | N7 | 1H:986 | 4.71 | 1.00 | 23.46 |  |
| 12 | G | N7 | 1H:984 | 4.78 | 1.00 | 36.75 |  |
| 13 | G | N1 | 1H:986 | 4.97 | 1.00 | 20.98 |  |
  |

  
**1^st^ shell ligand angles:**   
1-2: 66; 1-3: 81; 1-4: 66; 2-3: 96; 2-4: 78; 3-4: 146;   
  
 Contact(s): /O4/O6/O6/  
 Coded contact list: 3O_b_  
 Exclusion zone contact(s): /O6/  

### K:1H:3086

|  | Res. type | Atom | Residue | Distance | Occupancy | B-factor | Closest solute to 1^st^ shell water |
| --- | --- | --- | --- | --- | --- | --- | --- |
| **Ion** | | | |  | 1.00 | 66.80 || 1 | U | O4 | 1H:147 | 2.57 | 1.00 | 41.09 |
| 2 | G | O6 | 1H:129 | 2.80 | 1.00 | 28.83 |  |
| 3 | G | O6 | 1H:130 | 3.00 | 1.00 | 37.71 |  |
| 4 | G | O6 | 1H:146 | 3.37 | 1.00 | 44.29 |  |
| 5 | C | N4 | 1H:148 | 3.81 | 1.00 | 28.18 |  |
| 6 | U | N3 | 1H:147 | 3.97 | 1.00 | 35.30 |  |
| 7 | G | N1 | 1H:129 | 4.48 | 1.00 | 26.13 |  |
| 8 | C | N4 | 1H:131 | 4.71 | 1.00 | 30.15 |  |
| 9 | G | N7 | 1H:129 | 4.81 | 1.00 | 27.79 |  |
  |

  
**1^st^ shell ligand angles:**   
1-2: 107; 1-3: 76; 1-4: 65; 2-3: 78; 2-4: 138; 3-4: 60;   
  
 Contact(s): /O6/O6/O6/  
 Coded contact list: 3O_b_  
 Exclusion zone contact(s): /O4/  

### K:1H:3087

|  | Res. type | Atom | Residue | Distance | Occupancy | B-factor | Closest solute to 1^st^ shell water |
| --- | --- | --- | --- | --- | --- | --- | --- |
| **Ion** | | | |  | 1.00 | 31.55 || 1 | U | O4 | 1H:1444 | 2.81 | 1.00 | 29.64 |
| 2 | U | O2' | 1H:1441 | 2.81 | 1.00 | 31.34 |  |
| 3 | A | O5' | 1H:1442 | 2.83 | 1.00 | 29.15 |  |
| 4 | U | O2' | 1H:1649 | 2.84 | 1.00 | 28.74 |  |
| 5 | A | OP1 | 1H:1650 | 2.86 | 1.00 | 24.49 |  |
| 6 | U | O3' | 1H:1441 | 3.03 | 1.00 | 27.74 |  |
| 7 | A | OP1 | 1H:1442 | 3.17 | 1.00 | 24.15 |  |
| 8 | U | O3' | 1H:1649 | 3.20 | 1.00 | 24.22 |  |
| 9 | A | O5' | 1H:1650 | 4.29 | 1.00 | 26.99 |  |
| 10 | Water 1 | | 1H:4172 | 4.30 | 1.00 | 26.41 | 2.86 (A:OP1 1H:1442) |
| 11 | U | N3 | 1H:1444 | 4.51 | 1.00 | 27.79 |  |
| 12 | U | OP1 | 1H:1388 | 4.60 | 1.00 | 27.23 |  |
| 13 | U | OP2 | 1H:1388 | 4.61 | 1.00 | 27.10 |  |
| 14 | A | OP2 | 1H:1442 | 4.64 | 1.00 | 31.38 |  |
| 15 | C | OP2 | 1H:1651 | 4.83 | 1.00 | 24.25 |  |
| 16 | A | OP2 | 1H:1650 | 4.89 | 1.00 | 28.95 |  |
  |

  
**1^st^ shell ligand angles:**   
1-2: 92; 1-3: 91; 1-4: 83; 1-5: 150; 1-6: 134; 1-7: 133; 1-8: 107; 2-3: 72; 2-4: 83; 2-5: 117; 2-6: 55; 2-7: 100; 2-8: 127; 3-4: 154; 3-5: 107; 3-6: 51; 3-7: 52; 3-8: 152; 4-5: 90; 4-6: 119; 4-7: 143; 4-8: 52; 5-6: 75; 5-7: 55; 5-8: 49; 6-7: 46; 6-8: 119; 7-8: 102;   
  
 Contact(s): /O2'/O2'/O3'/O3'/O4/O5'/OP1/OP1/  
 Coded contact list: 2O_ph_5O_r_O_b_  

### K:1H:3088

|  | Res. type | Atom | Residue | Distance | Occupancy | B-factor | Closest solute to 1^st^ shell water |
| --- | --- | --- | --- | --- | --- | --- | --- |
| **Ion** | | | |  | 1.00 | 50.79 || 1 | U | OP2 | 1H:1347 | 2.74 | 1.00 | 29.43 |
| 2 | G | O6 | 1H:1674 | 2.80 | 1.00 | 35.28 |  |
| 3 | G | O6 | 1H:1675 | 3.06 | 1.00 | 40.37 |  |
| 4 | G | O3' | 1H:1346 | 3.19 | 1.00 | 31.66 |  |
| 5 | G | N7 | 1H:1674 | 3.31 | 1.00 | 33.47 |  |
| 6 | G | N7 | 1H:1675 | 3.75 | 1.00 | 37.50 |  |
| 7 | G | OP2 | 1H:1674 | 4.50 | 1.00 | 28.44 |  |
| 8 | U | OP1 | 1H:1347 | 4.50 | 1.00 | 23.40 |  |
| 9 | C | N4 | 1H:1686 | 4.56 | 1.00 | 31.96 |  |
| 10 | U | O4 | 1H:1687 | 4.58 | 1.00 | 28.74 |  |
| 11 | G | O2' | 1H:1346 | 4.61 | 1.00 | 31.35 |  |
| 12 | G | N1 | 1H:1674 | 4.63 | 1.00 | 30.79 |  |
| 13 | U | O5' | 1H:1347 | 4.70 | 1.00 | 23.05 |  |
| 14 | U | N3 | 1H:1687 | 4.80 | 1.00 | 33.14 |  |
  |

  
**1^st^ shell ligand angles:**   
1-2: 127; 1-3: 110; 1-4: 49; 1-5: 137; 2-3: 71; 2-4: 81; 2-5: 60; 3-4: 114; 3-5: 112; 4-5: 102;   
  
 Contact(s): /N7/O3'/O6/O6/OP2/  
 Coded contact list: O_ph_O_r_2O_b_N_b_  

### K:1H:3089

|  | Res. type | Atom | Residue | Distance | Occupancy | B-factor | Closest solute to 1^st^ shell water |
| --- | --- | --- | --- | --- | --- | --- | --- |
| **Ion** | | | |  | 1.00 | 36.46 || 1 | G | O6 | 1H:16 | 2.68 | 1.00 | 36.22 |
| 2 | G | O6 | 1H:15 | 2.87 | 1.00 | 32.36 |  |
| 3 | A | O2' | 1H:13 | 2.92 | 1.00 | 46.70 |  |
| 4 | U | O4 | 1H:551 | 3.23 | 1.00 | 41.55 |  |
| 5 | U | O4 | 1H:550 | 3.24 | 1.00 | 38.19 |  |
| 6 | U | N3 | 1H:551 | 3.76 | 1.00 | 35.78 |  |
| 7 | G | N7 | 1H:16 | 4.05 | 1.00 | 33.30 |  |
| 8 | U | N3 | 1H:550 | 4.11 | 1.00 | 38.65 |  |
| 9 | G | N7 | 1H:15 | 4.34 | 1.00 | 37.47 |  |
| 10 | G | N1 | 1H:15 | 4.82 | 1.00 | 32.86 |  |
| 11 | G | N1 | 1H:16 | 4.88 | 1.00 | 32.70 |  |
| 12 | A | N9 | 1H:13 | 4.88 | 1.00 | 35.37 |  |
  |

  
**1^st^ shell ligand angles:**   
1-2: 75; 1-3: 134; 1-4: 79; 1-5: 71; 2-3: 61; 2-4: 71; 2-5: 122; 3-4: 96; 3-5: 143; 4-5: 58;   
  
 Contact(s): /O2'/O4/O4/O6/O6/  
 Coded contact list: O_r_4O_b_  

### K:1H:3090

|  | Res. type | Atom | Residue | Distance | Occupancy | B-factor | Closest solute to 1^st^ shell water |
| --- | --- | --- | --- | --- | --- | --- | --- |
| **Ion** | | | |  | 1.00 | 55.06 || 1 | G | N1 | 1H:1973 | 2.66 | 1.00 | 39.42 |
| 2 | U | O4 | 1H:1974 | 3.02 | 1.00 | 39.36 |  |
| 3 | G | O6 | 1H:1973 | 3.09 | 1.00 | 39.70 |  |
| 4 | G | OP2 | 1H:1977 | 3.32 | 1.00 | 27.72 |  |
| 5 | G | N2 | 1H:1973 | 3.55 | 1.00 | 36.33 |  |
| 6 | U | N3 | 1H:1974 | 4.42 | 1.00 | 28.38 |  |
| 7 | G | OP1 | 1H:1977 | 4.61 | 1.00 | 31.95 |  |
| 8 | G | N7 | 1H:1972 | 4.81 | 1.00 | 30.20 |  |
| 9 | G | O6 | 1H:1972 | 4.93 | 1.00 | 29.73 |  |
| 10 | G | N3 | 1H:1973 | 4.95 | 1.00 | 27.07 |  |
  |

  
**1^st^ shell ligand angles:**   
1-2: 59; 1-3: 46; 1-4: 95; 2-3: 85; 2-4: 56; 3-4: 80;   
  
 Contact(s): /N1_G_/O4/O6/OP2/  
 Coded contact list: O_ph_2O_b_Anion  

### K:1H:3091

|  | Res. type | Atom | Residue | Distance | Occupancy | B-factor | Closest solute to 1^st^ shell water |
| --- | --- | --- | --- | --- | --- | --- | --- |
| **Ion** | | | |  | 1.00 | 64.49 || 1 | G | O6 | 1H:704 | 2.90 | 1.00 | 45.63 |
| 2 | U | O4 | 1H:705 | 2.95 | 1.00 | 41.26 |  |
| 3 | G | O6 | 1H:624 | 3.23 | 1.00 | 38.70 |  |
| 4 | G | O6 | 1H:623 | 3.29 | 1.00 | 40.77 |  |
| 5 | C | N4 | 1H:625 | 3.74 | 1.00 | 40.35 |  |
| 6 | U | N3 | 1H:705 | 4.18 | 1.00 | 42.09 |  |
| 7 | G | N7 | 1H:624 | 4.45 | 1.00 | 36.76 |  |
| 8 | C | N4 | 1H:706 | 4.47 | 1.00 | 40.19 |  |
| 9 | G | N7 | 1H:623 | 4.76 | 1.00 | 42.28 |  |
| 10 | G | N1 | 1H:704 | 4.78 | 1.00 | 38.72 |  |
  |

  
**1^st^ shell ligand angles:**   
1-2: 64; 1-3: 75; 1-4: 130; 2-3: 72; 2-4: 86; 3-4: 57;   
  
 Contact(s): /O4/O6/O6/O6/  
 Coded contact list: 4O_b_  

### K:1H:3092

|  | Res. type | Atom | Residue | Distance | Occupancy | B-factor | Closest solute to 1^st^ shell water |
| --- | --- | --- | --- | --- | --- | --- | --- |
| **Ion** | | | |  | 1.00 | 37.38 || 1 | A | OP2 | 1H:498 | 2.85 | 1.00 | 23.37 |
| 2 | C | OP1 | 1H:480 | 2.90 | 1.00 | 19.27 |  |
| 3 | G | O3' | 1H:479 | 2.90 | 1.00 | 23.38 |  |
| 4 | A | OP1 | 1H:498 | 3.06 | 1.00 | 21.71 |  |
| 5 | Water 1 | | 1H:4321 | 3.14 | 1.00 | 26.03 | 3.12 (C:OP1 1H:480) |
| 6 | A | O3' | 1H:497 | 3.59 | 1.00 | 24.84 |  |
| 7 | G | O2' | 1H:479 | 3.76 | 1.00 | 19.03 |  |
| 8 | Water 2 | | 1H:3975 | 4.22 | 1.00 | 20.47 | 2.62 (C:OP1 1H:480) |
| 9 | C | O5' | 1H:480 | 4.35 | 1.00 | 26.09 |  |
| 10 | A | O5' | 1H:498 | 4.74 | 1.00 | 24.53 |  |
| 11 | A | O5' | 1H:497 | 4.74 | 1.00 | 18.83 |  |
| 12 | C | OP2 | 1H:480 | 4.77 | 1.00 | 19.78 |  |
| 13 | TYR | OH | 31:59 | 4.83 | 1.00 | 31.66 |  |
| 14 | A | OP1 | 1H:497 | 4.86 | 1.00 | 21.89 |  |
  |

  
**1^st^ shell ligand angles:**   
1-2: 97; 1-3: 117; 1-4: 52; 1-5: 74; 2-3: 50; 2-4: 127; 2-5: 62; 3-4: 169; 3-5: 111; 4-5: 68;   
  
 Contact(s): /O3'/O_w_/OP1/OP1/OP2/  
 Coded contact list: 3O_ph_O_r_O_w_  

### K:1H:3093

|  | Res. type | Atom | Residue | Distance | Occupancy | B-factor | Closest solute to 1^st^ shell water |
| --- | --- | --- | --- | --- | --- | --- | --- |
| **Ion** | | | |  | 1.00 | 42.07 || 1 | A | O2' | 1H:2585 | 2.75 | 1.00 | 26.61 |
| 2 | C | OP2 | 1H:2586 | 2.81 | 1.00 | 27.39 |  |
| 3 | U | OP1 | 1H:2519 | 2.93 | 1.00 | 42.22 |  |
| 4 | ARG | NH2 | 21:144 | 3.52 | 1.00 | 22.93 |  |
| 5 | A | O3' | 1H:2585 | 3.58 | 1.00 | 19.72 |  |
| 6 | A | N3 | 1H:2585 | 3.62 | 1.00 | 21.08 |  |
| 7 | A | O4' | 1H:2466 | 3.67 | 1.00 | 19.45 |  |
| 8 | A | O2' | 1H:2466 | 3.78 | 1.00 | 24.67 |  |
| 9 | A | N9 | 1H:2585 | 4.43 | 1.00 | 19.50 |  |
| 10 | C | O2' | 1H:2465 | 4.57 | 1.00 | 28.83 |  |
| 11 | U | O5' | 1H:2519 | 4.61 | 1.00 | 31.29 |  |
| 12 | C | OP1 | 1H:2586 | 4.66 | 1.00 | 27.58 |  |
| 13 | U | OP2 | 1H:2519 | 4.71 | 1.00 | 30.31 |  |
| 14 | C | O5' | 1H:2586 | 4.92 | 1.00 | 27.80 |  |
  |

  
**1^st^ shell ligand angles:**   
1-2: 75; 1-3: 115; 2-3: 122;   
  
 Contact(s): /O2'/OP1/OP2/  
 Coded contact list: 2O_ph_O_r_  

### K:1H:3094

|  | Res. type | Atom | Residue | Distance | Occupancy | B-factor | Closest solute to 1^st^ shell water |
| --- | --- | --- | --- | --- | --- | --- | --- |
| **Ion** | | | |  | 1.00 | 51.99 || 1 | G | O6 | 1H:2722 | 2.76 | 1.00 | 32.76 |
| 2 | U | O4 | 1H:2711 | 2.91 | 1.00 | 37.46 |  |
| 3 | G | O6 | 1H:2721 | 3.17 | 1.00 | 31.38 |  |
| 4 | G | O6 | 1H:2710 | 3.29 | 1.00 | 30.76 |  |
| 5 | C | N4 | 1H:2723 | 3.82 | 1.00 | 34.81 |  |
| 6 | U | N3 | 1H:2711 | 3.89 | 1.00 | 29.14 |  |
| 7 | C | N4 | 1H:2712 | 4.28 | 1.00 | 27.32 |  |
| 8 | G | N7 | 1H:2722 | 4.37 | 1.00 | 28.52 |  |
| 9 | G | N7 | 1H:2721 | 4.66 | 1.00 | 27.78 |  |
| 10 | U | O4 | 1H:2709 | 4.75 | 1.00 | 34.70 |  |
| 11 | G | N1 | 1H:2721 | 4.87 | 1.00 | 25.32 |  |
  |

  
**1^st^ shell ligand angles:**   
1-2: 82; 1-3: 69; 1-4: 66; 2-3: 94; 2-4: 66; 3-4: 133;   
  
 Contact(s): /O4/O6/O6/O6/  
 Coded contact list: 4O_b_  

### K:1H:3095

|  | Res. type | Atom | Residue | Distance | Occupancy | B-factor | Closest solute to 1^st^ shell water |
| --- | --- | --- | --- | --- | --- | --- | --- |
| **Ion** | | | |  | 1.00 | 58.53 || 1 | G | O6 | 1H:2521 | 2.90 | 1.00 | 27.43 |
| 2 | U | O4 | 1H:2593 | 3.13 | 1.00 | 28.21 |  |
| 3 | G | N7 | 1H:2521 | 3.31 | 1.00 | 23.90 |  |
| 4 | G | O6 | 1H:2522 | 3.71 | 1.00 | 20.84 |  |
| 5 | C | N4 | 1H:2520 | 4.11 | 1.00 | 28.54 |  |
| 6 | G | N7 | 1H:2589 | 4.23 | 1.00 | 25.42 |  |
| 7 | U | N3 | 1H:2593 | 4.40 | 1.00 | 24.82 |  |
| 8 | C | N4 | 1H:2592 | 4.55 | 1.00 | 21.35 |  |
| 9 | C | OP2 | 1H:2520 | 4.72 | 1.00 | 42.63 |  |
| 10 | G | N9 | 1H:2589 | 4.89 | 1.00 | 24.28 |  |
| 11 | G | N7 | 1H:2522 | 4.98 | 1.00 | 22.05 |  |
| 12 | G | N1 | 1H:2521 | 4.99 | 1.00 | 20.88 |  |
  |

  
**1^st^ shell ligand angles:**   
1-2: 61; 1-3: 59; 2-3: 119;   
  
 Contact(s): /N7/O4/O6/  
 Coded contact list: 2O_b_N_b_  

### K:1H:3096

|  | Res. type | Atom | Residue | Distance | Occupancy | B-factor | Closest solute to 1^st^ shell water |
| --- | --- | --- | --- | --- | --- | --- | --- |
| **Ion** | | | |  | 1.00 | 52.15 || 1 | C | O2' | 1H:2477 | 2.61 | 1.00 | 38.45 |
| 2 | A | OP2 | 1H:1173 | 2.76 | 1.00 | 38.16 |  |
| 3 | A | OP1 | 1H:1173 | 2.89 | 1.00 | 39.55 |  |
| 4 | C | O2 | 1H:2477 | 3.34 | 1.00 | 34.00 |  |
| 5 | C | O4' | 1H:2478 | 3.61 | 1.00 | 40.10 |  |
| 6 | G | O3' | 1H:1172 | 3.92 | 1.00 | 37.96 |  |
| 7 | G | N2 | 1H:2499 | 3.94 | 1.00 | 32.56 |  |
| 8 | A | O5' | 1H:1173 | 4.60 | 1.00 | 41.43 |  |
| 9 | A | N6 | 1H:1075 | 4.66 | 1.00 | 32.55 |  |
| 10 | A | N1 | 1H:1075 | 4.76 | 1.00 | 31.27 |  |
| 11 | G | N2 | 1H:2500 | 4.78 | 1.00 | 34.94 |  |
| 12 | C | N1 | 1H:2477 | 4.81 | 1.00 | 31.29 |  |
| 13 | G | O2' | 1H:1172 | 4.86 | 1.00 | 27.71 |  |
  |

  
**1^st^ shell ligand angles:**   
1-2: 77; 1-3: 129; 1-4: 69; 2-3: 55; 2-4: 146; 3-4: 158;   
  
 Contact(s): /O2/O2'/OP1/OP2/  
 Coded contact list: 2O_ph_O_r_O_b_  

### K:1H:3097

|  | Res. type | Atom | Residue | Distance | Occupancy | B-factor | Closest solute to 1^st^ shell water |
| --- | --- | --- | --- | --- | --- | --- | --- |
| **Ion** | | | |  | 1.00 | 25.66 || 1 | A | OP2 | 1H:1959 | 2.64 | 1.00 | 26.97 |
| 2 | U | O4 | 1H:1963 | 2.77 | 1.00 | 38.08 |  |
| 3 | 5MC | N3 | 1H:1985 | 2.86 | 1.00 | 30.91 |  |
| 4 | A | N3 | 1H:1959 | 2.96 | 1.00 | 29.79 |  |
| 5 | 5MC | O2 | 1H:1985 | 3.04 | 1.00 | 31.52 |  |
| 6 | A | O4' | 1H:1959 | 3.15 | 1.00 | 27.76 |  |
| 7 | A | O5' | 1H:1959 | 3.60 | 1.00 | 22.94 |  |
| 8 | G | O2' | 1H:1958 | 3.77 | 1.00 | 27.09 |  |
| 9 | U | N3 | 1H:1963 | 3.82 | 1.00 | 27.13 |  |
| 10 | A | N9 | 1H:1959 | 4.20 | 1.00 | 28.39 |  |
| 11 | C | N4 | 1H:1984 | 4.25 | 1.00 | 26.17 |  |
| 12 | G | O3' | 1H:1958 | 4.34 | 1.00 | 26.00 |  |
| 13 | G | N3 | 1H:1958 | 4.52 | 1.00 | 29.71 |  |
| 14 | 5MC | N4 | 1H:1985 | 4.55 | 1.00 | 27.48 |  |
| 15 | A | N1 | 1H:1959 | 4.67 | 1.00 | 34.84 |  |
| 16 | G | N2 | 1H:1958 | 4.76 | 1.00 | 30.61 |  |
| 17 | 5MC | N1 | 1H:1985 | 4.84 | 1.00 | 31.46 |  |
| 18 | A | OP1 | 1H:1959 | 4.97 | 1.00 | 22.22 |  |
  |

  
**1^st^ shell ligand angles:**   
1-2: 136; 1-3: 63; 1-4: 106; 1-5: 109; 1-6: 102; 2-3: 115; 2-4: 115; 2-5: 87; 2-6: 81; 3-4: 109; 3-5: 47; 3-6: 163; 4-5: 90; 4-6: 66; 5-6: 145;   
  
 Contact(s): /N3/N3_A_/O2/O4/O4'/OP2/  
 Coded contact list: O_ph_O_r_2O_b_N_b_  

### K:1H:3098

|  | Res. type | Atom | Residue | Distance | Occupancy | B-factor | Closest solute to 1^st^ shell water |
| --- | --- | --- | --- | --- | --- | --- | --- |
| **Ion** | | | |  | 1.00 | 41.07 || 1 | U | O4 | 1H:1873 | 2.67 | 1.00 | 33.85 |
| 2 | U | O4 | 1H:1921 | 2.71 | 1.00 | 37.38 |  |
| 3 | G | O6 | 1H:1874 | 2.82 | 1.00 | 33.56 |  |
| 4 | C | O2 | 1H:1870 | 3.43 | 1.00 | 34.11 |  |
| 5 | G | O6 | 1H:1920 | 3.82 | 1.00 | 39.68 |  |
| 6 | U | N3 | 1H:1921 | 3.83 | 1.00 | 36.18 |  |
| 7 | G | N7 | 1H:1874 | 4.16 | 1.00 | 29.71 |  |
| 8 | A | N6 | 1H:1924 | 4.47 | 1.00 | 30.07 |  |
| 9 | U | N3 | 1H:1873 | 4.65 | 1.00 | 28.60 |  |
| 10 | G | OP2 | 1H:1872 | 4.70 | 1.00 | 35.36 |  |
| 11 | C | N4 | 1H:1875 | 4.72 | 1.00 | 24.92 |  |
| 12 | G | N1 | 1H:1874 | 4.92 | 1.00 | 26.75 |  |
  |

  
**1^st^ shell ligand angles:**   
1-2: 120; 1-3: 82; 2-3: 78;   
  
 Contact(s): /O4/O4/O6/  
 Coded contact list: 3O_b_  

### K:1H:3099

|  | Res. type | Atom | Residue | Distance | Occupancy | B-factor | Closest solute to 1^st^ shell water |
| --- | --- | --- | --- | --- | --- | --- | --- |
| **Ion** | | | |  | 1.00 | 59.79 || 1 | G | O6 | 1H:1998 | 3.12 | 1.00 | 23.37 |
| 2 | U | O4 | 1H:1999 | 3.50 | 1.00 | 27.72 |  |
| 3 | Water 1 | | 1H:4248 | 3.87 | 1.00 | 15.08 | 3.46 (G:OP2 1H:1804) |
| 4 | A | OP2 | 1H:1805 | 4.03 | 1.00 | 18.46 |  |
| 5 | C | N4 | 1H:1862 | 4.26 | 1.00 | 21.77 |  |
| 6 | G | N7 | 1H:1998 | 4.48 | 1.00 | 21.32 |  |
| 7 | C | N4 | 1H:1997 | 4.58 | 1.00 | 22.76 |  |
| 8 | Water 2 | | 1H:3807 | 4.73 | 1.00 | 16.26 | 2.86 (A:OP2 1H:1805) |
| 9 | A | O5' | 1H:1805 | 4.78 | 1.00 | 20.64 |  |
| 10 | A | OP2 | 1H:1861 | 4.87 | 1.00 | 20.58 |  |
| 11 | U | N3 | 1H:1999 | 4.92 | 1.00 | 22.85 |  |
| 12 | G | OP2 | 1H:1804 | 4.96 | 1.00 | 17.94 |  |
  |

  
 Contact(s): /O6/  
 Coded contact list: O_b_  

### K:1H:3100

|  | Res. type | Atom | Residue | Distance | Occupancy | B-factor | Closest solute to 1^st^ shell water |
| --- | --- | --- | --- | --- | --- | --- | --- |
| **Ion** | | | |  | 1.00 | 44.51 || 1 | G | O6 | 1H:2582 | 2.71 | 1.00 | 31.31 |
| 2 | G | O6 | 1H:2583 | 3.23 | 1.00 | 27.62 |  |
| 3 | U | O4 | 1H:2527 | 3.26 | 1.00 | 23.67 |  |
| 4 | G | O6 | 1H:2526 | 3.56 | 1.00 | 28.29 |  |
| 5 | C | N4 | 1H:2528 | 3.87 | 1.00 | 32.13 |  |
| 6 | G | N7 | 1H:2583 | 4.13 | 1.00 | 23.55 |  |
| 7 | C | N4 | 1H:2584 | 4.19 | 1.00 | 22.69 |  |
| 8 | G | N1 | 1H:2582 | 4.37 | 1.00 | 25.75 |  |
| 9 | U | N3 | 1H:2527 | 4.48 | 1.00 | 26.92 |  |
| 10 | G | N7 | 1H:2582 | 4.75 | 1.00 | 29.28 |  |
  |

  
**1^st^ shell ligand angles:**   
1-2: 86; 1-3: 104; 2-3: 63;   
  
 Contact(s): /O4/O6/O6/  
 Coded contact list: 3O_b_  

### K:1H:3101

|  | Res. type | Atom | Residue | Distance | Occupancy | B-factor | Closest solute to 1^st^ shell water |
| --- | --- | --- | --- | --- | --- | --- | --- |
| **Ion** | | | |  | 1.00 | 49.84 || 1 | U | OP2 | 1H:2734 | 2.90 | 1.00 | 36.98 |
| 2 | U | O4 | 1H:2700 | 3.16 | 1.00 | 34.16 |  |
| 3 | A | OP2 | 1H:2735 | 3.38 | 1.00 | 45.49 |  |
| 4 | A | N7 | 1H:2735 | 3.42 | 1.00 | 33.39 |  |
| 5 | G | O6 | 1H:2736 | 3.42 | 1.00 | 37.58 |  |
| 6 | U | O5' | 1H:2734 | 3.46 | 1.00 | 40.25 |  |
| 7 | U | N3 | 1H:2700 | 3.80 | 1.00 | 34.21 |  |
| 8 | U | OP1 | 1H:2734 | 4.26 | 1.00 | 32.48 |  |
| 9 | G | N7 | 1H:2736 | 4.30 | 1.00 | 38.47 |  |
| 10 | G | O6 | 1H:2699 | 4.49 | 1.00 | 36.34 |  |
| 11 | U | N1 | 1H:2701 | 4.66 | 1.00 | 35.05 |  |
| 12 | U | O4 | 1H:2701 | 4.73 | 1.00 | 39.64 |  |
| 13 | A | N6 | 1H:2735 | 4.96 | 1.00 | 38.50 |  |
  |

  
**1^st^ shell ligand angles:**   
1-2: 73; 1-3: 109; 2-3: 129;   
  
 Contact(s): /O4/OP2/OP2/  
 Coded contact list: 2O_ph_O_b_  

### K:1H:3102

|  | Res. type | Atom | Residue | Distance | Occupancy | B-factor | Closest solute to 1^st^ shell water |
| --- | --- | --- | --- | --- | --- | --- | --- |
| **Ion** | | | |  | 1.00 | 51.58 || 1 | U | O4 | 1H:772 | 2.78 | 1.00 | 41.35 |
| 2 | G | O6 | 1H:755 | 2.88 | 1.00 | 37.29 |  |
| 3 | G | O6 | 1H:771 | 3.07 | 1.00 | 41.82 |  |
| 4 | G | N2 | 1H:773 | 3.52 | 1.00 | 32.49 |  |
| 5 | G | N1 | 1H:773 | 4.03 | 1.00 | 33.02 |  |
| 6 | U | N3 | 1H:772 | 4.05 | 1.00 | 37.68 |  |
| 7 | C | N4 | 1H:756 | 4.21 | 1.00 | 43.49 |  |
| 8 | A | OP2 | 1H:754 | 4.28 | 1.00 | 28.10 |  |
| 9 | A | OP2 | 1H:753 | 4.43 | 1.00 | 29.75 |  |
| 10 | G | N3 | 1H:773 | 4.53 | 1.00 | 33.00 |  |
| 11 | G | N7 | 1H:755 | 4.67 | 1.00 | 38.81 |  |
| 12 | G | N1 | 1H:771 | 4.87 | 1.00 | 42.40 |  |
  |

  
**1^st^ shell ligand angles:**   
1-2: 75; 1-3: 74; 2-3: 75;   
  
 Contact(s): /O4/O6/O6/  
 Coded contact list: 3O_b_  

### K:1H:3103

|  | Res. type | Atom | Residue | Distance | Occupancy | B-factor | Closest solute to 1^st^ shell water |
| --- | --- | --- | --- | --- | --- | --- | --- |
| **Ion** | | | |  | 1.00 | 33.06 || 1 | G | OP1 | 1H:2092 | 2.77 | 1.00 | 20.36 |
| 2 | Water 1 | | 1H:3866 | 2.89 | 1.00 | 19.59 | 2.47 (G:OP2 1H:2092) |
| 3 | A | O3' | 1H:187 | 2.98 | 1.00 | 15.24 |  |
| 4 | U | OP1 | 1H:2257 | 3.12 | 1.00 | 18.89 |  |
| 5 | C | OP1 | 1H:188 | 3.30 | 1.00 | 20.69 |  |
| 6 | G | OP2 | 1H:2092 | 3.32 | 1.00 | 22.39 |  |
| 7 | G | OP1 | 1H:2093 | 3.33 | 1.00 | 23.23 |  |
| 8 | A | O2' | 1H:187 | 3.44 | 1.00 | 26.23 |  |
| 9 | G | OP2 | 1H:2093 | 3.65 | 1.00 | 24.52 |  |
| 10 | G | O5' | 1H:2092 | 4.08 | 1.00 | 24.81 |  |
| 11 | U | OP2 | 1H:2257 | 4.31 | 1.00 | 18.79 |  |
| 12 | C | O5' | 1H:188 | 4.64 | 1.00 | 16.92 |  |
| 13 | G | O3' | 1H:2092 | 4.71 | 1.00 | 27.34 |  |
| 14 | U | O3' | 1H:2091 | 4.78 | 1.00 | 21.97 |  |
| 15 | C | OP2 | 1H:188 | 4.99 | 1.00 | 17.01 |  |
  |

  
**1^st^ shell ligand angles:**   
1-2: 96; 1-3: 88; 1-4: 130; 1-5: 110; 1-6: 51; 1-7: 98; 2-3: 134; 2-4: 75; 2-5: 154; 2-6: 46; 2-7: 86; 3-4: 68; 3-5: 46; 3-6: 130; 3-7: 140; 4-5: 84; 4-6: 113; 4-7: 130; 5-6: 160; 5-7: 96; 6-7: 81;   
  
 Contact(s): /O3'/O_w_/OP1/OP1/OP1/OP1/OP2/  
 Coded contact list: 5O_ph_O_r_O_w_  

### K:1H:3104

|  | Res. type | Atom | Residue | Distance | Occupancy | B-factor | Closest solute to 1^st^ shell water |
| --- | --- | --- | --- | --- | --- | --- | --- |
| **Ion** | | | |  | 1.00 | 42.03 || 1 | Water 1 | | 1H:3672 | 2.56 | 1.00 | 16.27 | 2.14 (G:OP2 1H:2460) |
| 2 | A | O2' | 1H:591 | 2.79 | 1.00 | 16.20 |  |
| 3 | Water 2 | | 1H:4436 | 3.22 | 1.00 | 31.02 | 4.20 (G:OP2 1H:2460) |
| 4 | G | OP1 | 1H:2460 | 3.31 | 1.00 | 26.48 |  |
| 5 | Water 3 | | 1H:3916 | 4.06 | 1.00 | 13.34 | 3.74 (G:OP1 1H:2459) |
| 6 | G | OP2 | 1H:2460 | 4.16 | 1.00 | 22.10 |  |
| 7 | A | O4' | 1H:591 | 4.40 | 1.00 | 18.50 |  |
| 8 | G | OP1 | 1H:856 | 4.59 | 1.00 | 19.09 |  |
| 9 | A | N3 | 1H:591 | 4.69 | 1.00 | 20.75 |  |
| 10 | Water 4 | | 1H:3814 | 4.89 | 1.00 | 29.53 | 2.40 (G:OP2 1H:2460) |
| 11 | G | OP1 | 1H:2459 | 4.90 | 1.00 | 19.33 |  |
  |

  
**1^st^ shell ligand angles:**   
1-2: 130; 1-3: 66; 1-4: 60; 2-3: 164; 2-4: 90; 3-4: 97;   
  
 Contact(s): /O2'/O_w_/OP1/  
 Coded contact list: O_ph_O_r_O_w_  
 Exclusion zone contact(s): /O_w_/  
K hydration shell issue: water too close (< 2.4 Angs.)  

### K:1H:3105

|  | Res. type | Atom | Residue | Distance | Occupancy | B-factor | Closest solute to 1^st^ shell water |
| --- | --- | --- | --- | --- | --- | --- | --- |
| **Ion** | | | |  | 1.00 | 50.57 || 1 | U | O4 | 1H:1370 | 3.02 | 1.00 | 33.44 |
| 2 | G | O6 | 1H:1371 | 3.38 | 1.00 | 31.57 |  |
| 3 | C | O2 | 1H:1367 | 3.41 | 1.00 | 30.30 |  |
| 4 | C | O2' | 1H:1367 | 3.42 | 1.00 | 34.75 |  |
| 5 | A | N7 | 1H:1369 | 3.75 | 1.00 | 24.72 |  |
| 6 | Water 1 | | 1H:4416 | 3.94 | 1.00 | 32.64 | 3.35 (G:N7 1H:1371) |
| 7 | A | OP2 | 1H:1369 | 4.09 | 1.00 | 33.67 |  |
| 8 | C | N4 | 1H:1377 | 4.19 | 1.00 | 29.79 |  |
| 9 | C | O3' | 1H:1367 | 4.47 | 1.00 | 30.30 |  |
| 10 | C | N1 | 1H:1367 | 4.64 | 1.00 | 32.27 |  |
| 11 | G | N7 | 1H:1371 | 4.91 | 1.00 | 30.01 |  |
| 12 | A | N6 | 1H:1378 | 4.96 | 1.00 | 30.88 |  |
  |

  
**1^st^ shell ligand angles:**   
1-2: 53;   
  
 Contact(s): /O4/O6/  
 Coded contact list: 2O_b_  

### K:1H:3106

|  | Res. type | Atom | Residue | Distance | Occupancy | B-factor | Closest solute to 1^st^ shell water |
| --- | --- | --- | --- | --- | --- | --- | --- |
| **Ion** | | | |  | 1.00 | 59.65 || 1 | A | O2' | 1H:2222 | 2.93 | 1.00 | 43.35 |
| 2 | LYS | O | J8:48 | 3.03 | 1.00 | 36.25 |  |
| 3 | A | O3' | 1H:2222 | 3.15 | 1.00 | 36.40 |  |
| 4 | C | OP1 | 1H:2223 | 3.52 | 1.00 | 37.92 |  |
| 5 | VAL | N | J8:49 | 4.45 | 1.00 | 43.02 |  |
| 6 | U | OP2 | 1H:2115 | 4.59 | 1.00 | 50.16 |  |
| 7 | C | O5' | 1H:2223 | 4.59 | 1.00 | 37.51 |  |
| 8 | ARG | N | J8:50 | 4.64 | 1.00 | 44.09 |  |
| 9 | U | OP1 | 1H:2115 | 4.66 | 1.00 | 31.99 |  |
| 10 | ASN | OD1 | 61:28 | 4.87 | 1.00 | 47.88 |  |
  |

  
**1^st^ shell ligand angles:**   
1-2: 145; 1-3: 51; 2-3: 129;   
  
 Contact(s): /O2'/O3'/O_back_/  
 Coded contact list: 2O_r_O_back_  

### K:1H:3107

|  | Res. type | Atom | Residue | Distance | Occupancy | B-factor | Closest solute to 1^st^ shell water |
| --- | --- | --- | --- | --- | --- | --- | --- |
| **Ion** | | | |  | 1.00 | 44.66 || 1 | G | OP1 | 1H:857 | 2.71 | 1.00 | 25.58 |
| 2 | U | O2 | 1H:600 | 2.95 | 1.00 | 23.58 |  |
| 3 | C | O2 | 1H:589 | 2.97 | 1.00 | 20.21 |  |
| 4 | U | O4' | 1H:590 | 3.92 | 1.00 | 25.77 |  |
| 5 | G | N3 | 1H:601 | 3.95 | 1.00 | 21.55 |  |
| 6 | G | N9 | 1H:601 | 4.02 | 1.00 | 24.16 |  |
| 7 | G | O4' | 1H:601 | 4.04 | 1.00 | 16.89 |  |
| 8 | U | O2' | 1H:590 | 4.04 | 1.00 | 19.36 |  |
| 9 | G | O3' | 1H:856 | 4.16 | 1.00 | 19.71 |  |
| 10 | U | O2 | 1H:590 | 4.52 | 1.00 | 18.56 |  |
| 11 | A | O2' | 1H:1300 | 4.61 | 1.00 | 31.24 |  |
| 12 | U | N3 | 1H:600 | 4.64 | 1.00 | 21.19 |  |
| 13 | G | OP2 | 1H:857 | 4.68 | 1.00 | 20.73 |  |
| 14 | U | N1 | 1H:590 | 4.76 | 1.00 | 20.57 |  |
| 15 | C | O2' | 1H:589 | 4.84 | 1.00 | 24.47 |  |
| 16 | U | O2' | 1H:600 | 4.91 | 1.00 | 19.75 |  |
| 17 | C | N3 | 1H:589 | 4.92 | 1.00 | 22.00 |  |
  |

  
**1^st^ shell ligand angles:**   
1-2: 142; 1-3: 115; 2-3: 78;   
  
 Contact(s): /O2/O2/OP1/  
 Coded contact list: O_ph_2O_b_  

### K:1H:3108

|  | Res. type | Atom | Residue | Distance | Occupancy | B-factor | Closest solute to 1^st^ shell water |
| --- | --- | --- | --- | --- | --- | --- | --- |
| **Ion** | | | |  | 1.00 | 76.83 || 1 | U | O4 | 1H:1766 | 2.63 | 1.00 | 83.22 |
| 2 | G | O6 | 1H:1765 | 2.85 | 1.00 | 78.85 |  |
| 3 | G | O6 | 1H:1772 | 3.79 | 1.00 | 68.45 |  |
| 4 | C | N4 | 1H:1773 | 3.90 | 1.00 | 65.86 |  |
| 5 | U | N3 | 1H:1766 | 4.02 | 1.00 | 76.75 |  |
| 6 | G | OP2 | 1H:1770 | 4.24 | 1.00 | 96.19 |  |
| 7 | A | OP2 | 1H:1771 | 4.51 | 1.00 | 84.86 |  |
| 8 | G | N7 | 1H:1767 | 4.54 | 1.00 | 73.94 |  |
| 9 | G | O6 | 1H:1764 | 4.69 | 1.00 | 81.43 |  |
| 10 | G | N7 | 1H:1772 | 4.78 | 1.00 | 71.15 |  |
| 11 | G | O6 | 1H:1767 | 4.88 | 1.00 | 82.05 |  |
| 12 | G | N1 | 1H:1765 | 4.93 | 1.00 | 71.90 |  |
  |

  
**1^st^ shell ligand angles:**   
1-2: 66;   
  
 Contact(s): /O4/O6/  
 Coded contact list: 2O_b_  

### K:1H:3109

|  | Res. type | Atom | Residue | Distance | Occupancy | B-factor | Closest solute to 1^st^ shell water |
| --- | --- | --- | --- | --- | --- | --- | --- |
| **Ion** | | | |  | 1.00 | 57.54 || 1 | U | O4 | 1H:2114 | 3.14 | 1.00 | 33.48 |
| 2 | U | O4 | 1H:2115 | 3.21 | 1.00 | 39.83 |  |
| 3 | G | O6 | 1H:2241 | 3.21 | 1.00 | 34.64 |  |
| 4 | A | OP2 | 1H:2240 | 3.22 | 1.00 | 30.14 |  |
| 5 | C | OP2 | 1H:2239 | 3.83 | 1.00 | 37.80 |  |
| 6 | A | N7 | 1H:2240 | 3.93 | 1.00 | 37.26 |  |
| 7 | G | O6 | 1H:2113 | 3.96 | 1.00 | 27.18 |  |
| 8 | C | N4 | 1H:2242 | 3.98 | 1.00 | 30.85 |  |
| 9 | G | N7 | 1H:2241 | 4.08 | 1.00 | 33.53 |  |
| 10 | C | O5' | 1H:2239 | 4.23 | 1.00 | 37.42 |  |
| 11 | U | N3 | 1H:2114 | 4.36 | 1.00 | 27.71 |  |
| 12 | U | N3 | 1H:2115 | 4.49 | 1.00 | 38.13 |  |
| 13 | A | O5' | 1H:2240 | 4.92 | 1.00 | 36.84 |  |
| 14 | A | N6 | 1H:2240 | 4.98 | 1.00 | 32.18 |  |
  |

  
**1^st^ shell ligand angles:**   
1-2: 61; 1-3: 63; 1-4: 157; 2-3: 64; 2-4: 118; 3-4: 140;   
  
 Contact(s): /O4/O4/O6/OP2/  
 Coded contact list: O_ph_3O_b_  

### K:1H:3110

|  | Res. type | Atom | Residue | Distance | Occupancy | B-factor | Closest solute to 1^st^ shell water |
| --- | --- | --- | --- | --- | --- | --- | --- |
| **Ion** | | | |  | 1.00 | 60.31 || 1 | G | OP1 | 1H:1795 | 2.74 | 1.00 | 35.42 |
| 2 | G | O6 | 1H:1747 | 2.94 | 1.00 | 48.74 |  |
| 3 | G | O4' | 1H:1795 | 2.95 | 1.00 | 35.65 |  |
| 4 | A | O3' | 1H:1794 | 3.20 | 1.00 | 40.76 |  |
| 5 | G | OP1 | 1H:1730 | 3.39 | 1.00 | 38.37 |  |
| 6 | G | O5' | 1H:1795 | 3.88 | 1.00 | 41.97 |  |
| 7 | G | O3' | 1H:1729 | 4.54 | 1.00 | 36.06 |  |
| 8 | G | N1 | 1H:1747 | 4.58 | 1.00 | 40.66 |  |
| 9 | G | OP2 | 1H:1795 | 4.82 | 1.00 | 46.20 |  |
| 10 | A | O2' | 1H:1794 | 4.85 | 1.00 | 45.78 |  |
| 11 | G | N9 | 1H:1795 | 4.93 | 1.00 | 32.70 |  |
  |

  
**1^st^ shell ligand angles:**   
1-2: 132; 1-3: 84; 1-4: 51; 1-5: 153; 2-3: 59; 2-4: 155; 2-5: 65; 3-4: 100; 3-5: 91; 4-5: 105;   
  
 Contact(s): /O3'/O4'/O6/OP1/OP1/  
 Coded contact list: 2O_ph_2O_r_O_b_  

### K:1H:3111

|  | Res. type | Atom | Residue | Distance | Occupancy | B-factor | Closest solute to 1^st^ shell water |
| --- | --- | --- | --- | --- | --- | --- | --- |
| **Ion** | | | |  | 1.00 | 51.93 || 1 | G | O6 | 1H:1427 | 2.88 | 1.00 | 32.46 |
| 2 | Water 1 | | 1H:4216 | 3.22 | 1.00 | 50.00 | 2.91 (G:OP1 1H:1394) |
| 3 | A | O2' | 1H:1425 | 3.30 | 1.00 | 35.06 |  |
| 4 | G | O6 | 1H:1428 | 3.43 | 1.00 | 36.09 |  |
| 5 | C | N4 | 1H:1398 | 3.78 | 1.00 | 25.98 |  |
| 6 | Water 2 | | 1H:3896 | 3.99 | 1.00 | 32.03 | 3.51 (G:O6 1H:1428) |
| 7 | C | N4 | 1H:1397 | 4.15 | 1.00 | 35.48 |  |
| 8 | G | OP1 | 1H:1395 | 4.25 | 1.00 | 37.07 |  |
| 9 | A | OP2 | 1H:1426 | 4.27 | 1.00 | 45.64 |  |
| 10 | G | N7 | 1H:1427 | 4.39 | 1.00 | 26.75 |  |
| 11 | G | OP1 | 1H:1394 | 4.59 | 1.00 | 49.83 |  |
| 12 | Water 3 | | 1H:3708 | 4.87 | 1.00 | 42.18 | 3.27 (G:O6 1H:1428) |
| 13 | G | OP2 | 1H:1395 | 4.94 | 1.00 | 37.93 |  |
| 14 | G | N1 | 1H:1427 | 4.94 | 1.00 | 27.42 |  |
| 15 | G | O3' | 1H:1394 | 4.99 | 1.00 | 29.54 |  |
  |

  
**1^st^ shell ligand angles:**   
1-2: 115; 1-3: 69; 2-3: 93;   
  
 Contact(s): /O2'/O6/O_w_/  
 Coded contact list: O_r_O_b_O_w_  

### K:1H:3112

|  | Res. type | Atom | Residue | Distance | Occupancy | B-factor | Closest solute to 1^st^ shell water |
| --- | --- | --- | --- | --- | --- | --- | --- |
| **Ion** | | | |  | 1.00 | 48.71 || 1 | G | O6 | 1H:67 | 2.68 | 1.00 | 38.56 |
| 2 | U | O4 | 1H:58 | 3.09 | 1.00 | 39.25 |  |
| 3 | U | O4 | 1H:66 | 3.21 | 1.00 | 37.27 |  |
| 4 | G | O6 | 1H:57 | 3.25 | 1.00 | 36.03 |  |
| 5 | U | N3 | 1H:58 | 3.69 | 1.00 | 36.30 |  |
| 6 | C | N4 | 1H:68 | 4.44 | 1.00 | 31.13 |  |
| 7 | G | N7 | 1H:67 | 4.61 | 1.00 | 35.57 |  |
| 8 | G | N7 | 1H:57 | 4.87 | 1.00 | 29.56 |  |
| 9 | G | N1 | 1H:67 | 4.90 | 1.00 | 37.18 |  |
  |

  
**1^st^ shell ligand angles:**   
1-2: 94; 1-3: 72; 1-4: 64; 2-3: 118; 2-4: 71; 3-4: 136;   
  
 Contact(s): /O4/O4/O6/O6/  
 Coded contact list: 4O_b_  

### K:1H:3113

|  | Res. type | Atom | Residue | Distance | Occupancy | B-factor | Closest solute to 1^st^ shell water |
| --- | --- | --- | --- | --- | --- | --- | --- |
| **Ion** | | | |  | 1.00 | 30.67 || 1 | A | O2' | 1H:841 | 2.58 | 1.00 | 22.26 |
| 2 | G | OP2 | 1H:842 | 2.74 | 1.00 | 16.73 |  |
| 3 | G | N7 | 1H:824 | 2.87 | 1.00 | 24.87 |  |
| 4 | A | O3' | 1H:841 | 3.40 | 1.00 | 12.79 |  |
| 5 | G | O2' | 1H:823 | 3.46 | 1.00 | 18.09 |  |
| 6 | G | O3' | 1H:823 | 3.56 | 1.00 | 18.54 |  |
| 7 | G | OP2 | 1H:824 | 3.62 | 1.00 | 29.63 |  |
| 8 | G | N3 | 1H:823 | 4.18 | 1.00 | 15.94 |  |
| 9 | A | N3 | 1H:841 | 4.31 | 1.00 | 21.40 |  |
| 10 | G | O5' | 1H:842 | 4.57 | 1.00 | 17.54 |  |
| 11 | G | N9 | 1H:824 | 4.68 | 1.00 | 22.79 |  |
| 12 | A | N9 | 1H:841 | 4.68 | 1.00 | 21.80 |  |
| 13 | G | O5' | 1H:824 | 4.69 | 1.00 | 19.56 |  |
| 14 | G | OP1 | 1H:842 | 4.87 | 1.00 | 19.07 |  |
| 15 | G | O6 | 1H:824 | 4.89 | 1.00 | 22.19 |  |
  |

  
**1^st^ shell ligand angles:**   
1-2: 77; 1-3: 65; 1-4: 52; 2-3: 141; 2-4: 47; 3-4: 109;   
  
 Contact(s): /N7/O3'/OP2/  
 Coded contact list: O_ph_O_r_N_b_  
 Exclusion zone contact(s): /O2'/  

### K:1H:3114

|  | Res. type | Atom | Residue | Distance | Occupancy | B-factor | Closest solute to 1^st^ shell water |
| --- | --- | --- | --- | --- | --- | --- | --- |
| **Ion** | | | |  | 1.00 | 55.75 || 1 | G | O6 | 1H:366 | 2.65 | 1.00 | 43.25 |
| 2 | G | O6 | 1H:367 | 3.46 | 1.00 | 43.34 |  |
| 3 | G | N7 | 1H:366 | 4.00 | 1.00 | 42.10 |  |
| 4 | U | OP2 | 1H:364 | 4.09 | 1.00 | 42.98 |  |
| 5 | A | OP2 | 1H:365 | 4.10 | 1.00 | 41.47 |  |
| 6 | U | OP1 | 1H:364 | 4.22 | 1.00 | 41.26 |  |
| 7 | G | N7 | 1H:367 | 4.28 | 1.00 | 42.10 |  |
| 8 | C | N4 | 1H:322 | 4.28 | 1.00 | 39.30 |  |
| 9 | G | N2 | 1H:323 | 4.47 | 1.00 | 39.72 |  |
| 10 | C | N4 | 1H:321 | 4.69 | 1.00 | 47.04 |  |
| 11 | G | N1 | 1H:366 | 4.89 | 1.00 | 43.13 |  |
  |

  
 Contact(s): /O6/  
 Coded contact list: O_b_  

### K:1H:3115

|  | Res. type | Atom | Residue | Distance | Occupancy | B-factor | Closest solute to 1^st^ shell water |
| --- | --- | --- | --- | --- | --- | --- | --- |
| **Ion** | | | |  | 1.00 | 52.47 || 1 | A | O2' | 1H:836 | 2.86 | 1.00 | 28.43 |
| 2 | G | O6 | 1H:492 | 2.95 | 1.00 | 25.96 |  |
| 3 | U | O4 | 1H:491 | 3.35 | 1.00 | 20.47 |  |
| 4 | G | N1 | 1H:492 | 4.07 | 1.00 | 18.91 |  |
| 5 | A | OP2 | 1H:837 | 4.53 | 1.00 | 16.44 |  |
  |

  
**1^st^ shell ligand angles:**   
1-2: 115; 1-3: 63; 2-3: 66;   
  
 Contact(s): /O2'/O4/O6/  
 Coded contact list: O_r_2O_b_  

### K:1H:3116

|  | Res. type | Atom | Residue | Distance | Occupancy | B-factor | Closest solute to 1^st^ shell water |
| --- | --- | --- | --- | --- | --- | --- | --- |
| **Ion** | | | |  | 1.00 | 61.81 || 1 | G | OP1 | 1H:1283 | 2.77 | 1.00 | 32.47 |
| 2 | A | O5' | 1H:531 | 2.81 | 1.00 | 50.79 |  |
| 3 | U | OP1 | 1H:537 | 3.04 | 1.00 | 43.61 |  |
| 4 | G | OP2 | 1H:1283 | 3.25 | 1.00 | 32.50 |  |
| 5 | A | O4' | 1H:531 | 3.38 | 1.00 | 42.52 |  |
| 6 | A | OP2 | 1H:531 | 3.47 | 1.00 | 52.13 |  |
| 7 | C | O3' | 1H:536 | 3.63 | 1.00 | 30.57 |  |
| 8 | G | O3' | 1H:1282 | 4.51 | 1.00 | 32.92 |  |
| 9 | G | OP1 | 1H:1282 | 4.52 | 1.00 | 43.67 |  |
| 10 | U | O5' | 1H:537 | 4.63 | 1.00 | 30.65 |  |
| 11 | G | O5' | 1H:1283 | 4.64 | 1.00 | 34.31 |  |
| 12 | C | O2' | 1H:536 | 4.78 | 1.00 | 36.90 |  |
| 13 | U | O3' | 1H:530 | 4.83 | 1.00 | 59.65 |  |
| 14 | A | OP1 | 1H:531 | 4.91 | 1.00 | 61.08 |  |
  |

  
**1^st^ shell ligand angles:**   
1-2: 172; 1-3: 88; 1-4: 50; 1-5: 124; 2-3: 84; 2-4: 133; 2-5: 59; 3-4: 115; 3-5: 100; 4-5: 143;   
  
 Contact(s): /O4'/O5'/OP1/OP1/OP2/  
 Coded contact list: 3O_ph_2O_r_  

### K:1H:3117

|  | Res. type | Atom | Residue | Distance | Occupancy | B-factor | Closest solute to 1^st^ shell water |
| --- | --- | --- | --- | --- | --- | --- | --- |
| **Ion** | | | |  | 1.00 | 57.52 || 1 | G | O2' | 1H:332 | 2.92 | 1.00 | 50.05 |
| 2 | A | O2' | 1H:1256 | 3.04 | 1.00 | 46.19 |  |
| 3 | G | O6 | 1H:334 | 3.11 | 1.00 | 46.29 |  |
| 4 | U | OP2 | 1H:1257 | 3.43 | 1.00 | 42.34 |  |
| 5 | U | OP1 | 1H:1257 | 3.84 | 1.00 | 46.90 |  |
| 6 | G | N7 | 1H:333 | 4.39 | 1.00 | 53.47 |  |
| 7 | G | O3' | 1H:332 | 4.39 | 1.00 | 51.39 |  |
| 8 | A | N1 | 1H:355 | 4.39 | 1.00 | 45.45 |  |
| 9 | G | O6 | 1H:1258 | 4.42 | 1.00 | 46.37 |  |
| 10 | A | O3' | 1H:1256 | 4.45 | 1.00 | 45.34 |  |
| 11 | G | N7 | 1H:334 | 4.65 | 1.00 | 47.41 |  |
| 12 | G | O4' | 1H:332 | 4.78 | 1.00 | 51.13 |  |
| 13 | G | O6 | 1H:333 | 4.81 | 1.00 | 58.05 |  |
  |

  
**1^st^ shell ligand angles:**   
1-2: 115; 1-3: 78; 2-3: 74;   
  
 Contact(s): /O2'/O2'/O6/  
 Coded contact list: 2O_r_O_b_  

### K:1H:3118

|  | Res. type | Atom | Residue | Distance | Occupancy | B-factor | Closest solute to 1^st^ shell water |
| --- | --- | --- | --- | --- | --- | --- | --- |
| **Ion** | | | |  | 1.00 | 56.64 || 1 | G | O6 | 1H:1566 | 2.77 | 1.00 | 51.93 |
| 2 | U | O4 | 1H:1567 | 2.79 | 1.00 | 56.69 |  |
| 3 | G | O6 | 1H:1521 | 3.07 | 1.00 | 62.41 |  |
| 4 | C | N4 | 1H:1522 | 3.51 | 1.00 | 61.21 |  |
| 5 | G | N2 | 1H:1568 | 3.59 | 1.00 | 61.39 |  |
| 6 | Water 1 | | 1H:4042 | 3.66 | 1.00 | 47.02 | 2.68 (U:O4 1H:1567) |
| 7 | U | N3 | 1H:1567 | 4.02 | 1.00 | 57.08 |  |
| 8 | Water 2 | | 1H:3804 | 4.10 | 1.00 | 56.55 | 2.66 (G:N1 1H:1568) |
| 9 | G | N1 | 1H:1568 | 4.15 | 1.00 | 58.41 |  |
| 10 | Water 3 | | 1H:4580 | 4.33 | 1.00 | 53.55 | 4.60 (A:OP2 1H:1519) |
| 11 | C | N4 | 1H:1565 | 4.60 | 1.00 | 50.76 |  |
| 12 | A | OP2 | 1H:1519 | 4.67 | 1.00 | 51.48 |  |
| 13 | A | OP2 | 1H:1520 | 4.68 | 1.00 | 53.86 |  |
| 14 | G | N1 | 1H:1566 | 4.75 | 1.00 | 54.52 |  |
| 15 | G | N3 | 1H:1568 | 4.88 | 1.00 | 62.32 |  |
| 16 | G | N7 | 1H:1566 | 4.90 | 1.00 | 57.34 |  |
| 17 | G | N7 | 1H:1521 | 4.97 | 1.00 | 65.46 |  |
  |

  
**1^st^ shell ligand angles:**   
1-2: 70; 1-3: 84; 2-3: 75;   
  
 Contact(s): /O4/O6/O6/  
 Coded contact list: 3O_b_  

### K:1H:3119

|  | Res. type | Atom | Residue | Distance | Occupancy | B-factor | Closest solute to 1^st^ shell water |
| --- | --- | --- | --- | --- | --- | --- | --- |
| **Ion** | | | |  | 1.00 | 46.13 || 1 | G | OP2 | 1H:496 | 3.04 | 1.00 | 16.98 |
| 2 | G | O6 | 1H:485 | 3.07 | 1.00 | 21.12 |  |
| 3 | ALA | O | 31:58 | 3.26 | 1.00 | 25.63 |  |
| 4 | G | N1 | 1H:485 | 4.02 | 1.00 | 19.80 |  |
| 5 | G | O5' | 1H:496 | 4.42 | 1.00 | 15.74 |  |
| 6 | G | OP1 | 1H:495 | 4.45 | 1.00 | 18.00 |  |
| 7 | G | N7 | 1H:496 | 4.48 | 1.00 | 17.83 |  |
| 8 | G | OP2 | 1H:495 | 4.69 | 1.00 | 16.20 |  |
| 9 | G | OP1 | 1H:496 | 4.79 | 1.00 | 15.16 |  |
| 10 | TYR | N | 31:59 | 4.84 | 1.00 | 21.98 |  |
| 11 | A | OP2 | 1H:497 | 4.90 | 1.00 | 17.05 |  |
| 12 | LYS | NZ | P8:37 | 4.93 | 1.00 | 17.77 |  |
  |

  
**1^st^ shell ligand angles:**   
1-2: 144; 1-3: 98; 2-3: 102;   
  
 Contact(s): /O6/O_back_/OP2/  
 Coded contact list: O_ph_O_b_O_back_  

### K:1H:3120

|  | Res. type | Atom | Residue | Distance | Occupancy | B-factor | Closest solute to 1^st^ shell water |
| --- | --- | --- | --- | --- | --- | --- | --- |
| **Ion** | | | |  | 1.00 | 60.83 || 1 | G | O4' | 1H:1372 | 2.77 | 1.00 | 39.05 |
| 2 | G | O6 | 1H:1318 | 2.89 | 1.00 | 39.51 |  |
| 3 | G | O5' | 1H:1372 | 3.06 | 1.00 | 27.39 |  |
| 4 | G | N9 | 1H:1372 | 3.87 | 1.00 | 32.67 |  |
| 5 | G | OP2 | 1H:1372 | 4.00 | 1.00 | 33.97 |  |
| 6 | G | N7 | 1H:1372 | 4.35 | 1.00 | 37.85 |  |
| 7 | G | N7 | 1H:1318 | 4.37 | 1.00 | 39.27 |  |
| 8 | G | OP1 | 1H:1371 | 4.54 | 1.00 | 23.60 |  |
  |

  
**1^st^ shell ligand angles:**   
1-2: 132; 1-3: 59; 2-3: 126;   
  
 Contact(s): /O4'/O5'/O6/  
 Coded contact list: 2O_r_O_b_  

### K:1H:3121

|  | Res. type | Atom | Residue | Distance | Occupancy | B-factor | Closest solute to 1^st^ shell water |
| --- | --- | --- | --- | --- | --- | --- | --- |
| **Ion** | | | |  | 1.00 | 62.68 || 1 | Water 1 | | 1H:4397 | 2.73 | 1.00 | 33.58 | 3.29 (G:OP1 1H:1297) |
| 2 | G | OP2 | 1H:609 | 2.74 | 1.00 | 26.20 |  |
| 3 | THR | OG1 | C8:6 | 3.20 | 1.00 | 33.25 |  |
| 4 | Water 2 | | 1H:3998 | 3.58 | 1.00 | 25.35 | 2.75 (G:OP1 1H:1297) |
| 5 | G | OP1 | 1H:1297 | 3.92 | 1.00 | 23.29 |  |
| 6 | C | OP2 | 1H:608 | 4.22 | 1.00 | 27.83 |  |
| 7 | G | O5' | 1H:609 | 4.33 | 1.00 | 27.72 |  |
| 8 | ARG | NH2 | C8:10 | 4.37 | 1.00 | 24.91 |  |
| 9 | Water 3 | | 1H:3710 | 4.46 | 1.00 | 26.73 | 2.23 (G:OP1 1H:609) |
| 10 | G | N7 | 1H:609 | 4.59 | 1.00 | 26.45 |  |
| 11 | C | OP1 | 1H:608 | 4.67 | 1.00 | 31.69 |  |
| 12 | THR | N | C8:6 | 4.68 | 1.00 | 29.12 |  |
| 13 | G | OP1 | 1H:609 | 4.71 | 1.00 | 26.54 |  |
| 14 | U | O3' | 1H:1296 | 4.93 | 1.00 | 24.49 |  |
  |

  
**1^st^ shell ligand angles:**   
1-2: 79; 1-3: 130; 2-3: 107;   
  
 Contact(s): /O_w_/OG1_Thr_/OP2/  
 Coded contact list: O_ph_OH_Prot_O_w_  

### K:1H:3122

|  | Res. type | Atom | Residue | Distance | Occupancy | B-factor | Closest solute to 1^st^ shell water |
| --- | --- | --- | --- | --- | --- | --- | --- |
| **Ion** | | | |  | 1.00 | 65.63 || 1 | C | OP1 | 1H:403 | 2.85 | 1.00 | 31.51 |
| 2 | C | OP1 | 1H:421 | 3.01 | 1.00 | 36.99 |  |
| 3 | C | OP2 | 1H:421 | 3.23 | 1.00 | 31.98 |  |
| 4 | G | O2' | 1H:436 | 3.41 | 1.00 | 30.63 |  |
| 5 | G | O3' | 1H:436 | 3.77 | 1.00 | 35.90 |  |
| 6 | C | OP2 | 1H:403 | 3.85 | 1.00 | 37.08 |  |
| 7 | Water 1 | | 1H:4029 | 4.03 | 1.00 | 40.14 | 2.97 (C:OP2 1H:421) |
| 8 | A | O3' | 1H:402 | 4.11 | 1.00 | 41.33 |  |
| 9 | A | N6 | 1H:213 | 4.20 | 1.00 | 41.30 |  |
| 10 | Water 2 | | 1H:3799 | 4.39 | 1.00 | 39.57 | 2.38 (C:OP1 1H:403) |
| 11 | A | N7 | 1H:213 | 4.39 | 1.00 | 37.05 |  |
| 12 | C | O5' | 1H:421 | 4.40 | 1.00 | 34.89 |  |
| 13 | A | OP2 | 1H:422 | 4.69 | 1.00 | 33.89 |  |
| 14 | C | OP1 | 1H:437 | 4.84 | 1.00 | 36.34 |  |
| 15 | C | O3' | 1H:420 | 4.86 | 1.00 | 33.27 |  |
  |

  
**1^st^ shell ligand angles:**   
1-2: 146; 1-3: 112; 2-3: 49;   
  
 Contact(s): /OP1/OP1/OP2/  
 Coded contact list: 3O_ph_  

### K:1H:3123

|  | Res. type | Atom | Residue | Distance | Occupancy | B-factor | Closest solute to 1^st^ shell water |
| --- | --- | --- | --- | --- | --- | --- | --- |
| **Ion** | | | |  | 1.00 | 50.90 || 1 | C | OP2 | 1H:182 | 2.63 | 1.00 | 18.50 |
| 2 | Water 1 | | 1H:4307 | 2.74 | 1.00 | 17.50 | 3.08 (A:OP2 1H:189) |
| 3 | U | O4 | 1H:192 | 2.94 | 1.00 | 26.43 |  |
| 4 | U | O4 | 1H:183 | 3.11 | 1.00 | 23.58 |  |
| 5 | Water 2 | | 1H:3673 | 3.87 | 1.00 | 15.36 | 2.15 (C:OP1 1H:182) |
| 6 | U | N3 | 1H:192 | 4.23 | 1.00 | 19.53 |  |
| 7 | C | OP1 | 1H:182 | 4.51 | 1.00 | 20.96 |  |
| 8 | C | N4 | 1H:191 | 4.61 | 1.00 | 20.30 |  |
| 9 | A | O3' | 1H:181 | 4.79 | 1.00 | 20.50 |  |
| 10 | G | O6 | 1H:184 | 4.82 | 1.00 | 19.11 |  |
| 11 | A | OP2 | 1H:189 | 4.87 | 1.00 | 16.84 |  |
  |

  
**1^st^ shell ligand angles:**   
1-2: 79; 1-3: 135; 1-4: 80; 2-3: 119; 2-4: 75; 3-4: 68;   
  
 Contact(s): /O4/O4/O_w_/OP2/  
 Coded contact list: O_ph_2O_b_O_w_  
K hydration shell issue: water too close (< 2.4 Angs.)  

### K:1H:3124

|  | Res. type | Atom | Residue | Distance | Occupancy | B-factor | Closest solute to 1^st^ shell water |
| --- | --- | --- | --- | --- | --- | --- | --- |
| **Ion** | | | |  | 1.00 | 64.99 || 1 | A | O2' | 1H:1847 | 3.24 | 1.00 | 32.35 |
| 2 | A | O3' | 1H:1847 | 3.32 | 1.00 | 29.42 |  |
| 3 | G | O4' | 1H:1848 | 3.40 | 1.00 | 35.01 |  |
| 4 | G | OP2 | 1H:1848 | 3.48 | 1.00 | 33.09 |  |
| 5 | G | OP2 | 1H:1846 | 4.16 | 1.00 | 21.95 |  |
| 6 | G | OP1 | 1H:1845 | 4.18 | 1.00 | 20.46 |  |
| 7 | Water 1 | | 1H:4189 | 4.57 | 1.00 | 35.72 | 3.80 (A:N3 1H:1847) |
| 8 | G | N3 | 1H:1848 | 4.65 | 1.00 | 57.67 |  |
| 9 | G | O5' | 1H:1848 | 4.66 | 1.00 | 28.67 |  |
| 10 | A | N3 | 1H:1847 | 4.93 | 1.00 | 25.04 |  |
  |

  
**1^st^ shell ligand angles:**   
1-2: 50;   
  
 Contact(s): /O2'/O3'/  
 Coded contact list: 2O_r_  

### K:1H:3125

|  | Res. type | Atom | Residue | Distance | Occupancy | B-factor | Closest solute to 1^st^ shell water |
| --- | --- | --- | --- | --- | --- | --- | --- |
| **Ion** | | | |  | 1.00 | 64.60 || 1 | U | OP2 | 1H:1251 | 2.98 | 1.00 | 45.50 |
| 2 | G | OP2 | 1H:1249 | 3.86 | 1.00 | 38.36 |  |
| 3 | A | N7 | 1H:1250 | 3.92 | 1.00 | 46.30 |  |
| 4 | A | N9 | 1H:1250 | 4.04 | 1.00 | 49.31 |  |
| 5 | A | O2' | 1H:1250 | 4.29 | 1.00 | 55.14 |  |
| 6 | A | O3' | 1H:1250 | 4.37 | 1.00 | 42.97 |  |
| 7 | C | OP2 | 1H:1248 | 4.39 | 1.00 | 33.19 |  |
| 8 | C | OP1 | 1H:1248 | 4.61 | 1.00 | 36.11 |  |
  |

  
 Contact(s): /OP2/  
 Coded contact list: O_ph_  

### K:1H:3126

|  | Res. type | Atom | Residue | Distance | Occupancy | B-factor | Closest solute to 1^st^ shell water |
| --- | --- | --- | --- | --- | --- | --- | --- |
| **Ion** | | | |  | 1.00 | 86.66 || 1 | G | O6 | 1H:2111 | 4.16 | 1.00 | 29.12 |
| 2 | U | O4 | 1H:2112 | 4.43 | 1.00 | 37.47 |  |
| 3 | G | OP2 | 1H:2110 | 4.72 | 1.00 | 33.00 |  |
| 4 | G | N7 | 1H:2110 | 4.74 | 1.00 | 32.94 |  |
| 5 | U | O4 | 1H:2109 | 4.96 | 1.00 | 35.36 |  |
| 6 | G | N7 | 1H:2111 | 4.97 | 1.00 | 30.75 |  |
| 7 | ARG | NH2 | 11:263 | 4.99 | 1.00 | 48.78 |  |
  |

  
 Coded contact list:   
1^st^ K coordination shell missing (see Notes)  

### K:1H:3127

|  | Res. type | Atom | Residue | Distance | Occupancy | B-factor | Closest solute to 1^st^ shell water |
| --- | --- | --- | --- | --- | --- | --- | --- |
| **Ion** | | | |  | 1.00 | 47.09 || 1 | C | O2' | 1H:437 | 2.67 | 1.00 | 45.28 |
| 2 | G | N7 | 1H:216 | 2.76 | 1.00 | 40.39 |  |
| 3 | G | O6 | 1H:216 | 2.88 | 1.00 | 38.75 |  |
| 4 | G | O4' | 1H:438 | 3.21 | 1.00 | 34.35 |  |
| 5 | C | O2 | 1H:437 | 3.61 | 1.00 | 32.48 |  |
| 6 | A | N7 | 1H:215 | 4.09 | 1.00 | 33.73 |  |
| 7 | A | OP2 | 1H:215 | 4.27 | 1.00 | 45.71 |  |
| 8 | G | N2 | 1H:446 | 4.57 | 1.00 | 32.67 |  |
| 9 | G | N9 | 1H:438 | 4.67 | 1.00 | 29.62 |  |
| 10 | G | N1 | 1H:216 | 4.84 | 1.00 | 39.02 |  |
| 11 | C | O2 | 1H:447 | 4.91 | 1.00 | 36.68 |  |
| 12 | G | OP2 | 1H:214 | 4.91 | 1.00 | 27.89 |  |
| 13 | G | N9 | 1H:216 | 4.94 | 1.00 | 38.16 |  |
  |

  
**1^st^ shell ligand angles:**   
1-2: 141; 1-3: 134; 1-4: 73; 2-3: 66; 2-4: 78; 3-4: 85;   
  
 Contact(s): /N7/O2'/O4'/O6/  
 Coded contact list: 2O_r_O_b_N_b_  

### K:1H:3128

|  | Res. type | Atom | Residue | Distance | Occupancy | B-factor | Closest solute to 1^st^ shell water |
| --- | --- | --- | --- | --- | --- | --- | --- |
| **Ion** | | | |  | 1.00 | 64.23 || 1 | A | O3' | 1H:718 | 2.72 | 1.00 | 24.43 |
| 2 | A | O2' | 1H:718 | 2.89 | 1.00 | 35.46 |  |
| 3 | C | OP2 | 1H:719 | 3.09 | 1.00 | 30.59 |  |
| 4 | U | O4 | 1H:612 | 3.29 | 1.00 | 26.61 |  |
| 5 | C | OP1 | 1H:719 | 4.68 | 1.00 | 19.72 |  |
| 6 | C | O5' | 1H:719 | 4.69 | 1.00 | 18.63 |  |
| 7 | G | O6 | 1H:716 | 4.69 | 1.00 | 35.81 |  |
| 8 | C | N4 | 1H:613 | 4.93 | 1.00 | 29.35 |  |
  |

  
**1^st^ shell ligand angles:**   
1-2: 61; 1-3: 50; 1-4: 68; 2-3: 109; 2-4: 66; 3-4: 96;   
  
 Contact(s): /O2'/O3'/O4/OP2/  
 Coded contact list: O_ph_2O_r_O_b_  

### K:1H:3129

|  | Res. type | Atom | Residue | Distance | Occupancy | B-factor | Closest solute to 1^st^ shell water |
| --- | --- | --- | --- | --- | --- | --- | --- |
| **Ion** | | | |  | 1.00 | 60.33 || 1 | G | O6 | 1H:2732 | 3.23 | 1.00 | 43.42 |
| 2 | G | O6 | 1H:2731 | 3.53 | 1.00 | 39.17 |  |
| 3 | G | O6 | 1H:2733 | 3.90 | 1.00 | 46.45 |  |
| 4 | G | N7 | 1H:2732 | 4.16 | 1.00 | 38.65 |  |
| 5 | U | O4 | 1H:2730 | 4.44 | 1.00 | 35.74 |  |
| 6 | A | N1 | 1H:2727 | 4.69 | 1.00 | 47.13 |  |
| 7 | G | N7 | 1H:2731 | 4.77 | 1.00 | 37.05 |  |
| 8 | A | N3 | 1H:2727 | 4.97 | 1.00 | 32.91 |  |
  |

  
 Contact(s): /O6/  
 Coded contact list: O_b_  

### K:1H:3130

|  | Res. type | Atom | Residue | Distance | Occupancy | B-factor | Closest solute to 1^st^ shell water |
| --- | --- | --- | --- | --- | --- | --- | --- |
| **Ion** | | | |  | 1.00 | 61.29 || 1 | G | O6 | 1H:285 | 2.79 | 1.00 | 46.21 |
| 2 | G | O6 | 1H:284 | 2.81 | 1.00 | 53.24 |  |
| 3 | A | OP2 | 1H:260 | 3.23 | 1.00 | 46.70 |  |
| 4 | Water 1 | | 1H:4287 | 3.38 | 1.00 | 61.66 | 3.03 (G:O6 1H:284) |
| 5 | G | N1 | 1H:285 | 3.96 | 1.00 | 47.38 |  |
| 6 | A | OP1 | 1H:260 | 3.97 | 1.00 | 47.92 |  |
| 7 | U | O3' | 1H:259 | 4.16 | 1.00 | 42.25 |  |
| 8 | G | N7 | 1H:284 | 4.34 | 1.00 | 58.73 |  |
| 9 | G | OP2 | 1H:283 | 4.36 | 1.00 | 73.85 |  |
| 10 | U | OP1 | 1H:259 | 4.59 | 1.00 | 57.94 |  |
| 11 | Water 2 | | 1H:3836 | 4.71 | 1.00 | 61.85 | 2.43 (A:OP1 1H:260) |
| 12 | Water 3 | | 1H:3823 | 4.79 | 1.00 | 55.17 | 2.64 (G:O6 1H:284) |
  |

  
**1^st^ shell ligand angles:**   
1-2: 64; 1-3: 71; 1-4: 121; 2-3: 54; 2-4: 58; 3-4: 84;   
  
 Contact(s): /O6/O6/O_w_/OP2/  
 Coded contact list: O_ph_2O_b_O_w_  

### K:1H:3131

|  | Res. type | Atom | Residue | Distance | Occupancy | B-factor | Closest solute to 1^st^ shell water |
| --- | --- | --- | --- | --- | --- | --- | --- |
| **Ion** | | | |  | 1.00 | 63.18 || 1 | A | OP1 | 1H:841 | 3.38 | 1.00 | 21.19 |
| 2 | G | O2' | 1H:840 | 3.44 | 1.00 | 24.32 |  |
| 3 | C | O2' | 1H:2454 | 3.49 | 1.00 | 24.64 |  |
| 4 | C | O2 | 1H:2454 | 4.04 | 1.00 | 20.30 |  |
| 5 | G | O3' | 1H:840 | 4.15 | 1.00 | 18.57 |  |
| 6 | A | O2' | 1H:2094 | 4.26 | 1.00 | 20.59 |  |
| 7 | C | O4' | 1H:2455 | 4.39 | 1.00 | 14.98 |  |
| 8 | C | O2 | 1H:2453 | 4.54 | 1.00 | 19.95 |  |
| 9 | C | N3 | 1H:2453 | 4.56 | 1.00 | 19.04 |  |
| 10 | G | N2 | 1H:2093 | 4.57 | 1.00 | 13.35 |  |
| 11 | G | OP1 | 1H:840 | 4.77 | 1.00 | 22.30 |  |
  |

  
 Contact(s): /OP1/  
 Coded contact list: O_ph_  

### K:1H:3132

|  | Res. type | Atom | Residue | Distance | Occupancy | B-factor | Closest solute to 1^st^ shell water |
| --- | --- | --- | --- | --- | --- | --- | --- |
| **Ion** | | | |  | 1.00 | 87.33 || 1 | G | O6 | 1H:1889 | 2.65 | 1.00 | 71.96 |
| 2 | G | O6 | 1H:1890 | 2.80 | 1.00 | 70.12 |  |
| 3 | Water 1 | | 1H:4424 | 3.40 | 1.00 | 48.42 | 3.38 (G:O6 1H:1889) |
| 4 | G | OP2 | 1H:1888 | 4.13 | 1.00 | 48.17 |  |
| 5 | G | N1 | 1H:1890 | 4.26 | 1.00 | 68.97 |  |
| 6 | G | N7 | 1H:1889 | 4.40 | 1.00 | 68.05 |  |
| 7 | Water 2 | | 1H:4407 | 4.73 | 1.00 | 63.52 | 3.34 (G:O6 1H:1889) |
| 8 | G | N1 | 1H:1889 | 4.90 | 1.00 | 61.52 |  |
  |

  
**1^st^ shell ligand angles:**   
1-2: 68; 1-3: 67; 2-3: 134;   
  
 Contact(s): /O6/O6/O_w_/  
 Coded contact list: 2O_b_O_w_  

### K:1H:3133

|  | Res. type | Atom | Residue | Distance | Occupancy | B-factor | Closest solute to 1^st^ shell water |
| --- | --- | --- | --- | --- | --- | --- | --- |
| **Ion** | | | |  | 1.00 | 61.34 || 1 | U | O4 | 1H:1978 | 2.69 | 1.00 | 42.08 |
| 2 | C | O2 | 1H:2569 | 3.12 | 1.00 | 38.17 |  |
| 3 | U | O4 | 1H:2567 | 3.21 | 1.00 | 48.27 |  |
| 4 | C | N3 | 1H:2569 | 3.23 | 1.00 | 39.23 |  |
| 5 | U | O4 | 1H:2568 | 3.69 | 1.00 | 44.15 |  |
| 6 | U | N3 | 1H:1978 | 4.60 | 1.00 | 30.56 |  |
| 7 | C | N4 | 1H:2569 | 4.74 | 1.00 | 37.40 |  |
| 8 | C | N1 | 1H:2569 | 4.96 | 1.00 | 36.34 |  |
  |

  
**1^st^ shell ligand angles:**   
1-2: 115; 1-3: 69; 1-4: 73; 2-3: 104; 2-4: 42; 3-4: 92;   
  
 Contact(s): /N3_C_/O2/O4/O4/  
 Coded contact list: 3O_b_N_b_  

### K:1H:3134

|  | Res. type | Atom | Residue | Distance | Occupancy | B-factor | Closest solute to 1^st^ shell water |
| --- | --- | --- | --- | --- | --- | --- | --- |
| **Ion** | | | |  | 1.00 | 54.45 || 1 | G | O6 | 1H:1261 | 2.81 | 1.00 | 29.79 |
| 2 | G | O6 | 1H:1262 | 2.88 | 1.00 | 40.02 |  |
| 3 | G | N7 | 1H:1261 | 3.61 | 1.00 | 35.52 |  |
| 4 | U | O4 | 1H:1281 | 3.83 | 1.00 | 32.24 |  |
| 5 | G | N2 | 1H:1282 | 3.95 | 1.00 | 28.70 |  |
| 6 | G | N7 | 1H:1262 | 4.16 | 1.00 | 38.96 |  |
| 7 | G | N1 | 1H:1282 | 4.35 | 1.00 | 30.62 |  |
| 8 | A | OP2 | 1H:1260 | 4.39 | 1.00 | 30.29 |  |
| 9 | U | N3 | 1H:1281 | 4.53 | 1.00 | 33.24 |  |
| 10 | C | N4 | 1H:1280 | 4.57 | 1.00 | 39.31 |  |
| 11 | A | OP2 | 1H:1259 | 4.88 | 1.00 | 29.50 |  |
| 12 | G | N1 | 1H:1262 | 4.89 | 1.00 | 36.42 |  |
| 13 | G | N3 | 1H:1282 | 4.90 | 1.00 | 28.38 |  |
| 14 | G | N1 | 1H:1261 | 4.91 | 1.00 | 31.08 |  |
  |

  
**1^st^ shell ligand angles:**   
1-2: 72;   
  
 Contact(s): /O6/O6/  
 Coded contact list: 2O_b_  

### K:1H:3135

|  | Res. type | Atom | Residue | Distance | Occupancy | B-factor | Closest solute to 1^st^ shell water |
| --- | --- | --- | --- | --- | --- | --- | --- |
| **Ion** | | | |  | 1.00 | 82.62 || 1 | A | OP1 | 1H:349 | 2.91 | 1.00 | 46.22 |
| 2 | G | OP2 | 1H:348 | 3.36 | 1.00 | 36.69 |  |
| 3 | A | O4' | 1H:349 | 3.65 | 1.00 | 39.79 |  |
| 4 | U | O2 | 1H:639 | 3.69 | 1.00 | 88.13 |  |
| 5 | A | O5' | 1H:349 | 4.57 | 1.00 | 43.36 |  |
| 6 | G | O5' | 1H:348 | 4.58 | 1.00 | 36.89 |  |
  |

  
**1^st^ shell ligand angles:**   
1-2: 81;   
  
 Contact(s): /OP1/OP2/  
 Coded contact list: 2O_ph_  

### K:1H:3136

|  | Res. type | Atom | Residue | Distance | Occupancy | B-factor | Closest solute to 1^st^ shell water |
| --- | --- | --- | --- | --- | --- | --- | --- |
| **Ion** | | | |  | 1.00 | 54.76 || 1 | U | O4 | 1H:2245 | 2.53 | 1.00 | 30.63 |
| 2 | U | O4 | 1H:2109 | 2.89 | 1.00 | 35.36 |  |
| 3 | U | O4 | 1H:2246 | 3.31 | 1.00 | 28.86 |  |
| 4 | G | O6 | 1H:2110 | 3.39 | 1.00 | 30.62 |  |
| 5 | U | N3 | 1H:2109 | 3.64 | 1.00 | 24.03 |  |
| 6 | U | N3 | 1H:2245 | 4.00 | 1.00 | 24.83 |  |
| 7 | C | N4 | 1H:2244 | 4.33 | 1.00 | 26.28 |  |
| 8 | G | O6 | 1H:2111 | 4.42 | 1.00 | 29.12 |  |
| 9 | G | N7 | 1H:2110 | 4.90 | 1.00 | 32.94 |  |
| 10 | U | N3 | 1H:2246 | 4.98 | 1.00 | 26.40 |  |
  |

  
**1^st^ shell ligand angles:**   
1-2: 140; 1-3: 82; 1-4: 68; 2-3: 72; 2-4: 121; 3-4: 60;   
  
 Contact(s): /O4/O4/O6/  
 Coded contact list: 3O_b_  
 Exclusion zone contact(s): /O4/  

### K:1H:3137

|  | Res. type | Atom | Residue | Distance | Occupancy | B-factor | Closest solute to 1^st^ shell water |
| --- | --- | --- | --- | --- | --- | --- | --- |
| **Ion** | | | |  | 1.00 | 63.56 || 1 | G | O6 | 1H:168 | 3.24 | 1.00 | 27.65 |
| 2 | G | O6 | 1H:169 | 3.42 | 1.00 | 26.66 |  |
| 3 | G | N7 | 1H:168 | 3.56 | 1.00 | 32.34 |  |
| 4 | G | N7 | 1H:169 | 3.81 | 1.00 | 26.81 |  |
| 5 | A | OP2 | 1H:203 | 3.81 | 1.00 | 51.93 |  |
| 6 | G | OP1 | 1H:167 | 4.20 | 1.00 | 35.24 |  |
| 7 | G | O6 | 1H:170 | 4.65 | 1.00 | 31.87 |  |
| 8 | C | N4 | 1H:46 | 4.92 | 1.00 | 31.74 |  |
  |

  
 Contact(s): /O6/  
 Coded contact list: O_b_  

### K:1H:3138

|  | Res. type | Atom | Residue | Distance | Occupancy | B-factor | Closest solute to 1^st^ shell water |
| --- | --- | --- | --- | --- | --- | --- | --- |
| **Ion** | | | |  | 1.00 | 62.69 || 1 | G | OP2 | 1H:500 | 3.26 | 1.00 | 30.29 |
| 2 | A | O2' | 1H:474 | 3.27 | 1.00 | 29.47 |  |
| 3 | A | O3' | 1H:474 | 3.31 | 1.00 | 28.94 |  |
| 4 | A | OP2 | 1H:499 | 3.34 | 1.00 | 28.43 |  |
| 5 | U | OP2 | 1H:475 | 3.85 | 1.00 | 38.67 |  |
| 6 | A | OP1 | 1H:499 | 4.00 | 1.00 | 46.43 |  |
| 7 | A | O5' | 1H:499 | 4.06 | 1.00 | 25.81 |  |
| 8 | U | O5' | 1H:475 | 4.12 | 1.00 | 23.07 |  |
| 9 | Water 1 | | 1H:4130 | 4.26 | 1.00 | 28.81 | 3.90 (C:N4 1H:482) |
| 10 | G | OP1 | 1H:500 | 4.33 | 1.00 | 39.69 |  |
| 11 | C | N4 | 1H:482 | 4.47 | 1.00 | 32.67 |  |
| 12 | A | N3 | 1H:474 | 4.53 | 1.00 | 32.67 |  |
| 13 | G | N7 | 1H:500 | 4.59 | 1.00 | 35.92 |  |
| 14 | Water 2 | | 1H:3953 | 4.82 | 1.00 | 24.78 | 3.92 (U:O5' 1H:475) |
| 15 | A | O4' | 1H:474 | 4.99 | 1.00 | 27.18 |  |
  |

  
**1^st^ shell ligand angles:**   
1-2: 94; 1-3: 142; 1-4: 96; 2-3: 48; 2-4: 164; 3-4: 121;   
  
 Contact(s): /O2'/O3'/OP2/OP2/  
 Coded contact list: 2O_ph_2O_r_  

### K:1H:3139

|  | Res. type | Atom | Residue | Distance | Occupancy | B-factor | Closest solute to 1^st^ shell water |
| --- | --- | --- | --- | --- | --- | --- | --- |
| **Ion** | | | |  | 1.00 | 64.30 || 1 | G | O6 | 1H:748 | 2.53 | 1.00 | 35.94 |
| 2 | G | O6 | 1H:749 | 3.39 | 1.00 | 33.32 |  |
| 3 | G | N7 | 1H:748 | 4.08 | 1.00 | 33.03 |  |
| 4 | G | N7 | 1H:749 | 4.11 | 1.00 | 32.82 |  |
| 5 | C | N4 | 1H:780 | 4.31 | 1.00 | 28.44 |  |
| 6 | G | N1 | 1H:748 | 4.60 | 1.00 | 31.06 |  |
| 7 | U | OP2 | 1H:810 | 4.62 | 1.00 | 24.87 |  |
| 8 | C | N4 | 1H:779 | 4.78 | 1.00 | 21.96 |  |
| 9 | G | N2 | 1H:781 | 4.85 | 1.00 | 28.57 |  |
  |

  
**1^st^ shell ligand angles:**   
1-2: 77;   
  
 Contact(s): /O6/  
 Coded contact list: O_b_  
 Exclusion zone contact(s): /O6/  

### K:1H:3140

|  | Res. type | Atom | Residue | Distance | Occupancy | B-factor | Closest solute to 1^st^ shell water |
| --- | --- | --- | --- | --- | --- | --- | --- |
| **Ion** | | | |  | 1.00 | 62.74 || 1 | A | O2' | 1H:1312 | 2.87 | 1.00 | 36.54 |
| 2 | U | OP1 | 1H:1310 | 3.64 | 1.00 | 27.79 |  |
| 3 | A | N3 | 1H:1312 | 3.64 | 1.00 | 23.59 |  |
| 4 | ARG | NH1 | E8:15 | 3.69 | 1.00 | 29.74 |  |
| 5 | ARG | NH2 | N8:20 | 3.72 | 1.00 | 27.79 |  |
| 6 | G | OP2 | 1H:1311 | 3.76 | 1.00 | 24.56 |  |
| 7 | U | OP2 | 1H:1310 | 3.99 | 1.00 | 22.56 |  |
| 8 | A | N9 | 1H:1312 | 4.77 | 1.00 | 25.40 |  |
| 9 | U | O5' | 1H:1310 | 4.86 | 1.00 | 25.78 |  |
| 10 | G | N7 | 1H:1311 | 4.98 | 1.00 | 31.38 |  |
| 11 | A | N1 | 1H:1312 | 4.99 | 1.00 | 29.21 |  |
  |

  
 Contact(s): /O2'/  
 Coded contact list: O_r_  

### K:1H:3141

|  | Res. type | Atom | Residue | Distance | Occupancy | B-factor | Closest solute to 1^st^ shell water |
| --- | --- | --- | --- | --- | --- | --- | --- |
| **Ion** | | | |  | 1.00 | 41.63 || 1 | A | OP1 | 1H:2461 | 2.89 | 1.00 | 25.13 |
| 2 | A | O5' | 1H:2461 | 3.35 | 1.00 | 16.40 |  |
| 3 | G | OP1 | 1H:2460 | 3.35 | 1.00 | 26.48 |  |
| 4 | G | O5' | 1H:2460 | 3.54 | 1.00 | 20.05 |  |
| 5 | A | N6 | 1H:991 | 3.97 | 1.00 | 18.04 |  |
| 6 | G | O3' | 1H:2460 | 4.35 | 1.00 | 22.35 |  |
| 7 | G | O6 | 1H:594 | 4.40 | 1.00 | 24.35 |  |
| 8 | U | OP2 | 1H:2513 | 4.45 | 1.00 | 20.03 |  |
| 9 | U | O2' | 1H:592 | 4.50 | 1.00 | 24.12 |  |
| 10 | G | O3' | 1H:2459 | 4.51 | 1.00 | 18.40 |  |
| 11 | C | OP1 | 1H:2512 | 4.51 | 1.00 | 20.16 |  |
| 12 | A | O4' | 1H:2461 | 4.95 | 1.00 | 20.40 |  |
  |

  
**1^st^ shell ligand angles:**   
1-2: 47; 1-3: 135; 2-3: 104;   
  
 Contact(s): /O5'/OP1/OP1/  
 Coded contact list: 2O_ph_O_r_  

### K:1H:3142

|  | Res. type | Atom | Residue | Distance | Occupancy | B-factor | Closest solute to 1^st^ shell water |
| --- | --- | --- | --- | --- | --- | --- | --- |
| **Ion** | | | |  | 1.00 | 78.67 || 1 | U | OP2 | 1H:1320 | 2.91 | 1.00 | 29.76 |
| 2 | G | OP1 | 1H:1318 | 3.18 | 1.00 | 27.41 |  |
| 3 | G | OP2 | 1H:1318 | 3.35 | 1.00 | 34.09 |  |
| 4 | A | OP1 | 1H:1319 | 3.43 | 1.00 | 34.65 |  |
| 5 | U | OP1 | 1H:1320 | 4.21 | 1.00 | 24.68 |  |
| 6 | G | O5' | 1H:1318 | 4.43 | 1.00 | 29.85 |  |
| 7 | U | O5' | 1H:1320 | 4.80 | 1.00 | 27.49 |  |
  |

  
**1^st^ shell ligand angles:**   
1-2: 141; 1-3: 120; 2-3: 47;   
  
 Contact(s): /OP1/OP2/OP2/  
 Coded contact list: 3O_ph_  

### K:1H:3143

|  | Res. type | Atom | Residue | Distance | Occupancy | B-factor | Closest solute to 1^st^ shell water |
| --- | --- | --- | --- | --- | --- | --- | --- |
| **Ion** | | | |  | 1.00 | 71.81 || 1 | G | O6 | 1H:1468 | 2.60 | 1.00 | 54.54 |
| 2 | G | O6 | 1H:1469 | 2.99 | 1.00 | 44.25 |  |
| 3 | U | O4 | 1H:1624 | 3.84 | 1.00 | 46.18 |  |
| 4 | G | O6 | 1H:1470 | 3.90 | 1.00 | 44.97 |  |
| 5 | G | N7 | 1H:1469 | 3.91 | 1.00 | 50.05 |  |
| 6 | G | N7 | 1H:1468 | 3.91 | 1.00 | 66.86 |  |
| 7 | C | N4 | 1H:1625 | 4.10 | 1.00 | 47.98 |  |
| 8 | G | N1 | 1H:1468 | 4.46 | 1.00 | 55.35 |  |
| 9 | C | N4 | 1H:1623 | 4.60 | 1.00 | 39.90 |  |
  |

  
**1^st^ shell ligand angles:**   
1-2: 74;   
  
 Contact(s): /O6/O6/  
 Coded contact list: 2O_b_  

### K:1H:3144

|  | Res. type | Atom | Residue | Distance | Occupancy | B-factor | Closest solute to 1^st^ shell water |
| --- | --- | --- | --- | --- | --- | --- | --- |
| **Ion** | | | |  | 1.00 | 56.21 || 1 | U | O4 | 1H:751 | 2.80 | 1.00 | 32.16 |
| 2 | G | O6 | 1H:776 | 2.88 | 1.00 | 30.96 |  |
| 3 | G | O6 | 1H:750 | 3.40 | 1.00 | 32.64 |  |
| 4 | C | N4 | 1H:778 | 3.78 | 1.00 | 32.50 |  |
| 5 | G | N2 | 1H:752 | 3.80 | 1.00 | 31.43 |  |
| 6 | U | N3 | 1H:751 | 3.91 | 1.00 | 29.65 |  |
| 7 | G | N7 | 1H:776 | 4.08 | 1.00 | 25.85 |  |
| 8 | A | OP2 | 1H:775 | 4.27 | 1.00 | 25.73 |  |
| 9 | G | O2' | 1H:774 | 4.57 | 1.00 | 31.63 |  |
| 10 | A | N7 | 1H:775 | 4.59 | 1.00 | 28.32 |  |
| 11 | G | N1 | 1H:752 | 4.60 | 1.00 | 31.89 |  |
  |

  
**1^st^ shell ligand angles:**   
1-2: 74; 1-3: 58; 2-3: 81;   
  
 Contact(s): /O4/O6/O6/  
 Coded contact list: 3O_b_  

### MG:1H:3145

|  | Res. type | Atom | Residue | Distance | Occupancy | B-factor | Closest solute to 1^st^ shell water |
| --- | --- | --- | --- | --- | --- | --- | --- |
| **Ion** | | | |  | 1.00 | 17.62 || 1 | G | OP2 | 1H:1860 | 2.15 | 1.00 | 16.68 |
| 2 | A | OP2 | 1H:1995 | 3.79 | 1.00 | 23.13 |  |
| 3 | G | N7 | 1H:1860 | 3.98 | 1.00 | 18.17 |  |
| 4 | C | OP2 | 1H:1859 | 4.04 | 1.00 | 16.50 |  |
| 5 | C | O5' | 1H:1859 | 4.13 | 1.00 | 15.92 |  |
| 6 | Water 1 | | 1H:3630 | 4.23 | 1.00 | 17.42 | 1.97 (G:OP1 1H:1860) |
| 7 | C | O3' | 1H:1859 | 4.30 | 1.00 | 15.42 |  |
| 8 | C | OP1 | 1H:1859 | 4.36 | 1.00 | 16.87 |  |
| 9 | ARG | NH2 | 11:222 | 4.45 | 1.00 | 15.30 |  |
| 10 | G | O5' | 1H:1860 | 4.52 | 1.00 | 17.64 |  |
| 11 | G | OP1 | 1H:1860 | 4.52 | 1.00 | 15.93 |  |
| 12 | ARG | NE | 11:222 | 4.88 | 1.00 | 15.67 |  |
| 13 | ARG | NE | 11:239 | 4.89 | 1.00 | 24.33 |  |
| 14 | ARG | NH1 | 11:222 | 4.92 | 1.00 | 15.29 |  |
  |

  
 Contact(s): /OP2/  
 Coded contact list: O_ph_  

### MG:1H:3146

|  | Res. type | Atom | Residue | Distance | Occupancy | B-factor | Closest solute to 1^st^ shell water |
| --- | --- | --- | --- | --- | --- | --- | --- |
| **Ion** | | | |  | 1.00 | 38.43 || 1 | G | O6 | 1H:1681 | 2.02 | 1.00 | 39.19 |
| 2 | A | N7 | 1H:1680 | 2.67 | 1.00 | 37.40 |  |
| 3 | A | N6 | 1H:1680 | 3.39 | 1.00 | 40.36 |  |
| 4 | A | N1 | 1H:1794 | 3.81 | 1.00 | 49.26 |  |
| 5 | A | OP1 | 1H:1679 | 3.95 | 1.00 | 40.28 |  |
| 6 | A | OP2 | 1H:1680 | 4.01 | 1.00 | 36.60 |  |
| 7 | G | N2 | 1H:1683 | 4.02 | 1.00 | 30.01 |  |
| 8 | C | O2 | 1H:1678 | 4.10 | 1.00 | 36.19 |  |
| 9 | A | N6 | 1H:1794 | 4.15 | 1.00 | 44.04 |  |
| 10 | G | N1 | 1H:1681 | 4.17 | 1.00 | 33.19 |  |
| 11 | G | N7 | 1H:1681 | 4.31 | 1.00 | 37.29 |  |
| 12 | G | N1 | 1H:1683 | 4.40 | 1.00 | 28.09 |  |
| 13 | A | N7 | 1H:1679 | 4.51 | 1.00 | 46.93 |  |
| 14 | C | N3 | 1H:1678 | 4.55 | 1.00 | 34.24 |  |
| 15 | C | O2' | 1H:1678 | 4.71 | 1.00 | 44.05 |  |
| 16 | C | N1 | 1H:1678 | 4.79 | 1.00 | 40.08 |  |
| 17 | G | N3 | 1H:1683 | 4.82 | 1.00 | 30.79 |  |
| 18 | A | N9 | 1H:1680 | 4.85 | 1.00 | 32.70 |  |
| 19 | A | O5' | 1H:1679 | 4.95 | 1.00 | 51.87 |  |
  |

  
 Contact(s): /O6/  
 Coded contact list: O_b_  
 Exclusion zone contact(s): /N6/N7/  
Consider K in place of MG (2 valid contact(s) for K vs. 1 for MG)  

### MG:1H:3147

|  | Res. type | Atom | Residue | Distance | Occupancy | B-factor | Closest solute to 1^st^ shell water |
| --- | --- | --- | --- | --- | --- | --- | --- |
| **Ion** | | | |  | 1.00 | 54.70 || 1 | G | OP2 | 1H:1504 | 1.93 | 1.00 | 53.09 |
| 2 | G | OP2 | 1H:1503 | 2.32 | 1.00 | 48.93 |  |
| 3 | G | OP1 | 1H:1504 | 3.86 | 1.00 | 57.23 |  |
| 4 | A | OP2 | 1H:1501 | 4.10 | 1.00 | 61.45 |  |
| 5 | G | O5' | 1H:1504 | 4.28 | 1.00 | 52.52 |  |
| 6 | U | O3' | 1H:1502 | 4.31 | 1.00 | 61.04 |  |
| 7 | G | O3' | 1H:1503 | 4.33 | 1.00 | 45.44 |  |
| 8 | G | O5' | 1H:1503 | 4.69 | 1.00 | 53.46 |  |
| 9 | G | N7 | 1H:1504 | 4.73 | 1.00 | 51.45 |  |
| 10 | G | OP1 | 1H:1503 | 4.82 | 1.00 | 47.52 |  |
  |

  
**1^st^ shell ligand angles:**   
1-2: 82;   
  
 Contact(s): /OP2/OP2/  
 Coded contact list: 2O_ph_  

### MG:1H:3148

|  | Res. type | Atom | Residue | Distance | Occupancy | B-factor | Closest solute to 1^st^ shell water |
| --- | --- | --- | --- | --- | --- | --- | --- |
| **Ion** | | | |  | 1.00 | 35.60 || 1 | G | N7 | 1H:2441 | 3.32 | 1.00 | 25.17 |
| 2 | G | O6 | 1H:2441 | 3.81 | 1.00 | 26.46 |  |
| 3 | C | OP1 | 1H:873 | 3.85 | 1.00 | 29.83 |  |
| 4 | C | OP1 | 1H:2273 | 4.36 | 1.00 | 28.75 |  |
| 5 | G | OP2 | 1H:2441 | 4.57 | 1.00 | 23.82 |  |
| 6 | C | O5' | 1H:873 | 4.80 | 1.00 | 19.70 |  |
  |

  
 Coded contact list:   
 Exclusion zone contact(s): /N7/  
Consider K in place of MG (1 valid contact(s) for K vs. 0 for MG)  

### MG:1H:3149

|  | Res. type | Atom | Residue | Distance | Occupancy | B-factor | Closest solute to 1^st^ shell water |
| --- | --- | --- | --- | --- | --- | --- | --- |
| **Ion** | | | |  | 1.00 | 40.15 || 1 | A | OP2 | 1H:450 | 1.94 | 1.00 | 49.86 |
| 2 | G | OP2 | 1H:398 | 2.11 | 1.00 | 44.49 |  |
| 3 | A | O5' | 1H:450 | 3.92 | 1.00 | 38.59 |  |
| 4 | C | O3' | 1H:397 | 3.93 | 1.00 | 41.72 |  |
| 5 | G | N7 | 1H:398 | 4.23 | 1.00 | 39.89 |  |
| 6 | U | O3' | 1H:449 | 4.31 | 1.00 | 36.97 |  |
| 7 | G | O5' | 1H:398 | 4.31 | 1.00 | 38.27 |  |
| 8 | A | OP1 | 1H:450 | 4.40 | 1.00 | 41.34 |  |
| 9 | C | N4 | 1H:287 | 4.42 | 1.00 | 71.77 |  |
| 10 | C | O2' | 1H:397 | 4.52 | 1.00 | 43.66 |  |
| 11 | G | OP1 | 1H:398 | 4.57 | 1.00 | 42.04 |  |
| 12 | U | O2' | 1H:449 | 4.60 | 1.00 | 44.37 |  |
| 13 | A | OP2 | 1H:451 | 4.64 | 1.00 | 35.23 |  |
| 14 | A | N7 | 1H:450 | 4.76 | 1.00 | 38.23 |  |
| 15 | A | N7 | 1H:451 | 4.77 | 1.00 | 38.97 |  |
  |

  
**1^st^ shell ligand angles:**   
1-2: 157;   
  
 Contact(s): /OP2/OP2/  
 Coded contact list: 2O_ph_  

### MG:1H:3150

|  | Res. type | Atom | Residue | Distance | Occupancy | B-factor | Closest solute to 1^st^ shell water |
| --- | --- | --- | --- | --- | --- | --- | --- |
| **Ion** | | | |  | 1.00 | 25.48 || 1 | G | O6 | 1H:1233 | 1.82 | 1.00 | 29.27 |
| 2 | G | OP2 | 1H:866 | 2.40 | 1.00 | 27.92 |  |
| 3 | C | O5' | 1H:865 | 3.70 | 1.00 | 25.87 |  |
| 4 | C | OP1 | 1H:865 | 3.83 | 1.00 | 34.86 |  |
| 5 | Water 1 | | 1H:4244 | 3.98 | 1.00 | 32.31 | 2.96 (C:OP2 1H:865) |
| 6 | G | N1 | 1H:1233 | 3.99 | 1.00 | 28.43 |  |
| 7 | G | OP1 | 1H:1232 | 4.04 | 1.00 | 29.55 |  |
| 8 | G | N7 | 1H:1233 | 4.13 | 1.00 | 31.09 |  |
| 9 | C | OP2 | 1H:865 | 4.43 | 1.00 | 26.22 |  |
| 10 | G | N7 | 1H:1232 | 4.47 | 1.00 | 30.49 |  |
| 11 | G | O5' | 1H:866 | 4.50 | 1.00 | 30.21 |  |
| 12 | G | N7 | 1H:866 | 4.51 | 1.00 | 28.23 |  |
| 13 | G | OP1 | 1H:866 | 4.77 | 1.00 | 31.23 |  |
| 14 | C | O3' | 1H:865 | 4.85 | 1.00 | 28.13 |  |
  |

  
**1^st^ shell ligand angles:**   
1-2: 84;   
  
 Contact(s): /O6/OP2/  
 Coded contact list: O_ph_O_b_  

### MG:1H:3151

|  | Res. type | Atom | Residue | Distance | Occupancy | B-factor | Closest solute to 1^st^ shell water |
| --- | --- | --- | --- | --- | --- | --- | --- |
| **Ion** | | | |  | 1.00 | 41.59 || 1 | C | O2' | 1H:1213 | 3.62 | 1.00 | 49.35 |
| 2 | C | O2' | 1H:864 | 3.64 | 1.00 | 26.77 |  |
| 3 | G | N7 | 1H:978 | 3.93 | 1.00 | 37.32 |  |
| 4 | C | O3' | 1H:864 | 3.95 | 1.00 | 26.19 |  |
| 5 | C | O2 | 1H:1213 | 4.02 | 1.00 | 45.79 |  |
| 6 | G | N2 | 1H:1230 | 4.04 | 1.00 | 40.58 |  |
| 7 | G | N3 | 1H:1230 | 4.44 | 1.00 | 47.27 |  |
| 8 | U | O4' | 1H:1214 | 4.70 | 1.00 | 57.11 |  |
| 9 | G | O2' | 1H:1230 | 4.82 | 1.00 | 43.64 |  |
  |

  
 Coded contact list:   
1^st^ MG coordination shell missing (see Notes)  

### MG:1H:3152

|  | Res. type | Atom | Residue | Distance | Occupancy | B-factor | Closest solute to 1^st^ shell water |
| --- | --- | --- | --- | --- | --- | --- | --- |
| **Ion** | | | |  | 1.00 | 36.87 || 1 | Water 1 | | 1H:3714 | 2.30 | 1.00 | 41.28 | 2.23 (G:OP1 1H:1653) |
| 2 | G | OP1 | 1H:1653 | 2.54 | 1.00 | 35.64 |  |
| 3 | Water 2 | | 1H:4038 | 2.97 | 1.00 | 32.55 | 2.67 (G:OP2 1H:1653) |
| 4 | Water 3 | | 1H:4395 | 3.18 | 1.00 | 37.66 | 4.52 (G:OP1 1H:1653) |
| 5 | G | OP2 | 1H:1653 | 3.69 | 1.00 | 30.15 |  |
| 6 | C | O3' | 1H:1652 | 3.70 | 1.00 | 25.05 |  |
| 7 | G | O5' | 1H:1653 | 4.78 | 1.00 | 26.45 |  |
| 8 | Water 4 | | 1H:3626 | 4.80 | 1.00 | 32.53 | 3.34 (G:OP1 1H:1653) |
| 9 | Water 5 | | 1H:4059 | 4.96 | 1.00 | 39.56 | 4.91 (G:OP1 1H:1653) |
  |

  
 Contact(s): /O_w_/  
 Coded contact list: O_w_  
 Exclusion zone contact(s): /O_w_/O_w_/OP1/  
Consider NA in place of MG (1 valid contact(s) for NA vs. 0 for MG)  
MG hydration shell issue: water too close (< 2.4 Angs.)  

### MG:1H:3153

|  | Res. type | Atom | Residue | Distance | Occupancy | B-factor | Closest solute to 1^st^ shell water |
| --- | --- | --- | --- | --- | --- | --- | --- |
| **Ion** | | | |  | 1.00 | 45.53 || 1 | G | OP2 | 1H:1991 | 3.04 | 1.00 | 25.10 |
| 2 | Water 1 | | 1H:4233 | 3.19 | 1.00 | 26.06 | 2.94 (G:OP2 1H:1991) |
| 3 | A | OP2 | 1H:1992 | 4.09 | 1.00 | 28.15 |  |
| 4 | A | N6 | 1H:1994 | 4.11 | 1.00 | 27.49 |  |
| 5 | A | OP1 | 1H:1992 | 4.26 | 1.00 | 40.28 |  |
| 6 | G | O6 | 1H:1953 | 4.75 | 1.00 | 34.21 |  |
| 7 | G | O5' | 1H:1991 | 4.94 | 1.00 | 23.80 |  |
| 8 | Water 2 | | 1H:4425 | 4.98 | 1.00 | 34.22 | 3.39 (A:OP1 1H:1992) |
  |

  
 Coded contact list:   
 Exclusion zone contact(s): /O_w_/OP2/  
Consider K in place of MG (1 valid contact(s) for K vs. 0 for MG)  

### MG:1H:3154

|  | Res. type | Atom | Residue | Distance | Occupancy | B-factor | Closest solute to 1^st^ shell water |
| --- | --- | --- | --- | --- | --- | --- | --- |
| **Ion** | | | |  | 1.00 | 26.87 || 1 | G | OP2 | 1H:910 | 2.19 | 1.00 | 30.56 |
| 2 | Water 1 | | 1H:4022 | 2.86 | 1.00 | 32.67 | 2.66 (A:OP2 1H:909) |
| 3 | G | O5' | 1H:910 | 3.71 | 1.00 | 46.58 |  |
| 4 | G | OP1 | 1H:910 | 3.96 | 1.00 | 41.23 |  |
| 5 | G | N7 | 1H:910 | 4.04 | 1.00 | 35.56 |  |
| 6 | A | N7 | 1H:911 | 4.19 | 1.00 | 32.91 |  |
| 7 | A | OP2 | 1H:909 | 4.21 | 1.00 | 39.78 |  |
| 8 | A | OP1 | 1H:909 | 4.31 | 1.00 | 31.82 |  |
| 9 | A | OP2 | 1H:911 | 4.53 | 1.00 | 38.86 |  |
| 10 | A | OP1 | 1H:957 | 4.54 | 1.00 | 31.87 |  |
| 11 | A | O3' | 1H:909 | 4.58 | 1.00 | 26.36 |  |
| 12 | A | N6 | 1H:911 | 4.92 | 1.00 | 34.18 |  |
  |

  
 Contact(s): /OP2/  
 Coded contact list: O_ph_  
 Exclusion zone contact(s): /O_w_/  

### MG:1H:3155

|  | Res. type | Atom | Residue | Distance | Occupancy | B-factor | Closest solute to 1^st^ shell water |
| --- | --- | --- | --- | --- | --- | --- | --- |
| **Ion** | | | |  | 0.50 | 19.89 || 1 | C | OP2 | 1H:557 | 2.02 | 1.00 | 31.98 |
| 2 | C | OP1 | 1H:2059 | 2.70 | 1.00 | 31.62 |  |
| 3 | C | O5' | 1H:557 | 3.97 | 1.00 | 26.85 |  |
| 4 | C | OP1 | 1H:557 | 4.03 | 1.00 | 26.65 |  |
| 5 | G | N1 | 1H:585 | 4.15 | 1.00 | 30.32 |  |
| 6 | G | N3 | 1H:585 | 4.28 | 1.00 | 28.29 |  |
| 7 | G | N2 | 1H:585 | 4.40 | 1.00 | 24.93 |  |
| 8 | G | O3' | 1H:2058 | 4.53 | 1.00 | 31.92 |  |
| 9 | G | O3' | 1H:556 | 4.61 | 1.00 | 25.42 |  |
| 10 | G | O2' | 1H:556 | 4.69 | 1.00 | 34.46 |  |
| 11 | C | OP2 | 1H:2059 | 4.90 | 1.00 | 31.81 |  |
  |

  
 Contact(s): /OP2/  
 Coded contact list: O_ph_  
 Exclusion zone contact(s): /OP1/  

### MG:1H:3155

|  | Res. type | Atom | Residue | Distance | Occupancy | B-factor | Closest solute to 1^st^ shell water |
| --- | --- | --- | --- | --- | --- | --- | --- |
| **Ion** | | | |  | 0.50 | 22.49 || 1 | C | OP1 | 1H:2059 | 2.12 | 1.00 | 31.62 |
| 2 | C | OP2 | 1H:557 | 2.15 | 1.00 | 31.98 |  |
| 3 | G | O2' | 1H:556 | 3.16 | 1.00 | 34.46 |  |
| 4 | G | O3' | 1H:2058 | 3.32 | 1.00 | 31.92 |  |
| 5 | C | OP2 | 1H:2059 | 3.33 | 1.00 | 31.81 |  |
| 6 | G | O3' | 1H:556 | 4.17 | 1.00 | 25.42 |  |
| 7 | C | O5' | 1H:557 | 4.34 | 1.00 | 26.85 |  |
| 8 | C | O5' | 1H:2059 | 4.48 | 1.00 | 20.70 |  |
| 9 | C | OP1 | 1H:557 | 4.73 | 1.00 | 26.65 |  |
  |

  
**1^st^ shell ligand angles:**   
1-2: 133;   
  
 Contact(s): /OP1/OP2/  
 Coded contact list: 2O_ph_  
 Exclusion zone contact(s): /O2'/O3'/OP2/  
Consider K in place of MG (3 valid contact(s) for K vs. 2 for MG)  

### MG:1H:3156

|  | Res. type | Atom | Residue | Distance | Occupancy | B-factor | Closest solute to 1^st^ shell water |
| --- | --- | --- | --- | --- | --- | --- | --- |
| **Ion** | | | |  | 1.00 | 33.38 || 1 | G | O6 | 1H:2508 | 3.60 | 1.00 | 26.42 |
| 2 | Water 1 | | 1H:4470 | 3.99 | 1.00 | 25.18 | 3.71 (G:O6 1H:2468) |
| 3 | G | O6 | 1H:2507 | 4.10 | 1.00 | 29.14 |  |
| 4 | G | N7 | 1H:2507 | 4.12 | 1.00 | 29.93 |  |
| 5 | C | N4 | 1H:2469 | 4.12 | 1.00 | 20.40 |  |
| 6 | G | O6 | 1H:2468 | 4.39 | 1.00 | 22.74 |  |
| 7 | U | O4 | 1H:2470 | 4.48 | 1.00 | 28.78 |  |
  |

  
 Coded contact list:   
1^st^ MG coordination shell missing (see Notes)  

### MG:1H:3157

|  | Res. type | Atom | Residue | Distance | Occupancy | B-factor | Closest solute to 1^st^ shell water |
| --- | --- | --- | --- | --- | --- | --- | --- |
| **Ion** | | | |  | 1.00 | 49.73 || 1 | G | OP2 | 1H:168 | 2.70 | 1.00 | 32.87 |
| 2 | G | O5' | 1H:168 | 3.88 | 1.00 | 25.71 |  |
| 3 | G | N7 | 1H:168 | 3.95 | 1.00 | 32.34 |  |
| 4 | G | OP2 | 1H:202 | 4.35 | 1.00 | 31.03 |  |
| 5 | G | OP1 | 1H:202 | 4.38 | 1.00 | 37.89 |  |
| 6 | G | OP2 | 1H:169 | 4.54 | 1.00 | 20.94 |  |
| 7 | G | O5' | 1H:202 | 4.65 | 1.00 | 23.86 |  |
| 8 | G | O3' | 1H:167 | 4.79 | 1.00 | 32.03 |  |
| 9 | A | OP2 | 1H:203 | 4.92 | 1.00 | 51.93 |  |
| 10 | G | OP1 | 1H:168 | 4.99 | 1.00 | 27.11 |  |
  |

  
 Coded contact list:   
 Exclusion zone contact(s): /OP2/  
Consider K in place of MG (1 valid contact(s) for K vs. 0 for MG)  

### MG:1H:3158

|  | Res. type | Atom | Residue | Distance | Occupancy | B-factor | Closest solute to 1^st^ shell water |
| --- | --- | --- | --- | --- | --- | --- | --- |
| **Ion** | | | |  | 1.00 | 57.64 || 1 | Water 1 | | 1H:4451 | 2.81 | 1.00 | 77.21 | 4.07 (G:N7 1H:282) |
| 2 | G | OP2 | 1H:282 | 2.96 | 1.00 | 83.92 |  |
| 3 | Water 2 | | 1H:4287 | 3.19 | 1.00 | 61.66 | 4.04 (G:N7 1H:282) |
| 4 | Water 3 | | 1H:3836 | 3.28 | 1.00 | 61.85 | 2.43 (A:OP1 1H:260) |
| 5 | C | OP2 | 1H:281 | 4.05 | 1.00 | 84.07 |  |
| 6 | A | OP1 | 1H:260 | 4.35 | 1.00 | 47.92 |  |
| 7 | C | O5' | 1H:281 | 4.35 | 1.00 | 84.37 |  |
| 8 | G | N7 | 1H:282 | 4.71 | 1.00 | 82.31 |  |
| 9 | G | O5' | 1H:282 | 4.93 | 1.00 | 70.00 |  |
  |

  
 Coded contact list:   
 Exclusion zone contact(s): /O_w_/O_w_/O_w_/OP2/  
Consider K in place of MG (1 valid contact(s) for K vs. 0 for MG)  

### MG:1H:3159

|  | Res. type | Atom | Residue | Distance | Occupancy | B-factor | Closest solute to 1^st^ shell water |
| --- | --- | --- | --- | --- | --- | --- | --- |
| **Ion** | | | |  | 1.00 | 44.73 || 1 | G | O6 | 1H:1472 | 3.12 | 1.00 | 38.09 |
| 2 | G | O6 | 1H:1471 | 3.43 | 1.00 | 35.54 |  |
| 3 | G | N1 | 1H:1472 | 4.14 | 1.00 | 29.39 |  |
| 4 | G | O6 | 1H:1470 | 4.16 | 1.00 | 44.97 |  |
| 5 | A | OP2 | 1H:1620 | 4.23 | 1.00 | 27.07 |  |
| 6 | G | N7 | 1H:1471 | 4.37 | 1.00 | 36.09 |  |
| 7 | G | N7 | 1H:1470 | 4.43 | 1.00 | 47.32 |  |
| 8 | A | OP1 | 1H:1620 | 4.63 | 1.00 | 22.37 |  |
| 9 | C | N4 | 1H:1622 | 4.92 | 1.00 | 42.37 |  |
  |

  
 Coded contact list:   
 Exclusion zone contact(s): /O6/  
Consider K in place of MG (1 valid contact(s) for K vs. 0 for MG)  

### MG:1H:3160

|  | Res. type | Atom | Residue | Distance | Occupancy | B-factor | Closest solute to 1^st^ shell water |
| --- | --- | --- | --- | --- | --- | --- | --- |
| **Ion** | | | |  | 1.00 | 29.00 || 1 | G | O6 | 1H:1972 | 3.74 | 1.00 | 29.73 |
| 2 | G | O6 | 1H:1973 | 3.99 | 1.00 | 39.70 |  |
| 3 | U | OP2 | 1H:1978 | 4.13 | 1.00 | 17.43 |  |
| 4 | G | O6 | 1H:1971 | 4.29 | 1.00 | 30.02 |  |
| 5 | G | N7 | 1H:1971 | 4.61 | 1.00 | 24.88 |  |
| 6 | G | N7 | 1H:1972 | 4.63 | 1.00 | 30.20 |  |
| 7 | G | N1 | 1H:1973 | 4.79 | 1.00 | 39.42 |  |
| 8 | G | O2' | 1H:1977 | 4.91 | 1.00 | 23.43 |  |
  |

  
 Coded contact list:   
1^st^ MG coordination shell missing (see Notes)  

### MG:1H:3161

|  | Res. type | Atom | Residue | Distance | Occupancy | B-factor | Closest solute to 1^st^ shell water |
| --- | --- | --- | --- | --- | --- | --- | --- |
| **Ion** | | | |  | 1.00 | 55.17 || 1 | G | OP2 | 1H:1881 | 3.52 | 1.00 | 63.40 |
| 2 | U | O4 | 1H:1883 | 3.93 | 1.00 | 53.03 |  |
| 3 | G | OP2 | 1H:1882 | 4.12 | 1.00 | 60.28 |  |
| 4 | G | N7 | 1H:1881 | 4.50 | 1.00 | 55.32 |  |
| 5 | G | N7 | 1H:1882 | 4.76 | 1.00 | 51.55 |  |
| 6 | G | O5' | 1H:1881 | 4.84 | 1.00 | 47.09 |  |
  |

  
 Coded contact list:   
1^st^ MG coordination shell missing (see Notes)  

### MG:1H:3162

|  | Res. type | Atom | Residue | Distance | Occupancy | B-factor | Closest solute to 1^st^ shell water |
| --- | --- | --- | --- | --- | --- | --- | --- |
| **Ion** | | | |  | 1.00 | 58.12 || 1 | A | OP2 | 1H:1542 | 2.36 | 1.00 | 72.56 |
| 2 | A | OP2 | 1H:1541 | 3.50 | 1.00 | 73.05 |  |
| 3 | A | N1 | 1H:1543 | 3.73 | 1.00 | 74.12 |  |
| 4 | A | OP1 | 1H:1541 | 3.80 | 1.00 | 77.86 |  |
| 5 | A | N7 | 1H:1542 | 3.99 | 1.00 | 64.71 |  |
| 6 | A | O5' | 1H:1542 | 4.25 | 1.00 | 69.07 |  |
| 7 | A | O5' | 1H:1541 | 4.57 | 1.00 | 79.45 |  |
| 8 | A | O3' | 1H:1541 | 4.61 | 1.00 | 66.47 |  |
| 9 | A | OP1 | 1H:1542 | 4.92 | 1.00 | 68.14 |  |
  |

  
 Contact(s): /OP2/  
 Coded contact list: O_ph_  

### MG:1H:3163

|  | Res. type | Atom | Residue | Distance | Occupancy | B-factor | Closest solute to 1^st^ shell water |
| --- | --- | --- | --- | --- | --- | --- | --- |
| **Ion** | | | |  | 1.00 | 55.27 || 1 | G | O6 | 1H:1890 | 2.87 | 1.00 | 70.12 |
| 2 | G | N7 | 1H:1890 | 3.29 | 1.00 | 73.52 |  |
| 3 | G | N7 | 1H:1889 | 3.84 | 1.00 | 68.05 |  |
| 4 | G | OP2 | 1H:1889 | 4.38 | 1.00 | 55.83 |  |
| 5 | G | O6 | 1H:1889 | 4.46 | 1.00 | 71.96 |  |
| 6 | G | OP2 | 1H:1888 | 4.51 | 1.00 | 48.17 |  |
| 7 | G | O6 | 1H:1906 | 4.89 | 1.00 | 62.61 |  |
| 8 | G | OP1 | 1H:1888 | 4.92 | 1.00 | 46.78 |  |
| 9 | G | N1 | 1H:1890 | 4.93 | 1.00 | 68.97 |  |
  |

  
 Coded contact list:   
 Exclusion zone contact(s): /N7/O6/  
Consider K in place of MG (2 valid contact(s) for K vs. 0 for MG)  

### MG:1H:3164

|  | Res. type | Atom | Residue | Distance | Occupancy | B-factor | Closest solute to 1^st^ shell water |
| --- | --- | --- | --- | --- | --- | --- | --- |
| **Ion** | | | |  | 1.00 | 58.93 || 1 | G | OP2 | 1H:1919 | 3.79 | 1.00 | 51.51 |
| 2 | C | OP2 | 1H:1918 | 3.91 | 1.00 | 49.95 |  |
| 3 | G | N7 | 1H:1919 | 4.27 | 1.00 | 47.23 |  |
| 4 | C | O5' | 1H:1918 | 4.89 | 1.00 | 43.86 |  |
| 5 | G | OP2 | 1H:1871 | 4.94 | 1.00 | 40.08 |  |
| 6 | G | OP1 | 1H:1872 | 4.95 | 1.00 | 34.75 |  |
  |

  
 Coded contact list:   
1^st^ MG coordination shell missing (see Notes)  

### MG:1H:3165

|  | Res. type | Atom | Residue | Distance | Occupancy | B-factor | Closest solute to 1^st^ shell water |
| --- | --- | --- | --- | --- | --- | --- | --- |
| **Ion** | | | |  | 1.00 | 40.40 || 1 | U | OP1 | 1H:183 | 2.59 | 1.00 | 22.50 |
| 2 | U | O2 | 1H:851 | 3.82 | 1.00 | 18.06 |  |
| 3 | U | O2' | 1H:851 | 4.01 | 1.00 | 23.48 |  |
| 4 | A | N6 | 1H:724 | 4.11 | 1.00 | 16.58 |  |
| 5 | G | O2' | 1H:2092 | 4.18 | 1.00 | 16.71 |  |
| 6 | G | O3' | 1H:2092 | 4.21 | 1.00 | 27.34 |  |
| 7 | G | OP1 | 1H:2093 | 4.60 | 1.00 | 23.23 |  |
| 8 | U | O5' | 1H:183 | 4.62 | 1.00 | 20.80 |  |
| 9 | A | N6 | 1H:723 | 4.86 | 1.00 | 22.34 |  |
| 10 | U | OP2 | 1H:183 | 4.94 | 1.00 | 20.57 |  |
| 11 | C | O3' | 1H:182 | 4.96 | 1.00 | 15.16 |  |
| 12 | G | OP2 | 1H:184 | 4.98 | 1.00 | 19.80 |  |
  |

  
 Coded contact list:   
 Exclusion zone contact(s): /OP1/  
Consider NA in place of MG (1 valid contact(s) for NA vs. 0 for MG)  

### MG:1H:3166

|  | Res. type | Atom | Residue | Distance | Occupancy | B-factor | Closest solute to 1^st^ shell water |
| --- | --- | --- | --- | --- | --- | --- | --- |
| **Ion** | | | |  | 1.00 | 41.01 || 1 | Water 1 | | 1H:4485 | 2.37 | 1.00 | 53.50 | 3.63 (G:O6 1H:1436) |
| 2 | Water 2 | | 1H:4486 | 2.98 | 1.00 | 45.48 | 4.06 (G:N7 1H:1436) |
| 3 | U | O4 | 1H:1437 | 3.52 | 1.00 | 44.16 |  |
| 4 | G | N7 | 1H:1436 | 3.73 | 1.00 | 46.16 |  |
| 5 | Water 3 | | 1H:4573 | 4.03 | 1.00 | 55.13 | 6.46 (G:O6 1H:1436) |
| 6 | G | O6 | 1H:1436 | 4.24 | 1.00 | 52.43 |  |
| 7 | G | OP2 | 1H:1436 | 4.56 | 1.00 | 50.25 |  |
| 8 | Water 4 | | 1H:4364 | 4.61 | 1.00 | 42.31 | 3.36 (G:O6 1H:1436) |
  |

  
 Contact(s): /O_w_/  
 Coded contact list: O_w_  
 Exclusion zone contact(s): /O_w_/  

### MG:1H:3167

|  | Res. type | Atom | Residue | Distance | Occupancy | B-factor | Closest solute to 1^st^ shell water |
| --- | --- | --- | --- | --- | --- | --- | --- |
| **Ion** | | | |  | 1.00 | 20.65 || 1 | C | OP2 | 1H:2512 | 2.03 | 1.00 | 18.26 |
| 2 | C | OP2 | 1H:2511 | 2.20 | 1.00 | 22.83 |  |
| 3 | A | OP2 | 1H:2461 | 2.26 | 1.00 | 20.08 |  |
| 4 | G | O3' | 1H:2460 | 3.58 | 1.00 | 22.35 |  |
| 5 | A | O3' | 1H:2510 | 3.59 | 1.00 | 22.39 |  |
| 6 | A | OP1 | 1H:2461 | 4.10 | 1.00 | 25.13 |  |
| 7 | A | O2' | 1H:2510 | 4.26 | 1.00 | 21.26 |  |
| 8 | C | O3' | 1H:2511 | 4.32 | 1.00 | 19.87 |  |
| 9 | C | O5' | 1H:2511 | 4.32 | 1.00 | 13.13 |  |
| 10 | C | O5' | 1H:2512 | 4.36 | 1.00 | 17.43 |  |
| 11 | C | OP1 | 1H:2512 | 4.43 | 1.00 | 20.16 |  |
| 12 | G | O2' | 1H:2460 | 4.50 | 1.00 | 21.91 |  |
| 13 | C | OP1 | 1H:2511 | 4.62 | 1.00 | 18.77 |  |
| 14 | U | OP2 | 1H:2462 | 4.67 | 1.00 | 23.25 |  |
| 15 | A | O5' | 1H:2461 | 4.75 | 1.00 | 16.40 |  |
  |

  
**1^st^ shell ligand angles:**   
1-2: 76; 1-3: 84; 2-3: 79;   
  
 Contact(s): /OP2/OP2/OP2/  
 Coded contact list: 3O_ph_  

### MG:1H:3168

|  | Res. type | Atom | Residue | Distance | Occupancy | B-factor | Closest solute to 1^st^ shell water |
| --- | --- | --- | --- | --- | --- | --- | --- |
| **Ion** | | | |  | 1.00 | 15.12 || 1 | C | OP2 | 1H:188 | 2.27 | 1.00 | 17.01 |
| 2 | A | OP1 | 1H:187 | 2.31 | 1.00 | 16.05 |  |
| 3 | A | N7 | 1H:185 | 2.56 | 1.00 | 17.81 |  |
| 4 | G | OP2 | 1H:184 | 3.54 | 1.00 | 19.80 |  |
| 5 | A | OP2 | 1H:185 | 3.61 | 1.00 | 25.14 |  |
| 6 | G | N7 | 1H:184 | 3.68 | 1.00 | 17.96 |  |
| 7 | C | OP1 | 1H:188 | 3.97 | 1.00 | 20.69 |  |
| 8 | A | N6 | 1H:185 | 4.30 | 1.00 | 20.29 |  |
| 9 | G | O5' | 1H:184 | 4.34 | 1.00 | 20.73 |  |
| 10 | A | O5' | 1H:187 | 4.48 | 1.00 | 16.08 |  |
| 11 | A | O3' | 1H:186 | 4.53 | 1.00 | 13.95 |  |
| 12 | A | N9 | 1H:185 | 4.53 | 1.00 | 17.13 |  |
| 13 | A | O3' | 1H:187 | 4.70 | 1.00 | 15.24 |  |
| 14 | C | O5' | 1H:188 | 4.73 | 1.00 | 16.92 |  |
| 15 | G | N9 | 1H:184 | 4.74 | 1.00 | 20.57 |  |
| 16 | Water 1 | | 1H:4015 | 4.80 | 1.00 | 20.77 | 3.65 (C:OP1 1H:188) |
| 17 | A | OP2 | 1H:187 | 4.80 | 1.00 | 17.26 |  |
  |

  
**1^st^ shell ligand angles:**   
1-2: 72;   
  
 Contact(s): /OP1/OP2/  
 Coded contact list: 2O_ph_  
 Exclusion zone contact(s): /N7/  

### MG:1H:3169

|  | Res. type | Atom | Residue | Distance | Occupancy | B-factor | Closest solute to 1^st^ shell water |
| --- | --- | --- | --- | --- | --- | --- | --- |
| **Ion** | | | |  | 1.00 | 15.91 || 1 | G | OP2 | 1H:833 | 2.25 | 1.00 | 17.49 |
| 2 | G | OP2 | 1H:840 | 3.42 | 1.00 | 16.91 |  |
| 3 | C | O2' | 1H:2453 | 3.66 | 1.00 | 20.55 |  |
| 4 | A | O3' | 1H:832 | 4.18 | 1.00 | 16.04 |  |
| 5 | G | N7 | 1H:840 | 4.22 | 1.00 | 16.07 |  |
| 6 | G | OP1 | 1H:833 | 4.50 | 1.00 | 13.48 |  |
| 7 | G | O5' | 1H:833 | 4.68 | 1.00 | 15.06 |  |
| 8 | C | O2 | 1H:2453 | 4.79 | 1.00 | 19.95 |  |
| 9 | G | O5' | 1H:840 | 4.89 | 1.00 | 16.89 |  |
  |

  
 Contact(s): /OP2/  
 Coded contact list: O_ph_  

### MG:1H:3170

|  | Res. type | Atom | Residue | Distance | Occupancy | B-factor | Closest solute to 1^st^ shell water |
| --- | --- | --- | --- | --- | --- | --- | --- |
| **Ion** | | | |  | 1.00 | 18.52 || 1 | C | OP1 | 1H:2586 | 2.38 | 1.00 | 27.58 |
| 2 | C | OP2 | 1H:2588 | 2.46 | 1.00 | 25.07 |  |
| 3 | G | OP1 | 1H:2587 | 2.49 | 1.00 | 28.26 |  |
| 4 | C | OP1 | 1H:2520 | 3.04 | 1.00 | 35.58 |  |
| 5 | C | OP2 | 1H:2586 | 3.49 | 1.00 | 27.39 |  |
| 6 | C | O5' | 1H:2586 | 4.06 | 1.00 | 27.80 |  |
| 7 | C | OP1 | 1H:2588 | 4.30 | 1.00 | 19.30 |  |
| 8 | G | O3' | 1H:2587 | 4.47 | 1.00 | 24.00 |  |
| 9 | C | O3' | 1H:2586 | 4.60 | 1.00 | 32.52 |  |
| 10 | G | O5' | 1H:2587 | 4.74 | 1.00 | 26.10 |  |
| 11 | A | O3' | 1H:2585 | 4.77 | 1.00 | 19.72 |  |
| 12 | G | OP2 | 1H:2521 | 4.87 | 1.00 | 31.75 |  |
| 13 | A | O2' | 1H:2585 | 4.91 | 1.00 | 26.61 |  |
| 14 | C | O5' | 1H:2588 | 4.93 | 1.00 | 19.12 |  |
  |

  
 Contact(s): /OP1/  
 Coded contact list: O_ph_  
 Exclusion zone contact(s): /OP1/OP1/OP2/  
Consider NA in place of MG (2 valid contact(s) for NA vs. 1 for MG)  

### MG:1H:3171

|  | Res. type | Atom | Residue | Distance | Occupancy | B-factor | Closest solute to 1^st^ shell water |
| --- | --- | --- | --- | --- | --- | --- | --- |
| **Ion** | | | |  | 1.00 | 30.81 || 1 | G | OP1 | 1H:439 | 2.11 | 1.00 | 37.47 |
| 2 | G | OP1 | 1H:2420 | 2.34 | 1.00 | 42.92 |  |
| 3 | U | O3' | 1H:2419 | 2.94 | 1.00 | 42.10 |  |
| 4 | A | OP1 | 1H:440 | 3.59 | 1.00 | 30.30 |  |
| 5 | Water 1 | | 1H:3791 | 4.12 | 1.00 | 40.71 | 4.43 (U:O4 1H:2421) |
| 6 | G | OP2 | 1H:2420 | 4.28 | 1.00 | 38.60 |  |
| 7 | Water 2 | | 1H:3937 | 4.31 | 1.00 | 37.37 | 3.01 (G:O5' 1H:2420) |
| 8 | G | O3' | 1H:438 | 4.42 | 1.00 | 36.54 |  |
| 9 | G | O5' | 1H:2420 | 4.42 | 1.00 | 35.09 |  |
| 10 | G | O5' | 1H:439 | 4.46 | 1.00 | 29.66 |  |
| 11 | G | N7 | 1H:2420 | 4.49 | 1.00 | 38.13 |  |
| 12 | G | OP2 | 1H:439 | 4.51 | 1.00 | 36.29 |  |
| 13 | U | O4 | 1H:2421 | 4.66 | 1.00 | 38.99 |  |
  |

  
**1^st^ shell ligand angles:**   
1-2: 72;   
  
 Contact(s): /OP1/OP1/  
 Coded contact list: 2O_ph_  
 Exclusion zone contact(s): /O3'/  

### MG:1H:3172

|  | Res. type | Atom | Residue | Distance | Occupancy | B-factor | Closest solute to 1^st^ shell water |
| --- | --- | --- | --- | --- | --- | --- | --- |
| **Ion** | | | |  | 1.00 | 31.94 || 1 | C | OP2 | 1H:1060 | 2.47 | 1.00 | 43.97 |
| 2 | U | OP2 | 1H:1061 | 2.78 | 1.00 | 57.31 |  |
| 3 | C | OP1 | 1H:1060 | 3.89 | 1.00 | 56.19 |  |
| 4 | A | OP2 | 1H:1190 | 4.30 | 1.00 | 35.97 |  |
| 5 | U | O3' | 1H:1059 | 4.42 | 1.00 | 48.36 |  |
| 6 | A | OP1 | 1H:1189 | 4.48 | 1.00 | 41.08 |  |
| 7 | U | OP1 | 1H:1061 | 4.53 | 1.00 | 51.84 |  |
| 8 | C | O3' | 1H:1060 | 4.71 | 1.00 | 48.74 |  |
| 9 | C | O5' | 1H:1060 | 4.84 | 1.00 | 48.51 |  |
  |

  
 Coded contact list:   
 Exclusion zone contact(s): /OP2/OP2/  
Consider NA in place of MG (1 valid contact(s) for NA vs. 0 for MG)  

### MG:1H:3173

|  | Res. type | Atom | Residue | Distance | Occupancy | B-factor | Closest solute to 1^st^ shell water |
| --- | --- | --- | --- | --- | --- | --- | --- |
| **Ion** | | | |  | 1.00 | 28.18 || 1 | C | OP1 | 1H:785 | 3.48 | 1.00 | 23.51 |
| 2 | U | OP1 | 1H:1668 | 3.50 | 1.00 | 26.83 |  |
| 3 | G | OP1 | 1H:786 | 4.09 | 1.00 | 24.90 |  |
| 4 | G | O3' | 1H:1667 | 4.40 | 1.00 | 20.21 |  |
| 5 | C | O5' | 1H:785 | 4.51 | 1.00 | 17.66 |  |
| 6 | G | OP2 | 1H:786 | 4.62 | 1.00 | 18.56 |  |
| 7 | C | O3' | 1H:785 | 4.77 | 1.00 | 26.37 |  |
  |

  
 Coded contact list:   
1^st^ MG coordination shell missing (see Notes)  

### MG:1H:3174

|  | Res. type | Atom | Residue | Distance | Occupancy | B-factor | Closest solute to 1^st^ shell water |
| --- | --- | --- | --- | --- | --- | --- | --- |
| **Ion** | | | |  | 1.00 | 60.76 || 1 | G | N7 | 1H:366 | 3.53 | 1.00 | 42.10 |
| 2 | G | OP2 | 1H:366 | 3.88 | 1.00 | 40.55 |  |
| 3 | G | N7 | 1H:367 | 3.91 | 1.00 | 42.10 |  |
| 4 | G | O5' | 1H:366 | 4.84 | 1.00 | 42.91 |  |
| 5 | G | OP2 | 1H:367 | 4.97 | 1.00 | 46.65 |  |
  |

  
 Coded contact list:   
1^st^ MG coordination shell missing (see Notes)  

### MG:1H:3175

|  | Res. type | Atom | Residue | Distance | Occupancy | B-factor | Closest solute to 1^st^ shell water |
| --- | --- | --- | --- | --- | --- | --- | --- |
| **Ion** | | | |  | 1.00 | 31.11 || 1 | U | OP1 | 1H:2695 | 1.75 | 1.00 | 38.17 |
| 2 | A | OP1 | 1H:2735 | 1.76 | 1.00 | 37.33 |  |
| 3 | G | OP2 | 1H:2736 | 3.65 | 1.00 | 38.48 |  |
| 4 | A | OP2 | 1H:2735 | 3.76 | 1.00 | 45.49 |  |
| 5 | A | O5' | 1H:2735 | 3.86 | 1.00 | 32.85 |  |
| 6 | C | O2' | 1H:2694 | 3.92 | 1.00 | 36.14 |  |
| 7 | C | O3' | 1H:2694 | 3.93 | 1.00 | 29.79 |  |
| 8 | U | O5' | 1H:2695 | 4.13 | 1.00 | 32.56 |  |
| 9 | U | OP2 | 1H:2695 | 4.18 | 1.00 | 46.33 |  |
| 10 | U | O3' | 1H:2734 | 4.29 | 1.00 | 32.20 |  |
  |

  
**1^st^ shell ligand angles:**   
1-2: 168;   
  
 Coded contact list:   
 Exclusion zone contact(s): /OP1/OP1/  
1^st^ MG coordination shell missing (see Notes)  

### MG:1H:3176

|  | Res. type | Atom | Residue | Distance | Occupancy | B-factor | Closest solute to 1^st^ shell water |
| --- | --- | --- | --- | --- | --- | --- | --- |
| **Ion** | | | |  | 0.50 | 21.21 || 1 | G | OP2 | 1H:1318 | 1.93 | 1.00 | 34.09 |
| 2 | G | OP1 | 1H:1695 | 2.03 | 1.00 | 28.42 |  |
| 3 | C | OP1 | 1H:1696 | 2.72 | 1.00 | 32.04 |  |
| 4 | A | OP1 | 1H:1319 | 3.24 | 1.00 | 34.65 |  |
| 5 | C | O5' | 1H:1696 | 3.34 | 1.00 | 24.18 |  |
| 6 | A | OP2 | 1H:1319 | 3.54 | 1.00 | 26.16 |  |
| 7 | G | O5' | 1H:1695 | 3.74 | 1.00 | 28.29 |  |
| 8 | C | O3' | 1H:1317 | 3.89 | 1.00 | 26.45 |  |
| 9 | G | O5' | 1H:1318 | 4.09 | 1.00 | 29.85 |  |
| 10 | G | OP1 | 1H:1318 | 4.49 | 1.00 | 27.41 |  |
| 11 | C | O3' | 1H:1694 | 4.51 | 1.00 | 26.26 |  |
| 12 | G | OP2 | 1H:1695 | 4.53 | 1.00 | 26.21 |  |
| 13 | G | O3' | 1H:1695 | 4.54 | 1.00 | 28.09 |  |
| 14 | C | OP2 | 1H:1696 | 4.77 | 1.00 | 23.13 |  |
| 15 | Water 1 | | 1H:4224 | 4.83 | 1.00 | 34.02 | 2.92 (A:OP2 1H:1319) |
| 16 | G | O3' | 1H:1318 | 4.87 | 1.00 | 29.84 |  |
  |

  
**1^st^ shell ligand angles:**   
1-2: 142;   
  
 Contact(s): /OP1/OP2/  
 Coded contact list: 2O_ph_  
 Exclusion zone contact(s): /O5'/OP1/OP1/  
Consider K in place of MG (3 valid contact(s) for K vs. 2 for MG)  

### MG:1H:3176

|  | Res. type | Atom | Residue | Distance | Occupancy | B-factor | Closest solute to 1^st^ shell water |
| --- | --- | --- | --- | --- | --- | --- | --- |
| **Ion** | | | |  | 0.50 | 20.46 || 1 | G | OP1 | 1H:1695 | 2.00 | 1.00 | 28.42 |
| 2 | G | OP2 | 1H:1318 | 2.11 | 1.00 | 34.09 |  |
| 3 | C | OP1 | 1H:1696 | 2.27 | 1.00 | 32.04 |  |
| 4 | A | OP2 | 1H:1319 | 2.77 | 1.00 | 26.16 |  |
| 5 | A | OP1 | 1H:1319 | 3.26 | 1.00 | 34.65 |  |
| 6 | G | O5' | 1H:1318 | 3.38 | 1.00 | 29.85 |  |
| 7 | C | O3' | 1H:1317 | 3.59 | 1.00 | 26.45 |  |
| 8 | Water 1 | | 1H:4224 | 3.66 | 1.00 | 34.02 | 2.92 (A:OP2 1H:1319) |
| 9 | C | O5' | 1H:1696 | 3.89 | 1.00 | 24.18 |  |
| 10 | G | O3' | 1H:1318 | 4.06 | 1.00 | 29.84 |  |
| 11 | G | OP2 | 1H:1695 | 4.25 | 1.00 | 26.21 |  |
| 12 | G | O5' | 1H:1695 | 4.31 | 1.00 | 28.29 |  |
| 13 | C | OP2 | 1H:1696 | 4.50 | 1.00 | 23.13 |  |
| 14 | C | O3' | 1H:1694 | 4.51 | 1.00 | 26.26 |  |
| 15 | G | OP1 | 1H:1318 | 4.59 | 1.00 | 27.41 |  |
| 16 | G | O3' | 1H:1695 | 4.64 | 1.00 | 28.09 |  |
| 17 | A | O5' | 1H:1319 | 4.78 | 1.00 | 34.76 |  |
  |

  
**1^st^ shell ligand angles:**   
1-2: 131; 1-3: 78; 2-3: 103;   
  
 Contact(s): /OP1/OP1/OP2/  
 Coded contact list: 3O_ph_  
 Exclusion zone contact(s): /O5'/OP1/OP2/  

### MG:1H:3177

|  | Res. type | Atom | Residue | Distance | Occupancy | B-factor | Closest solute to 1^st^ shell water |
| --- | --- | --- | --- | --- | --- | --- | --- |
| **Ion** | | | |  | 1.00 | 29.20 || 1 | G | N7 | 1H:490 | 3.78 | 1.00 | 20.47 |
| 2 | G | OP1 | 1H:492 | 3.91 | 1.00 | 17.26 |  |
| 3 | G | O6 | 1H:490 | 4.26 | 1.00 | 28.68 |  |
| 4 | G | OP2 | 1H:492 | 4.75 | 1.00 | 17.94 |  |
  |

  
 Coded contact list:   
1^st^ MG coordination shell missing (see Notes)  

### MG:1H:3178

|  | Res. type | Atom | Residue | Distance | Occupancy | B-factor | Closest solute to 1^st^ shell water |
| --- | --- | --- | --- | --- | --- | --- | --- |
| **Ion** | | | |  | 1.00 | 17.25 || 1 | G | OP1 | 1H:853 | 2.18 | 1.00 | 18.82 |
| 2 | A | OP1 | 1H:852 | 2.31 | 1.00 | 27.50 |  |
| 3 | GLY | O | 78:44 | 3.88 | 1.00 | 21.21 |  |
| 4 | A | O5' | 1H:852 | 3.98 | 1.00 | 21.31 |  |
| 5 | GLY | N | 78:43 | 4.12 | 1.00 | 20.23 |  |
| 6 | G | O5' | 1H:853 | 4.13 | 1.00 | 17.99 |  |
| 7 | A | OP2 | 1H:852 | 4.38 | 1.00 | 20.17 |  |
| 8 | G | OP2 | 1H:853 | 4.43 | 1.00 | 18.17 |  |
| 9 | GLY | N | 78:44 | 4.60 | 1.00 | 21.07 |  |
| 10 | A | O3' | 1H:852 | 4.66 | 1.00 | 13.36 |  |
| 11 | U | O3' | 1H:851 | 4.84 | 1.00 | 20.38 |  |
  |

  
**1^st^ shell ligand angles:**   
1-2: 87;   
  
 Contact(s): /OP1/OP1/  
 Coded contact list: 2O_ph_  

### MG:1H:3179

|  | Res. type | Atom | Residue | Distance | Occupancy | B-factor | Closest solute to 1^st^ shell water |
| --- | --- | --- | --- | --- | --- | --- | --- |
| **Ion** | | | |  | 1.00 | 30.89 || 1 | G | O6 | 1H:1660 | 3.65 | 1.00 | 24.48 |
| 2 | G | N7 | 1H:1660 | 3.71 | 1.00 | 17.72 |  |
| 3 | C | OP2 | 1H:1664 | 3.85 | 1.00 | 29.87 |  |
| 4 | C | OP1 | 1H:1662 | 4.35 | 1.00 | 20.62 |  |
| 5 | A | OP1 | 1H:1657 | 4.57 | 1.00 | 23.92 |  |
| 6 | C | OP2 | 1H:1662 | 4.62 | 1.00 | 22.59 |  |
| 7 | C | O2' | 1H:1662 | 4.70 | 1.00 | 28.68 |  |
| 8 | Water 1 | | 1H:4257 | 4.79 | 1.00 | 26.28 | 2.98 (G:O6 1H:1660) |
| 9 | C | N4 | 1H:1659 | 4.87 | 1.00 | 23.62 |  |
  |

  
 Coded contact list:   
1^st^ MG coordination shell missing (see Notes)  

### MG:1H:3180

|  | Res. type | Atom | Residue | Distance | Occupancy | B-factor | Closest solute to 1^st^ shell water |
| --- | --- | --- | --- | --- | --- | --- | --- |
| **Ion** | | | |  | 1.00 | 43.95 || 1 | G | OP2 | 1H:1609 | 2.24 | 1.00 | 53.40 |
| 2 | G | OP1 | 1H:1608 | 2.30 | 1.00 | 49.49 |  |
| 3 | G | OP1 | 1H:1609 | 4.24 | 1.00 | 53.22 |  |
| 4 | G | O3' | 1H:1608 | 4.57 | 1.00 | 44.76 |  |
| 5 | G | O3' | 1H:1607 | 4.57 | 1.00 | 50.23 |  |
| 6 | G | OP2 | 1H:1608 | 4.60 | 1.00 | 55.51 |  |
| 7 | G | O5' | 1H:1608 | 4.61 | 1.00 | 49.56 |  |
| 8 | G | O5' | 1H:1609 | 4.65 | 1.00 | 39.98 |  |
| 9 | G | N7 | 1H:1609 | 4.96 | 1.00 | 43.97 |  |
  |

  
**1^st^ shell ligand angles:**   
1-2: 86;   
  
 Contact(s): /OP1/OP2/  
 Coded contact list: 2O_ph_  

### MG:1H:3181

|  | Res. type | Atom | Residue | Distance | Occupancy | B-factor | Closest solute to 1^st^ shell water |
| --- | --- | --- | --- | --- | --- | --- | --- |
| **Ion** | | | |  | 1.00 | 36.24 || 1 | G | N2 | 1H:1379 | 2.82 | 1.00 | 40.27 |
| 2 | G | N1 | 1H:1379 | 3.57 | 1.00 | 39.21 |  |
| 3 | Water 1 | | 1H:3714 | 3.84 | 1.00 | 41.28 | 2.23 (G:OP1 1H:1653) |
| 4 | G | OP1 | 1H:1653 | 3.86 | 1.00 | 35.64 |  |
| 5 | C | O3' | 1H:1652 | 4.07 | 1.00 | 25.05 |  |
| 6 | C | O2' | 1H:1652 | 4.43 | 1.00 | 27.52 |  |
| 7 | A | N7 | 1H:1657 | 4.68 | 1.00 | 29.50 |  |
| 8 | A | OP1 | 1H:1656 | 4.82 | 1.00 | 33.82 |  |
| 9 | A | OP2 | 1H:1657 | 4.88 | 1.00 | 27.61 |  |
  |

  
 Coded contact list:   
 Exclusion zone contact(s): /N2/  
Consider K in place of MG (1 valid contact(s) for K vs. 0 for MG)  
MG hydration shell issue: water too close (< 2.4 Angs.)  

### MG:1H:3182

|  | Res. type | Atom | Residue | Distance | Occupancy | B-factor | Closest solute to 1^st^ shell water |
| --- | --- | --- | --- | --- | --- | --- | --- |
| **Ion** | | | |  | 1.00 | 49.32 || 1 | A | OP1 | 1H:1497 | 2.42 | 1.00 | 78.80 |
| 2 | A | OP1 | 1H:1590 | 2.62 | 1.00 | 69.18 |  |
| 3 | A | O5' | 1H:1497 | 3.24 | 1.00 | 70.41 |  |
| 4 | A | OP2 | 1H:1590 | 3.85 | 1.00 | 80.12 |  |
| 5 | G | O3' | 1H:1496 | 4.24 | 1.00 | 69.29 |  |
| 6 | G | O3' | 1H:1589 | 4.25 | 1.00 | 75.47 |  |
| 7 | A | OP2 | 1H:1497 | 4.59 | 1.00 | 65.47 |  |
  |

  
 Coded contact list:   
 Exclusion zone contact(s): /O5'/OP1/OP1/  
Consider K in place of MG (2 valid contact(s) for K vs. 0 for MG)  

### MG:1H:3183

|  | Res. type | Atom | Residue | Distance | Occupancy | B-factor | Closest solute to 1^st^ shell water |
| --- | --- | --- | --- | --- | --- | --- | --- |
| **Ion** | | | |  | 1.00 | 47.98 || 1 | C | OP1 | 1H:2714 | 2.66 | 1.00 | 46.69 |
| 2 | U | O5' | 1H:2715 | 2.69 | 1.00 | 33.30 |  |
| 3 | U | OP2 | 1H:2715 | 3.52 | 1.00 | 47.49 |  |
| 4 | C | O5' | 1H:2714 | 3.59 | 1.00 | 34.38 |  |
| 5 | C | O3' | 1H:2714 | 3.66 | 1.00 | 40.26 |  |
| 6 | C | OP2 | 1H:2714 | 4.27 | 1.00 | 35.83 |  |
| 7 | U | OP1 | 1H:2715 | 4.84 | 1.00 | 46.87 |  |
| 8 | C | O3' | 1H:2713 | 4.93 | 1.00 | 41.38 |  |
  |

  
 Coded contact list:   
 Exclusion zone contact(s): /O5'/OP1/  
Consider K in place of MG (2 valid contact(s) for K vs. 0 for MG)  

### MG:1H:3184

|  | Res. type | Atom | Residue | Distance | Occupancy | B-factor | Closest solute to 1^st^ shell water |
| --- | --- | --- | --- | --- | --- | --- | --- |
| **Ion** | | | |  | 1.00 | 40.01 || 1 | A | OP2 | 1H:782 | 3.65 | 1.00 | 25.28 |
| 2 | G | O6 | 1H:808 | 3.83 | 1.00 | 21.61 |  |
| 3 | G | N7 | 1H:807 | 3.87 | 1.00 | 22.81 |  |
| 4 | G | O6 | 1H:807 | 3.93 | 1.00 | 22.72 |  |
| 5 | A | N6 | 1H:809 | 4.65 | 1.00 | 24.49 |  |
| 6 | G | N7 | 1H:808 | 4.72 | 1.00 | 20.53 |  |
  |

  
 Coded contact list:   
1^st^ MG coordination shell missing (see Notes)  

### MG:1H:3185

|  | Res. type | Atom | Residue | Distance | Occupancy | B-factor | Closest solute to 1^st^ shell water |
| --- | --- | --- | --- | --- | --- | --- | --- |
| **Ion** | | | |  | 1.00 | 40.73 || 1 | G | O6 | 1H:789 | 3.98 | 1.00 | 19.84 |
| 2 | C | N4 | 1H:803 | 4.18 | 1.00 | 19.71 |  |
| 3 | G | O6 | 1H:790 | 4.21 | 1.00 | 24.29 |  |
| 4 | C | N4 | 1H:804 | 4.64 | 1.00 | 19.24 |  |
| 5 | G | O2' | 1H:787 | 4.65 | 1.00 | 26.66 |  |
| 6 | G | N3 | 1H:787 | 4.75 | 1.00 | 22.93 |  |
| 7 | G | N2 | 1H:787 | 4.78 | 1.00 | 19.05 |  |
| 8 | U | O4 | 1H:788 | 4.82 | 1.00 | 19.15 |  |
| 9 | C | N4 | 1H:802 | 4.98 | 1.00 | 19.21 |  |
  |

  
 Coded contact list:   
1^st^ MG coordination shell missing (see Notes)  

### MG:1H:3186

|  | Res. type | Atom | Residue | Distance | Occupancy | B-factor | Closest solute to 1^st^ shell water |
| --- | --- | --- | --- | --- | --- | --- | --- |
| **Ion** | | | |  | 1.00 | 55.55 || 1 | G | O6 | 1H:1278 | 2.97 | 1.00 | 48.46 |
| 2 | G | N7 | 1H:1278 | 3.37 | 1.00 | 51.01 |  |
| 3 | G | O6 | 1H:1279 | 3.83 | 1.00 | 47.26 |  |
| 4 | G | N7 | 1H:1279 | 4.61 | 1.00 | 44.32 |  |
| 5 | G | N1 | 1H:1278 | 4.85 | 1.00 | 48.79 |  |
| 6 | C | N4 | 1H:1277 | 4.86 | 1.00 | 48.24 |  |
| 7 | C | N4 | 1H:1280 | 4.88 | 1.00 | 39.31 |  |
  |

  
 Coded contact list:   
 Exclusion zone contact(s): /N7/O6/  
Consider K in place of MG (2 valid contact(s) for K vs. 0 for MG)  

### MG:1H:3187

|  | Res. type | Atom | Residue | Distance | Occupancy | B-factor | Closest solute to 1^st^ shell water |
| --- | --- | --- | --- | --- | --- | --- | --- |
| **Ion** | | | |  | 1.00 | 30.77 || 1 | G | OP1 | 1H:238 | 2.58 | 1.00 | 16.59 |
| 2 | G | O2' | 1H:414 | 3.85 | 1.00 | 28.39 |  |
| 3 | U | OP1 | 1H:415 | 3.85 | 1.00 | 25.82 |  |
| 4 | G | O3' | 1H:414 | 4.03 | 1.00 | 23.41 |  |
| 5 | LYS | O | J8:32 | 4.33 | 1.00 | 34.16 |  |
| 6 | G | O3' | 1H:237 | 4.38 | 1.00 | 19.99 |  |
| 7 | G | OP2 | 1H:238 | 4.41 | 1.00 | 14.58 |  |
| 8 | G | N3 | 1H:414 | 4.78 | 1.00 | 20.81 |  |
  |

  
 Coded contact list:   
 Exclusion zone contact(s): /OP1/  
Consider NA in place of MG (1 valid contact(s) for NA vs. 0 for MG)  

### MG:1H:3188

|  | Res. type | Atom | Residue | Distance | Occupancy | B-factor | Closest solute to 1^st^ shell water |
| --- | --- | --- | --- | --- | --- | --- | --- |
| **Ion** | | | |  | 1.00 | 20.11 || 1 | U | OP1 | 1H:875 | 1.81 | 1.00 | 23.02 |
| 2 | A | OP2 | 1H:2443 | 1.98 | 1.00 | 28.39 |  |
| 3 | G | OP2 | 1H:2442 | 2.52 | 1.00 | 20.29 |  |
| 4 | U | O3' | 1H:874 | 3.27 | 1.00 | 21.12 |  |
| 5 | A | OP1 | 1H:2443 | 3.41 | 1.00 | 23.05 |  |
| 6 | A | O5' | 1H:2443 | 4.04 | 1.00 | 17.74 |  |
| 7 | U | OP2 | 1H:875 | 4.07 | 1.00 | 26.49 |  |
| 8 | U | O5' | 1H:875 | 4.15 | 1.00 | 21.76 |  |
| 9 | G | O3' | 1H:2442 | 4.50 | 1.00 | 25.04 |  |
| 10 | G | O5' | 1H:2442 | 4.54 | 1.00 | 18.50 |  |
| 11 | G | O3' | 1H:2441 | 4.62 | 1.00 | 21.53 |  |
| 12 | U | O4' | 1H:875 | 4.64 | 1.00 | 27.89 |  |
| 13 | G | OP1 | 1H:2442 | 4.89 | 1.00 | 19.80 |  |
  |

  
**1^st^ shell ligand angles:**   
1-2: 150;   
  
 Contact(s): /OP1/OP2/  
 Coded contact list: 2O_ph_  
 Exclusion zone contact(s): /O3'/OP2/  

### MG:1H:3189

|  | Res. type | Atom | Residue | Distance | Occupancy | B-factor | Closest solute to 1^st^ shell water |
| --- | --- | --- | --- | --- | --- | --- | --- |
| **Ion** | | | |  | 1.00 | 41.71 || 1 | C | OP2 | 1H:797 | 2.95 | 1.00 | 32.46 |
| 2 | G | OP2 | 1H:796 | 3.19 | 1.00 | 26.32 |  |
| 3 | A | OP2 | 1H:798 | 3.50 | 1.00 | 28.24 |  |
| 4 | A | OP1 | 1H:798 | 3.74 | 1.00 | 24.59 |  |
| 5 | G | O5' | 1H:796 | 3.79 | 1.00 | 25.13 |  |
| 6 | PRO | O | E8:87 | 4.21 | 1.00 | 28.65 |  |
| 7 | ALA | N | E8:89 | 4.62 | 1.00 | 31.03 |  |
| 8 | C | O5' | 1H:797 | 4.63 | 1.00 | 23.96 |  |
| 9 | ARG | NE | E8:88 | 4.72 | 1.00 | 30.01 |  |
| 10 | C | O3' | 1H:797 | 4.77 | 1.00 | 23.88 |  |
| 11 | G | O6 | 1H:1372 | 4.86 | 1.00 | 48.52 |  |
| 12 | G | OP1 | 1H:796 | 4.93 | 1.00 | 25.86 |  |
  |

  
 Coded contact list:   
 Exclusion zone contact(s): /OP2/OP2/  
Consider K in place of MG (2 valid contact(s) for K vs. 0 for MG)  

### MG:1H:3190

|  | Res. type | Atom | Residue | Distance | Occupancy | B-factor | Closest solute to 1^st^ shell water |
| --- | --- | --- | --- | --- | --- | --- | --- |
| **Ion** | | | |  | 1.00 | 32.41 || 1 | G | O6 | 1H:1670 | 3.66 | 1.00 | 29.77 |
| 2 | C | OP2 | 1H:1654 | 3.89 | 1.00 | 31.25 |  |
| 3 | G | O6 | 1H:1671 | 4.22 | 1.00 | 31.70 |  |
| 4 | C | N4 | 1H:1352 | 4.67 | 1.00 | 34.70 |  |
| 5 | G | N7 | 1H:1670 | 4.74 | 1.00 | 34.22 |  |
| 6 | G | O6 | 1H:1669 | 4.92 | 1.00 | 33.91 |  |
  |

  
 Coded contact list:   
1^st^ MG coordination shell missing (see Notes)  

### MG:1H:3191

|  | Res. type | Atom | Residue | Distance | Occupancy | B-factor | Closest solute to 1^st^ shell water |
| --- | --- | --- | --- | --- | --- | --- | --- |
| **Ion** | | | |  | 1.00 | 24.72 || 1 | C | OP2 | 1H:1362 | 1.94 | 1.00 | 27.71 |
| 2 | G | OP1 | 1H:1379 | 2.07 | 1.00 | 25.93 |  |
| 3 | C | OP1 | 1H:1361 | 2.34 | 1.00 | 30.94 |  |
| 4 | C | O5' | 1H:1361 | 3.32 | 1.00 | 26.02 |  |
| 5 | C | OP1 | 1H:1362 | 4.06 | 1.00 | 26.01 |  |
| 6 | C | O3' | 1H:1361 | 4.12 | 1.00 | 28.57 |  |
| 7 | G | O5' | 1H:1379 | 4.16 | 1.00 | 29.10 |  |
| 8 | G | OP2 | 1H:1379 | 4.40 | 1.00 | 24.49 |  |
| 9 | U | O3' | 1H:1360 | 4.45 | 1.00 | 26.56 |  |
| 10 | A | O3' | 1H:1378 | 4.45 | 1.00 | 27.07 |  |
| 11 | C | O5' | 1H:1362 | 4.48 | 1.00 | 25.74 |  |
| 12 | C | OP2 | 1H:1361 | 4.55 | 1.00 | 25.36 |  |
  |

  
**1^st^ shell ligand angles:**   
1-2: 164; 1-3: 95; 2-3: 83;   
  
 Contact(s): /OP1/OP1/OP2/  
 Coded contact list: 3O_ph_  
 Exclusion zone contact(s): /O5'/  

### MG:1H:3192

|  | Res. type | Atom | Residue | Distance | Occupancy | B-factor | Closest solute to 1^st^ shell water |
| --- | --- | --- | --- | --- | --- | --- | --- |
| **Ion** | | | |  | 1.00 | 37.96 || 1 | G | OP1 | 1H:2595 | 2.18 | 1.00 | 29.23 |
| 2 | Water 1 | | 1H:3675 | 3.26 | 1.00 | 24.68 | 2.15 (G:OP2 1H:2595) |
| 3 | G | OP2 | 1H:2621 | 3.31 | 1.00 | 17.03 |  |
| 4 | G | OP2 | 1H:2595 | 3.68 | 1.00 | 26.35 |  |
| 5 | G | OP1 | 1H:2621 | 4.06 | 1.00 | 15.51 |  |
| 6 | G | O5' | 1H:2621 | 4.12 | 1.00 | 15.72 |  |
| 7 | G | O5' | 1H:2595 | 4.35 | 1.00 | 20.82 |  |
| 8 | G | O3' | 1H:2594 | 4.46 | 1.00 | 23.56 |  |
| 9 | U | OP1 | 1H:2622 | 4.47 | 1.00 | 26.20 |  |
| 10 | Water 2 | | 1H:4205 | 4.85 | 1.00 | 42.21 | 4.16 (G:OP2 1H:2595) |
  |

  
 Contact(s): /OP1/  
 Coded contact list: O_ph_  
 Exclusion zone contact(s): /O_w_/OP2/  
MG hydration shell issue: water too close (< 2.4 Angs.)  

### MG:1H:3193

|  | Res. type | Atom | Residue | Distance | Occupancy | B-factor | Closest solute to 1^st^ shell water |
| --- | --- | --- | --- | --- | --- | --- | --- |
| **Ion** | | | |  | 1.00 | 40.69 || 1 | G | N7 | 1H:813 | 4.10 | 1.00 | 19.55 |
| 2 | G | OP2 | 1H:813 | 4.32 | 1.00 | 22.60 |  |
| 3 | G | N7 | 1H:811 | 4.79 | 1.00 | 22.97 |  |
| 4 | G | OP1 | 1H:813 | 4.80 | 1.00 | 23.50 |  |
  |

  
 Coded contact list:   
1^st^ MG coordination shell missing (see Notes)  

### MG:1H:3194

|  | Res. type | Atom | Residue | Distance | Occupancy | B-factor | Closest solute to 1^st^ shell water |
| --- | --- | --- | --- | --- | --- | --- | --- |
| **Ion** | | | |  | 1.00 | 14.28 || 1 | G | OP2 | 1H:1998 | 2.31 | 1.00 | 18.39 |
| 2 | C | OP1 | 1H:1803 | 3.36 | 1.00 | 16.43 |  |
| 3 | G | OP1 | 1H:1998 | 4.18 | 1.00 | 26.67 |  |
| 4 | C | OP2 | 1H:1803 | 4.40 | 1.00 | 17.72 |  |
| 5 | C | O3' | 1H:1997 | 4.50 | 1.00 | 20.11 |  |
| 6 | C | O5' | 1H:1997 | 4.59 | 1.00 | 20.14 |  |
| 7 | G | O5' | 1H:1998 | 4.74 | 1.00 | 17.25 |  |
| 8 | C | OP2 | 1H:1997 | 4.95 | 1.00 | 27.39 |  |
  |

  
 Contact(s): /OP2/  
 Coded contact list: O_ph_  
 Exclusion zone contact(s): /OP1/  

### MG:1H:3195

|  | Res. type | Atom | Residue | Distance | Occupancy | B-factor | Closest solute to 1^st^ shell water |
| --- | --- | --- | --- | --- | --- | --- | --- |
| **Ion** | | | |  | 1.00 | 37.41 || 1 | C | OP2 | 1H:1200 | 1.80 | 1.00 | 37.06 |
| 2 | U | OP2 | 1H:1046 | 1.92 | 1.00 | 39.06 |  |
| 3 | U | OP1 | 1H:1046 | 3.64 | 1.00 | 40.36 |  |
| 4 | C | O3' | 1H:1045 | 4.03 | 1.00 | 36.36 |  |
| 5 | C | OP1 | 1H:1200 | 4.03 | 1.00 | 32.41 |  |
| 6 | C | O5' | 1H:1200 | 4.05 | 1.00 | 34.61 |  |
| 7 | C | O3' | 1H:1199 | 4.25 | 1.00 | 37.84 |  |
| 8 | C | OP1 | 1H:1045 | 4.34 | 1.00 | 37.42 |  |
| 9 | U | O5' | 1H:1046 | 4.43 | 1.00 | 33.44 |  |
| 10 | G | O6 | 1H:1201 | 4.48 | 1.00 | 40.96 |  |
| 11 | C | OP2 | 1H:1045 | 4.51 | 1.00 | 33.97 |  |
| 12 | C | O5' | 1H:1045 | 4.60 | 1.00 | 31.64 |  |
| 13 | G | N7 | 1H:1201 | 4.76 | 1.00 | 37.47 |  |
  |

  
**1^st^ shell ligand angles:**   
1-2: 171;   
  
 Contact(s): /OP2/OP2/  
 Coded contact list: 2O_ph_  

### MG:1H:3196

|  | Res. type | Atom | Residue | Distance | Occupancy | B-factor | Closest solute to 1^st^ shell water |
| --- | --- | --- | --- | --- | --- | --- | --- |
| **Ion** | | | |  | 1.00 | 40.03 || 1 | C | OP1 | 1H:1045 | 3.51 | 1.00 | 37.42 |
| 2 | G | O3' | 1H:1044 | 3.84 | 1.00 | 30.16 |  |
| 3 | LYS | NZ | C8:93 | 4.03 | 1.00 | 28.38 |  |
| 4 | ILE | O | C8:88 | 4.41 | 1.00 | 50.14 |  |
  |

  
 Coded contact list:   
1^st^ MG coordination shell missing (see Notes)  

### MG:1H:3197

|  | Res. type | Atom | Residue | Distance | Occupancy | B-factor | Closest solute to 1^st^ shell water |
| --- | --- | --- | --- | --- | --- | --- | --- |
| **Ion** | | | |  | 1.00 | 30.64 || 1 | A | N3 | 1H:1737 | 3.37 | 1.00 | 36.61 |
| 2 | G | OP1 | 13:1475 | 3.52 | 1.00 | 45.84 |  |
| 3 | A | O2' | 1H:1737 | 3.69 | 1.00 | 37.86 |  |
| 4 | A | O4' | 1H:1738 | 3.77 | 1.00 | 30.18 |  |
| 5 | A | N9 | 1H:1738 | 4.70 | 1.00 | 32.31 |  |
| 6 | A | N9 | 1H:1737 | 4.72 | 1.00 | 36.88 |  |
  |

  
 Coded contact list:   
 Exclusion zone contact(s): /N3_A_/  
Consider K in place of MG (1 valid contact(s) for K vs. 0 for MG)  

### MG:1H:3198

|  | Res. type | Atom | Residue | Distance | Occupancy | B-factor | Closest solute to 1^st^ shell water |
| --- | --- | --- | --- | --- | --- | --- | --- |
| **Ion** | | | |  | 1.00 | 20.55 || 1 | U | O4 | 1H:1719 | 1.94 | 1.00 | 26.02 |
| 2 | G | OP1 | 1H:2563 | 2.35 | 1.00 | 26.91 |  |
| 3 | A | OP2 | 1H:1717 | 2.53 | 1.00 | 20.85 |  |
| 4 | C | OP2 | 1H:1718 | 2.64 | 1.00 | 21.37 |  |
| 5 | Water 1 | | 1H:4337 | 3.67 | 1.00 | 27.49 | 3.15 (A:OP1 1H:1717) |
| 6 | C | OP2 | 1H:2564 | 3.71 | 1.00 | 26.83 |  |
| 7 | G | OP2 | 1H:2563 | 3.88 | 1.00 | 27.73 |  |
| 8 | U | N3 | 1H:1719 | 4.03 | 1.00 | 22.37 |  |
| 9 | G | O5' | 1H:2563 | 4.14 | 1.00 | 25.87 |  |
| 10 | C | OP1 | 1H:1718 | 4.29 | 1.00 | 27.86 |  |
| 11 | A | OP1 | 1H:1717 | 4.70 | 1.00 | 20.71 |  |
| 12 | G | O3' | 1H:2562 | 4.74 | 1.00 | 26.30 |  |
| 13 | A | O5' | 1H:1717 | 4.80 | 1.00 | 20.72 |  |
| 14 | A | O3' | 1H:1717 | 4.88 | 1.00 | 22.18 |  |
| 15 | A | O3' | 1H:1716 | 4.88 | 1.00 | 18.64 |  |
| 16 | C | O5' | 1H:1718 | 4.88 | 1.00 | 24.37 |  |
| 17 | C | OP1 | 1H:2564 | 4.89 | 1.00 | 27.53 |  |
  |

  
**1^st^ shell ligand angles:**   
1-2: 139;   
  
 Contact(s): /O4/OP1/  
 Coded contact list: O_ph_O_b_  
 Exclusion zone contact(s): /OP2/OP2/  

### MG:1H:3199

|  | Res. type | Atom | Residue | Distance | Occupancy | B-factor | Closest solute to 1^st^ shell water |
| --- | --- | --- | --- | --- | --- | --- | --- |
| **Ion** | | | |  | 1.00 | 34.81 || 1 | U | OP1 | 1H:1399 | 2.36 | 1.00 | 35.11 |
| 2 | Water 1 | | 1H:4056 | 3.92 | 1.00 | 34.54 | 2.70 (C:OP2 1H:1422) |
| 3 | U | OP2 | 1H:1399 | 4.14 | 1.00 | 23.94 |  |
| 4 | C | O3' | 1H:1398 | 4.22 | 1.00 | 26.18 |  |
| 5 | A | O2' | 1H:1619 | 4.41 | 1.00 | 26.55 |  |
| 6 | C | OP2 | 1H:1422 | 4.55 | 1.00 | 35.58 |  |
| 7 | C | OP1 | 1H:1422 | 4.56 | 1.00 | 41.98 |  |
| 8 | U | O5' | 1H:1399 | 4.83 | 1.00 | 29.00 |  |
  |

  
 Contact(s): /OP1/  
 Coded contact list: O_ph_  

### MG:1H:3200

|  | Res. type | Atom | Residue | Distance | Occupancy | B-factor | Closest solute to 1^st^ shell water |
| --- | --- | --- | --- | --- | --- | --- | --- |
| **Ion** | | | |  | 1.00 | 36.98 || 1 | C | OP1 | 1H:1658 | 2.39 | 1.00 | 21.31 |
| 2 | C | O5' | 1H:1658 | 3.39 | 1.00 | 23.82 |  |
| 3 | G | O2' | 1H:1357 | 3.59 | 1.00 | 23.01 |  |
| 4 | C | OP2 | 1H:1659 | 3.63 | 1.00 | 19.05 |  |
| 5 | G | O3' | 1H:1357 | 4.13 | 1.00 | 25.10 |  |
| 6 | C | OP1 | 1H:1659 | 4.47 | 1.00 | 18.06 |  |
| 7 | C | OP2 | 1H:1658 | 4.57 | 1.00 | 22.77 |  |
| 8 | A | O3' | 1H:1657 | 4.62 | 1.00 | 27.95 |  |
| 9 | C | O3' | 1H:1658 | 4.69 | 1.00 | 22.16 |  |
| 10 | G | OP1 | 1H:1358 | 4.89 | 1.00 | 29.01 |  |
  |

  
 Contact(s): /OP1/  
 Coded contact list: O_ph_  
 Exclusion zone contact(s): /O5'/  

### MG:1H:3201

|  | Res. type | Atom | Residue | Distance | Occupancy | B-factor | Closest solute to 1^st^ shell water |
| --- | --- | --- | --- | --- | --- | --- | --- |
| **Ion** | | | |  | 1.00 | 45.84 || 1 | G | O6 | 1H:2235 | 3.23 | 1.00 | 39.55 |
| 2 | G | N7 | 1H:2235 | 3.80 | 1.00 | 37.40 |  |
| 3 | G | N7 | 1H:2236 | 4.02 | 1.00 | 39.33 |  |
| 4 | G | O6 | 1H:2237 | 4.05 | 1.00 | 48.09 |  |
| 5 | G | O6 | 1H:2236 | 4.10 | 1.00 | 36.23 |  |
| 6 | G | N1 | 1H:2237 | 4.24 | 1.00 | 44.75 |  |
| 7 | U | O2' | 1H:2220 | 4.56 | 1.00 | 43.76 |  |
| 8 | G | O6 | 1H:2234 | 4.80 | 1.00 | 44.25 |  |
  |

  
 Coded contact list:   
 Exclusion zone contact(s): /O6/  
Consider K in place of MG (1 valid contact(s) for K vs. 0 for MG)  

### MG:1H:3202

|  | Res. type | Atom | Residue | Distance | Occupancy | B-factor | Closest solute to 1^st^ shell water |
| --- | --- | --- | --- | --- | --- | --- | --- |
| **Ion** | | | |  | 1.00 | 45.21 || 1 | G | O6 | 1H:2638 | 3.26 | 1.00 | 24.88 |
| 2 | G | O6 | 1H:2637 | 3.36 | 1.00 | 19.74 |  |
| 3 | G | N7 | 1H:2637 | 3.44 | 1.00 | 23.82 |  |
| 4 | G | N7 | 1H:2638 | 4.17 | 1.00 | 23.96 |  |
| 5 | C | N4 | 1H:2066 | 4.64 | 1.00 | 20.95 |  |
| 6 | G | N7 | 1H:2636 | 4.72 | 1.00 | 22.64 |  |
| 7 | G | O6 | 1H:2636 | 4.91 | 1.00 | 19.93 |  |
| 8 | G | O6 | 1H:2791 | 4.95 | 1.00 | 26.81 |  |
  |

  
 Coded contact list:   
 Exclusion zone contact(s): /O6/O6/  
Consider K in place of MG (2 valid contact(s) for K vs. 0 for MG)  

### MG:1H:3203

|  | Res. type | Atom | Residue | Distance | Occupancy | B-factor | Closest solute to 1^st^ shell water |
| --- | --- | --- | --- | --- | --- | --- | --- |
| **Ion** | | | |  | 1.00 | 26.98 || 1 | U | OP1 | 1H:2045 | 2.14 | 1.00 | 38.62 |
| 2 | G | O6 | 1H:556 | 2.53 | 1.00 | 37.15 |  |
| 3 | G | N1 | 1H:556 | 2.79 | 1.00 | 34.84 |  |
| 4 | U | OP2 | 1H:2045 | 4.19 | 1.00 | 27.79 |  |
| 5 | G | N2 | 1H:556 | 4.39 | 1.00 | 45.97 |  |
| 6 | C | O3' | 1H:2044 | 4.48 | 1.00 | 31.60 |  |
| 7 | U | O5' | 1H:2045 | 4.56 | 1.00 | 32.94 |  |
| 8 | G | OP2 | 1H:2046 | 4.86 | 1.00 | 29.77 |  |
| 9 | A | N7 | 1H:555 | 4.93 | 1.00 | 39.22 |  |
  |

  
 Contact(s): /OP1/  
 Coded contact list: O_ph_  
 Exclusion zone contact(s): /N1_G_/O6/  

### MG:1H:3204

|  | Res. type | Atom | Residue | Distance | Occupancy | B-factor | Closest solute to 1^st^ shell water |
| --- | --- | --- | --- | --- | --- | --- | --- |
| **Ion** | | | |  | 1.00 | 29.06 || 1 | G | OP1 | 1H:807 | 2.50 | 1.00 | 31.95 |
| 2 | G | OP2 | 1H:781 | 3.69 | 1.00 | 25.37 |  |
| 3 | G | OP2 | 1H:808 | 3.80 | 1.00 | 17.18 |  |
| 4 | C | OP1 | 1H:780 | 3.82 | 1.00 | 27.62 |  |
| 5 | G | OP2 | 1H:807 | 4.26 | 1.00 | 24.52 |  |
| 6 | G | OP1 | 1H:781 | 4.35 | 1.00 | 32.15 |  |
| 7 | G | O5' | 1H:807 | 4.54 | 1.00 | 19.47 |  |
  |

  
 Coded contact list:   
 Exclusion zone contact(s): /OP1/  
Consider NA in place of MG (1 valid contact(s) for NA vs. 0 for MG)  

### MG:1H:3205

|  | Res. type | Atom | Residue | Distance | Occupancy | B-factor | Closest solute to 1^st^ shell water |
| --- | --- | --- | --- | --- | --- | --- | --- |
| **Ion** | | | |  | 1.00 | 33.34 || 1 | G | OP2 | 1H:2733 | 2.37 | 1.00 | 38.32 |
| 2 | U | OP2 | 1H:2700 | 3.50 | 1.00 | 34.42 |  |
| 3 | G | O5' | 1H:2733 | 4.57 | 1.00 | 36.63 |  |
| 4 | G | O3' | 1H:2732 | 4.72 | 1.00 | 35.61 |  |
| 5 | G | OP1 | 1H:2733 | 4.79 | 1.00 | 35.42 |  |
| 6 | G | OP2 | 1H:2699 | 4.97 | 1.00 | 37.15 |  |
  |

  
 Contact(s): /OP2/  
 Coded contact list: O_ph_  

### MG:1H:3206

|  | Res. type | Atom | Residue | Distance | Occupancy | B-factor | Closest solute to 1^st^ shell water |
| --- | --- | --- | --- | --- | --- | --- | --- |
| **Ion** | | | |  | 1.00 | 25.11 || 1 | G | OP2 | 1H:2833 | 2.14 | 1.00 | 22.92 |
| 2 | A | OP2 | 1H:2832 | 2.17 | 1.00 | 30.68 |  |
| 3 | HIS | NE2 | 98:3 | 2.28 | 1.00 | 22.02 |  |
| 4 | GLY | O | 21:110 | 2.61 | 1.00 | 27.72 |  |
| 5 | A | O3' | 1H:2831 | 3.41 | 1.00 | 26.34 |  |
| 6 | G | N7 | 1H:2833 | 3.50 | 1.00 | 27.03 |  |
| 7 | A | O5' | 1H:2832 | 3.85 | 1.00 | 20.06 |  |
| 8 | A | O3' | 1H:2832 | 4.19 | 1.00 | 30.00 |  |
| 9 | G | O5' | 1H:2833 | 4.28 | 1.00 | 21.78 |  |
| 10 | HIS | ND1 | 98:3 | 4.42 | 1.00 | 25.58 |  |
| 11 | GLY | N | 21:110 | 4.63 | 1.00 | 32.16 |  |
| 12 | A | OP1 | 1H:2832 | 4.66 | 1.00 | 26.56 |  |
| 13 | G | OP1 | 1H:2833 | 4.71 | 1.00 | 22.00 |  |
| 14 | G | O6 | 1H:2833 | 4.80 | 1.00 | 33.50 |  |
| 15 | ARG | N | 21:111 | 4.80 | 1.00 | 30.37 |  |
  |

  
**1^st^ shell ligand angles:**   
1-2: 82; 1-3: 138; 2-3: 79;   
  
 Contact(s): /NE2_His_/OP2/OP2/  
 Coded contact list: 2O_ph_His  
 Exclusion zone contact(s): /O_back_/  

### MG:1H:3207

|  | Res. type | Atom | Residue | Distance | Occupancy | B-factor | Closest solute to 1^st^ shell water |
| --- | --- | --- | --- | --- | --- | --- | --- |
| **Ion** | | | |  | 1.00 | 38.06 || 1 | G | O2' | 1H:878 | 3.05 | 1.00 | 30.93 |
| 2 | G | O4' | 1H:879 | 3.12 | 1.00 | 21.98 |  |
| 3 | A | OP1 | 1H:877 | 3.72 | 1.00 | 23.12 |  |
| 4 | G | O2' | 1H:853 | 3.80 | 1.00 | 26.93 |  |
| 5 | G | O3' | 1H:853 | 4.18 | 1.00 | 14.41 |  |
| 6 | G | O5' | 1H:879 | 4.29 | 1.00 | 19.41 |  |
| 7 | U | OP2 | 1H:2091 | 4.29 | 1.00 | 24.93 |  |
| 8 | G | O3' | 1H:878 | 4.56 | 1.00 | 15.34 |  |
| 9 | C | OP1 | 1H:854 | 4.59 | 1.00 | 19.24 |  |
| 10 | G | N3 | 1H:853 | 4.66 | 1.00 | 25.02 |  |
| 11 | G | N9 | 1H:879 | 4.76 | 1.00 | 17.00 |  |
  |

  
 Coded contact list:   
 Exclusion zone contact(s): /O2'/O4'/  
Consider K in place of MG (2 valid contact(s) for K vs. 0 for MG)  

### MG:1H:3208

|  | Res. type | Atom | Residue | Distance | Occupancy | B-factor | Closest solute to 1^st^ shell water |
| --- | --- | --- | --- | --- | --- | --- | --- |
| **Ion** | | | |  | 1.00 | 37.30 || 1 | C | OP2 | 1H:2276 | 2.58 | 1.00 | 31.63 |
| 2 | U | OP2 | 1H:2275 | 2.68 | 1.00 | 26.45 |  |
| 3 | Water 1 | | 1H:4196 | 2.89 | 1.00 | 41.60 | 2.89 (C:OP2 1H:2276) |
| 4 | SER | OG | I8:16 | 2.95 | 1.00 | 30.59 |  |
| 5 | Water 2 | | I8:202 | 3.19 | 1.00 | 29.44 | 3.69 (GLN:O I8:17) |
| 6 | GLN | O | I8:17 | 3.21 | 1.00 | 33.79 |  |
| 7 | U | O5' | 1H:2275 | 3.65 | 1.00 | 26.17 |  |
| 8 | U | OP1 | 1H:2275 | 4.10 | 1.00 | 25.13 |  |
| 9 | Water 3 | | I8:201 | 4.14 | 1.00 | 36.25 | 3.73 (C:OP2 1H:2276) |
| 10 | C | OP1 | 1H:2276 | 4.48 | 1.00 | 22.19 |  |
| 11 | U | O3' | 1H:2275 | 4.65 | 1.00 | 22.03 |  |
| 12 | LYS | N | I8:19 | 4.68 | 1.00 | 29.77 |  |
| 13 | GLN | N | I8:17 | 4.89 | 1.00 | 24.90 |  |
| 14 | ALA | N | I8:18 | 4.95 | 1.00 | 26.81 |  |
  |

  
 Coded contact list:   
 Exclusion zone contact(s): /O_back_/O_w_/O_w_/OG_Ser_/OP2/OP2/  
Consider K in place of MG (3 valid contact(s) for K vs. 0 for MG)  

### MG:1H:3209

|  | Res. type | Atom | Residue | Distance | Occupancy | B-factor | Closest solute to 1^st^ shell water |
| --- | --- | --- | --- | --- | --- | --- | --- |
| **Ion** | | | |  | 1.00 | 52.39 || 1 | G | O6 | 1H:677 | 3.51 | 1.00 | 67.05 |
| 2 | G | N7 | 1H:677 | 3.98 | 1.00 | 58.50 |  |
| 3 | C | N4 | 1H:676 | 4.09 | 1.00 | 51.29 |  |
| 4 | G | OP2 | 1H:675 | 4.57 | 1.00 | 42.59 |  |
  |

  
 Coded contact list:   
1^st^ MG coordination shell missing (see Notes)  

### MG:1H:3210

|  | Res. type | Atom | Residue | Distance | Occupancy | B-factor | Closest solute to 1^st^ shell water |
| --- | --- | --- | --- | --- | --- | --- | --- |
| **Ion** | | | |  | 1.00 | 38.45 || 1 | G | O6 | 1H:1790 | 3.35 | 1.00 | 51.26 |
| 2 | U | OP1 | 1H:2709 | 3.68 | 1.00 | 40.08 |  |
| 3 | G | N1 | 1H:1790 | 4.24 | 1.00 | 52.40 |  |
| 4 | C | O4' | 1H:1756 | 4.33 | 1.00 | 42.82 |  |
| 5 | G | O2' | 1H:1755 | 4.33 | 1.00 | 38.31 |  |
| 6 | G | N2 | 1H:1755 | 4.33 | 1.00 | 45.02 |  |
| 7 | G | N3 | 1H:1755 | 4.45 | 1.00 | 48.18 |  |
| 8 | C | O3' | 1H:2708 | 4.74 | 1.00 | 42.57 |  |
| 9 | G | N2 | 1H:1788 | 4.76 | 1.00 | 45.06 |  |
  |

  
 Coded contact list:   
 Exclusion zone contact(s): /O6/  
Consider K in place of MG (1 valid contact(s) for K vs. 0 for MG)  

### MG:1H:3211

|  | Res. type | Atom | Residue | Distance | Occupancy | B-factor | Closest solute to 1^st^ shell water |
| --- | --- | --- | --- | --- | --- | --- | --- |
| **Ion** | | | |  | 1.00 | 56.95 || 1 | G | O6 | 1H:2237 | 3.15 | 1.00 | 48.09 |
| 2 | G | OP2 | 1H:2235 | 3.50 | 1.00 | 44.04 |  |
| 3 | G | OP2 | 1H:2236 | 4.13 | 1.00 | 33.57 |  |
| 4 | U | OP2 | 1H:2220 | 4.27 | 1.00 | 53.24 |  |
| 5 | G | N7 | 1H:2235 | 4.75 | 1.00 | 37.40 |  |
| 6 | G | O5' | 1H:2235 | 4.80 | 1.00 | 45.94 |  |
| 7 | G | N7 | 1H:2237 | 4.81 | 1.00 | 36.92 |  |
  |

  
 Coded contact list:   
 Exclusion zone contact(s): /O6/  
Consider K in place of MG (1 valid contact(s) for K vs. 0 for MG)  

### MG:1H:3212

|  | Res. type | Atom | Residue | Distance | Occupancy | B-factor | Closest solute to 1^st^ shell water |
| --- | --- | --- | --- | --- | --- | --- | --- |
| **Ion** | | | |  | 1.00 | 44.46 || 1 | G | O6 | 1H:2471 | 2.79 | 1.00 | 29.83 |
| 2 | G | N7 | 1H:2471 | 3.63 | 1.00 | 35.56 |  |
| 3 | U | OP2 | 1H:2470 | 3.78 | 1.00 | 31.57 |  |
| 4 | G | O2' | 1H:2503 | 3.88 | 1.00 | 36.93 |  |
| 5 | G | N3 | 1H:2503 | 3.92 | 1.00 | 29.03 |  |
| 6 | G | N2 | 1H:2503 | 4.03 | 1.00 | 49.96 |  |
| 7 | G | N1 | 1H:2503 | 4.63 | 1.00 | 35.70 |  |
| 8 | C | OP2 | 1H:2469 | 4.70 | 1.00 | 26.25 |  |
| 9 | G | N1 | 1H:2471 | 4.94 | 1.00 | 29.45 |  |
  |

  
 Coded contact list:   
 Exclusion zone contact(s): /O6/  
Consider K in place of MG (1 valid contact(s) for K vs. 0 for MG)  

### MG:1H:3213

|  | Res. type | Atom | Residue | Distance | Occupancy | B-factor | Closest solute to 1^st^ shell water |
| --- | --- | --- | --- | --- | --- | --- | --- |
| **Ion** | | | |  | 1.00 | 25.72 || 1 | U | O2' | 1H:1811 | 3.35 | 1.00 | 18.50 |
| 2 | C | OP2 | 1H:1814 | 3.55 | 1.00 | 14.13 |  |
| 3 | A | OP2 | 1H:1812 | 3.85 | 1.00 | 16.90 |  |
| 4 | U | O3' | 1H:1811 | 4.24 | 1.00 | 17.62 |  |
| 5 | Water 1 | | 1H:3732 | 4.70 | 1.00 | 29.47 | 3.08 (U:O2' 1H:1811) |
| 6 | C | O2' | 1H:1813 | 4.87 | 1.00 | 25.24 |  |
| 7 | A | N6 | 1H:1815 | 4.96 | 1.00 | 11.47 |  |
  |

  
 Coded contact list:   
 Exclusion zone contact(s): /O2'/  
Consider K in place of MG (1 valid contact(s) for K vs. 0 for MG)  

### MG:1H:3214

|  | Res. type | Atom | Residue | Distance | Occupancy | B-factor | Closest solute to 1^st^ shell water |
| --- | --- | --- | --- | --- | --- | --- | --- |
| **Ion** | | | |  | 1.00 | 41.74 || 1 | ASN | ND2 | 98:11 | 3.57 | 1.00 | 35.25 |
| 2 | G | O6 | 1H:1699 | 4.06 | 1.00 | 28.53 |  |
| 3 | G | N7 | 1H:1699 | 4.07 | 1.00 | 30.39 |  |
| 4 | G | OP2 | 1H:2027 | 4.37 | 1.00 | 28.46 |  |
| 5 | G | O6 | 1H:1698 | 4.46 | 1.00 | 31.00 |  |
| 6 | G | OP1 | 1H:2027 | 4.52 | 1.00 | 30.66 |  |
| 7 | G | N7 | 1H:1698 | 4.63 | 1.00 | 28.98 |  |
| 8 | A | N6 | 1H:1700 | 4.80 | 1.00 | 29.86 |  |
| 9 | ASN | OD1 | 98:11 | 4.82 | 1.00 | 32.98 |  |
  |

  
 Coded contact list:   
1^st^ MG coordination shell missing (see Notes)  

### MG:1H:3215

|  | Res. type | Atom | Residue | Distance | Occupancy | B-factor | Closest solute to 1^st^ shell water |
| --- | --- | --- | --- | --- | --- | --- | --- |
| **Ion** | | | |  | 1.00 | 40.71 || 1 | G | OP2 | 1H:2796 | 1.80 | 1.00 | 41.61 |
| 2 | G | N7 | 1H:2796 | 3.66 | 1.00 | 39.97 |  |
| 3 | G | O5' | 1H:2796 | 3.76 | 1.00 | 39.54 |  |
| 4 | G | OP1 | 1H:2796 | 4.17 | 1.00 | 36.41 |  |
| 5 | A | O3' | 1H:2795 | 4.25 | 1.00 | 41.46 |  |
| 6 | G | N7 | 1H:2797 | 4.29 | 1.00 | 46.74 |  |
| 7 | A | O2' | 1H:2790 | 4.64 | 1.00 | 41.49 |  |
| 8 | G | OP2 | 1H:2797 | 4.78 | 1.00 | 45.61 |  |
| 9 | G | O6 | 1H:2796 | 4.93 | 1.00 | 40.11 |  |
| 10 | G | O6 | 1H:2797 | 4.95 | 1.00 | 54.00 |  |
  |

  
 Contact(s): /OP2/  
 Coded contact list: O_ph_  

### MG:1H:3216

|  | Res. type | Atom | Residue | Distance | Occupancy | B-factor | Closest solute to 1^st^ shell water |
| --- | --- | --- | --- | --- | --- | --- | --- |
| **Ion** | | | |  | 1.00 | 34.63 || 1 | U | O4 | 1H:1676 | 3.45 | 1.00 | 37.11 |
| 2 | G | N7 | 1H:1675 | 3.71 | 1.00 | 37.50 |  |
| 3 | G | O6 | 1H:1675 | 3.85 | 1.00 | 40.37 |  |
| 4 | G | OP2 | 1H:1674 | 4.32 | 1.00 | 28.44 |  |
| 5 | U | OP2 | 1H:1347 | 4.79 | 1.00 | 29.43 |  |
| 6 | C | N4 | 1H:1684 | 4.81 | 1.00 | 31.04 |  |
| 7 | A | N6 | 1H:1685 | 4.98 | 1.00 | 32.44 |  |
| 8 | G | OP2 | 1H:1675 | 4.98 | 1.00 | 36.61 |  |
  |

  
 Coded contact list:   
1^st^ MG coordination shell missing (see Notes)  

### MG:1H:3217

|  | Res. type | Atom | Residue | Distance | Occupancy | B-factor | Closest solute to 1^st^ shell water |
| --- | --- | --- | --- | --- | --- | --- | --- |
| **Ion** | | | |  | 1.00 | 31.09 || 1 | G | O6 | 1H:2841 | 3.68 | 1.00 | 39.16 |
| 2 | G | N7 | 1H:2841 | 3.96 | 1.00 | 35.39 |  |
| 3 | G | O6 | 1H:2842 | 3.98 | 1.00 | 38.30 |  |
| 4 | G | N7 | 1H:2842 | 4.14 | 1.00 | 40.61 |  |
  |

  
 Coded contact list:   
1^st^ MG coordination shell missing (see Notes)  

### MG:1H:3218

|  | Res. type | Atom | Residue | Distance | Occupancy | B-factor | Closest solute to 1^st^ shell water |
| --- | --- | --- | --- | --- | --- | --- | --- |
| **Ion** | | | |  | 1.00 | 28.01 || 1 | C | O2 | 1H:553 | 2.22 | 1.00 | 35.70 |
| 2 | C | N3 | 1H:553 | 3.53 | 1.00 | 35.13 |  |
| 3 | A | N6 | 1H:554 | 4.09 | 1.00 | 36.84 |  |
| 4 | G | OP1 | 1H:2794 | 4.18 | 1.00 | 57.71 |  |
| 5 | C | N1 | 1H:553 | 4.50 | 1.00 | 32.19 |  |
| 6 | U | OP1 | 1H:2793 | 4.56 | 1.00 | 42.44 |  |
| 7 | U | O2' | 1H:2793 | 4.66 | 1.00 | 39.84 |  |
| 8 | C | O2' | 1H:553 | 4.88 | 1.00 | 42.18 |  |
  |

  
 Contact(s): /O2/  
 Coded contact list: O_b_  

### MG:1H:3219

|  | Res. type | Atom | Residue | Distance | Occupancy | B-factor | Closest solute to 1^st^ shell water |
| --- | --- | --- | --- | --- | --- | --- | --- |
| **Ion** | | | |  | 1.00 | 31.25 || 1 | G | O6 | 1H:108 | 4.00 | 1.00 | 39.27 |
| 2 | G | N7 | 1H:107 | 4.05 | 1.00 | 32.67 |  |
| 3 | G | N7 | 1H:108 | 4.09 | 1.00 | 34.88 |  |
| 4 | G | O6 | 1H:107 | 4.22 | 1.00 | 32.25 |  |
  |

  
 Coded contact list:   
1^st^ MG coordination shell missing (see Notes)  

### MG:1H:3220

|  | Res. type | Atom | Residue | Distance | Occupancy | B-factor | Closest solute to 1^st^ shell water |
| --- | --- | --- | --- | --- | --- | --- | --- |
| **Ion** | | | |  | 1.00 | 50.05 || 1 | G | OP2 | 1H:209 | 3.43 | 1.00 | 26.87 |
| 2 | G | N7 | 1H:210 | 3.50 | 1.00 | 31.60 |  |
| 3 | G | O6 | 1H:210 | 3.84 | 1.00 | 33.41 |  |
| 4 | U | OP2 | 1H:455 | 4.10 | 1.00 | 35.30 |  |
| 5 | G | N7 | 1H:209 | 4.59 | 1.00 | 32.08 |  |
| 6 | G | OP2 | 1H:210 | 4.67 | 1.00 | 30.48 |  |
| 7 | G | O5' | 1H:209 | 4.67 | 1.00 | 32.31 |  |
| 8 | C | OP1 | 1H:454 | 4.71 | 1.00 | 34.96 |  |
| 9 | C | OP2 | 1H:454 | 4.73 | 1.00 | 35.18 |  |
  |

  
 Coded contact list:   
1^st^ MG coordination shell missing (see Notes)  

### MG:1H:3221

|  | Res. type | Atom | Residue | Distance | Occupancy | B-factor | Closest solute to 1^st^ shell water |
| --- | --- | --- | --- | --- | --- | --- | --- |
| **Ion** | | | |  | 1.00 | 34.18 || 1 | A | OP2 | 1H:2652 | 2.49 | 1.00 | 45.15 |
| 2 | G | N7 | 1H:2653 | 2.74 | 1.00 | 34.11 |  |
| 3 | G | O6 | 1H:2653 | 3.31 | 1.00 | 36.02 |  |
| 4 | G | O2' | 1H:2651 | 3.42 | 1.00 | 42.04 |  |
| 5 | A | O5' | 1H:2652 | 4.43 | 1.00 | 39.05 |  |
| 6 | G | O3' | 1H:2651 | 4.50 | 1.00 | 36.14 |  |
| 7 | G | N7 | 1H:2654 | 4.56 | 1.00 | 35.40 |  |
| 8 | G | O6 | 1H:2654 | 4.57 | 1.00 | 34.86 |  |
| 9 | A | N1 | 1H:2789 | 4.78 | 1.00 | 30.69 |  |
| 10 | G | N9 | 1H:2653 | 4.98 | 1.00 | 37.80 |  |
  |

  
 Coded contact list:   
 Exclusion zone contact(s): /N7/O6/OP2/  
Consider K in place of MG (2 valid contact(s) for K vs. 0 for MG)  

### MG:1H:3222

|  | Res. type | Atom | Residue | Distance | Occupancy | B-factor | Closest solute to 1^st^ shell water |
| --- | --- | --- | --- | --- | --- | --- | --- |
| **Ion** | | | |  | 1.00 | 32.11 || 1 | G | O6 | 1H:1878 | 3.69 | 1.00 | 34.14 |
| 2 | G | O6 | 1H:1877 | 3.77 | 1.00 | 42.03 |  |
| 3 | G | N7 | 1H:1877 | 3.81 | 1.00 | 40.82 |  |
| 4 | G | N7 | 1H:1878 | 4.11 | 1.00 | 42.27 |  |
  |

  
 Coded contact list:   
1^st^ MG coordination shell missing (see Notes)  

### MG:1H:3223

|  | Res. type | Atom | Residue | Distance | Occupancy | B-factor | Closest solute to 1^st^ shell water |
| --- | --- | --- | --- | --- | --- | --- | --- |
| **Ion** | | | |  | 1.00 | 30.47 || 1 | G | OP2 | 1H:1669 | 2.63 | 1.00 | 23.82 |
| 2 | G | OP1 | 1H:1669 | 3.44 | 1.00 | 33.97 |  |
| 3 | U | OP1 | 1H:1668 | 3.78 | 1.00 | 26.83 |  |
| 4 | C | OP1 | 1H:785 | 4.01 | 1.00 | 23.51 |  |
| 5 | G | O5' | 1H:1669 | 4.26 | 1.00 | 26.42 |  |
| 6 | U | O5' | 1H:1668 | 4.44 | 1.00 | 21.40 |  |
| 7 | C | O3' | 1H:784 | 4.46 | 1.00 | 20.51 |  |
| 8 | C | OP2 | 1H:785 | 4.47 | 1.00 | 15.79 |  |
| 9 | C | OP1 | 1H:784 | 4.53 | 1.00 | 17.71 |  |
| 10 | C | N4 | 1H:1654 | 4.65 | 1.00 | 27.86 |  |
| 11 | U | OP2 | 1H:1668 | 4.69 | 1.00 | 20.86 |  |
| 12 | U | O3' | 1H:1668 | 4.92 | 1.00 | 25.07 |  |
| 13 | C | O5' | 1H:784 | 4.95 | 1.00 | 17.69 |  |
| 14 | C | N3 | 1H:1654 | 4.95 | 1.00 | 28.19 |  |
  |

  
 Coded contact list:   
 Exclusion zone contact(s): /OP2/  
Consider K in place of MG (1 valid contact(s) for K vs. 0 for MG)  

### MG:1H:3224

|  | Res. type | Atom | Residue | Distance | Occupancy | B-factor | Closest solute to 1^st^ shell water |
| --- | --- | --- | --- | --- | --- | --- | --- |
| **Ion** | | | |  | 1.00 | 35.70 || 1 | G | O6 | 1H:984 | 3.42 | 1.00 | 30.95 |
| 2 | U | O4 | 1H:983 | 4.07 | 1.00 | 31.95 |  |
| 3 | G | N7 | 1H:984 | 4.33 | 1.00 | 36.75 |  |
| 4 | U | O4 | 1H:887 | 4.66 | 1.00 | 22.83 |  |
| 5 | C | N4 | 1H:888 | 4.70 | 1.00 | 28.52 |  |
  |

  
 Coded contact list:   
1^st^ MG coordination shell missing (see Notes)  

### MG:1H:3225

|  | Res. type | Atom | Residue | Distance | Occupancy | B-factor | Closest solute to 1^st^ shell water |
| --- | --- | --- | --- | --- | --- | --- | --- |
| **Ion** | | | |  | 1.00 | 37.96 || 1 | G | O6 | 1H:992 | 3.22 | 1.00 | 24.84 |
| 2 | G | O6 | 1H:993 | 3.68 | 1.00 | 32.34 |  |
| 3 | G | N7 | 1H:992 | 3.71 | 1.00 | 23.67 |  |
| 4 | A | OP2 | 1H:869 | 4.22 | 1.00 | 18.12 |  |
| 5 | G | N2 | 1H:1018 | 4.45 | 1.00 | 22.89 |  |
| 6 | C | N4 | 1H:1016 | 4.57 | 1.00 | 26.49 |  |
| 7 | U | O4 | 1H:1015 | 4.89 | 1.00 | 29.48 |  |
  |

  
 Coded contact list:   
 Exclusion zone contact(s): /O6/  
Consider K in place of MG (1 valid contact(s) for K vs. 0 for MG)  

### MG:1H:3226

|  | Res. type | Atom | Residue | Distance | Occupancy | B-factor | Closest solute to 1^st^ shell water |
| --- | --- | --- | --- | --- | --- | --- | --- |
| **Ion** | | | |  | 1.00 | 40.08 || 1 | A | OP2 | 1H:1048 | 3.43 | 1.00 | 35.32 |
| 2 | G | O6 | 1H:1049 | 3.59 | 1.00 | 38.36 |  |
| 3 | G | O6 | 1H:1050 | 4.03 | 1.00 | 39.48 |  |
| 4 | A | OP2 | 1H:1047 | 4.26 | 1.00 | 31.54 |  |
| 5 | G | N7 | 1H:1049 | 4.34 | 1.00 | 34.68 |  |
| 6 | A | OP1 | 1H:1047 | 4.41 | 1.00 | 29.54 |  |
| 7 | G | N7 | 1H:1050 | 4.68 | 1.00 | 37.08 |  |
| 8 | G | N2 | 1H:1201 | 4.97 | 1.00 | 35.76 |  |
| 9 | C | N4 | 1H:1200 | 4.97 | 1.00 | 36.75 |  |
  |

  
 Coded contact list:   
1^st^ MG coordination shell missing (see Notes)  

### MG:1H:3227

|  | Res. type | Atom | Residue | Distance | Occupancy | B-factor | Closest solute to 1^st^ shell water |
| --- | --- | --- | --- | --- | --- | --- | --- |
| **Ion** | | | |  | 1.00 | 29.99 || 1 | U | O4 | 1H:2102 | 3.24 | 1.00 | 19.38 |
| 2 | G | O6 | 1H:2103 | 3.41 | 1.00 | 19.13 |  |
| 3 | G | OP2 | 1H:2251 | 3.55 | 1.00 | 37.44 |  |
| 4 | G | N7 | 1H:2103 | 4.20 | 1.00 | 18.43 |  |
| 5 | G | N1 | 1H:2251 | 4.58 | 1.00 | 19.16 |  |
| 6 | G | O6 | 1H:2251 | 4.61 | 1.00 | 20.36 |  |
| 7 | C | N4 | 1H:2253 | 4.78 | 1.00 | 20.74 |  |
| 8 | A | N6 | 1H:2254 | 4.81 | 1.00 | 18.14 |  |
  |

  
 Coded contact list:   
 Exclusion zone contact(s): /O4/  
Consider K in place of MG (1 valid contact(s) for K vs. 0 for MG)  

### MG:1H:3228

|  | Res. type | Atom | Residue | Distance | Occupancy | B-factor | Closest solute to 1^st^ shell water |
| --- | --- | --- | --- | --- | --- | --- | --- |
| **Ion** | | | |  | 1.00 | 31.69 || 1 | G | N3 | 1H:1729 | 3.82 | 1.00 | 36.06 |
| 2 | G | O2' | 1H:1729 | 3.91 | 1.00 | 48.69 |  |
| 3 | G | N1 | 1H:1729 | 3.96 | 1.00 | 44.07 |  |
| 4 | U | O4 | 1H:1789 | 3.99 | 1.00 | 49.59 |  |
| 5 | G | N9 | 1H:1729 | 4.13 | 1.00 | 36.99 |  |
| 6 | A | O2' | 1H:1794 | 4.14 | 1.00 | 45.78 |  |
| 7 | G | N7 | 1H:1729 | 4.23 | 1.00 | 33.07 |  |
| 8 | G | O6 | 1H:1729 | 4.53 | 1.00 | 49.77 |  |
| 9 | G | N2 | 1H:1729 | 4.75 | 1.00 | 38.85 |  |
  |

  
 Coded contact list:   
1^st^ MG coordination shell missing (see Notes)  

### MG:1H:3229

|  | Res. type | Atom | Residue | Distance | Occupancy | B-factor | Closest solute to 1^st^ shell water |
| --- | --- | --- | --- | --- | --- | --- | --- |
| **Ion** | | | |  | 1.00 | 43.96 || 1 | G | O6 | 1H:410 | 3.26 | 1.00 | 32.70 |
| 2 | G | OP2 | 1H:414 | 3.54 | 1.00 | 43.37 |  |
| 3 | U | O4 | 1H:415 | 3.57 | 1.00 | 27.41 |  |
| 4 | G | O6 | 1H:416 | 4.10 | 1.00 | 39.77 |  |
| 5 | U | O4 | 1H:411 | 4.20 | 1.00 | 33.55 |  |
| 6 | G | O6 | 1H:409 | 4.26 | 1.00 | 35.46 |  |
| 7 | G | N7 | 1H:410 | 4.50 | 1.00 | 34.64 |  |
| 8 | G | N7 | 1H:409 | 4.76 | 1.00 | 28.58 |  |
  |

  
 Coded contact list:   
 Exclusion zone contact(s): /O6/  
Consider K in place of MG (1 valid contact(s) for K vs. 0 for MG)  

### MG:1H:3230

|  | Res. type | Atom | Residue | Distance | Occupancy | B-factor | Closest solute to 1^st^ shell water |
| --- | --- | --- | --- | --- | --- | --- | --- |
| **Ion** | | | |  | 1.00 | 39.68 || 1 | C | O2 | 1H:1021 | 2.60 | 1.00 | 30.16 |
| 2 | A | OP1 | 1H:1030 | 3.21 | 1.00 | 33.71 |  |
| 3 | C | N3 | 1H:1021 | 3.71 | 1.00 | 27.06 |  |
| 4 | C | O3' | 1H:1029 | 3.84 | 1.00 | 41.22 |  |
| 5 | C | N1 | 1H:1021 | 4.83 | 1.00 | 23.16 |  |
| 6 | C | O5' | 1H:1029 | 4.95 | 1.00 | 30.72 |  |
| 7 | A | OP2 | 1H:1030 | 4.96 | 1.00 | 40.06 |  |
  |

  
 Coded contact list:   
 Exclusion zone contact(s): /O2/OP1/  
Consider K in place of MG (2 valid contact(s) for K vs. 0 for MG)  

### MG:1H:3231

|  | Res. type | Atom | Residue | Distance | Occupancy | B-factor | Closest solute to 1^st^ shell water |
| --- | --- | --- | --- | --- | --- | --- | --- |
| **Ion** | | | |  | 1.00 | 43.42 || 1 | G | OP2 | 1H:410 | 3.06 | 1.00 | 31.77 |
| 2 | G | OP1 | 1H:414 | 3.76 | 1.00 | 34.00 |  |
| 3 | G | N7 | 1H:410 | 3.92 | 1.00 | 34.64 |  |
| 4 | G | OP2 | 1H:414 | 4.03 | 1.00 | 43.37 |  |
| 5 | G | OP2 | 1H:409 | 4.11 | 1.00 | 25.79 |  |
| 6 | G | O5' | 1H:409 | 4.54 | 1.00 | 30.59 |  |
| 7 | G | O5' | 1H:410 | 4.92 | 1.00 | 28.91 |  |
  |

  
 Coded contact list:   
 Exclusion zone contact(s): /OP2/  
Consider K in place of MG (1 valid contact(s) for K vs. 0 for MG)  

### MG:1H:3232

|  | Res. type | Atom | Residue | Distance | Occupancy | B-factor | Closest solute to 1^st^ shell water |
| --- | --- | --- | --- | --- | --- | --- | --- |
| **Ion** | | | |  | 1.00 | 43.75 || 1 | U | OP1 | 1H:1070 | 2.07 | 1.00 | 48.28 |
| 2 | G | OP2 | 1H:1071 | 2.11 | 1.00 | 40.29 |  |
| 3 | G | O2' | 1H:1072 | 2.44 | 1.00 | 45.44 |  |
| 4 | G | O3' | 1H:1072 | 3.44 | 1.00 | 43.11 |  |
| 5 | U | OP1 | 1H:1073 | 3.48 | 1.00 | 48.41 |  |
| 6 | U | O5' | 1H:1073 | 3.81 | 1.00 | 63.34 |  |
| 7 | U | OP2 | 1H:1070 | 3.98 | 1.00 | 42.78 |  |
| 8 | U | O5' | 1H:1070 | 4.29 | 1.00 | 45.09 |  |
| 9 | G | OP1 | 1H:1071 | 4.43 | 1.00 | 48.72 |  |
| 10 | U | O3' | 1H:1070 | 4.45 | 1.00 | 50.00 |  |
| 11 | G | O3' | 1H:1069 | 4.46 | 1.00 | 43.88 |  |
| 12 | G | O5' | 1H:1071 | 4.52 | 1.00 | 41.09 |  |
| 13 | G | N7 | 1H:1071 | 4.85 | 1.00 | 45.30 |  |
  |

  
**1^st^ shell ligand angles:**   
1-2: 90;   
  
 Contact(s): /OP1/OP2/  
 Coded contact list: 2O_ph_  
 Exclusion zone contact(s): /O2'/  

### MG:1H:3233

|  | Res. type | Atom | Residue | Distance | Occupancy | B-factor | Closest solute to 1^st^ shell water |
| --- | --- | --- | --- | --- | --- | --- | --- |
| **Ion** | | | |  | 1.00 | 51.32 || 1 | A | OP2 | 1H:1955 | 2.82 | 1.00 | 38.99 |
| 2 | C | N4 | 1H:1957 | 4.22 | 1.00 | 36.88 |  |
| 3 | U | OP2 | 1H:1954 | 4.26 | 1.00 | 39.32 |  |
| 4 | G | O6 | 1H:1956 | 4.50 | 1.00 | 32.80 |  |
| 5 | A | OP1 | 1H:1955 | 4.65 | 1.00 | 46.99 |  |
| 6 | G | O6 | 1H:1987 | 4.75 | 1.00 | 30.13 |  |
| 7 | G | N7 | 1H:1956 | 4.75 | 1.00 | 32.35 |  |
| 8 | U | O5' | 1H:1954 | 4.87 | 1.00 | 35.69 |  |
| 9 | G | N2 | 1H:1991 | 4.98 | 1.00 | 36.53 |  |
  |

  
 Coded contact list:   
 Exclusion zone contact(s): /OP2/  
Consider K in place of MG (1 valid contact(s) for K vs. 0 for MG)  

### MG:1H:3234

|  | Res. type | Atom | Residue | Distance | Occupancy | B-factor | Closest solute to 1^st^ shell water |
| --- | --- | --- | --- | --- | --- | --- | --- |
| **Ion** | | | |  | 1.00 | 39.11 || 1 | C | OP2 | 1H:2219 | 2.63 | 1.00 | 60.74 |
| 2 | G | OP2 | 1H:2237 | 3.79 | 1.00 | 45.84 |  |
| 3 | ARG | NH1 | 11:268 | 4.61 | 1.00 | 57.93 |  |
| 4 | C | O5' | 1H:2219 | 4.70 | 1.00 | 48.24 |  |
| 5 | C | O5' | 1H:2218 | 4.70 | 1.00 | 70.36 |  |
| 6 | C | OP1 | 1H:2219 | 4.76 | 1.00 | 52.78 |  |
  |

  
 Coded contact list:   
 Exclusion zone contact(s): /OP2/  
Consider K in place of MG (1 valid contact(s) for K vs. 0 for MG)  

### MG:1H:3235

|  | Res. type | Atom | Residue | Distance | Occupancy | B-factor | Closest solute to 1^st^ shell water |
| --- | --- | --- | --- | --- | --- | --- | --- |
| **Ion** | | | |  | 1.00 | 42.11 || 1 | G | O6 | 1H:2562 | 3.61 | 1.00 | 30.07 |
| 2 | G | N7 | 1H:2561 | 3.89 | 1.00 | 29.17 |  |
| 3 | G | N7 | 1H:2562 | 3.89 | 1.00 | 27.18 |  |
| 4 | G | O6 | 1H:2561 | 4.05 | 1.00 | 28.80 |  |
  |

  
 Coded contact list:   
1^st^ MG coordination shell missing (see Notes)  

### MG:1H:3236

|  | Res. type | Atom | Residue | Distance | Occupancy | B-factor | Closest solute to 1^st^ shell water |
| --- | --- | --- | --- | --- | --- | --- | --- |
| **Ion** | | | |  | 1.00 | 43.55 || 1 | G | OP1 | 1H:2362 | 2.99 | 1.00 | 51.00 |
| 2 | C | OP2 | 1H:2363 | 3.67 | 1.00 | 45.30 |  |
| 3 | A | OP1 | 1H:2431 | 4.06 | 1.00 | 41.00 |  |
| 4 | C | O3' | 1H:2430 | 4.42 | 1.00 | 40.32 |  |
| 5 | G | O5' | 1H:2362 | 4.66 | 1.00 | 39.51 |  |
  |

  
 Coded contact list:   
 Exclusion zone contact(s): /OP1/  
Consider K in place of MG (1 valid contact(s) for K vs. 0 for MG)  

### MG:1H:3237

|  | Res. type | Atom | Residue | Distance | Occupancy | B-factor | Closest solute to 1^st^ shell water |
| --- | --- | --- | --- | --- | --- | --- | --- |
| **Ion** | | | |  | 1.00 | 38.31 || 1 | C | OP1 | 1H:2465 | 2.68 | 1.00 | 29.01 |
| 2 | A | OP1 | 1K:76 | 3.68 | 1.00 | 92.58 |  |
| 3 | ALA | N | I8:2 | 3.74 | 1.00 | 60.51 |  |
| 4 | G | OP2 | 1H:2507 | 4.04 | 1.00 | 52.00 |  |
| 5 | A | O5' | 1K:76 | 4.05 | 1.00 | 83.77 |  |
| 6 | U | OP1 | 1H:2506 | 4.46 | 1.00 | 48.58 |  |
| 7 | A | OP2 | 1H:2466 | 4.63 | 1.00 | 22.53 |  |
| 8 | A | OP1 | 1H:2466 | 4.66 | 1.00 | 39.59 |  |
| 9 | C | O3' | 1K:75 | 4.81 | 1.00 | 74.25 |  |
| 10 | C | O5' | 1H:2465 | 4.94 | 1.00 | 24.61 |  |
| 11 | A | O3' | 1H:2464 | 4.96 | 1.00 | 29.46 |  |
  |

  
 Coded contact list:   
 Exclusion zone contact(s): /OP1/  
Consider K in place of MG (1 valid contact(s) for K vs. 0 for MG)  

### MG:1H:3238

|  | Res. type | Atom | Residue | Distance | Occupancy | B-factor | Closest solute to 1^st^ shell water |
| --- | --- | --- | --- | --- | --- | --- | --- |
| **Ion** | | | |  | 1.00 | 39.74 || 1 | Water 1 | | 1H:3717 | 2.48 | 1.00 | 25.44 | 2.24 (G:OP2 1H:195) |
| 2 | G | OP2 | 1H:195 | 2.68 | 1.00 | 34.51 |  |
| 3 | G | O5' | 1H:195 | 3.06 | 1.00 | 32.78 |  |
| 4 | A | O2' | 1H:194 | 3.12 | 1.00 | 24.94 |  |
| 5 | G | O4' | 1H:195 | 3.56 | 1.00 | 21.23 |  |
| 6 | A | O3' | 1H:194 | 3.76 | 1.00 | 24.45 |  |
| 7 | A | N3 | 1H:194 | 4.21 | 1.00 | 16.58 |  |
| 8 | Water 2 | | 1H:4413 | 4.60 | 1.00 | 27.75 | 3.35 (G:OP2 1H:195) |
| 9 | G | OP1 | 1H:195 | 4.87 | 1.00 | 33.49 |  |
| 10 | G | N9 | 1H:195 | 4.89 | 1.00 | 20.74 |  |
  |

  
 Coded contact list:   
 Exclusion zone contact(s): /O2'/O5'/O_w_/OP2/  
Consider K in place of MG (3 valid contact(s) for K vs. 0 for MG)  
MG hydration shell issue: water too close (< 2.4 Angs.)  

### MG:1H:3239

|  | Res. type | Atom | Residue | Distance | Occupancy | B-factor | Closest solute to 1^st^ shell water |
| --- | --- | --- | --- | --- | --- | --- | --- |
| **Ion** | | | |  | 1.00 | 26.33 || 1 | U | OP2 | 1H:1830 | 3.86 | 1.00 | 14.74 |
| 2 | ARG | O | 11:273 | 3.89 | 1.00 | 29.64 |  |
| 3 | C | OP2 | 1H:1829 | 4.62 | 1.00 | 16.10 |  |
| 4 | A | N1 | 1H:1851 | 4.89 | 1.00 | 19.47 |  |
| 5 | C | O5' | 1H:1829 | 4.97 | 1.00 | 15.79 |  |
  |

  
 Coded contact list:   
1^st^ MG coordination shell missing (see Notes)  

### MG:1H:3240

|  | Res. type | Atom | Residue | Distance | Occupancy | B-factor | Closest solute to 1^st^ shell water |
| --- | --- | --- | --- | --- | --- | --- | --- |
| **Ion** | | | |  | 1.00 | 39.45 || 1 | A | OP2 | 1H:1913 | 2.52 | 1.00 | 27.16 |
| 2 | G | O6 | 1H:1914 | 3.26 | 1.00 | 49.43 |  |
| 3 | G | N7 | 1H:1914 | 3.36 | 1.00 | 38.80 |  |
| 4 | A | N7 | 1H:1913 | 3.95 | 1.00 | 36.62 |  |
| 5 | A | O5' | 1H:1913 | 4.20 | 1.00 | 39.72 |  |
| 6 | C | N4 | 1H:1915 | 4.22 | 1.00 | 39.46 |  |
| 7 | U | O4 | 1H:1883 | 4.29 | 1.00 | 53.03 |  |
| 8 | C | O2 | 1H:1884 | 4.72 | 1.00 | 51.72 |  |
| 9 | G | O6 | 1H:1882 | 4.74 | 1.00 | 50.32 |  |
| 10 | A | O3' | 1H:1912 | 4.85 | 1.00 | 34.86 |  |
| 11 | U | N3 | 1H:1883 | 4.91 | 1.00 | 46.57 |  |
| 12 | A | OP1 | 1H:1913 | 4.93 | 1.00 | 36.52 |  |
| 13 | C | N3 | 1H:1884 | 4.95 | 1.00 | 58.84 |  |
  |

  
 Coded contact list:   
 Exclusion zone contact(s): /N7/O6/OP2/  
Consider K in place of MG (2 valid contact(s) for K vs. 0 for MG)  

### MG:1H:3241

|  | Res. type | Atom | Residue | Distance | Occupancy | B-factor | Closest solute to 1^st^ shell water |
| --- | --- | --- | --- | --- | --- | --- | --- |
| **Ion** | | | |  | 1.00 | 30.18 || 1 | C | OP1 | 1H:2454 | 4.00 | 1.00 | 16.76 |
| 2 | C | OP1 | 1H:2455 | 4.10 | 1.00 | 27.68 |  |
| 3 | C | OP2 | 1H:2454 | 4.52 | 1.00 | 20.89 |  |
| 4 | C | O2 | 1H:2599 | 4.80 | 1.00 | 22.11 |  |
  |

  
 Coded contact list:   
1^st^ MG coordination shell missing (see Notes)  

### MG:1H:3242

|  | Res. type | Atom | Residue | Distance | Occupancy | B-factor | Closest solute to 1^st^ shell water |
| --- | --- | --- | --- | --- | --- | --- | --- |
| **Ion** | | | |  | 1.00 | 37.37 || 1 | A | OP2 | 1H:1961 | 2.65 | 1.00 | 29.20 |
| 2 | C | OP1 | 1H:1980 | 3.38 | 1.00 | 27.91 |  |
| 3 | C | OP2 | 1H:1981 | 3.48 | 1.00 | 30.48 |  |
| 4 | C | OP1 | 1H:1981 | 3.74 | 1.00 | 28.85 |  |
| 5 | C | OP2 | 1H:1980 | 4.53 | 1.00 | 26.47 |  |
| 6 | A | O3' | 1H:1960 | 4.58 | 1.00 | 29.47 |  |
| 7 | C | O5' | 1H:1980 | 4.68 | 1.00 | 21.79 |  |
| 8 | C | O3' | 1H:1980 | 4.89 | 1.00 | 21.69 |  |
| 9 | A | OP1 | 1H:1961 | 4.90 | 1.00 | 30.12 |  |
  |

  
 Coded contact list:   
 Exclusion zone contact(s): /OP1/OP2/  
Consider K in place of MG (2 valid contact(s) for K vs. 0 for MG)  

### MG:1H:3243

|  | Res. type | Atom | Residue | Distance | Occupancy | B-factor | Closest solute to 1^st^ shell water |
| --- | --- | --- | --- | --- | --- | --- | --- |
| **Ion** | | | |  | 1.00 | 41.87 || 1 | A | OP2 | 1H:2365 | 3.70 | 1.00 | 43.21 |
| 2 | G | N7 | 1H:2366 | 3.71 | 1.00 | 39.80 |  |
| 3 | G | O6 | 1H:2366 | 4.10 | 1.00 | 34.05 |  |
| 4 | G | O6 | 1H:2367 | 4.46 | 1.00 | 28.19 |  |
| 5 | G | N7 | 1H:2367 | 4.53 | 1.00 | 32.51 |  |
| 6 | A | OP1 | 1H:2365 | 4.78 | 1.00 | 47.13 |  |
  |

  
 Coded contact list:   
1^st^ MG coordination shell missing (see Notes)  

### MG:1H:3244

|  | Res. type | Atom | Residue | Distance | Occupancy | B-factor | Closest solute to 1^st^ shell water |
| --- | --- | --- | --- | --- | --- | --- | --- |
| **Ion** | | | |  | 1.00 | 34.50 || 1 | U | OP1 | 1H:71 | 2.39 | 1.00 | 33.99 |
| 2 | A | OP2 | 1H:70 | 3.64 | 1.00 | 30.05 |  |
| 3 | G | OP1 | 1H:111 | 3.78 | 1.00 | 37.27 |  |
| 4 | U | O3' | 1H:110 | 4.10 | 1.00 | 29.22 |  |
| 5 | MET | N | F8:1 | 4.19 | 1.00 | 56.87 |  |
| 6 | A | O2' | 1H:70 | 4.29 | 1.00 | 21.95 |  |
| 7 | U | O2' | 1H:110 | 4.46 | 1.00 | 29.48 |  |
| 8 | A | O3' | 1H:70 | 4.53 | 1.00 | 25.82 |  |
| 9 | U | O5' | 1H:71 | 4.66 | 1.00 | 23.60 |  |
| 10 | U | OP2 | 1H:71 | 4.81 | 1.00 | 33.32 |  |
  |

  
 Contact(s): /OP1/  
 Coded contact list: O_ph_  

### MG:1H:3245

|  | Res. type | Atom | Residue | Distance | Occupancy | B-factor | Closest solute to 1^st^ shell water |
| --- | --- | --- | --- | --- | --- | --- | --- |
| **Ion** | | | |  | 1.00 | 50.49 || 1 | G | O6 | 1H:1182 | 3.96 | 1.00 | 41.50 |
| 2 | C | N4 | 1H:1054 | 4.51 | 1.00 | 34.05 |  |
| 3 | C | N3 | 1H:1055 | 4.66 | 1.00 | 37.98 |  |
| 4 | C | N4 | 1H:1055 | 4.85 | 1.00 | 35.79 |  |
| 5 | G | O3' | 1H:1050 | 4.95 | 1.00 | 36.24 |  |
  |

  
 Coded contact list:   
1^st^ MG coordination shell missing (see Notes)  

### MG:1H:3246

|  | Res. type | Atom | Residue | Distance | Occupancy | B-factor | Closest solute to 1^st^ shell water |
| --- | --- | --- | --- | --- | --- | --- | --- |
| **Ion** | | | |  | 1.00 | 35.22 || 1 | A | OP1 | 1H:2281 | 2.19 | 1.00 | 36.41 |
| 2 | A | OP1 | 1H:2282 | 2.82 | 1.00 | 35.67 |  |
| 3 | Water 1 | | 1H:4140 | 3.67 | 1.00 | 35.91 | 2.81 (G:O6 1H:2284) |
| 4 | A | N3 | 1H:2280 | 3.85 | 1.00 | 33.99 |  |
| 5 | G | O6 | 1H:2284 | 3.85 | 1.00 | 33.17 |  |
| 6 | A | O3' | 1H:2280 | 4.34 | 1.00 | 31.70 |  |
| 7 | A | OP2 | 1H:2280 | 4.42 | 1.00 | 33.91 |  |
| 8 | A | O5' | 1H:2281 | 4.56 | 1.00 | 27.32 |  |
| 9 | A | OP2 | 1H:2281 | 4.61 | 1.00 | 36.87 |  |
| 10 | A | O3' | 1H:2281 | 4.61 | 1.00 | 33.25 |  |
| 11 | A | O5' | 1H:2280 | 4.75 | 1.00 | 23.13 |  |
  |

  
 Contact(s): /OP1/  
 Coded contact list: O_ph_  
 Exclusion zone contact(s): /OP1/  

### MG:1H:3247

|  | Res. type | Atom | Residue | Distance | Occupancy | B-factor | Closest solute to 1^st^ shell water |
| --- | --- | --- | --- | --- | --- | --- | --- |
| **Ion** | | | |  | 1.00 | 21.58 || 1 | U | O4 | 1H:2432 | 2.14 | 1.00 | 45.46 |
| 2 | ARG | O | Q8:30 | 4.06 | 1.00 | 42.18 |  |
| 3 | U | N3 | 1H:2432 | 4.38 | 1.00 | 39.92 |  |
| 4 | A | N6 | 1H:2431 | 4.43 | 1.00 | 44.85 |  |
| 5 | A | N7 | 1H:2431 | 4.59 | 1.00 | 42.77 |  |
| 6 | C | N4 | 1H:2433 | 4.77 | 1.00 | 32.99 |  |
| 7 | ARG | NE | Q8:30 | 4.91 | 1.00 | 30.20 |  |
  |

  
 Contact(s): /O4/  
 Coded contact list: O_b_  

### MG:1H:3248

|  | Res. type | Atom | Residue | Distance | Occupancy | B-factor | Closest solute to 1^st^ shell water |
| --- | --- | --- | --- | --- | --- | --- | --- |
| **Ion** | | | |  | 1.00 | 34.62 || 1 | A | OP2 | 1H:1284 | 2.64 | 1.00 | 37.61 |
| 2 | A | OP1 | 1H:1284 | 2.97 | 1.00 | 39.82 |  |
| 3 | G | OP1 | 1H:503 | 3.56 | 1.00 | 27.42 |  |
| 4 | A | OP2 | 1H:504 | 3.78 | 1.00 | 45.58 |  |
| 5 | G | O3' | 1H:1283 | 4.29 | 1.00 | 38.39 |  |
| 6 | A | O5' | 1H:1284 | 4.59 | 1.00 | 31.77 |  |
  |

  
 Coded contact list:   
 Exclusion zone contact(s): /OP1/OP2/  
Consider K in place of MG (2 valid contact(s) for K vs. 0 for MG)  

### MG:1H:3249

|  | Res. type | Atom | Residue | Distance | Occupancy | B-factor | Closest solute to 1^st^ shell water |
| --- | --- | --- | --- | --- | --- | --- | --- |
| **Ion** | | | |  | 1.00 | 31.13 || 1 | C | OP1 | 1H:1029 | 2.81 | 1.00 | 43.62 |
| 2 | U | OP1 | 1H:586 | 3.06 | 1.00 | 32.89 |  |
| 3 | C | O2' | 1H:2059 | 3.08 | 1.00 | 24.80 |  |
| 4 | U | OP2 | 1H:586 | 3.30 | 1.00 | 23.09 |  |
| 5 | C | O3' | 1H:2059 | 3.50 | 1.00 | 20.12 |  |
| 6 | A | O3' | 1H:1028 | 3.90 | 1.00 | 31.50 |  |
| 7 | G | OP1 | 1H:2060 | 4.12 | 1.00 | 25.02 |  |
| 8 | G | OP1 | 1H:585 | 4.52 | 1.00 | 37.54 |  |
| 9 | U | O5' | 1H:586 | 4.65 | 1.00 | 27.08 |  |
| 10 | C | OP2 | 1H:1029 | 4.71 | 1.00 | 40.41 |  |
| 11 | G | O3' | 1H:585 | 4.80 | 1.00 | 36.36 |  |
| 12 | C | O4' | 1H:2059 | 4.84 | 1.00 | 24.61 |  |
  |

  
 Coded contact list:   
 Exclusion zone contact(s): /O2'/OP1/OP1/OP2/  
Consider K in place of MG (4 valid contact(s) for K vs. 0 for MG)  

### MG:1H:3250

|  | Res. type | Atom | Residue | Distance | Occupancy | B-factor | Closest solute to 1^st^ shell water |
| --- | --- | --- | --- | --- | --- | --- | --- |
| **Ion** | | | |  | 1.00 | 38.38 || 1 | G | OP2 | 1H:15 | 2.18 | 1.00 | 38.64 |
| 2 | A | OP2 | 1H:14 | 2.27 | 1.00 | 44.21 |  |
| 3 | A | O5' | 1H:14 | 4.10 | 1.00 | 39.56 |  |
| 4 | G | OP1 | 1H:15 | 4.37 | 1.00 | 34.32 |  |
| 5 | G | O5' | 1H:15 | 4.41 | 1.00 | 32.84 |  |
| 6 | A | OP1 | 1H:14 | 4.63 | 1.00 | 49.84 |  |
| 7 | A | O3' | 1H:13 | 4.70 | 1.00 | 34.94 |  |
| 8 | A | O3' | 1H:14 | 4.70 | 1.00 | 37.25 |  |
  |

  
**1^st^ shell ligand angles:**   
1-2: 92;   
  
 Contact(s): /OP2/OP2/  
 Coded contact list: 2O_ph_  

### MG:1H:3251

|  | Res. type | Atom | Residue | Distance | Occupancy | B-factor | Closest solute to 1^st^ shell water |
| --- | --- | --- | --- | --- | --- | --- | --- |
| **Ion** | | | |  | 1.00 | 39.20 || 1 | Water 1 | | 1H:3950 | 3.55 | 1.00 | 35.31 | 2.59 (C:OP2 1H:1011) |
| 2 | C | OP1 | 1H:1010 | 3.69 | 1.00 | 24.18 |  |
| 3 | Water 2 | | 1H:3801 | 3.88 | 1.00 | 31.09 | 2.38 (U:OP2 1H:870) |
| 4 | Water 3 | | 1H:3769 | 4.00 | 1.00 | 38.95 | 4.61 (C:OP2 1H:1011) |
| 5 | C | OP1 | 1H:1011 | 4.19 | 1.00 | 18.49 |  |
| 6 | Water 4 | | 1H:3767 | 4.41 | 1.00 | 26.74 | 2.32 (G:O5' 1H:992) |
| 7 | C | OP2 | 1H:1011 | 4.42 | 1.00 | 22.96 |  |
| 8 | G | O5' | 1H:992 | 4.54 | 1.00 | 19.98 |  |
| 9 | Water 5 | | 1H:3697 | 4.83 | 1.00 | 26.30 | 2.19 (G:OP2 1H:992) |
| 10 | U | OP2 | 1H:870 | 4.85 | 1.00 | 27.84 |  |
| 11 | C | O5' | 1H:1010 | 4.93 | 1.00 | 24.82 |  |
| 12 | G | OP2 | 1H:992 | 4.97 | 1.00 | 23.54 |  |
| 13 | C | OP2 | 1H:1010 | 4.99 | 1.00 | 27.85 |  |
| 14 | Water 6 | | 1H:4439 | 5.00 | 1.00 | 35.21 | 3.66 (U:OP2 1H:870) |
  |

  
 Coded contact list:   
1^st^ MG coordination shell missing (see Notes)  
MG hydration shell issue: water too close (< 2.4 Angs.)  

### MG:1H:3252

|  | Res. type | Atom | Residue | Distance | Occupancy | B-factor | Closest solute to 1^st^ shell water |
| --- | --- | --- | --- | --- | --- | --- | --- |
| **Ion** | | | |  | 1.00 | 42.85 || 1 | G | N7 | 1H:351 | 3.01 | 1.00 | 39.51 |
| 2 | G | OP2 | 1H:351 | 3.50 | 1.00 | 44.79 |  |
| 3 | G | N7 | 1H:352 | 4.49 | 1.00 | 41.93 |  |
| 4 | G | O6 | 1H:351 | 4.50 | 1.00 | 42.18 |  |
| 5 | G | O5' | 1H:351 | 4.86 | 1.00 | 41.31 |  |
| 6 | G | N9 | 1H:351 | 4.90 | 1.00 | 34.61 |  |
| 7 | G | N7 | 1H:350 | 4.95 | 1.00 | 40.89 |  |
  |

  
 Coded contact list:   
 Exclusion zone contact(s): /N7/  
Consider K in place of MG (1 valid contact(s) for K vs. 0 for MG)  

### MG:1H:3253

|  | Res. type | Atom | Residue | Distance | Occupancy | B-factor | Closest solute to 1^st^ shell water |
| --- | --- | --- | --- | --- | --- | --- | --- |
| **Ion** | | | |  | 1.00 | 41.75 || 1 | G | O6 | 1H:616 | 4.07 | 1.00 | 27.54 |
| 2 | G | O6 | 1H:617 | 4.17 | 1.00 | 27.11 |  |
| 3 | G | N7 | 1H:616 | 4.76 | 1.00 | 24.95 |  |
| 4 | G | N7 | 1H:617 | 4.84 | 1.00 | 25.52 |  |
  |

  
 Coded contact list:   
1^st^ MG coordination shell missing (see Notes)  

### MG:1H:3254

|  | Res. type | Atom | Residue | Distance | Occupancy | B-factor | Closest solute to 1^st^ shell water |
| --- | --- | --- | --- | --- | --- | --- | --- |
| **Ion** | | | |  | 1.00 | 29.20 || 1 | C | OP2 | 1H:1205 | 2.57 | 1.00 | 31.71 |
| 2 | G | OP1 | 1H:1204 | 3.02 | 1.00 | 34.29 |  |
| 3 | G | O5' | 1H:1204 | 3.34 | 1.00 | 27.30 |  |
| 4 | C | OP1 | 1H:1205 | 3.98 | 1.00 | 36.26 |  |
| 5 | G | O3' | 1H:1204 | 4.29 | 1.00 | 37.54 |  |
| 6 | A | OP1 | 1H:1037 | 4.33 | 1.00 | 36.88 |  |
| 7 | G | OP2 | 1H:1204 | 4.43 | 1.00 | 30.27 |  |
| 8 | A | O5' | 1H:1037 | 4.47 | 1.00 | 28.53 |  |
| 9 | C | O5' | 1H:1205 | 4.89 | 1.00 | 40.09 |  |
| 10 | A | O3' | 1H:1203 | 4.95 | 1.00 | 31.89 |  |
  |

  
 Coded contact list:   
 Exclusion zone contact(s): /O5'/OP1/OP2/  
Consider K in place of MG (2 valid contact(s) for K vs. 0 for MG)  

### MG:1H:3255

|  | Res. type | Atom | Residue | Distance | Occupancy | B-factor | Closest solute to 1^st^ shell water |
| --- | --- | --- | --- | --- | --- | --- | --- |
| **Ion** | | | |  | 1.00 | 47.48 || 1 | C | OP2 | 1H:2890 | 2.32 | 1.00 | 57.79 |
| 2 | U | OP1 | 1H:2889 | 3.14 | 1.00 | 62.74 |  |
| 3 | C | OP1 | 1H:2890 | 3.91 | 1.00 | 60.84 |  |
| 4 | U | O5' | 1H:2889 | 4.07 | 1.00 | 58.99 |  |
| 5 | U | OP2 | 1H:2889 | 4.26 | 1.00 | 62.19 |  |
| 6 | A | N7 | 1H:2846 | 4.60 | 1.00 | 50.48 |  |
| 7 | C | O5' | 1H:2890 | 4.63 | 1.00 | 56.05 |  |
| 8 | U | O3' | 1H:2889 | 4.65 | 1.00 | 55.00 |  |
  |

  
 Contact(s): /OP2/  
 Coded contact list: O_ph_  
 Exclusion zone contact(s): /OP1/  

### MG:1H:3256

|  | Res. type | Atom | Residue | Distance | Occupancy | B-factor | Closest solute to 1^st^ shell water |
| --- | --- | --- | --- | --- | --- | --- | --- |
| **Ion** | | | |  | 1.00 | 40.50 || 1 | U | O4 | 1H:2444 | 3.56 | 1.00 | 35.15 |
| 2 | A | OP2 | 1H:2447 | 3.76 | 1.00 | 21.55 |  |
| 3 | A | O5' | 1H:2447 | 3.83 | 1.00 | 25.99 |  |
| 4 | A | OP2 | 1H:2448 | 4.18 | 1.00 | 18.39 |  |
| 5 | A | OP1 | 1H:2447 | 4.49 | 1.00 | 29.86 |  |
| 6 | U | O4 | 1H:875 | 4.87 | 1.00 | 27.37 |  |
| 7 | U | N3 | 1H:2444 | 4.93 | 1.00 | 22.36 |  |
  |

  
 Coded contact list:   
1^st^ MG coordination shell missing (see Notes)  

### MG:1H:3257

|  | Res. type | Atom | Residue | Distance | Occupancy | B-factor | Closest solute to 1^st^ shell water |
| --- | --- | --- | --- | --- | --- | --- | --- |
| **Ion** | | | |  | 1.00 | 40.13 || 1 | A | OP2 | 1H:1288 | 2.19 | 1.00 | 48.28 |
| 2 | G | O6 | 1H:1249 | 2.33 | 1.00 | 50.25 |  |
| 3 | U | O3' | 1H:1287 | 3.20 | 1.00 | 40.22 |  |
| 4 | A | N6 | 1H:1250 | 3.31 | 1.00 | 49.28 |  |
| 5 | G | N1 | 1H:1249 | 3.71 | 1.00 | 47.38 |  |
| 6 | A | OP1 | 1H:1288 | 3.94 | 1.00 | 59.85 |  |
| 7 | U | O2' | 1H:1287 | 4.32 | 1.00 | 57.85 |  |
| 8 | A | O5' | 1H:1288 | 4.50 | 1.00 | 52.58 |  |
| 9 | U | N1 | 1H:1287 | 4.52 | 1.00 | 41.20 |  |
| 10 | U | O5' | 1H:1287 | 4.54 | 1.00 | 42.65 |  |
| 11 | A | N1 | 1H:1250 | 4.66 | 1.00 | 48.83 |  |
| 12 | U | O4' | 1H:1287 | 4.84 | 1.00 | 41.31 |  |
  |

  
**1^st^ shell ligand angles:**   
1-2: 71;   
  
 Contact(s): /O6/OP2/  
 Coded contact list: O_ph_O_b_  
 Exclusion zone contact(s): /N6/O3'/  

### MG:1H:3258

|  | Res. type | Atom | Residue | Distance | Occupancy | B-factor | Closest solute to 1^st^ shell water |
| --- | --- | --- | --- | --- | --- | --- | --- |
| **Ion** | | | |  | 1.00 | 52.48 || 1 | G | OP2 | 1H:1535 | 3.84 | 1.00 | 73.43 |
| 2 | U | O4 | 1H:1536 | 3.86 | 1.00 | 71.54 |  |
| 3 | G | N7 | 1H:1535 | 4.28 | 1.00 | 71.33 |  |
| 4 | G | OP2 | 1H:1534 | 4.96 | 1.00 | 93.99 |  |
  |

  
 Coded contact list:   
1^st^ MG coordination shell missing (see Notes)  

### MG:1H:3259

|  | Res. type | Atom | Residue | Distance | Occupancy | B-factor | Closest solute to 1^st^ shell water |
| --- | --- | --- | --- | --- | --- | --- | --- |
| **Ion** | | | |  | 1.00 | 46.30 || 1 | G | O6 | 1H:93 | 2.26 | 1.00 | 46.25 |
| 2 | G | O6 | 1H:94 | 3.67 | 1.00 | 37.12 |  |
| 3 | G | N7 | 1H:94 | 4.08 | 1.00 | 40.73 |  |
| 4 | G | N7 | 1H:93 | 4.33 | 1.00 | 58.82 |  |
| 5 | G | N1 | 1H:93 | 4.42 | 1.00 | 42.18 |  |
| 6 | C | OP2 | 1H:92 | 4.86 | 1.00 | 88.14 |  |
  |

  
 Contact(s): /O6/  
 Coded contact list: O_b_  

### MG:1H:3260

|  | Res. type | Atom | Residue | Distance | Occupancy | B-factor | Closest solute to 1^st^ shell water |
| --- | --- | --- | --- | --- | --- | --- | --- |
| **Ion** | | | |  | 1.00 | 24.57 || 1 | Water 1 | | 1H:3616 | 2.48 | 1.00 | 23.65 | 1.89 (G:OP1 1H:808) |
| 2 | G | OP1 | 1H:808 | 2.81 | 1.00 | 21.39 |  |
| 3 | Water 2 | | 1H:4355 | 2.91 | 1.00 | 30.03 | 3.18 (G:OP1 1H:808) |
| 4 | C | OP1 | 1H:780 | 2.94 | 1.00 | 27.62 |  |
| 5 | Water 3 | | 1H:3987 | 3.37 | 1.00 | 22.62 | 3.25 (C:O3' 1H:779) |
| 6 | G | OP2 | 1H:808 | 3.74 | 1.00 | 17.18 |  |
| 7 | C | OP2 | 1H:780 | 3.94 | 1.00 | 24.67 |  |
| 8 | G | O3' | 1H:807 | 4.35 | 1.00 | 17.46 |  |
| 9 | C | O3' | 1H:779 | 4.51 | 1.00 | 27.47 |  |
| 10 | Water 4 | | 1H:4171 | 4.75 | 1.00 | 29.67 | 2.86 (G:OP1 1H:808) |
| 11 | G | OP1 | 1H:2003 | 4.86 | 1.00 | 23.39 |  |
  |

  
 Coded contact list:   
 Exclusion zone contact(s): /O_w_/O_w_/O_w_/OP1/OP1/  
Consider K in place of MG (2 valid contact(s) for K vs. 0 for MG)  
MG hydration shell issue: water too close (< 2.4 Angs.)  

### MG:1H:3261

|  | Res. type | Atom | Residue | Distance | Occupancy | B-factor | Closest solute to 1^st^ shell water |
| --- | --- | --- | --- | --- | --- | --- | --- |
| **Ion** | | | |  | 1.00 | 44.15 || 1 | G | OP1 | 1H:1350 | 1.92 | 1.00 | 35.94 |
| 2 | Water 1 | | 1H:4059 | 2.09 | 1.00 | 39.56 | 2.70 (A:OP1 1H:1655) |
| 3 | Water 2 | | 1H:4395 | 2.17 | 1.00 | 37.66 | 3.28 (C:OP2 1H:1351) |
| 4 | G | O5' | 1H:1350 | 3.45 | 1.00 | 33.21 |  |
| 5 | G | OP2 | 1H:1350 | 3.53 | 1.00 | 30.35 |  |
| 6 | Water 3 | | 1H:3714 | 3.57 | 1.00 | 41.28 | 4.76 (A:OP1 1H:1655) |
| 7 | Water 4 | | 1H:3731 | 3.77 | 1.00 | 50.00 | 2.26 (C:OP2 1H:1351) |
| 8 | G | O2' | 1H:1691 | 4.09 | 1.00 | 31.76 |  |
| 9 | A | O3' | 1H:1349 | 4.28 | 1.00 | 32.49 |  |
| 10 | A | OP1 | 1H:1655 | 4.40 | 1.00 | 34.75 |  |
| 11 | C | OP2 | 1H:1351 | 4.62 | 1.00 | 33.28 |  |
| 12 | Water 5 | | 1H:3626 | 4.74 | 1.00 | 32.53 | 4.20 (A:OP1 1H:1655) |
| 13 | Water 6 | | 1H:4038 | 4.78 | 1.00 | 32.55 | 3.59 (C:OP2 1H:1351) |
  |

  
**1^st^ shell ligand angles:**   
1-2: 89; 1-3: 137; 2-3: 96;   
  
 Contact(s): /O_w_/O_w_/OP1/  
 Coded contact list: O_ph_2O_w_  

### MG:1H:3262

|  | Res. type | Atom | Residue | Distance | Occupancy | B-factor | Closest solute to 1^st^ shell water |
| --- | --- | --- | --- | --- | --- | --- | --- |
| **Ion** | | | |  | 1.00 | 46.03 || 1 | G | O6 | 1H:2830 | 3.77 | 1.00 | 28.40 |
| 2 | A | N1 | 1H:2832 | 4.12 | 1.00 | 24.80 |  |
| 3 | G | N7 | 1H:2830 | 4.52 | 1.00 | 28.65 |  |
  |

  
 Coded contact list:   
1^st^ MG coordination shell missing (see Notes)  

### MG:1H:3263

|  | Res. type | Atom | Residue | Distance | Occupancy | B-factor | Closest solute to 1^st^ shell water |
| --- | --- | --- | --- | --- | --- | --- | --- |
| **Ion** | | | |  | 1.00 | 36.17 || 1 | Water 1 | | 1H:3789 | 2.79 | 1.00 | 19.44 | 2.36 (C:OP1 1H:2029) |
| 2 | Water 2 | | 1H:3702 | 2.82 | 1.00 | 39.20 | 2.21 (A:OP2 1H:1315) |
| 3 | C | OP2 | 1H:2030 | 3.34 | 1.00 | 25.35 |  |
| 4 | C | OP2 | 1H:2029 | 3.37 | 1.00 | 23.46 |  |
| 5 | A | OP2 | 1H:1315 | 3.49 | 1.00 | 29.83 |  |
| 6 | A | OP1 | 1H:1315 | 3.64 | 1.00 | 25.82 |  |
| 7 | C | OP1 | 1H:2029 | 3.79 | 1.00 | 23.97 |  |
| 8 | Water 3 | | 1H:4465 | 3.94 | 1.00 | 40.22 | 3.56 (U:OP1 1H:1314) |
| 9 | U | O3' | 1H:1314 | 4.21 | 1.00 | 24.51 |  |
| 10 | U | O5' | 1H:1314 | 4.28 | 1.00 | 29.16 |  |
| 11 | C | O5' | 1H:2029 | 4.50 | 1.00 | 23.90 |  |
| 12 | Water 4 | | 1H:3837 | 4.81 | 1.00 | 34.77 | 3.66 (C:OP2 1H:2029) |
| 13 | U | OP1 | 1H:1314 | 4.84 | 1.00 | 36.95 |  |
| 14 | Water 5 | | 1H:3705 | 4.95 | 1.00 | 23.22 | 3.17 (C:OP2 1H:2029) |
  |

  
 Coded contact list:   
 Exclusion zone contact(s): /O_w_/O_w_/OP2/OP2/  
Consider K in place of MG (2 valid contact(s) for K vs. 0 for MG)  
MG hydration shell issue: water too close (< 2.4 Angs.)  

### MG:1H:3264

|  | Res. type | Atom | Residue | Distance | Occupancy | B-factor | Closest solute to 1^st^ shell water |
| --- | --- | --- | --- | --- | --- | --- | --- |
| **Ion** | | | |  | 1.00 | 55.11 || 1 | U | OP2 | 1H:1567 | 2.57 | 1.00 | 50.51 |
| 2 | G | N7 | 1H:1518 | 3.28 | 1.00 | 66.25 |  |
| 3 | G | O6 | 1H:1518 | 3.32 | 1.00 | 70.30 |  |
| 4 | Water 1 | | 1H:4381 | 3.48 | 1.00 | 49.67 | 3.25 (G:O6 1H:1568) |
| 5 | G | O6 | 1H:1568 | 4.43 | 1.00 | 62.70 |  |
| 6 | G | N7 | 1H:1568 | 4.44 | 1.00 | 63.17 |  |
| 7 | U | O5' | 1H:1567 | 4.53 | 1.00 | 62.16 |  |
| 8 | U | OP1 | 1H:1567 | 4.60 | 1.00 | 62.59 |  |
  |

  
 Coded contact list:   
 Exclusion zone contact(s): /N7/O6/OP2/  
Consider K in place of MG (2 valid contact(s) for K vs. 0 for MG)  

### MG:1H:3265

|  | Res. type | Atom | Residue | Distance | Occupancy | B-factor | Closest solute to 1^st^ shell water |
| --- | --- | --- | --- | --- | --- | --- | --- |
| **Ion** | | | |  | 1.00 | 48.30 || 1 | Water 1 | | 1H:4085 | 2.62 | 1.00 | 51.56 | 2.74 (A:OP2 1H:1517) |
| 2 | A | OP1 | 1H:1486 | 2.66 | 1.00 | 39.48 |  |
| 3 | G | OP1 | 1H:1566 | 2.97 | 1.00 | 40.67 |  |
| 4 | U | O3' | 1H:1485 | 3.28 | 1.00 | 43.24 |  |
| 5 | Water 2 | | 1H:4234 | 3.98 | 1.00 | 51.38 | 3.09 (A:OP1 1H:1486) |
| 6 | U | O2' | 1H:1485 | 4.09 | 1.00 | 42.00 |  |
| 7 | A | OP1 | 1H:1517 | 4.10 | 1.00 | 61.67 |  |
| 8 | A | OP2 | 1H:1517 | 4.28 | 1.00 | 53.39 |  |
| 9 | Water 3 | | 1H:4378 | 4.33 | 1.00 | 56.23 | 3.25 (A:OP1 1H:1486) |
| 10 | A | O5' | 1H:1486 | 4.36 | 1.00 | 47.78 |  |
| 11 | G | OP2 | 1H:1566 | 4.77 | 1.00 | 41.16 |  |
| 12 | A | OP2 | 1H:1486 | 4.85 | 1.00 | 39.34 |  |
| 13 | C | O3' | 1H:1565 | 4.97 | 1.00 | 40.97 |  |
  |

  
 Coded contact list:   
 Exclusion zone contact(s): /O3'/O_w_/OP1/OP1/  
Consider K in place of MG (3 valid contact(s) for K vs. 0 for MG)  

### MG:1H:3266

|  | Res. type | Atom | Residue | Distance | Occupancy | B-factor | Closest solute to 1^st^ shell water |
| --- | --- | --- | --- | --- | --- | --- | --- |
| **Ion** | | | |  | 1.00 | 39.52 || 1 | Water 1 | | 1H:4422 | 2.67 | 1.00 | 33.42 | 3.38 (A:O3' 1H:1334) |
| 2 | Water 2 | | 1H:4484 | 2.96 | 1.00 | 47.36 | 3.63 (G:O6 1H:1326) |
| 3 | Water 3 | | 1H:3685 | 3.74 | 1.00 | 49.11 | 3.77 (G:N7 1H:1326) |
| 4 | G | O6 | 1H:1327 | 3.78 | 1.00 | 34.70 |  |
| 5 | G | O6 | 1H:1326 | 4.08 | 1.00 | 40.68 |  |
| 6 | U | OP1 | 1H:1335 | 4.10 | 1.00 | 35.66 |  |
| 7 | A | O3' | 1H:1334 | 4.45 | 1.00 | 30.88 |  |
| 8 | G | N7 | 1H:1326 | 4.61 | 1.00 | 41.33 |  |
| 9 | Water 4 | | 1H:3951 | 4.99 | 1.00 | 46.72 | 3.73 (A:O3' 1H:1334) |
  |

  
 Coded contact list:   
 Exclusion zone contact(s): /O_w_/O_w_/  
1^st^ MG coordination shell missing (see Notes)  

### MG:1H:3267

|  | Res. type | Atom | Residue | Distance | Occupancy | B-factor | Closest solute to 1^st^ shell water |
| --- | --- | --- | --- | --- | --- | --- | --- |
| **Ion** | | | |  | 1.00 | 53.96 || 1 | A | OP1 | 1H:349 | 2.56 | 1.00 | 46.22 |
| 2 | A | OP2 | 1H:349 | 3.26 | 1.00 | 41.90 |  |
| 3 | U | O2 | 1H:639 | 3.34 | 1.00 | 88.13 |  |
| 4 | U | N3 | 1H:639 | 3.35 | 1.00 | 91.27 |  |
| 5 | U | O4 | 1H:1251 | 3.61 | 1.00 | 48.37 |  |
| 6 | ARG | O | 31:168 | 3.93 | 1.00 | 46.68 |  |
| 7 | G | O3' | 1H:348 | 4.02 | 1.00 | 44.56 |  |
| 8 | U | O4 | 1H:639 | 4.59 | 1.00 | 73.36 |  |
| 9 | A | O5' | 1H:349 | 4.69 | 1.00 | 43.36 |  |
| 10 | G | O2' | 1H:348 | 4.89 | 1.00 | 43.11 |  |
| 11 | ASN | OD1 | 31:169 | 4.92 | 1.00 | 45.31 |  |
  |

  
 Coded contact list:   
 Exclusion zone contact(s): /N3_U_/O2/OP1/OP2/  
Consider K in place of MG (3 valid contact(s) for K vs. 0 for MG)  

### MG:1H:3268

|  | Res. type | Atom | Residue | Distance | Occupancy | B-factor | Closest solute to 1^st^ shell water |
| --- | --- | --- | --- | --- | --- | --- | --- |
| **Ion** | | | |  | 1.00 | 34.84 || 1 | Water 1 | | 1H:4051 | 2.04 | 1.00 | 23.40 | 2.69 (G:N7 1H:606) |
| 2 | Water 2 | | 1H:4368 | 2.75 | 1.00 | 24.52 | 3.22 (A:OP1 1H:1300) |
| 3 | Water 3 | | 1H:4093 | 3.43 | 1.00 | 25.61 | 2.96 (A:OP1 1H:1300) |
| 4 | C | OP2 | 1H:605 | 3.90 | 1.00 | 29.78 |  |
| 5 | G | OP2 | 1H:606 | 3.96 | 1.00 | 20.68 |  |
| 6 | A | OP1 | 1H:1300 | 4.12 | 1.00 | 26.45 |  |
| 7 | Water 4 | | 1H:4206 | 4.26 | 1.00 | 18.47 | 2.90 (G:OP2 1H:606) |
| 8 | G | N7 | 1H:606 | 4.39 | 1.00 | 21.56 |  |
| 9 | C | O5' | 1H:605 | 4.54 | 1.00 | 24.31 |  |
| 10 | Water 5 | | 1H:3608 | 4.89 | 1.00 | 32.49 | 3.20 (G:N7 1H:606) |
  |

  
 Contact(s): /O_w_/  
 Coded contact list: O_w_  
 Exclusion zone contact(s): /O_w_/  

### MG:1H:3269

|  | Res. type | Atom | Residue | Distance | Occupancy | B-factor | Closest solute to 1^st^ shell water |
| --- | --- | --- | --- | --- | --- | --- | --- |
| **Ion** | | | |  | 1.00 | 33.07 || 1 | A | OP1 | 1H:1031 | 2.07 | 1.00 | 26.05 |
| 2 | A | O5' | 1H:1030 | 3.14 | 1.00 | 27.74 |  |
| 3 | A | OP2 | 1H:1030 | 3.23 | 1.00 | 40.06 |  |
| 4 | Water 1 | | 1H:4431 | 3.23 | 1.00 | 30.72 | 3.41 (A:OP1 1H:1031) |
| 5 | A | OP2 | 1H:1031 | 3.33 | 1.00 | 26.52 |  |
| 6 | C | N4 | 1H:1032 | 3.47 | 1.00 | 27.47 |  |
| 7 | G | N1 | 1H:1026 | 3.55 | 1.00 | 36.31 |  |
| 8 | A | O3' | 1H:1030 | 3.68 | 1.00 | 23.88 |  |
| 9 | G | O6 | 1H:1026 | 4.00 | 1.00 | 32.09 |  |
| 10 | Water 2 | | 1H:4537 | 4.02 | 1.00 | 35.08 | 3.89 (A:OP1 1H:1031) |
| 11 | G | O6 | 1H:1025 | 4.05 | 1.00 | 31.85 |  |
| 12 | A | O5' | 1H:1031 | 4.30 | 1.00 | 26.03 |  |
| 13 | G | N2 | 1H:1026 | 4.37 | 1.00 | 35.55 |  |
| 14 | C | O3' | 1H:1029 | 4.79 | 1.00 | 41.22 |  |
| 15 | Water 3 | | 1H:4366 | 4.96 | 1.00 | 24.30 | 3.37 (A:O3' 1H:1030) |
  |

  
 Contact(s): /OP1/  
 Coded contact list: O_ph_  
 Exclusion zone contact(s): /O5'/O_w_/OP2/OP2/  
Consider K in place of MG (3 valid contact(s) for K vs. 1 for MG)  

### MG:1H:3270

|  | Res. type | Atom | Residue | Distance | Occupancy | B-factor | Closest solute to 1^st^ shell water |
| --- | --- | --- | --- | --- | --- | --- | --- |
| **Ion** | | | |  | 1.00 | 42.51 || 1 | G | O6 | 1H:1745 | 3.51 | 1.00 | 41.18 |
| 2 | G | N7 | 1H:1744 | 3.95 | 1.00 | 36.72 |  |
| 3 | G | O6 | 1H:1744 | 4.62 | 1.00 | 31.19 |  |
| 4 | G | N1 | 1H:1745 | 4.95 | 1.00 | 33.95 |  |
  |

  
 Coded contact list:   
1^st^ MG coordination shell missing (see Notes)  

### MG:1H:3271

|  | Res. type | Atom | Residue | Distance | Occupancy | B-factor | Closest solute to 1^st^ shell water |
| --- | --- | --- | --- | --- | --- | --- | --- |
| **Ion** | | | |  | 1.00 | 47.86 || 1 | G | O6 | 1H:1745 | 3.69 | 1.00 | 41.18 |
| 2 | G | OP2 | 1H:1744 | 4.05 | 1.00 | 33.19 |  |
| 3 | G | N7 | 1H:1745 | 4.15 | 1.00 | 40.98 |  |
| 4 | G | N7 | 1H:1744 | 4.39 | 1.00 | 36.72 |  |
| 5 | A | N7 | 1H:1746 | 4.80 | 1.00 | 32.75 |  |
| 6 | G | OP1 | 1H:1744 | 4.99 | 1.00 | 38.73 |  |
  |

  
 Coded contact list:   
1^st^ MG coordination shell missing (see Notes)  

### MG:1H:3272

|  | Res. type | Atom | Residue | Distance | Occupancy | B-factor | Closest solute to 1^st^ shell water |
| --- | --- | --- | --- | --- | --- | --- | --- |
| **Ion** | | | |  | 1.00 | 55.18 || 1 | G | N7 | 1H:2388 | 3.58 | 1.00 | 37.51 |
| 2 | G | O6 | 1H:2388 | 3.59 | 1.00 | 46.50 |  |
| 3 | A | OP1 | 1H:2390 | 3.62 | 1.00 | 55.79 |  |
| 4 | A | OP2 | 1H:2390 | 3.72 | 1.00 | 41.64 |  |
| 5 | A | O3' | 1H:2389 | 4.58 | 1.00 | 53.34 |  |
  |

  
 Coded contact list:   
1^st^ MG coordination shell missing (see Notes)  

### MG:1H:3273

|  | Res. type | Atom | Residue | Distance | Occupancy | B-factor | Closest solute to 1^st^ shell water |
| --- | --- | --- | --- | --- | --- | --- | --- |
| **Ion** | | | |  | 1.00 | 61.62 || 1 | G | N7 | 1H:333 | 3.30 | 1.00 | 53.47 |
| 2 | G | O6 | 1H:1258 | 3.40 | 1.00 | 46.37 |  |
| 3 | G | O6 | 1H:333 | 4.15 | 1.00 | 58.05 |  |
| 4 | A | O2' | 1H:527 | 4.54 | 1.00 | 49.49 |  |
| 5 | G | O6 | 1H:334 | 4.93 | 1.00 | 46.29 |  |
  |

  
 Coded contact list:   
 Exclusion zone contact(s): /N7/O6/  
Consider K in place of MG (2 valid contact(s) for K vs. 0 for MG)  

### MG:1H:3274

|  | Res. type | Atom | Residue | Distance | Occupancy | B-factor | Closest solute to 1^st^ shell water |
| --- | --- | --- | --- | --- | --- | --- | --- |
| **Ion** | | | |  | 1.00 | 31.92 || 1 | C | OP2 | 1H:2059 | 2.81 | 1.00 | 31.81 |
| 2 | Water 1 | | 1H:4562 | 2.86 | 1.00 | 32.94 | 4.11 (C:OP2 1H:2059) |
| 3 | G | OP2 | 1H:2060 | 2.90 | 1.00 | 23.19 |  |
| 4 | Water 2 | | 1H:4288 | 2.97 | 1.00 | 22.52 | 3.63 (G:N7 1H:2060) |
| 5 | Water 3 | | 1H:4122 | 3.61 | 1.00 | 30.63 | 3.27 (G:N7 1H:2060) |
| 6 | C | O5' | 1H:2059 | 3.68 | 1.00 | 20.70 |  |
| 7 | C | O3' | 1H:2059 | 4.61 | 1.00 | 20.12 |  |
| 8 | G | N7 | 1H:2060 | 4.64 | 1.00 | 24.39 |  |
| 9 | C | OP1 | 1H:2059 | 4.67 | 1.00 | 31.62 |  |
| 10 | G | O5' | 1H:2060 | 4.72 | 1.00 | 23.42 |  |
| 11 | G | O3' | 1H:2058 | 4.95 | 1.00 | 31.92 |  |
  |

  
 Coded contact list:   
 Exclusion zone contact(s): /O_w_/O_w_/OP2/OP2/  
Consider K in place of MG (2 valid contact(s) for K vs. 0 for MG)  

### MG:1H:3275

|  | Res. type | Atom | Residue | Distance | Occupancy | B-factor | Closest solute to 1^st^ shell water |
| --- | --- | --- | --- | --- | --- | --- | --- |
| **Ion** | | | |  | 1.00 | 44.86 || 1 | U | OP1 | 1H:2505 | 2.71 | 1.00 | 36.63 |
| 2 | C | O4' | 1H:2586 | 3.83 | 1.00 | 36.40 |  |
| 3 | A | OP1 | 1H:2466 | 4.00 | 1.00 | 39.59 |  |
| 4 | G | OP2 | 1H:2467 | 4.22 | 1.00 | 21.39 |  |
| 5 | U | OP2 | 1H:2505 | 4.45 | 1.00 | 30.80 |  |
| 6 | U | OP2 | 1H:2506 | 4.47 | 1.00 | 36.81 |  |
| 7 | G | OP1 | 1H:2467 | 4.51 | 1.00 | 26.27 |  |
| 8 | C | N1 | 1H:2586 | 4.58 | 1.00 | 31.74 |  |
| 9 | U | O3' | 1H:2504 | 4.72 | 1.00 | 29.60 |  |
  |

  
 Coded contact list:   
 Exclusion zone contact(s): /OP1/  
Consider K in place of MG (1 valid contact(s) for K vs. 0 for MG)  

### MG:1H:3276

|  | Res. type | Atom | Residue | Distance | Occupancy | B-factor | Closest solute to 1^st^ shell water |
| --- | --- | --- | --- | --- | --- | --- | --- |
| **Ion** | | | |  | 1.00 | 44.72 || 1 | G | OP1 | 1H:2467 | 3.16 | 1.00 | 26.27 |
| 2 | U | OP1 | 1H:2505 | 3.69 | 1.00 | 36.63 |  |
| 3 | U | O3' | 1H:2504 | 3.72 | 1.00 | 29.60 |  |
| 4 | U | OP2 | 1H:2504 | 3.96 | 1.00 | 33.62 |  |
| 5 | C | OP1 | 1H:2584 | 3.97 | 1.00 | 16.58 |  |
| 6 | U | OP2 | 1H:2505 | 4.10 | 1.00 | 30.80 |  |
| 7 | C | O4' | 1H:2586 | 4.28 | 1.00 | 36.40 |  |
| 8 | G | OP2 | 1H:2467 | 4.78 | 1.00 | 21.39 |  |
| 9 | U | O5' | 1H:2504 | 4.84 | 1.00 | 32.77 |  |
| 10 | C | OP2 | 1H:2584 | 4.92 | 1.00 | 23.31 |  |
| 11 | G | OP1 | 1H:2583 | 4.98 | 1.00 | 31.32 |  |
  |

  
 Coded contact list:   
 Exclusion zone contact(s): /OP1/  
Consider K in place of MG (1 valid contact(s) for K vs. 0 for MG)  

### MG:1H:3277

|  | Res. type | Atom | Residue | Distance | Occupancy | B-factor | Closest solute to 1^st^ shell water |
| --- | --- | --- | --- | --- | --- | --- | --- |
| **Ion** | | | |  | 1.00 | 42.71 || 1 | A | OP1 | 1H:498 | 3.56 | 1.00 | 21.71 |
| 2 | A | O3' | 1H:497 | 3.60 | 1.00 | 24.84 |  |
| 3 | A | OP1 | 1H:497 | 3.97 | 1.00 | 21.89 |  |
| 4 | A | O2' | 1H:497 | 4.32 | 1.00 | 17.59 |  |
| 5 | VAL | O | 31:84 | 4.75 | 1.00 | 23.15 |  |
| 6 | TYR | OH | 31:59 | 4.76 | 1.00 | 31.66 |  |
  |

  
 Coded contact list:   
1^st^ MG coordination shell missing (see Notes)  

### MG:1H:3278

|  | Res. type | Atom | Residue | Distance | Occupancy | B-factor | Closest solute to 1^st^ shell water |
| --- | --- | --- | --- | --- | --- | --- | --- |
| **Ion** | | | |  | 1.00 | 28.55 || 1 | A | OP1 | 1H:2085 | 2.95 | 1.00 | 19.28 |
| 2 | Water 1 | | 1H:3642 | 3.13 | 1.00 | 17.48 | 2.02 (A:OP1 1H:2085) |
| 3 | G | O5' | 1H:2084 | 3.24 | 1.00 | 14.13 |  |
| 4 | A | N7 | 1H:2085 | 3.66 | 1.00 | 67.40 |  |
| 5 | A | O3' | 1H:2083 | 3.76 | 1.00 | 19.85 |  |
| 6 | G | O3' | 1H:2084 | 3.87 | 1.00 | 21.68 |  |
| 7 | HIS | ND1 | 31:69 | 4.01 | 1.00 | 20.47 |  |
| 8 | G | OP1 | 1H:2084 | 4.18 | 1.00 | 15.09 |  |
| 9 | A | O5' | 1H:2085 | 4.25 | 1.00 | 34.39 |  |
| 10 | HIS | NE2 | 31:69 | 4.26 | 1.00 | 16.86 |  |
| 11 | 2MA | N6 | 1H:2516 | 4.59 | 1.00 | 26.47 |  |
| 12 | 2MA | N1 | 1H:2516 | 4.65 | 1.00 | 24.72 |  |
| 13 | A | O2' | 1H:2082 | 4.69 | 1.00 | 33.39 |  |
| 14 | LYS | NZ | 31:68 | 4.82 | 1.00 | 13.89 |  |
  |

  
 Coded contact list:   
 Exclusion zone contact(s): /O5'/O_w_/OP1/  
Consider K in place of MG (2 valid contact(s) for K vs. 0 for MG)  
MG hydration shell issue: water too close (< 2.4 Angs.)  

### MG:1H:3279

|  | Res. type | Atom | Residue | Distance | Occupancy | B-factor | Closest solute to 1^st^ shell water |
| --- | --- | --- | --- | --- | --- | --- | --- |
| **Ion** | | | |  | 1.00 | 32.47 || 1 | A | OP2 | 1H:498 | 2.67 | 1.00 | 23.37 |
| 2 | Water 1 | | 1H:4130 | 2.95 | 1.00 | 28.81 | 2.80 (A:OP1 1H:481) |
| 3 | A | N6 | 1H:499 | 3.16 | 1.00 | 24.66 |  |
| 4 | A | N7 | 1H:499 | 3.20 | 1.00 | 23.74 |  |
| 5 | Water 2 | | 1H:4321 | 3.37 | 1.00 | 26.03 | 3.12 (C:OP1 1H:480) |
| 6 | A | N6 | 1H:484 | 3.83 | 1.00 | 21.30 |  |
| 7 | C | OP1 | 1H:480 | 3.95 | 1.00 | 19.27 |  |
| 8 | C | O3' | 1H:480 | 4.10 | 1.00 | 25.91 |  |
| 9 | A | O5' | 1H:498 | 4.19 | 1.00 | 24.53 |  |
| 10 | C | N4 | 1H:482 | 4.31 | 1.00 | 32.67 |  |
| 11 | A | OP1 | 1H:481 | 4.34 | 1.00 | 24.51 |  |
| 12 | A | OP2 | 1H:499 | 4.40 | 1.00 | 28.43 |  |
| 13 | A | N7 | 1H:498 | 4.47 | 1.00 | 24.71 |  |
| 14 | A | OP1 | 1H:498 | 4.56 | 1.00 | 21.71 |  |
| 15 | C | O5' | 1H:480 | 4.66 | 1.00 | 26.09 |  |
  |

  
 Coded contact list:   
 Exclusion zone contact(s): /N6/N7/O_w_/O_w_/OP2/  
Consider K in place of MG (3 valid contact(s) for K vs. 0 for MG)  

### MG:1H:3280

|  | Res. type | Atom | Residue | Distance | Occupancy | B-factor | Closest solute to 1^st^ shell water |
| --- | --- | --- | --- | --- | --- | --- | --- |
| **Ion** | | | |  | 1.00 | 86.30 || 1 | G | OP2 | 1H:2485 | 2.42 | 1.00 | 84.56 |
| 2 | C | OP1 | 1H:2488 | 3.00 | 1.00 | 105.77 |  |
| 3 | C | O3' | 1H:2487 | 3.39 | 1.00 | 124.12 |  |
| 4 | C | OP2 | 1H:2488 | 3.42 | 1.00 | 119.02 |  |
| 5 | C | OP2 | 1H:2487 | 3.70 | 1.00 | 114.59 |  |
| 6 | C | O5' | 1H:2487 | 4.22 | 1.00 | 119.37 |  |
| 7 | G | O5' | 1H:2485 | 4.32 | 1.00 | 91.50 |  |
| 8 | G | N7 | 1H:2485 | 4.33 | 1.00 | 88.43 |  |
| 9 | G | OP1 | 1H:2485 | 4.53 | 1.00 | 97.20 |  |
| 10 | C | OP2 | 1H:2484 | 4.56 | 1.00 | 75.08 |  |
| 11 | C | O5' | 1H:2488 | 4.98 | 1.00 | 122.76 |  |
  |

  
 Coded contact list:   
 Exclusion zone contact(s): /O3'/OP1/OP2/  
Consider K in place of MG (2 valid contact(s) for K vs. 0 for MG)  

### MG:1H:3281

|  | Res. type | Atom | Residue | Distance | Occupancy | B-factor | Closest solute to 1^st^ shell water |
| --- | --- | --- | --- | --- | --- | --- | --- |
| **Ion** | | | |  | 1.00 | 49.33 || 1 | G | OP2 | 1H:1290 | 2.93 | 1.00 | 51.49 |
| 2 | A | OP1 | 1H:1289 | 3.18 | 1.00 | 60.14 |  |
| 3 | A | O5' | 1H:1289 | 4.46 | 1.00 | 50.31 |  |
| 4 | G | OP1 | 1H:1290 | 4.65 | 1.00 | 50.07 |  |
| 5 | A | OP2 | 1H:1289 | 4.93 | 1.00 | 63.80 |  |
  |

  
 Coded contact list:   
 Exclusion zone contact(s): /OP1/OP2/  
Consider K in place of MG (2 valid contact(s) for K vs. 0 for MG)  

### MG:1H:3282

|  | Res. type | Atom | Residue | Distance | Occupancy | B-factor | Closest solute to 1^st^ shell water |
| --- | --- | --- | --- | --- | --- | --- | --- |
| **Ion** | | | |  | 1.00 | 49.16 || 1 | G | N7 | 1H:142 | 3.51 | 1.00 | 32.37 |
| 2 | C | N4 | 1H:141 | 3.58 | 1.00 | 35.11 |  |
| 3 | G | O6 | 1H:142 | 3.80 | 1.00 | 38.91 |  |
| 4 | A | OP1 | 1H:140 | 3.89 | 1.00 | 42.52 |  |
| 5 | C | N4 | 1H:143 | 4.07 | 1.00 | 39.50 |  |
| 6 | Water 1 | | 1H:4151 | 4.31 | 1.00 | 46.93 | 4.27 (G:O6 1H:142) |
| 7 | Water 2 | | 1H:4593 | 4.40 | 1.00 | 53.50 | 4.64 (A:OP1 1H:140) |
| 8 | A | OP2 | 1H:140 | 4.46 | 1.00 | 40.36 |  |
| 9 | Water 3 | | 1H:4414 | 4.56 | 1.00 | 51.70 | 4.32 (C:N4 1H:143) |
| 10 | C | N3 | 1H:141 | 4.85 | 1.00 | 30.45 |  |
  |

  
 Coded contact list:   
1^st^ MG coordination shell missing (see Notes)  

### MG:1H:3283

|  | Res. type | Atom | Residue | Distance | Occupancy | B-factor | Closest solute to 1^st^ shell water |
| --- | --- | --- | --- | --- | --- | --- | --- |
| **Ion** | | | |  | 1.00 | 44.22 || 1 | A | N1 | 1H:1474 | 3.97 | 1.00 | 25.55 |
| 2 | LYS | O | 11:31 | 3.97 | 1.00 | 35.40 |  |
| 3 | A | OP1 | 1H:1618 | 4.07 | 1.00 | 20.92 |  |
| 4 | VAL | O | 11:34 | 4.20 | 1.00 | 41.16 |  |
| 5 | G | O6 | 1H:1473 | 4.30 | 1.00 | 35.37 |  |
| 6 | A | OP2 | 1H:1619 | 4.45 | 1.00 | 21.29 |  |
  |

  
 Coded contact list:   
1^st^ MG coordination shell missing (see Notes)  

### MG:1H:3284

|  | Res. type | Atom | Residue | Distance | Occupancy | B-factor | Closest solute to 1^st^ shell water |
| --- | --- | --- | --- | --- | --- | --- | --- |
| **Ion** | | | |  | 1.00 | 36.02 || 1 | Water 1 | | 1H:3929 | 1.96 | 1.00 | 22.33 | 2.55 (C:OP1 1H:239) |
| 2 | G | OP1 | 1H:240 | 2.80 | 1.00 | 22.59 |  |
| 3 | G | O5' | 1H:240 | 3.22 | 1.00 | 19.35 |  |
| 4 | Water 2 | | 1H:4324 | 3.32 | 1.00 | 26.24 | 3.13 (G:O4' 1H:240) |
| 5 | Water 3 | | 1H:4449 | 3.57 | 1.00 | 25.01 | 4.18 (A:N3 1H:2406) |
| 6 | C | O4' | 1H:2407 | 3.87 | 1.00 | 30.27 |  |
| 7 | C | OP1 | 1H:239 | 4.12 | 1.00 | 20.96 |  |
| 8 | A | N3 | 1H:2406 | 4.28 | 1.00 | 27.69 |  |
| 9 | G | O4' | 1H:240 | 4.38 | 1.00 | 18.80 |  |
| 10 | C | O2' | 1H:2407 | 4.42 | 1.00 | 26.27 |  |
| 11 | C | O3' | 1H:239 | 4.57 | 1.00 | 30.58 |  |
| 12 | Water 4 | | 1H:4148 | 4.69 | 1.00 | 34.39 | 3.20 (C:OP1 1H:239) |
| 13 | G | OP2 | 1H:240 | 4.84 | 1.00 | 22.54 |  |
| 14 | C | O5' | 1H:239 | 5.00 | 1.00 | 22.61 |  |
  |

  
 Contact(s): /O_w_/  
 Coded contact list: O_w_  
 Exclusion zone contact(s): /O5'/O_w_/OP1/  
Consider K in place of MG (2 valid contact(s) for K vs. 0 for MG)  

### MG:1H:3285

|  | Res. type | Atom | Residue | Distance | Occupancy | B-factor | Closest solute to 1^st^ shell water |
| --- | --- | --- | --- | --- | --- | --- | --- |
| **Ion** | | | |  | 1.00 | 43.85 || 1 | A | OP2 | 1H:1030 | 3.04 | 1.00 | 40.06 |
| 2 | Water 1 | | 1H:3812 | 3.60 | 1.00 | 50.17 | 2.39 (G:O6 1H:1026) |
| 3 | C | OP1 | 1H:1029 | 4.07 | 1.00 | 43.62 |  |
| 4 | C | O5' | 1H:1029 | 4.11 | 1.00 | 30.72 |  |
| 5 | C | OP2 | 1H:1029 | 4.33 | 1.00 | 40.41 |  |
| 6 | A | OP1 | 1H:1030 | 4.43 | 1.00 | 33.71 |  |
| 7 | C | OP2 | 1H:1023 | 4.54 | 1.00 | 35.06 |  |
| 8 | G | O6 | 1H:1026 | 4.75 | 1.00 | 32.09 |  |
  |

  
 Coded contact list:   
 Exclusion zone contact(s): /OP2/  
Consider K in place of MG (1 valid contact(s) for K vs. 0 for MG)  
MG hydration shell issue: water too close (< 2.4 Angs.)  

### MG:1H:3286

|  | Res. type | Atom | Residue | Distance | Occupancy | B-factor | Closest solute to 1^st^ shell water |
| --- | --- | --- | --- | --- | --- | --- | --- |
| **Ion** | | | |  | 1.00 | 48.88 || 1 | G | O6 | 1H:2336 | 2.90 | 1.00 | 34.33 |
| 2 | A | OP2 | 1H:2335 | 3.30 | 1.00 | 51.50 |  |
| 3 | U | O4 | 1H:2345 | 3.39 | 1.00 | 32.99 |  |
| 4 | G | O6 | 1H:2344 | 3.84 | 1.00 | 42.60 |  |
| 5 | C | N4 | 1H:2337 | 3.86 | 1.00 | 35.73 |  |
| 6 | G | OP2 | 1H:2334 | 3.93 | 1.00 | 44.99 |  |
| 7 | G | N7 | 1H:2336 | 4.03 | 1.00 | 39.63 |  |
| 8 | G | O5' | 1H:2334 | 4.06 | 1.00 | 53.65 |  |
| 9 | U | N3 | 1H:2345 | 4.24 | 1.00 | 28.44 |  |
| 10 | A | N7 | 1H:2335 | 4.31 | 1.00 | 41.96 |  |
  |

  
 Coded contact list:   
 Exclusion zone contact(s): /O4/O6/OP2/  
Consider K in place of MG (3 valid contact(s) for K vs. 0 for MG)  

### MG:1H:3287

|  | Res. type | Atom | Residue | Distance | Occupancy | B-factor | Closest solute to 1^st^ shell water |
| --- | --- | --- | --- | --- | --- | --- | --- |
| **Ion** | | | |  | 1.00 | 37.67 || 1 | A | OP2 | 1H:852 | 3.41 | 1.00 | 20.17 |
| 2 | A | OP1 | 1H:718 | 3.60 | 1.00 | 26.91 |  |
| 3 | U | OP2 | 1H:851 | 3.86 | 1.00 | 27.39 |  |
| 4 | G | O3' | 1H:717 | 4.07 | 1.00 | 24.30 |  |
| 5 | U | O5' | 1H:851 | 4.65 | 1.00 | 19.74 |  |
| 6 | U | OP1 | 1H:851 | 4.93 | 1.00 | 23.83 |  |
  |

  
 Coded contact list:   
1^st^ MG coordination shell missing (see Notes)  

### MG:1H:3288

|  | Res. type | Atom | Residue | Distance | Occupancy | B-factor | Closest solute to 1^st^ shell water |
| --- | --- | --- | --- | --- | --- | --- | --- |
| **Ion** | | | |  | 1.00 | 44.82 || 1 | Water 1 | | 1H:3787 | 3.54 | 1.00 | 24.98 | 4.17 (A:N7 1H:852) |
| 2 | G | N7 | 1H:722 | 4.08 | 1.00 | 19.75 |  |
| 3 | A | N7 | 1H:852 | 4.13 | 1.00 | 19.17 |  |
| 4 | G | O6 | 1H:722 | 4.25 | 1.00 | 23.61 |  |
| 5 | Water 2 | | 1H:3634 | 4.57 | 1.00 | 23.70 | 5.20 (G:N7 1H:722) |
  |

  
 Coded contact list:   
1^st^ MG coordination shell missing (see Notes)  

### MG:1H:3289

|  | Res. type | Atom | Residue | Distance | Occupancy | B-factor | Closest solute to 1^st^ shell water |
| --- | --- | --- | --- | --- | --- | --- | --- |
| **Ion** | | | |  | 1.00 | 35.64 || 1 | C | OP2 | 1H:32 | 3.92 | 1.00 | 33.93 |
| 2 | Water 1 | | 1H:4548 | 3.97 | 1.00 | 33.87 | 5.50 (C:OP2 1H:32) |
| 3 | C | OP1 | 1H:32 | 4.27 | 1.00 | 28.72 |  |
| 4 | U | OP2 | 1H:33 | 4.67 | 1.00 | 40.81 |  |
| 5 | C | O5' | 1H:32 | 4.91 | 1.00 | 23.99 |  |
  |

  
 Coded contact list:   
1^st^ MG coordination shell missing (see Notes)  

### MG:1H:3290

|  | Res. type | Atom | Residue | Distance | Occupancy | B-factor | Closest solute to 1^st^ shell water |
| --- | --- | --- | --- | --- | --- | --- | --- |
| **Ion** | | | |  | 1.00 | 38.40 || 1 | A | OP2 | 1H:63 | 2.77 | 1.00 | 42.84 |
| 2 | U | O2' | 1H:62 | 3.12 | 1.00 | 35.78 |  |
| 3 | U | O3' | 1H:62 | 4.02 | 1.00 | 44.10 |  |
| 4 | U | OP2 | 1H:58 | 4.68 | 1.00 | 31.37 |  |
| 5 | A | O5' | 1H:63 | 4.91 | 1.00 | 44.20 |  |
| 6 | A | OP1 | 1H:63 | 4.98 | 1.00 | 45.52 |  |
  |

  
 Coded contact list:   
 Exclusion zone contact(s): /O2'/OP2/  
Consider K in place of MG (2 valid contact(s) for K vs. 0 for MG)  

### MG:1H:3291

|  | Res. type | Atom | Residue | Distance | Occupancy | B-factor | Closest solute to 1^st^ shell water |
| --- | --- | --- | --- | --- | --- | --- | --- |
| **Ion** | | | |  | 1.00 | 47.59 || 1 | A | OP2 | 1H:63 | 2.77 | 1.00 | 42.84 |
| 2 | C | OP2 | 1H:64 | 3.16 | 1.00 | 33.09 |  |
| 3 | LYS | O | F8:72 | 3.49 | 1.00 | 35.35 |  |
| 4 | A | O5' | 1H:63 | 3.83 | 1.00 | 44.20 |  |
| 5 | PRO | N | F8:74 | 4.48 | 1.00 | 27.05 |  |
| 6 | U | O3' | 1H:62 | 4.60 | 1.00 | 44.10 |  |
| 7 | C | OP1 | 1H:64 | 4.69 | 1.00 | 33.08 |  |
| 8 | U | O2' | 1H:62 | 4.70 | 1.00 | 35.78 |  |
| 9 | ARG | N | F8:73 | 4.84 | 1.00 | 31.32 |  |
| 10 | A | O3' | 1H:63 | 4.86 | 1.00 | 33.79 |  |
  |

  
 Coded contact list:   
 Exclusion zone contact(s): /OP2/OP2/  
Consider K in place of MG (2 valid contact(s) for K vs. 0 for MG)  

### MG:1H:3292

|  | Res. type | Atom | Residue | Distance | Occupancy | B-factor | Closest solute to 1^st^ shell water |
| --- | --- | --- | --- | --- | --- | --- | --- |
| **Ion** | | | |  | 1.00 | 32.65 || 1 | G | O6 | 1H:1008 | 2.75 | 1.00 | 35.79 |
| 2 | Water 1 | | 1H:4606 | 3.03 | 1.00 | 41.50 | 5.17 (G:O6 1H:1008) |
| 3 | Water 2 | | 1H:4230 | 3.42 | 1.00 | 31.55 | 2.94 (G:N7 1H:1008) |
| 4 | Water 3 | | 1H:4247 | 3.45 | 1.00 | 34.38 | 2.96 (U:O4 1H:1009) |
| 5 | U | O4 | 1H:1009 | 3.47 | 1.00 | 31.82 |  |
| 6 | C | N4 | 1H:1001 | 3.90 | 1.00 | 27.77 |  |
| 7 | G | O6 | 1H:1000 | 4.30 | 1.00 | 28.16 |  |
| 8 | G | N1 | 1H:1002 | 4.51 | 1.00 | 33.02 |  |
| 9 | G | N7 | 1H:1008 | 4.65 | 1.00 | 30.33 |  |
| 10 | G | O6 | 1H:1002 | 4.83 | 1.00 | 46.56 |  |
| 11 | G | N1 | 1H:1008 | 4.92 | 1.00 | 29.16 |  |
| 12 | U | N3 | 1H:1009 | 4.96 | 1.00 | 23.83 |  |
  |

  
 Coded contact list:   
 Exclusion zone contact(s): /O6/O_w_/  
Consider K in place of MG (1 valid contact(s) for K vs. 0 for MG)  

### MG:1H:3293

|  | Res. type | Atom | Residue | Distance | Occupancy | B-factor | Closest solute to 1^st^ shell water |
| --- | --- | --- | --- | --- | --- | --- | --- |
| **Ion** | | | |  | 1.00 | 40.91 || 1 | G | O6 | 1H:890 | 3.50 | 1.00 | 34.30 |
| 2 | G | N7 | 1H:890 | 3.59 | 1.00 | 31.30 |  |
| 3 | G | O6 | 1H:891 | 3.96 | 1.00 | 45.21 |  |
| 4 | G | N7 | 1H:891 | 4.27 | 1.00 | 39.19 |  |
| 5 | A | N7 | 1H:889 | 4.94 | 1.00 | 30.40 |  |
  |

  
 Coded contact list:   
1^st^ MG coordination shell missing (see Notes)  

### MG:1H:3294

|  | Res. type | Atom | Residue | Distance | Occupancy | B-factor | Closest solute to 1^st^ shell water |
| --- | --- | --- | --- | --- | --- | --- | --- |
| **Ion** | | | |  | 1.00 | 59.32 || 1 | Water 1 | | 1H:4508 | 3.18 | 1.00 | 61.37 | 3.74 (G:O6 1H:380) |
| 2 | Water 2 | | 1H:4523 | 3.42 | 1.00 | 66.21 | 5.58 (G:N7 1H:380) |
| 3 | Water 3 | | 1H:4225 | 3.52 | 1.00 | 72.52 | 3.49 (G:N7 1H:380) |
| 4 | G | O6 | 1H:380 | 3.63 | 1.00 | 67.56 |  |
| 5 | G | N7 | 1H:380 | 3.64 | 1.00 | 72.40 |  |
| 6 | G | N7 | 1H:381 | 3.88 | 1.00 | 70.19 |  |
| 7 | G | O6 | 1H:381 | 3.96 | 1.00 | 65.69 |  |
  |

  
 Coded contact list:   
 Exclusion zone contact(s): /O_w_/  
1^st^ MG coordination shell missing (see Notes)  

### MG:1H:3295

|  | Res. type | Atom | Residue | Distance | Occupancy | B-factor | Closest solute to 1^st^ shell water |
| --- | --- | --- | --- | --- | --- | --- | --- |
| **Ion** | | | |  | 1.00 | 29.53 || 1 | G | OP1 | 1H:27 | 2.78 | 1.00 | 26.31 |
| 2 | G | O2' | 1H:1306 | 3.52 | 1.00 | 24.39 |  |
| 3 | G | O5' | 1H:27 | 4.06 | 1.00 | 21.97 |  |
| 4 | G | O3' | 1H:1306 | 4.12 | 1.00 | 19.73 |  |
| 5 | G | OP2 | 1H:27 | 4.67 | 1.00 | 30.41 |  |
  |

  
 Coded contact list:   
 Exclusion zone contact(s): /OP1/  
Consider K in place of MG (1 valid contact(s) for K vs. 0 for MG)  

### MG:1H:3296

|  | Res. type | Atom | Residue | Distance | Occupancy | B-factor | Closest solute to 1^st^ shell water |
| --- | --- | --- | --- | --- | --- | --- | --- |
| **Ion** | | | |  | 1.00 | 32.27 || 1 | G | O3' | 1H:1306 | 3.77 | 1.00 | 19.73 |
| 2 | G | OP1 | 1H:1307 | 3.77 | 1.00 | 28.70 |  |
| 3 | G | OP1 | 1H:27 | 4.56 | 1.00 | 26.31 |  |
| 4 | A | O2' | 1H:498 | 4.71 | 1.00 | 28.12 |  |
  |

  
 Coded contact list:   
1^st^ MG coordination shell missing (see Notes)  

### MG:1H:3297

|  | Res. type | Atom | Residue | Distance | Occupancy | B-factor | Closest solute to 1^st^ shell water |
| --- | --- | --- | --- | --- | --- | --- | --- |
| **Ion** | | | |  | 1.00 | 40.33 || 1 | G | N7 | 1H:2265 | 2.95 | 1.00 | 30.26 |
| 2 | G | OP2 | 1H:2265 | 3.40 | 1.00 | 23.08 |  |
| 3 | OMG | OP1 | 1H:2264 | 3.74 | 1.00 | 24.67 |  |
| 4 | G | O6 | 1H:2266 | 4.14 | 1.00 | 37.10 |  |
| 5 | G | N7 | 1H:2266 | 4.22 | 1.00 | 29.79 |  |
| 6 | OMG | O5' | 1H:2264 | 4.48 | 1.00 | 20.47 |  |
| 7 | G | N9 | 1H:2265 | 4.55 | 1.00 | 21.44 |  |
| 8 | OMG | N7 | 1H:2264 | 4.61 | 1.00 | 24.44 |  |
| 9 | U | O2' | 1H:2262 | 4.62 | 1.00 | 26.89 |  |
| 10 | G | O5' | 1H:2265 | 4.63 | 1.00 | 23.51 |  |
  |

  
 Coded contact list:   
 Exclusion zone contact(s): /N7/OP2/  
Consider K in place of MG (2 valid contact(s) for K vs. 0 for MG)  

### MG:1H:3298

|  | Res. type | Atom | Residue | Distance | Occupancy | B-factor | Closest solute to 1^st^ shell water |
| --- | --- | --- | --- | --- | --- | --- | --- |
| **Ion** | | | |  | 1.00 | 28.41 || 1 | U | O2' | 1H:2262 | 2.47 | 1.00 | 26.89 |
| 2 | LYS | NZ | 88:85 | 3.16 | 1.00 | 30.77 |  |
| 3 | U | O3' | 1H:2262 | 3.31 | 1.00 | 20.39 |  |
| 4 | G | OP2 | 1H:2265 | 3.50 | 1.00 | 23.08 |  |
| 5 | OMG | OP1 | 1H:2264 | 3.88 | 1.00 | 24.67 |  |
| 6 | U | O2 | 1H:2262 | 3.97 | 1.00 | 29.46 |  |
| 7 | G | OP1 | 1H:2263 | 3.99 | 1.00 | 25.46 |  |
| 8 | G | O5' | 1H:2263 | 4.17 | 1.00 | 22.28 |  |
| 9 | OMG | O5' | 1H:2264 | 4.72 | 1.00 | 20.47 |  |
  |

  
 Coded contact list:   
 Exclusion zone contact(s): /NZ_Lys_/O2'/O3'/  
Consider K in place of MG (2 valid contact(s) for K vs. 0 for MG)  

### MG:1H:3299

|  | Res. type | Atom | Residue | Distance | Occupancy | B-factor | Closest solute to 1^st^ shell water |
| --- | --- | --- | --- | --- | --- | --- | --- |
| **Ion** | | | |  | 1.00 | 47.60 || 1 | G | OP2 | 1H:1034 | 3.05 | 1.00 | 33.00 |
| 2 | G | OP1 | 1H:1034 | 3.11 | 1.00 | 33.55 |  |
| 3 | LYS | NZ | L8:10 | 3.47 | 1.00 | 41.96 |  |
| 4 | TYR | OH | L8:15 | 3.82 | 1.00 | 36.96 |  |
| 5 | C | O3' | 1H:1033 | 4.06 | 1.00 | 32.54 |  |
| 6 | C | OP1 | 1H:1033 | 4.19 | 1.00 | 34.39 |  |
| 7 | Water 1 | | L8:202 | 4.20 | 1.00 | 38.02 | 3.48 (G:OP2 1H:1034) |
| 8 | Water 2 | | 1H:4185 | 4.28 | 1.00 | 37.35 | 3.28 (C:OP1 1H:1033) |
| 9 | C | OP2 | 16:84 | 4.40 | 1.00 | 43.97 |  |
| 10 | C | OP1 | 16:84 | 4.54 | 1.00 | 54.72 |  |
| 11 | G | O5' | 1H:1034 | 4.94 | 1.00 | 27.97 |  |
  |

  
 Coded contact list:   
 Exclusion zone contact(s): /OP1/OP2/  
Consider K in place of MG (2 valid contact(s) for K vs. 0 for MG)  

### MG:1H:3300

|  | Res. type | Atom | Residue | Distance | Occupancy | B-factor | Closest solute to 1^st^ shell water |
| --- | --- | --- | --- | --- | --- | --- | --- |
| **Ion** | | | |  | 1.00 | 59.51 || 1 | C | OP2 | 1H:1096 | 3.30 | 1.00 | 93.19 |
| 2 | A | N1 | 1H:1097 | 3.86 | 1.00 | 97.11 |  |
| 3 | G | O6 | 1H:1154 | 4.10 | 1.00 | 119.76 |  |
| 4 | G | O6 | 1H:1098 | 4.13 | 1.00 | 107.02 |  |
| 5 | G | N7 | 1H:1098 | 4.21 | 1.00 | 94.18 |  |
| 6 | A | N3 | 1H:1097 | 4.47 | 1.00 | 88.24 |  |
| 7 | U | O4 | 1H:1155 | 4.64 | 1.00 | 111.15 |  |
| 8 | A | OP2 | 1H:1097 | 4.79 | 1.00 | 100.81 |  |
| 9 | C | N4 | 1H:1099 | 4.99 | 1.00 | 103.87 |  |
  |

  
 Coded contact list:   
 Exclusion zone contact(s): /OP2/  
Consider K in place of MG (1 valid contact(s) for K vs. 0 for MG)  

### MG:1H:3301

|  | Res. type | Atom | Residue | Distance | Occupancy | B-factor | Closest solute to 1^st^ shell water |
| --- | --- | --- | --- | --- | --- | --- | --- |
| **Ion** | | | |  | 1.00 | 41.40 || 1 | A | OP2 | 1H:555 | 4.20 | 1.00 | 61.62 |
| 2 | A | O5' | 1H:555 | 4.23 | 1.00 | 56.59 |  |
| 3 | ARG | NH2 | 58:114 | 4.24 | 1.00 | 39.75 |  |
| 4 | A | O3' | 1H:554 | 4.32 | 1.00 | 53.40 |  |
| 5 | G | OP1 | 1H:556 | 4.44 | 1.00 | 52.14 |  |
| 6 | A | O2' | 1H:554 | 4.55 | 1.00 | 46.75 |  |
| 7 | A | O4' | 1H:555 | 4.87 | 1.00 | 38.83 |  |
| 8 | A | N1 | 1H:2065 | 4.93 | 1.00 | 34.28 |  |
| 9 | A | N9 | 1H:554 | 4.95 | 1.00 | 38.01 |  |
  |

  
 Coded contact list:   
1^st^ MG coordination shell missing (see Notes)  

### MG:1H:3302

|  | Res. type | Atom | Residue | Distance | Occupancy | B-factor | Closest solute to 1^st^ shell water |
| --- | --- | --- | --- | --- | --- | --- | --- |
| **Ion** | | | |  | 1.00 | 29.06 || 1 | Water 1 | | 1H:3713 | 2.47 | 1.00 | 16.03 | 2.23 (A:OP1 1H:1815) |
| 2 | A | OP1 | 1H:1815 | 2.64 | 1.00 | 17.36 |  |
| 3 | C | O3' | 1H:1814 | 2.75 | 1.00 | 15.30 |  |
| 4 | C | O2' | 1H:1814 | 3.12 | 1.00 | 19.23 |  |
| 5 | Water 2 | | 1H:3828 | 3.53 | 1.00 | 19.87 | 3.29 (A:OP1 1H:1815) |
| 6 | A | O5' | 1H:1815 | 4.22 | 1.00 | 21.16 |  |
| 7 | A | OP2 | 1H:1815 | 4.47 | 1.00 | 16.41 |  |
  |

  
 Coded contact list:   
 Exclusion zone contact(s): /O2'/O3'/O_w_/OP1/  
Consider K in place of MG (3 valid contact(s) for K vs. 0 for MG)  
MG hydration shell issue: water too close (< 2.4 Angs.)  

### MG:1H:3303

|  | Res. type | Atom | Residue | Distance | Occupancy | B-factor | Closest solute to 1^st^ shell water |
| --- | --- | --- | --- | --- | --- | --- | --- |
| **Ion** | | | |  | 1.00 | 46.67 || 1 | G | O6 | 1H:146 | 3.80 | 1.00 | 44.29 |
| 2 | G | O6 | 1H:145 | 3.82 | 1.00 | 34.58 |  |
| 3 | G | N7 | 1H:145 | 4.25 | 1.00 | 44.58 |  |
| 4 | C | N4 | 1H:144 | 4.40 | 1.00 | 40.70 |  |
| 5 | U | O4 | 1H:147 | 4.87 | 1.00 | 41.09 |  |
| 6 | G | N7 | 1H:146 | 4.97 | 1.00 | 37.69 |  |
  |

  
 Coded contact list:   
1^st^ MG coordination shell missing (see Notes)  

### MG:1H:3304

|  | Res. type | Atom | Residue | Distance | Occupancy | B-factor | Closest solute to 1^st^ shell water |
| --- | --- | --- | --- | --- | --- | --- | --- |
| **Ion** | | | |  | 1.00 | 38.84 || 1 | U | OP1 | 1H:102 | 3.89 | 1.00 | 44.59 |
| 2 | C | OP2 | 1H:76 | 3.99 | 1.00 | 37.84 |  |
| 3 | C | OP1 | 1H:76 | 4.67 | 1.00 | 37.46 |  |
| 4 | C | OP1 | 1H:75 | 4.90 | 1.00 | 27.88 |  |
  |

  
 Coded contact list:   
1^st^ MG coordination shell missing (see Notes)  

### MG:1H:3305

|  | Res. type | Atom | Residue | Distance | Occupancy | B-factor | Closest solute to 1^st^ shell water |
| --- | --- | --- | --- | --- | --- | --- | --- |
| **Ion** | | | |  | 1.00 | 41.66 || 1 | U | O4 | 1H:25 | 3.52 | 1.00 | 29.64 |
| 2 | G | N7 | 1H:24 | 3.70 | 1.00 | 25.33 |  |
| 3 | G | OP2 | 1H:24 | 4.12 | 1.00 | 30.28 |  |
| 4 | A | OP2 | 1H:539 | 4.15 | 1.00 | 28.03 |  |
| 5 | G | N7 | 1H:23 | 4.55 | 1.00 | 30.40 |  |
| 6 | G | O6 | 1H:24 | 4.87 | 1.00 | 31.26 |  |
  |

  
 Coded contact list:   
1^st^ MG coordination shell missing (see Notes)  

### MG:1H:3306

|  | Res. type | Atom | Residue | Distance | Occupancy | B-factor | Closest solute to 1^st^ shell water |
| --- | --- | --- | --- | --- | --- | --- | --- |
| **Ion** | | | |  | 1.00 | 34.50 || 1 | Water 1 | | 1H:3867 | 2.54 | 1.00 | 22.55 | 2.47 (A:OP2 1H:1203) |
| 2 | G | OP2 | 1H:1204 | 2.85 | 1.00 | 30.27 |  |
| 3 | A | OP2 | 1H:1203 | 2.97 | 1.00 | 34.25 |  |
| 4 | G | N7 | 1H:1204 | 3.22 | 1.00 | 33.40 |  |
| 5 | C | N4 | 1H:1205 | 3.35 | 1.00 | 33.51 |  |
| 6 | A | O5' | 1H:1203 | 3.53 | 1.00 | 31.06 |  |
| 7 | G | O6 | 1H:1204 | 3.72 | 1.00 | 34.50 |  |
| 8 | A | O3' | 1H:1203 | 4.42 | 1.00 | 31.89 |  |
| 9 | Water 2 | | 1H:4088 | 4.72 | 1.00 | 38.93 | 3.26 (A:OP2 1H:1203) |
| 10 | A | O4' | 1H:1203 | 4.72 | 1.00 | 35.59 |  |
| 11 | G | O6 | 1H:1044 | 4.85 | 1.00 | 36.33 |  |
| 12 | A | O3' | 1H:1202 | 4.86 | 1.00 | 35.88 |  |
| 13 | G | N9 | 1H:1204 | 4.90 | 1.00 | 32.96 |  |
| 14 | A | O2' | 1H:1202 | 4.90 | 1.00 | 32.88 |  |
| 15 | G | OP1 | 1H:1204 | 4.96 | 1.00 | 34.29 |  |
  |

  
 Coded contact list:   
 Exclusion zone contact(s): /N4/N7/O_w_/OP2/OP2/  
Consider K in place of MG (4 valid contact(s) for K vs. 0 for MG)  

### MG:1H:3307

|  | Res. type | Atom | Residue | Distance | Occupancy | B-factor | Closest solute to 1^st^ shell water |
| --- | --- | --- | --- | --- | --- | --- | --- |
| **Ion** | | | |  | 1.00 | 25.95 || 1 | Water 1 | | 1H:4064 | 2.56 | 1.00 | 22.25 | 2.71 (G:N7 1H:2071) |
| 2 | Water 2 | | 1H:4579 | 2.96 | 1.00 | 28.86 | 4.64 (U:O4 1H:2070) |
| 3 | U | O4 | 1H:2070 | 3.48 | 1.00 | 20.14 |  |
| 4 | Water 3 | | 1H:4318 | 3.54 | 1.00 | 31.37 | 3.12 (G:O6 1H:2072) |
| 5 | G | O6 | 1H:2072 | 3.63 | 1.00 | 21.16 |  |
| 6 | G | O6 | 1H:2071 | 3.72 | 1.00 | 19.35 |  |
| 7 | C | N4 | 1H:2632 | 3.81 | 1.00 | 19.64 |  |
| 8 | C | N4 | 1H:2633 | 4.34 | 1.00 | 22.15 |  |
| 9 | A | N6 | 1H:2634 | 4.54 | 1.00 | 16.90 |  |
| 10 | Water 4 | | 1H:4379 | 4.74 | 1.00 | 27.03 | 3.53 (G:N7 1H:2071) |
| 11 | G | N7 | 1H:2071 | 4.92 | 1.00 | 18.12 |  |
  |

  
 Coded contact list:   
 Exclusion zone contact(s): /O_w_/O_w_/  
1^st^ MG coordination shell missing (see Notes)  

### MG:1H:3308

|  | Res. type | Atom | Residue | Distance | Occupancy | B-factor | Closest solute to 1^st^ shell water |
| --- | --- | --- | --- | --- | --- | --- | --- |
| **Ion** | | | |  | 1.00 | 27.17 || 1 | G | OP1 | 1H:1379 | 2.91 | 1.00 | 25.93 |
| 2 | A | O3' | 1H:1378 | 3.36 | 1.00 | 27.07 |  |
| 3 | G | O5' | 1H:1379 | 4.53 | 1.00 | 29.10 |  |
| 4 | C | OP2 | 1H:1362 | 4.74 | 1.00 | 27.71 |  |
| 5 | A | O2' | 1H:1378 | 4.98 | 1.00 | 29.19 |  |
  |

  
 Coded contact list:   
 Exclusion zone contact(s): /O3'/OP1/  
Consider K in place of MG (2 valid contact(s) for K vs. 0 for MG)  

### MG:1H:3309

|  | Res. type | Atom | Residue | Distance | Occupancy | B-factor | Closest solute to 1^st^ shell water |
| --- | --- | --- | --- | --- | --- | --- | --- |
| **Ion** | | | |  | 1.00 | 52.87 || 1 | Water 1 | | 1H:4114 | 2.35 | 1.00 | 42.39 | 2.78 (G:N7 1H:1691) |
| 2 | Water 2 | | 1H:4594 | 2.93 | 1.00 | 62.26 | 4.70 (G:O6 1H:1691) |
| 3 | Water 3 | | 1H:4445 | 3.19 | 1.00 | 49.03 | 5.60 (G:N7 1H:1691) |
| 4 | G | N7 | 1H:1691 | 3.99 | 1.00 | 31.99 |  |
| 5 | G | O6 | 1H:1691 | 4.77 | 1.00 | 32.36 |  |
| 6 | Water 4 | | 1H:4345 | 4.78 | 1.00 | 28.85 | 3.37 (G:O6 1H:1691) |
| 7 | C | N4 | 1H:1692 | 4.92 | 1.00 | 34.25 |  |
| 8 | Water 5 | | 1H:3723 | 4.95 | 1.00 | 45.75 | 5.03 (G:N7 1H:1691) |
| 9 | G | OP2 | 1H:1691 | 4.97 | 1.00 | 34.48 |  |
  |

  
 Contact(s): /O_w_/  
 Coded contact list: O_w_  
 Exclusion zone contact(s): /O_w_/O_w_/  

### MG:1H:3310

|  | Res. type | Atom | Residue | Distance | Occupancy | B-factor | Closest solute to 1^st^ shell water |
| --- | --- | --- | --- | --- | --- | --- | --- |
| **Ion** | | | |  | 1.00 | 18.98 || 1 | C | OP2 | 1H:2629 | 2.15 | 1.00 | 27.64 |
| 2 | U | OP2 | 1H:2628 | 2.57 | 1.00 | 27.17 |  |
| 3 | Water 1 | | 1H:4317 | 2.85 | 1.00 | 39.90 | 3.11 (U:OP2 1H:2628) |
| 4 | Water 2 | | 1H:4318 | 3.57 | 1.00 | 31.37 | 4.46 (A:O2' 1H:2627) |
| 5 | Water 3 | | 1H:4579 | 3.59 | 1.00 | 28.86 | 4.65 (C:OP2 1H:2629) |
| 6 | U | O5' | 1H:2628 | 3.61 | 1.00 | 24.53 |  |
| 7 | A | O3' | 1H:2627 | 3.83 | 1.00 | 24.73 |  |
| 8 | U | O3' | 1H:2628 | 4.02 | 1.00 | 19.51 |  |
| 9 | C | OP1 | 1H:2629 | 4.17 | 1.00 | 25.74 |  |
| 10 | A | O2' | 1H:2627 | 4.50 | 1.00 | 22.78 |  |
| 11 | C | O5' | 1H:2629 | 4.67 | 1.00 | 27.01 |  |
| 12 | Water 4 | | 1H:3873 | 4.78 | 1.00 | 24.88 | 2.48 (U:OP2 1H:2628) |
| 13 | U | OP1 | 1H:2628 | 4.89 | 1.00 | 27.30 |  |
| 14 | Water 5 | | 1H:3625 | 4.96 | 1.00 | 22.15 | 2.79 (U:OP2 1H:2628) |
  |

  
 Contact(s): /OP2/  
 Coded contact list: O_ph_  
 Exclusion zone contact(s): /O_w_/OP2/  

### MG:1H:3311

|  | Res. type | Atom | Residue | Distance | Occupancy | B-factor | Closest solute to 1^st^ shell water |
| --- | --- | --- | --- | --- | --- | --- | --- |
| **Ion** | | | |  | 1.00 | 47.50 || 1 | G | O6 | 1H:1755 | 2.90 | 1.00 | 51.17 |
| 2 | G | O6 | 1H:1788 | 3.13 | 1.00 | 50.67 |  |
| 3 | Water 1 | | 1H:4272 | 3.14 | 1.00 | 47.72 | 3.01 (G:O6 1H:1788) |
| 4 | U | OP1 | 1H:1754 | 3.29 | 1.00 | 39.88 |  |
| 5 | Water 2 | | 1H:4191 | 3.30 | 1.00 | 54.14 | 2.88 (G:N7 1H:1788) |
| 6 | G | N7 | 1H:1755 | 3.67 | 1.00 | 45.44 |  |
| 7 | G | N7 | 1H:1788 | 4.17 | 1.00 | 47.42 |  |
| 8 | Water 3 | | 1H:4468 | 4.35 | 1.00 | 58.84 | 4.04 (U:OP1 1H:1754) |
| 9 | C | N4 | 1H:1783 | 4.36 | 1.00 | 53.36 |  |
| 10 | G | N1 | 1H:1755 | 4.78 | 1.00 | 45.56 |  |
| 11 | G | O6 | 1H:1782 | 4.99 | 1.00 | 69.10 |  |
  |

  
 Coded contact list:   
 Exclusion zone contact(s): /O6/O6/O_w_/O_w_/OP1/  
Consider K in place of MG (3 valid contact(s) for K vs. 0 for MG)  

### MG:1H:3312

|  | Res. type | Atom | Residue | Distance | Occupancy | B-factor | Closest solute to 1^st^ shell water |
| --- | --- | --- | --- | --- | --- | --- | --- |
| **Ion** | | | |  | 1.00 | 39.80 || 1 | U | OP2 | 1H:2039 | 3.38 | 1.00 | 20.97 |
| 2 | Water 1 | | 31:408 | 3.38 | 1.00 | 39.14 | 3.52 (U:OP1 1H:2039) |
| 3 | U | OP2 | 1H:2040 | 3.71 | 1.00 | 21.65 |  |
| 4 | U | OP1 | 1H:2039 | 3.75 | 1.00 | 26.41 |  |
| 5 | U | O5' | 1H:2039 | 4.22 | 1.00 | 20.73 |  |
| 6 | Water 2 | | 1H:3686 | 4.27 | 1.00 | 22.54 | 2.17 (U:OP2 1H:2040) |
| 7 | G | OP2 | 1H:603 | 4.45 | 1.00 | 18.30 |  |
| 8 | Water 3 | | 1H:4460 | 4.89 | 1.00 | 25.49 | 4.90 (U:OP2 1H:2040) |
  |

  
 Coded contact list:   
 Exclusion zone contact(s): /O_w_/OP2/  
Consider K in place of MG (1 valid contact(s) for K vs. 0 for MG)  

### MG:1H:3313

|  | Res. type | Atom | Residue | Distance | Occupancy | B-factor | Closest solute to 1^st^ shell water |
| --- | --- | --- | --- | --- | --- | --- | --- |
| **Ion** | | | |  | 1.00 | 44.89 || 1 | Water 1 | | 1H:3714 | 2.19 | 1.00 | 41.28 | 2.23 (G:OP1 1H:1653) |
| 2 | Water 2 | | 1H:4059 | 3.51 | 1.00 | 39.56 | 2.70 (A:OP1 1H:1655) |
| 3 | A | OP1 | 1H:1656 | 3.58 | 1.00 | 33.82 |  |
| 4 | Water 3 | | 1H:4395 | 4.04 | 1.00 | 37.66 | 4.52 (G:OP1 1H:1653) |
| 5 | G | OP1 | 1H:1653 | 4.14 | 1.00 | 35.64 |  |
| 6 | A | OP1 | 1H:1655 | 4.49 | 1.00 | 34.75 |  |
| 7 | A | O5' | 1H:1655 | 4.72 | 1.00 | 29.36 |  |
| 8 | Water 4 | | 1H:3626 | 4.85 | 1.00 | 32.53 | 3.34 (G:OP1 1H:1653) |
| 9 | A | O4' | 1H:1655 | 4.94 | 1.00 | 23.90 |  |
  |

  
 Contact(s): /O_w_/  
 Coded contact list: O_w_  
MG hydration shell issue: water too close (< 2.4 Angs.)  

### MG:1H:3314

|  | Res. type | Atom | Residue | Distance | Occupancy | B-factor | Closest solute to 1^st^ shell water |
| --- | --- | --- | --- | --- | --- | --- | --- |
| **Ion** | | | |  | 1.00 | 41.45 || 1 | Water 1 | | 1H:4549 | 2.84 | 1.00 | 45.60 | 4.08 (G:N7 1H:1394) |
| 2 | Water 2 | | 1H:4308 | 3.64 | 1.00 | 48.29 | 3.08 (C:O2 1H:1430) |
| 3 | Water 3 | | 1H:4169 | 3.73 | 1.00 | 46.15 | 2.85 (A:N6 1H:1431) |
| 4 | G | N7 | 1H:1395 | 3.97 | 1.00 | 33.40 |  |
| 5 | G | N7 | 1H:1394 | 4.26 | 1.00 | 35.21 |  |
| 6 | C | O2 | 1H:1430 | 4.70 | 1.00 | 42.27 |  |
| 7 | A | N6 | 1H:1431 | 4.86 | 1.00 | 38.40 |  |
  |

  
 Coded contact list:   
 Exclusion zone contact(s): /O_w_/  
1^st^ MG coordination shell missing (see Notes)  

### MG:1H:3315

|  | Res. type | Atom | Residue | Distance | Occupancy | B-factor | Closest solute to 1^st^ shell water |
| --- | --- | --- | --- | --- | --- | --- | --- |
| **Ion** | | | |  | 1.00 | 43.67 || 1 | A | N7 | 1H:1031 | 4.84 | 1.00 | 27.58 |
  |

  
 Coded contact list:   
1^st^ MG coordination shell missing (see Notes)  

### MG:1H:3316

|  | Res. type | Atom | Residue | Distance | Occupancy | B-factor | Closest solute to 1^st^ shell water |
| --- | --- | --- | --- | --- | --- | --- | --- |
| **Ion** | | | |  | 1.00 | 60.58 || 1 | Water 1 | | 1H:4588 | 2.75 | 1.00 | 54.35 | 4.55 (G:OP2 1H:1640) |
| 2 | G | OP2 | 1H:1640 | 3.62 | 1.00 | 60.16 |  |
| 3 | Water 2 | | 1H:4420 | 4.64 | 1.00 | 42.60 | 4.60 (G:OP2 1H:1640) |
| 4 | Water 3 | | 1H:4560 | 4.72 | 1.00 | 45.37 | 6.60 (G:OP2 1H:1640) |
| 5 | G | OP1 | 1H:1640 | 4.74 | 1.00 | 60.43 |  |
  |

  
 Coded contact list:   
 Exclusion zone contact(s): /O_w_/  
1^st^ MG coordination shell missing (see Notes)  

### MG:1H:3317

|  | Res. type | Atom | Residue | Distance | Occupancy | B-factor | Closest solute to 1^st^ shell water |
| --- | --- | --- | --- | --- | --- | --- | --- |
| **Ion** | | | |  | 1.00 | 52.18 || 1 | G | O6 | 1H:1514 | 3.56 | 1.00 | 60.60 |
| 2 | G | O6 | 1H:1513 | 3.81 | 1.00 | 57.60 |  |
| 3 | C | N4 | 1H:1512 | 3.87 | 1.00 | 55.32 |  |
| 4 | G | N7 | 1H:1513 | 4.00 | 1.00 | 54.44 |  |
| 5 | G | N7 | 1H:1514 | 4.83 | 1.00 | 56.79 |  |
| 6 | C | N4 | 1H:1493 | 4.84 | 1.00 | 64.88 |  |
| 7 | G | OP2 | 1H:1491 | 4.99 | 1.00 | 53.43 |  |
  |

  
 Coded contact list:   
1^st^ MG coordination shell missing (see Notes)  

### MG:1H:3318

|  | Res. type | Atom | Residue | Distance | Occupancy | B-factor | Closest solute to 1^st^ shell water |
| --- | --- | --- | --- | --- | --- | --- | --- |
| **Ion** | | | |  | 1.00 | 56.14 || 1 | U | O4 | 1H:375 | 2.34 | 1.00 | 54.23 |
| 2 | Water 1 | | 1H:4202 | 3.14 | 1.00 | 52.31 | 2.90 (G:O6 1H:374) |
| 3 | Water 2 | | 1H:4586 | 3.49 | 1.00 | 69.63 | 4.54 (G:O6 1H:374) |
| 4 | G | O6 | 1H:374 | 3.58 | 1.00 | 50.50 |  |
| 5 | G | O6 | 1H:376 | 3.63 | 1.00 | 56.30 |  |
| 6 | G | N7 | 1H:374 | 3.82 | 1.00 | 48.33 |  |
| 7 | G | N1 | 1H:376 | 4.44 | 1.00 | 55.26 |  |
| 8 | U | N3 | 1H:375 | 4.44 | 1.00 | 50.20 |  |
| 9 | Water 3 | | 1H:4469 | 4.86 | 1.00 | 55.41 | 5.91 (G:O6 1H:374) |
  |

  
 Contact(s): /O4/  
 Coded contact list: O_b_  
 Exclusion zone contact(s): /O_w_/  

### MG:1H:3319

|  | Res. type | Atom | Residue | Distance | Occupancy | B-factor | Closest solute to 1^st^ shell water |
| --- | --- | --- | --- | --- | --- | --- | --- |
| **Ion** | | | |  | 1.00 | 50.87 || 1 | Water 1 | | 1H:3865 | 2.82 | 1.00 | 43.24 | 2.47 (U:OP1 1H:1287) |
| 2 | U | OP1 | 1H:1287 | 3.37 | 1.00 | 46.04 |  |
| 3 | U | OP1 | 1H:1245 | 3.94 | 1.00 | 23.04 |  |
| 4 | Water 2 | | 1H:3995 | 3.97 | 1.00 | 32.19 | 4.55 (G:O3' 1H:1286) |
| 5 | G | OP1 | 1H:1261 | 4.09 | 1.00 | 31.99 |  |
| 6 | A | O3' | 1H:1260 | 4.13 | 1.00 | 30.48 |  |
| 7 | G | O3' | 1H:1286 | 4.19 | 1.00 | 33.10 |  |
| 8 | Water 3 | | 1H:3790 | 4.48 | 1.00 | 40.42 | 2.36 (U:OP1 1H:1245) |
| 9 | U | OP2 | 1H:1245 | 4.54 | 1.00 | 36.31 |  |
  |

  
 Coded contact list:   
 Exclusion zone contact(s): /O_w_/OP1/  
Consider K in place of MG (1 valid contact(s) for K vs. 0 for MG)  

### MG:1H:3320

|  | Res. type | Atom | Residue | Distance | Occupancy | B-factor | Closest solute to 1^st^ shell water |
| --- | --- | --- | --- | --- | --- | --- | --- |
| **Ion** | | | |  | 0.50 | 8.94 || 1 | G | O2' | 1H:176 | 2.94 | 1.00 | 24.07 |
| 2 | G | O4' | 1H:177 | 3.39 | 1.00 | 18.24 |  |
| 3 | G | N3 | 1H:176 | 4.02 | 1.00 | 18.19 |  |
| 4 | G | N2 | 1H:176 | 4.77 | 1.00 | 17.41 |  |
| 5 | G | O5' | 1H:177 | 4.85 | 1.00 | 13.79 |  |
| 6 | G | N9 | 1H:176 | 4.98 | 1.00 | 18.18 |  |
  |

  
 Coded contact list:   
 Exclusion zone contact(s): /O2'/O4'/  
Consider K in place of MG (2 valid contact(s) for K vs. 0 for MG)  

### MG:1H:3320

|  | Res. type | Atom | Residue | Distance | Occupancy | B-factor | Closest solute to 1^st^ shell water |
| --- | --- | --- | --- | --- | --- | --- | --- |
| **Ion** | | | |  | 0.50 | 9.02 || 1 | G | N2 | 1H:176 | 3.65 | 1.00 | 17.41 |
| 2 | G | O4' | 1H:177 | 3.78 | 1.00 | 18.24 |  |
| 3 | G | O2' | 1H:177 | 4.33 | 1.00 | 19.00 |  |
| 4 | A | O4' | 1H:1413 | 4.37 | 1.00 | 26.29 |  |
| 5 | G | N3 | 1H:176 | 4.40 | 1.00 | 18.19 |  |
| 6 | A | O2' | 1H:201 | 4.49 | 1.00 | 25.83 |  |
| 7 | G | N9 | 1H:177 | 4.65 | 1.00 | 14.81 |  |
| 8 | C | O2 | 1H:200 | 4.77 | 1.00 | 17.41 |  |
| 9 | A | O2' | 1H:1413 | 4.85 | 1.00 | 24.24 |  |
  |

  
 Coded contact list:   
1^st^ MG coordination shell missing (see Notes)  

### MG:1H:3321

|  | Res. type | Atom | Residue | Distance | Occupancy | B-factor | Closest solute to 1^st^ shell water |
| --- | --- | --- | --- | --- | --- | --- | --- |
| **Ion** | | | |  | 1.00 | 36.81 || 1 | C | O2 | 1H:611 | 3.60 | 1.00 | 29.19 |
| 2 | C | OP1 | 1H:712 | 4.16 | 1.00 | 34.03 |  |
| 3 | ARG | NH1 | 78:33 | 4.51 | 1.00 | 13.87 |  |
| 4 | C | O2' | 1H:611 | 4.63 | 1.00 | 22.44 |  |
| 5 | ALA | O | 78:31 | 4.67 | 1.00 | 23.47 |  |
| 6 | C | N3 | 1H:611 | 4.88 | 1.00 | 21.85 |  |
  |

  
 Coded contact list:   
1^st^ MG coordination shell missing (see Notes)  

### MG:1H:3322

|  | Res. type | Atom | Residue | Distance | Occupancy | B-factor | Closest solute to 1^st^ shell water |
| --- | --- | --- | --- | --- | --- | --- | --- |
| **Ion** | | | |  | 1.00 | 23.61 || 1 | U | OP2 | 1H:612 | 2.49 | 1.00 | 25.93 |
| 2 | U | OP1 | 1H:612 | 3.39 | 1.00 | 36.67 |  |
| 3 | C | O2' | 1H:611 | 3.63 | 1.00 | 22.44 |  |
| 4 | C | OP2 | 1H:712 | 3.69 | 1.00 | 28.67 |  |
| 5 | C | O3' | 1H:611 | 4.59 | 1.00 | 33.48 |  |
| 6 | U | O5' | 1H:612 | 4.66 | 1.00 | 26.14 |  |
  |

  
 Coded contact list:   
 Exclusion zone contact(s): /OP1/OP2/  
Consider NA in place of MG (1 valid contact(s) for NA vs. 0 for MG)  

### MG:1H:3323

|  | Res. type | Atom | Residue | Distance | Occupancy | B-factor | Closest solute to 1^st^ shell water |
| --- | --- | --- | --- | --- | --- | --- | --- |
| **Ion** | | | |  | 1.00 | 52.52 || 1 | G | O6 | 1H:2384 | 2.73 | 1.00 | 49.35 |
| 2 | G | N7 | 1H:2384 | 3.46 | 1.00 | 47.55 |  |
| 3 | G | N7 | 1H:2383 | 3.81 | 1.00 | 52.96 |  |
| 4 | G | OP2 | 1H:2383 | 4.09 | 1.00 | 59.53 |  |
| 5 | G | N1 | 1H:2384 | 4.95 | 1.00 | 44.81 |  |
  |

  
 Coded contact list:   
 Exclusion zone contact(s): /O6/  
Consider K in place of MG (1 valid contact(s) for K vs. 0 for MG)  

### MG:1H:3324

|  | Res. type | Atom | Residue | Distance | Occupancy | B-factor | Closest solute to 1^st^ shell water |
| --- | --- | --- | --- | --- | --- | --- | --- |
| **Ion** | | | |  | 1.00 | 12.36 || 1 | G | OP1 | 1H:2441 | 2.09 | 1.00 | 18.31 |
| 2 | U | OP1 | 1H:874 | 2.16 | 1.00 | 21.97 |  |
| 3 | G | OP1 | 1H:2442 | 2.26 | 1.00 | 19.80 |  |
| 4 | C | O5' | 1H:2440 | 3.10 | 1.00 | 19.58 |  |
| 5 | C | OP1 | 1H:2440 | 3.22 | 1.00 | 18.09 |  |
| 6 | U | OP2 | 1H:874 | 3.45 | 1.00 | 14.92 |  |
| 7 | C | OP2 | 1H:2440 | 3.82 | 1.00 | 17.35 |  |
| 8 | U | O5' | 1H:874 | 4.13 | 1.00 | 21.35 |  |
| 9 | G | OP2 | 1H:2441 | 4.14 | 1.00 | 23.82 |  |
| 10 | G | O3' | 1H:2441 | 4.21 | 1.00 | 21.53 |  |
| 11 | G | O5' | 1H:2441 | 4.21 | 1.00 | 24.47 |  |
| 12 | C | O3' | 1H:873 | 4.43 | 1.00 | 16.38 |  |
| 13 | U | OP2 | 1H:875 | 4.60 | 1.00 | 26.49 |  |
| 14 | G | OP2 | 1H:2442 | 4.61 | 1.00 | 20.29 |  |
| 15 | C | O3' | 1H:2440 | 4.66 | 1.00 | 18.51 |  |
| 16 | G | O5' | 1H:2442 | 4.79 | 1.00 | 18.50 |  |
| 17 | A | O3' | 1H:2439 | 4.95 | 1.00 | 24.04 |  |
  |

  
**1^st^ shell ligand angles:**   
1-2: 88; 1-3: 74; 2-3: 76;   
  
 Contact(s): /OP1/OP1/OP1/  
 Coded contact list: 3O_ph_  
 Exclusion zone contact(s): /O5'/OP1/  

### MG:1H:3325

|  | Res. type | Atom | Residue | Distance | Occupancy | B-factor | Closest solute to 1^st^ shell water |
| --- | --- | --- | --- | --- | --- | --- | --- |
| **Ion** | | | |  | 1.00 | 50.07 || 1 | C | OP2 | 1H:396 | 2.94 | 1.00 | 57.17 |
| 2 | C | OP2 | 1H:395 | 3.12 | 1.00 | 72.48 |  |
| 3 | C | OP1 | 1H:432 | 3.28 | 1.00 | 54.78 |  |
| 4 | A | O3' | 1H:394 | 3.50 | 1.00 | 76.55 |  |
| 5 | C | OP2 | 1H:432 | 3.51 | 1.00 | 55.74 |  |
| 6 | U | O3' | 1H:431 | 3.62 | 1.00 | 53.53 |  |
| 7 | A | O2' | 1H:394 | 4.02 | 1.00 | 76.40 |  |
| 8 | U | O2' | 1H:431 | 4.41 | 1.00 | 51.37 |  |
| 9 | C | O5' | 1H:395 | 4.59 | 1.00 | 68.26 |  |
  |

  
 Coded contact list:   
 Exclusion zone contact(s): /OP1/OP2/OP2/  
Consider K in place of MG (3 valid contact(s) for K vs. 0 for MG)  

### MG:1H:3326

|  | Res. type | Atom | Residue | Distance | Occupancy | B-factor | Closest solute to 1^st^ shell water |
| --- | --- | --- | --- | --- | --- | --- | --- |
| **Ion** | | | |  | 1.00 | 60.44 || 1 | G | OP2 | 1H:1261 | 3.60 | 1.00 | 37.38 |
| 2 | G | O6 | 1H:1262 | 4.51 | 1.00 | 40.02 |  |
| 3 | G | N7 | 1H:1261 | 4.57 | 1.00 | 35.52 |  |
| 4 | A | OP2 | 1H:1260 | 4.69 | 1.00 | 30.29 |  |
| 5 | G | N7 | 1H:1262 | 4.73 | 1.00 | 38.96 |  |
  |

  
 Coded contact list:   
1^st^ MG coordination shell missing (see Notes)  

### MG:1H:3327

|  | Res. type | Atom | Residue | Distance | Occupancy | B-factor | Closest solute to 1^st^ shell water |
| --- | --- | --- | --- | --- | --- | --- | --- |
| **Ion** | | | |  | 1.00 | 40.26 || 1 | U | O4 | 1H:2650 | 2.87 | 1.00 | 37.39 |
| 2 | A | OP2 | 1H:2790 | 3.22 | 1.00 | 34.36 |  |
| 3 | U | O4 | 1H:2649 | 3.36 | 1.00 | 45.03 |  |
| 4 | G | O6 | 1H:2651 | 3.41 | 1.00 | 33.28 |  |
| 5 | A | OP1 | 1H:2790 | 3.58 | 1.00 | 28.44 |  |
| 6 | G | O6 | 1H:2796 | 3.94 | 1.00 | 40.11 |  |
| 7 | G | N1 | 1H:2651 | 3.96 | 1.00 | 28.60 |  |
| 8 | U | N3 | 1H:2649 | 4.18 | 1.00 | 43.44 |  |
| 9 | A | O5' | 1H:2790 | 4.50 | 1.00 | 31.97 |  |
  |

  
 Coded contact list:   
 Exclusion zone contact(s): /O4/O4/OP2/  
Consider K in place of MG (3 valid contact(s) for K vs. 0 for MG)  

### MG:1H:3328

|  | Res. type | Atom | Residue | Distance | Occupancy | B-factor | Closest solute to 1^st^ shell water |
| --- | --- | --- | --- | --- | --- | --- | --- |
| **Ion** | | | |  | 0.50 | 22.52 || 1 | G | O6 | 1H:230 | 2.41 | 1.00 | 29.85 |
| 2 | U | O4 | 1H:229 | 2.80 | 1.00 | 33.26 |  |
| 3 | G | N1 | 1H:230 | 3.93 | 1.00 | 30.07 |  |
| 4 | A | OP2 | 1H:246 | 4.67 | 1.00 | 27.34 |  |
| 5 | U | N3 | 1H:229 | 4.94 | 1.00 | 29.44 |  |
  |

  
 Coded contact list:   
 Exclusion zone contact(s): /O4/O6/  
Consider NA in place of MG (1 valid contact(s) for NA vs. 0 for MG)  

### MG:1H:3328

|  | Res. type | Atom | Residue | Distance | Occupancy | B-factor | Closest solute to 1^st^ shell water |
| --- | --- | --- | --- | --- | --- | --- | --- |
| **Ion** | | | |  | 0.50 | 27.38 || 1 | G | O6 | 1H:230 | 2.90 | 1.00 | 29.85 |
| 2 | A | N1 | 1H:231 | 3.22 | 1.00 | 28.80 |  |
| 3 | A | OP2 | 1H:246 | 3.62 | 1.00 | 27.34 |  |
| 4 | A | OP1 | 1H:245 | 4.43 | 1.00 | 28.79 |  |
| 5 | G | N1 | 1H:230 | 4.50 | 1.00 | 30.07 |  |
| 6 | A | N6 | 1H:231 | 4.51 | 1.00 | 29.59 |  |
| 7 | A | N3 | 1H:231 | 4.84 | 1.00 | 30.17 |  |
  |

  
 Coded contact list:   
 Exclusion zone contact(s): /N1_A_/O6/  
Consider K in place of MG (2 valid contact(s) for K vs. 0 for MG)  

### MG:1H:3329

|  | Res. type | Atom | Residue | Distance | Occupancy | B-factor | Closest solute to 1^st^ shell water |
| --- | --- | --- | --- | --- | --- | --- | --- |
| **Ion** | | | |  | 1.00 | 63.10 || 1 | G | O6 | 1H:526 | 2.49 | 1.00 | 59.23 |
| 2 | Water 1 | | 1H:4385 | 2.92 | 1.00 | 61.40 | 3.26 (G:O6 1H:526) |
| 3 | U | O4 | 1H:525 | 3.76 | 1.00 | 50.63 |  |
| 4 | A | OP2 | 1H:529 | 4.25 | 1.00 | 49.84 |  |
| 5 | G | N1 | 1H:526 | 4.68 | 1.00 | 49.59 |  |
| 6 | G | N7 | 1H:526 | 4.80 | 1.00 | 52.30 |  |
  |

  
 Coded contact list:   
 Exclusion zone contact(s): /O6/O_w_/  
Consider NA in place of MG (1 valid contact(s) for NA vs. 0 for MG)  

### MG:1H:3330

|  | Res. type | Atom | Residue | Distance | Occupancy | B-factor | Closest solute to 1^st^ shell water |
| --- | --- | --- | --- | --- | --- | --- | --- |
| **Ion** | | | |  | 1.00 | 45.91 || 1 | G | N7 | 1H:146 | 2.44 | 1.00 | 37.69 |
| 2 | U | O4 | 1H:147 | 3.35 | 1.00 | 41.09 |  |
| 3 | G | N7 | 1H:145 | 3.62 | 1.00 | 44.58 |  |
| 4 | G | O6 | 1H:146 | 3.83 | 1.00 | 44.29 |  |
| 5 | G | OP2 | 1H:146 | 4.27 | 1.00 | 42.42 |  |
| 6 | G | N9 | 1H:146 | 4.54 | 1.00 | 36.70 |  |
| 7 | G | O6 | 1H:145 | 4.55 | 1.00 | 34.58 |  |
| 8 | G | N9 | 1H:145 | 4.77 | 1.00 | 40.24 |  |
  |

  
 Coded contact list:   
 Exclusion zone contact(s): /N7/O4/  
Consider NA in place of MG (1 valid contact(s) for NA vs. 0 for MG)  

### MG:1H:3331

|  | Res. type | Atom | Residue | Distance | Occupancy | B-factor | Closest solute to 1^st^ shell water |
| --- | --- | --- | --- | --- | --- | --- | --- |
| **Ion** | | | |  | 1.00 | 54.95 || 1 | Water 1 | | 1H:4534 | 2.78 | 1.00 | 42.34 | 3.87 (G:O6 1H:1751) |
| 2 | Water 2 | | 1H:3900 | 3.08 | 1.00 | 49.36 | 2.52 (G:N7 1H:1751) |
| 3 | G | O6 | 1H:1752 | 3.17 | 1.00 | 46.75 |  |
| 4 | G | O6 | 1H:1751 | 3.48 | 1.00 | 44.03 |  |
| 5 | Water 3 | | 1H:4376 | 3.74 | 1.00 | 50.36 | 5.55 (G:N7 1H:1751) |
| 6 | G | N7 | 1H:1751 | 4.00 | 1.00 | 45.77 |  |
| 7 | C | N4 | 1H:1732 | 4.03 | 1.00 | 41.99 |  |
| 8 | C | N4 | 1H:1731 | 4.76 | 1.00 | 35.03 |  |
| 9 | G | O6 | 1H:1753 | 4.77 | 1.00 | 42.47 |  |
| 10 | C | N4 | 1H:1733 | 4.82 | 1.00 | 45.76 |  |
  |

  
 Coded contact list:   
 Exclusion zone contact(s): /O6/O_w_/O_w_/  
Consider K in place of MG (1 valid contact(s) for K vs. 0 for MG)  

### MG:1H:3332

|  | Res. type | Atom | Residue | Distance | Occupancy | B-factor | Closest solute to 1^st^ shell water |
| --- | --- | --- | --- | --- | --- | --- | --- |
| **Ion** | | | |  | 1.00 | 48.00 || 1 | Water 1 | | 1H:3819 | 2.84 | 1.00 | 39.43 | 2.40 (G:O6 1H:1750) |
| 2 | Water 2 | | 1H:4270 | 2.95 | 1.00 | 46.80 | 3.00 (G:N3 1H:1747) |
| 3 | Water 3 | | 1H:4534 | 3.15 | 1.00 | 42.34 | 4.00 (G:N2 1H:1747) |
| 4 | A | OP1 | 1H:1748 | 3.41 | 1.00 | 37.28 |  |
| 5 | G | O3' | 1H:1747 | 3.91 | 1.00 | 32.95 |  |
| 6 | G | N3 | 1H:1747 | 4.54 | 1.00 | 45.35 |  |
| 7 | G | O4' | 1H:1747 | 4.64 | 1.00 | 30.08 |  |
| 8 | G | O6 | 1H:1750 | 4.72 | 1.00 | 43.01 |  |
| 9 | C | N4 | 1H:1734 | 4.76 | 1.00 | 43.33 |  |
| 10 | C | N4 | 1H:1733 | 4.85 | 1.00 | 45.76 |  |
| 11 | G | N1 | 1H:1735 | 4.93 | 1.00 | 34.16 |  |
| 12 | G | N2 | 1H:1747 | 4.98 | 1.00 | 43.19 |  |
  |

  
 Coded contact list:   
 Exclusion zone contact(s): /O_w_/O_w_/O_w_/  
1^st^ MG coordination shell missing (see Notes)  
MG hydration shell issue: water too close (< 2.4 Angs.)  

### MG:1H:3333

|  | Res. type | Atom | Residue | Distance | Occupancy | B-factor | Closest solute to 1^st^ shell water |
| --- | --- | --- | --- | --- | --- | --- | --- |
| **Ion** | | | |  | 1.00 | 50.85 || 1 | U | OP2 | 1H:1382 | 2.74 | 1.00 | 32.52 |
| 2 | Water 1 | | 1H:4298 | 3.11 | 1.00 | 50.20 | 4.17 (U:OP2 1H:1382) |
| 3 | Water 2 | | 1H:3920 | 3.74 | 1.00 | 31.94 | 4.93 (LYS:O F8:63) |
| 4 | LYS | O | F8:63 | 3.74 | 1.00 | 42.77 |  |
| 5 | U | OP1 | 1H:1382 | 3.97 | 1.00 | 42.78 |  |
| 6 | A | OP2 | 1H:1383 | 4.31 | 1.00 | 34.79 |  |
| 7 | U | O5' | 1H:1382 | 4.38 | 1.00 | 33.40 |  |
| 8 | Water 3 | | 1H:4453 | 4.79 | 1.00 | 45.52 | 3.70 (LYS:O F8:63) |
| 9 | ARG | NH1 | F8:65 | 4.92 | 1.00 | 51.75 |  |
| 10 | ARG | N | F8:65 | 4.97 | 1.00 | 45.42 |  |
  |

  
 Coded contact list:   
 Exclusion zone contact(s): /O_w_/OP2/  
Consider K in place of MG (1 valid contact(s) for K vs. 0 for MG)  

### MG:1H:3334

|  | Res. type | Atom | Residue | Distance | Occupancy | B-factor | Closest solute to 1^st^ shell water |
| --- | --- | --- | --- | --- | --- | --- | --- |
| **Ion** | | | |  | 1.00 | 43.98 || 1 | G | OP1 | 1H:2833 | 3.01 | 1.00 | 22.00 |
| 2 | HIS | ND1 | 21:159 | 3.35 | 1.00 | 40.86 |  |
| 3 | GLY | O | 21:161 | 3.54 | 1.00 | 36.97 |  |
| 4 | A | O3' | 1H:2832 | 4.45 | 1.00 | 30.00 |  |
| 5 | TYR | N | 21:160 | 4.68 | 1.00 | 25.09 |  |
| 6 | G | O5' | 1H:2833 | 4.68 | 1.00 | 21.78 |  |
| 7 | TYR | O | 21:160 | 4.78 | 1.00 | 31.94 |  |
| 8 | ALA | O | 21:162 | 4.85 | 1.00 | 36.76 |  |
| 9 | C | OP1 | 1H:2635 | 4.96 | 1.00 | 34.26 |  |
  |

  
 Coded contact list:   
 Exclusion zone contact(s): /ND1_His_/OP1/  
Consider K in place of MG (2 valid contact(s) for K vs. 0 for MG)  

### MG:1H:3335

|  | Res. type | Atom | Residue | Distance | Occupancy | B-factor | Closest solute to 1^st^ shell water |
| --- | --- | --- | --- | --- | --- | --- | --- |
| **Ion** | | | |  | 1.00 | 40.77 || 1 | G | O6 | 1H:2690 | 3.66 | 1.00 | 40.93 |
| 2 | G | O6 | 1H:2743 | 4.42 | 1.00 | 37.24 |  |
| 3 | G | N7 | 1H:2690 | 4.81 | 1.00 | 39.12 |  |
| 4 | C | N4 | 1H:2689 | 4.83 | 1.00 | 33.46 |  |
| 5 | C | N4 | 1H:2691 | 4.98 | 1.00 | 35.68 |  |
  |

  
 Coded contact list:   
1^st^ MG coordination shell missing (see Notes)  

### MG:1H:3336

|  | Res. type | Atom | Residue | Distance | Occupancy | B-factor | Closest solute to 1^st^ shell water |
| --- | --- | --- | --- | --- | --- | --- | --- |
| **Ion** | | | |  | 1.00 | 45.88 || 1 | G | N7 | 1H:2409 | 3.76 | 1.00 | 37.30 |
| 2 | G | OP2 | 1H:2409 | 3.99 | 1.00 | 36.56 |  |
| 3 | G | OP1 | 1H:2428 | 4.51 | 1.00 | 32.98 |  |
| 4 | C | OP2 | 1H:2408 | 4.55 | 1.00 | 34.24 |  |
  |

  
 Coded contact list:   
1^st^ MG coordination shell missing (see Notes)  

### MG:1H:3337

|  | Res. type | Atom | Residue | Distance | Occupancy | B-factor | Closest solute to 1^st^ shell water |
| --- | --- | --- | --- | --- | --- | --- | --- |
| **Ion** | | | |  | 1.00 | 23.20 || 1 | G | N7 | 1H:1856 | 3.47 | 1.00 | 18.41 |
| 2 | G | OP2 | 1H:1856 | 4.14 | 1.00 | 16.62 |  |
| 3 | G | N7 | 1H:1855 | 4.36 | 1.00 | 17.18 |  |
| 4 | G | OP2 | 1H:1855 | 4.47 | 1.00 | 19.15 |  |
| 5 | HIS | NE2 | 11:220 | 4.75 | 1.00 | 25.14 |  |
| 6 | G | O6 | 1H:1856 | 4.77 | 1.00 | 21.10 |  |
| 7 | G | O5' | 1H:1855 | 4.89 | 1.00 | 17.30 |  |
  |

  
 Coded contact list:   
1^st^ MG coordination shell missing (see Notes)  

### MG:1H:3338

|  | Res. type | Atom | Residue | Distance | Occupancy | B-factor | Closest solute to 1^st^ shell water |
| --- | --- | --- | --- | --- | --- | --- | --- |
| **Ion** | | | |  | 1.00 | 43.36 || 1 | U | OP1 | 1H:1485 | 4.76 | 1.00 | 42.96 |
| 2 | U | OP2 | 1H:1485 | 4.91 | 1.00 | 44.75 |  |
  |

  
 Coded contact list:   
1^st^ MG coordination shell missing (see Notes)  

### MG:1H:3339

|  | Res. type | Atom | Residue | Distance | Occupancy | B-factor | Closest solute to 1^st^ shell water |
| --- | --- | --- | --- | --- | --- | --- | --- |
| **Ion** | | | |  | 1.00 | 55.37 || 1 | G | O6 | 1H:2529 | 3.42 | 1.00 | 30.05 |
| 2 | C | N4 | 1H:2528 | 3.63 | 1.00 | 32.13 |  |
| 3 | C | N4 | 1H:2530 | 3.67 | 1.00 | 32.36 |  |
| 4 | G | O6 | 1H:2580 | 4.03 | 1.00 | 30.81 |  |
| 5 | G | N7 | 1H:2529 | 4.31 | 1.00 | 31.87 |  |
| 6 | U | O4 | 1H:2559 | 4.53 | 1.00 | 26.09 |  |
| 7 | C | N4 | 1H:2581 | 4.88 | 1.00 | 26.28 |  |
  |

  
 Coded contact list:   
1^st^ MG coordination shell missing (see Notes)  

### MG:1H:3340

|  | Res. type | Atom | Residue | Distance | Occupancy | B-factor | Closest solute to 1^st^ shell water |
| --- | --- | --- | --- | --- | --- | --- | --- |
| **Ion** | | | |  | 1.00 | 48.29 || 1 | G | N7 | 1H:2529 | 3.37 | 1.00 | 31.87 |
| 2 | C | N4 | 1H:2530 | 3.61 | 1.00 | 32.36 |  |
| 3 | G | OP2 | 1H:2529 | 3.74 | 1.00 | 32.35 |  |
| 4 | A | O2' | 1H:2555 | 4.35 | 1.00 | 38.16 |  |
| 5 | G | O6 | 1H:2529 | 4.66 | 1.00 | 30.05 |  |
  |

  
 Coded contact list:   
 Exclusion zone contact(s): /N7/  
Consider K in place of MG (1 valid contact(s) for K vs. 0 for MG)  

### MG:1H:3341

|  | Res. type | Atom | Residue | Distance | Occupancy | B-factor | Closest solute to 1^st^ shell water |
| --- | --- | --- | --- | --- | --- | --- | --- |
| **Ion** | | | |  | 1.00 | 24.92 || 1 | G | O6 | 1H:729 | 3.23 | 1.00 | 17.27 |
| 2 | G | O6 | 1H:730 | 3.55 | 1.00 | 15.16 |  |
| 3 | C | N4 | 1H:843 | 3.79 | 1.00 | 16.02 |  |
| 4 | G | N7 | 1H:729 | 4.47 | 1.00 | 18.26 |  |
| 5 | C | N4 | 1H:844 | 4.54 | 1.00 | 16.85 |  |
| 6 | G | N7 | 1H:728 | 4.63 | 1.00 | 17.63 |  |
| 7 | G | O6 | 1H:728 | 4.67 | 1.00 | 18.32 |  |
| 8 | G | O2' | 1H:823 | 4.87 | 1.00 | 18.09 |  |
  |

  
 Coded contact list:   
 Exclusion zone contact(s): /O6/  
Consider K in place of MG (1 valid contact(s) for K vs. 0 for MG)  

### MG:1H:3342

|  | Res. type | Atom | Residue | Distance | Occupancy | B-factor | Closest solute to 1^st^ shell water |
| --- | --- | --- | --- | --- | --- | --- | --- |
| **Ion** | | | |  | 1.00 | 27.95 || 1 | A | N3 | 1H:307 | 3.62 | 1.00 | 61.29 |
| 2 | A | O2' | 1H:308 | 3.96 | 1.00 | 41.81 |  |
| 3 | A | N3 | 1H:308 | 4.36 | 1.00 | 34.53 |  |
| 4 | C | O2' | 1H:257 | 4.54 | 1.00 | 42.18 |  |
| 5 | A | N9 | 1H:308 | 4.62 | 1.00 | 39.19 |  |
| 6 | A | N1 | 1H:307 | 4.69 | 1.00 | 66.07 |  |
| 7 | A | N1 | 1H:308 | 4.78 | 1.00 | 36.39 |  |
  |

  
 Coded contact list:   
1^st^ MG coordination shell missing (see Notes)  

### MG:1H:3343

|  | Res. type | Atom | Residue | Distance | Occupancy | B-factor | Closest solute to 1^st^ shell water |
| --- | --- | --- | --- | --- | --- | --- | --- |
| **Ion** | | | |  | 0.50 | 55.96 || 1 | U | OP1 | 1H:1588 | 3.48 | 1.00 | 104.25 |
| 2 | U | OP2 | 1H:1588 | 4.23 | 1.00 | 108.19 |  |
| 3 | U | O5' | 1H:1588 | 4.58 | 1.00 | 108.47 |  |
| 4 | G | OP2 | 1H:1589 | 4.59 | 1.00 | 89.50 |  |
  |

  
 Coded contact list:   
1^st^ MG coordination shell missing (see Notes)  

### MG:1H:3343

|  | Res. type | Atom | Residue | Distance | Occupancy | B-factor | Closest solute to 1^st^ shell water |
| --- | --- | --- | --- | --- | --- | --- | --- |
| **Ion** | | | |  | 0.50 | 68.21 || 1 | C | O5' | 1H:1591 | 3.68 | 1.00 | 84.14 |
| 2 | A | OP2 | 1H:1592 | 3.86 | 1.00 | 80.83 |  |
| 3 | A | O2' | 1H:1590 | 4.17 | 1.00 | 80.44 |  |
| 4 | C | OP2 | 1H:1591 | 4.81 | 1.00 | 89.24 |  |
  |

  
 Coded contact list:   
1^st^ MG coordination shell missing (see Notes)  

### MG:1H:3344

|  | Res. type | Atom | Residue | Distance | Occupancy | B-factor | Closest solute to 1^st^ shell water |
| --- | --- | --- | --- | --- | --- | --- | --- |
| **Ion** | | | |  | 1.00 | 39.26 || 1 | A | OP2 | 1H:2000 | 3.25 | 1.00 | 22.54 |
| 2 | U | OP2 | 1H:1999 | 4.82 | 1.00 | 26.45 |  |
| 3 | C | N4 | 1H:1803 | 4.84 | 1.00 | 17.99 |  |
| 4 | G | O6 | 1H:1804 | 4.85 | 1.00 | 20.61 |  |
  |

  
 Coded contact list:   
 Exclusion zone contact(s): /OP2/  
Consider K in place of MG (1 valid contact(s) for K vs. 0 for MG)  

### MG:1H:3345

|  | Res. type | Atom | Residue | Distance | Occupancy | B-factor | Closest solute to 1^st^ shell water |
| --- | --- | --- | --- | --- | --- | --- | --- |
| **Ion** | | | |  | 0.50 | 23.09 || 1 | G | OP2 | 1H:1653 | 3.34 | 1.00 | 30.15 |
| 2 | G | N7 | 1H:1653 | 3.36 | 1.00 | 33.05 |  |
| 3 | C | OP2 | 1H:1352 | 3.60 | 1.00 | 44.66 |  |
| 4 | C | OP2 | 1H:1652 | 3.77 | 1.00 | 24.97 |  |
| 5 | C | OP1 | 1H:1652 | 4.23 | 1.00 | 23.35 |  |
| 6 | C | O5' | 1H:1652 | 4.52 | 1.00 | 27.22 |  |
| 7 | G | O5' | 1H:1653 | 4.81 | 1.00 | 26.45 |  |
| 8 | Water 1 | | 1H:4038 | 4.84 | 1.00 | 32.55 | 2.67 (G:OP2 1H:1653) |
| 9 | G | N9 | 1H:1653 | 4.94 | 1.00 | 28.45 |  |
| 10 | C | OP2 | 1H:1353 | 4.99 | 1.00 | 44.62 |  |
  |

  
 Coded contact list:   
 Exclusion zone contact(s): /N7/OP2/  
Consider K in place of MG (2 valid contact(s) for K vs. 0 for MG)  

### MG:1H:3345

|  | Res. type | Atom | Residue | Distance | Occupancy | B-factor | Closest solute to 1^st^ shell water |
| --- | --- | --- | --- | --- | --- | --- | --- |
| **Ion** | | | |  | 0.50 | 24.06 || 1 | G | OP2 | 1H:1653 | 2.85 | 1.00 | 30.15 |
| 2 | Water 1 | | 1H:4038 | 3.29 | 1.00 | 32.55 | 2.67 (G:OP2 1H:1653) |
| 3 | C | OP2 | 1H:1352 | 3.32 | 1.00 | 44.66 |  |
| 4 | C | OP1 | 1H:1652 | 3.68 | 1.00 | 23.35 |  |
| 5 | C | OP2 | 1H:1351 | 4.77 | 1.00 | 33.28 |  |
| 6 | C | OP1 | 1H:1351 | 4.84 | 1.00 | 35.15 |  |
| 7 | C | OP2 | 1H:1652 | 4.88 | 1.00 | 24.97 |  |
| 8 | C | O5' | 1H:1652 | 4.89 | 1.00 | 27.22 |  |
| 9 | G | OP1 | 1H:1653 | 4.98 | 1.00 | 35.64 |  |
  |

  
 Coded contact list:   
 Exclusion zone contact(s): /O_w_/OP2/OP2/  
Consider K in place of MG (2 valid contact(s) for K vs. 0 for MG)  

### MG:1H:3346

|  | Res. type | Atom | Residue | Distance | Occupancy | B-factor | Closest solute to 1^st^ shell water |
| --- | --- | --- | --- | --- | --- | --- | --- |
| **Ion** | | | |  | 1.00 | 55.33 || 1 | G | N7 | 1H:2500 | 3.55 | 1.00 | 35.69 |
| 2 | G | O6 | 1H:2499 | 3.87 | 1.00 | 34.35 |  |
| 3 | G | O6 | 1H:2500 | 4.13 | 1.00 | 32.81 |  |
| 4 | G | N7 | 1H:2499 | 4.23 | 1.00 | 31.72 |  |
  |

  
 Coded contact list:   
1^st^ MG coordination shell missing (see Notes)  

### MG:1H:3347

|  | Res. type | Atom | Residue | Distance | Occupancy | B-factor | Closest solute to 1^st^ shell water |
| --- | --- | --- | --- | --- | --- | --- | --- |
| **Ion** | | | |  | 1.00 | 38.40 || 1 | G | N7 | 1H:1855 | 3.85 | 1.00 | 17.18 |
| 2 | G | OP2 | 1H:1855 | 4.16 | 1.00 | 19.15 |  |
| 3 | G | N7 | 1H:1854 | 4.54 | 1.00 | 17.79 |  |
| 4 | G | N7 | 1H:1856 | 4.99 | 1.00 | 18.41 |  |
  |

  
 Coded contact list:   
1^st^ MG coordination shell missing (see Notes)  

### MG:1H:3348

|  | Res. type | Atom | Residue | Distance | Occupancy | B-factor | Closest solute to 1^st^ shell water |
| --- | --- | --- | --- | --- | --- | --- | --- |
| **Ion** | | | |  | 1.00 | 30.67 || 1 | Water 1 | | 1H:4527 | 3.10 | 1.00 | 21.84 | 3.83 (G:N7 1H:819) |
| 2 | G | N7 | 1H:818 | 3.10 | 1.00 | 17.85 |  |
| 3 | Water 2 | | 1H:4120 | 3.48 | 1.00 | 24.02 | 2.78 (G:OP2 1H:818) |
| 4 | G | O6 | 1H:818 | 3.73 | 1.00 | 23.56 |  |
| 5 | G | N7 | 1H:817 | 3.78 | 1.00 | 22.62 |  |
| 6 | Water 3 | | 1H:3986 | 3.79 | 1.00 | 25.60 | 2.63 (G:N7 1H:817) |
| 7 | Water 4 | | 1H:3794 | 3.84 | 1.00 | 27.36 | 2.37 (G:OP2 1H:818) |
| 8 | Water 5 | | 1H:4404 | 4.03 | 1.00 | 21.36 | 3.31 (G:O6 1H:817) |
| 9 | G | O6 | 1H:817 | 4.37 | 1.00 | 23.47 |  |
| 10 | G | OP2 | 1H:818 | 4.56 | 1.00 | 24.96 |  |
| 11 | G | OP2 | 1H:817 | 4.58 | 1.00 | 21.76 |  |
| 12 | G | N7 | 1H:819 | 4.65 | 1.00 | 19.10 |  |
  |

  
 Coded contact list:   
 Exclusion zone contact(s): /N7/O_w_/  
Consider K in place of MG (1 valid contact(s) for K vs. 0 for MG)  

### MG:1H:3349

|  | Res. type | Atom | Residue | Distance | Occupancy | B-factor | Closest solute to 1^st^ shell water |
| --- | --- | --- | --- | --- | --- | --- | --- |
| **Ion** | | | |  | 1.00 | 48.49 || 1 | G | O6 | 1H:990 | 3.32 | 1.00 | 30.70 |
| 2 | U | OP2 | 1H:881 | 3.34 | 1.00 | 21.79 |  |
| 3 | U | O4 | 1H:989 | 3.37 | 1.00 | 26.03 |  |
| 4 | G | OP2 | 1H:988 | 4.60 | 1.00 | 23.22 |  |
| 5 | LYS | NZ | 78:39 | 4.63 | 1.00 | 25.91 |  |
  |

  
 Coded contact list:   
 Exclusion zone contact(s): /O4/O6/OP2/  
Consider K in place of MG (3 valid contact(s) for K vs. 0 for MG)  

### MG:1H:3350

|  | Res. type | Atom | Residue | Distance | Occupancy | B-factor | Closest solute to 1^st^ shell water |
| --- | --- | --- | --- | --- | --- | --- | --- |
| **Ion** | | | |  | 1.00 | 25.47 || 1 | A | OP1 | 1H:2461 | 1.87 | 1.00 | 25.13 |
| 2 | G | O6 | 1H:594 | 2.20 | 1.00 | 24.35 |  |
| 3 | C | OP1 | 1H:2512 | 2.29 | 1.00 | 20.16 |  |
| 4 | A | O5' | 1H:2461 | 3.60 | 1.00 | 16.40 |  |
| 5 | A | OP2 | 1H:2461 | 3.65 | 1.00 | 20.08 |  |
| 6 | U | O4 | 1H:593 | 3.66 | 1.00 | 27.30 |  |
| 7 | G | N1 | 1H:594 | 3.89 | 1.00 | 18.29 |  |
| 8 | C | OP2 | 1H:2512 | 3.98 | 1.00 | 18.26 |  |
| 9 | C | O3' | 1H:2511 | 4.15 | 1.00 | 19.87 |  |
| 10 | U | O2' | 1H:592 | 4.30 | 1.00 | 24.12 |  |
| 11 | G | O3' | 1H:2460 | 4.36 | 1.00 | 22.35 |  |
| 12 | U | OP2 | 1H:2513 | 4.56 | 1.00 | 20.03 |  |
| 13 | C | OP2 | 1H:2511 | 4.70 | 1.00 | 22.83 |  |
| 14 | C | O5' | 1H:2512 | 4.70 | 1.00 | 17.43 |  |
  |

  
**1^st^ shell ligand angles:**   
1-2: 138; 1-3: 71; 2-3: 73;   
  
 Contact(s): /O6/OP1/OP1/  
 Coded contact list: 2O_ph_O_b_  

### MG:1H:3351

|  | Res. type | Atom | Residue | Distance | Occupancy | B-factor | Closest solute to 1^st^ shell water |
| --- | --- | --- | --- | --- | --- | --- | --- |
| **Ion** | | | |  | 1.00 | 79.08 || 1 | U | O4 | 1H:1529 | 2.52 | 1.00 | 71.08 |
| 2 | G | O6 | 1H:1559 | 2.74 | 1.00 | 79.45 |  |
| 3 | G | O6 | 1H:1528 | 3.17 | 1.00 | 71.33 |  |
| 4 | A | OP2 | 1H:1558 | 3.74 | 1.00 | 123.57 |  |
| 5 | C | N4 | 1H:1560 | 4.12 | 1.00 | 82.45 |  |
| 6 | U | N3 | 1H:1529 | 4.25 | 1.00 | 71.94 |  |
| 7 | G | N7 | 1H:1559 | 4.48 | 1.00 | 91.34 |  |
| 8 | A | N7 | 1H:1558 | 4.49 | 1.00 | 116.17 |  |
  |

  
 Coded contact list:   
 Exclusion zone contact(s): /O4/O6/O6/  
Consider K in place of MG (2 valid contact(s) for K vs. 0 for MG)  

### MG:1H:3352

|  | Res. type | Atom | Residue | Distance | Occupancy | B-factor | Closest solute to 1^st^ shell water |
| --- | --- | --- | --- | --- | --- | --- | --- |
| **Ion** | | | |  | 1.00 | 43.03 || 1 | G | N7 | 1H:1081 | 2.88 | 1.00 | 46.69 |
| 2 | G | OP1 | 1H:1081 | 3.49 | 1.00 | 64.26 |  |
| 3 | A | N1 | 1H:1079 | 3.88 | 1.00 | 36.58 |  |
| 4 | U | O4 | 1H:1082 | 4.03 | 1.00 | 51.58 |  |
| 5 | G | O6 | 1H:1081 | 4.23 | 1.00 | 43.63 |  |
| 6 | A | N3 | 1H:1079 | 4.44 | 1.00 | 42.88 |  |
| 7 | A | N6 | 1H:1079 | 4.45 | 1.00 | 39.41 |  |
| 8 | G | OP2 | 1H:1081 | 4.72 | 1.00 | 59.77 |  |
| 9 | G | N9 | 1H:1081 | 4.94 | 1.00 | 42.68 |  |
  |

  
 Coded contact list:   
 Exclusion zone contact(s): /N7/  
Consider K in place of MG (1 valid contact(s) for K vs. 0 for MG)  

### MG:1H:3353

|  | Res. type | Atom | Residue | Distance | Occupancy | B-factor | Closest solute to 1^st^ shell water |
| --- | --- | --- | --- | --- | --- | --- | --- |
| **Ion** | | | |  | 1.00 | 33.41 || 1 | G | O6 | 1H:170 | 3.78 | 1.00 | 31.87 |
| 2 | G | N7 | 1H:169 | 3.97 | 1.00 | 26.81 |  |
| 3 | G | O6 | 1H:169 | 4.39 | 1.00 | 26.66 |  |
| 4 | A | OP2 | 1H:203 | 4.42 | 1.00 | 51.93 |  |
| 5 | G | N7 | 1H:170 | 4.77 | 1.00 | 25.64 |  |
| 6 | G | OP2 | 1H:169 | 4.94 | 1.00 | 20.94 |  |
  |

  
 Coded contact list:   
1^st^ MG coordination shell missing (see Notes)  

### MG:1H:3354

|  | Res. type | Atom | Residue | Distance | Occupancy | B-factor | Closest solute to 1^st^ shell water |
| --- | --- | --- | --- | --- | --- | --- | --- |
| **Ion** | | | |  | 1.00 | 29.95 || 1 | U | O2 | 1H:590 | 2.93 | 1.00 | 18.56 |
| 2 | U | O2' | 1H:590 | 3.09 | 1.00 | 19.36 |  |
| 3 | A | O4' | 1H:591 | 3.58 | 1.00 | 18.50 |  |
| 4 | U | O2 | 1H:600 | 3.70 | 1.00 | 23.58 |  |
| 5 | G | OP1 | 1H:856 | 3.83 | 1.00 | 19.09 |  |
| 6 | Water 1 | | 1H:3916 | 4.15 | 1.00 | 13.34 | 3.12 (G:O6 1H:2515) |
| 7 | U | N1 | 1H:590 | 4.51 | 1.00 | 20.57 |  |
| 8 | U | O4' | 1H:600 | 4.54 | 1.00 | 17.56 |  |
| 9 | G | O6 | 1H:2515 | 4.55 | 1.00 | 17.81 |  |
| 10 | U | N1 | 1H:600 | 4.61 | 1.00 | 20.01 |  |
| 11 | U | O2' | 1H:600 | 4.62 | 1.00 | 19.75 |  |
| 12 | Water 2 | | 1H:4436 | 4.73 | 1.00 | 31.02 | 3.42 (U:O4' 1H:600) |
| 13 | A | N9 | 1H:591 | 4.96 | 1.00 | 19.83 |  |
  |

  
 Coded contact list:   
 Exclusion zone contact(s): /O2/O2'/  
Consider K in place of MG (2 valid contact(s) for K vs. 0 for MG)  

### MG:1H:3355

|  | Res. type | Atom | Residue | Distance | Occupancy | B-factor | Closest solute to 1^st^ shell water |
| --- | --- | --- | --- | --- | --- | --- | --- |
| **Ion** | | | |  | 1.00 | 57.33 || 1 | Water 1 | | 1H:4106 | 2.51 | 1.00 | 57.89 | 3.43 (G:OP2 1H:637) |
| 2 | G | OP2 | 1H:637 | 2.64 | 1.00 | 46.16 |  |
| 3 | Water 2 | | 1H:4565 | 2.88 | 1.00 | 57.69 | 4.17 (C:OP2 1H:636) |
| 4 | Water 3 | | 1H:4531 | 3.13 | 1.00 | 65.85 | 3.87 (G:OP2 1H:637) |
| 5 | C | OP2 | 1H:636 | 3.24 | 1.00 | 43.27 |  |
| 6 | C | O5' | 1H:636 | 3.34 | 1.00 | 41.82 |  |
| 7 | C | OP1 | 1H:636 | 3.94 | 1.00 | 45.20 |  |
| 8 | C | O3' | 1H:636 | 4.55 | 1.00 | 47.94 |  |
| 9 | G | O5' | 1H:637 | 4.91 | 1.00 | 39.86 |  |
| 10 | Water 4 | | 1H:4103 | 4.97 | 1.00 | 46.43 | 5.90 (C:OP2 1H:636) |
  |

  
 Coded contact list:   
 Exclusion zone contact(s): /O5'/O_w_/O_w_/O_w_/OP2/OP2/  
Consider K in place of MG (3 valid contact(s) for K vs. 0 for MG)  

### MG:1H:3356

|  | Res. type | Atom | Residue | Distance | Occupancy | B-factor | Closest solute to 1^st^ shell water |
| --- | --- | --- | --- | --- | --- | --- | --- |
| **Ion** | | | |  | 1.00 | 43.71 || 1 | U | O2 | 1H:2370 | 3.08 | 1.00 | 35.16 |
| 2 | A | N6 | 1H:2374 | 3.32 | 1.00 | 36.69 |  |
| 3 | A | N7 | 1H:2374 | 3.42 | 1.00 | 33.92 |  |
| 4 | A | OP2 | 1H:2373 | 3.81 | 1.00 | 33.75 |  |
| 5 | A | N7 | 1H:2373 | 4.19 | 1.00 | 31.63 |  |
| 6 | C | OP2 | 1H:2372 | 4.36 | 1.00 | 26.37 |  |
| 7 | A | N6 | 1H:2373 | 4.74 | 1.00 | 28.07 |  |
| 8 | C | OP1 | 1H:2372 | 4.77 | 1.00 | 30.12 |  |
  |

  
 Coded contact list:   
 Exclusion zone contact(s): /N6/O2/  
Consider K in place of MG (2 valid contact(s) for K vs. 0 for MG)  

### MG:1H:3357

|  | Res. type | Atom | Residue | Distance | Occupancy | B-factor | Closest solute to 1^st^ shell water |
| --- | --- | --- | --- | --- | --- | --- | --- |
| **Ion** | | | |  | 1.00 | 28.78 || 1 | G | OP2 | 1H:734 | 2.42 | 1.00 | 21.32 |
| 2 | C | OP2 | 1H:735 | 3.05 | 1.00 | 20.17 |  |
| 3 | A | O3' | 1H:733 | 3.40 | 1.00 | 18.65 |  |
| 4 | U | O4 | 1H:736 | 3.61 | 1.00 | 21.95 |  |
| 5 | A | O2' | 1H:733 | 4.28 | 1.00 | 19.92 |  |
| 6 | G | OP1 | 1H:734 | 4.46 | 1.00 | 17.15 |  |
| 7 | Water 1 | | 1H:4527 | 4.57 | 1.00 | 21.84 | 3.92 (G:OP2 1H:734) |
| 8 | G | O5' | 1H:734 | 4.59 | 1.00 | 19.50 |  |
  |

  
 Coded contact list:   
 Exclusion zone contact(s): /OP2/OP2/  
Consider NA in place of MG (1 valid contact(s) for NA vs. 0 for MG)  

### MG:1H:3358

|  | Res. type | Atom | Residue | Distance | Occupancy | B-factor | Closest solute to 1^st^ shell water |
| --- | --- | --- | --- | --- | --- | --- | --- |
| **Ion** | | | |  | 1.00 | 41.64 || 1 | G | O6 | 1H:738 | 3.53 | 1.00 | 19.20 |
| 2 | G | O6 | 1H:819 | 4.06 | 1.00 | 23.99 |  |
| 3 | C | N4 | 1H:820 | 4.32 | 1.00 | 17.66 |  |
| 4 | C | N4 | 1H:739 | 4.33 | 1.00 | 18.69 |  |
| 5 | A | O2' | 1H:733 | 4.48 | 1.00 | 19.92 |  |
| 6 | U | O4 | 1H:736 | 4.48 | 1.00 | 21.95 |  |
| 7 | A | N6 | 1H:737 | 4.56 | 1.00 | 18.89 |  |
| 8 | G | N7 | 1H:738 | 4.65 | 1.00 | 17.14 |  |
| 9 | A | N7 | 1H:737 | 4.66 | 1.00 | 16.21 |  |
| 10 | G | OP2 | 1H:734 | 5.00 | 1.00 | 21.32 |  |
  |

  
 Coded contact list:   
1^st^ MG coordination shell missing (see Notes)  

### MG:1H:3359

|  | Res. type | Atom | Residue | Distance | Occupancy | B-factor | Closest solute to 1^st^ shell water |
| --- | --- | --- | --- | --- | --- | --- | --- |
| **Ion** | | | |  | 1.00 | 62.40 || 1 | G | O6 | 1H:377 | 3.23 | 1.00 | 73.68 |
| 2 | G | N7 | 1H:377 | 3.71 | 1.00 | 78.69 |  |
| 3 | G | OP1 | 1H:376 | 3.79 | 1.00 | 53.24 |  |
| 4 | A | OP2 | 1H:206 | 4.25 | 1.00 | 36.28 |  |
| 5 | A | OP1 | 1H:206 | 4.56 | 1.00 | 35.34 |  |
| 6 | G | N1 | 1H:377 | 4.97 | 1.00 | 76.78 |  |
  |

  
 Coded contact list:   
 Exclusion zone contact(s): /O6/  
Consider K in place of MG (1 valid contact(s) for K vs. 0 for MG)  

### MG:1H:3360

|  | Res. type | Atom | Residue | Distance | Occupancy | B-factor | Closest solute to 1^st^ shell water |
| --- | --- | --- | --- | --- | --- | --- | --- |
| **Ion** | | | |  | 1.00 | 28.95 || 1 | OMU | OP2 | 1H:2565 | 2.26 | 1.00 | 25.56 |
| 2 | U | OP1 | 1H:1721 | 2.76 | 1.00 | 29.89 |  |
| 3 | OMU | OP1 | 1H:2565 | 3.68 | 1.00 | 24.62 |  |
| 4 | OMU | O5' | 1H:2565 | 3.85 | 1.00 | 25.76 |  |
| 5 | C | O3' | 1H:1720 | 3.98 | 1.00 | 19.02 |  |
| 6 | C | OP1 | 1H:2564 | 4.28 | 1.00 | 27.53 |  |
| 7 | U | O5' | 1H:1721 | 4.32 | 1.00 | 20.12 |  |
| 8 | C | O5' | 1H:2564 | 4.60 | 1.00 | 25.75 |  |
| 9 | C | O3' | 1H:2564 | 4.65 | 1.00 | 23.69 |  |
| 10 | C | OP2 | 1H:2564 | 4.95 | 1.00 | 26.83 |  |
  |

  
 Contact(s): /OP2/  
 Coded contact list: O_ph_  
 Exclusion zone contact(s): /OP1/  

### MG:1H:3361

|  | Res. type | Atom | Residue | Distance | Occupancy | B-factor | Closest solute to 1^st^ shell water |
| --- | --- | --- | --- | --- | --- | --- | --- |
| **Ion** | | | |  | 1.00 | 57.26 || 1 | U | O2' | 1H:49 | 2.99 | 1.00 | 44.83 |
| 2 | U | O3' | 1H:49 | 3.39 | 1.00 | 37.17 |  |
| 3 | G | OP1 | 1H:50 | 3.65 | 1.00 | 42.62 |  |
| 4 | U | O2 | 1H:49 | 4.54 | 1.00 | 51.43 |  |
| 5 | A | OP1 | 1H:48 | 4.57 | 1.00 | 31.12 |  |
  |

  
 Coded contact list:   
 Exclusion zone contact(s): /O2'/O3'/  
Consider K in place of MG (2 valid contact(s) for K vs. 0 for MG)  

### MG:1H:3362

|  | Res. type | Atom | Residue | Distance | Occupancy | B-factor | Closest solute to 1^st^ shell water |
| --- | --- | --- | --- | --- | --- | --- | --- |
| **Ion** | | | |  | 1.00 | 64.67 || 1 | GLY | O | 98:46 | 3.89 | 1.00 | 44.14 |
| 2 | C | OP2 | 1H:2851 | 4.18 | 1.00 | 41.24 |  |
| 3 | C | OP1 | 1H:2851 | 4.19 | 1.00 | 45.96 |  |
| 4 | G | OP1 | 1H:2850 | 4.20 | 1.00 | 41.23 |  |
| 5 | G | OP2 | 1H:2883 | 4.23 | 1.00 | 62.17 |  |
| 6 | HIS | ND1 | 98:50 | 4.43 | 1.00 | 45.84 |  |
  |

  
 Coded contact list:   
1^st^ MG coordination shell missing (see Notes)  

### MG:1H:3363

|  | Res. type | Atom | Residue | Distance | Occupancy | B-factor | Closest solute to 1^st^ shell water |
| --- | --- | --- | --- | --- | --- | --- | --- |
| **Ion** | | | |  | 1.00 | 56.45 || 1 | G | O6 | 1H:750 | 3.09 | 1.00 | 32.64 |
| 2 | G | O6 | 1H:749 | 4.03 | 1.00 | 33.32 |  |
| 3 | G | N7 | 1H:750 | 4.26 | 1.00 | 30.84 |  |
| 4 | U | O4 | 1H:751 | 4.41 | 1.00 | 32.16 |  |
| 5 | C | N4 | 1H:778 | 4.52 | 1.00 | 32.50 |  |
| 6 | G | N7 | 1H:749 | 4.93 | 1.00 | 32.82 |  |
  |

  
 Coded contact list:   
 Exclusion zone contact(s): /O6/  
Consider K in place of MG (1 valid contact(s) for K vs. 0 for MG)  

### MG:1H:3364

|  | Res. type | Atom | Residue | Distance | Occupancy | B-factor | Closest solute to 1^st^ shell water |
| --- | --- | --- | --- | --- | --- | --- | --- |
| **Ion** | | | |  | 1.00 | 58.51 || 1 | G | N7 | 1H:1509 | 3.44 | 1.00 | 72.00 |
| 2 | G | O6 | 1H:1509 | 3.73 | 1.00 | 66.35 |  |
| 3 | G | N7 | 1H:1507 | 3.79 | 1.00 | 73.06 |  |
| 4 | G | O6 | 1H:1507 | 3.87 | 1.00 | 65.95 |  |
| 5 | G | OP2 | 1H:1509 | 4.92 | 1.00 | 90.40 |  |
  |

  
 Coded contact list:   
1^st^ MG coordination shell missing (see Notes)  

### MG:1H:3365

|  | Res. type | Atom | Residue | Distance | Occupancy | B-factor | Closest solute to 1^st^ shell water |
| --- | --- | --- | --- | --- | --- | --- | --- |
| **Ion** | | | |  | 1.00 | 58.96 || 1 | G | O6 | 1H:1507 | 3.24 | 1.00 | 65.95 |
| 2 | C | O2 | 1H:1506 | 4.02 | 1.00 | 63.75 |  |
| 3 | G | N7 | 1H:1507 | 4.26 | 1.00 | 73.06 |  |
| 4 | G | O6 | 1H:1509 | 4.76 | 1.00 | 66.35 |  |
| 5 | C | N4 | 1H:1500 | 4.88 | 1.00 | 63.57 |  |
  |

  
 Coded contact list:   
 Exclusion zone contact(s): /O6/  
Consider K in place of MG (1 valid contact(s) for K vs. 0 for MG)  

### MG:1H:3366

|  | Res. type | Atom | Residue | Distance | Occupancy | B-factor | Closest solute to 1^st^ shell water |
| --- | --- | --- | --- | --- | --- | --- | --- |
| **Ion** | | | |  | 1.00 | 40.91 || 1 | G | O4' | 1H:1489 | 2.77 | 1.00 | 42.69 |
| 2 | G | O2' | 1H:1488 | 2.82 | 1.00 | 37.26 |  |
| 3 | G | N3 | 1H:1488 | 3.55 | 1.00 | 44.52 |  |
| 4 | G | N2 | 1H:1488 | 3.71 | 1.00 | 44.68 |  |
| 5 | G | N9 | 1H:1489 | 4.21 | 1.00 | 43.54 |  |
| 6 | U | OP1 | 1H:1676 | 4.64 | 1.00 | 40.05 |  |
| 7 | G | O2' | 1H:1489 | 4.73 | 1.00 | 45.38 |  |
| 8 | G | OP1 | 1H:1675 | 4.93 | 1.00 | 27.88 |  |
  |

  
 Coded contact list:   
 Exclusion zone contact(s): /O2'/O4'/  
Consider K in place of MG (2 valid contact(s) for K vs. 0 for MG)  

### MG:1H:3367

|  | Res. type | Atom | Residue | Distance | Occupancy | B-factor | Closest solute to 1^st^ shell water |
| --- | --- | --- | --- | --- | --- | --- | --- |
| **Ion** | | | |  | 1.00 | 35.85 || 1 | U | O4 | 1H:1728 | 3.00 | 1.00 | 42.22 |
| 2 | G | OP2 | 1H:1796 | 3.10 | 1.00 | 36.03 |  |
| 3 | U | O4 | 1H:1727 | 3.95 | 1.00 | 32.93 |  |
| 4 | G | N2 | 1H:1729 | 3.96 | 1.00 | 38.85 |  |
| 5 | G | OP1 | 1H:1796 | 4.35 | 1.00 | 30.11 |  |
| 6 | U | N3 | 1H:1728 | 4.49 | 1.00 | 35.95 |  |
| 7 | G | N1 | 1H:1729 | 4.80 | 1.00 | 44.07 |  |
  |

  
 Coded contact list:   
 Exclusion zone contact(s): /O4/OP2/  
Consider K in place of MG (2 valid contact(s) for K vs. 0 for MG)  

### MG:1H:3368

|  | Res. type | Atom | Residue | Distance | Occupancy | B-factor | Closest solute to 1^st^ shell water |
| --- | --- | --- | --- | --- | --- | --- | --- |
| **Ion** | | | |  | 1.00 | 44.78 || 1 | G | OP2 | 1H:438 | 2.48 | 1.00 | 34.79 |
| 2 | Water 1 | | 1H:4089 | 3.62 | 1.00 | 34.96 | 2.75 (C:OP2 1H:437) |
| 3 | C | OP2 | 1H:437 | 3.82 | 1.00 | 37.23 |  |
| 4 | C | OP1 | 1H:437 | 3.86 | 1.00 | 36.34 |  |
| 5 | C | OP1 | 1H:420 | 4.16 | 1.00 | 34.42 |  |
| 6 | C | OP2 | 1H:441 | 4.24 | 1.00 | 36.04 |  |
| 7 | G | O3' | 1H:419 | 4.32 | 1.00 | 38.00 |  |
| 8 | C | O5' | 1H:437 | 4.36 | 1.00 | 38.68 |  |
| 9 | G | OP1 | 1H:438 | 4.50 | 1.00 | 32.61 |  |
| 10 | G | O2' | 1H:419 | 4.56 | 1.00 | 31.86 |  |
| 11 | C | OP1 | 1H:441 | 4.76 | 1.00 | 37.88 |  |
| 12 | C | O3' | 1H:437 | 4.77 | 1.00 | 35.66 |  |
| 13 | G | O5' | 1H:438 | 4.93 | 1.00 | 31.10 |  |
  |

  
 Coded contact list:   
 Exclusion zone contact(s): /OP2/  
Consider NA in place of MG (1 valid contact(s) for NA vs. 0 for MG)  

### MG:1H:3369

|  | Res. type | Atom | Residue | Distance | Occupancy | B-factor | Closest solute to 1^st^ shell water |
| --- | --- | --- | --- | --- | --- | --- | --- |
| **Ion** | | | |  | 1.00 | 46.41 || 1 | G | O6 | 1H:2316 | 3.26 | 1.00 | 59.65 |
| 2 | G | N7 | 1H:2316 | 3.67 | 1.00 | 61.21 |  |
| 3 | G | N7 | 1H:2315 | 3.79 | 1.00 | 63.95 |  |
| 4 | G | O6 | 1H:2315 | 3.94 | 1.00 | 61.53 |  |
| 5 | G | O6 | 1H:2317 | 4.27 | 1.00 | 65.59 |  |
  |

  
 Coded contact list:   
 Exclusion zone contact(s): /O6/  
Consider K in place of MG (1 valid contact(s) for K vs. 0 for MG)  

### MG:1H:3370

|  | Res. type | Atom | Residue | Distance | Occupancy | B-factor | Closest solute to 1^st^ shell water |
| --- | --- | --- | --- | --- | --- | --- | --- |
| **Ion** | | | |  | 1.00 | 61.73 || 1 | G | N7 | 1H:2316 | 3.78 | 1.00 | 61.21 |
| 2 | G | O6 | 1H:2317 | 3.93 | 1.00 | 65.59 |  |
| 3 | G | N7 | 1H:2317 | 4.10 | 1.00 | 61.35 |  |
| 4 | G | O6 | 1H:2316 | 4.41 | 1.00 | 59.65 |  |
| 5 | G | OP2 | 1H:2316 | 4.49 | 1.00 | 67.60 |  |
  |

  
 Coded contact list:   
1^st^ MG coordination shell missing (see Notes)  

### MG:1H:3371

|  | Res. type | Atom | Residue | Distance | Occupancy | B-factor | Closest solute to 1^st^ shell water |
| --- | --- | --- | --- | --- | --- | --- | --- |
| **Ion** | | | |  | 1.00 | 31.09 || 1 | G | N7 | 1H:1674 | 3.07 | 1.00 | 33.47 |
| 2 | G | O3' | 1H:1346 | 3.32 | 1.00 | 31.66 |  |
| 3 | G | O2' | 1H:1673 | 3.33 | 1.00 | 25.35 |  |
| 4 | U | OP2 | 1H:1347 | 3.44 | 1.00 | 29.43 |  |
| 5 | G | OP2 | 1H:1674 | 3.57 | 1.00 | 28.44 |  |
| 6 | G | O2' | 1H:1346 | 3.68 | 1.00 | 31.35 |  |
| 7 | G | O3' | 1H:1673 | 3.78 | 1.00 | 28.53 |  |
| 8 | A | N1 | 1H:1348 | 4.03 | 1.00 | 28.99 |  |
| 9 | U | O5' | 1H:1347 | 4.15 | 1.00 | 23.05 |  |
| 10 | G | O6 | 1H:1674 | 4.31 | 1.00 | 35.28 |  |
| 11 | U | O4' | 1H:1347 | 4.67 | 1.00 | 27.30 |  |
| 12 | A | N6 | 1H:1348 | 4.93 | 1.00 | 31.79 |  |
| 13 | C | N4 | 1H:1688 | 4.98 | 1.00 | 33.33 |  |
  |

  
 Coded contact list:   
 Exclusion zone contact(s): /N7/O2'/O3'/  
Consider K in place of MG (3 valid contact(s) for K vs. 0 for MG)  

### MG:1H:3372

|  | Res. type | Atom | Residue | Distance | Occupancy | B-factor | Closest solute to 1^st^ shell water |
| --- | --- | --- | --- | --- | --- | --- | --- |
| **Ion** | | | |  | 1.00 | 31.43 || 1 | C | O2' | 1H:1813 | 2.98 | 1.00 | 25.24 |
| 2 | C | OP2 | 1H:1814 | 3.71 | 1.00 | 14.13 |  |
| 3 | A | OP2 | 1H:1812 | 4.18 | 1.00 | 16.90 |  |
| 4 | C | O2 | 1H:1813 | 4.27 | 1.00 | 20.97 |  |
| 5 | C | O3' | 1H:1813 | 4.53 | 1.00 | 14.19 |  |
  |

  
 Coded contact list:   
 Exclusion zone contact(s): /O2'/  
Consider K in place of MG (1 valid contact(s) for K vs. 0 for MG)  

### MG:1H:3373

|  | Res. type | Atom | Residue | Distance | Occupancy | B-factor | Closest solute to 1^st^ shell water |
| --- | --- | --- | --- | --- | --- | --- | --- |
| **Ion** | | | |  | 1.00 | 43.93 || 1 | G | O6 | 1H:1490 | 3.39 | 1.00 | 58.59 |
| 2 | G | N7 | 1H:1490 | 3.44 | 1.00 | 47.99 |  |
| 3 | G | O6 | 1H:1491 | 4.05 | 1.00 | 62.60 |  |
| 4 | G | N7 | 1H:1491 | 4.53 | 1.00 | 57.04 |  |
| 5 | G | N7 | 1H:1514 | 4.80 | 1.00 | 56.79 |  |
  |

  
 Coded contact list:   
 Exclusion zone contact(s): /O6/  
Consider K in place of MG (1 valid contact(s) for K vs. 0 for MG)  

### MG:1H:3374

|  | Res. type | Atom | Residue | Distance | Occupancy | B-factor | Closest solute to 1^st^ shell water |
| --- | --- | --- | --- | --- | --- | --- | --- |
| **Ion** | | | |  | 1.00 | 28.06 || 1 | U | OP1 | 1H:2039 | 1.91 | 1.00 | 26.41 |
| 2 | Water 1 | | 31:408 | 2.31 | 1.00 | 39.14 | 3.52 (U:OP1 1H:2039) |
| 3 | A | OP1 | 1H:2081 | 3.14 | 1.00 | 19.11 |  |
| 4 | Water 2 | | 1H:4460 | 3.29 | 1.00 | 25.49 | 3.80 (A:OP1 1H:2081) |
| 5 | U | O5' | 1H:2039 | 3.64 | 1.00 | 20.73 |  |
| 6 | Water 3 | | 1H:3792 | 3.86 | 1.00 | 29.43 | 2.37 (A:OP1 1H:2082) |
| 7 | U | OP2 | 1H:2039 | 3.87 | 1.00 | 20.97 |  |
| 8 | A | OP1 | 1H:2082 | 3.94 | 1.00 | 26.02 |  |
| 9 | Water 4 | | 1H:3686 | 4.00 | 1.00 | 22.54 | 2.17 (U:OP2 1H:2040) |
| 10 | U | OP2 | 1H:2040 | 4.14 | 1.00 | 21.65 |  |
| 11 | A | OP2 | 1H:2082 | 4.26 | 1.00 | 17.54 |  |
| 12 | A | O3' | 1H:2038 | 4.35 | 1.00 | 22.58 |  |
| 13 | Water 5 | | 1H:3691 | 4.55 | 1.00 | 25.22 | 2.18 (A:OP2 1H:2082) |
| 14 | A | O5' | 1H:2081 | 4.72 | 1.00 | 18.59 |  |
  |

  
**1^st^ shell ligand angles:**   
1-2: 113;   
  
 Contact(s): /O_w_/OP1/  
 Coded contact list: O_ph_O_w_  
 Exclusion zone contact(s): /O_w_/OP1/  

### MG:1H:3375

|  | Res. type | Atom | Residue | Distance | Occupancy | B-factor | Closest solute to 1^st^ shell water |
| --- | --- | --- | --- | --- | --- | --- | --- |
| **Ion** | | | |  | 1.00 | 36.73 || 1 | G | OP2 | 1H:2023 | 3.34 | 1.00 | 21.57 |
| 2 | A | OP1 | 1H:1703 | 4.23 | 1.00 | 29.39 |  |
| 3 | Water 1 | | 1H:4545 | 4.29 | 1.00 | 25.19 | 3.97 (G:OP2 1H:2023) |
| 4 | A | OP2 | 1H:2024 | 4.39 | 1.00 | 30.02 |  |
| 5 | G | OP1 | 1H:2023 | 4.51 | 1.00 | 30.65 |  |
| 6 | A | OP1 | 1H:1702 | 4.61 | 1.00 | 32.56 |  |
| 7 | G | O5' | 1H:2023 | 4.67 | 1.00 | 22.90 |  |
| 8 | Water 2 | | 1H:4511 | 4.73 | 1.00 | 25.61 | 4.34 (A:OP2 1H:2024) |
| 9 | MET | O | 98:1 | 4.83 | 1.00 | 41.38 |  |
| 10 | Water 3 | | 1H:4515 | 4.84 | 1.00 | 32.20 | 5.74 (A:OP1 1H:1703) |
  |

  
 Coded contact list:   
 Exclusion zone contact(s): /OP2/  
Consider K in place of MG (1 valid contact(s) for K vs. 0 for MG)  

### MG:1H:3376

|  | Res. type | Atom | Residue | Distance | Occupancy | B-factor | Closest solute to 1^st^ shell water |
| --- | --- | --- | --- | --- | --- | --- | --- |
| **Ion** | | | |  | 1.00 | 54.75 || 1 | G | OP1 | 1H:1674 | 2.63 | 1.00 | 28.39 |
| 2 | G | O5' | 1H:1674 | 4.03 | 1.00 | 35.14 |  |
| 3 | G | OP2 | 1H:1674 | 4.15 | 1.00 | 28.44 |  |
| 4 | A | OP1 | 1H:1682 | 4.27 | 1.00 | 33.94 |  |
| 5 | G | OP1 | 1H:1673 | 4.81 | 1.00 | 44.93 |  |
| 6 | U | O4 | 1H:1347 | 4.99 | 1.00 | 36.43 |  |
  |

  
 Coded contact list:   
 Exclusion zone contact(s): /OP1/  
Consider K in place of MG (1 valid contact(s) for K vs. 0 for MG)  

### MG:1H:3377

|  | Res. type | Atom | Residue | Distance | Occupancy | B-factor | Closest solute to 1^st^ shell water |
| --- | --- | --- | --- | --- | --- | --- | --- |
| **Ion** | | | |  | 1.00 | 26.37 || 1 | U | O4 | 1H:2040 | 3.87 | 1.00 | 21.22 |
| 2 | G | N7 | 1H:603 | 3.96 | 1.00 | 27.43 |  |
| 3 | U | OP2 | 1H:2039 | 4.49 | 1.00 | 20.97 |  |
| 4 | G | OP1 | 1H:603 | 4.67 | 1.00 | 24.68 |  |
  |

  
 Coded contact list:   
1^st^ MG coordination shell missing (see Notes)  

### MG:1H:3378

|  | Res. type | Atom | Residue | Distance | Occupancy | B-factor | Closest solute to 1^st^ shell water |
| --- | --- | --- | --- | --- | --- | --- | --- |
| **Ion** | | | |  | 1.00 | 28.95 || 1 | G | OP2 | 1H:2092 | 2.65 | 1.00 | 22.39 |
| 2 | U | O2' | 1H:2091 | 2.72 | 1.00 | 27.64 |  |
| 3 | Water 1 | | 1H:4570 | 3.07 | 1.00 | 39.50 | 4.24 (G:OP2 1H:2092) |
| 4 | U | O3' | 1H:2091 | 3.37 | 1.00 | 21.97 |  |
| 5 | Water 2 | | 1H:4504 | 3.54 | 1.00 | 39.30 | 3.94 (G:OP2 1H:2093) |
| 6 | Water 3 | | 1H:3866 | 3.68 | 1.00 | 19.59 | 2.47 (G:OP2 1H:2092) |
| 7 | G | O2' | 1H:2090 | 4.04 | 1.00 | 16.56 |  |
| 8 | G | O5' | 1H:2092 | 4.54 | 1.00 | 24.81 |  |
| 9 | G | OP2 | 1H:2093 | 4.59 | 1.00 | 24.52 |  |
| 10 | G | OP1 | 1H:2092 | 4.87 | 1.00 | 20.36 |  |
| 11 | Water 4 | | 1H:4231 | 4.98 | 1.00 | 39.26 | 2.94 (G:OP2 1H:2093) |
  |

  
 Coded contact list:   
 Exclusion zone contact(s): /O2'/O3'/O_w_/OP2/  
Consider K in place of MG (3 valid contact(s) for K vs. 0 for MG)  

### MG:1H:3379

|  | Res. type | Atom | Residue | Distance | Occupancy | B-factor | Closest solute to 1^st^ shell water |
| --- | --- | --- | --- | --- | --- | --- | --- |
| **Ion** | | | |  | 1.00 | 47.89 || 1 | C | OP1 | 1H:535 | 3.07 | 1.00 | 49.11 |
| 2 | G | O6 | 1H:27 | 3.56 | 1.00 | 38.72 |  |
| 3 | G | N7 | 1H:27 | 3.73 | 1.00 | 29.51 |  |
| 4 | G | OP2 | 1H:26 | 4.55 | 1.00 | 20.95 |  |
| 5 | C | O5' | 1H:535 | 4.56 | 1.00 | 38.92 |  |
| 6 | G | N7 | 1H:26 | 4.73 | 1.00 | 26.99 |  |
| 7 | G | O3' | 1H:534 | 4.87 | 1.00 | 43.97 |  |
| 8 | Water 1 | | 1H:4012 | 4.89 | 1.00 | 39.02 | 4.00 (G:O6 1H:27) |
  |

  
 Coded contact list:   
 Exclusion zone contact(s): /OP1/  
Consider K in place of MG (1 valid contact(s) for K vs. 0 for MG)  

### MG:1H:3380

|  | Res. type | Atom | Residue | Distance | Occupancy | B-factor | Closest solute to 1^st^ shell water |
| --- | --- | --- | --- | --- | --- | --- | --- |
| **Ion** | | | |  | 1.00 | 44.48 || 1 | U | OP2 | 1H:954 | 3.34 | 1.00 | 48.62 |
| 2 | U | OP1 | 1H:954 | 3.72 | 1.00 | 55.79 |  |
| 3 | G | OP1 | 1H:912 | 3.81 | 1.00 | 47.27 |  |
| 4 | U | O5' | 1H:954 | 4.87 | 1.00 | 58.25 |  |
  |

  
 Coded contact list:   
 Exclusion zone contact(s): /OP2/  
Consider K in place of MG (1 valid contact(s) for K vs. 0 for MG)  

### MG:1H:3381

|  | Res. type | Atom | Residue | Distance | Occupancy | B-factor | Closest solute to 1^st^ shell water |
| --- | --- | --- | --- | --- | --- | --- | --- |
| **Ion** | | | |  | 1.00 | 39.99 || 1 | C | O2 | 1H:1694 | 3.24 | 1.00 | 39.48 |
| 2 | A | N6 | 1H:1321 | 3.73 | 1.00 | 31.59 |  |
| 3 | C | N3 | 1H:1694 | 3.88 | 1.00 | 32.96 |  |
| 4 | A | N7 | 1H:1321 | 3.91 | 1.00 | 30.52 |  |
| 5 | A | N6 | 1H:1349 | 4.82 | 1.00 | 31.51 |  |
| 6 | A | N7 | 1H:1349 | 4.95 | 1.00 | 28.68 |  |
  |

  
 Coded contact list:   
 Exclusion zone contact(s): /O2/  
Consider K in place of MG (1 valid contact(s) for K vs. 0 for MG)  

### MG:1H:3382

|  | Res. type | Atom | Residue | Distance | Occupancy | B-factor | Closest solute to 1^st^ shell water |
| --- | --- | --- | --- | --- | --- | --- | --- |
| **Ion** | | | |  | 1.00 | 13.33 || 1 | Water 1 | | 1H:4098 | 2.15 | 1.00 | 16.55 | 2.75 (G:OP2 1H:857) |
| 2 | Water 2 | | 1H:3726 | 2.17 | 1.00 | 19.14 | 2.26 (U:O4 1H:858) |
| 3 | Water 3 | | 1H:3985 | 2.18 | 1.00 | 18.14 | 2.63 (G:OP2 1H:856) |
| 4 | Water 4 | | 1H:4160 | 2.20 | 1.00 | 15.80 | 2.84 (G:N7 1H:857) |
| 5 | Water 5 | | 78:307 | 2.23 | 1.00 | 19.27 | 3.68 (A:OP1 1H:591) |
| 6 | A | OP1 | 1H:591 | 2.47 | 1.00 | 17.16 |  |
| 7 | U | O4 | 1H:858 | 3.82 | 1.00 | 16.69 |  |
| 8 | U | O3' | 1H:590 | 4.10 | 1.00 | 18.56 |  |
| 9 | G | OP2 | 1H:856 | 4.21 | 1.00 | 20.05 |  |
| 10 | A | O5' | 1H:591 | 4.27 | 1.00 | 17.20 |  |
| 11 | G | N7 | 1H:857 | 4.39 | 1.00 | 19.01 |  |
| 12 | G | OP2 | 1H:857 | 4.81 | 1.00 | 20.73 |  |
  |

  
**1^st^ shell ligand angles:**   
1-2: 91; 1-3: 82; 1-4: 88; 1-5: 176; 2-3: 170; 2-4: 87; 2-5: 93; 3-4: 99; 3-5: 94; 4-5: 91;   
  
 Contact(s): /O_w_/O_w_/O_w_/O_w_/O_w_/  
 Coded contact list: 5O_w_  
 Exclusion zone contact(s): /OP1/  
Consider NA in place of MG (1 valid contact(s) for NA vs. 0 for MG)  
MG hydration shell issue: water too close (< 2.4 Angs.)  

### MG:1H:3383

|  | Res. type | Atom | Residue | Distance | Occupancy | B-factor | Closest solute to 1^st^ shell water |
| --- | --- | --- | --- | --- | --- | --- | --- |
| **Ion** | | | |  | 1.00 | 18.26 || 1 | Water 1 | | 1H:3756 | 2.15 | 1.00 | 19.04 | 2.30 (G:OP2 1H:2079) |
| 2 | Water 2 | | 1H:3945 | 2.18 | 1.00 | 20.96 | 2.57 (2MA:O2' 1H:2516) |
| 3 | Water 3 | | 1H:4243 | 2.18 | 1.00 | 27.00 | 2.96 (A:N7 1H:2080) |
| 4 | Water 4 | | 1H:4111 | 2.19 | 1.00 | 18.51 | 2.77 (A:N7 1H:2081) |
| 5 | G | OP2 | 1H:2079 | 2.40 | 1.00 | 20.39 |  |
| 6 | A | OP2 | 1H:2080 | 2.46 | 1.00 | 20.35 |  |
| 7 | G | O5' | 1H:2079 | 3.42 | 1.00 | 16.23 |  |
| 8 | A | N6 | 1H:2081 | 3.91 | 1.00 | 20.23 |  |
| 9 | A | N7 | 1H:2080 | 4.25 | 1.00 | 23.14 |  |
| 10 | G | OP1 | 1H:2079 | 4.36 | 1.00 | 25.14 |  |
| 11 | Water 5 | | 1H:3813 | 4.45 | 1.00 | 18.26 | 3.91 (2MA:O3' 1H:2516) |
| 12 | G | OP1 | 1H:2518 | 4.56 | 1.00 | 24.64 |  |
| 13 | C | O3' | 1H:2078 | 4.57 | 1.00 | 21.80 |  |
| 14 | A | N7 | 1H:2081 | 4.65 | 1.00 | 25.47 |  |
| 15 | A | OP1 | 1H:2080 | 4.66 | 1.00 | 22.71 |  |
| 16 | A | O5' | 1H:2080 | 4.67 | 1.00 | 16.69 |  |
| 17 | 2MA | O2' | 1H:2516 | 4.69 | 1.00 | 24.00 |  |
| 18 | G | O3' | 1H:2079 | 4.76 | 1.00 | 22.88 |  |
| 19 | 2MA | O3' | 1H:2516 | 4.84 | 1.00 | 17.03 |  |
  |

  
**1^st^ shell ligand angles:**   
1-2: 88; 1-3: 175; 1-4: 82; 2-3: 97; 2-4: 80; 3-4: 100;   
  
 Contact(s): /O_w_/O_w_/O_w_/O_w_/  
 Coded contact list: 4O_w_  
 Exclusion zone contact(s): /OP2/OP2/  
Consider NA in place of MG (2 valid contact(s) for NA vs. 0 for MG)  
MG hydration shell issue: water too close (< 2.4 Angs.)  

### MG:1H:3384

|  | Res. type | Atom | Residue | Distance | Occupancy | B-factor | Closest solute to 1^st^ shell water |
| --- | --- | --- | --- | --- | --- | --- | --- |
| **Ion** | | | |  | 1.00 | 11.68 || 1 | Water 1 | | 1H:3635 | 2.13 | 1.00 | 18.98 | 2.01 (U:OP1 1H:600) |
| 2 | Water 2 | | 1H:3813 | 2.15 | 1.00 | 18.26 | 2.39 (2MA:OP2 1H:2516) |
| 3 | Water 3 | | 1H:3720 | 2.15 | 1.00 | 27.55 | 2.24 (U:OP1 1H:600) |
| 4 | Water 4 | | 1H:3973 | 2.18 | 1.00 | 17.20 | 2.61 (2MA:OP1 1H:2516) |
| 5 | Water 5 | | 1H:4295 | 2.19 | 1.00 | 33.55 | 3.06 (G:OP1 1H:2079) |
| 6 | Water 6 | | 1H:4007 | 2.20 | 1.00 | 30.42 | 2.65 (A:OP1 1H:2080) |
| 7 | U | OP1 | 1H:600 | 2.68 | 1.00 | 19.93 |  |
| 8 | 2MA | OP1 | 1H:2516 | 3.17 | 1.00 | 15.66 |  |
| 9 | 2MA | OP2 | 1H:2516 | 3.70 | 1.00 | 13.66 |  |
| 10 | Water 7 | | 1H:3756 | 3.89 | 1.00 | 19.04 | 3.45 (G:OP1 1H:2079) |
| 11 | U | OP2 | 1H:2517 | 4.06 | 1.00 | 19.86 |  |
| 12 | U | OP2 | 1H:600 | 4.38 | 1.00 | 20.11 |  |
| 13 | A | OP1 | 1H:2080 | 4.54 | 1.00 | 22.71 |  |
| 14 | G | O3' | 1H:2515 | 4.54 | 1.00 | 16.33 |  |
| 15 | A | O3' | 1H:599 | 4.55 | 1.00 | 19.41 |  |
| 16 | Water 8 | | 1H:4383 | 4.92 | 1.00 | 26.32 | 5.29 (U:OP1 1H:600) |
| 17 | G | OP1 | 1H:2079 | 4.95 | 1.00 | 25.14 |  |
  |

  
**1^st^ shell ligand angles:**   
1-2: 85; 1-3: 95; 1-4: 92; 1-5: 84; 1-6: 174; 2-3: 180; 2-4: 96; 2-5: 89; 2-6: 91; 3-4: 83; 3-5: 91; 3-6: 89; 4-5: 173; 4-6: 92; 5-6: 93;   
  
 Contact(s): /O_w_/O_w_/O_w_/O_w_/O_w_/O_w_/  
 Coded contact list: 6O_w_  
 Exclusion zone contact(s): /OP1/OP1/  
Consider K in place of MG (2 valid contact(s) for K vs. 0 for MG)  
MG hydration shell issue: water too close (< 2.4 Angs.)  

### MG:1H:3385

|  | Res. type | Atom | Residue | Distance | Occupancy | B-factor | Closest solute to 1^st^ shell water |
| --- | --- | --- | --- | --- | --- | --- | --- |
| **Ion** | | | |  | 1.00 | 20.15 || 1 | Water 1 | | 1H:3643 | 2.16 | 1.00 | 24.04 | 2.04 (G:OP2 1H:587) |
| 2 | Water 2 | | 1H:4025 | 2.16 | 1.00 | 22.05 | 2.66 (G:OP2 1H:587) |
| 3 | Water 3 | | 1H:4241 | 2.17 | 1.00 | 23.91 | 2.96 (C:O2' 1H:1021) |
| 4 | Water 4 | | 1H:3917 | 2.17 | 1.00 | 35.74 | 2.54 (G:OP2 1H:1022) |
| 5 | Water 5 | | 1H:3695 | 2.18 | 1.00 | 25.02 | 2.19 (G:OP2 1H:587) |
| 6 | Water 6 | | 1H:4212 | 2.19 | 1.00 | 31.14 | 2.91 (G:OP1 1H:1022) |
| 7 | G | OP2 | 1H:587 | 2.70 | 1.00 | 25.28 |  |
| 8 | G | OP1 | 1H:1022 | 4.08 | 1.00 | 33.48 |  |
| 9 | G | OP2 | 1H:1022 | 4.24 | 1.00 | 29.69 |  |
| 10 | C | O2' | 1H:1021 | 4.42 | 1.00 | 29.91 |  |
| 11 | C | OP1 | 1H:588 | 4.49 | 1.00 | 22.54 |  |
| 12 | C | O3' | 1H:1021 | 4.59 | 1.00 | 29.58 |  |
| 13 | U | O2' | 1H:586 | 4.60 | 1.00 | 26.30 |  |
| 14 | A | OP2 | 1H:596 | 4.60 | 1.00 | 24.33 |  |
| 15 | G | OP1 | 1H:587 | 4.61 | 1.00 | 26.98 |  |
| 16 | G | O5' | 1H:587 | 4.72 | 1.00 | 26.18 |  |
  |

  
**1^st^ shell ligand angles:**   
1-2: 85; 1-3: 88; 1-4: 91; 1-5: 92; 1-6: 179; 2-3: 85; 2-4: 177; 2-5: 91; 2-6: 96; 3-4: 94; 3-5: 176; 3-6: 92; 4-5: 89; 4-6: 87; 5-6: 88;   
  
 Contact(s): /O_w_/O_w_/O_w_/O_w_/O_w_/O_w_/  
 Coded contact list: 6O_w_  
 Exclusion zone contact(s): /OP2/  
Consider K in place of MG (1 valid contact(s) for K vs. 0 for MG)  
MG hydration shell issue: water too close (< 2.4 Angs.)  

### MG:1H:3386

|  | Res. type | Atom | Residue | Distance | Occupancy | B-factor | Closest solute to 1^st^ shell water |
| --- | --- | --- | --- | --- | --- | --- | --- |
| **Ion** | | | |  | 1.00 | 31.80 || 1 | Water 1 | | 1H:4245 | 2.17 | 1.00 | 24.34 | 2.96 (C:OP2 1H:1423) |
| 2 | Water 2 | | 1H:3822 | 2.18 | 1.00 | 32.01 | 2.40 (G:O6 1H:1424) |
| 3 | Water 3 | | 1H:4056 | 2.19 | 1.00 | 34.54 | 2.70 (C:OP2 1H:1422) |
| 4 | Water 4 | | 1H:4058 | 2.20 | 1.00 | 41.66 | 2.70 (C:OP1 1H:1398) |
| 5 | U | OP2 | 1H:1399 | 2.45 | 1.00 | 23.94 |  |
| 6 | C | OP2 | 1H:1423 | 2.49 | 1.00 | 30.17 |  |
| 7 | U | O5' | 1H:1399 | 3.29 | 1.00 | 29.00 |  |
| 8 | C | OP1 | 1H:1423 | 3.89 | 1.00 | 37.56 |  |
| 9 | U | OP1 | 1H:1399 | 4.08 | 1.00 | 35.11 |  |
| 10 | G | O6 | 1H:1424 | 4.12 | 1.00 | 34.79 |  |
| 11 | C | O5' | 1H:1422 | 4.59 | 1.00 | 34.55 |  |
| 12 | C | O5' | 1H:1423 | 4.59 | 1.00 | 30.31 |  |
| 13 | C | OP2 | 1H:1398 | 4.60 | 1.00 | 28.67 |  |
| 14 | C | OP1 | 1H:1398 | 4.63 | 1.00 | 25.78 |  |
| 15 | C | O3' | 1H:1398 | 4.65 | 1.00 | 26.18 |  |
| 16 | C | O3' | 1H:1422 | 4.67 | 1.00 | 30.35 |  |
| 17 | C | OP2 | 1H:1422 | 4.70 | 1.00 | 35.58 |  |
| 18 | C | O5' | 1H:1398 | 4.91 | 1.00 | 26.18 |  |
  |

  
**1^st^ shell ligand angles:**   
1-2: 90; 1-3: 178; 1-4: 93; 2-3: 90; 2-4: 173; 3-4: 87;   
  
 Contact(s): /O_w_/O_w_/O_w_/O_w_/  
 Coded contact list: 4O_w_  
 Exclusion zone contact(s): /O5'/OP2/OP2/  
Consider NA in place of MG (2 valid contact(s) for NA vs. 0 for MG)  

### MG:1H:3387

|  | Res. type | Atom | Residue | Distance | Occupancy | B-factor | Closest solute to 1^st^ shell water |
| --- | --- | --- | --- | --- | --- | --- | --- |
| **Ion** | | | |  | 1.00 | 16.11 || 1 | Water 1 | | 1H:3614 | 2.14 | 1.00 | 21.66 | 1.85 (G:OP2 1H:2601) |
| 2 | Water 2 | | 1H:3809 | 2.15 | 1.00 | 13.90 | 2.39 (A:N7 1H:2602) |
| 3 | Water 3 | | 1H:3654 | 2.17 | 1.00 | 18.81 | 2.08 (G:OP2 1H:2601) |
| 4 | Water 4 | | 1H:4256 | 2.19 | 1.00 | 21.72 | 2.98 (A:N6 1H:2602) |
| 5 | Water 5 | | 11:406 | 2.19 | 1.00 | 28.89 | 2.81 (GLU:OE1 11:237) |
| 6 | Water 6 | | 1H:4047 | 2.21 | 1.00 | 22.52 | 2.69 (A:OP1 1H:2600) |
| 7 | G | OP2 | 1H:2601 | 2.70 | 1.00 | 14.99 |  |
| 8 | A | OP1 | 1H:2600 | 3.62 | 1.00 | 20.12 |  |
| 9 | GLU | OE1 | 11:237 | 4.11 | 1.00 | 23.05 |  |
| 10 | A | N7 | 1H:2602 | 4.12 | 1.00 | 17.64 |  |
| 11 | A | N6 | 1H:2602 | 4.39 | 1.00 | 16.70 |  |
| 12 | G | N7 | 1H:2601 | 4.41 | 1.00 | 17.13 |  |
| 13 | G | O5' | 1H:2601 | 4.75 | 1.00 | 14.48 |  |
| 14 | A | OP2 | 1H:2602 | 4.86 | 1.00 | 16.66 |  |
| 15 | C | OP2 | 1H:2453 | 4.90 | 1.00 | 17.51 |  |
| 16 | A | O2' | 1H:2452 | 4.96 | 1.00 | 23.89 |  |
| 17 | G | OP1 | 1H:2601 | 5.00 | 1.00 | 16.92 |  |
  |

  
**1^st^ shell ligand angles:**   
1-2: 84; 1-3: 91; 1-4: 85; 1-5: 178; 1-6: 92; 2-3: 86; 2-4: 90; 2-5: 98; 2-6: 176; 3-4: 174; 3-5: 90; 3-6: 94; 4-5: 94; 4-6: 90; 5-6: 86;   
  
 Contact(s): /O_w_/O_w_/O_w_/O_w_/O_w_/O_w_/  
 Coded contact list: 6O_w_  
 Exclusion zone contact(s): /OP2/  
Consider K in place of MG (1 valid contact(s) for K vs. 0 for MG)  
MG hydration shell issue: water too close (< 2.4 Angs.)  

### MG:1H:3388

|  | Res. type | Atom | Residue | Distance | Occupancy | B-factor | Closest solute to 1^st^ shell water |
| --- | --- | --- | --- | --- | --- | --- | --- |
| **Ion** | | | |  | 1.00 | 15.28 || 1 | Water 1 | | 1H:4087 | 2.13 | 1.00 | 11.75 | 2.74 (C:OP2 1H:2604) |
| 2 | Water 2 | | 1H:4027 | 2.13 | 1.00 | 21.43 | 2.66 (G:N7 1H:2605) |
| 3 | Water 3 | | 1H:3829 | 2.18 | 1.00 | 28.09 | 2.42 (G:OP2 1H:2605) |
| 4 | Water 4 | | 1H:4387 | 2.18 | 1.00 | 17.12 | 3.26 (A:N6 1H:2613) |
| 5 | Water 5 | | 1H:4394 | 2.19 | 1.00 | 25.84 | 3.28 (A:N1 1H:1994) |
| 6 | U | O4 | 1H:2606 | 2.46 | 1.00 | 20.96 |  |
| 7 | GLY | O | 11:238 | 3.94 | 1.00 | 25.91 |  |
| 8 | G | N7 | 1H:2605 | 4.22 | 1.00 | 22.92 |  |
| 9 | G | OP2 | 1H:2605 | 4.30 | 1.00 | 19.90 |  |
| 10 | A | N6 | 1H:2613 | 4.39 | 1.00 | 17.60 |  |
| 11 | A | N1 | 1H:1994 | 4.41 | 1.00 | 21.96 |  |
| 12 | G | O6 | 1H:2605 | 4.41 | 1.00 | 17.89 |  |
| 13 | C | N4 | 1H:2607 | 4.58 | 1.00 | 20.33 |  |
| 14 | U | N3 | 1H:2606 | 4.63 | 1.00 | 20.25 |  |
| 15 | C | OP2 | 1H:2604 | 4.77 | 1.00 | 15.68 |  |
| 16 | G | O6 | 1H:2612 | 4.93 | 1.00 | 21.21 |  |
  |

  
**1^st^ shell ligand angles:**   
1-2: 86; 1-3: 86; 1-4: 93; 1-5: 101; 2-3: 80; 2-4: 97; 2-5: 172; 3-4: 176; 3-5: 96; 4-5: 88;   
  
 Contact(s): /O_w_/O_w_/O_w_/O_w_/O_w_/  
 Coded contact list: 5O_w_  
 Exclusion zone contact(s): /O4/  
Consider NA in place of MG (1 valid contact(s) for NA vs. 0 for MG)  

### MG:1H:3389

|  | Res. type | Atom | Residue | Distance | Occupancy | B-factor | Closest solute to 1^st^ shell water |
| --- | --- | --- | --- | --- | --- | --- | --- |
| **Ion** | | | |  | 1.00 | 13.51 || 1 | Water 1 | | 1H:4017 | 2.15 | 1.00 | 20.89 | 2.66 (U:OP2 1H:1707) |
| 2 | Water 2 | | 1H:3656 | 2.15 | 1.00 | 19.96 | 2.08 (G:OP1 1H:792) |
| 3 | Water 3 | | 21:406 | 2.15 | 1.00 | 15.33 | 2.91 (HIS:NE2 21:135) |
| 4 | Water 4 | | 1H:4074 | 2.20 | 1.00 | 21.32 | 2.73 (C:OP1 1H:1706) |
| 5 | C | OP1 | 1H:1706 | 2.49 | 1.00 | 19.86 |  |
| 6 | HIS | NE2 | 21:135 | 2.68 | 1.00 | 20.66 |  |
| 7 | C | O5' | 1H:1706 | 3.37 | 1.00 | 20.67 |  |
| 8 | G | OP1 | 1H:792 | 4.01 | 1.00 | 21.36 |  |
| 9 | U | OP2 | 1H:1707 | 4.13 | 1.00 | 17.93 |  |
| 10 | G | O3' | 1H:791 | 4.17 | 1.00 | 17.66 |  |
| 11 | C | OP2 | 1H:1706 | 4.33 | 1.00 | 23.68 |  |
| 12 | LYS | N | 21:133 | 4.37 | 1.00 | 18.72 |  |
| 13 | ALA | O | 21:131 | 4.39 | 1.00 | 23.37 |  |
| 14 | C | O3' | 1H:1705 | 4.44 | 1.00 | 22.99 |  |
| 15 | G | O2' | 1H:791 | 4.60 | 1.00 | 21.82 |  |
| 16 | HIS | ND1 | 21:135 | 4.74 | 1.00 | 22.18 |  |
  |

  
**1^st^ shell ligand angles:**   
1-2: 86; 1-3: 98; 1-4: 94; 2-3: 95; 2-4: 176; 3-4: 88;   
  
 Contact(s): /O_w_/O_w_/O_w_/O_w_/  
 Coded contact list: 4O_w_  
 Exclusion zone contact(s): /NE2_His_/O5'/OP1/  
Consider K in place of MG (2 valid contact(s) for K vs. 0 for MG)  
MG hydration shell issue: water too close (< 2.4 Angs.)  

### MG:1H:3390

|  | Res. type | Atom | Residue | Distance | Occupancy | B-factor | Closest solute to 1^st^ shell water |
| --- | --- | --- | --- | --- | --- | --- | --- |
| **Ion** | | | |  | 1.00 | 26.24 || 1 | Water 1 | | 1H:4137 | 2.15 | 1.00 | 24.56 | 2.81 (A:N6 1H:1725) |
| 2 | Water 2 | | 1H:3963 | 2.16 | 1.00 | 18.17 | 2.60 (A:OP2 1H:1724) |
| 3 | Water 3 | | 1H:3700 | 2.17 | 1.00 | 18.23 | 2.20 (A:OP2 1H:1724) |
| 4 | Water 4 | | 1H:3746 | 2.18 | 1.00 | 23.80 | 2.29 (A:OP2 1H:1724) |
| 5 | Water 5 | | 1H:4571 | 2.18 | 1.00 | 27.94 | 4.26 (C:OP2 1H:1723) |
| 6 | Water 6 | | 1H:4503 | 2.19 | 1.00 | 36.70 | 3.74 (G:O6 1H:1726) |
| 7 | A | OP2 | 1H:1724 | 2.78 | 1.00 | 19.23 |  |
| 8 | A | N7 | 1H:1725 | 4.50 | 1.00 | 24.25 |  |
| 9 | A | OP1 | 1H:1724 | 4.50 | 1.00 | 21.87 |  |
| 10 | C | OP2 | 1H:1723 | 4.53 | 1.00 | 25.65 |  |
| 11 | A | N7 | 1H:1724 | 4.62 | 1.00 | 21.58 |  |
| 12 | A | O5' | 1H:1724 | 4.66 | 1.00 | 21.07 |  |
| 13 | A | N6 | 1H:1725 | 4.73 | 1.00 | 23.45 |  |
| 14 | G | O6 | 1H:1726 | 4.76 | 1.00 | 38.93 |  |
  |

  
**1^st^ shell ligand angles:**   
1-2: 90; 1-3: 92; 1-4: 176; 1-5: 88; 1-6: 86; 2-3: 91; 2-4: 94; 2-5: 87; 2-6: 176; 3-4: 88; 3-5: 178; 3-6: 89; 4-5: 92; 4-6: 90; 5-6: 93;   
  
 Contact(s): /O_w_/O_w_/O_w_/O_w_/O_w_/O_w_/  
 Coded contact list: 6O_w_  
 Exclusion zone contact(s): /OP2/  
Consider K in place of MG (1 valid contact(s) for K vs. 0 for MG)  
MG hydration shell issue: water too close (< 2.4 Angs.)  

### MG:1H:3391

|  | Res. type | Atom | Residue | Distance | Occupancy | B-factor | Closest solute to 1^st^ shell water |
| --- | --- | --- | --- | --- | --- | --- | --- |
| **Ion** | | | |  | 1.00 | 20.00 || 1 | Water 1 | | 1H:3752 | 2.15 | 1.00 | 22.50 | 2.30 (A:OP1 1H:1712) |
| 2 | Water 2 | | 1H:3775 | 2.16 | 1.00 | 30.96 | 2.33 (A:OP2 1H:1713) |
| 3 | Water 3 | | 1H:3768 | 2.18 | 1.00 | 36.28 | 2.32 (A:OP1 1H:1712) |
| 4 | Water 4 | | 1H:4300 | 2.19 | 1.00 | 34.48 | 3.06 (G:OP2 1H:1714) |
| 5 | Water 5 | | 1H:4546 | 2.20 | 1.00 | 45.16 | 3.98 (G:O6 1H:1715) |
| 6 | A | OP2 | 1H:1713 | 2.37 | 1.00 | 26.36 |  |
| 7 | A | OP1 | 1H:1712 | 3.06 | 1.00 | 28.52 |  |
| 8 | A | O5' | 1H:1712 | 3.38 | 1.00 | 26.14 |  |
| 9 | A | OP2 | 1H:1712 | 3.94 | 1.00 | 26.83 |  |
| 10 | Water 6 | | 1H:4081 | 3.99 | 1.00 | 26.43 | 2.74 (G:O6 1H:1715) |
| 11 | A | O5' | 1H:1713 | 4.22 | 1.00 | 35.35 |  |
| 12 | Water 7 | | 1H:4336 | 4.32 | 1.00 | 27.44 | 3.15 (A:OP1 1H:1712) |
| 13 | G | O6 | 1H:1715 | 4.41 | 1.00 | 25.29 |  |
| 14 | A | OP1 | 1H:1713 | 4.45 | 1.00 | 31.93 |  |
| 15 | A | O3' | 1H:1712 | 4.72 | 1.00 | 28.92 |  |
| 16 | G | OP2 | 1H:1714 | 4.82 | 1.00 | 34.36 |  |
  |

  
**1^st^ shell ligand angles:**   
1-2: 90; 1-3: 93; 1-4: 178; 1-5: 87; 1-6: 89; 2-3: 177; 2-4: 89; 2-5: 93; 2-6: 62; 3-4: 88; 3-5: 88; 3-6: 118; 4-5: 92; 4-6: 91; 5-6: 154;   
  
 Contact(s): /O_w_/O_w_/O_w_/O_w_/O_w_/OP2/  
 Coded contact list: O_ph_5O_w_  
 Exclusion zone contact(s): /O5'/OP1/  
Consider K in place of MG (2 valid contact(s) for K vs. 1 for MG)  
MG hydration shell issue: water too close (< 2.4 Angs.)  

### MG:1H:3392

|  | Res. type | Atom | Residue | Distance | Occupancy | B-factor | Closest solute to 1^st^ shell water |
| --- | --- | --- | --- | --- | --- | --- | --- |
| **Ion** | | | |  | 1.00 | 51.19 || 1 | Water 1 | | 1H:3795 | 2.16 | 1.00 | 36.40 | 2.37 (C:OP1 1H:1246) |
| 2 | Water 2 | | 1H:3633 | 2.19 | 1.00 | 40.57 | 1.99 (C:OP1 1H:31) |
| 3 | Water 3 | | 1H:3995 | 2.19 | 1.00 | 32.19 | 2.63 (C:OP1 1H:31) |
| 4 | Water 4 | | 1H:3989 | 2.19 | 1.00 | 38.26 | 2.63 (G:OP1 1H:1286) |
| 5 | Water 5 | | 1H:3865 | 2.20 | 1.00 | 43.24 | 2.47 (U:OP1 1H:1287) |
| 6 | Water 6 | | 1H:3790 | 2.20 | 1.00 | 40.42 | 2.36 (U:OP1 1H:1245) |
| 7 | C | OP1 | 1H:31 | 2.96 | 1.00 | 32.22 |  |
| 8 | U | OP1 | 1H:1245 | 3.75 | 1.00 | 23.04 |  |
| 9 | C | OP1 | 1H:1246 | 4.03 | 1.00 | 21.55 |  |
| 10 | G | O3' | 1H:30 | 4.26 | 1.00 | 31.32 |  |
| 11 | G | OP1 | 1H:1286 | 4.26 | 1.00 | 38.80 |  |
| 12 | U | OP1 | 1H:1287 | 4.40 | 1.00 | 46.04 |  |
| 13 | C | OP2 | 1H:31 | 4.59 | 1.00 | 36.19 |  |
| 14 | U | OP2 | 1H:1287 | 4.64 | 1.00 | 34.41 |  |
| 15 | C | OP2 | 1H:1246 | 4.72 | 1.00 | 24.29 |  |
| 16 | U | O5' | 1H:1245 | 4.87 | 1.00 | 26.65 |  |
| 17 | G | O3' | 1H:1286 | 4.88 | 1.00 | 33.10 |  |
  |

  
**1^st^ shell ligand angles:**   
1-2: 85; 1-3: 176; 1-4: 91; 1-5: 85; 1-6: 91; 2-3: 99; 2-4: 87; 2-5: 170; 2-6: 89; 3-4: 91; 3-5: 91; 3-6: 87; 4-5: 90; 4-6: 176; 5-6: 94;   
  
 Contact(s): /O_w_/O_w_/O_w_/O_w_/O_w_/O_w_/  
 Coded contact list: 6O_w_  
 Exclusion zone contact(s): /OP1/  
Consider K in place of MG (1 valid contact(s) for K vs. 0 for MG)  
MG hydration shell issue: water too close (< 2.4 Angs.)  

### MG:1H:3393

|  | Res. type | Atom | Residue | Distance | Occupancy | B-factor | Closest solute to 1^st^ shell water |
| --- | --- | --- | --- | --- | --- | --- | --- |
| **Ion** | | | |  | 1.00 | 17.02 || 1 | Water 1 | | 1H:4028 | 2.15 | 1.00 | 20.15 | 2.66 (A:OP1 1H:1818) |
| 2 | Water 2 | | 1H:4014 | 2.16 | 1.00 | 16.86 | 2.65 (U:OP2 1H:1807) |
| 3 | Water 3 | | 1H:4403 | 2.20 | 1.00 | 19.72 | 3.31 (G:O6 1H:1808) |
| 4 | Water 4 | | 1H:4352 | 2.20 | 1.00 | 22.19 | 3.17 (G:O2' 1H:1804) |
| 5 | Water 5 | | 1H:4221 | 2.21 | 1.00 | 23.04 | 2.92 (C:OP1 1H:1806) |
| 6 | C | OP1 | 1H:1806 | 2.37 | 1.00 | 17.45 |  |
| 7 | G | O2' | 1H:1804 | 3.74 | 1.00 | 21.65 |  |
| 8 | A | OP2 | 1H:1818 | 4.04 | 1.00 | 15.86 |  |
| 9 | G | O6 | 1H:1808 | 4.11 | 1.00 | 16.60 |  |
| 10 | G | N3 | 1H:1804 | 4.18 | 1.00 | 19.70 |  |
| 11 | C | O5' | 1H:1806 | 4.19 | 1.00 | 18.10 |  |
| 12 | C | OP2 | 1H:1806 | 4.32 | 1.00 | 18.56 |  |
| 13 | U | OP2 | 1H:1807 | 4.35 | 1.00 | 17.60 |  |
| 14 | G | N2 | 1H:1804 | 4.46 | 1.00 | 19.35 |  |
| 15 | A | OP1 | 1H:1818 | 4.53 | 1.00 | 20.06 |  |
| 16 | A | O3' | 1H:1805 | 4.69 | 1.00 | 19.53 |  |
| 17 | C | N4 | 1H:1820 | 4.81 | 1.00 | 18.25 |  |
| 18 | A | N6 | 1H:1821 | 4.82 | 1.00 | 20.67 |  |
| 19 | Water 6 | | 1H:3988 | 4.91 | 1.00 | 16.02 | 2.63 (U:OP2 1H:1807) |
  |

  
**1^st^ shell ligand angles:**   
1-2: 89; 1-3: 98; 1-4: 87; 1-5: 171; 1-6: 92; 2-3: 85; 2-4: 176; 2-5: 90; 2-6: 75; 3-4: 94; 3-5: 90; 3-6: 158; 4-5: 94; 4-6: 106; 5-6: 79;   
  
 Contact(s): /O_w_/O_w_/O_w_/O_w_/O_w_/OP1/  
 Coded contact list: O_ph_5O_w_  

### MG:1H:3394

|  | Res. type | Atom | Residue | Distance | Occupancy | B-factor | Closest solute to 1^st^ shell water |
| --- | --- | --- | --- | --- | --- | --- | --- |
| **Ion** | | | |  | 1.00 | 11.72 || 1 | Water 1 | | 1H:4077 | 2.15 | 1.00 | 18.84 | 2.73 (G:OP2 1H:2457) |
| 2 | Water 2 | | 1H:3820 | 2.16 | 1.00 | 12.43 | 2.40 (A:OP1 1H:2085) |
| 3 | Water 3 | | 1H:3642 | 2.17 | 1.00 | 17.48 | 2.02 (A:OP1 1H:2085) |
| 4 | Water 4 | | 1H:4004 | 2.18 | 1.00 | 16.85 | 2.64 (G:N7 1H:2457) |
| 5 | Water 5 | | 1H:4326 | 2.18 | 1.00 | 24.00 | 3.13 (G:OP2 1H:2457) |
| 6 | Water 6 | | 1H:3895 | 2.19 | 1.00 | 21.76 | 2.51 (A:OP2 1H:2085) |
| 7 | A | OP1 | 1H:2085 | 2.76 | 1.00 | 19.28 |  |
| 8 | A | OP2 | 1H:2085 | 3.55 | 1.00 | 18.12 |  |
| 9 | Water 7 | | 1H:4123 | 3.72 | 1.00 | 21.98 | 2.79 (G:OP1 1H:2084) |
| 10 | G | OP1 | 1H:2084 | 3.89 | 1.00 | 15.09 |  |
| 11 | G | OP2 | 1H:2457 | 4.00 | 1.00 | 13.53 |  |
| 12 | G | N7 | 1H:2457 | 4.00 | 1.00 | 14.16 |  |
| 13 | G | N7 | 1H:2458 | 4.29 | 1.00 | 12.39 |  |
| 14 | LYS | NZ | 31:68 | 4.49 | 1.00 | 13.89 |  |
| 15 | A | O5' | 1H:2085 | 4.63 | 1.00 | 34.39 |  |
| 16 | G | O3' | 1H:2084 | 4.67 | 1.00 | 21.68 |  |
| 17 | G | O6 | 1H:2457 | 4.77 | 1.00 | 13.62 |  |
| 18 | C | OP2 | 1H:2456 | 4.84 | 1.00 | 14.38 |  |
  |

  
**1^st^ shell ligand angles:**   
1-2: 84; 1-3: 94; 1-4: 87; 1-5: 89; 1-6: 172; 2-3: 97; 2-4: 87; 2-5: 173; 2-6: 97; 3-4: 176; 3-5: 81; 3-6: 94; 4-5: 96; 4-6: 85; 5-6: 90;   
  
 Contact(s): /O_w_/O_w_/O_w_/O_w_/O_w_/O_w_/  
 Coded contact list: 6O_w_  
 Exclusion zone contact(s): /OP1/  
Consider K in place of MG (1 valid contact(s) for K vs. 0 for MG)  
MG hydration shell issue: water too close (< 2.4 Angs.)  

### MG:1H:3395

|  | Res. type | Atom | Residue | Distance | Occupancy | B-factor | Closest solute to 1^st^ shell water |
| --- | --- | --- | --- | --- | --- | --- | --- |
| **Ion** | | | |  | 1.00 | 23.11 || 1 | Water 1 | | 1H:3649 | 2.14 | 1.00 | 23.14 | 2.06 (A:N1 1H:2461) |
| 2 | Water 2 | | 1H:3716 | 2.15 | 1.00 | 18.98 | 2.24 (G:OP1 1H:879) |
| 3 | Water 3 | | 1H:4116 | 2.16 | 1.00 | 17.12 | 2.78 (U:OP1 1H:592) |
| 4 | Water 4 | | 78:306 | 2.16 | 1.00 | 18.47 | 3.67 (G:OP2 1H:990) |
| 5 | Water 5 | | 1H:4067 | 2.18 | 1.00 | 29.17 | 2.71 (A:N3 1H:991) |
| 6 | Water 6 | | 1H:4262 | 2.20 | 1.00 | 19.03 | 2.98 (G:OP1 1H:879) |
| 7 | G | OP1 | 1H:879 | 3.68 | 1.00 | 19.84 |  |
| 8 | A | N3 | 1H:991 | 3.86 | 1.00 | 18.12 |  |
| 9 | A | N1 | 1H:2461 | 4.03 | 1.00 | 22.20 |  |
| 10 | A | N6 | 1H:2461 | 4.06 | 1.00 | 19.25 |  |
| 11 | G | OP1 | 1H:990 | 4.28 | 1.00 | 19.16 |  |
| 12 | G | O3' | 1H:878 | 4.47 | 1.00 | 15.34 |  |
| 13 | G | OP2 | 1H:990 | 4.57 | 1.00 | 28.14 |  |
| 14 | U | OP1 | 1H:592 | 4.90 | 1.00 | 13.12 |  |
  |

  
**1^st^ shell ligand angles:**   
1-2: 96; 1-3: 93; 1-4: 177; 1-5: 83; 1-6: 85; 2-3: 96; 2-4: 83; 2-5: 178; 2-6: 86; 3-4: 84; 3-5: 86; 3-6: 177; 4-5: 97; 4-6: 98; 5-6: 92;   
  
 Contact(s): /O_w_/O_w_/O_w_/O_w_/O_w_/O_w_/  
 Coded contact list: 6O_w_  
MG hydration shell issue: water too close (< 2.4 Angs.)  

### MG:1H:3396

|  | Res. type | Atom | Residue | Distance | Occupancy | B-factor | Closest solute to 1^st^ shell water |
| --- | --- | --- | --- | --- | --- | --- | --- |
| **Ion** | | | |  | 1.00 | 16.42 || 1 | Water 1 | | 1H:4121 | 2.15 | 1.00 | 22.07 | 2.79 (G:O2' 1H:1018) |
| 2 | Water 2 | | 1H:4583 | 2.16 | 1.00 | 25.28 | 4.99 (A:OP2 1H:1019) |
| 3 | Water 3 | | 1H:3887 | 2.17 | 1.00 | 16.73 | 2.50 (A:OP2 1H:1019) |
| 4 | Water 4 | | 1H:4194 | 2.17 | 1.00 | 20.38 | 2.89 (A:N1 1H:868) |
| 5 | Water 5 | | 1H:4265 | 2.18 | 1.00 | 21.79 | 2.99 (A:OP1 1H:1019) |
| 6 | Water 6 | | 1H:4086 | 2.19 | 1.00 | 25.56 | 2.74 (U:OP2 1H:593) |
| 7 | A | OP2 | 1H:1019 | 4.32 | 1.00 | 20.65 |  |
| 8 | G | O2' | 1H:1018 | 4.36 | 1.00 | 20.85 |  |
| 9 | A | OP1 | 1H:1019 | 4.36 | 1.00 | 19.51 |  |
| 10 | G | OP1 | 1H:594 | 4.37 | 1.00 | 17.65 |  |
| 11 | A | N6 | 1H:868 | 4.40 | 1.00 | 18.21 |  |
| 12 | A | N1 | 1H:868 | 4.49 | 1.00 | 20.08 |  |
| 13 | A | N1 | 1H:869 | 4.51 | 1.00 | 16.38 |  |
| 14 | G | OP2 | 1H:594 | 4.75 | 1.00 | 22.18 |  |
| 15 | U | OP2 | 1H:593 | 4.77 | 1.00 | 25.23 |  |
| 16 | G | O3' | 1H:1018 | 4.89 | 1.00 | 20.25 |  |
  |

  
**1^st^ shell ligand angles:**   
1-2: 174; 1-3: 94; 1-4: 84; 1-5: 91; 1-6: 91; 2-3: 91; 2-4: 91; 2-5: 92; 2-6: 87; 3-4: 176; 3-5: 85; 3-6: 93; 4-5: 91; 4-6: 91; 5-6: 177;   
  
 Contact(s): /O_w_/O_w_/O_w_/O_w_/O_w_/O_w_/  
 Coded contact list: 6O_w_  

### MG:1H:3397

|  | Res. type | Atom | Residue | Distance | Occupancy | B-factor | Closest solute to 1^st^ shell water |
| --- | --- | --- | --- | --- | --- | --- | --- |
| **Ion** | | | |  | 1.00 | 17.48 || 1 | Water 1 | | 1H:3741 | 2.14 | 1.00 | 24.73 | 2.28 (G:OP2 1H:1233) |
| 2 | Water 2 | | 1H:3964 | 2.14 | 1.00 | 28.17 | 2.60 (G:OP2 1H:1233) |
| 3 | Water 3 | | 1H:3754 | 2.16 | 1.00 | 31.05 | 2.30 (C:OP2 1H:865) |
| 4 | Water 4 | | 1H:4001 | 2.18 | 1.00 | 24.67 | 2.64 (C:OP2 1H:864) |
| 5 | Water 5 | | 1H:4244 | 2.19 | 1.00 | 32.31 | 2.96 (C:OP2 1H:865) |
| 6 | G | OP2 | 1H:1233 | 2.48 | 1.00 | 24.11 |  |
| 7 | C | OP2 | 1H:865 | 3.43 | 1.00 | 26.22 |  |
| 8 | C | OP2 | 1H:864 | 3.91 | 1.00 | 24.47 |  |
| 9 | G | N7 | 1H:1233 | 4.07 | 1.00 | 31.09 |  |
| 10 | C | O2' | 1H:1231 | 4.30 | 1.00 | 40.88 |  |
| 11 | G | O5' | 1H:1233 | 4.32 | 1.00 | 28.39 |  |
| 12 | G | OP1 | 1H:1233 | 4.44 | 1.00 | 23.39 |  |
| 13 | C | O3' | 1H:1231 | 4.46 | 1.00 | 30.38 |  |
| 14 | G | O6 | 1H:866 | 4.47 | 1.00 | 26.35 |  |
| 15 | C | O5' | 1H:864 | 4.61 | 1.00 | 18.15 |  |
| 16 | C | OP1 | 1H:864 | 4.71 | 1.00 | 25.66 |  |
| 17 | G | O5' | 1H:1232 | 4.74 | 1.00 | 32.14 |  |
| 18 | G | O3' | 1H:1232 | 4.92 | 1.00 | 31.39 |  |
  |

  
**1^st^ shell ligand angles:**   
1-2: 86; 1-3: 177; 1-4: 95; 1-5: 90; 2-3: 93; 2-4: 88; 2-5: 176; 3-4: 88; 3-5: 91; 4-5: 91;   
  
 Contact(s): /O_w_/O_w_/O_w_/O_w_/O_w_/  
 Coded contact list: 5O_w_  
 Exclusion zone contact(s): /OP2/  
Consider NA in place of MG (1 valid contact(s) for NA vs. 0 for MG)  
MG hydration shell issue: water too close (< 2.4 Angs.)  

### MG:1H:3398

|  | Res. type | Atom | Residue | Distance | Occupancy | B-factor | Closest solute to 1^st^ shell water |
| --- | --- | --- | --- | --- | --- | --- | --- |
| **Ion** | | | |  | 1.00 | 52.68 || 1 | Water 1 | | 1H:3890 | 2.18 | 1.00 | 42.57 | 2.50 (G:OP2 1H:980) |
| 2 | Water 2 | | 1H:4159 | 2.18 | 1.00 | 27.71 | 2.84 (C:OP1 1H:886) |
| 3 | Water 3 | | 1H:4370 | 2.19 | 1.00 | 41.16 | 3.23 (C:OP1 1H:886) |
| 4 | Water 4 | | 1H:4406 | 2.19 | 1.00 | 36.80 | 3.33 (U:OP2 1H:887) |
| 5 | Water 5 | | 1H:4443 | 2.19 | 1.00 | 39.00 | 3.48 (A:OP2 1H:979) |
| 6 | Water 6 | | 1H:4514 | 2.20 | 1.00 | 43.59 | 3.83 (U:OP1 1H:887) |
| 7 | G | OP2 | 1H:980 | 4.15 | 1.00 | 38.51 |  |
| 8 | U | OP2 | 1H:887 | 4.21 | 1.00 | 23.37 |  |
| 9 | C | OP1 | 1H:886 | 4.23 | 1.00 | 35.08 |  |
| 10 | A | OP2 | 1H:979 | 4.34 | 1.00 | 47.86 |  |
| 11 | G | OP1 | 1H:980 | 4.36 | 1.00 | 30.28 |  |
| 12 | U | OP1 | 1H:887 | 4.63 | 1.00 | 29.81 |  |
  |

  
**1^st^ shell ligand angles:**   
1-2: 178; 1-3: 97; 1-4: 86; 1-5: 88; 1-6: 88; 2-3: 83; 2-4: 96; 2-5: 90; 2-6: 92; 3-4: 89; 3-5: 90; 3-6: 174; 4-5: 174; 4-6: 90; 5-6: 93;   
  
 Contact(s): /O_w_/O_w_/O_w_/O_w_/O_w_/O_w_/  
 Coded contact list: 6O_w_  

### MG:1H:3399

|  | Res. type | Atom | Residue | Distance | Occupancy | B-factor | Closest solute to 1^st^ shell water |
| --- | --- | --- | --- | --- | --- | --- | --- |
| **Ion** | | | |  | 1.00 | 24.11 || 1 | Water 1 | | 1H:3881 | 2.15 | 1.00 | 14.35 | 2.49 (U:OP1 1H:1296) |
| 2 | Water 2 | | 1H:3811 | 2.15 | 1.00 | 28.87 | 2.39 (U:OP1 1H:1296) |
| 3 | Water 3 | | 1H:3636 | 2.15 | 1.00 | 27.87 | 2.01 (U:OP1 1H:1296) |
| 4 | Water 4 | | 1H:3710 | 2.15 | 1.00 | 26.73 | 2.23 (G:OP1 1H:609) |
| 5 | Water 5 | | 1H:3998 | 2.21 | 1.00 | 25.35 | 2.64 (A:OP2 1H:610) |
| 6 | Water 6 | | 1H:4397 | 2.21 | 1.00 | 33.58 | 3.29 (G:OP1 1H:1297) |
| 7 | U | OP1 | 1H:1296 | 2.79 | 1.00 | 23.99 |  |
| 8 | A | OP2 | 1H:610 | 2.83 | 1.00 | 23.80 |  |
| 9 | G | OP1 | 1H:1297 | 3.65 | 1.00 | 23.29 |  |
| 10 | U | O5' | 1H:1296 | 3.98 | 1.00 | 19.72 |  |
| 11 | G | OP1 | 1H:609 | 3.99 | 1.00 | 26.54 |  |
| 12 | A | OP1 | 1H:610 | 4.19 | 1.00 | 17.96 |  |
| 13 | G | O3' | 1H:1295 | 4.37 | 1.00 | 16.21 |  |
| 14 | U | O4 | 1H:859 | 4.44 | 1.00 | 27.13 |  |
| 15 | G | O5' | 1H:609 | 4.46 | 1.00 | 27.72 |  |
| 16 | A | O5' | 1H:610 | 4.54 | 1.00 | 19.35 |  |
| 17 | U | O3' | 1H:1296 | 4.60 | 1.00 | 24.49 |  |
| 18 | G | OP2 | 1H:609 | 4.66 | 1.00 | 26.20 |  |
  |

  
**1^st^ shell ligand angles:**   
1-2: 87; 1-3: 77; 1-4: 91; 1-5: 103; 1-6: 173; 2-3: 96; 2-4: 178; 2-5: 87; 2-6: 91; 3-4: 82; 3-5: 176; 3-6: 97; 4-5: 94; 4-6: 90; 5-6: 84;   
  
 Contact(s): /O_w_/O_w_/O_w_/O_w_/O_w_/O_w_/  
 Coded contact list: 6O_w_  
 Exclusion zone contact(s): /OP1/OP2/  
Consider K in place of MG (2 valid contact(s) for K vs. 0 for MG)  
MG hydration shell issue: water too close (< 2.4 Angs.)  

### MG:1H:3400

|  | Res. type | Atom | Residue | Distance | Occupancy | B-factor | Closest solute to 1^st^ shell water |
| --- | --- | --- | --- | --- | --- | --- | --- |
| **Ion** | | | |  | 1.00 | 20.32 || 1 | Water 1 | | 1H:3953 | 2.14 | 1.00 | 24.78 | 2.59 (A:N7 1H:476) |
| 2 | Water 2 | | 1H:4321 | 2.16 | 1.00 | 26.03 | 3.12 (C:OP1 1H:480) |
| 3 | Water 3 | | 1H:3975 | 2.17 | 1.00 | 20.47 | 2.62 (C:OP1 1H:480) |
| 4 | Water 4 | | 1H:4130 | 2.18 | 1.00 | 28.81 | 2.80 (A:OP1 1H:481) |
| 5 | G | O6 | 1H:477 | 2.43 | 1.00 | 23.89 |  |
| 6 | C | OP1 | 1H:480 | 2.49 | 1.00 | 19.27 |  |
| 7 | G | N7 | 1H:477 | 4.01 | 1.00 | 19.82 |  |
| 8 | A | N6 | 1H:476 | 4.01 | 1.00 | 19.69 |  |
| 9 | A | OP1 | 1H:481 | 4.14 | 1.00 | 24.51 |  |
| 10 | A | N7 | 1H:476 | 4.16 | 1.00 | 20.56 |  |
| 11 | C | OP2 | 1H:480 | 4.43 | 1.00 | 19.78 |  |
| 12 | C | O5' | 1H:480 | 4.43 | 1.00 | 26.09 |  |
| 13 | G | N1 | 1H:477 | 4.53 | 1.00 | 22.28 |  |
| 14 | A | OP2 | 1H:476 | 4.85 | 1.00 | 17.46 |  |
| 15 | G | O3' | 1H:479 | 4.88 | 1.00 | 23.38 |  |
  |

  
**1^st^ shell ligand angles:**   
1-2: 88; 1-3: 83; 1-4: 94; 2-3: 93; 2-4: 95; 3-4: 171;   
  
 Contact(s): /O_w_/O_w_/O_w_/O_w_/  
 Coded contact list: 4O_w_  
 Exclusion zone contact(s): /O6/OP1/  
Consider NA in place of MG (2 valid contact(s) for NA vs. 0 for MG)  

### MG:1H:3401

|  | Res. type | Atom | Residue | Distance | Occupancy | B-factor | Closest solute to 1^st^ shell water |
| --- | --- | --- | --- | --- | --- | --- | --- |
| **Ion** | | | |  | 1.00 | 16.00 || 1 | Water 1 | | 1H:3788 | 2.15 | 1.00 | 16.52 | 2.36 (U:O4 1H:2097) |
| 2 | Water 2 | | 1H:4069 | 2.16 | 1.00 | 16.07 | 2.72 (U:O4 1H:2098) |
| 3 | Water 3 | | 1H:4040 | 2.16 | 1.00 | 15.85 | 2.68 (G:O6 1H:2255) |
| 4 | Water 4 | | 1H:4354 | 2.17 | 1.00 | 23.10 | 3.18 (G:N7 1H:2255) |
| 5 | Water 5 | | 1H:4170 | 2.17 | 1.00 | 19.28 | 2.85 (A:N7 1H:2254) |
| 6 | Water 6 | | 1H:4575 | 2.19 | 1.00 | 24.00 | 4.71 (U:O4 1H:2097) |
| 7 | U | O4 | 1H:2097 | 3.97 | 1.00 | 15.97 |  |
| 8 | G | N7 | 1H:2251 | 4.07 | 1.00 | 17.07 |  |
| 9 | U | O4 | 1H:2098 | 4.12 | 1.00 | 17.83 |  |
| 10 | U | O4 | 1H:2256 | 4.12 | 1.00 | 16.51 |  |
| 11 | A | N7 | 1H:2254 | 4.46 | 1.00 | 16.98 |  |
| 12 | G | O6 | 1H:2255 | 4.59 | 1.00 | 17.51 |  |
| 13 | G | N7 | 1H:2255 | 4.69 | 1.00 | 14.84 |  |
| 14 | A | N6 | 1H:2254 | 4.91 | 1.00 | 18.14 |  |
  |

  
**1^st^ shell ligand angles:**   
1-2: 87; 1-3: 90; 1-4: 88; 1-5: 175; 1-6: 92; 2-3: 91; 2-4: 174; 2-5: 89; 2-6: 94; 3-4: 92; 3-5: 85; 3-6: 175; 4-5: 95; 4-6: 83; 5-6: 92;   
  
 Contact(s): /O_w_/O_w_/O_w_/O_w_/O_w_/O_w_/  
 Coded contact list: 6O_w_  
MG hydration shell issue: water too close (< 2.4 Angs.)  

### MG:1H:3402

|  | Res. type | Atom | Residue | Distance | Occupancy | B-factor | Closest solute to 1^st^ shell water |
| --- | --- | --- | --- | --- | --- | --- | --- |
| **Ion** | | | |  | 1.00 | 23.67 || 1 | Water 1 | | 1H:3761 | 2.16 | 1.00 | 36.70 | 2.31 (A:OP2 1H:1656) |
| 2 | Water 2 | | 1H:4258 | 2.16 | 1.00 | 30.98 | 2.98 (G:OP2 1H:1666) |
| 3 | Water 3 | | 1H:3990 | 2.17 | 1.00 | 20.42 | 2.63 (C:O2' 1H:1664) |
| 4 | Water 4 | | 1H:4353 | 2.18 | 1.00 | 27.64 | 3.26 (G:O6 1H:1666) |
| 5 | Water 5 | | 1H:3665 | 2.18 | 1.00 | 31.77 | 2.11 (C:OP1 1H:1664) |
| 6 | Water 6 | | 1H:4257 | 2.18 | 1.00 | 26.28 | 2.98 (G:O6 1H:1660) |
| 7 | C | OP1 | 1H:1664 | 4.00 | 1.00 | 27.48 |  |
| 8 | A | OP2 | 1H:1656 | 4.10 | 1.00 | 28.85 |  |
| 9 | G | O6 | 1H:1666 | 4.23 | 1.00 | 20.10 |  |
| 10 | G | OP2 | 1H:1666 | 4.26 | 1.00 | 20.66 |  |
| 11 | G | O6 | 1H:1660 | 4.29 | 1.00 | 24.48 |  |
| 12 | G | N7 | 1H:1666 | 4.33 | 1.00 | 20.29 |  |
| 13 | C | O2' | 1H:1664 | 4.52 | 1.00 | 22.17 |  |
| 14 | A | O2' | 1H:1655 | 4.57 | 1.00 | 22.90 |  |
| 15 | C | OP2 | 1H:1664 | 4.81 | 1.00 | 29.87 |  |
  |

  
**1^st^ shell ligand angles:**   
1-2: 92; 1-3: 177; 1-4: 90; 1-5: 88; 1-6: 85; 2-3: 85; 2-4: 96; 2-5: 85; 2-6: 176; 3-4: 91; 3-5: 91; 3-6: 98; 4-5: 178; 4-6: 87; 5-6: 92;   
  
 Contact(s): /O_w_/O_w_/O_w_/O_w_/O_w_/O_w_/  
 Coded contact list: 6O_w_  
MG hydration shell issue: water too close (< 2.4 Angs.)  

### MG:1H:3403

|  | Res. type | Atom | Residue | Distance | Occupancy | B-factor | Closest solute to 1^st^ shell water |
| --- | --- | --- | --- | --- | --- | --- | --- |
| **Ion** | | | |  | 1.00 | 27.38 || 1 | Water 1 | | 1H:4269 | 2.15 | 1.00 | 30.89 | 3.00 (G:O6 1H:1381) |
| 2 | Water 2 | | 1H:4079 | 2.16 | 1.00 | 35.68 | 2.74 (U:O4 1H:1382) |
| 3 | Water 3 | | 1H:3875 | 2.17 | 1.00 | 31.13 | 2.49 (G:N7 1H:1381) |
| 4 | Water 4 | | 1H:3818 | 2.17 | 1.00 | 28.84 | 2.40 (C:OP2 1H:1380) |
| 5 | Water 5 | | 1H:4162 | 2.18 | 1.00 | 33.86 | 2.84 (C:OP2 1H:1380) |
| 6 | Water 6 | | 1H:4357 | 2.20 | 1.00 | 31.18 | 3.18 (U:OP2 1H:1360) |
| 7 | C | OP2 | 1H:1380 | 3.66 | 1.00 | 29.35 |  |
| 8 | U | OP2 | 1H:1360 | 4.12 | 1.00 | 28.38 |  |
| 9 | U | O4 | 1H:1382 | 4.17 | 1.00 | 30.58 |  |
| 10 | G | OP2 | 1H:1381 | 4.30 | 1.00 | 27.56 |  |
| 11 | G | N7 | 1H:1381 | 4.38 | 1.00 | 32.84 |  |
| 12 | G | O6 | 1H:1381 | 4.67 | 1.00 | 32.15 |  |
| 13 | G | OP2 | 1H:1379 | 4.75 | 1.00 | 24.49 |  |
| 14 | A | N6 | 1H:1383 | 4.92 | 1.00 | 34.48 |  |
| 15 | U | O4 | 1H:1363 | 4.94 | 1.00 | 34.72 |  |
  |

  
**1^st^ shell ligand angles:**   
1-2: 88; 1-3: 84; 1-4: 90; 1-5: 177; 1-6: 96; 2-3: 88; 2-4: 177; 2-5: 94; 2-6: 88; 3-4: 90; 3-5: 94; 3-6: 176; 4-5: 88; 4-6: 94; 5-6: 86;   
  
 Contact(s): /O_w_/O_w_/O_w_/O_w_/O_w_/O_w_/  
 Coded contact list: 6O_w_  
MG hydration shell issue: water too close (< 2.4 Angs.)  

### MG:1H:3404

|  | Res. type | Atom | Residue | Distance | Occupancy | B-factor | Closest solute to 1^st^ shell water |
| --- | --- | --- | --- | --- | --- | --- | --- |
| **Ion** | | | |  | 1.00 | 43.55 || 1 | Water 1 | | 1H:4331 | 2.17 | 1.00 | 52.38 | 3.14 (C:OP1 1H:1377) |
| 2 | Water 2 | | 1H:4390 | 2.18 | 1.00 | 52.55 | 3.50 (C:OP2 1H:1365) |
| 3 | Water 3 | | 1H:4152 | 2.18 | 1.00 | 37.43 | 2.83 (C:OP2 1H:1365) |
| 4 | Water 4 | | 1H:4400 | 2.19 | 1.00 | 42.38 | 3.49 (A:OP2 1H:1378) |
| 5 | Water 5 | | 1H:4330 | 2.19 | 1.00 | 37.72 | 3.14 (A:OP2 1H:1378) |
| 6 | Water 6 | | 1H:3852 | 2.20 | 1.00 | 32.04 | 2.45 (C:OP2 1H:1377) |
| 7 | C | OP1 | 1H:1377 | 3.82 | 1.00 | 30.17 |  |
| 8 | C | OP2 | 1H:1365 | 4.24 | 1.00 | 45.10 |  |
| 9 | A | OP2 | 1H:1378 | 4.26 | 1.00 | 29.95 |  |
| 10 | C | OP2 | 1H:1377 | 4.44 | 1.00 | 29.94 |  |
| 11 | G | N7 | 1H:1366 | 4.80 | 1.00 | 31.78 |  |
  |

  
**1^st^ shell ligand angles:**   
1-2: 89; 1-3: 86; 1-4: 177; 1-5: 89; 1-6: 92; 2-3: 87; 2-4: 92; 2-5: 90; 2-6: 178; 3-4: 91; 3-5: 175; 3-6: 94; 4-5: 93; 4-6: 86; 5-6: 89;   
  
 Contact(s): /O_w_/O_w_/O_w_/O_w_/O_w_/O_w_/  
 Coded contact list: 6O_w_  

### MG:1H:3405

|  | Res. type | Atom | Residue | Distance | Occupancy | B-factor | Closest solute to 1^st^ shell water |
| --- | --- | --- | --- | --- | --- | --- | --- |
| **Ion** | | | |  | 1.00 | 16.64 || 1 | Water 1 | | 1H:3807 | 2.14 | 1.00 | 16.26 | 2.39 (G:OP1 1H:1860) |
| 2 | Water 2 | | 1H:4021 | 2.15 | 1.00 | 13.73 | 2.66 (G:OP1 1H:1860) |
| 3 | Water 3 | | 1H:3630 | 2.16 | 1.00 | 17.42 | 1.97 (G:OP1 1H:1860) |
| 4 | Water 4 | | 1H:4248 | 2.19 | 1.00 | 15.08 | 2.96 (G:OP1 1H:1804) |
| 5 | Water 5 | | 1H:4176 | 2.20 | 1.00 | 21.56 | 2.87 (G:OP1 1H:1804) |
| 6 | Water 6 | | 1H:4036 | 2.20 | 1.00 | 26.63 | 2.67 (G:OP2 1H:1996) |
| 7 | G | OP1 | 1H:1860 | 2.71 | 1.00 | 15.93 |  |
| 8 | G | OP1 | 1H:1804 | 3.87 | 1.00 | 16.84 |  |
| 9 | G | OP2 | 1H:1996 | 4.01 | 1.00 | 21.74 |  |
| 10 | A | OP2 | 1H:1805 | 4.23 | 1.00 | 18.46 |  |
| 11 | A | O5' | 1H:1995 | 4.55 | 1.00 | 17.06 |  |
| 12 | A | OP2 | 1H:1995 | 4.58 | 1.00 | 23.13 |  |
| 13 | G | OP2 | 1H:1860 | 4.69 | 1.00 | 16.68 |  |
  |

  
**1^st^ shell ligand angles:**   
1-2: 85; 1-3: 96; 1-4: 85; 1-5: 91; 1-6: 179; 2-3: 92; 2-4: 91; 2-5: 174; 2-6: 94; 3-4: 177; 3-5: 85; 3-6: 84; 4-5: 93; 4-6: 95; 5-6: 90;   
  
 Contact(s): /O_w_/O_w_/O_w_/O_w_/O_w_/O_w_/  
 Coded contact list: 6O_w_  
 Exclusion zone contact(s): /OP1/  
Consider K in place of MG (1 valid contact(s) for K vs. 0 for MG)  
MG hydration shell issue: water too close (< 2.4 Angs.)  

### MG:1H:3406

|  | Res. type | Atom | Residue | Distance | Occupancy | B-factor | Closest solute to 1^st^ shell water |
| --- | --- | --- | --- | --- | --- | --- | --- |
| **Ion** | | | |  | 1.00 | 14.78 || 1 | Water 1 | | 1H:4275 | 2.16 | 1.00 | 31.35 | 3.01 (G:OP1 1H:1927) |
| 2 | Water 2 | | 1H:4388 | 2.16 | 1.00 | 31.18 | 3.26 (G:OP1 1H:1927) |
| 3 | Water 3 | | 1H:4060 | 2.16 | 1.00 | 13.87 | 2.71 (G:OP2 1H:2610) |
| 4 | Water 4 | | 1H:4499 | 2.17 | 1.00 | 23.00 | 3.85 (G:OP1 1H:2610) |
| 5 | Water 5 | | 1H:4351 | 2.17 | 1.00 | 27.83 | 3.17 (G:O6 1H:2608) |
| 6 | Water 6 | | 1H:4062 | 2.18 | 1.00 | 26.65 | 2.71 (G:N7 1H:2608) |
| 7 | G | N7 | 1H:2608 | 4.07 | 1.00 | 20.38 |  |
| 8 | G | OP1 | 1H:2610 | 4.13 | 1.00 | 15.08 |  |
| 9 | G | OP1 | 1H:1927 | 4.20 | 1.00 | 29.44 |  |
| 10 | G | O6 | 1H:2608 | 4.57 | 1.00 | 25.22 |  |
| 11 | G | OP2 | 1H:2610 | 4.85 | 1.00 | 17.67 |  |
  |

  
**1^st^ shell ligand angles:**   
1-2: 90; 1-3: 91; 1-4: 88; 1-5: 175; 1-6: 94; 2-3: 177; 2-4: 88; 2-5: 85; 2-6: 93; 3-4: 90; 3-5: 94; 3-6: 89; 4-5: 94; 4-6: 178; 5-6: 85;   
  
 Contact(s): /O_w_/O_w_/O_w_/O_w_/O_w_/O_w_/  
 Coded contact list: 6O_w_  

### MG:1H:3407

|  | Res. type | Atom | Residue | Distance | Occupancy | B-factor | Closest solute to 1^st^ shell water |
| --- | --- | --- | --- | --- | --- | --- | --- |
| **Ion** | | | |  | 1.00 | 16.59 || 1 | Water 1 | | 1H:3632 | 2.16 | 1.00 | 23.43 | 1.99 (G:O6 1H:601) |
| 2 | Water 2 | | 1H:4124 | 2.16 | 1.00 | 22.36 | 2.79 (G:O6 1H:587) |
| 3 | Water 3 | | 1H:3687 | 2.16 | 1.00 | 26.11 | 2.17 (C:OP2 1H:598) |
| 4 | Water 4 | | 1H:3835 | 2.16 | 1.00 | 18.76 | 2.43 (C:OP2 1H:598) |
| 5 | Water 5 | | 1H:3747 | 2.17 | 1.00 | 19.24 | 2.29 (C:OP2 1H:598) |
| 6 | Water 6 | | 1H:4302 | 2.20 | 1.00 | 28.62 | 3.06 (G:N7 1H:601) |
| 7 | C | OP2 | 1H:598 | 2.61 | 1.00 | 22.15 |  |
| 8 | G | O6 | 1H:601 | 3.45 | 1.00 | 23.01 |  |
| 9 | U | O4 | 1H:600 | 4.07 | 1.00 | 18.57 |  |
| 10 | G | O6 | 1H:587 | 4.19 | 1.00 | 25.10 |  |
| 11 | G | OP1 | 1H:597 | 4.32 | 1.00 | 23.98 |  |
| 12 | C | OP1 | 1H:598 | 4.32 | 1.00 | 16.42 |  |
| 13 | Water 7 | | 1H:3766 | 4.33 | 1.00 | 26.74 | 2.32 (C:OP1 1H:598) |
| 14 | A | N6 | 1H:602 | 4.38 | 1.00 | 21.07 |  |
| 15 | U | OP2 | 1H:600 | 4.42 | 1.00 | 20.11 |  |
| 16 | C | O5' | 1H:598 | 4.60 | 1.00 | 23.11 |  |
| 17 | G | N7 | 1H:601 | 4.67 | 1.00 | 21.80 |  |
| 18 | C | N4 | 1H:588 | 4.80 | 1.00 | 20.50 |  |
| 19 | G | O3' | 1H:597 | 4.94 | 1.00 | 19.86 |  |
  |

  
**1^st^ shell ligand angles:**   
1-2: 95; 1-3: 81; 1-4: 170; 1-5: 91; 1-6: 90; 2-3: 81; 2-4: 91; 2-5: 174; 2-6: 95; 3-4: 92; 3-5: 100; 3-6: 170; 4-5: 83; 4-6: 97; 5-6: 84;   
  
 Contact(s): /O_w_/O_w_/O_w_/O_w_/O_w_/O_w_/  
 Coded contact list: 6O_w_  
 Exclusion zone contact(s): /OP2/  
Consider K in place of MG (1 valid contact(s) for K vs. 0 for MG)  
MG hydration shell issue: water too close (< 2.4 Angs.)  

### MG:1H:3408

|  | Res. type | Atom | Residue | Distance | Occupancy | B-factor | Closest solute to 1^st^ shell water |
| --- | --- | --- | --- | --- | --- | --- | --- |
| **Ion** | | | |  | 1.00 | 29.29 || 1 | Water 1 | | 1H:4145 | 2.16 | 1.00 | 32.10 | 2.82 (G:N7 1H:1405) |
| 2 | Water 2 | | 1H:3729 | 2.17 | 1.00 | 31.77 | 2.26 (G:O6 1H:1418) |
| 3 | Water 3 | | 1H:3749 | 2.17 | 1.00 | 30.65 | 2.29 (G:OP2 1H:1405) |
| 4 | Water 4 | | 1H:3772 | 2.18 | 1.00 | 36.36 | 2.33 (U:OP2 1H:1404) |
| 5 | Water 5 | | 1H:4118 | 2.19 | 1.00 | 30.00 | 2.78 (C:OP2 1H:1417) |
| 6 | Water 6 | | 1H:4421 | 2.19 | 1.00 | 31.82 | 3.45 (G:N7 1H:1418) |
| 7 | U | OP2 | 1H:1404 | 3.76 | 1.00 | 25.86 |  |
| 8 | C | OP2 | 1H:1417 | 3.92 | 1.00 | 32.73 |  |
| 9 | G | OP2 | 1H:1405 | 3.97 | 1.00 | 36.94 |  |
| 10 | G | N7 | 1H:1405 | 4.02 | 1.00 | 38.04 |  |
| 11 | G | N7 | 1H:1418 | 4.07 | 1.00 | 35.11 |  |
| 12 | G | O6 | 1H:1418 | 4.36 | 1.00 | 40.82 |  |
| 13 | U | O5' | 1H:1404 | 4.39 | 1.00 | 33.15 |  |
| 14 | U | OP1 | 1H:1404 | 4.74 | 1.00 | 25.84 |  |
| 15 | C | O5' | 1H:1417 | 4.79 | 1.00 | 21.55 |  |
  |

  
**1^st^ shell ligand angles:**   
1-2: 94; 1-3: 92; 1-4: 86; 1-5: 179; 1-6: 85; 2-3: 88; 2-4: 179; 2-5: 87; 2-6: 94; 3-4: 91; 3-5: 87; 3-6: 177; 4-5: 93; 4-6: 87; 5-6: 95;   
  
 Contact(s): /O_w_/O_w_/O_w_/O_w_/O_w_/O_w_/  
 Coded contact list: 6O_w_  
MG hydration shell issue: water too close (< 2.4 Angs.)  

### MG:1H:3409

|  | Res. type | Atom | Residue | Distance | Occupancy | B-factor | Closest solute to 1^st^ shell water |
| --- | --- | --- | --- | --- | --- | --- | --- |
| **Ion** | | | |  | 1.00 | 14.54 || 1 | Water 1 | | 1H:3965 | 2.13 | 1.00 | 18.56 | 2.60 (A:N7 1H:809) |
| 2 | Water 2 | | 1H:4024 | 2.15 | 1.00 | 19.02 | 2.66 (G:N7 1H:781) |
| 3 | Water 3 | | 1H:3947 | 2.15 | 1.00 | 27.26 | 2.58 (C:OP2 1H:780) |
| 4 | Water 4 | | 1H:3750 | 2.17 | 1.00 | 24.41 | 2.29 (A:OP2 1H:809) |
| 5 | G | N7 | 1H:781 | 2.50 | 1.00 | 25.90 |  |
| 6 | A | N7 | 1H:809 | 2.53 | 1.00 | 19.82 |  |
| 7 | G | O6 | 1H:781 | 3.42 | 1.00 | 22.14 |  |
| 8 | A | OP2 | 1H:809 | 3.81 | 1.00 | 19.14 |  |
| 9 | C | OP2 | 1H:780 | 3.91 | 1.00 | 24.67 |  |
| 10 | G | OP2 | 1H:781 | 4.08 | 1.00 | 25.37 |  |
| 11 | A | N9 | 1H:809 | 4.30 | 1.00 | 20.94 |  |
| 12 | A | N6 | 1H:809 | 4.34 | 1.00 | 24.49 |  |
| 13 | G | N9 | 1H:781 | 4.62 | 1.00 | 24.91 |  |
| 14 | A | O5' | 1H:809 | 4.87 | 1.00 | 16.67 |  |
| 15 | C | O5' | 1H:780 | 4.88 | 1.00 | 21.92 |  |
| 16 | G | O5' | 1H:781 | 4.91 | 1.00 | 26.15 |  |
| 17 | G | OP2 | 1H:808 | 4.94 | 1.00 | 17.18 |  |
  |

  
**1^st^ shell ligand angles:**   
1-2: 77; 1-3: 97; 1-4: 94; 2-3: 81; 2-4: 169; 3-4: 95;   
  
 Contact(s): /O_w_/O_w_/O_w_/O_w_/  
 Coded contact list: 4O_w_  
 Exclusion zone contact(s): /N7/N7/  
Consider NA in place of MG (2 valid contact(s) for NA vs. 0 for MG)  
MG hydration shell issue: water too close (< 2.4 Angs.)  

### MG:1H:3410

|  | Res. type | Atom | Residue | Distance | Occupancy | B-factor | Closest solute to 1^st^ shell water |
| --- | --- | --- | --- | --- | --- | --- | --- |
| **Ion** | | | |  | 1.00 | 16.18 || 1 | Water 1 | | 1H:4084 | 2.16 | 1.00 | 23.05 | 2.74 (G:O6 1H:2467) |
| 2 | Water 2 | | 1H:3847 | 2.17 | 1.00 | 24.95 | 2.44 (C:OP2 1H:2465) |
| 3 | Water 3 | | 1H:4382 | 2.17 | 1.00 | 18.08 | 3.25 (A:OP2 1H:2466) |
| 4 | Water 4 | | 1H:4470 | 2.17 | 1.00 | 25.18 | 3.57 (C:N4 1H:2509) |
| 5 | Water 5 | | 1H:3954 | 2.18 | 1.00 | 18.84 | 2.59 (A:N7 1H:2466) |
| 6 | Water 6 | | 1H:3892 | 2.19 | 1.00 | 14.52 | 2.51 (A:N1 1H:2510) |
| 7 | A | N7 | 1H:2466 | 4.04 | 1.00 | 19.19 |  |
| 8 | G | N7 | 1H:2467 | 4.19 | 1.00 | 18.66 |  |
| 9 | C | OP2 | 1H:2465 | 4.33 | 1.00 | 27.76 |  |
| 10 | A | N1 | 1H:2510 | 4.36 | 1.00 | 16.99 |  |
| 11 | A | OP2 | 1H:2466 | 4.37 | 1.00 | 22.53 |  |
| 12 | G | O6 | 1H:2468 | 4.45 | 1.00 | 22.74 |  |
| 13 | C | N4 | 1H:2509 | 4.46 | 1.00 | 22.42 |  |
| 14 | G | O6 | 1H:2467 | 4.74 | 1.00 | 19.66 |  |
  |

  
**1^st^ shell ligand angles:**   
1-2: 175; 1-3: 90; 1-4: 89; 1-5: 87; 1-6: 94; 2-3: 86; 2-4: 94; 2-5: 90; 2-6: 91; 3-4: 90; 3-5: 92; 3-6: 176; 4-5: 176; 4-6: 88; 5-6: 90;   
  
 Contact(s): /O_w_/O_w_/O_w_/O_w_/O_w_/O_w_/  
 Coded contact list: 6O_w_  

### MG:1H:3411

|  | Res. type | Atom | Residue | Distance | Occupancy | B-factor | Closest solute to 1^st^ shell water |
| --- | --- | --- | --- | --- | --- | --- | --- |
| **Ion** | | | |  | 1.00 | 20.14 || 1 | Water 1 | | 1H:3615 | 2.12 | 1.00 | 23.65 | 1.87 (A:OP2 1H:991) |
| 2 | Water 2 | | 1H:4055 | 2.15 | 1.00 | 18.20 | 2.70 (A:OP2 1H:991) |
| 3 | Water 3 | | 1H:3697 | 2.16 | 1.00 | 26.30 | 2.19 (G:OP2 1H:992) |
| 4 | Water 4 | | 1H:3767 | 2.18 | 1.00 | 26.74 | 2.32 (G:O5' 1H:992) |
| 5 | Water 5 | | 1H:4181 | 2.19 | 1.00 | 25.66 | 2.87 (G:O3' 1H:990) |
| 6 | Water 6 | | 1H:4439 | 2.19 | 1.00 | 35.21 | 3.66 (U:OP2 1H:870) |
| 7 | G | OP2 | 1H:992 | 2.72 | 1.00 | 23.54 |  |
| 8 | A | OP2 | 1H:991 | 2.84 | 1.00 | 24.06 |  |
| 9 | G | O3' | 1H:990 | 3.74 | 1.00 | 23.73 |  |
| 10 | U | OP2 | 1H:870 | 3.97 | 1.00 | 27.84 |  |
| 11 | G | O5' | 1H:992 | 4.17 | 1.00 | 19.98 |  |
| 12 | A | O3' | 1H:869 | 4.20 | 1.00 | 18.79 |  |
| 13 | A | O3' | 1H:991 | 4.34 | 1.00 | 27.07 |  |
| 14 | A | O2' | 1H:869 | 4.41 | 1.00 | 23.02 |  |
| 15 | A | O5' | 1H:991 | 4.69 | 1.00 | 18.51 |  |
| 16 | G | O2' | 1H:990 | 4.92 | 1.00 | 30.47 |  |
  |

  
**1^st^ shell ligand angles:**   
1-2: 103; 1-3: 86; 1-4: 81; 1-5: 84; 1-6: 169; 2-3: 90; 2-4: 176; 2-5: 83; 2-6: 88; 3-4: 91; 3-5: 167; 3-6: 94; 4-5: 97; 4-6: 88; 5-6: 97;   
  
 Contact(s): /O_w_/O_w_/O_w_/O_w_/O_w_/O_w_/  
 Coded contact list: 6O_w_  
 Exclusion zone contact(s): /OP2/OP2/  
Consider K in place of MG (2 valid contact(s) for K vs. 0 for MG)  
MG hydration shell issue: water too close (< 2.4 Angs.)  

### MG:1H:3412

|  | Res. type | Atom | Residue | Distance | Occupancy | B-factor | Closest solute to 1^st^ shell water |
| --- | --- | --- | --- | --- | --- | --- | --- |
| **Ion** | | | |  | 1.00 | 16.25 || 1 | Water 1 | | 1H:3922 | 2.15 | 1.00 | 15.33 | 2.54 (A:OP1 1H:991) |
| 2 | Water 2 | | 1H:4020 | 2.17 | 1.00 | 20.86 | 2.66 (G:OP2 1H:879) |
| 3 | Water 3 | | 1H:4052 | 2.19 | 1.00 | 20.99 | 2.69 (G:O6 1H:879) |
| 4 | Water 4 | | 1H:4411 | 2.20 | 1.00 | 27.11 | 3.34 (G:O2' 1H:990) |
| 5 | Water 5 | | 1H:4039 | 2.21 | 1.00 | 21.68 | 2.68 (G:N3 1H:990) |
| 6 | A | OP1 | 1H:991 | 2.46 | 1.00 | 21.67 |  |
| 7 | G | O3' | 1H:990 | 3.69 | 1.00 | 23.73 |  |
| 8 | A | OP2 | 1H:991 | 4.12 | 1.00 | 24.06 |  |
| 9 | G | N7 | 1H:879 | 4.14 | 1.00 | 18.30 |  |
| 10 | Water 6 | | 1H:4055 | 4.27 | 1.00 | 18.20 | 2.70 (A:OP2 1H:991) |
| 11 | G | OP2 | 1H:879 | 4.33 | 1.00 | 18.44 |  |
| 12 | G | N3 | 1H:990 | 4.45 | 1.00 | 26.44 |  |
| 13 | G | O2' | 1H:990 | 4.48 | 1.00 | 30.47 |  |
| 14 | G | O6 | 1H:880 | 4.62 | 1.00 | 21.06 |  |
| 15 | G | N7 | 1H:880 | 4.65 | 1.00 | 17.53 |  |
| 16 | G | O6 | 1H:879 | 4.66 | 1.00 | 19.74 |  |
| 17 | G | N2 | 1H:990 | 4.80 | 1.00 | 25.29 |  |
| 18 | A | O5' | 1H:991 | 4.83 | 1.00 | 18.51 |  |
  |

  
**1^st^ shell ligand angles:**   
1-2: 81; 1-3: 91; 1-4: 100; 1-5: 168; 2-3: 82; 2-4: 169; 2-5: 94; 3-4: 87; 3-5: 99; 4-5: 87;   
  
 Contact(s): /O_w_/O_w_/O_w_/O_w_/O_w_/  
 Coded contact list: 5O_w_  
 Exclusion zone contact(s): /OP1/  
Consider NA in place of MG (1 valid contact(s) for NA vs. 0 for MG)  

### MG:1H:3413

|  | Res. type | Atom | Residue | Distance | Occupancy | B-factor | Closest solute to 1^st^ shell water |
| --- | --- | --- | --- | --- | --- | --- | --- |
| **Ion** | | | |  | 1.00 | 49.12 || 1 | Water 1 | | 1H:3934 | 2.16 | 1.00 | 38.03 | 2.55 (G:N7 1H:323) |
| 2 | Water 2 | | 1H:3786 | 2.17 | 1.00 | 47.14 | 2.36 (G:O6 1H:323) |
| 3 | Water 3 | | 1H:4481 | 2.18 | 1.00 | 55.21 | 3.63 (G:OP2 1H:323) |
| 4 | Water 4 | | 1H:3816 | 2.18 | 1.00 | 53.62 | 2.40 (G:N7 1H:323) |
| 5 | Water 5 | | 1H:4285 | 2.18 | 1.00 | 45.55 | 3.03 (G:OP2 1H:363) |
| 6 | Water 6 | | 1H:3898 | 2.20 | 1.00 | 50.28 | 2.52 (G:OP2 1H:363) |
| 7 | G | N7 | 1H:323 | 2.95 | 1.00 | 42.35 |  |
| 8 | G | OP2 | 1H:363 | 3.31 | 1.00 | 48.81 |  |
| 9 | G | O6 | 1H:323 | 4.27 | 1.00 | 40.12 |  |
| 10 | G | OP2 | 1H:323 | 4.37 | 1.00 | 44.96 |  |
| 11 | Water 7 | | 1H:4563 | 4.60 | 1.00 | 50.76 | 6.10 (G:OP2 1H:323) |
| 12 | G | N9 | 1H:323 | 4.76 | 1.00 | 41.19 |  |
| 13 | Water 8 | | 1H:4472 | 4.83 | 1.00 | 46.88 | 5.28 (G:N9 1H:323) |
  |

  
**1^st^ shell ligand angles:**   
1-2: 90; 1-3: 89; 1-4: 94; 1-5: 88; 1-6: 179; 2-3: 178; 2-4: 89; 2-5: 90; 2-6: 91; 3-4: 91; 3-5: 90; 3-6: 91; 4-5: 178; 4-6: 85; 5-6: 93;   
  
 Contact(s): /O_w_/O_w_/O_w_/O_w_/O_w_/O_w_/  
 Coded contact list: 6O_w_  
 Exclusion zone contact(s): /N7/OP2/  
Consider K in place of MG (2 valid contact(s) for K vs. 0 for MG)  
MG hydration shell issue: water too close (< 2.4 Angs.)  

### MG:1H:3414

|  | Res. type | Atom | Residue | Distance | Occupancy | B-factor | Closest solute to 1^st^ shell water |
| --- | --- | --- | --- | --- | --- | --- | --- |
| **Ion** | | | |  | 1.00 | 39.03 || 1 | Water 1 | | 1H:3849 | 2.15 | 1.00 | 34.49 | 2.44 (A:N6 1H:325) |
| 2 | Water 2 | | 1H:4405 | 2.17 | 1.00 | 38.23 | 3.33 (G:N7 1H:363) |
| 3 | Water 3 | | 1H:3871 | 2.18 | 1.00 | 46.35 | 2.48 (A:OP2 1H:324) |
| 4 | Water 4 | | 1H:4472 | 2.18 | 1.00 | 46.88 | 3.58 (A:OP2 1H:324) |
| 5 | Water 5 | | 1H:3843 | 2.19 | 1.00 | 50.56 | 2.43 (C:OP2 1H:361) |
| 6 | Water 6 | | 1H:4563 | 2.19 | 1.00 | 50.76 | 4.49 (C:OP2 1H:361) |
| 7 | A | OP2 | 1H:324 | 4.16 | 1.00 | 43.21 |  |
| 8 | C | OP2 | 1H:361 | 4.47 | 1.00 | 35.31 |  |
| 9 | A | N6 | 1H:325 | 4.54 | 1.00 | 50.00 |  |
| 10 | G | O6 | 1H:363 | 4.68 | 1.00 | 42.22 |  |
| 11 | Water 7 | | 1H:3816 | 4.82 | 1.00 | 53.62 | 4.97 (G:N7 1H:363) |
| 12 | A | N7 | 1H:324 | 4.83 | 1.00 | 46.56 |  |
| 13 | G | N7 | 1H:363 | 4.91 | 1.00 | 47.58 |  |
| 14 | Water 8 | | 1H:3898 | 4.92 | 1.00 | 50.28 | 4.41 (G:N7 1H:363) |
| 15 | Water 9 | | 1H:4481 | 4.93 | 1.00 | 55.21 | 5.99 (A:OP2 1H:324) |
| 16 | A | N7 | 1H:325 | 4.97 | 1.00 | 45.61 |  |
  |

  
**1^st^ shell ligand angles:**   
1-2: 89; 1-3: 90; 1-4: 90; 1-5: 90; 1-6: 178; 2-3: 179; 2-4: 92; 2-5: 87; 2-6: 93; 3-4: 88; 3-5: 93; 3-6: 88; 4-5: 179; 4-6: 90; 5-6: 90;   
  
 Contact(s): /O_w_/O_w_/O_w_/O_w_/O_w_/O_w_/  
 Coded contact list: 6O_w_  

### MG:1H:3415

|  | Res. type | Atom | Residue | Distance | Occupancy | B-factor | Closest solute to 1^st^ shell water |
| --- | --- | --- | --- | --- | --- | --- | --- |
| **Ion** | | | |  | 1.00 | 16.90 || 1 | Water 1 | | 1H:3957 | 2.16 | 1.00 | 20.54 | 2.59 (G:OP2 1H:847) |
| 2 | Water 2 | | 1H:4110 | 2.16 | 1.00 | 30.00 | 2.77 (G:OP2 1H:849) |
| 3 | Water 3 | | 31:404 | 2.17 | 1.00 | 30.65 | 2.63 (GLY:N 31:61) |
| 4 | Water 4 | | 31:406 | 2.17 | 1.00 | 23.34 | 3.21 (GLY:O 31:55) |
| 5 | Water 5 | | 31:407 | 2.18 | 1.00 | 29.21 | 3.24 (ARG:O 31:54) |
| 6 | G | OP2 | 1H:849 | 2.34 | 1.00 | 16.84 |  |
| 7 | A | O3' | 1H:848 | 3.64 | 1.00 | 18.09 |  |
| 8 | GLY | O | 31:55 | 4.25 | 1.00 | 16.29 |  |
| 9 | TYR | O | 31:59 | 4.42 | 1.00 | 23.98 |  |
| 10 | G | OP1 | 1H:849 | 4.45 | 1.00 | 23.01 |  |
| 11 | GLY | N | 31:61 | 4.49 | 1.00 | 20.49 |  |
| 12 | G | OP2 | 1H:847 | 4.54 | 1.00 | 12.24 |  |
| 13 | G | OP1 | 1H:846 | 4.56 | 1.00 | 20.05 |  |
| 14 | G | O5' | 1H:849 | 4.57 | 1.00 | 17.52 |  |
| 15 | ARG | O | 31:54 | 4.57 | 1.00 | 19.66 |  |
| 16 | ASP | OD2 | 31:77 | 5.00 | 1.00 | 30.98 |  |
  |

  
**1^st^ shell ligand angles:**   
1-2: 85; 1-3: 87; 1-4: 95; 1-5: 172; 1-6: 90; 2-3: 89; 2-4: 179; 2-5: 88; 2-6: 76; 3-4: 92; 3-5: 89; 3-6: 165; 4-5: 92; 4-6: 103; 5-6: 93;   
  
 Contact(s): /O_w_/O_w_/O_w_/O_w_/O_w_/OP2/  
 Coded contact list: O_ph_5O_w_  

### MG:1H:3416

|  | Res. type | Atom | Residue | Distance | Occupancy | B-factor | Closest solute to 1^st^ shell water |
| --- | --- | --- | --- | --- | --- | --- | --- |
| **Ion** | | | |  | 1.00 | 69.11 || 1 | Water 1 | | 1H:3671 | 2.18 | 1.00 | 45.01 | 2.14 (G:O6 1H:1785) |
| 2 | Water 2 | | 1H:4191 | 2.18 | 1.00 | 54.14 | 2.88 (G:N7 1H:1788) |
| 3 | Water 3 | | 1H:4272 | 2.19 | 1.00 | 47.72 | 3.01 (G:O6 1H:1788) |
| 4 | Water 4 | | B8:201 | 2.19 | 1.00 | 66.50 | 4.62 (G:O6 1H:1785) |
| 5 | Water 5 | | 1H:4468 | 2.19 | 1.00 | 58.84 | 3.57 (U:OP2 1H:1754) |
| 6 | Water 6 | | 1H:4518 | 2.19 | 1.00 | 50.52 | 3.79 (G:OP2 1H:1788) |
| 7 | G | O6 | 1H:1788 | 3.80 | 1.00 | 50.67 |  |
| 8 | G | O6 | 1H:1785 | 4.14 | 1.00 | 49.38 |  |
| 9 | G | OP2 | 1H:1788 | 4.34 | 1.00 | 42.68 |  |
| 10 | U | OP2 | 1H:1754 | 4.58 | 1.00 | 44.06 |  |
| 11 | G | N7 | 1H:1788 | 4.76 | 1.00 | 47.42 |  |
  |

  
**1^st^ shell ligand angles:**   
1-2: 91; 1-3: 91; 1-4: 88; 1-5: 178; 1-6: 89; 2-3: 91; 2-4: 179; 2-5: 87; 2-6: 91; 3-4: 90; 3-5: 88; 3-6: 179; 4-5: 94; 4-6: 89; 5-6: 92;   
  
 Contact(s): /O_w_/O_w_/O_w_/O_w_/O_w_/O_w_/  
 Coded contact list: 6O_w_  
MG hydration shell issue: water too close (< 2.4 Angs.)  

### MG:1H:3417

|  | Res. type | Atom | Residue | Distance | Occupancy | B-factor | Closest solute to 1^st^ shell water |
| --- | --- | --- | --- | --- | --- | --- | --- |
| **Ion** | | | |  | 1.00 | 61.68 || 1 | Water 1 | | 1H:3601 | 2.17 | 1.00 | 57.78 | 1.30 (G:N7 1H:332) |
| 2 | Water 2 | | 1H:4253 | 2.18 | 1.00 | 51.51 | 2.98 (G:OP2 1H:332) |
| 3 | Water 3 | | 1H:4444 | 2.18 | 1.00 | 66.31 | 3.48 (G:O6 1H:332) |
| 4 | Water 4 | | 1H:3936 | 2.19 | 1.00 | 70.47 | 2.56 (U:OP2 1H:331) |
| 5 | Water 5 | | 1H:4340 | 2.19 | 1.00 | 55.77 | 3.43 (G:N7 1H:332) |
| 6 | Water 6 | | 1H:4605 | 2.19 | 1.00 | 59.40 | 5.10 (G:N7 1H:332) |
| 7 | G | N7 | 1H:332 | 2.99 | 1.00 | 58.70 |  |
| 8 | G | O6 | 1H:332 | 4.04 | 1.00 | 54.64 |  |
| 9 | G | OP2 | 1H:332 | 4.26 | 1.00 | 54.32 |  |
| 10 | U | OP2 | 1H:331 | 4.43 | 1.00 | 56.34 |  |
| 11 | G | OP2 | 1H:334 | 4.87 | 1.00 | 46.86 |  |
  |

  
**1^st^ shell ligand angles:**   
1-2: 92; 1-3: 87; 1-4: 86; 1-5: 93; 1-6: 178; 2-3: 178; 2-4: 90; 2-5: 91; 2-6: 88; 3-4: 91; 3-5: 87; 3-6: 93; 4-5: 178; 4-6: 91; 5-6: 90;   
  
 Contact(s): /O_w_/O_w_/O_w_/O_w_/O_w_/O_w_/  
 Coded contact list: 6O_w_  
 Exclusion zone contact(s): /N7/  
Consider K in place of MG (1 valid contact(s) for K vs. 0 for MG)  
MG hydration shell issue: water too close (< 2.4 Angs.)  

### MG:1H:3418

|  | Res. type | Atom | Residue | Distance | Occupancy | B-factor | Closest solute to 1^st^ shell water |
| --- | --- | --- | --- | --- | --- | --- | --- |
| **Ion** | | | |  | 1.00 | 29.09 || 1 | Water 1 | | 1H:4278 | 2.16 | 1.00 | 23.84 | 3.01 (U:OP2 1H:1314) |
| 2 | Water 2 | | 1H:3939 | 2.16 | 1.00 | 27.45 | 2.57 (G:N7 1H:2033) |
| 3 | Water 3 | | 1H:4463 | 2.18 | 1.00 | 42.00 | 4.21 (U:OP1 1H:1314) |
| 4 | Water 4 | | 1H:4465 | 2.18 | 1.00 | 40.22 | 3.56 (U:OP1 1H:1314) |
| 5 | Water 5 | | 1H:3808 | 2.18 | 1.00 | 34.91 | 2.39 (G:O6 1H:2033) |
| 6 | Water 6 | | 1H:3966 | 2.18 | 1.00 | 35.76 | 2.60 (G:N7 1H:2032) |
| 7 | G | N7 | 1H:2032 | 3.96 | 1.00 | 27.77 |  |
| 8 | G | O6 | 1H:2033 | 4.02 | 1.00 | 30.24 |  |
| 9 | G | O6 | 1H:2032 | 4.29 | 1.00 | 31.80 |  |
| 10 | U | O4 | 1H:2034 | 4.47 | 1.00 | 25.17 |  |
| 11 | G | N7 | 1H:2033 | 4.48 | 1.00 | 32.08 |  |
| 12 | U | OP2 | 1H:1314 | 4.55 | 1.00 | 27.15 |  |
| 13 | U | OP1 | 1H:1314 | 4.66 | 1.00 | 36.95 |  |
  |

  
**1^st^ shell ligand angles:**   
1-2: 85; 1-3: 92; 1-4: 94; 1-5: 90; 1-6: 175; 2-3: 92; 2-4: 178; 2-5: 90; 2-6: 91; 3-4: 85; 3-5: 177; 3-6: 91; 4-5: 92; 4-6: 90; 5-6: 87;   
  
 Contact(s): /O_w_/O_w_/O_w_/O_w_/O_w_/O_w_/  
 Coded contact list: 6O_w_  
MG hydration shell issue: water too close (< 2.4 Angs.)  

### MG:1H:3419

|  | Res. type | Atom | Residue | Distance | Occupancy | B-factor | Closest solute to 1^st^ shell water |
| --- | --- | --- | --- | --- | --- | --- | --- |
| **Ion** | | | |  | 1.00 | 27.53 || 1 | Water 1 | | 1H:3789 | 2.16 | 1.00 | 19.44 | 2.36 (C:OP1 1H:2029) |
| 2 | Water 2 | | 1H:3666 | 2.17 | 1.00 | 27.23 | 2.11 (U:O3' 1H:1314) |
| 3 | Water 3 | | 1H:3972 | 2.17 | 1.00 | 22.58 | 2.61 (C:OP1 1H:2029) |
| 4 | Water 4 | | 1H:3651 | 2.18 | 1.00 | 23.71 | 2.07 (C:OP1 1H:2029) |
| 5 | Water 5 | | 1H:4483 | 2.21 | 1.00 | 31.64 | 4.20 (G:OP2 1H:2072) |
| 6 | Water 6 | | 1H:3914 | 2.21 | 1.00 | 35.48 | 2.53 (G:OP1 1H:2071) |
| 7 | C | OP1 | 1H:2029 | 2.87 | 1.00 | 23.97 |  |
| 8 | A | OP1 | 1H:1315 | 3.11 | 1.00 | 25.82 |  |
| 9 | U | O3' | 1H:1314 | 3.53 | 1.00 | 24.51 |  |
| 10 | G | OP1 | 1H:2071 | 3.60 | 1.00 | 22.11 |  |
| 11 | Water 7 | | 1H:4547 | 3.71 | 1.00 | 31.55 | 4.35 (G:OP1 1H:2071) |
| 12 | Water 8 | | 1H:3832 | 4.25 | 1.00 | 40.29 | 3.54 (U:O2' 1H:1314) |
| 13 | U | O2' | 1H:1314 | 4.67 | 1.00 | 29.76 |  |
| 14 | G | OP2 | 1H:2072 | 4.75 | 1.00 | 22.59 |  |
| 15 | A | O5' | 1H:1315 | 4.77 | 1.00 | 26.58 |  |
| 16 | A | O3' | 1H:2028 | 4.96 | 1.00 | 26.45 |  |
  |

  
**1^st^ shell ligand angles:**   
1-2: 91; 1-3: 92; 1-4: 85; 1-5: 176; 1-6: 92; 2-3: 176; 2-4: 92; 2-5: 90; 2-6: 86; 3-4: 86; 3-5: 87; 3-6: 96; 4-5: 98; 4-6: 177; 5-6: 84;   
  
 Contact(s): /O_w_/O_w_/O_w_/O_w_/O_w_/O_w_/  
 Coded contact list: 6O_w_  
 Exclusion zone contact(s): /OP1/OP1/  
Consider K in place of MG (2 valid contact(s) for K vs. 0 for MG)  
MG hydration shell issue: water too close (< 2.4 Angs.)  

### MG:1H:3420

|  | Res. type | Atom | Residue | Distance | Occupancy | B-factor | Closest solute to 1^st^ shell water |
| --- | --- | --- | --- | --- | --- | --- | --- |
| **Ion** | | | |  | 1.00 | 20.48 || 1 | Water 1 | | 1H:3784 | 2.14 | 1.00 | 26.31 | 2.35 (A:OP1 1H:2726) |
| 2 | Water 2 | | 1H:3991 | 2.17 | 1.00 | 26.03 | 2.63 (C:O3' 1H:2723) |
| 3 | Water 3 | | 1H:4528 | 2.18 | 1.00 | 21.74 | 3.84 (A:OP1 1H:2726) |
| 4 | Water 4 | | 1H:4003 | 2.21 | 1.00 | 36.63 | 2.64 (C:OP1 1H:1344) |
| 5 | A | OP1 | 1H:2724 | 2.46 | 1.00 | 23.77 |  |
| 6 | A | OP2 | 1H:2726 | 2.68 | 1.00 | 35.93 |  |
| 7 | A | OP1 | 1H:2726 | 2.83 | 1.00 | 25.43 |  |
| 8 | C | O3' | 1H:2723 | 3.61 | 1.00 | 31.40 |  |
| 9 | C | OP1 | 1H:1344 | 3.73 | 1.00 | 30.02 |  |
| 10 | Water 5 | | 1H:3815 | 3.81 | 1.00 | 27.58 | 2.40 (A:OP2 1H:2726) |
| 11 | U | O3' | 1H:2725 | 4.12 | 1.00 | 30.59 |  |
| 12 | A | OP2 | 1H:2724 | 4.53 | 1.00 | 28.54 |  |
| 13 | A | O5' | 1H:2726 | 4.65 | 1.00 | 27.97 |  |
| 14 | A | O5' | 1H:2724 | 4.69 | 1.00 | 23.06 |  |
| 15 | U | O2' | 1H:2725 | 4.74 | 1.00 | 39.87 |  |
  |

  
**1^st^ shell ligand angles:**   
1-2: 82; 1-3: 91; 1-4: 174; 2-3: 98; 2-4: 94; 3-4: 94;   
  
 Contact(s): /O_w_/O_w_/O_w_/O_w_/  
 Coded contact list: 4O_w_  
 Exclusion zone contact(s): /OP1/OP1/OP2/  
Consider K in place of MG (2 valid contact(s) for K vs. 0 for MG)  
MG hydration shell issue: water too close (< 2.4 Angs.)  

### MG:1H:3421

|  | Res. type | Atom | Residue | Distance | Occupancy | B-factor | Closest solute to 1^st^ shell water |
| --- | --- | --- | --- | --- | --- | --- | --- |
| **Ion** | | | |  | 1.00 | 36.78 || 1 | Water 1 | | 1H:3824 | 2.15 | 1.00 | 33.77 | 2.41 (G:O6 1H:1402) |
| 2 | Water 2 | | 1H:4128 | 2.17 | 1.00 | 27.21 | 2.80 (A:OP1 1H:1400) |
| 3 | Water 3 | | 1H:4446 | 2.19 | 1.00 | 40.42 | 3.48 (A:OP1 1H:1420) |
| 4 | Water 4 | | 1H:4519 | 2.19 | 1.00 | 44.58 | 3.79 (A:OP2 1H:1420) |
| 5 | Water 5 | | 1H:4293 | 2.20 | 1.00 | 32.04 | 3.12 (G:O6 1H:1402) |
| 6 | Water 6 | | 1H:4138 | 2.22 | 1.00 | 32.64 | 2.81 (G:O6 1H:1403) |
| 7 | G | O6 | 1H:1402 | 3.79 | 1.00 | 28.73 |  |
| 8 | G | O6 | 1H:1403 | 4.11 | 1.00 | 34.84 |  |
| 9 | C | N4 | 1H:1422 | 4.36 | 1.00 | 31.38 |  |
| 10 | A | OP1 | 1H:1400 | 4.55 | 1.00 | 24.41 |  |
| 11 | A | OP2 | 1H:1401 | 4.67 | 1.00 | 25.34 |  |
| 12 | A | OP2 | 1H:1400 | 4.76 | 1.00 | 25.23 |  |
| 13 | A | OP2 | 1H:1420 | 4.79 | 1.00 | 58.75 |  |
| 14 | A | OP1 | 1H:1401 | 4.89 | 1.00 | 24.06 |  |
| 15 | A | OP1 | 1H:1420 | 4.91 | 1.00 | 46.82 |  |
  |

  
**1^st^ shell ligand angles:**   
1-2: 85; 1-3: 90; 1-4: 179; 1-5: 89; 1-6: 93; 2-3: 86; 2-4: 95; 2-5: 88; 2-6: 176; 3-4: 90; 3-5: 174; 3-6: 90; 4-5: 90; 4-6: 86; 5-6: 96;   
  
 Contact(s): /O_w_/O_w_/O_w_/O_w_/O_w_/O_w_/  
 Coded contact list: 6O_w_  

### MG:1H:3422

|  | Res. type | Atom | Residue | Distance | Occupancy | B-factor | Closest solute to 1^st^ shell water |
| --- | --- | --- | --- | --- | --- | --- | --- |
| **Ion** | | | |  | 1.00 | 29.01 || 1 | Water 1 | | 1H:3904 | 2.15 | 1.00 | 28.82 | 2.52 (C:OP1 1H:553) |
| 2 | Water 2 | | 1H:4246 | 2.16 | 1.00 | 32.63 | 2.96 (C:OP1 1H:553) |
| 3 | Water 3 | | 1H:4488 | 2.18 | 1.00 | 39.34 | 3.64 (C:OP2 1H:553) |
| 4 | Water 4 | | 1H:4384 | 2.18 | 1.00 | 60.08 | 3.25 (A:OP1 1H:552) |
| 5 | C | OP1 | 1H:553 | 2.49 | 1.00 | 30.68 |  |
| 6 | A | OP1 | 1H:552 | 2.57 | 1.00 | 36.47 |  |
| 7 | A | O5' | 1H:552 | 3.53 | 1.00 | 32.96 |  |
| 8 | C | OP2 | 1H:553 | 3.71 | 1.00 | 24.98 |  |
| 9 | A | OP2 | 1H:552 | 4.07 | 1.00 | 41.49 |  |
| 10 | C | O5' | 1H:553 | 4.44 | 1.00 | 33.71 |  |
| 11 | A | O3' | 1H:552 | 4.59 | 1.00 | 25.68 |  |
| 12 | U | O3' | 1H:551 | 4.74 | 1.00 | 39.11 |  |
| 13 | A | OP1 | 1H:554 | 4.84 | 1.00 | 31.86 |  |
  |

  
**1^st^ shell ligand angles:**   
1-2: 85; 1-3: 174; 1-4: 89; 2-3: 92; 2-4: 95; 3-4: 97;   
  
 Contact(s): /O_w_/O_w_/O_w_/O_w_/  
 Coded contact list: 4O_w_  
 Exclusion zone contact(s): /OP1/OP1/  
Consider NA in place of MG (2 valid contact(s) for NA vs. 0 for MG)  

### MG:1H:3423

|  | Res. type | Atom | Residue | Distance | Occupancy | B-factor | Closest solute to 1^st^ shell water |
| --- | --- | --- | --- | --- | --- | --- | --- |
| **Ion** | | | |  | 1.00 | 35.39 || 1 | Water 1 | | 1H:4205 | 2.18 | 1.00 | 42.21 | 2.90 (G:N7 1H:2621) |
| 2 | Water 2 | | 1H:3776 | 2.19 | 1.00 | 26.93 | 2.34 (G:OP2 1H:2596) |
| 3 | Water 3 | | 1H:4127 | 2.19 | 1.00 | 36.47 | 2.80 (U:O4 1H:2597) |
| 4 | Water 4 | | 1H:3675 | 2.20 | 1.00 | 24.68 | 2.15 (G:OP2 1H:2595) |
| 5 | G | OP2 | 1H:2595 | 2.45 | 1.00 | 26.35 |  |
| 6 | G | OP2 | 1H:2596 | 2.65 | 1.00 | 26.26 |  |
| 7 | G | O3' | 1H:2594 | 4.05 | 1.00 | 23.56 |  |
| 8 | G | O6 | 1H:2621 | 4.19 | 1.00 | 31.15 |  |
| 9 | G | N7 | 1H:2621 | 4.26 | 1.00 | 23.22 |  |
| 10 | G | OP1 | 1H:2595 | 4.35 | 1.00 | 29.23 |  |
| 11 | G | O2' | 1H:2594 | 4.57 | 1.00 | 22.93 |  |
| 12 | G | N7 | 1H:2596 | 4.70 | 1.00 | 33.98 |  |
| 13 | G | O5' | 1H:2596 | 4.81 | 1.00 | 28.02 |  |
| 14 | G | O5' | 1H:2595 | 4.86 | 1.00 | 20.82 |  |
| 15 | U | N3 | 1H:2597 | 4.91 | 1.00 | 35.73 |  |
| 16 | U | O4 | 1H:2597 | 4.92 | 1.00 | 53.52 |  |
| 17 | G | OP1 | 1H:2596 | 4.95 | 1.00 | 29.52 |  |
| 18 | G | O3' | 1H:2595 | 4.96 | 1.00 | 23.90 |  |
  |

  
**1^st^ shell ligand angles:**   
1-2: 94; 1-3: 94; 1-4: 90; 2-3: 91; 2-4: 86; 3-4: 175;   
  
 Contact(s): /O_w_/O_w_/O_w_/O_w_/  
 Coded contact list: 4O_w_  
 Exclusion zone contact(s): /OP2/OP2/  
Consider NA in place of MG (1 valid contact(s) for NA vs. 0 for MG)  
MG hydration shell issue: water too close (< 2.4 Angs.)  

### MG:1H:3424

|  | Res. type | Atom | Residue | Distance | Occupancy | B-factor | Closest solute to 1^st^ shell water |
| --- | --- | --- | --- | --- | --- | --- | --- |
| **Ion** | | | |  | 1.00 | 42.85 || 1 | Water 1 | | 2K:203 | 2.16 | 1.00 | 28.04 | 2.35 (C:OP1 2K:75) |
| 2 | Water 2 | | 2K:204 | 2.18 | 1.00 | 43.50 | 2.57 (A:OP2 2K:76) |
| 3 | Water 3 | | 1H:4019 | 2.19 | 1.00 | 48.63 | 2.66 (U:OP1 1H:2598) |
| 4 | C | OP1 | 2K:75 | 2.56 | 1.00 | 35.95 |  |
| 5 | A | OP1 | 1H:2615 | 2.66 | 1.00 | 39.92 |  |
| 6 | A | OP2 | 2K:76 | 2.79 | 1.00 | 38.22 |  |
| 7 | A | O5' | 1H:2615 | 3.15 | 1.00 | 52.85 |  |
| 8 | A | O5' | 2K:76 | 3.49 | 1.00 | 44.68 |  |
| 9 | U | OP1 | 1H:2598 | 3.56 | 1.00 | 44.19 |  |
| 10 | C | O3' | 1H:2614 | 3.90 | 1.00 | 36.03 |  |
| 11 | C | O5' | 2K:75 | 4.25 | 1.00 | 34.05 |  |
| 12 | A | OP1 | 2K:76 | 4.39 | 1.00 | 52.04 |  |
| 13 | C | OP2 | 2K:75 | 4.43 | 1.00 | 47.06 |  |
| 14 | A | O3' | 2K:76 | 4.56 | 1.00 | 56.66 |  |
| 15 | A | OP2 | 1H:2615 | 4.80 | 1.00 | 43.34 |  |
  |

  
**1^st^ shell ligand angles:**   
1-2: 84; 1-3: 177; 2-3: 97;   
  
 Contact(s): /O_w_/O_w_/O_w_/  
 Coded contact list: 3O_w_  
 Exclusion zone contact(s): /O5'/OP1/OP1/OP2/  
Consider K in place of MG (3 valid contact(s) for K vs. 0 for MG)  
MG hydration shell issue: water too close (< 2.4 Angs.)  

### MG:1H:3425

|  | Res. type | Atom | Residue | Distance | Occupancy | B-factor | Closest solute to 1^st^ shell water |
| --- | --- | --- | --- | --- | --- | --- | --- |
| **Ion** | | | |  | 1.00 | 16.24 || 1 | Water 1 | | 1H:4260 | 2.16 | 1.00 | 14.90 | 2.98 (G:O6 1H:846) |
| 2 | Water 2 | | 1H:4526 | 2.17 | 1.00 | 30.22 | 3.83 (G:O6 1H:847) |
| 3 | Water 3 | | 1H:4371 | 2.18 | 1.00 | 24.59 | 3.23 (G:N7 1H:846) |
| 4 | Water 4 | | 31:403 | 2.19 | 1.00 | 22.71 | 2.36 (GLY:O 31:61) |
| 5 | Water 5 | | 31:409 | 2.19 | 1.00 | 14.27 | 3.72 (GLY:O 31:61) |
| 6 | Water 6 | | 1H:4464 | 2.21 | 1.00 | 26.55 | 3.56 (G:N7 1H:846) |
| 7 | GLY | O | 31:61 | 4.13 | 1.00 | 14.97 |  |
| 8 | G | N7 | 1H:846 | 4.31 | 1.00 | 17.63 |  |
| 9 | G | O6 | 1H:847 | 4.50 | 1.00 | 17.48 |  |
| 10 | G | O6 | 1H:846 | 4.58 | 1.00 | 17.16 |  |
  |

  
**1^st^ shell ligand angles:**   
1-2: 85; 1-3: 87; 1-4: 175; 1-5: 97; 1-6: 96; 2-3: 87; 2-4: 92; 2-5: 92; 2-6: 180; 3-4: 90; 3-5: 176; 3-6: 92; 4-5: 86; 4-6: 87; 5-6: 88;   
  
 Contact(s): /O_w_/O_w_/O_w_/O_w_/O_w_/O_w_/  
 Coded contact list: 6O_w_  
MG hydration shell issue: water too close (< 2.4 Angs.)  

### MG:1H:3426

|  | Res. type | Atom | Residue | Distance | Occupancy | B-factor | Closest solute to 1^st^ shell water |
| --- | --- | --- | --- | --- | --- | --- | --- |
| **Ion** | | | |  | 1.00 | 12.50 || 1 | Water 1 | | 1H:3623 | 2.14 | 1.00 | 13.68 | 1.95 (G:OP1 1H:2601) |
| 2 | Water 2 | | 11:408 | 2.16 | 1.00 | 14.59 | 2.97 (ASN:OD1 11:227) |
| 3 | Water 3 | | 1H:4310 | 2.18 | 1.00 | 30.87 | 3.34 (G:OP1 1H:2601) |
| 4 | Water 4 | | 11:410 | 2.20 | 1.00 | 35.54 | 3.61 (A:OP2 1H:2602) |
| 5 | G | OP1 | 1H:2601 | 2.47 | 1.00 | 16.92 |  |
| 6 | A | OP1 | 1H:832 | 2.64 | 1.00 | 15.56 |  |
| 7 | A | OP2 | 1H:2602 | 3.52 | 1.00 | 16.66 |  |
| 8 | Water 5 | | 1H:3638 | 3.68 | 1.00 | 16.91 | 2.01 (A:OP2 1H:832) |
| 9 | Water 6 | | 1H:3654 | 3.74 | 1.00 | 18.81 | 2.08 (G:OP2 1H:2601) |
| 10 | G | OP2 | 1H:2601 | 3.90 | 1.00 | 14.99 |  |
| 11 | ASN | OD1 | 11:227 | 4.32 | 1.00 | 13.22 |  |
| 12 | A | OP2 | 1H:832 | 4.38 | 1.00 | 16.68 |  |
| 13 | G | O6 | 1H:840 | 4.59 | 1.00 | 15.42 |  |
| 14 | G | O5' | 1H:2601 | 4.67 | 1.00 | 14.48 |  |
| 15 | A | OP1 | 1H:2602 | 4.67 | 1.00 | 21.28 |  |
| 16 | A | O3' | 1H:2600 | 4.74 | 1.00 | 17.37 |  |
| 17 | A | O5' | 1H:832 | 4.85 | 1.00 | 12.37 |  |
| 18 | G | O3' | 1H:2601 | 4.97 | 1.00 | 17.36 |  |
| 19 | A | O3' | 1H:831 | 4.98 | 1.00 | 19.90 |  |
  |

  
**1^st^ shell ligand angles:**   
1-2: 75; 1-3: 87; 1-4: 167; 2-3: 96; 2-4: 96; 3-4: 85;   
  
 Contact(s): /O_w_/O_w_/O_w_/O_w_/  
 Coded contact list: 4O_w_  
 Exclusion zone contact(s): /OP1/OP1/  
Consider NA in place of MG (1 valid contact(s) for NA vs. 0 for MG)  
MG hydration shell issue: water too close (< 2.4 Angs.)  

### MG:1H:3427

|  | Res. type | Atom | Residue | Distance | Occupancy | B-factor | Closest solute to 1^st^ shell water |
| --- | --- | --- | --- | --- | --- | --- | --- |
| **Ion** | | | |  | 1.00 | 36.51 || 1 | Water 1 | | 1H:3704 | 2.17 | 1.00 | 27.84 | 2.21 (G:OP1 1H:490) |
| 2 | Water 2 | | 1H:4487 | 2.17 | 1.00 | 30.15 | 3.64 (G:OP1 1H:490) |
| 3 | Water 3 | | 1H:4398 | 2.18 | 1.00 | 28.86 | 3.29 (U:OP1 1H:491) |
| 4 | Water 4 | | 1H:4377 | 2.19 | 1.00 | 30.25 | 3.25 (A:N6 1H:837) |
| 5 | Water 5 | | 1H:4441 | 2.20 | 1.00 | 29.70 | 3.45 (G:OP1 1H:490) |
| 6 | Water 6 | | 1H:4482 | 2.20 | 1.00 | 33.93 | 3.63 (U:OP1 1H:491) |
| 7 | G | OP1 | 1H:490 | 3.75 | 1.00 | 22.81 |  |
| 8 | U | OP1 | 1H:491 | 4.24 | 1.00 | 26.58 |  |
| 9 | A | N7 | 1H:837 | 4.69 | 1.00 | 23.14 |  |
| 10 | A | N6 | 1H:837 | 4.73 | 1.00 | 23.68 |  |
| 11 | U | OP2 | 1H:491 | 4.76 | 1.00 | 23.27 |  |
| 12 | G | OP1 | 1H:1660 | 4.80 | 1.00 | 16.76 |  |
  |

  
**1^st^ shell ligand angles:**   
1-2: 89; 1-3: 91; 1-4: 175; 1-5: 89; 1-6: 90; 2-3: 92; 2-4: 87; 2-5: 91; 2-6: 177; 3-4: 86; 3-5: 177; 3-6: 91; 4-5: 94; 4-6: 95; 5-6: 87;   
  
 Contact(s): /O_w_/O_w_/O_w_/O_w_/O_w_/O_w_/  
 Coded contact list: 6O_w_  
MG hydration shell issue: water too close (< 2.4 Angs.)  

### MG:1H:3428

|  | Res. type | Atom | Residue | Distance | Occupancy | B-factor | Closest solute to 1^st^ shell water |
| --- | --- | --- | --- | --- | --- | --- | --- |
| **Ion** | | | |  | 1.00 | 23.34 || 1 | Water 1 | | 1H:3981 | 2.16 | 1.00 | 26.07 | 2.62 (A:OP1 1H:1994) |
| 2 | Water 2 | | 1H:3930 | 2.17 | 1.00 | 27.04 | 2.55 (A:OP1 1H:1994) |
| 3 | Water 3 | | 1H:4489 | 2.18 | 1.00 | 37.58 | 3.65 (A:OP1 1H:1993) |
| 4 | Water 4 | | 1H:4425 | 2.19 | 1.00 | 34.22 | 3.47 (A:O5' 1H:1992) |
| 5 | A | OP1 | 1H:1994 | 2.63 | 1.00 | 23.61 |  |
| 6 | A | OP1 | 1H:1993 | 2.75 | 1.00 | 30.89 |  |
| 7 | A | O5' | 1H:1993 | 3.10 | 1.00 | 29.84 |  |
| 8 | A | O3' | 1H:1992 | 4.06 | 1.00 | 21.98 |  |
| 9 | Water 5 | | 1H:4320 | 4.59 | 1.00 | 39.83 | 3.12 (A:OP1 1H:1993) |
| 10 | A | O5' | 1H:1994 | 4.84 | 1.00 | 20.80 |  |
| 11 | A | O5' | 1H:1992 | 4.86 | 1.00 | 19.40 |  |
| 12 | A | OP2 | 1H:1994 | 4.86 | 1.00 | 21.67 |  |
| 13 | Water 6 | | 1H:3955 | 4.93 | 1.00 | 25.11 | 3.03 (A:OP1 1H:1993) |
| 14 | A | OP2 | 1H:1993 | 4.95 | 1.00 | 31.75 |  |
| 15 | A | O3' | 1H:1993 | 4.97 | 1.00 | 19.03 |  |
  |

  
**1^st^ shell ligand angles:**   
1-2: 84; 1-3: 175; 1-4: 90; 2-3: 92; 2-4: 86; 3-4: 87;   
  
 Contact(s): /O_w_/O_w_/O_w_/O_w_/  
 Coded contact list: 4O_w_  
 Exclusion zone contact(s): /O5'/OP1/OP1/  
Consider K in place of MG (3 valid contact(s) for K vs. 0 for MG)  

### MG:1H:3429

|  | Res. type | Atom | Residue | Distance | Occupancy | B-factor | Closest solute to 1^st^ shell water |
| --- | --- | --- | --- | --- | --- | --- | --- |
| **Ion** | | | |  | 1.00 | 29.95 || 1 | Water 1 | | 1H:3629 | 2.17 | 1.00 | 20.90 | 1.97 (U:O4 1H:1649) |
| 2 | Water 2 | | 1H:3758 | 2.19 | 1.00 | 30.84 | 2.31 (G:N7 1H:1648) |
| 3 | Water 3 | | 1H:4341 | 2.21 | 1.00 | 35.10 | 3.16 (C:OP2 1H:1647) |
| 4 | Water 4 | | F8:201 | 2.21 | 1.00 | 32.64 | 3.53 (THR:OG1 F8:56) |
| 5 | U | O4 | 1H:1649 | 2.62 | 1.00 | 24.59 |  |
| 6 | A | OP2 | 1H:1389 | 2.74 | 1.00 | 32.66 |  |
| 7 | THR | O | F8:56 | 3.54 | 1.00 | 29.02 |  |
| 8 | G | N7 | 1H:1648 | 4.08 | 1.00 | 28.78 |  |
| 9 | THR | OG1 | F8:56 | 4.16 | 1.00 | 30.77 |  |
| 10 | A | OP1 | 1H:1389 | 4.58 | 1.00 | 26.75 |  |
| 11 | C | OP2 | 1H:1647 | 4.71 | 1.00 | 31.12 |  |
| 12 | G | OP2 | 1H:1648 | 4.77 | 1.00 | 23.69 |  |
| 13 | U | N3 | 1H:1649 | 4.89 | 1.00 | 26.69 |  |
| 14 | G | O6 | 1H:1648 | 4.91 | 1.00 | 31.46 |  |
| 15 | A | O5' | 1H:1389 | 4.96 | 1.00 | 24.58 |  |
  |

  
**1^st^ shell ligand angles:**   
1-2: 86; 1-3: 170; 1-4: 89; 2-3: 84; 2-4: 93; 3-4: 89;   
  
 Contact(s): /O_w_/O_w_/O_w_/O_w_/  
 Coded contact list: 4O_w_  
 Exclusion zone contact(s): /O4/OP2/  
Consider K in place of MG (2 valid contact(s) for K vs. 0 for MG)  
MG hydration shell issue: water too close (< 2.4 Angs.)  

### MG:1H:3430

|  | Res. type | Atom | Residue | Distance | Occupancy | B-factor | Closest solute to 1^st^ shell water |
| --- | --- | --- | --- | --- | --- | --- | --- |
| **Ion** | | | |  | 1.00 | 54.74 || 1 | Water 1 | | 1H:3785 | 2.18 | 1.00 | 58.74 | 2.35 (C:OP1 1H:1464) |
| 2 | Water 2 | | 1H:3927 | 2.18 | 1.00 | 48.59 | 2.55 (G:O6 1H:1465) |
| 3 | Water 3 | | 1H:4054 | 2.18 | 1.00 | 53.11 | 2.69 (G:OP2 1H:1465) |
| 4 | Water 4 | | 1H:4033 | 2.19 | 1.00 | 57.25 | 2.67 (U:OP2 1H:1626) |
| 5 | Water 5 | | 1H:4043 | 2.19 | 1.00 | 55.15 | 2.68 (C:OP2 1H:1625) |
| 6 | C | OP2 | 1H:1464 | 2.52 | 1.00 | 55.03 |  |
| 7 | U | OP2 | 1H:1626 | 3.52 | 1.00 | 49.91 |  |
| 8 | C | OP1 | 1H:1464 | 3.75 | 1.00 | 58.65 |  |
| 9 | C | OP2 | 1H:1625 | 4.21 | 1.00 | 45.32 |  |
| 10 | G | N7 | 1H:1465 | 4.35 | 1.00 | 53.57 |  |
| 11 | C | O5' | 1H:1464 | 4.59 | 1.00 | 50.19 |  |
| 12 | G | O6 | 1H:1465 | 4.67 | 1.00 | 58.07 |  |
| 13 | G | O3' | 1H:1463 | 4.68 | 1.00 | 67.21 |  |
| 14 | G | OP2 | 1H:1465 | 4.69 | 1.00 | 34.35 |  |
| 15 | C | O5' | 1H:1625 | 4.71 | 1.00 | 50.24 |  |
| 16 | C | OP1 | 1H:1625 | 4.83 | 1.00 | 56.64 |  |
| 17 | G | O2' | 1H:1463 | 4.91 | 1.00 | 67.53 |  |
  |

  
**1^st^ shell ligand angles:**   
1-2: 177; 1-3: 90; 1-4: 89; 1-5: 88; 2-3: 87; 2-4: 93; 2-5: 94; 3-4: 178; 3-5: 93; 4-5: 89;   
  
 Contact(s): /O_w_/O_w_/O_w_/O_w_/O_w_/  
 Coded contact list: 5O_w_  
 Exclusion zone contact(s): /OP2/  
Consider NA in place of MG (1 valid contact(s) for NA vs. 0 for MG)  
MG hydration shell issue: water too close (< 2.4 Angs.)  

### MG:1H:3431

|  | Res. type | Atom | Residue | Distance | Occupancy | B-factor | Closest solute to 1^st^ shell water |
| --- | --- | --- | --- | --- | --- | --- | --- |
| **Ion** | | | |  | 1.00 | 46.69 || 1 | Water 1 | | 1H:4108 | 2.16 | 1.00 | 49.87 | 2.77 (G:O6 1H:2855) |
| 2 | Water 2 | | 1H:4274 | 2.17 | 1.00 | 46.08 | 3.01 (C:N4 1H:2882) |
| 3 | Water 3 | | 1H:4356 | 2.18 | 1.00 | 58.93 | 3.18 (G:O6 1H:2856) |
| 4 | Water 4 | | 1H:4209 | 2.18 | 1.00 | 42.06 | 2.90 (G:N7 1H:2854) |
| 5 | Water 5 | | 1H:4457 | 2.19 | 1.00 | 56.63 | 3.54 (G:OP2 1H:2854) |
| 6 | Water 6 | | 1H:4282 | 2.19 | 1.00 | 53.53 | 3.02 (G:N7 1H:2854) |
| 7 | G | N7 | 1H:2854 | 4.04 | 1.00 | 49.88 |  |
| 8 | C | N4 | 1H:2882 | 4.17 | 1.00 | 48.20 |  |
| 9 | G | O6 | 1H:2856 | 4.24 | 1.00 | 50.65 |  |
| 10 | C | N3 | 1H:2882 | 4.25 | 1.00 | 45.78 |  |
| 11 | G | N7 | 1H:2855 | 4.50 | 1.00 | 44.71 |  |
| 12 | G | O6 | 1H:2855 | 4.78 | 1.00 | 49.79 |  |
| 13 | G | OP2 | 1H:2854 | 4.82 | 1.00 | 54.57 |  |
| 14 | G | O6 | 1H:2854 | 4.99 | 1.00 | 48.92 |  |
  |

  
**1^st^ shell ligand angles:**   
1-2: 87; 1-3: 88; 1-4: 92; 1-5: 176; 1-6: 91; 2-3: 92; 2-4: 89; 2-5: 90; 2-6: 178; 3-4: 179; 3-5: 89; 3-6: 88; 4-5: 91; 4-6: 91; 5-6: 92;   
  
 Contact(s): /O_w_/O_w_/O_w_/O_w_/O_w_/O_w_/  
 Coded contact list: 6O_w_  

### MG:1H:3432

|  | Res. type | Atom | Residue | Distance | Occupancy | B-factor | Closest solute to 1^st^ shell water |
| --- | --- | --- | --- | --- | --- | --- | --- |
| **Ion** | | | |  | 1.00 | 55.09 || 1 | Water 1 | | 1H:3880 | 2.17 | 1.00 | 48.63 | 2.49 (G:O6 1H:133) |
| 2 | Water 2 | | 1H:4591 | 2.17 | 1.00 | 47.48 | 4.58 (G:N7 1H:133) |
| 3 | Water 3 | | 1H:4593 | 2.18 | 1.00 | 53.50 | 5.60 (G:O6 1H:134) |
| 4 | Water 4 | | 1H:4530 | 2.19 | 1.00 | 56.53 | 3.96 (G:N7 1H:133) |
| 5 | Water 5 | | 1H:4414 | 2.19 | 1.00 | 51.70 | 3.35 (G:O6 1H:133) |
| 6 | Water 6 | | 1H:4151 | 2.20 | 1.00 | 46.93 | 2.83 (G:O6 1H:134) |
| 7 | G | O6 | 1H:133 | 3.97 | 1.00 | 40.63 |  |
| 8 | G | O6 | 1H:134 | 4.16 | 1.00 | 39.88 |  |
| 9 | G | N7 | 1H:133 | 4.31 | 1.00 | 44.32 |  |
| 10 | G | N7 | 1H:134 | 4.66 | 1.00 | 42.21 |  |
  |

  
**1^st^ shell ligand angles:**   
1-2: 86; 1-3: 177; 1-4: 90; 1-5: 91; 1-6: 88; 2-3: 91; 2-4: 88; 2-5: 176; 2-6: 93; 3-4: 90; 3-5: 91; 3-6: 92; 4-5: 88; 4-6: 177; 5-6: 90;   
  
 Contact(s): /O_w_/O_w_/O_w_/O_w_/O_w_/O_w_/  
 Coded contact list: 6O_w_  

### MG:1H:3433

|  | Res. type | Atom | Residue | Distance | Occupancy | B-factor | Closest solute to 1^st^ shell water |
| --- | --- | --- | --- | --- | --- | --- | --- |
| **Ion** | | | |  | 1.00 | 51.44 || 1 | Water 1 | | 1H:4276 | 2.17 | 1.00 | 43.07 | 3.01 (A:OP2 1H:336) |
| 2 | Water 2 | | 1H:3621 | 2.17 | 1.00 | 54.99 | 1.95 (A:OP2 1H:336) |
| 3 | Water 3 | | 1H:3667 | 2.18 | 1.00 | 52.72 | 2.11 (U:O4 1H:330) |
| 4 | Water 4 | | 1H:4456 | 2.18 | 1.00 | 50.22 | 3.54 (U:O4 1H:330) |
| 5 | Water 5 | | 1H:3901 | 2.18 | 1.00 | 62.38 | 2.52 (G:N7 1H:329) |
| 6 | Water 6 | | 1H:3725 | 2.18 | 1.00 | 52.82 | 2.25 (A:OP2 1H:336) |
| 7 | A | OP2 | 1H:336 | 2.73 | 1.00 | 50.48 |  |
| 8 | U | O4 | 1H:330 | 3.58 | 1.00 | 53.28 |  |
| 9 | A | O2' | 1H:335 | 3.72 | 1.00 | 47.64 |  |
| 10 | G | O6 | 1H:329 | 3.95 | 1.00 | 55.16 |  |
| 11 | U | O4 | 1H:328 | 4.40 | 1.00 | 55.65 |  |
| 12 | G | N7 | 1H:329 | 4.52 | 1.00 | 57.95 |  |
| 13 | A | O5' | 1H:336 | 4.75 | 1.00 | 40.52 |  |
| 14 | G | O6 | 1H:337 | 4.82 | 1.00 | 55.03 |  |
| 15 | A | O3' | 1H:335 | 4.92 | 1.00 | 45.04 |  |
  |

  
**1^st^ shell ligand angles:**   
1-2: 90; 1-3: 174; 1-4: 92; 1-5: 91; 1-6: 88; 2-3: 89; 2-4: 85; 2-5: 175; 2-6: 95; 3-4: 95; 3-5: 91; 3-6: 85; 4-5: 89; 4-6: 180; 5-6: 90;   
  
 Contact(s): /O_w_/O_w_/O_w_/O_w_/O_w_/O_w_/  
 Coded contact list: 6O_w_  
 Exclusion zone contact(s): /OP2/  
Consider K in place of MG (1 valid contact(s) for K vs. 0 for MG)  
MG hydration shell issue: water too close (< 2.4 Angs.)  

### MG:1H:3434

|  | Res. type | Atom | Residue | Distance | Occupancy | B-factor | Closest solute to 1^st^ shell water |
| --- | --- | --- | --- | --- | --- | --- | --- |
| **Ion** | | | |  | 1.00 | 46.89 || 1 | Water 1 | | 1H:3773 | 2.17 | 1.00 | 36.43 | 2.33 (G:N7 1H:546) |
| 2 | Water 2 | | 1H:4532 | 2.18 | 1.00 | 35.00 | 3.87 (G:N7 1H:546) |
| 3 | Water 3 | | 1H:4607 | 2.19 | 1.00 | 37.28 | 5.76 (G:O6 1H:547) |
| 4 | Water 4 | | 1H:4492 | 2.19 | 1.00 | 28.96 | 3.66 (G:O6 1H:547) |
| 5 | Water 5 | | 1H:4349 | 2.20 | 1.00 | 30.08 | 3.17 (G:O6 1H:547) |
| 6 | Water 6 | | 1H:4399 | 2.20 | 1.00 | 50.39 | 3.43 (G:N7 1H:546) |
| 7 | G | N7 | 1H:546 | 3.90 | 1.00 | 36.42 |  |
| 8 | G | O6 | 1H:547 | 4.03 | 1.00 | 34.46 |  |
| 9 | G | N7 | 1H:547 | 4.18 | 1.00 | 36.43 |  |
| 10 | G | O6 | 1H:546 | 4.55 | 1.00 | 39.21 |  |
  |

  
**1^st^ shell ligand angles:**   
1-2: 86; 1-3: 177; 1-4: 88; 1-5: 94; 1-6: 89; 2-3: 91; 2-4: 93; 2-5: 179; 2-6: 87; 3-4: 94; 3-5: 89; 3-6: 89; 4-5: 86; 4-6: 177; 5-6: 94;   
  
 Contact(s): /O_w_/O_w_/O_w_/O_w_/O_w_/O_w_/  
 Coded contact list: 6O_w_  
MG hydration shell issue: water too close (< 2.4 Angs.)  

### MG:1H:3435

|  | Res. type | Atom | Residue | Distance | Occupancy | B-factor | Closest solute to 1^st^ shell water |
| --- | --- | --- | --- | --- | --- | --- | --- |
| **Ion** | | | |  | 1.00 | 46.29 || 1 | Water 1 | | 1H:4346 | 2.17 | 1.00 | 25.86 | 3.17 (A:OP1 1H:1006) |
| 2 | Water 2 | | 1H:3733 | 2.17 | 1.00 | 34.01 | 2.27 (C:OP2 1H:1007) |
| 3 | Water 3 | | 1H:4606 | 2.19 | 1.00 | 41.50 | 5.14 (A:OP1 1H:1006) |
| 4 | Water 4 | | 1H:4247 | 2.19 | 1.00 | 34.38 | 2.96 (U:O4 1H:1009) |
| 5 | Water 5 | | 1H:4230 | 2.19 | 1.00 | 31.55 | 2.94 (A:OP2 1H:1006) |
| 6 | Water 6 | | 1H:4529 | 2.19 | 1.00 | 40.96 | 4.70 (C:OP2 1H:1007) |
| 7 | C | OP2 | 1H:1007 | 4.02 | 1.00 | 32.21 |  |
| 8 | G | N7 | 1H:1008 | 4.28 | 1.00 | 30.33 |  |
| 9 | A | OP1 | 1H:1006 | 4.51 | 1.00 | 29.71 |  |
| 10 | A | OP2 | 1H:1006 | 4.85 | 1.00 | 26.85 |  |
| 11 | G | OP2 | 1H:1008 | 4.87 | 1.00 | 28.95 |  |
| 12 | A | O5' | 1H:1006 | 4.95 | 1.00 | 27.73 |  |
| 13 | U | O4 | 1H:1009 | 4.98 | 1.00 | 31.82 |  |
  |

  
**1^st^ shell ligand angles:**   
1-2: 84; 1-3: 90; 1-4: 179; 1-5: 93; 1-6: 92; 2-3: 173; 2-4: 97; 2-5: 90; 2-6: 89; 3-4: 90; 3-5: 91; 3-6: 90; 4-5: 87; 4-6: 89; 5-6: 175;   
  
 Contact(s): /O_w_/O_w_/O_w_/O_w_/O_w_/O_w_/  
 Coded contact list: 6O_w_  
MG hydration shell issue: water too close (< 2.4 Angs.)  

### MG:1H:3436

|  | Res. type | Atom | Residue | Distance | Occupancy | B-factor | Closest solute to 1^st^ shell water |
| --- | --- | --- | --- | --- | --- | --- | --- |
| **Ion** | | | |  | 1.00 | 57.23 || 1 | Water 1 | | 1H:4323 | 2.17 | 1.00 | 56.19 | 3.13 (G:OP1 1H:2870) |
| 2 | Water 2 | | 1H:4552 | 2.18 | 1.00 | 59.80 | 4.00 (G:OP1 1H:2870) |
| 3 | Water 3 | | 1H:4603 | 2.18 | 1.00 | 73.18 | 5.56 (G:N7 1H:2868) |
| 4 | Water 4 | | 1H:4314 | 2.19 | 1.00 | 63.62 | 3.10 (C:N4 1H:2867) |
| 5 | Water 5 | | 1H:4393 | 2.19 | 1.00 | 59.68 | 3.70 (G:N7 1H:2868) |
| 6 | Water 6 | | 1H:3605 | 2.20 | 1.00 | 71.98 | 1.30 (G:N7 1H:2868) |
| 7 | G | N7 | 1H:2868 | 3.41 | 1.00 | 65.32 |  |
| 8 | G | O6 | 1H:2868 | 4.26 | 1.00 | 57.94 |  |
| 9 | G | OP1 | 1H:2870 | 4.42 | 1.00 | 64.32 |  |
| 10 | C | N4 | 1H:2867 | 4.62 | 1.00 | 59.33 |  |
| 11 | G | OP2 | 1H:2870 | 4.62 | 1.00 | 63.13 |  |
  |

  
**1^st^ shell ligand angles:**   
1-2: 85; 1-3: 87; 1-4: 91; 1-5: 177; 1-6: 95; 2-3: 88; 2-4: 176; 2-5: 96; 2-6: 91; 3-4: 92; 3-5: 91; 3-6: 178; 4-5: 88; 4-6: 89; 5-6: 88;   
  
 Contact(s): /O_w_/O_w_/O_w_/O_w_/O_w_/O_w_/  
 Coded contact list: 6O_w_  

### MG:1H:3437

|  | Res. type | Atom | Residue | Distance | Occupancy | B-factor | Closest solute to 1^st^ shell water |
| --- | --- | --- | --- | --- | --- | --- | --- |
| **Ion** | | | |  | 1.00 | 51.30 || 1 | Water 1 | | 1H:4294 | 2.18 | 1.00 | 39.69 | 3.06 (C:N4 1H:1446) |
| 2 | Water 2 | | 1H:4218 | 2.18 | 1.00 | 43.83 | 2.92 (G:O6 1H:1448) |
| 3 | Water 3 | | 1H:4507 | 2.18 | 1.00 | 44.31 | 4.47 (C:OP2 1H:1445) |
| 4 | Water 4 | | 1H:4183 | 2.19 | 1.00 | 42.42 | 2.87 (U:OP1 1H:1444) |
| 5 | Water 5 | | 1H:4201 | 2.19 | 1.00 | 46.43 | 2.90 (G:O6 1H:1432) |
| 6 | Water 6 | | 1H:4392 | 2.20 | 1.00 | 52.17 | 3.28 (C:OP2 1H:1445) |
| 7 | U | OP1 | 1H:1444 | 4.08 | 1.00 | 35.95 |  |
| 8 | G | O6 | 1H:1448 | 4.22 | 1.00 | 52.03 |  |
| 9 | C | N4 | 1H:1446 | 4.22 | 1.00 | 41.42 |  |
| 10 | C | OP2 | 1H:1445 | 4.54 | 1.00 | 32.74 |  |
| 11 | G | O6 | 1H:1447 | 4.76 | 1.00 | 47.52 |  |
| 12 | G | O6 | 1H:1432 | 4.95 | 1.00 | 44.92 |  |
| 13 | C | N4 | 1H:1433 | 4.97 | 1.00 | 44.86 |  |
  |

  
**1^st^ shell ligand angles:**   
1-2: 92; 1-3: 87; 1-4: 92; 1-5: 177; 1-6: 86; 2-3: 92; 2-4: 85; 2-5: 90; 2-6: 178; 3-4: 177; 3-5: 92; 3-6: 89; 4-5: 89; 4-6: 94; 5-6: 92;   
  
 Contact(s): /O_w_/O_w_/O_w_/O_w_/O_w_/O_w_/  
 Coded contact list: 6O_w_  

### MG:1H:3438

|  | Res. type | Atom | Residue | Distance | Occupancy | B-factor | Closest solute to 1^st^ shell water |
| --- | --- | --- | --- | --- | --- | --- | --- |
| **Ion** | | | |  | 1.00 | 74.52 || 1 | Water 1 | | 1H:4551 | 2.17 | 1.00 | 71.99 | 4.00 (U:OP1 1H:2768) |
| 2 | Water 2 | | 1H:3748 | 2.18 | 1.00 | 69.41 | 2.29 (C:OP1 1H:2769) |
| 3 | Water 3 | | 1H:4329 | 2.19 | 1.00 | 74.05 | 3.13 (C:OP1 1H:2759) |
| 4 | Water 4 | | 1H:3764 | 2.19 | 1.00 | 56.21 | 2.32 (C:OP2 1H:2759) |
| 5 | Water 5 | | 1H:4166 | 2.20 | 1.00 | 66.80 | 2.85 (U:OP1 1H:2768) |
| 6 | Water 6 | | 1H:4479 | 2.20 | 1.00 | 67.48 | 3.62 (C:OP1 1H:2769) |
| 7 | C | OP1 | 1H:2769 | 3.40 | 1.00 | 66.05 |  |
| 8 | C | OP2 | 1H:2759 | 3.91 | 1.00 | 48.80 |  |
| 9 | C | OP2 | 1H:2769 | 4.01 | 1.00 | 54.75 |  |
| 10 | U | OP1 | 1H:2768 | 4.01 | 1.00 | 60.98 |  |
| 11 | C | OP1 | 1H:2759 | 4.63 | 1.00 | 52.46 |  |
| 12 | C | O5' | 1H:2769 | 4.91 | 1.00 | 54.33 |  |
  |

  
**1^st^ shell ligand angles:**   
1-2: 87; 1-3: 91; 1-4: 179; 1-5: 89; 1-6: 91; 2-3: 177; 2-4: 93; 2-5: 90; 2-6: 91; 3-4: 89; 3-5: 89; 3-6: 91; 4-5: 93; 4-6: 87; 5-6: 180;   
  
 Contact(s): /O_w_/O_w_/O_w_/O_w_/O_w_/O_w_/  
 Coded contact list: 6O_w_  
 Exclusion zone contact(s): /OP1/  
Consider K in place of MG (1 valid contact(s) for K vs. 0 for MG)  
MG hydration shell issue: water too close (< 2.4 Angs.)  

### MG:1H:3439

|  | Res. type | Atom | Residue | Distance | Occupancy | B-factor | Closest solute to 1^st^ shell water |
| --- | --- | --- | --- | --- | --- | --- | --- |
| **Ion** | | | |  | 1.00 | 53.02 || 1 | Water 1 | | 1H:3735 | 2.17 | 1.00 | 45.55 | 2.27 (G:N7 1H:1064) |
| 2 | Water 2 | | 1H:4496 | 2.17 | 1.00 | 48.83 | 3.67 (A:OP2 1H:1189) |
| 3 | Water 3 | | 1H:4044 | 2.18 | 1.00 | 52.63 | 2.68 (G:O6 1H:1063) |
| 4 | Water 4 | | 1H:4535 | 2.18 | 1.00 | 41.23 | 3.88 (U:O2' 1H:1188) |
| 5 | Water 5 | | 1H:3993 | 2.19 | 1.00 | 52.66 | 2.63 (G:N7 1H:1063) |
| 6 | Water 6 | | 1H:4254 | 2.19 | 1.00 | 51.86 | 2.98 (A:OP2 1H:1189) |
| 7 | G | O6 | 1H:1064 | 3.77 | 1.00 | 44.34 |  |
| 8 | G | N7 | 1H:1063 | 3.79 | 1.00 | 48.79 |  |
| 9 | G | O6 | 1H:1063 | 3.98 | 1.00 | 49.31 |  |
| 10 | G | N7 | 1H:1064 | 4.39 | 1.00 | 52.91 |  |
| 11 | A | OP2 | 1H:1189 | 4.47 | 1.00 | 54.86 |  |
| 12 | U | O3' | 1H:1188 | 4.78 | 1.00 | 50.93 |  |
| 13 | U | O2' | 1H:1188 | 4.87 | 1.00 | 60.76 |  |
  |

  
**1^st^ shell ligand angles:**   
1-2: 178; 1-3: 91; 1-4: 85; 1-5: 92; 1-6: 92; 2-3: 88; 2-4: 96; 2-5: 86; 2-6: 90; 3-4: 176; 3-5: 90; 3-6: 90; 4-5: 92; 4-6: 89; 5-6: 176;   
  
 Contact(s): /O_w_/O_w_/O_w_/O_w_/O_w_/O_w_/  
 Coded contact list: 6O_w_  
MG hydration shell issue: water too close (< 2.4 Angs.)  

### MG:1H:3440

|  | Res. type | Atom | Residue | Distance | Occupancy | B-factor | Closest solute to 1^st^ shell water |
| --- | --- | --- | --- | --- | --- | --- | --- |
| **Ion** | | | |  | 1.00 | 62.00 || 1 | Water 1 | | 1H:4386 | 2.18 | 1.00 | 58.91 | 3.26 (A:OP1 1H:528) |
| 2 | Water 2 | | 1H:4599 | 2.19 | 1.00 | 66.46 | 4.85 (U:OP2 1H:525) |
| 3 | Water 3 | | 1H:4385 | 2.19 | 1.00 | 61.40 | 3.40 (G:N7 1H:526) |
| 4 | Water 4 | | 1H:4595 | 2.19 | 1.00 | 42.73 | 5.16 (U:OP2 1H:525) |
| 5 | Water 5 | | 1H:4283 | 2.19 | 1.00 | 64.31 | 3.02 (G:N7 1H:526) |
| 6 | Water 6 | | 1H:3942 | 2.20 | 1.00 | 53.93 | 2.57 (U:OP2 1H:525) |
| 7 | G | N7 | 1H:526 | 4.14 | 1.00 | 52.30 |  |
| 8 | A | OP1 | 1H:528 | 4.56 | 1.00 | 52.73 |  |
| 9 | U | OP2 | 1H:525 | 4.69 | 1.00 | 51.44 |  |
  |

  
**1^st^ shell ligand angles:**   
1-2: 88; 1-3: 88; 1-4: 91; 1-5: 92; 1-6: 179; 2-3: 85; 2-4: 96; 2-5: 179; 2-6: 92; 3-4: 178; 3-5: 95; 3-6: 91; 4-5: 84; 4-6: 90; 5-6: 89;   
  
 Contact(s): /O_w_/O_w_/O_w_/O_w_/O_w_/O_w_/  
 Coded contact list: 6O_w_  

### MG:1H:3441

|  | Res. type | Atom | Residue | Distance | Occupancy | B-factor | Closest solute to 1^st^ shell water |
| --- | --- | --- | --- | --- | --- | --- | --- |
| **Ion** | | | |  | 1.00 | 61.50 || 1 | Water 1 | | 1H:4076 | 2.19 | 1.00 | 47.32 | 2.73 (C:O2 1H:254) |
| 2 | Water 2 | | 1H:4556 | 2.19 | 1.00 | 58.20 | 4.03 (A:OP2 1H:223) |
| 3 | Water 3 | | 1H:4327 | 2.19 | 1.00 | 37.90 | 3.13 (C:O2 1H:254) |
| 4 | Water 4 | | 1H:4500 | 2.20 | 1.00 | 44.63 | 3.71 (G:OP1 1H:256) |
| 5 | Water 5 | | 1H:4325 | 2.20 | 1.00 | 45.73 | 3.13 (A:OP2 1H:223) |
| 6 | Water 6 | | 1H:4374 | 2.20 | 1.00 | 49.44 | 3.24 (A:OP2 1H:223) |
| 7 | C | O2 | 1H:254 | 3.98 | 1.00 | 39.49 |  |
| 8 | A | OP2 | 1H:223 | 4.17 | 1.00 | 49.31 |  |
| 9 | G | OP1 | 1H:256 | 4.51 | 1.00 | 38.08 |  |
| 10 | C | N3 | 1H:254 | 4.85 | 1.00 | 39.86 |  |
  |

  
**1^st^ shell ligand angles:**   
1-2: 85; 1-3: 92; 1-4: 90; 1-5: 177; 1-6: 92; 2-3: 176; 2-4: 90; 2-5: 92; 2-6: 92; 3-4: 92; 3-5: 91; 3-6: 86; 4-5: 89; 4-6: 177; 5-6: 89;   
  
 Contact(s): /O_w_/O_w_/O_w_/O_w_/O_w_/O_w_/  
 Coded contact list: 6O_w_  

### MG:1H:3442

|  | Res. type | Atom | Residue | Distance | Occupancy | B-factor | Closest solute to 1^st^ shell water |
| --- | --- | --- | --- | --- | --- | --- | --- |
| **Ion** | | | |  | 1.00 | 56.17 || 1 | Water 1 | | 1H:3907 | 2.17 | 1.00 | 52.34 | 2.53 (G:O6 1H:1495) |
| 2 | Water 2 | | 1H:4576 | 2.18 | 1.00 | 42.65 | 4.69 (C:N4 1H:1494) |
| 3 | Water 3 | | 1H:3971 | 2.18 | 1.00 | 49.12 | 2.61 (G:N7 1H:1496) |
| 4 | Water 4 | | 1H:4342 | 2.18 | 1.00 | 52.93 | 3.16 (G:O6 1H:1496) |
| 5 | Water 5 | | 1H:4402 | 2.19 | 1.00 | 66.96 | 3.31 (G:N7 1H:1495) |
| 6 | Water 6 | | 1H:4609 | 2.19 | 1.00 | 58.78 | 5.50 (G:O6 1H:1496) |
| 7 | G | O6 | 1H:1496 | 3.91 | 1.00 | 58.55 |  |
| 8 | G | N7 | 1H:1495 | 4.16 | 1.00 | 61.78 |  |
| 9 | G | N7 | 1H:1496 | 4.41 | 1.00 | 62.95 |  |
| 10 | G | O6 | 1H:1495 | 4.51 | 1.00 | 58.64 |  |
| 11 | C | N4 | 1H:1494 | 4.94 | 1.00 | 59.46 |  |
  |

  
**1^st^ shell ligand angles:**   
1-2: 91; 1-3: 91; 1-4: 91; 1-5: 87; 1-6: 178; 2-3: 178; 2-4: 90; 2-5: 87; 2-6: 91; 3-4: 89; 3-5: 93; 3-6: 88; 4-5: 176; 4-6: 91; 5-6: 92;   
  
 Contact(s): /O_w_/O_w_/O_w_/O_w_/O_w_/O_w_/  
 Coded contact list: 6O_w_  

### MG:1H:3443

|  | Res. type | Atom | Residue | Distance | Occupancy | B-factor | Closest solute to 1^st^ shell water |
| --- | --- | --- | --- | --- | --- | --- | --- |
| **Ion** | | | |  | 1.00 | 42.31 || 1 | Water 1 | | 1H:4558 | 2.18 | 1.00 | 47.80 | 4.08 (G:OP1 1H:1681) |
| 2 | Water 2 | | 1H:4182 | 2.19 | 1.00 | 50.02 | 2.87 (U:OP2 1H:1676) |
| 3 | Water 3 | | 1H:3655 | 2.19 | 1.00 | 40.17 | 2.08 (G:OP2 1H:1681) |
| 4 | Water 4 | | 1H:3976 | 2.20 | 1.00 | 41.18 | 2.62 (G:N7 1H:1677) |
| 5 | Water 5 | | 1H:4490 | 2.20 | 1.00 | 40.86 | 3.65 (G:OP2 1H:1677) |
| 6 | Water 6 | | 1H:4577 | 2.20 | 1.00 | 39.47 | 4.36 (G:OP1 1H:1681) |
| 7 | G | OP2 | 1H:1681 | 4.18 | 1.00 | 33.22 |  |
| 8 | C | N4 | 1H:1678 | 4.29 | 1.00 | 33.49 |  |
| 9 | G | N7 | 1H:1677 | 4.47 | 1.00 | 38.81 |  |
| 10 | G | OP1 | 1H:1681 | 4.79 | 1.00 | 30.24 |  |
| 11 | G | OP2 | 1H:1677 | 4.88 | 1.00 | 42.69 |  |
| 12 | U | OP2 | 1H:1676 | 4.97 | 1.00 | 39.71 |  |
  |

  
**1^st^ shell ligand angles:**   
1-2: 96; 1-3: 84; 1-4: 179; 1-5: 89; 1-6: 90; 2-3: 178; 2-4: 85; 2-5: 90; 2-6: 88; 3-4: 95; 3-5: 89; 3-6: 94; 4-5: 91; 4-6: 89; 5-6: 177;   
  
 Contact(s): /O_w_/O_w_/O_w_/O_w_/O_w_/O_w_/  
 Coded contact list: 6O_w_  
MG hydration shell issue: water too close (< 2.4 Angs.)  

### MG:1H:3444

|  | Res. type | Atom | Residue | Distance | Occupancy | B-factor | Closest solute to 1^st^ shell water |
| --- | --- | --- | --- | --- | --- | --- | --- |
| **Ion** | | | |  | 1.00 | 23.03 || 1 | Water 1 | | 1H:4035 | 2.15 | 1.00 | 18.76 | 2.67 (A:OP2 1H:28) |
| 2 | Water 2 | | 1H:3706 | 2.16 | 1.00 | 28.29 | 2.22 (G:O6 1H:501) |
| 3 | Water 3 | | 1H:4299 | 2.17 | 1.00 | 43.42 | 3.06 (A:O2' 1H:474) |
| 4 | Water 4 | | 1H:4070 | 2.18 | 1.00 | 31.67 | 2.72 (G:O6 1H:500) |
| 5 | Water 5 | | 1H:4480 | 2.19 | 1.00 | 29.08 | 3.62 (A:OP2 1H:28) |
| 6 | Water 6 | | 1H:4478 | 2.19 | 1.00 | 40.08 | 3.61 (U:OP2 1H:29) |
| 7 | G | O6 | 1H:501 | 4.12 | 1.00 | 29.41 |  |
| 8 | G | O6 | 1H:500 | 4.21 | 1.00 | 31.32 |  |
| 9 | A | OP2 | 1H:28 | 4.22 | 1.00 | 22.46 |  |
| 10 | G | O6 | 1H:30 | 4.42 | 1.00 | 28.53 |  |
| 11 | A | OP2 | 1H:474 | 4.47 | 1.00 | 23.22 |  |
| 12 | A | O2' | 1H:474 | 4.57 | 1.00 | 29.47 |  |
| 13 | C | N4 | 1H:31 | 4.57 | 1.00 | 29.10 |  |
| 14 | U | O4 | 1H:29 | 4.68 | 1.00 | 30.52 |  |
| 15 | U | OP2 | 1H:29 | 4.72 | 1.00 | 25.07 |  |
  |

  
**1^st^ shell ligand angles:**   
1-2: 89; 1-3: 93; 1-4: 177; 1-5: 89; 1-6: 93; 2-3: 96; 2-4: 89; 2-5: 178; 2-6: 85; 3-4: 85; 3-5: 83; 3-6: 174; 4-5: 93; 4-6: 89; 5-6: 96;   
  
 Contact(s): /O_w_/O_w_/O_w_/O_w_/O_w_/O_w_/  
 Coded contact list: 6O_w_  
MG hydration shell issue: water too close (< 2.4 Angs.)  

### MG:1H:3445

|  | Res. type | Atom | Residue | Distance | Occupancy | B-factor | Closest solute to 1^st^ shell water |
| --- | --- | --- | --- | --- | --- | --- | --- |
| **Ion** | | | |  | 1.00 | 50.89 || 1 | Water 1 | | 1H:4389 | 2.17 | 1.00 | 40.10 | 3.27 (G:N7 1H:1210) |
| 2 | Water 2 | | 1H:3910 | 2.18 | 1.00 | 49.96 | 2.53 (G:O6 1H:1209) |
| 3 | Water 3 | | 1H:3969 | 2.18 | 1.00 | 39.09 | 2.60 (G:N7 1H:1209) |
| 4 | Water 4 | | 1H:4608 | 2.18 | 1.00 | 58.94 | 5.74 (G:N7 1H:1209) |
| 5 | Water 5 | | 1H:4566 | 2.19 | 1.00 | 46.85 | 4.18 (G:O6 1H:1210) |
| 6 | Water 6 | | 1H:4415 | 2.19 | 1.00 | 40.07 | 3.35 (C:N4 1H:1208) |
| 7 | G | N7 | 1H:1209 | 4.30 | 1.00 | 42.51 |  |
| 8 | G | O6 | 1H:1210 | 4.34 | 1.00 | 47.99 |  |
| 9 | G | N7 | 1H:1210 | 4.38 | 1.00 | 41.92 |  |
| 10 | G | O6 | 1H:1209 | 4.48 | 1.00 | 39.29 |  |
| 11 | C | N4 | 1H:1208 | 4.62 | 1.00 | 44.43 |  |
  |

  
**1^st^ shell ligand angles:**   
1-2: 89; 1-3: 87; 1-4: 91; 1-5: 91; 1-6: 178; 2-3: 87; 2-4: 179; 2-5: 94; 2-6: 89; 3-4: 93; 3-5: 178; 3-6: 92; 4-5: 87; 4-6: 91; 5-6: 89;   
  
 Contact(s): /O_w_/O_w_/O_w_/O_w_/O_w_/O_w_/  
 Coded contact list: 6O_w_  

### MG:1H:3446

|  | Res. type | Atom | Residue | Distance | Occupancy | B-factor | Closest solute to 1^st^ shell water |
| --- | --- | --- | --- | --- | --- | --- | --- |
| **Ion** | | | |  | 1.00 | 48.82 || 1 | Water 1 | | 1H:4168 | 2.17 | 1.00 | 49.75 | 2.85 (G:OP2 1H:2856) |
| 2 | Water 2 | | 1H:4597 | 2.18 | 1.00 | 68.97 | 4.79 (G:N7 1H:2856) |
| 3 | Water 3 | | B8:202 | 2.18 | 1.00 | 50.90 | 3.44 (ARG:NE B8:54) |
| 4 | Water 4 | | 1H:4049 | 2.19 | 1.00 | 41.99 | 2.69 (G:N7 1H:2857) |
| 5 | Water 5 | | 1H:4048 | 2.19 | 1.00 | 44.24 | 2.69 (G:N1 1H:2859) |
| 6 | Water 6 | | 1H:4242 | 2.19 | 1.00 | 51.41 | 2.96 (G:O6 1H:2859) |
| 7 | U | O4 | 1H:2858 | 3.51 | 1.00 | 45.17 |  |
| 8 | G | O6 | 1H:2859 | 4.20 | 1.00 | 49.19 |  |
| 9 | G | N7 | 1H:2857 | 4.51 | 1.00 | 45.65 |  |
| 10 | G | OP2 | 1H:2857 | 4.59 | 1.00 | 40.73 |  |
| 11 | G | N1 | 1H:2859 | 4.64 | 1.00 | 48.07 |  |
| 12 | ARG | NE | B8:54 | 4.70 | 1.00 | 45.48 |  |
| 13 | G | N7 | 1H:2856 | 4.86 | 1.00 | 49.13 |  |
| 14 | G | OP2 | 1H:2856 | 4.90 | 1.00 | 46.75 |  |
  |

  
**1^st^ shell ligand angles:**   
1-2: 88; 1-3: 90; 1-4: 84; 1-5: 178; 1-6: 94; 2-3: 89; 2-4: 89; 2-5: 90; 2-6: 178; 3-4: 174; 3-5: 89; 3-6: 90; 4-5: 96; 4-6: 92; 5-6: 88;   
  
 Contact(s): /O_w_/O_w_/O_w_/O_w_/O_w_/O_w_/  
 Coded contact list: 6O_w_  

### MG:1H:3447

|  | Res. type | Atom | Residue | Distance | Occupancy | B-factor | Closest solute to 1^st^ shell water |
| --- | --- | --- | --- | --- | --- | --- | --- |
| **Ion** | | | |  | 1.00 | 19.06 || 1 | Water 1 | | 1H:4578 | 2.15 | 1.00 | 22.81 | 5.48 (G:O6 1H:2023) |
| 2 | Water 2 | | 1H:4150 | 2.17 | 1.00 | 26.78 | 2.83 (G:O6 1H:2023) |
| 3 | Water 3 | | 1H:4502 | 2.17 | 1.00 | 31.80 | 3.73 (C:N4 1H:2022) |
| 4 | Water 4 | | 1H:4515 | 2.17 | 1.00 | 32.20 | 3.78 (G:N7 1H:2023) |
| 5 | Water 5 | | 1H:4545 | 2.20 | 1.00 | 25.19 | 3.97 (G:N7 1H:2023) |
| 6 | Water 6 | | 1H:3611 | 2.22 | 1.00 | 34.79 | 1.31 (G:N7 1H:2023) |
| 7 | G | N7 | 1H:2023 | 3.48 | 1.00 | 31.55 |  |
| 8 | G | O6 | 1H:2023 | 3.88 | 1.00 | 22.81 |  |
| 9 | C | N4 | 1H:2022 | 4.31 | 1.00 | 20.42 |  |
  |

  
**1^st^ shell ligand angles:**   
1-2: 86; 1-3: 83; 1-4: 90; 1-5: 94; 1-6: 179; 2-3: 84; 2-4: 91; 2-5: 177; 2-6: 94; 3-4: 172; 3-5: 93; 3-6: 98; 4-5: 92; 4-6: 89; 5-6: 85;   
  
 Contact(s): /O_w_/O_w_/O_w_/O_w_/O_w_/O_w_/  
 Coded contact list: 6O_w_  

### MG:1H:3448

|  | Res. type | Atom | Residue | Distance | Occupancy | B-factor | Closest solute to 1^st^ shell water |
| --- | --- | --- | --- | --- | --- | --- | --- |
| **Ion** | | | |  | 1.00 | 57.33 || 1 | Water 1 | | 1H:4164 | 2.17 | 1.00 | 55.16 | 2.84 (A:N1 1H:394) |
| 2 | Water 2 | | 1H:4590 | 2.17 | 1.00 | 52.11 | 5.02 (A:N1 1H:394) |
| 3 | Water 3 | | 1H:3932 | 2.18 | 1.00 | 54.73 | 2.55 (G:O6 1H:291) |
| 4 | Water 4 | | 1H:4311 | 2.18 | 1.00 | 56.11 | 3.10 (G:N7 1H:291) |
| 5 | Water 5 | | 1H:4380 | 2.18 | 1.00 | 69.37 | 3.60 (G:O6 1H:292) |
| 6 | Water 6 | | 1H:4432 | 2.19 | 1.00 | 56.33 | 3.41 (G:N7 1H:292) |
| 7 | G | N7 | 1H:291 | 3.95 | 1.00 | 50.81 |  |
| 8 | G | N7 | 1H:292 | 4.25 | 1.00 | 51.58 |  |
| 9 | G | O6 | 1H:292 | 4.57 | 1.00 | 56.80 |  |
| 10 | A | N1 | 1H:394 | 4.63 | 1.00 | 80.25 |  |
| 11 | G | O6 | 1H:291 | 4.65 | 1.00 | 45.96 |  |
  |

  
**1^st^ shell ligand angles:**   
1-2: 88; 1-3: 90; 1-4: 93; 1-5: 89; 1-6: 178; 2-3: 177; 2-4: 91; 2-5: 91; 2-6: 91; 3-4: 88; 3-5: 91; 3-6: 91; 4-5: 178; 4-6: 89; 5-6: 89;   
  
 Contact(s): /O_w_/O_w_/O_w_/O_w_/O_w_/O_w_/  
 Coded contact list: 6O_w_  

### MG:1H:3449

|  | Res. type | Atom | Residue | Distance | Occupancy | B-factor | Closest solute to 1^st^ shell water |
| --- | --- | --- | --- | --- | --- | --- | --- |
| **Ion** | | | |  | 1.00 | 15.11 || 1 | Water 1 | | 1H:3659 | 2.14 | 1.00 | 23.92 | 2.09 (U:OP2 1H:989) |
| 2 | Water 2 | | 1H:3797 | 2.15 | 1.00 | 23.40 | 2.37 (U:OP2 1H:989) |
| 3 | Water 3 | | 78:302 | 2.16 | 1.00 | 19.34 | 2.66 (GLY:N 78:37) |
| 4 | Water 4 | | 1H:3977 | 2.16 | 1.00 | 22.37 | 2.62 (U:OP2 1H:989) |
| 5 | Water 5 | | 1H:3663 | 2.17 | 1.00 | 19.88 | 2.10 (G:OP1 1H:880) |
| 6 | Water 6 | | 78:304 | 2.20 | 1.00 | 28.70 | 3.17 (LYS:O 78:36) |
| 7 | U | OP2 | 1H:989 | 2.70 | 1.00 | 20.15 |  |
| 8 | G | OP1 | 1H:880 | 3.91 | 1.00 | 17.18 |  |
| 9 | HIS | O | 78:35 | 4.05 | 1.00 | 18.38 |  |
| 10 | GLY | N | 78:37 | 4.19 | 1.00 | 21.83 |  |
| 11 | GLY | O | 78:34 | 4.33 | 1.00 | 23.11 |  |
| 12 | U | OP1 | 1H:989 | 4.33 | 1.00 | 19.21 |  |
| 13 | LYS | O | 78:36 | 4.61 | 1.00 | 23.30 |  |
| 14 | G | OP2 | 1H:880 | 4.61 | 1.00 | 21.21 |  |
| 15 | ARG | O | 78:33 | 4.83 | 1.00 | 18.66 |  |
| 16 | G | O3' | 1H:988 | 4.86 | 1.00 | 19.97 |  |
| 17 | U | O5' | 1H:989 | 4.99 | 1.00 | 18.01 |  |
  |

  
**1^st^ shell ligand angles:**   
1-2: 97; 1-3: 78; 1-4: 93; 1-5: 172; 1-6: 87; 2-3: 90; 2-4: 86; 2-5: 89; 2-6: 175; 3-4: 169; 3-5: 97; 3-6: 93; 4-5: 93; 4-6: 91; 5-6: 87;   
  
 Contact(s): /O_w_/O_w_/O_w_/O_w_/O_w_/O_w_/  
 Coded contact list: 6O_w_  
 Exclusion zone contact(s): /OP2/  
Consider K in place of MG (1 valid contact(s) for K vs. 0 for MG)  
MG hydration shell issue: water too close (< 2.4 Angs.)  

### MG:1H:3450

|  | Res. type | Atom | Residue | Distance | Occupancy | B-factor | Closest solute to 1^st^ shell water |
| --- | --- | --- | --- | --- | --- | --- | --- |
| **Ion** | | | |  | 1.00 | 52.62 || 1 | Water 1 | | 1H:3951 | 2.17 | 1.00 | 46.72 | 2.59 (A:O2' 1H:1334) |
| 2 | Water 2 | | 1H:4447 | 2.17 | 1.00 | 41.29 | 4.19 (A:N7 1H:1325) |
| 3 | Water 3 | | 1H:4422 | 2.18 | 1.00 | 33.42 | 3.38 (A:O3' 1H:1334) |
| 4 | Water 4 | | 1H:3685 | 2.19 | 1.00 | 49.11 | 2.17 (A:N7 1H:1325) |
| 5 | Water 5 | | 1H:4536 | 2.20 | 1.00 | 46.33 | 3.89 (A:N7 1H:1325) |
| 6 | Water 6 | | 1H:4484 | 2.20 | 1.00 | 47.36 | 3.66 (U:OP1 1H:1335) |
| 7 | A | N7 | 1H:1325 | 4.01 | 1.00 | 35.23 |  |
| 8 | A | O2' | 1H:1334 | 4.22 | 1.00 | 33.24 |  |
| 9 | A | O3' | 1H:1334 | 4.73 | 1.00 | 30.88 |  |
| 10 | U | OP1 | 1H:1335 | 4.77 | 1.00 | 35.66 |  |
| 11 | A | N6 | 1H:1325 | 4.96 | 1.00 | 39.46 |  |
  |

  
**1^st^ shell ligand angles:**   
1-2: 86; 1-3: 86; 1-4: 178; 1-5: 91; 1-6: 96; 2-3: 91; 2-4: 92; 2-5: 90; 2-6: 177; 3-4: 94; 3-5: 177; 3-6: 87; 4-5: 89; 4-6: 87; 5-6: 92;   
  
 Contact(s): /O_w_/O_w_/O_w_/O_w_/O_w_/O_w_/  
 Coded contact list: 6O_w_  
MG hydration shell issue: water too close (< 2.4 Angs.)  

### MG:1H:3451

|  | Res. type | Atom | Residue | Distance | Occupancy | B-factor | Closest solute to 1^st^ shell water |
| --- | --- | --- | --- | --- | --- | --- | --- |
| **Ion** | | | |  | 1.00 | 82.05 || 1 | Water 1 | | 1H:4523 | 2.18 | 1.00 | 66.21 | 3.81 (G:OP2 1H:378) |
| 2 | Water 2 | | 1H:4225 | 2.18 | 1.00 | 72.52 | 2.92 (G:N7 1H:379) |
| 3 | Water 3 | | 1H:4155 | 2.18 | 1.00 | 67.40 | 2.84 (G:OP2 1H:378) |
| 4 | Water 4 | | 1H:4494 | 2.19 | 1.00 | 61.44 | 3.67 (G:OP2 1H:378) |
| 5 | Water 5 | | 1H:4508 | 2.19 | 1.00 | 61.37 | 3.74 (G:O6 1H:380) |
| 6 | Water 6 | | 1H:4141 | 2.20 | 1.00 | 59.28 | 2.82 (G:OP2 1H:377) |
| 7 | G | OP2 | 1H:378 | 4.17 | 1.00 | 62.15 |  |
| 8 | G | OP2 | 1H:377 | 4.40 | 1.00 | 59.30 |  |
| 9 | G | O6 | 1H:379 | 4.60 | 1.00 | 67.42 |  |
| 10 | G | N7 | 1H:379 | 4.63 | 1.00 | 64.19 |  |
| 11 | G | N7 | 1H:378 | 4.81 | 1.00 | 68.74 |  |
| 12 | G | O6 | 1H:380 | 4.87 | 1.00 | 67.56 |  |
| 13 | G | O5' | 1H:377 | 4.98 | 1.00 | 66.74 |  |
  |

  
**1^st^ shell ligand angles:**   
1-2: 86; 1-3: 88; 1-4: 91; 1-5: 91; 1-6: 177; 2-3: 90; 2-4: 178; 2-5: 90; 2-6: 91; 3-4: 90; 3-5: 179; 3-6: 89; 4-5: 89; 4-6: 91; 5-6: 92;   
  
 Contact(s): /O_w_/O_w_/O_w_/O_w_/O_w_/O_w_/  
 Coded contact list: 6O_w_  

### MG:1H:3452

|  | Res. type | Atom | Residue | Distance | Occupancy | B-factor | Closest solute to 1^st^ shell water |
| --- | --- | --- | --- | --- | --- | --- | --- |
| **Ion** | | | |  | 1.00 | 68.79 || 1 | Water 1 | | 1H:4213 | 2.18 | 1.00 | 49.92 | 2.91 (G:N7 1H:373) |
| 2 | Water 2 | | 1H:4469 | 2.19 | 1.00 | 55.41 | 4.14 (G:N7 1H:373) |
| 3 | Water 3 | | 1H:4475 | 2.19 | 1.00 | 51.26 | 3.59 (G:O6 1H:373) |
| 4 | Water 4 | | 1H:4263 | 2.19 | 1.00 | 64.33 | 4.16 (A:O2' 1H:370) |
| 5 | Water 5 | | 1H:4202 | 2.19 | 1.00 | 52.31 | 2.90 (G:O6 1H:374) |
| 6 | Water 6 | | 1H:4586 | 2.20 | 1.00 | 69.63 | 4.54 (G:O6 1H:374) |
| 7 | G | N7 | 1H:373 | 4.04 | 1.00 | 46.98 |  |
| 8 | G | O6 | 1H:373 | 4.16 | 1.00 | 47.02 |  |
| 9 | G | O6 | 1H:374 | 4.37 | 1.00 | 50.50 |  |
| 10 | A | O2' | 1H:370 | 4.81 | 1.00 | 47.23 |  |
| 11 | A | N7 | 1H:372 | 4.99 | 1.00 | 44.72 |  |
  |

  
**1^st^ shell ligand angles:**   
1-2: 90; 1-3: 89; 1-4: 88; 1-5: 90; 1-6: 178; 2-3: 179; 2-4: 87; 2-5: 90; 2-6: 92; 3-4: 92; 3-5: 91; 3-6: 89; 4-5: 177; 4-6: 92; 5-6: 89;   
  
 Contact(s): /O_w_/O_w_/O_w_/O_w_/O_w_/O_w_/  
 Coded contact list: 6O_w_  

### MG:1H:3453

|  | Res. type | Atom | Residue | Distance | Occupancy | B-factor | Closest solute to 1^st^ shell water |
| --- | --- | --- | --- | --- | --- | --- | --- |
| **Ion** | | | |  | 1.00 | 20.70 || 1 | Water 1 | | 1H:3770 | 2.14 | 1.00 | 19.57 | 2.32 (C:OP2 1H:1684) |
| 2 | Water 2 | | 1H:3641 | 2.15 | 1.00 | 24.57 | 2.02 (C:OP2 1H:1684) |
| 3 | Water 3 | | 1H:4153 | 2.17 | 1.00 | 25.37 | 2.83 (A:OP2 1H:1685) |
| 4 | Water 4 | | 1H:3742 | 2.19 | 1.00 | 24.70 | 2.28 (G:OP1 1H:1683) |
| 5 | Water 5 | | 1H:3817 | 2.19 | 1.00 | 28.29 | 2.40 (U:OP1 1H:1347) |
| 6 | Water 6 | | 1H:4232 | 2.21 | 1.00 | 40.46 | 2.94 (A:OP1 1H:1348) |
| 7 | C | OP2 | 1H:1684 | 2.67 | 1.00 | 25.53 |  |
| 8 | A | OP1 | 1H:1348 | 3.81 | 1.00 | 19.82 |  |
| 9 | G | OP1 | 1H:1346 | 3.84 | 1.00 | 22.65 |  |
| 10 | G | OP1 | 1H:1683 | 4.11 | 1.00 | 29.52 |  |
| 11 | U | OP1 | 1H:1347 | 4.34 | 1.00 | 23.40 |  |
| 12 | C | OP1 | 1H:1684 | 4.38 | 1.00 | 25.22 |  |
| 13 | C | O5' | 1H:1684 | 4.38 | 1.00 | 22.89 |  |
| 14 | A | OP2 | 1H:1685 | 4.53 | 1.00 | 28.53 |  |
| 15 | G | OP2 | 1H:1683 | 4.62 | 1.00 | 27.46 |  |
| 16 | U | O5' | 1H:1347 | 4.83 | 1.00 | 23.05 |  |
  |

  
**1^st^ shell ligand angles:**   
1-2: 99; 1-3: 89; 1-4: 88; 1-5: 87; 1-6: 179; 2-3: 87; 2-4: 93; 2-5: 174; 2-6: 80; 3-4: 176; 3-5: 94; 3-6: 92; 4-5: 87; 4-6: 91; 5-6: 94;   
  
 Contact(s): /O_w_/O_w_/O_w_/O_w_/O_w_/O_w_/  
 Coded contact list: 6O_w_  
 Exclusion zone contact(s): /OP2/  
Consider K in place of MG (1 valid contact(s) for K vs. 0 for MG)  
MG hydration shell issue: water too close (< 2.4 Angs.)  

### MG:1H:3454

|  | Res. type | Atom | Residue | Distance | Occupancy | B-factor | Closest solute to 1^st^ shell water |
| --- | --- | --- | --- | --- | --- | --- | --- |
| **Ion** | | | |  | 1.00 | 23.38 || 1 | Water 1 | | 1H:3677 | 2.16 | 1.00 | 24.26 | 2.15 (A:OP2 1H:1312) |
| 2 | Water 2 | | 1H:3873 | 2.17 | 1.00 | 24.88 | 2.48 (U:OP2 1H:2628) |
| 3 | Water 3 | | 1H:4023 | 2.18 | 1.00 | 25.17 | 2.66 (A:O5' 1H:1312) |
| 4 | Water 4 | | 1H:3625 | 2.19 | 1.00 | 22.15 | 1.96 (A:OP2 1H:1312) |
| 5 | Water 5 | | 1H:4547 | 2.20 | 1.00 | 31.55 | 4.46 (A:OP2 1H:1312) |
| 6 | Water 6 | | 1H:3832 | 2.20 | 1.00 | 40.29 | 2.43 (A:OP1 1H:2627) |
| 7 | A | OP2 | 1H:1312 | 2.68 | 1.00 | 25.62 |  |
| 8 | U | OP1 | 1H:2628 | 3.03 | 1.00 | 27.30 |  |
| 9 | U | OP2 | 1H:2628 | 3.80 | 1.00 | 27.17 |  |
| 10 | A | OP1 | 1H:2627 | 3.85 | 1.00 | 22.76 |  |
| 11 | A | O5' | 1H:1312 | 3.90 | 1.00 | 22.48 |  |
| 12 | Water 7 | | 1H:4483 | 4.15 | 1.00 | 31.64 | 3.63 (A:OP1 1H:2627) |
| 13 | A | OP1 | 1H:1312 | 4.16 | 1.00 | 24.24 |  |
| 14 | U | O2' | 1H:1314 | 4.21 | 1.00 | 29.76 |  |
| 15 | Water 8 | | 1H:3914 | 4.27 | 1.00 | 35.48 | 4.73 (U:O2' 1H:1314) |
| 16 | Water 9 | | 1H:3666 | 4.41 | 1.00 | 27.23 | 2.54 (U:O2' 1H:1314) |
| 17 | A | O3' | 1H:2627 | 4.85 | 1.00 | 24.73 |  |
  |

  
**1^st^ shell ligand angles:**   
1-2: 93; 1-3: 88; 1-4: 92; 1-5: 173; 1-6: 88; 2-3: 179; 2-4: 84; 2-5: 91; 2-6: 90; 3-4: 95; 3-5: 88; 3-6: 91; 4-5: 94; 4-6: 174; 5-6: 87;   
  
 Contact(s): /O_w_/O_w_/O_w_/O_w_/O_w_/O_w_/  
 Coded contact list: 6O_w_  
 Exclusion zone contact(s): /OP1/OP2/  
Consider K in place of MG (2 valid contact(s) for K vs. 0 for MG)  
MG hydration shell issue: water too close (< 2.4 Angs.)  

### MG:1H:3455

|  | Res. type | Atom | Residue | Distance | Occupancy | B-factor | Closest solute to 1^st^ shell water |
| --- | --- | --- | --- | --- | --- | --- | --- |
| **Ion** | | | |  | 1.00 | 11.67 || 1 | Water 1 | | 1H:3740 | 2.12 | 1.00 | 13.25 | 2.28 (C:OP1 1H:1814) |
| 2 | Water 2 | | 1H:3732 | 2.18 | 1.00 | 29.47 | 2.26 (A:N7 1H:1815) |
| 3 | Water 3 | | 1H:4163 | 2.19 | 1.00 | 18.16 | 2.84 (A:OP2 1H:1815) |
| 4 | C | OP1 | 1H:1814 | 2.41 | 1.00 | 14.31 |  |
| 5 | A | OP2 | 1H:1815 | 2.57 | 1.00 | 16.41 |  |
| 6 | A | OP1 | 1H:1812 | 2.62 | 1.00 | 19.43 |  |
| 7 | A | OP2 | 1H:1812 | 3.32 | 1.00 | 16.90 |  |
| 8 | U | O3' | 1H:1811 | 4.00 | 1.00 | 17.62 |  |
| 9 | C | OP2 | 1H:1814 | 4.02 | 1.00 | 14.13 |  |
| 10 | A | N7 | 1H:1815 | 4.09 | 1.00 | 17.06 |  |
| 11 | A | O5' | 1H:1815 | 4.23 | 1.00 | 21.16 |  |
| 12 | U | O2' | 1H:1811 | 4.29 | 1.00 | 18.50 |  |
| 13 | U | O2 | 1H:1811 | 4.48 | 1.00 | 18.12 |  |
| 14 | C | O5' | 1H:1814 | 4.54 | 1.00 | 12.64 |  |
| 15 | C | O3' | 1H:1814 | 4.57 | 1.00 | 15.30 |  |
| 16 | C | O3' | 1H:1813 | 4.62 | 1.00 | 14.19 |  |
| 17 | A | O5' | 1H:1812 | 4.77 | 1.00 | 20.44 |  |
| 18 | Water 4 | | 1H:4031 | 4.83 | 1.00 | 17.88 | 3.06 (A:OP2 1H:1815) |
| 19 | C | OP2 | 1H:1813 | 4.85 | 1.00 | 17.88 |  |
| 20 | A | OP1 | 1H:1815 | 4.97 | 1.00 | 17.36 |  |
  |

  
**1^st^ shell ligand angles:**   
1-2: 172; 1-3: 92; 2-3: 94;   
  
 Contact(s): /O_w_/O_w_/O_w_/  
 Coded contact list: 3O_w_  
 Exclusion zone contact(s): /OP1/OP1/OP2/OP2/  
Consider NA in place of MG (2 valid contact(s) for NA vs. 0 for MG)  
MG hydration shell issue: water too close (< 2.4 Angs.)  

### MG:1H:3456

|  | Res. type | Atom | Residue | Distance | Occupancy | B-factor | Closest solute to 1^st^ shell water |
| --- | --- | --- | --- | --- | --- | --- | --- |
| **Ion** | | | |  | 1.00 | 14.32 || 1 | Water 1 | | 1H:4000 | 2.13 | 1.00 | 13.85 | 2.64 (A:N3 1H:2083) |
| 2 | Water 2 | | 1H:4177 | 2.15 | 1.00 | 11.92 | 2.87 (G:OP2 1H:2459) |
| 3 | Water 3 | | 1H:4123 | 2.16 | 1.00 | 21.98 | 2.79 (G:OP1 1H:2084) |
| 4 | Water 4 | | 1H:3854 | 2.16 | 1.00 | 18.83 | 2.46 (C:O2' 1H:2514) |
| 5 | G | OP2 | 1H:2515 | 2.29 | 1.00 | 13.96 |  |
| 6 | G | OP1 | 1H:2084 | 2.71 | 1.00 | 15.09 |  |
| 7 | G | OP2 | 1H:2084 | 3.33 | 1.00 | 19.63 |  |
| 8 | G | OP1 | 1H:2515 | 3.96 | 1.00 | 20.03 |  |
| 9 | Water 5 | | 1H:4326 | 4.06 | 1.00 | 24.00 | 3.30 (G:OP1 1H:2084) |
| 10 | C | O2' | 1H:2514 | 4.13 | 1.00 | 17.22 |  |
| 11 | C | O3' | 1H:2514 | 4.25 | 1.00 | 17.98 |  |
| 12 | G | N7 | 1H:2458 | 4.26 | 1.00 | 12.39 |  |
| 13 | Water 6 | | 1H:3895 | 4.29 | 1.00 | 21.76 | 3.54 (G:N7 1H:2458) |
| 14 | A | O2' | 1H:2083 | 4.38 | 1.00 | 20.10 |  |
| 15 | G | N7 | 1H:2459 | 4.41 | 1.00 | 17.75 |  |
| 16 | G | O5' | 1H:2515 | 4.44 | 1.00 | 13.96 |  |
| 17 | G | O5' | 1H:2084 | 4.48 | 1.00 | 14.13 |  |
| 18 | A | O3' | 1H:2083 | 4.59 | 1.00 | 19.85 |  |
| 19 | G | OP2 | 1H:2459 | 4.70 | 1.00 | 19.73 |  |
| 20 | A | N3 | 1H:2083 | 4.72 | 1.00 | 14.56 |  |
| 21 | G | O6 | 1H:2459 | 4.85 | 1.00 | 16.64 |  |
| 22 | Water 7 | | 1H:3642 | 4.94 | 1.00 | 17.48 | 2.60 (G:OP1 1H:2084) |
  |

  
**1^st^ shell ligand angles:**   
1-2: 90; 1-3: 83; 1-4: 177; 1-5: 92; 2-3: 91; 2-4: 90; 2-5: 102; 3-4: 100; 3-5: 167; 4-5: 85;   
  
 Contact(s): /O_w_/O_w_/O_w_/O_w_/OP2/  
 Coded contact list: O_ph_4O_w_  
 Exclusion zone contact(s): /OP1/OP2/  
Consider K in place of MG (2 valid contact(s) for K vs. 1 for MG)  

### MG:1H:3457

|  | Res. type | Atom | Residue | Distance | Occupancy | B-factor | Closest solute to 1^st^ shell water |
| --- | --- | --- | --- | --- | --- | --- | --- |
| **Ion** | | | |  | 1.00 | 25.00 || 1 | Water 1 | | 1H:4172 | 2.15 | 1.00 | 26.41 | 2.86 (A:OP1 1H:1442) |
| 2 | Water 2 | | 1H:3692 | 2.19 | 1.00 | 24.07 | 2.18 (C:OP2 1H:1651) |
| 3 | Water 3 | | 1H:4363 | 2.20 | 1.00 | 49.59 | 4.03 (C:OP2 1H:1651) |
| 4 | A | OP1 | 1H:1650 | 2.46 | 1.00 | 24.49 |  |
| 5 | C | OP2 | 1H:1651 | 2.49 | 1.00 | 24.25 |  |
| 6 | A | OP1 | 1H:1442 | 2.70 | 1.00 | 24.15 |  |
| 7 | A | OP2 | 1H:1650 | 3.22 | 1.00 | 28.95 |  |
| 8 | U | O3' | 1H:1649 | 4.26 | 1.00 | 24.22 |  |
| 9 | A | O5' | 1H:1650 | 4.60 | 1.00 | 26.99 |  |
| 10 | A | O3' | 1H:1650 | 4.67 | 1.00 | 25.36 |  |
| 11 | U | O3' | 1H:1441 | 4.78 | 1.00 | 27.74 |  |
| 12 | C | O5' | 1H:1651 | 4.79 | 1.00 | 24.01 |  |
| 13 | C | OP1 | 1H:1651 | 4.90 | 1.00 | 24.36 |  |
  |

  
**1^st^ shell ligand angles:**   
1-2: 98; 1-3: 93; 2-3: 89;   
  
 Contact(s): /O_w_/O_w_/O_w_/  
 Coded contact list: 3O_w_  
 Exclusion zone contact(s): /OP1/OP1/OP2/OP2/  
Consider NA in place of MG (2 valid contact(s) for NA vs. 0 for MG)  
MG hydration shell issue: water too close (< 2.4 Angs.)  

### MG:1H:3458

|  | Res. type | Atom | Residue | Distance | Occupancy | B-factor | Closest solute to 1^st^ shell water |
| --- | --- | --- | --- | --- | --- | --- | --- |
| **Ion** | | | |  | 1.00 | 21.71 || 1 | Water 1 | | 1H:3944 | 2.15 | 1.00 | 29.18 | 2.57 (G:O6 1H:1346) |
| 2 | Water 2 | | 1H:4065 | 2.15 | 1.00 | 32.12 | 2.71 (C:OP1 1H:1686) |
| 3 | Water 3 | | 1H:3803 | 2.18 | 1.00 | 23.58 | 2.38 (C:OP2 1H:1344) |
| 4 | Water 4 | | 1H:4401 | 2.18 | 1.00 | 25.17 | 3.31 (U:OP2 1H:1687) |
| 5 | C | OP2 | 1H:1345 | 2.43 | 1.00 | 25.64 |  |
| 6 | U | OP2 | 1H:1687 | 2.46 | 1.00 | 29.43 |  |
| 7 | C | OP1 | 1H:1345 | 3.88 | 1.00 | 24.95 |  |
| 8 | C | O5' | 1H:1345 | 4.04 | 1.00 | 24.71 |  |
| 9 | Water 5 | | 1H:3680 | 4.17 | 1.00 | 31.97 | 2.15 (U:OP1 1H:1687) |
| 10 | G | O6 | 1H:1346 | 4.34 | 1.00 | 28.66 |  |
| 11 | U | OP1 | 1H:1687 | 4.38 | 1.00 | 28.44 |  |
| 12 | U | O5' | 1H:1687 | 4.44 | 1.00 | 27.80 |  |
| 13 | C | OP2 | 1H:1344 | 4.48 | 1.00 | 30.90 |  |
| 14 | C | O5' | 1H:1686 | 4.50 | 1.00 | 29.08 |  |
| 15 | C | O3' | 1H:1344 | 4.54 | 1.00 | 26.02 |  |
| 16 | G | N7 | 1H:1346 | 4.58 | 1.00 | 29.45 |  |
| 17 | C | OP1 | 1H:1686 | 4.72 | 1.00 | 29.85 |  |
| 18 | C | OP2 | 1H:1686 | 4.78 | 1.00 | 29.10 |  |
| 19 | C | O3' | 1H:1686 | 4.84 | 1.00 | 27.45 |  |
| 20 | C | O5' | 1H:1344 | 4.86 | 1.00 | 29.67 |  |
  |

  
**1^st^ shell ligand angles:**   
1-2: 177; 1-3: 91; 1-4: 90; 2-3: 86; 2-4: 92; 3-4: 177;   
  
 Contact(s): /O_w_/O_w_/O_w_/O_w_/  
 Coded contact list: 4O_w_  
 Exclusion zone contact(s): /OP2/OP2/  
Consider NA in place of MG (2 valid contact(s) for NA vs. 0 for MG)  
MG hydration shell issue: water too close (< 2.4 Angs.)  

### MG:1H:3459

|  | Res. type | Atom | Residue | Distance | Occupancy | B-factor | Closest solute to 1^st^ shell water |
| --- | --- | --- | --- | --- | --- | --- | --- |
| **Ion** | | | |  | 1.00 | 15.67 || 1 | Water 1 | | 1H:3884 | 2.17 | 1.00 | 20.82 | 2.50 (C:O2 1H:182) |
| 2 | Water 2 | | 1H:3941 | 2.17 | 1.00 | 21.41 | 2.57 (A:OP1 1H:848) |
| 3 | Water 3 | | 1H:3753 | 2.17 | 1.00 | 22.95 | 2.30 (C:N3 1H:182) |
| 4 | Water 4 | | 1H:3628 | 2.17 | 1.00 | 15.24 | 1.97 (A:OP1 1H:848) |
| 5 | Water 5 | | 1H:3781 | 2.19 | 1.00 | 17.72 | 2.34 (A:N1 1H:181) |
| 6 | A | OP1 | 1H:848 | 2.38 | 1.00 | 20.41 |  |
| 7 | G | O3' | 1H:847 | 3.51 | 1.00 | 17.69 |  |
| 8 | C | N3 | 1H:182 | 3.62 | 1.00 | 18.45 |  |
| 9 | A | N1 | 1H:181 | 3.78 | 1.00 | 17.71 |  |
| 10 | G | O2' | 1H:847 | 3.91 | 1.00 | 19.33 |  |
| 11 | A | N7 | 1H:848 | 3.93 | 1.00 | 17.76 |  |
| 12 | C | O2 | 1H:182 | 4.27 | 1.00 | 16.98 |  |
| 13 | A | O5' | 1H:848 | 4.28 | 1.00 | 13.25 |  |
| 14 | C | N4 | 1H:182 | 4.38 | 1.00 | 17.57 |  |
| 15 | A | OP2 | 1H:848 | 4.63 | 1.00 | 21.52 |  |
  |

  
**1^st^ shell ligand angles:**   
1-2: 88; 1-3: 83; 1-4: 173; 1-5: 96; 1-6: 128; 2-3: 89; 2-4: 85; 2-5: 176; 2-6: 69; 3-4: 94; 3-5: 92; 3-6: 139; 4-5: 91; 4-6: 51; 5-6: 108;   
  
 Contact(s): /O_w_/O_w_/O_w_/O_w_/O_w_/OP1/  
 Coded contact list: O_ph_5O_w_  
Large octahedral topology discrepancy!!!  
MG hydration shell issue: water too close (< 2.4 Angs.)  

### MG:1H:3460

|  | Res. type | Atom | Residue | Distance | Occupancy | B-factor | Closest solute to 1^st^ shell water |
| --- | --- | --- | --- | --- | --- | --- | --- |
| **Ion** | | | |  | 1.00 | 11.21 || 1 | Water 1 | | 1H:3988 | 2.14 | 1.00 | 16.02 | 2.63 (U:OP2 1H:1807) |
| 2 | Water 2 | | 1H:4418 | 2.17 | 1.00 | 29.52 | 3.36 (G:OP1 1H:1808) |
| 3 | Water 3 | | 1H:4367 | 2.17 | 1.00 | 17.79 | 3.36 (G:N7 1H:1808) |
| 4 | Water 4 | | 1H:4251 | 2.17 | 1.00 | 16.70 | 2.97 (U:OP1 1H:1807) |
| 5 | Water 5 | | 1H:4333 | 2.18 | 1.00 | 24.07 | 3.14 (A:OP1 1H:2004) |
| 6 | G | OP2 | 1H:1808 | 2.60 | 1.00 | 19.80 |  |
| 7 | G | OP1 | 1H:1808 | 3.61 | 1.00 | 14.99 |  |
| 8 | G | N7 | 1H:1808 | 4.24 | 1.00 | 14.73 |  |
| 9 | U | OP1 | 1H:1807 | 4.24 | 1.00 | 13.05 |  |
| 10 | U | O3' | 1H:1807 | 4.29 | 1.00 | 17.40 |  |
| 11 | A | OP2 | 1H:2004 | 4.38 | 1.00 | 20.91 |  |
| 12 | A | OP1 | 1H:2004 | 4.43 | 1.00 | 24.73 |  |
| 13 | U | OP2 | 1H:1807 | 4.58 | 1.00 | 17.60 |  |
| 14 | U | O5' | 1H:1807 | 4.59 | 1.00 | 15.81 |  |
| 15 | Water 6 | | 1H:4403 | 4.70 | 1.00 | 19.72 | 4.27 (G:N7 1H:1808) |
| 16 | G | O5' | 1H:1808 | 4.71 | 1.00 | 15.04 |  |
  |

  
**1^st^ shell ligand angles:**   
1-2: 177; 1-3: 89; 1-4: 89; 1-5: 87; 2-3: 88; 2-4: 94; 2-5: 93; 3-4: 178; 3-5: 92; 4-5: 88;   
  
 Contact(s): /O_w_/O_w_/O_w_/O_w_/O_w_/  
 Coded contact list: 5O_w_  
 Exclusion zone contact(s): /OP2/  
Consider NA in place of MG (1 valid contact(s) for NA vs. 0 for MG)  

### MG:1H:3461

|  | Res. type | Atom | Residue | Distance | Occupancy | B-factor | Closest solute to 1^st^ shell water |
| --- | --- | --- | --- | --- | --- | --- | --- |
| **Ion** | | | |  | 1.00 | 20.17 || 1 | Water 1 | | 1H:3929 | 2.15 | 1.00 | 22.33 | 2.55 (C:OP1 1H:239) |
| 2 | Water 2 | | 1H:4449 | 2.17 | 1.00 | 25.01 | 3.51 (U:OP1 1H:2444) |
| 3 | Water 3 | | 1H:4324 | 2.17 | 1.00 | 26.24 | 3.13 (G:O4' 1H:240) |
| 4 | Water 4 | | 1H:4148 | 2.17 | 1.00 | 34.39 | 2.83 (A:OP1 1H:2445) |
| 5 | Water 5 | | 1H:3782 | 2.19 | 1.00 | 28.14 | 2.35 (A:OP1 1H:2445) |
| 6 | C | OP1 | 1H:239 | 2.39 | 1.00 | 20.96 |  |
| 7 | A | OP1 | 1H:2445 | 3.50 | 1.00 | 24.54 |  |
| 8 | U | OP1 | 1H:2444 | 3.93 | 1.00 | 18.14 |  |
| 9 | C | OP2 | 1H:239 | 4.02 | 1.00 | 27.47 |  |
| 10 | Water 6 | | 3K:101 | 4.55 | 1.00 | 35.75 | 3.47 (A:OP1 1H:2445) |
| 11 | G | O3' | 1H:238 | 4.55 | 1.00 | 25.83 |  |
| 12 | C | O2' | 1H:2407 | 4.59 | 1.00 | 26.27 |  |
| 13 | C | O5' | 1H:239 | 4.60 | 1.00 | 22.61 |  |
| 14 | G | O2' | 1H:238 | 4.64 | 1.00 | 25.75 |  |
| 15 | U | OP2 | 1H:2444 | 4.87 | 1.00 | 19.53 |  |
| 16 | G | O4' | 1H:240 | 4.97 | 1.00 | 18.80 |  |
  |

  
**1^st^ shell ligand angles:**   
1-2: 95; 1-3: 92; 1-4: 87; 1-5: 178; 1-6: 68; 2-3: 92; 2-4: 92; 2-5: 86; 2-6: 163; 3-4: 176; 3-5: 89; 3-6: 87; 4-5: 92; 4-6: 89; 5-6: 110;   
  
 Contact(s): /O_w_/O_w_/O_w_/O_w_/O_w_/OP1/  
 Coded contact list: O_ph_5O_w_  
MG hydration shell issue: water too close (< 2.4 Angs.)  

### MG:1H:3462

|  | Res. type | Atom | Residue | Distance | Occupancy | B-factor | Closest solute to 1^st^ shell water |
| --- | --- | --- | --- | --- | --- | --- | --- |
| **Ion** | | | |  | 1.00 | 27.03 || 1 | Water 1 | | 1H:3619 | 2.17 | 1.00 | 28.85 | 1.93 (U:OP1 1H:1687) |
| 2 | Water 2 | | 1H:3858 | 2.17 | 1.00 | 29.51 | 2.46 (G:O2' 1H:2722) |
| 3 | Water 3 | | 1H:4440 | 2.18 | 1.00 | 34.47 | 3.45 (U:OP1 1H:1687) |
| 4 | Water 4 | | 1H:3680 | 2.19 | 1.00 | 31.97 | 2.15 (U:OP1 1H:1687) |
| 5 | Water 5 | | 1H:4419 | 2.19 | 1.00 | 28.39 | 3.36 (G:OP2 1H:1343) |
| 6 | Water 6 | | 1H:4437 | 2.20 | 1.00 | 39.36 | 3.43 (G:O2' 1H:2722) |
| 7 | U | OP1 | 1H:1687 | 2.58 | 1.00 | 28.44 |  |
| 8 | G | O2' | 1H:2722 | 3.98 | 1.00 | 28.27 |  |
| 9 | U | O5' | 1H:1687 | 4.34 | 1.00 | 27.80 |  |
| 10 | C | OP2 | 1H:1344 | 4.37 | 1.00 | 30.90 |  |
| 11 | G | N3 | 1H:2722 | 4.38 | 1.00 | 30.39 |  |
| 12 | G | OP2 | 1H:1343 | 4.62 | 1.00 | 28.77 |  |
| 13 | C | O3' | 1H:1686 | 4.82 | 1.00 | 27.45 |  |
| 14 | U | OP2 | 1H:1687 | 4.83 | 1.00 | 29.43 |  |
| 15 | Water 7 | | 1H:3803 | 4.84 | 1.00 | 23.58 | 2.38 (C:OP2 1H:1344) |
| 16 | G | OP1 | 1H:1343 | 4.85 | 1.00 | 28.21 |  |
| 17 | G | N2 | 1H:2722 | 4.89 | 1.00 | 28.77 |  |
  |

  
**1^st^ shell ligand angles:**   
1-2: 86; 1-3: 88; 1-4: 99; 1-5: 173; 1-6: 83; 2-3: 172; 2-4: 93; 2-5: 88; 2-6: 92; 3-4: 83; 3-5: 98; 3-6: 92; 4-5: 82; 4-6: 175; 5-6: 96;   
  
 Contact(s): /O_w_/O_w_/O_w_/O_w_/O_w_/O_w_/  
 Coded contact list: 6O_w_  
 Exclusion zone contact(s): /OP1/  
Consider NA in place of MG (1 valid contact(s) for NA vs. 0 for MG)  
MG hydration shell issue: water too close (< 2.4 Angs.)  

### MG:1H:3463

|  | Res. type | Atom | Residue | Distance | Occupancy | B-factor | Closest solute to 1^st^ shell water |
| --- | --- | --- | --- | --- | --- | --- | --- |
| **Ion** | | | |  | 1.00 | 17.66 || 1 | Water 1 | | 11:403 | 2.14 | 1.00 | 16.55 | 2.63 (PRO:O 11:228) |
| 2 | Water 2 | | 11:405 | 2.17 | 1.00 | 31.00 | 2.71 (ARG:NH1 11:244) |
| 3 | Water 3 | | 1H:3846 | 2.17 | 1.00 | 20.49 | 2.44 (U:OP1 1H:2097) |
| 4 | Water 4 | | 1H:3862 | 2.17 | 1.00 | 17.38 | 2.46 (G:O3' 1H:2251) |
| 5 | Water 5 | | 1H:4369 | 2.18 | 1.00 | 21.95 | 3.23 (G:OP2 1H:2252) |
| 6 | U | OP1 | 1H:2097 | 2.37 | 1.00 | 14.34 |  |
| 7 | G | OP2 | 1H:2252 | 3.71 | 1.00 | 18.23 |  |
| 8 | PRO | O | 11:228 | 3.78 | 1.00 | 14.28 |  |
| 9 | C | OP2 | 1H:2253 | 4.23 | 1.00 | 18.30 |  |
| 10 | G | O3' | 1H:2251 | 4.25 | 1.00 | 19.49 |  |
| 11 | U | OP2 | 1H:2098 | 4.32 | 1.00 | 17.95 |  |
| 12 | U | OP2 | 1H:2097 | 4.42 | 1.00 | 17.18 |  |
| 13 | U | O5' | 1H:2097 | 4.49 | 1.00 | 19.09 |  |
| 14 | G | O2' | 1H:2251 | 4.59 | 1.00 | 18.86 |  |
| 15 | C | O3' | 1H:2096 | 4.73 | 1.00 | 9.05 |  |
| 16 | HIS | O | 11:231 | 4.76 | 1.00 | 18.56 |  |
| 17 | ARG | NH1 | 11:244 | 4.81 | 1.00 | 19.04 |  |
| 18 | PRO | N | 11:232 | 4.82 | 1.00 | 19.47 |  |
  |

  
**1^st^ shell ligand angles:**   
1-2: 87; 1-3: 89; 1-4: 178; 1-5: 91; 1-6: 74; 2-3: 175; 2-4: 93; 2-5: 90; 2-6: 111; 3-4: 91; 3-5: 94; 3-6: 65; 4-5: 86; 4-6: 108; 5-6: 154;   
  
 Contact(s): /O_w_/O_w_/O_w_/O_w_/O_w_/OP1/  
 Coded contact list: O_ph_5O_w_  

### MG:1H:3464

|  | Res. type | Atom | Residue | Distance | Occupancy | B-factor | Closest solute to 1^st^ shell water |
| --- | --- | --- | --- | --- | --- | --- | --- |
| **Ion** | | | |  | 1.00 | 23.36 || 1 | Water 1 | | 1H:3646 | 2.13 | 1.00 | 21.94 | 2.05 (C:OP2 1H:779) |
| 2 | Water 2 | | 1H:3664 | 2.18 | 1.00 | 15.47 | 2.10 (A:OP1 1H:809) |
| 3 | Water 3 | | 1H:3618 | 2.19 | 1.00 | 24.38 | 1.93 (G:OP1 1H:811) |
| 4 | A | OP1 | 1H:809 | 2.48 | 1.00 | 19.45 |  |
| 5 | C | OP2 | 1H:778 | 2.64 | 1.00 | 19.91 |  |
| 6 | C | OP2 | 1H:779 | 3.10 | 1.00 | 23.29 |  |
| 7 | G | OP1 | 1H:811 | 3.37 | 1.00 | 17.20 |  |
| 8 | G | O3' | 1H:777 | 3.64 | 1.00 | 18.11 |  |
| 9 | A | O5' | 1H:809 | 3.67 | 1.00 | 16.67 |  |
| 10 | U | OP1 | 1H:810 | 4.05 | 1.00 | 19.65 |  |
| 11 | G | O5' | 1H:811 | 4.23 | 1.00 | 20.41 |  |
| 12 | Water 4 | | 1H:3779 | 4.35 | 1.00 | 24.88 | 2.34 (A:OP1 1H:809) |
| 13 | G | O3' | 1H:808 | 4.43 | 1.00 | 19.17 |  |
| 14 | C | O5' | 1H:778 | 4.58 | 1.00 | 19.16 |  |
| 15 | A | OP2 | 1H:809 | 4.74 | 1.00 | 19.14 |  |
| 16 | C | OP1 | 1H:778 | 4.84 | 1.00 | 21.88 |  |
| 17 | C | O3' | 1H:778 | 4.92 | 1.00 | 23.08 |  |
  |

  
**1^st^ shell ligand angles:**   
1-2: 75; 1-3: 79; 2-3: 95;   
  
 Contact(s): /O_w_/O_w_/O_w_/  
 Coded contact list: 3O_w_  
 Exclusion zone contact(s): /OP1/OP1/OP2/OP2/  
Consider K in place of MG (3 valid contact(s) for K vs. 0 for MG)  
MG hydration shell issue: water too close (< 2.4 Angs.)  

### MG:1H:3465

|  | Res. type | Atom | Residue | Distance | Occupancy | B-factor | Closest solute to 1^st^ shell water |
| --- | --- | --- | --- | --- | --- | --- | --- |
| **Ion** | | | |  | 1.00 | 21.32 || 1 | Water 1 | | 1H:4006 | 2.17 | 1.00 | 23.53 | 2.65 (C:N3 1H:2078) |
| 2 | Water 2 | | 1H:3793 | 2.17 | 1.00 | 20.21 | 2.37 (C:O2 1H:2078) |
| 3 | Water 3 | | 1H:4100 | 2.18 | 1.00 | 19.32 | 2.76 (G:O6 1H:2055) |
| 4 | Water 4 | | 1H:4107 | 2.18 | 1.00 | 24.39 | 2.77 (G:OP1 1H:2055) |
| 5 | Water 5 | | 1H:3921 | 2.19 | 1.00 | 18.23 | 2.54 (C:N3 1H:2512) |
| 6 | G | N7 | 1H:2055 | 2.51 | 1.00 | 22.26 |  |
| 7 | C | N3 | 1H:2078 | 3.84 | 1.00 | 21.15 |  |
| 8 | C | O2 | 1H:2078 | 3.95 | 1.00 | 23.58 |  |
| 9 | C | N3 | 1H:2512 | 4.10 | 1.00 | 19.68 |  |
| 10 | G | OP1 | 1H:2055 | 4.15 | 1.00 | 28.86 |  |
| 11 | C | O2 | 1H:2512 | 4.23 | 1.00 | 19.07 |  |
| 12 | G | N9 | 1H:2055 | 4.35 | 1.00 | 24.31 |  |
| 13 | G | O6 | 1H:2055 | 4.38 | 1.00 | 23.59 |  |
| 14 | A | N1 | 1H:2585 | 4.38 | 1.00 | 22.22 |  |
| 15 | A | N3 | 1H:2466 | 4.42 | 1.00 | 18.88 |  |
| 16 | U | O2 | 1H:2513 | 4.50 | 1.00 | 18.12 |  |
| 17 | G | O2' | 1H:2467 | 4.66 | 1.00 | 24.04 |  |
| 18 | G | N3 | 1H:2467 | 4.82 | 1.00 | 17.41 |  |
| 19 | A | N1 | 1H:2466 | 4.93 | 1.00 | 18.61 |  |
  |

  
**1^st^ shell ligand angles:**   
1-2: 93; 1-3: 176; 1-4: 90; 1-5: 88; 2-3: 87; 2-4: 177; 2-5: 94; 3-4: 90; 3-5: 96; 4-5: 86;   
  
 Contact(s): /O_w_/O_w_/O_w_/O_w_/O_w_/  
 Coded contact list: 5O_w_  
 Exclusion zone contact(s): /N7/  
Consider NA in place of MG (1 valid contact(s) for NA vs. 0 for MG)  
MG hydration shell issue: water too close (< 2.4 Angs.)  

### MG:1H:3466

|  | Res. type | Atom | Residue | Distance | Occupancy | B-factor | Closest solute to 1^st^ shell water |
| --- | --- | --- | --- | --- | --- | --- | --- |
| **Ion** | | | |  | 1.00 | 22.91 || 1 | Water 1 | | 1H:3888 | 2.14 | 1.00 | 21.15 | 2.50 (C:OP1 1H:472) |
| 2 | Water 2 | | 1H:3622 | 2.16 | 1.00 | 21.60 | 1.95 (G:OP2 1H:473) |
| 3 | Water 3 | | 1H:3745 | 2.17 | 1.00 | 25.10 | 2.29 (G:OP2 1H:473) |
| 4 | Water 4 | | 1H:3709 | 2.18 | 1.00 | 26.43 | 2.22 (U:O3' 1H:1245) |
| 5 | Water 5 | | 1H:4303 | 2.18 | 1.00 | 33.40 | 3.07 (C:OP2 1H:472) |
| 6 | Water 6 | | 1H:4548 | 2.20 | 1.00 | 33.87 | 3.98 (G:OP2 1H:473) |
| 7 | G | OP2 | 1H:473 | 2.61 | 1.00 | 24.41 |  |
| 8 | C | OP1 | 1H:472 | 4.04 | 1.00 | 20.24 |  |
| 9 | U | O3' | 1H:1245 | 4.22 | 1.00 | 22.53 |  |
| 10 | C | OP2 | 1H:472 | 4.29 | 1.00 | 26.13 |  |
| 11 | G | O5' | 1H:473 | 4.47 | 1.00 | 22.92 |  |
| 12 | U | O2' | 1H:1245 | 4.62 | 1.00 | 20.17 |  |
| 13 | G | OP1 | 1H:473 | 4.65 | 1.00 | 23.47 |  |
| 14 | G | N7 | 1H:473 | 4.66 | 1.00 | 27.90 |  |
| 15 | C | OP1 | 1H:1246 | 4.67 | 1.00 | 21.55 |  |
| 16 | C | O5' | 1H:472 | 4.89 | 1.00 | 18.59 |  |
  |

  
**1^st^ shell ligand angles:**   
1-2: 87; 1-3: 95; 1-4: 87; 1-5: 88; 1-6: 176; 2-3: 94; 2-4: 174; 2-5: 82; 2-6: 94; 3-4: 89; 3-5: 175; 3-6: 82; 4-5: 95; 4-6: 92; 5-6: 96;   
  
 Contact(s): /O_w_/O_w_/O_w_/O_w_/O_w_/O_w_/  
 Coded contact list: 6O_w_  
 Exclusion zone contact(s): /OP2/  
Consider K in place of MG (1 valid contact(s) for K vs. 0 for MG)  
MG hydration shell issue: water too close (< 2.4 Angs.)  

### MG:1H:3467

|  | Res. type | Atom | Residue | Distance | Occupancy | B-factor | Closest solute to 1^st^ shell water |
| --- | --- | --- | --- | --- | --- | --- | --- |
| **Ion** | | | |  | 1.00 | 44.20 || 1 | Water 1 | | 1H:3679 | 2.16 | 1.00 | 51.56 | 2.15 (G:O6 1H:80) |
| 2 | Water 2 | | 1H:4197 | 2.18 | 1.00 | 52.19 | 2.89 (G:O6 1H:81) |
| 3 | Water 3 | | 1H:4306 | 2.18 | 1.00 | 56.11 | 3.07 (A:OP1 1H:101) |
| 4 | Water 4 | | 1H:4557 | 2.19 | 1.00 | 48.16 | 4.99 (G:N7 1H:79) |
| 5 | Water 5 | | 1H:4334 | 2.19 | 1.00 | 48.26 | 3.14 (G:N7 1H:80) |
| 6 | Water 6 | | 1H:4279 | 2.20 | 1.00 | 48.73 | 3.01 (G:O6 1H:79) |
| 7 | G | O6 | 1H:81 | 4.02 | 1.00 | 49.44 |  |
| 8 | G | N7 | 1H:80 | 4.27 | 1.00 | 41.34 |  |
| 9 | G | O6 | 1H:80 | 4.28 | 1.00 | 44.24 |  |
| 10 | A | OP1 | 1H:101 | 4.34 | 1.00 | 35.98 |  |
| 11 | G | O6 | 1H:79 | 4.48 | 1.00 | 41.12 |  |
| 12 | G | N7 | 1H:79 | 4.55 | 1.00 | 41.00 |  |
| 13 | G | N1 | 1H:81 | 4.87 | 1.00 | 40.52 |  |
  |

  
**1^st^ shell ligand angles:**   
1-2: 87; 1-3: 92; 1-4: 177; 1-5: 89; 1-6: 87; 2-3: 87; 2-4: 94; 2-5: 90; 2-6: 173; 3-4: 91; 3-5: 177; 3-6: 88; 4-5: 89; 4-6: 92; 5-6: 94;   
  
 Contact(s): /O_w_/O_w_/O_w_/O_w_/O_w_/O_w_/  
 Coded contact list: 6O_w_  
MG hydration shell issue: water too close (< 2.4 Angs.)  

### MG:1H:3468

|  | Res. type | Atom | Residue | Distance | Occupancy | B-factor | Closest solute to 1^st^ shell water |
| --- | --- | --- | --- | --- | --- | --- | --- |
| **Ion** | | | |  | 1.00 | 24.42 || 1 | Water 1 | | 1H:3688 | 2.16 | 1.00 | 29.29 | 2.17 (G:OP1 1H:2058) |
| 2 | Water 2 | | 1H:3912 | 2.17 | 1.00 | 27.99 | 2.53 (G:O6 1H:2060) |
| 3 | Water 3 | | 1H:3617 | 2.17 | 1.00 | 28.69 | 1.91 (G:O6 1H:2047) |
| 4 | Water 4 | | 1H:4288 | 2.17 | 1.00 | 22.52 | 3.04 (G:OP1 1H:2058) |
| 5 | Water 5 | | 1H:4562 | 2.19 | 1.00 | 32.94 | 4.64 (G:O5' 1H:2058) |
| 6 | Water 6 | | 1H:4122 | 2.20 | 1.00 | 30.63 | 2.79 (G:O6 1H:2061) |
| 7 | G | OP1 | 1H:2058 | 3.30 | 1.00 | 22.58 |  |
| 8 | G | O6 | 1H:2061 | 3.61 | 1.00 | 26.73 |  |
| 9 | G | O6 | 1H:2047 | 3.96 | 1.00 | 26.51 |  |
| 10 | G | O6 | 1H:2060 | 4.15 | 1.00 | 22.96 |  |
| 11 | C | N4 | 1H:2048 | 4.21 | 1.00 | 23.92 |  |
| 12 | G | O5' | 1H:2058 | 4.32 | 1.00 | 24.72 |  |
| 13 | G | N7 | 1H:2060 | 4.42 | 1.00 | 24.39 |  |
| 14 | C | N4 | 1H:2062 | 4.59 | 1.00 | 28.72 |  |
| 15 | C | N4 | 1H:2059 | 4.66 | 1.00 | 22.89 |  |
| 16 | U | O3' | 1H:2057 | 4.66 | 1.00 | 21.68 |  |
  |

  
**1^st^ shell ligand angles:**   
1-2: 87; 1-3: 89; 1-4: 94; 1-5: 89; 1-6: 175; 2-3: 90; 2-4: 87; 2-5: 176; 2-6: 95; 3-4: 176; 3-5: 91; 3-6: 87; 4-5: 92; 4-6: 91; 5-6: 89;   
  
 Contact(s): /O_w_/O_w_/O_w_/O_w_/O_w_/O_w_/  
 Coded contact list: 6O_w_  
 Exclusion zone contact(s): /OP1/  
Consider K in place of MG (1 valid contact(s) for K vs. 0 for MG)  
MG hydration shell issue: water too close (< 2.4 Angs.)  

### MG:1H:3469

|  | Res. type | Atom | Residue | Distance | Occupancy | B-factor | Closest solute to 1^st^ shell water |
| --- | --- | --- | --- | --- | --- | --- | --- |
| **Ion** | | | |  | 1.00 | 13.47 || 1 | Water 1 | | 1H:3724 | 2.13 | 1.00 | 21.84 | 2.25 (G:OP1 1H:2076) |
| 2 | Water 2 | | 1H:3997 | 2.16 | 1.00 | 24.15 | 2.64 (G:OP1 1H:2589) |
| 3 | Water 3 | | 1H:3805 | 2.16 | 1.00 | 25.18 | 2.38 (G:OP1 1H:2589) |
| 4 | Water 4 | | 1H:3850 | 2.16 | 1.00 | 19.10 | 2.45 (G:OP2 1H:2075) |
| 5 | Water 5 | | 1H:3639 | 2.16 | 1.00 | 29.59 | 2.02 (A:OP1 1H:2590) |
| 6 | A | OP1 | 1H:2590 | 2.45 | 1.00 | 20.08 |  |
| 7 | G | OP1 | 1H:2589 | 2.46 | 1.00 | 20.23 |  |
| 8 | C | O3' | 1H:2588 | 3.32 | 1.00 | 15.09 |  |
| 9 | G | OP2 | 1H:2075 | 3.99 | 1.00 | 14.15 |  |
| 10 | A | OP2 | 1H:2590 | 4.07 | 1.00 | 21.44 |  |
| 11 | G | OP1 | 1H:2076 | 4.09 | 1.00 | 24.21 |  |
| 12 | G | OP2 | 1H:2076 | 4.09 | 1.00 | 23.64 |  |
| 13 | G | OP2 | 1H:2589 | 4.41 | 1.00 | 19.26 |  |
| 14 | G | O5' | 1H:2589 | 4.45 | 1.00 | 22.33 |  |
| 15 | A | O5' | 1H:2590 | 4.64 | 1.00 | 20.02 |  |
| 16 | G | O3' | 1H:2589 | 4.66 | 1.00 | 21.23 |  |
| 17 | G | O5' | 1H:2075 | 4.71 | 1.00 | 17.97 |  |
| 18 | G | N2 | 1H:2589 | 4.79 | 1.00 | 27.60 |  |
| 19 | C | O2' | 1H:2588 | 5.00 | 1.00 | 17.92 |  |
  |

  
**1^st^ shell ligand angles:**   
1-2: 88; 1-3: 84; 1-4: 100; 1-5: 89; 2-3: 86; 2-4: 172; 2-5: 94; 3-4: 99; 3-5: 173; 4-5: 82;   
  
 Contact(s): /O_w_/O_w_/O_w_/O_w_/O_w_/  
 Coded contact list: 5O_w_  
 Exclusion zone contact(s): /O3'/OP1/OP1/  
Consider NA in place of MG (2 valid contact(s) for NA vs. 0 for MG)  
MG hydration shell issue: water too close (< 2.4 Angs.)  

### MG:1H:3470

|  | Res. type | Atom | Residue | Distance | Occupancy | B-factor | Closest solute to 1^st^ shell water |
| --- | --- | --- | --- | --- | --- | --- | --- |
| **Ion** | | | |  | 1.00 | 24.36 || 1 | Water 1 | | 1H:3867 | 2.16 | 1.00 | 22.55 | 2.47 (A:OP2 1H:1203) |
| 2 | Water 2 | | 1H:4266 | 2.17 | 1.00 | 32.76 | 3.00 (A:OP2 1H:1202) |
| 3 | Water 3 | | 1H:4088 | 2.17 | 1.00 | 38.93 | 2.75 (A:OP2 1H:1202) |
| 4 | Water 4 | | 1H:4434 | 2.18 | 1.00 | 49.06 | 3.42 (A:OP2 1H:1203) |
| 5 | Water 5 | | 1H:4516 | 2.19 | 1.00 | 44.92 | 3.78 (C:N4 1H:1045) |
| 6 | A | OP2 | 1H:1203 | 2.55 | 1.00 | 34.25 |  |
| 7 | A | OP1 | 1H:1203 | 3.86 | 1.00 | 31.14 |  |
| 8 | A | OP2 | 1H:1202 | 4.04 | 1.00 | 42.41 |  |
| 9 | A | O5' | 1H:1202 | 4.35 | 1.00 | 40.02 |  |
| 10 | A | O3' | 1H:1202 | 4.39 | 1.00 | 35.88 |  |
| 11 | G | O6 | 1H:1204 | 4.60 | 1.00 | 34.50 |  |
| 12 | C | N4 | 1H:1045 | 4.77 | 1.00 | 38.77 |  |
| 13 | A | O5' | 1H:1203 | 4.84 | 1.00 | 31.06 |  |
  |

  
**1^st^ shell ligand angles:**   
1-2: 177; 1-3: 94; 1-4: 87; 1-5: 88; 2-3: 86; 2-4: 92; 2-5: 95; 3-4: 178; 3-5: 89; 4-5: 92;   
  
 Contact(s): /O_w_/O_w_/O_w_/O_w_/O_w_/  
 Coded contact list: 5O_w_  
 Exclusion zone contact(s): /OP2/  
Consider NA in place of MG (1 valid contact(s) for NA vs. 0 for MG)  

### MG:1H:3471

|  | Res. type | Atom | Residue | Distance | Occupancy | B-factor | Closest solute to 1^st^ shell water |
| --- | --- | --- | --- | --- | --- | --- | --- |
| **Ion** | | | |  | 1.00 | 56.16 || 1 | Water 1 | | 1H:3838 | 2.18 | 1.00 | 45.34 | 2.43 (G:OP2 1H:1255) |
| 2 | Water 2 | | 1H:3959 | 2.18 | 1.00 | 54.94 | 2.59 (A:N1 1H:356) |
| 3 | Water 3 | | 1H:4344 | 2.19 | 1.00 | 42.11 | 3.16 (G:OP2 1H:1255) |
| 4 | Water 4 | | 1H:3891 | 2.19 | 1.00 | 52.48 | 2.51 (C:OP1 1H:338) |
| 5 | Water 5 | | 1H:4581 | 2.19 | 1.00 | 51.20 | 4.43 (G:O2' 1H:337) |
| 6 | Water 6 | | 1H:4167 | 2.20 | 1.00 | 57.09 | 2.85 (G:OP2 1H:1255) |
| 7 | G | OP2 | 1H:1255 | 3.43 | 1.00 | 37.42 |  |
| 8 | A | N1 | 1H:356 | 4.08 | 1.00 | 36.15 |  |
| 9 | G | O2' | 1H:337 | 4.45 | 1.00 | 39.93 |  |
| 10 | G | O3' | 1H:337 | 4.63 | 1.00 | 45.74 |  |
| 11 | C | OP1 | 1H:338 | 4.68 | 1.00 | 52.32 |  |
| 12 | G | O5' | 1H:1255 | 4.72 | 1.00 | 36.82 |  |
| 13 | A | OP2 | 1H:1256 | 4.78 | 1.00 | 37.19 |  |
  |

  
**1^st^ shell ligand angles:**   
1-2: 88; 1-3: 92; 1-4: 180; 1-5: 90; 1-6: 89; 2-3: 92; 2-4: 92; 2-5: 87; 2-6: 177; 3-4: 88; 3-5: 177; 3-6: 88; 4-5: 90; 4-6: 91; 5-6: 94;   
  
 Contact(s): /O_w_/O_w_/O_w_/O_w_/O_w_/O_w_/  
 Coded contact list: 6O_w_  

### MG:1H:3472

|  | Res. type | Atom | Residue | Distance | Occupancy | B-factor | Closest solute to 1^st^ shell water |
| --- | --- | --- | --- | --- | --- | --- | --- |
| **Ion** | | | |  | 1.00 | 17.01 || 1 | Water 1 | | 1H:4569 | 2.17 | 1.00 | 27.42 | 4.26 (C:OP1 1H:2511) |
| 2 | Water 2 | | 1H:4186 | 2.17 | 1.00 | 22.15 | 2.98 (U:OP1 1H:1009) |
| 3 | Water 3 | | 1H:3992 | 2.17 | 1.00 | 21.96 | 2.63 (C:OP1 1H:2511) |
| 4 | Water 4 | | 1H:4301 | 2.18 | 1.00 | 35.77 | 3.06 (C:OP2 1H:1010) |
| 5 | Water 5 | | 1H:4292 | 2.19 | 1.00 | 20.68 | 3.05 (G:OP1 1H:993) |
| 6 | U | OP1 | 1H:1009 | 2.50 | 1.00 | 20.46 |  |
| 7 | Water 6 | | 1H:3743 | 2.95 | 1.00 | 34.05 | 2.28 (U:OP1 1H:1009) |
| 8 | Water 7 | | 1H:4226 | 3.87 | 1.00 | 31.26 | 2.92 (U:OP1 1H:1009) |
| 9 | U | O5' | 1H:1009 | 4.13 | 1.00 | 20.03 |  |
| 10 | C | OP1 | 1H:2511 | 4.25 | 1.00 | 18.77 |  |
| 11 | G | OP1 | 1H:993 | 4.29 | 1.00 | 17.28 |  |
| 12 | G | O3' | 1H:1008 | 4.82 | 1.00 | 22.28 |  |
| 13 | C | OP2 | 1H:1010 | 4.92 | 1.00 | 27.85 |  |
| 14 | U | OP2 | 1H:1009 | 4.98 | 1.00 | 22.04 |  |
  |

  
**1^st^ shell ligand angles:**   
1-2: 84; 1-3: 105; 1-4: 86; 1-5: 87; 2-3: 88; 2-4: 90; 2-5: 170; 3-4: 169; 3-5: 89; 4-5: 95;   
  
 Contact(s): /O_w_/O_w_/O_w_/O_w_/O_w_/  
 Coded contact list: 5O_w_  
 Exclusion zone contact(s): /O_w_/OP1/  
Consider NA in place of MG (1 valid contact(s) for NA vs. 0 for MG)  

### MG:1H:3473

|  | Res. type | Atom | Residue | Distance | Occupancy | B-factor | Closest solute to 1^st^ shell water |
| --- | --- | --- | --- | --- | --- | --- | --- |
| **Ion** | | | |  | 1.00 | 19.40 || 1 | Water 1 | | 1H:3802 | 2.16 | 1.00 | 14.90 | 2.38 (A:OP2 1H:1019) |
| 2 | Water 2 | | 1H:4220 | 2.17 | 1.00 | 23.15 | 2.92 (U:O4 1H:592) |
| 3 | Water 3 | | D8:201 | 2.20 | 1.00 | 29.33 | 2.32 (LYS:O D8:78) |
| 4 | Water 4 | | 1H:4450 | 2.21 | 1.00 | 36.61 | 3.51 (A:O5' 1H:1019) |
| 5 | A | OP2 | 1H:1019 | 2.53 | 1.00 | 20.65 |  |
| 6 | U | O4 | 1H:592 | 2.56 | 1.00 | 21.19 |  |
| 7 | A | O5' | 1H:1019 | 3.42 | 1.00 | 21.26 |  |
| 8 | G | O3' | 1H:1018 | 4.07 | 1.00 | 20.25 |  |
| 9 | U | N3 | 1H:592 | 4.29 | 1.00 | 22.70 |  |
| 10 | LYS | O | D8:78 | 4.37 | 1.00 | 25.01 |  |
| 11 | Water 5 | | 1H:3887 | 4.47 | 1.00 | 16.73 | 2.50 (A:OP2 1H:1019) |
| 12 | A | OP2 | 1H:595 | 4.66 | 1.00 | 22.27 |  |
| 13 | A | OP1 | 1H:595 | 4.68 | 1.00 | 29.76 |  |
| 14 | A | OP1 | 1H:1019 | 4.84 | 1.00 | 19.51 |  |
  |

  
**1^st^ shell ligand angles:**   
1-2: 89; 1-3: 176; 1-4: 96; 2-3: 94; 2-4: 87; 3-4: 88;   
  
 Contact(s): /O_w_/O_w_/O_w_/O_w_/  
 Coded contact list: 4O_w_  
 Exclusion zone contact(s): /O4/OP2/  
Consider NA in place of MG (2 valid contact(s) for NA vs. 0 for MG)  
MG hydration shell issue: water too close (< 2.4 Angs.)  

### MG:1H:3474

|  | Res. type | Atom | Residue | Distance | Occupancy | B-factor | Closest solute to 1^st^ shell water |
| --- | --- | --- | --- | --- | --- | --- | --- |
| **Ion** | | | |  | 1.00 | 24.96 || 1 | Water 1 | | 1H:3604 | 2.18 | 1.00 | 27.13 | 1.30 (G:N7 1H:1236) |
| 2 | Water 2 | | 1H:3851 | 2.18 | 1.00 | 18.59 | 2.45 (A:OP2 1H:1235) |
| 3 | Water 3 | | 1H:4375 | 2.18 | 1.00 | 24.96 | 3.24 (A:OP2 1H:1235) |
| 4 | Water 4 | | 1H:4250 | 2.18 | 1.00 | 37.34 | 2.97 (G:OP2 1H:1236) |
| 5 | Water 5 | | 1H:4467 | 2.21 | 1.00 | 32.88 | 3.57 (G:O6 1H:1236) |
| 6 | A | OP2 | 1H:1235 | 2.35 | 1.00 | 18.78 |  |
| 7 | G | N7 | 1H:1236 | 3.06 | 1.00 | 25.65 |  |
| 8 | A | OP1 | 1H:1235 | 3.65 | 1.00 | 28.58 |  |
| 9 | G | OP2 | 1H:1236 | 3.81 | 1.00 | 25.11 |  |
| 10 | G | O6 | 1H:1236 | 3.95 | 1.00 | 20.88 |  |
| 11 | GLY | O | 78:26 | 4.19 | 1.00 | 37.70 |  |
| 12 | A | O5' | 1H:1235 | 4.35 | 1.00 | 22.34 |  |
| 13 | U | O3' | 1H:1234 | 4.57 | 1.00 | 22.89 |  |
  |

  
**1^st^ shell ligand angles:**   
1-2: 90; 1-3: 168; 1-4: 88; 1-5: 94; 1-6: 78; 2-3: 90; 2-4: 177; 2-5: 92; 2-6: 65; 3-4: 92; 3-5: 98; 3-6: 91; 4-5: 86; 4-6: 116; 5-6: 155;   
  
 Contact(s): /O_w_/O_w_/O_w_/O_w_/O_w_/OP2/  
 Coded contact list: O_ph_5O_w_  
 Exclusion zone contact(s): /N7/  
MG hydration shell issue: water too close (< 2.4 Angs.)  

### MG:1H:3475

|  | Res. type | Atom | Residue | Distance | Occupancy | B-factor | Closest solute to 1^st^ shell water |
| --- | --- | --- | --- | --- | --- | --- | --- |
| **Ion** | | | |  | 1.00 | 40.16 || 1 | Water 1 | | 1H:3826 | 2.18 | 1.00 | 40.86 | 2.41 (G:O6 1H:975) |
| 2 | Water 2 | | 1H:4045 | 2.18 | 1.00 | 37.36 | 2.69 (U:OP1 1H:895) |
| 3 | Water 3 | | 1H:4034 | 2.18 | 1.00 | 45.78 | 2.67 (G:O6 1H:975) |
| 4 | Water 4 | | 1H:4319 | 2.18 | 1.00 | 54.49 | 3.12 (A:OP2 1H:973) |
| 5 | Water 5 | | 1H:3728 | 2.19 | 1.00 | 58.12 | 2.26 (U:OP1 1H:895) |
| 6 | G | O6 | 1H:975 | 2.55 | 1.00 | 42.58 |  |
| 7 | U | OP2 | 1H:895 | 2.93 | 1.00 | 35.85 |  |
| 8 | U | OP1 | 1H:895 | 3.18 | 1.00 | 46.17 |  |
| 9 | U | O5' | 1H:895 | 4.10 | 1.00 | 50.17 |  |
| 10 | G | OP2 | 1H:974 | 4.24 | 1.00 | 50.71 |  |
| 11 | G | N7 | 1H:974 | 4.53 | 1.00 | 47.37 |  |
| 12 | G | N1 | 1H:975 | 4.59 | 1.00 | 37.49 |  |
| 13 | G | N7 | 1H:975 | 4.65 | 1.00 | 48.86 |  |
| 14 | A | N7 | 1H:973 | 4.67 | 1.00 | 51.64 |  |
| 15 | C | O3' | 1H:894 | 4.85 | 1.00 | 53.01 |  |
| 16 | G | OP2 | 1H:896 | 4.91 | 1.00 | 40.42 |  |
| 17 | A | OP2 | 1H:973 | 4.98 | 1.00 | 60.13 |  |
  |

  
**1^st^ shell ligand angles:**   
1-2: 176; 1-3: 91; 1-4: 93; 1-5: 85; 2-3: 87; 2-4: 91; 2-5: 96; 3-4: 86; 3-5: 172; 4-5: 87;   
  
 Contact(s): /O_w_/O_w_/O_w_/O_w_/O_w_/  
 Coded contact list: 5O_w_  
 Exclusion zone contact(s): /O6/OP1/OP2/  
Consider K in place of MG (2 valid contact(s) for K vs. 0 for MG)  
MG hydration shell issue: water too close (< 2.4 Angs.)  

### MG:1H:3476

|  | Res. type | Atom | Residue | Distance | Occupancy | B-factor | Closest solute to 1^st^ shell water |
| --- | --- | --- | --- | --- | --- | --- | --- |
| **Ion** | | | |  | 1.00 | 15.73 || 1 | Water 1 | | 1H:3853 | 2.15 | 1.00 | 26.47 | 2.45 (C:OP2 1H:478) |
| 2 | Water 2 | | 1H:3698 | 2.15 | 1.00 | 16.29 | 2.19 (G:OP2 1H:477) |
| 3 | Water 3 | | 1H:3699 | 2.16 | 1.00 | 13.40 | 2.19 (G:OP2 1H:477) |
| 4 | Water 4 | | 1H:3652 | 2.16 | 1.00 | 29.56 | 2.07 (G:OP2 1H:479) |
| 5 | Water 5 | | 31:401 | 2.17 | 1.00 | 20.56 | 2.24 (GLY:O 31:85) |
| 6 | Water 6 | | 1H:4180 | 2.19 | 1.00 | 22.26 | 2.87 (G:OP2 1H:479) |
| 7 | G | OP2 | 1H:477 | 2.70 | 1.00 | 18.18 |  |
| 8 | G | OP2 | 1H:479 | 3.46 | 1.00 | 22.87 |  |
| 9 | GLY | O | 31:86 | 4.05 | 1.00 | 15.94 |  |
| 10 | C | OP2 | 1H:478 | 4.23 | 1.00 | 24.81 |  |
| 11 | G | O5' | 1H:477 | 4.34 | 1.00 | 14.63 |  |
| 12 | GLY | O | 31:85 | 4.39 | 1.00 | 20.46 |  |
| 13 | A | OP1 | 1H:476 | 4.56 | 1.00 | 18.04 |  |
| 14 | A | OP2 | 1H:476 | 4.86 | 1.00 | 17.46 |  |
| 15 | G | OP1 | 1H:477 | 4.91 | 1.00 | 18.37 |  |
  |

  
**1^st^ shell ligand angles:**   
1-2: 84; 1-3: 96; 1-4: 88; 1-5: 174; 1-6: 90; 2-3: 90; 2-4: 89; 2-5: 96; 2-6: 174; 3-4: 176; 3-5: 90; 3-6: 92; 4-5: 87; 4-6: 90; 5-6: 90;   
  
 Contact(s): /O_w_/O_w_/O_w_/O_w_/O_w_/O_w_/  
 Coded contact list: 6O_w_  
 Exclusion zone contact(s): /OP2/  
Consider K in place of MG (1 valid contact(s) for K vs. 0 for MG)  
MG hydration shell issue: water too close (< 2.4 Angs.)  

### MG:1H:3477

|  | Res. type | Atom | Residue | Distance | Occupancy | B-factor | Closest solute to 1^st^ shell water |
| --- | --- | --- | --- | --- | --- | --- | --- |
| **Ion** | | | |  | 1.00 | 28.25 || 1 | Water 1 | | 1H:3660 | 2.16 | 1.00 | 14.36 | 2.09 (U:OP2 1H:2444) |
| 2 | Water 2 | | 1H:3926 | 2.19 | 1.00 | 32.55 | 2.55 (A:OP2 1H:2445) |
| 3 | Water 3 | | 1H:4448 | 2.19 | 1.00 | 42.47 | 4.01 (A:N1 1H:2406) |
| 4 | Water 4 | | 3K:101 | 2.20 | 1.00 | 35.75 | 3.18 (A:O2' 3K:76) |
| 5 | U | OP2 | 1H:2444 | 2.42 | 1.00 | 19.53 |  |
| 6 | A | OP2 | 1H:2445 | 2.63 | 1.00 | 22.72 |  |
| 7 | A | OP1 | 1H:2445 | 3.77 | 1.00 | 24.54 |  |
| 8 | Water 5 | | 1H:3782 | 4.28 | 1.00 | 28.14 | 2.35 (A:OP1 1H:2445) |
| 9 | Water 6 | | 1H:4148 | 4.35 | 1.00 | 34.39 | 2.83 (A:OP1 1H:2445) |
| 10 | Water 7 | | 1H:4449 | 4.40 | 1.00 | 25.01 | 3.51 (U:OP1 1H:2444) |
| 11 | A | N1 | 1H:2406 | 4.54 | 1.00 | 29.37 |  |
| 12 | A | O5' | 1H:2445 | 4.56 | 1.00 | 19.53 |  |
| 13 | A | O3' | 1H:2443 | 4.56 | 1.00 | 16.24 |  |
| 14 | U | OP1 | 1H:2444 | 4.64 | 1.00 | 18.14 |  |
| 15 | U | O5' | 1H:2444 | 4.80 | 1.00 | 18.20 |  |
| 16 | U | O3' | 1H:2444 | 4.80 | 1.00 | 20.65 |  |
| 17 | A | O2' | 3K:76 | 4.99 | 1.00 | 32.52 |  |
| 18 | A | OP2 | 1H:2446 | 5.00 | 1.00 | 22.85 |  |
  |

  
**1^st^ shell ligand angles:**   
1-2: 79; 1-3: 93; 1-4: 174; 2-3: 91; 2-4: 95; 3-4: 88;   
  
 Contact(s): /O_w_/O_w_/O_w_/O_w_/  
 Coded contact list: 4O_w_  
 Exclusion zone contact(s): /OP2/OP2/  
Consider NA in place of MG (1 valid contact(s) for NA vs. 0 for MG)  
MG hydration shell issue: water too close (< 2.4 Angs.)  

### MG:1H:3478

|  | Res. type | Atom | Residue | Distance | Occupancy | B-factor | Closest solute to 1^st^ shell water |
| --- | --- | --- | --- | --- | --- | --- | --- |
| **Ion** | | | |  | 1.00 | 44.77 || 1 | Water 1 | | 1H:4038 | 2.18 | 1.00 | 32.55 | 2.67 (G:OP2 1H:1653) |
| 2 | Water 2 | | 1H:3626 | 2.19 | 1.00 | 32.53 | 1.97 (C:OP1 1H:1654) |
| 3 | Water 3 | | 1H:4395 | 2.20 | 1.00 | 37.66 | 3.28 (C:OP2 1H:1351) |
| 4 | Water 4 | | 1H:3714 | 2.20 | 1.00 | 41.28 | 2.23 (G:OP1 1H:1653) |
| 5 | Water 5 | | 1H:3731 | 2.20 | 1.00 | 50.00 | 2.26 (C:OP2 1H:1351) |
| 6 | Water 6 | | 1H:4059 | 2.20 | 1.00 | 39.56 | 2.70 (A:OP1 1H:1655) |
| 7 | G | OP1 | 1H:1653 | 3.32 | 1.00 | 35.64 |  |
| 8 | C | OP2 | 1H:1351 | 3.66 | 1.00 | 33.28 |  |
| 9 | C | OP1 | 1H:1654 | 3.97 | 1.00 | 25.44 |  |
| 10 | G | OP2 | 1H:1350 | 4.13 | 1.00 | 30.35 |  |
| 11 | G | OP1 | 1H:1350 | 4.34 | 1.00 | 35.94 |  |
| 12 | A | OP1 | 1H:1655 | 4.55 | 1.00 | 34.75 |  |
| 13 | G | OP2 | 1H:1653 | 4.60 | 1.00 | 30.15 |  |
| 14 | G | O5' | 1H:1350 | 4.72 | 1.00 | 33.21 |  |
| 15 | G | O5' | 1H:1653 | 4.97 | 1.00 | 26.45 |  |
  |

  
**1^st^ shell ligand angles:**   
1-2: 88; 1-3: 90; 1-4: 90; 1-5: 90; 1-6: 177; 2-3: 178; 2-4: 92; 2-5: 89; 2-6: 89; 3-4: 89; 3-5: 91; 3-6: 92; 4-5: 179; 4-6: 90; 5-6: 90;   
  
 Contact(s): /O_w_/O_w_/O_w_/O_w_/O_w_/O_w_/  
 Coded contact list: 6O_w_  
 Exclusion zone contact(s): /OP1/  
Consider K in place of MG (1 valid contact(s) for K vs. 0 for MG)  
MG hydration shell issue: water too close (< 2.4 Angs.)  

### MG:1H:3479

|  | Res. type | Atom | Residue | Distance | Occupancy | B-factor | Closest solute to 1^st^ shell water |
| --- | --- | --- | --- | --- | --- | --- | --- |
| **Ion** | | | |  | 1.00 | 18.60 || 1 | Water 1 | | 1H:3689 | 2.16 | 1.00 | 19.70 | 2.17 (G:OP1 1H:2605) |
| 2 | Water 2 | | 1H:4233 | 2.18 | 1.00 | 26.06 | 2.94 (G:OP2 1H:1991) |
| 3 | Water 3 | | 1H:4187 | 2.20 | 1.00 | 23.58 | 2.88 (G:OP2 1H:2605) |
| 4 | Water 4 | | 1H:4190 | 2.21 | 1.00 | 30.59 | 2.88 (G:OP1 1H:1996) |
| 5 | G | OP1 | 1H:1991 | 2.29 | 1.00 | 24.94 |  |
| 6 | G | OP1 | 1H:2605 | 2.45 | 1.00 | 22.53 |  |
| 7 | G | OP2 | 1H:1991 | 3.63 | 1.00 | 25.10 |  |
| 8 | G | OP2 | 1H:2605 | 3.99 | 1.00 | 19.90 |  |
| 9 | C | O3' | 1H:1990 | 4.20 | 1.00 | 25.30 |  |
| 10 | C | O3' | 1H:2604 | 4.24 | 1.00 | 23.78 |  |
| 11 | G | O5' | 1H:1991 | 4.36 | 1.00 | 23.80 |  |
| 12 | G | OP1 | 1H:1996 | 4.64 | 1.00 | 23.88 |  |
| 13 | G | O5' | 1H:2605 | 4.72 | 1.00 | 25.84 |  |
  |

  
**1^st^ shell ligand angles:**   
1-2: 167; 1-3: 91; 1-4: 88; 1-5: 73; 2-3: 97; 2-4: 101; 2-5: 96; 3-4: 97; 3-5: 159; 4-5: 96;   
  
 Contact(s): /O_w_/O_w_/O_w_/O_w_/OP1/  
 Coded contact list: O_ph_4O_w_  
 Exclusion zone contact(s): /OP1/  
MG hydration shell issue: water too close (< 2.4 Angs.)  

### MG:1H:3480

|  | Res. type | Atom | Residue | Distance | Occupancy | B-factor | Closest solute to 1^st^ shell water |
| --- | --- | --- | --- | --- | --- | --- | --- |
| **Ion** | | | |  | 1.00 | 18.27 || 1 | Water 1 | | 1H:4185 | 2.16 | 1.00 | 37.35 | 2.88 (C:OP2 1H:1033) |
| 2 | Water 2 | | L8:202 | 2.16 | 1.00 | 38.02 | 2.52 (ILE:O L8:13) |
| 3 | Water 3 | | 1H:4203 | 2.16 | 1.00 | 27.09 | 2.90 (A:N7 1H:1035) |
| 4 | Water 4 | | L8:201 | 2.16 | 1.00 | 35.02 | 2.51 (ILE:O L8:13) |
| 5 | Water 5 | | 1H:4174 | 2.18 | 1.00 | 26.95 | 2.86 (G:OP2 1H:1034) |
| 6 | G | OP2 | 1H:1034 | 2.45 | 1.00 | 33.00 |  |
| 7 | ILE | O | L8:13 | 3.66 | 1.00 | 37.80 |  |
| 8 | G | O5' | 1H:1034 | 3.98 | 1.00 | 27.97 |  |
| 9 | C | OP2 | 1H:1033 | 4.12 | 1.00 | 32.76 |  |
| 10 | A | N7 | 1H:1035 | 4.58 | 1.00 | 32.88 |  |
| 11 | G | OP1 | 1H:1034 | 4.60 | 1.00 | 33.55 |  |
| 12 | A | OP2 | 1H:1035 | 4.66 | 1.00 | 32.09 |  |
| 13 | C | O3' | 1H:1033 | 4.76 | 1.00 | 32.54 |  |
| 14 | C | O5' | 1H:1033 | 4.80 | 1.00 | 31.03 |  |
| 15 | C | OP1 | 1H:1033 | 4.82 | 1.00 | 34.39 |  |
  |

  
**1^st^ shell ligand angles:**   
1-2: 88; 1-3: 174; 1-4: 89; 1-5: 88; 2-3: 95; 2-4: 84; 2-5: 173; 3-4: 86; 3-5: 89; 4-5: 102;   
  
 Contact(s): /O_w_/O_w_/O_w_/O_w_/O_w_/  
 Coded contact list: 5O_w_  
 Exclusion zone contact(s): /OP2/  
Consider NA in place of MG (1 valid contact(s) for NA vs. 0 for MG)  

### MG:1H:3481

|  | Res. type | Atom | Residue | Distance | Occupancy | B-factor | Closest solute to 1^st^ shell water |
| --- | --- | --- | --- | --- | --- | --- | --- |
| **Ion** | | | |  | 1.00 | 21.56 || 1 | Water 1 | | 1H:4255 | 2.15 | 1.00 | 24.41 | 2.98 (U:OP2 1H:2018) |
| 2 | Water 2 | | 21:404 | 2.16 | 1.00 | 24.36 | 2.53 (GLY:O 21:125) |
| 3 | Water 3 | | 1H:3949 | 2.16 | 1.00 | 24.72 | 2.59 (C:O3' 1H:2019) |
| 4 | Water 4 | | 21:407 | 2.17 | 1.00 | 35.52 | 2.97 (ASP:OD2 21:127) |
| 5 | Water 5 | | 1H:3863 | 2.18 | 1.00 | 25.09 | 2.46 (C:OP1 1H:2017) |
| 6 | G | OP2 | 1H:2020 | 2.36 | 1.00 | 22.31 |  |
| 7 | C | O3' | 1H:2019 | 3.55 | 1.00 | 23.95 |  |
| 8 | G | OP1 | 1H:2020 | 3.94 | 1.00 | 26.75 |  |
| 9 | ASP | OD2 | 21:127 | 3.99 | 1.00 | 30.03 |  |
| 10 | ALA | O | 21:123 | 3.99 | 1.00 | 27.62 |  |
| 11 | GLY | O | 21:125 | 4.13 | 1.00 | 23.61 |  |
| 12 | U | OP2 | 1H:2018 | 4.29 | 1.00 | 17.86 |  |
| 13 | C | OP1 | 1H:2017 | 4.39 | 1.00 | 16.91 |  |
| 14 | G | O5' | 1H:2020 | 4.48 | 1.00 | 27.61 |  |
| 15 | Water 6 | | 1H:4013 | 4.73 | 1.00 | 31.71 | 3.54 (ASP:OD2 21:127) |
| 16 | C | OP2 | 1H:2017 | 4.91 | 1.00 | 21.61 |  |
  |

  
**1^st^ shell ligand angles:**   
1-2: 92; 1-3: 87; 1-4: 175; 1-5: 99; 1-6: 90; 2-3: 176; 2-4: 86; 2-5: 95; 2-6: 96; 3-4: 95; 3-5: 88; 3-6: 80; 4-5: 86; 4-6: 86; 5-6: 165;   
  
 Contact(s): /O_w_/O_w_/O_w_/O_w_/O_w_/OP2/  
 Coded contact list: O_ph_5O_w_  

### MG:1H:3482

|  | Res. type | Atom | Residue | Distance | Occupancy | B-factor | Closest solute to 1^st^ shell water |
| --- | --- | --- | --- | --- | --- | --- | --- |
| **Ion** | | | |  | 1.00 | 45.14 || 1 | Water 1 | | 1H:3708 | 2.16 | 1.00 | 42.18 | 2.22 (G:N7 1H:1429) |
| 2 | Water 2 | | 1H:4216 | 2.17 | 1.00 | 50.00 | 2.91 (G:OP1 1H:1394) |
| 3 | Water 3 | | 1H:3896 | 2.17 | 1.00 | 32.03 | 2.51 (G:N7 1H:1428) |
| 4 | Water 4 | | 1H:3678 | 2.18 | 1.00 | 38.79 | 2.15 (G:OP2 1H:1427) |
| 5 | Water 5 | | 1H:4525 | 2.18 | 1.00 | 44.41 | 3.82 (C:N4 1H:1430) |
| 6 | Water 6 | | 1H:4154 | 2.20 | 1.00 | 46.57 | 2.83 (G:OP2 1H:1428) |
| 7 | G | OP2 | 1H:1427 | 3.64 | 1.00 | 41.43 |  |
| 8 | G | N7 | 1H:1428 | 4.13 | 1.00 | 30.58 |  |
| 9 | G | N7 | 1H:1429 | 4.14 | 1.00 | 33.27 |  |
| 10 | G | OP2 | 1H:1428 | 4.18 | 1.00 | 30.05 |  |
| 11 | G | O6 | 1H:1429 | 4.43 | 1.00 | 49.93 |  |
| 12 | G | O6 | 1H:1428 | 4.51 | 1.00 | 36.09 |  |
| 13 | C | N4 | 1H:1430 | 4.59 | 1.00 | 40.58 |  |
| 14 | G | OP1 | 1H:1394 | 4.67 | 1.00 | 49.83 |  |
  |

  
**1^st^ shell ligand angles:**   
1-2: 91; 1-3: 87; 1-4: 175; 1-5: 95; 1-6: 85; 2-3: 83; 2-4: 92; 2-5: 93; 2-6: 174; 3-4: 96; 3-5: 175; 3-6: 93; 4-5: 81; 4-6: 93; 5-6: 92;   
  
 Contact(s): /O_w_/O_w_/O_w_/O_w_/O_w_/O_w_/  
 Coded contact list: 6O_w_  
MG hydration shell issue: water too close (< 2.4 Angs.)  

### MG:1H:3483

|  | Res. type | Atom | Residue | Distance | Occupancy | B-factor | Closest solute to 1^st^ shell water |
| --- | --- | --- | --- | --- | --- | --- | --- |
| **Ion** | | | |  | 1.00 | 36.88 || 1 | Water 1 | | 1H:4102 | 2.17 | 1.00 | 33.06 | 2.76 (G:N7 1H:2072) |
| 2 | Water 2 | | 1H:4318 | 2.19 | 1.00 | 31.37 | 3.12 (G:O6 1H:2072) |
| 3 | Water 3 | | 1H:4064 | 2.19 | 1.00 | 22.25 | 2.71 (G:N7 1H:2071) |
| 4 | Water 4 | | 1H:4317 | 2.19 | 1.00 | 39.90 | 4.39 (G:OP2 1H:2071) |
| 5 | Water 5 | | 1H:4379 | 2.19 | 1.00 | 27.03 | 3.25 (G:OP2 1H:2071) |
| 6 | Water 6 | | 1H:4579 | 2.20 | 1.00 | 28.86 | 4.64 (U:O4 1H:2070) |
| 7 | G | N7 | 1H:2072 | 3.98 | 1.00 | 17.16 |  |
| 8 | G | N7 | 1H:2071 | 4.08 | 1.00 | 18.12 |  |
| 9 | G | O6 | 1H:2072 | 4.41 | 1.00 | 21.16 |  |
| 10 | G | OP2 | 1H:2071 | 4.85 | 1.00 | 27.06 |  |
| 11 | U | O4 | 1H:2070 | 4.99 | 1.00 | 20.14 |  |
  |

  
**1^st^ shell ligand angles:**   
1-2: 92; 1-3: 87; 1-4: 87; 1-5: 90; 1-6: 174; 2-3: 89; 2-4: 96; 2-5: 178; 2-6: 83; 3-4: 173; 3-5: 90; 3-6: 97; 4-5: 86; 4-6: 89; 5-6: 95;   
  
 Contact(s): /O_w_/O_w_/O_w_/O_w_/O_w_/O_w_/  
 Coded contact list: 6O_w_  

### MG:1H:3484

|  | Res. type | Atom | Residue | Distance | Occupancy | B-factor | Closest solute to 1^st^ shell water |
| --- | --- | --- | --- | --- | --- | --- | --- |
| **Ion** | | | |  | 1.00 | 16.51 || 1 | Water 1 | | 1H:3984 | 2.17 | 1.00 | 27.95 | 2.63 (G:N7 1H:2050) |
| 2 | Water 2 | | 1H:4112 | 2.17 | 1.00 | 27.54 | 2.77 (A:OP1 1H:2056) |
| 3 | Water 3 | | 1H:3780 | 2.18 | 1.00 | 21.09 | 2.34 (A:OP1 1H:2056) |
| 4 | Water 4 | | 1H:4158 | 2.19 | 1.00 | 33.37 | 2.84 (G:O6 1H:2050) |
| 5 | U | O4 | 1H:2051 | 2.35 | 1.00 | 22.66 |  |
| 6 | A | OP1 | 1H:2056 | 2.54 | 1.00 | 26.21 |  |
| 7 | G | O6 | 1H:2050 | 3.45 | 1.00 | 28.45 |  |
| 8 | G | N7 | 1H:2050 | 3.77 | 1.00 | 22.86 |  |
| 9 | A | O5' | 1H:2056 | 3.86 | 1.00 | 22.87 |  |
| 10 | G | O6 | 1H:2052 | 4.18 | 1.00 | 23.63 |  |
| 11 | U | N3 | 1H:2051 | 4.43 | 1.00 | 21.58 |  |
| 12 | A | N7 | 1H:2056 | 4.52 | 1.00 | 23.52 |  |
| 13 | G | O3' | 1H:2055 | 4.74 | 1.00 | 30.03 |  |
| 14 | G | N1 | 1H:2052 | 4.87 | 1.00 | 24.50 |  |
| 15 | A | OP2 | 1H:2056 | 4.89 | 1.00 | 24.98 |  |
| 16 | A | N9 | 1H:2056 | 4.96 | 1.00 | 22.57 |  |
  |

  
**1^st^ shell ligand angles:**   
1-2: 85; 1-3: 90; 1-4: 94; 1-5: 106; 2-3: 84; 2-4: 94; 2-5: 144; 3-4: 175; 3-5: 63; 4-5: 118;   
  
 Contact(s): /O4/O_w_/O_w_/O_w_/O_w_/  
 Coded contact list: O_b_4O_w_  
 Exclusion zone contact(s): /OP1/  
Large octahedral topology discrepancy!!!  
MG hydration shell issue: water too close (< 2.4 Angs.)  

### MG:1H:3485

|  | Res. type | Atom | Residue | Distance | Occupancy | B-factor | Closest solute to 1^st^ shell water |
| --- | --- | --- | --- | --- | --- | --- | --- |
| **Ion** | | | |  | 1.00 | 29.83 || 1 | Water 1 | | 1H:4005 | 2.18 | 1.00 | 29.16 | 2.64 (C:OP2 1H:2013) |
| 2 | Water 2 | | 1H:4286 | 2.19 | 1.00 | 32.75 | 3.03 (G:OP2 1H:2012) |
| 3 | Water 3 | | 1H:4497 | 2.19 | 1.00 | 28.74 | 3.67 (G:O5' 1H:1715) |
| 4 | Water 4 | | 1H:4113 | 2.19 | 1.00 | 34.14 | 2.77 (G:O2' 1H:1715) |
| 5 | Water 5 | | 1H:4165 | 2.19 | 1.00 | 30.55 | 2.84 (G:OP2 1H:1715) |
| 6 | Water 6 | | 1H:3693 | 2.20 | 1.00 | 40.03 | 2.18 (C:OP2 1H:2013) |
| 7 | C | OP2 | 1H:2013 | 3.42 | 1.00 | 28.13 |  |
| 8 | G | OP2 | 1H:2012 | 4.06 | 1.00 | 25.82 |  |
| 9 | G | O2' | 1H:1715 | 4.27 | 1.00 | 23.25 |  |
| 10 | G | OP2 | 1H:1715 | 4.73 | 1.00 | 25.96 |  |
| 11 | G | O5' | 1H:1715 | 4.89 | 1.00 | 28.38 |  |
  |

  
**1^st^ shell ligand angles:**   
1-2: 84; 1-3: 96; 1-4: 91; 1-5: 177; 1-6: 87; 2-3: 92; 2-4: 174; 2-5: 93; 2-6: 92; 3-4: 92; 3-5: 82; 3-6: 174; 4-5: 92; 4-6: 84; 5-6: 94;   
  
 Contact(s): /O_w_/O_w_/O_w_/O_w_/O_w_/O_w_/  
 Coded contact list: 6O_w_  

### MG:1H:3486

|  | Res. type | Atom | Residue | Distance | Occupancy | B-factor | Closest solute to 1^st^ shell water |
| --- | --- | --- | --- | --- | --- | --- | --- |
| **Ion** | | | |  | 1.00 | 36.26 || 1 | Water 1 | | 1H:3903 | 2.17 | 1.00 | 47.55 | 2.52 (A:OP2 1H:2718) |
| 2 | Water 2 | | 1H:3681 | 2.17 | 1.00 | 34.54 | 2.16 (G:O6 1H:2719) |
| 3 | Water 3 | | 1H:3913 | 2.17 | 1.00 | 38.13 | 2.53 (C:OP1 1H:2717) |
| 4 | Water 4 | | 1H:3983 | 2.18 | 1.00 | 41.75 | 2.63 (G:N7 1H:2719) |
| 5 | Water 5 | | 1H:3765 | 2.19 | 1.00 | 39.76 | 2.32 (A:N7 1H:2718) |
| 6 | A | OP2 | 1H:2718 | 2.58 | 1.00 | 41.42 |  |
| 7 | G | O6 | 1H:2719 | 3.63 | 1.00 | 38.02 |  |
| 8 | A | O5' | 1H:2718 | 3.85 | 1.00 | 32.33 |  |
| 9 | G | N7 | 1H:2719 | 3.91 | 1.00 | 33.77 |  |
| 10 | A | N7 | 1H:2718 | 3.92 | 1.00 | 41.81 |  |
| 11 | C | O5' | 1H:2717 | 4.09 | 1.00 | 43.18 |  |
| 12 | C | OP1 | 1H:2717 | 4.27 | 1.00 | 41.67 |  |
| 13 | C | OP2 | 1H:2717 | 4.32 | 1.00 | 51.43 |  |
| 14 | C | O3' | 1H:2717 | 4.47 | 1.00 | 44.65 |  |
| 15 | A | OP1 | 1H:2718 | 4.90 | 1.00 | 41.12 |  |
  |

  
**1^st^ shell ligand angles:**   
1-2: 91; 1-3: 89; 1-4: 87; 1-5: 177; 2-3: 92; 2-4: 93; 2-5: 92; 3-4: 174; 3-5: 92; 4-5: 92;   
  
 Contact(s): /O_w_/O_w_/O_w_/O_w_/O_w_/  
 Coded contact list: 5O_w_  
 Exclusion zone contact(s): /OP2/  
Consider NA in place of MG (1 valid contact(s) for NA vs. 0 for MG)  
MG hydration shell issue: water too close (< 2.4 Angs.)  

### MG:1H:3487

|  | Res. type | Atom | Residue | Distance | Occupancy | B-factor | Closest solute to 1^st^ shell water |
| --- | --- | --- | --- | --- | --- | --- | --- |
| **Ion** | | | |  | 1.00 | 41.77 || 1 | Water 1 | | 1H:3657 | 2.16 | 1.00 | 39.91 | 2.09 (G:OP2 1H:2268) |
| 2 | Water 2 | | 1H:4050 | 2.18 | 1.00 | 31.77 | 2.69 (G:N7 1H:2269) |
| 3 | Water 3 | | 1H:3974 | 2.18 | 1.00 | 36.66 | 2.61 (C:OP2 1H:2267) |
| 4 | Water 4 | | 1H:4072 | 2.18 | 1.00 | 51.22 | 2.73 (G:OP2 1H:2268) |
| 5 | Water 5 | | 1H:4335 | 2.19 | 1.00 | 34.70 | 3.14 (G:N7 1H:2268) |
| 6 | Water 6 | | 1H:4026 | 2.19 | 1.00 | 36.28 | 2.66 (U:O4 1H:2270) |
| 7 | G | OP2 | 1H:2268 | 3.10 | 1.00 | 31.68 |  |
| 8 | C | OP2 | 1H:2267 | 3.50 | 1.00 | 27.84 |  |
| 9 | G | O6 | 1H:2269 | 4.09 | 1.00 | 24.34 |  |
| 10 | G | N7 | 1H:2269 | 4.13 | 1.00 | 29.78 |  |
| 11 | G | N7 | 1H:2268 | 4.20 | 1.00 | 30.23 |  |
| 12 | C | O5' | 1H:2267 | 4.53 | 1.00 | 24.53 |  |
| 13 | G | O5' | 1H:2268 | 4.68 | 1.00 | 32.22 |  |
| 14 | U | O4 | 1H:2270 | 4.84 | 1.00 | 28.12 |  |
| 15 | C | OP1 | 1H:2267 | 4.90 | 1.00 | 28.30 |  |
  |

  
**1^st^ shell ligand angles:**   
1-2: 91; 1-3: 92; 1-4: 92; 1-5: 82; 1-6: 176; 2-3: 176; 2-4: 88; 2-5: 96; 2-6: 85; 3-4: 90; 3-5: 87; 3-6: 92; 4-5: 173; 4-6: 89; 5-6: 97;   
  
 Contact(s): /O_w_/O_w_/O_w_/O_w_/O_w_/O_w_/  
 Coded contact list: 6O_w_  
 Exclusion zone contact(s): /OP2/  
Consider K in place of MG (1 valid contact(s) for K vs. 0 for MG)  
MG hydration shell issue: water too close (< 2.4 Angs.)  

### MG:1H:3488

|  | Res. type | Atom | Residue | Distance | Occupancy | B-factor | Closest solute to 1^st^ shell water |
| --- | --- | --- | --- | --- | --- | --- | --- |
| **Ion** | | | |  | 1.00 | 35.94 || 1 | Water 1 | | 1H:4051 | 2.17 | 1.00 | 23.40 | 2.69 (G:N7 1H:606) |
| 2 | Water 2 | | 1H:4368 | 2.17 | 1.00 | 24.52 | 3.22 (A:OP1 1H:1300) |
| 3 | Water 3 | | 1H:4206 | 2.17 | 1.00 | 18.47 | 2.90 (G:OP2 1H:606) |
| 4 | Water 4 | | 1H:4252 | 2.21 | 1.00 | 22.31 | 2.97 (C:N4 1H:608) |
| 5 | Water 5 | | 1H:3608 | 2.22 | 1.00 | 32.49 | 1.30 (G:N7 1H:607) |
| 6 | Water 6 | | 1H:4093 | 2.25 | 1.00 | 25.61 | 2.75 (G:O6 1H:607) |
| 7 | G | N7 | 1H:607 | 3.47 | 1.00 | 26.48 |  |
| 8 | A | OP1 | 1H:1300 | 4.00 | 1.00 | 26.45 |  |
| 9 | G | N7 | 1H:606 | 4.00 | 1.00 | 21.56 |  |
| 10 | G | O6 | 1H:607 | 4.03 | 1.00 | 22.46 |  |
| 11 | C | N4 | 1H:608 | 4.14 | 1.00 | 24.93 |  |
| 12 | G | OP2 | 1H:606 | 4.21 | 1.00 | 20.68 |  |
| 13 | ARG | NH1 | C8:10 | 4.75 | 1.00 | 22.29 |  |
| 14 | G | OP2 | 1H:607 | 4.86 | 1.00 | 19.42 |  |
| 15 | C | O2' | 1H:1298 | 4.97 | 1.00 | 22.23 |  |
| 16 | G | O6 | 1H:609 | 4.99 | 1.00 | 26.57 |  |
  |

  
**1^st^ shell ligand angles:**   
1-2: 88; 1-3: 87; 1-4: 177; 1-5: 92; 1-6: 89; 2-3: 82; 2-4: 89; 2-5: 174; 2-6: 97; 3-4: 93; 3-5: 92; 3-6: 176; 4-5: 91; 4-6: 91; 5-6: 89;   
  
 Contact(s): /O_w_/O_w_/O_w_/O_w_/O_w_/O_w_/  
 Coded contact list: 6O_w_  
MG hydration shell issue: water too close (< 2.4 Angs.)  

### MG:1H:3489

|  | Res. type | Atom | Residue | Distance | Occupancy | B-factor | Closest solute to 1^st^ shell water |
| --- | --- | --- | --- | --- | --- | --- | --- |
| **Ion** | | | |  | 1.00 | 57.14 || 1 | Water 1 | | 1H:3859 | 2.17 | 1.00 | 56.82 | 2.46 (G:O6 1H:901) |
| 2 | Water 2 | | 1H:4223 | 2.18 | 1.00 | 38.75 | 2.92 (G:O6 1H:900) |
| 3 | Water 3 | | 1H:3980 | 2.18 | 1.00 | 55.52 | 2.62 (G:OP2 1H:900) |
| 4 | Water 4 | | 1H:4430 | 2.18 | 1.00 | 61.73 | 3.63 (G:OP2 1H:900) |
| 5 | Water 5 | | 1H:4587 | 2.19 | 1.00 | 58.13 | 4.55 (G:O6 1H:902) |
| 6 | Water 6 | | 1H:3609 | 2.20 | 1.00 | 63.70 | 1.30 (G:N7 1H:900) |
| 7 | G | N7 | 1H:900 | 3.26 | 1.00 | 59.21 |  |
| 8 | G | O6 | 1H:901 | 3.58 | 1.00 | 58.48 |  |
| 9 | G | N7 | 1H:901 | 3.78 | 1.00 | 58.15 |  |
| 10 | G | O6 | 1H:900 | 4.07 | 1.00 | 51.14 |  |
| 11 | G | OP2 | 1H:900 | 4.16 | 1.00 | 49.73 |  |
| 12 | G | O6 | 1H:902 | 4.85 | 1.00 | 50.93 |  |
  |

  
**1^st^ shell ligand angles:**   
1-2: 92; 1-3: 175; 1-4: 84; 1-5: 87; 1-6: 95; 2-3: 92; 2-4: 175; 2-5: 88; 2-6: 89; 3-4: 91; 3-5: 91; 3-6: 88; 4-5: 96; 4-6: 87; 5-6: 177;   
  
 Contact(s): /O_w_/O_w_/O_w_/O_w_/O_w_/O_w_/  
 Coded contact list: 6O_w_  
 Exclusion zone contact(s): /N7/  
Consider K in place of MG (1 valid contact(s) for K vs. 0 for MG)  

### MG:1H:3490

|  | Res. type | Atom | Residue | Distance | Occupancy | B-factor | Closest solute to 1^st^ shell water |
| --- | --- | --- | --- | --- | --- | --- | --- |
| **Ion** | | | |  | 1.00 | 32.37 || 1 | Water 1 | | 1H:3744 | 2.16 | 1.00 | 25.53 | 2.29 (A:O3' 1H:117) |
| 2 | Water 2 | | 1H:4011 | 2.17 | 1.00 | 22.70 | 2.65 (G:O6 1H:121) |
| 3 | Water 3 | | 1H:4002 | 2.18 | 1.00 | 22.94 | 2.64 (G:N7 1H:121) |
| 4 | Water 4 | | 1H:4210 | 2.19 | 1.00 | 36.66 | 2.91 (G:OP1 1H:119) |
| 5 | Water 5 | | 1H:3718 | 2.19 | 1.00 | 26.49 | 2.24 (G:N7 1H:120) |
| 6 | U | OP2 | 1H:118 | 2.47 | 1.00 | 30.69 |  |
| 7 | Water 6 | | 1H:3739 | 3.76 | 1.00 | 22.63 | 2.27 (G:O6 1H:122) |
| 8 | A | O3' | 1H:117 | 3.85 | 1.00 | 29.15 |  |
| 9 | G | N7 | 1H:120 | 4.09 | 1.00 | 18.95 |  |
| 10 | G | O6 | 1H:120 | 4.20 | 1.00 | 24.03 |  |
| 11 | G | O6 | 1H:122 | 4.21 | 1.00 | 22.92 |  |
| 12 | G | O6 | 1H:121 | 4.28 | 1.00 | 19.09 |  |
| 13 | G | OP1 | 1H:119 | 4.30 | 1.00 | 33.51 |  |
| 14 | G | N7 | 1H:121 | 4.33 | 1.00 | 21.75 |  |
| 15 | U | OP1 | 1H:118 | 4.66 | 1.00 | 31.29 |  |
| 16 | U | O5' | 1H:118 | 4.73 | 1.00 | 34.35 |  |
| 17 | A | O2' | 1H:117 | 4.84 | 1.00 | 33.90 |  |
| 18 | G | OP2 | 1H:115 | 4.88 | 1.00 | 25.77 |  |
  |

  
**1^st^ shell ligand angles:**   
1-2: 89; 1-3: 93; 1-4: 87; 1-5: 175; 2-3: 85; 2-4: 101; 2-5: 89; 3-4: 174; 3-5: 92; 4-5: 89;   
  
 Contact(s): /O_w_/O_w_/O_w_/O_w_/O_w_/  
 Coded contact list: 5O_w_  
 Exclusion zone contact(s): /OP2/  
Consider NA in place of MG (1 valid contact(s) for NA vs. 0 for MG)  
MG hydration shell issue: water too close (< 2.4 Angs.)  

### MG:1H:3491

|  | Res. type | Atom | Residue | Distance | Occupancy | B-factor | Closest solute to 1^st^ shell water |
| --- | --- | --- | --- | --- | --- | --- | --- |
| **Ion** | | | |  | 1.00 | 42.97 || 1 | Water 1 | | 1H:3650 | 2.18 | 1.00 | 31.27 | 2.06 (G:N7 1H:122) |
| 2 | Water 2 | | 1H:4471 | 2.18 | 1.00 | 30.86 | 3.57 (G:OP2 1H:121) |
| 3 | Water 3 | | 1H:3759 | 2.18 | 1.00 | 33.92 | 2.31 (U:OP1 1H:118) |
| 4 | Water 4 | | 1H:3739 | 2.19 | 1.00 | 22.63 | 2.27 (G:O6 1H:122) |
| 5 | Water 5 | | P8:201 | 2.20 | 1.00 | 45.84 | 2.83 (ARG:NH2 P8:19) |
| 6 | Water 6 | | 1H:3899 | 2.21 | 1.00 | 42.75 | 2.52 (G:OP2 1H:121) |
| 7 | U | OP1 | 1H:118 | 3.37 | 1.00 | 31.29 |  |
| 8 | G | O6 | 1H:122 | 3.72 | 1.00 | 22.92 |  |
| 9 | Water 7 | | 1H:4002 | 3.96 | 1.00 | 22.94 | 2.86 (G:O6 1H:122) |
| 10 | G | N7 | 1H:122 | 4.06 | 1.00 | 23.52 |  |
| 11 | G | OP2 | 1H:121 | 4.18 | 1.00 | 25.05 |  |
| 12 | ARG | NH2 | P8:19 | 4.25 | 1.00 | 18.18 |  |
| 13 | U | OP2 | 1H:118 | 4.32 | 1.00 | 30.69 |  |
  |

  
**1^st^ shell ligand angles:**   
1-2: 87; 1-3: 177; 1-4: 89; 1-5: 92; 1-6: 90; 2-3: 95; 2-4: 176; 2-5: 89; 2-6: 89; 3-4: 90; 3-5: 85; 3-6: 93; 4-5: 92; 4-6: 90; 5-6: 177;   
  
 Contact(s): /O_w_/O_w_/O_w_/O_w_/O_w_/O_w_/  
 Coded contact list: 6O_w_  
 Exclusion zone contact(s): /OP1/  
Consider K in place of MG (1 valid contact(s) for K vs. 0 for MG)  
MG hydration shell issue: water too close (< 2.4 Angs.)  

### MG:1H:3492

|  | Res. type | Atom | Residue | Distance | Occupancy | B-factor | Closest solute to 1^st^ shell water |
| --- | --- | --- | --- | --- | --- | --- | --- |
| **Ion** | | | |  | 1.00 | 63.87 || 1 | Water 1 | | 1H:4451 | 2.17 | 1.00 | 77.21 | 3.51 (C:N4 1H:281) |
| 2 | Water 2 | | 1H:3703 | 2.17 | 1.00 | 65.69 | 2.21 (G:O6 1H:282) |
| 3 | Water 3 | | 1H:3836 | 2.18 | 1.00 | 61.85 | 2.43 (A:OP1 1H:260) |
| 4 | Water 4 | | 1H:4501 | 2.18 | 1.00 | 54.15 | 3.72 (A:OP2 1H:261) |
| 5 | Water 5 | | 1H:3823 | 2.19 | 1.00 | 55.17 | 2.41 (G:N1 1H:284) |
| 6 | Water 6 | | 1H:4287 | 2.19 | 1.00 | 61.66 | 3.03 (G:O6 1H:284) |
| 7 | G | O6 | 1H:284 | 3.90 | 1.00 | 53.24 |  |
| 8 | G | O6 | 1H:283 | 4.17 | 1.00 | 63.56 |  |
| 9 | A | OP2 | 1H:261 | 4.24 | 1.00 | 52.41 |  |
| 10 | G | O6 | 1H:282 | 4.27 | 1.00 | 74.47 |  |
| 11 | A | OP1 | 1H:260 | 4.35 | 1.00 | 47.92 |  |
| 12 | G | N1 | 1H:284 | 4.37 | 1.00 | 53.41 |  |
| 13 | G | N7 | 1H:283 | 4.55 | 1.00 | 62.32 |  |
| 14 | A | OP2 | 1H:260 | 4.72 | 1.00 | 46.70 |  |
| 15 | G | N7 | 1H:282 | 4.83 | 1.00 | 82.31 |  |
| 16 | C | N4 | 1H:281 | 4.99 | 1.00 | 91.74 |  |
  |

  
**1^st^ shell ligand angles:**   
1-2: 90; 1-3: 89; 1-4: 87; 1-5: 175; 1-6: 93; 2-3: 177; 2-4: 89; 2-5: 94; 2-6: 88; 3-4: 94; 3-5: 88; 3-6: 89; 4-5: 89; 4-6: 177; 5-6: 91;   
  
 Contact(s): /O_w_/O_w_/O_w_/O_w_/O_w_/O_w_/  
 Coded contact list: 6O_w_  
MG hydration shell issue: water too close (< 2.4 Angs.)  

### MG:1H:3493

|  | Res. type | Atom | Residue | Distance | Occupancy | B-factor | Closest solute to 1^st^ shell water |
| --- | --- | --- | --- | --- | --- | --- | --- |
| **Ion** | | | |  | 1.00 | 27.17 || 1 | Water 1 | | 1H:3771 | 2.16 | 1.00 | 24.92 | 2.33 (C:OP1 1H:2564) |
| 2 | Water 2 | | 1H:4442 | 2.18 | 1.00 | 20.21 | 3.47 (A:OP1 1H:2009) |
| 3 | Water 3 | | 1H:3882 | 2.19 | 1.00 | 28.64 | 2.49 (G:OP1 1H:2010) |
| 4 | Water 4 | | 1H:4337 | 2.19 | 1.00 | 27.49 | 3.15 (A:OP1 1H:1717) |
| 5 | Water 5 | | 1H:4207 | 2.19 | 1.00 | 40.23 | 2.90 (A:OP1 1H:1717) |
| 6 | Water 6 | | 1H:4491 | 2.20 | 1.00 | 29.90 | 3.66 (G:O6 1H:1977) |
| 7 | C | OP1 | 1H:2564 | 3.98 | 1.00 | 27.53 |  |
| 8 | A | OP1 | 1H:2009 | 3.99 | 1.00 | 17.32 |  |
| 9 | G | O6 | 1H:1977 | 4.10 | 1.00 | 32.01 |  |
| 10 | A | OP1 | 1H:1717 | 4.13 | 1.00 | 20.71 |  |
| 11 | G | OP1 | 1H:2010 | 4.21 | 1.00 | 21.76 |  |
| 12 | G | OP2 | 1H:2010 | 4.36 | 1.00 | 21.82 |  |
| 13 | G | O3' | 1H:2563 | 4.51 | 1.00 | 27.81 |  |
| 14 | G | N1 | 1H:1977 | 4.62 | 1.00 | 27.50 |  |
  |

  
**1^st^ shell ligand angles:**   
1-2: 93; 1-3: 94; 1-4: 89; 1-5: 179; 1-6: 86; 2-3: 93; 2-4: 89; 2-5: 87; 2-6: 175; 3-4: 177; 3-5: 87; 3-6: 83; 4-5: 91; 4-6: 95; 5-6: 94;   
  
 Contact(s): /O_w_/O_w_/O_w_/O_w_/O_w_/O_w_/  
 Coded contact list: 6O_w_  
MG hydration shell issue: water too close (< 2.4 Angs.)  

### MG:1H:3494

|  | Res. type | Atom | Residue | Distance | Occupancy | B-factor | Closest solute to 1^st^ shell water |
| --- | --- | --- | --- | --- | --- | --- | --- |
| **Ion** | | | |  | 1.00 | 43.72 || 1 | Water 1 | | 1H:4350 | 2.17 | 1.00 | 42.73 | 3.17 (G:O6 1H:2418) |
| 2 | Water 2 | | 1H:3919 | 2.18 | 1.00 | 48.52 | 2.54 (G:OP2 1H:2418) |
| 3 | Water 3 | | 1H:3869 | 2.18 | 1.00 | 48.65 | 2.48 (G:OP2 1H:2423) |
| 4 | Water 4 | | 1H:4585 | 2.18 | 1.00 | 53.69 | 4.50 (G:OP2 1H:2423) |
| 5 | Water 5 | | 1H:4309 | 2.19 | 1.00 | 52.60 | 3.09 (G:OP2 1H:2418) |
| 6 | Water 6 | | 1H:3928 | 2.20 | 1.00 | 35.43 | 2.55 (G:N7 1H:2418) |
| 7 | G | N7 | 1H:2418 | 3.56 | 1.00 | 43.48 |  |
| 8 | G | OP2 | 1H:2418 | 3.82 | 1.00 | 43.25 |  |
| 9 | G | OP2 | 1H:2423 | 4.25 | 1.00 | 45.23 |  |
| 10 | C | OP2 | 1H:2417 | 4.52 | 1.00 | 41.40 |  |
| 11 | G | O6 | 1H:2418 | 4.61 | 1.00 | 46.05 |  |
  |

  
**1^st^ shell ligand angles:**   
1-2: 99; 1-3: 88; 1-4: 82; 1-5: 179; 1-6: 90; 2-3: 89; 2-4: 176; 2-5: 82; 2-6: 91; 3-4: 88; 3-5: 93; 3-6: 178; 4-5: 97; 4-6: 92; 5-6: 88;   
  
 Contact(s): /O_w_/O_w_/O_w_/O_w_/O_w_/O_w_/  
 Coded contact list: 6O_w_  

### MG:1H:3495

|  | Res. type | Atom | Residue | Distance | Occupancy | B-factor | Closest solute to 1^st^ shell water |
| --- | --- | --- | --- | --- | --- | --- | --- |
| **Ion** | | | |  | 1.00 | 48.17 || 1 | Water 1 | | 1H:3798 | 2.17 | 1.00 | 48.96 | 2.37 (A:OP1 1H:451) |
| 2 | Water 2 | | 1H:4101 | 2.19 | 1.00 | 46.48 | 2.76 (G:O6 1H:452) |
| 3 | Water 3 | | 1H:4132 | 2.19 | 1.00 | 52.99 | 2.80 (A:OP1 1H:450) |
| 4 | Water 4 | | 1H:4533 | 2.19 | 1.00 | 50.89 | 4.42 (A:OP1 1H:451) |
| 5 | Water 5 | | 1H:4458 | 2.20 | 1.00 | 43.54 | 3.54 (G:O6 1H:453) |
| 6 | Water 6 | | 1H:4157 | 2.20 | 1.00 | 37.53 | 2.84 (G:N7 1H:453) |
| 7 | G | N7 | 1H:452 | 4.29 | 1.00 | 35.41 |  |
| 8 | A | OP1 | 1H:451 | 4.35 | 1.00 | 32.04 |  |
| 9 | G | O6 | 1H:452 | 4.57 | 1.00 | 33.24 |  |
| 10 | G | N7 | 1H:453 | 4.66 | 1.00 | 37.70 |  |
| 11 | G | O6 | 1H:453 | 4.69 | 1.00 | 33.29 |  |
| 12 | A | OP1 | 1H:450 | 4.86 | 1.00 | 41.34 |  |
  |

  
**1^st^ shell ligand angles:**   
1-2: 88; 1-3: 82; 1-4: 94; 1-5: 174; 1-6: 92; 2-3: 87; 2-4: 177; 2-5: 89; 2-6: 95; 3-4: 95; 3-5: 93; 3-6: 174; 4-5: 89; 4-6: 83; 5-6: 93;   
  
 Contact(s): /O_w_/O_w_/O_w_/O_w_/O_w_/O_w_/  
 Coded contact list: 6O_w_  
MG hydration shell issue: water too close (< 2.4 Angs.)  

### MG:1H:3496

|  | Res. type | Atom | Residue | Distance | Occupancy | B-factor | Closest solute to 1^st^ shell water |
| --- | --- | --- | --- | --- | --- | --- | --- |
| **Ion** | | | |  | 1.00 | 48.29 || 1 | Water 1 | | 1H:4133 | 2.17 | 1.00 | 49.87 | 2.81 (C:OP2 1H:2783) |
| 2 | Water 2 | | 1H:4305 | 2.17 | 1.00 | 37.93 | 3.07 (G:O6 1H:2656) |
| 3 | Water 3 | | 1H:4240 | 2.19 | 1.00 | 48.21 | 2.95 (G:O6 1H:2657) |
| 4 | Water 4 | | 1H:4193 | 2.19 | 1.00 | 50.93 | 2.89 (G:N7 1H:2656) |
| 5 | Water 5 | | 1H:4289 | 2.19 | 1.00 | 50.06 | 3.05 (G:N7 1H:2655) |
| 6 | Water 6 | | 1H:4574 | 2.20 | 1.00 | 46.97 | 4.52 (LYS:NZ 58:83) |
| 7 | G | O6 | 1H:2656 | 3.92 | 1.00 | 37.94 |  |
| 8 | G | O6 | 1H:2657 | 4.25 | 1.00 | 46.60 |  |
| 9 | G | N7 | 1H:2655 | 4.26 | 1.00 | 34.52 |  |
| 10 | G | O6 | 1H:2655 | 4.60 | 1.00 | 37.64 |  |
| 11 | C | OP2 | 1H:2783 | 4.74 | 1.00 | 59.01 |  |
| 12 | G | N7 | 1H:2656 | 4.92 | 1.00 | 37.03 |  |
| 13 | LYS | NZ | 58:83 | 4.99 | 1.00 | 46.57 |  |
  |

  
**1^st^ shell ligand angles:**   
1-2: 86; 1-3: 93; 1-4: 179; 1-5: 85; 1-6: 93; 2-3: 94; 2-4: 93; 2-5: 85; 2-6: 178; 3-4: 87; 3-5: 178; 3-6: 87; 4-5: 95; 4-6: 88; 5-6: 93;   
  
 Contact(s): /O_w_/O_w_/O_w_/O_w_/O_w_/O_w_/  
 Coded contact list: 6O_w_  

### MG:1H:3497

|  | Res. type | Atom | Residue | Distance | Occupancy | B-factor | Closest solute to 1^st^ shell water |
| --- | --- | --- | --- | --- | --- | --- | --- |
| **Ion** | | | |  | 1.00 | 55.46 || 1 | Water 1 | | 1H:4270 | 2.18 | 1.00 | 46.80 | 3.00 (G:N3 1H:1747) |
| 2 | Water 2 | | 1H:4376 | 2.18 | 1.00 | 50.36 | 3.24 (G:O6 1H:1795) |
| 3 | Water 3 | | 1H:3819 | 2.18 | 1.00 | 39.43 | 2.40 (G:O6 1H:1750) |
| 4 | Water 4 | | 1H:4534 | 2.19 | 1.00 | 42.34 | 3.87 (G:O6 1H:1751) |
| 5 | Water 5 | | 1H:3908 | 2.20 | 1.00 | 33.93 | 2.53 (A:OP2 1H:1749) |
| 6 | Water 6 | | 1H:3900 | 2.20 | 1.00 | 49.36 | 2.52 (G:N7 1H:1751) |
| 7 | A | OP2 | 1H:1749 | 3.87 | 1.00 | 38.73 |  |
| 8 | G | O6 | 1H:1750 | 4.01 | 1.00 | 43.01 |  |
| 9 | G | N7 | 1H:1750 | 4.13 | 1.00 | 40.15 |  |
| 10 | G | N2 | 1H:1747 | 4.25 | 1.00 | 43.19 |  |
| 11 | G | O6 | 1H:1795 | 4.50 | 1.00 | 41.65 |  |
| 12 | G | N7 | 1H:1751 | 4.57 | 1.00 | 45.77 |  |
| 13 | G | O6 | 1H:1751 | 4.80 | 1.00 | 44.03 |  |
| 14 | G | N3 | 1H:1747 | 4.97 | 1.00 | 45.35 |  |
  |

  
**1^st^ shell ligand angles:**   
1-2: 90; 1-3: 86; 1-4: 90; 1-5: 92; 1-6: 179; 2-3: 177; 2-4: 87; 2-5: 90; 2-6: 91; 3-4: 92; 3-5: 91; 3-6: 93; 4-5: 177; 4-6: 90; 5-6: 89;   
  
 Contact(s): /O_w_/O_w_/O_w_/O_w_/O_w_/O_w_/  
 Coded contact list: 6O_w_  
MG hydration shell issue: water too close (< 2.4 Angs.)  

### MG:1H:3498

|  | Res. type | Atom | Residue | Distance | Occupancy | B-factor | Closest solute to 1^st^ shell water |
| --- | --- | --- | --- | --- | --- | --- | --- |
| **Ion** | | | |  | 1.00 | 24.71 || 1 | Water 1 | | 1H:4120 | 2.16 | 1.00 | 24.02 | 2.78 (G:OP2 1H:818) |
| 2 | Water 2 | | 1H:3794 | 2.17 | 1.00 | 27.36 | 2.37 (G:OP2 1H:818) |
| 3 | Water 3 | | 1H:4540 | 2.19 | 1.00 | 18.92 | 4.50 (ASN:ND2 P8:8) |
| 4 | Water 4 | | 1H:4527 | 2.19 | 1.00 | 21.84 | 4.56 (G:N7 1H:818) |
| 5 | Water 5 | | 1H:4105 | 2.19 | 1.00 | 24.28 | 2.76 (G:OP2 1H:819) |
| 6 | Water 6 | | 1H:3943 | 2.22 | 1.00 | 22.11 | 2.57 (G:OP1 1H:818) |
| 7 | G | OP2 | 1H:818 | 3.21 | 1.00 | 24.96 |  |
| 8 | G | OP2 | 1H:819 | 3.97 | 1.00 | 19.22 |  |
| 9 | G | O5' | 1H:818 | 4.13 | 1.00 | 18.48 |  |
| 10 | G | OP1 | 1H:818 | 4.29 | 1.00 | 22.22 |  |
| 11 | G | N7 | 1H:818 | 4.85 | 1.00 | 17.85 |  |
| 12 | ASN | ND2 | P8:8 | 4.86 | 1.00 | 17.32 |  |
  |

  
**1^st^ shell ligand angles:**   
1-2: 92; 1-3: 176; 1-4: 87; 1-5: 84; 1-6: 92; 2-3: 85; 2-4: 87; 2-5: 174; 2-6: 95; 3-4: 95; 3-5: 99; 3-6: 86; 4-5: 89; 4-6: 178; 5-6: 89;   
  
 Contact(s): /O_w_/O_w_/O_w_/O_w_/O_w_/O_w_/  
 Coded contact list: 6O_w_  
 Exclusion zone contact(s): /OP2/  
Consider K in place of MG (1 valid contact(s) for K vs. 0 for MG)  
MG hydration shell issue: water too close (< 2.4 Angs.)  

### MG:1H:3499

|  | Res. type | Atom | Residue | Distance | Occupancy | B-factor | Closest solute to 1^st^ shell water |
| --- | --- | --- | --- | --- | --- | --- | --- |
| **Ion** | | | |  | 1.00 | 32.74 || 1 | Water 1 | | 1H:4097 | 2.16 | 1.00 | 34.47 | 2.75 (G:OP1 1H:232) |
| 2 | Water 2 | | 1H:4372 | 2.17 | 1.00 | 40.55 | 3.24 (C:OP1 1H:650) |
| 3 | Water 3 | | 1H:4204 | 2.17 | 1.00 | 51.41 | 2.90 (C:OP1 1H:650) |
| 4 | Water 4 | | 1H:4473 | 2.18 | 1.00 | 33.96 | 3.58 (G:OP1 1H:232) |
| 5 | Water 5 | | Q8:201 | 2.19 | 1.00 | 32.94 | 2.60 (TYR:O Q8:64) |
| 6 | C | OP1 | 1H:650 | 2.34 | 1.00 | 27.19 |  |
| 7 | G | OP1 | 1H:232 | 3.96 | 1.00 | 26.47 |  |
| 8 | G | O3' | 1H:649 | 4.01 | 1.00 | 32.97 |  |
| 9 | TYR | O | Q8:64 | 4.37 | 1.00 | 50.97 |  |
| 10 | C | OP2 | 1H:650 | 4.48 | 1.00 | 28.84 |  |
| 11 | C | O5' | 1H:650 | 4.73 | 1.00 | 24.50 |  |
  |

  
**1^st^ shell ligand angles:**   
1-2: 86; 1-3: 176; 1-4: 94; 1-5: 94; 1-6: 100; 2-3: 90; 2-4: 99; 2-5: 172; 2-6: 92; 3-4: 87; 3-5: 90; 3-6: 80; 4-5: 89; 4-6: 164; 5-6: 81;   
  
 Contact(s): /O_w_/O_w_/O_w_/O_w_/O_w_/OP1/  
 Coded contact list: O_ph_5O_w_  

### MG:1H:3500

|  | Res. type | Atom | Residue | Distance | Occupancy | B-factor | Closest solute to 1^st^ shell water |
| --- | --- | --- | --- | --- | --- | --- | --- |
| **Ion** | | | |  | 1.00 | 63.92 || 1 | Water 1 | | 1H:3645 | 2.17 | 1.00 | 60.88 | 2.05 (G:OP2 1H:2773) |
| 2 | Water 2 | | 1H:3777 | 2.18 | 1.00 | 66.11 | 2.34 (U:O2' 1H:2770) |
| 3 | Water 3 | | 1H:4520 | 2.19 | 1.00 | 65.40 | 3.80 (A:OP2 1H:2772) |
| 4 | Water 4 | | 1H:4348 | 2.19 | 1.00 | 69.40 | 3.17 (G:OP2 1H:2773) |
| 5 | A | OP2 | 1H:2772 | 2.60 | 1.00 | 65.73 |  |
| 6 | G | OP2 | 1H:2773 | 2.87 | 1.00 | 56.95 |  |
| 7 | U | O2' | 1H:2770 | 3.78 | 1.00 | 69.04 |  |
| 8 | A | O5' | 1H:2772 | 4.18 | 1.00 | 57.95 |  |
| 9 | A | OP1 | 1H:2772 | 4.36 | 1.00 | 70.54 |  |
| 10 | G | N7 | 1H:2773 | 4.62 | 1.00 | 62.24 |  |
| 11 | G | O5' | 1H:2773 | 4.70 | 1.00 | 67.54 |  |
| 12 | U | O2 | 1H:2770 | 4.73 | 1.00 | 60.92 |  |
| 13 | G | OP1 | 1H:2773 | 4.91 | 1.00 | 71.87 |  |
| 14 | U | O3' | 1H:2770 | 4.93 | 1.00 | 69.22 |  |
| 15 | A | O3' | 1H:2771 | 4.99 | 1.00 | 64.54 |  |
  |

  
**1^st^ shell ligand angles:**   
1-2: 90; 1-3: 175; 1-4: 90; 2-3: 92; 2-4: 84; 3-4: 94;   
  
 Contact(s): /O_w_/O_w_/O_w_/O_w_/  
 Coded contact list: 4O_w_  
 Exclusion zone contact(s): /OP2/OP2/  
Consider K in place of MG (2 valid contact(s) for K vs. 0 for MG)  
MG hydration shell issue: water too close (< 2.4 Angs.)  

### MG:1H:3501

|  | Res. type | Atom | Residue | Distance | Occupancy | B-factor | Closest solute to 1^st^ shell water |
| --- | --- | --- | --- | --- | --- | --- | --- |
| **Ion** | | | |  | 1.00 | 57.41 || 1 | Water 1 | | 1H:4364 | 2.18 | 1.00 | 42.31 | 3.36 (G:O6 1H:1436) |
| 2 | Water 2 | | 1H:4485 | 2.18 | 1.00 | 53.50 | 3.63 (G:O6 1H:1436) |
| 3 | Water 3 | | 1H:4573 | 2.19 | 1.00 | 55.13 | 4.29 (U:OP2 1H:1443) |
| 4 | Water 4 | | 1H:4486 | 2.19 | 1.00 | 45.48 | 3.66 (G:N7 1H:1435) |
| 5 | Water 5 | | 1H:4538 | 2.20 | 1.00 | 38.69 | 3.96 (U:O4' 1H:1443) |
| 6 | Water 6 | | 1H:3821 | 2.22 | 1.00 | 56.23 | 2.40 (U:O5' 1H:1443) |
| 7 | U | OP2 | 1H:1443 | 4.39 | 1.00 | 34.69 |  |
| 8 | G | N7 | 1H:1435 | 4.49 | 1.00 | 50.49 |  |
| 9 | U | O5' | 1H:1443 | 4.61 | 1.00 | 32.41 |  |
| 10 | G | O6 | 1H:1436 | 4.71 | 1.00 | 52.43 |  |
| 11 | U | O4' | 1H:1443 | 4.88 | 1.00 | 48.88 |  |
  |

  
**1^st^ shell ligand angles:**   
1-2: 86; 1-3: 175; 1-4: 97; 1-5: 91; 1-6: 86; 2-3: 90; 2-4: 86; 2-5: 176; 2-6: 95; 3-4: 84; 3-5: 94; 3-6: 93; 4-5: 91; 4-6: 177; 5-6: 88;   
  
 Contact(s): /O_w_/O_w_/O_w_/O_w_/O_w_/O_w_/  
 Coded contact list: 6O_w_  

### MG:1H:3502

|  | Res. type | Atom | Residue | Distance | Occupancy | B-factor | Closest solute to 1^st^ shell water |
| --- | --- | --- | --- | --- | --- | --- | --- |
| **Ion** | | | |  | 1.00 | 25.62 || 1 | Water 1 | | 1H:3627 | 2.13 | 1.00 | 29.19 | 1.97 (C:OP2 1H:1374) |
| 2 | Water 2 | | 1H:4238 | 2.16 | 1.00 | 23.67 | 2.95 (U:OP2 1H:1373) |
| 3 | Water 3 | | 1H:3694 | 2.18 | 1.00 | 42.87 | 2.19 (C:OP2 1H:1374) |
| 4 | Water 4 | | 1H:4416 | 2.18 | 1.00 | 32.64 | 3.35 (G:N7 1H:1371) |
| 5 | Water 5 | | 1H:3889 | 2.19 | 1.00 | 39.12 | 2.50 (C:OP1 1H:1374) |
| 6 | Water 6 | | 1H:4239 | 2.20 | 1.00 | 40.18 | 2.95 (U:OP2 1H:1370) |
| 7 | C | OP2 | 1H:1374 | 2.75 | 1.00 | 26.44 |  |
| 8 | C | OP1 | 1H:1374 | 3.74 | 1.00 | 23.24 |  |
| 9 | U | OP1 | 1H:1373 | 4.02 | 1.00 | 21.31 |  |
| 10 | G | N7 | 1H:1371 | 4.34 | 1.00 | 30.01 |  |
| 11 | U | OP2 | 1H:1373 | 4.35 | 1.00 | 28.61 |  |
| 12 | C | O5' | 1H:1374 | 4.46 | 1.00 | 25.94 |  |
| 13 | U | OP2 | 1H:1370 | 4.71 | 1.00 | 22.69 |  |
| 14 | G | O6 | 1H:1375 | 4.83 | 1.00 | 29.07 |  |
| 15 | G | N7 | 1H:1375 | 4.83 | 1.00 | 30.92 |  |
| 16 | LYS | NZ | E8:98 | 4.84 | 1.00 | 32.11 |  |
| 17 | U | O3' | 1H:1373 | 4.89 | 1.00 | 28.06 |  |
| 18 | G | OP2 | 1H:1375 | 4.92 | 1.00 | 28.58 |  |
  |

  
**1^st^ shell ligand angles:**   
1-2: 74; 1-3: 94; 1-4: 90; 1-5: 96; 1-6: 177; 2-3: 89; 2-4: 89; 2-5: 170; 2-6: 106; 3-4: 175; 3-5: 89; 3-6: 82; 4-5: 94; 4-6: 93; 5-6: 84;   
  
 Contact(s): /O_w_/O_w_/O_w_/O_w_/O_w_/O_w_/  
 Coded contact list: 6O_w_  
 Exclusion zone contact(s): /OP2/  
Consider K in place of MG (1 valid contact(s) for K vs. 0 for MG)  
MG hydration shell issue: water too close (< 2.4 Angs.)  

### MG:1H:3503

|  | Res. type | Atom | Residue | Distance | Occupancy | B-factor | Closest solute to 1^st^ shell water |
| --- | --- | --- | --- | --- | --- | --- | --- |
| **Ion** | | | |  | 1.00 | 29.08 || 1 | Water 1 | | 1H:3962 | 2.16 | 1.00 | 19.93 | 2.60 (C:OP1 1H:1651) |
| 2 | Water 2 | | 1H:4391 | 2.18 | 1.00 | 48.73 | 3.27 (A:O3' 1H:1440) |
| 3 | Water 3 | | 1H:4542 | 2.18 | 1.00 | 43.08 | 3.93 (U:OP1 1H:1441) |
| 4 | Water 4 | | 1H:4267 | 2.19 | 1.00 | 32.27 | 3.00 (C:OP1 1H:1651) |
| 5 | C | OP1 | 1H:1651 | 2.59 | 1.00 | 24.36 |  |
| 6 | U | OP1 | 1H:1441 | 2.60 | 1.00 | 32.22 |  |
| 7 | A | O3' | 1H:1440 | 3.41 | 1.00 | 29.68 |  |
| 8 | U | OP2 | 1H:1441 | 3.89 | 1.00 | 33.55 |  |
| 9 | C | O5' | 1H:1651 | 4.66 | 1.00 | 24.01 |  |
| 10 | U | O5' | 1H:1441 | 4.73 | 1.00 | 29.09 |  |
| 11 | A | O3' | 1H:1650 | 4.90 | 1.00 | 25.36 |  |
| 12 | C | OP2 | 1H:1651 | 4.92 | 1.00 | 24.25 |  |
| 13 | A | OP2 | 1H:1442 | 4.95 | 1.00 | 31.38 |  |
  |

  
**1^st^ shell ligand angles:**   
1-2: 178; 1-3: 87; 1-4: 93; 2-3: 94; 2-4: 86; 3-4: 89;   
  
 Contact(s): /O_w_/O_w_/O_w_/O_w_/  
 Coded contact list: 4O_w_  
 Exclusion zone contact(s): /OP1/OP1/  
Consider NA in place of MG (1 valid contact(s) for NA vs. 0 for MG)  

### MG:1H:3504

|  | Res. type | Atom | Residue | Distance | Occupancy | B-factor | Closest solute to 1^st^ shell water |
| --- | --- | --- | --- | --- | --- | --- | --- |
| **Ion** | | | |  | 1.00 | 20.74 || 1 | Water 1 | | 1H:4073 | 2.15 | 1.00 | 24.38 | 2.73 (G:OP1 1H:2728) |
| 2 | Water 2 | | 1H:4010 | 2.16 | 1.00 | 28.00 | 2.65 (A:OP2 1H:2727) |
| 3 | Water 3 | | 1H:3815 | 2.16 | 1.00 | 27.58 | 2.40 (A:OP2 1H:2726) |
| 4 | G | OP2 | 1H:2728 | 2.33 | 1.00 | 30.06 |  |
| 5 | A | OP2 | 1H:2726 | 2.56 | 1.00 | 35.93 |  |
| 6 | A | OP2 | 1H:2724 | 2.57 | 1.00 | 28.54 |  |
| 7 | U | O3' | 1H:2725 | 2.96 | 1.00 | 30.59 |  |
| 8 | A | O5' | 1H:2726 | 3.65 | 1.00 | 27.97 |  |
| 9 | A | OP1 | 1H:2724 | 3.69 | 1.00 | 23.77 |  |
| 10 | A | OP2 | 1H:2727 | 3.70 | 1.00 | 27.41 |  |
| 11 | G | OP1 | 1H:2728 | 3.83 | 1.00 | 30.61 |  |
| 12 | SER | OG | 98:15 | 4.06 | 1.00 | 25.89 |  |
| 13 | A | O3' | 1H:2727 | 4.30 | 1.00 | 33.97 |  |
| 14 | C | OP1 | 1H:2723 | 4.44 | 1.00 | 30.64 |  |
| 15 | C | O3' | 1H:2723 | 4.48 | 1.00 | 31.40 |  |
| 16 | G | O5' | 1H:2728 | 4.52 | 1.00 | 35.07 |  |
| 17 | A | OP1 | 1H:2726 | 4.58 | 1.00 | 25.43 |  |
| 18 | A | O5' | 1H:2724 | 4.77 | 1.00 | 23.06 |  |
| 19 | C | O5' | 1H:2723 | 4.99 | 1.00 | 29.01 |  |
  |

  
**1^st^ shell ligand angles:**   
1-2: 103; 1-3: 79; 1-4: 85; 2-3: 93; 2-4: 118; 3-4: 148;   
  
 Contact(s): /O_w_/O_w_/O_w_/OP2/  
 Coded contact list: O_ph_3O_w_  
 Exclusion zone contact(s): /O3'/OP2/OP2/  
Large octahedral topology discrepancy!!!  
Consider NA in place of MG (2 valid contact(s) for NA vs. 1 for MG)  
MG hydration shell issue: water too close (< 2.4 Angs.)  

### MG:1H:3505

|  | Res. type | Atom | Residue | Distance | Occupancy | B-factor | Closest solute to 1^st^ shell water |
| --- | --- | --- | --- | --- | --- | --- | --- |
| **Ion** | | | |  | 1.00 | 16.57 || 1 | Water 1 | | 1H:3705 | 2.15 | 1.00 | 23.22 | 2.21 (A:OP1 1H:2028) |
| 2 | Water 2 | | 1H:3727 | 2.15 | 1.00 | 23.96 | 2.26 (A:OP2 1H:1316) |
| 3 | Water 3 | | 1H:3837 | 2.17 | 1.00 | 34.77 | 2.43 (A:OP1 1H:2028) |
| 4 | Water 4 | | 1H:3653 | 2.18 | 1.00 | 26.83 | 2.08 (A:OP1 1H:2028) |
| 5 | Water 5 | | 1H:3702 | 2.18 | 1.00 | 39.20 | 2.21 (A:OP2 1H:1315) |
| 6 | Water 6 | | 1H:4281 | 2.19 | 1.00 | 34.78 | 3.02 (C:N4 1H:1317) |
| 7 | A | OP1 | 1H:2028 | 2.62 | 1.00 | 23.40 |  |
| 8 | A | OP2 | 1H:1316 | 2.89 | 1.00 | 21.44 |  |
| 9 | A | OP2 | 1H:1315 | 3.76 | 1.00 | 29.83 |  |
| 10 | A | N7 | 1H:1316 | 4.12 | 1.00 | 26.43 |  |
| 11 | C | OP2 | 1H:2029 | 4.21 | 1.00 | 23.46 |  |
| 12 | A | OP2 | 1H:2028 | 4.42 | 1.00 | 28.71 |  |
| 13 | A | OP1 | 1H:1315 | 4.58 | 1.00 | 25.82 |  |
| 14 | A | O5' | 1H:2028 | 4.77 | 1.00 | 19.64 |  |
| 15 | A | O3' | 1H:1315 | 4.86 | 1.00 | 23.14 |  |
| 16 | A | O5' | 1H:1315 | 4.89 | 1.00 | 26.58 |  |
| 17 | A | OP1 | 1H:1316 | 4.95 | 1.00 | 24.09 |  |
| 18 | C | N4 | 1H:1317 | 4.99 | 1.00 | 33.47 |  |
  |

  
**1^st^ shell ligand angles:**   
1-2: 90; 1-3: 92; 1-4: 90; 1-5: 88; 1-6: 178; 2-3: 175; 2-4: 97; 2-5: 84; 2-6: 89; 3-4: 88; 3-5: 91; 3-6: 89; 4-5: 178; 4-6: 92; 5-6: 90;   
  
 Contact(s): /O_w_/O_w_/O_w_/O_w_/O_w_/O_w_/  
 Coded contact list: 6O_w_  
 Exclusion zone contact(s): /OP1/OP2/  
Consider K in place of MG (2 valid contact(s) for K vs. 0 for MG)  
MG hydration shell issue: water too close (< 2.4 Angs.)  

### MG:1H:3506

|  | Res. type | Atom | Residue | Distance | Occupancy | B-factor | Closest solute to 1^st^ shell water |
| --- | --- | --- | --- | --- | --- | --- | --- |
| **Ion** | | | |  | 1.00 | 15.19 || 1 | Water 1 | | 1H:3638 | 2.15 | 1.00 | 16.91 | 2.01 (A:OP2 1H:832) |
| 2 | Water 2 | | 1H:3999 | 2.17 | 1.00 | 20.00 | 2.64 (A:OP2 1H:832) |
| 3 | Water 3 | | 11:409 | 2.21 | 1.00 | 25.04 | 3.00 (MET:O 11:226) |
| 4 | A | OP2 | 1H:831 | 2.34 | 1.00 | 15.77 |  |
| 5 | A | OP1 | 1H:2602 | 2.59 | 1.00 | 21.28 |  |
| 6 | A | OP2 | 1H:832 | 2.92 | 1.00 | 16.68 |  |
| 7 | ALA | O | 11:225 | 3.59 | 1.00 | 13.19 |  |
| 8 | A | O3' | 1H:830 | 3.92 | 1.00 | 15.92 |  |
| 9 | ASN | OD1 | 11:227 | 4.29 | 1.00 | 13.22 |  |
| 10 | MET | O | 11:226 | 4.31 | 1.00 | 16.68 |  |
| 11 | A | OP1 | 1H:831 | 4.53 | 1.00 | 17.27 |  |
| 12 | A | O5' | 1H:831 | 4.63 | 1.00 | 20.28 |  |
| 13 | G | O3' | 1H:2601 | 4.73 | 1.00 | 17.36 |  |
| 14 | A | OP2 | 1H:2602 | 4.90 | 1.00 | 16.66 |  |
| 15 | A | O5' | 1H:2602 | 4.91 | 1.00 | 15.26 |  |
  |

  
**1^st^ shell ligand angles:**   
1-2: 86; 1-3: 95; 1-4: 100; 2-3: 93; 2-4: 91; 3-4: 164;   
  
 Contact(s): /O_w_/O_w_/O_w_/OP2/  
 Coded contact list: O_ph_3O_w_  
 Exclusion zone contact(s): /OP1/OP2/  
MG hydration shell issue: water too close (< 2.4 Angs.)  

### MG:1H:3507

|  | Res. type | Atom | Residue | Distance | Occupancy | B-factor | Closest solute to 1^st^ shell water |
| --- | --- | --- | --- | --- | --- | --- | --- |
| **Ion** | | | |  | 1.00 | 17.34 || 1 | Water 1 | | 1H:3673 | 2.16 | 1.00 | 15.36 | 2.15 (C:OP1 1H:182) |
| 2 | Water 2 | | 1H:3848 | 2.16 | 1.00 | 16.86 | 2.44 (U:OP1 1H:2256) |
| 3 | Water 3 | | 1H:3738 | 2.16 | 1.00 | 15.14 | 2.27 (U:OP1 1H:2256) |
| 4 | Water 4 | | 1H:4015 | 2.20 | 1.00 | 20.77 | 2.66 (U:OP2 1H:183) |
| 5 | Water 5 | | 1H:4307 | 2.21 | 1.00 | 17.50 | 3.08 (A:OP2 1H:189) |
| 6 | C | OP1 | 1H:182 | 2.52 | 1.00 | 20.96 |  |
| 7 | U | OP1 | 1H:2256 | 3.15 | 1.00 | 14.29 |  |
| 8 | C | OP2 | 1H:182 | 3.33 | 1.00 | 18.50 |  |
| 9 | C | O5' | 1H:182 | 3.79 | 1.00 | 17.94 |  |
| 10 | U | OP2 | 1H:183 | 3.79 | 1.00 | 20.57 |  |
| 11 | A | OP1 | 1H:189 | 4.21 | 1.00 | 14.99 |  |
| 12 | A | OP2 | 1H:189 | 4.38 | 1.00 | 16.84 |  |
| 13 | U | O5' | 1H:2256 | 4.48 | 1.00 | 14.06 |  |
| 14 | A | O3' | 1H:181 | 4.62 | 1.00 | 20.50 |  |
| 15 | G | O3' | 1H:2255 | 4.91 | 1.00 | 12.05 |  |
  |

  
**1^st^ shell ligand angles:**   
1-2: 75; 1-3: 89; 1-4: 174; 1-5: 93; 2-3: 81; 2-4: 100; 2-5: 103; 3-4: 88; 3-5: 176; 4-5: 89;   
  
 Contact(s): /O_w_/O_w_/O_w_/O_w_/O_w_/  
 Coded contact list: 5O_w_  
 Exclusion zone contact(s): /OP1/OP1/OP2/  
Consider K in place of MG (2 valid contact(s) for K vs. 0 for MG)  
MG hydration shell issue: water too close (< 2.4 Angs.)  

### MG:1H:3508

|  | Res. type | Atom | Residue | Distance | Occupancy | B-factor | Closest solute to 1^st^ shell water |
| --- | --- | --- | --- | --- | --- | --- | --- |
| **Ion** | | | |  | 1.00 | 22.54 || 1 | Water 1 | | 1H:3760 | 2.13 | 1.00 | 24.73 | 2.31 (U:OP1 1H:2624) |
| 2 | Water 2 | | 21:402 | 2.16 | 1.00 | 19.71 | 2.40 (HIS:NE2 21:132) |
| 3 | Water 3 | | 1H:3683 | 2.18 | 1.00 | 32.13 | 2.17 (A:N7 1H:794) |
| 4 | Water 4 | | 1H:3958 | 2.20 | 1.00 | 28.24 | 2.59 (C:OP1 1H:2625) |
| 5 | HIS | NE2 | 21:132 | 2.54 | 1.00 | 21.83 |  |
| 6 | C | OP1 | 1H:2625 | 2.71 | 1.00 | 29.05 |  |
| 7 | C | OP2 | 1H:2625 | 3.28 | 1.00 | 24.90 |  |
| 8 | A | N7 | 1H:794 | 3.56 | 1.00 | 27.09 |  |
| 9 | U | O3' | 1H:2624 | 4.01 | 1.00 | 23.74 |  |
| 10 | HIS | ND1 | 21:132 | 4.02 | 1.00 | 25.98 |  |
| 11 | G | OP2 | 1H:793 | 4.17 | 1.00 | 19.88 |  |
| 12 | U | OP1 | 1H:2624 | 4.25 | 1.00 | 30.31 |  |
| 13 | U | O5' | 1H:2624 | 4.62 | 1.00 | 25.39 |  |
| 14 | C | O5' | 1H:2625 | 4.69 | 1.00 | 23.34 |  |
| 15 | LYS | NZ | 21:133 | 4.99 | 1.00 | 9.06 |  |
  |

  
**1^st^ shell ligand angles:**   
1-2: 84; 1-3: 90; 1-4: 178; 2-3: 169; 2-4: 95; 3-4: 91;   
  
 Contact(s): /O_w_/O_w_/O_w_/O_w_/  
 Coded contact list: 4O_w_  
 Exclusion zone contact(s): /NE2_His_/OP1/OP2/  
Consider K in place of MG (2 valid contact(s) for K vs. 0 for MG)  
MG hydration shell issue: water too close (< 2.4 Angs.)  

### MG:1H:3509

|  | Res. type | Atom | Residue | Distance | Occupancy | B-factor | Closest solute to 1^st^ shell water |
| --- | --- | --- | --- | --- | --- | --- | --- |
| **Ion** | | | |  | 1.00 | 27.97 || 1 | Water 1 | | 1H:4126 | 2.16 | 1.00 | 28.21 | 2.80 (A:OP2 1H:457) |
| 2 | Water 2 | | 1H:4147 | 2.17 | 1.00 | 39.27 | 2.82 (A:O3' 1H:206) |
| 3 | Water 3 | | 1H:4143 | 2.18 | 1.00 | 30.28 | 2.82 (A:N7 1H:208) |
| 4 | Water 4 | | 1H:4338 | 2.18 | 1.00 | 35.68 | 3.15 (A:OP2 1H:457) |
| 5 | Water 5 | | 1H:3674 | 2.19 | 1.00 | 29.72 | 2.15 (A:OP1 1H:456) |
| 6 | G | OP2 | 1H:207 | 2.58 | 1.00 | 27.41 |  |
| 7 | A | O3' | 1H:206 | 3.37 | 1.00 | 26.91 |  |
| 8 | A | OP2 | 1H:457 | 3.98 | 1.00 | 29.26 |  |
| 9 | G | O5' | 1H:207 | 4.09 | 1.00 | 23.73 |  |
| 10 | A | OP1 | 1H:456 | 4.11 | 1.00 | 25.72 |  |
| 11 | G | O6 | 1H:209 | 4.13 | 1.00 | 39.75 |  |
| 12 | A | O5' | 1H:456 | 4.15 | 1.00 | 26.20 |  |
| 13 | G | N7 | 1H:207 | 4.61 | 1.00 | 25.70 |  |
| 14 | A | N6 | 1H:208 | 4.79 | 1.00 | 36.90 |  |
| 15 | A | N7 | 1H:208 | 4.80 | 1.00 | 33.08 |  |
| 16 | G | OP1 | 1H:207 | 4.84 | 1.00 | 32.38 |  |
| 17 | A | OP2 | 1H:456 | 4.88 | 1.00 | 30.57 |  |
  |

  
**1^st^ shell ligand angles:**   
1-2: 87; 1-3: 176; 1-4: 93; 1-5: 91; 2-3: 90; 2-4: 87; 2-5: 177; 3-4: 90; 3-5: 92; 4-5: 92;   
  
 Contact(s): /O_w_/O_w_/O_w_/O_w_/O_w_/  
 Coded contact list: 5O_w_  
 Exclusion zone contact(s): /O3'/OP2/  
Consider NA in place of MG (1 valid contact(s) for NA vs. 0 for MG)  
MG hydration shell issue: water too close (< 2.4 Angs.)  

### MG:1H:3510

|  | Res. type | Atom | Residue | Distance | Occupancy | B-factor | Closest solute to 1^st^ shell water |
| --- | --- | --- | --- | --- | --- | --- | --- |
| **Ion** | | | |  | 1.00 | 19.65 || 1 | Water 1 | | 1H:4412 | 2.16 | 1.00 | 33.14 | 3.35 (C:OP1 1H:598) |
| 2 | Water 2 | | 1H:3996 | 2.18 | 1.00 | 22.93 | 2.63 (G:O6 1H:2058) |
| 3 | Water 3 | | 1H:3766 | 2.18 | 1.00 | 26.74 | 2.32 (C:OP1 1H:598) |
| 4 | Water 4 | | 1H:4459 | 2.19 | 1.00 | 32.27 | 3.67 (C:OP2 1H:2078) |
| 5 | Water 5 | | 1H:4061 | 2.19 | 1.00 | 23.35 | 2.71 (C:OP1 1H:598) |
| 6 | C | OP1 | 1H:598 | 2.32 | 1.00 | 16.42 |  |
| 7 | Water 6 | | 1H:3835 | 4.33 | 1.00 | 18.76 | 2.43 (C:OP2 1H:598) |
| 8 | G | O3' | 1H:597 | 4.50 | 1.00 | 19.86 |  |
| 9 | C | O5' | 1H:598 | 4.51 | 1.00 | 23.11 |  |
| 10 | C | OP2 | 1H:2078 | 4.57 | 1.00 | 19.68 |  |
| 11 | C | OP2 | 1H:598 | 4.71 | 1.00 | 22.15 |  |
| 12 | Water 7 | | 1H:3687 | 4.79 | 1.00 | 26.11 | 2.17 (C:OP2 1H:598) |
| 13 | G | O6 | 1H:2058 | 4.80 | 1.00 | 28.93 |  |
| 14 | G | O6 | 1H:2041 | 4.88 | 1.00 | 26.57 |  |
  |

  
**1^st^ shell ligand angles:**   
1-2: 96; 1-3: 88; 1-4: 90; 1-5: 170; 1-6: 97; 2-3: 86; 2-4: 91; 2-5: 93; 2-6: 145; 3-4: 177; 3-5: 89; 3-6: 62; 4-5: 93; 4-6: 121; 5-6: 74;   
  
 Contact(s): /O_w_/O_w_/O_w_/O_w_/O_w_/OP1/  
 Coded contact list: O_ph_5O_w_  
Large octahedral topology discrepancy!!!  
MG hydration shell issue: water too close (< 2.4 Angs.)  

### MG:1H:3511

|  | Res. type | Atom | Residue | Distance | Occupancy | B-factor | Closest solute to 1^st^ shell water |
| --- | --- | --- | --- | --- | --- | --- | --- |
| **Ion** | | | |  | 1.00 | 26.78 || 1 | Water 1 | | 1H:3877 | 2.15 | 1.00 | 30.66 | 2.49 (U:OP1 1H:2040) |
| 2 | Water 2 | | 1H:3631 | 2.16 | 1.00 | 25.85 | 1.97 (U:OP1 1H:2040) |
| 3 | Water 3 | | 1H:3686 | 2.18 | 1.00 | 22.54 | 2.17 (U:OP2 1H:2040) |
| 4 | Water 4 | | 1H:3637 | 2.18 | 1.00 | 24.22 | 2.01 (A:OP2 1H:602) |
| 5 | Water 5 | | 1H:4383 | 2.19 | 1.00 | 26.32 | 3.25 (G:OP2 1H:601) |
| 6 | Water 6 | | 1H:4316 | 2.19 | 1.00 | 40.76 | 3.11 (G:OP1 1H:601) |
| 7 | U | OP1 | 1H:2040 | 2.80 | 1.00 | 21.44 |  |
| 8 | A | OP2 | 1H:602 | 3.54 | 1.00 | 21.50 |  |
| 9 | U | OP2 | 1H:2040 | 3.77 | 1.00 | 21.65 |  |
| 10 | Water 7 | | 1H:4460 | 4.25 | 1.00 | 25.49 | 3.80 (A:OP1 1H:2081) |
| 11 | G | OP2 | 1H:601 | 4.49 | 1.00 | 18.77 |  |
| 12 | G | OP2 | 1H:2041 | 4.60 | 1.00 | 23.18 |  |
| 13 | A | OP1 | 1H:2081 | 4.64 | 1.00 | 19.11 |  |
| 14 | U | O5' | 1H:2040 | 4.75 | 1.00 | 21.85 |  |
| 15 | U | O3' | 1H:2039 | 4.78 | 1.00 | 21.27 |  |
| 16 | G | OP1 | 1H:601 | 4.91 | 1.00 | 20.50 |  |
  |

  
**1^st^ shell ligand angles:**   
1-2: 98; 1-3: 87; 1-4: 176; 1-5: 87; 1-6: 85; 2-3: 94; 2-4: 79; 2-5: 89; 2-6: 175; 3-4: 89; 3-5: 174; 3-6: 90; 4-5: 97; 4-6: 98; 5-6: 87;   
  
 Contact(s): /O_w_/O_w_/O_w_/O_w_/O_w_/O_w_/  
 Coded contact list: 6O_w_  
 Exclusion zone contact(s): /OP1/  
Consider K in place of MG (1 valid contact(s) for K vs. 0 for MG)  
MG hydration shell issue: water too close (< 2.4 Angs.)  

### MG:1H:3512

|  | Res. type | Atom | Residue | Distance | Occupancy | B-factor | Closest solute to 1^st^ shell water |
| --- | --- | --- | --- | --- | --- | --- | --- |
| **Ion** | | | |  | 1.00 | 39.45 || 1 | Water 1 | | 1H:3602 | 2.16 | 1.00 | 36.80 | 1.30 (G:N7 1H:1026) |
| 2 | Water 2 | | 1H:3707 | 2.16 | 1.00 | 44.88 | 2.22 (A:OP1 1H:1028) |
| 3 | Water 3 | | 1H:4264 | 2.18 | 1.00 | 37.35 | 2.99 (G:N7 1H:1025) |
| 4 | Water 4 | | 1H:4506 | 2.19 | 1.00 | 45.37 | 3.74 (A:OP1 1H:1028) |
| 5 | Water 5 | | 1H:3812 | 2.20 | 1.00 | 50.17 | 2.39 (G:O6 1H:1026) |
| 6 | A | OP1 | 1H:1028 | 2.40 | 1.00 | 35.49 |  |
| 7 | G | N7 | 1H:1026 | 3.31 | 1.00 | 37.75 |  |
| 8 | G | O6 | 1H:1026 | 3.57 | 1.00 | 32.09 |  |
| 9 | C | OP2 | 1H:1029 | 3.59 | 1.00 | 40.41 |  |
| 10 | A | O5' | 1H:1028 | 4.14 | 1.00 | 31.45 |  |
| 11 | G | OP2 | 1H:1025 | 4.20 | 1.00 | 39.12 |  |
| 12 | A | OP2 | 1H:1028 | 4.46 | 1.00 | 33.62 |  |
| 13 | G | N7 | 1H:1025 | 4.77 | 1.00 | 36.12 |  |
| 14 | A | O3' | 1H:1027 | 4.79 | 1.00 | 26.63 |  |
| 15 | G | OP2 | 1H:1026 | 4.94 | 1.00 | 31.06 |  |
  |

  
**1^st^ shell ligand angles:**   
1-2: 86; 1-3: 92; 1-4: 173; 1-5: 89; 1-6: 64; 2-3: 92; 2-4: 89; 2-5: 174; 2-6: 58; 3-4: 93; 3-5: 91; 3-6: 141; 4-5: 96; 4-6: 109; 5-6: 117;   
  
 Contact(s): /O_w_/O_w_/O_w_/O_w_/O_w_/OP1/  
 Coded contact list: O_ph_5O_w_  
 Exclusion zone contact(s): /N7/  
Large octahedral topology discrepancy!!!  
MG hydration shell issue: water too close (< 2.4 Angs.)  

### MG:1H:3513

|  | Res. type | Atom | Residue | Distance | Occupancy | B-factor | Closest solute to 1^st^ shell water |
| --- | --- | --- | --- | --- | --- | --- | --- |
| **Ion** | | | |  | 1.00 | 30.51 || 1 | Water 1 | | 1H:4171 | 2.17 | 1.00 | 29.67 | 2.86 (G:OP1 1H:808) |
| 2 | Water 2 | | 1H:3987 | 2.18 | 1.00 | 22.62 | 2.63 (C:OP1 1H:779) |
| 3 | Water 3 | | 1H:3616 | 2.18 | 1.00 | 23.65 | 1.89 (G:OP1 1H:808) |
| 4 | Water 4 | | 1H:3779 | 2.18 | 1.00 | 24.88 | 2.34 (A:OP1 1H:809) |
| 5 | Water 5 | | 1H:4355 | 2.19 | 1.00 | 30.03 | 3.18 (G:OP1 1H:808) |
| 6 | Water 6 | | 1H:3755 | 2.21 | 1.00 | 26.47 | 2.30 (C:OP1 1H:779) |
| 7 | C | OP1 | 1H:779 | 2.80 | 1.00 | 22.75 |  |
| 8 | G | OP1 | 1H:808 | 3.06 | 1.00 | 21.39 |  |
| 9 | A | OP2 | 1H:809 | 4.27 | 1.00 | 19.14 |  |
| 10 | C | OP2 | 1H:780 | 4.38 | 1.00 | 24.67 |  |
| 11 | A | OP1 | 1H:809 | 4.47 | 1.00 | 19.45 |  |
| 12 | C | OP2 | 1H:779 | 4.54 | 1.00 | 23.29 |  |
| 13 | C | O5' | 1H:779 | 4.70 | 1.00 | 19.11 |  |
| 14 | G | O5' | 1H:808 | 4.78 | 1.00 | 14.42 |  |
| 15 | U | O3' | 1H:1807 | 4.93 | 1.00 | 17.40 |  |
  |

  
**1^st^ shell ligand angles:**   
1-2: 176; 1-3: 95; 1-4: 84; 1-5: 89; 1-6: 89; 2-3: 85; 2-4: 100; 2-5: 87; 2-6: 92; 3-4: 86; 3-5: 94; 3-6: 172; 4-5: 173; 4-6: 87; 5-6: 93;   
  
 Contact(s): /O_w_/O_w_/O_w_/O_w_/O_w_/O_w_/  
 Coded contact list: 6O_w_  
 Exclusion zone contact(s): /OP1/OP1/  
Consider K in place of MG (2 valid contact(s) for K vs. 0 for MG)  
MG hydration shell issue: water too close (< 2.4 Angs.)  

### MG:1H:3514

|  | Res. type | Atom | Residue | Distance | Occupancy | B-factor | Closest solute to 1^st^ shell water |
| --- | --- | --- | --- | --- | --- | --- | --- |
| **Ion** | | | |  | 1.00 | 53.97 || 1 | Water 1 | | 1H:3774 | 2.17 | 1.00 | 56.11 | 2.33 (A:OP2 1H:531) |
| 2 | Water 2 | | 1H:3839 | 2.18 | 1.00 | 37.79 | 2.43 (U:O3' 1H:1281) |
| 3 | Water 3 | | 1H:3911 | 2.18 | 1.00 | 58.63 | 2.53 (U:O2' 1H:530) |
| 4 | Water 4 | | 1H:3684 | 2.18 | 1.00 | 47.11 | 2.17 (G:OP2 1H:538) |
| 5 | A | OP2 | 1H:531 | 2.64 | 1.00 | 52.13 |  |
| 6 | G | OP1 | 1H:1282 | 2.74 | 1.00 | 43.67 |  |
| 7 | U | O3' | 1H:1281 | 3.52 | 1.00 | 34.25 |  |
| 8 | U | O2' | 1H:530 | 3.99 | 1.00 | 70.31 |  |
| 9 | G | OP2 | 1H:538 | 4.08 | 1.00 | 33.13 |  |
| 10 | U | O2' | 1H:1281 | 4.10 | 1.00 | 38.67 |  |
| 11 | U | OP1 | 1H:537 | 4.11 | 1.00 | 43.61 |  |
| 12 | U | O3' | 1H:530 | 4.17 | 1.00 | 59.65 |  |
| 13 | G | O5' | 1H:1282 | 4.19 | 1.00 | 39.08 |  |
| 14 | A | O5' | 1H:531 | 4.87 | 1.00 | 50.79 |  |
| 15 | A | OP1 | 1H:531 | 4.88 | 1.00 | 61.08 |  |
  |

  
**1^st^ shell ligand angles:**   
1-2: 92; 1-3: 88; 1-4: 177; 2-3: 90; 2-4: 90; 3-4: 91;   
  
 Contact(s): /O_w_/O_w_/O_w_/O_w_/  
 Coded contact list: 4O_w_  
 Exclusion zone contact(s): /OP1/OP2/  
Consider K in place of MG (2 valid contact(s) for K vs. 0 for MG)  
MG hydration shell issue: water too close (< 2.4 Angs.)  

### MG:1H:3515

|  | Res. type | Atom | Residue | Distance | Occupancy | B-factor | Closest solute to 1^st^ shell water |
| --- | --- | --- | --- | --- | --- | --- | --- |
| **Ion** | | | |  | 1.00 | 40.85 || 1 | Water 1 | | 1H:4169 | 2.17 | 1.00 | 46.15 | 2.85 (A:N6 1H:1431) |
| 2 | Water 2 | | 1H:3831 | 2.19 | 1.00 | 31.00 | 2.42 (C:OP2 1H:1392) |
| 3 | Water 3 | | 1H:4308 | 2.20 | 1.00 | 48.29 | 3.08 (C:O2 1H:1430) |
| 4 | Water 4 | | 1H:4549 | 2.20 | 1.00 | 45.60 | 3.99 (G:OP2 1H:1393) |
| 5 | C | OP2 | 1H:1392 | 2.60 | 1.00 | 41.55 |  |
| 6 | G | OP2 | 1H:1393 | 2.67 | 1.00 | 37.31 |  |
| 7 | C | O2 | 1H:1430 | 3.81 | 1.00 | 42.27 |  |
| 8 | C | N3 | 1H:1430 | 4.44 | 1.00 | 52.72 |  |
| 9 | G | O5' | 1H:1393 | 4.62 | 1.00 | 36.00 |  |
| 10 | A | N6 | 1H:1431 | 4.66 | 1.00 | 38.40 |  |
| 11 | A | N7 | 1H:1431 | 4.75 | 1.00 | 38.38 |  |
| 12 | C | O5' | 1H:1392 | 4.77 | 1.00 | 40.42 |  |
| 13 | G | O3' | 1H:1391 | 4.83 | 1.00 | 36.52 |  |
| 14 | C | OP1 | 1H:1392 | 4.90 | 1.00 | 37.71 |  |
  |

  
**1^st^ shell ligand angles:**   
1-2: 89; 1-3: 94; 1-4: 91; 2-3: 89; 2-4: 179; 3-4: 90;   
  
 Contact(s): /O_w_/O_w_/O_w_/O_w_/  
 Coded contact list: 4O_w_  
 Exclusion zone contact(s): /OP2/OP2/  
Consider NA in place of MG (1 valid contact(s) for NA vs. 0 for MG)  

### MG:1H:3516

|  | Res. type | Atom | Residue | Distance | Occupancy | B-factor | Closest solute to 1^st^ shell water |
| --- | --- | --- | --- | --- | --- | --- | --- |
| **Ion** | | | |  | 1.00 | 21.60 || 1 | Water 1 | | 1H:3670 | 2.16 | 1.00 | 14.28 | 2.14 (G:OP1 1H:994) |
| 2 | Water 2 | | 1H:3743 | 2.19 | 1.00 | 34.05 | 2.28 (U:OP1 1H:1009) |
| 3 | Water 3 | | 1H:4226 | 2.20 | 1.00 | 31.26 | 2.92 (U:OP1 1H:1009) |
| 4 | G | OP1 | 1H:994 | 2.47 | 1.00 | 20.91 |  |
| 5 | U | OP2 | 1H:1009 | 2.53 | 1.00 | 22.04 |  |
| 6 | G | OP1 | 1H:1008 | 2.63 | 1.00 | 23.35 |  |
| 7 | U | OP1 | 1H:1009 | 2.97 | 1.00 | 20.46 |  |
| 8 | G | OP2 | 1H:994 | 3.77 | 1.00 | 14.29 |  |
| 9 | G | O3' | 1H:993 | 3.91 | 1.00 | 25.93 |  |
| 10 | U | O5' | 1H:1009 | 4.27 | 1.00 | 20.03 |  |
| 11 | G | O3' | 1H:1008 | 4.36 | 1.00 | 22.28 |  |
| 12 | Water 4 | | 1H:4292 | 4.40 | 1.00 | 20.68 | 3.05 (G:OP1 1H:993) |
| 13 | G | OP1 | 1H:993 | 4.51 | 1.00 | 17.28 |  |
| 14 | Water 5 | | 1H:3992 | 4.61 | 1.00 | 21.96 | 2.82 (U:OP1 1H:1009) |
| 15 | G | O5' | 1H:994 | 4.81 | 1.00 | 18.60 |  |
| 16 | C | O3' | 1H:1007 | 4.83 | 1.00 | 23.54 |  |
| 17 | G | O5' | 1H:1008 | 4.89 | 1.00 | 24.80 |  |
| 18 | G | OP2 | 1H:1008 | 4.96 | 1.00 | 28.95 |  |
  |

  
**1^st^ shell ligand angles:**   
1-2: 169; 1-3: 91; 2-3: 89;   
  
 Contact(s): /O_w_/O_w_/O_w_/  
 Coded contact list: 3O_w_  
 Exclusion zone contact(s): /OP1/OP1/OP1/OP2/  
Consider NA in place of MG (2 valid contact(s) for NA vs. 0 for MG)  
MG hydration shell issue: water too close (< 2.4 Angs.)  

### MG:1H:3517

|  | Res. type | Atom | Residue | Distance | Occupancy | B-factor | Closest solute to 1^st^ shell water |
| --- | --- | --- | --- | --- | --- | --- | --- |
| **Ion** | | | |  | 1.00 | 26.04 || 1 | Water 1 | | 1H:3711 | 2.18 | 1.00 | 40.01 | 2.23 (C:OP2 1H:1231) |
| 2 | Water 2 | | 1H:4046 | 2.19 | 1.00 | 41.41 | 2.69 (C:OP1 1H:1231) |
| 3 | Water 3 | | 1H:3844 | 2.20 | 1.00 | 42.07 | 2.44 (A:OP1 1H:1037) |
| 4 | A | OP2 | 1H:1037 | 2.50 | 1.00 | 35.00 |  |
| 5 | G | OP2 | 1H:1232 | 2.57 | 1.00 | 30.29 |  |
| 6 | C | OP2 | 1H:1038 | 2.94 | 1.00 | 23.71 |  |
| 7 | A | OP1 | 1H:1037 | 3.48 | 1.00 | 36.88 |  |
| 8 | G | O2' | 1H:1036 | 3.65 | 1.00 | 36.35 |  |
| 9 | G | OP1 | 1H:1232 | 3.85 | 1.00 | 29.55 |  |
| 10 | C | OP2 | 1H:1231 | 3.89 | 1.00 | 28.22 |  |
| 11 | G | O3' | 1H:1036 | 4.00 | 1.00 | 31.96 |  |
| 12 | C | OP1 | 1H:1231 | 4.43 | 1.00 | 39.08 |  |
| 13 | C | O5' | 1H:1231 | 4.45 | 1.00 | 40.57 |  |
| 14 | G | O5' | 1H:1232 | 4.49 | 1.00 | 32.14 |  |
| 15 | A | O5' | 1H:1037 | 4.69 | 1.00 | 28.53 |  |
| 16 | C | O3' | 1H:1231 | 4.75 | 1.00 | 30.38 |  |
| 17 | C | O5' | 1H:1038 | 4.82 | 1.00 | 31.93 |  |
  |

  
**1^st^ shell ligand angles:**   
1-2: 88; 1-3: 88; 2-3: 84;   
  
 Contact(s): /O_w_/O_w_/O_w_/  
 Coded contact list: 3O_w_  
 Exclusion zone contact(s): /OP2/OP2/OP2/  
Consider NA in place of MG (2 valid contact(s) for NA vs. 0 for MG)  
MG hydration shell issue: water too close (< 2.4 Angs.)  

### MG:1H:3518

|  | Res. type | Atom | Residue | Distance | Occupancy | B-factor | Closest solute to 1^st^ shell water |
| --- | --- | --- | --- | --- | --- | --- | --- |
| **Ion** | | | |  | 1.00 | 37.63 || 1 | Water 1 | | 78:301 | 2.17 | 1.00 | 33.10 | 1.29 (HIS:NE2 78:27) |
| 2 | Water 2 | | 78:303 | 2.18 | 1.00 | 31.69 | 2.67 (SER:O 78:25) |
| 3 | Water 3 | | 1H:3658 | 2.18 | 1.00 | 28.39 | 2.09 (G:OP2 1H:1238) |
| 4 | Water 4 | | 1H:4091 | 2.20 | 1.00 | 34.76 | 2.75 (G:OP2 1H:1237) |
| 5 | Water 5 | | 1H:4284 | 2.20 | 1.00 | 43.90 | 3.02 (G:N7 1H:1239) |
| 6 | G | N7 | 1H:1238 | 2.68 | 1.00 | 31.15 |  |
| 7 | HIS | NE2 | 78:27 | 3.25 | 1.00 | 34.87 |  |
| 8 | G | OP2 | 1H:1238 | 4.15 | 1.00 | 25.37 |  |
| 9 | G | O6 | 1H:1238 | 4.25 | 1.00 | 32.80 |  |
| 10 | G | N7 | 1H:1237 | 4.42 | 1.00 | 28.44 |  |
| 11 | SER | O | 78:25 | 4.43 | 1.00 | 35.18 |  |
| 12 | ARG | NH1 | 78:18 | 4.43 | 1.00 | 32.49 |  |
| 13 | G | N9 | 1H:1238 | 4.58 | 1.00 | 31.92 |  |
| 14 | G | OP2 | 1H:1237 | 4.77 | 1.00 | 25.01 |  |
| 15 | G | N7 | 1H:1239 | 4.89 | 1.00 | 35.13 |  |
| 16 | HIS | N | 78:27 | 4.95 | 1.00 | 32.93 |  |
| 17 | HIS | ND1 | 78:27 | 4.98 | 1.00 | 36.01 |  |
  |

  
**1^st^ shell ligand angles:**   
1-2: 94; 1-3: 89; 1-4: 83; 1-5: 93; 2-3: 175; 2-4: 93; 2-5: 86; 3-4: 91; 3-5: 90; 4-5: 176;   
  
 Contact(s): /O_w_/O_w_/O_w_/O_w_/O_w_/  
 Coded contact list: 5O_w_  
 Exclusion zone contact(s): /N7/NE2_His_/  
Consider K in place of MG (2 valid contact(s) for K vs. 0 for MG)  
MG hydration shell issue: water too close (< 2.4 Angs.)  

### MG:1H:3519

|  | Res. type | Atom | Residue | Distance | Occupancy | B-factor | Closest solute to 1^st^ shell water |
| --- | --- | --- | --- | --- | --- | --- | --- |
| **Ion** | | | |  | 1.00 | 25.70 || 1 | Water 1 | | 1H:3757 | 2.15 | 1.00 | 29.05 | 2.31 (U:OP2 1H:537) |
| 2 | Water 2 | | 1H:4012 | 2.17 | 1.00 | 39.02 | 2.65 (G:N7 1H:538) |
| 3 | Water 3 | | 1H:4149 | 2.18 | 1.00 | 44.68 | 2.83 (C:O2' 1H:535) |
| 4 | U | OP2 | 1H:537 | 2.51 | 1.00 | 35.69 |  |
| 5 | C | OP1 | 1H:536 | 2.56 | 1.00 | 27.84 |  |
| 6 | C | O3' | 1H:535 | 2.76 | 1.00 | 29.20 |  |
| 7 | G | N7 | 1H:538 | 2.81 | 1.00 | 33.17 |  |
| 8 | C | O5' | 1H:536 | 3.84 | 1.00 | 31.85 |  |
| 9 | G | O6 | 1H:538 | 3.88 | 1.00 | 42.32 |  |
| 10 | C | O2' | 1H:535 | 4.05 | 1.00 | 37.83 |  |
| 11 | C | O3' | 1H:536 | 4.29 | 1.00 | 30.57 |  |
| 12 | C | OP2 | 1H:536 | 4.51 | 1.00 | 39.46 |  |
| 13 | U | OP1 | 1H:537 | 4.64 | 1.00 | 43.61 |  |
| 14 | U | O5' | 1H:537 | 4.90 | 1.00 | 30.65 |  |
| 15 | G | N9 | 1H:538 | 4.99 | 1.00 | 31.41 |  |
  |

  
**1^st^ shell ligand angles:**   
1-2: 83; 1-3: 84; 2-3: 98;   
  
 Contact(s): /O_w_/O_w_/O_w_/  
 Coded contact list: 3O_w_  
 Exclusion zone contact(s): /N7/O3'/OP1/OP2/  
Consider NA in place of MG (2 valid contact(s) for NA vs. 0 for MG)  
MG hydration shell issue: water too close (< 2.4 Angs.)  

### MG:1H:3520

|  | Res. type | Atom | Residue | Distance | Occupancy | B-factor | Closest solute to 1^st^ shell water |
| --- | --- | --- | --- | --- | --- | --- | --- |
| **Ion** | | | |  | 1.00 | 37.39 || 1 | Water 1 | | 1H:4413 | 2.17 | 1.00 | 27.75 | 3.35 (G:OP2 1H:195) |
| 2 | Water 2 | | 1H:4199 | 2.18 | 1.00 | 33.65 | 2.89 (C:O3' 1H:243) |
| 3 | Water 3 | | 1H:4339 | 2.18 | 1.00 | 41.13 | 3.15 (C:O2' 1H:243) |
| 4 | Water 4 | | 1H:3902 | 2.19 | 1.00 | 31.83 | 2.52 (C:OP2 1H:193) |
| 5 | Water 5 | | 1H:3717 | 2.20 | 1.00 | 25.44 | 2.24 (G:OP2 1H:195) |
| 6 | G | OP2 | 1H:195 | 2.46 | 1.00 | 34.51 |  |
| 7 | G | OP1 | 1H:195 | 3.47 | 1.00 | 33.49 |  |
| 8 | G | O5' | 1H:195 | 4.01 | 1.00 | 32.78 |  |
| 9 | C | O2' | 1H:243 | 4.06 | 1.00 | 25.31 |  |
| 10 | C | OP2 | 1H:193 | 4.15 | 1.00 | 23.75 |  |
| 11 | A | O2' | 1H:194 | 4.61 | 1.00 | 24.94 |  |
| 12 | U | OP1 | 1H:192 | 4.61 | 1.00 | 17.23 |  |
| 13 | A | O3' | 1H:194 | 4.63 | 1.00 | 24.45 |  |
| 14 | C | O3' | 1H:243 | 4.65 | 1.00 | 28.40 |  |
  |

  
**1^st^ shell ligand angles:**   
1-2: 85; 1-3: 96; 1-4: 177; 1-5: 95; 2-3: 95; 2-4: 93; 2-5: 174; 3-4: 83; 3-5: 91; 4-5: 87;   
  
 Contact(s): /O_w_/O_w_/O_w_/O_w_/O_w_/  
 Coded contact list: 5O_w_  
 Exclusion zone contact(s): /OP2/  
Consider NA in place of MG (1 valid contact(s) for NA vs. 0 for MG)  
MG hydration shell issue: water too close (< 2.4 Angs.)  

### MG:1H:3521

|  | Res. type | Atom | Residue | Distance | Occupancy | B-factor | Closest solute to 1^st^ shell water |
| --- | --- | --- | --- | --- | --- | --- | --- |
| **Ion** | | | |  | 1.00 | 14.49 || 1 | Water 1 | | 1H:4066 | 2.15 | 1.00 | 17.93 | 2.71 (U:OP1 1H:835) |
| 2 | Water 2 | | 1H:4297 | 2.16 | 1.00 | 23.24 | 3.06 (A:N6 1H:1812) |
| 3 | Water 3 | | 1H:3906 | 2.17 | 1.00 | 21.96 | 2.53 (U:OP2 1H:835) |
| 4 | Water 4 | | 1H:3842 | 2.18 | 1.00 | 24.83 | 2.43 (A:OP1 1H:837) |
| 5 | U | OP1 | 1H:835 | 2.30 | 1.00 | 19.37 |  |
| 6 | C | OP2 | 1H:838 | 2.44 | 1.00 | 19.75 |  |
| 7 | U | OP2 | 1H:835 | 3.53 | 1.00 | 15.97 |  |
| 8 | A | N6 | 1H:1812 | 4.01 | 1.00 | 20.15 |  |
| 9 | C | O3' | 1H:834 | 4.03 | 1.00 | 16.27 |  |
| 10 | A | OP1 | 1H:837 | 4.15 | 1.00 | 21.62 |  |
| 11 | C | OP2 | 1H:839 | 4.35 | 1.00 | 24.29 |  |
| 12 | A | O3' | 1H:837 | 4.35 | 1.00 | 22.96 |  |
| 13 | U | O5' | 1H:835 | 4.45 | 1.00 | 14.80 |  |
| 14 | C | OP1 | 1H:838 | 4.63 | 1.00 | 35.33 |  |
| 15 | A | N7 | 1H:1812 | 4.67 | 1.00 | 20.82 |  |
| 16 | C | O5' | 1H:838 | 4.80 | 1.00 | 28.97 |  |
  |

  
**1^st^ shell ligand angles:**   
1-2: 81; 1-3: 96; 1-4: 172; 1-5: 75; 2-3: 177; 2-4: 100; 2-5: 91; 3-4: 82; 3-5: 90; 4-5: 97;   
  
 Contact(s): /O_w_/O_w_/O_w_/O_w_/OP1/  
 Coded contact list: O_ph_4O_w_  
 Exclusion zone contact(s): /OP2/  

### MG:1H:3522

|  | Res. type | Atom | Residue | Distance | Occupancy | B-factor | Closest solute to 1^st^ shell water |
| --- | --- | --- | --- | --- | --- | --- | --- |
| **Ion** | | | |  | 1.00 | 49.85 || 1 | Water 1 | | 1H:4452 | 2.18 | 1.00 | 32.01 | 3.52 (G:O6 1H:1673) |
| 2 | Water 2 | | 1H:4109 | 2.18 | 1.00 | 40.37 | 2.77 (G:OP2 1H:1671) |
| 3 | Water 3 | | 1H:4134 | 2.19 | 1.00 | 31.49 | 2.81 (C:OP1 1H:746) |
| 4 | Water 4 | | 1H:4063 | 2.19 | 1.00 | 43.76 | 2.71 (G:OP2 1H:1671) |
| 5 | Water 5 | | 1H:4227 | 2.19 | 1.00 | 40.90 | 2.93 (A:N1 1H:1682) |
| 6 | Water 6 | | 1H:4195 | 2.20 | 1.00 | 46.91 | 2.89 (A:N1 1H:1682) |
| 7 | G | OP2 | 1H:1671 | 3.77 | 1.00 | 34.46 |  |
| 8 | A | N1 | 1H:1682 | 3.81 | 1.00 | 29.72 |  |
| 9 | G | O6 | 1H:1673 | 4.19 | 1.00 | 37.76 |  |
| 10 | G | OP1 | 1H:1671 | 4.75 | 1.00 | 33.50 |  |
| 11 | C | OP1 | 1H:746 | 4.79 | 1.00 | 32.80 |  |
| 12 | C | O3' | 1H:745 | 4.92 | 1.00 | 30.99 |  |
| 13 | A | N6 | 1H:1682 | 4.95 | 1.00 | 30.50 |  |
| 14 | A | O2' | 1H:783 | 4.95 | 1.00 | 30.78 |  |
  |

  
**1^st^ shell ligand angles:**   
1-2: 97; 1-3: 174; 1-4: 90; 1-5: 88; 1-6: 86; 2-3: 88; 2-4: 90; 2-5: 175; 2-6: 86; 3-4: 94; 3-5: 88; 3-6: 90; 4-5: 87; 4-6: 174; 5-6: 97;   
  
 Contact(s): /O_w_/O_w_/O_w_/O_w_/O_w_/O_w_/  
 Coded contact list: 6O_w_  

### MG:1H:3523

|  | Res. type | Atom | Residue | Distance | Occupancy | B-factor | Closest solute to 1^st^ shell water |
| --- | --- | --- | --- | --- | --- | --- | --- |
| **Ion** | | | |  | 1.00 | 32.94 || 1 | Water 1 | | 1H:4555 | 2.16 | 1.00 | 36.64 | 4.03 (U:OP1 1H:1360) |
| 2 | Water 2 | | 1H:4030 | 2.17 | 1.00 | 29.84 | 2.67 (G:OP1 1H:1381) |
| 3 | Water 3 | | 1H:4453 | 2.18 | 1.00 | 45.52 | 3.53 (U:OP1 1H:1360) |
| 4 | Water 4 | | 1H:3920 | 2.18 | 1.00 | 31.94 | 2.54 (U:OP1 1H:1360) |
| 5 | Water 5 | | 1H:4298 | 2.18 | 1.00 | 50.20 | 3.06 (G:OP1 1H:1381) |
| 6 | U | OP1 | 1H:1360 | 2.54 | 1.00 | 27.19 |  |
| 7 | G | OP1 | 1H:1381 | 3.93 | 1.00 | 30.96 |  |
| 8 | U | OP2 | 1H:1360 | 4.43 | 1.00 | 28.38 |  |
| 9 | U | O3' | 1H:1359 | 4.53 | 1.00 | 26.49 |  |
| 10 | C | OP1 | 1H:1380 | 4.60 | 1.00 | 23.82 |  |
| 11 | Water 6 | | 1H:4162 | 4.68 | 1.00 | 33.86 | 2.95 (U:OP2 1H:1360) |
| 12 | G | OP2 | 1H:1381 | 4.80 | 1.00 | 27.56 |  |
| 13 | U | O5' | 1H:1360 | 4.93 | 1.00 | 25.52 |  |
  |

  
**1^st^ shell ligand angles:**   
1-2: 93; 1-3: 88; 1-4: 177; 1-5: 87; 2-3: 177; 2-4: 89; 2-5: 89; 3-4: 91; 3-5: 94; 4-5: 91;   
  
 Contact(s): /O_w_/O_w_/O_w_/O_w_/O_w_/  
 Coded contact list: 5O_w_  
 Exclusion zone contact(s): /OP1/  
Consider NA in place of MG (1 valid contact(s) for NA vs. 0 for MG)  

### MG:1H:3524

|  | Res. type | Atom | Residue | Distance | Occupancy | B-factor | Closest solute to 1^st^ shell water |
| --- | --- | --- | --- | --- | --- | --- | --- |
| **Ion** | | | |  | 1.00 | 16.11 || 1 | Water 1 | | 1H:3713 | 2.17 | 1.00 | 16.03 | 2.23 (A:OP1 1H:1815) |
| 2 | Water 2 | | 1H:4031 | 2.17 | 1.00 | 17.88 | 2.67 (A:OP2 1H:1816) |
| 3 | Water 3 | | 1H:3828 | 2.18 | 1.00 | 19.87 | 2.42 (G:OP2 1H:789) |
| 4 | U | OP2 | 1H:788 | 2.55 | 1.00 | 21.63 |  |
| 5 | A | OP1 | 1H:1815 | 2.56 | 1.00 | 17.36 |  |
| 6 | A | OP2 | 1H:1816 | 2.58 | 1.00 | 20.33 |  |
| 7 | A | OP1 | 1H:1816 | 3.15 | 1.00 | 19.37 |  |
| 8 | U | OP1 | 1H:788 | 3.30 | 1.00 | 18.17 |  |
| 9 | G | OP2 | 1H:789 | 4.05 | 1.00 | 19.99 |  |
| 10 | A | OP2 | 1H:1815 | 4.12 | 1.00 | 16.41 |  |
| 11 | A | O3' | 1H:1815 | 4.20 | 1.00 | 19.12 |  |
| 12 | U | O5' | 1H:788 | 4.30 | 1.00 | 20.25 |  |
| 13 | G | O3' | 1H:787 | 4.40 | 1.00 | 27.36 |  |
| 14 | A | O5' | 1H:1816 | 4.41 | 1.00 | 18.09 |  |
| 15 | A | O5' | 1H:1815 | 4.68 | 1.00 | 21.16 |  |
| 16 | C | O3' | 1H:1814 | 4.82 | 1.00 | 15.30 |  |
  |

  
**1^st^ shell ligand angles:**   
1-2: 86; 1-3: 87; 2-3: 168;   
  
 Contact(s): /O_w_/O_w_/O_w_/  
 Coded contact list: 3O_w_  
 Exclusion zone contact(s): /OP1/OP1/OP1/OP2/OP2/  
Consider NA in place of MG (3 valid contact(s) for NA vs. 0 for MG)  
MG hydration shell issue: water too close (< 2.4 Angs.)  

### MG:1H:3525

|  | Res. type | Atom | Residue | Distance | Occupancy | B-factor | Closest solute to 1^st^ shell water |
| --- | --- | --- | --- | --- | --- | --- | --- |
| **Ion** | | | |  | 1.00 | 15.77 || 1 | Water 1 | | 1H:3690 | 2.15 | 1.00 | 19.68 | 2.17 (G:O3' 1H:2620) |
| 2 | Water 2 | | 1H:3734 | 2.16 | 1.00 | 27.99 | 2.27 (U:O3' 1H:788) |
| 3 | Water 3 | | 1H:4259 | 2.17 | 1.00 | 26.11 | 2.98 (G:OP1 1H:789) |
| 4 | Water 4 | | 1H:3868 | 2.17 | 1.00 | 16.35 | 2.47 (A:OP1 1H:1816) |
| 5 | Water 5 | | 1H:3640 | 2.18 | 1.00 | 14.15 | 2.02 (C:OP1 1H:2006) |
| 6 | G | OP1 | 1H:789 | 2.31 | 1.00 | 16.38 |  |
| 7 | U | O3' | 1H:788 | 3.63 | 1.00 | 15.20 |  |
| 8 | C | OP1 | 1H:2006 | 4.00 | 1.00 | 15.54 |  |
| 9 | A | OP1 | 1H:1816 | 4.00 | 1.00 | 19.37 |  |
| 10 | G | O3' | 1H:2620 | 4.02 | 1.00 | 16.63 |  |
| 11 | G | O5' | 1H:789 | 4.20 | 1.00 | 17.06 |  |
| 12 | G | OP1 | 1H:2621 | 4.34 | 1.00 | 15.51 |  |
| 13 | A | OP1 | 1H:1817 | 4.52 | 1.00 | 18.20 |  |
| 14 | G | OP2 | 1H:789 | 4.69 | 1.00 | 19.99 |  |
| 15 | G | O2' | 1H:2620 | 4.88 | 1.00 | 16.34 |  |
| 16 | Water 6 | | 1H:3828 | 4.95 | 1.00 | 19.87 | 2.42 (G:OP2 1H:789) |
  |

  
**1^st^ shell ligand angles:**   
1-2: 176; 1-3: 89; 1-4: 91; 1-5: 97; 1-6: 101; 2-3: 92; 2-4: 87; 2-5: 88; 2-6: 75; 3-4: 177; 3-5: 83; 3-6: 83; 4-5: 100; 4-6: 94; 5-6: 158;   
  
 Contact(s): /O_w_/O_w_/O_w_/O_w_/O_w_/OP1/  
 Coded contact list: O_ph_5O_w_  
MG hydration shell issue: water too close (< 2.4 Angs.)  

### MG:1H:3526

|  | Res. type | Atom | Residue | Distance | Occupancy | B-factor | Closest solute to 1^st^ shell water |
| --- | --- | --- | --- | --- | --- | --- | --- |
| **Ion** | | | |  | 1.00 | 21.22 || 1 | Water 1 | | 1H:3874 | 2.16 | 1.00 | 16.05 | 2.48 (G:OP1 1H:787) |
| 2 | Water 2 | | 1H:3737 | 2.17 | 1.00 | 23.77 | 2.27 (U:OP1 1H:1810) |
| 3 | Water 3 | | 1H:3948 | 2.17 | 1.00 | 22.00 | 2.58 (U:OP2 1H:1811) |
| 4 | Water 4 | | 1H:3669 | 2.18 | 1.00 | 26.14 | 2.13 (G:OP1 1H:787) |
| 5 | Water 5 | | 1H:4543 | 2.19 | 1.00 | 31.79 | 3.93 (G:OP1 1H:787) |
| 6 | Water 6 | | 1H:4144 | 2.20 | 1.00 | 20.60 | 2.82 (U:OP1 1H:1811) |
| 7 | G | OP1 | 1H:787 | 2.91 | 1.00 | 26.95 |  |
| 8 | U | OP2 | 1H:1811 | 4.12 | 1.00 | 17.61 |  |
| 9 | U | OP1 | 1H:1810 | 4.24 | 1.00 | 21.51 |  |
| 10 | G | OP2 | 1H:787 | 4.38 | 1.00 | 22.66 |  |
| 11 | U | OP1 | 1H:1811 | 4.39 | 1.00 | 16.87 |  |
| 12 | U | O5' | 1H:1810 | 4.57 | 1.00 | 16.30 |  |
| 13 | U | OP2 | 1H:1810 | 4.58 | 1.00 | 18.51 |  |
| 14 | G | O3' | 1H:786 | 4.61 | 1.00 | 20.45 |  |
| 15 | A | N1 | 1H:1816 | 4.98 | 1.00 | 18.62 |  |
  |

  
**1^st^ shell ligand angles:**   
1-2: 89; 1-3: 83; 1-4: 100; 1-5: 90; 1-6: 176; 2-3: 88; 2-4: 171; 2-5: 95; 2-6: 88; 3-4: 94; 3-5: 173; 3-6: 95; 4-5: 84; 4-6: 83; 5-6: 92;   
  
 Contact(s): /O_w_/O_w_/O_w_/O_w_/O_w_/O_w_/  
 Coded contact list: 6O_w_  
 Exclusion zone contact(s): /OP1/  
Consider K in place of MG (1 valid contact(s) for K vs. 0 for MG)  
MG hydration shell issue: water too close (< 2.4 Angs.)  

### MG:1H:3527

|  | Res. type | Atom | Residue | Distance | Occupancy | B-factor | Closest solute to 1^st^ shell water |
| --- | --- | --- | --- | --- | --- | --- | --- |
| **Ion** | | | |  | 1.00 | 26.02 || 1 | Water 1 | | 1H:4037 | 2.15 | 1.00 | 29.72 | 2.67 (U:O2' 1H:1866) |
| 2 | Water 2 | | 1H:4099 | 2.16 | 1.00 | 19.66 | 2.75 (A:OP2 1H:1993) |
| 3 | Water 3 | | 1H:4320 | 2.18 | 1.00 | 39.83 | 3.12 (A:OP1 1H:1993) |
| 4 | Water 4 | | 1H:3955 | 2.19 | 1.00 | 25.11 | 2.59 (G:O6 1H:1867) |
| 5 | Water 5 | | 1H:4427 | 2.20 | 1.00 | 34.41 | 3.40 (G:O2' 1H:1922) |
| 6 | A | OP1 | 1H:1993 | 2.66 | 1.00 | 30.89 |  |
| 7 | A | OP2 | 1H:1993 | 3.16 | 1.00 | 31.75 |  |
| 8 | G | OP2 | 1H:1867 | 4.13 | 1.00 | 23.10 |  |
| 9 | G | O6 | 1H:1867 | 4.19 | 1.00 | 29.97 |  |
| 10 | A | O5' | 1H:1993 | 4.31 | 1.00 | 29.84 |  |
| 11 | G | O2' | 1H:1922 | 4.43 | 1.00 | 37.22 |  |
| 12 | A | O3' | 1H:1992 | 4.44 | 1.00 | 21.98 |  |
| 13 | G | N7 | 1H:1867 | 4.45 | 1.00 | 30.40 |  |
| 14 | A | OP2 | 1H:1923 | 4.46 | 1.00 | 28.91 |  |
| 15 | U | O2' | 1H:1866 | 4.52 | 1.00 | 23.21 |  |
| 16 | C | N4 | 1H:1868 | 4.55 | 1.00 | 31.05 |  |
| 17 | A | O5' | 1H:1923 | 4.72 | 1.00 | 24.25 |  |
  |

  
**1^st^ shell ligand angles:**   
1-2: 87; 1-3: 177; 1-4: 89; 1-5: 95; 2-3: 93; 2-4: 176; 2-5: 89; 3-4: 90; 3-5: 88; 4-5: 92;   
  
 Contact(s): /O_w_/O_w_/O_w_/O_w_/O_w_/  
 Coded contact list: 5O_w_  
 Exclusion zone contact(s): /OP1/OP2/  
Consider K in place of MG (2 valid contact(s) for K vs. 0 for MG)  

### MG:1H:3528

|  | Res. type | Atom | Residue | Distance | Occupancy | B-factor | Closest solute to 1^st^ shell water |
| --- | --- | --- | --- | --- | --- | --- | --- |
| **Ion** | | | |  | 1.00 | 50.63 || 1 | Water 1 | | 1H:3644 | 2.17 | 1.00 | 29.72 | 2.04 (A:OP2 1H:957) |
| 2 | Water 2 | | 1H:3620 | 2.18 | 1.00 | 43.82 | 1.95 (U:O4 1H:960) |
| 3 | Water 3 | | 1H:4433 | 2.18 | 1.00 | 29.43 | 3.41 (A:OP1 1H:909) |
| 4 | Water 4 | | 1H:4396 | 2.19 | 1.00 | 32.52 | 3.28 (U:O4 1H:960) |
| 5 | Water 5 | | 1H:4022 | 2.21 | 1.00 | 32.67 | 2.66 (A:OP2 1H:909) |
| 6 | Water 6 | | 1H:3924 | 2.21 | 1.00 | 36.87 | 2.54 (A:OP2 1H:909) |
| 7 | U | O4 | 1H:960 | 3.38 | 1.00 | 33.23 |  |
| 8 | A | OP2 | 1H:909 | 3.44 | 1.00 | 39.78 |  |
| 9 | A | N6 | 1H:911 | 4.08 | 1.00 | 34.18 |  |
| 10 | A | OP2 | 1H:957 | 4.19 | 1.00 | 33.36 |  |
| 11 | A | OP1 | 1H:909 | 4.31 | 1.00 | 31.82 |  |
| 12 | U | O3' | 1H:908 | 4.39 | 1.00 | 33.19 |  |
| 13 | U | O5' | 1H:908 | 4.55 | 1.00 | 31.68 |  |
| 14 | U | OP2 | 1H:908 | 4.83 | 1.00 | 39.47 |  |
| 15 | C | N4 | 1H:959 | 4.95 | 1.00 | 32.53 |  |
| 16 | A | OP1 | 1H:957 | 4.99 | 1.00 | 31.87 |  |
  |

  
**1^st^ shell ligand angles:**   
1-2: 89; 1-3: 90; 1-4: 92; 1-5: 86; 1-6: 174; 2-3: 178; 2-4: 93; 2-5: 87; 2-6: 85; 3-4: 85; 3-5: 95; 3-6: 96; 4-5: 178; 4-6: 89; 5-6: 93;   
  
 Contact(s): /O_w_/O_w_/O_w_/O_w_/O_w_/O_w_/  
 Coded contact list: 6O_w_  
 Exclusion zone contact(s): /O4/  
Consider K in place of MG (1 valid contact(s) for K vs. 0 for MG)  
MG hydration shell issue: water too close (< 2.4 Angs.)  

### MG:1H:3529

|  | Res. type | Atom | Residue | Distance | Occupancy | B-factor | Closest solute to 1^st^ shell water |
| --- | --- | --- | --- | --- | --- | --- | --- |
| **Ion** | | | |  | 1.00 | 54.09 || 1 | Water 1 | | 1H:3701 | 2.18 | 1.00 | 37.15 | 2.20 (G:O6 1H:1846) |
| 2 | Water 2 | | 1H:4476 | 2.19 | 1.00 | 48.23 | 3.59 (A:OP2 1H:1844) |
| 3 | Water 3 | | 1H:3878 | 2.19 | 1.00 | 33.96 | 2.49 (G:O6 1H:1846) |
| 4 | Water 4 | | 1H:4189 | 2.20 | 1.00 | 35.72 | 2.88 (G:OP2 1H:1845) |
| 5 | Water 5 | | 1H:4184 | 2.20 | 1.00 | 33.23 | 2.88 (A:OP2 1H:1844) |
| 6 | Water 6 | | 1H:4602 | 2.20 | 1.00 | 50.58 | 4.93 (G:O6 1H:1846) |
| 7 | G | O6 | 1H:1846 | 3.24 | 1.00 | 37.47 |  |
| 8 | G | OP2 | 1H:1845 | 3.77 | 1.00 | 27.02 |  |
| 9 | A | OP2 | 1H:1844 | 4.36 | 1.00 | 24.40 |  |
| 10 | A | N1 | 1H:1847 | 4.85 | 1.00 | 26.77 |  |
| 11 | A | O5' | 1H:1844 | 4.95 | 1.00 | 29.88 |  |
  |

  
**1^st^ shell ligand angles:**   
1-2: 88; 1-3: 91; 1-4: 93; 1-5: 174; 1-6: 87; 2-3: 87; 2-4: 174; 2-5: 90; 2-6: 92; 3-4: 87; 3-5: 95; 3-6: 179; 4-5: 90; 4-6: 93; 5-6: 87;   
  
 Contact(s): /O_w_/O_w_/O_w_/O_w_/O_w_/O_w_/  
 Coded contact list: 6O_w_  
 Exclusion zone contact(s): /O6/  
Consider K in place of MG (1 valid contact(s) for K vs. 0 for MG)  
MG hydration shell issue: water too close (< 2.4 Angs.)  

### MG:1H:3530

|  | Res. type | Atom | Residue | Distance | Occupancy | B-factor | Closest solute to 1^st^ shell water |
| --- | --- | --- | --- | --- | --- | --- | --- |
| **Ion** | | | |  | 1.00 | 21.54 || 1 | Water 1 | | 1H:3909 | 2.18 | 1.00 | 16.94 | 2.53 (G:OP2 1H:826) |
| 2 | Water 2 | | 1H:3721 | 2.18 | 1.00 | 26.64 | 2.25 (G:OP2 1H:826) |
| 3 | Water 3 | | 11:411 | 2.19 | 1.00 | 32.08 | 4.34 (U:O4 1H:827) |
| 4 | Water 4 | | 1H:4156 | 2.45 | 1.00 | 17.21 | 2.84 (G:N7 1H:826) |
| 5 | Water 5 | | 1H:4435 | 2.81 | 1.00 | 27.04 | 5.25 (U:O4 1H:827) |
| 6 | Water 6 | | 1H:3905 | 2.94 | 1.00 | 22.69 | 2.52 (U:O4 1H:827) |
| 7 | G | OP2 | 1H:826 | 2.99 | 1.00 | 16.82 |  |
| 8 | G | N7 | 1H:826 | 4.04 | 1.00 | 16.09 |  |
| 9 | U | O4 | 1H:827 | 4.07 | 1.00 | 15.43 |  |
| 10 | Water 7 | | 1H:4409 | 4.81 | 1.00 | 21.27 | 5.38 (G:N7 1H:826) |
| 11 | Water 8 | | 1H:3613 | 4.89 | 1.00 | 21.74 | 3.68 (U:O4 1H:827) |
  |

  
**1^st^ shell ligand angles:**   
1-2: 85; 1-3: 83; 2-3: 94;   
  
 Contact(s): /O_w_/O_w_/O_w_/  
 Coded contact list: 3O_w_  
 Exclusion zone contact(s): /O_w_/O_w_/O_w_/OP2/  
Consider K in place of MG (1 valid contact(s) for K vs. 0 for MG)  
MG hydration shell issue: water too close (< 2.4 Angs.)  

### MG:1H:3531

|  | Res. type | Atom | Residue | Distance | Occupancy | B-factor | Closest solute to 1^st^ shell water |
| --- | --- | --- | --- | --- | --- | --- | --- |
| **Ion** | | | |  | 1.00 | 21.26 || 1 | Water 1 | | 1H:3613 | 2.15 | 1.00 | 21.74 | 1.84 (G:O6 1H:826) |
| 2 | Water 2 | | 1H:3905 | 2.17 | 1.00 | 22.69 | 2.52 (U:O4 1H:827) |
| 3 | Water 3 | | 1H:4156 | 2.17 | 1.00 | 17.21 | 2.84 (G:N7 1H:826) |
| 4 | Water 4 | | 1H:4435 | 2.17 | 1.00 | 27.04 | 3.42 (A:N1 1H:841) |
| 5 | Water 5 | | 1H:4455 | 2.18 | 1.00 | 19.50 | 4.21 (C:N4 1H:834) |
| 6 | Water 6 | | 1H:4409 | 2.20 | 1.00 | 21.27 | 3.44 (A:N6 1H:825) |
| 7 | G | O6 | 1H:826 | 3.70 | 1.00 | 18.84 |  |
| 8 | Water 7 | | 11:411 | 3.81 | 1.00 | 32.08 | 4.34 (U:O4 1H:827) |
| 9 | U | O4 | 1H:827 | 4.01 | 1.00 | 15.43 |  |
| 10 | G | N7 | 1H:826 | 4.32 | 1.00 | 16.09 |  |
| 11 | Water 8 | | 1H:3909 | 4.45 | 1.00 | 16.94 | 2.81 (U:O4 1H:827) |
| 12 | A | N6 | 1H:825 | 4.56 | 1.00 | 15.83 |  |
| 13 | A | N1 | 1H:841 | 4.64 | 1.00 | 18.70 |  |
| 14 | C | N4 | 1H:834 | 4.81 | 1.00 | 11.86 |  |
| 15 | G | O6 | 1H:833 | 4.96 | 1.00 | 16.35 |  |
  |

  
**1^st^ shell ligand angles:**   
1-2: 92; 1-3: 95; 1-4: 176; 1-5: 76; 1-6: 92; 2-3: 94; 2-4: 85; 2-5: 86; 2-6: 175; 3-4: 88; 3-5: 172; 3-6: 88; 4-5: 101; 4-6: 90; 5-6: 93;   
  
 Contact(s): /O_w_/O_w_/O_w_/O_w_/O_w_/O_w_/  
 Coded contact list: 6O_w_  
MG hydration shell issue: water too close (< 2.4 Angs.)  

### MG:1H:3532

|  | Res. type | Atom | Residue | Distance | Occupancy | B-factor | Closest solute to 1^st^ shell water |
| --- | --- | --- | --- | --- | --- | --- | --- |
| **Ion** | | | |  | 1.00 | 56.05 || 1 | Water 1 | | 1H:4106 | 2.18 | 1.00 | 57.89 | 2.77 (G:N7 1H:637) |
| 2 | Water 2 | | 1H:3783 | 2.18 | 1.00 | 48.65 | 2.35 (U:N3 1H:638) |
| 3 | Water 3 | | 1H:4531 | 2.19 | 1.00 | 65.85 | 3.87 (G:OP2 1H:637) |
| 4 | Water 4 | | 1H:4565 | 2.19 | 1.00 | 57.69 | 5.21 (G:OP2 1H:637) |
| 5 | Water 5 | | 1H:4498 | 2.19 | 1.00 | 45.95 | 3.68 (U:O4 1H:638) |
| 6 | Water 6 | | 1H:4103 | 2.20 | 1.00 | 46.43 | 2.76 (G:O6 1H:637) |
| 7 | U | O4 | 1H:638 | 3.90 | 1.00 | 54.47 |  |
| 8 | G | N7 | 1H:637 | 4.03 | 1.00 | 45.27 |  |
| 9 | G | O6 | 1H:637 | 4.17 | 1.00 | 51.52 |  |
| 10 | U | N3 | 1H:638 | 4.51 | 1.00 | 44.98 |  |
| 11 | G | OP2 | 1H:637 | 4.88 | 1.00 | 46.16 |  |
  |

  
**1^st^ shell ligand angles:**   
1-2: 93; 1-3: 88; 1-4: 86; 1-5: 178; 1-6: 89; 2-3: 88; 2-4: 178; 2-5: 89; 2-6: 90; 3-4: 90; 3-5: 91; 3-6: 176; 4-5: 92; 4-6: 92; 5-6: 92;   
  
 Contact(s): /O_w_/O_w_/O_w_/O_w_/O_w_/O_w_/  
 Coded contact list: 6O_w_  
MG hydration shell issue: water too close (< 2.4 Angs.)  

### MG:1H:3533

|  | Res. type | Atom | Residue | Distance | Occupancy | B-factor | Closest solute to 1^st^ shell water |
| --- | --- | --- | --- | --- | --- | --- | --- |
| **Ion** | | | |  | 1.00 | 49.62 || 1 | Water 1 | | 1H:3970 | 2.17 | 1.00 | 38.96 | 2.60 (U:O4 1H:2278) |
| 2 | Water 2 | | 1H:4568 | 2.18 | 1.00 | 44.20 | 5.33 (U:O4 1H:2278) |
| 3 | Water 3 | | I8:201 | 2.18 | 1.00 | 36.25 | 2.94 (ASP:OD1 I8:15) |
| 4 | Water 4 | | 1H:4196 | 2.18 | 1.00 | 41.60 | 2.89 (C:OP2 1H:2276) |
| 5 | Water 5 | | I8:202 | 2.19 | 1.00 | 29.44 | 3.80 (ASP:OD1 I8:15) |
| 6 | Water 6 | | 1H:4261 | 2.20 | 1.00 | 55.47 | 2.98 (C:OP2 1H:2277) |
| 7 | C | OP2 | 1H:2277 | 4.35 | 1.00 | 31.58 |  |
| 8 | ASP | OD1 | I8:15 | 4.40 | 1.00 | 29.24 |  |
| 9 | ASP | OD2 | I8:15 | 4.41 | 1.00 | 25.23 |  |
| 10 | C | OP2 | 1H:2276 | 4.50 | 1.00 | 31.63 |  |
| 11 | U | O4 | 1H:2278 | 4.58 | 1.00 | 33.11 |  |
  |

  
**1^st^ shell ligand angles:**   
1-2: 88; 1-3: 92; 1-4: 178; 1-5: 95; 1-6: 85; 2-3: 179; 2-4: 93; 2-5: 96; 2-6: 85; 3-4: 87; 3-5: 83; 3-6: 96; 4-5: 87; 4-6: 94; 5-6: 179;   
  
 Contact(s): /O_w_/O_w_/O_w_/O_w_/O_w_/O_w_/  
 Coded contact list: 6O_w_  

### MG:1H:3534

|  | Res. type | Atom | Residue | Distance | Occupancy | B-factor | Closest solute to 1^st^ shell water |
| --- | --- | --- | --- | --- | --- | --- | --- |
| **Ion** | | | |  | 1.00 | 27.16 || 1 | Water 1 | | 1H:3986 | 2.15 | 1.00 | 25.60 | 2.63 (G:N7 1H:817) |
| 2 | Water 2 | | 1H:4280 | 2.16 | 1.00 | 23.83 | 3.01 (G:N7 1H:816) |
| 3 | Water 3 | | 1H:4404 | 2.17 | 1.00 | 21.36 | 3.31 (G:O6 1H:817) |
| 4 | Water 4 | | 1H:4539 | 2.17 | 1.00 | 31.25 | 3.90 (U:O4 1H:815) |
| 5 | Water 5 | | 1H:4188 | 2.17 | 1.00 | 24.29 | 2.88 (G:O6 1H:816) |
| 6 | Water 6 | | 1H:3915 | 2.20 | 1.00 | 47.57 | 2.54 (G:OP2 1H:816) |
| 7 | G | N7 | 1H:816 | 3.75 | 1.00 | 24.05 |  |
| 8 | G | OP2 | 1H:816 | 3.87 | 1.00 | 20.58 |  |
| 9 | G | O6 | 1H:817 | 3.98 | 1.00 | 23.47 |  |
| 10 | G | N7 | 1H:817 | 4.58 | 1.00 | 22.62 |  |
| 11 | G | O6 | 1H:816 | 4.59 | 1.00 | 23.20 |  |
| 12 | U | O4 | 1H:815 | 4.90 | 1.00 | 29.24 |  |
| 13 | U | N3 | 1H:815 | 4.91 | 1.00 | 26.36 |  |
  |

  
**1^st^ shell ligand angles:**   
1-2: 97; 1-3: 81; 1-4: 176; 1-5: 94; 1-6: 86; 2-3: 177; 2-4: 85; 2-5: 89; 2-6: 90; 3-4: 98; 3-5: 90; 3-6: 91; 4-5: 90; 4-6: 90; 5-6: 179;   
  
 Contact(s): /O_w_/O_w_/O_w_/O_w_/O_w_/O_w_/  
 Coded contact list: 6O_w_  

### MG:1H:3535

|  | Res. type | Atom | Residue | Distance | Occupancy | B-factor | Closest solute to 1^st^ shell water |
| --- | --- | --- | --- | --- | --- | --- | --- |
| **Ion** | | | |  | 1.00 | 64.84 || 1 | Water 1 | | 1H:4541 | 2.17 | 1.00 | 64.67 | 3.92 (A:OP1 1H:357) |
| 2 | Water 2 | | 1H:4521 | 2.18 | 1.00 | 72.70 | 4.18 (A:OP1 1H:357) |
| 3 | Water 3 | | 1H:4304 | 2.19 | 1.00 | 40.96 | 3.59 (A:OP1 1H:357) |
| 4 | Water 4 | | 1H:4495 | 2.19 | 1.00 | 63.35 | 4.78 (G:O6 1H:1255) |
| 5 | Water 5 | | 1H:4365 | 2.19 | 1.00 | 52.74 | 3.41 (G:OP1 1H:1252) |
| 6 | Water 6 | | 1H:4146 | 2.20 | 1.00 | 57.57 | 2.82 (G:O6 1H:1255) |
| 7 | G | O6 | 1H:1255 | 4.31 | 1.00 | 40.70 |  |
| 8 | A | OP1 | 1H:357 | 4.74 | 1.00 | 37.74 |  |
| 9 | G | OP1 | 1H:1252 | 4.99 | 1.00 | 41.54 |  |
  |

  
**1^st^ shell ligand angles:**   
1-2: 87; 1-3: 89; 1-4: 95; 1-5: 87; 1-6: 176; 2-3: 91; 2-4: 177; 2-5: 90; 2-6: 91; 3-4: 87; 3-5: 176; 3-6: 94; 4-5: 92; 4-6: 88; 5-6: 90;   
  
 Contact(s): /O_w_/O_w_/O_w_/O_w_/O_w_/O_w_/  
 Coded contact list: 6O_w_  

### MG:1H:3536

|  | Res. type | Atom | Residue | Distance | Occupancy | B-factor | Closest solute to 1^st^ shell water |
| --- | --- | --- | --- | --- | --- | --- | --- |
| **Ion** | | | |  | 1.00 | 23.61 || 1 | Water 1 | | 1H:4366 | 2.16 | 1.00 | 24.30 | 3.20 (C:OP1 1H:1021) |
| 2 | Water 2 | | 1H:4219 | 2.17 | 1.00 | 31.49 | 2.92 (C:OP1 1H:1021) |
| 3 | Water 3 | | 1H:3834 | 2.18 | 1.00 | 30.32 | 2.43 (C:OP2 1H:1021) |
| 4 | Water 4 | | 1H:4537 | 2.19 | 1.00 | 35.08 | 3.89 (A:OP1 1H:1031) |
| 5 | Water 5 | | 1H:4431 | 2.19 | 1.00 | 30.72 | 3.41 (A:OP1 1H:1031) |
| 6 | C | OP1 | 1H:1021 | 2.57 | 1.00 | 30.04 |  |
| 7 | C | OP2 | 1H:1021 | 3.46 | 1.00 | 24.31 |  |
| 8 | G | O2' | 1H:1020 | 4.30 | 1.00 | 31.40 |  |
| 9 | A | OP1 | 1H:1031 | 4.46 | 1.00 | 26.05 |  |
| 10 | C | O5' | 1H:1021 | 4.49 | 1.00 | 26.74 |  |
| 11 | C | N4 | 1H:1023 | 4.57 | 1.00 | 34.50 |  |
| 12 | G | O3' | 1H:1020 | 4.65 | 1.00 | 28.27 |  |
| 13 | A | O3' | 1H:1030 | 4.66 | 1.00 | 23.88 |  |
| 14 | A | O2' | 1H:1030 | 4.71 | 1.00 | 25.48 |  |
  |

  
**1^st^ shell ligand angles:**   
1-2: 91; 1-3: 171; 1-4: 99; 1-5: 88; 2-3: 86; 2-4: 92; 2-5: 175; 3-4: 89; 3-5: 96; 4-5: 83;   
  
 Contact(s): /O_w_/O_w_/O_w_/O_w_/O_w_/  
 Coded contact list: 5O_w_  
 Exclusion zone contact(s): /OP1/  
Consider NA in place of MG (1 valid contact(s) for NA vs. 0 for MG)  

### MG:1H:3537

|  | Res. type | Atom | Residue | Distance | Occupancy | B-factor | Closest solute to 1^st^ shell water |
| --- | --- | --- | --- | --- | --- | --- | --- |
| **Ion** | | | |  | 1.00 | 54.14 || 1 | Water 1 | | 1H:4071 | 2.18 | 1.00 | 36.89 | 2.72 (G:N7 1H:54) |
| 2 | Water 2 | | 1H:4600 | 2.18 | 1.00 | 59.44 | 4.89 (A:OP2 1H:52) |
| 3 | Water 3 | | 1H:3668 | 2.19 | 1.00 | 36.78 | 2.13 (A:OP2 1H:52) |
| 4 | Water 4 | | 1H:3830 | 2.19 | 1.00 | 37.70 | 2.42 (G:N7 1H:53) |
| 5 | Water 5 | | 1H:4596 | 2.19 | 1.00 | 58.46 | 4.78 (A:OP2 1H:52) |
| 6 | Water 6 | | 1H:3833 | 2.19 | 1.00 | 33.46 | 2.43 (G:O6 1H:53) |
| 7 | A | OP2 | 1H:52 | 3.56 | 1.00 | 28.45 |  |
| 8 | G | O6 | 1H:53 | 3.82 | 1.00 | 29.57 |  |
| 9 | G | N7 | 1H:53 | 3.95 | 1.00 | 27.68 |  |
| 10 | G | N7 | 1H:54 | 4.43 | 1.00 | 23.72 |  |
| 11 | G | O6 | 1H:54 | 4.84 | 1.00 | 29.16 |  |
  |

  
**1^st^ shell ligand angles:**   
1-2: 90; 1-3: 178; 1-4: 92; 1-5: 88; 1-6: 90; 2-3: 89; 2-4: 86; 2-5: 92; 2-6: 179; 3-4: 91; 3-5: 90; 3-6: 90; 4-5: 178; 4-6: 95; 5-6: 87;   
  
 Contact(s): /O_w_/O_w_/O_w_/O_w_/O_w_/O_w_/  
 Coded contact list: 6O_w_  
MG hydration shell issue: water too close (< 2.4 Angs.)  

### MG:1H:3538

|  | Res. type | Atom | Residue | Distance | Occupancy | B-factor | Closest solute to 1^st^ shell water |
| --- | --- | --- | --- | --- | --- | --- | --- |
| **Ion** | | | |  | 1.00 | 27.15 || 1 | Water 1 | | 1H:3918 | 2.15 | 1.00 | 26.12 | 2.54 (G:OP2 1H:2025) |
| 2 | Water 2 | | 1H:4511 | 2.15 | 1.00 | 25.61 | 3.75 (G:N7 1H:2025) |
| 3 | Water 3 | | 1H:3607 | 2.16 | 1.00 | 26.72 | 1.30 (G:N7 1H:2025) |
| 4 | Water 4 | | 1H:4222 | 2.17 | 1.00 | 29.30 | 2.92 (G:O6 1H:2026) |
| 5 | Water 5 | | 1H:4510 | 2.19 | 1.00 | 37.46 | 3.75 (G:N2 1H:1701) |
| 6 | Water 6 | | 1H:4343 | 2.19 | 1.00 | 33.91 | 3.16 (G:N7 1H:2025) |
| 7 | G | N7 | 1H:2025 | 2.81 | 1.00 | 27.90 |  |
| 8 | G | O6 | 1H:2025 | 4.18 | 1.00 | 21.64 |  |
| 9 | G | OP2 | 1H:2025 | 4.31 | 1.00 | 24.23 |  |
| 10 | G | O6 | 1H:2026 | 4.39 | 1.00 | 25.08 |  |
| 11 | G | N7 | 1H:2026 | 4.45 | 1.00 | 25.09 |  |
| 12 | G | N9 | 1H:2025 | 4.61 | 1.00 | 23.12 |  |
| 13 | G | N2 | 1H:1701 | 4.87 | 1.00 | 31.11 |  |
  |

  
**1^st^ shell ligand angles:**   
1-2: 90; 1-3: 91; 1-4: 175; 1-5: 84; 1-6: 93; 2-3: 81; 2-4: 92; 2-5: 95; 2-6: 173; 3-4: 95; 3-5: 174; 3-6: 92; 4-5: 90; 4-6: 86; 5-6: 92;   
  
 Contact(s): /O_w_/O_w_/O_w_/O_w_/O_w_/O_w_/  
 Coded contact list: 6O_w_  
 Exclusion zone contact(s): /N7/  
Consider K in place of MG (1 valid contact(s) for K vs. 0 for MG)  
MG hydration shell issue: water too close (< 2.4 Angs.)  

### MG:1H:3539

|  | Res. type | Atom | Residue | Distance | Occupancy | B-factor | Closest solute to 1^st^ shell water |
| --- | --- | --- | --- | --- | --- | --- | --- |
| **Ion** | | | |  | 1.00 | 66.21 || 1 | Water 1 | | 1H:4582 | 2.17 | 1.00 | 46.82 | 4.44 (G:N7 1H:82) |
| 2 | Water 2 | | 1H:4068 | 2.18 | 1.00 | 54.01 | 2.72 (G:OP2 1H:81) |
| 3 | Water 3 | | 1H:3762 | 2.18 | 1.00 | 43.02 | 2.31 (G:O6 1H:82) |
| 4 | Water 4 | | 1H:4461 | 2.20 | 1.00 | 50.37 | 3.67 (G:OP2 1H:80) |
| 5 | Water 5 | | 1H:4512 | 2.20 | 1.00 | 54.55 | 3.75 (G:OP2 1H:80) |
| 6 | Water 6 | | 1H:3861 | 2.21 | 1.00 | 56.98 | 2.46 (G:OP2 1H:80) |
| 7 | G | N7 | 1H:81 | 3.97 | 1.00 | 42.01 |  |
| 8 | G | OP2 | 1H:80 | 4.01 | 1.00 | 46.82 |  |
| 9 | G | OP2 | 1H:81 | 4.07 | 1.00 | 41.67 |  |
| 10 | G | O6 | 1H:82 | 4.42 | 1.00 | 49.91 |  |
| 11 | G | N7 | 1H:82 | 4.72 | 1.00 | 42.93 |  |
| 12 | G | O6 | 1H:81 | 5.00 | 1.00 | 49.44 |  |
  |

  
**1^st^ shell ligand angles:**   
1-2: 83; 1-3: 88; 1-4: 94; 1-5: 93; 1-6: 178; 2-3: 91; 2-4: 87; 2-5: 176; 2-6: 96; 3-4: 177; 3-5: 88; 3-6: 90; 4-5: 93; 4-6: 88; 5-6: 88;   
  
 Contact(s): /O_w_/O_w_/O_w_/O_w_/O_w_/O_w_/  
 Coded contact list: 6O_w_  
MG hydration shell issue: water too close (< 2.4 Angs.)  

### MG:1H:3540

|  | Res. type | Atom | Residue | Distance | Occupancy | B-factor | Closest solute to 1^st^ shell water |
| --- | --- | --- | --- | --- | --- | --- | --- |
| **Ion** | | | |  | 1.00 | 50.95 || 1 | Water 1 | | 1H:3876 | 2.17 | 1.00 | 48.05 | 2.49 (G:O6 1H:2849) |
| 2 | Water 2 | | 1H:4115 | 2.17 | 1.00 | 46.30 | 2.78 (G:N7 1H:2848) |
| 3 | Water 3 | | 1H:4136 | 2.18 | 1.00 | 48.08 | 2.81 (G:OP2 1H:2848) |
| 4 | Water 4 | | 1H:4082 | 2.19 | 1.00 | 49.09 | 2.74 (A:O2' 1H:2846) |
| 5 | Water 5 | | 1H:3886 | 2.20 | 1.00 | 45.49 | 2.50 (G:OP2 1H:2848) |
| 6 | Water 6 | | 1H:3612 | 2.20 | 1.00 | 53.15 | 1.52 (G:N7 1H:2848) |
| 7 | G | N7 | 1H:2848 | 3.05 | 1.00 | 47.86 |  |
| 8 | G | OP2 | 1H:2848 | 3.65 | 1.00 | 43.92 |  |
| 9 | G | O6 | 1H:2849 | 3.96 | 1.00 | 45.83 |  |
| 10 | G | N7 | 1H:2849 | 3.98 | 1.00 | 47.30 |  |
| 11 | A | N3 | 1H:2846 | 4.21 | 1.00 | 51.33 |  |
| 12 | A | O2' | 1H:2846 | 4.39 | 1.00 | 51.19 |  |
| 13 | U | O4 | 1H:2889 | 4.54 | 1.00 | 57.67 |  |
| 14 | G | N9 | 1H:2848 | 4.61 | 1.00 | 45.39 |  |
| 15 | G | O6 | 1H:2848 | 4.71 | 1.00 | 52.95 |  |
| 16 | G | O5' | 1H:2848 | 4.96 | 1.00 | 42.00 |  |
  |

  
**1^st^ shell ligand angles:**   
1-2: 84; 1-3: 177; 1-4: 92; 1-5: 94; 1-6: 90; 2-3: 94; 2-4: 90; 2-5: 177; 2-6: 89; 3-4: 86; 3-5: 88; 3-6: 92; 4-5: 92; 4-6: 177; 5-6: 89;   
  
 Contact(s): /O_w_/O_w_/O_w_/O_w_/O_w_/O_w_/  
 Coded contact list: 6O_w_  
 Exclusion zone contact(s): /N7/  
Consider K in place of MG (1 valid contact(s) for K vs. 0 for MG)  

### MG:1H:3541

|  | Res. type | Atom | Residue | Distance | Occupancy | B-factor | Closest solute to 1^st^ shell water |
| --- | --- | --- | --- | --- | --- | --- | --- |
| **Ion** | | | |  | 1.00 | 66.38 || 1 | Water 1 | | 1H:4420 | 2.18 | 1.00 | 42.60 | 3.47 (G:O6 1H:1641) |
| 2 | Water 2 | | 1H:3712 | 2.18 | 1.00 | 42.01 | 2.23 (G:O6 1H:1641) |
| 3 | Water 3 | | 1H:4588 | 2.19 | 1.00 | 54.35 | 5.73 (G:N7 1H:1641) |
| 4 | Water 4 | | 1H:4560 | 2.19 | 1.00 | 45.37 | 4.10 (U:O4 1H:1453) |
| 5 | Water 5 | | 1H:4291 | 2.20 | 1.00 | 47.83 | 3.05 (G:N7 1H:1641) |
| 6 | Water 6 | | 1H:4513 | 2.21 | 1.00 | 51.47 | 3.77 (U:O4 1H:1453) |
| 7 | G | O6 | 1H:1641 | 3.95 | 1.00 | 44.05 |  |
| 8 | G | O6 | 1H:1642 | 4.18 | 1.00 | 40.70 |  |
| 9 | G | N7 | 1H:1641 | 4.42 | 1.00 | 47.76 |  |
| 10 | U | O4 | 1H:1453 | 4.94 | 1.00 | 35.16 |  |
  |

  
**1^st^ shell ligand angles:**   
1-2: 85; 1-3: 93; 1-4: 87; 1-5: 93; 1-6: 174; 2-3: 177; 2-4: 95; 2-5: 88; 2-6: 91; 3-4: 87; 3-5: 89; 3-6: 92; 4-5: 177; 4-6: 89; 5-6: 91;   
  
 Contact(s): /O_w_/O_w_/O_w_/O_w_/O_w_/O_w_/  
 Coded contact list: 6O_w_  
MG hydration shell issue: water too close (< 2.4 Angs.)  

### MG:1H:3542

|  | Res. type | Atom | Residue | Distance | Occupancy | B-factor | Closest solute to 1^st^ shell water |
| --- | --- | --- | --- | --- | --- | --- | --- |
| **Ion** | | | |  | 1.00 | 33.90 || 1 | Water 1 | | 1H:4322 | 2.18 | 1.00 | 31.12 | 3.12 (C:OP1 1H:1417) |
| 2 | Water 2 | | 1H:3800 | 2.18 | 1.00 | 24.66 | 2.38 (C:O3' 1H:820) |
| 3 | Water 3 | | 1H:4347 | 2.18 | 1.00 | 24.83 | 3.17 (U:OP1 1H:821) |
| 4 | Water 4 | | 1H:3872 | 2.19 | 1.00 | 31.42 | 2.48 (G:OP1 1H:1403) |
| 5 | Water 5 | | 1H:4217 | 2.20 | 1.00 | 34.30 | 2.92 (C:OP1 1H:1417) |
| 6 | Water 6 | | 1H:4229 | 2.21 | 1.00 | 33.57 | 2.93 (U:OP1 1H:821) |
| 7 | U | OP1 | 1H:821 | 3.55 | 1.00 | 20.41 |  |
| 8 | G | OP1 | 1H:1403 | 3.70 | 1.00 | 23.87 |  |
| 9 | C | OP1 | 1H:1417 | 4.01 | 1.00 | 26.16 |  |
| 10 | C | O3' | 1H:820 | 4.23 | 1.00 | 18.48 |  |
| 11 | C | OP2 | 1H:1417 | 4.96 | 1.00 | 32.73 |  |
  |

  
**1^st^ shell ligand angles:**   
1-2: 83; 1-3: 86; 1-4: 93; 1-5: 95; 1-6: 177; 2-3: 82; 2-4: 94; 2-5: 177; 2-6: 98; 3-4: 176; 3-5: 95; 3-6: 96; 4-5: 88; 4-6: 85; 5-6: 84;   
  
 Contact(s): /O_w_/O_w_/O_w_/O_w_/O_w_/O_w_/  
 Coded contact list: 6O_w_  
MG hydration shell issue: water too close (< 2.4 Angs.)  

### MG:1H:3543

|  | Res. type | Atom | Residue | Distance | Occupancy | B-factor | Closest solute to 1^st^ shell water |
| --- | --- | --- | --- | --- | --- | --- | --- |
| **Ion** | | | |  | 1.00 | 48.71 || 1 | Water 1 | | 1H:3933 | 2.17 | 1.00 | 56.73 | 2.55 (C:OP1 1H:1516) |
| 2 | Water 2 | | 1H:4378 | 2.18 | 1.00 | 56.23 | 3.25 (A:OP1 1H:1486) |
| 3 | Water 3 | | 1H:4085 | 2.19 | 1.00 | 51.56 | 2.74 (A:OP2 1H:1517) |
| 4 | Water 4 | | 1H:4234 | 2.19 | 1.00 | 51.38 | 2.94 (C:OP1 1H:1516) |
| 5 | A | OP1 | 1H:1486 | 2.43 | 1.00 | 39.48 |  |
| 6 | C | OP1 | 1H:1516 | 2.57 | 1.00 | 55.16 |  |
| 7 | C | OP2 | 1H:1516 | 3.21 | 1.00 | 58.39 |  |
| 8 | A | OP2 | 1H:1486 | 3.94 | 1.00 | 39.34 |  |
| 9 | C | O5' | 1H:1516 | 4.11 | 1.00 | 57.47 |  |
| 10 | G | OP2 | 1H:1487 | 4.30 | 1.00 | 48.47 |  |
| 11 | Water 5 | | 1H:4567 | 4.41 | 1.00 | 62.13 | 4.18 (C:OP1 1H:1516) |
| 12 | A | OP2 | 1H:1517 | 4.49 | 1.00 | 53.39 |  |
| 13 | A | O5' | 1H:1486 | 4.52 | 1.00 | 47.78 |  |
| 14 | U | O3' | 1H:1485 | 4.54 | 1.00 | 43.24 |  |
| 15 | C | O3' | 1H:1515 | 4.55 | 1.00 | 62.57 |  |
| 16 | Water 6 | | 1H:4016 | 4.95 | 1.00 | 49.14 | 2.66 (A:OP2 1H:1486) |
  |

  
**1^st^ shell ligand angles:**   
1-2: 89; 1-3: 179; 1-4: 86; 2-3: 92; 2-4: 172; 3-4: 93;   
  
 Contact(s): /O_w_/O_w_/O_w_/O_w_/  
 Coded contact list: 4O_w_  
 Exclusion zone contact(s): /OP1/OP1/OP2/  
Consider NA in place of MG (2 valid contact(s) for NA vs. 0 for MG)  

### MG:1H:3544

|  | Res. type | Atom | Residue | Distance | Occupancy | B-factor | Closest solute to 1^st^ shell water |
| --- | --- | --- | --- | --- | --- | --- | --- |
| **Ion** | | | |  | 1.00 | 63.80 || 1 | Water 1 | | 1H:4426 | 2.18 | 1.00 | 59.99 | 3.39 (A:OP2 1H:2546) |
| 2 | Water 2 | | 1H:3979 | 2.18 | 1.00 | 61.42 | 2.62 (G:OP1 1H:2545) |
| 3 | Water 3 | | 1H:3806 | 2.18 | 1.00 | 65.10 | 2.39 (A:OP2 1H:2547) |
| 4 | Water 4 | | 1H:3968 | 2.18 | 1.00 | 57.82 | 2.60 (A:N6 1H:2547) |
| 5 | Water 5 | | 1H:4428 | 2.18 | 1.00 | 73.38 | 3.40 (A:N6 1H:2547) |
| 6 | A | OP2 | 1H:2546 | 2.61 | 1.00 | 53.92 |  |
| 7 | A | N7 | 1H:2547 | 3.93 | 1.00 | 65.33 |  |
| 8 | A | O5' | 1H:2546 | 4.08 | 1.00 | 66.78 |  |
| 9 | A | N6 | 1H:2547 | 4.14 | 1.00 | 61.74 |  |
| 10 | A | N7 | 1H:2546 | 4.32 | 1.00 | 63.84 |  |
| 11 | A | OP2 | 1H:2547 | 4.40 | 1.00 | 56.94 |  |
| 12 | A | OP1 | 1H:2546 | 4.43 | 1.00 | 52.75 |  |
| 13 | G | O5' | 1H:2545 | 4.58 | 1.00 | 58.63 |  |
| 14 | G | OP1 | 1H:2545 | 4.62 | 1.00 | 54.90 |  |
| 15 | G | OP2 | 1H:2545 | 4.67 | 1.00 | 58.49 |  |
| 16 | A | O2' | 1H:2543 | 4.97 | 1.00 | 62.04 |  |
| 17 | G | O3' | 1H:2545 | 4.99 | 1.00 | 61.02 |  |
  |

  
**1^st^ shell ligand angles:**   
1-2: 86; 1-3: 92; 1-4: 175; 1-5: 90; 2-3: 176; 2-4: 90; 2-5: 95; 3-4: 92; 3-5: 89; 4-5: 88;   
  
 Contact(s): /O_w_/O_w_/O_w_/O_w_/O_w_/  
 Coded contact list: 5O_w_  
 Exclusion zone contact(s): /OP2/  
Consider K in place of MG (1 valid contact(s) for K vs. 0 for MG)  
MG hydration shell issue: water too close (< 2.4 Angs.)  

### MG:1H:3545

|  | Res. type | Atom | Residue | Distance | Occupancy | B-factor | Closest solute to 1^st^ shell water |
| --- | --- | --- | --- | --- | --- | --- | --- |
| **Ion** | | | |  | 1.00 | 45.83 || 1 | Water 1 | | 1H:3801 | 2.16 | 1.00 | 31.09 | 2.38 (U:OP2 1H:870) |
| 2 | Water 2 | | 1H:3769 | 2.17 | 1.00 | 38.95 | 2.32 (G:OP2 1H:993) |
| 3 | Water 3 | | 1H:4572 | 2.18 | 1.00 | 46.64 | 4.27 (C:OP2 1H:1011) |
| 4 | Water 4 | | 1H:4438 | 2.18 | 1.00 | 27.51 | 3.44 (U:OP1 1H:870) |
| 5 | Water 5 | | 1H:3950 | 2.19 | 1.00 | 35.31 | 2.59 (C:OP2 1H:1011) |
| 6 | Water 6 | | 1H:4429 | 2.19 | 1.00 | 41.00 | 3.40 (U:OP1 1H:870) |
| 7 | U | OP1 | 1H:870 | 3.78 | 1.00 | 19.06 |  |
| 8 | G | OP2 | 1H:993 | 3.91 | 1.00 | 25.52 |  |
| 9 | U | OP2 | 1H:870 | 4.11 | 1.00 | 27.84 |  |
| 10 | C | OP2 | 1H:1011 | 4.42 | 1.00 | 22.96 |  |
  |

  
**1^st^ shell ligand angles:**   
1-2: 87; 1-3: 178; 1-4: 92; 1-5: 85; 1-6: 92; 2-3: 90; 2-4: 180; 2-5: 90; 2-6: 91; 3-4: 90; 3-5: 94; 3-6: 89; 4-5: 90; 4-6: 89; 5-6: 177;   
  
 Contact(s): /O_w_/O_w_/O_w_/O_w_/O_w_/O_w_/  
 Coded contact list: 6O_w_  
MG hydration shell issue: water too close (< 2.4 Angs.)  

### MG:1H:3546

|  | Res. type | Atom | Residue | Distance | Occupancy | B-factor | Closest solute to 1^st^ shell water |
| --- | --- | --- | --- | --- | --- | --- | --- |
| **Ion** | | | |  | 1.00 | 61.23 || 1 | Water 1 | | 1H:4016 | 2.18 | 1.00 | 49.14 | 2.66 (A:OP2 1H:1486) |
| 2 | Water 2 | | 1H:3883 | 2.18 | 1.00 | 46.33 | 2.49 (G:O6 1H:1488) |
| 3 | Water 3 | | 1H:4567 | 2.19 | 1.00 | 62.13 | 4.33 (A:OP2 1H:1486) |
| 4 | Water 4 | | 1H:4544 | 2.19 | 1.00 | 55.17 | 3.96 (A:OP2 1H:1486) |
| 5 | Water 5 | | 1H:4408 | 2.19 | 1.00 | 46.68 | 3.34 (G:N7 1H:1488) |
| 6 | Water 6 | | 1H:4466 | 2.20 | 1.00 | 50.89 | 3.56 (G:O6 1H:1488) |
| 7 | G | N7 | 1H:1488 | 4.12 | 1.00 | 43.91 |  |
| 8 | G | O6 | 1H:1488 | 4.14 | 1.00 | 46.18 |  |
| 9 | G | N7 | 1H:1487 | 4.28 | 1.00 | 41.58 |  |
| 10 | Water 7 | | 1H:3933 | 4.30 | 1.00 | 56.73 | 3.49 (A:OP2 1H:1486) |
| 11 | A | OP2 | 1H:1486 | 4.40 | 1.00 | 39.34 |  |
| 12 | G | O6 | 1H:1487 | 4.41 | 1.00 | 45.84 |  |
  |

  
**1^st^ shell ligand angles:**   
1-2: 92; 1-3: 86; 1-4: 86; 1-5: 92; 1-6: 178; 2-3: 178; 2-4: 84; 2-5: 93; 2-6: 89; 3-4: 94; 3-5: 88; 3-6: 92; 4-5: 177; 4-6: 94; 5-6: 88;   
  
 Contact(s): /O_w_/O_w_/O_w_/O_w_/O_w_/O_w_/  
 Coded contact list: 6O_w_  

### MG:1H:3547

|  | Res. type | Atom | Residue | Distance | Occupancy | B-factor | Closest solute to 1^st^ shell water |
| --- | --- | --- | --- | --- | --- | --- | --- |
| **Ion** | | | |  | 1.00 | 52.95 || 1 | Water 1 | | 88:301 | 2.17 | 1.00 | 59.28 | 3.91 (LYS:NZ 88:128) |
| 2 | Water 2 | | 1H:3893 | 2.18 | 1.00 | 43.14 | 2.51 (G:O6 1H:1077) |
| 3 | Water 3 | | 1H:3894 | 2.18 | 1.00 | 34.80 | 2.51 (G:N7 1H:1077) |
| 4 | Water 4 | | 1H:4592 | 2.18 | 1.00 | 41.44 | 4.93 (G:O6 1H:1077) |
| 5 | Water 5 | | 1H:3885 | 2.19 | 1.00 | 45.42 | 2.50 (A:OP1 1H:1076) |
| 6 | Water 6 | | 1H:4041 | 2.21 | 1.00 | 34.49 | 2.68 (A:OP2 1H:1076) |
| 7 | A | OP2 | 1H:1076 | 3.46 | 1.00 | 36.56 |  |
| 8 | A | OP1 | 1H:1076 | 3.85 | 1.00 | 37.71 |  |
| 9 | G | N7 | 1H:1077 | 3.97 | 1.00 | 39.22 |  |
| 10 | G | O6 | 1H:1077 | 4.00 | 1.00 | 30.93 |  |
| 11 | LYS | NZ | 88:128 | 4.75 | 1.00 | 43.77 |  |
| 12 | G | O6 | 1H:1078 | 4.93 | 1.00 | 39.94 |  |
  |

  
**1^st^ shell ligand angles:**   
1-2: 87; 1-3: 81; 1-4: 94; 1-5: 91; 1-6: 178; 2-3: 92; 2-4: 86; 2-5: 177; 2-6: 93; 3-4: 174; 3-5: 90; 3-6: 98; 4-5: 92; 4-6: 88; 5-6: 89;   
  
 Contact(s): /O_w_/O_w_/O_w_/O_w_/O_w_/O_w_/  
 Coded contact list: 6O_w_  

### MG:1H:3548

|  | Res. type | Atom | Residue | Distance | Occupancy | B-factor | Closest solute to 1^st^ shell water |
| --- | --- | --- | --- | --- | --- | --- | --- |
| **Ion** | | | |  | 1.00 | 32.98 || 1 | Water 1 | | 1H:4008 | 2.17 | 1.00 | 20.79 | 2.65 (G:OP2 1H:2459) |
| 2 | Water 2 | | 1H:4139 | 2.18 | 1.00 | 20.34 | 2.81 (G:N7 1H:2515) |
| 3 | Water 3 | | 1H:3672 | 2.19 | 1.00 | 16.27 | 2.14 (G:OP2 1H:2460) |
| 4 | Water 4 | | 1H:3916 | 2.19 | 1.00 | 13.34 | 2.54 (G:N7 1H:2515) |
| 5 | Water 5 | | 1H:3814 | 2.20 | 1.00 | 29.53 | 2.40 (G:OP2 1H:2460) |
| 6 | Water 6 | | 1H:4436 | 2.22 | 1.00 | 31.02 | 3.42 (U:O4' 1H:600) |
| 7 | G | OP2 | 1H:2460 | 3.19 | 1.00 | 22.10 |  |
| 8 | G | N7 | 1H:2515 | 3.54 | 1.00 | 15.79 |  |
| 9 | G | OP1 | 1H:2459 | 3.93 | 1.00 | 19.33 |  |
| 10 | G | OP1 | 1H:2515 | 4.08 | 1.00 | 20.03 |  |
| 11 | U | O4' | 1H:600 | 4.56 | 1.00 | 17.56 |  |
| 12 | G | OP2 | 1H:2459 | 4.71 | 1.00 | 19.73 |  |
| 13 | G | OP1 | 1H:2460 | 4.81 | 1.00 | 26.48 |  |
| 14 | G | O5' | 1H:2459 | 4.85 | 1.00 | 14.18 |  |
  |

  
**1^st^ shell ligand angles:**   
1-2: 93; 1-3: 87; 1-4: 84; 1-5: 91; 1-6: 178; 2-3: 174; 2-4: 93; 2-5: 85; 2-6: 87; 3-4: 93; 3-5: 89; 3-6: 93; 4-5: 175; 4-6: 95; 5-6: 90;   
  
 Contact(s): /O_w_/O_w_/O_w_/O_w_/O_w_/O_w_/  
 Coded contact list: 6O_w_  
 Exclusion zone contact(s): /OP2/  
Consider K in place of MG (1 valid contact(s) for K vs. 0 for MG)  
MG hydration shell issue: water too close (< 2.4 Angs.)  

### MG:1H:3549

|  | Res. type | Atom | Residue | Distance | Occupancy | B-factor | Closest solute to 1^st^ shell water |
| --- | --- | --- | --- | --- | --- | --- | --- |
| **Ion** | | | |  | 1.00 | 60.73 || 1 | Water 1 | | 1H:4114 | 2.17 | 1.00 | 42.39 | 2.78 (G:N7 1H:1691) |
| 2 | Water 2 | | 1H:4345 | 2.18 | 1.00 | 28.85 | 3.17 (G:O6 1H:1690) |
| 3 | Water 3 | | 1H:4594 | 2.19 | 1.00 | 62.26 | 4.70 (G:O6 1H:1691) |
| 4 | Water 4 | | 1H:3723 | 2.19 | 1.00 | 45.75 | 2.25 (A:OP2 1H:1689) |
| 5 | Water 5 | | 1H:4445 | 2.20 | 1.00 | 49.03 | 3.48 (A:OP2 1H:1689) |
| 6 | Water 6 | | 1H:4080 | 2.22 | 1.00 | 47.91 | 2.74 (A:OP2 1H:1689) |
| 7 | A | OP2 | 1H:1689 | 3.35 | 1.00 | 35.77 |  |
| 8 | G | O6 | 1H:1690 | 3.95 | 1.00 | 29.96 |  |
| 9 | G | N7 | 1H:1690 | 4.02 | 1.00 | 36.16 |  |
| 10 | G | O6 | 1H:1691 | 4.52 | 1.00 | 32.36 |  |
| 11 | G | N7 | 1H:1691 | 4.87 | 1.00 | 31.99 |  |
| 12 | C | OP2 | 1H:1688 | 4.88 | 1.00 | 32.51 |  |
  |

  
**1^st^ shell ligand angles:**   
1-2: 87; 1-3: 88; 1-4: 90; 1-5: 90; 1-6: 178; 2-3: 91; 2-4: 86; 2-5: 176; 2-6: 93; 3-4: 177; 3-5: 92; 3-6: 90; 4-5: 90; 4-6: 92; 5-6: 90;   
  
 Contact(s): /O_w_/O_w_/O_w_/O_w_/O_w_/O_w_/  
 Coded contact list: 6O_w_  
 Exclusion zone contact(s): /OP2/  
Consider K in place of MG (1 valid contact(s) for K vs. 0 for MG)  
MG hydration shell issue: water too close (< 2.4 Angs.)  

### MG:1H:3550

|  | Res. type | Atom | Residue | Distance | Occupancy | B-factor | Closest solute to 1^st^ shell water |
| --- | --- | --- | --- | --- | --- | --- | --- |
| **Ion** | | | |  | 1.00 | 64.41 || 1 | Water 1 | | 1H:4179 | 2.18 | 1.00 | 47.71 | 2.87 (U:OP1 1H:502) |
| 2 | Water 2 | | 1H:4198 | 2.18 | 1.00 | 57.53 | 2.89 (A:OP1 1H:505) |
| 3 | Water 3 | | 1H:4589 | 2.19 | 1.00 | 71.26 | 4.57 (U:OP1 1H:502) |
| 4 | Water 4 | | 1H:3961 | 2.19 | 1.00 | 44.08 | 2.60 (U:OP1 1H:502) |
| 5 | Water 5 | | 1H:3603 | 2.19 | 1.00 | 59.79 | 1.30 (G:N7 1H:503) |
| 6 | Water 6 | | 1H:4373 | 2.21 | 1.00 | 39.43 | 3.24 (A:OP2 1H:505) |
| 7 | G | N7 | 1H:503 | 3.42 | 1.00 | 44.01 |  |
| 8 | U | OP1 | 1H:502 | 3.79 | 1.00 | 38.13 |  |
| 9 | A | OP2 | 1H:505 | 4.25 | 1.00 | 45.07 |  |
| 10 | A | OP1 | 1H:505 | 4.59 | 1.00 | 53.96 |  |
| 11 | G | OP2 | 1H:503 | 4.67 | 1.00 | 33.51 |  |
  |

  
**1^st^ shell ligand angles:**   
1-2: 85; 1-3: 93; 1-4: 90; 1-5: 87; 1-6: 178; 2-3: 88; 2-4: 175; 2-5: 91; 2-6: 96; 3-4: 92; 3-5: 179; 3-6: 90; 4-5: 89; 4-6: 90; 5-6: 90;   
  
 Contact(s): /O_w_/O_w_/O_w_/O_w_/O_w_/O_w_/  
 Coded contact list: 6O_w_  
MG hydration shell issue: water too close (< 2.4 Angs.)  

### MG:1H:3551

|  | Res. type | Atom | Residue | Distance | Occupancy | B-factor | Closest solute to 1^st^ shell water |
| --- | --- | --- | --- | --- | --- | --- | --- |
| **Ion** | | | |  | 1.00 | 45.39 || 1 | Water 1 | | 1H:3778 | 2.18 | 1.00 | 33.61 | 2.34 (G:OP1 1H:129) |
| 2 | Water 2 | | 1H:4554 | 2.19 | 1.00 | 45.91 | 4.02 (G:OP1 1H:129) |
| 3 | Water 3 | | 1H:4290 | 2.19 | 1.00 | 49.23 | 3.05 (G:OP1 1H:129) |
| 4 | Water 4 | | 1H:3982 | 2.19 | 1.00 | 48.88 | 2.62 (G:OP2 1H:142) |
| 5 | Water 5 | | 1H:3715 | 2.20 | 1.00 | 47.26 | 2.23 (C:OP1 1H:141) |
| 6 | G | OP1 | 1H:129 | 2.48 | 1.00 | 32.60 |  |
| 7 | G | O5' | 1H:129 | 3.96 | 1.00 | 36.20 |  |
| 8 | G | OP2 | 1H:142 | 4.25 | 1.00 | 25.60 |  |
| 9 | G | OP2 | 1H:129 | 4.35 | 1.00 | 27.46 |  |
| 10 | C | OP1 | 1H:141 | 4.39 | 1.00 | 32.49 |  |
| 11 | A | O2' | 1H:1396 | 4.66 | 1.00 | 30.99 |  |
| 12 | C | O3' | 1H:128 | 4.82 | 1.00 | 26.52 |  |
| 13 | G | OP1 | 1H:142 | 4.84 | 1.00 | 32.00 |  |
  |

  
**1^st^ shell ligand angles:**   
1-2: 178; 1-3: 91; 1-4: 88; 1-5: 88; 2-3: 91; 2-4: 91; 2-5: 93; 3-4: 178; 3-5: 88; 4-5: 93;   
  
 Contact(s): /O_w_/O_w_/O_w_/O_w_/O_w_/  
 Coded contact list: 5O_w_  
 Exclusion zone contact(s): /OP1/  
Consider NA in place of MG (1 valid contact(s) for NA vs. 0 for MG)  
MG hydration shell issue: water too close (< 2.4 Angs.)  

### MG:1H:3552

|  | Res. type | Atom | Residue | Distance | Occupancy | B-factor | Closest solute to 1^st^ shell water |
| --- | --- | --- | --- | --- | --- | --- | --- |
| **Ion** | | | |  | 1.00 | 85.39 || 1 | Water 1 | | 1H:4604 | 2.19 | 1.00 | 69.72 | 5.05 (C:OP2 1H:135) |
| 2 | Water 2 | | 1H:4125 | 2.19 | 1.00 | 53.65 | 2.80 (A:OP2 1H:139) |
| 3 | Water 3 | | 1H:4090 | 2.19 | 1.00 | 51.38 | 2.75 (G:OP1 1H:138) |
| 4 | Water 4 | | 1H:4135 | 2.20 | 1.00 | 55.92 | 2.81 (C:OP2 1H:135) |
| 5 | Water 5 | | 1H:4462 | 2.20 | 1.00 | 71.69 | 3.55 (G:OP1 1H:138) |
| 6 | Water 6 | | 1H:4524 | 2.20 | 1.00 | 62.79 | 3.82 (A:OP2 1H:139) |
| 7 | G | OP1 | 1H:138 | 4.21 | 1.00 | 42.83 |  |
| 8 | A | OP2 | 1H:139 | 4.37 | 1.00 | 48.05 |  |
| 9 | C | OP2 | 1H:135 | 4.47 | 1.00 | 47.49 |  |
| 10 | G | O5' | 1H:138 | 4.95 | 1.00 | 46.55 |  |
  |

  
**1^st^ shell ligand angles:**   
1-2: 89; 1-3: 87; 1-4: 92; 1-5: 91; 1-6: 179; 2-3: 92; 2-4: 88; 2-5: 179; 2-6: 91; 3-4: 179; 3-5: 88; 3-6: 93; 4-5: 93; 4-6: 87; 5-6: 88;   
  
 Contact(s): /O_w_/O_w_/O_w_/O_w_/O_w_/O_w_/  
 Coded contact list: 6O_w_  

### MG:1H:3553

|  | Res. type | Atom | Residue | Distance | Occupancy | B-factor | Closest solute to 1^st^ shell water |
| --- | --- | --- | --- | --- | --- | --- | --- |
| **Ion** | | | |  | 1.00 | 60.79 || 1 | Water 1 | | 1H:3870 | 2.17 | 1.00 | 49.42 | 2.48 (G:N7 1H:1482) |
| 2 | Water 2 | | 1H:3960 | 2.18 | 1.00 | 52.32 | 2.60 (G:O6 1H:1483) |
| 3 | Water 3 | | 1H:4477 | 2.18 | 1.00 | 42.65 | 4.10 (A:OP2 1H:1601) |
| 4 | Water 4 | | 1H:4315 | 2.19 | 1.00 | 57.06 | 3.81 (G:OP2 1H:1482) |
| 5 | Water 5 | | 1H:4423 | 2.19 | 1.00 | 36.88 | 3.38 (G:N7 1H:1483) |
| 6 | Water 6 | | 1H:4328 | 2.19 | 1.00 | 56.74 | 3.13 (G:OP2 1H:1482) |
| 7 | G | N7 | 1H:1482 | 3.93 | 1.00 | 52.82 |  |
| 8 | G | OP2 | 1H:1482 | 3.93 | 1.00 | 49.48 |  |
| 9 | G | O6 | 1H:1483 | 4.04 | 1.00 | 51.54 |  |
| 10 | G | N7 | 1H:1483 | 4.29 | 1.00 | 50.67 |  |
| 11 | A | OP2 | 1H:1601 | 4.85 | 1.00 | 45.73 |  |
| 12 | G | O6 | 1H:1482 | 4.99 | 1.00 | 51.21 |  |
  |

  
**1^st^ shell ligand angles:**   
1-2: 92; 1-3: 179; 1-4: 92; 1-5: 87; 1-6: 86; 2-3: 89; 2-4: 81; 2-5: 97; 2-6: 178; 3-4: 87; 3-5: 93; 3-6: 93; 4-5: 178; 4-6: 98; 5-6: 83;   
  
 Contact(s): /O_w_/O_w_/O_w_/O_w_/O_w_/O_w_/  
 Coded contact list: 6O_w_  

### MG:1H:3554

|  | Res. type | Atom | Residue | Distance | Occupancy | B-factor | Closest solute to 1^st^ shell water |
| --- | --- | --- | --- | --- | --- | --- | --- |
| **Ion** | | | |  | 1.00 | 77.78 || 1 | Water 1 | | 1H:4598 | 2.17 | 1.00 | 95.42 | 4.84 (G:N7 1H:1530) |
| 2 | Water 2 | | 1H:3935 | 2.18 | 1.00 | 92.70 | 2.56 (G:O6 1H:1531) |
| 3 | Water 3 | | 1H:3946 | 2.18 | 1.00 | 89.15 | 2.58 (G:N7 1H:1532) |
| 4 | Water 4 | | 1H:4192 | 2.18 | 1.00 | 87.28 | 2.88 (G:O6 1H:1532) |
| 5 | Water 5 | | 1H:4505 | 2.18 | 1.00 | 86.61 | 3.74 (G:N7 1H:1531) |
| 6 | Water 6 | | 1H:4601 | 2.18 | 1.00 | 87.52 | 4.91 (G:O6 1H:1532) |
| 7 | G | O6 | 1H:1532 | 3.96 | 1.00 | 89.87 |  |
| 8 | G | O6 | 1H:1531 | 4.02 | 1.00 | 102.10 |  |
| 9 | G | N7 | 1H:1531 | 4.38 | 1.00 | 97.78 |  |
| 10 | G | N7 | 1H:1532 | 4.66 | 1.00 | 97.15 |  |
| 11 | G | N7 | 1H:1530 | 4.84 | 1.00 | 101.49 |  |
  |

  
**1^st^ shell ligand angles:**   
1-2: 87; 1-3: 179; 1-4: 91; 1-5: 89; 1-6: 92; 2-3: 94; 2-4: 92; 2-5: 88; 2-6: 178; 3-4: 90; 3-5: 90; 3-6: 88; 4-5: 180; 4-6: 87; 5-6: 94;   
  
 Contact(s): /O_w_/O_w_/O_w_/O_w_/O_w_/O_w_/  
 Coded contact list: 6O_w_  

### MG:1H:3555

|  | Res. type | Atom | Residue | Distance | Occupancy | B-factor | Closest solute to 1^st^ shell water |
| --- | --- | --- | --- | --- | --- | --- | --- |
| **Ion** | | | |  | 1.00 | 56.62 || 1 | Water 1 | | 31:402 | 2.18 | 1.00 | 42.07 | 2.30 (SER:OG 31:98) |
| 2 | Water 2 | | 31:405 | 2.19 | 1.00 | 34.01 | 2.92 (SER:OG 31:98) |
| 3 | Water 3 | | 1H:4564 | 2.20 | 1.00 | 42.23 | 4.15 (A:OP2 1H:633) |
| 4 | Water 4 | | 1H:4057 | 2.20 | 1.00 | 48.87 | 2.70 (A:OP1 1H:633) |
| 5 | Water 5 | | 1H:4078 | 2.21 | 1.00 | 41.31 | 2.74 (A:OP1 1H:632) |
| 6 | Water 6 | | 31:410 | 2.22 | 1.00 | 36.56 | 4.16 (SER:OG 31:98) |
| 7 | SER | OG | 31:98 | 3.66 | 1.00 | 37.41 |  |
| 8 | A | OP1 | 1H:633 | 4.34 | 1.00 | 37.04 |  |
| 9 | THR | OG1 | 31:100 | 4.52 | 1.00 | 28.49 |  |
| 10 | A | OP1 | 1H:632 | 4.67 | 1.00 | 27.74 |  |
| 11 | A | OP2 | 1H:633 | 4.82 | 1.00 | 39.19 |  |
| 12 | ASP | OD2 | 31:96 | 4.85 | 1.00 | 40.06 |  |
  |

  
**1^st^ shell ligand angles:**   
1-2: 89; 1-3: 92; 1-4: 86; 1-5: 176; 1-6: 92; 2-3: 87; 2-4: 175; 2-5: 95; 2-6: 89; 3-4: 92; 3-5: 87; 3-6: 175; 4-5: 90; 4-6: 92; 5-6: 90;   
  
 Contact(s): /O_w_/O_w_/O_w_/O_w_/O_w_/O_w_/  
 Coded contact list: 6O_w_  
MG hydration shell issue: water too close (< 2.4 Angs.)  

### MG:1H:3556

|  | Res. type | Atom | Residue | Distance | Occupancy | B-factor | Closest solute to 1^st^ shell water |
| --- | --- | --- | --- | --- | --- | --- | --- |
| **Ion** | | | |  | 1.00 | 47.98 || 1 | Water 1 | | 1H:3879 | 2.18 | 1.00 | 29.58 | 2.49 (G:OP2 1H:792) |
| 2 | Water 2 | | 1H:4119 | 2.19 | 1.00 | 19.27 | 2.78 (G:N7 1H:792) |
| 3 | Water 3 | | 1H:4361 | 2.19 | 1.00 | 49.55 | 3.19 (G:OP2 1H:791) |
| 4 | Water 4 | | 1H:4550 | 2.20 | 1.00 | 37.75 | 3.99 (G:OP2 1H:792) |
| 5 | Water 5 | | 1H:4236 | 2.21 | 1.00 | 43.04 | 2.95 (G:O6 1H:793) |
| 6 | Water 6 | | 1H:4178 | 2.22 | 1.00 | 26.89 | 2.87 (G:N7 1H:793) |
| 7 | G | O6 | 1H:793 | 3.59 | 1.00 | 21.95 |  |
| 8 | G | N7 | 1H:792 | 4.10 | 1.00 | 22.54 |  |
| 9 | G | OP2 | 1H:792 | 4.13 | 1.00 | 21.38 |  |
| 10 | A | N6 | 1H:794 | 4.24 | 1.00 | 30.65 |  |
| 11 | G | OP2 | 1H:791 | 4.47 | 1.00 | 22.75 |  |
| 12 | G | N7 | 1H:793 | 4.60 | 1.00 | 23.17 |  |
| 13 | G | N7 | 1H:791 | 4.80 | 1.00 | 25.63 |  |
| 14 | C | OP1 | 1H:2623 | 4.94 | 1.00 | 29.86 |  |
  |

  
**1^st^ shell ligand angles:**   
1-2: 89; 1-3: 92; 1-4: 90; 1-5: 175; 1-6: 88; 2-3: 82; 2-4: 176; 2-5: 96; 2-6: 96; 3-4: 95; 3-5: 86; 3-6: 177; 4-5: 85; 4-6: 88; 5-6: 94;   
  
 Contact(s): /O_w_/O_w_/O_w_/O_w_/O_w_/O_w_/  
 Coded contact list: 6O_w_  

### MG:1H:3557

|  | Res. type | Atom | Residue | Distance | Occupancy | B-factor | Closest solute to 1^st^ shell water |
| --- | --- | --- | --- | --- | --- | --- | --- |
| **Ion** | | | |  | 1.00 | 22.13 || 1 | Water 1 | | 1H:4094 | 2.15 | 1.00 | 26.26 | 2.75 (G:N7 1H:2015) |
| 2 | Water 2 | | 1H:3952 | 2.16 | 1.00 | 23.72 | 2.59 (A:N6 1H:1713) |
| 3 | Water 3 | | 1H:4081 | 2.17 | 1.00 | 26.43 | 2.74 (G:O6 1H:1715) |
| 4 | Water 4 | | 1H:3696 | 2.17 | 1.00 | 20.60 | 2.19 (G:OP2 1H:2015) |
| 5 | Water 5 | | 1H:4336 | 2.19 | 1.00 | 27.44 | 3.15 (A:OP1 1H:1712) |
| 6 | A | OP2 | 1H:1712 | 2.43 | 1.00 | 26.83 |  |
| 7 | Water 6 | | 1H:3768 | 2.86 | 1.00 | 36.28 | 2.32 (A:OP1 1H:1712) |
| 8 | A | OP1 | 1H:1712 | 3.82 | 1.00 | 28.52 |  |
| 9 | G | O6 | 1H:1715 | 3.86 | 1.00 | 25.29 |  |
| 10 | G | OP2 | 1H:2015 | 3.87 | 1.00 | 25.17 |  |
| 11 | G | N7 | 1H:2015 | 3.94 | 1.00 | 22.61 |  |
| 12 | G | OP1 | 1H:2015 | 4.11 | 1.00 | 20.50 |  |
| 13 | A | N6 | 1H:1713 | 4.23 | 1.00 | 24.07 |  |
| 14 | A | N7 | 1H:1713 | 4.25 | 1.00 | 29.05 |  |
| 15 | G | O6 | 1H:2015 | 4.25 | 1.00 | 24.65 |  |
| 16 | C | O3' | 1H:1711 | 4.28 | 1.00 | 26.38 |  |
| 17 | G | O6 | 1H:1714 | 4.35 | 1.00 | 26.43 |  |
| 18 | G | N1 | 1H:1715 | 4.50 | 1.00 | 26.11 |  |
| 19 | A | N7 | 1H:1712 | 4.63 | 1.00 | 30.02 |  |
| 20 | A | O5' | 1H:1712 | 4.68 | 1.00 | 26.14 |  |
| 21 | G | N7 | 1H:1714 | 4.86 | 1.00 | 26.43 |  |
  |

  
**1^st^ shell ligand angles:**   
1-2: 89; 1-3: 176; 1-4: 92; 1-5: 89; 2-3: 87; 2-4: 94; 2-5: 175; 3-4: 90; 3-5: 94; 4-5: 91;   
  
 Contact(s): /O_w_/O_w_/O_w_/O_w_/O_w_/  
 Coded contact list: 5O_w_  
 Exclusion zone contact(s): /O_w_/OP2/  
Consider NA in place of MG (1 valid contact(s) for NA vs. 0 for MG)  
MG hydration shell issue: water too close (< 2.4 Angs.)  

### MG:1H:3558

|  | Res. type | Atom | Residue | Distance | Occupancy | B-factor | Closest solute to 1^st^ shell water |
| --- | --- | --- | --- | --- | --- | --- | --- |
| **Ion** | | | |  | 1.00 | 56.41 || 1 | Water 1 | | 1H:4228 | 2.17 | 1.00 | 46.09 | 2.93 (C:OP1 1H:2885) |
| 2 | Water 2 | | 1H:3855 | 2.17 | 1.00 | 38.12 | 2.46 (A:OP1 1H:2831) |
| 3 | Water 3 | | 1H:4417 | 2.18 | 1.00 | 41.35 | 3.36 (C:OP1 1H:2885) |
| 4 | Water 4 | | 1H:3624 | 2.18 | 1.00 | 40.15 | 1.96 (G:OP1 1H:2830) |
| 5 | Water 5 | | 1H:4362 | 2.19 | 1.00 | 47.70 | 3.19 (G:OP1 1H:2830) |
| 6 | Water 6 | | 1H:4313 | 2.20 | 1.00 | 59.81 | 3.10 (G:OP1 1H:2849) |
| 7 | G | OP1 | 1H:2830 | 3.43 | 1.00 | 31.11 |  |
| 8 | C | OP1 | 1H:2885 | 4.31 | 1.00 | 34.58 |  |
| 9 | A | OP1 | 1H:2831 | 4.62 | 1.00 | 22.51 |  |
| 10 | G | OP1 | 1H:2849 | 4.67 | 1.00 | 54.06 |  |
| 11 | LYS | O | 98:5 | 4.79 | 1.00 | 40.82 |  |
| 12 | SER | OG | 98:6 | 4.82 | 1.00 | 54.02 |  |
  |

  
**1^st^ shell ligand angles:**   
1-2: 90; 1-3: 86; 1-4: 179; 1-5: 93; 1-6: 89; 2-3: 86; 2-4: 88; 2-5: 94; 2-6: 179; 3-4: 93; 3-5: 179; 3-6: 92; 4-5: 88; 4-6: 92; 5-6: 87;   
  
 Contact(s): /O_w_/O_w_/O_w_/O_w_/O_w_/O_w_/  
 Coded contact list: 6O_w_  
MG hydration shell issue: water too close (< 2.4 Angs.)  

### MG:1H:3559

|  | Res. type | Atom | Residue | Distance | Occupancy | B-factor | Closest solute to 1^st^ shell water |
| --- | --- | --- | --- | --- | --- | --- | --- |
| **Ion** | | | |  | 1.00 | 15.75 || 1 | Water 1 | | 1H:3845 | 2.15 | 1.00 | 28.07 | 2.44 (C:O3' 1H:1694) |
| 2 | Water 2 | | 1H:4332 | 2.16 | 1.00 | 21.76 | 3.14 (G:OP2 1H:1695) |
| 3 | Water 3 | | 1H:4224 | 2.16 | 1.00 | 34.02 | 2.92 (A:OP2 1H:1319) |
| 4 | Water 4 | | 1H:3864 | 2.18 | 1.00 | 32.47 | 2.47 (G:OP1 1H:1372) |
| 5 | Water 5 | | 1H:4129 | 2.19 | 1.00 | 28.41 | 2.80 (A:O2' 1H:1663) |
| 6 | G | OP2 | 1H:1695 | 2.37 | 1.00 | 26.21 |  |
| 7 | C | O3' | 1H:1694 | 3.38 | 1.00 | 26.26 |  |
| 8 | A | O2' | 1H:1663 | 3.58 | 1.00 | 24.56 |  |
| 9 | G | OP1 | 1H:1695 | 4.00 | 1.00 | 28.42 |  |
| 10 | G | OP1 | 1H:1372 | 4.34 | 1.00 | 25.91 |  |
| 11 | A | OP2 | 1H:1319 | 4.62 | 1.00 | 26.16 |  |
| 12 | G | O5' | 1H:1695 | 4.68 | 1.00 | 28.29 |  |
| 13 | C | O2' | 1H:1694 | 4.72 | 1.00 | 28.88 |  |
  |

  
**1^st^ shell ligand angles:**   
1-2: 89; 1-3: 93; 1-4: 176; 1-5: 90; 1-6: 87; 2-3: 176; 2-4: 90; 2-5: 86; 2-6: 88; 3-4: 88; 3-5: 91; 3-6: 96; 4-5: 95; 4-6: 89; 5-6: 173;   
  
 Contact(s): /O_w_/O_w_/O_w_/O_w_/O_w_/OP2/  
 Coded contact list: O_ph_5O_w_  
 Exclusion zone contact(s): /O3'/  

### MG:1H:3560

|  | Res. type | Atom | Residue | Distance | Occupancy | B-factor | Closest solute to 1^st^ shell water |
| --- | --- | --- | --- | --- | --- | --- | --- |
| **Ion** | | | |  | 1.00 | 13.82 || 1 | Water 1 | | 1H:3825 | 2.14 | 1.00 | 17.00 | 2.41 (C:OP1 1H:2089) |
| 2 | Water 2 | | 1H:4214 | 2.15 | 1.00 | 26.16 | 2.91 (C:OP1 1H:2261) |
| 3 | Water 3 | | 1H:3978 | 2.15 | 1.00 | 24.42 | 2.62 (A:OP1 1H:2260) |
| 4 | Water 4 | | 1H:4009 | 2.17 | 1.00 | 21.49 | 2.65 (U:O4 1H:2262) |
| 5 | C | OP2 | 1H:2261 | 2.45 | 1.00 | 23.72 |  |
| 6 | U | O4 | 1H:2262 | 2.54 | 1.00 | 27.48 |  |
| 7 | C | OP1 | 1H:2261 | 3.74 | 1.00 | 25.12 |  |
| 8 | C | OP1 | 1H:2089 | 3.90 | 1.00 | 16.32 |  |
| 9 | A | O5' | 1H:2260 | 4.03 | 1.00 | 21.25 |  |
| 10 | G | OP1 | 1H:2266 | 4.18 | 1.00 | 16.30 |  |
| 11 | A | O3' | 1H:2260 | 4.43 | 1.00 | 22.31 |  |
| 12 | C | O5' | 1H:2261 | 4.55 | 1.00 | 21.43 |  |
| 13 | U | N3 | 1H:2262 | 4.60 | 1.00 | 22.13 |  |
| 14 | A | OP1 | 1H:2260 | 4.66 | 1.00 | 19.50 |  |
| 15 | A | OP2 | 1H:2260 | 4.85 | 1.00 | 24.54 |  |
  |

  
**1^st^ shell ligand angles:**   
1-2: 89; 1-3: 99; 1-4: 90; 2-3: 98; 2-4: 176; 3-4: 85;   
  
 Contact(s): /O_w_/O_w_/O_w_/O_w_/  
 Coded contact list: 4O_w_  
 Exclusion zone contact(s): /O4/OP2/  
Consider NA in place of MG (2 valid contact(s) for NA vs. 0 for MG)  

### MG:1H:3561

|  | Res. type | Atom | Residue | Distance | Occupancy | B-factor | Closest solute to 1^st^ shell water |
| --- | --- | --- | --- | --- | --- | --- | --- |
| **Ion** | | | |  | 1.00 | 38.91 || 1 | Water 1 | | 1H:4235 | 2.16 | 1.00 | 34.58 | 2.94 (G:O6 1H:1358) |
| 2 | Water 2 | | 1H:3722 | 2.17 | 1.00 | 34.94 | 2.25 (G:O6 1H:1357) |
| 3 | Water 3 | | 1H:3925 | 2.18 | 1.00 | 28.23 | 2.55 (A:OP2 1H:1355) |
| 4 | Water 4 | | 1H:4211 | 2.18 | 1.00 | 31.47 | 2.91 (G:O6 1H:1356) |
| 5 | Water 5 | | 1H:4559 | 2.18 | 1.00 | 41.28 | 4.09 (G:O6 1H:1358) |
| 6 | Water 6 | | 1H:3610 | 2.21 | 1.00 | 38.99 | 1.31 (G:N7 1H:1356) |
| 7 | G | N7 | 1H:1356 | 3.41 | 1.00 | 30.53 |  |
| 8 | G | O6 | 1H:1356 | 3.71 | 1.00 | 27.47 |  |
| 9 | G | O6 | 1H:1357 | 4.10 | 1.00 | 32.61 |  |
| 10 | G | O6 | 1H:1358 | 4.11 | 1.00 | 41.55 |  |
| 11 | G | N7 | 1H:1357 | 4.18 | 1.00 | 29.23 |  |
| 12 | A | OP2 | 1H:1355 | 4.46 | 1.00 | 25.77 |  |
  |

  
**1^st^ shell ligand angles:**   
1-2: 94; 1-3: 87; 1-4: 178; 1-5: 85; 1-6: 91; 2-3: 178; 2-4: 86; 2-5: 90; 2-6: 91; 3-4: 92; 3-5: 88; 3-6: 92; 4-5: 93; 4-6: 91; 5-6: 176;   
  
 Contact(s): /O_w_/O_w_/O_w_/O_w_/O_w_/O_w_/  
 Coded contact list: 6O_w_  
MG hydration shell issue: water too close (< 2.4 Angs.)  

### MG:1H:3562

|  | Res. type | Atom | Residue | Distance | Occupancy | B-factor | Closest solute to 1^st^ shell water |
| --- | --- | --- | --- | --- | --- | --- | --- |
| **Ion** | | | |  | 1.00 | 31.31 || 1 | Water 1 | | 1H:3736 | 2.15 | 1.00 | 36.11 | 2.27 (G:N7 1H:2284) |
| 2 | Water 2 | | 1H:3676 | 2.16 | 1.00 | 36.27 | 2.15 (U:O4 1H:2285) |
| 3 | Water 3 | | 1H:4360 | 2.18 | 1.00 | 40.57 | 3.19 (G:OP1 1H:2283) |
| 4 | Water 4 | | 1H:3751 | 2.18 | 1.00 | 39.10 | 2.30 (A:OP2 1H:2280) |
| 5 | Water 5 | | 1H:4140 | 2.19 | 1.00 | 35.91 | 2.81 (G:O6 1H:2284) |
| 6 | G | OP2 | 1H:2283 | 2.43 | 1.00 | 30.81 |  |
| 7 | A | OP2 | 1H:2280 | 3.78 | 1.00 | 33.91 |  |
| 8 | G | N7 | 1H:2284 | 3.94 | 1.00 | 29.23 |  |
| 9 | G | O5' | 1H:2283 | 3.98 | 1.00 | 29.47 |  |
| 10 | G | OP1 | 1H:2283 | 4.02 | 1.00 | 36.67 |  |
| 11 | G | O6 | 1H:2284 | 4.06 | 1.00 | 33.17 |  |
| 12 | U | O4 | 1H:2285 | 4.23 | 1.00 | 34.65 |  |
| 13 | G | OP2 | 1H:2284 | 4.58 | 1.00 | 26.99 |  |
| 14 | A | O3' | 1H:2282 | 4.73 | 1.00 | 41.77 |  |
| 15 | A | O5' | 1H:2282 | 4.82 | 1.00 | 34.85 |  |
| 16 | G | N7 | 1H:2283 | 4.86 | 1.00 | 30.17 |  |
| 17 | A | OP1 | 1H:2282 | 4.91 | 1.00 | 35.67 |  |
| 18 | A | O3' | 1H:2281 | 4.93 | 1.00 | 33.25 |  |
| 19 | A | O3' | 1H:2279 | 4.94 | 1.00 | 31.03 |  |
  |

  
**1^st^ shell ligand angles:**   
1-2: 86; 1-3: 87; 1-4: 177; 1-5: 90; 2-3: 92; 2-4: 94; 2-5: 90; 3-4: 89; 3-5: 176; 4-5: 93;   
  
 Contact(s): /O_w_/O_w_/O_w_/O_w_/O_w_/  
 Coded contact list: 5O_w_  
 Exclusion zone contact(s): /OP2/  
Consider NA in place of MG (1 valid contact(s) for NA vs. 0 for MG)  
MG hydration shell issue: water too close (< 2.4 Angs.)  

### MG:1H:3563

|  | Res. type | Atom | Residue | Distance | Occupancy | B-factor | Closest solute to 1^st^ shell water |
| --- | --- | --- | --- | --- | --- | --- | --- |
| **Ion** | | | |  | 1.00 | 75.64 || 1 | Water 1 | | 1H:3967 | 2.17 | 1.00 | 72.71 | 2.60 (G:OP2 1H:640) |
| 2 | Water 2 | | 1H:4131 | 2.18 | 1.00 | 68.61 | 2.80 (U:O3' 1H:639) |
| 3 | Water 3 | | 1H:4208 | 2.18 | 1.00 | 61.42 | 2.90 (U:OP2 1H:639) |
| 4 | Water 4 | | 1H:4474 | 2.18 | 1.00 | 64.80 | 3.59 (U:N1 1H:638) |
| 5 | Water 5 | | 1H:4553 | 2.19 | 1.00 | 71.66 | 4.02 (U:OP2 1H:639) |
| 6 | Water 6 | | 1H:4561 | 2.19 | 1.00 | 60.38 | 4.91 (U:N1 1H:638) |
| 7 | U | OP2 | 1H:639 | 3.95 | 1.00 | 56.76 |  |
| 8 | G | OP2 | 1H:640 | 4.37 | 1.00 | 78.91 |  |
| 9 | U | O3' | 1H:639 | 4.47 | 1.00 | 91.31 |  |
| 10 | U | N1 | 1H:638 | 4.68 | 1.00 | 49.04 |  |
  |

  
**1^st^ shell ligand angles:**   
1-2: 86; 1-3: 176; 1-4: 88; 1-5: 91; 1-6: 95; 2-3: 92; 2-4: 91; 2-5: 90; 2-6: 176; 3-4: 96; 3-5: 85; 3-6: 87; 4-5: 178; 4-6: 85; 5-6: 93;   
  
 Contact(s): /O_w_/O_w_/O_w_/O_w_/O_w_/O_w_/  
 Coded contact list: 6O_w_  

### MG:1H:3564

|  | Res. type | Atom | Residue | Distance | Occupancy | B-factor | Closest solute to 1^st^ shell water |
| --- | --- | --- | --- | --- | --- | --- | --- |
| **Ion** | | | |  | 1.00 | 25.47 || 1 | Water 1 | | 1H:3606 | 2.17 | 1.00 | 35.04 | 1.30 (A:N6 1H:1300) |
| 2 | Water 2 | | 1H:3938 | 2.18 | 1.00 | 22.63 | 2.56 (A:N7 1H:1300) |
| 3 | Water 3 | | 1H:3994 | 2.18 | 1.00 | 41.93 | 2.63 (U:OP1 1H:859) |
| 4 | Water 4 | | 1H:4075 | 2.19 | 1.00 | 22.31 | 2.73 (U:OP1 1H:590) |
| 5 | U | OP1 | 1H:858 | 2.49 | 1.00 | 23.01 |  |
| 6 | A | N7 | 1H:1300 | 2.76 | 1.00 | 24.08 |  |
| 7 | A | N6 | 1H:1300 | 3.40 | 1.00 | 38.03 |  |
| 8 | U | OP2 | 1H:858 | 3.46 | 1.00 | 22.11 |  |
| 9 | U | OP1 | 1H:859 | 3.83 | 1.00 | 23.97 |  |
| 10 | C | O3' | 1H:589 | 4.21 | 1.00 | 22.02 |  |
| 11 | U | O5' | 1H:858 | 4.22 | 1.00 | 16.41 |  |
| 12 | C | O2' | 1H:589 | 4.51 | 1.00 | 24.47 |  |
| 13 | GLN | OE1 | D8:80 | 4.55 | 1.00 | 29.40 |  |
| 14 | GLN | NE2 | D8:80 | 4.57 | 1.00 | 30.26 |  |
| 15 | G | O3' | 1H:857 | 4.59 | 1.00 | 18.82 |  |
| 16 | U | OP1 | 1H:590 | 4.81 | 1.00 | 16.39 |  |
| 17 | A | N9 | 1H:1300 | 4.94 | 1.00 | 25.26 |  |
  |

  
**1^st^ shell ligand angles:**   
1-2: 84; 1-3: 95; 1-4: 99; 2-3: 90; 2-4: 175; 3-4: 86;   
  
 Contact(s): /O_w_/O_w_/O_w_/O_w_/  
 Coded contact list: 4O_w_  
 Exclusion zone contact(s): /N7/OP1/  
Consider NA in place of MG (1 valid contact(s) for NA vs. 0 for MG)  
MG hydration shell issue: water too close (< 2.4 Angs.)  

### MG:1H:3565

|  | Res. type | Atom | Residue | Distance | Occupancy | B-factor | Closest solute to 1^st^ shell water |
| --- | --- | --- | --- | --- | --- | --- | --- |
| **Ion** | | | |  | 1.00 | 25.61 || 1 | Water 1 | | 1H:4095 | 2.15 | 1.00 | 34.96 | 2.75 (G:N7 1H:2423) |
| 2 | Water 2 | | 1H:3840 | 2.17 | 1.00 | 41.65 | 2.43 (U:OP2 1H:2421) |
| 3 | Water 3 | | 1H:3937 | 2.18 | 1.00 | 37.37 | 2.56 (U:OP2 1H:2421) |
| 4 | Water 4 | | 1H:3791 | 2.19 | 1.00 | 40.71 | 2.36 (U:OP1 1H:2419) |
| 5 | Water 5 | | 1H:3856 | 2.19 | 1.00 | 39.65 | 2.46 (G:N7 1H:2422) |
| 6 | U | OP1 | 1H:2419 | 2.56 | 1.00 | 35.03 |  |
| 7 | G | O3' | 1H:2418 | 3.39 | 1.00 | 42.96 |  |
| 8 | U | OP2 | 1H:2421 | 3.60 | 1.00 | 41.03 |  |
| 9 | U | O5' | 1H:2419 | 3.92 | 1.00 | 37.22 |  |
| 10 | G | O6 | 1H:2423 | 4.16 | 1.00 | 35.81 |  |
| 11 | G | N7 | 1H:2423 | 4.39 | 1.00 | 37.49 |  |
| 12 | G | N7 | 1H:2422 | 4.42 | 1.00 | 41.77 |  |
| 13 | U | O4 | 1H:2421 | 4.56 | 1.00 | 38.99 |  |
| 14 | U | OP2 | 1H:2419 | 4.70 | 1.00 | 43.70 |  |
| 15 | U | O3' | 1H:2419 | 4.83 | 1.00 | 42.10 |  |
| 16 | A | N6 | 1H:2424 | 4.88 | 1.00 | 32.96 |  |
| 17 | G | O6 | 1H:2422 | 4.94 | 1.00 | 42.55 |  |
| 18 | G | OP2 | 1H:2422 | 4.96 | 1.00 | 43.81 |  |
  |

  
**1^st^ shell ligand angles:**   
1-2: 97; 1-3: 179; 1-4: 84; 1-5: 84; 2-3: 84; 2-4: 179; 2-5: 94; 3-4: 95; 3-5: 96; 4-5: 85;   
  
 Contact(s): /O_w_/O_w_/O_w_/O_w_/O_w_/  
 Coded contact list: 5O_w_  
 Exclusion zone contact(s): /O3'/OP1/  
Consider NA in place of MG (1 valid contact(s) for NA vs. 0 for MG)  
MG hydration shell issue: water too close (< 2.4 Angs.)  

### MG:1H:3566

|  | Res. type | Atom | Residue | Distance | Occupancy | B-factor | Closest solute to 1^st^ shell water |
| --- | --- | --- | --- | --- | --- | --- | --- |
| **Ion** | | | |  | 1.00 | 37.50 || 1 | Water 1 | | 1H:4029 | 2.16 | 1.00 | 40.14 | 2.67 (C:OP1 1H:420) |
| 2 | Water 2 | | 1H:4089 | 2.18 | 1.00 | 34.96 | 2.75 (C:OP2 1H:437) |
| 3 | Water 3 | | 1H:3719 | 2.18 | 1.00 | 35.28 | 2.24 (C:OP2 1H:404) |
| 4 | Water 4 | | 1H:3799 | 2.20 | 1.00 | 39.57 | 2.38 (C:OP1 1H:403) |
| 5 | C | OP1 | 1H:420 | 2.38 | 1.00 | 34.42 |  |
| 6 | C | OP1 | 1H:437 | 2.46 | 1.00 | 36.34 |  |
| 7 | C | OP2 | 1H:437 | 3.56 | 1.00 | 37.23 |  |
| 8 | G | O3' | 1H:436 | 3.72 | 1.00 | 35.90 |  |
| 9 | C | OP1 | 1H:404 | 3.88 | 1.00 | 34.60 |  |
| 10 | C | OP1 | 1H:403 | 4.09 | 1.00 | 31.51 |  |
| 11 | C | OP2 | 1H:404 | 4.11 | 1.00 | 34.60 |  |
| 12 | C | OP2 | 1H:420 | 4.33 | 1.00 | 38.33 |  |
| 13 | C | O5' | 1H:420 | 4.53 | 1.00 | 31.49 |  |
| 14 | C | O5' | 1H:437 | 4.59 | 1.00 | 38.68 |  |
| 15 | G | O3' | 1H:419 | 4.66 | 1.00 | 38.00 |  |
| 16 | C | OP2 | 1H:421 | 4.88 | 1.00 | 31.98 |  |
  |

  
**1^st^ shell ligand angles:**   
1-2: 170; 1-3: 83; 1-4: 96; 1-5: 72; 2-3: 104; 2-4: 90; 2-5: 100; 3-4: 92; 3-5: 103; 4-5: 158;   
  
 Contact(s): /O_w_/O_w_/O_w_/O_w_/OP1/  
 Coded contact list: O_ph_4O_w_  
 Exclusion zone contact(s): /OP1/  
Large octahedral topology discrepancy!!!  
MG hydration shell issue: water too close (< 2.4 Angs.)  

### MG:1H:3567

|  | Res. type | Atom | Residue | Distance | Occupancy | B-factor | Closest solute to 1^st^ shell water |
| --- | --- | --- | --- | --- | --- | --- | --- |
| **Ion** | | | |  | 1.00 | 24.37 || 1 | Water 1 | | 1H:3787 | 2.17 | 1.00 | 24.98 | 2.36 (A:OP1 1H:850) |
| 2 | Water 2 | | 1H:3860 | 2.19 | 1.00 | 24.06 | 2.46 (A:OP2 1H:723) |
| 3 | Water 3 | | 1H:3634 | 2.19 | 1.00 | 23.70 | 1.99 (G:OP2 1H:722) |
| 4 | Water 4 | | 1H:4359 | 2.20 | 1.00 | 19.34 | 3.18 (U:O4 1H:851) |
| 5 | Water 5 | | 1H:4032 | 2.22 | 1.00 | 24.16 | 2.67 (A:OP2 1H:723) |
| 6 | A | OP1 | 1H:850 | 2.51 | 1.00 | 19.69 |  |
| 7 | G | OP2 | 1H:722 | 3.53 | 1.00 | 17.62 |  |
| 8 | A | OP2 | 1H:850 | 3.56 | 1.00 | 19.58 |  |
| 9 | A | OP2 | 1H:723 | 3.60 | 1.00 | 16.82 |  |
| 10 | U | O4 | 1H:851 | 4.49 | 1.00 | 18.47 |  |
| 11 | G | O3' | 1H:849 | 4.51 | 1.00 | 21.99 |  |
| 12 | A | O5' | 1H:723 | 4.51 | 1.00 | 18.75 |  |
| 13 | G | OP1 | 1H:722 | 4.59 | 1.00 | 22.86 |  |
| 14 | ARG | NH2 | 31:54 | 4.61 | 1.00 | 17.09 |  |
| 15 | G | O5' | 1H:722 | 4.69 | 1.00 | 17.25 |  |
| 16 | A | O5' | 1H:850 | 4.69 | 1.00 | 18.74 |  |
| 17 | A | N6 | 1H:852 | 4.85 | 1.00 | 18.22 |  |
  |

  
**1^st^ shell ligand angles:**   
1-2: 94; 1-3: 83; 1-4: 91; 1-5: 178; 2-3: 91; 2-4: 89; 2-5: 88; 3-4: 174; 3-5: 96; 4-5: 90;   
  
 Contact(s): /O_w_/O_w_/O_w_/O_w_/O_w_/  
 Coded contact list: 5O_w_  
 Exclusion zone contact(s): /OP1/  
Consider NA in place of MG (1 valid contact(s) for NA vs. 0 for MG)  
MG hydration shell issue: water too close (< 2.4 Angs.)  

### MG:1H:3568

|  | Res. type | Atom | Residue | Distance | Occupancy | B-factor | Closest solute to 1^st^ shell water |
| --- | --- | --- | --- | --- | --- | --- | --- |
| **Ion** | | | |  | 1.00 | 34.36 || 1 | Water 1 | | 1H:3940 | 2.16 | 1.00 | 48.41 | 2.57 (C:OP2 1H:2788) |
| 2 | Water 2 | | 1H:3730 | 2.17 | 1.00 | 38.20 | 2.26 (A:N7 1H:2789) |
| 3 | Water 3 | | 21:410 | 2.18 | 1.00 | 34.14 | 3.48 (ARG:NH1 21:164) |
| 4 | Water 4 | | 1H:4273 | 2.18 | 1.00 | 35.75 | 3.01 (A:OP2 1H:2789) |
| 5 | Water 5 | | 1H:4237 | 2.18 | 1.00 | 62.16 | 2.95 (C:OP2 1H:2788) |
| 6 | Water 6 | | 1H:4410 | 2.20 | 1.00 | 38.64 | 3.34 (A:N7 1H:2789) |
| 7 | C | OP2 | 1H:2788 | 3.59 | 1.00 | 39.88 |  |
| 8 | A | N7 | 1H:2789 | 3.80 | 1.00 | 33.81 |  |
| 9 | A | OP2 | 1H:2789 | 4.25 | 1.00 | 37.39 |  |
| 10 | ARG | NH1 | 21:164 | 4.58 | 1.00 | 49.10 |  |
| 11 | C | O5' | 1H:2788 | 4.63 | 1.00 | 29.34 |  |
| 12 | C | OP1 | 1H:2788 | 4.78 | 1.00 | 41.62 |  |
  |

  
**1^st^ shell ligand angles:**   
1-2: 93; 1-3: 86; 1-4: 86; 1-5: 90; 1-6: 178; 2-3: 179; 2-4: 88; 2-5: 88; 2-6: 89; 3-4: 93; 3-5: 91; 3-6: 92; 4-5: 174; 4-6: 95; 5-6: 89;   
  
 Contact(s): /O_w_/O_w_/O_w_/O_w_/O_w_/O_w_/  
 Coded contact list: 6O_w_  
MG hydration shell issue: water too close (< 2.4 Angs.)  

### MG:1H:3569

|  | Res. type | Atom | Residue | Distance | Occupancy | B-factor | Closest solute to 1^st^ shell water |
| --- | --- | --- | --- | --- | --- | --- | --- |
| **Ion** | | | |  | 1.00 | 62.12 || 1 | Water 1 | | 1H:4584 | 2.19 | 1.00 | 46.52 | 4.50 (G:OP1 1H:1069) |
| 2 | Water 2 | | 1H:3857 | 2.19 | 1.00 | 52.11 | 2.46 (A:O2' 1H:1068) |
| 3 | Water 3 | | 1H:4271 | 2.19 | 1.00 | 56.05 | 3.01 (C:OP1 1H:1171) |
| 4 | Water 4 | | 1H:4277 | 2.19 | 1.00 | 34.62 | 3.01 (C:OP2 1H:1171) |
| 5 | Water 5 | | 1H:4358 | 2.20 | 1.00 | 52.05 | 3.18 (C:OP1 1H:1171) |
| 6 | C | OP1 | 1H:1171 | 2.47 | 1.00 | 32.03 |  |
| 7 | C | OP2 | 1H:1171 | 3.89 | 1.00 | 41.55 |  |
| 8 | A | O2' | 1H:1068 | 4.03 | 1.00 | 48.08 |  |
| 9 | C | O3' | 1H:1170 | 4.46 | 1.00 | 44.17 |  |
| 10 | C | O5' | 1H:1171 | 4.62 | 1.00 | 34.80 |  |
| 11 | G | OP1 | 1H:1069 | 4.91 | 1.00 | 49.94 |  |
  |

  
**1^st^ shell ligand angles:**   
1-2: 94; 1-3: 91; 1-4: 94; 1-5: 89; 2-3: 175; 2-4: 89; 2-5: 91; 3-4: 90; 3-5: 89; 4-5: 177;   
  
 Contact(s): /O_w_/O_w_/O_w_/O_w_/O_w_/  
 Coded contact list: 5O_w_  
 Exclusion zone contact(s): /OP1/  
Consider NA in place of MG (1 valid contact(s) for NA vs. 0 for MG)  

### MG:1H:3570

|  | Res. type | Atom | Residue | Distance | Occupancy | B-factor | Closest solute to 1^st^ shell water |
| --- | --- | --- | --- | --- | --- | --- | --- |
| **Ion** | | | |  | 1.00 | 50.81 || 1 | Water 1 | | 1H:3763 | 2.17 | 1.00 | 49.73 | 2.32 (G:N7 1H:1887) |
| 2 | Water 2 | | 1H:4493 | 2.18 | 1.00 | 47.34 | 3.66 (A:OP2 1H:1886) |
| 3 | Water 3 | | 1H:3897 | 2.18 | 1.00 | 56.77 | 2.51 (G:O6 1H:1888) |
| 4 | Water 4 | | 1H:4407 | 2.18 | 1.00 | 63.52 | 3.34 (G:O6 1H:1889) |
| 5 | Water 5 | | 1H:4424 | 2.18 | 1.00 | 48.42 | 3.38 (G:O6 1H:1889) |
| 6 | Water 6 | | 1H:4509 | 2.19 | 1.00 | 50.62 | 3.75 (A:OP2 1H:1886) |
| 7 | G | N7 | 1H:1887 | 3.91 | 1.00 | 45.31 |  |
| 8 | G | O6 | 1H:1887 | 3.97 | 1.00 | 52.99 |  |
| 9 | G | O6 | 1H:1888 | 4.25 | 1.00 | 59.95 |  |
| 10 | A | OP2 | 1H:1886 | 4.31 | 1.00 | 37.82 |  |
| 11 | G | N7 | 1H:1888 | 4.32 | 1.00 | 60.05 |  |
| 12 | G | O6 | 1H:1889 | 4.49 | 1.00 | 71.96 |  |
| 13 | G | OP2 | 1H:1887 | 4.84 | 1.00 | 40.99 |  |
  |

  
**1^st^ shell ligand angles:**   
1-2: 85; 1-3: 91; 1-4: 178; 1-5: 93; 1-6: 89; 2-3: 175; 2-4: 94; 2-5: 94; 2-6: 88; 3-4: 91; 3-5: 87; 3-6: 92; 4-5: 87; 4-6: 90; 5-6: 177;   
  
 Contact(s): /O_w_/O_w_/O_w_/O_w_/O_w_/O_w_/  
 Coded contact list: 6O_w_  
MG hydration shell issue: water too close (< 2.4 Angs.)  

### MG:1H:3571

|  | Res. type | Atom | Residue | Distance | Occupancy | B-factor | Closest solute to 1^st^ shell water |
| --- | --- | --- | --- | --- | --- | --- | --- |
| **Ion** | | | |  | 1.00 | 26.33 || 1 | Water 1 | | 1H:3647 | 2.15 | 1.00 | 27.75 | 2.05 (A:O5' 1H:2531) |
| 2 | Water 2 | | 1H:4268 | 2.16 | 1.00 | 40.61 | 3.00 (A:OP2 1H:2531) |
| 3 | Water 3 | | 1H:4117 | 2.17 | 1.00 | 27.01 | 2.78 (A:OP2 1H:2554) |
| 4 | Water 4 | | 1H:4296 | 2.17 | 1.00 | 32.17 | 3.06 (A:N7 1H:2554) |
| 5 | Water 5 | | 1H:4175 | 2.17 | 1.00 | 33.50 | 2.87 (C:O2' 1H:2530) |
| 6 | A | OP2 | 1H:2531 | 2.50 | 1.00 | 37.77 |  |
| 7 | A | O5' | 1H:2531 | 3.07 | 1.00 | 28.73 |  |
| 8 | C | O3' | 1H:2530 | 3.89 | 1.00 | 29.98 |  |
| 9 | A | OP2 | 1H:2554 | 4.10 | 1.00 | 32.96 |  |
| 10 | C | OP2 | 1H:2553 | 4.19 | 1.00 | 33.36 |  |
| 11 | A | N7 | 1H:2554 | 4.40 | 1.00 | 32.84 |  |
| 12 | C | O2' | 1H:2530 | 4.42 | 1.00 | 32.37 |  |
| 13 | A | OP1 | 1H:2531 | 4.49 | 1.00 | 37.32 |  |
| 14 | U | OP2 | 1H:2532 | 4.57 | 1.00 | 33.70 |  |
| 15 | A | O3' | 1H:2531 | 4.66 | 1.00 | 35.80 |  |
  |

  
**1^st^ shell ligand angles:**   
1-2: 89; 1-3: 177; 1-4: 92; 1-5: 89; 2-3: 88; 2-4: 98; 2-5: 173; 3-4: 87; 3-5: 94; 4-5: 89;   
  
 Contact(s): /O_w_/O_w_/O_w_/O_w_/O_w_/  
 Coded contact list: 5O_w_  
 Exclusion zone contact(s): /O5'/OP2/  
Consider NA in place of MG (1 valid contact(s) for NA vs. 0 for MG)  
MG hydration shell issue: water too close (< 2.4 Angs.)  

### MG:1H:3572

|  | Res. type | Atom | Residue | Distance | Occupancy | B-factor | Closest solute to 1^st^ shell water |
| --- | --- | --- | --- | --- | --- | --- | --- |
| **Ion** | | | |  | 1.00 | 25.28 || 1 | Water 1 | | 1H:3923 | 2.15 | 1.00 | 21.86 | 2.54 (U:OP1 1H:2626) |
| 2 | Water 2 | | 1H:4454 | 2.16 | 1.00 | 16.75 | 3.53 (A:OP2 1H:2627) |
| 3 | Water 3 | | 1H:4092 | 2.19 | 1.00 | 32.70 | 2.75 (G:OP2 1H:2591) |
| 4 | Water 4 | | 1H:3841 | 2.19 | 1.00 | 26.08 | 2.43 (A:OP2 1H:2074) |
| 5 | Water 5 | | 1H:4517 | 2.20 | 1.00 | 18.21 | 3.78 (C:OP2 1H:2073) |
| 6 | Water 6 | | 1H:3956 | 2.20 | 1.00 | 20.00 | 2.59 (G:OP1 1H:2591) |
| 7 | G | OP1 | 1H:2591 | 3.91 | 1.00 | 19.86 |  |
| 8 | G | OP2 | 1H:2591 | 4.03 | 1.00 | 21.75 |  |
| 9 | U | OP1 | 1H:2626 | 4.21 | 1.00 | 24.77 |  |
| 10 | A | OP2 | 1H:2074 | 4.32 | 1.00 | 20.42 |  |
| 11 | A | N7 | 1H:2627 | 4.34 | 1.00 | 23.07 |  |
| 12 | A | N7 | 1H:2074 | 4.52 | 1.00 | 21.40 |  |
| 13 | A | OP2 | 1H:2627 | 4.78 | 1.00 | 21.62 |  |
| 14 | C | OP2 | 1H:2073 | 4.81 | 1.00 | 19.60 |  |
  |

  
**1^st^ shell ligand angles:**   
1-2: 87; 1-3: 96; 1-4: 175; 1-5: 96; 1-6: 82; 2-3: 176; 2-4: 92; 2-5: 91; 2-6: 92; 3-4: 86; 3-5: 85; 3-6: 91; 4-5: 89; 4-6: 93; 5-6: 176;   
  
 Contact(s): /O_w_/O_w_/O_w_/O_w_/O_w_/O_w_/  
 Coded contact list: 6O_w_  

### MG:1H:3573

|  | Res. type | Atom | Residue | Distance | Occupancy | B-factor | Closest solute to 1^st^ shell water |
| --- | --- | --- | --- | --- | --- | --- | --- |
| **Ion** | | | |  | 1.00 | 40.19 || 1 | Water 1 | | 1H:3792 | 2.16 | 1.00 | 29.43 | 2.37 (A:OP1 1H:2082) |
| 2 | Water 2 | | 1H:4460 | 2.19 | 1.00 | 25.49 | 3.55 (U:O2 1H:1302) |
| 3 | Water 3 | | 1H:3691 | 2.20 | 1.00 | 25.22 | 2.18 (A:OP2 1H:2082) |
| 4 | Water 4 | | 31:408 | 2.20 | 1.00 | 39.14 | 3.43 (GLY:O 31:71) |
| 5 | U | O2 | 1H:1302 | 2.82 | 1.00 | 22.61 |  |
| 6 | A | OP2 | 1H:2082 | 3.01 | 1.00 | 17.54 |  |
| 7 | A | OP1 | 1H:2082 | 3.64 | 1.00 | 26.02 |  |
| 8 | GLY | O | 31:71 | 3.72 | 1.00 | 27.50 |  |
| 9 | A | OP1 | 1H:2081 | 4.19 | 1.00 | 19.11 |  |
| 10 | U | OP1 | 1H:2039 | 4.38 | 1.00 | 26.41 |  |
| 11 | U | O2' | 1H:1302 | 4.55 | 1.00 | 21.54 |  |
| 12 | A | O5' | 1H:2082 | 4.62 | 1.00 | 21.62 |  |
| 13 | U | N3 | 1H:1302 | 4.85 | 1.00 | 22.58 |  |
| 14 | A | OP2 | 1H:2083 | 4.94 | 1.00 | 19.84 |  |
| 15 | Water 5 | | 1H:3686 | 4.99 | 1.00 | 22.54 | 3.84 (A:OP1 1H:2081) |
| 16 | U | N1 | 1H:1302 | 4.99 | 1.00 | 17.98 |  |
  |

  
**1^st^ shell ligand angles:**   
1-2: 174; 1-3: 92; 1-4: 81; 2-3: 92; 2-4: 94; 3-4: 173;   
  
 Contact(s): /O_w_/O_w_/O_w_/O_w_/  
 Coded contact list: 4O_w_  
 Exclusion zone contact(s): /O2/OP2/  
Consider K in place of MG (2 valid contact(s) for K vs. 0 for MG)  
MG hydration shell issue: water too close (< 2.4 Angs.)  

### MG:1H:3574

|  | Res. type | Atom | Residue | Distance | Occupancy | B-factor | Closest solute to 1^st^ shell water |
| --- | --- | --- | --- | --- | --- | --- | --- |
| **Ion** | | | |  | 1.00 | 24.51 || 1 | Water 1 | | 1H:4104 | 2.16 | 1.00 | 31.63 | 2.76 (C:OP2 1H:1054) |
| 2 | Water 2 | | 1H:4096 | 2.16 | 1.00 | 32.70 | 2.75 (A:OP2 1H:1057) |
| 3 | Water 3 | | 1H:4142 | 2.17 | 1.00 | 36.97 | 2.82 (C:O3' 1H:1055) |
| 4 | Water 4 | | 1H:3682 | 2.17 | 1.00 | 40.54 | 2.16 (A:OP1 1H:1057) |
| 5 | A | OP2 | 1H:1057 | 2.57 | 1.00 | 38.44 |  |
| 6 | A | OP2 | 1H:1056 | 2.68 | 1.00 | 34.38 |  |
| 7 | A | O5' | 1H:1056 | 3.11 | 1.00 | 38.14 |  |
| 8 | A | OP1 | 1H:1057 | 3.43 | 1.00 | 28.91 |  |
| 9 | C | O3' | 1H:1055 | 3.72 | 1.00 | 39.24 |  |
| 10 | LYS | NZ | 58:37 | 3.92 | 1.00 | 36.16 |  |
| 11 | C | OP2 | 1H:1054 | 4.09 | 1.00 | 32.72 |  |
| 12 | A | O3' | 1H:1056 | 4.16 | 1.00 | 43.73 |  |
| 13 | A | N7 | 1H:1057 | 4.21 | 1.00 | 36.59 |  |
| 14 | C | OP2 | 1H:1053 | 4.49 | 1.00 | 38.06 |  |
| 15 | A | OP1 | 1H:1056 | 4.70 | 1.00 | 43.73 |  |
| 16 | A | O5' | 1H:1057 | 4.70 | 1.00 | 33.88 |  |
| 17 | C | OP1 | 1H:1053 | 4.78 | 1.00 | 40.44 |  |
| 18 | C | O2' | 1H:1055 | 4.89 | 1.00 | 36.45 |  |
  |

  
**1^st^ shell ligand angles:**   
1-2: 79; 1-3: 95; 1-4: 90; 2-3: 92; 2-4: 92; 3-4: 174;   
  
 Contact(s): /O_w_/O_w_/O_w_/O_w_/  
 Coded contact list: 4O_w_  
 Exclusion zone contact(s): /O5'/OP2/OP2/  
Consider K in place of MG (2 valid contact(s) for K vs. 0 for MG)  
MG hydration shell issue: water too close (< 2.4 Angs.)  

### MG:1H:3575

|  | Res. type | Atom | Residue | Distance | Occupancy | B-factor | Closest solute to 1^st^ shell water |
| --- | --- | --- | --- | --- | --- | --- | --- |
| **Ion** | | | |  | 1.00 | 17.73 || 1 | Water 1 | | 1H:3931 | 2.17 | 1.00 | 21.09 | 2.55 (C:N3 1H:1664) |
| 2 | Water 2 | | 1H:4018 | 2.18 | 1.00 | 19.48 | 2.66 (A:OP1 1H:1661) |
| 3 | Water 3 | | 1H:4173 | 2.18 | 1.00 | 28.15 | 2.86 (A:N7 1H:799) |
| 4 | Water 4 | | 1H:4215 | 2.19 | 1.00 | 24.24 | 2.91 (C:N4 1H:1662) |
| 5 | A | OP1 | 1H:1661 | 2.30 | 1.00 | 13.52 |  |
| 6 | A | OP1 | 1H:799 | 2.37 | 1.00 | 19.58 |  |
| 7 | A | N7 | 1H:1661 | 3.55 | 1.00 | 25.26 |  |
| 8 | A | OP2 | 1H:799 | 3.71 | 1.00 | 24.71 |  |
| 9 | A | OP2 | 1H:1661 | 3.94 | 1.00 | 20.37 |  |
| 10 | C | N3 | 1H:1664 | 4.28 | 1.00 | 17.64 |  |
| 11 | G | O3' | 1H:1660 | 4.33 | 1.00 | 18.52 |  |
| 12 | A | N7 | 1H:799 | 4.42 | 1.00 | 24.40 |  |
| 13 | A | O3' | 1H:798 | 4.42 | 1.00 | 22.03 |  |
| 14 | C | N4 | 1H:1662 | 4.45 | 1.00 | 19.16 |  |
| 15 | A | O5' | 1H:799 | 4.46 | 1.00 | 25.02 |  |
| 16 | C | N4 | 1H:1664 | 4.54 | 1.00 | 17.03 |  |
| 17 | A | O5' | 1H:1661 | 4.55 | 1.00 | 19.16 |  |
| 18 | A | N6 | 1H:837 | 4.60 | 1.00 | 23.68 |  |
| 19 | G | O2' | 1H:1660 | 4.65 | 1.00 | 18.61 |  |
| 20 | GLY | N | E8:91 | 4.67 | 1.00 | 26.32 |  |
| 21 | A | N1 | 1H:837 | 4.68 | 1.00 | 23.72 |  |
  |

  
**1^st^ shell ligand angles:**   
1-2: 83; 1-3: 174; 1-4: 91; 1-5: 84; 1-6: 75; 2-3: 91; 2-4: 173; 2-5: 73; 2-6: 77; 3-4: 94; 3-5: 95; 3-6: 104; 4-5: 110; 4-6: 98; 5-6: 145;   
  
 Contact(s): /O_w_/O_w_/O_w_/O_w_/OP1/OP1/  
 Coded contact list: 2O_ph_4O_w_  
Large octahedral topology discrepancy!!!  

### MG:1H:3576

|  | Res. type | Atom | Residue | Distance | Occupancy | B-factor | Closest solute to 1^st^ shell water |
| --- | --- | --- | --- | --- | --- | --- | --- |
| **Ion** | | | |  | 1.00 | 69.76 || 1 | Water 1 | | 1H:3804 | 2.19 | 1.00 | 56.55 | 2.38 (G:O6 1H:1568) |
| 2 | Water 2 | | 1H:4042 | 2.19 | 1.00 | 47.02 | 2.68 (U:O4 1H:1567) |
| 3 | Water 3 | | 1H:4249 | 2.19 | 1.00 | 50.80 | 2.96 (G:OP2 1H:1566) |
| 4 | Water 4 | | 1H:4083 | 2.19 | 1.00 | 56.00 | 2.74 (G:OP1 1H:1518) |
| 5 | Water 5 | | 1H:4580 | 2.19 | 1.00 | 53.55 | 4.40 (A:OP1 1H:1519) |
| 6 | Water 6 | | 1H:4381 | 2.20 | 1.00 | 49.67 | 3.25 (G:O6 1H:1568) |
| 7 | G | O6 | 1H:1568 | 3.86 | 1.00 | 62.70 |  |
| 8 | U | O4 | 1H:1567 | 4.01 | 1.00 | 56.69 |  |
| 9 | G | OP2 | 1H:1566 | 4.39 | 1.00 | 41.16 |  |
| 10 | A | OP1 | 1H:1519 | 4.46 | 1.00 | 57.77 |  |
| 11 | G | OP2 | 1H:1518 | 4.50 | 1.00 | 53.13 |  |
| 12 | G | N1 | 1H:1568 | 4.70 | 1.00 | 58.41 |  |
| 13 | A | OP2 | 1H:1519 | 4.83 | 1.00 | 51.48 |  |
| 14 | G | OP1 | 1H:1518 | 4.93 | 1.00 | 50.78 |  |
  |

  
**1^st^ shell ligand angles:**   
1-2: 91; 1-3: 178; 1-4: 88; 1-5: 89; 1-6: 91; 2-3: 89; 2-4: 179; 2-5: 88; 2-6: 93; 3-4: 91; 3-5: 89; 3-6: 91; 4-5: 92; 4-6: 87; 5-6: 179;   
  
 Contact(s): /O_w_/O_w_/O_w_/O_w_/O_w_/O_w_/  
 Coded contact list: 6O_w_  
MG hydration shell issue: water too close (< 2.4 Angs.)  

### MG:1H:3577

|  | Res. type | Atom | Residue | Distance | Occupancy | B-factor | Closest solute to 1^st^ shell water |
| --- | --- | --- | --- | --- | --- | --- | --- |
| **Ion** | | | |  | 1.00 | 37.74 || 1 | Water 1 | | 1H:3866 | 2.17 | 1.00 | 19.59 | 2.47 (G:OP2 1H:2092) |
| 2 | Water 2 | | 1H:4570 | 2.19 | 1.00 | 39.50 | 4.24 (G:OP2 1H:2092) |
| 3 | Water 3 | | 1H:4231 | 2.20 | 1.00 | 39.26 | 2.94 (G:OP2 1H:2093) |
| 4 | Water 4 | | 1H:4504 | 2.20 | 1.00 | 39.30 | 3.94 (G:OP2 1H:2093) |
| 5 | G | OP2 | 1H:2093 | 2.60 | 1.00 | 24.52 |  |
| 6 | G | OP2 | 1H:2092 | 2.76 | 1.00 | 22.39 |  |
| 7 | G | O5' | 1H:2093 | 4.64 | 1.00 | 17.41 |  |
| 8 | G | O5' | 1H:2092 | 4.79 | 1.00 | 24.81 |  |
| 9 | G | O3' | 1H:2092 | 4.84 | 1.00 | 27.34 |  |
| 10 | U | OP2 | 1H:2257 | 4.91 | 1.00 | 18.79 |  |
| 11 | U | O2' | 1H:2091 | 4.94 | 1.00 | 27.64 |  |
| 12 | U | O3' | 1H:2091 | 4.97 | 1.00 | 21.97 |  |
  |

  
**1^st^ shell ligand angles:**   
1-2: 92; 1-3: 88; 1-4: 176; 2-3: 95; 2-4: 91; 3-4: 89;   
  
 Contact(s): /O_w_/O_w_/O_w_/O_w_/  
 Coded contact list: 4O_w_  
 Exclusion zone contact(s): /OP2/OP2/  
Consider K in place of MG (2 valid contact(s) for K vs. 0 for MG)  

### K:1J:201

|  | Res. type | Atom | Residue | Distance | Occupancy | B-factor | Closest solute to 1^st^ shell water |
| --- | --- | --- | --- | --- | --- | --- | --- |
| **Ion** | | | |  | 1.00 | 81.19 || 1 | G | O6 | 1J:82 | 2.59 | 1.00 | 79.09 |
| 2 | C | N4 | 1J:94 | 2.95 | 1.00 | 82.03 |  |
| 3 | U | O4 | 1J:95 | 3.06 | 1.00 | 81.18 |  |
| 4 | U | O4 | 1J:80 | 3.21 | 1.00 | 66.25 |  |
| 5 | G | N1 | 1J:81 | 3.25 | 1.00 | 73.88 |  |
| 6 | U | N3 | 1J:80 | 3.28 | 1.00 | 61.68 |  |
| 7 | G | N2 | 1J:81 | 3.34 | 1.00 | 72.16 |  |
| 8 | G | N1 | 1J:82 | 3.71 | 1.00 | 78.58 |  |
| 9 | G | O6 | 1J:96 | 3.81 | 1.00 | 70.74 |  |
| 10 | U | N3 | 1J:95 | 4.12 | 1.00 | 78.43 |  |
| 11 | G | O6 | 1J:81 | 4.42 | 1.00 | 74.29 |  |
| 12 | C | N3 | 1J:94 | 4.43 | 1.00 | 81.34 |  |
| 13 | U | O2 | 1J:80 | 4.59 | 1.00 | 66.57 |  |
| 14 | G | N1 | 1J:96 | 4.63 | 1.00 | 71.11 |  |
| 15 | U | N1 | 1J:80 | 4.77 | 1.00 | 64.32 |  |
| 16 | G | N7 | 1J:82 | 4.77 | 1.00 | 75.32 |  |
  |

  
**1^st^ shell ligand angles:**   
1-2: 75; 1-3: 121; 1-4: 161; 1-5: 79; 1-6: 147; 1-7: 81; 2-3: 67; 2-4: 97; 2-5: 129; 2-6: 137; 2-7: 156; 3-4: 69; 3-5: 92; 3-6: 85; 3-7: 127; 4-5: 118; 4-6: 40; 4-7: 106; 5-6: 81; 5-7: 41; 6-7: 67;   
  
 Contact(s): /N1_G_/N2/N3_U_/N4/O4/O4/  
 Coded contact list: 2O_b_4Anion  
 Exclusion zone contact(s): /O6/  

### MG:1J:202

|  | Res. type | Atom | Residue | Distance | Occupancy | B-factor | Closest solute to 1^st^ shell water |
| --- | --- | --- | --- | --- | --- | --- | --- |
| **Ion** | | | |  | 1.00 | 83.12 || 1 | A | OP2 | 1J:57 | 2.14 | 1.00 | 97.40 |
| 2 | A | OP2 | 1J:58 | 2.97 | 1.00 | 88.73 |  |
| 3 | G | O6 | 1J:23 | 2.99 | 1.00 | 95.70 |  |
| 4 | U | O4 | 1J:22 | 3.05 | 1.00 | 102.32 |  |
| 5 | A | OP1 | 1J:58 | 3.37 | 1.00 | 102.03 |  |
| 6 | G | O3' | 1J:56 | 3.64 | 1.00 | 84.32 |  |
| 7 | C | N4 | 1J:60 | 3.91 | 1.00 | 100.15 |  |
| 8 | A | N6 | 1J:59 | 4.02 | 1.00 | 99.81 |  |
| 9 | A | OP1 | 1J:57 | 4.38 | 1.00 | 93.41 |  |
| 10 | A | O5' | 1J:57 | 4.48 | 1.00 | 82.41 |  |
| 11 | G | O2' | 1J:56 | 4.73 | 1.00 | 90.43 |  |
| 12 | A | O5' | 1J:58 | 4.74 | 1.00 | 101.15 |  |
| 13 | A | O3' | 1J:57 | 4.88 | 1.00 | 90.95 |  |
  |

  
 Contact(s): /OP2/  
 Coded contact list: O_ph_  
 Exclusion zone contact(s): /O4/O6/OP1/OP2/  
Consider K in place of MG (4 valid contact(s) for K vs. 1 for MG)  

### MG:1J:203

|  | Res. type | Atom | Residue | Distance | Occupancy | B-factor | Closest solute to 1^st^ shell water |
| --- | --- | --- | --- | --- | --- | --- | --- |
| **Ion** | | | |  | 1.00 | 48.46 || 1 | G | N7 | 1J:102 | 3.30 | 1.00 | 68.23 |
| 2 | A | OP2 | 1J:101 | 3.66 | 1.00 | 67.18 |  |
| 3 | Water 1 | | 1J:303 | 3.66 | 1.00 | 77.12 | 2.34 (A:OP2 1J:73) |
| 4 | G | OP2 | 1J:102 | 3.86 | 1.00 | 64.27 |  |
| 5 | Water 2 | | 1J:301 | 4.24 | 1.00 | 66.08 | 3.07 (A:OP2 1J:101) |
| 6 | Water 3 | | 1J:304 | 4.27 | 1.00 | 68.34 | 3.26 (A:OP2 1J:101) |
| 7 | G | O6 | 1J:102 | 4.30 | 1.00 | 67.75 |  |
| 8 | U | O4 | 1J:103 | 4.33 | 1.00 | 79.29 |  |
| 9 | G | N1 | 1J:72 | 4.50 | 1.00 | 72.21 |  |
| 10 | G | O6 | 1J:72 | 4.61 | 1.00 | 87.40 |  |
| 11 | A | OP2 | 1J:73 | 4.65 | 1.00 | 79.02 |  |
| 12 | A | OP1 | 1J:101 | 4.79 | 1.00 | 64.70 |  |
| 13 | G | N7 | 1J:72 | 4.93 | 1.00 | 84.99 |  |
| 14 | G | N3 | 1J:72 | 4.97 | 1.00 | 77.52 |  |
  |

  
 Coded contact list:   
 Exclusion zone contact(s): /N7/  
Consider K in place of MG (1 valid contact(s) for K vs. 0 for MG)  
MG hydration shell issue: water too close (< 2.4 Angs.)  

### MG:1J:204

|  | Res. type | Atom | Residue | Distance | Occupancy | B-factor | Closest solute to 1^st^ shell water |
| --- | --- | --- | --- | --- | --- | --- | --- |
| **Ion** | | | |  | 1.00 | 64.73 || 1 | G | O6 | 1J:76 | 3.00 | 1.00 | 66.88 |
| 2 | G | N7 | 1J:75 | 3.63 | 1.00 | 80.84 |  |
| 3 | G | N7 | 1J:76 | 3.78 | 1.00 | 72.30 |  |
| 4 | U | OP2 | 1J:74 | 4.11 | 1.00 | 95.10 |  |
| 5 | A | O5' | 1J:99 | 4.15 | 1.00 | 66.47 |  |
| 6 | G | OP2 | 1J:75 | 4.72 | 1.00 | 95.85 |  |
| 7 | Water 1 | | 1J:305 | 4.78 | 1.00 | 67.82 | 4.06 (G:O6 1J:75) |
| 8 | G | O6 | 1J:75 | 5.00 | 1.00 | 75.05 |  |
  |

  
 Coded contact list:   
 Exclusion zone contact(s): /O6/  
Consider K in place of MG (1 valid contact(s) for K vs. 0 for MG)  

### MG:1J:205

|  | Res. type | Atom | Residue | Distance | Occupancy | B-factor | Closest solute to 1^st^ shell water |
| --- | --- | --- | --- | --- | --- | --- | --- |
| **Ion** | | | |  | 1.00 | 68.34 || 1 | Water 1 | | 1J:301 | 2.17 | 1.00 | 66.08 | 2.08 (A:N7 1J:101) |
| 2 | Water 2 | | 1J:302 | 2.18 | 1.00 | 70.38 | 2.22 (G:OP2 1J:100) |
| 3 | Water 3 | | 1J:303 | 2.18 | 1.00 | 77.12 | 2.34 (A:OP2 1J:73) |
| 4 | Water 4 | | 1J:304 | 2.18 | 1.00 | 68.34 | 2.64 (G:OP2 1J:100) |
| 5 | Water 5 | | 1J:305 | 2.19 | 1.00 | 67.82 | 3.33 (A:N6 1J:101) |
| 6 | G | OP2 | 1J:100 | 2.67 | 1.00 | 85.85 |  |
| 7 | G | O5' | 1J:100 | 3.91 | 1.00 | 64.73 |  |
| 8 | A | OP2 | 1J:73 | 3.94 | 1.00 | 79.02 |  |
| 9 | G | O6 | 1J:102 | 4.18 | 1.00 | 67.75 |  |
| 10 | A | N7 | 1J:101 | 4.20 | 1.00 | 63.31 |  |
| 11 | A | OP2 | 1J:101 | 4.32 | 1.00 | 67.18 |  |
| 12 | A | N6 | 1J:101 | 4.47 | 1.00 | 64.32 |  |
| 13 | A | O3' | 1J:99 | 4.84 | 1.00 | 78.05 |  |
| 14 | G | OP1 | 1J:100 | 4.96 | 1.00 | 78.46 |  |
  |

  
**1^st^ shell ligand angles:**   
1-2: 178; 1-3: 94; 1-4: 89; 1-5: 90; 2-3: 88; 2-4: 90; 2-5: 91; 3-4: 88; 3-5: 92; 4-5: 179;   
  
 Contact(s): /O_w_/O_w_/O_w_/O_w_/O_w_/  
 Coded contact list: 5O_w_  
 Exclusion zone contact(s): /OP2/  
Consider K in place of MG (1 valid contact(s) for K vs. 0 for MG)  
MG hydration shell issue: water too close (< 2.4 Angs.)  

### K:1K:101

|  | Res. type | Atom | Residue | Distance | Occupancy | B-factor | Closest solute to 1^st^ shell water |
| --- | --- | --- | --- | --- | --- | --- | --- |
| **Ion** | | | |  | 1.00 | 83.86 || 1 | U | O4 | 1K:33 | 3.16 | 1.00 | 71.42 |
| 2 | PSU | O2 | 1K:32 | 3.22 | 1.00 | 66.89 |  |
| 3 | A | OP2 | 1K:36 | 3.36 | 1.00 | 51.63 |  |
| 4 | A | OP1 | 1K:36 | 3.62 | 1.00 | 68.28 |  |
| 5 | A | OP2 | 1K:31 | 4.47 | 1.00 | 95.64 |  |
| 6 | U | N3 | 1K:33 | 4.72 | 1.00 | 55.45 |  |
| 7 | PSU | N1 | 1K:32 | 4.90 | 1.00 | 73.78 |  |
| 8 | A | O3' | 1K:35 | 5.00 | 1.00 | 57.16 |  |
  |

  
**1^st^ shell ligand angles:**   
1-2: 65; 1-3: 57; 2-3: 70;   
  
 Contact(s): /O2/O4/OP2/  
 Coded contact list: O_ph_2O_b_  

### K:21:301

|  | Res. type | Atom | Residue | Distance | Occupancy | B-factor | Closest solute to 1^st^ shell water |
| --- | --- | --- | --- | --- | --- | --- | --- |
| **Ion** | | | |  | 1.00 | 62.69 || 1 | LYS | O | 21:118 | 2.72 | 1.00 | 39.89 |
| 2 | TYR | OH | 21:160 | 3.04 | 1.00 | 42.63 |  |
| 3 | ARG | O | 21:119 | 3.08 | 1.00 | 35.27 |  |
| 4 | ASN | ND2 | 21:121 | 3.62 | 1.00 | 41.43 |  |
| 5 | C | OP1 | 1H:2691 | 3.62 | 1.00 | 45.91 |  |
| 6 | ASN | OD1 | 21:121 | 4.07 | 1.00 | 40.76 |  |
| 7 | C | OP2 | 1H:2691 | 4.33 | 1.00 | 30.42 |  |
| 8 | C | O5' | 1H:2691 | 4.44 | 1.00 | 39.68 |  |
| 9 | A | OP2 | 1H:2692 | 4.46 | 1.00 | 35.61 |  |
| 10 | ARG | N | 21:119 | 4.52 | 1.00 | 30.99 |  |
| 11 | TRP | N | 21:120 | 4.57 | 1.00 | 28.71 |  |
  |

  
**1^st^ shell ligand angles:**   
1-2: 84; 1-3: 67; 2-3: 106;   
  
 Contact(s): /O_back_/O_back_/OH_Tyr_/  
 Coded contact list: 2O_back_OH_Prot_  

### MG:21:302

|  | Res. type | Atom | Residue | Distance | Occupancy | B-factor | Closest solute to 1^st^ shell water |
| --- | --- | --- | --- | --- | --- | --- | --- |
| **Ion** | | | |  | 1.00 | 14.87 || 1 | Water 1 | | 21:409 | 2.14 | 1.00 | 25.28 | 3.13 (ASP:OD2 21:127) |
| 2 | Water 2 | | 1H:3648 | 2.16 | 1.00 | 26.55 | 2.05 (U:OP2 1H:2016) |
| 3 | Water 3 | | 1H:4013 | 2.16 | 1.00 | 31.71 | 2.65 (G:N7 1H:2020) |
| 4 | Water 4 | | 21:408 | 2.16 | 1.00 | 45.25 | 3.02 (ASP:OD1 21:127) |
| 5 | Water 5 | | 1H:4053 | 2.18 | 1.00 | 33.63 | 2.69 (G:O2' 1H:2015) |
| 6 | ASP | OD2 | 21:127 | 2.61 | 1.00 | 30.03 |  |
| 7 | ASP | OD1 | 21:127 | 3.78 | 1.00 | 30.96 |  |
| 8 | U | OP2 | 1H:2016 | 3.85 | 1.00 | 31.06 |  |
| 9 | G | O2' | 1H:2015 | 3.99 | 1.00 | 22.75 |  |
| 10 | C | OP2 | 1H:2017 | 4.18 | 1.00 | 21.61 |  |
| 11 | G | O6 | 1H:2020 | 4.34 | 1.00 | 25.84 |  |
| 12 | Water 6 | | 21:407 | 4.38 | 1.00 | 35.52 | 2.97 (ASP:OD2 21:127) |
| 13 | G | N7 | 1H:2020 | 4.44 | 1.00 | 27.29 |  |
| 14 | Water 7 | | 1H:3863 | 4.46 | 1.00 | 25.09 | 2.67 (ASP:OD2 21:127) |
  |

  
**1^st^ shell ligand angles:**   
1-2: 177; 1-3: 94; 1-4: 88; 1-5: 90; 2-3: 86; 2-4: 92; 2-5: 88; 3-4: 178; 3-5: 90; 4-5: 89;   
  
 Contact(s): /O_w_/O_w_/O_w_/O_w_/O_w_/  
 Coded contact list: 5O_w_  
 Exclusion zone contact(s): /OD2_Asp_/  
Consider K in place of MG (1 valid contact(s) for K vs. 0 for MG)  
MG hydration shell issue: water too close (< 2.4 Angs.)  

### MG:21:303

|  | Res. type | Atom | Residue | Distance | Occupancy | B-factor | Closest solute to 1^st^ shell water |
| --- | --- | --- | --- | --- | --- | --- | --- |
| **Ion** | | | |  | 1.00 | 33.81 || 1 | Water 1 | | 21:401 | 2.18 | 1.00 | 30.70 | 2.29 (PRO:O 21:147) |
| 2 | Water 2 | | 1H:4312 | 2.18 | 1.00 | 49.06 | 3.10 (U:OP2 1H:2057) |
| 3 | Water 3 | | 21:405 | 2.18 | 1.00 | 42.27 | 2.88 (PRO:O 21:147) |
| 4 | Water 4 | | 21:403 | 2.18 | 1.00 | 28.50 | 2.52 (PRO:O 21:147) |
| 5 | Water 5 | | 1H:3810 | 2.19 | 1.00 | 26.93 | 2.39 (C:OP2 1H:2048) |
| 6 | Water 6 | | 1H:4522 | 2.19 | 1.00 | 27.57 | 3.80 (U:OP2 1H:2057) |
| 7 | PRO | O | 21:147 | 2.90 | 1.00 | 30.88 |  |
| 8 | C | OP2 | 1H:2048 | 3.53 | 1.00 | 27.03 |  |
| 9 | C | OP1 | 1H:2048 | 3.94 | 1.00 | 29.99 |  |
| 10 | U | O4 | 1H:1177 | 4.25 | 1.00 | 32.44 |  |
| 11 | U | OP2 | 1H:2057 | 4.48 | 1.00 | 28.61 |  |
| 12 | LYS | O | 21:145 | 4.53 | 1.00 | 23.09 |  |
  |

  
**1^st^ shell ligand angles:**   
1-2: 86; 1-3: 92; 1-4: 96; 1-5: 86; 1-6: 176; 2-3: 83; 2-4: 178; 2-5: 96; 2-6: 91; 3-4: 97; 3-5: 178; 3-6: 86; 4-5: 84; 4-6: 87; 5-6: 96;   
  
 Contact(s): /O_w_/O_w_/O_w_/O_w_/O_w_/O_w_/  
 Coded contact list: 6O_w_  
 Exclusion zone contact(s): /O_back_/  
Consider K in place of MG (1 valid contact(s) for K vs. 0 for MG)  
MG hydration shell issue: water too close (< 2.4 Angs.)  

### K:29:301

|  | Res. type | Atom | Residue | Distance | Occupancy | B-factor | Closest solute to 1^st^ shell water |
| --- | --- | --- | --- | --- | --- | --- | --- |
| **Ion** | | | |  | 1.00 | 86.83 || 1 | LYS | O | 29:118 | 2.93 | 1.00 | 56.21 |
| 2 | TYR | OH | 29:160 | 3.30 | 1.00 | 64.42 |  |
| 3 | ASN | ND2 | 29:121 | 3.32 | 1.00 | 65.06 |  |
| 4 | A | OP2 | 14:2679 | 4.20 | 1.00 | 62.40 |  |
| 5 | C | OP2 | 14:2678 | 4.25 | 1.00 | 60.62 |  |
| 6 | ASN | OD1 | 29:121 | 4.30 | 1.00 | 55.88 |  |
| 7 | C | OP1 | 14:2678 | 4.30 | 1.00 | 73.82 |  |
| 8 | C | O5' | 14:2678 | 4.85 | 1.00 | 71.27 |  |
| 9 | ARG | O | 29:119 | 4.87 | 1.00 | 51.85 |  |
  |

  
**1^st^ shell ligand angles:**   
1-2: 79; 1-3: 98; 2-3: 175;   
  
 Contact(s): /ND2_Asn_/O_back_/OH_Tyr_/  
 Coded contact list: O_back_OH_Prot_Anion  

### MG:29:302

|  | Res. type | Atom | Residue | Distance | Occupancy | B-factor | Closest solute to 1^st^ shell water |
| --- | --- | --- | --- | --- | --- | --- | --- |
| **Ion** | | | |  | 1.00 | 38.35 || 1 | ASP | OD2 | 29:127 | 2.10 | 1.00 | 54.09 |
| 2 | G | O6 | 14:1997 | 3.03 | 1.00 | 38.70 |  |
| 3 | Water 1 | | 29:402 | 3.46 | 1.00 | 48.95 | 2.78 (ASP:OD1 29:127) |
| 4 | G | O2' | 14:1992 | 3.74 | 1.00 | 46.57 |  |
| 5 | G | N7 | 14:1997 | 3.90 | 1.00 | 43.12 |  |
| 6 | ASP | OD1 | 29:127 | 4.09 | 1.00 | 52.86 |  |
| 7 | U | OP2 | 14:1993 | 4.18 | 1.00 | 47.19 |  |
| 8 | G | N3 | 14:1992 | 4.55 | 1.00 | 42.96 |  |
| 9 | C | OP2 | 14:1994 | 4.74 | 1.00 | 47.40 |  |
| 10 | G | N9 | 14:1992 | 4.86 | 1.00 | 39.78 |  |
| 11 | C | N4 | 14:1663 | 4.97 | 1.00 | 37.89 |  |
  |

  
 Contact(s): /OD2_Asp_/  
 Coded contact list: O_coo_  
 Exclusion zone contact(s): /O6/  

### MG:29:303

|  | Res. type | Atom | Residue | Distance | Occupancy | B-factor | Closest solute to 1^st^ shell water |
| --- | --- | --- | --- | --- | --- | --- | --- |
| **Ion** | | | |  | 1.00 | 28.21 || 1 | Water 1 | | 14:3665 | 2.16 | 1.00 | 38.48 | 2.52 (G:OP1 14:744) |
| 2 | Water 2 | | 29:406 | 2.16 | 1.00 | 45.58 | 3.42 (HIS:NE2 29:135) |
| 3 | Water 3 | | 29:404 | 2.17 | 1.00 | 49.69 | 3.40 (HIS:NE2 29:135) |
| 4 | Water 4 | | 14:3761 | 2.17 | 1.00 | 41.74 | 2.68 (G:O2' 14:743) |
| 5 | HIS | NE2 | 29:135 | 2.56 | 1.00 | 43.32 |  |
| 6 | C | OP1 | 14:1658 | 2.59 | 1.00 | 41.12 |  |
| 7 | C | OP2 | 14:1658 | 3.99 | 1.00 | 36.73 |  |
| 8 | G | O3' | 14:743 | 4.20 | 1.00 | 33.91 |  |
| 9 | U | OP2 | 14:1659 | 4.21 | 1.00 | 42.37 |  |
| 10 | C | O5' | 14:1658 | 4.26 | 1.00 | 36.99 |  |
| 11 | ALA | O | 29:131 | 4.28 | 1.00 | 38.40 |  |
| 12 | G | OP1 | 14:744 | 4.33 | 1.00 | 34.85 |  |
| 13 | HIS | ND1 | 29:135 | 4.36 | 1.00 | 46.17 |  |
| 14 | LYS | N | 29:133 | 4.76 | 1.00 | 35.80 |  |
| 15 | G | O2' | 14:743 | 4.85 | 1.00 | 40.61 |  |
| 16 | C | O3' | 14:1657 | 4.87 | 1.00 | 35.77 |  |
  |

  
**1^st^ shell ligand angles:**   
1-2: 92; 1-3: 173; 1-4: 90; 2-3: 91; 2-4: 90; 3-4: 96;   
  
 Contact(s): /O_w_/O_w_/O_w_/O_w_/  
 Coded contact list: 4O_w_  
 Exclusion zone contact(s): /NE2_His_/OP1/  
Consider NA in place of MG (2 valid contact(s) for NA vs. 0 for MG)  

### K:2A:201

|  | Res. type | Atom | Residue | Distance | Occupancy | B-factor | Closest solute to 1^st^ shell water |
| --- | --- | --- | --- | --- | --- | --- | --- |
| **Ion** | | | |  | 1.00 | 70.49 || 1 | ASN | O | 2A:26 | 2.57 | 1.00 | 70.33 |
| 2 | U | O4 | 1G:692 | 2.69 | 1.00 | 69.51 |  |
| 3 | GLY | O | 2A:52 | 2.87 | 1.00 | 61.83 |  |
| 4 | G | O6 | 1G:691 | 3.15 | 1.00 | 65.16 |  |
| 5 | ASN | OD1 | 2A:26 | 3.40 | 1.00 | 67.73 |  |
| 6 | G | N7 | 1G:691 | 3.65 | 1.00 | 65.06 |  |
| 7 | SER | N | 2A:53 | 4.48 | 1.00 | 67.11 |  |
| 8 | GLY | N | 2A:56 | 4.49 | 1.00 | 65.81 |  |
| 9 | SER | O | 2A:53 | 4.54 | 1.00 | 70.28 |  |
| 10 | ASN | N | 2A:27 | 4.72 | 1.00 | 61.32 |  |
| 11 | G | N1 | 1G:691 | 4.78 | 1.00 | 63.28 |  |
| 12 | U | N3 | 1G:692 | 4.89 | 1.00 | 62.46 |  |
| 13 | LYS | N | 2A:55 | 4.98 | 1.00 | 64.68 |  |
  |

  
**1^st^ shell ligand angles:**   
1-2: 162; 1-3: 120; 1-4: 125; 1-5: 89; 2-3: 65; 2-4: 73; 2-5: 81; 3-4: 66; 3-5: 146; 4-5: 112;   
  
 Contact(s): /O4/O6/O_back_/OD1_Asn_/  
 Coded contact list: 2O_b_O_back_O_cno_  
 Exclusion zone contact(s): /O_back_/  

### K:2K:101

|  | Res. type | Atom | Residue | Distance | Occupancy | B-factor | Closest solute to 1^st^ shell water |
| --- | --- | --- | --- | --- | --- | --- | --- |
| **Ion** | | | |  | 1.00 | 62.35 || 1 | A | OP1 | 2K:14 | 2.83 | 1.00 | 61.40 |
| 2 | C | O3' | 1H:1947 | 3.02 | 1.00 | 48.63 |  |
| 3 | C | O3' | 2K:13 | 3.19 | 1.00 | 47.91 |  |
| 4 | C | O2' | 2K:13 | 3.27 | 1.00 | 50.39 |  |
| 5 | A | O5' | 2K:14 | 3.29 | 1.00 | 49.24 |  |
| 6 | C | O2' | 1H:1947 | 3.48 | 1.00 | 50.88 |  |
| 7 | C | OP1 | 1H:1948 | 3.49 | 1.00 | 42.18 |  |
| 8 | A | O4' | 2K:14 | 3.95 | 1.00 | 58.91 |  |
| 9 | C | O5' | 1H:1948 | 4.40 | 1.00 | 46.95 |  |
| 10 | A | OP2 | 2K:14 | 4.86 | 1.00 | 52.17 |  |
  |

  
**1^st^ shell ligand angles:**   
1-2: 133; 1-3: 47; 1-4: 96; 1-5: 47; 2-3: 100; 2-4: 76; 2-5: 138; 3-4: 52; 3-5: 46; 4-5: 64;   
  
 Contact(s): /O2'/O3'/O3'/O5'/OP1/  
 Coded contact list: O_ph_4O_r_  

### K:2K:102

|  | Res. type | Atom | Residue | Distance | Occupancy | B-factor | Closest solute to 1^st^ shell water |
| --- | --- | --- | --- | --- | --- | --- | --- |
| **Ion** | | | |  | 1.00 | 58.55 || 1 | G | N7 | 2K:24 | 2.76 | 1.00 | 47.24 |
| 2 | G | O6 | 2K:44 | 3.01 | 1.00 | 57.26 |  |
| 3 | G | O6 | 2K:24 | 3.01 | 1.00 | 47.19 |  |
| 4 | C | N4 | 2K:25 | 3.33 | 1.00 | 48.30 |  |
| 5 | A | N6 | 2K:9 | 3.40 | 1.00 | 54.54 |  |
| 6 | A | N7 | 2K:23 | 3.55 | 1.00 | 51.33 |  |
| 7 | A | N7 | 2K:9 | 3.59 | 1.00 | 54.82 |  |
| 8 | A | N6 | 2K:23 | 4.12 | 1.00 | 46.54 |  |
| 9 | G | N7 | 2K:44 | 4.31 | 1.00 | 70.07 |  |
| 10 | G | OP2 | 2K:24 | 4.35 | 1.00 | 59.35 |  |
| 11 | A | OP2 | 2K:23 | 4.42 | 1.00 | 68.34 |  |
| 12 | A | N9 | 2K:23 | 4.69 | 1.00 | 48.57 |  |
| 13 | G | O6 | 2K:10 | 4.88 | 1.00 | 48.69 |  |
| 14 | G | N9 | 2K:24 | 4.93 | 1.00 | 41.38 |  |
| 15 | A | N1 | 2K:23 | 4.95 | 1.00 | 44.61 |  |
| 16 | A | N1 | 2K:9 | 4.95 | 1.00 | 60.60 |  |
| 17 | G | N1 | 2K:24 | 4.96 | 1.00 | 42.36 |  |
  |

  
**1^st^ shell ligand angles:**   
1-2: 132; 1-3: 64; 1-4: 85; 1-5: 129; 2-3: 75; 2-4: 55; 2-5: 90; 3-4: 65; 3-5: 116; 4-5: 145;   
  
 Contact(s): /N4/N6/N7/O6/O6/  
 Coded contact list: 2O_b_N_b_2Anion  

### K:2K:103

|  | Res. type | Atom | Residue | Distance | Occupancy | B-factor | Closest solute to 1^st^ shell water |
| --- | --- | --- | --- | --- | --- | --- | --- |
| **Ion** | | | |  | 1.00 | 79.58 || 1 | C | N4 | 2K:62 | 3.08 | 1.00 | 69.34 |
| 2 | U | O4 | 2K:51 | 3.19 | 1.00 | 53.96 |  |
| 3 | G | O6 | 2K:63 | 3.27 | 1.00 | 67.98 |  |
| 4 | G | O6 | 2K:52 | 3.78 | 1.00 | 67.88 |  |
| 5 | U | OP1 | 2K:60 | 3.89 | 1.00 | 69.61 |  |
| 6 | U | O4 | 2K:50 | 4.06 | 1.00 | 62.35 |  |
| 7 | A | N6 | 2K:64 | 4.32 | 1.00 | 67.26 |  |
| 8 | C | N3 | 2K:62 | 4.61 | 1.00 | 59.71 |  |
| 9 | U | N3 | 2K:51 | 4.86 | 1.00 | 63.10 |  |
| 10 | G | N7 | 2K:63 | 4.97 | 1.00 | 68.21 |  |
  |

  
**1^st^ shell ligand angles:**   
1-2: 110; 1-3: 80; 2-3: 63;   
  
 Contact(s): /N4/O4/O6/  
 Coded contact list: 2O_b_Anion  

### MG:2K:104

|  | Res. type | Atom | Residue | Distance | Occupancy | B-factor | Closest solute to 1^st^ shell water |
| --- | --- | --- | --- | --- | --- | --- | --- |
| **Ion** | | | |  | 1.00 | 48.72 || 1 | G | O6 | 2K:53 | 3.87 | 1.00 | 70.70 |
| 2 | G | O6 | 2K:52 | 3.90 | 1.00 | 67.88 |  |
| 3 | G | N7 | 2K:52 | 4.09 | 1.00 | 65.39 |  |
| 4 | G | N7 | 2K:53 | 4.53 | 1.00 | 66.65 |  |
| 5 | A | O2' | 2K:58 | 4.55 | 1.00 | 59.17 |  |
| 6 | U | O4 | 2K:51 | 4.88 | 1.00 | 53.96 |  |
  |

  
 Coded contact list:   
1^st^ MG coordination shell missing (see Notes)  

### MG:2K:105

|  | Res. type | Atom | Residue | Distance | Occupancy | B-factor | Closest solute to 1^st^ shell water |
| --- | --- | --- | --- | --- | --- | --- | --- |
| **Ion** | | | |  | 1.00 | 45.40 || 1 | Water 1 | | 2K:205 | 2.17 | 1.00 | 47.04 | 2.65 (A:OP2 2K:9) |
| 2 | Water 2 | | 2K:206 | 2.17 | 1.00 | 51.46 | 2.85 (U:OP2 2K:12) |
| 3 | Water 3 | | 2K:201 | 2.17 | 1.00 | 55.44 | 2.06 (U:OP2 2K:12) |
| 4 | Water 4 | | 2K:202 | 2.18 | 1.00 | 59.29 | 2.23 (4SU:OP2 2K:8) |
| 5 | Water 5 | | 2K:208 | 2.19 | 1.00 | 44.66 | 3.58 (U:OP2 2K:12) |
| 6 | Water 6 | | 2K:207 | 2.19 | 1.00 | 43.98 | 3.13 (C:OP2 2K:11) |
| 7 | U | OP2 | 2K:12 | 3.36 | 1.00 | 42.13 |  |
| 8 | 4SU | OP2 | 2K:8 | 3.53 | 1.00 | 51.17 |  |
| 9 | 4SU | OP1 | 2K:8 | 4.29 | 1.00 | 42.63 |  |
| 10 | C | OP2 | 2K:11 | 4.56 | 1.00 | 43.26 |  |
| 11 | A | OP2 | 2K:9 | 4.61 | 1.00 | 47.64 |  |
| 12 | A | OP1 | 2K:9 | 4.86 | 1.00 | 43.92 |  |
  |

  
**1^st^ shell ligand angles:**   
1-2: 92; 1-3: 178; 1-4: 90; 1-5: 87; 1-6: 92; 2-3: 90; 2-4: 178; 2-5: 93; 2-6: 86; 3-4: 89; 3-5: 91; 3-6: 90; 4-5: 89; 4-6: 93; 5-6: 178;   
  
 Contact(s): /O_w_/O_w_/O_w_/O_w_/O_w_/O_w_/  
 Coded contact list: 6O_w_  
 Exclusion zone contact(s): /OP2/  
Consider K in place of MG (1 valid contact(s) for K vs. 0 for MG)  
MG hydration shell issue: water too close (< 2.4 Angs.)  

### MG:2L:101

|  | Res. type | Atom | Residue | Distance | Occupancy | B-factor | Closest solute to 1^st^ shell water |
| --- | --- | --- | --- | --- | --- | --- | --- |
| **Ion** | | | |  | 1.00 | 58.99 || 1 | A | OP2 | 2L:76 | 2.18 | 1.00 | 57.49 |
| 2 | C | OP1 | 2L:75 | 2.33 | 1.00 | 57.90 |  |
| 3 | A | OP1 | 14:2602 | 2.47 | 1.00 | 58.95 |  |
| 4 | C | OP1 | 14:2601 | 3.94 | 1.00 | 44.21 |  |
| 5 | A | OP2 | 14:2602 | 4.01 | 1.00 | 52.36 |  |
| 6 | C | O3' | 14:2601 | 4.03 | 1.00 | 50.31 |  |
| 7 | A | N1 | 14:2439 | 4.20 | 1.00 | 45.10 |  |
| 8 | A | O5' | 2L:76 | 4.21 | 1.00 | 69.82 |  |
| 9 | C | O5' | 2L:75 | 4.23 | 1.00 | 57.22 |  |
| 10 | A | N6 | 14:2439 | 4.24 | 1.00 | 55.35 |  |
| 11 | C | O5' | 14:2601 | 4.35 | 1.00 | 42.84 |  |
| 12 | A | OP1 | 2L:76 | 4.50 | 1.00 | 70.97 |  |
| 13 | U | OP1 | 14:2585 | 4.53 | 1.00 | 65.54 |  |
| 14 | C | OP2 | 2L:75 | 4.69 | 1.00 | 64.99 |  |
| 15 | C | O3' | 2L:75 | 4.71 | 1.00 | 56.68 |  |
| 16 | C | O3' | 2L:74 | 4.79 | 1.00 | 51.42 |  |
| 17 | A | O5' | 14:2602 | 4.86 | 1.00 | 63.34 |  |
| 18 | C | OP2 | 14:2601 | 4.96 | 1.00 | 39.87 |  |
  |

  
**1^st^ shell ligand angles:**   
1-2: 81;   
  
 Contact(s): /OP1/OP2/  
 Coded contact list: 2O_ph_  
 Exclusion zone contact(s): /OP1/  

### MG:2L:102

|  | Res. type | Atom | Residue | Distance | Occupancy | B-factor | Closest solute to 1^st^ shell water |
| --- | --- | --- | --- | --- | --- | --- | --- |
| **Ion** | | | |  | 1.00 | 56.51 || 1 | G | O6 | 2L:30 | 3.83 | 1.00 | 80.94 |
| 2 | G | N7 | 2L:30 | 4.08 | 1.00 | 76.75 |  |
| 3 | G | N7 | 2L:29 | 4.92 | 1.00 | 79.25 |  |
  |

  
 Coded contact list:   
1^st^ MG coordination shell missing (see Notes)  

### MG:2L:103

|  | Res. type | Atom | Residue | Distance | Occupancy | B-factor | Closest solute to 1^st^ shell water |
| --- | --- | --- | --- | --- | --- | --- | --- |
| **Ion** | | | |  | 1.00 | 45.41 || 1 | U | OP2 | 2L:12 | 3.49 | 1.00 | 64.87 |
| 2 | 4SU | OP1 | 2L:8 | 3.52 | 1.00 | 70.33 |  |
| 3 | A | OP2 | 2L:9 | 3.82 | 1.00 | 81.89 |  |
| 4 | 4SU | OP2 | 2L:8 | 3.91 | 1.00 | 72.99 |  |
| 5 | C | OP2 | 2L:11 | 4.21 | 1.00 | 74.05 |  |
| 6 | A | OP1 | 2L:9 | 4.55 | 1.00 | 79.25 |  |
  |

  
 Coded contact list:   
1^st^ MG coordination shell missing (see Notes)  

### K:31:301

|  | Res. type | Atom | Residue | Distance | Occupancy | B-factor | Closest solute to 1^st^ shell water |
| --- | --- | --- | --- | --- | --- | --- | --- |
| **Ion** | | | |  | 1.00 | 48.71 || 1 | VAL | O | 31:173 | 2.80 | 1.00 | 46.60 |
| 2 | ARG | O | 31:168 | 2.88 | 1.00 | 46.68 |  |
| 3 | LEU | O | 31:170 | 2.92 | 1.00 | 44.64 |  |
| 4 | U | O4 | 1H:1251 | 2.94 | 1.00 | 48.37 |  |
| 5 | U | O4 | 1H:639 | 3.88 | 1.00 | 73.36 |  |
| 6 | LEU | N | 31:170 | 4.39 | 1.00 | 42.71 |  |
| 7 | U | N3 | 1H:639 | 4.66 | 1.00 | 91.27 |  |
| 8 | VAL | N | 31:173 | 4.69 | 1.00 | 41.80 |  |
| 9 | ASN | N | 31:169 | 4.75 | 1.00 | 39.13 |  |
| 10 | VAL | N | 31:174 | 4.82 | 1.00 | 35.01 |  |
| 11 | PRO | N | 31:171 | 4.83 | 1.00 | 43.53 |  |
| 12 | U | N3 | 1H:1251 | 4.83 | 1.00 | 40.26 |  |
| 13 | ALA | O | 31:167 | 4.89 | 1.00 | 43.51 |  |
| 14 | G | O2' | 1H:348 | 4.90 | 1.00 | 43.11 |  |
| 15 | ASN | O | 31:169 | 4.93 | 1.00 | 43.08 |  |
  |

  
**1^st^ shell ligand angles:**   
1-2: 98; 1-3: 73; 1-4: 133; 2-3: 92; 2-4: 126; 3-4: 90;   
  
 Contact(s): /O4/O_back_/O_back_/O_back_/  
 Coded contact list: O_b_3O_back_  

### K:31:302

|  | Res. type | Atom | Residue | Distance | Occupancy | B-factor | Closest solute to 1^st^ shell water |
| --- | --- | --- | --- | --- | --- | --- | --- |
| **Ion** | | | |  | 1.00 | 45.34 || 1 | TYR | O | 31:59 | 2.75 | 1.00 | 23.98 |
| 2 | ALA | O | 31:58 | 3.17 | 1.00 | 25.63 |  |
| 3 | Water 1 | | 31:406 | 3.30 | 1.00 | 23.34 | 4.12 (TYR:O 31:59) |
| 4 | VAL | O | 31:57 | 3.34 | 1.00 | 21.17 |  |
| 5 | G | OP1 | 1H:846 | 3.78 | 1.00 | 20.05 |  |
| 6 | TYR | N | 31:59 | 4.12 | 1.00 | 21.98 |  |
| 7 | Water 2 | | 31:404 | 4.27 | 1.00 | 30.65 | 2.99 (G:OP1 1H:846) |
| 8 | SER | N | 31:60 | 4.69 | 1.00 | 21.81 |  |
| 9 | ALA | N | 31:58 | 4.72 | 1.00 | 21.07 |  |
| 10 | Water 3 | | 31:407 | 4.78 | 1.00 | 29.21 | 3.61 (TYR:O 31:59) |
  |

  
**1^st^ shell ligand angles:**   
1-2: 64; 1-3: 85; 1-4: 85; 2-3: 142; 2-4: 76; 3-4: 80;   
  
 Contact(s): /O_back_/O_back_/O_back_/O_w_/  
 Coded contact list: 3O_back_O_w_  

### MG:31:303

|  | Res. type | Atom | Residue | Distance | Occupancy | B-factor | Closest solute to 1^st^ shell water |
| --- | --- | --- | --- | --- | --- | --- | --- |
| **Ion** | | | |  | 1.00 | 37.64 || 1 | ARG | NH2 | 31:188 | 4.34 | 1.00 | 34.79 |
| 2 | ARG | NE | 31:188 | 4.96 | 1.00 | 36.69 |  |
  |

  
 Coded contact list:   
1^st^ MG coordination shell missing (see Notes)  

### K:32:301

|  | Res. type | Atom | Residue | Distance | Occupancy | B-factor | Closest solute to 1^st^ shell water |
| --- | --- | --- | --- | --- | --- | --- | --- |
| **Ion** | | | |  | 1.00 | 104.22 || 1 | SER | O | 32:83 | 2.73 | 1.00 | 77.38 |
| 2 | THR | OG1 | 32:89 | 2.98 | 1.00 | 82.80 |  |
| 3 | U | O2 | 1G:5 | 3.07 | 1.00 | 102.38 |  |
| 4 | LYS | O | 32:85 | 3.74 | 1.00 | 91.08 |  |
| 5 | ALA | O | 32:82 | 4.20 | 1.00 | 75.87 |  |
| 6 | G | OP2 | 1G:6 | 4.42 | 1.00 | 92.37 |  |
| 7 | LYS | O | 32:84 | 4.45 | 1.00 | 87.43 |  |
| 8 | LYS | N | 32:84 | 4.65 | 1.00 | 76.44 |  |
| 9 | GLY | N | 32:87 | 4.72 | 1.00 | 101.43 |  |
| 10 | SER | OG | 32:83 | 4.93 | 1.00 | 87.13 |  |
| 11 | U | N3 | 1G:5 | 4.97 | 1.00 | 103.94 |  |
  |

  
**1^st^ shell ligand angles:**   
1-2: 98; 1-3: 128; 2-3: 132;   
  
 Contact(s): /O2/O_back_/OG1_Thr_/  
 Coded contact list: O_b_O_back_OH_Prot_  

### MG:32:302

|  | Res. type | Atom | Residue | Distance | Occupancy | B-factor | Closest solute to 1^st^ shell water |
| --- | --- | --- | --- | --- | --- | --- | --- |
| **Ion** | | | |  | 1.00 | 79.58 || 1 | Water 1 | | 1G:1965 | 2.17 | 1.00 | 81.29 | 3.57 (G:OP2 1G:438) |
| 2 | Water 2 | | 1G:1849 | 2.17 | 1.00 | 85.17 | 2.42 (G:OP1 1G:438) |
| 3 | Water 3 | | 1G:1863 | 2.18 | 1.00 | 87.99 | 2.50 (G:OP2 1G:492) |
| 4 | Water 4 | | 1G:1862 | 2.18 | 1.00 | 88.06 | 2.49 (G:N7 1G:493) |
| 5 | Water 5 | | 32:401 | 2.18 | 1.00 | 90.52 | 1.30 (LYS:NZ 32:151) |
| 6 | Water 6 | | 32:403 | 2.19 | 1.00 | 83.22 | 3.00 (LYS:NZ 32:151) |
| 7 | LYS | NZ | 32:151 | 2.91 | 1.00 | 91.37 |  |
| 8 | G | OP2 | 1G:492 | 3.98 | 1.00 | 88.33 |  |
| 9 | G | OP1 | 1G:438 | 4.05 | 1.00 | 79.42 |  |
| 10 | G | O6 | 1G:493 | 4.40 | 1.00 | 84.55 |  |
| 11 | G | N7 | 1G:493 | 4.57 | 1.00 | 92.12 |  |
| 12 | G | OP2 | 1G:438 | 4.68 | 1.00 | 77.60 |  |
  |

  
**1^st^ shell ligand angles:**   
1-2: 90; 1-3: 88; 1-4: 89; 1-5: 91; 1-6: 180; 2-3: 178; 2-4: 90; 2-5: 90; 2-6: 90; 3-4: 89; 3-5: 91; 3-6: 92; 4-5: 180; 4-6: 91; 5-6: 89;   
  
 Contact(s): /O_w_/O_w_/O_w_/O_w_/O_w_/O_w_/  
 Coded contact list: 6O_w_  
 Exclusion zone contact(s): /NZ_Lys_/  
Consider K in place of MG (1 valid contact(s) for K vs. 0 for MG)  
MG hydration shell issue: water too close (< 2.4 Angs.)  

### K:39:301

|  | Res. type | Atom | Residue | Distance | Occupancy | B-factor | Closest solute to 1^st^ shell water |
| --- | --- | --- | --- | --- | --- | --- | --- |
| **Ion** | | | |  | 1.00 | 75.13 || 1 | LEU | O | 39:170 | 2.63 | 1.00 | 62.54 |
| 2 | U | O4 | 14:1205 | 2.68 | 1.00 | 66.92 |  |
| 3 | VAL | O | 39:173 | 2.68 | 1.00 | 59.31 |  |
| 4 | ARG | O | 39:168 | 3.20 | 1.00 | 63.38 |  |
| 5 | LEU | N | 39:170 | 4.13 | 1.00 | 56.72 |  |
| 6 | ASN | N | 39:169 | 4.48 | 1.00 | 67.87 |  |
| 7 | PRO | N | 39:171 | 4.61 | 1.00 | 60.04 |  |
| 8 | ALA | O | 39:167 | 4.64 | 1.00 | 61.93 |  |
| 9 | VAL | N | 39:173 | 4.71 | 1.00 | 57.12 |  |
| 10 | U | N3 | 14:1205 | 4.71 | 1.00 | 64.20 |  |
| 11 | VAL | N | 39:174 | 4.80 | 1.00 | 58.79 |  |
| 12 | ASN | O | 39:169 | 4.86 | 1.00 | 53.56 |  |
| 13 | G | O2' | 14:323 | 4.98 | 1.00 | 65.57 |  |
  |

  
**1^st^ shell ligand angles:**   
1-2: 85; 1-3: 74; 1-4: 107; 2-3: 138; 2-4: 109; 3-4: 112;   
  
 Contact(s): /O4/O_back_/O_back_/O_back_/  
 Coded contact list: O_b_3O_back_  

### K:39:302

|  | Res. type | Atom | Residue | Distance | Occupancy | B-factor | Closest solute to 1^st^ shell water |
| --- | --- | --- | --- | --- | --- | --- | --- |
| **Ion** | | | |  | 1.00 | 74.54 || 1 | TYR | O | 39:59 | 3.10 | 1.00 | 40.10 |
| 2 | VAL | O | 39:57 | 3.21 | 1.00 | 37.38 |  |
| 3 | Water 1 | | 39:401 | 3.25 | 1.00 | 38.32 | 2.72 (GLY:O 39:55) |
| 4 | Water 2 | | 14:3778 | 3.27 | 1.00 | 42.38 | 2.71 (G:OP1 14:798) |
| 5 | ALA | O | 39:58 | 3.40 | 1.00 | 41.57 |  |
| 6 | Water 3 | | 14:3911 | 3.54 | 1.00 | 39.33 | 3.79 (GLY:O 39:55) |
| 7 | G | OP1 | 14:798 | 3.99 | 1.00 | 40.53 |  |
| 8 | GLY | O | 39:55 | 4.29 | 1.00 | 43.00 |  |
| 9 | TYR | N | 39:59 | 4.58 | 1.00 | 37.15 |  |
| 10 | ALA | N | 39:58 | 4.91 | 1.00 | 39.74 |  |
  |

  
**1^st^ shell ligand angles:**   
1-2: 78; 1-3: 75; 1-4: 53; 2-3: 68; 2-4: 113; 3-4: 58;   
  
 Contact(s): /O_back_/O_back_/O_w_/O_w_/  
 Coded contact list: 2O_back_2O_w_  

### MG:39:303

|  | Res. type | Atom | Residue | Distance | Occupancy | B-factor | Closest solute to 1^st^ shell water |
| --- | --- | --- | --- | --- | --- | --- | --- |
| **Ion** | | | |  | 1.00 | 54.86 || 1 | ASP | OD2 | 39:185 | 2.54 | 1.00 | 63.10 |
| 2 | ASP | OD1 | 39:185 | 3.56 | 1.00 | 59.63 |  |
| 3 | A | OP2 | 14:616 | 3.67 | 1.00 | 83.28 |  |
| 4 | ARG | NH1 | 39:188 | 3.71 | 1.00 | 54.87 |  |
| 5 | A | O5' | 14:616 | 4.50 | 1.00 | 75.95 |  |
| 6 | THR | O | 39:175 | 4.63 | 1.00 | 69.59 |  |
| 7 | A | O4' | 14:616 | 4.99 | 1.00 | 59.97 |  |
  |

  
 Coded contact list:   
 Exclusion zone contact(s): /OD2_Asp_/  
Consider NA in place of MG (1 valid contact(s) for NA vs. 0 for MG)  

### MG:3E:301

|  | Res. type | Atom | Residue | Distance | Occupancy | B-factor | Closest solute to 1^st^ shell water |
| --- | --- | --- | --- | --- | --- | --- | --- |
| **Ion** | | | |  | 1.00 | 82.21 || 1 | Water 1 | | 13:1839 | 2.17 | 1.00 | 71.54 | 2.34 (A:OP1 13:621) |
| 2 | Water 2 | | 13:1828 | 2.17 | 1.00 | 81.62 | 2.26 (G:OP2 13:617) |
| 3 | Water 3 | | 13:1983 | 2.18 | 1.00 | 85.99 | 3.34 (G:N7 13:616) |
| 4 | Water 4 | | 3E:401 | 2.18 | 1.00 | 94.20 | 1.30 (ARG:NH1 3E:141) |
| 5 | Water 5 | | 13:1943 | 2.19 | 1.00 | 84.85 | 3.04 (G:N7 13:617) |
| 6 | Water 6 | | 13:1916 | 2.19 | 1.00 | 70.64 | 2.82 (C:OP1 13:620) |
| 7 | ARG | NH1 | 3E:141 | 3.37 | 1.00 | 95.86 |  |
| 8 | A | OP1 | 13:621 | 4.30 | 1.00 | 74.22 |  |
| 9 | G | OP2 | 13:617 | 4.38 | 1.00 | 83.37 |  |
| 10 | A | OP2 | 13:621 | 4.70 | 1.00 | 65.17 |  |
| 11 | G | N7 | 13:616 | 4.80 | 1.00 | 84.48 |  |
| 12 | ARG | NH2 | 3E:141 | 4.86 | 1.00 | 89.19 |  |
| 13 | G | N7 | 13:617 | 4.90 | 1.00 | 80.55 |  |
| 14 | C | OP1 | 13:620 | 4.99 | 1.00 | 72.11 |  |
  |

  
**1^st^ shell ligand angles:**   
1-2: 178; 1-3: 90; 1-4: 88; 1-5: 90; 1-6: 92; 2-3: 90; 2-4: 90; 2-5: 91; 2-6: 89; 3-4: 86; 3-5: 91; 3-6: 178; 4-5: 177; 4-6: 94; 5-6: 89;   
  
 Contact(s): /O_w_/O_w_/O_w_/O_w_/O_w_/O_w_/  
 Coded contact list: 6O_w_  
 Exclusion zone contact(s): /NH1_Arg_/  
Consider K in place of MG (1 valid contact(s) for K vs. 0 for MG)  
MG hydration shell issue: water too close (< 2.4 Angs.)  

### K:41:201

|  | Res. type | Atom | Residue | Distance | Occupancy | B-factor | Closest solute to 1^st^ shell water |
| --- | --- | --- | --- | --- | --- | --- | --- |
| **Ion** | | | |  | 1.00 | 67.82 || 1 | THR | O | 41:161 | 2.50 | 1.00 | 53.87 |
| 2 | ARG | O | 41:128 | 2.67 | 1.00 | 56.18 |  |
| 3 | ALA | O | 41:163 | 2.98 | 1.00 | 60.19 |  |
| 4 | ASN | ND2 | 41:130 | 3.58 | 1.00 | 56.19 |  |
| 5 | ASN | OD1 | 41:130 | 3.72 | 1.00 | 46.71 |  |
| 6 | GLY | O | 41:129 | 3.90 | 1.00 | 53.02 |  |
| 7 | THR | N | 41:161 | 4.29 | 1.00 | 53.01 |  |
| 8 | ALA | N | 41:163 | 4.49 | 1.00 | 47.40 |  |
| 9 | THR | N | 41:162 | 4.58 | 1.00 | 56.75 |  |
| 10 | GLY | N | 41:129 | 4.76 | 1.00 | 54.23 |  |
| 11 | THR | O | 41:162 | 4.81 | 1.00 | 50.43 |  |
| 12 | GLU | N | 41:164 | 4.93 | 1.00 | 55.03 |  |
  |

  
**1^st^ shell ligand angles:**   
1-2: 136; 1-3: 90; 2-3: 64;   
  
 Contact(s): /O_back_/O_back_/  
 Coded contact list: 2O_back_  
 Exclusion zone contact(s): /O_back_/  

### MG:41:202

|  | Res. type | Atom | Residue | Distance | Occupancy | B-factor | Closest solute to 1^st^ shell water |
| --- | --- | --- | --- | --- | --- | --- | --- |
| **Ion** | | | |  | 1.00 | 55.21 || 1 | PHE | O | 41:125 | 2.19 | 1.00 | 51.64 |
| 2 | ASN | O | 41:123 | 2.35 | 1.00 | 56.95 |  |
| 3 | PHE | N | 41:125 | 3.81 | 1.00 | 59.57 |  |
| 4 | ASP | OD2 | 41:166 | 4.03 | 1.00 | 59.19 |  |
| 5 | ASP | O | 41:126 | 4.31 | 1.00 | 73.40 |  |
| 6 | ASP | N | 41:126 | 4.45 | 1.00 | 56.20 |  |
| 7 | SER | N | 41:124 | 4.47 | 1.00 | 57.53 |  |
  |

  
**1^st^ shell ligand angles:**   
1-2: 88;   
  
 Contact(s): /O_back_/O_back_/  
 Coded contact list: 2O_back_  

### MG:42:201

|  | Res. type | Atom | Residue | Distance | Occupancy | B-factor | Closest solute to 1^st^ shell water |
| --- | --- | --- | --- | --- | --- | --- | --- |
| **Ion** | | | |  | 1.00 | 70.31 || 1 | ARG | NH2 | 42:27 | 2.07 | 1.00 | 94.36 |
| 2 | G | N7 | 1G:1082 | 2.91 | 1.00 | 77.50 |  |
| 3 | U | O4 | 1G:1083 | 3.19 | 1.00 | 85.28 |  |
| 4 | ARG | NH1 | 42:27 | 3.28 | 1.00 | 87.97 |  |
| 5 | G | O6 | 1G:1082 | 3.52 | 1.00 | 81.81 |  |
| 6 | ARG | NE | 42:27 | 4.24 | 1.00 | 95.81 |  |
| 7 | G | OP2 | 1G:1082 | 4.49 | 1.00 | 75.39 |  |
| 8 | G | N1 | 1G:1084 | 4.54 | 1.00 | 84.98 |  |
| 9 | U | N3 | 1G:1083 | 4.95 | 1.00 | 81.45 |  |
| 10 | G | N2 | 1G:1084 | 5.00 | 1.00 | 86.31 |  |
  |

  
 Contact(s): /NH2_Arg_/  
 Coded contact list: Anion  
 Exclusion zone contact(s): /N7/NH1_Arg_/O4/  
Consider K in place of MG (3 valid contact(s) for K vs. 1 for MG)  

### K:45:201

|  | Res. type | Atom | Residue | Distance | Occupancy | B-factor | Closest solute to 1^st^ shell water |
| --- | --- | --- | --- | --- | --- | --- | --- |
| **Ion** | | | |  | 1.00 | 90.05 || 1 | ILE | O | 45:127 | 2.72 | 1.00 | 81.67 |
| 2 | GLY | O | 45:122 | 2.89 | 1.00 | 79.25 |  |
| 3 | LEU | O | 45:125 | 3.04 | 1.00 | 77.52 |  |
| 4 | G | OP1 | 14:1030 | 3.10 | 1.00 | 92.58 |  |
| 5 | A | O3' | 14:1029 | 3.81 | 1.00 | 87.59 |  |
| 6 | HIS | O | 45:123 | 4.07 | 1.00 | 79.73 |  |
| 7 | HIS | N | 45:123 | 4.20 | 1.00 | 81.71 |  |
| 8 | A | O2' | 14:1029 | 4.26 | 1.00 | 80.62 |  |
| 9 | LYS | N | 45:128 | 4.33 | 1.00 | 81.21 |  |
| 10 | ILE | N | 45:127 | 4.53 | 1.00 | 78.44 |  |
| 11 | LEU | N | 45:125 | 4.67 | 1.00 | 76.75 |  |
| 12 | G | O2' | 14:2485 | 4.80 | 1.00 | 74.88 |  |
| 13 | G | O5' | 14:1030 | 4.90 | 1.00 | 72.20 |  |
  |

  
**1^st^ shell ligand angles:**   
1-2: 89; 1-3: 66; 1-4: 102; 2-3: 65; 2-4: 134; 3-4: 159;   
  
 Contact(s): /O_back_/O_back_/O_back_/OP1/  
 Coded contact list: O_ph_3O_back_  

### K:49:201

|  | Res. type | Atom | Residue | Distance | Occupancy | B-factor | Closest solute to 1^st^ shell water |
| --- | --- | --- | --- | --- | --- | --- | --- |
| **Ion** | | | |  | 1.00 | 108.83 || 1 | ARG | O | 49:128 | 2.88 | 1.00 | 89.75 |
| 2 | ALA | O | 49:163 | 3.30 | 1.00 | 98.12 |  |
| 3 | THR | O | 49:161 | 3.31 | 1.00 | 97.43 |  |
| 4 | ASN | ND2 | 49:130 | 4.19 | 1.00 | 92.83 |  |
| 5 | ALA | N | 49:163 | 4.31 | 1.00 | 85.48 |  |
| 6 | THR | O | 49:162 | 4.36 | 1.00 | 86.75 |  |
| 7 | ASN | OD1 | 49:130 | 4.43 | 1.00 | 91.31 |  |
| 8 | GLY | O | 49:129 | 4.53 | 1.00 | 95.16 |  |
| 9 | THR | N | 49:162 | 4.88 | 1.00 | 88.22 |  |
  |

  
**1^st^ shell ligand angles:**   
1-2: 57; 1-3: 98; 2-3: 95;   
  
 Contact(s): /O_back_/O_back_/O_back_/  
 Coded contact list: 3O_back_  

### K:4A:201

|  | Res. type | Atom | Residue | Distance | Occupancy | B-factor | Closest solute to 1^st^ shell water |
| --- | --- | --- | --- | --- | --- | --- | --- |
| **Ion** | | | |  | 1.00 | 96.06 || 1 | C | OP2 | 1G:1322 | 2.57 | 1.00 | 112.02 |
| 2 | GLN | O | 4A:101 | 3.02 | 1.00 | 88.99 |  |
| 3 | TYR | OH | 4A:87 | 3.38 | 1.00 | 113.83 |  |
| 4 | A | OP1 | 1G:1225 | 3.51 | 1.00 | 94.16 |  |
| 5 | GLN | N | 4A:101 | 3.88 | 1.00 | 101.67 |  |
| 6 | G | O3' | 1G:1224 | 4.05 | 1.00 | 98.23 |  |
| 7 | C | O3' | 1G:1321 | 4.07 | 1.00 | 104.32 |  |
| 8 | THR | OG1 | 4A:103 | 4.13 | 1.00 | 88.06 |  |
| 9 | C | OP2 | 1G:1226 | 4.16 | 1.00 | 90.98 |  |
| 10 | ARG | NH2 | 4A:91 | 4.19 | 1.00 | 99.70 |  |
| 11 | A | O5' | 1G:1225 | 4.44 | 1.00 | 86.45 |  |
| 12 | C | OP1 | 1G:1322 | 4.47 | 1.00 | 111.40 |  |
| 13 | G | O2' | 1G:1224 | 4.49 | 1.00 | 102.51 |  |
  |

  
**1^st^ shell ligand angles:**   
1-2: 114; 1-3: 97; 2-3: 129;   
  
 Contact(s): /O_back_/OH_Tyr_/  
 Coded contact list: O_back_OH_Prot_  
 Exclusion zone contact(s): /OP2/  

### K:4I:201

|  | Res. type | Atom | Residue | Distance | Occupancy | B-factor | Closest solute to 1^st^ shell water |
| --- | --- | --- | --- | --- | --- | --- | --- |
| **Ion** | | | |  | 1.00 | 71.46 || 1 | U | OP1 | 13:1330 | 2.51 | 1.00 | 85.15 |
| 2 | ILE | O | 4I:22 | 2.55 | 1.00 | 90.57 |  |
| 3 | THR | O | 4I:20 | 2.62 | 1.00 | 81.01 |  |
| 4 | ILE | O | 4I:25 | 2.87 | 1.00 | 86.61 |  |
| 5 | ILE | N | 4I:22 | 3.50 | 1.00 | 88.30 |  |
| 6 | ILE | N | 4I:25 | 3.64 | 1.00 | 85.52 |  |
| 7 | TYR | N | 4I:23 | 3.69 | 1.00 | 90.80 |  |
| 8 | U | OP2 | 13:1330 | 3.82 | 1.00 | 78.42 |  |
| 9 | GLY | N | 4I:24 | 3.85 | 1.00 | 89.51 |  |
| 10 | TYR | N | 4I:21 | 4.25 | 1.00 | 81.78 |  |
| 11 | LEU | O | 4I:19 | 4.29 | 1.00 | 77.56 |  |
| 12 | G | OP2 | 13:1331 | 4.45 | 1.00 | 91.17 |  |
| 13 | A | O3' | 13:1329 | 4.62 | 1.00 | 83.71 |  |
| 14 | U | O5' | 13:1330 | 4.65 | 1.00 | 87.96 |  |
| 15 | THR | N | 4I:20 | 4.89 | 1.00 | 85.64 |  |
| 16 | GLY | N | 4I:26 | 5.00 | 1.00 | 85.19 |  |
  |

  
**1^st^ shell ligand angles:**   
1-2: 88; 1-3: 154; 1-4: 90; 2-3: 118; 2-4: 89; 3-4: 93;   
  
 Contact(s): /O_back_/O_back_/  
 Coded contact list: 2O_back_  
 Exclusion zone contact(s): /O_back_/OP1/  

### MG:4L:101

|  | Res. type | Atom | Residue | Distance | Occupancy | B-factor | Closest solute to 1^st^ shell water |
| --- | --- | --- | --- | --- | --- | --- | --- |
| **Ion** | | | |  | 1.00 | 79.35 || 1 | U | OP1 | 4L:49 | 2.52 | 1.00 | 65.59 |
| 2 | 4OC | OP2 | 1G:1402 | 2.85 | 1.00 | 61.71 |  |
| 3 | U | OP2 | 4L:50 | 3.42 | 1.00 | 78.91 |  |
| 4 | G | OP1 | 1G:1401 | 3.60 | 1.00 | 67.76 |  |
| 5 | 4OC | OP1 | 1G:1402 | 3.97 | 1.00 | 71.55 |  |
| 6 | U | OP2 | 4L:49 | 4.30 | 1.00 | 71.64 |  |
| 7 | U | O5' | 4L:49 | 4.60 | 1.00 | 74.17 |  |
| 8 | U | OP1 | 4L:50 | 4.73 | 1.00 | 73.31 |  |
| 9 | U | O2' | 4L:48 | 4.88 | 1.00 | 62.91 |  |
| 10 | G | O3' | 1G:1401 | 4.93 | 1.00 | 65.88 |  |
  |

  
 Coded contact list:   
 Exclusion zone contact(s): /OP1/OP2/  
Consider NA in place of MG (1 valid contact(s) for NA vs. 0 for MG)  

### K:52:201

|  | Res. type | Atom | Residue | Distance | Occupancy | B-factor | Closest solute to 1^st^ shell water |
| --- | --- | --- | --- | --- | --- | --- | --- |
| **Ion** | | | |  | 1.00 | 76.72 || 1 | ARG | O | 52:80 | 2.96 | 1.00 | 60.90 |
| 2 | VAL | O | 52:85 | 3.17 | 1.00 | 65.95 |  |
| 3 | ARG | O | 52:82 | 3.28 | 1.00 | 61.75 |  |
| 4 | ASP | O | 52:83 | 3.94 | 1.00 | 71.80 |  |
| 5 | ILE | O | 52:81 | 4.34 | 1.00 | 63.90 |  |
| 6 | ASP | N | 52:83 | 4.39 | 1.00 | 68.91 |  |
| 7 | ASP | OD1 | 52:83 | 4.78 | 1.00 | 79.12 |  |
  |

  
**1^st^ shell ligand angles:**   
1-2: 81; 1-3: 57; 2-3: 58;   
  
 Contact(s): /O_back_/O_back_/O_back_/  
 Coded contact list: 3O_back_  

### K:5A:101

|  | Res. type | Atom | Residue | Distance | Occupancy | B-factor | Closest solute to 1^st^ shell water |
| --- | --- | --- | --- | --- | --- | --- | --- |
| **Ion** | | | |  | 1.00 | 82.78 || 1 | C | OP1 | 1G:1359 | 3.08 | 1.00 | 99.00 |
| 2 | VAL | N | 5A:33 | 3.31 | 1.00 | 91.81 |  |
| 3 | VAL | O | 5A:33 | 3.42 | 1.00 | 95.33 |  |
| 4 | THR | O | 5A:22 | 3.57 | 1.00 | 99.29 |  |
| 5 | THR | OG1 | 5A:22 | 3.67 | 1.00 | 99.03 |  |
| 6 | ALA | O | 5A:30 | 3.89 | 1.00 | 92.70 |  |
| 7 | G | OP2 | 1G:976 | 3.91 | 1.00 | 98.25 |  |
| 8 | SER | N | 5A:32 | 4.06 | 1.00 | 88.82 |  |
| 9 | U | O3' | 1G:1358 | 4.15 | 1.00 | 96.01 |  |
| 10 | THR | N | 5A:22 | 4.82 | 1.00 | 95.46 |  |
| 11 | ARG | N | 5A:23 | 4.85 | 1.00 | 95.08 |  |
| 12 | ARG | N | 5A:31 | 4.87 | 1.00 | 91.03 |  |
| 13 | ARG | O | 5A:31 | 4.90 | 1.00 | 85.09 |  |
| 14 | C | OP2 | 1G:1359 | 4.94 | 1.00 | 99.75 |  |
  |

  
**1^st^ shell ligand angles:**   
1-2: 122;   
  
 Contact(s): /N_back_/OP1/  
 Coded contact list: O_ph_Anion  

### ZN:5A:102

|  | Res. type | Atom | Residue | Distance | Occupancy | B-factor | Closest solute to 1^st^ shell water |
| --- | --- | --- | --- | --- | --- | --- | --- |
| **Ion** | | | |  | 1.00 | 82.96 || 1 | CYS | N | 5A:27 | 3.46 | 1.00 | 96.82 |
| 2 | CYS | N | 5A:40 | 3.78 | 1.00 | 91.36 |  |
| 3 | CYS | N | 5A:43 | 3.82 | 1.00 | 86.55 |  |
| 4 | G | N2 | 1G:1202 | 4.42 | 1.00 | 92.26 |  |
| 5 | CYS | O | 5A:27 | 4.67 | 1.00 | 91.26 |  |
| 6 | G | N3 | 1G:1202 | 4.70 | 1.00 | 93.88 |  |
| 7 | ARG | N | 5A:26 | 4.95 | 1.00 | 86.42 |  |
  |

  
 Coded contact list:   

### K:5E:201

|  | Res. type | Atom | Residue | Distance | Occupancy | B-factor | Closest solute to 1^st^ shell water |
| --- | --- | --- | --- | --- | --- | --- | --- |
| **Ion** | | | |  | 1.00 | 76.48 || 1 | VAL | O | 5E:85 | 2.80 | 1.00 | 70.27 |
| 2 | ARG | O | 5E:82 | 2.91 | 1.00 | 73.70 |  |
| 3 | ARG | O | 5E:80 | 3.09 | 1.00 | 61.45 |  |
| 4 | ILE | O | 5E:81 | 4.54 | 1.00 | 65.30 |  |
| 5 | VAL | N | 5E:85 | 4.62 | 1.00 | 67.06 |  |
| 6 | ASP | N | 5E:83 | 4.68 | 1.00 | 75.76 |  |
| 7 | ARG | N | 5E:86 | 4.74 | 1.00 | 57.70 |  |
| 8 | ARG | N | 5E:82 | 4.75 | 1.00 | 65.67 |  |
| 9 | ASP | O | 5E:83 | 4.76 | 1.00 | 73.50 |  |
| 10 | ILE | N | 5E:81 | 4.87 | 1.00 | 69.05 |  |
  |

  
**1^st^ shell ligand angles:**   
1-2: 73; 1-3: 122; 2-3: 86;   
  
 Contact(s): /O_back_/O_back_/O_back_/  
 Coded contact list: 3O_back_  

### ZN:5I:101

|  | Res. type | Atom | Residue | Distance | Occupancy | B-factor | Closest solute to 1^st^ shell water |
| --- | --- | --- | --- | --- | --- | --- | --- |
| **Ion** | | | |  | 1.00 | 70.91 || 1 | CYS | N | 5I:40 | 3.73 | 1.00 | 67.86 |
| 2 | CYS | N | 5I:43 | 3.92 | 1.00 | 68.07 |  |
| 3 | CYS | N | 5I:27 | 3.96 | 1.00 | 63.26 |  |
| 4 | G | N2 | 13:1202 | 4.68 | 1.00 | 66.01 |  |
| 5 | G | N3 | 13:1202 | 4.75 | 1.00 | 62.40 |  |
| 6 | GLY | N | 5I:28 | 4.87 | 1.00 | 68.23 |  |
| 7 | G | O2' | 13:1202 | 4.99 | 1.00 | 69.70 |  |
  |

  
 Coded contact list:   

### MG:6E:201

|  | Res. type | Atom | Residue | Distance | Occupancy | B-factor | Closest solute to 1^st^ shell water |
| --- | --- | --- | --- | --- | --- | --- | --- |
| **Ion** | | | |  | 1.00 | 62.30 || 1 | ARG | NH2 | 6E:32 | 3.45 | 1.00 | 96.66 |
| 2 | G | OP1 | 13:942 | 3.59 | 1.00 | 75.75 |  |
| 3 | C | N4 | 13:1237 | 4.02 | 1.00 | 73.45 |  |
| 4 | C | N3 | 13:1336 | 4.14 | 1.00 | 75.24 |  |
| 5 | C | O2 | 13:1336 | 4.39 | 1.00 | 79.15 |  |
| 6 | ARG | NH1 | 6E:32 | 4.53 | 1.00 | 90.18 |  |
  |

  
 Coded contact list:   
1^st^ MG coordination shell missing (see Notes)  

### MG:78:201

|  | Res. type | Atom | Residue | Distance | Occupancy | B-factor | Closest solute to 1^st^ shell water |
| --- | --- | --- | --- | --- | --- | --- | --- |
| **Ion** | | | |  | 1.00 | 38.55 || 1 | MET | O | 78:67 | 3.93 | 1.00 | 39.05 |
| 2 | GLN | O | 78:68 | 4.16 | 1.00 | 34.09 |  |
| 3 | G | OP2 | 1H:235 | 4.36 | 1.00 | 28.59 |  |
| 4 | A | OP1 | 1H:234 | 4.42 | 1.00 | 41.04 |  |
  |

  
 Coded contact list:   
1^st^ MG coordination shell missing (see Notes)  

### MG:78:202

|  | Res. type | Atom | Residue | Distance | Occupancy | B-factor | Closest solute to 1^st^ shell water |
| --- | --- | --- | --- | --- | --- | --- | --- |
| **Ion** | | | |  | 1.00 | 30.73 || 1 | Water 1 | | 1H:4161 | 2.14 | 1.00 | 32.14 | 2.84 (U:N3 1H:1296) |
| 2 | Water 2 | | 1H:4200 | 2.18 | 1.00 | 34.85 | 2.89 (G:N7 1H:1297) |
| 3 | Water 3 | | 78:305 | 2.18 | 1.00 | 34.51 | 3.26 (ARG:O 78:18) |
| 4 | Water 4 | | 1H:3827 | 2.19 | 1.00 | 22.98 | 2.42 (G:OP2 1H:1297) |
| 5 | Water 5 | | 78:308 | 2.19 | 1.00 | 46.81 | 3.91 (ARG:O 78:18) |
| 6 | Water 6 | | 78:309 | 2.20 | 1.00 | 51.38 | 4.16 (ARG:O 78:18) |
| 7 | U | N3 | 1H:1296 | 3.88 | 1.00 | 23.07 |  |
| 8 | ARG | NH1 | 78:21 | 4.34 | 1.00 | 24.29 |  |
| 9 | U | O4 | 1H:1296 | 4.39 | 1.00 | 22.13 |  |
| 10 | G | OP2 | 1H:1297 | 4.53 | 1.00 | 25.93 |  |
| 11 | U | O2 | 1H:1296 | 4.60 | 1.00 | 25.66 |  |
| 12 | U | N1 | 1H:1296 | 4.64 | 1.00 | 23.07 |  |
| 13 | ARG | O | 78:18 | 4.72 | 1.00 | 40.50 |  |
| 14 | G | N7 | 1H:1297 | 4.78 | 1.00 | 25.37 |  |
| 15 | ARG | NE | 78:21 | 4.81 | 1.00 | 30.92 |  |
  |

  
**1^st^ shell ligand angles:**   
1-2: 97; 1-3: 103; 1-4: 73; 1-5: 176; 1-6: 79; 2-3: 97; 2-4: 82; 2-5: 86; 2-6: 175; 3-4: 176; 3-5: 80; 3-6: 80; 4-5: 104; 4-6: 100; 5-6: 98;   
  
 Contact(s): /O_w_/O_w_/O_w_/O_w_/O_w_/O_w_/  
 Coded contact list: 6O_w_  

### MG:7I:101

|  | Res. type | Atom | Residue | Distance | Occupancy | B-factor | Closest solute to 1^st^ shell water |
| --- | --- | --- | --- | --- | --- | --- | --- |
| **Ion** | | | |  | 1.00 | 71.55 || 1 | ARG | NH1 | 7I:75 | 3.41 | 1.00 | 104.18 |
| 2 | G | O6 | 13:476 | 4.12 | 1.00 | 119.48 |  |
| 3 | ARG | NE | 7I:75 | 4.12 | 1.00 | 101.23 |  |
| 4 | G | O6 | 13:475 | 4.54 | 1.00 | 127.04 |  |
| 5 | C | OP2 | 13:454 | 4.62 | 1.00 | 98.00 |  |
| 6 | ARG | NH2 | 7I:75 | 4.85 | 1.00 | 102.78 |  |
| 7 | C | OP1 | 13:454 | 4.86 | 1.00 | 105.07 |  |
| 8 | G | N7 | 13:475 | 4.90 | 1.00 | 127.69 |  |
  |

  
 Coded contact list:   
1^st^ MG coordination shell missing (see Notes)  

### K:88:201

|  | Res. type | Atom | Residue | Distance | Occupancy | B-factor | Closest solute to 1^st^ shell water |
| --- | --- | --- | --- | --- | --- | --- | --- |
| **Ion** | | | |  | 1.00 | 57.95 || 1 | LEU | O | 88:125 | 2.75 | 1.00 | 36.08 |
| 2 | ILE | O | 88:127 | 2.82 | 1.00 | 43.80 |  |
| 3 | GLY | O | 88:122 | 2.90 | 1.00 | 45.05 |  |
| 4 | G | OP1 | 1H:1077 | 3.29 | 1.00 | 45.79 |  |
| 5 | A | O2' | 1H:1076 | 3.86 | 1.00 | 44.97 |  |
| 6 | A | O3' | 1H:1076 | 3.92 | 1.00 | 46.95 |  |
| 7 | ILE | N | 88:127 | 4.01 | 1.00 | 36.26 |  |
| 8 | HIS | O | 88:123 | 4.22 | 1.00 | 39.94 |  |
| 9 | G | O2' | 1H:2498 | 4.23 | 1.00 | 36.30 |  |
| 10 | HIS | N | 88:123 | 4.59 | 1.00 | 41.75 |  |
| 11 | LEU | N | 88:125 | 4.71 | 1.00 | 38.62 |  |
| 12 | LYS | N | 88:128 | 4.81 | 1.00 | 39.90 |  |
| 13 | PRO | N | 88:126 | 4.85 | 1.00 | 39.22 |  |
  |

  
**1^st^ shell ligand angles:**   
1-2: 82; 1-3: 73; 1-4: 163; 2-3: 92; 2-4: 82; 3-4: 113;   
  
 Contact(s): /O_back_/O_back_/O_back_/OP1/  
 Coded contact list: O_ph_3O_back_  

### K:BA:201

|  | Res. type | Atom | Residue | Distance | Occupancy | B-factor | Closest solute to 1^st^ shell water |
| --- | --- | --- | --- | --- | --- | --- | --- |
| **Ion** | | | |  | 1.00 | 72.97 || 1 | LEU | O | BA:72 | 2.87 | 1.00 | 79.67 |
| 2 | ALA | O | BA:67 | 2.88 | 1.00 | 92.65 |  |
| 3 | HIS | O | BA:73 | 3.09 | 1.00 | 77.05 |  |
| 4 | GLY | N | BA:69 | 3.36 | 1.00 | 93.19 |  |
| 5 | GLY | O | BA:69 | 3.71 | 1.00 | 79.28 |  |
| 6 | LYS | N | BA:74 | 4.06 | 1.00 | 74.19 |  |
| 7 | HIS | N | BA:73 | 4.40 | 1.00 | 71.21 |  |
| 8 | LYS | N | BA:68 | 4.54 | 1.00 | 86.85 |  |
  |

  
**1^st^ shell ligand angles:**   
1-2: 96; 1-3: 81; 1-4: 93; 2-3: 128; 2-4: 56; 3-4: 172;   
  
 Contact(s): /N_back_/O_back_/O_back_/O_back_/  
 Coded contact list: 3O_back_Anion  

### K:BI:201

|  | Res. type | Atom | Residue | Distance | Occupancy | B-factor | Closest solute to 1^st^ shell water |
| --- | --- | --- | --- | --- | --- | --- | --- |
| **Ion** | | | |  | 1.00 | 108.63 || 1 | GLY | O | BI:69 | 2.83 | 1.00 | 79.64 |
| 2 | ALA | O | BI:67 | 3.05 | 1.00 | 94.14 |  |
| 3 | LEU | O | BI:72 | 3.10 | 1.00 | 78.42 |  |
| 4 | LYS | O | BI:68 | 3.84 | 1.00 | 95.08 |  |
| 5 | SER | N | BI:70 | 4.65 | 1.00 | 88.04 |  |
| 6 | HIS | ND1 | BI:73 | 4.72 | 1.00 | 84.26 |  |
| 7 | GLY | N | BI:69 | 4.93 | 1.00 | 96.78 |  |
| 8 | HIS | N | BI:73 | 4.95 | 1.00 | 76.02 |  |
| 9 | THR | N | BI:71 | 4.97 | 1.00 | 86.08 |  |
  |

  
**1^st^ shell ligand angles:**   
1-2: 69; 1-3: 81; 2-3: 72;   
  
 Contact(s): /O_back_/O_back_/O_back_/  
 Coded contact list: 3O_back_  

### MG:C8:201

|  | Res. type | Atom | Residue | Distance | Occupancy | B-factor | Closest solute to 1^st^ shell water |
| --- | --- | --- | --- | --- | --- | --- | --- |
| **Ion** | | | |  | 1.00 | 44.75 || 1 | ARG | NH1 | C8:33 | 3.18 | 1.00 | 31.20 |
| 2 | ARG | NH2 | C8:33 | 3.24 | 1.00 | 38.40 |  |
| 3 | C | OP2 | 1H:605 | 3.36 | 1.00 | 29.78 |  |
| 4 | ARG | NE | C8:33 | 3.93 | 1.00 | 37.63 |  |
| 5 | A | O2' | 1H:602 | 4.47 | 1.00 | 27.23 |  |
| 6 | C | OP2 | 1H:604 | 4.53 | 1.00 | 26.71 |  |
| 7 | C | O5' | 1H:604 | 4.67 | 1.00 | 26.28 |  |
| 8 | A | OP1 | 1H:1300 | 4.78 | 1.00 | 26.45 |  |
| 9 | Water 1 | | 1H:4051 | 4.81 | 1.00 | 23.40 | 4.98 (A:OP1 1H:1300) |
  |

  
 Coded contact list:   
 Exclusion zone contact(s): /NH1_Arg_/NH2_Arg_/OP2/  
Consider K in place of MG (3 valid contact(s) for K vs. 0 for MG)  

### MG:F8:101

|  | Res. type | Atom | Residue | Distance | Occupancy | B-factor | Closest solute to 1^st^ shell water |
| --- | --- | --- | --- | --- | --- | --- | --- |
| **Ion** | | | |  | 1.00 | 32.92 || 1 | VAL | O | F8:54 | 2.83 | 1.00 | 29.91 |
| 2 | G | OP2 | 1H:1390 | 3.10 | 1.00 | 30.86 |  |
| 3 | ASN | OD1 | F8:55 | 3.38 | 1.00 | 31.44 |  |
| 4 | LYS | NZ | F8:36 | 3.72 | 1.00 | 30.14 |  |
| 5 | VAL | N | F8:54 | 3.94 | 1.00 | 34.04 |  |
| 6 | ASN | N | F8:55 | 4.40 | 1.00 | 29.07 |  |
| 7 | G | OP1 | 1H:1390 | 4.56 | 1.00 | 32.27 |  |
| 8 | A | OP1 | 1H:1389 | 4.81 | 1.00 | 26.75 |  |
  |

  
 Coded contact list:   
 Exclusion zone contact(s): /O_back_/OD1_Asn_/OP2/  
Consider K in place of MG (3 valid contact(s) for K vs. 0 for MG)  

### MG:I8:101

|  | Res. type | Atom | Residue | Distance | Occupancy | B-factor | Closest solute to 1^st^ shell water |
| --- | --- | --- | --- | --- | --- | --- | --- |
| **Ion** | | | |  | 1.00 | 26.71 || 1 | ARG | NH1 | I8:14 | 2.90 | 1.00 | 31.16 |
| 2 | G | O6 | 1H:2293 | 3.78 | 1.00 | 32.49 |  |
| 3 | C | N4 | 1H:2294 | 3.94 | 1.00 | 25.01 |  |
| 4 | G | N7 | 1H:2293 | 4.03 | 1.00 | 25.47 |  |
| 5 | ARG | NH2 | I8:14 | 4.82 | 1.00 | 22.87 |  |
| 6 | ARG | NE | I8:14 | 4.94 | 1.00 | 28.56 |  |
  |

  
 Coded contact list:   
 Exclusion zone contact(s): /NH1_Arg_/  
Consider K in place of MG (1 valid contact(s) for K vs. 0 for MG)  

### MG:J8:101

|  | Res. type | Atom | Residue | Distance | Occupancy | B-factor | Closest solute to 1^st^ shell water |
| --- | --- | --- | --- | --- | --- | --- | --- |
| **Ion** | | | |  | 1.00 | 32.85 || 1 | U | OP1 | 1H:408 | 3.59 | 1.00 | 33.78 |
| 2 | SER | OG | J8:17 | 3.89 | 1.00 | 18.75 |  |
| 3 | G | O3' | 1H:407 | 3.91 | 1.00 | 33.59 |  |
| 4 | ILE | N | J8:18 | 4.15 | 1.00 | 23.67 |  |
| 5 | U | OP2 | 1H:2246 | 4.16 | 1.00 | 24.36 |  |
| 6 | GLN | NE2 | J8:19 | 4.41 | 1.00 | 26.90 |  |
| 7 | ILE | O | J8:18 | 4.50 | 1.00 | 26.73 |  |
| 8 | G | O2' | 1H:407 | 4.83 | 1.00 | 31.25 |  |
| 9 | U | O5' | 1H:408 | 4.97 | 1.00 | 30.99 |  |
  |

  
 Coded contact list:   
1^st^ MG coordination shell missing (see Notes)  

### MG:L8:101

|  | Res. type | Atom | Residue | Distance | Occupancy | B-factor | Closest solute to 1^st^ shell water |
| --- | --- | --- | --- | --- | --- | --- | --- |
| **Ion** | | | |  | 1.00 | 38.38 || 1 | A | OP2 | 1H:1035 | 2.28 | 1.00 | 32.09 |
| 2 | Water 1 | | L8:202 | 2.71 | 1.00 | 38.02 | 2.52 (ILE:O L8:13) |
| 3 | SER | O | L8:11 | 2.71 | 1.00 | 31.55 |  |
| 4 | Water 2 | | 1H:4203 | 3.35 | 1.00 | 27.09 | 3.00 (A:OP2 1H:1035) |
| 5 | SER | N | L8:11 | 3.74 | 1.00 | 32.02 |  |
| 6 | LYS | NZ | L8:10 | 3.85 | 1.00 | 41.96 |  |
| 7 | G | OP1 | 1H:1034 | 3.98 | 1.00 | 33.55 |  |
| 8 | TYR | OH | L8:15 | 4.01 | 1.00 | 36.96 |  |
| 9 | ILE | O | L8:13 | 4.17 | 1.00 | 37.80 |  |
| 10 | G | O5' | 1H:1034 | 4.17 | 1.00 | 27.97 |  |
| 11 | A | OP1 | 1H:1035 | 4.18 | 1.00 | 37.68 |  |
| 12 | G | OP2 | 1H:1034 | 4.18 | 1.00 | 33.00 |  |
| 13 | G | O3' | 1H:1034 | 4.38 | 1.00 | 25.71 |  |
| 14 | A | O5' | 1H:1035 | 4.79 | 1.00 | 26.86 |  |
| 15 | Water 3 | | L8:201 | 4.82 | 1.00 | 35.02 | 2.51 (ILE:O L8:13) |
| 16 | PRO | N | L8:12 | 4.91 | 1.00 | 29.98 |  |
  |

  
 Contact(s): /OP2/  
 Coded contact list: O_ph_  
 Exclusion zone contact(s): /O_back_/O_w_/O_w_/  

### MG:M5:101

|  | Res. type | Atom | Residue | Distance | Occupancy | B-factor | Closest solute to 1^st^ shell water |
| --- | --- | --- | --- | --- | --- | --- | --- |
| **Ion** | | | |  | 1.00 | 84.23 || 1 | Water 1 | | M5:206 | 2.18 | 1.00 | 62.97 | 4.33 (LYS:NZ M5:15) |
| 2 | Water 2 | | M5:205 | 2.18 | 1.00 | 59.04 | 4.01 (LYS:NZ M5:15) |
| 3 | Water 3 | | 14:3849 | 2.18 | 1.00 | 77.27 | 2.88 (G:N7 14:630) |
| 4 | Water 4 | | M5:201 | 2.18 | 1.00 | 84.42 | 1.30 (LYS:NZ M5:15) |
| 5 | Water 5 | | 14:4031 | 2.19 | 1.00 | 67.84 | 3.55 (A:OP1 14:632) |
| 6 | Water 6 | | 14:4025 | 2.20 | 1.00 | 62.14 | 3.50 (G:N7 14:630) |
| 7 | LYS | NZ | M5:15 | 3.37 | 1.00 | 79.40 |  |
| 8 | G | N7 | 14:630 | 4.26 | 1.00 | 66.86 |  |
| 9 | A | OP1 | 14:632 | 4.45 | 1.00 | 63.78 |  |
| 10 | Water 7 | | 14:4070 | 4.82 | 1.00 | 67.77 | 5.23 (A:OP1 14:632) |
| 11 | G | O6 | 14:630 | 4.93 | 1.00 | 69.57 |  |
  |

  
**1^st^ shell ligand angles:**   
1-2: 89; 1-3: 86; 1-4: 89; 1-5: 93; 1-6: 177; 2-3: 175; 2-4: 91; 2-5: 91; 2-6: 92; 3-4: 90; 3-5: 88; 3-6: 93; 4-5: 177; 4-6: 88; 5-6: 90;   
  
 Contact(s): /O_w_/O_w_/O_w_/O_w_/O_w_/O_w_/  
 Coded contact list: 6O_w_  
 Exclusion zone contact(s): /NZ_Lys_/  
Consider K in place of MG (1 valid contact(s) for K vs. 0 for MG)  
MG hydration shell issue: water too close (< 2.4 Angs.)  

### MG:M5:102

|  | Res. type | Atom | Residue | Distance | Occupancy | B-factor | Closest solute to 1^st^ shell water |
| --- | --- | --- | --- | --- | --- | --- | --- |
| **Ion** | | | |  | 1.00 | 58.06 || 1 | Water 1 | | 14:3983 | 2.16 | 1.00 | 61.51 | 3.42 (A:OP2 14:631) |
| 2 | Water 2 | | 35:205 | 2.17 | 1.00 | 54.51 | 3.10 (ARG:O 35:65) |
| 3 | Water 3 | | M5:203 | 2.18 | 1.00 | 63.44 | 2.76 (LYS:NZ M5:47) |
| 4 | Water 4 | | M5:202 | 2.18 | 1.00 | 62.88 | 2.59 (VAL:O M5:14) |
| 5 | Water 5 | | M5:204 | 2.18 | 1.00 | 47.76 | 3.92 (LYS:N M5:15) |
| 6 | Water 6 | | 14:4070 | 2.19 | 1.00 | 67.77 | 4.00 (A:OP2 14:631) |
| 7 | LYS | NZ | M5:47 | 4.38 | 1.00 | 56.72 |  |
| 8 | A | OP2 | 14:631 | 4.41 | 1.00 | 54.56 |  |
| 9 | LYS | N | M5:15 | 4.44 | 1.00 | 45.20 |  |
| 10 | VAL | O | M5:14 | 4.60 | 1.00 | 44.51 |  |
| 11 | ARG | O | 35:65 | 4.77 | 1.00 | 48.71 |  |
| 12 | VAL | N | M5:14 | 4.89 | 1.00 | 47.94 |  |
| 13 | Water 7 | | M5:206 | 5.00 | 1.00 | 62.97 | 5.94 (A:OP2 14:631) |
  |

  
**1^st^ shell ligand angles:**   
1-2: 92; 1-3: 86; 1-4: 178; 1-5: 87; 1-6: 92; 2-3: 178; 2-4: 86; 2-5: 95; 2-6: 88; 3-4: 96; 3-5: 85; 3-6: 92; 4-5: 95; 4-6: 86; 5-6: 177;   
  
 Contact(s): /O_w_/O_w_/O_w_/O_w_/O_w_/O_w_/  
 Coded contact list: 6O_w_  

### MG:N8:101

|  | Res. type | Atom | Residue | Distance | Occupancy | B-factor | Closest solute to 1^st^ shell water |
| --- | --- | --- | --- | --- | --- | --- | --- |
| **Ion** | | | |  | 1.00 | 41.27 || 1 | ASP | OD2 | N8:17 | 2.78 | 1.00 | 30.78 |
| 2 | G | OP2 | 1H:17 | 3.59 | 1.00 | 31.97 |  |
| 3 | G | OP1 | 1H:16 | 3.98 | 1.00 | 28.00 |  |
| 4 | G | OP1 | 1H:17 | 4.02 | 1.00 | 28.11 |  |
| 5 | G | OP2 | 1H:544 | 4.47 | 1.00 | 27.29 |  |
| 6 | LYS | O | N8:13 | 4.73 | 1.00 | 28.64 |  |
| 7 | ASP | OD1 | N8:17 | 4.78 | 1.00 | 28.53 |  |
| 8 | ARG | NH1 | N8:16 | 4.80 | 1.00 | 24.66 |  |
| 9 | G | O3' | 1H:16 | 4.94 | 1.00 | 31.89 |  |
  |

  
 Coded contact list:   
 Exclusion zone contact(s): /OD2_Asp_/  
Consider K in place of MG (1 valid contact(s) for K vs. 0 for MG)  

### MG:O8:101

|  | Res. type | Atom | Residue | Distance | Occupancy | B-factor | Closest solute to 1^st^ shell water |
| --- | --- | --- | --- | --- | --- | --- | --- |
| **Ion** | | | |  | 1.00 | 60.52 || 1 | C | O2' | 1H:2360 | 3.74 | 1.00 | 55.34 |
| 2 | CYS | O | O8:40 | 3.96 | 1.00 | 94.59 |  |
| 3 | C | O3' | 1H:2360 | 4.22 | 1.00 | 52.55 |  |
| 4 | HIS | ND1 | O8:49 | 4.80 | 1.00 | 104.43 |  |
| 5 | CYS | N | O8:40 | 4.87 | 1.00 | 82.87 |  |
  |

  
 Coded contact list:   
1^st^ MG coordination shell missing (see Notes)  

### MG:P8:101

|  | Res. type | Atom | Residue | Distance | Occupancy | B-factor | Closest solute to 1^st^ shell water |
| --- | --- | --- | --- | --- | --- | --- | --- |
| **Ion** | | | |  | 1.00 | 35.75 || 1 | A | OP1 | 1H:55 | 4.53 | 1.00 | 20.20 |
| 2 | TRP | NE1 | P8:40 | 4.68 | 1.00 | 24.44 |  |
| 3 | C | OP1 | 1H:65 | 4.96 | 1.00 | 41.10 |  |
  |

  
 Coded contact list:   
1^st^ MG coordination shell missing (see Notes)  

### MG:Q8:101

|  | Res. type | Atom | Residue | Distance | Occupancy | B-factor | Closest solute to 1^st^ shell water |
| --- | --- | --- | --- | --- | --- | --- | --- |
| **Ion** | | | |  | 0.50 | 13.93 || 1 | ARG | O | 78:65 | 4.04 | 1.00 | 27.05 |
| 2 | VAL | N | Q8:14 | 4.13 | 1.00 | 25.42 |  |
| 3 | VAL | O | Q8:14 | 4.14 | 1.00 | 30.24 |  |
| 4 | LYS | O | Q8:12 | 4.47 | 1.00 | 31.76 |  |
| 5 | LYS | N | Q8:15 | 4.70 | 1.00 | 26.73 |  |
| 6 | LYS | O | Q8:11 | 4.78 | 1.00 | 27.08 |  |
| 7 | ARG | N | Q8:13 | 4.97 | 1.00 | 25.49 |  |
  |

  
 Coded contact list:   
1^st^ MG coordination shell missing (see Notes)  

### MG:Q8:101

|  | Res. type | Atom | Residue | Distance | Occupancy | B-factor | Closest solute to 1^st^ shell water |
| --- | --- | --- | --- | --- | --- | --- | --- |
| **Ion** | | | |  | 0.50 | 11.81 || 1 | LYS | NZ | Q8:47 | 3.65 | 1.00 | 28.63 |
| 2 | VAL | O | Q8:23 | 4.22 | 1.00 | 32.14 |  |
| 3 | LYS | N | Q8:15 | 4.28 | 1.00 | 26.73 |  |
| 4 | A | OP2 | 1H:657 | 4.56 | 1.00 | 23.13 |  |
| 5 | VAL | N | Q8:14 | 4.75 | 1.00 | 25.42 |  |
| 6 | ARG | O | Q8:13 | 4.92 | 1.00 | 30.74 |  |
  |

  
 Coded contact list:   
1^st^ MG coordination shell missing (see Notes)  
\_\_\_\_\_\_\_\_\_\_\_\_\_\_\_\_\_\_\_\_\_\_\_\_\_\_\_\_\_\_\_\_\_\_\_\_\_\_\_\_\_\_\_\_\_\_\_\_\_\_\_\_\_\_\_\_\_\_\_\_\_\_\_\_  

#### 5) Sorted coded contact list

##### Go to ToC

NOTES:

  
- N. --> estimate of the coordination number as deduced from the PDB structure;  
- Excl. --> existence of contacts in the ligand exclusion zone;  
- Bad hyd. --> existence of 2^nd^ shell hydration contacts shorter than 2.4 Angs.;  
- Lines in red mark stereochemical issues that should be addressed;  
- Lines in green mark ions without stereochemical issues - Please note that this does not always warrant the quality of the ion identification;  
  
(1) N=0 --> No 1^st^ shell coordination: impossible to assess ion identity;  
(2) Exclusion contacts --> Probable wrong ion identity - poor modelization of the ion binding site is also possible;  
(3) Bad 2^nd^ hydration shell contacts --> Poor modelization of the hydration shell and/or binding site;  
(4) Check for anion (Cl^-^, ...);  
(5) Mixed contacts with unprotonated O/N and protonated N, suggest poorly modeled binding site;  
(6) Currently the ZN (Zinc) and CD (Cadmium) coordination is not evaluated;  
(7) Check occupancies - partial occupancies are not reliable at medium to low resolutions, especially for solvent particles;  
(8) Check B-factors;  
  
  

|  |  |  |  |  |  |  |  |
| --- | --- | --- | --- | --- | --- | --- | --- |
| **Ion** | **Number** | **Code** | **N.** | **Excl.** | **Bad hyd.** | **Notes** | **Ion list** |
| MG: | 245 | No 1^st^ shell | >1 | Yes | No | (2) | 13:1655 13:1658 13:1662 13:1665 13:1666 13:1667 13:1672 13:1673 13:1676 13:1681 13:1686 13:1687 13:1688 13:1690 13:1691 13:1692 13:1697 13:1698 13:1700 13:1701 13:1702 13:1703 13:1704 13:1705 13:1706 13:1707 13:1708 13:1711 13:1712 13:1713 13:1714 13:1715 13:1717 13:1720 13:1721 13:1722 14:3109 14:3111 14:3112 14:3118 14:3120 14:3121 14:3137 14:3141 14:3143 14:3145 14:3146 14:3150 14:3166 14:3167 14:3168 14:3174 14:3176 14:3180 14:3181 14:3182 14:3185 14:3186 14:3187 14:3190 14:3193 14:3195 14:3197 14:3201 14:3202 14:3209 14:3210 14:3213 14:3215 14:3220 14:3221 14:3225 14:3226 14:3227 14:3228 14:3230 14:3231 14:3232 14:3234 14:3235 14:3236 14:3241 14:3242 14:3243 14:3244 14:3246 14:3248 14:3249 14:3250 14:3251 14:3254 14:3256 14:3259 14:3260 14:3261 14:3263 14:3264 14:3268 14:3269 14:3270 14:3274 14:3276 14:3278 16:204 16:205 16:206 16:207 16:208 16:209 1G:1640 1G:1651 1G:1653 1G:1657 1G:1659 1G:1667 1G:1670 1G:1674 1G:1675 1G:1677 1G:1678 1G:1679 1G:1680 1G:1681 1G:1682 1G:1685 1G:1686 1G:1687 1G:1688 1G:1690 1G:1691 1G:1694 1H:3148 1H:3153 1H:3157 1H:3158 1H:3159 1H:3163 1H:3165 1H:3172 1H:3175 1H:3182 1H:3183 1H:3186 1H:3187 1H:3189 1H:3197 1H:3201 1H:3202 1H:3204 1H:3207 1H:3208 1H:3210 1H:3211 1H:3212 1H:3213 1H:3221 1H:3223 1H:3225 1H:3227 1H:3229 1H:3230 1H:3231 1H:3233 1H:3234 1H:3236 1H:3237 1H:3240 1H:3242 1H:3248 1H:3249 1H:3252 1H:3254 1H:3264 1H:3265 1H:3266 1H:3267 1H:3273 1H:3274 1H:3275 1H:3276 1H:3279 1H:3280 1H:3281 1H:3286 1H:3290 1H:3291 1H:3292 1H:3294 1H:3295 1H:3297 1H:3298 1H:3299 1H:3300 1H:3306 1H:3307 1H:3308 1H:3311 1H:3312 1H:3314 1H:3316 1H:3319 1H:3322 1H:3323 1H:3325 1H:3327 1H:3329 1H:3330 1H:3331 1H:3333 1H:3334 1H:3340 1H:3341 1H:3344 1H:3348 1H:3349 1H:3351 1H:3352 1H:3354 1H:3355 1H:3356 1H:3357 1H:3359 1H:3361 1H:3363 1H:3365 1H:3366 1H:3367 1H:3368 1H:3369 1H:3371 1H:3372 1H:3373 1H:3375 1H:3376 1H:3378 1H:3379 1H:3380 1H:3381 1J:204 39:303 4L:101 C8:201 F8:101 I8:101 N8:101 || MG: | 144 | No 1^st^ shell | =0 | No | No | (1) | 13:1652 13:1654 13:1669 13:1671 13:1675 13:1679 13:1680 13:1682 13:1689 13:1696 13:1699 13:1710 13:1716 13:1718 13:1719 13:1723 13:1725 14:3113 14:3115 14:3116 14:3117 14:3122 14:3125 14:3142 14:3148 14:3149 14:3159 14:3164 14:3165 14:3173 14:3178 14:3183 14:3184 14:3188 14:3194 14:3200 14:3203 14:3204 14:3205 14:3207 14:3208 14:3219 14:3224 14:3233 14:3238 14:3239 14:3240 14:3252 14:3262 14:3271 14:3272 14:3277 1G:1649 1G:1656 1G:1658 1G:1660 1G:1661 1G:1664 1G:1665 1G:1666 1G:1669 1G:1671 1G:1689 1G:1692 1G:1696 1G:1697 1G:1698 1G:1700 1H:3151 1H:3156 1H:3160 1H:3161 1H:3164 1H:3173 1H:3174 1H:3177 1H:3179 1H:3184 1H:3185 1H:3190 1H:3193 1H:3196 1H:3209 1H:3214 1H:3216 1H:3217 1H:3219 1H:3220 1H:3222 1H:3224 1H:3226 1H:3228 1H:3235 1H:3239 1H:3241 1H:3243 1H:3245 1H:3253 1H:3256 1H:3258 1H:3262 1H:3270 1H:3271 1H:3272 1H:3277 1H:3282 1H:3283 1H:3287 1H:3288 1H:3289 1H:3293 1H:3296 1H:3301 1H:3303 1H:3304 1H:3305 1H:3315 1H:3317 1H:3321 1H:3326 1H:3335 1H:3336 1H:3337 1H:3338 1H:3339 1H:3342 1H:3346 1H:3347 1H:3353 1H:3358 1H:3362 1H:3364 1H:3370 1H:3377 2K:104 2L:102 2L:103 31:303 6E:201 78:201 7I:101 J8:101 O8:101 P8:101 || MG: | 95 | 6O_w_ | 6 | No | Yes | (3) | 13:1727 13:1729 13:1730 13:1732 13:1734 13:1736 13:1738 13:1739 13:1740 13:1741 13:1742 13:1743 13:1756 13:1759 13:1760 13:1762 13:1764 13:1765 13:1767 13:1768 14:3284 14:3287 14:3288 14:3290 14:3296 14:3297 14:3299 14:3304 14:3324 14:3328 14:3335 14:3338 14:3339 14:3349 14:3353 14:3379 14:3383 14:3384 14:3390 14:3391 14:3396 14:3398 14:3399 16:214 16:215 1G:1702 1G:1704 1G:1705 1G:1709 1G:1710 1G:1712 1G:1714 1G:1716 1G:1717 1G:1725 1G:1727 1G:1728 1G:1732 1G:1734 1H:3395 1H:3401 1H:3402 1H:3403 1H:3408 1H:3416 1H:3418 1H:3425 1H:3427 1H:3434 1H:3435 1H:3439 1H:3443 1H:3444 1H:3450 1H:3467 1H:3482 1H:3488 1H:3492 1H:3493 1H:3495 1H:3497 1H:3531 1H:3532 1H:3537 1H:3539 1H:3541 1H:3542 1H:3545 1H:3550 1H:3555 1H:3558 1H:3561 1H:3568 1H:3570 1H:3576 || MG: | 73 | 6O_w_ | 6 | No | No |  | 13:1728 13:1733 13:1755 13:1758 13:1766 14:3294 14:3300 14:3303 14:3307 14:3308 14:3333 14:3336 14:3337 14:3345 14:3352 14:3367 14:3373 14:3375 14:3380 14:3386 14:3387 14:3388 14:3392 14:3393 14:3394 16:213 1G:1701 1G:1706 1G:1707 1G:1708 1G:1729 1G:1731 1G:1735 1H:3396 1H:3398 1H:3404 1H:3406 1H:3410 1H:3414 1H:3421 1H:3431 1H:3432 1H:3436 1H:3437 1H:3440 1H:3441 1H:3442 1H:3445 1H:3446 1H:3447 1H:3448 1H:3451 1H:3452 1H:3471 1H:3483 1H:3485 1H:3494 1H:3496 1H:3501 1H:3522 1H:3533 1H:3534 1H:3535 1H:3546 1H:3547 1H:3552 1H:3553 1H:3554 1H:3556 1H:3563 1H:3572 78:202 M5:102 || MG: | 61 | 5O_w_ | >5 | Yes | Yes | (2,3) | 13:1744 13:1745 13:1746 13:1750 13:1752 13:1753 13:1754 14:3281 14:3282 14:3285 14:3306 14:3310 14:3315 14:3316 14:3318 14:3319 14:3322 14:3323 14:3327 14:3331 14:3332 14:3350 14:3351 14:3354 14:3357 14:3361 14:3369 14:3377 14:3381 14:3385 14:3389 14:3395 14:3397 16:211 16:212 1G:1715 1G:1718 1G:1720 1G:1721 1G:1722 1H:3382 1H:3397 1H:3430 1H:3465 1H:3469 1H:3475 1H:3486 1H:3490 1H:3507 1H:3509 1H:3518 1H:3520 1H:3544 1H:3551 1H:3557 1H:3562 1H:3565 1H:3567 1H:3571 1J:205 21:302 || MG: | 60 | 6O_w_ | >6 | Yes | Yes | (2,3) | 11:302 13:1735 13:1737 13:1757 13:1763 14:3280 14:3289 14:3291 14:3295 14:3298 14:3301 14:3355 14:3366 14:3382 1G:1703 1G:1711 1G:1724 1G:1730 1G:1733 1H:3384 1H:3385 1H:3387 1H:3390 1H:3392 1H:3394 1H:3399 1H:3405 1H:3407 1H:3411 1H:3413 1H:3417 1H:3419 1H:3433 1H:3438 1H:3449 1H:3453 1H:3454 1H:3462 1H:3466 1H:3468 1H:3476 1H:3478 1H:3487 1H:3491 1H:3498 1H:3502 1H:3505 1H:3511 1H:3513 1H:3526 1H:3528 1H:3529 1H:3538 1H:3548 1H:3549 21:303 2K:105 32:302 3E:301 M5:101 || MG: | 45 | O_ph_ | >1 | Yes | --- | (2) | 13:1653 13:1656 13:1657 13:1661 13:1674 13:1684 14:3114 14:3127 14:3133 14:3140 14:3144 14:3151 14:3152 14:3158 14:3177 14:3189 14:3191 14:3198 14:3211 14:3212 14:3216 14:3218 14:3247 14:3265 14:3273 1G:1642 1G:1643 1G:1646 1G:1655 1G:1668 1G:1684 1G:1695 1G:1699 1H:3154 1H:3170 1H:3194 1H:3200 1H:3203 1H:3246 1H:3255 1H:3269 1H:3310 1H:3360 1J:202 L8:101 || MG: | 34 | O_ph_ | 1 | No | --- |  | 13:1660 13:1663 13:1670 13:1678 13:1693 13:1724 14:3110 14:3119 14:3155 14:3156 14:3161 14:3169 14:3172 14:3192 14:3196 14:3206 14:3222 14:3223 14:3229 14:3257 1G:1638 1G:1647 1G:1650 1G:1652 1G:1654 1G:1662 1G:1673 1H:3145 1H:3162 1H:3169 1H:3199 1H:3205 1H:3215 1H:3244 || MG: | 31 | 4O_w_ | >4 | Yes | Yes | (2,3) | 13:1751 14:3279 14:3305 14:3317 14:3329 14:3341 14:3342 14:3343 14:3344 14:3348 14:3365 14:3370 14:3371 14:3372 1G:1723 1H:3383 1H:3389 1H:3409 1H:3420 1H:3423 1H:3426 1H:3429 1H:3458 1H:3473 1H:3477 1H:3500 1H:3508 1H:3514 1H:3564 1H:3573 1H:3574 || MG: | 25 | 5O_w_ | >5 | Yes | No | (2) | 13:1731 13:1770 14:3283 14:3293 14:3326 14:3330 14:3334 14:3346 14:3363 14:3364 14:3376 14:3378 14:3400 1G:1726 1G:1736 1H:3388 1H:3412 1H:3460 1H:3470 1H:3472 1H:3480 1H:3523 1H:3527 1H:3536 1H:3569 || K: | 23 | 2O_b_ | 2 | No | --- |  | 13:1613 13:1646 14:3006 14:3057 14:3061 14:3096 14:3101 14:3106 1G:1602 1G:1630 1G:1631 1H:3002 1H:3004 1H:3011 1H:3017 1H:3034 1H:3041 1H:3044 1H:3049 1H:3105 1H:3108 1H:3134 1H:3143 || K: | 23 | 3O_b_ | 3 | No | --- |  | 13:1618 13:1634 13:1638 13:1639 13:1645 14:3003 14:3020 14:3045 14:3095 1G:1626 1G:1632 1G:1633 1H:3012 1H:3051 1H:3058 1H:3067 1H:3081 1H:3083 1H:3098 1H:3100 1H:3102 1H:3118 1H:3144 || K: | 18 | 4O_b_ | 4 | No | --- |  | 13:1602 13:1607 13:1647 14:3029 14:3030 14:3037 14:3043 14:3047 14:3048 14:3056 14:3074 14:3098 1H:3003 1H:3042 1H:3075 1H:3091 1H:3094 1H:3112 || K: | 18 | O_b_ | 1 | No | --- |  | 13:1611 13:1620 13:1641 13:1648 14:3012 14:3053 14:3062 14:3064 14:3091 14:3105 1G:1635 1H:3021 1H:3029 1H:3046 1H:3099 1H:3114 1H:3129 1H:3137 || MG: | 18 | 3O_w_ | >3 | Yes | Yes | (2,3) | 13:1747 13:1749 13:1769 14:3292 14:3320 14:3358 14:3360 14:3368 1G:1713 1H:3424 1H:3455 1H:3457 1H:3464 1H:3516 1H:3517 1H:3519 1H:3524 1H:3530 || MG: | 17 | 2O_ph_ | >2 | Yes | --- | (2) | 14:3108 14:3124 14:3126 14:3128 14:3129 14:3139 14:3153 14:3162 14:3199 1G:1645 1G:1648 1G:1693 1H:3168 1H:3171 1H:3188 1H:3232 2L:101 || MG: | 14 | 2O_ph_ | 2 | No | --- |  | 13:1677 14:3107 14:3136 14:3147 14:3154 14:3171 14:3217 14:3258 1H:3147 1H:3149 1H:3178 1H:3180 1H:3195 1H:3250 || MG: | 12 | 4O_w_ | >4 | Yes | No | (2) | 14:3312 14:3359 1H:3386 1H:3400 1H:3422 1H:3428 1H:3503 1H:3515 1H:3543 1H:3560 1H:3577 29:303 || K: | 10 | No 1^st^ shell | =0 | No | No | (1) | 13:1614 14:3011 14:3079 14:3088 1G:1604 1H:3005 1H:3026 1H:3037 1H:3050 1H:3126 || K: | 10 | O_ph_ | 1 | No | --- |  | 13:1617 13:1642 14:3009 1G:1601 1G:1606 1G:1623 1G:1627 1H:3028 1H:3125 1H:3131 || MG: | 9 | No 1^st^ shell | >1 | Yes | Yes | (2) | 1H:3181 1H:3238 1H:3260 1H:3263 1H:3278 1H:3285 1H:3302 1H:3332 1J:203 || MG: | 8 | O_b_ | >1 | Yes | --- | (2) | 13:1659 13:1664 13:1685 13:1695 14:3163 14:3267 1H:3146 1H:3318 || K: | 8 | O_ph_O_b_ | 2 | No | --- |  | 13:1606 14:3001 14:3004 14:3058 14:3071 14:3081 1G:1611 1H:3048 || MG: | 7 | O_b_ | 1 | No | --- |  | 13:1694 14:3245 14:3275 1G:1672 1H:3218 1H:3247 1H:3259 || MG: | 7 | 3O_ph_ | >3 | Yes | --- | (2) | 13:1668 13:1726 14:3130 14:3160 1G:1637 1H:3191 1H:3324 || MG: | 7 | O_ph_O_b_ | 2 | No | --- |  | 13:1683 14:3132 14:3134 14:3179 1G:1639 1G:1676 1H:3150 || MG: | 6 | O_ph_5O_w_ | 6 | No | No |  | 14:3356 1H:3393 1H:3415 1H:3463 1H:3481 1H:3499 || K: | 6 | 2O_r_O_b_ | 3 | No | --- |  | 13:1603 14:3046 14:3082 1H:3014 1H:3117 1H:3120 || MG: | 6 | O_ph_5O_w_ | 6 | No | Yes | (3) | 14:3286 14:3313 1H:3459 1H:3461 1H:3510 1H:3525 || MG: | 6 | O_ph_4O_w_ | >5 | Yes | Yes | (2,3) | 13:1748 14:3309 14:3321 14:3325 1H:3479 1H:3566 || K: | 5 | 2O_ph_ | 2 | No | --- |  | 14:3097 1G:1605 1G:1607 1H:3032 1H:3135 || MG: | 5 | O_w_ | >1 | Yes | No | (2) | 14:3255 1H:3166 1H:3268 1H:3284 1H:3309 || K: | 5 | 3O_ph_2O_r_ | 5 | No | --- |  | 13:1637 1G:1620 1G:1621 1G:1634 1H:3116 || K: | 5 | O_ph_O_r_ | 2 | No | --- |  | 14:3010 14:3066 14:3087 14:3093 1H:3052 || K: | 5 | 2O_b_Anion | 3 | No | --- | (5) | 14:3052 14:3070 14:3080 1G:1622 2K:103 || K: | 4 | O_ph_O_r_ | >2 | Yes | --- | (2) | 13:1609 13:1644 1G:1603 1H:3063 || K: | 4 | 3O_b_ | >3 | Yes | --- | (2) | 1H:3043 1H:3085 1H:3086 1H:3136 || MG: | 4 | 6O_w_ | >6 | Yes | No | (2) | 13:1761 14:3362 1H:3489 1H:3540 || K: | 4 | O_ph_2O_r_ | 3 | No | --- |  | 14:3090 14:3100 1H:3025 1H:3031 || K: | 4 | No 1^st^ shell | >1 | Yes | No | (2) | 14:3008 1H:3033 1H:3035 1H:3038 || K: | 4 | 2O_ph_O_b_ | 3 | No | --- |  | 13:1650 14:3063 1H:3013 1H:3101 || K: | 4 | O_r_3O_b_ | 4 | No | --- |  | 14:3016 14:3051 1H:3001 1H:3054 || MG: | 4 | O_ph_4O_w_ | >5 | Yes | No | (2) | 14:3302 14:3311 1H:3456 1H:3521 || K: | 4 | O_ph_2O_b_ | 3 | No | --- |  | 13:1616 14:3044 1H:3107 1K:101 || K: | 4 | 4O_b_Anion | 5 | No | --- | (5) | 13:1624 16:202 1G:1612 1H:3068 || K: | 4 | 3O_back_ | 3 | No | --- |  | 49:201 52:201 5E:201 BI:201 || K: | 3 | O_ph_O_r_O_b_N_b_ | 4 | No | --- |  | 13:1626 14:3023 1H:3024 || K: | 3 | 2O_ph_O_r_ | 3 | No | --- |  | 13:1649 1H:3093 1H:3141 || K: | 3 | O_r_ | 1 | No | --- |  | 14:3013 14:3067 1H:3140 || K: | 3 | 3O_ph_ | 3 | No | --- |  | 1G:1636 1H:3122 1H:3142 || MG: | 3 | O_ph_O_b_ | >2 | Yes | --- | (2) | 14:3214 1H:3198 1H:3257 || MG: | 3 | 3O_w_ | >3 | Yes | No | (2) | 14:3314 14:3340 14:3347 || MG: | 3 | O_ph_5O_w_ | >6 | Yes | Yes | (2,3) | 1H:3391 1H:3474 1H:3512 || K: | 3 | O_b_ | >1 | Yes | --- | (2) | 1H:3008 1H:3030 1H:3139 || K: | 3 | N_b_ | 1 | No | --- |  | 13:1601 13:1632 1G:1616 || K: | 3 | 2O_b_ | >2 | Yes | --- | (2) | 13:1635 14:3069 1H:3009 || K: | 3 | 2O_b_N_b_ | 3 | No | --- |  | 13:1604 14:3099 1H:3095 || K: | 3 | O_r_2O_b_ | 3 | No | --- |  | 1H:3072 1H:3084 1H:3115 || K: | 2 | O_r_O_b_ | 2 | No | --- |  | 14:3034 14:3077 || K: | 2 | O_ph_O_r_O_b_ | 3 | No | --- |  | 14:3072 1H:3045 || K: | 2 | O_r_3O_b_N_b_ | >4 | Yes | --- | (2) | 14:3021 1H:3071 || MG: | 2 | O_ph_3O_w_ | >4 | Yes | Yes | (2,3) | 1H:3504 1H:3506 || MG: | 2 | O_w_ | >1 | Yes | Yes | (2,3) | 14:3253 1H:3152 || MG: | 2 | 3O_ph_ | 3 | No | --- |  | 1G:1644 1H:3167 || K: | 2 | 2O_r_O_b_ | >3 | Yes | --- | (2) | 14:3032 1H:3070 || MG: | 2 | O_ph_O_w_ | >2 | Yes | No | (2) | 14:3123 1H:3374 || K: | 2 | O_ph_ | >1 | Yes | --- | (2) | 14:3005 1G:1629 || K: | 2 | 2O_ph_N_b_ | 3 | No | --- |  | 14:3042 1H:3080 || K: | 2 | 3O_ph_O_r_ | 4 | No | --- |  | 13:1631 1G:1615 || K: | 2 | O_b_O_w_ | 2 | No | No |  | 14:3076 14:3092 || K: | 2 | 4O_r_ | 4 | No | --- |  | 14:3007 1H:3055 || K: | 2 | O_ph_OH_Prot_O_w_ | 3 | No | No |  | 14:3083 1H:3121 || K: | 2 | 2O_r_O_back_ | 3 | No | No |  | 14:3068 1H:3106 || MG: | 2 | 2O_ph_His | >3 | Yes | --- | (2) | 14:3131 1H:3206 || MG: | 2 | O_ph_ | >1 | Yes | --- | (2) | 16:210 1H:3192 || ZN: |  |  |  |  |  | (6) | 5A:102 5I:101 || K: | 2 | O_b_3O_back_ | 4 | No | --- |  | 31:301 39:301 || K: | 2 | O_ph_3O_back_ | 4 | No | No |  | 45:201 88:201 || K: | 2 | 2O_back_ | >2 | Yes | --- | (2) | 41:201 4I:201 || K: | 1 | O_ph_O_r_O_b_ | >3 | Yes | --- |  | 14:3036 || MG: | 1 | O_b_N_b_ | >2 | Yes | --- | (2) | 14:3135 || K: | 1 | O_ph_O_r_N_b_ | 3 | No | No |  | 14:3102 || K: | 1 | O_r_O_b_2N_b_Anion | >5 | Yes | --- | (2,5) | 1H:3078 || K: | 1 | 3O_b_OH_Prot_ | 4 | No | --- |  | 1G:1618 || MG: | 1 | O_ph_2O_w_ | 3 | No | No |  | 1H:3261 || MG: | 1 | 2O_ph_ | 2 | No | --- |  | 14:3138 || MG: | 1 | O_ph_O_r_ | >2 | Yes | --- | (2) | 14:3266 || K: | 1 | O_r_3O_b_ | >4 | Yes | --- |  | 14:3033 || K: | 1 | 3O_ph_O_r_O_w_ | 5 | No | No |  | 1H:3092 || MG: | 1 | N_b_ | 1 | No | --- |  | 1G:1641 || K: | 1 | 2O_b_O_w_ | 3 | No | No |  | 1H:3132 || K: | 1 | O_b_Anion | >2 | Yes | --- | (2,5) | 13:1608 || MG: | 1 | 2O_w_ | >2 | Yes | Yes | (2,3) | 14:3374 || K: | 1 | O_ph_O_w_ | 2 | No | No |  | 1H:3022 || K: | 1 | 2O_ph_O_r_ | >3 | Yes | --- | (2) | 1G:1628 || K: | 1 | 2O_ph_ | >2 | Yes | --- | (2) | 1H:3069 || MG: | 1 | 2O_ph_4O_w_ | 6 | No | No |  | 1H:3575 || K: | 1 | O_ph_O_r_O_w_ | >3 | Yes | Yes | (2,3) | 1H:3104 || K: | 1 | O_r_2O_w_ | 3 | No | Yes | (3) | 1H:3020 || MG: | 1 | N_b_ | >1 | Yes | --- | (2) | 14:3170 || MG: | 1 | O_ph_N_b_ | >2 | Yes | --- | (2) | 14:3157 || K: | 1 | O_ph_O_r_N_b_ | >3 | Yes | No |  | 1H:3113 || MG: | 1 | O_b_4O_w_ | >5 | Yes | Yes | (2,3) | 1H:3484 || K: | 1 | 2O_r_ | 2 | No | --- |  | 1H:3124 || K: | 1 | 2O_b_ | 2 | No | --- |  | 1H:3019 || K: | 1 | 2O_b_N_b_ | 3 | No | --- |  | 14:3059 || MG: | 1 | No 1^st^ shell | =0 | No | Yes | (1) | 1H:3251 || K: | 1 | 4O_b_O_w_ | 5 | No | No |  | 14:3060 || MG: | 1 | 2O_ph_O_b_ | 3 | No | --- |  | 1H:3350 || K: | 1 | O_r_O_b_O_w_ | 3 | No | No |  | 1H:3111 || K: | 1 | O_b_N_b_ | 2 | No | --- |  | 1H:3007 || K: | 1 | O_ph_2O_r_ | >3 | Yes | --- | (2) | 14:3078 || K: | 1 | O_ph_N_b_ | 2 | No | --- |  | 14:3027 || MG: | 1 | O_w_ | 1 | No | Yes | (3) | 1H:3313 || K: | 1 | 2O_b_O_w_ | >3 | Yes | No | (2) | 1H:3006 || MG: | 1 | O_ph_5O_w_ | >6 | Yes | No | (2) | 1H:3559 || K: | 1 | O_r_O_b_N_b_ | 3 | No | --- |  | 13:1605 || K: | 1 | 2Anion | 2 | No | --- | (4) | 1H:3039 || K: | 1 | O_ph_O_r_ | >2 | Yes | --- | (2) | 14:3054 || K: | 1 | O_b_N_b_ | >2 | Yes | --- | (2) | 13:1651 || K: | 1 | O_ph_2O_r_N_b_ | 4 | No | --- |  | 1H:3027 || K: | 1 | 4O_ph_ | 4 | No | --- |  | 14:3104 || K: | 1 | 3O_b_Anion | 4 | No | --- | (5) | 13:1610 || K: | 1 | 2N_b_O_w_ | 3 | No | Yes | (3) | 1H:3066 || MG: | 1 | 2O_w_ | >2 | Yes | No | (2) | 1G:1719 || K: | 1 | O_ph_2O_w_ | 3 | No | Yes | (3) | 14:3103 || K: | 1 | O_ph_O_back_ | 2 | No | --- |  | 13:1625 || K: | 1 | 3O_w_ | >3 | Yes | Yes | (2,3) | 14:3094 || MG: | 1 | O_b_O_coo_ | >2 | Yes | --- | (2) | 14:3237 || K: | 1 | 3O_ph_O_w_ | 4 | No | No |  | 14:3084 || MG: | 1 | Anion | >1 | Yes | --- | (2,4) | 42:201 || K: | 1 | 2O_back_2O_w_ | 4 | No | No |  | 39:302 || MG: | 1 | 2O_back_ | 2 | No | --- |  | 41:202 || K: | 1 | 2O_back_ | 2 | No | --- |  | 19:301 || MG: | 1 | O_coo_ | >1 | Yes | --- | (2) | 29:302 || K: | 1 | O_b_O_back_OH_Prot_ | 3 | No | --- |  | 32:301 ||  |  |  |  |  |  |  | List of ions with partial occupancies || MG: | 9 | Occup. < 1.0 | --- | Yes | No | (7) | 13:1709 14:3175 1G:1663 1G:1683 1H:3320 1H:3328 1H:3328 1H:3345 1H:3345 || MG: | 8 | Occup. < 1.0 | --- | No | No | (7) | 13:1709 1G:1663 1G:1683 1H:3320 1H:3343 1H:3343 Q8:101 Q8:101 || K: | 2 | Occup. < 1.0 | --- | No | No | (7) | 1H:3040 1H:3040 || MG: | 2 | Occup. < 1.0 | --- | Yes | No | (7) | 1H:3155 1H:3176 || MG: | 1 | Occup. < 1.0 | --- | Yes | Yes | (7) | 14:3175 || MG: | 1 | Occup. < 1.0 | --- | Yes | No | (7) | 1H:3176 || MG: | 1 | Occup. < 1.0 | --- | Yes | No | (7) | 1H:3155 ||  |  |  |  |  |  | Total: | 24 |

  
**Total number of Mg^2+^: 1005** **Total number of Mg^2+^ with stereochemical clashes: 853** **Total number of Mg^2+^ without stereochemical clashes: 152** **Total number of K^+^: 367** **Total number of K^+^ with stereochemical clashes: 164** **Total number of K^+^ without stereochemical clashes: 203** **Total number of Na^+^: 0** **Total number of Na^+^ with stereochemical clashes: 0** **Total number of Na^+^ without stereochemical clashes: 0** 

### ### END ###
