## Supplementary material for "Deflating the RNA Mg^2+^ bubble. Stereochemistry to the rescue!": Spupplemental IV: 6SJ6.ion_diagnosis.html

 Ion report: 6SJ6  

### Ion report: 6SJ6

#### date: 2020-04-09 12:50

Authors: Filip Leonarski and Pascal Auffinger  
\*\*\* beta version: unpublished \*\*\*  
-> reference 1: Leonarski F., D'Ascenzo L., Auffinger P. (2017) NAR, 45, 987-1004  
-> reference 2: Leonarski F., D'Ascenzo L., Auffinger P. (2019) RNA, 25, 173-192  
  
Density maps not assigned - No density checks made in the current version!  
  
\_\_\_\_\_\_\_\_\_\_\_\_\_\_\_\_\_\_\_\_\_\_\_\_\_\_\_\_\_\_\_\_\_\_\_\_\_\_\_\_\_\_\_\_\_\_\_\_\_\_\_\_\_\_\_\_\_\_\_\_\_\_\_\_\_\_\_\_\_\_\_\_\_\_\_\_\_\_\_\_\_  

#### Table of content (ToC):

1) Number of ions of each type and coordination distance ranges used for validation  
2) Ion validation table  
3) Potential issue list  
4) Ion/ligand tables  
5) Sorted coded contact list  
\_\_\_\_\_\_\_\_\_\_\_\_\_\_\_\_\_\_\_\_\_\_\_\_\_\_\_\_\_\_\_\_\_\_\_\_\_\_\_\_\_\_\_\_\_\_\_\_\_\_\_\_\_\_\_\_\_\_\_\_\_\_\_\_\_\_\_\_\_\_\_\_\_\_\_\_\_\_\_\_\_  
  
REMARKS:  
- 1^st^ shell: correct distance range for coordinating atoms;  
- Exclusion zone (excl. zone): a distance range where \*no\* coordinating atoms should be present;  
- Currently coordination distances with sulfur (S) atoms are not considered --> no CD/ZN (cadmium/zinc) validation;  
\_\_\_\_\_\_\_\_\_\_\_\_\_\_\_\_\_\_\_\_\_\_\_\_\_\_\_\_\_\_\_\_\_\_\_\_\_\_\_\_\_\_\_\_\_\_\_\_\_\_\_\_\_\_\_\_\_\_\_\_\_\_\_\_  

#### 1) Number of ions of each categorie and coordination distance ranges used for validation

##### Go to ToC

| Ion type | Number | Ion list | Short excl. zone [Angs] | 1^st^ shell [Angs.] | Long excl. zone [Angs] |
| --- | --- | --- | --- | --- | --- |
| Sodium (NA) | --- |  | <2.4 | 2.4 - 2.6 | 2.6 - 3.6 |
| Potassium (K) | 3 | A:3057 A:3058 A:3059 | <2.6 | 2.6 - 3.4 | --- |
| Caesium (CS) | --- |  | <2.9 | 2.9 - 3.6 | --- |
| Magnesium (MG) | 56 | A:3001 A:3002 A:3003 A:3004 A:3005 A:3006 A:3007 A:3008 A:3009 A:3010 A:3011 A:3012 A:3013 A:3014 A:3015 A:3016 A:3017 A:3018 A:3019 A:3020 A:3021 A:3022 A:3023 A:3024 A:3025 A:3026 A:3027 A:3028 A:3029 A:3030 A:3031 A:3032 A:3033 A:3034 A:3035 A:3036 A:3037 A:3038 A:3039 A:3040 A:3041 A:3042 A:3043 A:3044 A:3045 A:3046 A:3047 A:3048 A:3049 A:3050 A:3051 A:3052 A:3053 A:3054 A:3055 A:3056 | <1.8 | 1.8 - 2.4 | 2.4 - 3.4 |
| Calcium (CA) | --- |  | <2.2 | 2.2 - 2.8 | 2.8 - 3.6 |
| Strontium (SR) | --- |  | <2.4 | 2.4 - 2.8 | 2.8 - 4.0 |
| Barium (BA) | --- |  | <2.6 | 2.6 - 3.2 | --- |
| Manganese (MN) | --- |  | <2.0 | 2.0 - 2.3 | 2.3 - 3.6 |
| Zinc (ZN) | --- |  | <1.9 | 1.9 - 2.3 | 2.3 - 3.4 |
| Cadmium (CD) | --- |  | <2.2 | 2.2 - 2.5 | 2.5 - 3.6 |
| Cobalt (CO) | --- |  | <1.9 | 1.9 - 2.3 | 2.3 - 3.5 |
| Iridium (IRI) | --- |  | <2.0 | 2.0 - 2.3 | 2.3 - 3.6 |
| Anion (Anion) | --- |  | <2.8 | 2.8 - 3.4 | --- |
| Chloride (CL) | --- |  | <2.8 | 2.8 - 3.4 | --- |
| Sulphate (SO4) | --- |  | <2.8 | 2.8 - 3.4 | --- |
| Phosphate (PO4) | --- |  | <2.8 | 2.8 - 3.4 | --- |
| **Total** | **59** |  |  |  |  |

---

\_\_\_\_\_\_\_\_\_\_\_\_\_\_\_\_\_\_\_\_\_\_\_\_\_\_\_\_\_\_\_\_\_\_\_\_\_\_\_\_\_\_\_\_\_\_\_\_\_\_\_\_\_\_\_\_\_\_\_\_\_\_\_\_  

#### 2) Ion validation table

##### Go to ToC

NOTES:

  
- O_ph_: OP1 or OP2 anionic phosphate oxygens;  
- O_r_: O2', O3', O4' or O5' hydroxyl and ester oxygens;  
- O_b_: O2, O4 or O6 carbonyl oxygens ;  
- N_b_: N1_A_, N3_C_, N3_G_, N3_A_ or N7 non-protonated nitrogens;  
- O_back_: protein backbone carbonyl group;  
- O_coo_: ASP/GLU carboxyl group;  
- O_cno_: ASN/GLN carbonyl group;  
- OH_Prot_: SER/THR/TYR hydroxyl group;  
- O_w_: water molecule oxygen;  
- Anion: NH group (imino, amino and amonium groups) - suggests potential anion contact or poor binding site modeling;  
  
"Large octahedral discrepancy" means a cumulated ligand-ion-ligand angle deviation > 10 deg.;  
-> the ions that are currently checked for octahedral coordination are Mg^2+^, Mn^2+^ and Na^+^;  
-> could be a hint to the presence of a polyatomic ligand instead of a monoatomic ion;  
1^st^ coordination shell missing:   
1) if Mg^2+^ or Na^+^, consider adding a hydration shell and optimize the water to solute distances;   
2) if preceding step is not conclusive, consider the presence of another solvent molecule (check crystallisation conditions);   
Exclusion atoms:   
-> if O3'-O5' atoms are mentioned, this points to a defficient modelling of the RNA backbone;  
  

| Ion | Occup. | B-factor | Ligands (too close) | Ligands (1^st^ shell) | Ligands (excl. zone) | Contacts & potential issues |
| --- | --- | --- | --- | --- | --- | --- |
| MG:A:3001 | 1.00 | 26.13 |  | 1 | 1 | O_ph_ - Exclusion zone contact(s): /OP1/ |
| MG:A:3002 | 1.00 | 2.25 |  | 0 |  | 1^st^ MG coordination shell missing (see Notes) |
| MG:A:3003 | 1.00 | 26.13 |  | 0 | 2 | - Exclusion zone contact(s): /OP2/OP2/ Consider NA in place of MG (1 valid contact(s) for NA vs. 0 for MG) |
| MG:A:3004 | 1.00 | 26.13 |  | 0 |  | 1^st^ MG coordination shell missing (see Notes) |
| MG:A:3005 | 1.00 | 7.17 |  | 0 | 2 | - Exclusion zone contact(s): /N7/OP1/ Consider K in place of MG (2 valid contact(s) for K vs. 0 for MG) |
| MG:A:3006 | 1.00 | 26.13 |  | 3 |  | 3O_ph_ |
| MG:A:3007 | 1.00 | 8.87 |  | 1 |  | O_ph_ |
| MG:A:3008 | 1.00 | 28.38 |  | 1 |  | O_ph_ |
| MG:A:3009 | 1.00 | 11.52 |  | 0 |  | 1^st^ MG coordination shell missing (see Notes) |
| MG:A:3010 | 1.00 | 26.13 |  | 1 | 1 | N_b_ - Exclusion zone contact(s): /OP2/ |
| MG:A:3011 | 1.00 | 10.95 |  | 0 | 1 | - Exclusion zone contact(s): /OP1/ Consider K in place of MG (1 valid contact(s) for K vs. 0 for MG) |
| MG:A:3012 | 1.00 | 26.13 |  | 2 | 1 | O_ph_O_b_ - Exclusion zone contact(s): /OP1/ |
| MG:A:3013 | 1.00 | 26.13 |  | 0 | 2 | - Exclusion zone contact(s): /O2'/OP1/ Consider K in place of MG (2 valid contact(s) for K vs. 0 for MG) |
| MG:A:3014 | 1.00 | 15.03 |  | 0 | 1 | - Exclusion zone contact(s): /OP2/ Consider K in place of MG (1 valid contact(s) for K vs. 0 for MG) |
| MG:A:3015 | 1.00 | 11.75 |  | 0 |  | 1^st^ MG coordination shell missing (see Notes) |
| MG:A:3016 | 1.00 | 26.13 |  | 0 |  | 1^st^ MG coordination shell missing (see Notes) |
| MG:A:3017 | 1.00 | 26.13 |  | 1 | 2 | O_b_ - Exclusion zone contact(s): /N7/OP2/ |
| MG:A:3018 | 1.00 | 14.00 |  | 1 | 2 | O_ph_ - Exclusion zone contact(s): /O5'/OP2/ |
| MG:A:3019 | 1.00 | 26.13 |  | 1 |  | O_ph_ |
| MG:A:3020 | 1.00 | 26.13 |  | 0 | 2 | - Exclusion zone contact(s): /N7/OP2/ Consider NA in place of MG (1 valid contact(s) for NA vs. 0 for MG) |
| MG:A:3021 | 1.00 | 26.13 |  | 1 |  | O_ph_ |
| MG:A:3022 | 1.00 | 30.82 |  | 0 |  | 1^st^ MG coordination shell missing (see Notes) |
| MG:A:3023 | 1.00 | 26.13 |  | 1 | 1 | O_ph_ - Exclusion zone contact(s): /OP2/ |
| MG:A:3024 | 1.00 | 28.53 |  | 0 |  | 1^st^ MG coordination shell missing (see Notes) |
| MG:A:3025 | 1.00 | 26.13 |  | 1 |  | O_ph_ |
| MG:A:3026 | 1.00 | 8.03 |  | 1 | 1 | O_ph_ - Exclusion zone contact(s): /OP1/ |
| MG:A:3027 | 1.00 | 12.04 |  | 0 | 1 | - Exclusion zone contact(s): /N7/ Consider NA in place of MG (1 valid contact(s) for NA vs. 0 for MG) |
| MG:A:3028 | 1.00 | 18.26 |  | 0 |  | 1^st^ MG coordination shell missing (see Notes) |
| MG:A:3029 | 1.00 | 26.13 |  | 0 |  | 1^st^ MG coordination shell missing (see Notes) |
| MG:A:3030 | 1.00 | 26.13 |  | 1 | 1 | O_ph_ - Exclusion zone contact(s): /OP2/ |
| MG:A:3031 | 1.00 | 15.90 |  | 0 | 1 | - Exclusion zone contact(s): /OP2/ Consider K in place of MG (1 valid contact(s) for K vs. 0 for MG) |
| MG:A:3032 | 1.00 | 22.68 |  | 1 |  | O_b_ |
| MG:A:3033 | 1.00 | 13.65 |  | 1 | 1 | O_r_ - Exclusion zone contact(s): /O2/ |
| MG:A:3034 | 1.00 | 26.13 |  | 0 | 1 | - Exclusion zone contact(s): /N1_A_/ Consider NA in place of MG (1 valid contact(s) for NA vs. 0 for MG) |
| MG:A:3035 | 1.00 | 13.93 |  | 0 | 2 | - Exclusion zone contact(s): /O4/OP2/ Consider K in place of MG (2 valid contact(s) for K vs. 0 for MG) |
| MG:A:3036 | 1.00 | 13.32 |  | 0 |  | 1^st^ MG coordination shell missing (see Notes) |
| MG:A:3037 | 1.00 | 6.73 |  | 0 |  | 1^st^ MG coordination shell missing (see Notes) |
| MG:A:3038 | 1.00 | 13.51 |  | 1 |  | O_ph_ |
| MG:A:3039 | 1.00 | 26.13 |  | 0 | 1 | - Exclusion zone contact(s): /OP1/ Consider K in place of MG (1 valid contact(s) for K vs. 0 for MG) |
| MG:A:3040 | 1.00 | 26.13 |  | 0 | 1 | - Exclusion zone contact(s): /O6/ Consider K in place of MG (1 valid contact(s) for K vs. 0 for MG) |
| MG:A:3041 | 1.00 | 26.13 |  | 0 |  | 1^st^ MG coordination shell missing (see Notes) |
| MG:A:3042 | 1.00 | 14.14 |  | 0 |  | 1^st^ MG coordination shell missing (see Notes) |
| MG:A:3043 | 1.00 | 26.13 |  | 0 | 2 | - Exclusion zone contact(s): /OP1/OP2/ Consider K in place of MG (2 valid contact(s) for K vs. 0 for MG) |
| MG:A:3044 | 1.00 | 12.14 |  | 1 | 1 | O_ph_ - Exclusion zone contact(s): /OP2/ |
| MG:A:3045 | 1.00 | 11.01 |  | 1 |  | O_ph_ |
| MG:A:3046 | 1.00 | 14.10 |  | 0 | 1 | - Exclusion zone contact(s): /OP2/ Consider K in place of MG (1 valid contact(s) for K vs. 0 for MG) |
| MG:A:3047 | 1.00 | 16.96 |  | 1 |  | O_ph_ |
| MG:A:3048 | 1.00 | 26.13 |  | 1 |  | O_ph_ |
| MG:A:3049 | 1.00 | 26.13 |  | 1 |  | O_ph_ |
| MG:A:3050 | 1.00 | 10.36 |  | 1 |  | O_ph_ |
| MG:A:3051 | 1.00 | 26.13 |  | 1 |  | O_ph_ |
| MG:A:3052 | 1.00 | 17.62 |  | 1 | 1 | O_ph_ - Exclusion zone contact(s): /OP2/ |
| MG:A:3053 | 1.00 | 13.79 |  | 0 | 2 | - Exclusion zone contact(s): /O2'/O2'/ Consider K in place of MG (2 valid contact(s) for K vs. 0 for MG) |
| MG:A:3054 | 1.00 | 18.51 |  | 0 | 1 | - Exclusion zone contact(s): /OP1/ Consider K in place of MG (1 valid contact(s) for K vs. 0 for MG) |
| MG:A:3055 | 1.00 | 26.13 |  | 0 | 1 | - Exclusion zone contact(s): /OP1/ Consider K in place of MG (1 valid contact(s) for K vs. 0 for MG) |
| MG:A:3056 | 1.00 | 30.00 |  | 0 |  | 1^st^ MG coordination shell missing (see Notes) |
| K:A:3057 | 1.00 | 26.13 |  | 2 |  | 2O_ph_ |
| K:A:3058 | 1.00 | 9.19 |  | 0 |  | 1^st^ K coordination shell missing (see Notes) |
| K:A:3059 | 1.00 | 26.13 |  | 1 |  | O_ph_ |

---

\_\_\_\_\_\_\_\_\_\_\_\_\_\_\_\_\_\_\_\_\_\_\_\_\_\_\_\_\_\_\_\_\_\_\_\_\_\_\_\_\_\_\_\_\_\_\_\_\_\_\_\_\_\_\_\_\_\_\_\_\_\_\_\_  

#### 3) Potential issue list

##### Go to ToC

|  |  |  |
| --- | --- | --- |
| **Problem** | **Number** | **Affected ions** |
| **1) MG:  could be K** | 13 | A:3005  A:3011  A:3013  A:3014  A:3031  A:3035  A:3039  A:3040  A:3043  A:3046  A:3053  A:3054  A:3055 |
| **1) MG:  could be NA** | 4 | A:3003  A:3020  A:3027  A:3034 |
| **2) K: 1^st^ shell missing** | 1 | A:3058 |
| **2) MG: 1^st^ shell missing** | 14 | A:3002  A:3004  A:3009  A:3015  A:3016  A:3022  A:3024  A:3028  A:3029  A:3036  A:3037  A:3041  A:3042  A:3056 |
| **3) K/OP1 contacts** | 1 | A:3057 |
| **3) K/OP2 contacts** | 2 | A:3057  A:3059 |
| **3) MG/N7 contacts** | 1 | A:3010 |
| **3) MG/O2' contacts** | 1 | A:3033 |
| **3) MG/O4 contacts** | 1 | A:3032 |
| **3) MG/O6 contacts** | 2 | A:3012  A:3017 |
| **3) MG/OP1 contacts** | 10 | A:3001  A:3006  A:3006  A:3012  A:3023  A:3025  A:3026  A:3038  A:3047  A:3048 |
| **3) MG/OP2 contacts** | 13 | A:3006  A:3007  A:3008  A:3018  A:3019  A:3021  A:3030  A:3044  A:3045  A:3049  A:3050  A:3051  A:3052 |
| **3) MG/N1_A_ excl. contacts** | 1 | A:3034 |
| **3) MG/N7 excl. contacts** | 4 | A:3005  A:3017  A:3020  A:3027 |
| **3) MG/O2 excl. contacts** | 1 | A:3033 |
| **3) MG/O2' excl. contacts** | 3 | A:3013  A:3053  A:3053 |
| **3) MG/O4 excl. contacts** | 1 | A:3035 |
| **3) MG/O5' excl. contacts** | 1 | A:3018 |
| **3) MG/O6 excl. contacts** | 1 | A:3040 |
| **3) MG/OP1 excl. contacts** | 10 | A:3001  A:3005  A:3011  A:3012  A:3013  A:3026  A:3039  A:3043  A:3054  A:3055 |
| **3) MG/OP2 excl. contacts** | 15 | A:3003  A:3003  A:3010  A:3014  A:3017  A:3018  A:3020  A:3023  A:3030  A:3031  A:3035  A:3043  A:3044  A:3046  A:3052 |

---

\_\_\_\_\_\_\_\_\_\_\_\_\_\_\_\_\_\_\_\_\_\_\_\_\_\_\_\_\_\_\_\_\_\_\_\_\_\_\_\_\_\_\_\_\_\_\_\_\_\_\_\_\_\_\_\_\_\_\_\_\_\_\_\_  

#### 4) Ion/ligand tables: all contacts up to 5.0 Angs. are listed

##### Go to ToC

### MG:A:3001

|  | Res. type | Atom | Residue | Distance | Occupancy | B-factor | Closest solute to 1^st^ shell water |
| --- | --- | --- | --- | --- | --- | --- | --- |
| **Ion** | | | |  | 1.00 | 26.13 || 1 | G | OP1 | A:1288 | 2.02 | 1.00 | 10.58 |
| 2 | U | OP1 | A:1287 | 3.12 | 1.00 | 13.61 |  |
| 3 | U | O3' | A:1287 | 3.54 | 1.00 | 13.61 |  |
| 4 | G | OP2 | A:1288 | 4.22 | 1.00 | 10.58 |  |
| 5 | G | OP2 | A:628 | 4.23 | 1.00 | 12.49 |  |
| 6 | A | OP2 | A:629 | 4.38 | 1.00 | 9.12 |  |
| 7 | G | O5' | A:1288 | 4.41 | 1.00 | 10.58 |  |
| 8 | A | N1 | A:1289 | 4.58 | 1.00 | 9.93 |  |
| 9 | U | O5' | A:1287 | 4.83 | 1.00 | 13.61 |  |
| 10 | G | O3' | A:1286 | 4.86 | 1.00 | 13.83 |  |
| 11 | U | N3 | A:856 | 4.90 | 1.00 | 9.98 |  |
| 12 | U | O4 | A:856 | 4.94 | 1.00 | 9.98 |  |
  |

  
 Contact(s): /OP1/  
 Coded contact list: O_ph_  
 Exclusion zone contact(s): /OP1/  

### MG:A:3002

|  | Res. type | Atom | Residue | Distance | Occupancy | B-factor | Closest solute to 1^st^ shell water |
| --- | --- | --- | --- | --- | --- | --- | --- |
| **Ion** | | | |  | 1.00 | 2.25 || 1 | G | O6 | A:520 | 4.62 | 1.00 | 14.65 |
| 2 | U | OP2 | A:29 | 4.65 | 1.00 | 11.39 |  |
| 3 | G | O6 | A:30 | 4.74 | 1.00 | 13.86 |  |
| 4 | U | O4 | A:29 | 4.94 | 1.00 | 11.39 |  |
| 5 | A | O5' | A:493 | 4.96 | 1.00 | 13.30 |  |
  |

  
 Coded contact list:   
1^st^ MG coordination shell missing (see Notes)  

### MG:A:3003

|  | Res. type | Atom | Residue | Distance | Occupancy | B-factor | Closest solute to 1^st^ shell water |
| --- | --- | --- | --- | --- | --- | --- | --- |
| **Ion** | | | |  | 1.00 | 26.13 || 1 | G | OP2 | A:2084 | 2.51 | 1.00 | 11.67 |
| 2 | G | OP2 | A:2083 | 2.61 | 1.00 | 11.17 |  |
| 3 | G | OP1 | A:2083 | 3.84 | 1.00 | 11.17 |  |
| 4 | G | OP1 | A:2532 | 4.19 | 1.00 | 15.85 |  |
| 5 | G | O5' | A:2083 | 4.19 | 1.00 | 11.17 |  |
| 6 | G | N7 | A:2084 | 4.32 | 1.00 | 11.67 |  |
| 7 | G | O5' | A:2084 | 4.41 | 1.00 | 11.67 |  |
| 8 | G | OP1 | A:2084 | 4.55 | 1.00 | 11.67 |  |
| 9 | A | N6 | A:2085 | 4.96 | 1.00 | 11.44 |  |
| 10 | A | N7 | A:2085 | 4.99 | 1.00 | 11.44 |  |
  |

  
 Coded contact list:   
 Exclusion zone contact(s): /OP2/OP2/  
Consider NA in place of MG (1 valid contact(s) for NA vs. 0 for MG)  

### MG:A:3004

|  | Res. type | Atom | Residue | Distance | Occupancy | B-factor | Closest solute to 1^st^ shell water |
| --- | --- | --- | --- | --- | --- | --- | --- |
| **Ion** | | | |  | 1.00 | 26.13 || 1 | C | OP1 | A:988 | 3.96 | 1.00 | 9.74 |
| 2 | A | N7 | A:989 | 4.08 | 1.00 | 22.70 |  |
| 3 | G | OP1 | A:876 | 4.31 | 1.00 | 15.65 |  |
| 4 | A | N6 | A:2475 | 4.46 | 1.00 | 23.61 |  |
| 5 | C | OP2 | A:988 | 4.50 | 1.00 | 9.74 |  |
| 6 | A | N1 | A:2475 | 4.58 | 1.00 | 23.61 |  |
| 7 | U | OP1 | A:611 | 4.65 | 1.00 | 26.13 |  |
  |

  
 Coded contact list:   
1^st^ MG coordination shell missing (see Notes)  

### MG:A:3005

|  | Res. type | Atom | Residue | Distance | Occupancy | B-factor | Closest solute to 1^st^ shell water |
| --- | --- | --- | --- | --- | --- | --- | --- |
| **Ion** | | | |  | 1.00 | 7.17 || 1 | U | OP1 | A:610 | 2.87 | 1.00 | 28.90 |
| 2 | G | N7 | A:854 | 3.36 | 1.00 | 7.77 |  |
| 3 | G | OP2 | A:853 | 4.35 | 1.00 | 8.21 |  |
| 4 | G | N7 | A:853 | 4.49 | 1.00 | 8.21 |  |
| 5 | G | OP2 | A:854 | 4.51 | 1.00 | 7.77 |  |
| 6 | U | O5' | A:610 | 4.54 | 1.00 | 28.90 |  |
| 7 | U | O3' | A:609 | 4.64 | 1.00 | 20.17 |  |
| 8 | G | O5' | A:853 | 4.78 | 1.00 | 8.21 |  |
| 9 | G | O6 | A:854 | 4.80 | 1.00 | 7.77 |  |
| 10 | U | O4 | A:855 | 4.80 | 1.00 | 7.93 |  |
  |

  
 Coded contact list:   
 Exclusion zone contact(s): /N7/OP1/  
Consider K in place of MG (2 valid contact(s) for K vs. 0 for MG)  

### MG:A:3006

|  | Res. type | Atom | Residue | Distance | Occupancy | B-factor | Closest solute to 1^st^ shell water |
| --- | --- | --- | --- | --- | --- | --- | --- |
| **Ion** | | | |  | 1.00 | 26.13 || 1 | C | OP2 | A:1352 | 1.90 | 1.00 | 25.69 |
| 2 | G | OP1 | A:1369 | 1.96 | 1.00 | 15.52 |  |
| 3 | C | OP1 | A:1351 | 2.32 | 1.00 | 18.81 |  |
| 4 | C | OP1 | A:1352 | 3.74 | 1.00 | 25.69 |  |
| 5 | G | O5' | A:1369 | 3.77 | 1.00 | 15.52 |  |
| 6 | C | O3' | A:1351 | 4.20 | 1.00 | 18.81 |  |
| 7 | C | O5' | A:1352 | 4.28 | 1.00 | 25.69 |  |
| 8 | G | OP2 | A:1369 | 4.30 | 1.00 | 15.52 |  |
| 9 | C | O3' | A:1368 | 4.32 | 1.00 | 14.91 |  |
| 10 | C | OP2 | A:1351 | 4.37 | 1.00 | 18.81 |  |
| 11 | C | O5' | A:1351 | 4.39 | 1.00 | 18.81 |  |
| 12 | U | O3' | A:1350 | 4.81 | 1.00 | 16.36 |  |
| 13 | G | O4' | A:1369 | 4.82 | 1.00 | 15.52 |  |
  |

  
**1^st^ shell ligand angles:**   
1-2: 156; 1-3: 85; 2-3: 108;   
  
 Contact(s): /OP1/OP1/OP2/  
 Coded contact list: 3O_ph_  

### MG:A:3007

|  | Res. type | Atom | Residue | Distance | Occupancy | B-factor | Closest solute to 1^st^ shell water |
| --- | --- | --- | --- | --- | --- | --- | --- |
| **Ion** | | | |  | 1.00 | 8.87 || 1 | G | OP2 | A:2056 | 2.18 | 1.00 | 13.91 |
| 2 | G | O5' | A:2056 | 4.11 | 1.00 | 13.91 |  |
| 3 | G | OP1 | A:2056 | 4.42 | 1.00 | 13.91 |  |
| 4 | U | OP1 | A:2055 | 4.54 | 1.00 | 12.57 |  |
| 5 | U | O5' | A:2055 | 4.57 | 1.00 | 12.57 |  |
| 6 | U | O3' | A:2055 | 4.65 | 1.00 | 12.57 |  |
| 7 | G | N7 | A:2056 | 4.66 | 1.00 | 13.91 |  |
| 8 | U | OP2 | A:2055 | 4.99 | 1.00 | 12.57 |  |
  |

  
 Contact(s): /OP2/  
 Coded contact list: O_ph_  

### MG:A:3008

|  | Res. type | Atom | Residue | Distance | Occupancy | B-factor | Closest solute to 1^st^ shell water |
| --- | --- | --- | --- | --- | --- | --- | --- |
| **Ion** | | | |  | 1.00 | 28.38 || 1 | U | OP2 | A:2518 | 2.00 | 1.00 | 24.07 |
| 2 | G | OP2 | A:2482 | 3.46 | 1.00 | 19.37 |  |
| 3 | G | OP1 | A:2481 | 3.99 | 1.00 | 22.73 |  |
| 4 | G | O3' | A:2517 | 4.08 | 1.00 | 22.05 |  |
| 5 | U | OP1 | A:2518 | 4.12 | 1.00 | 24.07 |  |
| 6 | U | O5' | A:2518 | 4.45 | 1.00 | 24.07 |  |
| 7 | G | O2' | A:2517 | 4.46 | 1.00 | 22.05 |  |
| 8 | G | OP1 | A:2482 | 4.62 | 1.00 | 19.37 |  |
| 9 | U | OP1 | A:2598 | 4.90 | 1.00 | 17.75 |  |
  |

  
 Contact(s): /OP2/  
 Coded contact list: O_ph_  

### MG:A:3009

|  | Res. type | Atom | Residue | Distance | Occupancy | B-factor | Closest solute to 1^st^ shell water |
| --- | --- | --- | --- | --- | --- | --- | --- |
| **Ion** | | | |  | 1.00 | 11.52 || 1 | G | N7 | A:2521 | 3.62 | 1.00 | 15.89 |
| 2 | G | O6 | A:2521 | 3.88 | 1.00 | 15.89 |  |
| 3 | G | O6 | A:2522 | 4.15 | 1.00 | 13.74 |  |
  |

  
 Coded contact list:   
1^st^ MG coordination shell missing (see Notes)  

### MG:A:3010

|  | Res. type | Atom | Residue | Distance | Occupancy | B-factor | Closest solute to 1^st^ shell water |
| --- | --- | --- | --- | --- | --- | --- | --- |
| **Ion** | | | |  | 1.00 | 26.13 || 1 | G | N7 | A:2522 | 2.17 | 1.00 | 13.74 |
| 2 | G | OP2 | A:2522 | 3.18 | 1.00 | 13.74 |  |
| 3 | G | O5' | A:2522 | 3.97 | 1.00 | 13.74 |  |
| 4 | G | N7 | A:2521 | 3.98 | 1.00 | 15.89 |  |
| 5 | G | N9 | A:2522 | 4.07 | 1.00 | 13.74 |  |
| 6 | G | O6 | A:2522 | 4.20 | 1.00 | 13.74 |  |
| 7 | G | N9 | A:2521 | 4.25 | 1.00 | 15.89 |  |
| 8 | C | N4 | A:2523 | 4.52 | 1.00 | 15.90 |  |
| 9 | G | O3' | A:2521 | 4.66 | 1.00 | 15.89 |  |
  |

  
 Contact(s): /N7/  
 Coded contact list: N_b_  
 Exclusion zone contact(s): /OP2/  

### MG:A:3011

|  | Res. type | Atom | Residue | Distance | Occupancy | B-factor | Closest solute to 1^st^ shell water |
| --- | --- | --- | --- | --- | --- | --- | --- |
| **Ion** | | | |  | 1.00 | 10.95 || 1 | U | OP1 | A:1007 | 2.82 | 1.00 | 15.13 |
| 2 | C | OP1 | A:2525 | 3.81 | 1.00 | 26.13 |  |
| 3 | U | O5' | A:1007 | 4.49 | 1.00 | 15.13 |  |
| 4 | A | OP1 | A:2524 | 4.77 | 1.00 | 20.05 |  |
| 5 | U | OP2 | A:1007 | 4.79 | 1.00 | 15.13 |  |
  |

  
 Coded contact list:   
 Exclusion zone contact(s): /OP1/  
Consider K in place of MG (1 valid contact(s) for K vs. 0 for MG)  

### MG:A:3012

|  | Res. type | Atom | Residue | Distance | Occupancy | B-factor | Closest solute to 1^st^ shell water |
| --- | --- | --- | --- | --- | --- | --- | --- |
| **Ion** | | | |  | 1.00 | 26.13 || 1 | G | O6 | A:613 | 2.18 | 1.00 | 29.41 |
| 2 | A | OP1 | A:2475 | 2.31 | 1.00 | 23.61 |  |
| 3 | C | OP1 | A:2526 | 2.72 | 1.00 | 21.22 |  |
| 4 | A | OP2 | A:2475 | 3.79 | 1.00 | 23.61 |  |
| 5 | A | O5' | A:2475 | 3.80 | 1.00 | 23.61 |  |
| 6 | U | O4 | A:612 | 3.84 | 1.00 | 26.13 |  |
| 7 | G | N1 | A:613 | 4.07 | 1.00 | 29.41 |  |
| 8 | U | O2' | A:611 | 4.60 | 1.00 | 26.13 |  |
| 9 | C | O3' | A:2525 | 4.70 | 1.00 | 26.13 |  |
| 10 | G | O3' | A:2474 | 4.80 | 1.00 | 17.04 |  |
| 11 | G | N7 | A:613 | 4.90 | 1.00 | 29.41 |  |
| 12 | C | OP2 | A:2526 | 4.98 | 1.00 | 21.22 |  |
  |

  
**1^st^ shell ligand angles:**   
1-2: 142;   
  
 Contact(s): /O6/OP1/  
 Coded contact list: O_ph_O_b_  
 Exclusion zone contact(s): /OP1/  

### MG:A:3013

|  | Res. type | Atom | Residue | Distance | Occupancy | B-factor | Closest solute to 1^st^ shell water |
| --- | --- | --- | --- | --- | --- | --- | --- |
| **Ion** | | | |  | 1.00 | 26.13 || 1 | A | OP1 | A:618 | 2.70 | 1.00 | 26.13 |
| 2 | A | O2' | A:2057 | 3.16 | 1.00 | 23.16 |  |
| 3 | U | O2 | A:614 | 3.43 | 1.00 | 27.44 |  |
| 4 | A | OP2 | A:618 | 3.59 | 1.00 | 26.13 |  |
| 5 | C | O2' | A:2526 | 3.72 | 1.00 | 21.22 |  |
| 6 | A | N3 | A:2057 | 3.81 | 1.00 | 23.16 |  |
| 7 | G | O2' | A:616 | 3.96 | 1.00 | 14.53 |  |
| 8 | U | N3 | A:614 | 4.50 | 1.00 | 27.44 |  |
| 9 | G | N3 | A:616 | 4.57 | 1.00 | 14.53 |  |
| 10 | C | O3' | A:2526 | 4.64 | 1.00 | 21.22 |  |
| 11 | A | N9 | A:2057 | 4.68 | 1.00 | 23.16 |  |
| 12 | A | O5' | A:618 | 4.69 | 1.00 | 26.13 |  |
| 13 | A | O3' | A:617 | 4.79 | 1.00 | 10.35 |  |
| 14 | A | O4' | A:618 | 4.91 | 1.00 | 26.13 |  |
  |

  
 Coded contact list:   
 Exclusion zone contact(s): /O2'/OP1/  
Consider K in place of MG (2 valid contact(s) for K vs. 0 for MG)  

### MG:A:3014

|  | Res. type | Atom | Residue | Distance | Occupancy | B-factor | Closest solute to 1^st^ shell water |
| --- | --- | --- | --- | --- | --- | --- | --- |
| **Ion** | | | |  | 1.00 | 15.03 || 1 | A | OP2 | A:621 | 2.73 | 1.00 | 10.53 |
| 2 | G | OP2 | A:620 | 3.61 | 1.00 | 14.12 |  |
| 3 | A | O5' | A:621 | 4.57 | 1.00 | 10.53 |  |
| 4 | C | OP1 | A:2044 | 4.61 | 1.00 | 13.78 |  |
| 5 | G | O5' | A:620 | 4.76 | 1.00 | 14.12 |  |
| 6 | A | OP1 | A:2045 | 4.77 | 1.00 | 11.38 |  |
| 7 | A | N7 | A:621 | 4.80 | 1.00 | 10.53 |  |
| 8 | G | O3' | A:620 | 4.90 | 1.00 | 14.12 |  |
| 9 | A | OP2 | A:2045 | 4.96 | 1.00 | 11.38 |  |
| 10 | G | OP1 | A:620 | 5.00 | 1.00 | 14.12 |  |
  |

  
 Coded contact list:   
 Exclusion zone contact(s): /OP2/  
Consider K in place of MG (1 valid contact(s) for K vs. 0 for MG)  

### MG:A:3015

|  | Res. type | Atom | Residue | Distance | Occupancy | B-factor | Closest solute to 1^st^ shell water |
| --- | --- | --- | --- | --- | --- | --- | --- |
| **Ion** | | | |  | 1.00 | 11.75 || 1 | A | OP1 | A:1017 | 4.12 | 1.00 | 11.69 |
| 2 | G | OP1 | A:613 | 4.16 | 1.00 | 29.41 |  |
| 3 | G | OP2 | A:613 | 4.47 | 1.00 | 29.41 |  |
| 4 | A | OP2 | A:1017 | 4.53 | 1.00 | 11.69 |  |
| 5 | G | O2' | A:1016 | 4.61 | 1.00 | 10.51 |  |
| 6 | U | OP2 | A:612 | 4.83 | 1.00 | 26.13 |  |
| 7 | G | O3' | A:1016 | 4.86 | 1.00 | 10.51 |  |
| 8 | A | N6 | A:865 | 4.88 | 1.00 | 7.60 |  |
| 9 | A | N1 | A:866 | 4.94 | 1.00 | 9.85 |  |
  |

  
 Coded contact list:   
1^st^ MG coordination shell missing (see Notes)  

### MG:A:3016

|  | Res. type | Atom | Residue | Distance | Occupancy | B-factor | Closest solute to 1^st^ shell water |
| --- | --- | --- | --- | --- | --- | --- | --- |
| **Ion** | | | |  | 1.00 | 26.13 || 1 | ASN | ND2 | U:81 | 4.19 | 1.00 | 18.92 |
| 2 | U | OP1 | A:609 | 4.23 | 1.00 | 20.17 |  |
| 3 | C | O3' | A:608 | 4.28 | 1.00 | 14.73 |  |
| 4 | U | OP2 | A:855 | 4.44 | 1.00 | 7.93 |  |
| 5 | U | OP1 | A:856 | 4.44 | 1.00 | 9.98 |  |
| 6 | C | O2' | A:608 | 4.84 | 1.00 | 14.73 |  |
  |

  
 Coded contact list:   
1^st^ MG coordination shell missing (see Notes)  

### MG:A:3017

|  | Res. type | Atom | Residue | Distance | Occupancy | B-factor | Closest solute to 1^st^ shell water |
| --- | --- | --- | --- | --- | --- | --- | --- |
| **Ion** | | | |  | 1.00 | 26.13 || 1 | G | O6 | A:1226 | 2.04 | 1.00 | 10.30 |
| 2 | G | OP2 | A:863 | 2.51 | 1.00 | 8.12 |  |
| 3 | G | N7 | A:1226 | 3.39 | 1.00 | 10.30 |  |
| 4 | G | N7 | A:863 | 3.86 | 1.00 | 8.12 |  |
| 5 | G | N1 | A:1226 | 4.24 | 1.00 | 10.30 |  |
| 6 | G | O5' | A:863 | 4.39 | 1.00 | 8.12 |  |
| 7 | C | OP2 | A:862 | 4.45 | 1.00 | 9.46 |  |
| 8 | G | OP2 | A:1225 | 4.48 | 1.00 | 75.20 |  |
| 9 | C | OP1 | A:862 | 4.48 | 1.00 | 9.46 |  |
| 10 | C | O5' | A:862 | 4.71 | 1.00 | 9.46 |  |
| 11 | C | O3' | A:862 | 4.74 | 1.00 | 9.46 |  |
| 12 | G | N7 | A:1225 | 4.89 | 1.00 | 20.00 |  |
| 13 | G | OP1 | A:863 | 4.96 | 1.00 | 8.12 |  |
  |

  
 Contact(s): /O6/  
 Coded contact list: O_b_  
 Exclusion zone contact(s): /N7/OP2/  

### MG:A:3018

|  | Res. type | Atom | Residue | Distance | Occupancy | B-factor | Closest solute to 1^st^ shell water |
| --- | --- | --- | --- | --- | --- | --- | --- |
| **Ion** | | | |  | 1.00 | 14.00 || 1 | A | OP2 | A:1034 | 1.99 | 1.00 | 12.73 |
| 2 | C | OP2 | A:1035 | 2.59 | 1.00 | 16.47 |  |
| 3 | A | O5' | A:1034 | 3.38 | 1.00 | 12.73 |  |
| 4 | G | O5' | A:1225 | 3.75 | 1.00 | 78.76 |  |
| 5 | U | O3' | A:1224 | 3.87 | 1.00 | 77.17 |  |
| 6 | A | OP1 | A:1034 | 3.98 | 1.00 | 12.73 |  |
| 7 | G | O2' | A:1033 | 4.20 | 1.00 | 13.85 |  |
| 8 | U | O5' | A:1224 | 4.30 | 1.00 | 78.76 |  |
| 9 | G | O3' | A:1033 | 4.42 | 1.00 | 13.85 |  |
| 10 | U | OP1 | A:1224 | 4.58 | 1.00 | 78.59 |  |
| 11 | C | OP1 | A:1035 | 4.60 | 1.00 | 16.47 |  |
| 12 | U | OP2 | A:1224 | 4.77 | 1.00 | 75.20 |  |
| 13 | A | O3' | A:1034 | 4.89 | 1.00 | 12.73 |  |
| 14 | C | O5' | A:1035 | 4.92 | 1.00 | 16.47 |  |
| 15 | G | O4' | A:1225 | 4.92 | 1.00 | 79.49 |  |
  |

  
 Contact(s): /OP2/  
 Coded contact list: O_ph_  
 Exclusion zone contact(s): /O5'/OP2/  

### MG:A:3019

|  | Res. type | Atom | Residue | Distance | Occupancy | B-factor | Closest solute to 1^st^ shell water |
| --- | --- | --- | --- | --- | --- | --- | --- |
| **Ion** | | | |  | 1.00 | 26.13 || 1 | A | OP2 | A:1200 | 2.13 | 1.00 | 13.49 |
| 2 | A | O5' | A:1199 | 3.57 | 1.00 | 14.20 |  |
| 3 | G | O6 | A:1201 | 3.62 | 1.00 | 13.84 |  |
| 4 | A | OP2 | A:1199 | 3.73 | 1.00 | 14.20 |  |
| 5 | A | O3' | A:1199 | 4.28 | 1.00 | 14.20 |  |
| 6 | A | O5' | A:1200 | 4.35 | 1.00 | 13.49 |  |
| 7 | C | N4 | A:1042 | 4.35 | 1.00 | 18.00 |  |
| 8 | G | O6 | A:1041 | 4.49 | 1.00 | 18.23 |  |
| 9 | A | OP1 | A:1200 | 4.60 | 1.00 | 13.49 |  |
| 10 | A | OP1 | A:1199 | 4.62 | 1.00 | 14.20 |  |
  |

  
 Contact(s): /OP2/  
 Coded contact list: O_ph_  

### MG:A:3020

|  | Res. type | Atom | Residue | Distance | Occupancy | B-factor | Closest solute to 1^st^ shell water |
| --- | --- | --- | --- | --- | --- | --- | --- |
| **Ion** | | | |  | 1.00 | 26.13 || 1 | A | OP2 | A:1228 | 2.42 | 1.00 | 7.46 |
| 2 | G | N7 | A:1229 | 2.61 | 1.00 | 9.45 |  |
| 3 | G | O6 | A:1229 | 3.68 | 1.00 | 9.45 |  |
| 4 | GLY | O | O:26 | 4.33 | 1.00 | 26.13 |  |
| 5 | A | OP1 | A:1228 | 4.34 | 1.00 | 7.46 |  |
| 6 | A | O5' | A:1228 | 4.40 | 1.00 | 7.46 |  |
| 7 | G | OP2 | A:1229 | 4.66 | 1.00 | 9.45 |  |
| 8 | G | N2 | A:863 | 4.76 | 1.00 | 8.12 |  |
| 9 | G | N9 | A:1229 | 4.79 | 1.00 | 9.45 |  |
| 10 | U | OP1 | A:1227 | 4.80 | 1.00 | 9.33 |  |
| 11 | U | OP2 | A:1227 | 4.85 | 1.00 | 9.33 |  |
| 12 | U | O3' | A:1227 | 4.92 | 1.00 | 9.33 |  |
  |

  
 Coded contact list:   
 Exclusion zone contact(s): /N7/OP2/  
Consider NA in place of MG (1 valid contact(s) for NA vs. 0 for MG)  

### MG:A:3021

|  | Res. type | Atom | Residue | Distance | Occupancy | B-factor | Closest solute to 1^st^ shell water |
| --- | --- | --- | --- | --- | --- | --- | --- |
| **Ion** | | | |  | 1.00 | 26.13 || 1 | U | OP2 | A:987 | 2.21 | 1.00 | 8.73 |
| 2 | HIS | O | O:35 | 3.48 | 1.00 | 26.13 |  |
| 3 | U | OP1 | A:987 | 3.89 | 1.00 | 8.73 |  |
| 4 | G | OP1 | A:877 | 4.42 | 1.00 | 13.07 |  |
| 5 | U | O5' | A:987 | 4.43 | 1.00 | 8.73 |  |
| 6 | G | O3' | A:986 | 4.63 | 1.00 | 11.15 |  |
| 7 | GLY | N | O:37 | 4.79 | 1.00 | 26.13 |  |
| 8 | HIS | N | O:35 | 4.92 | 1.00 | 26.13 |  |
| 9 | LYS | O | O:36 | 5.00 | 1.00 | 22.86 |  |
  |

  
 Contact(s): /OP2/  
 Coded contact list: O_ph_  

### MG:A:3022

|  | Res. type | Atom | Residue | Distance | Occupancy | B-factor | Closest solute to 1^st^ shell water |
| --- | --- | --- | --- | --- | --- | --- | --- |
| **Ion** | | | |  | 1.00 | 30.82 || 1 | C | OP2 | A:789 | 3.56 | 1.00 | 12.84 |
| 2 | HIS | NE2 | E:146 | 4.15 | 1.00 | 15.84 |  |
| 3 | U | O4 | A:791 | 4.80 | 1.00 | 17.70 |  |
  |

  
 Coded contact list:   
1^st^ MG coordination shell missing (see Notes)  

### MG:A:3023

|  | Res. type | Atom | Residue | Distance | Occupancy | B-factor | Closest solute to 1^st^ shell water |
| --- | --- | --- | --- | --- | --- | --- | --- |
| **Ion** | | | |  | 1.00 | 26.13 || 1 | U | OP1 | A:2642 | 1.96 | 1.00 | 10.31 |
| 2 | A | OP2 | A:1303 | 2.76 | 1.00 | 9.47 |  |
| 3 | U | O5' | A:2642 | 3.84 | 1.00 | 10.31 |  |
| 4 | U | OP2 | A:2642 | 4.19 | 1.00 | 10.31 |  |
| 5 | A | OP1 | A:1303 | 4.35 | 1.00 | 9.47 |  |
| 6 | A | O3' | A:2641 | 4.48 | 1.00 | 12.07 |  |
| 7 | U | O2' | A:1305 | 4.59 | 1.00 | 8.19 |  |
| 8 | G | O3' | A:1302 | 4.95 | 1.00 | 10.87 |  |
  |

  
 Contact(s): /OP1/  
 Coded contact list: O_ph_  
 Exclusion zone contact(s): /OP2/  

### MG:A:3024

|  | Res. type | Atom | Residue | Distance | Occupancy | B-factor | Closest solute to 1^st^ shell water |
| --- | --- | --- | --- | --- | --- | --- | --- |
| **Ion** | | | |  | 1.00 | 28.53 || 1 | G | N7 | A:2036 | 4.30 | 1.00 | 10.91 |
| 2 | U | OP2 | A:1305 | 4.42 | 1.00 | 8.19 |  |
| 3 | C | N4 | A:2035 | 4.74 | 1.00 | 11.37 |  |
| 4 | G | O6 | A:2036 | 4.89 | 1.00 | 10.91 |  |
| 5 | U | OP1 | A:1305 | 4.96 | 1.00 | 8.19 |  |
  |

  
 Coded contact list:   
1^st^ MG coordination shell missing (see Notes)  

### MG:A:3025

|  | Res. type | Atom | Residue | Distance | Occupancy | B-factor | Closest solute to 1^st^ shell water |
| --- | --- | --- | --- | --- | --- | --- | --- |
| **Ion** | | | |  | 1.00 | 26.13 || 1 | U | OP1 | A:1305 | 2.31 | 1.00 | 8.19 |
| 2 | C | OP2 | A:2035 | 4.21 | 1.00 | 11.37 |  |
| 3 | U | O5' | A:1305 | 4.36 | 1.00 | 8.19 |  |
| 4 | U | OP2 | A:1305 | 4.46 | 1.00 | 8.19 |  |
| 5 | G | O3' | A:1304 | 4.78 | 1.00 | 8.70 |  |
  |

  
 Contact(s): /OP1/  
 Coded contact list: O_ph_  

### MG:A:3026

|  | Res. type | Atom | Residue | Distance | Occupancy | B-factor | Closest solute to 1^st^ shell water |
| --- | --- | --- | --- | --- | --- | --- | --- |
| **Ion** | | | |  | 1.00 | 8.03 || 1 | C | OP1 | A:2033 | 1.99 | 1.00 | 12.91 |
| 2 | A | OP1 | A:1306 | 3.07 | 1.00 | 6.99 |  |
| 3 | A | OP2 | A:2076 | 3.93 | 1.00 | 9.58 |  |
| 4 | A | O3' | A:2032 | 3.98 | 1.00 | 15.03 |  |
| 5 | C | O5' | A:2033 | 4.14 | 1.00 | 12.91 |  |
| 6 | G | OP1 | A:2075 | 4.31 | 1.00 | 8.20 |  |
| 7 | C | OP2 | A:2033 | 4.49 | 1.00 | 12.91 |  |
| 8 | U | O3' | A:1305 | 4.57 | 1.00 | 8.19 |  |
  |

  
 Contact(s): /OP1/  
 Coded contact list: O_ph_  
 Exclusion zone contact(s): /OP1/  

### MG:A:3027

|  | Res. type | Atom | Residue | Distance | Occupancy | B-factor | Closest solute to 1^st^ shell water |
| --- | --- | --- | --- | --- | --- | --- | --- |
| **Ion** | | | |  | 1.00 | 12.04 || 1 | G | N7 | A:2059 | 2.44 | 1.00 | 26.13 |
| 2 | G | N9 | A:2059 | 3.95 | 1.00 | 26.13 |  |
| 3 | C | N3 | A:2082 | 3.98 | 1.00 | 11.07 |  |
| 4 | C | O2 | A:2526 | 4.17 | 1.00 | 21.22 |  |
| 5 | C | N3 | A:2526 | 4.32 | 1.00 | 21.22 |  |
| 6 | C | O2 | A:2082 | 4.50 | 1.00 | 11.07 |  |
| 7 | C | N4 | A:2082 | 4.68 | 1.00 | 11.07 |  |
| 8 | G | O6 | A:2059 | 4.90 | 1.00 | 26.13 |  |
| 9 | G | O2' | A:2481 | 4.94 | 1.00 | 22.73 |  |
  |

  
 Coded contact list:   
 Exclusion zone contact(s): /N7/  
Consider NA in place of MG (1 valid contact(s) for NA vs. 0 for MG)  

### MG:A:3028

|  | Res. type | Atom | Residue | Distance | Occupancy | B-factor | Closest solute to 1^st^ shell water |
| --- | --- | --- | --- | --- | --- | --- | --- |
| **Ion** | | | |  | 1.00 | 18.26 || 1 | G | O6 | A:2054 | 4.26 | 1.00 | 10.81 |
| 2 | U | O4 | A:2053 | 4.43 | 1.00 | 11.87 |  |
| 3 | A | OP1 | A:2060 | 4.79 | 1.00 | 10.69 |  |
| 4 | C | OP2 | A:2052 | 4.89 | 1.00 | 11.36 |  |
  |

  
 Coded contact list:   
1^st^ MG coordination shell missing (see Notes)  

### MG:A:3029

|  | Res. type | Atom | Residue | Distance | Occupancy | B-factor | Closest solute to 1^st^ shell water |
| --- | --- | --- | --- | --- | --- | --- | --- |
| **Ion** | | | |  | 1.00 | 26.13 || 1 | G | O6 | A:2065 | 3.84 | 1.00 | 12.24 |
| 2 | G | N7 | A:2065 | 4.27 | 1.00 | 12.24 |  |
| 3 | C | OP2 | A:2063 | 4.30 | 1.00 | 9.64 |  |
| 4 | A | OP2 | A:2064 | 4.33 | 1.00 | 10.88 |  |
| 5 | A | N7 | A:2064 | 4.54 | 1.00 | 10.88 |  |
| 6 | G | O6 | A:2066 | 4.61 | 1.00 | 13.82 |  |
| 7 | G | OP2 | A:575 | 4.88 | 1.00 | 17.81 |  |
  |

  
 Coded contact list:   
1^st^ MG coordination shell missing (see Notes)  

### MG:A:3030

|  | Res. type | Atom | Residue | Distance | Occupancy | B-factor | Closest solute to 1^st^ shell water |
| --- | --- | --- | --- | --- | --- | --- | --- |
| **Ion** | | | |  | 1.00 | 26.13 || 1 | G | OP2 | A:2529 | 2.01 | 1.00 | 16.97 |
| 2 | G | OP2 | A:2088 | 2.73 | 1.00 | 17.88 |  |
| 3 | G | N7 | A:2473 | 3.95 | 1.00 | 26.13 |  |
| 4 | G | O5' | A:2529 | 4.26 | 1.00 | 16.97 |  |
| 5 | G | OP1 | A:2529 | 4.30 | 1.00 | 16.97 |  |
| 6 | C | O3' | A:2528 | 4.32 | 1.00 | 16.51 |  |
| 7 | G | N7 | A:2472 | 4.34 | 1.00 | 12.58 |  |
| 8 | G | OP2 | A:2473 | 4.42 | 1.00 | 26.13 |  |
| 9 | A | O2' | A:2087 | 4.68 | 1.00 | 18.70 |  |
| 10 | G | O6 | A:2473 | 4.70 | 1.00 | 26.13 |  |
| 11 | G | OP1 | A:2088 | 4.75 | 1.00 | 17.88 |  |
| 12 | C | O2' | A:2528 | 4.77 | 1.00 | 16.51 |  |
| 13 | A | N3 | A:2087 | 4.80 | 1.00 | 18.70 |  |
| 14 | A | O3' | A:2087 | 4.95 | 1.00 | 18.70 |  |
  |

  
 Contact(s): /OP2/  
 Coded contact list: O_ph_  
 Exclusion zone contact(s): /OP2/  

### MG:A:3031

|  | Res. type | Atom | Residue | Distance | Occupancy | B-factor | Closest solute to 1^st^ shell water |
| --- | --- | --- | --- | --- | --- | --- | --- |
| **Ion** | | | |  | 1.00 | 15.90 || 1 | A | OP2 | A:2089 | 2.79 | 1.00 | 26.13 |
| 2 | A | OP1 | A:2089 | 3.52 | 1.00 | 26.13 |  |
| 3 | G | O6 | A:2471 | 3.90 | 1.00 | 12.08 |  |
| 4 | G | N7 | A:2471 | 3.93 | 1.00 | 12.08 |  |
| 5 | G | N7 | A:2472 | 4.09 | 1.00 | 12.58 |  |
| 6 | A | O5' | A:2089 | 4.50 | 1.00 | 26.13 |  |
| 7 | G | O6 | A:2472 | 4.53 | 1.00 | 12.58 |  |
| 8 | G | O3' | A:2088 | 4.86 | 1.00 | 17.88 |  |
  |

  
 Coded contact list:   
 Exclusion zone contact(s): /OP2/  
Consider K in place of MG (1 valid contact(s) for K vs. 0 for MG)  

### MG:A:3032

|  | Res. type | Atom | Residue | Distance | Occupancy | B-factor | Closest solute to 1^st^ shell water |
| --- | --- | --- | --- | --- | --- | --- | --- |
| **Ion** | | | |  | 1.00 | 22.68 || 1 | U | O4 | A:1011 | 2.26 | 1.00 | 22.38 |
| 2 | G | O6 | A:1012 | 4.03 | 1.00 | 19.46 |  |
| 3 | G | O6 | A:1010 | 4.11 | 1.00 | 21.73 |  |
| 4 | G | N7 | A:1010 | 4.29 | 1.00 | 21.73 |  |
| 5 | U | N3 | A:1011 | 4.49 | 1.00 | 22.38 |  |
| 6 | A | N6 | A:994 | 4.93 | 1.00 | 23.34 |  |
  |

  
 Contact(s): /O4/  
 Coded contact list: O_b_  

### MG:A:3033

|  | Res. type | Atom | Residue | Distance | Occupancy | B-factor | Closest solute to 1^st^ shell water |
| --- | --- | --- | --- | --- | --- | --- | --- |
| **Ion** | | | |  | 1.00 | 13.65 || 1 | U | O2' | A:964 | 2.08 | 1.00 | 40.25 |
| 2 | U | O2 | A:964 | 3.11 | 1.00 | 40.25 |  |
| 3 | G | O4' | A:965 | 3.95 | 1.00 | 36.02 |  |
| 4 | U | N1 | A:964 | 4.07 | 1.00 | 40.25 |  |
| 5 | U | O4' | A:964 | 4.32 | 1.00 | 40.25 |  |
| 6 | C | O2' | B:79 | 4.79 | 1.00 | 58.81 |  |
| 7 | U | O3' | A:964 | 4.81 | 1.00 | 40.25 |  |
| 8 | A | N1 | A:963 | 4.83 | 1.00 | 43.13 |  |
| 9 | G | O5' | A:965 | 4.84 | 1.00 | 36.02 |  |
  |

  
 Contact(s): /O2'/  
 Coded contact list: O_r_  
 Exclusion zone contact(s): /O2/  

### MG:A:3034

|  | Res. type | Atom | Residue | Distance | Occupancy | B-factor | Closest solute to 1^st^ shell water |
| --- | --- | --- | --- | --- | --- | --- | --- |
| **Ion** | | | |  | 1.00 | 26.13 || 1 | A | N1 | A:963 | 2.55 | 1.00 | 43.13 |
| 2 | A | N6 | A:963 | 3.44 | 1.00 | 43.13 |  |
| 3 | A | OP2 | A:2295 | 4.44 | 1.00 | 30.94 |  |
| 4 | U | O2 | A:964 | 4.47 | 1.00 | 40.25 |  |
| 5 | A | N3 | A:963 | 4.71 | 1.00 | 43.13 |  |
| 6 | A | O2' | A:2294 | 4.94 | 1.00 | 23.93 |  |
  |

  
 Coded contact list:   
 Exclusion zone contact(s): /N1_A_/  
Consider NA in place of MG (1 valid contact(s) for NA vs. 0 for MG)  

### MG:A:3035

|  | Res. type | Atom | Residue | Distance | Occupancy | B-factor | Closest solute to 1^st^ shell water |
| --- | --- | --- | --- | --- | --- | --- | --- |
| **Ion** | | | |  | 1.00 | 13.93 || 1 | C | OP2 | A:2275 | 2.72 | 1.00 | 12.78 |
| 2 | U | O4 | A:2276 | 3.13 | 1.00 | 12.04 |  |
| 3 | C | OP1 | A:2093 | 3.74 | 1.00 | 10.78 |  |
| 4 | A | OP1 | A:2274 | 4.45 | 1.00 | 13.10 |  |
| 5 | C | OP1 | A:2275 | 4.50 | 1.00 | 12.78 |  |
| 6 | A | O5' | A:2274 | 4.64 | 1.00 | 13.10 |  |
| 7 | A | O3' | A:2274 | 4.89 | 1.00 | 13.10 |  |
| 8 | A | OP2 | A:2274 | 4.90 | 1.00 | 13.10 |  |
| 9 | G | OP1 | A:2280 | 4.94 | 1.00 | 15.84 |  |
  |

  
 Coded contact list:   
 Exclusion zone contact(s): /O4/OP2/  
Consider K in place of MG (2 valid contact(s) for K vs. 0 for MG)  

### MG:A:3036

|  | Res. type | Atom | Residue | Distance | Occupancy | B-factor | Closest solute to 1^st^ shell water |
| --- | --- | --- | --- | --- | --- | --- | --- |
| **Ion** | | | |  | 1.00 | 13.32 || 1 | U | O4 | A:2101 | 4.42 | 1.00 | 24.96 |
| 2 | U | O4 | A:2270 | 4.47 | 1.00 | 17.54 |  |
| 3 | G | O6 | A:2269 | 4.53 | 1.00 | 23.30 |  |
| 4 | G | N7 | A:2265 | 4.56 | 1.00 | 46.06 |  |
| 5 | A | N7 | A:2268 | 4.70 | 1.00 | 31.71 |  |
| 6 | U | O4 | A:2102 | 4.91 | 1.00 | 31.01 |  |
| 7 | G | N7 | A:2269 | 4.92 | 1.00 | 23.30 |  |
  |

  
 Coded contact list:   
1^st^ MG coordination shell missing (see Notes)  

### MG:A:3037

|  | Res. type | Atom | Residue | Distance | Occupancy | B-factor | Closest solute to 1^st^ shell water |
| --- | --- | --- | --- | --- | --- | --- | --- |
| **Ion** | | | |  | 1.00 | 6.73 || 1 | G | O6 | A:192 | 3.54 | 1.00 | 17.69 |
| 2 | G | N7 | A:208 | 4.28 | 1.00 | 12.20 |  |
| 3 | A | N7 | A:191 | 4.43 | 1.00 | 18.97 |  |
| 4 | G | OP2 | A:190 | 4.52 | 1.00 | 18.68 |  |
| 5 | A | OP2 | A:191 | 4.93 | 1.00 | 18.97 |  |
  |

  
 Coded contact list:   
1^st^ MG coordination shell missing (see Notes)  

### MG:A:3038

|  | Res. type | Atom | Residue | Distance | Occupancy | B-factor | Closest solute to 1^st^ shell water |
| --- | --- | --- | --- | --- | --- | --- | --- |
| **Ion** | | | |  | 1.00 | 13.51 || 1 | C | OP1 | A:31 | 2.06 | 1.00 | 16.62 |
| 2 | C | OP1 | A:1277 | 3.91 | 1.00 | 17.69 |  |
| 3 | C | OP2 | A:31 | 4.21 | 1.00 | 16.62 |  |
| 4 | C | OP1 | A:1239 | 4.26 | 1.00 | 13.54 |  |
| 5 | G | O3' | A:30 | 4.35 | 1.00 | 13.86 |  |
| 6 | C | O5' | A:31 | 4.40 | 1.00 | 16.62 |  |
| 7 | C | O3' | A:1277 | 4.76 | 1.00 | 17.69 |  |
| 8 | C | O5' | A:1277 | 4.86 | 1.00 | 17.69 |  |
  |

  
 Contact(s): /OP1/  
 Coded contact list: O_ph_  

### MG:A:3039

|  | Res. type | Atom | Residue | Distance | Occupancy | B-factor | Closest solute to 1^st^ shell water |
| --- | --- | --- | --- | --- | --- | --- | --- |
| **Ion** | | | |  | 1.00 | 26.13 || 1 | A | OP1 | A:1275 | 2.72 | 1.00 | 17.92 |
| 2 | A | OP2 | A:523 | 4.23 | 1.00 | 28.92 |  |
| 3 | G | OP1 | A:522 | 4.32 | 1.00 | 24.05 |  |
| 4 | A | O5' | A:1275 | 4.65 | 1.00 | 17.92 |  |
| 5 | G | OP2 | A:522 | 4.74 | 1.00 | 24.05 |  |
  |

  
 Coded contact list:   
 Exclusion zone contact(s): /OP1/  
Consider K in place of MG (1 valid contact(s) for K vs. 0 for MG)  

### MG:A:3040

|  | Res. type | Atom | Residue | Distance | Occupancy | B-factor | Closest solute to 1^st^ shell water |
| --- | --- | --- | --- | --- | --- | --- | --- |
| **Ion** | | | |  | 1.00 | 26.13 || 1 | G | O6 | A:381 | 3.38 | 1.00 | 28.33 |
| 2 | A | N6 | A:343 | 3.67 | 1.00 | 34.48 |  |
| 3 | A | N7 | A:342 | 4.15 | 1.00 | 35.16 |  |
| 4 | A | OP2 | A:342 | 4.16 | 1.00 | 35.16 |  |
| 5 | G | N7 | A:381 | 4.68 | 1.00 | 28.33 |  |
| 6 | A | N7 | A:343 | 4.88 | 1.00 | 34.48 |  |
  |

  
 Coded contact list:   
 Exclusion zone contact(s): /O6/  
Consider K in place of MG (1 valid contact(s) for K vs. 0 for MG)  

### MG:A:3041

|  | Res. type | Atom | Residue | Distance | Occupancy | B-factor | Closest solute to 1^st^ shell water |
| --- | --- | --- | --- | --- | --- | --- | --- |
| **Ion** | | | |  | 1.00 | 26.13 || 1 | G | O6 | A:539 | 4.21 | 1.00 | 22.88 |
| 2 | G | O6 | A:538 | 4.26 | 1.00 | 22.76 |  |
| 3 | A | OP2 | A:537 | 4.81 | 1.00 | 23.88 |  |
| 4 | G | N7 | A:538 | 4.82 | 1.00 | 22.76 |  |
  |

  
 Coded contact list:   
1^st^ MG coordination shell missing (see Notes)  

### MG:A:3042

|  | Res. type | Atom | Residue | Distance | Occupancy | B-factor | Closest solute to 1^st^ shell water |
| --- | --- | --- | --- | --- | --- | --- | --- |
| **Ion** | | | |  | 1.00 | 14.14 || 1 | G | N7 | A:1329 | 3.60 | 1.00 | 21.90 |
| 2 | G | O6 | A:1329 | 4.53 | 1.00 | 21.90 |  |
| 3 | U | O4 | A:1330 | 4.58 | 1.00 | 22.68 |  |
  |

  
 Coded contact list:   
1^st^ MG coordination shell missing (see Notes)  

### MG:A:3043

|  | Res. type | Atom | Residue | Distance | Occupancy | B-factor | Closest solute to 1^st^ shell water |
| --- | --- | --- | --- | --- | --- | --- | --- |
| **Ion** | | | |  | 1.00 | 26.13 || 1 | C | OP2 | A:1334 | 2.74 | 1.00 | 22.45 |
| 2 | U | OP1 | A:1683 | 3.07 | 1.00 | 23.47 |  |
| 3 | A | OP1 | A:1333 | 4.07 | 1.00 | 21.24 |  |
| 4 | C | OP1 | A:1334 | 4.37 | 1.00 | 22.45 |  |
| 5 | U | OP2 | A:1683 | 4.46 | 1.00 | 23.47 |  |
| 6 | A | O5' | A:1333 | 4.50 | 1.00 | 21.24 |  |
| 7 | G | N2 | A:2736 | 4.53 | 1.00 | 22.20 |  |
| 8 | G | O2' | A:2736 | 4.54 | 1.00 | 22.20 |  |
| 9 | G | N3 | A:2736 | 4.55 | 1.00 | 22.20 |  |
| 10 | A | OP2 | A:1333 | 4.69 | 1.00 | 21.24 |  |
| 11 | A | O3' | A:1333 | 4.72 | 1.00 | 21.24 |  |
  |

  
 Coded contact list:   
 Exclusion zone contact(s): /OP1/OP2/  
Consider K in place of MG (2 valid contact(s) for K vs. 0 for MG)  

### MG:A:3044

|  | Res. type | Atom | Residue | Distance | Occupancy | B-factor | Closest solute to 1^st^ shell water |
| --- | --- | --- | --- | --- | --- | --- | --- |
| **Ion** | | | |  | 1.00 | 12.14 || 1 | C | OP2 | A:1335 | 2.17 | 1.00 | 24.80 |
| 2 | U | OP2 | A:1683 | 3.17 | 1.00 | 23.47 |  |
| 3 | G | N7 | A:1336 | 3.78 | 1.00 | 26.97 |  |
| 4 | G | O6 | A:1336 | 4.04 | 1.00 | 26.97 |  |
| 5 | U | O5' | A:1683 | 4.10 | 1.00 | 23.47 |  |
| 6 | C | O5' | A:1335 | 4.33 | 1.00 | 24.80 |  |
| 7 | C | OP1 | A:1335 | 4.39 | 1.00 | 24.80 |  |
| 8 | C | O3' | A:1334 | 4.64 | 1.00 | 22.45 |  |
| 9 | C | O5' | A:1682 | 4.74 | 1.00 | 18.59 |  |
| 10 | C | OP2 | A:1682 | 4.76 | 1.00 | 18.59 |  |
| 11 | C | OP1 | A:1682 | 4.82 | 1.00 | 18.59 |  |
  |

  
 Contact(s): /OP2/  
 Coded contact list: O_ph_  
 Exclusion zone contact(s): /OP2/  

### MG:A:3045

|  | Res. type | Atom | Residue | Distance | Occupancy | B-factor | Closest solute to 1^st^ shell water |
| --- | --- | --- | --- | --- | --- | --- | --- |
| **Ion** | | | |  | 1.00 | 11.01 || 1 | U | OP2 | A:1680 | 1.97 | 1.00 | 17.96 |
| 2 | U | O5' | A:1680 | 3.87 | 1.00 | 17.96 |  |
| 3 | A | OP1 | A:1679 | 3.92 | 1.00 | 21.29 |  |
| 4 | U | OP1 | A:1680 | 3.93 | 1.00 | 17.96 |  |
| 5 | U | OP2 | A:1681 | 4.13 | 1.00 | 15.79 |  |
| 6 | A | OP1 | A:1337 | 4.26 | 1.00 | 32.54 |  |
| 7 | A | O3' | A:1679 | 4.46 | 1.00 | 21.29 |  |
  |

  
 Contact(s): /OP2/  
 Coded contact list: O_ph_  

### MG:A:3046

|  | Res. type | Atom | Residue | Distance | Occupancy | B-factor | Closest solute to 1^st^ shell water |
| --- | --- | --- | --- | --- | --- | --- | --- |
| **Ion** | | | |  | 1.00 | 14.10 || 1 | A | OP2 | A:1786 | 2.96 | 1.00 | 78.59 |
| 2 | A | N7 | A:1787 | 3.74 | 1.00 | 19.95 |  |
| 3 | A | N6 | A:1787 | 4.08 | 1.00 | 19.95 |  |
| 4 | G | O5' | A:1785 | 4.30 | 1.00 | 78.76 |  |
| 5 | U | N3 | A:1784 | 4.37 | 1.00 | 24.51 |  |
| 6 | A | O5' | A:1786 | 4.38 | 1.00 | 78.76 |  |
| 7 | U | O4 | A:1788 | 4.47 | 1.00 | 21.80 |  |
| 8 | U | O4 | A:1784 | 4.64 | 1.00 | 24.51 |  |
| 9 | U | N1 | A:1784 | 4.65 | 1.00 | 24.51 |  |
| 10 | A | OP2 | A:1787 | 4.69 | 1.00 | 19.95 |  |
| 11 | A | N7 | A:1786 | 4.95 | 1.00 | 20.00 |  |
  |

  
 Coded contact list:   
 Exclusion zone contact(s): /OP2/  
Consider K in place of MG (1 valid contact(s) for K vs. 0 for MG)  

### MG:A:3047

|  | Res. type | Atom | Residue | Distance | Occupancy | B-factor | Closest solute to 1^st^ shell water |
| --- | --- | --- | --- | --- | --- | --- | --- |
| **Ion** | | | |  | 1.00 | 16.96 || 1 | A | OP1 | A:847 | 2.22 | 1.00 | 14.11 |
| 2 | G | OP2 | A:719 | 4.22 | 1.00 | 10.39 |  |
| 3 | A | OP2 | A:720 | 4.30 | 1.00 | 11.53 |  |
| 4 | A | OP2 | A:847 | 4.40 | 1.00 | 14.11 |  |
| 5 | A | O5' | A:847 | 4.52 | 1.00 | 14.11 |  |
| 6 | G | O3' | A:846 | 4.53 | 1.00 | 17.01 |  |
| 7 | U | O4 | A:848 | 4.74 | 1.00 | 26.13 |  |
  |

  
 Contact(s): /OP1/  
 Coded contact list: O_ph_  

### MG:A:3048

|  | Res. type | Atom | Residue | Distance | Occupancy | B-factor | Closest solute to 1^st^ shell water |
| --- | --- | --- | --- | --- | --- | --- | --- |
| **Ion** | | | |  | 1.00 | 26.13 || 1 | A | OP1 | A:845 | 2.01 | 1.00 | 10.91 |
| 2 | A | OP2 | A:845 | 3.99 | 1.00 | 10.91 |  |
| 3 | G | O3' | A:844 | 4.26 | 1.00 | 9.13 |  |
| 4 | A | O5' | A:845 | 4.41 | 1.00 | 10.91 |  |
| 5 | C | N3 | A:195 | 4.51 | 1.00 | 8.54 |  |
| 6 | A | N7 | A:845 | 4.67 | 1.00 | 10.91 |  |
| 7 | C | N4 | A:195 | 4.80 | 1.00 | 8.54 |  |
| 8 | A | N1 | A:194 | 4.87 | 1.00 | 9.22 |  |
  |

  
 Contact(s): /OP1/  
 Coded contact list: O_ph_  

### MG:A:3049

|  | Res. type | Atom | Residue | Distance | Occupancy | B-factor | Closest solute to 1^st^ shell water |
| --- | --- | --- | --- | --- | --- | --- | --- |
| **Ion** | | | |  | 1.00 | 26.13 || 1 | G | OP2 | A:496 | 2.16 | 1.00 | 8.97 |
| 2 | G | OP2 | A:498 | 4.10 | 1.00 | 9.39 |  |
| 3 | GLY | O | F:86 | 4.20 | 1.00 | 27.49 |  |
| 4 | G | OP1 | A:496 | 4.27 | 1.00 | 8.97 |  |
| 5 | G | O5' | A:496 | 4.34 | 1.00 | 8.97 |  |
| 6 | A | OP1 | A:495 | 4.61 | 1.00 | 12.55 |  |
| 7 | A | O3' | A:495 | 4.63 | 1.00 | 12.55 |  |
| 8 | GLY | O | F:85 | 4.77 | 1.00 | 26.13 |  |
| 9 | A | O5' | A:495 | 4.94 | 1.00 | 12.55 |  |
  |

  
 Contact(s): /OP2/  
 Coded contact list: O_ph_  

### MG:A:3050

|  | Res. type | Atom | Residue | Distance | Occupancy | B-factor | Closest solute to 1^st^ shell water |
| --- | --- | --- | --- | --- | --- | --- | --- |
| **Ion** | | | |  | 1.00 | 10.36 || 1 | G | OP2 | A:1226 | 2.33 | 1.00 | 10.30 |
| 2 | C | OP2 | A:861 | 3.53 | 1.00 | 9.55 |  |
| 3 | G | OP2 | A:1225 | 3.64 | 1.00 | 75.20 |  |
| 4 | C | OP2 | A:862 | 3.66 | 1.00 | 9.46 |  |
| 5 | G | OP1 | A:1226 | 4.12 | 1.00 | 10.30 |  |
| 6 | G | O5' | A:1226 | 4.14 | 1.00 | 10.30 |  |
| 7 | U | O2' | A:1224 | 4.28 | 1.00 | 79.34 |  |
| 8 | G | OP1 | A:1225 | 4.35 | 1.00 | 78.59 |  |
| 9 | C | OP1 | A:861 | 4.46 | 1.00 | 9.55 |  |
| 10 | C | O5' | A:861 | 4.51 | 1.00 | 9.55 |  |
| 11 | G | O3' | A:1225 | 4.82 | 1.00 | 77.17 |  |
| 12 | G | O6 | A:863 | 4.83 | 1.00 | 8.12 |  |
  |

  
 Contact(s): /OP2/  
 Coded contact list: O_ph_  

### MG:A:3051

|  | Res. type | Atom | Residue | Distance | Occupancy | B-factor | Closest solute to 1^st^ shell water |
| --- | --- | --- | --- | --- | --- | --- | --- |
| **Ion** | | | |  | 1.00 | 26.13 || 1 | G | OP2 | A:1803 | 2.13 | 1.00 | 16.76 |
| 2 | G | OP1 | A:1803 | 4.05 | 1.00 | 16.76 |  |
| 3 | G | N7 | A:1803 | 4.13 | 1.00 | 16.76 |  |
| 4 | A | OP1 | A:2008 | 4.33 | 1.00 | 14.73 |  |
| 5 | G | O5' | A:1803 | 4.48 | 1.00 | 16.76 |  |
| 6 | U | O3' | A:1802 | 4.49 | 1.00 | 17.21 |  |
  |

  
 Contact(s): /OP2/  
 Coded contact list: O_ph_  

### MG:A:3052

|  | Res. type | Atom | Residue | Distance | Occupancy | B-factor | Closest solute to 1^st^ shell water |
| --- | --- | --- | --- | --- | --- | --- | --- |
| **Ion** | | | |  | 1.00 | 17.62 || 1 | C | OP2 | A:776 | 2.32 | 1.00 | 22.95 |
| 2 | A | OP2 | A:775 | 2.92 | 1.00 | 22.79 |  |
| 3 | A | OP1 | A:806 | 3.41 | 1.00 | 14.92 |  |
| 4 | C | O5' | A:776 | 4.25 | 1.00 | 22.95 |  |
| 5 | A | O3' | A:775 | 4.33 | 1.00 | 22.79 |  |
| 6 | G | OP1 | A:808 | 4.33 | 1.00 | 20.57 |  |
| 7 | A | O5' | A:806 | 4.36 | 1.00 | 14.92 |  |
| 8 | A | OP2 | A:806 | 4.43 | 1.00 | 14.92 |  |
| 9 | A | N3 | A:775 | 4.70 | 1.00 | 22.79 |  |
| 10 | A | O5' | A:775 | 4.72 | 1.00 | 22.79 |  |
| 11 | G | O3' | A:774 | 4.82 | 1.00 | 19.97 |  |
| 12 | C | OP1 | A:776 | 4.91 | 1.00 | 22.95 |  |
| 13 | U | OP1 | A:807 | 4.98 | 1.00 | 20.69 |  |
  |

  
 Contact(s): /OP2/  
 Coded contact list: O_ph_  
 Exclusion zone contact(s): /OP2/  

### MG:A:3053

|  | Res. type | Atom | Residue | Distance | Occupancy | B-factor | Closest solute to 1^st^ shell water |
| --- | --- | --- | --- | --- | --- | --- | --- |
| **Ion** | | | |  | 1.00 | 13.79 || 1 | A | O2' | A:660 | 2.81 | 1.00 | 31.89 |
| 2 | U | O2' | A:661 | 3.22 | 1.00 | 30.89 |  |
| 3 | ARG | NE | F:106 | 3.55 | 1.00 | 17.14 |  |
| 4 | A | O4' | A:660 | 4.06 | 1.00 | 31.89 |  |
| 5 | G | OP2 | A:662 | 4.82 | 1.00 | 32.03 |  |
  |

  
 Coded contact list:   
 Exclusion zone contact(s): /O2'/O2'/  
Consider K in place of MG (2 valid contact(s) for K vs. 0 for MG)  

### MG:A:3054

|  | Res. type | Atom | Residue | Distance | Occupancy | B-factor | Closest solute to 1^st^ shell water |
| --- | --- | --- | --- | --- | --- | --- | --- |
| **Ion** | | | |  | 1.00 | 18.51 || 1 | C | OP1 | A:2836 | 3.33 | 1.00 | 25.09 |
| 2 | LEU | O | 4:39 | 3.59 | 1.00 | 48.74 |  |
| 3 | C | O3' | A:2835 | 4.58 | 1.00 | 36.85 |  |
| 4 | GLU | OE2 | Q:115 | 4.66 | 1.00 | 16.11 |  |
| 5 | GLU | OE1 | Q:115 | 4.68 | 1.00 | 16.11 |  |
  |

  
 Coded contact list:   
 Exclusion zone contact(s): /OP1/  
Consider K in place of MG (1 valid contact(s) for K vs. 0 for MG)  

### MG:A:3055

|  | Res. type | Atom | Residue | Distance | Occupancy | B-factor | Closest solute to 1^st^ shell water |
| --- | --- | --- | --- | --- | --- | --- | --- |
| **Ion** | | | |  | 1.00 | 26.13 || 1 | C | OP1 | A:777 | 2.64 | 1.00 | 22.74 |
| 2 | G | OP1 | A:2007 | 3.97 | 1.00 | 20.17 |  |
| 3 | G | OP1 | A:805 | 4.39 | 1.00 | 12.56 |  |
| 4 | C | OP2 | A:777 | 4.54 | 1.00 | 22.74 |  |
| 5 | G | OP2 | A:805 | 4.71 | 1.00 | 12.56 |  |
| 6 | C | O3' | A:776 | 4.94 | 1.00 | 22.95 |  |
| 7 | C | O5' | A:777 | 4.94 | 1.00 | 22.74 |  |
  |

  
 Coded contact list:   
 Exclusion zone contact(s): /OP1/  
Consider K in place of MG (1 valid contact(s) for K vs. 0 for MG)  

### MG:A:3056

|  | Res. type | Atom | Residue | Distance | Occupancy | B-factor | Closest solute to 1^st^ shell water |
| --- | --- | --- | --- | --- | --- | --- | --- |
| **Ion** | | | |  | 1.00 | 30.00 || 1 | A | OP2 | A:2480 | 3.80 | 1.00 | 23.38 |
| 2 | A | N7 | A:2480 | 4.01 | 1.00 | 23.38 |  |
| 3 | G | N7 | A:2481 | 4.16 | 1.00 | 22.73 |  |
| 4 | G | O6 | A:2481 | 4.74 | 1.00 | 22.73 |  |
| 5 | C | OP2 | A:2479 | 4.75 | 1.00 | 26.13 |  |
  |

  
 Coded contact list:   
1^st^ MG coordination shell missing (see Notes)  

### K:A:3057

|  | Res. type | Atom | Residue | Distance | Occupancy | B-factor | Closest solute to 1^st^ shell water |
| --- | --- | --- | --- | --- | --- | --- | --- |
| **Ion** | | | |  | 1.00 | 26.13 || 1 | C | OP1 | A:1714 | 2.75 | 1.00 | 13.13 |
| 2 | U | OP2 | A:1715 | 2.78 | 1.00 | 14.86 |  |
| 3 | G | OP2 | A:2577 | 3.97 | 1.00 | 35.40 |  |
| 4 | G | OP2 | A:2576 | 4.11 | 1.00 | 26.76 |  |
| 5 | C | O3' | A:1714 | 4.51 | 1.00 | 13.13 |  |
| 6 | U | OP1 | A:1715 | 4.66 | 1.00 | 14.86 |  |
| 7 | A | O3' | A:1713 | 4.92 | 1.00 | 14.81 |  |
| 8 | C | O5' | A:1714 | 4.92 | 1.00 | 13.13 |  |
  |

  
**1^st^ shell ligand angles:**   
1-2: 75;   
  
 Contact(s): /OP1/OP2/  
 Coded contact list: 2O_ph_  

### K:A:3058

|  | Res. type | Atom | Residue | Distance | Occupancy | B-factor | Closest solute to 1^st^ shell water |
| --- | --- | --- | --- | --- | --- | --- | --- |
| **Ion** | | | |  | 1.00 | 9.19 || 1 | G | O6 | A:707 | 4.14 | 1.00 | 17.35 |
| 2 | G | N7 | A:707 | 4.24 | 1.00 | 17.35 |  |
| 3 | U | O4 | A:637 | 4.51 | 1.00 | 17.75 |  |
| 4 | A | N6 | A:636 | 4.94 | 1.00 | 15.72 |  |
| 5 | U | O4 | A:709 | 4.99 | 1.00 | 14.86 |  |
  |

  
 Coded contact list:   
1^st^ K coordination shell missing (see Notes)  

### K:A:3059

|  | Res. type | Atom | Residue | Distance | Occupancy | B-factor | Closest solute to 1^st^ shell water |
| --- | --- | --- | --- | --- | --- | --- | --- |
| **Ion** | | | |  | 1.00 | 26.13 || 1 | A | OP2 | A:880 | 3.19 | 1.00 | 14.20 |
| 2 | G | O6 | A:986 | 4.50 | 1.00 | 11.15 |  |
| 3 | A | N7 | A:880 | 4.59 | 1.00 | 14.20 |  |
| 4 | A | O5' | A:880 | 4.81 | 1.00 | 14.20 |  |
| 5 | U | OP2 | A:879 | 4.84 | 1.00 | 12.68 |  |
| 6 | G | O6 | A:881 | 4.96 | 1.00 | 14.18 |  |
| 7 | U | O5' | A:879 | 4.96 | 1.00 | 12.68 |  |
  |

  
 Contact(s): /OP2/  
 Coded contact list: O_ph_  
\_\_\_\_\_\_\_\_\_\_\_\_\_\_\_\_\_\_\_\_\_\_\_\_\_\_\_\_\_\_\_\_\_\_\_\_\_\_\_\_\_\_\_\_\_\_\_\_\_\_\_\_\_\_\_\_\_\_\_\_\_\_\_\_  

#### 5) Sorted coded contact list

##### Go to ToC

NOTES:

  
- N. --> estimate of the coordination number as deduced from the PDB structure;  
- Excl. --> existence of contacts in the ligand exclusion zone;  
- Bad hyd. --> existence of 2^nd^ shell hydration contacts shorter than 2.4 Angs.;  
- Lines in red mark stereochemical issues that should be addressed;  
- Lines in green mark ions without stereochemical issues - Please note that this does not always warrant the quality of the ion identification;  
  
(1) N=0 --> No 1^st^ shell coordination: impossible to assess ion identity;  
(2) Exclusion contacts --> Probable wrong ion identity - poor modelization of the ion binding site is also possible;  
(3) Bad 2^nd^ hydration shell contacts --> Poor modelization of the hydration shell and/or binding site;  
(4) Check for anion (Cl^-^, ...);  
(5) Mixed contacts with unprotonated O/N and protonated N, suggest poorly modeled binding site;  
(6) Currently the ZN (Zinc) and CD (Cadmium) coordination is not evaluated;  
(7) Check occupancies - partial occupancies are not reliable at medium to low resolutions, especially for solvent particles;  
(8) Check B-factors;  
  
  

|  |  |  |  |  |  |  |  |
| --- | --- | --- | --- | --- | --- | --- | --- |
| **Ion** | **Number** | **Code** | **N.** | **Excl.** | **Bad hyd.** | **Notes** | **Ion list** |
| MG: | 17 | No 1^st^ shell | >1 | Yes | No | (2) | A:3003 A:3005 A:3011 A:3013 A:3014 A:3020 A:3027 A:3031 A:3034 A:3035 A:3039 A:3040 A:3043 A:3046 A:3053 A:3054 A:3055 || MG: | 14 | No 1^st^ shell | =0 | No | No | (1) | A:3002 A:3004 A:3009 A:3015 A:3016 A:3022 A:3024 A:3028 A:3029 A:3036 A:3037 A:3041 A:3042 A:3056 || MG: | 12 | O_ph_ | 1 | No | --- |  | A:3007 A:3008 A:3019 A:3021 A:3025 A:3038 A:3045 A:3047 A:3048 A:3049 A:3050 A:3051 || MG: | 7 | O_ph_ | >1 | Yes | --- | (2) | A:3001 A:3018 A:3023 A:3026 A:3030 A:3044 A:3052 || MG: | 1 | N_b_ | >1 | Yes | --- | (2) | A:3010 || MG: | 1 | O_r_ | >1 | Yes | --- | (2) | A:3033 || MG: | 1 | O_ph_O_b_ | >2 | Yes | --- | (2) | A:3012 || K: | 1 | O_ph_ | 1 | No | --- |  | A:3059 || K: | 1 | 2O_ph_ | 2 | No | --- |  | A:3057 || MG: | 1 | O_b_ | >1 | Yes | --- | (2) | A:3017 || K: | 1 | No 1^st^ shell | =0 | No | No | (1) | A:3058 || MG: | 1 | O_b_ | 1 | No | --- |  | A:3032 || MG: | 1 | 3O_ph_ | 3 | No | --- |  | A:3006 ||  |  |  |  |  |  |  | List of ions with partial occupancies ||  |  |  |  |  |  | Total: | 0 |

  
**Total number of Mg^2+^: 56** **Total number of Mg^2+^ with stereochemical issues: 41** **Total number of Mg^2+^ without stereochemical issues: 15** **Total number of K^+^: 3** **Total number of K^+^ with stereochemical issues: 1** **Total number of K^+^ without stereochemical issues: 2** **Total number of Na^+^: 0** **Total number of Na^+^ with stereochemical issues: 0** **Total number of Na^+^ without stereochemical issues: 0** 

### ### END ###
